# Wild-type *Caenorhabditis elegans* Isolates Exhibit Distinct Gene Expression Profiles in Response to Microbial Infection

**DOI:** 10.1101/2021.10.16.464663

**Authors:** Patrick Lansdon, Maci Carlson, Brian D. Ackley

## Abstract

The soil-dwelling nematode *Caenorhabditis elegans* serves as a model system to study innate immunity against microbial pathogens. *C. elegans* have been collected from around the world, where they, presumably, adapted to regional microbial ecologies. Here we use survival assays and RNA- sequencing to better understand how two isolates from disparate climates respond to pathogenic bacteria. We found that, relative to N2 (originally isolated in Bristol, UK), CB4856 (isolated in Hawaii), was more susceptible to the Gram-positive pathogen, *Staphylococcus epidermidis*, but equally susceptible to *Staphylococcus aureus* as well as two Gram-negative pathogens, *Providencia rettgeri* and *Pseudomonas aeruginosa*. We performed transcriptome analysis of infected worms and found gene- expression profiles were considerably different in an isolate-specific and pathogen-specific manner. We performed GO term analysis to categorize differential gene expression in response to *S. epidermidis*. In N2, genes that encoded detoxification enzymes and extracellular matrix proteins were significantly enriched, while in CB4856, genes that encoded detoxification enzymes, C-type lectins, and lipid metabolism proteins were enriched, suggesting they have different responses to these bacterial pathogens, despite being the same species. Overall, discerning gene expression signatures in an isolate by pathogen manner can help us to understand the different possibilities for the evolution of immune responses within organisms.

## Introduction

Antimicrobial resistance is a looming public health crisis as microbial pathogens evade drugs of last resort. When pathogenic microbes bypass the physical barriers of the host (*e.g.,* an epidermis), an organism must rely on cellular and humoral defenses (*e.g.,* the innate immune system) to protect itself from infections that can compromise organismal fitness. While all organisms will inevitably become infected over their lifespan, genetic variation in individuals will influence the outcomes of infections. Thus, our ability to link genetic variation to infection susceptibility may enable more sophisticated strategies for deploying antibiotics or help us enhance host immunity to treat microbial infection.^1, 2^

Genetic variation in organisms can create tradeoffs in susceptibility to different pathogens. The presence of non-synonymous single nucleotide polymorphisms (SNPs) in the coding sequence of immune effector genes has been shown to alter the propensity of individuals to become infected and develop disease.^3^ For instance, a SNP in the *TLR5* gene introduces a premature stop codon, creating a truncated form of the protein and impairing the ability of TLR5 to recognize the flagellin protein in flagellated bacteria.^4^ Individuals carrying this SNP are more susceptible to infection by *Legionella pneumophila*, the vector for Legionnaires’ disease, but not *Salmonella enterica serotype Typhi,* the microbial vector for typhoid fever.^5, 6^

Animal models have been critical to our understanding of the innate immune system. However, the reliance on canonical laboratory strains often ignores genetic variation in innate immunity and susceptibility to pathogens.^7^ The soil-dwelling nematode *Caenorhabditis elegans* is a genetically tractable model organism to examine host-pathogen interactions.^8, 9^ *C. elegans* possesses a robust microbial defense system with evolutionarily conserved mechanisms in microbial pathogen detection, immune activation and pathogen killing (*e.g.* the induction of reactive oxygen species (ROS), deployment of antimicrobial and antifungal peptides, and expression of bacterial-binding C-type lectins).^10–14^ These evolutionarily conserved responses to infection make *C. elegans* an ideal model to study bacterial pathogenesis.^9^

*C. elegans* wild isolates collected from geographically discrete regions are apt to vary in their ability to fight infections either because of a history of different selective pressures or simply genetic drift. Many discoveries regarding *C. elegans* host-microbe interactions have come from studying wild isolates, including identifying virus that infect *C. elegans* and the characterization of the *C. elegans* gut microbiome.^15–17^ The genomic diversity of wild isolates has also provided a platform for quantitatively examining the genetic variation underlying complex traits, including behavioral differences in foraging as well as pathogen avoidance and susceptibility.^18–23^ However, less attention has been paid to the variation in the transcriptomic responses of wild isolates to multiple, discrete microbial pathogens.

In this study, we show that the *C. elegans* Hawaiian wild isolate CB4856 is significantly more susceptible to infection by the Gram-positive pathogen *Staphylococcus epidermidis* than the N2 lab-conditioned strain. We used high-throughput RNA sequencing (RNA-seq) to identify differentially expressed genes in staged N2 and CB4856 worms exposed to pathogenic bacteria. The differentially expressed genes identified in pathogen-infected animals provide insight into isolate-specific transcriptomic responses of *C. elegans* to pathogenic bacteria. We believe that this knowledge will provide a foundation for better understanding the role of natural genetic variation in the *C. elegans* innate immune response to microbial pathogens.

## Materials and Methods

### *C. elegans* and Bacterial Strains

*C. elegans* N2 and CB4856 wild-type strains were maintained at 20°C on nematode growth medium (NGM) agar plates seeded with *Escherichia coli* OP50.^24^ *Staphylococcus epidermidis* (EVL2000, this study), *Staphylococcus epidermidis* (1191; ATCC#700562), *Providencia rettgeri* (Dmel1)^25^, *Pseudomonas aeruginosa* (PA-14)^26^, and *Staphylococcus aureus* (Newman)^27^ were used as pathogens in this study. All bacteria strains were grown at 37°C with shaking in low-salt Luria Bertani (LB) medium (10 g Bacto-tryptone, 5 g yeast extract, 5 g NaCl).

### Identification of *S. epidermidis* Strain

DNA was extracted from 1mL of fresh culture using a QiaQuick miniprep column (Qiagen) and the 16S ribosomal region was amplified using universal primers 27F (5’- AGAGTTTGATCCTGGCTCAG-3’) and 1492R (5’-CGGTTACCTTGTTACGACTT-3’). PCR cycling conditions were as follows: 1) 92°C for 2 minutes; 2) 35 cycles at 92°C for 15-sec, 55°C for 15-sec, 68°C for 75-seconds; and 3) 68°C for 10 minutes. The amplified PCR product was purified, Sanger sequenced, and the NCBI BLAST tool was used to identify the species.

### Survival Assays

NGM plates were seeded with 40 µL of bacteria (*E. coli*, *S. epidermidis*, *P. rettgeri*, *P. aeruginosa*, *or S. aureus*) and incubated overnight at 37°C. Plates were acclimated to room temperature and 30 late-stage L4 worms were added. Each day, living worms were transferred to fresh plates and the number of dead worms was recorded.

### Pathogen Avoidance Assays

NGM plates were seeded with a 10 µL dot of pathogenic bacteria (*S. epidermidis*, *P. rettgeri*, *S. aureus*, or *P. aeruginosa*) on one half, a 10 µL dot of non-pathogenic *E. coli* OP50 on the other half and incubated overnight at 37°C. 20 late-stage L4 worms were placed on blank plates for 30 minutes to remove any external bacteria while pathogen avoidance plates acclimated to room temperature. Worms were placed at the midline and their location on the plate was recorded at 1-hour, 2-hour, 4-hour, 8-hour, and 24-hour time points. To avoid ambiguity in the number of animals associated with the specific bacteria, only animals residing within the spot of bacteria were assigned “preference” for either the control (OP50) or pathogen.

### Exposure of Nematodes to Pathogen

NGM plates were seeded with 250 µL of bacteria incubated overnight at 37°C. Approximately 2,000 L4 stage worms grown on *E. coli* OP50 were transferred to NGM plates seeded with bacteria and incubated at 20°C for 24 hours. After 24 hours, the worms were washed with M9 buffer three times to remove any residual bacteria. The worms were resuspended in 100 µL of M9 buffer and mechanically disrupted in liquid nitrogen using a ceramic mortar and pestle. Frozen tissue was transferred to a microcentrifuge tube and 1 mL TRIZOL reagent was added. The tubes were flash frozen in liquid nitrogen and stored at -80°C. Three biological replicates were collected for each worm and pathogen strain.

### RNA isolation and Sequencing

Frozen worm tissue was thawed, vortexed for 5 seconds and incubated at room temperature for 5 minutes. After the addition of 470 µL chloroform (mixed by inversion and phase separated for 2 minutes at room temperature), the samples were centrifuged at 15,000 RPM at 4°C for 15 minutes. The upper aqueous phase containing RNA (approximately 600 µL) was transferred to a new RNase-free Eppendorf tube. Total RNA extraction was carried out using the Monarch RNA Cleanup Kit (New England Biolabs) according to the manufacturer’s instructions. Total RNA was quantified using the Qubit (ThermoFisher Scientific) and RNA quality and integrity was assessed using the Agilent Tapestation 2200 (Agilent Technologies). Sequence libraries were prepared using the TruSeq Stranded mRNA Library Prep Kit (Illumina) and sequenced using single-end 1×75 bp sequencing on the Illumina NextSeq 550.

### RNA-seq Analysis and Quantification of Differentially Expressed Genes

The quality of the 75-bp single-end reads generated using the Illumina NextSeq 550 was assessed using FastQC^28^ (v.0.11.5). Illumina sequencing adapters and low-quality bases (Phred score <30) were trimmed from reads using Fastp^29^ (v.0.19.8). Reads were first aligned to the *E. coli*, *S. epidermidis*, *P. rettgeri*, *S. aureus*, or *P. aeruginosa* genomes to evaluate bacterial contamination, then aligned to the *C. elegans* genome assembly (Wormbase WS273 release) using HISAT2^30^ (v.2.1.0) with default settings. Aligned reads were mapped to the *C. elegans* annotated genome (Wormbase WS273 release) and transcript abundances were quantified using featureCounts^31^ (v.1.6.0). Data normalization and differential gene expression analysis was performed with DESeq2^32^ (v.3.1.0) in R (v.3.6.0) using a false discovery rate (FDR)-adjusted P-value <0.05 and fold change ≥2 as cutoffs. Separate design formulas were used to determine differential gene expression for infection (∼Infection), Gram-stain (∼Gram-stain), and individual pathogens (∼Pathogen). Geneset enrichment analysis was performed with WormCat^33^ using an FDR-adjusted P-value <0.05 as the significance cutoff.

### Statistical Analysis

All statistical analysis was carried out using SigmaPlot 14.0 (Systat Software, Inc.). Log rank (Mantel-Cox) analysis was used for pairwise comparison of survival curves and to calculate median survival (LT_50_). Chi-squared or Fisher’s Exact Test was used to assess statistical significance of the pathogen avoidance data. Spearman’s correlation was used to determine the correlation coefficient between log_2_ fold changes in gene expression. Statistically significant differences are defined in the figure legend and noted in the figure with asterisks.

## Results

### CB4856 and N2 exhibit differences in susceptibility to the pathogen *S. epidermidis*

During our studies on susceptibility to microbial pathogens we serendipitously cultured a bacterial contaminant that exhibited differences in a *C. elegans* isolate-specific manner. We identified this bacterium as *Staphylococcus epidermidis* by 16S sequencing (see Materials and Methods) and provided it with the strain designation EVL2000. Survival assays conducted with the EVL2000 *S. epidermidis* strain found CB4856 animals to be significantly more susceptible than N2, with a median survival (LT_50_) of 11.3 ± 1.6 days, whereas 50% of N2 animals were killed after an average of 18.8 ± 1.5 days (*p*<0.001) (Figure 1A, Table S1).

**Figure 1:**
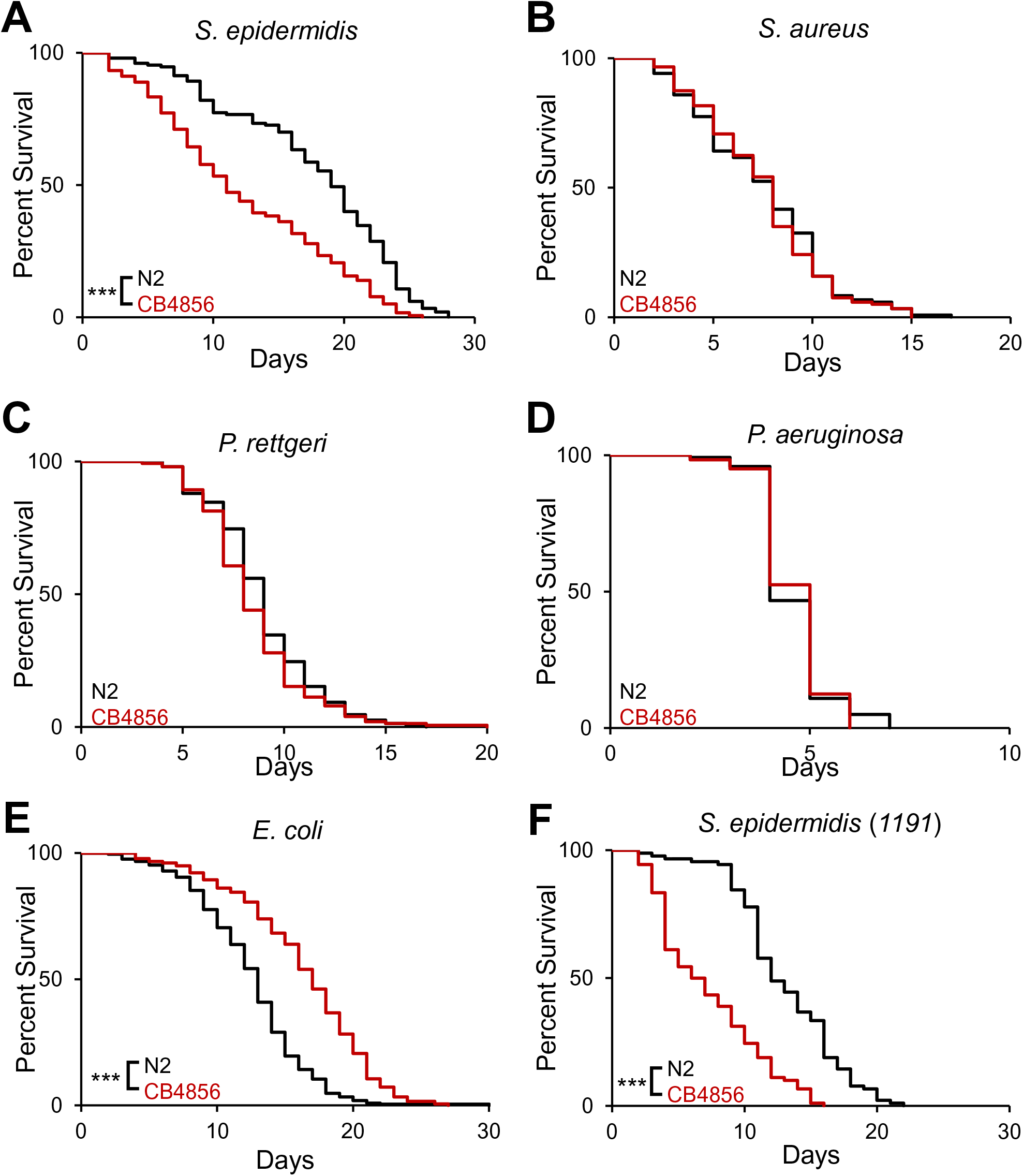
CB4856 is more susceptible to *S. epidermidis,* but not other bacterial pathogens, relative to N2. (A-E) Survival of the *C. elegans* Hawaiian wild isolate, CB4856, versus N2 on nematode growth medium (NGM) with a lawn of *S. epidermidis* (A)*, S. aureus* (B), *P. rettgeri* (C), *P. aeruginosa* (D), or *E. coli* (E). We tested a second *S. epidermidis* strain (1191) to confirm the differences in genotypes (F). In each experiment, 30 worms were placed on the bacterial lawn and transferred every day while scored for survival. Survival curves represent 3-7 independent experiments. For each pathogen, statistical analysis by Mantel-Cox (CB486 vs N2). ***, *p*<0.001.

We then compared N2 and CB4856 susceptibility to other known *C. elegans* pathogens, *Staphylococcus aureus, Providencia rettgeri*, and *Pseudomonas aeruginosa.* In each case, the susceptibility of N2 and CB4856 to these pathogens was not significantly different (Figures 1B-D, Table S1). We also grew animals on non-pathogenic *E. coli* OP50 to evaluate the possibility that the increased susceptibility of *S. epidermidis* was due to a general defect in lifespan. Interestingly, the lifespan of N2 worms was significantly shorter than CB4856, with 50% of animals dying at 12.0 ± 0.4 and 16.2 ± 0.6 days for N2 and CB4856 animals, respectively (*p*<0.001) (Figure 1E, Table S1). While there were differences in lifespan on *E. coli*, our data are consistent with the idea that the increased susceptibility of CB4856 to *S. epidermidis* is not due to a universal decrease in lifespan.

Relative to worms fed *E. coli*, infection with the EVL2000 *S. epidermidis* strain actually prolonged the median lifespan of N2 animals (*E. coli* OP50 LT_50_ = 12.0 ± 0.4 days, *S. epidermidis* EVL2000 LT_50_ = 18.8 ± 1.5 days; *p*<0.001). To determine if the differences in susceptibility were unique to the EVL2000 strain, we infected N2 and CB4856 animals with an additional *S. epidermidis* strain (1191). CB4856 animals infected with the 1191 strain had a median survival of 5.7 ± 1.2 days, whereas 50% of N2 animals were killed after an average of 12.3 ± 0.9 days (*p*<0.001) (Figure 1F, Table S1). Together, our results indicate that CB4856 is more susceptible to *S. epidermidis* infection and that this increase in susceptibility is observed across multiple strains of *S. epidermidis.* Since infection by the *S. epidermidis* EVL2000 strain provided the greatest difference in susceptibility between N2 and CB4856, follow-up experiments used this bacterial strain.

### Pathogen avoidance of N2 and CB4856 isolates depends on the pathogen and time of observation

We performed two-choice pathogen avoidance assays to determine if the difference in susceptibility could be explained in part by differences in animal behavior (Figure 2A). At the 1- hr time point there was a significant difference in *S. epidermidis* avoidance, with 100% of N2 and 70% of CB4586 animals on the OP50 lawn (Figure 2B) (*p*<0.001). By 24 hours the difference in pathogen avoidance was no longer significant and nearly 100% of worms from both isolates were found on the OP50 lawn. This suggests that both genotypes perceived *S. epidermidis* as pathogenic and avoided them accordingly.

**Figure 2:**
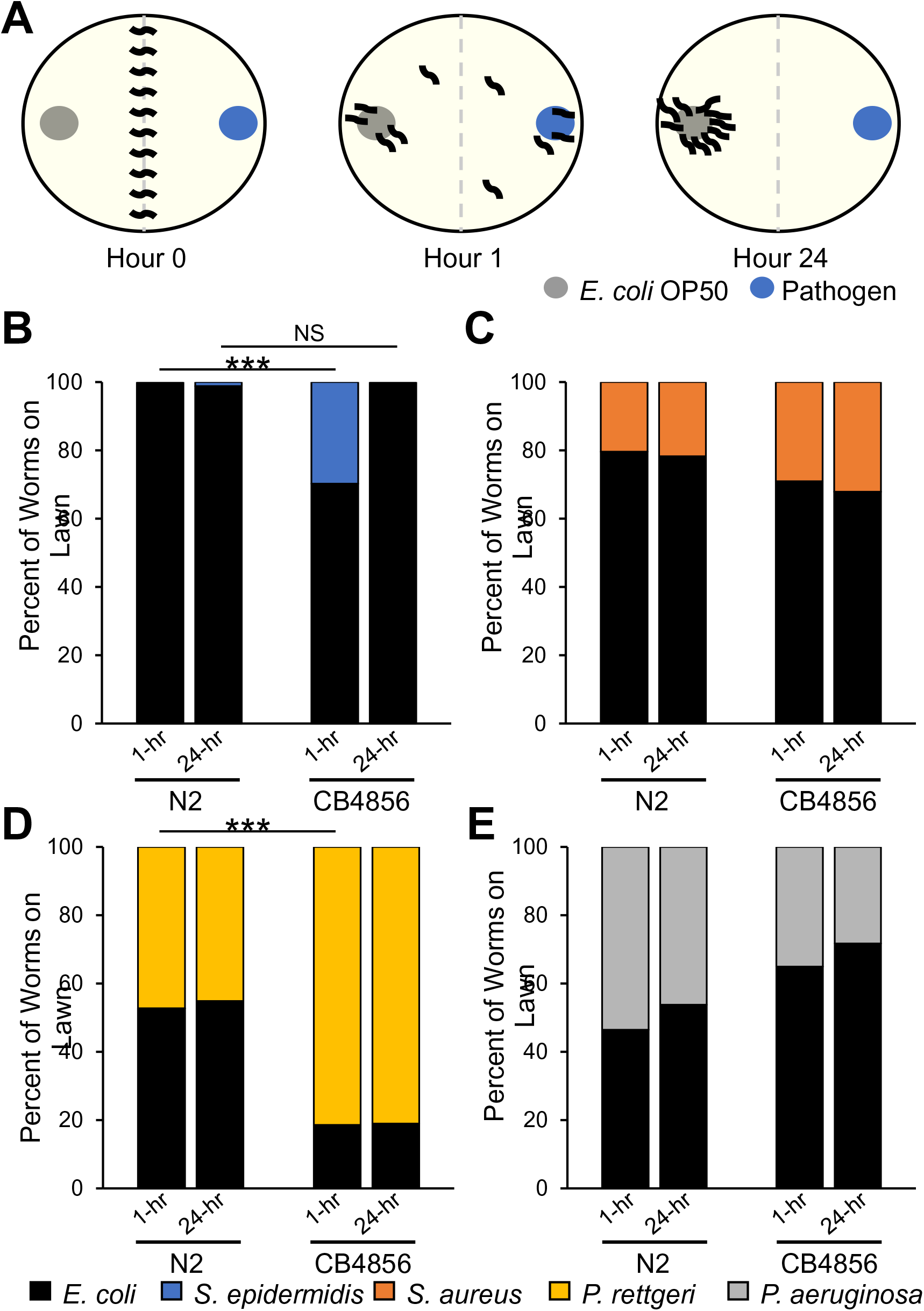
Pathogen avoidance of N2 and CB4856 isolates depends on the pathogen and time of observation. (A) Two-choice pathogen avoidance assay. L4 worms were washed and then placed at the midline of test plates containing two bacterial lawns, OP50 or pathogen. At 1-hr and 24-hr timepoints the percentage of worms on each lawn was calculated. (B-E) Percentage of N2 and CB4856 isolates on a bacterial lawn when given a choice of *E. coli* OP50 or. *S. epidermidis* (B), *S. aureus* (C), *P. rettgeri* (D) or *P. aeruginosa* (E). In each experiment, 20 worms were placed on NGM plates and the number of worms on the bacterial lawn was recorded at regular intervals. Data represent 5 independent experiments. ***, *p*<0.001; NS, *p>*0.05.

We also examined avoidance behavior in N2 and CB4856 animals exposed to *S. aureus*, *P. rettgeri* or *P. aeruginosa* and found differences in the initial avoidance of pathogenic bacteria. At the 1-hr time point avoidance of *S. aureus* was not significantly different between the two isolates, with 80% and 70% of animals observed on the OP50 lawn for N2 and CB4856, respectively (Figure 2C) (*p*=0.390). Avoidance of *P. rettgeri* at the 1-hr time point was significantly more likely in N2 worms, with 53% of N2 and 19% of CB4856 animals on the OP50 lawn (Figure 2D) (*p*<0.001). The avoidance of *P. aeruginosa* trended in the opposite direction with 46% of N2 and 65% of CB4856 animals on the OP50 lawn (Figure 2E) (*p*=0.051). Neither the N2 nor CB4856 genotypes had significant changes in pathogen avoidance between the 1-hr and 24-hr time points for these three pathogens (Figure 2C-E). Natural variation in olfactory preference may partially explain these differences in pathogen avoidance.^34^ While the results are interesting and warrant follow-up, they suggest that the avoidance behavior of N2 and CB4856 animals likely has little impact on susceptibility to *S. epidermidis*.

### Gene expression changes during infection cluster by genotype, rather than pathogen type

To identify gene expression changes due to pathogen we performed RNA sequencing comparing CB4856 and N2 animals fed pathogenic bacteria versus non-pathogenic *E. coli* OP50 for 24 hours. *E. coli* OP50 is a standardized food source for laboratory-bred *C. elegans*, and thus represents the nonpathogenic control diet. We analyzed gene expression in N2 and CB4856 animals exposed to either *S. epidermidis, S. aureus*, *P. rettgeri* or *P. aeruginosa*.

Although both *C. elegans* isolates showed similar susceptibility to three of these pathogens, we hypothesized geographical isolation might have resulted in variations in the immune response, which could be analyzed through transcriptional profiling. Principal component analysis (PCA) of gene expression found that N2 and CB4856 samples exposed to *E. coli*, the Gram-positive pathogen *S. epidermidis* or the pathogenic Gram-negative *P. rettgeri* clustered in an isolate-specific manner (Figure 3A). N2 samples exposed to the virulent Gram-negative *P. aeruginosa* were also part of this N2-specific cluster, whereas CB4856 samples exposed to *P. aeruginosa* formed their own distinct cluster. Further, both genotypes exposed to the moderately virulent Gram-positive *S. aureus* formed distinct groups that did not cluster with other samples from the same genotype, nor did they cluster with each other (Figure 3A).

**Figure 3:**
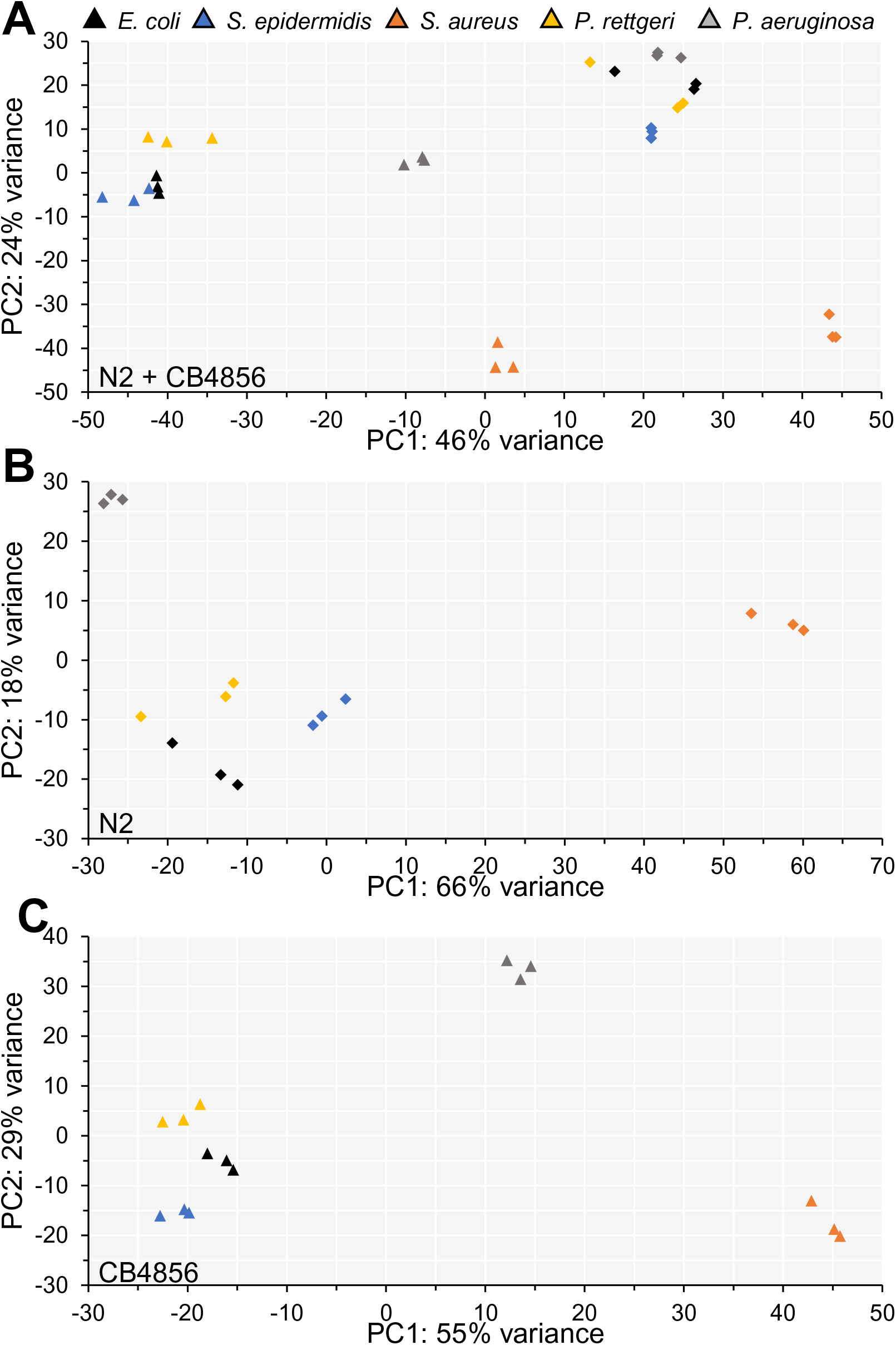
Principal component analysis clusters samples based on genotype, rather than pathogen exposure. (A-C) Principal component plots of expression data from the Hawaiian wild isolate, CB4856 (triangles) and N2 (diamonds) samples (A), only N2 samples (B), and only CB4856 samples (C).

We also performed PCA of gene expression using samples from a single genotype. For both *C. elegans* isolates *S. epidermidis* and *P. rettgeri* samples again clustered with *E. coli*, whereas. *S. aureus* samples formed a separate group along the first principal component axis (PC1). *P. aeruginosa* samples also formed individual clusters for both isolates, with separation along the second principal component axis (PC2) in N2 and separation along both the PC1 and PC2 axes in CB4856 (Figures 3B, 3C). Thus, it appears that both nematode genotype and microbial taxonomy contribute to the gene expression differences between N2 and CB4856.

### Differential Gene Expression is Largely Unique in N2 and CB4856 Animals After Infection with Pathogenic Bacteria

We first sought to identify how gene expression changes in these two isolates, as a function of infection. Thus, we identified genes differentially expressed in response to all pathogens (for all analyses, differentially expressed genes (DEGs) were defined as those with an FDR-adjusted P-value <0.05 and fold change ≥2, infected vs. control). We identified 273 and 117 genes that were differentially expressed in N2 and CB4856, respectively, as a function of infection. In either genotype the majority of DEGs were upregulated (N2, 236/273 = 86%; CB4856, 96/117 = 82%) (Table 1, Table 2, Table S2). Interestingly, only six out of the 384 total genes were differentially expressed in both N2 and CB4856 (N2-only, 267/273 = 98%; CB4856-only, 111/117 = 95%) (Figure 4A, Table S2).

**Table 1:** Differentially expressed genes (DEGs) in N2 animals following 24-hour exposure to microbial pathogens.

| Condition | <sup>1</sup>Total DEGs | Total Upregulated | Total Downregulated | <sup>2</sup>Total Genes Expressed |
| --- | --- | --- | --- | --- |
| All Pathogens | 273 | 236 | 37 | 23938 |
| Gram Positive | 3833 | 2964 | 869 | 22934 |
| Gram Negative | 628 | 254 | 374 | 21646 |
| *Staphylococcus epidermidis* | 2384 | 1754 | 630 | 21124 |
| *Staphylococcus aureus* | 8754 | 4901 | 3853 | 21433 |
| *Pseudomonas aeruginosa* | 5802 | 1804 | 3998 | 20304 |
| *Providencia rettgeri* | 1029 | 504 | 525 | 20295 |
| *Escherichia coli* | - | - | - | 19961 |

<sup>1</sup>Genes with a false discovery rate (FDR)-adjusted p-value of <0.05 and log2 fold change >1
<sup>2</sup>Total number of genes with >0 counts

**Figure 4:**
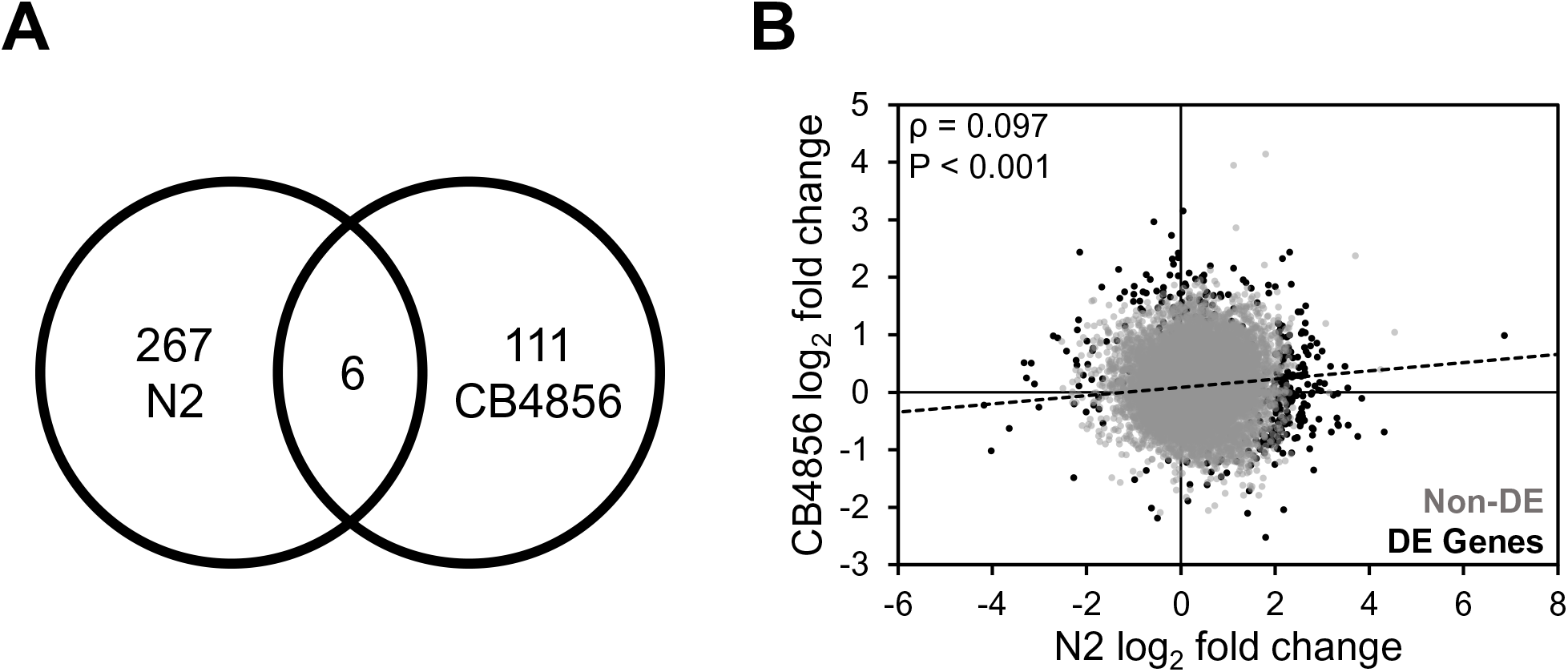
Expression differences in N2 and CB4856 animals after infection with pathogenic bacteria. (A) Intersection of differentially expressed genes and (B) correlation between log_2_ fold changes in N2 and CB4856 animals infected with pathogenic bacteria. Spearman’s rho value and P-values are presented in the upper left corner. Gray (Non-DE) and black dots (DE Genes) represent expressed and differentially expressed genes, respectively.

**Table 2:** Differentially expressed genes (DEGs) in CB4856 animals following 24-hour exposure to microbial pathogens.

| Condition | <sup>1</sup>Total DEGs | Total Upregulated | Total Downregulated | <sup>2</sup>Total Genes Expressed |
| --- | --- | --- | --- | --- |
| All Pathogens | 117 | 96 | 21 | 24333 |
| Gram Positive | 2565 | 1962 | 603 | 23297 |
| Gram Negative | 1720 | 456 | 1264 | 21847 |
| *Staphylococcus epidermidis* | 1776 | 1296 | 480 | 21252 |
| *Staphylococcus aureus* | 7033 | 3597 | 3436 | 21675 |
| *Pseudomonas aeruginosa* | 5171 | 1460 | 3711 | 20963 |
| *Providencia rettgeri* | 2253 | 694 | 1559 | 19731 |
| *Escherichia coli* | - | - | - | 20288 |

<sup>1</sup>Genes with a false discovery rate (FDR)-adjusted p-value of <0.05 and log2 fold change >1
<sup>2</sup>Total number of genes with >0 counts

We examined the correlation of gene expression using the log_2_ fold changes after infection. Expression levels were weakly, but positively, correlated between N2 and CB4856 (ρ = 0.097; *p*< 0.001) (Figure 4B). A correlation of baseline gene expression was determined using normalized gene counts from worms exposed to the control bacteria (OP50 *E. coli*). In that context N2 and CB4856 had similar patterns of expression (ρ = 0.826), suggesting that gene expression in response to infection diverges from a common baseline. Taken together, these data suggest that while there is some conservation of expression in response to infection, the pattern is relatively weak between these two isolates.

We further examined the DEGs identified in both isolates. Of them, only Y65B4BR.1 has any functional annotation relating to infection. Previous data suggests that RNAi knockdown increases susceptibility to infection by *S. aureus.*^35^ A single gene, *oac-14*, was identified as differentially expressed in opposing directions, *i.e.*, up in N2 and down in CB4856. RNAi knockdown delays or slows development, suggesting *oac-14* contributes to organismal growth.^36^ To take a more holistic approach, we used geneset enrichment analysis to analyze DEGs. The GO terms “Stress response: pathogen”, “Transmembrane transport: lipid”, and “Metabolism: lipid” were observed to be enriched in N2 (N2-only), whereas genes corresponding to the GO term “Metabolism: lipid” were overrepresented in CB4856 animals (CB4856-only) (Table 3, Table S3). Taken together, these results indicate that although there was little overlap in the transcriptomic response to microbial pathogen in N2 and CB4856 animals, both genotypes exhibit an enrichment of genes involved in lipid metabolism.

**Table 3:**
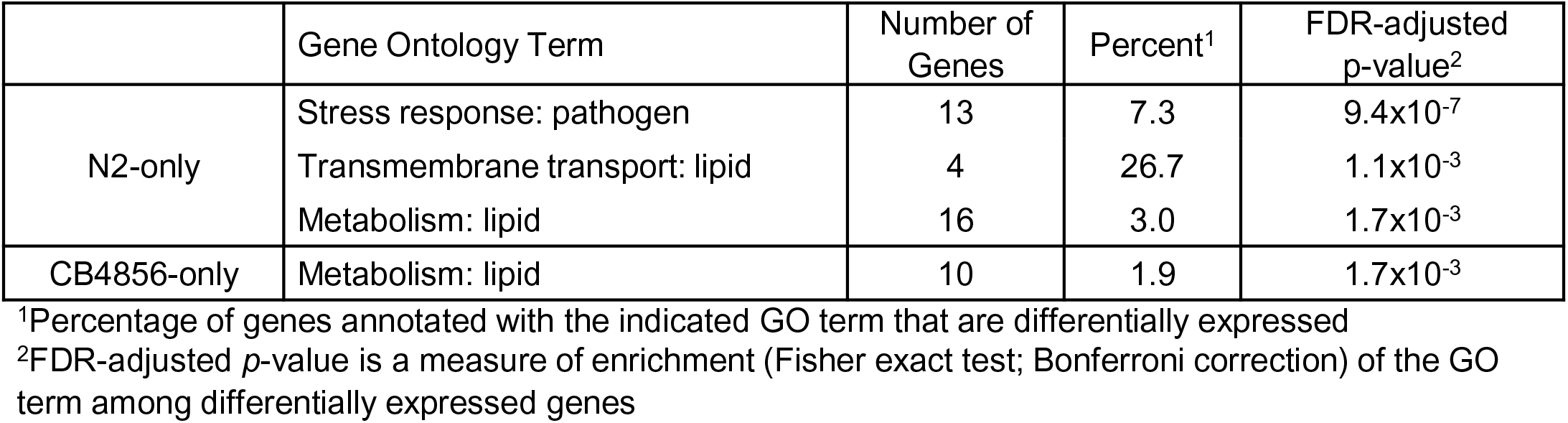
Enriched Gene Ontology Terms after Infection with Pathogenic Bacteria.

|  | Gene Ontology Term | Number of Genes | Percent<sup>1</sup> | FDR-adjusted p-value<sup>2</sup> |
| --- | --- | --- | --- | --- |
| N2-only | Stress response: pathogen | 13 | 7.3 | 9.4x10<sup>-7</sup> |
|  | Transmembrane transport: lipid | 4 | 26.7 | 1.1x10<sup>-3</sup> |
|  | Metabolism: lipid | 16 | 3.0 | 1.7x10<sup>-3</sup> |
| CB4856-only | Metabolism: lipid | 10 | 1.9 | 1.7x10<sup>-3</sup> |

<sup>1</sup>Percentage of genes annotated with the indicated GO term that are differentially expressed
<sup>2</sup>FDR-adjusted *p*-value is a measure of enrichment (Fisher exact test; Bonferroni correction) of the GO term among differentially expressed genes

Finally, to examine the change in transcriptional networks, we cross referenced our DEG lists with a table of known *C. elegans* transcription factors (File S1). We found that in N2, the transcription factors *cky-1, nhr-19, nhr-99, nhr-116, nhr-127, peb-1* and *unc-42* were all upregulated in response to infection by pathogens, whereas in CB4856-infected animals, *ham-1* expression was repressed and *klu-2* expression was induced (Table S2). These results are consistent with the divergence of gene expression profiles between N2 and CB4856, suggesting that, upon infection, these isolates rely on different immune pathways.

### N2 and CB4856 Transcriptomic Responses Exhibit More Overlap Following Infection with Gram-positive than Gram-negative Pathogens

We then asked how gene expression profiles differed when we grouped pathogens by Gram-stain. We identified 3833 and 2565 genes that were differentially expressed in response to Gram-positive bacteria in N2 and CB4856, respectively. In either genotype the majority of DEGs were upregulated (N2, 2964/3833 = 77%; CB4856, 1962/2565 = 76%) (Table 1, Table 2, Table S4). While most genes were unique to a genotype (N2-only, 2661/3833 = 69%; CB4856-only, 1393/2565 = 54%), 1172 genes were differentially expressed in both N2 and CB4856 worms (Figure 5A, Table S4). Further, the majority of DEGs identified in both genotypes were expressed in the same direction, while only 1% of genes were differentially expressed in opposing directions, *i.e.*, down in N2 and up in CB4856 (14/1172 = 1%) (Figure 5C, Table S4). These results were reflected in a strong, positive correlation between N2 and CB4856 gene expression levels (ρ = 0.613; *p*< 0.001) (Figure 5A).

**Figure 5:**
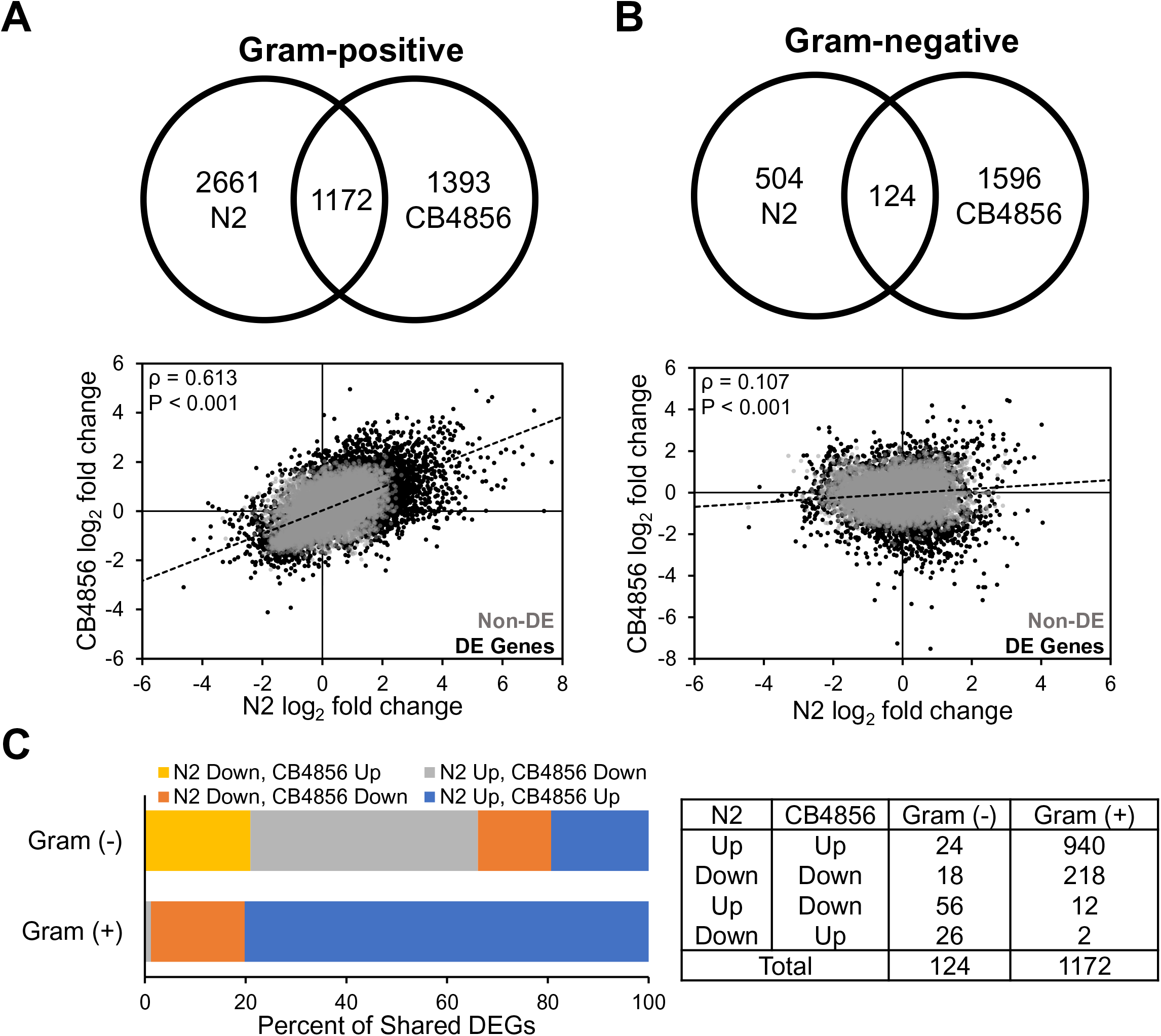
Expression differences in N2 and CB4856 animals infected with Gram-positive or Gram- negative pathogens. (A) Intersection of differentially expressed genes (top) and correlation between log_2_ fold changes (bottom) in N2 and CB4856 animals infected with Gram-positive pathogens (*S. epidermidis, S. aureus*) or (B) Gram-negative pathogens (*P. rettgeri*, *P. aeruginosa*). Spearman’s rho value and P-values are presented in the upper left corner. Gray (Non-DE) and black dots (DE Genes) represent expressed and differentially expressed genes, respectively. (C) The distribution of up- and down-regulated genes that are differentially expressed in both N2 and CB4856 animals infected with either Gram-negative or Gram-positive pathogens.

We again looked for transcription factors that were differentially expressed in both N2 and CB4856 as a function of infection by Gram-positive bacteria. In this list, there were 50 transcription factor genes that were significantly different from baseline (Table S4). As observed for the overall gene expression similarity, for each of the transcription factors, we observed a similar pattern for the direction of expression (*e.g.,* genes upregulated in N2 were upregulated in CB4856, *etc.*).

In response to Gram-negative bacteria, 628 and 1720 genes were differentially expressed in N2 and CB4856 animals, respectively. Interestingly, in both genotypes a minority of DEGs were upregulated (N2, 254/628 = 40%; CB4856, 456/1720 = 27%) (Table 1, Table 2, Table S5). Like the transcriptomic response when all pathogens were grouped together, most genes were unique to a genotype with a total of 124 genes differentially expressed in both N2 and CB4856 (N2-only, 504/628 = 80%; CB4856-only, 1596/1720 = 93%) (Figure 5B, Table S5). Further, the majority of DEGs identified in both genotypes were expressed in opposing directions (82/124 = 66%), leaving only 42 genes that were differentially expressed in both genotypes and in the same direction (Figure 5C, Table S5). Gene expression profiles were again weakly, but positively correlated (ρ = 0.107; *p*< 0.001) (Figure 5B). Finally, there were no transcription factors that were coordinately regulated in both isolates in response to Gram-negative bacteria (Table S5).

These results highlight the dramatic differences when we compare the transcriptomic response of N2 and CB4856 worms infected with Gram-positive or Gram-negative pathogens. Interestingly, exposure to Gram-positive pathogens resulted in differential gene expression profiles with considerable overlap between the two isolates, suggesting that there are conserved responses to some bacteria. In contrast, the transcriptomic responses of N2 and CB4856 animals to Gram-negative pathogens were relatively divergent, with few differentially expressed genes in common.

### Overlap in Differentially Expressed Genes between N2 and CB4856 Depends on the Microbial Pathogen

Finally, we analyzed gene expression by individual pathogen exposure. This identified 2384 (*S. epidermidis*), 8754 (*S. aureus*), 5802 (*P. aeruginosa*), and 1029 (*P. rettgeri*) differentially expressed genes in N2 and 1776 (*S. epidermidis*), 7033 (*S. aureus*), 5171 (*P. aeruginosa*), and 2253 (*P. rettgeri*) differentially expressed genes in CB4856 (Table 1, Table 2). In response to *S. epidermidis*, upregulated DEGs outnumbered downregulated DEGs at a ratio of 3:1 (N2, 1754/2384 = 74%; CB4856, 1296/1776 = 73%), whereas the response to *S. aureus* was split more evenly between upregulated and down regulated genes, although upregulated DEGs still represented a majority (N2, 4901/8754 = 56%; CB4856, 3597/7033 = 51%) (Table 1, Table 2, Table S6, Table S7).

Although both belong to the *Staphylococcus* genus, the overlap between N2 and CB4856 transcriptomic responses to *S. epidermidis* and *S. aureus* were considerably different. In *S. epidermidis-*infected animals the minority of DEGs, 511 in total, were differentially expressed in both N2 and CB4856 (N2-only, 1873/2384 = 79%; CB4856-only, 1265/1776 = 71%) (Figure 6A, Table S6). The correlation of gene expression in response to *S. epidermidis* was ρ = 0.294 (*p*< 0.001). In contrast, a majority of differentially expressed genes, 5039 in total, overlapped in N2 and CB4856 animals infected with *S. aureus* (N2-only, 3715/8754 = 42%; CB4856-only, 1994/7033 = 28%) (Figure 6B, Table S7), with a correlation of ρ = 0.721 (*p*< 0.001).

**Figure 6:**
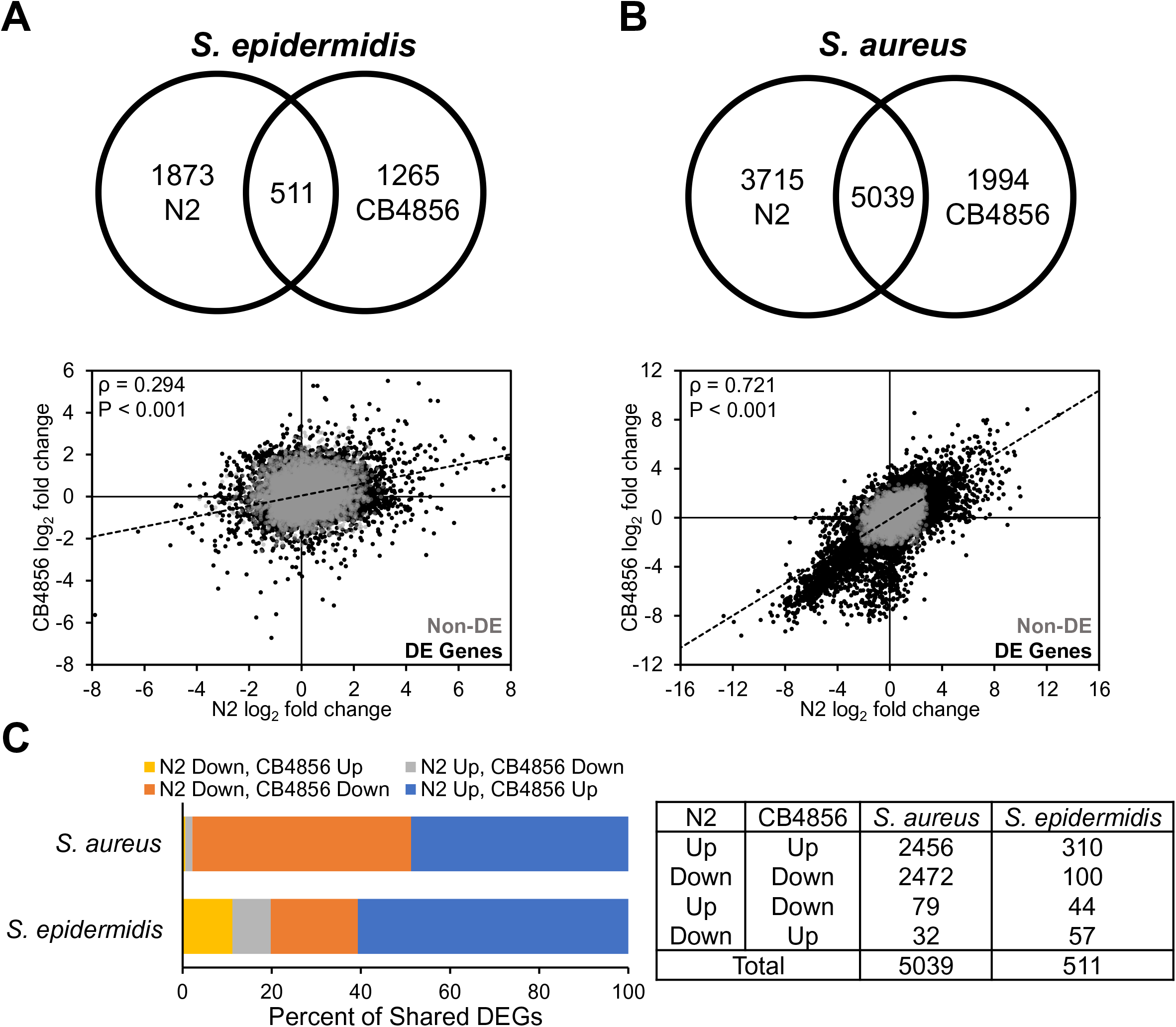
Expression differences in N2 and CB4856 animals infected with the gram-positive pathogens, *S. epidermidis* or *S. aureus*.

The majority of DEGs identified in both genotypes following either *S. epidermidis* or *S. aureus* infection were expressed in the same direction (*S. epidermidis*, 410/511 = 80%; *S. aureus*, 4928/5039 = 98%) (Figure 6C, Table S6, Table S7). Lastly, we only identified nine transcription factors that were coordinately regulated in *S. epidermidis*-infected animals, whereas 346 transcription factors genes were differentially expressed in both genotypes in the same direction after *S. aureus* infection (Table S6, Table S7). Together, these results suggest N2 and CB4856 worms exhibit considerable similarity in their transcriptomic response to *S. aureus*, whereas there is much less overlap in their response to *S. epidermidis* infection, which may perhaps explain some of the difference in susceptibility to infection by *S. epidermis*.

Exposure to either Gram-negative pathogen resulted in a minority of DEGs being upregulated in both genotypes. In *P. aeruginosa*-infected animals, upregulated genes represented less than one-third of all DEGs in both genotypes (N2, 1804/5802 = 31%; CB4856, 1460/5171 = 28%), whereas, after *P. rettgeri* infection, differential expression analysis found nearly equivalent numbers of upregulated and downregulated genes in N2 and upregulation of less than one-third of all DEGS in CB4856 (N2, 504/1029 =49%; CB4856, 694/2253 =31%) (Table 1, Table 2, Table S8, Table S9).

We observed considerable differences in the overlap of gene expression profiles between N2 and CB4856 worms infected with *P. aeruginosa* or *P. rettgeri*. In animals infected with *P. aeruginosa*, a majority of differentially expressed genes, 3092 in total, overlapped (N2-only, 2710/5802 = 47%; CB4856-only, 2079/5171 = 40%) (Figure 7A, Table S8), with a correlation of ρ = 0.539 (*p*< 0.001). Like infection with *S. epidermidis* or *S. aureus*, the majority of DEGs identified in both genotypes following *P. aeruginosa* infection were expressed in the same direction (2971/3092 = 96%) (Figure 7C, Table S8). In response to *P. rettgeri*, 581 genes were differentially expressed in both genotypes, representing a majority of DEGs in N2 but a minority in CB4856 (N2-only, 448/1029 = 44% CB4856-only, 1672/2253 = 74%) (Figure 7B, Table S9). Interestingly, the vast majority of DEGs found in both genotypes following *P. rettgeri* infection were expressed in opposing directions (564/581 = 97%), leaving only 17 genes that were coordinately expressed (Figure 7C, Table S9). Finally, we identified 114 transcription factor genes that were differentially expressed in both genotypes in the same direction after *P. aeruginosa* infection, whereas no transcription factors were coordinately regulated in *P. rettgeri*-infected animals (Table S8, Table S9).

**Figure 7:**
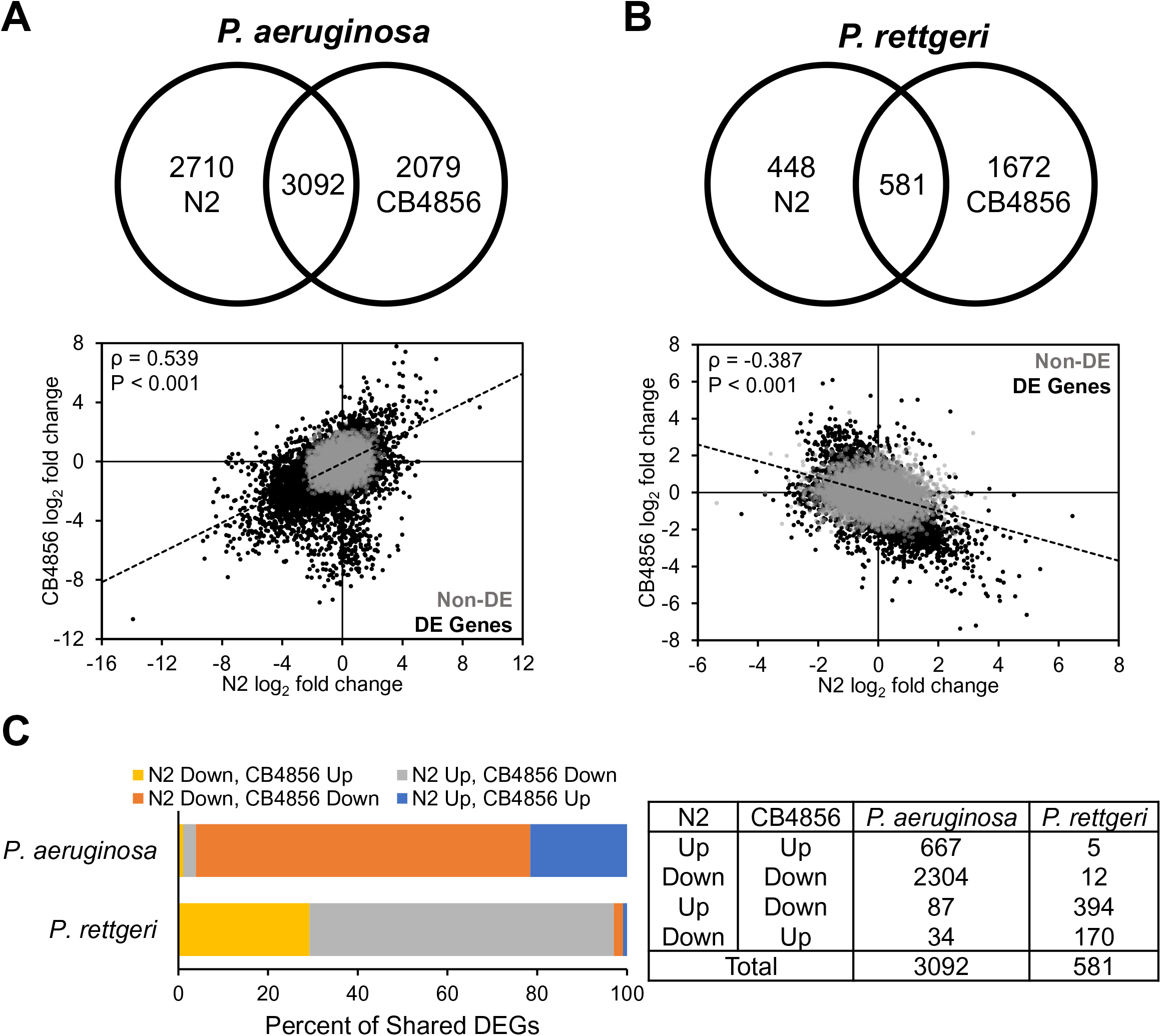
Expression differences in N2 and CB4856 animals infected with the gram-negative pathogens, *P. rettgeri* or *P. aeruginosa*.

Taken together, these results indicate that the N2 and CB4856 transcriptomic responses to *P. aeruginosa* have more in common, whereas little overlap in differential gene expression exists when N2 and C4856 are infected with *P. rettgeri*. In fact, of all the gene expression profiles compared, *P. rettgeri* infection was the only one to have a negative correlation (ρ = -0.387; *p*< 0.001). These results are consistent with the responses to infection being driven both by the isolate and the pathogen.

### Geneset Enrichment Analysis Identifies Unique GO Terms Enriched in N2 and CB4856 Worms Exposed to *S. epidermidis*

Since N2 and CB4856 exhibited survival differences after *S. epidermidis* infection but not *S. aureus* infection, we were interested in identifying biological processes that may be enriched in these animals. We cross-referenced DEGs identified in N2 following *S. epidermidis* and *S. aureus* infection and identified 556 genes that were either exclusively expressed in response to *S. epidermidis* or were expressed in response to both pathogens but in opposing directions (*e.g.,* upregulated in *S. epidermidis* and downregulated in *S. aureus*). The same analysis performed for DEGs in CB4856 animals identified 749 genes. Only 73 of these genes were differentially expressed in both N2 and CB4856 (N2-only, 483/556 = 87%; CB4856-only, 676/749 = 90%) (Table S10).

We performed geneset enrichment analysis on genes that were exclusive to N2 (N2-only), genes that were exclusive to CB4856 (CB4856-only), and genes that were differentially expressed in both N2 and CB4856 animals (N2 & CB4856). Biological functions pertaining to the extracellular matrix, stress response, and metabolism were enriched in N2 and CB4856 worms following *S. epidermidis* infection (Table 4, Table S11). The Gene Ontology (GO) terms “Extracellular material: collagen” and “Extracellular material: cuticlin” were enriched in N2-exclusive DEGs, whereas genes corresponding to the GO terms “Stress response: C-type Lectin” and “Metabolism: lipid” were overrepresented in DEGs exclusive to CB4856. Genes in the GO category “Stress response: detoxification” were overrepresented in both isolates, even if the individual genes were not necessarily the same. Genes that were differentially expressed in both isolates were enriched for a single GO term, “Stress response: pathogen”. A separate group of genes that was expressed exclusively in CB4856 animals was also enriched for this GO term. Notably, we did not find significant enrichment of the GO term “Stress response: pathogen” in DEGs exclusive to N2 animals. Thus, when only considering genes that are differentially expressed in an *S. epidermidis*- specific manner, our results indicate a common enrichment of biological processes related to stress response and innate immunity in N2 and CB4856 animals. At the same time, we also observe an N2-specific enrichment of extracellular matrix genes and a CB4856-specific enrichment of lipid metabolism genes, suggesting a degree of divergence in the transcriptomic response of these isolates to *S. epidermidis* infection.

**Table 4:** Enriched Gene Ontology Terms in N2 and CB4856 Worms Exposed to S. epidermidis.

|  | Gene Ontology Term | Number of Genes | Percent<sup>1</sup> | FDR-adjusted p-value<sup>2</sup> |
| --- | --- | --- | --- | --- |
| N2-only | Extracellular material: collagen | 55 | 29.7 | 4.5x10<sup>-44</sup> |
|  | Extracellular material: cuticlin | 7 | 20.0 | 1.5x10<sup>-4</sup> |
|  | Stress response: detoxification | 13 | 8.6 | 2.8x10<sup>-4</sup> |
| CB485-only | Stress response: pathogen | 22 | 12.4 | 4.6x10<sup>-8</sup> |
|  | Stress Response: C-type Lectin | 26 | 10.1 | 7.8x10<sup>-8</sup> |
|  | Stress response: detoxification | 20 | 13.2 | 1.0x10<sup>-7</sup> |
|  | Metabolism: lipid | 30 | 5.7 | 5.2x10<sup>-4</sup> |
| N2 & CB4856 | Stress response: pathogen | 7 | 4.0 | 1.4x10<sup>-5</sup> |

<sup>1</sup>Percentage of genes annotated with the indicated GO term that are differentially expressed
<sup>2</sup>FDR-adjusted *p*-value is a measure of enrichment (Fisher exact test; Bonferroni Correction) of the GO term among differentially expressed genes

### Comparing DEGs between N2 and CB4856 fed OP50 versus *S. epidermidis*

The genomes of N2 and CB4856 animals differ at approximately one polymorphism per kb and gene expression between these two isolates, including the expression of innate immune genes, is known to vary across similar developmental stages, even in the absence of infection.^37–39^ Since principal component analysis found that genotype is the greatest predictor of isolate-specific differences in gene expression following *S. epidermidis* infection (Figure 3), we performed an additional gene expression analysis to identify genes that were differentially expressed between N2 and CB4856 animals fed either the non-pathogenic *E. coli* OP50 or the pathogenic *S. epidermidis*.

Using N2 expression as a baseline, we found 3384 and 1892 genes to be differentially expressed in CB4856 animals fed *E. coli* OP50 and *S. epidermidis*, respectively. Upregulated genes outnumbered those that were downregulated at a ratio of 2:1, regardless of whether the worms were fed OP50 or *S. epidermidis* (OP50 Upregulated, 2121/3384 = 63%; OP50 Downregulated, 1263/3384 =37%) (*S. epidermidis* Upregulated, 1245/1892 = 66%; *S. epidermidis* Downregulated, 647/1892 =34%) (Table S12, Table S13).

Next, we cross-referenced genes that were differentially expressed between N2 and CB4856 animals in the OP50 and *S. epidermidis* conditions and classified them into four categories based on changes in gene expression, or lack thereof. Relative to N2, a total of 2280 genes were differentially expressed in CB4856 animals fed OP50 but were no longer differentially expressed when worms were fed *S. epidermidis*. This suggests that the final level of expression for these gene was similar in N2 and CB4856 animals following infection. A total of 1072 genes were differentially expressed between N2 and CB4856 animals fed OP50 and continued to be differentially expressed in the same direction in worms fed *S. epidermidis*, suggesting that they are likely uninvolved in the transcriptional response to the pathogen. A third group of genes, 788 in total, were exclusively differentially expressed between N2 and CB4856 animals fed *S. epidermidis*. Finally, we identified 32 genes that were differentially expressed between N2 and CB4856 animals fed either OP50 or *S. epidermidis* but in opposing directions, *i.e.*, up in CB4856 fed OP50 and down in CB4856 fed *S. epidermidis* (Table S14).

The isolate- and pathogen-specific expression pattern of genes from these latter two categories prompted us to perform geneset enrichment analysis to identify biological processes that may be differentially enriched in N2 and CB4856 animals following *S. epidermidis* infection. The 788 genes that were exclusively differentially expressed in CB4856 animals fed *S. epidermidis* were enriched for GO terms pertaining to immunity (Stress response: C-type lectin), cell signaling and transduction (Signaling: phosphatase; Signaling: tyrosine kinase), reproduction (Major sperm protein), posttranslational modification (Protein modification: glycoprotein), and cytoskeletal movement (Cytoskeleton: microtubule). From the 32 genes that were differentially expressed in opposing directions in CB4856 animals fed either OP50 or *S. epidermidis* geneset enrichment analysis identified GO terms corresponding to immunity (Stress response: pathogen) and the extracellular matrix (Extracellular material: collagen) (Table 5) (Table S15). Thus, by considering differences in gene expression between N2 and CB4856 animals fed non-pathogenic *E. coli* OP50, we have identified additional sets of genes that are enriched for GO terms pertaining to, among other categories, stress response, innate immunity, and the extracellular matrix.

**Table 5:**
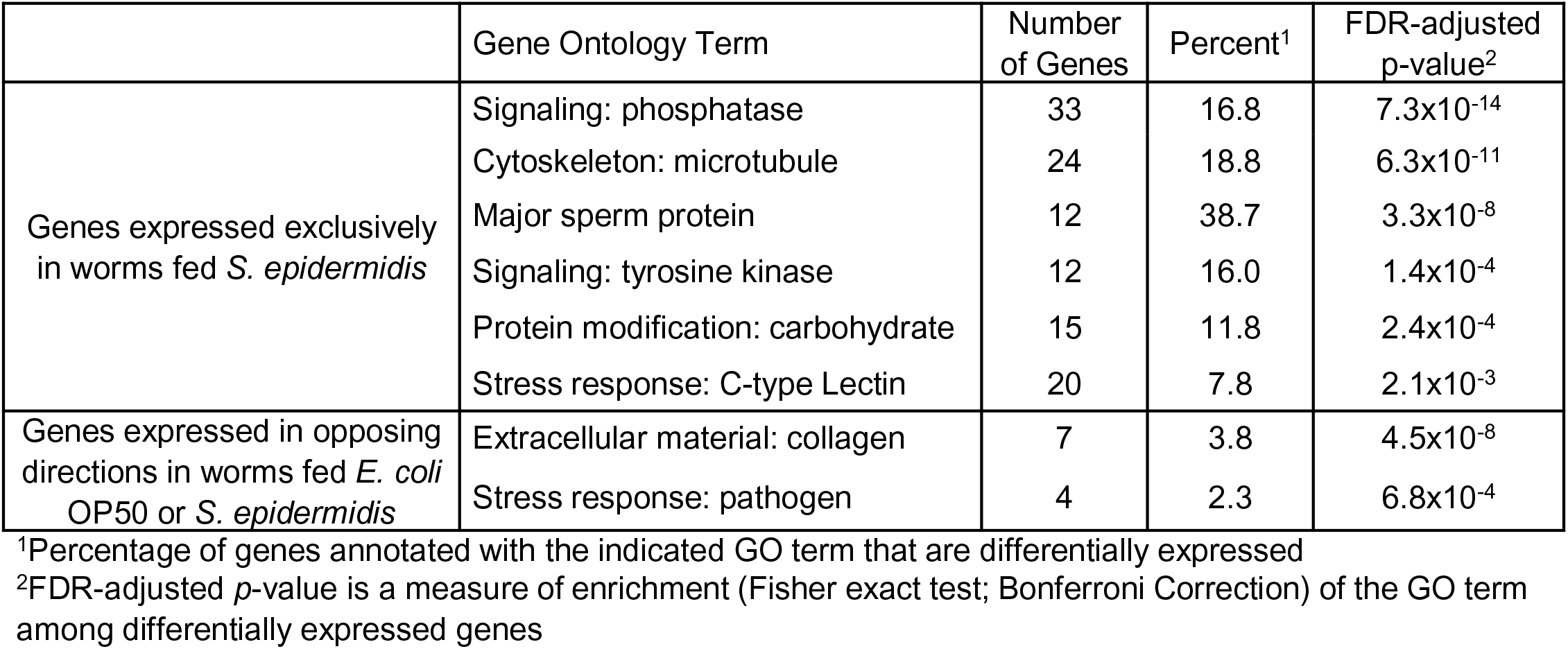
**Enriched Gene Ontology Terms when Comparing N2 and CB4856 Worms fed OP50 versus *S. epidermidis***

|  | Gene Ontology Term | Number of Genes | Percent<sup>1</sup> | FDR-adjusted p-value<sup>2</sup> |
| --- | --- | --- | --- | --- |
| Genes expressed exclusively in worms fed *S. epidermidis* | Signaling: phosphatase | 33 | 16.8 | 7.3x10<sup>-14</sup> |
|  | Cytoskeleton: microtubule | 24 | 18.8 | 6.3x10<sup>-11</sup> |
|  | Major sperm protein | 12 | 38.7 | 3.3x10<sup>-8</sup> |
|  | Signaling: tyrosine kinase | 12 | 16.0 | 1.4x10<sup>-4</sup> |
|  | Protein modification: carbohydrate | 15 | 11.8 | 2.4x10<sup>-4</sup> |
|  | Stress response: C-type Lectin | 20 | 7.8 | 2.1x10<sup>-3</sup> |
| Genes expressed in opposing directions in worms fed *E. coli* OP50 or *S. epidermidis* | Extracellular material: collagen | 7 | 3.8 | 4.5x10<sup>-8</sup> |
|  | Stress response: pathogen | 4 | 2.3 | 6.8x10<sup>-4</sup> |

<sup>1</sup>Percentage of genes annotated with the indicated GO term that are differentially expressed
<sup>2</sup>FDR-adjusted *p*-value is a measure of enrichment (Fisher exact test; Bonferroni Correction) of the GO term among differentially expressed genes

Finally, to better understand the transcriptional regulators that might be significantly different upon infection by *S. epidermidis,* we looked for transcription factors that were differentially expressed only upon infection. There were 22 protein coding genes (1 pseudogene), that fit these criteria. Interestingly, 13 of the 22 were members of the nuclear hormone receptor (*nhr*) family. There are 266 *nhr* genes in our list of 1077 transcription factors. The incidence of *nhr* genes in our DEG list was significantly enriched (P = 0.007, Fisher’s exact test, odds ratio 4.4).

## Discussion

We conducted survival, behavioral and transcriptional assays using the *C. elegans* strains N2 and CB4856 to identify differences in pathogen susceptibility between two wild-type isolates. CB4856 was significantly more susceptible than N2 to infection by the Gram-positive pathogen *S. epidermidis*. CB4856 and N2 were equally susceptible to a related species, *S. aureus,* and two non-*Staphylococcus* pathogens, *P. aeruginosa* and *P. rettgeri.* We confirmed the CB4856 susceptibility using a second isolate of *S. epidermidis*.

What might underlie the differences in susceptibility? Initially we hypothesized that, perhaps, the CB4856 strain lacked the ability to detect *S. epidermidis* as pathogenic. However, CB4856 worms avoided *S. epidermidis,* when given *E. coli* as an alternative, suggesting they were averse to the bacteria. Further, we found robust changes in gene expression in response to the *S. epidermis* infection. Unexpectedly, we found that, despite *P. rettgeri* being equally pathogenic to both isolates, CB4856 animals preferred *P. rettgeri* to *E. coli,* while N2 did not. This observation was also highlighted by the fact that changes in gene expression were negatively correlated in these two isolates in response to *P. rettgeri*.

### Isolates exhibit distinct transcriptomic responses to infection

The pathogens used in this study are diverse both in terms of their genome size and known virulence mechanisms.^40–43^ Principal component analysis of the sequencing dataset found that gene expression changes during infection clustered largely by genotype, rather than pathogen, even in instances where susceptibility was equivalent. At a glance, this suggests that the strategies these strains have adopted for dealing with microbial infection have diverged since they were separated from one another.

When we looked for genes that were differentially expressed in response to infection, *i.e.,* was identified in response to all pathogens, there were hundreds, compared to the lists for individual pathogen (thousands). Further, we found only six DEGs in common between N2 and CB4856, with those genes largely uncharacterized functionally. Overall, these results suggest that N2 and CB4856 respond to infection by altering gene transcription, but the core infection response represents only a small fraction of the total genes regulated, and that the core transcriptional response in each background has diverged. It also suggests there are still a number of genes involved in immunity that need to be characterized.

By categorizing the genes that were differentially expressed in response to all pathogens, we found an overrepresentation of genes involved in lipid metabolism in both N2- and CB4856- infected animals. Lipid metabolism has been shown to be required for proper innate immune system activation and pathogen defense.^44^ These genes could play similar roles in both isolates and thus, while they may reflect a divergence in specific gene activation, they could represent a similarity in how N2 and CB4856 animals utilize lipid metabolism enzymes to facilitate the innate immune response.

### Gram-stain level analysis suggests an absence of conserved master regulatory pathways

In many organisms, the Toll signaling pathway is activated by infection with Gram-positive bacteria and/or fungi, while the IMD pathway is activated in response to Gram-negative bacteria.

Thus, we examined the results grouping pathogens by Gram-stain, and then compared between isolates. Our results found that the majority of DEGs were strain-specific, consistent with a divergence of the responses to pathogens, but that, generally, there was a stronger conservation of responses to Gram-positive bacteria, compared to Gram-negative. Our results are consistent with the evidence that Toll and IMD signaling is diminished or absent in *C. elegans* as master regulators of immunity and suggests each isolate deals with Gram-negative and Gram-positive pathogens differently.

We examined the list of DEGs for transcription factors that could be contributing to the differential expression profiles in each of the isolates. There were hundreds of transcription factors regulated by infection with Gram-positive pathogens, with 50 overlapping between the N2 and CB4856 isolates. In contrast, for Gram-negative pathogens, N2 had only nine differentially expressed transcription factors, while CB4856 had 97, with none in common.

One possible explanation for the large overlap in N2 and CB4856 transcriptomic responses to Gram-positive bacteria is the identity of the microbial pathogens. *S. aureus* and *S. epidermidis* are both members of the *Staphylococcus* genus, share a number of virulence factors and invoke overlapping host immune responses following infection.^41, 45, 46^ Thus, the greater proportion of overlapping DEGs in N2 and CB4856 animals infected with Gram-positive bacteria may reflect a common response to *Staphylococcus* pathogens. Similar results for infection of N2 animals by *Enterococcus faecalis* and *Enterococcus faecium* suggest that, within species of pathogens, there are core transcriptional similarities.^47^

### Individual pathogens induce different responses in *C. elegans* isolates

Since N2 and CB4856 worms exhibited similar susceptibility to infection by *P. aeruginosa*, *S. aureus* or *P. rettgeri*, we posited that RNA sequencing would reveal considerable overlap in gene expression between the two isolates. Indeed, after infecting N2 and CB4856 worms with either *P. aeruginosa* or *S. aureus* we identified thousands of differentially expressed genes, a majority of which were coordinately regulated in both isolates, supporting the idea that infection by either of these pathogens invokes similar kinds of transcriptomic responses in N2 and CB4856 animals. However, animals infected with *P. rettgeri* exhibited very little overlap in gene expression with fewer than 20 genes differentially expressed in both isolates and expressed in the same direction.

It is possible that N2 and CB4856 worms have adopted two very different immune responses to *P. rettgeri* infection, with neither providing a perceivable survival advantage. But why do we not observe a similar lack of overlap in gene expression after infection with the Gram-negative pathogen, *P. aeruginosa*, which also showed similar lethality in N2 and CB4856 animals? Analysis of gene expression at regular intervals throughout an infection has demonstrated that the host immune response is a dynamic process which needs to be initiated and sustained to fight an infection.^48, 49^ At 24 hours post-infection, the timepoint at which RNA was collected for gene expression analysis, animals infected with *P. aeruginosa* were much closer to dying than animals infected with *P. rettgeri* (N2 *P. aeruginosa* LT_50_ = 3.8 ± 0.3 days, CB4856 *P. aeruginosa* LT_50_ = 3.5 ± 0.3 days; N2 *P. rettgeri* LT_50_ = 7.8 ± 0.7 days, CB4856 *P. rettgeri* LT_50_ = 7.2 ± 0.6 days). Perhaps animals exposed to *P. aeruginosa* or *P. rettgeri* are at two different stages in combatting their respective infections and the overlap in gene expression between N2 and CB4856 animals grows larger as animals progress from earlier to later stages of infection.

### Changes in gene expression by infection with *S. epidermidis*

We found differences in survival in response to *S. epidermidis.* For each isolate we narrowed down the list of differentially expressed genes to those that were expressed in a *S. epidermidis*- specific manner and performed geneset enrichment analysis on sets of genes that were either exclusive to N2, exclusive to CB4856, or expressed in both N2 and CB4856 animals. We observed an enrichment of genes encoding extracellular matrix (ECM) proteins, such as collagens and cuticlins, exclusively in N2 animals. Cuticular collagens are an integral component of the *C. elegans* epidermal barrier and have established roles in the host epidermal immune response to pathogens.^50–52^ Further, ECM proteins are likely involved in promoting longevity as changes in collagen gene expression have been used as *in vivo* biomarkers in *C. elegans* aging research.^53^

Together, our data reveal isolate-specific changes in ECM gene expression after *S. epidermidis* exposure, suggesting a role for collagen genes in the increased resistance of N2 worms to *S. epidermidis* infection. In DEGs exclusive to *S. epidermidis*-infected N2 animals as well as DEGs exclusive to *S. epidermidis*-infected CB4856 animals, our analysis detected an enrichment of genes encoding detoxification enzymes. These enzymes are primarily responsible for metabolizing xenobiotic substances and have demonstrated roles in the *C. elegans* immune response.^54, 55^ In both N2 and CB4856 animals, the DEGs were either members of the Cytochrome P450 (CYP) family or possessed UDP glucuronosyltransferase (UGT) activity. It is possible that these genes play similar roles in both isolates and thus may reflect a divergence in how N2 and CB4856 animals utilize detoxification enzymes to counteract the microbial infection.

Finally, we took advantage of the ability to look for genes with expression differences between N2 and CB4856 on control bacteria. This allowed us to sort differentially expressed genes in the *S. epidermidis* condition into those that started out different, but ended up being similar, and those that changed as a function of infection to end up at different levels of expression. *A priori* we might consider the former group as being less likely to underlie the differences in susceptibility. In those analyses, approximately 90% of the genes were upregulated in response to infection.

When we examined the transcription factors that were differentially expressed, we found an overrepresentation of nuclear hormone receptors. What is more, of the 13 *nhr* genes, seven were upregulated, while six were downregulated during infection. Nuclear hormone receptors are used in many contexts to systemically coordinate signaling. In *C. elegans* multiple *nhr* genes have been linked to innate immunity, including *nhr-8, nhr-23, nhr-25, nhr-45, nhr-49, nhr-57, nhr-80,* and *nhr-112.*^56–63^ Thus, this is an interesting family of genes to potentially contribute to differences in organismal response and/or susceptibility *in vivo*.

## Conclusions

In populations, genetic and genomic diversity drive phenotypes. Immune signaling is no exception, and yet, our understanding of *C. elegans* immunity has largely been driven by the analysis of the canonical laboratory strain, N2. We found that the N2 strain and CB4856 wild isolate were significantly different when we measured the susceptibility to *S. epidermidis.* It is possible that the relative abundance of *S. epidermidis* or related pathogens in their respective ecological niches resulted in the shift in immune capacity between the N2 and CB4856 lines. Unfortunately, without sufficient metagenomic sampling of the environment, this would be difficult to address. However, the robust differences observed suggests that isolating the animals and transferring them to the laboratory environment has fixed genetic variation that underlies pathogen-specific susceptibility.

Perhaps more surprisingly, we found significantly divergent transcriptomic responses between these isolates, even when pathogen susceptibility was equivalent. These differences could come from protein coding variants in the molecules that detect pathogens or the transcription factors that coordinate immune defenses, non-coding variants in regulatory regions, or, more likely, a combination of the two. By narrowing down the regions of the genome that are important for the observed differences, we will be able to better understand how genetic variation contributes to different immune outcomes.

## Declarations

### Ethics approval and consent to participate

Not Applicable

### Consent for publication

Not Applicable

### Availability of data and materials

The raw sequence files used for differential gene expression analysis are available in FASTQ format in the NCBI BioProject repository (BioProject Accession number: PRJNA770277; https://www.ncbi.nlm.nih.gov/sra/PRJNA770277).

### Authors’ contributions

PL and BDA designed and conceived the project; PL, MC and BDA acquired data; PL and BDA analyzed and interpreted data and wrote the manuscript. All authors have approved the manuscript.

## Acknowledgements

We thank Dr. Robert Unckless for his helpful advice in sequencing and identifying the *S. epidermidis* strain. We also thank Dr. Erik Lundquist and his laboratory for helpful discussions.

