## Supplemental Tables for "Wild-type *Caenorhabditis elegans* Isolates Exhibit Distinct Gene Expression Profiles in Response to Microbial Infection"

**Supplemental File 1**

**A list of all of the transcription factors identified to date in *C. elegans* used to identify differentially expressed TF genes**

Supplemental File 1 – *C. elegans* Transcription Factors

| WormBase Gene ID | Gene Name | Sequence Name |
|---|---|---|
| WBGene00000095 | aha-1 | C25A1.11 |
| WBGene00000096 | ahr-1 | C41G7.5 |
| WBGene00000120 | aly-1 | C01F6.5 |
| WBGene00000121 | aly-2 | F23B2.6 |
| WBGene00000220 | atf-2 | K08F8.2 |
| WBGene00000221 | atf-5 | T04C10.4 |
| WBGene00000222 | atf-6 | F45E6.2 |
| WBGene00000223 | atf-7 | C07G2.2 |
| WBGene00000271 | brf-1 | F45E12.2 |
| WBGene00000272 | bro-1 | F56A3.5 |
| WBGene00000366 | cbp-1 | R10E11.1 |
| WBGene00000383 | cdc-14 | C17G10.4 |
| WBGene00000408 | cdk-7 | Y39G10AL.3 |
| WBGene00000428 | ceh-1 | F16H11.4 |
| WBGene00000429 | ceh-2 | C27A12.5 |
| WBGene00000430 | ceh-5 | C16C2.1 |
| WBGene00000431 | ceh-6 | K02B12.1 |
| WBGene00000432 | ceh-7 | C34C6.8 |
| WBGene00000433 | ceh-8 | ZK265.4 |
| WBGene00000434 | ceh-9 | Y65B4BR.9 |
| WBGene00000435 | ceh-10 | W03A3.1 |
| WBGene00000436 | ceh-12 | F33D11.4 |
| WBGene00000437 | ceh-13 | R13A5.5 |
| WBGene00000438 | ceh-14 | F46C8.5 |
| WBGene00000439 | ceh-16 | C13G5.1 |
| WBGene00000440 | ceh-17 | D1007.1 |
| WBGene00000441 | ceh-18 | ZC64.3 |
| WBGene00000442 | ceh-19 | F20D12.6 |
| WBGene00000443 | ceh-20 | F31E3.1 |
| WBGene00000444 | ceh-21 | T26C11.6 |
| WBGene00000445 | ceh-22 | F29F11.5 |
| WBGene00000446 | ceh-23 | ZK652.5 |
| WBGene00000447 | ceh-24 | F55B12.1 |
| WBGene00000448 | pros-1 | K12H4.1 |
| WBGene00000449 | ceh-27 | F46F3.1 |
| WBGene00000450 | ceh-28 | K03A11.3 |
| WBGene00000451 | ceh-30 | C33D12.7 |
| WBGene00000452 | ceh-31 | C33D12.1 |
| WBGene00000453 | ceh-32 | W05E10.3 |
| WBGene00000454 | ceh-33 | C10G8.7 |
| WBGene00000455 | ceh-34 | C10G8.6 |

| | | |
|---|---|---|
| WBGene00000457 | ceh-36 | C37E2.4 |
| WBGene00000458 | ceh-37 | C37E2.5 |
| WBGene00000459 | ceh-38 | F22D3.1 |
| WBGene00000460 | ceh-39 | T26C11.7 |
| WBGene00000461 | ceh-40 | F17A2.5 |
| WBGene00000462 | ceh-41 | T26C11.5 |
| WBGene00000463 | ceh-43 | C28A5.4 |
| WBGene00000464 | ceh-44 | Y54F10AM.4 |
| WBGene00000467 | cep-1 | F52B5.5 |
| WBGene00000468 | ces-1 | F43G9.11 |
| WBGene00000469 | ces-2 | ZK909.4 |
| WBGene00000472 | cey-1 | F33A8.3 |
| WBGene00000473 | cey-2 | F46F11.2 |
| WBGene00000474 | cey-3 | M01E11.5 |
| WBGene00000475 | cey-4 | Y39A1C.3 |
| WBGene00000476 | cfi-1 | T23D8.8 |
| WBGene00000483 | che-1 | C55B7.12 |
| WBGene00000521 | cky-1 | C15C8.2 |
| WBGene00000561 | cnd-1 | C34E10.7 |
| WBGene00000584 | cog-1 | R03C1.3 |
| WBGene00000793 | crh-1 | Y41C4A.4 |
| WBGene00000895 | dac-1 | B0412.1 |
| WBGene00000899 | daf-3 | F25E2.5 |
| WBGene00000901 | daf-5 | W01G7.1 |
| WBGene00000904 | daf-8 | R05D11.1 |
| WBGene00000908 | daf-12 | F11A1.3 |
| WBGene00000910 | daf-14 | F01G10.8 |
| WBGene00000912 | daf-16 | R13H8.1 |
| WBGene00000914 | daf-19 | F33H1.1 |
| WBGene00000936 | dbl-1 | T25F10.2 |
| WBGene00000938 | dcp-66 | C26C6.5 |
| WBGene00000995 | die-1 | C18D1.1 |
| WBGene00001029 | dnj-11 | F38A5.13 |
| WBGene00001035 | dnj-17 | T03F6.2 |
| WBGene00001061 | dpl-1 | T23G7.1 |
| WBGene00001062 | dpr-1 | R10D12.2 |
| WBGene00001079 | dpy-20 | T22B3.1 |
| WBGene00001081 | dpy-22 | F47A4.2 |
| WBGene00001092 | dro-1 | F53A2.5 |
| WBGene00001096 | dsc-1 | C18B12.3 |
| WBGene00001159 | eff-1 | C26D10.5 |
| WBGene00001161 | efl-1 | Y102A5C.18 |
| WBGene00001162 | efl-2 | Y48C3A.17 |

| | | |
|---|---|---|
| WBGene00001174 | egl-5 | C08C3.1 |
| WBGene00001182 | egl-13 | T22B7.1 |
| WBGene00001186 | egl-18 | F55A8.1 |
| WBGene00001194 | egl-27 | C04A2.3 |
| WBGene00001204 | egl-38 | C04G2.7 |
| WBGene00001207 | egl-43 | R53.3 |
| WBGene00001208 | egl-44 | F28B12.2 |
| WBGene00001210 | egl-46 | K11G9.4 |
| WBGene00001223 | ehn-3 | ZK616.10 |
| WBGene00001235 | elb-1 | Y41C4A.10 |
| WBGene00001249 | elt-1 | W09C2.1 |
| WBGene00001250 | elt-2 | C33D3.1 |
| WBGene00001251 | elt-3 | K02B9.4 |
| WBGene00001252 | elt-4 | C39B10.6 |
| WBGene00001253 | elt-6 | F52C12.5 |
| WBGene00001259 | emb-5 | T04A8.14 |
| WBGene00001310 | end-1 | F58E10.2 |
| WBGene00001311 | end-3 | F58E10.5 |
| WBGene00001324 | eor-1 | R11E3.6 |
| WBGene00001345 | fos-1 | F29G9.4 |
| WBGene00001400 | fax-1 | F56E3.4 |
| WBGene00001434 | fkh-2 | T14G12.4 |
| WBGene00001435 | fkh-3 | C29F7.4 |
| WBGene00001436 | fkh-4 | C29F7.5 |
| WBGene00001437 | fkh-5 | F26A1.2 |
| WBGene00001438 | fkh-6 | B0286.5 |
| WBGene00001439 | fkh-7 | F26D12.1 |
| WBGene00001440 | fkh-8 | F40H3.4 |
| WBGene00001441 | fkh-9 | K03C7.2 |
| WBGene00001442 | fkh-10 | C25A1.2 |
| WBGene00001493 | frm-7 | C51F7.1 |
| WBGene00001514 | xnd-1 | C05D2.5 |
| WBGene00001560 | gei-3 | T22H6.6 |
| WBGene00001565 | gei-8 | C14B9.6 |
| WBGene00001568 | snpc-4 | F32H2.1 |
| WBGene00001570 | gei-13 | F58A4.11 |
| WBGene00001574 | gei-17 | W10D5.3 |
| WBGene00001707 | grh-1 | Y48G8AR.1 |
| WBGene00001820 | ham-1 | F53B2.6 |
| WBGene00001821 | ham-2 | C07A12.1 |
| WBGene00001824 | hbl-1 | F13D11.2 |
| WBGene00001827 | hcf-1 | C46A5.9 |
| WBGene00001851 | hif-1 | F38A6.3 |

| | | |
|---|---|---|
| WBGene00001867 | him-8 | T07G12.12 |
| WBGene00001948 | hlh-1 | B0304.1 |
| WBGene00001949 | hlh-2 | M05B5.5 |
| WBGene00001950 | hlh-3 | T24B8.6 |
| WBGene00001951 | hlh-4 | T05G5.2 |
| WBGene00001952 | hlh-6 | T15H9.3 |
| WBGene00001953 | hlh-8 | C02B8.4 |
| WBGene00001954 | hlh-10 | ZK682.4 |
| WBGene00001955 | hlh-11 | F58A4.7 |
| WBGene00001956 | hlh-12 | C28C12.8 |
| WBGene00001957 | hlh-13 | F48D6.3 |
| WBGene00001958 | hlh-14 | C18A3.8 |
| WBGene00001959 | hlh-15 | C43H6.8 |
| WBGene00001960 | hlh-16 | DY3.3 |
| WBGene00001961 | hlh-17 | F38C2.2 |
| WBGene00001962 | hlh-19 | F57C12.3 |
| WBGene00001964 | hlh-25 | C17C3.7 |
| WBGene00001965 | hlh-26 | C17C3.8 |
| WBGene00001966 | hlh-27 | C17C3.10 |
| WBGene00001967 | hlh-28 | F31A3.2 |
| WBGene00001968 | hlh-29 | F31A3.4 |
| WBGene00001971 | hmg-1.1 | Y48B6A.14 |
| WBGene00001972 | hmg-1.2 | F47D12.4 |
| WBGene00001973 | hmg-3 | C32F10.5 |
| WBGene00001974 | hmg-4 | T20B12.8 |
| WBGene00001975 | hmg-5 | F45E4.9 |
| WBGene00001976 | hmg-11 | T05A7.4 |
| WBGene00001977 | hmg-12 | Y17G7A.1 |
| WBGene00001981 | hnd-1 | C44C10.8 |
| WBGene00002002 | hsb-1 | K08E7.2 |
| WBGene00002004 | hsf-1 | Y53C10A.12 |
| WBGene00002025 | hsp-60 | Y22D7AL.5 |
| WBGene00002045 | icd-1 | C56C10.8 |
| WBGene00002169 | isw-1 | F37A4.8 |
| WBGene00002245 | lag-1 | K08B4.1 |
| WBGene00002261 | ldb-1 | F58A3.1 |
| WBGene00002601 | let-381 | F26B1.7 |
| WBGene00002717 | let-526 | C01G8.9 |
| WBGene00002783 | let-607 | F57B10.1 |
| WBGene00002987 | lim-4 | ZC64.4 |
| WBGene00002988 | lim-6 | K03E6.1 |
| WBGene00002989 | lim-7 | C04F1.3 |
| WBGene00002990 | lin-1 | C37F5.1 |

| | | |
|---|---|---|
| WBGene00003000 | lin-11 | ZC247.3 |
| WBGene00003002 | lin-13 | C03B8.4 |
| WBGene00003008 | lin-22 | Y54G2A.1 |
| WBGene00003012 | lin-26 | F18A1.2 |
| WBGene00003014 | lin-28 | F02E9.2 |
| WBGene00003015 | lin-29 | W03C9.4 |
| WBGene00003017 | lin-31 | K10G6.1 |
| WBGene00003018 | lin-32 | T14F9.5 |
| WBGene00003021 | lin-36 | F44B9.6 |
| WBGene00003024 | lin-39 | C07H6.7 |
| WBGene00003025 | lin-40 | T27C4.4 |
| WBGene00003033 | lin-48 | F34D10.5 |
| WBGene00003037 | lin-54 | JC8.6 |
| WBGene00003044 | lir-1 | F18A1.3 |
| WBGene00003045 | lir-2 | F18A1.4 |
| WBGene00003046 | lir-3 | F37H8.1 |
| WBGene00003059 | lpd-2 | C48E7.3 |
| WBGene00003087 | lsy-2 | F49H12.1 |
| WBGene00003100 | mab-3 | Y53C12B.5 |
| WBGene00003102 | mab-5 | C08C3.3 |
| WBGene00003106 | mab-9 | T27A1.6 |
| WBGene00003107 | mab-10 | R166.1 |
| WBGene00003114 | mab-23 | C32C4.5 |
| WBGene00003148 | mbf-1 | H21P03.1 |
| WBGene00003163 | mdl-1 | R03E9.1 |
| WBGene00003167 | mec-3 | F01D4.6 |
| WBGene00003180 | med-1 | T24D3.1 |
| WBGene00003181 | med-2 | K04C2.6 |
| WBGene00003182 | mef-2 | W10D5.1 |
| WBGene00003210 | mel-28 | C38D4.3 |
| WBGene00003218 | mep-1 | M04B2.1 |
| WBGene00003228 | mex-1 | W03C9.7 |
| WBGene00003230 | mex-5 | W02A2.7 |
| WBGene00003231 | mex-6 | AH6.5 |
| WBGene00003233 | mgl-2 | F45H11.4 |
| WBGene00003241 | mig-5 | T05C12.6 |
| WBGene00003376 | mls-1 | H14A12.4 |
| WBGene00003377 | mls-2 | C39E6.4 |
| WBGene00003378 | mml-1 | T20B12.6 |
| WBGene00003380 | mnm-2 | C10A4.8 |
| WBGene00003388 | moe-3 | F32A11.6 |
| WBGene00003406 | mrg-1 | Y37D8A.9 |
| WBGene00003480 | klf-3 | F54H5.4 |

| | | |
|---|---|---|
| WBGene00003509 | mxl-1 | T19B10.11 |
| WBGene00003510 | mxl-2 | F40G9.11 |
| WBGene00003511 | mxl-3 | F46G10.6 |
| WBGene00003592 | nfi-1 | ZK1290.4 |
| WBGene00003595 | ngn-1 | Y69A2AR.29 |
| WBGene00003600 | nhr-1 | R09G11.2 |
| WBGene00003601 | nhr-2 | C32F10.6 |
| WBGene00003602 | nhr-3 | H01A20.1 |
| WBGene00003603 | nhr-4 | F32B6.1 |
| WBGene00003604 | nhr-5 | Y73F8A.21 |
| WBGene00003605 | nhr-6 | C48D5.1 |
| WBGene00003606 | nhr-7 | F54D1.4 |
| WBGene00003607 | nhr-8 | F33D4.1 |
| WBGene00003608 | nhr-9 | ZK418.1 |
| WBGene00003609 | nhr-10 | B0280.8 |
| WBGene00003610 | nhr-11 | ZC410.1 |
| WBGene00003611 | nhr-12 | R04B5.4 |
| WBGene00003612 | nhr-13 | Y5H2B.2 |
| WBGene00003613 | nhr-14 | T01B10.4 |
| WBGene00003614 | nhr-15 | F33E11.1 |
| WBGene00003615 | nhr-16 | T12C9.6 |
| WBGene00003616 | nhr-17 | C02B4.2 |
| WBGene00003617 | nhr-18 | F44C8.3 |
| WBGene00003618 | nhr-19 | E02H1.7 |
| WBGene00003619 | nhr-20 | F43C1.4 |
| WBGene00003620 | nhr-21 | F21D12.1 |
| WBGene00003621 | nhr-22 | K06A1.4 |
| WBGene00003622 | nhr-23 | C01H6.5 |
| WBGene00003623 | nhr-25 | F11C1.6 |
| WBGene00003624 | nhr-28 | C11G6.4 |
| WBGene00003625 | nhr-31 | C26B2.3 |
| WBGene00003626 | nhr-32 | K08H2.8 |
| WBGene00003627 | nhr-34 | F58G6.5 |
| WBGene00003628 | nhr-35 | C07A12.3 |
| WBGene00003629 | nhr-38 | K01H12.3 |
| WBGene00003630 | nhr-40 | T03G6.2 |
| WBGene00003632 | nhr-42 | C33G8.6 |
| WBGene00003633 | nhr-43 | C29E6.5 |
| WBGene00003634 | nhr-44 | T19A5.4 |
| WBGene00003635 | nhr-45 | F16H11.5 |
| WBGene00003636 | nhr-46 | C45E5.6 |
| WBGene00003637 | nhr-47 | C24G6.4 |
| WBGene00003638 | nhr-48 | ZK662.3 |

| | | |
|---|---|---|
| WBGene00003639 | nhr-49 | K10C3.6 |
| WBGene00003640 | nhr-50 | C06C6.5 |
| WBGene00003641 | nhr-51 | K06B4.1 |
| WBGene00003642 | nhr-52 | K06B4.2 |
| WBGene00003643 | nhr-53 | K06B4.11 |
| WBGene00003644 | nhr-54 | F36D3.2 |
| WBGene00003645 | nhr-55 | T01G6.7 |
| WBGene00003646 | nhr-56 | F44C8.6 |
| WBGene00003647 | nhr-57 | T05B4.2 |
| WBGene00003648 | nhr-58 | R11G11.2 |
| WBGene00003649 | nhr-59 | T27B7.1 |
| WBGene00003650 | nhr-60 | F57A10.5 |
| WBGene00003651 | nhr-61 | W01D2.2 |
| WBGene00003652 | nhr-62 | Y67A6A.2 |
| WBGene00003653 | nhr-63 | C06C6.4 |
| WBGene00003654 | nhr-64 | C45E1.1 |
| WBGene00003655 | nhr-65 | Y17D7A.3 |
| WBGene00003656 | nhr-66 | T09A12.4 |
| WBGene00003657 | nhr-67 | C08F8.8 |
| WBGene00003658 | nhr-68 | H12C20.3 |
| WBGene00003659 | nhr-69 | T23H4.2 |
| WBGene00003660 | nhr-70 | Y51A2D.17 |
| WBGene00003661 | nhr-71 | K11E4.5 |
| WBGene00003662 | nhr-72 | C17A2.8 |
| WBGene00003663 | nhr-73 | C27C7.4 |
| WBGene00003664 | nhr-74 | C27C7.3 |
| WBGene00003665 | nhr-75 | C49D10.6 |
| WBGene00003667 | nhr-77 | T15D6.6 |
| WBGene00003668 | nhr-78 | F36A4.14 |
| WBGene00003669 | nhr-79 | T26H2.9 |
| WBGene00003670 | nhr-80 | H10E21.3 |
| WBGene00003671 | nhr-81 | C47F8.8 |
| WBGene00003672 | nhr-82 | F41D3.1 |
| WBGene00003673 | nhr-83 | F48G7.3 |
| WBGene00003674 | nhr-84 | T06C12.7 |
| WBGene00003675 | nhr-85 | W05B5.3 |
| WBGene00003676 | nhr-86 | Y40B10A.8 |
| WBGene00003677 | nhr-87 | Y41D4B.7 |
| WBGene00003678 | nhr-88 | K08A2.5 |
| WBGene00003679 | nhr-89 | E03H4.13 |
| WBGene00003680 | nhr-90 | ZK488.2 |
| WBGene00003681 | nhr-91 | Y15E3A.1 |
| WBGene00003682 | nhr-92 | Y41D4B.8 |

| | | |
|---|---|---|
| WBGene00003683 | nhr-93 | T24A6.9 |
| WBGene00003684 | nhr-94 | C12D5.8 |
| WBGene00003685 | nhr-95 | Y39B6A.17 |
| WBGene00003686 | nhr-96 | F44C8.11 |
| WBGene00003687 | nhr-97 | H27C11.1 |
| WBGene00003688 | nhr-98 | M02H5.6 |
| WBGene00003689 | nhr-99 | M02H5.1 |
| WBGene00003690 | nhr-100 | C28D4.1 |
| WBGene00003691 | nhr-101 | H12C20.6 |
| WBGene00003692 | nhr-102 | T06C12.6 |
| WBGene00003693 | nhr-103 | F44C8.4 |
| WBGene00003694 | nhr-104 | R11E3.5 |
| WBGene00003695 | nhr-105 | C06G3.1 |
| WBGene00003696 | nhr-106 | T01G6.4 |
| WBGene00003697 | nhr-107 | C33G8.11 |
| WBGene00003698 | nhr-108 | F35E8.12 |
| WBGene00003699 | nhr-109 | T12C9.5 |
| WBGene00003700 | nhr-110 | Y46H3D.5 |
| WBGene00003701 | nhr-111 | F44G3.9 |
| WBGene00003702 | nhr-112 | Y70C5C.6 |
| WBGene00003703 | nhr-113 | ZK1025.9 |
| WBGene00003704 | nhr-114 | Y45G5AM.1 |
| WBGene00003705 | nhr-115 | T27B7.4 |
| WBGene00003706 | nhr-116 | F09C6.9 |
| WBGene00003707 | nhr-117 | F16B4.12 |
| WBGene00003708 | nhr-118 | F13A2.8 |
| WBGene00003709 | nhr-119 | K12H6.1 |
| WBGene00003710 | nhr-120 | C25B8.6 |
| WBGene00003711 | nhr-121 | E02H9.8 |
| WBGene00003712 | nhr-122 | Y41D4B.9 |
| WBGene00003713 | nhr-123 | M02H5.7 |
| WBGene00003714 | nhr-124 | C17E7.8 |
| WBGene00003715 | nhr-125 | R02D1.1 |
| WBGene00003716 | nhr-126 | F44C8.10 |
| WBGene00003717 | nhr-127 | T13F3.3 |
| WBGene00003718 | nhr-128 | F44C8.5 |
| WBGene00003719 | nhr-129 | C50B6.14 |
| WBGene00003720 | nhr-130 | T01G6.8 |
| WBGene00003721 | nhr-131 | T01G6.2 |
| WBGene00003722 | nhr-132 | R11G11.1 |
| WBGene00003723 | nhr-133 | F44C8.8 |
| WBGene00003724 | nhr-134 | F44C8.2 |
| WBGene00003725 | nhr-135 | VC5.5 |

| | | |
|---|---|---|
| WBGene00003726 | nhr-136 | C13C4.3 |
| WBGene00003727 | nhr-137 | C56E10.4 |
| WBGene00003728 | nhr-138 | C28D4.9 |
| WBGene00003779 | nob-1 | Y75B8A.2 |
| WBGene00003834 | nxf-1 | C15H11.3 |
| WBGene00003845 | odd-1 | B0280.4 |
| WBGene00003846 | odd-2 | C34H3.2 |
| WBGene00003847 | blmp-1 | F25D7.3 |
| WBGene00003854 | odr-7 | T18D3.2 |
| WBGene00003864 | oma-1 | C09G9.6 |
| WBGene00003865 | oma-2 | ZC513.6 |
| WBGene00003909 | pag-3 | F45B8.4 |
| WBGene00003912 | pal-1 | C38D4.6 |
| WBGene00003933 | pat-9 | T27B1.2 |
| WBGene00003937 | pax-1 | K07C11.1 |
| WBGene00003938 | pax-2 | K06B9.5 |
| WBGene00003939 | pax-3 | F27E5.2 |
| WBGene00003968 | peb-1 | T14F9.4 |
| WBGene00003976 | pes-1 | T28H11.4 |
| WBGene00004011 | pha-2 | M6.3 |
| WBGene00004013 | pha-4 | F38A6.1 |
| WBGene00004024 | php-3 | Y75B8A.1 |
| WBGene00004027 | pie-1 | Y49E10.14 |
| WBGene00004046 | plp-1 | F45E4.2 |
| WBGene00004048 | plx-2 | K04B12.1 |
| WBGene00004077 | pop-1 | W10C8.2 |
| WBGene00004078 | pos-1 | F52E1.1 |
| WBGene00004096 | pqm-1 | F40F8.7 |
| WBGene00004106 | pqn-15 | C24A8.3 |
| WBGene00004111 | pqn-21 | C37A2.5 |
| WBGene00004134 | myrf-1 | F59B10.1 |
| WBGene00004136 | pqn-51 | K11D12.2 |
| WBGene00004157 | pqn-75 | W03D2.1 |
| WBGene00004203 | swsn-1 | Y113G7B.23 |
| WBGene00004204 | swsn-4 | F01G4.1 |
| WBGene00004319 | rbr-2 | ZK593.4 |
| WBGene00004334 | ref-1 | T01E8.2 |
| WBGene00004335 | ref-2 | C47C12.3 |
| WBGene00004393 | rnt-1 | B0414.2 |
| WBGene00004703 | rsp-6 | C33H5.12 |
| WBGene00004735 | sbp-1 | Y47D3B.7 |
| WBGene00004745 | sdc-1 | F52E10.1 |
| WBGene00004747 | sdc-3 | C25D7.3 |

| | | |
|---|---|---|
| WBGene00004750 | sea-1 | F19B10.9 |
| WBGene00004751 | sea-2 | K10G6.3 |
| WBGene00004771 | sem-2 | C32E12.5 |
| WBGene00004773 | sem-4 | F15C11.1 |
| WBGene00004786 | sex-1 | F44A6.2 |
| WBGene00004803 | sir-2.4 | C06A5.11 |
| WBGene00004804 | skn-1 | T19E7.2 |
| WBGene00004856 | sma-2 | ZK370.2 |
| WBGene00004857 | sma-3 | R13F6.9 |
| WBGene00004858 | sma-4 | R12B2.1 |
| WBGene00004860 | sma-6 | C32D5.2 |
| WBGene00004862 | sma-9 | T05A10.1 |
| WBGene00004911 | snf-12 | T25B6.7 |
| WBGene00004949 | sox-2 | K08A8.2 |
| WBGene00004950 | sox-3 | F40E10.2 |
| WBGene00004964 | spe-10 | AC3.10 |
| WBGene00005006 | spr-1 | D1014.8 |
| WBGene00005008 | spr-3 | C07A12.5 |
| WBGene00005009 | spr-4 | C09H6.1 |
| WBGene00005011 | | F26F4.8 |
| WBGene00005013 | jmjd-1.1 | F43G6.6 |
| WBGene00005014 | spt-4 | F54C4.2 |
| WBGene00005015 | spt-5 | K08E4.1 |
| WBGene00006342 | sup-37 | C01B7.1 |
| WBGene00006379 | sys-1 | T23D8.9 |
| WBGene00006380 | tab-1 | F31E8.3 |
| WBGene00006382 | taf-1 | W04A8.7 |
| WBGene00006383 | taf-2 | Y37E11B.4 |
| WBGene00006384 | taf-3 | C11G6.1 |
| WBGene00006385 | taf-4 | R119.6 |
| WBGene00006386 | taf-5 | F30F8.8 |
| WBGene00006387 | taf-6.1 | W09B6.2 |
| WBGene00006388 | taf-7.1 | F54F7.1 |
| WBGene00006389 | taf-7.2 | Y111B2A.16 |
| WBGene00006390 | taf-8 | ZK1320.12 |
| WBGene00006391 | taf-9 | T12D8.7 |
| WBGene00006392 | taf-10 | K03B4.3 |
| WBGene00006393 | taf-11.1 | F48D6.1 |
| WBGene00006394 | taf-11.2 | K10D3.3 |
| WBGene00006395 | taf-11.3 | F43D9.5 |
| WBGene00006396 | taf-12 | Y56A3A.4 |
| WBGene00006397 | taf-13 | C14A4.10 |
| WBGene00006424 | ctbp-1 | F49E10.5 |

| | | |
|---|---|---|
| WBGene00006445 | tag-68 | F37D6.6 |
| WBGene00006462 | tag-97 | C33A11.4 |
| WBGene00006471 | nhr-233 | Y32B12B.6 |
| WBGene00006474 | wdr-5.1 | C14B1.4 |
| WBGene00006476 | rhgf-2 | T08H4.1 |
| WBGene00006492 | let-391 | C27A12.3 |
| WBGene00006542 | tbp-1 | T20B12.2 |
| WBGene00006543 | tbx-2 | F21H11.3 |
| WBGene00006544 | tbx-7 | ZK328.8 |
| WBGene00006545 | tbx-8 | T07C4.2 |
| WBGene00006546 | tbx-9 | T07C4.6 |
| WBGene00006547 | tbx-11 | F40H6.4 |
| WBGene00006549 | tbx-30 | Y59E9AR.3 |
| WBGene00006550 | tbx-31 | C36C9.2 |
| WBGene00006551 | tbx-32 | ZK380.1 |
| WBGene00006552 | tbx-33 | Y66A7A.8 |
| WBGene00006553 | tbx-34 | Y47D3A.10 |
| WBGene00006554 | tbx-35 | ZK177.10 |
| WBGene00006555 | tbx-36 | ZK829.5 |
| WBGene00006556 | tbx-37 | Y47D3A.12 |
| WBGene00006557 | tbx-38 | C24H11.3 |
| WBGene00006558 | tbx-39 | Y73F8A.16 |
| WBGene00006559 | tbx-40 | Y73F8A.17 |
| WBGene00006560 | tbx-41 | T26C11.1 |
| WBGene00006577 | tlf-1 | F39H11.2 |
| WBGene00006580 | tlp-1 | T23G4.1 |
| WBGene00006604 | tra-1 | Y47D3A.6 |
| WBGene00006648 | ttb-1 | W03F9.5 |
| WBGene00006652 | ttx-1 | Y113G7A.6 |
| WBGene00006654 | ttx-3 | C40H5.5 |
| WBGene00006698 | uaf-2 | Y116A8C.35 |
| WBGene00006706 | ubc-9 | F29B9.6 |
| WBGene00006726 | ubl-5 | F46F11.4 |
| WBGene00006739 | ulp-4 | C41C4.6 |
| WBGene00006743 | unc-3 | Y16B4A.1 |
| WBGene00006744 | unc-4 | F26C11.2 |
| WBGene00006766 | unc-30 | B0564.10 |
| WBGene00006773 | unc-37 | W02D3.9 |
| WBGene00006775 | unc-39 | F56A12.1 |
| WBGene00006778 | unc-42 | F58E6.10 |
| WBGene00006790 | unc-55 | F55D12.4 |
| WBGene00006796 | unc-62 | T28F12.2 |
| WBGene00006818 | unc-86 | C30A5.7 |

| | | |
|---|---|---|
| WBGene00006827 | unc-98 | F08C6.7 |
| WBGene00006844 | unc-120 | D1081.2 |
| WBGene00006853 | unc-130 | C47G2.2 |
| WBGene00006870 | vab-3 | F14F3.1 |
| WBGene00006873 | vab-7 | M142.4 |
| WBGene00006881 | vab-15 | R07B1.1 |
| WBGene00006943 | wrm-1 | B0336.1 |
| WBGene00006959 | xbp-1 | R74.3 |
| WBGene00006970 | zag-1 | F28F9.1 |
| WBGene00006986 | zip-1 | Y75B8A.35 |
| WBGene00007042 | pbrm-1 | C26C6.1 |
| WBGene00007048 | nfx-1 | C16A3.7 |
| WBGene00007053 | chd-7 | T04D1.4 |
| WBGene00007058 | dmd-6 | F13G11.1 |
| WBGene00007063 | | 2L52.1 |
| WBGene00007094 | | B0019.2 |
| WBGene00007105 | | B0035.1 |
| WBGene00007121 | | B0250.4 |
| WBGene00007195 | thoc-7 | B0513.2 |
| WBGene00007223 | | C01F6.9 |
| WBGene00007242 | madf-5 | C01G12.1 |
| WBGene00007367 | nhr-150 | C06B8.1 |
| WBGene00007387 | jmjd-5 | C06H2.3 |
| WBGene00007416 | ceh-57 | C07E3.5 |
| WBGene00007417 | ceh-58 | C07E3.6 |
| WBGene00007433 | swsn-7 | C08B11.3 |
| WBGene00007496 | npax-4 | C09G9.7 |
| WBGene00007546 | nhr-153 | C13C4.1 |
| WBGene00007547 | nhr-154 | C13C4.2 |
| WBGene00007625 | | C16C10.4 |
| WBGene00007637 | dhhc-7 | C17D12.1 |
| WBGene00007645 | | C17E4.6 |
| WBGene00007732 | spe-44 | C25G4.4 |
| WBGene00007749 | ceh-79 | C26E1.3 |
| WBGene00007761 | rad-26 | C27B7.4 |
| WBGene00007772 | egrh-1 | C27C12.2 |
| WBGene00007776 | dmd-4 | C27C12.6 |
| WBGene00007813 | jmjd-3.3 | C29F7.6 |
| WBGene00007877 | nfki-1 | C33A11.1 |
| WBGene00007907 | | C34B4.2 |
| WBGene00007929 | dmd-10 | C34D1.1 |
| WBGene00007930 | dmd-11 | C34D1.2 |
| WBGene00007932 | zip-5 | C34D1.5 |

| | | |
|---|---|---|
| WBGene00007943 | | C34F6.9 |
| WBGene00007961 | | C35D6.4 |
| WBGene00007980 | | C36E8.1 |
| WBGene00007984 | irx-1 | C36F7.1 |
| WBGene00008007 | | C38D4.7 |
| WBGene00008056 | nhr-164 | C41G6.5 |
| WBGene00008081 | athp-1 | C44B9.4 |
| WBGene00008092 | gmeb-3 | C44F1.2 |
| WBGene00008118 | madf-8 | C46F11.3 |
| WBGene00008158 | nhr-165 | C47F8.2 |
| WBGene00008195 | ceh-88 | C49C3.5 |
| WBGene00008208 | nhr-167 | C49F5.4 |
| WBGene00008209 | | C49F5.5 |
| WBGene00008221 | nhr-168 | C50B6.8 |
| WBGene00008242 | ceh-75 | C50H2.6 |
| WBGene00008289 | nhr-169 | C54C8.1 |
| WBGene00008309 | nhr-170 | C54E10.5 |
| WBGene00008338 | pafo-1 | C55A6.9 |
| WBGene00008363 | | D1046.2 |
| WBGene00008386 | cdc-5L | D1081.8 |
| WBGene00008417 | | D2030.7 |
| WBGene00008419 | wdr-23 | D2030.9 |
| WBGene00008474 | nhr-174 | E03H4.6 |
| WBGene00008606 | dhhc-1 | F09B12.2 |
| WBGene00008619 | nhr-262 | F09C6.8 |
| WBGene00008630 | nhr-175 | F09F3.10 |
| WBGene00008640 | | F10B5.3 |
| WBGene00008683 | repo-1 | F11A10.2 |
| WBGene00008748 | yap-1 | F13E6.4 |
| WBGene00008762 | ztf-2 | F13G3.1 |
| WBGene00008778 | nhr-264 | F14A5.1 |
| WBGene00008830 | nhr-176 | F14H3.11 |
| WBGene00008882 | | F16B12.6 |
| WBGene00008884 | nhr-281 | F16B12.8 |
| WBGene00008901 | nhr-27 | F16H9.2 |
| WBGene00008976 | plp-2 | F20D1.4 |
| WBGene00008999 | myrf-2 | F21A10.2 |
| WBGene00009014 | | F21D5.9 |
| WBGene00009026 | | F21G4.5 |
| WBGene00009050 | | F22D6.2 |
| WBGene00009071 | | F23A7.6 |
| WBGene00009084 | | F23B12.7 |
| WBGene00009089 | jmjd-3.2 | F23D12.5 |

| | | |
|---|---|---|
| WBGene00009133 | bed-3 | F25H8.6 |
| WBGene00009174 | | F26H9.2 |
| WBGene00009180 | nurf-1 | F26H11.2 |
| WBGene00009188 | lsy-22 | F27D4.2 |
| WBGene00009189 | | F27D4.4 |
| WBGene00009190 | | F27D4.6 |
| WBGene00009202 | aptf-4 | F28C6.1 |
| WBGene00009203 | aptf-3 | F28C6.2 |
| WBGene00009224 | | F28F8.7 |
| WBGene00009231 | ceh-89 | F28H6.2 |
| WBGene00009341 | thoc-3 | F32H2.4 |
| WBGene00009365 | | F33H1.4 |
| WBGene00009448 | zfp-2 | F35H8.3 |
| WBGene00009461 | madf-2 | F36D1.1 |
| WBGene00009475 | | F36F2.1 |
| WBGene00009508 | row-1 | F37D6.2 |
| WBGene00009532 | ccch-1 | F38B7.1 |
| WBGene00009537 | ccch-2 | F38C2.5 |
| WBGene00009539 | | F38C2.7 |
| WBGene00009540 | hlh-31 | F38C2.8 |
| WBGene00009553 | hinf-1 | F39B2.1 |
| WBGene00009560 | psa-3 | F39D8.2 |
| WBGene00009584 | drap-1 | F40F9.7 |
| WBGene00009608 | nhr-265 | F41D3.3 |
| WBGene00009650 | ccnk-1 | F43D2.1 |
| WBGene00009672 | | F43G9.12 |
| WBGene00009687 | | F44D12.10 |
| WBGene00009743 | sptf-1 | F45H11.1 |
| WBGene00009772 | ztf-7 | F46B6.7 |
| WBGene00009827 | hmg-6 | F47G4.6 |
| WBGene00009834 | lsy-27 | F47H4.1 |
| WBGene00009899 | efl-3 | F49E12.6 |
| WBGene00009923 | | F52B5.7 |
| WBGene00009937 | lsl-1 | F52F12.4 |
| WBGene00009939 | ztf-11 | F52F12.6 |
| WBGene00009965 | frpr-9 | F53B7.2 |
| WBGene00009998 | klf-2 | F53F8.1 |
| WBGene00010010 | gmeb-2 | F53H4.5 |
| WBGene00010012 | saeg-1 | F53H10.2 |
| WBGene00010054 | gtf-2E2 | F54D5.11 |
| WBGene00010086 | | F55B11.4 |
| WBGene00010137 | ztf-26 | F55H12.6 |
| WBGene00010180 | nhr-192 | F57A8.5 |

| | | |
|---|---|---|
| WBGene00010186 | nhr-283 | F57A10.6 |
| WBGene00010215 | nhr-193 | F57G8.6 |
| WBGene00010251 | sta-2 | F58E6.1 |
| WBGene00010264 | | F58G1.2 |
| WBGene00010323 | dhhc-12 | F59C6.2 |
| WBGene00010401 | | H16D19.3 |
| WBGene00010406 | math-33 | H19N07.2 |
| WBGene00010410 | nhr-267 | H22D14.1 |
| WBGene00010453 | fozi-1 | K01B6.1 |
| WBGene00010476 | rnf-113 | K01G5.1 |
| WBGene00010553 | | K04C1.3 |
| WBGene00010600 | nhr-196 | K06B4.5 |
| WBGene00010601 | nhr-268 | K06B4.6 |
| WBGene00010602 | nhr-197 | K06B4.7 |
| WBGene00010603 | nhr-198 | K06B4.8 |
| WBGene00010604 | nhr-199 | K06B4.10 |
| WBGene00010704 | | K09A11.1 |
| WBGene00010770 | | K11D2.4 |
| WBGene00010781 | | K11H3.4 |
| WBGene00010868 | somi-1 | M04G12.4 |
| WBGene00010892 | dhhc-6 | M18.8 |
| WBGene00010936 | ztf-14 | M163.2 |
| WBGene00010990 | tceb-3 | R03D7.4 |
| WBGene00010995 | ceh-90 | R03E1.4 |
| WBGene00011002 | nhr-205 | R04B5.3 |
| WBGene00011066 | ztf-15 | R06C7.9 |
| WBGene00011069 | ceh-62 | R06F6.6 |
| WBGene00011097 | nhr-206 | R07B7.13 |
| WBGene00011098 | nhr-207 | R07B7.14 |
| WBGene00011099 | nhr-208 | R07B7.15 |
| WBGene00011100 | nhr-209 | R07B7.16 |
| WBGene00011113 | | R07E5.5 |
| WBGene00011130 | zip-6 | R07H5.10 |
| WBGene00011150 | nhr-269 | R08H2.9 |
| WBGene00011206 | | R10E4.11 |
| WBGene00011257 | npax-3 | R13.2 |
| WBGene00011259 | miz-1 | R13.4 |
| WBGene00011315 | mbr-1 | T01C1.2 |
| WBGene00011319 | | T01C3.2 |
| WBGene00011327 | hlh-34 | T01D3.2 |
| WBGene00011368 | tftc-5 | T02C12.3 |
| WBGene00011376 | gla-3 | T02E1.3 |
| WBGene00011396 | nhr-271 | T03E6.3 |

| | | |
|---|---|---|
| WBGene00011505 | pzf-1 | T05G11.1 |
| WBGene00011520 | nhr-213 | T06C12.13 |
| WBGene00011547 | | T06G6.5 |
| WBGene00011563 | jmjd-4 | T07C4.11 |
| WBGene00011565 | nhr-272 | T07C5.2 |
| WBGene00011566 | nhr-214 | T07C5.3 |
| WBGene00011567 | nhr-215 | T07C5.4 |
| WBGene00011568 | nhr-26 | T07C5.5 |
| WBGene00011575 | madf-4 | T07C12.11 |
| WBGene00011583 | | T07D10.3 |
| WBGene00011597 | zim-1 | T07G12.6 |
| WBGene00011600 | zim-2 | T07G12.10 |
| WBGene00011601 | zim-3 | T07G12.11 |
| WBGene00011614 | nfya-1 | T08D10.1 |
| WBGene00011615 | lsd-1 | T08D10.2 |
| WBGene00011626 | | T08G5.7 |
| WBGene00011639 | ztf-17 | T09A5.12 |
| WBGene00011651 | nhr-217 | T09E11.2 |
| WBGene00011661 | ztf-27 | T09F3.1 |
| WBGene00011696 | eea-1 | T10G3.5 |
| WBGene00011722 | | T11G6.8 |
| WBGene00011729 | set-16 | T12D8.1 |
| WBGene00011743 | | T13F2.2 |
| WBGene00011750 | nhr-218 | T13F3.2 |
| WBGene00011814 | gtf-2H2C | T16H12.4 |
| WBGene00011824 | | T18D3.7 |
| WBGene00011924 | | T22C8.3 |
| WBGene00011925 | | T22C8.4 |
| WBGene00011926 | sptf-2 | T22C8.5 |
| WBGene00011956 | | T23F11.4 |
| WBGene00011964 | saeg-2 | T23G5.6 |
| WBGene00012000 | | T24H10.1 |
| WBGene00012005 | jun-1 | T24H10.7 |
| WBGene00012050 | nhr-223 | T26E4.8 |
| WBGene00012056 | nhr-285 | T26E4.16 |
| WBGene00012074 | | T27A8.2 |
| WBGene00012101 | zip-10 | T27F2.4 |
| WBGene00012209 | hmg-20 | W02D9.3 |
| WBGene00012243 | | W04D2.4 |
| WBGene00012277 | ccch-3 | W05B10.2 |
| WBGene00012317 | ztf-6 | W06H12.1 |
| WBGene00012330 | zip-3 | W07G1.3 |
| WBGene00012385 | | Y5F2A.4 |

| | | |
|---|---|---|
| WBGene00012405 | ztf-25 | Y6G8.3 |
| WBGene00012435 | flh-1 | Y11D7A.12 |
| WBGene00012436 | flh-3 | Y11D7A.13 |
| WBGene00012446 | nhr-230 | Y17D7A.1 |
| WBGene00012449 | nhr-231 | Y17D7B.1 |
| WBGene00012473 | | Y17G7B.22 |
| WBGene00012474 | attf-6 | Y18D10A.1 |
| WBGene00012494 | nhr-232 | Y22F5A.1 |
| WBGene00012578 | ccct-1 | Y37H9A.3 |
| WBGene00012584 | ceh-100 | Y38E10A.6 |
| WBGene00012596 | nhr-234 | Y38E10A.18 |
| WBGene00012639 | fezf-1 | Y38H8A.5 |
| WBGene00012644 | rabx-5 | Y39A1A.5 |
| WBGene00012674 | bed-1 | Y39B6A.12 |
| WBGene00012694 | gtf-2F2 | Y39B6A.36 |
| WBGene00012702 | ztf-20 | Y39B6A.46 |
| WBGene00012703 | nhr-145 | Y39B6A.47 |
| WBGene00012715 | snpc-1.2 | Y39E4B.2 |
| WBGene00012718 | dhhc-8 | Y39E4B.7 |
| WBGene00012735 | sptf-3 | Y40B1A.4 |
| WBGene00012832 | dmd-3 | Y43F8C.10 |
| WBGene00012943 | bed-2 | Y47D3B.9 |
| WBGene00012948 | dhhc-2 | Y47H9C.2 |
| WBGene00012974 | | Y48A6C.1 |
| WBGene00012976 | sup-35 | Y48A6C.3 |
| WBGene00012988 | ztf-22 | Y48C3A.4 |
| WBGene00012994 | | Y48C3A.12 |
| WBGene00013006 | lin-38 | Y48E1B.7 |
| WBGene00013096 | mcd-1 | Y51H1A.6 |
| WBGene00013100 | zip-7 | Y51H4A.4 |
| WBGene00013111 | sta-1 | Y51H4A.17 |
| WBGene00013128 | dxbp-1 | Y52B11A.9 |
| WBGene00013134 | hsf-2 | Y53C10A.3 |
| WBGene00013147 | eyg-1 | Y53C12C.1 |
| WBGene00013150 | | Y53F4B.3 |
| WBGene00013152 | | Y53F4B.5 |
| WBGene00013178 | | Y53H1A.2 |
| WBGene00013219 | elof-1 | Y54G11A.11 |
| WBGene00013225 | | Y56A3A.2 |
| WBGene00013236 | | Y56A3A.18 |
| WBGene00013240 | | Y56A3A.28 |
| WBGene00013270 | | Y57A10A.31 |
| WBGene00013319 | ccch-5 | Y57G11C.25 |

| | | |
|---|---|---|
| WBGene00013350 | slr-2 | Y59A8B.13 |
| WBGene00013370 | | Y60A9.3 |
| WBGene00013383 | aptf-2 | Y62E10A.17 |
| WBGene00013425 | ceh-91 | Y66A7A.5 |
| WBGene00013438 | ztf-29 | Y66D12A.12 |
| WBGene00013441 | xpb-1 | Y66D12A.15 |
| WBGene00013465 | | Y67H2A.10 |
| WBGene00013483 | nhr-241 | Y69H2.8 |
| WBGene00013505 | | Y71A12B.8 |
| WBGene00013512 | nhr-276 | Y71A12C.1 |
| WBGene00013529 | gft-2H4 | Y73F8A.24 |
| WBGene00013537 | | Y73F8A.33 |
| WBGene00013560 | zip-12 | Y75B8A.29 |
| WBGene00013580 | | Y79H2A.3 |
| WBGene00013583 | ceh-51 | Y80D3A.3 |
| WBGene00013584 | nhr-243 | Y80D3A.4 |
| WBGene00013595 | atg-4.1 | Y87G2A.3 |
| WBGene00013632 | | Y105C5A.1 |
| WBGene00013639 | | Y105C5A.15 |
| WBGene00013665 | hlh-32 | Y105C5B.29 |
| WBGene00013676 | ekl-4 | Y105E8A.17 |
| WBGene00013717 | madf-10 | Y106G6H.4 |
| WBGene00013734 | | Y111B2A.10 |
| WBGene00013736 | gtf-2A2 | Y111B2A.13 |
| WBGene00013738 | elpc-2 | Y111B2A.17 |
| WBGene00013794 | dct-13 | Y116A8C.17 |
| WBGene00013795 | nhr-229 | Y116A8C.18 |
| WBGene00013796 | | Y116A8C.19 |
| WBGene00013797 | | Y116A8C.20 |
| WBGene00013799 | athp-3 | Y116A8C.22 |
| WBGene00013876 | ceh-74 | ZC376.4 |
| WBGene00013878 | atfs-1 | ZC376.7 |
| WBGene00013940 | nhr-248 | ZK218.6 |
| WBGene00013966 | ztf-9 | ZK287.6 |
| WBGene00013970 | klu-1 | ZK337.2 |
| WBGene00013976 | nhr-33 | ZK455.6 |
| WBGene00013998 | gtf-2E1 | ZK550.4 |
| WBGene00014060 | | ZK673.4 |
| WBGene00014068 | nhr-255 | ZK678.2 |
| WBGene00014075 | dhhc-4 | ZK757.4 |
| WBGene00014114 | tftc-3 | ZK856.13 |
| WBGene00014131 | sdz-38 | ZK892.7 |
| WBGene00014189 | nhr-245 | ZK1025.10 |

| | | |
|---|---|---|
| WBGene00014193 | nhr-247 | ZK1037.5 |
| WBGene00014208 | | ZK1067.2 |
| WBGene00014230 | gtf-2H3 | ZK1128.4 |
| WBGene00015091 | | B0261.1 |
| WBGene00015138 | | B0310.2 |
| WBGene00015143 | | B0336.3 |
| WBGene00015147 | | B0336.7 |
| WBGene00015150 | | B0336.13 |
| WBGene00015285 | gmeb-1 | C01B12.2 |
| WBGene00015296 | gtf-2F1 | C01F1.1 |
| WBGene00015352 | | C02F5.12 |
| WBGene00015395 | nhr-147 | C03G6.8 |
| WBGene00015396 | nhr-148 | C03G6.10 |
| WBGene00015397 | nhr-149 | C03G6.12 |
| WBGene00015447 | srab-2 | C04F5.5 |
| WBGene00015451 | | C04F5.9 |
| WBGene00015468 | madf-11 | C05D2.6 |
| WBGene00015477 | attf-4 | C05D10.1 |
| WBGene00015497 | nhr-76 | C05G6.1 |
| WBGene00015523 | ztf-30 | C06E1.8 |
| WBGene00015527 | | C06E2.1 |
| WBGene00015555 | ufl-1 | C06G3.9 |
| WBGene00015620 | | C08G9.2 |
| WBGene00015649 | | C09F5.3 |
| WBGene00015651 | ceh-53 | C09G12.1 |
| WBGene00015705 | nhr-152 | C12D5.2 |
| WBGene00015716 | sox-4 | C12D12.5 |
| WBGene00015758 | nhr-155 | C14C6.4 |
| WBGene00015806 | smrc-1 | C16A3.1 |
| WBGene00015809 | | C16A3.4 |
| WBGene00015813 | thoc-2 | C16A3.8 |
| WBGene00015869 | nhr-257 | C17A2.1 |
| WBGene00015897 | nhr-156 | C17E7.1 |
| WBGene00015900 | nhr-157 | C17E7.5 |
| WBGene00015901 | nhr-158 | C17E7.6 |
| WBGene00015902 | nhr-159 | C17E7.7 |
| WBGene00015934 | ceh-48 | C17H12.9 |
| WBGene00015971 | swsn-2.2 | C18E3.2 |
| WBGene00015981 | elt-7 | C18G1.2 |
| WBGene00016029 | elf-1 | C24A1.2 |
| WBGene00016091 | nhr-30 | C25E10.1 |
| WBGene00016126 | nhr-258 | C26B2.4 |
| WBGene00016138 | flh-2 | C26E6.2 |

| | | |
|---|---|---|
| WBGene00016154 | | C27A12.2 |
| WBGene00016162 | crh-2 | C27D6.4 |
| WBGene00016189 | | C28G1.4 |
| WBGene00016200 | dpff-1 | C28H8.9 |
| WBGene00016220 | | C29F9.5 |
| WBGene00016233 | srt-58 | C29G2.5 |
| WBGene00016364 | nhr-161 | C33G8.7 |
| WBGene00016365 | nhr-139 | C33G8.8 |
| WBGene00016366 | nhr-140 | C33G8.9 |
| WBGene00016367 | nhr-162 | C33G8.10 |
| WBGene00016368 | nhr-163 | C33G8.12 |
| WBGene00016383 | zgpa-1 | C33H5.17 |
| WBGene00016407 | | C34D10.2 |
| WBGene00016427 | | C34H4.5 |
| WBGene00016517 | nhr-260 | C38C3.9 |
| WBGene00016557 | ceh-84 | C40D2.4 |
| WBGene00016562 | | C41D11.3 |
| WBGene00016600 | ets-5 | C42D8.4 |
| WBGene00016620 | dhhc-9 | C43H6.7 |
| WBGene00016712 | | C46E10.8 |
| WBGene00016713 | | C46E10.9 |
| WBGene00016754 | cebp-2 | C48E7.11 |
| WBGene00016772 | nhr-166 | C49D10.2 |
| WBGene00016777 | nhr-261 | C49D10.9 |
| WBGene00016798 | ets-8 | C50A2.4 |
| WBGene00016865 | ets-9 | C52B9.2 |
| WBGene00016870 | ets-10 | C52B9.11 |
| WBGene00016888 | | C52E12.1 |
| WBGene00016890 | lst-5 | C52E12.6 |
| WBGene00016905 | ztf-3 | C53D5.4 |
| WBGene00016926 | nhr-171 | C54F6.8 |
| WBGene00016927 | nhr-172 | C54F6.9 |
| WBGene00016930 | madf-6 | C54G6.1 |
| WBGene00016948 | scrt-1 | C55C2.1 |
| WBGene00016975 | nhr-173 | C56E10.1 |
| WBGene00016997 | cebp-1 | D1005.3 |
| WBGene00017011 | eaf-1 | D1007.16 |
| WBGene00017075 | nap-1 | D2096.8 |
| WBGene00017198 | nhr-36 | F07C3.10 |
| WBGene00017317 | attf-2 | F09G2.9 |
| WBGene00017326 | dmd-5 | F10C1.5 |
| WBGene00017352 | | F10E7.11 |
| WBGene00017406 | sdz-12 | F12E12.5 |

| | | |
|---|---|---|
| WBGene00017423 | | F13C5.2 |
| WBGene00017430 | bcl-11 | F13H6.1 |
| WBGene00017482 | set-9 | F15E6.1 |
| WBGene00017503 | nhr-177 | F16B4.1 |
| WBGene00017510 | nhr-178 | F16B4.9 |
| WBGene00017512 | nhr-179 | F16B4.11 |
| WBGene00017535 | atf-8 | F17A9.3 |
| WBGene00017538 | ceh-49 | F17A9.6 |
| WBGene00017598 | | F19F10.1 |
| WBGene00017601 | ets-7 | F19F10.5 |
| WBGene00017606 | ets-6 | F19F10.10 |
| WBGene00017651 | | F21A9.2 |
| WBGene00017664 | npax-1 | F21D12.5 |
| WBGene00017687 | ets-4 | F22A3.1 |
| WBGene00017690 | ceh-60 | F22A3.5 |
| WBGene00017733 | ubxn-1 | F23C8.4 |
| WBGene00017742 | nfyc-1 | F23F1.1 |
| WBGene00017755 | zip-8 | F23F12.9 |
| WBGene00017758 | | F23H11.2 |
| WBGene00017787 | nhr-141 | F25E5.6 |
| WBGene00017814 | | F26A10.2 |
| WBGene00017961 | nhr-180 | F31F4.12 |
| WBGene00017967 | ada-2 | F32A5.1 |
| WBGene00018009 | dhhc-3 | F33D11.12 |
| WBGene00018022 | ceh-87 | F34D6.2 |
| WBGene00018099 | ztf-28 | F36F12.8 |
| WBGene00018140 | | F37B4.10 |
| WBGene00018189 | nhr-181 | F38H12.3 |
| WBGene00018248 | | F40G9.14 |
| WBGene00018265 | nhr-182 | F41B5.9 |
| WBGene00018266 | nhr-183 | F41B5.10 |
| WBGene00018355 | ceh-86 | F42G2.6 |
| WBGene00018391 | rpb-4 | F43E2.2 |
| WBGene00018404 | nhr-39 | F44A2.4 |
| WBGene00018412 | nhr-37 | F44C4.2 |
| WBGene00018415 | nhr-184 | F44C8.9 |
| WBGene00018420 | | F44E2.7 |
| WBGene00018430 | nhr-142 | F44E7.8 |
| WBGene00018432 | irld-33 | F45C12.1 |
| WBGene00018433 | ceh-82 | F45C12.2 |
| WBGene00018434 | ceh-81 | F45C12.3 |
| WBGene00018446 | ceh-83 | F45C12.15 |
| WBGene00018539 | nhr-185 | F47C10.1 |

| | | |
|---|---|---|
| WBGene00018541 | nhr-186 | F47C10.3 |
| WBGene00018542 | nhr-187 | F47C10.4 |
| WBGene00018544 | nhr-188 | F47C10.7 |
| WBGene00018545 | nhr-189 | F47C10.8 |
| WBGene00018567 | | F47E1.3 |
| WBGene00018591 | npax-2 | F48B9.5 |
| WBGene00018622 | nhr-190 | F48G7.11 |
| WBGene00018636 | | F49E8.2 |
| WBGene00018704 | ztf-13 | F52E4.8 |
| WBGene00018740 | tra-4 | F53B3.1 |
| WBGene00018769 | mnat-1 | F53G2.7 |
| WBGene00018786 | hmbx-1 | F54A5.1 |
| WBGene00018794 | attf-3 | F54C4.3 |
| WBGene00018833 | ztf-1 | F54F2.5 |
| WBGene00018836 | | F54F2.9 |
| WBGene00018892 | cir-1 | F55F8.4 |
| WBGene00018959 | | F56D1.1 |
| WBGene00018990 | klf-1 | F56F11.3 |
| WBGene00018993 | nhr-266 | F56H1.2 |
| WBGene00019011 | | F57C9.4 |
| WBGene00019115 | nhr-195 | F59E11.10 |
| WBGene00019116 | nhr-143 | F59E11.11 |
| WBGene00019137 | ceh-85 | F59H6.6 |
| WBGene00019217 | athp-2 | H20J04.2 |
| WBGene00019218 | madf-3 | H20J04.3 |
| WBGene00019257 | dhhc-13 | H32C10.3 |
| WBGene00019299 | snai-1 | K02D7.2 |
| WBGene00019327 | zip-2 | K02F3.4 |
| WBGene00019344 | dhhc-10 | K02G10.1 |
| WBGene00019347 | mbl-1 | K02H8.1 |
| WBGene00019380 | K04C2.2 | K04C2.2 |
| WBGene00019407 | | K05F1.5 |
| WBGene00019424 | aptf-1 | K06A1.1 |
| WBGene00019521 | dmd-7 | K08B12.2 |
| WBGene00019598 | | K09H9.7 |
| WBGene00019611 | | K10B3.5 |
| WBGene00019629 | cid-1 | K10D2.3 |
| WBGene00019651 | | K11D12.12 |
| WBGene00019691 | | K12H6.12 |
| WBGene00019741 | nhr-201 | M02H5.3 |
| WBGene00019742 | nhr-202 | M02H5.4 |
| WBGene00019743 | nhr-203 | M02H5.5 |
| WBGene00019751 | | M03D4.4 |

| | | |
|---|---|---|
| WBGene00019816 | nhr-204 | R02C2.4 |
| WBGene00019821 | gtf-2H1 | R02D3.3 |
| WBGene00019824 | | R02D3.7 |
| WBGene00019864 | ceh-93 | R04A9.5 |
| WBGene00019878 | | R05D3.3 |
| WBGene00019960 | ztf-16 | R08E3.4 |
| WBGene00020015 | nhr-210 | R11G11.12 |
| WBGene00020062 | nhr-270 | R13D11.8 |
| WBGene00020066 | dhhc-5 | R13F6.5 |
| WBGene00020093 | | R144.3 |
| WBGene00020111 | | R151.8 |
| WBGene00020152 | nhr-211 | T01G6.5 |
| WBGene00020153 | nhr-212 | T01G6.6 |
| WBGene00020172 | thoc-1 | T02H6.2 |
| WBGene00020189 | tfbm-1 | T03F1.7 |
| WBGene00020214 | attf-5 | T04G9.1 |
| WBGene00020251 | camt-1 | T05C1.4 |
| WBGene00020320 | | T07F8.4 |
| WBGene00020368 | ast-1 | T08H4.3 |
| WBGene00020385 | nhr-216 | T09D3.4 |
| WBGene00020399 | ztf-4 | T10B11.3 |
| WBGene00020408 | | T10D4.6 |
| WBGene00020460 | nhr-273 | T12C9.1 |
| WBGene00020480 | ssup-72 | T13C2.4 |
| WBGene00020485 | ceh-54 | T13C5.4 |
| WBGene00020555 | nhr-219 | T19A5.5 |
| WBGene00020591 | nhr-220 | T19H12.8 |
| WBGene00020630 | | T20F7.1 |
| WBGene00020635 | | T20H4.2 |
| WBGene00020694 | dhhc-11 | T22E7.2 |
| WBGene00020708 | dmd-8 | T22H9.4 |
| WBGene00020748 | nhr-221 | T24A6.8 |
| WBGene00020750 | nhr-222 | T24A6.11 |
| WBGene00020763 | ztf-18 | T24C4.7 |
| WBGene00020779 | | T24G10.2 |
| WBGene00020823 | | T26A5.8 |
| WBGene00020827 | | T26A8.4 |
| WBGene00020849 | nhr-225 | T27B7.2 |
| WBGene00020850 | nhr-226 | T27B7.3 |
| WBGene00020851 | nhr-227 | T27B7.5 |
| WBGene00020852 | nhr-228 | T27B7.6 |
| WBGene00020930 | hlh-30 | W02C12.3 |
| WBGene00020942 | | W02D7.6 |

| | | |
|---|---|---|
| WBGene00020961 | sknr-1 | W02H5.7 |
| WBGene00021000 | | W03F9.2 |
| WBGene00021019 | | W04B5.2 |
| WBGene00021047 | zfp-3 | W05H7.4 |
| WBGene00021082 | zip-11 | W08E12.1 |
| WBGene00021132 | nfyb-1 | W10D9.4 |
| WBGene00021163 | nhr-275 | Y5H2A.2 |
| WBGene00021237 | crtc-1 | Y20F4.2 |
| WBGene00021254 | | Y22D7AL.16 |
| WBGene00021281 | ell-1 | Y24D9A.1 |
| WBGene00021311 | thoc-5 | Y32H12A.2 |
| WBGene00021363 | taf-6.2 | Y37E11AL.8 |
| WBGene00021374 | | Y37E11B.1 |
| WBGene00021387 | | Y37F4.6 |
| WBGene00021417 | nhr-236 | Y38F2AL.5 |
| WBGene00021423 | tftc-1 | Y38F2AR.5 |
| WBGene00021446 | hlh-33 | Y39A3CR.6 |
| WBGene00021522 | nhr-274 | Y41D4B.21 |
| WBGene00021538 | rabs-5 | Y42H9AR.3 |
| WBGene00021552 | zip-4 | Y44E3B.1 |
| WBGene00021610 | nhr-237 | Y46H3D.6 |
| WBGene00021611 | nhr-238 | Y46H3D.7 |
| WBGene00021636 | pcaf-1 | Y47G6A.6 |
| WBGene00021637 | | Y47G6A.7 |
| WBGene00021689 | | Y48G8AL.10 |
| WBGene00021704 | | Y48G9A.11 |
| WBGene00021714 | cyh-1 | Y49F6B.1 |
| WBGene00021812 | | Y53G8AR.5 |
| WBGene00021816 | | Y53G8AR.9 |
| WBGene00021830 | | Y54E10A.10 |
| WBGene00021845 | rpb-7 | Y54E10BR.6 |
| WBGene00021846 | ztf-23 | Y54E10BR.8 |
| WBGene00021848 | nhr-239 | Y54F10AM.1 |
| WBGene00021885 | | Y54G2A.20 |
| WBGene00021904 | gtf-2H5 | Y55B1AL.2 |
| WBGene00021924 | | Y55F3AM.6 |
| WBGene00021931 | | Y55F3AM.14 |
| WBGene00021942 | madf-1 | Y55F3BR.5 |
| WBGene00022000 | tbx-42 | Y59E9AR.5 |
| WBGene00022060 | dmd-9 | Y67D8A.3 |
| WBGene00022097 | nhr-242 | Y69A2AR.26 |
| WBGene00022182 | swsn-3 | Y71H2AM.17 |
| WBGene00022207 | elk-2 | Y73B3A.5 |

| | | |
|---|---|---|
| WBGene00022278 | rcor-1 | Y74C9A.4 |
| WBGene00022334 | | Y82E9BR.1 |
| WBGene00022349 | | Y82E9BR.17 |
| WBGene00022374 | nhr-277 | Y94H6A.1 |
| WBGene00022423 | nhr-41 | Y104H12A.1 |
| WBGene00022500 | lfi-1 | ZC8.4 |
| WBGene00022518 | zfh-2 | ZC123.3 |
| WBGene00022554 | duxl-1 | ZC204.2 |
| WBGene00022592 | klu-2 | ZC328.2 |
| WBGene00022598 | ztf-8 | ZC395.8 |
| WBGene00022608 | madf-9 | ZC416.1 |
| WBGene00022636 | nhr-278 | ZK6.1 |
| WBGene00022637 | nhr-252 | ZK6.2 |
| WBGene00022639 | nhr-253 | ZK6.4 |
| WBGene00022640 | nhr-254 | ZK6.5 |
| WBGene00022671 | | ZK177.3 |
| WBGene00022681 | | ZK185.1 |
| WBGene00022695 | | ZK328.6 |
| WBGene00022755 | nhr-250 | ZK488.1 |
| WBGene00022756 | nhr-251 | ZK488.4 |
| WBGene00022762 | | ZK546.5 |
| WBGene00022785 | | ZK652.6 |
| WBGene00022794 | snu-23 | ZK686.4 |
| WBGene00022795 | | ZK686.5 |
| WBGene00022805 | nhr-256 | ZK697.2 |
| WBGene00022837 | ceh-45 | ZK993.1 |
| WBGene00022861 | dve-1 | ZK1193.5 |
| WBGene00023497 | lin-15B | ZK662.4 |
| WBGene00043056 | nfya-2 | Y53H1A.5 |
| WBGene00043705 | egrh-2 | Y55F3AM.7 |
| WBGene00044032 | ceh-99 | T21B4.17 |
| WBGene00044068 | syd-9 | ZK867.1 |
| WBGene00044071 | dhhc-14 | D2021.2 |
| WBGene00044072 | ham-3 | ZK1128.5 |
| WBGene00044078 | tag-243 | T04A8.4 |
| WBGene00044330 | alr-1 | R08B4.2 |
| WBGene00044386 | | C27A2.7 |
| WBGene00044508 | nsy-7 | C18F3.4 |
| WBGene00044699 | nhr-286 | VC5.6 |
| WBGene00044791 | bnc-1 | F55C5.11 |
| WBGene00044798 | tbx-43 | Y46E12A.4 |
| WBGene00044805 | | Y53C10A.15 |
| WBGene00045215 | ceh-63 | C02F12.10 |

| | | |
|---|---|---|
| WBGene00045255 | nhr-146 | Y41D4B.27 |
| WBGene00045515 | | ZK1037.13 |
| WBGene00077521 | maf-1 | F45H11.6 |
| WBGene00077761 | zip-9 | F17C11.17 |

**Supplemental Table 1**

**All trials of survival data for N2 and CB4856 isolates on control or pathogenic bacteria**

**Supplemental Table 1: Median survival of pathogen susceptibility experiments**

| Experiment | Trial | Total worms | Median survival (days) |
|---|---|---|---|
| *S. epidermidis* (EVL 2000) | | | |
| N2 | 1 | 30 | 22 |
| N2 | 2 | 30 | 22 |
| N2 | 3 | 30 | 19 |
| N2 | 4 | 30 | 15 |
| N2 | 5 | 30 | 16 |
| CB4856 | 1 | 30 | 8 |
| CB4856 | 2 | 30 | 16 |
| CB4856 | 3 | 30 | 13 |
| CB4856 | 4 | 30 | 10 |
| CB4856 | 5 | 30 | 6 |
| CB4856 | 6 | 30 | 15 |
| *S. epidermidis* (ATCC 700562) | | | |
| N2 | 1 | 30 | 11 |
| N2 | 2 | 30 | 12 |
| N2 | 3 | 30 | 14 |
| CB4856 | 1 | 30 | 4 |
| CB4856 | 2 | 30 | 5 |
| CB4856 | 3 | 30 | 8 |
| *P. aeruginosa* (PA-14) | | | |
| N2 | 1 | 30 | 4 |
| N2 | 2 | 30 | 4 |
| N2 | 3 | 30 | 4 |
| N2 | 4 | 30 | 3 |
| CB4856 | 1 | 30 | 3 |
| CB4856 | 2 | 30 | 4 |
| CB4856 | 3 | 30 | 4 |
| CB4856 | 4 | 30 | 3 |
| *P. rettgeri* (Dmel1) | | | |
| N2 | 1 | 30 | 10 |
| N2 | 2 | 30 | 8 |
| N2 | 3 | 30 | 8 |

| | | | |
|---|---|---|---|
| N2 | 4 | 30 | 6 |
| N2 | 5 | 30 | 7 |
| CB4856 | 1 | 30 | 7 |
| CB4856 | 2 | 30 | 8 |
| CB4856 | 3 | 30 | 8 |
| CB4856 | 4 | 30 | 5 |
| CB4856 | 5 | 30 | 8 |
| *S. aureus* (Newman) | | | |
| N2 | 1 | 30 | 8 |
| N2 | 2 | 30 | 9 |
| N2 | 3 | 30 | 3 |
| N2 | 4 | 30 | 8 |
| CB4856 | 1 | 30 | 9 |
| CB4856 | 2 | 30 | 7 |
| CB4856 | 3 | 30 | 4 |
| CB4856 | 4 | 30 | 8 |
| *E. coli* OP50 | | | |
| N2 | 1 | 30 | 11 |
| N2 | 2 | 30 | 11 |
| N2 | 3 | 30 | 11 |
| N2 | 4 | 30 | 13 |
| N2 | 5 | 30 | 13 |
| N2 | 6 | 30 | 12 |
| N2 | 7 | 30 | 13 |
| CB4856 | 1 | 30 | 16 |
| CB4856 | 2 | 30 | 15 |
| CB4856 | 3 | 30 | 16 |
| CB4856 | 4 | 30 | 16 |
| CB4856 | 5 | 30 | 15 |
| CB4856 | 6 | 30 | 19 |

**Supplemental Table 2**

**Lists of differentially expressed genes that are identified in either N2, CB4856, or both, after infection by all pathogens (compared to *E. coli*)**

**Supplmental Table 2 - Differentially Expressed Genes Identify as being Common To All Pathogens**

**N2**-exclusive DEGs found across all pathogens (*S. epidermidis* , *P. rettgeri* , *P. aeruginosa* , or *S. aureus* vs. OP50)

| Gene Name | Wormbase Gene ID | Type | $log_2FC$ | FC | Adj P-value |
|---|---|---|---|---|---|
| *ech-9* | WBGene00001158 | protein_coding_gene | 6.9 | 116.59 | 1.7E-04 |
| *T26H5.9* | WBGene00044206 | protein_coding_gene | 4.3 | 19.82 | 4.6E-10 |
| *F53A9.8* | WBGene00018731 | protein_coding_gene | 3.8 | 14.25 | 3.3E-09 |
| *cyp-37B1* | WBGene00009226 | protein_coding_gene | 3.8 | 13.49 | 8.2E-07 |
| *T26H5.14* | WBGene00235133 | protein_coding_gene | 3.6 | 11.71 | 3.2E-04 |
| *tts-1* | WBGene00006650 | ncRNA_gene | 3.5 | 11.61 | 4.8E-08 |
| *F47B8.10* | WBGene00009811 | protein_coding_gene | 3.5 | 11.17 | 5.1E-03 |
| *F46A8.13* | WBGene00219950 | ncRNA_gene | 3.3 | 10.13 | 2.0E-03 |
| *cyp-32B1* | WBGene00021167 | protein_coding_gene | 3.3 | 10.02 | 1.5E-05 |
| *C50F7.5* | WBGene00016845 | protein_coding_gene | 3.3 | 10.01 | 2.7E-07 |
| *F47B8.4* | WBGene00009805 | protein_coding_gene | 3.3 | 9.91 | 3.3E-04 |
| *T26H5.4* | WBGene00012069 | protein_coding_gene | 3.2 | 9.40 | 5.4E-04 |
| *T07G12.5* | WBGene00011596 | protein_coding_gene | 3.2 | 9.13 | 3.0E-05 |
| *fbxa-24* | WBGene00018803 | protein_coding_gene | 3.2 | 9.01 | 1.7E-06 |
| *T26H5.10* | WBGene00044783 | protein_coding_gene | 3.0 | 8.27 | 6.3E-03 |
| *F11C7.6* | WBGene00017378 | protein_coding_gene | 3.0 | 7.96 | 8.3E-04 |
| *F21C10.11* | WBGene00017660 | protein_coding_gene | 3.0 | 7.81 | 1.6E-04 |
| *Y53F4B.45* | WBGene00044728 | ncRNA_gene | 2.9 | 7.68 | 2.1E-04 |
| *wrt-2* | WBGene00006948 | protein_coding_gene | 2.9 | 7.62 | 5.1E-05 |
| *spp-2* | WBGene00004987 | protein_coding_gene | 2.9 | 7.58 | 2.6E-04 |
| *best-5* | WBGene00007404 | protein_coding_gene | 2.9 | 7.41 | 1.9E-04 |
| *clec-17* | WBGene00008477 | protein_coding_gene | 2.9 | 7.38 | 2.2E-03 |
| *lipl-2* | WBGene00009773 | protein_coding_gene | 2.8 | 7.18 | 1.6E-06 |
| *scl-2* | WBGene00009895 | protein_coding_gene | 2.8 | 7.07 | 6.2E-04 |
| *C18H7.1* | WBGene00015993 | protein_coding_gene | 2.8 | 7.00 | 2.9E-03 |
| *F45D3.4* | WBGene00009724 | protein_coding_gene | 2.8 | 6.93 | 4.7E-04 |
| *C25F9.11* | WBGene00045411 | protein_coding_gene | 2.8 | 6.88 | 7.9E-04 |
| *F55H12.7* | WBGene00219283 | protein_coding_gene | 2.8 | 6.87 | 1.5E-02 |
| *scl-14* | WBGene00008625 | protein_coding_gene | 2.8 | 6.83 | 1.7E-02 |
| *fbxa-48* | WBGene00021862 | protein_coding_gene | 2.8 | 6.79 | 3.9E-03 |
| *C07G3.8* | WBGene00015576 | protein_coding_gene | 2.8 | 6.73 | 3.6E-02 |
| *cky-1* | WBGene00000521 | protein_coding_gene | 2.7 | 6.65 | 1.6E-02 |
| *ttr-23* | WBGene00011894 | protein_coding_gene | 2.7 | 6.50 | 2.2E-03 |
| *mtl-1* | WBGene00003473 | protein_coding_gene | 2.7 | 6.44 | 2.0E-06 |
| *nhr-127* | WBGene00003717 | protein_coding_gene | 2.7 | 6.40 | 1.4E-02 |
| *math-34* | WBGene00019699 | protein_coding_gene | 2.7 | 6.30 | 2.2E-03 |
| *fbxa-66* | WBGene00021865 | protein_coding_gene | 2.6 | 6.22 | 4.2E-02 |
| *srz-99* | WBGene00012028 | protein_coding_gene | 2.6 | 6.21 | 2.2E-02 |
| *ZK105.13* | WBGene00235114 | protein_coding_gene | 2.6 | 6.19 | 2.2E-02 |
| *F46C5.1* | WBGene00009778 | protein_coding_gene | 2.6 | 6.18 | 1.7E-07 |
| *tsp-1* | WBGene00006627 | protein_coding_gene | 2.6 | 6.17 | 3.8E-03 |
| *lipl-1* | WBGene00010062 | protein_coding_gene | 2.6 | 6.16 | 6.7E-05 |
| *Y54G2A.76* | WBGene00235268 | protein_coding_gene | 2.6 | 6.14 | 2.5E-03 |
| *Y37A1B.7* | WBGene00012540 | protein_coding_gene | 2.6 | 6.12 | 3.2E-03 |
| *pals-39* | WBGene00008301 | protein_coding_gene | 2.6 | 6.02 | 4.4E-03 |
| *sodh-1* | WBGene00010790 | protein_coding_gene | 2.6 | 6.02 | 2.2E-03 |
| *F49H6.5* | WBGene00009908 | protein_coding_gene | 2.6 | 5.99 | 2.2E-03 |
| *F35E12.4* | WBGene00009428 | pseudogene | 2.6 | 5.98 | 3.9E-03 |
| *Y7A5A.6* | WBGene00012412 | protein_coding_gene | 2.6 | 5.97 | 2.3E-02 |
| *ZC395.5* | WBGene00022597 | protein_coding_gene | 2.6 | 5.95 | 1.6E-04 |
| *Y47H10A.3* | WBGene00012959 | protein_coding_gene | 2.6 | 5.95 | 2.3E-02 |
| *cup-16* | WBGene00007919 | protein_coding_gene | 2.6 | 5.89 | 2.0E-03 |
| *ugt-18* | WBGene00013900 | protein_coding_gene | 2.6 | 5.88 | 1.1E-05 |
| *Y26D4A.3* | WBGene00012501 | protein_coding_gene | 2.5 | 5.85 | 9.6E-03 |
| *srj-29* | WBGene00005616 | protein_coding_gene | 2.5 | 5.85 | 4.5E-02 |
| *sru-20* | WBGene00005683 | protein_coding_gene | 2.5 | 5.83 | 3.6E-02 |
| *F53A9.7* | WBGene00018730 | protein_coding_gene | 2.5 | 5.83 | 3.9E-03 |
| *C25H3.10* | WBGene00016119 | protein_coding_gene | 2.5 | 5.82 | 2.2E-03 |
| *unc-42* | WBGene00006778 | protein_coding_gene | 2.5 | 5.77 | 2.3E-03 |
| *T21D12.7* | WBGene00020648 | protein_coding_gene | 2.5 | 5.71 | 4.9E-02 |
| *ZC443.3* | WBGene00013898 | protein_coding_gene | 2.5 | 5.68 | 2.9E-03 |
| *kreg-1* | WBGene00018725 | protein_coding_gene | 2.5 | 5.65 | 9.3E-03 |
| *T12D8.5* | WBGene00011733 | protein_coding_gene | 2.5 | 5.63 | 1.7E-03 |
| *F37A8.5* | WBGene00009503 | protein_coding_gene | 2.5 | 5.63 | 4.8E-03 |
| *Y51H4A.5* | WBGene00013101 | protein_coding_gene | 2.5 | 5.61 | 9.6E-03 |
| *Y73B6BL.37* | WBGene00022256 | protein_coding_gene | 2.5 | 5.60 | 1.3E-02 |
| *pals-37* | WBGene00138721 | protein_coding_gene | 2.5 | 5.56 | 1.7E-02 |
| *C44H9.7* | WBGene00008102 | protein_coding_gene | 2.5 | 5.54 | 4.0E-03 |
| *T20B3.1* | WBGene00011850 | protein_coding_gene | 2.5 | 5.49 | 2.5E-03 |

| | | | | | |
|---|---|---|---|---|---|
| *F58F12.12* | WBGene00271812 | protein_coding_gene | 2.4 | 5.46 | 6.3E-03 |
| *ZC443.2* | WBGene00013897 | protein_coding_gene | 2.4 | 5.34 | 4.8E-02 |
| *peb-1* | WBGene00003968 | protein_coding_gene | 2.4 | 5.26 | 2.2E-03 |
| *nhr-99* | WBGene00003689 | protein_coding_gene | 2.4 | 5.19 | 9.6E-03 |
| *ZK666.11* | WBGene00014049 | protein_coding_gene | 2.4 | 5.18 | 4.4E-02 |
| *C25F9.12* | WBGene00045412 | protein_coding_gene | 2.4 | 5.13 | 3.9E-02 |
| *far-7* | WBGene00001391 | protein_coding_gene | 2.4 | 5.13 | 1.2E-02 |
| *dod-3* | WBGene00016052 | protein_coding_gene | 2.4 | 5.12 | 2.9E-03 |
| *F09F7.6* | WBGene00017303 | protein_coding_gene | 2.4 | 5.11 | 2.6E-04 |
| *C50F4.9* | WBGene00008234 | protein_coding_gene | 2.3 | 5.07 | 2.6E-04 |
| *dod-22* | WBGene00010125 | protein_coding_gene | 2.3 | 5.04 | 1.7E-02 |
| *T26H5.8* | WBGene00012070 | protein_coding_gene | 2.3 | 4.98 | 6.7E-03 |
| *EGAP1.1* | WBGene00017146 | protein_coding_gene | 2.3 | 4.96 | 3.0E-02 |
| *H42K12.3* | WBGene00019272 | protein_coding_gene | 2.3 | 4.94 | 7.6E-03 |
| *irld-11* | WBGene00019391 | protein_coding_gene | 2.3 | 4.94 | 4.8E-02 |
| *Y65B4BM.3* | WBGene00302974 | protein_coding_gene | 2.3 | 4.93 | 6.4E-03 |
| *C06G3.3* | WBGene00015549 | protein_coding_gene | 2.3 | 4.88 | 2.0E-02 |
| *clec-172* | WBGene00044719 | protein_coding_gene | 2.3 | 4.87 | 1.9E-02 |
| *C06B3.6* | WBGene00007365 | protein_coding_gene | 2.3 | 4.86 | 7.9E-04 |
| *W02A2.9* | WBGene00044280 | protein_coding_gene | 2.3 | 4.86 | 3.9E-03 |
| *nnt-1* | WBGene00003778 | protein_coding_gene | 2.3 | 4.85 | 2.0E-02 |
| *F11E6.9* | WBGene00008712 | protein_coding_gene | 2.3 | 4.84 | 2.8E-02 |
| *C36B1.13* | WBGene00044120 | protein_coding_gene | 2.3 | 4.80 | 3.9E-02 |
| *btb-16* | WBGene00018195 | protein_coding_gene | 2.3 | 4.79 | 2.9E-03 |
| *ZC204.12* | WBGene00022562 | protein_coding_gene | 2.3 | 4.78 | 2.2E-02 |
| *phat-5* | WBGene00020242 | protein_coding_gene | 2.3 | 4.77 | 2.6E-02 |
| *C35A5.11* | WBGene00044024 | protein_coding_gene | 2.2 | 4.75 | 3.1E-02 |
| *irg-1* | WBGene00015574 | protein_coding_gene | 2.2 | 4.75 | 2.4E-02 |
| *fil-1* | WBGene00011321 | protein_coding_gene | 2.2 | 4.74 | 3.6E-02 |
| *T05F1.11* | WBGene00011495 | protein_coding_gene | 2.2 | 4.73 | 1.7E-02 |
| *fbxc-20* | WBGene00018352 | protein_coding_gene | 2.2 | 4.69 | 2.5E-03 |
| *C37A5.3* | WBGene00014698 | protein_coding_gene | 2.2 | 4.67 | 3.4E-02 |
| *F56D2.3* | WBGene00018965 | protein_coding_gene | 2.2 | 4.66 | 7.5E-03 |
| *F58H1.2* | WBGene00010285 | protein_coding_gene | 2.2 | 4.66 | 1.8E-02 |
| *E02H4.4* | WBGene00008462 | protein_coding_gene | 2.2 | 4.66 | 2.3E-02 |
| *ftn-1* | WBGene00001500 | protein_coding_gene | 2.2 | 4.64 | 9.6E-03 |
| *K08B12.1* | WBGene00019520 | protein_coding_gene | 2.2 | 4.62 | 2.2E-03 |
| *best-1* | WBGene00007203 | protein_coding_gene | 2.2 | 4.59 | 2.1E-02 |
| *F15B9.6* | WBGene00008850 | protein_coding_gene | 2.2 | 4.57 | 1.7E-02 |
| *oac-31* | WBGene00018707 | protein_coding_gene | 2.2 | 4.57 | 3.8E-02 |
| *asp-11* | WBGene00017677 | pseudogene | 2.2 | 4.57 | 4.7E-02 |
| *C52E2.4* | WBGene00016883 | protein_coding_gene | 2.2 | 4.54 | 6.8E-03 |
| *ttr-29* | WBGene00011286 | protein_coding_gene | 2.2 | 4.53 | 2.8E-02 |
| *srw-85* | WBGene00005832 | protein_coding_gene | 2.2 | 4.53 | 3.9E-02 |
| *hil-1* | WBGene00001852 | protein_coding_gene | 2.2 | 4.53 | 3.4E-02 |
| *W09G12.7* | WBGene00021121 | protein_coding_gene | 2.2 | 4.52 | 9.3E-03 |
| *cyp-35B1* | WBGene00019472 | protein_coding_gene | 2.2 | 4.49 | 3.9E-02 |
| *C17B7.4* | WBGene00015878 | protein_coding_gene | 2.2 | 4.48 | 2.2E-02 |
| *cutl-28* | WBGene00018256 | protein_coding_gene | 2.2 | 4.48 | 6.3E-03 |
| *lys-3* | WBGene00003092 | protein_coding_gene | 2.2 | 4.48 | 4.5E-02 |
| *T09E11.11* | WBGene00011660 | pseudogene | 2.2 | 4.47 | 1.5E-02 |
| *srw-86* | WBGene00005833 | protein_coding_gene | 2.2 | 4.46 | 4.0E-02 |
| *T16G1.4* | WBGene00011798 | protein_coding_gene | 2.2 | 4.46 | 3.6E-02 |
| *C14A11.2* | WBGene00015748 | protein_coding_gene | 2.2 | 4.44 | 5.0E-02 |
| *clec-42* | WBGene00008891 | protein_coding_gene | 2.1 | 4.44 | 1.9E-02 |
| *Y68A4A.13* | WBGene00044212 | protein_coding_gene | 2.1 | 4.44 | 2.2E-03 |
| *lpr-4* | WBGene00012257 | protein_coding_gene | 2.1 | 4.43 | 5.4E-04 |
| *fbxa-21* | WBGene00022487 | protein_coding_gene | 2.1 | 4.39 | 1.9E-04 |
| *C35C5.8* | WBGene00007957 | protein_coding_gene | 2.1 | 4.39 | 3.9E-02 |
| *daao-1* | WBGene00022076 | protein_coding_gene | 2.1 | 4.38 | 3.2E-02 |
| *Y43F8B.9* | WBGene00012819 | protein_coding_gene | 2.1 | 4.37 | 1.7E-03 |
| *F43C11.7* | WBGene00018384 | protein_coding_gene | 2.1 | 4.35 | 5.4E-03 |
| *btb-21* | WBGene00018540 | pseudogene | 2.1 | 4.33 | 1.4E-03 |
| *C50B8.4* | WBGene00008226 | protein_coding_gene | 2.1 | 4.33 | 1.3E-02 |
| *ZK1290.10* | WBGene00022891 | protein_coding_gene | 2.1 | 4.31 | 5.0E-02 |
| *Y43F8B.23* | WBGene00194815 | protein_coding_gene | 2.1 | 4.30 | 9.5E-03 |
| *coel-1* | WBGene00016866 | protein_coding_gene | 2.1 | 4.29 | 1.9E-02 |
| *fmo-1* | WBGene00001476 | protein_coding_gene | 2.1 | 4.28 | 1.2E-02 |
| *tsp-2* | WBGene00006628 | protein_coding_gene | 2.1 | 4.27 | 4.7E-02 |
| *srr-6* | WBGene00005657 | protein_coding_gene | 2.1 | 4.27 | 4.2E-02 |
| *C06B3.7* | WBGene00007366 | protein_coding_gene | 2.1 | 4.26 | 5.0E-02 |
| *T02G5.15* | WBGene00206521 | protein_coding_gene | 2.1 | 4.26 | 7.7E-03 |
| *C08F11.13* | WBGene00007460 | protein_coding_gene | 2.1 | 4.25 | 2.1E-02 |

| | | | | | |
|---|---|---|---|---|---|
| *F49C12.7* | WBGene00009877 | protein_coding_gene | 2.1 | 4.24 | 4.9E-02 |
| *Y53C12B.7* | WBGene00013146 | protein_coding_gene | 2.1 | 4.22 | 4.8E-02 |
| *asah-1* | WBGene00010769 | protein_coding_gene | 2.1 | 4.21 | 4.7E-02 |
| *W03F9.4* | WBGene00021002 | protein_coding_gene | 2.1 | 4.19 | 4.7E-02 |
| *C06E4.8* | WBGene00015537 | protein_coding_gene | 2.1 | 4.18 | 2.6E-02 |
| *sdz-24* | WBGene00019495 | protein_coding_gene | 2.1 | 4.17 | 1.9E-02 |
| *R03H10.2* | WBGene00019855 | protein_coding_gene | 2.1 | 4.16 | 2.8E-03 |
| *F11E6.11* | WBGene00023501 | protein_coding_gene | 2.1 | 4.16 | 4.2E-02 |
| *T03G6.1* | WBGene00020194 | protein_coding_gene | 2.1 | 4.14 | 6.1E-03 |
| *K08F9.1* | WBGene00010684 | protein_coding_gene | 2.1 | 4.14 | 2.3E-02 |
| *hil-7* | WBGene00001858 | protein_coding_gene | 2.0 | 4.13 | 4.0E-02 |
| *C07E3.3* | WBGene00007414 | protein_coding_gene | 2.0 | 4.09 | 1.8E-02 |
| *F13H8.1* | WBGene00017434 | protein_coding_gene | 2.0 | 4.08 | 5.4E-03 |
| *ZC449.1* | WBGene00022611 | protein_coding_gene | 2.0 | 4.05 | 4.2E-02 |
| *gbh-2* | WBGene00001523 | protein_coding_gene | 2.0 | 4.04 | 9.3E-05 |
| *ilys-2* | WBGene00016669 | protein_coding_gene | 2.0 | 4.04 | 1.4E-02 |
| *Y54G2A.37* | WBGene00023484 | protein_coding_gene | 2.0 | 4.03 | 4.0E-03 |
| *lipl-3* | WBGene00020016 | protein_coding_gene | 2.0 | 3.99 | 1.8E-02 |
| *nhr-19* | WBGene00003618 | protein_coding_gene | 2.0 | 3.98 | 2.4E-02 |
| *math-38* | WBGene00020326 | protein_coding_gene | 2.0 | 3.96 | 7.6E-03 |
| *F18G5.6* | WBGene00017582 | protein_coding_gene | 2.0 | 3.96 | 1.3E-02 |
| *tsp-17* | WBGene00006643 | protein_coding_gene | 2.0 | 3.95 | 1.9E-02 |
| *Y17D7C.4* | WBGene00050903 | pseudogene | 2.0 | 3.92 | 1.4E-02 |
| *C32F10.4* | WBGene00016332 | protein_coding_gene | 2.0 | 3.92 | 2.8E-02 |
| *F10E9.12* | WBGene00017361 | protein_coding_gene | 2.0 | 3.87 | 8.3E-04 |
| *spp-13* | WBGene00004998 | protein_coding_gene | 1.9 | 3.85 | 3.6E-02 |
| *swt-1* | WBGene00019300 | protein_coding_gene | 1.9 | 3.85 | 4.4E-02 |
| *dpy-3* | WBGene00001065 | protein_coding_gene | 1.9 | 3.84 | 1.4E-02 |
| *ptr-10* | WBGene00004224 | protein_coding_gene | 1.9 | 3.81 | 4.2E-02 |
| *Y110A2AL.4* | WBGene00022441 | protein_coding_gene | 1.9 | 3.78 | 8.8E-03 |
| *F59B2.8* | WBGene00010310 | protein_coding_gene | 1.9 | 3.74 | 3.4E-03 |
| *K08F9.3* | WBGene00010686 | protein_coding_gene | 1.9 | 3.73 | 4.3E-02 |
| *lpr-5* | WBGene00012256 | protein_coding_gene | 1.9 | 3.69 | 1.3E-02 |
| *hhat-1* | WBGene00013852 | protein_coding_gene | 1.9 | 3.68 | 1.4E-02 |
| *T10G3.1* | WBGene00011692 | protein_coding_gene | 1.9 | 3.62 | 4.0E-02 |
| *dpy-7* | WBGene00001069 | protein_coding_gene | 1.8 | 3.60 | 9.6E-03 |
| *ZC123.1* | WBGene00022517 | protein_coding_gene | 1.8 | 3.59 | 2.9E-02 |
| *T04F8.7* | WBGene00011445 | protein_coding_gene | 1.8 | 3.55 | 1.3E-02 |
| *F53A9.6* | WBGene00018729 | protein_coding_gene | 1.8 | 3.51 | 2.3E-02 |
| *T20F5.4* | WBGene00020626 | protein_coding_gene | 1.8 | 3.48 | 1.8E-02 |
| *nmad-1* | WBGene00017304 | protein_coding_gene | 1.8 | 3.48 | 4.3E-02 |
| *C48B4.3* | WBGene00008168 | protein_coding_gene | 1.8 | 3.47 | 4.9E-02 |
| *C14F11.6* | WBGene00015781 | protein_coding_gene | 1.8 | 3.42 | 1.0E-02 |
| *mlt-11* | WBGene00012186 | protein_coding_gene | 1.8 | 3.39 | 4.6E-02 |
| *W04A8.4* | WBGene00012239 | protein_coding_gene | 1.7 | 3.35 | 4.4E-03 |
| *dao-2* | WBGene00000928 | protein_coding_gene | 1.7 | 3.35 | 8.8E-03 |
| *Y44A6C.1* | WBGene00012850 | protein_coding_gene | 1.7 | 3.33 | 4.2E-02 |
| *B0250.12* | WBGene00050927 | pseudogene | 1.7 | 3.31 | 1.7E-02 |
| *lpr-6* | WBGene00012255 | protein_coding_gene | 1.7 | 3.30 | 3.4E-02 |
| *glf-1* | WBGene00019154 | protein_coding_gene | 1.7 | 3.29 | 3.3E-02 |
| *lpr-3* | WBGene00012261 | protein_coding_gene | 1.7 | 3.28 | 1.4E-02 |
| *M03B6.3* | WBGene00010835 | protein_coding_gene | 1.7 | 3.26 | 3.0E-02 |
| *F31D4.5* | WBGene00009292 | protein_coding_gene | 1.7 | 3.21 | 4.5E-02 |
| *C18D4.8* | WBGene00007677 | pseudogene | 1.7 | 3.19 | 1.0E-02 |
| *Y69A2AL.2* | WBGene00044468 | protein_coding_gene | 1.7 | 3.19 | 2.0E-02 |
| *cest-29* | WBGene00017205 | pseudogene | 1.7 | 3.15 | 1.7E-03 |
| *F21A3.11* | WBGene00219316 | protein_coding_gene | 1.6 | 3.10 | 2.0E-02 |
| *T19C4.1* | WBGene00011838 | protein_coding_gene | 1.6 | 3.04 | 4.8E-02 |
| *T28H10.3* | WBGene00012144 | protein_coding_gene | 1.6 | 3.02 | 4.3E-02 |
| *nhr-116* | WBGene00003706 | protein_coding_gene | 1.6 | 3.01 | 2.5E-02 |
| *dpy-10* | WBGene00001072 | protein_coding_gene | 1.5 | 2.90 | 3.0E-02 |
| *sqt-3* | WBGene00005018 | protein_coding_gene | 1.5 | 2.89 | 1.7E-02 |
| *asm-3* | WBGene00000213 | protein_coding_gene | 1.5 | 2.84 | 1.4E-02 |
| *pals-23* | WBGene00016218 | protein_coding_gene | 1.5 | 2.82 | 1.2E-02 |
| *T02E9.5* | WBGene00011383 | protein_coding_gene | 1.5 | 2.79 | 1.8E-02 |
| *Y110A2AL.3* | WBGene00022440 | protein_coding_gene | 1.5 | 2.79 | 1.9E-02 |
| *cki-1* | WBGene00000516 | protein_coding_gene | 1.5 | 2.78 | 6.7E-03 |
| *F32D8.13* | WBGene00009335 | protein_coding_gene | 1.5 | 2.78 | 4.1E-02 |
| *mks-6* | WBGene00010642 | protein_coding_gene | 1.5 | 2.74 | 3.6E-02 |
| *EEED8.2* | WBGene00017133 | protein_coding_gene | 1.4 | 2.69 | 1.3E-03 |
| *tag-234* | WBGene00018874 | protein_coding_gene | 1.4 | 2.68 | 4.2E-02 |
| *aqp-2* | WBGene00000170 | protein_coding_gene | 1.4 | 2.66 | 4.4E-02 |
| *clec-76* | WBGene00021587 | protein_coding_gene | 1.4 | 2.62 | 1.8E-03 |

| | | | | | |
|---|---|---|---|---|---|
| *T28H10.1* | WBGene00012142 | protein_coding_gene | 1.4 | 2.62 | 2.2E-02 |
| *thn-1* | WBGene00009213 | protein_coding_gene | 1.4 | 2.57 | 2.0E-02 |
| *wrt-10* | WBGene00006956 | protein_coding_gene | 1.4 | 2.57 | 3.6E-02 |
| *clec-60* | WBGene00014046 | protein_coding_gene | 1.3 | 2.41 | 3.1E-04 |
| *fbxl-1* | WBGene00015350 | protein_coding_gene | 1.2 | 2.29 | 3.2E-02 |
| *fkb-7* | WBGene00001432 | protein_coding_gene | 1.2 | 2.27 | 2.8E-02 |
| *T25D10.4* | WBGene00020801 | protein_coding_gene | 1.2 | 2.24 | 2.6E-02 |
| *fbxa-108* | WBGene00010293 | protein_coding_gene | 1.1 | 2.22 | 1.3E-02 |
| *T22F3.11* | WBGene00020700 | protein_coding_gene | 1.1 | 2.19 | 6.7E-04 |
| *ilys-3* | WBGene00016670 | protein_coding_gene | 1.1 | 2.16 | 5.4E-04 |
| *C23G10.11* | WBGene00016017 | protein_coding_gene | 1.1 | 2.16 | 2.2E-03 |
| *acs-11* | WBGene00018269 | protein_coding_gene | 1.1 | 2.13 | 9.3E-03 |
| *tth-1* | WBGene00006649 | protein_coding_gene | 1.1 | 2.08 | 1.4E-02 |
| *F17B5.1* | WBGene00008905 | protein_coding_gene | 1.1 | 2.07 | 2.3E-02 |
| *F15D4.2* | WBGene00008859 | protein_coding_gene | 1.0 | 2.05 | 4.7E-02 |
| *pes-9* | WBGene00003982 | protein_coding_gene | -1.1 | 0.47 | 4.8E-02 |
| *trap-1* | WBGene00022122 | protein_coding_gene | -1.1 | 0.47 | 3.9E-02 |
| *ahcy-1* | WBGene00019322 | protein_coding_gene | -1.2 | 0.43 | 2.8E-02 |
| *alh-9* | WBGene00000115 | protein_coding_gene | -1.3 | 0.40 | 2.6E-02 |
| *col-158* | WBGene00000731 | protein_coding_gene | -1.4 | 0.39 | 8.0E-04 |
| *gst-4* | WBGene00001752 | protein_coding_gene | -1.6 | 0.34 | 4.5E-04 |
| *dop-4* | WBGene00016872 | protein_coding_gene | -1.7 | 0.32 | 1.3E-02 |
| *DC2.5* | WBGene00017082 | protein_coding_gene | -1.7 | 0.32 | 2.0E-02 |
| *K10C2.12* | WBGene00206386 | protein_coding_gene | -1.7 | 0.30 | 4.5E-02 |
| *tbb-6* | WBGene00006539 | protein_coding_gene | -1.8 | 0.30 | 3.2E-02 |
| *W06A7.4* | WBGene00012293 | protein_coding_gene | -1.8 | 0.28 | 2.2E-03 |
| *cest-7* | WBGene00007693 | protein_coding_gene | -1.9 | 0.27 | 2.8E-02 |
| *C45B2.2* | WBGene00016659 | protein_coding_gene | -1.9 | 0.27 | 3.9E-02 |
| *C52D10.3* | WBGene00016877 | protein_coding_gene | -1.9 | 0.27 | 4.5E-02 |
| *arf-1.1* | WBGene00000190 | protein_coding_gene | -2.0 | 0.25 | 2.8E-02 |
| *acs-1* | WBGene00018488 | protein_coding_gene | -2.0 | 0.25 | 3.6E-07 |
| *T20D4.7* | WBGene00020613 | protein_coding_gene | -2.0 | 0.25 | 3.9E-02 |
| *ZK6.6* | WBGene00022641 | protein_coding_gene | -2.1 | 0.24 | 1.6E-02 |
| *F46F2.3* | WBGene00009787 | protein_coding_gene | -2.1 | 0.23 | 1.0E-02 |
| *fbxa-72* | WBGene00017962 | protein_coding_gene | -2.2 | 0.22 | 7.1E-03 |
| *sur-5* | WBGene00006351 | protein_coding_gene | -2.2 | 0.22 | 1.8E-08 |
| *lbp-8* | WBGene00002260 | protein_coding_gene | -2.2 | 0.22 | 4.4E-02 |
| *F58G6.7* | WBGene00010277 | protein_coding_gene | -2.2 | 0.22 | 1.7E-02 |
| *F18E3.12* | WBGene00045275 | protein_coding_gene | -2.2 | 0.21 | 2.6E-04 |
| *ugt-63* | WBGene00015449 | protein_coding_gene | -2.3 | 0.21 | 1.3E-02 |
| *T10B5.8* | WBGene00020394 | protein_coding_gene | -2.3 | 0.21 | 7.4E-03 |
| *dhs-25* | WBGene00000988 | protein_coding_gene | -2.4 | 0.19 | 1.0E-02 |
| *F58G6.3* | WBGene00010274 | protein_coding_gene | -2.6 | 0.16 | 8.2E-07 |
| *Y48E1B.8* | WBGene00013007 | protein_coding_gene | -2.7 | 0.15 | 2.6E-04 |
| *nas-27* | WBGene00003545 | protein_coding_gene | -3.0 | 0.12 | 6.1E-04 |
| *C07G1.7* | WBGene00015573 | protein_coding_gene | -3.1 | 0.12 | 1.3E-06 |
| *C04C3.14* | WBGene00219410 | protein_coding_gene | -3.2 | 0.11 | 6.8E-03 |
| *F58G6.9* | WBGene00044107 | protein_coding_gene | -3.3 | 0.10 | 5.6E-11 |
| *R03G8.4* | WBGene00010999 | pseudogene | -3.3 | 0.10 | 3.0E-05 |
| *Y41C4A.32* | WBGene00270321 | protein_coding_gene | -3.6 | 0.08 | 3.6E-07 |
| *clec-210* | WBGene00022261 | protein_coding_gene | -4.0 | 0.06 | 1.6E-10 |
| *B0205.13* | WBGene00044644 | protein_coding_gene | -4.2 | 0.06 | 2.0E-10 |

**CB4856**-exclusive DEGs found across all pathogens (*S. epidermidis* , *P. rettgeri* , *P. aeruginosa* , or *S. aureus* vs. OP50)

| Gene Name | Wormbase Gene ID | Type | $log_2FC$ | FC | Adj P-value |
|---|---|---|---|---|---|
| *scl-6* | WBGene00008028 | protein_coding_gene | 3.2 | 8.94 | 2.1E-06 |
| *R193.2* | WBGene00020128 | protein_coding_gene | 3.0 | 7.82 | 1.7E-07 |
| *hrg-4* | WBGene00009493 | protein_coding_gene | 2.7 | 6.64 | 3.8E-05 |
| *Y51H7C.13* | WBGene00021791 | protein_coding_gene | 2.4 | 5.38 | 4.5E-02 |
| *pcp-2* | WBGene00003957 | protein_coding_gene | 2.3 | 5.09 | 3.4E-06 |
| *T16G12.1* | WBGene00011803 | protein_coding_gene | 2.3 | 5.01 | 3.4E-06 |
| *C02G6.1* | WBGene00015359 | pseudogene | 2.2 | 4.70 | 2.9E-02 |
| *spp-23* | WBGene00021335 | protein_coding_gene | 2.2 | 4.61 | 2.7E-04 |
| *F20G2.1* | WBGene00008985 | protein_coding_gene | 2.2 | 4.48 | 6.7E-04 |
| *pho-1* | WBGene00004020 | protein_coding_gene | 2.1 | 4.41 | 1.9E-04 |
| *C05D12.3* | WBGene00014666 | protein_coding_gene | 2.1 | 4.23 | 2.0E-03 |
| *zipt-2.3* | WBGene00013207 | protein_coding_gene | 2.1 | 4.15 | 7.2E-03 |
| *Y47G6A.15* | WBGene00021641 | protein_coding_gene | 2.0 | 4.13 | 3.7E-02 |
| *T20B6.1* | WBGene00020597 | protein_coding_gene | 2.0 | 4.13 | 2.9E-02 |
| *hsp-17* | WBGene00002021 | protein_coding_gene | 2.0 | 4.08 | 5.0E-05 |
| *fbxa-15* | WBGene00022327 | protein_coding_gene | 2.0 | 4.03 | 4.5E-02 |
| *Y16B4A.2* | WBGene00012445 | protein_coding_gene | 2.0 | 3.93 | 1.9E-03 |
| *ins-32* | WBGene00002115 | protein_coding_gene | 2.0 | 3.92 | 4.9E-02 |

| | | | | | |
|---|---|---|---|---|---|
| *F09C8.1* | WBGene00008621 | protein_coding_gene | 2.0 | 3.89 | 1.5E-03 |
| *H43E16.1* | WBGene00019273 | protein_coding_gene | 1.9 | 3.86 | 2.0E-16 |
| *K10C9.9* | WBGene00044515 | pseudogene | 1.9 | 3.71 | 4.7E-02 |
| *Y39G8B.9* | WBGene00012728 | protein_coding_gene | 1.9 | 3.65 | 3.7E-02 |
| *fbxa-84* | WBGene00011670 | protein_coding_gene | 1.9 | 3.62 | 2.7E-06 |
| *elo-2* | WBGene00001240 | protein_coding_gene | 1.9 | 3.61 | 6.8E-03 |
| *K03H6.2* | WBGene00019368 | protein_coding_gene | 1.8 | 3.59 | 2.2E-02 |
| *gcsh-1* | WBGene00008354 | protein_coding_gene | 1.8 | 3.58 | 5.5E-04 |
| *F57F4.4* | WBGene00019017 | protein_coding_gene | 1.8 | 3.56 | 7.2E-03 |
| *lips-5* | WBGene00011725 | protein_coding_gene | 1.8 | 3.52 | 2.9E-02 |
| *F53B3.6* | WBGene00018744 | protein_coding_gene | 1.8 | 3.40 | 3.8E-02 |
| *F09B12.3* | WBGene00008607 | protein_coding_gene | 1.8 | 3.37 | 2.1E-02 |
| *scav-4* | WBGene00008692 | protein_coding_gene | 1.7 | 3.36 | 7.0E-03 |
| *F42A8.1* | WBGene00009626 | protein_coding_gene | 1.7 | 3.36 | 2.5E-02 |
| *F14D7.6* | WBGene00008793 | protein_coding_gene | 1.7 | 3.36 | 1.2E-04 |
| *hrg-7* | WBGene00007605 | protein_coding_gene | 1.7 | 3.35 | 2.7E-02 |
| *sptl-2* | WBGene00018398 | protein_coding_gene | 1.7 | 3.32 | 4.0E-03 |
| *spin-3* | WBGene00008598 | protein_coding_gene | 1.7 | 3.32 | 5.4E-04 |
| *C41H7.1* | WBGene00016571 | protein_coding_gene | 1.7 | 3.30 | 2.4E-02 |
| *C06E1.7* | WBGene00015522 | protein_coding_gene | 1.7 | 3.28 | 3.8E-04 |
| *elo-5* | WBGene00001243 | protein_coding_gene | 1.7 | 3.28 | 2.8E-02 |
| *nmy-3* | WBGene00012437 | protein_coding_gene | 1.7 | 3.25 | 4.5E-02 |
| *R02C2.8* | WBGene00255728 | pseudogene | 1.7 | 3.24 | 2.9E-02 |
| *Y54G2A.45* | WBGene00044488 | protein_coding_gene | 1.7 | 3.22 | 4.9E-02 |
| *Y37H2A.14* | WBGene00045416 | protein_coding_gene | 1.7 | 3.22 | 4.9E-02 |
| *F10C1.8* | WBGene00017327 | protein_coding_gene | 1.7 | 3.17 | 3.1E-02 |
| *daf-28* | WBGene00000920 | protein_coding_gene | 1.7 | 3.17 | 7.7E-04 |
| *C35A11.4* | WBGene00016431 | protein_coding_gene | 1.6 | 3.12 | 4.3E-02 |
| *nep-22* | WBGene00020788 | protein_coding_gene | 1.6 | 3.12 | 2.8E-02 |
| *oac-48* | WBGene00011960 | protein_coding_gene | 1.6 | 3.06 | 4.9E-02 |
| *ZK228.4* | WBGene00013947 | protein_coding_gene | 1.6 | 3.02 | 4.3E-02 |
| *gst-26* | WBGene00001774 | protein_coding_gene | 1.6 | 3.00 | 4.1E-02 |
| *F19B6.3* | WBGene00008949 | protein_coding_gene | 1.6 | 3.00 | 4.5E-02 |
| *Y73B3A.18* | WBGene00022219 | pseudogene | 1.6 | 2.98 | 4.4E-03 |
| *F28H7.3* | WBGene00009237 | protein_coding_gene | 1.6 | 2.97 | 3.1E-03 |
| *cpr-3* | WBGene00000783 | protein_coding_gene | 1.6 | 2.96 | 4.3E-02 |
| *F19C7.4* | WBGene00017594 | protein_coding_gene | 1.6 | 2.95 | 2.8E-02 |
| *F01D5.3* | WBGene00008494 | protein_coding_gene | 1.6 | 2.94 | 2.7E-02 |
| *oac-58* | WBGene00019580 | protein_coding_gene | 1.5 | 2.89 | 1.5E-03 |
| *pqn-36* | WBGene00004123 | protein_coding_gene | 1.5 | 2.87 | 2.4E-02 |
| *ctsa-4.2* | WBGene00021503 | protein_coding_gene | 1.5 | 2.79 | 1.1E-04 |
| *F40E12.2* | WBGene00018235 | protein_coding_gene | 1.5 | 2.78 | 2.2E-02 |
| *F26A1.4* | WBGene00017803 | protein_coding_gene | 1.5 | 2.78 | 1.4E-02 |
| *mth-1* | WBGene00017948 | protein_coding_gene | 1.5 | 2.77 | 2.5E-02 |
| *F42A10.6* | WBGene00018342 | protein_coding_gene | 1.5 | 2.75 | 2.9E-02 |
| *math-2* | WBGene00015606 | protein_coding_gene | 1.5 | 2.74 | 2.8E-02 |
| *cln-3.3* | WBGene00000541 | protein_coding_gene | 1.5 | 2.73 | 2.9E-02 |
| *snf-4* | WBGene00004903 | pseudogene | 1.4 | 2.73 | 4.3E-02 |
| *C34H4.2* | WBGene00016425 | protein_coding_gene | 1.4 | 2.72 | 1.2E-02 |
| *F22E5.8* | WBGene00017707 | protein_coding_gene | 1.4 | 2.69 | 5.0E-03 |
| *act-5* | WBGene00000067 | protein_coding_gene | 1.4 | 2.69 | 4.3E-02 |
| *T19D12.1* | WBGene00020576 | protein_coding_gene | 1.4 | 2.61 | 2.4E-02 |
| *lec-8* | WBGene00002271 | protein_coding_gene | 1.4 | 2.58 | 9.9E-03 |
| *F39G3.5* | WBGene00018210 | protein_coding_gene | 1.4 | 2.57 | 2.9E-02 |
| *chk-2* | WBGene00000499 | protein_coding_gene | 1.4 | 2.57 | 7.9E-03 |
| *C12D12.1* | WBGene00015713 | protein_coding_gene | 1.3 | 2.52 | 8.2E-04 |
| *F15D4.5* | WBGene00008862 | protein_coding_gene | 1.3 | 2.50 | 3.7E-02 |
| *F43C1.5* | WBGene00009649 | protein_coding_gene | 1.3 | 2.49 | 4.5E-02 |
| *klu-2* | WBGene00022592 | protein_coding_gene | 1.3 | 2.48 | 1.2E-02 |
| *lec-9* | WBGene00002272 | protein_coding_gene | 1.3 | 2.47 | 3.8E-05 |
| *F28B4.3* | WBGene00017892 | protein_coding_gene | 1.3 | 2.47 | 4.5E-02 |
| *kri-1* | WBGene00013955 | protein_coding_gene | 1.3 | 2.44 | 1.7E-02 |
| *ZK856.5* | WBGene00014106 | protein_coding_gene | 1.2 | 2.37 | 4.5E-02 |
| *fbxa-190* | WBGene00018939 | protein_coding_gene | 1.2 | 2.37 | 4.0E-02 |
| *T25C12.3* | WBGene00012018 | protein_coding_gene | 1.2 | 2.34 | 4.5E-02 |
| *C55A6.4* | WBGene00008333 | protein_coding_gene | 1.2 | 2.24 | 5.5E-04 |
| *F42A10.7* | WBGene00018343 | protein_coding_gene | 1.2 | 2.23 | 2.9E-02 |
| *F19C7.1* | WBGene00017591 | protein_coding_gene | 1.1 | 2.21 | 1.5E-02 |
| *aqp-4* | WBGene00000172 | protein_coding_gene | 1.1 | 2.18 | 3.4E-06 |
| *sqrd-1* | WBGene00008538 | protein_coding_gene | 1.0 | 2.06 | 2.6E-02 |
| *W03F8.2* | WBGene00020992 | pseudogene | 1.0 | 2.06 | 2.6E-02 |
| *W04B5.3* | WBGene00021020 | protein_coding_gene | 1.0 | 2.02 | 4.3E-02 |
| *C17C3.15* | WBGene00015896 | protein_coding_gene | 1.0 | 2.02 | 3.7E-02 |

| | | | | | |
|---|---|---|---|---|---|
| *ham-1* | WBGene00001820 | protein_coding_gene | -1.0 | 0.49 | 1.1E-02 |
| *M7.8* | WBGene00010884 | protein_coding_gene | -1.2 | 0.45 | 1.5E-02 |
| *BE0003N10.3* | WBGene00015268 | protein_coding_gene | -1.2 | 0.44 | 6.9E-03 |
| *cpt-1* | WBGene00012907 | protein_coding_gene | -1.2 | 0.44 | 1.5E-04 |
| *ZK512.1* | WBGene00013982 | protein_coding_gene | -1.2 | 0.43 | 3.0E-02 |
| *cht-1* | WBGene00000503 | protein_coding_gene | -1.2 | 0.43 | 3.9E-08 |
| *frpr-11* | WBGene00019444 | protein_coding_gene | -1.2 | 0.43 | 1.6E-03 |
| *R09E12.9* | WBGene00044514 | protein_coding_gene | -1.4 | 0.39 | 4.3E-02 |
| *F40G9.20* | WBGene00195034 | ncRNA_gene | -1.4 | 0.38 | 3.7E-02 |
| *ins-7* | WBGene00002090 | protein_coding_gene | -1.4 | 0.38 | 1.5E-04 |
| *F22F7.3* | WBGene00017721 | protein_coding_gene | -1.5 | 0.36 | 2.9E-02 |
| *F36H1.12* | WBGene00009497 | protein_coding_gene | -1.5 | 0.35 | 2.8E-02 |
| *F31D5.6* | WBGene00017949 | protein_coding_gene | -1.6 | 0.33 | 4.5E-02 |
| *C42D4.1* | WBGene00016594 | protein_coding_gene | -1.6 | 0.33 | 2.5E-03 |
| *cpt-4* | WBGene00019644 | protein_coding_gene | -1.7 | 0.31 | 1.1E-04 |
| *ugt-4* | WBGene00013905 | protein_coding_gene | -1.9 | 0.27 | 3.7E-04 |
| *W10G11.2* | WBGene00021135 | protein_coding_gene | -2.0 | 0.25 | 2.7E-02 |
| *F17B5.6* | WBGene00044058 | protein_coding_gene | -2.1 | 0.23 | 2.5E-02 |
| *W10G11.4* | WBGene00021137 | protein_coding_gene | -2.2 | 0.22 | 2.4E-02 |
| *F15E6.3* | WBGene00017484 | protein_coding_gene | -2.5 | 0.17 | 1.4E-03 |

Shared DEGs (N2 & CB4856) found across all pathogens (*S. epidermidis* , *P. rettgeri* , *P. aeruginosa* , *or S. aureus* vs. OP50)

| Gene Name | Wormbase Gene ID | Type | $log_2$FC | FC | Adj P-value | $log_2$FC | FC | Adj P-value |
|---|---|---|---|---|---|---|---|---|
| *Y65B4BR.1* | WBGene00022040 | protein_coding_gene | 2.6 | 6.3 | 3.2E-05 | 1.5 | 2.8 | 9.1E-03 |
| *H02F09.3* | WBGene00019146 | protein_coding_gene | 2.3 | 5.1 | 1.5E-02 | 1.9 | 3.7 | 2.1E-05 |
| *Y41D4B.17* | WBGene00021519 | protein_coding_gene | 2.3 | 4.9 | 1.1E-02 | 2.4 | 5.4 | 1.1E-04 |
| ***oac-14*** | **WBGene00008602** | **protein_coding_gene** | **2.2** | **4.5** | **6.4E-03** | **-2.0** | **0.2** | **1.7E-02** |
| *Y47H9C.1* | WBGene00012947 | protein_coding_gene | 2.2 | 4.4 | 3.0E-03 | 2.3 | 5.0 | 1.6E-03 |
| *T07H8.11* | WBGene00304201 | protein_coding_gene | 1.4 | 2.6 | 2.9E-02 | 1.9 | 3.8 | 4.4E-03 |

*Genes in bold are differentially expressed in opposing directions

Transcription Factors Differentially Expressed in Rsponse to Pathogen Infection

| Gene Name | WormBase Gene ID | Sequence Name | $log_2$FC | $log_2$FC |
|---|---|---|---|---|
| | | N2-Specific | | |
| *cky-1* | WBGene00000521 | C15C8.2 | 2.7 | Not DE |
| *nhr-19* | WBGene00003618 | E02H1.7 | 2.0 | Not DE |
| *nhr-99* | WBGene00003689 | M02H5.1 | 2.4 | Not DE |
| *nhr-116* | WBGene00003706 | F09C6.9 | 1.6 | Not DE |
| *nhr-127* | WBGene00003717 | T13F3.3 | 2.7 | Not DE |
| *peb-1* | WBGene00003968 | T14F9.4 | 2.4 | Not DE |
| *unc-42* | WBGene00006778 | F58E6.10 | 2.5 | Not DE |
| | | CB4856-Specific | | |
| *ham-1* | WBGene00001820 | F53B2.6 | Not DE | -1.0 |
| *klu-2* | WBGene00022592 | ZC328.2 | Not DE | 1.3 |

**Supplemental Table 3**

**GO terms found to be significantly enriched after infection by all pathogens (compared to *E. coli*)**

**Supplemental Table 3 - GO Terms Enriched in Animals Infected (All Pathogens Combined)**

N2-Specific

| Category | Number of DEGs | Genes in Geneset | Downregulated Genes in Geneset | Upregulated Genes in Geneset | p-value | FDR-adjusted p-value | Differentially Expressed Genes within a Given Geneset |
|---|---|---|---|---|---|---|---|
| Stress response: pathogen | 13 | 177 | 0 | 13 | 1.5E-08 | 9.4E-07 | WBGene00004987,WBGene00004998,WBGene00006627,WBGene00008234,WBGene00009213,WBGene00009805,WBGene00010125,WBGene00015574,WBGene00016669,WBGene00017582,WBGene00018729,WBGene00018731,WBGene00020326 |
| Unknown | 114 | 8160 | 13 | 101 | 1.2E-05 | 7.2E-04 | WBGene00005657,WBGene00007366,WBGene00007414,WBGene00007460,WBGene00007693,WBGene00007957,WBGene00008102,WBGene00008168,WBGene00008226,WBGene00008301,WBGene00008462,WBGene00008712,WBGene00008850,WBGene00008905,WBGene00009292,WBGene00009503,WBGene00009724,WBGene00009778,WBGene00009787,WBGene00010285,WBGene00010686,WBGene00010835,WBGene00011286,WBGene00011383,WBGene00011445,WBGene00011692,WBGene00011798,WBGene00011838,WBGene00011894,WBGene00012069,WBGene00012186,WBGene00012412,WBGene00012501,WBGene00012540,WBGene00012819,WBGene00012850,WBGene00012959,WBGene00013007,WBGene00013101,WBGene00013146,WBGene00013897,WBGene00014049,WBGene00014698,WBGene00015537,WBGene00015549,WBGene00015573,WBGene00015576,WBGene00015748,WBGene00015878,WBGene00015993,WBGene00016017,WBGene00016052,WBGene00016119,WBGene00016218,WBGene00016332,WBGene00016659,WBGene00016845,WBGene00016877,WBGene00016883,WBGene00017133,WBGene00017146,WBGene00017205,WBGene00017303,WBGene00017361,WBGene00017378,WBGene00017434,WBGene00017660,WBGene00018195,WBGene00018730,WBGene00018874,WBGene00018965,WBGene00019272,WBGene00019391,WBGene00019495,WBGene00019520,WBGene00019699,WBGene00019855,WBGene00020194,WBGene00020242,WBGene00020394,WBGene00020613,WBGene00020626,WBGene00020700,WBGene00020801,WBGene00021121,WBGene00022440,WBGene00022441,WBGene00022517,WBGene00022562,WBGene00022597,WBGene00022611,WBGene00022891,WBGene00023501,WBGene00044024,WBGene00044120,WBGene00044206,WBGene00044212,WBGene00044280,WBGene00044644,WBGene00044783,WBGene00045275,WBGene00045411,WBGene00045412,WBGene00194815,WBGene00206386,WBGene00206521,WBGene00219283,WBGene00219410,WBGene00235114,WBGene00235133,WBGene00235268,WBGene00270321,WBGene00271812,WBGene00302974 |
| Transmembrane transport: lipid | 4 | 15 | 0 | 4 | 1.8E-05 | 1.1E-03 | WBGene00012255,WBGene00012256,WBGene00012257,WBGene00012261 |
| Metabolism: lipid | 16 | 525 | 4 | 12 | 2.7E-05 | 1.7E-03 | WBGene00000213,WBGene00000988,WBGene00001158,WBGene00001391,WBGene00002260,WBGene00006351,WBGene00007365,WBGene00009773,WBGene00010062,WBGene00010769,WBGene00011321,WBGene00018269,WBGene00018488,WBGene00018707,WBGene00020016,WBGene00021002 |

CB4856-Specific

| Category | Number of DEGs | Genes in Geneset | Downregulated Genes in Geneset | Upregulated Genes in Geneset | p-value | FDR-adjusted p-value | Differentially Expressed Genes within a Given Geneset |
|---|---|---|---|---|---|---|---|
| Metabolism: lipid | 10 | 525 | 2 | 8 | 3.7E-05 | 1.7E-03 | WBGene00001240,WBGene00001243,WBGene00008598,WBGene00011725,WBGene00011960,WBGene00012907,WBGene00014106,WBGene00018398,WBGene00019580,WBGene00019644 |

**Supplemental Table 4**

**Lists of differentially expressed genes that are identified in either N2, CB4856, or both, after infection Gram-positive pathogens (compared to *E. coli*)**

**Supplemental Table 4 - Differentially Expressed Genes in Gram-positive Infected Animals**

**N2**-exclusive DEGs in response to Gram-positive pathogens (*S. epidermidis* or *S. aureus* vs. OP50)

| Gene Name | Wormbase Gene ID | Type | $log_2FC$ | FC | Adj P-value |
|---|---|---|---|---|---|
| *clec-76* | WBGene00021587 | protein_coding_gene | 7.4 | 167.14 | 7.3E-15 |
| *thn-1* | WBGene00009213 | protein_coding_gene | 6.1 | 68.18 | 3.6E-13 |
| *tts-1* | WBGene00006650 | ncRNA_gene | 6.0 | 65.61 | 5.5E-20 |
| *Y68A4A.13* | WBGene00044212 | protein_coding_gene | 5.8 | 57.20 | 2.3E-10 |
| *lipl-2* | WBGene00009773 | protein_coding_gene | 5.6 | 49.60 | 2.6E-13 |
| *C50F7.5* | WBGene00016845 | protein_coding_gene | 5.4 | 43.70 | 9.5E-13 |
| *fbxa-24* | WBGene00018803 | protein_coding_gene | 5.4 | 43.70 | 2.9E-13 |
| *T26H5.9* | WBGene00044206 | protein_coding_gene | 5.0 | 31.70 | 7.1E-12 |
| *cest-29* | WBGene00017205 | pseudogene | 4.8 | 27.18 | 5.2E-07 |
| *clec-70* | WBGene00021581 | protein_coding_gene | 4.7 | 26.78 | 1.0E-09 |
| *cyp-37B1* | WBGene00009226 | protein_coding_gene | 4.7 | 26.32 | 5.3E-10 |
| *F49H6.5* | WBGene00009908 | protein_coding_gene | 4.7 | 25.61 | 3.4E-08 |
| *F14D7.10* | WBGene00008797 | protein_coding_gene | 4.6 | 24.31 | 4.2E-05 |
| *spp-2* | WBGene00004987 | protein_coding_gene | 4.6 | 23.80 | 9.7E-23 |
| *mtl-1* | WBGene00003473 | protein_coding_gene | 4.5 | 23.34 | 1.1E-09 |
| *T26H5.10* | WBGene00044783 | protein_coding_gene | 4.5 | 23.30 | 1.1E-05 |
| *F46A8.13* | WBGene00219950 | ncRNA_gene | 4.5 | 23.29 | 6.6E-06 |
| *F47B8.10* | WBGene00009811 | protein_coding_gene | 4.5 | 23.16 | 6.7E-06 |
| *sodh-1* | WBGene00010790 | protein_coding_gene | 4.5 | 23.07 | 5.0E-10 |
| *T20D4.12* | WBGene00020618 | protein_coding_gene | 4.5 | 22.22 | 1.6E-04 |
| *T26H5.4* | WBGene00012069 | protein_coding_gene | 4.4 | 21.57 | 4.2E-06 |
| *F11A5.17* | WBGene00045510 | protein_coding_gene | 4.4 | 21.56 | 3.6E-07 |
| *cyp-32B1* | WBGene00021167 | protein_coding_gene | 4.3 | 20.09 | 7.2E-10 |
| *ftn-1* | WBGene00001500 | protein_coding_gene | 4.3 | 20.05 | 5.5E-07 |
| *K03D3.2* | WBGene00010528 | protein_coding_gene | 4.3 | 19.70 | 1.7E-04 |
| *E02H4.4* | WBGene00008462 | protein_coding_gene | 4.3 | 19.28 | 7.2E-08 |
| *clec-75* | WBGene00021586 | protein_coding_gene | 4.2 | 18.90 | 3.5E-06 |
| *F53A9.8* | WBGene00018731 | protein_coding_gene | 4.2 | 18.39 | 3.8E-10 |
| *Y53F4B.45* | WBGene00044728 | ncRNA_gene | 4.2 | 18.38 | 7.4E-12 |
| *clec-77* | WBGene00044343 | pseudogene | 4.2 | 17.91 | 6.8E-05 |
| *T07G12.5* | WBGene00011596 | protein_coding_gene | 4.2 | 17.79 | 3.0E-07 |
| *T26H5.14* | WBGene00235133 | protein_coding_gene | 4.2 | 17.75 | 4.7E-06 |
| *pals-37* | WBGene00138721 | protein_coding_gene | 4.1 | 17.53 | 1.6E-05 |
| *ugt-18* | WBGene00013900 | protein_coding_gene | 4.1 | 17.49 | 2.9E-08 |
| *Y47D7A.12* | WBGene00021624 | protein_coding_gene | 4.1 | 17.23 | 1.9E-04 |
| *C18D11.6* | WBGene00007682 | protein_coding_gene | 4.1 | 16.93 | 2.3E-04 |
| *F47B8.4* | WBGene00009805 | protein_coding_gene | 4.1 | 16.70 | 9.1E-06 |
| *far-7* | WBGene00001391 | protein_coding_gene | 4.0 | 16.51 | 3.2E-07 |
| *clec-34* | WBGene00012023 | protein_coding_gene | 4.0 | 16.27 | 7.8E-04 |
| *oac-29* | WBGene00018295 | protein_coding_gene | 4.0 | 16.26 | 1.4E-04 |
| *fil-1* | WBGene00011321 | protein_coding_gene | 4.0 | 16.13 | 1.2E-05 |
| *fbxa-164* | WBGene00015599 | protein_coding_gene | 4.0 | 15.94 | 3.2E-04 |
| *M02E1.4* | WBGene00284848 | protein_coding_gene | 4.0 | 15.73 | 4.0E-06 |
| *F41C3.2* | WBGene00018268 | protein_coding_gene | 3.9 | 15.38 | 1.7E-05 |
| *Y47H10A.3* | WBGene00012959 | protein_coding_gene | 3.9 | 15.32 | 6.2E-05 |
| *oac-18* | WBGene00009230 | pseudogene | 3.9 | 15.06 | 2.0E-04 |
| *W02D7.11* | WBGene00020946 | protein_coding_gene | 3.9 | 15.03 | 9.8E-05 |
| *B0284.5* | WBGene00045188 | pseudogene | 3.9 | 14.79 | 3.2E-04 |
| *clec-71* | WBGene00021582 | protein_coding_gene | 3.9 | 14.72 | 4.7E-06 |
| *T20D4.10* | WBGene00020616 | protein_coding_gene | 3.9 | 14.64 | 7.8E-04 |
| *asp-16* | WBGene00012682 | protein_coding_gene | 3.9 | 14.58 | 7.4E-04 |
| *ilys-2* | WBGene00016669 | protein_coding_gene | 3.9 | 14.55 | 2.6E-05 |
| *fbxa-48* | WBGene00021862 | protein_coding_gene | 3.8 | 14.32 | 8.6E-07 |
| *sqst-4* | WBGene00012516 | protein_coding_gene | 3.8 | 14.23 | 2.4E-03 |

| | | | | | |
|---|---|---|---|---|---|
| *K09C4.4* | WBGene00019549 | protein_coding_gene | 3.8 | 14.13 | 7.0E-04 |
| *pals-32* | WBGene00016281 | protein_coding_gene | 3.8 | 14.12 | 1.5E-06 |
| *F54F3.2* | WBGene00014770 | pseudogene | 3.8 | 14.05 | 1.3E-03 |
| *ZC204.12* | WBGene00022562 | protein_coding_gene | 3.8 | 13.93 | 2.5E-10 |
| *Y46C8AM.1* | WBGene00271795 | protein_coding_gene | 3.8 | 13.90 | 1.4E-07 |
| *best-5* | WBGene00007404 | protein_coding_gene | 3.8 | 13.65 | 9.4E-10 |
| *ttr-23* | WBGene00011894 | protein_coding_gene | 3.8 | 13.54 | 3.4E-06 |
| *ins-20* | WBGene00002103 | protein_coding_gene | 3.8 | 13.54 | 3.6E-04 |
| *F55H12.7* | WBGene00219283 | protein_coding_gene | 3.7 | 13.44 | 1.5E-04 |
| *thn-2* | WBGene00009215 | protein_coding_gene | 3.7 | 13.34 | 3.5E-05 |
| *cky-1* | WBGene00000521 | protein_coding_gene | 3.7 | 13.21 | 2.0E-04 |
| *wrt-2* | WBGene00006948 | protein_coding_gene | 3.7 | 13.17 | 3.5E-11 |
| *Y105C5B.7* | WBGene00013648 | pseudogene | 3.7 | 13.13 | 4.0E-04 |
| *rgba-1* | WBGene00077766 | protein_coding_gene | 3.7 | 13.10 | 2.4E-03 |
| *T28A11.18* | WBGene00020880 | protein_coding_gene | 3.7 | 12.96 | 1.3E-03 |
| *fbxa-173* | WBGene00044960 | pseudogene | 3.7 | 12.79 | 1.3E-03 |
| *C17B7.4* | WBGene00015878 | protein_coding_gene | 3.6 | 12.48 | 1.5E-04 |
| *hil-1* | WBGene00001852 | protein_coding_gene | 3.6 | 12.40 | 3.6E-07 |
| *cyp-35B2* | WBGene00019471 | protein_coding_gene | 3.6 | 12.13 | 2.2E-03 |
| *F53B7.7* | WBGene00009969 | protein_coding_gene | 3.6 | 12.12 | 7.8E-05 |
| *T21D12.7* | WBGene00020648 | protein_coding_gene | 3.6 | 12.11 | 3.1E-04 |
| *F47H4.2* | WBGene00009835 | protein_coding_gene | 3.6 | 12.05 | 2.3E-07 |
| *Y65B4BL.1* | WBGene00022033 | protein_coding_gene | 3.6 | 11.95 | 4.9E-04 |
| *C35C5.8* | WBGene00007957 | protein_coding_gene | 3.6 | 11.93 | 1.2E-05 |
| *clec-17* | WBGene00008477 | protein_coding_gene | 3.6 | 11.79 | 1.1E-04 |
| *C18H7.1* | WBGene00015993 | protein_coding_gene | 3.6 | 11.75 | 6.9E-05 |
| *F49F1.5* | WBGene00018645 | protein_coding_gene | 3.5 | 11.71 | 4.7E-06 |
| *C17B7.10* | WBGene00015884 | protein_coding_gene | 3.5 | 11.65 | 8.2E-04 |
| *sru-40* | WBGene00005703 | protein_coding_gene | 3.5 | 11.57 | 2.1E-03 |
| *F53A9.7* | WBGene00018730 | protein_coding_gene | 3.5 | 11.57 | 1.1E-06 |
| *xtr-1* | WBGene00006965 | protein_coding_gene | 3.5 | 11.41 | 4.2E-09 |
| *cup-16* | WBGene00007919 | protein_coding_gene | 3.5 | 11.27 | 2.2E-08 |
| *F21C10.11* | WBGene00017660 | protein_coding_gene | 3.5 | 11.18 | 1.7E-06 |
| *T05F1.11* | WBGene00011495 | protein_coding_gene | 3.5 | 11.18 | 4.9E-11 |
| *T20D4.11* | WBGene00020617 | protein_coding_gene | 3.4 | 10.90 | 3.5E-04 |
| *chil-16* | WBGene00011158 | protein_coding_gene | 3.4 | 10.83 | 3.3E-03 |
| *F48G7.8* | WBGene00018619 | protein_coding_gene | 3.4 | 10.81 | 3.2E-03 |
| *R07C12.1* | WBGene00019931 | protein_coding_gene | 3.4 | 10.75 | 3.8E-04 |
| *srbc-20* | WBGene00016693 | protein_coding_gene | 3.4 | 10.74 | 2.7E-03 |
| *B0563.9* | WBGene00044687 | protein_coding_gene | 3.4 | 10.73 | 6.2E-03 |
| *F58F12.12* | WBGene00271812 | protein_coding_gene | 3.4 | 10.61 | 1.1E-08 |
| *srr-6* | WBGene00005657 | protein_coding_gene | 3.4 | 10.57 | 5.2E-04 |
| *T10C6.15* | WBGene00050899 | protein_coding_gene | 3.4 | 10.52 | 2.2E-04 |
| *thn-8* | WBGene00044285 | pseudogene | 3.4 | 10.51 | 4.5E-03 |
| *C25H3.10* | WBGene00016119 | protein_coding_gene | 3.4 | 10.50 | 2.8E-07 |
| *C18D4.4* | WBGene00007674 | protein_coding_gene | 3.4 | 10.43 | 4.7E-03 |
| *F59E11.7* | WBGene00019113 | protein_coding_gene | 3.4 | 10.36 | 6.8E-06 |
| *F45D3.4* | WBGene00009724 | protein_coding_gene | 3.4 | 10.34 | 3.1E-05 |
| *spp-6* | WBGene00004991 | protein_coding_gene | 3.4 | 10.22 | 2.6E-04 |
| *F26G1.15* | WBGene00270318 | protein_coding_gene | 3.4 | 10.22 | 1.2E-03 |
| *lim-4* | WBGene00002987 | protein_coding_gene | 3.3 | 10.11 | 2.9E-04 |
| *Y39A3B.7* | WBGene00206512 | protein_coding_gene | 3.3 | 10.09 | 5.0E-03 |
| *F37A8.5* | WBGene00009503 | protein_coding_gene | 3.3 | 9.90 | 7.0E-06 |
| *fbxa-138* | WBGene00022345 | protein_coding_gene | 3.3 | 9.84 | 1.8E-05 |
| *T03E6.9* | WBGene00045356 | protein_coding_gene | 3.3 | 9.83 | 1.8E-03 |
| *F45D11.1* | WBGene00018448 | pseudogene | 3.3 | 9.71 | 4.5E-05 |
| *Y65B4BM.3* | WBGene00302974 | protein_coding_gene | 3.3 | 9.68 | 1.5E-09 |
| *F16B4.7* | WBGene00017509 | protein_coding_gene | 3.3 | 9.61 | 1.4E-03 |

| | | | | | |
|---|---|---|---|---|---|
| *C25F9.2* | WBGene00007723 | protein_coding_gene | 3.3 | 9.58 | 2.5E-03 |
| *H42K12.3* | WBGene00019272 | protein_coding_gene | 3.3 | 9.58 | 2.1E-08 |
| *F57E7.2* | WBGene00010203 | protein_coding_gene | 3.3 | 9.57 | 3.0E-04 |
| *C14A11.2* | WBGene00015748 | protein_coding_gene | 3.2 | 9.50 | 9.4E-06 |
| *Y87G2A.16* | WBGene00013602 | protein_coding_gene | 3.2 | 9.45 | 5.3E-09 |
| *btb-16* | WBGene00018195 | protein_coding_gene | 3.2 | 9.43 | 2.3E-22 |
| *Y62H9A.11* | WBGene00013399 | protein_coding_gene | 3.2 | 9.43 | 3.3E-03 |
| *K01A6.7* | WBGene00045096 | protein_coding_gene | 3.2 | 9.39 | 5.1E-03 |
| *nhr-99* | WBGene00003689 | protein_coding_gene | 3.2 | 9.32 | 1.3E-05 |
| *T12D8.5* | WBGene00011733 | protein_coding_gene | 3.2 | 9.24 | 6.4E-07 |
| *C25F9.12* | WBGene00045412 | protein_coding_gene | 3.2 | 9.13 | 1.3E-03 |
| *ZC443.2* | WBGene00013897 | protein_coding_gene | 3.2 | 9.09 | 7.9E-04 |
| *oac-31* | WBGene00018707 | protein_coding_gene | 3.2 | 9.00 | 1.5E-04 |
| *ZC443.3* | WBGene00013898 | protein_coding_gene | 3.2 | 8.94 | 1.4E-05 |
| *R05A10.4* | WBGene00011021 | protein_coding_gene | 3.2 | 8.88 | 1.8E-04 |
| *unc-42* | WBGene00006778 | protein_coding_gene | 3.1 | 8.87 | 5.3E-07 |
| *peb-1* | WBGene00003968 | protein_coding_gene | 3.1 | 8.84 | 4.5E-08 |
| *fbxc-41* | WBGene00016304 | pseudogene | 3.1 | 8.79 | 2.2E-04 |
| *C07G3.15* | WBGene00269378 | protein_coding_gene | 3.1 | 8.78 | 9.9E-03 |
| *T24E12.5* | WBGene00020774 | protein_coding_gene | 3.1 | 8.77 | 3.8E-04 |
| *srw-85* | WBGene00005832 | protein_coding_gene | 3.1 | 8.73 | 3.2E-04 |
| *str-124* | WBGene00006175 | protein_coding_gene | 3.1 | 8.73 | 7.3E-03 |
| *C13A2.1* | WBGene00015718 | protein_coding_gene | 3.1 | 8.70 | 1.8E-02 |
| *asp-11* | WBGene00017677 | pseudogene | 3.1 | 8.63 | 9.5E-05 |
| *ZK993.5* | WBGene00236809 | protein_coding_gene | 3.1 | 8.62 | 5.0E-03 |
| *clec-37* | WBGene00013609 | protein_coding_gene | 3.1 | 8.61 | 4.6E-04 |
| *fbxa-44* | WBGene00008616 | protein_coding_gene | 3.1 | 8.58 | 8.1E-04 |
| *osm-8* | WBGene00003888 | protein_coding_gene | 3.1 | 8.55 | 2.1E-05 |
| *scl-13* | WBGene00019179 | protein_coding_gene | 3.1 | 8.41 | 2.3E-02 |
| *F23H12.5* | WBGene00009094 | protein_coding_gene | 3.1 | 8.40 | 7.8E-07 |
| *Y82E9BR.6* | WBGene00022339 | protein_coding_gene | 3.1 | 8.39 | 2.5E-03 |
| *F57F4.2* | WBGene00019016 | protein_coding_gene | 3.1 | 8.37 | 1.3E-03 |
| *Y60C6A.1* | WBGene00022013 | protein_coding_gene | 3.1 | 8.37 | 4.7E-03 |
| *T16G1.4* | WBGene00011798 | protein_coding_gene | 3.1 | 8.36 | 2.0E-04 |
| *K10D11.2* | WBGene00010746 | protein_coding_gene | 3.1 | 8.35 | 3.1E-03 |
| *pals-36* | WBGene00044787 | pseudogene | 3.1 | 8.33 | 7.6E-03 |
| *fbxa-7* | WBGene00020637 | protein_coding_gene | 3.1 | 8.32 | 1.2E-02 |
| *C44H9.7* | WBGene00008102 | protein_coding_gene | 3.0 | 8.27 | 1.3E-05 |
| *F09C6.14* | WBGene00044278 | protein_coding_gene | 3.0 | 8.27 | 4.6E-04 |
| *F59E11.6* | WBGene00019112 | protein_coding_gene | 3.0 | 8.21 | 5.5E-03 |
| *zmp-5* | WBGene00010423 | protein_coding_gene | 3.0 | 8.21 | 3.3E-03 |
| *F11E6.9* | WBGene00008712 | protein_coding_gene | 3.0 | 8.18 | 8.3E-05 |
| *F09C6.13* | WBGene00044183 | protein_coding_gene | 3.0 | 8.16 | 5.2E-07 |
| *srz-99* | WBGene00012028 | protein_coding_gene | 3.0 | 8.16 | 2.0E-03 |
| *B0205.4* | WBGene00015019 | protein_coding_gene | 3.0 | 8.15 | 9.8E-05 |
| *pals-31* | WBGene00018614 | protein_coding_gene | 3.0 | 8.10 | 8.6E-05 |
| *F56D2.3* | WBGene00018965 | protein_coding_gene | 3.0 | 8.08 | 4.1E-08 |
| *F58H1.2* | WBGene00010285 | protein_coding_gene | 3.0 | 8.08 | 1.2E-05 |
| *F55H12.2* | WBGene00010133 | protein_coding_gene | 3.0 | 8.00 | 2.1E-09 |
| *ZC395.5* | WBGene00022597 | protein_coding_gene | 3.0 | 7.99 | 6.7E-07 |
| *clec-2* | WBGene00015198 | protein_coding_gene | 3.0 | 7.97 | 1.1E-02 |
| *R11D1.12* | WBGene00045336 | protein_coding_gene | 3.0 | 7.94 | 1.1E-04 |
| *fbxa-137* | WBGene00020853 | protein_coding_gene | 3.0 | 7.93 | 4.3E-03 |
| *C25F9.11* | WBGene00045411 | protein_coding_gene | 3.0 | 7.89 | 2.1E-04 |
| *C06E4.8* | WBGene00015537 | protein_coding_gene | 3.0 | 7.88 | 7.6E-08 |
| *ttr-29* | WBGene00011286 | protein_coding_gene | 3.0 | 7.87 | 8.1E-05 |
| *spp-21* | WBGene00044284 | protein_coding_gene | 3.0 | 7.80 | 6.9E-03 |
| *kreg-1* | WBGene00018725 | protein_coding_gene | 3.0 | 7.78 | 4.3E-04 |

| | | | | | |
|---|---|---|---|---|---|
| *R09D1.12* | WBGene00011168 | protein_coding_gene | 3.0 | 7.77 | 2.8E-07 |
| *linc-84* | WBGene00219592 | lincRNA_gene | 3.0 | 7.75 | 1.9E-03 |
| *clec-72* | WBGene00021583 | protein_coding_gene | 3.0 | 7.75 | 6.6E-04 |
| *T28A11.20* | WBGene00020882 | protein_coding_gene | 2.9 | 7.72 | 1.5E-02 |
| *F35D2.6* | WBGene00194916 | protein_coding_gene | 2.9 | 7.71 | 1.0E-02 |
| *C25E10.16* | WBGene00202512 | protein_coding_gene | 2.9 | 7.69 | 9.4E-03 |
| *Y53C12B.7* | WBGene00013146 | protein_coding_gene | 2.9 | 7.67 | 2.5E-04 |
| *Y7A5A.6* | WBGene00012412 | protein_coding_gene | 2.9 | 7.66 | 1.1E-03 |
| *Y54G2A.76* | WBGene00235268 | protein_coding_gene | 2.9 | 7.61 | 1.0E-04 |
| *pals-7* | WBGene00007661 | protein_coding_gene | 2.9 | 7.60 | 1.1E-02 |
| *B0478.3* | WBGene00015202 | protein_coding_gene | 2.9 | 7.59 | 1.7E-03 |
| *Y110A7A.24* | WBGene00271632 | ncRNA_gene | 2.9 | 7.58 | 5.9E-03 |
| *clec-169* | WBGene00009526 | protein_coding_gene | 2.9 | 7.58 | 9.4E-03 |
| *nhr-127* | WBGene00003717 | protein_coding_gene | 2.9 | 7.57 | 8.9E-04 |
| *R11D1.7* | WBGene00011246 | protein_coding_gene | 2.9 | 7.56 | 1.4E-02 |
| *F21H7.12* | WBGene00009033 | protein_coding_gene | 2.9 | 7.55 | 2.5E-03 |
| *W03F9.4* | WBGene00021002 | protein_coding_gene | 2.9 | 7.49 | 2.1E-04 |
| *F10D7.4* | WBGene00017341 | protein_coding_gene | 2.9 | 7.48 | 1.0E-02 |
| *ZC190.6* | WBGene00022540 | protein_coding_gene | 2.9 | 7.42 | 1.1E-02 |
| *K04A8.21* | WBGene00271815 | protein_coding_gene | 2.9 | 7.41 | 7.1E-03 |
| *gpa-8* | WBGene00001670 | protein_coding_gene | 2.9 | 7.39 | 1.6E-03 |
| *cpr-1* | WBGene00000781 | protein_coding_gene | 2.9 | 7.37 | 1.3E-03 |
| *ins-11* | WBGene00002094 | protein_coding_gene | 2.9 | 7.36 | 5.3E-03 |
| *srm-4* | WBGene00009808 | protein_coding_gene | 2.9 | 7.35 | 1.6E-02 |
| *C40H1.9* | WBGene00008044 | protein_coding_gene | 2.9 | 7.33 | 1.6E-03 |
| *T26C5.2* | WBGene00012032 | protein_coding_gene | 2.9 | 7.29 | 1.2E-06 |
| *spp-7* | WBGene00004992 | protein_coding_gene | 2.9 | 7.28 | 5.5E-03 |
| *Y46E12BR.1* | WBGene00021598 | protein_coding_gene | 2.9 | 7.28 | 2.8E-03 |
| *cpr-4* | WBGene00000784 | protein_coding_gene | 2.9 | 7.24 | 1.1E-06 |
| *T20B3.1* | WBGene00011850 | protein_coding_gene | 2.9 | 7.24 | 9.4E-05 |
| *Y39A1A.27* | WBGene00194731 | protein_coding_gene | 2.9 | 7.24 | 1.0E-02 |
| *nhr-113* | WBGene00003703 | protein_coding_gene | 2.9 | 7.23 | 1.6E-02 |
| *cnc-6* | WBGene00000560 | protein_coding_gene | 2.9 | 7.22 | 2.2E-02 |
| *EGAP1.1* | WBGene00017146 | protein_coding_gene | 2.8 | 7.19 | 7.1E-04 |
| *nhr-37* | WBGene00018412 | protein_coding_gene | 2.8 | 7.17 | 1.7E-02 |
| *linc-69* | WBGene00219644 | lincRNA_gene | 2.8 | 7.16 | 7.9E-03 |
| *Y69H2.10* | WBGene00013485 | protein_coding_gene | 2.8 | 7.15 | 1.7E-03 |
| *W02A2.9* | WBGene00044280 | protein_coding_gene | 2.8 | 7.15 | 1.1E-05 |
| *thn-3* | WBGene00009214 | pseudogene | 2.8 | 7.15 | 2.4E-02 |
| *T19D12.3* | WBGene00020578 | protein_coding_gene | 2.8 | 7.15 | 9.3E-03 |
| *nlp-34* | WBGene00015046 | protein_coding_gene | 2.8 | 7.14 | 5.0E-03 |
| *M01B2.13* | WBGene00045338 | protein_coding_gene | 2.8 | 7.13 | 2.0E-03 |
| *Y32B12C.5* | WBGene00194769 | pseudogene | 2.8 | 7.11 | 4.4E-05 |
| *srg-25* | WBGene00005182 | protein_coding_gene | 2.8 | 7.10 | 1.6E-02 |
| *T08A9.13* | WBGene00044442 | protein_coding_gene | 2.8 | 7.09 | 1.9E-03 |
| *fbxa-12* | WBGene00016525 | protein_coding_gene | 2.8 | 7.08 | 8.6E-03 |
| *srw-86* | WBGene00005833 | protein_coding_gene | 2.8 | 7.06 | 2.2E-03 |
| *C14A6.3* | WBGene00007567 | protein_coding_gene | 2.8 | 7.05 | 1.9E-02 |
| *pitr-2* | WBGene00015054 | protein_coding_gene | 2.8 | 7.05 | 1.7E-02 |
| *F35F10.7* | WBGene00018057 | protein_coding_gene | 2.8 | 7.03 | 2.0E-02 |
| *Y77E11A.14* | WBGene00022314 | protein_coding_gene | 2.8 | 7.03 | 7.8E-05 |
| *C44H9.2* | WBGene00008098 | pseudogene | 2.8 | 6.97 | 1.3E-02 |
| *glr-3* | WBGene00001614 | protein_coding_gene | 2.8 | 6.96 | 6.9E-03 |
| *lys-10* | WBGene00003099 | protein_coding_gene | 2.8 | 6.96 | 1.0E-02 |
| *Y43C5A.3* | WBGene00012783 | protein_coding_gene | 2.8 | 6.94 | 2.7E-04 |
| *F38B6.3* | WBGene00018173 | protein_coding_gene | 2.8 | 6.94 | 8.1E-03 |
| *fbxa-185* | WBGene00009836 | protein_coding_gene | 2.8 | 6.94 | 1.7E-02 |
| *fbxa-127* | WBGene00012027 | protein_coding_gene | 2.8 | 6.93 | 2.4E-03 |

| | | | | | |
|---|---|---|---|---|---|
| *Y15E3A.5* | WBGene00012444 | protein_coding_gene | 2.8 | 6.93 | 6.4E-05 |
| *C32F10.4* | WBGene00016332 | protein_coding_gene | 2.8 | 6.86 | 2.1E-06 |
| *Y37A1B.7* | WBGene00012540 | protein_coding_gene | 2.8 | 6.85 | 4.7E-04 |
| *ptr-10* | WBGene00004224 | protein_coding_gene | 2.8 | 6.84 | 6.0E-07 |
| *scl-12* | WBGene00019178 | protein_coding_gene | 2.8 | 6.82 | 2.0E-02 |
| *ZC449.1* | WBGene00022611 | protein_coding_gene | 2.8 | 6.80 | 3.7E-05 |
| *clec-107* | WBGene00012502 | protein_coding_gene | 2.8 | 6.80 | 6.9E-03 |
| *C36B7.4* | WBGene00016467 | protein_coding_gene | 2.8 | 6.78 | 2.0E-02 |
| *M60.4* | WBGene00019780 | protein_coding_gene | 2.8 | 6.77 | 3.9E-04 |
| *fbxa-172* | WBGene00008014 | protein_coding_gene | 2.8 | 6.76 | 9.6E-06 |
| *poml-4* | WBGene00044697 | protein_coding_gene | 2.8 | 6.74 | 2.8E-04 |
| *sre-23* | WBGene00013957 | protein_coding_gene | 2.8 | 6.74 | 1.9E-03 |
| *F47G4.5* | WBGene00009826 | protein_coding_gene | 2.8 | 6.73 | 9.0E-03 |
| *twk-47* | WBGene00018002 | protein_coding_gene | 2.7 | 6.72 | 1.2E-03 |
| *lurp-3* | WBGene00009136 | protein_coding_gene | 2.7 | 6.69 | 1.2E-02 |
| *F13E9.9* | WBGene00008758 | protein_coding_gene | 2.7 | 6.67 | 1.3E-02 |
| *K08F9.1* | WBGene00010684 | protein_coding_gene | 2.7 | 6.65 | 2.0E-05 |
| *T24E12.12* | WBGene00044380 | protein_coding_gene | 2.7 | 6.61 | 2.3E-02 |
| *F56C4.4* | WBGene00044977 | protein_coding_gene | 2.7 | 6.60 | 1.0E-04 |
| *srh-282* | WBGene00005487 | protein_coding_gene | 2.7 | 6.60 | 1.5E-02 |
| *T23F6.2* | WBGene00011950 | protein_coding_gene | 2.7 | 6.59 | 3.0E-02 |
| *K08B12.1* | WBGene00019520 | protein_coding_gene | 2.7 | 6.57 | 1.1E-06 |
| *F21C10.19* | WBGene00271808 | protein_coding_gene | 2.7 | 6.55 | 1.3E-02 |
| *ZK1240.1* | WBGene00022866 | protein_coding_gene | 2.7 | 6.55 | 9.3E-04 |
| *pho-6* | WBGene00018696 | protein_coding_gene | 2.7 | 6.54 | 2.8E-05 |
| *str-112* | WBGene00006163 | protein_coding_gene | 2.7 | 6.54 | 9.6E-03 |
| *srz-24* | WBGene00017113 | protein_coding_gene | 2.7 | 6.53 | 1.3E-02 |
| *F09F7.6* | WBGene00017303 | protein_coding_gene | 2.7 | 6.53 | 2.1E-06 |
| *F09E10.10* | WBGene00022982 | ncRNA_gene | 2.7 | 6.52 | 1.5E-02 |
| *srv-4* | WBGene00005715 | protein_coding_gene | 2.7 | 6.51 | 1.9E-02 |
| *nhr-19* | WBGene00003618 | protein_coding_gene | 2.7 | 6.51 | 1.5E-05 |
| *srv-8* | WBGene00005719 | protein_coding_gene | 2.7 | 6.50 | 1.6E-02 |
| *C08E3.13* | WBGene00015605 | protein_coding_gene | 2.7 | 6.50 | 1.9E-02 |
| *C04A11.5* | WBGene00007288 | protein_coding_gene | 2.7 | 6.50 | 3.9E-03 |
| *srh-239* | WBGene00005446 | protein_coding_gene | 2.7 | 6.50 | 2.1E-02 |
| *C40C9.4* | WBGene00008037 | protein_coding_gene | 2.7 | 6.47 | 2.5E-02 |
| *ugt-15* | WBGene00008097 | protein_coding_gene | 2.7 | 6.45 | 5.1E-05 |
| *anr-51* | WBGene00219610 | antisense_lncRNA_gene | 2.7 | 6.44 | 1.5E-02 |
| *F19G12.3* | WBGene00017611 | protein_coding_gene | 2.7 | 6.43 | 2.3E-02 |
| *C02H6.3* | WBGene00220257 | protein_coding_gene | 2.7 | 6.42 | 3.0E-02 |
| *fozi-1* | WBGene00010453 | protein_coding_gene | 2.7 | 6.42 | 3.7E-05 |
| *Y43F8B.25* | WBGene00195177 | protein_coding_gene | 2.7 | 6.41 | 1.8E-02 |
| *cyp-33B1* | WBGene00016092 | protein_coding_gene | 2.7 | 6.40 | 2.4E-02 |
| *D1086.2* | WBGene00008389 | protein_coding_gene | 2.7 | 6.40 | 9.4E-06 |
| *abf-2* | WBGene00000013 | protein_coding_gene | 2.7 | 6.40 | 1.1E-03 |
| *clec-38* | WBGene00012025 | protein_coding_gene | 2.7 | 6.40 | 2.2E-02 |
| *K01A2.10* | WBGene00019284 | protein_coding_gene | 2.7 | 6.39 | 4.8E-04 |
| *F43C11.7* | WBGene00018384 | protein_coding_gene | 2.7 | 6.37 | 7.7E-06 |
| *F45D3.3* | WBGene00009723 | protein_coding_gene | 2.7 | 6.37 | 7.0E-04 |
| *T10H4.13* | WBGene00045339 | protein_coding_gene | 2.7 | 6.36 | 1.2E-02 |
| *phat-5* | WBGene00020242 | protein_coding_gene | 2.7 | 6.35 | 1.3E-03 |
| *ador-1* | WBGene00011878 | protein_coding_gene | 2.7 | 6.34 | 3.2E-03 |
| *nspc-16* | WBGene00008357 | protein_coding_gene | 2.7 | 6.33 | 2.6E-08 |
| *K02C4.2* | WBGene00010501 | protein_coding_gene | 2.7 | 6.32 | 1.1E-03 |
| *F47H4.12* | WBGene00043053 | pseudogene | 2.7 | 6.32 | 1.5E-03 |
| *gpa-9* | WBGene00001671 | protein_coding_gene | 2.7 | 6.32 | 3.4E-10 |
| *linc-47* | WBGene00219641 | lincRNA_gene | 2.7 | 6.31 | 7.2E-03 |
| *slc-36.3* | WBGene00012629 | protein_coding_gene | 2.7 | 6.31 | 9.6E-04 |

| | | | | | |
|---|---|---|---|---|---|
| *C36B1.13* | WBGene00044120 | protein_coding_gene | 2.7 | 6.30 | 2.0E-03 |
| *R03H10.2* | WBGene00019855 | protein_coding_gene | 2.7 | 6.29 | 1.4E-08 |
| *C50B8.4* | WBGene00008226 | protein_coding_gene | 2.7 | 6.29 | 5.4E-05 |
| *ZK355.3* | WBGene00022713 | protein_coding_gene | 2.7 | 6.29 | 3.5E-02 |
| *F33D4.6* | WBGene00017998 | protein_coding_gene | 2.7 | 6.28 | 1.9E-05 |
| *F56D6.9* | WBGene00044471 | protein_coding_gene | 2.7 | 6.28 | 1.5E-02 |
| *F53F4.1* | WBGene00009985 | protein_coding_gene | 2.7 | 6.28 | 2.0E-03 |
| *C06B3.6* | WBGene00007365 | protein_coding_gene | 2.7 | 6.28 | 9.0E-06 |
| *C33D9.10* | WBGene00044239 | protein_coding_gene | 2.7 | 6.28 | 1.2E-02 |
| *C31H5.1* | WBGene00007854 | protein_coding_gene | 2.7 | 6.28 | 9.9E-03 |
| *F56D6.8* | WBGene00044292 | protein_coding_gene | 2.6 | 6.27 | 1.7E-02 |
| *F53C3.6* | WBGene00018750 | protein_coding_gene | 2.6 | 6.26 | 7.6E-04 |
| *fbxa-154* | WBGene00007163 | protein_coding_gene | 2.6 | 6.26 | 1.4E-02 |
| *T28A11.5* | WBGene00020872 | pseudogene | 2.6 | 6.26 | 1.2E-02 |
| *mec-3* | WBGene00003167 | protein_coding_gene | 2.6 | 6.24 | 1.3E-07 |
| *str-97* | WBGene00006154 | protein_coding_gene | 2.6 | 6.23 | 3.5E-02 |
| *spp-13* | WBGene00004998 | protein_coding_gene | 2.6 | 6.22 | 2.2E-05 |
| *K08C9.5* | WBGene00010652 | protein_coding_gene | 2.6 | 6.20 | 1.8E-02 |
| *cnc-11* | WBGene00044900 | protein_coding_gene | 2.6 | 6.19 | 3.6E-04 |
| *W09G12.7* | WBGene00021121 | protein_coding_gene | 2.6 | 6.18 | 1.5E-04 |
| *C50F4.9* | WBGene00008234 | protein_coding_gene | 2.6 | 6.17 | 4.0E-06 |
| *H10D18.1* | WBGene00019177 | protein_coding_gene | 2.6 | 6.17 | 6.1E-03 |
| *glt-5* | WBGene00001623 | protein_coding_gene | 2.6 | 6.17 | 7.0E-04 |
| *fut-2* | WBGene00001506 | protein_coding_gene | 2.6 | 6.16 | 1.7E-03 |
| *F56H9.9* | WBGene00206460 | protein_coding_gene | 2.6 | 6.13 | 2.3E-02 |
| *F28G4.3* | WBGene00009228 | protein_coding_gene | 2.6 | 6.12 | 3.8E-02 |
| *linc-103* | WBGene00219743 | lincRNA_gene | 2.6 | 6.10 | 1.2E-02 |
| *F11A5.5* | WBGene00008673 | protein_coding_gene | 2.6 | 6.08 | 8.3E-03 |
| *lpr-4* | WBGene00012257 | protein_coding_gene | 2.6 | 6.07 | 7.6E-08 |
| *T10D4.15* | WBGene00077522 | protein_coding_gene | 2.6 | 6.07 | 3.5E-03 |
| *fbxc-20* | WBGene00018352 | protein_coding_gene | 2.6 | 6.06 | 7.9E-06 |
| *C54C8.8* | WBGene00014714 | pseudogene | 2.6 | 6.04 | 1.8E-02 |
| *Y102A11A.5* | WBGene00022415 | protein_coding_gene | 2.6 | 6.03 | 1.9E-04 |
| *T10G3.1* | WBGene00011692 | protein_coding_gene | 2.6 | 6.03 | 5.3E-06 |
| *F18G5.6* | WBGene00017582 | protein_coding_gene | 2.6 | 6.01 | 9.5E-06 |
| *hlh-13* | WBGene00001957 | protein_coding_gene | 2.6 | 6.01 | 1.9E-02 |
| *C54F6.18* | WBGene00235093 | protein_coding_gene | 2.6 | 5.98 | 1.2E-02 |
| *C42D4.13* | WBGene00016598 | protein_coding_gene | 2.6 | 5.97 | 3.8E-03 |
| *clec-67* | WBGene00018971 | protein_coding_gene | 2.6 | 5.95 | 3.7E-03 |
| *C07E3.3* | WBGene00007414 | protein_coding_gene | 2.6 | 5.94 | 1.3E-04 |
| *mlt-9* | WBGene00008605 | protein_coding_gene | 2.6 | 5.92 | 4.2E-06 |
| *R08A2.7* | WBGene00011137 | protein_coding_gene | 2.6 | 5.92 | 3.5E-04 |
| *cyp-34A1* | WBGene00011698 | protein_coding_gene | 2.6 | 5.91 | 1.1E-02 |
| *F49F1.14* | WBGene00044477 | protein_coding_gene | 2.6 | 5.91 | 2.2E-02 |
| *Y57G11C.1131* | WBGene00194661 | protein_coding_gene | 2.6 | 5.90 | 3.1E-02 |
| *F32A7.8* | WBGene00044245 | protein_coding_gene | 2.6 | 5.89 | 3.0E-02 |
| *T05A7.11* | WBGene00044383 | protein_coding_gene | 2.6 | 5.88 | 6.5E-03 |
| *C01F1.5* | WBGene00015300 | protein_coding_gene | 2.6 | 5.88 | 7.8E-04 |
| *srh-72* | WBGene00005293 | protein_coding_gene | 2.6 | 5.87 | 3.2E-02 |
| *F09F9.2* | WBGene00017307 | protein_coding_gene | 2.5 | 5.86 | 9.1E-04 |
| *W02D7.8* | WBGene00020943 | protein_coding_gene | 2.5 | 5.83 | 8.7E-05 |
| *nspe-6* | WBGene00012603 | protein_coding_gene | 2.5 | 5.83 | 1.6E-02 |
| *Y38H8A.1* | WBGene00012635 | protein_coding_gene | 2.5 | 5.82 | 1.5E-03 |
| *B0281.6* | WBGene00015113 | protein_coding_gene | 2.5 | 5.81 | 2.0E-03 |
| *ttr-26* | WBGene00013079 | protein_coding_gene | 2.5 | 5.81 | 2.9E-04 |
| *K03H6.4* | WBGene00019369 | protein_coding_gene | 2.5 | 5.80 | 1.6E-02 |
| *nhr-21* | WBGene00003620 | protein_coding_gene | 2.5 | 5.80 | 2.4E-08 |
| *F08H9.3* | WBGene00008591 | protein_coding_gene | 2.5 | 5.80 | 5.6E-03 |

| | | | | | |
|---|---|---|---|---|---|
| *oac-47* | WBGene00011933 | pseudogene | 2.5 | 5.80 | 1.2E-02 |
| *ZC123.1* | WBGene00022517 | protein_coding_gene | 2.5 | 5.80 | 1.1E-06 |
| *sre-1* | WBGene00005149 | protein_coding_gene | 2.5 | 5.79 | 2.0E-02 |
| *F22B7.9* | WBGene00017698 | protein_coding_gene | 2.5 | 5.79 | 2.7E-09 |
| *F11A6.3* | WBGene00077549 | pseudogene | 2.5 | 5.78 | 1.1E-02 |
| *T19H5.6* | WBGene00044807 | protein_coding_gene | 2.5 | 5.77 | 1.0E-02 |
| *K02F3.15* | WBGene00220001 | snoRNA_gene | 2.5 | 5.77 | 2.0E-02 |
| *linc-85* | WBGene00219754 | lincRNA_gene | 2.5 | 5.76 | 6.2E-04 |
| *math-21* | WBGene00016556 | protein_coding_gene | 2.5 | 5.75 | 9.3E-04 |
| *ZK105.11* | WBGene00194895 | protein_coding_gene | 2.5 | 5.75 | 2.8E-02 |
| *clec-55* | WBGene00008597 | protein_coding_gene | 2.5 | 5.75 | 3.2E-04 |
| *K01A6.8* | WBGene00194869 | protein_coding_gene | 2.5 | 5.73 | 2.1E-02 |
| *cutl-28* | WBGene00018256 | protein_coding_gene | 2.5 | 5.72 | 1.1E-04 |
| *K10C2.2* | WBGene00019618 | protein_coding_gene | 2.5 | 5.70 | 3.0E-02 |
| *cng-3* | WBGene00000563 | protein_coding_gene | 2.5 | 5.70 | 9.4E-03 |
| *C48B4.3* | WBGene00008168 | protein_coding_gene | 2.5 | 5.69 | 1.8E-06 |
| *B0462.1* | WBGene00007181 | pseudogene | 2.5 | 5.69 | 8.0E-03 |
| *lys-7* | WBGene00003096 | protein_coding_gene | 2.5 | 5.69 | 4.2E-04 |
| *C35E7.12* | WBGene00271779 | protein_coding_gene | 2.5 | 5.69 | 2.5E-02 |
| *F40H7.12* | WBGene00044379 | protein_coding_gene | 2.5 | 5.69 | 4.2E-02 |
| *K08D8.9* | WBGene00194918 | protein_coding_gene | 2.5 | 5.68 | 3.7E-02 |
| *C14A6.16* | WBGene00206536 | pseudogene | 2.5 | 5.68 | 4.3E-03 |
| *lys-3* | WBGene00003092 | protein_coding_gene | 2.5 | 5.68 | 4.9E-03 |
| *ZK1290.1* | WBGene00022886 | protein_coding_gene | 2.5 | 5.68 | 2.0E-02 |
| *skr-5* | WBGene00004811 | protein_coding_gene | 2.5 | 5.67 | 3.7E-03 |
| *hsd-3* | WBGene00022616 | protein_coding_gene | 2.5 | 5.67 | 3.5E-02 |
| *C08F11.3* | WBGene00007452 | protein_coding_gene | 2.5 | 5.66 | 7.9E-03 |
| *C56G3.2* | WBGene00016985 | protein_coding_gene | 2.5 | 5.65 | 1.3E-02 |
| *F28C6.9* | WBGene00009208 | protein_coding_gene | 2.5 | 5.65 | 3.4E-02 |
| *B0563.5* | WBGene00015262 | protein_coding_gene | 2.5 | 5.64 | 5.8E-05 |
| *lpr-5* | WBGene00012256 | protein_coding_gene | 2.5 | 5.63 | 7.4E-07 |
| *srt-65* | WBGene00015368 | protein_coding_gene | 2.5 | 5.63 | 2.9E-02 |
| *F09C6.11* | WBGene00044181 | protein_coding_gene | 2.5 | 5.62 | 3.0E-05 |
| *F38B2.2* | WBGene00009529 | protein_coding_gene | 2.5 | 5.62 | 1.6E-04 |
| *F13H8.1* | WBGene00017434 | protein_coding_gene | 2.5 | 5.62 | 1.4E-05 |
| *K10G6.4* | WBGene00019641 | protein_coding_gene | 2.5 | 5.62 | 7.9E-03 |
| *W04C9.9* | WBGene00235388 | protein_coding_gene | 2.5 | 5.61 | 4.4E-02 |
| *T03G6.1* | WBGene00020194 | protein_coding_gene | 2.5 | 5.60 | 2.2E-05 |
| *C12D5.3* | WBGene00015706 | protein_coding_gene | 2.5 | 5.60 | 8.2E-03 |
| *hhat-1* | WBGene00013852 | protein_coding_gene | 2.5 | 5.60 | 9.4E-07 |
| *C30G4.2* | WBGene00016268 | protein_coding_gene | 2.5 | 5.59 | 1.7E-02 |
| *M60.7* | WBGene00019782 | protein_coding_gene | 2.5 | 5.58 | 5.5E-03 |
| *nhr-178* | WBGene00017510 | protein_coding_gene | 2.5 | 5.58 | 2.2E-03 |
| *C23G10.10* | WBGene00016016 | protein_coding_gene | 2.5 | 5.58 | 4.5E-02 |
| *Y97E10AL.1* | WBGene00022392 | protein_coding_gene | 2.5 | 5.58 | 3.4E-03 |
| *nmad-1* | WBGene00017304 | protein_coding_gene | 2.5 | 5.57 | 1.9E-05 |
| *T13F3.7* | WBGene00011754 | protein_coding_gene | 2.5 | 5.57 | 6.2E-03 |
| *F36D1.7* | WBGene00009466 | protein_coding_gene | 2.5 | 5.55 | 2.6E-06 |
| *F48G7.10* | WBGene00018621 | protein_coding_gene | 2.5 | 5.55 | 1.2E-02 |
| *M7.10* | WBGene00010886 | protein_coding_gene | 2.5 | 5.53 | 1.3E-02 |
| *irld-24* | WBGene00007844 | protein_coding_gene | 2.5 | 5.53 | 2.6E-02 |
| *dpy-3* | WBGene00001065 | protein_coding_gene | 2.5 | 5.52 | 2.5E-05 |
| *F47B10.3* | WBGene00009814 | protein_coding_gene | 2.5 | 5.52 | 1.4E-02 |
| *K02E7.10* | WBGene00019314 | protein_coding_gene | 2.5 | 5.52 | 2.9E-06 |
| *gcy-34* | WBGene00001554 | protein_coding_gene | 2.5 | 5.52 | 1.2E-02 |
| *nlp-32* | WBGene00003770 | protein_coding_gene | 2.5 | 5.51 | 4.7E-02 |
| *F15E6.10* | WBGene00044479 | protein_coding_gene | 2.5 | 5.51 | 1.5E-02 |
| *Y7A5A.8* | WBGene00012414 | protein_coding_gene | 2.5 | 5.51 | 2.6E-02 |

| | | | | | |
|---|---|---|---|---|---|
| *T22F3.8* | WBGene00020698 | protein_coding_gene | 2.5 | 5.49 | 1.2E-05 |
| *C25F9.15* | WBGene00050967 | protein_coding_gene | 2.5 | 5.49 | 2.6E-03 |
| *pals-19* | WBGene00044761 | protein_coding_gene | 2.5 | 5.48 | 1.8E-05 |
| *fbxc-7* | WBGene00016302 | protein_coding_gene | 2.5 | 5.48 | 1.2E-02 |
| *F28F9.2* | WBGene00017906 | protein_coding_gene | 2.5 | 5.48 | 2.7E-02 |
| *C37A5.3* | WBGene00014698 | protein_coding_gene | 2.5 | 5.47 | 5.4E-03 |
| *rncs-1* | WBGene00023069 | ncRNA_gene | 2.5 | 5.47 | 1.5E-03 |
| *Y60C6A.3* | WBGene00271832 | protein_coding_gene | 2.5 | 5.47 | 2.3E-02 |
| *K08F9.3* | WBGene00010686 | protein_coding_gene | 2.5 | 5.47 | 1.4E-04 |
| *nhr-206* | WBGene00011097 | protein_coding_gene | 2.5 | 5.46 | 2.0E-05 |
| *AH9.4* | WBGene00015000 | protein_coding_gene | 2.4 | 5.44 | 1.9E-02 |
| *F11E6.11* | WBGene00023501 | protein_coding_gene | 2.4 | 5.43 | 2.6E-03 |
| *glr-8* | WBGene00001619 | protein_coding_gene | 2.4 | 5.41 | 5.2E-03 |
| *math-8* | WBGene00015837 | protein_coding_gene | 2.4 | 5.41 | 2.0E-02 |
| *F46F3.3* | WBGene00009791 | protein_coding_gene | 2.4 | 5.40 | 3.1E-03 |
| *F15B9.6* | WBGene00008850 | protein_coding_gene | 2.4 | 5.40 | 1.7E-03 |
| *C45E5.3* | WBGene00016666 | pseudogene | 2.4 | 5.39 | 7.0E-04 |
| *F40F9.10* | WBGene00009586 | protein_coding_gene | 2.4 | 5.39 | 1.4E-04 |
| *fmo-1* | WBGene00001476 | protein_coding_gene | 2.4 | 5.39 | 5.4E-04 |
| *F49E10.2* | WBGene00018640 | protein_coding_gene | 2.4 | 5.38 | 1.2E-03 |
| *aqp-5* | WBGene00000173 | protein_coding_gene | 2.4 | 5.37 | 1.3E-02 |
| *tsp-1* | WBGene00006627 | protein_coding_gene | 2.4 | 5.36 | 3.0E-03 |
| *sri-30* | WBGene00005542 | protein_coding_gene | 2.4 | 5.36 | 3.4E-02 |
| *T08G5.14* | WBGene00077572 | pseudogene | 2.4 | 5.35 | 3.4E-02 |
| *grd-2* | WBGene00001691 | protein_coding_gene | 2.4 | 5.34 | 6.1E-03 |
| *C08F11.13* | WBGene00007460 | protein_coding_gene | 2.4 | 5.34 | 1.5E-03 |
| *anr-45* | WBGene00219710 | antisense_lncRNA_gene | 2.4 | 5.34 | 2.9E-02 |
| *gstk-2* | WBGene00017054 | protein_coding_gene | 2.4 | 5.34 | 1.0E-04 |
| *F28G4.4* | WBGene00009229 | protein_coding_gene | 2.4 | 5.33 | 6.8E-03 |
| *col-56* | WBGene00000633 | protein_coding_gene | 2.4 | 5.33 | 2.9E-02 |
| *mboa-1* | WBGene00007174 | protein_coding_gene | 2.4 | 5.33 | 1.8E-03 |
| *tsp-17* | WBGene00006643 | protein_coding_gene | 2.4 | 5.33 | 1.5E-04 |
| *tpst-2* | WBGene00018365 | protein_coding_gene | 2.4 | 5.32 | 2.0E-02 |
| *B0348.9* | WBGene00235314 | ncRNA_gene | 2.4 | 5.32 | 2.5E-02 |
| *B0454.18* | WBGene00199082 | ncRNA_gene | 2.4 | 5.31 | 4.4E-02 |
| *C17F4.3* | WBGene00015911 | protein_coding_gene | 2.4 | 5.30 | 3.0E-02 |
| *fbxa-21* | WBGene00022487 | protein_coding_gene | 2.4 | 5.29 | 1.0E-06 |
| *bnc-1* | WBGene00044791 | protein_coding_gene | 2.4 | 5.29 | 2.9E-02 |
| *clec-25* | WBGene00044331 | protein_coding_gene | 2.4 | 5.28 | 1.1E-02 |
| *T09E11.11* | WBGene00011660 | pseudogene | 2.4 | 5.28 | 1.5E-03 |
| *F22F7.4* | WBGene00017722 | protein_coding_gene | 2.4 | 5.28 | 5.3E-03 |
| *F35F10.5* | WBGene00018055 | protein_coding_gene | 2.4 | 5.27 | 2.7E-02 |
| *T04F8.7* | WBGene00011445 | protein_coding_gene | 2.4 | 5.27 | 1.1E-06 |
| *str-9* | WBGene00006077 | protein_coding_gene | 2.4 | 5.26 | 3.0E-02 |
| *C33A12.4* | WBGene00007883 | protein_coding_gene | 2.4 | 5.23 | 1.6E-06 |
| *F11C1.9* | WBGene00195213 | protein_coding_gene | 2.4 | 5.23 | 4.8E-02 |
| *coel-1* | WBGene00016866 | protein_coding_gene | 2.4 | 5.23 | 1.6E-03 |
| *F02A9.7* | WBGene00044406 | protein_coding_gene | 2.4 | 5.23 | 6.8E-03 |
| *grl-2* | WBGene00001711 | protein_coding_gene | 2.4 | 5.21 | 3.0E-02 |
| *F13H10.1* | WBGene00008773 | protein_coding_gene | 2.4 | 5.21 | 6.9E-03 |
| *srab-14* | WBGene00019999 | protein_coding_gene | 2.4 | 5.20 | 1.0E-02 |
| *nlp-45* | WBGene00020136 | protein_coding_gene | 2.4 | 5.20 | 1.2E-02 |
| *R102.1* | WBGene00011288 | protein_coding_gene | 2.4 | 5.20 | 2.7E-02 |
| *T19C9.10* | WBGene00050934 | protein_coding_gene | 2.4 | 5.19 | 2.8E-02 |
| *srw-12* | WBGene00005759 | protein_coding_gene | 2.4 | 5.19 | 5.0E-02 |
| *gst-29* | WBGene00001777 | protein_coding_gene | 2.4 | 5.19 | 4.8E-04 |
| *R02D5.6* | WBGene00010979 | protein_coding_gene | 2.4 | 5.18 | 1.9E-02 |
| *nape-1* | WBGene00021371 | protein_coding_gene | 2.4 | 5.18 | 8.7E-04 |

| | | | | | |
|---|---|---|---|---|---|
| *cbn-1* | WBGene00019285 | protein_coding_gene | 2.4 | 5.18 | 3.2E-05 |
| *F11C1.10* | WBGene00195214 | protein_coding_gene | 2.4 | 5.18 | 3.6E-02 |
| *D1014.7* | WBGene00017020 | protein_coding_gene | 2.4 | 5.18 | 3.8E-02 |
| *D1053.3* | WBGene00008368 | protein_coding_gene | 2.4 | 5.17 | 3.1E-04 |
| *sru-20* | WBGene00005683 | protein_coding_gene | 2.4 | 5.16 | 8.9E-03 |
| *nhr-221* | WBGene00020748 | protein_coding_gene | 2.4 | 5.16 | 6.4E-03 |
| *C17B7.15* | WBGene00268213 | protein_coding_gene | 2.4 | 5.15 | 4.6E-02 |
| *F49C5.10* | WBGene00044162 | protein_coding_gene | 2.4 | 5.14 | 4.2E-02 |
| *T20D4.13* | WBGene00020619 | pseudogene | 2.4 | 5.14 | 1.6E-02 |
| *C49C8.8* | WBGene00235271 | protein_coding_gene | 2.4 | 5.13 | 1.5E-02 |
| *F47D2.11* | WBGene00206392 | protein_coding_gene | 2.4 | 5.12 | 4.8E-02 |
| *C05C9.2* | WBGene00007326 | protein_coding_gene | 2.4 | 5.11 | 4.5E-02 |
| *abu-13* | WBGene00004133 | protein_coding_gene | 2.3 | 5.09 | 2.1E-04 |
| *C14A4.9* | WBGene00007560 | protein_coding_gene | 2.3 | 5.08 | 9.9E-04 |
| *K01F9.2* | WBGene00010475 | protein_coding_gene | 2.3 | 5.08 | 1.9E-03 |
| *fbxa-142* | WBGene00044659 | protein_coding_gene | 2.3 | 5.08 | 2.8E-02 |
| *qua-1* | WBGene00004264 | protein_coding_gene | 2.3 | 5.07 | 1.2E-03 |
| *T08G5.15* | WBGene00077573 | protein_coding_gene | 2.3 | 5.07 | 1.5E-02 |
| *K01A6.4* | WBGene00010446 | protein_coding_gene | 2.3 | 5.07 | 2.7E-02 |
| *nstp-3* | WBGene00022577 | protein_coding_gene | 2.3 | 5.07 | 1.0E-03 |
| *H39E23.2* | WBGene00010432 | protein_coding_gene | 2.3 | 5.07 | 2.8E-02 |
| *linc-43* | WBGene00219579 | lincRNA_gene | 2.3 | 5.06 | 2.9E-02 |
| *srv-13* | WBGene00005724 | protein_coding_gene | 2.3 | 5.06 | 4.6E-02 |
| *glb-23* | WBGene00020045 | protein_coding_gene | 2.3 | 5.05 | 3.8E-03 |
| *D1014.5* | WBGene00017018 | protein_coding_gene | 2.3 | 5.05 | 3.8E-05 |
| *ceh-57* | WBGene00007416 | protein_coding_gene | 2.3 | 5.05 | 2.4E-03 |
| *str-133* | WBGene00006181 | pseudogene | 2.3 | 5.05 | 4.7E-02 |
| *fipr-23* | WBGene00007990 | protein_coding_gene | 2.3 | 5.05 | 2.1E-03 |
| *F12A10.9* | WBGene00077704 | protein_coding_gene | 2.3 | 5.05 | 4.9E-02 |
| *F07E5.9* | WBGene00017214 | protein_coding_gene | 2.3 | 5.04 | 3.8E-02 |
| *bath-25* | WBGene00020859 | protein_coding_gene | 2.3 | 5.03 | 2.7E-02 |
| *C02F4.4* | WBGene00007264 | protein_coding_gene | 2.3 | 5.02 | 3.7E-03 |
| *K10H10.10* | WBGene00010765 | protein_coding_gene | 2.3 | 5.02 | 1.2E-02 |
| *C25E10.13* | WBGene00023431 | protein_coding_gene | 2.3 | 5.01 | 1.1E-02 |
| *Y70G10A.2* | WBGene00013498 | protein_coding_gene | 2.3 | 5.01 | 9.9E-03 |
| *F59F5.4* | WBGene00010343 | protein_coding_gene | 2.3 | 5.01 | 3.9E-02 |
| *C17F4.12* | WBGene00044374 | protein_coding_gene | 2.3 | 5.00 | 2.4E-02 |
| *oac-56* | WBGene00013450 | protein_coding_gene | 2.3 | 4.98 | 4.2E-02 |
| *Y46H3A.5* | WBGene00021601 | protein_coding_gene | 2.3 | 4.98 | 3.1E-02 |
| *R11F4.1* | WBGene00020007 | protein_coding_gene | 2.3 | 4.98 | 7.9E-07 |
| *C12D5.10* | WBGene00015711 | protein_coding_gene | 2.3 | 4.98 | 1.8E-02 |
| *ZK757.2* | WBGene00014074 | protein_coding_gene | 2.3 | 4.98 | 1.4E-05 |
| *ttll-9* | WBGene00044187 | protein_coding_gene | 2.3 | 4.97 | 4.3E-02 |
| *Y17D7C.4* | WBGene00050903 | pseudogene | 2.3 | 4.97 | 1.5E-04 |
| *T20D4.9* | WBGene00020615 | pseudogene | 2.3 | 4.97 | 4.1E-02 |
| *Y71H2AM.14* | WBGene00022179 | protein_coding_gene | 2.3 | 4.97 | 1.4E-04 |
| *C01C4.2* | WBGene00015292 | protein_coding_gene | 2.3 | 4.97 | 3.5E-02 |
| *C07G1.15* | WBGene00235296 | protein_coding_gene | 2.3 | 4.97 | 1.2E-02 |
| *ptr-9* | WBGene00004223 | protein_coding_gene | 2.3 | 4.96 | 4.3E-02 |
| *T27A3.8* | WBGene00044611 | protein_coding_gene | 2.3 | 4.96 | 1.9E-02 |
| *ins-4* | WBGene00002087 | protein_coding_gene | 2.3 | 4.96 | 2.8E-03 |
| *arrd-11* | WBGene00011052 | protein_coding_gene | 2.3 | 4.95 | 3.2E-02 |
| *T14E8.4* | WBGene00043981 | protein_coding_gene | 2.3 | 4.94 | 8.6E-04 |
| *K10G4.3* | WBGene00010752 | protein_coding_gene | 2.3 | 4.92 | 5.9E-03 |
| *sups-1* | WBGene00017716 | protein_coding_gene | 2.3 | 4.92 | 3.1E-05 |
| *gbh-2* | WBGene00001523 | protein_coding_gene | 2.3 | 4.92 | 1.5E-07 |
| *srab-16* | WBGene00020060 | protein_coding_gene | 2.3 | 4.91 | 3.5E-02 |
| *zig-4* | WBGene00006981 | protein_coding_gene | 2.3 | 4.90 | 3.5E-04 |

| | | | | | |
|---|---|---|---|---|---|
| *R10F2.5* | WBGene00019996 | protein_coding_gene | 2.3 | 4.89 | 3.7E-02 |
| *C26E1.2* | WBGene00007748 | protein_coding_gene | 2.3 | 4.88 | 2.2E-04 |
| *osm-12* | WBGene00003892 | protein_coding_gene | 2.3 | 4.87 | 5.7E-05 |
| *spi-1* | WBGene00023416 | protein_coding_gene | 2.3 | 4.87 | 1.4E-03 |
| *T05C1.3* | WBGene00020250 | protein_coding_gene | 2.3 | 4.87 | 9.4E-03 |
| *F16C3.2* | WBGene00008886 | protein_coding_gene | 2.3 | 4.86 | 4.5E-03 |
| *nspc-19* | WBGene00008360 | protein_coding_gene | 2.3 | 4.86 | 7.7E-08 |
| *fat-3* | WBGene00001395 | protein_coding_gene | 2.3 | 4.85 | 3.2E-07 |
| *dpy-7* | WBGene00001069 | protein_coding_gene | 2.3 | 4.85 | 2.4E-05 |
| *uev-2* | WBGene00006731 | protein_coding_gene | 2.3 | 4.85 | 1.8E-02 |
| *F14D7.4* | WBGene00008791 | pseudogene | 2.3 | 4.83 | 7.6E-03 |
| *cil-7* | WBGene00021017 | protein_coding_gene | 2.3 | 4.83 | 8.2E-03 |
| *droe-8* | WBGene00022550 | protein_coding_gene | 2.3 | 4.81 | 1.3E-02 |
| *T12A7.7* | WBGene00011728 | pseudogene | 2.3 | 4.81 | 3.9E-02 |
| *gcy-22* | WBGene00001547 | protein_coding_gene | 2.3 | 4.81 | 6.0E-04 |
| *Y44A6C.1* | WBGene00012850 | protein_coding_gene | 2.3 | 4.80 | 1.6E-04 |
| *fbxa-162* | WBGene00015597 | pseudogene | 2.3 | 4.80 | 8.9E-03 |
| *srj-29* | WBGene00005616 | protein_coding_gene | 2.3 | 4.80 | 1.6E-02 |
| *F48G7.5* | WBGene00018616 | protein_coding_gene | 2.3 | 4.80 | 2.9E-02 |
| *K04H4.2* | WBGene00010573 | protein_coding_gene | 2.3 | 4.79 | 2.8E-05 |
| *Y5H2B.1* | WBGene00021164 | protein_coding_gene | 2.3 | 4.79 | 1.4E-02 |
| *bro-1* | WBGene00000272 | protein_coding_gene | 2.3 | 4.79 | 4.9E-04 |
| *med-2* | WBGene00003181 | protein_coding_gene | 2.3 | 4.79 | 3.3E-02 |
| *ikb-1* | WBGene00002069 | protein_coding_gene | 2.3 | 4.78 | 1.3E-03 |
| *pef-1* | WBGene00003969 | protein_coding_gene | 2.3 | 4.78 | 1.0E-03 |
| *unk-1* | WBGene00016407 | protein_coding_gene | 2.3 | 4.78 | 3.8E-03 |
| *asp-6* | WBGene00000219 | protein_coding_gene | 2.3 | 4.77 | 4.9E-05 |
| *F36F12.1* | WBGene00018092 | protein_coding_gene | 2.3 | 4.76 | 1.5E-02 |
| *F22B8.2* | WBGene00014741 | pseudogene | 2.2 | 4.75 | 3.4E-02 |
| *W04A8.4* | WBGene00012239 | protein_coding_gene | 2.2 | 4.75 | 9.4E-09 |
| *C06C3.9* | WBGene00014668 | protein_coding_gene | 2.2 | 4.75 | 4.6E-03 |
| *nhr-18* | WBGene00003617 | protein_coding_gene | 2.2 | 4.74 | 5.6E-04 |
| *ugt-43* | WBGene00008485 | protein_coding_gene | 2.2 | 4.74 | 4.8E-04 |
| *C07B5.4* | WBGene00007409 | protein_coding_gene | 2.2 | 4.73 | 2.0E-04 |
| *ZK1307.2* | WBGene00014245 | protein_coding_gene | 2.2 | 4.73 | 2.4E-02 |
| *twk-7* | WBGene00006662 | protein_coding_gene | 2.2 | 4.72 | 2.5E-03 |
| *C30F2.4* | WBGene00007818 | protein_coding_gene | 2.2 | 4.70 | 4.1E-03 |
| *B0250.12* | WBGene00050927 | pseudogene | 2.2 | 4.70 | 1.5E-06 |
| *F55A4.4* | WBGene00018856 | pseudogene | 2.2 | 4.69 | 9.4E-04 |
| *srg-32* | WBGene00005189 | protein_coding_gene | 2.2 | 4.68 | 4.5E-02 |
| *Y64H9A.2* | WBGene00022024 | protein_coding_gene | 2.2 | 4.68 | 1.3E-02 |
| *B0462.5* | WBGene00194695 | protein_coding_gene | 2.2 | 4.68 | 3.0E-03 |
| *Y43F8C.3* | WBGene00012825 | protein_coding_gene | 2.2 | 4.67 | 1.4E-04 |
| *srd-33* | WBGene00005111 | protein_coding_gene | 2.2 | 4.66 | 4.2E-02 |
| *F20G2.5* | WBGene00008988 | protein_coding_gene | 2.2 | 4.65 | 7.4E-03 |
| *K08D10.10* | WBGene00019533 | protein_coding_gene | 2.2 | 4.65 | 3.8E-02 |
| *nas-14* | WBGene00003533 | protein_coding_gene | 2.2 | 4.65 | 5.9E-03 |
| *F11A5.13* | WBGene00008679 | protein_coding_gene | 2.2 | 4.65 | 4.6E-02 |
| *T12A7.6* | WBGene00011727 | protein_coding_gene | 2.2 | 4.65 | 9.0E-04 |
| *F43D9.1* | WBGene00009653 | protein_coding_gene | 2.2 | 4.65 | 1.3E-03 |
| *mlt-11* | WBGene00012186 | protein_coding_gene | 2.2 | 4.64 | 6.9E-04 |
| *F08C6.5* | WBGene00017243 | protein_coding_gene | 2.2 | 4.64 | 6.4E-04 |
| *W05E10.5* | WBGene00012282 | protein_coding_gene | 2.2 | 4.64 | 2.8E-03 |
| *ptr-3* | WBGene00004218 | protein_coding_gene | 2.2 | 4.63 | 1.1E-02 |
| *Y26D4A.3* | WBGene00012501 | protein_coding_gene | 2.2 | 4.63 | 4.7E-03 |
| *dos-3* | WBGene00019331 | protein_coding_gene | 2.2 | 4.63 | 3.8E-02 |
| *R07B7.12* | WBGene00011096 | protein_coding_gene | 2.2 | 4.62 | 4.5E-02 |
| *M01A8.1* | WBGene00010795 | protein_coding_gene | 2.2 | 4.62 | 1.7E-03 |

| | | | | | |
|---|---|---|---|---|---|
| *F30A10.2* | WBGene00009261 | protein_coding_gene | 2.2 | 4.62 | 1.5E-03 |
| *glf-1* | WBGene00019154 | protein_coding_gene | 2.2 | 4.61 | 9.3E-05 |
| *F17C8.3* | WBGene00008910 | protein_coding_gene | 2.2 | 4.61 | 1.3E-03 |
| *ceh-9* | WBGene00000434 | protein_coding_gene | 2.2 | 4.61 | 2.6E-02 |
| *R07C12.3* | WBGene00019933 | protein_coding_gene | 2.2 | 4.61 | 3.9E-02 |
| *osr-1* | WBGene00016329 | protein_coding_gene | 2.2 | 4.61 | 7.1E-04 |
| *cyp-33C6* | WBGene00018264 | protein_coding_gene | 2.2 | 4.61 | 4.1E-03 |
| *droe-4* | WBGene00010262 | protein_coding_gene | 2.2 | 4.60 | 2.1E-03 |
| *C06A12.19* | WBGene00255702 | protein_coding_gene | 2.2 | 4.59 | 3.2E-02 |
| *Y11D7A.5* | WBGene00012429 | protein_coding_gene | 2.2 | 4.59 | 3.8E-03 |
| *nhr-290* | WBGene00016233 | protein_coding_gene | 2.2 | 4.59 | 4.5E-02 |
| *F26A1.6* | WBGene00017804 | protein_coding_gene | 2.2 | 4.59 | 3.0E-02 |
| *M03D4.3* | WBGene00019750 | protein_coding_gene | 2.2 | 4.59 | 3.0E-03 |
| *M04C3.5* | WBGene00206418 | protein_coding_gene | 2.2 | 4.59 | 4.3E-03 |
| *F28F5.4* | WBGene00043297 | protein_coding_gene | 2.2 | 4.59 | 2.9E-03 |
| *T04C12.32* | WBGene00235086 | ncRNA_gene | 2.2 | 4.59 | 3.7E-02 |
| *mks-3* | WBGene00018042 | protein_coding_gene | 2.2 | 4.58 | 6.4E-04 |
| *C34B4.3* | WBGene00007908 | protein_coding_gene | 2.2 | 4.58 | 1.6E-05 |
| *glb-5* | WBGene00015964 | protein_coding_gene | 2.2 | 4.58 | 3.3E-02 |
| *F59B2.11* | WBGene00010313 | pseudogene | 2.2 | 4.58 | 5.2E-04 |
| *F16H6.7* | WBGene00008897 | protein_coding_gene | 2.2 | 4.58 | 1.6E-02 |
| *tre-5* | WBGene00006611 | protein_coding_gene | 2.2 | 4.58 | 1.3E-02 |
| *ZC13.10* | WBGene00044557 | protein_coding_gene | 2.2 | 4.58 | 5.0E-02 |
| *C30A5.10* | WBGene00016242 | protein_coding_gene | 2.2 | 4.57 | 1.5E-02 |
| *F59F5.3* | WBGene00010342 | protein_coding_gene | 2.2 | 4.57 | 3.2E-04 |
| *W08G11.1* | WBGene00012346 | protein_coding_gene | 2.2 | 4.56 | 6.2E-03 |
| *T07D3.6* | WBGene00020309 | pseudogene | 2.2 | 4.56 | 5.1E-03 |
| *nhr-218* | WBGene00011750 | protein_coding_gene | 2.2 | 4.55 | 1.8E-02 |
| *M03B6.3* | WBGene00010835 | protein_coding_gene | 2.2 | 4.55 | 5.2E-05 |
| *F46B3.1* | WBGene00009756 | protein_coding_gene | 2.2 | 4.55 | 4.4E-02 |
| *K09C8.7* | WBGene00010717 | protein_coding_gene | 2.2 | 4.54 | 1.7E-02 |
| *D2024.4* | WBGene00017052 | protein_coding_gene | 2.2 | 4.54 | 2.9E-02 |
| *T05A7.13* | WBGene00271415 | protein_coding_gene | 2.2 | 4.53 | 1.2E-02 |
| *gcy-5* | WBGene00001532 | protein_coding_gene | 2.2 | 4.52 | 1.3E-02 |
| *C37A5.6* | WBGene00014700 | protein_coding_gene | 2.2 | 4.52 | 3.0E-02 |
| *ugt-52* | WBGene00018931 | protein_coding_gene | 2.2 | 4.52 | 1.5E-03 |
| *Y43F8B.23* | WBGene00194815 | protein_coding_gene | 2.2 | 4.52 | 1.4E-03 |
| *ZK593.2* | WBGene00014002 | protein_coding_gene | 2.2 | 4.51 | 2.4E-04 |
| *F21C10.10* | WBGene00017659 | protein_coding_gene | 2.2 | 4.51 | 9.8E-04 |
| *C08F1.8* | WBGene00015611 | protein_coding_gene | 2.2 | 4.51 | 4.0E-02 |
| *R10H1.1* | WBGene00019998 | protein_coding_gene | 2.2 | 4.51 | 2.4E-04 |
| *ZK381.2* | WBGene00022725 | protein_coding_gene | 2.2 | 4.50 | 4.4E-02 |
| *che-11* | WBGene00000490 | protein_coding_gene | 2.2 | 4.50 | 9.5E-06 |
| *T21D12.14* | WBGene00045177 | protein_coding_gene | 2.2 | 4.50 | 1.3E-02 |
| *Y116A8C.25* | WBGene00013802 | protein_coding_gene | 2.2 | 4.49 | 1.9E-03 |
| *ZK742.7* | WBGene00194921 | protein_coding_gene | 2.2 | 4.49 | 6.7E-04 |
| *K02A6.2* | WBGene00019293 | protein_coding_gene | 2.2 | 4.49 | 1.0E-02 |
| *pals-18* | WBGene00022353 | protein_coding_gene | 2.2 | 4.48 | 1.8E-04 |
| *ptr-4* | WBGene00004219 | protein_coding_gene | 2.2 | 4.48 | 1.4E-04 |
| *F09C6.3* | WBGene00008617 | protein_coding_gene | 2.2 | 4.48 | 5.9E-03 |
| *F16G10.15* | WBGene00017529 | pseudogene | 2.2 | 4.47 | 4.7E-03 |
| *srd-65* | WBGene00005142 | protein_coding_gene | 2.2 | 4.47 | 4.9E-02 |
| *F31F7.1* | WBGene00017964 | protein_coding_gene | 2.2 | 4.47 | 1.5E-04 |
| *chil-23* | WBGene00011167 | protein_coding_gene | 2.2 | 4.47 | 3.6E-03 |
| *sre-3* | WBGene00015568 | protein_coding_gene | 2.2 | 4.46 | 3.2E-02 |
| *swt-1* | WBGene00019300 | protein_coding_gene | 2.2 | 4.46 | 5.5E-03 |
| *T28A11.17* | WBGene00020879 | protein_coding_gene | 2.2 | 4.45 | 4.8E-02 |
| *lpr-6* | WBGene00012255 | protein_coding_gene | 2.2 | 4.45 | 2.6E-04 |

| | | | | | |
|---|---|---|---|---|---|
| *R05G6.10* | WBGene00019902 | protein_coding_gene | 2.2 | 4.45 | 1.5E-04 |
| *Y41D4B.6* | WBGene00021513 | protein_coding_gene | 2.2 | 4.45 | 1.8E-02 |
| *Y43F8B.9* | WBGene00012819 | protein_coding_gene | 2.2 | 4.45 | 4.1E-04 |
| *nep-16* | WBGene00018472 | protein_coding_gene | 2.2 | 4.45 | 5.2E-03 |
| *R06A10.1* | WBGene00019909 | protein_coding_gene | 2.2 | 4.44 | 8.3E-04 |
| *T02C12.5* | WBGene00194692 | protein_coding_gene | 2.2 | 4.44 | 4.1E-05 |
| *C25A6.1* | WBGene00016081 | protein_coding_gene | 2.2 | 4.44 | 2.4E-03 |
| *bbs-1* | WBGene00000241 | protein_coding_gene | 2.2 | 4.44 | 1.8E-03 |
| *ztf-30* | WBGene00015523 | protein_coding_gene | 2.1 | 4.43 | 1.4E-03 |
| *F46F5.15* | WBGene00018506 | protein_coding_gene | 2.1 | 4.43 | 4.2E-02 |
| *C39B10.7* | WBGene00164973 | protein_coding_gene | 2.1 | 4.43 | 9.0E-04 |
| *F13E9.12* | WBGene00008761 | protein_coding_gene | 2.1 | 4.43 | 6.4E-03 |
| *glc-1* | WBGene00001591 | protein_coding_gene | 2.1 | 4.43 | 3.4E-03 |
| *F53A9.6* | WBGene00018729 | protein_coding_gene | 2.1 | 4.42 | 7.5E-04 |
| *F09C6.12* | WBGene00044182 | protein_coding_gene | 2.1 | 4.42 | 7.3E-05 |
| *F21A3.11* | WBGene00219316 | protein_coding_gene | 2.1 | 4.42 | 1.8E-06 |
| *T07D1.3* | WBGene00020302 | protein_coding_gene | 2.1 | 4.41 | 1.9E-02 |
| *ZK1025.3* | WBGene00014183 | protein_coding_gene | 2.1 | 4.41 | 1.8E-02 |
| *fkb-4* | WBGene00001429 | protein_coding_gene | 2.1 | 4.41 | 3.7E-03 |
| *C25F6.7* | WBGene00016106 | protein_coding_gene | 2.1 | 4.40 | 4.5E-04 |
| *fbxa-222* | WBGene00045295 | protein_coding_gene | 2.1 | 4.40 | 3.5E-02 |
| *fezf-1* | WBGene00012639 | protein_coding_gene | 2.1 | 4.39 | 2.9E-02 |
| *C17H11.1* | WBGene00015921 | protein_coding_gene | 2.1 | 4.39 | 4.1E-03 |
| *F53F10.1* | WBGene00018763 | protein_coding_gene | 2.1 | 4.38 | 2.3E-03 |
| *T06G6.12* | WBGene00011551 | protein_coding_gene | 2.1 | 4.38 | 1.8E-02 |
| *dhhc-11* | WBGene00020694 | protein_coding_gene | 2.1 | 4.38 | 1.8E-02 |
| *atln-2* | WBGene00012763 | protein_coding_gene | 2.1 | 4.38 | 9.8E-05 |
| *K06A9.2* | WBGene00019436 | protein_coding_gene | 2.1 | 4.37 | 6.1E-03 |
| *nlp-66* | WBGene00011141 | protein_coding_gene | 2.1 | 4.36 | 3.3E-03 |
| *fbxa-174* | WBGene00008015 | protein_coding_gene | 2.1 | 4.36 | 2.6E-02 |
| *chil-20* | WBGene00011163 | protein_coding_gene | 2.1 | 4.36 | 4.7E-02 |
| *F59B2.8* | WBGene00010310 | protein_coding_gene | 2.1 | 4.36 | 4.0E-05 |
| *Y37F4.8* | WBGene00044289 | protein_coding_gene | 2.1 | 4.35 | 5.5E-03 |
| *E02H4.5* | WBGene00008463 | protein_coding_gene | 2.1 | 4.35 | 2.1E-02 |
| *fkh-10* | WBGene00001442 | protein_coding_gene | 2.1 | 4.34 | 9.4E-03 |
| *C18D4.8* | WBGene00007677 | pseudogene | 2.1 | 4.34 | 1.3E-06 |
| *F57E7.4* | WBGene00077616 | protein_coding_gene | 2.1 | 4.33 | 1.4E-03 |
| *Y75B12B.3* | WBGene00013568 | protein_coding_gene | 2.1 | 4.33 | 1.8E-04 |
| *C43C3.4* | WBGene00195084 | protein_coding_gene | 2.1 | 4.33 | 9.0E-06 |
| *F48G7.13* | WBGene00018624 | protein_coding_gene | 2.1 | 4.33 | 2.7E-02 |
| *C54E4.12* | WBGene00235274 | protein_coding_gene | 2.1 | 4.33 | 8.6E-03 |
| *T16G1.2* | WBGene00011796 | protein_coding_gene | 2.1 | 4.32 | 3.4E-03 |
| *fbxa-68* | WBGene00020641 | pseudogene | 2.1 | 4.32 | 3.9E-02 |
| *acs-2* | WBGene00009221 | protein_coding_gene | 2.1 | 4.32 | 4.6E-02 |
| *nhr-116* | WBGene00003706 | protein_coding_gene | 2.1 | 4.31 | 3.1E-07 |
| *R03H10.7* | WBGene00019859 | protein_coding_gene | 2.1 | 4.31 | 5.9E-03 |
| *C23H5.15* | WBGene00219423 | protein_coding_gene | 2.1 | 4.30 | 1.4E-02 |
| *R07B1.5* | WBGene00011077 | protein_coding_gene | 2.1 | 4.29 | 1.9E-02 |
| *K10D6.2* | WBGene00010742 | protein_coding_gene | 2.1 | 4.28 | 3.9E-06 |
| *lpr-3* | WBGene00012261 | protein_coding_gene | 2.1 | 4.28 | 7.5E-05 |
| *C25D7.12* | WBGene00007721 | pseudogene | 2.1 | 4.28 | 5.0E-02 |
| *T20F5.4* | WBGene00020626 | protein_coding_gene | 2.1 | 4.27 | 5.1E-04 |
| *F57C12.2* | WBGene00019014 | protein_coding_gene | 2.1 | 4.27 | 1.0E-04 |
| *clec-30* | WBGene00014842 | pseudogene | 2.1 | 4.27 | 4.0E-02 |
| *T16H12.9* | WBGene00011817 | protein_coding_gene | 2.1 | 4.27 | 3.5E-03 |
| *F53C11.9* | WBGene00044921 | protein_coding_gene | 2.1 | 4.27 | 7.1E-04 |
| *F30H5.3* | WBGene00017937 | protein_coding_gene | 2.1 | 4.27 | 1.8E-03 |
| *F16B3.2* | WBGene00017502 | protein_coding_gene | 2.1 | 4.27 | 1.5E-02 |

| | | | | | |
|---|---|---|---|---|---|
| *Y71G12B.42* | WBGene00220198 | ncRNA_gene | 2.1 | 4.26 | 1.0E-02 |
| *dsl-5* | WBGene00001107 | protein_coding_gene | 2.1 | 4.24 | 4.5E-02 |
| *T19C4.5* | WBGene00011839 | protein_coding_gene | 2.1 | 4.23 | 1.6E-04 |
| *dhs-16* | WBGene00000979 | protein_coding_gene | 2.1 | 4.21 | 1.1E-02 |
| *T23F4.2* | WBGene00020740 | protein_coding_gene | 2.1 | 4.21 | 2.0E-03 |
| *F14D2.19* | WBGene00206522 | protein_coding_gene | 2.1 | 4.21 | 3.6E-03 |
| *ttr-44* | WBGene00008341 | protein_coding_gene | 2.1 | 4.20 | 6.2E-03 |
| *zmp-1* | WBGene00006987 | protein_coding_gene | 2.1 | 4.19 | 5.8E-03 |
| *F59A7.2* | WBGene00019090 | protein_coding_gene | 2.1 | 4.19 | 4.2E-02 |
| *fbxa-157* | WBGene00007393 | protein_coding_gene | 2.1 | 4.18 | 1.2E-02 |
| *F10E9.12* | WBGene00017361 | protein_coding_gene | 2.1 | 4.18 | 1.0E-04 |
| *T19C4.1* | WBGene00011838 | protein_coding_gene | 2.1 | 4.18 | 1.8E-04 |
| *F35G2.5* | WBGene00009438 | protein_coding_gene | 2.1 | 4.18 | 1.8E-02 |
| *slc-36.5* | WBGene00016628 | protein_coding_gene | 2.1 | 4.18 | 1.7E-02 |
| *C24H10.3* | WBGene00016069 | protein_coding_gene | 2.1 | 4.17 | 3.9E-03 |
| *hil-7* | WBGene00001858 | protein_coding_gene | 2.1 | 4.17 | 1.1E-02 |
| *F49D11.6* | WBGene00018630 | protein_coding_gene | 2.1 | 4.17 | 6.9E-03 |
| *fbxa-30* | WBGene00013837 | protein_coding_gene | 2.1 | 4.16 | 1.5E-02 |
| *K08E7.6* | WBGene00010674 | protein_coding_gene | 2.1 | 4.16 | 1.0E-03 |
| *nspc-18* | WBGene00008358 | protein_coding_gene | 2.1 | 4.16 | 6.0E-05 |
| *grl-7* | WBGene00001716 | protein_coding_gene | 2.1 | 4.16 | 1.8E-03 |
| *math-45* | WBGene00021772 | protein_coding_gene | 2.1 | 4.16 | 3.8E-07 |
| *Y105C5B.25* | WBGene00013662 | protein_coding_gene | 2.1 | 4.15 | 1.7E-02 |
| *ZK287.1* | WBGene00013962 | protein_coding_gene | 2.1 | 4.15 | 3.1E-05 |
| *T02G5.15* | WBGene00206521 | protein_coding_gene | 2.1 | 4.15 | 1.9E-03 |
| *col-172* | WBGene00000745 | protein_coding_gene | 2.1 | 4.14 | 1.0E-02 |
| *M03A1.8* | WBGene00077490 | protein_coding_gene | 2.1 | 4.14 | 1.4E-02 |
| *chil-4* | WBGene00007469 | protein_coding_gene | 2.1 | 4.14 | 2.9E-02 |
| *C34F11.8* | WBGene00016417 | protein_coding_gene | 2.1 | 4.14 | 2.5E-02 |
| *Y50E8A.1* | WBGene00013046 | protein_coding_gene | 2.0 | 4.14 | 3.6E-05 |
| *Y43F8B.24* | WBGene00195176 | protein_coding_gene | 2.0 | 4.13 | 1.6E-03 |
| *F02C12.2* | WBGene00008516 | protein_coding_gene | 2.0 | 4.13 | 4.4E-02 |
| *unc-3* | WBGene00006743 | protein_coding_gene | 2.0 | 4.12 | 3.8E-04 |
| *btb-21* | WBGene00018540 | pseudogene | 2.0 | 4.12 | 5.7E-04 |
| *fbxa-87* | WBGene00044660 | pseudogene | 2.0 | 4.10 | 3.1E-03 |
| *W06G6.20* | WBGene00219308 | protein_coding_gene | 2.0 | 4.10 | 3.7E-02 |
| *sodh-2* | WBGene00010791 | protein_coding_gene | 2.0 | 4.10 | 1.9E-02 |
| *C11G10.1* | WBGene00007525 | protein_coding_gene | 2.0 | 4.10 | 2.4E-02 |
| *K03H1.5* | WBGene00010540 | protein_coding_gene | 2.0 | 4.10 | 9.1E-04 |
| *C07B5.3* | WBGene00007408 | protein_coding_gene | 2.0 | 4.09 | 1.4E-02 |
| *try-9* | WBGene00023425 | protein_coding_gene | 2.0 | 4.09 | 4.8E-03 |
| *nspb-7* | WBGene00007243 | protein_coding_gene | 2.0 | 4.08 | 4.7E-02 |
| *cyp-23A1* | WBGene00015135 | protein_coding_gene | 2.0 | 4.08 | 8.7E-03 |
| *Y45F3A.8* | WBGene00012863 | protein_coding_gene | 2.0 | 4.08 | 3.7E-02 |
| *ift-20* | WBGene00022465 | protein_coding_gene | 2.0 | 4.08 | 2.0E-02 |
| *suex-2* | WBGene00008271 | protein_coding_gene | 2.0 | 4.07 | 1.5E-02 |
| *C08G5.7* | WBGene00044418 | protein_coding_gene | 2.0 | 4.07 | 2.3E-04 |
| *nhr-35* | WBGene00003628 | protein_coding_gene | 2.0 | 4.06 | 2.6E-03 |
| *R01E6.2* | WBGene00010968 | protein_coding_gene | 2.0 | 4.05 | 3.1E-02 |
| *C04E12.2* | WBGene00015433 | protein_coding_gene | 2.0 | 4.05 | 6.7E-03 |
| *fbxa-29* | WBGene00013836 | protein_coding_gene | 2.0 | 4.04 | 1.8E-04 |
| *T01D3.6* | WBGene00011330 | protein_coding_gene | 2.0 | 4.04 | 3.2E-06 |
| *tts-2* | WBGene00006651 | ncRNA_gene | 2.0 | 4.03 | 5.3E-07 |
| *aagr-1* | WBGene00017071 | protein_coding_gene | 2.0 | 4.03 | 8.3E-04 |
| *R57.2* | WBGene00020083 | protein_coding_gene | 2.0 | 4.03 | 2.7E-03 |
| *cal-2* | WBGene00000286 | protein_coding_gene | 2.0 | 4.03 | 1.4E-03 |
| *F54C8.6* | WBGene00010039 | protein_coding_gene | 2.0 | 4.02 | 9.0E-06 |
| *Y17D7C.6* | WBGene00194680 | protein_coding_gene | 2.0 | 4.02 | 4.9E-02 |

| | | | | | |
|---|---|---|---|---|---|
| *K10D3.4* | WBGene00010738 | protein_coding_gene | 2.0 | 4.02 | 3.2E-04 |
| *C35A5.11* | WBGene00044024 | protein_coding_gene | 2.0 | 4.01 | 1.6E-02 |
| *acr-21* | WBGene00000060 | protein_coding_gene | 2.0 | 4.01 | 7.8E-03 |
| *F58B4.6* | WBGene00010239 | protein_coding_gene | 2.0 | 4.01 | 7.0E-03 |
| *Y43F8B.15* | WBGene00045415 | protein_coding_gene | 2.0 | 4.01 | 3.1E-02 |
| *nhr-150* | WBGene00007367 | protein_coding_gene | 2.0 | 4.00 | 6.5E-03 |
| *slc-28.2* | WBGene00017868 | protein_coding_gene | 2.0 | 4.00 | 4.0E-03 |
| *vab-23* | WBGene00014153 | protein_coding_gene | 2.0 | 4.00 | 1.4E-03 |
| *tba-6* | WBGene00006532 | protein_coding_gene | 2.0 | 4.00 | 3.5E-02 |
| *T23E7.6* | WBGene00044772 | protein_coding_gene | 2.0 | 4.00 | 2.9E-02 |
| *M04C3.2* | WBGene00010851 | protein_coding_gene | 2.0 | 4.00 | 1.5E-04 |
| *Y55D5A.1* | WBGene00021915 | protein_coding_gene | 2.0 | 3.99 | 2.0E-04 |
| *wrt-1* | WBGene00006947 | protein_coding_gene | 2.0 | 3.99 | 2.3E-04 |
| *C50H2.13* | WBGene00008246 | protein_coding_gene | 2.0 | 3.99 | 5.8E-08 |
| *C14F11.6* | WBGene00015781 | protein_coding_gene | 2.0 | 3.98 | 4.4E-04 |
| *W04C9.8* | WBGene00235387 | ncRNA_gene | 2.0 | 3.97 | 1.8E-03 |
| *F22F4.9* | WBGene00271606 | protein_coding_gene | 2.0 | 3.97 | 8.2E-04 |
| *fbxa-25* | WBGene00022321 | protein_coding_gene | 2.0 | 3.97 | 2.7E-02 |
| *C34C12.7* | WBGene00007926 | protein_coding_gene | 2.0 | 3.97 | 1.0E-04 |
| *oac-49* | WBGene00012068 | protein_coding_gene | 2.0 | 3.96 | 4.0E-02 |
| *F20C5.5* | WBGene00008970 | protein_coding_gene | 2.0 | 3.96 | 3.0E-04 |
| *M03D4.4* | WBGene00019751 | protein_coding_gene | 2.0 | 3.95 | 2.6E-04 |
| *R13H4.7* | WBGene00011265 | protein_coding_gene | 2.0 | 3.95 | 1.7E-02 |
| *C39D10.5* | WBGene00016532 | protein_coding_gene | 2.0 | 3.95 | 2.6E-02 |
| *dpy-2* | WBGene00001064 | protein_coding_gene | 2.0 | 3.95 | 8.7E-04 |
| *F26A10.2* | WBGene00017814 | protein_coding_gene | 2.0 | 3.94 | 1.2E-03 |
| *str-173* | WBGene00006217 | protein_coding_gene | 2.0 | 3.94 | 4.4E-02 |
| *Y110A2AL.4* | WBGene00022441 | protein_coding_gene | 2.0 | 3.94 | 1.7E-03 |
| *pqn-32* | WBGene00004120 | protein_coding_gene | 2.0 | 3.94 | 1.9E-03 |
| *C33D9.5* | WBGene00007899 | pseudogene | 2.0 | 3.94 | 7.8E-07 |
| *C36A4.10* | WBGene00007970 | protein_coding_gene | 2.0 | 3.94 | 2.1E-02 |
| *cwp-5* | WBGene00009844 | protein_coding_gene | 2.0 | 3.93 | 2.6E-03 |
| *F47B8.3* | WBGene00009804 | protein_coding_gene | 2.0 | 3.93 | 3.1E-04 |
| *F59E11.2* | WBGene00019109 | protein_coding_gene | 2.0 | 3.93 | 1.0E-03 |
| *F44G4.7* | WBGene00009716 | protein_coding_gene | 2.0 | 3.93 | 3.0E-02 |
| *col-131* | WBGene00000705 | protein_coding_gene | 2.0 | 3.92 | 2.7E-02 |
| *unc-86* | WBGene00006818 | protein_coding_gene | 2.0 | 3.92 | 1.8E-03 |
| *dod-22* | WBGene00010125 | protein_coding_gene | 2.0 | 3.90 | 1.6E-02 |
| *ugt-57* | WBGene00017154 | protein_coding_gene | 2.0 | 3.90 | 1.2E-03 |
| *K09H9.8* | WBGene00045387 | protein_coding_gene | 2.0 | 3.90 | 9.5E-04 |
| *ZK686.6* | WBGene00044799 | protein_coding_gene | 2.0 | 3.90 | 1.7E-02 |
| *M04B2.6* | WBGene00010848 | protein_coding_gene | 2.0 | 3.89 | 2.8E-03 |
| *mab-5* | WBGene00003102 | protein_coding_gene | 2.0 | 3.89 | 1.7E-03 |
| *mfb-1* | WBGene00008439 | protein_coding_gene | 2.0 | 3.89 | 3.3E-03 |
| *Y6G8.9* | WBGene00050974 | pseudogene | 2.0 | 3.89 | 1.5E-02 |
| *igdb-3* | WBGene00020691 | protein_coding_gene | 2.0 | 3.89 | 1.0E-02 |
| *lin-48* | WBGene00003033 | protein_coding_gene | 2.0 | 3.88 | 3.3E-03 |
| *fipr-22* | WBGene00007989 | protein_coding_gene | 2.0 | 3.88 | 9.3E-03 |
| *ZC168.2* | WBGene00013856 | protein_coding_gene | 2.0 | 3.88 | 4.0E-03 |
| *B0563.7* | WBGene00015264 | protein_coding_gene | 2.0 | 3.88 | 6.1E-03 |
| *ins-18* | WBGene00002101 | protein_coding_gene | 2.0 | 3.88 | 1.6E-03 |
| *F59A1.11* | WBGene00010297 | protein_coding_gene | 2.0 | 3.88 | 2.4E-02 |
| *Y54G2A.11* | WBGene00021876 | protein_coding_gene | 2.0 | 3.88 | 7.4E-03 |
| *srd-32* | WBGene00005110 | protein_coding_gene | 2.0 | 3.88 | 7.8E-03 |
| *ceh-22* | WBGene00000445 | protein_coding_gene | 2.0 | 3.87 | 8.5E-05 |
| *C27A2.8* | WBGene00044387 | protein_coding_gene | 2.0 | 3.87 | 4.6E-02 |
| *lgc-1* | WBGene00020236 | protein_coding_gene | 2.0 | 3.86 | 2.4E-02 |
| *B0310.3* | WBGene00015139 | protein_coding_gene | 2.0 | 3.86 | 1.0E-02 |

| | | | | | |
|---|---|---|---|---|---|
| *srr-2* | WBGene00005653 | protein_coding_gene | 1.9 | 3.86 | 2.6E-02 |
| *slc-28.1* | WBGene00017867 | protein_coding_gene | 1.9 | 3.86 | 1.4E-03 |
| *T23G7.3* | WBGene00011966 | protein_coding_gene | 1.9 | 3.86 | 2.4E-03 |
| *T05A8.3* | WBGene00020229 | protein_coding_gene | 1.9 | 3.85 | 1.9E-03 |
| *K01A2.4* | WBGene00019279 | protein_coding_gene | 1.9 | 3.85 | 4.7E-04 |
| *fmo-2* | WBGene00001477 | protein_coding_gene | 1.9 | 3.84 | 2.2E-04 |
| *F53F1.6* | WBGene00009984 | protein_coding_gene | 1.9 | 3.84 | 4.9E-03 |
| *srd-28* | WBGene00005106 | protein_coding_gene | 1.9 | 3.84 | 4.8E-02 |
| *elt-4* | WBGene00001252 | protein_coding_gene | 1.9 | 3.83 | 5.4E-03 |
| *nlp-48* | WBGene00008002 | protein_coding_gene | 1.9 | 3.83 | 7.1E-03 |
| *irld-13* | WBGene00019845 | pseudogene | 1.9 | 3.83 | 4.2E-02 |
| *trx-3* | WBGene00019717 | protein_coding_gene | 1.9 | 3.83 | 1.0E-02 |
| *clec-63* | WBGene00009394 | protein_coding_gene | 1.9 | 3.83 | 6.0E-03 |
| *F22E5.21* | WBGene00044363 | protein_coding_gene | 1.9 | 3.83 | 9.3E-03 |
| *nhr-275* | WBGene00021163 | protein_coding_gene | 1.9 | 3.82 | 1.4E-03 |
| *fbxa-52* | WBGene00017225 | protein_coding_gene | 1.9 | 3.82 | 4.4E-02 |
| *F10D7.10* | WBGene00271701 | protein_coding_gene | 1.9 | 3.82 | 9.4E-03 |
| *T09B9.3* | WBGene00011644 | protein_coding_gene | 1.9 | 3.82 | 7.1E-03 |
| *F08B12.4* | WBGene00008572 | protein_coding_gene | 1.9 | 3.81 | 1.8E-07 |
| *T08E11.1* | WBGene00020357 | protein_coding_gene | 1.9 | 3.81 | 2.7E-02 |
| *clec-78* | WBGene00018547 | protein_coding_gene | 1.9 | 3.81 | 2.1E-02 |
| *btb-17* | WBGene00018200 | protein_coding_gene | 1.9 | 3.81 | 1.2E-07 |
| *math-37* | WBGene00020080 | protein_coding_gene | 1.9 | 3.81 | 2.6E-04 |
| *ttm-2* | WBGene00017840 | protein_coding_gene | 1.9 | 3.80 | 4.7E-05 |
| *F49C12.6* | WBGene00009876 | protein_coding_gene | 1.9 | 3.80 | 3.0E-04 |
| *aakg-4* | WBGene00020135 | protein_coding_gene | 1.9 | 3.80 | 1.5E-02 |
| *C16D9.9* | WBGene00015864 | protein_coding_gene | 1.9 | 3.80 | 1.4E-02 |
| *gei-13* | WBGene00001570 | protein_coding_gene | 1.9 | 3.80 | 1.5E-04 |
| *F56D5.6* | WBGene00010150 | protein_coding_gene | 1.9 | 3.79 | 1.2E-04 |
| *lgc-27* | WBGene00022283 | protein_coding_gene | 1.9 | 3.79 | 8.5E-05 |
| *ttx-3* | WBGene00006654 | protein_coding_gene | 1.9 | 3.79 | 3.5E-02 |
| *unc-75* | WBGene00006807 | protein_coding_gene | 1.9 | 3.78 | 1.6E-02 |
| *del-5* | WBGene00010334 | protein_coding_gene | 1.9 | 3.78 | 6.2E-05 |
| *cki-1* | WBGene00000516 | protein_coding_gene | 1.9 | 3.78 | 2.4E-11 |
| *flp-19* | WBGene00001462 | protein_coding_gene | 1.9 | 3.77 | 3.1E-03 |
| *Y14H12A.2* | WBGene00044382 | protein_coding_gene | 1.9 | 3.77 | 2.1E-02 |
| *srj-14* | WBGene00005602 | protein_coding_gene | 1.9 | 3.77 | 4.7E-03 |
| *ins-17* | WBGene00002100 | protein_coding_gene | 1.9 | 3.77 | 5.3E-06 |
| *F41C3.6* | WBGene00018272 | protein_coding_gene | 1.9 | 3.77 | 1.0E-02 |
| *mltn-12* | WBGene00008277 | protein_coding_gene | 1.9 | 3.77 | 9.7E-03 |
| *F53C3.4* | WBGene00018748 | protein_coding_gene | 1.9 | 3.77 | 2.6E-04 |
| *ast-1* | WBGene00020368 | protein_coding_gene | 1.9 | 3.76 | 1.3E-03 |
| *sqt-3* | WBGene00005018 | protein_coding_gene | 1.9 | 3.76 | 2.0E-05 |
| *C36B1.6* | WBGene00007973 | protein_coding_gene | 1.9 | 3.76 | 2.6E-04 |
| *T28H10.3* | WBGene00012144 | protein_coding_gene | 1.9 | 3.76 | 1.3E-03 |
| *C30G4.4* | WBGene00016269 | protein_coding_gene | 1.9 | 3.76 | 8.3E-04 |
| *frpr-18* | WBGene00020586 | protein_coding_gene | 1.9 | 3.76 | 3.7E-03 |
| *zag-1* | WBGene00006970 | protein_coding_gene | 1.9 | 3.76 | 5.5E-04 |
| *tax-4* | WBGene00006526 | protein_coding_gene | 1.9 | 3.75 | 4.2E-02 |
| *Y73B6BL.44* | WBGene00044350 | protein_coding_gene | 1.9 | 3.74 | 3.6E-02 |
| *aff-1* | WBGene00016625 | protein_coding_gene | 1.9 | 3.74 | 9.8E-04 |
| *F27E5.9* | WBGene00206354 | protein_coding_gene | 1.9 | 3.74 | 1.7E-02 |
| *C17B7.5* | WBGene00015879 | protein_coding_gene | 1.9 | 3.74 | 2.1E-02 |
| *R05G9R.1* | WBGene00019906 | protein_coding_gene | 1.9 | 3.74 | 9.3E-04 |
| *T14A8.2* | WBGene00020497 | protein_coding_gene | 1.9 | 3.73 | 5.9E-03 |
| *ttr-21* | WBGene00008628 | protein_coding_gene | 1.9 | 3.73 | 1.3E-04 |
| *cex-1* | WBGene00023407 | protein_coding_gene | 1.9 | 3.73 | 1.1E-02 |
| *aqp-2* | WBGene00000170 | protein_coding_gene | 1.9 | 3.72 | 2.1E-06 |

| | | | | | |
|---|---|---|---|---|---|
| *clec-7* | WBGene00017364 | protein_coding_gene | 1.9 | 3.72 | 3.9E-02 |
| *Y39A3A.3* | WBGene00021434 | protein_coding_gene | 1.9 | 3.72 | 3.3E-02 |
| *fbxa-3* | WBGene00020363 | protein_coding_gene | 1.9 | 3.72 | 2.2E-03 |
| *cyp-13B1* | WBGene00008519 | protein_coding_gene | 1.9 | 3.71 | 3.3E-02 |
| *tag-196* | WBGene00007055 | protein_coding_gene | 1.9 | 3.71 | 3.8E-03 |
| *spp-1* | WBGene00004986 | protein_coding_gene | 1.9 | 3.71 | 8.2E-03 |
| *K09F6.13* | WBGene00235164 | protein_coding_gene | 1.9 | 3.70 | 3.0E-02 |
| *gst-20* | WBGene00001768 | protein_coding_gene | 1.9 | 3.70 | 2.9E-02 |
| *cutl-8* | WBGene00013960 | protein_coding_gene | 1.9 | 3.70 | 2.0E-02 |
| *F18C5.9* | WBGene00017562 | protein_coding_gene | 1.9 | 3.70 | 4.6E-02 |
| *pqn-73* | WBGene00004155 | protein_coding_gene | 1.9 | 3.69 | 3.5E-02 |
| *C43H6.6* | WBGene00016619 | protein_coding_gene | 1.9 | 3.69 | 2.5E-02 |
| *cutl-17* | WBGene00009541 | protein_coding_gene | 1.9 | 3.69 | 8.2E-03 |
| *F37A4.3* | WBGene00018133 | protein_coding_gene | 1.9 | 3.69 | 3.1E-03 |
| *ZK970.7* | WBGene00014173 | protein_coding_gene | 1.9 | 3.68 | 3.4E-02 |
| *F59B10.4* | WBGene00010320 | protein_coding_gene | 1.9 | 3.68 | 2.4E-02 |
| *clec-9* | WBGene00013493 | protein_coding_gene | 1.9 | 3.68 | 1.0E-02 |
| *B0507.3* | WBGene00015220 | protein_coding_gene | 1.9 | 3.67 | 6.5E-05 |
| *aat-8* | WBGene00000009 | protein_coding_gene | 1.9 | 3.67 | 6.9E-03 |
| *chpf-2* | WBGene00019108 | protein_coding_gene | 1.9 | 3.67 | 3.4E-05 |
| *C25F9.9* | WBGene00007728 | protein_coding_gene | 1.9 | 3.66 | 2.1E-05 |
| *Y51H7BR.7* | WBGene00021778 | protein_coding_gene | 1.9 | 3.65 | 4.5E-02 |
| *F16A11.5* | WBGene00023472 | ncRNA_gene | 1.9 | 3.65 | 4.8E-02 |
| *ZK930.2* | WBGene00014152 | protein_coding_gene | 1.9 | 3.65 | 7.6E-11 |
| *ncx-5* | WBGene00003570 | protein_coding_gene | 1.9 | 3.65 | 3.4E-02 |
| *W03F8.6* | WBGene00020995 | protein_coding_gene | 1.9 | 3.65 | 6.7E-03 |
| *hic-1* | WBGene00020887 | protein_coding_gene | 1.9 | 3.64 | 1.8E-02 |
| *ets-10* | WBGene00016870 | protein_coding_gene | 1.9 | 3.64 | 3.1E-02 |
| *F40D4.16* | WBGene00194681 | ncRNA_gene | 1.9 | 3.64 | 1.1E-03 |
| *grh-1* | WBGene00001707 | protein_coding_gene | 1.9 | 3.64 | 4.3E-04 |
| *prkl-1* | WBGene00022727 | protein_coding_gene | 1.9 | 3.64 | 9.4E-07 |
| *C52B9.19* | WBGene00304788 | protein_coding_gene | 1.9 | 3.63 | 6.1E-03 |
| *unc-129* | WBGene00006852 | protein_coding_gene | 1.9 | 3.63 | 4.2E-05 |
| *aat-1* | WBGene00000002 | protein_coding_gene | 1.9 | 3.63 | 4.1E-04 |
| *clec-180* | WBGene00017991 | protein_coding_gene | 1.9 | 3.63 | 3.3E-03 |
| *ZC239.16* | WBGene00022574 | protein_coding_gene | 1.9 | 3.63 | 1.8E-03 |
| *mps-1* | WBGene00003403 | protein_coding_gene | 1.9 | 3.62 | 1.1E-02 |
| *Y57G11C.44* | WBGene00013333 | protein_coding_gene | 1.9 | 3.62 | 3.3E-02 |
| *nlp-31* | WBGene00003769 | protein_coding_gene | 1.9 | 3.62 | 2.8E-02 |
| *F52A8.3* | WBGene00009916 | protein_coding_gene | 1.9 | 3.62 | 3.4E-03 |
| *F13B6.1* | WBGene00017416 | protein_coding_gene | 1.9 | 3.62 | 1.4E-02 |
| *math-27* | WBGene00018101 | pseudogene | 1.9 | 3.62 | 1.8E-04 |
| *C38C3.4* | WBGene00016513 | protein_coding_gene | 1.9 | 3.62 | 2.8E-03 |
| *F07C3.9* | WBGene00017197 | protein_coding_gene | 1.9 | 3.62 | 1.0E-04 |
| *gcy-15* | WBGene00001541 | protein_coding_gene | 1.9 | 3.62 | 1.8E-02 |
| *cpz-2* | WBGene00000789 | protein_coding_gene | 1.9 | 3.61 | 1.8E-03 |
| *T28A11.2* | WBGene00020869 | protein_coding_gene | 1.9 | 3.61 | 3.2E-02 |
| *fipr-24* | WBGene00007992 | protein_coding_gene | 1.9 | 3.61 | 2.5E-03 |
| *C15A11.7* | WBGene00007599 | protein_coding_gene | 1.9 | 3.61 | 1.4E-02 |
| *C01G6.9* | WBGene00007229 | protein_coding_gene | 1.9 | 3.61 | 1.6E-04 |
| *sre-4* | WBGene00011745 | protein_coding_gene | 1.9 | 3.61 | 3.8E-02 |
| *C54G6.3* | WBGene00016932 | protein_coding_gene | 1.9 | 3.61 | 2.7E-02 |
| *pho-4* | WBGene00004023 | protein_coding_gene | 1.9 | 3.61 | 3.8E-04 |
| *K08D10.14* | WBGene00219216 | protein_coding_gene | 1.8 | 3.60 | 3.7E-03 |
| *C30F8.3* | WBGene00016259 | protein_coding_gene | 1.8 | 3.60 | 1.7E-02 |
| *Y71G12B.2* | WBGene00022142 | protein_coding_gene | 1.8 | 3.60 | 3.1E-02 |
| *C28G1.10* | WBGene00219376 | protein_coding_gene | 1.8 | 3.60 | 7.6E-04 |
| *cutl-24* | WBGene00021396 | protein_coding_gene | 1.8 | 3.60 | 1.3E-03 |

| | | | | | |
|---|---|---|---|---|---|
| *F26F12.4* | WBGene00017835 | protein_coding_gene | 1.8 | 3.60 | 4.6E-04 |
| *T28F4.5* | WBGene00012140 | protein_coding_gene | 1.8 | 3.59 | 4.7E-03 |
| *F53B2.8* | WBGene00009957 | protein_coding_gene | 1.8 | 3.59 | 1.7E-02 |
| *F47B8.5* | WBGene00009806 | protein_coding_gene | 1.8 | 3.59 | 3.4E-02 |
| *T14B1.1* | WBGene00011763 | protein_coding_gene | 1.8 | 3.59 | 2.1E-04 |
| *C16B8.2* | WBGene00015822 | protein_coding_gene | 1.8 | 3.59 | 7.4E-04 |
| *lec-11* | WBGene00002274 | protein_coding_gene | 1.8 | 3.59 | 2.5E-04 |
| *T05E7.4* | WBGene00020260 | protein_coding_gene | 1.8 | 3.59 | 1.5E-02 |
| *sod-4* | WBGene00004933 | protein_coding_gene | 1.8 | 3.59 | 1.3E-03 |
| *endu-2* | WBGene00019779 | protein_coding_gene | 1.8 | 3.59 | 1.6E-02 |
| *F33D11.16* | WBGene00269382 | protein_coding_gene | 1.8 | 3.58 | 1.3E-02 |
| *mdl-1* | WBGene00003163 | protein_coding_gene | 1.8 | 3.58 | 1.2E-03 |
| *clec-264* | WBGene00009291 | protein_coding_gene | 1.8 | 3.58 | 2.5E-03 |
| *Y110A2AL.3* | WBGene00022440 | protein_coding_gene | 1.8 | 3.58 | 1.6E-05 |
| *F01G10.6* | WBGene00008509 | protein_coding_gene | 1.8 | 3.58 | 6.6E-03 |
| *ZK265.7* | WBGene00013959 | protein_coding_gene | 1.8 | 3.57 | 2.8E-04 |
| *ZK470.14* | WBGene00219362 | protein_coding_gene | 1.8 | 3.57 | 1.3E-02 |
| *Y60C6A.2* | WBGene00202498 | protein_coding_gene | 1.8 | 3.57 | 3.7E-02 |
| *M163.1* | WBGene00010935 | protein_coding_gene | 1.8 | 3.57 | 5.7E-03 |
| *pqn-42* | WBGene00004129 | protein_coding_gene | 1.8 | 3.57 | 9.5E-04 |
| *cdh-9* | WBGene00000401 | protein_coding_gene | 1.8 | 3.57 | 2.0E-04 |
| *M02D8.6* | WBGene00019732 | protein_coding_gene | 1.8 | 3.57 | 3.0E-03 |
| *osm-6* | WBGene00003886 | protein_coding_gene | 1.8 | 3.57 | 6.0E-03 |
| *cutl-21* | WBGene00018861 | protein_coding_gene | 1.8 | 3.56 | 3.8E-04 |
| *F42G4.6* | WBGene00009643 | protein_coding_gene | 1.8 | 3.56 | 2.9E-03 |
| *ubc-8* | WBGene00006705 | protein_coding_gene | 1.8 | 3.56 | 9.4E-03 |
| *srh-2* | WBGene00005228 | protein_coding_gene | 1.8 | 3.55 | 4.2E-02 |
| *egl-15* | WBGene00001184 | protein_coding_gene | 1.8 | 3.55 | 3.1E-02 |
| *F22B7.1* | WBGene00017692 | protein_coding_gene | 1.8 | 3.55 | 4.1E-03 |
| *F13A7.12* | WBGene00008727 | protein_coding_gene | 1.8 | 3.55 | 1.9E-02 |
| *C35C5.9* | WBGene00007958 | protein_coding_gene | 1.8 | 3.54 | 8.6E-04 |
| *Y8A9A.2* | WBGene00021171 | protein_coding_gene | 1.8 | 3.54 | 2.5E-02 |
| *F55A4.7* | WBGene00018859 | protein_coding_gene | 1.8 | 3.54 | 1.6E-02 |
| *ssu-2* | WBGene00013455 | protein_coding_gene | 1.8 | 3.54 | 1.4E-02 |
| *Y37D8A.16* | WBGene00012554 | protein_coding_gene | 1.8 | 3.54 | 8.3E-05 |
| *egl-46* | WBGene00001210 | protein_coding_gene | 1.8 | 3.54 | 1.4E-02 |
| *lury-1* | WBGene00013548 | protein_coding_gene | 1.8 | 3.54 | 2.7E-04 |
| *fbxa-163* | WBGene00015598 | protein_coding_gene | 1.8 | 3.53 | 4.8E-02 |
| *cyn-17* | WBGene00000893 | protein_coding_gene | 1.8 | 3.53 | 3.7E-03 |
| *ZK836.3* | WBGene00014099 | protein_coding_gene | 1.8 | 3.53 | 1.5E-03 |
| *C28H8.2* | WBGene00016193 | protein_coding_gene | 1.8 | 3.53 | 1.8E-03 |
| *Y41G9A.5* | WBGene00021529 | protein_coding_gene | 1.8 | 3.53 | 6.4E-03 |
| *che-2* | WBGene00000484 | protein_coding_gene | 1.8 | 3.52 | 4.4E-03 |
| *F10D7.3* | WBGene00017340 | protein_coding_gene | 1.8 | 3.52 | 3.4E-02 |
| *frpr-15* | WBGene00010735 | protein_coding_gene | 1.8 | 3.52 | 1.0E-02 |
| *ZK829.3* | WBGene00014094 | protein_coding_gene | 1.8 | 3.52 | 2.8E-02 |
| *F25E5.4* | WBGene00017785 | protein_coding_gene | 1.8 | 3.52 | 2.2E-04 |
| *F41G4.8* | WBGene00018311 | protein_coding_gene | 1.8 | 3.52 | 2.6E-03 |
| *osm-5* | WBGene00003885 | protein_coding_gene | 1.8 | 3.51 | 1.3E-03 |
| *sra-10* | WBGene00005036 | protein_coding_gene | 1.8 | 3.51 | 9.0E-03 |
| *cutl-16* | WBGene00019426 | protein_coding_gene | 1.8 | 3.51 | 2.5E-03 |
| *irld-34* | WBGene00010074 | protein_coding_gene | 1.8 | 3.51 | 4.8E-02 |
| *C49A9.1* | WBGene00016755 | protein_coding_gene | 1.8 | 3.51 | 5.9E-04 |
| *egl-43* | WBGene00001207 | protein_coding_gene | 1.8 | 3.51 | 9.5E-04 |
| *aptf-1* | WBGene00019424 | protein_coding_gene | 1.8 | 3.50 | 3.1E-02 |
| *inx-6* | WBGene00002128 | protein_coding_gene | 1.8 | 3.50 | 3.3E-04 |
| *hbl-1* | WBGene00001824 | protein_coding_gene | 1.8 | 3.50 | 9.2E-03 |
| *T01B6.1* | WBGene00020134 | protein_coding_gene | 1.8 | 3.49 | 2.6E-04 |

| | | | | | |
|---|---|---|---|---|---|
| *tsp-6* | WBGene00006632 | protein_coding_gene | 1.8 | 3.49 | 9.8E-03 |
| *C01G5.25* | WBGene00235291 | protein_coding_gene | 1.8 | 3.49 | 1.6E-02 |
| *hlh-6* | WBGene00001952 | protein_coding_gene | 1.8 | 3.49 | 2.5E-03 |
| *subs-4* | WBGene00012939 | protein_coding_gene | 1.8 | 3.49 | 7.2E-04 |
| *nlp-64* | WBGene00019734 | protein_coding_gene | 1.8 | 3.49 | 9.5E-04 |
| *Y67D8B.2* | WBGene00022062 | protein_coding_gene | 1.8 | 3.48 | 2.2E-02 |
| *F35B12.3* | WBGene00009383 | protein_coding_gene | 1.8 | 3.48 | 6.3E-04 |
| *Y34F4.5* | WBGene00021340 | protein_coding_gene | 1.8 | 3.47 | 1.4E-04 |
| *C49H3.12* | WBGene00016795 | protein_coding_gene | 1.8 | 3.47 | 2.0E-02 |
| *Y73E7A.8* | WBGene00022274 | protein_coding_gene | 1.8 | 3.47 | 1.2E-03 |
| *nsy-4* | WBGene00021415 | protein_coding_gene | 1.8 | 3.47 | 4.5E-05 |
| *R11G1.2* | WBGene00020011 | protein_coding_gene | 1.8 | 3.47 | 1.9E-03 |
| *K07H8.11* | WBGene00044510 | protein_coding_gene | 1.8 | 3.47 | 3.4E-03 |
| *ZK1290.13* | WBGene00022893 | protein_coding_gene | 1.8 | 3.46 | 9.3E-05 |
| *dhs-15* | WBGene00000978 | protein_coding_gene | 1.8 | 3.46 | 2.0E-02 |
| *grl-5* | WBGene00001714 | protein_coding_gene | 1.8 | 3.46 | 6.3E-03 |
| *F09F9.4* | WBGene00017309 | protein_coding_gene | 1.8 | 3.46 | 6.1E-03 |
| *B0348.5* | WBGene00015153 | protein_coding_gene | 1.8 | 3.46 | 1.2E-02 |
| *E02H9.6* | WBGene00017104 | protein_coding_gene | 1.8 | 3.45 | 4.9E-02 |
| *C08E8.4* | WBGene00007440 | protein_coding_gene | 1.8 | 3.45 | 1.7E-02 |
| *glr-2* | WBGene00001613 | protein_coding_gene | 1.8 | 3.45 | 1.1E-03 |
| *C13C12.2* | WBGene00007552 | protein_coding_gene | 1.8 | 3.45 | 1.3E-03 |
| *Y43F8A.1* | WBGene00012808 | protein_coding_gene | 1.8 | 3.45 | 3.0E-03 |
| *php-3* | WBGene00004024 | protein_coding_gene | 1.8 | 3.44 | 3.0E-05 |
| *daf-10* | WBGene00000906 | protein_coding_gene | 1.8 | 3.44 | 4.7E-05 |
| *F55G11.7* | WBGene00010127 | protein_coding_gene | 1.8 | 3.43 | 4.8E-02 |
| *C18A11.1* | WBGene00015947 | protein_coding_gene | 1.8 | 3.43 | 1.5E-02 |
| *fbxa-99* | WBGene00009490 | protein_coding_gene | 1.8 | 3.43 | 3.2E-02 |
| *C30G12.4* | WBGene00016276 | protein_coding_gene | 1.8 | 3.43 | 2.8E-02 |
| *W04B5.2* | WBGene00021019 | protein_coding_gene | 1.8 | 3.43 | 2.9E-05 |
| *vamp-8* | WBGene00007200 | protein_coding_gene | 1.8 | 3.43 | 2.3E-04 |
| *pha-4* | WBGene00004013 | protein_coding_gene | 1.8 | 3.43 | 3.5E-04 |
| *flp-32* | WBGene00010982 | protein_coding_gene | 1.8 | 3.43 | 9.3E-04 |
| *K07D4.4* | WBGene00019484 | protein_coding_gene | 1.8 | 3.43 | 4.7E-02 |
| *C03F11.2* | WBGene00015388 | protein_coding_gene | 1.8 | 3.43 | 3.9E-02 |
| *ZK1290.10* | WBGene00022891 | protein_coding_gene | 1.8 | 3.42 | 3.1E-02 |
| *E02A10.3* | WBGene00008453 | protein_coding_gene | 1.8 | 3.42 | 2.8E-03 |
| *twk-23* | WBGene00006676 | protein_coding_gene | 1.8 | 3.42 | 1.8E-02 |
| *F13D11.3* | WBGene00017428 | protein_coding_gene | 1.8 | 3.42 | 6.7E-03 |
| *sulp-3* | WBGene00018283 | protein_coding_gene | 1.8 | 3.42 | 1.6E-02 |
| *ZC455.1* | WBGene00013902 | protein_coding_gene | 1.8 | 3.41 | 4.6E-03 |
| *T24C2.5* | WBGene00011984 | protein_coding_gene | 1.8 | 3.41 | 9.1E-04 |
| *Y51B9A.9* | WBGene00013091 | protein_coding_gene | 1.8 | 3.41 | 3.7E-03 |
| *C07A9.5* | WBGene00007401 | protein_coding_gene | 1.8 | 3.41 | 4.2E-02 |
| *Y11D7A.8* | WBGene00012432 | protein_coding_gene | 1.8 | 3.41 | 1.7E-02 |
| *linc-56* | WBGene00219650 | lincRNA_gene | 1.8 | 3.41 | 4.6E-02 |
| *K03A11.5* | WBGene00010522 | protein_coding_gene | 1.8 | 3.41 | 5.2E-05 |
| *H04M03.12* | WBGene00019156 | protein_coding_gene | 1.8 | 3.41 | 1.2E-02 |
| *F13H6.5* | WBGene00017433 | protein_coding_gene | 1.8 | 3.41 | 2.1E-02 |
| *K08E7.5* | WBGene00010673 | protein_coding_gene | 1.8 | 3.41 | 2.1E-02 |
| *C50F4.1* | WBGene00008229 | protein_coding_gene | 1.8 | 3.41 | 5.6E-07 |
| *ZC84.6* | WBGene00013849 | protein_coding_gene | 1.8 | 3.41 | 2.1E-02 |
| *ZC239.17* | WBGene00022575 | protein_coding_gene | 1.8 | 3.40 | 1.3E-02 |
| *C54D2.1* | WBGene00016915 | protein_coding_gene | 1.8 | 3.40 | 1.1E-02 |
| *grl-14* | WBGene00001723 | protein_coding_gene | 1.8 | 3.40 | 1.2E-02 |
| *pros-1* | WBGene00000448 | protein_coding_gene | 1.8 | 3.40 | 1.0E-03 |
| *ZC132.3* | WBGene00022522 | protein_coding_gene | 1.8 | 3.39 | 2.9E-02 |
| *mks-6* | WBGene00010642 | protein_coding_gene | 1.8 | 3.39 | 1.5E-04 |

| | | | | | |
|---|---|---|---|---|---|
| *ptr-17* | WBGene00004231 | protein_coding_gene | 1.8 | 3.39 | 2.0E-02 |
| *F53B1.4* | WBGene00018737 | protein_coding_gene | 1.8 | 3.39 | 1.3E-03 |
| *pax-1* | WBGene00003937 | protein_coding_gene | 1.8 | 3.39 | 4.9E-02 |
| *K10G6.9* | WBGene00206374 | protein_coding_gene | 1.8 | 3.39 | 1.6E-02 |
| *igcm-2* | WBGene00020130 | protein_coding_gene | 1.8 | 3.38 | 2.3E-03 |
| *Y38C1AA.6* | WBGene00021397 | protein_coding_gene | 1.8 | 3.38 | 2.0E-03 |
| *flp-25* | WBGene00010575 | protein_coding_gene | 1.8 | 3.38 | 4.7E-03 |
| *F31D4.5* | WBGene00009292 | protein_coding_gene | 1.8 | 3.37 | 7.4E-03 |
| *C14C6.2* | WBGene00015756 | protein_coding_gene | 1.8 | 3.37 | 2.3E-02 |
| *lec-2* | WBGene00002265 | protein_coding_gene | 1.8 | 3.37 | 1.6E-02 |
| *F11C7.2* | WBGene00017376 | protein_coding_gene | 1.8 | 3.37 | 2.0E-03 |
| *C06E7.88* | WBGene00189995 | protein_coding_gene | 1.8 | 3.37 | 4.0E-02 |
| *K09F6.9* | WBGene00019592 | protein_coding_gene | 1.8 | 3.37 | 9.5E-04 |
| *ttx-1* | WBGene00006652 | protein_coding_gene | 1.8 | 3.37 | 7.6E-04 |
| *C13E3.1* | WBGene00015740 | protein_coding_gene | 1.8 | 3.37 | 3.0E-04 |
| *nspc-17* | WBGene00008359 | protein_coding_gene | 1.8 | 3.36 | 4.2E-03 |
| *clec-233* | WBGene00009030 | protein_coding_gene | 1.7 | 3.36 | 1.3E-03 |
| *Y7A5A.11* | WBGene00044155 | pseudogene | 1.7 | 3.36 | 1.6E-02 |
| *W10C8.4* | WBGene00021127 | protein_coding_gene | 1.7 | 3.36 | 4.1E-03 |
| *T26C5.4* | WBGene00012034 | protein_coding_gene | 1.7 | 3.35 | 3.0E-02 |
| *glb-25* | WBGene00020279 | protein_coding_gene | 1.7 | 3.35 | 1.8E-03 |
| *F59B10.5* | WBGene00010321 | protein_coding_gene | 1.7 | 3.35 | 1.1E-02 |
| *M6.4* | WBGene00019776 | protein_coding_gene | 1.7 | 3.35 | 3.4E-02 |
| *F45E12.6* | WBGene00018476 | protein_coding_gene | 1.7 | 3.35 | 3.0E-02 |
| *asp-5* | WBGene00000218 | protein_coding_gene | 1.7 | 3.35 | 3.0E-03 |
| *fax-1* | WBGene00001400 | protein_coding_gene | 1.7 | 3.35 | 2.8E-02 |
| *nhr-263* | WBGene00017367 | protein_coding_gene | 1.7 | 3.35 | 2.7E-02 |
| *F58E6.11* | WBGene00010258 | protein_coding_gene | 1.7 | 3.34 | 5.5E-03 |
| *C10C5.7* | WBGene00007511 | protein_coding_gene | 1.7 | 3.34 | 4.9E-03 |
| *Y2H9A.4* | WBGene00012381 | protein_coding_gene | 1.7 | 3.34 | 1.2E-02 |
| *pdl-1* | WBGene00003966 | protein_coding_gene | 1.7 | 3.34 | 9.6E-04 |
| *C05E11.7* | WBGene00015496 | protein_coding_gene | 1.7 | 3.34 | 4.8E-02 |
| *rcan-1* | WBGene00004321 | protein_coding_gene | 1.7 | 3.34 | 5.8E-03 |
| *C04E12.4* | WBGene00015434 | protein_coding_gene | 1.7 | 3.34 | 3.6E-02 |
| *F37C12.21* | WBGene00194742 | protein_coding_gene | 1.7 | 3.34 | 1.1E-02 |
| *Y69A2AR.11* | WBGene00022082 | pseudogene | 1.7 | 3.34 | 2.0E-02 |
| *Y4C6B.2* | WBGene00021156 | protein_coding_gene | 1.7 | 3.34 | 2.6E-02 |
| *ppm-1.H* | WBGene00016580 | protein_coding_gene | 1.7 | 3.33 | 4.6E-03 |
| *nhr-120* | WBGene00003710 | protein_coding_gene | 1.7 | 3.33 | 6.8E-03 |
| *R02E12.12* | WBGene00303425 | protein_coding_gene | 1.7 | 3.33 | 9.8E-03 |
| *T05B11.7* | WBGene00044342 | protein_coding_gene | 1.7 | 3.32 | 7.4E-04 |
| *nlp-42* | WBGene00013588 | protein_coding_gene | 1.7 | 3.32 | 1.4E-02 |
| *T05E11.9* | WBGene00014505 | ncRNA_gene | 1.7 | 3.32 | 4.0E-04 |
| *F41E6.5* | WBGene00018286 | protein_coding_gene | 1.7 | 3.32 | 2.7E-02 |
| *nhr-42* | WBGene00003632 | protein_coding_gene | 1.7 | 3.32 | 1.9E-05 |
| *nphp-4* | WBGene00011261 | protein_coding_gene | 1.7 | 3.31 | 2.0E-03 |
| *nhr-236* | WBGene00021417 | protein_coding_gene | 1.7 | 3.31 | 1.3E-02 |
| *F56H9.2* | WBGene00010174 | protein_coding_gene | 1.7 | 3.31 | 4.6E-08 |
| *C06G1.2* | WBGene00015546 | protein_coding_gene | 1.7 | 3.31 | 4.8E-03 |
| *mltn-1* | WBGene00009311 | protein_coding_gene | 1.7 | 3.31 | 4.1E-02 |
| *F18C5.5* | WBGene00017560 | protein_coding_gene | 1.7 | 3.30 | 2.4E-02 |
| *Y64G10A.7* | WBGene00013416 | protein_coding_gene | 1.7 | 3.30 | 5.4E-03 |
| *F09C3.2* | WBGene00008610 | protein_coding_gene | 1.7 | 3.30 | 5.2E-04 |
| *C36A4.11* | WBGene00044167 | protein_coding_gene | 1.7 | 3.30 | 1.4E-02 |
| *Y97E10C.1* | WBGene00022410 | protein_coding_gene | 1.7 | 3.30 | 2.7E-04 |
| *F44E2.4* | WBGene00018418 | protein_coding_gene | 1.7 | 3.30 | 2.0E-03 |
| *Y22D7AR.6* | WBGene00021260 | protein_coding_gene | 1.7 | 3.29 | 2.0E-03 |
| *C50F4.8* | WBGene00008233 | protein_coding_gene | 1.7 | 3.29 | 4.3E-02 |

| | | | | | |
|---|---|---|---|---|---|
| *grl-15* | WBGene00001724 | protein_coding_gene | 1.7 | 3.29 | 1.8E-02 |
| *F33A8.7* | WBGene00009355 | protein_coding_gene | 1.7 | 3.28 | 2.0E-02 |
| *F44F4.3* | WBGene00009702 | protein_coding_gene | 1.7 | 3.28 | 1.5E-02 |
| *F54D5.3* | WBGene00010049 | protein_coding_gene | 1.7 | 3.28 | 3.9E-05 |
| *C53B7.3* | WBGene00016894 | protein_coding_gene | 1.7 | 3.27 | 2.4E-03 |
| *eva-1* | WBGene00009304 | protein_coding_gene | 1.7 | 3.27 | 1.7E-04 |
| *nhr-173* | WBGene00016975 | protein_coding_gene | 1.7 | 3.26 | 2.5E-03 |
| *F13G3.3* | WBGene00008763 | protein_coding_gene | 1.7 | 3.26 | 1.6E-02 |
| *F13E9.11* | WBGene00008760 | protein_coding_gene | 1.7 | 3.26 | 3.4E-03 |
| *sqst-1* | WBGene00011737 | protein_coding_gene | 1.7 | 3.26 | 9.4E-03 |
| *K02E11.5* | WBGene00010514 | protein_coding_gene | 1.7 | 3.26 | 2.6E-03 |
| *atf-8* | WBGene00017535 | protein_coding_gene | 1.7 | 3.26 | 5.0E-03 |
| *fip-2* | WBGene00020862 | protein_coding_gene | 1.7 | 3.25 | 7.1E-10 |
| *nhr-44* | WBGene00003634 | protein_coding_gene | 1.7 | 3.25 | 2.5E-03 |
| *C05D12.2* | WBGene00007340 | protein_coding_gene | 1.7 | 3.25 | 4.7E-02 |
| *F23F12.8* | WBGene00017754 | protein_coding_gene | 1.7 | 3.24 | 1.6E-02 |
| *siah-1* | WBGene00021369 | protein_coding_gene | 1.7 | 3.24 | 1.8E-02 |
| *nas-25* | WBGene00003544 | protein_coding_gene | 1.7 | 3.24 | 1.2E-02 |
| *col-111* | WBGene00000685 | protein_coding_gene | 1.7 | 3.24 | 3.5E-02 |
| *sup-26* | WBGene00006331 | protein_coding_gene | 1.7 | 3.24 | 5.5E-04 |
| *R07G3.8* | WBGene00019944 | protein_coding_gene | 1.7 | 3.24 | 2.5E-03 |
| *cex-2* | WBGene00023408 | protein_coding_gene | 1.7 | 3.24 | 1.2E-02 |
| *F57A8.1* | WBGene00010177 | protein_coding_gene | 1.7 | 3.24 | 1.9E-04 |
| *glb-11* | WBGene00007981 | protein_coding_gene | 1.7 | 3.24 | 1.0E-02 |
| *ZC190.4* | WBGene00022538 | protein_coding_gene | 1.7 | 3.24 | 4.9E-03 |
| *C03D6.1* | WBGene00007275 | protein_coding_gene | 1.7 | 3.24 | 7.4E-03 |
| *abf-5* | WBGene00000016 | protein_coding_gene | 1.7 | 3.23 | 1.0E-04 |
| *maph-9* | WBGene00016397 | protein_coding_gene | 1.7 | 3.23 | 9.5E-03 |
| *dpy-10* | WBGene00001072 | protein_coding_gene | 1.7 | 3.22 | 2.6E-03 |
| *bus-12* | WBGene00010442 | protein_coding_gene | 1.7 | 3.22 | 2.4E-02 |
| *Y7A9C.1* | WBGene00012418 | pseudogene | 1.7 | 3.22 | 4.6E-08 |
| *C54D10.10* | WBGene00008304 | protein_coding_gene | 1.7 | 3.22 | 2.4E-02 |
| *sma-2* | WBGene00004856 | protein_coding_gene | 1.7 | 3.22 | 4.1E-03 |
| *Y79H2A.12* | WBGene00043997 | protein_coding_gene | 1.7 | 3.21 | 2.7E-02 |
| *C08B6.10* | WBGene00007430 | protein_coding_gene | 1.7 | 3.21 | 3.9E-02 |
| *Y38F1A.8* | WBGene00012610 | protein_coding_gene | 1.7 | 3.21 | 2.4E-02 |
| *mod-5* | WBGene00003387 | protein_coding_gene | 1.7 | 3.21 | 7.3E-03 |
| *Y51A2D.13* | WBGene00013080 | protein_coding_gene | 1.7 | 3.21 | 1.5E-02 |
| *C13C4.7* | WBGene00007551 | protein_coding_gene | 1.7 | 3.21 | 1.1E-03 |
| *F42C5.4* | WBGene00018346 | protein_coding_gene | 1.7 | 3.21 | 1.4E-03 |
| *F01D5.5* | WBGene00008495 | protein_coding_gene | 1.7 | 3.20 | 3.7E-02 |
| *dao-2* | WBGene00000928 | protein_coding_gene | 1.7 | 3.20 | 3.2E-03 |
| *ceh-45* | WBGene00022837 | protein_coding_gene | 1.7 | 3.19 | 7.9E-03 |
| *his-71* | WBGene00001945 | protein_coding_gene | 1.7 | 3.19 | 1.5E-03 |
| *nlp-17* | WBGene00003755 | protein_coding_gene | 1.7 | 3.19 | 1.3E-03 |
| *ptp-1* | WBGene00004213 | protein_coding_gene | 1.7 | 3.19 | 9.6E-04 |
| *marc-4* | WBGene00016903 | protein_coding_gene | 1.7 | 3.19 | 9.4E-03 |
| *msa-1* | WBGene00013882 | protein_coding_gene | 1.7 | 3.19 | 1.5E-02 |
| *wrt-6* | WBGene00006952 | protein_coding_gene | 1.7 | 3.19 | 1.9E-02 |
| *F52F10.2* | WBGene00018706 | protein_coding_gene | 1.7 | 3.19 | 3.7E-02 |
| *clec-65* | WBGene00009396 | protein_coding_gene | 1.7 | 3.18 | 3.0E-03 |
| *ceh-2* | WBGene00000429 | protein_coding_gene | 1.7 | 3.17 | 2.2E-02 |
| *F21C10.3* | WBGene00017653 | protein_coding_gene | 1.7 | 3.17 | 2.9E-03 |
| *F23F12.13* | WBGene00044455 | protein_coding_gene | 1.7 | 3.17 | 6.6E-03 |
| *nog-1* | WBGene00020297 | protein_coding_gene | 1.7 | 3.17 | 6.2E-03 |
| *tsp-16* | WBGene00006642 | protein_coding_gene | 1.7 | 3.17 | 3.2E-04 |
| *Y52B11A.4* | WBGene00013124 | protein_coding_gene | 1.7 | 3.17 | 4.8E-03 |
| *C32D5.12* | WBGene00016319 | protein_coding_gene | 1.7 | 3.16 | 6.5E-03 |

| | | | | | |
|---|---|---|---|---|---|
| *F54D1.6* | WBGene00010047 | protein_coding_gene | 1.7 | 3.16 | 1.0E-03 |
| *F07H5.4* | WBGene00014308 | pseudogene | 1.7 | 3.16 | 6.6E-04 |
| *atf-2* | WBGene00000220 | protein_coding_gene | 1.7 | 3.16 | 2.0E-02 |
| *ctsa-3.2* | WBGene00007462 | protein_coding_gene | 1.7 | 3.16 | 2.1E-02 |
| *C03G6.17* | WBGene00015402 | protein_coding_gene | 1.7 | 3.16 | 2.3E-02 |
| *lipl-6* | WBGene00021963 | protein_coding_gene | 1.7 | 3.16 | 1.2E-02 |
| *rom-2* | WBGene00004401 | protein_coding_gene | 1.7 | 3.16 | 1.6E-03 |
| *T17H7.7* | WBGene00020551 | protein_coding_gene | 1.7 | 3.15 | 2.6E-02 |
| *F47D12.6* | WBGene00018562 | protein_coding_gene | 1.7 | 3.15 | 1.4E-02 |
| *nhr-180* | WBGene00017961 | protein_coding_gene | 1.7 | 3.15 | 8.9E-03 |
| *ist-1* | WBGene00002163 | protein_coding_gene | 1.7 | 3.14 | 2.0E-02 |
| *F13H10.5* | WBGene00008776 | protein_coding_gene | 1.7 | 3.14 | 3.7E-03 |
| *Y38H6A.5* | WBGene00077669 | protein_coding_gene | 1.7 | 3.14 | 2.9E-02 |
| *nhr-258* | WBGene00016126 | protein_coding_gene | 1.7 | 3.14 | 2.7E-02 |
| *Y43E12A.2* | WBGene00012794 | protein_coding_gene | 1.7 | 3.14 | 3.0E-03 |
| *nhr-32* | WBGene00003626 | protein_coding_gene | 1.7 | 3.14 | 1.3E-03 |
| *fkh-8* | WBGene00001440 | protein_coding_gene | 1.7 | 3.14 | 2.9E-04 |
| *F35C11.7* | WBGene00009406 | protein_coding_gene | 1.7 | 3.14 | 5.1E-05 |
| *C15H11.11* | WBGene00007619 | protein_coding_gene | 1.6 | 3.14 | 1.1E-02 |
| *F26G1.2* | WBGene00017838 | protein_coding_gene | 1.6 | 3.13 | 2.1E-02 |
| *C33D9.6* | WBGene00007900 | protein_coding_gene | 1.6 | 3.13 | 3.4E-06 |
| *lgc-13* | WBGene00011354 | protein_coding_gene | 1.6 | 3.13 | 2.9E-02 |
| *F45H10.5* | WBGene00009742 | protein_coding_gene | 1.6 | 3.12 | 6.7E-03 |
| *ifta-2* | WBGene00012132 | protein_coding_gene | 1.6 | 3.12 | 1.1E-02 |
| *ZC449.4* | WBGene00022614 | protein_coding_gene | 1.6 | 3.12 | 1.1E-02 |
| *C54D10.13* | WBGene00045400 | protein_coding_gene | 1.6 | 3.12 | 1.4E-02 |
| *kel-1* | WBGene00002184 | protein_coding_gene | 1.6 | 3.12 | 8.8E-04 |
| *frpr-14* | WBGene00019496 | protein_coding_gene | 1.6 | 3.12 | 4.2E-02 |
| *nspc-20* | WBGene00008356 | protein_coding_gene | 1.6 | 3.12 | 1.9E-04 |
| *hlh-30* | WBGene00020930 | protein_coding_gene | 1.6 | 3.12 | 1.0E-02 |
| *C06E4.2* | WBGene00015531 | protein_coding_gene | 1.6 | 3.12 | 1.3E-02 |
| *exc-5* | WBGene00001366 | protein_coding_gene | 1.6 | 3.11 | 3.3E-03 |
| *tat-6* | WBGene00017174 | protein_coding_gene | 1.6 | 3.10 | 2.8E-02 |
| *bbs-8* | WBGene00000244 | protein_coding_gene | 1.6 | 3.10 | 1.2E-03 |
| *nac-2* | WBGene00003518 | protein_coding_gene | 1.6 | 3.10 | 9.4E-09 |
| *F08A7.1* | WBGene00045013 | protein_coding_gene | 1.6 | 3.10 | 2.3E-03 |
| *clec-62* | WBGene00009393 | protein_coding_gene | 1.6 | 3.10 | 1.3E-02 |
| *F59C6.8* | WBGene00010327 | protein_coding_gene | 1.6 | 3.10 | 2.2E-03 |
| *clec-40* | WBGene00011853 | protein_coding_gene | 1.6 | 3.10 | 3.5E-02 |
| *nhr-147* | WBGene00015395 | protein_coding_gene | 1.6 | 3.09 | 4.2E-02 |
| *C01G5.7* | WBGene00015309 | protein_coding_gene | 1.6 | 3.09 | 4.9E-02 |
| *ctbp-1* | WBGene00006424 | protein_coding_gene | 1.6 | 3.09 | 1.0E-02 |
| *R11D1.13* | WBGene00045517 | protein_coding_gene | 1.6 | 3.09 | 4.2E-02 |
| *inx-20* | WBGene00002142 | protein_coding_gene | 1.6 | 3.09 | 1.3E-02 |
| *F21D12.3* | WBGene00017663 | protein_coding_gene | 1.6 | 3.09 | 9.1E-03 |
| *Y55B1BL.1* | WBGene00021909 | protein_coding_gene | 1.6 | 3.09 | 3.6E-04 |
| *fkh-7* | WBGene00001439 | protein_coding_gene | 1.6 | 3.09 | 6.1E-03 |
| *Y41C4A.11* | WBGene00012757 | protein_coding_gene | 1.6 | 3.08 | 9.1E-03 |
| *lron-1* | WBGene00008093 | protein_coding_gene | 1.6 | 3.08 | 1.3E-02 |
| *M03C11.1* | WBGene00010838 | protein_coding_gene | 1.6 | 3.08 | 5.0E-04 |
| *K07A1.3* | WBGene00010610 | protein_coding_gene | 1.6 | 3.08 | 2.0E-04 |
| *dmsr-6* | WBGene00013222 | protein_coding_gene | 1.6 | 3.07 | 7.2E-03 |
| *fbxa-10* | WBGene00020450 | protein_coding_gene | 1.6 | 3.07 | 1.4E-03 |
| *grsp-3* | WBGene00016403 | protein_coding_gene | 1.6 | 3.07 | 3.0E-02 |
| *C16A3.6* | WBGene00015811 | protein_coding_gene | 1.6 | 3.07 | 1.7E-03 |
| *del-3* | WBGene00009144 | protein_coding_gene | 1.6 | 3.07 | 4.0E-02 |
| *K02E7.5* | WBGene00019310 | pseudogene | 1.6 | 3.07 | 7.5E-03 |
| *M03E7.4* | WBGene00019756 | protein_coding_gene | 1.6 | 3.07 | 4.1E-02 |

| | | | | | |
|---|---|---|---|---|---|
| *icmt-1* | WBGene00017673 | protein_coding_gene | 1.6 | 3.06 | 3.3E-02 |
| *C25F9.4* | WBGene00007724 | protein_coding_gene | 1.6 | 3.06 | 7.7E-05 |
| *ncx-10* | WBGene00003575 | protein_coding_gene | 1.6 | 3.06 | 1.2E-02 |
| *Y73B6BL.47* | WBGene00045250 | protein_coding_gene | 1.6 | 3.06 | 2.2E-02 |
| *dyf-5* | WBGene00001121 | protein_coding_gene | 1.6 | 3.06 | 1.1E-02 |
| *clec-83* | WBGene00021879 | protein_coding_gene | 1.6 | 3.05 | 2.8E-04 |
| *fipr-5* | WBGene00044174 | protein_coding_gene | 1.6 | 3.05 | 2.4E-04 |
| *K08H10.10* | WBGene00077491 | protein_coding_gene | 1.6 | 3.05 | 4.2E-02 |
| *nlp-69* | WBGene00013752 | protein_coding_gene | 1.6 | 3.05 | 2.8E-02 |
| *plx-2* | WBGene00004048 | protein_coding_gene | 1.6 | 3.05 | 8.8E-04 |
| *ceh-34* | WBGene00000455 | protein_coding_gene | 1.6 | 3.05 | 1.7E-04 |
| *F57G12.2* | WBGene00010222 | protein_coding_gene | 1.6 | 3.05 | 3.6E-02 |
| *H12D21.9* | WBGene00010385 | protein_coding_gene | 1.6 | 3.05 | 2.7E-02 |
| *lgc-50* | WBGene00020605 | protein_coding_gene | 1.6 | 3.04 | 2.0E-02 |
| *T10H9.8* | WBGene00044779 | protein_coding_gene | 1.6 | 3.04 | 2.7E-02 |
| *lrx-1* | WBGene00003075 | protein_coding_gene | 1.6 | 3.04 | 1.3E-02 |
| *dyf-18* | WBGene00010349 | protein_coding_gene | 1.6 | 3.04 | 2.5E-02 |
| *zig-10* | WBGene00020800 | protein_coding_gene | 1.6 | 3.04 | 8.5E-03 |
| *tsp-3* | WBGene00006629 | protein_coding_gene | 1.6 | 3.04 | 8.5E-03 |
| *F19B10.4* | WBGene00017586 | protein_coding_gene | 1.6 | 3.03 | 2.1E-02 |
| *Y22D7AL.15* | WBGene00021253 | protein_coding_gene | 1.6 | 3.03 | 2.8E-02 |
| *vab-3* | WBGene00006870 | protein_coding_gene | 1.6 | 3.03 | 2.1E-03 |
| *C32E8.6* | WBGene00016324 | protein_coding_gene | 1.6 | 3.02 | 4.9E-02 |
| *W05E10.2* | WBGene00012281 | protein_coding_gene | 1.6 | 3.02 | 2.7E-02 |
| *E04D5.2* | WBGene00008481 | protein_coding_gene | 1.6 | 3.02 | 3.9E-03 |
| *cfi-1* | WBGene00000476 | protein_coding_gene | 1.6 | 3.02 | 6.5E-03 |
| *ZC239.22* | WBGene00194717 | protein_coding_gene | 1.6 | 3.01 | 1.8E-02 |
| *nhr-87* | WBGene00003677 | protein_coding_gene | 1.6 | 3.01 | 1.8E-04 |
| *F35E12.4* | WBGene00009428 | pseudogene | 1.6 | 3.01 | 1.7E-02 |
| *E03A3.10* | WBGene00189944 | pseudogene | 1.6 | 3.01 | 1.7E-02 |
| *F53E10.1* | WBGene00018758 | protein_coding_gene | 1.6 | 3.01 | 1.2E-03 |
| *flp-22* | WBGene00009562 | protein_coding_gene | 1.6 | 3.01 | 7.2E-04 |
| *mig-18* | WBGene00003249 | protein_coding_gene | 1.6 | 3.01 | 1.2E-02 |
| *F09C6.1* | WBGene00008615 | protein_coding_gene | 1.6 | 3.00 | 2.0E-02 |
| *nhr-55* | WBGene00003645 | protein_coding_gene | 1.6 | 3.00 | 4.4E-04 |
| *C34E11.2* | WBGene00007935 | protein_coding_gene | 1.6 | 3.00 | 3.9E-02 |
| *alr-1* | WBGene00044330 | protein_coding_gene | 1.6 | 3.00 | 4.4E-04 |
| *Y39G10AR.11* | WBGene00021469 | protein_coding_gene | 1.6 | 3.00 | 2.3E-02 |
| *W03D2.9* | WBGene00020983 | protein_coding_gene | 1.6 | 3.00 | 4.7E-02 |
| *aqp-3* | WBGene00000171 | protein_coding_gene | 1.6 | 2.99 | 1.1E-02 |
| *sym-2* | WBGene00006367 | protein_coding_gene | 1.6 | 2.99 | 2.3E-03 |
| *F18E9.1* | WBGene00017568 | protein_coding_gene | 1.6 | 2.99 | 1.4E-02 |
| *T23B12.8* | WBGene00020722 | protein_coding_gene | 1.6 | 2.99 | 2.4E-03 |
| *slcf-2* | WBGene00006865 | protein_coding_gene | 1.6 | 2.99 | 1.7E-03 |
| *Y9C9A.8* | WBGene00021178 | protein_coding_gene | 1.6 | 2.98 | 2.9E-02 |
| *hlh-33* | WBGene00021446 | protein_coding_gene | 1.6 | 2.98 | 6.8E-03 |
| *K02E11.4* | WBGene00010513 | protein_coding_gene | 1.6 | 2.98 | 7.5E-04 |
| *bigr-1* | WBGene00009512 | protein_coding_gene | 1.6 | 2.98 | 8.1E-03 |
| *R02F11.1* | WBGene00019839 | protein_coding_gene | 1.6 | 2.98 | 3.3E-03 |
| *pag-3* | WBGene00003909 | protein_coding_gene | 1.6 | 2.98 | 3.3E-02 |
| *fkh-9* | WBGene00001441 | protein_coding_gene | 1.6 | 2.97 | 1.9E-02 |
| *Y34F4.3* | WBGene00021338 | protein_coding_gene | 1.6 | 2.97 | 1.4E-02 |
| *tpra-1* | WBGene00008366 | protein_coding_gene | 1.6 | 2.97 | 8.9E-03 |
| *W03B1.3* | WBGene00020973 | protein_coding_gene | 1.6 | 2.97 | 8.2E-03 |
| *tps-1* | WBGene00006602 | protein_coding_gene | 1.6 | 2.97 | 2.4E-02 |
| *Y62F5A.10* | WBGene00013388 | protein_coding_gene | 1.6 | 2.97 | 2.5E-03 |
| *cnc-10* | WBGene00012152 | protein_coding_gene | 1.6 | 2.97 | 4.4E-04 |
| *cutl-20* | WBGene00018787 | protein_coding_gene | 1.6 | 2.97 | 7.9E-04 |

| | | | | | |
|---|---|---|---|---|---|
| *cyp-13A1* | WBGene00011677 | protein_coding_gene | 1.6 | 2.97 | 1.5E-02 |
| *F36F2.2* | WBGene00009476 | protein_coding_gene | 1.6 | 2.96 | 2.4E-03 |
| *C52E2.5* | WBGene00016884 | protein_coding_gene | 1.6 | 2.96 | 7.9E-03 |
| *cbp-2* | WBGene00010545 | protein_coding_gene | 1.6 | 2.96 | 6.1E-03 |
| *nhr-94* | WBGene00003684 | protein_coding_gene | 1.6 | 2.96 | 7.1E-03 |
| *dhs-20* | WBGene00000983 | protein_coding_gene | 1.6 | 2.96 | 2.0E-03 |
| *B0302.5* | WBGene00015122 | protein_coding_gene | 1.6 | 2.95 | 5.0E-04 |
| *Y69A2AL.2* | WBGene00044468 | protein_coding_gene | 1.6 | 2.95 | 8.4E-03 |
| *ZK813.5* | WBGene00022824 | protein_coding_gene | 1.6 | 2.95 | 4.3E-02 |
| *oac-51* | WBGene00020976 | protein_coding_gene | 1.6 | 2.95 | 5.7E-03 |
| *lurp-1* | WBGene00009700 | protein_coding_gene | 1.6 | 2.95 | 3.9E-03 |
| *F08G2.8* | WBGene00008579 | protein_coding_gene | 1.6 | 2.95 | 2.7E-02 |
| *gcy-36* | WBGene00001556 | protein_coding_gene | 1.6 | 2.95 | 8.9E-03 |
| *C13C4.6* | WBGene00007550 | protein_coding_gene | 1.6 | 2.95 | 5.3E-03 |
| *myo-3* | WBGene00003515 | protein_coding_gene | 1.6 | 2.95 | 8.4E-03 |
| *ttr-17* | WBGene00012383 | protein_coding_gene | 1.6 | 2.95 | 2.6E-03 |
| *nhr-112* | WBGene00003702 | protein_coding_gene | 1.6 | 2.95 | 4.2E-02 |
| *ZK131.11* | WBGene00013934 | protein_coding_gene | 1.6 | 2.95 | 4.6E-04 |
| *moe-3* | WBGene00003388 | protein_coding_gene | 1.6 | 2.94 | 3.4E-02 |
| *C45B11.6* | WBGene00045379 | protein_coding_gene | 1.6 | 2.94 | 4.9E-04 |
| *col-76* | WBGene00000652 | protein_coding_gene | 1.6 | 2.94 | 1.6E-05 |
| *Y70C5A.3* | WBGene00050904 | protein_coding_gene | 1.6 | 2.94 | 4.7E-03 |
| *D2045.8* | WBGene00008425 | protein_coding_gene | 1.6 | 2.94 | 2.7E-02 |
| *cpna-5* | WBGene00007044 | protein_coding_gene | 1.6 | 2.94 | 2.3E-03 |
| *Y37H2B.1* | WBGene00012572 | pseudogene | 1.6 | 2.94 | 8.1E-03 |
| *F58E6.13* | WBGene00077697 | protein_coding_gene | 1.6 | 2.93 | 1.7E-02 |
| *glc-3* | WBGene00001593 | protein_coding_gene | 1.6 | 2.93 | 3.9E-02 |
| *nphp-1* | WBGene00010898 | protein_coding_gene | 1.6 | 2.93 | 1.4E-02 |
| *W03D2.6* | WBGene00020981 | pseudogene | 1.6 | 2.93 | 6.7E-05 |
| *nhr-107* | WBGene00003697 | protein_coding_gene | 1.5 | 2.93 | 2.4E-03 |
| *F56C9.8* | WBGene00018952 | protein_coding_gene | 1.5 | 2.93 | 1.5E-03 |
| *F26F12.3* | WBGene00017834 | protein_coding_gene | 1.5 | 2.93 | 2.4E-02 |
| *F35B12.9* | WBGene00009388 | protein_coding_gene | 1.5 | 2.93 | 3.7E-02 |
| *R06F6.14* | WBGene00269421 | protein_coding_gene | 1.5 | 2.93 | 3.2E-03 |
| *C35E7.2* | WBGene00016454 | protein_coding_gene | 1.5 | 2.92 | 1.8E-03 |
| *Y34F4.1* | WBGene00021336 | protein_coding_gene | 1.5 | 2.92 | 1.9E-04 |
| *cutl-29* | WBGene00017421 | protein_coding_gene | 1.5 | 2.92 | 2.8E-02 |
| *C40H5.2* | WBGene00008045 | protein_coding_gene | 1.5 | 2.92 | 2.5E-03 |
| *C31H5.7* | WBGene00007858 | protein_coding_gene | 1.5 | 2.92 | 7.4E-03 |
| *T28C6.10* | WBGene00045183 | protein_coding_gene | 1.5 | 2.92 | 1.6E-02 |
| *C05C12.4* | WBGene00007336 | protein_coding_gene | 1.5 | 2.92 | 2.0E-03 |
| *psa-3* | WBGene00009560 | protein_coding_gene | 1.5 | 2.92 | 5.0E-03 |
| *F42G2.2* | WBGene00018351 | protein_coding_gene | 1.5 | 2.92 | 4.5E-04 |
| *otpl-7* | WBGene00018480 | protein_coding_gene | 1.5 | 2.92 | 2.4E-02 |
| *linc-91* | WBGene00219746 | lincRNA_gene | 1.5 | 2.92 | 2.1E-02 |
| *cdf-2* | WBGene00011821 | protein_coding_gene | 1.5 | 2.91 | 1.3E-02 |
| *nhr-88* | WBGene00003678 | protein_coding_gene | 1.5 | 2.91 | 1.2E-02 |
| *Y32B12C.1* | WBGene00012525 | protein_coding_gene | 1.5 | 2.91 | 1.3E-02 |
| *C04E7.3* | WBGene00015430 | protein_coding_gene | 1.5 | 2.90 | 1.8E-02 |
| *F01E11.3* | WBGene00017156 | protein_coding_gene | 1.5 | 2.90 | 7.9E-03 |
| *bus-8* | WBGene00044623 | protein_coding_gene | 1.5 | 2.90 | 6.2E-04 |
| *ZC487.1* | WBGene00022623 | protein_coding_gene | 1.5 | 2.90 | 4.0E-03 |
| *snf-11* | WBGene00004910 | protein_coding_gene | 1.5 | 2.90 | 1.2E-02 |
| *chil-13* | WBGene00010945 | protein_coding_gene | 1.5 | 2.90 | 3.4E-02 |
| *chil-1* | WBGene00007465 | protein_coding_gene | 1.5 | 2.90 | 4.5E-02 |
| *K02A6.3* | WBGene00019294 | protein_coding_gene | 1.5 | 2.90 | 3.6E-03 |
| *egl-44* | WBGene00001208 | protein_coding_gene | 1.5 | 2.89 | 2.7E-03 |
| *fmi-1* | WBGene00001475 | protein_coding_gene | 1.5 | 2.89 | 1.8E-02 |

| | | | | | |
|---|---|---|---|---|---|
| *ugt-59* | WBGene00011238 | protein_coding_gene | 1.5 | 2.89 | 1.1E-02 |
| *K07C5.9* | WBGene00010632 | protein_coding_gene | 1.5 | 2.89 | 2.5E-03 |
| *Y37F4.1* | WBGene00021382 | protein_coding_gene | 1.5 | 2.89 | 3.1E-02 |
| *C33H5.13* | WBGene00016379 | protein_coding_gene | 1.5 | 2.89 | 3.0E-03 |
| *zip-10* | WBGene00012101 | protein_coding_gene | 1.5 | 2.89 | 4.1E-02 |
| *dmd-4* | WBGene00007776 | protein_coding_gene | 1.5 | 2.88 | 2.1E-02 |
| *nhr-103* | WBGene00003693 | protein_coding_gene | 1.5 | 2.88 | 1.1E-02 |
| *C10A4.3* | WBGene00015664 | protein_coding_gene | 1.5 | 2.88 | 3.0E-02 |
| *oig-4* | WBGene00043050 | protein_coding_gene | 1.5 | 2.88 | 5.6E-03 |
| *magu-2* | WBGene00006467 | protein_coding_gene | 1.5 | 2.87 | 2.8E-03 |
| *noah-2* | WBGene00009926 | protein_coding_gene | 1.5 | 2.87 | 1.8E-02 |
| *abu-12* | WBGene00004127 | protein_coding_gene | 1.5 | 2.87 | 4.2E-02 |
| *T26C5.5* | WBGene00012035 | protein_coding_gene | 1.5 | 2.87 | 3.2E-03 |
| *phg-1* | WBGene00004017 | protein_coding_gene | 1.5 | 2.87 | 4.0E-02 |
| *nhr-143* | WBGene00019116 | protein_coding_gene | 1.5 | 2.87 | 4.8E-03 |
| *Y53F4B.25* | WBGene00013171 | protein_coding_gene | 1.5 | 2.87 | 3.8E-02 |
| *cdr-1* | WBGene00000412 | protein_coding_gene | 1.5 | 2.87 | 1.9E-02 |
| *frpr-16* | WBGene00020023 | protein_coding_gene | 1.5 | 2.87 | 1.0E-02 |
| *F57E7.1* | WBGene00010202 | protein_coding_gene | 1.5 | 2.86 | 5.8E-04 |
| *bus-4* | WBGene00044620 | protein_coding_gene | 1.5 | 2.86 | 4.5E-02 |
| *sox-2* | WBGene00004949 | protein_coding_gene | 1.5 | 2.86 | 1.0E-02 |
| *Y47G6A.5* | WBGene00021635 | protein_coding_gene | 1.5 | 2.86 | 4.7E-03 |
| *F14H3.12* | WBGene00008831 | protein_coding_gene | 1.5 | 2.86 | 6.6E-04 |
| *F26F2.1* | WBGene00009166 | protein_coding_gene | 1.5 | 2.86 | 4.4E-02 |
| *C44H9.5* | WBGene00008100 | protein_coding_gene | 1.5 | 2.86 | 3.7E-03 |
| *R10E8.8* | WBGene00011213 | protein_coding_gene | 1.5 | 2.86 | 2.1E-02 |
| *rnt-1* | WBGene00004393 | protein_coding_gene | 1.5 | 2.85 | 4.0E-02 |
| *F53C3.3* | WBGene00018747 | protein_coding_gene | 1.5 | 2.85 | 4.5E-02 |
| *tbc-16* | WBGene00018639 | protein_coding_gene | 1.5 | 2.85 | 2.8E-02 |
| *F21C10.13* | WBGene00195086 | protein_coding_gene | 1.5 | 2.85 | 4.6E-02 |
| *Y17G7B.22* | WBGene00012473 | protein_coding_gene | 1.5 | 2.84 | 4.0E-02 |
| *pha-2* | WBGene00004011 | protein_coding_gene | 1.5 | 2.84 | 3.3E-02 |
| *T18D3.7* | WBGene00011824 | protein_coding_gene | 1.5 | 2.84 | 2.5E-02 |
| *K02C4.5* | WBGene00010503 | protein_coding_gene | 1.5 | 2.84 | 4.8E-03 |
| *C13A10.2* | WBGene00015729 | protein_coding_gene | 1.5 | 2.84 | 2.1E-02 |
| *ift-81* | WBGene00017973 | protein_coding_gene | 1.5 | 2.83 | 1.1E-02 |
| *T07G12.3* | WBGene00011594 | protein_coding_gene | 1.5 | 2.83 | 3.0E-02 |
| *F08G5.3* | WBGene00008582 | protein_coding_gene | 1.5 | 2.82 | 9.5E-04 |
| *C15B12.4* | WBGene00015786 | protein_coding_gene | 1.5 | 2.82 | 4.8E-03 |
| *T09F5.1* | WBGene00011665 | protein_coding_gene | 1.5 | 2.82 | 1.2E-02 |
| *eat-2* | WBGene00001133 | protein_coding_gene | 1.5 | 2.82 | 2.7E-02 |
| *T03F1.11* | WBGene00020192 | protein_coding_gene | 1.5 | 2.82 | 3.9E-05 |
| *C28G1.2* | WBGene00016187 | protein_coding_gene | 1.5 | 2.82 | 3.1E-02 |
| *nspc-14* | WBGene00009638 | protein_coding_gene | 1.5 | 2.82 | 2.9E-02 |
| *nhr-140* | WBGene00016366 | protein_coding_gene | 1.5 | 2.82 | 1.4E-02 |
| *F55C12.19* | WBGene00271817 | protein_coding_gene | 1.5 | 2.82 | 3.2E-02 |
| *Y41D4A.7* | WBGene00021509 | protein_coding_gene | 1.5 | 2.82 | 2.5E-02 |
| *ssr-2* | WBGene00006055 | protein_coding_gene | 1.5 | 2.82 | 1.2E-02 |
| *ZK54.3* | WBGene00022648 | protein_coding_gene | 1.5 | 2.81 | 1.1E-02 |
| *pals-23* | WBGene00016218 | protein_coding_gene | 1.5 | 2.81 | 2.7E-03 |
| *F54D5.15* | WBGene00010057 | protein_coding_gene | 1.5 | 2.81 | 1.2E-06 |
| *tag-234* | WBGene00018874 | protein_coding_gene | 1.5 | 2.81 | 6.4E-03 |
| *F17C11.4* | WBGene00008915 | protein_coding_gene | 1.5 | 2.81 | 2.8E-03 |
| *asp-14* | WBGene00019619 | protein_coding_gene | 1.5 | 2.80 | 3.6E-03 |
| *Y53F4B.12* | WBGene00013159 | protein_coding_gene | 1.5 | 2.80 | 9.5E-03 |
| *ncam-1* | WBGene00017184 | protein_coding_gene | 1.5 | 2.80 | 1.1E-03 |
| *ZC13.2* | WBGene00022503 | protein_coding_gene | 1.5 | 2.80 | 3.5E-02 |
| *pqn-13* | WBGene00004104 | protein_coding_gene | 1.5 | 2.80 | 5.5E-03 |

| | | | | | |
|---|---|---|---|---|---|
| *Y71G12B.31* | WBGene00044348 | protein_coding_gene | 1.5 | 2.80 | 2.3E-02 |
| *spp-5* | WBGene00004990 | protein_coding_gene | 1.5 | 2.80 | 2.2E-03 |
| *dve-1* | WBGene00022861 | protein_coding_gene | 1.5 | 2.80 | 3.0E-02 |
| *crtc-1* | WBGene00021237 | protein_coding_gene | 1.5 | 2.80 | 8.9E-03 |
| *K08D9.4* | WBGene00019525 | protein_coding_gene | 1.5 | 2.79 | 1.8E-02 |
| *K11E4.1* | WBGene00010773 | protein_coding_gene | 1.5 | 2.79 | 6.8E-03 |
| *tbx-7* | WBGene00006544 | protein_coding_gene | 1.5 | 2.79 | 2.0E-02 |
| *marc-5* | WBGene00013273 | protein_coding_gene | 1.5 | 2.79 | 4.5E-02 |
| *T20B3.14* | WBGene00011854 | protein_coding_gene | 1.5 | 2.79 | 1.2E-02 |
| *fbxa-6* | WBGene00008352 | protein_coding_gene | 1.5 | 2.79 | 1.0E-03 |
| *ceh-33* | WBGene00000454 | protein_coding_gene | 1.5 | 2.79 | 3.0E-02 |
| *klp-11* | WBGene00002222 | protein_coding_gene | 1.5 | 2.79 | 1.4E-02 |
| *cest-17* | WBGene00019477 | protein_coding_gene | 1.5 | 2.79 | 7.4E-04 |
| *K04C2.7* | WBGene00044311 | protein_coding_gene | 1.5 | 2.79 | 2.3E-02 |
| *pqn-62* | WBGene00004145 | protein_coding_gene | 1.5 | 2.79 | 2.0E-02 |
| *F46B6.9* | WBGene00009774 | protein_coding_gene | 1.5 | 2.79 | 2.6E-02 |
| *hhat-2* | WBGene00013312 | protein_coding_gene | 1.5 | 2.78 | 2.0E-03 |
| *F23A7.3* | WBGene00009068 | protein_coding_gene | 1.5 | 2.78 | 3.9E-02 |
| *eff-1* | WBGene00001159 | protein_coding_gene | 1.5 | 2.78 | 3.1E-02 |
| *ZC47.8* | WBGene00014978 | pseudogene | 1.5 | 2.78 | 3.5E-02 |
| *mef-2* | WBGene00003182 | protein_coding_gene | 1.5 | 2.78 | 3.8E-03 |
| *nhr-59* | WBGene00003649 | protein_coding_gene | 1.5 | 2.78 | 3.9E-03 |
| *B0244.5* | WBGene00015080 | protein_coding_gene | 1.5 | 2.77 | 1.1E-02 |
| *C05A9.2* | WBGene00007317 | protein_coding_gene | 1.5 | 2.77 | 2.6E-03 |
| *C29E4.10* | WBGene00016207 | protein_coding_gene | 1.5 | 2.77 | 8.3E-03 |
| *Y87G2A.19* | WBGene00044260 | protein_coding_gene | 1.5 | 2.77 | 1.1E-02 |
| *T22B7.8* | WBGene00044570 | protein_coding_gene | 1.5 | 2.77 | 2.5E-03 |
| *ttr-43* | WBGene00007590 | protein_coding_gene | 1.5 | 2.77 | 4.0E-03 |
| *F36F2.7* | WBGene00009480 | protein_coding_gene | 1.5 | 2.77 | 1.2E-02 |
| *F59B8.1* | WBGene00010316 | protein_coding_gene | 1.5 | 2.77 | 2.8E-02 |
| *tmed-13* | WBGene00022255 | protein_coding_gene | 1.5 | 2.76 | 1.1E-02 |
| *T10B11.5* | WBGene00020401 | protein_coding_gene | 1.5 | 2.76 | 1.3E-04 |
| *lpr-1* | WBGene00002393 | protein_coding_gene | 1.5 | 2.76 | 6.9E-03 |
| *mlt-8* | WBGene00021095 | protein_coding_gene | 1.5 | 2.76 | 3.8E-04 |
| *C09F9.2* | WBGene00007479 | protein_coding_gene | 1.5 | 2.75 | 4.1E-02 |
| *C29H12.6* | WBGene00016237 | protein_coding_gene | 1.5 | 2.75 | 9.2E-03 |
| *Y48A6B.7* | WBGene00012968 | protein_coding_gene | 1.5 | 2.75 | 6.9E-03 |
| *Y46G5A.20* | WBGene00012910 | protein_coding_gene | 1.5 | 2.75 | 1.2E-02 |
| *F15H10.8* | WBGene00008875 | protein_coding_gene | 1.5 | 2.75 | 2.3E-02 |
| *hot-4* | WBGene00001989 | protein_coding_gene | 1.5 | 2.75 | 8.6E-03 |
| *Y50E8A.5* | WBGene00013048 | protein_coding_gene | 1.5 | 2.75 | 1.2E-04 |
| *sek-6* | WBGene00012162 | protein_coding_gene | 1.5 | 2.75 | 1.2E-02 |
| *T13A10.1* | WBGene00020468 | protein_coding_gene | 1.5 | 2.75 | 3.2E-02 |
| *F54B11.10* | WBGene00010033 | protein_coding_gene | 1.5 | 2.75 | 4.8E-02 |
| *zip-2* | WBGene00019327 | protein_coding_gene | 1.5 | 2.75 | 3.9E-04 |
| *T01B10.5* | WBGene00020137 | protein_coding_gene | 1.5 | 2.75 | 1.1E-02 |
| *C26B9.3* | WBGene00016133 | protein_coding_gene | 1.5 | 2.75 | 5.7E-03 |
| *C03B1.5* | WBGene00015376 | protein_coding_gene | 1.5 | 2.75 | 3.0E-02 |
| *ras-1* | WBGene00004310 | protein_coding_gene | 1.5 | 2.75 | 3.9E-03 |
| *egl-38* | WBGene00001204 | protein_coding_gene | 1.5 | 2.74 | 2.5E-03 |
| *F08F3.1* | WBGene00017260 | protein_coding_gene | 1.5 | 2.74 | 3.2E-02 |
| *F13C5.1* | WBGene00017422 | protein_coding_gene | 1.5 | 2.74 | 8.4E-03 |
| *plpr-1* | WBGene00011524 | protein_coding_gene | 1.5 | 2.74 | 4.8E-03 |
| *F42G2.5* | WBGene00018354 | protein_coding_gene | 1.5 | 2.74 | 1.5E-04 |
| *magu-3* | WBGene00016841 | protein_coding_gene | 1.5 | 2.74 | 2.2E-03 |
| *F20D6.10* | WBGene00017639 | protein_coding_gene | 1.5 | 2.73 | 7.1E-03 |
| *nhr-58* | WBGene00003648 | protein_coding_gene | 1.4 | 2.73 | 3.3E-02 |
| *T25D10.4* | WBGene00020801 | protein_coding_gene | 1.4 | 2.73 | 1.0E-05 |

| | | | | | |
|---|---|---|---|---|---|
| *unc-24* | WBGene00006761 | protein_coding_gene | 1.4 | 2.73 | 4.7E-03 |
| *dhrs-4* | WBGene00010063 | protein_coding_gene | 1.4 | 2.73 | 8.4E-03 |
| *Y71H2AM.15* | WBGene00022180 | protein_coding_gene | 1.4 | 2.73 | 2.1E-03 |
| *M01H9.3* | WBGene00019719 | protein_coding_gene | 1.4 | 2.73 | 2.6E-03 |
| *smf-3* | WBGene00004878 | protein_coding_gene | 1.4 | 2.72 | 3.4E-02 |
| *R07C12.2* | WBGene00019932 | pseudogene | 1.4 | 2.72 | 1.1E-02 |
| *ZK682.7* | WBGene00022790 | protein_coding_gene | 1.4 | 2.72 | 2.6E-02 |
| *ceh-31* | WBGene00000452 | protein_coding_gene | 1.4 | 2.72 | 2.4E-02 |
| *cuti-1* | WBGene00022591 | protein_coding_gene | 1.4 | 2.72 | 5.0E-04 |
| *che-12* | WBGene00000491 | protein_coding_gene | 1.4 | 2.72 | 6.0E-03 |
| *T05E11.2* | WBGene00011479 | protein_coding_gene | 1.4 | 2.71 | 1.2E-02 |
| *F40E10.5* | WBGene00009573 | protein_coding_gene | 1.4 | 2.71 | 1.5E-02 |
| *mam-1* | WBGene00022504 | protein_coding_gene | 1.4 | 2.71 | 3.3E-02 |
| *F13E9.13* | WBGene00044797 | protein_coding_gene | 1.4 | 2.71 | 7.0E-04 |
| *H22K11.3* | WBGene00019226 | protein_coding_gene | 1.4 | 2.71 | 2.3E-02 |
| *nhr-163* | WBGene00016368 | protein_coding_gene | 1.4 | 2.71 | 9.4E-03 |
| *ubc-17* | WBGene00006712 | protein_coding_gene | 1.4 | 2.70 | 2.2E-02 |
| *acr-7* | WBGene00000046 | protein_coding_gene | 1.4 | 2.70 | 2.8E-02 |
| *jmjd-3.1* | WBGene00017571 | protein_coding_gene | 1.4 | 2.70 | 1.9E-04 |
| *daf-14* | WBGene00000910 | protein_coding_gene | 1.4 | 2.70 | 5.2E-03 |
| *W03F9.11* | WBGene00044512 | protein_coding_gene | 1.4 | 2.70 | 4.3E-02 |
| *Y105E8B.7* | WBGene00013692 | protein_coding_gene | 1.4 | 2.70 | 8.0E-04 |
| *lys-8* | WBGene00003097 | protein_coding_gene | 1.4 | 2.69 | 1.3E-02 |
| *F43G6.8* | WBGene00009660 | protein_coding_gene | 1.4 | 2.69 | 3.6E-02 |
| *ZK1128.7* | WBGene00014233 | protein_coding_gene | 1.4 | 2.69 | 1.2E-02 |
| *F46H5.5* | WBGene00018521 | protein_coding_gene | 1.4 | 2.69 | 5.0E-02 |
| *C18B10.6* | WBGene00015963 | protein_coding_gene | 1.4 | 2.69 | 4.4E-02 |
| *T19A5.3* | WBGene00020554 | protein_coding_gene | 1.4 | 2.69 | 2.0E-02 |
| *gar-1* | WBGene00001517 | protein_coding_gene | 1.4 | 2.69 | 3.8E-02 |
| *F59C6.11* | WBGene00010328 | protein_coding_gene | 1.4 | 2.69 | 2.6E-03 |
| *Y55F3BR.11* | WBGene00044757 | protein_coding_gene | 1.4 | 2.69 | 1.0E-02 |
| *C56E10.3* | WBGene00016976 | protein_coding_gene | 1.4 | 2.68 | 2.7E-02 |
| *acn-1* | WBGene00000039 | protein_coding_gene | 1.4 | 2.68 | 1.5E-02 |
| *B0393.5* | WBGene00007170 | protein_coding_gene | 1.4 | 2.68 | 1.7E-03 |
| *nspc-10* | WBGene00009640 | protein_coding_gene | 1.4 | 2.68 | 1.5E-03 |
| *lgx-1* | WBGene00002983 | protein_coding_gene | 1.4 | 2.68 | 1.1E-02 |
| *nlt-1* | WBGene00003773 | protein_coding_gene | 1.4 | 2.68 | 1.3E-03 |
| *R09A8.5* | WBGene00011154 | protein_coding_gene | 1.4 | 2.68 | 6.7E-03 |
| *C18G1.6* | WBGene00015984 | protein_coding_gene | 1.4 | 2.68 | 1.6E-02 |
| *F40F8.5* | WBGene00009578 | protein_coding_gene | 1.4 | 2.68 | 1.1E-02 |
| *ZK1010.8* | WBGene00014181 | protein_coding_gene | 1.4 | 2.68 | 2.7E-03 |
| *C35D10.12* | WBGene00016448 | protein_coding_gene | 1.4 | 2.68 | 2.4E-03 |
| *dma-1* | WBGene00011345 | protein_coding_gene | 1.4 | 2.68 | 7.0E-03 |
| *ttr-5* | WBGene00008040 | protein_coding_gene | 1.4 | 2.67 | 3.4E-03 |
| *Y54F10BM.3* | WBGene00021858 | protein_coding_gene | 1.4 | 2.67 | 1.5E-02 |
| *nfm-1* | WBGene00003593 | protein_coding_gene | 1.4 | 2.67 | 5.2E-03 |
| *lgc-55* | WBGene00013746 | protein_coding_gene | 1.4 | 2.67 | 3.5E-03 |
| *cest-35.1* | WBGene00015067 | protein_coding_gene | 1.4 | 2.67 | 4.5E-04 |
| *ilys-4* | WBGene00016958 | protein_coding_gene | 1.4 | 2.67 | 1.0E-02 |
| *syd-9* | WBGene00044068 | protein_coding_gene | 1.4 | 2.67 | 5.4E-03 |
| *ace-3* | WBGene00000037 | protein_coding_gene | 1.4 | 2.67 | 1.3E-03 |
| *F46G10.2* | WBGene00009797 | protein_coding_gene | 1.4 | 2.67 | 8.2E-03 |
| *C32D5.1* | WBGene00016310 | protein_coding_gene | 1.4 | 2.67 | 3.3E-02 |
| *ndx-1* | WBGene00003578 | protein_coding_gene | 1.4 | 2.67 | 2.3E-03 |
| *K07H8.12* | WBGene00044509 | protein_coding_gene | 1.4 | 2.66 | 4.8E-02 |
| *gyg-3* | WBGene00013535 | pseudogene | 1.4 | 2.66 | 2.6E-02 |
| *F18A11.2* | WBGene00008929 | protein_coding_gene | 1.4 | 2.66 | 1.2E-02 |
| *F41E7.2* | WBGene00009618 | protein_coding_gene | 1.4 | 2.65 | 1.8E-02 |

| | | | | | |
|---|---|---|---|---|---|
| *math-35* | WBGene00020075 | pseudogene | 1.4 | 2.65 | 6.2E-04 |
| *Y38H8A.8* | WBGene00045062 | pseudogene | 1.4 | 2.65 | 1.3E-02 |
| *pmt-1* | WBGene00022781 | protein_coding_gene | 1.4 | 2.65 | 1.6E-02 |
| *nlp-50* | WBGene00018298 | protein_coding_gene | 1.4 | 2.65 | 5.0E-04 |
| *ser-4* | WBGene00004779 | protein_coding_gene | 1.4 | 2.65 | 4.4E-03 |
| *C31C9.7* | WBGene00007841 | protein_coding_gene | 1.4 | 2.65 | 3.3E-02 |
| *B0284.3* | WBGene00007133 | protein_coding_gene | 1.4 | 2.65 | 2.6E-04 |
| *EEED8.2* | WBGene00017133 | protein_coding_gene | 1.4 | 2.65 | 2.3E-04 |
| *fbxa-85* | WBGene00044654 | protein_coding_gene | 1.4 | 2.65 | 1.6E-02 |
| *gtf-2A2* | WBGene00013736 | protein_coding_gene | 1.4 | 2.65 | 2.0E-03 |
| *acs-11* | WBGene00018269 | protein_coding_gene | 1.4 | 2.65 | 8.0E-13 |
| *W02B8.1* | WBGene00012197 | protein_coding_gene | 1.4 | 2.64 | 3.8E-02 |
| *F36H9.7* | WBGene00018115 | protein_coding_gene | 1.4 | 2.64 | 4.1E-02 |
| *nhr-126* | WBGene00003716 | protein_coding_gene | 1.4 | 2.64 | 8.0E-03 |
| *Y43B11AR.3* | WBGene00021544 | protein_coding_gene | 1.4 | 2.64 | 1.5E-02 |
| *feh-1* | WBGene00001410 | protein_coding_gene | 1.4 | 2.64 | 5.2E-03 |
| *hum-6* | WBGene00002039 | protein_coding_gene | 1.4 | 2.64 | 1.4E-05 |
| *math-24* | WBGene00016719 | protein_coding_gene | 1.4 | 2.64 | 1.1E-03 |
| *F28F9.3* | WBGene00017907 | protein_coding_gene | 1.4 | 2.64 | 4.3E-02 |
| *nhr-227* | WBGene00020851 | protein_coding_gene | 1.4 | 2.64 | 1.0E-02 |
| *flp-8* | WBGene00001451 | protein_coding_gene | 1.4 | 2.63 | 3.6E-02 |
| *F52D1.2* | WBGene00018683 | protein_coding_gene | 1.4 | 2.63 | 4.2E-02 |
| *Y54G2A.7* | WBGene00043702 | protein_coding_gene | 1.4 | 2.63 | 7.6E-04 |
| *rgs-4* | WBGene00004347 | protein_coding_gene | 1.4 | 2.63 | 6.4E-03 |
| *C39D10.11* | WBGene00043983 | protein_coding_gene | 1.4 | 2.63 | 2.0E-03 |
| *slt-1* | WBGene00004854 | protein_coding_gene | 1.4 | 2.63 | 9.4E-03 |
| *Y62E10A.19* | WBGene00013385 | protein_coding_gene | 1.4 | 2.63 | 2.4E-02 |
| *F23C8.14* | WBGene00255457 | protein_coding_gene | 1.4 | 2.63 | 8.3E-03 |
| *gmeb-2* | WBGene00010010 | protein_coding_gene | 1.4 | 2.63 | 3.5E-02 |
| *lido-17* | WBGene00007378 | protein_coding_gene | 1.4 | 2.63 | 3.4E-03 |
| *gcy-19* | WBGene00001544 | protein_coding_gene | 1.4 | 2.63 | 6.9E-03 |
| *fbxa-69* | WBGene00010018 | protein_coding_gene | 1.4 | 2.63 | 1.4E-02 |
| *olrn-1* | WBGene00007260 | protein_coding_gene | 1.4 | 2.63 | 3.0E-02 |
| *glct-6* | WBGene00019546 | protein_coding_gene | 1.4 | 2.62 | 8.4E-03 |
| *T11F9.1* | WBGene00011709 | protein_coding_gene | 1.4 | 2.62 | 1.8E-02 |
| *T05A10.6* | WBGene00011463 | protein_coding_gene | 1.4 | 2.62 | 2.4E-02 |
| *wrt-10* | WBGene00006956 | protein_coding_gene | 1.4 | 2.62 | 6.6E-03 |
| *noah-1* | WBGene00016422 | protein_coding_gene | 1.4 | 2.62 | 3.0E-02 |
| *Y52B11A.12* | WBGene00077539 | protein_coding_gene | 1.4 | 2.62 | 4.8E-02 |
| *jnk-1* | WBGene00002178 | protein_coding_gene | 1.4 | 2.62 | 2.5E-03 |
| *B0361.9* | WBGene00015163 | protein_coding_gene | 1.4 | 2.61 | 2.7E-02 |
| *yap-1* | WBGene00008748 | protein_coding_gene | 1.4 | 2.61 | 1.4E-02 |
| *aex-3* | WBGene00000086 | protein_coding_gene | 1.4 | 2.61 | 1.7E-02 |
| *ZC84.1* | WBGene00013846 | protein_coding_gene | 1.4 | 2.61 | 4.0E-02 |
| *dct-1* | WBGene00015776 | protein_coding_gene | 1.4 | 2.61 | 2.0E-03 |
| *T25B9.3* | WBGene00012009 | protein_coding_gene | 1.4 | 2.61 | 4.3E-02 |
| *ensh-1* | WBGene00017013 | protein_coding_gene | 1.4 | 2.61 | 1.4E-02 |
| *Y39E4B.6* | WBGene00012717 | protein_coding_gene | 1.4 | 2.61 | 1.1E-02 |
| *nhr-121* | WBGene00003711 | protein_coding_gene | 1.4 | 2.61 | 9.1E-03 |
| *C24B9.3* | WBGene00016048 | protein_coding_gene | 1.4 | 2.61 | 4.5E-02 |
| *F59C6.14* | WBGene00044792 | protein_coding_gene | 1.4 | 2.61 | 7.7E-03 |
| *R04E5.9* | WBGene00019872 | protein_coding_gene | 1.4 | 2.61 | 4.0E-03 |
| *K02E7.6* | WBGene00019311 | pseudogene | 1.4 | 2.60 | 7.0E-03 |
| *tnc-2* | WBGene00006583 | protein_coding_gene | 1.4 | 2.60 | 3.0E-03 |
| *F07A5.4* | WBGene00008543 | protein_coding_gene | 1.4 | 2.60 | 1.2E-02 |
| *sad-1* | WBGene00004719 | protein_coding_gene | 1.4 | 2.60 | 2.5E-02 |
| *Y23H5B.3* | WBGene00021274 | protein_coding_gene | 1.4 | 2.60 | 1.8E-02 |
| *clec-221* | WBGene00008916 | protein_coding_gene | 1.4 | 2.60 | 3.7E-02 |

| | | | | | |
|---|---|---|---|---|---|
| *F10D7.1* | WBGene00017338 | protein_coding_gene | 1.4 | 2.60 | 3.1E-02 |
| *F35B3.7* | WBGene00018033 | protein_coding_gene | 1.4 | 2.60 | 2.8E-02 |
| *exc-9* | WBGene00017644 | protein_coding_gene | 1.4 | 2.60 | 3.8E-04 |
| *mab-7* | WBGene00003104 | protein_coding_gene | 1.4 | 2.60 | 1.7E-02 |
| *R01E6.7* | WBGene00010971 | protein_coding_gene | 1.4 | 2.59 | 3.5E-02 |
| *twk-11* | WBGene00006666 | protein_coding_gene | 1.4 | 2.59 | 5.5E-03 |
| *C28G1.6* | WBGene00016191 | protein_coding_gene | 1.4 | 2.59 | 8.7E-04 |
| *cdh-5* | WBGene00000397 | protein_coding_gene | 1.4 | 2.59 | 1.0E-02 |
| *somi-1* | WBGene00010868 | protein_coding_gene | 1.4 | 2.58 | 2.0E-03 |
| *amt-2* | WBGene00000134 | protein_coding_gene | 1.4 | 2.58 | 4.7E-02 |
| *mam-8* | WBGene00020743 | protein_coding_gene | 1.4 | 2.58 | 2.2E-02 |
| *hda-5* | WBGene00009663 | protein_coding_gene | 1.4 | 2.58 | 7.3E-03 |
| *K08B5.2* | WBGene00019519 | protein_coding_gene | 1.4 | 2.58 | 1.2E-02 |
| *MTCE.7* | WBGene00014454 | rRNA_gene | 1.4 | 2.58 | 1.8E-02 |
| *F11D5.1* | WBGene00017380 | protein_coding_gene | 1.4 | 2.58 | 2.1E-04 |
| *C29F7.2* | WBGene00007811 | protein_coding_gene | 1.4 | 2.58 | 3.0E-02 |
| *iglr-1* | WBGene00008977 | protein_coding_gene | 1.4 | 2.58 | 3.6E-03 |
| *tag-241* | WBGene00044079 | protein_coding_gene | 1.4 | 2.58 | 2.9E-02 |
| *Y47A7.2* | WBGene00021614 | protein_coding_gene | 1.4 | 2.58 | 2.2E-02 |
| *linc-46* | WBGene00219748 | lincRNA_gene | 1.4 | 2.58 | 1.3E-02 |
| *pin-2* | WBGene00004030 | protein_coding_gene | 1.4 | 2.58 | 1.9E-04 |
| *C37C3.12* | WBGene00016504 | protein_coding_gene | 1.4 | 2.58 | 1.3E-02 |
| *bkip-1* | WBGene00014569 | protein_coding_gene | 1.4 | 2.57 | 2.1E-02 |
| *ZC262.13* | WBGene00284846 | protein_coding_gene | 1.4 | 2.57 | 4.0E-02 |
| *ceh-48* | WBGene00015934 | protein_coding_gene | 1.4 | 2.57 | 1.8E-04 |
| *rga-6* | WBGene00015303 | protein_coding_gene | 1.4 | 2.57 | 1.3E-02 |
| *arrd-14* | WBGene00011055 | protein_coding_gene | 1.4 | 2.57 | 3.1E-02 |
| *C25G4.2* | WBGene00007730 | protein_coding_gene | 1.4 | 2.57 | 4.8E-03 |
| *T27A10.6* | WBGene00020846 | protein_coding_gene | 1.4 | 2.57 | 3.9E-02 |
| *ZK593.9* | WBGene00014005 | pseudogene | 1.4 | 2.57 | 2.2E-02 |
| *F20C5.11* | WBGene00303015 | protein_coding_gene | 1.4 | 2.57 | 2.4E-05 |
| *mak-2* | WBGene00007007 | protein_coding_gene | 1.4 | 2.57 | 9.5E-03 |
| *atgp-1* | WBGene00000224 | protein_coding_gene | 1.4 | 2.57 | 1.4E-03 |
| *ZC123.4* | WBGene00022519 | protein_coding_gene | 1.4 | 2.57 | 1.1E-02 |
| *F53A2.1* | WBGene00009949 | protein_coding_gene | 1.4 | 2.57 | 1.2E-02 |
| *T19H5.4* | WBGene00011849 | protein_coding_gene | 1.4 | 2.56 | 1.4E-02 |
| *bus-18* | WBGene00044631 | protein_coding_gene | 1.4 | 2.56 | 1.1E-02 |
| *Y67A10A.7* | WBGene00013456 | protein_coding_gene | 1.4 | 2.56 | 2.1E-02 |
| *agr-1* | WBGene00018304 | protein_coding_gene | 1.4 | 2.56 | 3.5E-06 |
| *npr-20* | WBGene00011578 | protein_coding_gene | 1.4 | 2.56 | 4.9E-03 |
| *abf-6* | WBGene00000017 | protein_coding_gene | 1.4 | 2.56 | 2.8E-02 |
| *Y38H6C.20* | WBGene00012632 | protein_coding_gene | 1.4 | 2.56 | 1.1E-02 |
| *F39E9.10* | WBGene00023036 | pseudogene | 1.4 | 2.56 | 6.8E-05 |
| *ham-2* | WBGene00001821 | protein_coding_gene | 1.4 | 2.56 | 2.2E-02 |
| *ccep-290* | WBGene00021643 | protein_coding_gene | 1.4 | 2.55 | 2.2E-02 |
| *F10G8.8* | WBGene00008666 | protein_coding_gene | 1.4 | 2.55 | 2.2E-04 |
| *Y69A2AR.31* | WBGene00022100 | protein_coding_gene | 1.4 | 2.55 | 6.5E-03 |
| *sel-7* | WBGene00004764 | protein_coding_gene | 1.4 | 2.55 | 1.7E-02 |
| *C49H3.16* | WBGene00219436 | protein_coding_gene | 1.3 | 2.54 | 4.4E-02 |
| *akt-2* | WBGene00000103 | protein_coding_gene | 1.3 | 2.54 | 1.5E-02 |
| *crm-1* | WBGene00007103 | protein_coding_gene | 1.3 | 2.54 | 8.0E-03 |
| *lgc-47* | WBGene00009799 | protein_coding_gene | 1.3 | 2.54 | 1.9E-02 |
| *clec-84* | WBGene00021895 | protein_coding_gene | 1.3 | 2.54 | 1.9E-03 |
| *C37E2.2* | WBGene00007994 | protein_coding_gene | 1.3 | 2.54 | 3.8E-02 |
| *lin-42* | WBGene00018572 | protein_coding_gene | 1.3 | 2.54 | 2.2E-02 |
| *T28F3.9* | WBGene00012135 | protein_coding_gene | 1.3 | 2.53 | 4.6E-02 |
| *Y11D7A.9* | WBGene00012433 | protein_coding_gene | 1.3 | 2.53 | 2.8E-02 |
| *F22B5.4* | WBGene00009042 | protein_coding_gene | 1.3 | 2.53 | 2.4E-02 |

| | | | | | |
|---|---|---|---|---|---|
| *nhr-60* | WBGene00003650 | protein_coding_gene | 1.3 | 2.53 | 1.5E-02 |
| *F20A1.6* | WBGene00017618 | protein_coding_gene | 1.3 | 2.53 | 3.7E-02 |
| *mapk-15* | WBGene00015478 | protein_coding_gene | 1.3 | 2.52 | 5.6E-03 |
| *K01A12.3* | WBGene00019287 | protein_coding_gene | 1.3 | 2.52 | 4.3E-02 |
| *Y39G8B.5* | WBGene00012726 | protein_coding_gene | 1.3 | 2.52 | 3.3E-02 |
| *tsp-15* | WBGene00006641 | protein_coding_gene | 1.3 | 2.52 | 3.3E-02 |
| *egl-20* | WBGene00001188 | protein_coding_gene | 1.3 | 2.52 | 9.5E-04 |
| *Y97E10AR.1* | WBGene00022396 | protein_coding_gene | 1.3 | 2.52 | 9.3E-03 |
| *Y57G11C.32* | WBGene00013322 | protein_coding_gene | 1.3 | 2.51 | 3.0E-02 |
| *ltd-1* | WBGene00003089 | protein_coding_gene | 1.3 | 2.51 | 2.9E-03 |
| *MTCE.33* | WBGene00014472 | rRNA_gene | 1.3 | 2.51 | 2.6E-02 |
| *amt-3* | WBGene00000135 | protein_coding_gene | 1.3 | 2.51 | 8.3E-03 |
| *spp-15* | WBGene00005000 | protein_coding_gene | 1.3 | 2.51 | 8.7E-03 |
| *C17G1.5* | WBGene00007652 | protein_coding_gene | 1.3 | 2.51 | 1.8E-02 |
| *ZK1073.1* | WBGene00014213 | protein_coding_gene | 1.3 | 2.51 | 1.4E-02 |
| *F32D8.13* | WBGene00009335 | protein_coding_gene | 1.3 | 2.51 | 1.6E-02 |
| *lpd-7* | WBGene00003063 | protein_coding_gene | 1.3 | 2.51 | 1.7E-03 |
| *dyf-1* | WBGene00001117 | protein_coding_gene | 1.3 | 2.50 | 2.1E-02 |
| *F26A3.1* | WBGene00009140 | protein_coding_gene | 1.3 | 2.50 | 7.3E-03 |
| *F46C8.3* | WBGene00018485 | protein_coding_gene | 1.3 | 2.50 | 8.7E-03 |
| *grl-10* | WBGene00001719 | protein_coding_gene | 1.3 | 2.50 | 6.1E-03 |
| *dhc-3* | WBGene00007154 | protein_coding_gene | 1.3 | 2.50 | 3.7E-02 |
| *R03H10.1* | WBGene00019854 | protein_coding_gene | 1.3 | 2.50 | 1.4E-03 |
| *T19E7.6* | WBGene00044500 | protein_coding_gene | 1.3 | 2.50 | 8.7E-03 |
| *W05B10.4* | WBGene00012279 | protein_coding_gene | 1.3 | 2.50 | 6.8E-04 |
| *R05A10.2* | WBGene00011019 | protein_coding_gene | 1.3 | 2.50 | 2.7E-02 |
| *K09F6.6* | WBGene00019589 | protein_coding_gene | 1.3 | 2.50 | 2.7E-03 |
| *C53C7.5* | WBGene00014294 | pseudogene | 1.3 | 2.50 | 3.9E-02 |
| *C54F6.20* | WBGene00271796 | protein_coding_gene | 1.3 | 2.50 | 3.7E-02 |
| *fbxa-188* | WBGene00009839 | protein_coding_gene | 1.3 | 2.49 | 1.6E-02 |
| *T28H10.1* | WBGene00012142 | protein_coding_gene | 1.3 | 2.49 | 6.8E-03 |
| *F32D8.15* | WBGene00009325 | protein_coding_gene | 1.3 | 2.49 | 1.3E-02 |
| *F13B12.4* | WBGene00008732 | protein_coding_gene | 1.3 | 2.49 | 4.4E-02 |
| *F31E9.3* | WBGene00009296 | protein_coding_gene | 1.3 | 2.49 | 6.7E-03 |
| *clp-9* | WBGene00011705 | protein_coding_gene | 1.3 | 2.49 | 7.3E-03 |
| *Y56A3A.18* | WBGene00013236 | protein_coding_gene | 1.3 | 2.49 | 2.6E-03 |
| *C54G4.4* | WBGene00008314 | protein_coding_gene | 1.3 | 2.48 | 6.5E-03 |
| *oig-3* | WBGene00003861 | protein_coding_gene | 1.3 | 2.48 | 3.5E-03 |
| *F26F2.8* | WBGene00023504 | protein_coding_gene | 1.3 | 2.48 | 1.3E-02 |
| *nhr-139* | WBGene00016365 | protein_coding_gene | 1.3 | 2.48 | 1.3E-02 |
| *ift-74* | WBGene00016005 | protein_coding_gene | 1.3 | 2.48 | 2.4E-02 |
| *Y22D7AL.6* | WBGene00021245 | protein_coding_gene | 1.3 | 2.48 | 2.6E-02 |
| *lrch-1* | WBGene00015779 | protein_coding_gene | 1.3 | 2.48 | 2.9E-02 |
| *daam-1* | WBGene00018976 | protein_coding_gene | 1.3 | 2.48 | 4.2E-02 |
| *svh-5* | WBGene00006462 | protein_coding_gene | 1.3 | 2.48 | 4.1E-02 |
| *unc-53* | WBGene00006788 | protein_coding_gene | 1.3 | 2.48 | 2.1E-02 |
| *del-2* | WBGene00000953 | protein_coding_gene | 1.3 | 2.48 | 7.7E-03 |
| *F13H10.9* | WBGene00045056 | protein_coding_gene | 1.3 | 2.48 | 2.1E-02 |
| *cps-6* | WBGene00000787 | protein_coding_gene | 1.3 | 2.47 | 4.8E-03 |
| *C16H3.3* | WBGene00015868 | protein_coding_gene | 1.3 | 2.47 | 1.3E-04 |
| *ipla-7* | WBGene00011423 | protein_coding_gene | 1.3 | 2.47 | 5.0E-03 |
| *F15B9.8* | WBGene00008851 | protein_coding_gene | 1.3 | 2.47 | 2.5E-02 |
| *let-381* | WBGene00002601 | protein_coding_gene | 1.3 | 2.47 | 1.4E-02 |
| *arl-3* | WBGene00000188 | protein_coding_gene | 1.3 | 2.47 | 3.8E-03 |
| *T05C12.4* | WBGene00011468 | protein_coding_gene | 1.3 | 2.47 | 4.4E-02 |
| *acs-7* | WBGene00007228 | protein_coding_gene | 1.3 | 2.47 | 2.0E-02 |
| *B0286.1* | WBGene00015115 | protein_coding_gene | 1.3 | 2.47 | 3.1E-03 |
| *tub-1* | WBGene00006655 | protein_coding_gene | 1.3 | 2.47 | 1.1E-02 |

| | | | | | |
|---|---|---|---|---|---|
| *snt-1* | WBGene00004921 | protein_coding_gene | 1.3 | 2.47 | 4.0E-02 |
| *K06A4.2* | WBGene00010592 | protein_coding_gene | 1.3 | 2.47 | 4.1E-02 |
| *W02D7.3* | WBGene00020939 | protein_coding_gene | 1.3 | 2.47 | 4.2E-02 |
| *ZK1320.2* | WBGene00014252 | protein_coding_gene | 1.3 | 2.46 | 3.8E-04 |
| *C03A7.13* | WBGene00015371 | protein_coding_gene | 1.3 | 2.46 | 1.1E-02 |
| *srh-70* | WBGene00005291 | protein_coding_gene | 1.3 | 2.46 | 4.4E-02 |
| *F56H11.2* | WBGene00010175 | protein_coding_gene | 1.3 | 2.46 | 3.1E-02 |
| *Y39B6A.33* | WBGene00012692 | protein_coding_gene | 1.3 | 2.46 | 1.4E-02 |
| *arrd-21* | WBGene00015441 | pseudogene | 1.3 | 2.46 | 2.6E-02 |
| *M02G9.1* | WBGene00010830 | protein_coding_gene | 1.3 | 2.46 | 4.2E-02 |
| *Y75B8A.32* | WBGene00013562 | protein_coding_gene | 1.3 | 2.45 | 7.6E-03 |
| *C26B9.6* | WBGene00016135 | protein_coding_gene | 1.3 | 2.45 | 5.7E-03 |
| *pbo-4* | WBGene00003943 | protein_coding_gene | 1.3 | 2.45 | 5.2E-03 |
| *W10C4.1* | WBGene00021124 | protein_coding_gene | 1.3 | 2.45 | 3.8E-02 |
| *nhr-170* | WBGene00008309 | protein_coding_gene | 1.3 | 2.45 | 9.3E-03 |
| *cat-2* | WBGene00000296 | protein_coding_gene | 1.3 | 2.45 | 4.4E-02 |
| *glr-1* | WBGene00001612 | protein_coding_gene | 1.3 | 2.45 | 1.1E-02 |
| *Y37H9A.5* | WBGene00014871 | protein_coding_gene | 1.3 | 2.44 | 1.1E-02 |
| *D2023.3* | WBGene00008407 | protein_coding_gene | 1.3 | 2.44 | 1.4E-02 |
| *lin-1* | WBGene00002990 | protein_coding_gene | 1.3 | 2.44 | 2.3E-02 |
| *F19G12.9* | WBGene00195185 | protein_coding_gene | 1.3 | 2.44 | 2.8E-02 |
| *faah-1* | WBGene00015047 | protein_coding_gene | 1.3 | 2.44 | 1.8E-06 |
| *C40H1.2* | WBGene00008038 | protein_coding_gene | 1.3 | 2.44 | 2.3E-02 |
| *dlg-1* | WBGene00001006 | protein_coding_gene | 1.3 | 2.44 | 1.5E-03 |
| *C35E7.4* | WBGene00016456 | protein_coding_gene | 1.3 | 2.44 | 2.7E-04 |
| *calf-1* | WBGene00007119 | protein_coding_gene | 1.3 | 2.44 | 9.4E-03 |
| *F38E9.6* | WBGene00018188 | protein_coding_gene | 1.3 | 2.44 | 3.8E-02 |
| *dlc-3* | WBGene00012424 | protein_coding_gene | 1.3 | 2.43 | 2.3E-02 |
| *Y44E3A.1* | WBGene00021547 | protein_coding_gene | 1.3 | 2.43 | 2.6E-03 |
| *pqn-26* | WBGene00004115 | protein_coding_gene | 1.3 | 2.43 | 4.0E-02 |
| *C33G8.4* | WBGene00016362 | protein_coding_gene | 1.3 | 2.43 | 1.2E-02 |
| *K07E8.6* | WBGene00019497 | protein_coding_gene | 1.3 | 2.43 | 3.4E-02 |
| *Y34D9A.8* | WBGene00021333 | protein_coding_gene | 1.3 | 2.43 | 1.2E-02 |
| *let-522* | WBGene00010870 | protein_coding_gene | 1.3 | 2.43 | 6.6E-04 |
| *C32D5.6* | WBGene00016314 | protein_coding_gene | 1.3 | 2.43 | 4.0E-02 |
| *madd-4* | WBGene00009958 | protein_coding_gene | 1.3 | 2.42 | 1.9E-02 |
| *ZK154.1* | WBGene00022665 | protein_coding_gene | 1.3 | 2.42 | 3.2E-02 |
| *T28F3.8* | WBGene00012134 | protein_coding_gene | 1.3 | 2.42 | 3.9E-03 |
| *apl-1* | WBGene00000149 | protein_coding_gene | 1.3 | 2.42 | 4.3E-02 |
| *clc-2* | WBGene00000523 | protein_coding_gene | 1.3 | 2.42 | 8.6E-03 |
| *fbxa-209* | WBGene00012533 | protein_coding_gene | 1.3 | 2.42 | 2.6E-03 |
| *ugt-54* | WBGene00012013 | protein_coding_gene | 1.3 | 2.41 | 3.1E-02 |
| *T23B3.2* | WBGene00020710 | protein_coding_gene | 1.3 | 2.41 | 1.3E-05 |
| *rpi-2* | WBGene00019536 | protein_coding_gene | 1.3 | 2.41 | 5.0E-03 |
| *F41F3.8* | WBGene00044533 | protein_coding_gene | 1.3 | 2.41 | 2.6E-02 |
| *K07C10.3* | WBGene00194851 | ncRNA_gene | 1.3 | 2.41 | 2.4E-02 |
| *glb-27* | WBGene00012178 | protein_coding_gene | 1.3 | 2.41 | 1.9E-02 |
| *K09F5.6* | WBGene00044698 | protein_coding_gene | 1.3 | 2.41 | 2.3E-02 |
| *syd-1* | WBGene00006363 | protein_coding_gene | 1.3 | 2.41 | 1.7E-02 |
| *let-4* | WBGene00002282 | protein_coding_gene | 1.3 | 2.41 | 2.6E-02 |
| *K02D7.1* | WBGene00019298 | protein_coding_gene | 1.3 | 2.41 | 4.3E-02 |
| *lntl-1* | WBGene00016289 | protein_coding_gene | 1.3 | 2.41 | 1.7E-02 |
| *mig-10* | WBGene00003243 | protein_coding_gene | 1.3 | 2.41 | 5.5E-04 |
| *math-20* | WBGene00016555 | protein_coding_gene | 1.3 | 2.40 | 1.6E-03 |
| *gcy-35* | WBGene00001555 | protein_coding_gene | 1.3 | 2.40 | 1.4E-02 |
| *clc-5* | WBGene00000526 | protein_coding_gene | 1.3 | 2.40 | 6.8E-03 |
| *math-3* | WBGene00015608 | protein_coding_gene | 1.3 | 2.40 | 6.7E-03 |
| *sup-12* | WBGene00006321 | protein_coding_gene | 1.3 | 2.40 | 2.7E-03 |

| | | | | | |
|---|---|---|---|---|---|
| *R03G8.3* | WBGene00010998 | protein_coding_gene | 1.3 | 2.40 | 6.7E-03 |
| *mtl-2* | WBGene00003474 | protein_coding_gene | 1.3 | 2.40 | 4.6E-02 |
| *Y43C5A.2* | WBGene00012782 | protein_coding_gene | 1.3 | 2.40 | 1.3E-03 |
| *max-1* | WBGene00003143 | protein_coding_gene | 1.3 | 2.40 | 2.0E-03 |
| *Y48G9A.7* | WBGene00021701 | protein_coding_gene | 1.3 | 2.39 | 3.4E-02 |
| *F46H6.5* | WBGene00023460 | protein_coding_gene | 1.3 | 2.39 | 3.3E-02 |
| *C50B8.5* | WBGene00008227 | protein_coding_gene | 1.3 | 2.39 | 4.8E-03 |
| *C53C11.5* | WBGene00044770 | protein_coding_gene | 1.3 | 2.39 | 1.1E-03 |
| *lgc-20* | WBGene00007191 | protein_coding_gene | 1.3 | 2.39 | 1.6E-02 |
| *C09D4.2* | WBGene00015633 | protein_coding_gene | 1.3 | 2.39 | 4.4E-03 |
| *Y53G8AL.4* | WBGene00050917 | protein_coding_gene | 1.3 | 2.39 | 4.2E-02 |
| *Y52E8A.3* | WBGene00021796 | protein_coding_gene | 1.3 | 2.39 | 3.1E-03 |
| *F01D5.6* | WBGene00008496 | protein_coding_gene | 1.3 | 2.39 | 4.3E-02 |
| *sfrp-1* | WBGene00022242 | protein_coding_gene | 1.3 | 2.38 | 1.1E-02 |
| *nhr-50* | WBGene00003640 | protein_coding_gene | 1.3 | 2.38 | 1.0E-02 |
| *nekl-1* | WBGene00021461 | protein_coding_gene | 1.3 | 2.38 | 2.0E-02 |
| *Y97E10AR.2* | WBGene00022397 | protein_coding_gene | 1.3 | 2.38 | 3.0E-02 |
| *ceh-37* | WBGene00000458 | protein_coding_gene | 1.3 | 2.38 | 3.6E-02 |
| *nhr-193* | WBGene00010215 | protein_coding_gene | 1.3 | 2.38 | 5.2E-03 |
| *M03E7.1* | WBGene00019753 | protein_coding_gene | 1.3 | 2.38 | 1.7E-02 |
| *K10D6.4* | WBGene00010744 | protein_coding_gene | 1.3 | 2.38 | 1.6E-03 |
| *dyf-11* | WBGene00001127 | protein_coding_gene | 1.3 | 2.38 | 2.3E-02 |
| *pqn-89* | WBGene00004169 | protein_coding_gene | 1.2 | 2.38 | 1.1E-02 |
| *fbxl-1* | WBGene00015350 | protein_coding_gene | 1.2 | 2.38 | 4.1E-03 |
| *pes-4* | WBGene00003978 | protein_coding_gene | 1.2 | 2.37 | 1.4E-02 |
| *F19G12.8* | WBGene00195184 | protein_coding_gene | 1.2 | 2.37 | 2.7E-02 |
| *nhr-213* | WBGene00011520 | protein_coding_gene | 1.2 | 2.37 | 7.9E-03 |
| *K08D8.1* | WBGene00010655 | protein_coding_gene | 1.2 | 2.37 | 2.1E-02 |
| *math-39* | WBGene00020358 | protein_coding_gene | 1.2 | 2.37 | 3.9E-03 |
| *syx-2* | WBGene00006372 | protein_coding_gene | 1.2 | 2.37 | 2.5E-02 |
| *rrc-1* | WBGene00009800 | protein_coding_gene | 1.2 | 2.37 | 4.0E-02 |
| *ddr-2* | WBGene00017381 | protein_coding_gene | 1.2 | 2.36 | 7.3E-03 |
| *F33H2.8* | WBGene00009371 | protein_coding_gene | 1.2 | 2.36 | 1.1E-02 |
| *sek-4* | WBGene00018035 | protein_coding_gene | 1.2 | 2.36 | 1.3E-04 |
| *Y50D4A.10* | WBGene00235315 | protein_coding_gene | 1.2 | 2.36 | 2.1E-02 |
| *fbxa-14* | WBGene00022326 | protein_coding_gene | 1.2 | 2.36 | 3.3E-02 |
| *nab-1* | WBGene00003516 | protein_coding_gene | 1.2 | 2.36 | 2.4E-02 |
| *F40F8.4* | WBGene00009577 | protein_coding_gene | 1.2 | 2.36 | 3.1E-02 |
| *C47E12.9* | WBGene00008152 | protein_coding_gene | 1.2 | 2.36 | 7.0E-03 |
| *emre-1* | WBGene00008980 | protein_coding_gene | 1.2 | 2.36 | 7.8E-04 |
| *T28B8.1* | WBGene00012113 | protein_coding_gene | 1.2 | 2.36 | 2.9E-03 |
| *igcm-1* | WBGene00018215 | protein_coding_gene | 1.2 | 2.35 | 2.1E-02 |
| *hst-3.2* | WBGene00044303 | protein_coding_gene | 1.2 | 2.35 | 4.4E-03 |
| *skr-19* | WBGene00004825 | protein_coding_gene | 1.2 | 2.35 | 6.5E-03 |
| *F22E12.3* | WBGene00009060 | protein_coding_gene | 1.2 | 2.35 | 3.2E-02 |
| *mab-31* | WBGene00021834 | protein_coding_gene | 1.2 | 2.35 | 1.4E-02 |
| *cest-3* | WBGene00008451 | protein_coding_gene | 1.2 | 2.35 | 3.2E-02 |
| *F59G1.4* | WBGene00019128 | protein_coding_gene | 1.2 | 2.35 | 1.2E-02 |
| *E01G6.2* | WBGene00008450 | protein_coding_gene | 1.2 | 2.34 | 2.2E-02 |
| *T25G12.11* | WBGene00020817 | protein_coding_gene | 1.2 | 2.34 | 1.5E-02 |
| *hst-3.1* | WBGene00002030 | protein_coding_gene | 1.2 | 2.34 | 2.8E-03 |
| *Y42G9A.2* | WBGene00021532 | protein_coding_gene | 1.2 | 2.34 | 5.0E-03 |
| *F23A7.5* | WBGene00009070 | protein_coding_gene | 1.2 | 2.34 | 1.9E-02 |
| *F41C3.8* | WBGene00018274 | protein_coding_gene | 1.2 | 2.34 | 1.2E-02 |
| *pat-9* | WBGene00003933 | protein_coding_gene | 1.2 | 2.33 | 4.4E-03 |
| *tag-89* | WBGene00006459 | protein_coding_gene | 1.2 | 2.33 | 2.2E-02 |
| *madd-2* | WBGene00016539 | protein_coding_gene | 1.2 | 2.33 | 5.0E-03 |
| *fbxa-80* | WBGene00022330 | protein_coding_gene | 1.2 | 2.33 | 5.8E-03 |

| | | | | | |
|---|---|---|---|---|---|
| *madf-10* | WBGene00013717 | protein_coding_gene | 1.2 | 2.33 | 3.2E-03 |
| *memo-1* | WBGene00016500 | protein_coding_gene | 1.2 | 2.33 | 2.7E-02 |
| *Y71A12B.18* | WBGene00045077 | pseudogene | 1.2 | 2.32 | 3.6E-03 |
| *pqn-31* | WBGene00004119 | protein_coding_gene | 1.2 | 2.32 | 4.1E-02 |
| *trpl-1* | WBGene00018992 | pseudogene | 1.2 | 2.32 | 1.5E-02 |
| *F48F5.6* | WBGene00009848 | protein_coding_gene | 1.2 | 2.32 | 1.8E-02 |
| *ddo-1* | WBGene00008127 | protein_coding_gene | 1.2 | 2.32 | 2.2E-02 |
| *tba-5* | WBGene00006531 | protein_coding_gene | 1.2 | 2.32 | 3.3E-02 |
| *egl-36* | WBGene00001202 | protein_coding_gene | 1.2 | 2.31 | 9.4E-03 |
| *rrf-2* | WBGene00004509 | protein_coding_gene | 1.2 | 2.31 | 4.2E-02 |
| *sma-3* | WBGene00004857 | protein_coding_gene | 1.2 | 2.31 | 8.1E-03 |
| *lin-39* | WBGene00003024 | protein_coding_gene | 1.2 | 2.31 | 1.2E-02 |
| *F10E7.9* | WBGene00017350 | protein_coding_gene | 1.2 | 2.31 | 2.5E-03 |
| *K11E4.2* | WBGene00010774 | protein_coding_gene | 1.2 | 2.31 | 2.9E-04 |
| *ets-9* | WBGene00016865 | protein_coding_gene | 1.2 | 2.31 | 1.4E-02 |
| *nhr-69* | WBGene00003659 | protein_coding_gene | 1.2 | 2.31 | 1.7E-02 |
| *F35B12.10* | WBGene00009389 | protein_coding_gene | 1.2 | 2.30 | 2.2E-02 |
| *nlp-57* | WBGene00008589 | protein_coding_gene | 1.2 | 2.30 | 6.4E-03 |
| *F16B4.2* | WBGene00017504 | protein_coding_gene | 1.2 | 2.30 | 4.7E-02 |
| *rig-5* | WBGene00004372 | protein_coding_gene | 1.2 | 2.30 | 3.8E-02 |
| *C25D7.15* | WBGene00007722 | protein_coding_gene | 1.2 | 2.30 | 9.1E-03 |
| *K02D10.8* | WBGene00219329 | protein_coding_gene | 1.2 | 2.29 | 3.4E-02 |
| *mab-21* | WBGene00003112 | protein_coding_gene | 1.2 | 2.29 | 1.7E-02 |
| *C46F11.6* | WBGene00008121 | protein_coding_gene | 1.2 | 2.29 | 1.3E-03 |
| *ajm-1* | WBGene00000100 | protein_coding_gene | 1.2 | 2.29 | 2.8E-02 |
| *Y44A6D.3* | WBGene00012854 | protein_coding_gene | 1.2 | 2.29 | 1.9E-02 |
| *nlp-52* | WBGene00018363 | protein_coding_gene | 1.2 | 2.29 | 4.2E-02 |
| *nhr-136* | WBGene00003726 | protein_coding_gene | 1.2 | 2.29 | 1.0E-02 |
| *twk-31* | WBGene00006683 | protein_coding_gene | 1.2 | 2.29 | 3.5E-02 |
| *glb-33* | WBGene00022284 | protein_coding_gene | 1.2 | 2.29 | 1.8E-02 |
| *F09E5.14* | WBGene00017291 | protein_coding_gene | 1.2 | 2.28 | 1.4E-02 |
| *T15B7.10* | WBGene00020522 | pseudogene | 1.2 | 2.28 | 2.7E-03 |
| *cof-2* | WBGene00000583 | protein_coding_gene | 1.2 | 2.28 | 1.5E-02 |
| *cebp-1* | WBGene00016997 | protein_coding_gene | 1.2 | 2.28 | 1.8E-02 |
| *spv-1* | WBGene00014051 | protein_coding_gene | 1.2 | 2.28 | 4.7E-02 |
| *hpo-15* | WBGene00016061 | protein_coding_gene | 1.2 | 2.28 | 1.8E-03 |
| *lim-8* | WBGene00017904 | protein_coding_gene | 1.2 | 2.27 | 3.5E-03 |
| *sym-1* | WBGene00006366 | protein_coding_gene | 1.2 | 2.27 | 4.5E-02 |
| *uig-1* | WBGene00009337 | protein_coding_gene | 1.2 | 2.27 | 3.8E-04 |
| *F20C5.6* | WBGene00008971 | protein_coding_gene | 1.2 | 2.27 | 3.4E-03 |
| *myo-6* | WBGene00009730 | protein_coding_gene | 1.2 | 2.26 | 6.9E-03 |
| *Y54F10AM.6* | WBGene00021850 | protein_coding_gene | 1.2 | 2.26 | 1.4E-02 |
| *F32D1.3* | WBGene00017983 | protein_coding_gene | 1.2 | 2.26 | 9.6E-03 |
| *T10B5.7* | WBGene00020393 | protein_coding_gene | 1.2 | 2.26 | 1.1E-03 |
| *igeg-1* | WBGene00017901 | protein_coding_gene | 1.2 | 2.26 | 9.8E-03 |
| *tag-343* | WBGene00018374 | protein_coding_gene | 1.2 | 2.26 | 9.9E-03 |
| *che-13* | WBGene00000492 | protein_coding_gene | 1.2 | 2.26 | 1.4E-02 |
| *sta-1* | WBGene00013111 | protein_coding_gene | 1.2 | 2.26 | 2.3E-02 |
| *F23C8.3* | WBGene00017732 | protein_coding_gene | 1.2 | 2.26 | 2.6E-03 |
| *K10C9.1* | WBGene00019623 | protein_coding_gene | 1.2 | 2.26 | 4.9E-02 |
| *F23C8.13* | WBGene00017741 | protein_coding_gene | 1.2 | 2.26 | 4.7E-02 |
| *hda-11* | WBGene00007953 | protein_coding_gene | 1.2 | 2.25 | 1.1E-02 |
| *Y37E11B.1* | WBGene00021374 | protein_coding_gene | 1.2 | 2.25 | 1.0E-02 |
| *maa-1* | WBGene00007680 | protein_coding_gene | 1.2 | 2.25 | 1.6E-02 |
| *nhr-125* | WBGene00003715 | protein_coding_gene | 1.2 | 2.25 | 1.8E-02 |
| *F15E6.9* | WBGene00017489 | protein_coding_gene | 1.2 | 2.25 | 1.8E-02 |
| *spp-16* | WBGene00005001 | protein_coding_gene | 1.2 | 2.25 | 4.3E-03 |
| *nhr-96* | WBGene00003686 | protein_coding_gene | 1.2 | 2.25 | 7.8E-03 |

| | | | | | |
|---|---|---|---|---|---|
| *nhr-201* | WBGene00019741 | protein_coding_gene | 1.2 | 2.24 | 2.6E-02 |
| *C10B5.3* | WBGene00015671 | protein_coding_gene | 1.2 | 2.24 | 1.6E-02 |
| *Y42G9A.1* | WBGene00021531 | protein_coding_gene | 1.2 | 2.24 | 5.9E-03 |
| *T23D8.3* | WBGene00011944 | protein_coding_gene | 1.2 | 2.24 | 1.3E-02 |
| *pudl-1* | WBGene00044489 | protein_coding_gene | 1.2 | 2.23 | 1.3E-02 |
| *eri-12* | WBGene00007091 | protein_coding_gene | 1.2 | 2.23 | 5.4E-05 |
| *best-9* | WBGene00016613 | protein_coding_gene | 1.2 | 2.23 | 1.7E-02 |
| *Y57G11C.22* | WBGene00013317 | protein_coding_gene | 1.2 | 2.22 | 1.6E-02 |
| *dep-1* | WBGene00009717 | protein_coding_gene | 1.2 | 2.22 | 8.0E-04 |
| *nhr-128* | WBGene00003718 | protein_coding_gene | 1.2 | 2.22 | 2.6E-02 |
| *best-23* | WBGene00014989 | protein_coding_gene | 1.2 | 2.22 | 2.2E-02 |
| *klo-1* | WBGene00016848 | protein_coding_gene | 1.1 | 2.22 | 2.5E-03 |
| *F55G1.15* | WBGene00018907 | protein_coding_gene | 1.1 | 2.21 | 1.3E-02 |
| *F53F4.13* | WBGene00009995 | protein_coding_gene | 1.1 | 2.21 | 3.0E-02 |
| *B0024.15* | WBGene00007104 | protein_coding_gene | 1.1 | 2.21 | 3.5E-02 |
| *math-4* | WBGene00015609 | protein_coding_gene | 1.1 | 2.21 | 2.6E-03 |
| *fbxa-55* | WBGene00020453 | protein_coding_gene | 1.1 | 2.21 | 1.8E-02 |
| *tag-278* | WBGene00015356 | protein_coding_gene | 1.1 | 2.21 | 5.9E-03 |
| *C13A10.1* | WBGene00015728 | protein_coding_gene | 1.1 | 2.21 | 2.7E-03 |
| *H24G06.1* | WBGene00019237 | protein_coding_gene | 1.1 | 2.21 | 5.6E-03 |
| *fat-2* | WBGene00001394 | protein_coding_gene | 1.1 | 2.21 | 8.5E-04 |
| *ric-19* | WBGene00004368 | protein_coding_gene | 1.1 | 2.20 | 2.2E-03 |
| *T27E4.1* | WBGene00020861 | protein_coding_gene | 1.1 | 2.20 | 1.7E-02 |
| *B0252.8* | WBGene00044615 | protein_coding_gene | 1.1 | 2.20 | 1.5E-02 |
| *Y116A8B.4* | WBGene00013781 | protein_coding_gene | 1.1 | 2.20 | 2.7E-02 |
| *lagr-1* | WBGene00006505 | protein_coding_gene | 1.1 | 2.20 | 1.4E-02 |
| *hex-2* | WBGene00015767 | protein_coding_gene | 1.1 | 2.20 | 1.3E-04 |
| *fbxa-4* | WBGene00044672 | protein_coding_gene | 1.1 | 2.20 | 2.7E-02 |
| *T04A6.1* | WBGene00020200 | protein_coding_gene | 1.1 | 2.20 | 3.4E-02 |
| *C36E8.4* | WBGene00007983 | protein_coding_gene | 1.1 | 2.20 | 4.1E-03 |
| *C16B8.3* | WBGene00015823 | protein_coding_gene | 1.1 | 2.19 | 1.9E-03 |
| *F15D4.2* | WBGene00008859 | protein_coding_gene | 1.1 | 2.19 | 3.0E-03 |
| *fmil-1* | WBGene00021362 | protein_coding_gene | 1.1 | 2.19 | 1.3E-02 |
| *nhr-110* | WBGene00003700 | protein_coding_gene | 1.1 | 2.19 | 1.5E-03 |
| *Y53F4B.1* | WBGene00013149 | protein_coding_gene | 1.1 | 2.19 | 4.3E-02 |
| *C16B8.4* | WBGene00015824 | protein_coding_gene | 1.1 | 2.19 | 4.5E-02 |
| *C50C3.5* | WBGene00016801 | protein_coding_gene | 1.1 | 2.19 | 1.4E-02 |
| *C28C12.4* | WBGene00016180 | protein_coding_gene | 1.1 | 2.19 | 2.7E-02 |
| *elt-6* | WBGene00001253 | protein_coding_gene | 1.1 | 2.18 | 1.3E-03 |
| *vab-9* | WBGene00006875 | protein_coding_gene | 1.1 | 2.18 | 7.5E-03 |
| *twk-2* | WBGene00006657 | protein_coding_gene | 1.1 | 2.18 | 4.6E-02 |
| *K09F6.15* | WBGene00271744 | protein_coding_gene | 1.1 | 2.18 | 2.2E-03 |
| *best-18* | WBGene00011868 | protein_coding_gene | 1.1 | 2.18 | 1.1E-02 |
| *alh-3* | WBGene00000109 | protein_coding_gene | 1.1 | 2.18 | 5.0E-03 |
| *camt-1* | WBGene00020251 | protein_coding_gene | 1.1 | 2.18 | 1.4E-02 |
| *fbxa-105* | WBGene00015362 | protein_coding_gene | 1.1 | 2.17 | 3.8E-02 |
| *nhr-183* | WBGene00018266 | protein_coding_gene | 1.1 | 2.17 | 2.6E-02 |
| *F33G12.6* | WBGene00018018 | protein_coding_gene | 1.1 | 2.17 | 2.0E-02 |
| *sorb-1* | WBGene00012891 | protein_coding_gene | 1.1 | 2.17 | 3.7E-02 |
| *zip-5* | WBGene00007932 | protein_coding_gene | 1.1 | 2.17 | 2.9E-03 |
| *R07B7.6* | WBGene00011090 | protein_coding_gene | 1.1 | 2.17 | 2.1E-04 |
| *bre-2* | WBGene00000267 | protein_coding_gene | 1.1 | 2.16 | 1.5E-02 |
| *F27D9.2* | WBGene00017861 | protein_coding_gene | 1.1 | 2.16 | 4.0E-02 |
| *rei-2* | WBGene00019365 | protein_coding_gene | 1.1 | 2.16 | 3.7E-02 |
| *K02E2.6* | WBGene00010507 | protein_coding_gene | 1.1 | 2.16 | 1.4E-03 |
| *F22D6.15* | WBGene00009056 | protein_coding_gene | 1.1 | 2.16 | 1.1E-02 |
| *zip-1* | WBGene00006986 | protein_coding_gene | 1.1 | 2.16 | 1.7E-03 |
| *glt-3* | WBGene00001621 | protein_coding_gene | 1.1 | 2.15 | 4.4E-02 |

| | | | | | |
|---|---|---|---|---|---|
| *F36F2.1* | WBGene00009475 | protein_coding_gene | 1.1 | 2.15 | 4.2E-02 |
| *nlp-10* | WBGene00003748 | protein_coding_gene | 1.1 | 2.15 | 3.3E-02 |
| *frpr-5* | WBGene00008342 | protein_coding_gene | 1.1 | 2.15 | 4.7E-02 |
| *F21A9.2* | WBGene00017651 | protein_coding_gene | 1.1 | 2.15 | 8.7E-03 |
| *ceh-27* | WBGene00000449 | protein_coding_gene | 1.1 | 2.14 | 3.4E-02 |
| *nuc-1* | WBGene00003828 | protein_coding_gene | 1.1 | 2.14 | 3.2E-02 |
| *lin-17* | WBGene00003006 | protein_coding_gene | 1.1 | 2.14 | 4.3E-02 |
| *F38E11.6* | WBGene00009543 | protein_coding_gene | 1.1 | 2.14 | 4.4E-02 |
| *Y54E5A.8* | WBGene00013203 | protein_coding_gene | 1.1 | 2.14 | 1.2E-03 |
| *plc-2* | WBGene00004037 | protein_coding_gene | 1.1 | 2.14 | 1.5E-02 |
| *D2030.12* | WBGene00008421 | protein_coding_gene | 1.1 | 2.14 | 2.2E-02 |
| *abhd-14* | WBGene00021927 | protein_coding_gene | 1.1 | 2.14 | 1.8E-02 |
| *sma-4* | WBGene00004858 | protein_coding_gene | 1.1 | 2.14 | 3.6E-02 |
| *lron-15* | WBGene00016974 | protein_coding_gene | 1.1 | 2.14 | 3.5E-02 |
| *arrd-20* | WBGene00015440 | pseudogene | 1.1 | 2.14 | 1.7E-02 |
| *dmd-6* | WBGene00007058 | protein_coding_gene | 1.1 | 2.14 | 4.0E-02 |
| *rpl-24.2* | WBGene00004437 | protein_coding_gene | 1.1 | 2.14 | 3.9E-02 |
| *src-2* | WBGene00005078 | protein_coding_gene | 1.1 | 2.14 | 3.8E-02 |
| *alh-1* | WBGene00000107 | protein_coding_gene | 1.1 | 2.14 | 3.8E-03 |
| *C17H11.6* | WBGene00015926 | protein_coding_gene | 1.1 | 2.14 | 2.4E-02 |
| *F37C12.10* | WBGene00018153 | protein_coding_gene | 1.1 | 2.13 | 1.2E-02 |
| *dyf-2* | WBGene00001118 | protein_coding_gene | 1.1 | 2.13 | 1.2E-02 |
| *ugt-23* | WBGene00007650 | protein_coding_gene | 1.1 | 2.13 | 1.4E-02 |
| *C54E4.4* | WBGene00016919 | protein_coding_gene | 1.1 | 2.13 | 3.9E-02 |
| *acy-3* | WBGene00000070 | protein_coding_gene | 1.1 | 2.13 | 7.8E-03 |
| *T02B11.3* | WBGene00020155 | protein_coding_gene | 1.1 | 2.13 | 2.4E-02 |
| *M116.2* | WBGene00019789 | pseudogene | 1.1 | 2.13 | 9.6E-05 |
| *C44B11.6* | WBGene00016635 | protein_coding_gene | 1.1 | 2.12 | 1.5E-02 |
| *nhr-122* | WBGene00003712 | protein_coding_gene | 1.1 | 2.12 | 1.0E-02 |
| *F21G4.1* | WBGene00009023 | protein_coding_gene | 1.1 | 2.12 | 1.7E-03 |
| *ric-3* | WBGene00004363 | protein_coding_gene | 1.1 | 2.12 | 2.8E-02 |
| *Y54G2A.52* | WBGene00045397 | protein_coding_gene | 1.1 | 2.12 | 2.7E-03 |
| *nhr-90* | WBGene00003680 | protein_coding_gene | 1.1 | 2.12 | 3.5E-02 |
| *fbxa-146* | WBGene00018807 | pseudogene | 1.1 | 2.12 | 1.6E-02 |
| *C03A3.1* | WBGene00007266 | protein_coding_gene | 1.1 | 2.11 | 5.0E-02 |
| *Y46G5A.15* | WBGene00012906 | protein_coding_gene | 1.1 | 2.11 | 2.8E-02 |
| *frm-2* | WBGene00001489 | protein_coding_gene | 1.1 | 2.11 | 1.1E-02 |
| *C48D1.1* | WBGene00008177 | protein_coding_gene | 1.1 | 2.11 | 3.1E-02 |
| *F59B2.9* | WBGene00010311 | protein_coding_gene | 1.1 | 2.11 | 2.3E-02 |
| *nhr-195* | WBGene00019115 | protein_coding_gene | 1.1 | 2.11 | 4.5E-02 |
| *fhod-1* | WBGene00016735 | protein_coding_gene | 1.1 | 2.11 | 5.7E-03 |
| *K02E10.6* | WBGene00019320 | protein_coding_gene | 1.1 | 2.11 | 2.0E-02 |
| *fbxa-32* | WBGene00013839 | protein_coding_gene | 1.1 | 2.11 | 4.8E-02 |
| *K12B6.9* | WBGene00023432 | protein_coding_gene | 1.1 | 2.11 | 4.3E-02 |
| *egl-5* | WBGene00001174 | protein_coding_gene | 1.1 | 2.10 | 4.3E-02 |
| *C41D11.6* | WBGene00016565 | protein_coding_gene | 1.1 | 2.10 | 4.5E-02 |
| *pry-1* | WBGene00004202 | protein_coding_gene | 1.1 | 2.10 | 5.1E-03 |
| *nmtn-1* | WBGene00019434 | protein_coding_gene | 1.1 | 2.10 | 2.0E-03 |
| *nadk-1* | WBGene00022307 | protein_coding_gene | 1.1 | 2.10 | 7.7E-03 |
| *fbxa-219* | WBGene00044403 | protein_coding_gene | 1.1 | 2.10 | 1.8E-02 |
| *Y71H2AL.2* | WBGene00022165 | protein_coding_gene | 1.1 | 2.10 | 3.4E-02 |
| *nhr-133* | WBGene00003723 | protein_coding_gene | 1.1 | 2.09 | 4.1E-02 |
| *C49C3.8* | WBGene00008198 | protein_coding_gene | 1.1 | 2.09 | 2.1E-02 |
| *crb-3* | WBGene00044743 | protein_coding_gene | 1.1 | 2.09 | 3.3E-03 |
| *F26A3.4* | WBGene00009142 | protein_coding_gene | 1.1 | 2.09 | 7.9E-03 |
| *Y65A5A.1* | WBGene00013418 | protein_coding_gene | 1.1 | 2.09 | 2.1E-03 |
| *set-30* | WBGene00022499 | protein_coding_gene | 1.1 | 2.09 | 3.0E-02 |
| *nlp-7* | WBGene00003745 | protein_coding_gene | 1.1 | 2.08 | 1.0E-02 |

| | | | | | |
|---|---|---|---|---|---|
| *dohh-1* | WBGene00007555 | protein_coding_gene | 1.1 | 2.08 | 1.9E-02 |
| *F20C5.4* | WBGene00008969 | protein_coding_gene | 1.1 | 2.08 | 8.5E-03 |
| *T13H10.2* | WBGene00011762 | protein_coding_gene | 1.1 | 2.08 | 4.2E-02 |
| *F40F12.9* | WBGene00043995 | protein_coding_gene | 1.1 | 2.08 | 8.7E-03 |
| *nhr-64* | WBGene00003654 | protein_coding_gene | 1.1 | 2.08 | 1.3E-02 |
| *F38B6.6* | WBGene00018175 | protein_coding_gene | 1.1 | 2.07 | 3.7E-02 |
| *pdcd-2* | WBGene00011116 | protein_coding_gene | 1.0 | 2.07 | 3.1E-03 |
| *K06G5.1* | WBGene00010605 | protein_coding_gene | 1.0 | 2.07 | 1.8E-02 |
| *M176.5* | WBGene00010944 | protein_coding_gene | 1.0 | 2.07 | 1.0E-02 |
| *T14G11.1* | WBGene00020510 | protein_coding_gene | 1.0 | 2.07 | 5.7E-03 |
| *Y60A3A.16* | WBGene00013364 | protein_coding_gene | 1.0 | 2.07 | 2.5E-02 |
| *B0511.6* | WBGene00015232 | protein_coding_gene | 1.0 | 2.07 | 9.2E-03 |
| *Y75B8A.28* | WBGene00013559 | protein_coding_gene | 1.0 | 2.06 | 1.9E-03 |
| *M01F1.3* | WBGene00010809 | protein_coding_gene | 1.0 | 2.06 | 9.1E-03 |
| *unc-36* | WBGene00006772 | protein_coding_gene | 1.0 | 2.06 | 3.0E-02 |
| *ZK287.4* | WBGene00013965 | protein_coding_gene | 1.0 | 2.06 | 4.4E-02 |
| *R05A10.3* | WBGene00011020 | protein_coding_gene | 1.0 | 2.06 | 4.8E-02 |
| *F09F9.1* | WBGene00017306 | protein_coding_gene | 1.0 | 2.06 | 1.9E-02 |
| *efhd-1* | WBGene00012980 | protein_coding_gene | 1.0 | 2.05 | 1.9E-02 |
| *fbxa-108* | WBGene00010293 | protein_coding_gene | 1.0 | 2.05 | 4.1E-03 |
| *F39E9.28* | WBGene00271716 | pseudogene | 1.0 | 2.05 | 3.2E-03 |
| *byn-1* | WBGene00000276 | protein_coding_gene | 1.0 | 2.05 | 2.1E-02 |
| *cest-16* | WBGene00020330 | protein_coding_gene | 1.0 | 2.05 | 2.1E-02 |
| *W06F12.2* | WBGene00012306 | protein_coding_gene | 1.0 | 2.04 | 2.0E-03 |
| *tir-1* | WBGene00006575 | protein_coding_gene | 1.0 | 2.04 | 3.4E-02 |
| *T05E11.8* | WBGene00011484 | protein_coding_gene | 1.0 | 2.04 | 5.8E-03 |
| *unc-6* | WBGene00006746 | protein_coding_gene | 1.0 | 2.04 | 4.7E-02 |
| *F39E9.7* | WBGene00018199 | pseudogene | 1.0 | 2.04 | 1.8E-02 |
| *pqn-92* | WBGene00004172 | protein_coding_gene | 1.0 | 2.04 | 2.0E-02 |
| *ttr-10* | WBGene00017600 | protein_coding_gene | 1.0 | 2.04 | 3.7E-03 |
| *F46G10.1* | WBGene00009796 | protein_coding_gene | 1.0 | 2.04 | 1.1E-02 |
| *C30F2.5* | WBGene00007819 | protein_coding_gene | 1.0 | 2.03 | 1.5E-02 |
| *ptr-15* | WBGene00004229 | protein_coding_gene | 1.0 | 2.03 | 3.7E-02 |
| *cah-6* | WBGene00000284 | protein_coding_gene | 1.0 | 2.03 | 4.7E-03 |
| *bam-2* | WBGene00000237 | protein_coding_gene | 1.0 | 2.03 | 3.6E-02 |
| *F20A1.1* | WBGene00017613 | protein_coding_gene | 1.0 | 2.03 | 4.9E-02 |
| *hum-8* | WBGene00002041 | protein_coding_gene | 1.0 | 2.02 | 7.6E-03 |
| *ptp-4* | WBGene00010558 | protein_coding_gene | 1.0 | 2.02 | 2.9E-02 |
| *T28F4.4* | WBGene00012139 | protein_coding_gene | 1.0 | 2.02 | 3.0E-02 |
| *W02C12.2* | WBGene00020929 | protein_coding_gene | 1.0 | 2.02 | 4.0E-02 |
| *W10G11.19* | WBGene00021148 | protein_coding_gene | 1.0 | 2.02 | 1.5E-03 |
| *nhr-109* | WBGene00003699 | protein_coding_gene | 1.0 | 2.01 | 2.4E-02 |
| *nhr-105* | WBGene00003695 | protein_coding_gene | 1.0 | 2.01 | 2.9E-02 |
| *ZK384.7* | WBGene00269383 | ncRNA_gene | 1.0 | 2.00 | 4.9E-02 |
| *B0391.10* | WBGene00007165 | pseudogene | -1.0 | 0.50 | 5.0E-02 |
| *mig-22* | WBGene00003253 | protein_coding_gene | -1.0 | 0.50 | 6.8E-04 |
| *gad-1* | WBGene00001513 | protein_coding_gene | -1.0 | 0.50 | 4.2E-04 |
| *rbm-39* | WBGene00021921 | protein_coding_gene | -1.0 | 0.50 | 1.3E-03 |
| *C18E3.9* | WBGene00015976 | protein_coding_gene | -1.0 | 0.50 | 2.3E-02 |
| *W09C5.7* | WBGene00012353 | protein_coding_gene | -1.0 | 0.50 | 3.0E-03 |
| *swsn-9* | WBGene00007256 | protein_coding_gene | -1.0 | 0.50 | 8.3E-03 |
| *F26G1.1* | WBGene00017837 | protein_coding_gene | -1.0 | 0.50 | 1.3E-02 |
| *fkb-6* | WBGene00001431 | protein_coding_gene | -1.0 | 0.50 | 2.0E-04 |
| *smgl-1* | WBGene00008990 | protein_coding_gene | -1.0 | 0.50 | 7.9E-05 |
| *ZK945.4* | WBGene00014166 | protein_coding_gene | -1.0 | 0.50 | 4.6E-02 |
| *cchl-1* | WBGene00011527 | protein_coding_gene | -1.0 | 0.50 | 6.9E-03 |
| *nmt-1* | WBGene00020549 | protein_coding_gene | -1.0 | 0.50 | 2.4E-03 |
| *gcna-1* | WBGene00022694 | protein_coding_gene | -1.0 | 0.50 | 2.4E-02 |

| | | | | | |
|---|---|---|---|---|---|
| *Y4C6B.1* | WBGene00021155 | protein_coding_gene | -1.0 | 0.50 | 7.2E-03 |
| *tin-44* | WBGene00020383 | protein_coding_gene | -1.0 | 0.50 | 1.3E-04 |
| *sec-24.1* | WBGene00004755 | protein_coding_gene | -1.0 | 0.50 | 4.9E-04 |
| *mrpl-24* | WBGene00019076 | protein_coding_gene | -1.0 | 0.50 | 3.1E-04 |
| *Y50D4C.2* | WBGene00021747 | pseudogene | -1.0 | 0.50 | 2.2E-02 |
| *exo-3* | WBGene00001372 | protein_coding_gene | -1.0 | 0.50 | 4.2E-02 |
| *ect-2* | WBGene00002297 | protein_coding_gene | -1.0 | 0.50 | 2.9E-02 |
| *app-1* | WBGene00000155 | protein_coding_gene | -1.0 | 0.50 | 2.8E-03 |
| *cyn-3* | WBGene00000879 | protein_coding_gene | -1.0 | 0.50 | 6.6E-04 |
| *C28C12.12* | WBGene00016182 | protein_coding_gene | -1.0 | 0.50 | 3.7E-03 |
| *nfyc-1* | WBGene00017742 | protein_coding_gene | -1.0 | 0.50 | 1.4E-02 |
| *swsn-6* | WBGene00022774 | protein_coding_gene | -1.0 | 0.50 | 1.7E-02 |
| *Y48G10A.2* | WBGene00013019 | protein_coding_gene | -1.0 | 0.50 | 2.8E-02 |
| *lin-35* | WBGene00003020 | protein_coding_gene | -1.0 | 0.50 | 3.0E-03 |
| *such-1* | WBGene00013443 | protein_coding_gene | -1.0 | 0.50 | 6.3E-03 |
| *rpt-5* | WBGene00004505 | protein_coding_gene | -1.0 | 0.50 | 1.2E-04 |
| *Y56A3A.16* | WBGene00013235 | protein_coding_gene | -1.0 | 0.50 | 1.8E-02 |
| *pbs-1* | WBGene00003947 | protein_coding_gene | -1.0 | 0.50 | 1.6E-03 |
| *txdc-9* | WBGene00006515 | protein_coding_gene | -1.0 | 0.50 | 3.1E-02 |
| *ttr-53* | WBGene00011773 | protein_coding_gene | -1.0 | 0.50 | 2.6E-02 |
| *vps-53* | WBGene00011502 | protein_coding_gene | -1.0 | 0.49 | 5.0E-03 |
| *C52E12.1* | WBGene00016888 | protein_coding_gene | -1.0 | 0.49 | 2.9E-05 |
| *hal-3* | WBGene00011788 | protein_coding_gene | -1.0 | 0.49 | 2.8E-02 |
| *cct-2* | WBGene00000378 | protein_coding_gene | -1.0 | 0.49 | 3.3E-06 |
| *Y106G6H.14* | WBGene00013724 | protein_coding_gene | -1.0 | 0.49 | 1.1E-02 |
| *arle-14* | WBGene00015145 | protein_coding_gene | -1.0 | 0.49 | 3.0E-02 |
| *xnd-1* | WBGene00001514 | protein_coding_gene | -1.0 | 0.49 | 9.0E-03 |
| *F49C12.9* | WBGene00009878 | protein_coding_gene | -1.0 | 0.49 | 4.2E-02 |
| *rpt-2* | WBGene00004502 | protein_coding_gene | -1.0 | 0.49 | 4.2E-05 |
| *coa-5* | WBGene00012483 | protein_coding_gene | -1.0 | 0.49 | 1.2E-02 |
| *dro-1* | WBGene00001092 | protein_coding_gene | -1.0 | 0.49 | 1.8E-02 |
| *W03F9.1* | WBGene00020999 | protein_coding_gene | -1.0 | 0.49 | 1.2E-05 |
| *sun-1* | WBGene00006311 | protein_coding_gene | -1.0 | 0.49 | 1.9E-02 |
| *glh-4* | WBGene00001601 | protein_coding_gene | -1.0 | 0.49 | 3.9E-02 |
| *K07C5.6* | WBGene00010629 | protein_coding_gene | -1.0 | 0.49 | 2.9E-03 |
| *wdr-4* | WBGene00022420 | protein_coding_gene | -1.0 | 0.49 | 2.1E-02 |
| *duo-3* | WBGene00001112 | protein_coding_gene | -1.0 | 0.49 | 1.1E-02 |
| *rnh-2* | WBGene00004383 | protein_coding_gene | -1.0 | 0.49 | 9.2E-03 |
| *cra-1* | WBGene00020068 | protein_coding_gene | -1.0 | 0.49 | 4.0E-07 |
| *hst-1* | WBGene00002028 | protein_coding_gene | -1.0 | 0.49 | 3.6E-02 |
| *cua-1* | WBGene00000834 | protein_coding_gene | -1.0 | 0.49 | 2.3E-04 |
| *F07F6.4* | WBGene00017217 | protein_coding_gene | -1.0 | 0.49 | 6.2E-03 |
| *imp-2* | WBGene00011481 | protein_coding_gene | -1.0 | 0.49 | 4.8E-04 |
| *tra-2* | WBGene00006605 | protein_coding_gene | -1.0 | 0.49 | 6.9E-03 |
| *ufd-3* | WBGene00007333 | protein_coding_gene | -1.0 | 0.49 | 7.2E-03 |
| *haf-3* | WBGene00001813 | protein_coding_gene | -1.0 | 0.49 | 7.8E-05 |
| *txl-1* | WBGene00021826 | protein_coding_gene | -1.0 | 0.49 | 7.2E-03 |
| *Y73B3A.21* | WBGene00022222 | pseudogene | -1.0 | 0.49 | 1.7E-02 |
| *alg-5* | WBGene00011945 | protein_coding_gene | -1.0 | 0.49 | 4.8E-02 |
| *sep-1* | WBGene00004775 | protein_coding_gene | -1.0 | 0.49 | 1.8E-03 |
| *hcp-3* | WBGene00001831 | protein_coding_gene | -1.0 | 0.49 | 1.5E-02 |
| *aka-1* | WBGene00000101 | protein_coding_gene | -1.0 | 0.49 | 3.3E-02 |
| *pign-1* | WBGene00021840 | protein_coding_gene | -1.0 | 0.49 | 3.9E-02 |
| *zhp-2* | WBGene00008387 | protein_coding_gene | -1.0 | 0.49 | 8.6E-03 |
| *B0041.5* | WBGene00015009 | protein_coding_gene | -1.0 | 0.49 | 3.4E-02 |
| *F39H2.3* | WBGene00009563 | protein_coding_gene | -1.0 | 0.49 | 1.7E-02 |
| *hira-1* | WBGene00019627 | protein_coding_gene | -1.0 | 0.49 | 1.1E-04 |
| *pak-2* | WBGene00006443 | protein_coding_gene | -1.0 | 0.49 | 1.4E-02 |

| | | | | | |
|---|---|---|---|---|---|
| *tdpo-1* | WBGene00018678 | protein_coding_gene | -1.0 | 0.49 | 9.6E-03 |
| *cyb-1* | WBGene00000865 | protein_coding_gene | -1.0 | 0.49 | 4.3E-02 |
| *hil-4* | WBGene00001855 | protein_coding_gene | -1.0 | 0.49 | 2.1E-02 |
| *jmjd-2* | WBGene00012982 | protein_coding_gene | -1.0 | 0.49 | 1.3E-03 |
| *acaa-2* | WBGene00009952 | protein_coding_gene | -1.0 | 0.49 | 6.7E-03 |
| *B0001.2* | WBGene00007087 | protein_coding_gene | -1.0 | 0.49 | 3.7E-03 |
| *B0205.1* | WBGene00015017 | protein_coding_gene | -1.0 | 0.49 | 9.5E-05 |
| *spd-5* | WBGene00004955 | protein_coding_gene | -1.0 | 0.49 | 5.7E-05 |
| *T25D3.4* | WBGene00020798 | protein_coding_gene | -1.0 | 0.49 | 3.2E-02 |
| *K12D12.5* | WBGene00010787 | protein_coding_gene | -1.0 | 0.49 | 2.1E-03 |
| *Y71F9B.6* | WBGene00022128 | protein_coding_gene | -1.0 | 0.49 | 2.7E-04 |
| *snr-5* | WBGene00004918 | protein_coding_gene | -1.0 | 0.49 | 1.5E-02 |
| *cyn-7* | WBGene00000883 | protein_coding_gene | -1.0 | 0.49 | 5.7E-03 |
| *nars-1* | WBGene00003815 | protein_coding_gene | -1.0 | 0.49 | 6.7E-07 |
| *Y92H12BR.3* | WBGene00022369 | protein_coding_gene | -1.0 | 0.49 | 2.5E-02 |
| *C10G11.6* | WBGene00015686 | protein_coding_gene | -1.0 | 0.49 | 2.6E-03 |
| *glh-2* | WBGene00001599 | protein_coding_gene | -1.0 | 0.49 | 8.6E-03 |
| *rfc-2* | WBGene00004338 | protein_coding_gene | -1.0 | 0.49 | 2.8E-04 |
| *K08E3.5* | WBGene00010665 | protein_coding_gene | -1.0 | 0.49 | 2.2E-02 |
| *eif-2gamma* | WBGene00021466 | protein_coding_gene | -1.0 | 0.49 | 7.0E-09 |
| *pif-1* | WBGene00004028 | protein_coding_gene | -1.0 | 0.49 | 6.7E-06 |
| *R06C7.2* | WBGene00011062 | protein_coding_gene | -1.0 | 0.49 | 7.7E-03 |
| *C25E10.10* | WBGene00016099 | protein_coding_gene | -1.0 | 0.49 | 4.9E-02 |
| *dbt-1* | WBGene00014054 | protein_coding_gene | -1.0 | 0.48 | 6.3E-04 |
| *mom-5* | WBGene00003397 | protein_coding_gene | -1.0 | 0.48 | 2.2E-02 |
| *Y48G1C.8* | WBGene00021681 | protein_coding_gene | -1.0 | 0.48 | 4.9E-03 |
| *ccr-4* | WBGene00000376 | protein_coding_gene | -1.0 | 0.48 | 2.3E-02 |
| *npp-16* | WBGene00003802 | protein_coding_gene | -1.0 | 0.48 | 4.7E-03 |
| *Y54E10BR.2* | WBGene00021841 | protein_coding_gene | -1.0 | 0.48 | 2.9E-02 |
| *pdxk-1* | WBGene00019008 | protein_coding_gene | -1.0 | 0.48 | 2.3E-03 |
| *C56G2.15* | WBGene00016983 | protein_coding_gene | -1.0 | 0.48 | 2.4E-03 |
| *kat-1* | WBGene00002183 | protein_coding_gene | -1.0 | 0.48 | 8.5E-03 |
| *ints-1* | WBGene00015499 | protein_coding_gene | -1.1 | 0.48 | 1.4E-04 |
| *npp-9* | WBGene00003795 | protein_coding_gene | -1.1 | 0.48 | 8.8E-04 |
| *npp-25* | WBGene00012555 | protein_coding_gene | -1.1 | 0.48 | 3.5E-03 |
| *unc-40* | WBGene00006776 | protein_coding_gene | -1.1 | 0.48 | 1.4E-02 |
| *C50E3.5* | WBGene00016816 | protein_coding_gene | -1.1 | 0.48 | 3.0E-02 |
| *tars-1* | WBGene00006617 | protein_coding_gene | -1.1 | 0.48 | 2.8E-08 |
| *ptps-1* | WBGene00015010 | protein_coding_gene | -1.1 | 0.48 | 1.2E-02 |
| *Y50D4C.5* | WBGene00021749 | protein_coding_gene | -1.1 | 0.48 | 7.7E-04 |
| *F53H1.4* | WBGene00018778 | protein_coding_gene | -1.1 | 0.48 | 1.1E-02 |
| *clh-5* | WBGene00000532 | protein_coding_gene | -1.1 | 0.48 | 8.7E-04 |
| *Y73E7A.6* | WBGene00022273 | protein_coding_gene | -1.1 | 0.48 | 2.7E-04 |
| *phb-1* | WBGene00004014 | protein_coding_gene | -1.1 | 0.48 | 2.0E-12 |
| *T16H12.2* | WBGene00011812 | protein_coding_gene | -1.1 | 0.48 | 4.0E-02 |
| *W08E12.7* | WBGene00021088 | protein_coding_gene | -1.1 | 0.48 | 2.4E-03 |
| *pqn-20* | WBGene00004110 | protein_coding_gene | -1.1 | 0.48 | 4.8E-02 |
| *rpt-1* | WBGene00004501 | protein_coding_gene | -1.1 | 0.48 | 2.4E-04 |
| *snr-1* | WBGene00004914 | protein_coding_gene | -1.1 | 0.48 | 1.2E-03 |
| *eif-3.J* | WBGene00012738 | protein_coding_gene | -1.1 | 0.48 | 9.7E-08 |
| *rab-30* | WBGene00004282 | protein_coding_gene | -1.1 | 0.48 | 1.2E-06 |
| *B0511.7* | WBGene00015233 | protein_coding_gene | -1.1 | 0.48 | 1.3E-02 |
| *skr-17* | WBGene00004823 | protein_coding_gene | -1.1 | 0.48 | 2.8E-02 |
| *klp-18* | WBGene00002228 | protein_coding_gene | -1.1 | 0.48 | 3.0E-04 |
| *klp-10* | WBGene00002221 | pseudogene | -1.1 | 0.48 | 7.7E-03 |
| *R11.3* | WBGene00011228 | protein_coding_gene | -1.1 | 0.48 | 4.9E-02 |
| *lin-5* | WBGene00002994 | protein_coding_gene | -1.1 | 0.48 | 1.4E-04 |
| *T24A6.20* | WBGene00020756 | protein_coding_gene | -1.1 | 0.48 | 3.7E-02 |

| | | | | | |
|---|---|---|---|---|---|
| *F27D4.1* | WBGene00009187 | protein_coding_gene | -1.1 | 0.48 | 1.6E-02 |
| *pbs-3* | WBGene00003949 | protein_coding_gene | -1.1 | 0.48 | 6.1E-03 |
| *Y71G12B.13* | WBGene00022153 | protein_coding_gene | -1.1 | 0.48 | 7.3E-08 |
| *pbs-2* | WBGene00003948 | protein_coding_gene | -1.1 | 0.48 | 2.0E-03 |
| *knl-1* | WBGene00002231 | protein_coding_gene | -1.1 | 0.48 | 3.5E-04 |
| *C56E6.2* | WBGene00016970 | protein_coding_gene | -1.1 | 0.48 | 4.6E-02 |
| *sec-22* | WBGene00018853 | protein_coding_gene | -1.1 | 0.48 | 2.1E-04 |
| *F32B6.3* | WBGene00009320 | protein_coding_gene | -1.1 | 0.48 | 5.9E-03 |
| *C49H3.6* | WBGene00016792 | protein_coding_gene | -1.1 | 0.48 | 4.8E-02 |
| *pbs-5* | WBGene00003951 | protein_coding_gene | -1.1 | 0.48 | 1.7E-04 |
| *npp-7* | WBGene00003793 | protein_coding_gene | -1.1 | 0.48 | 9.6E-03 |
| *T10B11.8* | WBGene00020404 | protein_coding_gene | -1.1 | 0.48 | 7.9E-03 |
| *tram-1* | WBGene00007696 | protein_coding_gene | -1.1 | 0.48 | 2.1E-02 |
| *gex-2* | WBGene00001579 | protein_coding_gene | -1.1 | 0.48 | 8.6E-03 |
| *patr-1* | WBGene00009661 | protein_coding_gene | -1.1 | 0.48 | 4.8E-02 |
| *mpk-1* | WBGene00003401 | protein_coding_gene | -1.1 | 0.48 | 1.2E-03 |
| *C17C3.1* | WBGene00015887 | protein_coding_gene | -1.1 | 0.48 | 3.8E-02 |
| *T08A11.1* | WBGene00011604 | protein_coding_gene | -1.1 | 0.48 | 1.2E-02 |
| *rpn-13* | WBGene00016981 | protein_coding_gene | -1.1 | 0.48 | 1.5E-04 |
| *edc-3* | WBGene00011036 | protein_coding_gene | -1.1 | 0.48 | 1.4E-02 |
| *taf-6.1* | WBGene00006387 | protein_coding_gene | -1.1 | 0.48 | 8.7E-08 |
| *tat-3* | WBGene00012360 | protein_coding_gene | -1.1 | 0.48 | 3.4E-04 |
| *tag-153* | WBGene00006499 | protein_coding_gene | -1.1 | 0.48 | 3.1E-04 |
| *rsp-6* | WBGene00004703 | protein_coding_gene | -1.1 | 0.47 | 4.0E-03 |
| *Y51F10.11* | WBGene00044438 | protein_coding_gene | -1.1 | 0.47 | 4.2E-03 |
| *pfkb-1.2* | WBGene00019295 | protein_coding_gene | -1.1 | 0.47 | 9.6E-03 |
| *cdr-7* | WBGene00010472 | protein_coding_gene | -1.1 | 0.47 | 1.6E-02 |
| *C44B7.7* | WBGene00016629 | protein_coding_gene | -1.1 | 0.47 | 9.6E-03 |
| *pbs-6* | WBGene00003952 | protein_coding_gene | -1.1 | 0.47 | 1.3E-04 |
| *F55G1.6* | WBGene00018902 | protein_coding_gene | -1.1 | 0.47 | 2.5E-02 |
| *eif-3.L* | WBGene00015920 | protein_coding_gene | -1.1 | 0.47 | 7.0E-08 |
| *pas-5* | WBGene00003926 | protein_coding_gene | -1.1 | 0.47 | 1.9E-03 |
| *E01A2.2* | WBGene00017085 | protein_coding_gene | -1.1 | 0.47 | 1.8E-03 |
| *isw-1* | WBGene00002169 | protein_coding_gene | -1.1 | 0.47 | 3.1E-04 |
| *ran-1* | WBGene00004302 | protein_coding_gene | -1.1 | 0.47 | 1.0E-02 |
| *pqn-82* | WBGene00004163 | protein_coding_gene | -1.1 | 0.47 | 1.1E-02 |
| *natb-1* | WBGene00022394 | protein_coding_gene | -1.1 | 0.47 | 1.2E-02 |
| *C05C10.2* | WBGene00007329 | protein_coding_gene | -1.1 | 0.47 | 7.1E-04 |
| *Y53F4B.9* | WBGene00013156 | protein_coding_gene | -1.1 | 0.47 | 1.4E-04 |
| *gsy-1* | WBGene00001793 | protein_coding_gene | -1.1 | 0.47 | 1.9E-05 |
| *T14G8.3* | WBGene00011771 | protein_coding_gene | -1.1 | 0.47 | 2.0E-02 |
| *sel-10* | WBGene00004767 | protein_coding_gene | -1.1 | 0.47 | 1.2E-02 |
| *algn-9* | WBGene00007556 | protein_coding_gene | -1.1 | 0.47 | 2.0E-02 |
| *hasp-1* | WBGene00007258 | protein_coding_gene | -1.1 | 0.47 | 9.7E-03 |
| *ubc-7* | WBGene00006704 | protein_coding_gene | -1.1 | 0.47 | 6.7E-05 |
| *icp-1* | WBGene00002047 | protein_coding_gene | -1.1 | 0.47 | 4.1E-03 |
| *C08F8.3* | WBGene00007445 | protein_coding_gene | -1.1 | 0.47 | 3.7E-02 |
| *lex-1* | WBGene00008682 | protein_coding_gene | -1.1 | 0.47 | 5.0E-03 |
| *cyp-33C8* | WBGene00019967 | protein_coding_gene | -1.1 | 0.47 | 3.8E-02 |
| *dna-2* | WBGene00001016 | protein_coding_gene | -1.1 | 0.47 | 4.4E-04 |
| *tag-175* | WBGene00006520 | protein_coding_gene | -1.1 | 0.47 | 4.6E-04 |
| *sur-2* | WBGene00006349 | protein_coding_gene | -1.1 | 0.47 | 3.1E-02 |
| *pas-4* | WBGene00003925 | protein_coding_gene | -1.1 | 0.47 | 1.3E-03 |
| *C11D2.4* | WBGene00015702 | protein_coding_gene | -1.1 | 0.47 | 1.8E-02 |
| *B0001.7* | WBGene00007092 | protein_coding_gene | -1.1 | 0.47 | 1.0E-02 |
| *M142.5* | WBGene00010922 | protein_coding_gene | -1.1 | 0.47 | 1.1E-03 |
| *cul-1* | WBGene00000836 | protein_coding_gene | -1.1 | 0.47 | 2.0E-05 |
| *rnp-8* | WBGene00020091 | protein_coding_gene | -1.1 | 0.47 | 3.1E-02 |

| | | | | | |
|---|---|---|---|---|---|
| *T07C4.3* | WBGene00011560 | protein_coding_gene | -1.1 | 0.47 | 7.3E-04 |
| *D1007.5* | WBGene00017003 | protein_coding_gene | -1.1 | 0.47 | 1.8E-02 |
| *pbs-4* | WBGene00003950 | protein_coding_gene | -1.1 | 0.47 | 1.2E-06 |
| *acl-8* | WBGene00020264 | protein_coding_gene | -1.1 | 0.47 | 1.4E-03 |
| *nra-2* | WBGene00011488 | protein_coding_gene | -1.1 | 0.47 | 9.2E-03 |
| *trap-3* | WBGene00021420 | protein_coding_gene | -1.1 | 0.47 | 9.1E-04 |
| *C08F8.9* | WBGene00007449 | protein_coding_gene | -1.1 | 0.47 | 2.9E-02 |
| *R13A5.7* | WBGene00020050 | protein_coding_gene | -1.1 | 0.47 | 1.2E-02 |
| *apc-2* | WBGene00000143 | protein_coding_gene | -1.1 | 0.47 | 2.3E-05 |
| *eral-1* | WBGene00008456 | protein_coding_gene | -1.1 | 0.47 | 7.5E-04 |
| *fan-1* | WBGene00015310 | protein_coding_gene | -1.1 | 0.47 | 2.2E-02 |
| *sao-1* | WBGene00011195 | protein_coding_gene | -1.1 | 0.47 | 6.9E-03 |
| *F56C9.6* | WBGene00018950 | protein_coding_gene | -1.1 | 0.47 | 3.2E-02 |
| *cgt-3* | WBGene00019127 | protein_coding_gene | -1.1 | 0.47 | 3.0E-02 |
| *pas-7* | WBGene00003928 | protein_coding_gene | -1.1 | 0.47 | 2.6E-05 |
| *B0205.9* | WBGene00015023 | protein_coding_gene | -1.1 | 0.47 | 2.5E-03 |
| *rpn-12* | WBGene00004468 | protein_coding_gene | -1.1 | 0.47 | 2.5E-03 |
| *piga-1* | WBGene00008431 | protein_coding_gene | -1.1 | 0.47 | 2.9E-02 |
| *ostb-1* | WBGene00011638 | protein_coding_gene | -1.1 | 0.47 | 1.3E-03 |
| *tcc-1* | WBGene00013340 | protein_coding_gene | -1.1 | 0.47 | 4.2E-02 |
| *algn-12* | WBGene00022629 | protein_coding_gene | -1.1 | 0.47 | 3.0E-02 |
| *nhr-12* | WBGene00003611 | protein_coding_gene | -1.1 | 0.47 | 2.0E-02 |
| *emb-8* | WBGene00001262 | protein_coding_gene | -1.1 | 0.47 | 3.4E-04 |
| *rbg-3* | WBGene00007167 | protein_coding_gene | -1.1 | 0.47 | 4.5E-02 |
| *dpl-1* | WBGene00001061 | protein_coding_gene | -1.1 | 0.47 | 5.7E-03 |
| *F13H10.3* | WBGene00008774 | protein_coding_gene | -1.1 | 0.47 | 6.8E-03 |
| *sly-1* | WBGene00009654 | protein_coding_gene | -1.1 | 0.46 | 1.2E-06 |
| *lin-3* | WBGene00002992 | protein_coding_gene | -1.1 | 0.46 | 4.5E-02 |
| *ent-6* | WBGene00009499 | protein_coding_gene | -1.1 | 0.46 | 5.5E-03 |
| *flap-1* | WBGene00018998 | protein_coding_gene | -1.1 | 0.46 | 1.0E-03 |
| *M110.3* | WBGene00010913 | protein_coding_gene | -1.1 | 0.46 | 2.6E-02 |
| *aptf-3* | WBGene00009203 | protein_coding_gene | -1.1 | 0.46 | 4.1E-02 |
| *tpp-2* | WBGene00017686 | protein_coding_gene | -1.1 | 0.46 | 1.8E-03 |
| *usp-14* | WBGene00006856 | protein_coding_gene | -1.1 | 0.46 | 1.2E-03 |
| *Y82E9BR.2* | WBGene00022335 | protein_coding_gene | -1.1 | 0.46 | 6.0E-03 |
| *C41G7.3* | WBGene00008061 | protein_coding_gene | -1.1 | 0.46 | 1.4E-02 |
| *npp-10* | WBGene00003796 | protein_coding_gene | -1.1 | 0.46 | 3.3E-06 |
| *eif-3.H* | WBGene00001231 | protein_coding_gene | -1.1 | 0.46 | 5.9E-06 |
| *Y53C10A.6* | WBGene00013137 | protein_coding_gene | -1.1 | 0.46 | 3.5E-02 |
| *C47E12.2* | WBGene00008147 | protein_coding_gene | -1.1 | 0.46 | 6.0E-03 |
| *ced-5* | WBGene00000419 | protein_coding_gene | -1.1 | 0.46 | 1.7E-03 |
| *D1065.6* | WBGene00268212 | ncRNA_gene | -1.1 | 0.46 | 2.5E-03 |
| *C40A11.8* | WBGene00016551 | protein_coding_gene | -1.1 | 0.46 | 4.3E-02 |
| *arid-1* | WBGene00044689 | protein_coding_gene | -1.1 | 0.46 | 3.8E-02 |
| *mtrr-1* | WBGene00006510 | protein_coding_gene | -1.1 | 0.46 | 2.9E-02 |
| *C33E10.8* | WBGene00016349 | protein_coding_gene | -1.1 | 0.46 | 2.4E-02 |
| *K08E5.1* | WBGene00014801 | protein_coding_gene | -1.1 | 0.46 | 2.8E-02 |
| *usp-50* | WBGene00008441 | protein_coding_gene | -1.1 | 0.46 | 1.4E-04 |
| *ced-3* | WBGene00000417 | protein_coding_gene | -1.1 | 0.46 | 3.4E-02 |
| *zwl-1* | WBGene00021460 | protein_coding_gene | -1.1 | 0.46 | 3.9E-04 |
| *sec-61.G* | WBGene00001303 | protein_coding_gene | -1.1 | 0.46 | 5.2E-03 |
| *bckd-1B* | WBGene00006518 | protein_coding_gene | -1.1 | 0.46 | 1.2E-03 |
| *dab-1* | WBGene00000894 | protein_coding_gene | -1.1 | 0.46 | 2.0E-04 |
| *C53H9.3* | WBGene00044746 | protein_coding_gene | -1.1 | 0.46 | 4.2E-02 |
| *F41G3.21* | WBGene00044316 | protein_coding_gene | -1.1 | 0.46 | 2.1E-02 |
| *rrf-3* | WBGene00004510 | protein_coding_gene | -1.1 | 0.46 | 9.5E-05 |
| *vha-7* | WBGene00006916 | protein_coding_gene | -1.1 | 0.46 | 1.4E-02 |
| *dnj-9* | WBGene00001027 | protein_coding_gene | -1.1 | 0.46 | 2.2E-07 |

| | | | | | |
|---|---|---|---|---|---|
| *dnj-13* | WBGene00001031 | protein_coding_gene | -1.1 | 0.46 | 4.0E-05 |
| *ubr-5* | WBGene00009460 | protein_coding_gene | -1.1 | 0.46 | 1.5E-03 |
| *C47E12.3* | WBGene00008148 | protein_coding_gene | -1.1 | 0.46 | 3.8E-03 |
| *perm-5* | WBGene00016955 | protein_coding_gene | -1.1 | 0.46 | 4.2E-02 |
| *T25B9.9* | WBGene00012015 | protein_coding_gene | -1.1 | 0.46 | 2.7E-02 |
| *mbf-1* | WBGene00003148 | protein_coding_gene | -1.1 | 0.46 | 8.2E-08 |
| *hsr-9* | WBGene00002027 | protein_coding_gene | -1.1 | 0.46 | 3.1E-03 |
| *F56C9.11* | WBGene00018954 | protein_coding_gene | -1.1 | 0.46 | 4.8E-02 |
| *Y54G11A.3* | WBGene00013214 | protein_coding_gene | -1.1 | 0.46 | 2.6E-08 |
| *mdf-1* | WBGene00003160 | protein_coding_gene | -1.1 | 0.46 | 1.0E-04 |
| *pcyt-1* | WBGene00017241 | protein_coding_gene | -1.1 | 0.46 | 2.2E-02 |
| *Y66D12A.16* | WBGene00013442 | protein_coding_gene | -1.1 | 0.46 | 1.2E-02 |
| *vps-35* | WBGene00006933 | protein_coding_gene | -1.1 | 0.46 | 7.6E-04 |
| *zen-4* | WBGene00006974 | protein_coding_gene | -1.1 | 0.46 | 7.2E-03 |
| *tba-1* | WBGene00006528 | protein_coding_gene | -1.1 | 0.46 | 6.4E-03 |
| *mans-4* | WBGene00013881 | protein_coding_gene | -1.1 | 0.46 | 3.5E-02 |
| *cids-1* | WBGene00015347 | protein_coding_gene | -1.1 | 0.46 | 4.5E-03 |
| *D2030.8* | WBGene00008418 | protein_coding_gene | -1.1 | 0.45 | 2.1E-02 |
| *sdha-2* | WBGene00016392 | protein_coding_gene | -1.1 | 0.45 | 1.2E-04 |
| *his-68* | WBGene00001942 | protein_coding_gene | -1.1 | 0.45 | 4.9E-02 |
| *rsp-5* | WBGene00004702 | protein_coding_gene | -1.1 | 0.45 | 4.2E-03 |
| *nspc-4* | WBGene00044339 | protein_coding_gene | -1.1 | 0.45 | 4.3E-02 |
| *gex-3* | WBGene00001580 | protein_coding_gene | -1.1 | 0.45 | 1.3E-03 |
| *mig-14* | WBGene00003246 | protein_coding_gene | -1.1 | 0.45 | 3.4E-02 |
| *coq-8* | WBGene00000767 | protein_coding_gene | -1.1 | 0.45 | 5.3E-04 |
| *imb-2* | WBGene00002076 | protein_coding_gene | -1.1 | 0.45 | 1.0E-02 |
| *rsa-1* | WBGene00007710 | protein_coding_gene | -1.1 | 0.45 | 4.8E-02 |
| *spds-1* | WBGene00012909 | protein_coding_gene | -1.1 | 0.45 | 1.5E-02 |
| *coq-5* | WBGene00000765 | protein_coding_gene | -1.1 | 0.45 | 1.3E-02 |
| *hut-1* | WBGene00013740 | protein_coding_gene | -1.1 | 0.45 | 2.0E-02 |
| *tag-63* | WBGene00006440 | protein_coding_gene | -1.1 | 0.45 | 1.5E-03 |
| *sptl-3* | WBGene00011932 | protein_coding_gene | -1.1 | 0.45 | 2.9E-02 |
| *cdkr-3* | WBGene00013765 | protein_coding_gene | -1.1 | 0.45 | 1.0E-04 |
| *wdr-20* | WBGene00007428 | protein_coding_gene | -1.1 | 0.45 | 7.2E-03 |
| *erfa-1* | WBGene00020269 | protein_coding_gene | -1.1 | 0.45 | 6.1E-05 |
| *fars-1* | WBGene00001497 | protein_coding_gene | -1.1 | 0.45 | 1.3E-05 |
| *ani-2* | WBGene00019608 | protein_coding_gene | -1.1 | 0.45 | 4.4E-03 |
| *chc-1* | WBGene00011867 | protein_coding_gene | -1.1 | 0.45 | 1.4E-03 |
| *eif-3.C* | WBGene00001226 | protein_coding_gene | -1.1 | 0.45 | 1.3E-11 |
| *hsp-90* | WBGene00000915 | protein_coding_gene | -1.2 | 0.45 | 1.5E-05 |
| *adm-4* | WBGene00000075 | protein_coding_gene | -1.2 | 0.45 | 4.5E-02 |
| *bre-5* | WBGene00000270 | protein_coding_gene | -1.2 | 0.45 | 5.4E-03 |
| *Y57A10A.1* | WBGene00013246 | protein_coding_gene | -1.2 | 0.45 | 3.2E-03 |
| *ints-6* | WBGene00000994 | protein_coding_gene | -1.2 | 0.45 | 6.9E-04 |
| *C06C3.7* | WBGene00007377 | protein_coding_gene | -1.2 | 0.45 | 5.0E-02 |
| *plk-1* | WBGene00004042 | protein_coding_gene | -1.2 | 0.45 | 2.2E-02 |
| *T21B10.3* | WBGene00011885 | protein_coding_gene | -1.2 | 0.45 | 1.2E-06 |
| *yop-1* | WBGene00022127 | protein_coding_gene | -1.2 | 0.45 | 9.3E-04 |
| *erfa-3* | WBGene00010405 | protein_coding_gene | -1.2 | 0.45 | 2.4E-03 |
| *C14B1.7* | WBGene00007578 | pseudogene | -1.2 | 0.45 | 5.0E-02 |
| *C13B4.1* | WBGene00007545 | protein_coding_gene | -1.2 | 0.45 | 1.7E-03 |
| *tre-2* | WBGene00006608 | protein_coding_gene | -1.2 | 0.45 | 4.7E-03 |
| *F56D5.3* | WBGene00010148 | protein_coding_gene | -1.2 | 0.45 | 2.3E-02 |
| *asb-1* | WBGene00000206 | protein_coding_gene | -1.2 | 0.45 | 4.3E-03 |
| *Y53C10A.1* | WBGene00014915 | pseudogene | -1.2 | 0.45 | 1.3E-02 |
| *C31H2.4* | WBGene00016294 | protein_coding_gene | -1.2 | 0.45 | 4.6E-02 |
| *pid-3* | WBGene00021270 | protein_coding_gene | -1.2 | 0.45 | 2.3E-04 |
| *npp-15* | WBGene00003801 | protein_coding_gene | -1.2 | 0.45 | 4.2E-09 |

| | | | | | |
|---|---|---|---|---|---|
| *nspc-7* | WBGene00015392 | protein_coding_gene | -1.2 | 0.45 | 3.7E-02 |
| *uba-1* | WBGene00006699 | protein_coding_gene | -1.2 | 0.45 | 9.7E-04 |
| *rbm-12* | WBGene00013703 | protein_coding_gene | -1.2 | 0.44 | 2.1E-04 |
| *gls-1* | WBGene00007975 | protein_coding_gene | -1.2 | 0.44 | 1.7E-02 |
| *slc-30A5* | WBGene00013668 | protein_coding_gene | -1.2 | 0.44 | 1.8E-03 |
| *ZK809.5* | WBGene00014087 | protein_coding_gene | -1.2 | 0.44 | 1.4E-02 |
| *fbxa-129* | WBGene00010294 | pseudogene | -1.2 | 0.44 | 1.2E-02 |
| *cpt-6* | WBGene00020911 | protein_coding_gene | -1.2 | 0.44 | 4.2E-02 |
| *xpo-2* | WBGene00002079 | protein_coding_gene | -1.2 | 0.44 | 6.7E-03 |
| *rpn-7* | WBGene00004463 | protein_coding_gene | -1.2 | 0.44 | 3.0E-05 |
| *C38D4.4* | WBGene00008005 | protein_coding_gene | -1.2 | 0.44 | 1.4E-03 |
| *hinf-1* | WBGene00009553 | protein_coding_gene | -1.2 | 0.44 | 4.9E-02 |
| *haf-6* | WBGene00001816 | protein_coding_gene | -1.2 | 0.44 | 2.4E-04 |
| *klp-19* | WBGene00002229 | protein_coding_gene | -1.2 | 0.44 | 1.4E-02 |
| *wago-2* | WBGene00018862 | pseudogene | -1.2 | 0.44 | 1.4E-02 |
| *manf-1* | WBGene00021888 | protein_coding_gene | -1.2 | 0.44 | 8.7E-03 |
| *pqbp-1.2* | WBGene00020295 | pseudogene | -1.2 | 0.44 | 3.7E-03 |
| *C48D1.5* | WBGene00008178 | protein_coding_gene | -1.2 | 0.44 | 3.0E-02 |
| *Y75B8A.24* | WBGene00013557 | protein_coding_gene | -1.2 | 0.44 | 1.2E-03 |
| *npp-5* | WBGene00003791 | protein_coding_gene | -1.2 | 0.44 | 1.9E-05 |
| *EGAP4.1* | WBGene00017149 | protein_coding_gene | -1.2 | 0.44 | 3.1E-02 |
| *R01H10.7* | WBGene00010975 | protein_coding_gene | -1.2 | 0.44 | 2.8E-02 |
| *BE0003N10.1* | WBGene00015266 | protein_coding_gene | -1.2 | 0.44 | 1.7E-03 |
| *trap-4* | WBGene00013238 | protein_coding_gene | -1.2 | 0.44 | 1.1E-03 |
| *F52B5.2* | WBGene00009921 | protein_coding_gene | -1.2 | 0.44 | 3.4E-02 |
| *mppa-1* | WBGene00022159 | protein_coding_gene | -1.2 | 0.44 | 1.1E-08 |
| *spe-39* | WBGene00004975 | protein_coding_gene | -1.2 | 0.44 | 1.8E-02 |
| *gst-1* | WBGene00001749 | protein_coding_gene | -1.2 | 0.44 | 1.2E-02 |
| *R06F6.8* | WBGene00011071 | protein_coding_gene | -1.2 | 0.44 | 2.3E-04 |
| *Y47G7B.2* | WBGene00021654 | protein_coding_gene | -1.2 | 0.44 | 6.4E-03 |
| *pgap-1* | WBGene00011835 | protein_coding_gene | -1.2 | 0.44 | 2.3E-02 |
| *spg-7* | WBGene00004978 | protein_coding_gene | -1.2 | 0.44 | 8.0E-13 |
| *ile-1* | WBGene00002070 | protein_coding_gene | -1.2 | 0.44 | 3.9E-03 |
| *msh-2* | WBGene00003418 | protein_coding_gene | -1.2 | 0.44 | 1.8E-06 |
| *C52B11.5* | WBGene00016874 | protein_coding_gene | -1.2 | 0.44 | 4.7E-02 |
| *rga-3* | WBGene00019600 | protein_coding_gene | -1.2 | 0.43 | 4.8E-02 |
| *let-711* | WBGene00002845 | protein_coding_gene | -1.2 | 0.43 | 3.1E-03 |
| *pyk-1* | WBGene00009126 | protein_coding_gene | -1.2 | 0.43 | 3.2E-05 |
| *athp-1* | WBGene00008081 | protein_coding_gene | -1.2 | 0.43 | 2.5E-04 |
| *oxa-1* | WBGene00007215 | protein_coding_gene | -1.2 | 0.43 | 1.3E-04 |
| *ulp-2* | WBGene00006737 | protein_coding_gene | -1.2 | 0.43 | 1.1E-03 |
| *Y76B12C.6* | WBGene00022300 | protein_coding_gene | -1.2 | 0.43 | 1.3E-02 |
| *K01D12.6* | WBGene00010465 | protein_coding_gene | -1.2 | 0.43 | 1.9E-03 |
| *mrp-7* | WBGene00003413 | protein_coding_gene | -1.2 | 0.43 | 2.9E-04 |
| *pup-2* | WBGene00019628 | protein_coding_gene | -1.2 | 0.43 | 4.0E-02 |
| *sqv-3* | WBGene00005021 | protein_coding_gene | -1.2 | 0.43 | 3.3E-02 |
| *tag-266* | WBGene00044319 | protein_coding_gene | -1.2 | 0.43 | 2.9E-04 |
| *eif-2alpha* | WBGene00021351 | protein_coding_gene | -1.2 | 0.43 | 2.1E-04 |
| *sdz-15* | WBGene00017752 | protein_coding_gene | -1.2 | 0.43 | 3.9E-02 |
| *brd-1* | WBGene00000265 | protein_coding_gene | -1.2 | 0.43 | 3.8E-06 |
| *F14D2.5* | WBGene00017453 | protein_coding_gene | -1.2 | 0.43 | 4.8E-02 |
| *rpt-3* | WBGene00004503 | protein_coding_gene | -1.2 | 0.43 | 1.1E-04 |
| *cri-3* | WBGene00010303 | protein_coding_gene | -1.2 | 0.43 | 3.2E-04 |
| *F48A11.4* | WBGene00018585 | protein_coding_gene | -1.2 | 0.43 | 8.4E-03 |
| *T03F6.3* | WBGene00011399 | protein_coding_gene | -1.2 | 0.43 | 3.9E-02 |
| *C05C10.3* | WBGene00007330 | protein_coding_gene | -1.2 | 0.43 | 3.8E-08 |
| *Y75B8A.16* | WBGene00013551 | protein_coding_gene | -1.2 | 0.43 | 1.9E-03 |
| *dnj-11* | WBGene00001029 | protein_coding_gene | -1.2 | 0.43 | 1.8E-03 |

| | | | | | |
|---|---|---|---|---|---|
| *Y94H6A.10* | WBGene00022382 | protein_coding_gene | -1.2 | 0.43 | 1.4E-02 |
| *laf-1* | WBGene00002244 | protein_coding_gene | -1.2 | 0.43 | 2.1E-06 |
| *eif-2Bepsilon* | WBGene00008428 | protein_coding_gene | -1.2 | 0.43 | 3.1E-05 |
| *trap-1* | WBGene00022122 | protein_coding_gene | -1.2 | 0.43 | 1.7E-03 |
| *vps-54* | WBGene00006934 | protein_coding_gene | -1.2 | 0.43 | 5.0E-05 |
| *B0393.6* | WBGene00007171 | protein_coding_gene | -1.2 | 0.43 | 2.9E-02 |
| *catp-6* | WBGene00012341 | protein_coding_gene | -1.2 | 0.43 | 1.5E-05 |
| *D1014.2* | WBGene00017015 | protein_coding_gene | -1.2 | 0.43 | 4.4E-03 |
| *deps-1* | WBGene00022034 | protein_coding_gene | -1.2 | 0.42 | 2.9E-02 |
| *set-21* | WBGene00021282 | protein_coding_gene | -1.2 | 0.42 | 1.4E-03 |
| *ufd-2* | WBGene00006734 | protein_coding_gene | -1.2 | 0.42 | 1.1E-04 |
| *W08F4.3* | WBGene00021093 | protein_coding_gene | -1.2 | 0.42 | 2.9E-02 |
| *cct-8* | WBGene00021934 | protein_coding_gene | -1.2 | 0.42 | 1.9E-05 |
| *rpn-9* | WBGene00004465 | protein_coding_gene | -1.2 | 0.42 | 9.5E-04 |
| *tbb-1* | WBGene00006536 | protein_coding_gene | -1.2 | 0.42 | 1.1E-02 |
| *linc-7* | WBGene00219609 | lincRNA_gene | -1.2 | 0.42 | 1.5E-03 |
| *gon-14* | WBGene00002148 | protein_coding_gene | -1.2 | 0.42 | 9.3E-04 |
| *Y48G1A.7* | WBGene00235385 | pseudogene | -1.2 | 0.42 | 4.8E-04 |
| *ztf-4* | WBGene00020399 | protein_coding_gene | -1.3 | 0.42 | 1.7E-04 |
| *hsp-1* | WBGene00002005 | protein_coding_gene | -1.3 | 0.42 | 6.1E-08 |
| *rpn-5* | WBGene00004461 | protein_coding_gene | -1.3 | 0.42 | 1.2E-06 |
| *ZK637.6* | WBGene00014987 | pseudogene | -1.3 | 0.42 | 3.4E-02 |
| *top-2* | WBGene00010785 | protein_coding_gene | -1.3 | 0.42 | 2.8E-04 |
| *D2023.6* | WBGene00008410 | protein_coding_gene | -1.3 | 0.42 | 1.0E-02 |
| *toc-1* | WBGene00006590 | protein_coding_gene | -1.3 | 0.42 | 1.9E-03 |
| *pid-5* | WBGene00021555 | protein_coding_gene | -1.3 | 0.42 | 5.0E-05 |
| *K11H3.4* | WBGene00010781 | protein_coding_gene | -1.3 | 0.42 | 5.4E-03 |
| *pbs-7* | WBGene00003953 | protein_coding_gene | -1.3 | 0.42 | 4.4E-04 |
| *mtp-18* | WBGene00077690 | protein_coding_gene | -1.3 | 0.42 | 4.8E-03 |
| *H05L14.2* | WBGene00010367 | protein_coding_gene | -1.3 | 0.42 | 1.3E-05 |
| *bath-19* | WBGene00019132 | protein_coding_gene | -1.3 | 0.42 | 4.8E-02 |
| *drsh-1* | WBGene00009163 | protein_coding_gene | -1.3 | 0.42 | 4.4E-02 |
| *ilcr-2* | WBGene00018960 | protein_coding_gene | -1.3 | 0.41 | 1.5E-03 |
| *dgat-2* | WBGene00010296 | protein_coding_gene | -1.3 | 0.41 | 2.0E-02 |
| *pam-1* | WBGene00003914 | protein_coding_gene | -1.3 | 0.41 | 1.2E-04 |
| *mig-5* | WBGene00003241 | protein_coding_gene | -1.3 | 0.41 | 2.2E-02 |
| *tfg-1* | WBGene00006565 | protein_coding_gene | -1.3 | 0.41 | 4.7E-05 |
| *ugt-64* | WBGene00015577 | protein_coding_gene | -1.3 | 0.41 | 8.4E-04 |
| *pgl-3* | WBGene00003994 | protein_coding_gene | -1.3 | 0.41 | 1.2E-02 |
| *cdc-48.1* | WBGene00007352 | protein_coding_gene | -1.3 | 0.41 | 2.7E-04 |
| *F13H8.8* | WBGene00017441 | protein_coding_gene | -1.3 | 0.41 | 2.0E-04 |
| *sucg-1* | WBGene00016844 | protein_coding_gene | -1.3 | 0.41 | 7.6E-04 |
| *capg-2* | WBGene00010093 | protein_coding_gene | -1.3 | 0.41 | 1.6E-02 |
| *acdh-13* | WBGene00001566 | protein_coding_gene | -1.3 | 0.41 | 1.5E-06 |
| *hsp-6* | WBGene00002010 | protein_coding_gene | -1.3 | 0.41 | 4.4E-07 |
| *F44E7.4* | WBGene00018426 | protein_coding_gene | -1.3 | 0.41 | 1.8E-10 |
| *sgt-1* | WBGene00019893 | protein_coding_gene | -1.3 | 0.41 | 3.8E-08 |
| *C34C12.8* | WBGene00007927 | protein_coding_gene | -1.3 | 0.41 | 8.5E-05 |
| *cpi-1* | WBGene00000535 | protein_coding_gene | -1.3 | 0.41 | 2.7E-02 |
| *F52B11.2* | WBGene00009925 | protein_coding_gene | -1.3 | 0.41 | 3.8E-04 |
| *Y49E10.37* | WBGene00304219 | ncRNA_gene | -1.3 | 0.40 | 6.5E-04 |
| *gld-3* | WBGene00001597 | protein_coding_gene | -1.3 | 0.40 | 2.2E-02 |
| *F28B3.5* | WBGene00017886 | protein_coding_gene | -1.3 | 0.40 | 5.8E-03 |
| *dnj-2* | WBGene00001020 | protein_coding_gene | -1.3 | 0.40 | 3.4E-03 |
| *cogc-2* | WBGene00000585 | protein_coding_gene | -1.3 | 0.40 | 5.2E-04 |
| *ife-5* | WBGene00002063 | protein_coding_gene | -1.3 | 0.40 | 1.1E-04 |
| *ugt-28* | WBGene00015693 | protein_coding_gene | -1.3 | 0.40 | 1.2E-06 |
| *gld-4* | WBGene00014115 | protein_coding_gene | -1.3 | 0.40 | 5.7E-03 |

| | | | | | |
|---|---|---|---|---|---|
| *zipt-2.4* | WBGene00018897 | protein_coding_gene | -1.3 | 0.40 | 5.6E-03 |
| *cdc-48.2* | WBGene00008053 | protein_coding_gene | -1.3 | 0.40 | 1.2E-11 |
| *K02C4.3* | WBGene00010502 | protein_coding_gene | -1.3 | 0.40 | 8.3E-03 |
| *sucl-2* | WBGene00017759 | protein_coding_gene | -1.3 | 0.40 | 1.2E-05 |
| *sti-1* | WBGene00019983 | protein_coding_gene | -1.3 | 0.40 | 5.3E-06 |
| *Y57A10A.31* | WBGene00013270 | protein_coding_gene | -1.3 | 0.40 | 1.4E-04 |
| *gla-3* | WBGene00011376 | protein_coding_gene | -1.3 | 0.40 | 9.7E-03 |
| *imb-1* | WBGene00002075 | protein_coding_gene | -1.3 | 0.40 | 1.5E-04 |
| *ints-11* | WBGene00008642 | protein_coding_gene | -1.3 | 0.40 | 4.5E-04 |
| *mel-32* | WBGene00003214 | protein_coding_gene | -1.3 | 0.39 | 7.6E-03 |
| *C16C10.1* | WBGene00007622 | protein_coding_gene | -1.3 | 0.39 | 7.9E-03 |
| *R07E3.4* | WBGene00011105 | protein_coding_gene | -1.3 | 0.39 | 2.8E-02 |
| *ptc-1* | WBGene00004208 | protein_coding_gene | -1.4 | 0.39 | 1.6E-03 |
| *ttx-7* | WBGene00008765 | protein_coding_gene | -1.4 | 0.39 | 9.3E-04 |
| *cyp-13A2* | WBGene00011676 | protein_coding_gene | -1.4 | 0.39 | 3.7E-02 |
| *rod-1* | WBGene00018900 | protein_coding_gene | -1.4 | 0.39 | 1.5E-05 |
| *C48B4.9* | WBGene00008172 | protein_coding_gene | -1.4 | 0.39 | 3.7E-05 |
| *C29F7.3* | WBGene00007812 | protein_coding_gene | -1.4 | 0.39 | 2.1E-02 |
| *ugt-16* | WBGene00013901 | protein_coding_gene | -1.4 | 0.39 | 2.0E-02 |
| *F52D2.12* | WBGene00219451 | protein_coding_gene | -1.4 | 0.38 | 2.9E-02 |
| *F49D11.3* | WBGene00018627 | protein_coding_gene | -1.4 | 0.38 | 4.9E-02 |
| *F08F1.21* | WBGene00304789 | ncRNA_gene | -1.4 | 0.38 | 2.9E-02 |
| *rpn-1* | WBGene00004458 | protein_coding_gene | -1.4 | 0.38 | 4.7E-06 |
| *mltn-2* | WBGene00013126 | protein_coding_gene | -1.4 | 0.38 | 4.2E-02 |
| *R03D7.2* | WBGene00010989 | protein_coding_gene | -1.4 | 0.38 | 2.8E-03 |
| *hrde-1* | WBGene00007624 | protein_coding_gene | -1.4 | 0.38 | 1.3E-02 |
| *Y46H3C.5* | WBGene00021605 | protein_coding_gene | -1.4 | 0.38 | 2.8E-02 |
| *pes-9* | WBGene00003982 | protein_coding_gene | -1.4 | 0.38 | 5.2E-05 |
| *gst-16* | WBGene00001764 | protein_coding_gene | -1.4 | 0.38 | 1.1E-02 |
| *rhy-1* | WBGene00012324 | protein_coding_gene | -1.4 | 0.38 | 9.2E-05 |
| *tbb-6* | WBGene00006539 | protein_coding_gene | -1.4 | 0.38 | 1.5E-02 |
| *acbp-6* | WBGene00012457 | protein_coding_gene | -1.4 | 0.38 | 3.0E-02 |
| *Y105C5B.15* | WBGene00013654 | protein_coding_gene | -1.4 | 0.37 | 4.8E-02 |
| *dnj-28* | WBGene00001046 | protein_coding_gene | -1.4 | 0.37 | 1.7E-04 |
| *Y46G5A.8* | WBGene00012900 | protein_coding_gene | -1.4 | 0.37 | 2.5E-02 |
| *pup-3* | WBGene00019082 | protein_coding_gene | -1.4 | 0.37 | 4.0E-02 |
| *mnat-1* | WBGene00018769 | protein_coding_gene | -1.4 | 0.37 | 5.1E-05 |
| *hcp-1* | WBGene00001829 | protein_coding_gene | -1.4 | 0.37 | 9.4E-04 |
| *pygl-1* | WBGene00020696 | protein_coding_gene | -1.4 | 0.37 | 1.8E-02 |
| *sqv-6* | WBGene00005024 | protein_coding_gene | -1.4 | 0.37 | 2.4E-03 |
| *Y50D4B.2* | WBGene00021741 | protein_coding_gene | -1.4 | 0.37 | 1.5E-03 |
| *cuc-1* | WBGene00000835 | protein_coding_gene | -1.4 | 0.37 | 3.3E-06 |
| *C25A11.10* | WBGene00219374 | pseudogene | -1.4 | 0.37 | 2.5E-02 |
| *tric-1B.2* | WBGene00013268 | protein_coding_gene | -1.5 | 0.36 | 7.7E-03 |
| *DC2.5* | WBGene00017082 | protein_coding_gene | -1.5 | 0.36 | 1.0E-02 |
| *F53G2.12* | WBGene00302998 | protein_coding_gene | -1.5 | 0.36 | 9.9E-04 |
| *sqv-5* | WBGene00005023 | protein_coding_gene | -1.5 | 0.36 | 2.7E-02 |
| *wago-1* | WBGene00011061 | protein_coding_gene | -1.5 | 0.36 | 5.9E-03 |
| *tsn-1* | WBGene00006626 | protein_coding_gene | -1.5 | 0.36 | 1.3E-03 |
| *har-1* | WBGene00007630 | protein_coding_gene | -1.5 | 0.36 | 3.2E-06 |
| *pgph-2* | WBGene00018424 | protein_coding_gene | -1.5 | 0.36 | 4.7E-02 |
| *T15B7.1* | WBGene00020516 | protein_coding_gene | -1.5 | 0.36 | 1.9E-02 |
| *F35E12.10* | WBGene00009434 | protein_coding_gene | -1.5 | 0.35 | 3.0E-03 |
| *ZK380.t2* | WBGene00023401 | pseudogene | -1.5 | 0.35 | 4.9E-03 |
| *ugt-26* | WBGene00015695 | protein_coding_gene | -1.5 | 0.35 | 1.3E-02 |
| *sec-61.A* | WBGene00013311 | protein_coding_gene | -1.5 | 0.35 | 1.4E-03 |
| *F54D5.12* | WBGene00010055 | protein_coding_gene | -1.5 | 0.35 | 6.8E-07 |
| *T08A9.6* | WBGene00020338 | pseudogene | -1.5 | 0.35 | 1.7E-02 |

| | | | | | |
|---|---|---|---|---|---|
| *F25B4.4* | WBGene00017767 | protein_coding_gene | -1.5 | 0.35 | 1.4E-02 |
| *fbxb-68* | WBGene00008611 | protein_coding_gene | -1.5 | 0.35 | 3.3E-02 |
| *fbxa-153* | WBGene00007162 | protein_coding_gene | -1.5 | 0.34 | 3.2E-03 |
| *gale-1* | WBGene00008132 | protein_coding_gene | -1.5 | 0.34 | 1.8E-04 |
| *ZC434.8* | WBGene00013894 | protein_coding_gene | -1.5 | 0.34 | 2.0E-03 |
| *F59H6.2* | WBGene00019133 | pseudogene | -1.6 | 0.34 | 3.1E-03 |
| *F58G11.3* | WBGene00010281 | protein_coding_gene | -1.6 | 0.34 | 9.6E-05 |
| *lips-12* | WBGene00020473 | protein_coding_gene | -1.6 | 0.34 | 4.6E-02 |
| *inf-1* | WBGene00002083 | protein_coding_gene | -1.6 | 0.34 | 4.2E-09 |
| *zipt-7.1* | WBGene00044067 | protein_coding_gene | -1.6 | 0.34 | 3.7E-03 |
| *ech-6* | WBGene00001155 | protein_coding_gene | -1.6 | 0.34 | 3.0E-03 |
| *drd-1* | WBGene00009902 | protein_coding_gene | -1.6 | 0.34 | 3.0E-02 |
| *fitm-2* | WBGene00044094 | protein_coding_gene | -1.6 | 0.34 | 1.8E-03 |
| *ZK228.3* | WBGene00013946 | protein_coding_gene | -1.6 | 0.34 | 2.5E-02 |
| *K02F6.7* | WBGene00019341 | protein_coding_gene | -1.6 | 0.34 | 3.5E-02 |
| *Y53C12A.6* | WBGene00013141 | protein_coding_gene | -1.6 | 0.33 | 2.9E-03 |
| *hsp-4* | WBGene00002008 | protein_coding_gene | -1.6 | 0.33 | 3.8E-06 |
| *alh-9* | WBGene00000115 | protein_coding_gene | -1.6 | 0.33 | 4.6E-04 |
| *nlp-24* | WBGene00003762 | protein_coding_gene | -1.6 | 0.33 | 4.9E-02 |
| *F53C11.1* | WBGene00009971 | protein_coding_gene | -1.6 | 0.33 | 1.2E-03 |
| *C06A1.12* | WBGene00219803 | ncRNA_gene | -1.6 | 0.33 | 5.2E-03 |
| *pks-1* | WBGene00016558 | protein_coding_gene | -1.6 | 0.33 | 4.2E-03 |
| *ZC449.8* | WBGene00195189 | protein_coding_gene | -1.6 | 0.33 | 1.4E-02 |
| *C45B2.2* | WBGene00016659 | protein_coding_gene | -1.6 | 0.33 | 2.1E-02 |
| *taf-11.2* | WBGene00006394 | protein_coding_gene | -1.6 | 0.33 | 2.4E-02 |
| *H34C03.2* | WBGene00019259 | protein_coding_gene | -1.6 | 0.33 | 2.1E-04 |
| *C14C11.2* | WBGene00015766 | protein_coding_gene | -1.6 | 0.33 | 1.3E-03 |
| *F53F1.2* | WBGene00009980 | protein_coding_gene | -1.6 | 0.32 | 3.6E-04 |
| *C40A11.7* | WBGene00016550 | protein_coding_gene | -1.6 | 0.32 | 2.5E-02 |
| *mrp-4* | WBGene00003410 | protein_coding_gene | -1.6 | 0.32 | 4.4E-02 |
| *T05B11.4* | WBGene00020247 | protein_coding_gene | -1.6 | 0.32 | 6.3E-03 |
| *fbxa-206* | WBGene00013610 | protein_coding_gene | -1.6 | 0.32 | 9.5E-05 |
| *ahcy-1* | WBGene00019322 | protein_coding_gene | -1.6 | 0.32 | 1.2E-06 |
| *F23F12.12* | WBGene00017756 | protein_coding_gene | -1.6 | 0.32 | 4.3E-03 |
| *acly-1* | WBGene00016995 | protein_coding_gene | -1.7 | 0.32 | 4.4E-03 |
| *Y48B6A.10* | WBGene00012981 | protein_coding_gene | -1.7 | 0.31 | 4.5E-03 |
| *sdz-33* | WBGene00013233 | protein_coding_gene | -1.7 | 0.31 | 4.1E-02 |
| *K10C2.12* | WBGene00206386 | protein_coding_gene | -1.7 | 0.31 | 1.4E-02 |
| *F44E5.15* | WBGene00206377 | protein_coding_gene | -1.7 | 0.31 | 8.1E-03 |
| *T20D4.7* | WBGene00020613 | protein_coding_gene | -1.7 | 0.31 | 2.1E-02 |
| *ZK899.5* | WBGene00014143 | protein_coding_gene | -1.7 | 0.31 | 2.9E-02 |
| *atad-3* | WBGene00010015 | protein_coding_gene | -1.7 | 0.31 | 2.3E-22 |
| *C52D10.3* | WBGene00016877 | protein_coding_gene | -1.7 | 0.30 | 2.1E-02 |
| *Y47G6A.33* | WBGene00045399 | protein_coding_gene | -1.7 | 0.30 | 3.7E-02 |
| *str-227* | WBGene00006258 | protein_coding_gene | -1.7 | 0.30 | 2.5E-02 |
| *hsp-110* | WBGene00016250 | protein_coding_gene | -1.7 | 0.30 | 8.6E-08 |
| *clu-1* | WBGene00000550 | protein_coding_gene | -1.7 | 0.30 | 2.9E-14 |
| *nhr-74* | WBGene00003664 | protein_coding_gene | -1.7 | 0.30 | 3.9E-02 |
| *Y49F6A.11* | WBGene00303366 | protein_coding_gene | -1.7 | 0.30 | 2.9E-02 |
| *Y24F12A.3* | WBGene00012498 | pseudogene | -1.7 | 0.30 | 1.2E-02 |
| *clec-245* | WBGene00007851 | protein_coding_gene | -1.8 | 0.30 | 2.6E-02 |
| *F35E2.8* | WBGene00009415 | protein_coding_gene | -1.8 | 0.30 | 4.9E-02 |
| *T24D5.11* | WBGene00219279 | pseudogene | -1.8 | 0.29 | 9.7E-03 |
| *dhs-26* | WBGene00000989 | protein_coding_gene | -1.8 | 0.29 | 3.0E-02 |
| *F53H4.2* | WBGene00010007 | protein_coding_gene | -1.8 | 0.29 | 1.3E-02 |
| *fbxc-30* | WBGene00045302 | protein_coding_gene | -1.8 | 0.29 | 3.6E-02 |
| *T05E12.10* | WBGene00077787 | ncRNA_gene | -1.8 | 0.29 | 3.2E-02 |
| *C24D10.5* | WBGene00016056 | protein_coding_gene | -1.8 | 0.28 | 3.5E-02 |

| | | | | | |
|---|---|---|---|---|---|
| *str-146* | WBGene00006193 | protein_coding_gene | -1.8 | 0.28 | 3.7E-02 |
| *K08H2.2* | WBGene00010689 | protein_coding_gene | -1.8 | 0.28 | 4.8E-02 |
| *T24E12.11* | WBGene00020778 | protein_coding_gene | -1.8 | 0.28 | 1.7E-02 |
| *T24D5.5* | WBGene00011993 | pseudogene | -1.8 | 0.28 | 3.0E-02 |
| *Y46D2A.2* | WBGene00021590 | protein_coding_gene | -1.8 | 0.28 | 2.1E-03 |
| *F12E12.6* | WBGene00017407 | protein_coding_gene | -1.8 | 0.28 | 2.8E-02 |
| *D2023.8* | WBGene00050970 | pseudogene | -1.8 | 0.28 | 2.9E-02 |
| *sec-61.B* | WBGene00021427 | protein_coding_gene | -1.8 | 0.28 | 2.0E-04 |
| *str-179* | WBGene00006223 | pseudogene | -1.9 | 0.28 | 3.8E-02 |
| *K12B6.11* | WBGene00195164 | protein_coding_gene | -1.9 | 0.28 | 1.0E-02 |
| *sdz-19* | WBGene00009728 | protein_coding_gene | -1.9 | 0.27 | 2.8E-02 |
| *Y38E10A.3* | WBGene00012581 | protein_coding_gene | -1.9 | 0.27 | 4.2E-02 |
| *T26E3.11* | WBGene00044905 | pseudogene | -1.9 | 0.27 | 7.5E-03 |
| *ceh-82* | WBGene00018433 | protein_coding_gene | -1.9 | 0.27 | 3.0E-02 |
| *fbxc-32* | WBGene00016303 | protein_coding_gene | -1.9 | 0.27 | 3.0E-02 |
| *W06D11.2* | WBGene00012300 | protein_coding_gene | -1.9 | 0.27 | 3.0E-02 |
| *W08F4.7* | WBGene00021096 | protein_coding_gene | -1.9 | 0.27 | 2.1E-03 |
| *Y82E9BL.2* | WBGene00022319 | protein_coding_gene | -1.9 | 0.26 | 3.1E-02 |
| *dop-4* | WBGene00016872 | protein_coding_gene | -1.9 | 0.26 | 3.8E-04 |
| *K08H2.3* | WBGene00010690 | protein_coding_gene | -1.9 | 0.26 | 2.3E-02 |
| *lbp-8* | WBGene00002260 | protein_coding_gene | -1.9 | 0.26 | 2.1E-02 |
| *T10C6.16* | WBGene00077607 | protein_coding_gene | -1.9 | 0.26 | 8.5E-03 |
| *ugt-62* | WBGene00010904 | protein_coding_gene | -1.9 | 0.26 | 2.9E-02 |
| *ZK105.1* | WBGene00022653 | protein_coding_gene | -1.9 | 0.26 | 5.4E-04 |
| *F17C11.13* | WBGene00023503 | protein_coding_gene | -2.0 | 0.26 | 9.8E-03 |
| *F56F4.8* | WBGene00045498 | protein_coding_gene | -2.0 | 0.25 | 4.5E-02 |
| *F02E9.8* | WBGene00014731 | pseudogene | -2.0 | 0.25 | 1.8E-02 |
| *T22B7.7* | WBGene00020674 | protein_coding_gene | -2.0 | 0.25 | 4.6E-02 |
| *R03G8.6* | WBGene00011000 | protein_coding_gene | -2.0 | 0.25 | 2.2E-02 |
| *dot-1.5* | WBGene00022512 | protein_coding_gene | -2.0 | 0.25 | 2.4E-02 |
| *F25C8.1* | WBGene00009104 | pseudogene | -2.0 | 0.25 | 3.0E-02 |
| *hach-1* | WBGene00017301 | protein_coding_gene | -2.0 | 0.25 | 2.0E-03 |
| *W04A4.2* | WBGene00012231 | protein_coding_gene | -2.0 | 0.25 | 4.5E-02 |
| *C54G6.8* | WBGene00236808 | protein_coding_gene | -2.0 | 0.25 | 2.7E-02 |
| *Y50E8A.11* | WBGene00013054 | protein_coding_gene | -2.0 | 0.25 | 4.1E-02 |
| *EEED8.4* | WBGene00017135 | protein_coding_gene | -2.0 | 0.25 | 3.3E-02 |
| *his-38* | WBGene00001912 | protein_coding_gene | -2.0 | 0.25 | 3.7E-03 |
| *K03B8.14* | WBGene00194783 | protein_coding_gene | -2.0 | 0.24 | 4.7E-02 |
| *clec-206* | WBGene00019089 | protein_coding_gene | -2.0 | 0.24 | 3.5E-02 |
| *W06A7.4* | WBGene00012293 | protein_coding_gene | -2.1 | 0.24 | 9.8E-05 |
| *fbxc-35* | WBGene00019930 | pseudogene | -2.1 | 0.24 | 4.1E-02 |
| *sir-2.3* | WBGene00004802 | protein_coding_gene | -2.1 | 0.24 | 4.3E-05 |
| *Y39A1A.16* | WBGene00012654 | protein_coding_gene | -2.1 | 0.24 | 9.1E-03 |
| *C09B8.4* | WBGene00015623 | protein_coding_gene | -2.1 | 0.24 | 2.7E-02 |
| *F38C2.7* | WBGene00009539 | protein_coding_gene | -2.1 | 0.24 | 2.5E-02 |
| *hsp-60* | WBGene00002025 | protein_coding_gene | -2.1 | 0.24 | 1.1E-06 |
| *Y62H9A.3* | WBGene00013391 | protein_coding_gene | -2.1 | 0.23 | 1.5E-02 |
| *K10C2.7* | WBGene00019622 | protein_coding_gene | -2.1 | 0.23 | 7.1E-03 |
| *C38D9.8* | WBGene00008016 | protein_coding_gene | -2.1 | 0.23 | 9.8E-03 |
| *sdz-32* | WBGene00021717 | protein_coding_gene | -2.1 | 0.23 | 2.0E-02 |
| *K10C2.8* | WBGene00045265 | protein_coding_gene | -2.2 | 0.22 | 2.2E-02 |
| *F22E5.1* | WBGene00017703 | protein_coding_gene | -2.2 | 0.22 | 1.1E-03 |
| *F35E2.7* | WBGene00009414 | pseudogene | -2.2 | 0.22 | 2.1E-02 |
| *tbx-38* | WBGene00006557 | protein_coding_gene | -2.2 | 0.22 | 1.1E-02 |
| *Y73B6BL.14* | WBGene00022237 | protein_coding_gene | -2.2 | 0.22 | 2.0E-04 |
| *nhr-235* | WBGene00012597 | protein_coding_gene | -2.2 | 0.22 | 3.7E-02 |
| *end-1* | WBGene00001310 | protein_coding_gene | -2.2 | 0.21 | 2.9E-04 |
| *scav-5* | WBGene00011076 | protein_coding_gene | -2.2 | 0.21 | 1.3E-03 |

| | | | | | |
|---|---|---|---|---|---|
| *F54D8.10* | WBGene00284847 | protein_coding_gene | -2.2 | 0.21 | 4.4E-02 |
| *col-101* | WBGene00000675 | protein_coding_gene | -2.2 | 0.21 | 3.1E-02 |
| *vit-3* | WBGene00006927 | protein_coding_gene | -2.2 | 0.21 | 3.2E-02 |
| *F35E12.2* | WBGene00009426 | protein_coding_gene | -2.3 | 0.21 | 2.5E-02 |
| *ugt-41* | WBGene00017336 | protein_coding_gene | -2.3 | 0.21 | 1.0E-02 |
| *Y32B12C.3* | WBGene00012526 | pseudogene | -2.3 | 0.21 | 2.4E-02 |
| *cest-28* | WBGene00019654 | protein_coding_gene | -2.3 | 0.21 | 1.9E-02 |
| *K02F2.11* | WBGene00220000 | ncRNA_gene | -2.3 | 0.20 | 1.1E-02 |
| *F18E3.11* | WBGene00017567 | protein_coding_gene | -2.3 | 0.20 | 2.5E-02 |
| *Y60A9.2* | WBGene00013369 | pseudogene | -2.3 | 0.20 | 2.8E-03 |
| *F02H6.4* | WBGene00008537 | pseudogene | -2.3 | 0.20 | 8.0E-03 |
| *ZK177.9* | WBGene00022674 | protein_coding_gene | -2.3 | 0.20 | 2.8E-02 |
| *D1086.3* | WBGene00008390 | protein_coding_gene | -2.3 | 0.20 | 1.1E-02 |
| *sri-55* | WBGene00005567 | pseudogene | -2.3 | 0.20 | 1.9E-02 |
| *ZK6.6* | WBGene00022641 | protein_coding_gene | -2.3 | 0.20 | 1.6E-03 |
| *lips-14* | WBGene00019208 | pseudogene | -2.3 | 0.20 | 1.6E-02 |
| *acdh-2* | WBGene00015894 | protein_coding_gene | -2.3 | 0.20 | 3.4E-02 |
| *Y48A6C.10* | WBGene00220168 | snoRNA_gene | -2.3 | 0.20 | 2.3E-02 |
| *F54E2.1* | WBGene00018823 | protein_coding_gene | -2.3 | 0.20 | 4.7E-06 |
| *H17B01.2* | WBGene00000646 | protein_coding_gene | -2.3 | 0.20 | 2.9E-02 |
| *oac-32* | WBGene00018708 | protein_coding_gene | -2.4 | 0.20 | 2.2E-03 |
| *sdz-35* | WBGene00022570 | protein_coding_gene | -2.4 | 0.20 | 1.2E-02 |
| *F18E3.13* | WBGene00045276 | protein_coding_gene | -2.4 | 0.19 | 1.3E-02 |
| *Y62H9A.4* | WBGene00013392 | protein_coding_gene | -2.4 | 0.19 | 2.3E-03 |
| *plep-1* | WBGene00021797 | protein_coding_gene | -2.4 | 0.18 | 6.8E-03 |
| *tbx-39* | WBGene00006558 | protein_coding_gene | -2.4 | 0.18 | 1.2E-02 |
| *K01G12.3* | WBGene00010483 | protein_coding_gene | -2.5 | 0.18 | 5.6E-03 |
| *cest-10* | WBGene00016595 | protein_coding_gene | -2.5 | 0.18 | 2.3E-03 |
| *Y73B6BL.288* | WBGene00219218 | protein_coding_gene | -2.5 | 0.17 | 2.1E-05 |
| *cest-7* | WBGene00007693 | protein_coding_gene | -2.5 | 0.17 | 1.6E-04 |
| *vit-4* | WBGene00006928 | protein_coding_gene | -2.5 | 0.17 | 1.3E-02 |
| *W10G11.3* | WBGene00021136 | protein_coding_gene | -2.6 | 0.17 | 2.0E-03 |
| *clec-22* | WBGene00009855 | protein_coding_gene | -2.7 | 0.16 | 6.0E-03 |
| *C33H5.1* | WBGene00016370 | protein_coding_gene | -2.7 | 0.16 | 1.2E-02 |
| *T10C6.9* | WBGene00011690 | pseudogene | -2.7 | 0.15 | 5.0E-03 |
| *F49F1.7* | WBGene00018647 | protein_coding_gene | -2.7 | 0.15 | 2.5E-05 |
| *vit-1* | WBGene00006925 | protein_coding_gene | -2.7 | 0.15 | 1.2E-02 |
| *folt-2* | WBGene00018138 | protein_coding_gene | -2.8 | 0.15 | 6.8E-03 |
| *clec-190* | WBGene00010953 | protein_coding_gene | -2.8 | 0.14 | 5.0E-03 |
| *hlh-26* | WBGene00001965 | protein_coding_gene | -2.8 | 0.14 | 3.4E-03 |
| *grd-4* | WBGene00001693 | protein_coding_gene | -2.9 | 0.13 | 1.8E-03 |
| *C14E2.12* | WBGene00219816 | snoRNA_gene | -3.0 | 0.13 | 2.1E-06 |
| *dhs-25* | WBGene00000988 | protein_coding_gene | -3.0 | 0.13 | 1.8E-04 |
| *C05D9.12* | WBGene00219800 | ncRNA_gene | -3.0 | 0.12 | 3.9E-04 |
| *arf-1.1* | WBGene00000190 | protein_coding_gene | -3.1 | 0.12 | 4.7E-08 |
| *grl-27* | WBGene00001736 | protein_coding_gene | -3.1 | 0.12 | 2.3E-03 |
| *sdz-20* | WBGene00009950 | protein_coding_gene | -3.1 | 0.11 | 7.9E-04 |
| *F19G12.t1* | WBGene00023271 | pseudogene | -3.2 | 0.11 | 1.8E-03 |
| *nas-27* | WBGene00003545 | protein_coding_gene | -3.3 | 0.10 | 2.2E-04 |
| *nspc-5* | WBGene00044340 | protein_coding_gene | -3.5 | 0.09 | 3.2E-04 |
| *R03G8.4* | WBGene00010999 | pseudogene | -3.7 | 0.08 | 1.5E-05 |
| *C04C3.14* | WBGene00219410 | protein_coding_gene | -3.7 | 0.08 | 6.6E-04 |
| *Y41C4A.32* | WBGene00270321 | protein_coding_gene | -3.7 | 0.08 | 5.3E-07 |
| *C07G1.7* | WBGene00015573 | protein_coding_gene | -3.8 | 0.07 | 4.9E-10 |
| *B0205.13* | WBGene00044644 | protein_coding_gene | -4.3 | 0.05 | 7.1E-10 |

**CB4856**-exclusive DEGs in response to Gram-positive pathogens (*S. epidermidis* or *S. aureus* vs. OP50)

| Gene Name | Wormbase Gene ID | Type | $log_2FC$ | FC | Adj P-value |
|---|---|---|---|---|---|

| | | | | | |
|---|---|---|---|---|---|
| *scl-6* | WBGene00008028 | protein_coding_gene | 5.0 | 31.19 | 2.1E-17 |
| *R193.2* | WBGene00020128 | protein_coding_gene | 3.9 | 15.00 | 3.3E-11 |
| *tag-293* | WBGene00015398 | protein_coding_gene | 3.8 | 13.79 | 5.4E-05 |
| *hrg-4* | WBGene00009493 | protein_coding_gene | 3.8 | 13.56 | 7.8E-10 |
| *Y47G6A.15* | WBGene00021641 | protein_coding_gene | 3.6 | 11.99 | 3.6E-06 |
| *cdh-7* | WBGene00000399 | protein_coding_gene | 3.5 | 11.33 | 8.9E-05 |
| *T28A11.4* | WBGene00020871 | protein_coding_gene | 3.3 | 10.05 | 3.2E-03 |
| *ins-30* | WBGene00002113 | protein_coding_gene | 3.3 | 10.02 | 1.8E-05 |
| *F14F9.2* | WBGene00017465 | protein_coding_gene | 3.3 | 9.81 | 3.1E-03 |
| *dpy-6* | WBGene00001068 | protein_coding_gene | 3.3 | 9.74 | 4.8E-05 |
| *rol-6* | WBGene00004397 | protein_coding_gene | 3.3 | 9.69 | 9.6E-05 |
| *dpy-5* | WBGene00001067 | protein_coding_gene | 3.3 | 9.63 | 9.8E-05 |
| *sqt-1* | WBGene00005016 | protein_coding_gene | 3.2 | 9.31 | 1.8E-04 |
| *rol-8* | WBGene00004398 | protein_coding_gene | 3.2 | 9.19 | 2.5E-04 |
| *C29F3.3* | WBGene00007804 | protein_coding_gene | 3.2 | 8.99 | 2.4E-03 |
| *col-107* | WBGene00000681 | protein_coding_gene | 3.1 | 8.56 | 8.4E-09 |
| *col-49* | WBGene00000626 | protein_coding_gene | 3.1 | 8.31 | 3.8E-04 |
| *C05D12.3* | WBGene00014666 | protein_coding_gene | 3.0 | 8.23 | 2.5E-08 |
| *K02E11.10* | WBGene00044109 | protein_coding_gene | 3.0 | 8.02 | 8.8E-04 |
| *rol-1* | WBGene00004394 | protein_coding_gene | 3.0 | 8.00 | 6.6E-04 |
| *sqt-2* | WBGene00005017 | protein_coding_gene | 3.0 | 7.90 | 7.2E-04 |
| *T16G12.1* | WBGene00011803 | protein_coding_gene | 3.0 | 7.88 | 6.0E-15 |
| *col-79* | WBGene00000655 | protein_coding_gene | 3.0 | 7.78 | 6.8E-04 |
| *dpy-8* | WBGene00001070 | protein_coding_gene | 2.9 | 7.69 | 9.8E-04 |
| *col-168* | WBGene00000741 | protein_coding_gene | 2.9 | 7.60 | 5.6E-05 |
| *E01G4.6* | WBGene00008448 | protein_coding_gene | 2.9 | 7.58 | 9.3E-04 |
| *ZK180.6* | WBGene00022680 | protein_coding_gene | 2.9 | 7.50 | 6.2E-04 |
| *col-38* | WBGene00000615 | protein_coding_gene | 2.9 | 7.48 | 5.0E-04 |
| *dpf-6* | WBGene00001059 | protein_coding_gene | 2.9 | 7.48 | 1.1E-04 |
| *dpy-4* | WBGene00001066 | protein_coding_gene | 2.9 | 7.48 | 1.2E-03 |
| *srx-130* | WBGene00006021 | protein_coding_gene | 2.9 | 7.40 | 1.0E-02 |
| *C12D8.13* | WBGene00007540 | protein_coding_gene | 2.9 | 7.37 | 3.1E-03 |
| *srh-60* | WBGene00005282 | protein_coding_gene | 2.9 | 7.36 | 8.2E-03 |
| *C02G6.1* | WBGene00015359 | pseudogene | 2.9 | 7.28 | 8.5E-04 |
| *col-138* | WBGene00000711 | protein_coding_gene | 2.8 | 7.20 | 1.2E-03 |
| *F28A10.9* | WBGene00017877 | pseudogene | 2.8 | 7.19 | 3.0E-03 |
| *dpy-13* | WBGene00001074 | protein_coding_gene | 2.8 | 7.17 | 1.3E-03 |
| *col-173* | WBGene00000746 | protein_coding_gene | 2.8 | 7.11 | 1.5E-03 |
| *ZK84.1* | WBGene00022649 | protein_coding_gene | 2.8 | 7.11 | 8.8E-06 |
| *fbxa-147* | WBGene00011135 | protein_coding_gene | 2.8 | 6.84 | 1.5E-02 |
| *pqn-11* | WBGene00004103 | pseudogene | 2.8 | 6.80 | 2.3E-03 |
| *hrg-7* | WBGene00007605 | protein_coding_gene | 2.8 | 6.75 | 8.0E-09 |
| *F49C5.11* | WBGene00044163 | protein_coding_gene | 2.7 | 6.71 | 5.2E-03 |
| *col-63* | WBGene00000639 | protein_coding_gene | 2.7 | 6.68 | 2.0E-03 |
| *col-175* | WBGene00000748 | protein_coding_gene | 2.7 | 6.62 | 1.9E-03 |
| *col-170* | WBGene00000743 | protein_coding_gene | 2.7 | 6.57 | 1.5E-03 |
| *Y51H7C.13* | WBGene00021791 | protein_coding_gene | 2.7 | 6.55 | 5.6E-04 |
| *dot-1.3* | WBGene00012302 | protein_coding_gene | 2.7 | 6.53 | 2.3E-03 |
| *alh-2* | WBGene00000108 | protein_coding_gene | 2.7 | 6.46 | 4.3E-03 |
| *col-169* | WBGene00000742 | protein_coding_gene | 2.7 | 6.43 | 2.6E-03 |
| *col-71* | WBGene00000647 | protein_coding_gene | 2.7 | 6.37 | 1.5E-03 |
| *col-125* | WBGene00000699 | protein_coding_gene | 2.6 | 6.26 | 4.2E-06 |
| *col-50* | WBGene00000627 | protein_coding_gene | 2.6 | 6.25 | 1.5E-02 |
| *ram-2* | WBGene00004300 | protein_coding_gene | 2.6 | 6.20 | 1.2E-06 |
| *dpy-9* | WBGene00001071 | protein_coding_gene | 2.6 | 6.19 | 2.5E-03 |
| *col-17* | WBGene00000606 | protein_coding_gene | 2.6 | 6.17 | 4.1E-03 |
| *dpy-20* | WBGene00001079 | protein_coding_gene | 2.6 | 6.05 | 7.9E-03 |
| *col-176* | WBGene00000749 | protein_coding_gene | 2.6 | 6.01 | 4.8E-03 |

| | | | | | |
|---|---|---|---|---|---|
| *R08H2.8* | WBGene00011149 | pseudogene | 2.6 | 5.98 | 2.4E-02 |
| *str-131* | WBGene00006180 | protein_coding_gene | 2.6 | 5.92 | 2.7E-03 |
| *col-109* | WBGene00000683 | protein_coding_gene | 2.6 | 5.90 | 1.2E-03 |
| *npr-30* | WBGene00019183 | protein_coding_gene | 2.6 | 5.89 | 7.4E-03 |
| *F14F8.8* | WBGene00008816 | protein_coding_gene | 2.5 | 5.81 | 4.1E-03 |
| *F18F11.4* | WBGene00017577 | protein_coding_gene | 2.5 | 5.74 | 7.5E-03 |
| *Y48G8AL.12* | WBGene00021690 | protein_coding_gene | 2.5 | 5.72 | 8.1E-03 |
| *dmd-11* | WBGene00007930 | protein_coding_gene | 2.5 | 5.68 | 1.4E-02 |
| *C38D9.2* | WBGene00008010 | protein_coding_gene | 2.5 | 5.64 | 1.2E-05 |
| *Y55F3AM.14* | WBGene00021931 | protein_coding_gene | 2.5 | 5.57 | 3.7E-03 |
| *F53B3.6* | WBGene00018744 | protein_coding_gene | 2.5 | 5.55 | 1.3E-03 |
| *oac-52* | WBGene00020977 | protein_coding_gene | 2.5 | 5.53 | 6.6E-03 |
| *Y95B8A.2* | WBGene00022383 | protein_coding_gene | 2.5 | 5.51 | 7.8E-04 |
| *col-97* | WBGene00000672 | protein_coding_gene | 2.5 | 5.48 | 1.8E-03 |
| *nhr-219* | WBGene00020555 | protein_coding_gene | 2.4 | 5.46 | 2.4E-03 |
| *gst-34* | WBGene00001782 | protein_coding_gene | 2.4 | 5.43 | 2.5E-02 |
| *sra-6* | WBGene00005032 | protein_coding_gene | 2.4 | 5.41 | 3.0E-02 |
| *Y116F11B.9* | WBGene00013825 | protein_coding_gene | 2.4 | 5.41 | 1.6E-02 |
| *T17H7.1* | WBGene00020550 | protein_coding_gene | 2.4 | 5.41 | 1.5E-03 |
| *T25C12.5* | WBGene00197733 | ncRNA_gene | 2.4 | 5.39 | 2.8E-02 |
| *pals-3* | WBGene00007657 | protein_coding_gene | 2.4 | 5.36 | 2.0E-02 |
| *C27D9.2* | WBGene00016164 | protein_coding_gene | 2.4 | 5.35 | 1.1E-02 |
| *C33D3.3* | WBGene00007894 | protein_coding_gene | 2.4 | 5.35 | 1.4E-03 |
| *ZK1290.7* | WBGene00022889 | protein_coding_gene | 2.4 | 5.34 | 1.5E-04 |
| *nep-22* | WBGene00020788 | protein_coding_gene | 2.4 | 5.33 | 2.4E-07 |
| *K06H6.2* | WBGene00019450 | protein_coding_gene | 2.4 | 5.32 | 1.7E-02 |
| *stdh-3* | WBGene00007364 | protein_coding_gene | 2.4 | 5.26 | 1.6E-02 |
| *F35F10.13* | WBGene00018062 | protein_coding_gene | 2.4 | 5.26 | 4.3E-03 |
| *B0244.7* | WBGene00015082 | protein_coding_gene | 2.4 | 5.24 | 4.0E-03 |
| *clec-167* | WBGene00009517 | protein_coding_gene | 2.4 | 5.22 | 1.4E-02 |
| *srsx-21* | WBGene00011084 | protein_coding_gene | 2.4 | 5.20 | 3.9E-02 |
| *T19D12.9* | WBGene00020583 | protein_coding_gene | 2.4 | 5.20 | 4.2E-03 |
| *ins-32* | WBGene00002115 | protein_coding_gene | 2.4 | 5.15 | 4.2E-03 |
| *F21G4.3* | WBGene00009024 | protein_coding_gene | 2.4 | 5.12 | 1.5E-02 |
| *T05E11.7* | WBGene00011483 | pseudogene | 2.3 | 5.02 | 1.5E-02 |
| *F12E12.11* | WBGene00044811 | protein_coding_gene | 2.3 | 5.01 | 8.9E-08 |
| *mpst-7* | WBGene00011307 | protein_coding_gene | 2.3 | 4.99 | 2.6E-03 |
| *dao-4* | WBGene00000930 | protein_coding_gene | 2.3 | 4.93 | 1.5E-02 |
| *Y77E11A.16* | WBGene00044464 | protein_coding_gene | 2.3 | 4.93 | 1.8E-02 |
| *ferl-1* | WBGene00020262 | protein_coding_gene | 2.3 | 4.92 | 5.2E-03 |
| *K09E2.1* | WBGene00019570 | protein_coding_gene | 2.3 | 4.91 | 1.2E-02 |
| *bli-6* | WBGene00000256 | protein_coding_gene | 2.3 | 4.86 | 9.3E-03 |
| *gly-1* | WBGene00001626 | protein_coding_gene | 2.3 | 4.86 | 4.3E-03 |
| *gst-41* | WBGene00001789 | protein_coding_gene | 2.3 | 4.86 | 9.8E-04 |
| *abu-11* | WBGene00000034 | protein_coding_gene | 2.3 | 4.85 | 1.1E-03 |
| *C07D8.2* | WBGene00015562 | pseudogene | 2.3 | 4.84 | 1.9E-02 |
| *Y71G12B.18* | WBGene00022156 | protein_coding_gene | 2.3 | 4.83 | 1.9E-02 |
| *C38D9.13* | WBGene00235090 | ncRNA_gene | 2.3 | 4.80 | 1.6E-02 |
| *F09B12.3* | WBGene00008607 | protein_coding_gene | 2.3 | 4.78 | 2.0E-04 |
| *C41H7.1* | WBGene00016571 | protein_coding_gene | 2.3 | 4.77 | 3.8E-05 |
| *sptl-2* | WBGene00018398 | protein_coding_gene | 2.3 | 4.76 | 8.2E-07 |
| *T20B6.1* | WBGene00020597 | protein_coding_gene | 2.3 | 4.76 | 3.1E-03 |
| *ncx-8* | WBGene00003573 | protein_coding_gene | 2.2 | 4.75 | 1.3E-05 |
| *C14A6.5* | WBGene00007569 | protein_coding_gene | 2.2 | 4.73 | 7.9E-03 |
| *col-167* | WBGene00000740 | protein_coding_gene | 2.2 | 4.68 | 2.7E-04 |
| *H41C03.2* | WBGene00019269 | protein_coding_gene | 2.2 | 4.67 | 3.4E-02 |
| *col-117* | WBGene00000691 | protein_coding_gene | 2.2 | 4.64 | 3.5E-02 |
| *col-110* | WBGene00000684 | protein_coding_gene | 2.2 | 4.62 | 7.2E-03 |

| | | | | | |
|---|---|---|---|---|---|
| *pho-1* | WBGene00004020 | protein_coding_gene | 2.2 | 4.61 | 1.1E-04 |
| *R07B1.13* | WBGene00138711 | protein_coding_gene | 2.2 | 4.60 | 1.2E-02 |
| *pmp-1* | WBGene00004058 | protein_coding_gene | 2.2 | 4.60 | 1.2E-03 |
| *glb-22* | WBGene00011251 | protein_coding_gene | 2.2 | 4.59 | 2.2E-03 |
| *T10B5.10* | WBGene00020396 | protein_coding_gene | 2.2 | 4.57 | 3.6E-04 |
| *gst-27* | WBGene00001775 | protein_coding_gene | 2.2 | 4.54 | 1.3E-03 |
| *oac-48* | WBGene00011960 | protein_coding_gene | 2.2 | 4.52 | 7.0E-05 |
| *F56H6.7* | WBGene00010168 | protein_coding_gene | 2.2 | 4.51 | 2.7E-02 |
| *K06H6.1* | WBGene00019449 | protein_coding_gene | 2.2 | 4.51 | 2.6E-02 |
| *C18B2.1* | WBGene00015952 | protein_coding_gene | 2.2 | 4.49 | 2.2E-02 |
| *R08F11.4* | WBGene00019968 | protein_coding_gene | 2.2 | 4.48 | 1.1E-03 |
| *F55F3.4* | WBGene00010118 | protein_coding_gene | 2.2 | 4.48 | 4.4E-03 |
| *T10E9.3* | WBGene00020413 | protein_coding_gene | 2.2 | 4.47 | 5.3E-05 |
| *ZK262.19* | WBGene00206508 | protein_coding_gene | 2.2 | 4.44 | 9.9E-03 |
| *C14C6.3* | WBGene00015757 | protein_coding_gene | 2.1 | 4.42 | 4.5E-02 |
| *Y47D3A.34* | WBGene00206461 | protein_coding_gene | 2.1 | 4.42 | 2.6E-02 |
| *K01D12.5* | WBGene00010464 | protein_coding_gene | 2.1 | 4.40 | 2.5E-02 |
| *C26B9.2* | WBGene00016132 | protein_coding_gene | 2.1 | 4.39 | 2.8E-02 |
| *nhr-155* | WBGene00015758 | protein_coding_gene | 2.1 | 4.38 | 3.9E-02 |
| *H12D21.2* | WBGene00010378 | protein_coding_gene | 2.1 | 4.38 | 4.5E-02 |
| *F34H10.4* | WBGene00009381 | protein_coding_gene | 2.1 | 4.38 | 1.7E-02 |
| *pcp-2* | WBGene00003957 | protein_coding_gene | 2.1 | 4.36 | 1.4E-06 |
| *lon-3* | WBGene00003057 | protein_coding_gene | 2.1 | 4.36 | 1.5E-02 |
| *T25B9.2* | WBGene00012008 | protein_coding_gene | 2.1 | 4.33 | 9.2E-03 |
| *F08D12.12* | WBGene00017255 | protein_coding_gene | 2.1 | 4.33 | 9.7E-03 |
| *col-61* | WBGene00000637 | protein_coding_gene | 2.1 | 4.32 | 1.8E-02 |
| *col-73* | WBGene00000649 | protein_coding_gene | 2.1 | 4.31 | 3.8E-02 |
| *gsto-1* | WBGene00016204 | protein_coding_gene | 2.1 | 4.30 | 9.8E-06 |
| *B0294.1* | WBGene00015117 | protein_coding_gene | 2.1 | 4.30 | 1.6E-03 |
| *wago-10* | WBGene00020707 | protein_coding_gene | 2.1 | 4.29 | 1.8E-04 |
| *nhr-157* | WBGene00015900 | protein_coding_gene | 2.1 | 4.27 | 2.5E-02 |
| *R02C2.8* | WBGene00255728 | pseudogene | 2.1 | 4.25 | 4.2E-04 |
| *R10D12.6* | WBGene00011187 | protein_coding_gene | 2.1 | 4.25 | 4.7E-02 |
| *H01G02.4* | WBGene00045108 | protein_coding_gene | 2.1 | 4.24 | 5.7E-03 |
| *oig-5* | WBGene00023520 | protein_coding_gene | 2.1 | 4.23 | 9.6E-05 |
| *col-77* | WBGene00000653 | protein_coding_gene | 2.1 | 4.20 | 8.9E-03 |
| *AC3.5* | WBGene00007071 | protein_coding_gene | 2.1 | 4.18 | 5.3E-03 |
| *col-147* | WBGene00000720 | protein_coding_gene | 2.1 | 4.16 | 1.1E-02 |
| *K08D8.2* | WBGene00010656 | protein_coding_gene | 2.1 | 4.15 | 3.0E-02 |
| *T12B5.9* | WBGene00023345 | pseudogene | 2.1 | 4.15 | 3.6E-02 |
| *K04G2.7* | WBGene00010566 | protein_coding_gene | 2.0 | 4.14 | 8.1E-03 |
| *ZK488.5* | WBGene00022757 | protein_coding_gene | 2.0 | 4.14 | 2.3E-02 |
| *wrt-4* | WBGene00006950 | protein_coding_gene | 2.0 | 4.13 | 9.4E-03 |
| *irld-2* | WBGene00008518 | protein_coding_gene | 2.0 | 4.12 | 1.5E-02 |
| *K03D7.3* | WBGene00014796 | pseudogene | 2.0 | 4.11 | 3.1E-02 |
| *T05A8.2* | WBGene00020228 | protein_coding_gene | 2.0 | 4.10 | 3.0E-02 |
| *daf-28* | WBGene00000920 | protein_coding_gene | 2.0 | 4.10 | 5.8E-08 |
| *T04C12.33* | WBGene00271444 | protein_coding_gene | 2.0 | 4.10 | 3.9E-02 |
| *pes-8* | WBGene00003981 | protein_coding_gene | 2.0 | 4.10 | 3.1E-03 |
| *Y45F10B.8* | WBGene00012872 | protein_coding_gene | 2.0 | 4.09 | 3.2E-02 |
| *acr-25* | WBGene00043066 | protein_coding_gene | 2.0 | 4.08 | 2.5E-04 |
| *K10C9.9* | WBGene00044515 | pseudogene | 2.0 | 4.07 | 4.5E-03 |
| *clx-1* | WBGene00000551 | protein_coding_gene | 2.0 | 4.05 | 1.5E-05 |
| *Y46G5A.29* | WBGene00012917 | protein_coding_gene | 2.0 | 4.02 | 2.5E-02 |
| *T21C9.9* | WBGene00011895 | protein_coding_gene | 2.0 | 4.01 | 2.1E-03 |
| *srh-230* | WBGene00005438 | pseudogene | 2.0 | 4.01 | 4.3E-02 |
| *C28H8.5* | WBGene00016196 | protein_coding_gene | 2.0 | 4.00 | 2.7E-02 |
| *E02C12.6* | WBGene00017092 | protein_coding_gene | 2.0 | 4.00 | 5.0E-04 |

| | | | | | |
|---|---|---|---|---|---|
| *C47A4.4* | WBGene00014703 | pseudogene | 2.0 | 4.00 | 4.7E-03 |
| *F02D8.1* | WBGene00008520 | protein_coding_gene | 2.0 | 3.99 | 4.1E-02 |
| *scav-4* | WBGene00008692 | protein_coding_gene | 2.0 | 3.97 | 4.1E-04 |
| *sre-40* | WBGene00020225 | protein_coding_gene | 2.0 | 3.97 | 4.0E-02 |
| *C53D6.5* | WBGene00008282 | pseudogene | 2.0 | 3.96 | 1.2E-02 |
| *nhr-207* | WBGene00011098 | protein_coding_gene | 2.0 | 3.95 | 2.6E-02 |
| *col-144* | WBGene00000717 | protein_coding_gene | 2.0 | 3.95 | 9.3E-03 |
| *frpr-10* | WBGene00019019 | protein_coding_gene | 2.0 | 3.95 | 8.4E-03 |
| *piit-1* | WBGene00009384 | protein_coding_gene | 2.0 | 3.95 | 7.4E-08 |
| *col-182* | WBGene00000755 | pseudogene | 2.0 | 3.94 | 2.1E-02 |
| *ZC250.4* | WBGene00045305 | protein_coding_gene | 2.0 | 3.93 | 6.2E-03 |
| *C49F5.8* | WBGene00008212 | protein_coding_gene | 2.0 | 3.92 | 3.0E-02 |
| *F19B6.3* | WBGene00008949 | protein_coding_gene | 2.0 | 3.91 | 1.0E-03 |
| *F42A8.1* | WBGene00009626 | protein_coding_gene | 2.0 | 3.91 | 6.1E-03 |
| *fbxa-125* | WBGene00015229 | protein_coding_gene | 2.0 | 3.89 | 8.4E-03 |
| *K12D9.1* | WBGene00019675 | protein_coding_gene | 2.0 | 3.89 | 4.3E-02 |
| *ZK470.6* | WBGene00022747 | protein_coding_gene | 2.0 | 3.88 | 2.1E-02 |
| *clpr-2* | WBGene00023410 | protein_coding_gene | 2.0 | 3.88 | 5.7E-03 |
| *oac-19* | WBGene00009413 | protein_coding_gene | 2.0 | 3.88 | 1.9E-03 |
| *col-39* | WBGene00000616 | protein_coding_gene | 2.0 | 3.87 | 1.5E-02 |
| *C07E3.4* | WBGene00007415 | protein_coding_gene | 2.0 | 3.86 | 1.7E-05 |
| *C34F6.1* | WBGene00007938 | protein_coding_gene | 1.9 | 3.85 | 1.4E-02 |
| *ZK783.6* | WBGene00044448 | protein_coding_gene | 1.9 | 3.85 | 8.6E-04 |
| *T25E4.2* | WBGene00020804 | protein_coding_gene | 1.9 | 3.85 | 3.3E-02 |
| *lin-29* | WBGene00003015 | protein_coding_gene | 1.9 | 3.85 | 1.2E-04 |
| *gcy-32* | WBGene00001552 | protein_coding_gene | 1.9 | 3.84 | 1.7E-03 |
| *C38C6.5* | WBGene00007998 | protein_coding_gene | 1.9 | 3.84 | 4.0E-03 |
| *nceh-1* | WBGene00012810 | protein_coding_gene | 1.9 | 3.83 | 4.0E-03 |
| *Y51H4A.22* | WBGene00013116 | protein_coding_gene | 1.9 | 3.83 | 1.2E-02 |
| *F58A4.12* | WBGene00014777 | pseudogene | 1.9 | 3.83 | 2.7E-02 |
| *col-14* | WBGene00000603 | protein_coding_gene | 1.9 | 3.82 | 2.1E-02 |
| *nhr-198* | WBGene00010603 | protein_coding_gene | 1.9 | 3.80 | 2.9E-02 |
| *hrg-1* | WBGene00019830 | protein_coding_gene | 1.9 | 3.80 | 2.4E-05 |
| *zipt-2.3* | WBGene00013207 | protein_coding_gene | 1.9 | 3.79 | 2.7E-03 |
| *Y71F9AM.8* | WBGene00045398 | protein_coding_gene | 1.9 | 3.79 | 3.2E-02 |
| *col-115* | WBGene00000689 | protein_coding_gene | 1.9 | 3.79 | 1.8E-02 |
| *F28H7.3* | WBGene00009237 | protein_coding_gene | 1.9 | 3.78 | 5.4E-06 |
| *clec-152* | WBGene00008658 | pseudogene | 1.9 | 3.78 | 2.3E-03 |
| *C04C3.6* | WBGene00015416 | protein_coding_gene | 1.9 | 3.78 | 4.2E-02 |
| *pgp-9* | WBGene00004003 | protein_coding_gene | 1.9 | 3.78 | 3.8E-05 |
| *basl-2* | WBGene00017392 | pseudogene | 1.9 | 3.77 | 3.5E-03 |
| *T05C1.2* | WBGene00020249 | protein_coding_gene | 1.9 | 3.77 | 1.6E-02 |
| *M01E5.1* | WBGene00010804 | pseudogene | 1.9 | 3.75 | 7.2E-03 |
| *F26D2.3* | WBGene00009148 | protein_coding_gene | 1.9 | 3.74 | 2.4E-02 |
| *Y51A2B.5* | WBGene00013069 | protein_coding_gene | 1.9 | 3.74 | 1.4E-02 |
| *Y39A3B.1* | WBGene00021436 | protein_coding_gene | 1.9 | 3.74 | 4.3E-02 |
| *clec-204* | WBGene00021603 | protein_coding_gene | 1.9 | 3.73 | 4.0E-03 |
| *col-141* | WBGene00000714 | protein_coding_gene | 1.9 | 3.69 | 3.8E-02 |
| *C06B3.16* | WBGene00189947 | protein_coding_gene | 1.9 | 3.68 | 2.6E-02 |
| *T25B6.6* | WBGene00020792 | protein_coding_gene | 1.9 | 3.68 | 3.7E-02 |
| *col-104* | WBGene00000678 | protein_coding_gene | 1.9 | 3.67 | 3.4E-03 |
| *C25G6.4* | WBGene00016109 | protein_coding_gene | 1.9 | 3.66 | 2.5E-02 |
| *F35A5.4* | WBGene00018026 | protein_coding_gene | 1.9 | 3.66 | 3.2E-02 |
| *nhr-234* | WBGene00012596 | protein_coding_gene | 1.9 | 3.66 | 2.5E-02 |
| *fbxa-84* | WBGene00011670 | protein_coding_gene | 1.9 | 3.66 | 4.5E-07 |
| *R08F11.7* | WBGene00019970 | protein_coding_gene | 1.9 | 3.65 | 2.6E-02 |
| *C30H6.5* | WBGene00007823 | protein_coding_gene | 1.9 | 3.65 | 3.0E-02 |
| *scl-3* | WBGene00009896 | protein_coding_gene | 1.9 | 3.64 | 1.6E-10 |

| | | | | | |
|---|---|---|---|---|---|
| *C15F1.2* | WBGene00015795 | protein_coding_gene | 1.9 | 3.64 | 3.2E-02 |
| *B0034.1* | WBGene00015002 | protein_coding_gene | 1.9 | 3.63 | 7.8E-03 |
| *fbxc-48* | WBGene00015615 | protein_coding_gene | 1.9 | 3.63 | 2.7E-03 |
| *R11G11.6* | WBGene00020014 | protein_coding_gene | 1.9 | 3.62 | 4.7E-02 |
| *try-4* | WBGene00006622 | protein_coding_gene | 1.9 | 3.62 | 2.8E-02 |
| *clec-230* | WBGene00007806 | protein_coding_gene | 1.9 | 3.61 | 3.0E-02 |
| *C30G12.10* | WBGene00284859 | ncRNA_gene | 1.9 | 3.61 | 3.0E-02 |
| *nas-30* | WBGene00003548 | protein_coding_gene | 1.9 | 3.61 | 2.7E-03 |
| *Y102A11A.3* | WBGene00022413 | protein_coding_gene | 1.9 | 3.61 | 7.8E-06 |
| *K08E4.7* | WBGene00045036 | protein_coding_gene | 1.9 | 3.61 | 6.9E-04 |
| *C11E4.8* | WBGene00007522 | protein_coding_gene | 1.9 | 3.61 | 2.1E-03 |
| *spe-49* | WBGene00019667 | protein_coding_gene | 1.8 | 3.60 | 3.3E-03 |
| *pax-2* | WBGene00003938 | protein_coding_gene | 1.8 | 3.60 | 4.4E-03 |
| *T02C12.4* | WBGene00011369 | pseudogene | 1.8 | 3.59 | 6.7E-03 |
| *cln-3.3* | WBGene00000541 | protein_coding_gene | 1.8 | 3.59 | 1.4E-04 |
| *C28F5.4* | WBGene00016185 | protein_coding_gene | 1.8 | 3.59 | 2.0E-02 |
| *C34H4.2* | WBGene00016425 | protein_coding_gene | 1.8 | 3.58 | 3.6E-06 |
| *nspb-12* | WBGene00017305 | protein_coding_gene | 1.8 | 3.58 | 4.7E-02 |
| *F15B9.11* | WBGene00138724 | protein_coding_gene | 1.8 | 3.58 | 2.6E-02 |
| *R07E4.1* | WBGene00019935 | protein_coding_gene | 1.8 | 3.57 | 4.5E-04 |
| *nhr-31* | WBGene00003625 | protein_coding_gene | 1.8 | 3.57 | 9.0E-04 |
| *bus-1* | WBGene00044617 | protein_coding_gene | 1.8 | 3.56 | 2.0E-02 |
| *Y102A11A.7* | WBGene00022417 | protein_coding_gene | 1.8 | 3.56 | 4.1E-03 |
| *ins-27* | WBGene00002110 | protein_coding_gene | 1.8 | 3.55 | 3.2E-03 |
| *fipr-6* | WBGene00007544 | protein_coding_gene | 1.8 | 3.55 | 2.5E-02 |
| *W07A12.8* | WBGene00012325 | protein_coding_gene | 1.8 | 3.55 | 2.5E-02 |
| *Y106G6E.1* | WBGene00013705 | protein_coding_gene | 1.8 | 3.55 | 2.8E-02 |
| *col-130* | WBGene00000704 | protein_coding_gene | 1.8 | 3.55 | 2.2E-02 |
| *lec-8* | WBGene00002271 | protein_coding_gene | 1.8 | 3.54 | 2.8E-09 |
| *F15H9.1* | WBGene00008870 | protein_coding_gene | 1.8 | 3.53 | 1.1E-02 |
| *K09D9.11* | WBGene00019568 | pseudogene | 1.8 | 3.53 | 7.4E-03 |
| *pqn-29* | WBGene00004118 | protein_coding_gene | 1.8 | 3.51 | 3.3E-02 |
| *F46A8.9* | WBGene00009752 | pseudogene | 1.8 | 3.51 | 2.6E-02 |
| *fbxa-223* | WBGene00022630 | pseudogene | 1.8 | 3.51 | 7.0E-03 |
| *F23B2.10* | WBGene00009077 | protein_coding_gene | 1.8 | 3.50 | 2.6E-02 |
| *snf-8* | WBGene00004907 | pseudogene | 1.8 | 3.49 | 4.7E-03 |
| *gfi-1* | WBGene00001581 | protein_coding_gene | 1.8 | 3.49 | 3.1E-03 |
| *B0507.8* | WBGene00015225 | protein_coding_gene | 1.8 | 3.49 | 8.2E-03 |
| *dat-1* | WBGene00000934 | protein_coding_gene | 1.8 | 3.48 | 1.8E-03 |
| *cyp-34A5* | WBGene00015040 | protein_coding_gene | 1.8 | 3.48 | 3.1E-03 |
| *grd-10* | WBGene00001699 | protein_coding_gene | 1.8 | 3.47 | 2.6E-02 |
| *C49A9.5* | WBGene00016759 | protein_coding_gene | 1.8 | 3.47 | 2.0E-06 |
| *mec-4* | WBGene00003168 | protein_coding_gene | 1.8 | 3.46 | 8.8E-03 |
| *R12E2.7* | WBGene00020033 | protein_coding_gene | 1.8 | 3.45 | 2.9E-02 |
| *nhr-93* | WBGene00003683 | protein_coding_gene | 1.8 | 3.45 | 1.7E-02 |
| *C16E9.1* | WBGene00015865 | protein_coding_gene | 1.8 | 3.45 | 4.1E-02 |
| *elo-2* | WBGene00001240 | protein_coding_gene | 1.8 | 3.44 | 3.1E-03 |
| *K07A1.20* | WBGene00219285 | protein_coding_gene | 1.8 | 3.44 | 1.2E-02 |
| *ctsa-2* | WBGene00019617 | protein_coding_gene | 1.8 | 3.43 | 6.5E-03 |
| *lgc-40* | WBGene00020767 | protein_coding_gene | 1.8 | 3.43 | 2.0E-03 |
| *AH9.6* | WBGene00044701 | protein_coding_gene | 1.8 | 3.43 | 2.8E-04 |
| *clec-205* | WBGene00021602 | protein_coding_gene | 1.8 | 3.42 | 7.9E-03 |
| *str-44* | WBGene00006109 | protein_coding_gene | 1.8 | 3.41 | 3.5E-02 |
| *F52D10.4* | WBGene00009931 | protein_coding_gene | 1.8 | 3.41 | 8.9E-03 |
| *ZK896.1* | WBGene00014132 | protein_coding_gene | 1.8 | 3.41 | 5.7E-05 |
| *M195.2* | WBGene00010948 | protein_coding_gene | 1.8 | 3.41 | 3.6E-02 |
| *C39B10.5* | WBGene00008025 | protein_coding_gene | 1.8 | 3.40 | 4.0E-02 |
| *ZK1086.2* | WBGene00014216 | protein_coding_gene | 1.8 | 3.40 | 2.6E-02 |

| | | | | | |
|---|---|---|---|---|---|
| *gpa-6* | WBGene00001668 | protein_coding_gene | 1.8 | 3.40 | 4.4E-02 |
| *col-157* | WBGene00000730 | protein_coding_gene | 1.8 | 3.40 | 2.0E-02 |
| *fer-1* | WBGene00001414 | protein_coding_gene | 1.8 | 3.39 | 1.7E-04 |
| *C15C7.6* | WBGene00015792 | protein_coding_gene | 1.8 | 3.39 | 2.8E-02 |
| *F09B9.5* | WBGene00008604 | protein_coding_gene | 1.8 | 3.38 | 3.9E-03 |
| *cyp-35A4* | WBGene00016786 | protein_coding_gene | 1.8 | 3.38 | 4.4E-02 |
| *fat-5* | WBGene00001397 | protein_coding_gene | 1.8 | 3.37 | 5.0E-02 |
| *oac-3* | WBGene00007361 | protein_coding_gene | 1.8 | 3.37 | 3.4E-02 |
| *F22H10.1* | WBGene00017725 | protein_coding_gene | 1.8 | 3.37 | 4.8E-02 |
| *col-154* | WBGene00000727 | protein_coding_gene | 1.7 | 3.36 | 2.2E-02 |
| *slc-17.3* | WBGene00015333 | protein_coding_gene | 1.7 | 3.36 | 1.3E-03 |
| *F46F5.2* | WBGene00018494 | pseudogene | 1.7 | 3.36 | 2.1E-02 |
| *F57F4.4* | WBGene00019017 | protein_coding_gene | 1.7 | 3.35 | 3.4E-03 |
| *sprr-1* | WBGene00010986 | protein_coding_gene | 1.7 | 3.35 | 2.3E-03 |
| *mtd-1* | WBGene00003471 | protein_coding_gene | 1.7 | 3.35 | 6.7E-04 |
| *C14A6.10* | WBGene00045348 | pseudogene | 1.7 | 3.35 | 3.5E-02 |
| *Y47D3B.6* | WBGene00012942 | protein_coding_gene | 1.7 | 3.34 | 2.9E-02 |
| *snf-4* | WBGene00004903 | pseudogene | 1.7 | 3.33 | 9.9E-04 |
| *C27A2.5* | WBGene00016153 | protein_coding_gene | 1.7 | 3.33 | 4.0E-02 |
| *paic-1* | WBGene00015116 | protein_coding_gene | 1.7 | 3.33 | 6.6E-03 |
| *fbxa-196* | WBGene00020265 | protein_coding_gene | 1.7 | 3.32 | 7.0E-04 |
| *mpst-4* | WBGene00017387 | protein_coding_gene | 1.7 | 3.32 | 7.0E-03 |
| *F26C11.1* | WBGene00009146 | protein_coding_gene | 1.7 | 3.32 | 5.1E-03 |
| *C05C8.8* | WBGene00015465 | protein_coding_gene | 1.7 | 3.32 | 2.4E-02 |
| *lgc-49* | WBGene00010741 | protein_coding_gene | 1.7 | 3.32 | 2.1E-02 |
| *T23E1.1* | WBGene00020728 | protein_coding_gene | 1.7 | 3.31 | 2.5E-02 |
| *F38B6.4* | WBGene00018174 | protein_coding_gene | 1.7 | 3.31 | 8.7E-03 |
| *glb-7* | WBGene00016024 | protein_coding_gene | 1.7 | 3.31 | 5.3E-03 |
| *col-48* | WBGene00000625 | protein_coding_gene | 1.7 | 3.31 | 3.6E-02 |
| *fbxa-8* | WBGene00016526 | pseudogene | 1.7 | 3.31 | 4.2E-02 |
| *str-168* | WBGene00006212 | protein_coding_gene | 1.7 | 3.30 | 4.2E-02 |
| *C30A5.4* | WBGene00016239 | protein_coding_gene | 1.7 | 3.29 | 1.1E-02 |
| *F13H8.5* | WBGene00017438 | protein_coding_gene | 1.7 | 3.29 | 4.0E-03 |
| *nhr-30* | WBGene00016091 | protein_coding_gene | 1.7 | 3.28 | 1.9E-02 |
| *lido-16* | WBGene00020898 | protein_coding_gene | 1.7 | 3.28 | 1.4E-02 |
| *F55D10.4* | WBGene00018879 | protein_coding_gene | 1.7 | 3.28 | 2.2E-03 |
| *C08G5.6* | WBGene00015618 | protein_coding_gene | 1.7 | 3.28 | 2.3E-02 |
| *R07C3.13* | WBGene00019929 | protein_coding_gene | 1.7 | 3.28 | 3.9E-03 |
| *gcy-3* | WBGene00001530 | protein_coding_gene | 1.7 | 3.28 | 4.1E-02 |
| *T23B7.2* | WBGene00044298 | protein_coding_gene | 1.7 | 3.27 | 3.1E-02 |
| *C06C6.6* | WBGene00007380 | pseudogene | 1.7 | 3.27 | 2.1E-03 |
| *R02E4.2* | WBGene00044021 | protein_coding_gene | 1.7 | 3.27 | 2.0E-03 |
| *F58G1.9* | WBGene00010271 | pseudogene | 1.7 | 3.26 | 1.3E-02 |
| *elo-5* | WBGene00001243 | protein_coding_gene | 1.7 | 3.25 | 8.3E-03 |
| *Y73B3A.18* | WBGene00022219 | pseudogene | 1.7 | 3.25 | 3.9E-04 |
| *lev-9* | WBGene00002976 | protein_coding_gene | 1.7 | 3.24 | 7.3E-04 |
| *fbxa-195* | WBGene00019976 | pseudogene | 1.7 | 3.24 | 3.4E-02 |
| *math-2* | WBGene00015606 | protein_coding_gene | 1.7 | 3.24 | 7.1E-04 |
| *Y53G8AR.7* | WBGene00021814 | protein_coding_gene | 1.7 | 3.23 | 6.2E-03 |
| *pgp-12* | WBGene00004006 | protein_coding_gene | 1.7 | 3.23 | 1.9E-03 |
| *T23B12.5* | WBGene00020720 | protein_coding_gene | 1.7 | 3.22 | 2.6E-02 |
| *fbxa-15* | WBGene00022327 | protein_coding_gene | 1.7 | 3.21 | 2.9E-02 |
| *lin-31* | WBGene00003017 | protein_coding_gene | 1.7 | 3.21 | 5.0E-02 |
| *klu-2* | WBGene00022592 | protein_coding_gene | 1.7 | 3.20 | 4.5E-07 |
| *C47E8.10* | WBGene00044216 | pseudogene | 1.7 | 3.20 | 5.0E-03 |
| *hsp-16.41* | WBGene00002018 | protein_coding_gene | 1.7 | 3.19 | 3.4E-02 |
| *pcp-3* | WBGene00003958 | protein_coding_gene | 1.7 | 3.19 | 9.3E-04 |
| *msp-64* | WBGene00003457 | protein_coding_gene | 1.7 | 3.18 | 1.3E-02 |

| | | | | | |
|---|---|---|---|---|---|
| *scl-10* | WBGene00009891 | protein_coding_gene | 1.7 | 3.18 | 4.1E-02 |
| *ZC373.3* | WBGene00013868 | protein_coding_gene | 1.7 | 3.18 | 3.3E-02 |
| *F44E7.3* | WBGene00018425 | protein_coding_gene | 1.7 | 3.18 | 2.2E-02 |
| *cyp-33C5* | WBGene00018261 | protein_coding_gene | 1.7 | 3.18 | 3.3E-04 |
| *fbxa-60* | WBGene00020456 | protein_coding_gene | 1.7 | 3.18 | 2.0E-04 |
| *unc-47* | WBGene00006783 | protein_coding_gene | 1.7 | 3.18 | 1.8E-03 |
| *C02G6.2* | WBGene00015360 | protein_coding_gene | 1.7 | 3.17 | 9.2E-04 |
| *Y116A8A.4* | WBGene00013775 | protein_coding_gene | 1.7 | 3.17 | 1.3E-02 |
| *C01G12.7* | WBGene00007247 | protein_coding_gene | 1.7 | 3.16 | 3.2E-02 |
| *gcsh-1* | WBGene00008354 | protein_coding_gene | 1.7 | 3.16 | 3.9E-04 |
| *F33A8.10* | WBGene00009356 | protein_coding_gene | 1.7 | 3.16 | 2.8E-02 |
| *T05H4.7* | WBGene00020270 | protein_coding_gene | 1.7 | 3.16 | 3.8E-03 |
| *C09F12.2* | WBGene00007482 | protein_coding_gene | 1.7 | 3.15 | 1.1E-03 |
| *C49A9.4* | WBGene00016758 | protein_coding_gene | 1.7 | 3.15 | 1.4E-04 |
| *C47E8.9* | WBGene00008146 | protein_coding_gene | 1.7 | 3.15 | 3.3E-02 |
| *Y45F10D.2* | WBGene00012883 | protein_coding_gene | 1.7 | 3.15 | 3.5E-02 |
| *Y48G1BL.7* | WBGene00021666 | protein_coding_gene | 1.7 | 3.14 | 1.2E-02 |
| *T09B4.7* | WBGene00020381 | protein_coding_gene | 1.7 | 3.14 | 9.1E-03 |
| *Y50D4B.9* | WBGene00268189 | pseudogene | 1.6 | 3.14 | 4.3E-02 |
| *F31A3.3* | WBGene00017938 | protein_coding_gene | 1.6 | 3.14 | 3.9E-02 |
| *ZC449.7* | WBGene00045365 | protein_coding_gene | 1.6 | 3.13 | 5.3E-05 |
| *fbxa-183* | WBGene00018428 | protein_coding_gene | 1.6 | 3.13 | 2.5E-02 |
| *gmeb-3* | WBGene00008092 | protein_coding_gene | 1.6 | 3.12 | 1.1E-03 |
| *T05A1.5* | WBGene00011456 | protein_coding_gene | 1.6 | 3.12 | 1.4E-03 |
| *B0393.4* | WBGene00007169 | protein_coding_gene | 1.6 | 3.12 | 1.4E-03 |
| *C10A4.9* | WBGene00045456 | protein_coding_gene | 1.6 | 3.11 | 5.2E-03 |
| *W03A5.1* | WBGene00020966 | protein_coding_gene | 1.6 | 3.11 | 5.2E-04 |
| *T05F1.7* | WBGene00011492 | protein_coding_gene | 1.6 | 3.11 | 1.3E-02 |
| *Y37E11AL.1* | WBGene00021357 | protein_coding_gene | 1.6 | 3.11 | 1.4E-02 |
| *T20H9.6* | WBGene00020642 | protein_coding_gene | 1.6 | 3.11 | 3.4E-02 |
| *F14D7.12* | WBGene00045331 | protein_coding_gene | 1.6 | 3.10 | 3.9E-02 |
| *B0207.7* | WBGene00015030 | protein_coding_gene | 1.6 | 3.10 | 2.6E-05 |
| *elo-6* | WBGene00001244 | protein_coding_gene | 1.6 | 3.10 | 2.4E-02 |
| *ZK616.65* | WBGene00235383 | protein_coding_gene | 1.6 | 3.10 | 2.2E-04 |
| *R08C7.5* | WBGene00019949 | protein_coding_gene | 1.6 | 3.08 | 2.4E-05 |
| *ser-6* | WBGene00021897 | protein_coding_gene | 1.6 | 3.08 | 7.9E-03 |
| *grsp-4* | WBGene00017201 | protein_coding_gene | 1.6 | 3.07 | 2.0E-02 |
| *T03F6.8* | WBGene00044229 | pseudogene | 1.6 | 3.07 | 1.1E-02 |
| *D1065.3* | WBGene00017035 | pseudogene | 1.6 | 3.07 | 5.3E-03 |
| *B0457.3* | WBGene00014660 | pseudogene | 1.6 | 3.07 | 1.8E-02 |
| *F26A1.4* | WBGene00017803 | protein_coding_gene | 1.6 | 3.07 | 6.8E-04 |
| *T08D2.7* | WBGene00011612 | protein_coding_gene | 1.6 | 3.07 | 9.8E-03 |
| *T09F5.10* | WBGene00011669 | protein_coding_gene | 1.6 | 3.07 | 2.2E-02 |
| *ZK783.3* | WBGene00022818 | pseudogene | 1.6 | 3.06 | 2.4E-03 |
| *hsp-17* | WBGene00002021 | protein_coding_gene | 1.6 | 3.06 | 2.9E-08 |
| *acs-23* | WBGene00011174 | pseudogene | 1.6 | 3.06 | 9.3E-03 |
| *F22E5.8* | WBGene00017707 | protein_coding_gene | 1.6 | 3.06 | 7.2E-05 |
| *C45G9.8* | WBGene00016679 | protein_coding_gene | 1.6 | 3.06 | 7.6E-03 |
| *B0244.10* | WBGene00015085 | protein_coding_gene | 1.6 | 3.05 | 1.4E-03 |
| *col-120* | WBGene00000694 | protein_coding_gene | 1.6 | 3.05 | 4.3E-02 |
| *fbxa-9* | WBGene00021860 | protein_coding_gene | 1.6 | 3.04 | 3.7E-02 |
| *K02E10.7* | WBGene00019321 | protein_coding_gene | 1.6 | 3.04 | 9.8E-03 |
| *ZK1055.2* | WBGene00022843 | protein_coding_gene | 1.6 | 3.04 | 2.1E-02 |
| *F40E12.2* | WBGene00018235 | protein_coding_gene | 1.6 | 3.04 | 1.6E-03 |
| *F20G2.6* | WBGene00008989 | protein_coding_gene | 1.6 | 3.04 | 1.8E-02 |
| *Y50D4B.6* | WBGene00021745 | protein_coding_gene | 1.6 | 3.03 | 4.7E-02 |
| *T05B9.2* | WBGene00011465 | protein_coding_gene | 1.6 | 3.03 | 4.0E-02 |
| *glb-13* | WBGene00008957 | protein_coding_gene | 1.6 | 3.02 | 8.9E-03 |

| | | | | | |
|---|---|---|---|---|---|
| *F26B1.5* | WBGene00017817 | protein_coding_gene | 1.6 | 3.02 | 9.9E-04 |
| *pqn-74* | WBGene00004156 | protein_coding_gene | 1.6 | 3.02 | 1.2E-02 |
| *D1081.10* | WBGene00045508 | protein_coding_gene | 1.6 | 3.01 | 1.8E-02 |
| *M163.8* | WBGene00010940 | protein_coding_gene | 1.6 | 3.01 | 4.7E-03 |
| *mec-18* | WBGene00003179 | protein_coding_gene | 1.6 | 3.01 | 5.5E-03 |
| *T02G6.1* | WBGene00011384 | protein_coding_gene | 1.6 | 3.01 | 2.7E-02 |
| *ZK809.1* | WBGene00014085 | protein_coding_gene | 1.6 | 3.01 | 2.3E-02 |
| *ZK662.2* | WBGene00014039 | protein_coding_gene | 1.6 | 3.01 | 4.6E-02 |
| *act-5* | WBGene00000067 | protein_coding_gene | 1.6 | 3.00 | 3.8E-03 |
| *lgc-12* | WBGene00020048 | protein_coding_gene | 1.6 | 3.00 | 2.4E-02 |
| *F59F5.5* | WBGene00010344 | protein_coding_gene | 1.6 | 3.00 | 6.1E-03 |
| *tag-180* | WBGene00007041 | protein_coding_gene | 1.6 | 2.99 | 3.7E-04 |
| *F11D5.7* | WBGene00043980 | protein_coding_gene | 1.6 | 2.98 | 1.9E-04 |
| *Y62E10A.3* | WBGene00013374 | protein_coding_gene | 1.6 | 2.98 | 2.2E-02 |
| *ugt-10* | WBGene00020594 | protein_coding_gene | 1.6 | 2.98 | 1.8E-02 |
| *T22F7.4* | WBGene00020703 | protein_coding_gene | 1.6 | 2.98 | 1.1E-02 |
| *unc-8* | WBGene00006748 | protein_coding_gene | 1.6 | 2.98 | 1.4E-02 |
| *T06D8.10* | WBGene00011530 | protein_coding_gene | 1.6 | 2.97 | 1.8E-02 |
| *unc-17* | WBGene00006756 | protein_coding_gene | 1.6 | 2.97 | 3.2E-04 |
| *C30E1.6* | WBGene00016255 | pseudogene | 1.6 | 2.97 | 1.6E-02 |
| *F46C5.2* | WBGene00009779 | protein_coding_gene | 1.6 | 2.97 | 3.9E-04 |
| *nlp-16* | WBGene00003754 | protein_coding_gene | 1.6 | 2.97 | 5.3E-05 |
| *K09C4.5* | WBGene00019550 | protein_coding_gene | 1.6 | 2.97 | 4.0E-02 |
| *T10B5.2* | WBGene00020388 | pseudogene | 1.6 | 2.97 | 3.1E-03 |
| *T24C12.4* | WBGene00020766 | protein_coding_gene | 1.6 | 2.96 | 3.2E-02 |
| *F44B9.9* | WBGene00018410 | protein_coding_gene | 1.6 | 2.96 | 7.8E-03 |
| *F23H11.6* | WBGene00017761 | protein_coding_gene | 1.6 | 2.95 | 3.1E-02 |
| *fbxa-201* | WBGene00020342 | protein_coding_gene | 1.6 | 2.95 | 2.2E-02 |
| *gst-26* | WBGene00001774 | protein_coding_gene | 1.6 | 2.95 | 1.1E-02 |
| *Y116A8A.6* | WBGene00013776 | protein_coding_gene | 1.6 | 2.94 | 4.1E-03 |
| *F48F5.2* | WBGene00009846 | pseudogene | 1.6 | 2.94 | 2.3E-03 |
| *pdfr-1* | WBGene00015735 | protein_coding_gene | 1.6 | 2.94 | 5.5E-06 |
| *nas-1* | WBGene00003520 | protein_coding_gene | 1.6 | 2.94 | 3.7E-02 |
| *F58H7.1* | WBGene00019067 | protein_coding_gene | 1.6 | 2.94 | 8.1E-03 |
| *C23H5.11* | WBGene00044649 | protein_coding_gene | 1.6 | 2.94 | 3.7E-02 |
| *gur-4* | WBGene00001805 | protein_coding_gene | 1.6 | 2.93 | 3.4E-02 |
| *Y40C5A.4* | WBGene00021497 | protein_coding_gene | 1.6 | 2.93 | 4.8E-04 |
| *F43C1.5* | WBGene00009649 | protein_coding_gene | 1.5 | 2.93 | 9.9E-04 |
| *T06A4.3* | WBGene00020283 | protein_coding_gene | 1.5 | 2.92 | 2.4E-02 |
| *clp-3* | WBGene00000544 | protein_coding_gene | 1.5 | 2.92 | 8.4E-03 |
| *ppw-2* | WBGene00004094 | protein_coding_gene | 1.5 | 2.92 | 9.2E-03 |
| *clp-6* | WBGene00000546 | protein_coding_gene | 1.5 | 2.92 | 1.2E-02 |
| *math-18* | WBGene00016544 | protein_coding_gene | 1.5 | 2.92 | 4.0E-03 |
| *flr-4* | WBGene00001468 | protein_coding_gene | 1.5 | 2.90 | 3.6E-03 |
| *gab-1* | WBGene00001512 | protein_coding_gene | 1.5 | 2.90 | 1.6E-02 |
| *hrg-2* | WBGene00010473 | protein_coding_gene | 1.5 | 2.90 | 4.1E-02 |
| *Y22D7AL.11* | WBGene00021249 | protein_coding_gene | 1.5 | 2.90 | 4.4E-02 |
| *Y53F4B.51* | WBGene00194905 | protein_coding_gene | 1.5 | 2.89 | 7.5E-03 |
| *F35H10.3* | WBGene00018069 | protein_coding_gene | 1.5 | 2.89 | 1.2E-02 |
| *H03A11.2* | WBGene00010357 | protein_coding_gene | 1.5 | 2.89 | 1.4E-04 |
| *npax-2* | WBGene00018591 | protein_coding_gene | 1.5 | 2.89 | 3.6E-02 |
| *C25E10.17* | WBGene00235313 | protein_coding_gene | 1.5 | 2.89 | 2.7E-02 |
| *F01D4.3* | WBGene00008487 | pseudogene | 1.5 | 2.89 | 4.9E-03 |
| *irld-6* | WBGene00009239 | protein_coding_gene | 1.5 | 2.89 | 7.2E-03 |
| *nep-5* | WBGene00017393 | protein_coding_gene | 1.5 | 2.89 | 1.4E-02 |
| *C46G7.5* | WBGene00044502 | protein_coding_gene | 1.5 | 2.88 | 3.1E-02 |
| *ZC155.2* | WBGene00022530 | protein_coding_gene | 1.5 | 2.87 | 3.8E-03 |
| *moa-1* | WBGene00020116 | protein_coding_gene | 1.5 | 2.87 | 5.8E-04 |

| | | | | | |
|---|---|---|---|---|---|
| *ZC132.9* | WBGene00022527 | protein_coding_gene | 1.5 | 2.87 | 3.3E-02 |
| *ZK673.1* | WBGene00014057 | protein_coding_gene | 1.5 | 2.86 | 8.9E-07 |
| *C08F11.14* | WBGene00007461 | pseudogene | 1.5 | 2.86 | 2.1E-03 |
| *glr-4* | WBGene00001615 | protein_coding_gene | 1.5 | 2.85 | 6.7E-04 |
| *C35A11.4* | WBGene00016431 | protein_coding_gene | 1.5 | 2.85 | 1.7E-02 |
| *C30F2.2* | WBGene00007816 | protein_coding_gene | 1.5 | 2.85 | 3.2E-02 |
| *acox-1.4* | WBGene00008567 | protein_coding_gene | 1.5 | 2.85 | 2.8E-02 |
| *clp-10* | WBGene00021041 | protein_coding_gene | 1.5 | 2.84 | 4.3E-02 |
| *twk-48* | WBGene00010784 | protein_coding_gene | 1.5 | 2.84 | 2.6E-03 |
| *Y54G2A.45* | WBGene00044488 | protein_coding_gene | 1.5 | 2.84 | 1.8E-02 |
| *C09B9.7* | WBGene00015630 | protein_coding_gene | 1.5 | 2.84 | 5.4E-05 |
| *ZK1010.6* | WBGene00014180 | pseudogene | 1.5 | 2.84 | 3.4E-02 |
| *klp-13* | WBGene00002224 | protein_coding_gene | 1.5 | 2.84 | 1.3E-02 |
| *F22F1.3* | WBGene00017715 | protein_coding_gene | 1.5 | 2.83 | 8.9E-03 |
| *phat-4* | WBGene00020237 | protein_coding_gene | 1.5 | 2.82 | 2.2E-02 |
| *F39G3.5* | WBGene00018210 | protein_coding_gene | 1.5 | 2.82 | 2.1E-03 |
| *ser-5* | WBGene00008890 | protein_coding_gene | 1.5 | 2.82 | 9.2E-04 |
| *snpc-1.3* | WBGene00017265 | protein_coding_gene | 1.5 | 2.82 | 6.2E-04 |
| *gst-6* | WBGene00001754 | protein_coding_gene | 1.5 | 2.82 | 1.6E-02 |
| *K02E11.7* | WBGene00010516 | protein_coding_gene | 1.5 | 2.82 | 4.0E-03 |
| *R02F2.5* | WBGene00019834 | protein_coding_gene | 1.5 | 2.82 | 2.7E-03 |
| *F36H12.14* | WBGene00018127 | pseudogene | 1.5 | 2.82 | 1.3E-03 |
| *twk-40* | WBGene00006691 | protein_coding_gene | 1.5 | 2.81 | 4.5E-03 |
| *mab-10* | WBGene00003107 | protein_coding_gene | 1.5 | 2.81 | 5.9E-03 |
| *Y65B4A.9* | WBGene00022032 | protein_coding_gene | 1.5 | 2.81 | 6.8E-03 |
| *ctsa-4.2* | WBGene00021503 | protein_coding_gene | 1.5 | 2.81 | 1.7E-05 |
| *F28A10.1* | WBGene00017869 | protein_coding_gene | 1.5 | 2.81 | 2.7E-03 |
| *C55C2.3* | WBGene00016949 | protein_coding_gene | 1.5 | 2.80 | 4.1E-02 |
| *C04G2.10* | WBGene00007309 | protein_coding_gene | 1.5 | 2.80 | 2.4E-02 |
| *F29C6.1* | WBGene00009243 | protein_coding_gene | 1.5 | 2.80 | 3.1E-03 |
| *gbb-2* | WBGene00022675 | protein_coding_gene | 1.5 | 2.80 | 5.7E-03 |
| *W03F9.3* | WBGene00021001 | pseudogene | 1.5 | 2.80 | 2.8E-02 |
| *pck-1* | WBGene00021043 | protein_coding_gene | 1.5 | 2.80 | 4.9E-02 |
| *gcy-23* | WBGene00001548 | protein_coding_gene | 1.5 | 2.80 | 1.8E-02 |
| *Y73B6A.2* | WBGene00022229 | pseudogene | 1.5 | 2.80 | 1.3E-02 |
| *exp-1* | WBGene00001373 | protein_coding_gene | 1.5 | 2.79 | 2.2E-02 |
| *fbxa-202* | WBGene00020890 | protein_coding_gene | 1.5 | 2.79 | 8.1E-04 |
| *C28D4.5* | WBGene00007793 | protein_coding_gene | 1.5 | 2.79 | 1.5E-02 |
| *irk-1* | WBGene00002149 | protein_coding_gene | 1.5 | 2.79 | 4.5E-03 |
| *M03F8.5* | WBGene00019764 | protein_coding_gene | 1.5 | 2.79 | 9.0E-03 |
| *kcnl-1* | WBGene00007176 | protein_coding_gene | 1.5 | 2.79 | 5.4E-03 |
| *ZK250.14* | WBGene00235354 | protein_coding_gene | 1.5 | 2.79 | 4.1E-02 |
| *nhr-266* | WBGene00018993 | pseudogene | 1.5 | 2.79 | 3.3E-02 |
| *C08G5.3* | WBGene00015616 | pseudogene | 1.5 | 2.78 | 3.7E-02 |
| *slc-36.4* | WBGene00010421 | protein_coding_gene | 1.5 | 2.78 | 1.2E-03 |
| *F08F8.6* | WBGene00017271 | protein_coding_gene | 1.5 | 2.78 | 1.6E-03 |
| *F59A1.15* | WBGene00010300 | protein_coding_gene | 1.5 | 2.78 | 2.5E-03 |
| *hot-3* | WBGene00001988 | protein_coding_gene | 1.5 | 2.77 | 1.4E-02 |
| *Y4C6A.1* | WBGene00021151 | pseudogene | 1.5 | 2.77 | 7.5E-03 |
| *M05B5.3* | WBGene00010871 | protein_coding_gene | 1.5 | 2.76 | 2.5E-05 |
| *srp-10* | WBGene00005651 | pseudogene | 1.5 | 2.76 | 3.9E-02 |
| *ZK742.3* | WBGene00022813 | protein_coding_gene | 1.5 | 2.76 | 6.6E-04 |
| *tph-1* | WBGene00006600 | protein_coding_gene | 1.5 | 2.76 | 2.8E-02 |
| *K02B12.9* | WBGene00044679 | protein_coding_gene | 1.5 | 2.76 | 2.4E-02 |
| *tbc-8* | WBGene00008018 | protein_coding_gene | 1.5 | 2.76 | 4.2E-06 |
| *ugt-19* | WBGene00019515 | protein_coding_gene | 1.5 | 2.75 | 2.9E-02 |
| *molo-1* | WBGene00017299 | protein_coding_gene | 1.5 | 2.75 | 3.0E-02 |
| *Y4C6B.4* | WBGene00021158 | protein_coding_gene | 1.5 | 2.75 | 2.2E-02 |

| | | | | | |
|---|---|---|---|---|---|
| *che-10* | WBGene00018051 | protein_coding_gene | 1.5 | 2.75 | 2.8E-03 |
| *T10B10.4* | WBGene00011681 | protein_coding_gene | 1.5 | 2.75 | 2.6E-03 |
| *C14B9.3* | WBGene00015753 | protein_coding_gene | 1.5 | 2.75 | 9.5E-03 |
| *R09H10.5* | WBGene00011182 | protein_coding_gene | 1.5 | 2.75 | 1.9E-04 |
| *pitr-5* | WBGene00017312 | protein_coding_gene | 1.5 | 2.75 | 7.2E-03 |
| *nas-38* | WBGene00003554 | protein_coding_gene | 1.5 | 2.74 | 2.5E-02 |
| *F36D4.4* | WBGene00018089 | protein_coding_gene | 1.5 | 2.74 | 1.2E-02 |
| *W06H8.4* | WBGene00021067 | pseudogene | 1.5 | 2.74 | 1.1E-02 |
| *ZC262.4* | WBGene00022581 | protein_coding_gene | 1.5 | 2.74 | 2.1E-02 |
| *M04G7.2* | WBGene00019773 | protein_coding_gene | 1.5 | 2.73 | 3.3E-03 |
| *best-12* | WBGene00008821 | protein_coding_gene | 1.5 | 2.73 | 7.4E-03 |
| *C50C3.2* | WBGene00016800 | protein_coding_gene | 1.4 | 2.73 | 9.6E-04 |
| *epac-1* | WBGene00004255 | protein_coding_gene | 1.4 | 2.73 | 7.2E-03 |
| *ZK596.1* | WBGene00014006 | protein_coding_gene | 1.4 | 2.72 | 3.2E-02 |
| *glb-9* | WBGene00016184 | protein_coding_gene | 1.4 | 2.72 | 1.3E-02 |
| *ugt-6* | WBGene00013904 | protein_coding_gene | 1.4 | 2.72 | 3.3E-02 |
| *C09G1.5* | WBGene00007487 | protein_coding_gene | 1.4 | 2.72 | 3.2E-04 |
| *F09E10.7* | WBGene00017297 | protein_coding_gene | 1.4 | 2.72 | 1.7E-02 |
| *hrg-6* | WBGene00009495 | protein_coding_gene | 1.4 | 2.72 | 5.9E-03 |
| *lact-1* | WBGene00018523 | protein_coding_gene | 1.4 | 2.72 | 3.2E-03 |
| *C12D12.1* | WBGene00015713 | protein_coding_gene | 1.4 | 2.72 | 4.5E-05 |
| *F56B6.6* | WBGene00018937 | protein_coding_gene | 1.4 | 2.71 | 5.2E-03 |
| *scrm-4* | WBGene00008681 | protein_coding_gene | 1.4 | 2.71 | 2.3E-02 |
| *F49E2.5* | WBGene00009888 | protein_coding_gene | 1.4 | 2.71 | 1.1E-02 |
| *K09H9.9* | WBGene00219460 | protein_coding_gene | 1.4 | 2.71 | 1.5E-02 |
| *lgc-53* | WBGene00020657 | protein_coding_gene | 1.4 | 2.71 | 1.5E-02 |
| *acr-24* | WBGene00013534 | protein_coding_gene | 1.4 | 2.70 | 3.4E-03 |
| *Y116F11B.8* | WBGene00013824 | protein_coding_gene | 1.4 | 2.70 | 1.2E-02 |
| *lact-7* | WBGene00008047 | protein_coding_gene | 1.4 | 2.70 | 4.7E-03 |
| *math-23* | WBGene00016718 | protein_coding_gene | 1.4 | 2.70 | 2.1E-02 |
| *C04E6.4* | WBGene00015421 | protein_coding_gene | 1.4 | 2.70 | 1.3E-02 |
| *nhr-260* | WBGene00016517 | protein_coding_gene | 1.4 | 2.70 | 2.3E-02 |
| *F41B5.11* | WBGene00023039 | pseudogene | 1.4 | 2.70 | 4.5E-02 |
| *T10E10.3* | WBGene00020420 | protein_coding_gene | 1.4 | 2.69 | 3.5E-02 |
| *shl-1* | WBGene00022240 | protein_coding_gene | 1.4 | 2.69 | 4.8E-03 |
| *cdkl-1* | WBGene00012779 | protein_coding_gene | 1.4 | 2.69 | 8.6E-03 |
| *D1007.19* | WBGene00045048 | protein_coding_gene | 1.4 | 2.69 | 2.9E-02 |
| *anoh-1* | WBGene00010138 | protein_coding_gene | 1.4 | 2.68 | 4.1E-02 |
| *T25G12.6* | WBGene00020813 | protein_coding_gene | 1.4 | 2.68 | 6.2E-03 |
| *C01H6.4* | WBGene00007254 | protein_coding_gene | 1.4 | 2.68 | 6.3E-05 |
| *F58D5.6* | WBGene00010245 | protein_coding_gene | 1.4 | 2.68 | 1.1E-02 |
| *H20J04.1* | WBGene00019216 | protein_coding_gene | 1.4 | 2.68 | 3.2E-03 |
| *ocr-2* | WBGene00003839 | protein_coding_gene | 1.4 | 2.68 | 2.6E-03 |
| *F47G3.1* | WBGene00018576 | protein_coding_gene | 1.4 | 2.67 | 4.0E-02 |
| *F01G10.5* | WBGene00008508 | protein_coding_gene | 1.4 | 2.67 | 7.1E-05 |
| *F56D6.18* | WBGene00235285 | protein_coding_gene | 1.4 | 2.67 | 4.5E-02 |
| *C14A11.6* | WBGene00015751 | protein_coding_gene | 1.4 | 2.67 | 1.8E-02 |
| *gst-5* | WBGene00001753 | protein_coding_gene | 1.4 | 2.67 | 8.6E-03 |
| *nspb-6* | WBGene00019152 | protein_coding_gene | 1.4 | 2.67 | 4.1E-02 |
| *ZK105.3* | WBGene00022654 | protein_coding_gene | 1.4 | 2.67 | 1.4E-02 |
| *cng-2* | WBGene00022295 | protein_coding_gene | 1.4 | 2.66 | 2.3E-02 |
| *K09C6.2* | WBGene00019556 | protein_coding_gene | 1.4 | 2.66 | 5.0E-03 |
| *Y54G2A.41* | WBGene00044485 | protein_coding_gene | 1.4 | 2.66 | 1.5E-02 |
| *Y37F4.7* | WBGene00021388 | protein_coding_gene | 1.4 | 2.66 | 1.8E-02 |
| *cln-3.1* | WBGene00000539 | protein_coding_gene | 1.4 | 2.66 | 3.8E-02 |
| *clec-174* | WBGene00021580 | protein_coding_gene | 1.4 | 2.66 | 4.2E-02 |
| *T20B6.2* | WBGene00020598 | protein_coding_gene | 1.4 | 2.66 | 9.0E-03 |
| *C33H5.16* | WBGene00016382 | protein_coding_gene | 1.4 | 2.65 | 1.7E-02 |

| | | | | | |
|---|---|---|---|---|---|
| *dmsr-2* | WBGene00021275 | protein_coding_gene | 1.4 | 2.65 | 4.0E-03 |
| *nlp-63* | WBGene00044028 | protein_coding_gene | 1.4 | 2.65 | 2.0E-02 |
| *F57H12.5* | WBGene00019020 | protein_coding_gene | 1.4 | 2.65 | 3.3E-03 |
| *C06E2.1* | WBGene00015527 | protein_coding_gene | 1.4 | 2.64 | 3.5E-02 |
| *F35E2.5* | WBGene00009412 | protein_coding_gene | 1.4 | 2.63 | 5.7E-03 |
| *tsp-19* | WBGene00006645 | protein_coding_gene | 1.4 | 2.63 | 3.1E-03 |
| *R155.3* | WBGene00020117 | protein_coding_gene | 1.4 | 2.62 | 2.5E-03 |
| *C13G3.1* | WBGene00007553 | protein_coding_gene | 1.4 | 2.62 | 1.9E-02 |
| *ncx-6* | WBGene00003571 | protein_coding_gene | 1.4 | 2.62 | 1.6E-06 |
| *eat-4* | WBGene00001135 | protein_coding_gene | 1.4 | 2.62 | 2.4E-02 |
| *fbxa-155* | WBGene00007323 | protein_coding_gene | 1.4 | 2.62 | 3.8E-02 |
| *H04J21.1* | WBGene00019150 | protein_coding_gene | 1.4 | 2.62 | 1.7E-02 |
| *C55A6.4* | WBGene00008333 | protein_coding_gene | 1.4 | 2.61 | 1.6E-09 |
| *acr-9* | WBGene00000048 | protein_coding_gene | 1.4 | 2.61 | 2.3E-02 |
| *rimb-1* | WBGene00006513 | protein_coding_gene | 1.4 | 2.61 | 7.6E-03 |
| *gly-18* | WBGene00001643 | protein_coding_gene | 1.4 | 2.61 | 6.0E-03 |
| *anc-1* | WBGene00000140 | protein_coding_gene | 1.4 | 2.61 | 4.6E-03 |
| *nspc-1* | WBGene00023477 | protein_coding_gene | 1.4 | 2.61 | 8.3E-03 |
| *C15H7.2* | WBGene00014678 | pseudogene | 1.4 | 2.61 | 1.4E-02 |
| *Y39B6A.25* | WBGene00012684 | protein_coding_gene | 1.4 | 2.61 | 4.1E-03 |
| *asic-1* | WBGene00022815 | protein_coding_gene | 1.4 | 2.60 | 2.2E-02 |
| *K09B11.5* | WBGene00010712 | protein_coding_gene | 1.4 | 2.60 | 6.6E-03 |
| *K03H6.5* | WBGene00019370 | protein_coding_gene | 1.4 | 2.60 | 5.2E-03 |
| *D2023.1* | WBGene00014300 | protein_coding_gene | 1.4 | 2.60 | 1.8E-03 |
| *C09B9.85* | WBGene00220250 | protein_coding_gene | 1.4 | 2.60 | 1.2E-02 |
| *ZK105.12* | WBGene00235113 | protein_coding_gene | 1.4 | 2.60 | 2.4E-02 |
| *cut-4* | WBGene00008482 | protein_coding_gene | 1.4 | 2.59 | 4.7E-02 |
| *Y56A3A.6* | WBGene00013227 | protein_coding_gene | 1.4 | 2.59 | 1.5E-04 |
| *Y18H1A.15* | WBGene00044434 | protein_coding_gene | 1.4 | 2.59 | 4.7E-03 |
| *ser-1* | WBGene00004776 | protein_coding_gene | 1.4 | 2.59 | 1.8E-02 |
| *Y44A6D.2* | WBGene00012853 | protein_coding_gene | 1.4 | 2.58 | 4.1E-03 |
| *F28B4.3* | WBGene00017892 | protein_coding_gene | 1.4 | 2.58 | 6.1E-03 |
| *K10D2.8* | WBGene00189952 | protein_coding_gene | 1.4 | 2.58 | 6.5E-03 |
| *C50D2.2* | WBGene00016806 | protein_coding_gene | 1.4 | 2.58 | 7.0E-04 |
| *gyg-1* | WBGene00006863 | protein_coding_gene | 1.4 | 2.58 | 1.1E-02 |
| *ocr-1* | WBGene00003838 | protein_coding_gene | 1.4 | 2.58 | 3.4E-02 |
| *nas-8* | WBGene00003527 | protein_coding_gene | 1.4 | 2.58 | 5.7E-03 |
| *R05D7.2* | WBGene00011027 | protein_coding_gene | 1.4 | 2.57 | 7.6E-03 |
| *C50H2.7* | WBGene00008243 | protein_coding_gene | 1.4 | 2.57 | 3.7E-02 |
| *C18A11.3* | WBGene00015949 | protein_coding_gene | 1.4 | 2.57 | 1.4E-02 |
| *ins-6* | WBGene00002089 | protein_coding_gene | 1.4 | 2.57 | 1.1E-03 |
| *tnt-4* | WBGene00006589 | protein_coding_gene | 1.4 | 2.57 | 1.3E-03 |
| *F17B5.8* | WBGene00045458 | protein_coding_gene | 1.4 | 2.56 | 1.9E-02 |
| *F46C8.8* | WBGene00018487 | protein_coding_gene | 1.4 | 2.56 | 4.4E-02 |
| *B0361.4* | WBGene00015158 | protein_coding_gene | 1.4 | 2.56 | 5.5E-03 |
| *T06D8.2* | WBGene00011523 | protein_coding_gene | 1.4 | 2.56 | 3.8E-02 |
| *apt-9* | WBGene00000166 | protein_coding_gene | 1.4 | 2.56 | 1.3E-03 |
| *F29B9.12* | WBGene00044482 | protein_coding_gene | 1.4 | 2.56 | 3.3E-02 |
| *lips-15* | WBGene00012585 | protein_coding_gene | 1.4 | 2.55 | 6.8E-04 |
| *M02D8.7* | WBGene00044313 | protein_coding_gene | 1.4 | 2.55 | 3.4E-02 |
| *nhr-7* | WBGene00003606 | protein_coding_gene | 1.4 | 2.55 | 3.8E-04 |
| *B0244.17* | WBGene00271715 | protein_coding_gene | 1.4 | 2.55 | 4.3E-02 |
| *F35C11.2* | WBGene00009401 | protein_coding_gene | 1.3 | 2.55 | 1.8E-03 |
| *K12C11.5* | WBGene00044896 | protein_coding_gene | 1.3 | 2.54 | 8.8E-03 |
| *F07H5.7* | WBGene00008558 | protein_coding_gene | 1.3 | 2.54 | 2.3E-03 |
| *F45E1.5* | WBGene00018466 | protein_coding_gene | 1.3 | 2.54 | 3.1E-02 |
| *dpy-31* | WBGene00006592 | protein_coding_gene | 1.3 | 2.53 | 4.0E-02 |
| *F07D3.3* | WBGene00008556 | protein_coding_gene | 1.3 | 2.53 | 1.9E-02 |

| | | | | | |
|---|---|---|---|---|---|
| *unc-4* | WBGene00006744 | protein_coding_gene | 1.3 | 2.53 | 4.4E-02 |
| *Y69E1A.3* | WBGene00013475 | protein_coding_gene | 1.3 | 2.53 | 8.4E-03 |
| *fipr-11* | WBGene00008244 | protein_coding_gene | 1.3 | 2.52 | 3.1E-03 |
| *T25E12.16* | WBGene00194715 | protein_coding_gene | 1.3 | 2.52 | 2.3E-02 |
| *F08D12.7* | WBGene00017250 | protein_coding_gene | 1.3 | 2.52 | 3.4E-02 |
| *cest-19* | WBGene00020125 | protein_coding_gene | 1.3 | 2.52 | 2.7E-03 |
| *stg-2* | WBGene00017400 | protein_coding_gene | 1.3 | 2.51 | 4.1E-02 |
| *F10D11.6* | WBGene00008652 | protein_coding_gene | 1.3 | 2.51 | 2.9E-02 |
| *M05D6.3* | WBGene00010876 | protein_coding_gene | 1.3 | 2.51 | 1.3E-02 |
| *lon-8* | WBGene00013352 | protein_coding_gene | 1.3 | 2.51 | 3.4E-03 |
| *C49F8.3* | WBGene00008215 | protein_coding_gene | 1.3 | 2.51 | 2.0E-03 |
| *best-26* | WBGene00014103 | protein_coding_gene | 1.3 | 2.51 | 5.7E-03 |
| *F59E12.3* | WBGene00019119 | protein_coding_gene | 1.3 | 2.51 | 3.7E-02 |
| *lys-5* | WBGene00003094 | protein_coding_gene | 1.3 | 2.51 | 1.8E-02 |
| *C25G4.7* | WBGene00007734 | protein_coding_gene | 1.3 | 2.51 | 1.3E-02 |
| *K08H10.5* | WBGene00014805 | pseudogene | 1.3 | 2.51 | 4.3E-02 |
| *fbxa-178* | WBGene00017533 | pseudogene | 1.3 | 2.51 | 3.3E-02 |
| *Y105E8A.1* | WBGene00013666 | protein_coding_gene | 1.3 | 2.50 | 2.0E-02 |
| *frpr-17* | WBGene00011765 | protein_coding_gene | 1.3 | 2.50 | 2.5E-03 |
| *Y34B4A.10* | WBGene00021326 | protein_coding_gene | 1.3 | 2.50 | 5.8E-03 |
| *oac-58* | WBGene00019580 | protein_coding_gene | 1.3 | 2.50 | 3.8E-04 |
| *F35H10.10* | WBGene00018073 | protein_coding_gene | 1.3 | 2.50 | 3.2E-02 |
| *F23H11.7* | WBGene00017762 | protein_coding_gene | 1.3 | 2.50 | 9.0E-03 |
| *let-805* | WBGene00002915 | protein_coding_gene | 1.3 | 2.50 | 3.5E-02 |
| *glr-5* | WBGene00001616 | protein_coding_gene | 1.3 | 2.49 | 2.1E-04 |
| *oig-8* | WBGene00020582 | protein_coding_gene | 1.3 | 2.49 | 6.3E-03 |
| *aat-4* | WBGene00000005 | protein_coding_gene | 1.3 | 2.49 | 4.0E-02 |
| *C17C3.11* | WBGene00015893 | protein_coding_gene | 1.3 | 2.49 | 2.4E-02 |
| *ifd-2* | WBGene00002058 | protein_coding_gene | 1.3 | 2.49 | 3.2E-03 |
| *C15H7.4* | WBGene00007611 | protein_coding_gene | 1.3 | 2.48 | 1.8E-02 |
| *F26F12.2* | WBGene00017833 | protein_coding_gene | 1.3 | 2.48 | 2.6E-02 |
| *F46F5.11* | WBGene00018502 | protein_coding_gene | 1.3 | 2.48 | 1.0E-02 |
| *F22B3.10* | WBGene00045122 | protein_coding_gene | 1.3 | 2.48 | 4.7E-02 |
| *ZC581.9* | WBGene00022635 | protein_coding_gene | 1.3 | 2.47 | 3.3E-03 |
| *pept-1* | WBGene00003877 | protein_coding_gene | 1.3 | 2.47 | 2.9E-02 |
| *catp-5* | WBGene00019493 | protein_coding_gene | 1.3 | 2.47 | 3.1E-03 |
| *best-6* | WBGene00015628 | protein_coding_gene | 1.3 | 2.47 | 2.4E-02 |
| *T02D1.4* | WBGene00011371 | protein_coding_gene | 1.3 | 2.47 | 3.1E-02 |
| *F14H12.3* | WBGene00017471 | protein_coding_gene | 1.3 | 2.47 | 7.6E-03 |
| *Y52D5A.1* | WBGene00021792 | pseudogene | 1.3 | 2.47 | 1.1E-02 |
| *C49C3.2* | WBGene00008192 | pseudogene | 1.3 | 2.47 | 1.8E-02 |
| *C29F5.3* | WBGene00016212 | protein_coding_gene | 1.3 | 2.47 | 1.9E-03 |
| *gst-2* | WBGene00001750 | protein_coding_gene | 1.3 | 2.46 | 3.7E-02 |
| *Y22D7AR.12* | WBGene00021265 | pseudogene | 1.3 | 2.46 | 1.1E-02 |
| *ipla-5* | WBGene00019229 | protein_coding_gene | 1.3 | 2.46 | 1.9E-03 |
| *ceh-18* | WBGene00000441 | protein_coding_gene | 1.3 | 2.46 | 3.3E-03 |
| *C38D4.7* | WBGene00008007 | protein_coding_gene | 1.3 | 2.46 | 3.0E-02 |
| *Y53G8B.3* | WBGene00021819 | pseudogene | 1.3 | 2.46 | 2.9E-02 |
| *Y41C4A.12* | WBGene00012758 | protein_coding_gene | 1.3 | 2.45 | 5.3E-05 |
| *otpl-3* | WBGene00017449 | protein_coding_gene | 1.3 | 2.45 | 2.0E-02 |
| *fut-5* | WBGene00043986 | protein_coding_gene | 1.3 | 2.45 | 7.0E-03 |
| *Y55D5A.4* | WBGene00021918 | protein_coding_gene | 1.3 | 2.45 | 7.2E-03 |
| *nlp-67* | WBGene00011227 | protein_coding_gene | 1.3 | 2.45 | 2.6E-02 |
| *F19C7.8* | WBGene00017597 | protein_coding_gene | 1.3 | 2.45 | 1.3E-03 |
| *srx-45* | WBGene00005936 | protein_coding_gene | 1.3 | 2.45 | 2.8E-02 |
| *kvs-5* | WBGene00021948 | protein_coding_gene | 1.3 | 2.45 | 1.2E-02 |
| *F20D6.1* | WBGene00017633 | protein_coding_gene | 1.3 | 2.45 | 4.1E-02 |
| *T15H9.4* | WBGene00011789 | protein_coding_gene | 1.3 | 2.44 | 4.2E-02 |

| | | | | | |
|---|---|---|---|---|---|
| *F54G2.1* | WBGene00018837 | protein_coding_gene | 1.3 | 2.44 | 4.1E-04 |
| *ent-4* | WBGene00008123 | protein_coding_gene | 1.3 | 2.44 | 3.7E-03 |
| *rgs-2* | WBGene00004345 | protein_coding_gene | 1.3 | 2.44 | 5.8E-04 |
| *F35F10.1* | WBGene00018052 | protein_coding_gene | 1.3 | 2.44 | 1.5E-02 |
| *sipa-1* | WBGene00012100 | protein_coding_gene | 1.3 | 2.44 | 8.5E-04 |
| *tsp-18* | WBGene00006644 | protein_coding_gene | 1.3 | 2.43 | 1.4E-02 |
| *F56D5.2* | WBGene00010147 | protein_coding_gene | 1.3 | 2.43 | 1.4E-02 |
| *F41H10.5* | WBGene00018318 | protein_coding_gene | 1.3 | 2.43 | 1.5E-02 |
| *T25C12.3* | WBGene00012018 | protein_coding_gene | 1.3 | 2.43 | 5.5E-03 |
| *Y4C6A.68* | WBGene00235281 | pseudogene | 1.3 | 2.43 | 4.8E-02 |
| *Y71F9AL.2* | WBGene00022108 | protein_coding_gene | 1.3 | 2.43 | 5.6E-03 |
| *del-1* | WBGene00000952 | protein_coding_gene | 1.3 | 2.42 | 4.4E-02 |
| *F58D5.8* | WBGene00010247 | protein_coding_gene | 1.3 | 2.42 | 1.0E-02 |
| *glb-10* | WBGene00016215 | protein_coding_gene | 1.3 | 2.42 | 1.9E-03 |
| *nhr-14* | WBGene00003613 | protein_coding_gene | 1.3 | 2.42 | 1.1E-02 |
| *oac-23* | WBGene00018143 | protein_coding_gene | 1.3 | 2.41 | 3.7E-03 |
| *ndnf-1* | WBGene00021893 | protein_coding_gene | 1.3 | 2.41 | 2.5E-02 |
| *add-1* | WBGene00000072 | protein_coding_gene | 1.3 | 2.41 | 5.2E-03 |
| *F14D12.1* | WBGene00017463 | protein_coding_gene | 1.3 | 2.41 | 1.1E-02 |
| *hmit-1.1* | WBGene00013073 | protein_coding_gene | 1.3 | 2.41 | 3.7E-02 |
| *D2024.1* | WBGene00017050 | protein_coding_gene | 1.3 | 2.41 | 2.3E-02 |
| *ZK686.5* | WBGene00022795 | protein_coding_gene | 1.3 | 2.41 | 9.9E-03 |
| *R05C11.4* | WBGene00044467 | protein_coding_gene | 1.3 | 2.41 | 1.5E-02 |
| *Y26E6A.3* | WBGene00012514 | protein_coding_gene | 1.3 | 2.41 | 3.7E-03 |
| *fbxa-51* | WBGene00020448 | protein_coding_gene | 1.3 | 2.41 | 5.9E-03 |
| *nspd-7* | WBGene00012781 | protein_coding_gene | 1.3 | 2.41 | 2.2E-02 |
| *K03H1.9* | WBGene00010544 | protein_coding_gene | 1.3 | 2.40 | 2.5E-02 |
| *qui-1* | WBGene00004265 | protein_coding_gene | 1.3 | 2.40 | 2.5E-02 |
| *fkh-6* | WBGene00001438 | protein_coding_gene | 1.3 | 2.40 | 4.0E-02 |
| *F59B10.3* | WBGene00010319 | protein_coding_gene | 1.3 | 2.40 | 1.2E-02 |
| *C54E4.5* | WBGene00016920 | protein_coding_gene | 1.3 | 2.39 | 2.3E-02 |
| *glb-6* | WBGene00015969 | protein_coding_gene | 1.3 | 2.39 | 3.2E-02 |
| *exc-1* | WBGene00001362 | protein_coding_gene | 1.3 | 2.39 | 1.8E-02 |
| *gst-36* | WBGene00001784 | protein_coding_gene | 1.3 | 2.39 | 1.2E-02 |
| *nep-9* | WBGene00017554 | protein_coding_gene | 1.3 | 2.38 | 8.1E-03 |
| *spe-17* | WBGene00004971 | protein_coding_gene | 1.2 | 2.38 | 1.9E-02 |
| *ZK849.1* | WBGene00014100 | protein_coding_gene | 1.2 | 2.37 | 3.2E-03 |
| *pqn-75* | WBGene00004157 | protein_coding_gene | 1.2 | 2.37 | 1.2E-02 |
| *twk-17* | WBGene00006671 | protein_coding_gene | 1.2 | 2.37 | 2.9E-02 |
| *H38K22.9* | WBGene00284844 | protein_coding_gene | 1.2 | 2.36 | 3.1E-02 |
| *C14A4.13* | WBGene00007563 | protein_coding_gene | 1.2 | 2.36 | 6.9E-03 |
| *F16B4.5* | WBGene00017507 | protein_coding_gene | 1.2 | 2.36 | 2.1E-03 |
| *R07C3.17* | WBGene00269346 | ncRNA_gene | 1.2 | 2.36 | 3.2E-02 |
| *chil-12* | WBGene00010799 | protein_coding_gene | 1.2 | 2.36 | 3.2E-02 |
| *ZK666.8* | WBGene00014048 | protein_coding_gene | 1.2 | 2.36 | 1.3E-02 |
| *aman-3* | WBGene00018594 | protein_coding_gene | 1.2 | 2.36 | 2.6E-03 |
| *F29D10.1* | WBGene00009250 | protein_coding_gene | 1.2 | 2.35 | 1.2E-02 |
| *W04B5.3* | WBGene00021020 | protein_coding_gene | 1.2 | 2.35 | 2.1E-04 |
| *B0554.5* | WBGene00015258 | protein_coding_gene | 1.2 | 2.35 | 4.3E-02 |
| *B0034.4* | WBGene00015004 | protein_coding_gene | 1.2 | 2.35 | 8.4E-03 |
| *ser-2* | WBGene00004777 | protein_coding_gene | 1.2 | 2.35 | 1.5E-02 |
| *twk-14* | WBGene00006669 | protein_coding_gene | 1.2 | 2.35 | 9.7E-03 |
| *pde-6* | WBGene00022389 | protein_coding_gene | 1.2 | 2.35 | 5.1E-03 |
| *ZC404.1* | WBGene00022601 | protein_coding_gene | 1.2 | 2.34 | 1.8E-02 |
| *F54H5.2* | WBGene00018839 | protein_coding_gene | 1.2 | 2.34 | 3.5E-02 |
| *spe-43* | WBGene00008623 | protein_coding_gene | 1.2 | 2.34 | 1.6E-02 |
| *F56A12.2* | WBGene00010143 | protein_coding_gene | 1.2 | 2.34 | 2.7E-02 |
| *cpx-1* | WBGene00022271 | protein_coding_gene | 1.2 | 2.34 | 2.6E-03 |

| | | | | | |
|---|---|---|---|---|---|
| *Y105C5B.14* | WBGene00013653 | protein_coding_gene | 1.2 | 2.33 | 4.8E-02 |
| *col-34* | WBGene00000611 | protein_coding_gene | 1.2 | 2.33 | 4.7E-02 |
| *C55A6.11* | WBGene00008340 | protein_coding_gene | 1.2 | 2.33 | 1.3E-02 |
| *T06C10.3* | WBGene00020289 | protein_coding_gene | 1.2 | 2.33 | 3.3E-03 |
| *F27C1.11* | WBGene00017858 | protein_coding_gene | 1.2 | 2.33 | 6.1E-04 |
| *C53C11.2* | WBGene00016900 | protein_coding_gene | 1.2 | 2.33 | 5.0E-02 |
| *mth-1* | WBGene00017948 | protein_coding_gene | 1.2 | 2.33 | 9.5E-03 |
| *ncx-2* | WBGene00003567 | protein_coding_gene | 1.2 | 2.33 | 2.6E-02 |
| *cdc-25.4* | WBGene00000389 | protein_coding_gene | 1.2 | 2.33 | 4.3E-03 |
| *F28E10.5* | WBGene00044302 | pseudogene | 1.2 | 2.33 | 1.4E-02 |
| *amx-2* | WBGene00000138 | protein_coding_gene | 1.2 | 2.33 | 2.7E-02 |
| *tre-4* | WBGene00006610 | protein_coding_gene | 1.2 | 2.32 | 5.5E-03 |
| *fbxa-58* | WBGene00016225 | pseudogene | 1.2 | 2.32 | 1.6E-06 |
| *T06G6.3* | WBGene00011545 | protein_coding_gene | 1.2 | 2.32 | 4.7E-02 |
| *acd-3* | WBGene00007775 | protein_coding_gene | 1.2 | 2.32 | 1.8E-03 |
| *snn-1* | WBGene00004913 | protein_coding_gene | 1.2 | 2.32 | 2.5E-03 |
| *F47F2.1* | WBGene00018569 | protein_coding_gene | 1.2 | 2.32 | 1.3E-02 |
| *linc-36* | WBGene00219694 | lincRNA_gene | 1.2 | 2.32 | 3.3E-02 |
| *Y57G11C.23* | WBGene00013318 | protein_coding_gene | 1.2 | 2.32 | 3.6E-02 |
| *lec-9* | WBGene00002272 | protein_coding_gene | 1.2 | 2.32 | 1.6E-05 |
| *B0222.11* | WBGene00044582 | protein_coding_gene | 1.2 | 2.32 | 2.6E-02 |
| *hog-1* | WBGene00001984 | protein_coding_gene | 1.2 | 2.31 | 2.1E-02 |
| *ZK721.3* | WBGene00022810 | protein_coding_gene | 1.2 | 2.31 | 1.2E-02 |
| *F58F9.4* | WBGene00019058 | protein_coding_gene | 1.2 | 2.31 | 7.1E-05 |
| *K06A1.2* | WBGene00019425 | protein_coding_gene | 1.2 | 2.31 | 7.4E-03 |
| *dhhc-13* | WBGene00019257 | protein_coding_gene | 1.2 | 2.31 | 8.4E-03 |
| *B0403.5* | WBGene00015169 | protein_coding_gene | 1.2 | 2.31 | 2.9E-02 |
| *F28A10.5* | WBGene00017873 | protein_coding_gene | 1.2 | 2.31 | 6.1E-03 |
| *Y38H6C.15* | WBGene00012627 | protein_coding_gene | 1.2 | 2.31 | 3.3E-02 |
| *F46F5.4* | WBGene00018495 | pseudogene | 1.2 | 2.31 | 2.1E-02 |
| *T20B5.2* | WBGene00020595 | protein_coding_gene | 1.2 | 2.30 | 2.2E-04 |
| *npr-4* | WBGene00007635 | protein_coding_gene | 1.2 | 2.30 | 5.1E-03 |
| *npr-22* | WBGene00022004 | protein_coding_gene | 1.2 | 2.30 | 1.3E-03 |
| *F57F5.1* | WBGene00010204 | protein_coding_gene | 1.2 | 2.30 | 7.5E-03 |
| *rnp-9* | WBGene00007396 | protein_coding_gene | 1.2 | 2.30 | 1.6E-02 |
| *dhhc-12* | WBGene00010323 | protein_coding_gene | 1.2 | 2.29 | 7.0E-03 |
| *oac-50* | WBGene00020864 | protein_coding_gene | 1.2 | 2.29 | 4.8E-02 |
| *Y39F10A.3* | WBGene00021454 | protein_coding_gene | 1.2 | 2.28 | 2.3E-03 |
| *pde-1* | WBGene00011433 | protein_coding_gene | 1.2 | 2.28 | 1.5E-03 |
| *glb-16* | WBGene00018486 | protein_coding_gene | 1.2 | 2.28 | 3.5E-02 |
| *Y37D8A.5* | WBGene00012547 | protein_coding_gene | 1.2 | 2.28 | 8.5E-03 |
| *ZK1067.4* | WBGene00014210 | protein_coding_gene | 1.2 | 2.28 | 4.2E-03 |
| *C10E2.5* | WBGene00015675 | protein_coding_gene | 1.2 | 2.28 | 1.9E-02 |
| *C01G10.1* | WBGene00007230 | protein_coding_gene | 1.2 | 2.27 | 4.3E-02 |
| *B0250.4* | WBGene00007121 | pseudogene | 1.2 | 2.27 | 6.7E-03 |
| *cpb-2* | WBGene00000771 | protein_coding_gene | 1.2 | 2.27 | 1.3E-02 |
| *nhr-141* | WBGene00017787 | protein_coding_gene | 1.2 | 2.27 | 5.2E-03 |
| *alg-4* | WBGene00006449 | protein_coding_gene | 1.2 | 2.27 | 2.5E-03 |
| *K10H10.7* | WBGene00010763 | protein_coding_gene | 1.2 | 2.27 | 2.5E-02 |
| *T20H4.2* | WBGene00020635 | protein_coding_gene | 1.2 | 2.26 | 2.1E-02 |
| *avr-14* | WBGene00000232 | protein_coding_gene | 1.2 | 2.26 | 1.7E-03 |
| *C01G5.3* | WBGene00015305 | protein_coding_gene | 1.2 | 2.26 | 1.2E-02 |
| *C44C10.11* | WBGene00043985 | protein_coding_gene | 1.2 | 2.26 | 8.9E-03 |
| *T27A3.5* | WBGene00020841 | protein_coding_gene | 1.2 | 2.26 | 1.6E-03 |
| *Y32F6A.4* | WBGene00012529 | protein_coding_gene | 1.2 | 2.26 | 1.0E-02 |
| *ifa-1* | WBGene00002050 | protein_coding_gene | 1.2 | 2.26 | 7.4E-03 |
| *Y47D9A.5* | WBGene00021632 | protein_coding_gene | 1.2 | 2.26 | 7.8E-03 |
| *lips-1* | WBGene00009727 | protein_coding_gene | 1.2 | 2.26 | 4.2E-02 |

| | | | | | |
|---|---|---|---|---|---|
| *ttn-1* | WBGene00006436 | protein_coding_gene | 1.2 | 2.26 | 1.3E-02 |
| *daf-37* | WBGene00016246 | protein_coding_gene | 1.2 | 2.25 | 1.7E-02 |
| *plc-1* | WBGene00004036 | protein_coding_gene | 1.2 | 2.25 | 9.6E-03 |
| *aqp-4* | WBGene00000172 | protein_coding_gene | 1.2 | 2.25 | 5.7E-08 |
| *ZK688.12* | WBGene00044763 | protein_coding_gene | 1.2 | 2.25 | 8.0E-03 |
| *F38A5.6* | WBGene00018163 | protein_coding_gene | 1.2 | 2.25 | 2.8E-03 |
| *F23C8.7* | WBGene00017736 | protein_coding_gene | 1.2 | 2.24 | 1.1E-04 |
| *pll-1* | WBGene00004045 | protein_coding_gene | 1.2 | 2.24 | 9.1E-04 |
| *C17H12.10* | WBGene00015935 | protein_coding_gene | 1.2 | 2.24 | 1.3E-02 |
| *nekl-4* | WBGene00017033 | protein_coding_gene | 1.2 | 2.24 | 1.2E-02 |
| *C14B4.2* | WBGene00007583 | protein_coding_gene | 1.2 | 2.24 | 2.9E-02 |
| *C26C6.10* | WBGene00077553 | protein_coding_gene | 1.2 | 2.23 | 4.2E-02 |
| *F22E5.13* | WBGene00017711 | protein_coding_gene | 1.2 | 2.23 | 4.7E-02 |
| *F16F9.3* | WBGene00017514 | protein_coding_gene | 1.2 | 2.23 | 2.6E-02 |
| *csp-2* | WBGene00000820 | protein_coding_gene | 1.2 | 2.23 | 1.9E-02 |
| *clec-155* | WBGene00011405 | pseudogene | 1.2 | 2.23 | 2.8E-02 |
| *pfas-1* | WBGene00008654 | protein_coding_gene | 1.2 | 2.23 | 3.7E-02 |
| *mlt-2* | WBGene00007252 | protein_coding_gene | 1.2 | 2.23 | 3.9E-02 |
| *F18A1.1* | WBGene00017545 | protein_coding_gene | 1.2 | 2.23 | 4.5E-02 |
| *C35A5.4* | WBGene00007948 | protein_coding_gene | 1.2 | 2.23 | 2.9E-03 |
| *F59A3.8* | WBGene00019081 | protein_coding_gene | 1.2 | 2.22 | 5.5E-03 |
| *F44G3.7* | WBGene00009708 | protein_coding_gene | 1.2 | 2.22 | 3.0E-03 |
| *dig-1* | WBGene00000998 | protein_coding_gene | 1.2 | 2.22 | 3.7E-02 |
| *Y102A11A.2* | WBGene00022412 | protein_coding_gene | 1.2 | 2.22 | 3.4E-03 |
| *clec-236* | WBGene00013495 | pseudogene | 1.2 | 2.22 | 1.3E-02 |
| *F21A3.4* | WBGene00008994 | protein_coding_gene | 1.1 | 2.22 | 9.6E-03 |
| *T23F2.2* | WBGene00020735 | protein_coding_gene | 1.1 | 2.22 | 9.6E-03 |
| *lev-1* | WBGene00002974 | protein_coding_gene | 1.1 | 2.22 | 3.6E-02 |
| *W03D8.10* | WBGene00020991 | pseudogene | 1.1 | 2.21 | 3.9E-03 |
| *F55C12.4* | WBGene00018871 | protein_coding_gene | 1.1 | 2.21 | 3.4E-02 |
| *unc-41* | WBGene00006777 | protein_coding_gene | 1.1 | 2.21 | 1.4E-03 |
| *npr-6* | WBGene00009619 | protein_coding_gene | 1.1 | 2.21 | 6.4E-03 |
| *Y40B10A.4* | WBGene00021489 | pseudogene | 1.1 | 2.21 | 3.1E-02 |
| *F20H11.4* | WBGene00017647 | protein_coding_gene | 1.1 | 2.21 | 4.8E-03 |
| *K09C6.8* | WBGene00019562 | protein_coding_gene | 1.1 | 2.21 | 1.4E-02 |
| *C38D9.5* | WBGene00008013 | pseudogene | 1.1 | 2.20 | 2.6E-02 |
| *T01H3.5* | WBGene00011351 | protein_coding_gene | 1.1 | 2.20 | 8.4E-04 |
| *C01G5.4* | WBGene00015306 | protein_coding_gene | 1.1 | 2.20 | 8.0E-03 |
| *C44B9.2* | WBGene00008079 | protein_coding_gene | 1.1 | 2.20 | 1.5E-05 |
| *M28.4* | WBGene00010895 | protein_coding_gene | 1.1 | 2.20 | 4.2E-02 |
| *slo-2* | WBGene00004831 | protein_coding_gene | 1.1 | 2.20 | 8.5E-07 |
| *uggt-2* | WBGene00009178 | protein_coding_gene | 1.1 | 2.20 | 9.9E-03 |
| *F15D4.5* | WBGene00008862 | protein_coding_gene | 1.1 | 2.20 | 1.2E-02 |
| *gst-23* | WBGene00001771 | protein_coding_gene | 1.1 | 2.19 | 1.8E-02 |
| *Y66A7A.4* | WBGene00013424 | protein_coding_gene | 1.1 | 2.19 | 7.8E-03 |
| *math-44* | WBGene00021193 | protein_coding_gene | 1.1 | 2.19 | 3.3E-02 |
| *abts-3* | WBGene00019018 | protein_coding_gene | 1.1 | 2.19 | 2.6E-02 |
| *unc-55* | WBGene00006790 | protein_coding_gene | 1.1 | 2.19 | 4.7E-02 |
| *F13D12.9* | WBGene00008743 | protein_coding_gene | 1.1 | 2.19 | 3.6E-04 |
| *Y113G7C.1* | WBGene00013771 | protein_coding_gene | 1.1 | 2.19 | 1.2E-05 |
| *gst-40* | WBGene00001788 | protein_coding_gene | 1.1 | 2.19 | 1.6E-02 |
| *arrd-7* | WBGene00012464 | protein_coding_gene | 1.1 | 2.19 | 2.7E-02 |
| *Y23H5B.2* | WBGene00021273 | pseudogene | 1.1 | 2.19 | 4.3E-02 |
| *R08E3.1* | WBGene00019957 | protein_coding_gene | 1.1 | 2.19 | 1.3E-06 |
| *daf-12* | WBGene00000908 | protein_coding_gene | 1.1 | 2.18 | 4.1E-03 |
| *T08G3.4* | WBGene00011617 | pseudogene | 1.1 | 2.18 | 2.7E-02 |
| *acr-8* | WBGene00000047 | protein_coding_gene | 1.1 | 2.18 | 2.0E-02 |
| *kcnl-4* | WBGene00008265 | protein_coding_gene | 1.1 | 2.18 | 8.8E-03 |

| | | | | | |
|---|---|---|---|---|---|
| *bgnt-1.4* | WBGene00015982 | protein_coding_gene | 1.1 | 2.17 | 3.8E-02 |
| *clh-4* | WBGene00000531 | protein_coding_gene | 1.1 | 2.17 | 3.8E-02 |
| *F27E5.3* | WBGene00009193 | protein_coding_gene | 1.1 | 2.17 | 3.8E-03 |
| *K03A1.4* | WBGene00019352 | protein_coding_gene | 1.1 | 2.17 | 3.4E-02 |
| *gst-39* | WBGene00001787 | protein_coding_gene | 1.1 | 2.17 | 4.0E-02 |
| *ZK1251.3* | WBGene00014241 | protein_coding_gene | 1.1 | 2.17 | 7.8E-03 |
| *Y50E8A.6* | WBGene00013049 | protein_coding_gene | 1.1 | 2.16 | 2.7E-02 |
| *lron-4* | WBGene00008051 | protein_coding_gene | 1.1 | 2.16 | 1.0E-02 |
| *W03F8.2* | WBGene00020992 | pseudogene | 1.1 | 2.16 | 1.4E-03 |
| *C31G12.4* | WBGene00007853 | protein_coding_gene | 1.1 | 2.16 | 2.8E-02 |
| *F49E2.2* | WBGene00009886 | protein_coding_gene | 1.1 | 2.16 | 2.0E-03 |
| *W03G1.2* | WBGene00021010 | protein_coding_gene | 1.1 | 2.16 | 1.5E-02 |
| *unc-105* | WBGene00006832 | protein_coding_gene | 1.1 | 2.16 | 2.3E-02 |
| *F25B3.4* | WBGene00009101 | protein_coding_gene | 1.1 | 2.16 | 1.1E-02 |
| *F07F6.1* | WBGene00017215 | protein_coding_gene | 1.1 | 2.16 | 2.8E-03 |
| *unc-104* | WBGene00006831 | protein_coding_gene | 1.1 | 2.15 | 6.0E-03 |
| *Y76B12C.8* | WBGene00022302 | protein_coding_gene | 1.1 | 2.15 | 2.3E-02 |
| *D1086.9* | WBGene00008396 | protein_coding_gene | 1.1 | 2.15 | 1.1E-02 |
| *F38A1.9* | WBGene00009522 | protein_coding_gene | 1.1 | 2.15 | 8.0E-03 |
| *C18B2.5* | WBGene00015956 | protein_coding_gene | 1.1 | 2.15 | 1.2E-04 |
| *Y106G6H.1* | WBGene00013716 | protein_coding_gene | 1.1 | 2.15 | 1.7E-02 |
| *abt-1* | WBGene00000019 | protein_coding_gene | 1.1 | 2.14 | 4.0E-02 |
| *kin-24* | WBGene00002206 | protein_coding_gene | 1.1 | 2.14 | 3.9E-03 |
| *svh-1* | WBGene00006620 | protein_coding_gene | 1.1 | 2.14 | 3.9E-02 |
| *C34D4.2* | WBGene00016398 | protein_coding_gene | 1.1 | 2.14 | 7.8E-04 |
| *spe-47* | WBGene00021271 | protein_coding_gene | 1.1 | 2.14 | 3.4E-03 |
| *C05C12.1* | WBGene00007335 | protein_coding_gene | 1.1 | 2.14 | 2.7E-03 |
| *sqrd-1* | WBGene00008538 | protein_coding_gene | 1.1 | 2.13 | 2.1E-03 |
| *chk-2* | WBGene00000499 | protein_coding_gene | 1.1 | 2.13 | 2.4E-03 |
| *D2089.3* | WBGene00008434 | protein_coding_gene | 1.1 | 2.13 | 2.6E-02 |
| *C33F10.8* | WBGene00016356 | protein_coding_gene | 1.1 | 2.13 | 4.3E-03 |
| *dhhc-5* | WBGene00020066 | protein_coding_gene | 1.1 | 2.13 | 2.3E-02 |
| *dmsr-3* | WBGene00012993 | protein_coding_gene | 1.1 | 2.13 | 2.6E-02 |
| *Y69E1A.8* | WBGene00013478 | protein_coding_gene | 1.1 | 2.13 | 1.3E-02 |
| *F32B4.2* | WBGene00009313 | protein_coding_gene | 1.1 | 2.13 | 3.9E-03 |
| *F35E2.9* | WBGene00009416 | protein_coding_gene | 1.1 | 2.12 | 5.9E-03 |
| *bus-2* | WBGene00044618 | protein_coding_gene | 1.1 | 2.12 | 3.2E-02 |
| *csp-1* | WBGene00000819 | protein_coding_gene | 1.1 | 2.12 | 3.6E-02 |
| *ZK896.3* | WBGene00014134 | pseudogene | 1.1 | 2.12 | 1.7E-02 |
| *nas-31* | WBGene00003549 | protein_coding_gene | 1.1 | 2.12 | 4.3E-02 |
| *ceh-60* | WBGene00017690 | protein_coding_gene | 1.1 | 2.12 | 3.5E-09 |
| *twk-30* | WBGene00006682 | protein_coding_gene | 1.1 | 2.12 | 5.1E-04 |
| *mfsd-6* | WBGene00020051 | protein_coding_gene | 1.1 | 2.12 | 4.3E-04 |
| *srb-16* | WBGene00019025 | protein_coding_gene | 1.1 | 2.12 | 3.5E-02 |
| *C54G4.2* | WBGene00008312 | protein_coding_gene | 1.1 | 2.12 | 4.7E-03 |
| *T15B12.2* | WBGene00020531 | protein_coding_gene | 1.1 | 2.12 | 7.2E-03 |
| *unc-103* | WBGene00006830 | protein_coding_gene | 1.1 | 2.12 | 1.0E-03 |
| *Y20F4.8* | WBGene00044435 | protein_coding_gene | 1.1 | 2.12 | 3.9E-02 |
| *sulp-6* | WBGene00020914 | protein_coding_gene | 1.1 | 2.12 | 2.6E-03 |
| *Y39G8C.2* | WBGene00012731 | protein_coding_gene | 1.1 | 2.11 | 4.4E-02 |
| *unc-25* | WBGene00006762 | protein_coding_gene | 1.1 | 2.11 | 3.8E-04 |
| *F40G12.10* | WBGene00009605 | protein_coding_gene | 1.1 | 2.11 | 1.7E-02 |
| *R04B5.11* | WBGene00011008 | protein_coding_gene | 1.1 | 2.11 | 5.0E-02 |
| *Y37E3.19* | WBGene00044810 | protein_coding_gene | 1.1 | 2.11 | 2.1E-02 |
| *frpr-6* | WBGene00017661 | protein_coding_gene | 1.1 | 2.11 | 3.4E-02 |
| *F46F5.14* | WBGene00018505 | protein_coding_gene | 1.1 | 2.11 | 1.5E-02 |
| *F23B12.1* | WBGene00009079 | protein_coding_gene | 1.1 | 2.11 | 1.1E-02 |
| *Y69A2AR.14* | WBGene00022085 | pseudogene | 1.1 | 2.11 | 4.6E-02 |

| | | | | | |
|---|---|---|---|---|---|
| *ptr-11* | WBGene00004225 | protein_coding_gene | 1.1 | 2.10 | 2.2E-02 |
| *K10C9.7* | WBGene00019626 | protein_coding_gene | 1.1 | 2.10 | 6.8E-04 |
| *C01B10.44* | WBGene00194839 | protein_coding_gene | 1.1 | 2.10 | 3.6E-02 |
| *F58B6.1* | WBGene00019029 | protein_coding_gene | 1.1 | 2.10 | 1.6E-02 |
| *M04D8.7* | WBGene00010865 | protein_coding_gene | 1.1 | 2.10 | 1.6E-02 |
| *K06H7.8* | WBGene00019459 | protein_coding_gene | 1.1 | 2.10 | 4.1E-02 |
| *fbxa-50* | WBGene00013844 | protein_coding_gene | 1.1 | 2.10 | 3.2E-02 |
| *B0496.1* | WBGene00015210 | protein_coding_gene | 1.1 | 2.10 | 6.7E-03 |
| *T01C8.3* | WBGene00020144 | protein_coding_gene | 1.1 | 2.10 | 3.3E-02 |
| *Y53C10A.15* | WBGene00044805 | protein_coding_gene | 1.1 | 2.09 | 3.5E-02 |
| *unc-49* | WBGene00006784 | protein_coding_gene | 1.1 | 2.09 | 1.3E-03 |
| *C50D2.3* | WBGene00016807 | protein_coding_gene | 1.1 | 2.09 | 5.6E-03 |
| *kin-1* | WBGene00002189 | protein_coding_gene | 1.1 | 2.09 | 9.2E-03 |
| *unc-82* | WBGene00006814 | protein_coding_gene | 1.1 | 2.09 | 3.6E-02 |
| *B0285.6* | WBGene00007138 | protein_coding_gene | 1.1 | 2.08 | 3.2E-02 |
| *Y105C5B.11* | WBGene00013651 | protein_coding_gene | 1.1 | 2.08 | 1.5E-02 |
| *K08D8.12* | WBGene00194926 | protein_coding_gene | 1.1 | 2.08 | 3.9E-02 |
| *ZK1053.6* | WBGene00014201 | protein_coding_gene | 1.1 | 2.07 | 1.3E-02 |
| *T04A8.13* | WBGene00011414 | protein_coding_gene | 1.1 | 2.07 | 2.2E-02 |
| *C18E3.1* | WBGene00015970 | protein_coding_gene | 1.1 | 2.07 | 1.3E-02 |
| *Y60A3A.24* | WBGene00023519 | protein_coding_gene | 1.1 | 2.07 | 4.3E-03 |
| *osta-1* | WBGene00015287 | protein_coding_gene | 1.1 | 2.07 | 6.5E-03 |
| *dlk-1* | WBGene00001008 | protein_coding_gene | 1.0 | 2.07 | 9.7E-03 |
| *AH6.3* | WBGene00007081 | protein_coding_gene | 1.0 | 2.07 | 1.2E-02 |
| *cutl-15* | WBGene00011888 | protein_coding_gene | 1.0 | 2.07 | 2.1E-03 |
| *C51E3.9* | WBGene00008254 | protein_coding_gene | 1.0 | 2.07 | 4.1E-02 |
| *nhr-132* | WBGene00003722 | protein_coding_gene | 1.0 | 2.07 | 3.8E-02 |
| *F54F12.1* | WBGene00010072 | protein_coding_gene | 1.0 | 2.07 | 1.3E-02 |
| *clec-187* | WBGene00014137 | protein_coding_gene | 1.0 | 2.07 | 1.4E-02 |
| *unc-77* | WBGene00006809 | protein_coding_gene | 1.0 | 2.07 | 4.2E-02 |
| *C14A6.13* | WBGene00050956 | protein_coding_gene | 1.0 | 2.06 | 2.5E-02 |
| *snpc-3.1* | WBGene00015098 | protein_coding_gene | 1.0 | 2.06 | 2.1E-02 |
| *K03B8.6* | WBGene00010524 | protein_coding_gene | 1.0 | 2.06 | 1.4E-02 |
| *Y102A5C.4* | WBGene00013613 | protein_coding_gene | 1.0 | 2.06 | 8.7E-03 |
| *mpz-4* | WBGene00019165 | protein_coding_gene | 1.0 | 2.06 | 9.2E-03 |
| *tag-191* | WBGene00007049 | protein_coding_gene | 1.0 | 2.06 | 5.1E-03 |
| *Y67A10A.3* | WBGene00013452 | protein_coding_gene | 1.0 | 2.06 | 2.1E-02 |
| *haf-9* | WBGene00001819 | protein_coding_gene | 1.0 | 2.06 | 3.6E-02 |
| *Y43C5B.2* | WBGene00012785 | protein_coding_gene | 1.0 | 2.06 | 4.1E-03 |
| *F37A4.5* | WBGene00018135 | protein_coding_gene | 1.0 | 2.06 | 8.0E-04 |
| *mxt-1* | WBGene00012478 | protein_coding_gene | 1.0 | 2.06 | 2.0E-03 |
| *tra-1* | WBGene00006604 | protein_coding_gene | 1.0 | 2.06 | 5.1E-04 |
| *T11F8.4* | WBGene00020435 | protein_coding_gene | 1.0 | 2.06 | 1.4E-04 |
| *F46A9.2* | WBGene00009755 | protein_coding_gene | 1.0 | 2.05 | 1.6E-02 |
| *C31H1.1* | WBGene00016286 | protein_coding_gene | 1.0 | 2.05 | 3.4E-03 |
| *T22D1.8* | WBGene00020686 | protein_coding_gene | 1.0 | 2.05 | 2.4E-02 |
| *nep-17* | WBGene00010070 | protein_coding_gene | 1.0 | 2.05 | 1.9E-02 |
| *R03D7.8* | WBGene00010992 | protein_coding_gene | 1.0 | 2.05 | 4.9E-03 |
| *R07E5.6* | WBGene00011114 | protein_coding_gene | 1.0 | 2.05 | 5.1E-03 |
| *Y2H9A.6* | WBGene00086568 | protein_coding_gene | 1.0 | 2.05 | 1.3E-02 |
| *alh-12* | WBGene00000118 | protein_coding_gene | 1.0 | 2.05 | 4.0E-02 |
| *sox-4* | WBGene00015716 | protein_coding_gene | 1.0 | 2.05 | 2.9E-03 |
| *F59A6.2* | WBGene00019084 | protein_coding_gene | 1.0 | 2.04 | 9.5E-03 |
| *inx-1* | WBGene00002123 | protein_coding_gene | 1.0 | 2.04 | 4.5E-04 |
| *nrfl-1* | WBGene00006438 | protein_coding_gene | 1.0 | 2.04 | 5.2E-03 |
| *unc-2* | WBGene00006742 | protein_coding_gene | 1.0 | 2.04 | 4.9E-03 |
| *F23B2.7* | WBGene00009075 | protein_coding_gene | 1.0 | 2.04 | 1.2E-02 |
| *Y34B4A.4* | WBGene00021320 | protein_coding_gene | 1.0 | 2.03 | 8.9E-03 |

| | | | | | |
|---|---|---|---|---|---|
| *F10G8.1* | WBGene00008661 | protein_coding_gene | 1.0 | 2.03 | 1.6E-02 |
| *egl-13* | WBGene00001182 | protein_coding_gene | 1.0 | 2.03 | 3.3E-02 |
| *elo-7* | WBGene00001245 | protein_coding_gene | 1.0 | 2.03 | 8.4E-03 |
| *C15A11.2* | WBGene00007597 | protein_coding_gene | 1.0 | 2.03 | 1.4E-02 |
| *Y1A5A.1* | WBGene00012379 | protein_coding_gene | 1.0 | 2.03 | 2.6E-03 |
| *pbo-5* | WBGene00012857 | protein_coding_gene | 1.0 | 2.03 | 1.8E-02 |
| *trpl-5* | WBGene00004149 | protein_coding_gene | 1.0 | 2.03 | 4.0E-02 |
| *glb-14* | WBGene00008996 | protein_coding_gene | 1.0 | 2.02 | 4.2E-02 |
| *Y18H1A.11* | WBGene00021215 | protein_coding_gene | 1.0 | 2.02 | 8.6E-03 |
| *R102.10* | WBGene00011297 | protein_coding_gene | 1.0 | 2.02 | 9.5E-03 |
| *linc-8* | WBGene00219707 | lincRNA_gene | 1.0 | 2.02 | 7.9E-04 |
| *Y71H2AM.12* | WBGene00022177 | protein_coding_gene | 1.0 | 2.02 | 1.1E-03 |
| *cest-12* | WBGene00013540 | protein_coding_gene | 1.0 | 2.02 | 7.9E-04 |
| *F46C5.4* | WBGene00009780 | protein_coding_gene | 1.0 | 2.02 | 3.7E-02 |
| *W05F2.4* | WBGene00021036 | protein_coding_gene | 1.0 | 2.02 | 4.3E-02 |
| *F46A9.1* | WBGene00009754 | protein_coding_gene | 1.0 | 2.02 | 3.0E-02 |
| *B0280.2* | WBGene00015100 | protein_coding_gene | 1.0 | 2.02 | 1.3E-02 |
| *C55C3.3* | WBGene00016953 | protein_coding_gene | 1.0 | 2.02 | 3.0E-03 |
| *ZC513.3* | WBGene00022628 | protein_coding_gene | 1.0 | 2.02 | 2.8E-02 |
| *R13A5.11* | WBGene00020053 | protein_coding_gene | 1.0 | 2.02 | 2.6E-02 |
| *pmp-4* | WBGene00004061 | protein_coding_gene | 1.0 | 2.01 | 2.2E-02 |
| *ZK228.4* | WBGene00013947 | protein_coding_gene | 1.0 | 2.01 | 3.2E-02 |
| *alg-3* | WBGene00011910 | protein_coding_gene | 1.0 | 2.01 | 8.9E-03 |
| *F57A8.6* | WBGene00010181 | protein_coding_gene | 1.0 | 2.01 | 1.6E-02 |
| *aat-6* | WBGene00000007 | protein_coding_gene | 1.0 | 2.01 | 2.2E-02 |
| *F56D6.17* | WBGene00219307 | pseudogene | 1.0 | 2.01 | 7.9E-03 |
| *nkb-3* | WBGene00010117 | protein_coding_gene | 1.0 | 2.01 | 1.8E-02 |
| *wht-5* | WBGene00008950 | protein_coding_gene | 1.0 | 2.01 | 4.7E-03 |
| *F54F12.2* | WBGene00010073 | protein_coding_gene | 1.0 | 2.01 | 2.3E-02 |
| *nca-2* | WBGene00003558 | protein_coding_gene | 1.0 | 2.01 | 3.3E-02 |
| *F44D12.11* | WBGene00014375 | pseudogene | 1.0 | 2.01 | 2.7E-02 |
| *T25B9.6* | WBGene00012012 | protein_coding_gene | 1.0 | 2.01 | 3.2E-03 |
| *clhm-1* | WBGene00016626 | protein_coding_gene | 1.0 | 2.00 | 2.3E-02 |
| *C49A9.9* | WBGene00016763 | protein_coding_gene | 1.0 | 2.00 | 9.9E-03 |
| *fbxa-212* | WBGene00012568 | pseudogene | -1.0 | 0.50 | 3.9E-02 |
| *C05D9.7* | WBGene00015474 | protein_coding_gene | -1.0 | 0.50 | 3.4E-03 |
| *pri-1* | WBGene00004180 | protein_coding_gene | -1.0 | 0.50 | 3.2E-03 |
| *F10E9.7* | WBGene00017358 | protein_coding_gene | -1.0 | 0.50 | 1.2E-02 |
| *rfc-4* | WBGene00004340 | protein_coding_gene | -1.0 | 0.50 | 2.8E-04 |
| *C10B5.1* | WBGene00015670 | protein_coding_gene | -1.0 | 0.50 | 9.8E-04 |
| *dhs-27* | WBGene00000990 | protein_coding_gene | -1.0 | 0.50 | 3.1E-03 |
| *mop-25.3* | WBGene00020858 | protein_coding_gene | -1.0 | 0.50 | 7.9E-03 |
| *ham-1* | WBGene00001820 | protein_coding_gene | -1.0 | 0.50 | 1.9E-03 |
| *T01C3.2* | WBGene00011319 | protein_coding_gene | -1.0 | 0.50 | 3.9E-02 |
| *R13A5.15* | WBGene00194726 | protein_coding_gene | -1.0 | 0.50 | 3.2E-02 |
| *C50F4.12* | WBGene00008236 | protein_coding_gene | -1.0 | 0.50 | 1.0E-04 |
| *M01E11.1* | WBGene00019710 | protein_coding_gene | -1.0 | 0.50 | 4.3E-03 |
| *rnp-2* | WBGene00004385 | protein_coding_gene | -1.0 | 0.50 | 1.3E-04 |
| *cdh-4* | WBGene00000396 | protein_coding_gene | -1.0 | 0.50 | 3.4E-02 |
| *sld-2* | WBGene00020466 | protein_coding_gene | -1.0 | 0.50 | 2.2E-02 |
| *gst-7* | WBGene00001755 | protein_coding_gene | -1.0 | 0.50 | 4.9E-02 |
| *smo-1* | WBGene00004888 | protein_coding_gene | -1.0 | 0.50 | 2.6E-03 |
| *taf-9* | WBGene00006391 | protein_coding_gene | -1.0 | 0.50 | 4.4E-05 |
| *C01G12.12* | WBGene00044053 | pseudogene | -1.0 | 0.50 | 1.6E-03 |
| *C32E8.5* | WBGene00016323 | protein_coding_gene | -1.0 | 0.50 | 1.8E-04 |
| *bmy-1* | WBGene00017293 | protein_coding_gene | -1.0 | 0.50 | 9.8E-03 |
| *M117.6* | WBGene00044253 | protein_coding_gene | -1.0 | 0.50 | 9.9E-04 |
| *ceh-58* | WBGene00007417 | protein_coding_gene | -1.0 | 0.49 | 2.5E-02 |

| | | | | | |
|---|---|---|---|---|---|
| *ttr-37* | WBGene00017667 | protein_coding_gene | -1.0 | 0.49 | 4.9E-02 |
| *B0336.13* | WBGene00015150 | protein_coding_gene | -1.0 | 0.49 | 8.6E-03 |
| *kbp-4* | WBGene00022357 | protein_coding_gene | -1.0 | 0.49 | 4.9E-02 |
| *W04A8.2* | WBGene00012237 | protein_coding_gene | -1.0 | 0.49 | 4.1E-03 |
| *aph-1* | WBGene00000147 | protein_coding_gene | -1.0 | 0.49 | 6.6E-05 |
| *mdt-8* | WBGene00007013 | protein_coding_gene | -1.0 | 0.49 | 1.6E-02 |
| *kbp-2* | WBGene00017832 | protein_coding_gene | -1.0 | 0.49 | 2.5E-02 |
| *hrpf-1* | WBGene00020936 | protein_coding_gene | -1.0 | 0.49 | 2.6E-07 |
| *odd-1* | WBGene00003845 | protein_coding_gene | -1.0 | 0.49 | 3.0E-02 |
| *ccnk-1* | WBGene00009650 | protein_coding_gene | -1.0 | 0.49 | 6.3E-04 |
| *ooc-5* | WBGene00003870 | protein_coding_gene | -1.0 | 0.49 | 1.4E-02 |
| *mob-1* | WBGene00009551 | protein_coding_gene | -1.0 | 0.49 | 8.2E-03 |
| *erg-28* | WBGene00007589 | protein_coding_gene | -1.0 | 0.49 | 9.4E-03 |
| *mdt-11* | WBGene00007015 | protein_coding_gene | -1.0 | 0.49 | 1.9E-02 |
| *T26A5.8* | WBGene00020823 | protein_coding_gene | -1.0 | 0.49 | 2.1E-02 |
| *ssup-72* | WBGene00020480 | protein_coding_gene | -1.0 | 0.49 | 2.9E-04 |
| *fsn-1* | WBGene00001499 | protein_coding_gene | -1.0 | 0.49 | 1.2E-02 |
| *gut-2* | WBGene00001808 | protein_coding_gene | -1.0 | 0.49 | 2.9E-03 |
| *Y6D1A.1* | WBGene00012395 | protein_coding_gene | -1.0 | 0.49 | 6.8E-06 |
| *F58H1.8* | WBGene00138720 | protein_coding_gene | -1.0 | 0.49 | 2.4E-03 |
| *fubl-4* | WBGene00019692 | protein_coding_gene | -1.0 | 0.49 | 1.3E-04 |
| *mdt-6* | WBGene00003164 | protein_coding_gene | -1.0 | 0.49 | 2.1E-02 |
| *zipt-1* | WBGene00009286 | protein_coding_gene | -1.0 | 0.49 | 2.4E-02 |
| *dlc-5* | WBGene00043067 | protein_coding_gene | -1.0 | 0.49 | 1.7E-02 |
| *set-10* | WBGene00009370 | protein_coding_gene | -1.0 | 0.49 | 1.4E-05 |
| *mut-2* | WBGene00003499 | protein_coding_gene | -1.0 | 0.49 | 3.2E-03 |
| *Y105C5B.19* | WBGene00013658 | protein_coding_gene | -1.0 | 0.49 | 2.1E-02 |
| *cya-1* | WBGene00000863 | protein_coding_gene | -1.0 | 0.49 | 2.4E-02 |
| *fut-3* | WBGene00006402 | protein_coding_gene | -1.0 | 0.49 | 2.6E-03 |
| *metl-6* | WBGene00014205 | protein_coding_gene | -1.0 | 0.49 | 7.6E-04 |
| *lsm-3* | WBGene00003077 | protein_coding_gene | -1.0 | 0.49 | 1.0E-03 |
| *K08E4.2* | WBGene00010666 | protein_coding_gene | -1.0 | 0.49 | 1.9E-02 |
| *blos-8* | WBGene00010981 | protein_coding_gene | -1.0 | 0.48 | 1.6E-02 |
| *D2007.1* | WBGene00017041 | protein_coding_gene | -1.0 | 0.48 | 1.4E-04 |
| *Y41D4B.14* | WBGene00021516 | protein_coding_gene | -1.0 | 0.48 | 7.2E-03 |
| *C17E4.20* | WBGene00219313 | protein_coding_gene | -1.1 | 0.48 | 7.2E-03 |
| *D2089.8* | WBGene00271706 | protein_coding_gene | -1.1 | 0.48 | 1.3E-04 |
| *K01A2.3* | WBGene00019278 | pseudogene | -1.1 | 0.48 | 3.7E-02 |
| *C29F3.7* | WBGene00007807 | protein_coding_gene | -1.1 | 0.48 | 3.3E-02 |
| *lir-3* | WBGene00003046 | protein_coding_gene | -1.1 | 0.48 | 1.2E-02 |
| *C10C6.9* | WBGene00045175 | pseudogene | -1.1 | 0.48 | 3.8E-02 |
| *C14B1.12* | WBGene00184990 | protein_coding_gene | -1.1 | 0.48 | 2.6E-03 |
| *lbp-4* | WBGene00002256 | protein_coding_gene | -1.1 | 0.48 | 1.6E-04 |
| *cdc-7* | WBGene00016421 | protein_coding_gene | -1.1 | 0.48 | 2.9E-02 |
| *Y62F5A.12* | WBGene00077453 | protein_coding_gene | -1.1 | 0.48 | 7.3E-04 |
| *Y71F9AR.2* | WBGene00022123 | protein_coding_gene | -1.1 | 0.48 | 1.4E-03 |
| *R09B3.6* | WBGene00014820 | pseudogene | -1.1 | 0.48 | 2.1E-02 |
| *Y62E10A.2* | WBGene00013373 | protein_coding_gene | -1.1 | 0.48 | 2.3E-02 |
| *T18D3.1* | WBGene00011820 | protein_coding_gene | -1.1 | 0.48 | 3.9E-02 |
| *K03H1.7* | WBGene00010542 | protein_coding_gene | -1.1 | 0.48 | 3.6E-02 |
| *F23H12.10* | WBGene00044218 | protein_coding_gene | -1.1 | 0.48 | 3.7E-02 |
| *btb-19* | WBGene00010194 | protein_coding_gene | -1.1 | 0.48 | 7.1E-03 |
| *T06A1.1* | WBGene00020277 | protein_coding_gene | -1.1 | 0.48 | 4.6E-02 |
| *Y54G9A.5* | WBGene00013208 | protein_coding_gene | -1.1 | 0.48 | 3.1E-02 |
| *Y66D12A.7* | WBGene00013433 | protein_coding_gene | -1.1 | 0.47 | 6.2E-03 |
| *C05E11.3* | WBGene00015494 | protein_coding_gene | -1.1 | 0.47 | 2.1E-02 |
| *F01D4.5* | WBGene00008488 | protein_coding_gene | -1.1 | 0.47 | 7.2E-03 |
| *set-13* | WBGene00019690 | protein_coding_gene | -1.1 | 0.47 | 1.7E-02 |

| | | | | | |
|---|---|---|---|---|---|
| *tag-342* | WBGene00007187 | protein_coding_gene | -1.1 | 0.47 | 4.3E-03 |
| *frpr-11* | WBGene00019444 | protein_coding_gene | -1.1 | 0.47 | 3.3E-04 |
| *cdk-4* | WBGene00000406 | protein_coding_gene | -1.1 | 0.47 | 7.4E-03 |
| *sqv-4* | WBGene00005022 | protein_coding_gene | -1.1 | 0.47 | 1.7E-02 |
| *F42H10.2* | WBGene00018366 | protein_coding_gene | -1.1 | 0.47 | 3.3E-03 |
| *T03F1.12* | WBGene00020193 | protein_coding_gene | -1.1 | 0.47 | 3.7E-05 |
| *Y52E8A.2* | WBGene00021795 | protein_coding_gene | -1.1 | 0.47 | 6.6E-03 |
| *thk-1* | WBGene00006568 | protein_coding_gene | -1.1 | 0.47 | 8.6E-04 |
| *F47G3.3* | WBGene00018578 | protein_coding_gene | -1.1 | 0.47 | 5.0E-02 |
| *ced-9* | WBGene00000423 | protein_coding_gene | -1.1 | 0.47 | 3.5E-05 |
| *F40D4.17* | WBGene00194682 | protein_coding_gene | -1.1 | 0.47 | 4.7E-02 |
| *hpo-9* | WBGene00015463 | protein_coding_gene | -1.1 | 0.47 | 3.5E-03 |
| *C41D11.9* | WBGene00016567 | protein_coding_gene | -1.1 | 0.47 | 1.6E-03 |
| *kbp-3* | WBGene00009179 | protein_coding_gene | -1.1 | 0.47 | 8.3E-03 |
| *F52B5.7* | WBGene00009923 | protein_coding_gene | -1.1 | 0.47 | 1.9E-02 |
| *apr-1* | WBGene00000156 | protein_coding_gene | -1.1 | 0.47 | 7.5E-03 |
| *T05B9.1* | WBGene00011464 | protein_coding_gene | -1.1 | 0.47 | 4.3E-03 |
| *ska-1* | WBGene00013725 | protein_coding_gene | -1.1 | 0.47 | 1.4E-04 |
| *C06A5.12* | WBGene00044414 | protein_coding_gene | -1.1 | 0.47 | 4.6E-02 |
| *pie-1* | WBGene00004027 | protein_coding_gene | -1.1 | 0.47 | 1.5E-02 |
| *klc-1* | WBGene00002214 | protein_coding_gene | -1.1 | 0.47 | 4.0E-02 |
| *F10E9.11* | WBGene00017360 | protein_coding_gene | -1.1 | 0.47 | 2.6E-03 |
| *2RSSE.3* | WBGene00044951 | pseudogene | -1.1 | 0.47 | 4.8E-02 |
| *F47B10.9* | WBGene00009820 | protein_coding_gene | -1.1 | 0.47 | 1.8E-05 |
| *F37D6.4* | WBGene00009510 | protein_coding_gene | -1.1 | 0.46 | 2.1E-02 |
| *Y54G2A.12* | WBGene00021877 | protein_coding_gene | -1.1 | 0.46 | 1.7E-02 |
| *sas-5* | WBGene00009385 | protein_coding_gene | -1.1 | 0.46 | 1.3E-02 |
| *cgh-1* | WBGene00000479 | protein_coding_gene | -1.1 | 0.46 | 2.5E-02 |
| *Y69A2AR.21* | WBGene00022092 | protein_coding_gene | -1.1 | 0.46 | 5.7E-03 |
| *ceh-74* | WBGene00013876 | protein_coding_gene | -1.1 | 0.46 | 3.4E-03 |
| *fbxb-7* | WBGene00012492 | protein_coding_gene | -1.1 | 0.46 | 4.4E-05 |
| *T07D4.5* | WBGene00045407 | protein_coding_gene | -1.1 | 0.46 | 1.1E-02 |
| *cmt-1* | WBGene00016564 | protein_coding_gene | -1.1 | 0.46 | 3.5E-02 |
| *C50F4.4* | WBGene00008232 | protein_coding_gene | -1.1 | 0.46 | 2.9E-04 |
| *mei-1* | WBGene00003183 | protein_coding_gene | -1.1 | 0.46 | 3.1E-02 |
| *efl-3* | WBGene00009899 | protein_coding_gene | -1.1 | 0.46 | 1.5E-02 |
| *R11G1.7* | WBGene00044725 | protein_coding_gene | -1.1 | 0.46 | 1.7E-02 |
| *rfc-3* | WBGene00004339 | protein_coding_gene | -1.1 | 0.46 | 1.5E-02 |
| *orc-4* | WBGene00012651 | protein_coding_gene | -1.1 | 0.46 | 2.7E-04 |
| *C48B6.9* | WBGene00016745 | protein_coding_gene | -1.1 | 0.46 | 2.9E-02 |
| *M199.2* | WBGene00010951 | protein_coding_gene | -1.1 | 0.45 | 9.4E-03 |
| *C06E1.1* | WBGene00015518 | protein_coding_gene | -1.1 | 0.45 | 3.5E-02 |
| *vang-1* | WBGene00015171 | protein_coding_gene | -1.1 | 0.45 | 2.7E-02 |
| *madf-8* | WBGene00008118 | protein_coding_gene | -1.1 | 0.45 | 4.9E-03 |
| *F20D12.12* | WBGene00249815 | protein_coding_gene | -1.1 | 0.45 | 1.6E-02 |
| *mom-1* | WBGene00003394 | protein_coding_gene | -1.1 | 0.45 | 3.6E-02 |
| *phf-31* | WBGene00008747 | protein_coding_gene | -1.1 | 0.45 | 7.0E-03 |
| *Y71A12B.12* | WBGene00013509 | protein_coding_gene | -1.1 | 0.45 | 1.7E-02 |
| *dscc-1* | WBGene00019595 | protein_coding_gene | -1.2 | 0.45 | 1.6E-06 |
| *lsm-7* | WBGene00003081 | protein_coding_gene | -1.2 | 0.45 | 8.5E-07 |
| *dsh-2* | WBGene00001102 | protein_coding_gene | -1.2 | 0.45 | 2.8E-02 |
| *M03B6.4* | WBGene00010836 | protein_coding_gene | -1.2 | 0.45 | 2.7E-02 |
| *bcl-7* | WBGene00016192 | protein_coding_gene | -1.2 | 0.45 | 2.5E-02 |
| *K07A1.15* | WBGene00010619 | protein_coding_gene | -1.2 | 0.45 | 3.9E-03 |
| *F35D2.3* | WBGene00018041 | protein_coding_gene | -1.2 | 0.45 | 3.0E-02 |
| *T16H12.1* | WBGene00011811 | protein_coding_gene | -1.2 | 0.45 | 1.1E-04 |
| *Y58A7A.2* | WBGene00021976 | protein_coding_gene | -1.2 | 0.45 | 2.8E-02 |
| *srm-1* | WBGene00005638 | protein_coding_gene | -1.2 | 0.44 | 3.6E-02 |

| | | | | | |
|---|---|---|---|---|---|
| *cht-1* | WBGene00000503 | protein_coding_gene | -1.2 | 0.44 | 8.7E-09 |
| *cup-15* | WBGene00009627 | protein_coding_gene | -1.2 | 0.44 | 1.9E-02 |
| *cutl-11* | WBGene00011443 | protein_coding_gene | -1.2 | 0.44 | 4.0E-02 |
| *T09B4.2* | WBGene00020376 | protein_coding_gene | -1.2 | 0.44 | 4.3E-03 |
| *fbxa-211* | WBGene00012566 | protein_coding_gene | -1.2 | 0.44 | 1.4E-03 |
| *F28H1.1* | WBGene00017908 | protein_coding_gene | -1.2 | 0.44 | 8.2E-03 |
| *B0511.19* | WBGene00255714 | protein_coding_gene | -1.2 | 0.44 | 4.2E-04 |
| *ZK1010.10* | WBGene00164985 | protein_coding_gene | -1.2 | 0.44 | 3.0E-04 |
| *polk-1* | WBGene00017696 | protein_coding_gene | -1.2 | 0.44 | 2.1E-02 |
| *mdf-2* | WBGene00003161 | protein_coding_gene | -1.2 | 0.44 | 1.0E-03 |
| *Y39G8B.2* | WBGene00012723 | protein_coding_gene | -1.2 | 0.44 | 1.4E-04 |
| *ssna-1* | WBGene00020300 | protein_coding_gene | -1.2 | 0.44 | 2.6E-03 |
| *M01D1.12* | WBGene00269373 | protein_coding_gene | -1.2 | 0.44 | 2.3E-02 |
| *C25A11.2* | WBGene00016087 | protein_coding_gene | -1.2 | 0.44 | 4.3E-02 |
| *Y38H6C.23* | WBGene00012634 | protein_coding_gene | -1.2 | 0.44 | 3.4E-03 |
| *cey-3* | WBGene00000474 | protein_coding_gene | -1.2 | 0.44 | 1.5E-02 |
| *pph-4.2* | WBGene00004086 | protein_coding_gene | -1.2 | 0.44 | 4.4E-02 |
| *F42A9.8* | WBGene00018337 | protein_coding_gene | -1.2 | 0.43 | 3.5E-03 |
| *fbxb-85* | WBGene00011267 | protein_coding_gene | -1.2 | 0.43 | 2.7E-02 |
| *gmn-1* | WBGene00013552 | protein_coding_gene | -1.2 | 0.43 | 6.2E-03 |
| *Y75B8A.18* | WBGene00013553 | protein_coding_gene | -1.2 | 0.43 | 1.1E-02 |
| *C42C1.12* | WBGene00016590 | protein_coding_gene | -1.2 | 0.43 | 2.3E-03 |
| *C07E3.9* | WBGene00007419 | protein_coding_gene | -1.2 | 0.43 | 8.4E-03 |
| *K05C4.4* | WBGene00010581 | protein_coding_gene | -1.2 | 0.43 | 4.2E-03 |
| *lsm-1* | WBGene00003076 | protein_coding_gene | -1.2 | 0.43 | 1.5E-02 |
| *F40G9.20* | WBGene00195034 | ncRNA_gene | -1.2 | 0.43 | 1.3E-02 |
| *ZK643.2* | WBGene00014034 | protein_coding_gene | -1.2 | 0.43 | 1.4E-02 |
| *blos-9* | WBGene00010767 | protein_coding_gene | -1.2 | 0.43 | 2.0E-07 |
| *H31G24.3* | WBGene00019254 | protein_coding_gene | -1.2 | 0.42 | 1.9E-04 |
| *R07H5.11* | WBGene00011131 | protein_coding_gene | -1.2 | 0.42 | 2.0E-04 |
| *C36B1.14* | WBGene00044614 | protein_coding_gene | -1.2 | 0.42 | 3.8E-04 |
| *oac-6* | WBGene00007833 | protein_coding_gene | -1.2 | 0.42 | 1.3E-02 |
| *Y13C8A.4* | WBGene00219303 | pseudogene | -1.3 | 0.42 | 1.6E-02 |
| *Y74C9A.9* | WBGene00235381 | pseudogene | -1.3 | 0.42 | 4.4E-03 |
| *Y42H9B.3* | WBGene00021541 | protein_coding_gene | -1.3 | 0.42 | 1.4E-02 |
| *str-163* | WBGene00006208 | protein_coding_gene | -1.3 | 0.42 | 7.6E-03 |
| *C50F4.6* | WBGene00001910 | protein_coding_gene | -1.3 | 0.42 | 6.9E-03 |
| *cpt-1* | WBGene00012907 | protein_coding_gene | -1.3 | 0.42 | 8.9E-06 |
| *T02C1.1* | WBGene00011365 | protein_coding_gene | -1.3 | 0.42 | 1.8E-02 |
| *ins-7* | WBGene00002090 | protein_coding_gene | -1.3 | 0.42 | 4.6E-05 |
| *B0511.2* | WBGene00015228 | protein_coding_gene | -1.3 | 0.42 | 5.2E-03 |
| *duxl-1* | WBGene00022554 | protein_coding_gene | -1.3 | 0.42 | 4.5E-03 |
| *Y7A9D.1* | WBGene00012420 | protein_coding_gene | -1.3 | 0.42 | 3.0E-02 |
| *eya-1* | WBGene00001377 | protein_coding_gene | -1.3 | 0.41 | 3.4E-02 |
| *lido-11* | WBGene00016574 | protein_coding_gene | -1.3 | 0.41 | 3.7E-05 |
| *E01G4.7* | WBGene00045410 | pseudogene | -1.3 | 0.41 | 1.0E-02 |
| *BE0003N10.3* | WBGene00015268 | protein_coding_gene | -1.3 | 0.41 | 3.1E-04 |
| *crn-4* | WBGene00000797 | protein_coding_gene | -1.3 | 0.41 | 1.3E-03 |
| *ref-1* | WBGene00004334 | protein_coding_gene | -1.3 | 0.41 | 3.4E-02 |
| *ZK686.1* | WBGene00022791 | protein_coding_gene | -1.3 | 0.41 | 1.1E-05 |
| *linc-16* | WBGene00219728 | lincRNA_gene | -1.3 | 0.41 | 2.6E-02 |
| *ceh-86* | WBGene00018355 | protein_coding_gene | -1.3 | 0.41 | 2.6E-02 |
| *Y38H6A.4* | WBGene00014874 | pseudogene | -1.3 | 0.41 | 3.3E-02 |
| *E04A4.3* | WBGene00017118 | pseudogene | -1.3 | 0.41 | 2.3E-02 |
| *gadr-5* | WBGene00045058 | protein_coding_gene | -1.3 | 0.41 | 2.6E-03 |
| *ttr-52* | WBGene00008719 | protein_coding_gene | -1.3 | 0.40 | 4.3E-02 |
| *C14B1.9* | WBGene00007580 | protein_coding_gene | -1.3 | 0.40 | 4.7E-02 |
| *ceh-43* | WBGene00000463 | protein_coding_gene | -1.3 | 0.40 | 6.5E-03 |

| | | | | | |
|---|---|---|---|---|---|
| *ceh-53* | WBGene00015651 | protein_coding_gene | -1.3 | 0.40 | 3.8E-02 |
| *F40H3.6* | WBGene00018252 | protein_coding_gene | -1.3 | 0.40 | 3.6E-06 |
| *btb-11* | WBGene00019700 | protein_coding_gene | -1.3 | 0.40 | 2.9E-02 |
| *ugt-9* | WBGene00020587 | protein_coding_gene | -1.3 | 0.40 | 4.9E-02 |
| *C16A3.2* | WBGene00015807 | protein_coding_gene | -1.3 | 0.40 | 2.8E-04 |
| *blos-4* | WBGene00020783 | protein_coding_gene | -1.3 | 0.40 | 8.3E-03 |
| *Y47D3A.1* | WBGene00012922 | protein_coding_gene | -1.3 | 0.40 | 1.2E-02 |
| *epg-2* | WBGene00021468 | protein_coding_gene | -1.3 | 0.40 | 2.1E-02 |
| *lido-9* | WBGene00016576 | protein_coding_gene | -1.3 | 0.40 | 4.9E-02 |
| *Y53G8B.2* | WBGene00021818 | protein_coding_gene | -1.3 | 0.40 | 3.4E-06 |
| *tig-3* | WBGene00021594 | protein_coding_gene | -1.3 | 0.40 | 2.6E-02 |
| *ZK512.1* | WBGene00013982 | protein_coding_gene | -1.3 | 0.39 | 1.6E-03 |
| *Y9C9A.13* | WBGene00021180 | protein_coding_gene | -1.3 | 0.39 | 2.4E-02 |
| *C18H9.6* | WBGene00016004 | protein_coding_gene | -1.4 | 0.39 | 4.1E-04 |
| *memi-1* | WBGene00003216 | protein_coding_gene | -1.4 | 0.39 | 5.2E-03 |
| *F22F7.3* | WBGene00017721 | protein_coding_gene | -1.4 | 0.39 | 9.8E-03 |
| *col-83* | WBGene00000658 | protein_coding_gene | -1.4 | 0.39 | 3.1E-02 |
| *skr-21* | WBGene00004827 | protein_coding_gene | -1.4 | 0.39 | 1.1E-02 |
| *F19B2.12* | WBGene00194812 | pseudogene | -1.4 | 0.39 | 1.6E-02 |
| *pot-2* | WBGene00010195 | protein_coding_gene | -1.4 | 0.38 | 1.3E-07 |
| *F23D12.10* | WBGene00045291 | protein_coding_gene | -1.4 | 0.38 | 4.9E-02 |
| *Y18D10A.30* | WBGene00271598 | pseudogene | -1.4 | 0.38 | 1.6E-02 |
| *C06E2.5* | WBGene00015529 | protein_coding_gene | -1.4 | 0.38 | 2.5E-02 |
| *C52E12.11* | WBGene00304189 | pseudogene | -1.4 | 0.38 | 2.6E-02 |
| *R09E12.9* | WBGene00044514 | protein_coding_gene | -1.4 | 0.38 | 6.9E-03 |
| *F35C8.8* | WBGene00018038 | protein_coding_gene | -1.4 | 0.38 | 3.3E-02 |
| *fbxa-83* | WBGene00013453 | protein_coding_gene | -1.4 | 0.38 | 6.1E-03 |
| *C09G1.7* | WBGene00284856 | ncRNA_gene | -1.4 | 0.38 | 4.2E-02 |
| *fbxa-169* | WBGene00045289 | protein_coding_gene | -1.4 | 0.38 | 5.1E-03 |
| *fbxa-104* | WBGene00011542 | protein_coding_gene | -1.4 | 0.37 | 1.4E-02 |
| *dmd-5* | WBGene00017326 | protein_coding_gene | -1.4 | 0.37 | 2.4E-02 |
| *rnr-2* | WBGene00004392 | protein_coding_gene | -1.4 | 0.37 | 4.5E-02 |
| *C42D4.1* | WBGene00016594 | protein_coding_gene | -1.4 | 0.37 | 9.4E-04 |
| *F07H5.13* | WBGene00008563 | protein_coding_gene | -1.4 | 0.37 | 2.3E-02 |
| *T22D1.5* | WBGene00020684 | protein_coding_gene | -1.4 | 0.37 | 1.8E-03 |
| *lido-3* | WBGene00015588 | pseudogene | -1.4 | 0.37 | 4.6E-03 |
| *Y116A8C.40* | WBGene00013812 | protein_coding_gene | -1.5 | 0.36 | 2.6E-02 |
| *EEED8.13* | WBGene00017141 | protein_coding_gene | -1.5 | 0.36 | 2.9E-05 |
| *W04H10.6* | WBGene00206417 | protein_coding_gene | -1.5 | 0.36 | 4.7E-02 |
| *Y71A12B.19* | WBGene00077443 | pseudogene | -1.5 | 0.36 | 1.1E-03 |
| *K07H8.7* | WBGene00019507 | protein_coding_gene | -1.5 | 0.36 | 2.8E-03 |
| *T24D1.3* | WBGene00011986 | protein_coding_gene | -1.5 | 0.36 | 2.4E-02 |
| *F36H1.12* | WBGene00009497 | protein_coding_gene | -1.5 | 0.36 | 7.2E-03 |
| *bath-27* | WBGene00017450 | protein_coding_gene | -1.5 | 0.36 | 3.8E-02 |
| *cpt-4* | WBGene00019644 | protein_coding_gene | -1.5 | 0.36 | 3.5E-06 |
| *F48F7.3* | WBGene00009849 | protein_coding_gene | -1.5 | 0.36 | 2.6E-02 |
| *psf-1* | WBGene00011275 | protein_coding_gene | -1.5 | 0.36 | 9.9E-03 |
| *W04H10.5* | WBGene00206416 | protein_coding_gene | -1.5 | 0.35 | 4.3E-02 |
| *W04A8.9* | WBGene00195155 | protein_coding_gene | -1.5 | 0.35 | 2.0E-02 |
| *ugt-34* | WBGene00009255 | protein_coding_gene | -1.5 | 0.35 | 2.7E-02 |
| *Y105C5A.1268* | WBGene00185102 | protein_coding_gene | -1.5 | 0.35 | 1.1E-02 |
| *Y54G2A.36* | WBGene00021898 | protein_coding_gene | -1.5 | 0.35 | 2.0E-03 |
| *D1086.19* | WBGene00045406 | protein_coding_gene | -1.5 | 0.35 | 1.3E-02 |
| *C14B1.2* | WBGene00007573 | protein_coding_gene | -1.5 | 0.35 | 9.3E-03 |
| *ctf-8* | WBGene00011915 | protein_coding_gene | -1.5 | 0.35 | 9.2E-04 |
| *kin-33* | WBGene00017083 | protein_coding_gene | -1.5 | 0.35 | 6.1E-03 |
| *F54D5.4* | WBGene00010050 | protein_coding_gene | -1.5 | 0.35 | 1.1E-07 |
| *fbxb-93* | WBGene00013408 | protein_coding_gene | -1.5 | 0.35 | 1.2E-02 |

| | | | | | |
|---|---|---|---|---|---|
| *Y105C5A.1271* | WBGene00194667 | pseudogene | -1.5 | 0.35 | 4.0E-02 |
| *linc-6* | WBGene00219700 | lincRNA_gene | -1.5 | 0.34 | 5.0E-04 |
| *T11F8.1* | WBGene00020433 | protein_coding_gene | -1.5 | 0.34 | 2.7E-02 |
| *C52A10.3* | WBGene00016864 | protein_coding_gene | -1.5 | 0.34 | 4.9E-02 |
| *F44F1.5* | WBGene00009697 | pseudogene | -1.5 | 0.34 | 3.5E-02 |
| *C08F1.11* | WBGene00044417 | protein_coding_gene | -1.6 | 0.34 | 3.8E-02 |
| *C31E10.11* | WBGene00304783 | protein_coding_gene | -1.6 | 0.34 | 5.5E-03 |
| *R03H10.11* | WBGene00271824 | protein_coding_gene | -1.6 | 0.34 | 3.1E-02 |
| *gadr-4* | WBGene00021053 | protein_coding_gene | -1.6 | 0.34 | 2.8E-02 |
| *F55C5.10* | WBGene00010098 | protein_coding_gene | -1.6 | 0.34 | 3.5E-02 |
| *Y81B9A.5* | WBGene00304906 | ncRNA_gene | -1.6 | 0.33 | 3.4E-02 |
| *F54F7.6* | WBGene00010066 | protein_coding_gene | -1.6 | 0.33 | 4.2E-02 |
| *ugt-4* | WBGene00013905 | protein_coding_gene | -1.6 | 0.33 | 1.4E-10 |
| *Y82E9BL.6* | WBGene00022323 | protein_coding_gene | -1.6 | 0.33 | 3.8E-02 |
| *F40G9.9* | WBGene00018245 | protein_coding_gene | -1.6 | 0.33 | 4.1E-02 |
| *R08C7.4* | WBGene00019948 | protein_coding_gene | -1.6 | 0.33 | 4.4E-02 |
| *acd-1* | WBGene00016064 | protein_coding_gene | -1.6 | 0.32 | 2.0E-03 |
| *C10A4.1* | WBGene00015662 | protein_coding_gene | -1.6 | 0.32 | 4.3E-03 |
| *M151.9* | WBGene00219238 | pseudogene | -1.6 | 0.32 | 2.6E-02 |
| *C08F1.10* | WBGene00015613 | protein_coding_gene | -1.6 | 0.32 | 2.2E-02 |
| *fbxb-30* | WBGene00018458 | protein_coding_gene | -1.6 | 0.32 | 4.3E-02 |
| *F29G9.7* | WBGene00017933 | pseudogene | -1.6 | 0.32 | 2.6E-02 |
| *F58D2.3* | WBGene00044978 | protein_coding_gene | -1.7 | 0.32 | 4.8E-02 |
| *F12E12.3* | WBGene00017404 | pseudogene | -1.7 | 0.32 | 4.0E-02 |
| *Y71F9AL.6* | WBGene00022111 | protein_coding_gene | -1.7 | 0.32 | 3.6E-03 |
| *fbxb-99* | WBGene00021174 | protein_coding_gene | -1.7 | 0.32 | 2.9E-02 |
| *lido-4* | WBGene00015589 | protein_coding_gene | -1.7 | 0.32 | 6.0E-03 |
| *clec-3* | WBGene00016577 | protein_coding_gene | -1.7 | 0.32 | 9.8E-03 |
| *Y116F11A.1* | WBGene00013816 | protein_coding_gene | -1.7 | 0.31 | 5.1E-03 |
| *sdz-1* | WBGene00015129 | protein_coding_gene | -1.7 | 0.31 | 7.0E-03 |
| *F01G12.18* | WBGene00206382 | protein_coding_gene | -1.7 | 0.31 | 2.2E-02 |
| *F35C8.9* | WBGene00077784 | pseudogene | -1.7 | 0.31 | 4.8E-02 |
| *C39B5.10* | WBGene00016528 | protein_coding_gene | -1.7 | 0.31 | 1.2E-02 |
| *C09G5.14* | WBGene00271803 | protein_coding_gene | -1.7 | 0.31 | 3.1E-02 |
| *clec-218* | WBGene00020938 | protein_coding_gene | -1.7 | 0.30 | 2.0E-02 |
| *fbxb-31* | WBGene00019797 | protein_coding_gene | -1.7 | 0.30 | 3.0E-02 |
| *his-64* | WBGene00001938 | protein_coding_gene | -1.7 | 0.30 | 4.0E-02 |
| *F43C11.10* | WBGene00018387 | protein_coding_gene | -1.7 | 0.30 | 2.3E-02 |
| *Y27F2A.8* | WBGene00021299 | protein_coding_gene | -1.7 | 0.30 | 4.3E-02 |
| *bgnt-1.3* | WBGene00009032 | protein_coding_gene | -1.7 | 0.30 | 2.7E-02 |
| *W03D2.15* | WBGene00271791 | ncRNA_gene | -1.7 | 0.30 | 1.7E-02 |
| *Y20C6A.4* | WBGene00077495 | protein_coding_gene | -1.7 | 0.30 | 4.4E-02 |
| *dod-23* | WBGene00009898 | protein_coding_gene | -1.7 | 0.30 | 3.3E-02 |
| *fbxb-107* | WBGene00017912 | protein_coding_gene | -1.8 | 0.30 | 1.5E-02 |
| *K07D4.2* | WBGene00019483 | protein_coding_gene | -1.8 | 0.30 | 2.8E-02 |
| *Y51A2D.23* | WBGene00077653 | pseudogene | -1.8 | 0.29 | 4.4E-02 |
| *F55C9.15* | WBGene00206500 | protein_coding_gene | -1.8 | 0.29 | 1.9E-02 |
| *C01F6.16* | WBGene00235286 | ncRNA_gene | -1.8 | 0.29 | 4.2E-02 |
| *C11G6.2* | WBGene00007523 | protein_coding_gene | -1.8 | 0.29 | 3.7E-02 |
| *F40G9.12* | WBGene00018247 | protein_coding_gene | -1.8 | 0.29 | 2.6E-02 |
| *F45D11.5* | WBGene00018452 | protein_coding_gene | -1.8 | 0.29 | 3.8E-02 |
| *fbxb-22* | WBGene00013230 | protein_coding_gene | -1.8 | 0.29 | 6.0E-03 |
| *rrn-3.1* | WBGene00004622 | rRNA_gene | -1.8 | 0.29 | 4.6E-02 |
| *K01D12.1* | WBGene00010461 | protein_coding_gene | -1.8 | 0.29 | 1.9E-02 |
| *T22B2.2* | WBGene00020666 | protein_coding_gene | -1.8 | 0.29 | 3.1E-02 |
| *C27H5.12* | WBGene00271810 | protein_coding_gene | -1.8 | 0.29 | 4.0E-02 |
| *W10G6.1* | WBGene00012378 | protein_coding_gene | -1.8 | 0.29 | 2.3E-02 |
| *rrn-3.56* | WBGene00004677 | pseudogene | -1.8 | 0.29 | 4.4E-02 |

| | | | | | |
|---|---|---|---|---|---|
| *T12A2.3* | WBGene00020438 | protein_coding_gene | -1.8 | 0.29 | 2.6E-02 |
| *Y17G9B.2* | WBGene00021199 | protein_coding_gene | -1.8 | 0.29 | 1.3E-02 |
| *fbxb-105* | WBGene00017251 | protein_coding_gene | -1.8 | 0.29 | 1.5E-02 |
| *F36H5.4* | WBGene00018103 | protein_coding_gene | -1.8 | 0.28 | 3.2E-02 |
| *H25K10.4* | WBGene00010413 | protein_coding_gene | -1.8 | 0.28 | 1.9E-02 |
| *Y116A8B.1* | WBGene00013779 | protein_coding_gene | -1.8 | 0.28 | 2.6E-03 |
| *dhs-8* | WBGene00000972 | protein_coding_gene | -1.8 | 0.28 | 4.8E-05 |
| *fbxb-111* | WBGene00017254 | protein_coding_gene | -1.8 | 0.28 | 6.2E-03 |
| *sdz-29* | WBGene00020081 | pseudogene | -1.8 | 0.28 | 1.0E-02 |
| *C35D6.4* | WBGene00007961 | protein_coding_gene | -1.8 | 0.28 | 1.7E-02 |
| *sdz-37* | WBGene00014064 | protein_coding_gene | -1.9 | 0.27 | 9.4E-03 |
| *rrn-1.2* | WBGene00004513 | rRNA_gene | -1.9 | 0.27 | 3.8E-02 |
| *his-32* | WBGene00001906 | protein_coding_gene | -1.9 | 0.27 | 2.1E-03 |
| *W08F4.20* | WBGene00268207 | pseudogene | -1.9 | 0.27 | 1.8E-02 |
| *ZK185.9* | WBGene00220238 | snoRNA_gene | -1.9 | 0.27 | 3.0E-02 |
| *C32B5.18* | WBGene00235351 | protein_coding_gene | -1.9 | 0.27 | 1.3E-02 |
| *fbxb-39* | WBGene00019702 | protein_coding_gene | -1.9 | 0.27 | 1.5E-02 |
| *K08E3.2* | WBGene00010662 | protein_coding_gene | -1.9 | 0.26 | 1.1E-02 |
| *his-31* | WBGene00001905 | protein_coding_gene | -1.9 | 0.26 | 7.7E-03 |
| *srx-98* | WBGene00005989 | protein_coding_gene | -1.9 | 0.26 | 2.5E-02 |
| *C52D10.10* | WBGene00023259 | pseudogene | -1.9 | 0.26 | 3.7E-02 |
| *fbxb-9* | WBGene00007447 | protein_coding_gene | -2.0 | 0.26 | 9.7E-03 |
| *fbxb-60* | WBGene00010104 | protein_coding_gene | -2.0 | 0.26 | 1.3E-02 |
| *Y17G9A.4* | WBGene00021197 | protein_coding_gene | -2.0 | 0.26 | 2.3E-02 |
| *B0563.1* | WBGene00015261 | protein_coding_gene | -2.0 | 0.25 | 3.9E-03 |
| *Y116A8C.20* | WBGene00013797 | protein_coding_gene | -2.0 | 0.25 | 7.9E-03 |
| *fbxa-200* | WBGene00020304 | pseudogene | -2.0 | 0.25 | 2.2E-02 |
| *ugt-14* | WBGene00019233 | protein_coding_gene | -2.0 | 0.25 | 1.0E-02 |
| *M151.2* | WBGene00019794 | protein_coding_gene | -2.0 | 0.25 | 8.4E-03 |
| *fbxb-45* | WBGene00019705 | protein_coding_gene | -2.0 | 0.25 | 9.5E-03 |
| *T11F8.2* | WBGene00020434 | pseudogene | -2.0 | 0.24 | 8.6E-03 |
| *F46C3.5* | WBGene00044703 | pseudogene | -2.0 | 0.24 | 2.0E-02 |
| *his-63* | WBGene00001937 | protein_coding_gene | -2.0 | 0.24 | 1.3E-02 |
| *T16A1.10* | WBGene00269431 | protein_coding_gene | -2.1 | 0.24 | 2.4E-04 |
| *H16D19.4* | WBGene00010402 | pseudogene | -2.1 | 0.24 | 1.2E-02 |
| *his-5* | WBGene00001879 | protein_coding_gene | -2.1 | 0.24 | 1.6E-02 |
| *fbxb-25* | WBGene00019034 | protein_coding_gene | -2.1 | 0.24 | 2.4E-03 |
| *F02E11.3* | WBGene00017181 | protein_coding_gene | -2.1 | 0.24 | 6.8E-03 |
| *C24H12.8* | WBGene00016077 | protein_coding_gene | -2.1 | 0.23 | 2.7E-03 |
| *ceh-5* | WBGene00000430 | protein_coding_gene | -2.1 | 0.23 | 4.8E-03 |
| *B0302.2* | WBGene00015119 | protein_coding_gene | -2.2 | 0.22 | 1.4E-02 |
| *C49G7.13* | WBGene00269425 | protein_coding_gene | -2.2 | 0.22 | 2.1E-04 |
| *srt-63* | WBGene00020877 | protein_coding_gene | -2.2 | 0.22 | 4.0E-03 |
| *ZK666.1* | WBGene00014041 | protein_coding_gene | -2.2 | 0.22 | 9.9E-03 |
| *F17B5.6* | WBGene00044058 | protein_coding_gene | -2.2 | 0.21 | 7.6E-03 |
| *W10G11.4* | WBGene00021137 | protein_coding_gene | -2.2 | 0.21 | 9.2E-03 |
| *F31D5.6* | WBGene00017949 | protein_coding_gene | -2.2 | 0.21 | 1.2E-04 |
| *ins-8* | WBGene00002091 | protein_coding_gene | -2.3 | 0.21 | 6.0E-03 |
| *Y49F6A.5* | WBGene00021713 | protein_coding_gene | -2.3 | 0.21 | 2.3E-03 |
| *fbxc-37* | WBGene00017942 | pseudogene | -2.3 | 0.20 | 2.2E-04 |
| *B0563.8* | WBGene00015265 | protein_coding_gene | -2.3 | 0.20 | 8.9E-04 |
| *F15H9.8* | WBGene00194652 | pseudogene | -2.4 | 0.19 | 1.0E-04 |
| *dct-19* | WBGene00007876 | protein_coding_gene | -2.4 | 0.19 | 3.9E-03 |
| *fbxb-16* | WBGene00022560 | protein_coding_gene | -2.6 | 0.17 | 7.3E-04 |
| *Y53H1B.4* | WBGene00013182 | protein_coding_gene | -2.6 | 0.17 | 5.5E-05 |
| *T08B6.1* | WBGene00020351 | protein_coding_gene | -2.7 | 0.16 | 1.7E-03 |
| *F59A7.12* | WBGene00044520 | protein_coding_gene | -2.8 | 0.15 | 9.3E-05 |
| *fbxc-26* | WBGene00019043 | protein_coding_gene | -2.9 | 0.13 | 1.7E-05 |

| | | | | | |
|---|---|---|---|---|---|
| *dod-24* | WBGene00007875 | protein_coding_gene | -4.1 | 0.06 | 1.0E-16 |

Shared (N2&CB4856) DEGs in response to Gram-positive pathogens (*S. epidermidis* or *S. aureus* vs. OP50)

| | | | N2 | | | CB4856 | | |
|---|---|---|---|---|---|---|---|---|
| Gene Name | Wormbase Gene ID | Type | $log_2FC$ | FC | Adj P-value | $log_2FC$ | FC | Adj P-value |
| *C23G10.11* | WBGene00016017 | protein_coding_gene | 7.6 | 199.40 | 1.1E-12 | 2.0 | 4.00 | 4.2E-02 |
| *clec-60* | WBGene00014046 | protein_coding_gene | 7.1 | 132.87 | 1.4E-18 | 4.1 | 17.12 | 2.9E-08 |
| *chil-28* | WBGene00008842 | pseudogene | 7.0 | 128.36 | 6.7E-11 | 2.3 | 4.92 | 3.5E-05 |
| *T22F3.11* | WBGene00020700 | protein_coding_gene | 6.7 | 102.88 | 1.5E-14 | 1.9 | 3.77 | 3.8E-02 |
| *F17B5.1* | WBGene00008905 | protein_coding_gene | 6.7 | 100.52 | 2.7E-15 | 2.6 | 6.19 | 2.4E-03 |
| *Y54G2A.37* | WBGene00023484 | protein_coding_gene | 6.4 | 83.51 | 9.4E-09 | 2.4 | 5.29 | 4.4E-07 |
| *lipl-1* | WBGene00010062 | protein_coding_gene | 6.1 | 69.38 | 5.1E-20 | 2.5 | 5.71 | 7.3E-03 |
| *clec-61* | WBGene00014047 | protein_coding_gene | 5.9 | 61.76 | 1.3E-09 | 2.4 | 5.19 | 2.3E-05 |
| *T26H5.8* | WBGene00012070 | protein_coding_gene | 5.8 | 56.43 | 8.7E-22 | 1.1 | 2.07 | 1.4E-04 |
| *K05F6.10* | WBGene00019421 | protein_coding_gene | 5.7 | 52.63 | 2.1E-11 | 2.0 | 3.92 | 4.4E-03 |
| *T07H8.11* | WBGene00304201 | protein_coding_gene | 5.7 | 50.43 | 5.0E-10 | 4.6 | 24.92 | 5.3E-25 |
| *F40G12.5* | WBGene00009600 | protein_coding_gene | 5.5 | 46.09 | 1.3E-06 | 4.5 | 22.24 | 1.2E-06 |
| *clec-69* | WBGene00044642 | pseudogene | 5.5 | 45.52 | 4.0E-08 | 2.2 | 4.53 | 8.4E-03 |
| *F46C5.1* | WBGene00009778 | protein_coding_gene | 5.4 | 41.94 | 1.1E-12 | 1.7 | 3.14 | 3.1E-03 |
| *math-38* | WBGene00020326 | protein_coding_gene | 5.3 | 40.25 | 8.1E-12 | 1.2 | 2.31 | 5.9E-03 |
| *sdz-24* | WBGene00019495 | protein_coding_gene | 5.3 | 38.67 | 8.5E-29 | 2.2 | 4.58 | 1.6E-02 |
| *cpr-2* | WBGene00000782 | protein_coding_gene | 5.2 | 37.58 | 5.2E-10 | 3.0 | 7.81 | 5.2E-04 |
| *asm-3* | WBGene00000213 | protein_coding_gene | 5.1 | 35.03 | 7.7E-09 | 4.9 | 29.82 | 3.9E-10 |
| *dod-3* | WBGene00016052 | protein_coding_gene | 5.1 | 33.33 | 2.3E-10 | 1.6 | 3.03 | 8.8E-04 |
| *ech-9* | WBGene00001158 | protein_coding_gene | 5.0 | 32.10 | 3.7E-10 | 2.7 | 6.59 | 6.2E-03 |
| *clec-82* | WBGene00021873 | protein_coding_gene | 4.9 | 30.43 | 5.1E-09 | 1.8 | 3.47 | 1.2E-02 |
| *ZK666.15* | WBGene00195145 | protein_coding_gene | 4.8 | 28.69 | 2.8E-07 | 2.8 | 6.78 | 1.8E-05 |
| *math-34* | WBGene00019699 | protein_coding_gene | 4.8 | 28.54 | 1.1E-12 | 2.1 | 4.30 | 3.5E-04 |
| *ZK105.13* | WBGene00235114 | protein_coding_gene | 4.8 | 28.50 | 1.6E-06 | 1.4 | 2.58 | 7.6E-04 |
| *hsp-12.6* | WBGene00002013 | protein_coding_gene | 4.7 | 25.73 | 4.8E-07 | 3.4 | 10.42 | 2.5E-04 |
| *Y73B6BL.37* | WBGene00022256 | protein_coding_gene | 4.6 | 24.31 | 1.5E-05 | 3.5 | 11.22 | 6.3E-05 |
| *best-1* | WBGene00007203 | protein_coding_gene | 4.6 | 23.69 | 6.9E-13 | 2.0 | 3.90 | 1.2E-04 |
| *F53H2.1* | WBGene00010004 | protein_coding_gene | 4.5 | 23.40 | 4.6E-08 | 1.9 | 3.62 | 7.4E-05 |
| *ilys-3* | WBGene00016670 | protein_coding_gene | 4.5 | 23.37 | 3.7E-09 | 2.4 | 5.22 | 3.5E-04 |
| *C39B5.14* | WBGene00189950 | protein_coding_gene | 4.5 | 22.56 | 4.4E-06 | 3.6 | 11.92 | 8.9E-05 |
| *clec-172* | WBGene00044719 | protein_coding_gene | 4.5 | 22.15 | 1.0E-10 | 1.8 | 3.55 | 1.3E-03 |
| *T23F2.4* | WBGene00020737 | protein_coding_gene | 4.4 | 21.43 | 3.3E-05 | 3.3 | 9.58 | 5.5E-04 |
| *pals-39* | WBGene00008301 | protein_coding_gene | 4.4 | 21.24 | 1.6E-07 | 2.4 | 5.39 | 2.5E-02 |
| *F10B5.3* | WBGene00008640 | protein_coding_gene | 4.4 | 20.82 | 2.5E-05 | 3.1 | 8.40 | 2.1E-04 |
| *F42A10.7* | WBGene00018343 | protein_coding_gene | 4.4 | 20.62 | 9.0E-07 | 4.2 | 17.92 | 1.7E-07 |
| *K02E2.7* | WBGene00010508 | protein_coding_gene | 4.3 | 20.32 | 1.3E-04 | 2.3 | 4.78 | 9.2E-04 |
| *W07B8.4* | WBGene00021072 | protein_coding_gene | 4.3 | 20.07 | 2.3E-07 | 2.9 | 7.48 | 5.8E-05 |
| *cest-1.1* | WBGene00011362 | protein_coding_gene | 4.3 | 19.88 | 6.5E-10 | 2.9 | 7.47 | 6.2E-04 |
| *F14D7.7* | WBGene00008794 | protein_coding_gene | 4.3 | 19.62 | 6.5E-07 | 1.8 | 3.54 | 9.4E-05 |
| *asah-1* | WBGene00010769 | protein_coding_gene | 4.3 | 19.16 | 6.4E-18 | 2.9 | 7.57 | 4.8E-08 |
| *R05A10.8* | WBGene00011025 | protein_coding_gene | 4.3 | 19.05 | 6.2E-04 | 3.1 | 8.84 | 1.9E-03 |
| *F08H9.4* | WBGene00008592 | protein_coding_gene | 4.2 | 18.92 | 9.2E-05 | 3.6 | 12.18 | 4.8E-05 |
| *Y6E2A.4* | WBGene00012398 | protein_coding_gene | 4.2 | 18.79 | 1.0E-05 | 2.4 | 5.34 | 9.0E-03 |
| *C06G3.3* | WBGene00015549 | protein_coding_gene | 4.2 | 18.63 | 4.7E-08 | 2.1 | 4.36 | 4.6E-03 |
| *W05G11.8* | WBGene00271636 | ncRNA_gene | 4.2 | 18.12 | 2.2E-05 | 2.4 | 5.17 | 2.5E-03 |
| *Y65B4BR.1* | WBGene00022040 | protein_coding_gene | 4.2 | 18.12 | 7.6E-08 | 3.3 | 9.76 | 1.5E-05 |
| *F41C3.7* | WBGene00018273 | protein_coding_gene | 4.1 | 16.78 | 1.2E-05 | 3.0 | 7.88 | 1.5E-03 |
| *Y9C9A.1* | WBGene00021176 | protein_coding_gene | 4.1 | 16.70 | 7.9E-05 | 2.6 | 5.95 | 2.1E-03 |
| *Y110A2AL.2* | WBGene00022439 | protein_coding_gene | 4.1 | 16.68 | 1.3E-04 | 3.4 | 10.78 | 2.5E-04 |
| *Y51H4A.5* | WBGene00013101 | protein_coding_gene | 4.0 | 16.51 | 7.8E-09 | 1.3 | 2.48 | 1.4E-02 |
| *D1054.18* | WBGene00077783 | protein_coding_gene | 4.0 | 16.46 | 1.8E-05 | 2.1 | 4.34 | 2.1E-03 |
| *C54F6.12* | WBGene00016929 | protein_coding_gene | 4.0 | 16.34 | 2.5E-09 | 1.6 | 3.11 | 1.9E-02 |
| *Y57E12B.4* | WBGene00021964 | protein_coding_gene | 4.0 | 16.13 | 1.5E-03 | 3.2 | 9.03 | 2.9E-03 |

| | | | | | | | | |
|---|---|---|---|---|---|---|---|---|
| *lact-6* | WBGene00007358 | protein_coding_gene | 4.0 | 16.06 | 2.9E-05 | 3.8 | 14.11 | 1.4E-08 |
| *Y46G5A.36* | WBGene00012921 | protein_coding_gene | 4.0 | 15.97 | 2.9E-04 | 1.8 | 3.55 | 3.9E-02 |
| *T28A11.6* | WBGene00020873 | pseudogene | 4.0 | 15.83 | 2.7E-04 | 2.8 | 6.97 | 2.2E-03 |
| *F11C7.6* | WBGene00017378 | protein_coding_gene | 4.0 | 15.62 | 1.6E-07 | 1.5 | 2.73 | 9.7E-03 |
| *Y57E12B.11* | WBGene00235311 | protein_coding_gene | 3.9 | 15.26 | 5.5E-04 | 3.6 | 12.24 | 2.9E-04 |
| *sqst-3* | WBGene00012067 | protein_coding_gene | 3.9 | 15.14 | 2.6E-04 | 3.4 | 10.26 | 2.7E-04 |
| *F55B12.10* | WBGene00077439 | protein_coding_gene | 3.9 | 14.98 | 1.5E-04 | 3.3 | 9.56 | 5.7E-04 |
| *ZK666.11* | WBGene00014049 | protein_coding_gene | 3.9 | 14.93 | 4.0E-04 | 1.6 | 3.03 | 4.2E-02 |
| *F07C4.6* | WBGene00017200 | protein_coding_gene | 3.9 | 14.57 | 5.0E-07 | 3.2 | 9.27 | 3.4E-03 |
| *ZC116.5* | WBGene00119204 | protein_coding_gene | 3.9 | 14.55 | 6.7E-11 | 2.7 | 6.31 | 5.7E-08 |
| *cyp-35B1* | WBGene00019472 | protein_coding_gene | 3.8 | 14.16 | 8.5E-10 | 1.9 | 3.75 | 3.3E-04 |
| ***lipl-3*** | **WBGene00020016** | **protein_coding_gene** | **3.8** | **14.05** | **3.5E-05** | **-1.4** | **0.37** | **5.0E-03** |
| *irld-11* | WBGene00019391 | protein_coding_gene | 3.8 | 14.03 | 2.9E-04 | 1.9 | 3.80 | 2.6E-02 |
| *fbxa-136* | WBGene00007672 | protein_coding_gene | 3.8 | 13.90 | 5.9E-04 | 2.0 | 4.03 | 6.2E-04 |
| *F55H12.9* | WBGene00219976 | snoRNA_gene | 3.8 | 13.71 | 7.9E-04 | 1.8 | 3.55 | 1.1E-02 |
| *C45B11.7* | WBGene00050945 | pseudogene | 3.8 | 13.69 | 4.1E-05 | 2.4 | 5.21 | 3.4E-03 |
| *egas-4* | WBGene00018906 | protein_coding_gene | 3.8 | 13.48 | 1.2E-05 | 2.0 | 3.88 | 1.7E-02 |
| *dao-6* | WBGene00000932 | protein_coding_gene | 3.7 | 13.33 | 4.5E-04 | 2.5 | 5.74 | 1.6E-02 |
| *F49C12.7* | WBGene00009877 | protein_coding_gene | 3.7 | 13.29 | 8.2E-08 | 2.7 | 6.67 | 5.2E-06 |
| *ins-35* | WBGene00002118 | protein_coding_gene | 3.7 | 13.27 | 2.8E-05 | 2.1 | 4.43 | 9.2E-03 |
| *djr-1.2* | WBGene00016789 | protein_coding_gene | 3.7 | 13.04 | 2.8E-04 | 3.4 | 10.85 | 1.4E-04 |
| *lys-9* | WBGene00003098 | protein_coding_gene | 3.7 | 13.04 | 2.4E-03 | 2.5 | 5.74 | 1.8E-02 |
| *F46A8.1* | WBGene00014759 | pseudogene | 3.7 | 13.04 | 6.5E-08 | 3.0 | 8.23 | 4.5E-07 |
| *F09C12.2* | WBGene00017277 | protein_coding_gene | 3.7 | 12.65 | 1.4E-04 | 3.7 | 12.71 | 7.5E-05 |
| *sod-5* | WBGene00007036 | protein_coding_gene | 3.7 | 12.59 | 2.9E-04 | 3.8 | 14.15 | 7.5E-05 |
| *fbxa-66* | WBGene00021865 | protein_coding_gene | 3.6 | 12.52 | 5.0E-04 | 1.7 | 3.28 | 2.7E-02 |
| *ZK993.2* | WBGene00022838 | protein_coding_gene | 3.6 | 12.49 | 1.0E-03 | 1.5 | 2.88 | 4.8E-02 |
| *Y39G8B.10* | WBGene00012729 | protein_coding_gene | 3.6 | 12.44 | 4.2E-03 | 2.3 | 4.87 | 1.4E-02 |
| *T28A11.19* | WBGene00020881 | protein_coding_gene | 3.6 | 12.42 | 1.3E-04 | 2.2 | 4.72 | 1.4E-02 |
| *F14F9.8* | WBGene00044536 | protein_coding_gene | 3.6 | 12.40 | 1.9E-05 | 2.8 | 7.07 | 1.4E-03 |
| *clec-42* | WBGene00008891 | protein_coding_gene | 3.6 | 12.30 | 2.4E-04 | 2.0 | 4.03 | 2.2E-03 |
| *pals-26* | WBGene00007131 | protein_coding_gene | 3.6 | 12.18 | 9.6E-05 | 2.3 | 4.98 | 4.0E-03 |
| *ant-1.2* | WBGene00007057 | protein_coding_gene | 3.6 | 11.89 | 9.1E-04 | 3.4 | 10.33 | 2.5E-04 |
| *cyp-33D3* | WBGene00012448 | protein_coding_gene | 3.6 | 11.76 | 7.0E-04 | 2.9 | 7.59 | 3.0E-03 |
| *W01C8.1* | WBGene00020918 | protein_coding_gene | 3.5 | 11.68 | 1.1E-03 | 1.7 | 3.14 | 1.7E-02 |
| *exc-13* | WBGene00015393 | protein_coding_gene | 3.5 | 11.40 | 9.0E-05 | 1.9 | 3.68 | 1.9E-03 |
| *R09E10.13* | WBGene00194658 | protein_coding_gene | 3.5 | 11.36 | 8.5E-05 | 2.2 | 4.63 | 4.5E-02 |
| *nnt-1* | WBGene00003778 | protein_coding_gene | 3.5 | 11.36 | 2.1E-05 | 1.0 | 2.04 | 4.9E-02 |
| *arrd-8* | WBGene00012467 | protein_coding_gene | 3.5 | 11.34 | 8.6E-04 | 2.5 | 5.66 | 1.5E-03 |
| *T01D3.3* | WBGene00011328 | protein_coding_gene | 3.5 | 11.23 | 8.4E-07 | 1.4 | 2.56 | 3.7E-02 |
| *F01G10.4* | WBGene00008507 | protein_coding_gene | 3.5 | 11.03 | 5.3E-03 | 2.6 | 5.97 | 1.8E-02 |
| *C30G12.11* | WBGene00284861 | ncRNA_gene | 3.5 | 11.01 | 1.5E-03 | 1.9 | 3.80 | 4.3E-02 |
| ***scl-2*** | **WBGene00009895** | **protein_coding_gene** | **3.5** | **10.95** | **9.5E-06** | **-1.2** | **0.45** | **3.5E-05** |
| *pals-15* | WBGene00044707 | protein_coding_gene | 3.4 | 10.79 | 6.2E-03 | 3.0 | 8.12 | 6.9E-03 |
| *ZK971.1* | WBGene00014175 | protein_coding_gene | 3.4 | 10.79 | 7.0E-06 | 2.1 | 4.18 | 9.4E-03 |
| *pals-27* | WBGene00007132 | protein_coding_gene | 3.4 | 10.78 | 4.1E-04 | 2.4 | 5.38 | 6.0E-03 |
| *T28A11.16* | WBGene00020878 | protein_coding_gene | 3.4 | 10.75 | 7.0E-04 | 2.4 | 5.44 | 3.9E-03 |
| *T13F2.4* | WBGene00011744 | protein_coding_gene | 3.4 | 10.75 | 2.9E-03 | 2.3 | 5.09 | 2.4E-02 |
| *F35D2.1* | WBGene00018039 | protein_coding_gene | 3.4 | 10.67 | 4.5E-03 | 2.2 | 4.63 | 3.9E-02 |
| *R05H11.2* | WBGene00019908 | protein_coding_gene | 3.4 | 10.50 | 8.5E-04 | 2.2 | 4.59 | 1.2E-02 |
| *col-69* | WBGene00000645 | protein_coding_gene | 3.4 | 10.43 | 2.5E-04 | 1.8 | 3.48 | 6.7E-03 |
| *gst-8* | WBGene00001756 | protein_coding_gene | 3.4 | 10.39 | 3.1E-04 | 2.0 | 4.03 | 7.1E-03 |
| *spp-8* | WBGene00004993 | protein_coding_gene | 3.4 | 10.31 | 2.2E-10 | 1.3 | 2.40 | 1.2E-03 |
| *C54F6.17* | WBGene00235092 | protein_coding_gene | 3.4 | 10.31 | 3.1E-03 | 1.9 | 3.76 | 5.0E-02 |
| *C27H5.6* | WBGene00016175 | protein_coding_gene | 3.4 | 10.24 | 4.7E-03 | 2.1 | 4.35 | 5.0E-02 |
| *Y39G8B.9* | WBGene00012728 | protein_coding_gene | 3.3 | 10.11 | 1.8E-04 | 2.8 | 7.03 | 3.1E-04 |
| *math-42* | WBGene00020532 | protein_coding_gene | 3.3 | 10.00 | 3.3E-05 | 1.2 | 2.29 | 3.5E-02 |
| *sre-6* | WBGene00009242 | protein_coding_gene | 3.3 | 9.93 | 4.0E-03 | 2.8 | 6.90 | 1.8E-04 |

| | | | | | | | | |
|---|---|---|---|---|---|---|---|---|
| *clec-64* | WBGene00009395 | protein_coding_gene | 3.3 | 9.69 | 1.5E-03 | 1.7 | 3.18 | 2.0E-02 |
| *B0238.13* | WBGene00015077 | pseudogene | 3.3 | 9.67 | 4.0E-03 | 1.6 | 3.04 | 2.0E-02 |
| *fil-2* | WBGene00019668 | protein_coding_gene | 3.3 | 9.54 | 2.2E-03 | 2.4 | 5.13 | 1.3E-02 |
| *C54F6.5* | WBGene00016923 | protein_coding_gene | 3.2 | 9.48 | 3.9E-04 | 1.8 | 3.56 | 4.3E-02 |
| *C50D2.1* | WBGene00016805 | protein_coding_gene | 3.2 | 9.39 | 3.3E-06 | 1.3 | 2.41 | 1.6E-02 |
| *nlp-2* | WBGene00003740 | protein_coding_gene | 3.2 | 9.39 | 4.6E-06 | 1.4 | 2.65 | 3.8E-02 |
| *K03H6.2* | WBGene00019368 | protein_coding_gene | 3.2 | 9.29 | 2.3E-07 | 2.9 | 7.64 | 3.5E-07 |
| *daao-1* | WBGene00022076 | protein_coding_gene | 3.2 | 9.27 | 1.4E-06 | 1.4 | 2.66 | 3.1E-04 |
| *zip-6* | WBGene00011130 | protein_coding_gene | 3.2 | 9.27 | 3.3E-03 | 2.2 | 4.69 | 1.3E-02 |
| *F56F11.1* | WBGene00018988 | protein_coding_gene | 3.2 | 9.24 | 3.5E-03 | 1.7 | 3.22 | 3.6E-02 |
| *W04C9.6* | WBGene00021028 | protein_coding_gene | 3.2 | 9.20 | 4.6E-06 | 3.2 | 9.28 | 6.1E-07 |
| *Y39A1A.9* | WBGene00012648 | protein_coding_gene | 3.2 | 9.19 | 7.8E-04 | 2.8 | 6.82 | 1.6E-03 |
| *R09D1.4* | WBGene00011160 | pseudogene | 3.2 | 9.16 | 8.4E-03 | 1.8 | 3.51 | 1.5E-02 |
| *F57G8.7* | WBGene00010216 | protein_coding_gene | 3.2 | 9.16 | 1.1E-02 | 3.6 | 12.55 | 9.4E-04 |
| *odr-10* | WBGene00003856 | protein_coding_gene | 3.2 | 9.16 | 2.2E-06 | 1.0 | 2.03 | 4.3E-02 |
| *W02G9.4* | WBGene00020954 | protein_coding_gene | 3.2 | 9.16 | 1.6E-33 | 2.7 | 6.43 | 7.7E-20 |
| *scl-14* | WBGene00008625 | protein_coding_gene | 3.2 | 9.10 | 8.8E-04 | 2.3 | 4.88 | 5.6E-03 |
| *ZC443.4* | WBGene00013899 | protein_coding_gene | 3.2 | 9.02 | 3.9E-06 | 1.3 | 2.45 | 2.5E-02 |
| *Y41D4B.17* | WBGene00021519 | protein_coding_gene | 3.2 | 8.97 | 2.1E-06 | 2.9 | 7.22 | 5.1E-06 |
| *F01D5.7* | WBGene00008497 | protein_coding_gene | 3.2 | 8.92 | 2.5E-03 | 1.5 | 2.82 | 2.0E-02 |
| *pals-2* | WBGene00007656 | protein_coding_gene | 3.1 | 8.79 | 9.2E-03 | 2.0 | 4.10 | 2.5E-02 |
| *fipr-26* | WBGene00009963 | protein_coding_gene | 3.1 | 8.71 | 2.8E-03 | 2.3 | 4.87 | 4.9E-02 |
| *fbxa-158* | WBGene00015602 | protein_coding_gene | 3.1 | 8.70 | 1.4E-03 | 2.8 | 6.91 | 1.1E-03 |
| *Y105C5A.1270* | WBGene00194646 | protein_coding_gene | 3.1 | 8.64 | 1.2E-02 | 2.2 | 4.55 | 3.9E-02 |
| *T22F3.10* | WBGene00020699 | protein_coding_gene | 3.1 | 8.52 | 4.4E-04 | 1.4 | 2.63 | 1.3E-02 |
| *K10D3.6* | WBGene00010740 | protein_coding_gene | 3.1 | 8.48 | 4.9E-07 | 1.3 | 2.42 | 5.4E-05 |
| *F54B11.9* | WBGene00010032 | protein_coding_gene | 3.1 | 8.47 | 3.1E-03 | 1.9 | 3.72 | 1.3E-02 |
| *zmp-2* | WBGene00019212 | protein_coding_gene | 3.1 | 8.39 | 1.9E-04 | 2.0 | 4.01 | 1.1E-02 |
| *F20B6.4* | WBGene00017628 | protein_coding_gene | 3.1 | 8.38 | 5.0E-03 | 1.5 | 2.83 | 2.4E-02 |
| *F40E3.6* | WBGene00018233 | protein_coding_gene | 3.1 | 8.34 | 4.1E-03 | 2.7 | 6.53 | 9.9E-03 |
| *F58H10.1* | WBGene00010291 | protein_coding_gene | 3.0 | 8.25 | 8.7E-05 | 1.8 | 3.38 | 2.1E-02 |
| *gst-15* | WBGene00001763 | protein_coding_gene | 3.0 | 8.24 | 3.0E-04 | 2.4 | 5.38 | 5.3E-03 |
| *gcy-7* | WBGene00001534 | protein_coding_gene | 3.0 | 8.18 | 7.3E-03 | 2.7 | 6.31 | 5.2E-03 |
| *ZK418.11* | WBGene00044354 | protein_coding_gene | 3.0 | 8.11 | 5.7E-03 | 1.7 | 3.20 | 2.4E-02 |
| *fbxa-165* | WBGene00015600 | protein_coding_gene | 3.0 | 8.09 | 1.5E-03 | 1.6 | 3.10 | 2.7E-02 |
| *C11H1.5* | WBGene00007530 | protein_coding_gene | 3.0 | 8.05 | 4.1E-03 | 3.5 | 11.44 | 8.0E-05 |
| *chil-18* | WBGene00011161 | protein_coding_gene | 3.0 | 8.03 | 6.7E-04 | 1.7 | 3.25 | 3.5E-03 |
| *spp-12* | WBGene00004997 | protein_coding_gene | 3.0 | 8.02 | 2.4E-03 | 2.5 | 5.55 | 1.2E-02 |
| *T23B3.6* | WBGene00020714 | protein_coding_gene | 3.0 | 8.01 | 1.5E-05 | 1.6 | 3.06 | 3.6E-03 |
| *hsp-12.3* | WBGene00002012 | protein_coding_gene | 3.0 | 7.97 | 1.1E-05 | 2.6 | 6.00 | 1.0E-04 |
| *R12G8.1* | WBGene00011254 | protein_coding_gene | 3.0 | 7.95 | 9.0E-03 | 1.8 | 3.57 | 4.1E-02 |
| *R02F11.2* | WBGene00019840 | protein_coding_gene | 3.0 | 7.92 | 1.2E-05 | 2.0 | 3.99 | 5.2E-04 |
| *clec-52* | WBGene00015052 | protein_coding_gene | 3.0 | 7.84 | 6.4E-05 | 2.0 | 4.09 | 1.3E-02 |
| *B0348.2* | WBGene00015152 | protein_coding_gene | 3.0 | 7.81 | 2.9E-03 | 1.6 | 2.93 | 1.8E-02 |
| *tag-243* | WBGene00044078 | protein_coding_gene | 3.0 | 7.79 | 1.0E-04 | 1.6 | 2.96 | 1.2E-02 |
| *lips-5* | WBGene00011725 | protein_coding_gene | 3.0 | 7.77 | 3.5E-06 | 3.3 | 9.54 | 8.3E-16 |
| *C34C6.3* | WBGene00007916 | protein_coding_gene | 3.0 | 7.76 | 2.0E-04 | 1.3 | 2.46 | 1.6E-03 |
| *Y39H10A.1* | WBGene00021478 | protein_coding_gene | 3.0 | 7.74 | 7.3E-03 | 3.0 | 8.04 | 6.4E-03 |
| *C07G3.8* | WBGene00015576 | protein_coding_gene | 3.0 | 7.74 | 3.7E-03 | 2.3 | 4.80 | 8.2E-03 |
| *C40H1.3* | WBGene00008039 | protein_coding_gene | 2.9 | 7.65 | 3.2E-03 | 1.6 | 3.02 | 1.9E-02 |
| *bcmo-2* | WBGene00018755 | protein_coding_gene | 2.9 | 7.65 | 1.5E-08 | 3.2 | 8.92 | 2.9E-07 |
| *C15B12.8* | WBGene00015788 | protein_coding_gene | 2.9 | 7.58 | 1.8E-05 | 2.3 | 4.85 | 6.3E-04 |
| *B0336.12* | WBGene00015149 | protein_coding_gene | 2.9 | 7.56 | 2.2E-02 | 2.3 | 4.77 | 3.6E-02 |
| *cav-2* | WBGene00000302 | protein_coding_gene | 2.9 | 7.53 | 2.1E-06 | 2.0 | 4.12 | 3.5E-07 |
| *H02F09.3* | WBGene00019146 | protein_coding_gene | 2.9 | 7.48 | 8.9E-04 | 2.8 | 6.97 | 2.4E-06 |
| *Y39B6A.29* | WBGene00012688 | protein_coding_gene | 2.9 | 7.42 | 3.7E-04 | 1.6 | 3.11 | 2.3E-02 |
| *Y54G2A.57* | WBGene00189961 | protein_coding_gene | 2.9 | 7.36 | 2.0E-02 | 2.7 | 6.71 | 1.3E-02 |
| *R05A10.6* | WBGene00011023 | protein_coding_gene | 2.9 | 7.35 | 1.5E-02 | 2.1 | 4.27 | 4.3E-02 |

| | | | | | | | | |
|---|---|---|---|---|---|---|---|---|
| *C17B7.2* | WBGene00015876 | pseudogene | 2.9 | 7.33 | 2.7E-02 | 1.9 | 3.69 | 3.8E-02 |
| *H17B01.5* | WBGene00044643 | protein_coding_gene | 2.9 | 7.26 | 7.5E-04 | 2.1 | 4.15 | 1.1E-03 |
| *K06A9.1* | WBGene00019435 | protein_coding_gene | 2.9 | 7.26 | 4.6E-08 | 2.4 | 5.29 | 2.7E-09 |
| *gcy-29* | WBGene00007314 | protein_coding_gene | 2.9 | 7.25 | 4.7E-03 | 2.8 | 7.04 | 3.3E-03 |
| *asah-2* | WBGene00009192 | protein_coding_gene | 2.8 | 7.16 | 9.9E-06 | 1.5 | 2.78 | 2.7E-03 |
| *spin-3* | WBGene00008598 | protein_coding_gene | 2.8 | 7.14 | 2.1E-08 | 2.0 | 4.09 | 1.9E-06 |
| *Y40B10B.1* | WBGene00021494 | protein_coding_gene | 2.8 | 7.13 | 2.2E-02 | 2.5 | 5.55 | 1.1E-02 |
| *F56D3.1* | WBGene00018969 | protein_coding_gene | 2.8 | 7.07 | 2.5E-04 | 1.7 | 3.27 | 2.6E-03 |
| *dmd-10* | WBGene00007929 | protein_coding_gene | 2.8 | 7.04 | 6.4E-03 | 1.4 | 2.70 | 2.6E-02 |
| *lmd-4* | WBGene00017233 | protein_coding_gene | 2.8 | 6.99 | 1.2E-02 | 3.8 | 13.88 | 9.2E-05 |
| *nlp-81* | WBGene00022063 | protein_coding_gene | 2.8 | 6.98 | 7.5E-03 | 1.7 | 3.32 | 3.4E-02 |
| *nlp-51* | WBGene00011997 | protein_coding_gene | 2.8 | 6.89 | 8.6E-07 | 2.0 | 3.98 | 1.4E-06 |
| *W03F11.5* | WBGene00021008 | protein_coding_gene | 2.8 | 6.87 | 1.6E-07 | 1.5 | 2.85 | 2.6E-02 |
| *snt-7* | WBGene00011682 | protein_coding_gene | 2.8 | 6.83 | 5.6E-05 | 1.1 | 2.19 | 2.3E-02 |
| *tyr-3* | WBGene00009001 | protein_coding_gene | 2.8 | 6.75 | 1.1E-02 | 2.7 | 6.67 | 2.8E-03 |
| *K09H9.5* | WBGene00019597 | protein_coding_gene | 2.8 | 6.74 | 1.7E-03 | 2.3 | 4.94 | 4.7E-03 |
| *asp-7* | WBGene00019104 | protein_coding_gene | 2.8 | 6.73 | 2.4E-02 | 2.3 | 4.84 | 1.3E-02 |
| *sod-3* | WBGene00004932 | protein_coding_gene | 2.7 | 6.71 | 2.9E-04 | 2.7 | 6.58 | 1.4E-04 |
| *C39B10.1* | WBGene00008021 | protein_coding_gene | 2.7 | 6.64 | 1.7E-06 | 2.0 | 4.07 | 8.4E-06 |
| *C09B9.1* | WBGene00015626 | protein_coding_gene | 2.7 | 6.64 | 1.4E-03 | 1.9 | 3.83 | 1.8E-02 |
| *C15H9.2* | WBGene00015798 | protein_coding_gene | 2.7 | 6.63 | 8.6E-03 | 1.7 | 3.35 | 3.9E-02 |
| *msrp-7* | WBGene00011070 | protein_coding_gene | 2.7 | 6.63 | 1.0E-05 | 2.2 | 4.75 | 4.9E-05 |
| *C52E2.4* | WBGene00016883 | protein_coding_gene | 2.7 | 6.61 | 1.7E-05 | 1.4 | 2.63 | 3.2E-04 |
| *asic-2* | WBGene00012137 | protein_coding_gene | 2.7 | 6.56 | 4.1E-04 | 1.8 | 3.57 | 4.2E-03 |
| *nlp-76* | WBGene00045386 | protein_coding_gene | 2.7 | 6.53 | 9.8E-04 | 1.4 | 2.70 | 3.2E-02 |
| *C31H2.3* | WBGene00016293 | protein_coding_gene | 2.7 | 6.53 | 2.1E-03 | 1.5 | 2.89 | 4.2E-02 |
| *T03E6.8* | WBGene00011397 | protein_coding_gene | 2.7 | 6.52 | 6.9E-07 | 1.5 | 2.79 | 2.6E-02 |
| *Y94H6A.2* | WBGene00022375 | protein_coding_gene | 2.7 | 6.51 | 7.9E-04 | 1.4 | 2.61 | 5.0E-02 |
| *ZC434.3* | WBGene00013891 | protein_coding_gene | 2.7 | 6.50 | 4.6E-04 | 1.7 | 3.19 | 4.3E-02 |
| *F20A1.4* | WBGene00017616 | protein_coding_gene | 2.7 | 6.49 | 1.8E-03 | 1.4 | 2.67 | 2.8E-02 |
| *F13E9.14* | WBGene00045207 | protein_coding_gene | 2.7 | 6.46 | 3.3E-02 | 3.2 | 9.17 | 1.1E-03 |
| *T10G3.4* | WBGene00011695 | protein_coding_gene | 2.7 | 6.45 | 3.7E-02 | 2.0 | 3.88 | 4.4E-02 |
| *dlhd-1* | WBGene00020741 | protein_coding_gene | 2.7 | 6.35 | 1.8E-08 | 2.6 | 5.94 | 1.2E-06 |
| *hsp-70* | WBGene00002026 | protein_coding_gene | 2.7 | 6.31 | 3.7E-03 | 2.3 | 4.78 | 4.1E-03 |
| *F56C3.9* | WBGene00018946 | protein_coding_gene | 2.7 | 6.30 | 1.8E-05 | 1.2 | 2.31 | 1.7E-02 |
| *F45D3.2* | WBGene00009722 | protein_coding_gene | 2.7 | 6.29 | 2.9E-04 | 2.1 | 4.24 | 3.1E-03 |
| *F14H12.2* | WBGene00017470 | protein_coding_gene | 2.7 | 6.28 | 9.4E-03 | 1.7 | 3.30 | 3.9E-02 |
| *K02A2.5* | WBGene00019290 | protein_coding_gene | 2.6 | 6.26 | 8.2E-04 | 1.5 | 2.76 | 4.1E-02 |
| *ZC196.4* | WBGene00022548 | protein_coding_gene | 2.6 | 6.25 | 6.6E-04 | 2.0 | 3.91 | 1.9E-03 |
| *spp-18* | WBGene00005003 | protein_coding_gene | 2.6 | 6.24 | 2.8E-04 | 1.4 | 2.56 | 2.5E-02 |
| *F09C8.1* | WBGene00008621 | protein_coding_gene | 2.6 | 6.23 | 1.7E-08 | 2.7 | 6.38 | 1.4E-08 |
| *nhr-277* | WBGene00022374 | protein_coding_gene | 2.6 | 6.22 | 3.7E-04 | 1.9 | 3.80 | 1.2E-02 |
| *M04C9.2* | WBGene00010857 | protein_coding_gene | 2.6 | 6.20 | 1.7E-02 | 2.2 | 4.46 | 4.7E-02 |
| *T27C4.1* | WBGene00020854 | protein_coding_gene | 2.6 | 6.18 | 2.3E-06 | 1.9 | 3.62 | 1.0E-05 |
| *gmd-2* | WBGene00010166 | protein_coding_gene | 2.6 | 6.17 | 2.1E-02 | 3.1 | 8.47 | 2.3E-04 |
| *nhr-117* | WBGene00003707 | protein_coding_gene | 2.6 | 6.15 | 8.5E-07 | 1.2 | 2.26 | 8.7E-03 |
| *aqp-1* | WBGene00000169 | protein_coding_gene | 2.6 | 6.15 | 4.1E-03 | 2.2 | 4.48 | 1.3E-02 |
| *F56A6.5* | WBGene00044427 | protein_coding_gene | 2.6 | 6.13 | 3.9E-02 | 2.5 | 5.53 | 2.9E-02 |
| *bath-26* | WBGene00020329 | protein_coding_gene | 2.6 | 6.10 | 3.3E-09 | 1.3 | 2.51 | 2.4E-04 |
| *T05F1.9* | WBGene00011494 | protein_coding_gene | 2.6 | 6.09 | 2.3E-03 | 2.1 | 4.36 | 5.0E-03 |
| *pqm-1* | WBGene00004096 | protein_coding_gene | 2.6 | 6.06 | 2.4E-08 | 1.3 | 2.39 | 7.3E-06 |
| *W03G1.5* | WBGene00021011 | protein_coding_gene | 2.6 | 6.04 | 9.1E-04 | 1.7 | 3.27 | 2.8E-02 |
| *glb-34* | WBGene00077763 | protein_coding_gene | 2.6 | 6.04 | 1.8E-02 | 1.8 | 3.45 | 3.6E-02 |
| *T06G6.6* | WBGene00011548 | protein_coding_gene | 2.6 | 6.03 | 1.9E-02 | 2.3 | 4.77 | 1.9E-02 |
| *ZK863.8* | WBGene00014125 | protein_coding_gene | 2.6 | 6.01 | 2.9E-06 | 1.4 | 2.60 | 1.5E-02 |
| *C09E7.10* | WBGene00044295 | protein_coding_gene | 2.6 | 5.98 | 9.4E-03 | 2.1 | 4.38 | 2.0E-02 |
| *fipr-1* | WBGene00009096 | protein_coding_gene | 2.6 | 5.98 | 9.4E-08 | 1.3 | 2.49 | 2.7E-05 |
| *F13E9.15* | WBGene00045208 | protein_coding_gene | 2.6 | 5.98 | 9.6E-06 | 1.3 | 2.46 | 1.8E-04 |

| | | | | | | | | |
|---|---|---|---|---|---|---|---|---|
| *T09D3.8* | WBGene00044538 | protein_coding_gene | 2.6 | 5.95 | 4.2E-02 | 2.6 | 6.14 | 5.4E-03 |
| *ZK250.13* | WBGene00235352 | protein_coding_gene | 2.6 | 5.88 | 2.6E-06 | 2.0 | 3.94 | 1.1E-03 |
| *ubc-21* | WBGene00006716 | protein_coding_gene | 2.6 | 5.87 | 2.3E-02 | 2.2 | 4.60 | 3.9E-02 |
| *lmd-5* | WBGene00020141 | protein_coding_gene | 2.6 | 5.86 | 2.4E-03 | 2.1 | 4.26 | 1.7E-03 |
| *ZK355.8* | WBGene00086560 | protein_coding_gene | 2.6 | 5.86 | 2.9E-02 | 2.1 | 4.37 | 4.8E-02 |
| *EGAP9.3* | WBGene00017151 | protein_coding_gene | 2.5 | 5.85 | 1.4E-10 | 1.3 | 2.52 | 3.9E-05 |
| *W02F12.8* | WBGene00175027 | protein_coding_gene | 2.5 | 5.83 | 7.6E-03 | 2.2 | 4.63 | 1.0E-02 |
| *flp-21* | WBGene00001464 | protein_coding_gene | 2.5 | 5.83 | 9.2E-05 | 1.6 | 2.96 | 1.4E-03 |
| *F02A9.1* | WBGene00008513 | protein_coding_gene | 2.5 | 5.83 | 2.9E-04 | 1.2 | 2.28 | 3.3E-02 |
| *fipr-2* | WBGene00009097 | protein_coding_gene | 2.5 | 5.82 | 5.0E-07 | 1.4 | 2.61 | 1.7E-07 |
| *F47B8.2* | WBGene00009803 | protein_coding_gene | 2.5 | 5.78 | 1.1E-05 | 1.3 | 2.54 | 4.4E-02 |
| *pals-14* | WBGene00009061 | protein_coding_gene | 2.5 | 5.77 | 2.9E-03 | 1.4 | 2.64 | 2.6E-02 |
| *srg-15* | WBGene00005172 | protein_coding_gene | 2.5 | 5.74 | 3.5E-02 | 2.0 | 4.08 | 2.9E-02 |
| *bli-1* | WBGene00000251 | protein_coding_gene | 2.5 | 5.74 | 1.9E-02 | 3.2 | 9.30 | 2.8E-04 |
| *F26A1.19* | WBGene00270319 | protein_coding_gene | 2.5 | 5.71 | 7.6E-05 | 1.0 | 2.03 | 4.3E-02 |
| *K10D3.8* | WBGene00219281 | protein_coding_gene | 2.5 | 5.68 | 3.8E-02 | 2.6 | 6.12 | 1.1E-02 |
| *C14C11.4* | WBGene00015768 | protein_coding_gene | 2.5 | 5.67 | 2.1E-04 | 1.9 | 3.69 | 2.3E-04 |
| *T28A11.3* | WBGene00020870 | protein_coding_gene | 2.5 | 5.66 | 3.3E-02 | 2.0 | 3.99 | 2.7E-02 |
| *dur-1* | WBGene00001113 | protein_coding_gene | 2.5 | 5.63 | 2.6E-03 | 2.6 | 6.15 | 7.9E-04 |
| *cdh-10* | WBGene00000402 | protein_coding_gene | 2.5 | 5.63 | 1.0E-02 | 2.8 | 6.75 | 4.6E-04 |
| *gur-3* | WBGene00001804 | protein_coding_gene | 2.5 | 5.62 | 3.1E-02 | 2.4 | 5.16 | 3.6E-02 |
| *T19H12.15* | WBGene00271827 | protein_coding_gene | 2.5 | 5.62 | 7.8E-05 | 3.9 | 15.12 | 2.1E-33 |
| *pdf-2* | WBGene00018590 | protein_coding_gene | 2.5 | 5.62 | 4.1E-04 | 1.2 | 2.25 | 2.3E-02 |
| *pals-6* | WBGene00007660 | protein_coding_gene | 2.5 | 5.60 | 4.7E-03 | 2.6 | 5.93 | 1.1E-06 |
| *cyp-33C1* | WBGene00016686 | protein_coding_gene | 2.5 | 5.58 | 3.4E-02 | 2.4 | 5.32 | 3.0E-02 |
| *C50B6.7* | WBGene00008220 | protein_coding_gene | 2.5 | 5.58 | 1.1E-06 | 2.5 | 5.79 | 8.0E-07 |
| *nlp-71* | WBGene00022254 | protein_coding_gene | 2.5 | 5.56 | 2.9E-05 | 1.5 | 2.77 | 3.6E-03 |
| *srw-71* | WBGene00005818 | protein_coding_gene | 2.5 | 5.54 | 2.6E-02 | 2.1 | 4.41 | 3.0E-02 |
| *nlp-8* | WBGene00003746 | protein_coding_gene | 2.5 | 5.52 | 1.1E-08 | 1.6 | 3.02 | 1.9E-07 |
| *K08B5.1* | WBGene00019518 | protein_coding_gene | 2.5 | 5.51 | 4.9E-03 | 1.5 | 2.78 | 8.4E-03 |
| *C01G10.15* | WBGene00007240 | protein_coding_gene | 2.5 | 5.50 | 6.5E-05 | 1.2 | 2.36 | 2.4E-02 |
| *F28H7.2* | WBGene00009236 | protein_coding_gene | 2.5 | 5.50 | 2.8E-03 | 1.7 | 3.19 | 1.2E-02 |
| *Y54G2A.32* | WBGene00021894 | protein_coding_gene | 2.4 | 5.40 | 2.7E-05 | 1.1 | 2.14 | 5.2E-03 |
| *F42C5.3* | WBGene00018345 | protein_coding_gene | 2.4 | 5.39 | 3.7E-03 | 1.6 | 3.11 | 2.6E-02 |
| *xbx-3* | WBGene00010864 | protein_coding_gene | 2.4 | 5.38 | 6.2E-03 | 1.3 | 2.40 | 4.3E-02 |
| *F21H12.7* | WBGene00044287 | protein_coding_gene | 2.4 | 5.37 | 7.4E-05 | 2.6 | 5.93 | 8.8E-05 |
| *scl-5* | WBGene00008027 | protein_coding_gene | 2.4 | 5.37 | 3.5E-06 | 1.3 | 2.48 | 1.2E-05 |
| *fbxa-73* | WBGene00020640 | protein_coding_gene | 2.4 | 5.36 | 2.3E-02 | 1.5 | 2.82 | 1.6E-02 |
| *C05G5.7* | WBGene00050939 | protein_coding_gene | 2.4 | 5.35 | 1.4E-02 | 2.1 | 4.40 | 1.5E-02 |
| *gopc-1* | WBGene00014101 | protein_coding_gene | 2.4 | 5.33 | 2.7E-04 | 1.5 | 2.91 | 7.1E-03 |
| *Y75B8A.44* | WBGene00194855 | protein_coding_gene | 2.4 | 5.32 | 2.0E-05 | 2.2 | 4.48 | 3.3E-08 |
| *C15C7.4* | WBGene00015790 | protein_coding_gene | 2.4 | 5.29 | 7.2E-04 | 1.9 | 3.86 | 6.2E-07 |
| *W01A11.7* | WBGene00020912 | protein_coding_gene | 2.4 | 5.28 | 2.0E-03 | 1.6 | 3.02 | 3.9E-02 |
| *trh-1* | WBGene00007827 | protein_coding_gene | 2.4 | 5.27 | 5.8E-06 | 1.2 | 2.37 | 1.4E-03 |
| *T16G1.7* | WBGene00011801 | protein_coding_gene | 2.4 | 5.25 | 6.1E-03 | 1.1 | 2.17 | 1.3E-02 |
| *chil-3* | WBGene00007467 | protein_coding_gene | 2.4 | 5.21 | 1.8E-04 | 3.2 | 9.30 | 8.3E-05 |
| *ceh-23* | WBGene00000446 | protein_coding_gene | 2.4 | 5.21 | 9.5E-03 | 2.2 | 4.64 | 2.1E-03 |
| *C30E1.9* | WBGene00022951 | ncRNA_gene | 2.4 | 5.20 | 5.9E-06 | 1.3 | 2.53 | 1.5E-03 |
| *exc-14* | WBGene00019649 | protein_coding_gene | 2.4 | 5.20 | 3.2E-05 | 1.6 | 2.97 | 1.1E-03 |
| *R06C1.6* | WBGene00011060 | protein_coding_gene | 2.4 | 5.20 | 3.4E-04 | 1.6 | 3.07 | 7.4E-03 |
| *flp-9* | WBGene00001452 | protein_coding_gene | 2.4 | 5.19 | 1.2E-05 | 1.4 | 2.59 | 6.3E-04 |
| *ZK662.6* | WBGene00045494 | protein_coding_gene | 2.4 | 5.18 | 2.9E-02 | 3.2 | 8.95 | 7.1E-04 |
| *R07A4.3* | WBGene00011074 | protein_coding_gene | 2.4 | 5.14 | 1.7E-02 | 1.8 | 3.47 | 2.7E-02 |
| *F46A8.11* | WBGene00014761 | protein_coding_gene | 2.4 | 5.13 | 1.4E-04 | 2.4 | 5.26 | 8.1E-08 |
| *nhr-208* | WBGene00011099 | protein_coding_gene | 2.4 | 5.13 | 4.3E-04 | 1.5 | 2.80 | 1.0E-02 |
| *F59A7.5* | WBGene00019092 | protein_coding_gene | 2.4 | 5.13 | 1.0E-04 | 1.2 | 2.33 | 2.2E-03 |
| *nhr-168* | WBGene00008221 | protein_coding_gene | 2.4 | 5.12 | 1.3E-12 | 2.2 | 4.74 | 3.4E-05 |
| *C38C3.10* | WBGene00045390 | protein_coding_gene | 2.4 | 5.12 | 9.1E-04 | 1.5 | 2.86 | 1.6E-03 |

| | | | | | | | | |
|---|---|---|---|---|---|---|---|---|
| *nex-4* | WBGene00003591 | protein_coding_gene | 2.3 | 5.10 | 9.6E-04 | 1.6 | 3.05 | 1.4E-02 |
| *ubc-22* | WBGene00006717 | protein_coding_gene | 2.3 | 5.09 | 5.0E-03 | 1.9 | 3.67 | 1.5E-02 |
| *Y67D8B.1* | WBGene00022061 | protein_coding_gene | 2.3 | 5.06 | 8.9E-03 | 2.0 | 3.91 | 7.3E-03 |
| *twnk-1* | WBGene00018514 | protein_coding_gene | 2.3 | 5.03 | 4.1E-03 | 1.6 | 2.97 | 4.5E-03 |
| *R10F2.4* | WBGene00019995 | protein_coding_gene | 2.3 | 5.02 | 2.7E-03 | 2.3 | 5.06 | 2.3E-03 |
| *osm-9* | WBGene00003889 | protein_coding_gene | 2.3 | 5.01 | 1.2E-02 | 1.5 | 2.86 | 4.1E-02 |
| *col-3* | WBGene00000594 | protein_coding_gene | 2.3 | 5.01 | 2.8E-02 | 2.1 | 4.19 | 1.8E-02 |
| *pho-11* | WBGene00007331 | protein_coding_gene | 2.3 | 5.01 | 9.1E-04 | 1.8 | 3.46 | 5.0E-06 |
| *frpr-2* | WBGene00007346 | protein_coding_gene | 2.3 | 4.99 | 1.5E-02 | 1.6 | 3.07 | 4.7E-02 |
| *npr-8* | WBGene00016984 | protein_coding_gene | 2.3 | 4.99 | 6.1E-05 | 1.2 | 2.28 | 4.1E-03 |
| *clec-140* | WBGene00020220 | protein_coding_gene | 2.3 | 4.99 | 2.2E-02 | 1.7 | 3.29 | 2.4E-02 |
| *B0238.12* | WBGene00015076 | protein_coding_gene | 2.3 | 4.98 | 2.9E-03 | 2.5 | 5.54 | 2.9E-04 |
| *ocr-4* | WBGene00003841 | protein_coding_gene | 2.3 | 4.98 | 3.1E-02 | 2.4 | 5.39 | 9.3E-03 |
| *Y48A6B.8* | WBGene00012969 | protein_coding_gene | 2.3 | 4.95 | 1.5E-02 | 1.9 | 3.86 | 2.6E-02 |
| *nlp-35* | WBGene00007881 | protein_coding_gene | 2.3 | 4.95 | 2.4E-04 | 1.8 | 3.41 | 6.7E-04 |
| *T01H10.4* | WBGene00011357 | pseudogene | 2.3 | 4.94 | 4.4E-02 | 1.5 | 2.91 | 3.7E-02 |
| *F39H12.2* | WBGene00018213 | protein_coding_gene | 2.3 | 4.94 | 6.4E-03 | 2.4 | 5.35 | 8.0E-04 |
| *sri-74* | WBGene00005586 | protein_coding_gene | 2.3 | 4.93 | 4.4E-02 | 1.4 | 2.55 | 1.3E-02 |
| *gcy-37* | WBGene00001557 | protein_coding_gene | 2.3 | 4.93 | 2.7E-03 | 1.8 | 3.58 | 4.1E-03 |
| *klu-1* | WBGene00013970 | protein_coding_gene | 2.3 | 4.92 | 4.5E-04 | 2.0 | 3.95 | 1.1E-03 |
| *gcy-4* | WBGene00001531 | protein_coding_gene | 2.3 | 4.92 | 2.2E-02 | 1.8 | 3.46 | 3.9E-02 |
| *Y6G8.2* | WBGene00012404 | protein_coding_gene | 2.3 | 4.92 | 3.8E-05 | 1.6 | 3.03 | 2.4E-05 |
| *E04D5.4* | WBGene00008483 | protein_coding_gene | 2.3 | 4.91 | 6.7E-03 | 1.6 | 2.99 | 2.8E-02 |
| *flp-15* | WBGene00001458 | protein_coding_gene | 2.3 | 4.91 | 1.2E-05 | 1.5 | 2.76 | 5.8E-04 |
| *ddn-2* | WBGene00009399 | protein_coding_gene | 2.3 | 4.90 | 2.0E-03 | 1.7 | 3.26 | 3.5E-03 |
| *C46H11.2* | WBGene00016728 | protein_coding_gene | 2.3 | 4.90 | 8.7E-07 | 1.5 | 2.76 | 3.7E-05 |
| *H02F09.2* | WBGene00019145 | protein_coding_gene | 2.3 | 4.90 | 9.5E-03 | 1.9 | 3.61 | 1.5E-02 |
| *flp-27* | WBGene00016114 | protein_coding_gene | 2.3 | 4.88 | 7.6E-05 | 1.1 | 2.09 | 7.4E-03 |
| *clec-227* | WBGene00008593 | protein_coding_gene | 2.3 | 4.87 | 5.3E-03 | 2.2 | 4.58 | 3.2E-03 |
| *daf-9* | WBGene00000905 | protein_coding_gene | 2.3 | 4.86 | 1.5E-02 | 3.2 | 9.26 | 3.7E-04 |
| *F20G2.1* | WBGene00008985 | protein_coding_gene | 2.3 | 4.85 | 1.1E-03 | 3.0 | 8.09 | 2.3E-10 |
| *C30B5.6* | WBGene00016247 | protein_coding_gene | 2.3 | 4.85 | 7.1E-03 | 1.5 | 2.75 | 1.6E-02 |
| *Y42A5A.1* | WBGene00012776 | protein_coding_gene | 2.3 | 4.83 | 1.9E-03 | 1.6 | 3.07 | 5.5E-03 |
| *nlp-68* | WBGene00011899 | protein_coding_gene | 2.3 | 4.79 | 5.6E-04 | 1.4 | 2.64 | 1.4E-03 |
| *bli-2* | WBGene00000252 | protein_coding_gene | 2.3 | 4.79 | 3.8E-02 | 3.0 | 7.99 | 6.7E-04 |
| *C04E6.13* | WBGene00015427 | protein_coding_gene | 2.3 | 4.78 | 1.2E-05 | 1.1 | 2.09 | 4.5E-02 |
| *F13D11.4* | WBGene00017429 | protein_coding_gene | 2.3 | 4.77 | 6.1E-04 | 1.3 | 2.51 | 9.3E-03 |
| *ntc-1* | WBGene00044568 | protein_coding_gene | 2.3 | 4.77 | 2.0E-03 | 1.2 | 2.33 | 2.9E-02 |
| *ins-10* | WBGene00002093 | protein_coding_gene | 2.2 | 4.76 | 1.0E-02 | 1.7 | 3.30 | 1.7E-02 |
| *nhr-6* | WBGene00003605 | protein_coding_gene | 2.2 | 4.75 | 6.9E-04 | 1.5 | 2.92 | 6.3E-03 |
| *ins-1* | WBGene00002084 | protein_coding_gene | 2.2 | 4.74 | 8.6E-04 | 1.1 | 2.19 | 1.6E-02 |
| *gba-1* | WBGene00016335 | protein_coding_gene | 2.2 | 4.74 | 4.7E-03 | 1.1 | 2.18 | 2.0E-02 |
| *F40E3.5* | WBGene00018232 | protein_coding_gene | 2.2 | 4.74 | 1.5E-04 | 1.3 | 2.41 | 1.4E-02 |
| *acr-19* | WBGene00000058 | protein_coding_gene | 2.2 | 4.73 | 7.6E-04 | 1.5 | 2.90 | 4.2E-03 |
| *pals-33* | WBGene00021081 | protein_coding_gene | 2.2 | 4.72 | 2.5E-04 | 2.0 | 4.05 | 1.3E-04 |
| *Y51H4A.8* | WBGene00013104 | protein_coding_gene | 2.2 | 4.71 | 9.8E-06 | 1.5 | 2.84 | 2.6E-03 |
| *pals-38* | WBGene00008302 | protein_coding_gene | 2.2 | 4.71 | 6.2E-03 | 1.9 | 3.62 | 6.8E-03 |
| *col-166* | WBGene00000739 | protein_coding_gene | 2.2 | 4.70 | 9.4E-03 | 2.3 | 5.05 | 4.8E-03 |
| *asm-2* | WBGene00000212 | protein_coding_gene | 2.2 | 4.67 | 1.8E-03 | 2.1 | 4.42 | 2.3E-04 |
| *lact-3* | WBGene00010897 | protein_coding_gene | 2.2 | 4.66 | 3.0E-05 | 1.2 | 2.30 | 9.9E-03 |
| *nlp-38* | WBGene00007219 | protein_coding_gene | 2.2 | 4.66 | 9.8E-04 | 1.4 | 2.67 | 1.4E-02 |
| *efhc-1* | WBGene00013028 | protein_coding_gene | 2.2 | 4.65 | 2.8E-02 | 1.8 | 3.48 | 1.3E-02 |
| *gst-3* | WBGene00001751 | protein_coding_gene | 2.2 | 4.64 | 1.3E-03 | 1.7 | 3.22 | 1.2E-02 |
| *F42A10.6* | WBGene00018342 | protein_coding_gene | 2.2 | 4.63 | 3.5E-06 | 2.2 | 4.68 | 1.7E-24 |
| *C29F9.6* | WBGene00016221 | protein_coding_gene | 2.2 | 4.61 | 2.7E-02 | 1.8 | 3.58 | 2.8E-02 |
| *eol-1* | WBGene00045401 | protein_coding_gene | 2.2 | 4.60 | 1.1E-03 | 1.6 | 3.06 | 7.2E-03 |
| *F52H2.3* | WBGene00018715 | protein_coding_gene | 2.2 | 4.58 | 2.4E-04 | 1.6 | 2.96 | 3.9E-03 |
| *aat-7* | WBGene00000008 | protein_coding_gene | 2.2 | 4.58 | 1.8E-02 | 3.6 | 12.48 | 9.2E-06 |

| | | | | | | | | |
|---|---|---|---|---|---|---|---|---|
| ***oac-14*** | **WBGene00008602** | **protein_coding_gene** | **2.2** | **4.58** | **2.0E-03** | **-1.9** | **0.26** | **6.2E-03** |
| *Y43D4A.5* | WBGene00012791 | protein_coding_gene | 2.2 | 4.58 | 3.7E-02 | 2.1 | 4.14 | 3.0E-02 |
| *C18B2.2* | WBGene00015953 | protein_coding_gene | 2.2 | 4.57 | 2.2E-02 | 2.1 | 4.22 | 1.2E-02 |
| *degt-1* | WBGene00009109 | protein_coding_gene | 2.2 | 4.57 | 2.7E-04 | 2.6 | 6.06 | 5.7E-06 |
| *K12B6.4* | WBGene00019669 | protein_coding_gene | 2.2 | 4.55 | 2.5E-03 | 2.7 | 6.28 | 4.0E-05 |
| *nlp-21* | WBGene00003759 | protein_coding_gene | 2.2 | 4.54 | 6.4E-04 | 1.8 | 3.38 | 2.6E-04 |
| *pqn-48* | WBGene00004135 | protein_coding_gene | 2.2 | 4.54 | 1.8E-03 | 1.9 | 3.69 | 1.8E-03 |
| *bath-47* | WBGene00006464 | protein_coding_gene | 2.2 | 4.53 | 6.6E-04 | 1.1 | 2.18 | 1.8E-02 |
| *comt-1* | WBGene00018741 | protein_coding_gene | 2.2 | 4.52 | 4.8E-02 | 2.1 | 4.34 | 4.4E-02 |
| *ghi-1* | WBGene00013923 | protein_coding_gene | 2.2 | 4.52 | 5.6E-03 | 2.3 | 5.00 | 2.2E-04 |
| *B0432.14* | WBGene00044640 | protein_coding_gene | 2.2 | 4.51 | 5.3E-04 | 1.8 | 3.60 | 1.0E-03 |
| *T12A7.2* | WBGene00011723 | protein_coding_gene | 2.2 | 4.50 | 2.9E-04 | 1.4 | 2.66 | 3.7E-10 |
| *cdh-12* | WBGene00022103 | protein_coding_gene | 2.2 | 4.50 | 1.5E-03 | 1.5 | 2.75 | 4.2E-02 |
| *nas-6* | WBGene00003525 | protein_coding_gene | 2.2 | 4.49 | 6.8E-03 | 2.0 | 3.97 | 1.0E-02 |
| *oac-1* | WBGene00007177 | protein_coding_gene | 2.2 | 4.48 | 2.1E-02 | 2.9 | 7.63 | 6.8E-03 |
| *F56C4.1* | WBGene00010144 | protein_coding_gene | 2.2 | 4.48 | 8.2E-03 | 1.4 | 2.66 | 4.4E-02 |
| *F02E11.7* | WBGene00195073 | protein_coding_gene | 2.2 | 4.47 | 5.2E-04 | 1.2 | 2.36 | 2.3E-02 |
| *Y48G8AR.2* | WBGene00021693 | protein_coding_gene | 2.2 | 4.45 | 4.2E-03 | 1.3 | 2.52 | 7.0E-03 |
| *C45G9.11* | WBGene00016682 | protein_coding_gene | 2.2 | 4.45 | 1.0E-02 | 1.4 | 2.73 | 2.2E-02 |
| *far-4* | WBGene00001388 | protein_coding_gene | 2.2 | 4.44 | 4.0E-02 | 1.9 | 3.81 | 2.7E-02 |
| *T05H4.15* | WBGene00020276 | protein_coding_gene | 2.1 | 4.44 | 4.4E-04 | 1.4 | 2.69 | 7.9E-03 |
| *nlp-13* | WBGene00003751 | protein_coding_gene | 2.1 | 4.43 | 2.0E-04 | 1.6 | 3.06 | 8.2E-07 |
| *mboa-5* | WBGene00013995 | pseudogene | 2.1 | 4.42 | 3.3E-02 | 1.6 | 3.05 | 3.4E-02 |
| *flp-18* | WBGene00001461 | protein_coding_gene | 2.1 | 4.41 | 5.7E-04 | 1.4 | 2.60 | 2.2E-05 |
| *acr-23* | WBGene00000062 | protein_coding_gene | 2.1 | 4.40 | 1.5E-02 | 2.5 | 5.47 | 7.5E-03 |
| *capa-1* | WBGene00021268 | protein_coding_gene | 2.1 | 4.38 | 1.1E-03 | 1.2 | 2.25 | 3.8E-02 |
| *col-46* | WBGene00000623 | protein_coding_gene | 2.1 | 4.37 | 3.6E-02 | 2.7 | 6.50 | 7.9E-03 |
| *ugt-55* | WBGene00011452 | protein_coding_gene | 2.1 | 4.37 | 2.4E-04 | 1.6 | 3.01 | 2.7E-03 |
| *gpx-3* | WBGene00007517 | protein_coding_gene | 2.1 | 4.36 | 2.5E-05 | 2.1 | 4.15 | 5.6E-06 |
| *clec-80* | WBGene00043062 | protein_coding_gene | 2.1 | 4.35 | 6.0E-06 | 2.4 | 5.46 | 1.6E-06 |
| *R13H7.2* | WBGene00020069 | protein_coding_gene | 2.1 | 4.34 | 8.6E-04 | 1.6 | 2.94 | 2.3E-03 |
| *C47E8.3* | WBGene00008142 | protein_coding_gene | 2.1 | 4.34 | 7.9E-03 | 1.7 | 3.19 | 1.4E-03 |
| *tsp-10* | WBGene00006636 | protein_coding_gene | 2.1 | 4.34 | 1.1E-05 | 1.6 | 3.09 | 5.6E-04 |
| *F58B4.3* | WBGene00010236 | protein_coding_gene | 2.1 | 4.34 | 6.9E-04 | 1.7 | 3.14 | 4.0E-03 |
| ***cutl-9*** | **WBGene00010893** | **protein_coding_gene** | **2.1** | **4.33** | **5.2E-03** | **-1.3** | **0.41** | **2.9E-02** |
| *ZC21.3* | WBGene00022505 | protein_coding_gene | 2.1 | 4.32 | 1.1E-02 | 1.7 | 3.32 | 3.3E-05 |
| *Y49E10.18* | WBGene00013037 | protein_coding_gene | 2.1 | 4.32 | 8.5E-05 | 1.9 | 3.76 | 1.3E-07 |
| ***F15E6.3*** | **WBGene00017484** | **protein_coding_gene** | **2.1** | **4.32** | **1.5E-02** | **-2.4** | **0.19** | **1.5E-05** |
| *D1079.1* | WBGene00017040 | protein_coding_gene | 2.1 | 4.31 | 2.8E-03 | 1.2 | 2.34 | 1.1E-02 |
| *F11A5.9* | WBGene00008677 | protein_coding_gene | 2.1 | 4.30 | 1.3E-04 | 1.4 | 2.62 | 2.6E-02 |
| *ida-1* | WBGene00002048 | protein_coding_gene | 2.1 | 4.30 | 8.8E-04 | 1.6 | 3.05 | 4.6E-05 |
| *flp-16* | WBGene00001459 | protein_coding_gene | 2.1 | 4.29 | 7.8E-04 | 1.5 | 2.78 | 5.5E-04 |
| *pqn-94* | WBGene00004173 | protein_coding_gene | 2.1 | 4.29 | 2.1E-04 | 1.3 | 2.44 | 6.5E-03 |
| *dgk-3* | WBGene00000960 | protein_coding_gene | 2.1 | 4.28 | 2.1E-03 | 1.5 | 2.78 | 5.7E-03 |
| *F01D4.8* | WBGene00008490 | protein_coding_gene | 2.1 | 4.27 | 1.6E-02 | 2.5 | 5.70 | 1.2E-03 |
| *lgc-38* | WBGene00017389 | protein_coding_gene | 2.1 | 4.25 | 1.5E-03 | 2.1 | 4.43 | 1.6E-04 |
| *W10C8.6* | WBGene00044433 | protein_coding_gene | 2.1 | 4.25 | 1.6E-04 | 1.3 | 2.47 | 4.3E-02 |
| *nas-15* | WBGene00003534 | protein_coding_gene | 2.1 | 4.25 | 2.6E-02 | 1.9 | 3.85 | 3.1E-02 |
| *mpk-2* | WBGene00003402 | protein_coding_gene | 2.1 | 4.24 | 1.6E-07 | 1.5 | 2.87 | 8.1E-08 |
| *sto-6* | WBGene00006068 | protein_coding_gene | 2.1 | 4.24 | 1.8E-03 | 1.3 | 2.50 | 1.1E-02 |
| *F58A6.2* | WBGene00019023 | protein_coding_gene | 2.1 | 4.23 | 1.5E-02 | 1.9 | 3.81 | 1.2E-02 |
| *nlp-9* | WBGene00003747 | protein_coding_gene | 2.1 | 4.23 | 2.6E-03 | 1.5 | 2.92 | 1.4E-03 |
| *comt-2* | WBGene00012518 | protein_coding_gene | 2.1 | 4.23 | 3.1E-02 | 1.8 | 3.51 | 2.3E-02 |
| *Y18D10A.22* | WBGene00014864 | protein_coding_gene | 2.1 | 4.21 | 3.3E-05 | 1.8 | 3.38 | 3.7E-03 |
| *F58H1.7* | WBGene00010290 | protein_coding_gene | 2.1 | 4.20 | 1.8E-05 | 1.1 | 2.11 | 1.2E-03 |
| *B0222.1* | WBGene00015053 | protein_coding_gene | 2.1 | 4.19 | 2.0E-03 | 1.4 | 2.56 | 2.6E-02 |
| *nmy-3* | WBGene00012437 | protein_coding_gene | 2.1 | 4.18 | 6.2E-03 | 2.9 | 7.22 | 5.2E-08 |
| *egl-3* | WBGene00001172 | protein_coding_gene | 2.1 | 4.17 | 2.0E-03 | 2.0 | 4.07 | 2.8E-07 |

| | | | | | | | | |
|---|---|---|---|---|---|---|---|---|
| *Y22D7AL.16* | WBGene00021254 | protein_coding_gene | 2.1 | 4.16 | 4.1E-03 | 2.1 | 4.32 | 2.0E-03 |
| *lin-11* | WBGene00003000 | protein_coding_gene | 2.1 | 4.16 | 5.5E-03 | 1.3 | 2.45 | 4.8E-02 |
| *tag-244* | WBGene00044073 | protein_coding_gene | 2.1 | 4.15 | 3.3E-03 | 1.5 | 2.76 | 2.9E-02 |
| *pghm-1* | WBGene00022144 | protein_coding_gene | 2.1 | 4.15 | 1.3E-04 | 1.6 | 3.04 | 3.0E-06 |
| *acox-1.5* | WBGene00008167 | protein_coding_gene | 2.1 | 4.14 | 5.0E-03 | 1.5 | 2.90 | 1.8E-02 |
| *R10E8.6* | WBGene00011212 | protein_coding_gene | 2.0 | 4.14 | 1.7E-06 | 1.2 | 2.30 | 9.8E-03 |
| *C09G9.5* | WBGene00007495 | protein_coding_gene | 2.0 | 4.14 | 3.1E-03 | 1.8 | 3.41 | 7.3E-03 |
| *F10E9.1* | WBGene00017353 | protein_coding_gene | 2.0 | 4.13 | 2.6E-03 | 1.6 | 3.02 | 7.2E-03 |
| *Y9C9A.16* | WBGene00021183 | protein_coding_gene | 2.0 | 4.13 | 3.8E-02 | 2.2 | 4.50 | 1.1E-02 |
| *nlp-55* | WBGene00008820 | protein_coding_gene | 2.0 | 4.11 | 2.0E-03 | 1.3 | 2.54 | 4.5E-03 |
| *F41G4.7* | WBGene00018310 | protein_coding_gene | 2.0 | 4.11 | 7.2E-03 | 1.6 | 3.01 | 2.2E-02 |
| *pdf-1* | WBGene00020317 | protein_coding_gene | 2.0 | 4.11 | 9.8E-05 | 1.2 | 2.28 | 1.2E-04 |
| *dod-6* | WBGene00011869 | protein_coding_gene | 2.0 | 4.10 | 2.9E-05 | 1.3 | 2.46 | 1.8E-08 |
| *F02E11.2* | WBGene00017180 | protein_coding_gene | 2.0 | 4.09 | 1.1E-02 | 1.3 | 2.40 | 3.6E-03 |
| *W07A12.4* | WBGene00012322 | protein_coding_gene | 2.0 | 4.08 | 1.0E-02 | 1.2 | 2.32 | 1.1E-02 |
| *cwp-4* | WBGene00000862 | protein_coding_gene | 2.0 | 4.08 | 8.4E-03 | 1.2 | 2.37 | 1.6E-02 |
| *F53A9.1* | WBGene00018724 | protein_coding_gene | 2.0 | 4.08 | 1.5E-03 | 1.3 | 2.51 | 3.4E-02 |
| *F31E3.2* | WBGene00017950 | protein_coding_gene | 2.0 | 4.08 | 2.7E-04 | 1.5 | 2.78 | 2.4E-03 |
| *flp-2* | WBGene00001445 | protein_coding_gene | 2.0 | 4.07 | 2.2E-03 | 1.3 | 2.42 | 3.4E-02 |
| *ctg-2* | WBGene00011756 | protein_coding_gene | 2.0 | 4.06 | 2.6E-04 | 1.9 | 3.83 | 1.7E-05 |
| *npr-1* | WBGene00003807 | protein_coding_gene | 2.0 | 4.05 | 2.4E-02 | 2.0 | 4.09 | 9.6E-03 |
| *linc-125* | WBGene00219686 | lincRNA_gene | 2.0 | 4.05 | 1.1E-02 | 2.1 | 4.27 | 1.4E-02 |
| *T03F6.9* | WBGene00045482 | protein_coding_gene | 2.0 | 4.02 | 8.4E-03 | 1.2 | 2.34 | 4.0E-02 |
| *F42A9.18* | WBGene00249826 | ncRNA_gene | 2.0 | 4.01 | 3.2E-02 | 1.7 | 3.19 | 1.1E-02 |
| *W02D9.10* | WBGene00012216 | protein_coding_gene | 2.0 | 4.01 | 1.4E-03 | 1.8 | 3.52 | 5.1E-03 |
| *flp-5* | WBGene00001448 | protein_coding_gene | 2.0 | 4.00 | 4.3E-04 | 1.5 | 2.79 | 5.5E-04 |
| *nmr-2* | WBGene00003775 | protein_coding_gene | 2.0 | 4.00 | 2.4E-03 | 1.1 | 2.17 | 3.0E-02 |
| *nlp-14* | WBGene00003752 | protein_coding_gene | 2.0 | 4.00 | 4.8E-03 | 1.6 | 3.06 | 4.1E-04 |
| *sma-10* | WBGene00020649 | protein_coding_gene | 2.0 | 4.00 | 1.6E-03 | 1.1 | 2.12 | 1.7E-02 |
| *ttr-39* | WBGene00007304 | protein_coding_gene | 2.0 | 4.00 | 1.9E-04 | 1.5 | 2.87 | 2.8E-03 |
| *D1054.9* | WBGene00008376 | protein_coding_gene | 2.0 | 3.99 | 1.5E-03 | 1.5 | 2.80 | 4.1E-02 |
| *flp-14* | WBGene00001457 | protein_coding_gene | 2.0 | 3.99 | 7.4E-05 | 1.3 | 2.42 | 5.2E-05 |
| *tkr-2* | WBGene00016761 | protein_coding_gene | 2.0 | 3.99 | 3.3E-02 | 1.7 | 3.19 | 2.6E-02 |
| *F27B10.1* | WBGene00017850 | protein_coding_gene | 2.0 | 3.98 | 4.5E-03 | 1.2 | 2.29 | 1.0E-02 |
| *ZK856.5* | WBGene00014106 | protein_coding_gene | 2.0 | 3.98 | 1.0E-04 | 1.7 | 3.31 | 1.1E-07 |
| *lite-1* | WBGene00001803 | protein_coding_gene | 2.0 | 3.97 | 1.1E-03 | 1.5 | 2.76 | 3.4E-03 |
| *ins-24* | WBGene00002107 | protein_coding_gene | 2.0 | 3.97 | 8.4E-03 | 1.6 | 3.11 | 1.4E-02 |
| *nlp-1* | WBGene00003739 | protein_coding_gene | 2.0 | 3.97 | 1.8E-03 | 1.0 | 2.05 | 1.9E-02 |
| *F28B12.1* | WBGene00017894 | protein_coding_gene | 2.0 | 3.97 | 1.6E-03 | 1.5 | 2.78 | 2.5E-02 |
| *bath-46* | WBGene00020325 | protein_coding_gene | 2.0 | 3.97 | 3.6E-02 | 2.0 | 4.10 | 3.6E-03 |
| *Y6E2A.5* | WBGene00012399 | protein_coding_gene | 2.0 | 3.97 | 7.4E-03 | 1.0 | 2.06 | 4.1E-02 |
| *ZK669.5* | WBGene00014055 | protein_coding_gene | 2.0 | 3.96 | 5.1E-05 | 1.5 | 2.85 | 1.5E-04 |
| *kvs-2* | WBGene00007862 | protein_coding_gene | 2.0 | 3.96 | 1.0E-03 | 1.3 | 2.52 | 3.6E-03 |
| *F10D2.8* | WBGene00017334 | protein_coding_gene | 2.0 | 3.96 | 3.7E-04 | 1.6 | 2.98 | 8.3E-09 |
| *T22B11.4* | WBGene00020678 | protein_coding_gene | 2.0 | 3.95 | 3.1E-03 | 1.7 | 3.27 | 4.3E-04 |
| *ZK353.10* | WBGene00044677 | protein_coding_gene | 2.0 | 3.95 | 9.9E-03 | 1.2 | 2.33 | 4.8E-02 |
| *F11E6.6* | WBGene00008709 | protein_coding_gene | 2.0 | 3.94 | 1.5E-02 | 2.0 | 4.11 | 5.1E-03 |
| *cla-1* | WBGene00018468 | protein_coding_gene | 2.0 | 3.94 | 8.7E-04 | 1.8 | 3.59 | 3.5E-06 |
| *bcmo-1* | WBGene00012914 | protein_coding_gene | 2.0 | 3.93 | 2.2E-02 | 2.0 | 3.96 | 8.1E-03 |
| *vap-1* | WBGene00006886 | protein_coding_gene | 2.0 | 3.93 | 2.1E-04 | 1.1 | 2.18 | 5.0E-04 |
| *K09E4.4* | WBGene00010722 | protein_coding_gene | 2.0 | 3.93 | 5.1E-05 | 1.9 | 3.69 | 8.6E-05 |
| *C47D2.1* | WBGene00016738 | protein_coding_gene | 2.0 | 3.93 | 3.7E-03 | 1.9 | 3.71 | 5.4E-04 |
| *gst-14* | WBGene00001762 | protein_coding_gene | 2.0 | 3.93 | 4.1E-02 | 2.1 | 4.15 | 9.8E-03 |
| *B0507.6* | WBGene00015223 | protein_coding_gene | 2.0 | 3.92 | 6.8E-03 | 1.4 | 2.58 | 1.9E-03 |
| *aex-4* | WBGene00006454 | protein_coding_gene | 2.0 | 3.91 | 3.0E-04 | 1.7 | 3.26 | 7.6E-04 |
| *flp-33* | WBGene00011586 | protein_coding_gene | 2.0 | 3.91 | 1.1E-03 | 1.4 | 2.59 | 4.2E-03 |
| *R07B1.8* | WBGene00001608 | protein_coding_gene | 2.0 | 3.91 | 2.9E-02 | 2.2 | 4.60 | 1.7E-03 |
| *K09E9.7* | WBGene00194664 | protein_coding_gene | 2.0 | 3.91 | 2.1E-02 | 2.2 | 4.46 | 2.4E-02 |

| | | | | | | | | |
|---|---|---|---|---|---|---|---|---|
| *T27E4.7* | WBGene00020865 | protein_coding_gene | 2.0 | 3.91 | 5.9E-04 | 1.1 | 2.08 | 1.2E-02 |
| *Y119C1B.6* | WBGene00022472 | protein_coding_gene | 2.0 | 3.90 | 2.8E-03 | 1.5 | 2.85 | 1.2E-03 |
| *flp-1* | WBGene00001444 | protein_coding_gene | 2.0 | 3.90 | 2.2E-03 | 1.4 | 2.65 | 3.9E-03 |
| *ZK1290.14* | WBGene00044396 | protein_coding_gene | 2.0 | 3.89 | 6.2E-13 | 1.4 | 2.57 | 3.0E-03 |
| *flp-12* | WBGene00001455 | protein_coding_gene | 2.0 | 3.89 | 6.1E-05 | 1.1 | 2.20 | 1.1E-03 |
| *sul-2* | WBGene00006309 | protein_coding_gene | 2.0 | 3.88 | 1.5E-03 | 1.4 | 2.65 | 1.7E-03 |
| *ztf-2* | WBGene00008762 | protein_coding_gene | 2.0 | 3.88 | 1.6E-02 | 2.1 | 4.33 | 2.5E-03 |
| *nlp-80* | WBGene00011428 | protein_coding_gene | 2.0 | 3.88 | 2.0E-03 | 1.5 | 2.82 | 5.5E-04 |
| ***mir-238*** | **WBGene00003331** | **miRNA_gene** | **2.0** | **3.87** | **3.4E-02** | **-1.8** | **0.28** | **2.1E-02** |
| *nhr-179* | WBGene00017512 | protein_coding_gene | 2.0 | 3.87 | 8.7E-04 | 2.3 | 4.96 | 2.0E-05 |
| *gpa-11* | WBGene00001673 | protein_coding_gene | 1.9 | 3.86 | 3.4E-02 | 2.6 | 6.20 | 6.2E-03 |
| *T05C1.1* | WBGene00020248 | protein_coding_gene | 1.9 | 3.85 | 4.4E-04 | 1.6 | 3.06 | 2.0E-04 |
| *C44B11.4* | WBGene00016634 | protein_coding_gene | 1.9 | 3.84 | 9.5E-04 | 1.2 | 2.31 | 1.3E-02 |
| *nlp-15* | WBGene00003753 | protein_coding_gene | 1.9 | 3.84 | 2.3E-03 | 1.3 | 2.39 | 5.2E-03 |
| *C46H11.10* | WBGene00016734 | protein_coding_gene | 1.9 | 3.84 | 8.6E-04 | 1.5 | 2.80 | 2.3E-03 |
| *C54D2.2* | WBGene00016916 | protein_coding_gene | 1.9 | 3.84 | 2.4E-02 | 1.9 | 3.66 | 2.8E-03 |
| *Y20C6A.1* | WBGene00012491 | protein_coding_gene | 1.9 | 3.84 | 1.3E-06 | 1.1 | 2.21 | 2.1E-05 |
| *kcnl-3* | WBGene00015387 | protein_coding_gene | 1.9 | 3.83 | 5.4E-04 | 1.0 | 2.05 | 4.6E-02 |
| *glb-2* | WBGene00015536 | protein_coding_gene | 1.9 | 3.83 | 9.4E-03 | 1.4 | 2.69 | 1.8E-02 |
| *C30F12.3* | WBGene00016262 | protein_coding_gene | 1.9 | 3.83 | 3.2E-03 | 1.3 | 2.54 | 1.9E-02 |
| *C30B5.7* | WBGene00016248 | protein_coding_gene | 1.9 | 3.82 | 1.3E-02 | 1.7 | 3.23 | 9.4E-04 |
| *T10B9.9* | WBGene00011678 | protein_coding_gene | 1.9 | 3.82 | 2.8E-03 | 1.2 | 2.26 | 4.6E-03 |
| *B0035.13* | WBGene00007112 | protein_coding_gene | 1.9 | 3.81 | 5.9E-06 | 1.1 | 2.12 | 3.3E-04 |
| *EGAP5.1* | WBGene00017150 | protein_coding_gene | 1.9 | 3.81 | 2.1E-02 | 1.8 | 3.58 | 2.5E-02 |
| *F14B8.5* | WBGene00017447 | protein_coding_gene | 1.9 | 3.80 | 8.0E-04 | 1.1 | 2.22 | 1.4E-02 |
| *R155.4* | WBGene00020118 | protein_coding_gene | 1.9 | 3.80 | 3.1E-02 | 1.6 | 2.94 | 9.6E-03 |
| *npr-25* | WBGene00011381 | protein_coding_gene | 1.9 | 3.80 | 1.6E-03 | 1.1 | 2.16 | 4.4E-02 |
| *lec-7* | WBGene00002270 | protein_coding_gene | 1.9 | 3.80 | 8.8E-04 | 1.4 | 2.59 | 4.8E-02 |
| *T19D12.2* | WBGene00020577 | protein_coding_gene | 1.9 | 3.79 | 2.5E-10 | 1.3 | 2.54 | 6.2E-05 |
| *lpr-2* | WBGene00011726 | protein_coding_gene | 1.9 | 3.79 | 7.6E-03 | 1.2 | 2.31 | 4.4E-02 |
| *npr-3* | WBGene00007006 | protein_coding_gene | 1.9 | 3.78 | 1.3E-03 | 1.7 | 3.18 | 8.7E-04 |
| *flr-2* | WBGene00001466 | protein_coding_gene | 1.9 | 3.78 | 1.8E-03 | 1.7 | 3.32 | 4.9E-04 |
| *C12D5.5* | WBGene00015708 | protein_coding_gene | 1.9 | 3.77 | 2.0E-02 | 1.8 | 3.47 | 3.9E-02 |
| *tat-2* | WBGene00019166 | protein_coding_gene | 1.9 | 3.77 | 2.8E-04 | 1.6 | 3.13 | 5.4E-05 |
| *clec-199* | WBGene00007820 | protein_coding_gene | 1.9 | 3.77 | 6.1E-03 | 1.3 | 2.44 | 1.7E-03 |
| *lgc-11* | WBGene00000061 | protein_coding_gene | 1.9 | 3.76 | 4.4E-03 | 1.6 | 3.03 | 7.0E-03 |
| *tos-1* | WBGene00019466 | protein_coding_gene | 1.9 | 3.76 | 4.2E-04 | 1.5 | 2.77 | 7.8E-03 |
| *K07D4.9* | WBGene00019488 | protein_coding_gene | 1.9 | 3.75 | 1.5E-04 | 1.6 | 3.13 | 1.1E-05 |
| *dop-5* | WBGene00011382 | protein_coding_gene | 1.9 | 3.75 | 6.8E-03 | 1.1 | 2.15 | 1.2E-02 |
| *ZK856.6* | WBGene00014107 | protein_coding_gene | 1.9 | 3.74 | 2.0E-02 | 1.8 | 3.42 | 1.8E-02 |
| *lips-6* | WBGene00014009 | protein_coding_gene | 1.9 | 3.74 | 4.8E-02 | 3.1 | 8.52 | 1.9E-04 |
| *deg-1* | WBGene00000950 | protein_coding_gene | 1.9 | 3.74 | 3.5E-06 | 1.2 | 2.25 | 1.5E-04 |
| *C25E10.11* | WBGene00016100 | protein_coding_gene | 1.9 | 3.73 | 4.1E-02 | 1.7 | 3.19 | 2.2E-02 |
| *D1069.1* | WBGene00017036 | protein_coding_gene | 1.9 | 3.73 | 8.4E-03 | 1.0 | 2.02 | 4.0E-02 |
| *ace-4* | WBGene00000038 | protein_coding_gene | 1.9 | 3.72 | 8.9E-03 | 1.3 | 2.50 | 3.9E-02 |
| *snet-1* | WBGene00015354 | protein_coding_gene | 1.9 | 3.71 | 3.8E-03 | 1.6 | 3.06 | 3.4E-04 |
| *C04G2.14* | WBGene00255426 | ncRNA_gene | 1.9 | 3.71 | 2.8E-02 | 3.0 | 7.76 | 5.1E-04 |
| *nlp-12* | WBGene00003750 | protein_coding_gene | 1.9 | 3.70 | 4.6E-03 | 1.5 | 2.92 | 3.9E-03 |
| *odr-3* | WBGene00003850 | protein_coding_gene | 1.9 | 3.70 | 5.3E-03 | 1.4 | 2.61 | 5.7E-03 |
| *F10A3.4* | WBGene00008634 | protein_coding_gene | 1.9 | 3.69 | 2.2E-07 | 3.1 | 8.56 | 2.7E-14 |
| *C15A7.2* | WBGene00007595 | protein_coding_gene | 1.9 | 3.68 | 2.1E-02 | 1.3 | 2.44 | 4.7E-02 |
| *ckr-1* | WBGene00020712 | protein_coding_gene | 1.9 | 3.66 | 1.6E-02 | 2.0 | 3.97 | 9.4E-04 |
| *cyp-13A6* | WBGene00011673 | protein_coding_gene | 1.9 | 3.66 | 1.2E-02 | 2.0 | 4.03 | 4.0E-07 |
| *C35B1.7* | WBGene00016436 | protein_coding_gene | 1.9 | 3.66 | 8.1E-03 | 1.7 | 3.23 | 2.8E-02 |
| *clec-198* | WBGene00008203 | protein_coding_gene | 1.9 | 3.66 | 5.3E-03 | 1.7 | 3.22 | 2.7E-04 |
| *ZC196.1* | WBGene00022545 | pseudogene | 1.9 | 3.65 | 4.7E-03 | 1.5 | 2.91 | 5.7E-04 |
| *sbt-1* | WBGene00011392 | protein_coding_gene | 1.9 | 3.65 | 1.6E-03 | 1.3 | 2.54 | 1.7E-04 |
| *C27D6.12* | WBGene00044389 | protein_coding_gene | 1.9 | 3.65 | 2.1E-03 | 1.2 | 2.36 | 3.1E-03 |

| | | | | | | | | |
|---|---|---|---|---|---|---|---|---|
| *ifp-1* | WBGene00002067 | protein_coding_gene | 1.9 | 3.64 | 3.6E-07 | 2.1 | 4.19 | 3.9E-15 |
| *srr-1* | WBGene00005652 | protein_coding_gene | 1.9 | 3.64 | 2.6E-03 | 1.3 | 2.54 | 9.4E-04 |
| *glb-4* | WBGene00007502 | protein_coding_gene | 1.9 | 3.62 | 1.5E-02 | 1.3 | 2.47 | 3.8E-02 |
| *ZC513.2* | WBGene00022627 | protein_coding_gene | 1.9 | 3.62 | 1.0E-02 | 1.4 | 2.64 | 3.6E-02 |
| *cpna-4* | WBGene00006441 | protein_coding_gene | 1.9 | 3.62 | 6.5E-03 | 1.5 | 2.73 | 1.3E-03 |
| *nlp-49* | WBGene00019160 | protein_coding_gene | 1.9 | 3.62 | 2.7E-03 | 1.6 | 3.08 | 2.4E-05 |
| *T08B6.2* | WBGene00020352 | pseudogene | 1.9 | 3.61 | 1.2E-02 | 1.9 | 3.86 | 4.8E-03 |
| *cal-8* | WBGene00017394 | protein_coding_gene | 1.8 | 3.60 | 2.5E-03 | 1.1 | 2.21 | 3.3E-02 |
| *asns-2* | WBGene00019730 | protein_coding_gene | 1.8 | 3.59 | 3.4E-02 | 1.6 | 3.01 | 1.3E-02 |
| *rbf-1* | WBGene00004316 | protein_coding_gene | 1.8 | 3.59 | 3.4E-03 | 1.3 | 2.41 | 8.4E-03 |
| *Y54G2A.10* | WBGene00021875 | protein_coding_gene | 1.8 | 3.59 | 1.1E-03 | 1.3 | 2.44 | 2.2E-02 |
| *flp-20* | WBGene00001463 | protein_coding_gene | 1.8 | 3.59 | 2.8E-03 | 1.2 | 2.25 | 3.3E-02 |
| *npr-15* | WBGene00012084 | protein_coding_gene | 1.8 | 3.59 | 1.8E-03 | 1.1 | 2.09 | 2.4E-02 |
| *M04G7.1* | WBGene00019772 | protein_coding_gene | 1.8 | 3.58 | 2.4E-03 | 1.4 | 2.67 | 2.9E-03 |
| *dop-1* | WBGene00001052 | protein_coding_gene | 1.8 | 3.57 | 4.8E-03 | 2.0 | 4.13 | 3.6E-04 |
| *wrt-8* | WBGene00006954 | protein_coding_gene | 1.8 | 3.57 | 3.5E-02 | 2.3 | 5.00 | 1.1E-02 |
| *Y47H9C.1* | WBGene00012947 | protein_coding_gene | 1.8 | 3.56 | 2.6E-03 | 2.1 | 4.20 | 1.4E-03 |
| *T05C12.8* | WBGene00011471 | protein_coding_gene | 1.8 | 3.56 | 2.8E-02 | 1.6 | 3.07 | 7.5E-03 |
| *F17C11.2* | WBGene00008914 | protein_coding_gene | 1.8 | 3.56 | 3.6E-04 | 1.2 | 2.28 | 2.6E-05 |
| *H39E23.3* | WBGene00044201 | protein_coding_gene | 1.8 | 3.56 | 3.7E-02 | 2.2 | 4.64 | 3.2E-02 |
| *ggr-1* | WBGene00001586 | protein_coding_gene | 1.8 | 3.56 | 7.6E-03 | 1.4 | 2.57 | 1.3E-02 |
| *flp-3* | WBGene00001446 | protein_coding_gene | 1.8 | 3.55 | 3.4E-04 | 1.1 | 2.15 | 4.5E-03 |
| *Y110A2AL.9* | WBGene00022445 | protein_coding_gene | 1.8 | 3.54 | 2.5E-04 | 1.6 | 3.04 | 3.9E-02 |
| *npr-26* | WBGene00011372 | protein_coding_gene | 1.8 | 3.54 | 8.0E-03 | 1.7 | 3.28 | 1.4E-04 |
| *F36G3.3* | WBGene00009484 | protein_coding_gene | 1.8 | 3.53 | 4.5E-02 | 1.3 | 2.44 | 2.1E-02 |
| *fog-3* | WBGene00001483 | protein_coding_gene | 1.8 | 3.53 | 2.7E-02 | 2.0 | 4.04 | 3.0E-04 |
| *trp-1* | WBGene00006614 | protein_coding_gene | 1.8 | 3.53 | 3.6E-03 | 1.5 | 2.76 | 7.3E-03 |
| *fbxc-6* | WBGene00016301 | pseudogene | 1.8 | 3.53 | 3.0E-02 | 1.7 | 3.18 | 3.2E-02 |
| *C39D10.2* | WBGene00016530 | protein_coding_gene | 1.8 | 3.52 | 3.0E-03 | 1.3 | 2.39 | 4.8E-03 |
| *phat-3* | WBGene00016782 | protein_coding_gene | 1.8 | 3.52 | 1.7E-03 | 1.7 | 3.24 | 1.2E-02 |
| *lgc-46* | WBGene00022106 | protein_coding_gene | 1.8 | 3.51 | 1.7E-02 | 1.5 | 2.80 | 3.7E-03 |
| *H09F14.1* | WBGene00010375 | protein_coding_gene | 1.8 | 3.51 | 1.2E-02 | 1.8 | 3.51 | 2.9E-03 |
| *C27H5.2* | WBGene00016172 | protein_coding_gene | 1.8 | 3.51 | 1.3E-04 | 1.5 | 2.90 | 2.9E-05 |
| *C29F4.3* | WBGene00007809 | protein_coding_gene | 1.8 | 3.50 | 3.3E-02 | 1.5 | 2.81 | 2.2E-02 |
| *dmd-8* | WBGene00020708 | protein_coding_gene | 1.8 | 3.50 | 2.9E-02 | 2.2 | 4.52 | 9.0E-03 |
| *nlp-18* | WBGene00003756 | protein_coding_gene | 1.8 | 3.50 | 1.4E-02 | 1.7 | 3.21 | 1.2E-03 |
| *glb-12* | WBGene00008256 | protein_coding_gene | 1.8 | 3.49 | 1.2E-02 | 1.8 | 3.59 | 6.6E-03 |
| *F59F3.6* | WBGene00010335 | protein_coding_gene | 1.8 | 3.49 | 2.6E-03 | 1.2 | 2.35 | 2.4E-03 |
| *F10C2.3* | WBGene00008644 | protein_coding_gene | 1.8 | 3.49 | 1.7E-04 | 1.5 | 2.81 | 1.4E-03 |
| *F58G11.4* | WBGene00010282 | protein_coding_gene | 1.8 | 3.49 | 1.7E-03 | 1.3 | 2.44 | 1.5E-02 |
| *nlp-61* | WBGene00017813 | protein_coding_gene | 1.8 | 3.49 | 1.4E-03 | 1.3 | 2.41 | 1.2E-03 |
| *asp-1* | WBGene00000214 | protein_coding_gene | 1.8 | 3.48 | 4.6E-04 | 1.3 | 2.49 | 3.1E-03 |
| *ttr-28* | WBGene00011285 | protein_coding_gene | 1.8 | 3.47 | 9.4E-03 | 1.4 | 2.55 | 8.6E-04 |
| *aat-5* | WBGene00000006 | protein_coding_gene | 1.8 | 3.46 | 1.1E-03 | 1.5 | 2.91 | 9.6E-05 |
| *C18F10.2* | WBGene00015977 | protein_coding_gene | 1.8 | 3.46 | 8.3E-03 | 1.3 | 2.40 | 2.9E-02 |
| *snt-6* | WBGene00004926 | protein_coding_gene | 1.8 | 3.46 | 8.1E-04 | 1.3 | 2.38 | 7.2E-03 |
| *nas-29* | WBGene00003547 | protein_coding_gene | 1.8 | 3.46 | 3.7E-03 | 1.6 | 2.98 | 1.6E-02 |
| *K09H11.6* | WBGene00019603 | protein_coding_gene | 1.8 | 3.45 | 4.6E-02 | 1.9 | 3.65 | 1.0E-02 |
| *D1007.18* | WBGene00044421 | protein_coding_gene | 1.8 | 3.45 | 5.0E-03 | 1.4 | 2.64 | 7.6E-03 |
| ***tmem-231*** | **WBGene00020825** | **protein_coding_gene** | **1.8** | **3.45** | **1.6E-02** | **-1.1** | **0.48** | **4.6E-02** |
| *T21B4.15* | WBGene00043054 | protein_coding_gene | 1.8 | 3.45 | 1.8E-02 | 1.4 | 2.63 | 4.9E-02 |
| *ZK1055.4* | WBGene00022845 | protein_coding_gene | 1.8 | 3.44 | 3.7E-02 | 1.7 | 3.29 | 3.5E-02 |
| *cah-2* | WBGene00000280 | protein_coding_gene | 1.8 | 3.43 | 2.7E-04 | 1.2 | 2.33 | 1.7E-03 |
| *sox-3* | WBGene00004950 | protein_coding_gene | 1.8 | 3.43 | 1.3E-03 | 1.2 | 2.28 | 7.5E-03 |
| *cal-4* | WBGene00000288 | protein_coding_gene | 1.8 | 3.43 | 3.4E-03 | 1.2 | 2.23 | 9.8E-04 |
| *nlp-3* | WBGene00003741 | protein_coding_gene | 1.8 | 3.42 | 5.6E-04 | 1.1 | 2.22 | 1.1E-04 |
| *gpa-17* | WBGene00022196 | protein_coding_gene | 1.8 | 3.42 | 5.5E-04 | 1.4 | 2.64 | 1.9E-02 |
| *dct-15* | WBGene00012340 | protein_coding_gene | 1.8 | 3.42 | 1.0E-02 | 1.5 | 2.82 | 1.7E-02 |

| | | | | | | | | |
|---|---|---|---|---|---|---|---|---|
| *E02A10.4* | WBGene00008454 | protein_coding_gene | 1.8 | 3.41 | 5.5E-03 | 1.6 | 3.06 | 1.0E-03 |
| *ptl-1* | WBGene00004212 | protein_coding_gene | 1.8 | 3.40 | 4.8E-04 | 1.5 | 2.77 | 1.2E-05 |
| *R74.10* | WBGene00206525 | protein_coding_gene | 1.8 | 3.40 | 3.0E-04 | 1.2 | 2.35 | 4.4E-03 |
| *zig-5* | WBGene00006982 | protein_coding_gene | 1.8 | 3.40 | 1.0E-04 | 1.3 | 2.48 | 1.6E-03 |
| *scd-2* | WBGene00004740 | protein_coding_gene | 1.8 | 3.40 | 5.7E-03 | 1.7 | 3.28 | 1.3E-04 |
| *glb-32* | WBGene00021980 | protein_coding_gene | 1.8 | 3.40 | 1.0E-02 | 1.1 | 2.18 | 4.4E-02 |
| *cal-3* | WBGene00000287 | protein_coding_gene | 1.8 | 3.40 | 1.4E-03 | 1.2 | 2.31 | 3.0E-03 |
| *C25G6.1* | WBGene00016107 | protein_coding_gene | 1.8 | 3.39 | 1.1E-02 | 1.9 | 3.61 | 9.8E-03 |
| *C33D9.3* | WBGene00007898 | protein_coding_gene | 1.8 | 3.39 | 1.3E-02 | 1.9 | 3.62 | 2.3E-03 |
| *snt-4* | WBGene00004924 | protein_coding_gene | 1.8 | 3.39 | 4.7E-03 | 1.4 | 2.68 | 3.6E-04 |
| *marg-1* | WBGene00018538 | protein_coding_gene | 1.8 | 3.39 | 8.9E-03 | 1.8 | 3.51 | 5.6E-07 |
| *ZC116.1* | WBGene00013853 | protein_coding_gene | 1.8 | 3.39 | 6.5E-03 | 1.1 | 2.19 | 5.9E-03 |
| *nlp-58* | WBGene00077775 | protein_coding_gene | 1.8 | 3.38 | 1.4E-02 | 1.2 | 2.37 | 6.8E-03 |
| *cab-1* | WBGene00000277 | protein_coding_gene | 1.8 | 3.37 | 3.9E-03 | 1.6 | 3.02 | 2.0E-04 |
| *asp-8* | WBGene00019105 | protein_coding_gene | 1.8 | 3.37 | 7.9E-04 | 2.8 | 6.87 | 1.7E-10 |
| *shn-1* | WBGene00006444 | protein_coding_gene | 1.8 | 3.37 | 2.0E-03 | 1.3 | 2.45 | 1.4E-02 |
| *DH11.5* | WBGene00008438 | protein_coding_gene | 1.8 | 3.37 | 2.9E-03 | 1.3 | 2.49 | 2.0E-04 |
| *spp-23* | WBGene00021335 | protein_coding_gene | 1.8 | 3.37 | 2.3E-02 | 3.6 | 12.23 | 2.6E-07 |
| *T07F12.4* | WBGene00020324 | protein_coding_gene | 1.8 | 3.36 | 3.3E-04 | 1.4 | 2.57 | 6.8E-04 |
| *des-2* | WBGene00000955 | protein_coding_gene | 1.7 | 3.36 | 2.2E-02 | 1.4 | 2.59 | 6.2E-03 |
| *C24B5.1* | WBGene00016044 | protein_coding_gene | 1.7 | 3.36 | 6.7E-03 | 1.5 | 2.78 | 2.6E-03 |
| *T27A10.2* | WBGene00020844 | protein_coding_gene | 1.7 | 3.35 | 4.4E-03 | 1.6 | 3.08 | 8.1E-03 |
| *gpa-2* | WBGene00001664 | protein_coding_gene | 1.7 | 3.35 | 1.6E-02 | 1.4 | 2.65 | 1.9E-03 |
| *M28.8* | WBGene00010899 | protein_coding_gene | 1.7 | 3.35 | 5.5E-04 | 1.5 | 2.74 | 4.1E-06 |
| *fbxa-59* | WBGene00020455 | protein_coding_gene | 1.7 | 3.34 | 2.0E-03 | 1.6 | 3.00 | 3.7E-05 |
| ***T02E9.5*** | **WBGene00011383** | **protein_coding_gene** | **1.7** | **3.34** | **2.8E-04** | **-1.2** | **0.44** | **2.7E-02** |
| *ggr-2* | WBGene00001587 | protein_coding_gene | 1.7 | 3.34 | 1.2E-02 | 1.5 | 2.93 | 2.0E-03 |
| *gcy-11* | WBGene00001537 | protein_coding_gene | 1.7 | 3.33 | 4.4E-04 | 1.4 | 2.68 | 5.3E-06 |
| *mth-2* | WBGene00017947 | protein_coding_gene | 1.7 | 3.33 | 8.1E-04 | 1.4 | 2.61 | 4.3E-04 |
| *K04D7.6* | WBGene00010559 | protein_coding_gene | 1.7 | 3.33 | 4.6E-03 | 1.4 | 2.73 | 1.1E-02 |
| *hpo-28* | WBGene00015148 | protein_coding_gene | 1.7 | 3.33 | 3.5E-04 | 1.7 | 3.21 | 8.8E-06 |
| *kin-9* | WBGene00002195 | protein_coding_gene | 1.7 | 3.33 | 5.3E-03 | 1.5 | 2.74 | 9.6E-03 |
| *mgl-3* | WBGene00021152 | protein_coding_gene | 1.7 | 3.33 | 4.6E-03 | 1.1 | 2.09 | 9.0E-03 |
| *nlp-11* | WBGene00003749 | protein_coding_gene | 1.7 | 3.33 | 1.0E-03 | 1.6 | 2.97 | 1.4E-04 |
| *C49G7.3* | WBGene00016781 | protein_coding_gene | 1.7 | 3.33 | 5.9E-05 | 1.6 | 2.94 | 2.2E-12 |
| *C54D10.3* | WBGene00008298 | protein_coding_gene | 1.7 | 3.33 | 4.0E-08 | 1.1 | 2.17 | 9.2E-05 |
| *Y75B7AR.1* | WBGene00022287 | protein_coding_gene | 1.7 | 3.33 | 1.1E-02 | 2.0 | 3.94 | 1.0E-02 |
| *C07A12.2* | WBGene00015560 | protein_coding_gene | 1.7 | 3.32 | 2.6E-03 | 1.4 | 2.59 | 2.7E-03 |
| *arrd-6* | WBGene00009579 | protein_coding_gene | 1.7 | 3.32 | 2.0E-02 | 1.3 | 2.45 | 4.8E-03 |
| *R08C7.12* | WBGene00019955 | protein_coding_gene | 1.7 | 3.31 | 2.7E-03 | 1.7 | 3.20 | 1.8E-05 |
| *rege-1* | WBGene00016260 | protein_coding_gene | 1.7 | 3.31 | 1.7E-02 | 1.4 | 2.56 | 1.9E-02 |
| *glb-24* | WBGene00011287 | protein_coding_gene | 1.7 | 3.31 | 5.6E-03 | 1.3 | 2.50 | 2.2E-02 |
| *dhs-4* | WBGene00000968 | protein_coding_gene | 1.7 | 3.30 | 4.5E-06 | 1.3 | 2.45 | 1.5E-05 |
| *C25G6.3* | WBGene00016108 | protein_coding_gene | 1.7 | 3.30 | 2.5E-02 | 1.6 | 2.94 | 1.1E-02 |
| *nlp-43* | WBGene00016684 | protein_coding_gene | 1.7 | 3.30 | 3.5E-04 | 1.4 | 2.66 | 2.2E-08 |
| *gpa-3* | WBGene00001665 | protein_coding_gene | 1.7 | 3.30 | 7.8E-03 | 1.3 | 2.41 | 4.3E-03 |
| *F54C9.7* | WBGene00010043 | protein_coding_gene | 1.7 | 3.30 | 4.4E-03 | 1.5 | 2.84 | 6.1E-03 |
| *T06A4.1* | WBGene00020281 | protein_coding_gene | 1.7 | 3.30 | 1.8E-02 | 1.7 | 3.27 | 8.9E-03 |
| *dbl-1* | WBGene00000936 | protein_coding_gene | 1.7 | 3.29 | 2.2E-02 | 1.7 | 3.18 | 2.7E-03 |
| *C05E7.3* | WBGene00007345 | protein_coding_gene | 1.7 | 3.29 | 7.9E-04 | 1.3 | 2.54 | 3.4E-02 |
| *cpr-3* | WBGene00000783 | protein_coding_gene | 1.7 | 3.29 | 2.5E-02 | 1.8 | 3.46 | 3.4E-03 |
| *cdka-1* | WBGene00011955 | protein_coding_gene | 1.7 | 3.28 | 5.6E-04 | 1.4 | 2.64 | 3.4E-05 |
| *C18A11.2* | WBGene00015948 | protein_coding_gene | 1.7 | 3.28 | 1.2E-02 | 1.0 | 2.03 | 3.5E-02 |
| *ipla-2* | WBGene00009801 | protein_coding_gene | 1.7 | 3.28 | 9.2E-03 | 1.4 | 2.63 | 3.5E-02 |
| *F36H2.3* | WBGene00009500 | protein_coding_gene | 1.7 | 3.28 | 1.1E-02 | 1.6 | 2.94 | 1.8E-03 |
| *C45E1.4* | WBGene00016663 | protein_coding_gene | 1.7 | 3.28 | 7.9E-06 | 2.0 | 4.10 | 2.8E-07 |
| *deg-3* | WBGene00000951 | protein_coding_gene | 1.7 | 3.28 | 2.5E-02 | 1.1 | 2.17 | 2.7E-02 |
| *nlp-53* | WBGene00012440 | protein_coding_gene | 1.7 | 3.27 | 4.1E-03 | 1.8 | 3.48 | 1.0E-04 |

| | | | | | | | | |
|---|---|---|---|---|---|---|---|---|
| *flp-4* | WBGene00001447 | protein_coding_gene | 1.7 | 3.27 | 9.6E-04 | 1.2 | 2.29 | 4.4E-04 |
| *egl-21* | WBGene00001189 | protein_coding_gene | 1.7 | 3.27 | 4.2E-03 | 1.4 | 2.71 | 2.1E-03 |
| *K11H3.5* | WBGene00010782 | protein_coding_gene | 1.7 | 3.26 | 2.3E-03 | 1.8 | 3.46 | 4.0E-03 |
| *npr-29* | WBGene00013848 | protein_coding_gene | 1.7 | 3.26 | 1.0E-02 | 1.1 | 2.20 | 2.3E-02 |
| *ztf-14* | WBGene00010936 | protein_coding_gene | 1.7 | 3.25 | 1.6E-02 | 1.4 | 2.68 | 3.4E-02 |
| *sms-5* | WBGene00021075 | protein_coding_gene | 1.7 | 3.25 | 1.5E-03 | 1.4 | 2.69 | 5.6E-04 |
| *C06C3.12* | WBGene00175031 | protein_coding_gene | 1.7 | 3.25 | 1.3E-02 | 1.5 | 2.87 | 2.3E-03 |
| *myo-2* | WBGene00003514 | protein_coding_gene | 1.7 | 3.25 | 8.0E-03 | 1.8 | 3.47 | 1.1E-04 |
| *Y105C5A.24* | WBGene00013643 | protein_coding_gene | 1.7 | 3.24 | 5.2E-05 | 1.2 | 2.25 | 2.3E-07 |
| *R11D1.3* | WBGene00011244 | protein_coding_gene | 1.7 | 3.24 | 3.0E-02 | 1.9 | 3.78 | 3.2E-03 |
| *rgef-1* | WBGene00009100 | protein_coding_gene | 1.7 | 3.24 | 1.2E-02 | 1.2 | 2.24 | 3.5E-02 |
| *F15A4.6* | WBGene00008841 | protein_coding_gene | 1.7 | 3.23 | 3.0E-04 | 1.4 | 2.58 | 3.4E-08 |
| *flp-17* | WBGene00001460 | protein_coding_gene | 1.7 | 3.23 | 2.3E-02 | 1.1 | 2.18 | 3.2E-02 |
| *H10D18.5* | WBGene00019180 | protein_coding_gene | 1.7 | 3.22 | 9.0E-03 | 1.3 | 2.40 | 6.3E-03 |
| *sulp-1* | WBGene00016945 | protein_coding_gene | 1.7 | 3.22 | 2.3E-02 | 1.8 | 3.37 | 1.6E-03 |
| *fipr-10* | WBGene00008245 | protein_coding_gene | 1.7 | 3.22 | 4.3E-02 | 1.8 | 3.46 | 1.6E-03 |
| *linc-61* | WBGene00195170 | lincRNA_gene | 1.7 | 3.22 | 4.4E-02 | 1.6 | 2.93 | 2.0E-02 |
| *T26G10.5* | WBGene00012062 | protein_coding_gene | 1.7 | 3.22 | 3.6E-03 | 1.4 | 2.67 | 6.3E-03 |
| *npr-34* | WBGene00013187 | protein_coding_gene | 1.7 | 3.22 | 6.1E-03 | 1.6 | 3.06 | 1.2E-02 |
| *T09F5.12* | WBGene00014836 | lincRNA_gene | 1.7 | 3.21 | 2.5E-02 | 1.6 | 2.93 | 8.1E-03 |
| *T10G3.8* | WBGene00045340 | protein_coding_gene | 1.7 | 3.21 | 1.8E-02 | 1.5 | 2.79 | 1.4E-03 |
| *flp-13* | WBGene00001456 | protein_coding_gene | 1.7 | 3.20 | 3.0E-03 | 1.6 | 2.97 | 2.4E-04 |
| *F35F10.6* | WBGene00018056 | protein_coding_gene | 1.7 | 3.20 | 4.9E-02 | 2.6 | 6.05 | 2.5E-03 |
| *flp-7* | WBGene00001450 | protein_coding_gene | 1.7 | 3.20 | 7.2E-03 | 1.1 | 2.16 | 3.2E-03 |
| *Y105C5A.25* | WBGene00013644 | pseudogene | 1.7 | 3.19 | 2.2E-03 | 1.5 | 2.74 | 1.9E-03 |
| *lin-14* | WBGene00003003 | protein_coding_gene | 1.7 | 3.19 | 5.0E-03 | 1.3 | 2.54 | 8.7E-05 |
| *T08B1.4* | WBGene00020341 | protein_coding_gene | 1.7 | 3.19 | 2.4E-03 | 1.1 | 2.22 | 8.9E-03 |
| *ptr-6* | WBGene00004221 | protein_coding_gene | 1.7 | 3.19 | 2.8E-02 | 1.1 | 2.11 | 4.4E-02 |
| *F57C7.4* | WBGene00010200 | protein_coding_gene | 1.7 | 3.19 | 1.5E-02 | 1.6 | 3.01 | 1.9E-03 |
| *T03G11.3* | WBGene00020196 | protein_coding_gene | 1.7 | 3.18 | 1.3E-02 | 1.2 | 2.32 | 2.2E-02 |
| *C18H7.6* | WBGene00015996 | protein_coding_gene | 1.7 | 3.18 | 2.9E-02 | 2.0 | 4.05 | 2.4E-03 |
| *asp-3* | WBGene00000216 | protein_coding_gene | 1.7 | 3.18 | 1.9E-02 | 1.2 | 2.36 | 3.3E-02 |
| *F22F7.7* | WBGene00017724 | protein_coding_gene | 1.7 | 3.17 | 1.5E-02 | 1.5 | 2.85 | 1.6E-04 |
| *sptf-2* | WBGene00011926 | protein_coding_gene | 1.7 | 3.17 | 1.9E-02 | 1.5 | 2.80 | 1.6E-02 |
| *sol-1* | WBGene00004944 | protein_coding_gene | 1.7 | 3.16 | 5.8E-03 | 1.2 | 2.27 | 1.0E-02 |
| *nlp-70* | WBGene00044513 | protein_coding_gene | 1.7 | 3.16 | 1.8E-02 | 1.3 | 2.50 | 1.0E-02 |
| *glb-3* | WBGene00007389 | protein_coding_gene | 1.7 | 3.16 | 1.0E-02 | 1.4 | 2.69 | 9.7E-03 |
| *C01C4.3* | WBGene00015293 | protein_coding_gene | 1.7 | 3.16 | 2.7E-02 | 1.3 | 2.47 | 4.7E-03 |
| *C54D10.5* | WBGene00008300 | protein_coding_gene | 1.7 | 3.16 | 2.4E-02 | 2.1 | 4.15 | 2.3E-03 |
| *H10E21.5* | WBGene00019185 | protein_coding_gene | 1.7 | 3.15 | 6.0E-03 | 1.3 | 2.42 | 8.9E-03 |
| *xbx-5* | WBGene00011974 | protein_coding_gene | 1.7 | 3.15 | 1.4E-02 | 1.5 | 2.74 | 6.5E-03 |
| *cho-1* | WBGene00000501 | protein_coding_gene | 1.7 | 3.15 | 2.7E-03 | 1.6 | 2.93 | 8.1E-04 |
| *C03H5.6* | WBGene00015408 | protein_coding_gene | 1.7 | 3.15 | 2.8E-03 | 1.7 | 3.21 | 5.8E-03 |
| *clik-2* | WBGene00016898 | protein_coding_gene | 1.7 | 3.14 | 1.0E-02 | 1.4 | 2.60 | 4.0E-04 |
| *T14G8.6* | WBGene00304822 | protein_coding_gene | 1.7 | 3.14 | 3.9E-02 | 1.8 | 3.57 | 3.6E-02 |
| *twk-35* | WBGene00006687 | protein_coding_gene | 1.6 | 3.14 | 8.6E-03 | 1.6 | 2.93 | 1.3E-02 |
| *W05F2.7* | WBGene00021039 | protein_coding_gene | 1.6 | 3.14 | 4.0E-03 | 1.2 | 2.32 | 6.4E-04 |
| *pqn-35* | WBGene00004122 | protein_coding_gene | 1.6 | 3.13 | 4.1E-03 | 1.5 | 2.92 | 9.1E-04 |
| *xbx-9* | WBGene00007604 | protein_coding_gene | 1.6 | 3.13 | 2.7E-02 | 1.6 | 3.12 | 2.7E-04 |
| *C31C9.6* | WBGene00007840 | protein_coding_gene | 1.6 | 3.13 | 3.4E-03 | 1.1 | 2.16 | 9.2E-03 |
| *F14H12.8* | WBGene00017475 | protein_coding_gene | 1.6 | 3.12 | 1.4E-02 | 1.5 | 2.75 | 1.7E-02 |
| *nlp-6* | WBGene00003744 | protein_coding_gene | 1.6 | 3.12 | 1.4E-03 | 1.1 | 2.17 | 4.1E-05 |
| *T28B8.3* | WBGene00012114 | protein_coding_gene | 1.6 | 3.11 | 1.9E-02 | 2.0 | 4.10 | 2.1E-03 |
| *cmk-1* | WBGene00000553 | protein_coding_gene | 1.6 | 3.11 | 2.2E-03 | 1.2 | 2.36 | 2.6E-03 |
| *exp-2* | WBGene00001374 | protein_coding_gene | 1.6 | 3.11 | 6.8E-03 | 1.2 | 2.31 | 2.7E-03 |
| *dop-3* | WBGene00020506 | protein_coding_gene | 1.6 | 3.11 | 1.7E-02 | 1.7 | 3.27 | 8.6E-04 |
| *Y61A9LA.12* | WBGene00044586 | protein_coding_gene | 1.6 | 3.09 | 2.3E-02 | 1.5 | 2.80 | 7.9E-03 |
| *ZK822.6* | WBGene00044009 | protein_coding_gene | 1.6 | 3.09 | 2.0E-02 | 1.2 | 2.22 | 2.5E-02 |

| | | | | | | | | |
|---|---|---|---|---|---|---|---|---|
| *C16D9.1* | WBGene00015857 | protein_coding_gene | 1.6 | 3.09 | 2.1E-02 | 1.7 | 3.21 | 3.7E-02 |
| *cest-13* | WBGene00010116 | protein_coding_gene | 1.6 | 3.08 | 1.5E-02 | 2.4 | 5.40 | 6.7E-05 |
| ***F31E3.12*** | **WBGene00219317** | **protein_coding_gene** | **1.6** | **3.08** | **2.6E-02** | **-1.4** | **0.39** | **4.3E-02** |
| *C05D2.11* | WBGene00022910 | ncRNA_gene | 1.6 | 3.08 | 8.8E-04 | 1.2 | 2.23 | 5.9E-03 |
| *F13B6.3* | WBGene00017418 | protein_coding_gene | 1.6 | 3.07 | 5.9E-04 | 1.7 | 3.28 | 6.3E-04 |
| *F13B12.3* | WBGene00008731 | protein_coding_gene | 1.6 | 3.06 | 2.6E-04 | 1.0 | 2.04 | 1.2E-02 |
| *Y62H9A.15* | WBGene00045252 | protein_coding_gene | 1.6 | 3.06 | 1.1E-02 | 1.1 | 2.13 | 3.2E-02 |
| *C49A9.3* | WBGene00016757 | protein_coding_gene | 1.6 | 3.05 | 5.3E-03 | 1.8 | 3.44 | 3.8E-05 |
| *nhr-216* | WBGene00020385 | protein_coding_gene | 1.6 | 3.05 | 3.6E-02 | 2.0 | 3.93 | 1.2E-02 |
| *del-4* | WBGene00012116 | protein_coding_gene | 1.6 | 3.05 | 3.5E-02 | 1.1 | 2.11 | 3.7E-02 |
| *F32A5.4* | WBGene00017970 | protein_coding_gene | 1.6 | 3.05 | 6.7E-04 | 1.2 | 2.27 | 9.0E-04 |
| *flp-11* | WBGene00001454 | protein_coding_gene | 1.6 | 3.05 | 1.7E-03 | 1.6 | 3.00 | 1.4E-04 |
| *acs-21* | WBGene00016849 | protein_coding_gene | 1.6 | 3.04 | 7.9E-03 | 1.5 | 2.76 | 2.8E-02 |
| *C45E5.4* | WBGene00016667 | protein_coding_gene | 1.6 | 3.04 | 4.3E-03 | 2.1 | 4.29 | 5.8E-05 |
| *W06H8.6* | WBGene00021068 | protein_coding_gene | 1.6 | 3.03 | 2.8E-02 | 1.0 | 2.00 | 2.7E-02 |
| *flp-24* | WBGene00016028 | protein_coding_gene | 1.6 | 3.03 | 5.1E-04 | 1.3 | 2.50 | 7.0E-05 |
| *pgal-1* | WBGene00017671 | protein_coding_gene | 1.6 | 3.03 | 4.0E-03 | 1.2 | 2.26 | 1.9E-05 |
| *drn-1* | WBGene00016911 | protein_coding_gene | 1.6 | 3.03 | 1.9E-02 | 1.1 | 2.08 | 5.0E-04 |
| *cal-1* | WBGene00000285 | protein_coding_gene | 1.6 | 3.02 | 5.4E-03 | 1.1 | 2.11 | 3.3E-03 |
| *F35F10.4* | WBGene00018054 | pseudogene | 1.6 | 3.02 | 4.7E-02 | 2.7 | 6.57 | 2.4E-04 |
| *R05D7.1* | WBGene00011026 | protein_coding_gene | 1.6 | 3.02 | 6.1E-03 | 1.2 | 2.26 | 1.8E-02 |
| *F53F4.17* | WBGene00045195 | protein_coding_gene | 1.6 | 3.02 | 2.2E-02 | 1.5 | 2.76 | 1.0E-02 |
| *Y48C3A.5* | WBGene00012989 | protein_coding_gene | 1.6 | 2.99 | 1.0E-04 | 1.1 | 2.11 | 4.5E-03 |
| *sul-3* | WBGene00006310 | protein_coding_gene | 1.6 | 2.99 | 4.0E-02 | 1.6 | 2.95 | 2.3E-02 |
| *Y55B1BR.6* | WBGene00044401 | protein_coding_gene | 1.6 | 2.99 | 4.7E-03 | 1.1 | 2.12 | 2.9E-02 |
| *nep-11* | WBGene00017557 | protein_coding_gene | 1.6 | 2.98 | 2.6E-02 | 1.4 | 2.60 | 3.0E-03 |
| *gpa-14* | WBGene00001676 | protein_coding_gene | 1.6 | 2.98 | 4.0E-03 | 1.4 | 2.70 | 4.2E-03 |
| *Y9C2UA.1* | WBGene00012421 | protein_coding_gene | 1.6 | 2.97 | 2.8E-03 | 1.1 | 2.10 | 2.7E-03 |
| *cyp-33C3* | WBGene00018262 | protein_coding_gene | 1.6 | 2.97 | 3.2E-03 | 1.6 | 3.02 | 3.6E-04 |
| *osm-10* | WBGene00003890 | protein_coding_gene | 1.6 | 2.96 | 1.1E-02 | 1.3 | 2.51 | 3.4E-03 |
| *hacd-1* | WBGene00019978 | protein_coding_gene | 1.6 | 2.96 | 4.4E-02 | 2.2 | 4.61 | 7.9E-03 |
| *C18G1.1* | WBGene00015980 | protein_coding_gene | 1.6 | 2.96 | 2.1E-02 | 1.1 | 2.17 | 4.4E-02 |
| *ckk-1* | WBGene00000518 | protein_coding_gene | 1.6 | 2.96 | 2.0E-03 | 1.0 | 2.02 | 8.8E-03 |
| *dop-2* | WBGene00001053 | protein_coding_gene | 1.6 | 2.95 | 1.8E-03 | 1.1 | 2.14 | 1.6E-03 |
| *clh-2* | WBGene00000529 | protein_coding_gene | 1.6 | 2.95 | 1.9E-02 | 1.1 | 2.09 | 1.2E-02 |
| *flp-26* | WBGene00020126 | protein_coding_gene | 1.6 | 2.95 | 9.0E-04 | 1.2 | 2.28 | 1.8E-03 |
| *dyf-6* | WBGene00001122 | protein_coding_gene | 1.6 | 2.95 | 2.3E-03 | 1.1 | 2.10 | 9.4E-03 |
| *lgc-41* | WBGene00008022 | protein_coding_gene | 1.6 | 2.95 | 4.9E-02 | 1.5 | 2.83 | 3.8E-02 |
| *nspe-4* | WBGene00012589 | protein_coding_gene | 1.6 | 2.95 | 3.0E-02 | 2.8 | 7.14 | 9.8E-04 |
| *T03F6.4* | WBGene00011400 | protein_coding_gene | 1.6 | 2.94 | 3.3E-02 | 1.1 | 2.19 | 8.3E-03 |
| *cgt-1* | WBGene00011517 | protein_coding_gene | 1.6 | 2.94 | 1.5E-05 | 1.1 | 2.22 | 8.4E-03 |
| *pqn-18* | WBGene00004108 | protein_coding_gene | 1.6 | 2.93 | 9.0E-03 | 1.0 | 2.03 | 3.0E-03 |
| *T02E1.8* | WBGene00011380 | protein_coding_gene | 1.5 | 2.92 | 2.1E-02 | 1.1 | 2.11 | 1.5E-02 |
| *sro-1* | WBGene00005641 | protein_coding_gene | 1.5 | 2.91 | 1.2E-02 | 1.6 | 3.00 | 1.3E-02 |
| *phat-6* | WBGene00011683 | protein_coding_gene | 1.5 | 2.91 | 1.2E-02 | 1.1 | 2.07 | 4.0E-02 |
| *old-1* | WBGene00003862 | protein_coding_gene | 1.5 | 2.90 | 2.4E-02 | 1.5 | 2.78 | 4.1E-02 |
| *hpo-26* | WBGene00018492 | protein_coding_gene | 1.5 | 2.90 | 3.8E-04 | 1.6 | 3.12 | 1.4E-03 |
| *T14G8.4* | WBGene00011772 | protein_coding_gene | 1.5 | 2.90 | 4.0E-02 | 1.4 | 2.60 | 2.1E-02 |
| *ets-4* | WBGene00017687 | protein_coding_gene | 1.5 | 2.89 | 4.9E-03 | 1.8 | 3.49 | 5.7E-08 |
| *igeg-2* | WBGene00009842 | protein_coding_gene | 1.5 | 2.89 | 7.3E-03 | 1.2 | 2.37 | 2.3E-02 |
| ***F59C6.16*** | **WBGene00077764** | **protein_coding_gene** | **1.5** | **2.89** | **4.5E-02** | **-1.1** | **0.48** | **2.5E-03** |
| *fbxa-61* | WBGene00022486 | protein_coding_gene | 1.5 | 2.88 | 2.5E-03 | 1.3 | 2.44 | 2.8E-03 |
| *C25H3.1* | WBGene00016111 | protein_coding_gene | 1.5 | 2.88 | 2.8E-02 | 1.2 | 2.26 | 3.8E-02 |
| *npr-18* | WBGene00008065 | protein_coding_gene | 1.5 | 2.88 | 2.3E-02 | 1.3 | 2.44 | 4.8E-02 |
| *acr-20* | WBGene00000059 | protein_coding_gene | 1.5 | 2.87 | 9.4E-03 | 1.6 | 2.93 | 1.6E-03 |
| *F34D10.6* | WBGene00009375 | protein_coding_gene | 1.5 | 2.87 | 2.8E-03 | 1.4 | 2.67 | 2.5E-05 |
| *maf-1* | WBGene00077521 | protein_coding_gene | 1.5 | 2.87 | 6.5E-05 | 1.7 | 3.17 | 3.3E-05 |
| *rpl-25.1* | WBGene00004438 | protein_coding_gene | 1.5 | 2.87 | 3.8E-03 | 1.4 | 2.70 | 1.3E-02 |

| | | | | | | | | |
|---|---|---|---|---|---|---|---|---|
| *sup-1* | WBGene00012759 | protein_coding_gene | 1.5 | 2.86 | 2.3E-02 | 1.3 | 2.41 | 3.9E-02 |
| *F29G6.1* | WBGene00009257 | protein_coding_gene | 1.5 | 2.85 | 4.3E-02 | 2.7 | 6.51 | 6.9E-05 |
| *fipr-3* | WBGene00009090 | protein_coding_gene | 1.5 | 2.85 | 1.1E-02 | 1.5 | 2.85 | 8.0E-05 |
| *nhr-129* | WBGene00003719 | protein_coding_gene | 1.5 | 2.84 | 3.9E-03 | 1.4 | 2.60 | 2.6E-03 |
| *daf-7* | WBGene00000903 | protein_coding_gene | 1.5 | 2.84 | 5.5E-03 | 1.5 | 2.87 | 4.8E-05 |
| *sue-1* | WBGene00006306 | protein_coding_gene | 1.5 | 2.83 | 9.4E-03 | 1.5 | 2.83 | 5.3E-04 |
| *bcl-11* | WBGene00017430 | protein_coding_gene | 1.5 | 2.82 | 1.0E-02 | 1.4 | 2.69 | 8.9E-06 |
| *nrx-1* | WBGene00007799 | protein_coding_gene | 1.5 | 2.82 | 1.9E-03 | 1.3 | 2.50 | 2.0E-04 |
| *frpr-4* | WBGene00016909 | protein_coding_gene | 1.5 | 2.82 | 1.0E-02 | 1.3 | 2.40 | 6.5E-03 |
| *Y43F8B.3* | WBGene00012814 | protein_coding_gene | 1.5 | 2.82 | 2.2E-02 | 1.2 | 2.38 | 2.8E-02 |
| *pepm-1* | WBGene00009807 | protein_coding_gene | 1.5 | 2.82 | 9.2E-03 | 1.8 | 3.58 | 2.6E-02 |
| *C05E7.1* | WBGene00007343 | protein_coding_gene | 1.5 | 2.82 | 6.8E-03 | 1.1 | 2.12 | 1.7E-02 |
| *Y42G9A.3* | WBGene00021533 | protein_coding_gene | 1.5 | 2.82 | 1.6E-02 | 1.0 | 2.06 | 3.8E-02 |
| *lea-1* | WBGene00002263 | protein_coding_gene | 1.5 | 2.81 | 5.6E-03 | 1.3 | 2.41 | 3.2E-02 |
| *K02F2.5* | WBGene00019324 | protein_coding_gene | 1.5 | 2.81 | 2.8E-02 | 1.3 | 2.54 | 1.0E-03 |
| *F15D4.4* | WBGene00008861 | protein_coding_gene | 1.5 | 2.81 | 1.8E-02 | 1.6 | 2.93 | 3.9E-03 |
| *mec-2* | WBGene00003166 | protein_coding_gene | 1.5 | 2.80 | 9.4E-03 | 1.2 | 2.31 | 4.1E-03 |
| *hpo-30* | WBGene00011014 | protein_coding_gene | 1.5 | 2.79 | 2.0E-02 | 1.3 | 2.50 | 9.1E-03 |
| *T20B6.3* | WBGene00020599 | protein_coding_gene | 1.5 | 2.79 | 3.3E-02 | 1.3 | 2.45 | 4.2E-02 |
| *nep-26* | WBGene00014171 | protein_coding_gene | 1.5 | 2.78 | 8.0E-03 | 1.0 | 2.05 | 4.4E-04 |
| *tag-209* | WBGene00011072 | protein_coding_gene | 1.5 | 2.78 | 1.3E-02 | 1.0 | 2.05 | 1.6E-02 |
| *fbxa-39* | WBGene00021859 | protein_coding_gene | 1.5 | 2.78 | 3.3E-02 | 1.2 | 2.24 | 2.8E-02 |
| *fipr-9* | WBGene00044175 | protein_coding_gene | 1.5 | 2.78 | 3.8E-03 | 1.7 | 3.22 | 3.0E-05 |
| *myo-1* | WBGene00002348 | protein_coding_gene | 1.5 | 2.78 | 8.3E-03 | 1.7 | 3.15 | 4.5E-06 |
| *C30G7.2* | WBGene00044176 | protein_coding_gene | 1.5 | 2.78 | 2.0E-02 | 1.1 | 2.08 | 1.9E-03 |
| *nlp-5* | WBGene00003743 | protein_coding_gene | 1.5 | 2.77 | 3.0E-03 | 1.1 | 2.15 | 4.7E-03 |
| *C05D12.7* | WBGene00044029 | protein_coding_gene | 1.5 | 2.77 | 1.7E-02 | 1.2 | 2.24 | 1.6E-02 |
| *pqn-36* | WBGene00004123 | protein_coding_gene | 1.5 | 2.77 | 5.0E-02 | 2.0 | 3.96 | 1.8E-05 |
| *klf-1* | WBGene00018990 | protein_coding_gene | 1.5 | 2.77 | 2.0E-03 | 1.2 | 2.24 | 2.1E-02 |
| *CD4.1* | WBGene00016988 | protein_coding_gene | 1.5 | 2.76 | 2.0E-03 | 1.0 | 2.05 | 6.5E-03 |
| *K09E9.4* | WBGene00044143 | protein_coding_gene | 1.5 | 2.76 | 1.3E-02 | 1.2 | 2.32 | 7.5E-03 |
| *madf-9* | WBGene00022608 | protein_coding_gene | 1.5 | 2.76 | 1.7E-02 | 1.7 | 3.22 | 1.1E-03 |
| *Y39B6A.1* | WBGene00012664 | protein_coding_gene | 1.5 | 2.76 | 3.1E-04 | 1.2 | 2.31 | 3.0E-02 |
| *T01C1.4* | WBGene00011317 | protein_coding_gene | 1.5 | 2.76 | 2.5E-02 | 1.4 | 2.64 | 1.7E-02 |
| *kvs-3* | WBGene00018402 | protein_coding_gene | 1.5 | 2.76 | 2.3E-02 | 1.2 | 2.38 | 3.5E-02 |
| *cha-1* | WBGene00000481 | protein_coding_gene | 1.5 | 2.75 | 3.8E-03 | 1.7 | 3.26 | 2.1E-05 |
| *Y54G2A.49* | WBGene00044492 | protein_coding_gene | 1.5 | 2.75 | 1.5E-02 | 1.1 | 2.18 | 1.3E-02 |
| *ttll-11* | WBGene00019230 | protein_coding_gene | 1.5 | 2.75 | 2.2E-02 | 1.3 | 2.41 | 1.0E-02 |
| *C01B10.6* | WBGene00015280 | protein_coding_gene | 1.5 | 2.75 | 3.7E-03 | 1.1 | 2.12 | 3.6E-03 |
| *D2096.9* | WBGene00017076 | protein_coding_gene | 1.5 | 2.75 | 9.8E-03 | 1.3 | 2.40 | 2.8E-03 |
| *iff-2* | WBGene00002065 | protein_coding_gene | 1.5 | 2.74 | 5.2E-03 | 1.6 | 3.03 | 6.7E-03 |
| *C02B8.3* | WBGene00015322 | protein_coding_gene | 1.5 | 2.74 | 4.6E-02 | 1.8 | 3.49 | 5.9E-04 |
| *lgc-37* | WBGene00013914 | protein_coding_gene | 1.5 | 2.73 | 2.8E-02 | 1.8 | 3.40 | 2.1E-02 |
| *nlp-19* | WBGene00003757 | protein_coding_gene | 1.4 | 2.73 | 4.1E-02 | 1.3 | 2.43 | 1.2E-02 |
| *npr-27* | WBGene00018344 | protein_coding_gene | 1.4 | 2.73 | 3.0E-02 | 1.2 | 2.32 | 2.6E-02 |
| *snt-3* | WBGene00004923 | protein_coding_gene | 1.4 | 2.72 | 4.9E-03 | 1.1 | 2.08 | 2.0E-03 |
| *C45G7.13* | WBGene00235340 | protein_coding_gene | 1.4 | 2.72 | 4.9E-02 | 1.3 | 2.54 | 7.5E-03 |
| *flp-28* | WBGene00044686 | protein_coding_gene | 1.4 | 2.72 | 5.8E-04 | 1.4 | 2.72 | 1.4E-06 |
| *C01B9.1* | WBGene00007220 | protein_coding_gene | 1.4 | 2.72 | 1.2E-02 | 2.5 | 5.75 | 6.9E-06 |
| *PDB1.1* | WBGene00019803 | protein_coding_gene | 1.4 | 2.72 | 1.7E-03 | 1.9 | 3.75 | 3.1E-07 |
| *F17C8.8* | WBGene00044224 | pseudogene | 1.4 | 2.72 | 2.2E-02 | 2.0 | 3.96 | 3.5E-03 |
| *C39F7.1* | WBGene00016538 | protein_coding_gene | 1.4 | 2.72 | 2.8E-02 | 1.3 | 2.50 | 2.0E-02 |
| *W04E12.7* | WBGene00012252 | protein_coding_gene | 1.4 | 2.71 | 4.6E-02 | 1.9 | 3.81 | 2.7E-03 |
| *npr-7* | WBGene00018067 | protein_coding_gene | 1.4 | 2.71 | 4.2E-02 | 1.8 | 3.40 | 5.8E-03 |
| *Y49E10.16* | WBGene00013035 | protein_coding_gene | 1.4 | 2.71 | 6.9E-04 | 1.0 | 2.05 | 1.5E-02 |
| *fpn-1.2* | WBGene00019977 | protein_coding_gene | 1.4 | 2.70 | 2.8E-05 | 1.7 | 3.27 | 1.8E-05 |
| *T01G1.2* | WBGene00011337 | protein_coding_gene | 1.4 | 2.69 | 3.0E-02 | 1.1 | 2.12 | 3.2E-02 |
| *C06E1.7* | WBGene00015522 | protein_coding_gene | 1.4 | 2.69 | 1.0E-02 | 1.9 | 3.64 | 1.2E-05 |

| | | | | | | | | |
|---|---|---|---|---|---|---|---|---|
| *C26D10.4* | WBGene00007745 | protein_coding_gene | 1.4 | 2.68 | 1.2E-03 | 1.1 | 2.08 | 2.5E-02 |
| *nlp-59* | WBGene00008522 | protein_coding_gene | 1.4 | 2.68 | 1.4E-02 | 1.7 | 3.34 | 6.3E-04 |
| *F10C1.8* | WBGene00017327 | protein_coding_gene | 1.4 | 2.67 | 5.8E-04 | 2.0 | 4.14 | 3.4E-04 |
| *T28F3.5* | WBGene00012131 | protein_coding_gene | 1.4 | 2.66 | 2.6E-02 | 1.2 | 2.38 | 3.2E-02 |
| *F53A9.9* | WBGene00018732 | protein_coding_gene | 1.4 | 2.66 | 1.5E-03 | 1.0 | 2.04 | 1.4E-02 |
| *ugt-24* | WBGene00016762 | protein_coding_gene | 1.4 | 2.66 | 9.5E-04 | 1.1 | 2.14 | 6.9E-03 |
| *sem-4* | WBGene00004773 | protein_coding_gene | 1.4 | 2.65 | 6.5E-03 | 1.0 | 2.01 | 1.2E-02 |
| *F58E2.3* | WBGene00019047 | protein_coding_gene | 1.4 | 2.64 | 9.8E-03 | 1.5 | 2.78 | 8.4E-03 |
| *ttr-25* | WBGene00013078 | protein_coding_gene | 1.4 | 2.64 | 4.0E-02 | 1.6 | 2.99 | 2.2E-02 |
| *flp-6* | WBGene00001449 | protein_coding_gene | 1.4 | 2.64 | 4.1E-02 | 1.1 | 2.17 | 5.2E-06 |
| *ggr-3* | WBGene00001588 | protein_coding_gene | 1.4 | 2.63 | 2.1E-02 | 1.4 | 2.63 | 1.4E-02 |
| *kvs-4* | WBGene00012967 | protein_coding_gene | 1.4 | 2.62 | 1.5E-02 | 1.6 | 3.03 | 6.4E-04 |
| *galt-1* | WBGene00019763 | protein_coding_gene | 1.4 | 2.62 | 1.5E-02 | 1.7 | 3.31 | 1.9E-06 |
| *nhr-144* | WBGene00019117 | pseudogene | 1.4 | 2.62 | 8.7E-03 | 1.3 | 2.47 | 6.0E-03 |
| *F28E10.1* | WBGene00017900 | protein_coding_gene | 1.4 | 2.62 | 7.9E-03 | 1.2 | 2.34 | 4.4E-05 |
| *nhr-231* | WBGene00012449 | protein_coding_gene | 1.4 | 2.62 | 1.3E-02 | 1.4 | 2.59 | 1.6E-02 |
| *D1086.18* | WBGene00045405 | protein_coding_gene | 1.4 | 2.62 | 5.3E-03 | 1.2 | 2.33 | 1.4E-02 |
| *VM106R.1* | WBGene00014848 | protein_coding_gene | 1.4 | 2.61 | 4.6E-03 | 1.7 | 3.30 | 7.8E-05 |
| *sto-2* | WBGene00006064 | protein_coding_gene | 1.4 | 2.59 | 6.5E-03 | 1.6 | 3.07 | 6.3E-04 |
| *glb-19* | WBGene00010146 | protein_coding_gene | 1.4 | 2.59 | 2.3E-02 | 1.3 | 2.48 | 1.1E-02 |
| *pqn-25* | WBGene00004114 | protein_coding_gene | 1.4 | 2.58 | 8.1E-03 | 1.1 | 2.14 | 7.0E-03 |
| *C30F2.3* | WBGene00007817 | protein_coding_gene | 1.4 | 2.57 | 2.7E-02 | 2.0 | 4.04 | 3.3E-06 |
| *myo-5* | WBGene00019064 | protein_coding_gene | 1.4 | 2.57 | 1.9E-02 | 1.1 | 2.20 | 5.8E-05 |
| *axl-1* | WBGene00010497 | protein_coding_gene | 1.4 | 2.56 | 1.1E-02 | 1.0 | 2.03 | 1.8E-03 |
| *acc-1* | WBGene00010275 | protein_coding_gene | 1.4 | 2.56 | 4.0E-02 | 1.8 | 3.40 | 1.6E-04 |
| *lge-1* | WBGene00010716 | protein_coding_gene | 1.4 | 2.56 | 4.1E-02 | 1.7 | 3.33 | 2.7E-03 |
| *elt-2* | WBGene00001250 | protein_coding_gene | 1.4 | 2.56 | 1.0E-02 | 1.1 | 2.19 | 1.7E-05 |
| *T21H3.1* | WBGene00020662 | protein_coding_gene | 1.4 | 2.55 | 5.5E-03 | 1.1 | 2.17 | 6.1E-03 |
| *nas-4* | WBGene00003523 | protein_coding_gene | 1.3 | 2.55 | 2.7E-02 | 1.4 | 2.72 | 2.1E-02 |
| *haf-4* | WBGene00001814 | protein_coding_gene | 1.3 | 2.55 | 1.5E-02 | 1.7 | 3.27 | 1.2E-03 |
| *lat-2* | WBGene00002252 | protein_coding_gene | 1.3 | 2.55 | 2.1E-02 | 1.7 | 3.28 | 6.9E-04 |
| *ifd-1* | WBGene00002057 | protein_coding_gene | 1.3 | 2.55 | 6.6E-03 | 1.9 | 3.68 | 5.2E-10 |
| *Y73F8A.35* | WBGene00013539 | protein_coding_gene | 1.3 | 2.55 | 4.4E-03 | 1.1 | 2.16 | 4.8E-03 |
| *nfki-1* | WBGene00007877 | protein_coding_gene | 1.3 | 2.54 | 2.8E-02 | 1.1 | 2.12 | 1.0E-02 |
| *lim-6* | WBGene00002988 | protein_coding_gene | 1.3 | 2.54 | 2.3E-02 | 1.0 | 2.05 | 9.5E-03 |
| *F14B6.2* | WBGene00008783 | protein_coding_gene | 1.3 | 2.54 | 5.3E-03 | 1.2 | 2.32 | 8.9E-04 |
| *gcy-21* | WBGene00001546 | protein_coding_gene | 1.3 | 2.53 | 1.3E-02 | 1.3 | 2.43 | 6.1E-03 |
| *zig-2* | WBGene00006979 | protein_coding_gene | 1.3 | 2.53 | 4.1E-02 | 1.1 | 2.21 | 4.5E-02 |
| *R05D7.3* | WBGene00011028 | protein_coding_gene | 1.3 | 2.53 | 2.3E-02 | 1.1 | 2.20 | 9.5E-03 |
| *H34I24.2* | WBGene00019261 | protein_coding_gene | 1.3 | 2.53 | 3.1E-03 | 1.3 | 2.43 | 2.2E-03 |
| *hlh-11* | WBGene00001955 | protein_coding_gene | 1.3 | 2.53 | 2.1E-02 | 1.3 | 2.41 | 1.2E-02 |
| *casy-1* | WBGene00000403 | protein_coding_gene | 1.3 | 2.53 | 1.7E-02 | 1.2 | 2.29 | 6.8E-04 |
| *unc-10* | WBGene00006750 | protein_coding_gene | 1.3 | 2.52 | 2.4E-02 | 1.4 | 2.66 | 9.1E-05 |
| *ric-4* | WBGene00004364 | protein_coding_gene | 1.3 | 2.52 | 6.5E-03 | 1.0 | 2.03 | 2.8E-09 |
| *R151.1* | WBGene00020106 | protein_coding_gene | 1.3 | 2.52 | 1.7E-02 | 1.4 | 2.68 | 8.6E-03 |
| *seb-3* | WBGene00007664 | protein_coding_gene | 1.3 | 2.51 | 2.7E-02 | 1.0 | 2.03 | 2.5E-02 |
| *sqst-5* | WBGene00269434 | protein_coding_gene | 1.3 | 2.51 | 6.4E-03 | 1.8 | 3.47 | 1.4E-02 |
| *twk-21* | WBGene00006674 | protein_coding_gene | 1.3 | 2.51 | 4.8E-02 | 1.6 | 2.96 | 6.3E-03 |
| *F18A11.5* | WBGene00008932 | protein_coding_gene | 1.3 | 2.51 | 2.7E-03 | 1.2 | 2.27 | 4.7E-03 |
| *drh-2* | WBGene00001091 | pseudogene | 1.3 | 2.50 | 7.0E-03 | 1.4 | 2.62 | 2.0E-03 |
| *gpc-1* | WBGene00001681 | protein_coding_gene | 1.3 | 2.49 | 2.9E-02 | 1.1 | 2.12 | 1.3E-02 |
| *F15A4.5* | WBGene00008840 | protein_coding_gene | 1.3 | 2.49 | 3.0E-02 | 1.4 | 2.58 | 3.1E-04 |
| *C54F6.15* | WBGene00044113 | protein_coding_gene | 1.3 | 2.49 | 4.3E-02 | 2.4 | 5.38 | 1.1E-02 |
| *ifb-2* | WBGene00002054 | protein_coding_gene | 1.3 | 2.49 | 4.0E-04 | 1.3 | 2.53 | 2.2E-05 |
| *vhp-1* | WBGene00006923 | protein_coding_gene | 1.3 | 2.49 | 2.3E-02 | 1.0 | 2.04 | 3.4E-02 |
| *tre-3* | WBGene00006609 | protein_coding_gene | 1.3 | 2.48 | 1.1E-02 | 1.2 | 2.29 | 6.8E-04 |
| *dmsr-8* | WBGene00007951 | protein_coding_gene | 1.3 | 2.48 | 1.4E-02 | 1.3 | 2.54 | 3.6E-03 |
| *rgs-6* | WBGene00004349 | protein_coding_gene | 1.3 | 2.48 | 4.2E-02 | 1.2 | 2.36 | 3.5E-03 |

| | | | | | | | | |
|---|---|---|---|---|---|---|---|---|
| *F14D7.6* | WBGene00008793 | protein_coding_gene | 1.3 | 2.48 | 1.3E-03 | 2.1 | 4.22 | 3.6E-08 |
| *C34E7.3* | WBGene00007933 | protein_coding_gene | 1.3 | 2.48 | 4.2E-02 | 1.8 | 3.53 | 5.2E-03 |
| *ZC196.5* | WBGene00022549 | protein_coding_gene | 1.3 | 2.47 | 1.9E-03 | 1.1 | 2.07 | 1.9E-05 |
| *F14B6.4* | WBGene00008785 | protein_coding_gene | 1.3 | 2.47 | 4.6E-02 | 1.2 | 2.23 | 2.9E-02 |
| ***fkb-7*** | **WBGene00001432** | **protein_coding_gene** | **1.3** | **2.47** | **1.4E-03** | **-1.0** | **0.49** | **3.3E-02** |
| *cri-1* | WBGene00010614 | protein_coding_gene | 1.3 | 2.46 | 2.7E-02 | 1.8 | 3.58 | 4.2E-04 |
| *F41E6.7* | WBGene00018288 | protein_coding_gene | 1.3 | 2.46 | 3.8E-02 | 1.1 | 2.19 | 4.4E-02 |
| ***Y51H4A.1*** | **WBGene00013098** | **protein_coding_gene** | **1.3** | **2.45** | **3.4E-02** | **-1.1** | **0.48** | **1.6E-02** |
| *rgs-3* | WBGene00004346 | protein_coding_gene | 1.3 | 2.45 | 4.7E-03 | 1.2 | 2.25 | 3.1E-04 |
| *mab-9* | WBGene00003106 | protein_coding_gene | 1.3 | 2.45 | 2.9E-02 | 1.3 | 2.42 | 7.3E-03 |
| *T19D12.1* | WBGene00020576 | protein_coding_gene | 1.3 | 2.45 | 2.7E-03 | 2.0 | 3.88 | 2.5E-11 |
| *F19C6.3* | WBGene00008952 | protein_coding_gene | 1.3 | 2.45 | 2.4E-02 | 1.3 | 2.44 | 1.1E-03 |
| *nlp-82* | WBGene00044006 | protein_coding_gene | 1.3 | 2.45 | 1.9E-02 | 1.3 | 2.43 | 6.0E-03 |
| *npr-2* | WBGene00003808 | protein_coding_gene | 1.3 | 2.44 | 4.0E-02 | 1.0 | 2.06 | 2.1E-02 |
| *Y54E2A.10* | WBGene00013195 | protein_coding_gene | 1.3 | 2.43 | 4.7E-02 | 1.6 | 2.96 | 1.5E-04 |
| *prhg-1* | WBGene00022391 | protein_coding_gene | 1.3 | 2.43 | 4.6E-03 | 1.5 | 2.73 | 2.7E-04 |
| *odr-2* | WBGene00003849 | protein_coding_gene | 1.3 | 2.42 | 2.0E-02 | 1.0 | 2.06 | 1.4E-02 |
| *mec-9* | WBGene00003173 | protein_coding_gene | 1.3 | 2.42 | 4.1E-02 | 1.4 | 2.63 | 7.9E-03 |
| *gpx-6* | WBGene00020373 | protein_coding_gene | 1.3 | 2.42 | 2.7E-02 | 1.8 | 3.47 | 1.4E-02 |
| *kri-1* | WBGene00013955 | protein_coding_gene | 1.3 | 2.42 | 1.7E-03 | 1.5 | 2.79 | 1.4E-04 |
| *F38H4.3* | WBGene00009547 | protein_coding_gene | 1.3 | 2.39 | 2.1E-03 | 1.3 | 2.42 | 2.3E-04 |
| *lgc-31* | WBGene00008997 | protein_coding_gene | 1.3 | 2.39 | 2.2E-02 | 1.1 | 2.10 | 7.2E-03 |
| *tom-1* | WBGene00006594 | protein_coding_gene | 1.3 | 2.39 | 2.5E-03 | 1.1 | 2.14 | 2.9E-03 |
| *F49B2.6* | WBGene00009865 | protein_coding_gene | 1.3 | 2.39 | 9.4E-03 | 1.4 | 2.72 | 2.0E-04 |
| *C39H7.4* | WBGene00016542 | protein_coding_gene | 1.3 | 2.39 | 3.6E-05 | 1.2 | 2.34 | 3.3E-11 |
| *fbxa-190* | WBGene00018939 | protein_coding_gene | 1.3 | 2.38 | 4.3E-02 | 1.3 | 2.49 | 2.9E-03 |
| *K05F1.10* | WBGene00019411 | protein_coding_gene | 1.2 | 2.37 | 1.1E-05 | 1.3 | 2.43 | 2.7E-03 |
| *cyp-33A1* | WBGene00015709 | protein_coding_gene | 1.2 | 2.37 | 2.5E-04 | 1.3 | 2.49 | 2.0E-03 |
| *R10E11.5* | WBGene00011217 | protein_coding_gene | 1.2 | 2.36 | 5.1E-03 | 1.3 | 2.54 | 8.5E-05 |
| *W05H12.1* | WBGene00012289 | protein_coding_gene | 1.2 | 2.35 | 2.8E-02 | 1.5 | 2.82 | 8.2E-07 |
| *F25E5.16* | WBGene00017792 | protein_coding_gene | 1.2 | 2.34 | 3.8E-02 | 1.1 | 2.19 | 1.6E-02 |
| *ifc-2* | WBGene00002056 | protein_coding_gene | 1.2 | 2.32 | 4.6E-04 | 1.5 | 2.74 | 5.7E-05 |
| *unc-31* | WBGene00006767 | protein_coding_gene | 1.2 | 2.32 | 1.1E-02 | 1.4 | 2.57 | 1.7E-03 |
| *C54F6.6* | WBGene00016924 | protein_coding_gene | 1.2 | 2.31 | 2.9E-02 | 2.5 | 5.83 | 1.7E-02 |
| *Y38C1AA.9* | WBGene00021400 | protein_coding_gene | 1.2 | 2.31 | 1.1E-02 | 1.1 | 2.11 | 5.1E-04 |
| *F54C1.1* | WBGene00018789 | protein_coding_gene | 1.2 | 2.30 | 1.3E-02 | 2.1 | 4.19 | 3.3E-04 |
| *D2085.7* | WBGene00008432 | protein_coding_gene | 1.2 | 2.28 | 1.7E-02 | 1.2 | 2.29 | 8.7E-03 |
| *F58E2.4* | WBGene00019048 | protein_coding_gene | 1.2 | 2.28 | 1.0E-02 | 1.4 | 2.66 | 3.1E-03 |
| *Y16B4A.2* | WBGene00012445 | protein_coding_gene | 1.2 | 2.27 | 4.2E-02 | 2.8 | 6.74 | 1.3E-08 |
| *C55A6.7* | WBGene00008336 | protein_coding_gene | 1.2 | 2.27 | 6.2E-03 | 1.1 | 2.21 | 2.8E-04 |
| *mec-1* | WBGene00003165 | protein_coding_gene | 1.2 | 2.27 | 4.8E-02 | 1.4 | 2.58 | 6.6E-03 |
| *ddn-1* | WBGene00015227 | protein_coding_gene | 1.2 | 2.26 | 4.7E-02 | 2.0 | 3.98 | 5.2E-04 |
| *unc-79* | WBGene00006811 | protein_coding_gene | 1.2 | 2.26 | 4.6E-02 | 1.3 | 2.52 | 4.0E-03 |
| *F14B8.6* | WBGene00017448 | protein_coding_gene | 1.2 | 2.26 | 2.8E-03 | 1.3 | 2.49 | 1.9E-05 |
| *D1054.5* | WBGene00008372 | protein_coding_gene | 1.2 | 2.25 | 5.7E-03 | 1.2 | 2.34 | 8.5E-04 |
| *klo-2* | WBGene00017103 | protein_coding_gene | 1.2 | 2.24 | 2.8E-03 | 1.3 | 2.40 | 5.3E-04 |
| *ilcr-1* | WBGene00013415 | protein_coding_gene | 1.2 | 2.24 | 2.3E-02 | 1.4 | 2.65 | 2.0E-05 |
| *ifo-1* | WBGene00018350 | protein_coding_gene | 1.2 | 2.23 | 3.8E-02 | 1.4 | 2.56 | 1.3E-04 |
| *gbb-1* | WBGene00021528 | protein_coding_gene | 1.2 | 2.23 | 4.5E-02 | 1.3 | 2.48 | 9.5E-04 |
| *C02B8.12* | WBGene00219375 | protein_coding_gene | 1.2 | 2.23 | 6.9E-03 | 1.2 | 2.32 | 5.6E-03 |
| *F19C7.4* | WBGene00017594 | protein_coding_gene | 1.1 | 2.21 | 1.3E-02 | 2.1 | 4.16 | 3.3E-05 |
| *T11B7.5* | WBGene00011708 | protein_coding_gene | 1.1 | 2.21 | 2.6E-02 | 1.1 | 2.18 | 2.8E-03 |
| *ZK622.4* | WBGene00022782 | protein_coding_gene | 1.1 | 2.21 | 1.1E-02 | 1.1 | 2.14 | 6.1E-04 |
| *let-756* | WBGene00002881 | protein_coding_gene | 1.1 | 2.20 | 4.4E-03 | 1.0 | 2.01 | 6.1E-05 |
| *nhr-146* | WBGene00045255 | protein_coding_gene | 1.1 | 2.19 | 4.9E-02 | 1.2 | 2.25 | 3.3E-02 |
| *twk-49* | WBGene00019904 | protein_coding_gene | 1.1 | 2.19 | 2.1E-02 | 1.4 | 2.65 | 7.2E-03 |
| *kcc-2* | WBGene00019205 | protein_coding_gene | 1.1 | 2.18 | 3.5E-02 | 1.0 | 2.00 | 1.5E-03 |
| *nlp-79* | WBGene00008626 | protein_coding_gene | 1.1 | 2.18 | 4.4E-02 | 1.1 | 2.10 | 2.8E-02 |

| | | | | | | | | |
|---|---|---|---|---|---|---|---|---|
| *W07G4.5* | WBGene00012339 | protein_coding_gene | 1.1 | 2.18 | 5.3E-03 | 1.4 | 2.57 | 1.8E-06 |
| *T04A8.18* | WBGene00119203 | protein_coding_gene | 1.1 | 2.17 | 1.3E-02 | 1.6 | 2.95 | 2.2E-02 |
| *ncx-4* | WBGene00003569 | protein_coding_gene | 1.1 | 2.17 | 3.2E-02 | 1.7 | 3.26 | 1.1E-04 |
| *H43E16.1* | WBGene00019273 | protein_coding_gene | 1.1 | 2.17 | 4.9E-02 | 2.1 | 4.28 | 1.0E-22 |
| *tmc-1* | WBGene00020490 | protein_coding_gene | 1.1 | 2.17 | 2.6E-02 | 1.2 | 2.34 | 2.5E-03 |
| *arrd-18* | WBGene00018060 | protein_coding_gene | 1.1 | 2.17 | 2.7E-02 | 1.7 | 3.18 | 2.1E-03 |
| *lgc-39* | WBGene00017314 | protein_coding_gene | 1.1 | 2.16 | 3.4E-02 | 1.2 | 2.35 | 3.6E-04 |
| *cutl-23* | WBGene00021343 | protein_coding_gene | 1.1 | 2.15 | 3.9E-02 | 1.3 | 2.50 | 9.2E-04 |
| *F40A3.7* | WBGene00018222 | protein_coding_gene | 1.1 | 2.15 | 2.1E-02 | 1.0 | 2.04 | 1.9E-02 |
| *inpp-1* | WBGene00012016 | protein_coding_gene | 1.1 | 2.14 | 2.4E-02 | 1.1 | 2.10 | 1.3E-02 |
| *lmp-2* | WBGene00015471 | protein_coding_gene | 1.1 | 2.13 | 1.8E-02 | 1.7 | 3.16 | 3.6E-05 |
| *H01G02.3* | WBGene00010350 | protein_coding_gene | 1.1 | 2.12 | 3.7E-02 | 1.0 | 2.06 | 2.0E-02 |
| *fbxa-54* | WBGene00020449 | protein_coding_gene | 1.1 | 2.12 | 7.6E-03 | 1.4 | 2.55 | 2.5E-02 |
| *acc-4* | WBGene00012099 | protein_coding_gene | 1.1 | 2.11 | 2.4E-02 | 1.0 | 2.04 | 2.7E-03 |
| *trp-2* | WBGene00006615 | protein_coding_gene | 1.1 | 2.10 | 2.7E-02 | 1.0 | 2.01 | 2.3E-02 |
| *arrd-26* | WBGene00019873 | protein_coding_gene | 1.1 | 2.10 | 2.7E-02 | 1.7 | 3.14 | 1.0E-03 |
| *ncs-3* | WBGene00003565 | protein_coding_gene | 1.1 | 2.09 | 4.1E-02 | 1.1 | 2.11 | 9.1E-03 |
| *K04F10.1* | WBGene00019394 | protein_coding_gene | 1.0 | 2.07 | 1.5E-02 | 1.1 | 2.10 | 3.7E-03 |
| *bus-19* | WBGene00011590 | protein_coding_gene | 1.0 | 2.05 | 3.4E-02 | 1.2 | 2.36 | 2.2E-04 |
| *ZC247.1* | WBGene00013859 | protein_coding_gene | 1.0 | 2.05 | 4.3E-02 | 1.3 | 2.47 | 1.6E-02 |
| *rpl-11.2* | WBGene00004423 | protein_coding_gene | 1.0 | 2.05 | 2.4E-02 | 1.2 | 2.27 | 3.0E-02 |
| *ZK792.4* | WBGene00014077 | protein_coding_gene | 1.0 | 2.03 | 4.4E-02 | 1.2 | 2.33 | 8.9E-05 |
| *F09F7.5* | WBGene00017302 | protein_coding_gene | 1.0 | 2.03 | 1.5E-02 | 1.1 | 2.20 | 5.2E-03 |
| *E02H9.2* | WBGene00017100 | protein_coding_gene | 1.0 | 2.03 | 4.3E-02 | 1.4 | 2.59 | 1.9E-03 |
| *gar-2* | WBGene00001518 | protein_coding_gene | 1.0 | 2.01 | 4.3E-02 | 1.5 | 2.85 | 1.2E-05 |
| *unc-80* | WBGene00006812 | protein_coding_gene | 1.0 | 2.01 | 4.3E-02 | 1.2 | 2.22 | 1.2E-03 |
| *cest-15* | WBGene00012202 | pseudogene | 1.0 | 2.01 | 1.6E-03 | 1.6 | 2.98 | 3.4E-07 |
| *bgal-2* | WBGene00019225 | protein_coding_gene | 1.0 | 2.01 | 1.2E-02 | 1.2 | 2.23 | 2.9E-05 |
| *F41G3.10* | WBGene00018303 | protein_coding_gene | 1.0 | 2.00 | 4.9E-02 | 1.1 | 2.10 | 2.0E-03 |
| *tsp-12* | WBGene00006638 | protein_coding_gene | -1.0 | 0.50 | 4.3E-02 | -1.0 | 0.50 | 2.4E-02 |
| *tag-115* | WBGene00006466 | protein_coding_gene | -1.0 | 0.50 | 3.1E-02 | -1.0 | 0.50 | 4.6E-03 |
| *rbm-3.1* | WBGene00011155 | protein_coding_gene | -1.0 | 0.50 | 2.2E-02 | -1.0 | 0.49 | 4.2E-06 |
| *E01A2.1* | WBGene00017084 | protein_coding_gene | -1.0 | 0.50 | 7.1E-03 | -1.1 | 0.47 | 5.8E-04 |
| *T12F5.2* | WBGene00020467 | protein_coding_gene | -1.0 | 0.49 | 4.8E-02 | -1.3 | 0.41 | 6.9E-04 |
| *fog-2* | WBGene00001482 | protein_coding_gene | -1.0 | 0.49 | 1.1E-02 | -1.1 | 0.47 | 5.6E-05 |
| *lin-7* | WBGene00002996 | protein_coding_gene | -1.0 | 0.49 | 3.5E-03 | -1.1 | 0.48 | 4.8E-04 |
| *F33G12.3* | WBGene00018015 | protein_coding_gene | -1.0 | 0.49 | 2.2E-02 | -1.0 | 0.49 | 1.3E-02 |
| *bath-15* | WBGene00015590 | protein_coding_gene | -1.0 | 0.49 | 4.7E-02 | -1.1 | 0.47 | 2.1E-02 |
| *spr-3* | WBGene00005008 | protein_coding_gene | -1.0 | 0.49 | 2.8E-02 | -1.0 | 0.49 | 6.8E-03 |
| *C45G9.2* | WBGene00016674 | protein_coding_gene | -1.0 | 0.49 | 8.1E-03 | -1.1 | 0.47 | 5.0E-04 |
| *slx-1* | WBGene00018909 | protein_coding_gene | -1.0 | 0.49 | 6.2E-03 | -1.0 | 0.49 | 1.8E-04 |
| *pes-5* | WBGene00003979 | protein_coding_gene | -1.0 | 0.49 | 1.8E-02 | -1.0 | 0.48 | 1.5E-02 |
| *K04G7.1* | WBGene00019400 | protein_coding_gene | -1.0 | 0.49 | 2.2E-02 | -1.1 | 0.46 | 1.5E-03 |
| *R06F6.12* | WBGene00023418 | protein_coding_gene | -1.0 | 0.49 | 1.6E-03 | -1.1 | 0.48 | 3.7E-05 |
| *F54D10.5* | WBGene00018806 | protein_coding_gene | -1.0 | 0.49 | 3.4E-02 | -1.2 | 0.44 | 1.2E-03 |
| *czw-1* | WBGene00017643 | protein_coding_gene | -1.0 | 0.48 | 3.8E-03 | -1.0 | 0.50 | 1.1E-03 |
| *tbcd-1* | WBGene00008887 | protein_coding_gene | -1.1 | 0.48 | 3.2E-03 | -1.1 | 0.48 | 1.9E-04 |
| *air-2* | WBGene00000099 | protein_coding_gene | -1.1 | 0.48 | 4.4E-02 | -1.0 | 0.49 | 2.6E-02 |
| *exos-2* | WBGene00022232 | protein_coding_gene | -1.1 | 0.48 | 2.2E-03 | -1.0 | 0.50 | 8.9E-05 |
| *F55G11.4* | WBGene00010124 | protein_coding_gene | -1.1 | 0.48 | 1.3E-02 | -3.9 | 0.07 | 1.0E-12 |
| *R06B9.5* | WBGene00011056 | protein_coding_gene | -1.1 | 0.48 | 2.4E-02 | -1.3 | 0.40 | 1.1E-04 |
| *C36A4.4* | WBGene00007965 | protein_coding_gene | -1.1 | 0.48 | 2.2E-02 | -1.1 | 0.48 | 1.2E-02 |
| *rpb-3* | WBGene00007971 | protein_coding_gene | -1.1 | 0.48 | 7.0E-03 | -1.1 | 0.48 | 5.3E-06 |
| *mrpl-20* | WBGene00012992 | protein_coding_gene | -1.1 | 0.48 | 2.1E-03 | -1.0 | 0.49 | 2.0E-06 |
| *C53D6.4* | WBGene00008281 | protein_coding_gene | -1.1 | 0.48 | 3.2E-02 | -1.0 | 0.48 | 2.0E-03 |
| *gon-4* | WBGene00001653 | protein_coding_gene | -1.1 | 0.47 | 6.3E-04 | -1.0 | 0.50 | 9.1E-04 |
| *W04A8.1* | WBGene00012236 | protein_coding_gene | -1.1 | 0.47 | 4.9E-03 | -1.2 | 0.45 | 1.8E-03 |
| *let-391* | WBGene00006492 | protein_coding_gene | -1.1 | 0.47 | 2.2E-02 | -1.1 | 0.45 | 2.2E-03 |

| | | | | | | | | |
|---|---|---|---|---|---|---|---|---|
| *zhit-1* | WBGene00016992 | protein_coding_gene | -1.1 | 0.47 | 3.3E-02 | -1.3 | 0.40 | 3.0E-04 |
| *pch-2* | WBGene00008641 | protein_coding_gene | -1.1 | 0.47 | 4.6E-02 | -1.1 | 0.46 | 1.0E-02 |
| *flh-1* | WBGene00012435 | protein_coding_gene | -1.1 | 0.47 | 1.6E-02 | -1.1 | 0.48 | 1.4E-02 |
| *Y43F4B.10* | WBGene00012807 | protein_coding_gene | -1.1 | 0.47 | 7.8E-03 | -1.2 | 0.44 | 7.4E-04 |
| *mrps-23* | WBGene00014224 | protein_coding_gene | -1.1 | 0.47 | 3.5E-04 | -1.1 | 0.46 | 8.4E-07 |
| *F35D11.4* | WBGene00018045 | protein_coding_gene | -1.1 | 0.47 | 5.0E-03 | -1.3 | 0.42 | 2.8E-09 |
| *ima-1* | WBGene00002072 | protein_coding_gene | -1.1 | 0.47 | 4.5E-02 | -1.1 | 0.47 | 3.6E-03 |
| *F32H2.10* | WBGene00009346 | protein_coding_gene | -1.1 | 0.47 | 2.0E-02 | -1.5 | 0.36 | 1.1E-06 |
| *mec-15* | WBGene00003177 | protein_coding_gene | -1.1 | 0.46 | 1.9E-02 | -1.0 | 0.48 | 6.2E-03 |
| *lin-54* | WBGene00003037 | protein_coding_gene | -1.1 | 0.46 | 1.4E-02 | -1.0 | 0.48 | 5.2E-03 |
| *otub-3* | WBGene00009007 | protein_coding_gene | -1.1 | 0.46 | 4.5E-02 | -1.0 | 0.48 | 2.3E-02 |
| *bath-41* | WBGene00018137 | protein_coding_gene | -1.1 | 0.46 | 4.1E-02 | -1.0 | 0.49 | 1.2E-02 |
| *npp-23* | WBGene00007493 | protein_coding_gene | -1.1 | 0.46 | 1.2E-02 | -1.0 | 0.48 | 1.7E-03 |
| *psf-2* | WBGene00009287 | protein_coding_gene | -1.1 | 0.46 | 1.2E-02 | -1.3 | 0.39 | 6.1E-04 |
| *rpb-8* | WBGene00017830 | protein_coding_gene | -1.1 | 0.46 | 5.6E-03 | -1.0 | 0.49 | 1.7E-04 |
| *unc-59* | WBGene00006793 | protein_coding_gene | -1.1 | 0.46 | 2.5E-02 | -1.1 | 0.47 | 1.4E-02 |
| *skpt-1* | WBGene00018613 | protein_coding_gene | -1.1 | 0.46 | 8.1E-03 | -1.1 | 0.48 | 1.3E-04 |
| *ska-3* | WBGene00018830 | protein_coding_gene | -1.1 | 0.46 | 1.4E-02 | -1.2 | 0.43 | 1.6E-03 |
| *sygl-1* | WBGene00012104 | protein_coding_gene | -1.1 | 0.46 | 3.4E-02 | -1.1 | 0.48 | 2.1E-05 |
| *R07E5.11* | WBGene00011117 | protein_coding_gene | -1.1 | 0.46 | 1.6E-02 | -1.2 | 0.45 | 2.2E-04 |
| *evl-18* | WBGene00009372 | protein_coding_gene | -1.1 | 0.46 | 1.3E-02 | -1.1 | 0.47 | 1.4E-02 |
| *ubc-9* | WBGene00006706 | protein_coding_gene | -1.1 | 0.45 | 1.0E-02 | -1.0 | 0.49 | 3.6E-03 |
| *F43G6.3* | WBGene00009656 | pseudogene | -1.1 | 0.45 | 4.8E-02 | -1.4 | 0.37 | 8.8E-03 |
| *sys-1* | WBGene00006379 | protein_coding_gene | -1.1 | 0.45 | 7.8E-03 | -1.1 | 0.45 | 2.1E-02 |
| *him-14* | WBGene00001872 | protein_coding_gene | -1.1 | 0.45 | 1.3E-04 | -1.0 | 0.49 | 5.4E-05 |
| *snr-6* | WBGene00004919 | protein_coding_gene | -1.1 | 0.45 | 1.3E-03 | -1.2 | 0.43 | 6.8E-06 |
| *R02D3.7* | WBGene00019824 | protein_coding_gene | -1.1 | 0.45 | 3.0E-03 | -1.1 | 0.46 | 8.8E-04 |
| *M04B2.4* | WBGene00010847 | protein_coding_gene | -1.1 | 0.45 | 4.0E-06 | -1.0 | 0.49 | 1.0E-05 |
| *acly-2* | WBGene00007150 | protein_coding_gene | -1.1 | 0.45 | 1.0E-02 | -1.0 | 0.48 | 6.0E-03 |
| *F33H2.2* | WBGene00009366 | protein_coding_gene | -1.2 | 0.45 | 3.6E-02 | -1.0 | 0.49 | 1.9E-02 |
| *mzt-1* | WBGene00044431 | protein_coding_gene | -1.2 | 0.45 | 8.2E-03 | -1.2 | 0.45 | 6.3E-04 |
| *C06H2.7* | WBGene00007391 | protein_coding_gene | -1.2 | 0.45 | 6.4E-03 | -1.2 | 0.43 | 2.7E-02 |
| *F53A2.3* | WBGene00009951 | protein_coding_gene | -1.2 | 0.45 | 7.9E-03 | -1.3 | 0.39 | 1.6E-03 |
| *cdc-25.1* | WBGene00000386 | protein_coding_gene | -1.2 | 0.45 | 7.2E-03 | -1.1 | 0.48 | 1.8E-03 |
| *mcm-4* | WBGene00003156 | protein_coding_gene | -1.2 | 0.45 | 1.1E-05 | -1.2 | 0.44 | 2.3E-03 |
| *npp-1* | WBGene00003787 | protein_coding_gene | -1.2 | 0.45 | 6.4E-04 | -1.0 | 0.49 | 1.2E-04 |
| *Y59A8B.12* | WBGene00013349 | protein_coding_gene | -1.2 | 0.45 | 5.0E-02 | -1.2 | 0.42 | 1.2E-02 |
| *acl-3* | WBGene00006491 | protein_coding_gene | -1.2 | 0.45 | 5.4E-03 | -1.1 | 0.47 | 3.4E-05 |
| *ctf-4* | WBGene00008921 | protein_coding_gene | -1.2 | 0.45 | 5.3E-03 | -1.0 | 0.49 | 8.0E-03 |
| *ztf-15* | WBGene00011066 | protein_coding_gene | -1.2 | 0.45 | 4.0E-03 | -1.2 | 0.43 | 3.3E-04 |
| *R107.5* | WBGene00011300 | protein_coding_gene | -1.2 | 0.44 | 8.3E-03 | -1.0 | 0.49 | 7.0E-04 |
| *Y37H2A.12* | WBGene00044758 | protein_coding_gene | -1.2 | 0.44 | 3.1E-03 | -1.1 | 0.45 | 3.4E-02 |
| *dod-17* | WBGene00010745 | protein_coding_gene | -1.2 | 0.44 | 3.5E-02 | -1.5 | 0.36 | 1.0E-02 |
| *bag-1* | WBGene00000236 | protein_coding_gene | -1.2 | 0.44 | 1.6E-05 | -1.2 | 0.44 | 4.6E-06 |
| *npp-22* | WBGene00007118 | protein_coding_gene | -1.2 | 0.44 | 1.0E-02 | -1.1 | 0.45 | 2.3E-03 |
| *K11H3.8* | WBGene00185002 | protein_coding_gene | -1.2 | 0.44 | 6.6E-03 | -1.1 | 0.46 | 7.6E-03 |
| *oef-1* | WBGene00018636 | protein_coding_gene | -1.2 | 0.44 | 1.5E-03 | -1.1 | 0.47 | 5.9E-05 |
| *tofu-6* | WBGene00017132 | protein_coding_gene | -1.2 | 0.44 | 6.8E-04 | -1.3 | 0.40 | 1.7E-07 |
| *C27F2.10* | WBGene00016171 | protein_coding_gene | -1.2 | 0.44 | 1.9E-03 | -1.1 | 0.47 | 4.5E-04 |
| *emb-27* | WBGene00001281 | protein_coding_gene | -1.2 | 0.44 | 1.6E-05 | -1.0 | 0.49 | 1.9E-05 |
| *Y48B6A.13* | WBGene00012984 | protein_coding_gene | -1.2 | 0.43 | 1.7E-03 | -1.0 | 0.50 | 1.2E-05 |
| *snr-7* | WBGene00004920 | protein_coding_gene | -1.2 | 0.43 | 1.3E-02 | -1.2 | 0.45 | 2.5E-03 |
| *Y106G6H.6* | WBGene00013719 | protein_coding_gene | -1.2 | 0.43 | 1.3E-02 | -1.1 | 0.46 | 4.2E-03 |
| *har-2* | WBGene00011988 | protein_coding_gene | -1.2 | 0.43 | 1.4E-02 | -1.2 | 0.44 | 2.8E-02 |
| *apc-11* | WBGene00000145 | protein_coding_gene | -1.2 | 0.43 | 8.7E-03 | -1.5 | 0.36 | 5.0E-05 |
| *Y47G6A.25* | WBGene00021649 | protein_coding_gene | -1.2 | 0.42 | 6.7E-03 | -1.2 | 0.42 | 2.6E-03 |
| *C50B6.3* | WBGene00008219 | protein_coding_gene | -1.2 | 0.42 | 2.4E-02 | -1.2 | 0.45 | 1.9E-02 |
| *D1081.6* | WBGene00008384 | protein_coding_gene | -1.2 | 0.42 | 2.5E-03 | -1.1 | 0.48 | 1.6E-05 |

| | | | | | | | | |
|---|---|---|---|---|---|---|---|---|
| *C33F10.4* | WBGene00016353 | protein_coding_gene | -1.2 | 0.42 | 1.4E-02 | -1.3 | 0.41 | 1.7E-03 |
| *nud-1* | WBGene00003829 | protein_coding_gene | -1.2 | 0.42 | 3.5E-03 | -1.0 | 0.49 | 8.5E-03 |
| *rpl-41.1* | WBGene00023067 | protein_coding_gene | -1.2 | 0.42 | 2.1E-03 | -1.2 | 0.42 | 2.5E-08 |
| *F32D8.5* | WBGene00009330 | protein_coding_gene | -1.3 | 0.42 | 9.2E-03 | -1.0 | 0.49 | 5.9E-04 |
| *ZC155.4* | WBGene00022532 | protein_coding_gene | -1.3 | 0.42 | 2.9E-02 | -1.2 | 0.43 | 1.6E-02 |
| *snr-3* | WBGene00004916 | protein_coding_gene | -1.3 | 0.42 | 1.5E-06 | -1.1 | 0.46 | 8.3E-10 |
| *fzy-1* | WBGene00001511 | protein_coding_gene | -1.3 | 0.42 | 2.8E-02 | -1.0 | 0.49 | 2.2E-02 |
| *F42G8.19* | WBGene00249817 | protein_coding_gene | -1.3 | 0.42 | 1.9E-02 | -1.6 | 0.32 | 1.7E-02 |
| *C31H1.8* | WBGene00016291 | protein_coding_gene | -1.3 | 0.42 | 4.8E-02 | -1.1 | 0.45 | 1.8E-02 |
| *erh-2* | WBGene00009444 | protein_coding_gene | -1.3 | 0.41 | 4.4E-02 | -1.3 | 0.40 | 9.7E-03 |
| *F52E1.14* | WBGene00044696 | protein_coding_gene | -1.3 | 0.41 | 8.6E-04 | -1.5 | 0.36 | 7.5E-05 |
| *F44B9.8* | WBGene00018409 | protein_coding_gene | -1.3 | 0.41 | 3.5E-03 | -1.0 | 0.50 | 8.6E-04 |
| *Y39G10AR.9* | WBGene00021467 | protein_coding_gene | -1.3 | 0.41 | 2.9E-02 | -1.1 | 0.47 | 1.5E-02 |
| *C46C2.7* | WBGene00045078 | protein_coding_gene | -1.3 | 0.41 | 6.8E-03 | -1.1 | 0.48 | 2.0E-03 |
| *D1046.2* | WBGene00008363 | protein_coding_gene | -1.3 | 0.41 | 2.5E-03 | -1.0 | 0.50 | 6.8E-03 |
| *R05H5.3* | WBGene00011038 | protein_coding_gene | -1.3 | 0.41 | 1.5E-02 | -1.3 | 0.41 | 1.1E-02 |
| *orc-1* | WBGene00012650 | protein_coding_gene | -1.3 | 0.41 | 9.3E-03 | -1.1 | 0.47 | 9.1E-03 |
| *memb-1* | WBGene00007128 | protein_coding_gene | -1.3 | 0.41 | 5.0E-03 | -1.1 | 0.46 | 9.6E-03 |
| *daf-41* | WBGene00022599 | protein_coding_gene | -1.3 | 0.40 | 2.8E-04 | -1.1 | 0.47 | 2.1E-05 |
| *R07G3.7* | WBGene00019943 | protein_coding_gene | -1.3 | 0.40 | 3.8E-03 | -1.0 | 0.50 | 3.7E-03 |
| *C42C1.8* | WBGene00016586 | protein_coding_gene | -1.3 | 0.40 | 1.3E-02 | -1.2 | 0.44 | 2.3E-03 |
| *ify-1* | WBGene00002068 | protein_coding_gene | -1.3 | 0.40 | 3.2E-02 | -1.1 | 0.48 | 2.1E-02 |
| *rmh-2* | WBGene00011576 | protein_coding_gene | -1.3 | 0.40 | 5.7E-03 | -1.1 | 0.47 | 8.5E-03 |
| *B0041.8* | WBGene00015011 | protein_coding_gene | -1.3 | 0.40 | 8.1E-03 | -1.2 | 0.43 | 1.2E-03 |
| *snr-4* | WBGene00004917 | protein_coding_gene | -1.3 | 0.40 | 1.8E-05 | -1.1 | 0.46 | 2.2E-05 |
| *T24C4.5* | WBGene00020761 | protein_coding_gene | -1.3 | 0.40 | 7.9E-03 | -1.2 | 0.44 | 2.5E-03 |
| *spd-3* | WBGene00004954 | protein_coding_gene | -1.3 | 0.40 | 1.3E-04 | -1.1 | 0.48 | 2.0E-04 |
| *zif-1* | WBGene00006977 | protein_coding_gene | -1.3 | 0.40 | 1.1E-02 | -1.0 | 0.49 | 2.6E-02 |
| *F56F11.4* | WBGene00018991 | protein_coding_gene | -1.3 | 0.40 | 1.6E-02 | -1.3 | 0.41 | 1.4E-02 |
| *Y53C12A.3* | WBGene00013139 | protein_coding_gene | -1.3 | 0.40 | 5.2E-03 | -1.1 | 0.47 | 9.7E-05 |
| *dvc-1* | WBGene00011834 | protein_coding_gene | -1.3 | 0.40 | 4.1E-02 | -1.1 | 0.47 | 4.2E-02 |
| *C02F12.8* | WBGene00015357 | protein_coding_gene | -1.3 | 0.40 | 1.8E-03 | -1.0 | 0.49 | 1.1E-02 |
| *ubc-12* | WBGene00006707 | protein_coding_gene | -1.3 | 0.40 | 1.3E-03 | -1.2 | 0.45 | 3.6E-04 |
| *F11E6.7* | WBGene00008710 | protein_coding_gene | -1.3 | 0.40 | 1.4E-02 | -1.1 | 0.46 | 3.5E-02 |
| *F35G2.3* | WBGene00009437 | protein_coding_gene | -1.3 | 0.40 | 4.6E-02 | -1.3 | 0.41 | 4.8E-02 |
| *F54E12.2* | WBGene00010061 | protein_coding_gene | -1.3 | 0.39 | 4.2E-03 | -1.3 | 0.40 | 4.3E-04 |
| *vha-18* | WBGene00018698 | protein_coding_gene | -1.3 | 0.39 | 3.5E-03 | -1.2 | 0.45 | 2.3E-04 |
| *car-1* | WBGene00012484 | protein_coding_gene | -1.3 | 0.39 | 1.0E-02 | -1.2 | 0.43 | 9.5E-04 |
| *K07F5.12* | WBGene00010637 | protein_coding_gene | -1.3 | 0.39 | 3.3E-02 | -1.2 | 0.42 | 4.3E-02 |
| *pas-2* | WBGene00003923 | protein_coding_gene | -1.4 | 0.39 | 9.5E-05 | -1.1 | 0.47 | 3.1E-05 |
| *cpg-4* | WBGene00015677 | protein_coding_gene | -1.4 | 0.39 | 1.6E-02 | -1.2 | 0.44 | 1.8E-02 |
| *sup-17* | WBGene00006324 | protein_coding_gene | -1.4 | 0.39 | 1.0E-02 | -1.1 | 0.48 | 1.5E-02 |
| *xol-1* | WBGene00006962 | protein_coding_gene | -1.4 | 0.39 | 7.9E-03 | -1.3 | 0.41 | 4.0E-02 |
| *cpna-3* | WBGene00007221 | protein_coding_gene | -1.4 | 0.39 | 2.1E-02 | -1.0 | 0.49 | 3.8E-02 |
| *snr-2* | WBGene00004915 | protein_coding_gene | -1.4 | 0.38 | 2.5E-03 | -1.0 | 0.49 | 1.5E-03 |
| *lbp-9* | WBGene00021486 | protein_coding_gene | -1.4 | 0.38 | 2.6E-03 | -1.3 | 0.41 | 5.9E-08 |
| *D1081.7* | WBGene00008385 | protein_coding_gene | -1.4 | 0.38 | 1.5E-02 | -1.0 | 0.50 | 3.7E-03 |
| *mcm-5* | WBGene00003157 | protein_coding_gene | -1.4 | 0.38 | 1.1E-03 | -1.1 | 0.47 | 1.2E-02 |
| *B0393.3* | WBGene00007168 | protein_coding_gene | -1.4 | 0.38 | 1.5E-02 | -1.1 | 0.45 | 1.0E-02 |
| *mcm-7* | WBGene00003159 | protein_coding_gene | -1.4 | 0.38 | 1.2E-02 | -1.1 | 0.46 | 4.2E-02 |
| *linc-4* | WBGene00219306 | lincRNA_gene | -1.4 | 0.38 | 1.3E-04 | -1.2 | 0.42 | 2.1E-03 |
| *dut-1* | WBGene00010609 | protein_coding_gene | -1.4 | 0.38 | 9.4E-03 | -1.1 | 0.47 | 6.5E-03 |
| *mcm-6* | WBGene00003158 | protein_coding_gene | -1.4 | 0.38 | 1.0E-04 | -1.2 | 0.44 | 1.5E-03 |
| *rpt-4* | WBGene00004504 | protein_coding_gene | -1.4 | 0.38 | 1.9E-06 | -1.1 | 0.46 | 9.1E-08 |
| *cdd-2* | WBGene00000392 | protein_coding_gene | -1.4 | 0.38 | 1.9E-03 | -1.3 | 0.40 | 3.3E-04 |
| *ima-2* | WBGene00002073 | protein_coding_gene | -1.4 | 0.38 | 2.4E-02 | -1.0 | 0.50 | 4.0E-02 |
| *F12F6.8* | WBGene00008723 | pseudogene | -1.4 | 0.38 | 4.6E-02 | -1.8 | 0.28 | 5.9E-03 |
| ***T15B7.14*** | **WBGene00020526** | **protein_coding_gene** | **-1.4** | **0.37** | **3.6E-02** | **1.9** | **3.75** | **3.4E-03** |

| | | | | | | | | |
|---|---|---|---|---|---|---|---|---|
| *pfn-1* | WBGene00003989 | protein_coding_gene | -1.4 | 0.37 | 2.6E-03 | -1.0 | 0.49 | 2.4E-03 |
| *K08D8.6* | WBGene00010660 | protein_coding_gene | -1.4 | 0.37 | 4.1E-03 | -1.1 | 0.47 | 2.5E-02 |
| *F23A7.8* | WBGene00044638 | protein_coding_gene | -1.4 | 0.37 | 7.5E-03 | -1.9 | 0.27 | 3.0E-05 |
| *sup-35* | WBGene00012976 | protein_coding_gene | -1.4 | 0.37 | 1.9E-06 | -1.1 | 0.45 | 8.4E-09 |
| *F54H12.5* | WBGene00018845 | protein_coding_gene | -1.5 | 0.36 | 3.1E-02 | -1.5 | 0.35 | 3.1E-03 |
| *T04A8.7* | WBGene00011409 | protein_coding_gene | -1.5 | 0.36 | 3.4E-06 | -1.2 | 0.44 | 1.8E-05 |
| *wdr-5.3* | WBGene00013862 | protein_coding_gene | -1.5 | 0.36 | 2.1E-02 | -1.4 | 0.37 | 1.1E-02 |
| *ifet-1* | WBGene00004132 | protein_coding_gene | -1.5 | 0.36 | 2.0E-03 | -1.2 | 0.44 | 5.8E-04 |
| *F25G6.9* | WBGene00017800 | protein_coding_gene | -1.5 | 0.36 | 2.1E-06 | -1.1 | 0.45 | 4.8E-05 |
| *F13A2.4* | WBGene00017413 | protein_coding_gene | -1.5 | 0.36 | 4.4E-02 | -1.1 | 0.48 | 1.2E-02 |
| *F45D3.1* | WBGene00009721 | protein_coding_gene | -1.5 | 0.36 | 1.4E-02 | -1.9 | 0.27 | 1.2E-02 |
| *Y53C12A.11* | WBGene00194906 | protein_coding_gene | -1.5 | 0.35 | 9.4E-03 | -1.3 | 0.40 | 6.8E-04 |
| *rskn-1* | WBGene00011352 | protein_coding_gene | -1.5 | 0.35 | 6.5E-03 | -1.1 | 0.48 | 3.6E-02 |
| *Y105C5B.20* | WBGene00013659 | protein_coding_gene | -1.5 | 0.35 | 3.9E-02 | -2.0 | 0.26 | 1.5E-04 |
| *Y39G10AR.32* | WBGene00219275 | protein_coding_gene | -1.5 | 0.35 | 6.2E-12 | -1.8 | 0.28 | 4.9E-28 |
| *Y40A1A.6* | WBGene00304815 | ncRNA_gene | -1.5 | 0.35 | 1.0E-02 | -1.5 | 0.34 | 4.6E-03 |
| *rnr-1* | WBGene00004391 | protein_coding_gene | -1.5 | 0.34 | 2.9E-03 | -1.1 | 0.48 | 2.5E-02 |
| *mcm-3* | WBGene00003155 | protein_coding_gene | -1.5 | 0.34 | 1.6E-03 | -1.1 | 0.45 | 1.8E-02 |
| *perm-1* | WBGene00011350 | protein_coding_gene | -1.5 | 0.34 | 1.9E-02 | -1.5 | 0.36 | 7.9E-03 |
| *Y18D10A.11* | WBGene00012481 | protein_coding_gene | -1.6 | 0.34 | 4.1E-02 | -1.4 | 0.37 | 2.1E-02 |
| *cey-2* | WBGene00000473 | protein_coding_gene | -1.6 | 0.33 | 2.1E-02 | -1.5 | 0.35 | 3.7E-03 |
| *gei-14* | WBGene00001571 | protein_coding_gene | -1.6 | 0.33 | 1.5E-03 | -1.1 | 0.46 | 1.7E-02 |
| *K09D9.12* | WBGene00019569 | pseudogene | -1.6 | 0.33 | 4.9E-02 | -1.7 | 0.30 | 1.2E-02 |
| *F59A6.5* | WBGene00019087 | protein_coding_gene | -1.6 | 0.33 | 9.7E-04 | -1.5 | 0.37 | 5.3E-03 |
| *F23A7.4* | WBGene00009069 | protein_coding_gene | -1.6 | 0.33 | 1.4E-02 | -1.4 | 0.38 | 1.8E-03 |
| *aptf-2* | WBGene00013383 | protein_coding_gene | -1.6 | 0.32 | 2.9E-02 | -1.6 | 0.32 | 1.6E-02 |
| *fbxb-24* | WBGene00013234 | protein_coding_gene | -1.6 | 0.32 | 3.0E-02 | -1.8 | 0.29 | 1.4E-02 |
| *fbxb-118* | WBGene00021054 | protein_coding_gene | -1.6 | 0.32 | 4.7E-02 | -1.7 | 0.31 | 9.5E-03 |
| *gpd-1* | WBGene00001683 | protein_coding_gene | -1.6 | 0.32 | 1.3E-02 | -1.3 | 0.40 | 1.6E-02 |
| *scl-27* | WBGene00022638 | protein_coding_gene | -1.6 | 0.32 | 5.0E-02 | -2.2 | 0.22 | 1.7E-02 |
| *lin-41* | WBGene00003026 | protein_coding_gene | -1.6 | 0.32 | 5.2E-03 | -1.4 | 0.38 | 1.1E-02 |
| ***abhd-3.2*** | **WBGene00016644** | **protein_coding_gene** | **-1.7** | **0.31** | **2.8E-02** | **1.6** | **2.96** | **2.2E-02** |
| *mcm-2* | WBGene00003154 | protein_coding_gene | -1.7 | 0.31 | 4.7E-05 | -1.3 | 0.40 | 3.2E-04 |
| *F27B3.6* | WBGene00017848 | protein_coding_gene | -1.7 | 0.31 | 1.3E-02 | -1.5 | 0.35 | 3.9E-02 |
| *ZK673.5* | WBGene00014061 | protein_coding_gene | -1.7 | 0.31 | 2.1E-02 | -2.2 | 0.22 | 7.0E-04 |
| *W10G11.2* | WBGene00021135 | protein_coding_gene | -1.7 | 0.31 | 4.2E-02 | -2.0 | 0.26 | 9.1E-03 |
| *B0212.3* | WBGene00015037 | protein_coding_gene | -1.7 | 0.30 | 7.0E-10 | -1.2 | 0.43 | 1.3E-04 |
| *M117.7* | WBGene00044768 | pseudogene | -1.7 | 0.30 | 6.2E-03 | -1.5 | 0.35 | 2.6E-02 |
| *act-2* | WBGene00000064 | protein_coding_gene | -1.7 | 0.30 | 1.7E-05 | -1.1 | 0.46 | 7.4E-04 |
| *Y49E10.30* | WBGene00194981 | protein_coding_gene | -1.7 | 0.30 | 4.2E-02 | -1.4 | 0.39 | 2.6E-02 |
| *gfat-1* | WBGene00008546 | protein_coding_gene | -1.7 | 0.30 | 8.8E-03 | -1.0 | 0.49 | 4.2E-02 |
| *T05E12.8* | WBGene00045382 | protein_coding_gene | -1.8 | 0.29 | 1.5E-03 | -1.6 | 0.34 | 1.2E-02 |
| *wee-1.1* | WBGene00006938 | protein_coding_gene | -1.8 | 0.29 | 3.7E-03 | -1.5 | 0.36 | 1.7E-02 |
| *B0334.9* | WBGene00007147 | pseudogene | -1.8 | 0.29 | 3.8E-02 | -1.9 | 0.27 | 9.7E-03 |
| *F36F12.3* | WBGene00018094 | protein_coding_gene | -1.8 | 0.29 | 1.4E-02 | -2.1 | 0.24 | 2.2E-03 |
| *F21D5.1* | WBGene00009006 | protein_coding_gene | -1.8 | 0.28 | 2.5E-03 | -1.6 | 0.34 | 2.0E-03 |
| *his-67* | WBGene00001941 | protein_coding_gene | -1.8 | 0.28 | 2.9E-03 | -1.9 | 0.28 | 1.0E-02 |
| *C24H12.9* | WBGene00016078 | pseudogene | -1.9 | 0.27 | 9.4E-03 | -1.9 | 0.26 | 1.0E-02 |
| *gpd-4* | WBGene00001686 | protein_coding_gene | -1.9 | 0.27 | 2.0E-04 | -1.3 | 0.41 | 5.0E-03 |
| *T19H5.7* | WBGene00045414 | protein_coding_gene | -1.9 | 0.26 | 3.4E-03 | -1.6 | 0.34 | 3.9E-02 |
| *B0416.10* | WBGene00206482 | protein_coding_gene | -2.0 | 0.25 | 1.2E-02 | -1.5 | 0.35 | 1.0E-02 |
| *Y22D7AL.10* | WBGene00021248 | protein_coding_gene | -2.0 | 0.25 | 2.1E-08 | -1.3 | 0.41 | 1.6E-07 |
| *Y43D4A.6* | WBGene00012792 | protein_coding_gene | -2.0 | 0.25 | 1.7E-02 | -1.8 | 0.28 | 2.8E-02 |
| *Y44A6C.2* | WBGene00012851 | protein_coding_gene | -2.0 | 0.25 | 2.0E-02 | -1.8 | 0.28 | 1.2E-02 |
| *C06A1.4* | WBGene00014667 | pseudogene | -2.0 | 0.25 | 2.5E-03 | -1.2 | 0.43 | 9.4E-04 |
| *W04E12.2* | WBGene00012247 | protein_coding_gene | -2.0 | 0.24 | 1.1E-02 | -2.1 | 0.23 | 7.8E-04 |
| *F02H6.1* | WBGene00008534 | pseudogene | -2.1 | 0.24 | 1.6E-02 | -2.0 | 0.25 | 9.0E-03 |
| *F54D11.3* | WBGene00018813 | pseudogene | -2.1 | 0.24 | 1.6E-03 | -1.7 | 0.31 | 3.5E-03 |

| | | | | | | | | |
|---|---|---|---|---|---|---|---|---|
| *F55G11.2* | WBGene00010123 | protein_coding_gene | -2.2 | 0.22 | 5.0E-04 | -2.1 | 0.24 | 8.3E-04 |
| *ZK666.4* | WBGene00014044 | protein_coding_gene | -2.2 | 0.21 | 4.0E-03 | -2.2 | 0.22 | 7.3E-03 |
| *clec-74* | WBGene00021585 | protein_coding_gene | -2.2 | 0.21 | 9.8E-03 | -2.5 | 0.18 | 1.7E-05 |
| *F44E7.5* | WBGene00018427 | protein_coding_gene | -2.3 | 0.21 | 1.5E-05 | -1.4 | 0.39 | 9.5E-07 |
| *ckb-2* | WBGene00000512 | protein_coding_gene | -2.3 | 0.20 | 1.4E-08 | -1.1 | 0.45 | 1.1E-04 |
| *Y60A9.3* | WBGene00013370 | protein_coding_gene | -2.3 | 0.20 | 7.0E-03 | -2.1 | 0.24 | 4.3E-03 |
| *F07E5.12* | WBGene00219315 | protein_coding_gene | -2.4 | 0.19 | 1.5E-03 | -2.7 | 0.15 | 1.4E-03 |
| *M04D5.1* | WBGene00014807 | protein_coding_gene | -2.4 | 0.19 | 2.4E-03 | -1.8 | 0.29 | 2.4E-02 |
| *R11H6.6* | WBGene00014822 | pseudogene | -2.5 | 0.17 | 6.7E-05 | -1.7 | 0.32 | 1.1E-02 |
| *T10B5.8* | WBGene00020394 | protein_coding_gene | -2.6 | 0.17 | 5.7E-04 | -1.7 | 0.30 | 1.4E-02 |
| *drd-50* | WBGene00018643 | protein_coding_gene | -2.8 | 0.14 | 1.7E-03 | -1.9 | 0.27 | 5.4E-05 |
| *ZK6.11* | WBGene00022645 | protein_coding_gene | -2.8 | 0.14 | 1.2E-08 | -1.1 | 0.46 | 4.6E-02 |
| *C17F4.7* | WBGene00015913 | protein_coding_gene | -2.9 | 0.13 | 1.5E-05 | -2.0 | 0.26 | 1.4E-04 |
| *B0238.15* | WBGene00044537 | protein_coding_gene | -3.1 | 0.12 | 5.8E-05 | -2.1 | 0.24 | 1.3E-02 |
| *T24B8.5* | WBGene00011979 | protein_coding_gene | -3.3 | 0.10 | 4.3E-04 | -2.1 | 0.24 | 1.3E-02 |
| *clec-210* | WBGene00022261 | protein_coding_gene | -3.8 | 0.07 | 1.1E-08 | -1.6 | 0.33 | 5.8E-05 |
| *spp-17* | WBGene00005002 | protein_coding_gene | -4.6 | 0.04 | 1.2E-07 | -3.1 | 0.12 | 1.9E-04 |

*bold denotes genes that are differentially expressed in opposing directions

Transcription Factors Enriched in Gram-positive Infected Animals

| Gene Name | WormBase Gene ID | Sequence Name | N2 $log_2$FC | CB4856 $log_2$FC |
|---|---|---|---|---|
| *ceh-23* | WBGene00000446 | ZK652.5 | 2.4 | 2.2 |
| *cey-2* | WBGene00000473 | F46F11.2 | -1.6 | -1.5 |
| *dbl-1* | WBGene00000936 | T25F10.2 | 1.7 | 1.7 |
| *elt-2* | WBGene00001250 | C33D3.1 | 1.4 | 1.1 |
| *hlh-11* | WBGene00001955 | F58A4.7 | 1.3 | 1.3 |
| *lim-6* | WBGene00002988 | K03E6.1 | 1.3 | 1.0 |
| *lin-11* | WBGene00003000 | ZC247.3 | 2.1 | 1.3 |
| *lin-54* | WBGene00003037 | JC8.6 | -1.1 | -1.0 |
| *mab-9* | WBGene00003106 | T27A1.6 | 1.3 | 1.3 |
| *nhr-6* | WBGene00003605 | C48D5.1 | 2.2 | 1.5 |
| *nhr-117* | WBGene00003707 | F16B4.12 | 2.6 | 1.2 |
| *nhr-129* | WBGene00003719 | C50B6.14 | 1.5 | 1.4 |
| *pqm-1* | WBGene00004096 | F40F8.7 | 2.6 | 1.3 |
| *sem-4* | WBGene00004773 | F15C11.1 | 1.4 | 1.0 |
| *sox-3* | WBGene00004950 | F40E10.2 | 1.8 | 1.2 |
| *spr-3* | WBGene00005008 | C07A12.5 | -1.0 | -1.0 |
| *sys-1* | WBGene00006379 | T23D8.9 | -1.1 | -1.1 |
| *let-391* | WBGene00006492 | C27A12.3 | -1.1 | -1.1 |
| *ubc-9* | WBGene00006706 | F29B9.6 | -1.1 | -1.0 |
| *nfki-1* | WBGene00007877 | C33A11.1 | 1.3 | 1.1 |
| *dmd-10* | WBGene00007929 | C34D1.1 | 2.8 | 1.4 |
| *nhr-168* | WBGene00008221 | C50B6.8 | 2.4 | 2.2 |
| *D1046.2* | WBGene00008363 | D1046.2 | -1.3 | -1.0 |
| *F10B5.3* | WBGene00008640 | F10B5.3 | 4.4 | 3.1 |
| *ztf-2* | WBGene00008762 | F13G3.1 | 2.0 | 2.1 |
| *ztf-14* | WBGene00010936 | M163.2 | 1.7 | 1.4 |
| *ztf-15* | WBGene00011066 | R06C7.9 | -1.2 | -1.2 |
| *nhr-208* | WBGene00011099 | R07B7.15 | 2.4 | 1.5 |
| *zip-6* | WBGene00011130 | R07H5.10 | 3.2 | 2.2 |
| *sptf-2* | WBGene00011926 | T22C8.5 | 1.7 | 1.5 |
| *flh-1* | WBGene00012435 | Y11D7A.12 | -1.1 | -1.1 |
| *nhr-231* | WBGene00012449 | Y17D7B.1 | 1.4 | 1.4 |
| *sup-35* | WBGene00012976 | Y48A6C.3 | -1.4 | -1.1 |
| *Y60A9.3* | WBGene00013370 | Y60A9.3 | -2.3 | -2.1 |
| *aptf-2* | WBGene00013383 | Y62E10A.17 | -1.6 | -1.6 |
| *klu-1* | WBGene00013970 | ZK337.2 | 2.3 | 2.0 |
| *bcl-11* | WBGene00017430 | F13H6.1 | 1.5 | 1.4 |
| *nhr-179* | WBGene00017512 | F16B4.11 | 2.0 | 2.3 |

| | | | | |
|---|---|---|---|---|
| *ets-4* | WBGene00017687 | F22A3.1 | 1.5 | 1.8 |
| *oef-1* | WBGene00018636 | F49E8.2 | -1.2 | -1.1 |
| *klf-1* | WBGene00018990 | F56F11.3 | 1.5 | 1.2 |
| *R02D3.7* | WBGene00019824 | R02D3.7 | -1.1 | -1.1 |
| *nhr-216* | WBGene00020385 | T09D3.4 | 1.6 | 2.0 |
| *dmd-8* | WBGene00020708 | T22H9.4 | 1.8 | 2.2 |
| *Y22D7AL.16* | WBGene00021254 | Y22D7AL.16 | 2.1 | 2.1 |
| *nhr-277* | WBGene00022374 | Y94H6A.1 | 2.6 | 1.9 |
| *madf-9* | WBGene00022608 | ZC416.1 | 1.5 | 1.7 |
| *tag-243* | WBGene00044078 | T04A8.4 | 3.0 | 1.6 |
| *nhr-146* | WBGene00045255 | Y41D4B.27 | 1.1 | 1.2 |
| *maf-1* | WBGene00077521 | F45H11.6 | 1.5 | 1.7 |
| | | N2-Specific | | |
| *atf-2* | WBGene00000220 | K08F8.2 | 1.7 | Not DE |
| *bro-1* | WBGene00000272 | F56A3.5 | 2.3 | Not DE |
| *ceh-2* | WBGene00000429 | C27A12.5 | 1.7 | Not DE |
| *ceh-9* | WBGene00000434 | Y65B4BR.9 | 2.2 | Not DE |
| *ceh-22* | WBGene00000445 | F29F11.5 | 2.0 | Not DE |
| *pros-1* | WBGene00000448 | K12H4.1 | 1.8 | Not DE |
| *ceh-27* | WBGene00000449 | F46F3.1 | 1.1 | Not DE |
| *ceh-31* | WBGene00000452 | C33D12.1 | 1.4 | Not DE |
| *ceh-33* | WBGene00000454 | C10G8.7 | 1.5 | Not DE |
| *ceh-34* | WBGene00000455 | C10G8.6 | 1.6 | Not DE |
| *ceh-37* | WBGene00000458 | C37E2.5 | 1.3 | Not DE |
| *cfi-1* | WBGene00000476 | T23D8.8 | 1.6 | Not DE |
| *cky-1* | WBGene00000521 | C15C8.2 | 3.7 | Not DE |
| *daf-14* | WBGene00000910 | F01G10.8 | 1.4 | Not DE |
| *dnj-11* | WBGene00001029 | F38A5.13 | -1.2 | Not DE |
| *dpl-1* | WBGene00001061 | T23G7.1 | -1.1 | Not DE |
| *dro-1* | WBGene00001092 | F53A2.5 | -1.0 | Not DE |
| *eff-1* | WBGene00001159 | C26D10.5 | 1.5 | Not DE |
| *egl-5* | WBGene00001174 | C08C3.1 | 1.1 | Not DE |
| *egl-38* | WBGene00001204 | C04G2.7 | 1.5 | Not DE |
| *egl-43* | WBGene00001207 | R53.3 | 1.8 | Not DE |
| *egl-44* | WBGene00001208 | F28B12.2 | 1.5 | Not DE |
| *egl-46* | WBGene00001210 | K11G9.4 | 1.8 | Not DE |
| *elt-4* | WBGene00001252 | C39B10.6 | 1.9 | Not DE |
| *elt-6* | WBGene00001253 | F52C12.5 | 1.1 | Not DE |
| *end-1* | WBGene00001310 | F58E10.2 | -2.2 | Not DE |
| *fax-1* | WBGene00001400 | F56E3.4 | 1.7 | Not DE |
| *fkh-7* | WBGene00001439 | F26D12.1 | 1.6 | Not DE |
| *fkh-8* | WBGene00001440 | F40H3.4 | 1.7 | Not DE |
| *fkh-9* | WBGene00001441 | K03C7.2 | 1.6 | Not DE |
| *fkh-10* | WBGene00001442 | C25A1.2 | 2.1 | Not DE |
| *xnd-1* | WBGene00001514 | C05D2.5 | -1.0 | Not DE |
| *gei-13* | WBGene00001570 | F58A4.11 | 1.9 | Not DE |
| *grh-1* | WBGene00001707 | Y48G8AR.1 | 1.9 | Not DE |
| *ham-2* | WBGene00001821 | C07A12.1 | 1.4 | Not DE |
| *hbl-1* | WBGene00001824 | F13D11.2 | 1.8 | Not DE |
| *hlh-6* | WBGene00001952 | T15H9.3 | 1.8 | Not DE |
| *hlh-13* | WBGene00001957 | F48D6.3 | 2.6 | Not DE |
| *hsp-60* | WBGene00002025 | Y22D7AL.5 | -2.1 | Not DE |
| *isw-1* | WBGene00002169 | F37A4.8 | -1.1 | Not DE |
| *let-381* | WBGene00002601 | F26B1.7 | 1.3 | Not DE |
| *lim-4* | WBGene00002987 | ZC64.4 | 3.3 | Not DE |
| *lin-1* | WBGene00002990 | C37F5.1 | 1.3 | Not DE |
| *lin-39* | WBGene00003024 | C07H6.7 | 1.2 | Not DE |
| *lin-48* | WBGene00003033 | F34D10.5 | 2.0 | Not DE |

| | | | | |
|---|---|---|---|---|
| *mab-5* | WBGene00003102 | C08C3.3 | 2.0 | Not DE |
| *mbf-1* | WBGene00003148 | H21P03.1 | -1.1 | Not DE |
| *mdl-1* | WBGene00003163 | R03E9.1 | 1.8 | Not DE |
| *mec-3* | WBGene00003167 | F01D4.6 | 2.6 | Not DE |
| *med-2* | WBGene00003181 | K04C2.6 | 2.3 | Not DE |
| *mef-2* | WBGene00003182 | W10D5.1 | 1.5 | Not DE |
| *mig-5* | WBGene00003241 | T05C12.6 | -1.3 | Not DE |
| *moe-3* | WBGene00003388 | F32A11.6 | 1.6 | Not DE |
| *nhr-12* | WBGene00003611 | R04B5.4 | -1.1 | Not DE |
| *nhr-18* | WBGene00003617 | F44C8.3 | 2.2 | Not DE |
| *nhr-19* | WBGene00003618 | E02H1.7 | 2.7 | Not DE |
| *nhr-21* | WBGene00003620 | F21D12.1 | 2.5 | Not DE |
| *nhr-32* | WBGene00003626 | K08H2.8 | 1.7 | Not DE |
| *nhr-35* | WBGene00003628 | C07A12.3 | 2.0 | Not DE |
| *nhr-42* | WBGene00003632 | C33G8.6 | 1.7 | Not DE |
| *nhr-44* | WBGene00003634 | T19A5.4 | 1.7 | Not DE |
| *nhr-50* | WBGene00003640 | C06C6.5 | 1.3 | Not DE |
| *nhr-55* | WBGene00003645 | T01G6.7 | 1.6 | Not DE |
| *nhr-58* | WBGene00003648 | R11G11.2 | 1.4 | Not DE |
| *nhr-59* | WBGene00003649 | T27B7.1 | 1.5 | Not DE |
| *nhr-60* | WBGene00003650 | F57A10.5 | 1.3 | Not DE |
| *nhr-64* | WBGene00003654 | C45E1.1 | 1.1 | Not DE |
| *nhr-69* | WBGene00003659 | T23H4.2 | 1.2 | Not DE |
| *nhr-74* | WBGene00003664 | C27C7.3 | -1.7 | Not DE |
| *nhr-87* | WBGene00003677 | Y41D4B.7 | 1.6 | Not DE |
| *nhr-88* | WBGene00003678 | K08A2.5 | 1.5 | Not DE |
| *nhr-90* | WBGene00003680 | ZK488.2 | 1.1 | Not DE |
| *nhr-94* | WBGene00003684 | C12D5.8 | 1.6 | Not DE |
| *nhr-96* | WBGene00003686 | F44C8.11 | 1.2 | Not DE |
| *nhr-99* | WBGene00003689 | M02H5.1 | 3.2 | Not DE |
| *nhr-103* | WBGene00003693 | F44C8.4 | 1.5 | Not DE |
| *nhr-105* | WBGene00003695 | C06G3.1 | 1.0 | Not DE |
| *nhr-107* | WBGene00003697 | C33G8.11 | 1.5 | Not DE |
| *nhr-109* | WBGene00003699 | T12C9.5 | 1.0 | Not DE |
| *nhr-110* | WBGene00003700 | Y46H3D.5 | 1.1 | Not DE |
| *nhr-112* | WBGene00003702 | Y70C5C.6 | 1.6 | Not DE |
| *nhr-113* | WBGene00003703 | ZK1025.9 | 2.9 | Not DE |
| *nhr-116* | WBGene00003706 | F09C6.9 | 2.1 | Not DE |
| *nhr-120* | WBGene00003710 | C25B8.6 | 1.7 | Not DE |
| *nhr-121* | WBGene00003711 | E02H9.8 | 1.4 | Not DE |
| *nhr-122* | WBGene00003712 | Y41D4B.9 | 1.1 | Not DE |
| *nhr-125* | WBGene00003715 | R02D1.1 | 1.2 | Not DE |
| *nhr-126* | WBGene00003716 | F44C8.10 | 1.4 | Not DE |
| *nhr-127* | WBGene00003717 | T13F3.3 | 2.9 | Not DE |
| *nhr-128* | WBGene00003718 | F44C8.5 | 1.2 | Not DE |
| *nhr-133* | WBGene00003723 | F44C8.8 | 1.1 | Not DE |
| *nhr-136* | WBGene00003726 | C13C4.3 | 1.2 | Not DE |
| *pag-3* | WBGene00003909 | F45B8.4 | 1.6 | Not DE |
| *pat-9* | WBGene00003933 | T27B1.2 | 1.2 | Not DE |
| *pax-1* | WBGene00003937 | K07C11.1 | 1.8 | Not DE |
| *peb-1* | WBGene00003968 | T14F9.4 | 3.1 | Not DE |
| *pha-2* | WBGene00004011 | M6.3 | 1.5 | Not DE |
| *pha-4* | WBGene00004013 | F38A6.1 | 1.8 | Not DE |
| *php-3* | WBGene00004024 | Y75B8A.1 | 1.8 | Not DE |
| *plx-2* | WBGene00004048 | K04B12.1 | 1.6 | Not DE |
| *rnt-1* | WBGene00004393 | B0414.2 | 1.5 | Not DE |
| *rsp-6* | WBGene00004703 | C33H5.12 | -1.1 | Not DE |
| *sma-2* | WBGene00004856 | ZK370.2 | 1.7 | Not DE |

| | | | | |
|---|---|---|---|---|
| *sma-3* | WBGene00004857 | R13F6.9 | 1.2 | Not DE |
| *sma-4* | WBGene00004858 | R12B2.1 | 1.1 | Not DE |
| *sox-2* | WBGene00004949 | K08A8.2 | 1.5 | Not DE |
| *taf-6.1* | WBGene00006387 | W09B6.2 | -1.1 | Not DE |
| *taf-11.2* | WBGene00006394 | K10D3.3 | -1.6 | Not DE |
| *ctbp-1* | WBGene00006424 | F49E10.5 | 1.6 | Not DE |
| *svh-5* | WBGene00006462 | C33A11.4 | 1.3 | Not DE |
| *tbx-7* | WBGene00006544 | ZK328.8 | 1.5 | Not DE |
| *tbx-38* | WBGene00006557 | C24H11.3 | -2.2 | Not DE |
| *tbx-39* | WBGene00006558 | Y73F8A.16 | -2.4 | Not DE |
| *ttx-1* | WBGene00006652 | Y113G7A.6 | 1.8 | Not DE |
| *ttx-3* | WBGene00006654 | C40H5.5 | 1.9 | Not DE |
| *unc-3* | WBGene00006743 | Y16B4A.1 | 2.0 | Not DE |
| *unc-42* | WBGene00006778 | F58E6.10 | 3.1 | Not DE |
| *unc-86* | WBGene00006818 | C30A5.7 | 2.0 | Not DE |
| *vab-3* | WBGene00006870 | F14F3.1 | 1.6 | Not DE |
| *zag-1* | WBGene00006970 | F28F9.1 | 1.9 | Not DE |
| *zip-1* | WBGene00006986 | Y75B8A.35 | 1.1 | Not DE |
| *dmd-6* | WBGene00007058 | F13G11.1 | 1.1 | Not DE |
| *nhr-150* | WBGene00007367 | C06B8.1 | 2.0 | Not DE |
| *ceh-57* | WBGene00007416 | C07E3.5 | 2.3 | Not DE |
| *dmd-4* | WBGene00007776 | C27C12.6 | 1.5 | Not DE |
| *zip-5* | WBGene00007932 | C34D1.5 | 1.1 | Not DE |
| *athp-1* | WBGene00008081 | C44B9.4 | -1.2 | Not DE |
| *nhr-170* | WBGene00008309 | C54E10.5 | 1.3 | Not DE |
| *yap-1* | WBGene00008748 | F13E6.4 | 1.4 | Not DE |
| *aptf-3* | WBGene00009203 | F28C6.2 | -1.1 | Not DE |
| *F36F2.1* | WBGene00009475 | F36F2.1 | 1.1 | Not DE |
| *F38C2.7* | WBGene00009539 | F38C2.7 | -2.1 | Not DE |
| *hinf-1* | WBGene00009553 | F39B2.1 | -1.2 | Not DE |
| *psa-3* | WBGene00009560 | F39D8.2 | 1.5 | Not DE |
| *gmeb-2* | WBGene00010010 | F53H4.5 | 1.4 | Not DE |
| *nhr-193* | WBGene00010215 | F57G8.6 | 1.3 | Not DE |
| *fozi-1* | WBGene00010453 | K01B6.1 | 2.7 | Not DE |
| *K11H3.4* | WBGene00010781 | K11H3.4 | -1.3 | Not DE |
| *somi-1* | WBGene00010868 | M04G12.4 | 1.4 | Not DE |
| *nhr-206* | WBGene00011097 | R07B7.13 | 2.5 | Not DE |
| *gla-3* | WBGene00011376 | T02E1.3 | -1.3 | Not DE |
| *nhr-213* | WBGene00011520 | T06C12.13 | 1.2 | Not DE |
| *nhr-218* | WBGene00011750 | T13F3.2 | 2.2 | Not DE |
| *tsct-1* | WBGene00011824 | T18D3.7 | 1.5 | Not DE |
| *zip-10* | WBGene00012101 | T27F2.4 | 1.5 | Not DE |
| *Y17G7B.22* | WBGene00012473 | Y17G7B.22 | 1.5 | Not DE |
| *fezf-1* | WBGene00012639 | Y38H8A.5 | 2.1 | Not DE |
| *sta-1* | WBGene00013111 | Y51H4A.17 | 1.2 | Not DE |
| *znf-593* | WBGene00013236 | Y56A3A.18 | 1.3 | Not DE |
| *Y57A10A.31* | WBGene00013270 | Y57A10A.31 | -1.3 | Not DE |
| *madf-10* | WBGene00013717 | Y106G6H.4 | 1.2 | Not DE |
| *gtf-2A2* | WBGene00013736 | Y111B2A.13 | 1.4 | Not DE |
| *nhr-147* | WBGene00015395 | C03G6.8 | 1.6 | Not DE |
| *ztf-30* | WBGene00015523 | C06E1.8 | 2.1 | Not DE |
| *ceh-48* | WBGene00015934 | C17H12.9 | 1.4 | Not DE |
| *nhr-258* | WBGene00016126 | C26B2.4 | 1.7 | Not DE |
| *nhr-290* | WBGene00016233 | C29G2.5 | 2.2 | Not DE |
| *nhr-139* | WBGene00016365 | C33G8.8 | 1.3 | Not DE |
| *nhr-140* | WBGene00016366 | C33G8.9 | 1.5 | Not DE |
| *nhr-163* | WBGene00016368 | C33G8.12 | 1.4 | Not DE |
| *unk-1* | WBGene00016407 | C34D10.2 | 2.3 | Not DE |

| | | | | |
|---|---|---|---|---|
| *ets-9* | WBGene00016865 | C52B9.2 | 1.2 | Not DE |
| *ets-10* | WBGene00016870 | C52B9.11 | 1.9 | Not DE |
| *znf-598* | WBGene00016888 | C52E12.1 | -1.0 | Not DE |
| *nhr-173* | WBGene00016975 | C56E10.1 | 1.7 | Not DE |
| *cebp-1* | WBGene00016997 | D1005.3 | 1.2 | Not DE |
| *nhr-178* | WBGene00017510 | F16B4.9 | 2.5 | Not DE |
| *atf-8* | WBGene00017535 | F17A9.3 | 1.7 | Not DE |
| *F21A9.2* | WBGene00017651 | F21A9.2 | 1.1 | Not DE |
| *nfyc-1* | WBGene00017742 | F23F1.1 | -1.0 | Not DE |
| *F26A10.2* | WBGene00017814 | F26A10.2 | 2.0 | Not DE |
| *nhr-180* | WBGene00017961 | F31F4.12 | 1.7 | Not DE |
| *nhr-183* | WBGene00018266 | F41B5.10 | 1.1 | Not DE |
| *nhr-37* | WBGene00018412 | F44C4.2 | 2.8 | Not DE |
| *ceh-82* | WBGene00018433 | F45C12.2 | -1.9 | Not DE |
| *mnat-1* | WBGene00018769 | F53G2.7 | -1.4 | Not DE |
| *nhr-195* | WBGene00019115 | F59E11.10 | 1.1 | Not DE |
| *nhr-143* | WBGene00019116 | F59E11.11 | 1.5 | Not DE |
| *zip-2* | WBGene00019327 | K02F3.4 | 1.5 | Not DE |
| *aptf-1* | WBGene00019424 | K06A1.1 | 1.8 | Not DE |
| *nhr-201* | WBGene00019741 | M02H5.3 | 1.2 | Not DE |
| *M03D4.4* | WBGene00019751 | M03D4.4 | 2.0 | Not DE |
| *camt-1* | WBGene00020251 | T05C1.4 | 1.1 | Not DE |
| *ast-1* | WBGene00020368 | T08H4.3 | 1.9 | Not DE |
| *ztf-4* | WBGene00020399 | T10B11.3 | -1.3 | Not DE |
| *dhhc-11* | WBGene00020694 | T22E7.2 | 2.1 | Not DE |
| *nhr-221* | WBGene00020748 | T24A6.8 | 2.4 | Not DE |
| *nhr-227* | WBGene00020851 | T27B7.5 | 1.4 | Not DE |
| *hlh-30* | WBGene00020930 | W02C12.3 | 1.6 | Not DE |
| *W04B5.2* | WBGene00021019 | W04B5.2 | 1.8 | Not DE |
| *nhr-275* | WBGene00021163 | Y5H2A.2 | 1.9 | Not DE |
| *crtc-1* | WBGene00021237 | Y20F4.2 | 1.5 | Not DE |
| *Y37E11B.1* | WBGene00021374 | Y37E11B.1 | 1.2 | Not DE |
| *nhr-236* | WBGene00021417 | Y38F2AL.5 | 1.7 | Not DE |
| *hlh-33* | WBGene00021446 | Y39A3CR.6 | 1.6 | Not DE |
| *ceh-45* | WBGene00022837 | ZK993.1 | 1.7 | Not DE |
| *dve-1* | WBGene00022861 | ZK1193.5 | 1.5 | Not DE |
| *syd-9* | WBGene00044068 | ZK867.1 | 1.4 | Not DE |
| *alr-1* | WBGene00044330 | R08B4.2 | 1.6 | Not DE |
| *bnc-1* | WBGene00044791 | F55C5.11 | 2.4 | Not DE |
| *hlh-26* | WBGene00001965 | C17C3.8 | -2.8 | Not DE |
| | | CB4856-Specific | | |
| *ceh-5* | WBGene00000430 | C16C2.1 | Not DE | -2.1 |
| *ceh-18* | WBGene00000441 | ZC64.3 | Not DE | 1.3 |
| *ceh-43* | WBGene00000463 | C28A5.4 | Not DE | -1.3 |
| *cey-3* | WBGene00000474 | M01E11.5 | Not DE | -1.2 |
| *daf-12* | WBGene00000908 | F11A1.3 | Not DE | 1.1 |
| *dpy-20* | WBGene00001079 | T22B3.1 | Not DE | 2.6 |
| *egl-13* | WBGene00001182 | T22B7.1 | Not DE | 1.0 |
| *fkh-6* | WBGene00001438 | B0286.5 | Not DE | 1.3 |
| *ham-1* | WBGene00001820 | F53B2.6 | Not DE | -1.0 |
| *lin-29* | WBGene00003015 | W03C9.4 | Not DE | 1.9 |
| *lin-31* | WBGene00003017 | K10G6.1 | Not DE | 1.7 |
| *lir-3* | WBGene00003046 | F37H8.1 | Not DE | -1.1 |
| *mab-10* | WBGene00003107 | R166.1 | Not DE | 1.5 |
| *nhr-7* | WBGene00003606 | F54D1.4 | Not DE | 1.4 |
| *nhr-14* | WBGene00003613 | T01B10.4 | Not DE | 1.3 |
| *nhr-31* | WBGene00003625 | C26B2.3 | Not DE | 1.8 |
| *nhr-93* | WBGene00003683 | T24A6.9 | Not DE | 1.8 |

| | | | | |
|---|---|---|---|---|
| *nhr-132* | WBGene00003722 | R11G11.1 | Not DE | 1.0 |
| *odd-1* | WBGene00003845 | B0280.4 | Not DE | -1.0 |
| *pax-2* | WBGene00003938 | K06B9.5 | Not DE | 1.8 |
| *pie-1* | WBGene00004027 | Y49E10.14 | Not DE | -1.1 |
| *pqn-75* | WBGene00004157 | W03D2.1 | Not DE | 1.2 |
| *ref-1* | WBGene00004334 | T01E8.2 | Not DE | -1.3 |
| *taf-9* | WBGene00006391 | T12D8.7 | Not DE | -1.0 |
| *tra-1* | WBGene00006604 | Y47D3A.6 | Not DE | 1.0 |
| *unc-4* | WBGene00006744 | F26C11.2 | Not DE | 1.3 |
| *unc-55* | WBGene00006790 | F55D12.4 | Not DE | 1.1 |
| *B0250.4* | WBGene00007121 | B0250.4 | Not DE | 1.2 |
| *ceh-58* | WBGene00007417 | C07E3.6 | Not DE | -1.0 |
| *C35D6.4* | WBGene00007961 | C35D6.4 | Not DE | -1.8 |
| *C38D4.7* | WBGene00008007 | C38D4.7 | Not DE | 1.3 |
| *gmeb-3* | WBGene00008092 | C44F1.2 | Not DE | 1.6 |
| *madf-8* | WBGene00008118 | C46F11.3 | Not DE | -1.1 |
| *ccnk-1* | WBGene00009650 | F43D2.1 | Not DE | -1.0 |
| *efl-3* | WBGene00009899 | F49E12.6 | Not DE | -1.1 |
| *F52B5.7* | WBGene00009923 | F52B5.7 | Not DE | -1.1 |
| *dhhc-12* | WBGene00010323 | F59C6.2 | Not DE | 1.2 |
| *nhr-198* | WBGene00010603 | K06B4.8 | Not DE | 1.9 |
| *nhr-207* | WBGene00011098 | R07B7.14 | Not DE | 2.0 |
| *T01C3.2* | WBGene00011319 | T01C3.2 | Not DE | -1.0 |
| *nhr-234* | WBGene00012596 | Y38E10A.18 | Not DE | 1.9 |
| *Y116A8C.20* | WBGene00013797 | Y116A8C.20 | Not DE | -2.0 |
| *ceh-74* | WBGene00013876 | ZC376.4 | Not DE | -1.1 |
| *B0336.13* | WBGene00015150 | B0336.13 | Not DE | -1.0 |
| *C06E2.1* | WBGene00015527 | C06E2.1 | Not DE | 1.4 |
| *ceh-53* | WBGene00015651 | C09G12.1 | Not DE | -1.3 |
| *sox-4* | WBGene00015716 | C12D12.5 | Not DE | 1.0 |
| *nhr-155* | WBGene00015758 | C14C6.4 | Not DE | 2.1 |
| *nhr-157* | WBGene00015900 | C17E7.5 | Not DE | 2.1 |
| *nhr-30* | WBGene00016091 | C25E10.1 | Not DE | 1.7 |
| *nhr-260* | WBGene00016517 | C38C3.9 | Not DE | 1.4 |
| *dmd-5* | WBGene00017326 | F10C1.5 | Not DE | -1.4 |
| *ceh-60* | WBGene00017690 | F22A3.5 | Not DE | 1.1 |
| *nhr-141* | WBGene00017787 | F25E5.6 | Not DE | 1.2 |
| *ceh-86* | WBGene00018355 | F42G2.6 | Not DE | -1.3 |
| *npax-2* | WBGene00018591 | F48B9.5 | Not DE | 1.5 |
| *nhr-266* | WBGene00018993 | F56H1.2 | Not DE | 1.5 |
| *dhhc-13* | WBGene00019257 | H32C10.3 | Not DE | 1.2 |
| *dhhc-5* | WBGene00020066 | R13F6.5 | Not DE | 1.1 |
| *ssup-72* | WBGene00020480 | T13C2.4 | Not DE | -1.0 |
| *nhr-219* | WBGene00020555 | T19A5.5 | Not DE | 2.4 |
| *ztf-31* | WBGene00020635 | T20H4.2 | Not DE | 1.2 |
| *T26A5.8* | WBGene00020823 | T26A5.8 | Not DE | -1.0 |
| *znf-782* | WBGene00021931 | Y55F3AM.14 | Not DE | 2.5 |
| *duxl-1* | WBGene00022554 | ZC204.2 | Not DE | -1.3 |
| *klu-2* | WBGene00022592 | ZC328.2 | Not DE | 1.7 |
| *ZK686.5* | WBGene00022795 | ZK686.5 | Not DE | 1.3 |
| *Y53C10A.15* | WBGene00044805 | Y53C10A.15 | Not DE | 1.1 |

**Supplemental Table 5**

**Lists of differentially expressed genes that are identified in either N2, CB4856, or both, after infection Gram-negative pathogens (compared to *E. coli*)**

**Supplemental Table 5 - Genes Differentially Expressed in Gram-negative Infected Animals**

**N2**-exclusive DEGs in response to Gram-negative pathogens (*P. rettgeri* or *P. aeruginosa* vs. OP50)

| Gene Name | Wormbase Gene ID | Type | $log_2FC$ | FC | Adj P-value |
|---|---|---|---|---|---|
| *T26H5.9* | WBGene00044206 | protein_coding_gene | 4.0 | 16.40 | 3.8E-07 |
| *F53A9.8* | WBGene00018731 | protein_coding_gene | 3.6 | 12.14 | 1.2E-06 |
| *C06B3.7* | WBGene00007366 | protein_coding_gene | 3.5 | 11.48 | 2.0E-04 |
| *T24C4.4* | WBGene00020760 | protein_coding_gene | 3.4 | 10.20 | 1.6E-03 |
| *linc-37* | WBGene00219709 | lincRNA_gene | 3.2 | 9.43 | 7.8E-03 |
| *T26H5.14* | WBGene00235133 | protein_coding_gene | 3.2 | 9.42 | 2.3E-03 |
| *F08G2.5* | WBGene00008577 | protein_coding_gene | 3.1 | 8.59 | 6.3E-03 |
| *Y26D4A.3* | WBGene00012501 | protein_coding_gene | 3.0 | 7.78 | 6.6E-03 |
| *tsp-1* | WBGene00006627 | protein_coding_gene | 2.9 | 7.32 | 5.0E-03 |
| *T26H5.4* | WBGene00012069 | protein_coding_gene | 2.8 | 6.95 | 5.4E-03 |
| *dod-22* | WBGene00010125 | protein_coding_gene | 2.7 | 6.72 | 9.8E-03 |
| *tsp-2* | WBGene00006628 | protein_coding_gene | 2.7 | 6.67 | 7.9E-03 |
| *cyp-32B1* | WBGene00021167 | protein_coding_gene | 2.7 | 6.56 | 1.3E-03 |
| *irg-2* | WBGene00016783 | protein_coding_gene | 2.7 | 6.56 | 1.7E-02 |
| *fbxa-24* | WBGene00018803 | protein_coding_gene | 2.7 | 6.28 | 1.6E-03 |
| *C25F9.11* | WBGene00045411 | protein_coding_gene | 2.6 | 6.11 | 5.8E-03 |
| *F21C10.11* | WBGene00017660 | protein_coding_gene | 2.6 | 5.94 | 3.2E-03 |
| *C35A5.11* | WBGene00044024 | protein_coding_gene | 2.5 | 5.77 | 1.9E-02 |
| *Y37A1B.7* | WBGene00012540 | protein_coding_gene | 2.5 | 5.69 | 9.8E-03 |
| *nlp-39* | WBGene00008295 | protein_coding_gene | 2.5 | 5.64 | 4.8E-02 |
| *F11C7.6* | WBGene00017378 | protein_coding_gene | 2.5 | 5.57 | 6.1E-03 |
| *C45B11.2* | WBGene00008105 | protein_coding_gene | 2.5 | 5.54 | 2.0E-02 |
| *clec-17* | WBGene00008477 | protein_coding_gene | 2.5 | 5.53 | 1.6E-02 |
| *ZK1290.10* | WBGene00022891 | protein_coding_gene | 2.5 | 5.52 | 2.2E-02 |
| *lipl-2* | WBGene00009773 | protein_coding_gene | 2.4 | 5.43 | 2.4E-03 |
| *T26H5.10* | WBGene00044783 | protein_coding_gene | 2.4 | 5.30 | 3.4E-02 |
| *K05B2.4* | WBGene00019404 | protein_coding_gene | 2.4 | 5.28 | 3.4E-02 |
| *Y54G2A.76* | WBGene00235268 | protein_coding_gene | 2.4 | 5.25 | 9.3E-03 |
| *wrt-2* | WBGene00006948 | protein_coding_gene | 2.4 | 5.17 | 4.8E-04 |
| *srz-99* | WBGene00012028 | protein_coding_gene | 2.3 | 5.00 | 3.8E-02 |
| *T04C12.1* | WBGene00011427 | protein_coding_gene | 2.3 | 4.83 | 4.8E-02 |
| *C04G6.5* | WBGene00015455 | protein_coding_gene | 2.3 | 4.82 | 4.2E-02 |
| *K10G6.4* | WBGene00019641 | protein_coding_gene | 2.2 | 4.75 | 5.0E-02 |
| *Y58A7A.4* | WBGene00021978 | protein_coding_gene | 2.2 | 4.74 | 3.4E-02 |
| *Y39A3A.3* | WBGene00021434 | protein_coding_gene | 2.2 | 4.71 | 5.0E-02 |
| *btb-21* | WBGene00018540 | pseudogene | 2.2 | 4.70 | 1.9E-03 |
| *C06E7.88* | WBGene00189995 | protein_coding_gene | 2.2 | 4.69 | 4.6E-02 |
| *best-5* | WBGene00007404 | protein_coding_gene | 2.2 | 4.68 | 3.2E-03 |
| *Y71A12B.19* | WBGene00077443 | pseudogene | 2.2 | 4.66 | 8.5E-03 |
| *ZC395.5* | WBGene00022597 | protein_coding_gene | 2.2 | 4.66 | 2.5E-03 |
| *T02G5.15* | WBGene00206521 | protein_coding_gene | 2.2 | 4.57 | 6.8E-03 |
| *fbxa-128* | WBGene00012923 | protein_coding_gene | 2.2 | 4.56 | 5.7E-03 |
| *F31E3.12* | WBGene00219317 | protein_coding_gene | 2.2 | 4.50 | 2.0E-02 |
| *C50F4.9* | WBGene00008234 | protein_coding_gene | 2.1 | 4.32 | 2.4E-03 |
| *kreg-1* | WBGene00018725 | protein_coding_gene | 2.1 | 4.31 | 3.9E-02 |
| *Y43F8B.23* | WBGene00194815 | protein_coding_gene | 2.1 | 4.24 | 1.4E-02 |
| *parg-2* | WBGene00004052 | protein_coding_gene | 2.1 | 4.20 | 2.9E-02 |
| *Y39A3A.4* | WBGene00021435 | protein_coding_gene | 2.1 | 4.19 | 2.0E-02 |
| *hil-7* | WBGene00001858 | protein_coding_gene | 2.1 | 4.16 | 4.3E-02 |
| *Y53F4B.45* | WBGene00044728 | ncRNA_gene | 2.1 | 4.15 | 7.6E-03 |
| *ugt-18* | WBGene00013900 | protein_coding_gene | 2.0 | 4.14 | 3.2E-03 |
| *fbxa-48* | WBGene00021862 | protein_coding_gene | 2.0 | 4.12 | 2.9E-02 |
| *F15B9.6* | WBGene00008850 | protein_coding_gene | 2.0 | 3.99 | 3.8E-02 |

| | | | | | |
|---|---|---|---|---|---|
| *T09E11.11* | WBGene00011660 | pseudogene | 2.0 | 3.88 | 3.8E-02 |
| *C06B3.6* | WBGene00007365 | protein_coding_gene | 2.0 | 3.86 | 8.6E-03 |
| *fbxc-20* | WBGene00018352 | protein_coding_gene | 1.9 | 3.83 | 7.1E-03 |
| *ttr-23* | WBGene00011894 | protein_coding_gene | 1.9 | 3.81 | 4.2E-02 |
| *valv-1* | WBGene00015216 | protein_coding_gene | 1.9 | 3.77 | 4.4E-02 |
| *fbxa-21* | WBGene00022487 | protein_coding_gene | 1.9 | 3.75 | 1.6E-03 |
| *Y110A2AL.4* | WBGene00022441 | protein_coding_gene | 1.9 | 3.71 | 1.6E-02 |
| *F10E9.12* | WBGene00017361 | protein_coding_gene | 1.9 | 3.66 | 3.6E-03 |
| *F11E6.9* | WBGene00008712 | protein_coding_gene | 1.9 | 3.64 | 4.5E-02 |
| *Y49E10.7* | WBGene00013031 | pseudogene | 1.9 | 3.62 | 4.2E-02 |
| *dao-2* | WBGene00000928 | protein_coding_gene | 1.8 | 3.57 | 8.8E-03 |
| *Y69A2AL.2* | WBGene00044468 | protein_coding_gene | 1.8 | 3.50 | 1.4E-02 |
| *F54B8.4* | WBGene00010019 | protein_coding_gene | 1.8 | 3.48 | 4.9E-02 |
| *E03H4.8* | WBGene00008476 | protein_coding_gene | 1.8 | 3.48 | 1.6E-03 |
| *fmo-1* | WBGene00001476 | protein_coding_gene | 1.8 | 3.47 | 4.1E-02 |
| *Y22D7AL.15* | WBGene00021253 | protein_coding_gene | 1.8 | 3.46 | 5.0E-02 |
| *F59A7.7* | WBGene00019094 | protein_coding_gene | 1.8 | 3.44 | 5.0E-02 |
| *W02A2.9* | WBGene00044280 | protein_coding_gene | 1.8 | 3.42 | 2.9E-02 |
| *gbh-2* | WBGene00001523 | protein_coding_gene | 1.8 | 3.40 | 1.0E-03 |
| *K09H11.16* | WBGene00269384 | ncRNA_gene | 1.7 | 3.29 | 3.0E-02 |
| *Y54F10BM.12* | WBGene00021866 | protein_coding_gene | 1.7 | 3.28 | 1.1E-02 |
| *lipl-1* | WBGene00010062 | protein_coding_gene | 1.7 | 3.27 | 3.4E-02 |
| *Y48G9A.7* | WBGene00021701 | protein_coding_gene | 1.7 | 3.25 | 2.2E-02 |
| *Y17D7C.4* | WBGene00050903 | pseudogene | 1.7 | 3.24 | 2.7E-02 |
| *srv-7* | WBGene00005718 | protein_coding_gene | 1.7 | 3.23 | 4.3E-02 |
| *Y54G2A.36* | WBGene00021898 | protein_coding_gene | 1.7 | 3.23 | 3.2E-02 |
| *cest-29* | WBGene00017205 | pseudogene | 1.7 | 3.22 | 2.1E-02 |
| *Y111B2A.21* | WBGene00013741 | protein_coding_gene | 1.7 | 3.21 | 2.0E-03 |
| *C52E2.4* | WBGene00016883 | protein_coding_gene | 1.7 | 3.20 | 3.6E-02 |
| *gly-14* | WBGene00001639 | protein_coding_gene | 1.7 | 3.18 | 3.5E-02 |
| *C06E1.11* | WBGene00015526 | protein_coding_gene | 1.7 | 3.18 | 2.0E-02 |
| *F14D2.14* | WBGene00017462 | protein_coding_gene | 1.7 | 3.17 | 2.0E-02 |
| *fbxa-198* | WBGene00011514 | pseudogene | 1.7 | 3.16 | 6.1E-03 |
| *C16C4.18* | WBGene00194998 | protein_coding_gene | 1.7 | 3.15 | 6.1E-03 |
| *F58F12.12* | WBGene00271812 | protein_coding_gene | 1.6 | 3.14 | 3.1E-02 |
| *F32D8.13* | WBGene00009335 | protein_coding_gene | 1.6 | 3.13 | 1.7E-02 |
| *F13H8.1* | WBGene00017434 | protein_coding_gene | 1.6 | 3.05 | 2.7E-02 |
| *C50B8.4* | WBGene00008226 | protein_coding_gene | 1.6 | 3.05 | 5.0E-02 |
| *fbxc-46* | WBGene00012569 | pseudogene | 1.6 | 3.04 | 2.4E-03 |
| *fbxa-217* | WBGene00012954 | pseudogene | 1.6 | 3.02 | 1.4E-02 |
| *C14F11.6* | WBGene00015781 | protein_coding_gene | 1.6 | 3.01 | 2.6E-02 |
| *Y32G9A.13* | WBGene00044517 | pseudogene | 1.6 | 2.96 | 2.9E-02 |
| *ugt-31* | WBGene00021464 | protein_coding_gene | 1.6 | 2.95 | 1.2E-02 |
| *BE0003N10.6* | WBGene00255598 | protein_coding_gene | 1.6 | 2.95 | 4.4E-02 |
| *Y69A2AR.12* | WBGene00022083 | protein_coding_gene | 1.5 | 2.90 | 2.6E-02 |
| *F54D10.4* | WBGene00018805 | protein_coding_gene | 1.5 | 2.89 | 1.3E-02 |
| *M162.9* | WBGene00014811 | pseudogene | 1.5 | 2.88 | 3.9E-02 |
| *pals-23* | WBGene00016218 | protein_coding_gene | 1.5 | 2.88 | 1.4E-02 |
| *T16G1.5* | WBGene00011799 | protein_coding_gene | 1.5 | 2.84 | 4.2E-02 |
| *F56D2.3* | WBGene00018965 | protein_coding_gene | 1.5 | 2.83 | 3.4E-02 |
| *bath-35* | WBGene00012196 | protein_coding_gene | 1.5 | 2.82 | 4.7E-02 |
| *F40D4.17* | WBGene00194682 | protein_coding_gene | 1.5 | 2.79 | 2.7E-02 |
| *T28H10.1* | WBGene00012142 | protein_coding_gene | 1.5 | 2.79 | 1.4E-02 |
| *F52C6.3* | WBGene00018660 | protein_coding_gene | 1.5 | 2.78 | 3.3E-03 |
| *fbxa-212* | WBGene00012568 | pseudogene | 1.5 | 2.75 | 1.4E-02 |
| *F52C6.4* | WBGene00018661 | protein_coding_gene | 1.5 | 2.75 | 1.7E-02 |
| *F14D2.11* | WBGene00017459 | protein_coding_gene | 1.4 | 2.71 | 4.7E-02 |
| *C52E12.11* | WBGene00304189 | pseudogene | 1.4 | 2.65 | 1.4E-02 |

| | | | | | |
|---|---|---|---|---|---|
| *pals-24* | WBGene00016219 | protein_coding_gene | 1.4 | 2.64 | 9.3E-03 |
| *btb-1* | WBGene00001963 | pseudogene | 1.4 | 2.61 | 4.3E-02 |
| *Y55F3AM.11* | WBGene00021928 | protein_coding_gene | 1.4 | 2.60 | 4.6E-02 |
| *tag-234* | WBGene00018874 | protein_coding_gene | 1.4 | 2.60 | 4.4E-02 |
| *Y50D4A.5* | WBGene00021739 | protein_coding_gene | 1.4 | 2.60 | 1.2E-02 |
| *fbxa-140* | WBGene00008130 | protein_coding_gene | 1.4 | 2.58 | 3.6E-02 |
| *btb-16* | WBGene00018195 | protein_coding_gene | 1.4 | 2.57 | 2.0E-03 |
| *fbxa-105* | WBGene00015362 | protein_coding_gene | 1.4 | 2.57 | 4.4E-02 |
| *wrt-10* | WBGene00006956 | protein_coding_gene | 1.4 | 2.57 | 3.4E-02 |
| *anr-8* | WBGene00219575 | antisense_lncRNA_gene | 1.4 | 2.55 | 1.7E-02 |
| *Y37H2A.18* | WBGene00219210 | protein_coding_gene | 1.3 | 2.54 | 4.2E-02 |
| *D2096.10* | WBGene00017077 | protein_coding_gene | 1.3 | 2.54 | 4.6E-02 |
| *crn-2* | WBGene00000795 | protein_coding_gene | 1.3 | 2.53 | 2.1E-02 |
| *Y37H9A.5* | WBGene00014871 | protein_coding_gene | 1.3 | 2.53 | 3.5E-02 |
| *Y17G9B.4* | WBGene00021201 | protein_coding_gene | 1.3 | 2.52 | 4.5E-02 |
| *F14D2.8* | WBGene00017456 | protein_coding_gene | 1.3 | 2.51 | 2.2E-02 |
| *fbxa-211* | WBGene00012566 | protein_coding_gene | 1.3 | 2.50 | 9.0E-03 |
| *F54C4.4* | WBGene00018795 | protein_coding_gene | 1.3 | 2.50 | 7.2E-03 |
| *C17E4.11* | WBGene00044236 | protein_coding_gene | 1.3 | 2.49 | 3.3E-02 |
| *ift-43* | WBGene00016121 | protein_coding_gene | 1.3 | 2.47 | 5.4E-03 |
| *fbxa-124* | WBGene00014916 | pseudogene | 1.3 | 2.45 | 1.6E-02 |
| *bath-34* | WBGene00012195 | protein_coding_gene | 1.3 | 2.44 | 3.2E-02 |
| *fbxa-108* | WBGene00010293 | protein_coding_gene | 1.3 | 2.42 | 3.2E-03 |
| *fbxa-191* | WBGene00010209 | pseudogene | 1.3 | 2.39 | 4.0E-02 |
| *T04F8.7* | WBGene00011445 | protein_coding_gene | 1.3 | 2.39 | 4.8E-02 |
| *T09F3.4* | WBGene00011663 | protein_coding_gene | 1.2 | 2.37 | 5.1E-03 |
| *B0250.12* | WBGene00050927 | pseudogene | 1.2 | 2.36 | 4.0E-02 |
| *sws-1* | WBGene00012698 | protein_coding_gene | 1.2 | 2.31 | 4.9E-02 |
| *C08F11.6* | WBGene00014672 | pseudogene | 1.2 | 2.31 | 2.1E-02 |
| *mks-6* | WBGene00010642 | protein_coding_gene | 1.2 | 2.28 | 4.5E-02 |
| *C46G7.5* | WBGene00044502 | protein_coding_gene | 1.2 | 2.27 | 2.1E-02 |
| *spe-5* | WBGene00004959 | protein_coding_gene | 1.2 | 2.27 | 5.0E-02 |
| *Y110A2AL.3* | WBGene00022440 | protein_coding_gene | 1.2 | 2.25 | 3.4E-02 |
| *C24F3.2* | WBGene00007697 | protein_coding_gene | 1.2 | 2.24 | 3.4E-02 |
| *fbxl-1* | WBGene00015350 | protein_coding_gene | 1.2 | 2.23 | 3.4E-02 |
| *F13G3.6* | WBGene00008766 | protein_coding_gene | 1.1 | 2.22 | 2.0E-02 |
| *fbxa-122* | WBGene00014917 | pseudogene | 1.1 | 2.22 | 4.9E-02 |
| *Y71F9AL.8* | WBGene00022113 | protein_coding_gene | 1.1 | 2.20 | 2.1E-02 |
| *F08B4.3* | WBGene00017235 | pseudogene | 1.1 | 2.19 | 3.6E-02 |
| *F52C6.12* | WBGene00018669 | protein_coding_gene | 1.1 | 2.19 | 1.4E-02 |
| *F26F4.8* | WBGene00005011 | protein_coding_gene | 1.1 | 2.17 | 4.3E-02 |
| *mtcu-1* | WBGene00009557 | protein_coding_gene | 1.1 | 2.16 | 3.4E-02 |
| *R05D3.3* | WBGene00019878 | protein_coding_gene | 1.1 | 2.14 | 1.6E-02 |
| *JC8.4* | WBGene00010436 | protein_coding_gene | 1.1 | 2.11 | 4.9E-02 |
| *F35G12.5* | WBGene00009442 | protein_coding_gene | 1.1 | 2.11 | 4.9E-02 |
| *ZK1127.12* | WBGene00022857 | protein_coding_gene | 1.1 | 2.10 | 3.2E-02 |
| *R119.1* | WBGene00020087 | pseudogene | 1.1 | 2.09 | 4.2E-02 |
| *nhr-116* | WBGene00003706 | protein_coding_gene | 1.1 | 2.08 | 5.0E-02 |
| *R02D3.8* | WBGene00019825 | protein_coding_gene | 1.0 | 2.07 | 4.7E-02 |
| *fbxc-45* | WBGene00021887 | protein_coding_gene | 1.0 | 2.06 | 3.3E-02 |
| *T25B9.8* | WBGene00012014 | protein_coding_gene | 1.0 | 2.06 | 5.0E-02 |
| *fbxa-114* | WBGene00013758 | protein_coding_gene | 1.0 | 2.06 | 4.2E-02 |
| *C46A5.6* | WBGene00016702 | protein_coding_gene | 1.0 | 2.05 | 4.3E-02 |
| *Y57A10A.7* | WBGene00013252 | protein_coding_gene | 1.0 | 2.04 | 7.7E-03 |
| *tsen-34* | WBGene00019535 | protein_coding_gene | 1.0 | 2.03 | 4.1E-02 |
| *W07E6.5* | WBGene00021076 | protein_coding_gene | 1.0 | 2.02 | 3.3E-02 |
| *Y105E8B.6* | WBGene00013691 | protein_coding_gene | 1.0 | 2.02 | 4.6E-02 |
| *R11G1.7* | WBGene00044725 | protein_coding_gene | 1.0 | 2.02 | 3.9E-02 |

| | | | | | |
|---|---|---|---|---|---|
| *C08F11.7* | WBGene00007454 | protein_coding_gene | 1.0 | 2.02 | 4.6E-02 |
| *Y54G2A.50* | WBGene00044493 | protein_coding_gene | 1.0 | 2.01 | 3.2E-02 |
| *T16H12.1* | WBGene00011811 | protein_coding_gene | 1.0 | 2.01 | 4.2E-02 |
| *tsen-2* | WBGene00013231 | protein_coding_gene | 1.0 | 2.01 | 3.9E-02 |
| *F52C12.6* | WBGene00138717 | protein_coding_gene | 1.0 | 2.00 | 3.3E-02 |
| *Y34B4A.5* | WBGene00021321 | protein_coding_gene | -1.0 | 0.50 | 2.5E-02 |
| *cest-2.2* | WBGene00013874 | protein_coding_gene | -1.0 | 0.50 | 4.3E-02 |
| *F42A10.9* | WBGene00023489 | protein_coding_gene | -1.0 | 0.50 | 2.8E-02 |
| *vha-20* | WBGene00010993 | protein_coding_gene | -1.0 | 0.49 | 3.3E-02 |
| *F07A11.4* | WBGene00008547 | protein_coding_gene | -1.0 | 0.49 | 4.3E-02 |
| *C49A9.2* | WBGene00016756 | protein_coding_gene | -1.0 | 0.49 | 4.3E-02 |
| *T07F12.1* | WBGene00020321 | protein_coding_gene | -1.0 | 0.49 | 3.8E-02 |
| *M04C9.3* | WBGene00010858 | protein_coding_gene | -1.0 | 0.48 | 2.2E-02 |
| *C10G8.8* | WBGene00015684 | protein_coding_gene | -1.0 | 0.48 | 9.3E-03 |
| *F17A9.4* | WBGene00017536 | protein_coding_gene | -1.1 | 0.48 | 4.1E-02 |
| *F57B1.5* | WBGene00010188 | protein_coding_gene | -1.1 | 0.48 | 4.8E-02 |
| *ccm-2* | WBGene00019462 | protein_coding_gene | -1.1 | 0.48 | 3.1E-02 |
| *C44B7.7* | WBGene00016629 | protein_coding_gene | -1.1 | 0.47 | 3.9E-02 |
| *C08B6.11* | WBGene00007431 | protein_coding_gene | -1.1 | 0.47 | 3.5E-02 |
| *ucr-2.2* | WBGene00011679 | protein_coding_gene | -1.1 | 0.47 | 3.1E-02 |
| *F13D12.8* | WBGene00008742 | protein_coding_gene | -1.1 | 0.46 | 3.9E-02 |
| *K11C4.2* | WBGene00019643 | protein_coding_gene | -1.1 | 0.46 | 3.0E-03 |
| *C49A9.4* | WBGene00016758 | protein_coding_gene | -1.1 | 0.46 | 2.0E-02 |
| *R10E8.7* | WBGene00014821 | pseudogene | -1.1 | 0.46 | 1.9E-02 |
| *slc-25A18.1* | WBGene00008979 | protein_coding_gene | -1.1 | 0.45 | 3.9E-02 |
| *dyla-1* | WBGene00018307 | protein_coding_gene | -1.1 | 0.45 | 4.8E-02 |
| *C44C1.1* | WBGene00016641 | protein_coding_gene | -1.1 | 0.45 | 5.0E-02 |
| *cgr-1* | WBGene00020847 | protein_coding_gene | -1.2 | 0.45 | 3.9E-02 |
| *C18E9.5* | WBGene00007685 | protein_coding_gene | -1.2 | 0.45 | 4.5E-02 |
| *alh-9* | WBGene00000115 | protein_coding_gene | -1.2 | 0.45 | 4.3E-02 |
| *unc-105* | WBGene00006832 | protein_coding_gene | -1.2 | 0.45 | 2.1E-02 |
| *F07C3.2* | WBGene00017193 | protein_coding_gene | -1.2 | 0.44 | 3.5E-02 |
| *rimb-1* | WBGene00006513 | protein_coding_gene | -1.2 | 0.44 | 2.0E-02 |
| *Y48C3A.3* | WBGene00012987 | protein_coding_gene | -1.2 | 0.44 | 5.4E-03 |
| *nep-12* | WBGene00017842 | protein_coding_gene | -1.2 | 0.44 | 4.8E-02 |
| *chw-1* | WBGene00009059 | protein_coding_gene | -1.2 | 0.44 | 1.9E-02 |
| *C01H6.8* | WBGene00007257 | protein_coding_gene | -1.2 | 0.44 | 2.5E-02 |
| *T11F9.12* | WBGene00011713 | protein_coding_gene | -1.2 | 0.43 | 2.5E-02 |
| *F26C11.1* | WBGene00009146 | protein_coding_gene | -1.2 | 0.43 | 4.9E-02 |
| *ttr-35* | WBGene00017688 | protein_coding_gene | -1.2 | 0.43 | 4.7E-02 |
| *C23H3.9* | WBGene00016022 | protein_coding_gene | -1.2 | 0.43 | 1.5E-02 |
| *C26C6.9* | WBGene00007743 | protein_coding_gene | -1.2 | 0.43 | 2.5E-02 |
| *snt-5* | WBGene00004925 | protein_coding_gene | -1.2 | 0.43 | 4.3E-03 |
| *calu-1* | WBGene00019760 | protein_coding_gene | -1.2 | 0.43 | 1.8E-02 |
| *mxt-1* | WBGene00012478 | protein_coding_gene | -1.2 | 0.42 | 4.3E-02 |
| *fah-1* | WBGene00019620 | protein_coding_gene | -1.2 | 0.42 | 4.3E-02 |
| *B0252.1* | WBGene00015087 | protein_coding_gene | -1.2 | 0.42 | 2.8E-02 |
| *ugt-61* | WBGene00018206 | protein_coding_gene | -1.2 | 0.42 | 3.8E-03 |
| *nex-3* | WBGene00003590 | protein_coding_gene | -1.2 | 0.42 | 4.9E-02 |
| *T05H10.8* | WBGene00011512 | protein_coding_gene | -1.2 | 0.42 | 3.4E-02 |
| *arrd-18* | WBGene00018060 | protein_coding_gene | -1.3 | 0.42 | 5.0E-02 |
| *ZC581.9* | WBGene00022635 | protein_coding_gene | -1.3 | 0.42 | 4.5E-02 |
| *F15E6.6* | WBGene00017487 | protein_coding_gene | -1.3 | 0.42 | 2.8E-03 |
| *clec-80* | WBGene00043062 | protein_coding_gene | -1.3 | 0.42 | 4.3E-02 |
| *dhc-4* | WBGene00012272 | protein_coding_gene | -1.3 | 0.42 | 1.4E-02 |
| *inx-5* | WBGene00002127 | protein_coding_gene | -1.3 | 0.41 | 4.9E-02 |
| *phy-2* | WBGene00004025 | protein_coding_gene | -1.3 | 0.41 | 3.0E-02 |
| *Y54G11A.7* | WBGene00013216 | protein_coding_gene | -1.3 | 0.41 | 2.0E-03 |

| | | | | | |
|---|---|---|---|---|---|
| *C50D2.6* | WBGene00016809 | protein_coding_gene | -1.3 | 0.41 | 1.7E-02 |
| *Y54F10AM.8* | WBGene00021852 | protein_coding_gene | -1.3 | 0.40 | 4.9E-02 |
| *ckc-1* | WBGene00000515 | protein_coding_gene | -1.3 | 0.40 | 4.6E-02 |
| *F37H8.2* | WBGene00009511 | protein_coding_gene | -1.3 | 0.40 | 3.2E-02 |
| *K04G2.9* | WBGene00010567 | protein_coding_gene | -1.3 | 0.40 | 7.7E-03 |
| *C17H12.10* | WBGene00015935 | protein_coding_gene | -1.3 | 0.40 | 4.7E-02 |
| *C53D6.11* | WBGene00044241 | protein_coding_gene | -1.3 | 0.40 | 2.7E-02 |
| *atgp-2* | WBGene00000225 | protein_coding_gene | -1.3 | 0.40 | 4.7E-03 |
| *rft-1* | WBGene00044637 | protein_coding_gene | -1.3 | 0.39 | 2.9E-02 |
| *R08F11.4* | WBGene00019968 | protein_coding_gene | -1.3 | 0.39 | 3.3E-02 |
| *nhx-2* | WBGene00003730 | protein_coding_gene | -1.4 | 0.39 | 3.4E-02 |
| *fut-8* | WBGene00001508 | protein_coding_gene | -1.4 | 0.39 | 2.6E-02 |
| *cest-4* | WBGene00015930 | protein_coding_gene | -1.4 | 0.39 | 5.0E-02 |
| *F56D6.16* | WBGene00219304 | pseudogene | -1.4 | 0.39 | 1.7E-02 |
| *acsd-1* | WBGene00022104 | protein_coding_gene | -1.4 | 0.39 | 3.2E-02 |
| *mans-2* | WBGene00008258 | protein_coding_gene | -1.4 | 0.39 | 3.4E-02 |
| *W01C8.5* | WBGene00020921 | protein_coding_gene | -1.4 | 0.39 | 4.3E-02 |
| *C29G2.6* | WBGene00016234 | protein_coding_gene | -1.4 | 0.39 | 3.2E-02 |
| *fkb-3* | WBGene00001428 | protein_coding_gene | -1.4 | 0.39 | 3.0E-02 |
| *T10B5.10* | WBGene00020396 | protein_coding_gene | -1.4 | 0.39 | 4.1E-02 |
| *ZK185.3* | WBGene00022683 | protein_coding_gene | -1.4 | 0.39 | 5.0E-02 |
| *arrd-26* | WBGene00019873 | protein_coding_gene | -1.4 | 0.38 | 3.2E-02 |
| *R13A5.10* | WBGene00020052 | protein_coding_gene | -1.4 | 0.38 | 2.0E-02 |
| *paqr-3* | WBGene00013457 | protein_coding_gene | -1.4 | 0.38 | 2.0E-02 |
| *nrfl-1* | WBGene00006438 | protein_coding_gene | -1.4 | 0.38 | 3.8E-02 |
| *bca-1* | WBGene00000245 | protein_coding_gene | -1.4 | 0.38 | 3.9E-02 |
| *acp-1* | WBGene00022770 | protein_coding_gene | -1.4 | 0.38 | 4.9E-02 |
| *cat-1* | WBGene00000295 | protein_coding_gene | -1.4 | 0.38 | 5.0E-02 |
| *C51E3.9* | WBGene00008254 | protein_coding_gene | -1.4 | 0.38 | 5.0E-02 |
| *C14B4.2* | WBGene00007583 | protein_coding_gene | -1.4 | 0.38 | 1.3E-02 |
| *ZK328.6* | WBGene00022695 | protein_coding_gene | -1.4 | 0.38 | 3.0E-02 |
| *add-1* | WBGene00000072 | protein_coding_gene | -1.4 | 0.38 | 8.6E-03 |
| *sptl-3* | WBGene00011932 | protein_coding_gene | -1.4 | 0.38 | 3.4E-02 |
| *slc-17.5* | WBGene00017681 | protein_coding_gene | -1.4 | 0.37 | 1.6E-02 |
| *gbb-2* | WBGene00022675 | protein_coding_gene | -1.4 | 0.37 | 3.5E-02 |
| *F57E7.1* | WBGene00010202 | protein_coding_gene | -1.4 | 0.37 | 2.2E-02 |
| *F49C12.14* | WBGene00009883 | protein_coding_gene | -1.4 | 0.37 | 5.0E-02 |
| *pyk-2* | WBGene00014001 | protein_coding_gene | -1.4 | 0.37 | 3.1E-02 |
| *C55A6.12* | WBGene00044019 | protein_coding_gene | -1.4 | 0.37 | 2.4E-02 |
| *F09B9.5* | WBGene00008604 | protein_coding_gene | -1.4 | 0.37 | 4.0E-02 |
| *ugt-47* | WBGene00011006 | protein_coding_gene | -1.4 | 0.37 | 4.3E-02 |
| *F22F4.4* | WBGene00017717 | protein_coding_gene | -1.4 | 0.37 | 3.9E-02 |
| *K08C7.7* | WBGene00010649 | protein_coding_gene | -1.5 | 0.37 | 4.8E-02 |
| *spin-1* | WBGene00007549 | protein_coding_gene | -1.5 | 0.36 | 1.0E-02 |
| *nhr-137* | WBGene00003727 | protein_coding_gene | -1.5 | 0.36 | 4.4E-02 |
| *C06H5.6* | WBGene00007394 | protein_coding_gene | -1.5 | 0.36 | 5.7E-03 |
| *C31H2.4* | WBGene00016294 | protein_coding_gene | -1.5 | 0.36 | 4.7E-02 |
| *gsto-1* | WBGene00016204 | protein_coding_gene | -1.5 | 0.36 | 4.9E-02 |
| *cal-5* | WBGene00006861 | protein_coding_gene | -1.5 | 0.36 | 2.0E-02 |
| *clec-36* | WBGene00044332 | protein_coding_gene | -1.5 | 0.36 | 2.4E-02 |
| *dop-4* | WBGene00016872 | protein_coding_gene | -1.5 | 0.36 | 3.0E-02 |
| *K03B8.6* | WBGene00010524 | protein_coding_gene | -1.5 | 0.36 | 3.7E-02 |
| *aat-4* | WBGene00000005 | protein_coding_gene | -1.5 | 0.36 | 5.0E-02 |
| *dac-1* | WBGene00000895 | protein_coding_gene | -1.5 | 0.36 | 5.0E-02 |
| *C18D11.1* | WBGene00007679 | protein_coding_gene | -1.5 | 0.36 | 4.9E-02 |
| *C37H5.14* | WBGene00045308 | protein_coding_gene | -1.5 | 0.35 | 4.9E-02 |
| *sup-10* | WBGene00006319 | protein_coding_gene | -1.5 | 0.35 | 2.5E-02 |
| *F28A12.3* | WBGene00017880 | protein_coding_gene | -1.5 | 0.35 | 3.9E-02 |

| | | | | | |
|---|---|---|---|---|---|
| *afmd-1* | WBGene00017051 | protein_coding_gene | -1.5 | 0.35 | 3.9E-02 |
| *ugt-48* | WBGene00015965 | protein_coding_gene | -1.5 | 0.35 | 1.8E-02 |
| *F44B9.2* | WBGene00018407 | protein_coding_gene | -1.5 | 0.35 | 4.3E-02 |
| *Y73E7A.3* | WBGene00022270 | protein_coding_gene | -1.5 | 0.35 | 1.1E-02 |
| *gei-18* | WBGene00001575 | protein_coding_gene | -1.5 | 0.35 | 2.0E-02 |
| *comt-3* | WBGene00021487 | protein_coding_gene | -1.5 | 0.35 | 1.2E-02 |
| *fpn-1.2* | WBGene00019977 | protein_coding_gene | -1.5 | 0.35 | 4.8E-04 |
| *C52B11.5* | WBGene00016874 | protein_coding_gene | -1.5 | 0.34 | 4.3E-02 |
| *C01H6.4* | WBGene00007254 | protein_coding_gene | -1.5 | 0.34 | 2.3E-03 |
| *piit-1* | WBGene00009384 | protein_coding_gene | -1.6 | 0.34 | 4.2E-02 |
| *shl-1* | WBGene00022240 | protein_coding_gene | -1.6 | 0.34 | 7.5E-03 |
| *T04B2.3* | WBGene00011422 | protein_coding_gene | -1.6 | 0.34 | 2.0E-03 |
| *C52G5.2* | WBGene00008260 | protein_coding_gene | -1.6 | 0.34 | 3.9E-02 |
| *clec-56* | WBGene00008595 | protein_coding_gene | -1.6 | 0.34 | 4.3E-02 |
| *C01B9.1* | WBGene00007220 | protein_coding_gene | -1.6 | 0.34 | 3.2E-02 |
| *B0416.7* | WBGene00015182 | protein_coding_gene | -1.6 | 0.33 | 4.8E-02 |
| *clec-205* | WBGene00021602 | protein_coding_gene | -1.6 | 0.33 | 9.1E-03 |
| *lron-8* | WBGene00020693 | protein_coding_gene | -1.6 | 0.33 | 1.2E-02 |
| *gst-9* | WBGene00001757 | protein_coding_gene | -1.6 | 0.33 | 1.8E-02 |
| *C55C3.1* | WBGene00016951 | protein_coding_gene | -1.6 | 0.33 | 3.4E-02 |
| *F58B4.5* | WBGene00010238 | protein_coding_gene | -1.6 | 0.33 | 2.4E-02 |
| *Y37E11B.7* | WBGene00021379 | protein_coding_gene | -1.6 | 0.33 | 5.0E-02 |
| *sto-2* | WBGene00006064 | protein_coding_gene | -1.6 | 0.33 | 2.1E-02 |
| *pfn-3* | WBGene00003991 | protein_coding_gene | -1.6 | 0.33 | 1.1E-02 |
| *inx-18* | WBGene00002140 | protein_coding_gene | -1.6 | 0.32 | 2.3E-02 |
| *zip-3* | WBGene00012330 | protein_coding_gene | -1.6 | 0.32 | 3.4E-02 |
| *T18D3.5* | WBGene00011822 | protein_coding_gene | -1.6 | 0.32 | 4.6E-02 |
| *R13D11.4* | WBGene00020058 | protein_coding_gene | -1.7 | 0.32 | 6.9E-03 |
| *rap-3* | WBGene00004309 | protein_coding_gene | -1.7 | 0.32 | 1.4E-02 |
| *ugt-6* | WBGene00013904 | protein_coding_gene | -1.7 | 0.32 | 4.7E-02 |
| *ZK896.1* | WBGene00014132 | protein_coding_gene | -1.7 | 0.32 | 6.2E-03 |
| *F31D4.8* | WBGene00009293 | protein_coding_gene | -1.7 | 0.31 | 4.1E-02 |
| *tbc-19* | WBGene00007849 | protein_coding_gene | -1.7 | 0.31 | 3.8E-02 |
| *gon-1* | WBGene00001650 | protein_coding_gene | -1.7 | 0.31 | 2.0E-02 |
| *EGAP4.1* | WBGene00017149 | protein_coding_gene | -1.7 | 0.31 | 1.4E-02 |
| *W06A7.4* | WBGene00012293 | protein_coding_gene | -1.7 | 0.31 | 8.6E-03 |
| *cima-1* | WBGene00018473 | protein_coding_gene | -1.7 | 0.31 | 4.9E-02 |
| *D1005.4* | WBGene00016998 | protein_coding_gene | -1.7 | 0.31 | 2.3E-03 |
| *twk-42* | WBGene00006692 | protein_coding_gene | -1.7 | 0.30 | 1.6E-02 |
| *pept-1* | WBGene00003877 | protein_coding_gene | -1.7 | 0.30 | 1.8E-02 |
| *gldc-1* | WBGene00020022 | protein_coding_gene | -1.7 | 0.30 | 6.1E-03 |
| *W02H3.1* | WBGene00012217 | protein_coding_gene | -1.7 | 0.30 | 4.9E-02 |
| *C33E10.8* | WBGene00016349 | protein_coding_gene | -1.7 | 0.30 | 4.1E-03 |
| *K01B6.3* | WBGene00010454 | protein_coding_gene | -1.8 | 0.30 | 9.6E-04 |
| *let-805* | WBGene00002915 | protein_coding_gene | -1.8 | 0.30 | 3.1E-02 |
| *T22E5.1* | WBGene00020690 | protein_coding_gene | -1.8 | 0.30 | 2.0E-03 |
| *hrg-2* | WBGene00010473 | protein_coding_gene | -1.8 | 0.30 | 1.0E-02 |
| *acr-8* | WBGene00000047 | protein_coding_gene | -1.8 | 0.29 | 2.0E-03 |
| *lgc-12* | WBGene00020048 | protein_coding_gene | -1.8 | 0.29 | 3.6E-02 |
| *F10A3.4* | WBGene00008634 | protein_coding_gene | -1.8 | 0.29 | 7.8E-05 |
| *slc-17.8* | WBGene00017798 | protein_coding_gene | -1.8 | 0.29 | 3.0E-02 |
| *mpst-7* | WBGene00011307 | protein_coding_gene | -1.8 | 0.29 | 9.6E-03 |
| *tba-9* | WBGene00006535 | protein_coding_gene | -1.8 | 0.29 | 1.9E-02 |
| *adt-1* | WBGene00000082 | protein_coding_gene | -1.8 | 0.29 | 4.4E-02 |
| *nas-7* | WBGene00003526 | protein_coding_gene | -1.8 | 0.29 | 1.9E-02 |
| *C11H1.9* | WBGene00023500 | protein_coding_gene | -1.8 | 0.29 | 2.7E-02 |
| *unc-52* | WBGene00006787 | protein_coding_gene | -1.8 | 0.29 | 2.0E-02 |
| *gst-39* | WBGene00001787 | protein_coding_gene | -1.8 | 0.28 | 5.9E-03 |

| | | | | | |
|---|---|---|---|---|---|
| *K03B8.8* | WBGene00010525 | protein_coding_gene | -1.8 | 0.28 | 4.9E-02 |
| *C54D2.1* | WBGene00016915 | protein_coding_gene | -1.8 | 0.28 | 4.3E-02 |
| *K10C2.12* | WBGene00206386 | protein_coding_gene | -1.8 | 0.28 | 3.4E-02 |
| *ctsa-4.1* | WBGene00019605 | protein_coding_gene | -1.8 | 0.28 | 9.0E-03 |
| *C55A6.4* | WBGene00008333 | protein_coding_gene | -1.8 | 0.28 | 1.3E-03 |
| *cyp-33C8* | WBGene00019967 | protein_coding_gene | -1.8 | 0.28 | 3.5E-03 |
| *pgp-9* | WBGene00004003 | protein_coding_gene | -1.8 | 0.28 | 3.6E-02 |
| *W01A8.6* | WBGene00012168 | protein_coding_gene | -1.9 | 0.28 | 8.8E-03 |
| *T25G12.6* | WBGene00020813 | protein_coding_gene | -1.9 | 0.28 | 1.3E-03 |
| *old-1* | WBGene00003862 | protein_coding_gene | -1.9 | 0.27 | 3.4E-02 |
| *F54C1.1* | WBGene00018789 | protein_coding_gene | -1.9 | 0.27 | 2.0E-03 |
| *ttr-7* | WBGene00020566 | protein_coding_gene | -1.9 | 0.27 | 3.8E-02 |
| *F11F1.1* | WBGene00008714 | protein_coding_gene | -1.9 | 0.27 | 2.2E-02 |
| *T20D4.5* | WBGene00020611 | protein_coding_gene | -1.9 | 0.27 | 4.9E-02 |
| *C45E5.4* | WBGene00016667 | protein_coding_gene | -1.9 | 0.27 | 6.1E-03 |
| *elo-9* | WBGene00001247 | protein_coding_gene | -1.9 | 0.27 | 7.5E-03 |
| *F33E2.4* | WBGene00009360 | protein_coding_gene | -1.9 | 0.27 | 4.1E-02 |
| *best-17* | WBGene00020559 | protein_coding_gene | -1.9 | 0.27 | 3.2E-02 |
| *srd-53* | WBGene00005131 | protein_coding_gene | -1.9 | 0.27 | 2.2E-02 |
| *cln-3.1* | WBGene00000539 | protein_coding_gene | -1.9 | 0.27 | 3.3E-02 |
| *chil-5* | WBGene00007470 | protein_coding_gene | -1.9 | 0.27 | 4.6E-02 |
| *tkr-3* | WBGene00006428 | protein_coding_gene | -1.9 | 0.27 | 3.2E-03 |
| *F37B12.1* | WBGene00009504 | protein_coding_gene | -1.9 | 0.27 | 7.7E-03 |
| *ZK6.6* | WBGene00022641 | protein_coding_gene | -1.9 | 0.27 | 3.8E-02 |
| *ugt-60* | WBGene00007402 | protein_coding_gene | -1.9 | 0.27 | 3.0E-02 |
| *R07E3.4* | WBGene00011105 | protein_coding_gene | -1.9 | 0.27 | 1.2E-02 |
| *F41D9.2* | WBGene00018282 | protein_coding_gene | -1.9 | 0.26 | 1.7E-02 |
| *C09B8.8* | WBGene00015625 | protein_coding_gene | -1.9 | 0.26 | 3.2E-02 |
| *F31D5.1* | WBGene00017944 | protein_coding_gene | -1.9 | 0.26 | 2.3E-02 |
| *doxa-1* | WBGene00015519 | protein_coding_gene | -1.9 | 0.26 | 3.9E-02 |
| *Y39F10A.1* | WBGene00021452 | protein_coding_gene | -1.9 | 0.26 | 3.0E-02 |
| *F56D5.3* | WBGene00010148 | protein_coding_gene | -1.9 | 0.26 | 1.5E-03 |
| *T06A4.3* | WBGene00020283 | protein_coding_gene | -1.9 | 0.26 | 3.4E-02 |
| *C05C8.8* | WBGene00015465 | protein_coding_gene | -1.9 | 0.26 | 2.8E-02 |
| *mup-4* | WBGene00003497 | protein_coding_gene | -2.0 | 0.26 | 4.3E-02 |
| *nhx-1* | WBGene00003729 | protein_coding_gene | -2.0 | 0.26 | 6.1E-03 |
| *arrd-10* | WBGene00010268 | protein_coding_gene | -2.0 | 0.26 | 2.7E-02 |
| *C54E4.5* | WBGene00016920 | protein_coding_gene | -2.0 | 0.26 | 3.8E-02 |
| *F38A6.4* | WBGene00009528 | protein_coding_gene | -2.0 | 0.26 | 2.0E-02 |
| *DC2.5* | WBGene00017082 | protein_coding_gene | -2.0 | 0.26 | 4.4E-03 |
| *F26G1.5* | WBGene00017841 | protein_coding_gene | -2.0 | 0.26 | 7.8E-04 |
| *cut-4* | WBGene00008482 | protein_coding_gene | -2.0 | 0.26 | 3.4E-02 |
| *Y22D7AL.11* | WBGene00021249 | protein_coding_gene | -2.0 | 0.26 | 1.6E-02 |
| *B0034.1* | WBGene00015002 | protein_coding_gene | -2.0 | 0.25 | 2.3E-02 |
| *Y66C5A.1* | WBGene00044156 | protein_coding_gene | -2.0 | 0.25 | 2.5E-03 |
| *tag-10* | WBGene00006404 | protein_coding_gene | -2.0 | 0.25 | 5.0E-02 |
| *ZK455.5* | WBGene00013975 | protein_coding_gene | -2.0 | 0.25 | 5.0E-02 |
| *elo-6* | WBGene00001244 | protein_coding_gene | -2.0 | 0.25 | 4.2E-02 |
| *fipr-7* | WBGene00007543 | protein_coding_gene | -2.0 | 0.25 | 4.2E-02 |
| *T28C12.2* | WBGene00020889 | protein_coding_gene | -2.0 | 0.25 | 5.7E-03 |
| *clec-170* | WBGene00009515 | protein_coding_gene | -2.0 | 0.24 | 2.4E-02 |
| *ZK180.5* | WBGene00022679 | protein_coding_gene | -2.0 | 0.24 | 3.2E-02 |
| *D1022.3* | WBGene00017022 | protein_coding_gene | -2.1 | 0.24 | 2.0E-02 |
| *ugt-13* | WBGene00019232 | protein_coding_gene | -2.1 | 0.24 | 5.7E-03 |
| *clec-204* | WBGene00021603 | protein_coding_gene | -2.1 | 0.24 | 3.0E-03 |
| *nas-30* | WBGene00003548 | protein_coding_gene | -2.1 | 0.24 | 3.6E-02 |
| *F10D11.6* | WBGene00008652 | protein_coding_gene | -2.1 | 0.24 | 1.8E-02 |
| *nas-38* | WBGene00003554 | protein_coding_gene | -2.1 | 0.24 | 5.0E-02 |

| | | | | | |
|---|---|---|---|---|---|
| *T10B5.8* | WBGene00020394 | protein_coding_gene | -2.1 | 0.24 | 2.6E-02 |
| *cpg-9* | WBGene00022072 | protein_coding_gene | -2.1 | 0.24 | 3.0E-02 |
| *T12B5.14* | WBGene00050913 | protein_coding_gene | -2.1 | 0.23 | 3.0E-02 |
| *wrt-4* | WBGene00006950 | protein_coding_gene | -2.1 | 0.23 | 1.3E-02 |
| *Y47G6A.19* | WBGene00021645 | protein_coding_gene | -2.1 | 0.23 | 4.3E-02 |
| *adt-3* | WBGene00020567 | protein_coding_gene | -2.1 | 0.23 | 1.3E-02 |
| *T25B9.1* | WBGene00012007 | protein_coding_gene | -2.1 | 0.23 | 5.8E-03 |
| *ZC434.9* | WBGene00013895 | protein_coding_gene | -2.1 | 0.23 | 2.7E-02 |
| *Y7A9A.79* | WBGene00185076 | ncRNA_gene | -2.1 | 0.23 | 1.1E-02 |
| *dhs-25* | WBGene00000988 | protein_coding_gene | -2.1 | 0.23 | 3.5E-02 |
| *col-58* | WBGene00000634 | protein_coding_gene | -2.1 | 0.23 | 3.4E-02 |
| *VB0395L.1* | WBGene00044150 | protein_coding_gene | -2.1 | 0.23 | 2.4E-02 |
| *T19D12.9* | WBGene00020583 | protein_coding_gene | -2.1 | 0.23 | 3.7E-02 |
| *decr-1.1* | WBGene00018380 | protein_coding_gene | -2.1 | 0.23 | 4.3E-02 |
| *vha-3* | WBGene00006912 | protein_coding_gene | -2.1 | 0.23 | 4.2E-04 |
| *C52D10.3* | WBGene00016877 | protein_coding_gene | -2.2 | 0.23 | 2.1E-02 |
| *col-147* | WBGene00000720 | protein_coding_gene | -2.2 | 0.22 | 3.9E-02 |
| *C33D3.3* | WBGene00007894 | protein_coding_gene | -2.2 | 0.22 | 4.4E-02 |
| *mua-6* | WBGene00003485 | protein_coding_gene | -2.2 | 0.22 | 4.3E-02 |
| *ZK84.1* | WBGene00022649 | protein_coding_gene | -2.2 | 0.22 | 3.4E-02 |
| *clec-43* | WBGene00019917 | protein_coding_gene | -2.2 | 0.22 | 6.9E-03 |
| *ugt-30* | WBGene00011340 | protein_coding_gene | -2.2 | 0.22 | 2.1E-02 |
| *col-107* | WBGene00000681 | protein_coding_gene | -2.2 | 0.22 | 4.5E-02 |
| *C17B7.5* | WBGene00015879 | protein_coding_gene | -2.2 | 0.22 | 3.4E-02 |
| *pqn-74* | WBGene00004156 | protein_coding_gene | -2.2 | 0.22 | 1.6E-02 |
| *pcp-3* | WBGene00003958 | protein_coding_gene | -2.2 | 0.22 | 1.1E-04 |
| *pqn-63* | WBGene00004146 | protein_coding_gene | -2.2 | 0.22 | 4.9E-02 |
| *col-61* | WBGene00000637 | protein_coding_gene | -2.3 | 0.21 | 5.0E-02 |
| *C15F1.1* | WBGene00015794 | protein_coding_gene | -2.3 | 0.21 | 2.0E-03 |
| *Y17G9A.2* | WBGene00021195 | protein_coding_gene | -2.3 | 0.21 | 1.7E-02 |
| *ech-7* | WBGene00001156 | protein_coding_gene | -2.3 | 0.21 | 2.5E-03 |
| *ram-2* | WBGene00004300 | protein_coding_gene | -2.3 | 0.21 | 3.2E-02 |
| *col-113* | WBGene00000687 | protein_coding_gene | -2.3 | 0.21 | 2.4E-02 |
| *col-144* | WBGene00000717 | protein_coding_gene | -2.3 | 0.21 | 4.7E-02 |
| *nhr-115* | WBGene00003705 | protein_coding_gene | -2.3 | 0.21 | 1.1E-05 |
| *F40H3.2* | WBGene00018250 | protein_coding_gene | -2.3 | 0.21 | 4.2E-02 |
| *F38B6.4* | WBGene00018174 | protein_coding_gene | -2.3 | 0.20 | 2.0E-03 |
| *col-14* | WBGene00000603 | protein_coding_gene | -2.3 | 0.20 | 4.2E-02 |
| *ugt-40* | WBGene00017331 | protein_coding_gene | -2.3 | 0.20 | 2.2E-02 |
| *col-104* | WBGene00000678 | protein_coding_gene | -2.3 | 0.20 | 3.0E-02 |
| *Y50D4B.6* | WBGene00021745 | protein_coding_gene | -2.3 | 0.20 | 1.1E-02 |
| *C45B2.2* | WBGene00016659 | protein_coding_gene | -2.3 | 0.20 | 6.5E-03 |
| *cdr-2* | WBGene00008296 | protein_coding_gene | -2.3 | 0.20 | 2.0E-03 |
| *ZK180.6* | WBGene00022680 | protein_coding_gene | -2.3 | 0.20 | 4.6E-02 |
| *F57B1.1* | WBGene00010187 | protein_coding_gene | -2.4 | 0.20 | 2.0E-02 |
| *C39B5.5* | WBGene00016523 | protein_coding_gene | -2.4 | 0.20 | 2.6E-02 |
| *K10H10.4* | WBGene00010760 | protein_coding_gene | -2.4 | 0.19 | 4.1E-02 |
| *cyp-14A1* | WBGene00010705 | protein_coding_gene | -2.4 | 0.19 | 2.9E-02 |
| *col-97* | WBGene00000672 | protein_coding_gene | -2.4 | 0.19 | 2.5E-02 |
| *Y46G5A.29* | WBGene00012917 | protein_coding_gene | -2.4 | 0.19 | 3.0E-02 |
| *tbb-6* | WBGene00006539 | protein_coding_gene | -2.4 | 0.19 | 2.7E-04 |
| *C42D4.3* | WBGene00016596 | protein_coding_gene | -2.4 | 0.18 | 3.1E-02 |
| *Y95B8A.2* | WBGene00022383 | protein_coding_gene | -2.4 | 0.18 | 2.0E-02 |
| *fipr-10* | WBGene00008245 | protein_coding_gene | -2.5 | 0.18 | 1.3E-02 |
| *col-167* | WBGene00000740 | protein_coding_gene | -2.5 | 0.18 | 3.9E-02 |
| *grd-10* | WBGene00001699 | protein_coding_gene | -2.5 | 0.18 | 5.0E-03 |
| *T25G12.13* | WBGene00219274 | protein_coding_gene | -2.5 | 0.18 | 3.3E-02 |
| *T06E4.9* | WBGene00011536 | protein_coding_gene | -2.5 | 0.18 | 4.3E-02 |

| | | | | | |
|---|---|---|---|---|---|
| *C04E12.10* | WBGene00015439 | pseudogene | -2.5 | 0.17 | 4.9E-02 |
| *F57B1.9* | WBGene00045488 | protein_coding_gene | -2.5 | 0.17 | 3.0E-03 |
| *C06E8.5* | WBGene00015544 | protein_coding_gene | -2.5 | 0.17 | 2.1E-02 |
| *col-125* | WBGene00000699 | protein_coding_gene | -2.5 | 0.17 | 3.4E-02 |
| *pqn-2* | WBGene00004098 | protein_coding_gene | -2.5 | 0.17 | 3.7E-02 |
| *col-157* | WBGene00000730 | protein_coding_gene | -2.5 | 0.17 | 2.9E-02 |
| *T21C9.9* | WBGene00011895 | protein_coding_gene | -2.5 | 0.17 | 3.3E-02 |
| *F11G11.15* | WBGene00271822 | protein_coding_gene | -2.5 | 0.17 | 7.8E-03 |
| *asp-8* | WBGene00019105 | protein_coding_gene | -2.6 | 0.17 | 5.2E-06 |
| *cdr-4* | WBGene00010470 | protein_coding_gene | -2.6 | 0.17 | 4.1E-07 |
| *T01G5.1* | WBGene00011339 | protein_coding_gene | -2.6 | 0.17 | 3.7E-05 |
| *grd-13* | WBGene00001702 | protein_coding_gene | -2.6 | 0.17 | 5.1E-03 |
| *dct-5* | WBGene00017218 | protein_coding_gene | -2.6 | 0.17 | 2.4E-02 |
| *grl-16* | WBGene00001725 | protein_coding_gene | -2.6 | 0.17 | 1.4E-02 |
| *asns-1* | WBGene00003816 | protein_coding_gene | -2.6 | 0.16 | 2.0E-03 |
| *W07A12.8* | WBGene00012325 | protein_coding_gene | -2.6 | 0.16 | 1.6E-05 |
| *col-162* | WBGene00000735 | protein_coding_gene | -2.6 | 0.16 | 3.5E-02 |
| *F35B3.4* | WBGene00018031 | protein_coding_gene | -2.6 | 0.16 | 2.8E-02 |
| *col-152* | WBGene00000725 | protein_coding_gene | -2.7 | 0.16 | 2.9E-02 |
| *C05E7.2* | WBGene00007344 | protein_coding_gene | -2.7 | 0.16 | 1.4E-02 |
| *Y47D7A.13* | WBGene00021625 | protein_coding_gene | -2.7 | 0.16 | 8.0E-03 |
| *R12E2.7* | WBGene00020033 | protein_coding_gene | -2.7 | 0.16 | 1.1E-02 |
| *F53F1.4* | WBGene00009982 | protein_coding_gene | -2.7 | 0.16 | 9.3E-03 |
| *C30G12.2* | WBGene00016274 | protein_coding_gene | -2.7 | 0.15 | 2.7E-02 |
| *abu-6* | WBGene00000029 | protein_coding_gene | -2.7 | 0.15 | 3.4E-02 |
| *grd-14* | WBGene00001703 | protein_coding_gene | -2.8 | 0.15 | 1.9E-02 |
| *col-88* | WBGene00000663 | protein_coding_gene | -2.8 | 0.15 | 3.4E-02 |
| *col-60* | WBGene00000636 | protein_coding_gene | -2.8 | 0.15 | 2.0E-02 |
| *ins-33* | WBGene00002116 | protein_coding_gene | -2.8 | 0.15 | 4.2E-06 |
| *col-77* | WBGene00000653 | protein_coding_gene | -2.8 | 0.14 | 2.0E-02 |
| *col-180* | WBGene00000753 | protein_coding_gene | -2.8 | 0.14 | 2.1E-02 |
| *col-130* | WBGene00000704 | protein_coding_gene | -2.8 | 0.14 | 1.2E-02 |
| *grd-6* | WBGene00001695 | protein_coding_gene | -2.8 | 0.14 | 2.0E-02 |
| *col-145* | WBGene00000718 | protein_coding_gene | -2.8 | 0.14 | 2.4E-02 |
| *cyp-13A12* | WBGene00008810 | protein_coding_gene | -2.8 | 0.14 | 1.2E-02 |
| *C07G1.7* | WBGene00015573 | protein_coding_gene | -2.9 | 0.14 | 6.0E-06 |
| *T22B7.7* | WBGene00020674 | protein_coding_gene | -2.9 | 0.13 | 2.0E-02 |
| *col-120* | WBGene00000694 | protein_coding_gene | -2.9 | 0.13 | 1.2E-02 |
| *col-154* | WBGene00000727 | protein_coding_gene | -3.0 | 0.12 | 1.1E-02 |
| *Y48G8AL.12* | WBGene00021690 | protein_coding_gene | -3.1 | 0.12 | 9.3E-03 |
| *dhs-26* | WBGene00000989 | protein_coding_gene | -3.1 | 0.12 | 2.0E-03 |
| *col-137* | WBGene00000710 | protein_coding_gene | -3.2 | 0.11 | 7.7E-03 |
| *nhr-74* | WBGene00003664 | protein_coding_gene | -3.5 | 0.09 | 1.0E-03 |
| *Y41C4A.32* | WBGene00270321 | protein_coding_gene | -3.6 | 0.08 | 1.6E-05 |
| *B0205.13* | WBGene00044644 | protein_coding_gene | -4.1 | 0.06 | 8.8E-08 |

**CB4856**-exclusive DEGs in response to Gram-negative pathogens (*P. rettgeri* or *P. aeruginosa* vs. OP50)

| Gene Name | Wormbase Gene ID | Type | $log_2FC$ | FC | Adj P-value |
|---|---|---|---|---|---|
| *F01D5.3* | WBGene00008494 | protein_coding_gene | 4.2 | 18.40 | 7.0E-09 |
| *fbxa-15* | WBGene00022327 | protein_coding_gene | 3.6 | 12.48 | 1.7E-04 |
| *pqn-98* | WBGene00004177 | protein_coding_gene | 3.5 | 11.22 | 7.9E-04 |
| *F01D5.2* | WBGene00008493 | protein_coding_gene | 3.4 | 10.80 | 1.3E-03 |
| *drd-50* | WBGene00018643 | protein_coding_gene | 3.4 | 10.69 | 5.1E-26 |
| *C17H12.6* | WBGene00015932 | protein_coding_gene | 3.4 | 10.65 | 8.3E-04 |
| *gst-38* | WBGene00001786 | protein_coding_gene | 3.4 | 10.64 | 7.9E-05 |
| *C01G10.5* | WBGene00007232 | protein_coding_gene | 3.2 | 9.02 | 3.2E-04 |
| *Y48E1B.8* | WBGene00013007 | protein_coding_gene | 3.1 | 8.83 | 8.5E-04 |

| | | | | | |
|---|---|---|---|---|---|
| *C01G10.4* | WBGene00007231 | protein_coding_gene | 3.0 | 8.10 | 9.3E-04 |
| *pcp-2* | WBGene00003957 | protein_coding_gene | 2.9 | 7.49 | 3.3E-12 |
| *pud-2.1* | WBGene00017490 | protein_coding_gene | 2.9 | 7.38 | 3.5E-03 |
| *fbxa-88* | WBGene00008632 | protein_coding_gene | 2.8 | 7.14 | 2.1E-04 |
| *F55G11.2* | WBGene00010123 | protein_coding_gene | 2.8 | 7.05 | 1.1E-06 |
| *Y37H2A.14* | WBGene00045416 | protein_coding_gene | 2.8 | 6.89 | 1.1E-04 |
| *F55G11.6* | WBGene00010126 | protein_coding_gene | 2.8 | 6.84 | 5.2E-03 |
| *hsp-17* | WBGene00002021 | protein_coding_gene | 2.7 | 6.61 | 1.6E-24 |
| *fbxa-18* | WBGene00008633 | protein_coding_gene | 2.7 | 6.55 | 2.5E-03 |
| *F11D11.3* | WBGene00008698 | protein_coding_gene | 2.6 | 6.15 | 7.8E-03 |
| *F55G11.7* | WBGene00010127 | protein_coding_gene | 2.6 | 6.09 | 2.1E-03 |
| *F46F2.3* | WBGene00009787 | protein_coding_gene | 2.6 | 6.03 | 2.5E-03 |
| *Y51B9A.8* | WBGene00013090 | protein_coding_gene | 2.6 | 6.02 | 2.5E-03 |
| *cld-9* | WBGene00009432 | protein_coding_gene | 2.5 | 5.82 | 1.1E-04 |
| *clec-174* | WBGene00021580 | protein_coding_gene | 2.5 | 5.75 | 9.3E-04 |
| *zipt-2.3* | WBGene00013207 | protein_coding_gene | 2.5 | 5.71 | 1.1E-04 |
| *C01G10.17* | WBGene00044658 | protein_coding_gene | 2.5 | 5.68 | 4.2E-05 |
| *hrg-4* | WBGene00009493 | protein_coding_gene | 2.5 | 5.67 | 3.2E-05 |
| *linc-61* | WBGene00195170 | lincRNA_gene | 2.5 | 5.64 | 4.0E-04 |
| *F48G7.13* | WBGene00018624 | protein_coding_gene | 2.5 | 5.62 | 4.0E-03 |
| *C49G7.13* | WBGene00269425 | protein_coding_gene | 2.5 | 5.60 | 1.6E-05 |
| *F55A3.7* | WBGene00018852 | pseudogene | 2.5 | 5.56 | 3.1E-04 |
| *H20E11.2* | WBGene00019214 | protein_coding_gene | 2.5 | 5.56 | 2.1E-02 |
| *F17C11.14* | WBGene00045151 | snoRNA_gene | 2.5 | 5.54 | 1.8E-02 |
| *irg-3* | WBGene00018760 | protein_coding_gene | 2.5 | 5.52 | 4.1E-04 |
| *ZK228.4* | WBGene00013947 | protein_coding_gene | 2.4 | 5.42 | 8.9E-08 |
| *Y41D4B.17* | WBGene00021519 | protein_coding_gene | 2.4 | 5.41 | 7.9E-05 |
| *CD4.18* | WBGene00235160 | ncRNA_gene | 2.4 | 5.37 | 1.5E-02 |
| *C18C4.17* | WBGene00219824 | ncRNA_gene | 2.4 | 5.27 | 3.3E-02 |
| *F35E12.2* | WBGene00009426 | protein_coding_gene | 2.4 | 5.24 | 2.8E-02 |
| *argk-1* | WBGene00009706 | protein_coding_gene | 2.4 | 5.23 | 1.1E-02 |
| *Y53F4B.16* | WBGene00013162 | protein_coding_gene | 2.4 | 5.22 | 2.3E-03 |
| *dct-17* | WBGene00009431 | protein_coding_gene | 2.4 | 5.18 | 5.6E-04 |
| *T16G12.3* | WBGene00011804 | protein_coding_gene | 2.4 | 5.10 | 1.3E-03 |
| *sdz-35* | WBGene00022570 | protein_coding_gene | 2.3 | 5.03 | 2.9E-02 |
| *F01D5.1* | WBGene00008492 | protein_coding_gene | 2.3 | 4.82 | 1.5E-03 |
| *arrd-3* | WBGene00006429 | protein_coding_gene | 2.3 | 4.81 | 9.3E-03 |
| *Y6G8.5* | WBGene00044207 | protein_coding_gene | 2.3 | 4.81 | 3.4E-02 |
| *fbxa-20* | WBGene00022332 | protein_coding_gene | 2.3 | 4.79 | 1.9E-02 |
| *Y54G11A.14* | WBGene00013221 | protein_coding_gene | 2.3 | 4.78 | 1.6E-04 |
| *rab-11.2* | WBGene00004275 | protein_coding_gene | 2.3 | 4.77 | 1.6E-02 |
| *bath-7* | WBGene00018667 | protein_coding_gene | 2.2 | 4.69 | 3.8E-02 |
| *T20B6.1* | WBGene00020597 | protein_coding_gene | 2.2 | 4.65 | 4.3E-03 |
| *H02F09.3* | WBGene00019146 | protein_coding_gene | 2.2 | 4.65 | 2.6E-05 |
| *B0391.10* | WBGene00007165 | pseudogene | 2.2 | 4.64 | 8.4E-06 |
| *Y82E9BL.18* | WBGene00044402 | protein_coding_gene | 2.2 | 4.62 | 1.3E-02 |
| *ins-32* | WBGene00002115 | protein_coding_gene | 2.2 | 4.60 | 7.7E-03 |
| *gcsh-1* | WBGene00008354 | protein_coding_gene | 2.2 | 4.59 | 1.2E-06 |
| *Y46H3C.6* | WBGene00021606 | pseudogene | 2.2 | 4.58 | 9.9E-03 |
| *elo-2* | WBGene00001240 | protein_coding_gene | 2.2 | 4.56 | 3.0E-04 |
| *Y75B12B.12* | WBGene00045129 | snoRNA_gene | 2.2 | 4.55 | 4.3E-02 |
| *F57F4.4* | WBGene00019017 | protein_coding_gene | 2.2 | 4.53 | 2.7E-04 |
| *F58G1.9* | WBGene00010271 | pseudogene | 2.2 | 4.53 | 2.5E-03 |
| *M28.2* | WBGene00010894 | protein_coding_gene | 2.2 | 4.52 | 2.7E-02 |
| *F42A8.1* | WBGene00009626 | protein_coding_gene | 2.2 | 4.52 | 3.7E-03 |
| *Y54G2A.45* | WBGene00044488 | protein_coding_gene | 2.2 | 4.51 | 1.0E-03 |
| *rrn-1.1* | WBGene00004512 | rRNA_gene | 2.2 | 4.49 | 3.1E-02 |
| *F22E5.6* | WBGene00017705 | protein_coding_gene | 2.1 | 4.43 | 1.8E-03 |

| | | | | | |
|---|---|---|---|---|---|
| *hpo-6* | WBGene00021518 | protein_coding_gene | 2.1 | 4.39 | 1.2E-03 |
| *T24A6.7* | WBGene00020747 | protein_coding_gene | 2.1 | 4.36 | 2.7E-02 |
| *oac-34* | WBGene00010157 | protein_coding_gene | 2.1 | 4.34 | 1.7E-04 |
| *F10A3.1* | WBGene00008631 | protein_coding_gene | 2.1 | 4.34 | 9.3E-03 |
| *trp-4* | WBGene00006616 | protein_coding_gene | 2.1 | 4.32 | 7.8E-04 |
| *H25K10.1* | WBGene00010411 | protein_coding_gene | 2.1 | 4.30 | 1.2E-02 |
| *ZK185.9* | WBGene00220238 | snoRNA_gene | 2.1 | 4.29 | 3.6E-02 |
| *Y39D8A.1* | WBGene00021448 | protein_coding_gene | 2.1 | 4.22 | 2.1E-06 |
| *dod-19* | WBGene00022644 | protein_coding_gene | 2.1 | 4.18 | 1.9E-06 |
| *K10C9.9* | WBGene00044515 | pseudogene | 2.1 | 4.16 | 4.9E-03 |
| *R07E5.4* | WBGene00011112 | protein_coding_gene | 2.1 | 4.15 | 1.8E-02 |
| *lrg-1* | WBGene00003067 | pseudogene | 2.0 | 4.08 | 6.5E-05 |
| *sur-5* | WBGene00006351 | protein_coding_gene | 2.0 | 4.03 | 3.4E-03 |
| *fbxa-14* | WBGene00022326 | protein_coding_gene | 2.0 | 3.97 | 8.9E-03 |
| *ZK250.14* | WBGene00235354 | protein_coding_gene | 2.0 | 3.94 | 1.2E-02 |
| *otpl-5* | WBGene00018478 | protein_coding_gene | 2.0 | 3.93 | 4.0E-02 |
| *B0303.16* | WBGene00044678 | protein_coding_gene | 2.0 | 3.91 | 4.0E-02 |
| *K11H12.12* | WBGene00044948 | snoRNA_gene | 2.0 | 3.91 | 4.2E-02 |
| *C30E1.6* | WBGene00016255 | pseudogene | 2.0 | 3.91 | 1.0E-02 |
| *fbxa-84* | WBGene00011670 | protein_coding_gene | 2.0 | 3.86 | 7.5E-08 |
| *col-158* | WBGene00000731 | protein_coding_gene | 1.9 | 3.85 | 3.2E-02 |
| *str-131* | WBGene00006180 | protein_coding_gene | 1.9 | 3.85 | 1.5E-02 |
| *Y57G11C.51* | WBGene00044258 | protein_coding_gene | 1.9 | 3.84 | 1.3E-04 |
| *F55G11.8* | WBGene00010128 | protein_coding_gene | 1.9 | 3.82 | 4.6E-06 |
| *Y53F4B.17* | WBGene00013163 | protein_coding_gene | 1.9 | 3.81 | 3.0E-02 |
| *math-47* | WBGene00022688 | pseudogene | 1.9 | 3.81 | 7.1E-03 |
| *anr-32* | WBGene00219683 | antisense_lncRNA_gene | 1.9 | 3.80 | 1.4E-02 |
| *Y24F12A.4* | WBGene00012499 | pseudogene | 1.9 | 3.79 | 2.0E-03 |
| *Y105C5B.15* | WBGene00013654 | protein_coding_gene | 1.9 | 3.78 | 2.4E-04 |
| *clec-265* | WBGene00019738 | protein_coding_gene | 1.9 | 3.77 | 1.2E-21 |
| *Y53C10A.1* | WBGene00014915 | pseudogene | 1.9 | 3.77 | 1.1E-04 |
| *C04G6.6* | WBGene00015456 | protein_coding_gene | 1.9 | 3.77 | 5.2E-03 |
| *F58G6.3* | WBGene00010274 | protein_coding_gene | 1.9 | 3.75 | 3.6E-02 |
| *pqn-97* | WBGene00004176 | protein_coding_gene | 1.9 | 3.75 | 4.4E-02 |
| *Y106G6E.1* | WBGene00013705 | protein_coding_gene | 1.9 | 3.74 | 3.3E-02 |
| *clp-6* | WBGene00000546 | protein_coding_gene | 1.9 | 3.73 | 3.0E-03 |
| *F43C1.7* | WBGene00023423 | protein_coding_gene | 1.9 | 3.73 | 3.4E-02 |
| *hsp-16.41* | WBGene00002018 | protein_coding_gene | 1.9 | 3.69 | 2.8E-02 |
| *Y58A7A.5* | WBGene00021979 | protein_coding_gene | 1.9 | 3.67 | 1.0E-02 |
| *clec-67* | WBGene00018971 | protein_coding_gene | 1.9 | 3.65 | 2.1E-02 |
| *H43E16.1* | WBGene00019273 | protein_coding_gene | 1.9 | 3.63 | 3.4E-18 |
| *mth-1* | WBGene00017948 | protein_coding_gene | 1.9 | 3.63 | 6.3E-05 |
| *sid-1* | WBGene00004795 | protein_coding_gene | 1.9 | 3.62 | 7.0E-05 |
| *K09D9.11* | WBGene00019568 | pseudogene | 1.9 | 3.62 | 8.6E-03 |
| *oac-58* | WBGene00019580 | protein_coding_gene | 1.9 | 3.62 | 1.6E-07 |
| *F17B5.8* | WBGene00045458 | protein_coding_gene | 1.8 | 3.60 | 2.0E-03 |
| *C55B6.5* | WBGene00016941 | protein_coding_gene | 1.8 | 3.59 | 5.0E-02 |
| *linc-40* | WBGene00219770 | lincRNA_gene | 1.8 | 3.54 | 7.7E-03 |
| *prg-2* | WBGene00004179 | pseudogene | 1.8 | 3.54 | 7.1E-05 |
| *Y43B11AR.7* | WBGene00045086 | snoRNA_gene | 1.8 | 3.52 | 3.1E-04 |
| *T05F1.9* | WBGene00011494 | protein_coding_gene | 1.8 | 3.51 | 1.9E-02 |
| *C08G5.3* | WBGene00015616 | pseudogene | 1.8 | 3.50 | 1.7E-02 |
| *D1086.3* | WBGene00008390 | protein_coding_gene | 1.8 | 3.47 | 4.8E-02 |
| *lbp-8* | WBGene00002260 | protein_coding_gene | 1.8 | 3.46 | 4.7E-02 |
| *R11A8.1* | WBGene00011236 | protein_coding_gene | 1.8 | 3.46 | 2.3E-04 |
| *cyp-35D2* | WBGene00014735 | pseudogene | 1.8 | 3.45 | 4.4E-02 |
| *dct-18* | WBGene00010266 | protein_coding_gene | 1.8 | 3.44 | 8.9E-11 |
| *ZC15.3* | WBGene00013832 | protein_coding_gene | 1.8 | 3.43 | 4.9E-02 |

| | | | | | |
|---|---|---|---|---|---|
| *K11H12.3* | WBGene00019659 | protein_coding_gene | 1.8 | 3.41 | 4.7E-02 |
| *enu-3.5* | WBGene00012552 | protein_coding_gene | 1.8 | 3.41 | 2.3E-03 |
| *M03B6.1* | WBGene00010833 | protein_coding_gene | 1.8 | 3.41 | 9.4E-03 |
| *F25A2.1* | WBGene00017764 | protein_coding_gene | 1.8 | 3.39 | 1.9E-02 |
| *fbxa-178* | WBGene00017533 | pseudogene | 1.7 | 3.36 | 7.7E-03 |
| *clpr-2* | WBGene00023410 | protein_coding_gene | 1.7 | 3.36 | 1.7E-02 |
| *F22H10.6* | WBGene00017730 | protein_coding_gene | 1.7 | 3.36 | 1.4E-02 |
| *folt-3* | WBGene00044738 | protein_coding_gene | 1.7 | 3.36 | 3.3E-02 |
| *hsd-2* | WBGene00022498 | protein_coding_gene | 1.7 | 3.34 | 4.5E-02 |
| *C50H2.7* | WBGene00008243 | protein_coding_gene | 1.7 | 3.34 | 1.2E-02 |
| *K08D8.4* | WBGene00010658 | protein_coding_gene | 1.7 | 3.34 | 5.6E-03 |
| *C17H12.8* | WBGene00015933 | protein_coding_gene | 1.7 | 3.33 | 1.1E-02 |
| *C06A1.12* | WBGene00219803 | ncRNA_gene | 1.7 | 3.33 | 4.8E-08 |
| *F20G2.1* | WBGene00008985 | protein_coding_gene | 1.7 | 3.32 | 4.2E-04 |
| *T08D2.7* | WBGene00011612 | protein_coding_gene | 1.7 | 3.32 | 7.7E-03 |
| *F21D9.2* | WBGene00009016 | protein_coding_gene | 1.7 | 3.31 | 1.5E-02 |
| *T02G5.4* | WBGene00020164 | pseudogene | 1.7 | 3.30 | 3.2E-03 |
| *chk-2* | WBGene00000499 | protein_coding_gene | 1.7 | 3.29 | 5.3E-07 |
| *linc-29* | WBGene00189951 | lincRNA_gene | 1.7 | 3.28 | 2.4E-03 |
| *H02F09.2* | WBGene00019145 | protein_coding_gene | 1.7 | 3.27 | 2.7E-02 |
| *dod-20* | WBGene00015259 | protein_coding_gene | 1.7 | 3.25 | 4.7E-02 |
| *C30F2.4* | WBGene00007818 | protein_coding_gene | 1.7 | 3.25 | 2.2E-02 |
| *B0250.7* | WBGene00007124 | protein_coding_gene | 1.7 | 3.23 | 1.3E-04 |
| *Y9C9A.8* | WBGene00021178 | protein_coding_gene | 1.7 | 3.23 | 2.7E-02 |
| *H06I04.9* | WBGene00045092 | snoRNA_gene | 1.7 | 3.20 | 4.0E-02 |
| *C06E1.7* | WBGene00015522 | protein_coding_gene | 1.7 | 3.19 | 8.7E-05 |
| *F28A10.9* | WBGene00017877 | pseudogene | 1.7 | 3.18 | 4.6E-02 |
| *bath-46* | WBGene00020325 | protein_coding_gene | 1.7 | 3.18 | 2.0E-02 |
| *T23B12.14* | WBGene00220084 | ncRNA_gene | 1.7 | 3.18 | 2.7E-02 |
| *F53C11.1* | WBGene00009971 | protein_coding_gene | 1.7 | 3.17 | 1.9E-03 |
| *C05D12.3* | WBGene00014666 | protein_coding_gene | 1.7 | 3.15 | 3.0E-03 |
| *enu-3.6* | WBGene00010477 | protein_coding_gene | 1.7 | 3.15 | 4.6E-04 |
| *wago-2* | WBGene00018862 | pseudogene | 1.7 | 3.14 | 1.8E-07 |
| *C46C2.3* | WBGene00008111 | protein_coding_gene | 1.6 | 3.11 | 4.8E-03 |
| *Y48G1BM.8* | WBGene00021673 | pseudogene | 1.6 | 3.09 | 7.0E-03 |
| *F15D4.5* | WBGene00008862 | protein_coding_gene | 1.6 | 3.08 | 3.4E-04 |
| *T10G3.3* | WBGene00011694 | protein_coding_gene | 1.6 | 3.07 | 4.5E-02 |
| *F53F4.4* | WBGene00009988 | protein_coding_gene | 1.6 | 3.06 | 3.3E-02 |
| *Y43F11A.6* | WBGene00014576 | pseudogene | 1.6 | 3.05 | 7.1E-05 |
| *arl-7* | WBGene00000191 | pseudogene | 1.6 | 3.05 | 3.9E-02 |
| *F21C10.18* | WBGene00235330 | protein_coding_gene | 1.6 | 3.05 | 3.4E-02 |
| *Y39G8B.9* | WBGene00012728 | protein_coding_gene | 1.6 | 3.04 | 3.2E-02 |
| *Y53G8AM.5* | WBGene00021805 | protein_coding_gene | 1.6 | 3.04 | 1.6E-02 |
| *F41G4.7* | WBGene00018310 | protein_coding_gene | 1.6 | 3.04 | 3.0E-02 |
| *nduo-2* | WBGene00010961 | protein_coding_gene | 1.6 | 3.04 | 1.0E-02 |
| *his-62* | WBGene00001936 | protein_coding_gene | 1.6 | 3.04 | 2.7E-02 |
| *Y34B4A.6* | WBGene00021322 | protein_coding_gene | 1.6 | 3.04 | 2.8E-07 |
| *R03D7.2* | WBGene00010989 | protein_coding_gene | 1.6 | 3.04 | 1.7E-03 |
| *MTCE.7* | WBGene00014454 | rRNA_gene | 1.6 | 3.01 | 4.4E-02 |
| *Y34B4A.9* | WBGene00021325 | protein_coding_gene | 1.6 | 3.00 | 2.1E-04 |
| *dod-17* | WBGene00010745 | protein_coding_gene | 1.6 | 2.99 | 7.2E-03 |
| *spin-3* | WBGene00008598 | protein_coding_gene | 1.6 | 2.98 | 2.7E-04 |
| *Y73B3A.18* | WBGene00022219 | pseudogene | 1.6 | 2.97 | 1.2E-03 |
| *F09C8.1* | WBGene00008621 | protein_coding_gene | 1.6 | 2.97 | 1.3E-03 |
| *clec-84* | WBGene00021895 | protein_coding_gene | 1.6 | 2.96 | 4.9E-03 |
| *miro-3* | WBGene00021651 | pseudogene | 1.6 | 2.94 | 9.9E-03 |
| *F14D7.6* | WBGene00008793 | protein_coding_gene | 1.6 | 2.93 | 5.2E-05 |
| *cpr-3* | WBGene00000783 | protein_coding_gene | 1.5 | 2.92 | 1.5E-02 |

| | | | | | |
|---|---|---|---|---|---|
| *F27C1.18* | WBGene00219918 | ncRNA_gene | 1.5 | 2.92 | 1.9E-02 |
| *ZC250.4* | WBGene00045305 | protein_coding_gene | 1.5 | 2.91 | 3.8E-02 |
| *ctsa-4.2* | WBGene00021503 | protein_coding_gene | 1.5 | 2.91 | 6.6E-06 |
| *F59E12.3* | WBGene00019119 | protein_coding_gene | 1.5 | 2.90 | 2.4E-02 |
| *twk-28* | WBGene00006680 | protein_coding_gene | 1.5 | 2.89 | 7.2E-03 |
| *F09B12.3* | WBGene00008607 | protein_coding_gene | 1.5 | 2.88 | 1.4E-02 |
| *gst-6* | WBGene00001754 | protein_coding_gene | 1.5 | 2.88 | 2.0E-02 |
| *bath-12* | WBGene00019131 | pseudogene | 1.5 | 2.88 | 2.3E-03 |
| *enu-3.4* | WBGene00021034 | protein_coding_gene | 1.5 | 2.88 | 7.1E-03 |
| *R02C2.8* | WBGene00255728 | pseudogene | 1.5 | 2.87 | 1.2E-02 |
| *C15H9.11* | WBGene00015805 | protein_coding_gene | 1.5 | 2.86 | 2.5E-02 |
| *Y17G7B.8* | WBGene00012462 | protein_coding_gene | 1.5 | 2.85 | 2.9E-02 |
| *Y116F11B.10* | WBGene00013826 | pseudogene | 1.5 | 2.84 | 6.6E-03 |
| *let-383* | WBGene00044898 | protein_coding_gene | 1.5 | 2.84 | 1.3E-03 |
| *C18H2.2* | WBGene00015989 | pseudogene | 1.5 | 2.82 | 7.6E-04 |
| *H11L12.1* | WBGene00019189 | protein_coding_gene | 1.5 | 2.82 | 4.5E-02 |
| *B0457.3* | WBGene00014660 | pseudogene | 1.5 | 2.82 | 4.0E-02 |
| *F10C1.8* | WBGene00017327 | protein_coding_gene | 1.5 | 2.80 | 1.1E-02 |
| *K06B9.4* | WBGene00019441 | pseudogene | 1.5 | 2.80 | 2.3E-03 |
| *H20E11.3* | WBGene00019215 | protein_coding_gene | 1.5 | 2.80 | 6.5E-04 |
| *nhr-236* | WBGene00021417 | protein_coding_gene | 1.5 | 2.79 | 4.8E-02 |
| *fbxa-89* | WBGene00013555 | protein_coding_gene | 1.5 | 2.78 | 4.6E-02 |
| *fbxa-184* | WBGene00018439 | protein_coding_gene | 1.5 | 2.78 | 1.3E-02 |
| *hpo-38* | WBGene00021755 | protein_coding_gene | 1.5 | 2.78 | 2.9E-03 |
| *F40E12.2* | WBGene00018235 | protein_coding_gene | 1.5 | 2.78 | 4.6E-03 |
| *F19C7.1* | WBGene00017591 | protein_coding_gene | 1.4 | 2.73 | 3.0E-07 |
| *F26A1.4* | WBGene00017803 | protein_coding_gene | 1.4 | 2.73 | 3.0E-03 |
| *F40D4.13* | WBGene00009572 | protein_coding_gene | 1.4 | 2.71 | 2.4E-02 |
| *M01E5.1* | WBGene00010804 | pseudogene | 1.4 | 2.71 | 5.0E-02 |
| *lec-9* | WBGene00002272 | protein_coding_gene | 1.4 | 2.71 | 1.0E-07 |
| *Y71A12B.10* | WBGene00013507 | protein_coding_gene | 1.4 | 2.70 | 1.9E-02 |
| *T20D3.2* | WBGene00011856 | protein_coding_gene | 1.4 | 2.70 | 7.0E-05 |
| *C34E7.3* | WBGene00007933 | protein_coding_gene | 1.4 | 2.69 | 3.5E-02 |
| *K08D8.12* | WBGene00194926 | protein_coding_gene | 1.4 | 2.68 | 7.6E-03 |
| *acs-1* | WBGene00018488 | protein_coding_gene | 1.4 | 2.68 | 3.9E-02 |
| *C28A5.1* | WBGene00007788 | protein_coding_gene | 1.4 | 2.67 | 7.3E-04 |
| *pes-8* | WBGene00003981 | protein_coding_gene | 1.4 | 2.67 | 4.6E-02 |
| *F19B6.3* | WBGene00008949 | protein_coding_gene | 1.4 | 2.67 | 2.3E-02 |
| *clec-186* | WBGene00014138 | protein_coding_gene | 1.4 | 2.67 | 3.5E-06 |
| *gst-27* | WBGene00001775 | protein_coding_gene | 1.4 | 2.67 | 4.1E-02 |
| *nspd-7* | WBGene00012781 | protein_coding_gene | 1.4 | 2.66 | 1.5E-02 |
| *ZK218.8* | WBGene00013942 | pseudogene | 1.4 | 2.64 | 2.6E-03 |
| *T27E7.10* | WBGene00044268 | pseudogene | 1.4 | 2.64 | 2.2E-02 |
| *fbxa-203* | WBGene00021037 | pseudogene | 1.4 | 2.63 | 1.2E-02 |
| *clec-74* | WBGene00021585 | protein_coding_gene | 1.4 | 2.63 | 2.4E-02 |
| *math-46* | WBGene00021175 | protein_coding_gene | 1.4 | 2.61 | 4.5E-02 |
| *lurp-4* | WBGene00077698 | protein_coding_gene | 1.4 | 2.60 | 1.0E-02 |
| *Y48G1BM.6* | WBGene00021671 | pseudogene | 1.4 | 2.59 | 5.1E-03 |
| *T05C3.2* | WBGene00020254 | protein_coding_gene | 1.4 | 2.58 | 6.6E-03 |
| *gfi-1* | WBGene00001581 | protein_coding_gene | 1.4 | 2.58 | 3.1E-02 |
| *enu-3.3* | WBGene00021015 | protein_coding_gene | 1.4 | 2.58 | 3.0E-03 |
| *math-2* | WBGene00015606 | protein_coding_gene | 1.4 | 2.55 | 9.2E-03 |
| *Y54G11A.19* | WBGene00235256 | protein_coding_gene | 1.3 | 2.54 | 4.3E-03 |
| *T19D12.4* | WBGene00020579 | protein_coding_gene | 1.3 | 2.54 | 4.8E-02 |
| *F28H7.3* | WBGene00009237 | protein_coding_gene | 1.3 | 2.53 | 2.1E-03 |
| *F22E5.1* | WBGene00017703 | protein_coding_gene | 1.3 | 2.53 | 8.2E-03 |
| *T07H8.11* | WBGene00304201 | protein_coding_gene | 1.3 | 2.52 | 6.3E-03 |
| *F52C6.2* | WBGene00018659 | protein_coding_gene | 1.3 | 2.52 | 7.0E-03 |

| | | | | | |
|---|---|---|---|---|---|
| *F39G3.5* | WBGene00018210 | protein_coding_gene | 1.3 | 2.52 | 8.0E-03 |
| *Y50D4B.2* | WBGene00021741 | protein_coding_gene | 1.3 | 2.51 | 9.7E-03 |
| *oac-57* | WBGene00019579 | protein_coding_gene | 1.3 | 2.51 | 3.6E-02 |
| *F14H12.3* | WBGene00017471 | protein_coding_gene | 1.3 | 2.51 | 8.7E-03 |
| *fbxc-47* | WBGene00019723 | protein_coding_gene | 1.3 | 2.50 | 2.1E-02 |
| *hsf-2* | WBGene00013134 | pseudogene | 1.3 | 2.49 | 7.0E-03 |
| *Y57A10A.3* | WBGene00013248 | protein_coding_gene | 1.3 | 2.48 | 8.5E-03 |
| *T05E12.8* | WBGene00045382 | protein_coding_gene | 1.3 | 2.48 | 2.5E-02 |
| *lys-1* | WBGene00003090 | protein_coding_gene | 1.3 | 2.47 | 9.9E-03 |
| *Y66D12A.6* | WBGene00013432 | protein_coding_gene | 1.3 | 2.47 | 1.1E-02 |
| *snf-4* | WBGene00004903 | pseudogene | 1.3 | 2.47 | 1.8E-02 |
| *B0001.5* | WBGene00007090 | protein_coding_gene | 1.3 | 2.45 | 9.3E-04 |
| *K03H6.2* | WBGene00019368 | protein_coding_gene | 1.3 | 2.45 | 3.1E-02 |
| *T01C8.3* | WBGene00020144 | protein_coding_gene | 1.3 | 2.44 | 1.4E-02 |
| *C12D12.1* | WBGene00015713 | protein_coding_gene | 1.3 | 2.44 | 3.0E-04 |
| *linc-76* | WBGene00219619 | lincRNA_gene | 1.3 | 2.44 | 3.8E-02 |
| *Y53C12A.6* | WBGene00013141 | protein_coding_gene | 1.3 | 2.44 | 1.2E-03 |
| *Y105E8A.28* | WBGene00013685 | protein_coding_gene | 1.3 | 2.43 | 1.7E-02 |
| *C15H7.2* | WBGene00014678 | pseudogene | 1.3 | 2.43 | 3.1E-02 |
| *C47E8.10* | WBGene00044216 | pseudogene | 1.3 | 2.42 | 4.2E-02 |
| *F27B3.7* | WBGene00017849 | pseudogene | 1.3 | 2.41 | 3.3E-05 |
| *C38D9.5* | WBGene00008013 | pseudogene | 1.3 | 2.40 | 1.9E-02 |
| *snpc-3.2* | WBGene00021667 | protein_coding_gene | 1.3 | 2.40 | 2.7E-02 |
| *T07A9.14* | WBGene00044789 | protein_coding_gene | 1.3 | 2.39 | 8.4E-06 |
| *sri-74* | WBGene00005586 | protein_coding_gene | 1.3 | 2.39 | 2.1E-02 |
| *fbxa-190* | WBGene00018939 | protein_coding_gene | 1.3 | 2.38 | 6.0E-03 |
| *Y111B2A.1* | WBGene00013727 | protein_coding_gene | 1.3 | 2.38 | 1.0E-03 |
| *T25C12.3* | WBGene00012018 | protein_coding_gene | 1.3 | 2.38 | 8.7E-03 |
| *C14B1.7* | WBGene00007578 | pseudogene | 1.3 | 2.38 | 3.5E-03 |
| *ZK637.6* | WBGene00014987 | pseudogene | 1.2 | 2.37 | 1.9E-02 |
| *F19C7.4* | WBGene00017594 | protein_coding_gene | 1.2 | 2.37 | 1.7E-02 |
| *oac-48* | WBGene00011960 | protein_coding_gene | 1.2 | 2.37 | 3.0E-02 |
| *C34B4.2* | WBGene00007907 | protein_coding_gene | 1.2 | 2.37 | 4.1E-04 |
| *C49C8.5* | WBGene00016769 | protein_coding_gene | 1.2 | 2.36 | 1.7E-03 |
| *nos-1* | WBGene00003783 | protein_coding_gene | 1.2 | 2.36 | 3.6E-02 |
| *T06A1.5* | WBGene00020280 | protein_coding_gene | 1.2 | 2.36 | 9.3E-03 |
| *parn-1* | WBGene00010734 | protein_coding_gene | 1.2 | 2.36 | 1.1E-02 |
| *linc-7* | WBGene00219609 | lincRNA_gene | 1.2 | 2.35 | 1.6E-05 |
| *ctc-3* | WBGene00010962 | protein_coding_gene | 1.2 | 2.34 | 1.4E-02 |
| *asp-14* | WBGene00019619 | protein_coding_gene | 1.2 | 2.33 | 8.7E-08 |
| *K09H9.7* | WBGene00019598 | protein_coding_gene | 1.2 | 2.32 | 1.4E-02 |
| *T08D2.3* | WBGene00011608 | pseudogene | 1.2 | 2.32 | 1.9E-05 |
| *R05F9.11* | WBGene00019894 | protein_coding_gene | 1.2 | 2.32 | 8.3E-03 |
| *nbs-1* | WBGene00007504 | protein_coding_gene | 1.2 | 2.32 | 2.3E-05 |
| *F22B3.5* | WBGene00009036 | protein_coding_gene | 1.2 | 2.31 | 2.1E-02 |
| *C24H12.2* | WBGene00016072 | protein_coding_gene | 1.2 | 2.31 | 1.1E-02 |
| *pgph-2* | WBGene00018424 | protein_coding_gene | 1.2 | 2.31 | 2.2E-02 |
| *C14E2.12* | WBGene00219816 | snoRNA_gene | 1.2 | 2.30 | 4.7E-02 |
| *pqn-36* | WBGene00004123 | protein_coding_gene | 1.2 | 2.30 | 1.4E-02 |
| *W06A11.4* | WBGene00021055 | protein_coding_gene | 1.2 | 2.29 | 9.7E-03 |
| *C09E7.8* | WBGene00015642 | protein_coding_gene | 1.2 | 2.29 | 3.8E-04 |
| *cln-3.3* | WBGene00000541 | protein_coding_gene | 1.2 | 2.29 | 1.9E-02 |
| *Y65B4BL.4* | WBGene00022036 | protein_coding_gene | 1.2 | 2.28 | 4.6E-03 |
| *timm-17B.2* | WBGene00017069 | protein_coding_gene | 1.2 | 2.28 | 3.8E-03 |
| *F43C1.5* | WBGene00009649 | protein_coding_gene | 1.2 | 2.28 | 1.6E-02 |
| *nep-7* | WBGene00017552 | protein_coding_gene | 1.2 | 2.27 | 4.3E-02 |
| *ctc-1* | WBGene00010964 | protein_coding_gene | 1.2 | 2.26 | 8.5E-03 |
| *E02H9.3* | WBGene00017101 | protein_coding_gene | 1.2 | 2.25 | 1.2E-07 |

| | | | | | |
|---|---|---|---|---|---|
| *C28A5.2* | WBGene00007789 | protein_coding_gene | 1.2 | 2.24 | 6.3E-05 |
| *snpc-3.3* | WBGene00021662 | protein_coding_gene | 1.2 | 2.24 | 2.4E-03 |
| *rmh-1* | WBGene00019712 | protein_coding_gene | 1.2 | 2.24 | 1.0E-04 |
| *cdr-7* | WBGene00010472 | protein_coding_gene | 1.2 | 2.24 | 3.6E-04 |
| *C34H4.2* | WBGene00016425 | protein_coding_gene | 1.2 | 2.24 | 5.1E-03 |
| *ptps-1* | WBGene00015010 | protein_coding_gene | 1.2 | 2.23 | 7.9E-03 |
| *T20B6.2* | WBGene00020598 | protein_coding_gene | 1.2 | 2.22 | 4.4E-02 |
| *kri-1* | WBGene00013955 | protein_coding_gene | 1.1 | 2.22 | 4.5E-03 |
| *F18A11.6* | WBGene00008933 | pseudogene | 1.1 | 2.21 | 9.2E-03 |
| *Y67D8C.23* | WBGene00235353 | pseudogene | 1.1 | 2.21 | 3.7E-02 |
| *bir-2* | WBGene00000250 | protein_coding_gene | 1.1 | 2.21 | 1.9E-02 |
| *C18D11.3* | WBGene00007681 | protein_coding_gene | 1.1 | 2.20 | 8.9E-04 |
| *C06A5.6* | WBGene00015504 | protein_coding_gene | 1.1 | 2.20 | 8.4E-03 |
| *nduo-6* | WBGene00010957 | protein_coding_gene | 1.1 | 2.19 | 3.3E-02 |
| *C45G9.6* | WBGene00016677 | protein_coding_gene | 1.1 | 2.19 | 1.2E-03 |
| *R06C1.4* | WBGene00011059 | protein_coding_gene | 1.1 | 2.19 | 3.5E-25 |
| *tag-51* | WBGene00006430 | protein_coding_gene | 1.1 | 2.18 | 6.1E-05 |
| *pid-5* | WBGene00021555 | protein_coding_gene | 1.1 | 2.18 | 3.6E-15 |
| *CD4.10* | WBGene00022970 | ncRNA_gene | 1.1 | 2.17 | 2.5E-02 |
| *B0334.5* | WBGene00007145 | protein_coding_gene | 1.1 | 2.17 | 1.6E-05 |
| *bath-4* | WBGene00018665 | protein_coding_gene | 1.1 | 2.17 | 3.9E-02 |
| *C17C3.15* | WBGene00015896 | protein_coding_gene | 1.1 | 2.16 | 1.8E-03 |
| *W05F2.8* | WBGene00195045 | protein_coding_gene | 1.1 | 2.16 | 3.4E-02 |
| *hrg-7* | WBGene00007605 | protein_coding_gene | 1.1 | 2.16 | 2.9E-02 |
| *R144.10* | WBGene00020098 | protein_coding_gene | 1.1 | 2.16 | 2.1E-04 |
| *fbxc-44* | WBGene00015819 | protein_coding_gene | 1.1 | 2.16 | 3.7E-02 |
| *Y50D4C.6* | WBGene00021750 | protein_coding_gene | 1.1 | 2.16 | 1.3E-02 |
| *M04F3.2* | WBGene00019768 | protein_coding_gene | 1.1 | 2.15 | 4.2E-03 |
| *aqp-4* | WBGene00000172 | protein_coding_gene | 1.1 | 2.15 | 2.2E-07 |
| *tipn-1* | WBGene00017738 | protein_coding_gene | 1.1 | 2.15 | 1.2E-02 |
| *set-9* | WBGene00017482 | protein_coding_gene | 1.1 | 2.15 | 6.5E-19 |
| *ppw-1* | WBGene00004093 | protein_coding_gene | 1.1 | 2.14 | 1.7E-04 |
| *Y50D4A.6* | WBGene00195169 | protein_coding_gene | 1.1 | 2.14 | 3.0E-02 |
| *F55D12.5* | WBGene00010113 | protein_coding_gene | 1.1 | 2.14 | 4.6E-05 |
| *T25D10.1* | WBGene00020799 | protein_coding_gene | 1.1 | 2.13 | 3.0E-02 |
| *C18H2.4* | WBGene00015991 | pseudogene | 1.1 | 2.13 | 3.7E-02 |
| *K08F11.6* | WBGene00023323 | pseudogene | 1.1 | 2.12 | 4.8E-03 |
| *C48B4.7* | WBGene00008170 | protein_coding_gene | 1.1 | 2.12 | 2.2E-02 |
| *C08H9.16* | WBGene00086566 | protein_coding_gene | 1.1 | 2.12 | 8.5E-03 |
| *F27D4.3* | WBGene00014746 | pseudogene | 1.1 | 2.12 | 4.4E-02 |
| *rec-1* | WBGene00004332 | protein_coding_gene | 1.1 | 2.12 | 3.4E-03 |
| *enu-3.1* | WBGene00010362 | protein_coding_gene | 1.1 | 2.12 | 3.6E-03 |
| *Y22D7AR.7* | WBGene00021261 | protein_coding_gene | 1.1 | 2.11 | 3.3E-02 |
| *T04A8.13* | WBGene00011414 | protein_coding_gene | 1.1 | 2.11 | 2.7E-02 |
| *C16A11.5* | WBGene00015818 | protein_coding_gene | 1.1 | 2.10 | 9.9E-03 |
| *lip-1* | WBGene00003043 | protein_coding_gene | 1.1 | 2.10 | 6.4E-05 |
| *Y66D12A.16* | WBGene00013442 | protein_coding_gene | 1.1 | 2.10 | 1.0E-02 |
| *nduo-1* | WBGene00010959 | protein_coding_gene | 1.1 | 2.10 | 4.2E-02 |
| *selt-1.2* | WBGene00009238 | protein_coding_gene | 1.1 | 2.10 | 4.3E-02 |
| *ctb-1* | WBGene00000829 | protein_coding_gene | 1.1 | 2.09 | 2.4E-02 |
| *tsen-54* | WBGene00018680 | protein_coding_gene | 1.1 | 2.09 | 2.3E-03 |
| *snpc-3.1* | WBGene00015098 | protein_coding_gene | 1.1 | 2.08 | 2.7E-02 |
| *cpb-1* | WBGene00000770 | protein_coding_gene | 1.1 | 2.08 | 6.3E-05 |
| *F01G4.4* | WBGene00008503 | protein_coding_gene | 1.1 | 2.08 | 1.7E-10 |
| *T28D9.4* | WBGene00020896 | protein_coding_gene | 1.1 | 2.08 | 4.9E-04 |
| *ent-4* | WBGene00008123 | protein_coding_gene | 1.1 | 2.08 | 2.4E-02 |
| *enu-3.2* | WBGene00008003 | protein_coding_gene | 1.1 | 2.07 | 1.4E-03 |
| *C09G4.4* | WBGene00044001 | protein_coding_gene | 1.1 | 2.07 | 1.1E-02 |

| | | | | | |
|---|---|---|---|---|---|
| *Y73B3A.3* | WBGene00022205 | pseudogene | 1.1 | 2.07 | 2.7E-02 |
| *set-24* | WBGene00012845 | protein_coding_gene | 1.1 | 2.07 | 2.1E-04 |
| *ZK945.4* | WBGene00014166 | protein_coding_gene | 1.0 | 2.07 | 3.2E-02 |
| *VH15N14R.1* | WBGene00012151 | protein_coding_gene | 1.0 | 2.07 | 3.9E-02 |
| *K07D4.1* | WBGene00023318 | pseudogene | 1.0 | 2.07 | 9.9E-03 |
| *T07F10.5* | WBGene00011591 | protein_coding_gene | 1.0 | 2.07 | 4.3E-02 |
| *Y52E8A.3* | WBGene00021796 | protein_coding_gene | 1.0 | 2.06 | 1.7E-02 |
| *cye-1* | WBGene00000871 | protein_coding_gene | 1.0 | 2.06 | 2.5E-03 |
| *Y82E9BR.2* | WBGene00022335 | protein_coding_gene | 1.0 | 2.06 | 1.9E-02 |
| *rde-4* | WBGene00004326 | protein_coding_gene | 1.0 | 2.06 | 2.0E-02 |
| *sqrd-1* | WBGene00008538 | protein_coding_gene | 1.0 | 2.06 | 4.3E-03 |
| *pks-1* | WBGene00016558 | protein_coding_gene | 1.0 | 2.06 | 1.9E-02 |
| *Y54G2A.73* | WBGene00219425 | protein_coding_gene | 1.0 | 2.05 | 5.3E-05 |
| *msh-4* | WBGene00003420 | pseudogene | 1.0 | 2.05 | 4.0E-02 |
| *Y54E2A.8* | WBGene00013193 | protein_coding_gene | 1.0 | 2.05 | 2.9E-03 |
| *wago-4* | WBGene00010263 | protein_coding_gene | 1.0 | 2.05 | 2.4E-04 |
| *ani-3* | WBGene00012835 | protein_coding_gene | 1.0 | 2.05 | 1.8E-03 |
| *W03F8.2* | WBGene00020992 | pseudogene | 1.0 | 2.04 | 3.9E-03 |
| *agt-2* | WBGene00000094 | protein_coding_gene | 1.0 | 2.04 | 2.0E-02 |
| *wip-1* | WBGene00020094 | protein_coding_gene | 1.0 | 2.04 | 9.4E-06 |
| *F59A3.2* | WBGene00019075 | protein_coding_gene | 1.0 | 2.04 | 3.1E-03 |
| *Y73B3B.5* | WBGene00022227 | pseudogene | 1.0 | 2.04 | 7.6E-03 |
| *F43E2.1* | WBGene00018390 | protein_coding_gene | 1.0 | 2.04 | 8.7E-03 |
| *fbxa-31* | WBGene00013838 | protein_coding_gene | 1.0 | 2.04 | 2.4E-02 |
| *T12E12.3* | WBGene00020464 | protein_coding_gene | 1.0 | 2.04 | 5.4E-10 |
| *K12H6.2* | WBGene00019683 | protein_coding_gene | 1.0 | 2.03 | 7.5E-04 |
| *C06A1.4* | WBGene00014667 | pseudogene | 1.0 | 2.03 | 8.0E-03 |
| *hpr-9* | WBGene00001997 | protein_coding_gene | 1.0 | 2.03 | 1.7E-03 |
| *clec-173* | WBGene00044469 | protein_coding_gene | 1.0 | 2.03 | 9.6E-04 |
| *set-32* | WBGene00008062 | protein_coding_gene | 1.0 | 2.03 | 5.7E-04 |
| *R04F11.5* | WBGene00011017 | protein_coding_gene | 1.0 | 2.02 | 1.2E-04 |
| *Y54G2A.21* | WBGene00021886 | pseudogene | 1.0 | 2.02 | 1.8E-02 |
| *rbm-17* | WBGene00010233 | protein_coding_gene | 1.0 | 2.02 | 3.7E-03 |
| *R155.3* | WBGene00020117 | protein_coding_gene | 1.0 | 2.02 | 3.7E-02 |
| *Y73B3A.1* | WBGene00022203 | protein_coding_gene | 1.0 | 2.02 | 5.9E-06 |
| *F53G12.4* | WBGene00018772 | protein_coding_gene | 1.0 | 2.02 | 3.7E-02 |
| *Y71A12B.14* | WBGene00014947 | pseudogene | 1.0 | 2.02 | 2.2E-02 |
| *F19F10.11* | WBGene00017607 | protein_coding_gene | 1.0 | 2.02 | 4.7E-04 |
| *F56D5.4* | WBGene00014772 | pseudogene | 1.0 | 2.01 | 6.6E-03 |
| *cin-4* | WBGene00022854 | protein_coding_gene | 1.0 | 2.01 | 1.3E-02 |
| *F56F10.1* | WBGene00018984 | protein_coding_gene | 1.0 | 2.01 | 1.5E-03 |
| *Y53C10A.6* | WBGene00013137 | protein_coding_gene | 1.0 | 2.01 | 1.9E-02 |
| *R06C7.2* | WBGene00011062 | protein_coding_gene | 1.0 | 2.01 | 1.1E-02 |
| *cep-1* | WBGene00000467 | protein_coding_gene | 1.0 | 2.01 | 9.2E-04 |
| *klu-2* | WBGene00022592 | protein_coding_gene | 1.0 | 2.01 | 4.4E-03 |
| *eri-9* | WBGene00016143 | protein_coding_gene | 1.0 | 2.01 | 2.4E-02 |
| *T24H10.4* | WBGene00012002 | protein_coding_gene | 1.0 | 2.01 | 3.5E-02 |
| *heri-1* | WBGene00016236 | protein_coding_gene | 1.0 | 2.00 | 2.7E-02 |
| *lec-8* | WBGene00002271 | protein_coding_gene | 1.0 | 2.00 | 2.1E-03 |
| *ttr-20* | WBGene00018085 | protein_coding_gene | -1.0 | 0.50 | 1.4E-02 |
| *rig-3* | WBGene00004370 | protein_coding_gene | -1.0 | 0.50 | 4.4E-02 |
| *Y34D9A.8* | WBGene00021333 | protein_coding_gene | -1.0 | 0.50 | 7.6E-03 |
| *K12C11.7* | WBGene00045053 | protein_coding_gene | -1.0 | 0.50 | 1.4E-02 |
| *acdh-10* | WBGene00020366 | protein_coding_gene | -1.0 | 0.50 | 1.7E-06 |
| *sad-1* | WBGene00004719 | protein_coding_gene | -1.0 | 0.50 | 9.2E-03 |
| *ubc-17* | WBGene00006712 | protein_coding_gene | -1.0 | 0.50 | 2.1E-02 |
| *T19D7.6* | WBGene00020574 | protein_coding_gene | -1.0 | 0.50 | 1.2E-02 |
| *bbs-8* | WBGene00000244 | protein_coding_gene | -1.0 | 0.50 | 3.6E-02 |

| | | | | | |
|---|---|---|---|---|---|
| *acl-12* | WBGene00015295 | protein_coding_gene | -1.0 | 0.50 | 2.2E-03 |
| *tag-278* | WBGene00015356 | protein_coding_gene | -1.0 | 0.50 | 1.7E-05 |
| *ceh-10* | WBGene00000435 | protein_coding_gene | -1.0 | 0.50 | 2.5E-02 |
| *R03H10.6* | WBGene00019858 | protein_coding_gene | -1.0 | 0.50 | 4.0E-02 |
| *cdh-9* | WBGene00000401 | protein_coding_gene | -1.0 | 0.50 | 4.5E-02 |
| *ajm-1* | WBGene00000100 | protein_coding_gene | -1.0 | 0.50 | 4.5E-04 |
| *F53E10.1* | WBGene00018758 | protein_coding_gene | -1.0 | 0.50 | 3.6E-02 |
| *F55D12.1* | WBGene00010110 | protein_coding_gene | -1.0 | 0.50 | 2.2E-02 |
| *tag-343* | WBGene00018374 | protein_coding_gene | -1.0 | 0.50 | 1.6E-02 |
| *C33G3.4* | WBGene00007904 | protein_coding_gene | -1.0 | 0.50 | 5.0E-05 |
| *hpo-15* | WBGene00016061 | protein_coding_gene | -1.0 | 0.50 | 1.2E-07 |
| *gsto-3* | WBGene00019636 | protein_coding_gene | -1.0 | 0.50 | 1.3E-02 |
| *K10B4.3* | WBGene00019615 | protein_coding_gene | -1.0 | 0.50 | 6.8E-07 |
| *ceh-43* | WBGene00000463 | protein_coding_gene | -1.0 | 0.49 | 4.7E-02 |
| *dhs-5* | WBGene00000969 | protein_coding_gene | -1.0 | 0.49 | 3.9E-04 |
| *lgg-1* | WBGene00002980 | protein_coding_gene | -1.0 | 0.49 | 3.8E-05 |
| *tag-304* | WBGene00018757 | protein_coding_gene | -1.0 | 0.49 | 1.9E-08 |
| *prx-12* | WBGene00004197 | protein_coding_gene | -1.0 | 0.49 | 5.0E-04 |
| *F52B10.3* | WBGene00044304 | protein_coding_gene | -1.0 | 0.49 | 7.3E-04 |
| *tbx-2* | WBGene00006543 | protein_coding_gene | -1.0 | 0.49 | 3.5E-11 |
| *frm-2* | WBGene00001489 | protein_coding_gene | -1.0 | 0.49 | 2.6E-05 |
| *prx-3* | WBGene00004193 | protein_coding_gene | -1.0 | 0.49 | 6.9E-04 |
| *K02E10.1* | WBGene00019317 | protein_coding_gene | -1.0 | 0.49 | 7.0E-07 |
| *ppm-1.H* | WBGene00016580 | protein_coding_gene | -1.0 | 0.49 | 3.5E-02 |
| *cbp-3* | WBGene00009595 | protein_coding_gene | -1.0 | 0.49 | 1.5E-02 |
| *lin-17* | WBGene00003006 | protein_coding_gene | -1.0 | 0.49 | 4.9E-04 |
| *ZC190.4* | WBGene00022538 | protein_coding_gene | -1.0 | 0.49 | 2.5E-02 |
| *odd-1* | WBGene00003845 | protein_coding_gene | -1.0 | 0.49 | 4.2E-02 |
| *pqn-89* | WBGene00004169 | protein_coding_gene | -1.0 | 0.49 | 2.6E-03 |
| *ugt-49* | WBGene00007070 | protein_coding_gene | -1.0 | 0.49 | 1.7E-02 |
| *F27C8.2* | WBGene00009183 | protein_coding_gene | -1.0 | 0.49 | 3.2E-02 |
| *Y43F8B.2* | WBGene00012813 | protein_coding_gene | -1.0 | 0.49 | 1.3E-02 |
| *nsy-1* | WBGene00003822 | protein_coding_gene | -1.0 | 0.49 | 2.4E-05 |
| *exc-7* | WBGene00001368 | protein_coding_gene | -1.0 | 0.49 | 1.7E-03 |
| *C30G4.4* | WBGene00016269 | protein_coding_gene | -1.0 | 0.49 | 2.5E-02 |
| *spv-1* | WBGene00014051 | protein_coding_gene | -1.0 | 0.49 | 9.1E-03 |
| *slt-1* | WBGene00004854 | protein_coding_gene | -1.0 | 0.49 | 1.7E-03 |
| *fipr-1* | WBGene00009096 | protein_coding_gene | -1.0 | 0.49 | 1.5E-03 |
| *ttr-43* | WBGene00007590 | protein_coding_gene | -1.0 | 0.49 | 3.9E-02 |
| *linc-15* | WBGene00219571 | lincRNA_gene | -1.0 | 0.49 | 1.1E-04 |
| *C01F1.3* | WBGene00015298 | protein_coding_gene | -1.0 | 0.49 | 2.5E-03 |
| *pals-34* | WBGene00017823 | protein_coding_gene | -1.0 | 0.49 | 9.2E-03 |
| *wht-1* | WBGene00015479 | protein_coding_gene | -1.0 | 0.49 | 5.1E-04 |
| *F59C6.8* | WBGene00010327 | protein_coding_gene | -1.0 | 0.49 | 3.5E-02 |
| *mab-21* | WBGene00003112 | protein_coding_gene | -1.0 | 0.49 | 7.2E-03 |
| *W07A12.4* | WBGene00012322 | protein_coding_gene | -1.0 | 0.49 | 4.9E-02 |
| *F57A8.1* | WBGene00010177 | protein_coding_gene | -1.0 | 0.49 | 5.8E-04 |
| *ugt-20* | WBGene00019516 | protein_coding_gene | -1.0 | 0.49 | 3.3E-02 |
| *ugt-5* | WBGene00013906 | protein_coding_gene | -1.0 | 0.49 | 8.0E-03 |
| *che-12* | WBGene00000491 | protein_coding_gene | -1.0 | 0.48 | 8.5E-03 |
| *Y116A8C.3* | WBGene00013784 | protein_coding_gene | -1.0 | 0.48 | 4.7E-02 |
| *C17H11.6* | WBGene00015926 | protein_coding_gene | -1.0 | 0.48 | 3.9E-04 |
| *try-3* | WBGene00006621 | protein_coding_gene | -1.0 | 0.48 | 1.5E-02 |
| *nfm-1* | WBGene00003593 | protein_coding_gene | -1.0 | 0.48 | 8.2E-05 |
| *R09F10.5* | WBGene00019988 | protein_coding_gene | -1.0 | 0.48 | 2.5E-02 |
| *T25B9.3* | WBGene00012009 | protein_coding_gene | -1.1 | 0.48 | 3.4E-02 |
| *apl-1* | WBGene00000149 | protein_coding_gene | -1.1 | 0.48 | 4.7E-03 |
| *C32E12.4* | WBGene00016330 | protein_coding_gene | -1.1 | 0.48 | 7.9E-07 |

| | | | | | |
|---|---|---|---|---|---|
| *F18A11.2* | WBGene00008929 | protein_coding_gene | -1.1 | 0.48 | 1.4E-02 |
| *C34E11.2* | WBGene00007935 | protein_coding_gene | -1.1 | 0.48 | 1.3E-02 |
| *ceh-27* | WBGene00000449 | protein_coding_gene | -1.1 | 0.48 | 5.0E-03 |
| *lron-15* | WBGene00016974 | protein_coding_gene | -1.1 | 0.48 | 1.8E-03 |
| *C01B10.11* | WBGene00044294 | protein_coding_gene | -1.1 | 0.48 | 2.8E-02 |
| *C04E6.7* | WBGene00015423 | protein_coding_gene | -1.1 | 0.48 | 3.2E-05 |
| *rig-5* | WBGene00004372 | protein_coding_gene | -1.1 | 0.48 | 9.4E-03 |
| *nhr-88* | WBGene00003678 | protein_coding_gene | -1.1 | 0.48 | 2.8E-02 |
| *maoc-1* | WBGene00017123 | protein_coding_gene | -1.1 | 0.48 | 1.3E-03 |
| *E01G6.2* | WBGene00008450 | protein_coding_gene | -1.1 | 0.48 | 4.7E-04 |
| *cof-2* | WBGene00000583 | protein_coding_gene | -1.1 | 0.48 | 1.5E-02 |
| *F52H2.4* | WBGene00018716 | protein_coding_gene | -1.1 | 0.48 | 1.1E-02 |
| *unc-53* | WBGene00006788 | protein_coding_gene | -1.1 | 0.48 | 2.1E-04 |
| *Y48G8AL.13* | WBGene00021691 | protein_coding_gene | -1.1 | 0.48 | 2.2E-03 |
| *F16G10.15* | WBGene00017529 | pseudogene | -1.1 | 0.48 | 4.7E-02 |
| *R10E8.6* | WBGene00011212 | protein_coding_gene | -1.1 | 0.48 | 3.3E-02 |
| *T19C4.5* | WBGene00011839 | protein_coding_gene | -1.1 | 0.48 | 1.6E-03 |
| *K02D10.4* | WBGene00019304 | protein_coding_gene | -1.1 | 0.48 | 3.6E-02 |
| *K02C4.5* | WBGene00010503 | protein_coding_gene | -1.1 | 0.48 | 5.2E-04 |
| *T19B10.2* | WBGene00011831 | protein_coding_gene | -1.1 | 0.48 | 4.3E-02 |
| *T07D1.2* | WBGene00020301 | protein_coding_gene | -1.1 | 0.48 | 1.5E-08 |
| *Y42G9A.1* | WBGene00021531 | protein_coding_gene | -1.1 | 0.48 | 1.7E-02 |
| *gmeb-2* | WBGene00010010 | protein_coding_gene | -1.1 | 0.48 | 1.5E-02 |
| *E02H4.4* | WBGene00008462 | protein_coding_gene | -1.1 | 0.48 | 1.9E-04 |
| *igcm-3* | WBGene00020160 | protein_coding_gene | -1.1 | 0.48 | 8.2E-03 |
| *C54C6.7* | WBGene00044166 | protein_coding_gene | -1.1 | 0.48 | 1.9E-02 |
| *K10B3.5* | WBGene00019611 | protein_coding_gene | -1.1 | 0.47 | 2.6E-05 |
| *Y64G10A.7* | WBGene00013416 | protein_coding_gene | -1.1 | 0.47 | 1.1E-02 |
| *jmjd-3.1* | WBGene00017571 | protein_coding_gene | -1.1 | 0.47 | 1.5E-08 |
| *ads-1* | WBGene00000081 | protein_coding_gene | -1.1 | 0.47 | 4.7E-02 |
| *crm-1* | WBGene00007103 | protein_coding_gene | -1.1 | 0.47 | 5.4E-03 |
| *ncx-3* | WBGene00003568 | protein_coding_gene | -1.1 | 0.47 | 1.1E-05 |
| *rsf-1* | WBGene00011995 | protein_coding_gene | -1.1 | 0.47 | 7.7E-08 |
| *gana-1* | WBGene00011095 | protein_coding_gene | -1.1 | 0.47 | 6.4E-06 |
| *aex-3* | WBGene00000086 | protein_coding_gene | -1.1 | 0.47 | 2.0E-03 |
| *F53A2.1* | WBGene00009949 | protein_coding_gene | -1.1 | 0.47 | 6.3E-03 |
| *sek-6* | WBGene00012162 | protein_coding_gene | -1.1 | 0.47 | 9.2E-03 |
| *R09D1.9* | WBGene00011165 | pseudogene | -1.1 | 0.47 | 1.9E-02 |
| *K10D6.4* | WBGene00010744 | protein_coding_gene | -1.1 | 0.47 | 1.2E-09 |
| *C15B12.1* | WBGene00015783 | protein_coding_gene | -1.1 | 0.47 | 2.2E-03 |
| *ttx-1* | WBGene00006652 | protein_coding_gene | -1.1 | 0.47 | 1.7E-05 |
| *F57C9.6* | WBGene00019012 | protein_coding_gene | -1.1 | 0.47 | 4.8E-02 |
| *F25F8.1* | WBGene00017795 | protein_coding_gene | -1.1 | 0.47 | 2.5E-03 |
| *klo-1* | WBGene00016848 | protein_coding_gene | -1.1 | 0.47 | 2.1E-04 |
| *lfe-2* | WBGene00002979 | protein_coding_gene | -1.1 | 0.47 | 9.2E-04 |
| *ZK512.11* | WBGene00013989 | protein_coding_gene | -1.1 | 0.47 | 1.5E-03 |
| *viro-2* | WBGene00015107 | protein_coding_gene | -1.1 | 0.47 | 2.0E-02 |
| *F23F12.13* | WBGene00044455 | protein_coding_gene | -1.1 | 0.47 | 2.2E-02 |
| *fard-1* | WBGene00022200 | protein_coding_gene | -1.1 | 0.47 | 1.4E-03 |
| *ceh-34* | WBGene00000455 | protein_coding_gene | -1.1 | 0.47 | 9.1E-03 |
| *C45G7.13* | WBGene00235340 | protein_coding_gene | -1.1 | 0.47 | 4.9E-02 |
| *F08B12.4* | WBGene00008572 | protein_coding_gene | -1.1 | 0.47 | 9.0E-05 |
| *R03E9.2* | WBGene00019843 | protein_coding_gene | -1.1 | 0.47 | 3.9E-03 |
| *W06H8.6* | WBGene00021068 | protein_coding_gene | -1.1 | 0.47 | 2.1E-02 |
| *fmil-1* | WBGene00021362 | protein_coding_gene | -1.1 | 0.47 | 5.2E-03 |
| *C31H5.4* | WBGene00007855 | protein_coding_gene | -1.1 | 0.47 | 1.2E-08 |
| *T05E11.9* | WBGene00014505 | ncRNA_gene | -1.1 | 0.47 | 1.3E-03 |
| *M04D8.7* | WBGene00010865 | protein_coding_gene | -1.1 | 0.47 | 2.7E-02 |

| | | | | | |
|---|---|---|---|---|---|
| *daao-1* | WBGene00022076 | protein_coding_gene | -1.1 | 0.47 | 9.5E-03 |
| *D2045.8* | WBGene00008425 | protein_coding_gene | -1.1 | 0.46 | 2.3E-02 |
| *cyp-13A6* | WBGene00011673 | protein_coding_gene | -1.1 | 0.46 | 2.1E-02 |
| *Y18H1A.9* | WBGene00021213 | protein_coding_gene | -1.1 | 0.46 | 2.8E-02 |
| *Y54G2A.7* | WBGene00043702 | protein_coding_gene | -1.1 | 0.46 | 2.8E-02 |
| *ZK563.2* | WBGene00022767 | protein_coding_gene | -1.1 | 0.46 | 3.0E-02 |
| *glt-4* | WBGene00001622 | protein_coding_gene | -1.1 | 0.46 | 2.9E-02 |
| *cdh-8* | WBGene00017576 | protein_coding_gene | -1.1 | 0.46 | 4.1E-03 |
| *K07H8.12* | WBGene00044509 | protein_coding_gene | -1.1 | 0.46 | 1.3E-02 |
| *zig-7* | WBGene00006984 | protein_coding_gene | -1.1 | 0.46 | 3.1E-03 |
| *C16B8.4* | WBGene00015824 | protein_coding_gene | -1.1 | 0.46 | 5.3E-05 |
| *tep-1* | WBGene00013969 | protein_coding_gene | -1.1 | 0.46 | 3.5E-02 |
| *T01B6.1* | WBGene00020134 | protein_coding_gene | -1.1 | 0.46 | 3.0E-02 |
| *Y66D12A.13* | WBGene00013439 | protein_coding_gene | -1.1 | 0.46 | 4.9E-02 |
| *B0272.4* | WBGene00007130 | protein_coding_gene | -1.1 | 0.46 | 1.3E-03 |
| *ham-1* | WBGene00001820 | protein_coding_gene | -1.1 | 0.46 | 6.0E-04 |
| *pho-6* | WBGene00018696 | protein_coding_gene | -1.1 | 0.46 | 1.9E-02 |
| *fbxa-175* | WBGene00045293 | pseudogene | -1.1 | 0.46 | 2.2E-03 |
| *pdf-2* | WBGene00018590 | protein_coding_gene | -1.1 | 0.46 | 4.4E-02 |
| *C08G9.1* | WBGene00015619 | protein_coding_gene | -1.1 | 0.46 | 1.5E-03 |
| *vab-19* | WBGene00006882 | protein_coding_gene | -1.1 | 0.46 | 5.8E-03 |
| *acl-7* | WBGene00012911 | protein_coding_gene | -1.1 | 0.46 | 1.0E-03 |
| *C53C11.5* | WBGene00044770 | protein_coding_gene | -1.1 | 0.46 | 1.7E-04 |
| *Y57A10A.14* | WBGene00013257 | protein_coding_gene | -1.1 | 0.46 | 8.4E-05 |
| *C13C4.7* | WBGene00007551 | protein_coding_gene | -1.1 | 0.46 | 2.5E-03 |
| *C29H12.6* | WBGene00016237 | protein_coding_gene | -1.1 | 0.46 | 2.0E-03 |
| *C33D9.6* | WBGene00007900 | protein_coding_gene | -1.1 | 0.46 | 2.5E-04 |
| *let-23* | WBGene00002299 | protein_coding_gene | -1.1 | 0.46 | 6.0E-05 |
| *acox-1.6* | WBGene00010336 | protein_coding_gene | -1.1 | 0.46 | 1.2E-04 |
| *D1086.1* | WBGene00008388 | protein_coding_gene | -1.1 | 0.46 | 1.6E-03 |
| *fipr-11* | WBGene00008244 | protein_coding_gene | -1.1 | 0.46 | 3.2E-02 |
| *unc-130* | WBGene00006853 | protein_coding_gene | -1.1 | 0.46 | 5.6E-04 |
| *syd-1* | WBGene00006363 | protein_coding_gene | -1.1 | 0.46 | 1.8E-03 |
| *cft-1* | WBGene00000477 | protein_coding_gene | -1.1 | 0.46 | 2.9E-02 |
| *grdn-1* | WBGene00013082 | protein_coding_gene | -1.1 | 0.46 | 1.4E-08 |
| *pes-4* | WBGene00003978 | protein_coding_gene | -1.1 | 0.46 | 4.8E-05 |
| *E03A3.10* | WBGene00189944 | pseudogene | -1.1 | 0.46 | 2.3E-02 |
| *Y67A10A.9* | WBGene00013458 | protein_coding_gene | -1.1 | 0.46 | 3.3E-02 |
| *Y57A10A.26* | WBGene00013266 | protein_coding_gene | -1.1 | 0.46 | 6.2E-03 |
| *ctl-2* | WBGene00000831 | protein_coding_gene | -1.1 | 0.46 | 2.2E-02 |
| *sma-2* | WBGene00004856 | protein_coding_gene | -1.1 | 0.46 | 3.7E-02 |
| *ugt-11* | WBGene00020593 | protein_coding_gene | -1.1 | 0.46 | 2.6E-02 |
| *F17C11.6* | WBGene00008917 | protein_coding_gene | -1.1 | 0.46 | 1.5E-02 |
| *smp-2* | WBGene00004890 | protein_coding_gene | -1.1 | 0.46 | 8.5E-03 |
| *ndx-8* | WBGene00003585 | protein_coding_gene | -1.1 | 0.46 | 1.1E-04 |
| *let-522* | WBGene00010870 | protein_coding_gene | -1.1 | 0.46 | 1.8E-03 |
| *pha-2* | WBGene00004011 | protein_coding_gene | -1.1 | 0.46 | 1.3E-02 |
| *cest-35.2* | WBGene00016863 | protein_coding_gene | -1.1 | 0.45 | 2.9E-03 |
| *ace-3* | WBGene00000037 | protein_coding_gene | -1.1 | 0.45 | 9.0E-03 |
| *jmjd-3.3* | WBGene00007813 | protein_coding_gene | -1.1 | 0.45 | 2.0E-03 |
| *ceeh-1* | WBGene00019329 | protein_coding_gene | -1.1 | 0.45 | 1.2E-08 |
| *plpp-1.2* | WBGene00020895 | protein_coding_gene | -1.1 | 0.45 | 9.6E-03 |
| *C14A4.6* | WBGene00007557 | protein_coding_gene | -1.1 | 0.45 | 2.7E-03 |
| *BE0003N10.3* | WBGene00015268 | protein_coding_gene | -1.1 | 0.45 | 1.7E-03 |
| *nhr-110* | WBGene00003700 | protein_coding_gene | -1.1 | 0.45 | 1.8E-06 |
| *fln-2* | WBGene00016006 | protein_coding_gene | -1.1 | 0.45 | 1.2E-02 |
| *jkk-1* | WBGene00002177 | protein_coding_gene | -1.1 | 0.45 | 1.9E-02 |
| *bam-2* | WBGene00000237 | protein_coding_gene | -1.1 | 0.45 | 6.5E-04 |

| | | | | | |
|---|---|---|---|---|---|
| *nas-32* | WBGene00003550 | protein_coding_gene | -1.1 | 0.45 | 3.6E-02 |
| *unc-13* | WBGene00006752 | protein_coding_gene | -1.1 | 0.45 | 2.7E-02 |
| *lin-44* | WBGene00003029 | protein_coding_gene | -1.1 | 0.45 | 7.7E-03 |
| *zig-4* | WBGene00006981 | protein_coding_gene | -1.1 | 0.45 | 3.6E-03 |
| *clp-4* | WBGene00000545 | protein_coding_gene | -1.1 | 0.45 | 1.9E-03 |
| *cutl-21* | WBGene00018861 | protein_coding_gene | -1.1 | 0.45 | 9.7E-04 |
| *ZK512.1* | WBGene00013982 | protein_coding_gene | -1.1 | 0.45 | 9.2E-03 |
| *sgk-1* | WBGene00004789 | protein_coding_gene | -1.1 | 0.45 | 7.5E-04 |
| *F26F2.8* | WBGene00023504 | protein_coding_gene | -1.1 | 0.45 | 1.6E-02 |
| *ZC581.3* | WBGene00022633 | protein_coding_gene | -1.1 | 0.45 | 3.8E-03 |
| *C06G3.6* | WBGene00015552 | protein_coding_gene | -1.1 | 0.45 | 1.7E-07 |
| *che-3* | WBGene00000485 | protein_coding_gene | -1.1 | 0.45 | 5.2E-04 |
| *F41E7.2* | WBGene00009618 | protein_coding_gene | -1.1 | 0.45 | 8.6E-03 |
| *rei-2* | WBGene00019365 | protein_coding_gene | -1.2 | 0.45 | 5.6E-03 |
| *sulp-8* | WBGene00013963 | protein_coding_gene | -1.2 | 0.45 | 2.7E-04 |
| *R06F6.14* | WBGene00269421 | protein_coding_gene | -1.2 | 0.45 | 1.4E-02 |
| *bbs-9* | WBGene00016744 | protein_coding_gene | -1.2 | 0.45 | 3.3E-02 |
| *kel-1* | WBGene00002184 | protein_coding_gene | -1.2 | 0.45 | 4.5E-02 |
| *K09F5.6* | WBGene00044698 | protein_coding_gene | -1.2 | 0.45 | 8.8E-05 |
| *efl-3* | WBGene00009899 | protein_coding_gene | -1.2 | 0.45 | 1.8E-02 |
| *lgc-20* | WBGene00007191 | protein_coding_gene | -1.2 | 0.45 | 2.1E-02 |
| *F58A6.1* | WBGene00019022 | protein_coding_gene | -1.2 | 0.45 | 4.4E-03 |
| *hum-6* | WBGene00002039 | protein_coding_gene | -1.2 | 0.45 | 2.7E-05 |
| *glt-3* | WBGene00001621 | protein_coding_gene | -1.2 | 0.45 | 2.5E-02 |
| *col-89* | WBGene00000664 | protein_coding_gene | -1.2 | 0.45 | 2.2E-02 |
| *mec-3* | WBGene00003167 | protein_coding_gene | -1.2 | 0.45 | 1.6E-02 |
| *ssu-2* | WBGene00013455 | protein_coding_gene | -1.2 | 0.45 | 4.9E-02 |
| *frpr-18* | WBGene00020586 | protein_coding_gene | -1.2 | 0.45 | 4.7E-02 |
| *K09E2.3* | WBGene00019572 | protein_coding_gene | -1.2 | 0.45 | 2.9E-02 |
| *Y57G11C.22* | WBGene00013317 | protein_coding_gene | -1.2 | 0.45 | 2.4E-03 |
| *cpt-1* | WBGene00012907 | protein_coding_gene | -1.2 | 0.45 | 3.9E-05 |
| *Y55D5A.1* | WBGene00021915 | protein_coding_gene | -1.2 | 0.45 | 2.1E-05 |
| *nhr-171* | WBGene00016926 | protein_coding_gene | -1.2 | 0.45 | 3.1E-02 |
| *Y40B10A.9* | WBGene00021493 | protein_coding_gene | -1.2 | 0.45 | 1.5E-02 |
| *Y73F4A.2* | WBGene00013515 | protein_coding_gene | -1.2 | 0.45 | 4.9E-02 |
| *tbx-7* | WBGene00006544 | protein_coding_gene | -1.2 | 0.45 | 5.3E-03 |
| *epg-9* | WBGene00022078 | protein_coding_gene | -1.2 | 0.45 | 1.4E-04 |
| *abts-4* | WBGene00019844 | protein_coding_gene | -1.2 | 0.45 | 5.6E-05 |
| *C02B10.3* | WBGene00015328 | protein_coding_gene | -1.2 | 0.45 | 1.3E-02 |
| *glna-1* | WBGene00007480 | protein_coding_gene | -1.2 | 0.45 | 2.3E-02 |
| *F08F1.3* | WBGene00017257 | protein_coding_gene | -1.2 | 0.45 | 9.2E-04 |
| *nhr-36* | WBGene00017198 | protein_coding_gene | -1.2 | 0.45 | 9.7E-03 |
| *C12D5.3* | WBGene00015706 | protein_coding_gene | -1.2 | 0.45 | 3.4E-02 |
| *F23A7.3* | WBGene00009068 | protein_coding_gene | -1.2 | 0.45 | 2.8E-02 |
| *cpx-2* | WBGene00010520 | protein_coding_gene | -1.2 | 0.45 | 5.2E-03 |
| *R05G6.10* | WBGene00019902 | protein_coding_gene | -1.2 | 0.44 | 3.6E-02 |
| *srab-4* | WBGene00016363 | protein_coding_gene | -1.2 | 0.44 | 4.9E-02 |
| *R13A5.15* | WBGene00194726 | protein_coding_gene | -1.2 | 0.44 | 1.8E-02 |
| *hst-3.2* | WBGene00044303 | protein_coding_gene | -1.2 | 0.44 | 3.8E-04 |
| *cdh-1* | WBGene00019994 | protein_coding_gene | -1.2 | 0.44 | 3.6E-02 |
| *Y9C12A.1* | WBGene00021185 | protein_coding_gene | -1.2 | 0.44 | 3.2E-03 |
| *F10G8.8* | WBGene00008666 | protein_coding_gene | -1.2 | 0.44 | 1.7E-09 |
| *lips-15* | WBGene00012585 | protein_coding_gene | -1.2 | 0.44 | 4.8E-03 |
| *C07H4.1* | WBGene00007421 | protein_coding_gene | -1.2 | 0.44 | 2.1E-04 |
| *ttr-5* | WBGene00008040 | protein_coding_gene | -1.2 | 0.44 | 2.5E-02 |
| *hlh-30* | WBGene00020930 | protein_coding_gene | -1.2 | 0.44 | 2.4E-03 |
| *rdy-2* | WBGene00004328 | protein_coding_gene | -1.2 | 0.44 | 4.8E-02 |
| *ttr-54* | WBGene00011774 | protein_coding_gene | -1.2 | 0.44 | 4.4E-04 |

| | | | | | |
|---|---|---|---|---|---|
| *sym-2* | WBGene00006367 | protein_coding_gene | -1.2 | 0.44 | 2.9E-05 |
| *ptr-5* | WBGene00004220 | protein_coding_gene | -1.2 | 0.44 | 4.2E-03 |
| *tax-6* | WBGene00006527 | protein_coding_gene | -1.2 | 0.44 | 1.7E-03 |
| *R08D7.7* | WBGene00011147 | protein_coding_gene | -1.2 | 0.44 | 1.3E-03 |
| *nep-21* | WBGene00011794 | protein_coding_gene | -1.2 | 0.44 | 3.9E-02 |
| *ceh-31* | WBGene00000452 | protein_coding_gene | -1.2 | 0.44 | 9.7E-03 |
| *ctbp-1* | WBGene00006424 | protein_coding_gene | -1.2 | 0.44 | 9.6E-05 |
| *F59F5.7* | WBGene00010345 | protein_coding_gene | -1.2 | 0.44 | 2.6E-02 |
| *C28G1.6* | WBGene00016191 | protein_coding_gene | -1.2 | 0.44 | 2.7E-03 |
| *C09D4.1* | WBGene00015632 | protein_coding_gene | -1.2 | 0.44 | 4.0E-02 |
| *R07D5.2* | WBGene00011101 | protein_coding_gene | -1.2 | 0.44 | 9.1E-03 |
| *lin-22* | WBGene00003008 | protein_coding_gene | -1.2 | 0.44 | 2.5E-02 |
| *his-71* | WBGene00001945 | protein_coding_gene | -1.2 | 0.44 | 1.1E-02 |
| *unc-120* | WBGene00006844 | protein_coding_gene | -1.2 | 0.44 | 1.8E-06 |
| *acox-3* | WBGene00019060 | protein_coding_gene | -1.2 | 0.44 | 1.3E-04 |
| *R07B1.9* | WBGene00011081 | protein_coding_gene | -1.2 | 0.44 | 1.9E-02 |
| *aagr-1* | WBGene00017071 | protein_coding_gene | -1.2 | 0.44 | 6.2E-05 |
| *F15E6.9* | WBGene00017489 | protein_coding_gene | -1.2 | 0.44 | 2.7E-03 |
| *Y54G2A.52* | WBGene00045397 | protein_coding_gene | -1.2 | 0.44 | 4.2E-03 |
| *C02D5.4* | WBGene00043097 | protein_coding_gene | -1.2 | 0.44 | 1.8E-18 |
| *nlr-1* | WBGene00003772 | protein_coding_gene | -1.2 | 0.44 | 2.9E-03 |
| *unc-3* | WBGene00006743 | protein_coding_gene | -1.2 | 0.44 | 6.8E-04 |
| *F56H9.2* | WBGene00010174 | protein_coding_gene | -1.2 | 0.44 | 1.2E-04 |
| *nhr-67* | WBGene00003657 | protein_coding_gene | -1.2 | 0.44 | 1.7E-02 |
| *acs-14* | WBGene00008669 | protein_coding_gene | -1.2 | 0.44 | 4.1E-03 |
| *K02D7.1* | WBGene00019298 | protein_coding_gene | -1.2 | 0.44 | 5.7E-03 |
| *ceh-6* | WBGene00000431 | protein_coding_gene | -1.2 | 0.44 | 2.3E-04 |
| *T28B8.1* | WBGene00012113 | protein_coding_gene | -1.2 | 0.44 | 3.1E-05 |
| *D1005.2* | WBGene00016996 | protein_coding_gene | -1.2 | 0.44 | 4.7E-02 |
| *seb-2* | WBGene00014035 | protein_coding_gene | -1.2 | 0.44 | 4.1E-02 |
| *arrd-24* | WBGene00009852 | protein_coding_gene | -1.2 | 0.44 | 4.1E-02 |
| *F09F9.1* | WBGene00017306 | protein_coding_gene | -1.2 | 0.44 | 4.0E-03 |
| *F56C3.9* | WBGene00018946 | protein_coding_gene | -1.2 | 0.44 | 3.6E-02 |
| *C34C12.6* | WBGene00007925 | protein_coding_gene | -1.2 | 0.44 | 9.5E-03 |
| *C30G7.4* | WBGene00044178 | protein_coding_gene | -1.2 | 0.43 | 3.9E-02 |
| *K01A2.5* | WBGene00019280 | protein_coding_gene | -1.2 | 0.43 | 1.6E-03 |
| *F13C5.1* | WBGene00017422 | protein_coding_gene | -1.2 | 0.43 | 3.0E-02 |
| *ceeh-2* | WBGene00010628 | protein_coding_gene | -1.2 | 0.43 | 1.0E-02 |
| *tps-1* | WBGene00006602 | protein_coding_gene | -1.2 | 0.43 | 8.8E-03 |
| *Y39H10A.6* | WBGene00021482 | protein_coding_gene | -1.2 | 0.43 | 2.9E-03 |
| *ensh-1* | WBGene00017013 | protein_coding_gene | -1.2 | 0.43 | 2.2E-02 |
| *R10H1.1* | WBGene00019998 | protein_coding_gene | -1.2 | 0.43 | 2.8E-02 |
| *endu-1* | WBGene00010488 | protein_coding_gene | -1.2 | 0.43 | 2.2E-04 |
| *spp-22* | WBGene00044283 | protein_coding_gene | -1.2 | 0.43 | 1.1E-02 |
| *ptr-23* | WBGene00004237 | protein_coding_gene | -1.2 | 0.43 | 8.9E-03 |
| *tsp-20* | WBGene00006646 | protein_coding_gene | -1.2 | 0.43 | 2.6E-04 |
| *che-14* | WBGene00000493 | protein_coding_gene | -1.2 | 0.43 | 5.2E-03 |
| *K12B6.9* | WBGene00023432 | protein_coding_gene | -1.2 | 0.43 | 4.6E-04 |
| *twk-11* | WBGene00006666 | protein_coding_gene | -1.2 | 0.43 | 1.3E-02 |
| *shn-1* | WBGene00006444 | protein_coding_gene | -1.2 | 0.43 | 3.0E-02 |
| *nhr-206* | WBGene00011097 | protein_coding_gene | -1.2 | 0.43 | 3.9E-02 |
| *K01A2.3* | WBGene00019278 | pseudogene | -1.2 | 0.43 | 2.3E-02 |
| *F49H12.3* | WBGene00018654 | protein_coding_gene | -1.2 | 0.43 | 1.4E-02 |
| *nhr-32* | WBGene00003626 | protein_coding_gene | -1.2 | 0.43 | 1.3E-03 |
| *ZC443.4* | WBGene00013899 | protein_coding_gene | -1.2 | 0.43 | 4.9E-02 |
| *T19B10.5* | WBGene00011833 | protein_coding_gene | -1.2 | 0.43 | 9.8E-03 |
| *F32D1.11* | WBGene00044530 | protein_coding_gene | -1.2 | 0.43 | 2.4E-02 |
| *siah-1* | WBGene00021369 | protein_coding_gene | -1.2 | 0.43 | 4.9E-03 |

| | | | | | |
|---|---|---|---|---|---|
| *fbxa-42* | WBGene00044676 | protein_coding_gene | -1.2 | 0.43 | 8.3E-03 |
| *C13E3.1* | WBGene00015740 | protein_coding_gene | -1.2 | 0.43 | 2.0E-04 |
| *nhr-58* | WBGene00003648 | protein_coding_gene | -1.2 | 0.43 | 4.1E-02 |
| *che-13* | WBGene00000492 | protein_coding_gene | -1.2 | 0.43 | 1.8E-02 |
| *ckr-2* | WBGene00021439 | protein_coding_gene | -1.2 | 0.43 | 2.3E-02 |
| *ttr-17* | WBGene00012383 | protein_coding_gene | -1.2 | 0.43 | 2.3E-06 |
| *Y54F10BM.3* | WBGene00021858 | protein_coding_gene | -1.2 | 0.43 | 1.3E-02 |
| *Y105E8A.34* | WBGene00077689 | pseudogene | -1.2 | 0.43 | 2.4E-02 |
| *egl-38* | WBGene00001204 | protein_coding_gene | -1.2 | 0.43 | 1.5E-02 |
| *ttr-45* | WBGene00044737 | protein_coding_gene | -1.2 | 0.43 | 1.3E-02 |
| *K01A2.4* | WBGene00019279 | protein_coding_gene | -1.2 | 0.43 | 2.1E-05 |
| *nlp-82* | WBGene00044006 | protein_coding_gene | -1.2 | 0.43 | 2.1E-02 |
| *tag-273* | WBGene00013289 | protein_coding_gene | -1.2 | 0.43 | 2.6E-06 |
| *pho-4* | WBGene00004023 | protein_coding_gene | -1.2 | 0.43 | 9.5E-03 |
| *C05A9.2* | WBGene00007317 | protein_coding_gene | -1.2 | 0.43 | 7.6E-07 |
| *F53B3.3* | WBGene00018742 | protein_coding_gene | -1.2 | 0.43 | 2.0E-02 |
| *vab-15* | WBGene00006881 | protein_coding_gene | -1.2 | 0.43 | 3.6E-02 |
| *H41C03.1* | WBGene00019268 | protein_coding_gene | -1.2 | 0.43 | 1.4E-02 |
| *phlp-2* | WBGene00009647 | protein_coding_gene | -1.2 | 0.43 | 1.4E-02 |
| *C05B5.8* | WBGene00007324 | protein_coding_gene | -1.2 | 0.42 | 2.0E-02 |
| *mksr-1* | WBGene00019364 | protein_coding_gene | -1.2 | 0.42 | 2.2E-02 |
| *F57C12.2* | WBGene00019014 | protein_coding_gene | -1.2 | 0.42 | 9.0E-03 |
| *dgk-2* | WBGene00000959 | protein_coding_gene | -1.2 | 0.42 | 2.1E-04 |
| *lev-8* | WBGene00002975 | protein_coding_gene | -1.2 | 0.42 | 3.5E-03 |
| *F45E4.5* | WBGene00018470 | protein_coding_gene | -1.2 | 0.42 | 2.9E-03 |
| *dlg-1* | WBGene00001006 | protein_coding_gene | -1.2 | 0.42 | 8.3E-12 |
| *C44E12.1* | WBGene00016657 | protein_coding_gene | -1.2 | 0.42 | 3.6E-02 |
| *ZC262.13* | WBGene00284846 | protein_coding_gene | -1.2 | 0.42 | 2.6E-02 |
| *clc-2* | WBGene00000523 | protein_coding_gene | -1.2 | 0.42 | 3.5E-03 |
| *nhr-134* | WBGene00003724 | protein_coding_gene | -1.2 | 0.42 | 7.6E-04 |
| *ceh-37* | WBGene00000458 | protein_coding_gene | -1.2 | 0.42 | 6.7E-06 |
| *F58F12.4* | WBGene00044756 | protein_coding_gene | -1.2 | 0.42 | 5.6E-03 |
| *C25E10.12* | WBGene00016101 | protein_coding_gene | -1.2 | 0.42 | 2.8E-02 |
| *F38B6.6* | WBGene00018175 | protein_coding_gene | -1.2 | 0.42 | 4.2E-04 |
| *marc-4* | WBGene00016903 | protein_coding_gene | -1.3 | 0.42 | 5.9E-05 |
| *F32D1.3* | WBGene00017983 | protein_coding_gene | -1.3 | 0.42 | 3.4E-06 |
| *F53C3.8* | WBGene00018752 | protein_coding_gene | -1.3 | 0.42 | 1.3E-02 |
| *C24B5.4* | WBGene00016046 | protein_coding_gene | -1.3 | 0.42 | 1.3E-03 |
| *fipr-4* | WBGene00007541 | protein_coding_gene | -1.3 | 0.42 | 4.7E-02 |
| *T02C5.1* | WBGene00020158 | protein_coding_gene | -1.3 | 0.42 | 2.7E-03 |
| *sulp-4* | WBGene00010788 | protein_coding_gene | -1.3 | 0.42 | 1.7E-02 |
| *aat-1* | WBGene00000002 | protein_coding_gene | -1.3 | 0.42 | 3.9E-03 |
| *F22B7.9* | WBGene00017698 | protein_coding_gene | -1.3 | 0.42 | 4.7E-05 |
| *nhr-120* | WBGene00003710 | protein_coding_gene | -1.3 | 0.42 | 1.0E-02 |
| *gem-4* | WBGene00001577 | protein_coding_gene | -1.3 | 0.42 | 8.1E-06 |
| *C16H3.3* | WBGene00015868 | protein_coding_gene | -1.3 | 0.42 | 1.3E-07 |
| *VC5.2* | WBGene00020908 | protein_coding_gene | -1.3 | 0.42 | 5.8E-03 |
| *M01H9.4* | WBGene00019720 | protein_coding_gene | -1.3 | 0.42 | 6.0E-03 |
| *unc-29* | WBGene00006765 | protein_coding_gene | -1.3 | 0.42 | 1.3E-04 |
| *rab-28* | WBGene00004281 | protein_coding_gene | -1.3 | 0.42 | 1.4E-02 |
| *fbxa-157* | WBGene00007393 | protein_coding_gene | -1.3 | 0.42 | 4.4E-02 |
| *nex-2* | WBGene00003589 | protein_coding_gene | -1.3 | 0.42 | 8.9E-06 |
| *dhs-18* | WBGene00000981 | protein_coding_gene | -1.3 | 0.42 | 8.6E-09 |
| *pitr-3* | WBGene00015055 | protein_coding_gene | -1.3 | 0.42 | 1.8E-02 |
| *cest-33* | WBGene00017431 | protein_coding_gene | -1.3 | 0.42 | 2.8E-03 |
| *F49C12.6* | WBGene00009876 | protein_coding_gene | -1.3 | 0.42 | 1.8E-02 |
| *blmp-1* | WBGene00003847 | protein_coding_gene | -1.3 | 0.42 | 7.6E-03 |
| *C27D6.12* | WBGene00044389 | protein_coding_gene | -1.3 | 0.42 | 4.1E-03 |

| | | | | | |
|---|---|---|---|---|---|
| *ceh-32* | WBGene00000453 | protein_coding_gene | -1.3 | 0.42 | 2.6E-06 |
| *rskn-2* | WBGene00008311 | protein_coding_gene | -1.3 | 0.42 | 1.7E-02 |
| *mltn-9* | WBGene00008964 | protein_coding_gene | -1.3 | 0.42 | 4.3E-02 |
| *H35B03.1* | WBGene00019263 | protein_coding_gene | -1.3 | 0.42 | 3.5E-02 |
| *W03G11.4* | WBGene00012226 | protein_coding_gene | -1.3 | 0.41 | 2.2E-03 |
| *Y4C6B.3* | WBGene00021157 | protein_coding_gene | -1.3 | 0.41 | 1.9E-02 |
| *oac-6* | WBGene00007833 | protein_coding_gene | -1.3 | 0.41 | 1.6E-02 |
| *max-1* | WBGene00003143 | protein_coding_gene | -1.3 | 0.41 | 3.2E-05 |
| *nhr-41* | WBGene00022423 | protein_coding_gene | -1.3 | 0.41 | 4.9E-04 |
| *C07B5.4* | WBGene00007409 | protein_coding_gene | -1.3 | 0.41 | 1.6E-04 |
| *glna-2* | WBGene00008435 | protein_coding_gene | -1.3 | 0.41 | 4.7E-02 |
| *C05E7.1* | WBGene00007343 | protein_coding_gene | -1.3 | 0.41 | 7.6E-03 |
| *tub-1* | WBGene00006655 | protein_coding_gene | -1.3 | 0.41 | 4.7E-03 |
| *clec-54* | WBGene00008596 | protein_coding_gene | -1.3 | 0.41 | 7.1E-04 |
| *T24C12.1* | WBGene00020764 | protein_coding_gene | -1.3 | 0.41 | 4.5E-02 |
| *B0365.9* | WBGene00219208 | protein_coding_gene | -1.3 | 0.41 | 2.9E-02 |
| *M01D1.12* | WBGene00269373 | protein_coding_gene | -1.3 | 0.41 | 2.2E-02 |
| *T01D1.8* | WBGene00044641 | protein_coding_gene | -1.3 | 0.41 | 1.7E-02 |
| *F46H6.5* | WBGene00023460 | protein_coding_gene | -1.3 | 0.41 | 1.0E-02 |
| *ZK593.2* | WBGene00014002 | protein_coding_gene | -1.3 | 0.41 | 3.8E-02 |
| *atgp-1* | WBGene00000224 | protein_coding_gene | -1.3 | 0.41 | 1.6E-06 |
| *tmed-13* | WBGene00022255 | protein_coding_gene | -1.3 | 0.41 | 4.3E-03 |
| *Y55B1BL.1* | WBGene00021909 | protein_coding_gene | -1.3 | 0.41 | 1.1E-02 |
| *clec-25* | WBGene00044331 | protein_coding_gene | -1.3 | 0.41 | 1.1E-02 |
| *let-653* | WBGene00002827 | protein_coding_gene | -1.3 | 0.41 | 2.4E-03 |
| *nhr-131* | WBGene00003721 | protein_coding_gene | -1.3 | 0.41 | 3.6E-03 |
| *memo-1* | WBGene00016500 | protein_coding_gene | -1.3 | 0.41 | 1.1E-05 |
| *F38E9.1* | WBGene00018184 | protein_coding_gene | -1.3 | 0.41 | 1.0E-02 |
| *aff-1* | WBGene00016625 | protein_coding_gene | -1.3 | 0.41 | 2.6E-02 |
| *F20G2.3* | WBGene00008987 | protein_coding_gene | -1.3 | 0.41 | 3.8E-03 |
| *cht-1* | WBGene00000503 | protein_coding_gene | -1.3 | 0.41 | 4.5E-11 |
| *sox-2* | WBGene00004949 | protein_coding_gene | -1.3 | 0.41 | 3.4E-14 |
| *agr-1* | WBGene00018304 | protein_coding_gene | -1.3 | 0.41 | 6.6E-03 |
| *C34C6.3* | WBGene00007916 | protein_coding_gene | -1.3 | 0.41 | 3.5E-03 |
| *cest-27* | WBGene00018958 | protein_coding_gene | -1.3 | 0.41 | 5.4E-06 |
| *F56C9.8* | WBGene00018952 | protein_coding_gene | -1.3 | 0.41 | 4.0E-10 |
| *T27D12.6* | WBGene00194848 | protein_coding_gene | -1.3 | 0.41 | 2.1E-02 |
| *chil-23* | WBGene00011167 | protein_coding_gene | -1.3 | 0.41 | 3.0E-03 |
| *cest-3* | WBGene00008451 | protein_coding_gene | -1.3 | 0.41 | 7.4E-04 |
| *tsp-16* | WBGene00006642 | protein_coding_gene | -1.3 | 0.41 | 6.1E-06 |
| *C34C12.7* | WBGene00007926 | protein_coding_gene | -1.3 | 0.41 | 1.7E-02 |
| *nphp-4* | WBGene00011261 | protein_coding_gene | -1.3 | 0.41 | 6.5E-04 |
| *T25F10.3* | WBGene00020806 | protein_coding_gene | -1.3 | 0.41 | 2.6E-02 |
| *C24H10.3* | WBGene00016069 | protein_coding_gene | -1.3 | 0.41 | 2.8E-02 |
| *ccep-290* | WBGene00021643 | protein_coding_gene | -1.3 | 0.41 | 1.9E-03 |
| *C18G1.1* | WBGene00015980 | protein_coding_gene | -1.3 | 0.40 | 3.2E-02 |
| *E04F6.4* | WBGene00017124 | protein_coding_gene | -1.3 | 0.40 | 1.0E-02 |
| *T14B1.1* | WBGene00011763 | protein_coding_gene | -1.3 | 0.40 | 1.9E-06 |
| *F26A10.2* | WBGene00017814 | protein_coding_gene | -1.3 | 0.40 | 4.3E-03 |
| *Y47G6A.5* | WBGene00021635 | protein_coding_gene | -1.3 | 0.40 | 6.4E-04 |
| *ast-1* | WBGene00020368 | protein_coding_gene | -1.3 | 0.40 | 4.4E-03 |
| *T05A7.7* | WBGene00020224 | protein_coding_gene | -1.3 | 0.40 | 5.5E-03 |
| *T21D12.7* | WBGene00020648 | protein_coding_gene | -1.3 | 0.40 | 4.4E-02 |
| *magu-3* | WBGene00016841 | protein_coding_gene | -1.3 | 0.40 | 4.0E-03 |
| *lipl-6* | WBGene00021963 | protein_coding_gene | -1.3 | 0.40 | 1.7E-02 |
| *ric-3* | WBGene00004363 | protein_coding_gene | -1.3 | 0.40 | 2.6E-04 |
| *F57F5.3* | WBGene00010205 | protein_coding_gene | -1.3 | 0.40 | 3.1E-02 |
| *zipt-2.1* | WBGene00014669 | protein_coding_gene | -1.3 | 0.40 | 8.1E-03 |

| | | | | | |
|---|---|---|---|---|---|
| *mec-6* | WBGene00003170 | protein_coding_gene | -1.3 | 0.40 | 1.0E-02 |
| *C10C5.7* | WBGene00007511 | protein_coding_gene | -1.3 | 0.40 | 1.2E-03 |
| *C38C3.4* | WBGene00016513 | protein_coding_gene | -1.3 | 0.40 | 4.8E-03 |
| *W01B11.6* | WBGene00020917 | protein_coding_gene | -1.3 | 0.40 | 4.1E-02 |
| *D1086.5* | WBGene00008392 | protein_coding_gene | -1.3 | 0.40 | 1.7E-05 |
| *M79.2* | WBGene00010902 | protein_coding_gene | -1.3 | 0.40 | 4.1E-03 |
| *prmn-1* | WBGene00008571 | protein_coding_gene | -1.3 | 0.40 | 2.8E-03 |
| *C27B7.7* | WBGene00007764 | protein_coding_gene | -1.3 | 0.40 | 1.2E-03 |
| *bus-17* | WBGene00044630 | protein_coding_gene | -1.3 | 0.40 | 1.2E-02 |
| *tig-3* | WBGene00021594 | protein_coding_gene | -1.3 | 0.40 | 3.9E-02 |
| *C28G1.2* | WBGene00016187 | protein_coding_gene | -1.3 | 0.40 | 4.9E-02 |
| *math-34* | WBGene00019699 | protein_coding_gene | -1.3 | 0.40 | 4.5E-02 |
| *C31H5.7* | WBGene00007858 | protein_coding_gene | -1.3 | 0.40 | 2.2E-02 |
| *F53F10.1* | WBGene00018763 | protein_coding_gene | -1.3 | 0.40 | 3.1E-02 |
| *cest-16* | WBGene00020330 | protein_coding_gene | -1.3 | 0.40 | 2.3E-03 |
| *dyf-13* | WBGene00001129 | protein_coding_gene | -1.3 | 0.40 | 4.3E-02 |
| *K03H1.5* | WBGene00010540 | protein_coding_gene | -1.3 | 0.40 | 1.6E-02 |
| *ncx-10* | WBGene00003575 | protein_coding_gene | -1.3 | 0.40 | 6.0E-03 |
| *ncam-1* | WBGene00017184 | protein_coding_gene | -1.3 | 0.40 | 9.6E-09 |
| *cbn-1* | WBGene00019285 | protein_coding_gene | -1.3 | 0.40 | 2.6E-02 |
| *R05D8.7* | WBGene00019885 | protein_coding_gene | -1.3 | 0.40 | 7.3E-03 |
| *srp-2* | WBGene00005643 | protein_coding_gene | -1.3 | 0.40 | 3.3E-02 |
| *rhgf-2* | WBGene00006476 | protein_coding_gene | -1.3 | 0.40 | 4.1E-04 |
| *fkh-8* | WBGene00001440 | protein_coding_gene | -1.3 | 0.40 | 6.9E-04 |
| *C24H10.1* | WBGene00016067 | protein_coding_gene | -1.3 | 0.40 | 4.2E-04 |
| *unc-129* | WBGene00006852 | protein_coding_gene | -1.3 | 0.40 | 3.2E-04 |
| *F48D6.4* | WBGene00018602 | protein_coding_gene | -1.3 | 0.40 | 2.4E-02 |
| *C27A7.1* | WBGene00007753 | protein_coding_gene | -1.3 | 0.39 | 4.8E-02 |
| *clc-5* | WBGene00000526 | protein_coding_gene | -1.3 | 0.39 | 3.1E-03 |
| *cest-19* | WBGene00020125 | protein_coding_gene | -1.3 | 0.39 | 9.8E-03 |
| *B0403.6* | WBGene00043982 | protein_coding_gene | -1.3 | 0.39 | 7.4E-03 |
| *tag-196* | WBGene00007055 | protein_coding_gene | -1.3 | 0.39 | 4.5E-03 |
| *Y75B8A.6* | WBGene00013543 | protein_coding_gene | -1.3 | 0.39 | 3.3E-08 |
| *oac-46* | WBGene00011768 | protein_coding_gene | -1.3 | 0.39 | 1.4E-03 |
| *exc-5* | WBGene00001366 | protein_coding_gene | -1.3 | 0.39 | 2.9E-03 |
| *F01G10.9* | WBGene00008511 | protein_coding_gene | -1.3 | 0.39 | 3.6E-02 |
| *F20C5.6* | WBGene00008971 | protein_coding_gene | -1.4 | 0.39 | 2.4E-03 |
| *T13C2.3* | WBGene00020479 | protein_coding_gene | -1.4 | 0.39 | 2.8E-05 |
| *hpo-36* | WBGene00020855 | protein_coding_gene | -1.4 | 0.39 | 3.3E-02 |
| *ptr-6* | WBGene00004221 | protein_coding_gene | -1.4 | 0.39 | 1.5E-02 |
| *C25F6.7* | WBGene00016106 | protein_coding_gene | -1.4 | 0.39 | 2.5E-02 |
| *C10A4.3* | WBGene00015664 | protein_coding_gene | -1.4 | 0.39 | 2.3E-02 |
| *hhat-2* | WBGene00013312 | protein_coding_gene | -1.4 | 0.39 | 1.4E-04 |
| *hsp-12.1* | WBGene00011906 | protein_coding_gene | -1.4 | 0.39 | 2.3E-02 |
| *dep-1* | WBGene00009717 | protein_coding_gene | -1.4 | 0.39 | 6.5E-10 |
| *frpr-8* | WBGene00018728 | protein_coding_gene | -1.4 | 0.39 | 1.6E-02 |
| *Y34F4.1* | WBGene00021336 | protein_coding_gene | -1.4 | 0.39 | 7.2E-04 |
| *Y82E9BL.19* | WBGene00194735 | protein_coding_gene | -1.4 | 0.39 | 2.5E-02 |
| *C06G1.1* | WBGene00015545 | protein_coding_gene | -1.4 | 0.39 | 1.3E-03 |
| *F08F1.4* | WBGene00017258 | protein_coding_gene | -1.4 | 0.39 | 9.9E-03 |
| *C01A2.2* | WBGene00007214 | protein_coding_gene | -1.4 | 0.39 | 2.9E-03 |
| *ptr-17* | WBGene00004231 | protein_coding_gene | -1.4 | 0.39 | 4.5E-02 |
| *let-381* | WBGene00002601 | protein_coding_gene | -1.4 | 0.39 | 8.4E-03 |
| *osg-1* | WBGene00019832 | protein_coding_gene | -1.4 | 0.39 | 3.9E-03 |
| *egl-43* | WBGene00001207 | protein_coding_gene | -1.4 | 0.39 | 1.9E-07 |
| *tbc-16* | WBGene00018639 | protein_coding_gene | -1.4 | 0.39 | 1.3E-02 |
| *cwn-2* | WBGene00000858 | protein_coding_gene | -1.4 | 0.39 | 4.1E-04 |
| *oac-51* | WBGene00020976 | protein_coding_gene | -1.4 | 0.39 | 2.9E-03 |

| | | | | | |
|---|---|---|---|---|---|
| *F25B3.5* | WBGene00009102 | protein_coding_gene | -1.4 | 0.39 | 3.7E-02 |
| *sma-4* | WBGene00004858 | protein_coding_gene | -1.4 | 0.39 | 1.9E-03 |
| *cutl-17* | WBGene00009541 | protein_coding_gene | -1.4 | 0.39 | 1.4E-02 |
| *C06G3.5* | WBGene00015551 | protein_coding_gene | -1.4 | 0.39 | 5.8E-03 |
| *C26B9.6* | WBGene00016135 | protein_coding_gene | -1.4 | 0.39 | 1.4E-05 |
| *R148.5* | WBGene00020104 | protein_coding_gene | -1.4 | 0.39 | 2.5E-18 |
| *F11E6.8* | WBGene00008711 | protein_coding_gene | -1.4 | 0.39 | 1.2E-05 |
| *F53A2.9* | WBGene00009953 | protein_coding_gene | -1.4 | 0.39 | 7.9E-03 |
| *F58E6.13* | WBGene00077697 | protein_coding_gene | -1.4 | 0.39 | 4.7E-02 |
| *igdb-3* | WBGene00020691 | protein_coding_gene | -1.4 | 0.39 | 2.9E-03 |
| *fkb-4* | WBGene00001429 | protein_coding_gene | -1.4 | 0.39 | 1.1E-02 |
| *F25D1.5* | WBGene00009110 | protein_coding_gene | -1.4 | 0.39 | 4.3E-02 |
| *cfz-2* | WBGene00000478 | protein_coding_gene | -1.4 | 0.39 | 2.8E-02 |
| *F35D2.3* | WBGene00018041 | protein_coding_gene | -1.4 | 0.39 | 1.4E-02 |
| *Y116F11B.6* | WBGene00013822 | protein_coding_gene | -1.4 | 0.39 | 2.4E-02 |
| *Y57G11B.2* | WBGene00013295 | protein_coding_gene | -1.4 | 0.39 | 1.0E-02 |
| *hda-5* | WBGene00009663 | protein_coding_gene | -1.4 | 0.38 | 1.1E-03 |
| *vab-3* | WBGene00006870 | protein_coding_gene | -1.4 | 0.38 | 1.2E-09 |
| *F31D5.6* | WBGene00017949 | protein_coding_gene | -1.4 | 0.38 | 1.9E-02 |
| *pmt-1* | WBGene00022781 | protein_coding_gene | -1.4 | 0.38 | 1.6E-03 |
| *rrc-1* | WBGene00009800 | protein_coding_gene | -1.4 | 0.38 | 5.8E-05 |
| *cest-17* | WBGene00019477 | protein_coding_gene | -1.4 | 0.38 | 2.9E-05 |
| *lim-4* | WBGene00002987 | protein_coding_gene | -1.4 | 0.38 | 6.0E-03 |
| *F13B12.4* | WBGene00008732 | protein_coding_gene | -1.4 | 0.38 | 3.8E-02 |
| *ZC487.1* | WBGene00022623 | protein_coding_gene | -1.4 | 0.38 | 5.3E-04 |
| *T19A6.4* | WBGene00011829 | protein_coding_gene | -1.4 | 0.38 | 1.4E-04 |
| *Y48A6B.7* | WBGene00012968 | protein_coding_gene | -1.4 | 0.38 | 4.5E-04 |
| *D1069.1* | WBGene00017036 | protein_coding_gene | -1.4 | 0.38 | 1.2E-02 |
| *vglu-2* | WBGene00010755 | protein_coding_gene | -1.4 | 0.38 | 5.0E-03 |
| *col-153* | WBGene00000726 | protein_coding_gene | -1.4 | 0.38 | 2.0E-02 |
| *F28G4.2* | WBGene00009227 | protein_coding_gene | -1.4 | 0.38 | 1.9E-02 |
| *prkl-1* | WBGene00022727 | protein_coding_gene | -1.4 | 0.38 | 5.9E-05 |
| *lgc-28* | WBGene00044301 | protein_coding_gene | -1.4 | 0.38 | 4.0E-02 |
| *Y6E2A.5* | WBGene00012399 | protein_coding_gene | -1.4 | 0.38 | 9.1E-03 |
| *clh-4* | WBGene00000531 | protein_coding_gene | -1.4 | 0.38 | 1.6E-02 |
| *ceh-22* | WBGene00000445 | protein_coding_gene | -1.4 | 0.38 | 8.1E-04 |
| *dyf-1* | WBGene00001117 | protein_coding_gene | -1.4 | 0.38 | 1.3E-03 |
| *nmur-1* | WBGene00016747 | protein_coding_gene | -1.4 | 0.38 | 1.8E-02 |
| *ltd-1* | WBGene00003089 | protein_coding_gene | -1.4 | 0.38 | 2.1E-05 |
| *Y6B3B.7* | WBGene00012391 | protein_coding_gene | -1.4 | 0.38 | 3.4E-03 |
| *plpr-1* | WBGene00011524 | protein_coding_gene | -1.4 | 0.38 | 2.1E-02 |
| *flp-10* | WBGene00001453 | protein_coding_gene | -1.4 | 0.38 | 7.9E-03 |
| *jbts-14* | WBGene00018727 | protein_coding_gene | -1.4 | 0.38 | 2.9E-02 |
| *C40H1.2* | WBGene00008038 | protein_coding_gene | -1.4 | 0.38 | 5.2E-04 |
| *C02F4.4* | WBGene00007264 | protein_coding_gene | -1.4 | 0.38 | 3.5E-02 |
| *ZC196.3* | WBGene00022547 | protein_coding_gene | -1.4 | 0.38 | 1.3E-02 |
| *C03A7.2* | WBGene00015366 | protein_coding_gene | -1.4 | 0.38 | 3.6E-02 |
| *fkh-7* | WBGene00001439 | protein_coding_gene | -1.4 | 0.38 | 9.4E-03 |
| *ZK822.1* | WBGene00014089 | protein_coding_gene | -1.4 | 0.38 | 2.2E-03 |
| *glt-6* | WBGene00001624 | protein_coding_gene | -1.4 | 0.38 | 3.1E-02 |
| *grh-1* | WBGene00001707 | protein_coding_gene | -1.4 | 0.38 | 1.7E-04 |
| *unc-51* | WBGene00006786 | protein_coding_gene | -1.4 | 0.38 | 5.8E-04 |
| *aqp-2* | WBGene00000170 | protein_coding_gene | -1.4 | 0.38 | 1.2E-04 |
| *C24A8.5* | WBGene00016039 | protein_coding_gene | -1.4 | 0.38 | 3.3E-02 |
| *hot-6* | WBGene00001991 | protein_coding_gene | -1.4 | 0.38 | 5.8E-05 |
| *R09E12.9* | WBGene00044514 | protein_coding_gene | -1.4 | 0.38 | 8.6E-03 |
| *sym-1* | WBGene00006366 | protein_coding_gene | -1.4 | 0.38 | 2.4E-02 |
| *nhr-92* | WBGene00003682 | protein_coding_gene | -1.4 | 0.38 | 5.0E-03 |

| | | | | | |
|---|---|---|---|---|---|
| *ttc-36* | WBGene00009947 | protein_coding_gene | -1.4 | 0.38 | 1.8E-02 |
| *nhr-25* | WBGene00003623 | protein_coding_gene | -1.4 | 0.38 | 6.0E-12 |
| *cex-1* | WBGene00023407 | protein_coding_gene | -1.4 | 0.37 | 2.5E-02 |
| *F57C12.6* | WBGene00194687 | protein_coding_gene | -1.4 | 0.37 | 3.2E-02 |
| *F32D8.15* | WBGene00009325 | protein_coding_gene | -1.4 | 0.37 | 8.2E-03 |
| *F13B12.2* | WBGene00008730 | protein_coding_gene | -1.4 | 0.37 | 2.6E-02 |
| *M176.4* | WBGene00010943 | protein_coding_gene | -1.4 | 0.37 | 7.2E-03 |
| *asm-1* | WBGene00000211 | protein_coding_gene | -1.4 | 0.37 | 1.1E-05 |
| *C56G2.4* | WBGene00016979 | protein_coding_gene | -1.4 | 0.37 | 4.9E-03 |
| *ceh-48* | WBGene00015934 | protein_coding_gene | -1.4 | 0.37 | 2.6E-04 |
| *col-99* | WBGene00000674 | protein_coding_gene | -1.4 | 0.37 | 1.6E-03 |
| *nhr-119* | WBGene00003709 | protein_coding_gene | -1.4 | 0.37 | 1.9E-02 |
| *F19C7.2* | WBGene00017592 | protein_coding_gene | -1.4 | 0.37 | 4.3E-02 |
| *tpra-1* | WBGene00008366 | protein_coding_gene | -1.4 | 0.37 | 7.6E-03 |
| *ZC455.1* | WBGene00013902 | protein_coding_gene | -1.4 | 0.37 | 4.0E-05 |
| *syd-9* | WBGene00044068 | protein_coding_gene | -1.4 | 0.37 | 1.4E-04 |
| *K12C11.6* | WBGene00045052 | protein_coding_gene | -1.4 | 0.37 | 5.5E-05 |
| *aexr-1* | WBGene00016089 | protein_coding_gene | -1.4 | 0.37 | 1.7E-02 |
| *cgt-1* | WBGene00011517 | protein_coding_gene | -1.4 | 0.37 | 1.6E-03 |
| *F18G5.6* | WBGene00017582 | protein_coding_gene | -1.4 | 0.37 | 1.5E-02 |
| *nhr-69* | WBGene00003659 | protein_coding_gene | -1.4 | 0.37 | 2.8E-06 |
| *nhr-18* | WBGene00003617 | protein_coding_gene | -1.4 | 0.37 | 4.0E-02 |
| *dyf-5* | WBGene00001121 | protein_coding_gene | -1.4 | 0.37 | 8.8E-04 |
| *F10D7.10* | WBGene00271701 | protein_coding_gene | -1.4 | 0.37 | 3.8E-03 |
| *sod-4* | WBGene00004933 | protein_coding_gene | -1.4 | 0.37 | 1.7E-03 |
| *ham-2* | WBGene00001821 | protein_coding_gene | -1.4 | 0.37 | 5.5E-16 |
| *dlc-3* | WBGene00012424 | protein_coding_gene | -1.4 | 0.37 | 6.3E-05 |
| *dex-1* | WBGene00017028 | protein_coding_gene | -1.4 | 0.37 | 8.9E-03 |
| *crtc-1* | WBGene00021237 | protein_coding_gene | -1.4 | 0.37 | 3.5E-04 |
| *ttr-51* | WBGene00010440 | protein_coding_gene | -1.4 | 0.37 | 9.5E-04 |
| *ceh-79* | WBGene00007749 | protein_coding_gene | -1.4 | 0.37 | 5.8E-04 |
| *M03D4.3* | WBGene00019750 | protein_coding_gene | -1.4 | 0.37 | 2.0E-02 |
| *unc-30* | WBGene00006766 | protein_coding_gene | -1.4 | 0.37 | 3.3E-05 |
| *akt-2* | WBGene00000103 | protein_coding_gene | -1.5 | 0.37 | 5.0E-03 |
| *Y70G10A.3* | WBGene00013499 | protein_coding_gene | -1.5 | 0.37 | 3.8E-02 |
| *W03F9.4* | WBGene00021002 | protein_coding_gene | -1.5 | 0.37 | 1.6E-03 |
| *pef-1* | WBGene00003969 | protein_coding_gene | -1.5 | 0.36 | 4.3E-04 |
| *K01F9.2* | WBGene00010475 | protein_coding_gene | -1.5 | 0.36 | 6.7E-03 |
| *ugt-57* | WBGene00017154 | protein_coding_gene | -1.5 | 0.36 | 6.4E-03 |
| *Y50E8A.5* | WBGene00013048 | protein_coding_gene | -1.5 | 0.36 | 1.3E-03 |
| *C55A1.6* | WBGene00008323 | protein_coding_gene | -1.5 | 0.36 | 4.0E-02 |
| *Y59H11AR.4* | WBGene00022012 | protein_coding_gene | -1.5 | 0.36 | 2.5E-02 |
| *cyp-34A7* | WBGene00015042 | protein_coding_gene | -1.5 | 0.36 | 3.4E-02 |
| *F58E6.11* | WBGene00010258 | protein_coding_gene | -1.5 | 0.36 | 4.1E-03 |
| *mrp-3* | WBGene00003409 | protein_coding_gene | -1.5 | 0.36 | 9.7E-04 |
| *lam-3* | WBGene00002248 | protein_coding_gene | -1.5 | 0.36 | 1.3E-04 |
| *skr-19* | WBGene00004825 | protein_coding_gene | -1.5 | 0.36 | 1.8E-06 |
| *pals-33* | WBGene00021081 | protein_coding_gene | -1.5 | 0.36 | 1.3E-02 |
| *Y58A7A.2* | WBGene00021976 | protein_coding_gene | -1.5 | 0.36 | 9.0E-03 |
| *F59F5.3* | WBGene00010342 | protein_coding_gene | -1.5 | 0.36 | 4.5E-03 |
| *dhs-15* | WBGene00000978 | protein_coding_gene | -1.5 | 0.36 | 1.1E-02 |
| *T01D3.3* | WBGene00011328 | protein_coding_gene | -1.5 | 0.36 | 3.0E-02 |
| *K09E2.2* | WBGene00019571 | protein_coding_gene | -1.5 | 0.36 | 4.2E-02 |
| *afmd-2* | WBGene00012866 | protein_coding_gene | -1.5 | 0.36 | 3.4E-06 |
| *F17C8.3* | WBGene00008910 | protein_coding_gene | -1.5 | 0.36 | 1.6E-04 |
| *F32B4.5* | WBGene00009315 | protein_coding_gene | -1.5 | 0.36 | 9.4E-03 |
| *Y71H2AM.15* | WBGene00022180 | protein_coding_gene | -1.5 | 0.36 | 7.7E-05 |
| *droe-4* | WBGene00010262 | protein_coding_gene | -1.5 | 0.36 | 9.1E-03 |

| | | | | | |
|---|---|---|---|---|---|
| *K08B5.1* | WBGene00019518 | protein_coding_gene | -1.5 | 0.36 | 1.4E-02 |
| *T19H12.15* | WBGene00271827 | protein_coding_gene | -1.5 | 0.36 | 5.3E-04 |
| *cbp-2* | WBGene00010545 | protein_coding_gene | -1.5 | 0.36 | 7.6E-06 |
| *sulp-3* | WBGene00018283 | protein_coding_gene | -1.5 | 0.36 | 6.5E-03 |
| *R10D12.1* | WBGene00011185 | protein_coding_gene | -1.5 | 0.36 | 6.1E-03 |
| *F54C8.6* | WBGene00010039 | protein_coding_gene | -1.5 | 0.36 | 1.3E-05 |
| *Y22D7AL.6* | WBGene00021245 | protein_coding_gene | -1.5 | 0.36 | 1.6E-02 |
| *F28H7.2* | WBGene00009236 | protein_coding_gene | -1.5 | 0.36 | 4.9E-02 |
| *E02H4.5* | WBGene00008463 | protein_coding_gene | -1.5 | 0.36 | 3.7E-03 |
| *F41G4.8* | WBGene00018311 | protein_coding_gene | -1.5 | 0.36 | 2.9E-03 |
| *C36E8.4* | WBGene00007983 | protein_coding_gene | -1.5 | 0.36 | 3.7E-10 |
| *mks-3* | WBGene00018042 | protein_coding_gene | -1.5 | 0.36 | 1.3E-02 |
| *osm-5* | WBGene00003885 | protein_coding_gene | -1.5 | 0.36 | 6.1E-04 |
| *F07A5.4* | WBGene00008543 | protein_coding_gene | -1.5 | 0.36 | 2.6E-02 |
| *frpr-11* | WBGene00019444 | protein_coding_gene | -1.5 | 0.36 | 4.5E-07 |
| *egl-46* | WBGene00001210 | protein_coding_gene | -1.5 | 0.36 | 4.7E-02 |
| *F16C3.3* | WBGene00023415 | protein_coding_gene | -1.5 | 0.35 | 8.9E-03 |
| *nlp-45* | WBGene00020136 | protein_coding_gene | -1.5 | 0.35 | 9.7E-05 |
| *glb-8* | WBGene00007741 | protein_coding_gene | -1.5 | 0.35 | 5.8E-05 |
| *T28F4.5* | WBGene00012140 | protein_coding_gene | -1.5 | 0.35 | 3.7E-03 |
| *K09C8.8* | WBGene00010718 | protein_coding_gene | -1.5 | 0.35 | 3.0E-02 |
| *cyp-25A3* | WBGene00014697 | protein_coding_gene | -1.5 | 0.35 | 2.1E-05 |
| *pgp-11* | WBGene00004005 | protein_coding_gene | -1.5 | 0.35 | 2.7E-02 |
| *cex-2* | WBGene00023408 | protein_coding_gene | -1.5 | 0.35 | 5.8E-05 |
| *nhr-238* | WBGene00021611 | protein_coding_gene | -1.5 | 0.35 | 2.2E-02 |
| *eva-1* | WBGene00009304 | protein_coding_gene | -1.5 | 0.35 | 7.2E-05 |
| *unc-75* | WBGene00006807 | protein_coding_gene | -1.5 | 0.35 | 3.0E-03 |
| *ZK673.1* | WBGene00014057 | protein_coding_gene | -1.5 | 0.35 | 2.1E-06 |
| *Y116A8C.40* | WBGene00013812 | protein_coding_gene | -1.5 | 0.35 | 3.1E-02 |
| *mlt-8* | WBGene00021095 | protein_coding_gene | -1.5 | 0.35 | 2.9E-02 |
| *ptr-1* | WBGene00004216 | protein_coding_gene | -1.5 | 0.35 | 6.3E-03 |
| *ZK470.14* | WBGene00219362 | protein_coding_gene | -1.5 | 0.35 | 7.2E-03 |
| *Y42A5A.3* | WBGene00012778 | protein_coding_gene | -1.5 | 0.35 | 3.4E-02 |
| *T27A10.6* | WBGene00020846 | protein_coding_gene | -1.5 | 0.35 | 1.2E-03 |
| *nhr-205* | WBGene00011002 | protein_coding_gene | -1.5 | 0.35 | 5.5E-03 |
| *chpf-2* | WBGene00019108 | protein_coding_gene | -1.5 | 0.35 | 6.3E-06 |
| *T12A7.6* | WBGene00011727 | protein_coding_gene | -1.5 | 0.35 | 1.2E-03 |
| *dhs-16* | WBGene00000979 | protein_coding_gene | -1.5 | 0.35 | 1.8E-02 |
| *K08E7.6* | WBGene00010674 | protein_coding_gene | -1.5 | 0.35 | 3.8E-04 |
| *dhrs-4* | WBGene00010063 | protein_coding_gene | -1.5 | 0.35 | 1.3E-18 |
| *glc-3* | WBGene00001593 | protein_coding_gene | -1.5 | 0.35 | 1.4E-02 |
| *T22B7.3* | WBGene00020672 | protein_coding_gene | -1.5 | 0.35 | 2.0E-02 |
| *F47G3.4* | WBGene00194641 | protein_coding_gene | -1.5 | 0.35 | 2.2E-02 |
| *Y4C6B.2* | WBGene00021156 | protein_coding_gene | -1.5 | 0.35 | 1.9E-02 |
| *prk-2* | WBGene00004183 | protein_coding_gene | -1.5 | 0.35 | 5.2E-04 |
| *cyp-23A1* | WBGene00015135 | protein_coding_gene | -1.5 | 0.35 | 1.3E-02 |
| *insc-1* | WBGene00018392 | protein_coding_gene | -1.5 | 0.35 | 4.8E-05 |
| *cfi-1* | WBGene00000476 | protein_coding_gene | -1.5 | 0.35 | 7.9E-08 |
| *subs-4* | WBGene00012939 | protein_coding_gene | -1.5 | 0.35 | 1.2E-02 |
| *C14A11.2* | WBGene00015748 | protein_coding_gene | -1.5 | 0.35 | 2.5E-03 |
| *T01E8.1* | WBGene00011331 | protein_coding_gene | -1.5 | 0.35 | 6.8E-07 |
| *F52D1.2* | WBGene00018683 | protein_coding_gene | -1.5 | 0.35 | 8.0E-03 |
| *fipr-23* | WBGene00007990 | protein_coding_gene | -1.5 | 0.34 | 4.3E-02 |
| *ugt-2* | WBGene00007073 | protein_coding_gene | -1.5 | 0.34 | 4.9E-16 |
| *catp-7* | WBGene00022010 | protein_coding_gene | -1.5 | 0.34 | 7.0E-07 |
| *cyp-33C12* | WBGene00021168 | protein_coding_gene | -1.5 | 0.34 | 2.2E-02 |
| *Y52B11A.4* | WBGene00013124 | protein_coding_gene | -1.5 | 0.34 | 1.6E-07 |
| *R57.2* | WBGene00020083 | protein_coding_gene | -1.5 | 0.34 | 2.5E-02 |

| | | | | | |
|---|---|---|---|---|---|
| *mam-1* | WBGene00022504 | protein_coding_gene | -1.5 | 0.34 | 1.4E-02 |
| *vap-2* | WBGene00011462 | protein_coding_gene | -1.5 | 0.34 | 1.3E-02 |
| *F43G6.17* | WBGene00219378 | protein_coding_gene | -1.5 | 0.34 | 2.5E-03 |
| *bgnt-1.2* | WBGene00017723 | protein_coding_gene | -1.5 | 0.34 | 2.5E-02 |
| *F20D6.10* | WBGene00017639 | protein_coding_gene | -1.6 | 0.34 | 3.4E-10 |
| *Y43F8C.3* | WBGene00012825 | protein_coding_gene | -1.6 | 0.34 | 1.1E-05 |
| *ist-1* | WBGene00002163 | protein_coding_gene | -1.6 | 0.34 | 2.2E-02 |
| *lpr-1* | WBGene00002393 | protein_coding_gene | -1.6 | 0.34 | 8.9E-03 |
| *nhx-9* | WBGene00003736 | protein_coding_gene | -1.6 | 0.34 | 3.4E-03 |
| *K04H4.2* | WBGene00010573 | protein_coding_gene | -1.6 | 0.34 | 1.7E-02 |
| *sma-10* | WBGene00020649 | protein_coding_gene | -1.6 | 0.34 | 5.1E-04 |
| *dma-1* | WBGene00011345 | protein_coding_gene | -1.6 | 0.34 | 1.4E-06 |
| *C44H9.5* | WBGene00008100 | protein_coding_gene | -1.6 | 0.34 | 9.4E-06 |
| *Y6G8.2* | WBGene00012404 | protein_coding_gene | -1.6 | 0.34 | 1.2E-04 |
| *unc-86* | WBGene00006818 | protein_coding_gene | -1.6 | 0.34 | 3.3E-05 |
| *bkip-1* | WBGene00014569 | protein_coding_gene | -1.6 | 0.34 | 1.3E-04 |
| *C07A4.3* | WBGene00007398 | protein_coding_gene | -1.6 | 0.34 | 1.3E-03 |
| *dyf-7* | WBGene00001123 | protein_coding_gene | -1.6 | 0.33 | 4.2E-02 |
| *F13D11.3* | WBGene00017428 | protein_coding_gene | -1.6 | 0.33 | 2.2E-15 |
| *F46G11.6* | WBGene00045269 | protein_coding_gene | -1.6 | 0.33 | 4.7E-02 |
| *plx-2* | WBGene00004048 | protein_coding_gene | -1.6 | 0.33 | 2.8E-07 |
| *C26B9.3* | WBGene00016133 | protein_coding_gene | -1.6 | 0.33 | 7.7E-03 |
| *JC8.16* | WBGene00194915 | protein_coding_gene | -1.6 | 0.33 | 1.7E-02 |
| *F32H2.11* | WBGene00043049 | protein_coding_gene | -1.6 | 0.33 | 4.3E-02 |
| *W05B2.2* | WBGene00012271 | protein_coding_gene | -1.6 | 0.33 | 6.8E-03 |
| *M03F4.6* | WBGene00019759 | protein_coding_gene | -1.6 | 0.33 | 3.1E-04 |
| *hbl-1* | WBGene00001824 | protein_coding_gene | -1.6 | 0.33 | 2.4E-02 |
| *F12E12.11* | WBGene00044811 | protein_coding_gene | -1.6 | 0.33 | 1.1E-03 |
| *osm-6* | WBGene00003886 | protein_coding_gene | -1.6 | 0.33 | 1.1E-04 |
| *pgp-10* | WBGene00004004 | protein_coding_gene | -1.6 | 0.33 | 2.7E-02 |
| *C35E7.4* | WBGene00016456 | protein_coding_gene | -1.6 | 0.33 | 5.9E-06 |
| *F36F12.1* | WBGene00018092 | protein_coding_gene | -1.6 | 0.33 | 1.8E-02 |
| *F48E3.8* | WBGene00018607 | protein_coding_gene | -1.6 | 0.33 | 1.1E-11 |
| *lido-15* | WBGene00018804 | protein_coding_gene | -1.6 | 0.33 | 4.4E-05 |
| *lgc-27* | WBGene00022283 | protein_coding_gene | -1.6 | 0.33 | 1.3E-03 |
| *dcar-1* | WBGene00007395 | protein_coding_gene | -1.6 | 0.33 | 1.8E-02 |
| *ZK287.1* | WBGene00013962 | protein_coding_gene | -1.6 | 0.33 | 6.9E-04 |
| *W01C8.1* | WBGene00020918 | protein_coding_gene | -1.6 | 0.33 | 4.7E-02 |
| *fozi-1* | WBGene00010453 | protein_coding_gene | -1.6 | 0.33 | 2.5E-07 |
| *nlp-31* | WBGene00003769 | protein_coding_gene | -1.6 | 0.33 | 4.6E-02 |
| *glo-3* | WBGene00010341 | protein_coding_gene | -1.6 | 0.33 | 6.0E-04 |
| *C35E7.6* | WBGene00016458 | protein_coding_gene | -1.6 | 0.33 | 4.6E-08 |
| *K06A4.2* | WBGene00010592 | protein_coding_gene | -1.6 | 0.33 | 9.4E-03 |
| *igcm-1* | WBGene00018215 | protein_coding_gene | -1.6 | 0.33 | 2.9E-05 |
| *kpc-1* | WBGene00002232 | protein_coding_gene | -1.6 | 0.33 | 5.3E-14 |
| *ceh-5* | WBGene00000430 | protein_coding_gene | -1.6 | 0.33 | 4.3E-02 |
| *C17C3.3* | WBGene00015889 | protein_coding_gene | -1.6 | 0.33 | 3.1E-02 |
| *gln-3* | WBGene00001604 | protein_coding_gene | -1.6 | 0.33 | 2.4E-04 |
| *cyp-13A3* | WBGene00011675 | protein_coding_gene | -1.6 | 0.33 | 1.1E-02 |
| *W05B10.4* | WBGene00012279 | protein_coding_gene | -1.6 | 0.33 | 8.7E-05 |
| *T04A6.1* | WBGene00020200 | protein_coding_gene | -1.6 | 0.33 | 4.0E-12 |
| *ZK1128.7* | WBGene00014233 | protein_coding_gene | -1.6 | 0.33 | 4.1E-11 |
| *gcl-1* | WBGene00013382 | protein_coding_gene | -1.6 | 0.33 | 1.1E-04 |
| *M03D4.4* | WBGene00019751 | protein_coding_gene | -1.6 | 0.33 | 5.7E-06 |
| *clec-44* | WBGene00019928 | protein_coding_gene | -1.6 | 0.33 | 1.1E-02 |
| *F34H10.3* | WBGene00009380 | protein_coding_gene | -1.6 | 0.33 | 7.5E-05 |
| *nhr-172* | WBGene00016927 | protein_coding_gene | -1.6 | 0.33 | 4.7E-02 |
| *R07G3.8* | WBGene00019944 | protein_coding_gene | -1.6 | 0.33 | 1.6E-05 |

| | | | | | |
|---|---|---|---|---|---|
| *bro-1* | WBGene00000272 | protein_coding_gene | -1.6 | 0.33 | 4.0E-03 |
| *Y73E7A.8* | WBGene00022274 | protein_coding_gene | -1.6 | 0.33 | 1.7E-02 |
| *ZC449.2* | WBGene00022612 | protein_coding_gene | -1.6 | 0.32 | 2.6E-02 |
| *ddr-2* | WBGene00017381 | protein_coding_gene | -1.6 | 0.32 | 3.6E-06 |
| *F31C3.6* | WBGene00009288 | protein_coding_gene | -1.6 | 0.32 | 1.2E-07 |
| *F09F9.4* | WBGene00017309 | protein_coding_gene | -1.6 | 0.32 | 1.2E-03 |
| *F41G3.3* | WBGene00018300 | protein_coding_gene | -1.6 | 0.32 | 2.1E-02 |
| *faah-6* | WBGene00012750 | protein_coding_gene | -1.6 | 0.32 | 4.0E-02 |
| *zig-10* | WBGene00020800 | protein_coding_gene | -1.6 | 0.32 | 3.0E-03 |
| *T22C8.6* | WBGene00011927 | protein_coding_gene | -1.6 | 0.32 | 2.6E-02 |
| *cnc-11* | WBGene00044900 | protein_coding_gene | -1.6 | 0.32 | 2.1E-02 |
| *srt-63* | WBGene00020877 | protein_coding_gene | -1.6 | 0.32 | 3.7E-02 |
| *gba-1* | WBGene00016335 | protein_coding_gene | -1.6 | 0.32 | 6.5E-04 |
| *Y51B9A.9* | WBGene00013091 | protein_coding_gene | -1.6 | 0.32 | 6.2E-05 |
| *zag-1* | WBGene00006970 | protein_coding_gene | -1.6 | 0.32 | 5.6E-11 |
| *ZK105.13* | WBGene00235114 | protein_coding_gene | -1.6 | 0.32 | 2.6E-04 |
| *trx-3* | WBGene00019717 | protein_coding_gene | -1.6 | 0.32 | 1.1E-03 |
| *slc-28.2* | WBGene00017868 | protein_coding_gene | -1.6 | 0.32 | 6.2E-03 |
| *Y47H10A.5* | WBGene00012961 | protein_coding_gene | -1.6 | 0.32 | 3.3E-03 |
| *Y87G2A.25* | WBGene00219288 | protein_coding_gene | -1.6 | 0.32 | 1.7E-04 |
| *F26A1.9* | WBGene00017807 | protein_coding_gene | -1.6 | 0.32 | 1.1E-02 |
| *cuti-1* | WBGene00022591 | protein_coding_gene | -1.6 | 0.32 | 9.0E-03 |
| *bus-12* | WBGene00010442 | protein_coding_gene | -1.6 | 0.32 | 1.6E-02 |
| *cyp-33E1* | WBGene00016768 | protein_coding_gene | -1.6 | 0.32 | 7.0E-04 |
| *ugt-1* | WBGene00007072 | protein_coding_gene | -1.6 | 0.32 | 1.8E-02 |
| *F01G12.18* | WBGene00206382 | protein_coding_gene | -1.6 | 0.32 | 3.3E-02 |
| *ins-7* | WBGene00002090 | protein_coding_gene | -1.6 | 0.32 | 5.2E-08 |
| *F46C3.6* | WBGene00045253 | protein_coding_gene | -1.7 | 0.32 | 2.1E-04 |
| *Y2H9A.4* | WBGene00012381 | protein_coding_gene | -1.7 | 0.32 | 2.6E-03 |
| *F36D1.7* | WBGene00009466 | protein_coding_gene | -1.7 | 0.32 | 1.2E-07 |
| *acs-3* | WBGene00020343 | protein_coding_gene | -1.7 | 0.32 | 1.4E-13 |
| *igcm-2* | WBGene00020130 | protein_coding_gene | -1.7 | 0.32 | 1.6E-04 |
| *F53H2.1* | WBGene00010004 | protein_coding_gene | -1.7 | 0.32 | 2.2E-03 |
| *ZK402.5* | WBGene00022731 | protein_coding_gene | -1.7 | 0.32 | 2.3E-02 |
| *nlp-48* | WBGene00008002 | protein_coding_gene | -1.7 | 0.32 | 1.5E-02 |
| *T16G1.2* | WBGene00011796 | protein_coding_gene | -1.7 | 0.32 | 5.4E-03 |
| *F42C5.4* | WBGene00018346 | protein_coding_gene | -1.7 | 0.32 | 4.8E-05 |
| *nhr-180* | WBGene00017961 | protein_coding_gene | -1.7 | 0.32 | 4.1E-03 |
| *T17H7.7* | WBGene00020551 | protein_coding_gene | -1.7 | 0.32 | 9.2E-03 |
| *unk-1* | WBGene00016407 | protein_coding_gene | -1.7 | 0.32 | 2.8E-03 |
| *T01C3.11* | WBGene00011325 | protein_coding_gene | -1.7 | 0.32 | 2.3E-02 |
| *F49D11.6* | WBGene00018630 | protein_coding_gene | -1.7 | 0.31 | 2.3E-02 |
| *clec-40* | WBGene00011853 | protein_coding_gene | -1.7 | 0.31 | 1.7E-02 |
| *zmp-5* | WBGene00010423 | protein_coding_gene | -1.7 | 0.31 | 6.2E-04 |
| *twk-7* | WBGene00006662 | protein_coding_gene | -1.7 | 0.31 | 1.7E-03 |
| *mks-2* | WBGene00194710 | protein_coding_gene | -1.7 | 0.31 | 3.1E-02 |
| *C29F7.2* | WBGene00007811 | protein_coding_gene | -1.7 | 0.31 | 1.8E-31 |
| *nsy-4* | WBGene00021415 | protein_coding_gene | -1.7 | 0.31 | 2.1E-09 |
| *F17C11.4* | WBGene00008915 | protein_coding_gene | -1.7 | 0.31 | 1.1E-04 |
| *F01D5.6* | WBGene00008496 | protein_coding_gene | -1.7 | 0.31 | 1.2E-02 |
| *F58A3.3* | WBGene00010223 | protein_coding_gene | -1.7 | 0.31 | 1.1E-04 |
| *F07C3.9* | WBGene00017197 | protein_coding_gene | -1.7 | 0.31 | 3.2E-08 |
| *bath-25* | WBGene00020859 | protein_coding_gene | -1.7 | 0.31 | 3.2E-02 |
| *Y95B8A.6* | WBGene00022386 | protein_coding_gene | -1.7 | 0.31 | 1.2E-03 |
| *fbxa-163* | WBGene00015598 | protein_coding_gene | -1.7 | 0.31 | 2.5E-02 |
| *dct-1* | WBGene00015776 | protein_coding_gene | -1.7 | 0.31 | 4.5E-09 |
| *chil-28* | WBGene00008842 | pseudogene | -1.7 | 0.31 | 8.1E-03 |
| *mks-5* | WBGene00007490 | protein_coding_gene | -1.7 | 0.31 | 1.4E-03 |

| | | | | | |
|---|---|---|---|---|---|
| *Y77E11A.14* | WBGene00022314 | protein_coding_gene | -1.7 | 0.31 | 1.8E-02 |
| *Y53F4B.20* | WBGene00013166 | protein_coding_gene | -1.7 | 0.31 | 9.2E-03 |
| *F23H12.5* | WBGene00009094 | protein_coding_gene | -1.7 | 0.31 | 7.4E-03 |
| *tmem-218* | WBGene00044771 | protein_coding_gene | -1.7 | 0.31 | 3.2E-02 |
| *F22B7.1* | WBGene00017692 | protein_coding_gene | -1.7 | 0.31 | 7.3E-09 |
| *nas-33* | WBGene00003551 | protein_coding_gene | -1.7 | 0.31 | 1.3E-02 |
| *bus-8* | WBGene00044623 | protein_coding_gene | -1.7 | 0.31 | 2.9E-02 |
| *ZC239.14* | WBGene00022572 | protein_coding_gene | -1.7 | 0.31 | 4.8E-03 |
| *C45E5.3* | WBGene00016666 | pseudogene | -1.7 | 0.31 | 8.3E-04 |
| *aqp-5* | WBGene00000173 | protein_coding_gene | -1.7 | 0.31 | 1.0E-02 |
| *linc-103* | WBGene00219743 | lincRNA_gene | -1.7 | 0.31 | 3.9E-02 |
| *R06B10.2* | WBGene00019913 | protein_coding_gene | -1.7 | 0.31 | 1.6E-02 |
| *ZC84.6* | WBGene00013849 | protein_coding_gene | -1.7 | 0.31 | 1.3E-02 |
| *F40G9.20* | WBGene00195034 | ncRNA_gene | -1.7 | 0.31 | 9.1E-04 |
| *linc-43* | WBGene00219579 | lincRNA_gene | -1.7 | 0.31 | 4.1E-02 |
| *M7.8* | WBGene00010884 | protein_coding_gene | -1.7 | 0.31 | 8.6E-20 |
| *best-1* | WBGene00007203 | protein_coding_gene | -1.7 | 0.31 | 1.4E-03 |
| *gei-13* | WBGene00001570 | protein_coding_gene | -1.7 | 0.31 | 2.7E-05 |
| *daam-1* | WBGene00018976 | protein_coding_gene | -1.7 | 0.31 | 2.0E-06 |
| *fbxa-62* | WBGene00016521 | protein_coding_gene | -1.7 | 0.31 | 4.0E-02 |
| *W01F3.2* | WBGene00012185 | protein_coding_gene | -1.7 | 0.30 | 9.4E-04 |
| *Y37F4.8* | WBGene00044289 | protein_coding_gene | -1.7 | 0.30 | 1.5E-02 |
| *C06G3.3* | WBGene00015549 | protein_coding_gene | -1.7 | 0.30 | 3.1E-02 |
| *F36H1.12* | WBGene00009497 | protein_coding_gene | -1.7 | 0.30 | 2.4E-03 |
| *C03A3.1* | WBGene00007266 | protein_coding_gene | -1.7 | 0.30 | 3.6E-05 |
| *lgc-22* | WBGene00017483 | protein_coding_gene | -1.7 | 0.30 | 7.3E-03 |
| *Y39H10A.1* | WBGene00021478 | protein_coding_gene | -1.7 | 0.30 | 3.2E-02 |
| *lips-10* | WBGene00008803 | protein_coding_gene | -1.7 | 0.30 | 2.4E-02 |
| *Y71G12B.31* | WBGene00044348 | protein_coding_gene | -1.7 | 0.30 | 4.4E-03 |
| *C26E1.2* | WBGene00007748 | protein_coding_gene | -1.7 | 0.30 | 2.4E-03 |
| *ugt-51* | WBGene00015369 | protein_coding_gene | -1.7 | 0.30 | 1.7E-09 |
| *spp-8* | WBGene00004993 | protein_coding_gene | -1.7 | 0.30 | 3.5E-06 |
| *F43G6.8* | WBGene00009660 | protein_coding_gene | -1.7 | 0.30 | 3.9E-06 |
| *C07A9.12* | WBGene00043994 | protein_coding_gene | -1.7 | 0.30 | 2.4E-04 |
| *cyp-42A1* | WBGene00013585 | protein_coding_gene | -1.7 | 0.30 | 1.0E-06 |
| *F22F7.3* | WBGene00017721 | protein_coding_gene | -1.7 | 0.30 | 1.6E-03 |
| *Y57A10A.24* | WBGene00013264 | protein_coding_gene | -1.7 | 0.30 | 3.2E-02 |
| *F41C3.2* | WBGene00018268 | protein_coding_gene | -1.7 | 0.30 | 4.6E-02 |
| *gstk-2* | WBGene00017054 | protein_coding_gene | -1.7 | 0.30 | 1.2E-02 |
| *bgnt-1.3* | WBGene00009032 | protein_coding_gene | -1.7 | 0.30 | 3.8E-02 |
| *ttm-2* | WBGene00017840 | protein_coding_gene | -1.7 | 0.30 | 4.2E-10 |
| *moe-3* | WBGene00003388 | protein_coding_gene | -1.7 | 0.30 | 3.3E-03 |
| *pals-19* | WBGene00044761 | protein_coding_gene | -1.7 | 0.30 | 2.8E-03 |
| *F55H12.2* | WBGene00010133 | protein_coding_gene | -1.7 | 0.30 | 1.0E-08 |
| *C17H12.11* | WBGene00015936 | protein_coding_gene | -1.7 | 0.30 | 9.4E-06 |
| *cyp-14A3* | WBGene00010707 | protein_coding_gene | -1.7 | 0.30 | 2.8E-02 |
| *fbxa-27* | WBGene00022333 | protein_coding_gene | -1.7 | 0.30 | 1.6E-03 |
| *W05H9.3* | WBGene00021050 | ncRNA_gene | -1.7 | 0.30 | 3.2E-05 |
| *ZK836.3* | WBGene00014099 | protein_coding_gene | -1.7 | 0.30 | 5.4E-06 |
| *rga-2* | WBGene00013135 | protein_coding_gene | -1.8 | 0.30 | 2.2E-03 |
| *K10D3.4* | WBGene00010738 | protein_coding_gene | -1.8 | 0.30 | 1.2E-03 |
| *F57B1.6* | WBGene00010189 | protein_coding_gene | -1.8 | 0.30 | 1.9E-02 |
| *npr-20* | WBGene00011578 | protein_coding_gene | -1.8 | 0.30 | 9.9E-05 |
| *Y51H4A.5* | WBGene00013101 | protein_coding_gene | -1.8 | 0.30 | 1.2E-03 |
| *abhd-5.1* | WBGene00016506 | protein_coding_gene | -1.8 | 0.30 | 1.3E-03 |
| *nhr-91* | WBGene00003681 | protein_coding_gene | -1.8 | 0.29 | 7.2E-06 |
| *C34D1.4* | WBGene00007931 | protein_coding_gene | -1.8 | 0.29 | 2.8E-03 |
| *C32D5.12* | WBGene00016319 | protein_coding_gene | -1.8 | 0.29 | 1.9E-06 |

| | | | | | |
|---|---|---|---|---|---|
| *F55H12.7* | WBGene00219283 | protein_coding_gene | -1.8 | 0.29 | 1.5E-04 |
| *C08B6.2* | WBGene00007423 | protein_coding_gene | -1.8 | 0.29 | 5.6E-03 |
| *best-3* | WBGene00015286 | protein_coding_gene | -1.8 | 0.29 | 1.4E-03 |
| *ptr-9* | WBGene00004223 | protein_coding_gene | -1.8 | 0.29 | 2.4E-02 |
| *F30H5.3* | WBGene00017937 | protein_coding_gene | -1.8 | 0.29 | 2.2E-03 |
| *clec-145* | WBGene00013013 | protein_coding_gene | -1.8 | 0.29 | 2.0E-02 |
| *nlp-28* | WBGene00003766 | protein_coding_gene | -1.8 | 0.29 | 2.5E-02 |
| *F54B11.10* | WBGene00010033 | protein_coding_gene | -1.8 | 0.29 | 5.6E-03 |
| *D1086.2* | WBGene00008389 | protein_coding_gene | -1.8 | 0.29 | 3.6E-03 |
| *swip-10* | WBGene00018738 | protein_coding_gene | -1.8 | 0.29 | 8.5E-05 |
| *cutl-16* | WBGene00019426 | protein_coding_gene | -1.8 | 0.29 | 7.6E-07 |
| *nlp-34* | WBGene00015046 | protein_coding_gene | -1.8 | 0.29 | 4.7E-02 |
| *pxd-1* | WBGene00007982 | protein_coding_gene | -1.8 | 0.29 | 1.0E-17 |
| *fmi-1* | WBGene00001475 | protein_coding_gene | -1.8 | 0.29 | 5.7E-04 |
| *M60.4* | WBGene00019780 | protein_coding_gene | -1.8 | 0.29 | 2.5E-02 |
| *ptr-19* | WBGene00004233 | protein_coding_gene | -1.8 | 0.29 | 8.4E-03 |
| *dlc-5* | WBGene00043067 | protein_coding_gene | -1.8 | 0.29 | 6.2E-05 |
| *T18D3.7* | WBGene00011824 | protein_coding_gene | -1.8 | 0.29 | 1.6E-07 |
| *nab-1* | WBGene00003516 | protein_coding_gene | -1.8 | 0.29 | 6.3E-05 |
| *acs-22* | WBGene00017012 | protein_coding_gene | -1.8 | 0.29 | 2.0E-12 |
| *ZK550.2* | WBGene00013996 | protein_coding_gene | -1.8 | 0.29 | 1.4E-03 |
| *C25B8.8* | WBGene00044297 | protein_coding_gene | -1.8 | 0.29 | 2.5E-03 |
| *Y87G2A.16* | WBGene00013602 | protein_coding_gene | -1.8 | 0.29 | 4.8E-03 |
| *Y43F8A.1* | WBGene00012808 | protein_coding_gene | -1.8 | 0.29 | 7.5E-04 |
| *gcy-34* | WBGene00001554 | protein_coding_gene | -1.8 | 0.29 | 9.2E-03 |
| *pals-26* | WBGene00007131 | protein_coding_gene | -1.8 | 0.29 | 3.6E-02 |
| *ttr-37* | WBGene00017667 | protein_coding_gene | -1.8 | 0.29 | 6.2E-04 |
| *C06E4.8* | WBGene00015537 | protein_coding_gene | -1.8 | 0.29 | 2.7E-07 |
| *pho-11* | WBGene00007331 | protein_coding_gene | -1.8 | 0.29 | 3.1E-06 |
| *F56H11.2* | WBGene00010175 | protein_coding_gene | -1.8 | 0.29 | 3.1E-07 |
| *F59B10.4* | WBGene00010320 | protein_coding_gene | -1.8 | 0.29 | 1.7E-02 |
| *noah-1* | WBGene00016422 | protein_coding_gene | -1.8 | 0.28 | 2.3E-03 |
| *tmem-231* | WBGene00020825 | protein_coding_gene | -1.8 | 0.28 | 1.1E-03 |
| *cyp-33C7* | WBGene00018260 | protein_coding_gene | -1.8 | 0.28 | 5.8E-05 |
| *T04C12.8* | WBGene00011430 | protein_coding_gene | -1.8 | 0.28 | 4.9E-04 |
| *Y43C5A.3* | WBGene00012783 | protein_coding_gene | -1.8 | 0.28 | 1.6E-02 |
| *R04B5.5* | WBGene00011003 | protein_coding_gene | -1.8 | 0.28 | 1.4E-03 |
| *ptr-15* | WBGene00004229 | protein_coding_gene | -1.8 | 0.28 | 1.6E-04 |
| *Y97E10AR.1* | WBGene00022396 | protein_coding_gene | -1.8 | 0.28 | 4.9E-06 |
| *qua-1* | WBGene00004264 | protein_coding_gene | -1.8 | 0.28 | 1.3E-03 |
| *T05F1.11* | WBGene00011495 | protein_coding_gene | -1.8 | 0.28 | 8.2E-04 |
| *ZC84.1* | WBGene00013846 | protein_coding_gene | -1.8 | 0.28 | 3.6E-02 |
| *cky-1* | WBGene00000521 | protein_coding_gene | -1.8 | 0.28 | 1.9E-02 |
| *noah-2* | WBGene00009926 | protein_coding_gene | -1.8 | 0.28 | 4.9E-03 |
| *F36F2.2* | WBGene00009476 | protein_coding_gene | -1.8 | 0.28 | 2.1E-05 |
| *C27A7.8* | WBGene00007758 | protein_coding_gene | -1.8 | 0.28 | 3.4E-04 |
| *B0393.5* | WBGene00007170 | protein_coding_gene | -1.8 | 0.28 | 2.8E-03 |
| *B0507.8* | WBGene00015225 | protein_coding_gene | -1.8 | 0.28 | 1.2E-02 |
| *R07E3.6* | WBGene00011107 | protein_coding_gene | -1.8 | 0.28 | 1.8E-02 |
| *sri-23* | WBGene00005535 | pseudogene | -1.8 | 0.28 | 2.8E-02 |
| *F10D7.4* | WBGene00017341 | protein_coding_gene | -1.8 | 0.28 | 2.5E-03 |
| *T05E7.4* | WBGene00020260 | protein_coding_gene | -1.8 | 0.28 | 1.1E-02 |
| *F55G7.4* | WBGene00045493 | protein_coding_gene | -1.9 | 0.28 | 2.6E-02 |
| *pros-1* | WBGene00000448 | protein_coding_gene | -1.9 | 0.28 | 3.3E-08 |
| *F18C5.10* | WBGene00017563 | protein_coding_gene | -1.9 | 0.28 | 4.4E-10 |
| *ttr-44* | WBGene00008341 | protein_coding_gene | -1.9 | 0.28 | 8.5E-03 |
| *tbb-4* | WBGene00006538 | protein_coding_gene | -1.9 | 0.28 | 1.3E-03 |
| *Y65B4BL.1* | WBGene00022033 | protein_coding_gene | -1.9 | 0.28 | 1.9E-03 |

| | | | | | |
|---|---|---|---|---|---|
| *Y57E12B.10* | WBGene00202501 | protein_coding_gene | -1.9 | 0.28 | 1.4E-02 |
| *C27B7.9* | WBGene00007765 | protein_coding_gene | -1.9 | 0.28 | 3.7E-02 |
| *W08G11.1* | WBGene00012346 | protein_coding_gene | -1.9 | 0.28 | 2.6E-02 |
| *C35A5.6* | WBGene00007950 | protein_coding_gene | -1.9 | 0.27 | 9.8E-05 |
| *F33H2.8* | WBGene00009371 | protein_coding_gene | -1.9 | 0.27 | 1.3E-04 |
| *C09F5.1* | WBGene00015647 | protein_coding_gene | -1.9 | 0.27 | 2.8E-15 |
| *cdf-2* | WBGene00011821 | protein_coding_gene | -1.9 | 0.27 | 5.8E-06 |
| *pals-5* | WBGene00007659 | protein_coding_gene | -1.9 | 0.27 | 4.0E-02 |
| *cyp-35B1* | WBGene00019472 | protein_coding_gene | -1.9 | 0.27 | 8.6E-04 |
| *mab-7* | WBGene00003104 | protein_coding_gene | -1.9 | 0.27 | 1.0E-04 |
| *mod-5* | WBGene00003387 | protein_coding_gene | -1.9 | 0.27 | 3.1E-05 |
| *F20D6.8* | WBGene00017637 | protein_coding_gene | -1.9 | 0.27 | 9.8E-03 |
| *cyp-35A3* | WBGene00019565 | protein_coding_gene | -1.9 | 0.27 | 3.6E-02 |
| *chil-22* | WBGene00011166 | protein_coding_gene | -1.9 | 0.27 | 2.3E-03 |
| *fax-1* | WBGene00001400 | protein_coding_gene | -1.9 | 0.27 | 1.2E-03 |
| *nlp-30* | WBGene00003768 | protein_coding_gene | -1.9 | 0.27 | 2.0E-02 |
| *T14B4.19* | WBGene00302993 | protein_coding_gene | -1.9 | 0.27 | 7.3E-03 |
| *clec-78* | WBGene00018547 | protein_coding_gene | -1.9 | 0.27 | 1.0E-02 |
| *Y69H2.10* | WBGene00013485 | protein_coding_gene | -1.9 | 0.27 | 3.4E-02 |
| *C13C12.2* | WBGene00007552 | protein_coding_gene | -1.9 | 0.27 | 1.2E-03 |
| *T19C4.1* | WBGene00011838 | protein_coding_gene | -1.9 | 0.27 | 5.8E-05 |
| *glt-5* | WBGene00001623 | protein_coding_gene | -1.9 | 0.27 | 8.4E-10 |
| *T10H4.13* | WBGene00045339 | protein_coding_gene | -1.9 | 0.27 | 1.8E-03 |
| *nphp-1* | WBGene00010898 | protein_coding_gene | -1.9 | 0.27 | 1.7E-03 |
| *hmit-1.3* | WBGene00010811 | protein_coding_gene | -1.9 | 0.27 | 6.3E-08 |
| *bbs-1* | WBGene00000241 | protein_coding_gene | -1.9 | 0.27 | 2.3E-04 |
| *C01C4.2* | WBGene00015292 | protein_coding_gene | -1.9 | 0.27 | 1.2E-02 |
| *btb-15* | WBGene00015594 | pseudogene | -1.9 | 0.27 | 3.0E-02 |
| *C10C5.3* | WBGene00007507 | protein_coding_gene | -1.9 | 0.27 | 1.1E-02 |
| *ugt-52* | WBGene00018931 | protein_coding_gene | -1.9 | 0.27 | 8.8E-07 |
| *clec-75* | WBGene00021586 | protein_coding_gene | -1.9 | 0.27 | 9.6E-03 |
| *R05H10.1* | WBGene00011042 | protein_coding_gene | -1.9 | 0.27 | 8.4E-03 |
| *C48B4.3* | WBGene00008168 | protein_coding_gene | -1.9 | 0.27 | 3.2E-05 |
| *C05C9.1* | WBGene00007325 | protein_coding_gene | -1.9 | 0.27 | 1.1E-02 |
| *sta-2* | WBGene00010251 | protein_coding_gene | -1.9 | 0.27 | 1.2E-03 |
| *acn-1* | WBGene00000039 | protein_coding_gene | -1.9 | 0.27 | 4.2E-03 |
| *klp-11* | WBGene00002222 | protein_coding_gene | -1.9 | 0.27 | 1.1E-11 |
| *mlt-11* | WBGene00012186 | protein_coding_gene | -1.9 | 0.26 | 7.2E-03 |
| *fat-3* | WBGene00001395 | protein_coding_gene | -1.9 | 0.26 | 4.9E-10 |
| *T01B10.5* | WBGene00020137 | protein_coding_gene | -1.9 | 0.26 | 6.1E-06 |
| *wrt-1* | WBGene00006947 | protein_coding_gene | -1.9 | 0.26 | 1.3E-02 |
| *Y6E2A.4* | WBGene00012398 | protein_coding_gene | -1.9 | 0.26 | 3.8E-02 |
| *cutl-11* | WBGene00011443 | protein_coding_gene | -1.9 | 0.26 | 1.3E-03 |
| *C39D10.5* | WBGene00016532 | protein_coding_gene | -1.9 | 0.26 | 2.3E-03 |
| *T14E8.4* | WBGene00043981 | protein_coding_gene | -1.9 | 0.26 | 8.1E-04 |
| *F21A3.11* | WBGene00219316 | protein_coding_gene | -1.9 | 0.26 | 9.5E-08 |
| *ZK180.7* | WBGene00045248 | protein_coding_gene | -1.9 | 0.26 | 1.4E-03 |
| *lec-2* | WBGene00002265 | protein_coding_gene | -1.9 | 0.26 | 2.5E-04 |
| *W03F8.6* | WBGene00020995 | protein_coding_gene | -1.9 | 0.26 | 1.6E-06 |
| *C54F6.5* | WBGene00016923 | protein_coding_gene | -1.9 | 0.26 | 3.0E-02 |
| *dhs-2* | WBGene00000966 | protein_coding_gene | -1.9 | 0.26 | 9.2E-04 |
| *glf-1* | WBGene00019154 | protein_coding_gene | -1.9 | 0.26 | 3.1E-02 |
| *cut-5* | WBGene00011104 | protein_coding_gene | -1.9 | 0.26 | 2.6E-02 |
| *lact-6* | WBGene00007358 | protein_coding_gene | -1.9 | 0.26 | 1.3E-02 |
| *F13D11.4* | WBGene00017429 | protein_coding_gene | -1.9 | 0.26 | 9.0E-05 |
| *B0310.3* | WBGene00015139 | protein_coding_gene | -2.0 | 0.26 | 1.6E-08 |
| *glb-15* | WBGene00009387 | protein_coding_gene | -2.0 | 0.26 | 2.4E-05 |
| *slc-36.3* | WBGene00012629 | protein_coding_gene | -2.0 | 0.26 | 1.4E-03 |

| | | | | | |
|---|---|---|---|---|---|
| *clec-55* | WBGene00008597 | protein_coding_gene | -2.0 | 0.26 | 7.7E-04 |
| *F13E9.11* | WBGene00008760 | protein_coding_gene | -2.0 | 0.26 | 6.6E-05 |
| *pag-3* | WBGene00003909 | protein_coding_gene | -2.0 | 0.26 | 1.4E-04 |
| *R160.4* | WBGene00020121 | protein_coding_gene | -2.0 | 0.26 | 3.7E-03 |
| *ugt-36* | WBGene00017315 | protein_coding_gene | -2.0 | 0.26 | 3.5E-02 |
| *C09F9.2* | WBGene00007479 | protein_coding_gene | -2.0 | 0.26 | 1.1E-05 |
| *H42K12.3* | WBGene00019272 | protein_coding_gene | -2.0 | 0.25 | 1.1E-02 |
| *Y2H9A.6* | WBGene00086568 | protein_coding_gene | -2.0 | 0.25 | 9.6E-06 |
| *K10D6.2* | WBGene00010742 | protein_coding_gene | -2.0 | 0.25 | 2.7E-09 |
| *T28H10.3* | WBGene00012144 | protein_coding_gene | -2.0 | 0.25 | 1.3E-06 |
| *T03E6.8* | WBGene00011397 | protein_coding_gene | -2.0 | 0.25 | 3.9E-03 |
| *srr-6* | WBGene00005657 | protein_coding_gene | -2.0 | 0.25 | 3.2E-02 |
| *ZK669.3* | WBGene00014053 | protein_coding_gene | -2.0 | 0.25 | 7.0E-03 |
| *arl-6* | WBGene00000193 | protein_coding_gene | -2.0 | 0.25 | 4.6E-04 |
| *grl-2* | WBGene00001711 | protein_coding_gene | -2.0 | 0.25 | 2.2E-02 |
| *R09E10.5* | WBGene00011175 | protein_coding_gene | -2.0 | 0.25 | 3.4E-02 |
| *spp-1* | WBGene00004986 | protein_coding_gene | -2.0 | 0.25 | 4.5E-04 |
| *C54F6.12* | WBGene00016929 | protein_coding_gene | -2.0 | 0.25 | 5.8E-03 |
| *sulp-2* | WBGene00017464 | protein_coding_gene | -2.0 | 0.25 | 1.1E-02 |
| *ubc-8* | WBGene00006705 | protein_coding_gene | -2.0 | 0.25 | 3.2E-08 |
| *K01A6.7* | WBGene00045096 | protein_coding_gene | -2.0 | 0.25 | 2.9E-02 |
| *C37A5.3* | WBGene00014698 | protein_coding_gene | -2.0 | 0.25 | 3.4E-02 |
| *Y32F6B.1* | WBGene00012531 | protein_coding_gene | -2.0 | 0.25 | 6.1E-06 |
| *F15E6.10* | WBGene00044479 | protein_coding_gene | -2.0 | 0.25 | 1.7E-02 |
| *ZC443.2* | WBGene00013897 | protein_coding_gene | -2.0 | 0.25 | 1.8E-04 |
| *lpr-6* | WBGene00012255 | protein_coding_gene | -2.0 | 0.25 | 5.6E-03 |
| *Y71H2B.8* | WBGene00022197 | protein_coding_gene | -2.0 | 0.25 | 1.6E-05 |
| *clec-69* | WBGene00044642 | pseudogene | -2.0 | 0.25 | 1.6E-02 |
| *F53F1.6* | WBGene00009984 | protein_coding_gene | -2.0 | 0.25 | 3.7E-16 |
| *K10H10.10* | WBGene00010765 | protein_coding_gene | -2.0 | 0.25 | 1.9E-02 |
| *C42D4.1* | WBGene00016594 | protein_coding_gene | -2.0 | 0.25 | 1.3E-06 |
| *F48C1.11* | WBGene00195179 | protein_coding_gene | -2.0 | 0.25 | 3.8E-02 |
| *cyp-37A1* | WBGene00008499 | protein_coding_gene | -2.0 | 0.25 | 3.5E-05 |
| *gcy-8* | WBGene00001535 | protein_coding_gene | -2.0 | 0.25 | 4.1E-03 |
| *ptr-10* | WBGene00004224 | protein_coding_gene | -2.0 | 0.25 | 3.5E-04 |
| *fbn-1* | WBGene00022816 | protein_coding_gene | -2.0 | 0.25 | 4.0E-07 |
| *T19E7.6* | WBGene00044500 | protein_coding_gene | -2.0 | 0.25 | 1.2E-15 |
| *cutl-9* | WBGene00010893 | protein_coding_gene | -2.0 | 0.25 | 7.0E-04 |
| *F13H10.5* | WBGene00008776 | protein_coding_gene | -2.0 | 0.25 | 2.6E-03 |
| *dpy-2* | WBGene00001064 | protein_coding_gene | -2.0 | 0.24 | 3.3E-03 |
| *nlp-25* | WBGene00003763 | protein_coding_gene | -2.0 | 0.24 | 3.3E-02 |
| *K06A4.10* | WBGene00077497 | protein_coding_gene | -2.0 | 0.24 | 1.1E-02 |
| *F52E1.5* | WBGene00018694 | protein_coding_gene | -2.0 | 0.24 | 2.7E-08 |
| *W02D7.8* | WBGene00020943 | protein_coding_gene | -2.0 | 0.24 | 3.0E-07 |
| *F48F7.3* | WBGene00009849 | protein_coding_gene | -2.0 | 0.24 | 3.7E-03 |
| *ZK550.6* | WBGene00014000 | protein_coding_gene | -2.1 | 0.24 | 5.5E-09 |
| *F52C9.5* | WBGene00018675 | protein_coding_gene | -2.1 | 0.24 | 1.1E-04 |
| *T24C2.5* | WBGene00011984 | protein_coding_gene | -2.1 | 0.24 | 4.3E-10 |
| *C08D8.1* | WBGene00015592 | protein_coding_gene | -2.1 | 0.24 | 6.4E-05 |
| *R10E8.8* | WBGene00011213 | protein_coding_gene | -2.1 | 0.24 | 1.0E-05 |
| *srr-4* | WBGene00005655 | protein_coding_gene | -2.1 | 0.24 | 4.8E-12 |
| *C49H3.12* | WBGene00016795 | protein_coding_gene | -2.1 | 0.24 | 1.7E-03 |
| *ptr-22* | WBGene00004236 | protein_coding_gene | -2.1 | 0.24 | 1.4E-02 |
| *faah-2* | WBGene00015048 | protein_coding_gene | -2.1 | 0.24 | 4.4E-05 |
| *T02B11.6* | WBGene00020157 | protein_coding_gene | -2.1 | 0.24 | 7.5E-06 |
| *twk-36* | WBGene00006688 | protein_coding_gene | -2.1 | 0.23 | 1.1E-02 |
| *M03A1.8* | WBGene00077490 | protein_coding_gene | -2.1 | 0.23 | 1.7E-04 |
| *F41E6.5* | WBGene00018286 | protein_coding_gene | -2.1 | 0.23 | 6.1E-06 |

| | | | | | |
|---|---|---|---|---|---|
| *E04A4.3* | WBGene00017118 | pseudogene | -2.1 | 0.23 | 6.5E-04 |
| *ptr-3* | WBGene00004218 | protein_coding_gene | -2.1 | 0.23 | 5.6E-03 |
| *F52E1.12* | WBGene00018699 | pseudogene | -2.1 | 0.23 | 1.6E-02 |
| *F43D9.1* | WBGene00009653 | protein_coding_gene | -2.1 | 0.23 | 2.1E-04 |
| *rncs-1* | WBGene00023069 | ncRNA_gene | -2.1 | 0.23 | 5.9E-04 |
| *M03B6.3* | WBGene00010835 | protein_coding_gene | -2.1 | 0.23 | 1.2E-03 |
| *C36B1.6* | WBGene00007973 | protein_coding_gene | -2.1 | 0.23 | 2.1E-09 |
| *C35C5.9* | WBGene00007958 | protein_coding_gene | -2.1 | 0.23 | 1.7E-07 |
| *acs-17* | WBGene00016716 | protein_coding_gene | -2.1 | 0.23 | 5.2E-08 |
| *F41E7.20* | WBGene00219546 | ncRNA_gene | -2.1 | 0.23 | 8.5E-03 |
| *F21C10.10* | WBGene00017659 | protein_coding_gene | -2.1 | 0.23 | 9.3E-04 |
| *T04C12.32* | WBGene00235086 | ncRNA_gene | -2.1 | 0.23 | 1.7E-02 |
| *T10H9.8* | WBGene00044779 | protein_coding_gene | -2.2 | 0.22 | 7.8E-06 |
| *C43H6.6* | WBGene00016619 | protein_coding_gene | -2.2 | 0.22 | 5.1E-04 |
| *T09A12.8* | WBGene00235275 | protein_coding_gene | -2.2 | 0.22 | 2.5E-04 |
| *C14C6.2* | WBGene00015756 | protein_coding_gene | -2.2 | 0.22 | 5.5E-03 |
| *srg-34* | WBGene00005191 | protein_coding_gene | -2.2 | 0.22 | 2.7E-03 |
| *R05D8.9* | WBGene00019886 | protein_coding_gene | -2.2 | 0.22 | 1.1E-02 |
| *tag-297* | WBGene00007999 | protein_coding_gene | -2.2 | 0.22 | 3.7E-03 |
| *math-38* | WBGene00020326 | protein_coding_gene | -2.2 | 0.22 | 1.5E-07 |
| *Y53C12B.7* | WBGene00013146 | protein_coding_gene | -2.2 | 0.22 | 2.8E-07 |
| *gad-3* | WBGene00015057 | protein_coding_gene | -2.2 | 0.22 | 2.0E-08 |
| *Y46E12BR.1* | WBGene00021598 | protein_coding_gene | -2.2 | 0.22 | 2.9E-03 |
| *cpt-4* | WBGene00019644 | protein_coding_gene | -2.2 | 0.22 | 3.2E-13 |
| *icl-1* | WBGene00001564 | protein_coding_gene | -2.2 | 0.22 | 3.8E-04 |
| *M01A8.1* | WBGene00010795 | protein_coding_gene | -2.2 | 0.22 | 4.5E-10 |
| *F25E2.3* | WBGene00017781 | protein_coding_gene | -2.2 | 0.21 | 5.6E-07 |
| *F09F3.8* | WBGene00014732 | pseudogene | -2.2 | 0.21 | 2.7E-04 |
| *pals-38* | WBGene00008302 | protein_coding_gene | -2.2 | 0.21 | 2.2E-03 |
| *F44A6.5* | WBGene00009677 | protein_coding_gene | -2.2 | 0.21 | 2.2E-03 |
| *slc-17.9* | WBGene00011643 | protein_coding_gene | -2.2 | 0.21 | 9.3E-04 |
| *egl-15* | WBGene00001184 | protein_coding_gene | -2.2 | 0.21 | 6.7E-06 |
| *Y54G2A.11* | WBGene00021876 | protein_coding_gene | -2.2 | 0.21 | 3.8E-18 |
| *F43H9.4* | WBGene00018400 | protein_coding_gene | -2.2 | 0.21 | 3.1E-05 |
| *K09C4.4* | WBGene00019549 | protein_coding_gene | -2.2 | 0.21 | 1.5E-02 |
| *Y32B12C.5* | WBGene00194769 | pseudogene | -2.2 | 0.21 | 8.4E-05 |
| *F44E2.4* | WBGene00018418 | protein_coding_gene | -2.2 | 0.21 | 3.2E-05 |
| *F26F12.3* | WBGene00017834 | protein_coding_gene | -2.2 | 0.21 | 6.5E-27 |
| *ZK1307.2* | WBGene00014245 | protein_coding_gene | -2.2 | 0.21 | 7.3E-03 |
| *F13H6.5* | WBGene00017433 | protein_coding_gene | -2.2 | 0.21 | 5.2E-08 |
| *ztf-30* | WBGene00015523 | protein_coding_gene | -2.3 | 0.21 | 1.3E-07 |
| *F09E10.6* | WBGene00017296 | protein_coding_gene | -2.3 | 0.21 | 5.6E-04 |
| *F59E11.7* | WBGene00019113 | protein_coding_gene | -2.3 | 0.21 | 5.9E-04 |
| *K01A2.10* | WBGene00019284 | protein_coding_gene | -2.3 | 0.21 | 1.2E-05 |
| *C10C5.4* | WBGene00007508 | protein_coding_gene | -2.3 | 0.21 | 1.9E-13 |
| *C44C1.6* | WBGene00016645 | protein_coding_gene | -2.3 | 0.21 | 1.1E-03 |
| *R04B5.6* | WBGene00011004 | protein_coding_gene | -2.3 | 0.21 | 8.2E-04 |
| *Y39G10AR.11* | WBGene00021469 | protein_coding_gene | -2.3 | 0.21 | 3.2E-19 |
| *K10D3.6* | WBGene00010740 | protein_coding_gene | -2.3 | 0.21 | 6.8E-14 |
| *F49H6.5* | WBGene00009908 | protein_coding_gene | -2.3 | 0.21 | 1.8E-03 |
| *B0462.1* | WBGene00007181 | pseudogene | -2.3 | 0.20 | 1.6E-03 |
| *F17B5.6* | WBGene00044058 | protein_coding_gene | -2.3 | 0.20 | 7.9E-03 |
| *K09C8.7* | WBGene00010717 | protein_coding_gene | -2.3 | 0.20 | 1.8E-03 |
| *swt-1* | WBGene00019300 | protein_coding_gene | -2.3 | 0.20 | 6.3E-05 |
| *F17B5.1* | WBGene00008905 | protein_coding_gene | -2.3 | 0.20 | 6.7E-03 |
| *clec-218* | WBGene00020938 | protein_coding_gene | -2.3 | 0.20 | 2.4E-03 |
| *T19H5.6* | WBGene00044807 | protein_coding_gene | -2.3 | 0.20 | 2.6E-05 |
| *srbc-20* | WBGene00016693 | protein_coding_gene | -2.3 | 0.20 | 9.0E-03 |

| | | | | | |
|---|---|---|---|---|---|
| *ugt-54* | WBGene00012013 | protein_coding_gene | -2.3 | 0.20 | 2.6E-06 |
| *F42C5.3* | WBGene00018345 | protein_coding_gene | -2.3 | 0.20 | 1.4E-03 |
| *W08A12.2* | WBGene00021079 | protein_coding_gene | -2.3 | 0.20 | 2.3E-03 |
| *tsp-6* | WBGene00006632 | protein_coding_gene | -2.3 | 0.20 | 9.3E-07 |
| *Y71F9B.9* | WBGene00022130 | protein_coding_gene | -2.3 | 0.20 | 1.5E-23 |
| *cyp-13A9* | WBGene00014838 | pseudogene | -2.3 | 0.20 | 7.6E-03 |
| *clec-60* | WBGene00014046 | protein_coding_gene | -2.3 | 0.20 | 3.5E-03 |
| *Y32B12C.1* | WBGene00012525 | protein_coding_gene | -2.3 | 0.20 | 1.8E-03 |
| *T19A5.3* | WBGene00020554 | protein_coding_gene | -2.3 | 0.20 | 2.9E-04 |
| *mboa-1* | WBGene00007174 | protein_coding_gene | -2.3 | 0.20 | 1.2E-04 |
| *aakg-4* | WBGene00020135 | protein_coding_gene | -2.3 | 0.20 | 2.7E-04 |
| *clec-23* | WBGene00011851 | pseudogene | -2.4 | 0.20 | 5.3E-04 |
| *dct-8* | WBGene00044472 | protein_coding_gene | -2.4 | 0.19 | 1.4E-03 |
| *ugt-8* | WBGene00019234 | protein_coding_gene | -2.4 | 0.19 | 3.6E-04 |
| *pals-7* | WBGene00007661 | protein_coding_gene | -2.4 | 0.19 | 3.1E-03 |
| *cpg-7* | WBGene00010723 | protein_coding_gene | -2.4 | 0.19 | 8.3E-05 |
| *rom-2* | WBGene00004401 | protein_coding_gene | -2.4 | 0.19 | 2.4E-08 |
| *ins-20* | WBGene00002103 | protein_coding_gene | -2.4 | 0.19 | 1.1E-03 |
| *W05H9.1* | WBGene00021048 | protein_coding_gene | -2.4 | 0.19 | 1.6E-04 |
| *Y50E8A.1* | WBGene00013046 | protein_coding_gene | -2.4 | 0.19 | 3.0E-12 |
| *Y54G2A.37* | WBGene00023484 | protein_coding_gene | -2.4 | 0.19 | 6.7E-06 |
| *C04A11.5* | WBGene00007288 | protein_coding_gene | -2.4 | 0.19 | 7.8E-05 |
| *cpr-1* | WBGene00000781 | protein_coding_gene | -2.4 | 0.19 | 2.8E-04 |
| *nspe-6* | WBGene00012603 | protein_coding_gene | -2.4 | 0.19 | 9.0E-03 |
| *C54C8.4* | WBGene00008292 | protein_coding_gene | -2.4 | 0.19 | 1.9E-03 |
| *F49F1.5* | WBGene00018645 | protein_coding_gene | -2.4 | 0.19 | 2.6E-06 |
| *K09C4.1* | WBGene00019547 | protein_coding_gene | -2.4 | 0.19 | 3.2E-15 |
| *F10D7.3* | WBGene00017340 | protein_coding_gene | -2.4 | 0.19 | 5.3E-04 |
| *cyp-13A8* | WBGene00011674 | protein_coding_gene | -2.4 | 0.19 | 1.9E-03 |
| *W10G11.2* | WBGene00021135 | protein_coding_gene | -2.4 | 0.19 | 1.7E-03 |
| *Y70C5A.3* | WBGene00050904 | protein_coding_gene | -2.4 | 0.19 | 2.2E-08 |
| *clec-7* | WBGene00017364 | protein_coding_gene | -2.5 | 0.18 | 3.6E-04 |
| *K06A9.2* | WBGene00019436 | protein_coding_gene | -2.5 | 0.18 | 6.3E-05 |
| *F11D5.5* | WBGene00017382 | protein_coding_gene | -2.5 | 0.18 | 8.0E-08 |
| *ugt-9* | WBGene00020587 | protein_coding_gene | -2.5 | 0.18 | 2.9E-04 |
| *M01G12.14* | WBGene00010825 | protein_coding_gene | -2.5 | 0.18 | 7.8E-05 |
| *ptr-4* | WBGene00004219 | protein_coding_gene | -2.5 | 0.18 | 5.5E-06 |
| *T08G5.1* | WBGene00011622 | protein_coding_gene | -2.5 | 0.18 | 1.7E-04 |
| *R09D1.12* | WBGene00011168 | protein_coding_gene | -2.5 | 0.18 | 6.2E-13 |
| *W10G11.4* | WBGene00021137 | protein_coding_gene | -2.5 | 0.18 | 3.9E-03 |
| *C17G10.6* | WBGene00015918 | protein_coding_gene | -2.5 | 0.17 | 3.9E-03 |
| *C54C8.12* | WBGene00044242 | protein_coding_gene | -2.5 | 0.17 | 6.9E-03 |
| *aat-8* | WBGene00000009 | protein_coding_gene | -2.5 | 0.17 | 1.3E-06 |
| *gst-33* | WBGene00001781 | protein_coding_gene | -2.5 | 0.17 | 9.7E-08 |
| *F41C3.1* | WBGene00018267 | protein_coding_gene | -2.6 | 0.17 | 2.0E-04 |
| *F56D6.9* | WBGene00044471 | protein_coding_gene | -2.6 | 0.17 | 3.6E-03 |
| *clec-24* | WBGene00009856 | protein_coding_gene | -2.6 | 0.17 | 3.5E-05 |
| *C05D9.9* | WBGene00015476 | protein_coding_gene | -2.6 | 0.16 | 1.4E-05 |
| *oac-29* | WBGene00018295 | protein_coding_gene | -2.6 | 0.16 | 5.2E-05 |
| *hil-1* | WBGene00001852 | protein_coding_gene | -2.6 | 0.16 | 1.9E-04 |
| *F56H9.9* | WBGene00206460 | protein_coding_gene | -2.6 | 0.16 | 4.6E-04 |
| *M01B2.13* | WBGene00045338 | protein_coding_gene | -2.6 | 0.16 | 1.7E-06 |
| *ins-23* | WBGene00002106 | protein_coding_gene | -2.6 | 0.16 | 1.4E-05 |
| *ZK6.8* | WBGene00022643 | protein_coding_gene | -2.6 | 0.16 | 2.5E-04 |
| *mtl-2* | WBGene00003474 | protein_coding_gene | -2.7 | 0.16 | 1.0E-07 |
| *thn-2* | WBGene00009215 | protein_coding_gene | -2.7 | 0.16 | 6.3E-04 |
| *dlhd-1* | WBGene00020741 | protein_coding_gene | -2.7 | 0.16 | 1.2E-05 |
| *slc-36.5* | WBGene00016628 | protein_coding_gene | -2.7 | 0.16 | 1.3E-04 |

| | | | | | |
|---|---|---|---|---|---|
| *T26H5.8* | WBGene00012070 | protein_coding_gene | -2.7 | 0.16 | 1.0E-22 |
| *T21D12.14* | WBGene00045177 | protein_coding_gene | -2.7 | 0.15 | 1.8E-06 |
| *T05E12.3* | WBGene00011486 | protein_coding_gene | -2.7 | 0.15 | 8.4E-10 |
| *clec-8* | WBGene00012582 | protein_coding_gene | -2.7 | 0.15 | 1.5E-07 |
| *C18H9.5* | WBGene00016003 | protein_coding_gene | -2.7 | 0.15 | 1.2E-05 |
| *R160.10* | WBGene00219369 | protein_coding_gene | -2.7 | 0.15 | 1.0E-05 |
| *F16B4.7* | WBGene00017509 | protein_coding_gene | -2.7 | 0.15 | 2.2E-04 |
| *bigr-1* | WBGene00009512 | protein_coding_gene | -2.8 | 0.14 | 2.1E-06 |
| *pals-32* | WBGene00016281 | protein_coding_gene | -2.8 | 0.14 | 1.9E-07 |
| *Y17G9A.4* | WBGene00021197 | protein_coding_gene | -2.8 | 0.14 | 1.5E-03 |
| *chil-18* | WBGene00011161 | protein_coding_gene | -2.8 | 0.14 | 2.4E-05 |
| *T28C12.15* | WBGene00271821 | protein_coding_gene | -2.8 | 0.14 | 6.5E-04 |
| *H32K16.2* | WBGene00010422 | protein_coding_gene | -2.8 | 0.14 | 6.2E-07 |
| *ugt-4* | WBGene00013905 | protein_coding_gene | -2.9 | 0.14 | 3.7E-31 |
| *srbc-82* | WBGene00007832 | protein_coding_gene | -2.9 | 0.14 | 3.9E-04 |
| *C11G10.1* | WBGene00007525 | protein_coding_gene | -2.9 | 0.14 | 7.9E-07 |
| *K01A6.4* | WBGene00010446 | protein_coding_gene | -2.9 | 0.14 | 1.1E-06 |
| *thn-1* | WBGene00009213 | protein_coding_gene | -2.9 | 0.14 | 3.8E-05 |
| *smf-3* | WBGene00004878 | protein_coding_gene | -2.9 | 0.13 | 3.2E-07 |
| *T16G1.4* | WBGene00011798 | protein_coding_gene | -2.9 | 0.13 | 5.7E-06 |
| *clec-31* | WBGene00009854 | protein_coding_gene | -2.9 | 0.13 | 2.3E-05 |
| *F16C3.2* | WBGene00008886 | protein_coding_gene | -2.9 | 0.13 | 9.7E-08 |
| *nspe-7* | WBGene00012593 | protein_coding_gene | -2.9 | 0.13 | 1.1E-04 |
| *sodh-1* | WBGene00010790 | protein_coding_gene | -2.9 | 0.13 | 3.4E-13 |
| *fil-1* | WBGene00011321 | protein_coding_gene | -2.9 | 0.13 | 3.7E-04 |
| *T16G1.6* | WBGene00011800 | protein_coding_gene | -2.9 | 0.13 | 1.3E-12 |
| *F57A8.7* | WBGene00010182 | protein_coding_gene | -2.9 | 0.13 | 2.0E-04 |
| *dach-1* | WBGene00020386 | protein_coding_gene | -2.9 | 0.13 | 8.4E-07 |
| *nnt-1* | WBGene00003778 | protein_coding_gene | -3.0 | 0.13 | 9.7E-11 |
| *tre-5* | WBGene00006611 | protein_coding_gene | -3.0 | 0.13 | 6.0E-06 |
| *W10G11.1* | WBGene00021134 | protein_coding_gene | -3.0 | 0.13 | 4.4E-04 |
| *clec-77* | WBGene00044343 | pseudogene | -3.0 | 0.13 | 6.0E-42 |
| *clec-82* | WBGene00021873 | protein_coding_gene | -3.0 | 0.13 | 4.7E-06 |
| *F21H7.12* | WBGene00009033 | protein_coding_gene | -3.0 | 0.12 | 5.4E-04 |
| *Y45F10D.6* | WBGene00012886 | protein_coding_gene | -3.0 | 0.12 | 1.0E-04 |
| *asp-11* | WBGene00017677 | pseudogene | -3.0 | 0.12 | 6.8E-07 |
| *C35C5.8* | WBGene00007957 | protein_coding_gene | -3.0 | 0.12 | 2.2E-09 |
| *clec-32* | WBGene00009860 | protein_coding_gene | -3.0 | 0.12 | 5.8E-07 |
| *T04C12.7* | WBGene00011429 | protein_coding_gene | -3.1 | 0.12 | 1.5E-05 |
| *srh-2* | WBGene00005228 | protein_coding_gene | -3.1 | 0.12 | 4.6E-07 |
| *ftn-1* | WBGene00001500 | protein_coding_gene | -3.1 | 0.12 | 1.1E-05 |
| *C08F11.3* | WBGene00007452 | protein_coding_gene | -3.1 | 0.12 | 5.9E-05 |
| *dhs-23* | WBGene00000986 | protein_coding_gene | -3.1 | 0.12 | 8.0E-08 |
| *F59C6.16* | WBGene00077764 | protein_coding_gene | -3.1 | 0.11 | 2.5E-23 |
| *pitr-4* | WBGene00007141 | protein_coding_gene | -3.2 | 0.11 | 6.3E-05 |
| *H25K10.4* | WBGene00010413 | protein_coding_gene | -3.2 | 0.11 | 5.3E-05 |
| *cyp-13A5* | WBGene00011672 | protein_coding_gene | -3.2 | 0.11 | 1.3E-09 |
| *F56D6.8* | WBGene00044292 | protein_coding_gene | -3.3 | 0.10 | 3.5E-04 |
| *cpt-3* | WBGene00021703 | protein_coding_gene | -3.3 | 0.10 | 1.2E-04 |
| *cyp-34A2* | WBGene00011699 | protein_coding_gene | -3.3 | 0.10 | 9.1E-14 |
| *str-163* | WBGene00006208 | protein_coding_gene | -3.3 | 0.10 | 6.1E-10 |
| *F13H6.4* | WBGene00017432 | pseudogene | -3.3 | 0.10 | 1.1E-10 |
| *dct-7* | WBGene00017488 | protein_coding_gene | -3.3 | 0.10 | 2.4E-04 |
| *F11E6.11* | WBGene00023501 | protein_coding_gene | -3.3 | 0.10 | 3.8E-07 |
| *ilys-2* | WBGene00016669 | protein_coding_gene | -3.4 | 0.10 | 8.4E-06 |
| *far-7* | WBGene00001391 | protein_coding_gene | -3.4 | 0.10 | 3.2E-10 |
| *T23E7.6* | WBGene00044772 | protein_coding_gene | -3.4 | 0.10 | 1.0E-06 |
| *ugt-33* | WBGene00007946 | protein_coding_gene | -3.4 | 0.09 | 2.7E-09 |

| | | | | | |
|---|---|---|---|---|---|
| *C08E8.3* | WBGene00007439 | protein_coding_gene | -3.4 | 0.09 | 3.2E-05 |
| *E02C12.10* | WBGene00017095 | protein_coding_gene | -3.4 | 0.09 | 1.6E-05 |
| *W07B8.4* | WBGene00021072 | protein_coding_gene | -3.4 | 0.09 | 8.4E-07 |
| *F53F4.8* | WBGene00009991 | pseudogene | -3.6 | 0.08 | 1.2E-19 |
| *dod-3* | WBGene00016052 | protein_coding_gene | -3.7 | 0.08 | 5.1E-16 |
| *C23G10.11* | WBGene00016017 | protein_coding_gene | -3.7 | 0.08 | 3.0E-06 |
| *cyp-34A9* | WBGene00015044 | protein_coding_gene | -3.8 | 0.07 | 1.3E-11 |
| *M02E1.4* | WBGene00284848 | protein_coding_gene | -3.8 | 0.07 | 9.7E-09 |
| *cest-1.2* | WBGene00011364 | protein_coding_gene | -3.8 | 0.07 | 1.2E-54 |
| *ugt-34* | WBGene00009255 | protein_coding_gene | -3.8 | 0.07 | 3.2E-06 |
| *cyp-13A7* | WBGene00000372 | protein_coding_gene | -3.8 | 0.07 | 2.0E-08 |
| *alh-5* | WBGene00000111 | protein_coding_gene | -3.9 | 0.07 | 1.5E-23 |
| *glc-1* | WBGene00001591 | protein_coding_gene | -4.1 | 0.06 | 1.9E-22 |
| *F15E6.3* | WBGene00017484 | protein_coding_gene | -4.4 | 0.05 | 4.8E-17 |
| *ins-8* | WBGene00002091 | protein_coding_gene | -4.4 | 0.05 | 2.1E-07 |
| *T16G1.7* | WBGene00011801 | protein_coding_gene | -4.6 | 0.04 | 2.3E-33 |
| *clec-61* | WBGene00014047 | protein_coding_gene | -5.2 | 0.03 | 1.6E-18 |
| *T22F3.11* | WBGene00020700 | protein_coding_gene | -5.4 | 0.02 | 9.6E-15 |
| *cyp-34A10* | WBGene00015045 | protein_coding_gene | -5.5 | 0.02 | 6.7E-34 |
| *ilys-3* | WBGene00016670 | protein_coding_gene | -7.3 | 0.01 | 5.9E-37 |
| *lipl-3* | WBGene00020016 | protein_coding_gene | -7.5 | 0.01 | 5.3E-49 |

Shared (N2&CB4856) DEGs in response to Gram-negative pathogens (*P. rettgeri* or *P. aeruginosa* vs. OP50)

| | | | N2 | | | CB4856 | | |
|---|---|---|---|---|---|---|---|---|
| Gene Name | Wormbase Gene ID | Type | $log_2FC$ | FC | Adj P-value | $log_2FC$ | FC | Adj P-value |
| *C49G7.12* | WBGene00077593 | protein_coding_gene | 4.0 | 16.09 | 3.1E-05 | 3.3 | 9.64 | 1.6E-04 |
| *F35E12.4* | WBGene00009428 | pseudogene | 3.8 | 13.78 | 3.8E-07 | 1.7 | 3.26 | 1.6E-02 |
| *B0024.4* | WBGene00007097 | protein_coding_gene | 3.3 | 10.02 | 2.9E-03 | 2.3 | 4.96 | 5.6E-03 |
| ***cyp-37B1*** | **WBGene00009226** | **protein_coding_gene** | **3.3** | **9.85** | **1.1E-04** | **-2.5** | **0.18** | **9.3E-04** |
| *irg-1* | WBGene00015574 | protein_coding_gene | 3.2 | 9.47 | 3.0E-04 | 2.1 | 4.14 | 2.8E-02 |
| ***tts-1*** | **WBGene00006650** | **ncRNA_gene** | **3.2** | **8.99** | **1.6E-05** | **-2.2** | **0.22** | **6.7E-03** |
| *T24B8.5* | WBGene00011979 | protein_coding_gene | 3.1 | 8.45 | 2.3E-02 | 4.4 | 21.29 | 2.4E-09 |
| *irg-6* | WBGene00007864 | protein_coding_gene | 3.0 | 8.23 | 1.6E-02 | 1.4 | 2.70 | 3.5E-03 |
| ***C50F7.5*** | **WBGene00016845** | **protein_coding_gene** | **3.0** | **8.06** | **2.0E-04** | **-3.5** | **0.09** | **1.8E-06** |
| *irg-5* | WBGene00009429 | protein_coding_gene | 3.0 | 8.05 | 1.5E-03 | 4.5 | 22.02 | 5.1E-08 |
| *pgp-5* | WBGene00003999 | protein_coding_gene | 3.0 | 7.86 | 7.3E-06 | 2.0 | 4.06 | 1.0E-02 |
| ***F47B8.4*** | **WBGene00009805** | **protein_coding_gene** | **2.9** | **7.66** | **4.1E-03** | **-2.2** | **0.22** | **1.7E-02** |
| *srj-29* | WBGene00005616 | protein_coding_gene | 2.9 | 7.48 | 2.2E-02 | 2.8 | 7.06 | 3.2E-04 |
| *F44G3.10* | WBGene00009710 | protein_coding_gene | 2.9 | 7.43 | 4.2E-02 | 2.6 | 5.95 | 6.7E-04 |
| ***F46A8.13*** | **WBGene00219950** | **ncRNA_gene** | **2.8** | **6.79** | **1.5E-02** | **-1.8** | **0.29** | **4.8E-03** |
| ***T07G12.5*** | **WBGene00011596** | **protein_coding_gene** | **2.7** | **6.60** | **2.4E-03** | **-4.4** | **0.05** | **1.3E-29** |
| *F49H6.13* | WBGene00009914 | protein_coding_gene | 2.7 | 6.31 | 2.2E-02 | 1.6 | 3.13 | 4.5E-02 |
| *dod-24* | WBGene00007875 | protein_coding_gene | 2.6 | 6.25 | 2.6E-02 | 2.2 | 4.63 | 6.8E-05 |
| ***nhr-127*** | **WBGene00003717** | **protein_coding_gene** | **2.5** | **5.85** | **1.9E-02** | **-1.4** | **0.38** | **2.3E-02** |
| *Y47H9C.1* | WBGene00012947 | protein_coding_gene | 2.5 | 5.75 | 4.2E-04 | 3.4 | 10.90 | 1.4E-07 |
| ***F48C1.9*** | **WBGene00018601** | **protein_coding_gene** | **2.4** | **5.27** | **5.0E-02** | **-2.4** | **0.19** | **1.1E-02** |
| ***F45D3.4*** | **WBGene00009724** | **protein_coding_gene** | **2.4** | **5.20** | **1.1E-02** | **-4.6** | **0.04** | **1.1E-15** |
| ***Y7A5A.6*** | **WBGene00012412** | **protein_coding_gene** | **2.4** | **5.10** | **3.2E-02** | **-1.4** | **0.37** | **3.1E-02** |
| ***F46C5.1*** | **WBGene00009778** | **protein_coding_gene** | **2.3** | **5.06** | **5.5E-04** | **-3.9** | **0.07** | **2.2E-15** |
| ***scl-2*** | **WBGene00009895** | **protein_coding_gene** | **2.3** | **5.01** | **1.4E-02** | **-4.5** | **0.04** | **3.0E-75** |
| ***C18H7.1*** | **WBGene00015993** | **protein_coding_gene** | **2.3** | **4.93** | **2.4E-02** | **-5.2** | **0.03** | **8.2E-16** |
| *lys-2* | WBGene00003091 | protein_coding_gene | 2.3 | 4.91 | 2.0E-02 | 2.3 | 4.89 | 8.3E-04 |
| ***oac-14*** | **WBGene00008602** | **protein_coding_gene** | **2.2** | **4.59** | **1.2E-02** | **-2.5** | **0.17** | **4.1E-04** |
| *C34H4.1* | WBGene00016424 | protein_coding_gene | 2.2 | 4.55 | 7.5E-04 | 2.2 | 4.71 | 1.1E-03 |
| ***mtl-1*** | **WBGene00003473** | **protein_coding_gene** | **2.2** | **4.53** | **2.2E-03** | **-4.5** | **0.04** | **3.8E-10** |
| *E02C12.8* | WBGene00017093 | protein_coding_gene | 2.2 | 4.46 | 2.5E-03 | 1.5 | 2.89 | 3.2E-02 |
| ***Y43F8B.9*** | **WBGene00012819** | **protein_coding_gene** | **2.1** | **4.41** | **3.7E-03** | **-2.2** | **0.21** | **5.4E-05** |

| | | | | | | | | |
|---|---|---|---|---|---|---|---|---|
| T20B3.1 | WBGene00011850 | protein_coding_gene | 2.1 | 4.31 | 2.0E-02 | -2.2 | 0.21 | 1.3E-09 |
| Y65B4BR.1 | WBGene00022040 | protein_coding_gene | 2.1 | 4.16 | 7.5E-03 | 1.2 | 2.29 | 4.2E-02 |
| F09F7.6 | WBGene00017303 | protein_coding_gene | 2.0 | 4.13 | 3.5E-03 | -2.2 | 0.22 | 2.1E-16 |
| unc-42 | WBGene00006778 | protein_coding_gene | 2.0 | 4.06 | 9.3E-03 | -1.4 | 0.38 | 2.2E-03 |
| C44H9.7 | WBGene00008102 | protein_coding_gene | 2.0 | 4.05 | 2.0E-02 | -1.9 | 0.27 | 1.4E-03 |
| F47B8.10 | WBGene00009811 | protein_coding_gene | 2.0 | 4.03 | 3.3E-02 | -1.8 | 0.28 | 4.0E-02 |
| C32H11.4 | WBGene00007867 | protein_coding_gene | 2.0 | 3.96 | 4.7E-02 | 2.5 | 5.65 | 2.9E-03 |
| ZC443.3 | WBGene00013898 | protein_coding_gene | 1.9 | 3.81 | 3.3E-02 | -2.6 | 0.17 | 4.3E-06 |
| T12D8.5 | WBGene00011733 | protein_coding_gene | 1.9 | 3.63 | 2.2E-02 | -1.9 | 0.27 | 9.2E-03 |
| cutl-28 | WBGene00018256 | protein_coding_gene | 1.9 | 3.63 | 2.2E-02 | -1.7 | 0.31 | 2.3E-02 |
| coel-1 | WBGene00016866 | protein_coding_gene | 1.8 | 3.59 | 4.9E-02 | -2.6 | 0.17 | 1.2E-10 |
| spp-2 | WBGene00004987 | protein_coding_gene | 1.8 | 3.50 | 2.5E-03 | -1.4 | 0.39 | 3.3E-02 |
| C25H3.10 | WBGene00016119 | protein_coding_gene | 1.8 | 3.49 | 3.0E-02 | -2.8 | 0.14 | 7.2E-20 |
| K08B12.1 | WBGene00019520 | protein_coding_gene | 1.8 | 3.46 | 1.1E-02 | -1.6 | 0.32 | 4.8E-02 |
| F37A8.5 | WBGene00009503 | protein_coding_gene | 1.8 | 3.43 | 4.9E-02 | -2.1 | 0.23 | 1.9E-03 |
| W09G12.7 | WBGene00021121 | protein_coding_gene | 1.8 | 3.39 | 4.2E-02 | -1.5 | 0.34 | 2.5E-02 |
| irg-4 | WBGene00008584 | protein_coding_gene | 1.8 | 3.38 | 2.6E-02 | 4.1 | 17.50 | 1.2E-10 |
| lpr-4 | WBGene00012257 | protein_coding_gene | 1.8 | 3.38 | 3.5E-03 | -1.7 | 0.30 | 4.4E-02 |
| cup-16 | WBGene00007919 | protein_coding_gene | 1.7 | 3.36 | 2.8E-02 | -2.2 | 0.22 | 1.7E-05 |
| F59B2.8 | WBGene00010310 | protein_coding_gene | 1.7 | 3.36 | 6.1E-03 | -1.1 | 0.45 | 1.0E-02 |
| peb-1 | WBGene00003968 | protein_coding_gene | 1.7 | 3.30 | 2.0E-02 | -1.5 | 0.36 | 1.8E-02 |
| T03G6.1 | WBGene00020194 | protein_coding_gene | 1.7 | 3.21 | 2.3E-02 | -1.9 | 0.28 | 3.6E-02 |
| F31D4.5 | WBGene00009292 | protein_coding_gene | 1.6 | 3.12 | 4.6E-02 | -1.8 | 0.29 | 4.1E-02 |
| tsp-17 | WBGene00006643 | protein_coding_gene | 1.6 | 3.07 | 4.2E-02 | -1.7 | 0.31 | 3.7E-05 |
| F43C11.7 | WBGene00018384 | protein_coding_gene | 1.6 | 3.02 | 3.6E-02 | -2.4 | 0.19 | 3.6E-05 |
| T20F5.4 | WBGene00020626 | protein_coding_gene | 1.5 | 2.92 | 4.2E-02 | -1.6 | 0.33 | 2.5E-02 |
| R03H10.2 | WBGene00019855 | protein_coding_gene | 1.5 | 2.86 | 1.2E-02 | -1.4 | 0.38 | 3.9E-04 |
| EEED8.2 | WBGene00017133 | protein_coding_gene | 1.5 | 2.78 | 1.3E-03 | -1.2 | 0.44 | 3.4E-02 |
| dpy-3 | WBGene00001065 | protein_coding_gene | 1.5 | 2.74 | 4.9E-02 | -1.6 | 0.33 | 4.2E-02 |
| dpy-7 | WBGene00001069 | protein_coding_gene | 1.4 | 2.72 | 3.6E-02 | -1.9 | 0.27 | 7.9E-03 |
| Y71H2B.11 | WBGene00022198 | protein_coding_gene | 1.4 | 2.69 | 5.0E-02 | 1.5 | 2.73 | 1.1E-02 |
| F41B5.1 | WBGene00023284 | protein_coding_gene | 1.4 | 2.66 | 4.5E-02 | 1.2 | 2.33 | 4.4E-02 |
| dpy-10 | WBGene00001072 | protein_coding_gene | 1.4 | 2.65 | 4.6E-02 | -1.6 | 0.34 | 3.1E-02 |
| Y65B4BM.3 | WBGene00302974 | protein_coding_gene | 1.4 | 2.64 | 4.6E-02 | -1.4 | 0.37 | 1.0E-02 |
| lpr-3 | WBGene00012261 | protein_coding_gene | 1.3 | 2.54 | 4.6E-02 | -1.8 | 0.29 | 1.3E-02 |
| H04D03.3 | WBGene00010364 | protein_coding_gene | 1.3 | 2.53 | 3.9E-02 | 1.9 | 3.71 | 2.4E-03 |
| hhat-1 | WBGene00013852 | protein_coding_gene | 1.3 | 2.49 | 4.4E-02 | -2.1 | 0.23 | 3.1E-04 |
| lpr-5 | WBGene00012256 | protein_coding_gene | 1.3 | 2.43 | 4.9E-02 | -2.0 | 0.25 | 6.1E-03 |
| W04A8.4 | WBGene00012239 | protein_coding_gene | 1.2 | 2.37 | 1.5E-02 | -1.6 | 0.32 | 5.3E-05 |
| C18D4.8 | WBGene00007677 | pseudogene | 1.2 | 2.36 | 3.0E-02 | -1.0 | 0.50 | 2.5E-02 |
| T02E9.5 | WBGene00011383 | protein_coding_gene | 1.2 | 2.36 | 4.2E-02 | -1.5 | 0.36 | 7.2E-03 |
| sqt-3 | WBGene00005018 | protein_coding_gene | 1.1 | 2.21 | 4.7E-02 | -1.4 | 0.37 | 4.3E-02 |
| tth-1 | WBGene00006649 | protein_coding_gene | 1.1 | 2.20 | 5.9E-03 | -1.0 | 0.49 | 4.1E-02 |
| D2030.2 | WBGene00008412 | protein_coding_gene | 1.1 | 2.18 | 2.6E-02 | -1.4 | 0.39 | 3.8E-27 |
| F28F8.10 | WBGene00044219 | pseudogene | 1.1 | 2.17 | 3.8E-02 | 1.2 | 2.28 | 2.7E-03 |
| fkb-7 | WBGene00001432 | protein_coding_gene | 1.1 | 2.11 | 3.7E-02 | -1.5 | 0.35 | 2.6E-03 |
| Y57A10A.4 | WBGene00013249 | protein_coding_gene | 1.0 | 2.04 | 4.5E-02 | 1.1 | 2.17 | 3.3E-02 |
| cki-1 | WBGene00000516 | protein_coding_gene | 1.0 | 2.04 | 6.1E-03 | -1.0 | 0.49 | 7.8E-03 |
| C41H7.1 | WBGene00016571 | protein_coding_gene | -1.0 | 0.48 | 3.9E-02 | 1.4 | 2.71 | 1.1E-02 |
| R02C2.7 | WBGene00019818 | protein_coding_gene | -1.1 | 0.46 | 2.3E-02 | -1.2 | 0.43 | 2.6E-08 |
| F28B4.3 | WBGene00017892 | protein_coding_gene | -1.1 | 0.46 | 3.7E-02 | 1.3 | 2.54 | 9.2E-03 |
| T15B7.10 | WBGene00020522 | pseudogene | -1.3 | 0.40 | 1.2E-02 | -2.2 | 0.22 | 1.7E-03 |
| Y53G8B.1 | WBGene00021817 | protein_coding_gene | -1.3 | 0.40 | 4.1E-02 | -1.1 | 0.46 | 1.5E-02 |
| ctsa-3.1 | WBGene00017969 | protein_coding_gene | -1.4 | 0.39 | 3.5E-02 | 1.4 | 2.73 | 9.9E-03 |
| scav-4 | WBGene00008692 | protein_coding_gene | -1.5 | 0.36 | 2.6E-02 | 1.7 | 3.26 | 2.8E-03 |
| C02G6.1 | WBGene00015359 | pseudogene | -1.6 | 0.33 | 1.4E-02 | 2.2 | 4.50 | 9.2E-03 |
| arf-1.1 | WBGene00000190 | protein_coding_gene | -1.6 | 0.33 | 2.5E-02 | 1.2 | 2.26 | 4.3E-02 |
| zig-2 | WBGene00006979 | protein_coding_gene | -1.6 | 0.33 | 5.0E-02 | -1.4 | 0.38 | 2.2E-02 |

| | | | | | | | | |
|---|---|---|---|---|---|---|---|---|
| ***T16G12.1*** | **WBGene00011803** | **protein_coding_gene** | **-1.6** | **0.33** | **1.8E-02** | **2.0** | **4.02** | **3.0E-07** |
| ***C35A11.4*** | **WBGene00016431** | **protein_coding_gene** | **-1.6** | **0.32** | **2.9E-02** | **2.0** | **4.11** | **1.9E-03** |
| ***Y16B4A.2*** | **WBGene00012445** | **protein_coding_gene** | **-1.6** | **0.32** | **2.1E-02** | **1.6** | **2.94** | **1.9E-03** |
| *D2023.1* | WBGene00014300 | protein_coding_gene | -1.7 | 0.32 | 1.7E-02 | -1.0 | 0.49 | 3.2E-02 |
| ***T15B7.14*** | **WBGene00020526** | **protein_coding_gene** | **-1.7** | **0.31** | **4.5E-02** | **1.4** | **2.58** | **4.3E-02** |
| ***gst-26*** | **WBGene00001774** | **protein_coding_gene** | **-1.7** | **0.31** | **4.4E-02** | **1.8** | **3.56** | **3.7E-03** |
| ***elo-5*** | **WBGene00001243** | **protein_coding_gene** | **-1.7** | **0.30** | **3.7E-02** | **2.0** | **4.02** | **2.5E-03** |
| *pgp-14* | WBGene00004008 | protein_coding_gene | -1.7 | 0.30 | 3.7E-02 | -1.1 | 0.46 | 6.7E-03 |
| *lys-5* | WBGene00003094 | protein_coding_gene | -1.8 | 0.30 | 8.8E-05 | -2.0 | 0.25 | 3.4E-04 |
| *ddn-1* | WBGene00015227 | protein_coding_gene | -1.8 | 0.30 | 1.8E-02 | -1.5 | 0.36 | 2.0E-02 |
| ***nep-22*** | **WBGene00020788** | **protein_coding_gene** | **-1.8** | **0.29** | **8.5E-03** | **1.1** | **2.17** | **2.5E-02** |
| ***pho-1*** | **WBGene00004020** | **protein_coding_gene** | **-1.8** | **0.29** | **9.3E-03** | **2.3** | **5.07** | **3.2E-05** |
| ***act-5*** | **WBGene00000067** | **protein_coding_gene** | **-1.8** | **0.29** | **2.6E-03** | **1.4** | **2.67** | **1.3E-02** |
| *F54B11.11* | WBGene00010034 | protein_coding_gene | -1.8 | 0.28 | 3.4E-02 | -2.4 | 0.19 | 3.2E-05 |
| ***daf-28*** | **WBGene00000920** | **protein_coding_gene** | **-1.8** | **0.28** | **1.4E-02** | **1.4** | **2.66** | **2.8E-04** |
| *ugt-66* | WBGene00016013 | protein_coding_gene | -1.9 | 0.27 | 2.3E-02 | -1.9 | 0.26 | 2.2E-02 |
| ***paic-1*** | **WBGene00015116** | **protein_coding_gene** | **-1.9** | **0.27** | **2.0E-02** | **1.4** | **2.67** | **3.4E-02** |
| ***gst-10*** | **WBGene00001758** | **protein_coding_gene** | **-2.0** | **0.25** | **1.2E-02** | **1.7** | **3.28** | **4.6E-03** |
| ***sptl-2*** | **WBGene00018398** | **protein_coding_gene** | **-2.1** | **0.24** | **7.0E-04** | **1.4** | **2.65** | **3.0E-03** |
| ***F22E5.8*** | **WBGene00017707** | **protein_coding_gene** | **-2.1** | **0.24** | **2.4E-03** | **1.3** | **2.53** | **1.3E-03** |
| ***spp-23*** | **WBGene00021335** | **protein_coding_gene** | **-2.1** | **0.23** | **2.6E-02** | **2.1** | **4.39** | **1.2E-03** |
| ***abhd-3.2*** | **WBGene00016644** | **protein_coding_gene** | **-2.1** | **0.23** | **2.8E-02** | **2.0** | **3.91** | **6.6E-03** |
| ***R193.2*** | **WBGene00020128** | **protein_coding_gene** | **-2.1** | **0.23** | **1.9E-02** | **2.9** | **7.55** | **4.5E-07** |
| ***F53B3.6*** | **WBGene00018744** | **protein_coding_gene** | **-2.2** | **0.22** | **4.9E-02** | **1.7** | **3.16** | **2.7E-02** |
| ***F41E6.12*** | **WBGene00018293** | **protein_coding_gene** | **-2.2** | **0.22** | **3.5E-02** | **1.8** | **3.41** | **3.9E-02** |
| *C40H1.8* | WBGene00008043 | protein_coding_gene | -2.2 | 0.22 | 2.5E-03 | -2.5 | 0.17 | 4.5E-05 |
| *Y34F4.6* | WBGene00189956 | protein_coding_gene | -2.4 | 0.19 | 4.8E-04 | -2.0 | 0.24 | 1.2E-02 |
| ***scl-6*** | **WBGene00008028** | **protein_coding_gene** | **-2.4** | **0.19** | **4.1E-07** | **3.1** | **8.56** | **6.9E-08** |
| *clec-169* | WBGene00009526 | protein_coding_gene | -2.5 | 0.18 | 5.0E-02 | -1.9 | 0.27 | 4.4E-02 |
| *col-91* | WBGene00000666 | protein_coding_gene | -2.6 | 0.16 | 3.3E-02 | -2.0 | 0.25 | 3.3E-03 |
| *ugt-15* | WBGene00008097 | protein_coding_gene | -2.8 | 0.14 | 9.0E-04 | -2.6 | 0.17 | 4.7E-06 |
| *col-156* | WBGene00000729 | protein_coding_gene | -2.9 | 0.13 | 2.1E-02 | -1.7 | 0.31 | 3.7E-02 |
| *clec-70* | WBGene00021581 | protein_coding_gene | -3.1 | 0.12 | 2.0E-03 | -2.3 | 0.20 | 1.9E-08 |
| *pgp-1* | WBGene00003995 | protein_coding_gene | -4.4 | 0.05 | 4.1E-10 | -1.7 | 0.32 | 4.6E-02 |

*bold denotes genes that are differentially expressed in opposing directions

Transcription Factors Differentially Expressed in Gram-Negative Infected Animals

| Gene Name | WormBase Gene ID | Sequence Name | N2 $log_2$FC | CB4856 $log_2$FC |
|---|---|---|---|---|
| | | N2 Specific | | |
| *dac-1* | WBGene00000895 | B0412.1 | -1.5 | Not DE |
| *nhr-74* | WBGene00003664 | C27C7.3 | -3.5 | Not DE |
| *nhr-115* | WBGene00003705 | T27B7.4 | -2.3 | Not DE |
| *nhr-116* | WBGene00003706 | F09C6.9 | 1.1 | Not DE |
| *nhr-137* | WBGene00003727 | C56E10.4 | -1.5 | Not DE |
| *F26F4.8* | WBGene00005011 | F26F4.8 | 1.1 | Not DE |
| *zip-3* | WBGene00012330 | W07G1.3 | -1.6 | Not DE |
| *R05D3.3* | WBGene00019878 | R05D3.3 | 1.1 | Not DE |
| *ZK328.6* | WBGene00022695 | ZK328.6 | -1.4 | Not DE |
| | | CB4856 Specific | | |
| *bro-1* | WBGene00000272 | F56A3.5 | Not DE | -1.6 |
| *ceh-5* | WBGene00000430 | C16C2.1 | Not DE | -1.6 |
| *ceh-6* | WBGene00000431 | K02B12.1 | Not DE | -1.2 |
| *ceh-10* | WBGene00000435 | W03A3.1 | Not DE | -1.0 |
| *ceh-22* | WBGene00000445 | F29F11.5 | Not DE | -1.4 |
| *pros-1* | WBGene00000448 | K12H4.1 | Not DE | -1.9 |
| *ceh-27* | WBGene00000449 | F46F3.1 | Not DE | -1.1 |
| *ceh-31* | WBGene00000452 | C33D12.1 | Not DE | -1.2 |
| *ceh-32* | WBGene00000453 | W05E10.3 | Not DE | -1.3 |

| | | | | |
|---|---|---|---|---|
| *ceh-34* | WBGene00000455 | C10G8.6 | Not DE | -1.1 |
| *ceh-37* | WBGene00000458 | C37E2.5 | Not DE | -1.2 |
| *ceh-43* | WBGene00000463 | C28A5.4 | Not DE | -1.0 |
| *cep-1* | WBGene00000467 | F52B5.5 | Not DE | 1.0 |
| *cfi-1* | WBGene00000476 | T23D8.8 | Not DE | -1.5 |
| *cky-1* | WBGene00000521 | C15C8.2 | Not DE | -1.8 |
| *egl-38* | WBGene00001204 | C04G2.7 | Not DE | -1.2 |
| *egl-43* | WBGene00001207 | R53.3 | Not DE | -1.4 |
| *egl-46* | WBGene00001210 | K11G9.4 | Not DE | -1.5 |
| *fax-1* | WBGene00001400 | F56E3.4 | Not DE | -1.9 |
| *fkh-7* | WBGene00001439 | F26D12.1 | Not DE | -1.4 |
| *fkh-8* | WBGene00001440 | F40H3.4 | Not DE | -1.3 |
| *gei-13* | WBGene00001570 | F58A4.11 | Not DE | -1.7 |
| *grh-1* | WBGene00001707 | Y48G8AR.1 | Not DE | -1.4 |
| *ham-1* | WBGene00001820 | F53B2.6 | Not DE | -1.1 |
| *ham-2* | WBGene00001821 | C07A12.1 | Not DE | -1.4 |
| *hbl-1* | WBGene00001824 | F13D11.2 | Not DE | -1.6 |
| *let-381* | WBGene00002601 | F26B1.7 | Not DE | -1.4 |
| *lim-4* | WBGene00002987 | ZC64.4 | Not DE | -1.4 |
| *lin-22* | WBGene00003008 | Y54G2A.1 | Not DE | -1.2 |
| *mec-3* | WBGene00003167 | F01D4.6 | Not DE | -1.2 |
| *moe-3* | WBGene00003388 | F32A11.6 | Not DE | -1.7 |
| *nhr-18* | WBGene00003617 | F44C8.3 | Not DE | -1.4 |
| *nhr-25* | WBGene00003623 | F11C1.6 | Not DE | -1.4 |
| *nhr-32* | WBGene00003626 | K08H2.8 | Not DE | -1.2 |
| *nhr-58* | WBGene00003648 | R11G11.2 | Not DE | -1.2 |
| *nhr-67* | WBGene00003657 | C08F8.8 | Not DE | -1.2 |
| *nhr-69* | WBGene00003659 | T23H4.2 | Not DE | -1.4 |
| *nhr-88* | WBGene00003678 | K08A2.5 | Not DE | -1.1 |
| *nhr-91* | WBGene00003681 | Y15E3A.1 | Not DE | -1.8 |
| *nhr-92* | WBGene00003682 | Y41D4B.8 | Not DE | -1.4 |
| *nhr-110* | WBGene00003700 | Y46H3D.5 | Not DE | -1.1 |
| *nhr-119* | WBGene00003709 | K12H6.1 | Not DE | -1.4 |
| *nhr-120* | WBGene00003710 | C25B8.6 | Not DE | -1.3 |
| *nhr-131* | WBGene00003721 | T01G6.2 | Not DE | -1.3 |
| *nhr-134* | WBGene00003724 | F44C8.2 | Not DE | -1.2 |
| *odd-1* | WBGene00003845 | B0280.4 | Not DE | -1.0 |
| *blmp-1* | WBGene00003847 | F25D7.3 | Not DE | -1.3 |
| *pag-3* | WBGene00003909 | F45B8.4 | Not DE | -2.0 |
| *pha-2* | WBGene00004011 | M6.3 | Not DE | -1.1 |
| *plx-2* | WBGene00004048 | K04B12.1 | Not DE | -1.6 |
| *sma-2* | WBGene00004856 | ZK370.2 | Not DE | -1.1 |
| *sma-4* | WBGene00004858 | R12B2.1 | Not DE | -1.4 |
| *sox-2* | WBGene00004949 | K08A8.2 | Not DE | -1.3 |
| *ctbp-1* | WBGene00006424 | F49E10.5 | Not DE | -1.2 |
| *rhgf-2* | WBGene00006476 | T08H4.1 | Not DE | -1.3 |
| *tbx-2* | WBGene00006543 | F21H11.3 | Not DE | -1.0 |
| *tbx-7* | WBGene00006544 | ZK328.8 | Not DE | -1.2 |
| *ttx-1* | WBGene00006652 | Y113G7A.6 | Not DE | -1.1 |
| *unc-3* | WBGene00006743 | Y16B4A.1 | Not DE | -1.2 |
| *unc-30* | WBGene00006766 | B0564.10 | Not DE | -1.4 |
| *unc-86* | WBGene00006818 | C30A5.7 | Not DE | -1.6 |
| *unc-120* | WBGene00006844 | D1081.2 | Not DE | -1.2 |
| *unc-130* | WBGene00006853 | C47G2.2 | Not DE | -1.1 |
| *vab-3* | WBGene00006870 | F14F3.1 | Not DE | -1.4 |
| *vab-15* | WBGene00006881 | R07B1.1 | Not DE | -1.2 |
| *zag-1* | WBGene00006970 | F28F9.1 | Not DE | -1.6 |
| *ceh-79* | WBGene00007749 | C26E1.3 | Not DE | -1.4 |

| | | | | |
|---|---|---|---|---|
| *jmjd-3.3* | WBGene00007813 | C29F7.6 | Not DE | -1.1 |
| *C34B4.2* | WBGene00007907 | C34B4.2 | Not DE | 1.2 |
| *efl-3* | WBGene00009899 | F49E12.6 | Not DE | -1.2 |
| *gmeb-2* | WBGene00010010 | F53H4.5 | Not DE | -1.1 |
| *sta-2* | WBGene00010251 | F58E6.1 | Not DE | -1.9 |
| *fozi-1* | WBGene00010453 | K01B6.1 | Not DE | -1.6 |
| *nhr-205* | WBGene00011002 | R04B5.3 | Not DE | -1.5 |
| *nhr-206* | WBGene00011097 | R07B7.13 | Not DE | -1.2 |
| *tsct-1* | WBGene00011824 | T18D3.7 | Not DE | -1.8 |
| *hsf-2* | WBGene00013134 | Y53C10A.3 | Not DE | 1.3 |
| *ztf-30* | WBGene00015523 | C06E1.8 | Not DE | -2.3 |
| *ceh-48* | WBGene00015934 | C17H12.9 | Not DE | -1.4 |
| *unk-1* | WBGene00016407 | C34D10.2 | Not DE | -1.7 |
| *nhr-171* | WBGene00016926 | C54F6.8 | Not DE | -1.2 |
| *nhr-172* | WBGene00016927 | C54F6.9 | Not DE | -1.6 |
| *nhr-36* | WBGene00017198 | F07C3.10 | Not DE | -1.2 |
| *set-9* | WBGene00017482 | F15E6.1 | Not DE | 1.1 |
| *F26A10.2* | WBGene00017814 | F26A10.2 | Not DE | -1.3 |
| *nhr-180* | WBGene00017961 | F31F4.12 | Not DE | -1.7 |
| *K09H9.7* | WBGene00019598 | K09H9.7 | Not DE | 1.2 |
| *K10B3.5* | WBGene00019611 | K10B3.5 | Not DE | -1.1 |
| *M03D4.4* | WBGene00019751 | M03D4.4 | Not DE | -1.6 |
| *ast-1* | WBGene00020368 | T08H4.3 | Not DE | -1.3 |
| *hlh-30* | WBGene00020930 | W02C12.3 | Not DE | -1.2 |
| *crtc-1* | WBGene00021237 | Y20F4.2 | Not DE | -1.4 |
| *nhr-236* | WBGene00021417 | Y38F2AL.5 | Not DE | 1.5 |
| *nhr-238* | WBGene00021611 | Y46H3D.7 | Not DE | -1.5 |
| *nhr-41* | WBGene00022423 | Y104H12A.1 | Not DE | -1.3 |
| *klu-2* | WBGene00022592 | ZC328.2 | Not DE | 1.0 |
| *syd-9* | WBGene00044068 | ZK867.1 | Not DE | -1.4 |

**Supplemental Table 6**

**Lists of differentially expressed genes that are identified in either N2, CB4856, or both, after infection by *S. epidermidis* (compared to *E. coli*)**

**Supplemental Table 6 - Differentially Expressed Genes in *S. epidermidis* Infected Animals**

**N2**-exclusive DEGs in response to *S. epidermidis* (vs. OP50)

| Gene Name | Wormbase Gene ID | Type | $log_2FC$ | FC | Adj P-value |
|---|---|---|---|---|---|
| *C23G10.11* | WBGene00016017 | protein_coding_gene | 7.9 | 239.54 | 3.3E-15 |
| *thn-1* | WBGene00009213 | protein_coding_gene | 7.7 | 210.58 | 2.3E-33 |
| *clec-76* | WBGene00021587 | protein_coding_gene | 7.3 | 160.20 | 8.8E-17 |
| *math-38* | WBGene00020326 | protein_coding_gene | 5.4 | 41.16 | 2.2E-26 |
| *tts-1* | WBGene00006650 | ncRNA_gene | 5.3 | 40.30 | 2.3E-13 |
| *T07G12.5* | WBGene00011596 | protein_coding_gene | 5.2 | 36.50 | 6.0E-08 |
| *Y68A4A.13* | WBGene00044212 | protein_coding_gene | 5.1 | 34.23 | 3.4E-10 |
| *F14D7.10* | WBGene00008797 | protein_coding_gene | 4.9 | 30.02 | 4.0E-05 |
| *T26H5.9* | WBGene00044206 | protein_coding_gene | 4.8 | 27.13 | 2.3E-07 |
| *C14C6.8* | WBGene00015762 | protein_coding_gene | 4.6 | 25.07 | 1.2E-03 |
| *lipl-2* | WBGene00009773 | protein_coding_gene | 4.6 | 24.03 | 2.4E-07 |
| *cky-1* | WBGene00000521 | protein_coding_gene | 4.6 | 23.62 | 5.7E-05 |
| *clec-82* | WBGene00021873 | protein_coding_gene | 4.5 | 23.12 | 9.6E-20 |
| *cyp-35B3* | WBGene00019470 | protein_coding_gene | 4.5 | 23.02 | 2.0E-03 |
| *sodh-1* | WBGene00010790 | protein_coding_gene | 4.5 | 22.53 | 5.4E-24 |
| *clec-2* | WBGene00015198 | protein_coding_gene | 4.3 | 19.19 | 2.8E-04 |
| *fbxa-24* | WBGene00018803 | protein_coding_gene | 4.2 | 18.85 | 7.8E-10 |
| *spp-2* | WBGene00004987 | protein_coding_gene | 4.2 | 18.73 | 1.2E-23 |
| *oac-18* | WBGene00009230 | pseudogene | 4.2 | 18.17 | 7.4E-05 |
| *M02E1.4* | WBGene00284848 | protein_coding_gene | 4.2 | 17.82 | 2.4E-09 |
| *fbxa-164* | WBGene00015599 | protein_coding_gene | 4.1 | 17.68 | 2.3E-04 |
| *T26H5.14* | WBGene00235133 | protein_coding_gene | 4.1 | 17.46 | 2.8E-06 |
| *F55H12.7* | WBGene00219283 | protein_coding_gene | 4.1 | 17.26 | 1.4E-04 |
| *E02H4.4* | WBGene00008462 | protein_coding_gene | 4.1 | 17.13 | 2.4E-14 |
| *T26H5.4* | WBGene00012069 | protein_coding_gene | 4.1 | 16.69 | 5.3E-05 |
| *clec-69* | WBGene00044642 | pseudogene | 4.1 | 16.59 | 1.7E-08 |
| *Y41D4B.6* | WBGene00021513 | protein_coding_gene | 4.0 | 16.47 | 1.3E-06 |
| *M7.10* | WBGene00010886 | protein_coding_gene | 4.0 | 15.85 | 1.4E-03 |
| *thn-8* | WBGene00044285 | pseudogene | 4.0 | 15.70 | 2.4E-03 |
| *cyp-37B1* | WBGene00009226 | protein_coding_gene | 4.0 | 15.69 | 4.1E-06 |
| *cpr-2* | WBGene00000782 | protein_coding_gene | 3.9 | 15.28 | 8.6E-05 |
| *clec-17* | WBGene00008477 | protein_coding_gene | 3.8 | 14.01 | 1.7E-05 |
| *F54E2.2* | WBGene00018824 | protein_coding_gene | 3.8 | 13.93 | 1.8E-03 |
| *T05A7.11* | WBGene00044383 | protein_coding_gene | 3.8 | 13.86 | 8.5E-05 |
| *wrt-2* | WBGene00006948 | protein_coding_gene | 3.8 | 13.54 | 4.9E-10 |
| *F53A9.8* | WBGene00018731 | protein_coding_gene | 3.7 | 13.07 | 1.5E-07 |
| *clec-75* | WBGene00021586 | protein_coding_gene | 3.7 | 12.67 | 7.7E-07 |
| *Y47H10A.3* | WBGene00012959 | protein_coding_gene | 3.6 | 12.24 | 2.4E-04 |
| *oac-29* | WBGene00018295 | protein_coding_gene | 3.6 | 11.83 | 1.7E-04 |
| *cyp-32B1* | WBGene00021167 | protein_coding_gene | 3.6 | 11.78 | 4.4E-06 |
| *irld-11* | WBGene00019391 | protein_coding_gene | 3.6 | 11.72 | 1.4E-03 |
| *T21D12.7* | WBGene00020648 | protein_coding_gene | 3.5 | 11.70 | 7.8E-04 |
| *F11A5.5* | WBGene00008673 | protein_coding_gene | 3.5 | 11.59 | 2.9E-03 |
| *srbc-20* | WBGene00016693 | protein_coding_gene | 3.5 | 11.52 | 6.0E-03 |
| *clec-77* | WBGene00044343 | pseudogene | 3.5 | 11.41 | 3.5E-04 |
| *pals-39* | WBGene00008301 | protein_coding_gene | 3.5 | 11.40 | 3.1E-06 |
| *fbxa-48* | WBGene00021862 | protein_coding_gene | 3.5 | 11.14 | 1.2E-05 |
| *nhr-127* | WBGene00003717 | protein_coding_gene | 3.5 | 11.10 | 2.2E-04 |
| *pals-37* | WBGene00138721 | protein_coding_gene | 3.5 | 11.09 | 1.8E-04 |
| *C35A5.11* | WBGene00044024 | protein_coding_gene | 3.5 | 11.05 | 2.4E-07 |
| *fbxa-137* | WBGene00020853 | protein_coding_gene | 3.5 | 11.02 | 3.2E-03 |
| *lim-4* | WBGene00002987 | protein_coding_gene | 3.4 | 10.88 | 5.8E-04 |
| *C25H3.10* | WBGene00016119 | protein_coding_gene | 3.4 | 10.80 | 1.3E-11 |
| *K10G6.4* | WBGene00019641 | protein_coding_gene | 3.4 | 10.78 | 6.2E-04 |

| | | | | | |
|---|---|---|---|---|---|
| *nnt-1* | WBGene00003778 | protein_coding_gene | 3.4 | 10.69 | 9.1E-13 |
| *T05A7.13* | WBGene00271415 | protein_coding_gene | 3.4 | 10.64 | 2.3E-05 |
| *F58F12.12* | WBGene00271812 | protein_coding_gene | 3.4 | 10.59 | 9.5E-09 |
| *rgba-1* | WBGene00077766 | protein_coding_gene | 3.4 | 10.47 | 1.1E-02 |
| *F18C5.9* | WBGene00017562 | protein_coding_gene | 3.4 | 10.37 | 4.3E-03 |
| *ZK675.4* | WBGene00014067 | protein_coding_gene | 3.4 | 10.22 | 3.8E-04 |
| *dsl-6* | WBGene00001108 | protein_coding_gene | 3.3 | 10.04 | 7.6E-05 |
| *Y53F4B.45* | WBGene00044728 | ncRNA_gene | 3.3 | 10.01 | 3.4E-11 |
| *btb-16* | WBGene00018195 | protein_coding_gene | 3.3 | 10.01 | 9.4E-29 |
| *clec-70* | WBGene00021581 | protein_coding_gene | 3.3 | 9.97 | 2.0E-16 |
| *F56H9.9* | WBGene00206460 | protein_coding_gene | 3.3 | 9.86 | 1.2E-02 |
| *sru-20* | WBGene00005683 | protein_coding_gene | 3.3 | 9.84 | 3.6E-03 |
| *Y65B4BL.1* | WBGene00022033 | protein_coding_gene | 3.3 | 9.75 | 1.8E-03 |
| *srh-2* | WBGene00005228 | protein_coding_gene | 3.3 | 9.59 | 5.0E-06 |
| *B0284.5* | WBGene00045188 | pseudogene | 3.3 | 9.59 | 2.8E-03 |
| *T05F1.11* | WBGene00011495 | protein_coding_gene | 3.2 | 9.45 | 1.5E-13 |
| *pqn-73* | WBGene00004155 | protein_coding_gene | 3.2 | 9.31 | 6.4E-05 |
| *scl-14* | WBGene00008625 | protein_coding_gene | 3.2 | 9.27 | 3.0E-03 |
| *cutl-28* | WBGene00018256 | protein_coding_gene | 3.2 | 9.23 | 9.8E-08 |
| *clec-71* | WBGene00021582 | protein_coding_gene | 3.2 | 9.20 | 5.8E-11 |
| *pax-1* | WBGene00003937 | protein_coding_gene | 3.2 | 9.16 | 7.5E-03 |
| *hil-7* | WBGene00001858 | protein_coding_gene | 3.2 | 9.03 | 1.3E-04 |
| *best-5* | WBGene00007404 | protein_coding_gene | 3.2 | 8.90 | 1.8E-09 |
| *str-124* | WBGene00006175 | protein_coding_gene | 3.1 | 8.86 | 1.5E-02 |
| *F47B7.6* | WBGene00018537 | pseudogene | 3.1 | 8.84 | 1.2E-02 |
| *clec-78* | WBGene00018547 | protein_coding_gene | 3.1 | 8.81 | 2.7E-07 |
| *nhr-218* | WBGene00011750 | protein_coding_gene | 3.1 | 8.79 | 7.2E-03 |
| *C06G3.3* | WBGene00015549 | protein_coding_gene | 3.1 | 8.75 | 4.2E-09 |
| *T03E6.9* | WBGene00045356 | protein_coding_gene | 3.1 | 8.68 | 5.0E-03 |
| *T10D4.15* | WBGene00077522 | protein_coding_gene | 3.1 | 8.57 | 1.5E-03 |
| *dpy-14* | WBGene00001075 | protein_coding_gene | 3.1 | 8.57 | 1.1E-05 |
| *spp-7* | WBGene00004992 | protein_coding_gene | 3.1 | 8.55 | 6.6E-03 |
| *F21C10.10* | WBGene00017659 | protein_coding_gene | 3.1 | 8.46 | 2.0E-21 |
| *ZK993.2* | WBGene00022838 | protein_coding_gene | 3.1 | 8.39 | 6.7E-03 |
| *clec-34* | WBGene00012023 | protein_coding_gene | 3.1 | 8.35 | 7.1E-03 |
| *unc-42* | WBGene00006778 | protein_coding_gene | 3.1 | 8.34 | 2.0E-06 |
| *clec-55* | WBGene00008597 | protein_coding_gene | 3.1 | 8.32 | 2.1E-08 |
| *Y54G2A.76* | WBGene00235268 | protein_coding_gene | 3.1 | 8.29 | 8.0E-05 |
| *C06E7.88* | WBGene00189995 | protein_coding_gene | 3.0 | 8.28 | 1.9E-04 |
| *F49H6.5* | WBGene00009908 | protein_coding_gene | 3.0 | 8.26 | 2.6E-07 |
| *F26G1.15* | WBGene00270318 | protein_coding_gene | 3.0 | 8.24 | 4.8E-03 |
| *ZK381.2* | WBGene00022725 | protein_coding_gene | 3.0 | 8.19 | 1.9E-02 |
| *xtr-1* | WBGene00006965 | protein_coding_gene | 3.0 | 8.11 | 1.9E-08 |
| *cyp-35B2* | WBGene00019471 | protein_coding_gene | 3.0 | 7.95 | 6.4E-03 |
| *ZK813.6* | WBGene00044564 | protein_coding_gene | 3.0 | 7.95 | 1.9E-02 |
| *T02G6.7* | WBGene00011390 | protein_coding_gene | 3.0 | 7.94 | 1.8E-02 |
| *C27H5.6* | WBGene00016175 | protein_coding_gene | 3.0 | 7.94 | 1.8E-02 |
| *C18D4.4* | WBGene00007674 | protein_coding_gene | 3.0 | 7.92 | 2.4E-02 |
| *F43D9.1* | WBGene00009653 | protein_coding_gene | 3.0 | 7.89 | 1.2E-07 |
| *C14A6.16* | WBGene00206536 | pseudogene | 3.0 | 7.86 | 1.4E-03 |
| *F15B9.6* | WBGene00008850 | protein_coding_gene | 3.0 | 7.83 | 9.1E-08 |
| *R09E10.5* | WBGene00011175 | protein_coding_gene | 3.0 | 7.83 | 1.3E-05 |
| *F17B5.6* | WBGene00044058 | protein_coding_gene | 3.0 | 7.81 | 2.1E-02 |
| *cutl-9* | WBGene00010893 | protein_coding_gene | 3.0 | 7.81 | 5.6E-06 |
| *F38B6.3* | WBGene00018173 | protein_coding_gene | 3.0 | 7.79 | 1.3E-02 |
| *linc-69* | WBGene00219644 | lincRNA_gene | 3.0 | 7.77 | 1.2E-02 |
| *dpy-3* | WBGene00001065 | protein_coding_gene | 3.0 | 7.73 | 9.5E-08 |
| *Y37A1B.7* | WBGene00012540 | protein_coding_gene | 2.9 | 7.69 | 1.2E-05 |

| | | | | | |
|---|---|---|---|---|---|
| *H42K12.3* | WBGene00019272 | protein_coding_gene | 2.9 | 7.67 | 4.6E-09 |
| *R09D1.12* | WBGene00011168 | protein_coding_gene | 2.9 | 7.63 | 1.3E-06 |
| *T03G6.1* | WBGene00020194 | protein_coding_gene | 2.9 | 7.60 | 6.0E-07 |
| *far-7* | WBGene00001391 | protein_coding_gene | 2.9 | 7.56 | 6.6E-08 |
| *F47G4.5* | WBGene00009826 | protein_coding_gene | 2.9 | 7.53 | 1.3E-02 |
| *scl-15* | WBGene00017183 | protein_coding_gene | 2.9 | 7.50 | 2.0E-02 |
| *ZC449.1* | WBGene00022611 | protein_coding_gene | 2.9 | 7.49 | 7.2E-05 |
| *srz-99* | WBGene00012028 | protein_coding_gene | 2.9 | 7.48 | 6.0E-03 |
| *col-121* | WBGene00000695 | protein_coding_gene | 2.9 | 7.41 | 4.9E-03 |
| *F46B6.13* | WBGene00044192 | protein_coding_gene | 2.9 | 7.37 | 1.9E-02 |
| *egl-46* | WBGene00001210 | protein_coding_gene | 2.9 | 7.31 | 1.8E-08 |
| *clec-263* | WBGene00009171 | protein_coding_gene | 2.9 | 7.29 | 3.6E-02 |
| *nlp-81* | WBGene00022063 | protein_coding_gene | 2.9 | 7.26 | 1.3E-02 |
| *lpr-4* | WBGene00012257 | protein_coding_gene | 2.9 | 7.26 | 1.6E-08 |
| *dpy-17* | WBGene00001076 | protein_coding_gene | 2.9 | 7.25 | 2.1E-09 |
| *F11E6.9* | WBGene00008712 | protein_coding_gene | 2.8 | 7.19 | 5.8E-04 |
| *F09F7.6* | WBGene00017303 | protein_coding_gene | 2.8 | 7.11 | 5.4E-10 |
| *ZC395.5* | WBGene00022597 | protein_coding_gene | 2.8 | 7.10 | 9.8E-10 |
| *K08B12.1* | WBGene00019520 | protein_coding_gene | 2.8 | 7.09 | 3.6E-06 |
| *ZC204.12* | WBGene00022562 | protein_coding_gene | 2.8 | 7.09 | 2.8E-27 |
| *Y26D4A.3* | WBGene00012501 | protein_coding_gene | 2.8 | 7.08 | 4.0E-03 |
| *R05A10.8* | WBGene00011025 | protein_coding_gene | 2.8 | 7.06 | 1.4E-02 |
| *pals-32* | WBGene00016281 | protein_coding_gene | 2.8 | 7.02 | 1.1E-05 |
| *poml-4* | WBGene00044697 | protein_coding_gene | 2.8 | 7.01 | 5.8E-08 |
| *M176.10* | WBGene00010946 | protein_coding_gene | 2.8 | 6.95 | 1.9E-02 |
| *ZK666.11* | WBGene00014049 | protein_coding_gene | 2.8 | 6.93 | 9.8E-03 |
| *Y46C8AM.1* | WBGene00271795 | protein_coding_gene | 2.8 | 6.90 | 3.4E-07 |
| *srm-4* | WBGene00009808 | protein_coding_gene | 2.8 | 6.88 | 3.3E-02 |
| *bgnt-1.3* | WBGene00009032 | protein_coding_gene | 2.8 | 6.84 | 3.5E-02 |
| *R05A10.6* | WBGene00011023 | protein_coding_gene | 2.8 | 6.84 | 3.4E-02 |
| *ifa-3* | WBGene00002051 | protein_coding_gene | 2.8 | 6.84 | 2.8E-05 |
| *mlt-11* | WBGene00012186 | protein_coding_gene | 2.8 | 6.83 | 1.2E-07 |
| *T04C12.1* | WBGene00011427 | protein_coding_gene | 2.8 | 6.83 | 1.1E-03 |
| *C14A11.2* | WBGene00015748 | protein_coding_gene | 2.8 | 6.78 | 1.2E-04 |
| *hlh-32* | WBGene00013665 | protein_coding_gene | 2.8 | 6.75 | 3.6E-02 |
| *C01C4.2* | WBGene00015292 | protein_coding_gene | 2.8 | 6.75 | 2.4E-02 |
| *fbxc-20* | WBGene00018352 | protein_coding_gene | 2.7 | 6.71 | 9.9E-07 |
| *T20B3.1* | WBGene00011850 | protein_coding_gene | 2.7 | 6.70 | 6.0E-07 |
| *peb-1* | WBGene00003968 | protein_coding_gene | 2.7 | 6.69 | 8.5E-07 |
| *F47H4.2* | WBGene00009835 | protein_coding_gene | 2.7 | 6.67 | 8.1E-08 |
| *cup-16* | WBGene00007919 | protein_coding_gene | 2.7 | 6.67 | 1.2E-07 |
| *T09D3.3* | WBGene00020384 | protein_coding_gene | 2.7 | 6.60 | 3.8E-02 |
| *math-10* | WBGene00015839 | protein_coding_gene | 2.7 | 6.58 | 1.3E-05 |
| *F36F12.1* | WBGene00018092 | protein_coding_gene | 2.7 | 6.58 | 8.0E-03 |
| *dpy-7* | WBGene00001069 | protein_coding_gene | 2.7 | 6.56 | 8.1E-08 |
| *T20F5.4* | WBGene00020626 | protein_coding_gene | 2.7 | 6.54 | 4.1E-07 |
| *lpr-5* | WBGene00012256 | protein_coding_gene | 2.7 | 6.51 | 1.7E-08 |
| *C56G3.2* | WBGene00016985 | protein_coding_gene | 2.7 | 6.49 | 2.3E-02 |
| *lys-9* | WBGene00003098 | protein_coding_gene | 2.7 | 6.49 | 3.1E-02 |
| *hch-1* | WBGene00001828 | protein_coding_gene | 2.7 | 6.47 | 2.8E-05 |
| *Y32B12C.5* | WBGene00194769 | pseudogene | 2.7 | 6.46 | 3.7E-07 |
| *M03B6.3* | WBGene00010835 | protein_coding_gene | 2.7 | 6.46 | 4.8E-08 |
| *T16G1.4* | WBGene00011798 | protein_coding_gene | 2.7 | 6.45 | 1.9E-03 |
| *scl-8* | WBGene00008030 | protein_coding_gene | 2.7 | 6.43 | 1.8E-02 |
| *Y60C6A.1* | WBGene00022013 | protein_coding_gene | 2.7 | 6.40 | 2.4E-02 |
| *F58H1.2* | WBGene00010285 | protein_coding_gene | 2.7 | 6.33 | 7.8E-05 |
| *F54F3.2* | WBGene00014770 | pseudogene | 2.7 | 6.30 | 2.7E-02 |
| *ZK1240.1* | WBGene00022866 | protein_coding_gene | 2.7 | 6.29 | 4.9E-04 |

| | | | | | |
|---|---|---|---|---|---|
| *W07G4.1* | WBGene00012335 | protein_coding_gene | 2.7 | 6.28 | 3.3E-02 |
| *F22F7.4* | WBGene00017722 | protein_coding_gene | 2.6 | 6.27 | 5.5E-03 |
| *fbxa-66* | WBGene00021865 | protein_coding_gene | 2.6 | 6.24 | 9.6E-03 |
| *nhr-277* | WBGene00022374 | protein_coding_gene | 2.6 | 6.24 | 7.4E-04 |
| *mec-3* | WBGene00003167 | protein_coding_gene | 2.6 | 6.22 | 8.6E-07 |
| *slc-36.3* | WBGene00012629 | protein_coding_gene | 2.6 | 6.22 | 1.0E-05 |
| *Y110A2AL.4* | WBGene00022441 | protein_coding_gene | 2.6 | 6.22 | 1.0E-06 |
| *F55H12.9* | WBGene00219976 | snoRNA_gene | 2.6 | 6.20 | 1.6E-02 |
| *fbxc-41* | WBGene00016304 | pseudogene | 2.6 | 6.20 | 2.5E-03 |
| *F56D2.3* | WBGene00018965 | protein_coding_gene | 2.6 | 6.20 | 1.3E-06 |
| *C03C10.5* | WBGene00007271 | protein_coding_gene | 2.6 | 6.18 | 5.2E-04 |
| *lpr-6* | WBGene00012255 | protein_coding_gene | 2.6 | 6.17 | 6.0E-06 |
| *hil-1* | WBGene00001852 | protein_coding_gene | 2.6 | 6.09 | 2.0E-08 |
| *C44H9.7* | WBGene00008102 | protein_coding_gene | 2.6 | 6.08 | 3.0E-04 |
| *Y7A5A.6* | WBGene00012412 | protein_coding_gene | 2.6 | 6.06 | 6.4E-03 |
| *osm-8* | WBGene00003888 | protein_coding_gene | 2.6 | 6.06 | 3.9E-04 |
| *R160.4* | WBGene00020121 | protein_coding_gene | 2.6 | 6.05 | 1.2E-02 |
| *ptr-10* | WBGene00004224 | protein_coding_gene | 2.6 | 6.05 | 2.2E-06 |
| *C35E7.12* | WBGene00271779 | protein_coding_gene | 2.6 | 6.04 | 4.2E-02 |
| *cutl-13* | WBGene00021025 | protein_coding_gene | 2.6 | 6.02 | 4.3E-03 |
| *Y7A5A.8* | WBGene00012414 | protein_coding_gene | 2.6 | 5.99 | 4.0E-02 |
| *math-21* | WBGene00016556 | protein_coding_gene | 2.6 | 5.99 | 1.6E-03 |
| *fbxa-21* | WBGene00022487 | protein_coding_gene | 2.6 | 5.96 | 7.9E-07 |
| *F20B6.4* | WBGene00017628 | protein_coding_gene | 2.6 | 5.95 | 2.6E-02 |
| *F59E11.7* | WBGene00019113 | protein_coding_gene | 2.6 | 5.93 | 4.1E-07 |
| *C27A2.8* | WBGene00044387 | protein_coding_gene | 2.6 | 5.92 | 3.0E-02 |
| *F13H6.5* | WBGene00017433 | protein_coding_gene | 2.6 | 5.90 | 9.9E-08 |
| *math-42* | WBGene00020532 | protein_coding_gene | 2.6 | 5.90 | 5.0E-04 |
| *F55H12.2* | WBGene00010133 | protein_coding_gene | 2.6 | 5.89 | 1.4E-11 |
| *atf-2* | WBGene00000220 | protein_coding_gene | 2.6 | 5.87 | 1.9E-06 |
| *lpr-3* | WBGene00012261 | protein_coding_gene | 2.5 | 5.85 | 1.7E-08 |
| *F31D4.5* | WBGene00009292 | protein_coding_gene | 2.5 | 5.85 | 2.3E-06 |
| *F57E7.2* | WBGene00010203 | protein_coding_gene | 2.5 | 5.85 | 3.1E-03 |
| *glf-1* | WBGene00019154 | protein_coding_gene | 2.5 | 5.84 | 1.3E-05 |
| *hbl-1* | WBGene00001824 | protein_coding_gene | 2.5 | 5.80 | 5.6E-06 |
| *C06E4.8* | WBGene00015537 | protein_coding_gene | 2.5 | 5.78 | 1.3E-08 |
| *Y43F8B.9* | WBGene00012819 | protein_coding_gene | 2.5 | 5.77 | 9.8E-07 |
| *ZC443.3* | WBGene00013898 | protein_coding_gene | 2.5 | 5.77 | 7.1E-06 |
| *C48B4.3* | WBGene00008168 | protein_coding_gene | 2.5 | 5.76 | 1.4E-06 |
| *zmp-5* | WBGene00010423 | protein_coding_gene | 2.5 | 5.74 | 2.0E-02 |
| *sqst-3* | WBGene00012067 | protein_coding_gene | 2.5 | 5.73 | 1.6E-02 |
| *Y65B4BM.3* | WBGene00302974 | protein_coding_gene | 2.5 | 5.71 | 2.6E-09 |
| *str-173* | WBGene00006217 | protein_coding_gene | 2.5 | 5.70 | 3.2E-02 |
| *D1054.18* | WBGene00077783 | protein_coding_gene | 2.5 | 5.68 | 8.9E-04 |
| *nlp-66* | WBGene00011141 | protein_coding_gene | 2.5 | 5.67 | 4.8E-04 |
| *clec-208* | WBGene00018097 | protein_coding_gene | 2.5 | 5.66 | 1.3E-02 |
| *spp-8* | WBGene00004993 | protein_coding_gene | 2.5 | 5.64 | 9.9E-16 |
| *twk-47* | WBGene00018002 | protein_coding_gene | 2.5 | 5.64 | 5.1E-03 |
| *dpy-2* | WBGene00001064 | protein_coding_gene | 2.5 | 5.64 | 4.8E-07 |
| *K02E7.10* | WBGene00019314 | protein_coding_gene | 2.5 | 5.62 | 5.9E-06 |
| *F09C6.14* | WBGene00044278 | protein_coding_gene | 2.5 | 5.61 | 4.2E-03 |
| *F45D3.3* | WBGene00009723 | protein_coding_gene | 2.5 | 5.58 | 4.4E-21 |
| *F57F4.2* | WBGene00019016 | protein_coding_gene | 2.5 | 5.58 | 1.6E-02 |
| *lurp-3* | WBGene00009136 | protein_coding_gene | 2.5 | 5.55 | 4.0E-02 |
| *linc-85* | WBGene00219754 | lincRNA_gene | 2.5 | 5.54 | 2.0E-03 |
| *bro-1* | WBGene00000272 | protein_coding_gene | 2.5 | 5.54 | 4.3E-04 |
| *F45D11.1* | WBGene00018448 | pseudogene | 2.5 | 5.51 | 3.3E-04 |
| *F09C6.13* | WBGene00044183 | protein_coding_gene | 2.5 | 5.50 | 2.0E-07 |

| | | | | | |
|---|---|---|---|---|---|
| *ugt-15* | WBGene00008097 | protein_coding_gene | 2.5 | 5.48 | 6.9E-08 |
| *odr-10* | WBGene00003856 | protein_coding_gene | 2.5 | 5.48 | 6.4E-06 |
| *hhat-1* | WBGene00013852 | protein_coding_gene | 2.5 | 5.47 | 5.6E-06 |
| *W02D7.8* | WBGene00020943 | protein_coding_gene | 2.4 | 5.46 | 2.2E-06 |
| *K10D3.4* | WBGene00010738 | protein_coding_gene | 2.4 | 5.46 | 2.9E-08 |
| *E02H9.6* | WBGene00017104 | protein_coding_gene | 2.4 | 5.46 | 9.0E-03 |
| *ZC123.1* | WBGene00022517 | protein_coding_gene | 2.4 | 5.44 | 1.3E-05 |
| *K09C4.4* | WBGene00019549 | protein_coding_gene | 2.4 | 5.41 | 2.0E-02 |
| *linc-103* | WBGene00219743 | lincRNA_gene | 2.4 | 5.41 | 3.6E-02 |
| *nhr-21* | WBGene00003620 | protein_coding_gene | 2.4 | 5.39 | 2.4E-13 |
| *ugt-57* | WBGene00017154 | protein_coding_gene | 2.4 | 5.38 | 4.9E-05 |
| *F22B7.9* | WBGene00017698 | protein_coding_gene | 2.4 | 5.36 | 8.2E-40 |
| *C07A9.5* | WBGene00007401 | protein_coding_gene | 2.4 | 5.33 | 3.6E-02 |
| *T01D3.3* | WBGene00011328 | protein_coding_gene | 2.4 | 5.32 | 5.5E-06 |
| *linc-47* | WBGene00219641 | lincRNA_gene | 2.4 | 5.32 | 3.6E-02 |
| *dpy-10* | WBGene00001072 | protein_coding_gene | 2.4 | 5.32 | 1.4E-10 |
| *clec-7* | WBGene00017364 | protein_coding_gene | 2.4 | 5.32 | 2.6E-06 |
| *clec-37* | WBGene00013609 | protein_coding_gene | 2.4 | 5.28 | 1.9E-03 |
| *nstp-3* | WBGene00022577 | protein_coding_gene | 2.4 | 5.26 | 2.5E-03 |
| *T15D6.12* | WBGene00011786 | protein_coding_gene | 2.4 | 5.25 | 2.3E-02 |
| *C29A12.6* | WBGene00007800 | protein_coding_gene | 2.4 | 5.25 | 2.8E-03 |
| *Y71G12B.42* | WBGene00220198 | ncRNA_gene | 2.4 | 5.24 | 5.9E-03 |
| *R03H10.2* | WBGene00019855 | protein_coding_gene | 2.4 | 5.23 | 3.0E-07 |
| *ces-2* | WBGene00000469 | protein_coding_gene | 2.4 | 5.23 | 1.4E-06 |
| *K04H4.2* | WBGene00010573 | protein_coding_gene | 2.4 | 5.22 | 3.9E-07 |
| *EGAP1.1* | WBGene00017146 | protein_coding_gene | 2.4 | 5.22 | 3.7E-03 |
| *C08F1.11* | WBGene00044417 | protein_coding_gene | 2.4 | 5.20 | 4.6E-04 |
| *ZK287.1* | WBGene00013962 | protein_coding_gene | 2.4 | 5.20 | 4.8E-08 |
| *C40H1.3* | WBGene00008039 | protein_coding_gene | 2.4 | 5.20 | 1.8E-02 |
| *K11H12.11* | WBGene00044723 | protein_coding_gene | 2.4 | 5.19 | 6.5E-03 |
| *C07D10.6* | WBGene00044673 | protein_coding_gene | 2.4 | 5.17 | 1.6E-02 |
| *snt-7* | WBGene00011682 | protein_coding_gene | 2.4 | 5.17 | 4.7E-04 |
| *F56F11.1* | WBGene00018988 | protein_coding_gene | 2.4 | 5.16 | 3.4E-02 |
| *R05A10.4* | WBGene00011021 | protein_coding_gene | 2.4 | 5.15 | 1.8E-03 |
| *ZC84.6* | WBGene00013849 | protein_coding_gene | 2.4 | 5.14 | 2.0E-03 |
| *rocf-1* | WBGene00011948 | protein_coding_gene | 2.4 | 5.14 | 1.2E-02 |
| *C30G12.11* | WBGene00284861 | ncRNA_gene | 2.4 | 5.10 | 2.9E-02 |
| *F47B10.3* | WBGene00009814 | protein_coding_gene | 2.3 | 5.09 | 3.7E-02 |
| *spp-13* | WBGene00004998 | protein_coding_gene | 2.3 | 5.07 | 7.2E-05 |
| *unc-75* | WBGene00006807 | protein_coding_gene | 2.3 | 5.06 | 3.1E-04 |
| *dao-2* | WBGene00000928 | protein_coding_gene | 2.3 | 5.05 | 3.7E-10 |
| *F36D1.7* | WBGene00009466 | protein_coding_gene | 2.3 | 5.05 | 6.4E-06 |
| *nhr-99* | WBGene00003689 | protein_coding_gene | 2.3 | 5.00 | 3.9E-03 |
| *ins-35* | WBGene00002118 | protein_coding_gene | 2.3 | 5.00 | 1.1E-03 |
| *F55B11.5* | WBGene00010087 | protein_coding_gene | 2.3 | 4.99 | 4.2E-03 |
| *tmem-231* | WBGene00020825 | protein_coding_gene | 2.3 | 4.97 | 2.7E-03 |
| *cut-5* | WBGene00011104 | protein_coding_gene | 2.3 | 4.97 | 1.6E-03 |
| *gpa-9* | WBGene00001671 | protein_coding_gene | 2.3 | 4.96 | 2.6E-12 |
| *T21D12.14* | WBGene00045177 | protein_coding_gene | 2.3 | 4.95 | 1.8E-04 |
| *fezf-1* | WBGene00012639 | protein_coding_gene | 2.3 | 4.93 | 4.4E-02 |
| *T19D12.3* | WBGene00020578 | protein_coding_gene | 2.3 | 4.93 | 3.9E-02 |
| *Y17D7C.4* | WBGene00050903 | pseudogene | 2.3 | 4.93 | 5.6E-04 |
| *R11D1.12* | WBGene00045336 | protein_coding_gene | 2.3 | 4.92 | 9.5E-04 |
| *asp-16* | WBGene00012682 | protein_coding_gene | 2.3 | 4.92 | 4.0E-02 |
| *W02A2.9* | WBGene00044280 | protein_coding_gene | 2.3 | 4.92 | 1.6E-08 |
| *sqt-3* | WBGene00005018 | protein_coding_gene | 2.3 | 4.91 | 2.1E-08 |
| *coel-1* | WBGene00016866 | protein_coding_gene | 2.3 | 4.91 | 1.8E-04 |
| *T02G5.15* | WBGene00206521 | protein_coding_gene | 2.3 | 4.90 | 5.1E-04 |

| | | | | | |
|---|---|---|---|---|---|
| *rncs-1* | WBGene00023069 | ncRNA_gene | 2.3 | 4.90 | 1.9E-07 |
| *glt-5* | WBGene00001623 | protein_coding_gene | 2.3 | 4.89 | 5.5E-06 |
| *T03E6.8* | WBGene00011397 | protein_coding_gene | 2.3 | 4.88 | 9.5E-08 |
| *fipr-17* | WBGene00012965 | protein_coding_gene | 2.3 | 4.88 | 2.4E-02 |
| *F30H5.3* | WBGene00017937 | protein_coding_gene | 2.3 | 4.86 | 8.9E-05 |
| *C43C3.4* | WBGene00195084 | protein_coding_gene | 2.3 | 4.86 | 5.9E-06 |
| *ttr-44* | WBGene00008341 | protein_coding_gene | 2.3 | 4.85 | 3.0E-15 |
| *Y82E9BR.6* | WBGene00022339 | protein_coding_gene | 2.3 | 4.85 | 3.9E-02 |
| *Y46G5A.36* | WBGene00012921 | protein_coding_gene | 2.3 | 4.85 | 2.9E-02 |
| *F13E9.11* | WBGene00008760 | protein_coding_gene | 2.3 | 4.84 | 4.8E-08 |
| *M04C3.5* | WBGene00206418 | protein_coding_gene | 2.3 | 4.83 | 2.2E-08 |
| *dsl-3* | WBGene00001105 | protein_coding_gene | 2.3 | 4.83 | 9.0E-07 |
| *ces-1* | WBGene00000468 | protein_coding_gene | 2.3 | 4.82 | 7.5E-03 |
| *fbxa-173* | WBGene00044960 | pseudogene | 2.3 | 4.81 | 4.7E-02 |
| *spp-6* | WBGene00004991 | protein_coding_gene | 2.3 | 4.81 | 8.2E-03 |
| *icl-1* | WBGene00001564 | protein_coding_gene | 2.3 | 4.80 | 4.0E-07 |
| *B0205.4* | WBGene00015019 | protein_coding_gene | 2.3 | 4.80 | 9.5E-04 |
| *nep-16* | WBGene00018472 | protein_coding_gene | 2.2 | 4.75 | 9.5E-03 |
| *F14F9.8* | WBGene00044536 | protein_coding_gene | 2.2 | 4.75 | 4.0E-04 |
| *T19C4.5* | WBGene00011839 | protein_coding_gene | 2.2 | 4.73 | 1.9E-08 |
| *F23F12.8* | WBGene00017754 | protein_coding_gene | 2.2 | 4.73 | 9.6E-04 |
| *F49E10.2* | WBGene00018640 | protein_coding_gene | 2.2 | 4.71 | 1.5E-03 |
| *fat-3* | WBGene00001395 | protein_coding_gene | 2.2 | 4.70 | 5.5E-15 |
| *col-69* | WBGene00000645 | protein_coding_gene | 2.2 | 4.70 | 9.1E-03 |
| *T08G5.15* | WBGene00077573 | protein_coding_gene | 2.2 | 4.67 | 2.3E-02 |
| *T23B3.6* | WBGene00020714 | protein_coding_gene | 2.2 | 4.67 | 3.7E-05 |
| *B0281.6* | WBGene00015113 | protein_coding_gene | 2.2 | 4.65 | 1.0E-02 |
| *C14C6.5* | WBGene00015759 | protein_coding_gene | 2.2 | 4.64 | 3.1E-03 |
| *T26C5.2* | WBGene00012032 | protein_coding_gene | 2.2 | 4.63 | 2.4E-05 |
| *cutl-2* | WBGene00013145 | protein_coding_gene | 2.2 | 4.63 | 8.1E-05 |
| *Y39B6A.29* | WBGene00012688 | protein_coding_gene | 2.2 | 4.63 | 3.8E-03 |
| *noah-2* | WBGene00009926 | protein_coding_gene | 2.2 | 4.62 | 3.6E-07 |
| *K08B4.2* | WBGene00019514 | protein_coding_gene | 2.2 | 4.60 | 9.1E-04 |
| *K11H12.6* | WBGene00019662 | protein_coding_gene | 2.2 | 4.59 | 1.2E-02 |
| *K08F9.3* | WBGene00010686 | protein_coding_gene | 2.2 | 4.58 | 8.5E-04 |
| *F31E3.12* | WBGene00219317 | protein_coding_gene | 2.2 | 4.57 | 4.5E-03 |
| *bbs-1* | WBGene00000241 | protein_coding_gene | 2.2 | 4.55 | 3.8E-03 |
| *ZC443.2* | WBGene00013897 | protein_coding_gene | 2.2 | 4.54 | 1.9E-02 |
| *clec-232* | WBGene00009487 | protein_coding_gene | 2.2 | 4.52 | 5.0E-02 |
| *F59E11.6* | WBGene00019112 | protein_coding_gene | 2.2 | 4.52 | 4.8E-02 |
| *E02H4.5* | WBGene00008463 | protein_coding_gene | 2.2 | 4.51 | 5.9E-03 |
| *clec-25* | WBGene00044331 | protein_coding_gene | 2.2 | 4.50 | 9.3E-03 |
| *T22F3.10* | WBGene00020699 | protein_coding_gene | 2.2 | 4.50 | 5.0E-03 |
| *K03D3.2* | WBGene00010528 | protein_coding_gene | 2.2 | 4.49 | 2.5E-02 |
| *cyp-34A10* | WBGene00015045 | protein_coding_gene | 2.2 | 4.49 | 9.3E-03 |
| *nspc-16* | WBGene00008357 | protein_coding_gene | 2.2 | 4.49 | 1.3E-07 |
| *F11A6.3* | WBGene00077549 | pseudogene | 2.2 | 4.48 | 3.8E-02 |
| *K02C4.2* | WBGene00010501 | protein_coding_gene | 2.2 | 4.48 | 1.1E-02 |
| *C27A7.8* | WBGene00007758 | protein_coding_gene | 2.2 | 4.47 | 1.0E-08 |
| *sre-23* | WBGene00013957 | protein_coding_gene | 2.2 | 4.47 | 1.8E-02 |
| *tsp-17* | WBGene00006643 | protein_coding_gene | 2.2 | 4.47 | 8.8E-04 |
| *nspe-2* | WBGene00012604 | protein_coding_gene | 2.2 | 4.46 | 3.0E-02 |
| *fil-2* | WBGene00019668 | protein_coding_gene | 2.2 | 4.46 | 1.2E-02 |
| *Y41D4B.26* | WBGene00021525 | protein_coding_gene | 2.2 | 4.45 | 2.7E-04 |
| *fmi-1* | WBGene00001475 | protein_coding_gene | 2.2 | 4.44 | 4.8E-06 |
| *F49D11.6* | WBGene00018630 | protein_coding_gene | 2.1 | 4.42 | 1.4E-02 |
| *gcy-34* | WBGene00001554 | protein_coding_gene | 2.1 | 4.42 | 4.3E-02 |
| *C36B1.13* | WBGene00044120 | protein_coding_gene | 2.1 | 4.41 | 1.9E-02 |

| | | | | | |
|---|---|---|---|---|---|
| *fozi-1* | WBGene00010453 | protein_coding_gene | 2.1 | 4.40 | 2.3E-05 |
| *F57E7.4* | WBGene00077616 | protein_coding_gene | 2.1 | 4.40 | 8.8E-04 |
| *Y54G2A.11* | WBGene00021876 | protein_coding_gene | 2.1 | 4.39 | 1.8E-05 |
| *cutl-8* | WBGene00013960 | protein_coding_gene | 2.1 | 4.38 | 2.1E-02 |
| *ptr-4* | WBGene00004219 | protein_coding_gene | 2.1 | 4.37 | 4.1E-05 |
| *C24H10.3* | WBGene00016069 | protein_coding_gene | 2.1 | 4.37 | 2.5E-03 |
| *atf-8* | WBGene00017535 | protein_coding_gene | 2.1 | 4.36 | 5.1E-04 |
| *fbxa-127* | WBGene00012027 | protein_coding_gene | 2.1 | 4.36 | 7.7E-03 |
| *nlp-2* | WBGene00003740 | protein_coding_gene | 2.1 | 4.35 | 1.2E-04 |
| *T14B4.19* | WBGene00302993 | protein_coding_gene | 2.1 | 4.34 | 1.2E-04 |
| *F54C8.6* | WBGene00010039 | protein_coding_gene | 2.1 | 4.33 | 9.2E-07 |
| *C12D5.3* | WBGene00015706 | protein_coding_gene | 2.1 | 4.33 | 1.5E-03 |
| *osm-12* | WBGene00003892 | protein_coding_gene | 2.1 | 4.32 | 4.4E-04 |
| *C01G10.15* | WBGene00007240 | protein_coding_gene | 2.1 | 4.32 | 8.4E-04 |
| *T02E9.5* | WBGene00011383 | protein_coding_gene | 2.1 | 4.31 | 2.6E-06 |
| *clec-161* | WBGene00010228 | protein_coding_gene | 2.1 | 4.30 | 4.6E-02 |
| *gbh-2* | WBGene00001523 | protein_coding_gene | 2.1 | 4.30 | 1.4E-06 |
| *C14F11.6* | WBGene00015781 | protein_coding_gene | 2.1 | 4.30 | 2.5E-04 |
| *C34B4.3* | WBGene00007908 | protein_coding_gene | 2.1 | 4.30 | 7.7E-05 |
| *nphp-1* | WBGene00010898 | protein_coding_gene | 2.1 | 4.29 | 3.3E-04 |
| *try-9* | WBGene00023425 | protein_coding_gene | 2.1 | 4.29 | 1.5E-03 |
| *scl-5* | WBGene00008027 | protein_coding_gene | 2.1 | 4.29 | 3.9E-07 |
| *T22F3.8* | WBGene00020698 | protein_coding_gene | 2.1 | 4.28 | 2.6E-04 |
| *cyp-33C6* | WBGene00018264 | protein_coding_gene | 2.1 | 4.28 | 1.3E-02 |
| *pqm-1* | WBGene00004096 | protein_coding_gene | 2.1 | 4.28 | 4.9E-10 |
| *C03A7.2* | WBGene00015366 | protein_coding_gene | 2.1 | 4.26 | 2.7E-02 |
| *F59B2.8* | WBGene00010310 | protein_coding_gene | 2.1 | 4.26 | 2.5E-04 |
| *cbn-1* | WBGene00019285 | protein_coding_gene | 2.1 | 4.26 | 4.2E-04 |
| *cng-3* | WBGene00000563 | protein_coding_gene | 2.1 | 4.25 | 4.6E-02 |
| *gei-13* | WBGene00001570 | protein_coding_gene | 2.1 | 4.24 | 4.6E-05 |
| *F57B1.6* | WBGene00010189 | protein_coding_gene | 2.1 | 4.23 | 1.7E-03 |
| *F09F9.4* | WBGene00017309 | protein_coding_gene | 2.1 | 4.23 | 1.9E-04 |
| *fbxa-44* | WBGene00008616 | protein_coding_gene | 2.1 | 4.23 | 2.0E-02 |
| *fbxa-138* | WBGene00022345 | protein_coding_gene | 2.1 | 4.22 | 1.7E-04 |
| *C17B7.4* | WBGene00015878 | protein_coding_gene | 2.1 | 4.22 | 5.5E-03 |
| *mks-3* | WBGene00018042 | protein_coding_gene | 2.1 | 4.21 | 2.3E-03 |
| *noah-1* | WBGene00016422 | protein_coding_gene | 2.1 | 4.21 | 1.9E-07 |
| *zig-10* | WBGene00020800 | protein_coding_gene | 2.1 | 4.21 | 4.3E-04 |
| *F30A10.2* | WBGene00009261 | protein_coding_gene | 2.1 | 4.20 | 6.7E-03 |
| *fmo-1* | WBGene00001476 | protein_coding_gene | 2.1 | 4.19 | 6.3E-03 |
| *F23H12.5* | WBGene00009094 | protein_coding_gene | 2.1 | 4.19 | 6.1E-06 |
| *F53C3.3* | WBGene00018747 | protein_coding_gene | 2.1 | 4.19 | 7.7E-03 |
| *F09C6.11* | WBGene00044181 | protein_coding_gene | 2.1 | 4.19 | 5.8E-04 |
| *Y53C12B.7* | WBGene00013146 | protein_coding_gene | 2.1 | 4.17 | 1.0E-03 |
| *nlp-76* | WBGene00045386 | protein_coding_gene | 2.1 | 4.15 | 7.6E-03 |
| *C14A4.9* | WBGene00007560 | protein_coding_gene | 2.1 | 4.15 | 2.5E-03 |
| *cut-6* | WBGene00000853 | protein_coding_gene | 2.1 | 4.15 | 2.0E-03 |
| *aqp-2* | WBGene00000170 | protein_coding_gene | 2.1 | 4.14 | 2.9E-08 |
| *suex-2* | WBGene00008271 | protein_coding_gene | 2.0 | 4.14 | 1.6E-02 |
| *F13E9.12* | WBGene00008761 | protein_coding_gene | 2.0 | 4.13 | 2.1E-02 |
| *pals-31* | WBGene00018614 | protein_coding_gene | 2.0 | 4.11 | 3.9E-04 |
| *ador-1* | WBGene00011878 | protein_coding_gene | 2.0 | 4.11 | 2.7E-02 |
| *Y69H2.10* | WBGene00013485 | protein_coding_gene | 2.0 | 4.11 | 6.0E-03 |
| *hlh-16* | WBGene00001960 | protein_coding_gene | 2.0 | 4.08 | 4.1E-03 |
| *F20A1.4* | WBGene00017616 | protein_coding_gene | 2.0 | 4.08 | 2.0E-02 |
| *gcy-22* | WBGene00001547 | protein_coding_gene | 2.0 | 4.08 | 9.6E-05 |
| *T13F3.7* | WBGene00011754 | protein_coding_gene | 2.0 | 4.07 | 1.9E-02 |
| *C45B11.7* | WBGene00050945 | pseudogene | 2.0 | 4.05 | 3.8E-03 |

| | | | | | |
|---|---|---|---|---|---|
| *C09F9.2* | WBGene00007479 | protein_coding_gene | 2.0 | 4.05 | 8.3E-06 |
| *ZK930.2* | WBGene00014152 | protein_coding_gene | 2.0 | 4.04 | 7.6E-13 |
| *F13H10.5* | WBGene00008776 | protein_coding_gene | 2.0 | 4.04 | 4.4E-04 |
| *ech-9* | WBGene00001158 | protein_coding_gene | 2.0 | 4.02 | 5.8E-03 |
| *F21A3.11* | WBGene00219316 | protein_coding_gene | 2.0 | 4.01 | 1.7E-08 |
| *gpa-8* | WBGene00001670 | protein_coding_gene | 2.0 | 4.01 | 2.3E-02 |
| *hpo-2* | WBGene00012500 | protein_coding_gene | 2.0 | 4.01 | 4.3E-02 |
| *igcm-2* | WBGene00020130 | protein_coding_gene | 2.0 | 4.00 | 2.5E-04 |
| *Y64H9A.2* | WBGene00022024 | protein_coding_gene | 2.0 | 4.00 | 2.6E-02 |
| *T24E12.5* | WBGene00020774 | protein_coding_gene | 2.0 | 4.00 | 4.4E-02 |
| *ceh-57* | WBGene00007416 | protein_coding_gene | 2.0 | 3.99 | 1.7E-02 |
| *T05H10.3* | WBGene00011508 | protein_coding_gene | 2.0 | 3.99 | 4.0E-03 |
| *T19C4.1* | WBGene00011838 | protein_coding_gene | 2.0 | 3.98 | 4.7E-04 |
| *nhr-117* | WBGene00003707 | protein_coding_gene | 2.0 | 3.97 | 2.8E-06 |
| *F53C3.6* | WBGene00018750 | protein_coding_gene | 2.0 | 3.97 | 1.2E-02 |
| *ZC168.2* | WBGene00013856 | protein_coding_gene | 2.0 | 3.96 | 5.9E-03 |
| *C50F4.9* | WBGene00008234 | protein_coding_gene | 2.0 | 3.96 | 1.3E-06 |
| *C43H6.6* | WBGene00016619 | protein_coding_gene | 2.0 | 3.95 | 3.3E-02 |
| *wrt-10* | WBGene00006956 | protein_coding_gene | 2.0 | 3.95 | 1.7E-07 |
| *R08A2.7* | WBGene00011137 | protein_coding_gene | 2.0 | 3.95 | 4.0E-03 |
| *K08F9.1* | WBGene00010684 | protein_coding_gene | 2.0 | 3.93 | 3.4E-04 |
| *C04G6.5* | WBGene00015455 | protein_coding_gene | 2.0 | 3.91 | 6.6E-03 |
| *C01G6.9* | WBGene00007229 | protein_coding_gene | 2.0 | 3.91 | 1.5E-04 |
| *fax-1* | WBGene00001400 | protein_coding_gene | 2.0 | 3.91 | 1.9E-02 |
| *ceh-22* | WBGene00000445 | protein_coding_gene | 2.0 | 3.90 | 3.7E-05 |
| *math-14* | WBGene00015828 | protein_coding_gene | 2.0 | 3.90 | 7.7E-15 |
| *M01A8.1* | WBGene00010795 | protein_coding_gene | 2.0 | 3.89 | 9.0E-04 |
| *F13H8.1* | WBGene00017434 | protein_coding_gene | 2.0 | 3.89 | 3.6E-05 |
| *prkl-1* | WBGene00022727 | protein_coding_gene | 2.0 | 3.87 | 8.5E-07 |
| *wrt-1* | WBGene00006947 | protein_coding_gene | 2.0 | 3.87 | 8.7E-04 |
| *ttm-2* | WBGene00017840 | protein_coding_gene | 2.0 | 3.87 | 2.7E-07 |
| *unc-3* | WBGene00006743 | protein_coding_gene | 1.9 | 3.85 | 2.3E-04 |
| *ceh-45* | WBGene00022837 | protein_coding_gene | 1.9 | 3.84 | 4.3E-03 |
| *F37A4.3* | WBGene00018133 | protein_coding_gene | 1.9 | 3.84 | 3.1E-03 |
| *F02A9.1* | WBGene00008513 | protein_coding_gene | 1.9 | 3.83 | 2.3E-04 |
| *F41E6.5* | WBGene00018286 | protein_coding_gene | 1.9 | 3.83 | 5.1E-06 |
| *F43C11.7* | WBGene00018384 | protein_coding_gene | 1.9 | 3.82 | 4.1E-05 |
| *F41C3.1* | WBGene00018267 | protein_coding_gene | 1.9 | 3.82 | 2.0E-02 |
| *mlt-9* | WBGene00008605 | protein_coding_gene | 1.9 | 3.81 | 1.8E-04 |
| *C04A11.5* | WBGene00007288 | protein_coding_gene | 1.9 | 3.80 | 8.0E-03 |
| *mam-5* | WBGene00011572 | protein_coding_gene | 1.9 | 3.80 | 4.7E-02 |
| *C18D4.8* | WBGene00007677 | pseudogene | 1.9 | 3.80 | 2.3E-05 |
| *vap-2* | WBGene00011462 | protein_coding_gene | 1.9 | 3.80 | 7.6E-04 |
| *ins-23* | WBGene00002106 | protein_coding_gene | 1.9 | 3.79 | 1.0E-02 |
| *Y43F8B.23* | WBGene00194815 | protein_coding_gene | 1.9 | 3.79 | 6.6E-03 |
| *nhr-116* | WBGene00003706 | protein_coding_gene | 1.9 | 3.79 | 5.7E-07 |
| *cki-1* | WBGene00000516 | protein_coding_gene | 1.9 | 3.79 | 4.0E-10 |
| *cutl-21* | WBGene00018861 | protein_coding_gene | 1.9 | 3.77 | 4.3E-04 |
| *R04B3.1* | WBGene00019866 | protein_coding_gene | 1.9 | 3.77 | 4.6E-02 |
| *F28G4.4* | WBGene00009229 | protein_coding_gene | 1.9 | 3.76 | 4.9E-02 |
| *W04A8.4* | WBGene00012239 | protein_coding_gene | 1.9 | 3.75 | 1.5E-07 |
| *F58E6.11* | WBGene00010258 | protein_coding_gene | 1.9 | 3.75 | 6.6E-03 |
| *ceh-8* | WBGene00000433 | protein_coding_gene | 1.9 | 3.74 | 2.3E-02 |
| *Y46E12BR.1* | WBGene00021598 | protein_coding_gene | 1.9 | 3.74 | 4.2E-02 |
| *nhr-19* | WBGene00003618 | protein_coding_gene | 1.9 | 3.74 | 2.3E-05 |
| *F53A9.7* | WBGene00018730 | protein_coding_gene | 1.9 | 3.74 | 1.3E-05 |
| *F18C5.5* | WBGene00017560 | protein_coding_gene | 1.9 | 3.73 | 5.6E-03 |
| *oac-51* | WBGene00020976 | protein_coding_gene | 1.9 | 3.72 | 9.0E-04 |

| | | | | | |
|---|---|---|---|---|---|
| *sups-1* | WBGene00017716 | protein_coding_gene | 1.9 | 3.71 | 1.8E-04 |
| *M03D4.3* | WBGene00019750 | protein_coding_gene | 1.9 | 3.71 | 5.5E-04 |
| *zig-4* | WBGene00006981 | protein_coding_gene | 1.9 | 3.71 | 5.9E-05 |
| *ast-1* | WBGene00020368 | protein_coding_gene | 1.9 | 3.70 | 1.9E-03 |
| *C30F2.4* | WBGene00007818 | protein_coding_gene | 1.9 | 3.70 | 1.8E-03 |
| *slc-28.2* | WBGene00017868 | protein_coding_gene | 1.9 | 3.69 | 1.1E-04 |
| *K08B5.1* | WBGene00019518 | protein_coding_gene | 1.9 | 3.69 | 7.8E-05 |
| *W09G12.7* | WBGene00021121 | protein_coding_gene | 1.9 | 3.69 | 1.4E-03 |
| *M03D4.4* | WBGene00019751 | protein_coding_gene | 1.9 | 3.68 | 1.5E-05 |
| *B0250.12* | WBGene00050927 | pseudogene | 1.9 | 3.68 | 3.4E-05 |
| *zag-1* | WBGene00006970 | protein_coding_gene | 1.9 | 3.68 | 1.3E-04 |
| *F42G4.6* | WBGene00009643 | protein_coding_gene | 1.9 | 3.67 | 6.0E-03 |
| *math-45* | WBGene00021772 | protein_coding_gene | 1.9 | 3.67 | 2.2E-06 |
| *F22F7.3* | WBGene00017721 | protein_coding_gene | 1.9 | 3.66 | 7.9E-03 |
| *Y38H8A.1* | WBGene00012635 | protein_coding_gene | 1.9 | 3.65 | 2.2E-02 |
| *cutl-16* | WBGene00019426 | protein_coding_gene | 1.9 | 3.65 | 4.6E-04 |
| *F54B11.10* | WBGene00010033 | protein_coding_gene | 1.9 | 3.64 | 2.3E-02 |
| *cdh-9* | WBGene00000401 | protein_coding_gene | 1.9 | 3.63 | 2.1E-05 |
| *ZK757.2* | WBGene00014074 | protein_coding_gene | 1.9 | 3.63 | 5.7E-05 |
| *T10G3.1* | WBGene00011692 | protein_coding_gene | 1.9 | 3.62 | 7.0E-05 |
| *bus-12* | WBGene00010442 | protein_coding_gene | 1.9 | 3.62 | 2.4E-02 |
| *F14D7.4* | WBGene00008791 | pseudogene | 1.9 | 3.62 | 4.6E-02 |
| *F59E11.5* | WBGene00019111 | protein_coding_gene | 1.9 | 3.62 | 2.4E-10 |
| *ZC84.1* | WBGene00013846 | protein_coding_gene | 1.9 | 3.61 | 3.4E-03 |
| *F33D4.6* | WBGene00017998 | protein_coding_gene | 1.9 | 3.61 | 4.5E-04 |
| *del-3* | WBGene00009144 | protein_coding_gene | 1.9 | 3.61 | 4.4E-02 |
| *eva-1* | WBGene00009304 | protein_coding_gene | 1.8 | 3.60 | 3.6E-06 |
| *ztf-30* | WBGene00015523 | protein_coding_gene | 1.8 | 3.60 | 3.6E-03 |
| *F16G10.15* | WBGene00017529 | pseudogene | 1.8 | 3.60 | 1.8E-02 |
| *R160.10* | WBGene00219369 | protein_coding_gene | 1.8 | 3.59 | 1.9E-02 |
| *Y43F8C.18* | WBGene00012839 | ncRNA_gene | 1.8 | 3.59 | 4.1E-02 |
| *K07H8.11* | WBGene00044510 | protein_coding_gene | 1.8 | 3.58 | 4.1E-03 |
| *H17B01.5* | WBGene00044643 | protein_coding_gene | 1.8 | 3.58 | 3.5E-03 |
| *nhr-206* | WBGene00011097 | protein_coding_gene | 1.8 | 3.57 | 2.2E-04 |
| *C40H1.2* | WBGene00008038 | protein_coding_gene | 1.8 | 3.57 | 2.2E-04 |
| *C54E4.12* | WBGene00235274 | protein_coding_gene | 1.8 | 3.56 | 3.9E-02 |
| *asic-2* | WBGene00012137 | protein_coding_gene | 1.8 | 3.56 | 6.5E-03 |
| *Y71H2AM.14* | WBGene00022179 | protein_coding_gene | 1.8 | 3.56 | 3.3E-04 |
| *F53F1.6* | WBGene00009984 | protein_coding_gene | 1.8 | 3.55 | 1.1E-03 |
| *C25A6.1* | WBGene00016081 | protein_coding_gene | 1.8 | 3.55 | 1.9E-02 |
| *mks-6* | WBGene00010642 | protein_coding_gene | 1.8 | 3.55 | 3.0E-04 |
| *let-4* | WBGene00002282 | protein_coding_gene | 1.8 | 3.55 | 1.1E-04 |
| *plx-2* | WBGene00004048 | protein_coding_gene | 1.8 | 3.54 | 8.6E-06 |
| *Y20C6A.1* | WBGene00012491 | protein_coding_gene | 1.8 | 3.54 | 2.0E-06 |
| *Y23H5B.8* | WBGene00021279 | protein_coding_gene | 1.8 | 3.54 | 2.4E-04 |
| *pals-26* | WBGene00007131 | protein_coding_gene | 1.8 | 3.54 | 4.0E-03 |
| *R11F4.1* | WBGene00020007 | protein_coding_gene | 1.8 | 3.53 | 1.0E-09 |
| *Y73E7A.8* | WBGene00022274 | protein_coding_gene | 1.8 | 3.53 | 3.1E-03 |
| *M01D1.12* | WBGene00269373 | protein_coding_gene | 1.8 | 3.53 | 8.9E-03 |
| *ptr-19* | WBGene00004233 | protein_coding_gene | 1.8 | 3.53 | 1.8E-03 |
| *nlp-71* | WBGene00022254 | protein_coding_gene | 1.8 | 3.53 | 1.7E-07 |
| *sym-1* | WBGene00006366 | protein_coding_gene | 1.8 | 3.52 | 2.4E-06 |
| *nspc-19* | WBGene00008360 | protein_coding_gene | 1.8 | 3.52 | 6.4E-11 |
| *C35A5.6* | WBGene00007950 | protein_coding_gene | 1.8 | 3.52 | 2.8E-07 |
| *C09B9.1* | WBGene00015626 | protein_coding_gene | 1.8 | 3.52 | 2.4E-02 |
| *C14C6.2* | WBGene00015756 | protein_coding_gene | 1.8 | 3.52 | 3.4E-16 |
| *tsp-6* | WBGene00006632 | protein_coding_gene | 1.8 | 3.51 | 1.1E-03 |
| *pho-6* | WBGene00018696 | protein_coding_gene | 1.8 | 3.50 | 4.0E-05 |

| | | | | | |
|---|---|---|---|---|---|
| *che-11* | WBGene00000490 | protein_coding_gene | 1.8 | 3.50 | 1.1E-04 |
| *C07B5.4* | WBGene00007409 | protein_coding_gene | 1.8 | 3.49 | 9.1E-06 |
| *gst-29* | WBGene00001777 | protein_coding_gene | 1.8 | 3.49 | 7.1E-03 |
| *acr-19* | WBGene00000058 | protein_coding_gene | 1.8 | 3.49 | 2.2E-03 |
| *fkh-10* | WBGene00001442 | protein_coding_gene | 1.8 | 3.48 | 4.8E-02 |
| *osr-1* | WBGene00016329 | protein_coding_gene | 1.8 | 3.48 | 9.1E-03 |
| *F49F1.5* | WBGene00018645 | protein_coding_gene | 1.8 | 3.48 | 1.3E-04 |
| *Y110A2AL.3* | WBGene00022440 | protein_coding_gene | 1.8 | 3.47 | 1.1E-04 |
| *ceh-16* | WBGene00000439 | protein_coding_gene | 1.8 | 3.46 | 3.6E-03 |
| *C26C6.4* | WBGene00007739 | protein_coding_gene | 1.8 | 3.44 | 2.1E-02 |
| *E03A3.10* | WBGene00189944 | pseudogene | 1.8 | 3.44 | 1.7E-02 |
| *F10E9.12* | WBGene00017361 | protein_coding_gene | 1.8 | 3.44 | 9.5E-08 |
| *qua-1* | WBGene00004264 | protein_coding_gene | 1.8 | 3.43 | 3.1E-05 |
| *ugt-52* | WBGene00018931 | protein_coding_gene | 1.8 | 3.43 | 2.3E-03 |
| *F59F5.3* | WBGene00010342 | protein_coding_gene | 1.8 | 3.43 | 7.7E-05 |
| *T04F8.7* | WBGene00011445 | protein_coding_gene | 1.8 | 3.42 | 1.0E-05 |
| *M01H9.2* | WBGene00019718 | protein_coding_gene | 1.8 | 3.42 | 6.5E-03 |
| *C45B11.6* | WBGene00045379 | protein_coding_gene | 1.8 | 3.42 | 9.1E-05 |
| *cdh-5* | WBGene00000397 | protein_coding_gene | 1.8 | 3.41 | 1.6E-04 |
| *ZC443.4* | WBGene00013899 | protein_coding_gene | 1.8 | 3.41 | 5.2E-06 |
| *F53C3.4* | WBGene00018748 | protein_coding_gene | 1.8 | 3.41 | 9.0E-04 |
| *ceh-2* | WBGene00000429 | protein_coding_gene | 1.8 | 3.40 | 3.6E-02 |
| *F46F3.3* | WBGene00009791 | protein_coding_gene | 1.8 | 3.40 | 3.3E-02 |
| *dhs-16* | WBGene00000979 | protein_coding_gene | 1.8 | 3.40 | 3.1E-02 |
| *C45E5.3* | WBGene00016666 | pseudogene | 1.8 | 3.39 | 4.2E-03 |
| *F08B12.4* | WBGene00008572 | protein_coding_gene | 1.8 | 3.39 | 7.8E-07 |
| *trh-1* | WBGene00007827 | protein_coding_gene | 1.8 | 3.39 | 2.7E-06 |
| *F26A1.19* | WBGene00270319 | protein_coding_gene | 1.8 | 3.39 | 2.4E-04 |
| *W01C9.1* | WBGene00012175 | protein_coding_gene | 1.8 | 3.39 | 2.7E-02 |
| *gopc-1* | WBGene00014101 | protein_coding_gene | 1.8 | 3.39 | 2.9E-04 |
| *srd-32* | WBGene00005110 | protein_coding_gene | 1.8 | 3.37 | 2.8E-02 |
| *kcnl-3* | WBGene00015387 | protein_coding_gene | 1.8 | 3.37 | 3.5E-03 |
| *F09F9.2* | WBGene00017307 | protein_coding_gene | 1.7 | 3.36 | 7.1E-03 |
| *F38E9.1* | WBGene00018184 | protein_coding_gene | 1.7 | 3.36 | 6.8E-09 |
| *ZK971.1* | WBGene00014175 | protein_coding_gene | 1.7 | 3.36 | 2.3E-04 |
| *T01D3.6* | WBGene00011330 | protein_coding_gene | 1.7 | 3.35 | 6.1E-07 |
| *ZK822.1* | WBGene00014089 | protein_coding_gene | 1.7 | 3.34 | 1.2E-03 |
| *Y11D7A.5* | WBGene00012429 | protein_coding_gene | 1.7 | 3.34 | 2.1E-02 |
| *Y77E11A.14* | WBGene00022314 | protein_coding_gene | 1.7 | 3.34 | 1.7E-03 |
| *cutl-17* | WBGene00009541 | protein_coding_gene | 1.7 | 3.34 | 2.9E-02 |
| *T19H5.6* | WBGene00044807 | protein_coding_gene | 1.7 | 3.34 | 4.4E-02 |
| *T16G1.7* | WBGene00011801 | protein_coding_gene | 1.7 | 3.32 | 3.1E-02 |
| *C39B10.7* | WBGene00164973 | protein_coding_gene | 1.7 | 3.32 | 1.0E-02 |
| *acn-1* | WBGene00000039 | protein_coding_gene | 1.7 | 3.31 | 9.7E-05 |
| *F18E9.1* | WBGene00017568 | protein_coding_gene | 1.7 | 3.31 | 1.3E-02 |
| *Y43C5A.3* | WBGene00012783 | protein_coding_gene | 1.7 | 3.31 | 2.5E-07 |
| *Y44E3A.1* | WBGene00021547 | protein_coding_gene | 1.7 | 3.30 | 4.9E-08 |
| *ZK593.9* | WBGene00014005 | pseudogene | 1.7 | 3.30 | 6.4E-03 |
| *unk-1* | WBGene00016407 | protein_coding_gene | 1.7 | 3.30 | 2.2E-03 |
| *T05E12.3* | WBGene00011486 | protein_coding_gene | 1.7 | 3.29 | 2.2E-07 |
| *Y82E9BL.3* | WBGene00022320 | protein_coding_gene | 1.7 | 3.29 | 4.6E-02 |
| *D1014.5* | WBGene00017018 | protein_coding_gene | 1.7 | 3.29 | 7.8E-04 |
| *tts-2* | WBGene00006651 | ncRNA_gene | 1.7 | 3.29 | 2.6E-06 |
| *F18G5.6* | WBGene00017582 | protein_coding_gene | 1.7 | 3.29 | 2.7E-04 |
| *unc-86* | WBGene00006818 | protein_coding_gene | 1.7 | 3.28 | 4.8E-03 |
| *lite-1* | WBGene00001803 | protein_coding_gene | 1.7 | 3.28 | 9.9E-06 |
| *che-2* | WBGene00000484 | protein_coding_gene | 1.7 | 3.28 | 1.2E-02 |
| *nsy-4* | WBGene00021415 | protein_coding_gene | 1.7 | 3.27 | 1.5E-05 |

| | | | | | |
|---|---|---|---|---|---|
| *clec-233* | WBGene00009030 | protein_coding_gene | 1.7 | 3.27 | 4.8E-03 |
| *pqn-13* | WBGene00004104 | protein_coding_gene | 1.7 | 3.26 | 1.4E-03 |
| *ttx-1* | WBGene00006652 | protein_coding_gene | 1.7 | 3.25 | 1.5E-04 |
| *nac-2* | WBGene00003518 | protein_coding_gene | 1.7 | 3.25 | 5.1E-16 |
| *R10E8.6* | WBGene00011212 | protein_coding_gene | 1.7 | 3.25 | 9.0E-07 |
| *K02E7.5* | WBGene00019310 | pseudogene | 1.7 | 3.24 | 9.5E-03 |
| *ppm-1.H* | WBGene00016580 | protein_coding_gene | 1.7 | 3.24 | 1.5E-02 |
| *bus-18* | WBGene00044631 | protein_coding_gene | 1.7 | 3.24 | 2.2E-03 |
| *F22B7.1* | WBGene00017692 | protein_coding_gene | 1.7 | 3.24 | 1.3E-05 |
| *flp-27* | WBGene00016114 | protein_coding_gene | 1.7 | 3.24 | 1.6E-05 |
| *ZK250.13* | WBGene00235352 | protein_coding_gene | 1.7 | 3.24 | 1.2E-05 |
| *F20G2.5* | WBGene00008988 | protein_coding_gene | 1.7 | 3.23 | 6.4E-03 |
| *spp-1* | WBGene00004986 | protein_coding_gene | 1.7 | 3.23 | 2.5E-06 |
| *subs-4* | WBGene00012939 | protein_coding_gene | 1.7 | 3.23 | 3.9E-03 |
| *K08E7.6* | WBGene00010674 | protein_coding_gene | 1.7 | 3.22 | 4.3E-03 |
| *nlp-45* | WBGene00020136 | protein_coding_gene | 1.7 | 3.21 | 3.7E-02 |
| *F40F9.10* | WBGene00009586 | protein_coding_gene | 1.7 | 3.21 | 2.3E-03 |
| *C28G1.2* | WBGene00016187 | protein_coding_gene | 1.7 | 3.21 | 2.4E-02 |
| *C03A7.13* | WBGene00015371 | protein_coding_gene | 1.7 | 3.20 | 4.9E-04 |
| *ZK863.8* | WBGene00014125 | protein_coding_gene | 1.7 | 3.20 | 2.4E-06 |
| *T28A11.19* | WBGene00020881 | protein_coding_gene | 1.7 | 3.20 | 1.6E-02 |
| *K09C4.1* | WBGene00019547 | protein_coding_gene | 1.7 | 3.19 | 1.7E-03 |
| *F02E11.7* | WBGene00195073 | protein_coding_gene | 1.7 | 3.19 | 4.4E-03 |
| *F47H4.12* | WBGene00043053 | pseudogene | 1.7 | 3.19 | 9.1E-03 |
| *tsp-1* | WBGene00006627 | protein_coding_gene | 1.7 | 3.18 | 3.1E-03 |
| *K01A2.10* | WBGene00019284 | protein_coding_gene | 1.7 | 3.18 | 1.3E-03 |
| *rig-5* | WBGene00004372 | protein_coding_gene | 1.7 | 3.18 | 1.5E-03 |
| *F09C6.3* | WBGene00008617 | protein_coding_gene | 1.7 | 3.17 | 4.4E-02 |
| *lgc-27* | WBGene00022283 | protein_coding_gene | 1.7 | 3.17 | 9.0E-05 |
| *T07D3.6* | WBGene00020309 | pseudogene | 1.7 | 3.16 | 3.2E-02 |
| *btb-21* | WBGene00018540 | pseudogene | 1.7 | 3.16 | 2.6E-04 |
| *D1086.2* | WBGene00008389 | protein_coding_gene | 1.7 | 3.16 | 2.0E-06 |
| *dma-1* | WBGene00011345 | protein_coding_gene | 1.7 | 3.16 | 1.4E-03 |
| *syd-9* | WBGene00044068 | protein_coding_gene | 1.7 | 3.15 | 5.3E-05 |
| *cyp-23A1* | WBGene00015135 | protein_coding_gene | 1.7 | 3.15 | 2.6E-02 |
| *C33A12.4* | WBGene00007883 | protein_coding_gene | 1.7 | 3.15 | 1.2E-05 |
| *tps-1* | WBGene00006602 | protein_coding_gene | 1.7 | 3.15 | 8.7E-09 |
| *T02C12.5* | WBGene00194692 | protein_coding_gene | 1.7 | 3.15 | 7.4E-04 |
| *ubc-22* | WBGene00006717 | protein_coding_gene | 1.7 | 3.14 | 3.7E-02 |
| *Y46H3C.6* | WBGene00021606 | pseudogene | 1.7 | 3.14 | 2.1E-02 |
| *T14B1.1* | WBGene00011763 | protein_coding_gene | 1.6 | 3.14 | 1.5E-04 |
| *F02A9.7* | WBGene00044406 | protein_coding_gene | 1.6 | 3.14 | 2.8E-02 |
| *ZC449.2* | WBGene00022612 | protein_coding_gene | 1.6 | 3.12 | 7.4E-03 |
| *nspe-4* | WBGene00012589 | protein_coding_gene | 1.6 | 3.12 | 7.3E-03 |
| *C11G10.1* | WBGene00007525 | protein_coding_gene | 1.6 | 3.12 | 5.2E-04 |
| *btb-17* | WBGene00018200 | protein_coding_gene | 1.6 | 3.12 | 1.0E-06 |
| *nlp-38* | WBGene00007219 | protein_coding_gene | 1.6 | 3.11 | 1.4E-07 |
| *pef-1* | WBGene00003969 | protein_coding_gene | 1.6 | 3.11 | 1.3E-03 |
| *Y43F8C.3* | WBGene00012825 | protein_coding_gene | 1.6 | 3.10 | 7.4E-05 |
| *lron-1* | WBGene00008093 | protein_coding_gene | 1.6 | 3.10 | 3.1E-02 |
| *C32F10.4* | WBGene00016332 | protein_coding_gene | 1.6 | 3.10 | 1.1E-06 |
| *daf-10* | WBGene00000906 | protein_coding_gene | 1.6 | 3.10 | 5.5E-04 |
| *C50B8.4* | WBGene00008226 | protein_coding_gene | 1.6 | 3.09 | 2.3E-04 |
| *ceh-34* | WBGene00000455 | protein_coding_gene | 1.6 | 3.09 | 1.4E-04 |
| *pdf-2* | WBGene00018590 | protein_coding_gene | 1.6 | 3.09 | 1.6E-05 |
| *F38B2.2* | WBGene00009529 | protein_coding_gene | 1.6 | 3.09 | 1.6E-03 |
| *F14B4.1* | WBGene00008779 | protein_coding_gene | 1.6 | 3.09 | 2.3E-05 |
| *C50H2.13* | WBGene00008246 | protein_coding_gene | 1.6 | 3.08 | 6.4E-07 |

| | | | | | |
|---|---|---|---|---|---|
| *flp-15* | WBGene00001458 | protein_coding_gene | 1.6 | 3.08 | 3.9E-09 |
| *icmt-1* | WBGene00017673 | protein_coding_gene | 1.6 | 3.08 | 7.5E-03 |
| *cutl-29* | WBGene00017421 | protein_coding_gene | 1.6 | 3.07 | 2.2E-02 |
| *C13E3.1* | WBGene00015740 | protein_coding_gene | 1.6 | 3.07 | 5.5E-07 |
| *nspc-18* | WBGene00008358 | protein_coding_gene | 1.6 | 3.07 | 8.9E-06 |
| *F53F4.1* | WBGene00009985 | protein_coding_gene | 1.6 | 3.06 | 4.9E-02 |
| *F22F4.9* | WBGene00271606 | protein_coding_gene | 1.6 | 3.06 | 7.6E-04 |
| *F08C6.5* | WBGene00017243 | protein_coding_gene | 1.6 | 3.06 | 9.7E-05 |
| *C28C12.3* | WBGene00016179 | protein_coding_gene | 1.6 | 3.05 | 5.0E-02 |
| *B0393.5* | WBGene00007170 | protein_coding_gene | 1.6 | 3.05 | 2.1E-04 |
| *C33D9.5* | WBGene00007899 | pseudogene | 1.6 | 3.05 | 2.0E-06 |
| *Y51H7C.12* | WBGene00021790 | protein_coding_gene | 1.6 | 3.04 | 3.6E-02 |
| *hhat-2* | WBGene00013312 | protein_coding_gene | 1.6 | 3.04 | 2.0E-03 |
| *pals-19* | WBGene00044761 | protein_coding_gene | 1.6 | 3.04 | 2.8E-04 |
| *C25F9.9* | WBGene00007728 | protein_coding_gene | 1.6 | 3.04 | 4.1E-04 |
| *C26E1.2* | WBGene00007748 | protein_coding_gene | 1.6 | 3.04 | 3.4E-03 |
| *Y75B12B.3* | WBGene00013568 | protein_coding_gene | 1.6 | 3.04 | 2.2E-05 |
| *glb-15* | WBGene00009387 | protein_coding_gene | 1.6 | 3.04 | 2.1E-03 |
| *clec-147* | WBGene00007313 | protein_coding_gene | 1.6 | 3.04 | 2.9E-11 |
| *nlp-9* | WBGene00003747 | protein_coding_gene | 1.6 | 3.03 | 3.6E-05 |
| *nab-1* | WBGene00003516 | protein_coding_gene | 1.6 | 3.03 | 7.2E-05 |
| *F10E9.1* | WBGene00017353 | protein_coding_gene | 1.6 | 3.03 | 2.3E-02 |
| *F42G2.5* | WBGene00018354 | protein_coding_gene | 1.6 | 3.02 | 4.8E-05 |
| *T12A7.6* | WBGene00011727 | protein_coding_gene | 1.6 | 3.02 | 3.8E-03 |
| *spp-18* | WBGene00005003 | protein_coding_gene | 1.6 | 3.02 | 3.8E-08 |
| *tbb-4* | WBGene00006538 | protein_coding_gene | 1.6 | 3.02 | 6.4E-04 |
| *pros-1* | WBGene00000448 | protein_coding_gene | 1.6 | 3.01 | 6.2E-04 |
| *nhr-173* | WBGene00016975 | protein_coding_gene | 1.6 | 3.01 | 5.7E-05 |
| *ins-17* | WBGene00002100 | protein_coding_gene | 1.6 | 3.01 | 6.8E-09 |
| *bath-47* | WBGene00006464 | protein_coding_gene | 1.6 | 3.01 | 9.0E-04 |
| *T09F5.1* | WBGene00011665 | protein_coding_gene | 1.6 | 3.00 | 2.1E-02 |
| *sul-2* | WBGene00006309 | protein_coding_gene | 1.6 | 3.00 | 6.5E-04 |
| *F26F12.4* | WBGene00017835 | protein_coding_gene | 1.6 | 3.00 | 5.5E-03 |
| *F53F4.8* | WBGene00009991 | pseudogene | 1.6 | 2.99 | 3.1E-02 |
| *T24C2.5* | WBGene00011984 | protein_coding_gene | 1.6 | 2.99 | 1.1E-05 |
| *Y18D10A.22* | WBGene00014864 | protein_coding_gene | 1.6 | 2.99 | 2.3E-04 |
| *nmr-2* | WBGene00003775 | protein_coding_gene | 1.6 | 2.98 | 2.4E-02 |
| *fbn-1* | WBGene00022816 | protein_coding_gene | 1.6 | 2.98 | 1.2E-03 |
| *C03B1.5* | WBGene00015376 | protein_coding_gene | 1.6 | 2.98 | 2.8E-02 |
| *F58H1.7* | WBGene00010290 | protein_coding_gene | 1.6 | 2.98 | 8.7E-08 |
| *Y97E10AR.1* | WBGene00022396 | protein_coding_gene | 1.6 | 2.97 | 3.3E-04 |
| *C23H5.15* | WBGene00219423 | protein_coding_gene | 1.6 | 2.97 | 2.5E-07 |
| *K08D9.4* | WBGene00019525 | protein_coding_gene | 1.6 | 2.97 | 6.1E-03 |
| *cutl-25* | WBGene00021919 | protein_coding_gene | 1.6 | 2.97 | 2.2E-03 |
| *phat-3* | WBGene00016782 | protein_coding_gene | 1.6 | 2.97 | 1.9E-07 |
| *gstk-2* | WBGene00017054 | protein_coding_gene | 1.6 | 2.96 | 1.2E-05 |
| *F26A10.2* | WBGene00017814 | protein_coding_gene | 1.6 | 2.96 | 2.5E-03 |
| *ZK470.14* | WBGene00219362 | protein_coding_gene | 1.6 | 2.96 | 3.5E-02 |
| *spi-1* | WBGene00023416 | protein_coding_gene | 1.6 | 2.96 | 1.4E-02 |
| *C30E1.9* | WBGene00022951 | ncRNA_gene | 1.6 | 2.96 | 7.8E-05 |
| *magu-3* | WBGene00016841 | protein_coding_gene | 1.6 | 2.95 | 8.9E-04 |
| *agr-1* | WBGene00018304 | protein_coding_gene | 1.6 | 2.95 | 5.6E-09 |
| *W10C8.6* | WBGene00044433 | protein_coding_gene | 1.6 | 2.95 | 1.3E-03 |
| *sma-10* | WBGene00020649 | protein_coding_gene | 1.6 | 2.94 | 1.3E-05 |
| *ZK682.7* | WBGene00022790 | protein_coding_gene | 1.6 | 2.94 | 3.3E-02 |
| *nhr-171* | WBGene00016926 | protein_coding_gene | 1.6 | 2.94 | 1.9E-02 |
| *C25G6.1* | WBGene00016107 | protein_coding_gene | 1.6 | 2.93 | 4.9E-02 |
| *C38C3.10* | WBGene00045390 | protein_coding_gene | 1.6 | 2.93 | 7.8E-04 |

| | | | | | |
|---|---|---|---|---|---|
| *rom-2* | WBGene00004401 | protein_coding_gene | 1.6 | 2.93 | 5.9E-03 |
| *C49A9.1* | WBGene00016755 | protein_coding_gene | 1.6 | 2.93 | 6.2E-03 |
| *sra-10* | WBGene00005036 | protein_coding_gene | 1.5 | 2.93 | 2.5E-02 |
| *Y6E2A.4* | WBGene00012398 | protein_coding_gene | 1.5 | 2.92 | 1.3E-02 |
| *let-381* | WBGene00002601 | protein_coding_gene | 1.5 | 2.92 | 6.5E-03 |
| *tsp-16* | WBGene00006642 | protein_coding_gene | 1.5 | 2.92 | 2.7E-04 |
| *klp-11* | WBGene00002222 | protein_coding_gene | 1.5 | 2.92 | 2.2E-04 |
| *fbxb-101* | WBGene00013404 | protein_coding_gene | 1.5 | 2.92 | 3.5E-02 |
| *F09C6.12* | WBGene00044182 | protein_coding_gene | 1.5 | 2.92 | 1.6E-04 |
| *E01G4.7* | WBGene00045410 | pseudogene | 1.5 | 2.92 | 1.2E-09 |
| *egl-43* | WBGene00001207 | protein_coding_gene | 1.5 | 2.91 | 5.1E-05 |
| *pag-3* | WBGene00003909 | protein_coding_gene | 1.5 | 2.91 | 4.0E-02 |
| *egl-38* | WBGene00001204 | protein_coding_gene | 1.5 | 2.91 | 3.0E-03 |
| *nhr-55* | WBGene00003645 | protein_coding_gene | 1.5 | 2.91 | 9.9E-05 |
| *Y43F8A.1* | WBGene00012808 | protein_coding_gene | 1.5 | 2.91 | 1.3E-02 |
| *mod-5* | WBGene00003387 | protein_coding_gene | 1.5 | 2.91 | 4.0E-04 |
| *K09E9.7* | WBGene00194664 | protein_coding_gene | 1.5 | 2.91 | 4.7E-02 |
| *bus-8* | WBGene00044623 | protein_coding_gene | 1.5 | 2.91 | 1.3E-03 |
| *lury-1* | WBGene00013548 | protein_coding_gene | 1.5 | 2.91 | 2.0E-03 |
| *W03D2.6* | WBGene00020981 | pseudogene | 1.5 | 2.90 | 1.7E-08 |
| *ttr-29* | WBGene00011286 | protein_coding_gene | 1.5 | 2.90 | 2.6E-03 |
| *tag-243* | WBGene00044078 | protein_coding_gene | 1.5 | 2.90 | 9.5E-04 |
| *ceh-30* | WBGene00000451 | protein_coding_gene | 1.5 | 2.90 | 2.1E-02 |
| *M04C3.2* | WBGene00010851 | protein_coding_gene | 1.5 | 2.89 | 1.5E-05 |
| *nphp-4* | WBGene00011261 | protein_coding_gene | 1.5 | 2.89 | 3.0E-03 |
| *W05G11.8* | WBGene00271636 | ncRNA_gene | 1.5 | 2.89 | 5.7E-03 |
| *ZC434.3* | WBGene00013891 | protein_coding_gene | 1.5 | 2.88 | 3.1E-02 |
| *egas-4* | WBGene00018906 | protein_coding_gene | 1.5 | 2.88 | 3.0E-03 |
| *R02E12.12* | WBGene00303425 | protein_coding_gene | 1.5 | 2.88 | 4.5E-02 |
| *hsp-12.1* | WBGene00011906 | protein_coding_gene | 1.5 | 2.88 | 2.6E-03 |
| *dyf-7* | WBGene00001123 | protein_coding_gene | 1.5 | 2.88 | 2.3E-03 |
| *F52A8.3* | WBGene00009916 | protein_coding_gene | 1.5 | 2.88 | 2.2E-02 |
| *F44E2.4* | WBGene00018418 | protein_coding_gene | 1.5 | 2.88 | 2.7E-04 |
| *dpy-1* | WBGene00001063 | protein_coding_gene | 1.5 | 2.87 | 2.6E-02 |
| *hlh-6* | WBGene00001952 | protein_coding_gene | 1.5 | 2.87 | 2.1E-02 |
| *Y34F4.1* | WBGene00021336 | protein_coding_gene | 1.5 | 2.87 | 1.3E-05 |
| *F56C4.4* | WBGene00044977 | protein_coding_gene | 1.5 | 2.87 | 4.2E-03 |
| *Y116A8C.25* | WBGene00013802 | protein_coding_gene | 1.5 | 2.87 | 1.0E-03 |
| *K10D6.2* | WBGene00010742 | protein_coding_gene | 1.5 | 2.87 | 8.0E-06 |
| *T14E8.4* | WBGene00043981 | protein_coding_gene | 1.5 | 2.87 | 1.3E-02 |
| *T19D12.2* | WBGene00020577 | protein_coding_gene | 1.5 | 2.86 | 2.5E-12 |
| *F52E1.5* | WBGene00018694 | protein_coding_gene | 1.5 | 2.86 | 4.8E-03 |
| *Y51H4A.8* | WBGene00013104 | protein_coding_gene | 1.5 | 2.86 | 2.5E-07 |
| *C13C12.2* | WBGene00007552 | protein_coding_gene | 1.5 | 2.86 | 7.1E-03 |
| *T09E11.11* | WBGene00011660 | pseudogene | 1.5 | 2.86 | 5.1E-03 |
| *F41C3.7* | WBGene00018273 | protein_coding_gene | 1.5 | 2.86 | 2.1E-02 |
| *F53B7.7* | WBGene00009969 | protein_coding_gene | 1.5 | 2.86 | 1.6E-02 |
| *flp-9* | WBGene00001452 | protein_coding_gene | 1.5 | 2.86 | 4.6E-08 |
| *R05G9R.1* | WBGene00019906 | protein_coding_gene | 1.5 | 2.86 | 1.2E-02 |
| *Y70G10A.3* | WBGene00013499 | protein_coding_gene | 1.5 | 2.86 | 6.4E-04 |
| *K09F6.9* | WBGene00019592 | protein_coding_gene | 1.5 | 2.85 | 1.6E-03 |
| *chpf-2* | WBGene00019108 | protein_coding_gene | 1.5 | 2.85 | 1.2E-04 |
| *Y50E8A.1* | WBGene00013046 | protein_coding_gene | 1.5 | 2.85 | 4.9E-04 |
| *twk-7* | WBGene00006662 | protein_coding_gene | 1.5 | 2.85 | 1.5E-03 |
| *acs-11* | WBGene00018269 | protein_coding_gene | 1.5 | 2.85 | 2.2E-17 |
| *T27A10.2* | WBGene00020844 | protein_coding_gene | 1.5 | 2.84 | 5.4E-03 |
| *C17E4.19* | WBGene00219310 | protein_coding_gene | 1.5 | 2.84 | 4.8E-02 |
| *dhrs-4* | WBGene00010063 | protein_coding_gene | 1.5 | 2.84 | 6.4E-05 |

| | | | | | |
|---|---|---|---|---|---|
| *F01E11.3* | WBGene00017156 | protein_coding_gene | 1.5 | 2.83 | 2.5E-02 |
| *EEED8.2* | WBGene00017133 | protein_coding_gene | 1.5 | 2.83 | 2.7E-05 |
| *flp-18* | WBGene00001461 | protein_coding_gene | 1.5 | 2.83 | 1.6E-07 |
| *E04D5.2* | WBGene00008481 | protein_coding_gene | 1.5 | 2.82 | 1.0E-02 |
| *fkb-7* | WBGene00001432 | protein_coding_gene | 1.5 | 2.82 | 1.9E-04 |
| *nhr-275* | WBGene00021163 | protein_coding_gene | 1.5 | 2.82 | 1.8E-02 |
| *F26F12.3* | WBGene00017834 | protein_coding_gene | 1.5 | 2.82 | 4.9E-04 |
| *otpl-7* | WBGene00018480 | protein_coding_gene | 1.5 | 2.82 | 4.7E-02 |
| *C40H1.7* | WBGene00008042 | protein_coding_gene | 1.5 | 2.81 | 8.0E-08 |
| *T01B10.5* | WBGene00020137 | protein_coding_gene | 1.5 | 2.81 | 1.6E-02 |
| *twk-11* | WBGene00006666 | protein_coding_gene | 1.5 | 2.81 | 4.9E-03 |
| *bbs-8* | WBGene00000244 | protein_coding_gene | 1.5 | 2.81 | 4.4E-03 |
| *F15B9.8* | WBGene00008851 | protein_coding_gene | 1.5 | 2.81 | 1.4E-02 |
| *bigr-1* | WBGene00009512 | protein_coding_gene | 1.5 | 2.81 | 5.0E-05 |
| *Y110A2AL.9* | WBGene00022445 | protein_coding_gene | 1.5 | 2.81 | 1.5E-03 |
| *maph-9* | WBGene00016397 | protein_coding_gene | 1.5 | 2.81 | 4.6E-02 |
| *C31H5.7* | WBGene00007858 | protein_coding_gene | 1.5 | 2.81 | 2.3E-02 |
| *flp-32* | WBGene00010982 | protein_coding_gene | 1.5 | 2.81 | 1.8E-05 |
| *srx-125* | WBGene00006016 | protein_coding_gene | 1.5 | 2.80 | 2.1E-02 |
| *R05G6.10* | WBGene00019902 | protein_coding_gene | 1.5 | 2.80 | 5.3E-04 |
| *ccep-290* | WBGene00021643 | protein_coding_gene | 1.5 | 2.80 | 6.0E-03 |
| *flp-12* | WBGene00001455 | protein_coding_gene | 1.5 | 2.80 | 5.7E-06 |
| *ZC487.1* | WBGene00022623 | protein_coding_gene | 1.5 | 2.80 | 9.2E-03 |
| *F07C3.9* | WBGene00017197 | protein_coding_gene | 1.5 | 2.80 | 1.0E-03 |
| *C39D10.2* | WBGene00016530 | protein_coding_gene | 1.5 | 2.80 | 2.2E-03 |
| *pqn-94* | WBGene00004173 | protein_coding_gene | 1.5 | 2.80 | 3.2E-05 |
| *B0507.3* | WBGene00015220 | protein_coding_gene | 1.5 | 2.79 | 1.6E-04 |
| *F55G1.7* | WBGene00018903 | protein_coding_gene | 1.5 | 2.78 | 4.8E-07 |
| *Y39G10AR.11* | WBGene00021469 | protein_coding_gene | 1.5 | 2.78 | 8.5E-06 |
| *ncam-1* | WBGene00017184 | protein_coding_gene | 1.5 | 2.78 | 6.7E-07 |
| *pqn-42* | WBGene00004129 | protein_coding_gene | 1.5 | 2.77 | 3.4E-03 |
| *C34C12.7* | WBGene00007926 | protein_coding_gene | 1.5 | 2.77 | 5.3E-06 |
| *flp-21* | WBGene00001464 | protein_coding_gene | 1.5 | 2.77 | 1.2E-04 |
| *dyf-5* | WBGene00001121 | protein_coding_gene | 1.5 | 2.77 | 2.1E-02 |
| *C04E6.13* | WBGene00015427 | protein_coding_gene | 1.5 | 2.77 | 2.9E-06 |
| *B0563.5* | WBGene00015262 | protein_coding_gene | 1.5 | 2.77 | 8.1E-04 |
| *ins-1* | WBGene00002084 | protein_coding_gene | 1.5 | 2.76 | 2.1E-03 |
| *Y51H4A.1* | WBGene00013098 | protein_coding_gene | 1.5 | 2.76 | 3.6E-02 |
| *F47B8.2* | WBGene00009803 | protein_coding_gene | 1.5 | 2.76 | 4.8E-05 |
| *Y102A11A.5* | WBGene00022415 | protein_coding_gene | 1.5 | 2.75 | 6.0E-03 |
| *hsp-12.6* | WBGene00002013 | protein_coding_gene | 1.5 | 2.75 | 2.9E-03 |
| *C07E3.3* | WBGene00007414 | protein_coding_gene | 1.5 | 2.75 | 2.1E-03 |
| *W03F8.6* | WBGene00020995 | protein_coding_gene | 1.5 | 2.75 | 1.6E-03 |
| *acs-7* | WBGene00007228 | protein_coding_gene | 1.5 | 2.74 | 3.9E-03 |
| *K01A2.4* | WBGene00019279 | protein_coding_gene | 1.5 | 2.74 | 1.8E-03 |
| *fip-2* | WBGene00020862 | protein_coding_gene | 1.5 | 2.74 | 6.9E-16 |
| *atgp-1* | WBGene00000224 | protein_coding_gene | 1.5 | 2.74 | 6.7E-10 |
| *lido-17* | WBGene00007378 | protein_coding_gene | 1.5 | 2.74 | 5.9E-03 |
| *F35D2.3* | WBGene00018041 | protein_coding_gene | 1.5 | 2.74 | 1.8E-05 |
| *pitr-3* | WBGene00015055 | protein_coding_gene | 1.5 | 2.73 | 2.9E-04 |
| *Y52B11A.4* | WBGene00013124 | protein_coding_gene | 1.4 | 2.73 | 2.1E-03 |
| *Y70C5A.3* | WBGene00050904 | protein_coding_gene | 1.4 | 2.72 | 2.5E-03 |
| *nhr-150* | WBGene00007367 | protein_coding_gene | 1.4 | 2.72 | 4.6E-02 |
| *T19C9.8* | WBGene00011844 | protein_coding_gene | 1.4 | 2.72 | 3.3E-02 |
| *ceh-48* | WBGene00015934 | protein_coding_gene | 1.4 | 2.72 | 6.2E-05 |
| *M02H5.8* | WBGene00019744 | protein_coding_gene | 1.4 | 2.72 | 1.2E-07 |
| *capa-1* | WBGene00021268 | protein_coding_gene | 1.4 | 2.72 | 1.7E-04 |
| *dmd-5* | WBGene00017326 | protein_coding_gene | 1.4 | 2.72 | 1.9E-02 |

| | | | | | |
|---|---|---|---|---|---|
| *nmad-1* | WBGene00017304 | protein_coding_gene | 1.4 | 2.72 | 3.5E-09 |
| *npr-15* | WBGene00012084 | protein_coding_gene | 1.4 | 2.71 | 1.6E-02 |
| *H04M03.12* | WBGene00019156 | protein_coding_gene | 1.4 | 2.71 | 1.9E-03 |
| *twk-35* | WBGene00006687 | protein_coding_gene | 1.4 | 2.71 | 3.4E-02 |
| *mlt-8* | WBGene00021095 | protein_coding_gene | 1.4 | 2.70 | 9.7E-04 |
| *ZK742.7* | WBGene00194921 | protein_coding_gene | 1.4 | 2.70 | 5.3E-03 |
| *vab-3* | WBGene00006870 | protein_coding_gene | 1.4 | 2.70 | 1.5E-05 |
| *marc-4* | WBGene00016903 | protein_coding_gene | 1.4 | 2.70 | 1.3E-02 |
| *ins-24* | WBGene00002107 | protein_coding_gene | 1.4 | 2.70 | 1.5E-03 |
| *Y55D5A.1* | WBGene00021915 | protein_coding_gene | 1.4 | 2.69 | 1.5E-03 |
| *fkh-8* | WBGene00001440 | protein_coding_gene | 1.4 | 2.69 | 1.8E-03 |
| *F13D11.3* | WBGene00017428 | protein_coding_gene | 1.4 | 2.69 | 5.0E-03 |
| *C34F11.8* | WBGene00016417 | protein_coding_gene | 1.4 | 2.69 | 1.4E-04 |
| *R06A10.1* | WBGene00019909 | protein_coding_gene | 1.4 | 2.69 | 2.1E-02 |
| *W04C9.8* | WBGene00235387 | ncRNA_gene | 1.4 | 2.69 | 2.7E-02 |
| *F45H10.5* | WBGene00009742 | protein_coding_gene | 1.4 | 2.69 | 2.3E-02 |
| *inx-20* | WBGene00002142 | protein_coding_gene | 1.4 | 2.69 | 2.2E-02 |
| *sod-4* | WBGene00004933 | protein_coding_gene | 1.4 | 2.68 | 1.7E-07 |
| *asah-2* | WBGene00009192 | protein_coding_gene | 1.4 | 2.68 | 8.7E-06 |
| *D1079.1* | WBGene00017040 | protein_coding_gene | 1.4 | 2.68 | 2.9E-02 |
| *grh-1* | WBGene00001707 | protein_coding_gene | 1.4 | 2.68 | 2.9E-03 |
| *math-37* | WBGene00020080 | protein_coding_gene | 1.4 | 2.68 | 6.2E-04 |
| *flp-16* | WBGene00001459 | protein_coding_gene | 1.4 | 2.68 | 2.5E-11 |
| *nlp-48* | WBGene00008002 | protein_coding_gene | 1.4 | 2.68 | 4.8E-02 |
| *nlp-3* | WBGene00003741 | protein_coding_gene | 1.4 | 2.68 | 5.7E-07 |
| *R13H7.2* | WBGene00020069 | protein_coding_gene | 1.4 | 2.67 | 4.2E-03 |
| *M04G7.1* | WBGene00019772 | protein_coding_gene | 1.4 | 2.67 | 1.6E-05 |
| *D1053.3* | WBGene00008368 | protein_coding_gene | 1.4 | 2.67 | 1.3E-03 |
| *nlp-64* | WBGene00019734 | protein_coding_gene | 1.4 | 2.67 | 3.2E-03 |
| *osm-5* | WBGene00003885 | protein_coding_gene | 1.4 | 2.67 | 8.7E-03 |
| *F59A7.5* | WBGene00019092 | protein_coding_gene | 1.4 | 2.67 | 7.8E-05 |
| *flp-1* | WBGene00001444 | protein_coding_gene | 1.4 | 2.67 | 8.3E-05 |
| *F23F12.13* | WBGene00044455 | protein_coding_gene | 1.4 | 2.66 | 5.5E-04 |
| *zip-7* | WBGene00013100 | protein_coding_gene | 1.4 | 2.66 | 3.3E-03 |
| *ida-1* | WBGene00002048 | protein_coding_gene | 1.4 | 2.66 | 7.1E-05 |
| *igcm-1* | WBGene00018215 | protein_coding_gene | 1.4 | 2.66 | 1.0E-04 |
| *C35C5.9* | WBGene00007958 | protein_coding_gene | 1.4 | 2.65 | 1.9E-04 |
| *Y7A9D.1* | WBGene00012420 | protein_coding_gene | 1.4 | 2.65 | 1.0E-03 |
| *T26C12.3* | WBGene00020833 | protein_coding_gene | 1.4 | 2.65 | 4.0E-03 |
| *F02E11.2* | WBGene00017180 | protein_coding_gene | 1.4 | 2.65 | 1.5E-02 |
| *Y42A5A.1* | WBGene00012776 | protein_coding_gene | 1.4 | 2.64 | 1.3E-04 |
| *F53B3.5* | WBGene00018743 | protein_coding_gene | 1.4 | 2.64 | 2.6E-04 |
| *clec-52* | WBGene00015052 | protein_coding_gene | 1.4 | 2.63 | 2.2E-15 |
| *C27D6.12* | WBGene00044389 | protein_coding_gene | 1.4 | 2.63 | 1.2E-06 |
| *pdf-1* | WBGene00020317 | protein_coding_gene | 1.4 | 2.63 | 5.8E-08 |
| *T05B11.7* | WBGene00044342 | protein_coding_gene | 1.4 | 2.63 | 6.7E-03 |
| *abu-13* | WBGene00004133 | protein_coding_gene | 1.4 | 2.63 | 6.1E-03 |
| *clik-2* | WBGene00016898 | protein_coding_gene | 1.4 | 2.62 | 1.3E-05 |
| *F07A5.4* | WBGene00008543 | protein_coding_gene | 1.4 | 2.62 | 2.5E-02 |
| *C33A12.19* | WBGene00007890 | protein_coding_gene | 1.4 | 2.62 | 1.4E-08 |
| *C35D10.3* | WBGene00016441 | protein_coding_gene | 1.4 | 2.62 | 3.1E-02 |
| *dex-1* | WBGene00017028 | protein_coding_gene | 1.4 | 2.62 | 9.6E-04 |
| *T28H10.3* | WBGene00012144 | protein_coding_gene | 1.4 | 2.62 | 7.1E-03 |
| *W04B5.2* | WBGene00021019 | protein_coding_gene | 1.4 | 2.62 | 5.5E-04 |
| *C01F1.5* | WBGene00015300 | protein_coding_gene | 1.4 | 2.61 | 3.8E-02 |
| *Y71A12B.18* | WBGene00045077 | pseudogene | 1.4 | 2.61 | 1.6E-03 |
| *C08G5.7* | WBGene00044418 | protein_coding_gene | 1.4 | 2.61 | 3.9E-04 |
| *K07E3.4* | WBGene00019492 | protein_coding_gene | 1.4 | 2.61 | 2.8E-09 |

| | | | | | |
|---|---|---|---|---|---|
| *C14C11.4* | WBGene00015768 | protein_coding_gene | 1.4 | 2.61 | 2.0E-05 |
| *flp-2* | WBGene00001445 | protein_coding_gene | 1.4 | 2.61 | 2.0E-05 |
| *Y7A9C.1* | WBGene00012418 | pseudogene | 1.4 | 2.61 | 7.8E-09 |
| *C38C3.4* | WBGene00016513 | protein_coding_gene | 1.4 | 2.61 | 9.1E-03 |
| *osm-6* | WBGene00003886 | protein_coding_gene | 1.4 | 2.61 | 4.2E-02 |
| *flp-25* | WBGene00010575 | protein_coding_gene | 1.4 | 2.60 | 1.4E-03 |
| *best-16* | WBGene00020046 | protein_coding_gene | 1.4 | 2.60 | 1.5E-02 |
| *flp-19* | WBGene00001462 | protein_coding_gene | 1.4 | 2.60 | 1.7E-08 |
| *gcy-15* | WBGene00001541 | protein_coding_gene | 1.4 | 2.60 | 3.1E-02 |
| *Y15E3A.5* | WBGene00012444 | protein_coding_gene | 1.4 | 2.59 | 1.1E-03 |
| *snf-11* | WBGene00004910 | protein_coding_gene | 1.4 | 2.59 | 6.1E-04 |
| *plpr-1* | WBGene00011524 | protein_coding_gene | 1.4 | 2.59 | 1.1E-02 |
| *ddr-2* | WBGene00017381 | protein_coding_gene | 1.4 | 2.59 | 1.8E-04 |
| *H10E21.5* | WBGene00019185 | protein_coding_gene | 1.4 | 2.59 | 8.8E-05 |
| *M60.4* | WBGene00019780 | protein_coding_gene | 1.4 | 2.59 | 1.2E-03 |
| *F41G4.8* | WBGene00018311 | protein_coding_gene | 1.4 | 2.58 | 2.7E-04 |
| *F17C11.2* | WBGene00008914 | protein_coding_gene | 1.4 | 2.58 | 2.1E-09 |
| *cgt-1* | WBGene00011517 | protein_coding_gene | 1.4 | 2.58 | 1.3E-04 |
| *F56H9.2* | WBGene00010174 | protein_coding_gene | 1.4 | 2.58 | 1.1E-09 |
| *K11H3.5* | WBGene00010782 | protein_coding_gene | 1.4 | 2.57 | 8.8E-03 |
| *gba-1* | WBGene00016335 | protein_coding_gene | 1.4 | 2.56 | 1.9E-03 |
| *Y54G2A.7* | WBGene00043702 | protein_coding_gene | 1.4 | 2.56 | 4.7E-05 |
| *sel-7* | WBGene00004764 | protein_coding_gene | 1.4 | 2.56 | 2.8E-02 |
| *F28F5.4* | WBGene00043297 | protein_coding_gene | 1.4 | 2.56 | 4.3E-02 |
| *F48E3.8* | WBGene00018607 | protein_coding_gene | 1.4 | 2.55 | 1.8E-03 |
| *lgc-22* | WBGene00017483 | protein_coding_gene | 1.4 | 2.55 | 3.3E-04 |
| *F35C11.7* | WBGene00009406 | protein_coding_gene | 1.4 | 2.55 | 1.0E-03 |
| *ntc-1* | WBGene00044568 | protein_coding_gene | 1.4 | 2.55 | 2.1E-02 |
| *C05E7.3* | WBGene00007345 | protein_coding_gene | 1.3 | 2.55 | 1.0E-02 |
| *Y71H2AM.15* | WBGene00022180 | protein_coding_gene | 1.3 | 2.54 | 2.2E-03 |
| *ubc-8* | WBGene00006705 | protein_coding_gene | 1.3 | 2.54 | 3.1E-03 |
| *R74.10* | WBGene00206525 | protein_coding_gene | 1.3 | 2.54 | 2.2E-03 |
| *gcy-36* | WBGene00001556 | protein_coding_gene | 1.3 | 2.54 | 4.2E-02 |
| *cyn-17* | WBGene00000893 | protein_coding_gene | 1.3 | 2.54 | 3.7E-02 |
| *lpr-1* | WBGene00002393 | protein_coding_gene | 1.3 | 2.54 | 2.6E-02 |
| *twk-31* | WBGene00006683 | protein_coding_gene | 1.3 | 2.54 | 2.0E-03 |
| *del-2* | WBGene00000953 | protein_coding_gene | 1.3 | 2.53 | 1.6E-02 |
| *T01B6.1* | WBGene00020134 | protein_coding_gene | 1.3 | 2.53 | 4.3E-03 |
| *F14H12.8* | WBGene00017475 | protein_coding_gene | 1.3 | 2.53 | 1.1E-02 |
| *ZK265.7* | WBGene00013959 | protein_coding_gene | 1.3 | 2.53 | 7.0E-05 |
| *skr-5* | WBGene00004811 | protein_coding_gene | 1.3 | 2.53 | 3.9E-02 |
| *C10C5.7* | WBGene00007511 | protein_coding_gene | 1.3 | 2.52 | 3.0E-03 |
| *Y50E8A.5* | WBGene00013048 | protein_coding_gene | 1.3 | 2.52 | 1.7E-04 |
| *ace-3* | WBGene00000037 | protein_coding_gene | 1.3 | 2.52 | 5.0E-04 |
| *npr-29* | WBGene00013848 | protein_coding_gene | 1.3 | 2.52 | 2.9E-03 |
| *hst-3.2* | WBGene00044303 | protein_coding_gene | 1.3 | 2.52 | 3.7E-03 |
| *fbxa-29* | WBGene00013836 | protein_coding_gene | 1.3 | 2.51 | 3.8E-03 |
| *T05A8.3* | WBGene00020229 | protein_coding_gene | 1.3 | 2.51 | 3.8E-04 |
| *C28G1.10* | WBGene00219376 | protein_coding_gene | 1.3 | 2.51 | 1.3E-05 |
| *snt-6* | WBGene00004926 | protein_coding_gene | 1.3 | 2.51 | 1.4E-03 |
| *ZK54.3* | WBGene00022648 | protein_coding_gene | 1.3 | 2.51 | 4.5E-02 |
| *srap-1* | WBGene00011522 | protein_coding_gene | 1.3 | 2.51 | 1.2E-02 |
| *F11C7.2* | WBGene00017376 | protein_coding_gene | 1.3 | 2.51 | 1.5E-03 |
| *cfi-1* | WBGene00000476 | protein_coding_gene | 1.3 | 2.50 | 2.0E-04 |
| *nlp-58* | WBGene00077775 | protein_coding_gene | 1.3 | 2.50 | 1.2E-04 |
| *F46E10.2* | WBGene00018489 | protein_coding_gene | 1.3 | 2.50 | 1.2E-03 |
| *Y38H6C.21* | WBGene00012633 | protein_coding_gene | 1.3 | 2.50 | 1.0E-03 |
| *Y38C1AA.6* | WBGene00021397 | protein_coding_gene | 1.3 | 2.50 | 1.3E-02 |

| | | | | | |
|---|---|---|---|---|---|
| *droe-4* | WBGene00010262 | protein_coding_gene | 1.3 | 2.50 | 1.8E-04 |
| *lpr-2* | WBGene00011726 | protein_coding_gene | 1.3 | 2.50 | 6.7E-03 |
| *mboa-1* | WBGene00007174 | protein_coding_gene | 1.3 | 2.50 | 4.9E-02 |
| *C13C4.7* | WBGene00007551 | protein_coding_gene | 1.3 | 2.50 | 1.4E-03 |
| *ugt-55* | WBGene00011452 | protein_coding_gene | 1.3 | 2.49 | 1.7E-04 |
| *flp-14* | WBGene00001457 | protein_coding_gene | 1.3 | 2.49 | 2.1E-08 |
| *kvs-2* | WBGene00007862 | protein_coding_gene | 1.3 | 2.49 | 3.1E-03 |
| *T10B9.9* | WBGene00011678 | protein_coding_gene | 1.3 | 2.49 | 3.6E-02 |
| *Y44A6C.1* | WBGene00012850 | protein_coding_gene | 1.3 | 2.49 | 3.4E-04 |
| *egl-15* | WBGene00001184 | protein_coding_gene | 1.3 | 2.49 | 5.0E-03 |
| *C29H12.6* | WBGene00016237 | protein_coding_gene | 1.3 | 2.49 | 7.1E-04 |
| *flp-5* | WBGene00001448 | protein_coding_gene | 1.3 | 2.49 | 3.9E-07 |
| *tag-234* | WBGene00018874 | protein_coding_gene | 1.3 | 2.48 | 1.4E-04 |
| *ZK550.6* | WBGene00014000 | protein_coding_gene | 1.3 | 2.48 | 5.4E-03 |
| *nas-33* | WBGene00003551 | protein_coding_gene | 1.3 | 2.48 | 2.7E-02 |
| *clec-83* | WBGene00021879 | protein_coding_gene | 1.3 | 2.48 | 1.3E-03 |
| *C49H3.12* | WBGene00016795 | protein_coding_gene | 1.3 | 2.48 | 8.5E-03 |
| *F55A4.4* | WBGene00018856 | pseudogene | 1.3 | 2.48 | 2.5E-02 |
| *flp-34* | WBGene00019975 | protein_coding_gene | 1.3 | 2.48 | 7.6E-03 |
| *T28B8.1* | WBGene00012113 | protein_coding_gene | 1.3 | 2.48 | 5.8E-05 |
| *K04D7.6* | WBGene00010559 | protein_coding_gene | 1.3 | 2.47 | 3.6E-02 |
| *B0222.1* | WBGene00015053 | protein_coding_gene | 1.3 | 2.47 | 1.4E-02 |
| *rga-2* | WBGene00013135 | protein_coding_gene | 1.3 | 2.47 | 6.0E-03 |
| *ham-2* | WBGene00001821 | protein_coding_gene | 1.3 | 2.47 | 1.5E-03 |
| *ZC449.5* | WBGene00022615 | protein_coding_gene | 1.3 | 2.47 | 3.8E-04 |
| *pudl-1* | WBGene00044489 | protein_coding_gene | 1.3 | 2.47 | 3.8E-04 |
| *rpi-2* | WBGene00019536 | protein_coding_gene | 1.3 | 2.47 | 1.1E-02 |
| *E02A10.4* | WBGene00008454 | protein_coding_gene | 1.3 | 2.47 | 3.8E-03 |
| *F40E3.5* | WBGene00018232 | protein_coding_gene | 1.3 | 2.47 | 4.2E-04 |
| *F54D5.15* | WBGene00010057 | protein_coding_gene | 1.3 | 2.46 | 3.1E-05 |
| *Y54G2A.32* | WBGene00021894 | protein_coding_gene | 1.3 | 2.46 | 1.4E-07 |
| *sbt-1* | WBGene00011392 | protein_coding_gene | 1.3 | 2.46 | 1.2E-07 |
| *Y95B8A.6* | WBGene00022386 | protein_coding_gene | 1.3 | 2.46 | 3.8E-03 |
| *C46C11.4* | WBGene00044567 | protein_coding_gene | 1.3 | 2.46 | 8.4E-03 |
| *C50F4.1* | WBGene00008229 | protein_coding_gene | 1.3 | 2.46 | 1.4E-08 |
| *F17C8.3* | WBGene00008910 | protein_coding_gene | 1.3 | 2.45 | 8.4E-03 |
| *F59B2.9* | WBGene00010311 | protein_coding_gene | 1.3 | 2.45 | 3.7E-03 |
| *ncx-10* | WBGene00003575 | protein_coding_gene | 1.3 | 2.45 | 1.1E-02 |
| *F23C8.14* | WBGene00255457 | protein_coding_gene | 1.3 | 2.45 | 3.0E-02 |
| *F13D11.4* | WBGene00017429 | protein_coding_gene | 1.3 | 2.44 | 3.2E-04 |
| *mltn-1* | WBGene00009311 | protein_coding_gene | 1.3 | 2.44 | 1.5E-02 |
| *nape-1* | WBGene00021371 | protein_coding_gene | 1.3 | 2.44 | 1.2E-02 |
| *C33D9.6* | WBGene00007900 | protein_coding_gene | 1.3 | 2.44 | 6.0E-06 |
| *Y75B8A.6* | WBGene00013543 | protein_coding_gene | 1.3 | 2.44 | 3.9E-03 |
| *F01G10.6* | WBGene00008509 | protein_coding_gene | 1.3 | 2.44 | 2.8E-02 |
| *C44B11.4* | WBGene00016634 | protein_coding_gene | 1.3 | 2.44 | 1.2E-04 |
| *K03A11.5* | WBGene00010522 | protein_coding_gene | 1.3 | 2.44 | 4.5E-04 |
| *R07B7.6* | WBGene00011090 | protein_coding_gene | 1.3 | 2.43 | 9.0E-06 |
| *flp-7* | WBGene00001450 | protein_coding_gene | 1.3 | 2.43 | 1.3E-06 |
| *F16C3.2* | WBGene00008886 | protein_coding_gene | 1.3 | 2.43 | 4.5E-03 |
| *dlg-1* | WBGene00001006 | protein_coding_gene | 1.3 | 2.43 | 7.9E-05 |
| *rgef-1* | WBGene00009100 | protein_coding_gene | 1.3 | 2.43 | 6.0E-04 |
| *grl-7* | WBGene00001716 | protein_coding_gene | 1.3 | 2.43 | 3.2E-02 |
| *C33H5.13* | WBGene00016379 | protein_coding_gene | 1.3 | 2.43 | 3.2E-07 |
| *T05E11.2* | WBGene00011479 | protein_coding_gene | 1.3 | 2.43 | 2.7E-03 |
| *jmjd-3.1* | WBGene00017571 | protein_coding_gene | 1.3 | 2.43 | 9.6E-05 |
| *Y51B9A.9* | WBGene00013091 | protein_coding_gene | 1.3 | 2.42 | 6.0E-03 |
| *pha-4* | WBGene00004013 | protein_coding_gene | 1.3 | 2.42 | 3.5E-04 |

| | | | | | |
|---|---|---|---|---|---|
| *iglr-1* | WBGene00008977 | protein_coding_gene | 1.3 | 2.42 | 4.6E-03 |
| *slc-36.5* | WBGene00016628 | protein_coding_gene | 1.3 | 2.42 | 9.9E-03 |
| *clec-66* | WBGene00009397 | protein_coding_gene | 1.3 | 2.42 | 8.2E-04 |
| *cdf-2* | WBGene00011821 | protein_coding_gene | 1.3 | 2.42 | 1.8E-02 |
| *math-27* | WBGene00018101 | pseudogene | 1.3 | 2.42 | 2.4E-03 |
| *ceh-27* | WBGene00000449 | protein_coding_gene | 1.3 | 2.41 | 4.3E-03 |
| *clh-2* | WBGene00000529 | protein_coding_gene | 1.3 | 2.41 | 2.6E-03 |
| *Y57A10A.26* | WBGene00013266 | protein_coding_gene | 1.3 | 2.41 | 1.3E-05 |
| *F09C3.2* | WBGene00008610 | protein_coding_gene | 1.3 | 2.41 | 2.8E-04 |
| *mgl-3* | WBGene00021152 | protein_coding_gene | 1.3 | 2.41 | 8.5E-03 |
| *fat-1* | WBGene00001393 | protein_coding_gene | 1.3 | 2.41 | 6.7E-10 |
| *T14G8.4* | WBGene00011772 | protein_coding_gene | 1.3 | 2.40 | 3.2E-02 |
| *smf-3* | WBGene00004878 | protein_coding_gene | 1.3 | 2.40 | 1.3E-02 |
| *nex-4* | WBGene00003591 | protein_coding_gene | 1.3 | 2.40 | 1.8E-02 |
| *T03F1.11* | WBGene00020192 | protein_coding_gene | 1.3 | 2.40 | 3.3E-04 |
| *C54D10.3* | WBGene00008298 | protein_coding_gene | 1.3 | 2.39 | 5.6E-14 |
| *aat-8* | WBGene00000009 | protein_coding_gene | 1.3 | 2.39 | 2.1E-02 |
| *F57E7.1* | WBGene00010202 | protein_coding_gene | 1.3 | 2.39 | 6.5E-03 |
| *Y73F4A.3* | WBGene00013516 | protein_coding_gene | 1.3 | 2.39 | 2.0E-02 |
| *ins-18* | WBGene00002101 | protein_coding_gene | 1.3 | 2.39 | 3.6E-03 |
| *F25E2.3* | WBGene00017781 | protein_coding_gene | 1.3 | 2.39 | 5.2E-03 |
| *Y55B1BL.1* | WBGene00021909 | protein_coding_gene | 1.3 | 2.39 | 4.7E-10 |
| *C32D5.12* | WBGene00016319 | protein_coding_gene | 1.3 | 2.39 | 3.0E-02 |
| *K09H9.8* | WBGene00045387 | protein_coding_gene | 1.3 | 2.38 | 2.8E-06 |
| *nlp-42* | WBGene00013588 | protein_coding_gene | 1.3 | 2.38 | 1.3E-03 |
| *C35D10.12* | WBGene00016448 | protein_coding_gene | 1.3 | 2.38 | 1.3E-02 |
| *F56C9.8* | WBGene00018952 | protein_coding_gene | 1.3 | 2.38 | 2.3E-04 |
| *F45D3.2* | WBGene00009722 | protein_coding_gene | 1.3 | 2.38 | 2.0E-03 |
| *glb-24* | WBGene00011287 | protein_coding_gene | 1.3 | 2.38 | 2.0E-03 |
| *F58G11.4* | WBGene00010282 | protein_coding_gene | 1.2 | 2.38 | 8.9E-03 |
| *D2005.6* | WBGene00008401 | protein_coding_gene | 1.2 | 2.38 | 2.6E-02 |
| *pgp-11* | WBGene00004005 | protein_coding_gene | 1.2 | 2.37 | 4.9E-08 |
| *zig-5* | WBGene00006982 | protein_coding_gene | 1.2 | 2.37 | 1.8E-03 |
| *lec-2* | WBGene00002265 | protein_coding_gene | 1.2 | 2.37 | 7.2E-03 |
| *C16H3.3* | WBGene00015868 | protein_coding_gene | 1.2 | 2.37 | 2.4E-06 |
| *F58B4.6* | WBGene00010239 | protein_coding_gene | 1.2 | 2.37 | 1.2E-02 |
| *tsp-10* | WBGene00006636 | protein_coding_gene | 1.2 | 2.37 | 2.8E-06 |
| *C30G7.4* | WBGene00044178 | protein_coding_gene | 1.2 | 2.37 | 1.9E-03 |
| *ikb-1* | WBGene00002069 | protein_coding_gene | 1.2 | 2.37 | 2.7E-02 |
| *slcf-2* | WBGene00006865 | protein_coding_gene | 1.2 | 2.37 | 5.1E-04 |
| *T18D3.7* | WBGene00011824 | protein_coding_gene | 1.2 | 2.36 | 7.2E-03 |
| *C03G6.17* | WBGene00015402 | protein_coding_gene | 1.2 | 2.36 | 7.0E-04 |
| *K09F5.6* | WBGene00044698 | protein_coding_gene | 1.2 | 2.36 | 4.2E-03 |
| *lgc-11* | WBGene00000061 | protein_coding_gene | 1.2 | 2.36 | 4.8E-02 |
| *nlp-35* | WBGene00007881 | protein_coding_gene | 1.2 | 2.36 | 4.5E-03 |
| *atln-2* | WBGene00012763 | protein_coding_gene | 1.2 | 2.36 | 1.6E-04 |
| *C44H9.5* | WBGene00008100 | protein_coding_gene | 1.2 | 2.35 | 6.2E-05 |
| *F49C12.6* | WBGene00009876 | protein_coding_gene | 1.2 | 2.35 | 9.5E-04 |
| *K02E10.4* | WBGene00019318 | protein_coding_gene | 1.2 | 2.35 | 1.5E-02 |
| *ZC116.1* | WBGene00013853 | protein_coding_gene | 1.2 | 2.35 | 2.5E-03 |
| *lad-2* | WBGene00002243 | protein_coding_gene | 1.2 | 2.35 | 2.8E-03 |
| *lgc-55* | WBGene00013746 | protein_coding_gene | 1.2 | 2.35 | 1.1E-02 |
| *nhr-42* | WBGene00003632 | protein_coding_gene | 1.2 | 2.35 | 1.5E-04 |
| *C16B8.2* | WBGene00015822 | protein_coding_gene | 1.2 | 2.35 | 1.3E-02 |
| *F17C11.4* | WBGene00008915 | protein_coding_gene | 1.2 | 2.35 | 3.2E-03 |
| *C54D10.13* | WBGene00045400 | protein_coding_gene | 1.2 | 2.35 | 1.8E-02 |
| *ZK836.3* | WBGene00014099 | protein_coding_gene | 1.2 | 2.34 | 2.5E-02 |
| *glb-25* | WBGene00020279 | protein_coding_gene | 1.2 | 2.34 | 1.9E-03 |

| | | | | | |
|---|---|---|---|---|---|
| *acox-3* | WBGene00019060 | protein_coding_gene | 1.2 | 2.34 | 7.3E-03 |
| *hum-6* | WBGene00002039 | protein_coding_gene | 1.2 | 2.34 | 7.9E-05 |
| *slt-1* | WBGene00004854 | protein_coding_gene | 1.2 | 2.34 | 1.1E-02 |
| *aagr-1* | WBGene00017071 | protein_coding_gene | 1.2 | 2.34 | 5.1E-03 |
| *nas-37* | WBGene00003553 | protein_coding_gene | 1.2 | 2.34 | 3.6E-03 |
| *T15B7.10* | WBGene00020522 | pseudogene | 1.2 | 2.34 | 4.3E-03 |
| *K08B5.2* | WBGene00019519 | protein_coding_gene | 1.2 | 2.34 | 1.9E-03 |
| *K01F9.2* | WBGene00010475 | protein_coding_gene | 1.2 | 2.33 | 2.8E-02 |
| *F15H10.7* | WBGene00008874 | protein_coding_gene | 1.2 | 2.33 | 1.1E-02 |
| *B0454.6* | WBGene00015197 | protein_coding_gene | 1.2 | 2.33 | 1.1E-04 |
| *unc-53* | WBGene00006788 | protein_coding_gene | 1.2 | 2.33 | 2.9E-03 |
| *C05A9.2* | WBGene00007317 | protein_coding_gene | 1.2 | 2.33 | 3.3E-04 |
| *nspc-17* | WBGene00008359 | protein_coding_gene | 1.2 | 2.33 | 7.8E-04 |
| *C25F6.7* | WBGene00016106 | protein_coding_gene | 1.2 | 2.32 | 8.6E-03 |
| *R06F6.14* | WBGene00269421 | protein_coding_gene | 1.2 | 2.32 | 2.5E-05 |
| *dyf-6* | WBGene00001122 | protein_coding_gene | 1.2 | 2.32 | 1.7E-02 |
| *K07C10.3* | WBGene00194851 | ncRNA_gene | 1.2 | 2.32 | 4.7E-02 |
| *ptp-4* | WBGene00010558 | protein_coding_gene | 1.2 | 2.32 | 7.7E-03 |
| *nhr-87* | WBGene00003677 | protein_coding_gene | 1.2 | 2.32 | 8.6E-05 |
| *F31E3.2* | WBGene00017950 | protein_coding_gene | 1.2 | 2.32 | 5.9E-03 |
| *F14H3.12* | WBGene00008831 | protein_coding_gene | 1.2 | 2.32 | 2.0E-03 |
| *Y52E8A.3* | WBGene00021796 | protein_coding_gene | 1.2 | 2.31 | 2.6E-03 |
| *F57C12.2* | WBGene00019014 | protein_coding_gene | 1.2 | 2.31 | 1.5E-03 |
| *slc-28.1* | WBGene00017867 | protein_coding_gene | 1.2 | 2.31 | 2.2E-02 |
| *DH11.5* | WBGene00008438 | protein_coding_gene | 1.2 | 2.31 | 2.4E-06 |
| *Y64G10A.7* | WBGene00013416 | protein_coding_gene | 1.2 | 2.31 | 6.5E-04 |
| *flp-3* | WBGene00001446 | protein_coding_gene | 1.2 | 2.31 | 3.8E-06 |
| *E02H4.7* | WBGene00077757 | protein_coding_gene | 1.2 | 2.31 | 1.1E-05 |
| *hda-5* | WBGene00009663 | protein_coding_gene | 1.2 | 2.31 | 1.4E-03 |
| *Y69A2AR.31* | WBGene00022100 | protein_coding_gene | 1.2 | 2.31 | 6.0E-03 |
| *dhs-20* | WBGene00000983 | protein_coding_gene | 1.2 | 2.31 | 5.8E-03 |
| *alr-1* | WBGene00044330 | protein_coding_gene | 1.2 | 2.31 | 2.1E-04 |
| *Y6D1A.2* | WBGene00012396 | protein_coding_gene | 1.2 | 2.31 | 1.9E-02 |
| *mtl-2* | WBGene00003474 | protein_coding_gene | 1.2 | 2.31 | 3.7E-04 |
| *C03A3.1* | WBGene00007266 | protein_coding_gene | 1.2 | 2.30 | 2.3E-02 |
| *pqn-32* | WBGene00004120 | protein_coding_gene | 1.2 | 2.30 | 3.7E-02 |
| *C40H5.2* | WBGene00008045 | protein_coding_gene | 1.2 | 2.30 | 1.8E-02 |
| *inx-6* | WBGene00002128 | protein_coding_gene | 1.2 | 2.30 | 2.5E-03 |
| *T05E11.9* | WBGene00014505 | ncRNA_gene | 1.2 | 2.30 | 5.4E-10 |
| *F20C5.6* | WBGene00008971 | protein_coding_gene | 1.2 | 2.30 | 3.0E-04 |
| *Y32B12C.1* | WBGene00012525 | protein_coding_gene | 1.2 | 2.30 | 3.3E-02 |
| *T02G5.11* | WBGene00020167 | pseudogene | 1.2 | 2.29 | 9.6E-03 |
| *F20C5.11* | WBGene00303015 | protein_coding_gene | 1.2 | 2.29 | 2.6E-04 |
| *cdh-8* | WBGene00017576 | protein_coding_gene | 1.2 | 2.29 | 1.2E-03 |
| *F20C5.5* | WBGene00008970 | protein_coding_gene | 1.2 | 2.29 | 7.0E-03 |
| *F58B4.3* | WBGene00010236 | protein_coding_gene | 1.2 | 2.29 | 8.7E-03 |
| *C41G11.1* | WBGene00016568 | protein_coding_gene | 1.2 | 2.29 | 1.6E-04 |
| *D1054.9* | WBGene00008376 | protein_coding_gene | 1.2 | 2.29 | 1.8E-02 |
| *F27D9.2* | WBGene00017861 | protein_coding_gene | 1.2 | 2.29 | 2.4E-06 |
| *T26C5.5* | WBGene00012035 | protein_coding_gene | 1.2 | 2.28 | 2.9E-02 |
| *T07F12.4* | WBGene00020324 | protein_coding_gene | 1.2 | 2.28 | 1.6E-04 |
| *C28H8.2* | WBGene00016193 | protein_coding_gene | 1.2 | 2.28 | 1.4E-02 |
| *srw-85* | WBGene00005832 | protein_coding_gene | 1.2 | 2.28 | 4.0E-03 |
| *cab-1* | WBGene00000277 | protein_coding_gene | 1.2 | 2.28 | 1.0E-04 |
| *sox-2* | WBGene00004949 | protein_coding_gene | 1.2 | 2.28 | 1.6E-03 |
| *C30G4.4* | WBGene00016269 | protein_coding_gene | 1.2 | 2.28 | 1.8E-03 |
| *F18A11.2* | WBGene00008929 | protein_coding_gene | 1.2 | 2.28 | 2.1E-02 |
| *nlp-12* | WBGene00003750 | protein_coding_gene | 1.2 | 2.28 | 7.6E-04 |

| | | | | | |
|---|---|---|---|---|---|
| *cal-8* | WBGene00017394 | protein_coding_gene | 1.2 | 2.28 | 3.0E-02 |
| *ZK593.2* | WBGene00014002 | protein_coding_gene | 1.2 | 2.28 | 3.8E-03 |
| *sulp-3* | WBGene00018283 | protein_coding_gene | 1.2 | 2.28 | 2.7E-02 |
| *C44E12.1* | WBGene00016657 | protein_coding_gene | 1.2 | 2.28 | 1.3E-05 |
| *frpr-8* | WBGene00018728 | protein_coding_gene | 1.2 | 2.28 | 3.2E-02 |
| *F42G2.2* | WBGene00018351 | protein_coding_gene | 1.2 | 2.28 | 1.1E-03 |
| *jmjd-3.3* | WBGene00007813 | protein_coding_gene | 1.2 | 2.28 | 1.2E-02 |
| *T25B2.2* | WBGene00020786 | protein_coding_gene | 1.2 | 2.28 | 4.0E-04 |
| *nep-21* | WBGene00011794 | protein_coding_gene | 1.2 | 2.28 | 2.4E-04 |
| *ZK287.4* | WBGene00013965 | protein_coding_gene | 1.2 | 2.27 | 3.4E-02 |
| *myo-5* | WBGene00019064 | protein_coding_gene | 1.2 | 2.27 | 7.5E-05 |
| *cmk-1* | WBGene00000553 | protein_coding_gene | 1.2 | 2.27 | 4.0E-03 |
| *C35E7.4* | WBGene00016456 | protein_coding_gene | 1.2 | 2.27 | 1.1E-03 |
| *F20D6.10* | WBGene00017639 | protein_coding_gene | 1.2 | 2.27 | 1.0E-02 |
| *F42C5.4* | WBGene00018346 | protein_coding_gene | 1.2 | 2.27 | 7.9E-03 |
| *col-76* | WBGene00000652 | protein_coding_gene | 1.2 | 2.27 | 1.3E-04 |
| *nlp-1* | WBGene00003739 | protein_coding_gene | 1.2 | 2.26 | 5.7E-05 |
| *Y37D8A.16* | WBGene00012554 | protein_coding_gene | 1.2 | 2.26 | 6.5E-05 |
| *cal-2* | WBGene00000286 | protein_coding_gene | 1.2 | 2.26 | 2.2E-03 |
| *mdl-1* | WBGene00003163 | protein_coding_gene | 1.2 | 2.26 | 6.3E-03 |
| *gcy-21* | WBGene00001546 | protein_coding_gene | 1.2 | 2.26 | 2.9E-03 |
| *F10C2.3* | WBGene00008644 | protein_coding_gene | 1.2 | 2.26 | 1.7E-03 |
| *C04E7.3* | WBGene00015430 | protein_coding_gene | 1.2 | 2.25 | 2.0E-03 |
| *dct-1* | WBGene00015776 | protein_coding_gene | 1.2 | 2.25 | 3.0E-05 |
| *T27E4.1* | WBGene00020861 | protein_coding_gene | 1.2 | 2.25 | 1.1E-02 |
| *F57A8.1* | WBGene00010177 | protein_coding_gene | 1.2 | 2.25 | 2.1E-05 |
| *ensh-1* | WBGene00017013 | protein_coding_gene | 1.2 | 2.25 | 1.4E-02 |
| *E02A10.3* | WBGene00008453 | protein_coding_gene | 1.2 | 2.25 | 6.1E-03 |
| *hex-1* | WBGene00020509 | protein_coding_gene | 1.2 | 2.25 | 2.1E-05 |
| *T26G10.5* | WBGene00012062 | protein_coding_gene | 1.2 | 2.25 | 4.1E-02 |
| *K01A2.3* | WBGene00019278 | pseudogene | 1.2 | 2.25 | 1.9E-02 |
| *M176.9* | WBGene00014813 | protein_coding_gene | 1.2 | 2.25 | 2.4E-02 |
| *sad-1* | WBGene00004719 | protein_coding_gene | 1.2 | 2.25 | 3.4E-04 |
| *B0035.13* | WBGene00007112 | protein_coding_gene | 1.2 | 2.25 | 8.6E-08 |
| *pqn-18* | WBGene00004108 | protein_coding_gene | 1.2 | 2.25 | 3.2E-03 |
| *asp-14* | WBGene00019619 | protein_coding_gene | 1.2 | 2.24 | 2.6E-03 |
| *B0228.6* | WBGene00015063 | protein_coding_gene | 1.2 | 2.24 | 4.3E-05 |
| *sup-26* | WBGene00006331 | protein_coding_gene | 1.2 | 2.24 | 3.2E-03 |
| *glb-19* | WBGene00010146 | protein_coding_gene | 1.2 | 2.24 | 5.9E-03 |
| *pals-18* | WBGene00022353 | protein_coding_gene | 1.2 | 2.24 | 2.3E-03 |
| *pdl-1* | WBGene00003966 | protein_coding_gene | 1.2 | 2.24 | 1.6E-02 |
| *T27E4.7* | WBGene00020865 | protein_coding_gene | 1.2 | 2.24 | 1.2E-04 |
| *F23C8.3* | WBGene00017732 | protein_coding_gene | 1.2 | 2.23 | 6.6E-03 |
| *F53B1.4* | WBGene00018737 | protein_coding_gene | 1.2 | 2.23 | 1.1E-02 |
| *rbf-1* | WBGene00004316 | protein_coding_gene | 1.2 | 2.23 | 1.5E-03 |
| *asp-6* | WBGene00000219 | protein_coding_gene | 1.2 | 2.23 | 4.0E-05 |
| *ceh-32* | WBGene00000453 | protein_coding_gene | 1.2 | 2.23 | 1.3E-03 |
| *Y59H11AR.4* | WBGene00022012 | protein_coding_gene | 1.2 | 2.23 | 1.2E-03 |
| *K03H1.5* | WBGene00010540 | protein_coding_gene | 1.2 | 2.23 | 1.2E-02 |
| *cdka-1* | WBGene00011955 | protein_coding_gene | 1.2 | 2.23 | 1.5E-03 |
| *nlp-5* | WBGene00003743 | protein_coding_gene | 1.2 | 2.23 | 2.7E-02 |
| *F10G8.8* | WBGene00008666 | protein_coding_gene | 1.2 | 2.22 | 4.3E-06 |
| *T16H12.9* | WBGene00011817 | protein_coding_gene | 1.2 | 2.22 | 3.1E-02 |
| *zmp-2* | WBGene00019212 | protein_coding_gene | 1.2 | 2.22 | 2.2E-02 |
| *pat-9* | WBGene00003933 | protein_coding_gene | 1.2 | 2.22 | 3.6E-03 |
| *ech-1.1* | WBGene00001150 | protein_coding_gene | 1.1 | 2.22 | 6.1E-03 |
| *nlp-50* | WBGene00018298 | protein_coding_gene | 1.1 | 2.22 | 8.4E-04 |
| *F31F7.1* | WBGene00017964 | protein_coding_gene | 1.1 | 2.22 | 1.5E-04 |

| | | | | | |
|---|---|---|---|---|---|
| *lact-3* | WBGene00010897 | protein_coding_gene | 1.1 | 2.22 | 6.4E-06 |
| *F39E9.10* | WBGene00023036 | pseudogene | 1.1 | 2.22 | 3.2E-04 |
| *cal-3* | WBGene00000287 | protein_coding_gene | 1.1 | 2.22 | 1.5E-03 |
| *M04B2.6* | WBGene00010848 | protein_coding_gene | 1.1 | 2.21 | 7.2E-03 |
| *tag-209* | WBGene00011072 | protein_coding_gene | 1.1 | 2.21 | 4.5E-03 |
| *php-3* | WBGene00004024 | protein_coding_gene | 1.1 | 2.21 | 4.1E-05 |
| *ttr-21* | WBGene00008628 | protein_coding_gene | 1.1 | 2.21 | 3.6E-04 |
| *C02B8.1* | WBGene00015320 | protein_coding_gene | 1.1 | 2.21 | 2.8E-02 |
| *C04E12.4* | WBGene00015434 | protein_coding_gene | 1.1 | 2.21 | 2.6E-05 |
| *cutl-24* | WBGene00021396 | protein_coding_gene | 1.1 | 2.21 | 1.0E-02 |
| *daam-1* | WBGene00018976 | protein_coding_gene | 1.1 | 2.21 | 3.5E-03 |
| *abf-5* | WBGene00000016 | protein_coding_gene | 1.1 | 2.21 | 3.7E-04 |
| *nhr-121* | WBGene00003711 | protein_coding_gene | 1.1 | 2.21 | 6.8E-04 |
| *nlp-55* | WBGene00008820 | protein_coding_gene | 1.1 | 2.20 | 1.1E-03 |
| *F47E1.4* | WBGene00018568 | protein_coding_gene | 1.1 | 2.20 | 4.9E-03 |
| *tbc-16* | WBGene00018639 | protein_coding_gene | 1.1 | 2.20 | 2.1E-02 |
| *Y97E10C.1* | WBGene00022410 | protein_coding_gene | 1.1 | 2.20 | 4.1E-03 |
| *nspc-14* | WBGene00009638 | protein_coding_gene | 1.1 | 2.20 | 6.0E-03 |
| *C40A11.3* | WBGene00016546 | protein_coding_gene | 1.1 | 2.20 | 4.0E-02 |
| *Y65A5A.1* | WBGene00013418 | protein_coding_gene | 1.1 | 2.20 | 1.3E-03 |
| *del-5* | WBGene00010334 | protein_coding_gene | 1.1 | 2.20 | 2.0E-04 |
| *pudl-2* | WBGene00044490 | protein_coding_gene | 1.1 | 2.20 | 2.2E-02 |
| *cuti-1* | WBGene00022591 | protein_coding_gene | 1.1 | 2.20 | 6.7E-03 |
| *cex-2* | WBGene00023408 | protein_coding_gene | 1.1 | 2.20 | 6.8E-03 |
| *mks-5* | WBGene00007490 | protein_coding_gene | 1.1 | 2.20 | 3.0E-02 |
| *F27B10.1* | WBGene00017850 | protein_coding_gene | 1.1 | 2.20 | 1.4E-02 |
| *ddo-1* | WBGene00008127 | protein_coding_gene | 1.1 | 2.19 | 2.7E-03 |
| *nlp-14* | WBGene00003752 | protein_coding_gene | 1.1 | 2.19 | 2.0E-03 |
| *nlp-80* | WBGene00011428 | protein_coding_gene | 1.1 | 2.19 | 1.5E-04 |
| *flp-20* | WBGene00001463 | protein_coding_gene | 1.1 | 2.19 | 4.2E-03 |
| *klf-1* | WBGene00018990 | protein_coding_gene | 1.1 | 2.19 | 6.4E-05 |
| *sox-3* | WBGene00004950 | protein_coding_gene | 1.1 | 2.19 | 1.5E-02 |
| *Y51A2D.13* | WBGene00013080 | protein_coding_gene | 1.1 | 2.19 | 3.7E-02 |
| *ptp-1* | WBGene00004213 | protein_coding_gene | 1.1 | 2.19 | 2.5E-03 |
| *B0284.3* | WBGene00007133 | protein_coding_gene | 1.1 | 2.19 | 4.7E-03 |
| *unc-129* | WBGene00006852 | protein_coding_gene | 1.1 | 2.19 | 7.5E-05 |
| *col-99* | WBGene00000674 | protein_coding_gene | 1.1 | 2.19 | 1.1E-02 |
| *cest-16* | WBGene00020330 | protein_coding_gene | 1.1 | 2.18 | 1.1E-02 |
| *kin-9* | WBGene00002195 | protein_coding_gene | 1.1 | 2.18 | 3.2E-03 |
| *F14D2.19* | WBGene00206522 | protein_coding_gene | 1.1 | 2.18 | 8.0E-03 |
| *fbxa-3* | WBGene00020363 | protein_coding_gene | 1.1 | 2.18 | 2.8E-02 |
| *spin-2* | WBGene00008033 | protein_coding_gene | 1.1 | 2.18 | 3.5E-03 |
| *F54D1.6* | WBGene00010047 | protein_coding_gene | 1.1 | 2.18 | 6.8E-03 |
| *F22B5.4* | WBGene00009042 | protein_coding_gene | 1.1 | 2.18 | 4.3E-02 |
| *F58H10.1* | WBGene00010291 | protein_coding_gene | 1.1 | 2.18 | 1.1E-02 |
| *ZC455.1* | WBGene00013902 | protein_coding_gene | 1.1 | 2.18 | 3.3E-03 |
| *flp-8* | WBGene00001451 | protein_coding_gene | 1.1 | 2.18 | 3.7E-04 |
| *trpl-1* | WBGene00018992 | pseudogene | 1.1 | 2.18 | 4.8E-02 |
| *flp-26* | WBGene00020126 | protein_coding_gene | 1.1 | 2.18 | 3.2E-03 |
| *lipl-4* | WBGene00019376 | protein_coding_gene | 1.1 | 2.18 | 3.7E-02 |
| *trp-1* | WBGene00006614 | protein_coding_gene | 1.1 | 2.17 | 2.9E-02 |
| *bam-2* | WBGene00000237 | protein_coding_gene | 1.1 | 2.17 | 1.4E-04 |
| *W02D7.4* | WBGene00020940 | protein_coding_gene | 1.1 | 2.17 | 1.1E-06 |
| *C05D2.11* | WBGene00022910 | ncRNA_gene | 1.1 | 2.17 | 1.5E-03 |
| *C35E7.2* | WBGene00016454 | protein_coding_gene | 1.1 | 2.17 | 3.4E-03 |
| *Y87G2A.19* | WBGene00044260 | protein_coding_gene | 1.1 | 2.17 | 3.9E-03 |
| *sup-12* | WBGene00006321 | protein_coding_gene | 1.1 | 2.17 | 2.6E-03 |
| *snet-1* | WBGene00015354 | protein_coding_gene | 1.1 | 2.17 | 2.1E-03 |

| | | | | | |
|---|---|---|---|---|---|
| *lin-11* | WBGene00003000 | protein_coding_gene | 1.1 | 2.17 | 1.7E-02 |
| *nlp-15* | WBGene00003753 | protein_coding_gene | 1.1 | 2.16 | 2.3E-03 |
| *R06C1.6* | WBGene00011060 | protein_coding_gene | 1.1 | 2.16 | 1.1E-03 |
| *C29F7.2* | WBGene00007811 | protein_coding_gene | 1.1 | 2.16 | 9.6E-03 |
| *arrd-4* | WBGene00014159 | protein_coding_gene | 1.1 | 2.16 | 1.2E-02 |
| *che-13* | WBGene00000492 | protein_coding_gene | 1.1 | 2.16 | 3.9E-02 |
| *nspc-20* | WBGene00008356 | protein_coding_gene | 1.1 | 2.16 | 1.2E-03 |
| *C47E12.9* | WBGene00008152 | protein_coding_gene | 1.1 | 2.16 | 2.8E-02 |
| *oig-3* | WBGene00003861 | protein_coding_gene | 1.1 | 2.16 | 1.6E-04 |
| *F53A2.1* | WBGene00009949 | protein_coding_gene | 1.1 | 2.16 | 3.8E-02 |
| *F54D5.3* | WBGene00010049 | protein_coding_gene | 1.1 | 2.16 | 3.9E-14 |
| *mig-13* | WBGene00003245 | protein_coding_gene | 1.1 | 2.16 | 5.7E-04 |
| *nlp-68* | WBGene00011899 | protein_coding_gene | 1.1 | 2.15 | 2.9E-03 |
| *cest-17* | WBGene00019477 | protein_coding_gene | 1.1 | 2.15 | 5.9E-05 |
| *nhr-88* | WBGene00003678 | protein_coding_gene | 1.1 | 2.15 | 9.3E-03 |
| *T19E7.6* | WBGene00044500 | protein_coding_gene | 1.1 | 2.15 | 6.5E-03 |
| *tnc-2* | WBGene00006583 | protein_coding_gene | 1.1 | 2.15 | 3.8E-04 |
| *Y62F5A.9* | WBGene00013387 | protein_coding_gene | 1.1 | 2.15 | 2.3E-02 |
| *hlh-14* | WBGene00001958 | protein_coding_gene | 1.1 | 2.15 | 1.1E-02 |
| *tag-196* | WBGene00007055 | protein_coding_gene | 1.1 | 2.15 | 5.5E-04 |
| *nhr-91* | WBGene00003681 | protein_coding_gene | 1.1 | 2.15 | 7.3E-03 |
| *F37C12.10* | WBGene00018153 | protein_coding_gene | 1.1 | 2.15 | 2.1E-02 |
| *lec-11* | WBGene00002274 | protein_coding_gene | 1.1 | 2.15 | 5.7E-04 |
| *C25F9.4* | WBGene00007724 | protein_coding_gene | 1.1 | 2.15 | 2.8E-04 |
| *ZC449.4* | WBGene00022614 | protein_coding_gene | 1.1 | 2.14 | 5.9E-03 |
| *C28G1.6* | WBGene00016191 | protein_coding_gene | 1.1 | 2.14 | 7.8E-03 |
| *sqst-5* | WBGene00269434 | protein_coding_gene | 1.1 | 2.14 | 6.1E-03 |
| *F26G1.2* | WBGene00017838 | protein_coding_gene | 1.1 | 2.14 | 1.2E-04 |
| *lim-6* | WBGene00002988 | protein_coding_gene | 1.1 | 2.14 | 1.8E-03 |
| *T10B11.5* | WBGene00020401 | protein_coding_gene | 1.1 | 2.14 | 8.4E-04 |
| *cng-1* | WBGene00000562 | protein_coding_gene | 1.1 | 2.14 | 2.2E-03 |
| *F38E9.6* | WBGene00018188 | protein_coding_gene | 1.1 | 2.14 | 5.3E-03 |
| *C30G7.2* | WBGene00044176 | protein_coding_gene | 1.1 | 2.14 | 3.2E-03 |
| *fbxa-175* | WBGene00045293 | pseudogene | 1.1 | 2.14 | 2.5E-02 |
| *nmtn-1* | WBGene00019434 | protein_coding_gene | 1.1 | 2.13 | 1.6E-03 |
| *Y62F5A.10* | WBGene00013388 | protein_coding_gene | 1.1 | 2.13 | 2.8E-02 |
| *ZK1290.13* | WBGene00022893 | protein_coding_gene | 1.1 | 2.13 | 6.4E-04 |
| *vab-23* | WBGene00014153 | protein_coding_gene | 1.1 | 2.13 | 3.5E-02 |
| *nhr-32* | WBGene00003626 | protein_coding_gene | 1.1 | 2.13 | 3.4E-04 |
| *F25E5.4* | WBGene00017785 | protein_coding_gene | 1.1 | 2.13 | 3.4E-03 |
| *cest-3* | WBGene00008451 | protein_coding_gene | 1.1 | 2.13 | 4.0E-06 |
| *F23F1.6* | WBGene00017747 | protein_coding_gene | 1.1 | 2.13 | 9.1E-03 |
| *R02F11.1* | WBGene00019839 | protein_coding_gene | 1.1 | 2.13 | 2.4E-02 |
| *F39E9.28* | WBGene00271716 | pseudogene | 1.1 | 2.13 | 2.4E-03 |
| *cdh-12* | WBGene00022103 | protein_coding_gene | 1.1 | 2.13 | 2.6E-02 |
| *clec-54* | WBGene00008596 | protein_coding_gene | 1.1 | 2.12 | 9.3E-05 |
| *ptr-6* | WBGene00004221 | protein_coding_gene | 1.1 | 2.12 | 3.4E-04 |
| *F08A7.1* | WBGene00045013 | protein_coding_gene | 1.1 | 2.12 | 2.0E-02 |
| *kcc-2* | WBGene00019205 | protein_coding_gene | 1.1 | 2.12 | 7.7E-05 |
| *C23G10.14* | WBGene00271647 | ncRNA_gene | 1.1 | 2.12 | 4.3E-02 |
| *C46H11.10* | WBGene00016734 | protein_coding_gene | 1.1 | 2.12 | 1.8E-02 |
| *F52B10.3* | WBGene00044304 | protein_coding_gene | 1.1 | 2.12 | 1.0E-02 |
| *F21D12.3* | WBGene00017663 | protein_coding_gene | 1.1 | 2.12 | 1.0E-02 |
| *flp-22* | WBGene00009562 | protein_coding_gene | 1.1 | 2.12 | 2.3E-05 |
| *egl-21* | WBGene00001189 | protein_coding_gene | 1.1 | 2.11 | 1.1E-06 |
| *snt-4* | WBGene00004924 | protein_coding_gene | 1.1 | 2.11 | 8.0E-04 |
| *skr-21* | WBGene00004827 | protein_coding_gene | 1.1 | 2.11 | 1.1E-03 |
| *klo-1* | WBGene00016848 | protein_coding_gene | 1.1 | 2.11 | 1.0E-11 |

| | | | | | |
|---|---|---|---|---|---|
| *unc-24* | WBGene00006761 | protein_coding_gene | 1.1 | 2.11 | 4.5E-02 |
| *F40D4.16* | WBGene00194681 | ncRNA_gene | 1.1 | 2.11 | 2.4E-02 |
| *marc-5* | WBGene00013273 | protein_coding_gene | 1.1 | 2.11 | 6.2E-03 |
| *fbxa-55* | WBGene00020453 | protein_coding_gene | 1.1 | 2.11 | 2.4E-03 |
| *ttr-26* | WBGene00013079 | protein_coding_gene | 1.1 | 2.10 | 2.0E-02 |
| *F56H11.2* | WBGene00010175 | protein_coding_gene | 1.1 | 2.10 | 2.1E-02 |
| *mab-21* | WBGene00003112 | protein_coding_gene | 1.1 | 2.10 | 6.5E-03 |
| *F13B12.3* | WBGene00008731 | protein_coding_gene | 1.1 | 2.10 | 1.1E-03 |
| *gpa-3* | WBGene00001665 | protein_coding_gene | 1.1 | 2.10 | 3.8E-02 |
| *nhr-6* | WBGene00003605 | protein_coding_gene | 1.1 | 2.10 | 1.2E-02 |
| *dop-5* | WBGene00011382 | protein_coding_gene | 1.1 | 2.10 | 1.4E-02 |
| *Y48C3A.5* | WBGene00012989 | protein_coding_gene | 1.1 | 2.10 | 3.8E-04 |
| *C11H1.7* | WBGene00007531 | protein_coding_gene | 1.1 | 2.10 | 4.1E-02 |
| *R10H1.1* | WBGene00019998 | protein_coding_gene | 1.1 | 2.10 | 6.9E-04 |
| *dhs-2* | WBGene00000966 | protein_coding_gene | 1.1 | 2.10 | 1.3E-02 |
| *T05H4.15* | WBGene00020276 | protein_coding_gene | 1.1 | 2.09 | 9.6E-03 |
| *T05E11.8* | WBGene00011484 | protein_coding_gene | 1.1 | 2.09 | 8.1E-03 |
| *nlr-1* | WBGene00003772 | protein_coding_gene | 1.1 | 2.09 | 8.4E-05 |
| *odr-3* | WBGene00003850 | protein_coding_gene | 1.1 | 2.09 | 8.9E-03 |
| *F40F12.9* | WBGene00043995 | protein_coding_gene | 1.1 | 2.09 | 1.4E-02 |
| *nlp-73* | WBGene00013964 | protein_coding_gene | 1.1 | 2.09 | 1.8E-03 |
| *ins-22* | WBGene00002105 | protein_coding_gene | 1.1 | 2.09 | 3.8E-02 |
| *syg-1* | WBGene00006365 | protein_coding_gene | 1.1 | 2.09 | 8.3E-03 |
| *F13C5.5* | WBGene00017426 | protein_coding_gene | 1.1 | 2.09 | 9.0E-03 |
| *D1044.7* | WBGene00017032 | protein_coding_gene | 1.1 | 2.09 | 3.4E-02 |
| *pals-23* | WBGene00016218 | protein_coding_gene | 1.1 | 2.09 | 2.0E-02 |
| *C50E3.6* | WBGene00016817 | protein_coding_gene | 1.1 | 2.08 | 8.8E-03 |
| *sek-6* | WBGene00012162 | protein_coding_gene | 1.1 | 2.08 | 7.1E-03 |
| *B0252.8* | WBGene00044615 | protein_coding_gene | 1.1 | 2.08 | 1.4E-02 |
| *F53E10.1* | WBGene00018758 | protein_coding_gene | 1.1 | 2.08 | 1.0E-06 |
| *madd-2* | WBGene00016539 | protein_coding_gene | 1.1 | 2.08 | 4.0E-05 |
| *mab-5* | WBGene00003102 | protein_coding_gene | 1.1 | 2.08 | 2.0E-02 |
| *F15E6.9* | WBGene00017489 | protein_coding_gene | 1.1 | 2.08 | 1.0E-02 |
| *ttr-10* | WBGene00017600 | protein_coding_gene | 1.1 | 2.08 | 9.1E-05 |
| *syd-1* | WBGene00006363 | protein_coding_gene | 1.1 | 2.08 | 3.2E-02 |
| *egl-36* | WBGene00001202 | protein_coding_gene | 1.1 | 2.08 | 4.3E-03 |
| *C25G4.2* | WBGene00007730 | protein_coding_gene | 1.1 | 2.08 | 5.7E-05 |
| *vamp-8* | WBGene00007200 | protein_coding_gene | 1.1 | 2.08 | 3.9E-04 |
| *T10H9.8* | WBGene00044779 | protein_coding_gene | 1.1 | 2.07 | 1.4E-02 |
| *F21A9.2* | WBGene00017651 | protein_coding_gene | 1.1 | 2.07 | 1.5E-02 |
| *F53A9.6* | WBGene00018729 | protein_coding_gene | 1.1 | 2.07 | 3.0E-02 |
| *W03G11.4* | WBGene00012226 | protein_coding_gene | 1.1 | 2.07 | 3.2E-03 |
| *hst-3.1* | WBGene00002030 | protein_coding_gene | 1.1 | 2.07 | 2.1E-02 |
| *pgal-1* | WBGene00017671 | protein_coding_gene | 1.1 | 2.07 | 2.4E-05 |
| *ncx-2* | WBGene00003567 | protein_coding_gene | 1.1 | 2.07 | 6.8E-03 |
| *ugt-24* | WBGene00016762 | protein_coding_gene | 1.0 | 2.07 | 8.7E-03 |
| *srj-14* | WBGene00005602 | protein_coding_gene | 1.0 | 2.07 | 2.7E-02 |
| *R07G3.8* | WBGene00019944 | protein_coding_gene | 1.0 | 2.07 | 1.6E-03 |
| *F28E10.5* | WBGene00044302 | pseudogene | 1.0 | 2.07 | 4.9E-02 |
| *M116.2* | WBGene00019789 | pseudogene | 1.0 | 2.07 | 1.1E-05 |
| *dgk-1* | WBGene00000958 | protein_coding_gene | 1.0 | 2.07 | 7.4E-03 |
| *B0403.6* | WBGene00043982 | protein_coding_gene | 1.0 | 2.06 | 2.0E-02 |
| *nlp-6* | WBGene00003744 | protein_coding_gene | 1.0 | 2.06 | 1.4E-05 |
| *F59C6.8* | WBGene00010327 | protein_coding_gene | 1.0 | 2.06 | 3.4E-02 |
| *F38B6.6* | WBGene00018175 | protein_coding_gene | 1.0 | 2.06 | 1.7E-02 |
| *jnk-1* | WBGene00002178 | protein_coding_gene | 1.0 | 2.06 | 8.4E-04 |
| *Y22D7AL.15* | WBGene00021253 | protein_coding_gene | 1.0 | 2.06 | 1.4E-03 |
| *hlh-30* | WBGene00020930 | protein_coding_gene | 1.0 | 2.06 | 1.8E-02 |

| | | | | | |
|---|---|---|---|---|---|
| *glt-6* | WBGene00001624 | protein_coding_gene | 1.0 | 2.06 | 3.8E-03 |
| *K07C5.9* | WBGene00010632 | protein_coding_gene | 1.0 | 2.06 | 1.9E-02 |
| *his-71* | WBGene00001945 | protein_coding_gene | 1.0 | 2.06 | 2.1E-04 |
| *ges-1* | WBGene00001578 | protein_coding_gene | 1.0 | 2.05 | 1.4E-04 |
| *lec-7* | WBGene00002270 | protein_coding_gene | 1.0 | 2.05 | 1.2E-02 |
| *sem-4* | WBGene00004773 | protein_coding_gene | 1.0 | 2.05 | 2.2E-03 |
| *fbxa-6* | WBGene00008352 | protein_coding_gene | 1.0 | 2.05 | 2.8E-03 |
| *lron-15* | WBGene00016974 | protein_coding_gene | 1.0 | 2.05 | 1.9E-02 |
| *C53C11.5* | WBGene00044770 | protein_coding_gene | 1.0 | 2.05 | 1.5E-03 |
| *R04B5.5* | WBGene00011003 | protein_coding_gene | 1.0 | 2.05 | 1.0E-02 |
| *fmil-1* | WBGene00021362 | protein_coding_gene | 1.0 | 2.05 | 9.0E-03 |
| *fat-4* | WBGene00001396 | protein_coding_gene | 1.0 | 2.05 | 2.4E-07 |
| *aff-1* | WBGene00016625 | protein_coding_gene | 1.0 | 2.05 | 2.0E-02 |
| *zip-2* | WBGene00019327 | protein_coding_gene | 1.0 | 2.04 | 2.4E-03 |
| *crtc-1* | WBGene00021237 | protein_coding_gene | 1.0 | 2.04 | 9.3E-03 |
| *MTCE.7* | WBGene00014454 | rRNA_gene | 1.0 | 2.04 | 6.0E-03 |
| *F56D5.6* | WBGene00010150 | protein_coding_gene | 1.0 | 2.04 | 6.5E-04 |
| *sol-1* | WBGene00004944 | protein_coding_gene | 1.0 | 2.04 | 3.6E-02 |
| *syg-2* | WBGene00007750 | protein_coding_gene | 1.0 | 2.04 | 5.9E-03 |
| *R09A1.3* | WBGene00019973 | protein_coding_gene | 1.0 | 2.04 | 1.2E-04 |
| *D1007.18* | WBGene00044421 | protein_coding_gene | 1.0 | 2.04 | 3.8E-02 |
| *lgx-1* | WBGene00002983 | protein_coding_gene | 1.0 | 2.04 | 7.8E-03 |
| *K02A6.3* | WBGene00019294 | protein_coding_gene | 1.0 | 2.04 | 1.0E-02 |
| *F58A6.1* | WBGene00019022 | protein_coding_gene | 1.0 | 2.04 | 8.7E-03 |
| *F08D12.3* | WBGene00017247 | protein_coding_gene | 1.0 | 2.04 | 2.3E-02 |
| *elt-6* | WBGene00001253 | protein_coding_gene | 1.0 | 2.04 | 4.9E-03 |
| *T23F4.2* | WBGene00020740 | protein_coding_gene | 1.0 | 2.04 | 2.4E-02 |
| *lea-1* | WBGene00002263 | protein_coding_gene | 1.0 | 2.04 | 3.2E-04 |
| *abts-4* | WBGene00019844 | protein_coding_gene | 1.0 | 2.04 | 4.1E-05 |
| *exc-13* | WBGene00015393 | protein_coding_gene | 1.0 | 2.04 | 8.5E-03 |
| *cbp-2* | WBGene00010545 | protein_coding_gene | 1.0 | 2.04 | 9.5E-03 |
| *Y37H9A.5* | WBGene00014871 | protein_coding_gene | 1.0 | 2.03 | 4.0E-02 |
| *T09B4.5* | WBGene00020379 | protein_coding_gene | 1.0 | 2.03 | 1.0E-02 |
| *Y34F4.5* | WBGene00021340 | protein_coding_gene | 1.0 | 2.03 | 1.3E-05 |
| *max-1* | WBGene00003143 | protein_coding_gene | 1.0 | 2.03 | 1.3E-03 |
| *C05D12.2* | WBGene00007340 | protein_coding_gene | 1.0 | 2.03 | 2.3E-04 |
| *hsp-12.2* | WBGene00002011 | protein_coding_gene | 1.0 | 2.03 | 1.0E-03 |
| *cutl-20* | WBGene00018787 | protein_coding_gene | 1.0 | 2.03 | 1.0E-02 |
| *kal-1* | WBGene00002181 | protein_coding_gene | 1.0 | 2.03 | 4.7E-04 |
| *glr-2* | WBGene00001613 | protein_coding_gene | 1.0 | 2.03 | 2.5E-02 |
| *akt-2* | WBGene00000103 | protein_coding_gene | 1.0 | 2.03 | 2.4E-03 |
| *K02E11.5* | WBGene00010514 | protein_coding_gene | 1.0 | 2.03 | 2.4E-03 |
| *F31C3.6* | WBGene00009288 | protein_coding_gene | 1.0 | 2.03 | 7.4E-03 |
| *W10C8.4* | WBGene00021127 | protein_coding_gene | 1.0 | 2.03 | 4.2E-02 |
| *nspc-10* | WBGene00009640 | protein_coding_gene | 1.0 | 2.03 | 5.4E-04 |
| *exc-5* | WBGene00001366 | protein_coding_gene | 1.0 | 2.03 | 1.9E-03 |
| *C01C4.3* | WBGene00015293 | protein_coding_gene | 1.0 | 2.03 | 2.9E-02 |
| *flp-17* | WBGene00001460 | protein_coding_gene | 1.0 | 2.03 | 8.5E-03 |
| *catp-7* | WBGene00022010 | protein_coding_gene | 1.0 | 2.02 | 8.3E-03 |
| *T25D10.4* | WBGene00020801 | protein_coding_gene | 1.0 | 2.02 | 8.6E-06 |
| *ets-9* | WBGene00016865 | protein_coding_gene | 1.0 | 2.02 | 1.4E-03 |
| *T04A6.1* | WBGene00020200 | protein_coding_gene | 1.0 | 2.02 | 2.7E-04 |
| *sym-2* | WBGene00006367 | protein_coding_gene | 1.0 | 2.02 | 4.7E-03 |
| *pll-1* | WBGene00004045 | protein_coding_gene | 1.0 | 2.02 | 2.5E-02 |
| *dep-1* | WBGene00009717 | protein_coding_gene | 1.0 | 2.02 | 2.7E-05 |
| *DH11.2* | WBGene00008436 | protein_coding_gene | 1.0 | 2.01 | 3.8E-03 |
| *C13C4.6* | WBGene00007550 | protein_coding_gene | 1.0 | 2.01 | 2.8E-02 |
| *sto-1* | WBGene00006063 | protein_coding_gene | 1.0 | 2.01 | 1.4E-03 |

| | | | | | |
|---|---|---|---|---|---|
| *Y73F8A.5* | WBGene00013518 | protein_coding_gene | 1.0 | 2.01 | 1.6E-02 |
| *R09A8.5* | WBGene00011154 | protein_coding_gene | 1.0 | 2.01 | 1.2E-02 |
| *C18G1.1* | WBGene00015980 | protein_coding_gene | 1.0 | 2.01 | 5.0E-02 |
| *lgc-50* | WBGene00020605 | protein_coding_gene | 1.0 | 2.01 | 1.9E-02 |
| *Y43F8B.24* | WBGene00195176 | protein_coding_gene | 1.0 | 2.01 | 3.8E-02 |
| *grdn-1* | WBGene00013082 | protein_coding_gene | 1.0 | 2.01 | 1.7E-03 |
| *R11G1.2* | WBGene00020011 | protein_coding_gene | 1.0 | 2.01 | 4.2E-02 |
| *tos-1* | WBGene00019466 | protein_coding_gene | 1.0 | 2.00 | 9.0E-03 |
| *ddo-2* | WBGene00017565 | protein_coding_gene | 1.0 | 2.00 | 3.2E-04 |
| *F32D1.3* | WBGene00017983 | protein_coding_gene | 1.0 | 2.00 | 3.5E-02 |
| *lam-3* | WBGene00002248 | protein_coding_gene | 1.0 | 2.00 | 5.2E-03 |
| *mnp-1* | WBGene00007139 | protein_coding_gene | 1.0 | 2.00 | 8.1E-04 |
| *gst-39* | WBGene00001787 | protein_coding_gene | -1.0 | 0.50 | 1.1E-02 |
| *tyr-2* | WBGene00010661 | protein_coding_gene | -1.0 | 0.50 | 1.6E-02 |
| *ifet-1* | WBGene00004132 | protein_coding_gene | -1.0 | 0.50 | 2.1E-03 |
| *F49D11.10* | WBGene00045433 | protein_coding_gene | -1.0 | 0.50 | 3.0E-05 |
| *Y49E10.29* | WBGene00045239 | protein_coding_gene | -1.0 | 0.50 | 6.2E-04 |
| *ints-11* | WBGene00008642 | protein_coding_gene | -1.0 | 0.50 | 3.3E-13 |
| *catp-6* | WBGene00012341 | protein_coding_gene | -1.0 | 0.50 | 3.6E-05 |
| *mrp-4* | WBGene00003410 | protein_coding_gene | -1.0 | 0.50 | 9.4E-04 |
| *cct-5* | WBGene00000380 | protein_coding_gene | -1.0 | 0.50 | 2.7E-05 |
| *cct-3* | WBGene00018782 | protein_coding_gene | -1.0 | 0.49 | 5.6E-08 |
| *gpd-1* | WBGene00001683 | protein_coding_gene | -1.0 | 0.49 | 2.1E-05 |
| *C52B11.5* | WBGene00016874 | protein_coding_gene | -1.0 | 0.49 | 6.3E-04 |
| *C48B4.9* | WBGene00008172 | protein_coding_gene | -1.0 | 0.49 | 5.8E-09 |
| *nit-1* | WBGene00014206 | protein_coding_gene | -1.0 | 0.49 | 1.2E-03 |
| *cest-4* | WBGene00015930 | protein_coding_gene | -1.0 | 0.49 | 1.8E-04 |
| *aspm-1* | WBGene00008107 | protein_coding_gene | -1.0 | 0.49 | 2.4E-09 |
| *pas-2* | WBGene00003923 | protein_coding_gene | -1.0 | 0.49 | 5.9E-05 |
| *T04D3.1* | WBGene00011431 | protein_coding_gene | -1.0 | 0.49 | 1.8E-03 |
| *Y116F11B.10* | WBGene00013826 | pseudogene | -1.0 | 0.49 | 7.0E-03 |
| *W02H5.8* | WBGene00020962 | protein_coding_gene | -1.0 | 0.49 | 3.0E-04 |
| *sptl-3* | WBGene00011932 | protein_coding_gene | -1.0 | 0.49 | 3.6E-04 |
| *mcm-3* | WBGene00003155 | protein_coding_gene | -1.0 | 0.49 | 4.1E-05 |
| *fah-1* | WBGene00019620 | protein_coding_gene | -1.0 | 0.49 | 3.0E-04 |
| *rft-1* | WBGene00044637 | protein_coding_gene | -1.0 | 0.49 | 3.1E-02 |
| *F31A3.3* | WBGene00017938 | protein_coding_gene | -1.0 | 0.49 | 3.0E-02 |
| *duo-1* | WBGene00001110 | protein_coding_gene | -1.0 | 0.49 | 7.5E-07 |
| *Y73B6BL.31* | WBGene00022252 | protein_coding_gene | -1.0 | 0.49 | 3.9E-04 |
| *C29F7.3* | WBGene00007812 | protein_coding_gene | -1.0 | 0.49 | 1.3E-04 |
| *acl-1* | WBGene00010339 | protein_coding_gene | -1.0 | 0.49 | 1.7E-06 |
| *M04B2.4* | WBGene00010847 | protein_coding_gene | -1.0 | 0.49 | 3.9E-10 |
| *brd-1* | WBGene00000265 | protein_coding_gene | -1.0 | 0.49 | 1.1E-09 |
| *Y49E10.37* | WBGene00304219 | ncRNA_gene | -1.0 | 0.49 | 2.0E-03 |
| *mcm-6* | WBGene00003158 | protein_coding_gene | -1.0 | 0.49 | 1.1E-05 |
| *Y48G1A.7* | WBGene00235385 | pseudogene | -1.0 | 0.49 | 6.3E-03 |
| *sucl-2* | WBGene00017759 | protein_coding_gene | -1.0 | 0.49 | 5.5E-06 |
| *elpc-1* | WBGene00022463 | protein_coding_gene | -1.0 | 0.49 | 3.5E-17 |
| *twk-42* | WBGene00006692 | protein_coding_gene | -1.0 | 0.49 | 3.4E-03 |
| *C48D1.5* | WBGene00008178 | protein_coding_gene | -1.0 | 0.49 | 6.7E-03 |
| *F14H3.5* | WBGene00008824 | protein_coding_gene | -1.0 | 0.49 | 1.3E-04 |
| *sdz-30* | WBGene00011432 | protein_coding_gene | -1.0 | 0.49 | 4.0E-02 |
| *Y54G11A.3* | WBGene00013214 | protein_coding_gene | -1.0 | 0.48 | 4.3E-13 |
| *spg-7* | WBGene00004978 | protein_coding_gene | -1.0 | 0.48 | 1.1E-14 |
| *dnj-28* | WBGene00001046 | protein_coding_gene | -1.1 | 0.48 | 3.3E-09 |
| *D1014.2* | WBGene00017015 | protein_coding_gene | -1.1 | 0.48 | 1.1E-02 |
| *C44B7.7* | WBGene00016629 | protein_coding_gene | -1.1 | 0.48 | 1.2E-02 |
| *Y57A10A.1* | WBGene00013246 | protein_coding_gene | -1.1 | 0.48 | 1.4E-06 |

| | | | | | |
|---|---|---|---|---|---|
| *sec-61.A* | WBGene00013311 | protein_coding_gene | -1.1 | 0.48 | 1.6E-04 |
| *C34C12.8* | WBGene00007927 | protein_coding_gene | -1.1 | 0.48 | 3.3E-09 |
| *atfs-1* | WBGene00013878 | protein_coding_gene | -1.1 | 0.48 | 2.5E-13 |
| *nlp-77* | WBGene00015514 | protein_coding_gene | -1.1 | 0.48 | 3.0E-03 |
| *chs-1* | WBGene00000496 | protein_coding_gene | -1.1 | 0.48 | 2.1E-05 |
| *F27B3.6* | WBGene00017848 | protein_coding_gene | -1.1 | 0.48 | 3.5E-02 |
| *haly-1* | WBGene00009813 | protein_coding_gene | -1.1 | 0.48 | 3.2E-04 |
| *cima-1* | WBGene00018473 | protein_coding_gene | -1.1 | 0.48 | 1.3E-02 |
| *C10C5.5* | WBGene00007509 | protein_coding_gene | -1.1 | 0.48 | 1.5E-02 |
| *Y71H10B.1* | WBGene00022201 | protein_coding_gene | -1.1 | 0.48 | 4.5E-05 |
| *phb-1* | WBGene00004014 | protein_coding_gene | -1.1 | 0.48 | 3.4E-12 |
| *T06D4.1* | WBGene00020290 | protein_coding_gene | -1.1 | 0.48 | 4.4E-10 |
| *rod-1* | WBGene00018900 | protein_coding_gene | -1.1 | 0.48 | 6.2E-07 |
| *Y39E4B.5* | WBGene00012716 | protein_coding_gene | -1.1 | 0.48 | 6.5E-04 |
| *mrpl-24* | WBGene00019076 | protein_coding_gene | -1.1 | 0.48 | 6.3E-06 |
| *gpa-4* | WBGene00001666 | protein_coding_gene | -1.1 | 0.48 | 4.9E-02 |
| *sdz-33* | WBGene00013233 | protein_coding_gene | -1.1 | 0.48 | 1.8E-02 |
| *ugt-28* | WBGene00015693 | protein_coding_gene | -1.1 | 0.48 | 3.3E-06 |
| *C06A1.12* | WBGene00219803 | ncRNA_gene | -1.1 | 0.48 | 2.2E-03 |
| *rmd-1* | WBGene00011501 | protein_coding_gene | -1.1 | 0.48 | 6.0E-04 |
| *cra-1* | WBGene00020068 | protein_coding_gene | -1.1 | 0.48 | 2.0E-07 |
| *linc-4* | WBGene00219306 | lincRNA_gene | -1.1 | 0.48 | 5.3E-03 |
| *H05L14.2* | WBGene00010367 | protein_coding_gene | -1.1 | 0.47 | 2.6E-05 |
| *C10C6.7* | WBGene00007515 | protein_coding_gene | -1.1 | 0.47 | 1.9E-02 |
| *gst-20* | WBGene00001768 | protein_coding_gene | -1.1 | 0.47 | 2.0E-03 |
| *sgt-1* | WBGene00019893 | protein_coding_gene | -1.1 | 0.47 | 3.1E-09 |
| *gei-14* | WBGene00001571 | protein_coding_gene | -1.1 | 0.47 | 6.0E-09 |
| *T28A11.2* | WBGene00020869 | protein_coding_gene | -1.1 | 0.47 | 2.0E-05 |
| *tofu-4* | WBGene00019013 | protein_coding_gene | -1.1 | 0.47 | 2.3E-07 |
| *R03D7.2* | WBGene00010989 | protein_coding_gene | -1.1 | 0.47 | 7.7E-06 |
| *imb-1* | WBGene00002075 | protein_coding_gene | -1.1 | 0.47 | 2.9E-04 |
| *Y54E10A.17* | WBGene00021835 | protein_coding_gene | -1.1 | 0.47 | 6.0E-03 |
| *B0391.10* | WBGene00007165 | pseudogene | -1.1 | 0.47 | 2.6E-04 |
| *Y50D4B.2* | WBGene00021741 | protein_coding_gene | -1.1 | 0.47 | 6.3E-03 |
| *cept-1* | WBGene00009057 | protein_coding_gene | -1.1 | 0.47 | 1.5E-02 |
| *C14C11.2* | WBGene00015766 | protein_coding_gene | -1.1 | 0.47 | 6.3E-09 |
| *rpn-1* | WBGene00004458 | protein_coding_gene | -1.1 | 0.47 | 8.0E-07 |
| *cogc-2* | WBGene00000585 | protein_coding_gene | -1.1 | 0.47 | 4.2E-11 |
| *lin-41* | WBGene00003026 | protein_coding_gene | -1.1 | 0.47 | 1.7E-03 |
| *F59A6.5* | WBGene00019087 | protein_coding_gene | -1.1 | 0.47 | 2.9E-08 |
| *F52B11.2* | WBGene00009925 | protein_coding_gene | -1.1 | 0.47 | 3.3E-04 |
| *C46E10.8* | WBGene00016712 | protein_coding_gene | -1.1 | 0.47 | 5.4E-03 |
| *eif-3.C* | WBGene00001226 | protein_coding_gene | -1.1 | 0.47 | 8.7E-12 |
| *athp-3* | WBGene00013799 | protein_coding_gene | -1.1 | 0.47 | 3.2E-03 |
| *ZC434.8* | WBGene00013894 | protein_coding_gene | -1.1 | 0.47 | 8.4E-04 |
| *F44E7.4* | WBGene00018426 | protein_coding_gene | -1.1 | 0.47 | 6.7E-13 |
| *heri-1* | WBGene00016236 | protein_coding_gene | -1.1 | 0.47 | 1.1E-12 |
| *F28H7.8* | WBGene00009241 | protein_coding_gene | -1.1 | 0.47 | 1.0E-02 |
| *K12H4.7* | WBGene00019682 | protein_coding_gene | -1.1 | 0.47 | 1.9E-06 |
| *C31G12.1* | WBGene00007850 | protein_coding_gene | -1.1 | 0.47 | 1.9E-04 |
| *F14H3.4* | WBGene00008823 | protein_coding_gene | -1.1 | 0.46 | 3.9E-07 |
| *sti-1* | WBGene00019983 | protein_coding_gene | -1.1 | 0.46 | 2.3E-08 |
| *oig-2* | WBGene00003860 | protein_coding_gene | -1.1 | 0.46 | 4.2E-03 |
| *fitm-2* | WBGene00044094 | protein_coding_gene | -1.1 | 0.46 | 2.6E-03 |
| *T05E12.10* | WBGene00077787 | ncRNA_gene | -1.1 | 0.46 | 4.0E-02 |
| *hsp-6* | WBGene00002010 | protein_coding_gene | -1.1 | 0.46 | 2.1E-06 |
| *K01C8.1* | WBGene00010456 | protein_coding_gene | -1.1 | 0.46 | 6.5E-04 |
| *ets-7* | WBGene00017601 | protein_coding_gene | -1.1 | 0.46 | 1.7E-02 |

| | | | | | |
|---|---|---|---|---|---|
| *F25G6.9* | WBGene00017800 | protein_coding_gene | -1.1 | 0.46 | 4.9E-10 |
| *F14D2.5* | WBGene00017453 | protein_coding_gene | -1.1 | 0.46 | 3.0E-02 |
| *col-38* | WBGene00000615 | protein_coding_gene | -1.1 | 0.46 | 2.0E-03 |
| *clec-265* | WBGene00019738 | protein_coding_gene | -1.1 | 0.46 | 2.1E-02 |
| *cth-2* | WBGene00022856 | protein_coding_gene | -1.1 | 0.46 | 1.4E-03 |
| *bath-14* | WBGene00015016 | pseudogene | -1.1 | 0.46 | 6.0E-03 |
| *D1044.1* | WBGene00017027 | protein_coding_gene | -1.1 | 0.46 | 1.4E-03 |
| *F18A12.7* | WBGene00017556 | protein_coding_gene | -1.1 | 0.46 | 1.1E-02 |
| *acly-1* | WBGene00016995 | protein_coding_gene | -1.1 | 0.46 | 1.2E-05 |
| *C54E4.5* | WBGene00016920 | protein_coding_gene | -1.1 | 0.46 | 8.9E-03 |
| *gfat-1* | WBGene00008546 | protein_coding_gene | -1.1 | 0.46 | 2.2E-05 |
| *har-1* | WBGene00007630 | protein_coding_gene | -1.1 | 0.46 | 2.2E-09 |
| *fbxa-153* | WBGene00007162 | protein_coding_gene | -1.1 | 0.45 | 2.4E-03 |
| *hphd-1* | WBGene00012608 | protein_coding_gene | -1.1 | 0.45 | 5.3E-04 |
| *ugt-40* | WBGene00017331 | protein_coding_gene | -1.1 | 0.45 | 9.5E-03 |
| *T27A8.2* | WBGene00012074 | protein_coding_gene | -1.1 | 0.45 | 2.0E-03 |
| *ttr-42* | WBGene00045268 | protein_coding_gene | -1.1 | 0.45 | 1.3E-02 |
| *Y48B6A.10* | WBGene00012981 | protein_coding_gene | -1.1 | 0.45 | 2.1E-02 |
| *F25B4.4* | WBGene00017767 | protein_coding_gene | -1.1 | 0.45 | 2.5E-02 |
| *Y46G5A.28* | WBGene00012916 | protein_coding_gene | -1.1 | 0.45 | 4.3E-02 |
| *clec-57* | WBGene00008594 | protein_coding_gene | -1.1 | 0.45 | 1.0E-02 |
| *T02G6.5* | WBGene00011388 | protein_coding_gene | -1.1 | 0.45 | 2.0E-05 |
| *Y75D11A.3* | WBGene00022292 | protein_coding_gene | -1.1 | 0.45 | 4.9E-05 |
| *rpt-4* | WBGene00004504 | protein_coding_gene | -1.2 | 0.45 | 1.9E-06 |
| *F12E12.6* | WBGene00017407 | protein_coding_gene | -1.2 | 0.45 | 2.0E-02 |
| *eif-2gamma* | WBGene00021466 | protein_coding_gene | -1.2 | 0.45 | 2.3E-11 |
| *grl-21* | WBGene00001730 | protein_coding_gene | -1.2 | 0.45 | 5.0E-02 |
| *F59H6.2* | WBGene00019133 | pseudogene | -1.2 | 0.45 | 4.1E-02 |
| *acox-1.3* | WBGene00008566 | protein_coding_gene | -1.2 | 0.45 | 9.1E-04 |
| *mua-3* | WBGene00003482 | protein_coding_gene | -1.2 | 0.45 | 5.7E-03 |
| *T05B11.4* | WBGene00020247 | protein_coding_gene | -1.2 | 0.45 | 3.0E-05 |
| *sprr-2* | WBGene00009629 | protein_coding_gene | -1.2 | 0.45 | 1.6E-02 |
| *fbxc-42* | WBGene00018455 | protein_coding_gene | -1.2 | 0.44 | 2.5E-06 |
| *F25E2.2* | WBGene00017780 | protein_coding_gene | -1.2 | 0.44 | 1.6E-10 |
| *bath-31* | WBGene00017405 | protein_coding_gene | -1.2 | 0.44 | 3.5E-02 |
| *F13H8.8* | WBGene00017441 | protein_coding_gene | -1.2 | 0.44 | 1.0E-03 |
| *F58G11.3* | WBGene00010281 | protein_coding_gene | -1.2 | 0.44 | 6.7E-06 |
| *ZC53.1* | WBGene00022508 | protein_coding_gene | -1.2 | 0.44 | 1.6E-04 |
| *cct-2* | WBGene00000378 | protein_coding_gene | -1.2 | 0.44 | 7.1E-10 |
| *rme-2* | WBGene00004374 | protein_coding_gene | -1.2 | 0.44 | 1.2E-04 |
| *H34C03.2* | WBGene00019259 | protein_coding_gene | -1.2 | 0.44 | 6.5E-06 |
| *fbxb-17* | WBGene00019037 | protein_coding_gene | -1.2 | 0.44 | 1.4E-02 |
| *hsp-90* | WBGene00000915 | protein_coding_gene | -1.2 | 0.44 | 1.6E-05 |
| *npp-15* | WBGene00003801 | protein_coding_gene | -1.2 | 0.44 | 5.3E-11 |
| *cdr-2* | WBGene00008296 | protein_coding_gene | -1.2 | 0.44 | 8.4E-03 |
| *T24E12.11* | WBGene00020778 | protein_coding_gene | -1.2 | 0.44 | 1.4E-04 |
| *vwa-8* | WBGene00008694 | protein_coding_gene | -1.2 | 0.44 | 1.2E-12 |
| *R160.3* | WBGene00020120 | protein_coding_gene | -1.2 | 0.44 | 3.4E-02 |
| *mnat-1* | WBGene00018769 | protein_coding_gene | -1.2 | 0.44 | 1.4E-07 |
| *dgat-2* | WBGene00010296 | protein_coding_gene | -1.2 | 0.44 | 5.3E-03 |
| *pes-9* | WBGene00003982 | protein_coding_gene | -1.2 | 0.44 | 3.0E-04 |
| *F31D4.8* | WBGene00009293 | protein_coding_gene | -1.2 | 0.44 | 1.5E-02 |
| *hsp-1* | WBGene00002005 | protein_coding_gene | -1.2 | 0.44 | 6.5E-07 |
| *C44F1.1* | WBGene00008091 | protein_coding_gene | -1.2 | 0.44 | 2.4E-03 |
| *cdc-48.2* | WBGene00008053 | protein_coding_gene | -1.2 | 0.43 | 2.2E-10 |
| *nas-36* | WBGene00003552 | protein_coding_gene | -1.2 | 0.43 | 2.1E-02 |
| *Y47D3B.6* | WBGene00012942 | protein_coding_gene | -1.2 | 0.43 | 2.8E-02 |
| *col-17* | WBGene00000606 | protein_coding_gene | -1.2 | 0.43 | 1.8E-02 |

| | | | | | |
|---|---|---|---|---|---|
| *acbp-6* | WBGene00012457 | protein_coding_gene | -1.2 | 0.43 | 2.5E-02 |
| *Y51A2D.18* | WBGene00013083 | protein_coding_gene | -1.2 | 0.43 | 2.6E-02 |
| *T04A8.7* | WBGene00011409 | protein_coding_gene | -1.2 | 0.43 | 4.9E-07 |
| *Y53C12A.6* | WBGene00013141 | protein_coding_gene | -1.2 | 0.43 | 2.7E-03 |
| *col-12* | WBGene00000601 | protein_coding_gene | -1.2 | 0.43 | 4.8E-02 |
| *K08H2.2* | WBGene00010689 | protein_coding_gene | -1.2 | 0.43 | 2.0E-02 |
| *eif-2Bepsilon* | WBGene00008428 | protein_coding_gene | -1.2 | 0.43 | 2.7E-11 |
| *nhr-115* | WBGene00003705 | protein_coding_gene | -1.2 | 0.43 | 2.2E-03 |
| *C39E9.8* | WBGene00008032 | protein_coding_gene | -1.2 | 0.43 | 6.6E-08 |
| *col-176* | WBGene00000749 | protein_coding_gene | -1.2 | 0.43 | 1.2E-02 |
| *nmat-1* | WBGene00012295 | protein_coding_gene | -1.2 | 0.43 | 1.5E-03 |
| *T28C12.2* | WBGene00020889 | protein_coding_gene | -1.2 | 0.42 | 3.8E-02 |
| *col-124* | WBGene00000698 | protein_coding_gene | -1.2 | 0.42 | 3.4E-04 |
| *H10E21.4* | WBGene00019184 | protein_coding_gene | -1.2 | 0.42 | 3.2E-02 |
| *ech-6* | WBGene00001155 | protein_coding_gene | -1.2 | 0.42 | 1.2E-06 |
| *laf-1* | WBGene00002244 | protein_coding_gene | -1.2 | 0.42 | 4.4E-07 |
| *mcm-2* | WBGene00003154 | protein_coding_gene | -1.2 | 0.42 | 4.8E-08 |
| *col-63* | WBGene00000639 | protein_coding_gene | -1.2 | 0.42 | 1.2E-02 |
| *ugt-42* | WBGene00017959 | protein_coding_gene | -1.2 | 0.42 | 4.4E-02 |
| *acdh-9* | WBGene00017874 | protein_coding_gene | -1.2 | 0.42 | 2.3E-09 |
| *F13H8.3* | WBGene00017436 | protein_coding_gene | -1.2 | 0.42 | 4.4E-03 |
| *T10C6.7* | WBGene00011689 | protein_coding_gene | -1.2 | 0.42 | 2.3E-11 |
| *cld-9* | WBGene00009432 | protein_coding_gene | -1.2 | 0.42 | 1.6E-03 |
| *cyp-13A9* | WBGene00014838 | pseudogene | -1.3 | 0.42 | 3.2E-02 |
| *C49C3.9* | WBGene00008199 | protein_coding_gene | -1.3 | 0.42 | 1.3E-02 |
| *B0554.5* | WBGene00015258 | protein_coding_gene | -1.3 | 0.42 | 2.8E-02 |
| *ceh-82* | WBGene00018433 | protein_coding_gene | -1.3 | 0.42 | 1.0E-02 |
| *clec-97* | WBGene00013931 | protein_coding_gene | -1.3 | 0.41 | 2.2E-04 |
| *slc-17.4* | WBGene00012130 | protein_coding_gene | -1.3 | 0.41 | 7.6E-06 |
| *F53F1.2* | WBGene00009980 | protein_coding_gene | -1.3 | 0.41 | 1.6E-03 |
| *ZC449.8* | WBGene00195189 | protein_coding_gene | -1.3 | 0.41 | 2.6E-04 |
| *R02E4.3* | WBGene00044724 | protein_coding_gene | -1.3 | 0.41 | 2.2E-02 |
| *cutl-18* | WBGene00009830 | protein_coding_gene | -1.3 | 0.41 | 3.4E-02 |
| *W06D11.2* | WBGene00012300 | protein_coding_gene | -1.3 | 0.41 | 4.0E-02 |
| *F53G2.12* | WBGene00302998 | protein_coding_gene | -1.3 | 0.41 | 2.1E-03 |
| *gst-24* | WBGene00001772 | protein_coding_gene | -1.3 | 0.41 | 6.0E-04 |
| *R05G9.3* | WBGene00019905 | protein_coding_gene | -1.3 | 0.41 | 2.7E-05 |
| *T22B2.1* | WBGene00020665 | protein_coding_gene | -1.3 | 0.41 | 1.6E-03 |
| *ugt-26* | WBGene00015695 | protein_coding_gene | -1.3 | 0.41 | 1.0E-04 |
| *cyp-13A2* | WBGene00011676 | protein_coding_gene | -1.3 | 0.41 | 2.9E-03 |
| *C45B2.1* | WBGene00016658 | protein_coding_gene | -1.3 | 0.41 | 1.6E-02 |
| *mppa-1* | WBGene00022159 | protein_coding_gene | -1.3 | 0.41 | 6.2E-16 |
| *Y38E10A.3* | WBGene00012581 | protein_coding_gene | -1.3 | 0.40 | 7.4E-03 |
| *mig-21* | WBGene00003252 | protein_coding_gene | -1.3 | 0.40 | 5.3E-03 |
| *sdz-8* | WBGene00008334 | protein_coding_gene | -1.3 | 0.40 | 5.3E-04 |
| *F53H4.2* | WBGene00010007 | protein_coding_gene | -1.3 | 0.40 | 6.8E-03 |
| *Y5H2B.3* | WBGene00021165 | protein_coding_gene | -1.3 | 0.40 | 1.3E-02 |
| *sdz-16* | WBGene00009297 | protein_coding_gene | -1.3 | 0.40 | 1.7E-02 |
| *B0416.10* | WBGene00206482 | protein_coding_gene | -1.3 | 0.40 | 1.3E-02 |
| *cpi-1* | WBGene00000535 | protein_coding_gene | -1.3 | 0.40 | 3.4E-04 |
| *act-2* | WBGene00000064 | protein_coding_gene | -1.3 | 0.40 | 8.8E-07 |
| *inf-1* | WBGene00002083 | protein_coding_gene | -1.3 | 0.40 | 3.4E-10 |
| *col-110* | WBGene00000684 | protein_coding_gene | -1.3 | 0.40 | 2.5E-02 |
| *K10C2.12* | WBGene00206386 | protein_coding_gene | -1.3 | 0.40 | 4.4E-02 |
| *cyp-35C1* | WBGene00007362 | protein_coding_gene | -1.3 | 0.40 | 2.1E-04 |
| *suro-1* | WBGene00011235 | protein_coding_gene | -1.3 | 0.39 | 1.1E-02 |
| *Y55F3AM.14* | WBGene00021931 | protein_coding_gene | -1.3 | 0.39 | 4.5E-02 |
| *hsp-4* | WBGene00002008 | protein_coding_gene | -1.3 | 0.39 | 2.1E-05 |

| | | | | | |
|---|---|---|---|---|---|
| *Y47G7B.2* | WBGene00021654 | protein_coding_gene | -1.4 | 0.39 | 3.8E-03 |
| *col-14* | WBGene00000603 | protein_coding_gene | -1.4 | 0.39 | 7.0E-03 |
| *F19B2.5* | WBGene00008944 | protein_coding_gene | -1.4 | 0.39 | 1.0E-02 |
| *C25E10.10* | WBGene00016099 | protein_coding_gene | -1.4 | 0.39 | 1.6E-04 |
| *ZK218.1* | WBGene00013935 | protein_coding_gene | -1.4 | 0.39 | 4.6E-02 |
| *F54D5.12* | WBGene00010055 | protein_coding_gene | -1.4 | 0.39 | 1.8E-08 |
| *cbl-1* | WBGene00007533 | protein_coding_gene | -1.4 | 0.39 | 2.4E-03 |
| *vit-5* | WBGene00006929 | protein_coding_gene | -1.4 | 0.39 | 3.3E-03 |
| *clec-186* | WBGene00014138 | protein_coding_gene | -1.4 | 0.39 | 3.3E-04 |
| *col-8* | WBGene00000597 | protein_coding_gene | -1.4 | 0.39 | 2.0E-04 |
| *ZK180.6* | WBGene00022680 | protein_coding_gene | -1.4 | 0.39 | 2.6E-03 |
| *K08H2.3* | WBGene00010690 | protein_coding_gene | -1.4 | 0.39 | 2.1E-05 |
| *F41B4.3* | WBGene00018259 | protein_coding_gene | -1.4 | 0.39 | 2.8E-04 |
| *gpd-4* | WBGene00001686 | protein_coding_gene | -1.4 | 0.38 | 1.9E-10 |
| *msra-1* | WBGene00018393 | protein_coding_gene | -1.4 | 0.38 | 4.8E-03 |
| *col-7* | WBGene00000596 | protein_coding_gene | -1.4 | 0.38 | 3.5E-02 |
| *dot-1.5* | WBGene00022512 | protein_coding_gene | -1.4 | 0.38 | 8.0E-03 |
| *nspc-4* | WBGene00044339 | protein_coding_gene | -1.4 | 0.38 | 5.8E-04 |
| *Y39A1A.16* | WBGene00012654 | protein_coding_gene | -1.4 | 0.38 | 3.0E-02 |
| *fbxa-206* | WBGene00013610 | protein_coding_gene | -1.4 | 0.38 | 5.3E-14 |
| *W08F4.7* | WBGene00021096 | protein_coding_gene | -1.4 | 0.38 | 4.4E-02 |
| *K01D12.9* | WBGene00010468 | protein_coding_gene | -1.4 | 0.37 | 3.7E-02 |
| *miro-3* | WBGene00021651 | pseudogene | -1.4 | 0.37 | 8.8E-03 |
| *C05C10.3* | WBGene00007330 | protein_coding_gene | -1.4 | 0.37 | 6.6E-32 |
| *hsp-110* | WBGene00016250 | protein_coding_gene | -1.4 | 0.37 | 8.8E-09 |
| *C36C5.5* | WBGene00016474 | protein_coding_gene | -1.4 | 0.37 | 9.4E-08 |
| *sec-61.B* | WBGene00021427 | protein_coding_gene | -1.4 | 0.37 | 6.4E-05 |
| *dpy-13* | WBGene00001074 | protein_coding_gene | -1.4 | 0.37 | 7.5E-03 |
| *Y47G6A.33* | WBGene00045399 | protein_coding_gene | -1.4 | 0.37 | 7.1E-05 |
| *cdr-4* | WBGene00010470 | protein_coding_gene | -1.4 | 0.37 | 8.8E-04 |
| *hpd-1* | WBGene00001993 | protein_coding_gene | -1.4 | 0.37 | 4.3E-03 |
| *gst-12* | WBGene00001760 | protein_coding_gene | -1.4 | 0.37 | 1.4E-03 |
| *pho-12* | WBGene00016152 | protein_coding_gene | -1.4 | 0.37 | 1.9E-02 |
| *arl-7* | WBGene00000191 | pseudogene | -1.4 | 0.37 | 1.3E-02 |
| *T15B7.1* | WBGene00020516 | protein_coding_gene | -1.4 | 0.37 | 5.5E-06 |
| *Y47D7A.13* | WBGene00021625 | protein_coding_gene | -1.4 | 0.37 | 1.9E-02 |
| *Y119D3B.21* | WBGene00022497 | protein_coding_gene | -1.4 | 0.37 | 1.8E-04 |
| *C06A1.4* | WBGene00014667 | pseudogene | -1.4 | 0.37 | 8.6E-07 |
| *dct-18* | WBGene00010266 | protein_coding_gene | -1.4 | 0.37 | 1.0E-03 |
| *T05E12.8* | WBGene00045382 | protein_coding_gene | -1.4 | 0.37 | 2.5E-02 |
| *gale-1* | WBGene00008132 | protein_coding_gene | -1.4 | 0.37 | 4.4E-04 |
| *cuc-1* | WBGene00000835 | protein_coding_gene | -1.4 | 0.37 | 2.4E-06 |
| *pks-1* | WBGene00016558 | protein_coding_gene | -1.4 | 0.37 | 1.7E-03 |
| *C17C3.1* | WBGene00015887 | protein_coding_gene | -1.5 | 0.36 | 3.0E-14 |
| *cyp-35A5* | WBGene00019473 | protein_coding_gene | -1.5 | 0.36 | 2.6E-04 |
| *C07A9.9* | WBGene00007405 | protein_coding_gene | -1.5 | 0.36 | 5.2E-06 |
| *his-40* | WBGene00001914 | protein_coding_gene | -1.5 | 0.36 | 1.7E-02 |
| *F35E12.10* | WBGene00009434 | protein_coding_gene | -1.5 | 0.36 | 4.3E-04 |
| *Y57G11B.5* | WBGene00013297 | protein_coding_gene | -1.5 | 0.36 | 4.2E-08 |
| *wee-1.1* | WBGene00006938 | protein_coding_gene | -1.5 | 0.36 | 1.1E-02 |
| *T01D1.3* | WBGene00020148 | protein_coding_gene | -1.5 | 0.36 | 1.4E-02 |
| *gly-8* | WBGene00001633 | protein_coding_gene | -1.5 | 0.35 | 2.9E-03 |
| *F56D5.3* | WBGene00010148 | protein_coding_gene | -1.5 | 0.35 | 3.8E-05 |
| *C17H12.6* | WBGene00015932 | protein_coding_gene | -1.5 | 0.35 | 1.6E-02 |
| *C23H5.8* | WBGene00016027 | protein_coding_gene | -1.5 | 0.35 | 8.7E-05 |
| *B0212.3* | WBGene00015037 | protein_coding_gene | -1.5 | 0.35 | 3.6E-15 |
| *C33E10.8* | WBGene00016349 | protein_coding_gene | -1.5 | 0.35 | 1.2E-02 |
| *ule-1* | WBGene00021005 | protein_coding_gene | -1.5 | 0.35 | 1.6E-05 |

| | | | | | |
|---|---|---|---|---|---|
| *ZK228.3* | WBGene00013946 | protein_coding_gene | -1.5 | 0.35 | 4.1E-05 |
| *col-71* | WBGene00000647 | protein_coding_gene | -1.5 | 0.35 | 2.3E-03 |
| *ahcy-1* | WBGene00019322 | protein_coding_gene | -1.5 | 0.35 | 1.5E-07 |
| *ZK105.1* | WBGene00022653 | protein_coding_gene | -1.5 | 0.35 | 4.9E-07 |
| *Y51B9A.6* | WBGene00013088 | protein_coding_gene | -1.5 | 0.35 | 3.4E-02 |
| *F45E1.4* | WBGene00018465 | protein_coding_gene | -1.5 | 0.35 | 6.7E-04 |
| *C26B2.8* | WBGene00016130 | protein_coding_gene | -1.5 | 0.35 | 2.0E-02 |
| *R02D1.2* | WBGene00235310 | protein_coding_gene | -1.5 | 0.34 | 2.8E-03 |
| *dop-4* | WBGene00016872 | protein_coding_gene | -1.5 | 0.34 | 1.6E-03 |
| *EEED8.4* | WBGene00017135 | protein_coding_gene | -1.5 | 0.34 | 7.1E-03 |
| *sir-2.3* | WBGene00004802 | protein_coding_gene | -1.6 | 0.34 | 5.5E-06 |
| *F08F3.4* | WBGene00017262 | protein_coding_gene | -1.6 | 0.34 | 9.4E-04 |
| *sri-40* | WBGene00005552 | protein_coding_gene | -1.6 | 0.34 | 5.4E-13 |
| *pcp-4* | WBGene00003959 | protein_coding_gene | -1.6 | 0.34 | 8.0E-06 |
| *kynu-1* | WBGene00015802 | protein_coding_gene | -1.6 | 0.34 | 1.0E-03 |
| *F38C2.7* | WBGene00009539 | protein_coding_gene | -1.6 | 0.34 | 2.5E-02 |
| *K08D8.12* | WBGene00194926 | protein_coding_gene | -1.6 | 0.34 | 2.4E-03 |
| *F46F2.5* | WBGene00009789 | protein_coding_gene | -1.6 | 0.33 | 1.7E-02 |
| *tbx-38* | WBGene00006557 | protein_coding_gene | -1.6 | 0.33 | 2.7E-02 |
| *R12E2.7* | WBGene00020033 | protein_coding_gene | -1.6 | 0.33 | 2.2E-02 |
| *dpy-4* | WBGene00001066 | protein_coding_gene | -1.6 | 0.33 | 3.6E-04 |
| *Y43C5A.7* | WBGene00044922 | protein_coding_gene | -1.6 | 0.33 | 1.1E-02 |
| *col-104* | WBGene00000678 | protein_coding_gene | -1.6 | 0.33 | 3.2E-03 |
| *col-175* | WBGene00000748 | protein_coding_gene | -1.6 | 0.33 | 1.8E-03 |
| *pgp-8* | WBGene00004002 | protein_coding_gene | -1.6 | 0.33 | 4.1E-04 |
| *col-130* | WBGene00000704 | protein_coding_gene | -1.6 | 0.33 | 5.9E-03 |
| *ugt-37* | WBGene00017332 | protein_coding_gene | -1.6 | 0.33 | 4.9E-03 |
| *Y62H9A.3* | WBGene00013391 | protein_coding_gene | -1.6 | 0.33 | 3.2E-06 |
| *F17C11.13* | WBGene00023503 | protein_coding_gene | -1.6 | 0.32 | 2.8E-02 |
| *C30G12.2* | WBGene00016274 | protein_coding_gene | -1.6 | 0.32 | 2.3E-04 |
| *clu-1* | WBGene00000550 | protein_coding_gene | -1.6 | 0.32 | 3.3E-13 |
| *C50E3.12* | WBGene00016823 | protein_coding_gene | -1.6 | 0.32 | 9.1E-13 |
| *cyp-33C8* | WBGene00019967 | protein_coding_gene | -1.7 | 0.32 | 5.0E-04 |
| *Y39B6A.21* | WBGene00012680 | pseudogene | -1.7 | 0.32 | 2.1E-02 |
| *grl-16* | WBGene00001725 | protein_coding_gene | -1.7 | 0.32 | 2.1E-03 |
| *scav-5* | WBGene00011076 | protein_coding_gene | -1.7 | 0.31 | 2.1E-04 |
| *col-73* | WBGene00000649 | protein_coding_gene | -1.7 | 0.31 | 6.4E-03 |
| *col-90* | WBGene00000665 | protein_coding_gene | -1.7 | 0.31 | 7.4E-03 |
| *ttr-31* | WBGene00010225 | protein_coding_gene | -1.7 | 0.31 | 1.4E-03 |
| *nhr-114* | WBGene00003704 | protein_coding_gene | -1.7 | 0.31 | 2.9E-04 |
| *H23N18.5* | WBGene00019236 | protein_coding_gene | -1.7 | 0.31 | 1.6E-02 |
| *alh-9* | WBGene00000115 | protein_coding_gene | -1.7 | 0.31 | 1.5E-06 |
| *pgph-3* | WBGene00016892 | protein_coding_gene | -1.7 | 0.31 | 1.0E-02 |
| *hsp-60* | WBGene00002025 | protein_coding_gene | -1.7 | 0.31 | 1.9E-10 |
| *C50A2.3* | WBGene00016797 | protein_coding_gene | -1.7 | 0.31 | 1.3E-03 |
| *col-129* | WBGene00000703 | protein_coding_gene | -1.7 | 0.31 | 1.6E-03 |
| *grsp-1* | WBGene00012840 | protein_coding_gene | -1.7 | 0.31 | 3.4E-03 |
| *col-139* | WBGene00000712 | protein_coding_gene | -1.7 | 0.31 | 1.8E-02 |
| *Y60A9.2* | WBGene00013369 | pseudogene | -1.7 | 0.31 | 3.3E-02 |
| *rhy-1* | WBGene00012324 | protein_coding_gene | -1.7 | 0.31 | 9.4E-08 |
| *F23F12.12* | WBGene00017756 | protein_coding_gene | -1.7 | 0.30 | 5.1E-03 |
| *F41G3.21* | WBGene00044316 | protein_coding_gene | -1.7 | 0.30 | 3.9E-04 |
| *rol-1* | WBGene00004394 | protein_coding_gene | -1.7 | 0.30 | 4.1E-04 |
| *R03G8.6* | WBGene00011000 | protein_coding_gene | -1.7 | 0.30 | 1.2E-05 |
| *Y51H7C.1* | WBGene00021779 | protein_coding_gene | -1.7 | 0.30 | 1.6E-05 |
| *T20D4.3* | WBGene00020609 | protein_coding_gene | -1.8 | 0.30 | 5.7E-03 |
| *col-138* | WBGene00000711 | protein_coding_gene | -1.8 | 0.30 | 2.8E-04 |
| *mec-19* | WBGene00008216 | protein_coding_gene | -1.8 | 0.30 | 6.8E-03 |

| | | | | | |
|---|---|---|---|---|---|
| *Y73B6BL.14* | WBGene00022237 | protein_coding_gene | -1.8 | 0.29 | 6.2E-04 |
| *dhs-26* | WBGene00000989 | protein_coding_gene | -1.8 | 0.29 | 1.6E-04 |
| *ugt-46* | WBGene00015141 | protein_coding_gene | -1.8 | 0.29 | 1.8E-05 |
| *C42D4.3* | WBGene00016596 | protein_coding_gene | -1.8 | 0.29 | 5.8E-03 |
| *lact-1* | WBGene00018523 | protein_coding_gene | -1.8 | 0.29 | 2.5E-03 |
| *T06E4.9* | WBGene00011536 | protein_coding_gene | -1.8 | 0.29 | 3.9E-02 |
| *Y94H6A.10* | WBGene00022382 | protein_coding_gene | -1.8 | 0.29 | 1.8E-06 |
| *W04A4.2* | WBGene00012231 | protein_coding_gene | -1.8 | 0.28 | 3.2E-07 |
| *gst-16* | WBGene00001764 | protein_coding_gene | -1.8 | 0.28 | 7.0E-07 |
| *F02H6.4* | WBGene00008537 | pseudogene | -1.8 | 0.28 | 1.1E-21 |
| *col-77* | WBGene00000653 | protein_coding_gene | -1.8 | 0.28 | 1.0E-03 |
| *dod-20* | WBGene00015259 | protein_coding_gene | -1.8 | 0.28 | 4.7E-02 |
| *T22B7.7* | WBGene00020674 | protein_coding_gene | -1.9 | 0.28 | 3.3E-04 |
| *DC2.5* | WBGene00017082 | protein_coding_gene | -1.9 | 0.28 | 2.4E-06 |
| *nlp-24* | WBGene00003762 | protein_coding_gene | -1.9 | 0.28 | 3.1E-05 |
| *T25G12.13* | WBGene00219274 | protein_coding_gene | -1.9 | 0.27 | 9.5E-03 |
| *Y46G5A.29* | WBGene00012917 | protein_coding_gene | -1.9 | 0.27 | 1.2E-03 |
| *bli-6* | WBGene00000256 | protein_coding_gene | -1.9 | 0.27 | 3.2E-04 |
| *F16H6.4* | WBGene00008894 | protein_coding_gene | -1.9 | 0.27 | 1.0E-02 |
| *gst-4* | WBGene00001752 | protein_coding_gene | -1.9 | 0.27 | 1.2E-03 |
| *end-1* | WBGene00001310 | protein_coding_gene | -1.9 | 0.27 | 3.3E-03 |
| *F23D12.11* | WBGene00194674 | protein_coding_gene | -1.9 | 0.27 | 1.9E-04 |
| *wago-11* | WBGene00021711 | protein_coding_gene | -1.9 | 0.26 | 6.9E-05 |
| *hmit-1.2* | WBGene00013074 | protein_coding_gene | -1.9 | 0.26 | 1.9E-07 |
| *Y51H7C.13* | WBGene00021791 | protein_coding_gene | -1.9 | 0.26 | 2.4E-05 |
| *R04B5.6* | WBGene00011004 | protein_coding_gene | -2.0 | 0.26 | 3.2E-04 |
| *oac-20* | WBGene00009488 | protein_coding_gene | -2.0 | 0.26 | 2.1E-03 |
| *col-60* | WBGene00000636 | protein_coding_gene | -2.0 | 0.25 | 8.6E-03 |
| *col-81* | WBGene00000657 | protein_coding_gene | -2.0 | 0.25 | 5.9E-03 |
| *col-161* | WBGene00000734 | protein_coding_gene | -2.0 | 0.25 | 1.8E-02 |
| *swt-6* | WBGene00011190 | protein_coding_gene | -2.0 | 0.25 | 3.1E-05 |
| *Y39B6A.27* | WBGene00012686 | protein_coding_gene | -2.0 | 0.25 | 5.2E-06 |
| *K10C2.8* | WBGene00045265 | protein_coding_gene | -2.0 | 0.25 | 7.9E-12 |
| *C52D10.3* | WBGene00016877 | protein_coding_gene | -2.0 | 0.25 | 5.0E-03 |
| *col-152* | WBGene00000725 | protein_coding_gene | -2.0 | 0.25 | 3.2E-02 |
| *R03H10.12* | WBGene00303363 | pseudogene | -2.0 | 0.25 | 2.6E-02 |
| *Y62H9A.4* | WBGene00013392 | protein_coding_gene | -2.0 | 0.25 | 5.3E-08 |
| *C04E12.5* | WBGene00015435 | protein_coding_gene | -2.0 | 0.25 | 2.1E-02 |
| *R08E5.3* | WBGene00019963 | protein_coding_gene | -2.0 | 0.24 | 1.3E-05 |
| *clec-53* | WBGene00020191 | protein_coding_gene | -2.0 | 0.24 | 3.2E-06 |
| *K01G12.3* | WBGene00010483 | protein_coding_gene | -2.0 | 0.24 | 2.5E-13 |
| *grd-3* | WBGene00001692 | protein_coding_gene | -2.1 | 0.24 | 1.6E-08 |
| *tbb-6* | WBGene00006539 | protein_coding_gene | -2.1 | 0.24 | 2.5E-04 |
| *W06A7.4* | WBGene00012293 | protein_coding_gene | -2.1 | 0.24 | 1.6E-08 |
| *col-120* | WBGene00000694 | protein_coding_gene | -2.1 | 0.24 | 1.0E-03 |
| *linc-107* | WBGene00219655 | lincRNA_gene | -2.1 | 0.23 | 1.6E-02 |
| *ugt-62* | WBGene00010904 | protein_coding_gene | -2.1 | 0.23 | 7.5E-08 |
| *Y73B6BL.288* | WBGene00219218 | protein_coding_gene | -2.1 | 0.23 | 4.6E-10 |
| *cyp-29A3* | WBGene00021412 | protein_coding_gene | -2.1 | 0.23 | 5.0E-06 |
| *F52C6.14* | WBGene00018671 | protein_coding_gene | -2.1 | 0.23 | 3.3E-02 |
| *cyp-13A12* | WBGene00008810 | protein_coding_gene | -2.2 | 0.22 | 7.1E-03 |
| *ugt-63* | WBGene00015449 | protein_coding_gene | -2.2 | 0.22 | 1.1E-02 |
| *F58G6.7* | WBGene00010277 | protein_coding_gene | -2.2 | 0.21 | 2.9E-04 |
| *col-133* | WBGene00000707 | protein_coding_gene | -2.2 | 0.21 | 9.0E-03 |
| *F57B9.3* | WBGene00018997 | protein_coding_gene | -2.2 | 0.21 | 1.4E-02 |
| *his-38* | WBGene00001912 | protein_coding_gene | -2.2 | 0.21 | 2.2E-03 |
| *ugt-41* | WBGene00017336 | protein_coding_gene | -2.3 | 0.21 | 1.5E-13 |
| *C45B2.2* | WBGene00016659 | protein_coding_gene | -2.3 | 0.21 | 2.0E-04 |

| | | | | | |
|---|---|---|---|---|---|
| *gba-4* | WBGene00021160 | protein_coding_gene | -2.3 | 0.21 | 5.9E-06 |
| *grd-5* | WBGene00001694 | protein_coding_gene | -2.3 | 0.20 | 3.0E-07 |
| *col-101* | WBGene00000675 | protein_coding_gene | -2.3 | 0.20 | 6.8E-07 |
| *F35F10.13* | WBGene00018062 | protein_coding_gene | -2.3 | 0.20 | 3.1E-02 |
| *K10C2.7* | WBGene00019622 | protein_coding_gene | -2.3 | 0.20 | 5.5E-05 |
| *F22E5.1* | WBGene00017703 | protein_coding_gene | -2.3 | 0.20 | 1.6E-03 |
| *mth-1* | WBGene00017948 | protein_coding_gene | -2.3 | 0.20 | 8.9E-08 |
| *F53C11.1* | WBGene00009971 | protein_coding_gene | -2.3 | 0.20 | 2.6E-11 |
| *F52G3.3* | WBGene00018711 | pseudogene | -2.4 | 0.20 | 3.8E-02 |
| *acs-1* | WBGene00018488 | protein_coding_gene | -2.4 | 0.20 | 3.1E-07 |
| *ugt-30* | WBGene00011340 | protein_coding_gene | -2.4 | 0.19 | 1.4E-11 |
| *F18E3.11* | WBGene00017567 | protein_coding_gene | -2.4 | 0.19 | 1.3E-02 |
| *srm-6* | WBGene00009810 | protein_coding_gene | -2.4 | 0.19 | 3.3E-02 |
| *cest-7* | WBGene00007693 | protein_coding_gene | -2.4 | 0.19 | 1.1E-03 |
| *ugt-16* | WBGene00013901 | protein_coding_gene | -2.4 | 0.19 | 1.9E-14 |
| *col-162* | WBGene00000735 | protein_coding_gene | -2.4 | 0.19 | 3.6E-03 |
| *cest-28* | WBGene00019654 | protein_coding_gene | -2.4 | 0.19 | 1.6E-03 |
| *lbp-8* | WBGene00002260 | protein_coding_gene | -2.4 | 0.19 | 1.5E-02 |
| *T20D4.7* | WBGene00020613 | protein_coding_gene | -2.4 | 0.19 | 1.7E-02 |
| *clec-206* | WBGene00019089 | protein_coding_gene | -2.5 | 0.18 | 1.3E-02 |
| *Y45G12C.1* | WBGene00021565 | protein_coding_gene | -2.5 | 0.18 | 1.0E-04 |
| *C35A5.3* | WBGene00007947 | protein_coding_gene | -2.5 | 0.18 | 3.9E-14 |
| *R07E5.4* | WBGene00011112 | protein_coding_gene | -2.5 | 0.18 | 1.9E-02 |
| *far-3* | WBGene00001387 | protein_coding_gene | -2.5 | 0.18 | 3.2E-08 |
| *fbxa-72* | WBGene00017962 | protein_coding_gene | -2.5 | 0.18 | 1.3E-05 |
| *F58G6.3* | WBGene00010274 | protein_coding_gene | -2.5 | 0.18 | 8.8E-04 |
| *drd-1* | WBGene00009902 | protein_coding_gene | -2.5 | 0.17 | 3.1E-10 |
| *K11H12.3* | WBGene00019659 | protein_coding_gene | -2.5 | 0.17 | 5.5E-04 |
| *C55C3.7* | WBGene00043065 | protein_coding_gene | -2.5 | 0.17 | 3.7E-02 |
| *F18E3.13* | WBGene00045276 | protein_coding_gene | -2.5 | 0.17 | 8.9E-04 |
| *Y46D2A.2* | WBGene00021590 | protein_coding_gene | -2.6 | 0.16 | 8.3E-06 |
| *B0238.15* | WBGene00044537 | protein_coding_gene | -2.6 | 0.16 | 4.2E-03 |
| *cest-10* | WBGene00016595 | protein_coding_gene | -2.6 | 0.16 | 1.1E-13 |
| *B0205.14* | WBGene00044646 | protein_coding_gene | -2.7 | 0.16 | 6.6E-03 |
| *F49F1.7* | WBGene00018647 | protein_coding_gene | -2.7 | 0.16 | 3.2E-05 |
| *C33H5.1* | WBGene00016370 | protein_coding_gene | -2.7 | 0.15 | 4.4E-02 |
| *grd-14* | WBGene00001703 | protein_coding_gene | -2.7 | 0.15 | 1.0E-03 |
| *Y47D7A.15* | WBGene00021627 | protein_coding_gene | -2.7 | 0.15 | 8.0E-04 |
| *arf-1.1* | WBGene00000190 | protein_coding_gene | -2.7 | 0.15 | 5.6E-08 |
| *clec-190* | WBGene00010953 | protein_coding_gene | -2.7 | 0.15 | 4.1E-04 |
| *Y48G8AL.12* | WBGene00021690 | protein_coding_gene | -2.8 | 0.15 | 2.5E-04 |
| *Y75B8A.39* | WBGene00023483 | protein_coding_gene | -2.8 | 0.15 | 1.4E-02 |
| *col-159* | WBGene00000732 | protein_coding_gene | -2.8 | 0.15 | 2.3E-10 |
| *C04E12.10* | WBGene00015439 | pseudogene | -2.8 | 0.15 | 1.9E-03 |
| *ZK6.11* | WBGene00022645 | protein_coding_gene | -2.8 | 0.15 | 4.6E-08 |
| *C05D9.12* | WBGene00219800 | ncRNA_gene | -2.8 | 0.14 | 1.1E-05 |
| *col-88* | WBGene00000663 | protein_coding_gene | -2.8 | 0.14 | 6.2E-04 |
| *lips-14* | WBGene00019208 | pseudogene | -2.9 | 0.14 | 5.4E-03 |
| *C14E2.12* | WBGene00219816 | snoRNA_gene | -3.0 | 0.13 | 1.5E-07 |
| *oac-32* | WBGene00018708 | protein_coding_gene | -3.0 | 0.13 | 2.7E-03 |
| *F53F1.4* | WBGene00009982 | protein_coding_gene | -3.0 | 0.13 | 2.1E-05 |
| *pgp-1* | WBGene00003995 | protein_coding_gene | -3.0 | 0.13 | 6.6E-15 |
| *sdz-35* | WBGene00022570 | protein_coding_gene | -3.0 | 0.12 | 6.0E-03 |
| *nhr-74* | WBGene00003664 | protein_coding_gene | -3.0 | 0.12 | 1.5E-04 |
| *W10G11.3* | WBGene00021136 | protein_coding_gene | -3.0 | 0.12 | 2.6E-09 |
| *ZK6.6* | WBGene00022641 | protein_coding_gene | -3.1 | 0.12 | 3.7E-07 |
| *col-145* | WBGene00000718 | protein_coding_gene | -3.1 | 0.12 | 4.0E-05 |
| *col-156* | WBGene00000729 | protein_coding_gene | -3.1 | 0.12 | 3.6E-06 |

| | | | | | |
|---|---|---|---|---|---|
| *col-180* | WBGene00000753 | protein_coding_gene | -3.1 | 0.12 | 6.9E-08 |
| *F35E12.2* | WBGene00009426 | protein_coding_gene | -3.1 | 0.12 | 9.5E-03 |
| *Y48E1B.8* | WBGene00013007 | protein_coding_gene | -3.1 | 0.11 | 1.3E-02 |
| *plep-1* | WBGene00021797 | protein_coding_gene | -3.2 | 0.11 | 3.1E-06 |
| *col-137* | WBGene00000710 | protein_coding_gene | -3.2 | 0.11 | 6.2E-04 |
| *gpdh-1* | WBGene00009824 | protein_coding_gene | -3.3 | 0.10 | 2.0E-04 |
| *dhs-25* | WBGene00000988 | protein_coding_gene | -3.4 | 0.09 | 9.2E-20 |
| *Y41C4A.32* | WBGene00270321 | protein_coding_gene | -3.4 | 0.09 | 3.1E-08 |
| *folt-2* | WBGene00018138 | protein_coding_gene | -3.5 | 0.09 | 1.4E-18 |
| *vit-1* | WBGene00006925 | protein_coding_gene | -3.5 | 0.09 | 8.0E-09 |
| *grd-4* | WBGene00001693 | protein_coding_gene | -3.7 | 0.08 | 5.2E-05 |
| *nspc-5* | WBGene00044340 | protein_coding_gene | -3.7 | 0.08 | 2.0E-03 |
| *vit-3* | WBGene00006927 | protein_coding_gene | -3.8 | 0.07 | 8.8E-10 |
| *C07G1.7* | WBGene00015573 | protein_coding_gene | -3.9 | 0.07 | 2.2E-07 |
| *F19G12.t1* | WBGene00023271 | pseudogene | -3.9 | 0.07 | 2.0E-03 |
| *F18E3.12* | WBGene00045275 | protein_coding_gene | -4.1 | 0.06 | 1.1E-06 |
| *R03G8.4* | WBGene00010999 | pseudogene | -4.3 | 0.05 | 5.0E-06 |
| *vit-4* | WBGene00006928 | protein_coding_gene | -4.3 | 0.05 | 4.0E-08 |
| *C04C3.14* | WBGene00219410 | protein_coding_gene | -4.6 | 0.04 | 1.2E-03 |
| *nas-27* | WBGene00003545 | protein_coding_gene | -4.8 | 0.04 | 1.5E-04 |
| *D1086.3* | WBGene00008390 | protein_coding_gene | -4.8 | 0.04 | 4.3E-13 |

**CB4856**-exclusive DEGs in response to *S. epidermidis* (vs. OP50)

| Gene Name | Wormbase Gene ID | Type | $log_2FC$ | FC | Adj P-value |
|---|---|---|---|---|---|
| *R193.2* | WBGene00020128 | protein_coding_gene | 5.3 | 39.16 | 2.0E-95 |
| *scl-6* | WBGene00008028 | protein_coding_gene | 5.3 | 39.09 | 2.5E-20 |
| *hrg-4* | WBGene00009493 | protein_coding_gene | 4.7 | 26.06 | 1.2E-20 |
| *spp-23* | WBGene00021335 | protein_coding_gene | 4.6 | 24.77 | 2.5E-18 |
| *F56H6.7* | WBGene00010168 | protein_coding_gene | 4.3 | 19.76 | 1.3E-06 |
| *F42A8.1* | WBGene00009626 | protein_coding_gene | 3.9 | 15.00 | 2.5E-99 |
| *oac-35* | WBGene00010171 | protein_coding_gene | 3.6 | 12.36 | 7.2E-04 |
| *F08H9.4* | WBGene00008592 | protein_coding_gene | 3.4 | 10.55 | 1.2E-03 |
| *C02G6.1* | WBGene00015359 | pseudogene | 3.4 | 10.29 | 1.2E-04 |
| *F56H6.9* | WBGene00010170 | protein_coding_gene | 3.3 | 9.92 | 3.3E-03 |
| *C47F8.6* | WBGene00008162 | protein_coding_gene | 3.3 | 9.88 | 2.6E-03 |
| *T16G12.1* | WBGene00011803 | protein_coding_gene | 3.3 | 9.82 | 7.3E-24 |
| *hsd-2* | WBGene00022498 | protein_coding_gene | 3.2 | 9.40 | 3.1E-03 |
| *gmd-2* | WBGene00010166 | protein_coding_gene | 3.2 | 9.38 | 7.4E-04 |
| *col-94* | WBGene00000669 | protein_coding_gene | 3.2 | 9.18 | 8.9E-03 |
| *F57G8.7* | WBGene00010216 | protein_coding_gene | 3.1 | 8.76 | 4.2E-03 |
| *H25K10.1* | WBGene00010411 | protein_coding_gene | 3.0 | 8.16 | 9.3E-07 |
| *Y106G6E.1* | WBGene00013705 | protein_coding_gene | 3.0 | 8.11 | 1.2E-03 |
| *Y16B4A.2* | WBGene00012445 | protein_coding_gene | 3.0 | 8.00 | 6.3E-28 |
| *F53B3.6* | WBGene00018744 | protein_coding_gene | 3.0 | 7.81 | 4.1E-11 |
| *elo-6* | WBGene00001244 | protein_coding_gene | 2.9 | 7.56 | 6.1E-16 |
| *F18F11.4* | WBGene00017577 | protein_coding_gene | 2.9 | 7.38 | 4.2E-03 |
| *C49F5.8* | WBGene00008212 | protein_coding_gene | 2.9 | 7.29 | 6.4E-03 |
| *oac-1* | WBGene00007177 | protein_coding_gene | 2.8 | 7.18 | 1.2E-02 |
| *elo-5* | WBGene00001243 | protein_coding_gene | 2.8 | 7.12 | 6.8E-75 |
| *K09C4.5* | WBGene00019550 | protein_coding_gene | 2.8 | 7.11 | 6.9E-13 |
| *elo-2* | WBGene00001240 | protein_coding_gene | 2.8 | 6.98 | 4.2E-26 |
| *ins-30* | WBGene00002113 | protein_coding_gene | 2.8 | 6.85 | 1.8E-04 |
| *clec-230* | WBGene00007806 | protein_coding_gene | 2.7 | 6.56 | 2.5E-03 |
| *T28D6.3* | WBGene00012123 | protein_coding_gene | 2.7 | 6.28 | 2.7E-33 |
| *abu-11* | WBGene00000034 | protein_coding_gene | 2.6 | 6.12 | 1.9E-04 |
| *clec-174* | WBGene00021580 | protein_coding_gene | 2.6 | 6.07 | 5.1E-06 |
| *C05D12.3* | WBGene00014666 | protein_coding_gene | 2.6 | 5.86 | 1.1E-41 |
| *sptl-2* | WBGene00018398 | protein_coding_gene | 2.5 | 5.79 | 3.0E-39 |

| | | | | | |
|---|---|---|---|---|---|
| *oac-45* | WBGene00011656 | protein_coding_gene | 2.5 | 5.71 | 7.5E-03 |
| *tag-10* | WBGene00006404 | protein_coding_gene | 2.5 | 5.69 | 1.2E-48 |
| *K01D12.5* | WBGene00010464 | protein_coding_gene | 2.5 | 5.54 | 2.4E-02 |
| *H06A10.1* | WBGene00010368 | protein_coding_gene | 2.5 | 5.53 | 4.2E-11 |
| *clec-100* | WBGene00013179 | protein_coding_gene | 2.5 | 5.52 | 6.8E-08 |
| *acox-1.4* | WBGene00008567 | protein_coding_gene | 2.5 | 5.49 | 2.1E-12 |
| *gst-27* | WBGene00001775 | protein_coding_gene | 2.4 | 5.38 | 4.1E-04 |
| *F49D11.3* | WBGene00018627 | protein_coding_gene | 2.4 | 5.32 | 4.4E-02 |
| *tag-293* | WBGene00015398 | protein_coding_gene | 2.4 | 5.29 | 4.2E-03 |
| *ins-32* | WBGene00002115 | protein_coding_gene | 2.4 | 5.26 | 1.2E-02 |
| *T05E11.7* | WBGene00011483 | pseudogene | 2.4 | 5.19 | 2.7E-02 |
| *C54D2.1* | WBGene00016915 | protein_coding_gene | 2.4 | 5.15 | 1.6E-03 |
| *mua-6* | WBGene00003485 | protein_coding_gene | 2.4 | 5.12 | 2.7E-26 |
| *oac-48* | WBGene00011960 | protein_coding_gene | 2.3 | 5.07 | 8.9E-05 |
| *Y54G2A.45* | WBGene00044488 | protein_coding_gene | 2.3 | 5.00 | 2.2E-05 |
| *nep-22* | WBGene00020788 | protein_coding_gene | 2.3 | 4.92 | 3.1E-35 |
| *npr-30* | WBGene00019183 | protein_coding_gene | 2.3 | 4.91 | 2.7E-02 |
| *col-147* | WBGene00000720 | protein_coding_gene | 2.3 | 4.83 | 3.3E-19 |
| *F46B3.14* | WBGene00009765 | protein_coding_gene | 2.3 | 4.81 | 6.8E-04 |
| *F21G4.3* | WBGene00009024 | protein_coding_gene | 2.3 | 4.81 | 3.6E-02 |
| *F58F9.9* | WBGene00044307 | protein_coding_gene | 2.3 | 4.79 | 3.2E-04 |
| *C55B6.5* | WBGene00016941 | protein_coding_gene | 2.3 | 4.79 | 3.1E-02 |
| *T20B6.1* | WBGene00020597 | protein_coding_gene | 2.2 | 4.75 | 4.3E-03 |
| *gcy-7* | WBGene00001534 | protein_coding_gene | 2.2 | 4.74 | 2.5E-02 |
| *Y53F4B.23* | WBGene00013169 | protein_coding_gene | 2.2 | 4.74 | 2.0E-06 |
| *abu-1* | WBGene00000024 | protein_coding_gene | 2.2 | 4.72 | 4.0E-02 |
| *K02E11.10* | WBGene00044109 | protein_coding_gene | 2.2 | 4.68 | 3.7E-02 |
| *try-4* | WBGene00006622 | protein_coding_gene | 2.2 | 4.67 | 5.7E-03 |
| *H43E16.1* | WBGene00019273 | protein_coding_gene | 2.2 | 4.65 | 5.8E-27 |
| *piit-1* | WBGene00009384 | protein_coding_gene | 2.2 | 4.64 | 2.8E-13 |
| *oac-44* | WBGene00011654 | protein_coding_gene | 2.2 | 4.64 | 2.4E-02 |
| *C41H7.1* | WBGene00016571 | protein_coding_gene | 2.2 | 4.63 | 9.9E-05 |
| *C28F5.4* | WBGene00016185 | protein_coding_gene | 2.2 | 4.59 | 1.7E-02 |
| *Y39G8B.9* | WBGene00012728 | protein_coding_gene | 2.2 | 4.55 | 4.6E-04 |
| *F28A10.9* | WBGene00017877 | pseudogene | 2.2 | 4.52 | 2.6E-02 |
| *T28A11.4* | WBGene00020871 | protein_coding_gene | 2.2 | 4.51 | 4.2E-02 |
| *paic-1* | WBGene00015116 | protein_coding_gene | 2.2 | 4.45 | 3.2E-53 |
| *decr-1.1* | WBGene00018380 | protein_coding_gene | 2.1 | 4.43 | 2.2E-17 |
| *Y50D4B.4* | WBGene00021743 | protein_coding_gene | 2.1 | 4.43 | 3.5E-22 |
| *ZK84.1* | WBGene00022649 | protein_coding_gene | 2.1 | 4.41 | 4.7E-07 |
| *F29A7.3* | WBGene00017913 | protein_coding_gene | 2.1 | 4.41 | 7.1E-03 |
| *oig-5* | WBGene00023520 | protein_coding_gene | 2.1 | 4.41 | 1.2E-04 |
| *pqn-71* | WBGene00004153 | protein_coding_gene | 2.1 | 4.41 | 3.0E-02 |
| *C01B9.1* | WBGene00007220 | protein_coding_gene | 2.1 | 4.40 | 2.6E-13 |
| *tor-2* | WBGene00006598 | protein_coding_gene | 2.1 | 4.39 | 1.5E-02 |
| *gst-26* | WBGene00001774 | protein_coding_gene | 2.1 | 4.38 | 2.0E-21 |
| *F46B3.15* | WBGene00009766 | protein_coding_gene | 2.1 | 4.34 | 2.0E-03 |
| *F35A5.4* | WBGene00018026 | protein_coding_gene | 2.1 | 4.33 | 3.3E-02 |
| *wrt-8* | WBGene00006954 | protein_coding_gene | 2.1 | 4.31 | 4.7E-02 |
| *F38B6.4* | WBGene00018174 | protein_coding_gene | 2.1 | 4.31 | 3.1E-21 |
| *B0244.7* | WBGene00015082 | protein_coding_gene | 2.1 | 4.28 | 6.1E-03 |
| *gpa-11* | WBGene00001673 | protein_coding_gene | 2.1 | 4.28 | 3.7E-02 |
| *AC3.5* | WBGene00007071 | protein_coding_gene | 2.1 | 4.28 | 4.8E-37 |
| *D2024.4* | WBGene00017052 | protein_coding_gene | 2.1 | 4.24 | 3.4E-02 |
| *ugt-6* | WBGene00013904 | protein_coding_gene | 2.1 | 4.23 | 8.4E-18 |
| *M195.2* | WBGene00010948 | protein_coding_gene | 2.1 | 4.23 | 2.0E-02 |
| *Y50D4B.7* | WBGene00021746 | pseudogene | 2.1 | 4.21 | 4.2E-03 |
| *T10B5.10* | WBGene00020396 | protein_coding_gene | 2.1 | 4.18 | 7.2E-04 |

| | | | | | |
|---|---|---|---|---|---|
| *F26C11.1* | WBGene00009146 | protein_coding_gene | 2.1 | 4.15 | 2.3E-05 |
| *F59A6.3* | WBGene00019085 | protein_coding_gene | 2.1 | 4.15 | 3.8E-03 |
| *rol-6* | WBGene00004397 | protein_coding_gene | 2.0 | 4.14 | 3.2E-02 |
| *F58G1.9* | WBGene00010271 | pseudogene | 2.0 | 4.14 | 8.3E-03 |
| *nspb-12* | WBGene00017305 | protein_coding_gene | 2.0 | 4.13 | 4.9E-02 |
| *F40H6.5* | WBGene00018255 | protein_coding_gene | 2.0 | 4.12 | 2.5E-02 |
| *C53D6.5* | WBGene00008282 | pseudogene | 2.0 | 4.10 | 1.8E-02 |
| *ets-5* | WBGene00016600 | protein_coding_gene | 2.0 | 4.09 | 1.7E-02 |
| *ferl-1* | WBGene00020262 | protein_coding_gene | 2.0 | 4.03 | 1.3E-02 |
| *cyp-34A5* | WBGene00015040 | protein_coding_gene | 2.0 | 4.02 | 3.7E-03 |
| *gcsh-1* | WBGene00008354 | protein_coding_gene | 2.0 | 4.01 | 5.2E-12 |
| *K10C9.9* | WBGene00044515 | pseudogene | 2.0 | 4.00 | 1.4E-02 |
| *T25E4.1* | WBGene00020803 | protein_coding_gene | 2.0 | 3.99 | 1.2E-02 |
| *col-58* | WBGene00000634 | protein_coding_gene | 2.0 | 3.99 | 9.6E-03 |
| *lgc-40* | WBGene00020767 | protein_coding_gene | 2.0 | 3.98 | 9.1E-05 |
| *acr-23* | WBGene00000062 | protein_coding_gene | 2.0 | 3.97 | 3.6E-02 |
| *F59B1.4* | WBGene00019098 | protein_coding_gene | 2.0 | 3.96 | 3.2E-02 |
| *F37A4.6* | WBGene00018136 | protein_coding_gene | 2.0 | 3.96 | 1.3E-02 |
| *R03C1.1* | WBGene00010987 | protein_coding_gene | 2.0 | 3.96 | 2.2E-02 |
| *F34H10.4* | WBGene00009381 | protein_coding_gene | 2.0 | 3.94 | 2.0E-02 |
| *clec-1* | WBGene00017772 | protein_coding_gene | 2.0 | 3.94 | 1.0E-24 |
| *K02D3.2* | WBGene00010505 | protein_coding_gene | 2.0 | 3.92 | 2.9E-03 |
| *pals-6* | WBGene00007660 | protein_coding_gene | 2.0 | 3.88 | 1.9E-06 |
| *clec-245* | WBGene00007851 | protein_coding_gene | 2.0 | 3.88 | 3.1E-02 |
| *Y47D7A.2* | WBGene00021616 | protein_coding_gene | 2.0 | 3.88 | 4.5E-02 |
| *F46F2.3* | WBGene00009787 | protein_coding_gene | 2.0 | 3.87 | 6.6E-06 |
| *abu-8* | WBGene00000031 | protein_coding_gene | 2.0 | 3.87 | 1.3E-02 |
| *ech-7* | WBGene00001156 | protein_coding_gene | 1.9 | 3.85 | 4.0E-28 |
| *nmy-3* | WBGene00012437 | protein_coding_gene | 1.9 | 3.84 | 6.2E-12 |
| *T06E4.14* | WBGene00077691 | protein_coding_gene | 1.9 | 3.83 | 3.2E-02 |
| *fipr-6* | WBGene00007544 | protein_coding_gene | 1.9 | 3.81 | 3.9E-02 |
| *drd-5* | WBGene00018885 | protein_coding_gene | 1.9 | 3.77 | 3.4E-12 |
| *fbxa-73* | WBGene00020640 | protein_coding_gene | 1.9 | 3.77 | 2.3E-03 |
| *C18B2.2* | WBGene00015953 | protein_coding_gene | 1.9 | 3.77 | 4.1E-02 |
| *T06E4.8* | WBGene00011535 | protein_coding_gene | 1.9 | 3.74 | 4.7E-02 |
| *Y39F10A.1* | WBGene00021452 | protein_coding_gene | 1.9 | 3.74 | 1.3E-03 |
| *far-4* | WBGene00001388 | protein_coding_gene | 1.9 | 3.73 | 2.3E-02 |
| *ZK1290.7* | WBGene00022889 | protein_coding_gene | 1.9 | 3.73 | 3.1E-03 |
| *D1054.6* | WBGene00008373 | pseudogene | 1.9 | 3.72 | 3.3E-02 |
| *F22H10.6* | WBGene00017730 | protein_coding_gene | 1.9 | 3.67 | 6.8E-06 |
| *C16E9.1* | WBGene00015865 | protein_coding_gene | 1.9 | 3.66 | 1.4E-08 |
| *C50E3.9* | WBGene00016820 | protein_coding_gene | 1.9 | 3.65 | 8.0E-04 |
| *unc-8* | WBGene00006748 | protein_coding_gene | 1.9 | 3.65 | 4.1E-03 |
| *nlp-16* | WBGene00003754 | protein_coding_gene | 1.9 | 3.64 | 3.1E-28 |
| *Y102A11A.7* | WBGene00022417 | protein_coding_gene | 1.9 | 3.63 | 1.3E-02 |
| *lin-31* | WBGene00003017 | protein_coding_gene | 1.9 | 3.63 | 1.5E-03 |
| *clx-1* | WBGene00000551 | protein_coding_gene | 1.9 | 3.62 | 1.9E-02 |
| *clec-205* | WBGene00021602 | protein_coding_gene | 1.9 | 3.62 | 4.9E-06 |
| *F52D10.4* | WBGene00009931 | protein_coding_gene | 1.9 | 3.62 | 2.2E-02 |
| *acr-25* | WBGene00043066 | protein_coding_gene | 1.9 | 3.61 | 1.1E-03 |
| *scav-4* | WBGene00008692 | protein_coding_gene | 1.8 | 3.60 | 1.7E-10 |
| *Y38F1A.4* | WBGene00012607 | protein_coding_gene | 1.8 | 3.60 | 4.2E-02 |
| *ncx-8* | WBGene00003573 | protein_coding_gene | 1.8 | 3.59 | 3.9E-05 |
| *C54D10.5* | WBGene00008300 | protein_coding_gene | 1.8 | 3.59 | 9.3E-03 |
| *M03E7.4* | WBGene00019756 | protein_coding_gene | 1.8 | 3.59 | 4.1E-02 |
| *F25C8.1* | WBGene00009104 | pseudogene | 1.8 | 3.59 | 4.3E-05 |
| *mup-4* | WBGene00003497 | protein_coding_gene | 1.8 | 3.58 | 9.8E-35 |
| *wrt-4* | WBGene00006950 | protein_coding_gene | 1.8 | 3.57 | 4.5E-02 |

| | | | | | |
|---|---|---|---|---|---|
| *C39D10.8* | WBGene00016535 | protein_coding_gene | 1.8 | 3.57 | 6.5E-19 |
| *Y67D8C.7* | WBGene00022071 | protein_coding_gene | 1.8 | 3.56 | 3.7E-02 |
| *C10F3.7* | WBGene00015679 | protein_coding_gene | 1.8 | 3.54 | 7.7E-04 |
| *scl-3* | WBGene00009896 | protein_coding_gene | 1.8 | 3.54 | 3.5E-19 |
| *oac-52* | WBGene00020977 | protein_coding_gene | 1.8 | 3.54 | 4.4E-02 |
| *C08G5.3* | WBGene00015616 | pseudogene | 1.8 | 3.53 | 2.6E-02 |
| *F22E5.8* | WBGene00017707 | protein_coding_gene | 1.8 | 3.53 | 1.9E-05 |
| *gly-1* | WBGene00001626 | protein_coding_gene | 1.8 | 3.52 | 3.6E-02 |
| *arrd-10* | WBGene00010268 | protein_coding_gene | 1.8 | 3.52 | 5.7E-07 |
| *clpr-2* | WBGene00023410 | protein_coding_gene | 1.8 | 3.51 | 1.4E-02 |
| *pcp-3* | WBGene00003958 | protein_coding_gene | 1.8 | 3.49 | 8.0E-28 |
| *inx-18* | WBGene00002140 | protein_coding_gene | 1.8 | 3.49 | 6.3E-04 |
| *abu-10* | WBGene00000033 | protein_coding_gene | 1.8 | 3.48 | 4.8E-02 |
| *ser-6* | WBGene00021897 | protein_coding_gene | 1.8 | 3.48 | 2.2E-03 |
| *Y59E9AL.4* | WBGene00021994 | protein_coding_gene | 1.8 | 3.48 | 3.2E-07 |
| *R12A1.3* | WBGene00020017 | protein_coding_gene | 1.8 | 3.48 | 1.7E-02 |
| *comt-2* | WBGene00012518 | protein_coding_gene | 1.8 | 3.46 | 1.7E-02 |
| *ugt-21* | WBGene00007885 | protein_coding_gene | 1.8 | 3.45 | 7.8E-09 |
| *F33E2.6* | WBGene00009362 | protein_coding_gene | 1.8 | 3.45 | 1.5E-02 |
| *act-5* | WBGene00000067 | protein_coding_gene | 1.8 | 3.44 | 1.0E-04 |
| *C05C8.8* | WBGene00015465 | protein_coding_gene | 1.8 | 3.44 | 2.8E-02 |
| *C35A5.10* | WBGene00044016 | protein_coding_gene | 1.8 | 3.41 | 2.7E-02 |
| *lgc-54* | WBGene00020528 | protein_coding_gene | 1.8 | 3.40 | 4.5E-07 |
| *F10C1.8* | WBGene00017327 | protein_coding_gene | 1.8 | 3.39 | 4.8E-03 |
| *T17H7.1* | WBGene00020550 | protein_coding_gene | 1.8 | 3.39 | 2.8E-02 |
| *ZC434.9* | WBGene00013895 | protein_coding_gene | 1.8 | 3.38 | 5.0E-24 |
| *Y50D4B.9* | WBGene00268189 | pseudogene | 1.8 | 3.38 | 3.7E-02 |
| *frpr-10* | WBGene00019019 | protein_coding_gene | 1.8 | 3.38 | 3.9E-02 |
| *W07A12.8* | WBGene00012325 | protein_coding_gene | 1.8 | 3.38 | 3.5E-02 |
| *exp-1* | WBGene00001373 | protein_coding_gene | 1.8 | 3.36 | 3.8E-03 |
| *dmsr-2* | WBGene00021275 | protein_coding_gene | 1.7 | 3.35 | 5.8E-06 |
| *F29G6.1* | WBGene00009257 | protein_coding_gene | 1.7 | 3.35 | 1.5E-03 |
| *ncx-7* | WBGene00003572 | protein_coding_gene | 1.7 | 3.35 | 5.9E-17 |
| *Y57G11C.42* | WBGene00013331 | protein_coding_gene | 1.7 | 3.34 | 4.9E-03 |
| *ctsa-2* | WBGene00019617 | protein_coding_gene | 1.7 | 3.34 | 7.8E-05 |
| *pqn-74* | WBGene00004156 | protein_coding_gene | 1.7 | 3.33 | 2.1E-02 |
| *K09D9.11* | WBGene00019568 | pseudogene | 1.7 | 3.33 | 1.1E-02 |
| *nas-6* | WBGene00003525 | protein_coding_gene | 1.7 | 3.31 | 3.6E-03 |
| *col-166* | WBGene00000739 | protein_coding_gene | 1.7 | 3.31 | 1.4E-15 |
| *C14A6.5* | WBGene00007569 | protein_coding_gene | 1.7 | 3.31 | 4.9E-02 |
| *F57B1.9* | WBGene00045488 | protein_coding_gene | 1.7 | 3.31 | 6.9E-04 |
| *C30A5.10* | WBGene00016242 | protein_coding_gene | 1.7 | 3.30 | 2.9E-02 |
| *cdh-7* | WBGene00000399 | protein_coding_gene | 1.7 | 3.29 | 9.6E-03 |
| *F54E4.4* | WBGene00010060 | protein_coding_gene | 1.7 | 3.29 | 3.8E-02 |
| *srh-254* | WBGene00005460 | pseudogene | 1.7 | 3.28 | 1.3E-09 |
| *ZK662.6* | WBGene00045494 | protein_coding_gene | 1.7 | 3.28 | 4.2E-02 |
| *daf-28* | WBGene00000920 | protein_coding_gene | 1.7 | 3.28 | 3.5E-06 |
| *nhr-219* | WBGene00020555 | protein_coding_gene | 1.7 | 3.27 | 2.5E-02 |
| *cah-5* | WBGene00000283 | protein_coding_gene | 1.7 | 3.27 | 9.6E-17 |
| *F19H6.6* | WBGene00008961 | protein_coding_gene | 1.7 | 3.26 | 4.1E-03 |
| *F28H7.3* | WBGene00009237 | protein_coding_gene | 1.7 | 3.26 | 6.7E-06 |
| *ncx-4* | WBGene00003569 | protein_coding_gene | 1.7 | 3.25 | 1.6E-07 |
| *ZK228.4* | WBGene00013947 | protein_coding_gene | 1.7 | 3.25 | 3.7E-14 |
| *lge-1* | WBGene00010716 | protein_coding_gene | 1.7 | 3.25 | 1.3E-02 |
| *pho-13* | WBGene00008560 | protein_coding_gene | 1.7 | 3.24 | 3.9E-03 |
| *F14D7.6* | WBGene00008793 | protein_coding_gene | 1.7 | 3.24 | 1.7E-10 |
| *atic-1* | WBGene00016957 | protein_coding_gene | 1.7 | 3.23 | 2.9E-19 |
| *asns-1* | WBGene00003816 | protein_coding_gene | 1.7 | 3.23 | 1.5E-03 |

| | | | | | |
|---|---|---|---|---|---|
| *H14N18.4* | WBGene00019204 | protein_coding_gene | 1.7 | 3.23 | 1.1E-23 |
| *mct-6* | WBGene00015676 | protein_coding_gene | 1.7 | 3.22 | 8.6E-44 |
| *C06C6.6* | WBGene00007380 | pseudogene | 1.7 | 3.22 | 4.0E-03 |
| *Y73C8C.10* | WBGene00022267 | protein_coding_gene | 1.7 | 3.22 | 2.0E-03 |
| *lec-8* | WBGene00002271 | protein_coding_gene | 1.7 | 3.22 | 1.7E-07 |
| *mtd-1* | WBGene00003471 | protein_coding_gene | 1.7 | 3.22 | 2.9E-04 |
| *cyp-33C5* | WBGene00018261 | protein_coding_gene | 1.7 | 3.21 | 1.3E-03 |
| *frpr-3* | WBGene00016149 | protein_coding_gene | 1.7 | 3.21 | 4.4E-02 |
| *C34E7.3* | WBGene00007933 | protein_coding_gene | 1.7 | 3.21 | 2.4E-02 |
| *R05H10.7* | WBGene00011046 | protein_coding_gene | 1.7 | 3.20 | 2.7E-15 |
| *galt-1* | WBGene00019763 | protein_coding_gene | 1.7 | 3.19 | 3.0E-05 |
| *Y22D7AL.11* | WBGene00021249 | protein_coding_gene | 1.7 | 3.19 | 4.7E-05 |
| *F19H6.5* | WBGene00008960 | protein_coding_gene | 1.7 | 3.18 | 5.2E-03 |
| *F07H5.8* | WBGene00008559 | protein_coding_gene | 1.7 | 3.18 | 1.9E-02 |
| *nhr-31* | WBGene00003625 | protein_coding_gene | 1.7 | 3.18 | 3.7E-18 |
| *C49A9.4* | WBGene00016758 | protein_coding_gene | 1.7 | 3.18 | 3.3E-08 |
| *daf-36* | WBGene00007536 | protein_coding_gene | 1.7 | 3.17 | 2.7E-13 |
| *smd-1* | WBGene00004875 | protein_coding_gene | 1.7 | 3.17 | 5.1E-09 |
| *ldp-1* | WBGene00017719 | protein_coding_gene | 1.7 | 3.16 | 6.9E-58 |
| *haao-1* | WBGene00010595 | protein_coding_gene | 1.7 | 3.16 | 5.8E-13 |
| *ZK742.3* | WBGene00022813 | protein_coding_gene | 1.7 | 3.15 | 3.0E-08 |
| *T07G12.3* | WBGene00011594 | protein_coding_gene | 1.7 | 3.15 | 1.5E-02 |
| *phat-4* | WBGene00020237 | protein_coding_gene | 1.7 | 3.14 | 1.8E-03 |
| *C26D10.7* | WBGene00023420 | protein_coding_gene | 1.7 | 3.14 | 1.7E-02 |
| *T10B11.4* | WBGene00020400 | protein_coding_gene | 1.6 | 3.13 | 1.7E-02 |
| *K08D8.3* | WBGene00010657 | protein_coding_gene | 1.6 | 3.13 | 1.3E-11 |
| *twk-16* | WBGene00006670 | protein_coding_gene | 1.6 | 3.12 | 1.4E-03 |
| *C06E2.1* | WBGene00015527 | protein_coding_gene | 1.6 | 3.11 | 5.4E-04 |
| *gfi-1* | WBGene00001581 | protein_coding_gene | 1.6 | 3.11 | 5.7E-03 |
| *unc-47* | WBGene00006783 | protein_coding_gene | 1.6 | 3.10 | 6.2E-03 |
| *F26G1.5* | WBGene00017841 | protein_coding_gene | 1.6 | 3.10 | 3.4E-02 |
| *pes-8* | WBGene00003981 | protein_coding_gene | 1.6 | 3.10 | 4.3E-02 |
| *cpt-5* | WBGene00008629 | protein_coding_gene | 1.6 | 3.10 | 2.0E-14 |
| *C09F12.2* | WBGene00007482 | protein_coding_gene | 1.6 | 3.09 | 1.2E-03 |
| *C18H7.6* | WBGene00015996 | protein_coding_gene | 1.6 | 3.09 | 2.8E-02 |
| *C01G10.17* | WBGene00044658 | protein_coding_gene | 1.6 | 3.09 | 1.8E-03 |
| *B0507.8* | WBGene00015225 | protein_coding_gene | 1.6 | 3.09 | 1.1E-02 |
| *ins-33* | WBGene00002116 | protein_coding_gene | 1.6 | 3.07 | 1.5E-04 |
| *C07E3.4* | WBGene00007415 | protein_coding_gene | 1.6 | 3.07 | 5.0E-04 |
| *clp-6* | WBGene00000546 | protein_coding_gene | 1.6 | 3.07 | 6.6E-03 |
| *Y23H5B.12* | WBGene00194701 | protein_coding_gene | 1.6 | 3.06 | 3.5E-02 |
| *K04G2.7* | WBGene00010566 | protein_coding_gene | 1.6 | 3.06 | 3.8E-02 |
| *R07E4.1* | WBGene00019935 | protein_coding_gene | 1.6 | 3.06 | 7.9E-15 |
| *Y55D5A.4* | WBGene00021918 | protein_coding_gene | 1.6 | 3.05 | 1.0E-03 |
| *F56D6.12* | WBGene00044474 | protein_coding_gene | 1.6 | 3.05 | 3.4E-03 |
| *mgl-2* | WBGene00003233 | protein_coding_gene | 1.6 | 3.05 | 3.7E-09 |
| *hrg-2* | WBGene00010473 | protein_coding_gene | 1.6 | 3.05 | 2.6E-02 |
| *fbxa-84* | WBGene00011670 | protein_coding_gene | 1.6 | 3.04 | 4.5E-06 |
| *VM106R.1* | WBGene00014848 | protein_coding_gene | 1.6 | 3.04 | 1.4E-06 |
| *F41G4.7* | WBGene00018310 | protein_coding_gene | 1.6 | 3.04 | 4.8E-02 |
| *Y41D4B.1* | WBGene00021511 | protein_coding_gene | 1.6 | 3.03 | 2.5E-02 |
| *lon-1* | WBGene00003055 | protein_coding_gene | 1.6 | 3.03 | 3.4E-05 |
| *F10D11.6* | WBGene00008652 | protein_coding_gene | 1.6 | 3.03 | 5.4E-03 |
| *hrg-6* | WBGene00009495 | protein_coding_gene | 1.6 | 3.02 | 8.9E-04 |
| *lgc-4* | WBGene00017580 | protein_coding_gene | 1.6 | 3.02 | 6.0E-04 |
| *C29F4.3* | WBGene00007809 | protein_coding_gene | 1.6 | 3.02 | 4.6E-02 |
| *W02H3.1* | WBGene00012217 | protein_coding_gene | 1.6 | 3.02 | 1.2E-05 |
| *ifd-1* | WBGene00002057 | protein_coding_gene | 1.6 | 3.02 | 3.2E-12 |

| | | | | | |
|---|---|---|---|---|---|
| *C15F1.1* | WBGene00015794 | protein_coding_gene | 1.6 | 3.02 | 6.3E-11 |
| *bath-46* | WBGene00020325 | protein_coding_gene | 1.6 | 3.02 | 5.6E-04 |
| *pkd-2* | WBGene00004035 | protein_coding_gene | 1.6 | 3.02 | 1.4E-04 |
| *tph-1* | WBGene00006600 | protein_coding_gene | 1.6 | 3.01 | 3.0E-02 |
| *Y48G1BL.7* | WBGene00021666 | protein_coding_gene | 1.6 | 3.01 | 4.2E-02 |
| *C05C9.3* | WBGene00007327 | protein_coding_gene | 1.6 | 3.00 | 4.2E-06 |
| *C39B5.5* | WBGene00016523 | protein_coding_gene | 1.6 | 3.00 | 1.9E-02 |
| *C34H4.2* | WBGene00016425 | protein_coding_gene | 1.6 | 3.00 | 1.4E-07 |
| *mec-18* | WBGene00003179 | protein_coding_gene | 1.6 | 3.00 | 2.2E-04 |
| *F09B9.5* | WBGene00008604 | protein_coding_gene | 1.6 | 3.00 | 2.0E-02 |
| *ugt-10* | WBGene00020594 | protein_coding_gene | 1.6 | 2.99 | 4.0E-02 |
| *ckr-1* | WBGene00020712 | protein_coding_gene | 1.6 | 2.99 | 1.7E-02 |
| *pgp-9* | WBGene00004003 | protein_coding_gene | 1.6 | 2.99 | 5.0E-10 |
| *adt-2* | WBGene00000083 | protein_coding_gene | 1.6 | 2.98 | 2.7E-22 |
| *aat-4* | WBGene00000005 | protein_coding_gene | 1.6 | 2.98 | 1.4E-05 |
| *F55A3.7* | WBGene00018852 | pseudogene | 1.6 | 2.98 | 7.4E-04 |
| *ZC449.7* | WBGene00045365 | protein_coding_gene | 1.6 | 2.98 | 2.4E-04 |
| *acc-1* | WBGene00010275 | protein_coding_gene | 1.6 | 2.97 | 4.3E-03 |
| *ceh-23* | WBGene00000446 | protein_coding_gene | 1.6 | 2.97 | 3.3E-02 |
| *ppat-1* | WBGene00011407 | protein_coding_gene | 1.6 | 2.96 | 1.5E-10 |
| *C45G9.8* | WBGene00016679 | protein_coding_gene | 1.6 | 2.96 | 3.1E-02 |
| *T06A4.1* | WBGene00020281 | protein_coding_gene | 1.6 | 2.96 | 2.2E-02 |
| *col-165* | WBGene00000738 | protein_coding_gene | 1.6 | 2.96 | 7.4E-07 |
| *dmd-8* | WBGene00020708 | protein_coding_gene | 1.6 | 2.95 | 4.6E-02 |
| *C11E4.7* | WBGene00007521 | protein_coding_gene | 1.6 | 2.95 | 3.0E-10 |
| *T10G3.3* | WBGene00011694 | protein_coding_gene | 1.6 | 2.95 | 1.3E-07 |
| *col-143* | WBGene00000716 | protein_coding_gene | 1.6 | 2.94 | 5.8E-06 |
| *R02F2.8* | WBGene00019837 | protein_coding_gene | 1.6 | 2.94 | 5.7E-10 |
| *col-160* | WBGene00000733 | protein_coding_gene | 1.6 | 2.94 | 2.6E-30 |
| *Y43F8C.16* | WBGene00012837 | protein_coding_gene | 1.6 | 2.93 | 4.0E-03 |
| *dat-1* | WBGene00000934 | protein_coding_gene | 1.6 | 2.93 | 3.2E-03 |
| *cyp-36A1* | WBGene00007913 | protein_coding_gene | 1.5 | 2.91 | 1.7E-09 |
| *cest-15* | WBGene00012202 | pseudogene | 1.5 | 2.90 | 7.9E-08 |
| *D2096.6* | WBGene00017073 | protein_coding_gene | 1.5 | 2.89 | 4.5E-03 |
| *C46F2.1* | WBGene00016714 | protein_coding_gene | 1.5 | 2.88 | 1.0E-06 |
| *R02E4.2* | WBGene00044021 | protein_coding_gene | 1.5 | 2.88 | 1.6E-02 |
| *F55D10.4* | WBGene00018879 | protein_coding_gene | 1.5 | 2.88 | 4.3E-03 |
| *Y95B8A.2* | WBGene00022383 | protein_coding_gene | 1.5 | 2.88 | 1.8E-03 |
| *gbb-2* | WBGene00022675 | protein_coding_gene | 1.5 | 2.88 | 1.2E-09 |
| *T02D1.7* | WBGene00011373 | protein_coding_gene | 1.5 | 2.86 | 2.8E-02 |
| *lov-1* | WBGene00003058 | protein_coding_gene | 1.5 | 2.86 | 2.1E-02 |
| *F28B4.3* | WBGene00017892 | protein_coding_gene | 1.5 | 2.86 | 8.1E-03 |
| *R02C2.8* | WBGene00255728 | pseudogene | 1.5 | 2.85 | 1.2E-02 |
| *C06G1.2* | WBGene00015546 | protein_coding_gene | 1.5 | 2.85 | 4.3E-02 |
| *T22F3.7* | WBGene00020697 | protein_coding_gene | 1.5 | 2.83 | 3.0E-02 |
| *inpp-1* | WBGene00012016 | protein_coding_gene | 1.5 | 2.83 | 4.4E-08 |
| *F54H5.3* | WBGene00018840 | protein_coding_gene | 1.5 | 2.82 | 2.0E-04 |
| *F13H8.5* | WBGene00017438 | protein_coding_gene | 1.5 | 2.82 | 2.0E-02 |
| *R13A5.10* | WBGene00020052 | protein_coding_gene | 1.5 | 2.82 | 6.5E-06 |
| *K12H6.5* | WBGene00019685 | protein_coding_gene | 1.5 | 2.82 | 8.7E-05 |
| *ZK105.3* | WBGene00022654 | protein_coding_gene | 1.5 | 2.82 | 7.0E-03 |
| *nas-30* | WBGene00003548 | protein_coding_gene | 1.5 | 2.82 | 2.3E-02 |
| *ctsa-4.2* | WBGene00021503 | protein_coding_gene | 1.5 | 2.81 | 1.9E-05 |
| *fbxa-196* | WBGene00020265 | protein_coding_gene | 1.5 | 2.81 | 3.4E-05 |
| *Y61A9LA.12* | WBGene00044586 | protein_coding_gene | 1.5 | 2.79 | 2.6E-02 |
| *Y47D7A.7* | WBGene00021620 | protein_coding_gene | 1.5 | 2.79 | 2.4E-03 |
| *ins-27* | WBGene00002110 | protein_coding_gene | 1.5 | 2.79 | 1.1E-04 |
| *W05H12.1* | WBGene00012289 | protein_coding_gene | 1.5 | 2.78 | 3.4E-10 |

| | | | | | |
|---|---|---|---|---|---|
| *F36D4.4* | WBGene00018089 | protein_coding_gene | 1.5 | 2.78 | 1.2E-02 |
| *AH9.6* | WBGene00044701 | protein_coding_gene | 1.5 | 2.78 | 8.4E-04 |
| *Y6B3B.1* | WBGene00012387 | protein_coding_gene | 1.5 | 2.78 | 3.2E-04 |
| *sulp-1* | WBGene00016945 | protein_coding_gene | 1.5 | 2.78 | 1.2E-02 |
| *pqn-36* | WBGene00004123 | protein_coding_gene | 1.5 | 2.77 | 6.7E-05 |
| *C11E4.8* | WBGene00007522 | protein_coding_gene | 1.5 | 2.77 | 2.4E-02 |
| *T05A1.5* | WBGene00011456 | protein_coding_gene | 1.5 | 2.77 | 7.1E-03 |
| *cut-4* | WBGene00008482 | protein_coding_gene | 1.5 | 2.77 | 8.8E-03 |
| *B0207.7* | WBGene00015030 | protein_coding_gene | 1.5 | 2.77 | 1.1E-05 |
| *T25G12.6* | WBGene00020813 | protein_coding_gene | 1.5 | 2.77 | 1.9E-05 |
| *F44B9.9* | WBGene00018410 | protein_coding_gene | 1.5 | 2.76 | 2.7E-02 |
| *lgc-21* | WBGene00007903 | protein_coding_gene | 1.5 | 2.76 | 2.0E-02 |
| *C47A10.12* | WBGene00008129 | protein_coding_gene | 1.5 | 2.76 | 2.6E-08 |
| *nas-22* | WBGene00003541 | protein_coding_gene | 1.5 | 2.76 | 1.9E-02 |
| *acr-8* | WBGene00000047 | protein_coding_gene | 1.5 | 2.76 | 4.1E-04 |
| *pqn-22* | WBGene00004112 | protein_coding_gene | 1.5 | 2.76 | 8.8E-15 |
| *C38D9.2* | WBGene00008010 | protein_coding_gene | 1.5 | 2.76 | 8.6E-06 |
| *C01H6.4* | WBGene00007254 | protein_coding_gene | 1.5 | 2.75 | 1.2E-09 |
| *T04A8.13* | WBGene00011414 | protein_coding_gene | 1.5 | 2.75 | 9.8E-05 |
| *clec-155* | WBGene00011405 | pseudogene | 1.5 | 2.75 | 9.6E-03 |
| *glb-7* | WBGene00016024 | protein_coding_gene | 1.5 | 2.75 | 3.1E-02 |
| *Y54E2A.10* | WBGene00013195 | protein_coding_gene | 1.5 | 2.75 | 3.6E-03 |
| *F13H8.6* | WBGene00017439 | protein_coding_gene | 1.5 | 2.74 | 3.3E-02 |
| *T01C8.3* | WBGene00020144 | protein_coding_gene | 1.5 | 2.74 | 2.7E-03 |
| *fbxa-60* | WBGene00020456 | protein_coding_gene | 1.5 | 2.74 | 3.9E-03 |
| *ugt-32* | WBGene00018543 | protein_coding_gene | 1.5 | 2.74 | 1.1E-02 |
| *tor-1* | WBGene00006597 | protein_coding_gene | 1.5 | 2.74 | 4.9E-02 |
| *W06H8.2* | WBGene00021066 | protein_coding_gene | 1.5 | 2.74 | 2.3E-04 |
| *C30F2.3* | WBGene00007817 | protein_coding_gene | 1.5 | 2.73 | 6.2E-05 |
| *gst-41* | WBGene00001789 | protein_coding_gene | 1.5 | 2.73 | 2.3E-03 |
| *col-39* | WBGene00000616 | protein_coding_gene | 1.5 | 2.73 | 2.1E-14 |
| *F26A1.4* | WBGene00017803 | protein_coding_gene | 1.4 | 2.73 | 1.7E-03 |
| *T10B10.4* | WBGene00011681 | protein_coding_gene | 1.4 | 2.73 | 1.2E-13 |
| *sod-3* | WBGene00004932 | protein_coding_gene | 1.4 | 2.73 | 1.4E-06 |
| *T19D12.9* | WBGene00020583 | protein_coding_gene | 1.4 | 2.72 | 3.3E-02 |
| *Y40H7A.11* | WBGene00012748 | protein_coding_gene | 1.4 | 2.72 | 4.9E-02 |
| *unc-17* | WBGene00006756 | protein_coding_gene | 1.4 | 2.72 | 8.0E-04 |
| *F59F5.5* | WBGene00010344 | protein_coding_gene | 1.4 | 2.72 | 1.7E-02 |
| *adt-3* | WBGene00020567 | protein_coding_gene | 1.4 | 2.72 | 2.0E-05 |
| *lec-9* | WBGene00002272 | protein_coding_gene | 1.4 | 2.72 | 1.2E-13 |
| *F09E10.7* | WBGene00017297 | protein_coding_gene | 1.4 | 2.71 | 1.3E-04 |
| *tag-180* | WBGene00007041 | protein_coding_gene | 1.4 | 2.71 | 4.5E-07 |
| *H01G02.4* | WBGene00045108 | protein_coding_gene | 1.4 | 2.71 | 4.9E-02 |
| *clec-132* | WBGene00021143 | protein_coding_gene | 1.4 | 2.71 | 1.2E-05 |
| *T12A2.5* | WBGene00020439 | protein_coding_gene | 1.4 | 2.71 | 3.5E-07 |
| *ifc-1* | WBGene00002055 | protein_coding_gene | 1.4 | 2.71 | 4.5E-32 |
| *F25D7.5* | WBGene00009114 | protein_coding_gene | 1.4 | 2.70 | 1.1E-02 |
| *nas-38* | WBGene00003554 | protein_coding_gene | 1.4 | 2.70 | 4.3E-02 |
| *ugt-3* | WBGene00013903 | protein_coding_gene | 1.4 | 2.70 | 3.2E-02 |
| *ZK250.14* | WBGene00235354 | protein_coding_gene | 1.4 | 2.70 | 3.7E-02 |
| *K02B12.9* | WBGene00044679 | protein_coding_gene | 1.4 | 2.69 | 4.7E-02 |
| *F01D5.3* | WBGene00008494 | protein_coding_gene | 1.4 | 2.69 | 1.6E-02 |
| *doxa-1* | WBGene00015519 | protein_coding_gene | 1.4 | 2.69 | 1.7E-06 |
| *lev-9* | WBGene00002976 | protein_coding_gene | 1.4 | 2.68 | 1.1E-04 |
| *F15H9.7* | WBGene00077535 | pseudogene | 1.4 | 2.68 | 3.7E-02 |
| *F47F2.1* | WBGene00018569 | protein_coding_gene | 1.4 | 2.68 | 6.3E-08 |
| *unc-52* | WBGene00006787 | protein_coding_gene | 1.4 | 2.68 | 1.9E-24 |
| *egl-47* | WBGene00001211 | protein_coding_gene | 1.4 | 2.67 | 4.8E-02 |

| | | | | | |
|---|---|---|---|---|---|
| *ctsa-3.1* | WBGene00017969 | protein_coding_gene | 1.4 | 2.67 | 7.8E-03 |
| *dpf-6* | WBGene00001059 | protein_coding_gene | 1.4 | 2.67 | 1.3E-05 |
| *sro-1* | WBGene00005641 | protein_coding_gene | 1.4 | 2.67 | 1.2E-02 |
| *T06A4.3* | WBGene00020283 | protein_coding_gene | 1.4 | 2.66 | 2.0E-03 |
| *pde-5* | WBGene00016328 | protein_coding_gene | 1.4 | 2.66 | 3.6E-02 |
| *cest-19* | WBGene00020125 | protein_coding_gene | 1.4 | 2.66 | 7.5E-03 |
| *col-98* | WBGene00000673 | protein_coding_gene | 1.4 | 2.66 | 5.3E-24 |
| *C47D2.1* | WBGene00016738 | protein_coding_gene | 1.4 | 2.65 | 5.5E-03 |
| *math-2* | WBGene00015606 | protein_coding_gene | 1.4 | 2.65 | 1.7E-03 |
| *C02B8.3* | WBGene00015322 | protein_coding_gene | 1.4 | 2.65 | 5.0E-04 |
| *T25C12.3* | WBGene00012018 | protein_coding_gene | 1.4 | 2.65 | 8.6E-04 |
| *F11F1.1* | WBGene00008714 | protein_coding_gene | 1.4 | 2.64 | 4.0E-07 |
| *clec-105* | WBGene00012488 | protein_coding_gene | 1.4 | 2.64 | 6.3E-04 |
| *F35F10.4* | WBGene00018054 | pseudogene | 1.4 | 2.64 | 5.0E-03 |
| *hsp-17* | WBGene00002021 | protein_coding_gene | 1.4 | 2.64 | 2.3E-10 |
| *clec-204* | WBGene00021603 | protein_coding_gene | 1.4 | 2.64 | 3.6E-04 |
| *K12C11.5* | WBGene00044896 | protein_coding_gene | 1.4 | 2.63 | 1.4E-02 |
| *T04F3.1* | WBGene00011436 | protein_coding_gene | 1.4 | 2.63 | 3.3E-27 |
| *T10H9.1* | WBGene00020424 | protein_coding_gene | 1.4 | 2.63 | 1.1E-02 |
| *ZK402.5* | WBGene00022731 | protein_coding_gene | 1.4 | 2.63 | 2.1E-02 |
| *C31B8.12* | WBGene00016285 | protein_coding_gene | 1.4 | 2.62 | 3.0E-03 |
| *flr-4* | WBGene00001468 | protein_coding_gene | 1.4 | 2.62 | 6.9E-05 |
| *C17H12.37* | WBGene00236786 | protein_coding_gene | 1.4 | 2.62 | 4.4E-02 |
| *kvs-4* | WBGene00012967 | protein_coding_gene | 1.4 | 2.62 | 7.1E-03 |
| *add-1* | WBGene00000072 | protein_coding_gene | 1.4 | 2.62 | 3.3E-10 |
| *ZK783.6* | WBGene00044448 | protein_coding_gene | 1.4 | 2.61 | 4.9E-03 |
| *Y51A2D.1* | WBGene00013072 | protein_coding_gene | 1.4 | 2.61 | 3.8E-04 |
| *cha-1* | WBGene00000481 | protein_coding_gene | 1.4 | 2.61 | 2.2E-04 |
| *M03B6.1* | WBGene00010833 | protein_coding_gene | 1.4 | 2.61 | 7.7E-04 |
| *gly-18* | WBGene00001643 | protein_coding_gene | 1.4 | 2.61 | 2.0E-02 |
| *clec-108* | WBGene00012504 | protein_coding_gene | 1.4 | 2.60 | 4.2E-03 |
| *lev-1* | WBGene00002974 | protein_coding_gene | 1.4 | 2.60 | 4.0E-06 |
| *M163.8* | WBGene00010940 | protein_coding_gene | 1.4 | 2.60 | 8.8E-07 |
| *lgc-32* | WBGene00020569 | protein_coding_gene | 1.4 | 2.59 | 1.4E-06 |
| *B0457.3* | WBGene00014660 | pseudogene | 1.4 | 2.59 | 4.3E-02 |
| *R09H10.5* | WBGene00011182 | protein_coding_gene | 1.4 | 2.59 | 3.8E-05 |
| *D2023.1* | WBGene00014300 | protein_coding_gene | 1.4 | 2.59 | 3.6E-07 |
| *cyp-35D1* | WBGene00008829 | protein_coding_gene | 1.4 | 2.59 | 5.4E-03 |
| *twk-22* | WBGene00006675 | protein_coding_gene | 1.4 | 2.59 | 1.2E-06 |
| *glr-4* | WBGene00001615 | protein_coding_gene | 1.4 | 2.58 | 2.7E-06 |
| *C14B9.3* | WBGene00015753 | protein_coding_gene | 1.4 | 2.58 | 2.2E-04 |
| *T02G5.7* | WBGene00020166 | protein_coding_gene | 1.4 | 2.58 | 6.8E-23 |
| *lgc-45* | WBGene00021146 | protein_coding_gene | 1.4 | 2.57 | 4.6E-10 |
| *col-96* | WBGene00000671 | protein_coding_gene | 1.4 | 2.57 | 6.8E-08 |
| *W10C8.5* | WBGene00021128 | protein_coding_gene | 1.4 | 2.57 | 4.4E-16 |
| *Y43F8B.3* | WBGene00012814 | protein_coding_gene | 1.4 | 2.56 | 1.8E-03 |
| *nspd-7* | WBGene00012781 | protein_coding_gene | 1.4 | 2.56 | 4.2E-02 |
| *T21D12.12* | WBGene00020651 | protein_coding_gene | 1.4 | 2.56 | 6.5E-11 |
| *col-103* | WBGene00000677 | protein_coding_gene | 1.4 | 2.55 | 5.4E-23 |
| *F58H7.1* | WBGene00019067 | protein_coding_gene | 1.4 | 2.55 | 3.9E-02 |
| *Y43F8C.17* | WBGene00012838 | protein_coding_gene | 1.4 | 2.55 | 1.0E-10 |
| *vha-5* | WBGene00006914 | protein_coding_gene | 1.4 | 2.55 | 3.6E-23 |
| *clec-165* | WBGene00009523 | protein_coding_gene | 1.4 | 2.55 | 6.7E-04 |
| *unc-41* | WBGene00006777 | protein_coding_gene | 1.4 | 2.55 | 7.2E-07 |
| *F13B6.3* | WBGene00017418 | protein_coding_gene | 1.4 | 2.55 | 6.8E-03 |
| *C34F11.1* | WBGene00016413 | protein_coding_gene | 1.4 | 2.55 | 8.9E-03 |
| *let-805* | WBGene00002915 | protein_coding_gene | 1.3 | 2.55 | 1.2E-15 |
| *cpr-3* | WBGene00000783 | protein_coding_gene | 1.3 | 2.54 | 4.7E-10 |

| | | | | | |
|---|---|---|---|---|---|
| *F09E5.16* | WBGene00017292 | protein_coding_gene | 1.3 | 2.53 | 4.0E-07 |
| *F57F4.4* | WBGene00019017 | protein_coding_gene | 1.3 | 2.53 | 3.1E-02 |
| *skpo-3* | WBGene00017968 | protein_coding_gene | 1.3 | 2.53 | 3.0E-24 |
| *C50D2.2* | WBGene00016806 | protein_coding_gene | 1.3 | 2.53 | 3.0E-03 |
| *cwp-1* | WBGene00000859 | protein_coding_gene | 1.3 | 2.53 | 2.3E-04 |
| *shl-1* | WBGene00022240 | protein_coding_gene | 1.3 | 2.52 | 2.3E-04 |
| *hpo-28* | WBGene00015148 | protein_coding_gene | 1.3 | 2.52 | 3.3E-06 |
| *B0416.2* | WBGene00015178 | protein_coding_gene | 1.3 | 2.52 | 1.8E-02 |
| *pmp-1* | WBGene00004058 | protein_coding_gene | 1.3 | 2.52 | 4.1E-15 |
| *cwp-2* | WBGene00000860 | protein_coding_gene | 1.3 | 2.52 | 4.7E-02 |
| *pat-12* | WBGene00003936 | protein_coding_gene | 1.3 | 2.52 | 5.4E-21 |
| *bgnt-1.6* | WBGene00010167 | protein_coding_gene | 1.3 | 2.52 | 2.6E-02 |
| *nlp-26* | WBGene00003764 | protein_coding_gene | 1.3 | 2.51 | 3.0E-05 |
| *gar-2* | WBGene00001518 | protein_coding_gene | 1.3 | 2.51 | 7.8E-06 |
| *gst-36* | WBGene00001784 | protein_coding_gene | 1.3 | 2.51 | 1.5E-12 |
| *glb-13* | WBGene00008957 | protein_coding_gene | 1.3 | 2.51 | 6.2E-03 |
| *pals-38* | WBGene00008302 | protein_coding_gene | 1.3 | 2.51 | 1.7E-02 |
| *F23C8.7* | WBGene00017736 | protein_coding_gene | 1.3 | 2.51 | 2.9E-07 |
| *C12D12.1* | WBGene00015713 | protein_coding_gene | 1.3 | 2.51 | 2.8E-05 |
| *snf-4* | WBGene00004903 | pseudogene | 1.3 | 2.50 | 4.2E-03 |
| *pqn-72* | WBGene00004154 | protein_coding_gene | 1.3 | 2.50 | 7.3E-04 |
| *F56B6.6* | WBGene00018937 | protein_coding_gene | 1.3 | 2.49 | 3.0E-02 |
| *C35B1.4* | WBGene00016434 | protein_coding_gene | 1.3 | 2.49 | 1.1E-10 |
| *C14A11.6* | WBGene00015751 | protein_coding_gene | 1.3 | 2.49 | 2.5E-03 |
| *nhr-179* | WBGene00017512 | protein_coding_gene | 1.3 | 2.49 | 1.1E-05 |
| *VB0395L.1* | WBGene00044150 | protein_coding_gene | 1.3 | 2.48 | 3.8E-03 |
| *R08C7.5* | WBGene00019949 | protein_coding_gene | 1.3 | 2.48 | 1.0E-06 |
| *arrd-8* | WBGene00012467 | protein_coding_gene | 1.3 | 2.48 | 2.0E-02 |
| *T10G3.8* | WBGene00045340 | protein_coding_gene | 1.3 | 2.48 | 1.9E-02 |
| *pho-7* | WBGene00011879 | protein_coding_gene | 1.3 | 2.48 | 2.4E-04 |
| *amt-4* | WBGene00000136 | protein_coding_gene | 1.3 | 2.48 | 1.3E-11 |
| *slc-36.4* | WBGene00010421 | protein_coding_gene | 1.3 | 2.48 | 1.8E-06 |
| *lact-7* | WBGene00008047 | protein_coding_gene | 1.3 | 2.48 | 2.6E-02 |
| *cyp-25A1* | WBGene00007963 | protein_coding_gene | 1.3 | 2.48 | 1.5E-04 |
| *gcy-32* | WBGene00001552 | protein_coding_gene | 1.3 | 2.47 | 3.2E-02 |
| *ets-4* | WBGene00017687 | protein_coding_gene | 1.3 | 2.47 | 4.0E-17 |
| *pqn-35* | WBGene00004122 | protein_coding_gene | 1.3 | 2.47 | 2.7E-10 |
| *Y116F11A.3* | WBGene00013817 | protein_coding_gene | 1.3 | 2.47 | 2.7E-02 |
| *mab-3* | WBGene00003100 | protein_coding_gene | 1.3 | 2.47 | 5.9E-06 |
| *Y51H4A.935* | WBGene00194647 | protein_coding_gene | 1.3 | 2.47 | 2.4E-02 |
| *dmsr-3* | WBGene00012993 | protein_coding_gene | 1.3 | 2.47 | 1.5E-02 |
| *Y48G8AR.2* | WBGene00021693 | protein_coding_gene | 1.3 | 2.47 | 3.3E-02 |
| *C28D4.8* | WBGene00007795 | protein_coding_gene | 1.3 | 2.47 | 1.3E-02 |
| *C18B2.3* | WBGene00015954 | protein_coding_gene | 1.3 | 2.47 | 1.1E-07 |
| *F43C11.2* | WBGene00018379 | protein_coding_gene | 1.3 | 2.46 | 9.4E-08 |
| *twk-40* | WBGene00006691 | protein_coding_gene | 1.3 | 2.46 | 2.9E-02 |
| *gst-35* | WBGene00001783 | protein_coding_gene | 1.3 | 2.46 | 8.9E-04 |
| *F46A9.2* | WBGene00009755 | protein_coding_gene | 1.3 | 2.46 | 4.4E-06 |
| *Y53C10A.15* | WBGene00044805 | protein_coding_gene | 1.3 | 2.45 | 2.4E-02 |
| *clec-135* | WBGene00015821 | protein_coding_gene | 1.3 | 2.45 | 1.7E-06 |
| *frpr-17* | WBGene00011765 | protein_coding_gene | 1.3 | 2.45 | 6.3E-03 |
| *ceh-18* | WBGene00000441 | protein_coding_gene | 1.3 | 2.45 | 1.0E-10 |
| *kcc-3* | WBGene00019289 | protein_coding_gene | 1.3 | 2.45 | 2.1E-06 |
| *D1054.8* | WBGene00008375 | protein_coding_gene | 1.3 | 2.45 | 3.6E-13 |
| *che-10* | WBGene00018051 | protein_coding_gene | 1.3 | 2.45 | 3.6E-14 |
| *aqp-4* | WBGene00000172 | protein_coding_gene | 1.3 | 2.45 | 3.6E-09 |
| *T01C1.4* | WBGene00011317 | protein_coding_gene | 1.3 | 2.44 | 4.1E-02 |
| *prhg-1* | WBGene00022391 | protein_coding_gene | 1.3 | 2.44 | 6.1E-04 |

| | | | | | |
|---|---|---|---|---|---|
| *sur-5* | WBGene00006351 | protein_coding_gene | 1.3 | 2.44 | 1.3E-16 |
| *T05H4.7* | WBGene00020270 | protein_coding_gene | 1.3 | 2.44 | 1.9E-02 |
| *F15D4.5* | WBGene00008862 | protein_coding_gene | 1.3 | 2.44 | 9.2E-05 |
| *scl-26* | WBGene00016821 | protein_coding_gene | 1.3 | 2.44 | 8.5E-03 |
| *B0222.11* | WBGene00044582 | protein_coding_gene | 1.3 | 2.44 | 2.2E-02 |
| *ZC404.1* | WBGene00022601 | protein_coding_gene | 1.3 | 2.44 | 3.1E-02 |
| *ifc-2* | WBGene00002056 | protein_coding_gene | 1.3 | 2.43 | 9.4E-06 |
| *ggr-2* | WBGene00001587 | protein_coding_gene | 1.3 | 2.43 | 2.2E-02 |
| *amx-2* | WBGene00000138 | protein_coding_gene | 1.3 | 2.43 | 1.9E-12 |
| *ncx-6* | WBGene00003571 | protein_coding_gene | 1.3 | 2.43 | 1.4E-04 |
| *aqp-8* | WBGene00000176 | protein_coding_gene | 1.3 | 2.43 | 7.2E-22 |
| *grl-4* | WBGene00001713 | protein_coding_gene | 1.3 | 2.42 | 3.3E-06 |
| *B0238.12* | WBGene00015076 | protein_coding_gene | 1.3 | 2.42 | 5.9E-04 |
| *Y66C5A.1* | WBGene00044156 | protein_coding_gene | 1.3 | 2.42 | 3.4E-02 |
| *F59A6.11* | WBGene00044664 | protein_coding_gene | 1.3 | 2.42 | 9.2E-03 |
| *C49F8.3* | WBGene00008215 | protein_coding_gene | 1.3 | 2.42 | 7.2E-04 |
| *C47E8.10* | WBGene00044216 | pseudogene | 1.3 | 2.42 | 1.7E-02 |
| *Y116A8A.6* | WBGene00013776 | protein_coding_gene | 1.3 | 2.42 | 2.8E-02 |
| *twk-17* | WBGene00006671 | protein_coding_gene | 1.3 | 2.41 | 1.4E-03 |
| *exc-1* | WBGene00001362 | protein_coding_gene | 1.3 | 2.41 | 4.5E-02 |
| *pfas-1* | WBGene00008654 | protein_coding_gene | 1.3 | 2.40 | 1.8E-15 |
| *ketn-1* | WBGene00004130 | protein_coding_gene | 1.3 | 2.40 | 9.1E-14 |
| *F57A8.6* | WBGene00010181 | protein_coding_gene | 1.3 | 2.40 | 2.6E-03 |
| *eat-4* | WBGene00001135 | protein_coding_gene | 1.3 | 2.40 | 8.3E-04 |
| *F52H2.6* | WBGene00018718 | protein_coding_gene | 1.3 | 2.40 | 6.7E-10 |
| *F07H5.7* | WBGene00008558 | protein_coding_gene | 1.3 | 2.40 | 1.6E-02 |
| *ZK1193.2* | WBGene00022858 | protein_coding_gene | 1.3 | 2.39 | 6.0E-08 |
| *sprr-1* | WBGene00010986 | protein_coding_gene | 1.3 | 2.39 | 3.4E-02 |
| *sams-1* | WBGene00008205 | protein_coding_gene | 1.3 | 2.39 | 5.5E-23 |
| *frpr-6* | WBGene00017661 | protein_coding_gene | 1.3 | 2.39 | 2.8E-02 |
| *aqp-7* | WBGene00000175 | protein_coding_gene | 1.3 | 2.39 | 2.5E-13 |
| *C45E5.4* | WBGene00016667 | protein_coding_gene | 1.3 | 2.39 | 1.2E-05 |
| *acr-16* | WBGene00000055 | protein_coding_gene | 1.3 | 2.39 | 2.6E-03 |
| *F40F9.3* | WBGene00009581 | protein_coding_gene | 1.3 | 2.38 | 2.4E-05 |
| *T02E9.6* | WBGene00044203 | protein_coding_gene | 1.2 | 2.38 | 4.1E-02 |
| *glr-5* | WBGene00001616 | protein_coding_gene | 1.2 | 2.38 | 1.2E-04 |
| *R02F2.5* | WBGene00019834 | protein_coding_gene | 1.2 | 2.38 | 2.9E-02 |
| *F29C6.1* | WBGene00009243 | protein_coding_gene | 1.2 | 2.38 | 8.1E-10 |
| *F38A1.9* | WBGene00009522 | protein_coding_gene | 1.2 | 2.37 | 4.4E-04 |
| *C28D4.5* | WBGene00007793 | protein_coding_gene | 1.2 | 2.37 | 4.2E-02 |
| *mec-9* | WBGene00003173 | protein_coding_gene | 1.2 | 2.37 | 3.0E-03 |
| *T28B8.3* | WBGene00012114 | protein_coding_gene | 1.2 | 2.37 | 3.7E-02 |
| *ttn-1* | WBGene00006436 | protein_coding_gene | 1.2 | 2.37 | 1.1E-03 |
| *F14H12.3* | WBGene00017471 | protein_coding_gene | 1.2 | 2.36 | 1.7E-05 |
| *C55A6.4* | WBGene00008333 | protein_coding_gene | 1.2 | 2.36 | 1.9E-07 |
| *lys-5* | WBGene00003094 | protein_coding_gene | 1.2 | 2.36 | 3.2E-04 |
| *kvs-5* | WBGene00021948 | protein_coding_gene | 1.2 | 2.36 | 8.5E-08 |
| *dpf-2* | WBGene00001055 | protein_coding_gene | 1.2 | 2.36 | 2.8E-15 |
| *arrd-26* | WBGene00019873 | protein_coding_gene | 1.2 | 2.36 | 9.5E-03 |
| *fipr-11* | WBGene00008244 | protein_coding_gene | 1.2 | 2.36 | 1.5E-02 |
| *kcnl-1* | WBGene00007176 | protein_coding_gene | 1.2 | 2.35 | 1.6E-04 |
| *tbc-8* | WBGene00008018 | protein_coding_gene | 1.2 | 2.35 | 1.6E-04 |
| *oac-19* | WBGene00009413 | protein_coding_gene | 1.2 | 2.35 | 1.8E-02 |
| *epac-1* | WBGene00004255 | protein_coding_gene | 1.2 | 2.35 | 4.9E-02 |
| *ZK688.12* | WBGene00044763 | protein_coding_gene | 1.2 | 2.35 | 3.7E-03 |
| *cpn-4* | WBGene00000780 | protein_coding_gene | 1.2 | 2.35 | 2.6E-17 |
| *tnt-4* | WBGene00006589 | protein_coding_gene | 1.2 | 2.35 | 1.9E-15 |
| *rimb-1* | WBGene00006513 | protein_coding_gene | 1.2 | 2.35 | 5.5E-06 |

| | | | | | |
|---|---|---|---|---|---|
| *C09G1.5* | WBGene00007487 | protein_coding_gene | 1.2 | 2.35 | 1.9E-03 |
| *unc-23* | WBGene00006760 | protein_coding_gene | 1.2 | 2.34 | 6.0E-09 |
| *lat-2* | WBGene00002252 | protein_coding_gene | 1.2 | 2.34 | 7.4E-05 |
| *fbxa-54* | WBGene00020449 | protein_coding_gene | 1.2 | 2.34 | 3.1E-02 |
| *ilcr-1* | WBGene00013415 | protein_coding_gene | 1.2 | 2.34 | 3.6E-08 |
| *lin-29* | WBGene00003015 | protein_coding_gene | 1.2 | 2.34 | 1.5E-08 |
| *Y44A6D.2* | WBGene00012853 | protein_coding_gene | 1.2 | 2.33 | 1.2E-06 |
| *M03F8.5* | WBGene00019764 | protein_coding_gene | 1.2 | 2.33 | 5.0E-02 |
| *clec-92* | WBGene00013125 | protein_coding_gene | 1.2 | 2.33 | 9.0E-06 |
| *W07G4.5* | WBGene00012339 | protein_coding_gene | 1.2 | 2.33 | 4.6E-10 |
| *T20B6.2* | WBGene00020598 | protein_coding_gene | 1.2 | 2.33 | 2.5E-02 |
| *W03D8.5* | WBGene00020987 | protein_coding_gene | 1.2 | 2.33 | 5.4E-04 |
| *F46C5.2* | WBGene00009779 | protein_coding_gene | 1.2 | 2.33 | 8.2E-03 |
| *T22F7.4* | WBGene00020703 | protein_coding_gene | 1.2 | 2.33 | 4.7E-03 |
| *C01B10.44* | WBGene00194839 | protein_coding_gene | 1.2 | 2.33 | 3.5E-02 |
| *C49A9.3* | WBGene00016757 | protein_coding_gene | 1.2 | 2.33 | 5.2E-06 |
| *Y65B4A.7* | WBGene00022030 | protein_coding_gene | 1.2 | 2.32 | 2.5E-02 |
| *aco-1* | WBGene00000040 | protein_coding_gene | 1.2 | 2.32 | 1.9E-17 |
| *unc-46* | WBGene00006782 | protein_coding_gene | 1.2 | 2.32 | 5.0E-02 |
| *cat-1* | WBGene00000295 | protein_coding_gene | 1.2 | 2.32 | 6.8E-03 |
| *ZK622.4* | WBGene00022782 | protein_coding_gene | 1.2 | 2.31 | 1.2E-06 |
| *hex-5* | WBGene00013497 | protein_coding_gene | 1.2 | 2.31 | 2.8E-05 |
| *C01B10.3* | WBGene00015278 | protein_coding_gene | 1.2 | 2.31 | 2.0E-12 |
| *C30B5.7* | WBGene00016248 | protein_coding_gene | 1.2 | 2.31 | 2.1E-02 |
| *C09B9.7* | WBGene00015630 | protein_coding_gene | 1.2 | 2.31 | 1.1E-04 |
| *cdkl-1* | WBGene00012779 | protein_coding_gene | 1.2 | 2.31 | 4.7E-02 |
| *F37C4.6* | WBGene00018146 | protein_coding_gene | 1.2 | 2.31 | 1.5E-11 |
| *klu-2* | WBGene00022592 | protein_coding_gene | 1.2 | 2.31 | 1.4E-07 |
| *Y110A2AL.6* | WBGene00022443 | protein_coding_gene | 1.2 | 2.30 | 5.9E-04 |
| *F33C8.4* | WBGene00009358 | protein_coding_gene | 1.2 | 2.30 | 2.1E-07 |
| *lgc-53* | WBGene00020657 | protein_coding_gene | 1.2 | 2.30 | 3.8E-02 |
| *cal-6* | WBGene00022214 | pseudogene | 1.2 | 2.30 | 4.7E-02 |
| *sqrd-1* | WBGene00008538 | protein_coding_gene | 1.2 | 2.29 | 6.9E-09 |
| *B0034.1* | WBGene00015002 | protein_coding_gene | 1.2 | 2.29 | 1.8E-03 |
| *clec-99* | WBGene00013933 | protein_coding_gene | 1.2 | 2.29 | 2.3E-09 |
| *unc-25* | WBGene00006762 | protein_coding_gene | 1.2 | 2.29 | 5.6E-09 |
| *idh-1* | WBGene00010317 | protein_coding_gene | 1.2 | 2.29 | 4.6E-15 |
| *mec-1* | WBGene00003165 | protein_coding_gene | 1.2 | 2.29 | 1.8E-05 |
| *C12D5.9* | WBGene00015710 | protein_coding_gene | 1.2 | 2.29 | 4.6E-05 |
| *C49A9.5* | WBGene00016759 | protein_coding_gene | 1.2 | 2.29 | 1.4E-09 |
| *bca-1* | WBGene00000245 | protein_coding_gene | 1.2 | 2.29 | 7.8E-09 |
| *gst-11* | WBGene00001759 | protein_coding_gene | 1.2 | 2.29 | 3.5E-02 |
| *slc-17.3* | WBGene00015333 | protein_coding_gene | 1.2 | 2.28 | 4.2E-05 |
| *M02D8.5* | WBGene00019731 | protein_coding_gene | 1.2 | 2.28 | 1.5E-05 |
| *F26B1.5* | WBGene00017817 | protein_coding_gene | 1.2 | 2.28 | 1.3E-02 |
| *F19B6.3* | WBGene00008949 | protein_coding_gene | 1.2 | 2.28 | 1.6E-02 |
| *Y39B6A.41* | WBGene00012699 | protein_coding_gene | 1.2 | 2.28 | 4.2E-03 |
| *ocr-2* | WBGene00003839 | protein_coding_gene | 1.2 | 2.28 | 1.7E-02 |
| *fbxa-223* | WBGene00022630 | pseudogene | 1.2 | 2.28 | 3.3E-02 |
| *F35F10.1* | WBGene00018052 | protein_coding_gene | 1.2 | 2.28 | 8.4E-04 |
| *acr-24* | WBGene00013534 | protein_coding_gene | 1.2 | 2.27 | 3.2E-02 |
| *ifa-1* | WBGene00002050 | protein_coding_gene | 1.2 | 2.27 | 3.9E-16 |
| *flp-23* | WBGene00017693 | protein_coding_gene | 1.2 | 2.27 | 2.2E-02 |
| *ttll-11* | WBGene00019230 | protein_coding_gene | 1.2 | 2.27 | 3.6E-02 |
| *ins-28* | WBGene00002111 | protein_coding_gene | 1.2 | 2.27 | 8.4E-03 |
| *clec-130* | WBGene00021144 | protein_coding_gene | 1.2 | 2.27 | 1.0E-02 |
| *F58B6.1* | WBGene00019029 | protein_coding_gene | 1.2 | 2.27 | 1.4E-02 |
| *pqn-26* | WBGene00004115 | protein_coding_gene | 1.2 | 2.27 | 3.8E-02 |

| | | | | | |
|---|---|---|---|---|---|
| *Y46D2A.3* | WBGene00021591 | protein_coding_gene | 1.2 | 2.27 | 4.8E-03 |
| *T09B4.3* | WBGene00020377 | protein_coding_gene | 1.2 | 2.26 | 1.1E-02 |
| *clec-56* | WBGene00008595 | protein_coding_gene | 1.2 | 2.26 | 6.5E-11 |
| *Y6G8.16* | WBGene00194801 | protein_coding_gene | 1.2 | 2.26 | 3.3E-02 |
| *F44G3.10* | WBGene00009710 | protein_coding_gene | 1.2 | 2.26 | 2.9E-03 |
| *mct-3* | WBGene00010834 | protein_coding_gene | 1.2 | 2.26 | 3.7E-20 |
| *T05F1.9* | WBGene00011494 | protein_coding_gene | 1.2 | 2.26 | 3.3E-02 |
| *sng-1* | WBGene00004912 | protein_coding_gene | 1.2 | 2.25 | 2.4E-03 |
| *Y6G8.15* | WBGene00189937 | protein_coding_gene | 1.2 | 2.25 | 5.2E-05 |
| *gyg-1* | WBGene00006863 | protein_coding_gene | 1.2 | 2.25 | 6.4E-13 |
| *tmc-1* | WBGene00020490 | protein_coding_gene | 1.2 | 2.25 | 7.6E-05 |
| *Y71H2AM.12* | WBGene00022177 | protein_coding_gene | 1.2 | 2.25 | 1.3E-04 |
| *F46A9.1* | WBGene00009754 | protein_coding_gene | 1.2 | 2.25 | 8.8E-07 |
| *F38E1.10* | WBGene00018182 | protein_coding_gene | 1.2 | 2.25 | 8.8E-04 |
| *wago-10* | WBGene00020707 | protein_coding_gene | 1.2 | 2.25 | 1.7E-06 |
| *dop-3* | WBGene00020506 | protein_coding_gene | 1.2 | 2.25 | 1.4E-02 |
| *lgc-39* | WBGene00017314 | protein_coding_gene | 1.2 | 2.25 | 1.3E-04 |
| *fipr-3* | WBGene00009090 | protein_coding_gene | 1.2 | 2.25 | 1.4E-03 |
| *lgc-46* | WBGene00022106 | protein_coding_gene | 1.2 | 2.25 | 1.4E-02 |
| *W04E12.7* | WBGene00012252 | protein_coding_gene | 1.2 | 2.24 | 1.9E-04 |
| *fut-5* | WBGene00043986 | protein_coding_gene | 1.2 | 2.24 | 4.6E-02 |
| *Y69E1A.3* | WBGene00013475 | protein_coding_gene | 1.2 | 2.24 | 8.1E-03 |
| *qui-1* | WBGene00004265 | protein_coding_gene | 1.2 | 2.24 | 9.8E-07 |
| *F42G10.1* | WBGene00009645 | protein_coding_gene | 1.2 | 2.24 | 6.7E-06 |
| *B0294.1* | WBGene00015117 | protein_coding_gene | 1.2 | 2.24 | 4.7E-02 |
| *ggr-3* | WBGene00001588 | protein_coding_gene | 1.2 | 2.24 | 1.5E-02 |
| *Y43F8B.19* | WBGene00077688 | protein_coding_gene | 1.2 | 2.24 | 1.1E-02 |
| *Y77E11A.12* | WBGene00022313 | protein_coding_gene | 1.2 | 2.24 | 1.2E-02 |
| *ser-1* | WBGene00004776 | protein_coding_gene | 1.2 | 2.24 | 2.9E-03 |
| *K08E4.7* | WBGene00045036 | protein_coding_gene | 1.2 | 2.24 | 2.1E-03 |
| *inx-1* | WBGene00002123 | protein_coding_gene | 1.2 | 2.24 | 9.4E-06 |
| *R155.4* | WBGene00020118 | protein_coding_gene | 1.2 | 2.24 | 5.2E-03 |
| *W01G7.2* | WBGene00014851 | pseudogene | 1.2 | 2.24 | 2.9E-02 |
| *K01B6.3* | WBGene00010454 | protein_coding_gene | 1.2 | 2.23 | 1.2E-03 |
| *F39H12.2* | WBGene00018213 | protein_coding_gene | 1.2 | 2.23 | 1.9E-02 |
| *T05F1.5* | WBGene00011491 | protein_coding_gene | 1.2 | 2.23 | 2.4E-02 |
| *gon-1* | WBGene00001650 | protein_coding_gene | 1.2 | 2.23 | 2.6E-10 |
| *Y64G10A.2* | WBGene00013412 | protein_coding_gene | 1.2 | 2.23 | 3.6E-02 |
| *mxt-1* | WBGene00012478 | protein_coding_gene | 1.2 | 2.23 | 1.6E-08 |
| *clec-138* | WBGene00018049 | protein_coding_gene | 1.2 | 2.23 | 3.7E-06 |
| *F01D4.8* | WBGene00008490 | protein_coding_gene | 1.2 | 2.22 | 4.9E-03 |
| *Y102A11A.3* | WBGene00022413 | protein_coding_gene | 1.2 | 2.22 | 1.2E-11 |
| *gur-4* | WBGene00001805 | protein_coding_gene | 1.2 | 2.22 | 4.9E-02 |
| *npr-6* | WBGene00009619 | protein_coding_gene | 1.2 | 2.22 | 2.6E-02 |
| *Y37F4.7* | WBGene00021388 | protein_coding_gene | 1.2 | 2.22 | 9.8E-03 |
| *Y53C10A.5* | WBGene00013136 | protein_coding_gene | 1.1 | 2.22 | 6.5E-12 |
| *fbxa-58* | WBGene00016225 | pseudogene | 1.1 | 2.22 | 6.1E-06 |
| *gst-40* | WBGene00001788 | protein_coding_gene | 1.1 | 2.22 | 4.2E-02 |
| *frpr-4* | WBGene00016909 | protein_coding_gene | 1.1 | 2.22 | 2.5E-02 |
| *clec-110* | WBGene00008909 | protein_coding_gene | 1.1 | 2.21 | 1.3E-08 |
| *F46F5.1* | WBGene00018493 | protein_coding_gene | 1.1 | 2.21 | 3.2E-03 |
| *F37B12.1* | WBGene00009504 | protein_coding_gene | 1.1 | 2.21 | 3.2E-03 |
| *F31D5.2* | WBGene00017945 | protein_coding_gene | 1.1 | 2.21 | 6.2E-06 |
| *csp-2* | WBGene00000820 | protein_coding_gene | 1.1 | 2.21 | 2.1E-02 |
| *C08G5.1* | WBGene00015614 | pseudogene | 1.1 | 2.21 | 5.8E-03 |
| *alh-12* | WBGene00000118 | protein_coding_gene | 1.1 | 2.21 | 2.0E-18 |
| *ugt-48* | WBGene00015965 | protein_coding_gene | 1.1 | 2.20 | 8.9E-07 |
| *F58F9.8* | WBGene00044306 | protein_coding_gene | 1.1 | 2.20 | 2.6E-03 |

| | | | | | |
|---|---|---|---|---|---|
| *C36B7.5* | WBGene00016468 | protein_coding_gene | 1.1 | 2.20 | 4.0E-02 |
| *R08E3.1* | WBGene00019957 | protein_coding_gene | 1.1 | 2.20 | 1.2E-12 |
| *ctg-2* | WBGene00011756 | protein_coding_gene | 1.1 | 2.20 | 3.2E-09 |
| *M05B5.3* | WBGene00010871 | protein_coding_gene | 1.1 | 2.20 | 8.0E-05 |
| *C02G6.2* | WBGene00015360 | protein_coding_gene | 1.1 | 2.20 | 9.3E-03 |
| *W03G11.2* | WBGene00012224 | protein_coding_gene | 1.1 | 2.20 | 1.3E-02 |
| *frpr-5* | WBGene00008342 | protein_coding_gene | 1.1 | 2.20 | 1.8E-03 |
| *math-18* | WBGene00016544 | protein_coding_gene | 1.1 | 2.20 | 1.5E-02 |
| *M02D8.2* | WBGene00019728 | protein_coding_gene | 1.1 | 2.20 | 3.9E-02 |
| *F10C1.9* | WBGene00044296 | protein_coding_gene | 1.1 | 2.20 | 6.5E-11 |
| *nspc-1* | WBGene00023477 | protein_coding_gene | 1.1 | 2.19 | 2.1E-02 |
| *cutl-23* | WBGene00021343 | protein_coding_gene | 1.1 | 2.19 | 6.4E-03 |
| *C35A5.4* | WBGene00007948 | protein_coding_gene | 1.1 | 2.19 | 9.3E-04 |
| *nlp-53* | WBGene00012440 | protein_coding_gene | 1.1 | 2.19 | 8.0E-04 |
| *acr-20* | WBGene00000059 | protein_coding_gene | 1.1 | 2.19 | 3.6E-02 |
| *argk-1* | WBGene00009706 | protein_coding_gene | 1.1 | 2.19 | 1.6E-02 |
| *unc-10* | WBGene00006750 | protein_coding_gene | 1.1 | 2.19 | 3.7E-06 |
| *aldo-1* | WBGene00011474 | protein_coding_gene | 1.1 | 2.19 | 1.3E-13 |
| *ZK721.3* | WBGene00022810 | protein_coding_gene | 1.1 | 2.19 | 3.4E-02 |
| *F19C6.3* | WBGene00008952 | protein_coding_gene | 1.1 | 2.19 | 4.2E-03 |
| *best-12* | WBGene00008821 | protein_coding_gene | 1.1 | 2.19 | 3.1E-02 |
| *madd-4* | WBGene00009958 | protein_coding_gene | 1.1 | 2.19 | 6.3E-06 |
| *fipr-29* | WBGene00019171 | protein_coding_gene | 1.1 | 2.19 | 3.6E-02 |
| *Y51F10.7* | WBGene00021768 | protein_coding_gene | 1.1 | 2.18 | 2.8E-09 |
| *plg-1* | WBGene00004041 | pseudogene | 1.1 | 2.18 | 3.2E-11 |
| *twnk-1* | WBGene00018514 | protein_coding_gene | 1.1 | 2.18 | 4.7E-02 |
| *mec-12* | WBGene00003175 | protein_coding_gene | 1.1 | 2.18 | 7.2E-11 |
| *R07C3.13* | WBGene00019929 | protein_coding_gene | 1.1 | 2.18 | 1.4E-02 |
| *mec-5* | WBGene00003169 | protein_coding_gene | 1.1 | 2.18 | 3.8E-05 |
| *F16H6.9* | WBGene00008899 | protein_coding_gene | 1.1 | 2.18 | 6.5E-04 |
| *twk-48* | WBGene00010784 | protein_coding_gene | 1.1 | 2.18 | 1.2E-02 |
| *Y40C5A.4* | WBGene00021497 | protein_coding_gene | 1.1 | 2.17 | 9.9E-03 |
| *fer-1* | WBGene00001414 | protein_coding_gene | 1.1 | 2.17 | 3.4E-05 |
| *apt-9* | WBGene00000166 | protein_coding_gene | 1.1 | 2.17 | 2.1E-02 |
| *sdha-1* | WBGene00015391 | protein_coding_gene | 1.1 | 2.17 | 3.1E-15 |
| *F29D10.1* | WBGene00009250 | protein_coding_gene | 1.1 | 2.17 | 2.8E-02 |
| *F58A4.1* | WBGene00010226 | protein_coding_gene | 1.1 | 2.17 | 2.0E-06 |
| *avr-14* | WBGene00000232 | protein_coding_gene | 1.1 | 2.17 | 2.1E-05 |
| *K05F1.9* | WBGene00019410 | protein_coding_gene | 1.1 | 2.17 | 8.9E-08 |
| *T14A8.2* | WBGene00020497 | protein_coding_gene | 1.1 | 2.17 | 6.4E-04 |
| *C51E3.9* | WBGene00008254 | protein_coding_gene | 1.1 | 2.17 | 6.8E-09 |
| *pept-2* | WBGene00003876 | protein_coding_gene | 1.1 | 2.17 | 4.4E-08 |
| *R08C7.12* | WBGene00019955 | protein_coding_gene | 1.1 | 2.17 | 3.9E-07 |
| *Y59E9AL.5* | WBGene00021995 | pseudogene | 1.1 | 2.17 | 4.1E-02 |
| *Y34B4A.4* | WBGene00021320 | protein_coding_gene | 1.1 | 2.17 | 5.6E-06 |
| *C54G4.2* | WBGene00008312 | protein_coding_gene | 1.1 | 2.17 | 1.6E-03 |
| *glb-10* | WBGene00016215 | protein_coding_gene | 1.1 | 2.17 | 8.9E-03 |
| *C05C12.1* | WBGene00007335 | protein_coding_gene | 1.1 | 2.17 | 1.4E-04 |
| *F43C1.5* | WBGene00009649 | protein_coding_gene | 1.1 | 2.17 | 1.8E-02 |
| *R07B1.8* | WBGene00001608 | protein_coding_gene | 1.1 | 2.17 | 4.7E-02 |
| *xbx-9* | WBGene00007604 | protein_coding_gene | 1.1 | 2.17 | 4.8E-03 |
| *spin-1* | WBGene00007549 | protein_coding_gene | 1.1 | 2.16 | 5.4E-16 |
| *nas-20* | WBGene00003539 | protein_coding_gene | 1.1 | 2.16 | 1.1E-07 |
| *B0513.90* | WBGene00235301 | protein_coding_gene | 1.1 | 2.16 | 8.6E-03 |
| *F08H9.2* | WBGene00008590 | protein_coding_gene | 1.1 | 2.16 | 1.3E-06 |
| *R08F11.4* | WBGene00019968 | protein_coding_gene | 1.1 | 2.16 | 8.4E-03 |
| *C55A6.7* | WBGene00008336 | protein_coding_gene | 1.1 | 2.16 | 1.9E-04 |
| *Y57G11A.4* | WBGene00013292 | protein_coding_gene | 1.1 | 2.16 | 1.2E-07 |

| | | | | | |
|---|---|---|---|---|---|
| *nekl-4* | WBGene00017033 | protein_coding_gene | 1.1 | 2.16 | 2.9E-02 |
| *B0348.2* | WBGene00015152 | protein_coding_gene | 1.1 | 2.15 | 5.4E-03 |
| *clec-95* | WBGene00013927 | protein_coding_gene | 1.1 | 2.15 | 1.2E-08 |
| *daf-9* | WBGene00000905 | protein_coding_gene | 1.1 | 2.15 | 1.9E-02 |
| *H11E01.2* | WBGene00019187 | protein_coding_gene | 1.1 | 2.15 | 2.1E-04 |
| *ZC412.3* | WBGene00013884 | protein_coding_gene | 1.1 | 2.15 | 9.7E-12 |
| *Y67H2A.9* | WBGene00014941 | pseudogene | 1.1 | 2.15 | 4.1E-08 |
| *H01G02.3* | WBGene00010350 | protein_coding_gene | 1.1 | 2.15 | 2.3E-02 |
| *unc-9* | WBGene00006749 | protein_coding_gene | 1.1 | 2.15 | 9.5E-07 |
| *fipr-10* | WBGene00008245 | protein_coding_gene | 1.1 | 2.15 | 5.1E-03 |
| *C45G9.12* | WBGene00016683 | protein_coding_gene | 1.1 | 2.15 | 4.0E-02 |
| *C14C11.1* | WBGene00015765 | protein_coding_gene | 1.1 | 2.14 | 4.4E-04 |
| *F02H6.6* | WBGene00008539 | protein_coding_gene | 1.1 | 2.14 | 2.0E-03 |
| *C33F10.11* | WBGene00016357 | protein_coding_gene | 1.1 | 2.14 | 1.7E-05 |
| *K09A9.6* | WBGene00010703 | protein_coding_gene | 1.1 | 2.14 | 4.6E-08 |
| *gpa-2* | WBGene00001664 | protein_coding_gene | 1.1 | 2.14 | 3.2E-02 |
| *Y45F10C.4* | WBGene00012880 | protein_coding_gene | 1.1 | 2.14 | 3.5E-06 |
| *D2045.5* | WBGene00008423 | protein_coding_gene | 1.1 | 2.14 | 3.3E-03 |
| *slo-2* | WBGene00004831 | protein_coding_gene | 1.1 | 2.14 | 1.2E-10 |
| *F46F5.14* | WBGene00018505 | protein_coding_gene | 1.1 | 2.14 | 2.0E-02 |
| *T16G1.5* | WBGene00011799 | protein_coding_gene | 1.1 | 2.14 | 1.7E-03 |
| *cyp-35A2* | WBGene00015400 | protein_coding_gene | 1.1 | 2.13 | 1.2E-03 |
| *col-113* | WBGene00000687 | protein_coding_gene | 1.1 | 2.13 | 1.3E-02 |
| *mpst-4* | WBGene00017387 | protein_coding_gene | 1.1 | 2.13 | 1.9E-02 |
| *Y2H9A.6* | WBGene00086568 | protein_coding_gene | 1.1 | 2.13 | 1.5E-02 |
| *tbc-19* | WBGene00007849 | protein_coding_gene | 1.1 | 2.13 | 1.1E-05 |
| *C04G6.6* | WBGene00015456 | protein_coding_gene | 1.1 | 2.13 | 4.3E-02 |
| *F40G9.7* | WBGene00018243 | protein_coding_gene | 1.1 | 2.13 | 1.1E-02 |
| *ser-5* | WBGene00008890 | protein_coding_gene | 1.1 | 2.13 | 2.8E-02 |
| *C03F11.4* | WBGene00015390 | protein_coding_gene | 1.1 | 2.13 | 4.1E-06 |
| *C02B8.12* | WBGene00219375 | protein_coding_gene | 1.1 | 2.13 | 2.2E-02 |
| *F35H10.10* | WBGene00018073 | protein_coding_gene | 1.1 | 2.13 | 2.1E-06 |
| *cdc-25.4* | WBGene00000389 | protein_coding_gene | 1.1 | 2.13 | 2.8E-03 |
| *F38B2.4* | WBGene00009531 | protein_coding_gene | 1.1 | 2.13 | 4.6E-11 |
| *K05F1.1* | WBGene00019405 | protein_coding_gene | 1.1 | 2.12 | 4.5E-02 |
| *twk-39* | WBGene00006690 | protein_coding_gene | 1.1 | 2.12 | 1.0E-02 |
| *F36G3.1* | WBGene00009482 | protein_coding_gene | 1.1 | 2.12 | 1.8E-08 |
| *C17H12.12* | WBGene00015937 | protein_coding_gene | 1.1 | 2.12 | 1.7E-06 |
| *B0244.10* | WBGene00015085 | protein_coding_gene | 1.1 | 2.12 | 4.4E-03 |
| *daf-7* | WBGene00000903 | protein_coding_gene | 1.1 | 2.12 | 2.1E-04 |
| *pod-2* | WBGene00004076 | protein_coding_gene | 1.1 | 2.12 | 5.1E-21 |
| *W03F8.2* | WBGene00020992 | pseudogene | 1.1 | 2.12 | 2.3E-03 |
| *prmt-6* | WBGene00016023 | protein_coding_gene | 1.1 | 2.11 | 1.4E-03 |
| *R10E11.5* | WBGene00011217 | protein_coding_gene | 1.1 | 2.11 | 1.4E-03 |
| *F22G12.8* | WBGene00044712 | pseudogene | 1.1 | 2.11 | 1.6E-02 |
| *F39G3.5* | WBGene00018210 | protein_coding_gene | 1.1 | 2.11 | 6.5E-05 |
| *F28E10.1* | WBGene00017900 | protein_coding_gene | 1.1 | 2.11 | 3.9E-07 |
| *ifb-1* | WBGene00002053 | protein_coding_gene | 1.1 | 2.11 | 1.1E-12 |
| *ZK1067.4* | WBGene00014210 | protein_coding_gene | 1.1 | 2.11 | 4.5E-03 |
| *ivns-1* | WBGene00006493 | protein_coding_gene | 1.1 | 2.11 | 3.0E-17 |
| *Y49F6B.8* | WBGene00021720 | protein_coding_gene | 1.1 | 2.11 | 1.9E-02 |
| *Y69H2.15* | WBGene00050884 | protein_coding_gene | 1.1 | 2.11 | 4.9E-02 |
| *C17F3.3* | WBGene00015908 | protein_coding_gene | 1.1 | 2.11 | 4.6E-04 |
| *oig-8* | WBGene00020582 | protein_coding_gene | 1.1 | 2.11 | 3.0E-02 |
| *C33G3.5* | WBGene00007905 | protein_coding_gene | 1.1 | 2.11 | 1.5E-06 |
| *kvs-1* | WBGene00002242 | protein_coding_gene | 1.1 | 2.11 | 1.5E-02 |
| *F28A10.1* | WBGene00017869 | protein_coding_gene | 1.1 | 2.11 | 1.3E-03 |
| *F35A5.1* | WBGene00018024 | protein_coding_gene | 1.1 | 2.11 | 9.4E-11 |

| | | | | | |
|---|---|---|---|---|---|
| *fmo-3* | WBGene00001478 | protein_coding_gene | 1.1 | 2.11 | 1.3E-05 |
| *C47E8.3* | WBGene00008142 | protein_coding_gene | 1.1 | 2.10 | 2.9E-02 |
| *ZC250.4* | WBGene00045305 | protein_coding_gene | 1.1 | 2.10 | 6.0E-05 |
| *B0222.10* | WBGene00015058 | ncRNA_gene | 1.1 | 2.10 | 3.8E-07 |
| *cpr-5* | WBGene00000785 | protein_coding_gene | 1.1 | 2.10 | 5.3E-03 |
| *pept-1* | WBGene00003877 | protein_coding_gene | 1.1 | 2.10 | 1.8E-09 |
| *tsp-18* | WBGene00006644 | protein_coding_gene | 1.1 | 2.10 | 3.7E-02 |
| *Y71G12B.5* | WBGene00022145 | protein_coding_gene | 1.1 | 2.10 | 1.9E-02 |
| *pdfr-1* | WBGene00015735 | protein_coding_gene | 1.1 | 2.10 | 3.5E-07 |
| *scav-6* | WBGene00008542 | protein_coding_gene | 1.1 | 2.10 | 5.4E-06 |
| *H09F14.1* | WBGene00010375 | protein_coding_gene | 1.1 | 2.10 | 2.3E-02 |
| *col-186* | WBGene00000759 | protein_coding_gene | 1.1 | 2.10 | 3.9E-02 |
| *R155.3* | WBGene00020117 | protein_coding_gene | 1.1 | 2.10 | 9.6E-04 |
| *sipa-1* | WBGene00012100 | protein_coding_gene | 1.1 | 2.10 | 4.3E-09 |
| *F54C9.7* | WBGene00010043 | protein_coding_gene | 1.1 | 2.10 | 4.0E-02 |
| *E02C12.6* | WBGene00017092 | protein_coding_gene | 1.1 | 2.10 | 2.9E-03 |
| *lmp-2* | WBGene00015471 | protein_coding_gene | 1.1 | 2.10 | 2.7E-06 |
| *T14G12.11* | WBGene00219411 | protein_coding_gene | 1.1 | 2.10 | 7.8E-04 |
| *Y116A8C.44* | WBGene00013815 | protein_coding_gene | 1.1 | 2.09 | 2.1E-02 |
| *C45E5.1* | WBGene00016664 | protein_coding_gene | 1.1 | 2.09 | 4.1E-05 |
| *T08B2.12* | WBGene00020350 | protein_coding_gene | 1.1 | 2.09 | 3.7E-03 |
| *F57F5.1* | WBGene00010204 | protein_coding_gene | 1.1 | 2.09 | 2.4E-03 |
| *tba-5* | WBGene00006531 | protein_coding_gene | 1.1 | 2.09 | 2.8E-02 |
| *odr-2* | WBGene00003849 | protein_coding_gene | 1.1 | 2.09 | 9.6E-03 |
| *C49C3.11* | WBGene00008201 | protein_coding_gene | 1.1 | 2.09 | 1.5E-09 |
| *B0285.6* | WBGene00007138 | protein_coding_gene | 1.1 | 2.08 | 2.1E-02 |
| *gpc-1* | WBGene00001681 | protein_coding_gene | 1.1 | 2.08 | 4.0E-02 |
| *mrp-5* | WBGene00003411 | protein_coding_gene | 1.1 | 2.08 | 7.2E-22 |
| *T11F9.12* | WBGene00011713 | protein_coding_gene | 1.1 | 2.08 | 2.0E-12 |
| *Y7A5A.1* | WBGene00012407 | protein_coding_gene | 1.1 | 2.08 | 4.8E-08 |
| *klo-2* | WBGene00017103 | protein_coding_gene | 1.1 | 2.08 | 3.8E-04 |
| *cpr-8* | WBGene00021070 | protein_coding_gene | 1.1 | 2.08 | 1.1E-03 |
| *R07E5.6* | WBGene00011114 | protein_coding_gene | 1.1 | 2.08 | 2.3E-04 |
| *T13F3.8* | WBGene00044205 | protein_coding_gene | 1.1 | 2.08 | 1.7E-03 |
| *C06C3.12* | WBGene00175031 | protein_coding_gene | 1.1 | 2.07 | 4.3E-02 |
| *F46A8.5* | WBGene00009748 | protein_coding_gene | 1.1 | 2.07 | 1.0E-04 |
| *C17G10.7* | WBGene00015919 | protein_coding_gene | 1.1 | 2.07 | 1.1E-02 |
| *F32D8.12* | WBGene00009334 | protein_coding_gene | 1.0 | 2.07 | 2.0E-07 |
| *math-47* | WBGene00022688 | pseudogene | 1.0 | 2.07 | 2.8E-03 |
| *afmd-1* | WBGene00017051 | protein_coding_gene | 1.0 | 2.07 | 1.2E-02 |
| *Y41C4A.12* | WBGene00012758 | protein_coding_gene | 1.0 | 2.07 | 3.3E-04 |
| *col-142* | WBGene00000715 | protein_coding_gene | 1.0 | 2.07 | 6.0E-05 |
| *unc-77* | WBGene00006809 | protein_coding_gene | 1.0 | 2.07 | 3.2E-04 |
| *F58F9.4* | WBGene00019058 | protein_coding_gene | 1.0 | 2.06 | 7.3E-04 |
| *pxn-1* | WBGene00004256 | protein_coding_gene | 1.0 | 2.06 | 1.4E-09 |
| *gbb-1* | WBGene00021528 | protein_coding_gene | 1.0 | 2.06 | 5.1E-05 |
| *E02H9.2* | WBGene00017100 | protein_coding_gene | 1.0 | 2.06 | 6.4E-03 |
| *rig-6* | WBGene00016354 | protein_coding_gene | 1.0 | 2.06 | 7.0E-09 |
| *T11F8.4* | WBGene00020435 | protein_coding_gene | 1.0 | 2.06 | 4.4E-04 |
| *rab-3* | WBGene00004267 | protein_coding_gene | 1.0 | 2.06 | 2.6E-09 |
| *acs-18* | WBGene00011173 | protein_coding_gene | 1.0 | 2.06 | 3.3E-07 |
| *rhr-1* | WBGene00004358 | protein_coding_gene | 1.0 | 2.06 | 4.6E-18 |
| *F25E5.3* | WBGene00017784 | protein_coding_gene | 1.0 | 2.06 | 3.9E-07 |
| *ugt-47* | WBGene00011006 | protein_coding_gene | 1.0 | 2.06 | 5.9E-14 |
| *T01B4.3* | WBGene00011310 | protein_coding_gene | 1.0 | 2.05 | 7.3E-05 |
| *Y43F8C.15* | WBGene00012836 | protein_coding_gene | 1.0 | 2.05 | 1.6E-02 |
| *unc-104* | WBGene00006831 | protein_coding_gene | 1.0 | 2.05 | 3.7E-10 |
| *F38A5.6* | WBGene00018163 | protein_coding_gene | 1.0 | 2.05 | 5.3E-03 |

| | | | | | |
|---|---|---|---|---|---|
| *basl-2* | WBGene00017392 | pseudogene | 1.0 | 2.05 | 3.7E-02 |
| *try-8* | WBGene00017791 | protein_coding_gene | 1.0 | 2.05 | 3.8E-03 |
| *F15A4.5* | WBGene00008840 | protein_coding_gene | 1.0 | 2.05 | 5.6E-03 |
| *R102.3* | WBGene00011290 | protein_coding_gene | 1.0 | 2.05 | 2.2E-02 |
| *wht-4* | WBGene00017179 | protein_coding_gene | 1.0 | 2.05 | 1.9E-03 |
| *rgs-3* | WBGene00004346 | protein_coding_gene | 1.0 | 2.04 | 7.1E-03 |
| *Y18D10A.2* | WBGene00012475 | protein_coding_gene | 1.0 | 2.04 | 5.5E-04 |
| *del-10* | WBGene00020897 | protein_coding_gene | 1.0 | 2.04 | 4.2E-04 |
| *R05G6.1* | WBGene00019896 | protein_coding_gene | 1.0 | 2.04 | 2.6E-02 |
| *C41A3.2* | WBGene00016559 | protein_coding_gene | 1.0 | 2.04 | 5.8E-06 |
| *C15A11.2* | WBGene00007597 | protein_coding_gene | 1.0 | 2.04 | 3.1E-03 |
| *F11E6.3* | WBGene00008707 | protein_coding_gene | 1.0 | 2.04 | 6.4E-11 |
| *pde-1* | WBGene00011433 | protein_coding_gene | 1.0 | 2.04 | 9.9E-04 |
| *F57H12.5* | WBGene00019020 | protein_coding_gene | 1.0 | 2.04 | 7.7E-03 |
| *F49B2.6* | WBGene00009865 | protein_coding_gene | 1.0 | 2.04 | 1.0E-03 |
| *gmeb-3* | WBGene00008092 | protein_coding_gene | 1.0 | 2.04 | 5.2E-03 |
| *rgs-2* | WBGene00004345 | protein_coding_gene | 1.0 | 2.04 | 3.0E-05 |
| *cln-3.3* | WBGene00000541 | protein_coding_gene | 1.0 | 2.04 | 5.4E-04 |
| *clec-106* | WBGene00012482 | protein_coding_gene | 1.0 | 2.04 | 9.1E-04 |
| *C44C10.3* | WBGene00008084 | protein_coding_gene | 1.0 | 2.04 | 1.2E-03 |
| *unc-22* | WBGene00006759 | protein_coding_gene | 1.0 | 2.04 | 1.5E-04 |
| *C43F9.5* | WBGene00008073 | protein_coding_gene | 1.0 | 2.04 | 2.3E-02 |
| *ver-4* | WBGene00006897 | protein_coding_gene | 1.0 | 2.04 | 2.1E-02 |
| *T21B6.3* | WBGene00011880 | protein_coding_gene | 1.0 | 2.04 | 2.2E-09 |
| *Y6G8.14* | WBGene00138715 | protein_coding_gene | 1.0 | 2.04 | 2.5E-03 |
| *F07C3.2* | WBGene00017193 | protein_coding_gene | 1.0 | 2.04 | 2.0E-03 |
| *cal-5* | WBGene00006861 | protein_coding_gene | 1.0 | 2.04 | 2.6E-13 |
| *cest-13* | WBGene00010116 | protein_coding_gene | 1.0 | 2.03 | 9.7E-06 |
| *Y43F8C.13* | WBGene00012834 | protein_coding_gene | 1.0 | 2.03 | 2.1E-08 |
| *tyra-3* | WBGene00006475 | protein_coding_gene | 1.0 | 2.03 | 2.5E-02 |
| *clec-152* | WBGene00008658 | pseudogene | 1.0 | 2.03 | 1.2E-02 |
| *C35B1.8* | WBGene00016437 | protein_coding_gene | 1.0 | 2.03 | 1.7E-02 |
| *npr-22* | WBGene00022004 | protein_coding_gene | 1.0 | 2.03 | 5.8E-03 |
| *poml-2* | WBGene00017089 | protein_coding_gene | 1.0 | 2.03 | 8.9E-04 |
| *F35C5.3* | WBGene00009391 | protein_coding_gene | 1.0 | 2.03 | 2.6E-02 |
| *alg-4* | WBGene00006449 | protein_coding_gene | 1.0 | 2.03 | 7.1E-08 |
| *C35B1.3* | WBGene00016433 | protein_coding_gene | 1.0 | 2.03 | 1.2E-02 |
| *ZK616.61* | WBGene00194816 | protein_coding_gene | 1.0 | 2.03 | 5.0E-04 |
| *alg-3* | WBGene00011910 | protein_coding_gene | 1.0 | 2.03 | 2.5E-07 |
| *F19C7.8* | WBGene00017597 | protein_coding_gene | 1.0 | 2.03 | 1.7E-03 |
| *C49C3.4* | WBGene00008194 | protein_coding_gene | 1.0 | 2.03 | 1.7E-08 |
| *F18E9.7* | WBGene00017573 | protein_coding_gene | 1.0 | 2.03 | 2.2E-05 |
| *C04C11.1* | WBGene00007295 | protein_coding_gene | 1.0 | 2.03 | 2.6E-02 |
| *T01H3.5* | WBGene00011351 | protein_coding_gene | 1.0 | 2.03 | 4.8E-03 |
| *dyc-1* | WBGene00001116 | protein_coding_gene | 1.0 | 2.03 | 4.0E-05 |
| *acd-3* | WBGene00007775 | protein_coding_gene | 1.0 | 2.02 | 2.0E-03 |
| *gst-6* | WBGene00001754 | protein_coding_gene | 1.0 | 2.02 | 1.3E-05 |
| *D2085.7* | WBGene00008432 | protein_coding_gene | 1.0 | 2.02 | 1.5E-03 |
| *Y51A2D.8* | WBGene00013076 | protein_coding_gene | 1.0 | 2.02 | 9.7E-07 |
| *spe-12* | WBGene00004966 | protein_coding_gene | 1.0 | 2.02 | 9.2E-03 |
| *ZC196.5* | WBGene00022549 | protein_coding_gene | 1.0 | 2.02 | 1.0E-05 |
| *nlp-82* | WBGene00044006 | protein_coding_gene | 1.0 | 2.02 | 4.0E-02 |
| *Y39B6A.18* | WBGene00012679 | pseudogene | 1.0 | 2.02 | 3.0E-02 |
| *slc-9B.2* | WBGene00010214 | protein_coding_gene | 1.0 | 2.02 | 2.3E-02 |
| *ceh-60* | WBGene00017690 | protein_coding_gene | 1.0 | 2.02 | 7.1E-07 |
| *F59A6.2* | WBGene00019084 | protein_coding_gene | 1.0 | 2.02 | 4.7E-03 |
| *acr-18* | WBGene00000057 | protein_coding_gene | 1.0 | 2.02 | 4.2E-03 |
| *crn-6* | WBGene00000799 | protein_coding_gene | 1.0 | 2.02 | 5.8E-07 |

| | | | | | |
|---|---|---|---|---|---|
| *C25D7.15* | WBGene00007722 | protein_coding_gene | 1.0 | 2.01 | 3.6E-02 |
| *F49E11.7* | WBGene00009893 | protein_coding_gene | 1.0 | 2.01 | 6.5E-03 |
| *oac-59* | WBGene00016585 | protein_coding_gene | 1.0 | 2.01 | 1.3E-06 |
| *ins-31* | WBGene00002114 | protein_coding_gene | 1.0 | 2.01 | 3.2E-03 |
| *fbxa-202* | WBGene00020890 | protein_coding_gene | 1.0 | 2.01 | 4.3E-04 |
| *haf-4* | WBGene00001814 | protein_coding_gene | 1.0 | 2.01 | 9.1E-11 |
| *F36A2.10* | WBGene00009457 | protein_coding_gene | 1.0 | 2.01 | 5.1E-06 |
| *clec-124* | WBGene00015909 | protein_coding_gene | 1.0 | 2.01 | 1.4E-06 |
| *C14A6.6* | WBGene00007570 | protein_coding_gene | 1.0 | 2.01 | 1.2E-03 |
| *bus-19* | WBGene00011590 | protein_coding_gene | 1.0 | 2.01 | 1.0E-03 |
| *C48B4.12* | WBGene00008175 | protein_coding_gene | 1.0 | 2.01 | 4.9E-07 |
| *C45G9.10* | WBGene00016681 | protein_coding_gene | 1.0 | 2.01 | 2.4E-04 |
| *rnh-1.1* | WBGene00022888 | protein_coding_gene | 1.0 | 2.00 | 7.2E-05 |
| *ent-7* | WBGene00009686 | protein_coding_gene | 1.0 | 2.00 | 4.9E-08 |
| *exc-12* | WBGene00011476 | protein_coding_gene | 1.0 | 2.00 | 3.3E-02 |
| *F08F8.6* | WBGene00017271 | protein_coding_gene | 1.0 | 2.00 | 6.5E-03 |
| *glb-3* | WBGene00007389 | protein_coding_gene | 1.0 | 2.00 | 1.5E-02 |
| *fbp-1* | WBGene00001404 | protein_coding_gene | 1.0 | 2.00 | 1.0E-15 |
| *dhc-3* | WBGene00007154 | protein_coding_gene | 1.0 | 2.00 | 4.4E-02 |
| *col-42* | WBGene00013489 | protein_coding_gene | 1.0 | 2.00 | 1.2E-08 |
| *Y37E11AL.4* | WBGene00021360 | pseudogene | 1.0 | 2.00 | 5.0E-05 |
| *nkb-3* | WBGene00010117 | protein_coding_gene | 1.0 | 2.00 | 4.6E-15 |
| *F59B8.1* | WBGene00010316 | protein_coding_gene | 1.0 | 2.00 | 3.5E-02 |
| *ptl-1* | WBGene00004212 | protein_coding_gene | 1.0 | 2.00 | 1.8E-05 |
| *clec-162* | WBGene00007399 | protein_coding_gene | 1.0 | 2.00 | 4.4E-02 |
| *pxd-1* | WBGene00007982 | protein_coding_gene | -1.0 | 0.50 | 1.1E-10 |
| *vet-6* | WBGene00006902 | protein_coding_gene | -1.0 | 0.50 | 2.2E-06 |
| *dscc-1* | WBGene00019595 | protein_coding_gene | -1.0 | 0.50 | 1.3E-11 |
| *C16C8.13* | WBGene00015851 | protein_coding_gene | -1.0 | 0.50 | 2.3E-03 |
| *mrps-23* | WBGene00014224 | protein_coding_gene | -1.0 | 0.50 | 5.4E-10 |
| *F44E2.9* | WBGene00018422 | protein_coding_gene | -1.0 | 0.50 | 3.3E-08 |
| *mrps-17* | WBGene00015487 | protein_coding_gene | -1.0 | 0.50 | 2.2E-10 |
| *C52A10.3* | WBGene00016864 | protein_coding_gene | -1.0 | 0.50 | 2.5E-02 |
| *cpg-8* | WBGene00019357 | protein_coding_gene | -1.0 | 0.50 | 1.6E-13 |
| *glo-3* | WBGene00010341 | protein_coding_gene | -1.0 | 0.50 | 6.7E-04 |
| *fbxb-104* | WBGene00021719 | protein_coding_gene | -1.0 | 0.50 | 1.3E-02 |
| *F53F10.1* | WBGene00018763 | protein_coding_gene | -1.0 | 0.50 | 7.3E-05 |
| *R07H5.11* | WBGene00011131 | protein_coding_gene | -1.0 | 0.50 | 8.4E-05 |
| *C06G3.5* | WBGene00015551 | protein_coding_gene | -1.0 | 0.50 | 4.1E-08 |
| *anr-8* | WBGene00219575 | antisense_lncRNA_gene | -1.0 | 0.50 | 1.2E-03 |
| *tsen-34* | WBGene00019535 | protein_coding_gene | -1.0 | 0.50 | 4.2E-05 |
| *Y45G5AM.5* | WBGene00021557 | protein_coding_gene | -1.0 | 0.49 | 1.3E-03 |
| *ZK380.6* | WBGene00304812 | protein_coding_gene | -1.0 | 0.49 | 6.6E-13 |
| *fbxb-40* | WBGene00019704 | protein_coding_gene | -1.0 | 0.49 | 3.5E-02 |
| *mfb-1* | WBGene00008439 | protein_coding_gene | -1.0 | 0.49 | 2.2E-06 |
| *F44F1.4* | WBGene00009696 | protein_coding_gene | -1.0 | 0.49 | 1.3E-03 |
| *snr-3* | WBGene00004916 | protein_coding_gene | -1.0 | 0.49 | 2.3E-17 |
| *fbxb-7* | WBGene00012492 | protein_coding_gene | -1.0 | 0.49 | 2.0E-05 |
| *bath-35* | WBGene00012196 | protein_coding_gene | -1.0 | 0.49 | 3.1E-02 |
| *F45C12.17* | WBGene00269388 | protein_coding_gene | -1.0 | 0.49 | 4.9E-03 |
| *F33H2.8* | WBGene00009371 | protein_coding_gene | -1.0 | 0.49 | 3.7E-02 |
| *metl-6* | WBGene00014205 | protein_coding_gene | -1.0 | 0.49 | 2.1E-10 |
| *C50F4.4* | WBGene00008232 | protein_coding_gene | -1.0 | 0.49 | 5.7E-03 |
| *M162.5* | WBGene00010931 | protein_coding_gene | -1.0 | 0.49 | 9.3E-04 |
| *R05D3.3* | WBGene00019878 | protein_coding_gene | -1.0 | 0.49 | 2.9E-07 |
| *rpl-41.1* | WBGene00023067 | protein_coding_gene | -1.0 | 0.49 | 3.3E-13 |
| *C16A3.2* | WBGene00015807 | protein_coding_gene | -1.0 | 0.49 | 9.9E-10 |
| *Y71F9AL.25* | WBGene00304792 | protein_coding_gene | -1.0 | 0.49 | 7.5E-03 |

| | | | | | |
|---|---|---|---|---|---|
| *ttr-52* | WBGene00008719 | protein_coding_gene | -1.0 | 0.49 | 4.2E-02 |
| *fbxb-116* | WBGene00012066 | protein_coding_gene | -1.0 | 0.48 | 2.2E-02 |
| *ctf-8* | WBGene00011915 | protein_coding_gene | -1.0 | 0.48 | 2.1E-05 |
| *Y71F9AR.2* | WBGene00022123 | protein_coding_gene | -1.0 | 0.48 | 9.8E-03 |
| *F48E3.6* | WBGene00018606 | protein_coding_gene | -1.0 | 0.48 | 3.7E-06 |
| *rgs-9* | WBGene00004352 | protein_coding_gene | -1.0 | 0.48 | 2.5E-03 |
| *K12C11.6* | WBGene00045052 | protein_coding_gene | -1.1 | 0.48 | 4.2E-05 |
| *F52C12.2* | WBGene00018679 | protein_coding_gene | -1.1 | 0.48 | 1.5E-09 |
| *C07E3.9* | WBGene00007419 | protein_coding_gene | -1.1 | 0.48 | 6.4E-08 |
| *aptf-2* | WBGene00013383 | protein_coding_gene | -1.1 | 0.48 | 2.5E-08 |
| *F56G4.6* | WBGene00010161 | protein_coding_gene | -1.1 | 0.48 | 2.8E-10 |
| *fbxb-57* | WBGene00010101 | protein_coding_gene | -1.1 | 0.48 | 9.3E-03 |
| *Y111B2A.21* | WBGene00013741 | protein_coding_gene | -1.1 | 0.48 | 3.7E-02 |
| *dyf-17* | WBGene00012673 | protein_coding_gene | -1.1 | 0.48 | 3.1E-03 |
| *pes-10* | WBGene00003983 | protein_coding_gene | -1.1 | 0.48 | 3.2E-04 |
| *F46C5.10* | WBGene00009785 | protein_coding_gene | -1.1 | 0.48 | 6.4E-05 |
| *ins-20* | WBGene00002103 | protein_coding_gene | -1.1 | 0.48 | 7.1E-03 |
| *T08A9.13* | WBGene00044442 | protein_coding_gene | -1.1 | 0.48 | 4.6E-02 |
| *fbxa-157* | WBGene00007393 | protein_coding_gene | -1.1 | 0.48 | 1.7E-02 |
| *fbxb-26* | WBGene00019035 | protein_coding_gene | -1.1 | 0.48 | 8.3E-03 |
| *zip-11* | WBGene00021082 | protein_coding_gene | -1.1 | 0.48 | 9.4E-05 |
| *tyr-1* | WBGene00015332 | protein_coding_gene | -1.1 | 0.48 | 2.2E-06 |
| *F54F7.6* | WBGene00010066 | protein_coding_gene | -1.1 | 0.48 | 6.2E-08 |
| *clec-3* | WBGene00016577 | protein_coding_gene | -1.1 | 0.48 | 4.7E-03 |
| *fbxc-45* | WBGene00021887 | protein_coding_gene | -1.1 | 0.48 | 1.2E-06 |
| *apc-11* | WBGene00000145 | protein_coding_gene | -1.1 | 0.48 | 3.6E-12 |
| *K06H7.7* | WBGene00019458 | protein_coding_gene | -1.1 | 0.47 | 4.6E-11 |
| *BE0003N10.3* | WBGene00015268 | protein_coding_gene | -1.1 | 0.47 | 2.4E-03 |
| *F13B12.2* | WBGene00008730 | protein_coding_gene | -1.1 | 0.47 | 4.4E-05 |
| *F19B2.12* | WBGene00194812 | pseudogene | -1.1 | 0.47 | 1.1E-04 |
| *btb-7* | WBGene00018443 | protein_coding_gene | -1.1 | 0.47 | 2.0E-04 |
| *ZK1127.12* | WBGene00022857 | protein_coding_gene | -1.1 | 0.47 | 7.8E-08 |
| *K11D12.12* | WBGene00019651 | protein_coding_gene | -1.1 | 0.47 | 1.6E-10 |
| *Y40A1A.6* | WBGene00304815 | ncRNA_gene | -1.1 | 0.47 | 3.8E-10 |
| *K05C4.4* | WBGene00010581 | protein_coding_gene | -1.1 | 0.47 | 3.3E-05 |
| *M01G12.14* | WBGene00010825 | protein_coding_gene | -1.1 | 0.47 | 3.0E-02 |
| *csa-1* | WBGene00008403 | protein_coding_gene | -1.1 | 0.47 | 1.4E-04 |
| *ZK185.5* | WBGene00044481 | protein_coding_gene | -1.1 | 0.47 | 4.0E-06 |
| *F29G9.7* | WBGene00017933 | pseudogene | -1.1 | 0.46 | 2.5E-04 |
| *blos-9* | WBGene00010767 | protein_coding_gene | -1.1 | 0.46 | 1.7E-11 |
| *ccch-2* | WBGene00009537 | protein_coding_gene | -1.1 | 0.46 | 2.2E-03 |
| *C08F11.13* | WBGene00007460 | protein_coding_gene | -1.1 | 0.46 | 3.8E-10 |
| *btb-12* | WBGene00022555 | protein_coding_gene | -1.1 | 0.46 | 2.4E-02 |
| *abf-2* | WBGene00000013 | protein_coding_gene | -1.1 | 0.46 | 2.0E-03 |
| *slc-17.9* | WBGene00011643 | protein_coding_gene | -1.1 | 0.46 | 1.7E-02 |
| *B0478.3* | WBGene00015202 | protein_coding_gene | -1.1 | 0.46 | 3.2E-02 |
| *F40H3.6* | WBGene00018252 | protein_coding_gene | -1.1 | 0.46 | 5.9E-04 |
| *T04B8.2* | WBGene00020204 | pseudogene | -1.1 | 0.46 | 3.9E-02 |
| *clec-146* | WBGene00013008 | protein_coding_gene | -1.1 | 0.46 | 2.2E-07 |
| *F59B1.8* | WBGene00019100 | protein_coding_gene | -1.1 | 0.46 | 1.7E-08 |
| *Y49A3A.3* | WBGene00013026 | protein_coding_gene | -1.1 | 0.46 | 7.4E-13 |
| *F40G9.17* | WBGene00185078 | protein_coding_gene | -1.1 | 0.46 | 1.3E-11 |
| *Y49F6C.8* | WBGene00021730 | protein_coding_gene | -1.1 | 0.45 | 7.2E-05 |
| *F59C12.4* | WBGene00045272 | protein_coding_gene | -1.1 | 0.45 | 2.3E-08 |
| *dlc-5* | WBGene00043067 | protein_coding_gene | -1.1 | 0.45 | 3.1E-02 |
| *R11.4* | WBGene00011229 | protein_coding_gene | -1.1 | 0.45 | 4.7E-04 |
| *fbxa-212* | WBGene00012568 | pseudogene | -1.1 | 0.45 | 1.3E-05 |
| *F35E12.9* | WBGene00009433 | protein_coding_gene | -1.1 | 0.45 | 4.6E-04 |

| | | | | | |
|---|---|---|---|---|---|
| *R10E8.8* | WBGene00011213 | protein_coding_gene | -1.1 | 0.45 | 2.3E-05 |
| *fbxa-145* | WBGene00018595 | protein_coding_gene | -1.1 | 0.45 | 6.8E-04 |
| *sdz-21* | WBGene00018828 | protein_coding_gene | -1.1 | 0.45 | 4.5E-03 |
| *K04A8.20* | WBGene00235332 | protein_coding_gene | -1.1 | 0.45 | 1.7E-02 |
| *Y48A6B.7* | WBGene00012968 | protein_coding_gene | -1.2 | 0.45 | 1.0E-14 |
| *fbxb-111* | WBGene00017254 | protein_coding_gene | -1.2 | 0.45 | 4.9E-02 |
| *W04A8.2* | WBGene00012237 | protein_coding_gene | -1.2 | 0.45 | 9.5E-06 |
| *C43D7.8* | WBGene00008070 | protein_coding_gene | -1.2 | 0.45 | 4.7E-02 |
| *fbxb-22* | WBGene00013230 | protein_coding_gene | -1.2 | 0.45 | 8.4E-03 |
| *F52D1.2* | WBGene00018683 | protein_coding_gene | -1.2 | 0.44 | 7.3E-04 |
| *tsp-2* | WBGene00006628 | protein_coding_gene | -1.2 | 0.44 | 4.2E-04 |
| *fbxa-198* | WBGene00011514 | pseudogene | -1.2 | 0.44 | 6.7E-06 |
| *F22E5.20* | WBGene00017713 | protein_coding_gene | -1.2 | 0.44 | 8.9E-03 |
| *F32H2.10* | WBGene00009346 | protein_coding_gene | -1.2 | 0.44 | 9.1E-14 |
| *Y57G11B.2* | WBGene00013295 | protein_coding_gene | -1.2 | 0.44 | 4.3E-06 |
| *Y38H6C.23* | WBGene00012634 | protein_coding_gene | -1.2 | 0.44 | 2.5E-02 |
| *C05C10.7* | WBGene00007334 | protein_coding_gene | -1.2 | 0.44 | 2.4E-10 |
| *M03A1.8* | WBGene00077490 | protein_coding_gene | -1.2 | 0.44 | 1.1E-02 |
| *fbxb-107* | WBGene00017912 | protein_coding_gene | -1.2 | 0.44 | 2.1E-02 |
| *D2007.1* | WBGene00017041 | protein_coding_gene | -1.2 | 0.44 | 4.3E-05 |
| *ZK688.7* | WBGene00022802 | protein_coding_gene | -1.2 | 0.44 | 6.3E-07 |
| *T07G12.8* | WBGene00011599 | pseudogene | -1.2 | 0.44 | 8.4E-06 |
| *K09D9.12* | WBGene00019569 | pseudogene | -1.2 | 0.44 | 5.1E-04 |
| *Y57A10C.1* | WBGene00013280 | protein_coding_gene | -1.2 | 0.44 | 1.3E-02 |
| *K07C5.13* | WBGene00050947 | protein_coding_gene | -1.2 | 0.43 | 2.5E-02 |
| *tofu-6* | WBGene00017132 | protein_coding_gene | -1.2 | 0.43 | 4.0E-19 |
| *D2089.8* | WBGene00271706 | protein_coding_gene | -1.2 | 0.43 | 1.3E-06 |
| *Y74C9A.9* | WBGene00235381 | pseudogene | -1.2 | 0.43 | 1.9E-02 |
| *C05D9.9* | WBGene00015476 | protein_coding_gene | -1.2 | 0.43 | 6.3E-09 |
| *fbxb-24* | WBGene00013234 | protein_coding_gene | -1.2 | 0.43 | 1.3E-03 |
| *F42H10.6* | WBGene00018370 | protein_coding_gene | -1.2 | 0.43 | 4.3E-11 |
| *cyp-13A3* | WBGene00011675 | protein_coding_gene | -1.2 | 0.43 | 1.7E-02 |
| *F13C5.1* | WBGene00017422 | protein_coding_gene | -1.2 | 0.43 | 8.6E-07 |
| *K02A11.4* | WBGene00044607 | protein_coding_gene | -1.2 | 0.43 | 9.5E-06 |
| *K01A6.4* | WBGene00010446 | protein_coding_gene | -1.2 | 0.42 | 3.4E-02 |
| *Y73B3A.21* | WBGene00022222 | pseudogene | -1.2 | 0.42 | 7.3E-03 |
| *sdz-14* | WBGene00008782 | protein_coding_gene | -1.2 | 0.42 | 4.1E-06 |
| *F36F2.2* | WBGene00009476 | protein_coding_gene | -1.2 | 0.42 | 7.8E-08 |
| *faah-2* | WBGene00015048 | protein_coding_gene | -1.2 | 0.42 | 2.0E-04 |
| *F08B4.3* | WBGene00017235 | pseudogene | -1.2 | 0.42 | 2.2E-09 |
| *F12F6.8* | WBGene00008723 | pseudogene | -1.3 | 0.42 | 2.3E-03 |
| *T06A1.1* | WBGene00020277 | protein_coding_gene | -1.3 | 0.42 | 1.2E-02 |
| *C18A11.1* | WBGene00015947 | protein_coding_gene | -1.3 | 0.42 | 2.8E-09 |
| *C11D2.3* | WBGene00015701 | protein_coding_gene | -1.3 | 0.42 | 1.8E-02 |
| *Y71F9AL.6* | WBGene00022111 | protein_coding_gene | -1.3 | 0.42 | 1.1E-02 |
| *fbxa-122* | WBGene00014917 | pseudogene | -1.3 | 0.42 | 5.8E-06 |
| *F18C5.10* | WBGene00017563 | protein_coding_gene | -1.3 | 0.42 | 9.3E-25 |
| *sdz-37* | WBGene00014064 | protein_coding_gene | -1.3 | 0.42 | 4.6E-02 |
| *F01D5.5* | WBGene00008495 | protein_coding_gene | -1.3 | 0.42 | 7.1E-04 |
| *alh-5* | WBGene00000111 | protein_coding_gene | -1.3 | 0.42 | 2.3E-12 |
| *Y53G8B.2* | WBGene00021818 | protein_coding_gene | -1.3 | 0.42 | 4.1E-15 |
| *ins-7* | WBGene00002090 | protein_coding_gene | -1.3 | 0.42 | 4.5E-04 |
| *fbxa-95* | WBGene00009222 | protein_coding_gene | -1.3 | 0.41 | 1.1E-02 |
| *ZK550.2* | WBGene00013996 | protein_coding_gene | -1.3 | 0.41 | 6.6E-06 |
| *ZK896.4* | WBGene00014135 | protein_coding_gene | -1.3 | 0.41 | 1.1E-03 |
| *Y17G9B.2* | WBGene00021199 | protein_coding_gene | -1.3 | 0.41 | 8.4E-03 |
| *T12A2.3* | WBGene00020438 | protein_coding_gene | -1.3 | 0.41 | 8.4E-03 |
| *C54F6.5* | WBGene00016923 | protein_coding_gene | -1.3 | 0.41 | 9.9E-06 |

| | | | | | |
|---|---|---|---|---|---|
| *F21D9.5* | WBGene00009019 | pseudogene | -1.3 | 0.41 | 3.5E-06 |
| *cpt-4* | WBGene00019644 | protein_coding_gene | -1.3 | 0.41 | 8.4E-08 |
| *ugt-8* | WBGene00019234 | protein_coding_gene | -1.3 | 0.41 | 3.1E-07 |
| *ugt-17* | WBGene00007422 | protein_coding_gene | -1.3 | 0.40 | 4.8E-10 |
| *jmjd-4* | WBGene00011563 | protein_coding_gene | -1.3 | 0.40 | 5.4E-06 |
| *pot-2* | WBGene00010195 | protein_coding_gene | -1.3 | 0.40 | 2.5E-13 |
| *K10G6.9* | WBGene00206374 | protein_coding_gene | -1.3 | 0.40 | 1.6E-02 |
| *cyp-13A5* | WBGene00011672 | protein_coding_gene | -1.3 | 0.40 | 3.7E-14 |
| *F59A7.2* | WBGene00019090 | protein_coding_gene | -1.3 | 0.40 | 1.7E-06 |
| *W05H9.3* | WBGene00021050 | ncRNA_gene | -1.3 | 0.40 | 2.6E-13 |
| *Y116F11A.1* | WBGene00013816 | protein_coding_gene | -1.3 | 0.40 | 2.4E-02 |
| *F45D3.1* | WBGene00009721 | protein_coding_gene | -1.3 | 0.40 | 9.6E-03 |
| *D2045.8* | WBGene00008425 | protein_coding_gene | -1.3 | 0.40 | 5.0E-07 |
| *Y58A7A.3* | WBGene00021977 | protein_coding_gene | -1.3 | 0.40 | 1.0E-10 |
| *Y27F2A.8* | WBGene00021299 | protein_coding_gene | -1.3 | 0.40 | 1.7E-03 |
| *pot-3* | WBGene00007065 | protein_coding_gene | -1.3 | 0.40 | 3.1E-11 |
| *W01F3.2* | WBGene00012185 | protein_coding_gene | -1.3 | 0.40 | 3.0E-16 |
| *C08E8.10* | WBGene00077692 | protein_coding_gene | -1.3 | 0.39 | 2.4E-02 |
| *F45E10.2* | WBGene00009729 | protein_coding_gene | -1.3 | 0.39 | 3.8E-07 |
| *Y116A8C.20* | WBGene00013797 | protein_coding_gene | -1.3 | 0.39 | 3.2E-02 |
| *C18H9.5* | WBGene00016003 | protein_coding_gene | -1.3 | 0.39 | 6.6E-06 |
| *dct-17* | WBGene00009431 | protein_coding_gene | -1.3 | 0.39 | 3.7E-02 |
| *dach-1* | WBGene00020386 | protein_coding_gene | -1.4 | 0.39 | 1.2E-04 |
| *spp-22* | WBGene00044283 | protein_coding_gene | -1.4 | 0.39 | 1.9E-05 |
| *F09F3.8* | WBGene00014732 | pseudogene | -1.4 | 0.39 | 3.0E-02 |
| *lido-4* | WBGene00015589 | protein_coding_gene | -1.4 | 0.39 | 1.9E-02 |
| *frpr-11* | WBGene00019444 | protein_coding_gene | -1.4 | 0.39 | 1.1E-05 |
| *fbxa-140* | WBGene00008130 | protein_coding_gene | -1.4 | 0.39 | 8.0E-09 |
| *C36C9.10* | WBGene00219849 | ncRNA_gene | -1.4 | 0.38 | 2.2E-03 |
| *fbxa-211* | WBGene00012566 | protein_coding_gene | -1.4 | 0.38 | 3.6E-14 |
| *W01B11.6* | WBGene00020917 | protein_coding_gene | -1.4 | 0.38 | 4.8E-11 |
| *fbxa-162* | WBGene00015597 | pseudogene | -1.4 | 0.38 | 1.3E-04 |
| *C51E3.10* | WBGene00008255 | protein_coding_gene | -1.4 | 0.38 | 4.1E-04 |
| *dod-23* | WBGene00009898 | protein_coding_gene | -1.4 | 0.38 | 1.2E-04 |
| *F48D6.4* | WBGene00018602 | protein_coding_gene | -1.4 | 0.38 | 7.3E-20 |
| *EEED8.13* | WBGene00017141 | protein_coding_gene | -1.4 | 0.38 | 9.9E-06 |
| *R09B3.6* | WBGene00014820 | pseudogene | -1.4 | 0.38 | 6.8E-08 |
| *M151.2* | WBGene00019794 | protein_coding_gene | -1.4 | 0.38 | 9.1E-03 |
| *Y20C6A.4* | WBGene00077495 | protein_coding_gene | -1.4 | 0.37 | 8.9E-04 |
| *his-32* | WBGene00001906 | protein_coding_gene | -1.4 | 0.37 | 8.0E-03 |
| *M03B6.5* | WBGene00010837 | protein_coding_gene | -1.4 | 0.37 | 2.7E-02 |
| *ugt-29* | WBGene00021709 | protein_coding_gene | -1.4 | 0.37 | 2.6E-05 |
| *F36H5.4* | WBGene00018103 | protein_coding_gene | -1.4 | 0.37 | 3.4E-03 |
| *fbxa-99* | WBGene00009490 | protein_coding_gene | -1.4 | 0.37 | 2.1E-02 |
| *R09E12.9* | WBGene00044514 | protein_coding_gene | -1.5 | 0.37 | 5.1E-04 |
| *fbxa-163* | WBGene00015598 | protein_coding_gene | -1.5 | 0.37 | 3.8E-05 |
| *fbxb-60* | WBGene00010104 | protein_coding_gene | -1.5 | 0.36 | 1.3E-02 |
| *cpt-1* | WBGene00012907 | protein_coding_gene | -1.5 | 0.36 | 1.8E-20 |
| *C49G7.12* | WBGene00077593 | protein_coding_gene | -1.5 | 0.36 | 4.5E-04 |
| *F15E6.10* | WBGene00044479 | protein_coding_gene | -1.5 | 0.36 | 2.4E-02 |
| *oac-6* | WBGene00007833 | protein_coding_gene | -1.5 | 0.36 | 5.2E-04 |
| *lips-10* | WBGene00008803 | protein_coding_gene | -1.5 | 0.36 | 5.1E-08 |
| *Y105C5B.20* | WBGene00013659 | protein_coding_gene | -1.5 | 0.36 | 5.1E-10 |
| *F13A7.11* | WBGene00008726 | protein_coding_gene | -1.5 | 0.36 | 1.3E-05 |
| *M163.1* | WBGene00010935 | protein_coding_gene | -1.5 | 0.36 | 4.6E-15 |
| *F08H9.3* | WBGene00008591 | protein_coding_gene | -1.5 | 0.36 | 7.5E-04 |
| *Y71A12B.19* | WBGene00077443 | pseudogene | -1.5 | 0.35 | 8.6E-10 |
| *M60.7* | WBGene00019782 | protein_coding_gene | -1.5 | 0.35 | 8.4E-05 |

| | | | | | |
|---|---|---|---|---|---|
| *T22B7.3* | WBGene00020672 | protein_coding_gene | -1.5 | 0.35 | 2.2E-05 |
| *fbxb-45* | WBGene00019705 | protein_coding_gene | -1.5 | 0.35 | 7.2E-03 |
| *C52E12.11* | WBGene00304189 | pseudogene | -1.5 | 0.35 | 2.9E-02 |
| *tbc-6* | WBGene00015410 | protein_coding_gene | -1.5 | 0.35 | 4.0E-14 |
| *F23A7.8* | WBGene00044638 | protein_coding_gene | -1.5 | 0.35 | 4.6E-24 |
| *clec-40* | WBGene00011853 | protein_coding_gene | -1.5 | 0.34 | 9.2E-03 |
| *Y47G6A.14* | WBGene00021640 | protein_coding_gene | -1.5 | 0.34 | 2.4E-06 |
| *cnc-7* | WBGene00010005 | protein_coding_gene | -1.6 | 0.34 | 7.1E-04 |
| *fipr-26* | WBGene00009963 | protein_coding_gene | -1.6 | 0.34 | 8.3E-03 |
| *F40G9.20* | WBGene00195034 | ncRNA_gene | -1.6 | 0.34 | 9.3E-03 |
| *F36F12.3* | WBGene00018094 | protein_coding_gene | -1.6 | 0.34 | 3.5E-02 |
| *ugt-33* | WBGene00007946 | protein_coding_gene | -1.6 | 0.34 | 1.5E-03 |
| *T23F11.6* | WBGene00011957 | protein_coding_gene | -1.6 | 0.33 | 1.3E-03 |
| *F02E11.3* | WBGene00017181 | protein_coding_gene | -1.6 | 0.33 | 4.8E-02 |
| *his-63* | WBGene00001937 | protein_coding_gene | -1.6 | 0.33 | 4.7E-02 |
| *T11F8.2* | WBGene00020434 | pseudogene | -1.6 | 0.33 | 4.2E-07 |
| *fbxc-37* | WBGene00017942 | pseudogene | -1.6 | 0.33 | 3.4E-02 |
| *T04C12.8* | WBGene00011430 | protein_coding_gene | -1.6 | 0.33 | 1.0E-05 |
| *C06B3.7* | WBGene00007366 | protein_coding_gene | -1.6 | 0.33 | 2.0E-03 |
| *Y49F6A.5* | WBGene00021713 | protein_coding_gene | -1.6 | 0.33 | 4.2E-02 |
| *C04C11.25* | WBGene00284855 | protein_coding_gene | -1.6 | 0.32 | 1.3E-03 |
| *Y53F4B.38* | WBGene00014920 | pseudogene | -1.6 | 0.32 | 4.2E-02 |
| *clec-218* | WBGene00020938 | protein_coding_gene | -1.6 | 0.32 | 9.4E-21 |
| *D2024.10* | WBGene00044505 | pseudogene | -1.6 | 0.32 | 3.2E-02 |
| *dhs-8* | WBGene00000972 | protein_coding_gene | -1.7 | 0.32 | 1.2E-03 |
| *K08E3.2* | WBGene00010662 | protein_coding_gene | -1.7 | 0.31 | 1.3E-02 |
| *F11D5.5* | WBGene00017382 | protein_coding_gene | -1.7 | 0.31 | 2.2E-05 |
| *srt-63* | WBGene00020877 | protein_coding_gene | -1.7 | 0.31 | 4.6E-02 |
| *B0563.8* | WBGene00015265 | protein_coding_gene | -1.7 | 0.31 | 5.0E-02 |
| *Y37H2A.12* | WBGene00044758 | protein_coding_gene | -1.7 | 0.31 | 2.7E-05 |
| *ceh-5* | WBGene00000430 | protein_coding_gene | -1.7 | 0.31 | 9.5E-04 |
| *H25K10.4* | WBGene00010413 | protein_coding_gene | -1.7 | 0.31 | 2.4E-05 |
| *Y8A9A.2* | WBGene00021171 | protein_coding_gene | -1.7 | 0.31 | 1.5E-08 |
| *F31D5.6* | WBGene00017949 | protein_coding_gene | -1.7 | 0.30 | 1.7E-03 |
| *dao-6* | WBGene00000932 | protein_coding_gene | -1.7 | 0.30 | 9.0E-03 |
| *tre-5* | WBGene00006611 | protein_coding_gene | -1.7 | 0.30 | 2.6E-12 |
| *clec-23* | WBGene00011851 | pseudogene | -1.7 | 0.30 | 5.1E-03 |
| *T16A1.10* | WBGene00269431 | protein_coding_gene | -1.7 | 0.30 | 1.1E-02 |
| *T10C6.15* | WBGene00050899 | protein_coding_gene | -1.7 | 0.30 | 3.5E-07 |
| *Y37F4.8* | WBGene00044289 | protein_coding_gene | -1.7 | 0.30 | 1.7E-04 |
| *drd-50* | WBGene00018643 | protein_coding_gene | -1.8 | 0.30 | 6.3E-04 |
| *H32K16.2* | WBGene00010422 | protein_coding_gene | -1.8 | 0.30 | 6.4E-06 |
| *F15H9.8* | WBGene00194652 | pseudogene | -1.8 | 0.29 | 6.0E-03 |
| *pqn-16* | WBGene00004107 | protein_coding_gene | -1.8 | 0.29 | 4.0E-02 |
| *fbxa-200* | WBGene00020304 | pseudogene | -1.8 | 0.29 | 1.1E-06 |
| *irg-5* | WBGene00009429 | protein_coding_gene | -1.8 | 0.29 | 1.9E-04 |
| *W05H9.1* | WBGene00021048 | protein_coding_gene | -1.8 | 0.29 | 2.7E-37 |
| *oac-54* | WBGene00012323 | protein_coding_gene | -1.8 | 0.28 | 2.5E-03 |
| *K10D11.2* | WBGene00010746 | protein_coding_gene | -1.8 | 0.28 | 4.3E-02 |
| *C18H9.6* | WBGene00016004 | protein_coding_gene | -1.8 | 0.28 | 9.4E-10 |
| *F10D7.3* | WBGene00017340 | protein_coding_gene | -1.8 | 0.28 | 5.6E-17 |
| *clec-145* | WBGene00013013 | protein_coding_gene | -1.8 | 0.28 | 1.3E-02 |
| *pho-9* | WBGene00022181 | protein_coding_gene | -1.8 | 0.28 | 8.0E-11 |
| *ttr-37* | WBGene00017667 | protein_coding_gene | -1.8 | 0.28 | 9.2E-14 |
| *ins-19* | WBGene00002102 | protein_coding_gene | -1.9 | 0.28 | 2.3E-03 |
| *Y17G9A.4* | WBGene00021197 | protein_coding_gene | -1.9 | 0.27 | 1.8E-09 |
| *F16B4.7* | WBGene00017509 | protein_coding_gene | -1.9 | 0.27 | 3.3E-04 |
| *fbxb-16* | WBGene00022560 | protein_coding_gene | -1.9 | 0.27 | 3.9E-02 |

| | | | | | |
|---|---|---|---|---|---|
| *nlp-31* | WBGene00003769 | protein_coding_gene | -1.9 | 0.26 | 1.9E-23 |
| *F53B2.8* | WBGene00009957 | protein_coding_gene | -1.9 | 0.26 | 1.1E-06 |
| *C42D4.1* | WBGene00016594 | protein_coding_gene | -1.9 | 0.26 | 1.3E-35 |
| *K01A6.7* | WBGene00045096 | protein_coding_gene | -2.0 | 0.26 | 4.4E-03 |
| *irg-2* | WBGene00016783 | protein_coding_gene | -2.0 | 0.26 | 1.4E-03 |
| *Y81B9A.5* | WBGene00304906 | ncRNA_gene | -2.0 | 0.26 | 8.6E-03 |
| *B0563.1* | WBGene00015261 | protein_coding_gene | -2.0 | 0.25 | 2.7E-02 |
| *clec-47* | WBGene00011668 | protein_coding_gene | -2.0 | 0.25 | 1.0E-11 |
| *nspe-6* | WBGene00012603 | protein_coding_gene | -2.0 | 0.25 | 9.2E-03 |
| *ugt-54* | WBGene00012013 | protein_coding_gene | -2.0 | 0.25 | 4.0E-20 |
| *ZK673.5* | WBGene00014061 | protein_coding_gene | -2.0 | 0.25 | 5.5E-03 |
| *srd-28* | WBGene00005106 | protein_coding_gene | -2.0 | 0.24 | 4.6E-02 |
| *K09D9.1* | WBGene00019564 | protein_coding_gene | -2.1 | 0.24 | 1.1E-04 |
| *ZK1307.2* | WBGene00014245 | protein_coding_gene | -2.1 | 0.24 | 3.4E-05 |
| *C54F6.18* | WBGene00235093 | protein_coding_gene | -2.1 | 0.23 | 4.4E-03 |
| *Y54G2A.36* | WBGene00021898 | protein_coding_gene | -2.1 | 0.23 | 1.4E-32 |
| *ZK6.8* | WBGene00022643 | protein_coding_gene | -2.1 | 0.23 | 2.5E-08 |
| *nlp-34* | WBGene00015046 | protein_coding_gene | -2.1 | 0.23 | 1.0E-06 |
| *sri-36* | WBGene00005548 | protein_coding_gene | -2.1 | 0.23 | 1.9E-02 |
| *M04C9.2* | WBGene00010857 | protein_coding_gene | -2.1 | 0.23 | 8.6E-03 |
| *Y45F10D.6* | WBGene00012886 | protein_coding_gene | -2.1 | 0.23 | 6.8E-08 |
| *C31E10.11* | WBGene00304783 | protein_coding_gene | -2.2 | 0.22 | 8.0E-03 |
| *Y53H1B.4* | WBGene00013182 | protein_coding_gene | -2.2 | 0.22 | 9.3E-03 |
| *btb-15* | WBGene00015594 | pseudogene | -2.2 | 0.22 | 8.8E-03 |
| *ugt-34* | WBGene00009255 | protein_coding_gene | -2.2 | 0.22 | 1.5E-02 |
| *nlp-30* | WBGene00003768 | protein_coding_gene | -2.2 | 0.22 | 1.8E-10 |
| *ptr-22* | WBGene00004236 | protein_coding_gene | -2.2 | 0.22 | 7.6E-28 |
| *clec-24* | WBGene00009856 | protein_coding_gene | -2.2 | 0.21 | 2.0E-14 |
| *T04C12.32* | WBGene00235086 | ncRNA_gene | -2.3 | 0.21 | 8.9E-03 |
| *acd-1* | WBGene00016064 | protein_coding_gene | -2.3 | 0.20 | 3.0E-04 |
| *Y24D9A.9* | WBGene00044952 | snoRNA_gene | -2.3 | 0.20 | 1.3E-02 |
| *clec-32* | WBGene00009860 | protein_coding_gene | -2.3 | 0.20 | 1.9E-06 |
| *linc-6* | WBGene00219700 | lincRNA_gene | -2.3 | 0.20 | 2.7E-09 |
| *C27B7.9* | WBGene00007765 | protein_coding_gene | -2.3 | 0.20 | 6.8E-67 |
| *Y46H3A.5* | WBGene00021601 | protein_coding_gene | -2.4 | 0.19 | 2.5E-03 |
| *W08A12.2* | WBGene00021079 | protein_coding_gene | -2.4 | 0.19 | 7.6E-28 |
| *Y71F9AM.9* | WBGene00271781 | lincRNA_gene | -2.4 | 0.19 | 7.3E-04 |
| *fbxc-26* | WBGene00019043 | protein_coding_gene | -2.4 | 0.19 | 3.1E-05 |
| *dod-22* | WBGene00010125 | protein_coding_gene | -2.4 | 0.19 | 7.6E-24 |
| *pitr-4* | WBGene00007141 | protein_coding_gene | -2.5 | 0.18 | 1.4E-03 |
| *nlp-28* | WBGene00003766 | protein_coding_gene | -2.5 | 0.18 | 8.0E-39 |
| *cnc-6* | WBGene00000560 | protein_coding_gene | -2.5 | 0.18 | 1.6E-02 |
| *F59A7.12* | WBGene00044520 | protein_coding_gene | -2.5 | 0.18 | 6.8E-03 |
| *fipr-23* | WBGene00007990 | protein_coding_gene | -2.5 | 0.17 | 1.6E-03 |
| *W10G11.1* | WBGene00021134 | protein_coding_gene | -2.6 | 0.17 | 3.3E-03 |
| *C49G7.7* | WBGene00016785 | protein_coding_gene | -2.6 | 0.17 | 4.8E-06 |
| *C39B5.10* | WBGene00016528 | protein_coding_gene | -2.6 | 0.16 | 1.5E-33 |
| *C37A5.3* | WBGene00014698 | protein_coding_gene | -2.6 | 0.16 | 8.4E-03 |
| *T23E7.6* | WBGene00044772 | protein_coding_gene | -2.7 | 0.16 | 8.6E-21 |
| *srr-6* | WBGene00005657 | protein_coding_gene | -2.7 | 0.15 | 5.4E-11 |
| *W10G11.4* | WBGene00021137 | protein_coding_gene | -2.9 | 0.13 | 7.0E-04 |
| *cnc-2* | WBGene00000556 | protein_coding_gene | -2.9 | 0.13 | 2.4E-07 |
| *dct-19* | WBGene00007876 | protein_coding_gene | -2.9 | 0.13 | 2.1E-03 |
| *fipr-24* | WBGene00007992 | protein_coding_gene | -3.0 | 0.13 | 1.2E-05 |
| *C18D11.6* | WBGene00007682 | protein_coding_gene | -3.0 | 0.12 | 6.2E-06 |
| *C08E8.3* | WBGene00007439 | protein_coding_gene | -3.1 | 0.12 | 1.1E-05 |
| *C37A5.6* | WBGene00014700 | protein_coding_gene | -3.1 | 0.12 | 4.2E-03 |
| *T24B8.5* | WBGene00011979 | protein_coding_gene | -3.3 | 0.10 | 5.4E-09 |

| | | | | | |
|---|---|---|---|---|---|
| *cnc-11* | WBGene00044900 | protein_coding_gene | -3.3 | 0.10 | 7.1E-10 |
| *nlp-29* | WBGene00003767 | protein_coding_gene | -3.3 | 0.10 | 3.3E-62 |
| *cpt-3* | WBGene00021703 | protein_coding_gene | -3.3 | 0.10 | 8.6E-15 |
| *F56D6.9* | WBGene00044471 | protein_coding_gene | -3.4 | 0.10 | 5.3E-04 |
| *ins-8* | WBGene00002091 | protein_coding_gene | -3.7 | 0.08 | 6.3E-04 |
| *F48C1.11* | WBGene00195179 | protein_coding_gene | -3.8 | 0.07 | 7.3E-04 |
| *W02D7.11* | WBGene00020946 | protein_coding_gene | -3.9 | 0.07 | 4.9E-13 |
| *fmo-2* | WBGene00001477 | protein_coding_gene | -3.9 | 0.07 | 2.0E-07 |
| *dod-24* | WBGene00007875 | protein_coding_gene | -4.5 | 0.04 | 3.0E-136 |
| *F56D6.8* | WBGene00044292 | protein_coding_gene | -4.9 | 0.03 | 4.1E-06 |
| *dct-7* | WBGene00017488 | protein_coding_gene | -5.2 | 0.03 | 4.0E-08 |
| *F48C1.9* | WBGene00018601 | protein_coding_gene | -5.6 | 0.02 | 5.0E-22 |
| *dct-8* | WBGene00044472 | protein_coding_gene | -6.7 | 0.01 | 1.0E-09 |

Shared (N2&CB4856) DEGs in response to *S. epidermidis* (vs. OP50)

| | | | N2 | | | CB4856 | | |
|---|---|---|---|---|---|---|---|---|
| Gene Name | Wormbase Gene ID | Type | $log_2FC$ | FC | Adj P-value | log2FC | FC | Adj P-value |
| *ilys-3* | WBGene00016670 | protein_coding_gene | 7.8 | 227.922 | 1.8E-16 | 1.8 | 3.58 | 2.2E-12 |
| *T22F3.11* | WBGene00020700 | protein_coding_gene | 7.7 | 211.199 | 2.4E-17 | 2.3 | 4.99 | 1.1E-02 |
| *chil-28* | WBGene00008842 | pseudogene | 7.4 | 168.420 | 3.9E-12 | 2.7 | 6.67 | 3.6E-11 |
| *F46C5.1* | WBGene00009778 | protein_coding_gene | 7.1 | 135.390 | 3.9E-15 | 1.6 | 3.04 | 9.0E-03 |
| *F17B5.1* | WBGene00008905 | protein_coding_gene | 6.8 | 113.321 | 2.3E-33 | 1.7 | 3.16 | 1.8E-02 |
| *lys-10* | WBGene00003099 | protein_coding_gene | 6.5 | 93.165 | 3.8E-31 | 2.8 | 6.91 | 1.1E-15 |
| *clec-60* | WBGene00014046 | protein_coding_gene | 6.5 | 87.749 | 7.2E-16 | 2.9 | 7.35 | 2.3E-12 |
| *ilys-2* | WBGene00016669 | protein_coding_gene | 6.3 | 76.604 | 8.3E-18 | 1.4 | 2.56 | 3.7E-03 |
| *Y54G2A.37* | WBGene00023484 | protein_coding_gene | 6.1 | 70.775 | 1.9E-08 | 2.0 | 3.98 | 4.3E-09 |
| *T26H5.8* | WBGene00012070 | protein_coding_gene | 5.9 | 61.655 | 1.3E-22 | 1.1 | 2.21 | 7.4E-08 |
| *clec-61* | WBGene00014047 | protein_coding_gene | 5.8 | 56.786 | 5.3E-21 | 1.7 | 3.29 | 4.0E-12 |
| ***cest-29*** | **WBGene00017205** | **pseudogene** | **5.8** | **56.553** | **1.3E-06** | **-1.2** | **0.42** | **3.2E-05** |
| *thn-2* | WBGene00009215 | protein_coding_gene | 5.5 | 44.261 | 4.4E-29 | 2.7 | 6.31 | 1.3E-53 |
| *lipl-1* | WBGene00010062 | protein_coding_gene | 5.3 | 40.691 | 3.9E-13 | 2.9 | 7.36 | 3.7E-09 |
| *cest-1.1* | WBGene00011362 | protein_coding_gene | 5.2 | 36.972 | 6.3E-31 | 4.6 | 23.65 | 5.9E-24 |
| *K05F6.10* | WBGene00019421 | protein_coding_gene | 5.2 | 35.991 | 8.2E-13 | 1.7 | 3.36 | 2.7E-02 |
| *dod-3* | WBGene00016052 | protein_coding_gene | 5.1 | 35.269 | 5.7E-09 | 1.8 | 3.37 | 8.5E-05 |
| ***lipl-3*** | **WBGene00020016** | **protein_coding_gene** | **5.0** | **33.116** | **5.0E-12** | **-1.7** | **0.30** | **8.6E-03** |
| *ZK105.13* | WBGene00235114 | protein_coding_gene | 5.0 | 31.011 | 1.7E-06 | 1.3 | 2.50 | 2.0E-03 |
| *W01C8.1* | WBGene00020918 | protein_coding_gene | 4.9 | 30.495 | 2.6E-05 | 1.9 | 3.70 | 1.3E-02 |
| *T07H8.11* | WBGene00304201 | protein_coding_gene | 4.9 | 30.243 | 1.8E-14 | 4.6 | 23.95 | 6.1E-44 |
| *F11A5.17* | WBGene00045510 | protein_coding_gene | 4.9 | 29.645 | 1.6E-08 | 2.5 | 5.48 | 8.9E-03 |
| ***C50F7.5*** | **WBGene00016845** | **protein_coding_gene** | **4.8** | **27.131** | **4.4E-07** | **-2.8** | **0.15** | **6.3E-68** |
| *best-1* | WBGene00007203 | protein_coding_gene | 4.7 | 26.293 | 2.7E-48 | 2.2 | 4.48 | 9.5E-05 |
| *T26H5.10* | WBGene00044783 | protein_coding_gene | 4.7 | 25.897 | 2.6E-06 | 1.4 | 2.56 | 1.3E-03 |
| ***mtl-1*** | **WBGene00003473** | **protein_coding_gene** | **4.7** | **25.368** | **2.8E-07** | **-1.6** | **0.34** | **1.1E-04** |
| *sdz-24* | WBGene00019495 | protein_coding_gene | 4.5 | 23.239 | 1.3E-73 | 2.5 | 5.48 | 1.8E-02 |
| *asm-3* | WBGene00000213 | protein_coding_gene | 4.5 | 22.184 | 3.5E-09 | 5.4 | 42.52 | 7.6E-47 |
| *C54F6.12* | WBGene00016929 | protein_coding_gene | 4.5 | 21.890 | 6.9E-15 | 2.3 | 5.10 | 4.8E-09 |
| *F53H2.1* | WBGene00010004 | protein_coding_gene | 4.4 | 21.323 | 9.5E-08 | 2.0 | 3.98 | 2.6E-05 |
| *F14D7.7* | WBGene00008794 | protein_coding_gene | 4.3 | 20.036 | 8.7E-07 | 1.8 | 3.50 | 1.3E-03 |
| *ZK666.15* | WBGene00195145 | protein_coding_gene | 4.3 | 20.036 | 1.2E-06 | 2.3 | 4.87 | 1.7E-05 |
| *C35C5.8* | WBGene00007957 | protein_coding_gene | 4.2 | 18.491 | 5.4E-24 | 1.2 | 2.29 | 7.9E-04 |
| ***F46A8.13*** | **WBGene00219950** | **ncRNA_gene** | **4.2** | **18.115** | **7.6E-05** | **-1.1** | **0.47** | **7.3E-03** |
| *math-34* | WBGene00019699 | protein_coding_gene | 4.1 | 17.528 | 9.1E-14 | 1.7 | 3.17 | 3.3E-08 |
| *F07C4.6* | WBGene00017200 | protein_coding_gene | 4.0 | 16.059 | 6.1E-07 | 3.5 | 11.09 | 3.1E-03 |
| *aqp-1* | WBGene00000169 | protein_coding_gene | 4.0 | 15.482 | 5.7E-39 | 3.6 | 12.10 | 5.0E-21 |
| ***scl-2*** | **WBGene00009895** | **protein_coding_gene** | **3.9** | **15.229** | **3.3E-06** | **-1.2** | **0.43** | **3.0E-12** |
| *F11C7.6* | WBGene00017378 | protein_coding_gene | 3.9 | 15.109 | 4.4E-09 | 2.1 | 4.32 | 2.6E-13 |
| *K02E2.7* | WBGene00010508 | protein_coding_gene | 3.9 | 14.854 | 6.2E-04 | 1.6 | 3.10 | 1.4E-02 |
| *Y87G2A.16* | WBGene00013602 | protein_coding_gene | 3.8 | 14.398 | 4.7E-24 | 1.2 | 2.33 | 4.6E-02 |
| *acs-2* | WBGene00009221 | protein_coding_gene | 3.8 | 14.028 | 5.0E-21 | 2.5 | 5.77 | 7.6E-10 |

| | | | | | | | | |
|---|---|---|---|---|---|---|---|---|
| *daao-1* | WBGene00022076 | protein_coding_gene | 3.7 | 13.198 | 8.8E-19 | 1.8 | 3.46 | 6.4E-11 |
| *Y41D4B.17* | WBGene00021519 | protein_coding_gene | 3.7 | 12.846 | 5.0E-23 | 3.6 | 12.40 | 2.6E-17 |
| *F40G12.5* | WBGene00009600 | protein_coding_gene | 3.7 | 12.768 | 5.9E-04 | 2.1 | 4.28 | 1.2E-05 |
| ***ftn-1*** | **WBGene00001500** | **protein_coding_gene** | **3.6** | **12.342** | **4.5E-06** | **-1.4** | **0.39** | **3.2E-03** |
| *Y105C5B.7* | WBGene00013648 | pseudogene | 3.6 | 11.903 | 1.5E-03 | 1.6 | 2.98 | 2.3E-03 |
| *cpr-1* | WBGene00000781 | protein_coding_gene | 3.6 | 11.811 | 1.1E-27 | 1.6 | 3.02 | 9.8E-28 |
| ***F45D3.4*** | **WBGene00009724** | **protein_coding_gene** | **3.5** | **11.570** | **2.2E-06** | **-1.6** | **0.32** | **1.5E-07** |
| *Y65B4BR.1* | WBGene00022040 | protein_coding_gene | 3.5 | 11.486 | 5.1E-16 | 3.1 | 8.81 | 1.2E-09 |
| *asah-1* | WBGene00010769 | protein_coding_gene | 3.5 | 11.319 | 4.2E-23 | 2.7 | 6.49 | 2.2E-19 |
| ***F47B8.4*** | **WBGene00009805** | **protein_coding_gene** | **3.5** | **11.012** | **1.4E-03** | **-2.2** | **0.22** | **1.1E-09** |
| ***F47B8.10*** | **WBGene00009811** | **protein_coding_gene** | **3.4** | **10.923** | **6.3E-04** | **-1.0** | **0.50** | **3.2E-02** |
| *clec-172* | WBGene00044719 | protein_coding_gene | 3.4 | 10.922 | 2.3E-14 | 1.2 | 2.36 | 7.5E-05 |
| *lys-7* | WBGene00003096 | protein_coding_gene | 3.4 | 10.743 | 7.7E-22 | 2.3 | 5.01 | 3.2E-15 |
| *W07B8.4* | WBGene00021072 | protein_coding_gene | 3.4 | 10.590 | 1.3E-18 | 1.4 | 2.72 | 1.0E-08 |
| *ZC116.5* | WBGene00119204 | protein_coding_gene | 3.3 | 10.077 | 7.5E-11 | 2.2 | 4.70 | 2.4E-07 |
| *gst-8* | WBGene00001756 | protein_coding_gene | 3.3 | 9.987 | 8.5E-04 | 2.5 | 5.74 | 8.9E-04 |
| *aat-7* | WBGene00000008 | protein_coding_gene | 3.3 | 9.821 | 3.3E-23 | 5.5 | 46.14 | 2.7E-26 |
| *nlp-51* | WBGene00011997 | protein_coding_gene | 3.3 | 9.609 | 8.0E-29 | 2.6 | 5.94 | 1.1E-56 |
| *Y51H4A.5* | WBGene00013101 | protein_coding_gene | 3.2 | 8.953 | 4.9E-08 | 1.1 | 2.14 | 2.3E-03 |
| *cyp-35B1* | WBGene00019472 | protein_coding_gene | 3.2 | 8.917 | 2.7E-17 | 1.8 | 3.51 | 3.4E-07 |
| *zip-6* | WBGene00011130 | protein_coding_gene | 3.1 | 8.851 | 7.7E-03 | 2.1 | 4.20 | 4.3E-02 |
| *F56C3.9* | WBGene00018946 | protein_coding_gene | 3.1 | 8.782 | 6.8E-08 | 1.6 | 3.12 | 3.3E-04 |
| *K10D3.6* | WBGene00010740 | protein_coding_gene | 3.1 | 8.773 | 1.2E-30 | 1.4 | 2.68 | 3.4E-08 |
| *W02G9.4* | WBGene00020954 | protein_coding_gene | 3.1 | 8.572 | 1.0E-58 | 3.0 | 7.92 | 1.2E-41 |
| ***F21C10.11*** | **WBGene00017660** | **protein_coding_gene** | **3.1** | **8.348** | **9.3E-13** | **-1.2** | **0.43** | **2.2E-02** |
| *cpr-4* | WBGene00000784 | protein_coding_gene | 3.1 | 8.322 | 6.0E-21 | 1.1 | 2.09 | 2.6E-04 |
| *ugt-43* | WBGene00008485 | protein_coding_gene | 3.1 | 8.321 | 7.0E-13 | 1.3 | 2.40 | 1.4E-05 |
| *asp-11* | WBGene00017677 | pseudogene | 3.0 | 8.213 | 1.2E-04 | 1.2 | 2.23 | 2.0E-02 |
| *cyp-13A7* | WBGene00000372 | protein_coding_gene | 3.0 | 8.169 | 4.4E-03 | 2.0 | 4.05 | 3.4E-12 |
| *R09D1.4* | WBGene00011160 | pseudogene | 3.0 | 8.035 | 2.2E-02 | 1.8 | 3.60 | 3.4E-02 |
| ***C18H7.1*** | **WBGene00015993** | **protein_coding_gene** | **3.0** | **7.925** | **6.4E-04** | **-1.8** | **0.29** | **3.8E-08** |
| ***F15E6.3*** | **WBGene00017484** | **protein_coding_gene** | **3.0** | **7.861** | **6.4E-08** | **-2.8** | **0.14** | **6.2E-24** |
| *lact-6* | WBGene00007358 | protein_coding_gene | 3.0 | 7.825 | 3.6E-06 | 3.2 | 9.07 | 9.9E-12 |
| *W03F11.5* | WBGene00021008 | protein_coding_gene | 3.0 | 7.784 | 1.5E-07 | 2.1 | 4.34 | 1.4E-04 |
| *lips-6* | WBGene00014009 | protein_coding_gene | 2.9 | 7.641 | 8.5E-04 | 4.1 | 17.68 | 2.8E-08 |
| ***ugt-18*** | **WBGene00013900** | **protein_coding_gene** | **2.9** | **7.584** | **3.1E-03** | **-1.6** | **0.34** | **7.8E-04** |
| ***nlp-39*** | **WBGene00008295** | **protein_coding_gene** | **2.9** | **7.568** | **1.1E-03** | **-1.7** | **0.32** | **4.1E-03** |
| *Y73B6BL.37* | WBGene00022256 | protein_coding_gene | 2.9 | 7.531 | 4.0E-03 | 1.8 | 3.39 | 8.0E-03 |
| *clec-158* | WBGene00020067 | protein_coding_gene | 2.9 | 7.348 | 4.1E-03 | 1.4 | 2.63 | 8.1E-03 |
| *C07G3.8* | WBGene00015576 | protein_coding_gene | 2.9 | 7.306 | 9.4E-03 | 1.9 | 3.78 | 4.3E-02 |
| *C35D6.5* | WBGene00007962 | pseudogene | 2.8 | 7.183 | 1.4E-03 | 1.3 | 2.49 | 3.9E-03 |
| *chil-18* | WBGene00011161 | protein_coding_gene | 2.8 | 7.006 | 6.0E-04 | 1.2 | 2.30 | 2.9E-02 |
| *F11A5.9* | WBGene00008677 | protein_coding_gene | 2.8 | 6.890 | 1.5E-20 | 2.1 | 4.39 | 3.2E-23 |
| *K03H6.2* | WBGene00019368 | protein_coding_gene | 2.7 | 6.702 | 2.8E-11 | 3.2 | 9.15 | 4.2E-16 |
| *F01D5.7* | WBGene00008497 | protein_coding_gene | 2.7 | 6.610 | 1.1E-02 | 1.9 | 3.63 | 6.5E-03 |
| ***mir-238*** | **WBGene00003331** | **miRNA_gene** | **2.7** | **6.577** | **2.6E-02** | **-2.3** | **0.21** | **2.6E-02** |
| *W03F9.4* | WBGene00021002 | protein_coding_gene | 2.7 | 6.417 | 1.3E-11 | 1.0 | 2.05 | 1.7E-02 |
| *hsp-12.3* | WBGene00002012 | protein_coding_gene | 2.7 | 6.376 | 7.4E-22 | 1.9 | 3.81 | 2.0E-16 |
| *Y38H6C.19* | WBGene00012631 | protein_coding_gene | 2.7 | 6.335 | 9.6E-18 | 2.1 | 4.29 | 1.1E-02 |
| *K06A9.1* | WBGene00019435 | protein_coding_gene | 2.6 | 6.232 | 4.6E-09 | 2.5 | 5.76 | 2.3E-24 |
| *ZK1290.1* | WBGene00022886 | protein_coding_gene | 2.6 | 6.212 | 3.6E-02 | 1.0 | 2.00 | 1.3E-05 |
| *F09C12.2* | WBGene00017277 | protein_coding_gene | 2.6 | 6.164 | 5.5E-04 | 3.1 | 8.37 | 4.8E-04 |
| *F46A8.1* | WBGene00014759 | pseudogene | 2.6 | 6.142 | 4.1E-09 | 2.6 | 6.18 | 8.9E-20 |
| *clec-42* | WBGene00008891 | protein_coding_gene | 2.6 | 6.117 | 8.1E-03 | 1.2 | 2.22 | 1.1E-02 |
| *F42A10.7* | WBGene00018343 | protein_coding_gene | 2.6 | 6.015 | 2.9E-34 | 4.4 | 21.51 | 2.8E-37 |
| ***C08F11.3*** | **WBGene00007452** | **protein_coding_gene** | **2.6** | **5.899** | **1.2E-04** | **-1.3** | **0.40** | **1.3E-02** |
| *sre-6* | WBGene00009242 | protein_coding_gene | 2.6 | 5.894 | 2.9E-02 | 2.8 | 7.10 | 1.3E-04 |
| *Y94H6A.2* | WBGene00022375 | protein_coding_gene | 2.6 | 5.886 | 5.0E-05 | 1.8 | 3.60 | 1.3E-04 |

| | | | | | | | | |
|---|---|---|---|---|---|---|---|---|
| ***T12D8.5*** | **WBGene00011733** | **protein_coding_gene** | **2.5** | **5.808** | **1.6E-06** | **-1.2** | **0.43** | **2.3E-04** |
| *F22F7.7* | WBGene00017724 | protein_coding_gene | 2.5 | 5.652 | 2.4E-24 | 2.0 | 4.05 | 1.8E-19 |
| *fbxa-136* | WBGene00007672 | protein_coding_gene | 2.5 | 5.635 | 2.2E-02 | 1.4 | 2.73 | 2.6E-04 |
| *hacd-1* | WBGene00019978 | protein_coding_gene | 2.5 | 5.633 | 1.4E-11 | 3.7 | 12.98 | 3.0E-13 |
| *scl-7* | WBGene00008029 | protein_coding_gene | 2.5 | 5.571 | 4.6E-02 | 1.2 | 2.26 | 2.8E-06 |
| *Y9C9A.1* | WBGene00021176 | protein_coding_gene | 2.5 | 5.557 | 4.8E-03 | 1.4 | 2.56 | 3.5E-02 |
| *phat-5* | WBGene00020242 | protein_coding_gene | 2.5 | 5.551 | 8.4E-04 | 1.9 | 3.65 | 3.1E-04 |
| *asns-2* | WBGene00019730 | protein_coding_gene | 2.5 | 5.521 | 2.8E-06 | 1.9 | 3.77 | 4.0E-12 |
| *F49F1.15* | WBGene00235365 | protein_coding_gene | 2.5 | 5.494 | 3.3E-02 | 1.1 | 2.11 | 4.2E-05 |
| ***M01B2.13*** | **WBGene00045338** | **protein_coding_gene** | **2.4** | **5.459** | **2.6E-03** | **-1.1** | **0.48** | **6.9E-03** |
| *dlhd-1* | WBGene00020741 | protein_coding_gene | 2.4 | 5.439 | 4.4E-14 | 2.4 | 5.23 | 1.2E-07 |
| *C15B12.8* | WBGene00015788 | protein_coding_gene | 2.4 | 5.239 | 9.7E-06 | 2.3 | 4.80 | 1.2E-09 |
| *F54B11.9* | WBGene00010032 | protein_coding_gene | 2.4 | 5.225 | 2.7E-02 | 1.9 | 3.83 | 3.0E-02 |
| *fbxa-172* | WBGene00008014 | protein_coding_gene | 2.4 | 5.184 | 1.2E-04 | 1.3 | 2.42 | 1.4E-04 |
| *C46A5.4* | WBGene00016700 | protein_coding_gene | 2.4 | 5.165 | 1.8E-04 | 1.1 | 2.15 | 8.9E-03 |
| *Y47D7A.12* | WBGene00021624 | protein_coding_gene | 2.4 | 5.121 | 1.5E-02 | 1.4 | 2.57 | 2.0E-03 |
| *F13E9.15* | WBGene00045208 | protein_coding_gene | 2.3 | 5.089 | 1.4E-07 | 1.0 | 2.01 | 2.3E-07 |
| *cut-3* | WBGene00009041 | protein_coding_gene | 2.3 | 5.080 | 2.4E-02 | 1.4 | 2.67 | 1.1E-03 |
| *F43C9.1* | WBGene00018375 | protein_coding_gene | 2.3 | 5.075 | 4.7E-05 | 1.0 | 2.02 | 8.2E-03 |
| *col-74* | WBGene00000650 | protein_coding_gene | 2.3 | 5.075 | 2.0E-02 | 1.8 | 3.55 | 9.3E-11 |
| *aat-1* | WBGene00000002 | protein_coding_gene | 2.3 | 4.939 | 2.1E-23 | 1.1 | 2.13 | 1.0E-06 |
| ***F37A8.5*** | **WBGene00009503** | **protein_coding_gene** | **2.3** | **4.908** | **2.3E-04** | **-1.7** | **0.31** | **5.7E-22** |
| *dmsr-16* | WBGene00194821 | protein_coding_gene | 2.3 | 4.897 | 5.4E-03 | 1.5 | 2.87 | 2.7E-02 |
| *C39B10.1* | WBGene00008021 | protein_coding_gene | 2.3 | 4.844 | 7.5E-10 | 1.8 | 3.38 | 1.7E-06 |
| *Y6G8.2* | WBGene00012404 | protein_coding_gene | 2.3 | 4.800 | 2.1E-05 | 1.3 | 2.46 | 2.1E-05 |
| ***R09E10.13*** | **WBGene00194658** | **protein_coding_gene** | **2.3** | **4.757** | **2.1E-02** | **-2.0** | **0.26** | **4.9E-04** |
| *Y57E12B.11* | WBGene00235311 | protein_coding_gene | 2.2 | 4.749 | 3.9E-02 | 2.1 | 4.32 | 2.6E-02 |
| *F09C8.1* | WBGene00008621 | protein_coding_gene | 2.2 | 4.710 | 1.3E-07 | 2.6 | 6.01 | 4.6E-10 |
| *M02D8.6* | WBGene00019732 | protein_coding_gene | 2.2 | 4.699 | 1.5E-12 | 1.2 | 2.25 | 2.7E-08 |
| ***ins-2*** | **WBGene00002085** | **protein_coding_gene** | **2.2** | **4.685** | **3.0E-03** | **-1.0** | **0.49** | **2.3E-02** |
| *F59B2.11* | WBGene00010313 | pseudogene | 2.2 | 4.683 | 9.0E-04 | 1.0 | 2.01 | 3.0E-02 |
| *linc-125* | WBGene00219686 | lincRNA_gene | 2.2 | 4.618 | 9.5E-03 | 2.0 | 4.12 | 2.4E-02 |
| *fbxa-50* | WBGene00013844 | protein_coding_gene | 2.2 | 4.570 | 3.2E-04 | 1.5 | 2.82 | 3.2E-03 |
| *bath-26* | WBGene00020329 | protein_coding_gene | 2.2 | 4.498 | 9.0E-11 | 1.1 | 2.11 | 9.1E-04 |
| *EGAP9.3* | WBGene00017151 | protein_coding_gene | 2.1 | 4.425 | 3.1E-11 | 1.1 | 2.09 | 3.5E-04 |
| *C08B6.4* | WBGene00007425 | protein_coding_gene | 2.1 | 4.395 | 1.4E-03 | 1.4 | 2.67 | 6.4E-03 |
| *lgc-13* | WBGene00011354 | protein_coding_gene | 2.1 | 4.322 | 4.4E-03 | 2.1 | 4.37 | 1.2E-03 |
| *fipr-2* | WBGene00009097 | protein_coding_gene | 2.1 | 4.309 | 8.3E-13 | 1.3 | 2.45 | 3.8E-16 |
| *R02F11.2* | WBGene00019840 | protein_coding_gene | 2.1 | 4.297 | 8.5E-06 | 1.2 | 2.22 | 5.7E-03 |
| ***Y34F4.3*** | **WBGene00021338** | **protein_coding_gene** | **2.1** | **4.283** | **3.1E-04** | **-1.3** | **0.40** | **1.6E-03** |
| *C50D2.1* | WBGene00016805 | protein_coding_gene | 2.1 | 4.264 | 1.7E-05 | 1.1 | 2.09 | 1.8E-02 |
| *ttr-28* | WBGene00011285 | protein_coding_gene | 2.1 | 4.247 | 1.3E-05 | 1.5 | 2.76 | 5.5E-04 |
| *clec-227* | WBGene00008593 | protein_coding_gene | 2.1 | 4.232 | 7.0E-08 | 2.3 | 4.93 | 1.3E-13 |
| ***F21H7.12*** | **WBGene00009033** | **protein_coding_gene** | **2.1** | **4.167** | **8.5E-03** | **-1.5** | **0.35** | **3.8E-02** |
| *spin-3* | WBGene00008598 | protein_coding_gene | 2.0 | 4.116 | 1.6E-12 | 2.0 | 4.02 | 2.5E-08 |
| *C39B5.14* | WBGene00189950 | protein_coding_gene | 2.0 | 4.101 | 1.2E-02 | 1.1 | 2.17 | 1.8E-02 |
| ***oac-14*** | **WBGene00008602** | **protein_coding_gene** | **2.0** | **4.091** | **1.9E-02** | **-2.8** | **0.15** | **2.3E-21** |
| ***fil-1*** | **WBGene00011321** | **protein_coding_gene** | **2.0** | **4.088** | **3.9E-03** | **-2.6** | **0.17** | **5.7E-25** |
| *nhr-168* | WBGene00008221 | protein_coding_gene | 2.0 | 4.084 | 3.9E-16 | 2.1 | 4.19 | 1.1E-09 |
| *T19H12.15* | WBGene00271827 | protein_coding_gene | 2.0 | 4.066 | 4.7E-08 | 3.7 | 12.97 | 4.7E-49 |
| *T10H10.2* | WBGene00020426 | protein_coding_gene | 2.0 | 4.045 | 2.5E-07 | 2.4 | 5.18 | 3.4E-22 |
| *chil-3* | WBGene00007467 | protein_coding_gene | 2.0 | 4.040 | 2.3E-04 | 3.9 | 14.84 | 1.0E-06 |
| *fbxa-158* | WBGene00015602 | protein_coding_gene | 2.0 | 4.027 | 1.3E-02 | 1.9 | 3.64 | 9.2E-03 |
| *gpx-3* | WBGene00007517 | protein_coding_gene | 2.0 | 3.995 | 2.3E-15 | 1.8 | 3.57 | 1.0E-13 |
| *Y75B8A.44* | WBGene00194855 | protein_coding_gene | 2.0 | 3.976 | 2.2E-04 | 2.0 | 3.99 | 1.0E-06 |
| *Y70G10A.2* | WBGene00013498 | protein_coding_gene | 2.0 | 3.972 | 1.6E-02 | 1.7 | 3.29 | 1.0E-02 |
| *Y42G9A.3* | WBGene00021533 | protein_coding_gene | 2.0 | 3.943 | 2.1E-07 | 1.2 | 2.35 | 8.4E-04 |
| ***ttr-23*** | **WBGene00011894** | **protein_coding_gene** | **2.0** | **3.899** | **1.1E-03** | **-1.1** | **0.46** | **4.4E-06** |

| | | | | | | | | |
|---|---|---|---|---|---|---|---|---|
| *Y7A5A.11* | WBGene00044155 | pseudogene | 2.0 | 3.883 | 1.5E-02 | 1.6 | 3.11 | 1.7E-03 |
| *C52E2.4* | WBGene00016883 | protein_coding_gene | 1.9 | 3.856 | 4.9E-04 | 1.1 | 2.16 | 7.3E-04 |
| *asp-8* | WBGene00019105 | protein_coding_gene | 1.9 | 3.845 | 3.8E-13 | 2.9 | 7.34 | 1.2E-10 |
| ***K06A9.2*** | **WBGene00019436** | **protein_coding_gene** | **1.9** | **3.843** | **8.7E-04** | **-1.3** | **0.41** | **2.1E-07** |
| *pals-33* | WBGene00021081 | protein_coding_gene | 1.9 | 3.840 | 1.2E-03 | 1.5 | 2.79 | 3.0E-04 |
| ***cyp-34A9*** | **WBGene00015044** | **protein_coding_gene** | **1.9** | **3.804** | **2.0E-02** | **-1.2** | **0.45** | **3.8E-18** |
| *F37C12.21* | WBGene00194742 | protein_coding_gene | 1.9 | 3.792 | 4.2E-03 | 1.6 | 3.01 | 1.7E-03 |
| *cav-2* | WBGene00000302 | protein_coding_gene | 1.9 | 3.777 | 9.8E-08 | 1.7 | 3.21 | 4.0E-08 |
| *F49C12.7* | WBGene00009877 | protein_coding_gene | 1.9 | 3.759 | 1.1E-17 | 1.5 | 2.86 | 1.7E-05 |
| *F10A3.4* | WBGene00008634 | protein_coding_gene | 1.9 | 3.749 | 1.5E-07 | 3.1 | 8.57 | 9.6E-17 |
| ***swt-1*** | **WBGene00019300** | **protein_coding_gene** | **1.9** | **3.728** | **4.1E-04** | **-1.6** | **0.33** | **4.4E-10** |
| *bcmo-2* | WBGene00018755 | protein_coding_gene | 1.9 | 3.720 | 1.4E-24 | 2.3 | 5.04 | 2.2E-19 |
| *ZK1290.10* | WBGene00022891 | protein_coding_gene | 1.9 | 3.717 | 9.8E-05 | 1.1 | 2.15 | 5.4E-03 |
| *fipr-1* | WBGene00009096 | protein_coding_gene | 1.9 | 3.709 | 8.6E-18 | 1.2 | 2.25 | 2.1E-10 |
| *best-24* | WBGene00022797 | protein_coding_gene | 1.9 | 3.705 | 5.4E-07 | 1.3 | 2.39 | 8.8E-19 |
| ***K01D12.1*** | **WBGene00010461** | **protein_coding_gene** | **1.9** | **3.667** | **4.6E-02** | **-1.4** | **0.39** | **3.1E-02** |
| *F10D7.10* | WBGene00271701 | protein_coding_gene | 1.9 | 3.661 | 1.7E-03 | 1.0 | 2.04 | 2.2E-02 |
| *T27C4.1* | WBGene00020854 | protein_coding_gene | 1.9 | 3.650 | 1.6E-07 | 1.1 | 2.09 | 2.2E-14 |
| *nhr-208* | WBGene00011099 | protein_coding_gene | 1.9 | 3.634 | 3.1E-03 | 1.1 | 2.18 | 6.6E-03 |
| *F21H12.7* | WBGene00044287 | protein_coding_gene | 1.9 | 3.610 | 1.4E-03 | 2.6 | 5.95 | 1.1E-04 |
| *T12A7.2* | WBGene00011723 | protein_coding_gene | 1.8 | 3.598 | 1.1E-07 | 1.2 | 2.24 | 4.8E-10 |
| *hrg-1* | WBGene00019830 | protein_coding_gene | 1.8 | 3.585 | 7.0E-06 | 2.6 | 5.99 | 8.7E-35 |
| *lips-5* | WBGene00011725 | protein_coding_gene | 1.8 | 3.549 | 2.3E-07 | 3.1 | 8.64 | 4.1E-17 |
| *npr-8* | WBGene00016984 | protein_coding_gene | 1.8 | 3.536 | 3.8E-04 | 1.2 | 2.34 | 1.4E-02 |
| ***C06B3.6*** | **WBGene00007365** | **protein_coding_gene** | **1.8** | **3.529** | **1.2E-04** | **-1.6** | **0.34** | **4.6E-14** |
| *aat-5* | WBGene00000006 | protein_coding_gene | 1.8 | 3.515 | 4.1E-06 | 1.4 | 2.56 | 2.0E-08 |
| ***glc-1*** | **WBGene00001591** | **protein_coding_gene** | **1.8** | **3.514** | **1.4E-05** | **-1.0** | **0.50** | **2.1E-09** |
| *nlp-8* | WBGene00003746 | protein_coding_gene | 1.8 | 3.507 | 5.6E-16 | 1.3 | 2.52 | 7.3E-13 |
| *gcy-37* | WBGene00001557 | protein_coding_gene | 1.8 | 3.488 | 1.9E-02 | 1.5 | 2.88 | 1.4E-02 |
| *lmd-5* | WBGene00020141 | protein_coding_gene | 1.8 | 3.473 | 3.4E-02 | 1.7 | 3.18 | 3.0E-03 |
| *cwp-5* | WBGene00009844 | protein_coding_gene | 1.8 | 3.461 | 1.3E-02 | 2.0 | 3.97 | 6.8E-04 |
| *pho-11* | WBGene00007331 | protein_coding_gene | 1.8 | 3.460 | 1.3E-06 | 1.5 | 2.75 | 1.4E-07 |
| *gsto-1* | WBGene00016204 | protein_coding_gene | 1.8 | 3.450 | 1.3E-08 | 2.8 | 6.89 | 7.8E-26 |
| *W04C9.6* | WBGene00021028 | protein_coding_gene | 1.8 | 3.425 | 5.9E-07 | 2.3 | 5.09 | 2.3E-11 |
| ***F59C6.16*** | **WBGene00077764** | **protein_coding_gene** | **1.8** | **3.410** | **2.4E-03** | **-1.1** | **0.46** | **8.8E-07** |
| *eol-1* | WBGene00045401 | protein_coding_gene | 1.8 | 3.388 | 1.2E-02 | 1.2 | 2.27 | 1.8E-03 |
| *H02F09.3* | WBGene00019146 | protein_coding_gene | 1.8 | 3.377 | 3.0E-04 | 2.8 | 6.88 | 2.4E-06 |
| *pqn-84* | WBGene00004165 | protein_coding_gene | 1.8 | 3.376 | 1.1E-02 | 1.2 | 2.33 | 1.9E-10 |
| *deg-1* | WBGene00000950 | protein_coding_gene | 1.8 | 3.369 | 2.7E-06 | 1.1 | 2.20 | 9.2E-04 |
| *ZC21.3* | WBGene00022505 | protein_coding_gene | 1.7 | 3.363 | 7.8E-04 | 1.4 | 2.70 | 5.7E-25 |
| *F59E11.2* | WBGene00019109 | protein_coding_gene | 1.7 | 3.362 | 4.0E-03 | 1.1 | 2.09 | 8.6E-03 |
| *Y45F10D.2* | WBGene00012883 | protein_coding_gene | 1.7 | 3.306 | 1.6E-03 | 2.2 | 4.65 | 1.4E-03 |
| *ace-4* | WBGene00000038 | protein_coding_gene | 1.7 | 3.297 | 9.9E-03 | 1.2 | 2.38 | 3.3E-02 |
| *C50B6.7* | WBGene00008220 | protein_coding_gene | 1.7 | 3.292 | 8.5E-06 | 1.9 | 3.70 | 3.5E-21 |
| *msrp-7* | WBGene00011070 | protein_coding_gene | 1.7 | 3.256 | 2.0E-08 | 1.4 | 2.64 | 3.2E-11 |
| *R10F2.4* | WBGene00019995 | protein_coding_gene | 1.7 | 3.253 | 2.3E-02 | 1.6 | 3.11 | 3.3E-02 |
| *K12B6.4* | WBGene00019669 | protein_coding_gene | 1.7 | 3.240 | 2.4E-02 | 2.2 | 4.57 | 7.0E-04 |
| *Y47D7A.6* | WBGene00021619 | protein_coding_gene | 1.7 | 3.233 | 6.7E-03 | 2.2 | 4.63 | 1.7E-07 |
| *aat-2* | WBGene00000003 | protein_coding_gene | 1.7 | 3.233 | 1.5E-07 | 1.7 | 3.24 | 2.7E-12 |
| ***aakg-4*** | **WBGene00020135** | **protein_coding_gene** | **1.7** | **3.230** | **1.1E-04** | **-1.1** | **0.48** | **3.2E-07** |
| *T28A11.6* | WBGene00020873 | pseudogene | 1.7 | 3.211 | 2.7E-02 | 2.2 | 4.73 | 2.2E-02 |
| ***F11E6.11*** | **WBGene00023501** | **protein_coding_gene** | **1.7** | **3.206** | **9.1E-03** | **-2.4** | **0.18** | **7.2E-22** |
| *F42A10.6* | WBGene00018342 | protein_coding_gene | 1.7 | 3.168 | 1.7E-05 | 2.1 | 4.19 | 3.0E-30 |
| *ZK1290.14* | WBGene00044396 | protein_coding_gene | 1.6 | 3.125 | 7.9E-16 | 1.4 | 2.57 | 6.7E-09 |
| *clec-80* | WBGene00043062 | protein_coding_gene | 1.6 | 3.081 | 2.8E-11 | 2.3 | 5.03 | 2.1E-18 |
| *C46H11.2* | WBGene00016728 | protein_coding_gene | 1.6 | 3.065 | 3.7E-09 | 1.1 | 2.12 | 5.5E-08 |
| *Y47H9C.1* | WBGene00012947 | protein_coding_gene | 1.6 | 3.063 | 1.5E-02 | 1.4 | 2.68 | 1.6E-02 |
| *mpk-2* | WBGene00003402 | protein_coding_gene | 1.6 | 3.063 | 5.7E-15 | 1.5 | 2.88 | 1.8E-13 |

| | | | | | | | | |
|---|---|---|---|---|---|---|---|---|
| *poml-3* | WBGene00077701 | protein_coding_gene | 1.6 | 3.060 | 3.4E-24 | 1.2 | 2.33 | 9.7E-08 |
| *F46A8.11* | WBGene00014761 | protein_coding_gene | 1.6 | 3.052 | 1.4E-04 | 2.2 | 4.49 | 2.3E-08 |
| *C18F10.2* | WBGene00015977 | protein_coding_gene | 1.6 | 3.035 | 1.9E-03 | 1.8 | 3.40 | 4.1E-05 |
| *F10D2.8* | WBGene00017334 | protein_coding_gene | 1.6 | 3.034 | 5.3E-03 | 1.7 | 3.16 | 3.0E-09 |
| ***C36B1.6*** | **WBGene00007973** | **protein_coding_gene** | **1.6** | **3.023** | **4.5E-04** | **-1.0** | **0.49** | **1.9E-05** |
| *gst-34* | WBGene00001782 | protein_coding_gene | 1.6 | 2.992 | 2.9E-02 | 3.0 | 7.75 | 1.1E-02 |
| *ZK673.1* | WBGene00014057 | protein_coding_gene | 1.6 | 2.986 | 1.2E-10 | 1.6 | 2.99 | 4.2E-17 |
| *ddn-2* | WBGene00009399 | protein_coding_gene | 1.6 | 2.978 | 3.5E-02 | 1.4 | 2.67 | 4.1E-02 |
| *dgk-3* | WBGene00000960 | protein_coding_gene | 1.6 | 2.964 | 1.8E-02 | 1.3 | 2.48 | 3.7E-02 |
| *ZC196.1* | WBGene00022545 | pseudogene | 1.6 | 2.962 | 2.3E-02 | 1.6 | 3.02 | 4.9E-04 |
| *Y119C1B.6* | WBGene00022472 | protein_coding_gene | 1.6 | 2.931 | 1.7E-04 | 1.0 | 2.00 | 9.0E-03 |
| *acox-1.5* | WBGene00008167 | protein_coding_gene | 1.5 | 2.916 | 1.1E-07 | 1.2 | 2.29 | 7.7E-05 |
| *ifp-1* | WBGene00002067 | protein_coding_gene | 1.5 | 2.891 | 5.0E-06 | 2.0 | 4.03 | 3.8E-18 |
| *fat-2* | WBGene00001394 | protein_coding_gene | 1.5 | 2.888 | 8.0E-15 | 1.2 | 2.35 | 1.5E-21 |
| *srr-1* | WBGene00005652 | protein_coding_gene | 1.5 | 2.862 | 1.3E-03 | 1.3 | 2.53 | 3.6E-03 |
| *marg-1* | WBGene00018538 | protein_coding_gene | 1.5 | 2.856 | 2.9E-04 | 1.6 | 3.03 | 5.0E-15 |
| ***ZK970.7*** | **WBGene00014173** | **protein_coding_gene** | **1.5** | **2.833** | **1.1E-02** | **-3.2** | **0.11** | **1.3E-11** |
| *exc-14* | WBGene00019649 | protein_coding_gene | 1.5 | 2.799 | 4.1E-04 | 1.1 | 2.10 | 8.7E-03 |
| *K09E4.4* | WBGene00010722 | protein_coding_gene | 1.5 | 2.798 | 2.8E-04 | 1.3 | 2.49 | 1.0E-05 |
| *Y4C6B.4* | WBGene00021158 | protein_coding_gene | 1.5 | 2.798 | 2.6E-06 | 1.9 | 3.79 | 2.4E-09 |
| *fbxa-59* | WBGene00020455 | protein_coding_gene | 1.5 | 2.792 | 2.1E-05 | 1.4 | 2.64 | 3.1E-04 |
| *dhs-4* | WBGene00000968 | protein_coding_gene | 1.5 | 2.787 | 2.4E-06 | 1.3 | 2.52 | 5.4E-06 |
| ***T04C12.7*** | **WBGene00011429** | **protein_coding_gene** | **1.5** | **2.785** | **2.8E-02** | **-1.8** | **0.29** | **2.5E-04** |
| *pho-8* | WBGene00011262 | protein_coding_gene | 1.5 | 2.779 | 2.4E-02 | 1.9 | 3.71 | 1.9E-06 |
| *egl-3* | WBGene00001172 | protein_coding_gene | 1.5 | 2.753 | 2.8E-05 | 1.4 | 2.56 | 1.8E-23 |
| *Y105C5A.24* | WBGene00013643 | protein_coding_gene | 1.5 | 2.751 | 1.0E-06 | 1.1 | 2.09 | 2.4E-08 |
| *pck-1* | WBGene00021043 | protein_coding_gene | 1.5 | 2.743 | 5.2E-09 | 2.1 | 4.24 | 4.3E-64 |
| *degt-1* | WBGene00009109 | protein_coding_gene | 1.5 | 2.732 | 2.3E-03 | 1.9 | 3.83 | 3.9E-05 |
| *ZK669.5* | WBGene00014055 | protein_coding_gene | 1.4 | 2.727 | 1.0E-04 | 1.2 | 2.35 | 4.7E-04 |
| *nlp-21* | WBGene00003759 | protein_coding_gene | 1.4 | 2.720 | 1.6E-04 | 1.1 | 2.18 | 3.0E-10 |
| *nlp-13* | WBGene00003751 | protein_coding_gene | 1.4 | 2.712 | 1.7E-08 | 1.3 | 2.51 | 1.2E-16 |
| *dod-6* | WBGene00011869 | protein_coding_gene | 1.4 | 2.712 | 9.8E-05 | 1.0 | 2.01 | 4.4E-12 |
| *T08B6.2* | WBGene00020352 | pseudogene | 1.4 | 2.706 | 1.7E-02 | 1.1 | 2.21 | 4.4E-02 |
| *cah-4* | WBGene00000282 | protein_coding_gene | 1.4 | 2.699 | 4.1E-07 | 1.3 | 2.54 | 3.8E-18 |
| *acer-1* | WBGene00016630 | protein_coding_gene | 1.4 | 2.694 | 3.1E-10 | 1.1 | 2.15 | 5.3E-14 |
| *sto-6* | WBGene00006068 | protein_coding_gene | 1.4 | 2.678 | 1.4E-02 | 1.1 | 2.20 | 4.6E-02 |
| *F20G2.1* | WBGene00008985 | protein_coding_gene | 1.4 | 2.667 | 6.0E-03 | 2.9 | 7.35 | 1.3E-11 |
| *lgc-38* | WBGene00017389 | protein_coding_gene | 1.4 | 2.658 | 1.1E-02 | 1.6 | 2.97 | 2.5E-03 |
| *klu-1* | WBGene00013970 | protein_coding_gene | 1.4 | 2.658 | 2.0E-03 | 1.2 | 2.37 | 7.0E-04 |
| *pitr-5* | WBGene00017312 | protein_coding_gene | 1.4 | 2.658 | 1.5E-05 | 2.0 | 3.99 | 1.1E-09 |
| *pghm-1* | WBGene00022144 | protein_coding_gene | 1.4 | 2.651 | 4.1E-08 | 1.0 | 2.02 | 1.7E-14 |
| *C15C7.4* | WBGene00015790 | protein_coding_gene | 1.4 | 2.644 | 7.3E-03 | 1.4 | 2.66 | 7.4E-08 |
| *F12E12.11* | WBGene00044811 | protein_coding_gene | 1.4 | 2.639 | 1.7E-02 | 1.9 | 3.78 | 8.9E-09 |
| *ddn-1* | WBGene00015227 | protein_coding_gene | 1.4 | 2.633 | 2.9E-03 | 1.8 | 3.53 | 2.9E-04 |
| *Y17D7B.7* | WBGene00014562 | pseudogene | 1.4 | 2.632 | 1.2E-02 | 1.7 | 3.22 | 3.1E-03 |
| *C49G7.3* | WBGene00016781 | protein_coding_gene | 1.4 | 2.631 | 6.0E-07 | 1.4 | 2.56 | 3.8E-31 |
| *F19C7.4* | WBGene00017594 | protein_coding_gene | 1.4 | 2.626 | 1.0E-04 | 2.5 | 5.69 | 1.3E-09 |
| *cla-1* | WBGene00018468 | protein_coding_gene | 1.4 | 2.620 | 4.2E-03 | 1.4 | 2.60 | 2.2E-09 |
| *F41B5.11* | WBGene00023039 | pseudogene | 1.4 | 2.617 | 4.9E-02 | 1.3 | 2.42 | 1.7E-03 |
| *vap-1* | WBGene00006886 | protein_coding_gene | 1.4 | 2.612 | 2.4E-07 | 1.2 | 2.22 | 3.9E-10 |
| *K07D4.9* | WBGene00019488 | protein_coding_gene | 1.4 | 2.612 | 6.7E-04 | 1.1 | 2.13 | 1.3E-06 |
| ***C44C1.6*** | **WBGene00016645** | **protein_coding_gene** | **1.4** | **2.611** | **1.4E-02** | **-1.0** | **0.50** | **6.8E-04** |
| *Y22D7AL.16* | WBGene00021254 | protein_coding_gene | 1.4 | 2.611 | 4.2E-02 | 1.5 | 2.74 | 3.6E-02 |
| *fipr-5* | WBGene00044174 | protein_coding_gene | 1.4 | 2.599 | 2.1E-04 | 1.1 | 2.15 | 3.0E-05 |
| *PDB1.1* | WBGene00019803 | protein_coding_gene | 1.4 | 2.598 | 5.6E-05 | 2.0 | 4.02 | 8.6E-22 |
| *Y54G2A.49* | WBGene00044492 | protein_coding_gene | 1.4 | 2.595 | 1.7E-03 | 1.0 | 2.04 | 2.9E-02 |
| *sul-3* | WBGene00006310 | protein_coding_gene | 1.4 | 2.594 | 1.5E-05 | 2.0 | 4.05 | 3.4E-10 |
| *acs-19* | WBGene00007969 | protein_coding_gene | 1.4 | 2.583 | 8.9E-08 | 1.2 | 2.26 | 3.0E-17 |

| | | | | | | | | |
|---|---|---|---|---|---|---|---|---|
| *cpna-4* | WBGene00006441 | protein_coding_gene | 1.4 | 2.574 | 9.7E-03 | 1.1 | 2.09 | 9.4E-03 |
| *C45E1.4* | WBGene00016663 | protein_coding_gene | 1.3 | 2.547 | 5.7E-05 | 1.5 | 2.88 | 2.3E-08 |
| *fpn-1.2* | WBGene00019977 | protein_coding_gene | 1.3 | 2.528 | 1.0E-04 | 1.8 | 3.47 | 1.2E-05 |
| *nlp-49* | WBGene00019160 | protein_coding_gene | 1.3 | 2.527 | 1.7E-05 | 1.0 | 2.03 | 5.4E-10 |
| *nlp-43* | WBGene00016684 | protein_coding_gene | 1.3 | 2.525 | 3.9E-06 | 1.3 | 2.41 | 7.7E-07 |
| *myo-2* | WBGene00003514 | protein_coding_gene | 1.3 | 2.523 | 4.0E-04 | 1.2 | 2.36 | 1.5E-11 |
| *cyp-13A6* | WBGene00011673 | protein_coding_gene | 1.3 | 2.519 | 4.9E-02 | 1.7 | 3.30 | 6.5E-06 |
| *ttr-39* | WBGene00007304 | protein_coding_gene | 1.3 | 2.498 | 4.4E-04 | 1.0 | 2.05 | 7.0E-03 |
| *T26C5.4* | WBGene00012034 | protein_coding_gene | 1.3 | 2.495 | 7.1E-04 | 1.1 | 2.10 | 4.8E-04 |
| *cho-1* | WBGene00000501 | protein_coding_gene | 1.3 | 2.487 | 1.4E-02 | 1.1 | 2.19 | 5.3E-03 |
| *nep-11* | WBGene00017557 | protein_coding_gene | 1.3 | 2.487 | 6.8E-04 | 1.1 | 2.07 | 1.2E-07 |
| *ghi-1* | WBGene00013923 | protein_coding_gene | 1.3 | 2.476 | 1.2E-02 | 1.4 | 2.71 | 1.3E-04 |
| *T22B11.4* | WBGene00020678 | protein_coding_gene | 1.3 | 2.464 | 8.6E-06 | 1.1 | 2.10 | 7.7E-13 |
| *F59F3.6* | WBGene00010335 | protein_coding_gene | 1.3 | 2.451 | 3.0E-02 | 1.3 | 2.40 | 1.1E-02 |
| *hrg-7* | WBGene00007605 | protein_coding_gene | 1.3 | 2.451 | 1.4E-02 | 3.4 | 10.82 | 1.0E-19 |
| *gcy-11* | WBGene00001537 | protein_coding_gene | 1.3 | 2.441 | 4.0E-06 | 1.0 | 2.06 | 3.9E-06 |
| *ZC196.4* | WBGene00022548 | protein_coding_gene | 1.3 | 2.437 | 8.7E-04 | 1.5 | 2.80 | 4.2E-05 |
| *C06E1.7* | WBGene00015522 | protein_coding_gene | 1.3 | 2.427 | 1.4E-02 | 2.1 | 4.17 | 1.8E-06 |
| *gpa-14* | WBGene00001676 | protein_coding_gene | 1.3 | 2.425 | 3.3E-02 | 1.3 | 2.51 | 2.9E-02 |
| *clec-199* | WBGene00007820 | protein_coding_gene | 1.3 | 2.419 | 3.7E-02 | 1.2 | 2.34 | 2.4E-03 |
| *dop-1* | WBGene00001052 | protein_coding_gene | 1.3 | 2.418 | 3.9E-03 | 1.2 | 2.28 | 6.5E-03 |
| *Y37A1B.5* | WBGene00012538 | protein_coding_gene | 1.3 | 2.416 | 5.2E-08 | 1.1 | 2.21 | 8.6E-09 |
| *ctsa-3.2* | WBGene00007462 | protein_coding_gene | 1.3 | 2.414 | 2.3E-02 | 1.4 | 2.58 | 5.9E-03 |
| *sms-5* | WBGene00021075 | protein_coding_gene | 1.3 | 2.412 | 5.2E-04 | 1.0 | 2.01 | 3.8E-03 |
| *C34C6.3* | WBGene00007916 | protein_coding_gene | 1.3 | 2.408 | 1.7E-03 | 1.4 | 2.68 | 7.5E-05 |
| *maf-1* | WBGene00077521 | protein_coding_gene | 1.2 | 2.361 | 1.8E-04 | 1.3 | 2.50 | 2.5E-04 |
| *T05C1.1* | WBGene00020248 | protein_coding_gene | 1.2 | 2.348 | 7.4E-03 | 1.1 | 2.16 | 6.3E-03 |
| *lys-4* | WBGene00003093 | protein_coding_gene | 1.2 | 2.346 | 1.4E-07 | 1.3 | 2.50 | 2.4E-12 |
| *F56D3.1* | WBGene00018969 | protein_coding_gene | 1.2 | 2.342 | 2.8E-02 | 1.1 | 2.20 | 3.5E-02 |
| *C07A12.2* | WBGene00015560 | protein_coding_gene | 1.2 | 2.338 | 1.5E-02 | 1.1 | 2.19 | 2.7E-03 |
| ***F41C3.2*** | **WBGene00018268** | **protein_coding_gene** | **1.2** | **2.338** | **2.2E-02** | **-1.2** | **0.43** | **2.0E-06** |
| *C03H5.6* | WBGene00015408 | protein_coding_gene | 1.2 | 2.319 | 2.1E-02 | 1.6 | 3.09 | 1.9E-03 |
| *ZK896.1* | WBGene00014132 | protein_coding_gene | 1.2 | 2.309 | 1.0E-02 | 1.8 | 3.50 | 4.0E-04 |
| *npr-3* | WBGene00007006 | protein_coding_gene | 1.2 | 2.305 | 2.5E-02 | 1.2 | 2.33 | 1.8E-02 |
| *myo-1* | WBGene00002348 | protein_coding_gene | 1.2 | 2.302 | 1.2E-03 | 1.2 | 2.32 | 8.6E-11 |
| *frpr-16* | WBGene00020023 | protein_coding_gene | 1.2 | 2.302 | 1.6E-02 | 1.0 | 2.05 | 4.7E-02 |
| *F17B5.8* | WBGene00045458 | protein_coding_gene | 1.2 | 2.298 | 4.3E-02 | 1.7 | 3.14 | 6.6E-03 |
| *scd-2* | WBGene00004740 | protein_coding_gene | 1.2 | 2.291 | 6.0E-03 | 1.1 | 2.13 | 6.1E-05 |
| *flp-11* | WBGene00001454 | protein_coding_gene | 1.2 | 2.290 | 3.3E-04 | 1.0 | 2.03 | 2.4E-06 |
| *irg-3* | WBGene00018760 | protein_coding_gene | 1.2 | 2.289 | 6.0E-04 | 1.6 | 2.99 | 2.0E-09 |
| *clec-198* | WBGene00008203 | protein_coding_gene | 1.2 | 2.284 | 3.5E-02 | 1.5 | 2.84 | 4.8E-03 |
| *cah-2* | WBGene00000280 | protein_coding_gene | 1.2 | 2.276 | 4.8E-03 | 1.1 | 2.13 | 1.0E-02 |
| *ZK856.5* | WBGene00014106 | protein_coding_gene | 1.2 | 2.263 | 1.3E-03 | 1.3 | 2.48 | 3.0E-10 |
| *pepm-1* | WBGene00009807 | protein_coding_gene | 1.2 | 2.257 | 4.3E-02 | 1.7 | 3.23 | 4.6E-02 |
| *cyp-33C3* | WBGene00018262 | protein_coding_gene | 1.2 | 2.257 | 9.8E-04 | 1.3 | 2.44 | 4.9E-07 |
| *M28.8* | WBGene00010899 | protein_coding_gene | 1.2 | 2.251 | 3.5E-03 | 1.2 | 2.26 | 4.4E-09 |
| *glr-1* | WBGene00001612 | protein_coding_gene | 1.2 | 2.251 | 2.6E-02 | 1.3 | 2.48 | 2.2E-03 |
| *D1086.18* | WBGene00045405 | protein_coding_gene | 1.2 | 2.249 | 2.8E-02 | 1.5 | 2.78 | 3.6E-04 |
| *kri-1* | WBGene00013955 | protein_coding_gene | 1.2 | 2.249 | 3.2E-03 | 1.8 | 3.39 | 6.3E-06 |
| *hpo-26* | WBGene00018492 | protein_coding_gene | 1.2 | 2.240 | 2.6E-03 | 1.2 | 2.33 | 1.2E-04 |
| *F58E2.3* | WBGene00019047 | protein_coding_gene | 1.2 | 2.231 | 3.0E-03 | 1.0 | 2.00 | 2.0E-02 |
| *F54C1.1* | WBGene00018789 | protein_coding_gene | 1.2 | 2.222 | 1.4E-03 | 1.9 | 3.70 | 5.0E-06 |
| *F57C7.4* | WBGene00010200 | protein_coding_gene | 1.1 | 2.213 | 4.4E-02 | 1.1 | 2.10 | 3.1E-02 |
| *Y49E10.18* | WBGene00013037 | protein_coding_gene | 1.1 | 2.213 | 2.7E-04 | 1.4 | 2.62 | 8.7E-12 |
| *sto-2* | WBGene00006064 | protein_coding_gene | 1.1 | 2.212 | 3.9E-02 | 1.5 | 2.89 | 2.9E-03 |
| ***T28F4.5*** | **WBGene00012140** | **protein_coding_gene** | **1.1** | **2.210** | **6.8E-03** | **-1.2** | **0.42** | **2.1E-11** |
| *T19D12.1* | WBGene00020576 | protein_coding_gene | 1.1 | 2.204 | 7.7E-03 | 2.1 | 4.31 | 8.2E-32 |
| *aman-3* | WBGene00018594 | protein_coding_gene | 1.1 | 2.188 | 1.1E-04 | 1.8 | 3.40 | 1.5E-20 |

| | | | | | | | | |
|---|---|---|---|---|---|---|---|---|
| *tat-2* | WBGene00019166 | protein_coding_gene | 1.1 | 2.182 | 5.4E-03 | 1.1 | 2.13 | 6.0E-11 |
| *fipr-9* | WBGene00044175 | protein_coding_gene | 1.1 | 2.170 | 1.6E-03 | 1.6 | 3.12 | 3.5E-05 |
| *tag-290* | WBGene00009386 | protein_coding_gene | 1.1 | 2.167 | 2.3E-04 | 1.3 | 2.45 | 1.3E-11 |
| *Y105C5A.25* | WBGene00013644 | pseudogene | 1.1 | 2.154 | 2.1E-02 | 1.1 | 2.16 | 3.3E-02 |
| ***C08F1.10*** | **WBGene00015613** | **protein_coding_gene** | **1.1** | **2.135** | **3.2E-03** | **-1.1** | **0.46** | **3.8E-03** |
| *npr-26* | WBGene00011372 | protein_coding_gene | 1.1 | 2.133 | 3.9E-02 | 1.5 | 2.75 | 2.2E-03 |
| *F36H2.3* | WBGene00009500 | protein_coding_gene | 1.1 | 2.113 | 8.2E-03 | 1.0 | 2.03 | 4.9E-04 |
| *F41E6.15* | WBGene00044554 | protein_coding_gene | 1.1 | 2.113 | 8.2E-06 | 1.4 | 2.70 | 2.7E-08 |
| *F41G3.10* | WBGene00018303 | protein_coding_gene | 1.1 | 2.111 | 9.6E-05 | 1.2 | 2.30 | 1.0E-16 |
| ***F59B10.4*** | **WBGene00010320** | **protein_coding_gene** | **1.1** | **2.100** | **1.5E-02** | **-1.1** | **0.46** | **1.9E-04** |
| *clec-166* | WBGene00009518 | protein_coding_gene | 1.1 | 2.088 | 3.1E-04 | 1.1 | 2.07 | 5.5E-10 |
| *F11D5.7* | WBGene00043980 | protein_coding_gene | 1.1 | 2.087 | 7.2E-03 | 1.9 | 3.70 | 7.9E-08 |
| *pgp-13* | WBGene00004007 | protein_coding_gene | 1.1 | 2.085 | 7.3E-05 | 1.6 | 3.00 | 4.5E-13 |
| *F15A4.6* | WBGene00008841 | protein_coding_gene | 1.1 | 2.078 | 6.8E-06 | 1.1 | 2.08 | 4.1E-10 |
| *F34D10.6* | WBGene00009375 | protein_coding_gene | 1.0 | 2.066 | 6.6E-03 | 1.2 | 2.34 | 6.7E-04 |
| *C39H7.4* | WBGene00016542 | protein_coding_gene | 1.0 | 2.050 | 3.3E-04 | 1.1 | 2.11 | 2.2E-12 |
| *nlp-18* | WBGene00003756 | protein_coding_gene | 1.0 | 2.041 | 5.7E-04 | 1.1 | 2.11 | 2.8E-08 |
| *C27H5.2* | WBGene00016172 | protein_coding_gene | 1.0 | 2.025 | 1.3E-05 | 1.0 | 2.03 | 5.6E-07 |
| *nrx-1* | WBGene00007799 | protein_coding_gene | 1.0 | 2.015 | 3.0E-03 | 1.2 | 2.22 | 1.8E-06 |
| *cyp-33A1* | WBGene00015709 | protein_coding_gene | 1.0 | 2.015 | 1.3E-07 | 1.2 | 2.29 | 4.5E-08 |
| *oac-58* | WBGene00019580 | protein_coding_gene | 1.0 | 2.014 | 2.2E-02 | 1.6 | 3.07 | 6.0E-06 |
| *fat-5* | WBGene00001397 | protein_coding_gene | 1.0 | 2.007 | 1.4E-03 | 3.5 | 11.71 | 9.0E-21 |
| *F55G11.4* | WBGene00010124 | protein_coding_gene | -1.0 | 0.499 | 6.8E-03 | -4.1 | 0.06 | 7.6E-14 |
| ***oat-1*** | **WBGene00003837** | **protein_coding_gene** | **-1.0** | **0.498** | **1.2E-02** | **1.5** | **2.74** | **6.3E-07** |
| *bag-1* | WBGene00000236 | protein_coding_gene | -1.0 | 0.495 | 4.9E-10 | -1.1 | 0.48 | 3.0E-15 |
| ***col-109*** | **WBGene00000683** | **protein_coding_gene** | **-1.0** | **0.491** | **3.9E-02** | **2.0** | **3.92** | **3.4E-02** |
| *gstk-1* | WBGene00014251 | protein_coding_gene | -1.0 | 0.490 | 2.7E-07 | -1.4 | 0.38 | 3.4E-14 |
| ***cln-3.1*** | **WBGene00000539** | **protein_coding_gene** | **-1.0** | **0.488** | **9.6E-03** | **1.1** | **2.08** | **8.3E-04** |
| *trd-1* | WBGene00020600 | protein_coding_gene | -1.0 | 0.488 | 1.7E-13 | -1.1 | 0.48 | 8.9E-14 |
| *F21F3.4* | WBGene00017674 | protein_coding_gene | -1.0 | 0.485 | 8.9E-05 | -1.2 | 0.43 | 1.6E-10 |
| *lbp-9* | WBGene00021486 | protein_coding_gene | -1.0 | 0.485 | 1.8E-03 | -1.1 | 0.48 | 8.4E-17 |
| *F35D11.4* | WBGene00018045 | protein_coding_gene | -1.0 | 0.485 | 2.1E-03 | -1.3 | 0.41 | 1.3E-12 |
| ***F21A3.3*** | **WBGene00008993** | **protein_coding_gene** | **-1.1** | **0.478** | **1.1E-02** | **1.2** | **2.31** | **6.6E-04** |
| *clec-150* | WBGene00019914 | protein_coding_gene | -1.1 | 0.478 | 1.0E-04 | -1.3 | 0.41 | 7.0E-24 |
| *acdh-13* | WBGene00001566 | protein_coding_gene | -1.1 | 0.474 | 2.9E-16 | -1.0 | 0.49 | 1.1E-14 |
| *tag-276* | WBGene00013254 | protein_coding_gene | -1.1 | 0.473 | 3.0E-03 | -1.2 | 0.45 | 2.3E-04 |
| *C17E7.12* | WBGene00015905 | protein_coding_gene | -1.1 | 0.469 | 1.4E-06 | -1.2 | 0.44 | 7.1E-07 |
| *mrps-22* | WBGene00007564 | protein_coding_gene | -1.1 | 0.469 | 4.4E-18 | -1.1 | 0.47 | 7.7E-18 |
| *eif-3.H* | WBGene00001231 | protein_coding_gene | -1.1 | 0.468 | 1.2E-06 | -1.1 | 0.48 | 4.1E-17 |
| *C29F3.7* | WBGene00007807 | protein_coding_gene | -1.1 | 0.467 | 3.8E-03 | -1.6 | 0.33 | 3.3E-12 |
| *R06C7.2* | WBGene00011062 | protein_coding_gene | -1.1 | 0.467 | 1.1E-06 | -1.1 | 0.47 | 1.4E-07 |
| *ZK1307.1* | WBGene00014244 | protein_coding_gene | -1.1 | 0.467 | 2.2E-03 | -1.1 | 0.46 | 5.1E-08 |
| ***F58B4.5*** | **WBGene00010238** | **protein_coding_gene** | **-1.1** | **0.466** | **1.1E-04** | **1.2** | **2.36** | **8.2E-11** |
| *F12A10.1* | WBGene00017390 | protein_coding_gene | -1.1 | 0.464 | 4.6E-02 | -2.6 | 0.17 | 2.2E-19 |
| *him-14* | WBGene00001872 | protein_coding_gene | -1.1 | 0.460 | 1.7E-12 | -1.1 | 0.45 | 2.7E-13 |
| ***pqn-37*** | **WBGene00004124** | **protein_coding_gene** | **-1.1** | **0.460** | **6.8E-03** | **1.0** | **2.03** | **1.6E-02** |
| *eel-1* | WBGene00022069 | protein_coding_gene | -1.1 | 0.459 | 1.4E-03 | -1.0 | 0.49 | 1.3E-13 |
| ***Y50D4B.6*** | **WBGene00021745** | **protein_coding_gene** | **-1.1** | **0.457** | **3.6E-02** | **2.1** | **4.21** | **8.9E-04** |
| *D2063.1* | WBGene00017060 | protein_coding_gene | -1.1 | 0.457 | 1.6E-02 | -1.2 | 0.45 | 2.5E-05 |
| ***R07E3.4*** | **WBGene00011105** | **protein_coding_gene** | **-1.1** | **0.456** | **1.4E-02** | **1.3** | **2.39** | **7.6E-07** |
| *ugt-4* | WBGene00013905 | protein_coding_gene | -1.1 | 0.452 | 3.2E-02 | -2.0 | 0.26 | 2.3E-15 |
| *dnj-9* | WBGene00001027 | protein_coding_gene | -1.1 | 0.452 | 1.0E-13 | -1.1 | 0.47 | 1.8E-17 |
| ***acp-6*** | **WBGene00022245** | **protein_coding_gene** | **-1.1** | **0.452** | **3.2E-02** | **1.6** | **3.03** | **9.9E-24** |
| *F23A7.4* | WBGene00009069 | protein_coding_gene | -1.1 | 0.452 | 2.8E-02 | -1.1 | 0.47 | 7.2E-11 |
| ***grd-10*** | **WBGene00001699** | **protein_coding_gene** | **-1.1** | **0.451** | **2.2E-02** | **1.3** | **2.55** | **9.3E-05** |
| ***grsp-4*** | **WBGene00017201** | **protein_coding_gene** | **-1.1** | **0.451** | **1.6E-02** | **1.0** | **2.00** | **2.8E-02** |
| *irg-4* | WBGene00008584 | protein_coding_gene | -1.2 | 0.447 | 1.7E-02 | -1.5 | 0.35 | 1.0E-04 |
| *eif-2alpha* | WBGene00021351 | protein_coding_gene | -1.2 | 0.447 | 3.1E-13 | -1.0 | 0.50 | 3.6E-18 |

| | | | | | | | | |
|---|---|---|---|---|---|---|---|---|
| *ZK899.6* | WBGene00014144 | protein_coding_gene | -1.2 | 0.443 | 2.8E-02 | -1.1 | 0.46 | 8.1E-03 |
| *sgnh-1* | WBGene00020354 | protein_coding_gene | -1.2 | 0.442 | 3.6E-02 | -1.3 | 0.40 | 2.6E-03 |
| *C24D10.5* | WBGene00016056 | protein_coding_gene | -1.2 | 0.436 | 6.1E-03 | -1.1 | 0.47 | 1.7E-05 |
| ***pcp-2*** | **WBGene00003957** | **protein_coding_gene** | **-1.2** | **0.428** | **2.1E-02** | **2.0** | **3.91** | **4.9E-05** |
| *F21D5.1* | WBGene00009006 | protein_coding_gene | -1.2 | 0.425 | 3.9E-06 | -1.0 | 0.49 | 4.1E-16 |
| ***col-107*** | **WBGene00000681** | **protein_coding_gene** | **-1.2** | **0.424** | **1.3E-02** | **2.1** | **4.19** | **1.2E-20** |
| *his-67* | WBGene00001941 | protein_coding_gene | -1.2 | 0.423 | 3.2E-03 | -1.4 | 0.39 | 1.2E-02 |
| *Y73C8C.8* | WBGene00022265 | protein_coding_gene | -1.2 | 0.421 | 4.3E-04 | -1.1 | 0.47 | 2.5E-04 |
| *sup-35* | WBGene00012976 | protein_coding_gene | -1.3 | 0.415 | 3.9E-07 | -1.1 | 0.47 | 2.6E-07 |
| ***ugt-22*** | **WBGene00007455** | **protein_coding_gene** | **-1.3** | **0.411** | **2.2E-03** | **1.0** | **2.02** | **7.7E-03** |
| ***pho-1*** | **WBGene00004020** | **protein_coding_gene** | **-1.3** | **0.401** | **1.1E-03** | **2.1** | **4.35** | **5.0E-07** |
| *F55G11.8* | WBGene00010128 | protein_coding_gene | -1.3 | 0.401 | 1.2E-02 | -1.2 | 0.43 | 7.0E-05 |
| *Y116A8C.19* | WBGene00013796 | protein_coding_gene | -1.3 | 0.398 | 4.3E-02 | -1.1 | 0.46 | 9.1E-03 |
| ***col-115*** | **WBGene00000689** | **protein_coding_gene** | **-1.3** | **0.396** | **4.6E-02** | **2.1** | **4.17** | **8.1E-03** |
| ***mpst-7*** | **WBGene00011307** | **protein_coding_gene** | **-1.3** | **0.396** | **7.0E-06** | **1.4** | **2.58** | **3.9E-03** |
| *F42G8.19* | WBGene00249817 | protein_coding_gene | -1.3 | 0.394 | 1.6E-02 | -1.6 | 0.32 | 4.6E-03 |
| *Y39G10AR.32* | WBGene00219275 | protein_coding_gene | -1.3 | 0.393 | 7.6E-13 | -1.7 | 0.31 | 6.8E-26 |
| *fbxc-32* | WBGene00016303 | protein_coding_gene | -1.4 | 0.391 | 1.3E-07 | -1.1 | 0.47 | 2.8E-08 |
| *F49E12.10* | WBGene00009903 | protein_coding_gene | -1.4 | 0.380 | 5.8E-04 | -1.3 | 0.41 | 7.6E-04 |
| *fbxc-58* | WBGene00007506 | protein_coding_gene | -1.4 | 0.378 | 4.6E-02 | -1.1 | 0.45 | 1.0E-02 |
| ***abhd-3.2*** | **WBGene00016644** | **protein_coding_gene** | **-1.4** | **0.378** | **2.8E-02** | **2.5** | **5.81** | **7.4E-20** |
| ***C33D3.3*** | **WBGene00007894** | **protein_coding_gene** | **-1.4** | **0.377** | **1.3E-02** | **1.3** | **2.41** | **3.9E-02** |
| ***Y47G6A.15*** | **WBGene00021641** | **protein_coding_gene** | **-1.4** | **0.376** | **9.4E-04** | **2.0** | **4.00** | **3.1E-03** |
| *clec-86* | WBGene00016912 | protein_coding_gene | -1.4 | 0.375 | 1.0E-05 | -1.9 | 0.26 | 1.4E-16 |
| *W04E12.2* | WBGene00012247 | protein_coding_gene | -1.4 | 0.372 | 1.3E-03 | -1.6 | 0.33 | 6.7E-05 |
| *C40H1.8* | WBGene00008043 | protein_coding_gene | -1.4 | 0.368 | 1.3E-02 | -1.1 | 0.47 | 3.2E-02 |
| *ZK380.t2* | WBGene00023401 | pseudogene | -1.4 | 0.368 | 2.0E-03 | -1.4 | 0.37 | 1.7E-05 |
| ***F09B12.3*** | **WBGene00008607** | **protein_coding_gene** | **-1.4** | **0.367** | **2.2E-03** | **1.1** | **2.12** | **1.1E-08** |
| *C18H7.11* | WBGene00044457 | protein_coding_gene | -1.4 | 0.366 | 1.3E-02 | -1.4 | 0.37 | 1.8E-03 |
| *F07G6.11* | WBGene00304813 | ncRNA_gene | -1.5 | 0.366 | 2.2E-02 | -1.8 | 0.29 | 2.0E-04 |
| ***dpy-5*** | **WBGene00001067** | **protein_coding_gene** | **-1.5** | **0.365** | **3.3E-03** | **2.1** | **4.22** | **3.3E-02** |
| ***T21C9.9*** | **WBGene00011895** | **protein_coding_gene** | **-1.5** | **0.364** | **1.5E-02** | **1.4** | **2.66** | **1.6E-02** |
| ***col-97*** | **WBGene00000672** | **protein_coding_gene** | **-1.5** | **0.364** | **1.1E-03** | **2.0** | **3.99** | **3.2E-02** |
| *Y37A1A.2* | WBGene00012535 | protein_coding_gene | -1.5 | 0.362 | 2.2E-05 | -1.5 | 0.36 | 6.8E-05 |
| ***col-144*** | **WBGene00000717** | **protein_coding_gene** | **-1.5** | **0.361** | **9.6E-03** | **1.7** | **3.26** | **3.8E-03** |
| *F54D11.3* | WBGene00018813 | pseudogene | -1.5 | 0.361 | 4.2E-09 | -1.2 | 0.45 | 2.6E-05 |
| ***Y58A7A.1*** | **WBGene00021975** | **protein_coding_gene** | **-1.5** | **0.360** | **1.1E-02** | **1.1** | **2.13** | **1.3E-05** |
| *Y44A6C.2* | WBGene00012851 | protein_coding_gene | -1.5 | 0.358 | 1.8E-03 | -1.3 | 0.41 | 8.7E-05 |
| ***cyp-35A4*** | **WBGene00016786** | **protein_coding_gene** | **-1.5** | **0.357** | **4.4E-02** | **2.0** | **4.06** | **8.1E-03** |
| ***T10E9.3*** | **WBGene00020413** | **protein_coding_gene** | **-1.5** | **0.355** | **8.4E-04** | **1.6** | **3.03** | **2.3E-05** |
| *C32H11.4* | WBGene00007867 | protein_coding_gene | -1.5 | 0.354 | 1.4E-05 | -2.0 | 0.25 | 1.7E-07 |
| *F54D5.4* | WBGene00010050 | protein_coding_gene | -1.5 | 0.353 | 1.1E-10 | -1.7 | 0.31 | 2.8E-09 |
| *F02H6.1* | WBGene00008534 | pseudogene | -1.5 | 0.353 | 1.9E-06 | -1.5 | 0.34 | 7.3E-04 |
| ***C30H6.5*** | **WBGene00007823** | **protein_coding_gene** | **-1.5** | **0.347** | **1.3E-02** | **2.2** | **4.46** | **1.5E-02** |
| *W10G11.2* | WBGene00021135 | protein_coding_gene | -1.5 | 0.346 | 2.8E-02 | -2.5 | 0.18 | 3.8E-05 |
| *F52E1.14* | WBGene00044696 | protein_coding_gene | -1.5 | 0.346 | 2.1E-08 | -2.0 | 0.25 | 1.2E-25 |
| *F21D5.3* | WBGene00009008 | protein_coding_gene | -1.5 | 0.345 | 2.4E-06 | -1.0 | 0.49 | 3.0E-03 |
| ***kin-15*** | **WBGene00002199** | **protein_coding_gene** | **-1.5** | **0.344** | **1.0E-02** | **1.7** | **3.32** | **1.3E-07** |
| ***C35A11.4*** | **WBGene00016431** | **protein_coding_gene** | **-1.5** | **0.342** | **1.1E-03** | **1.9** | **3.61** | **3.6E-12** |
| *spd-3* | WBGene00004954 | protein_coding_gene | -1.6 | 0.341 | 4.1E-16 | -1.2 | 0.42 | 3.2E-13 |
| *F56F4.8* | WBGene00045498 | protein_coding_gene | -1.6 | 0.336 | 3.9E-03 | -1.2 | 0.43 | 2.5E-04 |
| ***col-169*** | **WBGene00000742** | **protein_coding_gene** | **-1.6** | **0.336** | **3.3E-03** | **1.2** | **2.28** | **3.7E-02** |
| *K08D8.6* | WBGene00010660 | protein_coding_gene | -1.6 | 0.334 | 1.1E-03 | -1.1 | 0.48 | 8.0E-03 |
| *atad-3* | WBGene00010015 | protein_coding_gene | -1.6 | 0.332 | 3.2E-26 | -1.1 | 0.45 | 1.7E-18 |
| ***F40H3.2*** | **WBGene00018250** | **protein_coding_gene** | **-1.6** | **0.331** | **2.3E-02** | **1.2** | **2.26** | **2.8E-03** |
| *C38D9.8* | WBGene00008016 | protein_coding_gene | -1.6 | 0.327 | 5.7E-03 | -1.3 | 0.41 | 2.3E-04 |
| ***T23F6.1*** | **WBGene00011949** | **protein_coding_gene** | **-1.6** | **0.323** | **3.0E-02** | **1.7** | **3.17** | **4.9E-02** |
| ***col-125*** | **WBGene00000699** | **protein_coding_gene** | **-1.6** | **0.323** | **2.4E-02** | **2.0** | **3.96** | **5.7E-05** |

| | | | | | | | | |
|---|---|---|---|---|---|---|---|---|
| ***Y47G6A.19*** | **WBGene00021645** | **protein_coding_gene** | **-1.6** | **0.321** | **4.5E-05** | **1.1** | **2.15** | **6.7E-07** |
| *Y22D7AL.10* | WBGene00021248 | protein_coding_gene | -1.6 | 0.320 | 2.2E-11 | -1.0 | 0.49 | 1.0E-18 |
| *nlp-33* | WBGene00003771 | protein_coding_gene | -1.6 | 0.320 | 8.0E-06 | -1.3 | 0.40 | 3.8E-03 |
| *F07E5.12* | WBGene00219315 | protein_coding_gene | -1.6 | 0.320 | 4.2E-02 | -2.2 | 0.22 | 3.3E-02 |
| *C49G7.10* | WBGene00016788 | protein_coding_gene | -1.7 | 0.314 | 1.3E-03 | -1.5 | 0.36 | 9.4E-04 |
| ***ram-2*** | **WBGene00004300** | **protein_coding_gene** | **-1.7** | **0.306** | **1.8E-04** | **1.9** | **3.76** | **1.9E-17** |
| *ugt-14* | WBGene00019233 | protein_coding_gene | -1.7 | 0.303 | 7.2E-03 | -2.2 | 0.21 | 1.6E-06 |
| ***dpy-6*** | **WBGene00001068** | **protein_coding_gene** | **-1.7** | **0.303** | **7.5E-03** | **2.1** | **4.22** | **1.6E-03** |
| ***col-49*** | **WBGene00000626** | **protein_coding_gene** | **-1.7** | **0.301** | **1.2E-03** | **2.0** | **4.13** | **4.5E-02** |
| *cnc-4* | WBGene00000558 | protein_coding_gene | -1.7 | 0.299 | 6.7E-03 | -5.7 | 0.02 | 5.9E-17 |
| *M04D5.1* | WBGene00014807 | protein_coding_gene | -1.8 | 0.292 | 5.1E-04 | -1.2 | 0.44 | 3.4E-02 |
| ***ugt-19*** | **WBGene00019515** | **protein_coding_gene** | **-1.8** | **0.292** | **1.4E-03** | **2.1** | **4.41** | **8.7E-21** |
| *Y60A9.3* | WBGene00013370 | protein_coding_gene | -1.8 | 0.289 | 4.8E-03 | -1.5 | 0.36 | 1.5E-02 |
| *R09H10.7* | WBGene00011184 | protein_coding_gene | -1.8 | 0.287 | 8.7E-03 | -1.3 | 0.40 | 1.0E-02 |
| *hach-1* | WBGene00017301 | protein_coding_gene | -1.8 | 0.278 | 1.3E-35 | -1.2 | 0.43 | 4.8E-03 |
| *clec-74* | WBGene00021585 | protein_coding_gene | -1.9 | 0.277 | 2.0E-02 | -2.1 | 0.23 | 3.9E-07 |
| ***Y82E9BL.18*** | **WBGene00044402** | **protein_coding_gene** | **-2.0** | **0.256** | **6.0E-04** | **1.9** | **3.63** | **4.5E-02** |
| *ugt-9* | WBGene00020587 | protein_coding_gene | -2.0 | 0.256 | 1.2E-07 | -1.8 | 0.28 | 7.2E-11 |
| *dod-17* | WBGene00010745 | protein_coding_gene | -2.0 | 0.255 | 8.3E-05 | -1.6 | 0.33 | 1.7E-02 |
| *R11H6.6* | WBGene00014822 | pseudogene | -2.0 | 0.254 | 3.9E-10 | -1.1 | 0.47 | 5.7E-03 |
| *F44E7.5* | WBGene00018427 | protein_coding_gene | -2.0 | 0.251 | 2.8E-09 | -1.1 | 0.48 | 1.7E-11 |
| ***gst-10*** | **WBGene00001758** | **protein_coding_gene** | **-2.0** | **0.244** | **2.3E-05** | **1.0** | **2.02** | **3.6E-04** |
| *F54E4.2* | WBGene00010058 | protein_coding_gene | -2.1 | 0.232 | 1.9E-02 | -3.7 | 0.08 | 1.9E-05 |
| ***C05E7.2*** | **WBGene00007344** | **protein_coding_gene** | **-2.1** | **0.231** | **2.0E-03** | **1.8** | **3.39** | **5.3E-03** |
| ***grd-6*** | **WBGene00001695** | **protein_coding_gene** | **-2.1** | **0.230** | **1.6E-03** | **1.9** | **3.84** | **1.3E-02** |
| ***col-167*** | **WBGene00000740** | **protein_coding_gene** | **-2.1** | **0.228** | **3.3E-05** | **1.7** | **3.34** | **1.9E-03** |
| ***B0334.13*** | **WBGene00007149** | **protein_coding_gene** | **-2.2** | **0.221** | **8.4E-03** | **2.1** | **4.18** | **2.7E-02** |
| ***col-168*** | **WBGene00000741** | **protein_coding_gene** | **-2.2** | **0.214** | **2.5E-05** | **2.1** | **4.14** | **3.3E-04** |
| ***col-170*** | **WBGene00000743** | **protein_coding_gene** | **-2.2** | **0.212** | **6.0E-04** | **1.9** | **3.80** | **2.7E-03** |
| ***col-91*** | **WBGene00000666** | **protein_coding_gene** | **-2.3** | **0.206** | **4.5E-04** | **1.0** | **2.00** | **4.5E-02** |
| ***pgp-12*** | **WBGene00004006** | **protein_coding_gene** | **-2.3** | **0.205** | **2.6E-09** | **1.3** | **2.42** | **5.7E-03** |
| *asp-12* | WBGene00017678 | protein_coding_gene | -2.3 | 0.199 | 2.2E-08 | -2.2 | 0.21 | 3.0E-13 |
| ***str-146*** | **WBGene00006193** | **protein_coding_gene** | **-2.4** | **0.194** | **8.4E-03** | **2.1** | **4.39** | **2.9E-02** |
| *ckb-2* | WBGene00000512 | protein_coding_gene | -2.4 | 0.192 | 2.0E-09 | -1.2 | 0.43 | 4.5E-08 |
| *C10C5.3* | WBGene00007507 | protein_coding_gene | -2.4 | 0.190 | 3.7E-07 | -2.2 | 0.21 | 6.7E-06 |
| ***F41E6.12*** | **WBGene00018293** | **protein_coding_gene** | **-2.4** | **0.186** | **2.2E-04** | **1.4** | **2.58** | **1.3E-04** |
| *C17F4.7* | WBGene00015913 | protein_coding_gene | -2.4 | 0.184 | 5.4E-13 | -1.5 | 0.36 | 7.3E-05 |
| *F23F12.3* | WBGene00017751 | protein_coding_gene | -2.5 | 0.178 | 9.5E-07 | -1.3 | 0.40 | 1.2E-02 |
| *F55G11.2* | WBGene00010123 | protein_coding_gene | -2.5 | 0.177 | 8.7E-05 | -1.6 | 0.33 | 1.2E-02 |
| *K12B6.11* | WBGene00195164 | protein_coding_gene | -2.5 | 0.173 | 3.5E-03 | -1.9 | 0.27 | 4.2E-02 |
| ***col-157*** | **WBGene00000730** | **protein_coding_gene** | **-2.6** | **0.167** | **2.6E-04** | **1.7** | **3.19** | **3.5E-02** |
| *catp-3* | WBGene00015660 | protein_coding_gene | -2.7 | 0.158 | 7.9E-09 | -1.6 | 0.33 | 3.4E-06 |
| ***grd-13*** | **WBGene00001702** | **protein_coding_gene** | **-2.8** | **0.148** | **4.6E-06** | **1.0** | **2.07** | **4.0E-02** |
| *T10B5.8* | WBGene00020394 | protein_coding_gene | -2.8 | 0.147 | 3.1E-15 | -1.7 | 0.31 | 2.6E-06 |
| *C17H12.8* | WBGene00015933 | protein_coding_gene | -2.8 | 0.143 | 5.2E-13 | -2.0 | 0.25 | 5.6E-21 |
| *nlp-25* | WBGene00003763 | protein_coding_gene | -2.8 | 0.140 | 1.1E-03 | -3.6 | 0.08 | 1.1E-07 |
| ***zipt-2.3*** | **WBGene00013207** | **protein_coding_gene** | **-2.9** | **0.138** | **8.6E-04** | **1.7** | **3.15** | **6.1E-03** |
| ***col-154*** | **WBGene00000727** | **protein_coding_gene** | **-2.9** | **0.134** | **2.9E-06** | **1.3** | **2.41** | **1.6E-03** |
| *C49G7.13* | WBGene00269425 | protein_coding_gene | -2.9 | 0.132 | 1.3E-02 | -2.3 | 0.20 | 3.8E-08 |
| ***col-149*** | **WBGene00000722** | **protein_coding_gene** | **-3.0** | **0.122** | **8.1E-10** | **2.1** | **4.16** | **5.4E-05** |
| *C09B8.4* | WBGene00015623 | protein_coding_gene | -3.1 | 0.117 | 2.0E-22 | -1.9 | 0.27 | 3.2E-08 |
| *F33H12.7* | WBGene00045457 | protein_coding_gene | -3.2 | 0.110 | 7.5E-04 | -2.6 | 0.16 | 3.6E-04 |
| *F15E6.4* | WBGene00017485 | protein_coding_gene | -3.2 | 0.110 | 2.2E-09 | -4.4 | 0.05 | 3.8E-11 |
| *F54E2.1* | WBGene00018823 | protein_coding_gene | -3.3 | 0.103 | 9.9E-18 | -2.5 | 0.17 | 6.7E-07 |
| *cnc-3* | WBGene00000557 | protein_coding_gene | -3.3 | 0.101 | 1.3E-11 | -2.3 | 0.20 | 2.4E-10 |
| *K11H12.4* | WBGene00019660 | protein_coding_gene | -3.3 | 0.100 | 1.3E-11 | -2.1 | 0.24 | 1.4E-03 |
| *fip-6* | WBGene00009964 | protein_coding_gene | -3.8 | 0.071 | 4.9E-08 | -2.1 | 0.24 | 2.0E-04 |
| *acdh-2* | WBGene00015894 | protein_coding_gene | -4.2 | 0.056 | 1.5E-18 | -1.7 | 0.31 | 1.5E-03 |

| | | | | | | | | |
|---|---|---|---|---|---|---|---|---|
| *clec-210* | WBGene00022261 | protein_coding_gene | -4.2 | 0.054 | 3.2E-06 | -1.4 | 0.37 | 1.2E-03 |
| *amt-1* | WBGene00000133 | protein_coding_gene | -4.6 | 0.041 | 6.3E-09 | -1.3 | 0.42 | 4.2E-02 |
| *F54D8.10* | WBGene00284847 | protein_coding_gene | -5.0 | 0.031 | 3.8E-06 | -2.3 | 0.20 | 5.0E-03 |
| *B0205.13* | WBGene00044644 | protein_coding_gene | -6.3 | 0.013 | 4.1E-28 | -1.7 | 0.32 | 6.2E-04 |
| *spp-17* | WBGene00005002 | protein_coding_gene | -7.9 | 0.004 | 3.8E-93 | -5.6 | 0.02 | 1.5E-171 |

*Genes in bold are differentially expressed in opposing directions

Transcription Factors Differentially Expressed in *S. epidermidis*

| Gene Name | WormBase Gene ID | Sequence Name | N2 $log_2$FC | CB4856 $log_2$FC |
|---|---|---|---|---|
| *nhr-168* | WBGene00008221 | C50B6.8 | 2.0 | 2.1 |
| *nhr-208* | WBGene00011099 | R07B7.15 | 1.9 | 1.1 |
| *zip-6* | WBGene00011130 | R07H5.10 | 3.1 | 2.1 |
| *sup-35* | WBGene00012976 | Y48A6C.3 | -1.3 | -1.1 |
| *Y60A9.3* | WBGene00013370 | Y60A9.3 | -1.8 | -1.5 |
| *Y116A8C.19* | WBGene00013796 | Y116A8C.19 | -1.3 | -1.1 |
| *klu-1* | WBGene00013970 | ZK337.2 | 1.4 | 1.2 |
| *Y22D7AL.16* | WBGene00021254 | Y22D7AL.16 | 1.4 | 1.5 |
| *maf-1* | WBGene00077521 | F45H11.6 | 1.2 | 1.3 |
| | | N2 Specific | | |
| *atf-2* | WBGene00000220 | K08F8.2 | 2.6 | Not DE |
| *bro-1* | WBGene00000272 | F56A3.5 | 2.5 | Not DE |
| *ceh-2* | WBGene00000429 | C27A12.5 | 1.8 | Not DE |
| *ceh-8* | WBGene00000433 | ZK265.4 | 1.9 | Not DE |
| *ceh-16* | WBGene00000439 | C13G5.1 | 1.8 | Not DE |
| *ceh-22* | WBGene00000445 | F29F11.5 | 2.0 | Not DE |
| *pros-1* | WBGene00000448 | K12H4.1 | 1.6 | Not DE |
| *ceh-27* | WBGene00000449 | F46F3.1 | 1.3 | Not DE |
| *ceh-30* | WBGene00000451 | C33D12.7 | 1.5 | Not DE |
| *ceh-32* | WBGene00000453 | W05E10.3 | 1.2 | Not DE |
| *ceh-34* | WBGene00000455 | C10G8.6 | 1.6 | Not DE |
| *ces-1* | WBGene00000468 | F43G9.11 | 2.3 | Not DE |
| *ces-2* | WBGene00000469 | ZK909.4 | 2.4 | Not DE |
| *cfi-1* | WBGene00000476 | T23D8.8 | 1.3 | Not DE |
| *cky-1* | WBGene00000521 | C15C8.2 | 4.6 | Not DE |
| *egl-38* | WBGene00001204 | C04G2.7 | 1.5 | Not DE |
| *egl-43* | WBGene00001207 | R53.3 | 1.5 | Not DE |
| *egl-46* | WBGene00001210 | K11G9.4 | 2.9 | Not DE |
| *elt-6* | WBGene00001253 | F52C12.5 | 1.0 | Not DE |
| *end-1* | WBGene00001310 | F58E10.2 | -1.9 | Not DE |
| *fax-1* | WBGene00001400 | F56E3.4 | 2.0 | Not DE |
| *fkh-8* | WBGene00001440 | F40H3.4 | 1.4 | Not DE |
| *fkh-10* | WBGene00001442 | C25A1.2 | 1.8 | Not DE |
| *gei-13* | WBGene00001570 | F58A4.11 | 2.1 | Not DE |
| *grh-1* | WBGene00001707 | Y48G8AR.1 | 1.4 | Not DE |
| *ham-2* | WBGene00001821 | C07A12.1 | 1.3 | Not DE |
| *hbl-1* | WBGene00001824 | F13D11.2 | 2.5 | Not DE |
| *hlh-6* | WBGene00001952 | T15H9.3 | 1.5 | Not DE |
| *hlh-14* | WBGene00001958 | C18A3.8 | 1.1 | Not DE |
| *hlh-16* | WBGene00001960 | DY3.3 | 2.0 | Not DE |
| *hsp-60* | WBGene00002025 | Y22D7AL.5 | -1.7 | Not DE |
| *let-381* | WBGene00002601 | F26B1.7 | 1.5 | Not DE |
| *lim-4* | WBGene00002987 | ZC64.4 | 3.4 | Not DE |
| *lim-6* | WBGene00002988 | K03E6.1 | 1.1 | Not DE |
| *lin-11* | WBGene00003000 | ZC247.3 | 1.1 | Not DE |
| *mab-5* | WBGene00003102 | C08C3.3 | 1.1 | Not DE |
| *mdl-1* | WBGene00003163 | R03E9.1 | 1.2 | Not DE |
| *mec-3* | WBGene00003167 | F01D4.6 | 2.6 | Not DE |
| *nhr-6* | WBGene00003605 | C48D5.1 | 1.1 | Not DE |

| | | | | |
|---|---|---|---|---|
| *nhr-19* | WBGene00003618 | E02H1.7 | 1.9 | Not DE |
| *nhr-21* | WBGene00003620 | F21D12.1 | 2.4 | Not DE |
| *nhr-32* | WBGene00003626 | K08H2.8 | 1.1 | Not DE |
| *nhr-42* | WBGene00003632 | C33G8.6 | 1.2 | Not DE |
| *nhr-55* | WBGene00003645 | T01G6.7 | 1.5 | Not DE |
| *nhr-74* | WBGene00003664 | C27C7.3 | -3.0 | Not DE |
| *nhr-87* | WBGene00003677 | Y41D4B.7 | 1.2 | Not DE |
| *nhr-88* | WBGene00003678 | K08A2.5 | 1.1 | Not DE |
| *nhr-91* | WBGene00003681 | Y15E3A.1 | 1.1 | Not DE |
| *nhr-99* | WBGene00003689 | M02H5.1 | 2.3 | Not DE |
| *nhr-114* | WBGene00003704 | Y45G5AM.1 | -1.7 | Not DE |
| *nhr-115* | WBGene00003705 | T27B7.4 | -1.2 | Not DE |
| *nhr-116* | WBGene00003706 | F09C6.9 | 1.9 | Not DE |
| *nhr-117* | WBGene00003707 | F16B4.12 | 2.0 | Not DE |
| *nhr-121* | WBGene00003711 | E02H9.8 | 1.1 | Not DE |
| *nhr-127* | WBGene00003717 | T13F3.3 | 3.5 | Not DE |
| *pag-3* | WBGene00003909 | F45B8.4 | 1.5 | Not DE |
| *pat-9* | WBGene00003933 | T27B1.2 | 1.2 | Not DE |
| *pax-1* | WBGene00003937 | K07C11.1 | 3.2 | Not DE |
| *peb-1* | WBGene00003968 | T14F9.4 | 2.7 | Not DE |
| *pha-4* | WBGene00004013 | F38A6.1 | 1.3 | Not DE |
| *php-3* | WBGene00004024 | Y75B8A.1 | 1.1 | Not DE |
| *plx-2* | WBGene00004048 | K04B12.1 | 1.8 | Not DE |
| *pqm-1* | WBGene00004096 | F40F8.7 | 2.1 | Not DE |
| *sem-4* | WBGene00004773 | F15C11.1 | 1.0 | Not DE |
| *sox-2* | WBGene00004949 | K08A8.2 | 1.2 | Not DE |
| *sox-3* | WBGene00004950 | F40E10.2 | 1.1 | Not DE |
| *tbx-38* | WBGene00006557 | C24H11.3 | -1.6 | Not DE |
| *ttx-1* | WBGene00006652 | Y113G7A.6 | 1.7 | Not DE |
| *unc-3* | WBGene00006743 | Y16B4A.1 | 1.9 | Not DE |
| *unc-42* | WBGene00006778 | F58E6.10 | 3.1 | Not DE |
| *unc-86* | WBGene00006818 | C30A5.7 | 1.7 | Not DE |
| *vab-3* | WBGene00006870 | F14F3.1 | 1.4 | Not DE |
| *zag-1* | WBGene00006970 | F28F9.1 | 1.9 | Not DE |
| *nhr-150* | WBGene00007367 | C06B8.1 | 1.4 | Not DE |
| *ceh-57* | WBGene00007416 | C07E3.5 | 2.0 | Not DE |
| *jmjd-3.3* | WBGene00007813 | C29F7.6 | 1.2 | Not DE |
| *F38C2.7* | WBGene00009539 | F38C2.7 | -1.6 | Not DE |
| *fozi-1* | WBGene00010453 | K01B6.1 | 2.1 | Not DE |
| *nhr-206* | WBGene00011097 | R07B7.13 | 1.8 | Not DE |
| *nhr-218* | WBGene00011750 | T13F3.2 | 3.1 | Not DE |
| *tsct-1* | WBGene00011824 | T18D3.7 | 1.2 | Not DE |
| *T27A8.2* | WBGene00012074 | T27A8.2 | -1.1 | Not DE |
| *fezf-1* | WBGene00012639 | Y38H8A.5 | 2.3 | Not DE |
| *zip-7* | WBGene00013100 | Y51H4A.4 | 1.4 | Not DE |
| *hlh-32* | WBGene00013665 | Y105C5B.29 | 2.8 | Not DE |
| *athp-3* | WBGene00013799 | Y116A8C.22 | -1.1 | Not DE |
| *atfs-1* | WBGene00013878 | ZC376.7 | -1.1 | Not DE |
| *ztf-30* | WBGene00015523 | C06E1.8 | 1.8 | Not DE |
| *ceh-48* | WBGene00015934 | C17H12.9 | 1.4 | Not DE |
| *unk-1* | WBGene00016407 | C34D10.2 | 1.7 | Not DE |
| *C46E10.8* | WBGene00016712 | C46E10.8 | -1.1 | Not DE |
| *ets-9* | WBGene00016865 | C52B9.2 | 1.0 | Not DE |
| *nhr-171* | WBGene00016926 | C54F6.8 | 1.6 | Not DE |
| *nhr-173* | WBGene00016975 | C56E10.1 | 1.6 | Not DE |
| *dmd-5* | WBGene00017326 | F10C1.5 | 1.4 | Not DE |
| *atf-8* | WBGene00017535 | F17A9.3 | 2.1 | Not DE |
| *ets-7* | WBGene00017601 | F19F10.5 | -1.1 | Not DE |

| | | | | |
|---|---|---|---|---|
| *F21A9.2* | WBGene00017651 | F21A9.2 | 1.1 | Not DE |
| *F26A10.2* | WBGene00017814 | F26A10.2 | 1.6 | Not DE |
| *ceh-82* | WBGene00018433 | F45C12.2 | -1.3 | Not DE |
| *mnat-1* | WBGene00018769 | F53G2.7 | -1.2 | Not DE |
| *klf-1* | WBGene00018990 | F56F11.3 | 1.1 | Not DE |
| *zip-2* | WBGene00019327 | K02F3.4 | 1.0 | Not DE |
| *M03D4.4* | WBGene00019751 | M03D4.4 | 1.9 | Not DE |
| *ast-1* | WBGene00020368 | T08H4.3 | 1.9 | Not DE |
| *hlh-30* | WBGene00020930 | W02C12.3 | 1.0 | Not DE |
| *W04B5.2* | WBGene00021019 | W04B5.2 | 1.4 | Not DE |
| *nhr-275* | WBGene00021163 | Y5H2A.2 | 1.5 | Not DE |
| *crtc-1* | WBGene00021237 | Y20F4.2 | 1.0 | Not DE |
| *znf-782* | WBGene00021931 | Y55F3AM.14 | -1.3 | Not DE |
| *nhr-277* | WBGene00022374 | Y94H6A.1 | 2.6 | Not DE |
| *ceh-45* | WBGene00022837 | ZK993.1 | 1.9 | Not DE |
| *syd-9* | WBGene00044068 | ZK867.1 | 1.7 | Not DE |
| *tag-243* | WBGene00044078 | T04A8.4 | 1.5 | Not DE |
| *alr-1* | WBGene00044330 | R08B4.2 | 1.2 | Not DE |
| | | CB4856 Specific | | |
| *ceh-5* | WBGene00000430 | C16C2.1 | Not DE | -1.7 |
| *ceh-18* | WBGene00000441 | ZC64.3 | Not DE | 1.3 |
| *ceh-23* | WBGene00000446 | ZK652.5 | Not DE | 1.6 |
| *lin-29* | WBGene00003015 | W03C9.4 | Not DE | 1.2 |
| *lin-31* | WBGene00003017 | K10G6.1 | Not DE | 1.9 |
| *mab-3* | WBGene00003100 | Y53C12B.5 | Not DE | 1.3 |
| *mgl-2* | WBGene00003233 | F45H11.4 | Not DE | 1.6 |
| *nhr-31* | WBGene00003625 | C26B2.3 | Not DE | 1.7 |
| *gmeb-3* | WBGene00008092 | C44F1.2 | Not DE | 1.0 |
| *ccch-2* | WBGene00009537 | F38C2.5 | Not DE | -1.1 |
| *jmjd-4* | WBGene00011563 | T07C4.11 | Not DE | -1.3 |
| *aptf-2* | WBGene00013383 | Y62E10A.17 | Not DE | -1.1 |
| *Y116A8C.20* | WBGene00013797 | Y116A8C.20 | Not DE | -1.3 |
| *C06E2.1* | WBGene00015527 | C06E2.1 | Not DE | 1.6 |
| *ets-5* | WBGene00016600 | C42D8.4 | Not DE | 2.0 |
| *nhr-179* | WBGene00017512 | F16B4.11 | Not DE | 1.3 |
| *ets-4* | WBGene00017687 | F22A3.1 | Not DE | 1.3 |
| *ceh-60* | WBGene00017690 | F22A3.5 | Not DE | 1.0 |
| *K11D12.12* | WBGene00019651 | K11D12.12 | Not DE | -1.1 |
| *R05D3.3* | WBGene00019878 | R05D3.3 | Not DE | -1.0 |
| *nhr-219* | WBGene00020555 | T19A5.5 | Not DE | 1.7 |
| *dmd-8* | WBGene00020708 | T22H9.4 | Not DE | 1.6 |
| *zip-11* | WBGene00021082 | W08E12.1 | Not DE | -1.1 |
| *klu-2* | WBGene00022592 | ZC328.2 | Not DE | 1.2 |
| *Y53C10A.15* | WBGene00044805 | Y53C10A.15 | Not DE | 1.3 |

**Supplemental Table 7**

**Lists of differentially expressed genes that are identified in either N2, CB4856, or both, after infection by *S. aureus* (compared to *E. coli*)**

**Supplemental Table 7 - Differentially Expressed Genes in *S. aureus* Infected Animals**

**N2**-exclusive DEGs in response to *S. aureus* (vs. OP50)

| Gene Name | Wormbase Gene ID | Type | $log_2$ FC | FC | Adj P-value |
|---|---|---|---|---|---|
| *C23G10.11* | WBGene00016017 | protein_coding_gene | 10.0 | 1003.431 | 1.8E-24 |
| *T22F3.11* | WBGene00020700 | protein_coding_gene | 9.2 | 585.939 | 1.7E-25 |
| *clec-76* | WBGene00021587 | protein_coding_gene | 9.0 | 508.077 | 6.2E-26 |
| *T20D4.12* | WBGene00020618 | protein_coding_gene | 7.8 | 219.129 | 1.2E-19 |
| *Y68A4A.13* | WBGene00044212 | protein_coding_gene | 7.6 | 198.465 | 7.3E-23 |
| *Y60C6A.2* | WBGene00202498 | protein_coding_gene | 7.3 | 157.620 | 2.4E-09 |
| *C50F7.5* | WBGene00016845 | protein_coding_gene | 6.9 | 116.634 | 1.8E-14 |
| *T20D4.10* | WBGene00020616 | protein_coding_gene | 6.4 | 85.993 | 7.6E-40 |
| *K03D3.2* | WBGene00010528 | protein_coding_gene | 6.4 | 84.358 | 3.1E-13 |
| *C17B7.4* | WBGene00015878 | protein_coding_gene | 6.3 | 80.632 | 7.0E-21 |
| *sqst-4* | WBGene00012516 | protein_coding_gene | 6.1 | 69.200 | 2.4E-07 |
| *dmd-3* | WBGene00012832 | protein_coding_gene | 6.1 | 68.874 | 5.8E-07 |
| *C17B7.10* | WBGene00015884 | protein_coding_gene | 6.1 | 67.991 | 1.1E-09 |
| *asp-16* | WBGene00012682 | protein_coding_gene | 6.1 | 66.647 | 5.3E-08 |
| *clec-70* | WBGene00021581 | protein_coding_gene | 6.0 | 65.129 | 8.5E-57 |
| *T28A11.5* | WBGene00020872 | pseudogene | 6.0 | 64.062 | 1.8E-07 |
| *spp-21* | WBGene00044284 | protein_coding_gene | 6.0 | 63.517 | 4.3E-07 |
| *T26H5.8* | WBGene00012070 | protein_coding_gene | 6.0 | 63.292 | 2.5E-24 |
| *cyp-37B1* | WBGene00009226 | protein_coding_gene | 5.9 | 60.653 | 1.6E-13 |
| *T26H5.9* | WBGene00044206 | protein_coding_gene | 5.8 | 56.806 | 1.4E-11 |
| *K10D11.2* | WBGene00010746 | protein_coding_gene | 5.8 | 56.484 | 1.4E-07 |
| *F54F3.2* | WBGene00014770 | pseudogene | 5.8 | 54.693 | 1.0E-06 |
| *K09C4.4* | WBGene00019549 | protein_coding_gene | 5.8 | 54.457 | 7.7E-09 |
| *sodh-1* | WBGene00010790 | protein_coding_gene | 5.8 | 54.222 | 3.7E-41 |
| *F46A8.13* | WBGene00219950 | ncRNA_gene | 5.8 | 54.106 | 9.0E-09 |
| *clec-75* | WBGene00021586 | protein_coding_gene | 5.7 | 51.968 | 8.3E-17 |
| *fipr-22* | WBGene00007989 | protein_coding_gene | 5.7 | 51.193 | 1.3E-10 |
| *B0563.9* | WBGene00044687 | protein_coding_gene | 5.6 | 49.778 | 2.0E-06 |
| *F35F10.7* | WBGene00018057 | protein_coding_gene | 5.6 | 48.500 | 3.1E-06 |
| *C25F9.11* | WBGene00045411 | protein_coding_gene | 5.6 | 48.435 | 2.1E-12 |
| *F45D3.4* | WBGene00009724 | protein_coding_gene | 5.6 | 47.851 | 3.8E-16 |
| *cyp-35B2* | WBGene00019471 | protein_coding_gene | 5.6 | 47.578 | 5.3E-08 |
| *pals-37* | WBGene00138721 | protein_coding_gene | 5.5 | 46.440 | 1.2E-10 |
| *C18H7.1* | WBGene00015993 | protein_coding_gene | 5.4 | 43.654 | 2.1E-11 |
| *E02H4.4* | WBGene00008462 | protein_coding_gene | 5.4 | 42.878 | 1.7E-26 |
| *srw-86* | WBGene00005833 | protein_coding_gene | 5.4 | 42.110 | 4.7E-14 |
| *F47B8.4* | WBGene00009805 | protein_coding_gene | 5.4 | 42.049 | 2.0E-07 |
| *Y60C6A.3* | WBGene00271832 | protein_coding_gene | 5.3 | 40.280 | 2.9E-05 |
| *C25F9.12* | WBGene00045412 | protein_coding_gene | 5.3 | 39.870 | 6.7E-09 |
| *T26H5.14* | WBGene00235133 | protein_coding_gene | 5.3 | 39.676 | 1.4E-10 |
| *T07G12.5* | WBGene00011596 | protein_coding_gene | 5.3 | 39.541 | 2.7E-09 |
| *clec-71* | WBGene00021582 | protein_coding_gene | 5.3 | 39.409 | 1.7E-31 |
| *cyp-32B1* | WBGene00021167 | protein_coding_gene | 5.3 | 39.330 | 1.1E-13 |
| *F53A9.8* | WBGene00018731 | protein_coding_gene | 5.2 | 37.631 | 1.1E-15 |
| *fbxa-164* | WBGene00015599 | protein_coding_gene | 5.1 | 35.439 | 8.4E-07 |
| *clec-77* | WBGene00044343 | pseudogene | 5.1 | 35.245 | 1.2E-08 |
| *spp-2* | WBGene00004987 | protein_coding_gene | 5.1 | 34.627 | 2.8E-36 |
| *Y47H10A.3* | WBGene00012959 | protein_coding_gene | 5.0 | 32.691 | 3.8E-08 |
| *F48G7.8* | WBGene00018619 | protein_coding_gene | 5.0 | 32.273 | 1.9E-07 |
| *Y32B12A.5* | WBGene00012520 | protein_coding_gene | 5.0 | 31.499 | 3.7E-04 |
| *R07C12.1* | WBGene00019931 | protein_coding_gene | 5.0 | 31.111 | 7.8E-11 |
| *srw-85* | WBGene00005832 | protein_coding_gene | 5.0 | 31.041 | 1.1E-49 |
| *nhr-113* | WBGene00003703 | protein_coding_gene | 4.9 | 30.580 | 5.6E-05 |
| *F11C7.6* | WBGene00017378 | protein_coding_gene | 4.9 | 29.694 | 5.2E-15 |
| *F14D7.10* | WBGene00008797 | protein_coding_gene | 4.9 | 29.050 | 8.9E-06 |
| *ZK993.2* | WBGene00022838 | protein_coding_gene | 4.8 | 28.512 | 8.3E-06 |
| *Y105C5B.7* | WBGene00013648 | pseudogene | 4.8 | 28.424 | 5.6E-06 |
| *ZC204.12* | WBGene00022562 | protein_coding_gene | 4.8 | 28.022 | 4.3E-83 |
| *oac-18* | WBGene00009230 | pseudogene | 4.8 | 27.974 | 8.5E-07 |
| *F16B4.7* | WBGene00017509 | protein_coding_gene | 4.8 | 27.626 | 3.7E-07 |
| *F01D5.7* | WBGene00008497 | protein_coding_gene | 4.8 | 27.611 | 2.5E-06 |
| *C37A5.3* | WBGene00014698 | protein_coding_gene | 4.7 | 25.673 | 2.5E-10 |
| *cnc-6* | WBGene00000560 | protein_coding_gene | 4.7 | 25.369 | 4.7E-07 |
| *F56D6.8* | WBGene00044292 | protein_coding_gene | 4.7 | 25.223 | 6.4E-05 |
| *C25F9.2* | WBGene00007723 | protein_coding_gene | 4.6 | 24.672 | 1.6E-06 |
| *nlp-34* | WBGene00015046 | protein_coding_gene | 4.6 | 24.388 | 2.9E-20 |
| *Y65B4BL.1* | WBGene00022033 | protein_coding_gene | 4.6 | 23.622 | 3.4E-06 |
| *cdr-3* | WBGene00008297 | protein_coding_gene | 4.6 | 23.536 | 1.7E-04 |
| *spp-6* | WBGene00004991 | protein_coding_gene | 4.5 | 23.379 | 4.8E-09 |
| *C17B7.15* | WBGene00268213 | protein_coding_gene | 4.5 | 23.364 | 2.2E-05 |
| *Y57G11C.1131* | WBGene00194661 | protein_coding_gene | 4.5 | 23.211 | 1.9E-06 |
| *clec-72* | WBGene00021583 | protein_coding_gene | 4.5 | 23.030 | 3.6E-64 |
| *fbxa-138* | WBGene00022345 | protein_coding_gene | 4.5 | 23.008 | 8.6E-21 |
| *scl-13* | WBGene00019179 | protein_coding_gene | 4.5 | 22.036 | 3.4E-04 |
| *F47H4.2* | WBGene00009835 | protein_coding_gene | 4.5 | 22.011 | 1.2E-21 |

| | | | | | |
|---|---|---|---|---|---|
| *C35C5.8* | WBGene00007957 | protein_coding_gene | 4.4 | 21.549 | 4.2E-28 |
| *F37A8.5* | WBGene00009503 | protein_coding_gene | 4.4 | 21.539 | 2.0E-15 |
| *C08E3.13* | WBGene00015605 | protein_coding_gene | 4.4 | 21.399 | 3.9E-04 |
| *fbxa-7* | WBGene00020637 | protein_coding_gene | 4.4 | 21.335 | 3.1E-04 |
| *ZK993.5* | WBGene00236809 | protein_coding_gene | 4.4 | 20.928 | 1.6E-04 |
| *cup-16* | WBGene00007919 | protein_coding_gene | 4.4 | 20.626 | 3.3E-20 |
| *F15E6.10* | WBGene00044479 | protein_coding_gene | 4.4 | 20.618 | 6.9E-05 |
| *Y82E9BR.6* | WBGene00022339 | protein_coding_gene | 4.4 | 20.487 | 5.7E-05 |
| *ZC443.2* | WBGene00013897 | protein_coding_gene | 4.4 | 20.435 | 3.9E-07 |
| *Y60C6A.1* | WBGene00022013 | protein_coding_gene | 4.3 | 20.088 | 1.2E-04 |
| *F35D2.6* | WBGene00194916 | protein_coding_gene | 4.3 | 19.931 | 2.3E-04 |
| *Y46E12BR.1* | WBGene00021598 | protein_coding_gene | 4.3 | 19.883 | 1.7E-07 |
| *F55H12.7* | WBGene00219283 | protein_coding_gene | 4.3 | 19.708 | 1.5E-05 |
| *Y46H3A.5* | WBGene00021601 | protein_coding_gene | 4.3 | 19.688 | 1.4E-04 |
| *suex-1* | WBGene00017396 | protein_coding_gene | 4.3 | 19.568 | 3.9E-04 |
| *F49C5.9* | WBGene00044031 | protein_coding_gene | 4.3 | 19.499 | 1.6E-03 |
| *pals-36* | WBGene00044787 | pseudogene | 4.2 | 19.023 | 2.7E-04 |
| *C25E10.16* | WBGene00202512 | protein_coding_gene | 4.2 | 18.767 | 4.0E-04 |
| *clec-37* | WBGene00013609 | protein_coding_gene | 4.2 | 18.589 | 9.4E-10 |
| *ZC443.3* | WBGene00013898 | protein_coding_gene | 4.2 | 18.530 | 1.8E-16 |
| *T24E12.5* | WBGene00020774 | protein_coding_gene | 4.2 | 18.420 | 1.5E-06 |
| *srx-111* | WBGene00006002 | protein_coding_gene | 4.2 | 18.365 | 2.9E-04 |
| *F11A5.17* | WBGene00045510 | protein_coding_gene | 4.2 | 18.095 | 2.1E-07 |
| *F57E7.2* | WBGene00010203 | protein_coding_gene | 4.1 | 17.483 | 9.0E-08 |
| *Y53C12B.7* | WBGene00013146 | protein_coding_gene | 4.1 | 17.096 | 1.9E-13 |
| *C54F6.12* | WBGene00016929 | protein_coding_gene | 4.1 | 17.073 | 7.3E-14 |
| *F09E10.10* | WBGene00022982 | ncRNA_gene | 4.0 | 16.513 | 1.1E-03 |
| *Y65B4BM.3* | WBGene00302974 | protein_coding_gene | 4.0 | 16.490 | 2.3E-25 |
| *T21D12.7* | WBGene00020648 | protein_coding_gene | 4.0 | 16.377 | 3.3E-05 |
| *srj-29* | WBGene00005616 | protein_coding_gene | 4.0 | 16.334 | 2.7E-05 |
| *C18D4.4* | WBGene00007674 | protein_coding_gene | 4.0 | 16.299 | 1.1E-03 |
| *Y43F8B.25* | WBGene00195177 | protein_coding_gene | 4.0 | 16.297 | 4.0E-04 |
| *C25H3.10* | WBGene00016119 | protein_coding_gene | 4.0 | 16.067 | 4.5E-17 |
| *C31H5.1* | WBGene00007854 | protein_coding_gene | 4.0 | 16.053 | 1.0E-05 |
| *F12A10.9* | WBGene00077704 | protein_coding_gene | 4.0 | 15.838 | 1.5E-03 |
| *F26G1.15* | WBGene00270318 | protein_coding_gene | 4.0 | 15.812 | 6.4E-05 |
| *T24E12.12* | WBGene00044380 | protein_coding_gene | 4.0 | 15.801 | 1.7E-03 |
| *dod-22* | WBGene00010125 | protein_coding_gene | 4.0 | 15.763 | 6.3E-41 |
| *clec-38* | WBGene00012025 | protein_coding_gene | 4.0 | 15.650 | 2.8E-04 |
| *F55G11.7* | WBGene00010127 | protein_coding_gene | 4.0 | 15.626 | 8.9E-05 |
| *F23H12.5* | WBGene00009094 | protein_coding_gene | 4.0 | 15.476 | 4.3E-22 |
| *W03F9.4* | WBGene00021002 | protein_coding_gene | 4.0 | 15.464 | 1.1E-26 |
| *M01B2.13* | WBGene00045338 | protein_coding_gene | 3.9 | 15.336 | 6.1E-08 |
| *C14A11.2* | WBGene00015748 | protein_coding_gene | 3.9 | 15.322 | 1.4E-09 |
| *wrt-2* | WBGene00006948 | protein_coding_gene | 3.9 | 15.029 | 5.3E-12 |
| *fbxa-161* | WBGene00015596 | pseudogene | 3.9 | 15.019 | 2.9E-04 |
| *srd-27* | WBGene00005105 | protein_coding_gene | 3.9 | 14.910 | 1.5E-03 |
| *nhr-37* | WBGene00018412 | protein_coding_gene | 3.9 | 14.838 | 1.4E-03 |
| *fbxa-163* | WBGene00015598 | protein_coding_gene | 3.9 | 14.717 | 3.0E-10 |
| *C07G3.15* | WBGene00269378 | protein_coding_gene | 3.9 | 14.691 | 1.1E-03 |
| *F35F10.5* | WBGene00018055 | protein_coding_gene | 3.9 | 14.603 | 8.2E-20 |
| *T19H5.6* | WBGene00044807 | protein_coding_gene | 3.8 | 14.417 | 2.3E-07 |
| *C33D9.10* | WBGene00044239 | protein_coding_gene | 3.8 | 14.393 | 6.9E-04 |
| *H42K12.3* | WBGene00019272 | protein_coding_gene | 3.8 | 14.288 | 1.5E-16 |
| *C23G10.10* | WBGene00016016 | protein_coding_gene | 3.8 | 14.141 | 2.5E-04 |
| *T03E6.9* | WBGene00045356 | protein_coding_gene | 3.8 | 14.134 | 1.7E-04 |
| *fbxa-154* | WBGene00007163 | protein_coding_gene | 3.8 | 14.092 | 9.8E-04 |
| *K08C9.5* | WBGene00010652 | protein_coding_gene | 3.8 | 14.059 | 7.1E-04 |
| *zmp-5* | WBGene00010423 | protein_coding_gene | 3.8 | 14.037 | 1.3E-04 |
| *abf-2* | WBGene00000013 | protein_coding_gene | 3.8 | 14.011 | 6.9E-11 |
| *F09C6.14* | WBGene00044278 | protein_coding_gene | 3.8 | 13.865 | 1.3E-06 |
| *nspe-6* | WBGene00012603 | protein_coding_gene | 3.8 | 13.725 | 2.4E-04 |
| *R102.1* | WBGene00011288 | protein_coding_gene | 3.8 | 13.690 | 8.6E-04 |
| *F10D7.4* | WBGene00017341 | protein_coding_gene | 3.8 | 13.653 | 9.5E-04 |
| *C17F4.3* | WBGene00015911 | protein_coding_gene | 3.8 | 13.582 | 3.9E-04 |
| *ins-25* | WBGene00002108 | protein_coding_gene | 3.7 | 13.412 | 3.4E-03 |
| *C29F7.11* | WBGene00206531 | ncRNA_gene | 3.7 | 13.400 | 1.8E-03 |
| *F57F4.2* | WBGene00019016 | protein_coding_gene | 3.7 | 13.342 | 5.4E-05 |
| *W01C8.1* | WBGene00020918 | protein_coding_gene | 3.7 | 13.339 | 3.1E-04 |
| *osm-8* | WBGene00003888 | protein_coding_gene | 3.7 | 13.288 | 1.6E-08 |
| *T04C12.32* | WBGene00235086 | ncRNA_gene | 3.7 | 13.280 | 1.7E-03 |
| *cyp-34A1* | WBGene00011698 | protein_coding_gene | 3.7 | 13.249 | 5.0E-05 |
| *skr-5* | WBGene00004811 | protein_coding_gene | 3.7 | 13.131 | 8.1E-12 |
| *EGAP1.1* | WBGene00017146 | protein_coding_gene | 3.7 | 13.038 | 5.6E-07 |
| *C44H9.7* | WBGene00008102 | protein_coding_gene | 3.7 | 12.903 | 1.4E-08 |
| *K01A6.4* | WBGene00010446 | protein_coding_gene | 3.7 | 12.888 | 2.4E-03 |
| *K02F3.15* | WBGene00220001 | snoRNA_gene | 3.7 | 12.874 | 1.9E-03 |
| *T20B3.1* | WBGene00011850 | protein_coding_gene | 3.7 | 12.871 | 2.9E-13 |
| *W02A2.9* | WBGene00044280 | protein_coding_gene | 3.7 | 12.851 | 3.4E-23 |
| *C17B7.12* | WBGene00015886 | protein_coding_gene | 3.7 | 12.804 | 9.1E-03 |

| | | | | | |
|---|---|---|---|---|---|
| *C17F4.12* | WBGene00044374 | protein_coding_gene | 3.7 | 12.736 | 1.2E-05 |
| *T23E7.6* | WBGene00044772 | protein_coding_gene | 3.7 | 12.699 | 5.8E-08 |
| *C13A2.3* | WBGene00015720 | protein_coding_gene | 3.7 | 12.599 | 1.9E-03 |
| *nlp-32* | WBGene00003770 | protein_coding_gene | 3.7 | 12.598 | 5.9E-03 |
| *F58F12.12* | WBGene00271812 | protein_coding_gene | 3.7 | 12.589 | 3.7E-11 |
| *F40H7.12* | WBGene00044379 | protein_coding_gene | 3.6 | 12.503 | 3.7E-03 |
| *F58H1.2* | WBGene00010285 | protein_coding_gene | 3.6 | 12.452 | 2.6E-09 |
| *fbxa-162* | WBGene00015597 | pseudogene | 3.6 | 12.436 | 2.0E-10 |
| *R11D1.7* | WBGene00011246 | protein_coding_gene | 3.6 | 12.415 | 3.1E-03 |
| *asp-11* | WBGene00017677 | pseudogene | 3.6 | 12.352 | 4.5E-07 |
| *irld-24* | WBGene00007844 | protein_coding_gene | 3.6 | 12.340 | 8.1E-04 |
| *glr-3* | WBGene00001614 | protein_coding_gene | 3.6 | 12.275 | 2.1E-04 |
| *Y37A1B.7* | WBGene00012540 | protein_coding_gene | 3.6 | 12.251 | 4.6E-09 |
| *Y7A5A.6* | WBGene00012412 | protein_coding_gene | 3.6 | 12.202 | 3.6E-05 |
| *scl-12* | WBGene00019178 | protein_coding_gene | 3.6 | 12.140 | 9.1E-04 |
| *ZC395.5* | WBGene00022597 | protein_coding_gene | 3.6 | 12.084 | 6.8E-17 |
| *B0393.8* | WBGene00007173 | protein_coding_gene | 3.6 | 12.045 | 1.4E-03 |
| *C08F11.13* | WBGene00007460 | protein_coding_gene | 3.6 | 12.020 | 4.3E-12 |
| *hlh-13* | WBGene00001957 | protein_coding_gene | 3.6 | 11.962 | 2.0E-03 |
| *srbc-20* | WBGene00016693 | protein_coding_gene | 3.6 | 11.915 | 2.1E-03 |
| *F11E6.11* | WBGene00023501 | protein_coding_gene | 3.6 | 11.885 | 2.3E-10 |
| *C06E4.8* | WBGene00015537 | protein_coding_gene | 3.6 | 11.798 | 2.2E-18 |
| *srh-282* | WBGene00005487 | protein_coding_gene | 3.6 | 11.743 | 3.4E-03 |
| *W09B7.2* | WBGene00021107 | protein_coding_gene | 3.5 | 11.694 | 1.0E-02 |
| *lim-4* | WBGene00002987 | protein_coding_gene | 3.5 | 11.628 | 1.1E-04 |
| *F55H12.2* | WBGene00010133 | protein_coding_gene | 3.5 | 11.570 | 2.2E-23 |
| *F09F9.2* | WBGene00017307 | protein_coding_gene | 3.5 | 11.553 | 5.3E-10 |
| *irld-15* | WBGene00021232 | protein_coding_gene | 3.5 | 11.547 | 2.1E-04 |
| *Y102A11A.5* | WBGene00022415 | protein_coding_gene | 3.5 | 11.452 | 1.9E-14 |
| *T26C5.2* | WBGene00012032 | protein_coding_gene | 3.5 | 11.446 | 7.2E-14 |
| *grd-2* | WBGene00001691 | protein_coding_gene | 3.5 | 11.440 | 6.8E-10 |
| *K03H6.4* | WBGene00019369 | protein_coding_gene | 3.5 | 11.356 | 2.7E-03 |
| *F46B3.1* | WBGene00009756 | protein_coding_gene | 3.5 | 11.352 | 2.3E-03 |
| *W09G12.7* | WBGene00021121 | protein_coding_gene | 3.5 | 11.282 | 1.8E-11 |
| *oac-5* | WBGene00007829 | protein_coding_gene | 3.5 | 11.257 | 1.5E-03 |
| *T27C5.8* | WBGene00012081 | protein_coding_gene | 3.5 | 11.231 | 1.1E-03 |
| *F56D2.3* | WBGene00018965 | protein_coding_gene | 3.5 | 11.226 | 2.1E-12 |
| *fip-1* | WBGene00017695 | protein_coding_gene | 3.5 | 11.188 | 3.7E-03 |
| *fbxa-25* | WBGene00022321 | protein_coding_gene | 3.5 | 11.170 | 9.1E-04 |
| *nlp-45* | WBGene00020136 | protein_coding_gene | 3.5 | 11.164 | 4.8E-07 |
| *fbxa-142* | WBGene00044659 | protein_coding_gene | 3.5 | 11.163 | 3.0E-03 |
| *F11E6.9* | WBGene00008712 | protein_coding_gene | 3.5 | 11.159 | 3.9E-06 |
| *ZK285.2* | WBGene00013961 | protein_coding_gene | 3.5 | 11.153 | 1.5E-02 |
| *K08F9.1* | WBGene00010684 | protein_coding_gene | 3.5 | 11.057 | 1.4E-12 |
| *linc-47* | WBGene00219641 | lincRNA_gene | 3.5 | 10.930 | 1.3E-03 |
| *C06B3.6* | WBGene00007365 | protein_coding_gene | 3.4 | 10.915 | 3.6E-16 |
| *unc-42* | WBGene00006778 | protein_coding_gene | 3.4 | 10.910 | 6.1E-09 |
| *R07C12.3* | WBGene00019933 | protein_coding_gene | 3.4 | 10.891 | 2.6E-03 |
| *glt-5* | WBGene00001623 | protein_coding_gene | 3.4 | 10.818 | 6.1E-14 |
| *C25F9.16* | WBGene00194803 | protein_coding_gene | 3.4 | 10.812 | 1.0E-03 |
| *C04A11.t1* | WBGene00000308 | tRNA_gene | 3.4 | 10.762 | 9.4E-03 |
| *C07G1.15* | WBGene00235296 | protein_coding_gene | 3.4 | 10.758 | 1.1E-06 |
| *K01A6.8* | WBGene00194869 | protein_coding_gene | 3.4 | 10.751 | 2.8E-03 |
| *Y43F8B.15* | WBGene00045415 | protein_coding_gene | 3.4 | 10.750 | 3.5E-07 |
| *srv-36* | WBGene00005747 | pseudogene | 3.4 | 10.742 | 1.8E-02 |
| *F43C11.7* | WBGene00018384 | protein_coding_gene | 3.4 | 10.742 | 7.1E-16 |
| *str-97* | WBGene00006154 | protein_coding_gene | 3.4 | 10.719 | 9.5E-03 |
| *T19C9.10* | WBGene00050934 | protein_coding_gene | 3.4 | 10.700 | 5.4E-04 |
| *F39G3.4* | WBGene00018209 | protein_coding_gene | 3.4 | 10.686 | 5.0E-03 |
| *nspb-3* | WBGene00018162 | protein_coding_gene | 3.4 | 10.666 | 4.7E-03 |
| *clec-168* | WBGene00009520 | protein_coding_gene | 3.4 | 10.596 | 8.6E-04 |
| *nhr-178* | WBGene00017510 | protein_coding_gene | 3.4 | 10.592 | 2.8E-08 |
| *F48C1.11* | WBGene00195179 | protein_coding_gene | 3.4 | 10.499 | 6.5E-04 |
| *clec-25* | WBGene00044331 | protein_coding_gene | 3.4 | 10.463 | 4.8E-06 |
| *K01F9.2* | WBGene00010475 | protein_coding_gene | 3.4 | 10.462 | 9.8E-13 |
| *srg-25* | WBGene00005182 | protein_coding_gene | 3.4 | 10.448 | 6.8E-03 |
| *str-137* | WBGene00006185 | pseudogene | 3.4 | 10.369 | 1.4E-02 |
| *clec-101* | WBGene00008853 | protein_coding_gene | 3.4 | 10.352 | 4.8E-03 |
| *daao-1* | WBGene00022076 | protein_coding_gene | 3.4 | 10.351 | 9.1E-17 |
| *srv-8* | WBGene00005719 | protein_coding_gene | 3.4 | 10.336 | 3.1E-03 |
| *F33D4.6* | WBGene00017998 | protein_coding_gene | 3.4 | 10.314 | 7.5E-13 |
| *F11C1.10* | WBGene00195214 | protein_coding_gene | 3.4 | 10.295 | 9.0E-04 |
| *F16C3.2* | WBGene00008886 | protein_coding_gene | 3.4 | 10.271 | 1.0E-17 |
| *str-124* | WBGene00006175 | protein_coding_gene | 3.4 | 10.234 | 4.1E-03 |
| *F18G5.6* | WBGene00017582 | protein_coding_gene | 3.4 | 10.200 | 1.4E-15 |
| *C50F4.9* | WBGene00008234 | protein_coding_gene | 3.3 | 10.183 | 8.4E-20 |
| *C40C9.4* | WBGene00008037 | protein_coding_gene | 3.3 | 10.145 | 5.4E-03 |
| *anr-51* | WBGene00219610 | antisense_lncRNA_gene | 3.3 | 10.080 | 4.9E-03 |
| *fbxc-7* | WBGene00016302 | protein_coding_gene | 3.3 | 10.006 | 1.4E-04 |
| *ZK1025.3* | WBGene00014183 | protein_coding_gene | 3.3 | 9.908 | 1.2E-11 |

| | | | | | |
|---|---|---|---|---|---|
| *T28A11.17* | WBGene00020879 | protein_coding_gene | 3.3 | 9.880 | 9.1E-05 |
| *linc-103* | WBGene00219743 | lincRNA_gene | 3.3 | 9.843 | 1.9E-03 |
| *clec-235* | WBGene00009474 | pseudogene | 3.3 | 9.837 | 2.6E-02 |
| *F32A7.8* | WBGene00044245 | protein_coding_gene | 3.3 | 9.814 | 9.2E-03 |
| *Y54G2A.76* | WBGene00235268 | protein_coding_gene | 3.3 | 9.813 | 3.2E-06 |
| *F16H6.7* | WBGene00008897 | protein_coding_gene | 3.3 | 9.779 | 7.0E-13 |
| *T10H4.13* | WBGene00045339 | protein_coding_gene | 3.3 | 9.770 | 5.8E-04 |
| *T20D4.9* | WBGene00020615 | pseudogene | 3.3 | 9.740 | 5.8E-04 |
| *T10G3.1* | WBGene00011692 | protein_coding_gene | 3.3 | 9.701 | 3.6E-15 |
| *C02F4.4* | WBGene00007264 | protein_coding_gene | 3.3 | 9.691 | 1.5E-07 |
| *C14A6.3* | WBGene00007567 | protein_coding_gene | 3.3 | 9.667 | 7.0E-03 |
| *F28C6.9* | WBGene00009208 | protein_coding_gene | 3.3 | 9.580 | 6.9E-03 |
| *H39E23.2* | WBGene00010432 | protein_coding_gene | 3.3 | 9.574 | 2.1E-03 |
| *Y38H8A.1* | WBGene00012635 | protein_coding_gene | 3.3 | 9.568 | 5.7E-06 |
| *fbxa-185* | WBGene00009836 | protein_coding_gene | 3.3 | 9.551 | 4.2E-03 |
| *clec-30* | WBGene00014842 | pseudogene | 3.3 | 9.548 | 2.8E-04 |
| *K01A12.2* | WBGene00019286 | protein_coding_gene | 3.2 | 9.510 | 4.5E-03 |
| *Y97E10AL.1* | WBGene00022392 | protein_coding_gene | 3.2 | 9.476 | 4.6E-06 |
| *twk-47* | WBGene00018002 | protein_coding_gene | 3.2 | 9.447 | 5.8E-05 |
| *btb-16* | WBGene00018195 | protein_coding_gene | 3.2 | 9.447 | 8.0E-29 |
| *B0454.18* | WBGene00199082 | ncRNA_gene | 3.2 | 9.411 | 1.4E-02 |
| *F46F3.3* | WBGene00009791 | protein_coding_gene | 3.2 | 9.392 | 7.9E-06 |
| *cyp-33B1* | WBGene00016092 | protein_coding_gene | 3.2 | 9.383 | 7.4E-03 |
| *C25F9.15* | WBGene00050967 | protein_coding_gene | 3.2 | 9.367 | 1.2E-06 |
| *F20G2.5* | WBGene00008988 | protein_coding_gene | 3.2 | 9.352 | 2.2E-09 |
| *anr-45* | WBGene00219710 | antisense_lncRNA_gene | 3.2 | 9.338 | 4.6E-03 |
| *glr-8* | WBGene00001619 | protein_coding_gene | 3.2 | 9.271 | 2.0E-05 |
| *poml-4* | WBGene00044697 | protein_coding_gene | 3.2 | 9.269 | 1.9E-11 |
| *pqn-16* | WBGene00004107 | protein_coding_gene | 3.2 | 9.250 | 1.0E-02 |
| *lurp-3* | WBGene00009136 | protein_coding_gene | 3.2 | 9.221 | 3.2E-03 |
| *F07E5.9* | WBGene00017214 | protein_coding_gene | 3.2 | 9.215 | 4.1E-03 |
| *K08D8.10* | WBGene00194919 | protein_coding_gene | 3.2 | 9.198 | 3.9E-02 |
| *C08F1.8* | WBGene00015611 | protein_coding_gene | 3.2 | 9.189 | 2.7E-03 |
| *fbxa-68* | WBGene00020641 | pseudogene | 3.2 | 9.184 | 1.0E-02 |
| *F47D2.11* | WBGene00206392 | protein_coding_gene | 3.2 | 9.170 | 1.4E-02 |
| *str-9* | WBGene00006077 | protein_coding_gene | 3.2 | 9.169 | 7.9E-03 |
| *aqp-5* | WBGene00000173 | protein_coding_gene | 3.2 | 9.151 | 1.1E-04 |
| *F11A5.13* | WBGene00008679 | protein_coding_gene | 3.2 | 9.133 | 1.7E-03 |
| *rncs-1* | WBGene00023069 | ncRNA_gene | 3.2 | 9.121 | 3.2E-15 |
| *tpst-2* | WBGene00018365 | protein_coding_gene | 3.2 | 9.060 | 1.4E-03 |
| *ZK105.11* | WBGene00194895 | protein_coding_gene | 3.2 | 8.963 | 4.7E-03 |
| *F38B6.3* | WBGene00018173 | protein_coding_gene | 3.2 | 8.949 | 3.4E-03 |
| *srg-17* | WBGene00005174 | protein_coding_gene | 3.2 | 8.918 | 2.8E-02 |
| *F31B9.4* | WBGene00009281 | protein_coding_gene | 3.2 | 8.916 | 2.6E-02 |
| *pals-19* | WBGene00044761 | protein_coding_gene | 3.2 | 8.911 | 4.7E-16 |
| *fbxa-172* | WBGene00008014 | protein_coding_gene | 3.2 | 8.890 | 1.3E-08 |
| *fbxa-73* | WBGene00020640 | protein_coding_gene | 3.1 | 8.875 | 2.2E-03 |
| *C45E5.3* | WBGene00016666 | pseudogene | 3.1 | 8.874 | 4.2E-09 |
| *Y105C5A.13* | WBGene00013637 | protein_coding_gene | 3.1 | 8.873 | 1.5E-02 |
| *T13F3.7* | WBGene00011754 | protein_coding_gene | 3.1 | 8.861 | 3.1E-05 |
| *T12E12.6* | WBGene00020465 | protein_coding_gene | 3.1 | 8.849 | 1.2E-02 |
| *R08A2.7* | WBGene00011137 | protein_coding_gene | 3.1 | 8.848 | 2.0E-07 |
| *mlt-9* | WBGene00008605 | protein_coding_gene | 3.1 | 8.847 | 7.2E-12 |
| *F13H10.1* | WBGene00008773 | protein_coding_gene | 3.1 | 8.846 | 8.8E-05 |
| *21ur-9997* | WBGene00168467 | piRNA_gene | 3.1 | 8.839 | 2.3E-02 |
| *slc-36.5* | WBGene00016628 | protein_coding_gene | 3.1 | 8.838 | 1.5E-13 |
| *M03A1.8* | WBGene00077490 | protein_coding_gene | 3.1 | 8.829 | 2.0E-11 |
| *ugt-15* | WBGene00008097 | protein_coding_gene | 3.1 | 8.825 | 5.6E-14 |
| *spp-13* | WBGene00004998 | protein_coding_gene | 3.1 | 8.751 | 3.9E-09 |
| *nspc-16* | WBGene00008357 | protein_coding_gene | 3.1 | 8.722 | 4.8E-17 |
| *F36G9.7* | WBGene00009486 | protein_coding_gene | 3.1 | 8.710 | 5.7E-03 |
| *F48G7.5* | WBGene00018616 | protein_coding_gene | 3.1 | 8.663 | 1.0E-03 |
| *F53F10.1* | WBGene00018763 | protein_coding_gene | 3.1 | 8.662 | 9.2E-29 |
| *srab-16* | WBGene00020060 | protein_coding_gene | 3.1 | 8.621 | 9.2E-03 |
| *B0348.9* | WBGene00235314 | ncRNA_gene | 3.1 | 8.617 | 6.2E-03 |
| *ptr-10* | WBGene00004224 | protein_coding_gene | 3.1 | 8.604 | 6.3E-10 |
| *dct-7* | WBGene00017488 | protein_coding_gene | 3.1 | 8.573 | 3.5E-03 |
| *abu-13* | WBGene00004133 | protein_coding_gene | 3.1 | 8.565 | 2.2E-12 |
| *qua-1* | WBGene00004264 | protein_coding_gene | 3.1 | 8.562 | 6.5E-16 |
| *thn-8* | WBGene00044285 | pseudogene | 3.1 | 8.554 | 7.6E-03 |
| *col-108* | WBGene00000682 | protein_coding_gene | 3.1 | 8.542 | 1.3E-02 |
| *coel-1* | WBGene00016866 | protein_coding_gene | 3.1 | 8.529 | 2.4E-08 |
| *T05C1.3* | WBGene00020250 | protein_coding_gene | 3.1 | 8.494 | 1.6E-04 |
| *fbxa-222* | WBGene00045295 | protein_coding_gene | 3.1 | 8.483 | 7.8E-03 |
| *Y44A6C.1* | WBGene00012850 | protein_coding_gene | 3.1 | 8.476 | 1.5E-21 |
| *C06C3.9* | WBGene00014668 | protein_coding_gene | 3.1 | 8.469 | 9.3E-08 |
| *cpr-4* | WBGene00000784 | protein_coding_gene | 3.1 | 8.466 | 1.1E-22 |
| *srh-239* | WBGene00005446 | protein_coding_gene | 3.1 | 8.440 | 8.7E-03 |
| *F49C5.10* | WBGene00044162 | protein_coding_gene | 3.1 | 8.426 | 1.5E-02 |
| *R09D1.12* | WBGene00011168 | protein_coding_gene | 3.1 | 8.403 | 4.0E-08 |

| | | | | | |
|---|---|---|---|---|---|
| *med-2* | WBGene00003181 | protein_coding_gene | 3.1 | 8.385 | 8.4E-03 |
| *C11G10.1* | WBGene00007525 | protein_coding_gene | 3.1 | 8.359 | 2.0E-13 |
| *F56H9.9* | WBGene00206460 | protein_coding_gene | 3.1 | 8.343 | 9.1E-03 |
| *chil-23* | WBGene00011167 | protein_coding_gene | 3.1 | 8.323 | 4.1E-12 |
| *T20D4.20* | WBGene00044718 | pseudogene | 3.0 | 8.281 | 4.0E-02 |
| *cyp-34A2* | WBGene00011699 | protein_coding_gene | 3.0 | 8.279 | 5.0E-07 |
| *fkb-4* | WBGene00001429 | protein_coding_gene | 3.0 | 8.248 | 1.1E-12 |
| *gpa-9* | WBGene00001671 | protein_coding_gene | 3.0 | 8.231 | 3.6E-23 |
| *AH9.4* | WBGene00015000 | protein_coding_gene | 3.0 | 8.230 | 3.1E-03 |
| *cil-7* | WBGene00021017 | protein_coding_gene | 3.0 | 8.217 | 2.7E-05 |
| *C12D5.10* | WBGene00015711 | protein_coding_gene | 3.0 | 8.213 | 7.8E-04 |
| *ptr-22* | WBGene00004236 | protein_coding_gene | 3.0 | 8.210 | 3.2E-06 |
| *cng-3* | WBGene00000563 | protein_coding_gene | 3.0 | 8.193 | 1.2E-03 |
| *F26A1.6* | WBGene00017804 | protein_coding_gene | 3.0 | 8.177 | 7.6E-03 |
| *Y82E9BL.9* | WBGene00023395 | pseudogene | 3.0 | 8.140 | 5.0E-04 |
| *F47G4.5* | WBGene00009826 | protein_coding_gene | 3.0 | 8.138 | 4.4E-03 |
| *sre-1* | WBGene00005149 | protein_coding_gene | 3.0 | 8.118 | 1.3E-02 |
| *F49F1.14* | WBGene00044477 | protein_coding_gene | 3.0 | 8.085 | 8.2E-03 |
| *B0281.6* | WBGene00015113 | protein_coding_gene | 3.0 | 8.054 | 9.8E-05 |
| *K10G4.5* | WBGene00010754 | protein_coding_gene | 3.0 | 8.039 | 1.1E-02 |
| *R03H10.2* | WBGene00019855 | protein_coding_gene | 3.0 | 8.019 | 2.3E-12 |
| *ilys-2* | WBGene00016669 | protein_coding_gene | 3.0 | 8.008 | 2.1E-05 |
| *swt-1* | WBGene00019300 | protein_coding_gene | 3.0 | 8.007 | 4.2E-10 |
| *Y46G5A.38* | WBGene00044043 | protein_coding_gene | 3.0 | 7.989 | 2.0E-03 |
| *F17C8.3* | WBGene00008910 | protein_coding_gene | 3.0 | 7.970 | 1.4E-12 |
| *nspb-7* | WBGene00007243 | protein_coding_gene | 3.0 | 7.961 | 2.7E-06 |
| *F23H12.21* | WBGene00219416 | ncRNA_gene | 3.0 | 7.949 | 2.0E-02 |
| *arrd-27* | WBGene00009853 | pseudogene | 3.0 | 7.945 | 2.0E-02 |
| *ttll-9* | WBGene00044187 | protein_coding_gene | 3.0 | 7.940 | 1.8E-02 |
| *C29F7.8* | WBGene00044405 | pseudogene | 3.0 | 7.929 | 2.4E-02 |
| *fmo-1* | WBGene00001476 | protein_coding_gene | 3.0 | 7.922 | 9.8E-06 |
| *ZK20.10* | WBGene00220237 | ncRNA_gene | 3.0 | 7.918 | 3.7E-02 |
| *T14E8.4* | WBGene00043981 | protein_coding_gene | 3.0 | 7.915 | 1.1E-08 |
| *D1014.7* | WBGene00017020 | protein_coding_gene | 3.0 | 7.894 | 1.2E-02 |
| *spi-1* | WBGene00023416 | protein_coding_gene | 3.0 | 7.892 | 5.5E-08 |
| *fbxa-137* | WBGene00020853 | protein_coding_gene | 3.0 | 7.883 | 4.3E-03 |
| *F28G4.4* | WBGene00009229 | protein_coding_gene | 3.0 | 7.879 | 4.8E-04 |
| *T08G5.15* | WBGene00077573 | protein_coding_gene | 3.0 | 7.876 | 6.7E-04 |
| *clec-107* | WBGene00012502 | protein_coding_gene | 3.0 | 7.868 | 5.6E-03 |
| *T20D4.13* | WBGene00020619 | pseudogene | 3.0 | 7.867 | 1.1E-03 |
| *F53C11.9* | WBGene00044921 | protein_coding_gene | 3.0 | 7.856 | 8.5E-40 |
| *C04E12.2* | WBGene00015433 | protein_coding_gene | 3.0 | 7.845 | 3.9E-30 |
| *F09F7.6* | WBGene00017303 | protein_coding_gene | 3.0 | 7.803 | 4.0E-12 |
| *R03H10.7* | WBGene00019859 | protein_coding_gene | 3.0 | 7.767 | 6.5E-08 |
| *srh-20* | WBGene00005245 | protein_coding_gene | 3.0 | 7.758 | 3.8E-02 |
| *linc-69* | WBGene00219644 | lincRNA_gene | 3.0 | 7.742 | 5.2E-03 |
| *F31F7.1* | WBGene00017964 | protein_coding_gene | 3.0 | 7.733 | 4.4E-28 |
| *spp-7* | WBGene00004992 | protein_coding_gene | 2.9 | 7.720 | 4.1E-03 |
| *F46F5.15* | WBGene00018506 | protein_coding_gene | 2.9 | 7.665 | 3.5E-03 |
| *K03B4.8* | WBGene00219235 | protein_coding_gene | 2.9 | 7.650 | 5.2E-03 |
| *col-172* | WBGene00000745 | protein_coding_gene | 2.9 | 7.636 | 4.8E-07 |
| *C03F11.2* | WBGene00015388 | protein_coding_gene | 2.9 | 7.599 | 1.0E-06 |
| *C14A4.9* | WBGene00007560 | protein_coding_gene | 2.9 | 7.593 | 1.4E-06 |
| *W02D7.8* | WBGene00020943 | protein_coding_gene | 2.9 | 7.547 | 7.1E-10 |
| *gcy-5* | WBGene00001532 | protein_coding_gene | 2.9 | 7.547 | 1.4E-04 |
| *grl-2* | WBGene00001711 | protein_coding_gene | 2.9 | 7.544 | 7.6E-03 |
| *F41E7.20* | WBGene00219546 | ncRNA_gene | 2.9 | 7.528 | 9.0E-05 |
| *D1014.5* | WBGene00017018 | protein_coding_gene | 2.9 | 7.525 | 8.2E-11 |
| *T12A7.6* | WBGene00011727 | protein_coding_gene | 2.9 | 7.523 | 1.9E-09 |
| *F14D2.19* | WBGene00206522 | protein_coding_gene | 2.9 | 7.522 | 1.0E-15 |
| *Y37F4.8* | WBGene00044289 | protein_coding_gene | 2.9 | 7.521 | 9.8E-07 |
| *Y5H2B.1* | WBGene00021164 | protein_coding_gene | 2.9 | 7.521 | 1.6E-03 |
| *Y43F8B.14* | WBGene00012824 | protein_coding_gene | 2.9 | 7.507 | 6.4E-09 |
| *sek-5* | WBGene00018034 | protein_coding_gene | 2.9 | 7.505 | 2.7E-02 |
| *K02A6.2* | WBGene00019293 | protein_coding_gene | 2.9 | 7.485 | 1.4E-05 |
| *igdb-3* | WBGene00020691 | protein_coding_gene | 2.9 | 7.480 | 1.5E-10 |
| *pals-18* | WBGene00022353 | protein_coding_gene | 2.9 | 7.472 | 1.3E-18 |
| *lgc-24* | WBGene00045385 | protein_coding_gene | 2.9 | 7.468 | 1.6E-02 |
| *F09C6.11* | WBGene00044181 | protein_coding_gene | 2.9 | 7.464 | 5.4E-08 |
| *C17H11.1* | WBGene00015921 | protein_coding_gene | 2.9 | 7.453 | 4.3E-07 |
| *C07B5.3* | WBGene00007408 | protein_coding_gene | 2.9 | 7.450 | 6.1E-07 |
| *C25E10.13* | WBGene00023431 | protein_coding_gene | 2.9 | 7.443 | 9.3E-04 |
| *T08G5.1* | WBGene00011622 | protein_coding_gene | 2.9 | 7.430 | 5.8E-04 |
| *fbxa-30* | WBGene00013837 | protein_coding_gene | 2.9 | 7.412 | 2.8E-04 |
| *srt-65* | WBGene00015368 | protein_coding_gene | 2.9 | 7.395 | 1.9E-02 |
| *nspb-8* | WBGene00007250 | protein_coding_gene | 2.9 | 7.388 | 1.2E-05 |
| *F28F9.2* | WBGene00017906 | protein_coding_gene | 2.9 | 7.385 | 1.2E-02 |
| *F49E10.2* | WBGene00018640 | protein_coding_gene | 2.9 | 7.383 | 4.7E-06 |
| *T12A7.7* | WBGene00011728 | pseudogene | 2.9 | 7.352 | 9.4E-03 |
| *Y38H6C.8* | WBGene00012621 | protein_coding_gene | 2.9 | 7.323 | 4.2E-03 |

| | | | | | |
|---|---|---|---|---|---|
| *linc-43* | WBGene00219579 | lincRNA_gene | 2.9 | 7.317 | 1.6E-02 |
| *irld-32* | WBGene00017497 | protein_coding_gene | 2.9 | 7.284 | 3.2E-02 |
| *glc-1* | WBGene00001591 | protein_coding_gene | 2.9 | 7.281 | 3.5E-14 |
| *B0310.3* | WBGene00015139 | protein_coding_gene | 2.9 | 7.277 | 9.2E-10 |
| *T16G1.2* | WBGene00011796 | protein_coding_gene | 2.9 | 7.248 | 7.4E-08 |
| *C34D10.4* | WBGene00201065 | ncRNA_gene | 2.9 | 7.233 | 3.0E-02 |
| *fbxa-174* | WBGene00008015 | protein_coding_gene | 2.9 | 7.231 | 1.4E-03 |
| *K07E8.4* | WBGene00023320 | pseudogene | 2.9 | 7.214 | 4.2E-02 |
| *C08E8.3* | WBGene00007439 | protein_coding_gene | 2.8 | 7.197 | 2.7E-03 |
| *F32F2.2* | WBGene00194792 | antisense_lncRNA_gene | 2.8 | 7.190 | 2.6E-02 |
| *fbxa-157* | WBGene00007393 | protein_coding_gene | 2.8 | 7.184 | 1.0E-04 |
| *T08G5.14* | WBGene00077572 | pseudogene | 2.8 | 7.169 | 2.1E-02 |
| *srd-23* | WBGene00005101 | protein_coding_gene | 2.8 | 7.167 | 2.1E-04 |
| *gcy-17* | WBGene00001542 | protein_coding_gene | 2.8 | 7.165 | 8.7E-03 |
| *C25F6.7* | WBGene00016106 | protein_coding_gene | 2.8 | 7.164 | 8.1E-13 |
| *srsx-32* | WBGene00017615 | protein_coding_gene | 2.8 | 7.155 | 3.1E-02 |
| *Y43F8C.3* | WBGene00012825 | protein_coding_gene | 2.8 | 7.151 | 1.6E-14 |
| *K08F9.3* | WBGene00010686 | protein_coding_gene | 2.8 | 7.141 | 1.5E-06 |
| *C35E7.12* | WBGene00271779 | protein_coding_gene | 2.8 | 7.139 | 1.4E-02 |
| *F59F5.4* | WBGene00010343 | protein_coding_gene | 2.8 | 7.130 | 1.1E-02 |
| *math-8* | WBGene00015837 | protein_coding_gene | 2.8 | 7.129 | 1.4E-02 |
| *clec-39* | WBGene00012022 | protein_coding_gene | 2.8 | 7.120 | 1.2E-02 |
| *zig-4* | WBGene00006981 | protein_coding_gene | 2.8 | 7.120 | 1.6E-11 |
| *tsp-17* | WBGene00006643 | protein_coding_gene | 2.8 | 7.099 | 1.2E-06 |
| *dhhc-11* | WBGene00020694 | protein_coding_gene | 2.8 | 7.096 | 6.6E-04 |
| *T14A8.2* | WBGene00020497 | protein_coding_gene | 2.8 | 7.064 | 1.2E-17 |
| *F07E5.7* | WBGene00017212 | protein_coding_gene | 2.8 | 7.054 | 1.8E-02 |
| *W08G11.1* | WBGene00012346 | protein_coding_gene | 2.8 | 7.054 | 7.9E-05 |
| *Y43F8B.24* | WBGene00195176 | protein_coding_gene | 2.8 | 7.047 | 3.8E-12 |
| *R01E6.2* | WBGene00010968 | protein_coding_gene | 2.8 | 7.038 | 7.4E-04 |
| *trx-3* | WBGene00019717 | protein_coding_gene | 2.8 | 7.026 | 2.0E-06 |
| *C07B5.4* | WBGene00007409 | protein_coding_gene | 2.8 | 7.024 | 1.8E-14 |
| *srd-12* | WBGene00005090 | protein_coding_gene | 2.8 | 7.021 | 3.3E-02 |
| *Y11D7A.5* | WBGene00012429 | protein_coding_gene | 2.8 | 7.016 | 2.1E-05 |
| *srh-271* | WBGene00005477 | protein_coding_gene | 2.8 | 6.997 | 1.8E-02 |
| *C17B7.5* | WBGene00015879 | protein_coding_gene | 2.8 | 6.979 | 9.8E-11 |
| *nhr-21* | WBGene00003620 | protein_coding_gene | 2.8 | 6.958 | 4.5E-19 |
| *T07D3.6* | WBGene00020309 | pseudogene | 2.8 | 6.954 | 3.2E-05 |
| *R10F2.5* | WBGene00019996 | protein_coding_gene | 2.8 | 6.936 | 1.1E-02 |
| *ceh-9* | WBGene00000434 | protein_coding_gene | 2.8 | 6.932 | 4.3E-03 |
| *ZC449.1* | WBGene00022611 | protein_coding_gene | 2.8 | 6.908 | 3.2E-05 |
| *T20D4.3* | WBGene00020609 | protein_coding_gene | 2.8 | 6.880 | 3.1E-10 |
| *ift-20* | WBGene00022465 | protein_coding_gene | 2.8 | 6.876 | 7.5E-04 |
| *srh-72* | WBGene00005293 | protein_coding_gene | 2.8 | 6.865 | 3.1E-02 |
| *F09C6.3* | WBGene00008617 | protein_coding_gene | 2.8 | 6.825 | 1.2E-04 |
| *C30A5.10* | WBGene00016242 | protein_coding_gene | 2.8 | 6.824 | 1.7E-03 |
| *T21B10.9* | WBGene00196442 | ncRNA_gene | 2.8 | 6.794 | 5.0E-02 |
| *F47B10.3* | WBGene00009814 | protein_coding_gene | 2.8 | 6.794 | 6.2E-03 |
| *F21C10.19* | WBGene00271808 | protein_coding_gene | 2.8 | 6.791 | 1.7E-02 |
| *ugt-52* | WBGene00018931 | protein_coding_gene | 2.8 | 6.775 | 9.3E-08 |
| *grd-7* | WBGene00001696 | protein_coding_gene | 2.8 | 6.764 | 3.8E-02 |
| *M04C3.5* | WBGene00206418 | protein_coding_gene | 2.8 | 6.760 | 1.7E-13 |
| *K08B12.1* | WBGene00019520 | protein_coding_gene | 2.8 | 6.759 | 1.0E-06 |
| *C05C9.2* | WBGene00007326 | protein_coding_gene | 2.8 | 6.757 | 2.2E-02 |
| *thn-2* | WBGene00009215 | protein_coding_gene | 2.8 | 6.749 | 1.2E-08 |
| *fbxa-99* | WBGene00009490 | protein_coding_gene | 2.8 | 6.744 | 2.6E-05 |
| *sups-1* | WBGene00017716 | protein_coding_gene | 2.8 | 6.737 | 1.1E-09 |
| *F38H4.15* | WBGene00200459 | ncRNA_gene | 2.8 | 6.736 | 2.9E-02 |
| *srd-36* | WBGene00005114 | protein_coding_gene | 2.8 | 6.728 | 3.1E-02 |
| *F10D2.16* | WBGene00271437 | protein_coding_gene | 2.7 | 6.725 | 2.5E-02 |
| *F26F2.10* | WBGene00138719 | protein_coding_gene | 2.7 | 6.721 | 6.6E-04 |
| *C54G7.1* | WBGene00016933 | protein_coding_gene | 2.7 | 6.707 | 7.4E-03 |
| *mltn-12* | WBGene00008277 | protein_coding_gene | 2.7 | 6.702 | 2.1E-08 |
| *nhr-290* | WBGene00016233 | protein_coding_gene | 2.7 | 6.687 | 2.6E-02 |
| *F16A11.5* | WBGene00023472 | ncRNA_gene | 2.7 | 6.668 | 1.3E-03 |
| *grl-7* | WBGene00001716 | protein_coding_gene | 2.7 | 6.666 | 9.4E-08 |
| *cbn-1* | WBGene00019285 | protein_coding_gene | 2.7 | 6.663 | 2.9E-07 |
| *bah-1* | WBGene00050875 | protein_coding_gene | 2.7 | 6.658 | 1.4E-04 |
| *M01A8.1* | WBGene00010795 | protein_coding_gene | 2.7 | 6.657 | 2.4E-07 |
| *F14D7.4* | WBGene00008791 | pseudogene | 2.7 | 6.633 | 8.1E-04 |
| *cyp-34A9* | WBGene00015044 | protein_coding_gene | 2.7 | 6.624 | 1.9E-04 |
| *F28G4.3* | WBGene00009228 | protein_coding_gene | 2.7 | 6.624 | 4.5E-02 |
| *ZC123.1* | WBGene00022517 | protein_coding_gene | 2.7 | 6.607 | 1.1E-07 |
| *T13C5.2* | WBGene00020483 | protein_coding_gene | 2.7 | 6.603 | 4.4E-02 |
| *ttr-22* | WBGene00011450 | protein_coding_gene | 2.7 | 6.595 | 1.6E-02 |
| *nspc-19* | WBGene00008360 | protein_coding_gene | 2.7 | 6.589 | 1.0E-26 |
| *oac-56* | WBGene00013450 | protein_coding_gene | 2.7 | 6.579 | 1.8E-02 |
| *K10G4.13* | WBGene00077629 | pseudogene | 2.7 | 6.578 | 2.9E-02 |
| *sre-22* | WBGene00009462 | protein_coding_gene | 2.7 | 6.576 | 3.6E-02 |
| *F56C4.1* | WBGene00010144 | protein_coding_gene | 2.7 | 6.563 | 3.5E-04 |

| | | | | | |
|---|---|---|---|---|---|
| *F17C8.12* | WBGene00200212 | ncRNA_gene | 2.7 | 6.558 | 4.0E-02 |
| *srd-33* | WBGene00005111 | protein_coding_gene | 2.7 | 6.553 | 1.2E-02 |
| *mec-3* | WBGene00003167 | protein_coding_gene | 2.7 | 6.539 | 4.6E-08 |
| *C26B9.7* | WBGene00016136 | protein_coding_gene | 2.7 | 6.534 | 3.0E-03 |
| *F53B2.8* | WBGene00009957 | protein_coding_gene | 2.7 | 6.523 | 2.3E-04 |
| *K08D12.8* | WBGene00195004 | protein_coding_gene | 2.7 | 6.508 | 1.3E-03 |
| *F59F5.3* | WBGene00010342 | protein_coding_gene | 2.7 | 6.500 | 1.9E-11 |
| *srx-1* | WBGene00005892 | protein_coding_gene | 2.7 | 6.495 | 2.8E-02 |
| *nhr-127* | WBGene00003717 | protein_coding_gene | 2.7 | 6.485 | 1.5E-03 |
| *F35F10.14* | WBGene00018063 | protein_coding_gene | 2.7 | 6.474 | 4.4E-02 |
| *srv-13* | WBGene00005724 | protein_coding_gene | 2.7 | 6.469 | 3.4E-02 |
| *ZK1320.5* | WBGene00014255 | protein_coding_gene | 2.7 | 6.469 | 1.7E-02 |
| *F14H8.4* | WBGene00008834 | protein_coding_gene | 2.7 | 6.464 | 4.6E-03 |
| *Y45F3A.8* | WBGene00012863 | protein_coding_gene | 2.7 | 6.463 | 3.5E-03 |
| *F36D1.7* | WBGene00009466 | protein_coding_gene | 2.7 | 6.461 | 1.3E-08 |
| *F16G10.15* | WBGene00017529 | pseudogene | 2.7 | 6.444 | 9.3E-05 |
| *F22B8.2* | WBGene00014741 | pseudogene | 2.7 | 6.444 | 1.6E-02 |
| *CC8.3* | WBGene00195049 | protein_coding_gene | 2.7 | 6.439 | 3.2E-02 |
| *Y87G2A.16* | WBGene00013602 | protein_coding_gene | 2.7 | 6.421 | 7.7E-13 |
| *aagr-1* | WBGene00017071 | protein_coding_gene | 2.7 | 6.406 | 2.8E-12 |
| *irld-38* | WBGene00019106 | protein_coding_gene | 2.7 | 6.404 | 3.7E-02 |
| *T06G6.12* | WBGene00011551 | protein_coding_gene | 2.7 | 6.400 | 2.0E-03 |
| *dhs-16* | WBGene00000979 | protein_coding_gene | 2.7 | 6.379 | 2.0E-04 |
| *oac-49* | WBGene00012068 | protein_coding_gene | 2.7 | 6.378 | 3.4E-03 |
| *W03F11.5* | WBGene00021008 | protein_coding_gene | 2.7 | 6.372 | 3.9E-07 |
| *Y75B12B.3* | WBGene00013568 | protein_coding_gene | 2.7 | 6.348 | 1.4E-15 |
| *F20D6.2* | WBGene00017634 | protein_coding_gene | 2.7 | 6.344 | 2.4E-03 |
| *elt-4* | WBGene00001252 | protein_coding_gene | 2.7 | 6.333 | 8.5E-07 |
| *osr-1* | WBGene00016329 | protein_coding_gene | 2.7 | 6.326 | 1.3E-05 |
| *ZC13.10* | WBGene00044557 | protein_coding_gene | 2.7 | 6.322 | 1.9E-02 |
| *math-21* | WBGene00016556 | protein_coding_gene | 2.7 | 6.310 | 3.3E-04 |
| *F09C6.12* | WBGene00044182 | protein_coding_gene | 2.7 | 6.305 | 1.6E-13 |
| *K08E7.5* | WBGene00010673 | protein_coding_gene | 2.7 | 6.298 | 1.8E-06 |
| *Y105C5B.25* | WBGene00013662 | protein_coding_gene | 2.7 | 6.294 | 6.7E-04 |
| *srb-18* | WBGene00016925 | protein_coding_gene | 2.7 | 6.294 | 3.7E-02 |
| *C06A12.19* | WBGene00255702 | protein_coding_gene | 2.7 | 6.291 | 1.4E-02 |
| *C36C9.5* | WBGene00016488 | protein_coding_gene | 2.7 | 6.285 | 9.7E-03 |
| *ZK970.7* | WBGene00014173 | protein_coding_gene | 2.6 | 6.274 | 1.8E-07 |
| *mab-5* | WBGene00003102 | protein_coding_gene | 2.6 | 6.273 | 3.3E-12 |
| *btb-21* | WBGene00018540 | pseudogene | 2.6 | 6.268 | 5.0E-11 |
| *ZC84.7* | WBGene00013850 | protein_coding_gene | 2.6 | 6.258 | 2.4E-02 |
| *K10C2.2* | WBGene00019618 | protein_coding_gene | 2.6 | 6.238 | 2.0E-02 |
| *Y43F8B.23* | WBGene00194815 | protein_coding_gene | 2.6 | 6.237 | 2.8E-05 |
| *fezf-1* | WBGene00012639 | protein_coding_gene | 2.6 | 6.233 | 1.1E-02 |
| *F32H2.8* | WBGene00009345 | protein_coding_gene | 2.6 | 6.227 | 2.6E-02 |
| *F47B8.5* | WBGene00009806 | protein_coding_gene | 2.6 | 6.222 | 8.2E-05 |
| *clec-180* | WBGene00017991 | protein_coding_gene | 2.6 | 6.215 | 5.5E-14 |
| *fbxc-20* | WBGene00018352 | protein_coding_gene | 2.6 | 6.211 | 4.4E-07 |
| *ZC132.11* | WBGene00269432 | protein_coding_gene | 2.6 | 6.193 | 8.0E-03 |
| *hhat-1* | WBGene00013852 | protein_coding_gene | 2.6 | 6.188 | 1.2E-07 |
| *F55C10.4* | WBGene00010108 | protein_coding_gene | 2.6 | 6.187 | 1.7E-02 |
| *Y39H10B.3* | WBGene00194902 | protein_coding_gene | 2.6 | 6.173 | 5.4E-04 |
| *T23G4.2* | WBGene00011958 | protein_coding_gene | 2.6 | 6.166 | 1.3E-02 |
| *Y40B10A.5* | WBGene00021490 | protein_coding_gene | 2.6 | 6.165 | 3.4E-02 |
| *F55A4.3* | WBGene00018855 | protein_coding_gene | 2.6 | 6.145 | 1.2E-02 |
| *C48B4.3* | WBGene00008168 | protein_coding_gene | 2.6 | 6.144 | 6.3E-08 |
| *chil-4* | WBGene00007469 | protein_coding_gene | 2.6 | 6.143 | 1.4E-03 |
| *ZK686.6* | WBGene00044799 | protein_coding_gene | 2.6 | 6.136 | 6.6E-04 |
| *Y7A5A.8* | WBGene00012414 | protein_coding_gene | 2.6 | 6.134 | 2.0E-02 |
| *H43I07.4* | WBGene00269429 | protein_coding_gene | 2.6 | 6.126 | 2.4E-02 |
| *srh-28* | WBGene00005251 | protein_coding_gene | 2.6 | 6.122 | 2.8E-02 |
| *K02F6.4* | WBGene00019338 | protein_coding_gene | 2.6 | 6.122 | 5.0E-02 |
| *K10D6.2* | WBGene00010742 | protein_coding_gene | 2.6 | 6.110 | 1.7E-17 |
| *C38C3.6* | WBGene00016514 | protein_coding_gene | 2.6 | 6.103 | 1.3E-02 |
| *W04A8.4* | WBGene00012239 | protein_coding_gene | 2.6 | 6.093 | 5.9E-15 |
| *F12E12.12* | WBGene00086558 | protein_coding_gene | 2.6 | 6.079 | 4.8E-03 |
| *B0250.12* | WBGene00050927 | pseudogene | 2.6 | 6.072 | 1.5E-10 |
| *F20C5.5* | WBGene00008970 | protein_coding_gene | 2.6 | 6.066 | 5.4E-12 |
| *fbxa-29* | WBGene00013836 | protein_coding_gene | 2.6 | 6.054 | 8.9E-11 |
| *ztf-30* | WBGene00015523 | protein_coding_gene | 2.6 | 6.039 | 3.9E-06 |
| *pqn-32* | WBGene00004120 | protein_coding_gene | 2.6 | 6.028 | 1.7E-07 |
| *nspb-11* | WBGene00009398 | protein_coding_gene | 2.6 | 6.025 | 1.2E-04 |
| *nhr-150* | WBGene00007367 | protein_coding_gene | 2.6 | 5.999 | 3.1E-05 |
| *slc-28.1* | WBGene00017867 | protein_coding_gene | 2.6 | 5.993 | 1.1E-08 |
| *srz-24* | WBGene00017113 | protein_coding_gene | 2.6 | 5.991 | 2.3E-02 |
| *ZK994.6* | WBGene00022842 | protein_coding_gene | 2.6 | 5.967 | 7.8E-04 |
| *F21C10.17* | WBGene00235309 | pseudogene | 2.6 | 5.966 | 1.4E-02 |
| *F25D1.2* | WBGene00009107 | protein_coding_gene | 2.6 | 5.962 | 2.4E-02 |
| *nlp-30* | WBGene00003768 | protein_coding_gene | 2.6 | 5.962 | 1.2E-06 |
| *nhr-112* | WBGene00003702 | protein_coding_gene | 2.6 | 5.950 | 3.1E-13 |

| | | | | | |
|---|---|---|---|---|---|
| *C45G9.14* | WBGene00044409 | protein_coding_gene | 2.6 | 5.947 | 4.0E-02 |
| *siah-1* | WBGene00021369 | protein_coding_gene | 2.6 | 5.939 | 2.0E-09 |
| *gbh-2* | WBGene00001523 | protein_coding_gene | 2.6 | 5.938 | 1.4E-10 |
| *clec-24* | WBGene00009856 | protein_coding_gene | 2.6 | 5.929 | 4.4E-03 |
| *ZK177.3* | WBGene00022671 | protein_coding_gene | 2.6 | 5.921 | 3.4E-02 |
| *C39D10.5* | WBGene00016532 | protein_coding_gene | 2.6 | 5.919 | 3.9E-03 |
| *tsp-2* | WBGene00006628 | protein_coding_gene | 2.6 | 5.904 | 6.9E-08 |
| *Y14H12A.2* | WBGene00044382 | protein_coding_gene | 2.6 | 5.902 | 2.5E-04 |
| *T28F4.5* | WBGene00012140 | protein_coding_gene | 2.6 | 5.894 | 4.9E-12 |
| *fbxa-52* | WBGene00017225 | protein_coding_gene | 2.6 | 5.890 | 6.2E-03 |
| *F33A8.7* | WBGene00009355 | protein_coding_gene | 2.6 | 5.885 | 4.4E-09 |
| *fbxa-3* | WBGene00020363 | protein_coding_gene | 2.6 | 5.885 | 2.9E-09 |
| *C39B10.7* | WBGene00164973 | protein_coding_gene | 2.6 | 5.874 | 1.6E-05 |
| *M04C7.4* | WBGene00010855 | protein_coding_gene | 2.6 | 5.872 | 3.8E-02 |
| *E02H4.5* | WBGene00008463 | protein_coding_gene | 2.6 | 5.871 | 2.9E-04 |
| *chil-20* | WBGene00011163 | protein_coding_gene | 2.5 | 5.854 | 1.8E-02 |
| *K08D10.14* | WBGene00219216 | protein_coding_gene | 2.5 | 5.841 | 5.1E-20 |
| *che-11* | WBGene00000490 | protein_coding_gene | 2.5 | 5.838 | 9.4E-10 |
| *col-131* | WBGene00000705 | protein_coding_gene | 2.5 | 5.836 | 3.1E-03 |
| *T21D12.14* | WBGene00045177 | protein_coding_gene | 2.5 | 5.828 | 5.7E-06 |
| *srt-40* | WBGene00021169 | protein_coding_gene | 2.5 | 5.819 | 4.5E-02 |
| *linc-38* | WBGene00219415 | lincRNA_gene | 2.5 | 5.804 | 2.6E-02 |
| *frpr-15* | WBGene00010735 | protein_coding_gene | 2.5 | 5.795 | 1.2E-06 |
| *cyp-23A1* | WBGene00015135 | protein_coding_gene | 2.5 | 5.777 | 9.5E-05 |
| *K02E7.10* | WBGene00019314 | protein_coding_gene | 2.5 | 5.762 | 6.0E-07 |
| *ubc-8* | WBGene00006705 | protein_coding_gene | 2.5 | 5.761 | 3.3E-10 |
| *str-130* | WBGene00006179 | protein_coding_gene | 2.5 | 5.759 | 4.7E-02 |
| *ZK856.14* | WBGene00044020 | protein_coding_gene | 2.5 | 5.754 | 1.9E-02 |
| *clec-33* | WBGene00009859 | protein_coding_gene | 2.5 | 5.752 | 3.9E-03 |
| *F10E9.12* | WBGene00017361 | protein_coding_gene | 2.5 | 5.749 | 1.2E-16 |
| *Y55D5A.1* | WBGene00021915 | protein_coding_gene | 2.5 | 5.747 | 1.7E-10 |
| *W04C9.8* | WBGene00235387 | ncRNA_gene | 2.5 | 5.745 | 6.0E-06 |
| *C06G4.6* | WBGene00044446 | protein_coding_gene | 2.5 | 5.737 | 1.3E-02 |
| *nhr-263* | WBGene00017367 | protein_coding_gene | 2.5 | 5.735 | 2.7E-06 |
| *osm-12* | WBGene00003892 | protein_coding_gene | 2.5 | 5.733 | 3.3E-06 |
| *sodh-2* | WBGene00010791 | protein_coding_gene | 2.5 | 5.732 | 3.7E-03 |
| *ZC434.10* | WBGene00220261 | protein_coding_gene | 2.5 | 5.720 | 4.3E-03 |
| *tax-4* | WBGene00006526 | protein_coding_gene | 2.5 | 5.718 | 3.4E-03 |
| *Y50E8A.1* | WBGene00013046 | protein_coding_gene | 2.5 | 5.713 | 3.8E-11 |
| *sri-30* | WBGene00005542 | protein_coding_gene | 2.5 | 5.707 | 4.3E-02 |
| *R07B7.12* | WBGene00011096 | protein_coding_gene | 2.5 | 5.680 | 2.9E-02 |
| *F58G6.8* | WBGene00010278 | protein_coding_gene | 2.5 | 5.675 | 2.2E-02 |
| *F47D12.6* | WBGene00018562 | protein_coding_gene | 2.5 | 5.662 | 1.4E-11 |
| *clec-62* | WBGene00009393 | protein_coding_gene | 2.5 | 5.661 | 1.0E-13 |
| *C01G10.17* | WBGene00044658 | protein_coding_gene | 2.5 | 5.651 | 1.2E-02 |
| *T04C4.3* | WBGene00235277 | protein_coding_gene | 2.5 | 5.622 | 5.5E-03 |
| *nspc-18* | WBGene00008358 | protein_coding_gene | 2.5 | 5.618 | 4.2E-14 |
| *fkh-10* | WBGene00001442 | protein_coding_gene | 2.5 | 5.592 | 1.9E-03 |
| *F30A10.2* | WBGene00009261 | protein_coding_gene | 2.5 | 5.584 | 2.9E-04 |
| *fbxa-86* | WBGene00021179 | protein_coding_gene | 2.5 | 5.584 | 4.8E-02 |
| *srh-30* | WBGene00005253 | protein_coding_gene | 2.5 | 5.574 | 3.9E-02 |
| *clec-28* | WBGene00009857 | protein_coding_gene | 2.5 | 5.568 | 1.4E-04 |
| *T08E11.1* | WBGene00020357 | protein_coding_gene | 2.5 | 5.565 | 3.6E-03 |
| *F11A5.5* | WBGene00008673 | protein_coding_gene | 2.5 | 5.563 | 1.6E-02 |
| *C34C12.7* | WBGene00007926 | protein_coding_gene | 2.5 | 5.557 | 1.6E-17 |
| *M03B6.5* | WBGene00010837 | protein_coding_gene | 2.5 | 5.539 | 2.2E-04 |
| *sre-4* | WBGene00011745 | protein_coding_gene | 2.5 | 5.538 | 2.4E-03 |
| *K08E7.6* | WBGene00010674 | protein_coding_gene | 2.5 | 5.531 | 2.1E-06 |
| *R13H4.7* | WBGene00011265 | protein_coding_gene | 2.5 | 5.530 | 1.4E-03 |
| *T11F9.10* | WBGene00011712 | protein_coding_gene | 2.5 | 5.528 | 8.1E-04 |
| *C52A11.3* | WBGene00008257 | protein_coding_gene | 2.5 | 5.523 | 4.2E-02 |
| *T28H10.3* | WBGene00012144 | protein_coding_gene | 2.5 | 5.511 | 6.2E-08 |
| *Y11D7A.8* | WBGene00012432 | protein_coding_gene | 2.5 | 5.509 | 1.5E-04 |
| *Y17D7C.4* | WBGene00050903 | pseudogene | 2.5 | 5.501 | 4.9E-05 |
| *grl-15* | WBGene00001724 | protein_coding_gene | 2.5 | 5.499 | 2.1E-05 |
| *irld-30* | WBGene00017493 | protein_coding_gene | 2.5 | 5.498 | 2.7E-02 |
| *F01D5.4* | WBGene00014730 | pseudogene | 2.5 | 5.498 | 3.9E-02 |
| *M04C3.2* | WBGene00010851 | protein_coding_gene | 2.5 | 5.497 | 9.9E-15 |
| *dhs-15* | WBGene00000978 | protein_coding_gene | 2.5 | 5.497 | 7.7E-06 |
| *nlp-48* | WBGene00008002 | protein_coding_gene | 2.5 | 5.495 | 6.8E-05 |
| *fat-3* | WBGene00001395 | protein_coding_gene | 2.5 | 5.494 | 1.6E-19 |
| *lpr-4* | WBGene00012257 | protein_coding_gene | 2.5 | 5.488 | 2.3E-07 |
| *F26A10.2* | WBGene00017814 | protein_coding_gene | 2.5 | 5.487 | 9.1E-08 |
| *Y113G7A.12* | WBGene00013749 | protein_coding_gene | 2.5 | 5.465 | 1.1E-03 |
| *linc-113* | WBGene00219598 | lincRNA_gene | 2.4 | 5.460 | 3.5E-02 |
| *W05H9.1* | WBGene00021048 | protein_coding_gene | 2.4 | 5.457 | 1.1E-08 |
| *C15H11.11* | WBGene00007619 | protein_coding_gene | 2.4 | 5.452 | 1.9E-10 |
| *lpr-5* | WBGene00012256 | protein_coding_gene | 2.4 | 5.438 | 4.9E-08 |
| *C54E4.12* | WBGene00235274 | protein_coding_gene | 2.4 | 5.433 | 1.8E-03 |
| *Y6G8.9* | WBGene00050974 | pseudogene | 2.4 | 5.431 | 9.8E-04 |

| | | | | | |
|---|---|---|---|---|---|
| *srh-21* | WBGene00005246 | protein_coding_gene | 2.4 | 5.425 | 3.7E-02 |
| *C03A3.3* | WBGene00007268 | protein_coding_gene | 2.4 | 5.417 | 3.8E-02 |
| *linc-56* | WBGene00219650 | lincRNA_gene | 2.4 | 5.411 | 2.7E-04 |
| *unc-129* | WBGene00006852 | protein_coding_gene | 2.4 | 5.410 | 2.1E-22 |
| *wrt-6* | WBGene00006952 | protein_coding_gene | 2.4 | 5.410 | 1.7E-07 |
| *C30G12.4* | WBGene00016276 | protein_coding_gene | 2.4 | 5.408 | 9.1E-04 |
| *F38B6.7* | WBGene00018176 | protein_coding_gene | 2.4 | 5.404 | 4.0E-02 |
| *F44A2.7* | WBGene00018406 | protein_coding_gene | 2.4 | 5.401 | 4.1E-03 |
| *sra-11* | WBGene00005037 | protein_coding_gene | 2.4 | 5.396 | 4.0E-02 |
| *K01A2.4* | WBGene00019279 | protein_coding_gene | 2.4 | 5.392 | 3.3E-09 |
| *D2045.8* | WBGene00008425 | protein_coding_gene | 2.4 | 5.375 | 7.8E-10 |
| *C54D2.1* | WBGene00016915 | protein_coding_gene | 2.4 | 5.367 | 2.4E-08 |
| *srd-65* | WBGene00005142 | protein_coding_gene | 2.4 | 5.359 | 2.9E-02 |
| *nstp-3* | WBGene00022577 | protein_coding_gene | 2.4 | 5.354 | 7.6E-04 |
| *T26C12.2* | WBGene00020832 | protein_coding_gene | 2.4 | 5.352 | 6.9E-04 |
| *mks-3* | WBGene00018042 | protein_coding_gene | 2.4 | 5.352 | 7.8E-05 |
| *C04E12.4* | WBGene00015434 | protein_coding_gene | 2.4 | 5.351 | 1.6E-23 |
| *F54E7.6* | WBGene00018829 | protein_coding_gene | 2.4 | 5.345 | 5.6E-03 |
| *col-65* | WBGene00000641 | protein_coding_gene | 2.4 | 5.340 | 1.6E-06 |
| *grl-3* | WBGene00001712 | protein_coding_gene | 2.4 | 5.340 | 1.3E-02 |
| *C49H3.12* | WBGene00016795 | protein_coding_gene | 2.4 | 5.336 | 1.4E-08 |
| *T04C12.7* | WBGene00011429 | protein_coding_gene | 2.4 | 5.325 | 3.6E-05 |
| *F45E12.6* | WBGene00018476 | protein_coding_gene | 2.4 | 5.321 | 1.0E-03 |
| *R11G1.2* | WBGene00020011 | protein_coding_gene | 2.4 | 5.311 | 5.8E-09 |
| *nhr-147* | WBGene00015395 | protein_coding_gene | 2.4 | 5.307 | 3.5E-06 |
| *F35G2.5* | WBGene00009438 | protein_coding_gene | 2.4 | 5.305 | 6.6E-03 |
| *W03F8.6* | WBGene00020995 | protein_coding_gene | 2.4 | 5.298 | 3.5E-09 |
| *F22F4.9* | WBGene00271606 | protein_coding_gene | 2.4 | 5.292 | 1.2E-08 |
| *grl-5* | WBGene00001714 | protein_coding_gene | 2.4 | 5.279 | 4.6E-06 |
| *F01G10.6* | WBGene00008509 | protein_coding_gene | 2.4 | 5.278 | 1.5E-06 |
| *F21A3.11* | WBGene00219316 | protein_coding_gene | 2.4 | 5.277 | 3.2E-13 |
| *C36A4.10* | WBGene00007970 | protein_coding_gene | 2.4 | 5.273 | 5.7E-03 |
| *glr-2* | WBGene00001613 | protein_coding_gene | 2.4 | 5.269 | 3.3E-10 |
| *C28G1.10* | WBGene00219376 | protein_coding_gene | 2.4 | 5.261 | 7.2E-19 |
| *K10G6.9* | WBGene00206374 | protein_coding_gene | 2.4 | 5.261 | 2.8E-04 |
| *math-37* | WBGene00020080 | protein_coding_gene | 2.4 | 5.245 | 3.6E-11 |
| *C01G10.6* | WBGene00007233 | protein_coding_gene | 2.4 | 5.239 | 3.6E-03 |
| *srd-49* | WBGene00005127 | protein_coding_gene | 2.4 | 5.236 | 1.7E-02 |
| *gcy-13* | WBGene00001539 | protein_coding_gene | 2.4 | 5.232 | 3.7E-02 |
| *C30F8.3* | WBGene00016259 | protein_coding_gene | 2.4 | 5.222 | 7.1E-04 |
| *best-2* | WBGene00007204 | protein_coding_gene | 2.4 | 5.219 | 4.0E-02 |
| *R102.7* | WBGene00011294 | pseudogene | 2.4 | 5.215 | 2.5E-03 |
| *F22E5.21* | WBGene00044363 | protein_coding_gene | 2.4 | 5.214 | 4.7E-04 |
| *Y67D8B.2* | WBGene00022062 | protein_coding_gene | 2.4 | 5.212 | 8.9E-04 |
| *clec-23* | WBGene00011851 | pseudogene | 2.4 | 5.210 | 9.9E-03 |
| *mig-18* | WBGene00003249 | protein_coding_gene | 2.4 | 5.198 | 5.9E-24 |
| *B0207.10* | WBGene00015033 | protein_coding_gene | 2.4 | 5.198 | 2.8E-02 |
| *Y62H9A.16* | WBGene00304791 | protein_coding_gene | 2.4 | 5.190 | 1.1E-03 |
| *cyp-33C6* | WBGene00018264 | protein_coding_gene | 2.4 | 5.188 | 1.6E-03 |
| *F41E6.11* | WBGene00018292 | protein_coding_gene | 2.4 | 5.184 | 6.5E-05 |
| *F22F7.4* | WBGene00017722 | protein_coding_gene | 2.4 | 5.177 | 6.1E-03 |
| *F13G3.3* | WBGene00008763 | protein_coding_gene | 2.4 | 5.167 | 2.9E-05 |
| *ptr-4* | WBGene00004219 | protein_coding_gene | 2.4 | 5.161 | 5.7E-07 |
| *T10D4.15* | WBGene00077522 | protein_coding_gene | 2.4 | 5.152 | 7.1E-03 |
| *C38C3.4* | WBGene00016513 | protein_coding_gene | 2.4 | 5.150 | 3.9E-07 |
| *math-27* | WBGene00018101 | pseudogene | 2.4 | 5.145 | 1.7E-10 |
| *ZC239.17* | WBGene00022575 | protein_coding_gene | 2.4 | 5.132 | 1.9E-04 |
| *T19D7.5* | WBGene00020573 | protein_coding_gene | 2.4 | 5.129 | 4.6E-03 |
| *C18D4.8* | WBGene00007677 | pseudogene | 2.4 | 5.121 | 1.2E-08 |
| *C33D9.5* | WBGene00007899 | pseudogene | 2.4 | 5.119 | 1.5E-14 |
| *F19H6.6* | WBGene00008961 | protein_coding_gene | 2.4 | 5.110 | 3.0E-02 |
| *K08D10.10* | WBGene00019533 | protein_coding_gene | 2.4 | 5.109 | 4.5E-02 |
| *K06B9.6* | WBGene00044478 | protein_coding_gene | 2.4 | 5.103 | 5.8E-05 |
| *R04B5.5* | WBGene00011003 | protein_coding_gene | 2.4 | 5.103 | 1.6E-11 |
| *nhr-116* | WBGene00003706 | protein_coding_gene | 2.4 | 5.101 | 2.1E-11 |
| *spp-1* | WBGene00004986 | protein_coding_gene | 2.4 | 5.101 | 8.1E-13 |
| *F13E9.12* | WBGene00008761 | protein_coding_gene | 2.4 | 5.100 | 2.9E-03 |
| *ZK836.3* | WBGene00014099 | protein_coding_gene | 2.3 | 5.081 | 6.1E-07 |
| *C49C3.3* | WBGene00008193 | pseudogene | 2.3 | 5.080 | 4.8E-02 |
| *slc-28.2* | WBGene00017868 | protein_coding_gene | 2.3 | 5.076 | 9.2E-08 |
| *M03E7.4* | WBGene00019756 | protein_coding_gene | 2.3 | 5.073 | 4.3E-05 |
| *vamp-8* | WBGene00007200 | protein_coding_gene | 2.3 | 5.068 | 1.6E-19 |
| *fbxa-21* | WBGene00022487 | protein_coding_gene | 2.3 | 5.067 | 1.4E-06 |
| *unc-86* | WBGene00006818 | protein_coding_gene | 2.3 | 5.055 | 1.5E-05 |
| *C08E8.4* | WBGene00007440 | protein_coding_gene | 2.3 | 5.053 | 1.5E-03 |
| *C32H11.5* | WBGene00007868 | protein_coding_gene | 2.3 | 5.051 | 2.6E-03 |
| *T01D3.6* | WBGene00011330 | protein_coding_gene | 2.3 | 5.038 | 3.7E-13 |
| *tts-2* | WBGene00006651 | ncRNA_gene | 2.3 | 5.031 | 2.7E-12 |
| *C34E11.2* | WBGene00007935 | protein_coding_gene | 2.3 | 5.017 | 1.4E-07 |
| *Y102A5C.11* | WBGene00014956 | pseudogene | 2.3 | 5.013 | 3.7E-02 |

| | | | | | |
|---|---|---|---|---|---|
| *F55G11.6* | WBGene00010126 | protein_coding_gene | 2.3 | 5.012 | 2.8E-02 |
| *Y17D7C.2* | WBGene00012456 | protein_coding_gene | 2.3 | 5.011 | 4.3E-02 |
| *clec-40* | WBGene00011853 | protein_coding_gene | 2.3 | 5.010 | 3.0E-04 |
| *clec-7* | WBGene00017364 | protein_coding_gene | 2.3 | 5.009 | 9.4E-07 |
| *lgc-18* | WBGene00011360 | protein_coding_gene | 2.3 | 5.003 | 3.7E-02 |
| *T20D4.15* | WBGene00020620 | protein_coding_gene | 2.3 | 5.002 | 9.6E-03 |
| *C49F8.15* | WBGene00304787 | protein_coding_gene | 2.3 | 5.000 | 7.0E-03 |
| *F13A7.12* | WBGene00008727 | protein_coding_gene | 2.3 | 4.998 | 1.6E-03 |
| *ZK697.14* | WBGene00022809 | protein_coding_gene | 2.3 | 4.994 | 2.3E-02 |
| *nhr-287* | WBGene00021521 | pseudogene | 2.3 | 4.992 | 4.6E-02 |
| *str-116* | WBGene00006167 | protein_coding_gene | 2.3 | 4.989 | 4.3E-02 |
| *mec-14* | WBGene00003176 | protein_coding_gene | 2.3 | 4.988 | 4.3E-02 |
| *F42G2.7* | WBGene00044362 | protein_coding_gene | 2.3 | 4.969 | 2.1E-02 |
| *K09B3.1* | WBGene00019545 | protein_coding_gene | 2.3 | 4.963 | 1.1E-02 |
| *K04H4.2* | WBGene00010573 | protein_coding_gene | 2.3 | 4.963 | 1.2E-07 |
| *ttr-44* | WBGene00008341 | protein_coding_gene | 2.3 | 4.963 | 5.0E-17 |
| *osm-6* | WBGene00003886 | protein_coding_gene | 2.3 | 4.959 | 7.9E-05 |
| *R02E4.3* | WBGene00044724 | protein_coding_gene | 2.3 | 4.956 | 2.7E-07 |
| *R13H4.8* | WBGene00011266 | protein_coding_gene | 2.3 | 4.955 | 1.7E-02 |
| *inx-6* | WBGene00002128 | protein_coding_gene | 2.3 | 4.941 | 3.8E-11 |
| *abu-1* | WBGene00000024 | protein_coding_gene | 2.3 | 4.937 | 2.7E-04 |
| *cex-2* | WBGene00023408 | protein_coding_gene | 2.3 | 4.936 | 3.5E-10 |
| *C12D5.4* | WBGene00015707 | protein_coding_gene | 2.3 | 4.932 | 4.3E-03 |
| *C36B1.6* | WBGene00007973 | protein_coding_gene | 2.3 | 4.929 | 1.6E-08 |
| *unc-3* | WBGene00006743 | protein_coding_gene | 2.3 | 4.929 | 1.3E-06 |
| *mam-2* | WBGene00008150 | protein_coding_gene | 2.3 | 4.922 | 2.7E-04 |
| *C35C5.9* | WBGene00007958 | protein_coding_gene | 2.3 | 4.921 | 8.2E-12 |
| *clec-27* | WBGene00013607 | protein_coding_gene | 2.3 | 4.915 | 6.7E-03 |
| *F59B2.11* | WBGene00010313 | pseudogene | 2.3 | 4.908 | 1.6E-04 |
| *ptr-9* | WBGene00004223 | protein_coding_gene | 2.3 | 4.905 | 4.9E-02 |
| *F53B1.4* | WBGene00018737 | protein_coding_gene | 2.3 | 4.903 | 6.2E-09 |
| *clec-55* | WBGene00008597 | protein_coding_gene | 2.3 | 4.901 | 8.7E-06 |
| *ZK1055.4* | WBGene00022845 | protein_coding_gene | 2.3 | 4.900 | 6.0E-03 |
| *K02E11.5* | WBGene00010514 | protein_coding_gene | 2.3 | 4.898 | 1.6E-15 |
| *R07B1.6* | WBGene00011078 | protein_coding_gene | 2.3 | 4.893 | 2.5E-02 |
| *K09F6.13* | WBGene00235164 | protein_coding_gene | 2.3 | 4.881 | 3.1E-03 |
| *R07G3.8* | WBGene00019944 | protein_coding_gene | 2.3 | 4.880 | 2.2E-15 |
| *abu-12* | WBGene00004127 | protein_coding_gene | 2.3 | 4.879 | 3.1E-06 |
| *grh-1* | WBGene00001707 | protein_coding_gene | 2.3 | 4.876 | 6.3E-08 |
| *C30H6.12* | WBGene00044238 | protein_coding_gene | 2.3 | 4.874 | 3.5E-02 |
| *Y43E12A.2* | WBGene00012794 | protein_coding_gene | 2.3 | 4.873 | 2.7E-15 |
| *Y64G10A.7* | WBGene00013416 | protein_coding_gene | 2.3 | 4.872 | 2.5E-13 |
| *F10D7.10* | WBGene00271701 | protein_coding_gene | 2.3 | 4.871 | 1.9E-05 |
| *C07E3.8* | WBGene00007418 | protein_coding_gene | 2.3 | 4.869 | 2.6E-02 |
| *srh-210* | WBGene00005420 | pseudogene | 2.3 | 4.864 | 4.4E-02 |
| *irld-13* | WBGene00019845 | pseudogene | 2.3 | 4.850 | 1.6E-02 |
| *B0348.5* | WBGene00015153 | protein_coding_gene | 2.3 | 4.839 | 7.6E-04 |
| *F26G1.2* | WBGene00017838 | protein_coding_gene | 2.3 | 4.836 | 3.1E-20 |
| *F10G2.2* | WBGene00017363 | protein_coding_gene | 2.3 | 4.836 | 7.6E-03 |
| *F49D11.2* | WBGene00018626 | protein_coding_gene | 2.3 | 4.834 | 1.7E-02 |
| *cnc-7* | WBGene00010005 | protein_coding_gene | 2.3 | 4.832 | 2.6E-05 |
| *Y54G2A.11* | WBGene00021876 | protein_coding_gene | 2.3 | 4.825 | 6.5E-07 |
| *R12C12.10* | WBGene00044395 | protein_coding_gene | 2.3 | 4.818 | 7.7E-03 |
| *Y51B9A.9* | WBGene00013091 | protein_coding_gene | 2.3 | 4.796 | 2.7E-08 |
| *cyn-17* | WBGene00000893 | protein_coding_gene | 2.3 | 4.794 | 4.9E-05 |
| *T10H9.8* | WBGene00044779 | protein_coding_gene | 2.3 | 4.789 | 9.0E-10 |
| *srw-145* | WBGene00011197 | protein_coding_gene | 2.3 | 4.787 | 2.8E-02 |
| *nspc-17* | WBGene00008359 | protein_coding_gene | 2.3 | 4.786 | 1.4E-12 |
| *pha-4* | WBGene00004013 | protein_coding_gene | 2.3 | 4.783 | 1.2E-12 |
| *ncx-5* | WBGene00003570 | protein_coding_gene | 2.3 | 4.780 | 8.9E-03 |
| *F59E11.2* | WBGene00019109 | protein_coding_gene | 2.3 | 4.773 | 3.1E-05 |
| *R07C12.4* | WBGene00019934 | pseudogene | 2.3 | 4.772 | 5.4E-03 |
| *Y69A2AR.11* | WBGene00022082 | pseudogene | 2.3 | 4.758 | 6.7E-04 |
| *Y17D7B.2* | WBGene00012450 | protein_coding_gene | 2.2 | 4.751 | 1.7E-02 |
| *D1086.21* | WBGene00304203 | protein_coding_gene | 2.2 | 4.751 | 3.0E-02 |
| *F54B3.4* | WBGene00175034 | protein_coding_gene | 2.2 | 4.750 | 4.4E-02 |
| *C29F4.3* | WBGene00007809 | protein_coding_gene | 2.2 | 4.750 | 6.5E-03 |
| *F57E7.4* | WBGene00077616 | protein_coding_gene | 2.2 | 4.740 | 1.2E-04 |
| *M03D4.4* | WBGene00019751 | protein_coding_gene | 2.2 | 4.738 | 1.5E-08 |
| *F58E6.13* | WBGene00077697 | protein_coding_gene | 2.2 | 4.735 | 8.5E-07 |
| *F59B2.8* | WBGene00010310 | protein_coding_gene | 2.2 | 4.735 | 1.5E-05 |
| *T01B6.1* | WBGene00020134 | protein_coding_gene | 2.2 | 4.732 | 4.6E-08 |
| *srd-32* | WBGene00005110 | protein_coding_gene | 2.2 | 4.731 | 1.4E-03 |
| *math-45* | WBGene00021772 | protein_coding_gene | 2.2 | 4.731 | 6.5E-10 |
| *fipr-4* | WBGene00007541 | protein_coding_gene | 2.2 | 4.728 | 2.0E-03 |
| *pals-30* | WBGene00013294 | protein_coding_gene | 2.2 | 4.721 | 4.5E-02 |
| *comt-4* | WBGene00021491 | protein_coding_gene | 2.2 | 4.720 | 1.9E-02 |
| *C03B1.2* | WBGene00015373 | protein_coding_gene | 2.2 | 4.718 | 1.8E-02 |
| *nhr-94* | WBGene00003684 | protein_coding_gene | 2.2 | 4.718 | 6.0E-17 |
| *cyp-35A1* | WBGene00015399 | protein_coding_gene | 2.2 | 4.716 | 5.0E-03 |

| | | | | | |
|---|---|---|---|---|---|
| *T05E11.9* | WBGene00014505 | ncRNA_gene | 2.2 | 4.712 | 6.9E-36 |
| *acr-14* | WBGene00000053 | protein_coding_gene | 2.2 | 4.710 | 3.2E-02 |
| *cyp-34A10* | WBGene00015045 | protein_coding_gene | 2.2 | 4.707 | 2.9E-03 |
| *chpf-2* | WBGene00019108 | protein_coding_gene | 2.2 | 4.706 | 1.7E-10 |
| *C32E8.6* | WBGene00016324 | protein_coding_gene | 2.2 | 4.704 | 1.2E-03 |
| *fbxa-10* | WBGene00020450 | protein_coding_gene | 2.2 | 4.701 | 2.8E-16 |
| *mab-3* | WBGene00003100 | protein_coding_gene | 2.2 | 4.699 | 4.6E-06 |
| *ZK829.3* | WBGene00014094 | protein_coding_gene | 2.2 | 4.691 | 6.2E-03 |
| *nmur-1* | WBGene00016747 | protein_coding_gene | 2.2 | 4.687 | 3.3E-04 |
| *pqn-42* | WBGene00004129 | protein_coding_gene | 2.2 | 4.682 | 5.1E-07 |
| *F13D11.3* | WBGene00017428 | protein_coding_gene | 2.2 | 4.678 | 7.6E-07 |
| *T05A8.8* | WBGene00044367 | protein_coding_gene | 2.2 | 4.674 | 3.1E-04 |
| *C55A1.6* | WBGene00008323 | protein_coding_gene | 2.2 | 4.656 | 1.6E-03 |
| *lgc-27* | WBGene00022283 | protein_coding_gene | 2.2 | 4.654 | 6.5E-09 |
| *nspe-2* | WBGene00012604 | protein_coding_gene | 2.2 | 4.654 | 1.3E-02 |
| *osm-5* | WBGene00003885 | protein_coding_gene | 2.2 | 4.652 | 2.2E-06 |
| *cnc-4* | WBGene00000558 | protein_coding_gene | 2.2 | 4.651 | 8.6E-06 |
| *col-151* | WBGene00000724 | protein_coding_gene | 2.2 | 4.651 | 1.9E-02 |
| *nspb-12* | WBGene00017305 | protein_coding_gene | 2.2 | 4.646 | 1.1E-04 |
| *gcy-27* | WBGene00001550 | protein_coding_gene | 2.2 | 4.645 | 3.9E-02 |
| *F53F1.6* | WBGene00009984 | protein_coding_gene | 2.2 | 4.644 | 1.1E-05 |
| *flp-25* | WBGene00010575 | protein_coding_gene | 2.2 | 4.642 | 5.0E-09 |
| *F07C3.9* | WBGene00017197 | protein_coding_gene | 2.2 | 4.640 | 3.5E-08 |
| *ctbp-1* | WBGene00006424 | protein_coding_gene | 2.2 | 4.628 | 2.0E-09 |
| *dsc-1* | WBGene00001096 | protein_coding_gene | 2.2 | 4.627 | 2.5E-02 |
| *Y17D7C.6* | WBGene00194680 | protein_coding_gene | 2.2 | 4.623 | 3.7E-02 |
| *pdl-1* | WBGene00003966 | protein_coding_gene | 2.2 | 4.621 | 1.1E-07 |
| *F56C3.9* | WBGene00018946 | protein_coding_gene | 2.2 | 4.613 | 6.6E-05 |
| *btb-17* | WBGene00018200 | protein_coding_gene | 2.2 | 4.611 | 5.8E-13 |
| *F14F7.5* | WBGene00008812 | protein_coding_gene | 2.2 | 4.602 | 7.9E-03 |
| *C54D10.13* | WBGene00045400 | protein_coding_gene | 2.2 | 4.600 | 9.9E-07 |
| *F21D12.3* | WBGene00017663 | protein_coding_gene | 2.2 | 4.598 | 2.2E-09 |
| *lgc-50* | WBGene00020605 | protein_coding_gene | 2.2 | 4.592 | 2.1E-09 |
| *tba-6* | WBGene00006532 | protein_coding_gene | 2.2 | 4.590 | 2.7E-02 |
| *R07E3.2* | WBGene00011103 | protein_coding_gene | 2.2 | 4.587 | 1.2E-04 |
| *C06E4.2* | WBGene00015531 | protein_coding_gene | 2.2 | 4.566 | 3.6E-05 |
| *ZK470.14* | WBGene00219362 | protein_coding_gene | 2.2 | 4.565 | 7.0E-04 |
| *F11C7.2* | WBGene00017376 | protein_coding_gene | 2.2 | 4.563 | 2.8E-09 |
| *gad-2* | WBGene00015666 | protein_coding_gene | 2.2 | 4.554 | 3.4E-06 |
| *F22B7.1* | WBGene00017692 | protein_coding_gene | 2.2 | 4.551 | 4.9E-10 |
| *F08G2.5* | WBGene00008577 | protein_coding_gene | 2.2 | 4.549 | 1.3E-08 |
| *C50F4.1* | WBGene00008229 | protein_coding_gene | 2.2 | 4.540 | 2.3E-25 |
| *bbs-1* | WBGene00000241 | protein_coding_gene | 2.2 | 4.539 | 1.4E-03 |
| *C05E11.7* | WBGene00015496 | protein_coding_gene | 2.2 | 4.536 | 9.5E-03 |
| *F52A8.3* | WBGene00009916 | protein_coding_gene | 2.2 | 4.535 | 1.7E-04 |
| *sup-26* | WBGene00006331 | protein_coding_gene | 2.2 | 4.531 | 4.5E-10 |
| *W03B1.3* | WBGene00020973 | protein_coding_gene | 2.2 | 4.530 | 5.2E-10 |
| *T03G6.1* | WBGene00020194 | protein_coding_gene | 2.2 | 4.530 | 7.7E-05 |
| *nhr-32* | WBGene00003626 | protein_coding_gene | 2.2 | 4.530 | 6.2E-16 |
| *Y38C1AA.6* | WBGene00021397 | protein_coding_gene | 2.2 | 4.525 | 3.0E-06 |
| *R11D1.13* | WBGene00045517 | protein_coding_gene | 2.2 | 4.525 | 4.8E-04 |
| *del-4* | WBGene00012116 | protein_coding_gene | 2.2 | 4.519 | 3.8E-04 |
| *T03F7.5* | WBGene00011402 | protein_coding_gene | 2.2 | 4.519 | 2.8E-02 |
| *egl-43* | WBGene00001207 | protein_coding_gene | 2.2 | 4.516 | 2.1E-10 |
| *R10E8.8* | WBGene00011213 | protein_coding_gene | 2.2 | 4.514 | 1.6E-05 |
| *twk-23* | WBGene00006676 | protein_coding_gene | 2.2 | 4.502 | 2.0E-03 |
| *Y71G12B.42* | WBGene00220198 | ncRNA_gene | 2.2 | 4.490 | 6.0E-03 |
| *Y69A2AL.2* | WBGene00044468 | protein_coding_gene | 2.2 | 4.488 | 1.6E-09 |
| *F57A8.1* | WBGene00010177 | protein_coding_gene | 2.2 | 4.485 | 1.9E-18 |
| *B0554.7* | WBGene00015260 | protein_coding_gene | 2.2 | 4.483 | 8.3E-03 |
| *nhr-42* | WBGene00003632 | protein_coding_gene | 2.2 | 4.477 | 6.1E-14 |
| *fbxa-23* | WBGene00020459 | protein_coding_gene | 2.2 | 4.468 | 1.2E-02 |
| *nlp-29* | WBGene00003767 | protein_coding_gene | 2.2 | 4.465 | 7.5E-06 |
| *M03E7.2* | WBGene00019754 | protein_coding_gene | 2.2 | 4.462 | 7.2E-04 |
| *F09C3.2* | WBGene00008610 | protein_coding_gene | 2.2 | 4.461 | 3.1E-12 |
| *C01G10.16* | WBGene00007241 | protein_coding_gene | 2.2 | 4.459 | 6.2E-03 |
| *aqp-3* | WBGene00000171 | protein_coding_gene | 2.2 | 4.457 | 1.8E-05 |
| *F30H5.3* | WBGene00017937 | protein_coding_gene | 2.2 | 4.455 | 5.5E-05 |
| *bus-4* | WBGene00044620 | protein_coding_gene | 2.2 | 4.453 | 1.6E-04 |
| *psa-3* | WBGene00009560 | protein_coding_gene | 2.2 | 4.451 | 2.4E-09 |
| *F36F2.2* | WBGene00009476 | protein_coding_gene | 2.2 | 4.450 | 1.9E-13 |
| *Y62E10A.19* | WBGene00013385 | protein_coding_gene | 2.2 | 4.447 | 8.0E-12 |
| *ptp-1* | WBGene00004213 | protein_coding_gene | 2.1 | 4.438 | 6.3E-11 |
| *F47B10.8* | WBGene00009819 | protein_coding_gene | 2.1 | 4.438 | 8.7E-04 |
| *T25B6.4* | WBGene00020790 | protein_coding_gene | 2.1 | 4.434 | 1.5E-02 |
| *T02G5.15* | WBGene00206521 | protein_coding_gene | 2.1 | 4.431 | 3.7E-04 |
| *Y54G2A.44* | WBGene00044487 | protein_coding_gene | 2.1 | 4.427 | 1.7E-02 |
| *R03C1.1* | WBGene00010987 | protein_coding_gene | 2.1 | 4.425 | 1.4E-03 |
| *lipl-6* | WBGene00021963 | protein_coding_gene | 2.1 | 4.425 | 1.9E-04 |
| *cyp-33C12* | WBGene00021168 | protein_coding_gene | 2.1 | 4.422 | 3.0E-02 |

| | | | | | |
|---|---|---|---|---|---|
| *ttr-17* | WBGene00012383 | protein_coding_gene | 2.1 | 4.415 | 5.3E-19 |
| *cpna-5* | WBGene00007044 | protein_coding_gene | 2.1 | 4.410 | 5.4E-21 |
| *suex-2* | WBGene00008271 | protein_coding_gene | 2.1 | 4.409 | 5.2E-03 |
| *W04B5.2* | WBGene00021019 | protein_coding_gene | 2.1 | 4.409 | 2.1E-09 |
| *F42C5.4* | WBGene00018346 | protein_coding_gene | 2.1 | 4.407 | 4.0E-08 |
| *Y17G7B.22* | WBGene00012473 | protein_coding_gene | 2.1 | 4.407 | 6.6E-04 |
| *K08D10.9* | WBGene00019532 | pseudogene | 2.1 | 4.406 | 2.5E-04 |
| *F13A7.11* | WBGene00008726 | protein_coding_gene | 2.1 | 4.399 | 1.1E-08 |
| *Y65B4BL.6* | WBGene00022038 | protein_coding_gene | 2.1 | 4.399 | 6.7E-04 |
| *cnc-10* | WBGene00012152 | protein_coding_gene | 2.1 | 4.393 | 1.0E-24 |
| *R07C12.2* | WBGene00019932 | pseudogene | 2.1 | 4.389 | 2.3E-12 |
| *cwp-5* | WBGene00009844 | protein_coding_gene | 2.1 | 4.383 | 9.3E-04 |
| *Y75B8A.33* | WBGene00013563 | protein_coding_gene | 2.1 | 4.374 | 3.6E-02 |
| *linc-91* | WBGene00219746 | lincRNA_gene | 2.1 | 4.374 | 1.8E-04 |
| *Y73B6BL.47* | WBGene00045250 | protein_coding_gene | 2.1 | 4.363 | 1.1E-03 |
| *magu-2* | WBGene00006467 | protein_coding_gene | 2.1 | 4.363 | 1.1E-12 |
| *C18F10.2* | WBGene00015977 | protein_coding_gene | 2.1 | 4.362 | 3.4E-06 |
| *wrt-1* | WBGene00006947 | protein_coding_gene | 2.1 | 4.360 | 6.3E-05 |
| *ctsa-3.2* | WBGene00007462 | protein_coding_gene | 2.1 | 4.360 | 1.0E-05 |
| *T19C4.5* | WBGene00011839 | protein_coding_gene | 2.1 | 4.359 | 1.2E-08 |
| *hlh-6* | WBGene00001952 | protein_coding_gene | 2.1 | 4.357 | 2.4E-04 |
| *C05D9.9* | WBGene00015476 | protein_coding_gene | 2.1 | 4.356 | 2.2E-05 |
| *dpy-3* | WBGene00001065 | protein_coding_gene | 2.1 | 4.351 | 4.8E-05 |
| *C25F9.9* | WBGene00007728 | protein_coding_gene | 2.1 | 4.349 | 1.6E-07 |
| *C14A6.16* | WBGene00206536 | pseudogene | 2.1 | 4.340 | 1.4E-02 |
| *F53C3.4* | WBGene00018748 | protein_coding_gene | 2.1 | 4.338 | 9.4E-06 |
| *glb-15* | WBGene00009387 | protein_coding_gene | 2.1 | 4.333 | 4.9E-06 |
| *nhr-140* | WBGene00016366 | protein_coding_gene | 2.1 | 4.331 | 1.3E-08 |
| *asp-9* | WBGene00015700 | protein_coding_gene | 2.1 | 4.321 | 1.8E-03 |
| *lgc-21* | WBGene00007903 | protein_coding_gene | 2.1 | 4.321 | 4.2E-06 |
| *F08A7.1* | WBGene00045013 | protein_coding_gene | 2.1 | 4.314 | 1.9E-07 |
| *srh-46* | WBGene00005269 | protein_coding_gene | 2.1 | 4.313 | 2.5E-02 |
| *cutl-17* | WBGene00009541 | protein_coding_gene | 2.1 | 4.312 | 2.8E-03 |
| *C05C9.1* | WBGene00007325 | protein_coding_gene | 2.1 | 4.310 | 2.3E-02 |
| *Y61A9LA.4* | WBGene00022016 | protein_coding_gene | 2.1 | 4.306 | 3.0E-02 |
| *F13C5.1* | WBGene00017422 | protein_coding_gene | 2.1 | 4.302 | 2.2E-10 |
| *otpl-2* | WBGene00017034 | protein_coding_gene | 2.1 | 4.297 | 2.8E-02 |
| *aptf-1* | WBGene00019424 | protein_coding_gene | 2.1 | 4.293 | 1.2E-02 |
| *F52D1.2* | WBGene00018683 | protein_coding_gene | 2.1 | 4.292 | 6.8E-06 |
| *C13C12.2* | WBGene00007552 | protein_coding_gene | 2.1 | 4.292 | 2.3E-05 |
| *gyg-3* | WBGene00013535 | pseudogene | 2.1 | 4.291 | 1.6E-06 |
| *T22F3.12* | WBGene00044705 | protein_coding_gene | 2.1 | 4.289 | 1.0E-02 |
| *C32D5.12* | WBGene00016319 | protein_coding_gene | 2.1 | 4.285 | 2.9E-05 |
| *Y73C8C.4* | WBGene00022263 | protein_coding_gene | 2.1 | 4.284 | 1.9E-02 |
| *T14B1.1* | WBGene00011763 | protein_coding_gene | 2.1 | 4.282 | 9.7E-08 |
| *sra-10* | WBGene00005036 | protein_coding_gene | 2.1 | 4.279 | 5.0E-04 |
| *Y44A6D.7* | WBGene00077681 | protein_coding_gene | 2.1 | 4.279 | 4.9E-02 |
| *srh-2* | WBGene00005228 | protein_coding_gene | 2.1 | 4.275 | 1.8E-03 |
| *K09C8.8* | WBGene00010718 | protein_coding_gene | 2.1 | 4.274 | 8.5E-03 |
| *F43G6.8* | WBGene00009660 | protein_coding_gene | 2.1 | 4.268 | 1.8E-07 |
| *lips-7* | WBGene00015645 | protein_coding_gene | 2.1 | 4.267 | 2.8E-04 |
| *F59C6.8* | WBGene00010327 | protein_coding_gene | 2.1 | 4.262 | 3.8E-07 |
| *cbp-2* | WBGene00010545 | protein_coding_gene | 2.1 | 4.258 | 1.4E-09 |
| *Y47G6A.30* | WBGene00021653 | protein_coding_gene | 2.1 | 4.257 | 1.4E-02 |
| *ilys-4* | WBGene00016958 | protein_coding_gene | 2.1 | 4.257 | 6.1E-13 |
| *F35E2.3* | WBGene00009411 | pseudogene | 2.1 | 4.256 | 9.9E-03 |
| *ZC168.2* | WBGene00013856 | protein_coding_gene | 2.1 | 4.254 | 1.3E-03 |
| *zag-1* | WBGene00006970 | protein_coding_gene | 2.1 | 4.242 | 3.0E-06 |
| *sym-2* | WBGene00006367 | protein_coding_gene | 2.1 | 4.241 | 2.6E-11 |
| *T24C2.5* | WBGene00011984 | protein_coding_gene | 2.1 | 4.241 | 1.8E-10 |
| *Y43F8B.9* | WBGene00012819 | protein_coding_gene | 2.1 | 4.240 | 1.6E-05 |
| *T19H5.4* | WBGene00011849 | protein_coding_gene | 2.1 | 4.235 | 1.5E-18 |
| *T23B12.8* | WBGene00020722 | protein_coding_gene | 2.1 | 4.233 | 1.2E-10 |
| *nspc-20* | WBGene00008356 | protein_coding_gene | 2.1 | 4.230 | 6.5E-12 |
| *nspb-9* | WBGene00007251 | protein_coding_gene | 2.1 | 4.229 | 4.2E-02 |
| *C14C6.2* | WBGene00015756 | protein_coding_gene | 2.1 | 4.223 | 9.3E-23 |
| *dyf-18* | WBGene00010349 | protein_coding_gene | 2.1 | 4.222 | 2.3E-03 |
| *T05B11.7* | WBGene00044342 | protein_coding_gene | 2.1 | 4.218 | 3.8E-06 |
| *Y105E8A.34* | WBGene00077689 | pseudogene | 2.1 | 4.215 | 1.6E-02 |
| *ast-1* | WBGene00020368 | protein_coding_gene | 2.1 | 4.214 | 1.5E-04 |
| *pqn-63* | WBGene00004146 | protein_coding_gene | 2.1 | 4.214 | 3.6E-05 |
| *daf-14* | WBGene00000910 | protein_coding_gene | 2.1 | 4.205 | 7.4E-20 |
| *spp-5* | WBGene00004990 | protein_coding_gene | 2.1 | 4.202 | 3.5E-20 |
| *C13C4.7* | WBGene00007551 | protein_coding_gene | 2.1 | 4.193 | 1.0E-08 |
| *B0244.16* | WBGene00271577 | protein_coding_gene | 2.1 | 4.187 | 7.8E-03 |
| *nhr-107* | WBGene00003697 | protein_coding_gene | 2.1 | 4.186 | 7.2E-10 |
| *egl-44* | WBGene00001208 | protein_coding_gene | 2.1 | 4.184 | 1.3E-11 |
| *F41E6.5* | WBGene00018286 | protein_coding_gene | 2.1 | 4.183 | 1.2E-07 |
| *F56H9.2* | WBGene00010174 | protein_coding_gene | 2.1 | 4.167 | 3.8E-23 |
| *H12D21.9* | WBGene00010385 | protein_coding_gene | 2.1 | 4.163 | 5.8E-04 |

| | | | | | |
|---|---|---|---|---|---|
| *F55C12.19* | WBGene00271817 | protein_coding_gene | 2.1 | 4.161 | 1.3E-04 |
| *tsp-3* | WBGene00006629 | protein_coding_gene | 2.1 | 4.148 | 6.6E-05 |
| *C13C4.6* | WBGene00007550 | protein_coding_gene | 2.1 | 4.142 | 1.9E-07 |
| *F23F12.13* | WBGene00044455 | protein_coding_gene | 2.1 | 4.142 | 1.6E-08 |
| *gst-31* | WBGene00001779 | protein_coding_gene | 2.0 | 4.140 | 6.6E-03 |
| *C52E2.5* | WBGene00016884 | protein_coding_gene | 2.0 | 4.139 | 4.8E-05 |
| *cyp-13A1* | WBGene00011677 | protein_coding_gene | 2.0 | 4.133 | 2.5E-04 |
| *F09C6.1* | WBGene00008615 | protein_coding_gene | 2.0 | 4.132 | 1.2E-03 |
| *F21C10.13* | WBGene00195086 | protein_coding_gene | 2.0 | 4.131 | 3.2E-03 |
| *Y48A6B.7* | WBGene00012968 | protein_coding_gene | 2.0 | 4.129 | 1.2E-13 |
| *sri-23* | WBGene00005535 | pseudogene | 2.0 | 4.127 | 2.3E-02 |
| *Y68A4A.12* | WBGene00014944 | pseudogene | 2.0 | 4.124 | 4.6E-02 |
| *pros-1* | WBGene00000448 | protein_coding_gene | 2.0 | 4.121 | 1.0E-06 |
| *sru-20* | WBGene00005683 | protein_coding_gene | 2.0 | 4.107 | 3.3E-02 |
| *ZK180.7* | WBGene00045248 | protein_coding_gene | 2.0 | 4.104 | 7.4E-03 |
| *tsp-6* | WBGene00006632 | protein_coding_gene | 2.0 | 4.102 | 4.5E-05 |
| *T24F1.7* | WBGene00011999 | protein_coding_gene | 2.0 | 4.100 | 2.5E-04 |
| *tat-6* | WBGene00017174 | protein_coding_gene | 2.0 | 4.099 | 3.5E-03 |
| *glct-6* | WBGene00019546 | protein_coding_gene | 2.0 | 4.095 | 2.6E-14 |
| *flp-10* | WBGene00001453 | protein_coding_gene | 2.0 | 4.095 | 3.5E-06 |
| *F44E2.4* | WBGene00018418 | protein_coding_gene | 2.0 | 4.093 | 6.5E-08 |
| *oig-4* | WBGene00043050 | protein_coding_gene | 2.0 | 4.092 | 3.6E-06 |
| *T07G12.3* | WBGene00011594 | protein_coding_gene | 2.0 | 4.085 | 3.8E-04 |
| *cutl-22* | WBGene00020663 | protein_coding_gene | 2.0 | 4.079 | 5.7E-03 |
| *K09F6.9* | WBGene00019592 | protein_coding_gene | 2.0 | 4.075 | 2.5E-06 |
| *Y43F8A.1* | WBGene00012808 | protein_coding_gene | 2.0 | 4.074 | 2.0E-04 |
| *Y52B11A.12* | WBGene00077539 | protein_coding_gene | 2.0 | 4.073 | 3.6E-05 |
| *ttm-2* | WBGene00017840 | protein_coding_gene | 2.0 | 4.071 | 8.4E-09 |
| *C56C10.4* | WBGene00016962 | protein_coding_gene | 2.0 | 4.068 | 8.1E-03 |
| *C25F9.4* | WBGene00007724 | protein_coding_gene | 2.0 | 4.066 | 3.7E-14 |
| *snai-1* | WBGene00019299 | protein_coding_gene | 2.0 | 4.064 | 3.5E-02 |
| *Y11D7A.9* | WBGene00012433 | protein_coding_gene | 2.0 | 4.061 | 1.2E-08 |
| *marc-4* | WBGene00016903 | protein_coding_gene | 2.0 | 4.055 | 6.9E-05 |
| *cutl-20* | WBGene00018787 | protein_coding_gene | 2.0 | 4.046 | 3.0E-09 |
| *Y55B1BL.1* | WBGene00021909 | protein_coding_gene | 2.0 | 4.038 | 2.1E-27 |
| *lys-8* | WBGene00003097 | protein_coding_gene | 2.0 | 4.035 | 4.1E-25 |
| *ceh-22* | WBGene00000445 | protein_coding_gene | 2.0 | 4.023 | 4.2E-06 |
| *R07A4.4* | WBGene00011075 | protein_coding_gene | 2.0 | 4.021 | 1.2E-02 |
| *K02C4.5* | WBGene00010503 | protein_coding_gene | 2.0 | 4.020 | 1.2E-09 |
| *ifta-2* | WBGene00012132 | protein_coding_gene | 2.0 | 4.020 | 1.2E-03 |
| *F37A4.3* | WBGene00018133 | protein_coding_gene | 2.0 | 4.018 | 7.2E-04 |
| *R02F11.1* | WBGene00019839 | protein_coding_gene | 2.0 | 4.001 | 1.7E-06 |
| *F41G3.3* | WBGene00018300 | protein_coding_gene | 2.0 | 3.999 | 1.0E-03 |
| *Y52B11A.4* | WBGene00013124 | protein_coding_gene | 2.0 | 3.999 | 1.7E-06 |
| *Y39G10AR.11* | WBGene00021469 | protein_coding_gene | 2.0 | 3.998 | 3.5E-11 |
| *ZK112.5* | WBGene00022661 | protein_coding_gene | 2.0 | 3.993 | 2.3E-02 |
| *ceh-24* | WBGene00000447 | protein_coding_gene | 2.0 | 3.992 | 6.7E-03 |
| *C06C3.10* | WBGene00044169 | protein_coding_gene | 2.0 | 3.991 | 7.1E-05 |
| *Y37H2B.1* | WBGene00012572 | pseudogene | 2.0 | 3.989 | 1.2E-04 |
| *K07H8.12* | WBGene00044509 | protein_coding_gene | 2.0 | 3.988 | 2.2E-09 |
| *nphp-4* | WBGene00011261 | protein_coding_gene | 2.0 | 3.985 | 1.3E-05 |
| *lin-42* | WBGene00018572 | protein_coding_gene | 2.0 | 3.984 | 4.5E-10 |
| *C33D9.6* | WBGene00007900 | protein_coding_gene | 2.0 | 3.979 | 8.1E-15 |
| *Y71G12B.31* | WBGene00044348 | protein_coding_gene | 2.0 | 3.978 | 9.7E-05 |
| *F54C8.6* | WBGene00010039 | protein_coding_gene | 2.0 | 3.976 | 6.8E-07 |
| *hlh-8* | WBGene00001953 | protein_coding_gene | 2.0 | 3.976 | 6.6E-05 |
| *F40E10.5* | WBGene00009573 | protein_coding_gene | 2.0 | 3.973 | 5.0E-06 |
| *ZK757.1* | WBGene00014073 | protein_coding_gene | 2.0 | 3.967 | 6.1E-03 |
| *che-2* | WBGene00000484 | protein_coding_gene | 2.0 | 3.960 | 1.1E-03 |
| *T28C6.10* | WBGene00045183 | protein_coding_gene | 2.0 | 3.960 | 8.6E-04 |
| *nhr-58* | WBGene00003648 | protein_coding_gene | 2.0 | 3.959 | 6.7E-05 |
| *gtf-2A2* | WBGene00013736 | protein_coding_gene | 2.0 | 3.951 | 8.1E-17 |
| *Y57G11C.44* | WBGene00013333 | protein_coding_gene | 2.0 | 3.948 | 3.1E-02 |
| *mod-5* | WBGene00003387 | protein_coding_gene | 2.0 | 3.947 | 3.8E-07 |
| *cyp-13A3* | WBGene00011675 | protein_coding_gene | 2.0 | 3.945 | 8.1E-05 |
| *Y7A9C.1* | WBGene00012418 | pseudogene | 2.0 | 3.940 | 1.3E-19 |
| *C43C3.4* | WBGene00195084 | protein_coding_gene | 2.0 | 3.940 | 2.6E-05 |
| *Y45F10D.6* | WBGene00012886 | protein_coding_gene | 2.0 | 3.929 | 6.7E-05 |
| *subs-4* | WBGene00012939 | protein_coding_gene | 2.0 | 3.928 | 1.7E-04 |
| *ZC262.13* | WBGene00284846 | protein_coding_gene | 2.0 | 3.928 | 4.0E-05 |
| *ZC239.22* | WBGene00194717 | protein_coding_gene | 2.0 | 3.923 | 1.9E-03 |
| *cht-2* | WBGene00011757 | protein_coding_gene | 2.0 | 3.923 | 2.8E-04 |
| *inx-20* | WBGene00002142 | protein_coding_gene | 2.0 | 3.919 | 3.1E-04 |
| *K02E2.1* | WBGene00010506 | protein_coding_gene | 2.0 | 3.910 | 3.1E-02 |
| *alr-1* | WBGene00044330 | protein_coding_gene | 2.0 | 3.902 | 9.2E-12 |
| *cfi-1* | WBGene00000476 | protein_coding_gene | 2.0 | 3.902 | 5.3E-10 |
| *ZK863.1* | WBGene00014122 | protein_coding_gene | 2.0 | 3.899 | 4.4E-02 |
| *arrd-22* | WBGene00020612 | protein_coding_gene | 2.0 | 3.898 | 1.5E-09 |
| *C35E7.2* | WBGene00016454 | protein_coding_gene | 2.0 | 3.894 | 3.2E-09 |
| *C34D1.4* | WBGene00007931 | protein_coding_gene | 2.0 | 3.894 | 1.5E-10 |

| | | | | | |
|---|---|---|---|---|---|
| *F26F2.1* | WBGene00009166 | protein_coding_gene | 2.0 | 3.889 | 4.2E-03 |
| *cyp-37A1* | WBGene00008499 | protein_coding_gene | 2.0 | 3.882 | 9.9E-07 |
| *poml-3* | WBGene00077701 | protein_coding_gene | 2.0 | 3.882 | 1.1E-37 |
| *F41E7.2* | WBGene00009618 | protein_coding_gene | 2.0 | 3.881 | 3.4E-10 |
| *T27A10.6* | WBGene00020846 | protein_coding_gene | 2.0 | 3.878 | 1.0E-08 |
| *F09B12.7* | WBGene00195212 | protein_coding_gene | 2.0 | 3.877 | 1.9E-03 |
| *C18B10.6* | WBGene00015963 | protein_coding_gene | 2.0 | 3.876 | 7.4E-04 |
| *rrf-2* | WBGene00004509 | protein_coding_gene | 2.0 | 3.874 | 1.0E-13 |
| *cdf-2* | WBGene00011821 | protein_coding_gene | 2.0 | 3.874 | 3.4E-05 |
| *crtc-1* | WBGene00021237 | protein_coding_gene | 2.0 | 3.874 | 1.9E-08 |
| *nsy-4* | WBGene00021415 | protein_coding_gene | 2.0 | 3.870 | 6.5E-08 |
| *R02E12.12* | WBGene00303425 | protein_coding_gene | 2.0 | 3.870 | 3.4E-03 |
| *Y70D2A.1* | WBGene00013496 | protein_coding_gene | 2.0 | 3.866 | 1.1E-03 |
| *Y32B12C.1* | WBGene00012525 | protein_coding_gene | 2.0 | 3.866 | 6.3E-05 |
| *daf-10* | WBGene00000906 | protein_coding_gene | 2.0 | 3.865 | 4.1E-06 |
| *F35B3.7* | WBGene00018033 | protein_coding_gene | 1.9 | 3.863 | 5.2E-07 |
| *ssr-2* | WBGene00006055 | protein_coding_gene | 1.9 | 3.861 | 4.2E-05 |
| *glf-1* | WBGene00019154 | protein_coding_gene | 1.9 | 3.860 | 3.4E-04 |
| *W09C2.7* | WBGene00185095 | protein_coding_gene | 1.9 | 3.858 | 2.9E-02 |
| *F26A1.9* | WBGene00017807 | protein_coding_gene | 1.9 | 3.855 | 4.0E-05 |
| *F46A9.1* | WBGene00009754 | protein_coding_gene | 1.9 | 3.855 | 7.2E-04 |
| *nhr-173* | WBGene00016975 | protein_coding_gene | 1.9 | 3.854 | 6.0E-08 |
| *oac-30* | WBGene00018573 | protein_coding_gene | 1.9 | 3.853 | 1.1E-02 |
| *F36H9.2* | WBGene00018111 | protein_coding_gene | 1.9 | 3.848 | 2.2E-02 |
| *cki-1* | WBGene00000516 | protein_coding_gene | 1.9 | 3.848 | 1.6E-11 |
| *K07C5.9* | WBGene00010632 | protein_coding_gene | 1.9 | 3.841 | 3.2E-07 |
| *F18C5.5* | WBGene00017560 | protein_coding_gene | 1.9 | 3.841 | 1.7E-03 |
| *grsp-3* | WBGene00016403 | protein_coding_gene | 1.9 | 3.841 | 3.2E-03 |
| *fbxa-38* | WBGene00016522 | protein_coding_gene | 1.9 | 3.833 | 1.6E-08 |
| *dpy-7* | WBGene00001069 | protein_coding_gene | 1.9 | 3.823 | 5.1E-05 |
| *K08H10.10* | WBGene00077491 | protein_coding_gene | 1.9 | 3.822 | 1.2E-02 |
| *F08F3.1* | WBGene00017260 | protein_coding_gene | 1.9 | 3.820 | 1.1E-03 |
| *sox-2* | WBGene00004949 | protein_coding_gene | 1.9 | 3.820 | 7.3E-09 |
| *nhr-163* | WBGene00016368 | protein_coding_gene | 1.9 | 3.819 | 9.7E-06 |
| *F54B11.8* | WBGene00010031 | protein_coding_gene | 1.9 | 3.819 | 3.8E-03 |
| *linc-24* | WBGene00219771 | lincRNA_gene | 1.9 | 3.819 | 3.5E-03 |
| *dhs-20* | WBGene00000983 | protein_coding_gene | 1.9 | 3.818 | 5.6E-07 |
| *T18D3.7* | WBGene00011824 | protein_coding_gene | 1.9 | 3.816 | 2.2E-06 |
| *R06F6.14* | WBGene00269421 | protein_coding_gene | 1.9 | 3.814 | 8.8E-14 |
| *ift-81* | WBGene00017973 | protein_coding_gene | 1.9 | 3.814 | 9.2E-05 |
| *F53F4.2* | WBGene00009986 | protein_coding_gene | 1.9 | 3.810 | 4.6E-02 |
| *Y54F10BM.3* | WBGene00021858 | protein_coding_gene | 1.9 | 3.809 | 7.3E-06 |
| *cdh-9* | WBGene00000401 | protein_coding_gene | 1.9 | 3.798 | 1.6E-06 |
| *yap-1* | WBGene00008748 | protein_coding_gene | 1.9 | 3.798 | 7.6E-09 |
| *F36F12.1* | WBGene00018092 | protein_coding_gene | 1.9 | 3.796 | 3.7E-02 |
| *maph-9* | WBGene00016397 | protein_coding_gene | 1.9 | 3.788 | 3.3E-03 |
| *ZK1128.7* | WBGene00014233 | protein_coding_gene | 1.9 | 3.783 | 5.8E-09 |
| *C04G6.5* | WBGene00015455 | protein_coding_gene | 1.9 | 3.781 | 3.4E-03 |
| *pals-23* | WBGene00016218 | protein_coding_gene | 1.9 | 3.777 | 1.3E-06 |
| *ttx-3* | WBGene00006654 | protein_coding_gene | 1.9 | 3.776 | 4.1E-02 |
| *ZK813.5* | WBGene00022824 | protein_coding_gene | 1.9 | 3.776 | 1.0E-02 |
| *unc-75* | WBGene00006807 | protein_coding_gene | 1.9 | 3.774 | 1.4E-03 |
| *F32D8.13* | WBGene00009335 | protein_coding_gene | 1.9 | 3.768 | 8.4E-09 |
| *T19A5.3* | WBGene00020554 | protein_coding_gene | 1.9 | 3.768 | 3.3E-08 |
| *MTCE.33* | WBGene00014472 | rRNA_gene | 1.9 | 3.762 | 1.2E-14 |
| *nfm-1* | WBGene00003593 | protein_coding_gene | 1.9 | 3.755 | 7.1E-09 |
| *C37E2.2* | WBGene00007994 | protein_coding_gene | 1.9 | 3.751 | 7.2E-05 |
| *nhr-258* | WBGene00016126 | protein_coding_gene | 1.9 | 3.751 | 8.8E-03 |
| *W02B8.1* | WBGene00012197 | protein_coding_gene | 1.9 | 3.750 | 3.9E-06 |
| *C44H9.9* | WBGene00077564 | pseudogene | 1.9 | 3.750 | 2.0E-02 |
| *F41F3.3* | WBGene00018297 | protein_coding_gene | 1.9 | 3.749 | 1.2E-04 |
| *clec-83* | WBGene00021879 | protein_coding_gene | 1.9 | 3.745 | 1.7E-07 |
| *best-3* | WBGene00015286 | protein_coding_gene | 1.9 | 3.745 | 9.6E-14 |
| *F57G12.2* | WBGene00010222 | protein_coding_gene | 1.9 | 3.743 | 1.4E-02 |
| *hlh-17* | WBGene00001961 | protein_coding_gene | 1.9 | 3.735 | 4.6E-02 |
| *T07D10.10* | WBGene00194657 | pseudogene | 1.9 | 3.734 | 2.3E-02 |
| *C33H5.13* | WBGene00016379 | protein_coding_gene | 1.9 | 3.733 | 8.0E-17 |
| *F26F12.3* | WBGene00017834 | protein_coding_gene | 1.9 | 3.731 | 8.9E-07 |
| *T01B7.13* | WBGene00206502 | protein_coding_gene | 1.9 | 3.730 | 2.6E-03 |
| *bigr-1* | WBGene00009512 | protein_coding_gene | 1.9 | 3.727 | 1.0E-08 |
| *icmt-1* | WBGene00017673 | protein_coding_gene | 1.9 | 3.727 | 4.8E-04 |
| *F42G2.2* | WBGene00018351 | protein_coding_gene | 1.9 | 3.725 | 4.0E-09 |
| *ttx-1* | WBGene00006652 | protein_coding_gene | 1.9 | 3.719 | 3.3E-06 |
| *nlp-66* | WBGene00011141 | protein_coding_gene | 1.9 | 3.717 | 4.4E-03 |
| *ras-1* | WBGene00004310 | protein_coding_gene | 1.9 | 3.713 | 8.7E-10 |
| *F32H2.11* | WBGene00043049 | protein_coding_gene | 1.9 | 3.712 | 9.1E-03 |
| *col-76* | WBGene00000652 | protein_coding_gene | 1.9 | 3.711 | 6.7E-12 |
| *F54E4.4* | WBGene00010060 | protein_coding_gene | 1.9 | 3.709 | 1.3E-03 |
| *fkh-8* | WBGene00001440 | protein_coding_gene | 1.9 | 3.709 | 3.3E-06 |
| *gana-1* | WBGene00011095 | protein_coding_gene | 1.9 | 3.703 | 1.9E-10 |

| | | | | | |
|---|---|---|---|---|---|
| *Y105E8B.7* | WBGene00013692 | protein_coding_gene | 1.9 | 3.702 | 1.1E-12 |
| *F32D8.1* | WBGene00009326 | protein_coding_gene | 1.9 | 3.696 | 2.2E-02 |
| *F56C9.8* | WBGene00018952 | protein_coding_gene | 1.9 | 3.695 | 5.8E-10 |
| *cdr-1* | WBGene00000412 | protein_coding_gene | 1.9 | 3.692 | 3.3E-03 |
| *ZC13.2* | WBGene00022503 | protein_coding_gene | 1.9 | 3.691 | 2.7E-03 |
| *mlt-11* | WBGene00012186 | protein_coding_gene | 1.9 | 3.691 | 1.4E-04 |
| *dhs-2* | WBGene00000966 | protein_coding_gene | 1.9 | 3.691 | 5.5E-07 |
| *F53F8.4* | WBGene00010001 | protein_coding_gene | 1.9 | 3.690 | 4.3E-10 |
| *K03H9.3* | WBGene00019372 | protein_coding_gene | 1.9 | 3.690 | 4.8E-03 |
| *C40H5.2* | WBGene00008045 | protein_coding_gene | 1.9 | 3.682 | 2.0E-05 |
| *clec-184* | WBGene00023502 | protein_coding_gene | 1.9 | 3.681 | 5.3E-05 |
| *sulp-2* | WBGene00017464 | protein_coding_gene | 1.9 | 3.681 | 2.4E-15 |
| *che-12* | WBGene00000491 | protein_coding_gene | 1.9 | 3.680 | 3.2E-06 |
| *npr-19* | WBGene00015364 | protein_coding_gene | 1.9 | 3.679 | 1.6E-02 |
| *F28F9.3* | WBGene00017907 | protein_coding_gene | 1.9 | 3.673 | 6.9E-04 |
| *fbxa-6* | WBGene00008352 | protein_coding_gene | 1.9 | 3.672 | 3.6E-10 |
| *C44H9.5* | WBGene00008100 | protein_coding_gene | 1.9 | 3.667 | 1.4E-11 |
| *nhr-53* | WBGene00003643 | protein_coding_gene | 1.9 | 3.666 | 1.2E-09 |
| *sek-6* | WBGene00012162 | protein_coding_gene | 1.9 | 3.666 | 3.6E-08 |
| *F42G4.6* | WBGene00009643 | protein_coding_gene | 1.9 | 3.666 | 2.4E-03 |
| *ugt-59* | WBGene00011238 | protein_coding_gene | 1.9 | 3.664 | 1.2E-03 |
| *pqn-72* | WBGene00004154 | protein_coding_gene | 1.9 | 3.662 | 3.1E-07 |
| *srab-12* | WBGene00008128 | protein_coding_gene | 1.9 | 3.659 | 3.7E-02 |
| *T22B7.3* | WBGene00020672 | protein_coding_gene | 1.9 | 3.656 | 3.9E-06 |
| *C26B9.3* | WBGene00016133 | protein_coding_gene | 1.9 | 3.656 | 2.9E-05 |
| *F13E9.13* | WBGene00044797 | protein_coding_gene | 1.9 | 3.656 | 2.8E-10 |
| *R01E6.7* | WBGene00010971 | protein_coding_gene | 1.9 | 3.653 | 1.4E-03 |
| *T05A10.6* | WBGene00011463 | protein_coding_gene | 1.9 | 3.651 | 2.8E-05 |
| *T25E4.1* | WBGene00020803 | protein_coding_gene | 1.9 | 3.651 | 7.0E-04 |
| *F26D11.2* | WBGene00017819 | protein_coding_gene | 1.9 | 3.649 | 3.5E-02 |
| *cyp-33E1* | WBGene00016768 | protein_coding_gene | 1.9 | 3.649 | 4.7E-13 |
| *C44C1.6* | WBGene00016645 | protein_coding_gene | 1.9 | 3.647 | 1.9E-04 |
| *abf-6* | WBGene00000017 | protein_coding_gene | 1.9 | 3.641 | 1.7E-08 |
| *fbxa-20* | WBGene00022332 | protein_coding_gene | 1.9 | 3.638 | 6.4E-03 |
| *Y55F3BR.11* | WBGene00044757 | protein_coding_gene | 1.9 | 3.638 | 8.2E-05 |
| *C39B10.3* | WBGene00008023 | protein_coding_gene | 1.9 | 3.638 | 3.1E-03 |
| *R04E5.9* | WBGene00019872 | protein_coding_gene | 1.9 | 3.637 | 1.1E-07 |
| *F36G9.4* | WBGene00014751 | pseudogene | 1.9 | 3.634 | 6.9E-03 |
| *F13B9.2* | WBGene00017420 | protein_coding_gene | 1.9 | 3.633 | 3.7E-03 |
| *Y70G10A.3* | WBGene00013499 | protein_coding_gene | 1.9 | 3.632 | 3.3E-06 |
| *vab-3* | WBGene00006870 | protein_coding_gene | 1.9 | 3.629 | 6.2E-10 |
| *Y87G2A.19* | WBGene00044260 | protein_coding_gene | 1.9 | 3.624 | 5.7E-08 |
| *K06A4.15* | WBGene00220007 | ncRNA_gene | 1.9 | 3.624 | 1.5E-02 |
| *chil-22* | WBGene00011166 | protein_coding_gene | 1.9 | 3.621 | 4.9E-18 |
| *T19D12.10* | WBGene00020584 | protein_coding_gene | 1.9 | 3.619 | 1.5E-02 |
| *asp-14* | WBGene00019619 | protein_coding_gene | 1.9 | 3.618 | 6.8E-08 |
| *gei-13* | WBGene00001570 | protein_coding_gene | 1.9 | 3.618 | 8.3E-05 |
| *Y67D8C.6* | WBGene00022070 | protein_coding_gene | 1.9 | 3.612 | 4.2E-02 |
| *fipr-5* | WBGene00044174 | protein_coding_gene | 1.9 | 3.608 | 2.9E-08 |
| *K02D7.1* | WBGene00019298 | protein_coding_gene | 1.8 | 3.604 | 6.9E-08 |
| *F46B6.9* | WBGene00009774 | protein_coding_gene | 1.8 | 3.603 | 7.7E-04 |
| *amt-2* | WBGene00000134 | protein_coding_gene | 1.8 | 3.602 | 3.8E-03 |
| *cest-17* | WBGene00019477 | protein_coding_gene | 1.8 | 3.594 | 7.4E-14 |
| *Y57G11C.32* | WBGene00013322 | protein_coding_gene | 1.8 | 3.585 | 1.2E-04 |
| *pqn-2* | WBGene00004098 | protein_coding_gene | 1.8 | 3.585 | 4.3E-03 |
| *fbxa-69* | WBGene00010018 | protein_coding_gene | 1.8 | 3.573 | 4.8E-07 |
| *linc-11* | WBGene00219758 | lincRNA_gene | 1.8 | 3.565 | 1.3E-05 |
| *srd-70* | WBGene00005147 | protein_coding_gene | 1.8 | 3.556 | 4.6E-02 |
| *npr-28* | WBGene00018886 | protein_coding_gene | 1.8 | 3.556 | 2.8E-09 |
| *ugt-54* | WBGene00012013 | protein_coding_gene | 1.8 | 3.555 | 8.4E-05 |
| *Y32H12A.1* | WBGene00021310 | protein_coding_gene | 1.8 | 3.554 | 2.2E-02 |
| *ppm-1.H* | WBGene00016580 | protein_coding_gene | 1.8 | 3.552 | 3.4E-03 |
| *Y38H6A.5* | WBGene00077669 | protein_coding_gene | 1.8 | 3.551 | 2.0E-02 |
| *C28D4.10* | WBGene00007796 | protein_coding_gene | 1.8 | 3.549 | 3.7E-02 |
| *dyf-5* | WBGene00001121 | protein_coding_gene | 1.8 | 3.549 | 1.1E-03 |
| *Y73E7A.8* | WBGene00022274 | protein_coding_gene | 1.8 | 3.548 | 1.0E-03 |
| *C43H6.6* | WBGene00016619 | protein_coding_gene | 1.8 | 3.547 | 2.9E-02 |
| *F13B12.4* | WBGene00008732 | protein_coding_gene | 1.8 | 3.546 | 1.2E-03 |
| *fbxa-1* | WBGene00021864 | protein_coding_gene | 1.8 | 3.546 | 2.3E-02 |
| *D1081.4* | WBGene00008382 | protein_coding_gene | 1.8 | 3.539 | 1.6E-02 |
| *lgx-1* | WBGene00002983 | protein_coding_gene | 1.8 | 3.539 | 6.2E-08 |
| *F14H3.12* | WBGene00008831 | protein_coding_gene | 1.8 | 3.535 | 1.6E-07 |
| *cyp-33C7* | WBGene00018260 | protein_coding_gene | 1.8 | 3.534 | 3.0E-05 |
| *nhr-227* | WBGene00020851 | protein_coding_gene | 1.8 | 3.532 | 1.2E-04 |
| *tag-234* | WBGene00018874 | protein_coding_gene | 1.8 | 3.528 | 4.8E-09 |
| *T25D10.4* | WBGene00020801 | protein_coding_gene | 1.8 | 3.528 | 2.5E-19 |
| *C17F4.2* | WBGene00015910 | protein_coding_gene | 1.8 | 3.523 | 7.1E-04 |
| *T26C5.5* | WBGene00012035 | protein_coding_gene | 1.8 | 3.523 | 1.3E-04 |
| *F23F1.2* | WBGene00017743 | protein_coding_gene | 1.8 | 3.523 | 3.2E-02 |
| *frpr-14* | WBGene00019496 | protein_coding_gene | 1.8 | 3.523 | 2.1E-02 |

| | | | | | |
|---|---|---|---|---|---|
| *col-58* | WBGene00000634 | protein_coding_gene | 1.8 | 3.523 | 4.1E-04 |
| *acr-5* | WBGene00000044 | protein_coding_gene | 1.8 | 3.523 | 7.8E-04 |
| *MTCE.7* | WBGene00014454 | rRNA_gene | 1.8 | 3.522 | 4.0E-08 |
| *Y23H5B.3* | WBGene00021274 | protein_coding_gene | 1.8 | 3.522 | 2.6E-04 |
| *ZK1010.8* | WBGene00014181 | protein_coding_gene | 1.8 | 3.513 | 1.4E-06 |
| *abu-15* | WBGene00004099 | protein_coding_gene | 1.8 | 3.512 | 1.5E-03 |
| *T20D4.5* | WBGene00020611 | protein_coding_gene | 1.8 | 3.509 | 2.5E-06 |
| *cutl-21* | WBGene00018861 | protein_coding_gene | 1.8 | 3.509 | 2.8E-04 |
| *tsp-15* | WBGene00006641 | protein_coding_gene | 1.8 | 3.509 | 5.1E-09 |
| *K09E2.2* | WBGene00019571 | protein_coding_gene | 1.8 | 3.505 | 1.7E-02 |
| *aqp-2* | WBGene00000170 | protein_coding_gene | 1.8 | 3.505 | 1.6E-07 |
| *arrd-16* | WBGene00013043 | protein_coding_gene | 1.8 | 3.502 | 4.0E-02 |
| *nspb-6* | WBGene00019152 | protein_coding_gene | 1.8 | 3.502 | 2.6E-02 |
| *Y43F8B.13* | WBGene00012823 | protein_coding_gene | 1.8 | 3.499 | 1.6E-02 |
| *bbs-8* | WBGene00000244 | protein_coding_gene | 1.8 | 3.496 | 1.0E-04 |
| *F38C2.4* | WBGene00009536 | protein_coding_gene | 1.8 | 3.496 | 2.5E-02 |
| *tsp-16* | WBGene00006642 | protein_coding_gene | 1.8 | 3.496 | 2.5E-06 |
| *clec-233* | WBGene00009030 | protein_coding_gene | 1.8 | 3.492 | 9.5E-04 |
| *tps-1* | WBGene00006602 | protein_coding_gene | 1.8 | 3.487 | 1.7E-11 |
| *T10B11.5* | WBGene00020401 | protein_coding_gene | 1.8 | 3.484 | 5.0E-10 |
| *mab-7* | WBGene00003104 | protein_coding_gene | 1.8 | 3.483 | 3.5E-05 |
| *C38C6.3* | WBGene00007996 | protein_coding_gene | 1.8 | 3.481 | 7.9E-03 |
| *ltd-1* | WBGene00003089 | protein_coding_gene | 1.8 | 3.479 | 2.8E-11 |
| *F17C11.4* | WBGene00008915 | protein_coding_gene | 1.8 | 3.477 | 1.3E-06 |
| *F37C12.21* | WBGene00194742 | protein_coding_gene | 1.8 | 3.474 | 3.3E-03 |
| *ZC123.4* | WBGene00022519 | protein_coding_gene | 1.8 | 3.473 | 3.0E-10 |
| *ndx-1* | WBGene00003578 | protein_coding_gene | 1.8 | 3.471 | 7.6E-10 |
| *T20B3.14* | WBGene00011854 | protein_coding_gene | 1.8 | 3.469 | 6.4E-04 |
| *F36F2.7* | WBGene00009480 | protein_coding_gene | 1.8 | 3.468 | 1.4E-03 |
| *F10G2.1* | WBGene00017362 | protein_coding_gene | 1.8 | 3.466 | 5.6E-30 |
| *F13H10.9* | WBGene00045056 | protein_coding_gene | 1.8 | 3.461 | 2.3E-04 |
| *rom-2* | WBGene00004401 | protein_coding_gene | 1.8 | 3.461 | 3.8E-04 |
| *tmed-13* | WBGene00022255 | protein_coding_gene | 1.8 | 3.459 | 4.1E-04 |
| *feh-1* | WBGene00001410 | protein_coding_gene | 1.8 | 3.457 | 4.6E-12 |
| *C43F9.7* | WBGene00008075 | protein_coding_gene | 1.8 | 3.454 | 1.3E-02 |
| *lips-9* | WBGene00010267 | protein_coding_gene | 1.8 | 3.453 | 4.6E-03 |
| *Y54G2A.28* | WBGene00021892 | protein_coding_gene | 1.8 | 3.449 | 2.2E-03 |
| *ser-4* | WBGene00004779 | protein_coding_gene | 1.8 | 3.448 | 1.2E-05 |
| *math-24* | WBGene00016719 | protein_coding_gene | 1.8 | 3.447 | 1.0E-07 |
| *lgc-47* | WBGene00009799 | protein_coding_gene | 1.8 | 3.441 | 3.0E-05 |
| *T28H10.1* | WBGene00012142 | protein_coding_gene | 1.8 | 3.441 | 1.1E-14 |
| *K05B2.4* | WBGene00019404 | protein_coding_gene | 1.8 | 3.436 | 1.7E-02 |
| *F11A5.15* | WBGene00008680 | protein_coding_gene | 1.8 | 3.434 | 9.7E-04 |
| *ceh-33* | WBGene00000454 | protein_coding_gene | 1.8 | 3.432 | 6.3E-03 |
| *Y47A7.2* | WBGene00021614 | protein_coding_gene | 1.8 | 3.432 | 3.3E-09 |
| *fipr-8* | WBGene00007537 | protein_coding_gene | 1.8 | 3.431 | 4.4E-02 |
| *unc-24* | WBGene00006761 | protein_coding_gene | 1.8 | 3.429 | 1.1E-04 |
| *K07H8.11* | WBGene00044510 | protein_coding_gene | 1.8 | 3.427 | 2.3E-03 |
| *pag-3* | WBGene00003909 | protein_coding_gene | 1.8 | 3.425 | 7.3E-03 |
| *dyf-1* | WBGene00001117 | protein_coding_gene | 1.8 | 3.424 | 3.7E-07 |
| *F08G12.11* | WBGene00044135 | protein_coding_gene | 1.8 | 3.423 | 2.9E-02 |
| *C05A9.2* | WBGene00007317 | protein_coding_gene | 1.8 | 3.422 | 5.1E-09 |
| *F25E2.3* | WBGene00017781 | protein_coding_gene | 1.8 | 3.422 | 9.0E-06 |
| *smf-3* | WBGene00004878 | protein_coding_gene | 1.8 | 3.420 | 7.3E-05 |
| *arrd-21* | WBGene00015441 | pseudogene | 1.8 | 3.419 | 1.2E-04 |
| *C29F7.2* | WBGene00007811 | protein_coding_gene | 1.8 | 3.417 | 3.0E-06 |
| *F46C8.3* | WBGene00018485 | protein_coding_gene | 1.8 | 3.415 | 2.3E-11 |
| *F28H7.7* | WBGene00009240 | protein_coding_gene | 1.8 | 3.412 | 4.6E-02 |
| *linc-46* | WBGene00219748 | lincRNA_gene | 1.8 | 3.411 | 1.8E-04 |
| *mapk-15* | WBGene00015478 | protein_coding_gene | 1.8 | 3.408 | 5.6E-08 |
| *F20D6.10* | WBGene00017639 | protein_coding_gene | 1.8 | 3.408 | 1.2E-05 |
| *clc-2* | WBGene00000523 | protein_coding_gene | 1.8 | 3.407 | 3.5E-07 |
| *gcy-35* | WBGene00001555 | protein_coding_gene | 1.8 | 3.406 | 4.5E-06 |
| *amt-3* | WBGene00000135 | protein_coding_gene | 1.8 | 3.404 | 6.1E-06 |
| *Y75B8A.32* | WBGene00013562 | protein_coding_gene | 1.8 | 3.399 | 1.4E-07 |
| *pqn-71* | WBGene00004153 | protein_coding_gene | 1.8 | 3.396 | 1.3E-03 |
| *Y19D10B.6* | WBGene00021235 | protein_coding_gene | 1.8 | 3.394 | 2.7E-08 |
| *exc-9* | WBGene00017644 | protein_coding_gene | 1.8 | 3.393 | 2.2E-11 |
| *gcy-36* | WBGene00001556 | protein_coding_gene | 1.8 | 3.389 | 2.2E-03 |
| *C05B5.5* | WBGene00007322 | protein_coding_gene | 1.8 | 3.387 | 4.1E-04 |
| *cutl-8* | WBGene00013960 | protein_coding_gene | 1.8 | 3.387 | 3.5E-02 |
| *tbx-7* | WBGene00006544 | protein_coding_gene | 1.8 | 3.382 | 4.8E-03 |
| *F49E10.4* | WBGene00018641 | protein_coding_gene | 1.8 | 3.381 | 1.5E-04 |
| *F26D11.12* | WBGene00045392 | protein_coding_gene | 1.8 | 3.380 | 1.7E-08 |
| *sta-2* | WBGene00010251 | protein_coding_gene | 1.8 | 3.380 | 4.5E-09 |
| *C33G8.4* | WBGene00016362 | protein_coding_gene | 1.8 | 3.379 | 5.3E-13 |
| *C53C7.5* | WBGene00014294 | pseudogene | 1.8 | 3.377 | 1.6E-08 |
| *abu-8* | WBGene00000031 | protein_coding_gene | 1.8 | 3.376 | 1.5E-03 |
| *W02D7.3* | WBGene00020939 | protein_coding_gene | 1.8 | 3.376 | 7.1E-04 |
| *F40F8.4* | WBGene00009577 | protein_coding_gene | 1.8 | 3.374 | 1.3E-13 |

| | | | | | |
|---|---|---|---|---|---|
| *mks-6* | WBGene00010642 | protein_coding_gene | 1.8 | 3.373 | 1.5E-04 |
| *snt-1* | WBGene00004921 | protein_coding_gene | 1.8 | 3.373 | 3.5E-10 |
| *lpr-6* | WBGene00012255 | protein_coding_gene | 1.8 | 3.364 | 1.3E-03 |
| *C54G4.4* | WBGene00008314 | protein_coding_gene | 1.7 | 3.361 | 4.7E-06 |
| *decr-1.1* | WBGene00018380 | protein_coding_gene | 1.7 | 3.357 | 8.3E-06 |
| *jnk-1* | WBGene00002178 | protein_coding_gene | 1.7 | 3.351 | 1.5E-10 |
| *grd-6* | WBGene00001695 | protein_coding_gene | 1.7 | 3.346 | 3.9E-03 |
| *T02C5.1* | WBGene00020158 | protein_coding_gene | 1.7 | 3.345 | 3.8E-08 |
| *M02G9.2* | WBGene00010831 | protein_coding_gene | 1.7 | 3.344 | 8.1E-03 |
| *cuti-1* | WBGene00022591 | protein_coding_gene | 1.7 | 3.344 | 2.2E-06 |
| *F57E7.1* | WBGene00010202 | protein_coding_gene | 1.7 | 3.340 | 1.8E-05 |
| *F20A1.6* | WBGene00017618 | protein_coding_gene | 1.7 | 3.340 | 1.7E-08 |
| *ZC21.6* | WBGene00022506 | protein_coding_gene | 1.7 | 3.337 | 3.3E-02 |
| *C27A2.5* | WBGene00016153 | protein_coding_gene | 1.7 | 3.337 | 4.7E-02 |
| *F40A3.4* | WBGene00018219 | protein_coding_gene | 1.7 | 3.334 | 3.6E-02 |
| *abu-14* | WBGene00004174 | protein_coding_gene | 1.7 | 3.331 | 9.1E-05 |
| *ZK930.2* | WBGene00014152 | protein_coding_gene | 1.7 | 3.329 | 1.0E-10 |
| *pin-2* | WBGene00004030 | protein_coding_gene | 1.7 | 3.328 | 3.6E-11 |
| *dmsr-11* | WBGene00022605 | protein_coding_gene | 1.7 | 3.325 | 4.4E-02 |
| *T25B9.3* | WBGene00012009 | protein_coding_gene | 1.7 | 3.324 | 7.2E-03 |
| *C10A4.3* | WBGene00015664 | protein_coding_gene | 1.7 | 3.323 | 1.5E-02 |
| *K09F6.10* | WBGene00019593 | protein_coding_gene | 1.7 | 3.321 | 1.0E-03 |
| *Y39A3A.4* | WBGene00021435 | protein_coding_gene | 1.7 | 3.317 | 4.0E-03 |
| *dlc-3* | WBGene00012424 | protein_coding_gene | 1.7 | 3.317 | 1.8E-06 |
| *lpr-3* | WBGene00012261 | protein_coding_gene | 1.7 | 3.313 | 5.5E-05 |
| *F11D5.1* | WBGene00017380 | protein_coding_gene | 1.7 | 3.313 | 7.2E-12 |
| *oac-22* | WBGene00018142 | protein_coding_gene | 1.7 | 3.312 | 4.5E-02 |
| *nlp-62* | WBGene00018005 | protein_coding_gene | 1.7 | 3.309 | 5.7E-04 |
| *fbxa-144* | WBGene00009709 | protein_coding_gene | 1.7 | 3.308 | 2.7E-03 |
| *F23C8.13* | WBGene00017741 | protein_coding_gene | 1.7 | 3.308 | 1.0E-04 |
| *Y70C5A.3* | WBGene00050904 | protein_coding_gene | 1.7 | 3.308 | 5.7E-05 |
| *spp-15* | WBGene00005000 | protein_coding_gene | 1.7 | 3.304 | 2.6E-10 |
| *D1054.19* | WBGene00194677 | protein_coding_gene | 1.7 | 3.304 | 2.7E-02 |
| *lin-44* | WBGene00003029 | protein_coding_gene | 1.7 | 3.303 | 4.8E-05 |
| *col-113* | WBGene00000687 | protein_coding_gene | 1.7 | 3.303 | 3.0E-04 |
| *somi-1* | WBGene00010868 | protein_coding_gene | 1.7 | 3.301 | 7.7E-10 |
| *H22K11.3* | WBGene00019226 | protein_coding_gene | 1.7 | 3.297 | 2.5E-03 |
| *C29H12.6* | WBGene00016237 | protein_coding_gene | 1.7 | 3.297 | 7.0E-07 |
| *T24A6.7* | WBGene00020747 | protein_coding_gene | 1.7 | 3.297 | 3.7E-02 |
| *R10D12.7* | WBGene00011188 | protein_coding_gene | 1.7 | 3.296 | 1.8E-02 |
| *F10D7.1* | WBGene00017338 | protein_coding_gene | 1.7 | 3.294 | 4.4E-03 |
| *ZK154.1* | WBGene00022665 | protein_coding_gene | 1.7 | 3.291 | 1.1E-04 |
| *klp-6* | WBGene00002218 | protein_coding_gene | 1.7 | 3.289 | 1.2E-02 |
| *F52E1.5* | WBGene00018694 | protein_coding_gene | 1.7 | 3.287 | 3.7E-04 |
| *abu-10* | WBGene00000033 | protein_coding_gene | 1.7 | 3.287 | 8.3E-04 |
| *npr-20* | WBGene00011578 | protein_coding_gene | 1.7 | 3.286 | 6.8E-06 |
| *tub-1* | WBGene00006655 | protein_coding_gene | 1.7 | 3.286 | 9.4E-06 |
| *dmd-4* | WBGene00007776 | protein_coding_gene | 1.7 | 3.285 | 1.2E-02 |
| *T03F1.11* | WBGene00020192 | protein_coding_gene | 1.7 | 3.285 | 5.4E-08 |
| *C54F6.20* | WBGene00271796 | protein_coding_gene | 1.7 | 3.285 | 2.0E-03 |
| *C17H12.6* | WBGene00015932 | protein_coding_gene | 1.7 | 3.283 | 3.8E-05 |
| *T13C2.2* | WBGene00020478 | protein_coding_gene | 1.7 | 3.283 | 1.1E-03 |
| *fax-1* | WBGene00001400 | protein_coding_gene | 1.7 | 3.280 | 2.4E-02 |
| *math-39* | WBGene00020358 | protein_coding_gene | 1.7 | 3.280 | 3.8E-10 |
| *F32D8.15* | WBGene00009325 | protein_coding_gene | 1.7 | 3.280 | 1.1E-04 |
| *F32B4.5* | WBGene00009315 | protein_coding_gene | 1.7 | 3.277 | 1.9E-04 |
| *ZK180.5* | WBGene00022679 | protein_coding_gene | 1.7 | 3.274 | 1.6E-04 |
| *swt-5* | WBGene00010594 | protein_coding_gene | 1.7 | 3.272 | 1.8E-03 |
| *pqn-89* | WBGene00004169 | protein_coding_gene | 1.7 | 3.271 | 1.9E-06 |
| *hmit-1.3* | WBGene00010811 | protein_coding_gene | 1.7 | 3.271 | 3.4E-05 |
| *rrc-1* | WBGene00009800 | protein_coding_gene | 1.7 | 3.269 | 2.5E-08 |
| *fbxa-188* | WBGene00009839 | protein_coding_gene | 1.7 | 3.268 | 2.4E-05 |
| *E04D5.2* | WBGene00008481 | protein_coding_gene | 1.7 | 3.267 | 1.0E-03 |
| *lrch-1* | WBGene00015779 | protein_coding_gene | 1.7 | 3.266 | 1.3E-06 |
| *alh-5* | WBGene00000111 | protein_coding_gene | 1.7 | 3.263 | 1.5E-04 |
| *fbxl-1* | WBGene00015350 | protein_coding_gene | 1.7 | 3.259 | 6.0E-08 |
| *C32H11.3* | WBGene00007866 | protein_coding_gene | 1.7 | 3.258 | 3.2E-04 |
| *clec-84* | WBGene00021895 | protein_coding_gene | 1.7 | 3.256 | 5.6E-15 |
| *ccch-1* | WBGene00009532 | protein_coding_gene | 1.7 | 3.255 | 2.3E-05 |
| *Y22D7AL.6* | WBGene00021245 | protein_coding_gene | 1.7 | 3.253 | 1.3E-03 |
| *ZK1320.2* | WBGene00014252 | protein_coding_gene | 1.7 | 3.252 | 3.9E-13 |
| *apl-1* | WBGene00000149 | protein_coding_gene | 1.7 | 3.251 | 6.9E-08 |
| *C25E10.7* | WBGene00016096 | protein_coding_gene | 1.7 | 3.250 | 3.0E-02 |
| *C15C8.6* | WBGene00007608 | protein_coding_gene | 1.7 | 3.249 | 4.8E-02 |
| *pbo-4* | WBGene00003943 | protein_coding_gene | 1.7 | 3.248 | 6.6E-15 |
| *pgp-10* | WBGene00004004 | protein_coding_gene | 1.7 | 3.243 | 1.2E-06 |
| *nspb-10* | WBGene00007246 | protein_coding_gene | 1.7 | 3.243 | 1.0E-02 |
| *math-20* | WBGene00016555 | protein_coding_gene | 1.7 | 3.241 | 7.8E-12 |
| *E01G6.1* | WBGene00008449 | protein_coding_gene | 1.7 | 3.239 | 7.9E-04 |
| *C09D4.2* | WBGene00015633 | protein_coding_gene | 1.7 | 3.239 | 1.9E-15 |

| | | | | | |
|---|---|---|---|---|---|
| *zig-13* | WBGene00020546 | protein_coding_gene | 1.7 | 3.238 | 3.0E-02 |
| *F45E4.5* | WBGene00018470 | protein_coding_gene | 1.7 | 3.238 | 1.1E-06 |
| *ZK673.11* | WBGene00014065 | protein_coding_gene | 1.7 | 3.237 | 1.5E-02 |
| *Y38H6C.20* | WBGene00012632 | protein_coding_gene | 1.7 | 3.236 | 4.3E-04 |
| *clec-35* | WBGene00013608 | protein_coding_gene | 1.7 | 3.233 | 1.8E-02 |
| *nhr-55* | WBGene00003645 | protein_coding_gene | 1.7 | 3.232 | 2.6E-06 |
| *pha-2* | WBGene00004011 | protein_coding_gene | 1.7 | 3.231 | 1.7E-02 |
| *T19C3.2* | WBGene00020560 | protein_coding_gene | 1.7 | 3.231 | 1.5E-04 |
| *D1007.13* | WBGene00017009 | protein_coding_gene | 1.7 | 3.230 | 4.4E-03 |
| *sad-1* | WBGene00004719 | protein_coding_gene | 1.7 | 3.228 | 6.0E-09 |
| *Y38H6C.19* | WBGene00012631 | protein_coding_gene | 1.7 | 3.226 | 3.3E-08 |
| *bkip-1* | WBGene00014569 | protein_coding_gene | 1.7 | 3.226 | 3.1E-03 |
| *Y50D4A.10* | WBGene00235315 | protein_coding_gene | 1.7 | 3.226 | 1.8E-05 |
| *nmur-3* | WBGene00017176 | protein_coding_gene | 1.7 | 3.225 | 3.3E-03 |
| *grl-21* | WBGene00001730 | protein_coding_gene | 1.7 | 3.221 | 6.7E-04 |
| *B0286.1* | WBGene00015115 | protein_coding_gene | 1.7 | 3.221 | 5.4E-08 |
| *F39F10.4* | WBGene00018204 | protein_coding_gene | 1.7 | 3.220 | 5.8E-07 |
| *unc-7* | WBGene00006747 | protein_coding_gene | 1.7 | 3.215 | 1.5E-05 |
| *cutl-29* | WBGene00017421 | protein_coding_gene | 1.7 | 3.208 | 8.1E-03 |
| *F18A11.2* | WBGene00008929 | protein_coding_gene | 1.7 | 3.206 | 1.8E-04 |
| *R03G8.3* | WBGene00010998 | protein_coding_gene | 1.7 | 3.205 | 3.8E-06 |
| *K10H10.4* | WBGene00010760 | protein_coding_gene | 1.7 | 3.205 | 5.2E-03 |
| *pgp-11* | WBGene00004005 | protein_coding_gene | 1.7 | 3.202 | 1.3E-15 |
| *K10D3.4* | WBGene00010738 | protein_coding_gene | 1.7 | 3.201 | 6.0E-05 |
| *Y67A10A.7* | WBGene00013456 | protein_coding_gene | 1.7 | 3.201 | 7.2E-04 |
| *cutl-14* | WBGene00015231 | protein_coding_gene | 1.7 | 3.200 | 4.3E-04 |
| *bus-12* | WBGene00010442 | protein_coding_gene | 1.7 | 3.200 | 2.4E-02 |
| *M176.4* | WBGene00010943 | protein_coding_gene | 1.7 | 3.200 | 6.7E-42 |
| *gmeb-2* | WBGene00010010 | protein_coding_gene | 1.7 | 3.198 | 1.2E-03 |
| *F54D5.15* | WBGene00010057 | protein_coding_gene | 1.7 | 3.197 | 2.4E-09 |
| *C26B9.6* | WBGene00016135 | protein_coding_gene | 1.7 | 3.195 | 5.5E-08 |
| *T24C4.4* | WBGene00020760 | protein_coding_gene | 1.7 | 3.195 | 1.7E-04 |
| *T01B7.8* | WBGene00011313 | protein_coding_gene | 1.7 | 3.194 | 2.3E-02 |
| *best-24* | WBGene00022797 | protein_coding_gene | 1.7 | 3.193 | 1.8E-06 |
| *M02G9.1* | WBGene00010830 | protein_coding_gene | 1.7 | 3.193 | 2.2E-03 |
| *T05E11.2* | WBGene00011479 | protein_coding_gene | 1.7 | 3.190 | 1.0E-05 |
| *Y48G9A.7* | WBGene00021701 | protein_coding_gene | 1.7 | 3.189 | 6.7E-04 |
| *clc-5* | WBGene00000526 | protein_coding_gene | 1.7 | 3.188 | 1.5E-08 |
| *cyp-33D1* | WBGene00010589 | protein_coding_gene | 1.7 | 3.185 | 3.1E-03 |
| *emre-1* | WBGene00008980 | protein_coding_gene | 1.7 | 3.185 | 1.0E-16 |
| *R05H10.1* | WBGene00011042 | protein_coding_gene | 1.7 | 3.185 | 1.4E-03 |
| *cyp-32A1* | WBGene00016147 | protein_coding_gene | 1.7 | 3.180 | 6.8E-08 |
| *acp-4* | WBGene00008804 | protein_coding_gene | 1.7 | 3.180 | 3.8E-02 |
| *dct-1* | WBGene00015776 | protein_coding_gene | 1.7 | 3.180 | 5.4E-11 |
| *dct-5* | WBGene00017218 | protein_coding_gene | 1.7 | 3.179 | 2.7E-02 |
| *B0284.3* | WBGene00007133 | protein_coding_gene | 1.7 | 3.179 | 1.7E-06 |
| *F46H5.5* | WBGene00018521 | protein_coding_gene | 1.7 | 3.177 | 9.9E-03 |
| *F26F2.8* | WBGene00023504 | protein_coding_gene | 1.7 | 3.175 | 3.4E-05 |
| *C04G6.2* | WBGene00015453 | protein_coding_gene | 1.7 | 3.174 | 2.3E-03 |
| *F55G1.15* | WBGene00018907 | protein_coding_gene | 1.7 | 3.174 | 4.7E-16 |
| *tnc-2* | WBGene00006583 | protein_coding_gene | 1.7 | 3.174 | 2.3E-09 |
| *mks-1* | WBGene00020100 | protein_coding_gene | 1.7 | 3.171 | 1.4E-03 |
| *C15H9.11* | WBGene00015805 | protein_coding_gene | 1.7 | 3.164 | 1.8E-15 |
| *mul-1* | WBGene00018646 | protein_coding_gene | 1.7 | 3.162 | 5.6E-04 |
| *egl-20* | WBGene00001188 | protein_coding_gene | 1.7 | 3.159 | 1.4E-08 |
| *K02E7.5* | WBGene00019310 | pseudogene | 1.7 | 3.158 | 5.1E-03 |
| *ceh-2* | WBGene00000429 | protein_coding_gene | 1.7 | 3.158 | 2.8E-02 |
| *fbxa-80* | WBGene00022330 | protein_coding_gene | 1.7 | 3.156 | 4.1E-08 |
| *M02D8.6* | WBGene00019732 | protein_coding_gene | 1.7 | 3.156 | 5.2E-08 |
| *gasr-8* | WBGene00008144 | protein_coding_gene | 1.7 | 3.152 | 4.5E-02 |
| *H23N18.6* | WBGene00044534 | protein_coding_gene | 1.7 | 3.148 | 1.9E-02 |
| *fbxa-85* | WBGene00044654 | protein_coding_gene | 1.7 | 3.145 | 4.3E-03 |
| *C02E7.7* | WBGene00015340 | protein_coding_gene | 1.7 | 3.145 | 3.8E-02 |
| *eva-1* | WBGene00009304 | protein_coding_gene | 1.7 | 3.145 | 8.2E-06 |
| *let-522* | WBGene00010870 | protein_coding_gene | 1.7 | 3.143 | 9.8E-10 |
| *mab-31* | WBGene00021834 | protein_coding_gene | 1.7 | 3.138 | 1.4E-08 |
| *F31E9.3* | WBGene00009296 | protein_coding_gene | 1.6 | 3.134 | 1.4E-04 |
| *ensh-1* | WBGene00017013 | protein_coding_gene | 1.6 | 3.132 | 7.9E-05 |
| *lpr-1* | WBGene00002393 | protein_coding_gene | 1.6 | 3.129 | 2.0E-03 |
| *scrm-2* | WBGene00014200 | protein_coding_gene | 1.6 | 3.129 | 5.7E-03 |
| *W08E12.8* | WBGene00021089 | protein_coding_gene | 1.6 | 3.128 | 4.4E-03 |
| *tax-2* | WBGene00006525 | protein_coding_gene | 1.6 | 3.128 | 1.5E-02 |
| *tag-343* | WBGene00018374 | protein_coding_gene | 1.6 | 3.127 | 2.7E-08 |
| *memo-1* | WBGene00016500 | protein_coding_gene | 1.6 | 3.126 | 1.8E-07 |
| *lron-1* | WBGene00008093 | protein_coding_gene | 1.6 | 3.125 | 1.5E-02 |
| *C28G1.6* | WBGene00016191 | protein_coding_gene | 1.6 | 3.121 | 6.4E-06 |
| *ZC317.2* | WBGene00022588 | protein_coding_gene | 1.6 | 3.121 | 4.9E-02 |
| *ZK669.3* | WBGene00014053 | protein_coding_gene | 1.6 | 3.117 | 1.8E-05 |
| *ceh-37* | WBGene00000458 | protein_coding_gene | 1.6 | 3.117 | 7.6E-06 |
| *F58E6.11* | WBGene00010258 | protein_coding_gene | 1.6 | 3.116 | 1.1E-02 |

| | | | | | |
|---|---|---|---|---|---|
| *egas-1* | WBGene00013486 | protein_coding_gene | 1.6 | 3.112 | 4.9E-02 |
| *madd-4* | WBGene00009958 | protein_coding_gene | 1.6 | 3.111 | 1.2E-09 |
| *Y47H10A.4* | WBGene00012960 | protein_coding_gene | 1.6 | 3.109 | 1.2E-05 |
| *C13F10.1* | WBGene00015741 | protein_coding_gene | 1.6 | 3.104 | 4.5E-04 |
| *Y53G8AL.4* | WBGene00050917 | protein_coding_gene | 1.6 | 3.104 | 3.7E-03 |
| *ZC487.1* | WBGene00022623 | protein_coding_gene | 1.6 | 3.101 | 1.4E-03 |
| *ceh-34* | WBGene00000455 | protein_coding_gene | 1.6 | 3.101 | 2.9E-05 |
| *T05H4.4* | WBGene00020267 | protein_coding_gene | 1.6 | 3.098 | 1.4E-03 |
| *ugt-44* | WBGene00008486 | protein_coding_gene | 1.6 | 3.098 | 9.3E-08 |
| *C49C3.8* | WBGene00008198 | protein_coding_gene | 1.6 | 3.096 | 7.0E-15 |
| *T16G1.6* | WBGene00011800 | protein_coding_gene | 1.6 | 3.095 | 1.1E-03 |
| *faah-1* | WBGene00015047 | protein_coding_gene | 1.6 | 3.093 | 6.0E-36 |
| *ZC15.5* | WBGene00013833 | protein_coding_gene | 1.6 | 3.092 | 4.0E-02 |
| *C31H5.7* | WBGene00007858 | protein_coding_gene | 1.6 | 3.088 | 5.5E-03 |
| *W03D2.6* | WBGene00020981 | pseudogene | 1.6 | 3.085 | 1.5E-10 |
| *grl-16* | WBGene00001725 | protein_coding_gene | 1.6 | 3.084 | 8.8E-04 |
| *acc-3* | WBGene00018880 | protein_coding_gene | 1.6 | 3.083 | 1.2E-02 |
| *anr-20* | WBGene00219581 | antisense_lncRNA_gene | 1.6 | 3.079 | 3.0E-02 |
| *F09A5.3* | WBGene00008600 | protein_coding_gene | 1.6 | 3.079 | 2.1E-02 |
| *C34B2.11* | WBGene00016396 | protein_coding_gene | 1.6 | 3.077 | 1.8E-10 |
| *F53A2.1* | WBGene00009949 | protein_coding_gene | 1.6 | 3.073 | 4.5E-04 |
| *Y53F4B.25* | WBGene00013171 | protein_coding_gene | 1.6 | 3.071 | 3.4E-02 |
| *tag-89* | WBGene00006459 | protein_coding_gene | 1.6 | 3.070 | 6.8E-05 |
| *jmjd-3.1* | WBGene00017571 | protein_coding_gene | 1.6 | 3.070 | 4.9E-08 |
| *slt-1* | WBGene00004854 | protein_coding_gene | 1.6 | 3.070 | 1.5E-04 |
| *srh-70* | WBGene00005291 | protein_coding_gene | 1.6 | 3.066 | 1.2E-02 |
| *Y34F4.1* | WBGene00021336 | protein_coding_gene | 1.6 | 3.060 | 4.8E-07 |
| *nhr-122* | WBGene00003712 | protein_coding_gene | 1.6 | 3.060 | 1.0E-19 |
| *T23B3.2* | WBGene00020710 | protein_coding_gene | 1.6 | 3.060 | 2.1E-22 |
| *W05B10.4* | WBGene00012279 | protein_coding_gene | 1.6 | 3.059 | 9.4E-07 |
| *F01E11.3* | WBGene00017156 | protein_coding_gene | 1.6 | 3.058 | 7.1E-03 |
| *K08B5.2* | WBGene00019519 | protein_coding_gene | 1.6 | 3.057 | 4.5E-06 |
| *pes-4* | WBGene00003978 | protein_coding_gene | 1.6 | 3.056 | 1.3E-08 |
| *T10H10.2* | WBGene00020426 | protein_coding_gene | 1.6 | 3.056 | 1.1E-05 |
| *madf-10* | WBGene00013717 | protein_coding_gene | 1.6 | 3.055 | 3.3E-07 |
| *nekl-1* | WBGene00021461 | protein_coding_gene | 1.6 | 3.055 | 7.0E-04 |
| *C49H3.16* | WBGene00219436 | protein_coding_gene | 1.6 | 3.053 | 1.7E-02 |
| *col-90* | WBGene00000665 | protein_coding_gene | 1.6 | 3.053 | 2.7E-03 |
| *Y38H8A.8* | WBGene00045062 | pseudogene | 1.6 | 3.048 | 3.8E-03 |
| *hda-5* | WBGene00009663 | protein_coding_gene | 1.6 | 3.047 | 1.8E-06 |
| *K01A12.3* | WBGene00019287 | protein_coding_gene | 1.6 | 3.046 | 1.1E-02 |
| *K09F6.6* | WBGene00019589 | protein_coding_gene | 1.6 | 3.045 | 6.4E-06 |
| *nmr-1* | WBGene00003774 | protein_coding_gene | 1.6 | 3.045 | 1.8E-02 |
| *C35D10.12* | WBGene00016448 | protein_coding_gene | 1.6 | 3.044 | 2.9E-04 |
| *set-30* | WBGene00022499 | protein_coding_gene | 1.6 | 3.044 | 1.7E-10 |
| *ceh-31* | WBGene00000452 | protein_coding_gene | 1.6 | 3.044 | 8.6E-03 |
| *lgc-55* | WBGene00013746 | protein_coding_gene | 1.6 | 3.043 | 1.7E-04 |
| *skpo-2* | WBGene00015841 | protein_coding_gene | 1.6 | 3.039 | 2.0E-03 |
| *nac-2* | WBGene00003518 | protein_coding_gene | 1.6 | 3.039 | 1.5E-15 |
| *R12E2.14* | WBGene00020039 | protein_coding_gene | 1.6 | 3.039 | 9.4E-03 |
| *klp-11* | WBGene00002222 | protein_coding_gene | 1.6 | 3.038 | 2.6E-05 |
| *daam-1* | WBGene00018976 | protein_coding_gene | 1.6 | 3.035 | 3.4E-06 |
| *mam-1* | WBGene00022504 | protein_coding_gene | 1.6 | 3.032 | 1.0E-02 |
| *F36H9.7* | WBGene00018115 | protein_coding_gene | 1.6 | 3.031 | 1.5E-02 |
| *F41F3.8* | WBGene00044533 | protein_coding_gene | 1.6 | 3.031 | 2.5E-03 |
| *math-3* | WBGene00015608 | protein_coding_gene | 1.6 | 3.031 | 2.8E-10 |
| *Y92H12BR.4* | WBGene00022370 | pseudogene | 1.6 | 3.030 | 2.2E-05 |
| *Y50E8A.5* | WBGene00013048 | protein_coding_gene | 1.6 | 3.028 | 5.8E-07 |
| *F56A11.4* | WBGene00018924 | protein_coding_gene | 1.6 | 3.027 | 4.9E-02 |
| *4R79.2* | WBGene00007067 | protein_coding_gene | 1.6 | 3.025 | 2.8E-03 |
| *F09E5.14* | WBGene00017291 | protein_coding_gene | 1.6 | 3.024 | 7.5E-08 |
| *F35H12.1* | WBGene00018074 | protein_coding_gene | 1.6 | 3.023 | 3.9E-04 |
| *F56H11.2* | WBGene00010175 | protein_coding_gene | 1.6 | 3.023 | 8.7E-05 |
| *T25G12.11* | WBGene00020817 | protein_coding_gene | 1.6 | 3.023 | 9.4E-05 |
| *Y71H2AM.15* | WBGene00022180 | protein_coding_gene | 1.6 | 3.021 | 5.3E-05 |
| *Y37H9A.5* | WBGene00014871 | protein_coding_gene | 1.6 | 3.021 | 2.0E-04 |
| *mig-10* | WBGene00003243 | protein_coding_gene | 1.6 | 3.020 | 5.4E-11 |
| *clec-174* | WBGene00021580 | protein_coding_gene | 1.6 | 3.020 | 1.4E-02 |
| *alh-1* | WBGene00000107 | protein_coding_gene | 1.6 | 3.017 | 4.6E-42 |
| *F33H2.8* | WBGene00009371 | protein_coding_gene | 1.6 | 3.017 | 2.6E-04 |
| *Y34D9A.8* | WBGene00021333 | protein_coding_gene | 1.6 | 3.017 | 1.2E-04 |
| *C06G1.1* | WBGene00015545 | protein_coding_gene | 1.6 | 3.017 | 4.8E-04 |
| *C48D1.1* | WBGene00008177 | protein_coding_gene | 1.6 | 3.013 | 2.7E-14 |
| *sfrp-1* | WBGene00022242 | protein_coding_gene | 1.6 | 3.012 | 1.3E-05 |
| *arl-3* | WBGene00000188 | protein_coding_gene | 1.6 | 3.012 | 9.2E-05 |
| *lin-39* | WBGene00003024 | protein_coding_gene | 1.6 | 3.011 | 1.3E-06 |
| *C50B8.5* | WBGene00008227 | protein_coding_gene | 1.6 | 3.011 | 2.8E-05 |
| *F15D4.2* | WBGene00008859 | protein_coding_gene | 1.6 | 3.008 | 7.6E-29 |
| *abch-1* | WBGene00016973 | protein_coding_gene | 1.6 | 3.008 | 1.8E-02 |
| *Y57G11C.22* | WBGene00013317 | protein_coding_gene | 1.6 | 3.007 | 2.3E-11 |

| | | | | | |
|---|---|---|---|---|---|
| *nhr-106* | WBGene00003696 | protein_coding_gene | 1.6 | 3.007 | 2.3E-05 |
| *F55G11.8* | WBGene00010128 | protein_coding_gene | 1.6 | 3.006 | 2.4E-04 |
| *Y57G11B.2* | WBGene00013295 | protein_coding_gene | 1.6 | 3.006 | 2.0E-06 |
| *T06E4.10* | WBGene00011537 | protein_coding_gene | 1.6 | 3.006 | 2.1E-02 |
| *ncx-9* | WBGene00003574 | protein_coding_gene | 1.6 | 3.006 | 4.6E-02 |
| *F01D5.6* | WBGene00008496 | protein_coding_gene | 1.6 | 3.005 | 6.9E-03 |
| *ncam-1* | WBGene00017184 | protein_coding_gene | 1.6 | 3.005 | 6.7E-09 |
| *math-4* | WBGene00015609 | protein_coding_gene | 1.6 | 2.999 | 3.2E-15 |
| *D2092.8* | WBGene00017068 | protein_coding_gene | 1.6 | 2.998 | 4.8E-02 |
| *ins-14* | WBGene00002097 | protein_coding_gene | 1.6 | 2.998 | 3.1E-02 |
| *calf-1* | WBGene00007119 | protein_coding_gene | 1.6 | 2.995 | 1.7E-04 |
| *nhr-36* | WBGene00017198 | protein_coding_gene | 1.6 | 2.994 | 1.1E-06 |
| *ahr-1* | WBGene00000096 | protein_coding_gene | 1.6 | 2.993 | 8.3E-03 |
| *cog-1* | WBGene00000584 | protein_coding_gene | 1.6 | 2.992 | 2.0E-03 |
| *F48F5.6* | WBGene00009848 | protein_coding_gene | 1.6 | 2.992 | 1.6E-04 |
| *E01G6.2* | WBGene00008450 | protein_coding_gene | 1.6 | 2.990 | 7.6E-09 |
| *nhr-193* | WBGene00010215 | protein_coding_gene | 1.6 | 2.988 | 9.4E-06 |
| *ceh-17* | WBGene00000440 | protein_coding_gene | 1.6 | 2.988 | 4.6E-02 |
| *nhr-69* | WBGene00003659 | protein_coding_gene | 1.6 | 2.985 | 1.9E-09 |
| *K09H11.4* | WBGene00019601 | protein_coding_gene | 1.6 | 2.984 | 2.3E-02 |
| *T01C3.11* | WBGene00011325 | protein_coding_gene | 1.6 | 2.983 | 3.5E-02 |
| *C29F9.2* | WBGene00016217 | protein_coding_gene | 1.6 | 2.981 | 8.6E-05 |
| *T19E7.6* | WBGene00044500 | protein_coding_gene | 1.6 | 2.981 | 1.2E-05 |
| *F32D8.7* | WBGene00009331 | protein_coding_gene | 1.6 | 2.978 | 1.7E-03 |
| *gad-3* | WBGene00015057 | protein_coding_gene | 1.6 | 2.976 | 1.1E-10 |
| *hum-6* | WBGene00002039 | protein_coding_gene | 1.6 | 2.976 | 2.2E-08 |
| *K08D9.4* | WBGene00019525 | protein_coding_gene | 1.6 | 2.974 | 2.4E-03 |
| *T28F4.4* | WBGene00012139 | protein_coding_gene | 1.6 | 2.973 | 5.2E-13 |
| *Y53F4B.27* | WBGene00013173 | protein_coding_gene | 1.6 | 2.972 | 5.7E-03 |
| *ajm-1* | WBGene00000100 | protein_coding_gene | 1.6 | 2.972 | 2.5E-07 |
| *K10D6.4* | WBGene00010744 | protein_coding_gene | 1.6 | 2.972 | 4.4E-09 |
| *spv-1* | WBGene00014051 | protein_coding_gene | 1.6 | 2.971 | 6.3E-09 |
| *glb-33* | WBGene00022284 | protein_coding_gene | 1.6 | 2.971 | 3.4E-12 |
| *F53F4.13* | WBGene00009995 | protein_coding_gene | 1.6 | 2.968 | 1.4E-10 |
| *R12E2.15* | WBGene00020040 | protein_coding_gene | 1.6 | 2.966 | 2.9E-03 |
| *ham-2* | WBGene00001821 | protein_coding_gene | 1.6 | 2.966 | 2.0E-05 |
| *F10G8.8* | WBGene00008666 | protein_coding_gene | 1.6 | 2.966 | 6.4E-12 |
| *bus-8* | WBGene00044623 | protein_coding_gene | 1.6 | 2.961 | 3.1E-04 |
| *ugt-57* | WBGene00017154 | protein_coding_gene | 1.6 | 2.960 | 5.3E-03 |
| *maa-1* | WBGene00007680 | protein_coding_gene | 1.6 | 2.959 | 2.9E-11 |
| *ZK550.6* | WBGene00014000 | protein_coding_gene | 1.6 | 2.958 | 2.1E-04 |
| *VZK822L.2* | WBGene00012163 | protein_coding_gene | 1.6 | 2.958 | 1.2E-02 |
| *F15H10.8* | WBGene00008875 | protein_coding_gene | 1.6 | 2.956 | 1.6E-02 |
| *nhr-96* | WBGene00003686 | protein_coding_gene | 1.6 | 2.954 | 3.2E-06 |
| *dpy-2* | WBGene00001064 | protein_coding_gene | 1.6 | 2.954 | 8.9E-04 |
| *his-60* | WBGene00001934 | protein_coding_gene | 1.6 | 2.953 | 1.5E-08 |
| *M01H9.10* | WBGene00271788 | protein_coding_gene | 1.6 | 2.953 | 8.1E-03 |
| *fbxa-4* | WBGene00044672 | protein_coding_gene | 1.6 | 2.952 | 6.7E-07 |
| *skr-19* | WBGene00004825 | protein_coding_gene | 1.6 | 2.952 | 1.1E-12 |
| *T01B11.1* | WBGene00020138 | protein_coding_gene | 1.6 | 2.951 | 2.9E-02 |
| *F34H10.3* | WBGene00009380 | protein_coding_gene | 1.6 | 2.949 | 2.1E-06 |
| *K06A4.2* | WBGene00010592 | protein_coding_gene | 1.6 | 2.946 | 3.9E-03 |
| *fipr-6* | WBGene00007544 | protein_coding_gene | 1.6 | 2.945 | 3.1E-02 |
| *oig-3* | WBGene00003861 | protein_coding_gene | 1.6 | 2.944 | 3.4E-09 |
| *C04A11.1* | WBGene00007285 | protein_coding_gene | 1.6 | 2.944 | 2.6E-03 |
| *nas-1* | WBGene00003520 | protein_coding_gene | 1.6 | 2.943 | 2.6E-02 |
| *F39E9.10* | WBGene00023036 | pseudogene | 1.6 | 2.940 | 5.3E-08 |
| *fbxa-105* | WBGene00015362 | protein_coding_gene | 1.6 | 2.935 | 2.8E-06 |
| *dhrs-4* | WBGene00010063 | protein_coding_gene | 1.6 | 2.934 | 6.4E-06 |
| *C07A9.12* | WBGene00043994 | protein_coding_gene | 1.6 | 2.931 | 3.6E-05 |
| *nhr-60* | WBGene00003650 | protein_coding_gene | 1.6 | 2.930 | 4.8E-03 |
| *Y54F10AM.6* | WBGene00021850 | protein_coding_gene | 1.6 | 2.929 | 8.9E-05 |
| *K07G5.4* | WBGene00010643 | protein_coding_gene | 1.5 | 2.926 | 2.8E-03 |
| *gpcp-1* | WBGene00019612 | protein_coding_gene | 1.5 | 2.926 | 2.4E-10 |
| *F33G12.6* | WBGene00018018 | protein_coding_gene | 1.5 | 2.921 | 2.0E-19 |
| *K10C9.1* | WBGene00019623 | protein_coding_gene | 1.5 | 2.919 | 5.2E-03 |
| *lgc-20* | WBGene00007191 | protein_coding_gene | 1.5 | 2.919 | 5.9E-04 |
| *mgl-1* | WBGene00003232 | protein_coding_gene | 1.5 | 2.917 | 4.4E-04 |
| *Y39G8B.5* | WBGene00012726 | protein_coding_gene | 1.5 | 2.913 | 7.3E-03 |
| *Y97E10AR.2* | WBGene00022397 | protein_coding_gene | 1.5 | 2.910 | 5.6E-04 |
| *lron-2* | WBGene00022789 | protein_coding_gene | 1.5 | 2.907 | 3.7E-03 |
| *nhr-158* | WBGene00015901 | protein_coding_gene | 1.5 | 2.907 | 1.4E-02 |
| *T05A7.7* | WBGene00020224 | protein_coding_gene | 1.5 | 2.904 | 4.2E-04 |
| *trf-2* | WBGene00022454 | protein_coding_gene | 1.5 | 2.903 | 2.5E-02 |
| *H24G06.1* | WBGene00019237 | protein_coding_gene | 1.5 | 2.900 | 3.8E-10 |
| *ace-3* | WBGene00000037 | protein_coding_gene | 1.5 | 2.899 | 8.9E-06 |
| *zip-5* | WBGene00007932 | protein_coding_gene | 1.5 | 2.899 | 8.5E-13 |
| *mlt-8* | WBGene00021095 | protein_coding_gene | 1.5 | 2.896 | 9.7E-05 |
| *Y69A2AR.31* | WBGene00022100 | protein_coding_gene | 1.5 | 2.893 | 9.0E-05 |
| *T28C12.6* | WBGene00020893 | protein_coding_gene | 1.5 | 2.889 | 1.9E-03 |

| | | | | | |
|---|---|---|---|---|---|
| *hot-5* | WBGene00001990 | protein_coding_gene | 1.5 | 2.888 | 2.4E-02 |
| *C50C3.5* | WBGene00016801 | protein_coding_gene | 1.5 | 2.888 | 1.8E-05 |
| *alh-3* | WBGene00000109 | protein_coding_gene | 1.5 | 2.888 | 3.1E-28 |
| *ric-19* | WBGene00004368 | protein_coding_gene | 1.5 | 2.883 | 1.3E-10 |
| *ztf-16* | WBGene00019960 | protein_coding_gene | 1.5 | 2.882 | 1.8E-06 |
| *pqn-92* | WBGene00004172 | protein_coding_gene | 1.5 | 2.880 | 1.0E-10 |
| *D2030.12* | WBGene00008421 | protein_coding_gene | 1.5 | 2.880 | 3.2E-06 |
| *F49E12.12* | WBGene00009904 | protein_coding_gene | 1.5 | 2.880 | 7.7E-03 |
| *C10B5.3* | WBGene00015671 | protein_coding_gene | 1.5 | 2.875 | 3.3E-11 |
| *T10E10.4* | WBGene00020421 | protein_coding_gene | 1.5 | 2.873 | 1.4E-02 |
| *ceh-79* | WBGene00007749 | protein_coding_gene | 1.5 | 2.871 | 3.2E-04 |
| *nlp-57* | WBGene00008589 | protein_coding_gene | 1.5 | 2.871 | 4.4E-05 |
| *Y7A5A.11* | WBGene00044155 | pseudogene | 1.5 | 2.871 | 4.1E-02 |
| *dhs-18* | WBGene00000981 | protein_coding_gene | 1.5 | 2.870 | 1.4E-04 |
| *nhr-45* | WBGene00003635 | protein_coding_gene | 1.5 | 2.870 | 1.4E-08 |
| *F20C5.11* | WBGene00303015 | protein_coding_gene | 1.5 | 2.870 | 2.5E-07 |
| *K02E10.6* | WBGene00019320 | protein_coding_gene | 1.5 | 2.866 | 6.7E-07 |
| *Y43F8B.2* | WBGene00012813 | protein_coding_gene | 1.5 | 2.866 | 4.9E-10 |
| *Y105C5A.15* | WBGene00013639 | protein_coding_gene | 1.5 | 2.865 | 1.2E-05 |
| *grl-1* | WBGene00001710 | protein_coding_gene | 1.5 | 2.864 | 3.5E-02 |
| *nhr-152* | WBGene00015705 | protein_coding_gene | 1.5 | 2.861 | 9.8E-07 |
| *Y74C9A.1* | WBGene00022275 | protein_coding_gene | 1.5 | 2.861 | 4.4E-02 |
| *syd-1* | WBGene00006363 | protein_coding_gene | 1.5 | 2.860 | 4.1E-04 |
| *F23C8.14* | WBGene00255457 | protein_coding_gene | 1.5 | 2.860 | 3.9E-03 |
| *F11C7.7* | WBGene00017379 | protein_coding_gene | 1.5 | 2.858 | 3.4E-02 |
| *C25D7.15* | WBGene00007722 | protein_coding_gene | 1.5 | 2.858 | 1.6E-04 |
| *sek-4* | WBGene00018035 | protein_coding_gene | 1.5 | 2.857 | 9.6E-10 |
| *tyr-1* | WBGene00015332 | protein_coding_gene | 1.5 | 2.857 | 8.2E-07 |
| *F57C12.6* | WBGene00194687 | protein_coding_gene | 1.5 | 2.856 | 4.3E-04 |
| *tba-5* | WBGene00006531 | protein_coding_gene | 1.5 | 2.855 | 4.1E-03 |
| *lpr-7* | WBGene00020571 | protein_coding_gene | 1.5 | 2.855 | 4.5E-03 |
| *ZK721.4* | WBGene00022811 | protein_coding_gene | 1.5 | 2.852 | 4.9E-02 |
| *nhr-90* | WBGene00003680 | protein_coding_gene | 1.5 | 2.851 | 7.7E-07 |
| *F59D12.3* | WBGene00010331 | protein_coding_gene | 1.5 | 2.851 | 4.4E-02 |
| *B0024.15* | WBGene00007104 | protein_coding_gene | 1.5 | 2.851 | 1.9E-03 |
| *cof-2* | WBGene00000583 | protein_coding_gene | 1.5 | 2.850 | 8.9E-06 |
| *F46C3.6* | WBGene00045253 | protein_coding_gene | 1.5 | 2.849 | 7.7E-06 |
| *max-1* | WBGene00003143 | protein_coding_gene | 1.5 | 2.844 | 8.6E-08 |
| *hpo-15* | WBGene00016061 | protein_coding_gene | 1.5 | 2.840 | 7.7E-11 |
| *K11D12.12* | WBGene00019651 | protein_coding_gene | 1.5 | 2.839 | 4.7E-13 |
| *C36E8.4* | WBGene00007983 | protein_coding_gene | 1.5 | 2.837 | 3.0E-08 |
| *ttr-14* | WBGene00011460 | protein_coding_gene | 1.5 | 2.837 | 1.4E-05 |
| *T06E4.9* | WBGene00011536 | protein_coding_gene | 1.5 | 2.833 | 4.7E-02 |
| *F18E9.1* | WBGene00017568 | protein_coding_gene | 1.5 | 2.833 | 1.8E-02 |
| *C18H9.5* | WBGene00016003 | protein_coding_gene | 1.5 | 2.830 | 6.7E-03 |
| *ZK121.2* | WBGene00022664 | protein_coding_gene | 1.5 | 2.829 | 7.7E-06 |
| *F16B3.3* | WBGene00044774 | protein_coding_gene | 1.5 | 2.827 | 2.9E-04 |
| *lagr-1* | WBGene00006505 | protein_coding_gene | 1.5 | 2.827 | 3.6E-05 |
| *F41C3.8* | WBGene00018274 | protein_coding_gene | 1.5 | 2.820 | 1.1E-04 |
| *iglr-1* | WBGene00008977 | protein_coding_gene | 1.5 | 2.819 | 2.0E-04 |
| *Y113G7A.13* | WBGene00013750 | protein_coding_gene | 1.5 | 2.818 | 1.3E-04 |
| *ceh-45* | WBGene00022837 | protein_coding_gene | 1.5 | 2.818 | 1.8E-02 |
| *pqn-67* | WBGene00004150 | protein_coding_gene | 1.5 | 2.816 | 6.8E-06 |
| *abhd-5.1* | WBGene00016506 | protein_coding_gene | 1.5 | 2.815 | 3.5E-04 |
| *acs-17* | WBGene00016716 | protein_coding_gene | 1.5 | 2.815 | 8.7E-04 |
| *unc-53* | WBGene00006788 | protein_coding_gene | 1.5 | 2.814 | 4.8E-05 |
| *fog-1* | WBGene00001481 | protein_coding_gene | 1.5 | 2.814 | 2.1E-11 |
| *vab-9* | WBGene00006875 | protein_coding_gene | 1.5 | 2.812 | 1.4E-07 |
| *AC7.3* | WBGene00014997 | protein_coding_gene | 1.5 | 2.812 | 1.5E-02 |
| *F36F2.1* | WBGene00009475 | protein_coding_gene | 1.5 | 2.812 | 7.9E-08 |
| *M176.5* | WBGene00010944 | protein_coding_gene | 1.5 | 2.809 | 2.2E-09 |
| *C53C11.5* | WBGene00044770 | protein_coding_gene | 1.5 | 2.809 | 2.4E-07 |
| *K10D11.3* | WBGene00010747 | protein_coding_gene | 1.5 | 2.809 | 1.4E-04 |
| *nuc-1* | WBGene00003828 | protein_coding_gene | 1.5 | 2.808 | 1.0E-10 |
| *T01B10.5* | WBGene00020137 | protein_coding_gene | 1.5 | 2.806 | 7.4E-03 |
| *F43G6.4* | WBGene00009657 | protein_coding_gene | 1.5 | 2.805 | 1.3E-03 |
| *F07E5.5* | WBGene00017210 | protein_coding_gene | 1.5 | 2.805 | 3.0E-10 |
| *K06A4.10* | WBGene00077497 | protein_coding_gene | 1.5 | 2.802 | 3.1E-02 |
| *E02D9.3* | WBGene00304811 | protein_coding_gene | 1.5 | 2.800 | 2.4E-10 |
| *ncr-1* | WBGene00003561 | protein_coding_gene | 1.5 | 2.799 | 1.6E-05 |
| *Y42G9A.1* | WBGene00021531 | protein_coding_gene | 1.5 | 2.798 | 4.1E-06 |
| *uig-1* | WBGene00009337 | protein_coding_gene | 1.5 | 2.798 | 6.3E-10 |
| *F11D5.5* | WBGene00017382 | protein_coding_gene | 1.5 | 2.798 | 1.6E-02 |
| *Y18H1A.9* | WBGene00021213 | protein_coding_gene | 1.5 | 2.797 | 1.5E-03 |
| *ceh-6* | WBGene00000431 | protein_coding_gene | 1.5 | 2.794 | 1.9E-03 |
| *clec-194* | WBGene00013778 | protein_coding_gene | 1.5 | 2.794 | 3.7E-02 |
| *hda-11* | WBGene00007953 | protein_coding_gene | 1.5 | 2.793 | 3.1E-04 |
| *F10E7.9* | WBGene00017350 | protein_coding_gene | 1.5 | 2.793 | 1.1E-07 |
| *clec-31* | WBGene00009854 | protein_coding_gene | 1.5 | 2.792 | 1.9E-04 |
| *bre-2* | WBGene00000267 | protein_coding_gene | 1.5 | 2.791 | 1.1E-05 |

| | | | | | |
|---|---|---|---|---|---|
| *hex-2* | WBGene00015767 | protein_coding_gene | 1.5 | 2.791 | 9.7E-21 |
| *C41D11.6* | WBGene00016565 | protein_coding_gene | 1.5 | 2.790 | 8.8E-04 |
| *dhp-2* | WBGene00000964 | protein_coding_gene | 1.5 | 2.790 | 6.5E-07 |
| *Y42G9A.2* | WBGene00021532 | protein_coding_gene | 1.5 | 2.787 | 1.5E-06 |
| *mam-8* | WBGene00020743 | protein_coding_gene | 1.5 | 2.787 | 1.7E-02 |
| *his-3* | WBGene00001877 | protein_coding_gene | 1.5 | 2.786 | 5.8E-03 |
| *Y71H2AM.25* | WBGene00044760 | protein_coding_gene | 1.5 | 2.784 | 3.4E-02 |
| *ptr-23* | WBGene00004237 | protein_coding_gene | 1.5 | 2.783 | 5.5E-10 |
| *dyf-2* | WBGene00001118 | protein_coding_gene | 1.5 | 2.782 | 8.9E-07 |
| *nhr-123* | WBGene00003713 | protein_coding_gene | 1.5 | 2.779 | 1.3E-03 |
| *prk-2* | WBGene00004183 | protein_coding_gene | 1.5 | 2.778 | 1.4E-10 |
| *ptd-2* | WBGene00004211 | protein_coding_gene | 1.5 | 2.778 | 2.7E-02 |
| *T09F5.1* | WBGene00011665 | protein_coding_gene | 1.5 | 2.771 | 1.8E-02 |
| *zip-1* | WBGene00006986 | protein_coding_gene | 1.5 | 2.770 | 5.7E-12 |
| *nhr-209* | WBGene00011100 | protein_coding_gene | 1.5 | 2.768 | 4.3E-05 |
| *C54E4.4* | WBGene00016919 | protein_coding_gene | 1.5 | 2.768 | 4.1E-07 |
| *lim-8* | WBGene00017904 | protein_coding_gene | 1.5 | 2.767 | 2.7E-08 |
| *acr-3* | WBGene00000043 | protein_coding_gene | 1.5 | 2.767 | 3.4E-02 |
| *dhc-3* | WBGene00007154 | protein_coding_gene | 1.5 | 2.765 | 2.2E-02 |
| *plx-2* | WBGene00004048 | protein_coding_gene | 1.5 | 2.764 | 1.2E-04 |
| *col-89* | WBGene00000664 | protein_coding_gene | 1.5 | 2.764 | 1.7E-03 |
| *C46F11.6* | WBGene00008121 | protein_coding_gene | 1.5 | 2.763 | 4.7E-06 |
| *cest-3* | WBGene00008451 | protein_coding_gene | 1.5 | 2.763 | 8.3E-12 |
| *Y54G2A.7* | WBGene00043702 | protein_coding_gene | 1.5 | 2.763 | 1.5E-06 |
| *C48E7.6* | WBGene00016751 | protein_coding_gene | 1.5 | 2.762 | 2.2E-05 |
| *ZK1236.9* | WBGene00044588 | protein_coding_gene | 1.5 | 2.761 | 1.3E-03 |
| *R13A1.10* | WBGene00044504 | protein_coding_gene | 1.5 | 2.759 | 3.5E-04 |
| *acy-1* | WBGene00000068 | protein_coding_gene | 1.5 | 2.758 | 2.2E-06 |
| *serp-1.1* | WBGene00010337 | protein_coding_gene | 1.5 | 2.753 | 3.8E-09 |
| *F21D9.4* | WBGene00009018 | protein_coding_gene | 1.5 | 2.752 | 1.1E-02 |
| *F38B7.2* | WBGene00009533 | protein_coding_gene | 1.5 | 2.750 | 9.5E-04 |
| *pdcd-2* | WBGene00011116 | protein_coding_gene | 1.5 | 2.748 | 7.3E-25 |
| *Y39B6A.7* | WBGene00012669 | protein_coding_gene | 1.5 | 2.748 | 5.0E-03 |
| *tir-1* | WBGene00006575 | protein_coding_gene | 1.5 | 2.748 | 3.5E-07 |
| *clec-49* | WBGene00012251 | protein_coding_gene | 1.5 | 2.745 | 4.0E-08 |
| *efhd-1* | WBGene00012980 | protein_coding_gene | 1.5 | 2.745 | 1.9E-07 |
| *F57F5.3* | WBGene00010205 | protein_coding_gene | 1.5 | 2.744 | 2.3E-02 |
| *cyp-33C4* | WBGene00018413 | protein_coding_gene | 1.5 | 2.742 | 3.4E-03 |
| *K08D8.1* | WBGene00010655 | protein_coding_gene | 1.5 | 2.742 | 6.4E-03 |
| *lin-22* | WBGene00003008 | protein_coding_gene | 1.5 | 2.742 | 1.3E-02 |
| *C06G3.5* | WBGene00015551 | protein_coding_gene | 1.5 | 2.741 | 1.3E-11 |
| *ctn-1* | WBGene00000832 | protein_coding_gene | 1.5 | 2.739 | 1.7E-22 |
| *dohh-1* | WBGene00007555 | protein_coding_gene | 1.5 | 2.738 | 5.0E-12 |
| *Y75B7AL.2* | WBGene00022285 | protein_coding_gene | 1.5 | 2.738 | 6.2E-04 |
| *F23A7.5* | WBGene00009070 | protein_coding_gene | 1.4 | 2.732 | 2.2E-03 |
| *F49E11.2* | WBGene00009889 | protein_coding_gene | 1.4 | 2.728 | 1.7E-04 |
| *asd-2* | WBGene00006423 | protein_coding_gene | 1.4 | 2.728 | 2.5E-05 |
| *aat-1* | WBGene00000002 | protein_coding_gene | 1.4 | 2.724 | 3.0E-10 |
| *gst-33* | WBGene00001781 | protein_coding_gene | 1.4 | 2.724 | 7.2E-03 |
| *F07H5.8* | WBGene00008559 | protein_coding_gene | 1.4 | 2.721 | 4.8E-03 |
| *R06A10.4* | WBGene00019911 | protein_coding_gene | 1.4 | 2.720 | 4.3E-07 |
| *ets-9* | WBGene00016865 | protein_coding_gene | 1.4 | 2.718 | 3.2E-07 |
| *Y95B8A.6* | WBGene00022386 | protein_coding_gene | 1.4 | 2.717 | 3.6E-04 |
| *K07F5.16* | WBGene00010640 | protein_coding_gene | 1.4 | 2.717 | 7.3E-11 |
| *tag-278* | WBGene00015356 | protein_coding_gene | 1.4 | 2.717 | 7.3E-08 |
| *fbxa-219* | WBGene00044403 | protein_coding_gene | 1.4 | 2.715 | 1.7E-04 |
| *frm-2* | WBGene00001489 | protein_coding_gene | 1.4 | 2.711 | 3.1E-07 |
| *sma-4* | WBGene00004858 | protein_coding_gene | 1.4 | 2.711 | 1.8E-09 |
| *Y41G9A.10* | WBGene00044901 | protein_coding_gene | 1.4 | 2.711 | 1.2E-07 |
| *madd-2* | WBGene00016539 | protein_coding_gene | 1.4 | 2.709 | 6.3E-10 |
| *sulp-4* | WBGene00010788 | protein_coding_gene | 1.4 | 2.707 | 1.5E-05 |
| *dyf-11* | WBGene00001127 | protein_coding_gene | 1.4 | 2.707 | 5.9E-03 |
| *ztf-22* | WBGene00012988 | protein_coding_gene | 1.4 | 2.705 | 1.2E-06 |
| *ges-1* | WBGene00001578 | protein_coding_gene | 1.4 | 2.704 | 5.0E-09 |
| *T05C12.4* | WBGene00011468 | protein_coding_gene | 1.4 | 2.704 | 3.6E-02 |
| *F31F7.2* | WBGene00017965 | protein_coding_gene | 1.4 | 2.704 | 1.4E-02 |
| *sup-12* | WBGene00006321 | protein_coding_gene | 1.4 | 2.703 | 1.6E-05 |
| *C16B8.4* | WBGene00015824 | protein_coding_gene | 1.4 | 2.703 | 9.7E-05 |
| *Y82E9BR.5* | WBGene00022338 | protein_coding_gene | 1.4 | 2.702 | 2.1E-05 |
| *Y49A3A.4* | WBGene00013027 | protein_coding_gene | 1.4 | 2.701 | 1.8E-03 |
| *plc-2* | WBGene00004037 | protein_coding_gene | 1.4 | 2.701 | 2.0E-08 |
| *upb-1* | WBGene00017440 | protein_coding_gene | 1.4 | 2.701 | 6.3E-22 |
| *C32E12.4* | WBGene00016330 | protein_coding_gene | 1.4 | 2.694 | 2.1E-04 |
| *K12B6.9* | WBGene00023432 | protein_coding_gene | 1.4 | 2.692 | 3.8E-05 |
| *K09C6.9* | WBGene00019563 | protein_coding_gene | 1.4 | 2.690 | 5.4E-05 |
| *valv-1* | WBGene00015216 | protein_coding_gene | 1.4 | 2.689 | 2.9E-05 |
| *T02B11.3* | WBGene00020155 | protein_coding_gene | 1.4 | 2.688 | 3.9E-13 |
| *R07E3.7* | WBGene00011108 | protein_coding_gene | 1.4 | 2.687 | 5.0E-04 |
| *nhr-64* | WBGene00003654 | protein_coding_gene | 1.4 | 2.686 | 1.3E-09 |
| *F49H12.4* | WBGene00018655 | protein_coding_gene | 1.4 | 2.686 | 2.9E-02 |

| | | | | | |
|---|---|---|---|---|---|
| *C50E3.6* | WBGene00016817 | protein_coding_gene | 1.4 | 2.685 | 6.8E-05 |
| *Y38A10A.2* | WBGene00021389 | protein_coding_gene | 1.4 | 2.685 | 1.9E-03 |
| *K06G5.1* | WBGene00010605 | protein_coding_gene | 1.4 | 2.683 | 1.3E-15 |
| *nhr-270* | WBGene00020062 | protein_coding_gene | 1.4 | 2.680 | 2.5E-02 |
| *F29A7.4* | WBGene00017914 | protein_coding_gene | 1.4 | 2.678 | 2.5E-02 |
| *col-60* | WBGene00000636 | protein_coding_gene | 1.4 | 2.676 | 4.0E-02 |
| *F59E11.5* | WBGene00019111 | protein_coding_gene | 1.4 | 2.674 | 3.4E-07 |
| *K02E2.8* | WBGene00010509 | protein_coding_gene | 1.4 | 2.673 | 4.3E-02 |
| *F58F9.1* | WBGene00019055 | protein_coding_gene | 1.4 | 2.673 | 3.0E-02 |
| *F18C5.10* | WBGene00017563 | protein_coding_gene | 1.4 | 2.672 | 3.9E-05 |
| *K09F5.6* | WBGene00044698 | protein_coding_gene | 1.4 | 2.669 | 2.8E-04 |
| *lin-17* | WBGene00003006 | protein_coding_gene | 1.4 | 2.668 | 3.4E-06 |
| *K11E4.2* | WBGene00010774 | protein_coding_gene | 1.4 | 2.667 | 2.3E-07 |
| *F46G10.1* | WBGene00009796 | protein_coding_gene | 1.4 | 2.665 | 6.0E-14 |
| *C30F12.5* | WBGene00016264 | protein_coding_gene | 1.4 | 2.664 | 7.4E-12 |
| *C17H11.6* | WBGene00015926 | protein_coding_gene | 1.4 | 2.663 | 9.6E-13 |
| *glr-1* | WBGene00001612 | protein_coding_gene | 1.4 | 2.663 | 2.3E-03 |
| *C35E7.4* | WBGene00016456 | protein_coding_gene | 1.4 | 2.663 | 1.5E-05 |
| *C25F9.5* | WBGene00007725 | protein_coding_gene | 1.4 | 2.661 | 9.9E-05 |
| *C16H3.3* | WBGene00015868 | protein_coding_gene | 1.4 | 2.660 | 5.9E-09 |
| *C42D8.1* | WBGene00016599 | protein_coding_gene | 1.4 | 2.658 | 1.8E-06 |
| *K09F6.15* | WBGene00271744 | protein_coding_gene | 1.4 | 2.658 | 3.3E-06 |
| *ace-2* | WBGene00000036 | protein_coding_gene | 1.4 | 2.657 | 2.5E-07 |
| *T19C3.4* | WBGene00020562 | protein_coding_gene | 1.4 | 2.655 | 4.6E-09 |
| *fbxa-11* | WBGene00020451 | protein_coding_gene | 1.4 | 2.655 | 2.2E-06 |
| *C28G1.2* | WBGene00016187 | protein_coding_gene | 1.4 | 2.649 | 3.9E-02 |
| *magu-3* | WBGene00016841 | protein_coding_gene | 1.4 | 2.645 | 1.1E-03 |
| *dcar-1* | WBGene00007395 | protein_coding_gene | 1.4 | 2.644 | 6.8E-05 |
| *T19D7.6* | WBGene00020574 | protein_coding_gene | 1.4 | 2.643 | 6.3E-04 |
| *F13H6.4* | WBGene00017432 | pseudogene | 1.4 | 2.642 | 6.9E-04 |
| *F39E9.7* | WBGene00018199 | pseudogene | 1.4 | 2.640 | 3.9E-07 |
| *C38H2.3* | WBGene00008020 | protein_coding_gene | 1.4 | 2.640 | 1.4E-03 |
| *K08C7.6* | WBGene00010648 | protein_coding_gene | 1.4 | 2.639 | 2.2E-07 |
| *hst-3.1* | WBGene00002030 | protein_coding_gene | 1.4 | 2.638 | 4.4E-04 |
| *F44A2.5* | WBGene00018405 | protein_coding_gene | 1.4 | 2.638 | 3.3E-06 |
| *lys-7* | WBGene00003096 | protein_coding_gene | 1.4 | 2.637 | 8.0E-05 |
| *Y71G12B.26* | WBGene00022161 | protein_coding_gene | 1.4 | 2.636 | 5.5E-03 |
| *F58D12.3* | WBGene00010250 | protein_coding_gene | 1.4 | 2.636 | 3.4E-02 |
| *ttr-8* | WBGene00020049 | protein_coding_gene | 1.4 | 2.636 | 7.5E-06 |
| *inpp-1* | WBGene00012016 | protein_coding_gene | 1.4 | 2.633 | 1.1E-05 |
| *gcl-1* | WBGene00013382 | protein_coding_gene | 1.4 | 2.632 | 2.8E-05 |
| *fbxa-33* | WBGene00013841 | pseudogene | 1.4 | 2.631 | 2.4E-03 |
| *nhr-134* | WBGene00003724 | protein_coding_gene | 1.4 | 2.630 | 2.9E-06 |
| *C49A1.10* | WBGene00008191 | protein_coding_gene | 1.4 | 2.628 | 1.2E-05 |
| *Y65B4BL.7* | WBGene00022039 | protein_coding_gene | 1.4 | 2.627 | 7.5E-03 |
| *dph-3* | WBGene00010484 | protein_coding_gene | 1.4 | 2.625 | 6.0E-12 |
| *egl-36* | WBGene00001202 | protein_coding_gene | 1.4 | 2.623 | 2.0E-05 |
| *frpr-9* | WBGene00009965 | protein_coding_gene | 1.4 | 2.619 | 1.0E-02 |
| *egl-38* | WBGene00001204 | protein_coding_gene | 1.4 | 2.618 | 3.6E-03 |
| *camt-1* | WBGene00020251 | protein_coding_gene | 1.4 | 2.618 | 2.8E-08 |
| *ttr-33* | WBGene00016505 | protein_coding_gene | 1.4 | 2.618 | 2.0E-07 |
| *fbxa-189* | WBGene00009840 | protein_coding_gene | 1.4 | 2.617 | 1.3E-04 |
| *mlt-10* | WBGene00015646 | protein_coding_gene | 1.4 | 2.616 | 1.2E-06 |
| *ZK682.7* | WBGene00022790 | protein_coding_gene | 1.4 | 2.615 | 3.6E-02 |
| *F53F4.8* | WBGene00009991 | pseudogene | 1.4 | 2.615 | 3.7E-02 |
| *sulp-8* | WBGene00013963 | protein_coding_gene | 1.4 | 2.614 | 2.3E-08 |
| *F35H12.6* | WBGene00018078 | protein_coding_gene | 1.4 | 2.614 | 2.6E-02 |
| *glo-3* | WBGene00010341 | protein_coding_gene | 1.4 | 2.613 | 1.1E-08 |
| *BE0003N10.6* | WBGene00255598 | protein_coding_gene | 1.4 | 2.610 | 1.8E-03 |
| *R05A10.3* | WBGene00011020 | protein_coding_gene | 1.4 | 2.610 | 5.1E-05 |
| *W10C4.1* | WBGene00021124 | protein_coding_gene | 1.4 | 2.609 | 2.8E-02 |
| *K01D12.9* | WBGene00010468 | protein_coding_gene | 1.4 | 2.606 | 2.0E-02 |
| *Y60A3A.16* | WBGene00013364 | protein_coding_gene | 1.4 | 2.606 | 1.5E-04 |
| *nhr-4* | WBGene00003603 | protein_coding_gene | 1.4 | 2.605 | 1.9E-18 |
| *C25F9.14* | WBGene00045461 | protein_coding_gene | 1.4 | 2.605 | 2.1E-04 |
| *M03A8.3* | WBGene00019749 | protein_coding_gene | 1.4 | 2.605 | 6.3E-09 |
| *F53B3.3* | WBGene00018742 | protein_coding_gene | 1.4 | 2.604 | 1.8E-03 |
| *ZK809.9* | WBGene00194934 | protein_coding_gene | 1.4 | 2.603 | 1.9E-03 |
| *vab-19* | WBGene00006882 | protein_coding_gene | 1.4 | 2.602 | 4.3E-07 |
| *ddo-1* | WBGene00008127 | protein_coding_gene | 1.4 | 2.602 | 4.8E-05 |
| *dsl-7* | WBGene00001109 | protein_coding_gene | 1.4 | 2.601 | 5.5E-03 |
| *nhr-110* | WBGene00003700 | protein_coding_gene | 1.4 | 2.600 | 4.4E-06 |
| *glb-8* | WBGene00007741 | protein_coding_gene | 1.4 | 2.598 | 1.9E-03 |
| *sel-7* | WBGene00004764 | protein_coding_gene | 1.4 | 2.597 | 1.3E-02 |
| *xbx-4* | WBGene00016025 | protein_coding_gene | 1.4 | 2.597 | 1.8E-02 |
| *fbxa-32* | WBGene00013839 | protein_coding_gene | 1.4 | 2.596 | 9.6E-05 |
| *phat-1* | WBGene00016732 | protein_coding_gene | 1.4 | 2.595 | 4.8E-02 |
| *arrd-20* | WBGene00015440 | pseudogene | 1.4 | 2.594 | 1.4E-03 |
| *nlg-1* | WBGene00006412 | protein_coding_gene | 1.4 | 2.594 | 3.7E-05 |
| *mab-21* | WBGene00003112 | protein_coding_gene | 1.4 | 2.593 | 8.6E-05 |

| | | | | | |
|---|---|---|---|---|---|
| *H10E21.4* | WBGene00019184 | protein_coding_gene | 1.4 | 2.592 | 6.0E-03 |
| *Y54E10BR.4* | WBGene00021843 | protein_coding_gene | 1.4 | 2.590 | 1.2E-25 |
| *nas-32* | WBGene00003550 | protein_coding_gene | 1.4 | 2.588 | 1.4E-04 |
| *ceh-19* | WBGene00000442 | protein_coding_gene | 1.4 | 2.588 | 1.0E-03 |
| *F30A10.9* | WBGene00009266 | protein_coding_gene | 1.4 | 2.588 | 4.7E-15 |
| *phat-2* | WBGene00016733 | protein_coding_gene | 1.4 | 2.588 | 8.1E-03 |
| *F32D1.3* | WBGene00017983 | protein_coding_gene | 1.4 | 2.586 | 9.2E-04 |
| *riok-1* | WBGene00019698 | protein_coding_gene | 1.4 | 2.586 | 5.5E-10 |
| *K11D12.8* | WBGene00019648 | protein_coding_gene | 1.4 | 2.579 | 1.1E-02 |
| *Y50D4B.1* | WBGene00021740 | protein_coding_gene | 1.4 | 2.576 | 5.2E-07 |
| *C47E12.9* | WBGene00008152 | protein_coding_gene | 1.4 | 2.568 | 2.3E-03 |
| *hhat-2* | WBGene00013312 | protein_coding_gene | 1.4 | 2.566 | 4.5E-03 |
| *nadk-1* | WBGene00022307 | protein_coding_gene | 1.4 | 2.565 | 1.8E-09 |
| *C45B11.6* | WBGene00045379 | protein_coding_gene | 1.4 | 2.565 | 1.4E-03 |
| *T14G11.1* | WBGene00020510 | protein_coding_gene | 1.4 | 2.565 | 8.6E-09 |
| *R12E2.7* | WBGene00020033 | protein_coding_gene | 1.4 | 2.565 | 3.0E-02 |
| *C49F8.1* | WBGene00008213 | protein_coding_gene | 1.4 | 2.563 | 2.0E-04 |
| *nhr-195* | WBGene00019115 | protein_coding_gene | 1.4 | 2.561 | 2.8E-03 |
| *dlg-1* | WBGene00001006 | protein_coding_gene | 1.4 | 2.557 | 4.7E-06 |
| *C25F9.6* | WBGene00007726 | protein_coding_gene | 1.4 | 2.557 | 7.8E-03 |
| *swip-10* | WBGene00018738 | protein_coding_gene | 1.4 | 2.556 | 6.4E-03 |
| *T04A6.1* | WBGene00020200 | protein_coding_gene | 1.4 | 2.556 | 6.4E-08 |
| *dop-6* | WBGene00016037 | protein_coding_gene | 1.4 | 2.555 | 4.0E-02 |
| *M176.9* | WBGene00014813 | protein_coding_gene | 1.4 | 2.555 | 3.3E-03 |
| *egal-1* | WBGene00015680 | protein_coding_gene | 1.3 | 2.547 | 6.6E-07 |
| *ZK563.2* | WBGene00022767 | protein_coding_gene | 1.3 | 2.546 | 1.3E-03 |
| *Y119C1B.3* | WBGene00022469 | protein_coding_gene | 1.3 | 2.545 | 1.8E-02 |
| *gba-3* | WBGene00008706 | protein_coding_gene | 1.3 | 2.542 | 5.7E-10 |
| *nhr-119* | WBGene00003709 | protein_coding_gene | 1.3 | 2.539 | 1.6E-05 |
| *M03F8.6* | WBGene00019765 | protein_coding_gene | 1.3 | 2.538 | 6.4E-03 |
| *ceh-62* | WBGene00011069 | protein_coding_gene | 1.3 | 2.538 | 1.2E-02 |
| *lips-3* | WBGene00008655 | protein_coding_gene | 1.3 | 2.534 | 7.9E-03 |
| *pph-2* | WBGene00004084 | protein_coding_gene | 1.3 | 2.532 | 3.5E-02 |
| *F15E6.9* | WBGene00017489 | protein_coding_gene | 1.3 | 2.530 | 2.4E-04 |
| *npr-12* | WBGene00020689 | protein_coding_gene | 1.3 | 2.529 | 1.2E-04 |
| *shw-3* | WBGene00004793 | protein_coding_gene | 1.3 | 2.529 | 2.3E-04 |
| *atgp-1* | WBGene00000224 | protein_coding_gene | 1.3 | 2.525 | 1.7E-09 |
| *F35E12.9* | WBGene00009433 | protein_coding_gene | 1.3 | 2.523 | 2.4E-06 |
| *F07H5.5* | WBGene00014309 | pseudogene | 1.3 | 2.521 | 7.8E-16 |
| *tag-260* | WBGene00010867 | protein_coding_gene | 1.3 | 2.518 | 6.6E-06 |
| *K07C10.3* | WBGene00194851 | ncRNA_gene | 1.3 | 2.517 | 1.4E-02 |
| *pat-9* | WBGene00003933 | protein_coding_gene | 1.3 | 2.517 | 1.8E-04 |
| *B0198.3* | WBGene00007116 | protein_coding_gene | 1.3 | 2.516 | 5.9E-05 |
| *F20A1.1* | WBGene00017613 | protein_coding_gene | 1.3 | 2.514 | 2.9E-07 |
| *scd-1* | WBGene00004739 | protein_coding_gene | 1.3 | 2.514 | 1.0E-05 |
| *B0281.3* | WBGene00015110 | protein_coding_gene | 1.3 | 2.513 | 6.2E-08 |
| *fbxa-108* | WBGene00010293 | protein_coding_gene | 1.3 | 2.512 | 1.1E-10 |
| *his-37* | WBGene00001911 | protein_coding_gene | 1.3 | 2.510 | 1.9E-10 |
| *atf-8* | WBGene00017535 | protein_coding_gene | 1.3 | 2.510 | 2.2E-02 |
| *nhl-3* | WBGene00003599 | protein_coding_gene | 1.3 | 2.509 | 1.1E-10 |
| *nhr-184* | WBGene00018415 | protein_coding_gene | 1.3 | 2.509 | 6.3E-05 |
| *W04A4.6* | WBGene00012235 | protein_coding_gene | 1.3 | 2.508 | 9.0E-07 |
| *Y54E5A.8* | WBGene00013203 | protein_coding_gene | 1.3 | 2.507 | 7.9E-09 |
| *F09E5.16* | WBGene00017292 | protein_coding_gene | 1.3 | 2.506 | 2.3E-05 |
| *dhs-29* | WBGene00000992 | protein_coding_gene | 1.3 | 2.506 | 3.6E-07 |
| *ttr-27* | WBGene00011284 | protein_coding_gene | 1.3 | 2.506 | 2.5E-03 |
| *nhr-225* | WBGene00020849 | protein_coding_gene | 1.3 | 2.505 | 1.1E-02 |
| *K08D8.4* | WBGene00010658 | protein_coding_gene | 1.3 | 2.504 | 2.0E-03 |
| *trpl-1* | WBGene00018992 | pseudogene | 1.3 | 2.501 | 8.0E-03 |
| *gcy-33* | WBGene00001553 | protein_coding_gene | 1.3 | 2.500 | 1.6E-02 |
| *best-11* | WBGene00008186 | protein_coding_gene | 1.3 | 2.499 | 1.3E-02 |
| *myrf-2* | WBGene00008999 | protein_coding_gene | 1.3 | 2.499 | 4.6E-06 |
| *Y52E8A.3* | WBGene00021796 | protein_coding_gene | 1.3 | 2.496 | 2.7E-04 |
| *F48E3.8* | WBGene00018607 | protein_coding_gene | 1.3 | 2.496 | 8.0E-04 |
| *gpa-10* | WBGene00001672 | protein_coding_gene | 1.3 | 2.495 | 4.3E-02 |
| *dep-1* | WBGene00009717 | protein_coding_gene | 1.3 | 2.492 | 1.6E-09 |
| *F11A5.9* | WBGene00008677 | protein_coding_gene | 1.3 | 2.489 | 9.1E-06 |
| *ccdc-149* | WBGene00009258 | protein_coding_gene | 1.3 | 2.489 | 3.9E-04 |
| *Y71H2AL.2* | WBGene00022165 | protein_coding_gene | 1.3 | 2.488 | 7.5E-03 |
| *F42G2.5* | WBGene00018354 | protein_coding_gene | 1.3 | 2.482 | 3.2E-04 |
| *ser-3* | WBGene00004778 | protein_coding_gene | 1.3 | 2.482 | 1.5E-03 |
| *eat-16* | WBGene00001145 | protein_coding_gene | 1.3 | 2.480 | 2.7E-06 |
| *ZK896.4* | WBGene00014135 | protein_coding_gene | 1.3 | 2.479 | 4.6E-02 |
| *R10E8.1* | WBGene00011207 | protein_coding_gene | 1.3 | 2.477 | 9.0E-11 |
| *F15H10.7* | WBGene00008874 | protein_coding_gene | 1.3 | 2.477 | 2.3E-03 |
| *C50F2.7* | WBGene00016840 | protein_coding_gene | 1.3 | 2.475 | 1.1E-03 |
| *best-18* | WBGene00011868 | protein_coding_gene | 1.3 | 2.474 | 1.4E-07 |
| *C03B1.5* | WBGene00015376 | protein_coding_gene | 1.3 | 2.474 | 4.7E-02 |
| *faah-2* | WBGene00015048 | protein_coding_gene | 1.3 | 2.474 | 1.3E-02 |
| *acs-11* | WBGene00018269 | protein_coding_gene | 1.3 | 2.473 | 2.5E-14 |

| | | | | | |
|---|---|---|---|---|---|
| *sax-1* | WBGene00004727 | protein_coding_gene | 1.3 | 2.472 | 7.2E-10 |
| *R07E3.6* | WBGene00011107 | protein_coding_gene | 1.3 | 2.471 | 2.5E-02 |
| *C14H10.2* | WBGene00007592 | protein_coding_gene | 1.3 | 2.471 | 7.2E-07 |
| *gsto-3* | WBGene00019636 | protein_coding_gene | 1.3 | 2.470 | 2.0E-05 |
| *lep-5* | WBGene00010424 | lincRNA_gene | 1.3 | 2.469 | 6.5E-17 |
| *F21G4.1* | WBGene00009023 | protein_coding_gene | 1.3 | 2.468 | 1.0E-05 |
| *F28F5.6* | WBGene00017905 | protein_coding_gene | 1.3 | 2.466 | 2.8E-04 |
| *shk-1* | WBGene00014261 | protein_coding_gene | 1.3 | 2.466 | 2.9E-06 |
| *tth-1* | WBGene00006649 | protein_coding_gene | 1.3 | 2.465 | 1.1E-07 |
| *del-2* | WBGene00000953 | protein_coding_gene | 1.3 | 2.464 | 9.9E-03 |
| *ceh-48* | WBGene00015934 | protein_coding_gene | 1.3 | 2.462 | 9.6E-05 |
| *Y37E11B.2* | WBGene00021375 | protein_coding_gene | 1.3 | 2.462 | 1.7E-05 |
| *F43D2.7* | WBGene00045516 | protein_coding_gene | 1.3 | 2.461 | 1.3E-03 |
| *pqn-13* | WBGene00004104 | protein_coding_gene | 1.3 | 2.458 | 8.5E-03 |
| *C04A2.15* | WBGene00206529 | protein_coding_gene | 1.3 | 2.458 | 4.2E-02 |
| *K12C11.6* | WBGene00045052 | protein_coding_gene | 1.3 | 2.457 | 3.5E-05 |
| *R07B1.9* | WBGene00011081 | protein_coding_gene | 1.3 | 2.454 | 5.1E-04 |
| *ZC15.10* | WBGene00045050 | protein_coding_gene | 1.3 | 2.454 | 2.1E-02 |
| *tag-297* | WBGene00007999 | protein_coding_gene | 1.3 | 2.453 | 2.1E-02 |
| *nhr-25* | WBGene00003623 | protein_coding_gene | 1.3 | 2.451 | 1.5E-03 |
| *K03B4.9* | WBGene00219236 | protein_coding_gene | 1.3 | 2.451 | 8.3E-03 |
| *M7.8* | WBGene00010884 | protein_coding_gene | 1.3 | 2.450 | 1.8E-03 |
| *egl-5* | WBGene00001174 | protein_coding_gene | 1.3 | 2.448 | 1.3E-03 |
| *R02D5.3* | WBGene00010977 | protein_coding_gene | 1.3 | 2.448 | 6.9E-03 |
| *F44G4.6* | WBGene00009715 | protein_coding_gene | 1.3 | 2.447 | 1.1E-02 |
| *lfe-2* | WBGene00002979 | protein_coding_gene | 1.3 | 2.446 | 3.4E-12 |
| *hlh-12* | WBGene00001956 | protein_coding_gene | 1.3 | 2.444 | 4.2E-02 |
| *clc-1* | WBGene00000522 | protein_coding_gene | 1.3 | 2.444 | 4.8E-04 |
| *E04F6.9* | WBGene00017128 | protein_coding_gene | 1.3 | 2.443 | 9.4E-10 |
| *fmil-1* | WBGene00021362 | protein_coding_gene | 1.3 | 2.443 | 2.4E-04 |
| *unc-30* | WBGene00006766 | protein_coding_gene | 1.3 | 2.443 | 5.1E-03 |
| *Y105C5B.9* | WBGene00013650 | protein_coding_gene | 1.3 | 2.442 | 3.0E-07 |
| *F19B2.6* | WBGene00008945 | protein_coding_gene | 1.3 | 2.440 | 8.2E-05 |
| *cytb-5.2* | WBGene00020931 | protein_coding_gene | 1.3 | 2.439 | 6.5E-08 |
| *fbxa-128* | WBGene00012923 | protein_coding_gene | 1.3 | 2.438 | 1.8E-04 |
| *C02E7.6* | WBGene00015339 | protein_coding_gene | 1.3 | 2.438 | 7.3E-03 |
| *C35E7.6* | WBGene00016458 | protein_coding_gene | 1.3 | 2.437 | 6.0E-08 |
| *acs-7* | WBGene00007228 | protein_coding_gene | 1.3 | 2.437 | 5.2E-03 |
| *fbxa-55* | WBGene00020453 | protein_coding_gene | 1.3 | 2.437 | 5.5E-05 |
| *aatf-1* | WBGene00022269 | protein_coding_gene | 1.3 | 2.436 | 3.5E-17 |
| *frpr-5* | WBGene00008342 | protein_coding_gene | 1.3 | 2.436 | 7.0E-04 |
| *C45G9.4* | WBGene00016675 | protein_coding_gene | 1.3 | 2.434 | 2.0E-02 |
| *F09F9.1* | WBGene00017306 | protein_coding_gene | 1.3 | 2.433 | 1.3E-04 |
| *islo-1* | WBGene00018356 | protein_coding_gene | 1.3 | 2.432 | 1.4E-06 |
| *exc-7* | WBGene00001368 | protein_coding_gene | 1.3 | 2.430 | 2.2E-06 |
| *try-3* | WBGene00006621 | protein_coding_gene | 1.3 | 2.428 | 1.8E-03 |
| *hpo-38* | WBGene00021755 | protein_coding_gene | 1.3 | 2.427 | 2.2E-03 |
| *fhod-1* | WBGene00016735 | protein_coding_gene | 1.3 | 2.427 | 1.4E-05 |
| *tpa-1* | WBGene00006599 | protein_coding_gene | 1.3 | 2.426 | 2.8E-07 |
| *F22D6.15* | WBGene00009056 | protein_coding_gene | 1.3 | 2.425 | 2.6E-03 |
| *ptr-18* | WBGene00004232 | protein_coding_gene | 1.3 | 2.424 | 1.8E-03 |
| *ric-3* | WBGene00004363 | protein_coding_gene | 1.3 | 2.424 | 3.8E-05 |
| *C27B7.7* | WBGene00007764 | protein_coding_gene | 1.3 | 2.421 | 1.0E-02 |
| *set-18* | WBGene00044070 | protein_coding_gene | 1.3 | 2.418 | 1.1E-09 |
| *fbxa-156* | WBGene00007392 | protein_coding_gene | 1.3 | 2.415 | 7.6E-03 |
| *T11B7.2* | WBGene00011707 | protein_coding_gene | 1.3 | 2.414 | 1.1E-02 |
| *unc-83* | WBGene00006815 | protein_coding_gene | 1.3 | 2.412 | 1.1E-13 |
| *sng-1* | WBGene00004912 | protein_coding_gene | 1.3 | 2.411 | 4.0E-05 |
| *F38E11.6* | WBGene00009543 | protein_coding_gene | 1.3 | 2.410 | 2.1E-02 |
| *klo-1* | WBGene00016848 | protein_coding_gene | 1.3 | 2.407 | 1.5E-17 |
| *gmeb-1* | WBGene00015285 | protein_coding_gene | 1.3 | 2.405 | 1.9E-08 |
| *twk-11* | WBGene00006666 | protein_coding_gene | 1.3 | 2.404 | 9.0E-03 |
| *dach-1* | WBGene00020386 | protein_coding_gene | 1.3 | 2.403 | 3.5E-03 |
| *Y43F4A.1* | WBGene00012796 | protein_coding_gene | 1.3 | 2.402 | 7.3E-08 |
| *F53H4.3* | WBGene00010008 | protein_coding_gene | 1.3 | 2.401 | 1.4E-03 |
| *Y41C4A.6* | WBGene00012753 | protein_coding_gene | 1.3 | 2.400 | 2.5E-03 |
| *R11G1.6* | WBGene00020012 | protein_coding_gene | 1.3 | 2.400 | 3.9E-06 |
| *lin-12* | WBGene00003001 | protein_coding_gene | 1.3 | 2.400 | 2.5E-07 |
| *ugt-33* | WBGene00007946 | protein_coding_gene | 1.3 | 2.398 | 2.9E-02 |
| *F31E9.11* | WBGene00195233 | ncRNA_gene | 1.3 | 2.397 | 5.0E-05 |
| *tmd-2* | WBGene00006582 | protein_coding_gene | 1.3 | 2.396 | 1.6E-03 |
| *crb-3* | WBGene00044743 | protein_coding_gene | 1.3 | 2.395 | 5.4E-05 |
| *unc-13* | WBGene00006752 | protein_coding_gene | 1.3 | 2.394 | 2.4E-04 |
| *rpi-2* | WBGene00019536 | protein_coding_gene | 1.3 | 2.393 | 6.5E-03 |
| *enu-3.6* | WBGene00010477 | protein_coding_gene | 1.3 | 2.393 | 6.4E-03 |
| *cest-27* | WBGene00018958 | protein_coding_gene | 1.3 | 2.392 | 1.7E-04 |
| *R05G6.1* | WBGene00019896 | protein_coding_gene | 1.3 | 2.391 | 3.8E-02 |
| *fbxa-94* | WBGene00009225 | protein_coding_gene | 1.3 | 2.389 | 1.6E-02 |
| *unc-64* | WBGene00006798 | protein_coding_gene | 1.3 | 2.389 | 4.1E-10 |
| *D2007.2* | WBGene00017042 | protein_coding_gene | 1.3 | 2.388 | 2.4E-03 |

| | | | | | |
|---|---|---|---|---|---|
| *syd-9* | WBGene00044068 | protein_coding_gene | 1.3 | 2.384 | 1.0E-03 |
| *M01H9.4* | WBGene00019720 | protein_coding_gene | 1.3 | 2.383 | 6.0E-04 |
| *Y57G11C.38* | WBGene00013327 | protein_coding_gene | 1.3 | 2.382 | 2.8E-05 |
| *Y54G2A.52* | WBGene00045397 | protein_coding_gene | 1.3 | 2.382 | 9.8E-05 |
| *elt-3* | WBGene00001251 | protein_coding_gene | 1.3 | 2.381 | 6.6E-05 |
| *fbxa-82* | WBGene00007716 | protein_coding_gene | 1.3 | 2.381 | 2.1E-02 |
| *nhr-151* | WBGene00077594 | pseudogene | 1.3 | 2.379 | 6.0E-04 |
| *F54C9.11* | WBGene00010045 | protein_coding_gene | 1.3 | 2.379 | 7.5E-06 |
| *fbxa-199* | WBGene00005931 | protein_coding_gene | 1.2 | 2.376 | 4.0E-02 |
| *F42C5.9* | WBGene00018349 | protein_coding_gene | 1.2 | 2.375 | 8.7E-16 |
| *ras-2* | WBGene00004311 | protein_coding_gene | 1.2 | 2.375 | 6.1E-12 |
| *K03B4.4* | WBGene00019355 | protein_coding_gene | 1.2 | 2.374 | 8.7E-04 |
| *T04C12.8* | WBGene00011430 | protein_coding_gene | 1.2 | 2.373 | 8.8E-03 |
| *cutl-3* | WBGene00008978 | protein_coding_gene | 1.2 | 2.372 | 1.5E-03 |
| *C04F12.8* | WBGene00007302 | protein_coding_gene | 1.2 | 2.371 | 2.3E-04 |
| *Y45F3A.4* | WBGene00012861 | protein_coding_gene | 1.2 | 2.371 | 6.5E-05 |
| *ZK822.4* | WBGene00014091 | protein_coding_gene | 1.2 | 2.371 | 2.8E-12 |
| *dyf-13* | WBGene00001129 | protein_coding_gene | 1.2 | 2.370 | 4.6E-02 |
| *tiam-1* | WBGene00015704 | protein_coding_gene | 1.2 | 2.366 | 8.3E-08 |
| *M02E1.3* | WBGene00023458 | protein_coding_gene | 1.2 | 2.366 | 2.8E-02 |
| *C37C3.7* | WBGene00016499 | protein_coding_gene | 1.2 | 2.364 | 5.4E-05 |
| *math-40* | WBGene00020359 | protein_coding_gene | 1.2 | 2.364 | 5.2E-06 |
| *chil-5* | WBGene00007470 | protein_coding_gene | 1.2 | 2.363 | 1.9E-02 |
| *C01A2.1* | WBGene00007213 | protein_coding_gene | 1.2 | 2.363 | 2.0E-02 |
| *F19G12.10* | WBGene00195186 | lincRNA_gene | 1.2 | 2.363 | 2.1E-04 |
| *F11E6.8* | WBGene00008711 | protein_coding_gene | 1.2 | 2.363 | 6.4E-05 |
| *lgc-25* | WBGene00010994 | protein_coding_gene | 1.2 | 2.360 | 3.1E-03 |
| *che-13* | WBGene00000492 | protein_coding_gene | 1.2 | 2.359 | 9.4E-03 |
| *elt-6* | WBGene00001253 | protein_coding_gene | 1.2 | 2.358 | 1.4E-04 |
| *nhr-101* | WBGene00003691 | protein_coding_gene | 1.2 | 2.358 | 5.0E-03 |
| *F35D11.3* | WBGene00018044 | protein_coding_gene | 1.2 | 2.357 | 5.9E-08 |
| *nhr-194* | WBGene00019114 | pseudogene | 1.2 | 2.357 | 8.6E-04 |
| *cah-6* | WBGene00000284 | protein_coding_gene | 1.2 | 2.352 | 1.0E-04 |
| *cyp-25A3* | WBGene00014697 | protein_coding_gene | 1.2 | 2.350 | 6.7E-04 |
| *Y73F4A.3* | WBGene00013516 | protein_coding_gene | 1.2 | 2.350 | 1.2E-02 |
| *twk-9* | WBGene00006664 | protein_coding_gene | 1.2 | 2.349 | 2.7E-02 |
| *T28B8.1* | WBGene00012113 | protein_coding_gene | 1.2 | 2.349 | 3.7E-05 |
| *neto-1* | WBGene00019360 | protein_coding_gene | 1.2 | 2.345 | 4.0E-03 |
| *col-120* | WBGene00000694 | protein_coding_gene | 1.2 | 2.345 | 3.7E-02 |
| *tni-4* | WBGene00006586 | protein_coding_gene | 1.2 | 2.341 | 2.6E-07 |
| *C01F1.4* | WBGene00015299 | protein_coding_gene | 1.2 | 2.340 | 1.9E-02 |
| *dma-1* | WBGene00011345 | protein_coding_gene | 1.2 | 2.339 | 1.1E-02 |
| *W06F12.2* | WBGene00012306 | protein_coding_gene | 1.2 | 2.338 | 5.2E-07 |
| *mam-4* | WBGene00011836 | protein_coding_gene | 1.2 | 2.338 | 1.1E-02 |
| *H03E18.1* | WBGene00019148 | protein_coding_gene | 1.2 | 2.338 | 1.5E-03 |
| *F52B11.5* | WBGene00009927 | protein_coding_gene | 1.2 | 2.338 | 2.3E-04 |
| *tax-6* | WBGene00006527 | protein_coding_gene | 1.2 | 2.338 | 1.5E-04 |
| *acn-1* | WBGene00000039 | protein_coding_gene | 1.2 | 2.333 | 3.1E-03 |
| *ZK688.1* | WBGene00022796 | protein_coding_gene | 1.2 | 2.333 | 3.6E-02 |
| *D1086.8* | WBGene00008395 | protein_coding_gene | 1.2 | 2.332 | 3.1E-09 |
| *nhr-153* | WBGene00007546 | protein_coding_gene | 1.2 | 2.330 | 2.1E-05 |
| *C45G9.6* | WBGene00016677 | protein_coding_gene | 1.2 | 2.328 | 5.6E-10 |
| *W03C9.6* | WBGene00012222 | protein_coding_gene | 1.2 | 2.327 | 2.5E-05 |
| *pals-24* | WBGene00016219 | protein_coding_gene | 1.2 | 2.326 | 1.2E-03 |
| *fbxa-62* | WBGene00016521 | protein_coding_gene | 1.2 | 2.325 | 4.5E-02 |
| *Y116A8B.4* | WBGene00013781 | protein_coding_gene | 1.2 | 2.325 | 2.3E-02 |
| *Y6D1A.2* | WBGene00012396 | protein_coding_gene | 1.2 | 2.325 | 8.5E-03 |
| *aakb-1* | WBGene00010115 | protein_coding_gene | 1.2 | 2.322 | 1.2E-11 |
| *C25F9.10* | WBGene00044291 | protein_coding_gene | 1.2 | 2.321 | 1.1E-05 |
| *F26A3.4* | WBGene00009142 | protein_coding_gene | 1.2 | 2.321 | 2.1E-09 |
| *unc-51* | WBGene00006786 | protein_coding_gene | 1.2 | 2.321 | 9.8E-05 |
| *lron-15* | WBGene00016974 | protein_coding_gene | 1.2 | 2.317 | 2.0E-03 |
| *C29F7.1* | WBGene00007810 | protein_coding_gene | 1.2 | 2.315 | 2.3E-06 |
| *agmo-1* | WBGene00007210 | protein_coding_gene | 1.2 | 2.314 | 1.5E-02 |
| *T05B11.1* | WBGene00020245 | protein_coding_gene | 1.2 | 2.314 | 2.9E-06 |
| *C18E9.9* | WBGene00007689 | protein_coding_gene | 1.2 | 2.313 | 8.5E-04 |
| *ZK384.7* | WBGene00269383 | ncRNA_gene | 1.2 | 2.313 | 3.3E-03 |
| *F14B6.6* | WBGene00008787 | protein_coding_gene | 1.2 | 2.312 | 7.6E-04 |
| *T07E3.2* | WBGene00020313 | protein_coding_gene | 1.2 | 2.312 | 2.2E-02 |
| *C39E9.7* | WBGene00008031 | protein_coding_gene | 1.2 | 2.311 | 6.5E-04 |
| *let-23* | WBGene00002299 | protein_coding_gene | 1.2 | 2.310 | 7.1E-05 |
| *ins-22* | WBGene00002105 | protein_coding_gene | 1.2 | 2.307 | 7.8E-03 |
| *nhr-56* | WBGene00003646 | protein_coding_gene | 1.2 | 2.305 | 2.7E-05 |
| *tsp-7* | WBGene00006633 | protein_coding_gene | 1.2 | 2.303 | 1.2E-04 |
| *gly-14* | WBGene00001639 | protein_coding_gene | 1.2 | 2.302 | 5.6E-04 |
| *F23C8.3* | WBGene00017732 | protein_coding_gene | 1.2 | 2.299 | 1.8E-03 |
| *C50B6.9* | WBGene00008222 | protein_coding_gene | 1.2 | 2.299 | 1.2E-03 |
| *dnj-24* | WBGene00001042 | protein_coding_gene | 1.2 | 2.298 | 4.3E-04 |
| *sma-9* | WBGene00004862 | protein_coding_gene | 1.2 | 2.297 | 4.5E-04 |
| *T19H12.3* | WBGene00020589 | protein_coding_gene | 1.2 | 2.292 | 9.6E-09 |

| | | | | | |
|---|---|---|---|---|---|
| *F27C8.2* | WBGene00009183 | protein_coding_gene | 1.2 | 2.291 | 2.0E-02 |
| *frm-7* | WBGene00001493 | protein_coding_gene | 1.2 | 2.290 | 2.7E-04 |
| *F13B12.2* | WBGene00008730 | protein_coding_gene | 1.2 | 2.289 | 6.0E-03 |
| *cccp-1* | WBGene00013042 | protein_coding_gene | 1.2 | 2.287 | 7.2E-07 |
| *C46G7.1* | WBGene00016721 | protein_coding_gene | 1.2 | 2.285 | 6.0E-11 |
| *C33A12.19* | WBGene00007890 | protein_coding_gene | 1.2 | 2.285 | 2.1E-07 |
| *icap-1* | WBGene00012889 | protein_coding_gene | 1.2 | 2.284 | 2.5E-07 |
| *C49C3.15* | WBGene00008204 | protein_coding_gene | 1.2 | 2.284 | 3.2E-06 |
| *Y53G8AL.1* | WBGene00021799 | protein_coding_gene | 1.2 | 2.283 | 6.9E-03 |
| *F16F9.1* | WBGene00017513 | protein_coding_gene | 1.2 | 2.280 | 8.7E-06 |
| *lec-3* | WBGene00002266 | protein_coding_gene | 1.2 | 2.278 | 3.0E-05 |
| *atf-6* | WBGene00000222 | protein_coding_gene | 1.2 | 2.277 | 4.6E-05 |
| *R13A1.5* | WBGene00020043 | protein_coding_gene | 1.2 | 2.277 | 4.1E-03 |
| *K10D11.6* | WBGene00010750 | protein_coding_gene | 1.2 | 2.273 | 4.4E-04 |
| *fbxa-67* | WBGene00020457 | protein_coding_gene | 1.2 | 2.272 | 1.5E-09 |
| *Y47D7A.13* | WBGene00021625 | protein_coding_gene | 1.2 | 2.272 | 3.4E-02 |
| *B0546.3* | WBGene00015250 | protein_coding_gene | 1.2 | 2.272 | 1.8E-07 |
| *tbc-18* | WBGene00018281 | protein_coding_gene | 1.2 | 2.271 | 1.1E-04 |
| *F21A9.2* | WBGene00017651 | protein_coding_gene | 1.2 | 2.271 | 2.1E-03 |
| *endu-1* | WBGene00010488 | protein_coding_gene | 1.2 | 2.268 | 1.0E-07 |
| *Y57A10A.14* | WBGene00013257 | protein_coding_gene | 1.2 | 2.264 | 1.2E-03 |
| *rig-1* | WBGene00004369 | protein_coding_gene | 1.2 | 2.264 | 7.8E-06 |
| *prx-2* | WBGene00004192 | protein_coding_gene | 1.2 | 2.263 | 9.0E-04 |
| *wdr-60* | WBGene00016165 | protein_coding_gene | 1.2 | 2.263 | 3.6E-03 |
| *nep-21* | WBGene00011794 | protein_coding_gene | 1.2 | 2.262 | 6.5E-05 |
| *K09C4.1* | WBGene00019547 | protein_coding_gene | 1.2 | 2.262 | 1.8E-02 |
| *F07F6.7* | WBGene00017219 | protein_coding_gene | 1.2 | 2.261 | 1.4E-04 |
| *ugt-5* | WBGene00013906 | protein_coding_gene | 1.2 | 2.260 | 3.4E-09 |
| *B0546.4* | WBGene00015251 | protein_coding_gene | 1.2 | 2.257 | 5.5E-08 |
| *Y75B8A.28* | WBGene00013559 | protein_coding_gene | 1.2 | 2.257 | 6.6E-06 |
| *M02H5.8* | WBGene00019744 | protein_coding_gene | 1.2 | 2.256 | 4.6E-06 |
| *Y40B1B.7* | WBGene00012739 | protein_coding_gene | 1.2 | 2.256 | 1.4E-13 |
| *aex-6* | WBGene00000089 | protein_coding_gene | 1.2 | 2.252 | 3.3E-06 |
| *skr-20* | WBGene00004826 | protein_coding_gene | 1.2 | 2.252 | 4.6E-14 |
| *unc-6* | WBGene00006746 | protein_coding_gene | 1.2 | 2.250 | 8.5E-05 |
| *C30F2.5* | WBGene00007819 | protein_coding_gene | 1.2 | 2.250 | 1.7E-04 |
| *che-3* | WBGene00000485 | protein_coding_gene | 1.2 | 2.249 | 5.3E-03 |
| *fbxa-167* | WBGene00007837 | protein_coding_gene | 1.2 | 2.247 | 5.1E-06 |
| *F41C3.11* | WBGene00018275 | protein_coding_gene | 1.2 | 2.243 | 3.7E-06 |
| *ZK512.11* | WBGene00013989 | protein_coding_gene | 1.2 | 2.243 | 1.7E-04 |
| *F36H5.14* | WBGene00219271 | protein_coding_gene | 1.2 | 2.242 | 9.4E-03 |
| *hst-3.2* | WBGene00044303 | protein_coding_gene | 1.2 | 2.240 | 5.8E-03 |
| *tag-273* | WBGene00013289 | protein_coding_gene | 1.2 | 2.239 | 5.8E-05 |
| *T22C8.3* | WBGene00011924 | protein_coding_gene | 1.2 | 2.238 | 1.9E-06 |
| *ulp-3* | WBGene00006738 | protein_coding_gene | 1.2 | 2.236 | 3.0E-12 |
| *pitr-3* | WBGene00015055 | protein_coding_gene | 1.2 | 2.236 | 1.7E-03 |
| *ZK1240.2* | WBGene00022867 | protein_coding_gene | 1.2 | 2.235 | 3.6E-08 |
| *Y34F4.3* | WBGene00021338 | protein_coding_gene | 1.2 | 2.232 | 3.8E-02 |
| *ctl-2* | WBGene00000831 | protein_coding_gene | 1.2 | 2.231 | 8.6E-06 |
| *eps-8* | WBGene00001330 | protein_coding_gene | 1.2 | 2.229 | 1.2E-05 |
| *C38H2.2* | WBGene00008019 | protein_coding_gene | 1.2 | 2.229 | 8.9E-13 |
| *K10H10.12* | WBGene00045403 | protein_coding_gene | 1.2 | 2.229 | 4.9E-02 |
| *npa-1* | WBGene00003786 | protein_coding_gene | 1.2 | 2.229 | 2.0E-06 |
| *F53A2.9* | WBGene00009953 | protein_coding_gene | 1.2 | 2.229 | 1.5E-05 |
| *F28D1.6* | WBGene00009216 | protein_coding_gene | 1.2 | 2.228 | 1.5E-03 |
| *ZC449.5* | WBGene00022615 | protein_coding_gene | 1.2 | 2.225 | 6.2E-04 |
| *pqn-44* | WBGene00004131 | protein_coding_gene | 1.2 | 2.225 | 4.0E-05 |
| *Y97E10AR.1* | WBGene00022396 | protein_coding_gene | 1.2 | 2.224 | 4.7E-03 |
| *ZK792.7* | WBGene00014079 | protein_coding_gene | 1.2 | 2.223 | 3.9E-06 |
| *F43C9.2* | WBGene00018376 | protein_coding_gene | 1.2 | 2.222 | 1.7E-04 |
| *W02B12.13* | WBGene00043534 | protein_coding_gene | 1.2 | 2.222 | 5.1E-03 |
| *F20A1.10* | WBGene00017621 | protein_coding_gene | 1.2 | 2.221 | 9.5E-06 |
| *M03F8.1* | WBGene00019761 | protein_coding_gene | 1.2 | 2.221 | 3.7E-02 |
| *blmp-1* | WBGene00003847 | protein_coding_gene | 1.2 | 2.220 | 2.0E-03 |
| *pmk-3* | WBGene00004057 | protein_coding_gene | 1.2 | 2.219 | 1.1E-06 |
| *M116.2* | WBGene00019789 | pseudogene | 1.1 | 2.219 | 1.4E-07 |
| *ZC247.2* | WBGene00013860 | protein_coding_gene | 1.1 | 2.219 | 1.2E-04 |
| *unc-14* | WBGene00006753 | protein_coding_gene | 1.1 | 2.216 | 7.9E-09 |
| *T27E4.1* | WBGene00020861 | protein_coding_gene | 1.1 | 2.216 | 6.1E-03 |
| *T13H10.2* | WBGene00011762 | protein_coding_gene | 1.1 | 2.215 | 2.7E-02 |
| *sgca-1* | WBGene00019227 | protein_coding_gene | 1.1 | 2.215 | 3.4E-07 |
| *F46G11.4* | WBGene00018517 | protein_coding_gene | 1.1 | 2.214 | 1.6E-02 |
| *Y57E12AR.1* | WBGene00044612 | protein_coding_gene | 1.1 | 2.210 | 2.3E-03 |
| *zfh-2* | WBGene00022518 | protein_coding_gene | 1.1 | 2.208 | 4.5E-04 |
| *linc-35* | WBGene00219680 | lincRNA_gene | 1.1 | 2.208 | 3.2E-02 |
| *F58A3.3* | WBGene00010223 | protein_coding_gene | 1.1 | 2.206 | 1.0E-03 |
| *unc-41* | WBGene00006777 | protein_coding_gene | 1.1 | 2.206 | 1.8E-04 |
| *C10E2.2* | WBGene00015673 | protein_coding_gene | 1.1 | 2.206 | 3.1E-04 |
| *fln-2* | WBGene00016006 | protein_coding_gene | 1.1 | 2.203 | 1.7E-03 |
| *chw-1* | WBGene00009059 | protein_coding_gene | 1.1 | 2.203 | 2.0E-05 |

| | | | | | |
|---|---|---|---|---|---|
| *F40A3.1* | WBGene00018216 | protein_coding_gene | 1.1 | 2.203 | 1.0E-10 |
| *K02D10.4* | WBGene00019304 | protein_coding_gene | 1.1 | 2.195 | 1.6E-03 |
| *stn-2* | WBGene00017866 | protein_coding_gene | 1.1 | 2.195 | 2.1E-04 |
| *acr-15* | WBGene00000054 | protein_coding_gene | 1.1 | 2.195 | 8.7E-04 |
| *M01G12.9* | WBGene00010822 | protein_coding_gene | 1.1 | 2.195 | 1.5E-02 |
| *dyf-19* | WBGene00012826 | protein_coding_gene | 1.1 | 2.194 | 4.2E-02 |
| *F38B6.6* | WBGene00018175 | protein_coding_gene | 1.1 | 2.193 | 3.7E-03 |
| *F49H12.3* | WBGene00018654 | protein_coding_gene | 1.1 | 2.193 | 3.4E-02 |
| *lgg-2* | WBGene00002981 | protein_coding_gene | 1.1 | 2.193 | 1.5E-05 |
| *R03A10.5* | WBGene00010985 | protein_coding_gene | 1.1 | 2.192 | 3.3E-08 |
| *Y56A3A.7* | WBGene00013228 | protein_coding_gene | 1.1 | 2.191 | 1.0E-10 |
| *Y102A11A.1* | WBGene00022411 | protein_coding_gene | 1.1 | 2.189 | 3.9E-02 |
| *nhr-5* | WBGene00003604 | protein_coding_gene | 1.1 | 2.188 | 9.3E-07 |
| *VW02B12L.2* | WBGene00012155 | protein_coding_gene | 1.1 | 2.188 | 2.9E-02 |
| *magu-4* | WBGene00009678 | protein_coding_gene | 1.1 | 2.188 | 1.7E-09 |
| *ugt-43* | WBGene00008485 | protein_coding_gene | 1.1 | 2.188 | 9.7E-03 |
| *nit-1* | WBGene00014206 | protein_coding_gene | 1.1 | 2.185 | 1.2E-05 |
| *linc-14* | WBGene00195060 | lincRNA_gene | 1.1 | 2.184 | 7.1E-03 |
| *K02A11.4* | WBGene00044607 | protein_coding_gene | 1.1 | 2.184 | 1.3E-03 |
| *bbs-5* | WBGene00010974 | protein_coding_gene | 1.1 | 2.180 | 6.5E-03 |
| *nhr-61* | WBGene00003651 | protein_coding_gene | 1.1 | 2.179 | 1.4E-08 |
| *math-26* | WBGene00018100 | protein_coding_gene | 1.1 | 2.179 | 4.5E-06 |
| *ZK899.1* | WBGene00014140 | protein_coding_gene | 1.1 | 2.179 | 1.5E-05 |
| *Y55F3AM.11* | WBGene00021928 | protein_coding_gene | 1.1 | 2.178 | 2.1E-06 |
| *C01H6.3* | WBGene00007253 | protein_coding_gene | 1.1 | 2.178 | 4.3E-02 |
| *col-153* | WBGene00000726 | protein_coding_gene | 1.1 | 2.174 | 2.2E-04 |
| *hmbx-1* | WBGene00018786 | protein_coding_gene | 1.1 | 2.174 | 1.4E-03 |
| *nob-1* | WBGene00003779 | protein_coding_gene | 1.1 | 2.173 | 5.0E-04 |
| *cyp-31A1* | WBGene00000374 | pseudogene | 1.1 | 2.172 | 5.4E-18 |
| *F47G9.4* | WBGene00009831 | protein_coding_gene | 1.1 | 2.170 | 5.2E-04 |
| *twk-31* | WBGene00006683 | protein_coding_gene | 1.1 | 2.170 | 5.1E-03 |
| *F07C6.3* | WBGene00008554 | protein_coding_gene | 1.1 | 2.169 | 1.9E-05 |
| *F27D9.7* | WBGene00017865 | protein_coding_gene | 1.1 | 2.169 | 2.0E-03 |
| *cyp-42A1* | WBGene00013585 | protein_coding_gene | 1.1 | 2.169 | 4.9E-03 |
| *spp-3* | WBGene00004988 | protein_coding_gene | 1.1 | 2.167 | 4.1E-08 |
| *C01F6.14* | WBGene00219207 | protein_coding_gene | 1.1 | 2.166 | 4.6E-04 |
| *T24H10.5* | WBGene00012003 | protein_coding_gene | 1.1 | 2.165 | 2.5E-02 |
| *C14A4.12* | WBGene00007562 | protein_coding_gene | 1.1 | 2.164 | 1.7E-08 |
| *T25F10.3* | WBGene00020806 | protein_coding_gene | 1.1 | 2.163 | 2.6E-04 |
| *mfn-1* | WBGene00012204 | protein_coding_gene | 1.1 | 2.163 | 6.3E-15 |
| *C09F9.2* | WBGene00007479 | protein_coding_gene | 1.1 | 2.161 | 9.7E-03 |
| *snx-17* | WBGene00008927 | protein_coding_gene | 1.1 | 2.158 | 6.0E-07 |
| *nhr-16* | WBGene00003615 | protein_coding_gene | 1.1 | 2.157 | 3.5E-03 |
| *mrps-11* | WBGene00012244 | protein_coding_gene | 1.1 | 2.157 | 9.0E-14 |
| *bath-38* | WBGene00009945 | protein_coding_gene | 1.1 | 2.157 | 6.3E-12 |
| *nhr-100* | WBGene00003690 | protein_coding_gene | 1.1 | 2.154 | 1.2E-09 |
| *frpr-8* | WBGene00018728 | protein_coding_gene | 1.1 | 2.153 | 2.7E-02 |
| *Y41C4A.8* | WBGene00012755 | protein_coding_gene | 1.1 | 2.152 | 5.1E-04 |
| *C36C5.12* | WBGene00016481 | protein_coding_gene | 1.1 | 2.152 | 1.6E-02 |
| *F27D9.2* | WBGene00017861 | protein_coding_gene | 1.1 | 2.152 | 2.5E-06 |
| *nhr-83* | WBGene00003673 | pseudogene | 1.1 | 2.150 | 5.0E-03 |
| *Y54F10BM.12* | WBGene00021866 | protein_coding_gene | 1.1 | 2.150 | 1.2E-02 |
| *dgk-5* | WBGene00019428 | protein_coding_gene | 1.1 | 2.150 | 7.9E-05 |
| *T23F2.3* | WBGene00020736 | protein_coding_gene | 1.1 | 2.149 | 2.9E-08 |
| *W06B4.2* | WBGene00021057 | protein_coding_gene | 1.1 | 2.148 | 3.9E-17 |
| *E02H1.1* | WBGene00008455 | protein_coding_gene | 1.1 | 2.148 | 4.9E-15 |
| *F37C12.10* | WBGene00018153 | protein_coding_gene | 1.1 | 2.148 | 1.0E-02 |
| *bar-1* | WBGene00000238 | protein_coding_gene | 1.1 | 2.146 | 2.3E-06 |
| *plp-2* | WBGene00008976 | protein_coding_gene | 1.1 | 2.146 | 4.3E-02 |
| *F52E1.2* | WBGene00018692 | protein_coding_gene | 1.1 | 2.143 | 1.1E-04 |
| *ncs-2* | WBGene00003564 | protein_coding_gene | 1.1 | 2.142 | 1.3E-07 |
| *kpc-1* | WBGene00002232 | protein_coding_gene | 1.1 | 2.138 | 1.5E-03 |
| *F48E8.3* | WBGene00018610 | protein_coding_gene | 1.1 | 2.137 | 1.7E-16 |
| *Y34B4A.4* | WBGene00021320 | protein_coding_gene | 1.1 | 2.137 | 7.7E-05 |
| *C09F5.1* | WBGene00015647 | protein_coding_gene | 1.1 | 2.134 | 2.8E-07 |
| *arl-13* | WBGene00021349 | protein_coding_gene | 1.1 | 2.133 | 3.3E-02 |
| *C17E4.11* | WBGene00044236 | protein_coding_gene | 1.1 | 2.133 | 3.5E-07 |
| *T01E8.1* | WBGene00011331 | protein_coding_gene | 1.1 | 2.131 | 1.8E-03 |
| *Y48G9A.9* | WBGene00021702 | protein_coding_gene | 1.1 | 2.131 | 9.2E-05 |
| *R02D5.1* | WBGene00010976 | protein_coding_gene | 1.1 | 2.127 | 9.7E-03 |
| *ZC239.15* | WBGene00022573 | protein_coding_gene | 1.1 | 2.127 | 1.2E-03 |
| *E02H9.4* | WBGene00017102 | protein_coding_gene | 1.1 | 2.127 | 8.9E-03 |
| *best-4* | WBGene00015288 | protein_coding_gene | 1.1 | 2.124 | 2.6E-02 |
| *F48F7.5* | WBGene00009850 | protein_coding_gene | 1.1 | 2.121 | 9.1E-05 |
| *H06I04.6* | WBGene00019170 | protein_coding_gene | 1.1 | 2.119 | 9.8E-04 |
| *F23H12.3* | WBGene00009093 | protein_coding_gene | 1.1 | 2.119 | 2.0E-06 |
| *ttr-30* | WBGene00020335 | protein_coding_gene | 1.1 | 2.119 | 4.4E-05 |
| *Y116A8C.29* | WBGene00013806 | protein_coding_gene | 1.1 | 2.118 | 1.1E-02 |
| *gpa-7* | WBGene00001669 | protein_coding_gene | 1.1 | 2.117 | 9.5E-06 |
| *Y45F10D.2* | WBGene00012883 | protein_coding_gene | 1.1 | 2.117 | 3.7E-02 |

| | | | | | |
|---|---|---|---|---|---|
| *cec-1* | WBGene00000414 | protein_coding_gene | 1.1 | 2.117 | 1.7E-08 |
| *Y59C2A.1* | WBGene00021984 | protein_coding_gene | 1.1 | 2.116 | 2.8E-04 |
| *C40H1.7* | WBGene00008042 | protein_coding_gene | 1.1 | 2.115 | 3.9E-05 |
| *F58A6.1* | WBGene00019022 | protein_coding_gene | 1.1 | 2.115 | 2.2E-03 |
| *nhr-135* | WBGene00003725 | protein_coding_gene | 1.1 | 2.113 | 9.2E-05 |
| *nhr-177* | WBGene00017503 | protein_coding_gene | 1.1 | 2.110 | 1.5E-04 |
| *gpn-1* | WBGene00001687 | protein_coding_gene | 1.1 | 2.110 | 2.0E-03 |
| *ZK1240.5* | WBGene00022870 | protein_coding_gene | 1.1 | 2.109 | 2.3E-03 |
| *M01G12.14* | WBGene00010825 | protein_coding_gene | 1.1 | 2.109 | 3.5E-02 |
| *msi-1* | WBGene00003423 | protein_coding_gene | 1.1 | 2.109 | 1.3E-03 |
| *C02D5.2* | WBGene00015336 | protein_coding_gene | 1.1 | 2.108 | 4.1E-04 |
| *col-150* | WBGene00000723 | protein_coding_gene | 1.1 | 2.107 | 1.9E-04 |
| *C36B7.5* | WBGene00016468 | protein_coding_gene | 1.1 | 2.105 | 2.3E-02 |
| *Y73F8A.5* | WBGene00013518 | protein_coding_gene | 1.1 | 2.104 | 4.2E-03 |
| *F40F12.9* | WBGene00043995 | protein_coding_gene | 1.1 | 2.103 | 5.8E-03 |
| *atat-2* | WBGene00021059 | protein_coding_gene | 1.1 | 2.100 | 2.3E-04 |
| *bbs-9* | WBGene00016744 | protein_coding_gene | 1.1 | 2.099 | 3.6E-02 |
| *C18E9.7* | WBGene00007687 | protein_coding_gene | 1.1 | 2.099 | 1.7E-02 |
| *D1086.5* | WBGene00008392 | protein_coding_gene | 1.1 | 2.099 | 1.3E-02 |
| *frm-3* | WBGene00001490 | protein_coding_gene | 1.1 | 2.099 | 2.6E-04 |
| *K08D8.7* | WBGene00185092 | protein_coding_gene | 1.1 | 2.097 | 7.0E-03 |
| *C01A2.2* | WBGene00007214 | protein_coding_gene | 1.1 | 2.096 | 4.3E-03 |
| *nhr-204* | WBGene00019816 | protein_coding_gene | 1.1 | 2.094 | 1.1E-04 |
| *T07D1.2* | WBGene00020301 | protein_coding_gene | 1.1 | 2.094 | 1.2E-04 |
| *lact-5* | WBGene00016355 | protein_coding_gene | 1.1 | 2.093 | 7.6E-04 |
| *nas-5* | WBGene00003524 | protein_coding_gene | 1.1 | 2.093 | 1.0E-02 |
| *M03F4.6* | WBGene00019759 | protein_coding_gene | 1.1 | 2.091 | 8.6E-03 |
| *F49E12.10* | WBGene00009903 | protein_coding_gene | 1.1 | 2.091 | 2.4E-03 |
| *mif-1* | WBGene00003234 | protein_coding_gene | 1.1 | 2.091 | 1.8E-07 |
| *Y39H10A.6* | WBGene00021482 | protein_coding_gene | 1.1 | 2.090 | 1.2E-05 |
| *C31H5.5* | WBGene00007856 | protein_coding_gene | 1.1 | 2.090 | 1.6E-04 |
| *Y71A12B.18* | WBGene00045077 | pseudogene | 1.1 | 2.089 | 9.1E-03 |
| *unc-120* | WBGene00006844 | protein_coding_gene | 1.1 | 2.088 | 1.4E-03 |
| *sup-37* | WBGene00006342 | protein_coding_gene | 1.1 | 2.088 | 1.4E-06 |
| *rrbs-1* | WBGene00007617 | protein_coding_gene | 1.1 | 2.087 | 5.9E-10 |
| *C28H8.14* | WBGene00219838 | ncRNA_gene | 1.1 | 2.086 | 6.7E-12 |
| *apm-1* | WBGene00000150 | protein_coding_gene | 1.1 | 2.086 | 5.5E-06 |
| *ace-1* | WBGene00000035 | protein_coding_gene | 1.1 | 2.083 | 1.9E-04 |
| *C46A5.6* | WBGene00016702 | protein_coding_gene | 1.1 | 2.083 | 1.3E-14 |
| *F35A5.2* | WBGene00018025 | protein_coding_gene | 1.1 | 2.081 | 2.1E-03 |
| *cah-4* | WBGene00000282 | protein_coding_gene | 1.1 | 2.078 | 7.2E-05 |
| *nmtn-1* | WBGene00019434 | protein_coding_gene | 1.1 | 2.077 | 8.7E-04 |
| *Y75B8A.6* | WBGene00013543 | protein_coding_gene | 1.1 | 2.076 | 9.9E-03 |
| *inso-1* | WBGene00016871 | protein_coding_gene | 1.1 | 2.075 | 1.3E-05 |
| *K02E11.3* | WBGene00010512 | protein_coding_gene | 1.1 | 2.074 | 4.3E-04 |
| *F35H8.2* | WBGene00009447 | protein_coding_gene | 1.1 | 2.073 | 1.2E-05 |
| *ctc-3* | WBGene00010962 | protein_coding_gene | 1.1 | 2.073 | 5.9E-06 |
| *Y106G6E.4* | WBGene00013708 | protein_coding_gene | 1.1 | 2.072 | 3.7E-08 |
| *F44A2.3* | WBGene00018403 | protein_coding_gene | 1.1 | 2.072 | 7.7E-07 |
| *F23F1.4* | WBGene00017745 | protein_coding_gene | 1.1 | 2.072 | 1.2E-11 |
| *elt-7* | WBGene00015981 | protein_coding_gene | 1.0 | 2.070 | 4.1E-04 |
| *ptr-15* | WBGene00004229 | protein_coding_gene | 1.0 | 2.070 | 3.6E-02 |
| *gpb-2* | WBGene00001680 | protein_coding_gene | 1.0 | 2.069 | 8.8E-12 |
| *nhr-176* | WBGene00008830 | protein_coding_gene | 1.0 | 2.069 | 3.5E-06 |
| *C46C11.3* | WBGene00016706 | protein_coding_gene | 1.0 | 2.069 | 2.1E-04 |
| *pbo-6* | WBGene00017375 | protein_coding_gene | 1.0 | 2.068 | 1.1E-02 |
| *fbxa-19* | WBGene00022331 | protein_coding_gene | 1.0 | 2.067 | 3.6E-04 |
| *Y32G9B.1* | WBGene00021309 | protein_coding_gene | 1.0 | 2.067 | 1.0E-02 |
| *ZK550.2* | WBGene00013996 | protein_coding_gene | 1.0 | 2.066 | 4.5E-02 |
| *dhs-23* | WBGene00000986 | protein_coding_gene | 1.0 | 2.066 | 2.4E-02 |
| *irg-7* | WBGene00018237 | protein_coding_gene | 1.0 | 2.065 | 2.8E-04 |
| *C06A1.2* | WBGene00007353 | protein_coding_gene | 1.0 | 2.064 | 2.0E-03 |
| *Y71H2AM.12* | WBGene00022177 | protein_coding_gene | 1.0 | 2.063 | 4.4E-05 |
| *F26E4.7* | WBGene00009162 | protein_coding_gene | 1.0 | 2.063 | 5.2E-09 |
| *rab-3* | WBGene00004267 | protein_coding_gene | 1.0 | 2.062 | 4.4E-06 |
| *sma-6* | WBGene00004860 | protein_coding_gene | 1.0 | 2.061 | 1.7E-07 |
| *nhr-131* | WBGene00003721 | protein_coding_gene | 1.0 | 2.061 | 9.8E-03 |
| *C35E7.11* | WBGene00016463 | protein_coding_gene | 1.0 | 2.060 | 3.4E-03 |
| *rdy-2* | WBGene00004328 | protein_coding_gene | 1.0 | 2.060 | 4.9E-02 |
| *T10B10.8* | WBGene00011684 | protein_coding_gene | 1.0 | 2.056 | 8.9E-05 |
| *lron-9* | WBGene00011971 | protein_coding_gene | 1.0 | 2.055 | 1.9E-04 |
| *nhr-43* | WBGene00003633 | protein_coding_gene | 1.0 | 2.055 | 2.7E-03 |
| *ZC239.6* | WBGene00022569 | protein_coding_gene | 1.0 | 2.054 | 7.9E-04 |
| *oac-46* | WBGene00011768 | protein_coding_gene | 1.0 | 2.054 | 2.0E-02 |
| *ptr-5* | WBGene00004220 | protein_coding_gene | 1.0 | 2.050 | 5.7E-03 |
| *K10G9.2* | WBGene00010756 | protein_coding_gene | 1.0 | 2.050 | 5.9E-04 |
| *C06A8.8* | WBGene00015517 | protein_coding_gene | 1.0 | 2.049 | 4.3E-06 |
| *glb-20* | WBGene00010970 | protein_coding_gene | 1.0 | 2.048 | 7.5E-03 |
| *obr-2* | WBGene00008832 | protein_coding_gene | 1.0 | 2.048 | 1.3E-06 |
| *F13H6.5* | WBGene00017433 | protein_coding_gene | 1.0 | 2.047 | 3.2E-02 |

| | | | | | |
|---|---|---|---|---|---|
| *F20G2.3* | WBGene00008987 | protein_coding_gene | 1.0 | 2.047 | 1.2E-04 |
| *H05C05.4* | WBGene00194747 | protein_coding_gene | 1.0 | 2.046 | 2.6E-02 |
| *ccb-1* | WBGene00000368 | protein_coding_gene | 1.0 | 2.046 | 2.9E-06 |
| *grdn-1* | WBGene00013082 | protein_coding_gene | 1.0 | 2.045 | 3.7E-04 |
| *ttr-10* | WBGene00017600 | protein_coding_gene | 1.0 | 2.045 | 3.0E-05 |
| *pudl-1* | WBGene00044489 | protein_coding_gene | 1.0 | 2.045 | 2.4E-03 |
| *B0353.1* | WBGene00015155 | protein_coding_gene | 1.0 | 2.043 | 3.4E-06 |
| *fbxc-54* | WBGene00017744 | protein_coding_gene | 1.0 | 2.042 | 8.4E-04 |
| *C35B8.3* | WBGene00016438 | protein_coding_gene | 1.0 | 2.042 | 1.2E-04 |
| *Y67D8C.9* | WBGene00022073 | protein_coding_gene | 1.0 | 2.040 | 2.7E-05 |
| *cest-5.2* | WBGene00015284 | protein_coding_gene | 1.0 | 2.039 | 1.4E-03 |
| *C18C4.7* | WBGene00015968 | protein_coding_gene | 1.0 | 2.039 | 1.7E-06 |
| *Y42G9A.3* | WBGene00021533 | protein_coding_gene | 1.0 | 2.038 | 5.0E-03 |
| *F52B10.3* | WBGene00044304 | protein_coding_gene | 1.0 | 2.037 | 7.3E-03 |
| *ZC239.14* | WBGene00022572 | protein_coding_gene | 1.0 | 2.037 | 2.3E-02 |
| *T06G6.4* | WBGene00011546 | protein_coding_gene | 1.0 | 2.036 | 2.4E-02 |
| *M153.4* | WBGene00044144 | protein_coding_gene | 1.0 | 2.035 | 5.6E-04 |
| *nape-2* | WBGene00021370 | protein_coding_gene | 1.0 | 2.034 | 8.7E-04 |
| *F21C10.4* | WBGene00017654 | protein_coding_gene | 1.0 | 2.034 | 2.5E-02 |
| *C16C10.8* | WBGene00007628 | protein_coding_gene | 1.0 | 2.032 | 8.2E-10 |
| *daf-5* | WBGene00000901 | protein_coding_gene | 1.0 | 2.024 | 3.4E-04 |
| *tsp-13* | WBGene00006639 | protein_coding_gene | 1.0 | 2.022 | 4.0E-03 |
| *tli-1* | WBGene00006578 | protein_coding_gene | 1.0 | 2.022 | 2.7E-04 |
| *dgk-2* | WBGene00000959 | protein_coding_gene | 1.0 | 2.022 | 3.1E-08 |
| *ceh-27* | WBGene00000449 | protein_coding_gene | 1.0 | 2.022 | 1.3E-02 |
| *F31C3.6* | WBGene00009288 | protein_coding_gene | 1.0 | 2.019 | 3.2E-03 |
| *B0507.1* | WBGene00015218 | protein_coding_gene | 1.0 | 2.019 | 6.5E-03 |
| *mig-39* | WBGene00018369 | protein_coding_gene | 1.0 | 2.019 | 1.9E-03 |
| *K07F5.22* | WBGene00303023 | protein_coding_gene | 1.0 | 2.016 | 5.0E-05 |
| *puf-12* | WBGene00014165 | protein_coding_gene | 1.0 | 2.016 | 2.0E-10 |
| *T13C2.3* | WBGene00020479 | protein_coding_gene | 1.0 | 2.016 | 3.8E-03 |
| *T01B7.9* | WBGene00011314 | protein_coding_gene | 1.0 | 2.014 | 2.6E-02 |
| *ppt-1* | WBGene00004092 | protein_coding_gene | 1.0 | 2.014 | 4.0E-03 |
| *F38A1.9* | WBGene00009522 | protein_coding_gene | 1.0 | 2.013 | 2.2E-02 |
| *E02H4.7* | WBGene00077757 | protein_coding_gene | 1.0 | 2.012 | 7.9E-05 |
| *T28D6.4* | WBGene00012124 | protein_coding_gene | 1.0 | 2.012 | 2.5E-03 |
| *lbp-1* | WBGene00002253 | protein_coding_gene | 1.0 | 2.011 | 2.0E-05 |
| *T01D3.1* | WBGene00011326 | protein_coding_gene | 1.0 | 2.011 | 3.1E-04 |
| *K04F10.2* | WBGene00019395 | protein_coding_gene | 1.0 | 2.010 | 4.7E-02 |
| *C05D10.4* | WBGene00015480 | protein_coding_gene | 1.0 | 2.009 | 7.4E-07 |
| *B0244.6* | WBGene00015081 | protein_coding_gene | 1.0 | 2.009 | 5.2E-03 |
| *pals-22* | WBGene00016216 | protein_coding_gene | 1.0 | 2.008 | 3.5E-04 |
| *C25E10.4* | WBGene00016094 | protein_coding_gene | 1.0 | 2.008 | 4.9E-04 |
| *C10A4.13* | WBGene00206485 | protein_coding_gene | 1.0 | 2.008 | 1.9E-02 |
| *F32D1.11* | WBGene00044530 | protein_coding_gene | 1.0 | 2.007 | 2.5E-02 |
| *nuaf-1* | WBGene00008225 | protein_coding_gene | 1.0 | 2.007 | 5.0E-09 |
| *T05E11.8* | WBGene00011484 | protein_coding_gene | 1.0 | 2.007 | 6.0E-03 |
| *nhr-108* | WBGene00003698 | protein_coding_gene | 1.0 | 2.006 | 1.4E-02 |
| *nhr-202* | WBGene00019742 | protein_coding_gene | 1.0 | 2.006 | 1.6E-03 |
| *shc-1* | WBGene00018788 | protein_coding_gene | 1.0 | 2.005 | 2.3E-05 |
| *F39E9.28* | WBGene00271716 | pseudogene | 1.0 | 2.004 | 2.3E-03 |
| *hda-6* | WBGene00018319 | protein_coding_gene | -1.0 | 0.500 | 1.6E-17 |
| *F44E2.7* | WBGene00018420 | protein_coding_gene | -1.0 | 0.500 | 3.8E-12 |
| *nstp-5* | WBGene00014139 | protein_coding_gene | -1.0 | 0.500 | 6.2E-09 |
| *paa-1* | WBGene00003901 | protein_coding_gene | -1.0 | 0.500 | 1.2E-05 |
| *tlf-1* | WBGene00006577 | protein_coding_gene | -1.0 | 0.500 | 1.2E-04 |
| *C10C5.5* | WBGene00007509 | protein_coding_gene | -1.0 | 0.500 | 1.1E-02 |
| *F10C2.5* | WBGene00008646 | protein_coding_gene | -1.0 | 0.500 | 8.9E-14 |
| *lst-6* | WBGene00016889 | protein_coding_gene | -1.0 | 0.500 | 4.5E-07 |
| *W02H5.19* | WBGene00302982 | protein_coding_gene | -1.0 | 0.499 | 1.7E-03 |
| *nmy-2* | WBGene00003777 | protein_coding_gene | -1.0 | 0.499 | 2.5E-06 |
| *bath-1* | WBGene00019139 | protein_coding_gene | -1.0 | 0.499 | 8.3E-04 |
| *set-9* | WBGene00017482 | protein_coding_gene | -1.0 | 0.499 | 4.6E-05 |
| *mdt-17* | WBGene00007017 | protein_coding_gene | -1.0 | 0.499 | 2.1E-09 |
| *fipr-13* | WBGene00010183 | protein_coding_gene | -1.0 | 0.499 | 4.3E-02 |
| *F55H2.7* | WBGene00010132 | protein_coding_gene | -1.0 | 0.499 | 2.0E-08 |
| *C24G6.8* | WBGene00016062 | protein_coding_gene | -1.0 | 0.499 | 1.1E-07 |
| *Y32H12A.7* | WBGene00021315 | protein_coding_gene | -1.0 | 0.499 | 1.1E-19 |
| *nsun-4* | WBGene00021476 | protein_coding_gene | -1.0 | 0.499 | 1.1E-11 |
| *xbx-6* | WBGene00009580 | protein_coding_gene | -1.0 | 0.499 | 9.7E-08 |
| *srf-3* | WBGene00005153 | protein_coding_gene | -1.0 | 0.499 | 8.6E-08 |
| *tpst-1* | WBGene00013737 | protein_coding_gene | -1.0 | 0.498 | 7.3E-10 |
| *ruvb-1* | WBGene00007784 | protein_coding_gene | -1.0 | 0.498 | 5.2E-15 |
| *F42A10.9* | WBGene00023489 | protein_coding_gene | -1.0 | 0.498 | 3.2E-03 |
| *ztf-18* | WBGene00020763 | protein_coding_gene | -1.0 | 0.498 | 3.9E-15 |
| *F14E5.2* | WBGene00008800 | protein_coding_gene | -1.0 | 0.498 | 5.2E-07 |
| *C34D4.4* | WBGene00016400 | protein_coding_gene | -1.0 | 0.498 | 2.8E-08 |
| *ebp-1* | WBGene00013344 | protein_coding_gene | -1.0 | 0.498 | 1.7E-04 |
| *usp-48* | WBGene00009267 | protein_coding_gene | -1.0 | 0.498 | 8.9E-08 |
| *ZK1321.4* | WBGene00014262 | protein_coding_gene | -1.0 | 0.498 | 8.3E-06 |

| | | | | | |
|---|---|---|---|---|---|
| *smr-1* | WBGene00004891 | protein_coding_gene | -1.0 | 0.498 | 3.2E-15 |
| *npp-18* | WBGene00003804 | protein_coding_gene | -1.0 | 0.498 | 1.8E-10 |
| *xpo-3* | WBGene00002080 | protein_coding_gene | -1.0 | 0.497 | 4.1E-06 |
| *Y38H8A.7* | WBGene00012640 | protein_coding_gene | -1.0 | 0.497 | 1.3E-07 |
| *Y48G1BL.5* | WBGene00021664 | pseudogene | -1.0 | 0.497 | 1.5E-04 |
| *mars-1* | WBGene00003415 | protein_coding_gene | -1.0 | 0.497 | 8.1E-09 |
| *fat-5* | WBGene00001397 | protein_coding_gene | -1.0 | 0.497 | 4.3E-04 |
| *glrx-5* | WBGene00013029 | protein_coding_gene | -1.0 | 0.496 | 9.0E-13 |
| *C18B12.6* | WBGene00007668 | protein_coding_gene | -1.0 | 0.496 | 1.5E-06 |
| *algn-8* | WBGene00007464 | protein_coding_gene | -1.0 | 0.496 | 4.8E-10 |
| *Y71F9AL.12* | WBGene00022117 | protein_coding_gene | -1.0 | 0.496 | 2.0E-08 |
| *R06C7.2* | WBGene00011062 | protein_coding_gene | -1.0 | 0.496 | 1.3E-06 |
| *rpac-19* | WBGene00010230 | protein_coding_gene | -1.0 | 0.495 | 7.8E-06 |
| *pdi-2* | WBGene00003963 | protein_coding_gene | -1.0 | 0.495 | 8.6E-05 |
| *sams-3* | WBGene00015538 | protein_coding_gene | -1.0 | 0.495 | 3.1E-08 |
| *H14E04.2* | WBGene00019199 | protein_coding_gene | -1.0 | 0.495 | 1.7E-09 |
| *cil-1* | WBGene00086546 | protein_coding_gene | -1.0 | 0.495 | 3.1E-08 |
| *F56D5.3* | WBGene00010148 | protein_coding_gene | -1.0 | 0.495 | 2.2E-03 |
| *hsf-1* | WBGene00002004 | protein_coding_gene | -1.0 | 0.495 | 1.5E-07 |
| *mppb-1* | WBGene00013880 | protein_coding_gene | -1.0 | 0.495 | 8.4E-15 |
| *nprl-3* | WBGene00018072 | protein_coding_gene | -1.0 | 0.495 | 3.9E-12 |
| *F55A12.2* | WBGene00018863 | protein_coding_gene | -1.0 | 0.494 | 2.2E-06 |
| *comp-1* | WBGene00018158 | protein_coding_gene | -1.0 | 0.494 | 3.5E-02 |
| *mel-28* | WBGene00003210 | protein_coding_gene | -1.0 | 0.494 | 9.9E-07 |
| *mrpl-39* | WBGene00044344 | protein_coding_gene | -1.0 | 0.493 | 1.8E-08 |
| *mask-1* | WBGene00011240 | protein_coding_gene | -1.0 | 0.493 | 1.8E-03 |
| *pdi-6* | WBGene00015168 | protein_coding_gene | -1.0 | 0.493 | 2.9E-03 |
| *W09C3.4* | WBGene00021112 | protein_coding_gene | -1.0 | 0.493 | 1.7E-09 |
| *tag-296* | WBGene00018632 | protein_coding_gene | -1.0 | 0.493 | 1.5E-11 |
| *Y73E7A.3* | WBGene00022270 | protein_coding_gene | -1.0 | 0.493 | 3.1E-03 |
| *nstp-1* | WBGene00008275 | protein_coding_gene | -1.0 | 0.493 | 2.7E-09 |
| *F23H11.4* | WBGene00017760 | protein_coding_gene | -1.0 | 0.493 | 5.2E-14 |
| *suco-1* | WBGene00020031 | protein_coding_gene | -1.0 | 0.493 | 2.5E-11 |
| *F14D2.15* | WBGene00023427 | protein_coding_gene | -1.0 | 0.493 | 6.4E-07 |
| *F31C3.3* | WBGene00009285 | protein_coding_gene | -1.0 | 0.493 | 7.9E-05 |
| *ben-1* | WBGene00000248 | protein_coding_gene | -1.0 | 0.493 | 3.1E-15 |
| *usp-5* | WBGene00020839 | protein_coding_gene | -1.0 | 0.492 | 4.2E-07 |
| *iffb-1* | WBGene00021857 | protein_coding_gene | -1.0 | 0.492 | 2.1E-08 |
| *efa-6* | WBGene00013223 | protein_coding_gene | -1.0 | 0.491 | 5.9E-09 |
| *ribo-1* | WBGene00020683 | protein_coding_gene | -1.0 | 0.491 | 6.1E-10 |
| *hmt-1* | WBGene00001815 | protein_coding_gene | -1.0 | 0.491 | 5.4E-13 |
| *mut-16* | WBGene00003508 | protein_coding_gene | -1.0 | 0.491 | 3.6E-04 |
| *lonp-1* | WBGene00016391 | protein_coding_gene | -1.0 | 0.491 | 9.0E-13 |
| *nspc-4* | WBGene00044339 | protein_coding_gene | -1.0 | 0.491 | 5.9E-03 |
| *acdh-5* | WBGene00016491 | protein_coding_gene | -1.0 | 0.491 | 4.0E-02 |
| *mboa-6* | WBGene00020115 | protein_coding_gene | -1.0 | 0.490 | 2.9E-11 |
| *Y92H12BR.2* | WBGene00022368 | protein_coding_gene | -1.0 | 0.490 | 7.7E-11 |
| *taf-10* | WBGene00006392 | protein_coding_gene | -1.0 | 0.490 | 1.4E-12 |
| *F46F11.10* | WBGene00018513 | protein_coding_gene | -1.0 | 0.490 | 9.8E-07 |
| *dnj-17* | WBGene00001035 | protein_coding_gene | -1.0 | 0.489 | 2.8E-07 |
| *pbrm-1* | WBGene00007042 | protein_coding_gene | -1.0 | 0.489 | 1.2E-04 |
| *F31D4.2* | WBGene00009290 | protein_coding_gene | -1.0 | 0.489 | 1.4E-09 |
| *hprt-1* | WBGene00013690 | protein_coding_gene | -1.0 | 0.489 | 1.1E-03 |
| *gsp-4* | WBGene00020187 | protein_coding_gene | -1.0 | 0.489 | 4.6E-02 |
| *Y71H10B.1* | WBGene00022201 | protein_coding_gene | -1.0 | 0.489 | 1.6E-05 |
| *hrpf-2* | WBGene00022253 | protein_coding_gene | -1.0 | 0.489 | 1.8E-07 |
| *F46B6.6* | WBGene00009771 | protein_coding_gene | -1.0 | 0.489 | 9.3E-16 |
| *Y39A3CL.1* | WBGene00021440 | protein_coding_gene | -1.0 | 0.489 | 5.1E-17 |
| *hcp-4* | WBGene00001832 | protein_coding_gene | -1.0 | 0.488 | 6.8E-13 |
| *C14C10.5* | WBGene00007588 | protein_coding_gene | -1.0 | 0.488 | 1.4E-04 |
| *C47E8.4* | WBGene00008143 | protein_coding_gene | -1.0 | 0.488 | 4.8E-08 |
| *eif-3.K* | WBGene00001233 | protein_coding_gene | -1.0 | 0.488 | 1.8E-08 |
| *F52B5.3* | WBGene00009922 | protein_coding_gene | -1.0 | 0.488 | 3.7E-06 |
| *fdps-1* | WBGene00011058 | protein_coding_gene | -1.0 | 0.488 | 2.3E-13 |
| *F27D4.4* | WBGene00009189 | protein_coding_gene | -1.0 | 0.488 | 1.4E-09 |
| *aos-1* | WBGene00000142 | protein_coding_gene | -1.0 | 0.488 | 3.2E-10 |
| *iars-2* | WBGene00002153 | protein_coding_gene | -1.0 | 0.488 | 2.4E-10 |
| *ntl-3* | WBGene00003826 | protein_coding_gene | -1.0 | 0.488 | 3.8E-07 |
| *ran-5* | WBGene00004306 | protein_coding_gene | -1.0 | 0.487 | 6.7E-17 |
| *ufd-1* | WBGene00006733 | protein_coding_gene | -1.0 | 0.487 | 5.8E-14 |
| *F01F1.15* | WBGene00017169 | protein_coding_gene | -1.0 | 0.487 | 8.4E-07 |
| *emb-30* | WBGene00001284 | protein_coding_gene | -1.0 | 0.487 | 7.3E-22 |
| *lin-9* | WBGene00002998 | protein_coding_gene | -1.0 | 0.487 | 5.3E-08 |
| *Y46G5A.7* | WBGene00012899 | protein_coding_gene | -1.0 | 0.486 | 1.6E-03 |
| *Y77E11A.7* | WBGene00022310 | protein_coding_gene | -1.0 | 0.486 | 1.1E-12 |
| *K07A1.5* | WBGene00010612 | protein_coding_gene | -1.0 | 0.486 | 4.2E-02 |
| *Y66D12A.21* | WBGene00013446 | protein_coding_gene | -1.0 | 0.486 | 3.1E-12 |
| *gly-7* | WBGene00001632 | protein_coding_gene | -1.0 | 0.486 | 2.2E-12 |
| *lgc-34* | WBGene00020836 | protein_coding_gene | -1.0 | 0.486 | 5.1E-05 |
| *ctps-1* | WBGene00012316 | protein_coding_gene | -1.0 | 0.486 | 3.6E-10 |

| | | | | | |
|---|---|---|---|---|---|
| *daf-31* | WBGene00000923 | protein_coding_gene | -1.0 | 0.486 | 4.6E-09 |
| *W07E11.1* | WBGene00012326 | protein_coding_gene | -1.0 | 0.486 | 4.3E-06 |
| *T26A5.6* | WBGene00020822 | protein_coding_gene | -1.0 | 0.485 | 6.0E-15 |
| *C53B4.4* | WBGene00008274 | protein_coding_gene | -1.0 | 0.485 | 3.9E-06 |
| *ptp-5.3* | WBGene00021207 | protein_coding_gene | -1.0 | 0.485 | 4.3E-02 |
| *W01G7.4* | WBGene00012188 | protein_coding_gene | -1.0 | 0.485 | 3.9E-14 |
| *catp-8* | WBGene00007514 | protein_coding_gene | -1.0 | 0.485 | 5.7E-10 |
| *Y48G1C.10* | WBGene00021683 | protein_coding_gene | -1.0 | 0.484 | 3.2E-05 |
| *moag-4* | WBGene00021348 | protein_coding_gene | -1.0 | 0.484 | 3.9E-05 |
| *tdpt-1* | WBGene00013405 | protein_coding_gene | -1.0 | 0.484 | 8.9E-12 |
| *romo-1* | WBGene00008860 | protein_coding_gene | -1.0 | 0.483 | 2.8E-09 |
| *dpf-5* | WBGene00001058 | protein_coding_gene | -1.0 | 0.483 | 5.6E-08 |
| *Y57G11A.2* | WBGene00013290 | protein_coding_gene | -1.0 | 0.483 | 3.0E-02 |
| *mrps-15* | WBGene00010624 | protein_coding_gene | -1.0 | 0.483 | 3.1E-09 |
| *ing-3* | WBGene00013095 | protein_coding_gene | -1.1 | 0.483 | 2.2E-19 |
| *F44B9.2* | WBGene00018407 | protein_coding_gene | -1.1 | 0.483 | 1.5E-03 |
| *Y82E9BR.19* | WBGene00022351 | protein_coding_gene | -1.1 | 0.483 | 1.7E-04 |
| *phb-1* | WBGene00004014 | protein_coding_gene | -1.1 | 0.482 | 3.7E-13 |
| *knl-3* | WBGene00020392 | protein_coding_gene | -1.1 | 0.482 | 8.1E-16 |
| *pqbp-1.1* | WBGene00020647 | protein_coding_gene | -1.1 | 0.482 | 2.2E-12 |
| *phf-10* | WBGene00018013 | protein_coding_gene | -1.1 | 0.482 | 3.9E-11 |
| *F19B6.1* | WBGene00008948 | protein_coding_gene | -1.1 | 0.482 | 8.2E-09 |
| *sec-10* | WBGene00016376 | protein_coding_gene | -1.1 | 0.481 | 2.0E-09 |
| *C28A5.2* | WBGene00007789 | protein_coding_gene | -1.1 | 0.481 | 1.1E-08 |
| *M02B7.2* | WBGene00019724 | protein_coding_gene | -1.1 | 0.481 | 7.9E-06 |
| *C04F5.9* | WBGene00015451 | protein_coding_gene | -1.1 | 0.481 | 4.3E-10 |
| *C26E6.12* | WBGene00016145 | protein_coding_gene | -1.1 | 0.481 | 3.4E-14 |
| *cah-5* | WBGene00000283 | protein_coding_gene | -1.1 | 0.481 | 7.6E-04 |
| *kdin-1* | WBGene00006490 | protein_coding_gene | -1.1 | 0.481 | 1.3E-07 |
| *Y37D8A.4* | WBGene00012546 | protein_coding_gene | -1.1 | 0.481 | 4.5E-08 |
| *F59E12.9* | WBGene00019124 | protein_coding_gene | -1.1 | 0.480 | 3.8E-04 |
| *C49H3.9* | WBGene00016794 | protein_coding_gene | -1.1 | 0.480 | 2.2E-07 |
| *mys-2* | WBGene00010537 | protein_coding_gene | -1.1 | 0.480 | 3.3E-07 |
| *cyp-35A5* | WBGene00019473 | protein_coding_gene | -1.1 | 0.480 | 3.9E-03 |
| *Y53H1A.2* | WBGene00013178 | protein_coding_gene | -1.1 | 0.480 | 1.0E-09 |
| *npl-4.1* | WBGene00019120 | protein_coding_gene | -1.1 | 0.480 | 1.8E-05 |
| *eat-6* | WBGene00001137 | protein_coding_gene | -1.1 | 0.479 | 8.7E-06 |
| *Y62E10A.20* | WBGene00044745 | protein_coding_gene | -1.1 | 0.479 | 6.7E-04 |
| *par-4* | WBGene00003919 | protein_coding_gene | -1.1 | 0.479 | 3.9E-07 |
| *W02B12.11* | WBGene00012206 | protein_coding_gene | -1.1 | 0.479 | 2.0E-13 |
| *cyy-1* | WBGene00022697 | protein_coding_gene | -1.1 | 0.479 | 1.7E-08 |
| *C04E6.11* | WBGene00015425 | protein_coding_gene | -1.1 | 0.479 | 6.7E-09 |
| *C01B10.9* | WBGene00015283 | protein_coding_gene | -1.1 | 0.478 | 1.0E-10 |
| *T24B8.7* | WBGene00011980 | protein_coding_gene | -1.1 | 0.478 | 4.2E-04 |
| *tceb-3* | WBGene00010990 | protein_coding_gene | -1.1 | 0.478 | 3.5E-08 |
| *R12C12.8* | WBGene00020028 | pseudogene | -1.1 | 0.478 | 8.7E-13 |
| *R186.3* | WBGene00011306 | protein_coding_gene | -1.1 | 0.478 | 6.5E-10 |
| *mep-1* | WBGene00003218 | protein_coding_gene | -1.1 | 0.478 | 3.1E-05 |
| *F37H8.5* | WBGene00009514 | protein_coding_gene | -1.1 | 0.478 | 2.7E-08 |
| *pdhb-1* | WBGene00015413 | protein_coding_gene | -1.1 | 0.477 | 1.5E-13 |
| *Y47G7B.2* | WBGene00021654 | protein_coding_gene | -1.1 | 0.477 | 1.1E-02 |
| *Y38H8A.4* | WBGene00012638 | protein_coding_gene | -1.1 | 0.477 | 4.4E-02 |
| *nbet-1* | WBGene00043064 | protein_coding_gene | -1.1 | 0.477 | 4.4E-09 |
| *F22G12.5* | WBGene00009065 | protein_coding_gene | -1.1 | 0.477 | 1.8E-04 |
| *eor-1* | WBGene00001324 | protein_coding_gene | -1.1 | 0.477 | 1.1E-08 |
| *unc-33* | WBGene00006769 | protein_coding_gene | -1.1 | 0.476 | 2.0E-12 |
| *acl-11* | WBGene00017888 | protein_coding_gene | -1.1 | 0.476 | 1.3E-11 |
| *sec-31* | WBGene00011338 | protein_coding_gene | -1.1 | 0.476 | 6.0E-06 |
| *rpn-8* | WBGene00004464 | protein_coding_gene | -1.1 | 0.476 | 2.5E-09 |
| *glp-4* | WBGene00006936 | protein_coding_gene | -1.1 | 0.476 | 3.7E-11 |
| *gldc-1* | WBGene00020022 | protein_coding_gene | -1.1 | 0.476 | 2.4E-03 |
| *mdt-21* | WBGene00007704 | protein_coding_gene | -1.1 | 0.476 | 7.4E-06 |
| *M28.5* | WBGene00010896 | protein_coding_gene | -1.1 | 0.475 | 2.6E-07 |
| *tbb-6* | WBGene00006539 | protein_coding_gene | -1.1 | 0.475 | 4.2E-02 |
| *aps-1* | WBGene00000159 | protein_coding_gene | -1.1 | 0.475 | 7.5E-05 |
| *stc-1* | WBGene00006059 | protein_coding_gene | -1.1 | 0.475 | 1.0E-21 |
| *Y67D8A.2* | WBGene00022059 | protein_coding_gene | -1.1 | 0.475 | 2.1E-06 |
| *C56C10.7* | WBGene00016964 | protein_coding_gene | -1.1 | 0.475 | 1.0E-07 |
| *wdfy-2* | WBGene00008402 | protein_coding_gene | -1.1 | 0.475 | 4.4E-08 |
| *zig-12* | WBGene00019727 | protein_coding_gene | -1.1 | 0.475 | 6.1E-07 |
| *R02D3.8* | WBGene00019825 | protein_coding_gene | -1.1 | 0.475 | 1.5E-19 |
| *C18E9.4* | WBGene00007684 | protein_coding_gene | -1.1 | 0.475 | 2.0E-05 |
| *ppk-3* | WBGene00004089 | protein_coding_gene | -1.1 | 0.475 | 1.4E-07 |
| *vps-15* | WBGene00014151 | protein_coding_gene | -1.1 | 0.474 | 7.3E-07 |
| *F25B5.3* | WBGene00017775 | protein_coding_gene | -1.1 | 0.474 | 1.0E-10 |
| *F20D1.1* | WBGene00008973 | protein_coding_gene | -1.1 | 0.474 | 4.4E-09 |
| *hrp-1* | WBGene00001999 | protein_coding_gene | -1.1 | 0.474 | 2.6E-12 |
| *C05C10.3* | WBGene00007330 | protein_coding_gene | -1.1 | 0.474 | 7.2E-20 |
| *stt-3* | WBGene00020437 | protein_coding_gene | -1.1 | 0.474 | 2.5E-12 |
| *sgcb-1* | WBGene00019277 | protein_coding_gene | -1.1 | 0.474 | 1.1E-03 |

| | | | | | |
|---|---|---|---|---|---|
| *hrde-4* | WBGene00020403 | protein_coding_gene | -1.1 | 0.474 | 2.4E-08 |
| *gpdh-3* | WBGene00012031 | protein_coding_gene | -1.1 | 0.474 | 2.0E-13 |
| *D2045.2* | WBGene00008422 | protein_coding_gene | -1.1 | 0.474 | 2.3E-06 |
| *C45G3.3* | WBGene00001590 | protein_coding_gene | -1.1 | 0.473 | 9.6E-09 |
| *mpst-1* | WBGene00008409 | protein_coding_gene | -1.1 | 0.473 | 1.4E-10 |
| *ostd-1* | WBGene00019693 | protein_coding_gene | -1.1 | 0.473 | 3.0E-09 |
| *prmt-6* | WBGene00016023 | protein_coding_gene | -1.1 | 0.473 | 2.0E-02 |
| *rde-10* | WBGene00021634 | protein_coding_gene | -1.1 | 0.472 | 2.6E-12 |
| *Y48C3A.12* | WBGene00012994 | protein_coding_gene | -1.1 | 0.472 | 3.6E-04 |
| *glt-1* | WBGene00001620 | protein_coding_gene | -1.1 | 0.472 | 4.3E-05 |
| *miro-1* | WBGene00019544 | protein_coding_gene | -1.1 | 0.472 | 5.1E-15 |
| *zip-3* | WBGene00012330 | protein_coding_gene | -1.1 | 0.472 | 3.2E-03 |
| *Y110A7A.15* | WBGene00022462 | protein_coding_gene | -1.1 | 0.472 | 4.9E-07 |
| *Y73B6BL.29* | WBGene00022250 | protein_coding_gene | -1.1 | 0.472 | 5.0E-06 |
| *W02A11.6* | WBGene00014852 | pseudogene | -1.1 | 0.472 | 4.4E-02 |
| *ire-1* | WBGene00002147 | protein_coding_gene | -1.1 | 0.472 | 8.2E-11 |
| *K12H6.2* | WBGene00019683 | protein_coding_gene | -1.1 | 0.471 | 1.8E-03 |
| *oxy-5* | WBGene00045483 | protein_coding_gene | -1.1 | 0.471 | 2.8E-08 |
| *lmn-1* | WBGene00003052 | protein_coding_gene | -1.1 | 0.471 | 2.2E-13 |
| *W02B3.4* | WBGene00020924 | protein_coding_gene | -1.1 | 0.471 | 1.2E-03 |
| *mon-2* | WBGene00008685 | protein_coding_gene | -1.1 | 0.471 | 8.4E-06 |
| *hmr-1* | WBGene00001980 | protein_coding_gene | -1.1 | 0.470 | 6.7E-05 |
| *skr-2* | WBGene00004808 | protein_coding_gene | -1.1 | 0.470 | 4.4E-07 |
| *fib-1* | WBGene00001423 | protein_coding_gene | -1.1 | 0.470 | 1.1E-09 |
| *pas-3* | WBGene00003924 | protein_coding_gene | -1.1 | 0.470 | 3.8E-08 |
| *cdt-2* | WBGene00011318 | protein_coding_gene | -1.1 | 0.470 | 1.9E-07 |
| *Y56A3A.19* | WBGene00013237 | protein_coding_gene | -1.1 | 0.470 | 3.7E-15 |
| *C30F12.2* | WBGene00016261 | protein_coding_gene | -1.1 | 0.470 | 8.2E-13 |
| *F39B2.5* | WBGene00009556 | protein_coding_gene | -1.1 | 0.470 | 4.4E-13 |
| *Y35H6.3* | WBGene00021342 | pseudogene | -1.1 | 0.470 | 1.9E-04 |
| *C26B2.7* | WBGene00016129 | protein_coding_gene | -1.1 | 0.470 | 2.2E-12 |
| *kle-2* | WBGene00016202 | protein_coding_gene | -1.1 | 0.469 | 5.6E-12 |
| *F57F4.1* | WBGene00019015 | protein_coding_gene | -1.1 | 0.469 | 2.4E-02 |
| *praf-3* | WBGene00017070 | protein_coding_gene | -1.1 | 0.469 | 7.2E-13 |
| *chn-1* | WBGene00000500 | protein_coding_gene | -1.1 | 0.469 | 2.4E-17 |
| *glrx-21* | WBGene00022663 | protein_coding_gene | -1.1 | 0.469 | 6.7E-10 |
| *C47D12.2* | WBGene00008136 | protein_coding_gene | -1.1 | 0.469 | 8.8E-09 |
| *F02E9.3* | WBGene00008529 | protein_coding_gene | -1.1 | 0.468 | 3.2E-02 |
| *Y54G11A.9* | WBGene00013218 | protein_coding_gene | -1.1 | 0.468 | 8.9E-11 |
| *skih-2* | WBGene00008502 | protein_coding_gene | -1.1 | 0.468 | 4.0E-10 |
| *bre-4* | WBGene00000269 | protein_coding_gene | -1.1 | 0.468 | 2.8E-14 |
| *hcp-2* | WBGene00001830 | protein_coding_gene | -1.1 | 0.467 | 1.0E-14 |
| *atln-1* | WBGene00021868 | protein_coding_gene | -1.1 | 0.467 | 1.1E-13 |
| *copb-1* | WBGene00021292 | protein_coding_gene | -1.1 | 0.467 | 4.7E-15 |
| *C14B9.8* | WBGene00015754 | protein_coding_gene | -1.1 | 0.467 | 1.7E-14 |
| *W04G3.5* | WBGene00012258 | protein_coding_gene | -1.1 | 0.467 | 6.1E-07 |
| *npl-4.2* | WBGene00019121 | protein_coding_gene | -1.1 | 0.467 | 2.7E-08 |
| *T08B2.11* | WBGene00020349 | protein_coding_gene | -1.1 | 0.467 | 8.2E-09 |
| *vps-8* | WBGene00016582 | protein_coding_gene | -1.1 | 0.467 | 2.2E-14 |
| *cogc-8* | WBGene00019820 | protein_coding_gene | -1.1 | 0.467 | 2.7E-22 |
| *rib-2* | WBGene00004361 | protein_coding_gene | -1.1 | 0.466 | 3.5E-11 |
| *ech-5* | WBGene00001154 | protein_coding_gene | -1.1 | 0.466 | 1.8E-13 |
| *ubh-4* | WBGene00006724 | protein_coding_gene | -1.1 | 0.466 | 2.2E-10 |
| *imb-3* | WBGene00002077 | protein_coding_gene | -1.1 | 0.466 | 1.8E-06 |
| *slc-17.4* | WBGene00012130 | protein_coding_gene | -1.1 | 0.466 | 2.7E-05 |
| *zipt-15* | WBGene00021936 | protein_coding_gene | -1.1 | 0.466 | 4.4E-13 |
| *mlcd-1* | WBGene00009439 | protein_coding_gene | -1.1 | 0.466 | 1.6E-16 |
| *mppa-1* | WBGene00022159 | protein_coding_gene | -1.1 | 0.466 | 7.8E-13 |
| *rabx-5* | WBGene00012644 | protein_coding_gene | -1.1 | 0.466 | 4.4E-10 |
| *lcmt-1* | WBGene00007137 | protein_coding_gene | -1.1 | 0.466 | 1.5E-06 |
| *pars-2* | WBGene00004190 | protein_coding_gene | -1.1 | 0.465 | 4.7E-14 |
| *daz-1* | WBGene00000935 | protein_coding_gene | -1.1 | 0.465 | 6.3E-04 |
| *gpa-16* | WBGene00001678 | protein_coding_gene | -1.1 | 0.464 | 1.1E-07 |
| *sec-8* | WBGene00004753 | protein_coding_gene | -1.1 | 0.464 | 1.8E-14 |
| *kars-1* | WBGene00002238 | protein_coding_gene | -1.1 | 0.464 | 7.8E-13 |
| *VB0395L.1* | WBGene00044150 | protein_coding_gene | -1.1 | 0.463 | 1.0E-02 |
| *F37F2.2* | WBGene00018159 | protein_coding_gene | -1.1 | 0.463 | 8.1E-10 |
| *rig-4* | WBGene00004371 | protein_coding_gene | -1.1 | 0.463 | 6.5E-07 |
| *hip-1* | WBGene00011735 | protein_coding_gene | -1.1 | 0.463 | 3.7E-10 |
| *csr-1* | WBGene00017641 | protein_coding_gene | -1.1 | 0.463 | 2.7E-06 |
| *slr-2* | WBGene00013350 | protein_coding_gene | -1.1 | 0.463 | 2.2E-07 |
| *lido-18* | WBGene00015988 | protein_coding_gene | -1.1 | 0.463 | 5.3E-05 |
| *dpt-1* | WBGene00008532 | protein_coding_gene | -1.1 | 0.463 | 1.8E-10 |
| *tufm-2* | WBGene00007001 | protein_coding_gene | -1.1 | 0.462 | 1.5E-15 |
| *cyn-11* | WBGene00000887 | protein_coding_gene | -1.1 | 0.462 | 2.4E-10 |
| *C34C12.2* | WBGene00007921 | protein_coding_gene | -1.1 | 0.462 | 7.9E-17 |
| *dnj-3* | WBGene00001021 | protein_coding_gene | -1.1 | 0.462 | 1.1E-07 |
| *Y71G12A.4* | WBGene00022140 | protein_coding_gene | -1.1 | 0.462 | 1.9E-02 |
| *hgap-1* | WBGene00021209 | protein_coding_gene | -1.1 | 0.462 | 2.0E-12 |
| *Y56A3A.2* | WBGene00013225 | protein_coding_gene | -1.1 | 0.462 | 1.7E-16 |

| | | | | | |
|---|---|---|---|---|---|
| *ekl-7* | WBGene00021465 | protein_coding_gene | -1.1 | 0.462 | 1.3E-08 |
| *F56A11.5* | WBGene00018925 | protein_coding_gene | -1.1 | 0.461 | 2.2E-07 |
| *pole-1* | WBGene00009368 | protein_coding_gene | -1.1 | 0.461 | 1.9E-04 |
| *F35G12.12* | WBGene00009445 | protein_coding_gene | -1.1 | 0.461 | 3.3E-08 |
| *eif-2Bbeta* | WBGene00012950 | protein_coding_gene | -1.1 | 0.461 | 1.5E-10 |
| *F38H4.5* | WBGene00009549 | protein_coding_gene | -1.1 | 0.461 | 1.6E-02 |
| *B0432.8* | WBGene00015189 | protein_coding_gene | -1.1 | 0.461 | 1.0E-09 |
| *ZK84.2* | WBGene00022650 | protein_coding_gene | -1.1 | 0.461 | 3.5E-02 |
| *let-92* | WBGene00002363 | protein_coding_gene | -1.1 | 0.461 | 4.1E-09 |
| *vps-51* | WBGene00015176 | protein_coding_gene | -1.1 | 0.461 | 3.3E-15 |
| *F02A9.10* | WBGene00303021 | protein_coding_gene | -1.1 | 0.461 | 1.3E-16 |
| *smu-1* | WBGene00004895 | protein_coding_gene | -1.1 | 0.460 | 2.8E-15 |
| *dnj-9* | WBGene00001027 | protein_coding_gene | -1.1 | 0.460 | 2.3E-14 |
| *Y116A8C.9* | WBGene00013789 | protein_coding_gene | -1.1 | 0.460 | 1.1E-08 |
| *Y69A2AR.1* | WBGene00022074 | protein_coding_gene | -1.1 | 0.460 | 4.3E-16 |
| *mop-25.2* | WBGene00013140 | protein_coding_gene | -1.1 | 0.460 | 7.4E-10 |
| *Y53G8AR.6* | WBGene00021813 | protein_coding_gene | -1.1 | 0.460 | 3.7E-17 |
| *timm-17B.1* | WBGene00017119 | protein_coding_gene | -1.1 | 0.460 | 1.7E-13 |
| *Y54E2A.4* | WBGene00013189 | protein_coding_gene | -1.1 | 0.460 | 4.2E-05 |
| *R11A8.1* | WBGene00011236 | protein_coding_gene | -1.1 | 0.459 | 2.1E-08 |
| *nduf-2.2* | WBGene00006463 | protein_coding_gene | -1.1 | 0.459 | 7.8E-12 |
| *ctsa-4.1* | WBGene00019605 | protein_coding_gene | -1.1 | 0.459 | 6.4E-03 |
| *bath-12* | WBGene00019131 | pseudogene | -1.1 | 0.459 | 1.0E-05 |
| *larp-5* | WBGene00004147 | protein_coding_gene | -1.1 | 0.459 | 5.0E-04 |
| *pdr-1* | WBGene00003967 | protein_coding_gene | -1.1 | 0.458 | 1.3E-12 |
| *gars-1* | WBGene00001744 | protein_coding_gene | -1.1 | 0.458 | 2.5E-10 |
| *syp-4* | WBGene00019247 | protein_coding_gene | -1.1 | 0.458 | 3.2E-14 |
| *rtcb-1* | WBGene00008877 | protein_coding_gene | -1.1 | 0.458 | 1.4E-20 |
| *R148.7* | WBGene00020105 | protein_coding_gene | -1.1 | 0.457 | 3.6E-02 |
| *Y66D12A.9* | WBGene00013435 | protein_coding_gene | -1.1 | 0.457 | 1.0E-07 |
| *Y82E9BL.18* | WBGene00044402 | protein_coding_gene | -1.1 | 0.457 | 2.0E-02 |
| *chl-1* | WBGene00010839 | protein_coding_gene | -1.1 | 0.457 | 2.9E-18 |
| *rab-33* | WBGene00004283 | protein_coding_gene | -1.1 | 0.457 | 8.7E-13 |
| *acdh-9* | WBGene00017874 | protein_coding_gene | -1.1 | 0.457 | 7.8E-09 |
| *eak-7* | WBGene00010671 | protein_coding_gene | -1.1 | 0.456 | 1.3E-13 |
| *tars-1* | WBGene00006617 | protein_coding_gene | -1.1 | 0.456 | 9.7E-14 |
| *ced-4* | WBGene00000418 | protein_coding_gene | -1.1 | 0.456 | 3.7E-09 |
| *tag-353* | WBGene00009112 | protein_coding_gene | -1.1 | 0.456 | 1.4E-07 |
| *spe-12* | WBGene00004966 | protein_coding_gene | -1.1 | 0.456 | 4.5E-02 |
| *ints-4* | WBGene00012234 | protein_coding_gene | -1.1 | 0.456 | 1.1E-14 |
| *R10E9.2* | WBGene00011214 | protein_coding_gene | -1.1 | 0.456 | 2.4E-02 |
| *cka-1* | WBGene00000509 | protein_coding_gene | -1.1 | 0.456 | 5.8E-25 |
| *cec-2* | WBGene00016796 | protein_coding_gene | -1.1 | 0.456 | 3.5E-05 |
| *Y73B6BL.31* | WBGene00022252 | protein_coding_gene | -1.1 | 0.455 | 1.9E-05 |
| *aps-2* | WBGene00000157 | protein_coding_gene | -1.1 | 0.455 | 6.1E-05 |
| *hsp-90* | WBGene00000915 | protein_coding_gene | -1.1 | 0.455 | 6.3E-06 |
| *B0361.3* | WBGene00015157 | protein_coding_gene | -1.1 | 0.455 | 7.1E-17 |
| *ZK1010.2* | WBGene00014176 | protein_coding_gene | -1.1 | 0.455 | 1.0E-15 |
| *H40L08.1* | WBGene00010433 | protein_coding_gene | -1.1 | 0.455 | 2.6E-10 |
| *lin-23* | WBGene00003009 | protein_coding_gene | -1.1 | 0.455 | 4.1E-09 |
| *eif-3.H* | WBGene00001231 | protein_coding_gene | -1.1 | 0.454 | 5.3E-08 |
| *Y51H7C.7* | WBGene00021785 | protein_coding_gene | -1.1 | 0.454 | 6.9E-10 |
| *hrpu-1* | WBGene00012769 | protein_coding_gene | -1.1 | 0.454 | 1.7E-04 |
| *prpf-4* | WBGene00004186 | protein_coding_gene | -1.1 | 0.454 | 7.0E-10 |
| *B0303.3* | WBGene00015125 | protein_coding_gene | -1.1 | 0.454 | 1.7E-16 |
| *sas-1* | WBGene00013742 | protein_coding_gene | -1.1 | 0.454 | 3.9E-13 |
| *pif-1* | WBGene00004028 | protein_coding_gene | -1.1 | 0.453 | 3.8E-09 |
| *F54B3.1* | WBGene00010013 | protein_coding_gene | -1.1 | 0.453 | 2.3E-07 |
| *agl-1* | WBGene00011050 | protein_coding_gene | -1.1 | 0.453 | 1.4E-10 |
| *M04C9.3* | WBGene00010858 | protein_coding_gene | -1.1 | 0.453 | 2.1E-04 |
| *sec-12* | WBGene00010496 | protein_coding_gene | -1.1 | 0.453 | 7.6E-16 |
| *edg-1* | WBGene00007109 | protein_coding_gene | -1.1 | 0.453 | 2.5E-15 |
| *cogc-4* | WBGene00021784 | protein_coding_gene | -1.1 | 0.453 | 4.2E-20 |
| *R148.3* | WBGene00020102 | protein_coding_gene | -1.1 | 0.453 | 3.9E-13 |
| *aco-1* | WBGene00000040 | protein_coding_gene | -1.1 | 0.453 | 1.4E-07 |
| *tofu-7* | WBGene00022737 | protein_coding_gene | -1.1 | 0.453 | 7.7E-11 |
| *cpr-6* | WBGene00000786 | protein_coding_gene | -1.1 | 0.452 | 1.4E-07 |
| *sptl-1* | WBGene00016020 | protein_coding_gene | -1.1 | 0.452 | 8.1E-08 |
| *mpdu-1* | WBGene00018181 | protein_coding_gene | -1.1 | 0.452 | 1.9E-09 |
| *Y76B12C.4* | WBGene00022298 | protein_coding_gene | -1.1 | 0.452 | 6.9E-03 |
| *C08B11.9* | WBGene00007436 | protein_coding_gene | -1.1 | 0.452 | 2.6E-09 |
| *C44B7.7* | WBGene00016629 | protein_coding_gene | -1.1 | 0.452 | 2.4E-03 |
| *mct-5* | WBGene00020168 | protein_coding_gene | -1.1 | 0.451 | 3.4E-12 |
| *F53F8.5* | WBGene00010002 | protein_coding_gene | -1.1 | 0.451 | 2.1E-09 |
| *Y14H12B.2* | WBGene00021192 | protein_coding_gene | -1.1 | 0.451 | 9.3E-12 |
| *Y25C1A.8* | WBGene00021295 | protein_coding_gene | -1.1 | 0.451 | 7.7E-13 |
| *ulp-5* | WBGene00006740 | protein_coding_gene | -1.1 | 0.451 | 2.4E-06 |
| *mog-3* | WBGene00003391 | protein_coding_gene | -1.1 | 0.451 | 5.5E-15 |
| *ZK688.5* | WBGene00022800 | protein_coding_gene | -1.1 | 0.451 | 4.7E-08 |
| *mat-3* | WBGene00003134 | protein_coding_gene | -1.1 | 0.451 | 6.2E-20 |

| | | | | | |
|---|---|---|---|---|---|
| *F56A8.3* | WBGene00010139 | protein_coding_gene | -1.2 | 0.451 | 5.2E-09 |
| *F52C6.2* | WBGene00018659 | protein_coding_gene | -1.2 | 0.450 | 4.4E-08 |
| *ZK686.2* | WBGene00022792 | protein_coding_gene | -1.2 | 0.450 | 7.4E-19 |
| *W09G3.6* | WBGene00012369 | protein_coding_gene | -1.2 | 0.450 | 2.0E-08 |
| *lnkn-1* | WBGene00014023 | protein_coding_gene | -1.2 | 0.450 | 6.3E-10 |
| *gst-9* | WBGene00001757 | protein_coding_gene | -1.2 | 0.450 | 8.5E-03 |
| *pabp-2* | WBGene00003904 | protein_coding_gene | -1.2 | 0.450 | 1.5E-09 |
| *C16C8.14* | WBGene00015852 | protein_coding_gene | -1.2 | 0.450 | 1.4E-09 |
| *pps-1* | WBGene00004091 | protein_coding_gene | -1.2 | 0.449 | 2.8E-09 |
| *ttc-1* | WBGene00016390 | protein_coding_gene | -1.2 | 0.449 | 1.7E-16 |
| *emc-5* | WBGene00195248 | protein_coding_gene | -1.2 | 0.449 | 1.8E-03 |
| *clec-147* | WBGene00007313 | protein_coding_gene | -1.2 | 0.449 | 1.7E-05 |
| *marc-6* | WBGene00018847 | protein_coding_gene | -1.2 | 0.449 | 2.3E-10 |
| *C17E4.6* | WBGene00007645 | protein_coding_gene | -1.2 | 0.449 | 1.9E-05 |
| *letm-1* | WBGene00010279 | protein_coding_gene | -1.2 | 0.449 | 6.9E-14 |
| *F26F4.5* | WBGene00017827 | protein_coding_gene | -1.2 | 0.449 | 2.8E-11 |
| *npp-15* | WBGene00003801 | protein_coding_gene | -1.2 | 0.449 | 8.4E-12 |
| *hpo-35* | WBGene00009744 | protein_coding_gene | -1.2 | 0.449 | 2.4E-06 |
| *rme-6* | WBGene00004377 | protein_coding_gene | -1.2 | 0.449 | 3.4E-07 |
| *C15C6.2* | WBGene00007601 | protein_coding_gene | -1.2 | 0.449 | 4.6E-02 |
| *glrx-10* | WBGene00021331 | protein_coding_gene | -1.2 | 0.448 | 8.0E-05 |
| *pas-1* | WBGene00003922 | protein_coding_gene | -1.2 | 0.448 | 4.4E-09 |
| *angl-1* | WBGene00020955 | protein_coding_gene | -1.2 | 0.448 | 1.0E-10 |
| *T10D4.6* | WBGene00020408 | protein_coding_gene | -1.2 | 0.448 | 2.9E-04 |
| *W03F8.4* | WBGene00020994 | protein_coding_gene | -1.2 | 0.448 | 3.7E-07 |
| *Y54G2A.73* | WBGene00219425 | protein_coding_gene | -1.2 | 0.448 | 4.8E-03 |
| *arp-1* | WBGene00013168 | protein_coding_gene | -1.2 | 0.448 | 2.7E-19 |
| *rpn-11* | WBGene00004467 | protein_coding_gene | -1.2 | 0.448 | 1.6E-14 |
| *rbm-22* | WBGene00011722 | protein_coding_gene | -1.2 | 0.448 | 5.7E-14 |
| *spd-3* | WBGene00004954 | protein_coding_gene | -1.2 | 0.448 | 1.5E-10 |
| *pigu-1* | WBGene00011914 | protein_coding_gene | -1.2 | 0.448 | 1.2E-13 |
| *akir-1* | WBGene00017088 | protein_coding_gene | -1.2 | 0.447 | 9.7E-11 |
| *unc-27* | WBGene00006764 | protein_coding_gene | -1.2 | 0.447 | 7.2E-09 |
| *atn-1* | WBGene00000228 | protein_coding_gene | -1.2 | 0.447 | 8.0E-08 |
| *rsr-2* | WBGene00013260 | protein_coding_gene | -1.2 | 0.446 | 2.5E-13 |
| *eas-1* | WBGene00018270 | protein_coding_gene | -1.2 | 0.446 | 1.0E-11 |
| *F08B4.7* | WBGene00017238 | protein_coding_gene | -1.2 | 0.446 | 1.4E-13 |
| *Y14H12A.1* | WBGene00021190 | protein_coding_gene | -1.2 | 0.446 | 2.6E-05 |
| *eif-3.J* | WBGene00012738 | protein_coding_gene | -1.2 | 0.445 | 3.2E-22 |
| *C32E8.9* | WBGene00016325 | protein_coding_gene | -1.2 | 0.445 | 1.9E-08 |
| *tpi-1* | WBGene00006601 | protein_coding_gene | -1.2 | 0.445 | 1.5E-22 |
| *ath-1* | WBGene00010564 | protein_coding_gene | -1.2 | 0.445 | 1.1E-09 |
| *nkat-3* | WBGene00010984 | protein_coding_gene | -1.2 | 0.444 | 5.6E-05 |
| *Y54F10AL.1* | WBGene00021847 | protein_coding_gene | -1.2 | 0.444 | 1.7E-12 |
| *poml-2* | WBGene00017089 | protein_coding_gene | -1.2 | 0.444 | 1.6E-04 |
| *Y54F10AR.1* | WBGene00021854 | protein_coding_gene | -1.2 | 0.444 | 2.3E-12 |
| *ifg-1* | WBGene00002066 | protein_coding_gene | -1.2 | 0.444 | 1.5E-06 |
| *F32B6.3* | WBGene00009320 | protein_coding_gene | -1.2 | 0.444 | 1.0E-06 |
| *tomm-20* | WBGene00009092 | protein_coding_gene | -1.2 | 0.444 | 3.4E-16 |
| *C16A11.2* | WBGene00015815 | protein_coding_gene | -1.2 | 0.444 | 9.6E-18 |
| *trap-2* | WBGene00020216 | protein_coding_gene | -1.2 | 0.443 | 3.1E-06 |
| *eif-1.A* | WBGene00019162 | protein_coding_gene | -1.2 | 0.443 | 4.9E-22 |
| *srpa-68* | WBGene00010097 | protein_coding_gene | -1.2 | 0.443 | 6.0E-17 |
| *F53C11.1* | WBGene00009971 | protein_coding_gene | -1.2 | 0.443 | 3.3E-04 |
| *Y57E12AL.6* | WBGene00021959 | protein_coding_gene | -1.2 | 0.443 | 1.4E-14 |
| *asd-1* | WBGene00011279 | protein_coding_gene | -1.2 | 0.442 | 1.0E-05 |
| *syx-18* | WBGene00020425 | protein_coding_gene | -1.2 | 0.442 | 1.3E-11 |
| *C27D8.4* | WBGene00007780 | protein_coding_gene | -1.2 | 0.442 | 2.1E-08 |
| *D1086.17* | WBGene00045355 | protein_coding_gene | -1.2 | 0.442 | 3.4E-02 |
| *ric-8* | WBGene00004367 | protein_coding_gene | -1.2 | 0.442 | 1.0E-10 |
| *taf-6.1* | WBGene00006387 | protein_coding_gene | -1.2 | 0.442 | 3.4E-16 |
| *ccpp-1* | WBGene00018995 | protein_coding_gene | -1.2 | 0.442 | 1.9E-12 |
| *D1069.3* | WBGene00017037 | protein_coding_gene | -1.2 | 0.442 | 1.9E-19 |
| *syp-5* | WBGene00021832 | protein_coding_gene | -1.2 | 0.441 | 4.0E-11 |
| *rhy-1* | WBGene00012324 | protein_coding_gene | -1.2 | 0.441 | 7.9E-05 |
| *pink-1* | WBGene00017137 | protein_coding_gene | -1.2 | 0.441 | 4.7E-14 |
| *rnp-1* | WBGene00004384 | protein_coding_gene | -1.2 | 0.440 | 3.8E-09 |
| *gspd-1* | WBGene00007108 | protein_coding_gene | -1.2 | 0.440 | 3.2E-09 |
| *pid-2* | WBGene00021676 | protein_coding_gene | -1.2 | 0.439 | 3.1E-10 |
| *F52H2.6* | WBGene00018718 | protein_coding_gene | -1.2 | 0.439 | 4.2E-03 |
| *Y52B11A.3* | WBGene00013123 | protein_coding_gene | -1.2 | 0.439 | 3.8E-16 |
| *ZK1307.8* | WBGene00014249 | protein_coding_gene | -1.2 | 0.439 | 5.1E-11 |
| *K06B9.4* | WBGene00019441 | pseudogene | -1.2 | 0.439 | 1.7E-04 |
| *K04G2.4* | WBGene00010563 | protein_coding_gene | -1.2 | 0.439 | 3.1E-02 |
| *him-14* | WBGene00001872 | protein_coding_gene | -1.2 | 0.439 | 3.6E-15 |
| *T08B2.12* | WBGene00020350 | protein_coding_gene | -1.2 | 0.439 | 4.5E-02 |
| *F41G3.5* | WBGene00018301 | protein_coding_gene | -1.2 | 0.438 | 4.1E-02 |
| *clp-8* | WBGene00009695 | protein_coding_gene | -1.2 | 0.438 | 3.7E-02 |
| *skr-18* | WBGene00018935 | protein_coding_gene | -1.2 | 0.438 | 4.3E-16 |
| *Y71H2AR.1* | WBGene00022188 | protein_coding_gene | -1.2 | 0.438 | 4.5E-08 |

| | | | | | |
|---|---|---|---|---|---|
| *npp-12* | WBGene00003798 | protein_coding_gene | -1.2 | 0.438 | 8.5E-08 |
| *sart-3* | WBGene00007111 | protein_coding_gene | -1.2 | 0.437 | 1.7E-10 |
| *C09E7.8* | WBGene00015642 | protein_coding_gene | -1.2 | 0.437 | 1.7E-11 |
| *C48B6.3* | WBGene00016741 | protein_coding_gene | -1.2 | 0.437 | 1.5E-13 |
| *R08B4.3* | WBGene00011139 | protein_coding_gene | -1.2 | 0.437 | 1.5E-07 |
| *cest-35.2* | WBGene00016863 | protein_coding_gene | -1.2 | 0.436 | 7.3E-03 |
| *tag-335* | WBGene00016583 | protein_coding_gene | -1.2 | 0.436 | 4.5E-10 |
| *cfim-1* | WBGene00009668 | protein_coding_gene | -1.2 | 0.436 | 2.0E-12 |
| *copa-1* | WBGene00022119 | protein_coding_gene | -1.2 | 0.436 | 1.0E-12 |
| *eif-2Bgamma* | WBGene00004090 | protein_coding_gene | -1.2 | 0.436 | 4.7E-13 |
| *gtbp-1* | WBGene00010677 | protein_coding_gene | -1.2 | 0.436 | 5.9E-10 |
| *C48B6.4* | WBGene00016742 | protein_coding_gene | -1.2 | 0.436 | 2.8E-02 |
| *clec-175* | WBGene00019950 | protein_coding_gene | -1.2 | 0.436 | 2.1E-02 |
| *elo-1* | WBGene00001239 | protein_coding_gene | -1.2 | 0.436 | 2.7E-13 |
| *unc-50* | WBGene00006785 | protein_coding_gene | -1.2 | 0.436 | 3.2E-14 |
| *mrps-16* | WBGene00018961 | protein_coding_gene | -1.2 | 0.436 | 1.4E-09 |
| *cul-3* | WBGene00000838 | protein_coding_gene | -1.2 | 0.436 | 4.5E-09 |
| *lin-36* | WBGene00003021 | protein_coding_gene | -1.2 | 0.435 | 8.5E-08 |
| *cest-2.2* | WBGene00013874 | protein_coding_gene | -1.2 | 0.435 | 3.0E-07 |
| *phb-2* | WBGene00004015 | protein_coding_gene | -1.2 | 0.435 | 8.3E-14 |
| *eif-3.L* | WBGene00015920 | protein_coding_gene | -1.2 | 0.434 | 5.3E-12 |
| *mtx-1* | WBGene00009559 | protein_coding_gene | -1.2 | 0.434 | 1.9E-16 |
| *cogc-3* | WBGene00022121 | protein_coding_gene | -1.2 | 0.434 | 1.4E-16 |
| *C24D10.4* | WBGene00016055 | protein_coding_gene | -1.2 | 0.434 | 2.7E-19 |
| *F10C2.4* | WBGene00008645 | protein_coding_gene | -1.2 | 0.434 | 2.0E-12 |
| *ucr-2.3* | WBGene00020757 | protein_coding_gene | -1.2 | 0.434 | 6.7E-10 |
| *W05H7.1* | WBGene00021044 | protein_coding_gene | -1.2 | 0.434 | 5.6E-05 |
| *msp-56* | WBGene00003449 | protein_coding_gene | -1.2 | 0.433 | 1.3E-02 |
| *C06A1.3* | WBGene00007354 | protein_coding_gene | -1.2 | 0.433 | 2.1E-02 |
| *misc-1* | WBGene00015186 | protein_coding_gene | -1.2 | 0.433 | 1.9E-08 |
| *bcs-1* | WBGene00010042 | protein_coding_gene | -1.2 | 0.433 | 3.0E-16 |
| *K08C7.1* | WBGene00010646 | protein_coding_gene | -1.2 | 0.433 | 5.1E-09 |
| *acdh-3* | WBGene00019433 | protein_coding_gene | -1.2 | 0.433 | 8.7E-12 |
| *ppfr-1* | WBGene00008878 | protein_coding_gene | -1.2 | 0.432 | 2.7E-13 |
| *taf-6.2* | WBGene00021363 | protein_coding_gene | -1.2 | 0.432 | 1.8E-11 |
| *C48E7.7* | WBGene00016752 | protein_coding_gene | -1.2 | 0.432 | 3.2E-02 |
| *B0273.1* | WBGene00015097 | protein_coding_gene | -1.2 | 0.432 | 3.2E-02 |
| *aars-2* | WBGene00000197 | protein_coding_gene | -1.2 | 0.432 | 2.4E-09 |
| *W04B5.5* | WBGene00021022 | protein_coding_gene | -1.2 | 0.432 | 5.5E-14 |
| *fkb-6* | WBGene00001431 | protein_coding_gene | -1.2 | 0.432 | 7.6E-10 |
| *let-721* | WBGene00002855 | protein_coding_gene | -1.2 | 0.432 | 3.2E-19 |
| *Y37A1A.2* | WBGene00012535 | protein_coding_gene | -1.2 | 0.432 | 1.2E-04 |
| *ints-2* | WBGene00013877 | protein_coding_gene | -1.2 | 0.432 | 2.9E-10 |
| *lrg-1* | WBGene00003067 | pseudogene | -1.2 | 0.431 | 1.5E-13 |
| *eif-3.C* | WBGene00001226 | protein_coding_gene | -1.2 | 0.431 | 9.6E-16 |
| *C27F2.9* | WBGene00043156 | protein_coding_gene | -1.2 | 0.431 | 4.3E-14 |
| *R05D3.2* | WBGene00019877 | protein_coding_gene | -1.2 | 0.431 | 6.4E-09 |
| *B0303.4* | WBGene00015126 | protein_coding_gene | -1.2 | 0.431 | 2.8E-13 |
| *ddp-1* | WBGene00000941 | protein_coding_gene | -1.2 | 0.431 | 2.5E-08 |
| *prp-19* | WBGene00020423 | protein_coding_gene | -1.2 | 0.431 | 1.9E-13 |
| *M70.4* | WBGene00019786 | protein_coding_gene | -1.2 | 0.431 | 9.3E-12 |
| *nra-4* | WBGene00015344 | protein_coding_gene | -1.2 | 0.431 | 2.0E-06 |
| *rnf-121* | WBGene00007626 | protein_coding_gene | -1.2 | 0.430 | 4.8E-12 |
| *rcor-1* | WBGene00022278 | protein_coding_gene | -1.2 | 0.430 | 2.8E-15 |
| *par-6* | WBGene00003921 | protein_coding_gene | -1.2 | 0.430 | 4.5E-08 |
| *vwa-8* | WBGene00008694 | protein_coding_gene | -1.2 | 0.430 | 1.2E-14 |
| *pole-2* | WBGene00017237 | protein_coding_gene | -1.2 | 0.430 | 3.5E-09 |
| *B0495.7* | WBGene00015206 | protein_coding_gene | -1.2 | 0.430 | 7.1E-16 |
| *ari-2* | WBGene00020462 | protein_coding_gene | -1.2 | 0.430 | 3.5E-12 |
| *B0379.7* | WBGene00007159 | protein_coding_gene | -1.2 | 0.429 | 1.8E-02 |
| *tbc-3* | WBGene00009322 | protein_coding_gene | -1.2 | 0.429 | 8.4E-21 |
| *Y32B12B.4* | WBGene00012524 | protein_coding_gene | -1.2 | 0.429 | 8.5E-13 |
| *F37B12.1* | WBGene00009504 | protein_coding_gene | -1.2 | 0.429 | 9.7E-03 |
| *mtss-1* | WBGene00019800 | protein_coding_gene | -1.2 | 0.429 | 5.1E-25 |
| *swsn-1* | WBGene00004203 | protein_coding_gene | -1.2 | 0.429 | 2.8E-05 |
| *prx-19* | WBGene00004201 | protein_coding_gene | -1.2 | 0.429 | 9.9E-12 |
| *ebp-2* | WBGene00012156 | protein_coding_gene | -1.2 | 0.429 | 8.4E-17 |
| *eipr-1* | WBGene00013599 | protein_coding_gene | -1.2 | 0.429 | 2.2E-12 |
| *ech-4* | WBGene00001153 | protein_coding_gene | -1.2 | 0.429 | 4.8E-21 |
| *Y106G6D.5* | WBGene00014965 | protein_coding_gene | -1.2 | 0.429 | 2.3E-17 |
| *ech-1.2* | WBGene00020347 | protein_coding_gene | -1.2 | 0.428 | 9.6E-17 |
| *rmh-1* | WBGene00019712 | protein_coding_gene | -1.2 | 0.428 | 2.4E-16 |
| *oxi-1* | WBGene00003898 | protein_coding_gene | -1.2 | 0.428 | 3.3E-20 |
| *lip-1* | WBGene00003043 | protein_coding_gene | -1.2 | 0.428 | 3.8E-04 |
| *rsp-2* | WBGene00004699 | protein_coding_gene | -1.2 | 0.427 | 8.5E-21 |
| *sqv-8* | WBGene00005026 | protein_coding_gene | -1.2 | 0.427 | 9.9E-24 |
| *tmem-131* | WBGene00016170 | protein_coding_gene | -1.2 | 0.427 | 1.2E-05 |
| *jmjd-5* | WBGene00007387 | protein_coding_gene | -1.2 | 0.427 | 2.0E-11 |
| *laf-1* | WBGene00002244 | protein_coding_gene | -1.2 | 0.427 | 7.8E-08 |
| *lem-4* | WBGene00021945 | protein_coding_gene | -1.2 | 0.427 | 1.5E-14 |

| | | | | | |
|---|---|---|---|---|---|
| *acs-4* | WBGene00018152 | protein_coding_gene | -1.2 | 0.427 | 9.9E-09 |
| *Y54G11A.3* | WBGene00013214 | protein_coding_gene | -1.2 | 0.426 | 3.2E-19 |
| *mcd-1* | WBGene00013096 | protein_coding_gene | -1.2 | 0.426 | 7.4E-15 |
| *bath-40* | WBGene00013689 | protein_coding_gene | -1.2 | 0.426 | 4.1E-09 |
| *ZK512.8* | WBGene00013987 | protein_coding_gene | -1.2 | 0.426 | 3.3E-02 |
| *Y43F8C.6* | WBGene00012828 | protein_coding_gene | -1.2 | 0.426 | 4.7E-08 |
| *H14A12.3* | WBGene00019196 | protein_coding_gene | -1.2 | 0.426 | 3.6E-15 |
| *swt-4* | WBGene00012647 | protein_coding_gene | -1.2 | 0.426 | 3.1E-06 |
| *Y97E10AR.4* | WBGene00022399 | protein_coding_gene | -1.2 | 0.426 | 6.3E-10 |
| *R12C12.7* | WBGene00020027 | protein_coding_gene | -1.2 | 0.425 | 6.8E-13 |
| *ZK616.5* | WBGene00022775 | protein_coding_gene | -1.2 | 0.425 | 8.5E-07 |
| *W02H5.2* | WBGene00020957 | protein_coding_gene | -1.2 | 0.425 | 1.7E-17 |
| *Y38E10A.14* | WBGene00012592 | protein_coding_gene | -1.2 | 0.425 | 1.5E-03 |
| *mrpl-12* | WBGene00012361 | protein_coding_gene | -1.2 | 0.425 | 3.9E-07 |
| *ham-3* | WBGene00044072 | protein_coding_gene | -1.2 | 0.425 | 6.3E-12 |
| *gly-10* | WBGene00001635 | protein_coding_gene | -1.2 | 0.425 | 3.4E-22 |
| *W02D9.4* | WBGene00012210 | protein_coding_gene | -1.2 | 0.425 | 5.1E-12 |
| *ints-1* | WBGene00015499 | protein_coding_gene | -1.2 | 0.424 | 2.5E-07 |
| *C33F10.11* | WBGene00016357 | protein_coding_gene | -1.2 | 0.424 | 2.5E-02 |
| *snf-10* | WBGene00004909 | protein_coding_gene | -1.2 | 0.424 | 8.2E-03 |
| *him-17* | WBGene00001874 | protein_coding_gene | -1.2 | 0.424 | 3.6E-19 |
| *ZK370.4* | WBGene00022718 | protein_coding_gene | -1.2 | 0.424 | 4.3E-13 |
| *W02D7.4* | WBGene00020940 | protein_coding_gene | -1.2 | 0.424 | 7.5E-07 |
| *H35N09.2* | WBGene00019266 | protein_coding_gene | -1.2 | 0.424 | 9.1E-05 |
| *ptp-2* | WBGene00004214 | protein_coding_gene | -1.2 | 0.423 | 1.1E-06 |
| *ubxn-3* | WBGene00018586 | protein_coding_gene | -1.2 | 0.423 | 1.3E-09 |
| *eea-1* | WBGene00011696 | protein_coding_gene | -1.2 | 0.423 | 6.8E-08 |
| *epg-3* | WBGene00012559 | protein_coding_gene | -1.2 | 0.423 | 5.2E-17 |
| *asna-2* | WBGene00018577 | pseudogene | -1.2 | 0.422 | 9.5E-06 |
| *ech-2* | WBGene00001151 | protein_coding_gene | -1.2 | 0.422 | 2.0E-09 |
| *saps-1* | WBGene00008166 | protein_coding_gene | -1.2 | 0.422 | 2.0E-08 |
| *pnk-4* | WBGene00006862 | protein_coding_gene | -1.2 | 0.422 | 3.3E-10 |
| *ehs-1* | WBGene00001224 | protein_coding_gene | -1.2 | 0.422 | 1.9E-07 |
| *arx-5* | WBGene00000203 | protein_coding_gene | -1.2 | 0.421 | 1.8E-16 |
| *mat-1* | WBGene00003132 | protein_coding_gene | -1.2 | 0.421 | 3.0E-11 |
| *K07A1.4* | WBGene00010611 | protein_coding_gene | -1.2 | 0.421 | 1.8E-02 |
| *slc-25A18.1* | WBGene00008979 | protein_coding_gene | -1.2 | 0.421 | 1.8E-09 |
| *acl-6* | WBGene00017261 | protein_coding_gene | -1.2 | 0.421 | 3.0E-23 |
| *pinn-1* | WBGene00022448 | protein_coding_gene | -1.2 | 0.421 | 1.2E-09 |
| *rip-1* | WBGene00010973 | protein_coding_gene | -1.2 | 0.421 | 5.6E-03 |
| *par-5* | WBGene00003920 | protein_coding_gene | -1.2 | 0.421 | 1.4E-10 |
| *Y66D12A.6* | WBGene00013432 | protein_coding_gene | -1.2 | 0.420 | 8.1E-08 |
| *M03C11.3* | WBGene00010840 | protein_coding_gene | -1.3 | 0.420 | 2.7E-05 |
| *gsy-1* | WBGene00001793 | protein_coding_gene | -1.3 | 0.420 | 3.9E-11 |
| *C55B7.11* | WBGene00016947 | protein_coding_gene | -1.3 | 0.420 | 6.3E-16 |
| *F42A10.5* | WBGene00018341 | protein_coding_gene | -1.3 | 0.420 | 2.4E-12 |
| *mspn-1* | WBGene00010557 | protein_coding_gene | -1.3 | 0.420 | 2.5E-15 |
| *F58B4.5* | WBGene00010238 | protein_coding_gene | -1.3 | 0.420 | 1.5E-06 |
| *cpr-8* | WBGene00021070 | protein_coding_gene | -1.3 | 0.420 | 1.0E-03 |
| *rft-2* | WBGene00021626 | protein_coding_gene | -1.3 | 0.420 | 8.7E-16 |
| *ssp-34* | WBGene00013858 | protein_coding_gene | -1.3 | 0.419 | 2.1E-02 |
| *C06A5.3* | WBGene00015501 | protein_coding_gene | -1.3 | 0.419 | 5.2E-16 |
| *Y23H5B.5* | WBGene00021276 | protein_coding_gene | -1.3 | 0.419 | 1.5E-19 |
| *calu-1* | WBGene00019760 | protein_coding_gene | -1.3 | 0.418 | 1.4E-06 |
| *rpn-3* | WBGene00004460 | protein_coding_gene | -1.3 | 0.418 | 1.6E-14 |
| *perm-3* | WBGene00022776 | protein_coding_gene | -1.3 | 0.418 | 6.9E-12 |
| *tsfm-1* | WBGene00010094 | protein_coding_gene | -1.3 | 0.418 | 1.6E-12 |
| *vdac-1* | WBGene00019900 | protein_coding_gene | -1.3 | 0.418 | 1.1E-11 |
| *rsp-7* | WBGene00004704 | protein_coding_gene | -1.3 | 0.418 | 9.3E-17 |
| *prp-17* | WBGene00018625 | protein_coding_gene | -1.3 | 0.418 | 4.6E-15 |
| *nep-8* | WBGene00017553 | protein_coding_gene | -1.3 | 0.417 | 1.5E-02 |
| *ZK938.1* | WBGene00014158 | protein_coding_gene | -1.3 | 0.417 | 2.5E-02 |
| *eif-2Bepsilon* | WBGene00008428 | protein_coding_gene | -1.3 | 0.417 | 3.3E-13 |
| *C33F10.1* | WBGene00016351 | protein_coding_gene | -1.3 | 0.417 | 3.0E-02 |
| *cyn-3* | WBGene00000879 | protein_coding_gene | -1.3 | 0.417 | 2.3E-07 |
| *Y47D7A.6* | WBGene00021619 | protein_coding_gene | -1.3 | 0.417 | 4.3E-02 |
| *Y38C1AA.12* | WBGene00021402 | protein_coding_gene | -1.3 | 0.416 | 6.8E-17 |
| *Y61A9LA.1* | WBGene00022014 | protein_coding_gene | -1.3 | 0.416 | 2.3E-16 |
| *T27E7.1* | WBGene00012087 | protein_coding_gene | -1.3 | 0.416 | 2.2E-02 |
| *rpn-10* | WBGene00004466 | protein_coding_gene | -1.3 | 0.416 | 2.1E-12 |
| *copg-1* | WBGene00011775 | protein_coding_gene | -1.3 | 0.416 | 4.3E-11 |
| *Y106G6H.13* | WBGene00013723 | protein_coding_gene | -1.3 | 0.415 | 1.6E-02 |
| *B0511.12* | WBGene00015237 | protein_coding_gene | -1.3 | 0.415 | 1.7E-05 |
| *cox-17* | WBGene00018240 | protein_coding_gene | -1.3 | 0.415 | 8.1E-09 |
| *emr-1* | WBGene00001309 | protein_coding_gene | -1.3 | 0.415 | 1.2E-20 |
| *mdh-1* | WBGene00018491 | protein_coding_gene | -1.3 | 0.414 | 2.9E-22 |
| *C37H5.13* | WBGene00016511 | protein_coding_gene | -1.3 | 0.414 | 2.2E-12 |
| *ZK1037.6* | WBGene00014194 | protein_coding_gene | -1.3 | 0.414 | 8.4E-04 |
| *syx-5* | WBGene00006373 | protein_coding_gene | -1.3 | 0.413 | 6.7E-13 |
| *papl-1* | WBGene00008936 | protein_coding_gene | -1.3 | 0.413 | 2.2E-03 |

| | | | | | |
|---|---|---|---|---|---|
| *pfk-1.1* | WBGene00022199 | protein_coding_gene | -1.3 | 0.413 | 2.2E-13 |
| *plst-1* | WBGene00022425 | protein_coding_gene | -1.3 | 0.412 | 9.1E-09 |
| *Y43F11A.6* | WBGene00014576 | pseudogene | -1.3 | 0.412 | 2.3E-09 |
| *spin-4* | WBGene00013739 | protein_coding_gene | -1.3 | 0.412 | 3.0E-07 |
| *fbxa-27* | WBGene00022333 | protein_coding_gene | -1.3 | 0.412 | 1.1E-03 |
| *sax-2* | WBGene00004728 | protein_coding_gene | -1.3 | 0.412 | 6.1E-08 |
| *Y105C5B.18* | WBGene00013657 | protein_coding_gene | -1.3 | 0.412 | 1.3E-02 |
| *F01G4.4* | WBGene00008503 | protein_coding_gene | -1.3 | 0.412 | 6.5E-07 |
| *Y59E9AL.2* | WBGene00043743 | protein_coding_gene | -1.3 | 0.412 | 2.0E-02 |
| *tbb-2* | WBGene00006537 | protein_coding_gene | -1.3 | 0.411 | 1.9E-09 |
| *F48E3.2* | WBGene00018603 | protein_coding_gene | -1.3 | 0.411 | 1.1E-02 |
| *dpf-3* | WBGene00001056 | protein_coding_gene | -1.3 | 0.411 | 1.1E-15 |
| *smgl-1* | WBGene00008990 | protein_coding_gene | -1.3 | 0.411 | 1.3E-21 |
| *cyh-1* | WBGene00021714 | protein_coding_gene | -1.3 | 0.410 | 1.4E-12 |
| *Y59E9AL.3* | WBGene00021993 | protein_coding_gene | -1.3 | 0.410 | 1.9E-02 |
| *arx-7* | WBGene00000205 | protein_coding_gene | -1.3 | 0.410 | 7.1E-12 |
| *nars-1* | WBGene00003815 | protein_coding_gene | -1.3 | 0.410 | 3.0E-20 |
| *R11.3* | WBGene00011228 | protein_coding_gene | -1.3 | 0.410 | 3.7E-03 |
| *xpf-1* | WBGene00008140 | protein_coding_gene | -1.3 | 0.409 | 3.1E-12 |
| *Y32F6A.6* | WBGene00044209 | pseudogene | -1.3 | 0.409 | 3.0E-05 |
| *C55B6.1* | WBGene00016939 | protein_coding_gene | -1.3 | 0.409 | 5.4E-14 |
| *tomm-7* | WBGene00022783 | protein_coding_gene | -1.3 | 0.409 | 3.4E-04 |
| *npp-11* | WBGene00003797 | protein_coding_gene | -1.3 | 0.409 | 8.6E-17 |
| *R05H5.4* | WBGene00011039 | protein_coding_gene | -1.3 | 0.409 | 5.2E-10 |
| *C10A4.4* | WBGene00015665 | protein_coding_gene | -1.3 | 0.408 | 5.1E-03 |
| *msp-59* | WBGene00003452 | protein_coding_gene | -1.3 | 0.408 | 1.2E-02 |
| *csn-4* | WBGene00000816 | protein_coding_gene | -1.3 | 0.408 | 4.7E-13 |
| *aly-3* | WBGene00000122 | protein_coding_gene | -1.3 | 0.408 | 5.6E-27 |
| *msp-142* | WBGene00003469 | protein_coding_gene | -1.3 | 0.407 | 8.6E-03 |
| *haf-3* | WBGene00001813 | protein_coding_gene | -1.3 | 0.407 | 1.8E-19 |
| *ced-6* | WBGene00000420 | protein_coding_gene | -1.3 | 0.407 | 1.2E-21 |
| *spcs-1* | WBGene00016395 | protein_coding_gene | -1.3 | 0.407 | 1.9E-12 |
| *plx-1* | WBGene00004047 | protein_coding_gene | -1.3 | 0.407 | 1.1E-08 |
| *zyg-1* | WBGene00006988 | protein_coding_gene | -1.3 | 0.407 | 6.8E-11 |
| *tmed-1* | WBGene00010670 | protein_coding_gene | -1.3 | 0.407 | 3.7E-15 |
| *fcho-1* | WBGene00018974 | protein_coding_gene | -1.3 | 0.407 | 9.2E-14 |
| *eif-2D* | WBGene00016113 | protein_coding_gene | -1.3 | 0.407 | 2.3E-16 |
| *gipc-2* | WBGene00009681 | protein_coding_gene | -1.3 | 0.407 | 4.3E-02 |
| *F36A2.9* | WBGene00009456 | protein_coding_gene | -1.3 | 0.407 | 1.9E-13 |
| *gad-1* | WBGene00001513 | protein_coding_gene | -1.3 | 0.406 | 4.7E-22 |
| *K10B2.4* | WBGene00019607 | protein_coding_gene | -1.3 | 0.406 | 9.0E-10 |
| *tmed-12* | WBGene00011606 | protein_coding_gene | -1.3 | 0.406 | 1.8E-02 |
| *lap-2* | WBGene00002250 | protein_coding_gene | -1.3 | 0.406 | 3.5E-12 |
| *Y67D2.7* | WBGene00022057 | protein_coding_gene | -1.3 | 0.406 | 4.5E-22 |
| *slc-25A46* | WBGene00012740 | protein_coding_gene | -1.3 | 0.406 | 2.0E-10 |
| *crls-1* | WBGene00017763 | protein_coding_gene | -1.3 | 0.406 | 4.5E-11 |
| *tomm-40* | WBGene00007686 | protein_coding_gene | -1.3 | 0.406 | 7.4E-19 |
| *rnp-5* | WBGene00004388 | protein_coding_gene | -1.3 | 0.406 | 5.0E-22 |
| *Y7A5A.1* | WBGene00012407 | protein_coding_gene | -1.3 | 0.405 | 2.7E-07 |
| *scav-2* | WBGene00013578 | protein_coding_gene | -1.3 | 0.405 | 3.8E-22 |
| *T27E9.2* | WBGene00012094 | protein_coding_gene | -1.3 | 0.405 | 1.3E-07 |
| *spch-1* | WBGene00007307 | protein_coding_gene | -1.3 | 0.404 | 1.3E-02 |
| *rom-1* | WBGene00004400 | protein_coding_gene | -1.3 | 0.404 | 4.3E-09 |
| *Y71G12B.13* | WBGene00022153 | protein_coding_gene | -1.3 | 0.404 | 6.4E-27 |
| *hpo-27* | WBGene00016418 | protein_coding_gene | -1.3 | 0.404 | 6.7E-17 |
| *Y66C5A.1* | WBGene00044156 | protein_coding_gene | -1.3 | 0.404 | 5.6E-03 |
| *pod-1* | WBGene00004075 | protein_coding_gene | -1.3 | 0.404 | 6.0E-11 |
| *met-1* | WBGene00016603 | protein_coding_gene | -1.3 | 0.404 | 9.8E-10 |
| *Y54G11A.1* | WBGene00013212 | protein_coding_gene | -1.3 | 0.403 | 1.3E-18 |
| *F26F2.7* | WBGene00009172 | protein_coding_gene | -1.3 | 0.403 | 4.3E-25 |
| *Y57A10A.1* | WBGene00013246 | protein_coding_gene | -1.3 | 0.403 | 9.9E-11 |
| *tin-44* | WBGene00020383 | protein_coding_gene | -1.3 | 0.403 | 3.6E-17 |
| *F25E5.1* | WBGene00017782 | protein_coding_gene | -1.3 | 0.402 | 1.1E-07 |
| *nra-1* | WBGene00012128 | protein_coding_gene | -1.3 | 0.402 | 1.3E-16 |
| *vpr-1* | WBGene00018008 | protein_coding_gene | -1.3 | 0.402 | 1.8E-19 |
| *tin-13* | WBGene00006574 | protein_coding_gene | -1.3 | 0.402 | 2.7E-04 |
| *ppat-1* | WBGene00011407 | protein_coding_gene | -1.3 | 0.402 | 2.2E-03 |
| *fbxa-120* | WBGene00016346 | protein_coding_gene | -1.3 | 0.402 | 5.9E-05 |
| *dhod-1* | WBGene00020932 | protein_coding_gene | -1.3 | 0.402 | 9.9E-21 |
| *ZK354.7* | WBGene00022708 | protein_coding_gene | -1.3 | 0.402 | 1.8E-02 |
| *C05D2.8* | WBGene00015469 | protein_coding_gene | -1.3 | 0.402 | 2.7E-09 |
| *T14G8.3* | WBGene00011771 | protein_coding_gene | -1.3 | 0.401 | 2.3E-05 |
| *R05H11.1* | WBGene00019907 | protein_coding_gene | -1.3 | 0.401 | 1.5E-14 |
| *Y51H7C.9* | WBGene00021787 | protein_coding_gene | -1.3 | 0.401 | 1.9E-02 |
| *C31H2.14* | WBGene00206484 | protein_coding_gene | -1.3 | 0.401 | 3.7E-03 |
| *M57.2* | WBGene00019778 | protein_coding_gene | -1.3 | 0.401 | 3.0E-21 |
| *lbp-2* | WBGene00002254 | protein_coding_gene | -1.3 | 0.400 | 6.4E-07 |
| *C10G11.6* | WBGene00015686 | protein_coding_gene | -1.3 | 0.400 | 8.1E-06 |
| *hsp-1* | WBGene00002005 | protein_coding_gene | -1.3 | 0.400 | 2.3E-09 |
| *ipgm-1* | WBGene00019001 | protein_coding_gene | -1.3 | 0.400 | 2.0E-12 |

| | | | | | |
|---|---|---|---|---|---|
| *ric-7* | WBGene00010259 | protein_coding_gene | -1.3 | 0.400 | 2.9E-15 |
| *F26B1.8* | WBGene00194703 | protein_coding_gene | -1.3 | 0.400 | 8.9E-03 |
| *C52E12.1* | WBGene00016888 | protein_coding_gene | -1.3 | 0.400 | 4.9E-18 |
| *cdk-2* | WBGene00019362 | protein_coding_gene | -1.3 | 0.399 | 3.0E-16 |
| *eif-2alpha* | WBGene00021351 | protein_coding_gene | -1.3 | 0.399 | 1.8E-18 |
| *sec-22* | WBGene00018853 | protein_coding_gene | -1.3 | 0.398 | 2.1E-09 |
| *cdr-7* | WBGene00010472 | protein_coding_gene | -1.3 | 0.398 | 7.2E-05 |
| *B0334.5* | WBGene00007145 | protein_coding_gene | -1.3 | 0.398 | 1.9E-10 |
| *Y111B2A.3* | WBGene00013729 | protein_coding_gene | -1.3 | 0.398 | 1.5E-18 |
| *sly-1* | WBGene00009654 | protein_coding_gene | -1.3 | 0.398 | 1.1E-13 |
| *C32E8.4* | WBGene00016322 | protein_coding_gene | -1.3 | 0.398 | 4.8E-02 |
| *ubxn-1* | WBGene00017733 | protein_coding_gene | -1.3 | 0.398 | 3.0E-21 |
| *F46B6.5* | WBGene00009770 | protein_coding_gene | -1.3 | 0.398 | 9.4E-09 |
| *R09E10.6* | WBGene00011176 | protein_coding_gene | -1.3 | 0.397 | 4.0E-02 |
| *C30B5.17* | WBGene00269379 | protein_coding_gene | -1.3 | 0.397 | 1.5E-02 |
| *kynu-1* | WBGene00015802 | protein_coding_gene | -1.3 | 0.397 | 2.2E-03 |
| *DC2.5* | WBGene00017082 | protein_coding_gene | -1.3 | 0.397 | 2.7E-04 |
| *pges-2* | WBGene00011239 | protein_coding_gene | -1.3 | 0.397 | 6.4E-22 |
| *W03F9.1* | WBGene00020999 | protein_coding_gene | -1.3 | 0.397 | 1.6E-24 |
| *rrf-3* | WBGene00004510 | protein_coding_gene | -1.3 | 0.396 | 4.1E-16 |
| *Y82E9BR.2* | WBGene00022335 | protein_coding_gene | -1.3 | 0.396 | 5.0E-15 |
| *EGAP4.1* | WBGene00017149 | protein_coding_gene | -1.3 | 0.396 | 3.6E-02 |
| *moc-1* | WBGene00003384 | protein_coding_gene | -1.3 | 0.396 | 4.6E-09 |
| *C16A11.5* | WBGene00015818 | protein_coding_gene | -1.3 | 0.395 | 3.0E-18 |
| *col-98* | WBGene00000673 | protein_coding_gene | -1.3 | 0.395 | 1.7E-04 |
| *F52H2.7* | WBGene00018719 | protein_coding_gene | -1.3 | 0.395 | 3.0E-10 |
| *ZK265.3* | WBGene00013956 | protein_coding_gene | -1.3 | 0.395 | 1.0E-02 |
| *parp-1* | WBGene00004049 | protein_coding_gene | -1.3 | 0.394 | 2.0E-14 |
| *dbn-1* | WBGene00010664 | protein_coding_gene | -1.3 | 0.394 | 1.8E-20 |
| *Y67D2.2* | WBGene00022052 | protein_coding_gene | -1.3 | 0.394 | 4.6E-21 |
| *C39H7.1* | WBGene00016541 | protein_coding_gene | -1.3 | 0.394 | 1.1E-02 |
| *acbp-5* | WBGene00011731 | protein_coding_gene | -1.3 | 0.393 | 2.9E-29 |
| *efr-3* | WBGene00016311 | protein_coding_gene | -1.3 | 0.393 | 2.0E-20 |
| *rsp-1* | WBGene00004698 | protein_coding_gene | -1.3 | 0.393 | 1.4E-18 |
| *Y41E3.1* | WBGene00012762 | protein_coding_gene | -1.3 | 0.393 | 6.6E-16 |
| *kbrl-1* | WBGene00009919 | protein_coding_gene | -1.3 | 0.392 | 1.4E-28 |
| *mbf-1* | WBGene00003148 | protein_coding_gene | -1.4 | 0.392 | 4.9E-22 |
| *K09C6.7* | WBGene00019561 | protein_coding_gene | -1.4 | 0.392 | 1.7E-02 |
| *R04F11.5* | WBGene00011017 | protein_coding_gene | -1.4 | 0.392 | 1.6E-08 |
| *npp-10* | WBGene00003796 | protein_coding_gene | -1.4 | 0.392 | 4.6E-13 |
| *pptr-2* | WBGene00007554 | protein_coding_gene | -1.4 | 0.391 | 1.8E-21 |
| *ZK380.6* | WBGene00304812 | protein_coding_gene | -1.4 | 0.391 | 4.5E-05 |
| *Y49F6B.9* | WBGene00021721 | protein_coding_gene | -1.4 | 0.391 | 7.8E-15 |
| *Y73E7A.6* | WBGene00022273 | protein_coding_gene | -1.4 | 0.390 | 2.1E-09 |
| *fipp-1* | WBGene00017988 | protein_coding_gene | -1.4 | 0.390 | 1.3E-21 |
| *bath-2* | WBGene00018668 | protein_coding_gene | -1.4 | 0.390 | 1.9E-06 |
| *cye-1* | WBGene00000871 | protein_coding_gene | -1.4 | 0.390 | 1.3E-11 |
| *C32D5.10* | WBGene00016317 | protein_coding_gene | -1.4 | 0.389 | 7.4E-28 |
| *F55A3.7* | WBGene00018852 | pseudogene | -1.4 | 0.389 | 2.2E-06 |
| *cdkr-3* | WBGene00013765 | protein_coding_gene | -1.4 | 0.388 | 6.9E-09 |
| *C46A5.1* | WBGene00016698 | protein_coding_gene | -1.4 | 0.388 | 5.5E-03 |
| *imp-2* | WBGene00011481 | protein_coding_gene | -1.4 | 0.388 | 2.4E-10 |
| *vglu-3* | WBGene00011556 | protein_coding_gene | -1.4 | 0.388 | 1.2E-06 |
| *tba-2* | WBGene00006529 | protein_coding_gene | -1.4 | 0.388 | 5.7E-15 |
| *C15H9.4* | WBGene00015800 | protein_coding_gene | -1.4 | 0.387 | 1.8E-15 |
| *ZC317.7* | WBGene00022590 | protein_coding_gene | -1.4 | 0.387 | 2.7E-04 |
| *ndg-4* | WBGene00003577 | protein_coding_gene | -1.4 | 0.387 | 3.5E-06 |
| *pbs-4* | WBGene00003950 | protein_coding_gene | -1.4 | 0.387 | 1.3E-23 |
| *rom-4* | WBGene00004403 | protein_coding_gene | -1.4 | 0.387 | 1.7E-07 |
| *C50D2.6* | WBGene00016809 | protein_coding_gene | -1.4 | 0.387 | 1.5E-04 |
| *inx-9* | WBGene00002131 | protein_coding_gene | -1.4 | 0.386 | 5.7E-08 |
| *rsd-6* | WBGene00004684 | protein_coding_gene | -1.4 | 0.386 | 3.0E-18 |
| *gpi-1* | WBGene00013597 | protein_coding_gene | -1.4 | 0.386 | 1.6E-20 |
| *ccm-3* | WBGene00007561 | protein_coding_gene | -1.4 | 0.386 | 4.1E-15 |
| *dnj-13* | WBGene00001031 | protein_coding_gene | -1.4 | 0.386 | 1.5E-13 |
| *exo-1* | WBGene00009731 | protein_coding_gene | -1.4 | 0.386 | 3.4E-26 |
| *cox-16* | WBGene00302991 | protein_coding_gene | -1.4 | 0.385 | 2.2E-05 |
| *C02B10.4* | WBGene00015329 | protein_coding_gene | -1.4 | 0.385 | 5.1E-19 |
| *spg-7* | WBGene00004978 | protein_coding_gene | -1.4 | 0.385 | 1.6E-26 |
| *F43H9.3* | WBGene00018399 | protein_coding_gene | -1.4 | 0.385 | 2.4E-09 |
| *sac-1* | WBGene00009264 | protein_coding_gene | -1.4 | 0.385 | 8.7E-26 |
| *iglr-2* | WBGene00022580 | protein_coding_gene | -1.4 | 0.385 | 9.1E-14 |
| *copb-2* | WBGene00009542 | protein_coding_gene | -1.4 | 0.385 | 1.2E-11 |
| *pbs-5* | WBGene00003951 | protein_coding_gene | -1.4 | 0.384 | 6.8E-12 |
| *rsd-2* | WBGene00004681 | protein_coding_gene | -1.4 | 0.384 | 4.3E-13 |
| *B0035.15* | WBGene00007113 | protein_coding_gene | -1.4 | 0.384 | 4.4E-20 |
| *pqn-22* | WBGene00004112 | protein_coding_gene | -1.4 | 0.384 | 5.6E-06 |
| *ZK512.10* | WBGene00013988 | protein_coding_gene | -1.4 | 0.384 | 3.1E-02 |
| *npp-14* | WBGene00003800 | protein_coding_gene | -1.4 | 0.384 | 2.0E-09 |
| *Y69H2.3* | WBGene00013481 | protein_coding_gene | -1.4 | 0.383 | 6.4E-10 |

| | | | | | |
|---|---|---|---|---|---|
| *syp-2* | WBGene00006376 | protein_coding_gene | -1.4 | 0.383 | 9.4E-13 |
| *attf-3* | WBGene00018794 | protein_coding_gene | -1.4 | 0.383 | 1.2E-07 |
| *sec-24.1* | WBGene00004755 | protein_coding_gene | -1.4 | 0.383 | 4.0E-13 |
| *acd-2* | WBGene00016066 | protein_coding_gene | -1.4 | 0.383 | 1.0E-06 |
| *dbt-1* | WBGene00014054 | protein_coding_gene | -1.4 | 0.383 | 3.6E-11 |
| *mig-22* | WBGene00003253 | protein_coding_gene | -1.4 | 0.383 | 6.7E-13 |
| *trap-3* | WBGene00021420 | protein_coding_gene | -1.4 | 0.382 | 9.5E-08 |
| *F22F1.2* | WBGene00017714 | protein_coding_gene | -1.4 | 0.382 | 4.0E-02 |
| *F14D2.5* | WBGene00017453 | protein_coding_gene | -1.4 | 0.382 | 3.6E-03 |
| *Y42H9AR.2* | WBGene00021537 | protein_coding_gene | -1.4 | 0.382 | 1.6E-02 |
| *C04F12.7* | WBGene00007301 | protein_coding_gene | -1.4 | 0.382 | 4.7E-02 |
| *K08H10.6* | WBGene00010698 | protein_coding_gene | -1.4 | 0.382 | 1.5E-02 |
| *F56C9.10* | WBGene00018953 | protein_coding_gene | -1.4 | 0.381 | 2.1E-09 |
| *eri-3* | WBGene00021103 | protein_coding_gene | -1.4 | 0.381 | 5.4E-28 |
| *mccc-1* | WBGene00009319 | protein_coding_gene | -1.4 | 0.381 | 4.9E-21 |
| *F19B10.10* | WBGene00017589 | protein_coding_gene | -1.4 | 0.381 | 1.4E-10 |
| *cogc-1* | WBGene00003258 | protein_coding_gene | -1.4 | 0.381 | 3.6E-24 |
| *Y66D12A.16* | WBGene00013442 | protein_coding_gene | -1.4 | 0.381 | 5.8E-15 |
| *C56G2.5* | WBGene00016980 | protein_coding_gene | -1.4 | 0.381 | 4.2E-10 |
| *ptps-1* | WBGene00015010 | protein_coding_gene | -1.4 | 0.380 | 2.4E-04 |
| *sam-10* | WBGene00012990 | protein_coding_gene | -1.4 | 0.380 | 5.2E-12 |
| *skat-1* | WBGene00010307 | protein_coding_gene | -1.4 | 0.380 | 1.4E-12 |
| *soap-1* | WBGene00015743 | protein_coding_gene | -1.4 | 0.380 | 4.5E-11 |
| *dna-2* | WBGene00001016 | protein_coding_gene | -1.4 | 0.380 | 1.9E-17 |
| *T24A6.20* | WBGene00020756 | protein_coding_gene | -1.4 | 0.379 | 7.6E-06 |
| *T05G5.4* | WBGene00011499 | protein_coding_gene | -1.4 | 0.379 | 1.1E-03 |
| *ugt-47* | WBGene00011006 | protein_coding_gene | -1.4 | 0.379 | 7.2E-04 |
| *fars-1* | WBGene00001497 | protein_coding_gene | -1.4 | 0.379 | 5.2E-15 |
| *bckd-1A* | WBGene00012713 | protein_coding_gene | -1.4 | 0.379 | 2.4E-19 |
| *B0432.1* | WBGene00015183 | protein_coding_gene | -1.4 | 0.379 | 3.9E-02 |
| *F55D12.6* | WBGene00010114 | protein_coding_gene | -1.4 | 0.378 | 1.3E-02 |
| *acs-15* | WBGene00019920 | protein_coding_gene | -1.4 | 0.378 | 2.7E-02 |
| *agef-1* | WBGene00012386 | protein_coding_gene | -1.4 | 0.378 | 2.1E-17 |
| *Y46D2A.2* | WBGene00021590 | protein_coding_gene | -1.4 | 0.377 | 8.2E-03 |
| *Y53F4B.9* | WBGene00013156 | protein_coding_gene | -1.4 | 0.377 | 3.2E-12 |
| *K12D12.5* | WBGene00010787 | protein_coding_gene | -1.4 | 0.376 | 4.0E-20 |
| *npp-25* | WBGene00012555 | protein_coding_gene | -1.4 | 0.376 | 5.7E-10 |
| *T21B10.4* | WBGene00011886 | protein_coding_gene | -1.4 | 0.376 | 2.5E-11 |
| *glo-2* | WBGene00019010 | protein_coding_gene | -1.4 | 0.375 | 1.4E-03 |
| *F22G12.4* | WBGene00009064 | protein_coding_gene | -1.4 | 0.374 | 4.7E-19 |
| *msp-55* | WBGene00003448 | protein_coding_gene | -1.4 | 0.374 | 6.1E-03 |
| *tat-3* | WBGene00012360 | protein_coding_gene | -1.4 | 0.373 | 1.6E-10 |
| *jmjd-2* | WBGene00012982 | protein_coding_gene | -1.4 | 0.373 | 1.1E-11 |
| *F25H8.2* | WBGene00009132 | protein_coding_gene | -1.4 | 0.373 | 3.1E-18 |
| *T16G12.6* | WBGene00011807 | protein_coding_gene | -1.4 | 0.372 | 4.6E-21 |
| *elc-1* | WBGene00001236 | protein_coding_gene | -1.4 | 0.372 | 7.3E-08 |
| *msrp-2* | WBGene00022751 | protein_coding_gene | -1.4 | 0.372 | 2.7E-02 |
| *par-3* | WBGene00003918 | protein_coding_gene | -1.4 | 0.372 | 1.5E-07 |
| *unc-101* | WBGene00006829 | protein_coding_gene | -1.4 | 0.371 | 5.4E-20 |
| *pbs-6* | WBGene00003952 | protein_coding_gene | -1.4 | 0.371 | 6.5E-14 |
| *hda-10* | WBGene00001838 | protein_coding_gene | -1.4 | 0.370 | 5.8E-19 |
| *Y47D9A.1* | WBGene00021628 | protein_coding_gene | -1.4 | 0.370 | 4.4E-18 |
| *F58D2.4* | WBGene00185077 | protein_coding_gene | -1.4 | 0.370 | 4.9E-02 |
| *B0250.5* | WBGene00007122 | protein_coding_gene | -1.4 | 0.370 | 7.6E-13 |
| *M28.9* | WBGene00010900 | protein_coding_gene | -1.4 | 0.369 | 4.9E-03 |
| *D1065.6* | WBGene00268212 | ncRNA_gene | -1.4 | 0.369 | 4.6E-05 |
| *enol-1* | WBGene00011884 | protein_coding_gene | -1.4 | 0.368 | 3.6E-14 |
| *B0454.5* | WBGene00015196 | protein_coding_gene | -1.4 | 0.368 | 1.4E-04 |
| *dad-1* | WBGene00000896 | protein_coding_gene | -1.4 | 0.368 | 2.3E-14 |
| *F13H8.8* | WBGene00017441 | protein_coding_gene | -1.4 | 0.368 | 9.9E-06 |
| *dpy-30* | WBGene00001088 | protein_coding_gene | -1.4 | 0.368 | 2.2E-17 |
| *fncm-1* | WBGene00018908 | protein_coding_gene | -1.4 | 0.368 | 3.1E-16 |
| *elo-3* | WBGene00001241 | protein_coding_gene | -1.4 | 0.367 | 1.6E-11 |
| *grd-5* | WBGene00001694 | protein_coding_gene | -1.4 | 0.367 | 6.0E-04 |
| *sdz-15* | WBGene00017752 | protein_coding_gene | -1.4 | 0.367 | 6.3E-03 |
| *C06G4.1* | WBGene00015557 | protein_coding_gene | -1.4 | 0.367 | 6.4E-18 |
| *rict-1* | WBGene00009245 | protein_coding_gene | -1.4 | 0.367 | 6.3E-10 |
| *cua-1* | WBGene00000834 | protein_coding_gene | -1.4 | 0.366 | 5.9E-22 |
| *hex-3* | WBGene00012663 | protein_coding_gene | -1.5 | 0.366 | 2.1E-10 |
| *gex-2* | WBGene00001579 | protein_coding_gene | -1.5 | 0.366 | 1.8E-11 |
| *nars-2* | WBGene00013447 | protein_coding_gene | -1.5 | 0.365 | 1.1E-10 |
| *sptl-3* | WBGene00011932 | protein_coding_gene | -1.5 | 0.365 | 2.0E-08 |
| *ZK858.6* | WBGene00014119 | protein_coding_gene | -1.5 | 0.365 | 4.2E-14 |
| *mys-4* | WBGene00007914 | protein_coding_gene | -1.5 | 0.365 | 4.7E-15 |
| *eif-2Bdelta* | WBGene00008670 | protein_coding_gene | -1.5 | 0.365 | 2.9E-27 |
| *Y41D4B.4* | WBGene00021512 | protein_coding_gene | -1.5 | 0.365 | 6.4E-12 |
| *zhp-4* | WBGene00012678 | protein_coding_gene | -1.5 | 0.364 | 1.0E-14 |
| *F27C1.1* | WBGene00017851 | protein_coding_gene | -1.5 | 0.364 | 8.4E-03 |
| *cdc-48.2* | WBGene00008053 | protein_coding_gene | -1.5 | 0.363 | 1.8E-16 |
| *M02G9.4* | WBGene00194935 | protein_coding_gene | -1.5 | 0.363 | 4.7E-02 |

| | | | | | |
|---|---|---|---|---|---|
| *hil-5* | WBGene00001856 | protein_coding_gene | -1.5 | 0.363 | 6.2E-25 |
| *F56C11.5* | WBGene00018957 | protein_coding_gene | -1.5 | 0.363 | 7.4E-14 |
| *T13C2.6* | WBGene00020481 | protein_coding_gene | -1.5 | 0.363 | 1.1E-16 |
| *R05H10.3* | WBGene00011044 | protein_coding_gene | -1.5 | 0.363 | 1.9E-19 |
| *hsf-2* | WBGene00013134 | pseudogene | -1.5 | 0.363 | 1.3E-03 |
| *F53B6.4* | WBGene00009959 | protein_coding_gene | -1.5 | 0.362 | 7.3E-03 |
| *ugt-13* | WBGene00019232 | protein_coding_gene | -1.5 | 0.362 | 4.0E-06 |
| *dhhc-7* | WBGene00007637 | protein_coding_gene | -1.5 | 0.362 | 1.4E-19 |
| *snx-14* | WBGene00013011 | protein_coding_gene | -1.5 | 0.362 | 1.3E-12 |
| *vnut-1* | WBGene00010758 | protein_coding_gene | -1.5 | 0.361 | 3.0E-23 |
| *ugt-12* | WBGene00020592 | protein_coding_gene | -1.5 | 0.361 | 3.0E-08 |
| *slc-30A9* | WBGene00022174 | protein_coding_gene | -1.5 | 0.361 | 8.3E-23 |
| *art-1* | WBGene00000198 | protein_coding_gene | -1.5 | 0.361 | 6.1E-12 |
| *cpi-1* | WBGene00000535 | protein_coding_gene | -1.5 | 0.360 | 1.1E-05 |
| *Y39A1A.22* | WBGene00012659 | protein_coding_gene | -1.5 | 0.360 | 3.2E-13 |
| *plpp-1.3* | WBGene00008749 | protein_coding_gene | -1.5 | 0.359 | 3.9E-11 |
| *F31D4.8* | WBGene00009293 | protein_coding_gene | -1.5 | 0.358 | 8.2E-04 |
| *rbc-2* | WBGene00004314 | protein_coding_gene | -1.5 | 0.358 | 1.7E-09 |
| *ssq-3* | WBGene00006052 | protein_coding_gene | -1.5 | 0.358 | 1.2E-02 |
| *rmd-3* | WBGene00007190 | protein_coding_gene | -1.5 | 0.358 | 2.8E-03 |
| *Y56A3A.22* | WBGene00013239 | protein_coding_gene | -1.5 | 0.357 | 2.9E-16 |
| *cpb-3* | WBGene00000772 | protein_coding_gene | -1.5 | 0.357 | 4.9E-14 |
| *atic-1* | WBGene00016957 | protein_coding_gene | -1.5 | 0.357 | 2.7E-03 |
| *Y69E1A.1* | WBGene00013473 | protein_coding_gene | -1.5 | 0.357 | 4.3E-03 |
| *Y73B3A.21* | WBGene00022222 | pseudogene | -1.5 | 0.357 | 1.0E-05 |
| *nlp-77* | WBGene00015514 | protein_coding_gene | -1.5 | 0.356 | 3.1E-06 |
| *ani-1* | WBGene00013038 | protein_coding_gene | -1.5 | 0.356 | 3.1E-10 |
| *tag-153* | WBGene00006499 | protein_coding_gene | -1.5 | 0.356 | 6.9E-16 |
| *oxa-1* | WBGene00007215 | protein_coding_gene | -1.5 | 0.355 | 8.7E-12 |
| *ubr-5* | WBGene00009460 | protein_coding_gene | -1.5 | 0.355 | 2.2E-08 |
| *T20F5.6* | WBGene00020628 | protein_coding_gene | -1.5 | 0.355 | 7.1E-15 |
| *T26A8.1* | WBGene00020824 | protein_coding_gene | -1.5 | 0.355 | 5.3E-16 |
| *csc-1* | WBGene00000804 | protein_coding_gene | -1.5 | 0.355 | 2.0E-25 |
| *F15D3.4* | WBGene00008854 | protein_coding_gene | -1.5 | 0.354 | 5.3E-04 |
| *pbs-1* | WBGene00003947 | protein_coding_gene | -1.5 | 0.354 | 4.9E-21 |
| *catp-3* | WBGene00015660 | protein_coding_gene | -1.5 | 0.353 | 4.9E-04 |
| *Y50D4C.2* | WBGene00021747 | pseudogene | -1.5 | 0.353 | 3.1E-06 |
| *H04D03.2* | WBGene00010363 | protein_coding_gene | -1.5 | 0.353 | 1.5E-21 |
| *Y17G7B.19* | WBGene00012470 | protein_coding_gene | -1.5 | 0.352 | 3.2E-02 |
| *Y50D4C.5* | WBGene00021749 | protein_coding_gene | -1.5 | 0.352 | 1.4E-14 |
| *coq-8* | WBGene00000767 | protein_coding_gene | -1.5 | 0.351 | 1.5E-30 |
| *W08E12.7* | WBGene00021088 | protein_coding_gene | -1.5 | 0.351 | 1.5E-11 |
| *Y48A6C.4* | WBGene00012977 | protein_coding_gene | -1.5 | 0.351 | 2.0E-14 |
| *C48B4.7* | WBGene00008170 | protein_coding_gene | -1.5 | 0.351 | 3.5E-10 |
| *H25K10.1* | WBGene00010411 | protein_coding_gene | -1.5 | 0.351 | 2.3E-02 |
| *nmt-1* | WBGene00020549 | protein_coding_gene | -1.5 | 0.351 | 1.5E-15 |
| *C13G5.2* | WBGene00015747 | protein_coding_gene | -1.5 | 0.351 | 2.8E-25 |
| *Y48G1A.7* | WBGene00235385 | pseudogene | -1.5 | 0.351 | 2.1E-05 |
| *ZK228.4* | WBGene00013947 | protein_coding_gene | -1.5 | 0.350 | 1.4E-08 |
| *cpg-9* | WBGene00022072 | protein_coding_gene | -1.5 | 0.350 | 4.3E-05 |
| *dab-1* | WBGene00000894 | protein_coding_gene | -1.5 | 0.350 | 9.4E-14 |
| *F27D4.3* | WBGene00014746 | pseudogene | -1.5 | 0.350 | 4.6E-05 |
| *rbm-39* | WBGene00021921 | protein_coding_gene | -1.5 | 0.350 | 2.6E-25 |
| *pigm-1* | WBGene00007189 | protein_coding_gene | -1.5 | 0.350 | 1.6E-19 |
| *hsp-6* | WBGene00002010 | protein_coding_gene | -1.5 | 0.349 | 2.0E-12 |
| *klp-10* | WBGene00002221 | pseudogene | -1.5 | 0.349 | 1.1E-13 |
| *folt-1* | WBGene00007388 | protein_coding_gene | -1.5 | 0.348 | 5.9E-07 |
| *sdha-2* | WBGene00016392 | protein_coding_gene | -1.5 | 0.348 | 4.9E-23 |
| *nhr-74* | WBGene00003664 | protein_coding_gene | -1.5 | 0.348 | 2.1E-02 |
| *catp-6* | WBGene00012341 | protein_coding_gene | -1.5 | 0.347 | 5.3E-12 |
| *F13A7.1* | WBGene00008724 | protein_coding_gene | -1.5 | 0.347 | 1.2E-02 |
| *F28E10.4* | WBGene00017902 | protein_coding_gene | -1.5 | 0.346 | 1.1E-02 |
| *T21E3.2* | WBGene00020653 | protein_coding_gene | -1.5 | 0.346 | 4.4E-02 |
| *dgat-2* | WBGene00010296 | protein_coding_gene | -1.5 | 0.346 | 6.9E-05 |
| *W09C5.7* | WBGene00012353 | protein_coding_gene | -1.5 | 0.346 | 2.4E-32 |
| *his-74* | WBGene00012276 | protein_coding_gene | -1.5 | 0.346 | 1.6E-14 |
| *K12B6.11* | WBGene00195164 | protein_coding_gene | -1.5 | 0.346 | 3.4E-02 |
| *hmg-12* | WBGene00001977 | protein_coding_gene | -1.5 | 0.346 | 1.8E-20 |
| *pgl-1* | WBGene00003992 | protein_coding_gene | -1.5 | 0.346 | 4.5E-27 |
| *F44E2.8* | WBGene00018421 | protein_coding_gene | -1.5 | 0.345 | 4.0E-15 |
| *ZK524.4* | WBGene00013994 | protein_coding_gene | -1.5 | 0.345 | 7.0E-11 |
| *app-1* | WBGene00000155 | protein_coding_gene | -1.5 | 0.345 | 9.8E-16 |
| *bckd-1B* | WBGene00006518 | protein_coding_gene | -1.5 | 0.345 | 3.1E-09 |
| *itsn-1* | WBGene00006405 | protein_coding_gene | -1.5 | 0.344 | 7.0E-13 |
| *uba-5* | WBGene00020184 | protein_coding_gene | -1.5 | 0.343 | 1.1E-22 |
| *F44E7.4* | WBGene00018426 | protein_coding_gene | -1.5 | 0.343 | 7.6E-27 |
| *smz-2* | WBGene00020661 | protein_coding_gene | -1.5 | 0.343 | 8.2E-04 |
| *ZK84.5* | WBGene00022652 | protein_coding_gene | -1.5 | 0.342 | 6.5E-03 |
| *dpf-7* | WBGene00001060 | protein_coding_gene | -1.5 | 0.342 | 2.2E-28 |
| *Y37H2A.12* | WBGene00044758 | protein_coding_gene | -1.5 | 0.342 | 3.3E-12 |

| | | | | | |
|---|---|---|---|---|---|
| *pyk-1* | WBGene00009126 | protein_coding_gene | -1.5 | 0.342 | 1.0E-14 |
| *fbxa-129* | WBGene00010294 | pseudogene | -1.6 | 0.341 | 1.4E-10 |
| *haao-1* | WBGene00010595 | protein_coding_gene | -1.6 | 0.341 | 6.1E-04 |
| *usp-50* | WBGene00008441 | protein_coding_gene | -1.6 | 0.341 | 7.4E-18 |
| *ugt-21* | WBGene00007885 | protein_coding_gene | -1.6 | 0.341 | 1.2E-05 |
| *Y63D3A.7* | WBGene00013406 | protein_coding_gene | -1.6 | 0.341 | 3.8E-10 |
| *sec-61.G* | WBGene00001303 | protein_coding_gene | -1.6 | 0.341 | 6.3E-09 |
| *erfa-1* | WBGene00020269 | protein_coding_gene | -1.6 | 0.340 | 1.9E-21 |
| *ZK930.6* | WBGene00014156 | protein_coding_gene | -1.6 | 0.340 | 3.5E-04 |
| *nspc-7* | WBGene00015392 | protein_coding_gene | -1.6 | 0.340 | 2.5E-03 |
| *F23C8.5* | WBGene00017734 | protein_coding_gene | -1.6 | 0.339 | 1.7E-22 |
| *T28A8.5* | WBGene00012111 | protein_coding_gene | -1.6 | 0.339 | 3.1E-19 |
| *sep-1* | WBGene00004775 | protein_coding_gene | -1.6 | 0.339 | 2.0E-24 |
| *Y38C1AA.7* | WBGene00021398 | protein_coding_gene | -1.6 | 0.339 | 3.1E-03 |
| *Y102A5C.2* | WBGene00013611 | protein_coding_gene | -1.6 | 0.339 | 2.4E-17 |
| *T10C6.6* | WBGene00011688 | protein_coding_gene | -1.6 | 0.338 | 1.5E-23 |
| *F17E9.5* | WBGene00017542 | protein_coding_gene | -1.6 | 0.338 | 3.5E-03 |
| *F20B6.9* | WBGene00017632 | protein_coding_gene | -1.6 | 0.338 | 6.0E-03 |
| *sel-11* | WBGene00004768 | protein_coding_gene | -1.6 | 0.337 | 3.4E-16 |
| *gst-30* | WBGene00001778 | protein_coding_gene | -1.6 | 0.336 | 3.1E-03 |
| *T07C4.3* | WBGene00011560 | protein_coding_gene | -1.6 | 0.335 | 6.6E-18 |
| *vrp-1* | WBGene00021869 | protein_coding_gene | -1.6 | 0.335 | 7.5E-07 |
| *pdxk-1* | WBGene00019008 | protein_coding_gene | -1.6 | 0.335 | 1.1E-17 |
| *dct-16* | WBGene00012615 | protein_coding_gene | -1.6 | 0.335 | 4.6E-09 |
| *acdh-13* | WBGene00001566 | protein_coding_gene | -1.6 | 0.335 | 3.5E-35 |
| *sgt-1* | WBGene00019893 | protein_coding_gene | -1.6 | 0.334 | 9.5E-21 |
| *C40A11.3* | WBGene00016546 | protein_coding_gene | -1.6 | 0.334 | 8.6E-03 |
| *Y65B4BL.3* | WBGene00022035 | protein_coding_gene | -1.6 | 0.334 | 3.6E-19 |
| *hpo-40* | WBGene00014113 | protein_coding_gene | -1.6 | 0.333 | 3.2E-15 |
| *alh-9* | WBGene00000115 | protein_coding_gene | -1.6 | 0.333 | 1.1E-06 |
| *K08D8.3* | WBGene00010657 | protein_coding_gene | -1.6 | 0.333 | 5.5E-08 |
| *C08F8.9* | WBGene00007449 | protein_coding_gene | -1.6 | 0.332 | 3.1E-08 |
| *ostb-1* | WBGene00011638 | protein_coding_gene | -1.6 | 0.331 | 1.9E-13 |
| *crml-1* | WBGene00010641 | protein_coding_gene | -1.6 | 0.331 | 3.4E-21 |
| *F52B11.2* | WBGene00009925 | protein_coding_gene | -1.6 | 0.330 | 6.4E-09 |
| *acdh-8* | WBGene00019406 | protein_coding_gene | -1.6 | 0.330 | 2.0E-03 |
| *F22F7.8* | WBGene00194908 | protein_coding_gene | -1.6 | 0.330 | 1.2E-07 |
| *xnd-1* | WBGene00001514 | protein_coding_gene | -1.6 | 0.329 | 2.3E-20 |
| *grld-1* | WBGene00017929 | protein_coding_gene | -1.6 | 0.329 | 4.6E-27 |
| *cyk-4* | WBGene00000875 | protein_coding_gene | -1.6 | 0.329 | 2.5E-17 |
| *T21C12.4* | WBGene00011900 | protein_coding_gene | -1.6 | 0.329 | 1.4E-03 |
| *cct-8* | WBGene00021934 | protein_coding_gene | -1.6 | 0.328 | 4.1E-16 |
| *sid-1* | WBGene00004795 | protein_coding_gene | -1.6 | 0.328 | 1.6E-11 |
| *dagl-2* | WBGene00018364 | protein_coding_gene | -1.6 | 0.328 | 4.0E-19 |
| *F57B1.9* | WBGene00045488 | protein_coding_gene | -1.6 | 0.327 | 6.3E-03 |
| *T06D8.9* | WBGene00011529 | protein_coding_gene | -1.6 | 0.327 | 1.7E-15 |
| *Y48G1C.8* | WBGene00021681 | protein_coding_gene | -1.6 | 0.326 | 1.1E-12 |
| *ttbk-2* | WBGene00018122 | protein_coding_gene | -1.6 | 0.326 | 1.9E-02 |
| *ced-5* | WBGene00000419 | protein_coding_gene | -1.6 | 0.325 | 1.7E-12 |
| *snf-6* | WBGene00004905 | protein_coding_gene | -1.6 | 0.324 | 9.4E-12 |
| *sti-1* | WBGene00019983 | protein_coding_gene | -1.6 | 0.324 | 8.8E-19 |
| *trap-4* | WBGene00013238 | protein_coding_gene | -1.6 | 0.324 | 5.5E-14 |
| *lem-3* | WBGene00002276 | protein_coding_gene | -1.6 | 0.324 | 2.0E-26 |
| *Y69E1A.2* | WBGene00013474 | protein_coding_gene | -1.6 | 0.324 | 1.1E-03 |
| *npp-16* | WBGene00003802 | protein_coding_gene | -1.6 | 0.324 | 2.7E-25 |
| *emb-8* | WBGene00001262 | protein_coding_gene | -1.6 | 0.323 | 3.1E-37 |
| *twk-16* | WBGene00006670 | protein_coding_gene | -1.6 | 0.323 | 4.2E-03 |
| *Y48G10A.1* | WBGene00013018 | protein_coding_gene | -1.6 | 0.323 | 8.2E-21 |
| *grd-3* | WBGene00001692 | protein_coding_gene | -1.6 | 0.323 | 1.6E-06 |
| *pgk-1* | WBGene00020185 | protein_coding_gene | -1.6 | 0.323 | 2.3E-08 |
| *prg-2* | WBGene00004179 | pseudogene | -1.6 | 0.323 | 1.2E-13 |
| *F44E2.10* | WBGene00018423 | protein_coding_gene | -1.6 | 0.323 | 4.2E-24 |
| *F41H10.3* | WBGene00018316 | protein_coding_gene | -1.6 | 0.323 | 3.1E-08 |
| *B0025.5* | WBGene00194700 | protein_coding_gene | -1.6 | 0.322 | 3.4E-03 |
| *C17F3.1* | WBGene00015907 | protein_coding_gene | -1.6 | 0.321 | 7.1E-03 |
| *Y43B11AR.7* | WBGene00045086 | snoRNA_gene | -1.6 | 0.321 | 1.1E-02 |
| *Y106G6A.4* | WBGene00013696 | protein_coding_gene | -1.6 | 0.320 | 2.1E-03 |
| *ugt-28* | WBGene00015693 | protein_coding_gene | -1.6 | 0.320 | 4.2E-14 |
| *cyp-14A1* | WBGene00010705 | protein_coding_gene | -1.6 | 0.320 | 2.5E-03 |
| *pbs-2* | WBGene00003948 | protein_coding_gene | -1.6 | 0.320 | 6.0E-22 |
| *ess-2* | WBGene00018371 | protein_coding_gene | -1.6 | 0.320 | 1.3E-26 |
| *npp-7* | WBGene00003793 | protein_coding_gene | -1.6 | 0.319 | 4.9E-09 |
| *F35E12.10* | WBGene00009434 | protein_coding_gene | -1.6 | 0.319 | 1.4E-05 |
| *cbl-1* | WBGene00007533 | protein_coding_gene | -1.7 | 0.319 | 4.2E-05 |
| *fbp-1* | WBGene00001404 | protein_coding_gene | -1.7 | 0.318 | 4.4E-12 |
| *C34C12.8* | WBGene00007927 | protein_coding_gene | -1.7 | 0.318 | 5.1E-23 |
| *F07F6.4* | WBGene00017217 | protein_coding_gene | -1.7 | 0.318 | 5.9E-17 |
| *tkt-1* | WBGene00008506 | protein_coding_gene | -1.7 | 0.317 | 3.1E-16 |
| *B0432.7* | WBGene00015188 | protein_coding_gene | -1.7 | 0.317 | 8.5E-13 |
| *kat-1* | WBGene00002183 | protein_coding_gene | -1.7 | 0.317 | 1.4E-24 |

| | | | | | |
|---|---|---|---|---|---|
| *let-754* | WBGene00002879 | protein_coding_gene | -1.7 | 0.317 | 1.6E-15 |
| *chc-1* | WBGene00011867 | protein_coding_gene | -1.7 | 0.317 | 1.3E-12 |
| *F21H7.5* | WBGene00009031 | protein_coding_gene | -1.7 | 0.317 | 3.0E-03 |
| *Y73F8A.14* | WBGene00013523 | protein_coding_gene | -1.7 | 0.316 | 2.6E-02 |
| *Y53C10A.1* | WBGene00014915 | pseudogene | -1.7 | 0.316 | 4.1E-16 |
| *haf-6* | WBGene00001816 | protein_coding_gene | -1.7 | 0.316 | 6.5E-17 |
| *asp-17* | WBGene00012683 | protein_coding_gene | -1.7 | 0.315 | 9.2E-03 |
| *F18A11.6* | WBGene00008933 | pseudogene | -1.7 | 0.315 | 1.0E-03 |
| *spd-2* | WBGene00004953 | protein_coding_gene | -1.7 | 0.314 | 6.1E-19 |
| *C17F4.8* | WBGene00015914 | protein_coding_gene | -1.7 | 0.314 | 1.4E-03 |
| *spds-1* | WBGene00012909 | protein_coding_gene | -1.7 | 0.313 | 8.5E-09 |
| *pes-9* | WBGene00003982 | protein_coding_gene | -1.7 | 0.313 | 1.5E-08 |
| *rbm-12* | WBGene00013703 | protein_coding_gene | -1.7 | 0.312 | 5.7E-24 |
| *F55C9.3* | WBGene00010100 | protein_coding_gene | -1.7 | 0.312 | 4.1E-02 |
| *pid-5* | WBGene00021555 | protein_coding_gene | -1.7 | 0.312 | 9.4E-26 |
| *W02D9.2* | WBGene00012208 | protein_coding_gene | -1.7 | 0.312 | 2.1E-24 |
| *msp-81* | WBGene00003467 | protein_coding_gene | -1.7 | 0.311 | 1.0E-03 |
| *lsl-1* | WBGene00009937 | protein_coding_gene | -1.7 | 0.311 | 2.2E-25 |
| *imb-1* | WBGene00002075 | protein_coding_gene | -1.7 | 0.311 | 3.8E-10 |
| *gale-1* | WBGene00008132 | protein_coding_gene | -1.7 | 0.311 | 6.2E-06 |
| *Y47H9B.2* | WBGene00012946 | protein_coding_gene | -1.7 | 0.311 | 6.1E-04 |
| *gex-3* | WBGene00001580 | protein_coding_gene | -1.7 | 0.310 | 4.2E-19 |
| *cyp-13A2* | WBGene00011676 | protein_coding_gene | -1.7 | 0.310 | 1.9E-05 |
| *pgph-3* | WBGene00016892 | protein_coding_gene | -1.7 | 0.310 | 4.3E-03 |
| *Y73B3A.1* | WBGene00022203 | protein_coding_gene | -1.7 | 0.309 | 6.9E-12 |
| *Y49E10.37* | WBGene00304219 | ncRNA_gene | -1.7 | 0.309 | 2.9E-08 |
| *top-2* | WBGene00010785 | protein_coding_gene | -1.7 | 0.309 | 8.7E-15 |
| *ZK380.t2* | WBGene00023401 | pseudogene | -1.7 | 0.309 | 6.5E-05 |
| *C52B11.5* | WBGene00016874 | protein_coding_gene | -1.7 | 0.308 | 2.6E-10 |
| *Y51F10.11* | WBGene00044438 | protein_coding_gene | -1.7 | 0.308 | 1.4E-14 |
| *F34D10.8* | WBGene00009377 | protein_coding_gene | -1.7 | 0.307 | 3.3E-04 |
| *C18D11.3* | WBGene00007681 | protein_coding_gene | -1.7 | 0.307 | 4.7E-35 |
| *ugt-65* | WBGene00008583 | protein_coding_gene | -1.7 | 0.306 | 2.3E-03 |
| *tre-2* | WBGene00006608 | protein_coding_gene | -1.7 | 0.305 | 3.4E-09 |
| *uba-1* | WBGene00006699 | protein_coding_gene | -1.7 | 0.305 | 1.2E-18 |
| *R06F6.8* | WBGene00011071 | protein_coding_gene | -1.7 | 0.305 | 2.4E-21 |
| *tfg-1* | WBGene00006565 | protein_coding_gene | -1.7 | 0.305 | 2.3E-15 |
| *C11E4.7* | WBGene00007521 | protein_coding_gene | -1.7 | 0.305 | 7.4E-06 |
| *linc-7* | WBGene00219609 | lincRNA_gene | -1.7 | 0.304 | 6.3E-12 |
| *algn-14* | WBGene00019725 | protein_coding_gene | -1.7 | 0.302 | 1.7E-21 |
| *Y75B8A.24* | WBGene00013557 | protein_coding_gene | -1.7 | 0.301 | 1.8E-14 |
| *sucl-2* | WBGene00017759 | protein_coding_gene | -1.7 | 0.301 | 9.5E-17 |
| *pam-1* | WBGene00003914 | protein_coding_gene | -1.7 | 0.301 | 1.9E-19 |
| *Y55F3AM.21* | WBGene00219399 | protein_coding_gene | -1.7 | 0.300 | 3.9E-04 |
| *K09H9.7* | WBGene00019598 | protein_coding_gene | -1.7 | 0.300 | 1.7E-11 |
| *klp-7* | WBGene00002219 | protein_coding_gene | -1.7 | 0.300 | 1.1E-17 |
| *F53G2.12* | WBGene00302998 | protein_coding_gene | -1.7 | 0.299 | 5.2E-06 |
| *F54D5.12* | WBGene00010055 | protein_coding_gene | -1.7 | 0.299 | 6.5E-15 |
| *C18A11.4* | WBGene00015950 | protein_coding_gene | -1.7 | 0.298 | 3.9E-03 |
| *emc-1* | WBGene00019209 | protein_coding_gene | -1.7 | 0.298 | 6.5E-26 |
| *T08A11.1* | WBGene00011604 | protein_coding_gene | -1.7 | 0.297 | 7.2E-11 |
| *ani-2* | WBGene00019608 | protein_coding_gene | -1.8 | 0.297 | 1.6E-12 |
| *col-160* | WBGene00000733 | protein_coding_gene | -1.8 | 0.296 | 1.5E-06 |
| *sss-1* | WBGene00006056 | protein_coding_gene | -1.8 | 0.295 | 6.0E-04 |
| *cri-3* | WBGene00010303 | protein_coding_gene | -1.8 | 0.294 | 1.4E-20 |
| *ldp-1* | WBGene00017719 | protein_coding_gene | -1.8 | 0.294 | 2.7E-11 |
| *H04D03.6* | WBGene00194861 | protein_coding_gene | -1.8 | 0.294 | 1.9E-04 |
| *F27D4.1* | WBGene00009187 | protein_coding_gene | -1.8 | 0.293 | 2.0E-10 |
| *Y7A5A.25* | WBGene00304824 | protein_coding_gene | -1.8 | 0.292 | 4.1E-02 |
| *ssq-4* | WBGene00006053 | protein_coding_gene | -1.8 | 0.292 | 6.0E-04 |
| *Y66A7A.9* | WBGene00194864 | protein_coding_gene | -1.8 | 0.291 | 2.0E-02 |
| *BE0003N10.1* | WBGene00015266 | protein_coding_gene | -1.8 | 0.291 | 3.6E-14 |
| *C52D10.3* | WBGene00016877 | protein_coding_gene | -1.8 | 0.290 | 5.7E-03 |
| *K08D12.6* | WBGene00019540 | protein_coding_gene | -1.8 | 0.289 | 1.9E-05 |
| *M02B1.4* | WBGene00010829 | protein_coding_gene | -1.8 | 0.288 | 1.9E-04 |
| *mnat-1* | WBGene00018769 | protein_coding_gene | -1.8 | 0.287 | 5.1E-18 |
| *F55G7.1* | WBGene00010119 | protein_coding_gene | -1.8 | 0.285 | 4.0E-03 |
| *lbp-6* | WBGene00002258 | protein_coding_gene | -1.8 | 0.284 | 3.3E-06 |
| *Y41E3.18* | WBGene00044003 | protein_coding_gene | -1.8 | 0.283 | 1.2E-02 |
| *C27C7.1* | WBGene00007766 | protein_coding_gene | -1.8 | 0.283 | 3.2E-24 |
| *Y51B9A.5* | WBGene00013087 | protein_coding_gene | -1.8 | 0.283 | 2.2E-03 |
| *ufm-1* | WBGene00044324 | protein_coding_gene | -1.8 | 0.282 | 2.2E-19 |
| *col-137* | WBGene00000710 | protein_coding_gene | -1.8 | 0.281 | 2.9E-02 |
| *srh-217* | WBGene00005427 | protein_coding_gene | -1.8 | 0.281 | 4.9E-02 |
| *F13H10.3* | WBGene00008774 | protein_coding_gene | -1.8 | 0.280 | 1.5E-19 |
| *gst-4* | WBGene00001752 | protein_coding_gene | -1.8 | 0.279 | 4.5E-04 |
| *set-21* | WBGene00021282 | protein_coding_gene | -1.8 | 0.278 | 2.4E-18 |
| *trap-1* | WBGene00022122 | protein_coding_gene | -1.8 | 0.278 | 2.0E-17 |
| *F56D6.19* | WBGene00235293 | protein_coding_gene | -1.9 | 0.277 | 1.2E-02 |
| *ahcy-1* | WBGene00019322 | protein_coding_gene | -1.9 | 0.277 | 2.9E-12 |

| | | | | | |
|---|---|---|---|---|---|
| *fbxb-112* | WBGene00021716 | protein_coding_gene | -1.9 | 0.276 | 3.4E-02 |
| *atad-3* | WBGene00010015 | protein_coding_gene | -1.9 | 0.275 | 2.1E-37 |
| *Y57A10A.31* | WBGene00013270 | protein_coding_gene | -1.9 | 0.275 | 3.0E-17 |
| *chil-25* | WBGene00011846 | protein_coding_gene | -1.9 | 0.275 | 2.9E-04 |
| *F08H9.2* | WBGene00008590 | protein_coding_gene | -1.9 | 0.273 | 1.0E-03 |
| *clu-1* | WBGene00000550 | protein_coding_gene | -1.9 | 0.273 | 1.2E-18 |
| *K07F5.17* | WBGene00045180 | protein_coding_gene | -1.9 | 0.272 | 3.7E-02 |
| *linc-63* | WBGene00219725 | lincRNA_gene | -1.9 | 0.272 | 1.6E-02 |
| *lbp-8* | WBGene00002260 | protein_coding_gene | -1.9 | 0.272 | 3.2E-02 |
| *inf-1* | WBGene00002083 | protein_coding_gene | -1.9 | 0.272 | 1.7E-21 |
| *Y82E9BL.3* | WBGene00022320 | protein_coding_gene | -1.9 | 0.270 | 4.4E-02 |
| *F36D3.4* | WBGene00009470 | protein_coding_gene | -1.9 | 0.270 | 1.7E-04 |
| *W02H3.1* | WBGene00012217 | protein_coding_gene | -1.9 | 0.270 | 1.5E-06 |
| *ife-5* | WBGene00002063 | protein_coding_gene | -1.9 | 0.270 | 5.1E-29 |
| *ech-7* | WBGene00001156 | protein_coding_gene | -1.9 | 0.270 | 3.0E-06 |
| *R03D7.2* | WBGene00010989 | protein_coding_gene | -1.9 | 0.269 | 2.2E-16 |
| *elo-6* | WBGene00001244 | protein_coding_gene | -1.9 | 0.269 | 2.8E-03 |
| *R07E3.4* | WBGene00011105 | protein_coding_gene | -1.9 | 0.267 | 3.2E-06 |
| *msp-33* | WBGene00003431 | protein_coding_gene | -1.9 | 0.267 | 1.3E-04 |
| *F54E2.1* | WBGene00018823 | protein_coding_gene | -1.9 | 0.267 | 2.4E-07 |
| *msp-77* | WBGene00003464 | protein_coding_gene | -1.9 | 0.266 | 1.6E-04 |
| *hsp-4* | WBGene00002008 | protein_coding_gene | -1.9 | 0.266 | 2.9E-11 |
| *snpc-3.3* | WBGene00021662 | protein_coding_gene | -1.9 | 0.265 | 1.4E-06 |
| *clec-57* | WBGene00008594 | protein_coding_gene | -1.9 | 0.264 | 1.7E-06 |
| *cpl-1* | WBGene00000776 | protein_coding_gene | -1.9 | 0.264 | 8.9E-13 |
| *R03D7.3* | WBGene00014814 | pseudogene | -1.9 | 0.263 | 3.1E-03 |
| *K11H3.4* | WBGene00010781 | protein_coding_gene | -1.9 | 0.263 | 8.5E-19 |
| *T12B5.15* | WBGene00050914 | protein_coding_gene | -1.9 | 0.262 | 1.3E-04 |
| *pks-1* | WBGene00016558 | protein_coding_gene | -1.9 | 0.260 | 3.3E-06 |
| *str-146* | WBGene00006193 | protein_coding_gene | -1.9 | 0.259 | 1.4E-02 |
| *str-227* | WBGene00006258 | protein_coding_gene | -1.9 | 0.259 | 3.6E-02 |
| *hmit-1.2* | WBGene00013074 | protein_coding_gene | -1.9 | 0.259 | 1.5E-08 |
| *acs-1* | WBGene00018488 | protein_coding_gene | -2.0 | 0.258 | 5.7E-06 |
| *mltn-2* | WBGene00013126 | protein_coding_gene | -2.0 | 0.258 | 1.9E-02 |
| *ile-1* | WBGene00002070 | protein_coding_gene | -2.0 | 0.258 | 3.1E-19 |
| *arrd-9* | WBGene00008843 | protein_coding_gene | -2.0 | 0.257 | 6.1E-03 |
| *Y102A5C.6* | WBGene00014955 | pseudogene | -2.0 | 0.254 | 4.6E-19 |
| *ugt-22* | WBGene00007455 | protein_coding_gene | -2.0 | 0.254 | 1.3E-07 |
| *nhr-172* | WBGene00016927 | protein_coding_gene | -2.0 | 0.254 | 2.2E-02 |
| *mxl-3* | WBGene00003511 | protein_coding_gene | -2.0 | 0.253 | 2.5E-04 |
| *ugt-26* | WBGene00015695 | protein_coding_gene | -2.0 | 0.250 | 6.2E-11 |
| *manf-1* | WBGene00021888 | protein_coding_gene | -2.0 | 0.249 | 2.0E-17 |
| *oac-32* | WBGene00018708 | protein_coding_gene | -2.0 | 0.249 | 1.4E-02 |
| *fbxa-206* | WBGene00013610 | protein_coding_gene | -2.0 | 0.247 | 3.9E-28 |
| *Y46B2A.4* | WBGene00284850 | protein_coding_gene | -2.0 | 0.247 | 3.9E-03 |
| *har-1* | WBGene00007630 | protein_coding_gene | -2.0 | 0.247 | 3.1E-30 |
| *his-38* | WBGene00001912 | protein_coding_gene | -2.0 | 0.246 | 2.1E-03 |
| *Y23H5B.7* | WBGene00021278 | pseudogene | -2.0 | 0.245 | 2.0E-02 |
| *Y50D4B.2* | WBGene00021741 | protein_coding_gene | -2.0 | 0.243 | 1.9E-07 |
| *wago-2* | WBGene00018862 | pseudogene | -2.0 | 0.243 | 1.3E-31 |
| *glh-4* | WBGene00001601 | protein_coding_gene | -2.0 | 0.243 | 2.3E-33 |
| *F40H3.2* | WBGene00018250 | protein_coding_gene | -2.0 | 0.243 | 1.1E-03 |
| *Y102A5C.5* | WBGene00014954 | pseudogene | -2.1 | 0.241 | 1.1E-18 |
| *C45E5.1* | WBGene00016664 | protein_coding_gene | -2.1 | 0.241 | 8.3E-05 |
| *K07D4.1* | WBGene00023318 | pseudogene | -2.1 | 0.240 | 8.2E-10 |
| *lon-1* | WBGene00003055 | protein_coding_gene | -2.1 | 0.240 | 1.9E-07 |
| *ZK228.3* | WBGene00013946 | protein_coding_gene | -2.1 | 0.239 | 1.3E-09 |
| *elo-2* | WBGene00001240 | protein_coding_gene | -2.1 | 0.238 | 5.5E-06 |
| *F20B6.7* | WBGene00017631 | pseudogene | -2.1 | 0.232 | 3.5E-03 |
| *F55B11.6* | WBGene00044249 | protein_coding_gene | -2.1 | 0.232 | 1.9E-14 |
| *cept-1* | WBGene00009057 | protein_coding_gene | -2.1 | 0.231 | 7.3E-08 |
| *AC8.1* | WBGene00004054 | pseudogene | -2.1 | 0.230 | 4.5E-03 |
| *T20F10.8* | WBGene00077765 | pseudogene | -2.1 | 0.230 | 2.0E-02 |
| *hsd-2* | WBGene00022498 | protein_coding_gene | -2.1 | 0.229 | 9.5E-04 |
| *daf-36* | WBGene00007536 | protein_coding_gene | -2.1 | 0.229 | 1.1E-06 |
| *F02E11.4* | WBGene00017182 | protein_coding_gene | -2.1 | 0.229 | 4.9E-02 |
| *F22E5.1* | WBGene00017703 | protein_coding_gene | -2.1 | 0.228 | 1.3E-03 |
| *srj-9* | WBGene00005597 | protein_coding_gene | -2.1 | 0.227 | 3.0E-02 |
| *T16H12.2* | WBGene00011812 | protein_coding_gene | -2.1 | 0.227 | 1.9E-10 |
| *T17A3.12* | WBGene00020548 | protein_coding_gene | -2.1 | 0.226 | 4.2E-02 |
| *ldh-1* | WBGene00002262 | protein_coding_gene | -2.1 | 0.226 | 9.4E-15 |
| *T25B9.9* | WBGene00012015 | protein_coding_gene | -2.2 | 0.225 | 8.3E-18 |
| *Y105C5B.15* | WBGene00013654 | protein_coding_gene | -2.2 | 0.224 | 2.7E-03 |
| *T28B8.4* | WBGene00012115 | protein_coding_gene | -2.2 | 0.224 | 7.1E-05 |
| *C08F1.12* | WBGene00269389 | protein_coding_gene | -2.2 | 0.224 | 3.6E-02 |
| *F59H6.2* | WBGene00019133 | pseudogene | -2.2 | 0.224 | 7.8E-05 |
| *osta-3* | WBGene00012182 | protein_coding_gene | -2.2 | 0.223 | 1.9E-19 |
| *fbxc-12* | WBGene00045298 | protein_coding_gene | -2.2 | 0.223 | 4.2E-02 |
| *Y17D7B.7* | WBGene00014562 | pseudogene | -2.2 | 0.222 | 5.6E-05 |
| *msp-53* | WBGene00003446 | protein_coding_gene | -2.2 | 0.221 | 6.6E-04 |

| | | | | | |
|---|---|---|---|---|---|
| *wago-4* | WBGene00010263 | protein_coding_gene | -2.2 | 0.221 | 4.0E-18 |
| *F44E5.16* | WBGene00219949 | ncRNA_gene | -2.2 | 0.221 | 3.5E-02 |
| *C35A5.3* | WBGene00007947 | protein_coding_gene | -2.2 | 0.221 | 2.3E-12 |
| *srsx-18* | WBGene00015419 | protein_coding_gene | -2.2 | 0.220 | 8.1E-03 |
| *C36C9.10* | WBGene00219849 | ncRNA_gene | -2.2 | 0.220 | 1.7E-06 |
| *21ur-14269* | WBGene00169029 | piRNA_gene | -2.2 | 0.219 | 3.3E-03 |
| *C32H11.7* | WBGene00007870 | pseudogene | -2.2 | 0.219 | 1.3E-03 |
| *hsp-110* | WBGene00016250 | protein_coding_gene | -2.2 | 0.219 | 8.0E-22 |
| *nspa-8* | WBGene00012294 | protein_coding_gene | -2.2 | 0.218 | 1.5E-04 |
| *nep-10* | WBGene00017555 | protein_coding_gene | -2.2 | 0.217 | 2.8E-04 |
| *ZK6.6* | WBGene00022641 | protein_coding_gene | -2.2 | 0.215 | 3.0E-05 |
| *F58D2.5* | WBGene00206469 | protein_coding_gene | -2.2 | 0.214 | 4.2E-02 |
| *F53G2.1* | WBGene00018766 | protein_coding_gene | -2.2 | 0.213 | 5.2E-05 |
| *oac-54* | WBGene00012323 | protein_coding_gene | -2.2 | 0.210 | 4.6E-06 |
| *ZK662.8* | WBGene00200998 | ncRNA_gene | -2.3 | 0.209 | 1.7E-02 |
| *srpr-2.2* | WBGene00195017 | ncRNA_gene | -2.3 | 0.208 | 2.3E-02 |
| *clec-139* | WBGene00018050 | protein_coding_gene | -2.3 | 0.207 | 1.0E-03 |
| *rmd-6* | WBGene00020070 | protein_coding_gene | -2.3 | 0.205 | 6.0E-03 |
| *tag-10* | WBGene00006404 | protein_coding_gene | -2.3 | 0.205 | 2.7E-14 |
| *F02E9.8* | WBGene00014731 | pseudogene | -2.3 | 0.205 | 2.4E-02 |
| *sdz-35* | WBGene00022570 | protein_coding_gene | -2.3 | 0.202 | 1.1E-02 |
| *linc-107* | WBGene00219655 | lincRNA_gene | -2.3 | 0.202 | 4.0E-03 |
| *C09B8.4* | WBGene00015623 | protein_coding_gene | -2.3 | 0.201 | 1.0E-14 |
| *sams-1* | WBGene00008205 | protein_coding_gene | -2.3 | 0.201 | 3.4E-09 |
| *C40A11.7* | WBGene00016550 | protein_coding_gene | -2.3 | 0.201 | 5.2E-03 |
| *T26C11.8* | WBGene00044563 | protein_coding_gene | -2.3 | 0.200 | 3.5E-02 |
| *H11L12.1* | WBGene00019189 | protein_coding_gene | -2.3 | 0.198 | 2.5E-10 |
| *nhr-249* | WBGene00022726 | protein_coding_gene | -2.4 | 0.195 | 2.9E-02 |
| *Y40C7B.4* | WBGene00021500 | protein_coding_gene | -2.4 | 0.195 | 3.2E-02 |
| *rrn-3.1* | WBGene00004622 | rRNA_gene | -2.4 | 0.195 | 9.2E-03 |
| *F11A10.9* | WBGene00045153 | snoRNA_gene | -2.4 | 0.194 | 3.0E-02 |
| *Y39A1A.17* | WBGene00012655 | protein_coding_gene | -2.4 | 0.193 | 2.1E-02 |
| *pgl-3* | WBGene00003994 | protein_coding_gene | -2.4 | 0.193 | 4.0E-31 |
| *sdz-11* | WBGene00017408 | protein_coding_gene | -2.4 | 0.192 | 4.7E-04 |
| *ttr-13* | WBGene00012849 | protein_coding_gene | -2.4 | 0.190 | 2.0E-02 |
| *AC8.5* | WBGene00007077 | pseudogene | -2.4 | 0.190 | 4.5E-02 |
| *sec-61.A* | WBGene00013311 | protein_coding_gene | -2.4 | 0.188 | 4.6E-22 |
| *F08D12.4* | WBGene00017248 | protein_coding_gene | -2.4 | 0.188 | 3.2E-02 |
| *clec-22* | WBGene00009855 | protein_coding_gene | -2.4 | 0.187 | 3.3E-02 |
| *bath-19* | WBGene00019132 | protein_coding_gene | -2.4 | 0.187 | 1.5E-18 |
| *clec-206* | WBGene00019089 | protein_coding_gene | -2.4 | 0.186 | 6.7E-03 |
| *Y57G11C.1143* | WBGene00219342 | pseudogene | -2.4 | 0.183 | 1.9E-02 |
| *clp-10* | WBGene00021041 | protein_coding_gene | -2.4 | 0.183 | 3.7E-02 |
| *ZC266.1* | WBGene00022584 | protein_coding_gene | -2.5 | 0.183 | 6.2E-14 |
| *C08F1.13* | WBGene00269390 | pseudogene | -2.5 | 0.183 | 3.7E-02 |
| *F55C9.11* | WBGene00010107 | protein_coding_gene | -2.5 | 0.182 | 3.2E-02 |
| *T04B8.6* | WBGene00268209 | pseudogene | -2.5 | 0.180 | 2.5E-02 |
| *D2023.8* | WBGene00050970 | pseudogene | -2.5 | 0.179 | 1.1E-02 |
| *rnh-1.3* | WBGene00007303 | protein_coding_gene | -2.5 | 0.178 | 2.9E-11 |
| *smd-1* | WBGene00004875 | protein_coding_gene | -2.5 | 0.176 | 1.0E-10 |
| *F54E7.11* | WBGene00044942 | snoRNA_gene | -2.5 | 0.176 | 2.3E-02 |
| *F22H10.6* | WBGene00017730 | protein_coding_gene | -2.5 | 0.171 | 7.0E-07 |
| *acox-1.4* | WBGene00008567 | protein_coding_gene | -2.5 | 0.171 | 2.2E-07 |
| *W03D2.11* | WBGene00219554 | pseudogene | -2.6 | 0.168 | 1.6E-02 |
| *ZK637.6* | WBGene00014987 | pseudogene | -2.6 | 0.168 | 7.1E-41 |
| *21ur-8696* | WBGene00173920 | piRNA_gene | -2.6 | 0.168 | 4.1E-02 |
| *pho-13* | WBGene00008560 | protein_coding_gene | -2.6 | 0.167 | 9.4E-07 |
| *F54D8.10* | WBGene00284847 | protein_coding_gene | -2.6 | 0.167 | 1.5E-03 |
| *T28D6.3* | WBGene00012123 | protein_coding_gene | -2.6 | 0.166 | 2.7E-09 |
| *tbx-35* | WBGene00006554 | protein_coding_gene | -2.6 | 0.165 | 1.3E-02 |
| *F35E2.7* | WBGene00009414 | pseudogene | -2.6 | 0.165 | 2.8E-02 |
| *R09A8.9* | WBGene00219278 | protein_coding_gene | -2.6 | 0.163 | 1.6E-03 |
| *str-55* | WBGene00006120 | protein_coding_gene | -2.6 | 0.162 | 3.2E-02 |
| *F53C3.8* | WBGene00018752 | protein_coding_gene | -2.6 | 0.162 | 1.3E-02 |
| *lips-14* | WBGene00019208 | pseudogene | -2.6 | 0.160 | 3.9E-03 |
| *Y67H2A.9* | WBGene00014941 | pseudogene | -2.6 | 0.160 | 2.8E-13 |
| *srh-77* | WBGene00005298 | pseudogene | -2.6 | 0.160 | 3.3E-02 |
| *Y50D4B.4* | WBGene00021743 | protein_coding_gene | -2.7 | 0.159 | 7.2E-12 |
| *Y49E10.5* | WBGene00014902 | pseudogene | -2.7 | 0.159 | 1.4E-02 |
| *fbxb-68* | WBGene00008611 | protein_coding_gene | -2.7 | 0.159 | 5.9E-05 |
| *sec-61.B* | WBGene00021427 | protein_coding_gene | -2.7 | 0.155 | 5.9E-17 |
| *dhs-26* | WBGene00000989 | protein_coding_gene | -2.7 | 0.153 | 2.3E-09 |
| *sdz-6* | WBGene00008068 | protein_coding_gene | -2.7 | 0.153 | 1.8E-02 |
| *ets-8* | WBGene00016798 | protein_coding_gene | -2.7 | 0.153 | 1.6E-02 |
| *Y24D9A.9* | WBGene00044952 | snoRNA_gene | -2.7 | 0.153 | 1.7E-02 |
| *H39E20.1* | WBGene00019267 | protein_coding_gene | -2.7 | 0.151 | 4.1E-03 |
| *ugt-62* | WBGene00010904 | protein_coding_gene | -2.7 | 0.151 | 3.5E-14 |
| *F25H2.14* | WBGene00045138 | snoRNA_gene | -2.7 | 0.151 | 7.9E-05 |
| *K02F2.11* | WBGene00220000 | ncRNA_gene | -2.8 | 0.148 | 1.6E-02 |
| *hpo-41* | WBGene00008048 | pseudogene | -2.8 | 0.146 | 6.6E-03 |

| | | | | | |
|---|---|---|---|---|---|
| *plep-1* | WBGene00021797 | protein_coding_gene | -2.8 | 0.146 | 2.6E-06 |
| *clec-245* | WBGene00007851 | protein_coding_gene | -2.8 | 0.144 | 3.8E-04 |
| *F36H5.15* | WBGene00270291 | pseudogene | -2.8 | 0.144 | 1.9E-02 |
| *Y48B6A.10* | WBGene00012981 | protein_coding_gene | -2.8 | 0.143 | 3.5E-07 |
| *T20D4.19* | WBGene00020624 | protein_coding_gene | -2.8 | 0.143 | 5.6E-03 |
| *zeel-1* | WBGene00021463 | protein_coding_gene | -2.8 | 0.143 | 9.9E-08 |
| *str-148* | WBGene00006195 | protein_coding_gene | -2.8 | 0.143 | 7.4E-03 |
| *Y50E8A.11* | WBGene00013054 | protein_coding_gene | -2.8 | 0.140 | 8.8E-03 |
| *hsp-60* | WBGene00002025 | protein_coding_gene | -2.8 | 0.139 | 1.4E-30 |
| *K05F6.14* | WBGene00219239 | protein_coding_gene | -2.9 | 0.137 | 1.1E-02 |
| *C27H5.12* | WBGene00271810 | protein_coding_gene | -2.9 | 0.136 | 4.4E-03 |
| *nas-27* | WBGene00003545 | protein_coding_gene | -2.9 | 0.136 | 1.8E-03 |
| *nhr-114* | WBGene00003704 | protein_coding_gene | -2.9 | 0.136 | 4.6E-12 |
| *acp-6* | WBGene00022245 | protein_coding_gene | -2.9 | 0.134 | 2.8E-10 |
| *srx-98* | WBGene00005989 | protein_coding_gene | -2.9 | 0.133 | 4.0E-03 |
| *F07C6.6* | WBGene00045067 | protein_coding_gene | -2.9 | 0.133 | 8.0E-03 |
| *F49F1.7* | WBGene00018647 | protein_coding_gene | -2.9 | 0.131 | 1.3E-06 |
| *C24H12.3* | WBGene00023244 | pseudogene | -2.9 | 0.131 | 2.5E-03 |
| *clec-238* | WBGene00013622 | protein_coding_gene | -2.9 | 0.131 | 7.2E-03 |
| *Y49E10.25* | WBGene00013044 | protein_coding_gene | -2.9 | 0.131 | 1.6E-02 |
| *sri-57* | WBGene00005569 | protein_coding_gene | -2.9 | 0.130 | 4.6E-03 |
| *cest-7* | WBGene00007693 | protein_coding_gene | -2.9 | 0.129 | 4.0E-05 |
| *fbxb-98* | WBGene00015830 | protein_coding_gene | -3.0 | 0.129 | 7.2E-05 |
| *irld-5* | WBGene00017454 | protein_coding_gene | -3.0 | 0.128 | 5.9E-03 |
| *F53B6.5* | WBGene00009960 | protein_coding_gene | -3.0 | 0.128 | 1.2E-02 |
| *Y73B6BL.14* | WBGene00022237 | protein_coding_gene | -3.0 | 0.128 | 3.0E-07 |
| *F58G6.7* | WBGene00010277 | protein_coding_gene | -3.0 | 0.127 | 9.2E-08 |
| *Y46G5A.8* | WBGene00012900 | protein_coding_gene | -3.0 | 0.126 | 3.3E-12 |
| *C29G2.2* | WBGene00016230 | pseudogene | -3.0 | 0.124 | 1.2E-18 |
| *C16C4.17* | WBGene00044359 | protein_coding_gene | -3.0 | 0.123 | 3.4E-03 |
| *K10G4.10* | WBGene00044202 | protein_coding_gene | -3.0 | 0.123 | 2.9E-03 |
| *bath-23* | WBGene00021727 | protein_coding_gene | -3.0 | 0.122 | 1.4E-03 |
| *W10G11.3* | WBGene00021136 | protein_coding_gene | -3.0 | 0.121 | 2.3E-10 |
| *fbxb-56* | WBGene00013409 | protein_coding_gene | -3.0 | 0.121 | 1.7E-03 |
| *sri-8* | WBGene00005520 | pseudogene | -3.0 | 0.121 | 5.5E-03 |
| *Y73F8A.22* | WBGene00013527 | protein_coding_gene | -3.0 | 0.121 | 7.5E-03 |
| *fbxc-27* | WBGene00019926 | protein_coding_gene | -3.1 | 0.117 | 3.0E-03 |
| *T26H2.10* | WBGene00044976 | pseudogene | -3.1 | 0.117 | 2.2E-04 |
| *fbxb-115* | WBGene00012064 | protein_coding_gene | -3.1 | 0.116 | 1.1E-03 |
| *btb-10* | WBGene00019313 | protein_coding_gene | -3.1 | 0.115 | 1.1E-09 |
| *Y51A2A.16* | WBGene00206507 | protein_coding_gene | -3.1 | 0.114 | 3.6E-03 |
| *hlh-26* | WBGene00001965 | protein_coding_gene | -3.2 | 0.112 | 5.5E-03 |
| *grl-27* | WBGene00001736 | protein_coding_gene | -3.2 | 0.112 | 1.3E-02 |
| *clec-118* | WBGene00021032 | protein_coding_gene | -3.2 | 0.112 | 5.2E-04 |
| *grd-4* | WBGene00001693 | protein_coding_gene | -3.2 | 0.107 | 6.3E-05 |
| *T10C6.9* | WBGene00011690 | pseudogene | -3.3 | 0.104 | 3.0E-03 |
| *Y110A2AL.1* | WBGene00022438 | protein_coding_gene | -3.3 | 0.103 | 1.9E-03 |
| *fbxb-21* | WBGene00022558 | protein_coding_gene | -3.3 | 0.102 | 5.5E-04 |
| *T26E3.11* | WBGene00044905 | pseudogene | -3.4 | 0.098 | 5.1E-04 |
| *fbxb-27* | WBGene00044695 | pseudogene | -3.4 | 0.097 | 7.2E-04 |
| *T02G6.11* | WBGene00086562 | protein_coding_gene | -3.4 | 0.094 | 7.1E-04 |
| *F19G12.t1* | WBGene00023271 | pseudogene | -3.4 | 0.093 | 2.3E-03 |
| *F46F2.3* | WBGene00009787 | protein_coding_gene | -3.5 | 0.091 | 1.9E-04 |
| *math-29* | WBGene00018441 | protein_coding_gene | -3.5 | 0.091 | 5.2E-04 |
| *C33H5.1* | WBGene00016370 | protein_coding_gene | -3.5 | 0.091 | 5.4E-03 |
| *cest-28* | WBGene00019654 | protein_coding_gene | -3.5 | 0.090 | 1.5E-06 |
| *T24D5.11* | WBGene00219279 | pseudogene | -3.5 | 0.090 | 2.2E-04 |
| *fbxa-103* | WBGene00008612 | protein_coding_gene | -3.5 | 0.089 | 2.8E-04 |
| *F59C6.10* | WBGene00014789 | pseudogene | -3.5 | 0.087 | 3.7E-05 |
| *T24D5.6* | WBGene00044149 | protein_coding_gene | -3.5 | 0.086 | 2.5E-04 |
| *Y48E1B.8* | WBGene00013007 | protein_coding_gene | -3.6 | 0.085 | 1.5E-03 |
| *C40H5.8* | WBGene00044585 | pseudogene | -3.6 | 0.084 | 1.8E-04 |
| *F20D1.22* | WBGene00304908 | protein_coding_gene | -3.6 | 0.084 | 2.3E-12 |
| *dhs-25* | WBGene00000988 | protein_coding_gene | -3.6 | 0.082 | 2.4E-23 |
| *F18E3.13* | WBGene00045276 | protein_coding_gene | -3.6 | 0.082 | 2.5E-07 |
| *fbxb-97* | WBGene00016885 | protein_coding_gene | -3.6 | 0.082 | 9.3E-05 |
| *C04C3.14* | WBGene00219410 | protein_coding_gene | -3.6 | 0.080 | 7.3E-03 |
| *acdh-2* | WBGene00015894 | protein_coding_gene | -3.7 | 0.078 | 6.0E-16 |
| *nspc-5* | WBGene00044340 | protein_coding_gene | -3.7 | 0.078 | 7.6E-04 |
| *Y73B6BL.288* | WBGene00219218 | protein_coding_gene | -3.7 | 0.077 | 8.7E-22 |
| *C53B7.6* | WBGene00016895 | protein_coding_gene | -3.8 | 0.071 | 1.1E-03 |
| *F18E3.11* | WBGene00017567 | protein_coding_gene | -3.8 | 0.071 | 7.1E-06 |
| *ZK177.9* | WBGene00022674 | protein_coding_gene | -3.9 | 0.069 | 1.6E-03 |
| *fbxb-20* | WBGene00022559 | protein_coding_gene | -3.9 | 0.066 | 2.2E-07 |
| *F58G6.3* | WBGene00010274 | protein_coding_gene | -4.0 | 0.064 | 7.8E-09 |
| *F36H5.8* | WBGene00018106 | protein_coding_gene | -4.0 | 0.064 | 7.1E-05 |
| *C07G1.7* | WBGene00015573 | protein_coding_gene | -4.0 | 0.063 | 2.1E-08 |
| *R03G8.4* | WBGene00010999 | pseudogene | -4.0 | 0.063 | 1.0E-06 |
| *arf-1.1* | WBGene00000190 | protein_coding_gene | -4.0 | 0.062 | 2.4E-14 |
| *fbxb-12* | WBGene00018109 | pseudogene | -4.0 | 0.061 | 1.4E-05 |

| | | | | | |
|---|---|---|---|---|---|
| *fbxb-51* | WBGene00019414 | protein_coding_gene | -4.0 | 0.061 | 1.4E-05 |
| *C30G12.2* | WBGene00016274 | protein_coding_gene | -4.1 | 0.059 | 5.4E-25 |
| *C40A11.6* | WBGene00016549 | protein_coding_gene | -4.1 | 0.057 | 1.8E-05 |
| *F36H5.17* | WBGene00270293 | protein_coding_gene | -4.1 | 0.057 | 1.1E-05 |
| *K05F6.12* | WBGene00045486 | protein_coding_gene | -4.1 | 0.057 | 7.5E-06 |
| *fbxb-116* | WBGene00012066 | protein_coding_gene | -4.2 | 0.056 | 2.3E-05 |
| *B0281.4* | WBGene00015111 | protein_coding_gene | -4.2 | 0.054 | 1.3E-05 |
| *scav-5* | WBGene00011076 | protein_coding_gene | -4.2 | 0.053 | 3.7E-19 |
| *folt-2* | WBGene00018138 | protein_coding_gene | -4.2 | 0.053 | 3.9E-28 |
| *C40A11.2* | WBGene00016545 | protein_coding_gene | -4.3 | 0.050 | 1.0E-06 |
| *fbxb-52* | WBGene00019417 | protein_coding_gene | -4.3 | 0.050 | 3.5E-06 |
| *fbxb-44* | WBGene00019416 | protein_coding_gene | -4.3 | 0.050 | 1.5E-06 |
| *C40A11.4* | WBGene00016547 | protein_coding_gene | -4.4 | 0.048 | 9.1E-07 |
| *Y32B12C.3* | WBGene00012526 | pseudogene | -4.6 | 0.042 | 4.6E-05 |
| *fbxb-50* | WBGene00019413 | protein_coding_gene | -4.6 | 0.042 | 5.9E-07 |
| *fbxb-95* | WBGene00016880 | protein_coding_gene | -4.6 | 0.041 | 7.6E-07 |
| *fbxb-15* | WBGene00022557 | protein_coding_gene | -4.7 | 0.039 | 6.7E-09 |
| *fbxb-96* | WBGene00016886 | protein_coding_gene | -4.7 | 0.038 | 4.0E-07 |
| *K05F6.4* | WBGene00019415 | protein_coding_gene | -4.7 | 0.038 | 8.5E-12 |
| *F39F10.3* | WBGene00018203 | protein_coding_gene | -4.8 | 0.036 | 1.5E-14 |
| *sdz-18* | WBGene00018442 | protein_coding_gene | -4.9 | 0.034 | 1.7E-07 |
| *fbxb-53* | WBGene00018104 | protein_coding_gene | -5.0 | 0.032 | 5.7E-08 |
| *C05D9.12* | WBGene00219800 | ncRNA_gene | -5.0 | 0.032 | 3.4E-07 |
| *ZC239.21* | WBGene00044372 | protein_coding_gene | -5.0 | 0.032 | 1.6E-07 |
| *fbxb-43* | WBGene00021774 | protein_coding_gene | -5.0 | 0.031 | 4.7E-08 |
| *fbxb-49* | WBGene00019412 | protein_coding_gene | -5.1 | 0.028 | 5.5E-08 |
| *F10E7.1* | WBGene00017343 | protein_coding_gene | -5.1 | 0.028 | 2.1E-08 |
| *fbxb-42* | WBGene00021773 | protein_coding_gene | -5.1 | 0.028 | 2.1E-08 |
| *T22B7.7* | WBGene00020674 | protein_coding_gene | -5.1 | 0.028 | 6.6E-26 |
| *fbxb-13* | WBGene00015607 | protein_coding_gene | -5.1 | 0.028 | 1.5E-08 |
| *T26E3.8* | WBGene00012041 | protein_coding_gene | -5.2 | 0.028 | 2.5E-08 |
| *K05F6.15* | WBGene00269419 | pseudogene | -5.2 | 0.027 | 9.2E-09 |
| *ZC239.20* | WBGene00044371 | protein_coding_gene | -5.3 | 0.025 | 1.4E-08 |
| *fbxb-46* | WBGene00019420 | protein_coding_gene | -5.3 | 0.025 | 1.6E-08 |
| *fbxb-54* | WBGene00019418 | protein_coding_gene | -5.4 | 0.023 | 3.4E-09 |
| *F18E3.12* | WBGene00045275 | protein_coding_gene | -5.7 | 0.020 | 8.1E-11 |
| *fbxb-91* | WBGene00044692 | protein_coding_gene | -5.8 | 0.017 | 1.3E-10 |
| *fbxb-37* | WBGene00020537 | protein_coding_gene | -6.1 | 0.015 | 3.8E-11 |
| *sri-40* | WBGene00005552 | protein_coding_gene | -6.1 | 0.015 | 9.6E-87 |
| *F54D10.7* | WBGene00018808 | protein_coding_gene | -7.2 | 0.007 | 4.1E-21 |

**CB4856**-exclusive DEGs in response to *S. aureus* (vs. OP50)

| Gene Name | Wormbase Gene ID | Type | $log_2$ FC | FC | Adj P-value |
|---|---|---|---|---|---|
| *alh-2* | WBGene00000108 | protein_coding_gene | 6.5 | 91.750 | 3.7E-09 |
| *C29F9.15* | WBGene00219840 | ncRNA_gene | 5.5 | 46.501 | 2.6E-06 |
| *C06B3.1* | WBGene00007360 | protein_coding_gene | 5.0 | 33.028 | 4.6E-05 |
| *T28A11.4* | WBGene00020871 | protein_coding_gene | 5.0 | 31.345 | 2.4E-06 |
| *stdh-3* | WBGene00007364 | protein_coding_gene | 4.9 | 29.632 | 3.6E-05 |
| *K06H6.2* | WBGene00019450 | protein_coding_gene | 4.3 | 19.332 | 2.9E-04 |
| *K02E11.10* | WBGene00044109 | protein_coding_gene | 4.2 | 18.440 | 1.1E-04 |
| *T02B11.10* | WBGene00235327 | protein_coding_gene | 4.2 | 17.992 | 1.8E-03 |
| *F28A10.9* | WBGene00017877 | pseudogene | 4.1 | 17.715 | 6.1E-05 |
| *nhr-155* | WBGene00015758 | protein_coding_gene | 4.1 | 17.199 | 2.3E-04 |
| *C14C6.3* | WBGene00015757 | protein_coding_gene | 4.1 | 17.066 | 9.6E-04 |
| *srh-60* | WBGene00005282 | protein_coding_gene | 4.0 | 15.647 | 2.3E-04 |
| *col-141* | WBGene00000714 | protein_coding_gene | 4.0 | 15.586 | 1.1E-08 |
| *F53B3.6* | WBGene00018744 | protein_coding_gene | 3.9 | 15.374 | 1.3E-20 |
| *F59A7.11* | WBGene00044519 | protein_coding_gene | 3.9 | 15.185 | 3.9E-03 |
| *T02B11.13* | WBGene00255733 | protein_coding_gene | 3.9 | 14.563 | 3.7E-03 |
| *mpst-7* | WBGene00011307 | protein_coding_gene | 3.7 | 12.882 | 1.2E-18 |
| *H12D21.2* | WBGene00010378 | protein_coding_gene | 3.6 | 12.164 | 4.6E-03 |
| *srsx-21* | WBGene00011084 | protein_coding_gene | 3.6 | 12.144 | 4.1E-03 |
| *gmd-2* | WBGene00010166 | protein_coding_gene | 3.6 | 11.802 | 9.7E-05 |
| *R193.2* | WBGene00020128 | protein_coding_gene | 3.5 | 11.646 | 1.8E-43 |
| *K09E2.1* | WBGene00019570 | protein_coding_gene | 3.5 | 11.525 | 1.4E-03 |
| *col-147* | WBGene00000720 | protein_coding_gene | 3.5 | 11.488 | 5.0E-49 |
| *oac-52* | WBGene00020977 | protein_coding_gene | 3.5 | 11.148 | 5.0E-05 |
| *K08H10.14* | WBGene00195600 | ncRNA_gene | 3.5 | 11.089 | 3.4E-02 |
| *str-131* | WBGene00006180 | protein_coding_gene | 3.5 | 11.045 | 5.0E-04 |
| *Y71G12B.18* | WBGene00022156 | protein_coding_gene | 3.4 | 10.610 | 1.8E-03 |
| *T17H7.1* | WBGene00020550 | protein_coding_gene | 3.4 | 10.518 | 5.3E-06 |
| *F21G4.3* | WBGene00009024 | protein_coding_gene | 3.4 | 10.483 | 2.1E-03 |
| *grd-10* | WBGene00001699 | protein_coding_gene | 3.4 | 10.385 | 2.0E-27 |
| *F09B12.3* | WBGene00008607 | protein_coding_gene | 3.4 | 10.337 | 8.3E-88 |
| *ZK488.5* | WBGene00022757 | protein_coding_gene | 3.4 | 10.306 | 3.2E-03 |
| *ferl-1* | WBGene00020262 | protein_coding_gene | 3.4 | 10.293 | 1.0E-05 |
| *hrg-4* | WBGene00009493 | protein_coding_gene | 3.3 | 10.178 | 1.0E-11 |
| *K06H6.1* | WBGene00019449 | protein_coding_gene | 3.3 | 10.163 | 8.9E-03 |
| *nhr-219* | WBGene00020555 | protein_coding_gene | 3.3 | 10.154 | 1.8E-06 |

| | | | | | |
|---|---|---|---|---|---|
| *R08F11.7* | WBGene00019970 | protein_coding_gene | 3.3 | 10.069 | 3.4E-07 |
| *C02G6.1* | WBGene00015359 | pseudogene | 3.3 | 10.063 | 6.5E-05 |
| *ZC250.4* | WBGene00045305 | protein_coding_gene | 3.3 | 10.009 | 2.2E-45 |
| *Y48G8AL.12* | WBGene00021690 | protein_coding_gene | 3.3 | 9.662 | 2.1E-03 |
| *ins-32* | WBGene00002115 | protein_coding_gene | 3.3 | 9.534 | 7.2E-04 |
| *F40F11.6* | WBGene00219941 | ncRNA_gene | 3.2 | 9.472 | 1.0E-02 |
| *clec-167* | WBGene00009517 | protein_coding_gene | 3.2 | 9.431 | 1.0E-03 |
| *C14A6.5* | WBGene00007569 | protein_coding_gene | 3.2 | 9.298 | 1.2E-04 |
| *T25C12.5* | WBGene00197733 | ncRNA_gene | 3.2 | 9.135 | 1.2E-02 |
| *sra-6* | WBGene00005032 | protein_coding_gene | 3.2 | 9.098 | 9.7E-03 |
| *dmd-11* | WBGene00007930 | protein_coding_gene | 3.2 | 9.095 | 6.4E-03 |
| *srg-27* | WBGene00005184 | protein_coding_gene | 3.2 | 8.961 | 1.5E-02 |
| *npr-30* | WBGene00019183 | protein_coding_gene | 3.2 | 8.944 | 2.3E-03 |
| *F35E8.9* | WBGene00009422 | protein_coding_gene | 3.2 | 8.916 | 2.4E-02 |
| *Y45F10B.8* | WBGene00012872 | protein_coding_gene | 3.2 | 8.887 | 2.4E-03 |
| *fbxa-125* | WBGene00015229 | protein_coding_gene | 3.1 | 8.657 | 2.0E-14 |
| *Y52B11B.1* | WBGene00013130 | protein_coding_gene | 3.1 | 8.592 | 1.1E-02 |
| *hsp-16.41* | WBGene00002018 | protein_coding_gene | 3.1 | 8.557 | 1.4E-04 |
| *T25B9.2* | WBGene00012008 | protein_coding_gene | 3.1 | 8.538 | 9.0E-05 |
| *C14C6.6* | WBGene00015760 | protein_coding_gene | 3.1 | 8.479 | 2.2E-02 |
| *srx-130* | WBGene00006021 | protein_coding_gene | 3.1 | 8.425 | 8.7E-03 |
| *F31F4.11* | WBGene00017960 | protein_coding_gene | 3.0 | 8.152 | 1.7E-02 |
| *R08F11.4* | WBGene00019968 | protein_coding_gene | 3.0 | 8.045 | 4.8E-16 |
| *str-113* | WBGene00006164 | protein_coding_gene | 3.0 | 8.043 | 1.5E-02 |
| *C12D8.13* | WBGene00007540 | protein_coding_gene | 3.0 | 7.978 | 5.1E-03 |
| *ZK1290.7* | WBGene00022889 | protein_coding_gene | 3.0 | 7.961 | 4.7E-07 |
| *F46A8.9* | WBGene00009752 | pseudogene | 3.0 | 7.942 | 1.3E-08 |
| *Y51H4A.22* | WBGene00013116 | protein_coding_gene | 3.0 | 7.910 | 1.2E-07 |
| *sre-40* | WBGene00020225 | protein_coding_gene | 3.0 | 7.843 | 1.3E-02 |
| *R03E9.6* | WBGene00196969 | ncRNA_gene | 2.9 | 7.678 | 3.1E-02 |
| *H01G02.4* | WBGene00045108 | protein_coding_gene | 2.9 | 7.629 | 1.1E-05 |
| *str-161* | WBGene00006206 | protein_coding_gene | 2.9 | 7.607 | 1.9E-02 |
| *gcy-14* | WBGene00001540 | protein_coding_gene | 2.9 | 7.600 | 1.7E-02 |
| *F58A4.12* | WBGene00014777 | pseudogene | 2.9 | 7.559 | 1.1E-03 |
| *smf-2* | WBGene00004877 | protein_coding_gene | 2.9 | 7.540 | 1.9E-02 |
| *pals-3* | WBGene00007657 | protein_coding_gene | 2.9 | 7.521 | 1.1E-02 |
| *Y39A3B.1* | WBGene00021436 | protein_coding_gene | 2.9 | 7.498 | 1.0E-02 |
| *srd-9* | WBGene00005087 | protein_coding_gene | 2.9 | 7.479 | 2.2E-02 |
| *C03B1.4* | WBGene00015375 | protein_coding_gene | 2.9 | 7.431 | 3.0E-02 |
| *T16G12.1* | WBGene00011803 | protein_coding_gene | 2.9 | 7.306 | 2.5E-19 |
| *C38C6.5* | WBGene00007998 | protein_coding_gene | 2.9 | 7.270 | 3.9E-08 |
| *fbxa-15* | WBGene00022327 | protein_coding_gene | 2.9 | 7.267 | 1.4E-03 |
| *C18B2.1* | WBGene00015952 | protein_coding_gene | 2.9 | 7.250 | 5.5E-03 |
| *Y51B9A.8* | WBGene00013090 | protein_coding_gene | 2.9 | 7.230 | 2.4E-03 |
| *T04C12.33* | WBGene00271444 | protein_coding_gene | 2.8 | 7.187 | 5.9E-03 |
| *T20B6.1* | WBGene00020597 | protein_coding_gene | 2.8 | 7.174 | 1.4E-04 |
| *gly-1* | WBGene00001626 | protein_coding_gene | 2.8 | 7.173 | 5.1E-04 |
| *basl-2* | WBGene00017392 | pseudogene | 2.8 | 7.033 | 7.8E-11 |
| *clec-152* | WBGene00008658 | pseudogene | 2.8 | 7.012 | 3.7E-15 |
| *R08H2.8* | WBGene00011149 | pseudogene | 2.8 | 6.963 | 2.2E-02 |
| *sri-4* | WBGene00005516 | protein_coding_gene | 2.8 | 6.917 | 1.3E-02 |
| *C06B3.16* | WBGene00189947 | protein_coding_gene | 2.8 | 6.861 | 1.1E-03 |
| *F19B6.3* | WBGene00008949 | protein_coding_gene | 2.8 | 6.828 | 2.3E-10 |
| *ZK105.5* | WBGene00022656 | protein_coding_gene | 2.8 | 6.804 | 3.7E-02 |
| *pho-1* | WBGene00004020 | protein_coding_gene | 2.8 | 6.789 | 3.6E-12 |
| *T15B7.14* | WBGene00020526 | protein_coding_gene | 2.8 | 6.753 | 1.3E-08 |
| *srab-17* | WBGene00011701 | protein_coding_gene | 2.8 | 6.749 | 4.8E-02 |
| *fbxc-48* | WBGene00015615 | protein_coding_gene | 2.7 | 6.726 | 1.8E-15 |
| *F02D8.1* | WBGene00008520 | protein_coding_gene | 2.7 | 6.684 | 1.1E-02 |
| *C36C5.4* | WBGene00016473 | protein_coding_gene | 2.7 | 6.678 | 2.0E-02 |
| *T25B6.6* | WBGene00020792 | protein_coding_gene | 2.7 | 6.659 | 1.1E-02 |
| *Y51A2B.8* | WBGene00013071 | protein_coding_gene | 2.7 | 6.631 | 4.8E-02 |
| *ncx-8* | WBGene00003573 | protein_coding_gene | 2.7 | 6.623 | 4.6E-11 |
| *ZK262.19* | WBGene00206508 | protein_coding_gene | 2.7 | 6.613 | 2.1E-03 |
| *lips-6* | WBGene00014009 | protein_coding_gene | 2.7 | 6.604 | 1.5E-04 |
| *sri-38* | WBGene00005550 | protein_coding_gene | 2.7 | 6.597 | 1.4E-02 |
| *W07A12.8* | WBGene00012325 | protein_coding_gene | 2.7 | 6.592 | 5.0E-04 |
| *F22H10.1* | WBGene00017725 | protein_coding_gene | 2.7 | 6.568 | 1.9E-02 |
| *oac-19* | WBGene00009413 | protein_coding_gene | 2.7 | 6.523 | 5.5E-09 |
| *nhr-234* | WBGene00012596 | protein_coding_gene | 2.7 | 6.467 | 2.0E-03 |
| *R02C2.8* | WBGene00255728 | pseudogene | 2.7 | 6.437 | 8.9E-07 |
| *F15H9.1* | WBGene00008870 | protein_coding_gene | 2.7 | 6.398 | 2.7E-06 |
| *F31A3.3* | WBGene00017938 | protein_coding_gene | 2.7 | 6.354 | 2.9E-07 |
| *fbxa-183* | WBGene00018428 | protein_coding_gene | 2.7 | 6.334 | 2.6E-07 |
| *clec-29* | WBGene00012024 | protein_coding_gene | 2.7 | 6.318 | 2.1E-02 |
| *C04C3.6* | WBGene00015416 | protein_coding_gene | 2.6 | 6.264 | 2.3E-02 |
| *col-154* | WBGene00000727 | protein_coding_gene | 2.6 | 6.256 | 2.0E-13 |
| *C30G12.10* | WBGene00284859 | ncRNA_gene | 2.6 | 6.237 | 1.4E-02 |
| *nas-3* | WBGene00003522 | protein_coding_gene | 2.6 | 6.219 | 1.5E-05 |
| *grd-13* | WBGene00001702 | protein_coding_gene | 2.6 | 6.194 | 3.8E-09 |

| | | | | | |
|---|---|---|---|---|---|
| *F33H12.7* | WBGene00045457 | protein_coding_gene | 2.6 | 6.100 | 2.9E-08 |
| *C14A6.10* | WBGene00045348 | pseudogene | 2.6 | 6.095 | 3.4E-04 |
| *ilc-17.1* | WBGene00011937 | protein_coding_gene | 2.6 | 6.094 | 2.6E-02 |
| *Y50D4B.6* | WBGene00021745 | protein_coding_gene | 2.6 | 6.088 | 8.2E-06 |
| *Y51A2B.5* | WBGene00013069 | protein_coding_gene | 2.6 | 6.073 | 1.2E-03 |
| *T23E1.1* | WBGene00020728 | protein_coding_gene | 2.6 | 6.025 | 7.3E-04 |
| *ZK783.6* | WBGene00044448 | protein_coding_gene | 2.6 | 5.971 | 7.0E-09 |
| *Y47D3A.34* | WBGene00206461 | protein_coding_gene | 2.6 | 5.950 | 1.4E-02 |
| *T02C12.4* | WBGene00011369 | pseudogene | 2.6 | 5.942 | 3.1E-05 |
| *snf-8* | WBGene00004907 | pseudogene | 2.6 | 5.934 | 2.4E-08 |
| *zipt-2.3* | WBGene00013207 | protein_coding_gene | 2.6 | 5.907 | 4.6E-06 |
| *clec-148* | WBGene00018246 | protein_coding_gene | 2.6 | 5.887 | 3.4E-02 |
| *T20H9.6* | WBGene00020642 | protein_coding_gene | 2.6 | 5.867 | 1.9E-04 |
| *F59B10.6* | WBGene00010322 | protein_coding_gene | 2.6 | 5.859 | 3.7E-02 |
| *fbxa-8* | WBGene00016526 | pseudogene | 2.5 | 5.838 | 1.2E-03 |
| *fbxa-195* | WBGene00019976 | pseudogene | 2.5 | 5.834 | 2.4E-03 |
| *K06H6.4* | WBGene00019452 | protein_coding_gene | 2.5 | 5.818 | 4.5E-02 |
| *T26E4.4* | WBGene00012046 | protein_coding_gene | 2.5 | 5.810 | 3.0E-02 |
| *T10B5.10* | WBGene00020396 | protein_coding_gene | 2.5 | 5.796 | 9.7E-06 |
| *K08E4.7* | WBGene00045036 | protein_coding_gene | 2.5 | 5.794 | 5.3E-14 |
| *srp-9* | WBGene00005650 | pseudogene | 2.5 | 5.755 | 2.7E-04 |
| *scav-4* | WBGene00008692 | protein_coding_gene | 2.5 | 5.748 | 1.3E-20 |
| *F42E8.1* | WBGene00009630 | protein_coding_gene | 2.5 | 5.728 | 2.5E-02 |
| *mpst-4* | WBGene00017387 | protein_coding_gene | 2.5 | 5.688 | 9.6E-10 |
| *C18G1.8* | WBGene00015986 | protein_coding_gene | 2.5 | 5.672 | 3.3E-02 |
| *K08D8.2* | WBGene00010656 | protein_coding_gene | 2.5 | 5.666 | 2.1E-02 |
| *fbxa-223* | WBGene00022630 | pseudogene | 2.5 | 5.649 | 5.5E-07 |
| *T05A8.2* | WBGene00020228 | protein_coding_gene | 2.5 | 5.648 | 2.8E-02 |
| *K08H2.5* | WBGene00010692 | protein_coding_gene | 2.5 | 5.646 | 1.3E-02 |
| *C41H7.1* | WBGene00016571 | protein_coding_gene | 2.5 | 5.614 | 3.5E-06 |
| *Y6E2A.7* | WBGene00012400 | protein_coding_gene | 2.5 | 5.604 | 4.9E-02 |
| *ZC373.3* | WBGene00013868 | protein_coding_gene | 2.5 | 5.583 | 8.1E-04 |
| *K07A1.20* | WBGene00219285 | protein_coding_gene | 2.5 | 5.579 | 2.2E-05 |
| *clpr-2* | WBGene00023410 | protein_coding_gene | 2.5 | 5.551 | 3.4E-04 |
| *T05E11.7* | WBGene00011483 | pseudogene | 2.5 | 5.547 | 1.6E-02 |
| *otpl-5* | WBGene00018478 | protein_coding_gene | 2.5 | 5.541 | 1.7E-02 |
| *F28B1.10* | WBGene00050958 | pseudogene | 2.5 | 5.529 | 2.8E-02 |
| *linc-49* | WBGene00219731 | lincRNA_gene | 2.5 | 5.496 | 7.9E-03 |
| *srx-56* | WBGene00005947 | protein_coding_gene | 2.5 | 5.474 | 3.1E-02 |
| *srd-51* | WBGene00005129 | protein_coding_gene | 2.4 | 5.446 | 3.0E-02 |
| *K02H11.4* | WBGene00019348 | protein_coding_gene | 2.4 | 5.423 | 4.6E-02 |
| *K10C9.4* | WBGene00019625 | protein_coding_gene | 2.4 | 5.419 | 2.9E-02 |
| *Y50D4B.7* | WBGene00021746 | pseudogene | 2.4 | 5.408 | 3.7E-04 |
| *srt-47* | WBGene00012770 | protein_coding_gene | 2.4 | 5.399 | 4.4E-02 |
| *hen-1* | WBGene00001841 | protein_coding_gene | 2.4 | 5.390 | 4.0E-03 |
| *R07C3.16* | WBGene00269338 | pseudogene | 2.4 | 5.369 | 2.2E-02 |
| *clec-144* | WBGene00012086 | protein_coding_gene | 2.4 | 5.352 | 3.5E-02 |
| *mec-4* | WBGene00003168 | protein_coding_gene | 2.4 | 5.341 | 2.2E-05 |
| *nhr-157* | WBGene00015900 | protein_coding_gene | 2.4 | 5.324 | 1.8E-02 |
| *T05F1.7* | WBGene00011492 | protein_coding_gene | 2.4 | 5.324 | 1.1E-06 |
| *T08D2.7* | WBGene00011612 | protein_coding_gene | 2.4 | 5.283 | 1.1E-08 |
| *M01E5.1* | WBGene00010804 | pseudogene | 2.4 | 5.274 | 1.2E-03 |
| *K02E7.4* | WBGene00019309 | protein_coding_gene | 2.4 | 5.267 | 4.7E-02 |
| *R07C3.13* | WBGene00019929 | protein_coding_gene | 2.4 | 5.247 | 3.7E-09 |
| *Y73B3A.18* | WBGene00022219 | pseudogene | 2.4 | 5.212 | 5.4E-18 |
| *Y105E8A.13* | WBGene00013673 | protein_coding_gene | 2.4 | 5.212 | 6.1E-05 |
| *Y47G6A.19* | WBGene00021645 | protein_coding_gene | 2.4 | 5.204 | 3.8E-32 |
| *C23H5.11* | WBGene00044649 | protein_coding_gene | 2.4 | 5.199 | 4.6E-05 |
| *daf-28* | WBGene00000920 | protein_coding_gene | 2.4 | 5.196 | 3.3E-12 |
| *frpr-10* | WBGene00019019 | protein_coding_gene | 2.4 | 5.182 | 3.2E-03 |
| *ceh-54* | WBGene00020485 | protein_coding_gene | 2.4 | 5.172 | 3.7E-02 |
| *K03D7.3* | WBGene00014796 | pseudogene | 2.4 | 5.149 | 2.2E-02 |
| *Y26D4A.22* | WBGene00077789 | pseudogene | 2.4 | 5.141 | 3.8E-02 |
| *F23B2.10* | WBGene00009077 | protein_coding_gene | 2.4 | 5.139 | 6.6E-03 |
| *C53D6.5* | WBGene00008282 | pseudogene | 2.4 | 5.122 | 3.8E-03 |
| *ZK1086.2* | WBGene00014216 | protein_coding_gene | 2.4 | 5.115 | 2.8E-03 |
| *T03F6.8* | WBGene00044229 | pseudogene | 2.4 | 5.105 | 5.5E-09 |
| *D1065.3* | WBGene00017035 | pseudogene | 2.3 | 5.077 | 1.4E-09 |
| *pgp-9* | WBGene00004003 | protein_coding_gene | 2.3 | 5.076 | 9.1E-23 |
| *C46G7.5* | WBGene00044502 | protein_coding_gene | 2.3 | 5.075 | 6.6E-05 |
| *fer-1* | WBGene00001414 | protein_coding_gene | 2.3 | 5.063 | 3.8E-21 |
| *F57F4.4* | WBGene00019017 | protein_coding_gene | 2.3 | 5.046 | 4.6E-05 |
| *K09C4.5* | WBGene00019550 | protein_coding_gene | 2.3 | 5.040 | 1.0E-09 |
| *T07D10.1* | WBGene00011581 | protein_coding_gene | 2.3 | 5.031 | 3.6E-02 |
| *hsp-16.2* | WBGene00002016 | protein_coding_gene | 2.3 | 5.021 | 4.8E-04 |
| *C55B6.5* | WBGene00016941 | protein_coding_gene | 2.3 | 5.020 | 2.0E-02 |
| *B0457.4* | WBGene00007179 | protein_coding_gene | 2.3 | 5.012 | 1.1E-02 |
| *srh-38* | WBGene00005261 | protein_coding_gene | 2.3 | 5.012 | 4.1E-02 |
| *Y75B8A.10* | WBGene00013547 | protein_coding_gene | 2.3 | 5.002 | 2.8E-02 |
| *B0001.11* | WBGene00197796 | ncRNA_gene | 2.3 | 4.971 | 4.6E-02 |

| | | | | | |
|---|---|---|---|---|---|
| *F46F5.2* | WBGene00018494 | pseudogene | 2.3 | 4.968 | 4.0E-03 |
| *K09D9.11* | WBGene00019568 | pseudogene | 2.3 | 4.968 | 2.7E-04 |
| *T09F5.10* | WBGene00011669 | protein_coding_gene | 2.3 | 4.967 | 3.0E-04 |
| *C30A5.4* | WBGene00016239 | protein_coding_gene | 2.3 | 4.965 | 4.9E-04 |
| *F14H3.8* | WBGene00008827 | pseudogene | 2.3 | 4.937 | 3.4E-02 |
| *T25G12.13* | WBGene00219274 | protein_coding_gene | 2.3 | 4.934 | 9.4E-03 |
| *R102.11* | WBGene00077714 | protein_coding_gene | 2.3 | 4.919 | 1.4E-02 |
| *C07E3.4* | WBGene00007415 | protein_coding_gene | 2.3 | 4.917 | 8.6E-08 |
| *irld-57* | WBGene00012401 | protein_coding_gene | 2.3 | 4.914 | 7.2E-03 |
| *srx-41* | WBGene00005932 | protein_coding_gene | 2.3 | 4.910 | 2.4E-02 |
| *sptl-2* | WBGene00018398 | protein_coding_gene | 2.3 | 4.908 | 1.5E-33 |
| *pqn-37* | WBGene00004124 | protein_coding_gene | 2.3 | 4.907 | 7.8E-10 |
| *irld-6* | WBGene00009239 | protein_coding_gene | 2.3 | 4.900 | 3.4E-14 |
| *sri-26* | WBGene00005538 | protein_coding_gene | 2.3 | 4.896 | 4.8E-02 |
| *Y57A10C.9* | WBGene00013287 | protein_coding_gene | 2.3 | 4.889 | 3.2E-02 |
| *H06I04.8* | WBGene00044940 | snoRNA_gene | 2.3 | 4.858 | 4.0E-02 |
| *srp-10* | WBGene00005651 | pseudogene | 2.3 | 4.845 | 1.8E-04 |
| *D1081.10* | WBGene00045508 | protein_coding_gene | 2.3 | 4.827 | 1.8E-04 |
| *gst-27* | WBGene00001775 | protein_coding_gene | 2.3 | 4.823 | 5.4E-04 |
| *K05F6.16* | WBGene00269420 | pseudogene | 2.3 | 4.823 | 2.1E-02 |
| *F36D4.6* | WBGene00018091 | protein_coding_gene | 2.3 | 4.789 | 2.7E-02 |
| *T10B5.2* | WBGene00020388 | pseudogene | 2.3 | 4.788 | 6.8E-13 |
| *C27A2.7* | WBGene00044386 | protein_coding_gene | 2.3 | 4.786 | 3.3E-02 |
| *Y62H9A.12* | WBGene00013400 | protein_coding_gene | 2.3 | 4.776 | 1.8E-05 |
| *ddn-1* | WBGene00015227 | protein_coding_gene | 2.3 | 4.776 | 1.7E-06 |
| *gpdh-1* | WBGene00009824 | protein_coding_gene | 2.3 | 4.770 | 4.0E-12 |
| *Y45F10B.9* | WBGene00012873 | protein_coding_gene | 2.3 | 4.768 | 4.6E-02 |
| *R08E5.4* | WBGene00019964 | protein_coding_gene | 2.3 | 4.762 | 8.1E-04 |
| *gfi-1* | WBGene00001581 | protein_coding_gene | 2.2 | 4.715 | 5.4E-05 |
| *F56D5.9* | WBGene00010152 | protein_coding_gene | 2.2 | 4.704 | 1.5E-03 |
| *T12B5.9* | WBGene00023345 | pseudogene | 2.2 | 4.703 | 3.5E-02 |
| *T07A5.1* | WBGene00011554 | protein_coding_gene | 2.2 | 4.697 | 2.3E-06 |
| *snf-4* | WBGene00004903 | pseudogene | 2.2 | 4.697 | 1.2E-07 |
| *C47E8.10* | WBGene00044216 | pseudogene | 2.2 | 4.688 | 3.5E-06 |
| *gst-38* | WBGene00001786 | protein_coding_gene | 2.2 | 4.685 | 7.5E-03 |
| *fbxa-9* | WBGene00021860 | protein_coding_gene | 2.2 | 4.668 | 3.6E-03 |
| *T01D1.3* | WBGene00020148 | protein_coding_gene | 2.2 | 4.666 | 8.5E-20 |
| *F40E12.2* | WBGene00018235 | protein_coding_gene | 2.2 | 4.660 | 1.1E-10 |
| *T09A5.14* | WBGene00011640 | protein_coding_gene | 2.2 | 4.637 | 3.3E-02 |
| *scrm-4* | WBGene00008681 | protein_coding_gene | 2.2 | 4.631 | 8.9E-14 |
| *Y46H3C.6* | WBGene00021606 | pseudogene | 2.2 | 4.620 | 2.7E-03 |
| *ZK809.1* | WBGene00014085 | protein_coding_gene | 2.2 | 4.612 | 5.0E-04 |
| *srh-125* | WBGene00005342 | protein_coding_gene | 2.2 | 4.608 | 4.3E-02 |
| *C18H2.3* | WBGene00015990 | pseudogene | 2.2 | 4.579 | 9.6E-07 |
| *acs-23* | WBGene00011174 | pseudogene | 2.2 | 4.575 | 3.8E-05 |
| *Y22D7AL.11* | WBGene00021249 | protein_coding_gene | 2.2 | 4.570 | 1.1E-08 |
| *C32A9.1* | WBGene00007861 | protein_coding_gene | 2.2 | 4.565 | 4.4E-02 |
| *nhr-198* | WBGene00010603 | protein_coding_gene | 2.2 | 4.559 | 2.5E-02 |
| *fbxa-84* | WBGene00011670 | protein_coding_gene | 2.2 | 4.559 | 1.6E-11 |
| *F48F5.2* | WBGene00009846 | pseudogene | 2.2 | 4.559 | 1.9E-17 |
| *cyp-35A4* | WBGene00016786 | protein_coding_gene | 2.2 | 4.546 | 2.5E-03 |
| *ZK250.14* | WBGene00235354 | protein_coding_gene | 2.2 | 4.544 | 5.9E-04 |
| *ZK616.65* | WBGene00235383 | protein_coding_gene | 2.2 | 4.537 | 2.6E-15 |
| *nhr-93* | WBGene00003683 | protein_coding_gene | 2.2 | 4.526 | 4.2E-03 |
| *B0457.3* | WBGene00014660 | pseudogene | 2.2 | 4.525 | 4.8E-04 |
| *C39B10.5* | WBGene00008025 | protein_coding_gene | 2.2 | 4.510 | 2.4E-02 |
| *C34H4.2* | WBGene00016425 | protein_coding_gene | 2.2 | 4.505 | 1.2E-14 |
| *gpa-6* | WBGene00001668 | protein_coding_gene | 2.2 | 4.499 | 2.2E-02 |
| *pgp-12* | WBGene00004006 | protein_coding_gene | 2.2 | 4.496 | 2.1E-07 |
| *M01H9.5* | WBGene00019721 | protein_coding_gene | 2.2 | 4.496 | 4.2E-02 |
| *F13E9.8* | WBGene00008757 | protein_coding_gene | 2.2 | 4.487 | 1.8E-02 |
| *Y62E10A.3* | WBGene00013374 | protein_coding_gene | 2.2 | 4.485 | 1.0E-03 |
| *nhr-266* | WBGene00018993 | pseudogene | 2.2 | 4.483 | 1.1E-04 |
| *fbxa-159* | WBGene00015603 | pseudogene | 2.2 | 4.479 | 2.2E-02 |
| *col-157* | WBGene00000730 | protein_coding_gene | 2.2 | 4.475 | 3.6E-03 |
| *C02G6.2* | WBGene00015360 | protein_coding_gene | 2.2 | 4.474 | 3.7E-08 |
| *col-41* | WBGene00000618 | protein_coding_gene | 2.2 | 4.447 | 1.0E-02 |
| *Y26D4A.21* | WBGene00077684 | protein_coding_gene | 2.2 | 4.442 | 1.5E-02 |
| *C05C8.8* | WBGene00015465 | protein_coding_gene | 2.2 | 4.439 | 5.0E-03 |
| *ugt-46* | WBGene00015141 | protein_coding_gene | 2.1 | 4.437 | 5.4E-31 |
| *C39E6.2* | WBGene00016537 | protein_coding_gene | 2.1 | 4.430 | 3.2E-02 |
| *T23B7.2* | WBGene00044298 | protein_coding_gene | 2.1 | 4.410 | 9.4E-03 |
| *srv-32* | WBGene00005743 | protein_coding_gene | 2.1 | 4.403 | 3.8E-02 |
| *fbxa-196* | WBGene00020265 | protein_coding_gene | 2.1 | 4.375 | 1.6E-10 |
| *T26H2.12* | WBGene00077662 | pseudogene | 2.1 | 4.345 | 4.1E-03 |
| *cln-3.1* | WBGene00000539 | protein_coding_gene | 2.1 | 4.330 | 1.8E-13 |
| *nep-5* | WBGene00017393 | protein_coding_gene | 2.1 | 4.328 | 1.2E-04 |
| *aat-7* | WBGene00000008 | protein_coding_gene | 2.1 | 4.303 | 6.5E-05 |
| *klu-2* | WBGene00022592 | protein_coding_gene | 2.1 | 4.292 | 6.5E-24 |
| *C33H5.16* | WBGene00016382 | protein_coding_gene | 2.1 | 4.286 | 8.0E-07 |

| | | | | | |
|---|---|---|---|---|---|
| *math-2* | WBGene00015606 | protein_coding_gene | 2.1 | 4.278 | 3.4E-07 |
| *F01D4.3* | WBGene00008487 | pseudogene | 2.1 | 4.272 | 1.0E-07 |
| *fbxa-201* | WBGene00020342 | protein_coding_gene | 2.1 | 4.270 | 1.8E-03 |
| *C33C12.12* | WBGene00219322 | protein_coding_gene | 2.1 | 4.259 | 4.7E-02 |
| *Y4C6A.1* | WBGene00021151 | pseudogene | 2.1 | 4.225 | 4.1E-08 |
| *C08F11.14* | WBGene00007461 | pseudogene | 2.1 | 4.222 | 7.7E-12 |
| *F21A3.3* | WBGene00008993 | protein_coding_gene | 2.1 | 4.220 | 1.2E-10 |
| *C08F11.7* | WBGene00007454 | protein_coding_gene | 2.1 | 4.216 | 1.2E-02 |
| *H04J21.1* | WBGene00019150 | protein_coding_gene | 2.1 | 4.209 | 1.2E-08 |
| *F15B9.11* | WBGene00138724 | protein_coding_gene | 2.1 | 4.196 | 2.4E-02 |
| *hrg-7* | WBGene00007605 | protein_coding_gene | 2.1 | 4.188 | 2.8E-08 |
| *str-7* | WBGene00006075 | protein_coding_gene | 2.1 | 4.184 | 6.1E-04 |
| *moa-1* | WBGene00020116 | protein_coding_gene | 2.1 | 4.182 | 9.9E-24 |
| *Y116A8A.4* | WBGene00013775 | protein_coding_gene | 2.1 | 4.174 | 3.8E-03 |
| *B0041.1* | WBGene00015006 | protein_coding_gene | 2.1 | 4.169 | 1.1E-02 |
| *clp-3* | WBGene00000544 | protein_coding_gene | 2.1 | 4.169 | 3.4E-05 |
| *Y37D8A.3* | WBGene00012545 | protein_coding_gene | 2.1 | 4.164 | 3.7E-26 |
| *W03F9.3* | WBGene00021001 | pseudogene | 2.1 | 4.142 | 8.4E-04 |
| *F38B6.4* | WBGene00018174 | protein_coding_gene | 2.0 | 4.130 | 3.0E-21 |
| *dos-2* | WBGene00019640 | protein_coding_gene | 2.0 | 4.114 | 2.3E-02 |
| *ttr-57* | WBGene00050892 | protein_coding_gene | 2.0 | 4.097 | 1.1E-03 |
| *lron-6* | WBGene00018157 | protein_coding_gene | 2.0 | 4.095 | 4.7E-02 |
| *ins-33* | WBGene00002116 | protein_coding_gene | 2.0 | 4.091 | 3.4E-07 |
| *B0507.8* | WBGene00015225 | protein_coding_gene | 2.0 | 4.084 | 6.6E-04 |
| *oac-50* | WBGene00020864 | protein_coding_gene | 2.0 | 4.073 | 7.3E-12 |
| *T15H9.4* | WBGene00011789 | protein_coding_gene | 2.0 | 4.066 | 7.5E-07 |
| *K08H10.5* | WBGene00014805 | pseudogene | 2.0 | 4.041 | 3.2E-04 |
| *F59A1.15* | WBGene00010300 | protein_coding_gene | 2.0 | 4.035 | 1.1E-10 |
| *Y73B6A.2* | WBGene00022229 | pseudogene | 2.0 | 4.033 | 6.1E-04 |
| *F07D3.3* | WBGene00008556 | protein_coding_gene | 2.0 | 4.020 | 9.2E-08 |
| *C04E6.4* | WBGene00015421 | protein_coding_gene | 2.0 | 4.017 | 4.6E-06 |
| *paic-1* | WBGene00015116 | protein_coding_gene | 2.0 | 4.017 | 1.6E-47 |
| *fbxa-178* | WBGene00017533 | pseudogene | 2.0 | 4.007 | 1.0E-04 |
| *F26B1.5* | WBGene00017817 | protein_coding_gene | 2.0 | 4.004 | 3.8E-06 |
| *F08F3.8* | WBGene00017264 | protein_coding_gene | 2.0 | 3.989 | 1.7E-03 |
| *C38D4.7* | WBGene00008007 | protein_coding_gene | 2.0 | 3.986 | 6.5E-08 |
| *F09B9.5* | WBGene00008604 | protein_coding_gene | 2.0 | 3.971 | 1.8E-03 |
| *str-44* | WBGene00006109 | protein_coding_gene | 2.0 | 3.960 | 3.6E-02 |
| *F09E10.15* | WBGene00219886 | ncRNA_gene | 2.0 | 3.956 | 6.0E-10 |
| *fbxb-72* | WBGene00019243 | protein_coding_gene | 2.0 | 3.955 | 8.3E-05 |
| *M04G7.2* | WBGene00019773 | protein_coding_gene | 2.0 | 3.951 | 6.3E-08 |
| *him-4* | WBGene00001863 | protein_coding_gene | 2.0 | 3.943 | 3.6E-28 |
| *pgp-5* | WBGene00003999 | protein_coding_gene | 2.0 | 3.938 | 1.9E-08 |
| *K04A8.2* | WBGene00019374 | pseudogene | 2.0 | 3.937 | 4.0E-02 |
| *hmit-1.1* | WBGene00013073 | protein_coding_gene | 2.0 | 3.929 | 8.0E-11 |
| *scl-3* | WBGene00009896 | protein_coding_gene | 2.0 | 3.923 | 9.0E-24 |
| *sre-14* | WBGene00016579 | protein_coding_gene | 2.0 | 3.920 | 3.2E-02 |
| *lact-1* | WBGene00018523 | protein_coding_gene | 2.0 | 3.911 | 8.9E-09 |
| *srr-5* | WBGene00005656 | pseudogene | 2.0 | 3.910 | 3.0E-02 |
| *Y65B4A.9* | WBGene00022032 | protein_coding_gene | 2.0 | 3.907 | 6.9E-05 |
| *R08C7.5* | WBGene00019949 | protein_coding_gene | 2.0 | 3.903 | 2.3E-15 |
| *F52G3.3* | WBGene00018711 | pseudogene | 2.0 | 3.899 | 3.9E-02 |
| *Y4C6A.68* | WBGene00235281 | pseudogene | 2.0 | 3.880 | 3.2E-04 |
| *Y39G10AR.16* | WBGene00021472 | protein_coding_gene | 2.0 | 3.875 | 5.2E-03 |
| *fbxa-202* | WBGene00020890 | protein_coding_gene | 2.0 | 3.868 | 5.3E-14 |
| *T02G6.1* | WBGene00011384 | protein_coding_gene | 2.0 | 3.866 | 8.3E-03 |
| *cyp-35D2* | WBGene00014735 | pseudogene | 1.9 | 3.859 | 3.6E-02 |
| *F08F8.6* | WBGene00017271 | protein_coding_gene | 1.9 | 3.858 | 3.4E-09 |
| *Y116F11B.8* | WBGene00013824 | protein_coding_gene | 1.9 | 3.856 | 6.9E-05 |
| *srsx-6* | WBGene00022405 | protein_coding_gene | 1.9 | 3.852 | 4.0E-02 |
| *F28A10.1* | WBGene00017869 | protein_coding_gene | 1.9 | 3.844 | 1.2E-10 |
| *T10E10.3* | WBGene00020420 | protein_coding_gene | 1.9 | 3.828 | 5.5E-03 |
| *F55H12.4* | WBGene00010135 | protein_coding_gene | 1.9 | 3.819 | 4.1E-45 |
| *F36D1.6* | WBGene00009465 | protein_coding_gene | 1.9 | 3.809 | 1.7E-03 |
| *K09C6.2* | WBGene00019556 | protein_coding_gene | 1.9 | 3.807 | 3.9E-09 |
| *C48D1.7* | WBGene00044909 | pseudogene | 1.9 | 3.801 | 3.9E-03 |
| *B0244.17* | WBGene00271715 | protein_coding_gene | 1.9 | 3.790 | 2.4E-05 |
| *C15H7.4* | WBGene00007611 | protein_coding_gene | 1.9 | 3.790 | 6.8E-08 |
| *Y43C5A.7* | WBGene00044922 | protein_coding_gene | 1.9 | 3.786 | 6.0E-06 |
| *clec-154* | WBGene00008657 | pseudogene | 1.9 | 3.767 | 4.2E-05 |
| *Y42H9AR.5* | WBGene00021540 | protein_coding_gene | 1.9 | 3.757 | 2.5E-05 |
| *ZK177.2* | WBGene00022670 | protein_coding_gene | 1.9 | 3.757 | 3.1E-02 |
| *F20D6.1* | WBGene00017633 | protein_coding_gene | 1.9 | 3.756 | 1.3E-04 |
| *ins-28* | WBGene00002111 | protein_coding_gene | 1.9 | 3.740 | 3.4E-06 |
| *K09H9.9* | WBGene00219460 | protein_coding_gene | 1.9 | 3.735 | 1.8E-03 |
| *adt-2* | WBGene00000083 | protein_coding_gene | 1.9 | 3.735 | 4.0E-34 |
| *best-6* | WBGene00015628 | protein_coding_gene | 1.9 | 3.734 | 9.6E-05 |
| *Y18H1A.15* | WBGene00044434 | protein_coding_gene | 1.9 | 3.733 | 2.6E-09 |
| *F26A1.4* | WBGene00017803 | protein_coding_gene | 1.9 | 3.720 | 9.9E-06 |
| *Y102A11A.7* | WBGene00022417 | protein_coding_gene | 1.9 | 3.703 | 7.8E-03 |

| | | | | | |
|---|---|---|---|---|---|
| *fbxa-160* | WBGene00015604 | protein_coding_gene | 1.9 | 3.702 | 2.4E-03 |
| *H06I04.9* | WBGene00045092 | snoRNA_gene | 1.9 | 3.689 | 2.2E-02 |
| *Y50D4B.9* | WBGene00268189 | pseudogene | 1.9 | 3.686 | 1.9E-02 |
| *W06H8.4* | WBGene00021067 | pseudogene | 1.9 | 3.679 | 1.0E-03 |
| *ZK1010.6* | WBGene00014180 | pseudogene | 1.9 | 3.678 | 1.2E-02 |
| *fbxa-60* | WBGene00020456 | protein_coding_gene | 1.9 | 3.675 | 6.1E-05 |
| *C55C2.3* | WBGene00016949 | protein_coding_gene | 1.9 | 3.669 | 1.9E-02 |
| *C49C3.2* | WBGene00008192 | pseudogene | 1.9 | 3.666 | 2.5E-05 |
| *hsp-17* | WBGene00002021 | protein_coding_gene | 1.9 | 3.661 | 1.3E-19 |
| *tyr-2* | WBGene00010661 | protein_coding_gene | 1.9 | 3.659 | 4.9E-12 |
| *tsp-19* | WBGene00006645 | protein_coding_gene | 1.9 | 3.659 | 4.7E-08 |
| *B0207.7* | WBGene00015030 | protein_coding_gene | 1.9 | 3.656 | 2.1E-09 |
| *C28D4.5* | WBGene00007793 | protein_coding_gene | 1.9 | 3.649 | 9.4E-04 |
| *best-26* | WBGene00014103 | protein_coding_gene | 1.9 | 3.648 | 3.8E-08 |
| *C25G4.7* | WBGene00007734 | protein_coding_gene | 1.9 | 3.636 | 5.2E-07 |
| *T05B9.2* | WBGene00011465 | protein_coding_gene | 1.9 | 3.632 | 3.4E-02 |
| *C06C6.6* | WBGene00007380 | pseudogene | 1.9 | 3.630 | 7.9E-04 |
| *ram-5* | WBGene00004301 | protein_coding_gene | 1.9 | 3.629 | 2.0E-02 |
| *F01G10.5* | WBGene00008508 | protein_coding_gene | 1.9 | 3.624 | 5.9E-22 |
| *K09B11.5* | WBGene00010712 | protein_coding_gene | 1.9 | 3.606 | 4.7E-07 |
| *ser-5* | WBGene00008890 | protein_coding_gene | 1.8 | 3.601 | 3.9E-05 |
| *best-12* | WBGene00008821 | protein_coding_gene | 1.8 | 3.597 | 1.0E-04 |
| *pqn-75* | WBGene00004157 | protein_coding_gene | 1.8 | 3.590 | 5.1E-16 |
| *gur-5* | WBGene00001806 | protein_coding_gene | 1.8 | 3.588 | 5.3E-03 |
| *piit-1* | WBGene00009384 | protein_coding_gene | 1.8 | 3.579 | 4.1E-10 |
| *F56D6.18* | WBGene00235285 | protein_coding_gene | 1.8 | 3.576 | 1.2E-02 |
| *M117.3* | WBGene00010919 | protein_coding_gene | 1.8 | 3.571 | 1.4E-02 |
| *C04G2.12* | WBGene00045104 | pseudogene | 1.8 | 3.566 | 2.3E-02 |
| *lgc-12* | WBGene00020048 | protein_coding_gene | 1.8 | 3.563 | 1.3E-02 |
| *oac-27* | WBGene00009609 | protein_coding_gene | 1.8 | 3.554 | 3.2E-03 |
| *nhr-30* | WBGene00016091 | protein_coding_gene | 1.8 | 3.548 | 2.7E-02 |
| *F33A8.10* | WBGene00009356 | protein_coding_gene | 1.8 | 3.545 | 2.9E-02 |
| *C35A11.4* | WBGene00016431 | protein_coding_gene | 1.8 | 3.545 | 8.3E-13 |
| *T07G12.2* | WBGene00011593 | protein_coding_gene | 1.8 | 3.539 | 1.5E-02 |
| *clp-6* | WBGene00000546 | protein_coding_gene | 1.8 | 3.534 | 1.1E-03 |
| *ttll-15* | WBGene00010630 | protein_coding_gene | 1.8 | 3.534 | 7.9E-12 |
| *clec-51* | WBGene00015050 | protein_coding_gene | 1.8 | 3.530 | 1.3E-25 |
| *spe-17* | WBGene00004971 | protein_coding_gene | 1.8 | 3.529 | 2.1E-06 |
| *M05D6.3* | WBGene00010876 | protein_coding_gene | 1.8 | 3.527 | 2.1E-04 |
| *mth-1* | WBGene00017948 | protein_coding_gene | 1.8 | 3.523 | 1.4E-09 |
| *sra-37* | WBGene00005063 | protein_coding_gene | 1.8 | 3.522 | 3.4E-02 |
| *C09B9.7* | WBGene00015630 | protein_coding_gene | 1.8 | 3.522 | 3.1E-10 |
| *F21D9.2* | WBGene00009016 | protein_coding_gene | 1.8 | 3.520 | 5.5E-03 |
| *Y38H6C.15* | WBGene00012627 | protein_coding_gene | 1.8 | 3.506 | 5.5E-09 |
| *pcp-3* | WBGene00003958 | protein_coding_gene | 1.8 | 3.504 | 2.1E-29 |
| *pept-1* | WBGene00003877 | protein_coding_gene | 1.8 | 3.500 | 3.5E-27 |
| *ZK686.5* | WBGene00022795 | protein_coding_gene | 1.8 | 3.494 | 1.6E-09 |
| *best-17* | WBGene00020559 | protein_coding_gene | 1.8 | 3.493 | 2.3E-09 |
| *C54E4.5* | WBGene00016920 | protein_coding_gene | 1.8 | 3.488 | 6.2E-20 |
| *C49A9.4* | WBGene00016758 | protein_coding_gene | 1.8 | 3.486 | 2.8E-10 |
| *Y22D7AR.12* | WBGene00021265 | pseudogene | 1.8 | 3.484 | 2.2E-08 |
| *C09F12.2* | WBGene00007482 | protein_coding_gene | 1.8 | 3.484 | 1.4E-04 |
| *R155.3* | WBGene00020117 | protein_coding_gene | 1.8 | 3.448 | 1.5E-09 |
| *F17C11.16* | WBGene00050890 | pseudogene | 1.8 | 3.436 | 4.6E-02 |
| *T20F7.5* | WBGene00020632 | pseudogene | 1.8 | 3.435 | 5.4E-09 |
| *H12I13.1* | WBGene00019190 | protein_coding_gene | 1.8 | 3.426 | 4.8E-05 |
| *T12B3.1* | WBGene00020444 | protein_coding_gene | 1.8 | 3.422 | 3.0E-03 |
| *F59F5.5* | WBGene00010344 | protein_coding_gene | 1.8 | 3.416 | 1.7E-03 |
| *F44B9.9* | WBGene00018410 | protein_coding_gene | 1.8 | 3.407 | 4.5E-03 |
| *F58H7.1* | WBGene00019067 | protein_coding_gene | 1.8 | 3.405 | 3.7E-03 |
| *glr-4* | WBGene00001615 | protein_coding_gene | 1.8 | 3.405 | 7.7E-11 |
| *fbxa-64* | WBGene00020883 | protein_coding_gene | 1.8 | 3.401 | 3.1E-06 |
| *T05C12.15* | WBGene00219289 | protein_coding_gene | 1.8 | 3.398 | 1.2E-02 |
| *hot-3* | WBGene00001988 | protein_coding_gene | 1.8 | 3.397 | 2.9E-03 |
| *ugt-30* | WBGene00011340 | protein_coding_gene | 1.8 | 3.397 | 6.4E-04 |
| *C25E10.17* | WBGene00235313 | protein_coding_gene | 1.8 | 3.384 | 2.2E-02 |
| *ZK896.1* | WBGene00014132 | protein_coding_gene | 1.8 | 3.382 | 3.0E-04 |
| *Y69H2.15* | WBGene00050884 | protein_coding_gene | 1.8 | 3.381 | 3.9E-04 |
| *R02F2.5* | WBGene00019834 | protein_coding_gene | 1.8 | 3.377 | 8.7E-04 |
| *ZK930.5* | WBGene00014155 | protein_coding_gene | 1.8 | 3.373 | 5.1E-08 |
| *Y55B1AR.3* | WBGene00021907 | protein_coding_gene | 1.8 | 3.366 | 3.4E-02 |
| *R05C11.4* | WBGene00044467 | protein_coding_gene | 1.7 | 3.358 | 1.5E-11 |
| *F26F12.2* | WBGene00017833 | protein_coding_gene | 1.7 | 3.348 | 2.2E-03 |
| *D2024.1* | WBGene00017050 | protein_coding_gene | 1.7 | 3.348 | 7.9E-04 |
| *Y26E6A.3* | WBGene00012514 | protein_coding_gene | 1.7 | 3.345 | 2.7E-13 |
| *nhr-260* | WBGene00016517 | protein_coding_gene | 1.7 | 3.344 | 1.1E-02 |
| *lgc-40* | WBGene00020767 | protein_coding_gene | 1.7 | 3.343 | 3.8E-04 |
| *F56D5.2* | WBGene00010147 | protein_coding_gene | 1.7 | 3.341 | 2.5E-06 |
| *M28.4* | WBGene00010895 | protein_coding_gene | 1.7 | 3.338 | 1.2E-04 |
| *T03G11.10* | WBGene00044732 | protein_coding_gene | 1.7 | 3.338 | 4.7E-02 |

| | | | | | |
|---|---|---|---|---|---|
| *ent-4* | WBGene00008123 | protein_coding_gene | 1.7 | 3.338 | 3.6E-13 |
| *ser-1* | WBGene00004776 | protein_coding_gene | 1.7 | 3.336 | 1.1E-06 |
| *anoh-1* | WBGene00010138 | protein_coding_gene | 1.7 | 3.335 | 2.2E-02 |
| *Y46H3D.1* | WBGene00021608 | protein_coding_gene | 1.7 | 3.334 | 2.2E-05 |
| *C14A11.5* | WBGene00015750 | protein_coding_gene | 1.7 | 3.333 | 4.6E-02 |
| *B0496.2* | WBGene00015211 | protein_coding_gene | 1.7 | 3.332 | 6.4E-03 |
| *K03H1.9* | WBGene00010544 | protein_coding_gene | 1.7 | 3.321 | 1.4E-06 |
| *Y23H5B.2* | WBGene00021273 | pseudogene | 1.7 | 3.319 | 1.1E-05 |
| *sre-43* | WBGene00012333 | protein_coding_gene | 1.7 | 3.319 | 2.3E-02 |
| *slc-36.4* | WBGene00010421 | protein_coding_gene | 1.7 | 3.314 | 1.2E-11 |
| *Y51B9A.7* | WBGene00013089 | protein_coding_gene | 1.7 | 3.313 | 3.6E-03 |
| *Y97E10AR.8* | WBGene00044552 | protein_coding_gene | 1.7 | 3.307 | 2.3E-03 |
| *ZK849.1* | WBGene00014100 | protein_coding_gene | 1.7 | 3.301 | 8.7E-11 |
| *F46F5.4* | WBGene00018495 | pseudogene | 1.7 | 3.298 | 2.0E-05 |
| *F08F3.4* | WBGene00017262 | protein_coding_gene | 1.7 | 3.297 | 4.4E-45 |
| *F58D5.8* | WBGene00010247 | protein_coding_gene | 1.7 | 3.294 | 2.2E-05 |
| *C17C3.11* | WBGene00015893 | protein_coding_gene | 1.7 | 3.293 | 3.0E-03 |
| *F35C11.2* | WBGene00009401 | protein_coding_gene | 1.7 | 3.292 | 1.3E-06 |
| *eat-4* | WBGene00001135 | protein_coding_gene | 1.7 | 3.284 | 8.8E-07 |
| *T20B6.2* | WBGene00020598 | protein_coding_gene | 1.7 | 3.278 | 6.5E-04 |
| *F09E10.7* | WBGene00017297 | protein_coding_gene | 1.7 | 3.278 | 1.1E-06 |
| *C15H7.2* | WBGene00014678 | pseudogene | 1.7 | 3.274 | 1.7E-03 |
| *Y41G9A.2* | WBGene00021526 | protein_coding_gene | 1.7 | 3.265 | 4.6E-02 |
| *gba-4* | WBGene00021160 | protein_coding_gene | 1.7 | 3.264 | 6.7E-19 |
| *F36G9.13* | WBGene00009489 | protein_coding_gene | 1.7 | 3.263 | 1.6E-06 |
| *F19C7.4* | WBGene00017594 | protein_coding_gene | 1.7 | 3.252 | 2.6E-05 |
| *C07A4.2* | WBGene00007397 | protein_coding_gene | 1.7 | 3.250 | 7.6E-04 |
| *Y7A9A.79* | WBGene00185076 | ncRNA_gene | 1.7 | 3.246 | 3.0E-03 |
| *F46F5.11* | WBGene00018502 | protein_coding_gene | 1.7 | 3.242 | 4.1E-06 |
| *dhhc-12* | WBGene00010323 | protein_coding_gene | 1.7 | 3.229 | 2.1E-09 |
| *rimb-1* | WBGene00006513 | protein_coding_gene | 1.7 | 3.221 | 2.3E-11 |
| *tbc-8* | WBGene00008018 | protein_coding_gene | 1.7 | 3.213 | 2.4E-08 |
| *math-44* | WBGene00021193 | protein_coding_gene | 1.7 | 3.213 | 1.2E-05 |
| *T08G3.4* | WBGene00011617 | pseudogene | 1.7 | 3.206 | 2.5E-08 |
| *T28C12.2* | WBGene00020889 | protein_coding_gene | 1.7 | 3.201 | 1.7E-06 |
| *oac-23* | WBGene00018143 | protein_coding_gene | 1.7 | 3.199 | 1.4E-06 |
| *ocr-2* | WBGene00003839 | protein_coding_gene | 1.7 | 3.179 | 2.9E-04 |
| *cpb-2* | WBGene00000771 | protein_coding_gene | 1.7 | 3.173 | 3.3E-15 |
| *ugt-10* | WBGene00020594 | protein_coding_gene | 1.7 | 3.170 | 2.3E-02 |
| *ZK666.8* | WBGene00014048 | protein_coding_gene | 1.7 | 3.169 | 3.0E-05 |
| *fbxa-51* | WBGene00020448 | protein_coding_gene | 1.7 | 3.164 | 1.0E-04 |
| *dhs-14* | WBGene00000977 | protein_coding_gene | 1.7 | 3.163 | 9.6E-26 |
| *C10E2.1* | WBGene00015672 | protein_coding_gene | 1.7 | 3.158 | 4.0E-02 |
| *T10B10.4* | WBGene00011681 | protein_coding_gene | 1.7 | 3.157 | 4.4E-19 |
| *act-5* | WBGene00000067 | protein_coding_gene | 1.7 | 3.154 | 1.6E-04 |
| *H25P06.5* | WBGene00219319 | protein_coding_gene | 1.7 | 3.152 | 1.2E-02 |
| *M02D8.7* | WBGene00044313 | protein_coding_gene | 1.7 | 3.147 | 2.1E-02 |
| *dhhc-13* | WBGene00019257 | protein_coding_gene | 1.7 | 3.146 | 1.0E-06 |
| *C36F7.5* | WBGene00007986 | protein_coding_gene | 1.7 | 3.143 | 3.4E-16 |
| *csp-1* | WBGene00000819 | protein_coding_gene | 1.7 | 3.142 | 1.7E-06 |
| *lgc-42* | WBGene00021437 | protein_coding_gene | 1.7 | 3.141 | 4.9E-03 |
| *C14A4.13* | WBGene00007563 | protein_coding_gene | 1.7 | 3.140 | 5.1E-11 |
| *F58G1.9* | WBGene00010271 | pseudogene | 1.6 | 3.137 | 2.7E-02 |
| *Y40B10A.4* | WBGene00021489 | pseudogene | 1.6 | 3.137 | 1.3E-04 |
| *gst-39* | WBGene00001787 | protein_coding_gene | 1.6 | 3.137 | 9.0E-23 |
| *H38K22.9* | WBGene00284844 | protein_coding_gene | 1.6 | 3.136 | 3.9E-03 |
| *R07C3.17* | WBGene00269346 | ncRNA_gene | 1.6 | 3.133 | 3.3E-03 |
| *M176.11* | WBGene00010947 | protein_coding_gene | 1.6 | 3.133 | 8.8E-20 |
| *Y57G11C.23* | WBGene00013318 | protein_coding_gene | 1.6 | 3.125 | 3.2E-03 |
| *Y59E9AL.4* | WBGene00021994 | protein_coding_gene | 1.6 | 3.123 | 1.1E-06 |
| *T02G6.2* | WBGene00011385 | protein_coding_gene | 1.6 | 3.123 | 4.3E-02 |
| *cyp-14A4* | WBGene00011009 | protein_coding_gene | 1.6 | 3.123 | 6.0E-03 |
| *K06A1.2* | WBGene00019425 | protein_coding_gene | 1.6 | 3.121 | 5.2E-07 |
| *C39D10.8* | WBGene00016535 | protein_coding_gene | 1.6 | 3.120 | 3.2E-16 |
| *shl-1* | WBGene00022240 | protein_coding_gene | 1.6 | 3.119 | 1.2E-06 |
| *Y44A6D.2* | WBGene00012853 | protein_coding_gene | 1.6 | 3.118 | 2.2E-12 |
| *ZC434.9* | WBGene00013895 | protein_coding_gene | 1.6 | 3.117 | 3.6E-22 |
| *lips-1* | WBGene00009727 | protein_coding_gene | 1.6 | 3.113 | 2.2E-03 |
| *T06C10.3* | WBGene00020289 | protein_coding_gene | 1.6 | 3.105 | 4.1E-11 |
| *ser-6* | WBGene00021897 | protein_coding_gene | 1.6 | 3.100 | 3.9E-03 |
| *C45G9.8* | WBGene00016679 | protein_coding_gene | 1.6 | 3.095 | 1.8E-02 |
| *F52D10.4* | WBGene00009931 | protein_coding_gene | 1.6 | 3.093 | 3.7E-02 |
| *F41E6.17* | WBGene00194909 | protein_coding_gene | 1.6 | 3.089 | 1.8E-02 |
| *let-805* | WBGene00002915 | protein_coding_gene | 1.6 | 3.089 | 6.5E-24 |
| *nhr-132* | WBGene00003722 | protein_coding_gene | 1.6 | 3.088 | 1.5E-06 |
| *R09H10.5* | WBGene00011182 | protein_coding_gene | 1.6 | 3.083 | 2.5E-07 |
| *rps-0* | WBGene00004469 | protein_coding_gene | 1.6 | 3.081 | 3.1E-31 |
| *B0034.4* | WBGene00015004 | protein_coding_gene | 1.6 | 3.079 | 2.7E-04 |
| *C12D12.1* | WBGene00015713 | protein_coding_gene | 1.6 | 3.071 | 6.0E-08 |
| *C14B4.2* | WBGene00007583 | protein_coding_gene | 1.6 | 3.069 | 7.2E-24 |

| | | | | | |
|---|---|---|---|---|---|
| *Y69E1A.3* | WBGene00013475 | protein_coding_gene | 1.6 | 3.067 | 6.5E-05 |
| *bath-13* | WBGene00018223 | protein_coding_gene | 1.6 | 3.061 | 1.6E-02 |
| *F26C11.1* | WBGene00009146 | protein_coding_gene | 1.6 | 3.060 | 6.0E-04 |
| *lgc-4* | WBGene00017580 | protein_coding_gene | 1.6 | 3.058 | 2.6E-04 |
| *K08D8.12* | WBGene00194926 | protein_coding_gene | 1.6 | 3.040 | 7.4E-05 |
| *pgp-1* | WBGene00003995 | protein_coding_gene | 1.6 | 3.036 | 2.2E-03 |
| *C44C10.11* | WBGene00043985 | protein_coding_gene | 1.6 | 3.035 | 3.7E-09 |
| *B0285.11* | WBGene00014641 | pseudogene | 1.6 | 3.026 | 3.7E-02 |
| *K10H10.7* | WBGene00010763 | protein_coding_gene | 1.6 | 3.022 | 1.2E-04 |
| *F46F5.7* | WBGene00018498 | protein_coding_gene | 1.6 | 3.015 | 9.8E-06 |
| *kmo-1* | WBGene00011089 | protein_coding_gene | 1.6 | 3.015 | 5.8E-22 |
| *col-180* | WBGene00000753 | protein_coding_gene | 1.6 | 3.014 | 6.5E-03 |
| *ipla-5* | WBGene00019229 | protein_coding_gene | 1.6 | 3.013 | 9.6E-05 |
| *K06H7.1* | WBGene00019455 | protein_coding_gene | 1.6 | 3.010 | 3.9E-02 |
| *Y69A2AR.32* | WBGene00022101 | protein_coding_gene | 1.6 | 3.006 | 5.1E-25 |
| *Y37D8A.5* | WBGene00012547 | protein_coding_gene | 1.6 | 3.002 | 1.1E-08 |
| *pde-6* | WBGene00022389 | protein_coding_gene | 1.6 | 3.002 | 7.1E-28 |
| *math-23* | WBGene00016718 | protein_coding_gene | 1.6 | 2.997 | 1.6E-02 |
| *hrg-2* | WBGene00010473 | protein_coding_gene | 1.6 | 2.994 | 2.1E-02 |
| *F08A8.8* | WBGene00045468 | protein_coding_gene | 1.6 | 2.992 | 5.2E-08 |
| *K09C6.8* | WBGene00019562 | protein_coding_gene | 1.6 | 2.970 | 1.1E-04 |
| *C01G5.3* | WBGene00015305 | protein_coding_gene | 1.6 | 2.968 | 1.9E-07 |
| *C01G10.1* | WBGene00007230 | protein_coding_gene | 1.6 | 2.968 | 2.6E-04 |
| *F10G2.11* | WBGene00271802 | pseudogene | 1.6 | 2.967 | 5.6E-03 |
| *T25G12.6* | WBGene00020813 | protein_coding_gene | 1.6 | 2.965 | 1.4E-06 |
| *F14D7.1* | WBGene00008788 | protein_coding_gene | 1.6 | 2.965 | 2.1E-02 |
| *T04B2.1* | WBGene00011421 | pseudogene | 1.6 | 2.961 | 9.3E-03 |
| *pgp-15* | WBGene00004009 | pseudogene | 1.6 | 2.960 | 5.0E-03 |
| *C50H2.7* | WBGene00008243 | protein_coding_gene | 1.6 | 2.958 | 2.9E-02 |
| *linc-36* | WBGene00219694 | lincRNA_gene | 1.6 | 2.957 | 1.4E-02 |
| *clec-236* | WBGene00013495 | pseudogene | 1.6 | 2.952 | 5.4E-05 |
| *nep-9* | WBGene00017554 | protein_coding_gene | 1.6 | 2.950 | 1.3E-04 |
| *W03D8.10* | WBGene00020991 | pseudogene | 1.6 | 2.950 | 1.9E-12 |
| *tsp-18* | WBGene00006644 | protein_coding_gene | 1.6 | 2.950 | 9.5E-04 |
| *uggt-2* | WBGene00009178 | protein_coding_gene | 1.6 | 2.938 | 3.5E-10 |
| *F46G11.2* | WBGene00018515 | protein_coding_gene | 1.6 | 2.938 | 6.4E-03 |
| *W04B5.3* | WBGene00021020 | protein_coding_gene | 1.6 | 2.934 | 1.7E-13 |
| *C15H11.13* | WBGene00077576 | protein_coding_gene | 1.6 | 2.933 | 1.2E-02 |
| *T25E12.16* | WBGene00194715 | protein_coding_gene | 1.6 | 2.929 | 1.2E-02 |
| *ctsa-4.2* | WBGene00021503 | protein_coding_gene | 1.5 | 2.925 | 2.8E-06 |
| *F14H12.3* | WBGene00017471 | protein_coding_gene | 1.5 | 2.921 | 9.7E-09 |
| *F23F12.3* | WBGene00017751 | protein_coding_gene | 1.5 | 2.920 | 1.7E-04 |
| *gcsh-1* | WBGene00008354 | protein_coding_gene | 1.5 | 2.919 | 4.8E-08 |
| *C01G5.4* | WBGene00015306 | protein_coding_gene | 1.5 | 2.917 | 1.5E-09 |
| *abt-1* | WBGene00000019 | protein_coding_gene | 1.5 | 2.917 | 4.1E-06 |
| *T25B9.1* | WBGene00012007 | protein_coding_gene | 1.5 | 2.917 | 5.3E-12 |
| *nas-38* | WBGene00003554 | protein_coding_gene | 1.5 | 2.915 | 2.2E-02 |
| *K02B12.9* | WBGene00044679 | protein_coding_gene | 1.5 | 2.914 | 2.4E-02 |
| *kvs-5* | WBGene00021948 | protein_coding_gene | 1.5 | 2.914 | 9.1E-13 |
| *snpc-3.1* | WBGene00015098 | protein_coding_gene | 1.5 | 2.913 | 2.0E-06 |
| *F23B12.1* | WBGene00009079 | protein_coding_gene | 1.5 | 2.908 | 9.7E-11 |
| *Y17D7B.10* | WBGene00194982 | protein_coding_gene | 1.5 | 2.904 | 3.0E-02 |
| *dig-1* | WBGene00000998 | protein_coding_gene | 1.5 | 2.902 | 3.1E-15 |
| *ZK896.3* | WBGene00014134 | pseudogene | 1.5 | 2.897 | 8.7E-05 |
| *Y57G7A.11* | WBGene00021974 | protein_coding_gene | 1.5 | 2.891 | 2.8E-02 |
| *anr-39* | WBGene00219696 | antisense_lncRNA_gene | 1.5 | 2.889 | 7.8E-03 |
| *nlp-63* | WBGene00044028 | protein_coding_gene | 1.5 | 2.884 | 2.4E-02 |
| *C45G9.15* | WBGene00050919 | protein_coding_gene | 1.5 | 2.879 | 9.9E-04 |
| *W03G1.2* | WBGene00021010 | protein_coding_gene | 1.5 | 2.878 | 3.6E-08 |
| *mlt-4* | WBGene00013835 | protein_coding_gene | 1.5 | 2.876 | 2.7E-04 |
| *Y69A2AR.14* | WBGene00022085 | pseudogene | 1.5 | 2.872 | 7.8E-05 |
| *F22B3.10* | WBGene00045122 | protein_coding_gene | 1.5 | 2.864 | 3.9E-02 |
| *Y105C5B.14* | WBGene00013653 | protein_coding_gene | 1.5 | 2.863 | 3.0E-02 |
| *Y39F10A.3* | WBGene00021454 | protein_coding_gene | 1.5 | 2.860 | 8.9E-06 |
| *C01C10.2* | WBGene00015294 | protein_coding_gene | 1.5 | 2.853 | 3.3E-05 |
| *ZK105.3* | WBGene00022654 | protein_coding_gene | 1.5 | 2.852 | 4.0E-03 |
| *R13A5.11* | WBGene00020053 | protein_coding_gene | 1.5 | 2.849 | 4.6E-09 |
| *F41G4.5* | WBGene00018309 | pseudogene | 1.5 | 2.848 | 4.8E-02 |
| *K08D10.5* | WBGene00019529 | pseudogene | 1.5 | 2.847 | 1.2E-02 |
| *R11G10.3* | WBGene00044045 | protein_coding_gene | 1.5 | 2.846 | 2.3E-02 |
| *snf-3* | WBGene00004902 | protein_coding_gene | 1.5 | 2.845 | 8.8E-17 |
| *mua-3* | WBGene00003482 | protein_coding_gene | 1.5 | 2.841 | 4.5E-05 |
| *Y66A7A.4* | WBGene00013424 | protein_coding_gene | 1.5 | 2.839 | 8.6E-06 |
| *C05G5.1* | WBGene00007347 | protein_coding_gene | 1.5 | 2.837 | 1.2E-15 |
| *C01G12.13* | WBGene00044333 | protein_coding_gene | 1.5 | 2.831 | 1.8E-04 |
| *C33F10.8* | WBGene00016356 | protein_coding_gene | 1.5 | 2.830 | 1.1E-09 |
| *Y50E8A.6* | WBGene00013049 | protein_coding_gene | 1.5 | 2.830 | 2.2E-03 |
| *Y39G8C.2* | WBGene00012731 | protein_coding_gene | 1.5 | 2.828 | 6.3E-05 |
| *trpl-5* | WBGene00004149 | protein_coding_gene | 1.5 | 2.826 | 4.1E-05 |
| *F32B4.2* | WBGene00009313 | protein_coding_gene | 1.5 | 2.821 | 2.4E-12 |

| | | | | | |
|---|---|---|---|---|---|
| *Y37E3.19* | WBGene00044810 | protein_coding_gene | 1.5 | 2.818 | 8.6E-05 |
| *F32H5.3* | WBGene00009349 | protein_coding_gene | 1.5 | 2.815 | 5.5E-09 |
| *C01H6.4* | WBGene00007254 | protein_coding_gene | 1.5 | 2.814 | 6.0E-11 |
| *ncx-6* | WBGene00003571 | protein_coding_gene | 1.5 | 2.813 | 2.2E-06 |
| *F35E2.9* | WBGene00009416 | protein_coding_gene | 1.5 | 2.813 | 9.8E-12 |
| *svh-1* | WBGene00006620 | protein_coding_gene | 1.5 | 2.811 | 5.0E-03 |
| *Y102A5C.4* | WBGene00013613 | protein_coding_gene | 1.5 | 2.810 | 4.3E-11 |
| *arrd-7* | WBGene00012464 | protein_coding_gene | 1.5 | 2.809 | 1.6E-06 |
| *W04E12.4* | WBGene00012249 | protein_coding_gene | 1.5 | 2.808 | 2.5E-02 |
| *F35F10.1* | WBGene00018052 | protein_coding_gene | 1.5 | 2.805 | 7.3E-06 |
| *Y38H6C.9* | WBGene00012622 | protein_coding_gene | 1.5 | 2.795 | 2.1E-02 |
| *Y82E9BL.12* | WBGene00022328 | protein_coding_gene | 1.5 | 2.795 | 1.9E-02 |
| *F40G12.10* | WBGene00009605 | protein_coding_gene | 1.5 | 2.794 | 1.2E-12 |
| *twk-13* | WBGene00006668 | protein_coding_gene | 1.5 | 2.794 | 2.7E-06 |
| *D2023.1* | WBGene00014300 | protein_coding_gene | 1.5 | 2.794 | 6.1E-09 |
| *Y47D9A.3* | WBGene00021630 | protein_coding_gene | 1.5 | 2.792 | 5.1E-03 |
| *C38D9.5* | WBGene00008013 | pseudogene | 1.5 | 2.790 | 4.2E-04 |
| *rpl-17* | WBGene00004429 | protein_coding_gene | 1.5 | 2.790 | 8.3E-33 |
| *F59E12.3* | WBGene00019119 | protein_coding_gene | 1.5 | 2.787 | 3.4E-02 |
| *F44G3.7* | WBGene00009708 | protein_coding_gene | 1.5 | 2.784 | 3.0E-11 |
| *K06H7.8* | WBGene00019459 | protein_coding_gene | 1.5 | 2.777 | 4.0E-07 |
| *T27A3.5* | WBGene00020841 | protein_coding_gene | 1.5 | 2.773 | 7.9E-11 |
| *skpo-3* | WBGene00017968 | protein_coding_gene | 1.5 | 2.772 | 7.0E-31 |
| *ZK1251.3* | WBGene00014241 | protein_coding_gene | 1.5 | 2.772 | 9.6E-05 |
| *F59A3.8* | WBGene00019081 | protein_coding_gene | 1.5 | 2.770 | 3.5E-07 |
| *F28A10.5* | WBGene00017873 | protein_coding_gene | 1.5 | 2.766 | 1.1E-05 |
| *C18E3.1* | WBGene00015970 | protein_coding_gene | 1.5 | 2.766 | 7.0E-07 |
| *lgc-32* | WBGene00020569 | protein_coding_gene | 1.5 | 2.759 | 5.4E-08 |
| *D1054.8* | WBGene00008375 | protein_coding_gene | 1.5 | 2.757 | 5.1E-18 |
| *Y52D5A.1* | WBGene00021792 | pseudogene | 1.5 | 2.756 | 1.1E-02 |
| *F22E5.8* | WBGene00017707 | protein_coding_gene | 1.5 | 2.755 | 3.8E-04 |
| *C06E2.1* | WBGene00015527 | protein_coding_gene | 1.5 | 2.754 | 1.3E-03 |
| *F07F6.1* | WBGene00017215 | protein_coding_gene | 1.5 | 2.752 | 1.6E-11 |
| *F13D12.9* | WBGene00008743 | protein_coding_gene | 1.5 | 2.750 | 4.3E-15 |
| *R06B10.1* | WBGene00019912 | pseudogene | 1.5 | 2.746 | 2.0E-03 |
| *T15B12.2* | WBGene00020531 | protein_coding_gene | 1.5 | 2.745 | 2.0E-08 |
| *F36F12.7* | WBGene00018098 | protein_coding_gene | 1.5 | 2.740 | 6.6E-04 |
| *dlk-1* | WBGene00001008 | protein_coding_gene | 1.5 | 2.737 | 2.2E-25 |
| *col-42* | WBGene00013489 | protein_coding_gene | 1.5 | 2.736 | 1.7E-18 |
| *ZK596.3* | WBGene00014008 | protein_coding_gene | 1.4 | 2.731 | 2.8E-03 |
| *ZK105.12* | WBGene00235113 | protein_coding_gene | 1.4 | 2.731 | 4.2E-02 |
| *C26C6.10* | WBGene00077553 | protein_coding_gene | 1.4 | 2.730 | 2.4E-02 |
| *hpd-1* | WBGene00001993 | protein_coding_gene | 1.4 | 2.729 | 1.1E-19 |
| *aat-6* | WBGene00000007 | protein_coding_gene | 1.4 | 2.728 | 1.2E-15 |
| *col-87* | WBGene00000662 | protein_coding_gene | 1.4 | 2.727 | 3.8E-02 |
| *mct-4* | WBGene00003986 | protein_coding_gene | 1.4 | 2.723 | 5.1E-23 |
| *acd-3* | WBGene00007775 | protein_coding_gene | 1.4 | 2.723 | 1.8E-06 |
| *Y39A3CL.4* | WBGene00021442 | protein_coding_gene | 1.4 | 2.719 | 6.7E-35 |
| *Y69E1A.8* | WBGene00013478 | protein_coding_gene | 1.4 | 2.710 | 2.3E-06 |
| *spe-43* | WBGene00008623 | protein_coding_gene | 1.4 | 2.710 | 9.2E-03 |
| *tdo-2* | WBGene00016201 | protein_coding_gene | 1.4 | 2.710 | 7.1E-12 |
| *F21H7.2* | WBGene00009028 | protein_coding_gene | 1.4 | 2.708 | 3.1E-05 |
| *cima-1* | WBGene00018473 | protein_coding_gene | 1.4 | 2.707 | 8.8E-06 |
| *F20H11.4* | WBGene00017647 | protein_coding_gene | 1.4 | 2.704 | 1.2E-13 |
| *F37H8.2* | WBGene00009511 | protein_coding_gene | 1.4 | 2.699 | 1.7E-06 |
| *snn-1* | WBGene00004913 | protein_coding_gene | 1.4 | 2.697 | 5.3E-04 |
| *nep-17* | WBGene00010070 | protein_coding_gene | 1.4 | 2.695 | 4.8E-34 |
| *kin-24* | WBGene00002206 | protein_coding_gene | 1.4 | 2.694 | 2.1E-06 |
| *hrg-6* | WBGene00009495 | protein_coding_gene | 1.4 | 2.693 | 1.9E-03 |
| *dsc-4* | WBGene00001099 | protein_coding_gene | 1.4 | 2.689 | 3.4E-23 |
| *Y22D7AR.2* | WBGene00021256 | protein_coding_gene | 1.4 | 2.688 | 1.1E-09 |
| *npr-23* | WBGene00021328 | protein_coding_gene | 1.4 | 2.687 | 4.3E-04 |
| *glb-16* | WBGene00018486 | protein_coding_gene | 1.4 | 2.686 | 2.4E-02 |
| *F49D11.10* | WBGene00045433 | protein_coding_gene | 1.4 | 2.682 | 5.1E-18 |
| *E04D5.5* | WBGene00008484 | protein_coding_gene | 1.4 | 2.682 | 4.1E-06 |
| *bgnt-1.4* | WBGene00015982 | protein_coding_gene | 1.4 | 2.680 | 4.4E-03 |
| *Y17G9A.3* | WBGene00021196 | protein_coding_gene | 1.4 | 2.678 | 3.7E-02 |
| *F54F12.1* | WBGene00010072 | protein_coding_gene | 1.4 | 2.676 | 1.8E-07 |
| *F27E5.3* | WBGene00009193 | protein_coding_gene | 1.4 | 2.675 | 2.7E-08 |
| *B0280.2* | WBGene00015100 | protein_coding_gene | 1.4 | 2.674 | 9.0E-06 |
| *F29D10.1* | WBGene00009250 | protein_coding_gene | 1.4 | 2.672 | 2.7E-03 |
| *fut-5* | WBGene00043986 | protein_coding_gene | 1.4 | 2.672 | 9.4E-03 |
| *cdc-25.4* | WBGene00000389 | protein_coding_gene | 1.4 | 2.666 | 3.0E-05 |
| *oat-1* | WBGene00003837 | protein_coding_gene | 1.4 | 2.666 | 3.7E-07 |
| *T10G3.3* | WBGene00011694 | protein_coding_gene | 1.4 | 2.664 | 5.9E-07 |
| *alg-4* | WBGene00006449 | protein_coding_gene | 1.4 | 2.663 | 2.2E-15 |
| *F44D12.11* | WBGene00014375 | pseudogene | 1.4 | 2.661 | 7.4E-04 |
| *pah-1* | WBGene00000240 | protein_coding_gene | 1.4 | 2.656 | 3.4E-31 |
| *clec-187* | WBGene00014137 | protein_coding_gene | 1.4 | 2.656 | 1.9E-09 |
| *Y54G2A.40* | WBGene00044495 | protein_coding_gene | 1.4 | 2.647 | 1.0E-02 |

| | | | | | |
|---|---|---|---|---|---|
| *slc-9B.2* | WBGene00010214 | protein_coding_gene | 1.4 | 2.644 | 6.1E-04 |
| *snpc-3.2* | WBGene00021667 | protein_coding_gene | 1.4 | 2.643 | 3.5E-03 |
| *trpl-2* | WBGene00020972 | protein_coding_gene | 1.4 | 2.642 | 2.8E-06 |
| *F28A10.4* | WBGene00017872 | protein_coding_gene | 1.4 | 2.642 | 3.7E-03 |
| *Y47D9A.5* | WBGene00021632 | protein_coding_gene | 1.4 | 2.639 | 3.6E-03 |
| *C49A9.9* | WBGene00016763 | protein_coding_gene | 1.4 | 2.628 | 6.6E-18 |
| *elo-7* | WBGene00001245 | protein_coding_gene | 1.4 | 2.626 | 1.6E-06 |
| *cyp-34A7* | WBGene00015042 | protein_coding_gene | 1.4 | 2.623 | 6.2E-03 |
| *twk-10* | WBGene00006665 | protein_coding_gene | 1.4 | 2.623 | 2.7E-02 |
| *gst-36* | WBGene00001784 | protein_coding_gene | 1.4 | 2.622 | 9.2E-15 |
| *twk-17* | WBGene00006671 | protein_coding_gene | 1.4 | 2.622 | 2.1E-04 |
| *ZK742.3* | WBGene00022813 | protein_coding_gene | 1.4 | 2.618 | 1.4E-06 |
| *Y59E9AL.8* | WBGene00044346 | protein_coding_gene | 1.4 | 2.616 | 4.9E-02 |
| *sulp-6* | WBGene00020914 | protein_coding_gene | 1.4 | 2.616 | 3.4E-07 |
| *Y76B12C.8* | WBGene00022302 | protein_coding_gene | 1.4 | 2.605 | 7.2E-03 |
| *del-6* | WBGene00011891 | protein_coding_gene | 1.4 | 2.605 | 9.0E-21 |
| *gsto-1* | WBGene00016204 | protein_coding_gene | 1.4 | 2.605 | 3.2E-07 |
| *T23B7.3* | WBGene00044767 | protein_coding_gene | 1.4 | 2.603 | 4.0E-04 |
| *Y54E10A.17* | WBGene00021835 | protein_coding_gene | 1.4 | 2.600 | 8.5E-14 |
| *ubl-1* | WBGene00006725 | protein_coding_gene | 1.4 | 2.599 | 4.8E-26 |
| *spe-19* | WBGene00013747 | protein_coding_gene | 1.4 | 2.596 | 1.4E-04 |
| *chk-2* | WBGene00000499 | protein_coding_gene | 1.4 | 2.596 | 1.0E-10 |
| *unc-105* | WBGene00006832 | protein_coding_gene | 1.4 | 2.587 | 8.8E-08 |
| *K12C11.5* | WBGene00044896 | protein_coding_gene | 1.4 | 2.582 | 1.2E-02 |
| *C48D1.9* | WBGene00194643 | protein_coding_gene | 1.4 | 2.582 | 1.6E-04 |
| *rpl-13* | WBGene00004425 | protein_coding_gene | 1.4 | 2.582 | 2.0E-22 |
| *spe-47* | WBGene00021271 | protein_coding_gene | 1.4 | 2.580 | 2.6E-05 |
| *ZK507.3* | WBGene00013979 | protein_coding_gene | 1.4 | 2.578 | 5.6E-04 |
| *dhhc-5* | WBGene00020066 | protein_coding_gene | 1.4 | 2.573 | 4.3E-03 |
| *arrd-10* | WBGene00010268 | protein_coding_gene | 1.4 | 2.573 | 1.1E-04 |
| *osta-1* | WBGene00015287 | protein_coding_gene | 1.4 | 2.572 | 4.2E-05 |
| *mec-5* | WBGene00003169 | protein_coding_gene | 1.4 | 2.568 | 9.7E-08 |
| *Y1A5A.1* | WBGene00012379 | protein_coding_gene | 1.4 | 2.567 | 2.9E-10 |
| *T07A5.7* | WBGene00045475 | protein_coding_gene | 1.4 | 2.565 | 4.8E-03 |
| *rps-10* | WBGene00004479 | protein_coding_gene | 1.4 | 2.563 | 7.2E-27 |
| *pho-7* | WBGene00011879 | protein_coding_gene | 1.4 | 2.561 | 6.2E-05 |
| *Y54G11A.7* | WBGene00013216 | protein_coding_gene | 1.4 | 2.557 | 1.7E-18 |
| *C44B9.2* | WBGene00008079 | protein_coding_gene | 1.4 | 2.554 | 7.8E-09 |
| *fbxa-166* | WBGene00015601 | protein_coding_gene | 1.4 | 2.552 | 2.2E-02 |
| *F38A5.6* | WBGene00018163 | protein_coding_gene | 1.3 | 2.537 | 9.9E-05 |
| *C34D4.10* | WBGene00016402 | protein_coding_gene | 1.3 | 2.535 | 1.0E-20 |
| *nhx-3* | WBGene00003731 | protein_coding_gene | 1.3 | 2.531 | 1.2E-06 |
| *Y45F10B.2* | WBGene00012870 | pseudogene | 1.3 | 2.531 | 1.9E-02 |
| *C31H1.1* | WBGene00016286 | protein_coding_gene | 1.3 | 2.529 | 2.5E-07 |
| *Y75B7B.1* | WBGene00022288 | protein_coding_gene | 1.3 | 2.528 | 2.2E-03 |
| *tag-191* | WBGene00007049 | protein_coding_gene | 1.3 | 2.526 | 2.8E-05 |
| *Y113G7C.1* | WBGene00013771 | protein_coding_gene | 1.3 | 2.522 | 4.9E-17 |
| *Y12A6A.1* | WBGene00012439 | protein_coding_gene | 1.3 | 2.518 | 1.2E-09 |
| *C34D4.2* | WBGene00016398 | protein_coding_gene | 1.3 | 2.515 | 9.7E-08 |
| *nhr-1* | WBGene00003600 | protein_coding_gene | 1.3 | 2.514 | 6.9E-22 |
| *rps-5* | WBGene00004474 | protein_coding_gene | 1.3 | 2.514 | 9.1E-22 |
| *Y67A10A.3* | WBGene00013452 | protein_coding_gene | 1.3 | 2.512 | 2.7E-05 |
| *ZC513.3* | WBGene00022628 | protein_coding_gene | 1.3 | 2.510 | 7.5E-05 |
| *T21D12.12* | WBGene00020651 | protein_coding_gene | 1.3 | 2.507 | 2.3E-11 |
| *pxn-1* | WBGene00004256 | protein_coding_gene | 1.3 | 2.504 | 3.0E-16 |
| *C44C10.3* | WBGene00008084 | protein_coding_gene | 1.3 | 2.502 | 7.5E-06 |
| *F10G8.1* | WBGene00008661 | protein_coding_gene | 1.3 | 2.496 | 2.2E-06 |
| *F28H7.8* | WBGene00009241 | protein_coding_gene | 1.3 | 2.496 | 1.4E-05 |
| *Y102A11A.9* | WBGene00022419 | protein_coding_gene | 1.3 | 2.495 | 1.2E-04 |
| *linc-8* | WBGene00219707 | lincRNA_gene | 1.3 | 2.495 | 9.8E-10 |
| *vha-6* | WBGene00006915 | protein_coding_gene | 1.3 | 2.494 | 1.3E-15 |
| *ZK39.10* | WBGene00050881 | protein_coding_gene | 1.3 | 2.493 | 1.0E-02 |
| *rpl-18* | WBGene00004430 | protein_coding_gene | 1.3 | 2.491 | 2.6E-23 |
| *F01D5.3* | WBGene00008494 | protein_coding_gene | 1.3 | 2.490 | 2.0E-02 |
| *T05F1.5* | WBGene00011491 | protein_coding_gene | 1.3 | 2.489 | 6.6E-03 |
| *Y54F10AM.8* | WBGene00021852 | protein_coding_gene | 1.3 | 2.486 | 1.6E-11 |
| *mca-2* | WBGene00019875 | protein_coding_gene | 1.3 | 2.485 | 5.0E-22 |
| *C29G2.6* | WBGene00016234 | protein_coding_gene | 1.3 | 2.479 | 3.9E-12 |
| *rpl-19* | WBGene00004431 | protein_coding_gene | 1.3 | 2.474 | 1.4E-24 |
| *fbxa-58* | WBGene00016225 | pseudogene | 1.3 | 2.474 | 4.6E-08 |
| *unc-132* | WBGene00021078 | protein_coding_gene | 1.3 | 2.473 | 2.2E-22 |
| *F46G10.4* | WBGene00009798 | protein_coding_gene | 1.3 | 2.472 | 5.9E-06 |
| *F28B4.3* | WBGene00017892 | protein_coding_gene | 1.3 | 2.472 | 1.8E-02 |
| *spe-4* | WBGene00004958 | protein_coding_gene | 1.3 | 2.468 | 5.4E-10 |
| *ZK1053.6* | WBGene00014201 | protein_coding_gene | 1.3 | 2.465 | 1.2E-03 |
| *Y58A7A.1* | WBGene00021975 | protein_coding_gene | 1.3 | 2.460 | 3.0E-08 |
| *C14A6.13* | WBGene00050956 | protein_coding_gene | 1.3 | 2.460 | 7.1E-07 |
| *lgc-45* | WBGene00021146 | protein_coding_gene | 1.3 | 2.458 | 5.5E-10 |
| *K10C9.7* | WBGene00019626 | protein_coding_gene | 1.3 | 2.456 | 1.7E-06 |
| *adt-3* | WBGene00020567 | protein_coding_gene | 1.3 | 2.453 | 6.5E-05 |

| | | | | | |
|---|---|---|---|---|---|
| *R04B5.11* | WBGene00011008 | protein_coding_gene | 1.3 | 2.450 | 2.2E-02 |
| *far-5* | WBGene00001389 | protein_coding_gene | 1.3 | 2.449 | 2.1E-05 |
| *cest-32* | WBGene00016862 | protein_coding_gene | 1.3 | 2.443 | 1.4E-16 |
| *rpl-4* | WBGene00004415 | protein_coding_gene | 1.3 | 2.441 | 2.9E-23 |
| *K02D3.1* | WBGene00010504 | protein_coding_gene | 1.3 | 2.438 | 2.9E-04 |
| *tag-344* | WBGene00015230 | protein_coding_gene | 1.3 | 2.436 | 5.8E-06 |
| *fbxa-89* | WBGene00013555 | protein_coding_gene | 1.3 | 2.435 | 1.1E-02 |
| *ifa-1* | WBGene00002050 | protein_coding_gene | 1.3 | 2.430 | 4.7E-20 |
| *T25C12.3* | WBGene00012018 | protein_coding_gene | 1.3 | 2.429 | 1.5E-03 |
| *add-1* | WBGene00000072 | protein_coding_gene | 1.3 | 2.427 | 1.5E-09 |
| *pfas-1* | WBGene00008654 | protein_coding_gene | 1.3 | 2.427 | 6.6E-17 |
| *C55C3.3* | WBGene00016953 | protein_coding_gene | 1.3 | 2.425 | 6.1E-08 |
| *R05G6.5* | WBGene00019899 | protein_coding_gene | 1.3 | 2.422 | 1.4E-03 |
| *T01H3.5* | WBGene00011351 | protein_coding_gene | 1.3 | 2.421 | 1.5E-04 |
| *cpr-5* | WBGene00000785 | protein_coding_gene | 1.3 | 2.418 | 4.5E-04 |
| *csp-2* | WBGene00000820 | protein_coding_gene | 1.3 | 2.417 | 6.2E-03 |
| *F37A4.5* | WBGene00018135 | protein_coding_gene | 1.3 | 2.416 | 1.5E-08 |
| *R102.10* | WBGene00011297 | protein_coding_gene | 1.3 | 2.413 | 3.6E-06 |
| *B0496.1* | WBGene00015210 | protein_coding_gene | 1.3 | 2.412 | 1.2E-03 |
| *T25B9.4* | WBGene00012010 | protein_coding_gene | 1.3 | 2.411 | 5.5E-04 |
| *gpa-13* | WBGene00001675 | protein_coding_gene | 1.3 | 2.408 | 3.9E-02 |
| *rhr-1* | WBGene00004358 | protein_coding_gene | 1.3 | 2.404 | 5.2E-28 |
| *ttr-31* | WBGene00010225 | protein_coding_gene | 1.3 | 2.404 | 6.9E-07 |
| *twk-12* | WBGene00006667 | protein_coding_gene | 1.3 | 2.403 | 8.5E-04 |
| *C37H5.14* | WBGene00045308 | protein_coding_gene | 1.3 | 2.403 | 5.5E-03 |
| *aakg-3* | WBGene00021527 | protein_coding_gene | 1.3 | 2.401 | 8.1E-04 |
| *unc-22* | WBGene00006759 | protein_coding_gene | 1.3 | 2.399 | 8.2E-07 |
| *F46A8.6* | WBGene00009749 | protein_coding_gene | 1.3 | 2.397 | 2.3E-02 |
| *maph-1.1* | WBGene00009306 | protein_coding_gene | 1.3 | 2.397 | 1.1E-22 |
| *srh-48* | WBGene00005271 | protein_coding_gene | 1.3 | 2.396 | 4.1E-03 |
| *W03D8.2* | WBGene00020985 | protein_coding_gene | 1.3 | 2.393 | 4.7E-07 |
| *F47G6.3* | WBGene00018580 | protein_coding_gene | 1.3 | 2.389 | 1.7E-03 |
| *rpl-5* | WBGene00004416 | protein_coding_gene | 1.3 | 2.389 | 6.6E-26 |
| *C10E2.5* | WBGene00015675 | protein_coding_gene | 1.3 | 2.387 | 3.4E-02 |
| *rpl-15* | WBGene00004427 | protein_coding_gene | 1.3 | 2.387 | 1.9E-21 |
| *zmp-3* | WBGene00016283 | protein_coding_gene | 1.3 | 2.386 | 4.6E-08 |
| *T02E1.7* | WBGene00011379 | protein_coding_gene | 1.3 | 2.386 | 3.0E-08 |
| *hpo-32* | WBGene00015313 | protein_coding_gene | 1.3 | 2.385 | 4.0E-25 |
| *ketn-1* | WBGene00004130 | protein_coding_gene | 1.3 | 2.385 | 1.4E-14 |
| *C17C3.15* | WBGene00015896 | protein_coding_gene | 1.3 | 2.384 | 6.7E-06 |
| *nca-2* | WBGene00003558 | protein_coding_gene | 1.3 | 2.384 | 1.5E-04 |
| *C50D2.3* | WBGene00016807 | protein_coding_gene | 1.3 | 2.382 | 8.0E-04 |
| *gnrr-1* | WBGene00018798 | protein_coding_gene | 1.3 | 2.379 | 4.7E-02 |
| *Y39B6A.21* | WBGene00012680 | pseudogene | 1.3 | 2.379 | 1.9E-02 |
| *M02B1.2* | WBGene00010827 | protein_coding_gene | 1.3 | 2.379 | 3.3E-10 |
| *doxa-1* | WBGene00015519 | protein_coding_gene | 1.2 | 2.378 | 1.3E-05 |
| *ZK1251.5* | WBGene00014996 | pseudogene | 1.2 | 2.375 | 1.8E-03 |
| *fbxa-63* | WBGene00022485 | protein_coding_gene | 1.2 | 2.374 | 1.7E-04 |
| *egl-23* | WBGene00001190 | protein_coding_gene | 1.2 | 2.370 | 5.8E-06 |
| *Y51H4A.13* | WBGene00013107 | protein_coding_gene | 1.2 | 2.369 | 4.5E-04 |
| *rpl-31* | WBGene00004445 | protein_coding_gene | 1.2 | 2.366 | 2.6E-22 |
| *C35A5.4* | WBGene00007948 | protein_coding_gene | 1.2 | 2.366 | 1.2E-04 |
| *C49A9.6* | WBGene00016760 | protein_coding_gene | 1.2 | 2.365 | 8.8E-13 |
| *F47E1.2* | WBGene00018566 | protein_coding_gene | 1.2 | 2.362 | 4.1E-05 |
| *F21G4.5* | WBGene00009026 | protein_coding_gene | 1.2 | 2.361 | 4.2E-03 |
| *mai-1* | WBGene00003124 | protein_coding_gene | 1.2 | 2.361 | 1.2E-06 |
| *B0222.11* | WBGene00044582 | protein_coding_gene | 1.2 | 2.361 | 2.1E-02 |
| *R11A5.6* | WBGene00011234 | protein_coding_gene | 1.2 | 2.360 | 3.5E-02 |
| *F23B2.7* | WBGene00009075 | protein_coding_gene | 1.2 | 2.359 | 9.8E-05 |
| *msp-42* | WBGene00003437 | pseudogene | 1.2 | 2.358 | 3.1E-03 |
| *his-73* | WBGene00001947 | protein_coding_gene | 1.2 | 2.355 | 3.7E-02 |
| *cysl-4* | WBGene00019096 | protein_coding_gene | 1.2 | 2.353 | 4.3E-03 |
| *pbo-1* | WBGene00003941 | protein_coding_gene | 1.2 | 2.353 | 5.2E-13 |
| *K03B8.8* | WBGene00010525 | protein_coding_gene | 1.2 | 2.350 | 9.3E-05 |
| *col-91* | WBGene00000666 | protein_coding_gene | 1.2 | 2.347 | 8.6E-03 |
| *slc-17.5* | WBGene00017681 | protein_coding_gene | 1.2 | 2.344 | 1.8E-07 |
| *irk-2* | WBGene00002150 | protein_coding_gene | 1.2 | 2.343 | 6.4E-03 |
| *ZC581.7* | WBGene00022634 | protein_coding_gene | 1.2 | 2.343 | 1.2E-06 |
| *R03D7.8* | WBGene00010992 | protein_coding_gene | 1.2 | 2.343 | 3.4E-05 |
| *ZK507.1* | WBGene00013978 | protein_coding_gene | 1.2 | 2.343 | 1.4E-04 |
| *ZK455.5* | WBGene00013975 | protein_coding_gene | 1.2 | 2.343 | 1.0E-02 |
| *mpz-4* | WBGene00019165 | protein_coding_gene | 1.2 | 2.342 | 2.2E-06 |
| *ZK470.2* | WBGene00022745 | protein_coding_gene | 1.2 | 2.342 | 6.6E-18 |
| *F35E2.10* | WBGene00044270 | pseudogene | 1.2 | 2.341 | 8.2E-04 |
| *nrfl-1* | WBGene00006438 | protein_coding_gene | 1.2 | 2.341 | 3.9E-16 |
| *wht-5* | WBGene00008950 | protein_coding_gene | 1.2 | 2.339 | 2.1E-08 |
| *F25A2.1* | WBGene00017764 | protein_coding_gene | 1.2 | 2.339 | 1.3E-02 |
| *npr-16* | WBGene00006864 | protein_coding_gene | 1.2 | 2.337 | 1.3E-02 |
| *rps-22* | WBGene00004491 | protein_coding_gene | 1.2 | 2.336 | 3.4E-16 |
| *rpl-43* | WBGene00004456 | protein_coding_gene | 1.2 | 2.333 | 5.4E-15 |

| | | | | | |
|---|---|---|---|---|---|
| *fbxa-100* | WBGene00008826 | protein_coding_gene | 1.2 | 2.331 | 8.6E-04 |
| *ttn-1* | WBGene00006436 | protein_coding_gene | 1.2 | 2.324 | 8.1E-04 |
| *C52E4.7* | WBGene00008259 | protein_coding_gene | 1.2 | 2.324 | 1.4E-06 |
| *F11F1.1* | WBGene00008714 | protein_coding_gene | 1.2 | 2.324 | 4.8E-06 |
| *ZC190.8* | WBGene00022542 | protein_coding_gene | 1.2 | 2.323 | 4.4E-03 |
| *glb-29* | WBGene00012461 | protein_coding_gene | 1.2 | 2.322 | 2.8E-02 |
| *C38C3.8* | WBGene00016516 | pseudogene | 1.2 | 2.321 | 2.1E-04 |
| *hrg-1* | WBGene00019830 | protein_coding_gene | 1.2 | 2.318 | 1.5E-08 |
| *abt-2* | WBGene00000020 | protein_coding_gene | 1.2 | 2.317 | 1.2E-18 |
| *ttr-9* | WBGene00007888 | protein_coding_gene | 1.2 | 2.315 | 1.0E-03 |
| *ndfl-4* | WBGene00010958 | protein_coding_gene | 1.2 | 2.314 | 2.2E-02 |
| *T25B9.6* | WBGene00012012 | protein_coding_gene | 1.2 | 2.314 | 4.6E-09 |
| *C07A9.9* | WBGene00007405 | protein_coding_gene | 1.2 | 2.314 | 3.1E-06 |
| *cnnm-5* | WBGene00011260 | protein_coding_gene | 1.2 | 2.313 | 7.6E-04 |
| *exc-1* | WBGene00001362 | protein_coding_gene | 1.2 | 2.310 | 4.6E-02 |
| *C09H10.9* | WBGene00007503 | protein_coding_gene | 1.2 | 2.310 | 1.1E-04 |
| *anr-35* | WBGene00219589 | antisense_lncRNA_gene | 1.2 | 2.310 | 8.5E-04 |
| *T16A1.2* | WBGene00020533 | protein_coding_gene | 1.2 | 2.309 | 4.6E-03 |
| *rpl-1* | WBGene00004412 | protein_coding_gene | 1.2 | 2.309 | 3.0E-19 |
| *acox-1.2* | WBGene00008565 | protein_coding_gene | 1.2 | 2.305 | 4.1E-05 |
| *scrm-7* | WBGene00013052 | protein_coding_gene | 1.2 | 2.304 | 5.8E-03 |
| *T04F3.1* | WBGene00011436 | protein_coding_gene | 1.2 | 2.303 | 1.5E-21 |
| *Y43C5B.2* | WBGene00012785 | protein_coding_gene | 1.2 | 2.302 | 2.0E-05 |
| *AH6.3* | WBGene00007081 | protein_coding_gene | 1.2 | 2.301 | 8.8E-05 |
| *K10D11.5* | WBGene00010749 | protein_coding_gene | 1.2 | 2.297 | 3.7E-03 |
| *B0403.3* | WBGene00015167 | protein_coding_gene | 1.2 | 2.294 | 9.5E-04 |
| *nhr-80* | WBGene00003670 | protein_coding_gene | 1.2 | 2.293 | 8.2E-08 |
| *C55A6.3* | WBGene00008332 | protein_coding_gene | 1.2 | 2.292 | 2.3E-06 |
| *C18D11.1* | WBGene00007679 | protein_coding_gene | 1.2 | 2.292 | 7.8E-08 |
| *clec-143* | WBGene00014063 | protein_coding_gene | 1.2 | 2.291 | 4.0E-02 |
| *F56H1.3* | WBGene00018994 | protein_coding_gene | 1.2 | 2.289 | 4.7E-03 |
| *R160.3* | WBGene00020120 | protein_coding_gene | 1.2 | 2.289 | 5.5E-03 |
| *fmo-5* | WBGene00001480 | protein_coding_gene | 1.2 | 2.287 | 2.7E-09 |
| *F36D3.8* | WBGene00009473 | protein_coding_gene | 1.2 | 2.287 | 1.5E-03 |
| *egl-9* | WBGene00001178 | protein_coding_gene | 1.2 | 2.285 | 1.0E-18 |
| *B0252.1* | WBGene00015087 | protein_coding_gene | 1.2 | 2.285 | 4.3E-05 |
| *Y50E8A.10* | WBGene00013053 | protein_coding_gene | 1.2 | 2.280 | 6.1E-04 |
| *unc-77* | WBGene00006809 | protein_coding_gene | 1.2 | 2.276 | 1.4E-05 |
| *BE10.1* | WBGene00007209 | protein_coding_gene | 1.2 | 2.275 | 4.1E-05 |
| *col-12* | WBGene00000601 | protein_coding_gene | 1.2 | 2.272 | 1.9E-02 |
| *ZK1055.6* | WBGene00022847 | protein_coding_gene | 1.2 | 2.271 | 1.7E-10 |
| *Y58A7A.5* | WBGene00021979 | protein_coding_gene | 1.2 | 2.269 | 4.1E-03 |
| *Y24D9B.1* | WBGene00021287 | protein_coding_gene | 1.2 | 2.269 | 5.5E-06 |
| *ZK688.12* | WBGene00044763 | protein_coding_gene | 1.2 | 2.268 | 3.6E-03 |
| *ugt-16* | WBGene00013901 | protein_coding_gene | 1.2 | 2.267 | 3.2E-06 |
| *F55F8.7* | WBGene00018895 | protein_coding_gene | 1.2 | 2.267 | 4.3E-07 |
| *F36A4.3* | WBGene00018082 | protein_coding_gene | 1.2 | 2.267 | 1.3E-03 |
| *rpl-7A* | WBGene00004419 | protein_coding_gene | 1.2 | 2.266 | 3.6E-20 |
| *W03F8.2* | WBGene00020992 | pseudogene | 1.2 | 2.265 | 4.2E-04 |
| *Y39E4B.13* | WBGene00012721 | protein_coding_gene | 1.2 | 2.261 | 6.1E-03 |
| *grl-4* | WBGene00001713 | protein_coding_gene | 1.2 | 2.260 | 7.3E-06 |
| *ZC404.1* | WBGene00022601 | protein_coding_gene | 1.2 | 2.252 | 4.1E-02 |
| *C09G5.7* | WBGene00007489 | protein_coding_gene | 1.2 | 2.252 | 2.8E-04 |
| *F40F11.4* | WBGene00009589 | protein_coding_gene | 1.2 | 2.251 | 2.4E-02 |
| *Y77E11A.12* | WBGene00022313 | protein_coding_gene | 1.2 | 2.251 | 8.1E-03 |
| *linc-76* | WBGene00219619 | lincRNA_gene | 1.2 | 2.243 | 4.1E-02 |
| *T18D3.6* | WBGene00011823 | protein_coding_gene | 1.2 | 2.243 | 6.3E-07 |
| *F40F12.3* | WBGene00009592 | protein_coding_gene | 1.2 | 2.243 | 5.6E-05 |
| *lgc-30* | WBGene00019069 | protein_coding_gene | 1.2 | 2.242 | 3.2E-02 |
| *tig-2* | WBGene00006570 | protein_coding_gene | 1.2 | 2.240 | 6.6E-07 |
| *Y56A3A.30* | WBGene00013242 | protein_coding_gene | 1.2 | 2.239 | 2.9E-09 |
| *col-146* | WBGene00000719 | protein_coding_gene | 1.2 | 2.238 | 3.8E-02 |
| *K07H8.5* | WBGene00019506 | protein_coding_gene | 1.2 | 2.235 | 5.4E-05 |
| *ugt-6* | WBGene00013904 | protein_coding_gene | 1.2 | 2.233 | 2.1E-06 |
| *Y80D3A.8* | WBGene00013586 | protein_coding_gene | 1.2 | 2.233 | 4.0E-03 |
| *nlp-40* | WBGene00022276 | protein_coding_gene | 1.2 | 2.233 | 1.5E-17 |
| *F14F7.4* | WBGene00008811 | protein_coding_gene | 1.2 | 2.231 | 4.5E-05 |
| *gly-8* | WBGene00001633 | protein_coding_gene | 1.2 | 2.231 | 5.5E-06 |
| *F22B3.5* | WBGene00009036 | protein_coding_gene | 1.2 | 2.230 | 6.3E-06 |
| *arl-7* | WBGene00000191 | pseudogene | 1.2 | 2.230 | 2.7E-02 |
| *F10D11.6* | WBGene00008652 | protein_coding_gene | 1.2 | 2.228 | 3.9E-02 |
| *W03D8.8* | WBGene00020989 | protein_coding_gene | 1.2 | 2.226 | 2.8E-04 |
| *F47F2.1* | WBGene00018569 | protein_coding_gene | 1.2 | 2.225 | 5.3E-06 |
| *C23G10.1* | WBGene00016010 | protein_coding_gene | 1.2 | 2.224 | 1.2E-04 |
| *C53A5.13* | WBGene00008270 | protein_coding_gene | 1.2 | 2.222 | 4.6E-08 |
| *ZC196.2* | WBGene00022546 | protein_coding_gene | 1.2 | 2.222 | 1.7E-02 |
| *enu-3.5* | WBGene00012552 | protein_coding_gene | 1.2 | 2.221 | 2.9E-02 |
| *W07E6.2* | WBGene00021074 | protein_coding_gene | 1.2 | 2.221 | 3.6E-12 |
| *math-47* | WBGene00022688 | pseudogene | 1.2 | 2.220 | 5.0E-04 |
| *dmsr-2* | WBGene00021275 | protein_coding_gene | 1.1 | 2.217 | 2.5E-03 |

| | | | | | |
|---|---|---|---|---|---|
| *H09G03.1* | WBGene00019174 | protein_coding_gene | 1.1 | 2.217 | 2.5E-02 |
| *T22D1.8* | WBGene00020686 | protein_coding_gene | 1.1 | 2.214 | 2.8E-02 |
| *ttr-49* | WBGene00023485 | protein_coding_gene | 1.1 | 2.214 | 4.5E-07 |
| *math-48* | WBGene00022689 | protein_coding_gene | 1.1 | 2.213 | 2.4E-03 |
| *slc-17.7* | WBGene00022788 | protein_coding_gene | 1.1 | 2.212 | 1.6E-11 |
| *rps-29* | WBGene00004498 | protein_coding_gene | 1.1 | 2.208 | 6.7E-16 |
| *btb-14* | WBGene00013275 | protein_coding_gene | 1.1 | 2.207 | 4.7E-02 |
| *C05C12.1* | WBGene00007335 | protein_coding_gene | 1.1 | 2.206 | 3.9E-05 |
| *C31G12.4* | WBGene00007853 | protein_coding_gene | 1.1 | 2.206 | 4.4E-02 |
| *best-21* | WBGene00013520 | protein_coding_gene | 1.1 | 2.205 | 8.5E-05 |
| *F45E4.6* | WBGene00018471 | protein_coding_gene | 1.1 | 2.204 | 3.3E-04 |
| *W01A8.6* | WBGene00012168 | protein_coding_gene | 1.1 | 2.199 | 2.2E-04 |
| *ZK930.7* | WBGene00014157 | protein_coding_gene | 1.1 | 2.198 | 4.7E-04 |
| *rps-20* | WBGene00004489 | protein_coding_gene | 1.1 | 2.198 | 3.2E-21 |
| *fbxa-90* | WBGene00013623 | protein_coding_gene | 1.1 | 2.196 | 8.5E-04 |
| *clpr-1* | WBGene00012233 | protein_coding_gene | 1.1 | 2.195 | 1.8E-02 |
| *ZK816.1* | WBGene00022825 | protein_coding_gene | 1.1 | 2.195 | 1.2E-02 |
| *F15D4.5* | WBGene00008862 | protein_coding_gene | 1.1 | 2.194 | 3.3E-04 |
| *T08B1.1* | WBGene00020340 | protein_coding_gene | 1.1 | 2.192 | 2.3E-03 |
| *rla-0* | WBGene00004408 | protein_coding_gene | 1.1 | 2.188 | 7.0E-20 |
| *Y71G12B.35* | WBGene00185098 | protein_coding_gene | 1.1 | 2.185 | 3.4E-07 |
| *nkb-3* | WBGene00010117 | protein_coding_gene | 1.1 | 2.185 | 3.0E-20 |
| *T16G1.5* | WBGene00011799 | protein_coding_gene | 1.1 | 2.183 | 6.4E-04 |
| *lev-1* | WBGene00002974 | protein_coding_gene | 1.1 | 2.182 | 9.5E-05 |
| *C51E3.9* | WBGene00008254 | protein_coding_gene | 1.1 | 2.181 | 8.3E-10 |
| *pmp-4* | WBGene00004061 | protein_coding_gene | 1.1 | 2.181 | 9.1E-09 |
| *cbs-2* | WBGene00018783 | protein_coding_gene | 1.1 | 2.180 | 6.0E-05 |
| *rps-21* | WBGene00004490 | protein_coding_gene | 1.1 | 2.177 | 3.4E-16 |
| *T16G12.7* | WBGene00011808 | protein_coding_gene | 1.1 | 2.173 | 3.5E-05 |
| *ZC21.8* | WBGene00022507 | protein_coding_gene | 1.1 | 2.172 | 9.6E-03 |
| *Y54E2A.7* | WBGene00013192 | protein_coding_gene | 1.1 | 2.170 | 3.7E-04 |
| *F55G11.1* | WBGene00010122 | protein_coding_gene | 1.1 | 2.169 | 1.4E-02 |
| *T05C3.2* | WBGene00020254 | protein_coding_gene | 1.1 | 2.165 | 3.4E-02 |
| *kin-26* | WBGene00002208 | protein_coding_gene | 1.1 | 2.163 | 7.0E-05 |
| *B0207.1* | WBGene00015026 | protein_coding_gene | 1.1 | 2.163 | 1.1E-03 |
| *nhr-130* | WBGene00003720 | protein_coding_gene | 1.1 | 2.161 | 4.7E-06 |
| *cyp-29A3* | WBGene00021412 | protein_coding_gene | 1.1 | 2.159 | 6.1E-04 |
| *F59A6.2* | WBGene00019084 | protein_coding_gene | 1.1 | 2.157 | 1.0E-03 |
| *acr-16* | WBGene00000055 | protein_coding_gene | 1.1 | 2.155 | 5.8E-03 |
| *B0285.6* | WBGene00007138 | protein_coding_gene | 1.1 | 2.151 | 1.1E-02 |
| *F58D5.7* | WBGene00010246 | protein_coding_gene | 1.1 | 2.150 | 4.4E-07 |
| *F43G6.17* | WBGene00219378 | protein_coding_gene | 1.1 | 2.147 | 7.3E-03 |
| *glc-2* | WBGene00001592 | protein_coding_gene | 1.1 | 2.146 | 1.2E-06 |
| *F07E5.8* | WBGene00017213 | protein_coding_gene | 1.1 | 2.146 | 4.2E-02 |
| *C54G4.2* | WBGene00008312 | protein_coding_gene | 1.1 | 2.143 | 1.1E-03 |
| *nhr-97* | WBGene00003687 | protein_coding_gene | 1.1 | 2.142 | 3.2E-22 |
| *F46F5.14* | WBGene00018505 | protein_coding_gene | 1.1 | 2.140 | 1.4E-02 |
| *acl-1* | WBGene00010339 | protein_coding_gene | 1.1 | 2.138 | 1.8E-12 |
| *Y69A2AR.8* | WBGene00022079 | pseudogene | 1.1 | 2.136 | 2.2E-02 |
| *unc-89* | WBGene00006820 | protein_coding_gene | 1.1 | 2.136 | 5.2E-05 |
| *R186.1* | WBGene00011305 | protein_coding_gene | 1.1 | 2.135 | 1.1E-03 |
| *ampd-1* | WBGene00016415 | protein_coding_gene | 1.1 | 2.135 | 6.8E-23 |
| *slc-9B.1* | WBGene00009617 | protein_coding_gene | 1.1 | 2.134 | 2.5E-05 |
| *rack-1* | WBGene00010556 | protein_coding_gene | 1.1 | 2.133 | 1.5E-16 |
| *trpa-1* | WBGene00007801 | protein_coding_gene | 1.1 | 2.133 | 8.8E-07 |
| *chil-14* | WBGene00019980 | protein_coding_gene | 1.1 | 2.133 | 2.3E-03 |
| *gst-10* | WBGene00001758 | protein_coding_gene | 1.1 | 2.128 | 4.9E-05 |
| *F15E6.6* | WBGene00017487 | protein_coding_gene | 1.1 | 2.126 | 8.1E-11 |
| *twk-5* | WBGene00006660 | protein_coding_gene | 1.1 | 2.125 | 2.6E-08 |
| *Y59E9AL.5* | WBGene00021995 | pseudogene | 1.1 | 2.125 | 3.7E-02 |
| *F42G4.2* | WBGene00009641 | protein_coding_gene | 1.1 | 2.123 | 1.2E-06 |
| *rps-1* | WBGene00004470 | protein_coding_gene | 1.1 | 2.122 | 1.3E-21 |
| *F54H12.8* | WBGene00045247 | protein_coding_gene | 1.1 | 2.121 | 1.6E-03 |
| *W06F12.3* | WBGene00012307 | protein_coding_gene | 1.1 | 2.121 | 2.4E-04 |
| *dct-18* | WBGene00010266 | protein_coding_gene | 1.1 | 2.119 | 2.1E-19 |
| *C15A11.2* | WBGene00007597 | protein_coding_gene | 1.1 | 2.119 | 1.0E-03 |
| *Y38F1A.1* | WBGene00012605 | protein_coding_gene | 1.1 | 2.119 | 5.2E-06 |
| *ZK930.4* | WBGene00014154 | protein_coding_gene | 1.1 | 2.118 | 1.3E-04 |
| *Y38F2AR.12* | WBGene00021430 | protein_coding_gene | 1.1 | 2.114 | 7.3E-10 |
| *elo-9* | WBGene00001247 | protein_coding_gene | 1.1 | 2.113 | 3.9E-05 |
| *ccm-2* | WBGene00019462 | protein_coding_gene | 1.1 | 2.109 | 4.3E-03 |
| *R07E5.6* | WBGene00011114 | protein_coding_gene | 1.1 | 2.108 | 7.6E-05 |
| *ncx-1* | WBGene00003566 | protein_coding_gene | 1.1 | 2.108 | 2.6E-18 |
| *rpl-26* | WBGene00004440 | protein_coding_gene | 1.1 | 2.107 | 2.1E-16 |
| *irld-4* | WBGene00008772 | protein_coding_gene | 1.1 | 2.107 | 1.2E-02 |
| *clec-43* | WBGene00019917 | protein_coding_gene | 1.1 | 2.104 | 2.8E-02 |
| *alg-3* | WBGene00011910 | protein_coding_gene | 1.1 | 2.104 | 1.1E-08 |
| *rdl-1* | WBGene00016703 | protein_coding_gene | 1.1 | 2.102 | 6.4E-03 |
| *rpl-6* | WBGene00004417 | protein_coding_gene | 1.1 | 2.102 | 1.5E-21 |
| *nep-13* | WBGene00018196 | protein_coding_gene | 1.1 | 2.102 | 4.0E-02 |

| | | | | | |
|---|---|---|---|---|---|
| *lgc-26* | WBGene00021981 | protein_coding_gene | 1.1 | 2.102 | 1.1E-03 |
| *rps-6* | WBGene00004475 | protein_coding_gene | 1.1 | 2.101 | 3.6E-18 |
| *F23B12.4* | WBGene00009081 | protein_coding_gene | 1.1 | 2.095 | 4.2E-12 |
| *spe-9* | WBGene00004963 | protein_coding_gene | 1.1 | 2.095 | 6.6E-03 |
| *C25H3.16* | WBGene00044392 | protein_coding_gene | 1.1 | 2.095 | 4.3E-02 |
| *twk-43* | WBGene00006693 | protein_coding_gene | 1.1 | 2.094 | 3.1E-04 |
| *clec-142* | WBGene00010460 | protein_coding_gene | 1.1 | 2.092 | 1.8E-02 |
| *ZK484.7* | WBGene00022753 | protein_coding_gene | 1.1 | 2.092 | 4.3E-06 |
| *R09E10.1* | WBGene00011171 | protein_coding_gene | 1.1 | 2.090 | 1.2E-04 |
| *aqp-4* | WBGene00000172 | protein_coding_gene | 1.1 | 2.090 | 4.6E-07 |
| *itr-1* | WBGene00002173 | protein_coding_gene | 1.1 | 2.089 | 9.6E-14 |
| *C14A6.6* | WBGene00007570 | protein_coding_gene | 1.1 | 2.089 | 2.9E-04 |
| *Y71H2AM.10* | WBGene00022175 | protein_coding_gene | 1.1 | 2.088 | 1.0E-03 |
| *H12D21.5* | WBGene00010381 | protein_coding_gene | 1.1 | 2.088 | 5.5E-03 |
| *hir-1* | WBGene00016059 | protein_coding_gene | 1.1 | 2.086 | 1.6E-11 |
| *best-22* | WBGene00013921 | protein_coding_gene | 1.1 | 2.086 | 1.1E-08 |
| *B0410.3* | WBGene00015172 | protein_coding_gene | 1.1 | 2.084 | 1.8E-09 |
| *frm-1* | WBGene00001488 | protein_coding_gene | 1.1 | 2.084 | 3.4E-24 |
| *tba-4* | WBGene00006530 | protein_coding_gene | 1.1 | 2.082 | 7.9E-09 |
| *rpl-16* | WBGene00004428 | protein_coding_gene | 1.1 | 2.081 | 4.3E-17 |
| *ckb-1* | WBGene00000511 | protein_coding_gene | 1.1 | 2.080 | 7.9E-05 |
| *ant-1.4* | WBGene00020140 | protein_coding_gene | 1.1 | 2.080 | 1.1E-05 |
| *rpl-36* | WBGene00004450 | protein_coding_gene | 1.1 | 2.080 | 2.6E-21 |
| *ZK550.5* | WBGene00013999 | protein_coding_gene | 1.1 | 2.079 | 8.9E-05 |
| *B0457.6* | WBGene00007180 | protein_coding_gene | 1.1 | 2.078 | 2.5E-05 |
| *fbxa-37* | WBGene00013845 | protein_coding_gene | 1.1 | 2.077 | 2.7E-04 |
| *utp-20* | WBGene00007050 | protein_coding_gene | 1.1 | 2.077 | 4.5E-14 |
| *W03C9.1* | WBGene00012219 | protein_coding_gene | 1.1 | 2.076 | 4.5E-03 |
| *fbl-1* | WBGene00001403 | protein_coding_gene | 1.1 | 2.076 | 4.6E-08 |
| *far-1* | WBGene00001385 | protein_coding_gene | 1.1 | 2.074 | 1.2E-16 |
| *C01G6.2* | WBGene00007224 | protein_coding_gene | 1.1 | 2.074 | 3.9E-03 |
| *C55C3.1* | WBGene00016951 | protein_coding_gene | 1.1 | 2.073 | 3.1E-02 |
| *trpa-2* | WBGene00010873 | protein_coding_gene | 1.1 | 2.073 | 7.3E-05 |
| *fbf-1* | WBGene00001401 | protein_coding_gene | 1.1 | 2.072 | 1.6E-08 |
| *T11F8.4* | WBGene00020435 | protein_coding_gene | 1.0 | 2.070 | 1.9E-04 |
| *C15C7.5* | WBGene00015791 | protein_coding_gene | 1.0 | 2.070 | 2.3E-11 |
| *nhr-230* | WBGene00012446 | protein_coding_gene | 1.0 | 2.068 | 2.8E-04 |
| *H11E01.2* | WBGene00019187 | protein_coding_gene | 1.0 | 2.065 | 2.4E-04 |
| *ZK1010.5* | WBGene00014179 | protein_coding_gene | 1.0 | 2.063 | 4.4E-08 |
| *F35A5.1* | WBGene00018024 | protein_coding_gene | 1.0 | 2.062 | 4.6E-11 |
| *sth-1* | WBGene00006060 | protein_coding_gene | 1.0 | 2.062 | 2.5E-03 |
| *C05C9.3* | WBGene00007327 | protein_coding_gene | 1.0 | 2.062 | 2.2E-03 |
| *ZK484.6* | WBGene00022752 | protein_coding_gene | 1.0 | 2.061 | 1.3E-02 |
| *abt-5* | WBGene00000023 | protein_coding_gene | 1.0 | 2.061 | 5.2E-06 |
| *R12B2.3* | WBGene00020019 | protein_coding_gene | 1.0 | 2.060 | 1.2E-02 |
| *F22H10.10* | WBGene00219912 | ncRNA_gene | 1.0 | 2.060 | 3.1E-02 |
| *F41B5.1* | WBGene00023284 | protein_coding_gene | 1.0 | 2.059 | 8.1E-04 |
| *zipt-2.2* | WBGene00017936 | protein_coding_gene | 1.0 | 2.058 | 6.3E-05 |
| *Y57G11C.36* | WBGene00013325 | protein_coding_gene | 1.0 | 2.058 | 6.0E-14 |
| *W01D2.1* | WBGene00012179 | protein_coding_gene | 1.0 | 2.058 | 2.4E-17 |
| *ZK1248.5* | WBGene00022876 | protein_coding_gene | 1.0 | 2.058 | 1.9E-05 |
| *unc-58* | WBGene00006792 | protein_coding_gene | 1.0 | 2.057 | 4.6E-05 |
| *T05H10.8* | WBGene00011512 | protein_coding_gene | 1.0 | 2.056 | 2.8E-02 |
| *rpl-9* | WBGene00004420 | protein_coding_gene | 1.0 | 2.056 | 1.5E-16 |
| *ZK370.6* | WBGene00022720 | protein_coding_gene | 1.0 | 2.055 | 4.0E-02 |
| *Y49E10.10* | WBGene00013033 | protein_coding_gene | 1.0 | 2.053 | 9.1E-04 |
| *F28A10.12* | WBGene00269337 | ncRNA_gene | 1.0 | 2.053 | 1.4E-02 |
| *spe-10* | WBGene00004964 | protein_coding_gene | 1.0 | 2.053 | 1.5E-03 |
| *B0303.11* | WBGene00015131 | protein_coding_gene | 1.0 | 2.051 | 7.8E-09 |
| *F35H8.1* | WBGene00009446 | protein_coding_gene | 1.0 | 2.049 | 1.7E-03 |
| *acl-7* | WBGene00012911 | protein_coding_gene | 1.0 | 2.048 | 2.4E-10 |
| *got-1.2* | WBGene00020146 | protein_coding_gene | 1.0 | 2.046 | 6.8E-18 |
| *mboa-3* | WBGene00016934 | protein_coding_gene | 1.0 | 2.045 | 1.1E-10 |
| *unc-122* | WBGene00006845 | protein_coding_gene | 1.0 | 2.045 | 4.3E-02 |
| *Y71G12B.3* | WBGene00022143 | protein_coding_gene | 1.0 | 2.044 | 5.3E-04 |
| *F23C8.7* | WBGene00017736 | protein_coding_gene | 1.0 | 2.042 | 3.9E-05 |
| *cest-25* | WBGene00022178 | protein_coding_gene | 1.0 | 2.042 | 2.4E-09 |
| *F48A9.2* | WBGene00018582 | protein_coding_gene | 1.0 | 2.042 | 7.3E-08 |
| *nep-2* | WBGene00020230 | protein_coding_gene | 1.0 | 2.042 | 9.9E-11 |
| *Y18D10A.23* | WBGene00012487 | protein_coding_gene | 1.0 | 2.042 | 6.6E-07 |
| *srv-1* | WBGene00005712 | protein_coding_gene | 1.0 | 2.040 | 8.3E-08 |
| *F26A1.3* | WBGene00017802 | protein_coding_gene | 1.0 | 2.040 | 5.9E-05 |
| *glr-6* | WBGene00001617 | protein_coding_gene | 1.0 | 2.038 | 6.3E-05 |
| *arrd-19* | WBGene00018061 | protein_coding_gene | 1.0 | 2.038 | 1.3E-04 |
| *T14B4.5* | WBGene00020501 | protein_coding_gene | 1.0 | 2.037 | 4.3E-03 |
| *R11E3.1* | WBGene00020003 | protein_coding_gene | 1.0 | 2.035 | 9.7E-06 |
| *pro-3* | WBGene00012676 | protein_coding_gene | 1.0 | 2.035 | 1.1E-10 |
| *best-25* | WBGene00014102 | protein_coding_gene | 1.0 | 2.033 | 2.5E-02 |
| *spe-8* | WBGene00004962 | protein_coding_gene | 1.0 | 2.033 | 2.0E-04 |
| *F59B2.15* | WBGene00185006 | protein_coding_gene | 1.0 | 2.032 | 2.1E-02 |

| | | | | | |
|---|---|---|---|---|---|
| *cnnm-3* | WBGene00016343 | protein_coding_gene | 1.0 | 2.031 | 3.0E-09 |
| *M03A1.3* | WBGene00019746 | protein_coding_gene | 1.0 | 2.031 | 1.0E-02 |
| *ncs-4* | WBGene00008998 | protein_coding_gene | 1.0 | 2.028 | 3.8E-02 |
| *fbxa-218* | WBGene00013036 | protein_coding_gene | 1.0 | 2.028 | 4.6E-02 |
| *rpl-7* | WBGene00004418 | protein_coding_gene | 1.0 | 2.027 | 6.4E-18 |
| *B0432.11* | WBGene00015192 | protein_coding_gene | 1.0 | 2.026 | 3.9E-02 |
| *fut-6* | WBGene00020222 | protein_coding_gene | 1.0 | 2.023 | 2.2E-04 |
| *F19C7.1* | WBGene00017591 | protein_coding_gene | 1.0 | 2.023 | 4.3E-04 |
| *nep-6* | WBGene00017550 | protein_coding_gene | 1.0 | 2.023 | 4.2E-04 |
| *F44D12.8* | WBGene00009685 | protein_coding_gene | 1.0 | 2.023 | 7.6E-06 |
| *B0361.11* | WBGene00015165 | protein_coding_gene | 1.0 | 2.020 | 6.7E-03 |
| *gnrr-5* | WBGene00019224 | protein_coding_gene | 1.0 | 2.020 | 2.2E-02 |
| *C45G9.12* | WBGene00016683 | protein_coding_gene | 1.0 | 2.019 | 4.9E-02 |
| *T02B11.9* | WBGene00044777 | protein_coding_gene | 1.0 | 2.018 | 2.9E-03 |
| *F32A5.8* | WBGene00017971 | protein_coding_gene | 1.0 | 2.018 | 1.5E-09 |
| *K08E7.8* | WBGene00010675 | protein_coding_gene | 1.0 | 2.016 | 3.5E-03 |
| *C43E11.5* | WBGene00016605 | protein_coding_gene | 1.0 | 2.015 | 1.3E-02 |
| *irld-14* | WBGene00011322 | protein_coding_gene | 1.0 | 2.015 | 1.2E-03 |
| *decr-1.3* | WBGene00011467 | protein_coding_gene | 1.0 | 2.015 | 2.1E-04 |
| *alh-8* | WBGene00000114 | protein_coding_gene | 1.0 | 2.015 | 2.4E-15 |
| *cest-9.2* | WBGene00007695 | protein_coding_gene | 1.0 | 2.014 | 4.6E-03 |
| *C49G7.12* | WBGene00077593 | protein_coding_gene | 1.0 | 2.012 | 9.5E-04 |
| *sdz-8* | WBGene00008334 | protein_coding_gene | 1.0 | 2.011 | 7.6E-03 |
| *F53C3.1* | WBGene00018745 | protein_coding_gene | 1.0 | 2.007 | 1.9E-03 |
| *lec-9* | WBGene00002272 | protein_coding_gene | 1.0 | 2.006 | 1.1E-07 |
| *rpl-37* | WBGene00004451 | protein_coding_gene | 1.0 | 2.005 | 1.1E-12 |
| *Y53F4B.21* | WBGene00013167 | protein_coding_gene | 1.0 | 2.005 | 3.0E-06 |
| *col-142* | WBGene00000715 | protein_coding_gene | 1.0 | 2.004 | 5.6E-05 |
| *F32H5.4* | WBGene00009350 | protein_coding_gene | 1.0 | 2.002 | 4.6E-03 |
| *col-103* | WBGene00000677 | protein_coding_gene | 1.0 | 2.001 | 9.4E-14 |
| *T20G5.8* | WBGene00011870 | protein_coding_gene | 1.0 | 2.001 | 2.3E-07 |
| *dyc-1* | WBGene00001116 | protein_coding_gene | 1.0 | 2.000 | 2.2E-05 |
| *ZK418.13* | WBGene00220259 | protein_coding_gene | 1.0 | 2.000 | 6.5E-03 |
| *peli-1* | WBGene00017766 | protein_coding_gene | -1.0 | 0.500 | 7.1E-10 |
| *K04C1.3* | WBGene00010553 | protein_coding_gene | -1.0 | 0.500 | 7.2E-07 |
| *K02B7.3* | WBGene00010491 | protein_coding_gene | -1.0 | 0.500 | 2.8E-11 |
| *aman-1* | WBGene00018877 | protein_coding_gene | -1.0 | 0.499 | 4.0E-05 |
| *umps-1* | WBGene00011559 | protein_coding_gene | -1.0 | 0.499 | 6.5E-15 |
| *mig-13* | WBGene00003245 | protein_coding_gene | -1.0 | 0.498 | 1.7E-05 |
| *mett-10* | WBGene00014228 | protein_coding_gene | -1.0 | 0.498 | 1.9E-11 |
| *magu-1* | WBGene00021914 | protein_coding_gene | -1.0 | 0.497 | 1.3E-10 |
| *algn-2* | WBGene00017282 | protein_coding_gene | -1.0 | 0.497 | 1.7E-15 |
| *dnj-26* | WBGene00001044 | protein_coding_gene | -1.0 | 0.497 | 1.5E-10 |
| *hlh-1* | WBGene00001948 | protein_coding_gene | -1.0 | 0.497 | 5.5E-04 |
| *lin-15A* | WBGene00023498 | protein_coding_gene | -1.0 | 0.497 | 1.3E-09 |
| *Y108G3AL.2* | WBGene00022434 | protein_coding_gene | -1.0 | 0.496 | 3.4E-14 |
| *W06A7.2* | WBGene00012291 | protein_coding_gene | -1.0 | 0.496 | 1.5E-05 |
| *Y38F2AL.12* | WBGene00219413 | protein_coding_gene | -1.0 | 0.495 | 1.1E-04 |
| *frg-1* | WBGene00014177 | protein_coding_gene | -1.0 | 0.495 | 1.6E-15 |
| *jtr-1* | WBGene00002180 | protein_coding_gene | -1.0 | 0.495 | 3.1E-11 |
| *R04B3.3* | WBGene00019868 | protein_coding_gene | -1.0 | 0.495 | 1.0E-02 |
| *F14H12.7* | WBGene00017474 | protein_coding_gene | -1.0 | 0.494 | 3.9E-02 |
| *aipr-1* | WBGene00016966 | protein_coding_gene | -1.0 | 0.494 | 1.9E-15 |
| *JC8.4* | WBGene00010436 | protein_coding_gene | -1.0 | 0.494 | 7.1E-07 |
| *C42D4.18* | WBGene00206380 | protein_coding_gene | -1.0 | 0.493 | 1.6E-03 |
| *insc-1* | WBGene00018392 | protein_coding_gene | -1.0 | 0.493 | 7.4E-06 |
| *ugt-66* | WBGene00016013 | protein_coding_gene | -1.0 | 0.493 | 1.8E-04 |
| *exoc-7* | WBGene00016606 | protein_coding_gene | -1.0 | 0.493 | 3.4E-14 |
| *apb-3* | WBGene00000163 | protein_coding_gene | -1.0 | 0.492 | 6.4E-24 |
| *C49C3.7* | WBGene00008197 | protein_coding_gene | -1.0 | 0.492 | 1.4E-19 |
| *lam-1* | WBGene00002247 | protein_coding_gene | -1.0 | 0.492 | 1.2E-13 |
| *Y48G1C.9* | WBGene00021682 | protein_coding_gene | -1.0 | 0.492 | 4.5E-10 |
| *C42C1.13* | WBGene00016591 | protein_coding_gene | -1.0 | 0.492 | 3.6E-09 |
| *cdk-7* | WBGene00000408 | protein_coding_gene | -1.0 | 0.492 | 5.7E-19 |
| *col-129* | WBGene00000703 | protein_coding_gene | -1.0 | 0.492 | 2.7E-03 |
| *ZK546.2* | WBGene00022759 | protein_coding_gene | -1.0 | 0.491 | 1.6E-21 |
| *cpd-2* | WBGene00012073 | protein_coding_gene | -1.0 | 0.491 | 8.7E-05 |
| *nhr-68* | WBGene00003658 | protein_coding_gene | -1.0 | 0.491 | 4.8E-15 |
| *W04E12.9* | WBGene00012254 | protein_coding_gene | -1.0 | 0.490 | 4.1E-02 |
| *cnt-2* | WBGene00000566 | protein_coding_gene | -1.0 | 0.490 | 2.0E-18 |
| *dct-10* | WBGene00012618 | pseudogene | -1.0 | 0.490 | 3.3E-02 |
| *T10E9.1* | WBGene00020411 | protein_coding_gene | -1.0 | 0.490 | 8.4E-15 |
| *cec-5* | WBGene00017993 | protein_coding_gene | -1.0 | 0.490 | 2.7E-26 |
| *T20B12.7* | WBGene00020604 | protein_coding_gene | -1.0 | 0.489 | 2.3E-20 |
| *btb-1* | WBGene00001963 | pseudogene | -1.0 | 0.489 | 4.6E-02 |
| *dpf-2* | WBGene00001055 | protein_coding_gene | -1.0 | 0.489 | 1.3E-10 |
| *ced-12* | WBGene00000426 | protein_coding_gene | -1.0 | 0.489 | 2.1E-18 |
| *lir-2* | WBGene00003045 | protein_coding_gene | -1.0 | 0.487 | 2.7E-11 |
| *chd-3* | WBGene00000482 | protein_coding_gene | -1.0 | 0.487 | 7.9E-14 |
| *Y47D3B.4* | WBGene00012941 | protein_coding_gene | -1.0 | 0.487 | 1.2E-02 |

| | | | | | |
|---|---|---|---|---|---|
| *ekl-4* | WBGene00013676 | protein_coding_gene | -1.0 | 0.486 | 5.5E-18 |
| *sax-3* | WBGene00004729 | protein_coding_gene | -1.0 | 0.486 | 3.7E-17 |
| *calm-1* | WBGene00009260 | protein_coding_gene | -1.0 | 0.485 | 5.2E-05 |
| *eri-1* | WBGene00001332 | protein_coding_gene | -1.0 | 0.485 | 4.9E-18 |
| *K09A11.1* | WBGene00010704 | protein_coding_gene | -1.0 | 0.485 | 4.9E-06 |
| *C56G2.4* | WBGene00016979 | protein_coding_gene | -1.0 | 0.485 | 2.4E-03 |
| *F35H10.6* | WBGene00018071 | protein_coding_gene | -1.0 | 0.485 | 2.6E-13 |
| *csn-5* | WBGene00000817 | protein_coding_gene | -1.0 | 0.484 | 4.8E-21 |
| *C06B8.2* | WBGene00007368 | protein_coding_gene | -1.0 | 0.484 | 5.0E-06 |
| *B0564.2* | WBGene00007202 | protein_coding_gene | -1.0 | 0.484 | 5.6E-08 |
| *T28A8.4* | WBGene00012110 | protein_coding_gene | -1.0 | 0.483 | 7.5E-15 |
| *Y23H5A.2* | WBGene00021269 | protein_coding_gene | -1.1 | 0.483 | 4.8E-08 |
| *ife-4* | WBGene00002062 | protein_coding_gene | -1.1 | 0.482 | 6.0E-07 |
| *H38K22.6* | WBGene00044911 | protein_coding_gene | -1.1 | 0.482 | 3.7E-02 |
| *coh-1* | WBGene00000591 | protein_coding_gene | -1.1 | 0.482 | 2.2E-15 |
| *nep-24* | WBGene00013786 | protein_coding_gene | -1.1 | 0.481 | 3.5E-02 |
| *mdt-30* | WBGene00004125 | protein_coding_gene | -1.1 | 0.481 | 3.2E-15 |
| *F36D4.5* | WBGene00018090 | protein_coding_gene | -1.1 | 0.481 | 2.2E-23 |
| *ZK1248.13* | WBGene00022882 | protein_coding_gene | -1.1 | 0.481 | 4.6E-18 |
| *R144.3* | WBGene00020093 | protein_coding_gene | -1.1 | 0.481 | 5.9E-14 |
| *ZC581.3* | WBGene00022633 | protein_coding_gene | -1.1 | 0.481 | 7.6E-11 |
| *apc-10* | WBGene00000144 | protein_coding_gene | -1.1 | 0.481 | 1.5E-12 |
| *K07F5.14* | WBGene00010638 | protein_coding_gene | -1.1 | 0.481 | 1.1E-17 |
| *snu-23* | WBGene00022794 | protein_coding_gene | -1.1 | 0.481 | 5.2E-21 |
| *ZC374.2* | WBGene00013872 | protein_coding_gene | -1.1 | 0.480 | 1.8E-03 |
| *syg-2* | WBGene00007750 | protein_coding_gene | -1.1 | 0.480 | 1.4E-06 |
| *Y69E1A.5* | WBGene00013477 | protein_coding_gene | -1.1 | 0.480 | 3.4E-09 |
| *C18H9.6* | WBGene00016004 | protein_coding_gene | -1.1 | 0.480 | 2.4E-04 |
| *R08D7.1* | WBGene00011142 | protein_coding_gene | -1.1 | 0.478 | 1.8E-19 |
| *hmg-11* | WBGene00001976 | protein_coding_gene | -1.1 | 0.477 | 1.4E-08 |
| *F09E5.11* | WBGene00017289 | protein_coding_gene | -1.1 | 0.477 | 5.2E-16 |
| *T04H1.2* | WBGene00011449 | protein_coding_gene | -1.1 | 0.477 | 2.0E-26 |
| *jac-1* | WBGene00002175 | protein_coding_gene | -1.1 | 0.476 | 4.2E-16 |
| *mfap-1* | WBGene00009671 | protein_coding_gene | -1.1 | 0.476 | 2.8E-23 |
| *snfc-5* | WBGene00011111 | protein_coding_gene | -1.1 | 0.476 | 9.1E-19 |
| *B0432.10* | WBGene00015191 | protein_coding_gene | -1.1 | 0.476 | 4.9E-10 |
| *F38B2.6* | WBGene00194663 | protein_coding_gene | -1.1 | 0.476 | 1.3E-02 |
| *C49H3.4* | WBGene00016791 | protein_coding_gene | -1.1 | 0.475 | 2.2E-09 |
| *T05E7.3* | WBGene00020259 | protein_coding_gene | -1.1 | 0.475 | 3.0E-20 |
| *fbxa-211* | WBGene00012566 | protein_coding_gene | -1.1 | 0.475 | 3.7E-10 |
| *pnn-1* | WBGene00011308 | protein_coding_gene | -1.1 | 0.474 | 1.8E-19 |
| *rab-10* | WBGene00004273 | protein_coding_gene | -1.1 | 0.473 | 2.5E-31 |
| *E02H1.5* | WBGene00008457 | protein_coding_gene | -1.1 | 0.473 | 2.2E-14 |
| *R13A5.15* | WBGene00194726 | protein_coding_gene | -1.1 | 0.471 | 3.2E-02 |
| *C27H6.9* | WBGene00077500 | protein_coding_gene | -1.1 | 0.471 | 8.7E-15 |
| *Y105E8B.9* | WBGene00013693 | protein_coding_gene | -1.1 | 0.470 | 1.6E-05 |
| *C10B5.1* | WBGene00015670 | protein_coding_gene | -1.1 | 0.470 | 1.5E-10 |
| *taf-13* | WBGene00006397 | protein_coding_gene | -1.1 | 0.470 | 8.9E-16 |
| *mat-2* | WBGene00003133 | protein_coding_gene | -1.1 | 0.470 | 5.8E-23 |
| *C33D9.13* | WBGene00194651 | protein_coding_gene | -1.1 | 0.469 | 1.0E-05 |
| *R10E12.2* | WBGene00011221 | protein_coding_gene | -1.1 | 0.469 | 1.4E-07 |
| *Y71F9AR.2* | WBGene00022123 | protein_coding_gene | -1.1 | 0.468 | 4.3E-03 |
| *R05D7.7* | WBGene00077559 | protein_coding_gene | -1.1 | 0.468 | 1.7E-04 |
| *ZK1307.3* | WBGene00014246 | protein_coding_gene | -1.1 | 0.468 | 1.1E-02 |
| *hus-1* | WBGene00002042 | protein_coding_gene | -1.1 | 0.468 | 1.8E-09 |
| *T28D9.4* | WBGene00020896 | protein_coding_gene | -1.1 | 0.467 | 7.0E-14 |
| *C05G5.5* | WBGene00007351 | protein_coding_gene | -1.1 | 0.467 | 2.4E-02 |
| *sym-3* | WBGene00006368 | protein_coding_gene | -1.1 | 0.467 | 6.9E-09 |
| *ttr-15* | WBGene00011561 | protein_coding_gene | -1.1 | 0.467 | 1.2E-20 |
| *ZK1058.9* | WBGene00014207 | protein_coding_gene | -1.1 | 0.466 | 3.5E-16 |
| *M02B7.7* | WBGene00044690 | protein_coding_gene | -1.1 | 0.466 | 2.5E-05 |
| *C01G5.9* | WBGene00015311 | protein_coding_gene | -1.1 | 0.465 | 1.0E-02 |
| *W03G11.4* | WBGene00012226 | protein_coding_gene | -1.1 | 0.464 | 9.4E-09 |
| *mlp-1* | WBGene00003375 | protein_coding_gene | -1.1 | 0.464 | 2.7E-10 |
| *act-3* | WBGene00000065 | protein_coding_gene | -1.1 | 0.463 | 1.2E-07 |
| *snpc-3.4* | WBGene00011367 | protein_coding_gene | -1.1 | 0.463 | 3.1E-17 |
| *nhr-246* | WBGene00014192 | protein_coding_gene | -1.1 | 0.462 | 4.6E-02 |
| *msp-10* | WBGene00003425 | protein_coding_gene | -1.1 | 0.461 | 6.0E-03 |
| *F26F4.6* | WBGene00017828 | protein_coding_gene | -1.1 | 0.461 | 2.2E-17 |
| *pqn-65* | WBGene00004148 | protein_coding_gene | -1.1 | 0.461 | 3.7E-20 |
| *F46F11.8* | WBGene00018511 | protein_coding_gene | -1.1 | 0.460 | 5.0E-19 |
| *sig-7* | WBGene00000890 | protein_coding_gene | -1.1 | 0.460 | 5.7E-23 |
| *lido-10* | WBGene00016575 | protein_coding_gene | -1.1 | 0.460 | 7.7E-04 |
| *linc-6* | WBGene00219700 | lincRNA_gene | -1.1 | 0.459 | 1.2E-03 |
| *Y51H4A.1* | WBGene00013098 | protein_coding_gene | -1.1 | 0.459 | 4.2E-02 |
| *Y6D1A.1* | WBGene00012395 | protein_coding_gene | -1.1 | 0.459 | 1.1E-12 |
| *C42C1.19* | WBGene00302978 | protein_coding_gene | -1.1 | 0.459 | 2.2E-14 |
| *T22C1.1* | WBGene00011912 | protein_coding_gene | -1.1 | 0.457 | 3.6E-22 |
| *nduo-6* | WBGene00010957 | protein_coding_gene | -1.1 | 0.457 | 1.1E-04 |
| *mdt-19* | WBGene00007019 | protein_coding_gene | -1.1 | 0.457 | 9.7E-23 |

| | | | | | |
|---|---|---|---|---|---|
| *set-23* | WBGene00021515 | protein_coding_gene | -1.1 | 0.457 | 8.9E-12 |
| *ZK822.1* | WBGene00014089 | protein_coding_gene | -1.1 | 0.456 | 6.7E-04 |
| *F46H5.2* | WBGene00018518 | protein_coding_gene | -1.1 | 0.456 | 6.9E-12 |
| *Y44E3A.1* | WBGene00021547 | protein_coding_gene | -1.1 | 0.456 | 1.0E-04 |
| *snf-9* | WBGene00004908 | protein_coding_gene | -1.1 | 0.456 | 5.0E-11 |
| *C16C8.13* | WBGene00015851 | protein_coding_gene | -1.1 | 0.455 | 2.8E-04 |
| *ZK1225.7* | WBGene00014994 | pseudogene | -1.1 | 0.455 | 3.8E-02 |
| *cpt-1* | WBGene00012907 | protein_coding_gene | -1.1 | 0.455 | 1.3E-13 |
| *ZK1248.11* | WBGene00022881 | protein_coding_gene | -1.1 | 0.454 | 1.0E-18 |
| *unc-85* | WBGene00006817 | protein_coding_gene | -1.1 | 0.453 | 1.2E-19 |
| *eat-20* | WBGene00001148 | protein_coding_gene | -1.1 | 0.453 | 8.9E-08 |
| *npp-2* | WBGene00003788 | protein_coding_gene | -1.2 | 0.451 | 1.6E-26 |
| *mig-2* | WBGene00003239 | protein_coding_gene | -1.2 | 0.450 | 2.3E-16 |
| *tag-275* | WBGene00016423 | protein_coding_gene | -1.2 | 0.449 | 3.6E-04 |
| *B0304.4* | WBGene00015136 | protein_coding_gene | -1.2 | 0.449 | 8.2E-11 |
| *unc-119* | WBGene00006843 | protein_coding_gene | -1.2 | 0.448 | 9.3E-09 |
| *F23A7.1* | WBGene00009067 | protein_coding_gene | -1.2 | 0.448 | 4.8E-03 |
| *spt-4* | WBGene00005014 | protein_coding_gene | -1.2 | 0.448 | 4.3E-16 |
| *mdt-10* | WBGene00007014 | protein_coding_gene | -1.2 | 0.448 | 1.5E-15 |
| *rbbp-5* | WBGene00017683 | protein_coding_gene | -1.2 | 0.446 | 3.3E-24 |
| *unc-84* | WBGene00006816 | protein_coding_gene | -1.2 | 0.446 | 1.1E-19 |
| *pals-34* | WBGene00017823 | protein_coding_gene | -1.2 | 0.446 | 1.8E-03 |
| *trul-1* | WBGene00015194 | protein_coding_gene | -1.2 | 0.446 | 1.6E-14 |
| *viln-1* | WBGene00015691 | protein_coding_gene | -1.2 | 0.445 | 1.4E-23 |
| *mec-6* | WBGene00003170 | protein_coding_gene | -1.2 | 0.444 | 7.7E-03 |
| *Y57G7A.8* | WBGene00021971 | pseudogene | -1.2 | 0.444 | 6.5E-04 |
| *cyn-8* | WBGene00000884 | protein_coding_gene | -1.2 | 0.443 | 2.4E-16 |
| *mdmh-35* | WBGene00011635 | protein_coding_gene | -1.2 | 0.442 | 2.2E-17 |
| *rap-1* | WBGene00004307 | protein_coding_gene | -1.2 | 0.442 | 8.3E-37 |
| *cwn-2* | WBGene00000858 | protein_coding_gene | -1.2 | 0.442 | 1.7E-07 |
| *T16G12.9* | WBGene00011810 | protein_coding_gene | -1.2 | 0.442 | 1.7E-02 |
| *D2030.11* | WBGene00008420 | protein_coding_gene | -1.2 | 0.442 | 1.9E-13 |
| *T25B2.2* | WBGene00020786 | protein_coding_gene | -1.2 | 0.442 | 7.9E-10 |
| *F26H11.4* | WBGene00009182 | protein_coding_gene | -1.2 | 0.441 | 2.4E-03 |
| *fbxa-123* | WBGene00014918 | pseudogene | -1.2 | 0.441 | 6.2E-08 |
| *lbp-5* | WBGene00002257 | protein_coding_gene | -1.2 | 0.441 | 8.2E-13 |
| *cir-1* | WBGene00018892 | protein_coding_gene | -1.2 | 0.440 | 6.5E-24 |
| *rabs-5* | WBGene00021538 | protein_coding_gene | -1.2 | 0.440 | 3.3E-14 |
| *epg-8* | WBGene00013695 | protein_coding_gene | -1.2 | 0.439 | 4.7E-25 |
| *aat-3* | WBGene00000004 | protein_coding_gene | -1.2 | 0.439 | 1.5E-12 |
| *C44C1.1* | WBGene00016641 | protein_coding_gene | -1.2 | 0.438 | 7.8E-06 |
| *F14E5.1* | WBGene00008799 | protein_coding_gene | -1.2 | 0.438 | 4.4E-03 |
| *T04D1.2* | WBGene00020212 | protein_coding_gene | -1.2 | 0.438 | 7.9E-03 |
| *madf-3* | WBGene00019218 | protein_coding_gene | -1.2 | 0.438 | 8.4E-23 |
| *anmt-3* | WBGene00011573 | protein_coding_gene | -1.2 | 0.436 | 7.2E-14 |
| *C11H1.7* | WBGene00007531 | protein_coding_gene | -1.2 | 0.436 | 5.4E-04 |
| *gkow-1* | WBGene00011237 | protein_coding_gene | -1.2 | 0.435 | 8.2E-26 |
| *C42D4.1* | WBGene00016594 | protein_coding_gene | -1.2 | 0.435 | 5.5E-15 |
| *smc-5* | WBGene00016151 | protein_coding_gene | -1.2 | 0.434 | 5.5E-33 |
| *ztf-29* | WBGene00013438 | protein_coding_gene | -1.2 | 0.433 | 1.6E-10 |
| *Y38H6C.23* | WBGene00012634 | protein_coding_gene | -1.2 | 0.433 | 1.6E-02 |
| *F37B4.10* | WBGene00018140 | protein_coding_gene | -1.2 | 0.433 | 3.6E-10 |
| *F15G9.1* | WBGene00008865 | protein_coding_gene | -1.2 | 0.433 | 6.2E-15 |
| *Y59H11AM.1* | WBGene00022005 | protein_coding_gene | -1.2 | 0.432 | 1.7E-06 |
| *C32E12.1* | WBGene00016327 | protein_coding_gene | -1.2 | 0.432 | 1.5E-06 |
| *cls-3* | WBGene00013847 | protein_coding_gene | -1.2 | 0.432 | 2.2E-16 |
| *T04H1.5* | WBGene00011451 | protein_coding_gene | -1.2 | 0.432 | 1.7E-19 |
| *Y47D3A.20* | WBGene00012931 | protein_coding_gene | -1.2 | 0.431 | 1.4E-08 |
| *Y87G2A.25* | WBGene00219288 | protein_coding_gene | -1.2 | 0.431 | 8.6E-03 |
| *bath-4* | WBGene00018665 | protein_coding_gene | -1.2 | 0.431 | 3.0E-03 |
| *T23B5.4* | WBGene00011941 | protein_coding_gene | -1.2 | 0.430 | 8.4E-13 |
| *nhr-67* | WBGene00003657 | protein_coding_gene | -1.2 | 0.429 | 1.5E-03 |
| *set-10* | WBGene00009370 | protein_coding_gene | -1.2 | 0.428 | 3.1E-08 |
| *ZC13.1* | WBGene00022502 | protein_coding_gene | -1.2 | 0.428 | 1.1E-13 |
| *rde-11* | WBGene00023421 | protein_coding_gene | -1.2 | 0.428 | 1.5E-23 |
| *vps-60* | WBGene00018290 | protein_coding_gene | -1.2 | 0.428 | 4.7E-26 |
| *tab-1* | WBGene00006380 | protein_coding_gene | -1.2 | 0.427 | 2.4E-02 |
| *F39H11.1* | WBGene00009565 | protein_coding_gene | -1.2 | 0.426 | 1.4E-23 |
| *mnm-2* | WBGene00003380 | protein_coding_gene | -1.2 | 0.425 | 6.1E-03 |
| *bra-1* | WBGene00000262 | protein_coding_gene | -1.2 | 0.425 | 2.9E-09 |
| *nlr-1* | WBGene00003772 | protein_coding_gene | -1.2 | 0.424 | 1.8E-08 |
| *crb-1* | WBGene00000792 | protein_coding_gene | -1.2 | 0.424 | 5.6E-12 |
| *acd-1* | WBGene00016064 | protein_coding_gene | -1.2 | 0.424 | 2.6E-02 |
| *gbh-1* | WBGene00001522 | protein_coding_gene | -1.2 | 0.423 | 9.6E-11 |
| *tag-294* | WBGene00016937 | protein_coding_gene | -1.2 | 0.423 | 4.6E-14 |
| *Y55F3C.9* | WBGene00044459 | protein_coding_gene | -1.2 | 0.423 | 3.2E-06 |
| *eyg-1* | WBGene00013147 | protein_coding_gene | -1.2 | 0.422 | 3.7E-02 |
| *F23F1.10* | WBGene00017750 | protein_coding_gene | -1.2 | 0.422 | 2.5E-12 |
| *hpo-7* | WBGene00016548 | protein_coding_gene | -1.2 | 0.422 | 4.7E-03 |
| *K07C5.3* | WBGene00010626 | protein_coding_gene | -1.2 | 0.422 | 2.8E-27 |

| | | | | | |
|---|---|---|---|---|---|
| *D1007.8* | WBGene00017005 | protein_coding_gene | -1.2 | 0.421 | 1.5E-16 |
| *F56D6.13* | WBGene00044475 | protein_coding_gene | -1.2 | 0.421 | 8.6E-05 |
| *dnj-27* | WBGene00001045 | protein_coding_gene | -1.3 | 0.420 | 2.2E-13 |
| *ZK1127.3* | WBGene00022850 | protein_coding_gene | -1.3 | 0.420 | 1.0E-21 |
| *Y119C1B.10* | WBGene00022475 | protein_coding_gene | -1.3 | 0.417 | 1.5E-08 |
| *EGAP2.1* | WBGene00017147 | protein_coding_gene | -1.3 | 0.417 | 1.0E-07 |
| *C04C11.1* | WBGene00007295 | protein_coding_gene | -1.3 | 0.417 | 9.6E-03 |
| *K12G11.6* | WBGene00010792 | protein_coding_gene | -1.3 | 0.416 | 2.8E-02 |
| *Y57A10A.26* | WBGene00013266 | protein_coding_gene | -1.3 | 0.415 | 1.2E-14 |
| *Y54G2A.36* | WBGene00021898 | protein_coding_gene | -1.3 | 0.415 | 2.6E-14 |
| *F44A6.5* | WBGene00009677 | protein_coding_gene | -1.3 | 0.415 | 2.4E-09 |
| *F58A4.6* | WBGene00010229 | protein_coding_gene | -1.3 | 0.415 | 3.5E-13 |
| *psmd-9* | WBGene00016623 | protein_coding_gene | -1.3 | 0.412 | 7.0E-22 |
| *ddi-1* | WBGene00015308 | protein_coding_gene | -1.3 | 0.412 | 6.4E-32 |
| *Y74C9A.9* | WBGene00235381 | pseudogene | -1.3 | 0.412 | 9.6E-03 |
| *gst-44* | WBGene00001792 | protein_coding_gene | -1.3 | 0.411 | 1.3E-02 |
| *set-20* | WBGene00020920 | protein_coding_gene | -1.3 | 0.410 | 5.1E-08 |
| *W03F9.2* | WBGene00021000 | protein_coding_gene | -1.3 | 0.410 | 8.2E-07 |
| *F13C5.2* | WBGene00017423 | protein_coding_gene | -1.3 | 0.409 | 1.3E-15 |
| *pat-10* | WBGene00003934 | protein_coding_gene | -1.3 | 0.409 | 2.7E-31 |
| *F52F12.9* | WBGene00009942 | protein_coding_gene | -1.3 | 0.408 | 3.0E-03 |
| *lect-2* | WBGene00019407 | protein_coding_gene | -1.3 | 0.407 | 2.4E-10 |
| *D1007.4* | WBGene00017002 | protein_coding_gene | -1.3 | 0.406 | 3.7E-14 |
| *dos-1* | WBGene00013980 | protein_coding_gene | -1.3 | 0.406 | 2.1E-07 |
| *clec-66* | WBGene00009397 | protein_coding_gene | -1.3 | 0.406 | 2.9E-09 |
| *lron-10* | WBGene00012646 | protein_coding_gene | -1.3 | 0.405 | 1.2E-19 |
| *Y45G5AM.7* | WBGene00021559 | protein_coding_gene | -1.3 | 0.405 | 7.0E-23 |
| *nep-4* | WBGene00016896 | protein_coding_gene | -1.3 | 0.403 | 2.1E-04 |
| *agt-2* | WBGene00000094 | protein_coding_gene | -1.3 | 0.403 | 1.9E-13 |
| *ugt-8* | WBGene00019234 | protein_coding_gene | -1.3 | 0.403 | 5.2E-08 |
| *F21D9.1* | WBGene00009015 | protein_coding_gene | -1.3 | 0.402 | 6.9E-03 |
| *cwn-1* | WBGene00000857 | protein_coding_gene | -1.3 | 0.402 | 1.0E-09 |
| *C53A3.1* | WBGene00016891 | protein_coding_gene | -1.3 | 0.400 | 4.8E-02 |
| *lig-1* | WBGene00002985 | protein_coding_gene | -1.3 | 0.400 | 1.0E-35 |
| *Y39B6A.37* | WBGene00012695 | protein_coding_gene | -1.3 | 0.399 | 1.1E-13 |
| *F23H12.10* | WBGene00044218 | protein_coding_gene | -1.3 | 0.399 | 4.8E-04 |
| *dod-17* | WBGene00010745 | protein_coding_gene | -1.3 | 0.397 | 3.7E-02 |
| *Y73F4A.1* | WBGene00013514 | protein_coding_gene | -1.3 | 0.396 | 1.9E-09 |
| *noah-1* | WBGene00016422 | protein_coding_gene | -1.3 | 0.394 | 2.7E-02 |
| *Y17G9B.8* | WBGene00021205 | protein_coding_gene | -1.3 | 0.394 | 4.8E-25 |
| *C13F10.7* | WBGene00015746 | protein_coding_gene | -1.3 | 0.393 | 4.8E-21 |
| *C08B6.4* | WBGene00007425 | protein_coding_gene | -1.3 | 0.392 | 9.8E-03 |
| *T04D3.8* | WBGene00011435 | protein_coding_gene | -1.4 | 0.392 | 4.7E-13 |
| *gpx-1* | WBGene00009165 | protein_coding_gene | -1.4 | 0.392 | 7.4E-08 |
| *got-2.2* | WBGene00015778 | protein_coding_gene | -1.4 | 0.392 | 2.6E-30 |
| *F28F8.7* | WBGene00009224 | protein_coding_gene | -1.4 | 0.391 | 4.8E-15 |
| *K12H4.2* | WBGene00019677 | protein_coding_gene | -1.4 | 0.391 | 1.1E-16 |
| *ubl-5* | WBGene00006726 | protein_coding_gene | -1.4 | 0.391 | 2.7E-18 |
| *F16B12.4* | WBGene00008880 | protein_coding_gene | -1.4 | 0.390 | 2.4E-13 |
| *rnst-2* | WBGene00019624 | protein_coding_gene | -1.4 | 0.389 | 1.1E-18 |
| *ugt-4* | WBGene00013905 | protein_coding_gene | -1.4 | 0.388 | 2.9E-09 |
| *Y41D4A.3* | WBGene00021505 | protein_coding_gene | -1.4 | 0.387 | 1.1E-05 |
| *ccch-3* | WBGene00012277 | protein_coding_gene | -1.4 | 0.386 | 3.0E-32 |
| *lmd-2* | WBGene00015008 | protein_coding_gene | -1.4 | 0.386 | 2.2E-10 |
| *Y49A10A.2* | WBGene00044949 | protein_coding_gene | -1.4 | 0.384 | 3.7E-10 |
| *pop-1* | WBGene00004077 | protein_coding_gene | -1.4 | 0.384 | 5.8E-26 |
| *cyd-1* | WBGene00000870 | protein_coding_gene | -1.4 | 0.383 | 6.4E-09 |
| *blos-9* | WBGene00010767 | protein_coding_gene | -1.4 | 0.382 | 1.8E-18 |
| *his-24* | WBGene00001898 | protein_coding_gene | -1.4 | 0.381 | 9.4E-25 |
| *sdn-1* | WBGene00004749 | protein_coding_gene | -1.4 | 0.381 | 1.8E-29 |
| *F07G6.10* | WBGene00045271 | protein_coding_gene | -1.4 | 0.381 | 1.6E-02 |
| *lam-3* | WBGene00002248 | protein_coding_gene | -1.4 | 0.380 | 9.0E-15 |
| *F54D5.4* | WBGene00010050 | protein_coding_gene | -1.4 | 0.380 | 3.6E-07 |
| *F20D12.12* | WBGene00249815 | protein_coding_gene | -1.4 | 0.380 | 4.4E-03 |
| *cht-1* | WBGene00000503 | protein_coding_gene | -1.4 | 0.379 | 1.9E-23 |
| *C52E12.11* | WBGene00304189 | pseudogene | -1.4 | 0.379 | 3.2E-02 |
| *F59D6.1* | WBGene00019103 | protein_coding_gene | -1.4 | 0.378 | 3.1E-02 |
| *mksr-2* | WBGene00021416 | protein_coding_gene | -1.4 | 0.376 | 2.7E-02 |
| *ugt-9* | WBGene00020587 | protein_coding_gene | -1.4 | 0.375 | 7.5E-08 |
| *F36D1.9* | WBGene00009468 | protein_coding_gene | -1.4 | 0.375 | 1.5E-04 |
| *Y81B9A.5* | WBGene00304906 | ncRNA_gene | -1.4 | 0.375 | 4.3E-02 |
| *mab-20* | WBGene00003111 | protein_coding_gene | -1.4 | 0.374 | 5.2E-22 |
| *T04A6.2* | WBGene00020201 | protein_coding_gene | -1.4 | 0.373 | 1.3E-02 |
| *Y47G6A.36* | WBGene00220165 | ncRNA_gene | -1.4 | 0.372 | 7.9E-05 |
| *lido-1* | WBGene00015586 | protein_coding_gene | -1.4 | 0.371 | 1.6E-10 |
| *C10H11.7* | WBGene00015696 | protein_coding_gene | -1.4 | 0.370 | 8.9E-05 |
| *fbxc-53* | WBGene00016711 | protein_coding_gene | -1.4 | 0.369 | 3.5E-02 |
| *arrd-4* | WBGene00014159 | protein_coding_gene | -1.4 | 0.368 | 3.3E-08 |
| *nlp-28* | WBGene00003766 | protein_coding_gene | -1.4 | 0.368 | 6.4E-15 |
| *Y53G8B.2* | WBGene00021818 | protein_coding_gene | -1.4 | 0.368 | 2.1E-20 |

| | | | | | |
|---|---|---|---|---|---|
| *ska-1* | WBGene00013725 | protein_coding_gene | -1.4 | 0.367 | 7.2E-27 |
| *F09D12.2* | WBGene00017281 | protein_coding_gene | -1.4 | 0.367 | 3.2E-02 |
| *rgs-7* | WBGene00004350 | protein_coding_gene | -1.4 | 0.366 | 1.3E-27 |
| *mlc-7* | WBGene00023451 | protein_coding_gene | -1.4 | 0.366 | 9.0E-16 |
| *DH11.2* | WBGene00008436 | protein_coding_gene | -1.5 | 0.365 | 1.2E-28 |
| *col-99* | WBGene00000674 | protein_coding_gene | -1.5 | 0.364 | 1.2E-09 |
| *noah-2* | WBGene00009926 | protein_coding_gene | -1.5 | 0.364 | 2.9E-02 |
| *F47B10.9* | WBGene00009820 | protein_coding_gene | -1.5 | 0.363 | 7.9E-13 |
| *F40D4.17* | WBGene00194682 | protein_coding_gene | -1.5 | 0.361 | 5.4E-07 |
| *K08D8.5* | WBGene00010659 | protein_coding_gene | -1.5 | 0.359 | 3.4E-05 |
| *tcl-2* | WBGene00006561 | protein_coding_gene | -1.5 | 0.359 | 1.7E-11 |
| *ZK1098.3* | WBGene00014220 | protein_coding_gene | -1.5 | 0.358 | 2.1E-10 |
| *C01G12.12* | WBGene00044053 | pseudogene | -1.5 | 0.358 | 1.2E-07 |
| *tipn-1* | WBGene00017738 | protein_coding_gene | -1.5 | 0.357 | 9.8E-24 |
| *irg-6* | WBGene00007864 | protein_coding_gene | -1.5 | 0.356 | 1.4E-04 |
| *osg-1* | WBGene00019832 | protein_coding_gene | -1.5 | 0.354 | 1.4E-07 |
| *grl-18* | WBGene00001727 | protein_coding_gene | -1.5 | 0.353 | 2.7E-02 |
| *nab-1* | WBGene00003516 | protein_coding_gene | -1.5 | 0.353 | 1.0E-10 |
| *F43C9.1* | WBGene00018375 | protein_coding_gene | -1.5 | 0.352 | 9.5E-05 |
| *AH9.3* | WBGene00014999 | protein_coding_gene | -1.5 | 0.351 | 1.5E-16 |
| *K08H2.4* | WBGene00010691 | protein_coding_gene | -1.5 | 0.350 | 2.9E-04 |
| *ham-1* | WBGene00001820 | protein_coding_gene | -1.5 | 0.349 | 1.0E-14 |
| *C27B7.9* | WBGene00007765 | protein_coding_gene | -1.5 | 0.348 | 3.4E-31 |
| *dhs-27* | WBGene00000990 | protein_coding_gene | -1.5 | 0.347 | 6.9E-14 |
| *ugt-53* | WBGene00020182 | protein_coding_gene | -1.5 | 0.347 | 4.3E-05 |
| *C53C11.1* | WBGene00016899 | protein_coding_gene | -1.5 | 0.345 | 4.7E-02 |
| *C39B5.10* | WBGene00016528 | protein_coding_gene | -1.5 | 0.345 | 4.7E-14 |
| *odd-1* | WBGene00003845 | protein_coding_gene | -1.5 | 0.344 | 6.0E-04 |
| *R09E12.9* | WBGene00044514 | protein_coding_gene | -1.5 | 0.342 | 9.3E-05 |
| *ttr-34* | WBGene00045245 | protein_coding_gene | -1.6 | 0.341 | 5.5E-11 |
| *Y38H6C.21* | WBGene00012633 | protein_coding_gene | -1.6 | 0.340 | 6.2E-06 |
| *phlp-2* | WBGene00009647 | protein_coding_gene | -1.6 | 0.340 | 1.4E-03 |
| *ZC84.6* | WBGene00013849 | protein_coding_gene | -1.6 | 0.340 | 3.6E-03 |
| *snb-5* | WBGene00004899 | protein_coding_gene | -1.6 | 0.339 | 2.9E-02 |
| *F20A1.8* | WBGene00017619 | protein_coding_gene | -1.6 | 0.339 | 3.3E-02 |
| *R07H5.11* | WBGene00011131 | protein_coding_gene | -1.6 | 0.339 | 5.9E-10 |
| *F10E9.11* | WBGene00017360 | protein_coding_gene | -1.6 | 0.338 | 1.1E-29 |
| *H01G02.1* | WBGene00010348 | protein_coding_gene | -1.6 | 0.337 | 1.9E-12 |
| *C27A7.8* | WBGene00007758 | protein_coding_gene | -1.6 | 0.337 | 4.6E-10 |
| *cut-2* | WBGene00009983 | protein_coding_gene | -1.6 | 0.335 | 4.0E-02 |
| *K07E12.2* | WBGene00019501 | protein_coding_gene | -1.6 | 0.334 | 5.9E-06 |
| *Y57E12B.10* | WBGene00202501 | protein_coding_gene | -1.6 | 0.333 | 4.3E-02 |
| *K11H12.11* | WBGene00044723 | protein_coding_gene | -1.6 | 0.333 | 1.3E-02 |
| *C09G1.7* | WBGene00284856 | ncRNA_gene | -1.6 | 0.331 | 3.3E-02 |
| *srx-125* | WBGene00006016 | protein_coding_gene | -1.6 | 0.331 | 1.4E-02 |
| *C33E10.3* | WBGene00023245 | pseudogene | -1.6 | 0.329 | 1.3E-02 |
| *Y71A12B.19* | WBGene00077443 | pseudogene | -1.6 | 0.329 | 5.3E-12 |
| *F48E3.9* | WBGene00018608 | protein_coding_gene | -1.6 | 0.327 | 2.9E-04 |
| *C05D9.7* | WBGene00015474 | protein_coding_gene | -1.6 | 0.326 | 3.4E-16 |
| *Y39B6A.31* | WBGene00012690 | pseudogene | -1.6 | 0.325 | 1.2E-08 |
| *B0228.6* | WBGene00015063 | protein_coding_gene | -1.6 | 0.324 | 1.5E-10 |
| *msp-79* | WBGene00003466 | protein_coding_gene | -1.6 | 0.324 | 3.7E-05 |
| *C39B5.2* | WBGene00016520 | protein_coding_gene | -1.6 | 0.324 | 1.5E-14 |
| *F08D12.2* | WBGene00017246 | protein_coding_gene | -1.6 | 0.319 | 1.0E-03 |
| *ttr-51* | WBGene00010440 | protein_coding_gene | -1.6 | 0.319 | 2.9E-26 |
| *2RSSE.3* | WBGene00044951 | pseudogene | -1.7 | 0.318 | 1.9E-02 |
| *dpy-10* | WBGene00001072 | protein_coding_gene | -1.7 | 0.318 | 1.8E-02 |
| *srh-254* | WBGene00005460 | pseudogene | -1.7 | 0.314 | 4.5E-07 |
| *D1086.10* | WBGene00044080 | protein_coding_gene | -1.7 | 0.313 | 2.1E-09 |
| *clec-183* | WBGene00011855 | protein_coding_gene | -1.7 | 0.312 | 1.4E-02 |
| *fmi-1* | WBGene00001475 | protein_coding_gene | -1.7 | 0.311 | 9.5E-12 |
| *K01A2.3* | WBGene00019278 | pseudogene | -1.7 | 0.311 | 3.1E-06 |
| *T05E12.3* | WBGene00011486 | protein_coding_gene | -1.7 | 0.310 | 1.5E-27 |
| *F38E9.1* | WBGene00018184 | protein_coding_gene | -1.7 | 0.310 | 3.2E-28 |
| *igcm-3* | WBGene00020160 | protein_coding_gene | -1.7 | 0.308 | 8.0E-31 |
| *Y41D4B.14* | WBGene00021516 | protein_coding_gene | -1.7 | 0.308 | 6.6E-07 |
| *F40F8.12* | WBGene00044917 | protein_coding_gene | -1.7 | 0.306 | 2.8E-02 |
| *C07A4.3* | WBGene00007398 | protein_coding_gene | -1.7 | 0.304 | 1.4E-05 |
| *cest-6* | WBGene00020688 | protein_coding_gene | -1.7 | 0.303 | 9.8E-09 |
| *str-163* | WBGene00006208 | protein_coding_gene | -1.8 | 0.297 | 2.1E-03 |
| *T24B8.5* | WBGene00011979 | protein_coding_gene | -1.8 | 0.294 | 1.9E-05 |
| *C44C10.5* | WBGene00008086 | protein_coding_gene | -1.8 | 0.293 | 9.9E-04 |
| *Y71G12B.5* | WBGene00022145 | protein_coding_gene | -1.8 | 0.292 | 3.2E-04 |
| *F55C9.5* | WBGene00010102 | protein_coding_gene | -1.8 | 0.292 | 5.8E-03 |
| *D2005.6* | WBGene00008401 | protein_coding_gene | -1.8 | 0.291 | 4.4E-07 |
| *T06G6.11* | WBGene00011550 | pseudogene | -1.8 | 0.289 | 2.7E-03 |
| *duxl-1* | WBGene00022554 | protein_coding_gene | -1.8 | 0.289 | 7.2E-04 |
| *tmem-231* | WBGene00020825 | protein_coding_gene | -1.8 | 0.289 | 1.1E-03 |
| *chk-1* | WBGene00000498 | protein_coding_gene | -1.8 | 0.287 | 1.4E-41 |
| *F27D9.13* | WBGene00270320 | protein_coding_gene | -1.8 | 0.287 | 2.4E-02 |

| | | | | | |
|---|---|---|---|---|---|
| *C32H11.4* | WBGene00007867 | protein_coding_gene | -1.8 | 0.285 | 5.0E-07 |
| *W03G11.2* | WBGene00012224 | protein_coding_gene | -1.8 | 0.284 | 3.0E-04 |
| *C35A5.5* | WBGene00007949 | protein_coding_gene | -1.8 | 0.284 | 6.6E-03 |
| *pax-3* | WBGene00003939 | protein_coding_gene | -1.8 | 0.281 | 2.3E-07 |
| *C17G10.10* | WBGene00195182 | protein_coding_gene | -1.8 | 0.280 | 7.5E-03 |
| *F58H1.5* | WBGene00010288 | protein_coding_gene | -1.8 | 0.278 | 1.5E-14 |
| *Y40H7A.147* | WBGene00077755 | pseudogene | -1.9 | 0.276 | 2.0E-07 |
| *C08D8.1* | WBGene00015592 | protein_coding_gene | -1.9 | 0.274 | 8.0E-07 |
| *ceh-53* | WBGene00015651 | protein_coding_gene | -1.9 | 0.273 | 1.3E-02 |
| *C54C8.4* | WBGene00008292 | protein_coding_gene | -1.9 | 0.271 | 2.1E-02 |
| *C26C6.4* | WBGene00007739 | protein_coding_gene | -1.9 | 0.271 | 6.6E-03 |
| *ttbk-3* | WBGene00009324 | protein_coding_gene | -1.9 | 0.271 | 1.2E-09 |
| *clec-68* | WBGene00018970 | protein_coding_gene | -1.9 | 0.270 | 3.4E-02 |
| *R11G1.7* | WBGene00044725 | protein_coding_gene | -1.9 | 0.270 | 2.1E-08 |
| *K02E10.4* | WBGene00019318 | protein_coding_gene | -1.9 | 0.266 | 3.2E-03 |
| *E04A4.3* | WBGene00017118 | pseudogene | -1.9 | 0.265 | 1.2E-02 |
| *cutl-11* | WBGene00011443 | protein_coding_gene | -1.9 | 0.264 | 8.9E-04 |
| *R01H2.8* | WBGene00045190 | protein_coding_gene | -1.9 | 0.264 | 1.6E-02 |
| *C06A5.12* | WBGene00044414 | protein_coding_gene | -1.9 | 0.263 | 1.5E-04 |
| *Y47D7A.2* | WBGene00021616 | protein_coding_gene | -1.9 | 0.263 | 4.7E-02 |
| *D1086.7* | WBGene00008394 | protein_coding_gene | -1.9 | 0.261 | 3.0E-28 |
| *Y48B6A.16* | WBGene00050948 | pseudogene | -2.0 | 0.258 | 2.3E-06 |
| *Y105C5A.1268* | WBGene00185102 | protein_coding_gene | -2.0 | 0.258 | 4.6E-03 |
| *F22B5.6* | WBGene00009044 | protein_coding_gene | -2.0 | 0.257 | 4.1E-05 |
| *efl-3* | WBGene00009899 | protein_coding_gene | -2.0 | 0.257 | 3.2E-20 |
| *ZK678.3* | WBGene00014069 | pseudogene | -2.0 | 0.256 | 3.5E-02 |
| *ZC84.1* | WBGene00013846 | protein_coding_gene | -2.0 | 0.256 | 3.6E-04 |
| *F22F7.3* | WBGene00017721 | protein_coding_gene | -2.0 | 0.256 | 8.0E-04 |
| *W08E12.6* | WBGene00021087 | protein_coding_gene | -2.0 | 0.256 | 4.3E-02 |
| *drd-50* | WBGene00018643 | protein_coding_gene | -2.0 | 0.256 | 9.2E-05 |
| *F21C10.10* | WBGene00017659 | protein_coding_gene | -2.0 | 0.255 | 1.1E-05 |
| *F25D7.5* | WBGene00009114 | protein_coding_gene | -2.0 | 0.255 | 2.5E-03 |
| *C47A4.1* | WBGene00008122 | protein_coding_gene | -2.0 | 0.255 | 5.6E-06 |
| *VH15N14R.1* | WBGene00012151 | protein_coding_gene | -2.0 | 0.255 | 7.6E-09 |
| *F01G12.18* | WBGene00206382 | protein_coding_gene | -2.0 | 0.254 | 3.8E-02 |
| *R04B3.1* | WBGene00019866 | protein_coding_gene | -2.0 | 0.254 | 1.4E-02 |
| *fbxb-87* | WBGene00043996 | protein_coding_gene | -2.0 | 0.254 | 3.0E-03 |
| *dex-1* | WBGene00017028 | protein_coding_gene | -2.0 | 0.253 | 4.7E-22 |
| *T25D1.3* | WBGene00020795 | protein_coding_gene | -2.0 | 0.253 | 4.0E-02 |
| *C55C3.7* | WBGene00043065 | protein_coding_gene | -2.0 | 0.252 | 3.4E-02 |
| *ttr-52* | WBGene00008719 | protein_coding_gene | -2.0 | 0.251 | 8.3E-05 |
| *M195.4* | WBGene00010949 | protein_coding_gene | -2.0 | 0.251 | 4.7E-03 |
| *T26A8.6* | WBGene00195245 | snoRNA_gene | -2.0 | 0.250 | 3.6E-02 |
| *Y43F8C.18* | WBGene00012839 | ncRNA_gene | -2.0 | 0.249 | 2.0E-02 |
| *ZK512.1* | WBGene00013982 | protein_coding_gene | -2.0 | 0.248 | 3.7E-12 |
| *mnp-1* | WBGene00007139 | protein_coding_gene | -2.0 | 0.248 | 2.0E-29 |
| *clec-47* | WBGene00011668 | protein_coding_gene | -2.0 | 0.247 | 3.9E-13 |
| *ssp-11* | WBGene00006040 | protein_coding_gene | -2.0 | 0.247 | 1.1E-04 |
| *col-83* | WBGene00000658 | protein_coding_gene | -2.0 | 0.246 | 5.9E-03 |
| *C30G7.4* | WBGene00044178 | protein_coding_gene | -2.0 | 0.246 | 8.5E-10 |
| *F58G6.9* | WBGene00044107 | protein_coding_gene | -2.0 | 0.245 | 6.0E-04 |
| *C18D4.6* | WBGene00007675 | protein_coding_gene | -2.0 | 0.245 | 1.8E-28 |
| *taf-7.1* | WBGene00006388 | protein_coding_gene | -2.0 | 0.244 | 9.0E-15 |
| *lips-16* | WBGene00012588 | pseudogene | -2.0 | 0.244 | 1.0E-15 |
| *C06E7.4* | WBGene00015541 | protein_coding_gene | -2.0 | 0.243 | 1.5E-05 |
| *T23C6.4* | WBGene00020726 | protein_coding_gene | -2.0 | 0.243 | 7.5E-21 |
| *his-64* | WBGene00001938 | protein_coding_gene | -2.1 | 0.241 | 2.1E-02 |
| *F19B2.12* | WBGene00194812 | pseudogene | -2.1 | 0.241 | 3.5E-13 |
| *C18D11.7* | WBGene00044232 | protein_coding_gene | -2.1 | 0.241 | 2.2E-02 |
| *nphp-1* | WBGene00010898 | protein_coding_gene | -2.1 | 0.240 | 3.9E-06 |
| *dhs-8* | WBGene00000972 | protein_coding_gene | -2.1 | 0.238 | 3.4E-05 |
| *R160.10* | WBGene00219369 | protein_coding_gene | -2.1 | 0.238 | 6.6E-04 |
| *W02B3.7* | WBGene00020927 | protein_coding_gene | -2.1 | 0.237 | 2.1E-02 |
| *F23F12.8* | WBGene00017754 | protein_coding_gene | -2.1 | 0.236 | 6.1E-04 |
| *tbb-4* | WBGene00006538 | protein_coding_gene | -2.1 | 0.235 | 4.0E-13 |
| *ins-8* | WBGene00002091 | protein_coding_gene | -2.1 | 0.235 | 2.1E-02 |
| *scrt-1* | WBGene00016948 | protein_coding_gene | -2.1 | 0.234 | 1.9E-02 |
| *Y59E9AR.1* | WBGene00021997 | protein_coding_gene | -2.1 | 0.234 | 3.0E-03 |
| *str-173* | WBGene00006217 | protein_coding_gene | -2.1 | 0.233 | 9.9E-03 |
| *hsp-12.1* | WBGene00011906 | protein_coding_gene | -2.1 | 0.233 | 2.2E-08 |
| *F54F7.9* | WBGene00045251 | protein_coding_gene | -2.1 | 0.232 | 4.0E-02 |
| *E01G4.7* | WBGene00045410 | pseudogene | -2.1 | 0.228 | 8.5E-05 |
| *C14C6.5* | WBGene00015759 | protein_coding_gene | -2.1 | 0.228 | 2.2E-07 |
| *Y75B8A.18* | WBGene00013553 | protein_coding_gene | -2.1 | 0.228 | 1.3E-46 |
| *C24H12.18* | WBGene00235333 | protein_coding_gene | -2.1 | 0.226 | 4.3E-02 |
| *F49F1.11* | WBGene00018651 | protein_coding_gene | -2.2 | 0.225 | 3.8E-02 |
| *C05E11.3* | WBGene00015494 | protein_coding_gene | -2.2 | 0.224 | 3.6E-22 |
| *T09E11.12* | WBGene00045103 | protein_coding_gene | -2.2 | 0.223 | 3.4E-02 |
| *linc-23* | WBGene00219692 | lincRNA_gene | -2.2 | 0.222 | 3.8E-02 |
| *rga-2* | WBGene00013135 | protein_coding_gene | -2.2 | 0.222 | 1.7E-15 |

| | | | | | |
|---|---|---|---|---|---|
| *F28H6.8* | WBGene00044137 | protein_coding_gene | -2.2 | 0.221 | 2.1E-02 |
| *fbxb-70* | WBGene00014149 | protein_coding_gene | -2.2 | 0.221 | 4.6E-03 |
| *ZK402.6* | WBGene00044748 | ncRNA_gene | -2.2 | 0.220 | 3.9E-02 |
| *K11H12.10* | WBGene00044458 | protein_coding_gene | -2.2 | 0.220 | 4.7E-02 |
| *W01B6.11* | WBGene00077750 | pseudogene | -2.2 | 0.220 | 2.5E-02 |
| *F28H1.1* | WBGene00017908 | protein_coding_gene | -2.2 | 0.219 | 5.6E-34 |
| *srm-1* | WBGene00005638 | protein_coding_gene | -2.2 | 0.219 | 8.2E-05 |
| *ceh-16* | WBGene00000439 | protein_coding_gene | -2.2 | 0.219 | 2.4E-07 |
| *Y116A8C.7* | WBGene00014976 | pseudogene | -2.2 | 0.219 | 1.9E-02 |
| *T15D6.10* | WBGene00011784 | protein_coding_gene | -2.2 | 0.218 | 2.8E-02 |
| *clec-259* | WBGene00010927 | pseudogene | -2.2 | 0.217 | 4.0E-02 |
| *C53A5.2* | WBGene00008263 | protein_coding_gene | -2.2 | 0.217 | 7.3E-06 |
| *F36H12.17* | WBGene00018130 | protein_coding_gene | -2.2 | 0.216 | 9.2E-04 |
| *Y53F4B.20* | WBGene00013166 | protein_coding_gene | -2.2 | 0.216 | 2.2E-04 |
| *C14E2.4* | WBGene00015773 | protein_coding_gene | -2.2 | 0.216 | 3.0E-02 |
| *Y53F4B.43* | WBGene00044726 | protein_coding_gene | -2.2 | 0.215 | 1.3E-02 |
| *C18D11.10* | WBGene00194727 | protein_coding_gene | -2.2 | 0.215 | 3.7E-02 |
| *T24H7.6* | WBGene00044936 | snoRNA_gene | -2.2 | 0.215 | 1.7E-02 |
| *C35D6.3* | WBGene00007960 | protein_coding_gene | -2.2 | 0.214 | 2.3E-02 |
| *W05B10.3* | WBGene00012278 | protein_coding_gene | -2.2 | 0.214 | 1.1E-02 |
| *Y47D7A.7* | WBGene00021620 | protein_coding_gene | -2.2 | 0.213 | 1.8E-04 |
| *tyr-5* | WBGene00021553 | protein_coding_gene | -2.2 | 0.212 | 2.4E-14 |
| *C49G7.13* | WBGene00269425 | protein_coding_gene | -2.2 | 0.212 | 3.4E-08 |
| *nspa-6* | WBGene00044273 | protein_coding_gene | -2.2 | 0.212 | 2.5E-02 |
| *unc-39* | WBGene00006775 | protein_coding_gene | -2.2 | 0.211 | 1.4E-07 |
| *R11E3.19* | WBGene00235374 | protein_coding_gene | -2.2 | 0.211 | 3.6E-04 |
| *tig-3* | WBGene00021594 | protein_coding_gene | -2.2 | 0.211 | 9.0E-06 |
| *nlp-23* | WBGene00003761 | protein_coding_gene | -2.3 | 0.209 | 2.8E-02 |
| *pes-23* | WBGene00003987 | protein_coding_gene | -2.3 | 0.208 | 1.1E-12 |
| *F40H6.2* | WBGene00018254 | protein_coding_gene | -2.3 | 0.206 | 3.7E-02 |
| *srh-132* | WBGene00005349 | protein_coding_gene | -2.3 | 0.205 | 3.1E-02 |
| *T05A7.13* | WBGene00271415 | protein_coding_gene | -2.3 | 0.205 | 1.5E-07 |
| *W08E3.4* | WBGene00012345 | protein_coding_gene | -2.3 | 0.204 | 1.3E-05 |
| *T26E3.6* | WBGene00012039 | protein_coding_gene | -2.3 | 0.204 | 3.4E-02 |
| *linc-16* | WBGene00219728 | lincRNA_gene | -2.3 | 0.204 | 1.9E-03 |
| *Y79H2A.2* | WBGene00013579 | protein_coding_gene | -2.3 | 0.203 | 3.4E-19 |
| *F55G7.5* | WBGene00195215 | protein_coding_gene | -2.3 | 0.202 | 3.6E-02 |
| *fbxb-77* | WBGene00021204 | protein_coding_gene | -2.3 | 0.202 | 2.4E-04 |
| *his-5* | WBGene00001879 | protein_coding_gene | -2.3 | 0.201 | 2.8E-02 |
| *grl-11* | WBGene00001720 | protein_coding_gene | -2.3 | 0.201 | 2.1E-02 |
| *fbxa-102* | WBGene00010534 | protein_coding_gene | -2.3 | 0.198 | 2.7E-02 |
| *hmg-6* | WBGene00009827 | protein_coding_gene | -2.3 | 0.197 | 9.6E-03 |
| *Y17G9B.11* | WBGene00194835 | protein_coding_gene | -2.3 | 0.197 | 4.3E-02 |
| *Y51H4A.937* | WBGene00194927 | protein_coding_gene | -2.3 | 0.197 | 3.8E-02 |
| *W03D2.9* | WBGene00020983 | protein_coding_gene | -2.3 | 0.197 | 9.4E-04 |
| *Y116F11B.17* | WBGene00189943 | protein_coding_gene | -2.3 | 0.197 | 3.8E-02 |
| *nhr-77* | WBGene00003667 | protein_coding_gene | -2.3 | 0.197 | 3.2E-02 |
| *ZK185.9* | WBGene00220238 | snoRNA_gene | -2.3 | 0.197 | 8.6E-03 |
| *T19B10.12* | WBGene00050898 | protein_coding_gene | -2.3 | 0.196 | 3.6E-02 |
| *C44C10.15* | WBGene00219301 | protein_coding_gene | -2.4 | 0.195 | 3.9E-02 |
| *ceh-43* | WBGene00000463 | protein_coding_gene | -2.4 | 0.194 | 2.4E-29 |
| *Y7A5A.5* | WBGene00012411 | protein_coding_gene | -2.4 | 0.194 | 4.4E-02 |
| *Y13C8A.4* | WBGene00219303 | pseudogene | -2.4 | 0.193 | 4.9E-13 |
| *K11H12.5* | WBGene00019661 | protein_coding_gene | -2.4 | 0.193 | 2.5E-13 |
| *F25H5.8* | WBGene00009130 | protein_coding_gene | -2.4 | 0.192 | 4.6E-22 |
| *F28D1.13* | WBGene00044962 | snoRNA_gene | -2.4 | 0.191 | 1.8E-02 |
| *T02C1.1* | WBGene00011365 | protein_coding_gene | -2.4 | 0.191 | 1.1E-10 |
| *F26D10.24* | WBGene00219917 | snoRNA_gene | -2.4 | 0.191 | 1.0E-02 |
| *K07E8.1* | WBGene00019494 | pseudogene | -2.4 | 0.190 | 1.7E-02 |
| *K07C5.10* | WBGene00023428 | snoRNA_gene | -2.4 | 0.190 | 9.6E-03 |
| *Y51A2D.23* | WBGene00077653 | pseudogene | -2.4 | 0.188 | 1.5E-02 |
| *skr-21* | WBGene00004827 | protein_coding_gene | -2.4 | 0.188 | 1.2E-40 |
| *Y71A12B.24* | WBGene00195228 | protein_coding_gene | -2.4 | 0.187 | 3.4E-04 |
| *ceh-8* | WBGene00000433 | protein_coding_gene | -2.4 | 0.187 | 2.5E-04 |
| *F48F7.3* | WBGene00009849 | protein_coding_gene | -2.4 | 0.186 | 8.0E-04 |
| *srz-56* | WBGene00016519 | protein_coding_gene | -2.4 | 0.185 | 8.3E-03 |
| *srg-58* | WBGene00005215 | protein_coding_gene | -2.4 | 0.185 | 3.5E-02 |
| *Y18D10A.30* | WBGene00271598 | pseudogene | -2.4 | 0.184 | 4.0E-04 |
| *Y60A9A.3* | WBGene00194675 | pseudogene | -2.4 | 0.184 | 1.4E-02 |
| *T07D3.9* | WBGene00020311 | protein_coding_gene | -2.4 | 0.184 | 5.6E-22 |
| *mls-2* | WBGene00003377 | protein_coding_gene | -2.5 | 0.183 | 4.7E-08 |
| *F59A1.18* | WBGene00206506 | protein_coding_gene | -2.5 | 0.183 | 1.9E-02 |
| *linc-68* | WBGene00219756 | lincRNA_gene | -2.5 | 0.183 | 2.0E-02 |
| *ehn-3* | WBGene00001223 | protein_coding_gene | -2.5 | 0.183 | 2.4E-02 |
| *clec-162* | WBGene00007399 | protein_coding_gene | -2.5 | 0.182 | 2.5E-05 |
| *C24H12.19* | WBGene00235338 | pseudogene | -2.5 | 0.182 | 1.1E-03 |
| *K05C4.3* | WBGene00010580 | protein_coding_gene | -2.5 | 0.182 | 2.7E-03 |
| *T04A8.17* | WBGene00045118 | snoRNA_gene | -2.5 | 0.181 | 1.2E-02 |
| *F18G5.5* | WBGene00017581 | pseudogene | -2.5 | 0.181 | 1.8E-02 |
| *F35D2.3* | WBGene00018041 | protein_coding_gene | -2.5 | 0.180 | 1.2E-25 |

| | | | | | |
|---|---|---|---|---|---|
| *kal-1* | WBGene00002181 | protein_coding_gene | -2.5 | 0.179 | 2.9E-28 |
| *oac-44* | WBGene00011654 | protein_coding_gene | -2.5 | 0.179 | 9.7E-03 |
| *C46A5.4* | WBGene00016700 | protein_coding_gene | -2.5 | 0.178 | 1.6E-07 |
| *oac-45* | WBGene00011656 | protein_coding_gene | -2.5 | 0.177 | 8.0E-03 |
| *W10D9.1* | WBGene00021129 | protein_coding_gene | -2.5 | 0.176 | 1.3E-02 |
| *cpn-2* | WBGene00000778 | protein_coding_gene | -2.5 | 0.174 | 1.5E-06 |
| *cfz-2* | WBGene00000478 | protein_coding_gene | -2.5 | 0.174 | 1.7E-15 |
| *F31E3.12* | WBGene00219317 | protein_coding_gene | -2.5 | 0.174 | 1.9E-04 |
| *rrn-4.2* | WBGene00077465 | rRNA_gene | -2.5 | 0.173 | 4.0E-03 |
| *ZC449.2* | WBGene00022612 | protein_coding_gene | -2.5 | 0.172 | 3.4E-08 |
| *sem-2* | WBGene00004771 | protein_coding_gene | -2.5 | 0.172 | 1.5E-09 |
| *F16G10.6* | WBGene00017521 | protein_coding_gene | -2.5 | 0.172 | 1.1E-02 |
| *lips-2* | WBGene00007291 | protein_coding_gene | -2.5 | 0.171 | 1.3E-02 |
| *Y45F10B.59* | WBGene00194917 | protein_coding_gene | -2.5 | 0.171 | 1.8E-02 |
| *F13E9.4* | WBGene00008753 | protein_coding_gene | -2.6 | 0.169 | 2.3E-03 |
| *C04F6.9* | WBGene00304901 | protein_coding_gene | -2.6 | 0.169 | 2.3E-02 |
| *W04H10.5* | WBGene00206416 | protein_coding_gene | -2.6 | 0.168 | 1.4E-03 |
| *F28B1.3* | WBGene00009197 | pseudogene | -2.6 | 0.168 | 1.1E-02 |
| *F25B3.2* | WBGene00009099 | protein_coding_gene | -2.6 | 0.168 | 3.3E-05 |
| *clec-171* | WBGene00022312 | protein_coding_gene | -2.6 | 0.167 | 8.5E-03 |
| *cut-6* | WBGene00000853 | protein_coding_gene | -2.6 | 0.167 | 1.5E-08 |
| *F46F11.12* | WBGene00045082 | snoRNA_gene | -2.6 | 0.166 | 2.0E-03 |
| *lhfp-4* | WBGene00009155 | protein_coding_gene | -2.6 | 0.165 | 4.5E-04 |
| *Y49F6C.2* | WBGene00021724 | protein_coding_gene | -2.6 | 0.165 | 3.1E-02 |
| *egl-1* | WBGene00001170 | protein_coding_gene | -2.6 | 0.164 | 3.4E-04 |
| *T04F8.9* | WBGene00011447 | protein_coding_gene | -2.6 | 0.163 | 2.4E-06 |
| *dct-19* | WBGene00007876 | protein_coding_gene | -2.6 | 0.163 | 2.7E-03 |
| *F17E9.16* | WBGene00195009 | protein_coding_gene | -2.6 | 0.161 | 5.9E-03 |
| *F17E9.15* | WBGene00194833 | protein_coding_gene | -2.6 | 0.161 | 8.3E-03 |
| *W10G11.4* | WBGene00021137 | protein_coding_gene | -2.6 | 0.161 | 8.8E-04 |
| *T09D3.3* | WBGene00020384 | protein_coding_gene | -2.6 | 0.160 | 1.3E-02 |
| *F32H2.6* | WBGene00009343 | protein_coding_gene | -2.6 | 0.160 | 7.3E-07 |
| *ceh-76* | WBGene00022395 | protein_coding_gene | -2.7 | 0.158 | 4.5E-03 |
| *his-31* | WBGene00001905 | protein_coding_gene | -2.7 | 0.158 | 2.2E-03 |
| *Y71A12B.15* | WBGene00013511 | protein_coding_gene | -2.7 | 0.157 | 8.0E-03 |
| *T27A8.7* | WBGene00206527 | protein_coding_gene | -2.7 | 0.155 | 4.4E-03 |
| *K07D4.2* | WBGene00019483 | protein_coding_gene | -2.7 | 0.154 | 6.8E-03 |
| *F59A6.3* | WBGene00019085 | protein_coding_gene | -2.7 | 0.153 | 5.9E-03 |
| *dgn-3* | WBGene00017221 | protein_coding_gene | -2.7 | 0.152 | 9.2E-03 |
| *R03H10.5* | WBGene00019857 | protein_coding_gene | -2.7 | 0.151 | 8.8E-03 |
| *F49B2.9* | WBGene00268216 | protein_coding_gene | -2.7 | 0.151 | 3.7E-03 |
| *nspa-1* | WBGene00010377 | protein_coding_gene | -2.7 | 0.150 | 6.3E-09 |
| *lov-1* | WBGene00003058 | protein_coding_gene | -2.7 | 0.150 | 8.3E-04 |
| *F54F7.3* | WBGene00010065 | protein_coding_gene | -2.7 | 0.149 | 1.1E-33 |
| *Y75B12B.13* | WBGene00045402 | protein_coding_gene | -2.7 | 0.149 | 1.1E-02 |
| *clec-93* | WBGene00013929 | protein_coding_gene | -2.8 | 0.148 | 5.6E-03 |
| *C17H12.37* | WBGene00236786 | protein_coding_gene | -2.8 | 0.148 | 1.5E-03 |
| *C54C8.12* | WBGene00044242 | protein_coding_gene | -2.8 | 0.148 | 1.3E-02 |
| *Y48G9A.6* | WBGene00021700 | protein_coding_gene | -2.8 | 0.148 | 5.5E-03 |
| *sdz-1* | WBGene00015129 | protein_coding_gene | -2.8 | 0.147 | 6.7E-05 |
| *ZK381.2* | WBGene00022725 | protein_coding_gene | -2.8 | 0.146 | 1.5E-03 |
| *C16C4.18* | WBGene00194998 | protein_coding_gene | -2.8 | 0.146 | 3.3E-03 |
| *F55G11.10* | WBGene00010129 | protein_coding_gene | -2.8 | 0.144 | 1.4E-02 |
| *ceh-30* | WBGene00000451 | protein_coding_gene | -2.8 | 0.144 | 3.0E-06 |
| *C35A11.3* | WBGene00016430 | protein_coding_gene | -2.8 | 0.144 | 5.8E-03 |
| *F21D5.9* | WBGene00009014 | protein_coding_gene | -2.8 | 0.143 | 5.7E-06 |
| *C47E12.13* | WBGene00008156 | protein_coding_gene | -2.8 | 0.143 | 6.0E-03 |
| *T04C12.1* | WBGene00011427 | protein_coding_gene | -2.8 | 0.142 | 2.0E-06 |
| *F47E1.4* | WBGene00018568 | protein_coding_gene | -2.8 | 0.140 | 4.9E-19 |
| *W09G10.9* | WBGene00271830 | protein_coding_gene | -2.8 | 0.139 | 7.8E-04 |
| *C05B5.1* | WBGene00007319 | protein_coding_gene | -2.9 | 0.137 | 4.6E-03 |
| *hbl-1* | WBGene00001824 | protein_coding_gene | -2.9 | 0.137 | 5.7E-24 |
| *Y7A9D.1* | WBGene00012420 | protein_coding_gene | -2.9 | 0.136 | 9.3E-35 |
| *Y62H9A.8* | WBGene00013396 | protein_coding_gene | -2.9 | 0.136 | 5.9E-03 |
| *fbxb-55* | WBGene00020979 | protein_coding_gene | -2.9 | 0.135 | 2.3E-03 |
| *Y74C9A.6* | WBGene00023193 | snoRNA_gene | -2.9 | 0.135 | 2.1E-03 |
| *Y77E11A.22* | WBGene00235266 | pseudogene | -2.9 | 0.134 | 3.1E-03 |
| *B0302.2* | WBGene00015119 | protein_coding_gene | -2.9 | 0.133 | 5.4E-03 |
| *Y19D10B.4* | WBGene00021233 | pseudogene | -2.9 | 0.132 | 7.3E-04 |
| *F16G10.9* | WBGene00017524 | protein_coding_gene | -2.9 | 0.132 | 5.2E-03 |
| *F35C8.9* | WBGene00077784 | pseudogene | -2.9 | 0.131 | 2.6E-03 |
| *W01C9.1* | WBGene00012175 | protein_coding_gene | -2.9 | 0.130 | 4.8E-05 |
| *dmsr-12* | WBGene00019262 | protein_coding_gene | -2.9 | 0.130 | 2.6E-12 |
| *E03H4.9* | WBGene00014729 | pseudogene | -2.9 | 0.130 | 2.2E-03 |
| *F02H6.7* | WBGene00008540 | protein_coding_gene | -3.0 | 0.129 | 3.3E-03 |
| *Y26G10.3* | WBGene00050973 | pseudogene | -3.0 | 0.129 | 4.2E-03 |
| *T08B6.64* | WBGene00206375 | snoRNA_gene | -3.0 | 0.128 | 1.3E-03 |
| *M199.9* | WBGene00044998 | protein_coding_gene | -3.0 | 0.128 | 1.7E-03 |
| *Y51H7C.12* | WBGene00021790 | protein_coding_gene | -3.0 | 0.126 | 2.4E-04 |
| *Y64G10A.10* | WBGene00045144 | protein_coding_gene | -3.0 | 0.126 | 4.0E-03 |

| | | | | | |
|---|---|---|---|---|---|
| *fbxb-114* | WBGene00016254 | protein_coding_gene | -3.0 | 0.122 | 5.0E-03 |
| *K08E3.2* | WBGene00010662 | protein_coding_gene | -3.0 | 0.122 | 2.8E-05 |
| *F59A7.12* | WBGene00044520 | protein_coding_gene | -3.0 | 0.122 | 2.1E-03 |
| *clc-4* | WBGene00000525 | protein_coding_gene | -3.0 | 0.122 | 2.8E-14 |
| *Y40A1A.2* | WBGene00021484 | protein_coding_gene | -3.0 | 0.122 | 1.1E-03 |
| *oac-43* | WBGene00011653 | protein_coding_gene | -3.0 | 0.121 | 2.7E-03 |
| *C54E10.10* | WBGene00206523 | protein_coding_gene | -3.1 | 0.121 | 1.7E-03 |
| *Y11D7A.19* | WBGene00077756 | protein_coding_gene | -3.1 | 0.120 | 8.6E-05 |
| *F18C5.9* | WBGene00017562 | protein_coding_gene | -3.1 | 0.120 | 8.3E-04 |
| *fbxb-38* | WBGene00016337 | protein_coding_gene | -3.1 | 0.118 | 3.6E-05 |
| *F53B3.5* | WBGene00018743 | protein_coding_gene | -3.1 | 0.118 | 1.4E-45 |
| *dmd-5* | WBGene00017326 | protein_coding_gene | -3.1 | 0.117 | 8.3E-11 |
| *fbxb-88* | WBGene00016359 | protein_coding_gene | -3.1 | 0.116 | 4.6E-24 |
| *F29A7.3* | WBGene00017913 | protein_coding_gene | -3.1 | 0.116 | 7.7E-04 |
| *C16C8.17* | WBGene00015854 | protein_coding_gene | -3.1 | 0.116 | 2.8E-05 |
| *fbxa-70* | WBGene00020458 | pseudogene | -3.1 | 0.115 | 1.9E-03 |
| *K08C9.10* | WBGene00086564 | protein_coding_gene | -3.1 | 0.114 | 1.8E-03 |
| *atf-2* | WBGene00000220 | protein_coding_gene | -3.1 | 0.113 | 2.2E-07 |
| *T10D4.7* | WBGene00020409 | protein_coding_gene | -3.2 | 0.110 | 2.3E-03 |
| *dyf-7* | WBGene00001123 | protein_coding_gene | -3.2 | 0.109 | 6.2E-09 |
| *C29A12.6* | WBGene00007800 | protein_coding_gene | -3.2 | 0.109 | 1.0E-08 |
| *Y6E2A.1* | WBGene00012397 | protein_coding_gene | -3.2 | 0.108 | 2.4E-03 |
| *W04H10.6* | WBGene00206417 | protein_coding_gene | -3.2 | 0.108 | 1.2E-03 |
| *Y47D7A.11* | WBGene00021623 | protein_coding_gene | -3.2 | 0.108 | 1.0E-03 |
| *T11F9.22* | WBGene00194784 | protein_coding_gene | -3.2 | 0.108 | 2.6E-05 |
| *Y53H1B.4* | WBGene00013182 | protein_coding_gene | -3.2 | 0.107 | 3.9E-04 |
| *C54E10.11* | WBGene00206524 | protein_coding_gene | -3.2 | 0.106 | 2.0E-03 |
| *F46B3.14* | WBGene00009765 | protein_coding_gene | -3.3 | 0.104 | 8.1E-04 |
| *C10F3.7* | WBGene00015679 | protein_coding_gene | -3.3 | 0.104 | 5.3E-05 |
| *clec-78* | WBGene00018547 | protein_coding_gene | -3.3 | 0.103 | 1.4E-32 |
| *cut-5* | WBGene00011104 | protein_coding_gene | -3.3 | 0.103 | 8.2E-09 |
| *K10G6.4* | WBGene00019641 | protein_coding_gene | -3.3 | 0.103 | 1.5E-08 |
| *C27D6.15* | WBGene00303362 | protein_coding_gene | -3.3 | 0.102 | 1.4E-03 |
| *linc-1* | WBGene00194659 | lincRNA_gene | -3.3 | 0.101 | 1.3E-30 |
| *ZC204.17* | WBGene00194714 | protein_coding_gene | -3.3 | 0.100 | 7.6E-04 |
| *C41G7.15* | WBGene00219284 | protein_coding_gene | -3.3 | 0.099 | 6.7E-04 |
| *nas-19* | WBGene00003538 | protein_coding_gene | -3.3 | 0.099 | 4.8E-04 |
| *Y57G11C.39* | WBGene00013328 | protein_coding_gene | -3.3 | 0.099 | 8.4E-04 |
| *T12A2.6* | WBGene00020440 | protein_coding_gene | -3.3 | 0.099 | 1.3E-03 |
| *Y43F8C.23* | WBGene00045344 | protein_coding_gene | -3.3 | 0.098 | 8.8E-04 |
| *F59A6.11* | WBGene00044664 | protein_coding_gene | -3.4 | 0.098 | 3.4E-07 |
| *F56D2.8* | WBGene00018968 | protein_coding_gene | -3.4 | 0.097 | 9.4E-12 |
| *egl-46* | WBGene00001210 | protein_coding_gene | -3.4 | 0.096 | 4.6E-33 |
| *F58F9.9* | WBGene00044307 | protein_coding_gene | -3.4 | 0.096 | 5.7E-04 |
| *F46B3.15* | WBGene00009766 | protein_coding_gene | -3.4 | 0.096 | 6.9E-04 |
| *cutl-13* | WBGene00021025 | protein_coding_gene | -3.4 | 0.092 | 2.8E-07 |
| *M01G12.10* | WBGene00010823 | protein_coding_gene | -3.5 | 0.089 | 2.9E-07 |
| *F31D4.5* | WBGene00009292 | protein_coding_gene | -3.5 | 0.088 | 2.2E-10 |
| *ZC178.2* | WBGene00022536 | protein_coding_gene | -3.5 | 0.088 | 4.4E-05 |
| *pax-1* | WBGene00003937 | protein_coding_gene | -3.5 | 0.087 | 4.9E-04 |
| *F28B1.2* | WBGene00009196 | protein_coding_gene | -3.5 | 0.087 | 1.9E-04 |
| *F55C9.12* | WBGene00044196 | pseudogene | -3.5 | 0.086 | 1.9E-04 |
| *C10G8.2* | WBGene00015681 | protein_coding_gene | -3.5 | 0.086 | 6.3E-04 |
| *F02E11.3* | WBGene00017181 | protein_coding_gene | -3.6 | 0.085 | 3.0E-04 |
| *cwp-2* | WBGene00000860 | protein_coding_gene | -3.6 | 0.085 | 3.0E-04 |
| *B0228.8* | WBGene00015065 | protein_coding_gene | -3.6 | 0.084 | 1.5E-04 |
| *F49C5.12* | WBGene00184956 | protein_coding_gene | -3.6 | 0.083 | 1.4E-04 |
| *ZK675.4* | WBGene00014067 | protein_coding_gene | -3.6 | 0.083 | 3.4E-07 |
| *C06A12.8* | WBGene00007359 | protein_coding_gene | -3.6 | 0.083 | 3.7E-04 |
| *H25K10.4* | WBGene00010413 | protein_coding_gene | -3.6 | 0.082 | 5.8E-16 |
| *C05C10.8* | WBGene00086567 | protein_coding_gene | -3.7 | 0.079 | 1.7E-04 |
| *clec-108* | WBGene00012504 | protein_coding_gene | -3.7 | 0.079 | 1.8E-08 |
| *cnd-1* | WBGene00000561 | protein_coding_gene | -3.7 | 0.077 | 4.7E-16 |
| *nep-25* | WBGene00022874 | protein_coding_gene | -3.7 | 0.076 | 5.5E-19 |
| *oac-4* | WBGene00015873 | protein_coding_gene | -3.7 | 0.075 | 1.2E-08 |
| *hil-7* | WBGene00001858 | protein_coding_gene | -3.8 | 0.074 | 5.1E-27 |
| *nas-24* | WBGene00003543 | protein_coding_gene | -3.8 | 0.074 | 1.7E-45 |
| *C44B12.9* | WBGene00044462 | protein_coding_gene | -3.8 | 0.073 | 2.7E-13 |
| *LLC1.121* | WBGene00235121 | pseudogene | -3.8 | 0.073 | 2.0E-04 |
| *clec-129* | WBGene00021145 | protein_coding_gene | -3.8 | 0.072 | 2.7E-04 |
| *fbxb-39* | WBGene00019702 | protein_coding_gene | -3.9 | 0.068 | 3.8E-05 |
| *F57B1.6* | WBGene00010189 | protein_coding_gene | -3.9 | 0.068 | 1.4E-14 |
| *Y116F11A.3* | WBGene00013817 | protein_coding_gene | -3.9 | 0.068 | 7.3E-05 |
| *F56C3.8* | WBGene00018945 | protein_coding_gene | -3.9 | 0.067 | 1.7E-19 |
| *bgnt-1.3* | WBGene00009032 | protein_coding_gene | -3.9 | 0.067 | 1.8E-04 |
| *K01D12.1* | WBGene00010461 | protein_coding_gene | -3.9 | 0.066 | 8.0E-08 |
| *hlh-16* | WBGene00001960 | protein_coding_gene | -3.9 | 0.066 | 6.9E-13 |
| *W07G4.1* | WBGene00012335 | protein_coding_gene | -3.9 | 0.065 | 2.8E-05 |
| *clec-100* | WBGene00013179 | protein_coding_gene | -4.0 | 0.065 | 2.5E-05 |
| *ZC204.6* | WBGene00022556 | protein_coding_gene | -4.0 | 0.065 | 1.1E-04 |

| | | | | | |
|---|---|---|---|---|---|
| *Y57G11C.1132* | WBGene00194666 | antisense_lncRNA_gene | -4.0 | 0.064 | 3.0E-05 |
| *flp-23* | WBGene00017693 | protein_coding_gene | -4.0 | 0.063 | 1.5E-06 |
| *F13E9.19* | WBGene00194922 | protein_coding_gene | -4.0 | 0.062 | 2.0E-11 |
| *srt-63* | WBGene00020877 | protein_coding_gene | -4.0 | 0.061 | 9.7E-05 |
| *klf-2* | WBGene00009998 | protein_coding_gene | -4.0 | 0.061 | 1.2E-05 |
| *C06E7.88* | WBGene00189995 | protein_coding_gene | -4.0 | 0.061 | 4.3E-13 |
| *C35B1.8* | WBGene00016437 | protein_coding_gene | -4.1 | 0.060 | 3.9E-10 |
| *rocf-1* | WBGene00011948 | protein_coding_gene | -4.1 | 0.059 | 4.2E-11 |
| *ZK813.6* | WBGene00044564 | protein_coding_gene | -4.1 | 0.059 | 2.6E-05 |
| *Y37F4.3* | WBGene00021384 | protein_coding_gene | -4.1 | 0.059 | 4.0E-05 |
| *B0391.14* | WBGene00206462 | protein_coding_gene | -4.1 | 0.059 | 4.1E-05 |
| *B0513.90* | WBGene00235301 | protein_coding_gene | -4.1 | 0.058 | 2.8E-09 |
| *Y17G7B.23* | WBGene00044030 | protein_coding_gene | -4.1 | 0.058 | 5.8E-07 |
| *F15H9.8* | WBGene00194652 | pseudogene | -4.1 | 0.057 | 1.1E-05 |
| *D2096.5* | WBGene00017072 | protein_coding_gene | -4.2 | 0.055 | 4.6E-20 |
| *ins-19* | WBGene00002102 | protein_coding_gene | -4.2 | 0.055 | 5.4E-06 |
| *E02H9.1* | WBGene00017099 | protein_coding_gene | -4.2 | 0.054 | 1.4E-18 |
| *F26C11.3* | WBGene00009147 | protein_coding_gene | -4.2 | 0.054 | 7.2E-06 |
| *clec-98* | WBGene00013932 | protein_coding_gene | -4.2 | 0.054 | 9.2E-06 |
| *F49B2.4* | WBGene00009864 | pseudogene | -4.2 | 0.054 | 2.8E-05 |
| *T02H6.9* | WBGene00020179 | protein_coding_gene | -4.2 | 0.054 | 2.2E-21 |
| *W09G12.6* | WBGene00021120 | protein_coding_gene | -4.2 | 0.054 | 2.8E-05 |
| *lys-10* | WBGene00003099 | protein_coding_gene | -4.3 | 0.052 | 2.2E-10 |
| *T02D1.7* | WBGene00011373 | protein_coding_gene | -4.3 | 0.051 | 9.5E-06 |
| *F49F1.2* | WBGene00235363 | protein_coding_gene | -4.3 | 0.050 | 5.4E-06 |
| *K12H6.8* | WBGene00019688 | protein_coding_gene | -4.3 | 0.050 | 1.7E-10 |
| *C08E3.14* | WBGene00044415 | protein_coding_gene | -4.3 | 0.050 | 5.8E-06 |
| *F17E9.18* | WBGene00235282 | protein_coding_gene | -4.4 | 0.049 | 7.3E-06 |
| *clec-232* | WBGene00009487 | protein_coding_gene | -4.4 | 0.049 | 8.6E-31 |
| *fip-7* | WBGene00013105 | protein_coding_gene | -4.4 | 0.048 | 3.3E-06 |
| *C16C8.10* | WBGene00015848 | protein_coding_gene | -4.4 | 0.048 | 4.5E-06 |
| *F17E9.19* | WBGene00249825 | pseudogene | -4.4 | 0.048 | 6.6E-06 |
| *nep-14* | WBGene00018225 | pseudogene | -4.4 | 0.048 | 5.3E-25 |
| *F58F9.8* | WBGene00044306 | protein_coding_gene | -4.4 | 0.047 | 4.6E-11 |
| *T02E9.6* | WBGene00044203 | protein_coding_gene | -4.4 | 0.047 | 1.5E-06 |
| *F46A8.4* | WBGene00009747 | protein_coding_gene | -4.4 | 0.046 | 7.9E-06 |
| *hpo-2* | WBGene00012500 | protein_coding_gene | -4.5 | 0.045 | 2.3E-55 |
| *pqn-73* | WBGene00004155 | protein_coding_gene | -4.5 | 0.044 | 2.0E-16 |
| *R05G6.3* | WBGene00043466 | protein_coding_gene | -4.5 | 0.044 | 2.7E-06 |
| *C47A10.12* | WBGene00008129 | protein_coding_gene | -4.5 | 0.044 | 3.3E-22 |
| *F17B5.6* | WBGene00044058 | protein_coding_gene | -4.5 | 0.043 | 1.2E-05 |
| *M01D1.11* | WBGene00220026 | snoRNA_gene | -4.6 | 0.042 | 3.3E-07 |
| *F59A1.16* | WBGene00010301 | protein_coding_gene | -4.6 | 0.042 | 8.6E-07 |
| *K08C9.6* | WBGene00010653 | protein_coding_gene | -4.6 | 0.042 | 1.3E-06 |
| *Y82E9BR.8* | WBGene00022341 | protein_coding_gene | -4.6 | 0.042 | 1.2E-06 |
| *Y43F8C.16* | WBGene00012837 | protein_coding_gene | -4.6 | 0.042 | 4.4E-07 |
| *nspf-1* | WBGene00044364 | protein_coding_gene | -4.6 | 0.041 | 1.7E-06 |
| *fipr-19* | WBGene00044451 | protein_coding_gene | -4.6 | 0.041 | 1.2E-06 |
| *T22C1.12* | WBGene00011921 | protein_coding_gene | -4.6 | 0.040 | 2.3E-10 |
| *F25E5.7* | WBGene00017788 | protein_coding_gene | -4.6 | 0.040 | 2.8E-59 |
| *F25E5.3* | WBGene00017784 | protein_coding_gene | -4.7 | 0.040 | 8.5E-41 |
| *F35C5.3* | WBGene00009391 | protein_coding_gene | -4.7 | 0.039 | 4.0E-07 |
| *Y64G10A.2* | WBGene00013412 | protein_coding_gene | -4.7 | 0.039 | 3.7E-07 |
| *Y57E12AL.4* | WBGene00021958 | protein_coding_gene | -4.7 | 0.039 | 6.5E-07 |
| *K12H6.5* | WBGene00019685 | protein_coding_gene | -4.7 | 0.038 | 2.1E-07 |
| *W07G1.7* | WBGene00012334 | protein_coding_gene | -4.7 | 0.037 | 3.9E-07 |
| *btb-13* | WBGene00022561 | protein_coding_gene | -4.8 | 0.036 | 6.1E-07 |
| *Y51A2D.1* | WBGene00013072 | protein_coding_gene | -4.8 | 0.036 | 1.5E-07 |
| *clec-106* | WBGene00012482 | protein_coding_gene | -4.8 | 0.036 | 1.3E-16 |
| *F02H6.6* | WBGene00008539 | protein_coding_gene | -4.8 | 0.036 | 1.3E-11 |
| *C28C12.3* | WBGene00016179 | protein_coding_gene | -4.8 | 0.036 | 2.2E-51 |
| *clec-126* | WBGene00021117 | protein_coding_gene | -4.8 | 0.035 | 1.3E-29 |
| *F47C12.8* | WBGene00018551 | protein_coding_gene | -4.8 | 0.035 | 5.5E-19 |
| *Y6G8.16* | WBGene00194801 | protein_coding_gene | -4.9 | 0.034 | 1.5E-07 |
| *C41D11.10* | WBGene00195180 | protein_coding_gene | -4.9 | 0.034 | 1.3E-07 |
| *B0222.10* | WBGene00015058 | ncRNA_gene | -4.9 | 0.034 | 1.7E-48 |
| *C35B1.3* | WBGene00016433 | protein_coding_gene | -4.9 | 0.033 | 1.6E-11 |
| *nspf-2* | WBGene00044365 | protein_coding_gene | -4.9 | 0.033 | 1.6E-07 |
| *Y73F8A.10* | WBGene00013519 | protein_coding_gene | -4.9 | 0.033 | 6.1E-08 |
| *F59A1.6* | WBGene00010292 | protein_coding_gene | -5.0 | 0.032 | 9.8E-08 |
| *Y46D2A.3* | WBGene00021591 | protein_coding_gene | -5.0 | 0.032 | 2.0E-07 |
| *C16C8.8* | WBGene00015846 | protein_coding_gene | -5.0 | 0.031 | 1.2E-07 |
| *C35A5.11* | WBGene00044024 | protein_coding_gene | -5.0 | 0.031 | 1.4E-17 |
| *D1022.2* | WBGene00017021 | protein_coding_gene | -5.1 | 0.030 | 9.5E-17 |
| *F26H11.8* | WBGene00195149 | protein_coding_gene | -5.1 | 0.030 | 3.7E-17 |
| *Y43F8B.19* | WBGene00077688 | protein_coding_gene | -5.1 | 0.029 | 1.9E-07 |
| *F55B11.5* | WBGene00010087 | protein_coding_gene | -5.1 | 0.029 | 7.2E-09 |
| *C03C10.5* | WBGene00007271 | protein_coding_gene | -5.1 | 0.029 | 4.3E-17 |
| *pkd-2* | WBGene00004035 | protein_coding_gene | -5.1 | 0.029 | 1.3E-08 |
| *F46A8.5* | WBGene00009748 | protein_coding_gene | -5.2 | 0.028 | 1.4E-21 |

| | | | | | |
|---|---|---|---|---|---|
| *ZK1290.11* | WBGene00022892 | protein_coding_gene | -5.2 | 0.028 | 3.1E-31 |
| *F54E2.2* | WBGene00018824 | protein_coding_gene | -5.2 | 0.027 | 3.7E-08 |
| *Y18D10A.2* | WBGene00012475 | protein_coding_gene | -5.2 | 0.027 | 9.9E-14 |
| *scl-26* | WBGene00016821 | protein_coding_gene | -5.2 | 0.027 | 7.7E-09 |
| *Y43F8C.24* | WBGene00045345 | pseudogene | -5.3 | 0.026 | 1.9E-08 |
| *fip-3* | WBGene00015520 | protein_coding_gene | -5.3 | 0.025 | 1.3E-08 |
| *clec-136* | WBGene00018047 | protein_coding_gene | -5.3 | 0.025 | 1.2E-11 |
| *C49C3.11* | WBGene00008201 | protein_coding_gene | -5.4 | 0.024 | 2.1E-87 |
| *F47C12.6* | WBGene00018549 | protein_coding_gene | -5.4 | 0.024 | 7.1E-09 |
| *T14G12.11* | WBGene00219411 | protein_coding_gene | -5.4 | 0.024 | 4.7E-12 |
| *E02H9.9* | WBGene00017106 | protein_coding_gene | -5.4 | 0.023 | 4.7E-26 |
| *R09E10.5* | WBGene00011175 | protein_coding_gene | -5.5 | 0.023 | 3.9E-21 |
| *F28B1.9* | WBGene00045381 | protein_coding_gene | -5.5 | 0.023 | 2.7E-08 |
| *F40G9.7* | WBGene00018243 | protein_coding_gene | -5.5 | 0.023 | 1.8E-09 |
| *clec-181* | WBGene00044511 | protein_coding_gene | -5.5 | 0.023 | 2.4E-08 |
| *clec-138* | WBGene00018049 | protein_coding_gene | -5.5 | 0.023 | 3.2E-20 |
| *clec-261* | WBGene00013834 | protein_coding_gene | -5.5 | 0.022 | 2.2E-09 |
| *F38E1.10* | WBGene00018182 | protein_coding_gene | -5.5 | 0.022 | 1.4E-08 |
| *F44F1.4* | WBGene00009696 | protein_coding_gene | -5.5 | 0.022 | 4.3E-20 |
| *Y49F6B.13* | WBGene00044377 | protein_coding_gene | -5.5 | 0.022 | 6.1E-09 |
| *F30A10.11* | WBGene00009268 | protein_coding_gene | -5.5 | 0.022 | 6.3E-26 |
| *T26A8.3* | WBGene00020826 | protein_coding_gene | -5.5 | 0.022 | 7.1E-25 |
| *clec-105* | WBGene00012488 | protein_coding_gene | -5.5 | 0.022 | 1.9E-09 |
| *C14F11.7* | WBGene00015782 | protein_coding_gene | -5.5 | 0.021 | 4.2E-33 |
| *ZK1225.10* | WBGene00044714 | pseudogene | -5.6 | 0.021 | 1.9E-09 |
| *T12A7.10* | WBGene00194933 | protein_coding_gene | -5.6 | 0.021 | 1.2E-09 |
| *C06E2.9* | WBGene00044579 | protein_coding_gene | -5.6 | 0.021 | 6.7E-09 |
| *F46A8.8* | WBGene00009751 | protein_coding_gene | -5.7 | 0.019 | 5.5E-09 |
| *clec-125* | WBGene00021118 | protein_coding_gene | -5.7 | 0.019 | 3.1E-63 |
| *cwp-1* | WBGene00000859 | protein_coding_gene | -5.7 | 0.019 | 3.2E-10 |
| *B0207.5* | WBGene00015028 | protein_coding_gene | -5.8 | 0.018 | 6.4E-59 |
| *F47C12.12* | WBGene00219377 | pseudogene | -5.8 | 0.018 | 1.3E-13 |
| *Y43F8C.15* | WBGene00012836 | protein_coding_gene | -5.8 | 0.018 | 2.9E-10 |
| *Y110A2AL.6* | WBGene00022443 | protein_coding_gene | -5.8 | 0.017 | 1.1E-10 |
| *F11A6.16* | WBGene00271780 | protein_coding_gene | -5.9 | 0.017 | 1.9E-09 |
| *clec-135* | WBGene00015821 | protein_coding_gene | -5.9 | 0.017 | 3.3E-14 |
| *W02D7.12* | WBGene00044356 | protein_coding_gene | -5.9 | 0.017 | 1.7E-09 |
| *F16H6.9* | WBGene00008899 | protein_coding_gene | -5.9 | 0.017 | 5.0E-11 |
| *Y51A2D.8* | WBGene00013076 | protein_coding_gene | -5.9 | 0.016 | 1.1E-66 |
| *F26C11.4* | WBGene00044159 | protein_coding_gene | -6.0 | 0.016 | 4.4E-11 |
| *clec-263* | WBGene00009171 | protein_coding_gene | -6.0 | 0.016 | 1.0E-10 |
| *clec-130* | WBGene00021144 | protein_coding_gene | -6.0 | 0.016 | 8.6E-11 |
| *K03B8.11* | WBGene00010526 | protein_coding_gene | -6.0 | 0.016 | 5.2E-11 |
| *K09C8.2* | WBGene00010715 | protein_coding_gene | -6.0 | 0.016 | 6.2E-11 |
| *C35E7.7* | WBGene00016459 | protein_coding_gene | -6.0 | 0.016 | 8.3E-10 |
| *Y49F6B.20* | WBGene00303369 | protein_coding_gene | -6.0 | 0.015 | 5.9E-11 |
| *T05A7.11* | WBGene00044383 | protein_coding_gene | -6.0 | 0.015 | 2.2E-14 |
| *scl-7* | WBGene00008029 | protein_coding_gene | -6.1 | 0.014 | 5.4E-22 |
| *scl-8* | WBGene00008030 | protein_coding_gene | -6.1 | 0.014 | 2.6E-10 |
| *clec-158* | WBGene00020067 | protein_coding_gene | -6.1 | 0.014 | 5.8E-19 |
| *R05H10.7* | WBGene00011046 | protein_coding_gene | -6.2 | 0.014 | 1.7E-19 |
| *T16G12.10* | WBGene00044222 | protein_coding_gene | -6.2 | 0.014 | 4.2E-43 |
| *ZC328.5* | WBGene00022594 | protein_coding_gene | -6.2 | 0.013 | 3.7E-12 |
| *T12A2.5* | WBGene00020439 | protein_coding_gene | -6.3 | 0.013 | 2.3E-12 |
| *F17E9.3* | WBGene00017540 | protein_coding_gene | -6.3 | 0.013 | 6.8E-12 |
| *F46A8.3* | WBGene00009746 | protein_coding_gene | -6.3 | 0.013 | 9.6E-11 |
| *clec-133* | WBGene00021142 | protein_coding_gene | -6.4 | 0.012 | 3.8E-12 |
| *C45G9.10* | WBGene00016681 | protein_coding_gene | -6.4 | 0.012 | 1.1E-12 |
| *tag-329* | WBGene00008231 | protein_coding_gene | -6.4 | 0.012 | 1.2E-48 |
| *W02B12.16* | WBGene00195148 | protein_coding_gene | -6.4 | 0.012 | 3.5E-12 |
| *F42A6.2* | WBGene00018324 | protein_coding_gene | -6.4 | 0.012 | 4.9E-12 |
| *clec-124* | WBGene00015909 | protein_coding_gene | -6.4 | 0.012 | 2.8E-28 |
| *Y6G8.14* | WBGene00138715 | protein_coding_gene | -6.4 | 0.012 | 2.0E-12 |
| *clec-161* | WBGene00010228 | protein_coding_gene | -6.4 | 0.011 | 1.4E-24 |
| *fipr-16* | WBGene00015521 | protein_coding_gene | -6.5 | 0.011 | 5.5E-13 |
| *Y49F6B.6* | WBGene00021718 | protein_coding_gene | -6.5 | 0.011 | 1.1E-12 |
| *fipr-20* | WBGene00044449 | protein_coding_gene | -6.6 | 0.011 | 4.4E-29 |
| *clec-132* | WBGene00021143 | protein_coding_gene | -6.6 | 0.010 | 4.5E-13 |
| *nspf-3* | WBGene00044366 | protein_coding_gene | -6.6 | 0.010 | 7.4E-13 |
| *F49F1.15* | WBGene00235365 | protein_coding_gene | -6.6 | 0.010 | 7.2E-12 |
| *T10B10.9* | WBGene00011685 | protein_coding_gene | -6.6 | 0.010 | 3.0E-13 |
| *Y6G8.15* | WBGene00189937 | protein_coding_gene | -6.6 | 0.010 | 8.3E-12 |
| *F07G6.8* | WBGene00017227 | protein_coding_gene | -6.7 | 0.010 | 2.6E-13 |
| *clec-92* | WBGene00013125 | protein_coding_gene | -6.7 | 0.010 | 5.5E-12 |
| *Y25C1A.2* | WBGene00021289 | protein_coding_gene | -6.7 | 0.010 | 1.5E-13 |
| *Y41D4B.6* | WBGene00021513 | protein_coding_gene | -6.7 | 0.010 | 7.5E-12 |
| *F16G10.10* | WBGene00017525 | protein_coding_gene | -6.7 | 0.009 | 2.0E-13 |
| *M7.10* | WBGene00010886 | protein_coding_gene | -6.8 | 0.009 | 9.3E-14 |
| *C16C8.19* | WBGene00015856 | protein_coding_gene | -6.8 | 0.009 | 9.6E-14 |
| *scl-9* | WBGene00009890 | protein_coding_gene | -6.8 | 0.009 | 1.8E-12 |

| | | | | | |
|---|---|---|---|---|---|
| *R53.8* | WBGene00011277 | protein_coding_gene | -6.8 | 0.009 | 2.7E-14 |
| *clec-96* | WBGene00013930 | protein_coding_gene | -6.9 | 0.009 | 1.4E-12 |
| *R09E10.8* | WBGene00011177 | protein_coding_gene | -6.9 | 0.008 | 2.7E-14 |
| *col-165* | WBGene00000738 | protein_coding_gene | -6.9 | 0.008 | 8.7E-15 |
| *F13E9.10* | WBGene00008759 | protein_coding_gene | -7.0 | 0.008 | 2.1E-14 |
| *C16C8.7* | WBGene00015845 | protein_coding_gene | -7.0 | 0.008 | 2.8E-14 |
| *clec-141* | WBGene00017139 | protein_coding_gene | -7.0 | 0.008 | 4.5E-13 |
| *clec-217* | WBGene00017824 | protein_coding_gene | -7.0 | 0.008 | 5.0E-13 |
| *T22G5.1* | WBGene00011929 | protein_coding_gene | -7.0 | 0.008 | 7.9E-15 |
| *C48B4.12* | WBGene00008175 | protein_coding_gene | -7.1 | 0.007 | 4.1E-30 |
| *F43C11.2* | WBGene00018379 | protein_coding_gene | -7.1 | 0.007 | 5.3E-15 |
| *M176.10* | WBGene00010946 | protein_coding_gene | -7.1 | 0.007 | 1.9E-15 |
| *F59B2.12* | WBGene00010314 | protein_coding_gene | -7.2 | 0.007 | 4.9E-19 |
| *clec-208* | WBGene00018097 | protein_coding_gene | -7.2 | 0.007 | 1.9E-56 |
| *C33G3.5* | WBGene00007905 | protein_coding_gene | -7.2 | 0.007 | 1.2E-13 |
| *Y74C10AR.2* | WBGene00022282 | protein_coding_gene | -7.2 | 0.007 | 5.3E-36 |
| *fipr-17* | WBGene00012965 | protein_coding_gene | -7.2 | 0.007 | 1.1E-30 |
| *T10D4.1* | WBGene00020406 | protein_coding_gene | -7.3 | 0.006 | 3.1E-16 |
| *Y110A2AL.7* | WBGene00022444 | protein_coding_gene | -7.4 | 0.006 | 3.2E-14 |
| *F46B6.13* | WBGene00044192 | protein_coding_gene | -7.4 | 0.006 | 8.5E-16 |
| *clec-110* | WBGene00008909 | protein_coding_gene | -7.5 | 0.006 | 9.8E-17 |
| *K12H6.9* | WBGene00019689 | protein_coding_gene | -7.6 | 0.005 | 2.2E-28 |
| *clec-137* | WBGene00018048 | protein_coding_gene | -7.6 | 0.005 | 5.4E-29 |
| *C16C8.9* | WBGene00015847 | protein_coding_gene | -7.6 | 0.005 | 2.6E-17 |
| *clec-99* | WBGene00013933 | protein_coding_gene | -7.6 | 0.005 | 1.6E-17 |
| *try-8* | WBGene00017791 | protein_coding_gene | -7.7 | 0.005 | 5.1E-27 |
| *fipr-28* | WBGene00013120 | protein_coding_gene | -7.7 | 0.005 | 4.4E-17 |
| *pqn-84* | WBGene00004165 | protein_coding_gene | -7.8 | 0.004 | 3.8E-18 |
| *F54D12.9* | WBGene00018822 | protein_coding_gene | -7.8 | 0.004 | 1.9E-17 |
| *clec-197* | WBGene00008202 | protein_coding_gene | -7.8 | 0.004 | 2.5E-69 |
| *F26D10.13* | WBGene00044244 | protein_coding_gene | -7.9 | 0.004 | 3.0E-16 |
| *T23G4.5* | WBGene00011961 | protein_coding_gene | -7.9 | 0.004 | 4.9E-18 |
| *clec-95* | WBGene00013927 | protein_coding_gene | -8.0 | 0.004 | 5.7E-19 |
| *fipr-18* | WBGene00044450 | protein_coding_gene | -8.1 | 0.004 | 1.4E-16 |
| *ZK1053.3* | WBGene00014198 | protein_coding_gene | -8.2 | 0.003 | 2.3E-20 |
| *K12H6.4* | WBGene00019684 | protein_coding_gene | -8.3 | 0.003 | 3.2E-19 |
| *ins-31* | WBGene00002114 | protein_coding_gene | -8.3 | 0.003 | 3.8E-26 |
| *clec-219* | WBGene00020945 | protein_coding_gene | -8.5 | 0.003 | 2.7E-49 |

Shared DEGs (N2 and CB4856) when exposed to *S. aureus* (vs. OP50)

| | | | | N2 | | | CB4856 | |
|---|---|---|---|---|---|---|---|---|
| Gene Name | Wormbase Gene ID | Type | $\log_2$ FC | FC | Adj P-value | $\log_2$ FC | FC | Adj P-value |
| *cpr-2* | WBGene00000782 | protein_coding_gene | 12.9 | 7490.0966 | 1.6E-36 | 8.4 | 336.174 | 8.2E-78 |
| *D1086.9* | WBGene00008396 | protein_coding_gene | 10.5 | 1450.0293 | 6.0E-21 | 8.9 | 469.160 | 4.2E-15 |
| *fmo-2* | WBGene00001477 | protein_coding_gene | 9.9 | 941.8149 | 4.6E-26 | 2.5 | 5.555 | 8.7E-04 |
| *ech-9* | WBGene00001158 | protein_coding_gene | 9.5 | 746.8991 | 4.0E-49 | 5.1 | 34.693 | 9.6E-16 |
| *clec-60* | WBGene00014046 | protein_coding_gene | 9.5 | 707.2606 | 1.2E-34 | 5.4 | 41.832 | 6.4E-44 |
| *R09E10.13* | WBGene00194658 | protein_coding_gene | 9.1 | 546.4793 | 1.2E-16 | 3.9 | 15.347 | 1.9E-25 |
| *C54F6.6* | WBGene00016924 | protein_coding_gene | 9.0 | 504.8586 | 7.9E-15 | 5.3 | 39.255 | 4.8E-13 |
| *F40G12.5* | WBGene00009600 | protein_coding_gene | 9.0 | 504.8527 | 2.2E-16 | 7.7 | 202.404 | 2.0E-71 |
| *clec-9* | WBGene00013493 | protein_coding_gene | 8.9 | 466.6411 | 2.2E-15 | 3.8 | 13.760 | 3.7E-03 |
| *chil-28* | WBGene00008842 | pseudogene | 8.7 | 411.1139 | 4.7E-17 | 1.9 | 3.785 | 2.1E-06 |
| *ilys-3* | WBGene00016670 | protein_coding_gene | 8.6 | 383.8322 | 6.3E-21 | 2.8 | 7.043 | 9.2E-30 |
| *fipr-23* | WBGene00007990 | protein_coding_gene | 8.6 | 381.5589 | 2.3E-15 | 4.1 | 16.903 | 1.1E-08 |
| *clec-61* | WBGene00014047 | protein_coding_gene | 8.4 | 348.1406 | 5.1E-45 | 2.9 | 7.579 | 2.1E-36 |
| *fbxa-165* | WBGene00015600 | protein_coding_gene | 8.3 | 324.7149 | 2.2E-14 | 2.4 | 5.380 | 1.5E-04 |
| *F46C5.1* | WBGene00009778 | protein_coding_gene | 8.3 | 317.3081 | 1.2E-21 | 1.6 | 2.993 | 7.1E-03 |
| *F17B5.1* | WBGene00008905 | protein_coding_gene | 8.3 | 313.6603 | 8.5E-51 | 3.5 | 11.287 | 5.4E-08 |
| *T23F2.4* | WBGene00020737 | protein_coding_gene | 8.2 | 299.2492 | 3.8E-22 | 7.1 | 135.499 | 3.9E-66 |
| *spp-12* | WBGene00004997 | protein_coding_gene | 8.2 | 288.4540 | 1.9E-12 | 4.7 | 25.209 | 1.7E-08 |
| *C54F6.15* | WBGene00044113 | protein_coding_gene | 8.0 | 261.4216 | 5.3E-12 | 5.6 | 49.907 | 1.5E-07 |
| *Y65B4BR.1* | WBGene00022040 | protein_coding_gene | 8.0 | 254.0424 | 6.1E-85 | 6.1 | 67.418 | 9.0E-36 |
| *fipr-24* | WBGene00007992 | protein_coding_gene | 7.9 | 238.7168 | 3.4E-11 | 3.6 | 12.331 | 3.2E-22 |
| *Y54G2A.37* | WBGene00023484 | protein_coding_gene | 7.8 | 223.6969 | 1.1E-13 | 2.8 | 7.079 | 3.7E-19 |
| *W02D7.11* | WBGene00020946 | protein_coding_gene | 7.8 | 217.0396 | 4.4E-15 | 3.9 | 15.411 | 1.4E-38 |
| *C37A5.5* | WBGene00014699 | protein_coding_gene | 7.6 | 200.5483 | 9.4E-11 | 7.2 | 145.826 | 1.9E-10 |
| *clec-69* | WBGene00044642 | pseudogene | 7.6 | 199.6483 | 2.7E-30 | 3.0 | 7.872 | 6.8E-06 |
| *cnc-11* | WBGene00044900 | protein_coding_gene | 7.6 | 197.9607 | 5.4E-10 | 4.5 | 22.192 | 3.2E-67 |
| *sod-5* | WBGene00007036 | protein_coding_gene | 7.6 | 196.7435 | 9.8E-23 | 6.8 | 111.809 | 5.0E-56 |
| *T07H8.11* | WBGene00304201 | protein_coding_gene | 7.6 | 189.6070 | 7.3E-35 | 5.1 | 35.420 | 3.0E-57 |
| *Y73B6BL.37* | WBGene00022256 | protein_coding_gene | 7.5 | 178.1350 | 2.2E-12 | 5.4 | 41.613 | 5.4E-19 |
| *C39B5.14* | WBGene00189950 | protein_coding_gene | 7.5 | 177.4187 | 8.3E-24 | 6.1 | 69.809 | 9.1E-54 |
| *lipl-2* | WBGene00009773 | protein_coding_gene | 7.5 | 177.1942 | 4.6E-19 | 1.5 | 2.890 | 2.4E-05 |
| *mtl-1* | WBGene00003473 | protein_coding_gene | 7.5 | 175.9932 | 3.8E-18 | 2.2 | 4.562 | 5.5E-09 |
| *fbxa-24* | WBGene00018803 | protein_coding_gene | 7.4 | 173.2206 | 8.8E-31 | 2.5 | 5.762 | 7.4E-16 |
| *F08H9.4* | WBGene00008592 | protein_coding_gene | 7.4 | 172.1009 | 7.6E-27 | 8.0 | 256.462 | 2.2E-12 |
| *asm-3* | WBGene00000213 | protein_coding_gene | 7.3 | 161.0578 | 3.7E-25 | 7.7 | 212.087 | 2.4E-97 |
| *H39E23.3* | WBGene00044201 | protein_coding_gene | 7.3 | 156.9103 | 2.4E-10 | 4.4 | 21.563 | 3.1E-04 |
| *tts-1* | WBGene00006650 | ncRNA_gene | 7.3 | 156.5966 | 4.8E-26 | 1.8 | 3.432 | 1.4E-11 |
| *T28A11.19* | WBGene00020881 | protein_coding_gene | 7.3 | 154.5272 | 2.0E-32 | 3.8 | 14.395 | 2.0E-09 |

| | | | | | | | | |
|---|---|---|---|---|---|---|---|---|
| *Y39H10A.1* | WBGene00021478 | protein_coding_gene | 7.1 | 141.7704 | 2.5E-09 | 5.5 | 44.887 | 6.2E-34 |
| *F10B5.3* | WBGene00008640 | protein_coding_gene | 7.1 | 141.4042 | 2.5E-31 | 7.9 | 246.870 | 2.2E-28 |
| *cest-29* | WBGene00017205 | pseudogene | 7.1 | 139.5820 | 6.8E-10 | 1.5 | 2.758 | 1.7E-08 |
| *lipl-1* | WBGene00010062 | protein_coding_gene | 7.1 | 139.5166 | 2.2E-24 | 3.6 | 12.174 | 6.6E-15 |
| *C37A5.6* | WBGene00014700 | protein_coding_gene | 7.1 | 133.9304 | 5.1E-09 | 3.9 | 14.842 | 4.3E-06 |
| *hsp-12.6* | WBGene00002013 | protein_coding_gene | 7.1 | 132.5586 | 6.3E-65 | 5.4 | 42.988 | 3.9E-107 |
| *fipr-26* | WBGene00009963 | protein_coding_gene | 7.0 | 127.5633 | 3.1E-13 | 4.7 | 26.877 | 2.2E-37 |
| *C18D11.6* | WBGene00007682 | protein_coding_gene | 7.0 | 125.4044 | 2.6E-14 | 3.2 | 8.930 | 1.4E-14 |
| *T28A11.18* | WBGene00020880 | protein_coding_gene | 6.9 | 123.3563 | 7.9E-10 | 3.1 | 8.472 | 2.3E-02 |
| *K05F6.10* | WBGene00019421 | protein_coding_gene | 6.9 | 118.6298 | 1.2E-23 | 2.1 | 4.351 | 4.3E-03 |
| *Y57E12B.4* | WBGene00021964 | protein_coding_gene | 6.9 | 117.7914 | 1.8E-09 | 5.8 | 55.060 | 1.9E-07 |
| *R05A10.8* | WBGene00011025 | protein_coding_gene | 6.9 | 116.8242 | 1.2E-09 | 5.3 | 38.406 | 9.4E-16 |
| *ugt-18* | WBGene00013900 | protein_coding_gene | 6.8 | 112.9217 | 1.1E-12 | 2.3 | 4.896 | 8.3E-08 |
| *F14F9.2* | WBGene00017465 | protein_coding_gene | 6.8 | 109.2747 | 1.0E-08 | 6.2 | 72.785 | 2.6E-09 |
| *Y6E2A.4* | WBGene00012398 | protein_coding_gene | 6.8 | 109.2328 | 5.0E-37 | 3.9 | 15.219 | 1.5E-73 |
| *F49H6.5* | WBGene00009908 | protein_coding_gene | 6.7 | 103.0572 | 1.0E-34 | 2.2 | 4.752 | 8.4E-18 |
| *W05G11.8* | WBGene00271636 | ncRNA_gene | 6.7 | 102.6458 | 9.3E-47 | 3.7 | 12.630 | 7.0E-33 |
| *math-38* | WBGene00020326 | protein_coding_gene | 6.7 | 101.8056 | 1.6E-42 | 1.7 | 3.203 | 3.5E-22 |
| *Y57E12B.11* | WBGene00235311 | protein_coding_gene | 6.7 | 101.0587 | 1.4E-09 | 6.6 | 97.405 | 6.1E-10 |
| *ins-20* | WBGene00002103 | protein_coding_gene | 6.6 | 100.2667 | 1.2E-09 | 1.3 | 2.525 | 1.5E-04 |
| *F47B8.10* | WBGene00009811 | protein_coding_gene | 6.6 | 96.2154 | 2.3E-10 | 2.7 | 6.439 | 1.5E-12 |
| *sqst-3* | WBGene00012067 | protein_coding_gene | 6.6 | 94.5407 | 1.2E-09 | 5.7 | 52.488 | 3.2E-25 |
| *Y110A2AL.2* | WBGene00022439 | protein_coding_gene | 6.5 | 89.7075 | 8.2E-18 | 6.7 | 100.746 | 1.5E-18 |
| *T26H5.4* | WBGene00012069 | protein_coding_gene | 6.4 | 87.0438 | 2.0E-10 | 1.2 | 2.355 | 8.0E-03 |
| *clec-82* | WBGene00021873 | protein_coding_gene | 6.4 | 86.1198 | 1.9E-41 | 2.4 | 5.245 | 6.3E-06 |
| *thn-1* | WBGene00009213 | protein_coding_gene | 6.4 | 85.8903 | 6.7E-25 | 1.9 | 3.608 | 9.1E-07 |
| *F55B12.10* | WBGene00077439 | protein_coding_gene | 6.4 | 84.5452 | 1.6E-18 | 5.3 | 40.329 | 2.0E-40 |
| *fil-1* | WBGene00011321 | protein_coding_gene | 6.4 | 83.0333 | 9.7E-25 | 1.4 | 2.628 | 5.8E-10 |
| *clec-42* | WBGene00008891 | protein_coding_gene | 6.3 | 80.0254 | 1.7E-11 | 2.9 | 7.512 | 6.6E-13 |
| *clec-17* | WBGene00008477 | protein_coding_gene | 6.3 | 78.8882 | 7.5E-14 | 1.9 | 3.826 | 1.8E-03 |
| *K01A6.7* | WBGene00045096 | protein_coding_gene | 6.3 | 77.0570 | 1.4E-07 | 3.4 | 10.411 | 1.3E-12 |
| *djr-1.2* | WBGene00016789 | protein_coding_gene | 6.2 | 73.6997 | 5.6E-14 | 5.6 | 47.965 | 3.0E-32 |
| *F41C3.7* | WBGene00018273 | protein_coding_gene | 6.2 | 72.3269 | 2.8E-28 | 5.3 | 38.748 | 5.7E-57 |
| *sdz-24* | WBGene00019495 | protein_coding_gene | 6.1 | 68.3490 | 2.8E-136 | 2.2 | 4.539 | 2.7E-02 |
| *Y46G5A.36* | WBGene00012921 | protein_coding_gene | 6.1 | 67.7519 | 1.8E-09 | 3.0 | 7.969 | 2.8E-07 |
| *F42A10.7* | WBGene00018343 | protein_coding_gene | 6.1 | 67.3532 | 4.3E-196 | 6.4 | 87.307 | 3.9E-81 |
| *dao-6* | WBGene00000932 | protein_coding_gene | 6.1 | 66.7066 | 5.6E-31 | 5.0 | 32.813 | 1.3E-19 |
| *R10D12.6* | WBGene00011187 | protein_coding_gene | 6.0 | 65.1282 | 5.7E-08 | 4.7 | 25.669 | 1.5E-04 |
| *F41C3.2* | WBGene00018268 | protein_coding_gene | 6.0 | 62.6091 | 7.1E-41 | 2.5 | 5.767 | 3.9E-29 |
| *K10D3.8* | WBGene00219281 | protein_coding_gene | 5.9 | 61.2769 | 3.1E-07 | 5.3 | 39.145 | 1.4E-10 |
| *T28A11.6* | WBGene00020873 | pseudogene | 5.9 | 60.8511 | 9.9E-21 | 5.7 | 52.436 | 9.4E-08 |
| *ftn-1* | WBGene00001500 | protein_coding_gene | 5.9 | 60.1130 | 3.8E-16 | 1.5 | 2.852 | 5.3E-04 |
| *math-34* | WBGene00019699 | protein_coding_gene | 5.9 | 59.2833 | 2.9E-29 | 2.7 | 6.475 | 6.5E-22 |
| *pals-39* | WBGene00008301 | protein_coding_gene | 5.9 | 59.0233 | 2.9E-17 | 4.2 | 18.937 | 5.5E-04 |
| *Y39G8B.10* | WBGene00012729 | protein_coding_gene | 5.9 | 57.8154 | 1.0E-07 | 3.4 | 10.521 | 2.7E-03 |
| *sru-40* | WBGene00005703 | protein_coding_gene | 5.9 | 57.7569 | 3.4E-10 | 2.6 | 6.169 | 2.0E-05 |
| *fbxa-173* | WBGene00044960 | pseudogene | 5.8 | 57.5106 | 2.0E-07 | 1.9 | 3.626 | 3.7E-02 |
| *ZK666.15* | WBGene00195145 | protein_coding_gene | 5.8 | 57.1618 | 2.4E-12 | 3.4 | 10.591 | 4.7E-12 |
| *Y47D7A.12* | WBGene00021624 | protein_coding_gene | 5.8 | 56.6174 | 2.7E-11 | 1.7 | 3.296 | 2.8E-05 |
| *F01G10.4* | WBGene00008507 | protein_coding_gene | 5.8 | 56.3321 | 3.4E-07 | 4.1 | 17.339 | 4.6E-04 |
| ***T26H5.10*** | **WBGene00044783** | **protein_coding_gene** | **5.8** | **56.2873** | **6.6E-10** | **-3.1** | **0.120** | **5.4E-07** |
| *srr-6* | WBGene00005657 | protein_coding_gene | 5.8 | 54.8701 | 4.2E-13 | 1.7 | 3.228 | 6.2E-08 |
| *fbxa-136* | WBGene00007672 | protein_coding_gene | 5.8 | 54.7960 | 1.5E-07 | 2.6 | 6.146 | 3.4E-13 |
| *dod-3* | WBGene00016052 | protein_coding_gene | 5.8 | 54.3193 | 3.0E-12 | 1.4 | 2.607 | 1.3E-03 |
| *arrd-8* | WBGene00012467 | protein_coding_gene | 5.7 | 52.5232 | 4.6E-18 | 3.5 | 11.518 | 1.1E-12 |
| *Y9C9A.1* | WBGene00021176 | protein_coding_gene | 5.7 | 52.4872 | 4.4E-13 | 3.8 | 14.136 | 1.8E-11 |
| *W07B8.4* | WBGene00021072 | protein_coding_gene | 5.7 | 52.4648 | 1.4E-55 | 3.9 | 15.391 | 2.8E-68 |
| *D1054.18* | WBGene00077783 | protein_coding_gene | 5.7 | 51.6038 | 3.0E-17 | 3.0 | 8.109 | 4.6E-15 |
| *T28A11.16* | WBGene00020878 | protein_coding_gene | 5.7 | 51.3028 | 9.5E-35 | 4.1 | 16.921 | 1.4E-17 |
| *T13F2.4* | WBGene00011744 | protein_coding_gene | 5.7 | 50.3758 | 3.8E-08 | 3.7 | 12.660 | 7.9E-06 |
| *C17B7.2* | WBGene00015876 | pseudogene | 5.6 | 50.0824 | 3.8E-06 | 3.2 | 9.248 | 1.8E-13 |
| *C54F6.17* | WBGene00235092 | protein_coding_gene | 5.6 | 49.7091 | 8.7E-07 | 3.5 | 11.638 | 3.6E-13 |
| *ZK666.11* | WBGene00014049 | protein_coding_gene | 5.6 | 49.6934 | 1.8E-07 | 2.4 | 5.341 | 1.6E-04 |
| *ant-1.2* | WBGene00007057 | protein_coding_gene | 5.6 | 49.5677 | 1.0E-28 | 5.6 | 47.326 | 7.1E-43 |
| *clec-172* | WBGene00044719 | protein_coding_gene | 5.6 | 49.0192 | 3.8E-40 | 2.5 | 5.476 | 4.0E-18 |
| *C06G3.3* | WBGene00015549 | protein_coding_gene | 5.6 | 48.8798 | 2.1E-30 | 3.2 | 9.125 | 2.8E-23 |
| *ttr-23* | WBGene00011894 | protein_coding_gene | 5.6 | 47.2693 | 5.9E-26 | 2.7 | 6.548 | 6.5E-45 |
| *far-7* | WBGene00001391 | protein_coding_gene | 5.6 | 46.8766 | 3.1E-29 | 1.4 | 2.587 | 1.2E-10 |
| ***lipl-3*** | **WBGene00020016** | **protein_coding_gene** | **5.6** | **46.8704** | **1.0E-15** | **-1.5** | **0.365** | **2.3E-02** |
| *F13E9.14* | WBGene00045207 | protein_coding_gene | 5.5 | 46.2804 | 3.3E-14 | 6.2 | 74.236 | 1.4E-44 |
| *oac-29* | WBGene00018295 | protein_coding_gene | 5.5 | 46.1555 | 3.1E-10 | 1.3 | 2.486 | 5.2E-03 |
| *lact-6* | WBGene00007358 | protein_coding_gene | 5.5 | 45.0396 | 2.0E-21 | 4.7 | 26.608 | 3.7E-27 |
| *pals-15* | WBGene00044707 | protein_coding_gene | 5.5 | 44.7990 | 2.6E-06 | 4.5 | 23.410 | 2.4E-05 |
| *cyp-33D3* | WBGene00012448 | protein_coding_gene | 5.5 | 44.2507 | 1.9E-18 | 4.8 | 27.137 | 1.4E-10 |
| *lys-9* | WBGene00003098 | protein_coding_gene | 5.4 | 43.3063 | 6.5E-06 | 4.8 | 27.782 | 7.5E-05 |
| *egas-4* | WBGene00018906 | protein_coding_gene | 5.4 | 43.1866 | 1.3E-36 | 3.1 | 8.873 | 7.4E-16 |
| *F57G8.7* | WBGene00010216 | protein_coding_gene | 5.4 | 42.6706 | 5.2E-06 | 4.7 | 26.182 | 1.8E-05 |
| *K02E2.7* | WBGene00010508 | protein_coding_gene | 5.4 | 42.5747 | 5.1E-07 | 2.8 | 7.048 | 2.9E-06 |
| *F55H12.9* | WBGene00219976 | snoRNA_gene | 5.4 | 42.4604 | 5.6E-07 | 2.4 | 5.336 | 3.0E-04 |
| *T20D4.11* | WBGene00020617 | protein_coding_gene | 5.4 | 42.2250 | 1.8E-80 | 3.3 | 10.042 | 1.3E-02 |

| | | | | | | | | |
|---|---|---|---|---|---|---|---|---|
| *Y39A3B.7* | WBGene00206512 | protein_coding_gene | 5.3 | 40.5461 | 4.3E-06 | 3.1 | 8.830 | 5.3E-03 |
| *C45B11.7* | WBGene00050945 | pseudogene | 5.3 | 40.4139 | 1.7E-18 | 3.6 | 12.535 | 3.0E-15 |
| *srz-99* | WBGene00012028 | protein_coding_gene | 5.3 | 40.2661 | 3.9E-07 | 2.5 | 5.675 | 3.0E-07 |
| *F09C12.2* | WBGene00017277 | protein_coding_gene | 5.3 | 40.2588 | 6.1E-15 | 4.9 | 30.382 | 8.4E-09 |
| *exc-13* | WBGene00015393 | protein_coding_gene | 5.3 | 40.0165 | 1.3E-66 | 2.8 | 6.898 | 7.5E-31 |
| *best-1* | WBGene00007203 | protein_coding_gene | 5.3 | 39.7355 | 2.0E-63 | 1.7 | 3.310 | 1.3E-03 |
| *clec-64* | WBGene00009395 | protein_coding_gene | 5.3 | 38.8613 | 2.9E-13 | 2.6 | 6.244 | 4.7E-07 |
| *C36B7.4* | WBGene00016467 | protein_coding_gene | 5.3 | 38.7460 | 1.2E-05 | 3.2 | 9.442 | 1.1E-02 |
| *M04C9.2* | WBGene00010857 | protein_coding_gene | 5.2 | 37.9640 | 2.7E-10 | 4.1 | 17.056 | 2.2E-16 |
| *K04A8.21* | WBGene00271815 | protein_coding_gene | 5.2 | 37.4576 | 6.5E-10 | 3.3 | 10.190 | 1.8E-10 |
| *clec-34* | WBGene00012023 | protein_coding_gene | 5.2 | 37.2427 | 6.3E-07 | 2.0 | 3.905 | 5.8E-03 |
| *Y53F4B.45* | WBGene00044728 | ncRNA_gene | 5.2 | 37.0114 | 1.0E-28 | 1.8 | 3.434 | 2.7E-14 |
| *H02F09.3* | WBGene00019146 | protein_coding_gene | 5.2 | 36.7490 | 7.6E-35 | 7.6 | 188.998 | 4.3E-38 |
| *ZK105.13* | WBGene00235114 | protein_coding_gene | 5.2 | 35.9926 | 7.4E-08 | 1.4 | 2.632 | 5.7E-04 |
| *R05H11.2* | WBGene00019908 | protein_coding_gene | 5.2 | 35.8149 | 2.3E-09 | 3.7 | 12.727 | 9.6E-10 |
| *chil-16* | WBGene00011158 | protein_coding_gene | 5.1 | 35.2763 | 1.1E-05 | 4.3 | 19.167 | 1.1E-05 |
| *C54F6.5* | WBGene00016923 | protein_coding_gene | 5.1 | 35.0923 | 5.2E-16 | 3.1 | 8.482 | 9.9E-40 |
| *T28A11.20* | WBGene00020882 | protein_coding_gene | 5.1 | 34.1542 | 1.8E-05 | 2.6 | 6.012 | 3.6E-02 |
| *pals-27* | WBGene00007132 | protein_coding_gene | 5.1 | 34.1123 | 3.2E-14 | 3.9 | 14.939 | 1.3E-16 |
| *F56D6.9* | WBGene00044471 | protein_coding_gene | 5.1 | 33.9735 | 4.7E-05 | 1.4 | 2.678 | 3.2E-02 |
| *T10C6.15* | WBGene00050899 | protein_coding_gene | 5.1 | 33.7178 | 7.7E-19 | 2.8 | 6.826 | 2.8E-25 |
| *Y51H4A.5* | WBGene00013101 | protein_coding_gene | 5.1 | 33.6592 | 1.8E-21 | 1.6 | 2.981 | 2.4E-06 |
| *F49C12.7* | WBGene00009877 | protein_coding_gene | 5.1 | 33.6409 | 2.0E-132 | 3.7 | 13.158 | 2.7E-30 |
| *pals-32* | WBGene00016281 | protein_coding_gene | 5.1 | 33.6069 | 2.0E-18 | 1.4 | 2.703 | 3.1E-09 |
| *pals-26* | WBGene00007131 | protein_coding_gene | 5.1 | 33.2847 | 1.7E-20 | 3.4 | 10.885 | 2.0E-21 |
| *B0336.12* | WBGene00015149 | protein_coding_gene | 5.1 | 33.2095 | 4.2E-05 | 4.2 | 18.668 | 4.7E-04 |
| *Y40B10B.1* | WBGene00021494 | protein_coding_gene | 5.0 | 33.0741 | 4.7E-06 | 4.1 | 16.786 | 7.6E-09 |
| *F53H2.1* | WBGene00010004 | protein_coding_gene | 5.0 | 32.9274 | 6.2E-11 | 1.7 | 3.297 | 1.6E-04 |
| *gcy-7* | WBGene00001534 | protein_coding_gene | 5.0 | 32.8394 | 1.4E-05 | 3.4 | 10.767 | 5.4E-04 |
| *C13A2.1* | WBGene00015718 | protein_coding_gene | 5.0 | 32.6199 | 5.4E-05 | 2.8 | 7.188 | 1.4E-02 |
| *F53B7.7* | WBGene00009969 | protein_coding_gene | 5.0 | 32.6063 | 8.8E-22 | 1.7 | 3.217 | 2.4E-02 |
| *ins-35* | WBGene00002118 | protein_coding_gene | 5.0 | 32.4788 | 1.5E-15 | 3.3 | 10.049 | 3.9E-11 |
| *H02F09.2* | WBGene00019145 | protein_coding_gene | 5.0 | 32.4688 | 2.2E-06 | 4.3 | 19.361 | 3.6E-05 |
| *asah-1* | WBGene00010769 | protein_coding_gene | 5.0 | 32.3049 | 2.5E-49 | 3.5 | 10.931 | 6.9E-33 |
| *hsd-3* | WBGene00022616 | protein_coding_gene | 5.0 | 32.0097 | 4.6E-05 | 4.5 | 22.184 | 7.9E-05 |
| *C30G12.11* | WBGene00284861 | ncRNA_gene | 5.0 | 31.2312 | 1.3E-06 | 2.9 | 7.491 | 9.7E-03 |
| *fbxa-66* | WBGene00021865 | protein_coding_gene | 5.0 | 31.0157 | 3.5E-07 | 2.7 | 6.333 | 1.4E-06 |
| *irld-11* | WBGene00019391 | protein_coding_gene | 5.0 | 31.0062 | 3.1E-06 | 2.3 | 5.080 | 1.2E-02 |
| *F14F9.8* | WBGene00044536 | protein_coding_gene | 4.9 | 30.4057 | 1.7E-18 | 4.4 | 20.715 | 1.4E-12 |
| *F53A9.7* | WBGene00018730 | protein_coding_gene | 4.9 | 30.3857 | 7.2E-37 | 1.9 | 3.619 | 1.2E-22 |
| *sri-39* | WBGene00005551 | protein_coding_gene | 4.9 | 30.3157 | 1.9E-04 | 3.6 | 11.888 | 2.9E-03 |
| *B0284.5* | WBGene00045188 | pseudogene | 4.9 | 30.2822 | 1.4E-06 | 2.6 | 6.134 | 3.3E-07 |
| *F49F1.5* | WBGene00018645 | protein_coding_gene | 4.9 | 30.0685 | 3.5E-34 | 1.0 | 2.047 | 4.2E-05 |
| *F35D2.1* | WBGene00018039 | protein_coding_gene | 4.9 | 29.0509 | 1.8E-05 | 3.1 | 8.297 | 1.2E-02 |
| *col-176* | WBGene00000749 | protein_coding_gene | 4.9 | 28.8824 | 1.3E-44 | 5.8 | 56.689 | 1.9E-18 |
| *nnt-1* | WBGene00003778 | protein_coding_gene | 4.9 | 28.8611 | 5.1E-27 | 1.6 | 3.114 | 3.0E-27 |
| *Y39A1A.9* | WBGene00012648 | protein_coding_gene | 4.8 | 28.6468 | 8.3E-22 | 4.6 | 24.163 | 1.3E-42 |
| ***Y46C8AM.1*** | **WBGene00271795** | **protein_coding_gene** | **4.8** | **28.3663** | **3.3E-22** | **-1.1** | **0.457** | **4.4E-02** |
| *ZK418.11* | WBGene00044354 | protein_coding_gene | 4.8 | 28.2180 | 2.3E-07 | 2.3 | 4.871 | 1.9E-03 |
| *Y39G8B.9* | WBGene00012728 | protein_coding_gene | 4.8 | 28.0757 | 3.5E-22 | 4.3 | 19.884 | 1.3E-13 |
| *B0348.2* | WBGene00015152 | protein_coding_gene | 4.8 | 27.7824 | 1.1E-12 | 2.3 | 4.770 | 2.4E-10 |
| *R12G8.1* | WBGene00011254 | protein_coding_gene | 4.8 | 27.6064 | 3.4E-08 | 3.0 | 8.131 | 1.5E-08 |
| *hil-1* | WBGene00001852 | protein_coding_gene | 4.8 | 27.5437 | 2.3E-29 | 2.2 | 4.595 | 1.6E-13 |
| *ins-11* | WBGene00002094 | protein_coding_gene | 4.8 | 27.4527 | 5.7E-08 | 2.1 | 4.335 | 1.4E-04 |
| *lmd-4* | WBGene00017233 | protein_coding_gene | 4.8 | 27.1737 | 1.6E-22 | 6.8 | 113.852 | 2.1E-42 |
| *W04C9.9* | WBGene00235388 | protein_coding_gene | 4.8 | 26.9804 | 2.4E-06 | 3.7 | 13.309 | 2.3E-03 |
| *Y26D4A.3* | WBGene00012501 | protein_coding_gene | 4.8 | 26.9216 | 1.2E-06 | 2.7 | 6.541 | 1.2E-02 |
| *catp-2* | WBGene00015338 | protein_coding_gene | 4.7 | 26.5169 | 8.9E-05 | 4.6 | 25.103 | 1.7E-04 |
| *cyp-35B1* | WBGene00019472 | protein_coding_gene | 4.7 | 26.5059 | 3.0E-41 | 2.1 | 4.435 | 1.2E-10 |
| *ZK971.1* | WBGene00014175 | protein_coding_gene | 4.7 | 26.3358 | 7.3E-31 | 3.3 | 10.036 | 1.0E-43 |
| *oac-31* | WBGene00018707 | protein_coding_gene | 4.7 | 26.1446 | 2.2E-20 | 1.1 | 2.185 | 3.1E-03 |
| *B0238.13* | WBGene00015077 | pseudogene | 4.7 | 25.9887 | 4.4E-07 | 2.4 | 5.114 | 3.5E-06 |
| *F21C10.11* | WBGene00017660 | protein_coding_gene | 4.7 | 25.8838 | 1.2E-31 | 1.1 | 2.217 | 2.2E-02 |
| *ZK355.8* | WBGene00086560 | protein_coding_gene | 4.7 | 25.8718 | 9.9E-05 | 4.1 | 16.583 | 7.6E-05 |
| *M02E1.4* | WBGene00284848 | protein_coding_gene | 4.7 | 25.8568 | 5.4E-13 | 1.4 | 2.702 | 1.4E-04 |
| *F46A8.1* | WBGene00014759 | pseudogene | 4.7 | 25.7747 | 8.4E-31 | 3.9 | 14.477 | 4.1E-45 |
| *T23F6.2* | WBGene00011950 | protein_coding_gene | 4.7 | 25.5779 | 4.1E-04 | 3.2 | 9.125 | 2.0E-02 |
| *C11H1.5* | WBGene00007530 | protein_coding_gene | 4.7 | 25.5566 | 2.5E-15 | 5.3 | 38.606 | 5.0E-10 |
| *thn-3* | WBGene00009214 | pseudogene | 4.7 | 25.4255 | 3.1E-04 | 5.3 | 39.035 | 2.1E-05 |
| *fil-2* | WBGene00019668 | protein_coding_gene | 4.7 | 25.3278 | 5.9E-10 | 3.8 | 14.183 | 7.1E-16 |
| *fbxa-158* | WBGene00015602 | protein_coding_gene | 4.7 | 25.2494 | 4.7E-11 | 4.0 | 15.825 | 1.0E-09 |
| *Y62H9A.11* | WBGene00013399 | protein_coding_gene | 4.7 | 25.1429 | 6.4E-06 | 1.9 | 3.743 | 1.8E-02 |
| *T28A11.3* | WBGene00020870 | protein_coding_gene | 4.6 | 25.1013 | 1.4E-04 | 4.4 | 21.709 | 1.1E-06 |
| *Y54G2A.57* | WBGene00189961 | protein_coding_gene | 4.6 | 24.9738 | 3.6E-04 | 4.8 | 27.996 | 6.5E-07 |
| *F14D7.7* | WBGene00008794 | protein_coding_gene | 4.6 | 24.7827 | 1.3E-08 | 1.8 | 3.459 | 8.1E-04 |
| *fbxa-48* | WBGene00021862 | protein_coding_gene | 4.6 | 24.5839 | 3.0E-10 | 1.3 | 2.405 | 6.3E-05 |
| *best-5* | WBGene00007404 | protein_coding_gene | 4.6 | 24.5711 | 1.9E-21 | 1.4 | 2.598 | 3.6E-09 |
| *ZC190.6* | WBGene00022540 | protein_coding_gene | 4.6 | 24.2067 | 8.6E-05 | 2.3 | 4.970 | 4.6E-02 |
| *T01D3.3* | WBGene00011328 | protein_coding_gene | 4.6 | 23.8502 | 1.3E-21 | 2.1 | 4.382 | 4.7E-14 |
| *gst-15* | WBGene00001763 | protein_coding_gene | 4.5 | 22.8234 | 9.9E-49 | 4.0 | 15.807 | 1.5E-69 |

| | | | | | | | | |
|---|---|---|---|---|---|---|---|---|
| *clec-67* | WBGene00018971 | protein_coding_gene | 4.5 | 22.7411 | 1.0E-21 | 2.2 | 4.711 | 5.0E-07 |
| *kreg-1* | WBGene00018725 | protein_coding_gene | 4.5 | 22.6410 | 7.6E-14 | 2.5 | 5.550 | 3.0E-06 |
| *pitr-2* | WBGene00015054 | protein_coding_gene | 4.5 | 22.4905 | 1.9E-04 | 3.3 | 10.086 | 2.7E-04 |
| *ZC116.5* | WBGene00119204 | protein_coding_gene | 4.5 | 22.1628 | 7.9E-21 | 3.1 | 8.840 | 5.2E-15 |
| *col-69* | WBGene00000645 | protein_coding_gene | 4.5 | 21.9645 | 4.3E-09 | 2.5 | 5.468 | 2.2E-06 |
| *B0478.3* | WBGene00015202 | protein_coding_gene | 4.4 | 21.8094 | 2.8E-11 | 1.7 | 3.355 | 2.4E-05 |
| *tag-293* | WBGene00015398 | protein_coding_gene | 4.4 | 21.7015 | 3.3E-06 | 5.8 | 56.633 | 3.6E-12 |
| *clec-169* | WBGene00009526 | protein_coding_gene | 4.4 | 21.1559 | 4.9E-17 | 2.5 | 5.594 | 5.5E-04 |
| *zmp-2* | WBGene00019212 | protein_coding_gene | 4.4 | 20.7913 | 1.5E-25 | 3.2 | 9.193 | 6.0E-26 |
| *T09D3.8* | WBGene00044538 | protein_coding_gene | 4.4 | 20.6533 | 1.4E-05 | 4.5 | 22.764 | 6.3E-16 |
| *F59E11.7* | WBGene00019113 | protein_coding_gene | 4.4 | 20.5473 | 3.0E-21 | 1.6 | 3.073 | 2.7E-07 |
| *sri-74* | WBGene00005586 | protein_coding_gene | 4.3 | 20.2578 | 5.6E-04 | 5.5 | 45.898 | 7.8E-08 |
| *C54F6.18* | WBGene00235093 | protein_coding_gene | 4.3 | 20.2213 | 4.4E-08 | 3.2 | 9.127 | 3.6E-07 |
| *C40H1.9* | WBGene00008044 | protein_coding_gene | 4.3 | 20.0603 | 1.7E-18 | 2.0 | 4.069 | 1.5E-07 |
| *F58H10.1* | WBGene00010291 | protein_coding_gene | 4.3 | 19.9336 | 6.1E-32 | 2.9 | 7.299 | 2.9E-20 |
| *gcy-29* | WBGene00007314 | protein_coding_gene | 4.3 | 19.6740 | 1.1E-08 | 5.1 | 34.772 | 4.9E-09 |
| *ZC443.4* | WBGene00013899 | protein_coding_gene | 4.3 | 19.5198 | 4.5E-35 | 2.0 | 4.135 | 1.5E-43 |
| *math-42* | WBGene00020532 | protein_coding_gene | 4.3 | 19.4551 | 1.0E-10 | 1.7 | 3.342 | 2.5E-05 |
| *W04C9.6* | WBGene00021028 | protein_coding_gene | 4.3 | 19.4169 | 7.8E-43 | 4.3 | 19.351 | 6.8E-40 |
| *Y53G8AM.5* | WBGene00021805 | protein_coding_gene | 4.3 | 19.3081 | 3.4E-05 | 3.8 | 14.409 | 3.3E-05 |
| *C50D2.1* | WBGene00016805 | protein_coding_gene | 4.3 | 19.2626 | 4.1E-23 | 1.5 | 2.866 | 2.8E-04 |
| *spp-11* | WBGene00004996 | protein_coding_gene | 4.3 | 19.1869 | 6.0E-04 | 2.4 | 5.270 | 4.3E-02 |
| *T08A9.13* | WBGene00044442 | protein_coding_gene | 4.2 | 18.9809 | 1.5E-11 | 3.0 | 8.280 | 3.3E-14 |
| *ZK355.3* | WBGene00022713 | protein_coding_gene | 4.2 | 18.9424 | 8.2E-04 | 2.8 | 7.153 | 2.6E-03 |
| *lys-3* | WBGene00003092 | protein_coding_gene | 4.2 | 18.9231 | 2.8E-09 | 1.5 | 2.851 | 3.7E-05 |
| *fbxa-127* | WBGene00012027 | protein_coding_gene | 4.2 | 18.8255 | 2.5E-09 | 1.8 | 3.398 | 2.1E-04 |
| *K08D8.9* | WBGene00194918 | protein_coding_gene | 4.2 | 18.7901 | 8.5E-04 | 2.6 | 5.882 | 1.3E-02 |
| *clec-52* | WBGene00015052 | protein_coding_gene | 4.2 | 18.7821 | 4.3E-148 | 3.2 | 8.968 | 4.3E-19 |
| *linc-84* | WBGene00219592 | lincRNA_gene | 4.2 | 18.7244 | 5.5E-08 | 1.6 | 3.035 | 9.7E-03 |
| *tsp-1* | WBGene00006627 | protein_coding_gene | 4.2 | 18.6899 | 8.6E-18 | 1.4 | 2.585 | 1.1E-02 |
| *nlp-2* | WBGene00003740 | protein_coding_gene | 4.2 | 18.5632 | 6.1E-18 | 2.3 | 4.844 | 3.7E-12 |
| *sre-6* | WBGene00009242 | protein_coding_gene | 4.2 | 18.4368 | 8.9E-05 | 3.0 | 7.919 | 2.2E-05 |
| *F21H7.12* | WBGene00009033 | protein_coding_gene | 4.2 | 18.2655 | 9.7E-10 | 1.2 | 2.352 | 4.9E-02 |
| *F56F11.1* | WBGene00018988 | protein_coding_gene | 4.2 | 18.0099 | 5.0E-05 | 2.5 | 5.755 | 2.7E-05 |
| *ttr-29* | WBGene00011286 | protein_coding_gene | 4.2 | 18.0079 | 7.1E-22 | 1.1 | 2.098 | 6.2E-06 |
| *spp-8* | WBGene00004993 | protein_coding_gene | 4.2 | 17.9095 | 1.5E-45 | 1.7 | 3.194 | 4.2E-12 |
| *C34C6.3* | WBGene00007916 | protein_coding_gene | 4.2 | 17.8522 | 2.1E-34 | 1.2 | 2.298 | 5.4E-04 |
| *T05F1.9* | WBGene00011494 | protein_coding_gene | 4.2 | 17.8206 | 4.3E-19 | 3.6 | 12.374 | 4.9E-14 |
| *rgba-1* | WBGene00077766 | protein_coding_gene | 4.2 | 17.8042 | 7.5E-04 | 2.3 | 4.784 | 1.4E-03 |
| *F40E3.6* | WBGene00018233 | protein_coding_gene | 4.1 | 17.6927 | 2.4E-05 | 4.1 | 17.198 | 1.0E-04 |
| *K09H9.5* | WBGene00019597 | protein_coding_gene | 4.1 | 17.6136 | 3.3E-56 | 3.8 | 14.208 | 8.6E-81 |
| *xtr-1* | WBGene00006965 | protein_coding_gene | 4.1 | 17.4295 | 7.9E-17 | 1.9 | 3.640 | 1.4E-06 |
| *cyp-33C1* | WBGene00016686 | protein_coding_gene | 4.1 | 17.3303 | 2.3E-04 | 3.4 | 10.746 | 2.8E-03 |
| *fbxa-44* | WBGene00008616 | protein_coding_gene | 4.1 | 17.2982 | 3.0E-07 | 1.6 | 3.089 | 1.8E-02 |
| *F45D11.1* | WBGene00018448 | pseudogene | 4.1 | 17.2970 | 1.5E-11 | 1.7 | 3.201 | 4.4E-07 |
| *T10G3.4* | WBGene00011695 | protein_coding_gene | 4.1 | 17.2680 | 1.0E-03 | 4.3 | 19.910 | 2.6E-04 |
| *pals-2* | WBGene00007656 | protein_coding_gene | 4.1 | 17.1150 | 8.1E-04 | 2.6 | 5.962 | 3.0E-03 |
| *str-112* | WBGene00006163 | protein_coding_gene | 4.1 | 17.0765 | 7.4E-07 | 2.4 | 5.136 | 1.3E-04 |
| *tag-243* | WBGene00044078 | protein_coding_gene | 4.1 | 17.0670 | 5.5E-24 | 2.4 | 5.408 | 6.5E-28 |
| *nhr-99* | WBGene00003689 | protein_coding_gene | 4.1 | 17.0159 | 1.2E-08 | 2.0 | 3.911 | 9.5E-20 |
| *Y105C5A.1270* | WBGene00194646 | protein_coding_gene | 4.1 | 17.0126 | 1.3E-03 | 2.8 | 6.852 | 1.6E-02 |
| *T22F3.10* | WBGene00020699 | protein_coding_gene | 4.1 | 16.9474 | 2.2E-09 | 1.7 | 3.216 | 1.4E-03 |
| *T19D12.3* | WBGene00020578 | protein_coding_gene | 4.1 | 16.9214 | 7.8E-05 | 1.5 | 2.919 | 2.4E-02 |
| ***Y39A1A.27*** | **WBGene00194731** | **protein_coding_gene** | **4.1** | **16.9062** | **2.6E-04** | **-1.7** | **0.306** | **1.3E-05** |
| *tyr-3* | WBGene00009001 | protein_coding_gene | 4.1 | 16.8569 | 2.6E-06 | 5.5 | 45.337 | 3.7E-07 |
| *T12D8.5* | WBGene00011733 | protein_coding_gene | 4.1 | 16.8080 | 3.4E-17 | 1.7 | 3.236 | 3.1E-08 |
| *F14F8.8* | WBGene00008816 | protein_coding_gene | 4.1 | 16.7665 | 1.4E-04 | 5.5 | 46.521 | 3.6E-07 |
| *fbxa-12* | WBGene00016525 | protein_coding_gene | 4.1 | 16.7387 | 2.3E-04 | 2.1 | 4.337 | 1.5E-02 |
| *col-50* | WBGene00000627 | protein_coding_gene | 4.1 | 16.6796 | 2.9E-03 | 5.6 | 47.614 | 1.9E-06 |
| *odr-10* | WBGene00003856 | protein_coding_gene | 4.1 | 16.6395 | 1.3E-16 | 1.5 | 2.805 | 4.2E-04 |
| *C04A11.5* | WBGene00007288 | protein_coding_gene | 4.1 | 16.5650 | 1.8E-10 | 1.1 | 2.173 | 2.7E-03 |
| *T06G6.6* | WBGene00011548 | protein_coding_gene | 4.0 | 16.5265 | 1.7E-10 | 3.4 | 10.265 | 3.6E-03 |
| *M60.7* | WBGene00019782 | protein_coding_gene | 4.0 | 16.4137 | 8.6E-13 | 3.1 | 8.502 | 1.1E-24 |
| *pals-31* | WBGene00018614 | protein_coding_gene | 4.0 | 16.4027 | 1.3E-15 | 2.1 | 4.205 | 2.0E-04 |
| *B0205.4* | WBGene00015019 | protein_coding_gene | 4.0 | 16.3251 | 3.8E-11 | 1.5 | 2.860 | 3.8E-05 |
| *comt-2* | WBGene00012518 | protein_coding_gene | 4.0 | 16.2407 | 3.7E-12 | 2.9 | 7.550 | 2.9E-05 |
| *C27H5.6* | WBGene00016175 | protein_coding_gene | 4.0 | 16.1632 | 5.7E-04 | 2.9 | 7.374 | 8.5E-03 |
| *hsp-70* | WBGene00002026 | protein_coding_gene | 4.0 | 16.1260 | 1.5E-13 | 3.9 | 14.731 | 7.2E-27 |
| *C15H9.2* | WBGene00015798 | protein_coding_gene | 4.0 | 16.1037 | 4.0E-06 | 2.5 | 5.624 | 1.4E-03 |
| *bli-1* | WBGene00000251 | protein_coding_gene | 4.0 | 16.0228 | 2.1E-07 | 4.5 | 22.599 | 7.9E-06 |
| *R05A10.4* | WBGene00011021 | protein_coding_gene | 4.0 | 16.0005 | 2.7E-09 | 2.2 | 4.563 | 4.5E-06 |
| *W02F12.8* | WBGene00175027 | protein_coding_gene | 4.0 | 15.7695 | 2.8E-18 | 3.6 | 12.147 | 6.8E-10 |
| *K03H6.2* | WBGene00019368 | protein_coding_gene | 4.0 | 15.6054 | 7.6E-25 | 3.3 | 9.990 | 1.7E-18 |
| *srv-4* | WBGene00005715 | protein_coding_gene | 4.0 | 15.5675 | 7.7E-04 | 3.2 | 9.384 | 2.9E-04 |
| *Y110A7A.24* | WBGene00271632 | ncRNA_gene | 4.0 | 15.5589 | 9.8E-05 | 1.8 | 3.564 | 1.5E-02 |
| *Y43C5A.3* | WBGene00012783 | protein_coding_gene | 4.0 | 15.5289 | 4.4E-39 | 2.0 | 4.014 | 2.6E-38 |
| *C31H2.3* | WBGene00016293 | protein_coding_gene | 4.0 | 15.4704 | 9.7E-11 | 2.5 | 5.599 | 2.8E-08 |
| *F08H9.3* | WBGene00008591 | protein_coding_gene | 3.9 | 15.3945 | 5.9E-17 | 3.4 | 10.227 | 7.1E-25 |
| *Y69H2.10* | WBGene00013485 | protein_coding_gene | 3.9 | 15.3827 | 1.4E-09 | 2.4 | 5.125 | 2.5E-08 |
| *T05F1.11* | WBGene00011495 | protein_coding_gene | 3.9 | 15.3731 | 1.8E-21 | 1.2 | 2.271 | 9.6E-07 |

| | | | | | | | | |
|---|---|---|---|---|---|---|---|---|
| *srd-28* | WBGene00005106 | protein_coding_gene | 3.9 | 15.1571 | 8.7E-04 | 2.0 | 3.900 | 1.3E-02 |
| ***scl-2*** | **WBGene00009895** | **protein_coding_gene** | **3.9** | **15.1137** | **5.1E-07** | **-1.2** | **0.440** | **1.2E-12** |
| *Y71F9AM.8* | WBGene00045398 | protein_coding_gene | 3.9 | 15.0839 | 1.9E-02 | 5.0 | 31.970 | 2.2E-05 |
| *T16G1.4* | WBGene00011798 | protein_coding_gene | 3.9 | 15.0418 | 4.8E-07 | 1.6 | 3.063 | 5.5E-06 |
| *M60.4* | WBGene00019780 | protein_coding_gene | 3.9 | 14.9686 | 9.3E-26 | 2.5 | 5.607 | 4.4E-75 |
| *dmd-10* | WBGene00007929 | protein_coding_gene | 3.9 | 14.9677 | 3.4E-05 | 2.0 | 3.998 | 9.9E-04 |
| *H17B01.5* | WBGene00044643 | protein_coding_gene | 3.9 | 14.8798 | 1.0E-12 | 2.9 | 7.486 | 1.2E-13 |
| *F56D3.1* | WBGene00018969 | protein_coding_gene | 3.9 | 14.8659 | 2.7E-16 | 2.2 | 4.541 | 8.7E-06 |
| *C54C8.8* | WBGene00014714 | pseudogene | 3.9 | 14.6945 | 1.6E-06 | 2.6 | 5.859 | 6.3E-06 |
| *C07G3.8* | WBGene00015576 | protein_coding_gene | 3.9 | 14.5723 | 2.2E-04 | 2.6 | 5.991 | 3.8E-03 |
| *F07C4.6* | WBGene00017200 | protein_coding_gene | 3.9 | 14.5627 | 2.1E-07 | 3.1 | 8.417 | 5.3E-03 |
| *Y15E3A.5* | WBGene00012444 | protein_coding_gene | 3.9 | 14.4923 | 1.1E-25 | 2.1 | 4.381 | 2.2E-31 |
| *scl-14* | WBGene00008625 | protein_coding_gene | 3.9 | 14.4406 | 1.4E-04 | 3.2 | 9.261 | 5.2E-06 |
| *C42D4.13* | WBGene00016598 | protein_coding_gene | 3.8 | 14.3984 | 1.5E-11 | 2.1 | 4.343 | 5.2E-03 |
| *F54B11.9* | WBGene00010032 | protein_coding_gene | 3.8 | 14.3726 | 7.6E-05 | 1.8 | 3.567 | 3.2E-02 |
| *fut-2* | WBGene00001506 | protein_coding_gene | 3.8 | 14.3153 | 5.1E-15 | 2.1 | 4.148 | 1.5E-08 |
| *F59E11.6* | WBGene00019112 | protein_coding_gene | 3.8 | 14.2397 | 7.9E-05 | 2.3 | 4.988 | 6.2E-03 |
| *F56A6.5* | WBGene00044427 | protein_coding_gene | 3.8 | 14.1867 | 3.4E-03 | 3.4 | 10.654 | 3.3E-03 |
| *R11D1.12* | WBGene00045336 | protein_coding_gene | 3.8 | 14.1683 | 6.7E-10 | 1.7 | 3.205 | 1.2E-05 |
| *fbxc-41* | WBGene00016304 | pseudogene | 3.8 | 14.1560 | 1.2E-06 | 2.0 | 3.898 | 3.1E-05 |
| *gpa-8* | WBGene00001670 | protein_coding_gene | 3.8 | 14.1373 | 7.8E-07 | 2.7 | 6.362 | 1.8E-04 |
| *sod-3* | WBGene00004932 | protein_coding_gene | 3.8 | 14.1203 | 1.4E-43 | 4.0 | 15.487 | 3.1E-49 |
| *F20B6.4* | WBGene00017628 | protein_coding_gene | 3.8 | 14.1135 | 3.3E-04 | 1.9 | 3.680 | 8.7E-03 |
| *bli-2* | WBGene00000252 | protein_coding_gene | 3.8 | 14.1005 | 1.1E-12 | 4.2 | 18.415 | 3.3E-05 |
| *R02F11.2* | WBGene00019840 | protein_coding_gene | 3.8 | 14.0964 | 1.1E-19 | 2.7 | 6.522 | 2.0E-13 |
| *lips-5* | WBGene00011725 | protein_coding_gene | 3.8 | 14.0931 | 1.3E-33 | 3.6 | 11.851 | 8.3E-24 |
| *T27A3.8* | WBGene00044611 | protein_coding_gene | 3.8 | 14.0593 | 6.8E-10 | 2.7 | 6.515 | 2.3E-08 |
| *cav-2* | WBGene00000302 | protein_coding_gene | 3.8 | 14.0465 | 2.7E-32 | 2.5 | 5.482 | 5.5E-18 |
| *asah-2* | WBGene00009192 | protein_coding_gene | 3.8 | 13.9729 | 3.5E-40 | 2.2 | 4.450 | 7.2E-45 |
| *gst-8* | WBGene00001756 | protein_coding_gene | 3.8 | 13.9470 | 3.0E-05 | 1.5 | 2.916 | 3.9E-02 |
| *Y48A6B.8* | WBGene00012969 | protein_coding_gene | 3.8 | 13.9153 | 1.2E-18 | 3.2 | 9.325 | 1.7E-23 |
| *T23B3.6* | WBGene00020714 | protein_coding_gene | 3.8 | 13.7895 | 2.2E-15 | 2.2 | 4.752 | 5.2E-09 |
| *chil-18* | WBGene00011161 | protein_coding_gene | 3.8 | 13.7095 | 3.6E-07 | 2.1 | 4.163 | 3.9E-05 |
| *C40H1.3* | WBGene00008039 | protein_coding_gene | 3.8 | 13.6922 | 3.1E-05 | 1.9 | 3.777 | 9.3E-03 |
| *zip-6* | WBGene00011130 | protein_coding_gene | 3.8 | 13.6614 | 5.4E-04 | 2.6 | 5.885 | 9.2E-03 |
| *phat-5* | WBGene00020242 | protein_coding_gene | 3.8 | 13.6249 | 1.7E-08 | 1.1 | 2.145 | 3.3E-02 |
| *C05G5.7* | WBGene00050939 | protein_coding_gene | 3.8 | 13.5447 | 4.1E-09 | 3.5 | 11.097 | 2.5E-04 |
| *K01A2.10* | WBGene00019284 | protein_coding_gene | 3.7 | 13.4491 | 2.1E-16 | 1.4 | 2.628 | 2.0E-14 |
| *Y116F11B.9* | WBGene00013825 | protein_coding_gene | 3.7 | 13.4247 | 3.5E-04 | 4.3 | 19.528 | 1.6E-04 |
| *cdh-10* | WBGene00000402 | protein_coding_gene | 3.7 | 13.4201 | 1.5E-15 | 4.2 | 18.829 | 2.4E-33 |
| *ZK1307.2* | WBGene00014245 | protein_coding_gene | 3.7 | 13.3317 | 7.8E-11 | 1.2 | 2.265 | 4.8E-03 |
| *bcmo-2* | WBGene00018755 | protein_coding_gene | 3.7 | 13.2870 | 1.3E-102 | 4.3 | 19.203 | 1.6E-68 |
| *sri-36* | WBGene00005548 | protein_coding_gene | 3.7 | 13.2830 | 4.9E-04 | 3.2 | 9.022 | 1.0E-07 |
| *W03G1.5* | WBGene00021011 | protein_coding_gene | 3.7 | 13.1589 | 9.3E-19 | 2.8 | 7.188 | 7.5E-21 |
| *dos-3* | WBGene00019331 | protein_coding_gene | 3.7 | 13.1451 | 1.0E-04 | 2.8 | 6.924 | 5.8E-04 |
| *F09C6.13* | WBGene00044183 | protein_coding_gene | 3.7 | 13.1274 | 5.6E-18 | 1.8 | 3.527 | 6.2E-03 |
| *tre-5* | WBGene00006611 | protein_coding_gene | 3.7 | 13.1244 | 5.0E-11 | 1.1 | 2.073 | 1.6E-06 |
| *srw-71* | WBGene00005818 | protein_coding_gene | 3.7 | 13.1231 | 1.1E-04 | 3.0 | 8.236 | 1.6E-03 |
| *spp-18* | WBGene00005003 | protein_coding_gene | 3.7 | 13.1179 | 3.3E-45 | 2.1 | 4.299 | 2.0E-53 |
| *Y77E11A.14* | WBGene00022314 | protein_coding_gene | 3.7 | 13.0890 | 1.3E-14 | 2.1 | 4.269 | 1.6E-12 |
| *dur-1* | WBGene00001113 | protein_coding_gene | 3.7 | 13.0697 | 7.7E-56 | 4.1 | 17.197 | 8.7E-132 |
| ***cky-1*** | **WBGene00000521** | **protein_coding_gene** | **3.7** | **13.0043** | **2.2E-04** | **-1.9** | **0.265** | **7.1E-03** |
| *F48G7.10* | WBGene00018621 | protein_coding_gene | 3.7 | 12.9491 | 1.3E-10 | 2.3 | 5.083 | 9.0E-04 |
| *C32F10.4* | WBGene00016332 | protein_coding_gene | 3.7 | 12.9175 | 3.9E-34 | 1.6 | 3.035 | 4.1E-08 |
| *H10D18.1* | WBGene00019177 | protein_coding_gene | 3.7 | 12.9023 | 1.5E-05 | 1.7 | 3.335 | 4.3E-04 |
| *F56C4.4* | WBGene00044977 | protein_coding_gene | 3.7 | 12.8894 | 8.0E-16 | 1.2 | 2.220 | 1.6E-03 |
| *peb-1* | WBGene00003968 | protein_coding_gene | 3.7 | 12.8856 | 5.1E-13 | 1.2 | 2.273 | 4.1E-08 |
| *ocr-4* | WBGene00003841 | protein_coding_gene | 3.7 | 12.8664 | 7.3E-07 | 4.1 | 16.672 | 3.3E-07 |
| *K02A2.5* | WBGene00019290 | protein_coding_gene | 3.7 | 12.8639 | 2.1E-14 | 2.3 | 4.927 | 1.0E-05 |
| *F45D3.2* | WBGene00009722 | protein_coding_gene | 3.7 | 12.7458 | 2.1E-26 | 3.1 | 8.811 | 1.0E-31 |
| *R09D1.4* | WBGene00011160 | pseudogene | 3.7 | 12.6140 | 2.3E-03 | 1.7 | 3.187 | 4.5E-02 |
| *ZC196.4* | WBGene00022548 | protein_coding_gene | 3.7 | 12.5910 | 1.6E-29 | 2.6 | 5.970 | 9.7E-15 |
| *hsp-12.3* | WBGene00002012 | protein_coding_gene | 3.7 | 12.5715 | 5.7E-43 | 3.5 | 11.223 | 5.4E-56 |
| *ZC434.3* | WBGene00013891 | protein_coding_gene | 3.7 | 12.5598 | 1.8E-09 | 2.3 | 4.843 | 1.1E-02 |
| *pals-14* | WBGene00009061 | protein_coding_gene | 3.6 | 12.5384 | 1.0E-09 | 2.0 | 4.008 | 3.3E-05 |
| *F47H4.12* | WBGene00043053 | pseudogene | 3.6 | 12.3903 | 6.1E-11 | 1.2 | 2.358 | 1.6E-03 |
| *ZK662.6* | WBGene00045494 | protein_coding_gene | 3.6 | 12.3276 | 1.1E-09 | 5.5 | 46.718 | 5.2E-11 |
| *arrd-11* | WBGene00011052 | protein_coding_gene | 3.6 | 12.3007 | 2.9E-06 | 1.7 | 3.262 | 2.3E-08 |
| *Y39B6A.29* | WBGene00012688 | protein_coding_gene | 3.6 | 12.2388 | 8.8E-08 | 2.1 | 4.198 | 4.0E-03 |
| *C15B12.8* | WBGene00015788 | protein_coding_gene | 3.6 | 12.2268 | 9.9E-14 | 2.9 | 7.332 | 1.9E-16 |
| *col-3* | WBGene00000594 | protein_coding_gene | 3.6 | 12.2125 | 8.5E-08 | 3.0 | 7.927 | 2.1E-04 |
| *pals-6* | WBGene00007660 | protein_coding_gene | 3.6 | 12.1435 | 1.4E-12 | 3.2 | 9.243 | 1.9E-17 |
| *Y43D4A.5* | WBGene00012791 | protein_coding_gene | 3.6 | 12.1055 | 1.2E-06 | 3.3 | 9.552 | 3.4E-05 |
| *C09B9.1* | WBGene00015626 | protein_coding_gene | 3.6 | 12.1024 | 2.4E-07 | 3.1 | 8.490 | 3.2E-06 |
| *K08B5.1* | WBGene00019518 | protein_coding_gene | 3.6 | 12.1008 | 9.2E-18 | 2.0 | 3.890 | 9.1E-09 |
| *nhr-247* | WBGene00014193 | protein_coding_gene | 3.6 | 12.1005 | 7.8E-03 | 1.9 | 3.727 | 3.3E-02 |
| *R07A4.3* | WBGene00011074 | protein_coding_gene | 3.6 | 12.0668 | 3.6E-06 | 2.8 | 7.140 | 4.3E-06 |
| *W01A11.7* | WBGene00020912 | protein_coding_gene | 3.6 | 12.0029 | 5.1E-19 | 2.8 | 6.784 | 6.0E-14 |
| *C50B8.4* | WBGene00008226 | protein_coding_gene | 3.6 | 11.9856 | 9.7E-21 | 1.4 | 2.570 | 2.8E-07 |
| *ttr-26* | WBGene00013079 | protein_coding_gene | 3.6 | 11.9657 | 8.1E-20 | 2.3 | 5.089 | 2.5E-44 |

| | | | | | | | | |
|---|---|---|---|---|---|---|---|---|
| *col-117* | WBGene00000691 | protein_coding_gene | 3.6 | 11.9152 | 4.7E-08 | 3.7 | 12.859 | 2.8E-04 |
| *msrp-7* | WBGene00011070 | protein_coding_gene | 3.6 | 11.8988 | 6.2E-39 | 3.0 | 8.178 | 2.7E-56 |
| *spin-3* | WBGene00008598 | protein_coding_gene | 3.6 | 11.8883 | 6.2E-42 | 2.2 | 4.688 | 6.1E-11 |
| *daf-9* | WBGene00000905 | protein_coding_gene | 3.6 | 11.8758 | 1.8E-32 | 5.4 | 41.552 | 3.5E-42 |
| *srg-32* | WBGene00005189 | protein_coding_gene | 3.6 | 11.7994 | 2.8E-03 | 1.8 | 3.557 | 4.9E-02 |
| *F45D3.3* | WBGene00009723 | protein_coding_gene | 3.5 | 11.6736 | 4.5E-45 | 1.2 | 2.255 | 4.9E-04 |
| *sre-23* | WBGene00013957 | protein_coding_gene | 3.5 | 11.6130 | 1.7E-05 | 1.2 | 2.311 | 4.8E-02 |
| *D1086.2* | WBGene00008389 | protein_coding_gene | 3.5 | 11.5734 | 8.3E-31 | 1.1 | 2.122 | 2.5E-03 |
| *C07E3.3* | WBGene00007414 | protein_coding_gene | 3.5 | 11.5707 | 1.6E-17 | 1.6 | 3.048 | 3.7E-23 |
| *B0462.1* | WBGene00007181 | pseudogene | 3.5 | 11.5649 | 2.4E-05 | 1.8 | 3.428 | 4.8E-07 |
| *pals-7* | WBGene00007661 | protein_coding_gene | 3.5 | 11.5132 | 3.1E-03 | 1.9 | 3.648 | 4.0E-03 |
| *F59A7.2* | WBGene00019090 | protein_coding_gene | 3.5 | 11.4730 | 3.4E-11 | 2.4 | 5.264 | 1.8E-27 |
| *F53F4.1* | WBGene00009985 | protein_coding_gene | 3.5 | 11.4708 | 4.4E-07 | 1.9 | 3.857 | 1.5E-02 |
| *nhr-19* | WBGene00003618 | protein_coding_gene | 3.5 | 11.3934 | 3.6E-18 | 1.5 | 2.916 | 8.5E-21 |
| *asic-2* | WBGene00012137 | protein_coding_gene | 3.5 | 11.3724 | 2.1E-09 | 2.5 | 5.707 | 7.9E-08 |
| *srg-15* | WBGene00005172 | protein_coding_gene | 3.5 | 11.3466 | 4.7E-03 | 2.9 | 7.238 | 8.5E-03 |
| *K10D3.6* | WBGene00010740 | protein_coding_gene | 3.5 | 11.3383 | 3.3E-40 | 1.1 | 2.167 | 7.4E-06 |
| *cest-1.1* | WBGene00011362 | protein_coding_gene | 3.5 | 11.2917 | 3.5E-15 | 1.0 | 2.030 | 3.7E-02 |
| *pho-6* | WBGene00018696 | protein_coding_gene | 3.5 | 11.2599 | 2.6E-19 | 1.3 | 2.534 | 1.6E-08 |
| *oac-47* | WBGene00011933 | pseudogene | 3.5 | 11.2561 | 1.8E-03 | 2.2 | 4.675 | 4.2E-02 |
| *C52E2.4* | WBGene00016883 | protein_coding_gene | 3.5 | 11.2450 | 1.9E-12 | 1.7 | 3.215 | 2.4E-08 |
| *C01F1.5* | WBGene00015300 | protein_coding_gene | 3.5 | 11.0949 | 6.4E-10 | 2.0 | 3.939 | 5.9E-03 |
| *E04D5.4* | WBGene00008483 | protein_coding_gene | 3.5 | 11.0923 | 5.7E-11 | 2.6 | 6.261 | 4.0E-07 |
| *C44H9.2* | WBGene00008098 | pseudogene | 3.5 | 11.0864 | 1.5E-03 | 1.7 | 3.344 | 4.9E-02 |
| *F14H12.2* | WBGene00017470 | protein_coding_gene | 3.5 | 11.0228 | 1.7E-04 | 2.3 | 4.861 | 3.4E-02 |
| *Y94H6A.2* | WBGene00022375 | protein_coding_gene | 3.5 | 11.0043 | 1.2E-09 | 1.2 | 2.345 | 9.9E-03 |
| *K10H10.10* | WBGene00010765 | protein_coding_gene | 3.5 | 10.9800 | 7.2E-08 | 2.3 | 4.957 | 1.2E-05 |
| *gcy-3* | WBGene00001530 | protein_coding_gene | 3.5 | 10.9759 | 2.9E-03 | 2.8 | 6.810 | 1.3E-04 |
| *C12D5.3* | WBGene00015706 | protein_coding_gene | 3.5 | 10.9696 | 3.3E-09 | 1.2 | 2.321 | 4.4E-03 |
| *C36B1.13* | WBGene00044120 | protein_coding_gene | 3.5 | 10.9646 | 2.4E-05 | 1.4 | 2.589 | 1.6E-03 |
| *cpr-1* | WBGene00000781 | protein_coding_gene | 3.4 | 10.9220 | 1.8E-27 | 1.4 | 2.588 | 6.9E-22 |
| *C14C11.4* | WBGene00015768 | protein_coding_gene | 3.4 | 10.8980 | 1.5E-33 | 2.6 | 6.008 | 6.4E-24 |
| *T09E11.11* | WBGene00011660 | pseudogene | 3.4 | 10.8049 | 3.0E-13 | 1.5 | 2.799 | 1.2E-09 |
| *nlp-76* | WBGene00045386 | protein_coding_gene | 3.4 | 10.8032 | 4.3E-07 | 2.2 | 4.442 | 6.0E-06 |
| *ptr-3* | WBGene00004218 | protein_coding_gene | 3.4 | 10.7440 | 1.2E-15 | 2.3 | 4.876 | 1.0E-17 |
| *F35F10.13* | WBGene00018062 | protein_coding_gene | 3.4 | 10.7276 | 1.1E-08 | 4.5 | 22.616 | 2.7E-56 |
| *F42C5.3* | WBGene00018345 | protein_coding_gene | 3.4 | 10.6938 | 2.9E-09 | 2.5 | 5.630 | 8.9E-13 |
| *C05B5.17* | WBGene00219314 | protein_coding_gene | 3.4 | 10.6615 | 5.6E-03 | 3.1 | 8.450 | 1.0E-02 |
| *F28H7.2* | WBGene00009236 | protein_coding_gene | 3.4 | 10.6590 | 4.6E-10 | 2.3 | 4.911 | 4.0E-06 |
| *glb-5* | WBGene00015964 | protein_coding_gene | 3.4 | 10.6459 | 9.5E-06 | 1.8 | 3.445 | 3.1E-05 |
| *ceh-23* | WBGene00000446 | protein_coding_gene | 3.4 | 10.6247 | 1.9E-07 | 2.9 | 7.652 | 1.5E-05 |
| *F43C11.8* | WBGene00018385 | protein_coding_gene | 3.4 | 10.5822 | 1.6E-03 | 2.6 | 6.145 | 6.9E-03 |
| ***F13E9.9*** | **WBGene00008758** | **protein_coding_gene** | **3.4** | **10.5674** | **3.6E-03** | **-1.3** | **0.409** | **7.3E-03** |
| *flp-21* | WBGene00001464 | protein_coding_gene | 3.4 | 10.5078 | 3.0E-24 | 2.2 | 4.718 | 4.3E-27 |
| *C30B5.6* | WBGene00016247 | protein_coding_gene | 3.4 | 10.4725 | 1.1E-12 | 1.9 | 3.832 | 1.1E-04 |
| *ador-1* | WBGene00011878 | protein_coding_gene | 3.4 | 10.4654 | 3.6E-05 | 1.9 | 3.764 | 1.0E-02 |
| *bath-25* | WBGene00020859 | protein_coding_gene | 3.4 | 10.4466 | 6.4E-04 | 1.6 | 3.126 | 1.7E-03 |
| *Y41D4B.17* | WBGene00021519 | protein_coding_gene | 3.4 | 10.3732 | 1.3E-20 | 2.9 | 7.481 | 3.9E-12 |
| *mboa-1* | WBGene00007174 | protein_coding_gene | 3.4 | 10.3496 | 2.9E-09 | 1.5 | 2.849 | 8.5E-07 |
| *F20A1.4* | WBGene00017616 | protein_coding_gene | 3.4 | 10.3286 | 1.1E-05 | 2.0 | 3.885 | 1.7E-03 |
| *fbxa-147* | WBGene00011135 | protein_coding_gene | 3.4 | 10.3275 | 1.6E-02 | 3.2 | 9.396 | 7.5E-03 |
| *F53C3.6* | WBGene00018750 | protein_coding_gene | 3.4 | 10.3145 | 1.5E-06 | 1.1 | 2.186 | 2.1E-02 |
| *fozi-1* | WBGene00010453 | protein_coding_gene | 3.4 | 10.2830 | 2.1E-13 | 1.2 | 2.295 | 5.0E-12 |
| *Y32B12C.5* | WBGene00194769 | pseudogene | 3.4 | 10.2426 | 6.0E-12 | 1.6 | 3.120 | 1.3E-06 |
| *K10G4.3* | WBGene00010752 | protein_coding_gene | 3.4 | 10.2090 | 8.8E-09 | 1.9 | 3.782 | 3.0E-02 |
| *nmad-1* | WBGene00017304 | protein_coding_gene | 3.4 | 10.2072 | 2.4E-51 | 1.8 | 3.397 | 3.8E-24 |
| *C29F9.6* | WBGene00016221 | protein_coding_gene | 3.4 | 10.2055 | 2.9E-08 | 2.7 | 6.675 | 1.2E-03 |
| *comt-1* | WBGene00018741 | protein_coding_gene | 3.3 | 10.1652 | 5.4E-05 | 3.8 | 13.745 | 1.7E-05 |
| *W02G9.4* | WBGene00020954 | protein_coding_gene | 3.3 | 10.1587 | 2.4E-70 | 2.5 | 5.465 | 2.8E-29 |
| *C09E7.10* | WBGene00044295 | protein_coding_gene | 3.3 | 10.1444 | 3.3E-04 | 2.8 | 6.883 | 4.4E-03 |
| *pdf-2* | WBGene00018590 | protein_coding_gene | 3.3 | 10.1006 | 2.9E-23 | 1.8 | 3.499 | 9.6E-24 |
| *F47B8.2* | WBGene00009803 | protein_coding_gene | 3.3 | 10.0929 | 1.8E-25 | 2.3 | 4.880 | 2.0E-39 |
| *C56G3.2* | WBGene00016985 | protein_coding_gene | 3.3 | 10.0838 | 2.3E-03 | 1.9 | 3.638 | 4.2E-02 |
| *ins-4* | WBGene00002087 | protein_coding_gene | 3.3 | 10.0778 | 1.4E-14 | 1.4 | 2.631 | 9.4E-06 |
| *ZK863.8* | WBGene00014125 | protein_coding_gene | 3.3 | 10.0399 | 3.2E-25 | 2.1 | 4.415 | 4.1E-34 |
| *C48D5.3* | WBGene00008180 | protein_coding_gene | 3.3 | 10.0354 | 6.6E-03 | 2.1 | 4.183 | 4.1E-02 |
| *col-166* | WBGene00000739 | protein_coding_gene | 3.3 | 10.0294 | 8.9E-26 | 3.8 | 13.659 | 3.1E-79 |
| *C02H6.3* | WBGene00220257 | protein_coding_gene | 3.3 | 10.0070 | 1.1E-02 | 2.6 | 5.955 | 2.1E-02 |
| *T27C4.1* | WBGene00020854 | protein_coding_gene | 3.3 | 9.9853 | 1.7E-24 | 2.5 | 5.621 | 3.3E-84 |
| *R06C1.6* | WBGene00011060 | protein_coding_gene | 3.3 | 9.8872 | 2.0E-29 | 2.5 | 5.552 | 6.9E-31 |
| *R05A10.6* | WBGene00011023 | protein_coding_gene | 3.3 | 9.8576 | 5.3E-03 | 2.6 | 5.876 | 2.7E-02 |
| *C39B10.1* | WBGene00008021 | protein_coding_gene | 3.3 | 9.8244 | 1.7E-22 | 2.4 | 5.144 | 4.6E-12 |
| *lmd-5* | WBGene00020141 | protein_coding_gene | 3.3 | 9.7499 | 7.7E-06 | 2.7 | 6.706 | 9.2E-08 |
| *mboa-5* | WBGene00013995 | pseudogene | 3.3 | 9.7451 | 5.0E-04 | 2.5 | 5.843 | 3.8E-06 |
| *F02A9.7* | WBGene00044406 | protein_coding_gene | 3.3 | 9.6999 | 3.4E-07 | 1.9 | 3.730 | 4.7E-04 |
| *ZK250.13* | WBGene00235352 | protein_coding_gene | 3.3 | 9.6973 | 2.4E-21 | 2.8 | 7.176 | 3.2E-29 |
| *nas-14* | WBGene00003533 | protein_coding_gene | 3.3 | 9.6916 | 1.0E-15 | 1.6 | 3.115 | 1.7E-02 |
| *asm-2* | WBGene00000212 | protein_coding_gene | 3.3 | 9.6759 | 4.2E-43 | 3.1 | 8.433 | 4.5E-91 |
| *nape-1* | WBGene00021371 | protein_coding_gene | 3.3 | 9.6635 | 1.7E-13 | 1.3 | 2.432 | 6.7E-10 |
| ***dct-8*** | **WBGene00044472** | **protein_coding_gene** | **3.3** | **9.6369** | **2.2E-03** | **-1.3** | **0.418** | **4.2E-03** |

| | | | | | | | | |
|---|---|---|---|---|---|---|---|---|
| *snt-7* | WBGene00011682 | protein_coding_gene | 3.3 | 9.6233 | 7.4E-08 | 1.5 | 2.850 | 1.1E-03 |
| *F38B2.2* | WBGene00009529 | protein_coding_gene | 3.3 | 9.6115 | 2.2E-13 | 1.3 | 2.536 | 1.7E-03 |
| *B0563.5* | WBGene00015262 | protein_coding_gene | 3.3 | 9.5909 | 5.1E-18 | 1.2 | 2.270 | 2.6E-03 |
| *Y54G2A.32* | WBGene00021894 | protein_coding_gene | 3.3 | 9.5542 | 8.8E-51 | 1.5 | 2.868 | 5.2E-15 |
| *K02C4.2* | WBGene00010501 | protein_coding_gene | 3.2 | 9.5087 | 2.0E-05 | 1.5 | 2.825 | 1.2E-02 |
| *Y67D8B.1* | WBGene00022061 | protein_coding_gene | 3.2 | 9.4695 | 1.6E-05 | 2.5 | 5.659 | 6.9E-04 |
| *Y9C9A.16* | WBGene00021183 | protein_coding_gene | 3.2 | 9.4543 | 1.0E-07 | 3.6 | 11.754 | 5.4E-14 |
| *Y105C5A.17* | WBGene00013640 | ncRNA_gene | 3.2 | 9.4525 | 7.8E-03 | 2.5 | 5.777 | 3.6E-02 |
| *glb-34* | WBGene00077763 | protein_coding_gene | 3.2 | 9.4430 | 3.6E-03 | 2.3 | 4.860 | 1.7E-02 |
| *slc-36.3* | WBGene00012629 | protein_coding_gene | 3.2 | 9.3976 | 3.0E-09 | 1.7 | 3.230 | 2.8E-08 |
| *R05A10.1* | WBGene00011018 | protein_coding_gene | 3.2 | 9.3806 | 5.8E-04 | 2.3 | 4.763 | 4.9E-02 |
| *K06A9.1* | WBGene00019435 | protein_coding_gene | 3.2 | 9.3521 | 1.3E-14 | 2.5 | 5.604 | 6.2E-25 |
| *C15C7.4* | WBGene00015790 | protein_coding_gene | 3.2 | 9.3476 | 1.4E-12 | 2.4 | 5.452 | 2.1E-24 |
| *F02A9.1* | WBGene00008513 | protein_coding_gene | 3.2 | 9.3447 | 3.2E-12 | 1.7 | 3.290 | 7.1E-06 |
| *F01D4.8* | WBGene00008490 | protein_coding_gene | 3.2 | 9.3444 | 7.6E-17 | 3.9 | 14.505 | 1.4E-27 |
| *nhr-117* | WBGene00003707 | protein_coding_gene | 3.2 | 9.3274 | 4.8E-17 | 1.6 | 2.969 | 5.7E-14 |
| *tsp-4* | WBGene00006630 | protein_coding_gene | 3.2 | 9.3245 | 1.1E-02 | 2.7 | 6.499 | 1.7E-02 |
| *nex-4* | WBGene00003591 | protein_coding_gene | 3.2 | 9.3163 | 3.8E-13 | 2.4 | 5.157 | 3.3E-07 |
| *bnc-1* | WBGene00044791 | protein_coding_gene | 3.2 | 9.2959 | 5.1E-03 | 2.0 | 4.098 | 4.1E-02 |
| *T03E6.8* | WBGene00011397 | protein_coding_gene | 3.2 | 9.2770 | 3.4E-16 | 2.2 | 4.541 | 3.8E-08 |
| *R02D5.6* | WBGene00010979 | protein_coding_gene | 3.2 | 9.2685 | 6.7E-04 | 1.8 | 3.571 | 1.8E-02 |
| *B0238.12* | WBGene00015076 | protein_coding_gene | 3.2 | 9.2530 | 1.8E-10 | 3.5 | 11.535 | 1.4E-26 |
| *gba-1* | WBGene00016335 | protein_coding_gene | 3.2 | 9.2523 | 7.2E-17 | 1.7 | 3.180 | 2.1E-15 |
| *nas-15* | WBGene00003534 | protein_coding_gene | 3.2 | 9.2450 | 3.9E-09 | 2.7 | 6.568 | 6.2E-03 |
| *F11A6.3* | WBGene00077549 | pseudogene | 3.2 | 9.2343 | 5.8E-04 | 1.9 | 3.721 | 1.0E-02 |
| *gst-14* | WBGene00001762 | protein_coding_gene | 3.2 | 9.2284 | 1.2E-13 | 3.2 | 9.286 | 1.9E-12 |
| *F26A1.19* | WBGene00270319 | protein_coding_gene | 3.2 | 9.2243 | 3.0E-14 | 1.5 | 2.833 | 3.2E-05 |
| *fipr-1* | WBGene00009096 | protein_coding_gene | 3.2 | 9.2219 | 9.0E-54 | 1.5 | 2.807 | 1.3E-17 |
| *B0462.5* | WBGene00194695 | protein_coding_gene | 3.2 | 9.2095 | 1.8E-12 | 1.2 | 2.300 | 3.8E-05 |
| *far-4* | WBGene00001388 | protein_coding_gene | 3.2 | 9.1707 | 1.2E-07 | 3.8 | 13.578 | 1.4E-06 |
| *C08F11.3* | WBGene00007452 | protein_coding_gene | 3.2 | 9.1608 | 1.1E-07 | 1.3 | 2.510 | 4.7E-03 |
| *C17G10.6* | WBGene00015918 | protein_coding_gene | 3.2 | 9.1483 | 6.8E-05 | 1.7 | 3.235 | 1.7E-02 |
| *glb-23* | WBGene00020045 | protein_coding_gene | 3.2 | 9.1360 | 1.7E-07 | 1.2 | 2.274 | 3.7E-02 |
| *T16G1.7* | WBGene00011801 | protein_coding_gene | 3.2 | 9.1346 | 5.3E-06 | 1.6 | 3.054 | 8.2E-33 |
| *F49H6.13* | WBGene00009914 | protein_coding_gene | 3.2 | 9.1223 | 2.0E-03 | 3.0 | 7.747 | 4.3E-03 |
| *D1053.3* | WBGene00008368 | protein_coding_gene | 3.2 | 9.1135 | 1.4E-16 | 1.5 | 2.767 | 1.7E-09 |
| *nhr-221* | WBGene00020748 | protein_coding_gene | 3.2 | 9.0741 | 4.4E-05 | 1.6 | 3.070 | 1.7E-02 |
| *oac-1* | WBGene00007177 | protein_coding_gene | 3.2 | 9.0630 | 3.3E-08 | 4.0 | 15.519 | 3.2E-04 |
| *Y64H9A.2* | WBGene00022024 | protein_coding_gene | 3.2 | 9.0534 | 7.6E-05 | 1.5 | 2.896 | 4.9E-02 |
| *xbx-3* | WBGene00010864 | protein_coding_gene | 3.2 | 9.0433 | 4.2E-05 | 1.8 | 3.511 | 1.5E-03 |
| *gst-20* | WBGene00001768 | protein_coding_gene | 3.2 | 9.0340 | 3.9E-28 | 2.3 | 5.016 | 8.2E-33 |
| *C30G4.2* | WBGene00016268 | protein_coding_gene | 3.2 | 9.0231 | 2.1E-03 | 1.6 | 2.965 | 2.0E-02 |
| *nhr-6* | WBGene00003605 | protein_coding_gene | 3.2 | 8.9936 | 3.4E-19 | 2.1 | 4.423 | 1.2E-07 |
| *nhr-18* | WBGene00003617 | protein_coding_gene | 3.2 | 8.9489 | 8.4E-21 | 1.5 | 2.877 | 1.5E-12 |
| *twnk-1* | WBGene00018514 | protein_coding_gene | 3.2 | 8.9408 | 8.7E-08 | 2.0 | 3.990 | 1.0E-04 |
| *ZK1240.1* | WBGene00022866 | protein_coding_gene | 3.2 | 8.8814 | 4.7E-06 | 2.8 | 6.790 | 1.4E-02 |
| *gstk-2* | WBGene00017054 | protein_coding_gene | 3.2 | 8.8806 | 1.3E-23 | 2.1 | 4.173 | 4.2E-17 |
| *srz-100* | WBGene00012026 | protein_coding_gene | 3.2 | 8.8775 | 7.5E-03 | 3.0 | 7.732 | 1.2E-05 |
| *Y42A5A.1* | WBGene00012776 | protein_coding_gene | 3.1 | 8.8501 | 5.5E-23 | 2.4 | 5.107 | 7.0E-22 |
| *gur-3* | WBGene00001804 | protein_coding_gene | 3.1 | 8.8497 | 8.7E-03 | 3.3 | 9.553 | 8.3E-03 |
| *sqt-1* | WBGene00005016 | protein_coding_gene | 3.1 | 8.8472 | 2.0E-16 | 4.5 | 22.060 | 5.1E-06 |
| *nlp-71* | WBGene00022254 | protein_coding_gene | 3.1 | 8.8346 | 1.5E-23 | 2.1 | 4.416 | 2.3E-20 |
| *droe-8* | WBGene00022550 | protein_coding_gene | 3.1 | 8.7984 | 2.7E-05 | 2.2 | 4.657 | 7.5E-06 |
| *asp-7* | WBGene00019104 | protein_coding_gene | 3.1 | 8.7816 | 1.5E-02 | 3.0 | 7.768 | 4.0E-04 |
| *F39H12.2* | WBGene00018213 | protein_coding_gene | 3.1 | 8.7715 | 8.5E-08 | 3.5 | 11.036 | 4.7E-16 |
| *srw-12* | WBGene00005759 | protein_coding_gene | 3.1 | 8.7711 | 1.1E-02 | 2.5 | 5.719 | 2.2E-02 |
| ***F48C1.9*** | **WBGene00018601** | **protein_coding_gene** | **3.1** | **8.7382** | **2.1E-03** | **-1.7** | **0.311** | **4.7E-05** |
| *ikb-1* | WBGene00002069 | protein_coding_gene | 3.1 | 8.7271 | 9.1E-11 | 1.5 | 2.862 | 5.7E-20 |
| *C49C8.8* | WBGene00235271 | protein_coding_gene | 3.1 | 8.7223 | 1.1E-03 | 1.7 | 3.325 | 1.5E-02 |
| *nlp-68* | WBGene00011899 | protein_coding_gene | 3.1 | 8.7018 | 9.0E-24 | 1.9 | 3.667 | 2.2E-09 |
| *F13H8.1* | WBGene00017434 | protein_coding_gene | 3.1 | 8.6942 | 2.4E-13 | 1.4 | 2.659 | 1.7E-10 |
| *F59A7.5* | WBGene00019092 | protein_coding_gene | 3.1 | 8.6768 | 4.8E-23 | 1.7 | 3.334 | 4.3E-18 |
| *C38C3.10* | WBGene00045390 | protein_coding_gene | 3.1 | 8.6705 | 8.3E-15 | 2.1 | 4.240 | 2.1E-11 |
| *pqm-1* | WBGene00004096 | protein_coding_gene | 3.1 | 8.6575 | 2.4E-23 | 1.6 | 3.012 | 2.5E-31 |
| *F48G7.13* | WBGene00018624 | protein_coding_gene | 3.1 | 8.6184 | 3.2E-04 | 2.1 | 4.265 | 1.1E-02 |
| *C34F11.8* | WBGene00016417 | protein_coding_gene | 3.1 | 8.5889 | 9.9E-21 | 1.5 | 2.733 | 4.0E-19 |
| *osm-9* | WBGene00003889 | protein_coding_gene | 3.1 | 8.5840 | 1.5E-04 | 2.3 | 4.897 | 7.7E-06 |
| *unk-1* | WBGene00016407 | protein_coding_gene | 3.1 | 8.5835 | 4.9E-10 | 1.6 | 3.037 | 5.0E-30 |
| *bath-46* | WBGene00020325 | protein_coding_gene | 3.1 | 8.5823 | 1.2E-09 | 3.0 | 8.140 | 4.0E-13 |
| *F19G12.3* | WBGene00017611 | protein_coding_gene | 3.1 | 8.5701 | 9.4E-03 | 1.7 | 3.361 | 9.9E-03 |
| *Y70G10A.2* | WBGene00013498 | protein_coding_gene | 3.1 | 8.5646 | 1.8E-05 | 1.7 | 3.325 | 6.5E-03 |
| *C47E8.3* | WBGene00008142 | protein_coding_gene | 3.1 | 8.5567 | 2.2E-12 | 2.2 | 4.508 | 6.8E-07 |
| *T28A11.2* | WBGene00020869 | protein_coding_gene | 3.1 | 8.5500 | 2.2E-54 | 2.3 | 4.929 | 1.2E-02 |
| *exc-14* | WBGene00019649 | protein_coding_gene | 3.1 | 8.5289 | 2.5E-17 | 2.0 | 4.068 | 3.9E-08 |
| *clec-140* | WBGene00020220 | protein_coding_gene | 3.1 | 8.5280 | 2.5E-03 | 2.1 | 4.341 | 1.1E-02 |
| *srm-4* | WBGene00009808 | protein_coding_gene | 3.1 | 8.5264 | 8.9E-03 | 2.1 | 4.211 | 2.9E-02 |
| *uev-2* | WBGene00006731 | protein_coding_gene | 3.1 | 8.5241 | 4.0E-04 | 2.1 | 4.213 | 1.6E-02 |
| *C50B6.7* | WBGene00008220 | protein_coding_gene | 3.1 | 8.5221 | 8.6E-19 | 3.2 | 9.347 | 4.6E-64 |
| *droe-4* | WBGene00010262 | protein_coding_gene | 3.1 | 8.5207 | 2.5E-23 | 1.3 | 2.462 | 7.7E-17 |
| *nas-6* | WBGene00003525 | protein_coding_gene | 3.1 | 8.5141 | 2.4E-08 | 2.9 | 7.312 | 1.4E-07 |

| | | | | | | | | |
|---|---|---|---|---|---|---|---|---|
| *flp-9* | WBGene00001452 | protein_coding_gene | 3.1 | 8.4835 | 1.8E-34 | 1.9 | 3.806 | 9.2E-50 |
| *F40F9.10* | WBGene00009586 | protein_coding_gene | 3.1 | 8.4831 | 1.1E-10 | 1.3 | 2.471 | 1.6E-03 |
| *pqn-48* | WBGene00004135 | protein_coding_gene | 3.1 | 8.4754 | 7.6E-32 | 2.7 | 6.639 | 7.3E-48 |
| *nlp-35* | WBGene00007881 | protein_coding_gene | 3.1 | 8.4630 | 1.3E-16 | 2.4 | 5.459 | 2.3E-20 |
| *gcy-34* | WBGene00001554 | protein_coding_gene | 3.1 | 8.4537 | 1.2E-03 | 1.3 | 2.410 | 2.3E-02 |
| *dpy-8* | WBGene00001070 | protein_coding_gene | 3.1 | 8.4509 | 5.8E-10 | 4.1 | 17.212 | 1.1E-05 |
| *col-46* | WBGene00000623 | protein_coding_gene | 3.1 | 8.4500 | 3.2E-04 | 4.5 | 22.472 | 3.4E-05 |
| *cyp-13B1* | WBGene00008519 | protein_coding_gene | 3.1 | 8.4366 | 1.2E-08 | 1.1 | 2.209 | 9.1E-05 |
| *ubc-22* | WBGene00006717 | protein_coding_gene | 3.1 | 8.4109 | 7.0E-06 | 2.8 | 6.982 | 2.7E-05 |
| *bcmo-1* | WBGene00012914 | protein_coding_gene | 3.1 | 8.4058 | 2.9E-19 | 3.0 | 8.010 | 1.8E-12 |
| *srab-14* | WBGene00019999 | protein_coding_gene | 3.1 | 8.3928 | 3.3E-04 | 1.7 | 3.175 | 3.4E-02 |
| *Y41D4B.15* | WBGene00021517 | protein_coding_gene | 3.1 | 8.3757 | 2.2E-03 | 2.7 | 6.538 | 3.6E-03 |
| *nhr-282* | WBGene00010017 | protein_coding_gene | 3.1 | 8.3524 | 4.8E-03 | 2.9 | 7.323 | 3.1E-02 |
| *gst-3* | WBGene00001751 | protein_coding_gene | 3.1 | 8.3447 | 1.1E-15 | 2.6 | 6.084 | 3.7E-13 |
| *F11C1.9* | WBGene00195213 | protein_coding_gene | 3.1 | 8.3437 | 1.5E-02 | 2.1 | 4.174 | 4.9E-02 |
| *F16B3.2* | WBGene00017502 | protein_coding_gene | 3.1 | 8.3390 | 1.1E-05 | 2.2 | 4.605 | 6.5E-08 |
| *dpy-6* | WBGene00001068 | protein_coding_gene | 3.1 | 8.3376 | 2.0E-08 | 6.6 | 99.454 | 5.4E-27 |
| *twk-7* | WBGene00006662 | protein_coding_gene | 3.1 | 8.3366 | 2.8E-13 | 1.3 | 2.378 | 3.0E-09 |
| *asp-6* | WBGene00000219 | protein_coding_gene | 3.1 | 8.3302 | 4.9E-34 | 1.5 | 2.879 | 9.5E-46 |
| *F26G1.10* | WBGene00044663 | protein_coding_gene | 3.1 | 8.3218 | 1.8E-04 | 2.2 | 4.516 | 1.4E-02 |
| *ntc-1* | WBGene00044568 | protein_coding_gene | 3.1 | 8.3014 | 8.8E-10 | 1.7 | 3.343 | 3.3E-05 |
| *Y73B6BL.44* | WBGene00044350 | protein_coding_gene | 3.1 | 8.2981 | 2.6E-27 | 1.5 | 2.835 | 8.8E-14 |
| *pho-11* | WBGene00007331 | protein_coding_gene | 3.1 | 8.2925 | 1.6E-19 | 2.1 | 4.322 | 5.2E-16 |
| *ghi-1* | WBGene00013923 | protein_coding_gene | 3.1 | 8.2838 | 9.1E-12 | 3.2 | 9.380 | 7.8E-22 |
| *C27D9.2* | WBGene00016164 | protein_coding_gene | 3.1 | 8.2838 | 1.3E-03 | 4.4 | 20.438 | 1.7E-05 |
| *T19H12.15* | WBGene00271827 | protein_coding_gene | 3.0 | 8.2789 | 1.6E-19 | 4.2 | 18.840 | 9.6E-67 |
| *gopc-1* | WBGene00014101 | protein_coding_gene | 3.0 | 8.2553 | 6.8E-13 | 2.2 | 4.548 | 2.8E-08 |
| *nhr-206* | WBGene00011097 | protein_coding_gene | 3.0 | 8.2420 | 5.6E-12 | 1.4 | 2.606 | 3.1E-06 |
| *col-169* | WBGene00000742 | protein_coding_gene | 3.0 | 8.2404 | 1.0E-13 | 4.4 | 20.874 | 3.0E-19 |
| *F09C8.1* | WBGene00008621 | protein_coding_gene | 3.0 | 8.2266 | 6.6E-15 | 3.0 | 8.091 | 2.1E-14 |
| *nlp-8* | WBGene00003746 | protein_coding_gene | 3.0 | 8.2153 | 4.3E-48 | 1.9 | 3.662 | 1.2E-26 |
| *fipr-2* | WBGene00009097 | protein_coding_gene | 3.0 | 8.2071 | 5.5E-28 | 1.5 | 2.841 | 3.6E-23 |
| *klu-1* | WBGene00013970 | protein_coding_gene | 3.0 | 8.1878 | 1.3E-14 | 2.7 | 6.705 | 6.0E-17 |
| *F46A8.11* | WBGene00014761 | protein_coding_gene | 3.0 | 8.1817 | 1.5E-16 | 2.7 | 6.669 | 4.4E-14 |
| *C30E1.9* | WBGene00022951 | ncRNA_gene | 3.0 | 8.1708 | 1.3E-17 | 1.9 | 3.641 | 4.1E-15 |
| *C54H2.4* | WBGene00016938 | protein_coding_gene | 3.0 | 8.1624 | 2.1E-04 | 2.3 | 5.074 | 2.1E-03 |
| *rab-11.2* | WBGene00004275 | protein_coding_gene | 3.0 | 8.1623 | 7.2E-04 | 3.0 | 8.215 | 2.8E-03 |
| *F42A9.18* | WBGene00249826 | ncRNA_gene | 3.0 | 8.1593 | 6.9E-05 | 2.4 | 5.412 | 2.5E-06 |
| *F13D11.4* | WBGene00017429 | protein_coding_gene | 3.0 | 8.1183 | 1.1E-21 | 2.0 | 3.942 | 2.7E-32 |
| *efhc-1* | WBGene00013028 | protein_coding_gene | 3.0 | 8.1128 | 1.7E-03 | 2.0 | 4.136 | 9.6E-03 |
| *R07B1.5* | WBGene00011077 | protein_coding_gene | 3.0 | 8.1126 | 6.1E-05 | 2.8 | 7.049 | 1.7E-03 |
| *C33A12.4* | WBGene00007883 | protein_coding_gene | 3.0 | 8.1005 | 5.2E-19 | 1.1 | 2.158 | 6.1E-06 |
| *W05E10.5* | WBGene00012282 | protein_coding_gene | 3.0 | 8.0955 | 4.0E-09 | 1.7 | 3.313 | 3.5E-04 |
| *pals-38* | WBGene00008302 | protein_coding_gene | 3.0 | 8.0920 | 4.6E-07 | 2.3 | 5.094 | 3.3E-06 |
| *Y71G12B.2* | WBGene00022142 | protein_coding_gene | 3.0 | 8.0911 | 1.6E-09 | 2.6 | 6.075 | 3.3E-06 |
| *cdh-12* | WBGene00022103 | protein_coding_gene | 3.0 | 8.0755 | 3.3E-13 | 2.2 | 4.514 | 2.8E-04 |
| *nmy-3* | WBGene00012437 | protein_coding_gene | 3.0 | 8.0712 | 1.6E-15 | 3.8 | 13.631 | 4.4E-48 |
| *R07B1.8* | WBGene00001608 | protein_coding_gene | 3.0 | 8.0678 | 2.2E-11 | 3.1 | 8.643 | 1.8E-10 |
| *F20G2.1* | WBGene00008985 | protein_coding_gene | 3.0 | 8.0626 | 1.5E-11 | 3.4 | 10.628 | 2.6E-17 |
| *W07A12.4* | WBGene00012322 | protein_coding_gene | 3.0 | 8.0616 | 9.5E-13 | 1.5 | 2.812 | 2.3E-06 |
| *W09C2.8* | WBGene00185116 | protein_coding_gene | 3.0 | 8.0560 | 3.3E-02 | 2.8 | 6.795 | 2.1E-02 |
| *lact-3* | WBGene00010897 | protein_coding_gene | 3.0 | 8.0544 | 6.7E-40 | 1.8 | 3.553 | 3.4E-44 |
| *bath-26* | WBGene00020329 | protein_coding_gene | 3.0 | 8.0285 | 2.1E-22 | 1.6 | 2.955 | 1.8E-07 |
| *F13E9.15* | WBGene00045208 | protein_coding_gene | 3.0 | 8.0260 | 2.7E-13 | 1.6 | 3.023 | 7.0E-19 |
| *F52H2.3* | WBGene00018715 | protein_coding_gene | 3.0 | 8.0237 | 1.1E-29 | 2.3 | 4.757 | 5.6E-12 |
| *D2024.4* | WBGene00017052 | protein_coding_gene | 3.0 | 8.0063 | 5.5E-04 | 2.1 | 4.180 | 2.8E-02 |
| *endu-2* | WBGene00019779 | protein_coding_gene | 3.0 | 8.0030 | 1.7E-26 | 1.9 | 3.724 | 6.0E-32 |
| *C04G2.14* | WBGene00255426 | ncRNA_gene | 3.0 | 8.0024 | 4.7E-18 | 4.8 | 27.788 | 8.5E-45 |
| *F53A9.6* | WBGene00018729 | protein_coding_gene | 3.0 | 7.9757 | 5.3E-13 | 1.0 | 2.061 | 5.8E-10 |
| *dlhd-1* | WBGene00020741 | protein_coding_gene | 3.0 | 7.9710 | 8.7E-23 | 2.8 | 7.192 | 1.9E-11 |
| *nlp-81* | WBGene00022063 | protein_coding_gene | 3.0 | 7.9505 | 3.9E-03 | 2.4 | 5.237 | 3.6E-03 |
| *R10H1.1* | WBGene00019998 | protein_coding_gene | 3.0 | 7.9111 | 1.1E-27 | 1.7 | 3.153 | 2.3E-22 |
| *B0432.14* | WBGene00044640 | protein_coding_gene | 3.0 | 7.9099 | 8.7E-20 | 2.6 | 6.207 | 4.8E-21 |
| *nhr-38* | WBGene00003629 | protein_coding_gene | 3.0 | 7.9081 | 9.0E-03 | 3.3 | 9.719 | 2.2E-03 |
| *F40E3.5* | WBGene00018232 | protein_coding_gene | 3.0 | 7.9077 | 1.8E-20 | 1.9 | 3.799 | 2.1E-13 |
| *F53A9.1* | WBGene00018724 | protein_coding_gene | 3.0 | 7.8941 | 3.0E-40 | 2.2 | 4.489 | 2.2E-22 |
| *C23H5.15* | WBGene00219423 | protein_coding_gene | 3.0 | 7.8878 | 1.1E-27 | 2.9 | 7.297 | 3.5E-48 |
| *dsl-5* | WBGene00001107 | protein_coding_gene | 3.0 | 7.8871 | 2.4E-03 | 2.7 | 6.536 | 6.4E-03 |
| *T04F8.7* | WBGene00011445 | protein_coding_gene | 3.0 | 7.8472 | 2.7E-16 | 1.1 | 2.095 | 9.0E-10 |
| *srh-17* | WBGene00005242 | protein_coding_gene | 3.0 | 7.8416 | 1.6E-02 | 3.0 | 7.781 | 3.9E-04 |
| *W06G6.20* | WBGene00219308 | protein_coding_gene | 3.0 | 7.8405 | 2.4E-03 | 1.5 | 2.756 | 4.5E-02 |
| *F10D7.3* | WBGene00017340 | protein_coding_gene | 3.0 | 7.8195 | 9.6E-09 | 1.2 | 2.344 | 1.0E-09 |
| *clec-63* | WBGene00009394 | protein_coding_gene | 3.0 | 7.8036 | 2.3E-23 | 1.5 | 2.869 | 5.7E-41 |
| *F28F5.4* | WBGene00043297 | protein_coding_gene | 3.0 | 7.8024 | 1.3E-07 | 1.3 | 2.462 | 1.3E-02 |
| *fbxa-152* | WBGene00007166 | protein_coding_gene | 3.0 | 7.7903 | 2.0E-02 | 2.2 | 4.478 | 1.9E-02 |
| *rol-6* | WBGene00004397 | protein_coding_gene | 3.0 | 7.7881 | 6.5E-14 | 4.4 | 21.730 | 1.0E-06 |
| *K09C8.7* | WBGene00010717 | protein_coding_gene | 3.0 | 7.7702 | 2.1E-04 | 1.4 | 2.705 | 8.1E-03 |
| *trh-1* | WBGene00007827 | protein_coding_gene | 3.0 | 7.7490 | 8.6E-19 | 1.6 | 3.107 | 4.1E-09 |
| *F21H12.7* | WBGene00044287 | protein_coding_gene | 3.0 | 7.7379 | 5.9E-09 | 2.8 | 7.207 | 6.1E-06 |
| *Y77E11A.16* | WBGene00044464 | protein_coding_gene | 3.0 | 7.7299 | 2.7E-03 | 3.2 | 9.502 | 2.8E-03 |

| | | | | | | | | |
|---|---|---|---|---|---|---|---|---|
| *ins-1* | WBGene00002084 | protein_coding_gene | 2.9 | 7.7226 | 1.1E-12 | 1.7 | 3.191 | 1.3E-08 |
| *pef-1* | WBGene00003969 | protein_coding_gene | 2.9 | 7.7197 | 4.7E-11 | 1.1 | 2.095 | 4.3E-06 |
| *dpy-20* | WBGene00001079 | protein_coding_gene | 2.9 | 7.7011 | 7.5E-07 | 4.0 | 15.671 | 4.7E-07 |
| *T07D1.3* | WBGene00020302 | protein_coding_gene | 2.9 | 7.6892 | 1.5E-05 | 2.7 | 6.726 | 3.7E-03 |
| *tag-244* | WBGene00044073 | protein_coding_gene | 2.9 | 7.6679 | 1.4E-10 | 2.5 | 5.509 | 2.8E-35 |
| *F55A4.4* | WBGene00018856 | pseudogene | 2.9 | 7.6611 | 2.3E-09 | 1.9 | 3.810 | 1.4E-04 |
| *hic-1* | WBGene00020887 | protein_coding_gene | 2.9 | 7.6562 | 2.9E-16 | 2.2 | 4.472 | 1.9E-18 |
| *srsx-24* | WBGene00015401 | protein_coding_gene | 2.9 | 7.6483 | 2.3E-02 | 2.9 | 7.484 | 9.6E-03 |
| *C33D3.4* | WBGene00007895 | protein_coding_gene | 2.9 | 7.6442 | 7.6E-03 | 2.4 | 5.257 | 4.1E-02 |
| *col-182* | WBGene00000755 | pseudogene | 2.9 | 7.6412 | 1.2E-10 | 2.5 | 5.541 | 1.2E-02 |
| *EGAP9.3* | WBGene00017151 | protein_coding_gene | 2.9 | 7.6394 | 7.5E-23 | 1.6 | 2.986 | 8.9E-09 |
| *C26E1.2* | WBGene00007748 | protein_coding_gene | 2.9 | 7.6190 | 1.5E-09 | 1.1 | 2.119 | 9.0E-12 |
| *zmp-1* | WBGene00006987 | protein_coding_gene | 2.9 | 7.5922 | 1.3E-07 | 2.3 | 5.040 | 1.1E-08 |
| *nlp-38* | WBGene00007219 | protein_coding_gene | 2.9 | 7.5884 | 9.7E-26 | 2.0 | 4.015 | 8.8E-16 |
| *ZK593.2* | WBGene00014002 | protein_coding_gene | 2.9 | 7.5867 | 1.1E-16 | 1.1 | 2.217 | 6.6E-05 |
| *T05H4.15* | WBGene00020276 | protein_coding_gene | 2.9 | 7.5821 | 3.5E-17 | 2.1 | 4.250 | 1.5E-17 |
| *ddn-2* | WBGene00009399 | protein_coding_gene | 2.9 | 7.5780 | 4.7E-06 | 1.9 | 3.776 | 2.8E-03 |
| *Y116A8C.25* | WBGene00013802 | protein_coding_gene | 2.9 | 7.5646 | 3.1E-13 | 1.9 | 3.643 | 1.2E-06 |
| *Y48G8AR.2* | WBGene00021693 | protein_coding_gene | 2.9 | 7.5640 | 2.7E-06 | 1.3 | 2.533 | 2.1E-02 |
| *srd-18* | WBGene00005096 | protein_coding_gene | 2.9 | 7.5170 | 2.5E-02 | 2.4 | 5.239 | 3.4E-02 |
| *nhr-208* | WBGene00011099 | protein_coding_gene | 2.9 | 7.5086 | 1.8E-07 | 1.9 | 3.748 | 3.8E-07 |
| *C04E6.13* | WBGene00015427 | protein_coding_gene | 2.9 | 7.4974 | 4.8E-26 | 1.6 | 2.967 | 1.7E-06 |
| *ztf-2* | WBGene00008762 | protein_coding_gene | 2.9 | 7.4802 | 4.7E-09 | 3.1 | 8.464 | 3.2E-13 |
| *ins-10* | WBGene00002093 | protein_coding_gene | 2.9 | 7.4758 | 2.3E-04 | 2.5 | 5.488 | 6.6E-05 |
| *R10F2.4* | WBGene00019995 | protein_coding_gene | 2.9 | 7.4628 | 7.4E-06 | 3.0 | 8.082 | 1.9E-05 |
| *rol-8* | WBGene00004398 | protein_coding_gene | 2.9 | 7.4581 | 1.1E-11 | 4.4 | 20.547 | 3.5E-06 |
| *flp-15* | WBGene00001458 | protein_coding_gene | 2.9 | 7.4533 | 3.3E-31 | 2.0 | 4.007 | 1.5E-19 |
| *nlp-31* | WBGene00003769 | protein_coding_gene | 2.9 | 7.4476 | 3.1E-17 | 1.0 | 2.019 | 1.4E-08 |
| *str-133* | WBGene00006181 | pseudogene | 2.9 | 7.4328 | 2.3E-02 | 2.3 | 5.003 | 3.5E-02 |
| *R13.5* | WBGene00014825 | pseudogene | 2.9 | 7.4288 | 4.3E-02 | 1.1 | 2.126 | 7.6E-04 |
| *nhr-35* | WBGene00003628 | protein_coding_gene | 2.9 | 7.4010 | 8.7E-19 | 1.1 | 2.139 | 2.4E-17 |
| *C09G9.5* | WBGene00007495 | protein_coding_gene | 2.9 | 7.3912 | 2.5E-18 | 2.7 | 6.367 | 8.8E-11 |
| *F11E6.6* | WBGene00008709 | protein_coding_gene | 2.9 | 7.3902 | 3.6E-10 | 3.1 | 8.743 | 4.1E-12 |
| *mfb-1* | WBGene00008439 | protein_coding_gene | 2.9 | 7.3667 | 7.1E-17 | 1.6 | 3.045 | 1.1E-16 |
| *T23F4.2* | WBGene00020740 | protein_coding_gene | 2.9 | 7.3618 | 1.2E-13 | 1.5 | 2.794 | 9.2E-16 |
| *T23G7.3* | WBGene00011966 | protein_coding_gene | 2.9 | 7.3545 | 1.5E-30 | 1.7 | 3.184 | 4.0E-47 |
| *C46H11.2* | WBGene00016728 | protein_coding_gene | 2.9 | 7.3531 | 1.6E-30 | 1.8 | 3.574 | 2.9E-23 |
| *C30F2.4* | WBGene00007818 | protein_coding_gene | 2.9 | 7.3501 | 6.6E-08 | 1.8 | 3.549 | 4.7E-05 |
| *F08C6.5* | WBGene00017243 | protein_coding_gene | 2.9 | 7.3498 | 2.2E-15 | 1.4 | 2.581 | 1.1E-04 |
| *T21D12.11* | WBGene00020650 | protein_coding_gene | 2.9 | 7.3427 | 1.9E-03 | 2.1 | 4.346 | 3.3E-02 |
| *gst-29* | WBGene00001777 | protein_coding_gene | 2.9 | 7.3409 | 8.3E-07 | 1.9 | 3.833 | 5.0E-06 |
| *ubc-21* | WBGene00006716 | protein_coding_gene | 2.9 | 7.3384 | 8.9E-03 | 2.9 | 7.524 | 9.5E-03 |
| *srw-78* | WBGene00005825 | pseudogene | 2.9 | 7.3067 | 5.5E-03 | 3.4 | 10.294 | 5.7E-04 |
| *nlp-21* | WBGene00003759 | protein_coding_gene | 2.9 | 7.2925 | 3.7E-17 | 2.4 | 5.120 | 2.5E-46 |
| *C18B2.2* | WBGene00015953 | protein_coding_gene | 2.9 | 7.2902 | 6.6E-04 | 2.2 | 4.493 | 1.5E-02 |
| *frpr-2* | WBGene00007346 | protein_coding_gene | 2.9 | 7.2675 | 1.7E-03 | 1.9 | 3.715 | 3.0E-02 |
| *C26B9.2* | WBGene00016132 | protein_coding_gene | 2.9 | 7.2640 | 5.6E-03 | 2.7 | 6.567 | 1.3E-02 |
| *scl-5* | WBGene00008027 | protein_coding_gene | 2.9 | 7.2496 | 5.4E-14 | 1.6 | 3.090 | 8.8E-23 |
| *F02C12.2* | WBGene00008516 | protein_coding_gene | 2.9 | 7.2383 | 2.0E-03 | 2.0 | 3.945 | 2.6E-02 |
| *T22F3.8* | WBGene00020698 | protein_coding_gene | 2.9 | 7.2364 | 2.5E-08 | 1.8 | 3.562 | 2.9E-05 |
| *flp-27* | WBGene00016114 | protein_coding_gene | 2.9 | 7.2297 | 7.2E-16 | 1.4 | 2.636 | 2.9E-10 |
| *Y51H4A.8* | WBGene00013104 | protein_coding_gene | 2.9 | 7.2288 | 1.8E-27 | 2.1 | 4.424 | 1.0E-28 |
| *ZK742.7* | WBGene00194921 | protein_coding_gene | 2.9 | 7.2131 | 9.3E-11 | 1.4 | 2.723 | 3.5E-04 |
| *clec-227* | WBGene00008593 | protein_coding_gene | 2.8 | 7.1846 | 9.2E-16 | 2.7 | 6.456 | 7.3E-20 |
| *T16H12.9* | WBGene00011817 | protein_coding_gene | 2.8 | 7.1775 | 3.0E-10 | 1.7 | 3.162 | 2.3E-10 |
| *T25E4.2* | WBGene00020804 | protein_coding_gene | 2.8 | 7.1713 | 3.2E-02 | 2.8 | 6.931 | 1.5E-02 |
| *F27E5.9* | WBGene00206354 | protein_coding_gene | 2.8 | 7.1662 | 6.0E-10 | 1.6 | 3.047 | 5.3E-06 |
| *R11F4.1* | WBGene00020007 | protein_coding_gene | 2.8 | 7.1579 | 6.3E-25 | 1.8 | 3.394 | 5.4E-23 |
| *lin-11* | WBGene00003000 | protein_coding_gene | 2.8 | 7.1544 | 1.3E-12 | 2.1 | 4.186 | 4.3E-12 |
| *pqn-29* | WBGene00004118 | protein_coding_gene | 2.8 | 7.1542 | 2.5E-06 | 3.0 | 7.776 | 1.3E-04 |
| *cwp-4* | WBGene00000862 | protein_coding_gene | 2.8 | 7.1490 | 8.9E-09 | 1.5 | 2.823 | 9.6E-04 |
| *sre-3* | WBGene00015568 | protein_coding_gene | 2.8 | 7.1488 | 2.4E-03 | 1.8 | 3.480 | 2.6E-02 |
| *gcy-6* | WBGene00001533 | protein_coding_gene | 2.8 | 7.1170 | 9.2E-03 | 1.8 | 3.382 | 3.9E-02 |
| *F58B4.3* | WBGene00010236 | protein_coding_gene | 2.8 | 7.1153 | 3.9E-13 | 2.4 | 5.236 | 9.6E-14 |
| *F54C1.6* | WBGene00018791 | protein_coding_gene | 2.8 | 7.1116 | 4.9E-03 | 1.8 | 3.486 | 1.7E-02 |
| *Y39A3A.3* | WBGene00021434 | protein_coding_gene | 2.8 | 7.0902 | 3.7E-04 | 1.7 | 3.156 | 1.4E-02 |
| *degt-1* | WBGene00009109 | protein_coding_gene | 2.8 | 7.0880 | 5.9E-12 | 3.3 | 10.015 | 1.4E-14 |
| *Y71H2AM.14* | WBGene00022179 | protein_coding_gene | 2.8 | 7.0760 | 4.1E-10 | 2.0 | 4.123 | 1.7E-05 |
| *F22B7.9* | WBGene00017698 | protein_coding_gene | 2.8 | 7.0741 | 1.3E-56 | 1.0 | 2.055 | 5.3E-08 |
| *Y75B8A.44* | WBGene00194855 | protein_coding_gene | 2.8 | 7.0709 | 4.5E-09 | 2.4 | 5.212 | 5.2E-10 |
| *W02D9.10* | WBGene00012216 | protein_coding_gene | 2.8 | 7.0701 | 6.6E-38 | 2.8 | 6.889 | 1.5E-36 |
| *npr-8* | WBGene00016984 | protein_coding_gene | 2.8 | 7.0666 | 4.1E-10 | 1.2 | 2.257 | 1.4E-02 |
| *gcy-37* | WBGene00001557 | protein_coding_gene | 2.8 | 7.0666 | 2.7E-05 | 2.2 | 4.693 | 9.9E-05 |
| *capa-1* | WBGene00021268 | protein_coding_gene | 2.8 | 7.0389 | 5.6E-17 | 1.9 | 3.647 | 7.6E-17 |
| *B0280.7* | WBGene00015103 | protein_coding_gene | 2.8 | 7.0388 | 2.2E-06 | 2.9 | 7.356 | 7.5E-04 |
| *C01G10.15* | WBGene00007240 | protein_coding_gene | 2.8 | 7.0348 | 6.4E-07 | 1.6 | 2.944 | 7.2E-04 |
| *C32H11.6* | WBGene00007869 | protein_coding_gene | 2.8 | 7.0306 | 4.8E-02 | 2.9 | 7.276 | 1.5E-02 |
| *F36G3.3* | WBGene00009484 | protein_coding_gene | 2.8 | 7.0180 | 1.4E-04 | 1.7 | 3.226 | 2.6E-03 |
| *H41C03.2* | WBGene00019269 | protein_coding_gene | 2.8 | 7.0159 | 2.7E-03 | 3.4 | 10.924 | 2.0E-03 |
| *cex-1* | WBGene00023407 | protein_coding_gene | 2.8 | 7.0134 | 1.1E-15 | 1.8 | 3.504 | 2.2E-26 |

| | | | | | | | | |
|---|---|---|---|---|---|---|---|---|
| *C18A11.1* | WBGene00015947 | protein_coding_gene | 2.8 | 6.9948 | 6.9E-12 | 1.3 | 2.476 | 1.0E-12 |
| *atln-2* | WBGene00012763 | protein_coding_gene | 2.8 | 6.9922 | 7.1E-23 | 1.1 | 2.159 | 1.7E-03 |
| *nlp-13* | WBGene00003751 | protein_coding_gene | 2.8 | 6.9866 | 9.8E-34 | 1.9 | 3.815 | 3.4E-37 |
| *Y49E10.18* | WBGene00013037 | protein_coding_gene | 2.8 | 6.9831 | 7.5E-24 | 2.4 | 5.191 | 2.2E-35 |
| *acr-23* | WBGene00000062 | protein_coding_gene | 2.8 | 6.9775 | 2.0E-04 | 3.4 | 10.257 | 2.4E-04 |
| *C16E9.1* | WBGene00015865 | protein_coding_gene | 2.8 | 6.9747 | 6.0E-26 | 3.2 | 9.047 | 8.3E-26 |
| *F28B12.1* | WBGene00017894 | protein_coding_gene | 2.8 | 6.9744 | 3.5E-22 | 2.4 | 5.187 | 2.0E-14 |
| *F14F3.4* | WBGene00008807 | protein_coding_gene | 2.8 | 6.9664 | 5.2E-04 | 2.0 | 3.996 | 3.8E-02 |
| *flp-18* | WBGene00001461 | protein_coding_gene | 2.8 | 6.9599 | 4.5E-27 | 1.8 | 3.408 | 1.3E-38 |
| *nlp-55* | WBGene00008820 | protein_coding_gene | 2.8 | 6.9587 | 1.8E-20 | 2.0 | 3.886 | 8.8E-25 |
| *lon-3* | WBGene00003057 | protein_coding_gene | 2.8 | 6.9553 | 8.4E-09 | 3.0 | 7.949 | 1.3E-03 |
| *nhr-277* | WBGene00022374 | protein_coding_gene | 2.8 | 6.9543 | 8.3E-05 | 2.5 | 5.782 | 4.5E-04 |
| *gcy-22* | WBGene00001547 | protein_coding_gene | 2.8 | 6.9534 | 1.8E-09 | 1.1 | 2.203 | 9.6E-04 |
| *fbxa-87* | WBGene00044660 | pseudogene | 2.8 | 6.9450 | 3.7E-07 | 1.1 | 2.091 | 1.0E-05 |
| *ugt-55* | WBGene00011452 | protein_coding_gene | 2.8 | 6.9303 | 1.5E-20 | 2.2 | 4.604 | 1.5E-10 |
| *ZC21.3* | WBGene00022505 | protein_coding_gene | 2.8 | 6.9287 | 1.9E-09 | 2.1 | 4.300 | 9.3E-57 |
| *lgc-1* | WBGene00020236 | protein_coding_gene | 2.8 | 6.9216 | 6.4E-05 | 1.3 | 2.394 | 3.0E-02 |
| *chil-3* | WBGene00007467 | protein_coding_gene | 2.8 | 6.9200 | 9.7E-09 | 2.9 | 7.221 | 1.8E-04 |
| *M01B2.12* | WBGene00045337 | protein_coding_gene | 2.8 | 6.9078 | 3.1E-02 | 2.4 | 5.307 | 4.2E-02 |
| *R05G6.10* | WBGene00019902 | protein_coding_gene | 2.8 | 6.9075 | 2.7E-13 | 1.2 | 2.323 | 2.1E-13 |
| *R06A10.1* | WBGene00019909 | protein_coding_gene | 2.8 | 6.9014 | 1.4E-07 | 1.7 | 3.200 | 1.6E-04 |
| *F44G4.7* | WBGene00009716 | protein_coding_gene | 2.8 | 6.8990 | 1.6E-04 | 1.7 | 3.321 | 2.6E-03 |
| *nlp-14* | WBGene00003752 | protein_coding_gene | 2.8 | 6.8914 | 4.0E-18 | 2.2 | 4.648 | 4.5E-39 |
| *F57C12.2* | WBGene00019014 | protein_coding_gene | 2.8 | 6.8904 | 1.6E-17 | 1.1 | 2.192 | 4.1E-05 |
| *ZK757.2* | WBGene00014074 | protein_coding_gene | 2.8 | 6.8827 | 1.2E-11 | 1.1 | 2.107 | 3.6E-03 |
| *ida-1* | WBGene00002048 | protein_coding_gene | 2.8 | 6.8697 | 2.0E-18 | 2.2 | 4.488 | 1.7E-78 |
| *acox-1.5* | WBGene00008167 | protein_coding_gene | 2.8 | 6.8635 | 1.6E-25 | 2.1 | 4.287 | 1.0E-13 |
| *fog-3* | WBGene00001483 | protein_coding_gene | 2.8 | 6.8541 | 1.3E-07 | 2.9 | 7.332 | 1.8E-37 |
| *tsp-10* | WBGene00006636 | protein_coding_gene | 2.8 | 6.8504 | 1.0E-31 | 2.3 | 4.912 | 4.6E-36 |
| *C54D2.2* | WBGene00016916 | protein_coding_gene | 2.8 | 6.8485 | 2.3E-08 | 2.6 | 6.181 | 9.1E-11 |
| *flp-16* | WBGene00001459 | protein_coding_gene | 2.8 | 6.8468 | 1.9E-46 | 2.0 | 4.078 | 2.4E-36 |
| *K06A9.2* | WBGene00019436 | protein_coding_gene | 2.8 | 6.8377 | 1.0E-07 | 1.0 | 2.066 | 2.2E-06 |
| *col-79* | WBGene00000655 | protein_coding_gene | 2.8 | 6.8372 | 1.4E-10 | 4.3 | 19.603 | 2.7E-05 |
| *srr-2* | WBGene00005653 | protein_coding_gene | 2.8 | 6.8232 | 2.2E-04 | 1.3 | 2.506 | 3.6E-02 |
| *F58B4.6* | WBGene00010239 | protein_coding_gene | 2.8 | 6.8027 | 6.5E-11 | 1.5 | 2.811 | 9.5E-07 |
| *ctg-2* | WBGene00011756 | protein_coding_gene | 2.8 | 6.7965 | 9.1E-35 | 2.6 | 6.070 | 8.4E-50 |
| *F59B10.4* | WBGene00010320 | protein_coding_gene | 2.8 | 6.7891 | 9.0E-14 | 1.3 | 2.382 | 1.5E-06 |
| *Y6E2A.5* | WBGene00012399 | protein_coding_gene | 2.8 | 6.7870 | 3.8E-08 | 1.6 | 3.063 | 4.1E-08 |
| *wrt-8* | WBGene00006954 | protein_coding_gene | 2.8 | 6.7840 | 9.6E-09 | 2.9 | 7.362 | 4.7E-03 |
| *cdh-7* | WBGene00000399 | protein_coding_gene | 2.8 | 6.7820 | 9.8E-08 | 5.4 | 43.167 | 1.7E-19 |
| *gcy-4* | WBGene00001531 | protein_coding_gene | 2.8 | 6.7699 | 6.7E-03 | 2.6 | 6.240 | 3.6E-03 |
| *D1079.1* | WBGene00017040 | protein_coding_gene | 2.8 | 6.7663 | 6.9E-07 | 1.5 | 2.852 | 1.1E-03 |
| *bath-47* | WBGene00006464 | protein_coding_gene | 2.8 | 6.7660 | 6.9E-11 | 1.7 | 3.139 | 4.2E-11 |
| *C30F12.3* | WBGene00016262 | protein_coding_gene | 2.8 | 6.7500 | 1.2E-17 | 2.1 | 4.197 | 9.3E-11 |
| *B0507.6* | WBGene00015223 | protein_coding_gene | 2.8 | 6.7424 | 1.8E-09 | 1.8 | 3.532 | 4.0E-10 |
| *cpr-3* | WBGene00000783 | protein_coding_gene | 2.8 | 6.7414 | 1.8E-16 | 2.6 | 5.954 | 2.0E-37 |
| *R57.2* | WBGene00020083 | protein_coding_gene | 2.8 | 6.7372 | 1.3E-09 | 1.2 | 2.231 | 2.6E-02 |
| *M163.1* | WBGene00010935 | protein_coding_gene | 2.8 | 6.7326 | 8.1E-14 | 1.3 | 2.427 | 7.9E-13 |
| *C07D8.2* | WBGene00015562 | pseudogene | 2.7 | 6.7208 | 4.0E-02 | 3.4 | 10.564 | 1.1E-03 |
| *col-111* | WBGene00000685 | protein_coding_gene | 2.7 | 6.7166 | 1.5E-13 | 1.6 | 3.123 | 5.8E-03 |
| *K03H1.5* | WBGene00010540 | protein_coding_gene | 2.7 | 6.7112 | 6.7E-12 | 1.7 | 3.148 | 2.0E-27 |
| *R13H7.2* | WBGene00020069 | protein_coding_gene | 2.7 | 6.7019 | 1.2E-10 | 2.1 | 4.194 | 7.4E-08 |
| *clec-146* | WBGene00013008 | protein_coding_gene | 2.7 | 6.6932 | 1.4E-26 | 1.5 | 2.766 | 1.3E-14 |
| *M03D4.3* | WBGene00019750 | protein_coding_gene | 2.7 | 6.6708 | 1.9E-08 | 1.5 | 2.800 | 1.3E-04 |
| *W01C9.2* | WBGene00012176 | protein_coding_gene | 2.7 | 6.6679 | 1.9E-06 | 1.7 | 3.209 | 6.0E-03 |
| *C50F4.8* | WBGene00008233 | protein_coding_gene | 2.7 | 6.6652 | 2.2E-17 | 1.4 | 2.709 | 1.4E-18 |
| *otpl-6* | WBGene00018479 | protein_coding_gene | 2.7 | 6.6531 | 2.1E-02 | 2.1 | 4.332 | 2.8E-02 |
| *linc-85* | WBGene00219754 | lincRNA_gene | 2.7 | 6.6522 | 1.5E-04 | 1.7 | 3.203 | 1.8E-02 |
| *T28A11.25* | WBGene00269355 | protein_coding_gene | 2.7 | 6.6518 | 3.8E-05 | 2.1 | 4.295 | 2.3E-04 |
| *col-17* | WBGene00000606 | protein_coding_gene | 2.7 | 6.6436 | 1.6E-10 | 3.7 | 12.926 | 2.3E-04 |
| *F27B10.1* | WBGene00017850 | protein_coding_gene | 2.7 | 6.6435 | 4.2E-12 | 1.7 | 3.306 | 1.8E-10 |
| *ZK1290.10* | WBGene00022891 | protein_coding_gene | 2.7 | 6.6330 | 2.7E-10 | 1.4 | 2.670 | 1.2E-04 |
| *cal-2* | WBGene00000286 | protein_coding_gene | 2.7 | 6.6152 | 3.3E-16 | 1.4 | 2.610 | 1.4E-14 |
| *Y38F1A.8* | WBGene00012610 | protein_coding_gene | 2.7 | 6.6120 | 7.8E-22 | 1.2 | 2.345 | 1.4E-06 |
| *nhr-179* | WBGene00017512 | protein_coding_gene | 2.7 | 6.6049 | 2.1E-24 | 3.2 | 9.071 | 5.9E-33 |
| *flp-33* | WBGene00011586 | protein_coding_gene | 2.7 | 6.6007 | 4.1E-25 | 1.9 | 3.609 | 2.6E-09 |
| *F47B8.3* | WBGene00009804 | protein_coding_gene | 2.7 | 6.5982 | 3.6E-26 | 1.1 | 2.209 | 8.1E-08 |
| *lgc-36* | WBGene00008551 | protein_coding_gene | 2.7 | 6.5937 | 3.9E-03 | 2.8 | 6.808 | 1.2E-02 |
| *B0222.1* | WBGene00015053 | protein_coding_gene | 2.7 | 6.5842 | 1.4E-09 | 2.0 | 4.009 | 9.0E-06 |
| *egl-3* | WBGene00001172 | protein_coding_gene | 2.7 | 6.5624 | 5.5E-18 | 2.6 | 6.109 | 1.9E-90 |
| *T09F5.12* | WBGene00014836 | lincRNA_gene | 2.7 | 6.5480 | 2.2E-21 | 2.5 | 5.486 | 2.3E-46 |
| *ceh-57* | WBGene00007416 | protein_coding_gene | 2.7 | 6.5350 | 2.4E-04 | 1.6 | 3.037 | 1.6E-02 |
| *vab-23* | WBGene00014153 | protein_coding_gene | 2.7 | 6.5336 | 8.9E-10 | 1.2 | 2.273 | 1.2E-06 |
| *C45G9.11* | WBGene00016682 | protein_coding_gene | 2.7 | 6.5299 | 4.2E-04 | 1.7 | 3.283 | 9.3E-03 |
| *npr-1* | WBGene00003807 | protein_coding_gene | 2.7 | 6.5270 | 2.6E-04 | 2.7 | 6.454 | 1.8E-04 |
| *acr-19* | WBGene00000058 | protein_coding_gene | 2.7 | 6.5248 | 1.7E-07 | 2.0 | 3.877 | 4.1E-05 |
| *clec-264* | WBGene00009291 | protein_coding_gene | 2.7 | 6.5136 | 3.0E-31 | 1.3 | 2.525 | 6.1E-09 |
| *ZK353.10* | WBGene00044677 | protein_coding_gene | 2.7 | 6.5076 | 3.0E-05 | 1.6 | 2.933 | 7.7E-03 |
| *aex-4* | WBGene00006454 | protein_coding_gene | 2.7 | 6.5067 | 1.2E-25 | 2.4 | 5.431 | 1.5E-25 |
| *ZC47.11* | WBGene00014979 | pseudogene | 2.7 | 6.5037 | 1.9E-02 | 2.6 | 6.068 | 2.3E-02 |

| | | | | | | | | |
|---|---|---|---|---|---|---|---|---|
| *cpz-2* | WBGene00000789 | protein_coding_gene | 2.7 | 6.5030 | 4.8E-37 | 1.7 | 3.282 | 1.1E-11 |
| *nlp-9* | WBGene00003747 | protein_coding_gene | 2.7 | 6.5025 | 8.7E-15 | 2.1 | 4.396 | 4.4E-33 |
| *lgc-38* | WBGene00017389 | protein_coding_gene | 2.7 | 6.4992 | 1.1E-08 | 2.7 | 6.477 | 1.3E-08 |
| ***ZK1290.1*** | **WBGene00022886** | **protein_coding_gene** | **2.7** | **6.4968** | **1.7E-02** | **-4.6** | **0.041** | **3.9E-40** |
| *flr-2* | WBGene00001466 | protein_coding_gene | 2.7 | 6.4938 | 3.6E-23 | 2.4 | 5.297 | 3.4E-18 |
| *F02E11.2* | WBGene00017180 | protein_coding_gene | 2.7 | 6.4933 | 5.6E-08 | 1.6 | 2.999 | 4.1E-05 |
| *nlp-1* | WBGene00003739 | protein_coding_gene | 2.7 | 6.4797 | 3.3E-26 | 1.6 | 2.929 | 8.8E-12 |
| *F13B6.1* | WBGene00017416 | protein_coding_gene | 2.7 | 6.4777 | 3.1E-08 | 1.2 | 2.319 | 1.2E-03 |
| *T01H10.4* | WBGene00011357 | pseudogene | 2.7 | 6.4774 | 3.8E-02 | 4.5 | 21.946 | 7.7E-05 |
| *ssu-2* | WBGene00013455 | protein_coding_gene | 2.7 | 6.4588 | 3.4E-11 | 1.0 | 2.005 | 1.7E-02 |
| *F42A10.6* | WBGene00018342 | protein_coding_gene | 2.7 | 6.4586 | 1.5E-14 | 2.4 | 5.338 | 4.3E-43 |
| *C30B5.7* | WBGene00016248 | protein_coding_gene | 2.7 | 6.4561 | 3.3E-06 | 2.1 | 4.298 | 8.8E-06 |
| *nhr-168* | WBGene00008221 | protein_coding_gene | 2.7 | 6.4043 | 2.3E-30 | 2.7 | 6.553 | 1.8E-17 |
| *C15A11.7* | WBGene00007599 | protein_coding_gene | 2.7 | 6.4023 | 2.0E-25 | 1.5 | 2.902 | 3.9E-13 |
| *C47D2.1* | WBGene00016738 | protein_coding_gene | 2.7 | 6.3865 | 4.0E-08 | 2.4 | 5.309 | 1.8E-07 |
| *dop-5* | WBGene00011382 | protein_coding_gene | 2.7 | 6.3858 | 3.2E-13 | 1.6 | 2.989 | 4.9E-08 |
| *T09E11.8* | WBGene00011657 | protein_coding_gene | 2.7 | 6.3857 | 4.8E-02 | 2.7 | 6.672 | 3.0E-02 |
| *C07C7.1* | WBGene00007411 | protein_coding_gene | 2.7 | 6.3739 | 1.2E-02 | 1.9 | 3.735 | 4.0E-02 |
| *egl-15* | WBGene00001184 | protein_coding_gene | 2.7 | 6.3699 | 8.2E-11 | 1.2 | 2.318 | 6.4E-15 |
| *M04B2.6* | WBGene00010848 | protein_coding_gene | 2.7 | 6.3686 | 1.5E-13 | 1.3 | 2.457 | 1.2E-04 |
| *fbxc-6* | WBGene00016301 | pseudogene | 2.7 | 6.3681 | 2.4E-05 | 2.3 | 4.860 | 4.5E-03 |
| *pqn-94* | WBGene00004173 | protein_coding_gene | 2.7 | 6.3595 | 7.3E-17 | 1.7 | 3.359 | 1.3E-20 |
| *T22B11.4* | WBGene00020678 | protein_coding_gene | 2.7 | 6.3558 | 3.4E-24 | 2.3 | 4.962 | 4.8E-62 |
| *flp-2* | WBGene00001445 | protein_coding_gene | 2.7 | 6.3551 | 1.2E-20 | 1.8 | 3.481 | 1.9E-09 |
| *F15B9.6* | WBGene00008850 | protein_coding_gene | 2.7 | 6.3482 | 2.8E-07 | 1.6 | 3.087 | 4.6E-03 |
| *T12A7.2* | WBGene00011723 | protein_coding_gene | 2.7 | 6.3446 | 5.6E-17 | 1.6 | 3.122 | 3.4E-21 |
| *F31E3.2* | WBGene00017950 | protein_coding_gene | 2.7 | 6.3368 | 3.0E-12 | 2.1 | 4.357 | 4.4E-24 |
| *R155.4* | WBGene00020118 | protein_coding_gene | 2.7 | 6.3331 | 7.1E-04 | 2.1 | 4.359 | 1.5E-08 |
| *zig-3* | WBGene00006980 | protein_coding_gene | 2.7 | 6.3281 | 8.3E-17 | 2.4 | 5.314 | 1.1E-20 |
| *slc-17.9* | WBGene00011643 | protein_coding_gene | 2.7 | 6.3271 | 2.7E-04 | 1.0 | 2.004 | 1.7E-02 |
| *ins-24* | WBGene00002107 | protein_coding_gene | 2.7 | 6.3220 | 7.5E-12 | 2.5 | 5.595 | 9.8E-11 |
| *sqt-2* | WBGene00005017 | protein_coding_gene | 2.7 | 6.3218 | 1.8E-12 | 4.2 | 17.757 | 1.5E-05 |
| *B0563.7* | WBGene00015264 | protein_coding_gene | 2.7 | 6.3155 | 9.6E-08 | 1.2 | 2.314 | 5.6E-03 |
| *C52B9.19* | WBGene00304788 | protein_coding_gene | 2.7 | 6.3031 | 1.9E-08 | 1.3 | 2.517 | 4.4E-03 |
| *eol-1* | WBGene00045401 | protein_coding_gene | 2.7 | 6.2975 | 1.6E-05 | 2.2 | 4.530 | 2.3E-10 |
| *R11G11.6* | WBGene00020014 | protein_coding_gene | 2.7 | 6.2925 | 6.4E-03 | 3.2 | 9.028 | 3.5E-03 |
| *sto-6* | WBGene00006068 | protein_coding_gene | 2.7 | 6.2917 | 8.2E-08 | 1.5 | 2.821 | 4.8E-03 |
| *F14B8.5* | WBGene00017447 | protein_coding_gene | 2.7 | 6.2908 | 5.8E-16 | 1.8 | 3.375 | 2.5E-20 |
| *K12B6.4* | WBGene00019669 | protein_coding_gene | 2.7 | 6.2856 | 4.8E-05 | 3.2 | 8.938 | 1.5E-07 |
| *pghm-1* | WBGene00022144 | protein_coding_gene | 2.6 | 6.2743 | 7.9E-30 | 2.1 | 4.284 | 1.4E-65 |
| *C16D9.9* | WBGene00015864 | protein_coding_gene | 2.6 | 6.2623 | 4.4E-05 | 1.8 | 3.459 | 2.6E-03 |
| *D1054.9* | WBGene00008376 | protein_coding_gene | 2.6 | 6.2611 | 6.8E-10 | 1.9 | 3.609 | 1.8E-02 |
| *T27E4.7* | WBGene00020865 | protein_coding_gene | 2.6 | 6.2568 | 2.0E-23 | 1.6 | 2.957 | 9.1E-20 |
| *ipla-2* | WBGene00009801 | protein_coding_gene | 2.6 | 6.2514 | 5.6E-46 | 2.3 | 5.091 | 2.3E-36 |
| *srj-14* | WBGene00005602 | protein_coding_gene | 2.6 | 6.2475 | 5.3E-11 | 1.4 | 2.622 | 1.1E-06 |
| *nlp-15* | WBGene00003753 | protein_coding_gene | 2.6 | 6.2405 | 9.7E-17 | 1.8 | 3.495 | 8.1E-25 |
| *Y22D7AL.16* | WBGene00021254 | protein_coding_gene | 2.6 | 6.2298 | 4.5E-06 | 2.8 | 6.738 | 1.3E-05 |
| *col-168* | WBGene00000741 | protein_coding_gene | 2.6 | 6.2257 | 6.2E-11 | 3.9 | 15.139 | 5.8E-14 |
| *nlp-80* | WBGene00011428 | protein_coding_gene | 2.6 | 6.2038 | 1.1E-23 | 2.0 | 4.046 | 2.4E-21 |
| *flp-5* | WBGene00001448 | protein_coding_gene | 2.6 | 6.1912 | 1.2E-29 | 2.1 | 4.155 | 8.4E-35 |
| *frpr-18* | WBGene00020586 | protein_coding_gene | 2.6 | 6.1776 | 1.7E-11 | 1.5 | 2.874 | 1.1E-04 |
| ***ZK666.14*** | **WBGene00194952** | **protein_coding_gene** | **2.6** | **6.1757** | **1.9E-02** | **-2.1** | **0.229** | **3.6E-03** |
| *R11D1.3* | WBGene00011244 | protein_coding_gene | 2.6 | 6.1757 | 1.2E-08 | 2.8 | 7.003 | 2.0E-23 |
| ***Y43F8C.17*** | **WBGene00012838** | **protein_coding_gene** | **2.6** | **6.1609** | **2.6E-02** | **-6.7** | **0.010** | **1.2E-13** |
| *pdf-1* | WBGene00020317 | protein_coding_gene | 2.6 | 6.1607 | 4.0E-29 | 1.6 | 3.008 | 1.1E-31 |
| *F41G4.7* | WBGene00018310 | protein_coding_gene | 2.6 | 6.1597 | 9.5E-05 | 1.8 | 3.499 | 1.9E-02 |
| *T05E7.4* | WBGene00020260 | protein_coding_gene | 2.6 | 6.1537 | 1.9E-08 | 1.3 | 2.434 | 6.4E-03 |
| *tag-196* | WBGene00007055 | protein_coding_gene | 2.6 | 6.1492 | 1.7E-20 | 1.5 | 2.832 | 4.9E-30 |
| *F02E11.7* | WBGene00195073 | protein_coding_gene | 2.6 | 6.1458 | 3.5E-07 | 1.7 | 3.208 | 8.2E-04 |
| *lin-48* | WBGene00003033 | protein_coding_gene | 2.6 | 6.1434 | 6.6E-07 | 1.3 | 2.459 | 1.3E-02 |
| *npr-25* | WBGene00011381 | protein_coding_gene | 2.6 | 6.1367 | 5.3E-11 | 1.7 | 3.217 | 8.4E-05 |
| *Y41G9A.5* | WBGene00021529 | protein_coding_gene | 2.6 | 6.1322 | 3.0E-10 | 1.3 | 2.464 | 2.1E-03 |
| *odr-3* | WBGene00003850 | protein_coding_gene | 2.6 | 6.1298 | 1.5E-14 | 1.9 | 3.639 | 2.0E-07 |
| *Y47G6A.15* | WBGene00021641 | protein_coding_gene | 2.6 | 6.1221 | 2.0E-12 | 5.6 | 49.344 | 2.9E-19 |
| *T17H7.7* | WBGene00020551 | protein_coding_gene | 2.6 | 6.1201 | 1.0E-15 | 1.4 | 2.679 | 1.4E-03 |
| *C07B5.2* | WBGene00007407 | protein_coding_gene | 2.6 | 6.1167 | 6.8E-03 | 2.5 | 5.753 | 4.1E-02 |
| *dgk-3* | WBGene00000960 | protein_coding_gene | 2.6 | 6.1164 | 5.3E-06 | 1.6 | 3.068 | 6.0E-03 |
| *srh-268* | WBGene00005474 | protein_coding_gene | 2.6 | 6.1125 | 3.8E-02 | 2.7 | 6.518 | 8.4E-03 |
| *K01D12.5* | WBGene00010464 | protein_coding_gene | 2.6 | 6.1024 | 4.3E-07 | 2.2 | 4.487 | 3.7E-02 |
| *T02C12.5* | WBGene00194692 | protein_coding_gene | 2.6 | 6.0960 | 1.6E-09 | 1.1 | 2.126 | 9.0E-03 |
| *asp-3* | WBGene00000216 | protein_coding_gene | 2.6 | 6.0882 | 4.4E-53 | 2.0 | 4.128 | 2.8E-75 |
| *F35F10.6* | WBGene00018056 | protein_coding_gene | 2.6 | 6.0791 | 8.2E-10 | 4.2 | 19.008 | 3.6E-15 |
| *E01G4.6* | WBGene00008448 | protein_coding_gene | 2.6 | 6.0686 | 1.7E-08 | 4.1 | 16.862 | 7.8E-05 |
| *tos-1* | WBGene00019466 | protein_coding_gene | 2.6 | 6.0599 | 7.0E-15 | 2.3 | 4.767 | 2.2E-43 |
| *ZK856.5* | WBGene00014106 | protein_coding_gene | 2.6 | 6.0585 | 5.4E-16 | 2.1 | 4.358 | 4.2E-28 |
| *ZC513.2* | WBGene00022627 | protein_coding_gene | 2.6 | 6.0256 | 1.5E-07 | 1.8 | 3.570 | 9.5E-04 |
| *lec-7* | WBGene00002270 | protein_coding_gene | 2.6 | 6.0255 | 4.2E-14 | 2.2 | 4.718 | 1.9E-08 |
| *clec-230* | WBGene00007806 | protein_coding_gene | 2.6 | 6.0224 | 7.7E-05 | 2.9 | 7.721 | 5.7E-04 |
| *C46H11.10* | WBGene00016734 | protein_coding_gene | 2.6 | 6.0162 | 2.1E-11 | 2.0 | 3.913 | 1.9E-08 |
| *ets-10* | WBGene00016870 | protein_coding_gene | 2.6 | 6.0110 | 1.0E-03 | 1.8 | 3.569 | 1.6E-02 |

| | | | | | | | | |
|---|---|---|---|---|---|---|---|---|
| *K09H9.8* | WBGene00045387 | protein_coding_gene | 2.6 | 6.0092 | 9.0E-28 | 1.7 | 3.256 | 2.4E-14 |
| *clec-80* | WBGene00043062 | protein_coding_gene | 2.6 | 5.9937 | 4.6E-30 | 2.9 | 7.235 | 4.8E-29 |
| ***C49C8.6*** | **WBGene00016770** | **protein_coding_gene** | **2.6** | **5.9936** | **4.1E-02** | **-2.0** | **0.248** | **2.7E-02** |
| *C08G5.7* | WBGene00044418 | protein_coding_gene | 2.6 | 5.9928 | 3.9E-14 | 1.4 | 2.727 | 8.4E-10 |
| *pals-33* | WBGene00021081 | protein_coding_gene | 2.6 | 5.9927 | 1.3E-06 | 2.5 | 5.779 | 1.3E-11 |
| *F20D6.8* | WBGene00017637 | protein_coding_gene | 2.6 | 5.9925 | 8.2E-05 | 1.1 | 2.089 | 1.9E-02 |
| *dbl-1* | WBGene00000936 | protein_coding_gene | 2.6 | 5.9901 | 9.1E-19 | 2.4 | 5.375 | 2.9E-22 |
| *lpr-2* | WBGene00011726 | protein_coding_gene | 2.6 | 5.9869 | 1.3E-09 | 1.8 | 3.533 | 2.2E-13 |
| *F56D5.6* | WBGene00010150 | protein_coding_gene | 2.6 | 5.9776 | 3.6E-22 | 1.5 | 2.849 | 9.4E-17 |
| *flp-14* | WBGene00001457 | protein_coding_gene | 2.6 | 5.9713 | 2.4E-33 | 1.7 | 3.203 | 8.6E-33 |
| *snet-1* | WBGene00015354 | protein_coding_gene | 2.6 | 5.9649 | 4.5E-16 | 2.2 | 4.518 | 5.1E-42 |
| *C30E1.6* | WBGene00016255 | pseudogene | 2.6 | 5.9640 | 4.0E-02 | 4.0 | 15.535 | 6.2E-05 |
| *C03D6.1* | WBGene00007275 | protein_coding_gene | 2.6 | 5.9513 | 8.6E-20 | 1.8 | 3.574 | 3.2E-10 |
| *sma-10* | WBGene00020649 | protein_coding_gene | 2.6 | 5.9472 | 1.8E-15 | 1.6 | 3.007 | 2.1E-24 |
| *Y8A9A.2* | WBGene00021171 | protein_coding_gene | 2.6 | 5.9363 | 1.7E-06 | 1.1 | 2.089 | 6.2E-05 |
| *F55A4.7* | WBGene00018859 | protein_coding_gene | 2.6 | 5.9268 | 4.3E-07 | 1.7 | 3.210 | 4.5E-04 |
| *C25A6.1* | WBGene00016081 | protein_coding_gene | 2.6 | 5.9157 | 1.9E-04 | 1.0 | 2.020 | 4.4E-04 |
| *W10C8.6* | WBGene00044433 | protein_coding_gene | 2.6 | 5.9092 | 2.1E-09 | 2.1 | 4.175 | 3.3E-12 |
| *F58H1.7* | WBGene00010290 | protein_coding_gene | 2.6 | 5.9027 | 6.6E-22 | 1.4 | 2.593 | 9.1E-11 |
| *Y75B7AR.1* | WBGene00022287 | protein_coding_gene | 2.6 | 5.9018 | 5.6E-14 | 2.8 | 6.971 | 2.8E-04 |
| *Y71G12B.32* | WBGene00044349 | protein_coding_gene | 2.6 | 5.9000 | 6.0E-07 | 1.6 | 3.006 | 2.7E-03 |
| *aakg-4* | WBGene00020135 | protein_coding_gene | 2.6 | 5.8980 | 7.3E-11 | 1.2 | 2.232 | 6.5E-10 |
| *T05A8.3* | WBGene00020229 | protein_coding_gene | 2.6 | 5.8898 | 1.5E-14 | 1.1 | 2.191 | 2.1E-12 |
| *rege-1* | WBGene00016260 | protein_coding_gene | 2.6 | 5.8869 | 9.8E-18 | 2.0 | 4.090 | 4.3E-25 |
| *C54D10.10* | WBGene00008304 | protein_coding_gene | 2.6 | 5.8827 | 2.1E-12 | 1.6 | 3.074 | 3.2E-04 |
| *ckr-1* | WBGene00020712 | protein_coding_gene | 2.6 | 5.8824 | 2.0E-05 | 2.3 | 5.054 | 1.3E-04 |
| *F58A6.2* | WBGene00019023 | protein_coding_gene | 2.6 | 5.8815 | 1.9E-03 | 2.4 | 5.151 | 3.9E-03 |
| *acr-21* | WBGene00000060 | protein_coding_gene | 2.6 | 5.8789 | 6.1E-05 | 2.0 | 4.114 | 1.5E-03 |
| *nlp-51* | WBGene00011997 | protein_coding_gene | 2.6 | 5.8773 | 6.1E-19 | 1.3 | 2.512 | 1.0E-15 |
| *linc-61* | WBGene00195170 | lincRNA_gene | 2.6 | 5.8611 | 3.1E-04 | 2.1 | 4.324 | 1.5E-03 |
| *kvs-2* | WBGene00007862 | protein_coding_gene | 2.6 | 5.8608 | 2.2E-11 | 1.7 | 3.170 | 6.3E-05 |
| *F01D5.5* | WBGene00008495 | protein_coding_gene | 2.6 | 5.8605 | 2.4E-06 | 1.7 | 3.140 | 9.9E-08 |
| *ins-18* | WBGene00002101 | protein_coding_gene | 2.6 | 5.8599 | 3.8E-12 | 1.2 | 2.331 | 2.0E-04 |
| *ttr-39* | WBGene00007304 | protein_coding_gene | 2.6 | 5.8565 | 8.3E-15 | 2.0 | 4.004 | 9.0E-09 |
| *C07D8.5* | WBGene00015564 | protein_coding_gene | 2.5 | 5.8510 | 4.7E-02 | 2.2 | 4.569 | 4.9E-02 |
| *dod-6* | WBGene00011869 | protein_coding_gene | 2.5 | 5.8485 | 1.4E-14 | 1.6 | 2.975 | 1.5E-30 |
| *col-10* | WBGene00000599 | protein_coding_gene | 2.5 | 5.8473 | 2.7E-02 | 2.6 | 6.257 | 4.3E-02 |
| *T21B4.15* | WBGene00043054 | protein_coding_gene | 2.5 | 5.8462 | 4.0E-07 | 2.0 | 4.132 | 1.4E-04 |
| *C33D9.3* | WBGene00007898 | protein_coding_gene | 2.5 | 5.8440 | 2.2E-17 | 2.7 | 6.507 | 1.2E-62 |
| *C30G4.4* | WBGene00016269 | protein_coding_gene | 2.5 | 5.8308 | 3.5E-14 | 1.1 | 2.079 | 5.2E-11 |
| ***oac-14*** | **WBGene00008602** | **protein_coding_gene** | **2.5** | **5.8193** | **1.0E-03** | **-2.1** | **0.239** | **1.2E-13** |
| *H09F14.1* | WBGene00010375 | protein_coding_gene | 2.5 | 5.8186 | 8.8E-07 | 2.5 | 5.541 | 3.7E-09 |
| *C44B11.4* | WBGene00016634 | protein_coding_gene | 2.5 | 5.8066 | 3.8E-18 | 1.7 | 3.279 | 2.1E-09 |
| *flp-1* | WBGene00001444 | protein_coding_gene | 2.5 | 5.8058 | 3.2E-15 | 1.9 | 3.789 | 7.4E-31 |
| *aff-1* | WBGene00016625 | protein_coding_gene | 2.5 | 5.8041 | 4.7E-12 | 1.2 | 2.359 | 1.3E-03 |
| *Y4C6B.2* | WBGene00021156 | protein_coding_gene | 2.5 | 5.8003 | 3.2E-09 | 1.4 | 2.653 | 3.9E-07 |
| *nog-1* | WBGene00020297 | protein_coding_gene | 2.5 | 5.7914 | 6.3E-43 | 2.0 | 3.878 | 4.8E-40 |
| *T03F6.9* | WBGene00045482 | protein_coding_gene | 2.5 | 5.7885 | 3.5E-04 | 1.7 | 3.332 | 3.8E-03 |
| *ZC239.16* | WBGene00022574 | protein_coding_gene | 2.5 | 5.7842 | 3.1E-11 | 1.6 | 3.018 | 7.4E-03 |
| *mpk-2* | WBGene00003402 | protein_coding_gene | 2.5 | 5.7789 | 2.9E-39 | 1.6 | 2.957 | 3.3E-15 |
| *nlp-12* | WBGene00003750 | protein_coding_gene | 2.5 | 5.7694 | 1.4E-16 | 2.2 | 4.682 | 6.3E-18 |
| *asp-5* | WBGene00000218 | protein_coding_gene | 2.5 | 5.7683 | 3.1E-50 | 1.6 | 3.095 | 5.5E-36 |
| *mps-1* | WBGene00003403 | protein_coding_gene | 2.5 | 5.7547 | 3.5E-06 | 1.8 | 3.580 | 1.1E-03 |
| *F08D12.12* | WBGene00017255 | protein_coding_gene | 2.5 | 5.7524 | 4.5E-03 | 2.7 | 6.438 | 1.3E-03 |
| *nlp-18* | WBGene00003756 | protein_coding_gene | 2.5 | 5.7435 | 3.4E-22 | 2.3 | 4.874 | 1.7E-38 |
| *del-5* | WBGene00010334 | protein_coding_gene | 2.5 | 5.7387 | 6.0E-21 | 1.2 | 2.333 | 2.0E-09 |
| *vap-1* | WBGene00006886 | protein_coding_gene | 2.5 | 5.7317 | 2.0E-25 | 1.2 | 2.219 | 5.5E-11 |
| *nhr-207* | WBGene00011098 | protein_coding_gene | 2.5 | 5.7316 | 1.0E-04 | 3.0 | 8.157 | 3.8E-04 |
| *asp-1* | WBGene00000214 | protein_coding_gene | 2.5 | 5.7268 | 7.8E-37 | 2.0 | 3.886 | 8.3E-54 |
| *Y18D10A.22* | WBGene00014864 | protein_coding_gene | 2.5 | 5.7262 | 3.1E-11 | 2.5 | 5.474 | 4.0E-08 |
| *T05C12.8* | WBGene00011471 | protein_coding_gene | 2.5 | 5.7259 | 2.6E-04 | 2.0 | 4.071 | 1.1E-03 |
| *C05D12.2* | WBGene00007340 | protein_coding_gene | 2.5 | 5.7236 | 2.2E-25 | 1.9 | 3.636 | 3.9E-14 |
| *ZK856.6* | WBGene00014107 | protein_coding_gene | 2.5 | 5.7225 | 1.2E-04 | 2.1 | 4.293 | 8.7E-03 |
| *B0035.13* | WBGene00007112 | protein_coding_gene | 2.5 | 5.7224 | 5.2E-37 | 1.4 | 2.711 | 1.9E-20 |
| *cla-1* | WBGene00018468 | protein_coding_gene | 2.5 | 5.7170 | 3.9E-09 | 2.3 | 4.887 | 8.6E-27 |
| *pax-2* | WBGene00003938 | protein_coding_gene | 2.5 | 5.7101 | 2.8E-02 | 2.5 | 5.487 | 3.0E-06 |
| *T05C1.1* | WBGene00020248 | protein_coding_gene | 2.5 | 5.7046 | 1.7E-10 | 2.0 | 4.093 | 2.4E-08 |
| *des-2* | WBGene00000955 | protein_coding_gene | 2.5 | 5.7013 | 3.7E-09 | 1.8 | 3.515 | 5.4E-10 |
| *dmd-8* | WBGene00020708 | protein_coding_gene | 2.5 | 5.6977 | 3.7E-06 | 2.9 | 7.594 | 4.3E-05 |
| *gst-12* | WBGene00001760 | protein_coding_gene | 2.5 | 5.6931 | 9.4E-13 | 2.1 | 4.383 | 5.0E-12 |
| *F10E9.1* | WBGene00017353 | protein_coding_gene | 2.5 | 5.6862 | 4.1E-05 | 2.1 | 4.320 | 9.4E-05 |
| *tat-2* | WBGene00019166 | protein_coding_gene | 2.5 | 5.6854 | 1.3E-12 | 2.2 | 4.440 | 7.5E-45 |
| *clec-199* | WBGene00007820 | protein_coding_gene | 2.5 | 5.6851 | 1.7E-06 | 1.4 | 2.599 | 3.0E-04 |
| *pho-4* | WBGene00004023 | protein_coding_gene | 2.5 | 5.6828 | 3.0E-48 | 1.2 | 2.231 | 6.8E-11 |
| *flp-19* | WBGene00001462 | protein_coding_gene | 2.5 | 5.6633 | 7.8E-30 | 1.6 | 3.054 | 1.5E-14 |
| ***F15E6.3*** | **WBGene00017484** | **protein_coding_gene** | **2.5** | **5.6604** | **1.2E-06** | **-2.7** | **0.159** | **5.7E-23** |
| *srsx-30* | WBGene00008251 | protein_coding_gene | 2.5 | 5.6542 | 1.1E-02 | 2.4 | 5.102 | 3.6E-02 |
| *B0024.4* | WBGene00007097 | protein_coding_gene | 2.5 | 5.6505 | 3.0E-06 | 2.0 | 4.097 | 5.1E-07 |
| *ggr-1* | WBGene00001586 | protein_coding_gene | 2.5 | 5.6475 | 7.4E-08 | 1.8 | 3.442 | 4.6E-04 |
| *Y119C1B.6* | WBGene00022472 | protein_coding_gene | 2.5 | 5.6443 | 6.5E-12 | 2.0 | 3.931 | 9.5E-09 |

| | | | | | | | | |
|---|---|---|---|---|---|---|---|---|
| *ttr-21* | WBGene00008628 | protein_coding_gene | 2.5 | 5.6374 | 9.1E-19 | 1.1 | 2.178 | 2.1E-11 |
| *F35F10.4* | WBGene00018054 | pseudogene | 2.5 | 5.6300 | 1.9E-08 | 4.0 | 15.738 | 1.2E-19 |
| *C12D5.5* | WBGene00015708 | protein_coding_gene | 2.5 | 5.6276 | 5.6E-04 | 2.3 | 5.073 | 1.2E-02 |
| *aat-8* | WBGene00000009 | protein_coding_gene | 2.5 | 5.6081 | 1.2E-07 | 1.3 | 2.544 | 1.3E-04 |
| *C35B1.7* | WBGene00016436 | protein_coding_gene | 2.5 | 5.6053 | 9.8E-07 | 2.5 | 5.757 | 2.0E-04 |
| *srh-230* | WBGene00005438 | pseudogene | 2.5 | 5.6039 | 4.7E-02 | 2.8 | 6.792 | 1.5E-02 |
| *dpy-5* | WBGene00001067 | protein_coding_gene | 2.5 | 5.6030 | 1.0E-08 | 4.3 | 20.306 | 3.2E-06 |
| *npr-3* | WBGene00007006 | protein_coding_gene | 2.5 | 5.6007 | 4.0E-08 | 2.1 | 4.203 | 9.3E-06 |
| *sqst-1* | WBGene00011737 | protein_coding_gene | 2.5 | 5.6002 | 8.3E-11 | 1.3 | 2.529 | 3.4E-39 |
| *Y54G2A.10* | WBGene00021875 | protein_coding_gene | 2.5 | 5.5992 | 1.5E-11 | 1.9 | 3.764 | 4.4E-12 |
| *F40D4.16* | WBGene00194681 | ncRNA_gene | 2.5 | 5.5990 | 4.0E-10 | 1.1 | 2.091 | 6.2E-03 |
| *sptf-2* | WBGene00011926 | protein_coding_gene | 2.5 | 5.5936 | 4.1E-08 | 2.2 | 4.607 | 1.9E-07 |
| *col-128* | WBGene00000702 | protein_coding_gene | 2.5 | 5.5917 | 1.0E-03 | 3.2 | 9.124 | 4.8E-04 |
| *cal-4* | WBGene00000288 | protein_coding_gene | 2.5 | 5.5889 | 2.8E-18 | 1.5 | 2.922 | 1.4E-11 |
| *R05G6.9* | WBGene00019901 | protein_coding_gene | 2.5 | 5.5748 | 4.1E-03 | 3.0 | 8.013 | 6.3E-03 |
| *F52F10.2* | WBGene00018706 | protein_coding_gene | 2.5 | 5.5657 | 3.7E-07 | 1.5 | 2.838 | 2.6E-02 |
| *C25E10.11* | WBGene00016100 | protein_coding_gene | 2.5 | 5.5594 | 4.6E-03 | 2.2 | 4.559 | 4.2E-03 |
| *K09E9.7* | WBGene00194664 | protein_coding_gene | 2.5 | 5.5545 | 1.8E-04 | 3.4 | 10.803 | 1.6E-04 |
| *F49C12.6* | WBGene00009876 | protein_coding_gene | 2.5 | 5.5510 | 1.6E-14 | 1.5 | 2.851 | 2.2E-06 |
| *gpa-2* | WBGene00001664 | protein_coding_gene | 2.5 | 5.5501 | 3.3E-08 | 1.7 | 3.175 | 3.6E-04 |
| *Y6G8.2* | WBGene00012404 | protein_coding_gene | 2.5 | 5.5459 | 3.8E-07 | 1.9 | 3.689 | 2.6E-11 |
| *clec-198* | WBGene00008203 | protein_coding_gene | 2.5 | 5.5420 | 2.6E-07 | 1.9 | 3.654 | 1.8E-04 |
| *ZK783.3* | WBGene00022818 | pseudogene | 2.5 | 5.5414 | 6.2E-03 | 2.3 | 4.888 | 1.3E-11 |
| *T10B9.9* | WBGene00011678 | protein_coding_gene | 2.5 | 5.5410 | 4.1E-06 | 1.4 | 2.679 | 1.3E-03 |
| *glb-4* | WBGene00007502 | protein_coding_gene | 2.5 | 5.5335 | 1.3E-04 | 1.8 | 3.540 | 1.5E-03 |
| *C01G5.25* | WBGene00235291 | protein_coding_gene | 2.5 | 5.5332 | 4.4E-08 | 1.2 | 2.319 | 5.3E-04 |
| *C08B6.10* | WBGene00007430 | protein_coding_gene | 2.5 | 5.5318 | 4.1E-16 | 2.1 | 4.231 | 4.0E-12 |
| *rbf-1* | WBGene00004316 | protein_coding_gene | 2.5 | 5.5281 | 1.2E-14 | 1.8 | 3.437 | 1.0E-21 |
| *C54G6.3* | WBGene00016932 | protein_coding_gene | 2.5 | 5.5256 | 1.7E-04 | 1.7 | 3.313 | 5.2E-03 |
| *lgc-46* | WBGene00022106 | protein_coding_gene | 2.5 | 5.5247 | 3.1E-06 | 1.8 | 3.523 | 2.5E-05 |
| *T08B6.2* | WBGene00020352 | pseudogene | 2.5 | 5.5244 | 1.8E-06 | 2.9 | 7.359 | 1.1E-08 |
| *cyp-13A6* | WBGene00011673 | protein_coding_gene | 2.5 | 5.5224 | 2.3E-05 | 2.3 | 4.872 | 1.3E-10 |
| *npr-26* | WBGene00011372 | protein_coding_gene | 2.5 | 5.5220 | 1.8E-08 | 2.0 | 3.902 | 8.6E-06 |
| *col-107* | WBGene00000681 | protein_coding_gene | 2.5 | 5.5218 | 1.1E-08 | 4.0 | 16.439 | 1.0E-83 |
| *mltn-1* | WBGene00009311 | protein_coding_gene | 2.5 | 5.5160 | 7.8E-08 | 1.8 | 3.427 | 2.7E-03 |
| *F44F4.3* | WBGene00009702 | protein_coding_gene | 2.5 | 5.5134 | 2.9E-06 | 1.9 | 3.702 | 5.9E-08 |
| *gcy-15* | WBGene00001541 | protein_coding_gene | 2.5 | 5.5058 | 8.9E-06 | 1.6 | 2.956 | 2.3E-02 |
| *glb-32* | WBGene00021980 | protein_coding_gene | 2.5 | 5.5042 | 6.5E-07 | 1.6 | 2.943 | 1.7E-03 |
| *ptr-17* | WBGene00004231 | protein_coding_gene | 2.5 | 5.4829 | 1.2E-05 | 1.2 | 2.374 | 1.7E-02 |
| *flp-20* | WBGene00001463 | protein_coding_gene | 2.5 | 5.4819 | 4.7E-13 | 1.6 | 2.964 | 6.5E-07 |
| *C36A4.11* | WBGene00044167 | protein_coding_gene | 2.5 | 5.4790 | 1.5E-05 | 1.1 | 2.164 | 3.7E-03 |
| *F35B12.3* | WBGene00009383 | protein_coding_gene | 2.5 | 5.4751 | 3.3E-18 | 1.3 | 2.443 | 1.5E-09 |
| *rcan-1* | WBGene00004321 | protein_coding_gene | 2.5 | 5.4729 | 3.3E-14 | 1.5 | 2.749 | 1.0E-20 |
| *F59B10.5* | WBGene00010321 | protein_coding_gene | 2.5 | 5.4726 | 5.6E-09 | 1.9 | 3.627 | 3.2E-07 |
| *arrd-6* | WBGene00009579 | protein_coding_gene | 2.5 | 5.4660 | 2.4E-08 | 1.7 | 3.230 | 8.5E-06 |
| *lec-11* | WBGene00002274 | protein_coding_gene | 2.4 | 5.4614 | 3.3E-18 | 1.1 | 2.120 | 7.6E-06 |
| *sbt-1* | WBGene00011392 | protein_coding_gene | 2.4 | 5.4576 | 7.5E-28 | 1.8 | 3.434 | 8.2E-36 |
| *T27D12.1* | WBGene00012085 | protein_coding_gene | 2.4 | 5.4538 | 8.6E-41 | 2.0 | 4.062 | 1.5E-37 |
| *nlp-61* | WBGene00017813 | protein_coding_gene | 2.4 | 5.4504 | 1.1E-15 | 1.7 | 3.204 | 4.8E-09 |
| *ZK669.5* | WBGene00014055 | protein_coding_gene | 2.4 | 5.4478 | 9.4E-14 | 1.8 | 3.487 | 3.0E-08 |
| *glb-2* | WBGene00015536 | protein_coding_gene | 2.4 | 5.4379 | 9.4E-05 | 1.9 | 3.756 | 1.1E-03 |
| *fipr-10* | WBGene00008245 | protein_coding_gene | 2.4 | 5.4269 | 9.5E-09 | 2.4 | 5.315 | 5.2E-12 |
| *T09B9.3* | WBGene00011644 | protein_coding_gene | 2.4 | 5.4252 | 7.9E-05 | 1.4 | 2.710 | 6.2E-03 |
| *Y51H7BR.7* | WBGene00021778 | protein_coding_gene | 2.4 | 5.4166 | 8.2E-03 | 2.3 | 5.096 | 2.8E-02 |
| *Y9C9A.8* | WBGene00021178 | protein_coding_gene | 2.4 | 5.4102 | 7.1E-10 | 2.5 | 5.512 | 6.0E-09 |
| *glb-11* | WBGene00007981 | protein_coding_gene | 2.4 | 5.4051 | 1.2E-17 | 1.7 | 3.184 | 4.9E-10 |
| *srap-1* | WBGene00011522 | protein_coding_gene | 2.4 | 5.3983 | 1.4E-07 | 1.9 | 3.634 | 7.0E-29 |
| *nhr-167* | WBGene00008208 | protein_coding_gene | 2.4 | 5.3973 | 7.3E-03 | 1.5 | 2.739 | 5.0E-02 |
| *Y51H4A.25* | WBGene00013119 | protein_coding_gene | 2.4 | 5.3922 | 1.4E-19 | 2.0 | 4.094 | 1.1E-18 |
| *T22C8.6* | WBGene00011927 | protein_coding_gene | 2.4 | 5.3910 | 2.0E-09 | 1.6 | 3.019 | 3.0E-08 |
| *lite-1* | WBGene00001803 | protein_coding_gene | 2.4 | 5.3889 | 3.4E-12 | 2.0 | 3.902 | 1.1E-11 |
| *trp-1* | WBGene00006614 | protein_coding_gene | 2.4 | 5.3662 | 2.3E-08 | 2.1 | 4.413 | 7.0E-13 |
| *acs-21* | WBGene00016849 | protein_coding_gene | 2.4 | 5.3655 | 2.7E-32 | 2.3 | 4.970 | 1.8E-13 |
| *lgc-11* | WBGene00000061 | protein_coding_gene | 2.4 | 5.3655 | 4.3E-06 | 2.1 | 4.432 | 7.6E-05 |
| *Y2H9A.4* | WBGene00012381 | protein_coding_gene | 2.4 | 5.3648 | 2.1E-11 | 1.4 | 2.690 | 7.1E-08 |
| *Y22D7AL.15* | WBGene00021253 | protein_coding_gene | 2.4 | 5.3611 | 2.4E-17 | 1.6 | 3.086 | 8.3E-16 |
| *deg-3* | WBGene00000951 | protein_coding_gene | 2.4 | 5.3588 | 1.4E-11 | 1.7 | 3.216 | 4.6E-08 |
| *F57H12.6* | WBGene00019021 | protein_coding_gene | 2.4 | 5.3587 | 6.1E-07 | 2.1 | 4.187 | 3.2E-06 |
| *moe-3* | WBGene00003388 | protein_coding_gene | 2.4 | 5.3576 | 8.7E-10 | 1.3 | 2.396 | 6.3E-12 |
| *F31B9.2* | WBGene00009279 | protein_coding_gene | 2.4 | 5.3504 | 4.8E-02 | 3.1 | 8.721 | 9.1E-03 |
| *grl-14* | WBGene00001723 | protein_coding_gene | 2.4 | 5.3496 | 2.1E-07 | 1.2 | 2.336 | 2.6E-02 |
| *K07D4.4* | WBGene00019484 | protein_coding_gene | 2.4 | 5.3490 | 3.4E-03 | 1.8 | 3.373 | 2.0E-02 |
| *ptl-1* | WBGene00004212 | protein_coding_gene | 2.4 | 5.3488 | 1.6E-25 | 1.9 | 3.674 | 7.9E-19 |
| *sulp-3* | WBGene00018283 | protein_coding_gene | 2.4 | 5.3478 | 1.1E-07 | 1.2 | 2.309 | 5.5E-04 |
| *cutl-24* | WBGene00021396 | protein_coding_gene | 2.4 | 5.3362 | 2.7E-10 | 1.1 | 2.139 | 4.4E-04 |
| *flp-12* | WBGene00001455 | protein_coding_gene | 2.4 | 5.3307 | 3.3E-16 | 1.5 | 2.811 | 1.4E-19 |
| *Y62H9A.15* | WBGene00045252 | protein_coding_gene | 2.4 | 5.3274 | 7.3E-13 | 1.6 | 3.111 | 9.4E-06 |
| *Y34F4.5* | WBGene00021340 | protein_coding_gene | 2.4 | 5.3248 | 3.8E-31 | 1.2 | 2.244 | 1.5E-15 |
| *Y61A9LA.12* | WBGene00044586 | protein_coding_gene | 2.4 | 5.3228 | 6.3E-05 | 1.5 | 2.883 | 1.6E-02 |
| *R10E8.6* | WBGene00011212 | protein_coding_gene | 2.4 | 5.3194 | 2.2E-14 | 1.8 | 3.447 | 1.9E-23 |

| | | | | | | | | |
|---|---|---|---|---|---|---|---|---|
| *mdl-1* | WBGene00003163 | protein_coding_gene | 2.4 | 5.3190 | 1.6E-10 | 1.3 | 2.540 | 6.6E-17 |
| *nlp-49* | WBGene00019160 | protein_coding_gene | 2.4 | 5.3160 | 7.8E-18 | 2.1 | 4.390 | 4.6E-44 |
| *K09E4.4* | WBGene00010722 | protein_coding_gene | 2.4 | 5.3152 | 2.6E-11 | 2.4 | 5.327 | 3.2E-19 |
| *nmr-2* | WBGene00003775 | protein_coding_gene | 2.4 | 5.3140 | 7.5E-05 | 1.2 | 2.323 | 3.6E-02 |
| *sul-2* | WBGene00006309 | protein_coding_gene | 2.4 | 5.3115 | 5.9E-09 | 1.9 | 3.725 | 6.9E-15 |
| *D1069.1* | WBGene00017036 | protein_coding_gene | 2.4 | 5.3112 | 1.4E-04 | 1.3 | 2.470 | 1.2E-02 |
| *C24B5.1* | WBGene00016044 | protein_coding_gene | 2.4 | 5.3086 | 7.5E-10 | 2.0 | 3.966 | 3.1E-08 |
| *C15A7.2* | WBGene00007595 | protein_coding_gene | 2.4 | 5.3084 | 1.4E-03 | 1.6 | 2.948 | 3.0E-02 |
| *gpx-3* | WBGene00007517 | protein_coding_gene | 2.4 | 5.3011 | 6.9E-24 | 2.4 | 5.428 | 1.8E-25 |
| *C27H5.2* | WBGene00016172 | protein_coding_gene | 2.4 | 5.3003 | 3.0E-31 | 2.0 | 3.965 | 4.5E-26 |
| *cest-13* | WBGene00010116 | protein_coding_gene | 2.4 | 5.2999 | 7.1E-16 | 3.5 | 11.293 | 2.5E-66 |
| *msa-1* | WBGene00013882 | protein_coding_gene | 2.4 | 5.2749 | 9.9E-22 | 1.7 | 3.198 | 5.2E-29 |
| *M6.4* | WBGene00019776 | protein_coding_gene | 2.4 | 5.2627 | 5.1E-05 | 1.5 | 2.800 | 1.2E-02 |
| *dop-1* | WBGene00001052 | protein_coding_gene | 2.4 | 5.2622 | 3.0E-10 | 2.8 | 6.962 | 5.8E-13 |
| *tkr-2* | WBGene00016761 | protein_coding_gene | 2.4 | 5.2620 | 9.7E-03 | 2.1 | 4.392 | 7.0E-03 |
| *K09H11.6* | WBGene00019603 | protein_coding_gene | 2.4 | 5.2617 | 3.0E-04 | 2.7 | 6.567 | 2.2E-05 |
| *C47A4.4* | WBGene00014703 | pseudogene | 2.4 | 5.2572 | 6.5E-03 | 2.8 | 6.870 | 5.9E-06 |
| *nhr-120* | WBGene00003710 | protein_coding_gene | 2.4 | 5.2540 | 1.2E-12 | 1.5 | 2.743 | 4.0E-12 |
| *ZC132.3* | WBGene00022522 | protein_coding_gene | 2.4 | 5.2530 | 8.0E-04 | 1.6 | 3.126 | 1.9E-02 |
| *snt-4* | WBGene00004924 | protein_coding_gene | 2.4 | 5.2418 | 3.8E-17 | 2.0 | 3.926 | 8.4E-48 |
| *C16D9.1* | WBGene00015857 | protein_coding_gene | 2.4 | 5.2409 | 2.9E-12 | 2.6 | 6.091 | 5.8E-12 |
| *C27D6.12* | WBGene00044389 | protein_coding_gene | 2.4 | 5.2347 | 3.3E-20 | 1.7 | 3.168 | 1.1E-11 |
| *irld-2* | WBGene00008518 | protein_coding_gene | 2.4 | 5.2271 | 4.8E-03 | 2.7 | 6.366 | 1.8E-03 |
| *T26C5.4* | WBGene00012034 | protein_coding_gene | 2.4 | 5.2241 | 2.8E-12 | 1.1 | 2.191 | 9.7E-05 |
| *dct-15* | WBGene00012340 | protein_coding_gene | 2.4 | 5.2222 | 7.0E-07 | 2.1 | 4.355 | 1.6E-05 |
| *kin-30* | WBGene00002211 | protein_coding_gene | 2.4 | 5.2162 | 2.9E-12 | 2.0 | 3.910 | 4.4E-07 |
| *D1007.18* | WBGene00044421 | protein_coding_gene | 2.4 | 5.2084 | 1.2E-08 | 1.9 | 3.806 | 7.8E-07 |
| *sma-2* | WBGene00004856 | protein_coding_gene | 2.4 | 5.2036 | 7.8E-16 | 1.4 | 2.565 | 2.0E-11 |
| *gpa-17* | WBGene00022196 | protein_coding_gene | 2.4 | 5.2019 | 8.5E-15 | 2.2 | 4.483 | 3.4E-37 |
| *cpna-4* | WBGene00006441 | protein_coding_gene | 2.4 | 5.1964 | 2.0E-07 | 1.8 | 3.503 | 1.1E-06 |
| *cal-8* | WBGene00017394 | protein_coding_gene | 2.4 | 5.1940 | 2.5E-07 | 1.6 | 3.020 | 3.2E-04 |
| *acr-24* | WBGene00013534 | protein_coding_gene | 2.4 | 5.1889 | 1.6E-06 | 1.7 | 3.150 | 1.2E-03 |
| *ZC455.1* | WBGene00013902 | protein_coding_gene | 2.4 | 5.1699 | 1.2E-12 | 1.3 | 2.405 | 5.5E-11 |
| *clec-65* | WBGene00009396 | protein_coding_gene | 2.4 | 5.1669 | 7.4E-58 | 1.3 | 2.525 | 4.7E-18 |
| *lec-2* | WBGene00002265 | protein_coding_gene | 2.4 | 5.1612 | 6.6E-09 | 1.2 | 2.344 | 3.4E-05 |
| *K07D4.9* | WBGene00019488 | protein_coding_gene | 2.4 | 5.1611 | 5.5E-11 | 2.1 | 4.366 | 3.7E-25 |
| *zip-10* | WBGene00012101 | protein_coding_gene | 2.4 | 5.1586 | 2.0E-06 | 2.0 | 3.904 | 3.0E-10 |
| *spp-23* | WBGene00021335 | protein_coding_gene | 2.4 | 5.1579 | 1.3E-06 | 4.9 | 30.570 | 5.9E-22 |
| *Y37D8A.16* | WBGene00012554 | protein_coding_gene | 2.4 | 5.1571 | 2.0E-19 | 1.2 | 2.234 | 3.5E-13 |
| *F10D2.8* | WBGene00017334 | protein_coding_gene | 2.4 | 5.1540 | 2.6E-06 | 1.5 | 2.834 | 2.1E-08 |
| *ptr-6* | WBGene00004221 | protein_coding_gene | 2.4 | 5.1537 | 4.1E-19 | 1.4 | 2.594 | 1.8E-08 |
| *shn-1* | WBGene00006444 | protein_coding_gene | 2.4 | 5.1532 | 5.5E-13 | 2.0 | 4.014 | 1.7E-36 |
| *ist-1* | WBGene00002163 | protein_coding_gene | 2.4 | 5.1523 | 3.3E-12 | 1.8 | 3.477 | 1.5E-21 |
| *C25G6.3* | WBGene00016108 | protein_coding_gene | 2.4 | 5.1517 | 4.0E-05 | 2.2 | 4.677 | 5.4E-07 |
| *mth-2* | WBGene00017947 | protein_coding_gene | 2.4 | 5.1470 | 1.7E-15 | 1.8 | 3.529 | 5.3E-12 |
| *flp-17* | WBGene00001460 | protein_coding_gene | 2.4 | 5.1421 | 2.0E-12 | 1.6 | 2.942 | 8.5E-18 |
| *C28H8.2* | WBGene00016193 | protein_coding_gene | 2.4 | 5.1407 | 1.3E-08 | 1.2 | 2.301 | 4.8E-03 |
| *flp-3* | WBGene00001446 | protein_coding_gene | 2.4 | 5.1400 | 5.9E-24 | 1.6 | 2.971 | 7.5E-19 |
| *T08B1.4* | WBGene00020341 | protein_coding_gene | 2.4 | 5.1398 | 5.4E-23 | 1.6 | 3.086 | 2.8E-08 |
| *nas-29* | WBGene00003547 | protein_coding_gene | 2.4 | 5.1377 | 2.6E-07 | 2.2 | 4.744 | 2.8E-05 |
| *clec-53* | WBGene00020191 | protein_coding_gene | 2.4 | 5.1373 | 2.0E-11 | 2.2 | 4.537 | 1.5E-09 |
| *F33D11.16* | WBGene00269382 | protein_coding_gene | 2.4 | 5.1355 | 8.4E-04 | 1.3 | 2.387 | 1.0E-02 |
| ***F25E5.4*** | **WBGene00017785** | **protein_coding_gene** | **2.4** | **5.1213** | **4.4E-13** | **-1.4** | **0.385** | **5.4E-17** |
| *dpy-4* | WBGene00001066 | protein_coding_gene | 2.4 | 5.1188 | 2.3E-09 | 3.9 | 15.205 | 1.4E-04 |
| *nhr-275* | WBGene00021163 | protein_coding_gene | 2.4 | 5.1177 | 1.7E-05 | 1.5 | 2.857 | 2.4E-02 |
| *gpa-3* | WBGene00001665 | protein_coding_gene | 2.4 | 5.1140 | 4.4E-08 | 1.6 | 3.107 | 6.2E-05 |
| *fkh-6* | WBGene00001438 | protein_coding_gene | 2.4 | 5.1111 | 2.2E-07 | 1.8 | 3.460 | 1.1E-07 |
| *C50H2.13* | WBGene00008246 | protein_coding_gene | 2.4 | 5.1046 | 1.6E-15 | 1.1 | 2.170 | 4.6E-05 |
| *irld-34* | WBGene00010074 | protein_coding_gene | 2.3 | 5.0941 | 5.1E-03 | 1.9 | 3.636 | 8.2E-03 |
| *nhr-236* | WBGene00021417 | protein_coding_gene | 2.3 | 5.0911 | 3.8E-05 | 1.5 | 2.916 | 4.6E-02 |
| *Y22D7AR.6* | WBGene00021260 | protein_coding_gene | 2.3 | 5.0891 | 3.1E-11 | 1.7 | 3.158 | 4.5E-18 |
| *R08C7.12* | WBGene00019955 | protein_coding_gene | 2.3 | 5.0872 | 1.4E-13 | 2.2 | 4.488 | 3.5E-27 |
| *EGAP5.1* | WBGene00017150 | protein_coding_gene | 2.3 | 5.0835 | 4.0E-03 | 2.5 | 5.488 | 4.5E-03 |
| *ZC190.4* | WBGene00022538 | protein_coding_gene | 2.3 | 5.0779 | 5.8E-16 | 1.6 | 2.956 | 1.3E-24 |
| *H10D18.5* | WBGene00019180 | protein_coding_gene | 2.3 | 5.0777 | 1.6E-14 | 1.7 | 3.241 | 1.7E-06 |
| *nhr-44* | WBGene00003634 | protein_coding_gene | 2.3 | 5.0762 | 7.7E-11 | 1.2 | 2.222 | 5.4E-05 |
| *sod-4* | WBGene00004933 | protein_coding_gene | 2.3 | 5.0759 | 4.1E-21 | 1.0 | 2.007 | 1.5E-08 |
| ***F36H1.11*** | **WBGene00009496** | **protein_coding_gene** | **2.3** | **5.0750** | **6.6E-03** | **-1.3** | **0.404** | **4.6E-02** |
| *C16A3.6* | WBGene00015811 | protein_coding_gene | 2.3 | 5.0719 | 1.4E-43 | 1.4 | 2.613 | 1.5E-21 |
| *glb-3* | WBGene00007389 | protein_coding_gene | 2.3 | 5.0705 | 7.8E-10 | 1.9 | 3.770 | 1.8E-07 |
| *M04G7.1* | WBGene00019772 | protein_coding_gene | 2.3 | 5.0692 | 2.6E-15 | 1.9 | 3.766 | 3.2E-27 |
| *ZK662.2* | WBGene00014039 | protein_coding_gene | 2.3 | 5.0689 | 3.6E-07 | 2.0 | 3.921 | 2.0E-02 |
| *nlp-58* | WBGene00077775 | protein_coding_gene | 2.3 | 5.0685 | 1.2E-14 | 1.7 | 3.189 | 1.5E-10 |
| *ZK1290.13* | WBGene00022893 | protein_coding_gene | 2.3 | 5.0634 | 1.0E-16 | 1.1 | 2.110 | 1.4E-09 |
| *nlp-53* | WBGene00012440 | protein_coding_gene | 2.3 | 5.0522 | 1.7E-09 | 2.4 | 5.146 | 4.3E-15 |
| *C01C4.3* | WBGene00015293 | protein_coding_gene | 2.3 | 5.0460 | 5.7E-09 | 1.9 | 3.664 | 1.1E-22 |
| *C34B4.3* | WBGene00007908 | protein_coding_gene | 2.3 | 5.0433 | 1.4E-06 | 1.0 | 2.021 | 3.5E-03 |
| *C11D2.3* | WBGene00015701 | protein_coding_gene | 2.3 | 5.0335 | 2.1E-04 | 2.1 | 4.268 | 2.5E-06 |
| *F08G2.8* | WBGene00008579 | protein_coding_gene | 2.3 | 5.0330 | 4.2E-07 | 1.4 | 2.641 | 6.4E-03 |
| *C16B8.2* | WBGene00015822 | protein_coding_gene | 2.3 | 5.0321 | 3.1E-08 | 1.2 | 2.251 | 1.3E-02 |

| | | | | | | | | |
|---|---|---|---|---|---|---|---|---|
| *srt-42* | WBGene00021599 | protein_coding_gene | 2.3 | 5.0303 | 1.8E-02 | 2.0 | 4.098 | 2.0E-02 |
| *nlp-11* | WBGene00003749 | protein_coding_gene | 2.3 | 5.0287 | 2.5E-14 | 2.1 | 4.426 | 5.0E-34 |
| *sulp-1* | WBGene00016945 | protein_coding_gene | 2.3 | 5.0266 | 1.7E-06 | 2.0 | 4.099 | 1.8E-04 |
| *nhr-216* | WBGene00020385 | protein_coding_gene | 2.3 | 5.0240 | 8.6E-05 | 3.0 | 8.009 | 5.0E-05 |
| *F36H2.3* | WBGene00009500 | protein_coding_gene | 2.3 | 5.0212 | 4.4E-11 | 2.1 | 4.150 | 1.4E-14 |
| *spl-2* | WBGene00006418 | protein_coding_gene | 2.3 | 5.0125 | 1.4E-06 | 1.3 | 2.390 | 3.3E-05 |
| *hpo-28* | WBGene00015148 | protein_coding_gene | 2.3 | 5.0104 | 6.8E-23 | 2.1 | 4.156 | 9.6E-15 |
| *gpa-11* | WBGene00001673 | protein_coding_gene | 2.3 | 5.0102 | 1.1E-02 | 3.4 | 10.902 | 5.9E-04 |
| *C53B7.3* | WBGene00016894 | protein_coding_gene | 2.3 | 5.0100 | 7.7E-23 | 1.6 | 2.969 | 4.4E-20 |
| *ZK265.7* | WBGene00013959 | protein_coding_gene | 2.3 | 5.0088 | 3.5E-15 | 1.2 | 2.279 | 6.8E-05 |
| *cab-1* | WBGene00000277 | protein_coding_gene | 2.3 | 4.9966 | 2.5E-17 | 2.1 | 4.382 | 1.1E-80 |
| *dpf-6* | WBGene00001059 | protein_coding_gene | 2.3 | 4.9921 | 7.7E-20 | 4.3 | 19.830 | 3.2E-50 |
| *xbx-9* | WBGene00007604 | protein_coding_gene | 2.3 | 4.9902 | 4.9E-09 | 2.1 | 4.243 | 4.2E-09 |
| *F10C2.3* | WBGene00008644 | protein_coding_gene | 2.3 | 4.9823 | 1.6E-12 | 2.0 | 3.996 | 2.6E-12 |
| *F41G4.8* | WBGene00018311 | protein_coding_gene | 2.3 | 4.9822 | 3.2E-12 | 1.2 | 2.335 | 3.8E-07 |
| *E02A10.3* | WBGene00008453 | protein_coding_gene | 2.3 | 4.9723 | 4.0E-10 | 1.1 | 2.086 | 9.5E-06 |
| *scd-2* | WBGene00004740 | protein_coding_gene | 2.3 | 4.9601 | 1.2E-09 | 2.3 | 4.840 | 1.5E-20 |
| *ztf-14* | WBGene00010936 | protein_coding_gene | 2.3 | 4.9576 | 7.1E-05 | 2.1 | 4.328 | 1.1E-04 |
| *Y51A2D.13* | WBGene00013080 | protein_coding_gene | 2.3 | 4.9556 | 8.3E-07 | 1.4 | 2.601 | 4.5E-26 |
| *W10C8.4* | WBGene00021127 | protein_coding_gene | 2.3 | 4.9539 | 3.5E-08 | 1.5 | 2.733 | 4.0E-04 |
| *F29G6.1* | WBGene00009257 | protein_coding_gene | 2.3 | 4.9472 | 2.4E-10 | 3.6 | 12.472 | 3.1E-13 |
| *nlp-42* | WBGene00013588 | protein_coding_gene | 2.3 | 4.9414 | 1.4E-11 | 1.2 | 2.297 | 1.9E-07 |
| *F54C9.7* | WBGene00010043 | protein_coding_gene | 2.3 | 4.9282 | 8.3E-07 | 1.9 | 3.811 | 3.8E-05 |
| *egl-21* | WBGene00001189 | protein_coding_gene | 2.3 | 4.9280 | 1.5E-30 | 2.0 | 3.910 | 3.5E-55 |
| *nhr-180* | WBGene00017961 | protein_coding_gene | 2.3 | 4.9271 | 2.0E-09 | 1.3 | 2.415 | 2.9E-06 |
| *F17C11.2* | WBGene00008914 | protein_coding_gene | 2.3 | 4.9270 | 1.4E-28 | 1.5 | 2.826 | 1.1E-17 |
| *flp-13* | WBGene00001456 | protein_coding_gene | 2.3 | 4.9265 | 5.3E-19 | 2.2 | 4.439 | 2.2E-29 |
| *F18F11.4* | WBGene00017577 | protein_coding_gene | 2.3 | 4.9199 | 6.8E-04 | 2.9 | 7.544 | 2.4E-03 |
| *DH11.5* | WBGene00008438 | protein_coding_gene | 2.3 | 4.9184 | 1.3E-23 | 1.7 | 3.252 | 2.4E-23 |
| *php-3* | WBGene00004024 | protein_coding_gene | 2.3 | 4.9177 | 2.0E-20 | 1.0 | 2.070 | 4.3E-10 |
| *F58G11.4* | WBGene00010282 | protein_coding_gene | 2.3 | 4.9173 | 1.8E-08 | 1.7 | 3.304 | 1.0E-04 |
| *T06A4.1* | WBGene00020281 | protein_coding_gene | 2.3 | 4.9142 | 1.2E-04 | 2.0 | 3.944 | 2.0E-03 |
| *dpy-13* | WBGene00001074 | protein_coding_gene | 2.3 | 4.9138 | 8.9E-07 | 3.9 | 14.725 | 7.9E-05 |
| ***R05G9R.1*** | **WBGene00019906** | **protein_coding_gene** | **2.3** | **4.9081** | **1.4E-05** | **-1.3** | **0.410** | **8.6E-03** |
| *H04M03.12* | WBGene00019156 | protein_coding_gene | 2.3 | 4.9078 | 1.9E-08 | 1.0 | 2.051 | 4.2E-04 |
| *sox-3* | WBGene00004950 | protein_coding_gene | 2.3 | 4.9054 | 6.5E-09 | 1.4 | 2.702 | 1.4E-04 |
| *hlh-33* | WBGene00021446 | protein_coding_gene | 2.3 | 4.9040 | 4.2E-18 | 1.6 | 2.933 | 1.8E-09 |
| *kin-9* | WBGene00002195 | protein_coding_gene | 2.3 | 4.9024 | 2.6E-12 | 2.1 | 4.411 | 3.0E-09 |
| *ggr-2* | WBGene00001587 | protein_coding_gene | 2.3 | 4.8979 | 7.7E-06 | 1.8 | 3.473 | 4.9E-04 |
| *T10G3.8* | WBGene00045340 | protein_coding_gene | 2.3 | 4.8957 | 5.0E-05 | 1.6 | 3.035 | 2.2E-03 |
| *ZC116.1* | WBGene00013853 | protein_coding_gene | 2.3 | 4.8778 | 2.3E-10 | 1.6 | 2.991 | 1.4E-19 |
| *flp-4* | WBGene00001447 | protein_coding_gene | 2.3 | 4.8695 | 1.1E-17 | 1.6 | 2.998 | 1.7E-12 |
| *tpra-1* | WBGene00008366 | protein_coding_gene | 2.3 | 4.8670 | 5.6E-16 | 1.0 | 2.015 | 9.1E-12 |
| *cal-3* | WBGene00000287 | protein_coding_gene | 2.3 | 4.8617 | 4.1E-13 | 1.6 | 2.952 | 2.5E-07 |
| *nlp-70* | WBGene00044513 | protein_coding_gene | 2.3 | 4.8583 | 1.7E-08 | 1.8 | 3.529 | 2.4E-05 |
| *C06C3.12* | WBGene00175031 | protein_coding_gene | 2.3 | 4.8567 | 2.2E-05 | 1.9 | 3.860 | 3.1E-05 |
| *fkh-7* | WBGene00001439 | protein_coding_gene | 2.3 | 4.8566 | 1.3E-11 | 1.0 | 2.043 | 3.9E-12 |
| *C10A4.9* | WBGene00045456 | protein_coding_gene | 2.3 | 4.8531 | 6.6E-07 | 2.3 | 4.839 | 9.3E-07 |
| *C06G1.2* | WBGene00015546 | protein_coding_gene | 2.3 | 4.8525 | 7.9E-07 | 1.9 | 3.634 | 8.0E-03 |
| *xbx-5* | WBGene00011974 | protein_coding_gene | 2.3 | 4.8481 | 4.6E-09 | 2.0 | 4.042 | 2.3E-07 |
| *F41C3.6* | WBGene00018272 | protein_coding_gene | 2.3 | 4.8445 | 1.4E-03 | 2.3 | 4.906 | 4.9E-04 |
| *srr-1* | WBGene00005652 | protein_coding_gene | 2.3 | 4.8380 | 3.5E-08 | 1.4 | 2.563 | 1.9E-03 |
| *lrx-1* | WBGene00003075 | protein_coding_gene | 2.3 | 4.8346 | 2.5E-06 | 1.3 | 2.400 | 3.7E-02 |
| *C49A9.3* | WBGene00016757 | protein_coding_gene | 2.3 | 4.8344 | 1.6E-21 | 2.3 | 4.976 | 1.7E-21 |
| *glb-12* | WBGene00008256 | protein_coding_gene | 2.3 | 4.8324 | 4.4E-04 | 2.5 | 5.763 | 2.3E-05 |
| *T19D12.2* | WBGene00020577 | protein_coding_gene | 2.3 | 4.8269 | 3.3E-29 | 1.7 | 3.254 | 5.5E-23 |
| *T03G11.3* | WBGene00020196 | protein_coding_gene | 2.3 | 4.8224 | 1.4E-05 | 1.6 | 3.083 | 1.1E-03 |
| *ora-1* | WBGene00003879 | protein_coding_gene | 2.3 | 4.8216 | 3.0E-04 | 2.3 | 5.076 | 7.6E-03 |
| *lin-14* | WBGene00003003 | protein_coding_gene | 2.3 | 4.8156 | 1.7E-10 | 1.8 | 3.403 | 5.3E-41 |
| *col-38* | WBGene00000615 | protein_coding_gene | 2.3 | 4.8139 | 7.6E-13 | 3.9 | 14.914 | 2.3E-05 |
| *B0507.3* | WBGene00015220 | protein_coding_gene | 2.3 | 4.8095 | 8.9E-11 | 1.3 | 2.388 | 4.6E-04 |
| ***T19C4.1*** | **WBGene00011838** | **protein_coding_gene** | **2.3** | **4.8068** | **1.1E-05** | **-1.2** | **0.436** | **4.6E-04** |
| *linc-96* | WBGene00219673 | lincRNA_gene | 2.3 | 4.8023 | 5.5E-03 | 2.2 | 4.752 | 1.9E-02 |
| *E02A10.4* | WBGene00008454 | protein_coding_gene | 2.3 | 4.8004 | 8.7E-09 | 2.2 | 4.600 | 1.7E-13 |
| ***C24H10.3*** | **WBGene00016069** | **protein_coding_gene** | **2.3** | **4.7959** | **3.7E-04** | **-1.4** | **0.373** | **5.2E-03** |
| *ins-17* | WBGene00002100 | protein_coding_gene | 2.3 | 4.7939 | 2.4E-19 | 1.3 | 2.382 | 3.0E-11 |
| *R05D7.1* | WBGene00011026 | protein_coding_gene | 2.3 | 4.7899 | 6.9E-10 | 1.8 | 3.510 | 4.8E-17 |
| *ZC47.8* | WBGene00014978 | pseudogene | 2.3 | 4.7806 | 2.3E-07 | 2.1 | 4.173 | 3.7E-08 |
| *F59F3.6* | WBGene00010335 | protein_coding_gene | 2.3 | 4.7800 | 9.8E-06 | 1.2 | 2.310 | 1.0E-02 |
| *madf-9* | WBGene00022608 | protein_coding_gene | 2.3 | 4.7763 | 2.0E-19 | 2.4 | 5.253 | 1.5E-25 |
| *dop-3* | WBGene00020506 | protein_coding_gene | 2.3 | 4.7596 | 5.9E-09 | 2.2 | 4.526 | 3.7E-07 |
| *pqn-35* | WBGene00004122 | protein_coding_gene | 2.2 | 4.7517 | 1.6E-18 | 1.9 | 3.749 | 4.1E-23 |
| *col-173* | WBGene00000746 | protein_coding_gene | 2.2 | 4.7507 | 5.2E-08 | 3.8 | 14.071 | 7.6E-05 |
| *npr-15* | WBGene00012084 | protein_coding_gene | 2.2 | 4.7485 | 1.4E-05 | 1.2 | 2.377 | 8.6E-03 |
| *nas-25* | WBGene00003544 | protein_coding_gene | 2.2 | 4.7480 | 3.1E-05 | 1.3 | 2.474 | 3.9E-03 |
| *cah-2* | WBGene00000280 | protein_coding_gene | 2.2 | 4.7475 | 3.8E-10 | 1.3 | 2.536 | 6.7E-04 |
| *F55C12.6* | WBGene00018873 | protein_coding_gene | 2.2 | 4.7471 | 5.9E-06 | 1.2 | 2.294 | 2.7E-02 |
| *ZK1290.14* | WBGene00044396 | protein_coding_gene | 2.2 | 4.7467 | 1.0E-31 | 1.5 | 2.879 | 6.4E-12 |
| *C31C9.6* | WBGene00007840 | protein_coding_gene | 2.2 | 4.7398 | 2.5E-09 | 1.6 | 3.085 | 1.0E-16 |
| *drn-1* | WBGene00016911 | protein_coding_gene | 2.2 | 4.7391 | 1.0E-15 | 1.3 | 2.547 | 1.1E-08 |

| | | | | | | | | |
|---|---|---|---|---|---|---|---|---|
| *F59A1.11* | WBGene00010297 | protein_coding_gene | 2.2 | 4.7320 | 9.6E-03 | 1.8 | 3.602 | 3.4E-02 |
| *sup-1* | WBGene00012759 | protein_coding_gene | 2.2 | 4.7264 | 2.5E-21 | 2.0 | 4.015 | 7.2E-34 |
| *F54B8.4* | WBGene00010019 | protein_coding_gene | 2.2 | 4.7261 | 9.8E-15 | 1.1 | 2.210 | 1.3E-05 |
| *Y37F4.1* | WBGene00021382 | protein_coding_gene | 2.2 | 4.7253 | 9.7E-06 | 1.5 | 2.869 | 1.5E-08 |
| *exp-2* | WBGene00001374 | protein_coding_gene | 2.2 | 4.7251 | 1.6E-12 | 1.7 | 3.157 | 1.6E-12 |
| *F11C1.4* | WBGene00008693 | protein_coding_gene | 2.2 | 4.7227 | 5.2E-05 | 1.8 | 3.595 | 5.4E-04 |
| *T07F12.4* | WBGene00020324 | protein_coding_gene | 2.2 | 4.7215 | 4.6E-16 | 1.8 | 3.424 | 2.8E-10 |
| *kel-1* | WBGene00002184 | protein_coding_gene | 2.2 | 4.7206 | 1.1E-15 | 1.1 | 2.097 | 1.1E-03 |
| *W04E12.7* | WBGene00012252 | protein_coding_gene | 2.2 | 4.7206 | 2.5E-23 | 2.7 | 6.727 | 3.3E-23 |
| *ocr-1* | WBGene00003838 | protein_coding_gene | 2.2 | 4.7179 | 7.6E-04 | 1.8 | 3.479 | 8.8E-03 |
| *F35B12.9* | WBGene00009388 | protein_coding_gene | 2.2 | 4.7142 | 1.0E-10 | 1.6 | 3.125 | 7.6E-14 |
| *F23G4.1* | WBGene00044574 | protein_coding_gene | 2.2 | 4.7114 | 1.7E-05 | 1.4 | 2.691 | 2.2E-03 |
| *pqn-36* | WBGene00004123 | protein_coding_gene | 2.2 | 4.7095 | 2.8E-06 | 2.5 | 5.712 | 9.4E-14 |
| *M28.8* | WBGene00010899 | protein_coding_gene | 2.2 | 4.7039 | 2.5E-10 | 1.7 | 3.355 | 9.9E-21 |
| *snt-6* | WBGene00004926 | protein_coding_gene | 2.2 | 4.6970 | 1.0E-09 | 1.7 | 3.363 | 3.1E-11 |
| *ZK896.5* | WBGene00014136 | protein_coding_gene | 2.2 | 4.6893 | 6.8E-11 | 1.5 | 2.838 | 8.4E-03 |
| *C18A11.2* | WBGene00015948 | protein_coding_gene | 2.2 | 4.6871 | 2.0E-04 | 1.4 | 2.685 | 2.0E-04 |
| *C36E6.8* | WBGene00044683 | protein_coding_gene | 2.2 | 4.6867 | 1.3E-03 | 1.7 | 3.341 | 1.4E-02 |
| *glb-24* | WBGene00011287 | protein_coding_gene | 2.2 | 4.6823 | 1.6E-10 | 1.7 | 3.332 | 5.8E-05 |
| *hlh-30* | WBGene00020930 | protein_coding_gene | 2.2 | 4.6822 | 5.8E-09 | 1.1 | 2.167 | 3.3E-21 |
| *cal-1* | WBGene00000285 | protein_coding_gene | 2.2 | 4.6815 | 5.0E-14 | 1.5 | 2.809 | 1.9E-12 |
| *glb-25* | WBGene00020279 | protein_coding_gene | 2.2 | 4.6811 | 8.0E-11 | 1.1 | 2.149 | 4.1E-04 |
| *F54D5.3* | WBGene00010049 | protein_coding_gene | 2.2 | 4.6779 | 4.2E-59 | 1.2 | 2.248 | 3.2E-26 |
| *Y41C4A.11* | WBGene00012757 | protein_coding_gene | 2.2 | 4.6771 | 2.0E-07 | 1.2 | 2.246 | 2.8E-03 |
| *rnt-1* | WBGene00004393 | protein_coding_gene | 2.2 | 4.6744 | 9.1E-06 | 1.4 | 2.704 | 2.1E-04 |
| *C03G6.17* | WBGene00015402 | protein_coding_gene | 2.2 | 4.6714 | 6.2E-12 | 2.0 | 4.018 | 1.8E-20 |
| *mgl-3* | WBGene00021152 | protein_coding_gene | 2.2 | 4.6703 | 7.9E-08 | 1.3 | 2.438 | 2.0E-03 |
| *rol-1* | WBGene00004394 | protein_coding_gene | 2.2 | 4.6697 | 3.4E-07 | 4.0 | 15.706 | 1.7E-04 |
| *his-71* | WBGene00001945 | protein_coding_gene | 2.2 | 4.6697 | 3.6E-19 | 1.4 | 2.673 | 3.2E-21 |
| *irg-1* | WBGene00015574 | protein_coding_gene | 2.2 | 4.6686 | 5.0E-04 | 1.7 | 3.174 | 8.2E-05 |
| *rpl-25.1* | WBGene00004438 | protein_coding_gene | 2.2 | 4.6681 | 1.6E-29 | 2.3 | 4.978 | 1.6E-82 |
| *ins-5* | WBGene00002088 | protein_coding_gene | 2.2 | 4.6637 | 3.9E-06 | 1.9 | 3.815 | 3.8E-04 |
| *C34F6.1* | WBGene00007938 | protein_coding_gene | 2.2 | 4.6609 | 1.4E-06 | 2.7 | 6.617 | 6.9E-04 |
| *F15A4.6* | WBGene00008841 | protein_coding_gene | 2.2 | 4.6584 | 5.2E-26 | 1.6 | 3.137 | 9.6E-26 |
| *F32A5.4* | WBGene00017970 | protein_coding_gene | 2.2 | 4.6579 | 5.7E-32 | 1.7 | 3.194 | 7.4E-41 |
| *Y20F4.8* | WBGene00044435 | protein_coding_gene | 2.2 | 4.6559 | 2.0E-02 | 1.6 | 2.936 | 2.9E-03 |
| *F07H5.4* | WBGene00014308 | pseudogene | 2.2 | 4.6551 | 3.9E-29 | 1.1 | 2.146 | 3.2E-13 |
| *fkh-9* | WBGene00001441 | protein_coding_gene | 2.2 | 4.6526 | 4.5E-08 | 1.3 | 2.489 | 7.4E-23 |
| *F19B10.4* | WBGene00017586 | protein_coding_gene | 2.2 | 4.6469 | 1.1E-04 | 1.6 | 2.951 | 3.0E-03 |
| *F21C10.3* | WBGene00017653 | protein_coding_gene | 2.2 | 4.6434 | 2.0E-13 | 1.4 | 2.622 | 1.1E-07 |
| *W06H8.6* | WBGene00021068 | protein_coding_gene | 2.2 | 4.6378 | 2.3E-06 | 1.5 | 2.804 | 1.7E-28 |
| *gcy-11* | WBGene00001537 | protein_coding_gene | 2.2 | 4.6358 | 8.6E-19 | 1.8 | 3.371 | 4.3E-17 |
| *C45E5.4* | WBGene00016667 | protein_coding_gene | 2.2 | 4.6358 | 1.4E-13 | 2.9 | 7.225 | 3.6E-28 |
| *C10C5.7* | WBGene00007511 | protein_coding_gene | 2.2 | 4.6352 | 1.6E-08 | 1.1 | 2.093 | 1.5E-03 |
| *cdka-1* | WBGene00011955 | protein_coding_gene | 2.2 | 4.6337 | 3.1E-12 | 1.8 | 3.433 | 2.2E-14 |
| *C54D10.5* | WBGene00008300 | protein_coding_gene | 2.2 | 4.6322 | 2.1E-04 | 2.4 | 5.233 | 3.2E-04 |
| *phg-1* | WBGene00004017 | protein_coding_gene | 2.2 | 4.6301 | 7.9E-05 | 1.2 | 2.319 | 3.3E-02 |
| *T02E1.8* | WBGene00011380 | protein_coding_gene | 2.2 | 4.6283 | 9.0E-08 | 1.4 | 2.729 | 8.1E-05 |
| *Y47H9C.1* | WBGene00012947 | protein_coding_gene | 2.2 | 4.6277 | 1.7E-04 | 3.1 | 8.531 | 9.4E-09 |
| *nlp-69* | WBGene00013752 | protein_coding_gene | 2.2 | 4.6259 | 9.2E-06 | 1.5 | 2.822 | 2.3E-03 |
| *T03F6.4* | WBGene00011400 | protein_coding_gene | 2.2 | 4.6257 | 1.6E-08 | 1.5 | 2.737 | 6.1E-06 |
| *lact-8* | WBGene00012777 | protein_coding_gene | 2.2 | 4.6206 | 1.8E-21 | 1.8 | 3.442 | 1.1E-12 |
| *chil-13* | WBGene00010945 | protein_coding_gene | 2.2 | 4.6195 | 3.0E-29 | 2.0 | 3.903 | 1.6E-20 |
| *F57C7.4* | WBGene00010200 | protein_coding_gene | 2.2 | 4.6169 | 4.5E-06 | 2.0 | 4.051 | 5.5E-06 |
| *K12D9.1* | WBGene00019675 | protein_coding_gene | 2.2 | 4.6103 | 4.9E-02 | 2.4 | 5.451 | 2.0E-02 |
| *C39D10.2* | WBGene00016530 | protein_coding_gene | 2.2 | 4.6069 | 2.7E-07 | 1.6 | 2.939 | 2.6E-06 |
| *Y55B1BR.6* | WBGene00044401 | protein_coding_gene | 2.2 | 4.6024 | 1.0E-09 | 1.5 | 2.895 | 2.1E-04 |
| *nlp-17* | WBGene00003755 | protein_coding_gene | 2.2 | 4.5996 | 1.8E-09 | 1.3 | 2.547 | 7.2E-04 |
| *pqn-54* | WBGene00004139 | protein_coding_gene | 2.2 | 4.5973 | 1.1E-04 | 1.5 | 2.902 | 1.9E-02 |
| *W05F2.7* | WBGene00021039 | protein_coding_gene | 2.2 | 4.5972 | 3.3E-09 | 1.6 | 3.023 | 7.2E-09 |
| *exc-5* | WBGene00001366 | protein_coding_gene | 2.2 | 4.5872 | 2.1E-14 | 1.5 | 2.750 | 5.0E-31 |
| *zig-5* | WBGene00006982 | protein_coding_gene | 2.2 | 4.5870 | 2.5E-10 | 1.7 | 3.265 | 1.4E-07 |
| *Y97E10C.1* | WBGene00022410 | protein_coding_gene | 2.2 | 4.5861 | 3.2E-10 | 1.2 | 2.289 | 4.1E-12 |
| *rgef-1* | WBGene00009100 | protein_coding_gene | 2.2 | 4.5851 | 2.1E-11 | 1.6 | 3.011 | 2.7E-08 |
| *ace-4* | WBGene00000038 | protein_coding_gene | 2.2 | 4.5808 | 2.0E-04 | 1.5 | 2.866 | 5.7E-03 |
| *phat-3* | WBGene00016782 | protein_coding_gene | 2.2 | 4.5770 | 1.9E-15 | 2.4 | 5.190 | 6.6E-06 |
| *ZC262.4* | WBGene00022581 | protein_coding_gene | 2.2 | 4.5768 | 1.2E-02 | 2.1 | 4.247 | 1.9E-04 |
| *sol-1* | WBGene00004944 | protein_coding_gene | 2.2 | 4.5767 | 1.3E-07 | 1.5 | 2.741 | 1.6E-04 |
| *col-49* | WBGene00000626 | protein_coding_gene | 2.2 | 4.5654 | 3.6E-06 | 4.0 | 15.685 | 5.5E-05 |
| *K03A11.5* | WBGene00010522 | protein_coding_gene | 2.2 | 4.5651 | 1.4E-11 | 1.6 | 2.977 | 7.5E-15 |
| *osm-10* | WBGene00003890 | protein_coding_gene | 2.2 | 4.5627 | 2.0E-09 | 1.8 | 3.478 | 1.9E-10 |
| *W05E10.2* | WBGene00012281 | protein_coding_gene | 2.2 | 4.5619 | 3.5E-04 | 2.3 | 4.849 | 2.7E-03 |
| *old-1* | WBGene00003862 | protein_coding_gene | 2.2 | 4.5606 | 1.6E-07 | 1.8 | 3.580 | 2.4E-03 |
| *col-48* | WBGene00000625 | protein_coding_gene | 2.2 | 4.5578 | 1.1E-07 | 2.0 | 4.067 | 3.0E-02 |
| *nlp-64* | WBGene00019734 | protein_coding_gene | 2.2 | 4.5568 | 2.2E-07 | 1.2 | 2.348 | 9.6E-04 |
| *F41C6.6* | WBGene00018279 | protein_coding_gene | 2.2 | 4.5516 | 3.9E-04 | 1.9 | 3.668 | 7.1E-03 |
| *C18H7.6* | WBGene00015996 | protein_coding_gene | 2.2 | 4.5499 | 9.8E-04 | 2.4 | 5.295 | 4.8E-04 |
| ***nep-16*** | **WBGene00018472** | **protein_coding_gene** | **2.2** | **4.5349** | **5.4E-03** | **-1.5** | **0.351** | **8.1E-03** |
| *M03C11.1* | WBGene00010838 | protein_coding_gene | 2.2 | 4.5289 | 2.0E-15 | 1.4 | 2.710 | 8.3E-09 |
| *lgc-41* | WBGene00008022 | protein_coding_gene | 2.2 | 4.5200 | 8.8E-04 | 1.9 | 3.721 | 1.2E-02 |

| | | | | | | | | |
|---|---|---|---|---|---|---|---|---|
| *C07A12.2* | WBGene00015560 | protein_coding_gene | 2.2 | 4.5181 | 4.9E-07 | 1.7 | 3.168 | 1.5E-06 |
| *npr-29* | WBGene00013848 | protein_coding_gene | 2.2 | 4.5180 | 2.1E-08 | 1.4 | 2.677 | 1.1E-03 |
| *R74.10* | WBGene00206525 | protein_coding_gene | 2.2 | 4.5147 | 1.7E-08 | 1.6 | 3.060 | 6.2E-07 |
| *fbxa-88* | WBGene00008632 | protein_coding_gene | 2.2 | 4.5139 | 6.4E-03 | 1.2 | 2.291 | 4.5E-02 |
| *nlp-3* | WBGene00003741 | protein_coding_gene | 2.2 | 4.5033 | 3.9E-17 | 1.5 | 2.756 | 1.1E-16 |
| *col-175* | WBGene00000748 | protein_coding_gene | 2.2 | 4.5032 | 1.6E-06 | 3.7 | 12.651 | 2.9E-04 |
| *Y110A2AL.9* | WBGene00022445 | protein_coding_gene | 2.2 | 4.4936 | 2.1E-07 | 2.6 | 5.926 | 1.9E-07 |
| *B0303.16* | WBGene00044678 | protein_coding_gene | 2.2 | 4.4920 | 3.3E-02 | 2.4 | 5.411 | 1.9E-02 |
| *Y55F3AM.14* | WBGene00021931 | protein_coding_gene | 2.2 | 4.4914 | 1.8E-05 | 4.3 | 19.067 | 4.7E-07 |
| ***bro-1*** | **WBGene00000272** | **protein_coding_gene** | **2.2** | **4.4853** | **7.9E-04** | **-1.2** | **0.439** | **2.0E-03** |
| *dmsr-6* | WBGene00013222 | protein_coding_gene | 2.2 | 4.4802 | 3.5E-06 | 1.5 | 2.756 | 5.7E-05 |
| *nlp-6* | WBGene00003744 | protein_coding_gene | 2.2 | 4.4792 | 4.6E-24 | 1.5 | 2.739 | 4.5E-20 |
| *ifp-1* | WBGene00002067 | protein_coding_gene | 2.2 | 4.4787 | 1.4E-12 | 2.2 | 4.515 | 1.7E-22 |
| *ttr-25* | WBGene00013078 | protein_coding_gene | 2.2 | 4.4785 | 3.5E-18 | 2.5 | 5.837 | 1.0E-71 |
| *C05D12.7* | WBGene00044029 | protein_coding_gene | 2.2 | 4.4728 | 1.0E-07 | 1.7 | 3.239 | 1.5E-07 |
| *F20G2.6* | WBGene00008989 | protein_coding_gene | 2.2 | 4.4722 | 8.8E-03 | 1.7 | 3.338 | 2.4E-02 |
| *marg-1* | WBGene00018538 | protein_coding_gene | 2.2 | 4.4722 | 8.4E-09 | 2.1 | 4.257 | 5.5E-27 |
| *K07A1.3* | WBGene00010610 | protein_coding_gene | 2.2 | 4.4721 | 5.4E-21 | 1.1 | 2.092 | 7.2E-11 |
| *srx-2* | WBGene00005893 | protein_coding_gene | 2.2 | 4.4690 | 3.6E-02 | 2.0 | 3.917 | 3.4E-02 |
| *lurp-1* | WBGene00009700 | protein_coding_gene | 2.2 | 4.4685 | 4.7E-10 | 1.0 | 2.029 | 3.6E-03 |
| *nep-15* | WBGene00018227 | protein_coding_gene | 2.2 | 4.4599 | 2.1E-03 | 2.5 | 5.641 | 1.4E-02 |
| *ZC449.4* | WBGene00022614 | protein_coding_gene | 2.2 | 4.4532 | 5.6E-10 | 1.2 | 2.374 | 4.7E-06 |
| *T01G1.2* | WBGene00011337 | protein_coding_gene | 2.2 | 4.4496 | 5.2E-16 | 1.6 | 3.137 | 8.8E-12 |
| *Y79H2A.12* | WBGene00043997 | protein_coding_gene | 2.2 | 4.4494 | 6.6E-04 | 1.3 | 2.470 | 2.5E-02 |
| *flp-7* | WBGene00001450 | protein_coding_gene | 2.2 | 4.4479 | 1.0E-19 | 1.4 | 2.667 | 1.8E-10 |
| *iff-2* | WBGene00002065 | protein_coding_gene | 2.2 | 4.4478 | 3.1E-41 | 2.5 | 5.742 | 7.6E-83 |
| *ZC196.1* | WBGene00022545 | pseudogene | 2.2 | 4.4469 | 3.7E-04 | 1.5 | 2.882 | 4.6E-04 |
| *C25G6.4* | WBGene00016109 | protein_coding_gene | 2.2 | 4.4465 | 2.1E-03 | 2.8 | 6.761 | 1.1E-04 |
| *cyp-13A10* | WBGene00014254 | protein_coding_gene | 2.2 | 4.4449 | 2.0E-05 | 1.5 | 2.749 | 9.7E-06 |
| *xdh-1* | WBGene00010083 | protein_coding_gene | 2.2 | 4.4438 | 3.3E-66 | 1.6 | 2.962 | 5.9E-27 |
| *T06D8.2* | WBGene00011523 | protein_coding_gene | 2.2 | 4.4412 | 1.3E-02 | 1.8 | 3.449 | 8.7E-03 |
| *nhr-129* | WBGene00003719 | protein_coding_gene | 2.2 | 4.4387 | 4.1E-16 | 1.8 | 3.544 | 5.4E-09 |
| *fbxa-59* | WBGene00020455 | protein_coding_gene | 2.1 | 4.4369 | 8.4E-12 | 1.8 | 3.472 | 7.0E-07 |
| *Y105C5A.25* | WBGene00013644 | pseudogene | 2.1 | 4.4354 | 1.3E-07 | 1.8 | 3.380 | 2.0E-04 |
| ***Y18H1A.8*** | **WBGene00021212** | **protein_coding_gene** | **2.1** | **4.4312** | **1.4E-03** | **-1.9** | **0.275** | **1.8E-06** |
| *C29F3.3* | WBGene00007804 | protein_coding_gene | 2.1 | 4.4266 | 4.5E-02 | 3.4 | 10.492 | 2.8E-03 |
| *T10E9.3* | WBGene00020413 | protein_coding_gene | 2.1 | 4.4213 | 2.7E-08 | 2.8 | 6.756 | 2.2E-15 |
| *abf-5* | WBGene00000016 | protein_coding_gene | 2.1 | 4.4186 | 2.2E-14 | 1.3 | 2.449 | 2.0E-07 |
| *eff-1* | WBGene00001159 | protein_coding_gene | 2.1 | 4.4184 | 2.6E-13 | 1.1 | 2.191 | 1.2E-03 |
| *kcnl-3* | WBGene00015387 | protein_coding_gene | 2.1 | 4.4183 | 6.3E-05 | 1.5 | 2.767 | 3.2E-03 |
| *myo-2* | WBGene00003514 | protein_coding_gene | 2.1 | 4.4170 | 2.3E-10 | 2.3 | 5.077 | 5.1E-42 |
| *ckk-1* | WBGene00000518 | protein_coding_gene | 2.1 | 4.4150 | 7.4E-12 | 1.4 | 2.634 | 2.0E-06 |
| *Y41D4A.7* | WBGene00021509 | protein_coding_gene | 2.1 | 4.4150 | 5.9E-10 | 1.5 | 2.800 | 6.2E-07 |
| *K02F2.5* | WBGene00019324 | protein_coding_gene | 2.1 | 4.4145 | 1.3E-16 | 1.8 | 3.540 | 1.2E-25 |
| *K07E1.1* | WBGene00019489 | protein_coding_gene | 2.1 | 4.4143 | 3.7E-13 | 2.1 | 4.207 | 3.0E-34 |
| *col-109* | WBGene00000683 | protein_coding_gene | 2.1 | 4.4124 | 3.1E-07 | 3.1 | 8.453 | 4.4E-04 |
| *dve-1* | WBGene00022861 | protein_coding_gene | 2.1 | 4.4061 | 1.4E-10 | 1.1 | 2.158 | 5.7E-12 |
| *F13B6.3* | WBGene00017418 | protein_coding_gene | 2.1 | 4.4027 | 4.5E-16 | 2.1 | 4.204 | 5.7E-06 |
| *ZK822.6* | WBGene00044009 | protein_coding_gene | 2.1 | 4.4011 | 2.0E-04 | 1.5 | 2.878 | 1.9E-03 |
| *C54D10.3* | WBGene00008298 | protein_coding_gene | 2.1 | 4.3997 | 3.2E-42 | 1.4 | 2.728 | 1.7E-37 |
| *nhr-103* | WBGene00003693 | protein_coding_gene | 2.1 | 4.3992 | 7.3E-08 | 1.1 | 2.103 | 4.0E-06 |
| *W02B12.1* | WBGene00012201 | protein_coding_gene | 2.1 | 4.3989 | 1.1E-32 | 1.4 | 2.556 | 2.0E-13 |
| *F53F4.17* | WBGene00045195 | protein_coding_gene | 2.1 | 4.3981 | 6.3E-05 | 1.9 | 3.834 | 1.4E-04 |
| *C56E10.3* | WBGene00016976 | protein_coding_gene | 2.1 | 4.3976 | 3.6E-08 | 1.1 | 2.170 | 1.1E-03 |
| *col-138* | WBGene00000711 | protein_coding_gene | 2.1 | 4.3890 | 8.0E-07 | 3.7 | 13.295 | 2.6E-04 |
| *col-144* | WBGene00000717 | protein_coding_gene | 2.1 | 4.3874 | 9.1E-07 | 2.6 | 6.250 | 1.0E-06 |
| *F54D1.6* | WBGene00010047 | protein_coding_gene | 2.1 | 4.3854 | 5.8E-09 | 1.2 | 2.242 | 1.6E-24 |
| *linc-53* | WBGene00219733 | lincRNA_gene | 2.1 | 4.3821 | 2.2E-02 | 2.0 | 4.118 | 2.5E-02 |
| *F08B12.4* | WBGene00008572 | protein_coding_gene | 2.1 | 4.3794 | 8.6E-11 | 1.2 | 2.310 | 5.5E-12 |
| ***F26F12.4*** | **WBGene00017835** | **protein_coding_gene** | **2.1** | **4.3790** | **2.4E-05** | **-1.3** | **0.420** | **1.1E-02** |
| *Y9C2UA.1* | WBGene00012421 | protein_coding_gene | 2.1 | 4.3724 | 6.3E-21 | 1.4 | 2.729 | 6.7E-14 |
| *F23A7.3* | WBGene00009068 | protein_coding_gene | 2.1 | 4.3698 | 8.4E-08 | 1.2 | 2.333 | 1.9E-05 |
| *C13A10.2* | WBGene00015729 | protein_coding_gene | 2.1 | 4.3689 | 4.8E-06 | 1.2 | 2.276 | 8.4E-03 |
| *col-73* | WBGene00000649 | protein_coding_gene | 2.1 | 4.3667 | 2.5E-05 | 3.0 | 7.763 | 1.5E-02 |
| *glb-22* | WBGene00011251 | protein_coding_gene | 2.1 | 4.3665 | 1.6E-04 | 3.0 | 7.973 | 3.8E-07 |
| *phat-6* | WBGene00011683 | protein_coding_gene | 2.1 | 4.3623 | 8.6E-11 | 1.5 | 2.759 | 1.8E-12 |
| *srtx-1* | WBGene00015214 | protein_coding_gene | 2.1 | 4.3620 | 1.5E-02 | 2.2 | 4.694 | 1.4E-02 |
| *col-39* | WBGene00000616 | protein_coding_gene | 2.1 | 4.3554 | 1.2E-05 | 3.0 | 8.062 | 4.7E-65 |
| *lury-1* | WBGene00013548 | protein_coding_gene | 2.1 | 4.3497 | 1.7E-06 | 1.1 | 2.118 | 4.6E-04 |
| *pgal-1* | WBGene00017671 | protein_coding_gene | 2.1 | 4.3439 | 2.3E-21 | 1.5 | 2.904 | 9.3E-43 |
| *col-71* | WBGene00000647 | protein_coding_gene | 2.1 | 4.3405 | 1.4E-06 | 3.6 | 12.409 | 4.9E-05 |
| *ets-4* | WBGene00017687 | protein_coding_gene | 2.1 | 4.3364 | 1.6E-14 | 2.2 | 4.755 | 1.1E-52 |
| *glc-3* | WBGene00001593 | protein_coding_gene | 2.1 | 4.3323 | 4.0E-04 | 1.3 | 2.520 | 2.1E-02 |
| *flp-32* | WBGene00010982 | protein_coding_gene | 2.1 | 4.3308 | 1.3E-11 | 1.3 | 2.428 | 6.3E-09 |
| *C18G1.1* | WBGene00015980 | protein_coding_gene | 2.1 | 4.3302 | 7.7E-07 | 1.6 | 2.943 | 2.7E-04 |
| *nlp-43* | WBGene00016684 | protein_coding_gene | 2.1 | 4.3269 | 3.3E-16 | 1.6 | 2.939 | 9.2E-11 |
| *flp-24* | WBGene00016028 | protein_coding_gene | 2.1 | 4.3246 | 2.3E-33 | 1.7 | 3.326 | 1.4E-24 |
| *F59B8.1* | WBGene00010316 | protein_coding_gene | 2.1 | 4.3245 | 4.8E-07 | 1.1 | 2.116 | 1.6E-02 |
| *nhr-250* | WBGene00022755 | pseudogene | 2.1 | 4.3232 | 1.5E-02 | 1.4 | 2.687 | 2.7E-02 |
| *myo-3* | WBGene00003515 | protein_coding_gene | 2.1 | 4.3139 | 9.0E-17 | 1.1 | 2.103 | 1.2E-13 |

| | | | | | | | | |
|---|---|---|---|---|---|---|---|---|
| *T26G10.5* | WBGene00012062 | protein_coding_gene | 2.1 | 4.3129 | 1.4E-05 | 1.8 | 3.589 | 1.9E-04 |
| *C15F1.2* | WBGene00015795 | protein_coding_gene | 2.1 | 4.2963 | 9.5E-06 | 2.9 | 7.416 | 5.8E-05 |
| *T27F6.8* | WBGene00012107 | protein_coding_gene | 2.1 | 4.2943 | 1.2E-09 | 1.9 | 3.731 | 7.1E-14 |
| *F53E10.1* | WBGene00018758 | protein_coding_gene | 2.1 | 4.2941 | 3.3E-27 | 1.3 | 2.395 | 2.0E-17 |
| *igeg-2* | WBGene00009842 | protein_coding_gene | 2.1 | 4.2908 | 2.7E-07 | 1.9 | 3.641 | 1.1E-06 |
| *F55F3.4* | WBGene00010118 | protein_coding_gene | 2.1 | 4.2855 | 2.8E-03 | 2.9 | 7.655 | 4.3E-06 |
| *chil-1* | WBGene00007465 | protein_coding_gene | 2.1 | 4.2831 | 2.5E-04 | 1.9 | 3.849 | 8.7E-03 |
| *K04D7.6* | WBGene00010559 | protein_coding_gene | 2.1 | 4.2824 | 8.3E-05 | 1.8 | 3.594 | 1.2E-03 |
| *nceh-1* | WBGene00012810 | protein_coding_gene | 2.1 | 4.2791 | 3.4E-10 | 2.9 | 7.375 | 5.9E-15 |
| *Y20C6A.1* | WBGene00012491 | protein_coding_gene | 2.1 | 4.2786 | 2.6E-09 | 1.4 | 2.639 | 6.1E-15 |
| *ZK899.7* | WBGene00014145 | protein_coding_gene | 2.1 | 4.2765 | 1.9E-03 | 1.4 | 2.727 | 3.6E-02 |
| *C05C12.4* | WBGene00007336 | protein_coding_gene | 2.1 | 4.2722 | 1.1E-11 | 1.2 | 2.283 | 2.7E-23 |
| *F49E2.5* | WBGene00009888 | protein_coding_gene | 2.1 | 4.2694 | 6.6E-08 | 2.1 | 4.185 | 8.3E-54 |
| ***F49D11.6*** | **WBGene00018630** | **protein_coding_gene** | **2.1** | **4.2672** | **7.9E-03** | **-1.9** | **0.266** | **1.1E-02** |
| *sms-5* | WBGene00021075 | protein_coding_gene | 2.1 | 4.2471 | 7.7E-11 | 1.8 | 3.439 | 1.7E-08 |
| *mef-2* | WBGene00003182 | protein_coding_gene | 2.1 | 4.2464 | 1.6E-19 | 1.5 | 2.754 | 4.3E-20 |
| *C45G7.13* | WBGene00235340 | protein_coding_gene | 2.1 | 4.2460 | 3.1E-09 | 1.9 | 3.796 | 5.6E-11 |
| *C31C9.7* | WBGene00007841 | protein_coding_gene | 2.1 | 4.2460 | 1.7E-08 | 1.8 | 3.368 | 4.8E-10 |
| *deg-1* | WBGene00000950 | protein_coding_gene | 2.1 | 4.2390 | 9.6E-10 | 1.2 | 2.300 | 2.1E-04 |
| *sue-1* | WBGene00006306 | protein_coding_gene | 2.1 | 4.2331 | 8.1E-10 | 2.1 | 4.164 | 1.0E-16 |
| *B0294.1* | WBGene00015117 | protein_coding_gene | 2.1 | 4.2291 | 7.0E-06 | 2.9 | 7.674 | 1.3E-08 |
| *daf-7* | WBGene00000903 | protein_coding_gene | 2.1 | 4.2281 | 8.3E-13 | 1.9 | 3.766 | 6.7E-13 |
| *unc-55* | WBGene00006790 | protein_coding_gene | 2.1 | 4.2266 | 5.7E-04 | 1.5 | 2.804 | 1.4E-02 |
| ***try-9*** | **WBGene00023425** | **protein_coding_gene** | **2.1** | **4.2242** | **5.4E-04** | **-2.8** | **0.144** | **2.7E-11** |
| *F13B12.3* | WBGene00008731 | protein_coding_gene | 2.1 | 4.2204 | 5.2E-13 | 1.5 | 2.735 | 3.8E-08 |
| *F35E12.4* | WBGene00009428 | pseudogene | 2.1 | 4.2199 | 6.2E-03 | 1.1 | 2.125 | 6.6E-04 |
| *Y95B8A.2* | WBGene00022383 | protein_coding_gene | 2.1 | 4.2181 | 5.1E-06 | 3.5 | 11.564 | 6.2E-16 |
| *K11H3.5* | WBGene00010782 | protein_coding_gene | 2.1 | 4.2123 | 4.4E-06 | 2.4 | 5.188 | 2.1E-05 |
| *C49G7.3* | WBGene00016781 | protein_coding_gene | 2.1 | 4.2112 | 5.2E-16 | 1.8 | 3.399 | 7.1E-55 |
| *ZK131.11* | WBGene00013934 | protein_coding_gene | 2.1 | 4.2097 | 6.9E-14 | 1.2 | 2.326 | 3.2E-21 |
| *dop-2* | WBGene00001053 | protein_coding_gene | 2.1 | 4.2078 | 8.7E-11 | 1.3 | 2.406 | 1.3E-05 |
| *C30F2.3* | WBGene00007817 | protein_coding_gene | 2.1 | 4.2036 | 1.6E-12 | 2.5 | 5.819 | 1.0E-14 |
| *C05D2.11* | WBGene00022910 | ncRNA_gene | 2.1 | 4.2029 | 2.8E-11 | 1.5 | 2.847 | 2.2E-18 |
| *D1005.2* | WBGene00016996 | protein_coding_gene | 2.1 | 4.2021 | 6.3E-08 | 1.3 | 2.426 | 4.1E-06 |
| *F01E11.18* | WBGene00219373 | protein_coding_gene | 2.1 | 4.1985 | 2.9E-06 | 1.2 | 2.286 | 1.9E-03 |
| *C32D5.1* | WBGene00016310 | protein_coding_gene | 2.1 | 4.1961 | 8.7E-12 | 1.2 | 2.291 | 3.3E-05 |
| *nhr-143* | WBGene00019116 | protein_coding_gene | 2.1 | 4.1956 | 2.0E-11 | 1.2 | 2.320 | 8.9E-10 |
| *Y46G5A.20* | WBGene00012910 | protein_coding_gene | 2.1 | 4.1943 | 1.1E-07 | 1.0 | 2.033 | 7.9E-03 |
| *T28B8.3* | WBGene00012114 | protein_coding_gene | 2.1 | 4.1936 | 9.8E-05 | 2.7 | 6.670 | 2.4E-07 |
| *K02E11.4* | WBGene00010513 | protein_coding_gene | 2.1 | 4.1878 | 2.9E-22 | 1.1 | 2.122 | 6.3E-07 |
| *fbxa-61* | WBGene00022486 | protein_coding_gene | 2.1 | 4.1835 | 3.9E-11 | 1.7 | 3.192 | 1.9E-13 |
| *Y43F8B.3* | WBGene00012814 | protein_coding_gene | 2.1 | 4.1833 | 7.6E-06 | 1.3 | 2.478 | 1.6E-03 |
| *F15D4.4* | WBGene00008861 | protein_coding_gene | 2.1 | 4.1831 | 1.1E-09 | 2.2 | 4.742 | 9.5E-15 |
| *mif-4* | WBGene00003237 | protein_coding_gene | 2.1 | 4.1785 | 2.7E-04 | 1.6 | 3.032 | 3.4E-03 |
| *cmk-1* | WBGene00000553 | protein_coding_gene | 2.1 | 4.1781 | 9.9E-09 | 1.6 | 3.120 | 3.9E-09 |
| *C05E7.3* | WBGene00007345 | protein_coding_gene | 2.1 | 4.1762 | 6.0E-06 | 2.0 | 3.992 | 5.9E-05 |
| *T27A10.2* | WBGene00020844 | protein_coding_gene | 2.1 | 4.1758 | 1.5E-05 | 2.0 | 4.049 | 1.7E-03 |
| *ins-30* | WBGene00002113 | protein_coding_gene | 2.1 | 4.1742 | 2.9E-07 | 4.3 | 19.425 | 1.4E-09 |
| *C49A9.1* | WBGene00016755 | protein_coding_gene | 2.1 | 4.1725 | 3.8E-05 | 1.3 | 2.495 | 9.4E-03 |
| *npr-7* | WBGene00018067 | protein_coding_gene | 2.1 | 4.1710 | 4.1E-05 | 2.3 | 4.968 | 9.9E-05 |
| *M01H9.3* | WBGene00019719 | protein_coding_gene | 2.1 | 4.1685 | 3.9E-24 | 1.8 | 3.513 | 6.7E-42 |
| *Y17G7B.8* | WBGene00012462 | protein_coding_gene | 2.1 | 4.1682 | 1.2E-15 | 1.1 | 2.118 | 8.3E-08 |
| ***C14F11.6*** | **WBGene00015781** | **protein_coding_gene** | **2.1** | **4.1674** | **8.8E-05** | **-1.6** | **0.326** | **7.2E-03** |
| *F23H11.6* | WBGene00017761 | protein_coding_gene | 2.1 | 4.1626 | 7.5E-03 | 2.0 | 3.921 | 1.2E-02 |
| *Y43F8C.11* | WBGene00012833 | protein_coding_gene | 2.1 | 4.1592 | 2.5E-04 | 1.7 | 3.341 | 1.5E-02 |
| *ZK795.6* | WBGene00194644 | pseudogene | 2.1 | 4.1574 | 1.3E-02 | 1.6 | 3.131 | 2.7E-02 |
| *ZC132.9* | WBGene00022527 | protein_coding_gene | 2.1 | 4.1485 | 1.6E-02 | 1.8 | 3.562 | 1.9E-02 |
| *C24B9.3* | WBGene00016048 | protein_coding_gene | 2.1 | 4.1477 | 1.1E-21 | 1.2 | 2.239 | 2.6E-05 |
| *C02B8.3* | WBGene00015322 | protein_coding_gene | 2.1 | 4.1461 | 3.9E-09 | 2.3 | 4.853 | 6.8E-10 |
| ***C31B8.12*** | **WBGene00016285** | **protein_coding_gene** | **2.1** | **4.1460** | **4.4E-02** | **-2.5** | **0.178** | **6.0E-06** |
| *nhr-59* | WBGene00003649 | protein_coding_gene | 2.0 | 4.1406 | 1.2E-11 | 1.2 | 2.361 | 1.6E-05 |
| *clik-2* | WBGene00016898 | protein_coding_gene | 2.0 | 4.1388 | 1.6E-12 | 1.8 | 3.386 | 4.2E-36 |
| *col-110* | WBGene00000684 | protein_coding_gene | 2.0 | 4.1382 | 1.0E-05 | 3.6 | 11.978 | 1.2E-04 |
| *C03H5.6* | WBGene00015408 | protein_coding_gene | 2.0 | 4.1365 | 5.0E-06 | 1.9 | 3.835 | 7.9E-05 |
| *C44H4.6* | WBGene00008095 | protein_coding_gene | 2.0 | 4.1347 | 1.2E-04 | 1.9 | 3.855 | 6.1E-04 |
| *F08G5.3* | WBGene00008582 | protein_coding_gene | 2.0 | 4.1293 | 4.6E-16 | 1.1 | 2.122 | 2.3E-06 |
| *bcl-11* | WBGene00017430 | protein_coding_gene | 2.0 | 4.1268 | 5.0E-09 | 1.8 | 3.599 | 6.2E-45 |
| *fipr-3* | WBGene00009090 | protein_coding_gene | 2.0 | 4.1261 | 6.4E-08 | 1.8 | 3.515 | 6.5E-08 |
| *C30F2.2* | WBGene00007816 | protein_coding_gene | 2.0 | 4.1227 | 2.0E-03 | 1.9 | 3.813 | 9.4E-03 |
| *npr-34* | WBGene00013187 | protein_coding_gene | 2.0 | 4.1178 | 4.4E-04 | 2.1 | 4.433 | 4.0E-04 |
| *gar-3* | WBGene00001519 | protein_coding_gene | 2.0 | 4.1142 | 5.0E-10 | 1.3 | 2.527 | 1.4E-07 |
| *gcy-23* | WBGene00001548 | protein_coding_gene | 2.0 | 4.1121 | 6.8E-05 | 2.1 | 4.363 | 1.6E-05 |
| *mak-2* | WBGene00007007 | protein_coding_gene | 2.0 | 4.1114 | 4.0E-38 | 1.6 | 3.107 | 9.4E-27 |
| *nlp-19* | WBGene00003757 | protein_coding_gene | 2.0 | 4.1100 | 2.3E-04 | 1.7 | 3.245 | 3.6E-04 |
| *flp-22* | WBGene00009562 | protein_coding_gene | 2.0 | 4.1081 | 4.3E-19 | 1.2 | 2.359 | 1.4E-17 |
| *nlp-67* | WBGene00011227 | protein_coding_gene | 2.0 | 4.1061 | 2.8E-05 | 2.0 | 3.973 | 5.2E-07 |
| *acs-16* | WBGene00018579 | protein_coding_gene | 2.0 | 4.1015 | 2.3E-08 | 1.0 | 2.057 | 1.5E-03 |
| *ubc-17* | WBGene00006712 | protein_coding_gene | 2.0 | 4.0972 | 5.0E-08 | 1.3 | 2.387 | 2.7E-04 |
| *dmsr-5* | WBGene00012962 | protein_coding_gene | 2.0 | 4.0939 | 1.0E-05 | 1.5 | 2.880 | 2.9E-03 |
| *B0302.5* | WBGene00015122 | protein_coding_gene | 2.0 | 4.0920 | 3.7E-13 | 1.0 | 2.054 | 5.2E-08 |

| | | | | | | | | |
|---|---|---|---|---|---|---|---|---|
| *H10E21.5* | WBGene00019185 | protein_coding_gene | 2.0 | 4.0898 | 6.7E-11 | 1.8 | 3.412 | 5.8E-10 |
| *C45E1.4* | WBGene00016663 | protein_coding_gene | 2.0 | 4.0830 | 1.1E-11 | 2.5 | 5.787 | 5.0E-24 |
| *K02E7.6* | WBGene00019311 | pseudogene | 2.0 | 4.0826 | 6.8E-15 | 1.3 | 2.433 | 1.8E-02 |
| *flp-6* | WBGene00001449 | protein_coding_gene | 2.0 | 4.0825 | 2.5E-17 | 1.4 | 2.648 | 2.1E-26 |
| *marc-5* | WBGene00013273 | protein_coding_gene | 2.0 | 4.0811 | 2.8E-09 | 1.5 | 2.750 | 4.6E-11 |
| *pmp-1* | WBGene00004058 | protein_coding_gene | 2.0 | 4.0799 | 1.5E-14 | 3.3 | 9.796 | 2.4E-97 |
| *pqn-62* | WBGene00004145 | protein_coding_gene | 2.0 | 4.0779 | 3.9E-12 | 1.0 | 2.015 | 9.8E-06 |
| *nlt-1* | WBGene00003773 | protein_coding_gene | 2.0 | 4.0706 | 2.1E-38 | 1.1 | 2.197 | 2.7E-20 |
| *nep-11* | WBGene00017557 | protein_coding_gene | 2.0 | 4.0700 | 4.3E-09 | 1.7 | 3.328 | 2.2E-21 |
| *T20B6.3* | WBGene00020599 | protein_coding_gene | 2.0 | 4.0638 | 1.1E-04 | 1.5 | 2.900 | 1.4E-02 |
| *haf-4* | WBGene00001814 | protein_coding_gene | 2.0 | 4.0615 | 4.3E-30 | 2.4 | 5.231 | 3.3E-62 |
| *nhr-88* | WBGene00003678 | protein_coding_gene | 2.0 | 4.0584 | 6.4E-08 | 1.4 | 2.568 | 1.3E-14 |
| *F46C8.8* | WBGene00018487 | protein_coding_gene | 2.0 | 4.0578 | 4.0E-09 | 1.6 | 2.977 | 3.3E-02 |
| *Y48C3A.5* | WBGene00012989 | protein_coding_gene | 2.0 | 4.0567 | 2.0E-14 | 1.5 | 2.743 | 2.8E-08 |
| *hot-4* | WBGene00001989 | protein_coding_gene | 2.0 | 4.0539 | 2.9E-07 | 1.0 | 2.029 | 1.3E-02 |
| *T02D1.4* | WBGene00011371 | protein_coding_gene | 2.0 | 4.0516 | 4.3E-03 | 1.6 | 3.013 | 2.0E-02 |
| *F14H12.8* | WBGene00017475 | protein_coding_gene | 2.0 | 4.0503 | 1.2E-05 | 2.1 | 4.148 | 1.6E-06 |
| *T22B7.8* | WBGene00044570 | protein_coding_gene | 2.0 | 4.0498 | 3.0E-10 | 1.3 | 2.523 | 4.8E-05 |
| *D2096.6* | WBGene00017073 | protein_coding_gene | 2.0 | 4.0449 | 1.7E-05 | 1.2 | 2.358 | 1.8E-02 |
| *dpy-9* | WBGene00001071 | protein_coding_gene | 2.0 | 4.0435 | 2.3E-07 | 3.9 | 15.273 | 7.1E-08 |
| *twk-8* | WBGene00006663 | protein_coding_gene | 2.0 | 4.0383 | 5.4E-13 | 1.7 | 3.300 | 1.4E-21 |
| *npr-18* | WBGene00008065 | protein_coding_gene | 2.0 | 4.0306 | 1.5E-04 | 1.7 | 3.350 | 1.2E-03 |
| ***C07A9.5*** | **WBGene00007401** | **protein_coding_gene** | **2.0** | **4.0277** | **4.7E-02** | **-2.1** | **0.238** | **4.8E-02** |
| *hpo-6* | WBGene00021518 | protein_coding_gene | 2.0 | 4.0265 | 3.0E-12 | 1.9 | 3.676 | 3.8E-05 |
| *gar-1* | WBGene00001517 | protein_coding_gene | 2.0 | 4.0261 | 6.5E-07 | 1.4 | 2.565 | 1.2E-05 |
| *K04C2.7* | WBGene00044311 | protein_coding_gene | 2.0 | 4.0256 | 4.0E-04 | 1.2 | 2.298 | 1.6E-02 |
| *flp-11* | WBGene00001454 | protein_coding_gene | 2.0 | 4.0256 | 1.2E-11 | 2.1 | 4.258 | 3.9E-26 |
| *nas-4* | WBGene00003523 | protein_coding_gene | 2.0 | 4.0253 | 4.8E-07 | 2.0 | 4.086 | 2.0E-04 |
| *clh-2* | WBGene00000529 | protein_coding_gene | 2.0 | 4.0252 | 5.6E-08 | 1.2 | 2.357 | 3.9E-04 |
| *C01B10.6* | WBGene00015280 | protein_coding_gene | 2.0 | 4.0252 | 3.5E-10 | 1.5 | 2.805 | 2.1E-20 |
| *W03F9.11* | WBGene00044512 | protein_coding_gene | 2.0 | 4.0234 | 1.8E-04 | 2.1 | 4.250 | 4.2E-04 |
| *asic-1* | WBGene00022815 | protein_coding_gene | 2.0 | 4.0230 | 7.7E-06 | 1.9 | 3.824 | 5.1E-05 |
| *col-115* | WBGene00000689 | protein_coding_gene | 2.0 | 4.0226 | 2.8E-04 | 2.0 | 3.929 | 8.0E-03 |
| *K02A6.3* | WBGene00019294 | protein_coding_gene | 2.0 | 4.0225 | 1.0E-08 | 1.1 | 2.072 | 1.6E-08 |
| *mec-2* | WBGene00003166 | protein_coding_gene | 2.0 | 4.0170 | 1.5E-10 | 1.5 | 2.784 | 2.0E-08 |
| *C04E7.3* | WBGene00015430 | protein_coding_gene | 2.0 | 4.0161 | 2.3E-09 | 1.2 | 2.230 | 4.0E-11 |
| *Y113G7A.16* | WBGene00013753 | protein_coding_gene | 2.0 | 4.0148 | 1.1E-02 | 2.5 | 5.558 | 4.0E-04 |
| *twk-29* | WBGene00006681 | protein_coding_gene | 2.0 | 4.0106 | 1.6E-05 | 1.1 | 2.161 | 4.4E-03 |
| *Y39B6A.33* | WBGene00012692 | protein_coding_gene | 2.0 | 4.0088 | 1.4E-22 | 1.9 | 3.697 | 1.2E-44 |
| *F59B10.3* | WBGene00010319 | protein_coding_gene | 2.0 | 4.0043 | 2.0E-04 | 1.5 | 2.873 | 3.5E-03 |
| *C47E8.9* | WBGene00008146 | protein_coding_gene | 2.0 | 4.0011 | 3.3E-02 | 2.5 | 5.551 | 2.5E-04 |
| *K09E9.4* | WBGene00044143 | protein_coding_gene | 2.0 | 4.0000 | 2.9E-07 | 1.7 | 3.333 | 6.0E-09 |
| *T14G8.6* | WBGene00304822 | protein_coding_gene | 2.0 | 3.9948 | 4.7E-03 | 2.6 | 6.160 | 4.7E-04 |
| *kvs-4* | WBGene00012967 | protein_coding_gene | 2.0 | 3.9939 | 1.9E-06 | 1.8 | 3.546 | 1.4E-04 |
| *lea-1* | WBGene00002263 | protein_coding_gene | 2.0 | 3.9922 | 4.2E-15 | 2.0 | 4.097 | 9.4E-52 |
| ***oac-6*** | **WBGene00007833** | **protein_coding_gene** | **2.0** | **3.9919** | **3.4E-06** | **-1.2** | **0.435** | **3.0E-03** |
| *Y47G6A.5* | WBGene00021635 | protein_coding_gene | 2.0 | 3.9914 | 1.4E-08 | 1.2 | 2.269 | 4.0E-16 |
| *Y62F5A.10* | WBGene00013388 | protein_coding_gene | 2.0 | 3.9872 | 3.4E-06 | 1.1 | 2.087 | 3.4E-04 |
| *C01B9.1* | WBGene00007220 | protein_coding_gene | 2.0 | 3.9850 | 2.1E-08 | 3.2 | 9.453 | 2.5E-32 |
| *gcy-32* | WBGene00001552 | protein_coding_gene | 2.0 | 3.9820 | 3.5E-04 | 2.5 | 5.645 | 4.9E-06 |
| *C39F7.1* | WBGene00016538 | protein_coding_gene | 2.0 | 3.9809 | 4.7E-04 | 1.6 | 2.957 | 1.0E-02 |
| *CD4.1* | WBGene00016988 | protein_coding_gene | 2.0 | 3.9803 | 4.5E-13 | 1.4 | 2.722 | 5.0E-07 |
| *Y39E4B.6* | WBGene00012717 | protein_coding_gene | 2.0 | 3.9799 | 4.7E-14 | 1.3 | 2.388 | 1.4E-18 |
| *R151.1* | WBGene00020106 | protein_coding_gene | 2.0 | 3.9786 | 7.0E-15 | 2.1 | 4.224 | 2.0E-12 |
| *Y44A6D.1* | WBGene00012852 | protein_coding_gene | 2.0 | 3.9783 | 3.7E-02 | 2.2 | 4.585 | 1.1E-02 |
| *C29E4.10* | WBGene00016207 | protein_coding_gene | 2.0 | 3.9767 | 6.2E-06 | 1.3 | 2.487 | 1.4E-03 |
| *dhs-4* | WBGene00000968 | protein_coding_gene | 2.0 | 3.9751 | 2.5E-12 | 1.3 | 2.426 | 4.7E-06 |
| *sul-3* | WBGene00006310 | protein_coding_gene | 2.0 | 3.9747 | 4.0E-12 | 1.5 | 2.778 | 2.6E-06 |
| *C01G12.7* | WBGene00007247 | protein_coding_gene | 2.0 | 3.9704 | 1.1E-03 | 2.5 | 5.715 | 2.9E-05 |
| *cho-1* | WBGene00000501 | protein_coding_gene | 2.0 | 3.9684 | 2.0E-05 | 2.0 | 3.905 | 8.2E-08 |
| *F41B5.11* | WBGene00023039 | pseudogene | 2.0 | 3.9672 | 1.2E-03 | 1.8 | 3.530 | 1.2E-06 |
| *W08A12.2* | WBGene00021079 | protein_coding_gene | 2.0 | 3.9636 | 5.8E-06 | 1.0 | 2.044 | 1.2E-07 |
| *cyp-33C3* | WBGene00018262 | protein_coding_gene | 2.0 | 3.9580 | 3.1E-10 | 1.9 | 3.827 | 3.7E-16 |
| *col-63* | WBGene00000639 | protein_coding_gene | 2.0 | 3.9554 | 3.9E-06 | 3.6 | 12.530 | 3.8E-04 |
| *C15B12.4* | WBGene00015786 | protein_coding_gene | 2.0 | 3.9550 | 2.4E-09 | 1.1 | 2.085 | 6.7E-06 |
| *C05E7.1* | WBGene00007343 | protein_coding_gene | 2.0 | 3.9513 | 9.3E-08 | 1.2 | 2.305 | 4.5E-04 |
| *nlp-46* | WBGene00012145 | protein_coding_gene | 2.0 | 3.9512 | 3.8E-02 | 1.4 | 2.616 | 1.3E-02 |
| *F19G12.9* | WBGene00195185 | protein_coding_gene | 2.0 | 3.9504 | 1.3E-11 | 1.2 | 2.243 | 1.6E-04 |
| *Y43B11AR.3* | WBGene00021544 | protein_coding_gene | 2.0 | 3.9474 | 6.2E-13 | 1.0 | 2.062 | 1.5E-09 |
| *C25G6.1* | WBGene00016107 | protein_coding_gene | 2.0 | 3.9468 | 4.0E-03 | 2.3 | 5.065 | 8.8E-04 |
| *nep-26* | WBGene00014171 | protein_coding_gene | 2.0 | 3.9390 | 1.5E-11 | 1.3 | 2.450 | 2.1E-09 |
| *pqn-18* | WBGene00004108 | protein_coding_gene | 2.0 | 3.9382 | 1.7E-08 | 1.4 | 2.615 | 5.0E-18 |
| *ncx-10* | WBGene00003575 | protein_coding_gene | 2.0 | 3.9320 | 9.6E-06 | 1.1 | 2.163 | 3.9E-03 |
| *Y49E10.16* | WBGene00013035 | protein_coding_gene | 2.0 | 3.9309 | 1.3E-26 | 1.6 | 3.067 | 4.6E-36 |
| *snt-3* | WBGene00004923 | protein_coding_gene | 2.0 | 3.9306 | 1.0E-09 | 1.3 | 2.524 | 3.6E-06 |
| *bus-2* | WBGene00044618 | protein_coding_gene | 2.0 | 3.9299 | 1.7E-04 | 1.3 | 2.444 | 1.9E-02 |
| *Y105C5A.24* | WBGene00013643 | protein_coding_gene | 2.0 | 3.9271 | 5.2E-13 | 1.3 | 2.424 | 8.7E-13 |
| *F59C6.11* | WBGene00010328 | protein_coding_gene | 2.0 | 3.9214 | 4.5E-11 | 1.3 | 2.432 | 6.2E-06 |
| *F47B8.8* | WBGene00009809 | protein_coding_gene | 2.0 | 3.9188 | 2.1E-12 | 1.5 | 2.910 | 8.7E-14 |
| *gcy-8* | WBGene00001535 | protein_coding_gene | 2.0 | 3.9183 | 5.2E-06 | 1.1 | 2.200 | 4.5E-02 |

| | | | | | | | | |
|---|---|---|---|---|---|---|---|---|
| *F34H10.4* | WBGene00009381 | protein_coding_gene | 2.0 | 3.9147 | 5.8E-04 | 3.2 | 9.207 | 6.7E-05 |
| *lin-29* | WBGene00003015 | protein_coding_gene | 2.0 | 3.9142 | 2.3E-08 | 2.6 | 6.220 | 1.1E-41 |
| *vhp-1* | WBGene00006923 | protein_coding_gene | 2.0 | 3.9077 | 1.8E-10 | 1.7 | 3.236 | 1.2E-38 |
| *F46G10.2* | WBGene00009797 | protein_coding_gene | 2.0 | 3.9017 | 1.4E-14 | 1.1 | 2.175 | 5.8E-12 |
| *Y53F4B.12* | WBGene00013159 | protein_coding_gene | 2.0 | 3.8983 | 4.2E-06 | 1.3 | 2.383 | 3.1E-04 |
| *ZC412.3* | WBGene00013884 | protein_coding_gene | 2.0 | 3.8968 | 1.9E-10 | 1.0 | 2.001 | 1.4E-10 |
| *acr-20* | WBGene00000059 | protein_coding_gene | 2.0 | 3.8939 | 8.6E-05 | 1.9 | 3.762 | 9.1E-05 |
| *flp-26* | WBGene00020126 | protein_coding_gene | 2.0 | 3.8932 | 3.8E-09 | 1.6 | 2.947 | 1.6E-09 |
| *T02B11.4* | WBGene00020156 | protein_coding_gene | 2.0 | 3.8925 | 1.6E-04 | 1.6 | 3.106 | 1.5E-10 |
| *C13E3.1* | WBGene00015740 | protein_coding_gene | 2.0 | 3.8925 | 3.7E-11 | 1.0 | 2.006 | 2.9E-05 |
| *F34D10.6* | WBGene00009375 | protein_coding_gene | 2.0 | 3.8888 | 3.3E-09 | 1.6 | 3.034 | 1.8E-06 |
| *nhr-162* | WBGene00016367 | protein_coding_gene | 2.0 | 3.8811 | 1.4E-05 | 1.4 | 2.558 | 1.1E-03 |
| *C25H3.1* | WBGene00016111 | protein_coding_gene | 2.0 | 3.8766 | 1.1E-03 | 1.5 | 2.741 | 1.1E-02 |
| *ttr-43* | WBGene00007590 | protein_coding_gene | 2.0 | 3.8763 | 7.4E-08 | 1.1 | 2.207 | 9.9E-04 |
| *F26A3.1* | WBGene00009140 | protein_coding_gene | 2.0 | 3.8755 | 2.4E-40 | 1.2 | 2.254 | 2.2E-14 |
| *T14G8.4* | WBGene00011772 | protein_coding_gene | 2.0 | 3.8753 | 1.2E-04 | 1.9 | 3.737 | 9.0E-05 |
| *M03F8.5* | WBGene00019764 | protein_coding_gene | 2.0 | 3.8738 | 2.2E-04 | 1.8 | 3.408 | 2.1E-03 |
| *snpc-1.3* | WBGene00017265 | protein_coding_gene | 2.0 | 3.8683 | 5.5E-09 | 2.0 | 3.983 | 4.0E-13 |
| *col-167* | WBGene00000740 | protein_coding_gene | 2.0 | 3.8683 | 1.9E-05 | 2.8 | 6.793 | 8.6E-08 |
| *C17G1.5* | WBGene00007652 | protein_coding_gene | 1.9 | 3.8624 | 2.9E-16 | 1.4 | 2.663 | 1.1E-09 |
| *fip-2* | WBGene00020862 | protein_coding_gene | 1.9 | 3.8568 | 3.6E-30 | 1.0 | 2.054 | 4.7E-16 |
| *ttll-11* | WBGene00019230 | protein_coding_gene | 1.9 | 3.8550 | 4.6E-04 | 1.4 | 2.597 | 9.4E-03 |
| *F17C8.8* | WBGene00044224 | pseudogene | 1.9 | 3.8549 | 1.4E-04 | 2.7 | 6.334 | 1.0E-05 |
| *hlh-11* | WBGene00001955 | protein_coding_gene | 1.9 | 3.8547 | 1.5E-10 | 2.0 | 3.912 | 8.3E-52 |
| *R07B1.13* | WBGene00138711 | protein_coding_gene | 1.9 | 3.8523 | 5.9E-05 | 3.2 | 9.157 | 6.8E-05 |
| *nspc-14* | WBGene00009638 | protein_coding_gene | 1.9 | 3.8514 | 1.0E-07 | 1.7 | 3.355 | 4.4E-03 |
| *K11E4.1* | WBGene00010773 | protein_coding_gene | 1.9 | 3.8508 | 8.6E-08 | 1.3 | 2.535 | 3.0E-05 |
| *tbc-16* | WBGene00018639 | protein_coding_gene | 1.9 | 3.8502 | 5.4E-06 | 1.1 | 2.099 | 1.5E-02 |
| *C54D1.7* | WBGene00016914 | protein_coding_gene | 1.9 | 3.8491 | 7.0E-09 | 1.1 | 2.178 | 1.0E-02 |
| *T16G12.12* | WBGene00269433 | protein_coding_gene | 1.9 | 3.8485 | 1.8E-05 | 1.2 | 2.275 | 1.9E-02 |
| *twk-14* | WBGene00006669 | protein_coding_gene | 1.9 | 3.8464 | 2.0E-03 | 1.5 | 2.850 | 1.4E-03 |
| *lys-6* | WBGene00003095 | protein_coding_gene | 1.9 | 3.8446 | 2.1E-06 | 1.8 | 3.516 | 2.2E-10 |
| *F44E7.3* | WBGene00018425 | protein_coding_gene | 1.9 | 3.8442 | 9.8E-03 | 2.6 | 6.070 | 1.9E-06 |
| *bus-1* | WBGene00044617 | protein_coding_gene | 1.9 | 3.8400 | 2.2E-02 | 2.3 | 5.025 | 4.4E-03 |
| *hog-1* | WBGene00001984 | protein_coding_gene | 1.9 | 3.8365 | 9.0E-05 | 1.7 | 3.156 | 4.4E-04 |
| *ncs-5* | WBGene00008307 | protein_coding_gene | 1.9 | 3.8363 | 2.5E-04 | 1.2 | 2.279 | 1.8E-02 |
| *C35A11.2* | WBGene00016429 | protein_coding_gene | 1.9 | 3.8342 | 8.1E-05 | 1.6 | 2.977 | 6.0E-05 |
| *Y110A2AL.3* | WBGene00022440 | protein_coding_gene | 1.9 | 3.8334 | 4.3E-06 | 2.3 | 4.808 | 5.8E-06 |
| ***cutl-16*** | **WBGene00019426** | **protein_coding_gene** | **1.9** | **3.8323** | **6.4E-05** | **-1.1** | **0.462** | **8.4E-05** |
| *acc-1* | WBGene00010275 | protein_coding_gene | 1.9 | 3.8272 | 1.3E-04 | 1.9 | 3.843 | 1.6E-04 |
| *F25B3.5* | WBGene00009102 | protein_coding_gene | 1.9 | 3.8262 | 5.4E-06 | 1.1 | 2.128 | 2.0E-04 |
| *B0361.9* | WBGene00015163 | protein_coding_gene | 1.9 | 3.8227 | 1.7E-30 | 1.2 | 2.256 | 9.5E-17 |
| *sprr-1* | WBGene00010986 | protein_coding_gene | 1.9 | 3.8218 | 1.3E-04 | 2.2 | 4.478 | 6.0E-05 |
| *Y48G1BL.7* | WBGene00021666 | protein_coding_gene | 1.9 | 3.8210 | 8.6E-03 | 1.8 | 3.491 | 1.5E-02 |
| *hpo-30* | WBGene00011014 | protein_coding_gene | 1.9 | 3.8192 | 3.6E-04 | 1.6 | 3.014 | 2.0E-03 |
| *ifd-1* | WBGene00002057 | protein_coding_gene | 1.9 | 3.8161 | 5.5E-18 | 2.2 | 4.566 | 2.1E-24 |
| *T06E4.14* | WBGene00077691 | protein_coding_gene | 1.9 | 3.8130 | 1.8E-03 | 1.8 | 3.595 | 3.3E-02 |
| *F35C11.7* | WBGene00009406 | protein_coding_gene | 1.9 | 3.8086 | 9.4E-08 | 1.0 | 2.039 | 3.4E-03 |
| *C06E8.5* | WBGene00015544 | protein_coding_gene | 1.9 | 3.8061 | 2.7E-05 | 1.9 | 3.800 | 5.1E-05 |
| *nhr-87* | WBGene00003677 | protein_coding_gene | 1.9 | 3.8047 | 2.1E-12 | 1.4 | 2.648 | 2.6E-09 |
| *F54H5.2* | WBGene00018839 | protein_coding_gene | 1.9 | 3.8045 | 1.8E-03 | 1.7 | 3.160 | 1.2E-05 |
| *T21H3.1* | WBGene00020662 | protein_coding_gene | 1.9 | 3.8043 | 4.7E-24 | 1.6 | 3.094 | 1.6E-31 |
| *srg-34* | WBGene00005191 | protein_coding_gene | 1.9 | 3.8031 | 2.0E-04 | 1.1 | 2.147 | 8.9E-03 |
| *F19H8.2* | WBGene00008963 | protein_coding_gene | 1.9 | 3.8028 | 1.7E-17 | 2.5 | 5.592 | 2.1E-18 |
| *arrd-14* | WBGene00011055 | protein_coding_gene | 1.9 | 3.7989 | 7.4E-33 | 1.7 | 3.239 | 1.5E-23 |
| *brp-1* | WBGene00000273 | protein_coding_gene | 1.9 | 3.7987 | 4.4E-14 | 1.9 | 3.848 | 1.5E-53 |
| *nhr-126* | WBGene00003716 | protein_coding_gene | 1.9 | 3.7966 | 1.2E-09 | 1.5 | 2.821 | 1.5E-10 |
| *galt-1* | WBGene00019763 | protein_coding_gene | 1.9 | 3.7798 | 1.6E-06 | 1.8 | 3.473 | 2.2E-06 |
| *slcf-2* | WBGene00006865 | protein_coding_gene | 1.9 | 3.7787 | 1.2E-09 | 1.1 | 2.171 | 2.7E-04 |
| *frpr-4* | WBGene00016909 | protein_coding_gene | 1.9 | 3.7780 | 3.9E-06 | 1.4 | 2.637 | 3.6E-03 |
| *K03A1.4* | WBGene00019352 | protein_coding_gene | 1.9 | 3.7771 | 2.5E-04 | 1.5 | 2.744 | 9.6E-03 |
| *nrx-1* | WBGene00007799 | protein_coding_gene | 1.9 | 3.7710 | 1.4E-10 | 1.5 | 2.888 | 1.0E-11 |
| *sulp-5* | WBGene00010789 | protein_coding_gene | 1.9 | 3.7699 | 7.7E-07 | 1.2 | 2.349 | 1.2E-02 |
| *F49E12.8* | WBGene00009901 | protein_coding_gene | 1.9 | 3.7697 | 2.2E-03 | 2.0 | 4.094 | 1.2E-03 |
| *C11E4.8* | WBGene00007522 | protein_coding_gene | 1.9 | 3.7658 | 4.2E-03 | 2.2 | 4.649 | 2.3E-04 |
| *gst-6* | WBGene00001754 | protein_coding_gene | 1.9 | 3.7647 | 3.7E-12 | 2.3 | 4.848 | 1.7E-27 |
| *F35A5.4* | WBGene00018026 | protein_coding_gene | 1.9 | 3.7646 | 8.7E-04 | 2.3 | 5.077 | 1.3E-02 |
| *wrt-4* | WBGene00006950 | protein_coding_gene | 1.9 | 3.7594 | 7.9E-07 | 2.3 | 4.928 | 7.9E-03 |
| *F40F8.5* | WBGene00009578 | protein_coding_gene | 1.9 | 3.7594 | 1.6E-10 | 1.1 | 2.124 | 4.7E-14 |
| *ZK1073.1* | WBGene00014213 | protein_coding_gene | 1.9 | 3.7520 | 4.9E-14 | 1.2 | 2.261 | 2.5E-16 |
| *T21G5.2* | WBGene00020660 | protein_coding_gene | 1.9 | 3.7504 | 5.4E-03 | 1.2 | 2.357 | 2.3E-02 |
| *fbxa-14* | WBGene00022326 | protein_coding_gene | 1.9 | 3.7490 | 1.7E-06 | 1.4 | 2.570 | 6.4E-04 |
| *acr-7* | WBGene00000046 | protein_coding_gene | 1.9 | 3.7476 | 9.8E-04 | 1.5 | 2.838 | 5.7E-03 |
| *olrn-1* | WBGene00007260 | protein_coding_gene | 1.9 | 3.7461 | 2.7E-07 | 1.3 | 2.546 | 3.1E-09 |
| *ttc-36* | WBGene00009947 | protein_coding_gene | 1.9 | 3.7427 | 3.8E-13 | 1.3 | 2.417 | 1.3E-13 |
| *Y51H7BR.4* | WBGene00021776 | protein_coding_gene | 1.9 | 3.7388 | 2.3E-03 | 2.0 | 3.988 | 7.0E-03 |
| *seb-3* | WBGene00007664 | protein_coding_gene | 1.9 | 3.7383 | 3.9E-08 | 1.4 | 2.687 | 3.4E-05 |
| *C53C11.2* | WBGene00016900 | protein_coding_gene | 1.9 | 3.7369 | 3.2E-06 | 2.0 | 4.104 | 2.7E-09 |
| *cest-35.1* | WBGene00015067 | protein_coding_gene | 1.9 | 3.7366 | 1.3E-17 | 1.4 | 2.638 | 1.1E-13 |
| *aat-5* | WBGene00000006 | protein_coding_gene | 1.9 | 3.7340 | 1.6E-07 | 1.8 | 3.464 | 1.7E-15 |

| | | | | | | | | |
|---|---|---|---|---|---|---|---|---|
| *gpx-6* | WBGene00020373 | protein_coding_gene | 1.9 | 3.7328 | 1.7E-10 | 2.8 | 6.924 | 3.1E-15 |
| *gnrr-2* | WBGene00007614 | protein_coding_gene | 1.9 | 3.7289 | 1.3E-02 | 1.8 | 3.541 | 2.7E-02 |
| *dot-1.3* | WBGene00012302 | protein_coding_gene | 1.9 | 3.7279 | 1.8E-02 | 4.6 | 23.874 | 1.5E-12 |
| *drh-2* | WBGene00001091 | pseudogene | 1.9 | 3.7259 | 2.7E-12 | 2.0 | 4.013 | 1.5E-28 |
| *M04D8.7* | WBGene00010865 | protein_coding_gene | 1.9 | 3.7184 | 6.1E-05 | 1.5 | 2.820 | 2.8E-05 |
| *fbxa-190* | WBGene00018939 | protein_coding_gene | 1.9 | 3.7157 | 7.7E-07 | 1.7 | 3.178 | 8.7E-05 |
| *C30G7.2* | WBGene00044176 | protein_coding_gene | 1.9 | 3.7140 | 6.0E-09 | 1.3 | 2.445 | 3.6E-07 |
| *dao-4* | WBGene00000930 | protein_coding_gene | 1.9 | 3.7126 | 8.1E-04 | 3.6 | 12.399 | 8.6E-04 |
| *T19D7.7* | WBGene00020575 | protein_coding_gene | 1.9 | 3.7125 | 2.4E-04 | 1.1 | 2.176 | 4.4E-02 |
| *C04G2.10* | WBGene00007309 | protein_coding_gene | 1.9 | 3.7096 | 2.7E-12 | 2.4 | 5.389 | 4.4E-16 |
| *R05A10.2* | WBGene00011019 | protein_coding_gene | 1.9 | 3.7089 | 1.5E-18 | 1.1 | 2.212 | 7.9E-08 |
| *F45H10.5* | WBGene00009742 | protein_coding_gene | 1.9 | 3.7082 | 5.8E-04 | 1.4 | 2.680 | 1.9E-02 |
| *D2096.9* | WBGene00017076 | protein_coding_gene | 1.9 | 3.7059 | 1.1E-04 | 1.5 | 2.828 | 4.0E-04 |
| *C26D10.4* | WBGene00007745 | protein_coding_gene | 1.9 | 3.6991 | 2.5E-13 | 1.6 | 3.006 | 3.6E-08 |
| *T24F1.4* | WBGene00011996 | protein_coding_gene | 1.9 | 3.6971 | 2.5E-03 | 1.4 | 2.700 | 3.7E-02 |
| *B0244.5* | WBGene00015080 | protein_coding_gene | 1.9 | 3.6946 | 9.6E-05 | 1.5 | 2.921 | 4.0E-04 |
| *F56B6.6* | WBGene00018937 | protein_coding_gene | 1.9 | 3.6916 | 6.6E-03 | 1.6 | 2.935 | 6.3E-03 |
| *F53A9.9* | WBGene00018732 | protein_coding_gene | 1.9 | 3.6897 | 1.2E-11 | 1.4 | 2.628 | 8.1E-05 |
| *B0361.4* | WBGene00015158 | protein_coding_gene | 1.9 | 3.6890 | 8.9E-05 | 1.7 | 3.336 | 3.1E-05 |
| *nhr-139* | WBGene00016365 | protein_coding_gene | 1.9 | 3.6845 | 4.4E-08 | 1.2 | 2.254 | 4.2E-04 |
| *Y18H1A.11* | WBGene00021215 | protein_coding_gene | 1.9 | 3.6824 | 7.0E-04 | 1.2 | 2.226 | 7.4E-03 |
| *Y39B6A.1* | WBGene00012664 | protein_coding_gene | 1.9 | 3.6808 | 1.8E-11 | 1.5 | 2.779 | 2.0E-02 |
| *C39D10.11* | WBGene00043983 | protein_coding_gene | 1.9 | 3.6802 | 6.9E-17 | 1.1 | 2.095 | 1.2E-07 |
| *Y37H2C.1* | WBGene00012573 | pseudogene | 1.9 | 3.6771 | 1.0E-04 | 2.3 | 4.957 | 1.9E-08 |
| *dyf-6* | WBGene00001122 | protein_coding_gene | 1.9 | 3.6769 | 2.1E-05 | 1.3 | 2.457 | 2.4E-03 |
| *npr-27* | WBGene00018344 | protein_coding_gene | 1.9 | 3.6748 | 9.0E-04 | 1.6 | 2.950 | 5.5E-03 |
| *nhr-144* | WBGene00019117 | pseudogene | 1.9 | 3.6740 | 1.4E-07 | 1.9 | 3.759 | 4.2E-15 |
| *lgc-28* | WBGene00044301 | protein_coding_gene | 1.9 | 3.6733 | 9.4E-06 | 1.2 | 2.334 | 1.1E-03 |
| *F16B4.2* | WBGene00017504 | protein_coding_gene | 1.9 | 3.6715 | 1.6E-07 | 1.4 | 2.672 | 5.1E-10 |
| *fbxa-39* | WBGene00021859 | protein_coding_gene | 1.9 | 3.6711 | 1.5E-04 | 1.6 | 3.050 | 6.8E-04 |
| *pmt-1* | WBGene00022781 | protein_coding_gene | 1.9 | 3.6698 | 6.7E-16 | 1.3 | 2.431 | 1.9E-29 |
| *kvs-3* | WBGene00018402 | protein_coding_gene | 1.9 | 3.6684 | 4.1E-05 | 1.7 | 3.262 | 9.2E-04 |
| *set-11* | WBGene00018023 | protein_coding_gene | 1.9 | 3.6675 | 2.1E-02 | 2.0 | 3.921 | 4.4E-02 |
| *C32D5.6* | WBGene00016314 | protein_coding_gene | 1.9 | 3.6636 | 4.1E-11 | 1.1 | 2.146 | 4.7E-06 |
| *asns-2* | WBGene00019730 | protein_coding_gene | 1.9 | 3.6633 | 1.4E-04 | 1.5 | 2.856 | 1.3E-08 |
| *C37C3.12* | WBGene00016504 | protein_coding_gene | 1.9 | 3.6632 | 3.4E-07 | 1.2 | 2.232 | 1.4E-04 |
| *T21C9.9* | WBGene00011895 | protein_coding_gene | 1.9 | 3.6631 | 2.0E-04 | 2.5 | 5.806 | 1.8E-06 |
| *clx-1* | WBGene00000551 | protein_coding_gene | 1.9 | 3.6629 | 1.1E-02 | 8.6 | 380.520 | 4.2E-12 |
| *F10A3.4* | WBGene00008634 | protein_coding_gene | 1.9 | 3.6617 | 3.0E-08 | 3.3 | 9.662 | 7.5E-20 |
| *R05D7.2* | WBGene00011027 | protein_coding_gene | 1.9 | 3.6592 | 6.8E-05 | 1.9 | 3.739 | 7.0E-06 |
| *rgs-4* | WBGene00004347 | protein_coding_gene | 1.9 | 3.6558 | 9.5E-17 | 1.5 | 2.891 | 4.8E-11 |
| *oac-3* | WBGene00007361 | protein_coding_gene | 1.9 | 3.6545 | 8.2E-03 | 3.2 | 9.013 | 9.7E-09 |
| *K08C7.4* | WBGene00010647 | protein_coding_gene | 1.9 | 3.6540 | 1.0E-03 | 1.0 | 2.022 | 5.2E-03 |
| *frpr-16* | WBGene00020023 | protein_coding_gene | 1.9 | 3.6537 | 1.5E-05 | 1.4 | 2.643 | 3.7E-03 |
| *T19D12.9* | WBGene00020583 | protein_coding_gene | 1.9 | 3.6536 | 1.9E-04 | 3.4 | 10.444 | 2.1E-08 |
| *ggr-3* | WBGene00001588 | protein_coding_gene | 1.9 | 3.6532 | 3.5E-06 | 1.7 | 3.246 | 1.1E-04 |
| *twk-35* | WBGene00006687 | protein_coding_gene | 1.9 | 3.6523 | 1.6E-03 | 1.9 | 3.701 | 4.8E-03 |
| *F45E1.5* | WBGene00018466 | protein_coding_gene | 1.9 | 3.6483 | 8.3E-04 | 1.9 | 3.663 | 1.9E-04 |
| *Y42A5A.3* | WBGene00012778 | protein_coding_gene | 1.9 | 3.6434 | 2.5E-02 | 1.3 | 2.451 | 1.1E-03 |
| *cha-1* | WBGene00000481 | protein_coding_gene | 1.9 | 3.6417 | 2.1E-09 | 2.0 | 4.069 | 4.1E-09 |
| *cdkl-1* | WBGene00012779 | protein_coding_gene | 1.9 | 3.6410 | 5.0E-04 | 1.6 | 3.120 | 3.5E-03 |
| *nfki-1* | WBGene00007877 | protein_coding_gene | 1.9 | 3.6322 | 7.9E-11 | 1.5 | 2.825 | 1.4E-28 |
| *unc-4* | WBGene00006744 | protein_coding_gene | 1.9 | 3.6319 | 5.9E-03 | 1.5 | 2.847 | 4.7E-02 |
| *snf-12* | WBGene00004911 | protein_coding_gene | 1.9 | 3.6307 | 1.1E-09 | 1.9 | 3.675 | 9.4E-10 |
| *ipla-7* | WBGene00011423 | protein_coding_gene | 1.9 | 3.6281 | 1.1E-30 | 1.4 | 2.636 | 7.5E-13 |
| *H34I24.2* | WBGene00019261 | protein_coding_gene | 1.9 | 3.6269 | 7.5E-13 | 1.9 | 3.653 | 8.8E-40 |
| *F59C6.14* | WBGene00044792 | protein_coding_gene | 1.9 | 3.6265 | 1.4E-09 | 1.1 | 2.182 | 5.7E-05 |
| *svh-5* | WBGene00006462 | protein_coding_gene | 1.9 | 3.6255 | 6.6E-13 | 1.4 | 2.593 | 1.7E-09 |
| *H23N18.4* | WBGene00019235 | protein_coding_gene | 1.9 | 3.6246 | 2.7E-02 | 1.5 | 2.780 | 1.6E-02 |
| ***cutl-28*** | **WBGene00018256** | **protein_coding_gene** | **1.9** | **3.6216** | **1.2E-03** | **-2.1** | **0.240** | **6.6E-04** |
| *gpa-14* | WBGene00001676 | protein_coding_gene | 1.9 | 3.6182 | 3.5E-04 | 1.5 | 2.883 | 7.5E-03 |
| *sro-1* | WBGene00005641 | protein_coding_gene | 1.9 | 3.6181 | 1.5E-03 | 1.9 | 3.828 | 2.1E-04 |
| *T28F3.5* | WBGene00012131 | protein_coding_gene | 1.9 | 3.6162 | 5.0E-07 | 1.9 | 3.606 | 7.0E-08 |
| *K02E11.7* | WBGene00010516 | protein_coding_gene | 1.9 | 3.6134 | 5.4E-11 | 2.0 | 3.886 | 3.8E-09 |
| *gly-18* | WBGene00001643 | protein_coding_gene | 1.9 | 3.6096 | 8.5E-04 | 1.3 | 2.544 | 1.8E-02 |
| *hpo-26* | WBGene00018492 | protein_coding_gene | 1.9 | 3.6092 | 5.3E-08 | 2.2 | 4.501 | 4.2E-14 |
| *lpd-7* | WBGene00003063 | protein_coding_gene | 1.9 | 3.6091 | 3.6E-40 | 1.5 | 2.864 | 8.6E-30 |
| *nas-9* | WBGene00003528 | protein_coding_gene | 1.8 | 3.6045 | 7.4E-10 | 1.5 | 2.907 | 3.2E-16 |
| *tag-209* | WBGene00011072 | protein_coding_gene | 1.8 | 3.6031 | 2.2E-07 | 1.4 | 2.648 | 9.6E-17 |
| *flp-28* | WBGene00044686 | protein_coding_gene | 1.8 | 3.5983 | 2.0E-17 | 1.8 | 3.578 | 8.0E-36 |
| *Y56A3A.18* | WBGene00013236 | protein_coding_gene | 1.8 | 3.5956 | 1.0E-27 | 1.7 | 3.268 | 8.1E-22 |
| *fipr-11* | WBGene00008244 | protein_coding_gene | 1.8 | 3.5922 | 1.2E-04 | 1.4 | 2.653 | 3.3E-03 |
| *unc-10* | WBGene00006750 | protein_coding_gene | 1.8 | 3.5890 | 2.3E-07 | 1.7 | 3.223 | 8.0E-14 |
| *math-35* | WBGene00020075 | pseudogene | 1.8 | 3.5847 | 1.4E-13 | 1.3 | 2.505 | 4.9E-10 |
| *R02E4.2* | WBGene00044021 | protein_coding_gene | 1.8 | 3.5842 | 6.7E-05 | 1.9 | 3.721 | 1.3E-03 |
| *T13A10.1* | WBGene00020468 | protein_coding_gene | 1.8 | 3.5830 | 1.9E-03 | 1.7 | 3.179 | 1.8E-02 |
| *AH9.6* | WBGene00044701 | protein_coding_gene | 1.8 | 3.5789 | 1.2E-05 | 2.1 | 4.362 | 1.8E-07 |
| *trk-1* | WBGene00017039 | protein_coding_gene | 1.8 | 3.5778 | 6.1E-09 | 1.6 | 3.125 | 1.9E-06 |
| *pqn-25* | WBGene00004114 | protein_coding_gene | 1.8 | 3.5778 | 1.0E-09 | 1.6 | 3.056 | 4.0E-14 |
| *ttr-5* | WBGene00008040 | protein_coding_gene | 1.8 | 3.5773 | 4.1E-20 | 1.1 | 2.158 | 9.9E-09 |

| | | | | | | | | |
|---|---|---|---|---|---|---|---|---|
| *ins-27* | WBGene00002110 | protein_coding_gene | 1.8 | 3.5762 | 4.9E-10 | 2.4 | 5.350 | 4.9E-12 |
| *F19G12.8* | WBGene00195184 | protein_coding_gene | 1.8 | 3.5727 | 7.9E-07 | 1.3 | 2.513 | 6.3E-04 |
| *zip-2* | WBGene00019327 | protein_coding_gene | 1.8 | 3.5719 | 1.1E-09 | 1.1 | 2.195 | 3.2E-21 |
| *lgc-31* | WBGene00008997 | protein_coding_gene | 1.8 | 3.5664 | 4.3E-10 | 1.3 | 2.439 | 2.9E-04 |
| *pals-17* | WBGene00022346 | protein_coding_gene | 1.8 | 3.5662 | 4.6E-11 | 1.3 | 2.388 | 3.8E-04 |
| *C09B9.85* | WBGene00220250 | protein_coding_gene | 1.8 | 3.5654 | 9.2E-03 | 1.9 | 3.748 | 4.5E-06 |
| *F35H10.3* | WBGene00018069 | protein_coding_gene | 1.8 | 3.5638 | 5.0E-07 | 2.3 | 4.847 | 9.0E-09 |
| *mltn-13* | WBGene00017794 | protein_coding_gene | 1.8 | 3.5631 | 2.2E-02 | 2.0 | 3.880 | 4.6E-02 |
| *F52E10.4* | WBGene00009935 | protein_coding_gene | 1.8 | 3.5613 | 3.6E-07 | 1.6 | 2.989 | 6.5E-06 |
| *nhr-50* | WBGene00003640 | protein_coding_gene | 1.8 | 3.5590 | 1.4E-12 | 1.2 | 2.291 | 7.5E-07 |
| *T09B4.7* | WBGene00020381 | protein_coding_gene | 1.8 | 3.5577 | 2.1E-03 | 2.3 | 4.936 | 1.5E-05 |
| *R09A8.5* | WBGene00011154 | protein_coding_gene | 1.8 | 3.5574 | 1.0E-07 | 1.1 | 2.140 | 4.9E-04 |
| *nlp-59* | WBGene00008522 | protein_coding_gene | 1.8 | 3.5548 | 3.3E-11 | 2.4 | 5.353 | 4.6E-27 |
| *lin-46* | WBGene00003031 | protein_coding_gene | 1.8 | 3.5543 | 4.1E-07 | 2.0 | 3.875 | 3.6E-06 |
| *T01C1.4* | WBGene00011317 | protein_coding_gene | 1.8 | 3.5519 | 2.2E-03 | 1.6 | 3.031 | 6.7E-03 |
| *klf-1* | WBGene00018990 | protein_coding_gene | 1.8 | 3.5439 | 6.5E-13 | 1.8 | 3.392 | 2.7E-29 |
| *syx-2* | WBGene00006372 | protein_coding_gene | 1.8 | 3.5432 | 1.6E-06 | 1.0 | 2.040 | 2.1E-06 |
| *molo-1* | WBGene00017299 | protein_coding_gene | 1.8 | 3.5431 | 9.3E-05 | 2.1 | 4.153 | 3.9E-06 |
| ***C01G5.7*** | **WBGene00015309** | **protein_coding_gene** | **1.8** | **3.5392** | **3.2E-02** | **-1.1** | **0.471** | **3.7E-03** |
| *nhr-63* | WBGene00003653 | protein_coding_gene | 1.8 | 3.5390 | 9.8E-05 | 1.3 | 2.485 | 6.9E-05 |
| *F46C5.2* | WBGene00009779 | protein_coding_gene | 1.8 | 3.5390 | 2.1E-05 | 1.9 | 3.714 | 7.0E-06 |
| *gst-5* | WBGene00001753 | protein_coding_gene | 1.8 | 3.5339 | 1.9E-10 | 2.2 | 4.626 | 3.6E-37 |
| *dmsr-8* | WBGene00007951 | protein_coding_gene | 1.8 | 3.5335 | 2.7E-06 | 1.8 | 3.413 | 6.8E-06 |
| *R12A1.3* | WBGene00020017 | protein_coding_gene | 1.8 | 3.5324 | 2.4E-03 | 1.7 | 3.218 | 2.0E-02 |
| *F10C1.8* | WBGene00017327 | protein_coding_gene | 1.8 | 3.5303 | 4.8E-10 | 2.4 | 5.359 | 3.2E-05 |
| *C49C8.5* | WBGene00016769 | protein_coding_gene | 1.8 | 3.5298 | 6.9E-11 | 1.3 | 2.487 | 4.4E-13 |
| *elt-2* | WBGene00001250 | protein_coding_gene | 1.8 | 3.5161 | 2.7E-12 | 1.4 | 2.596 | 8.4E-15 |
| *tag-241* | WBGene00044079 | protein_coding_gene | 1.8 | 3.5145 | 6.4E-07 | 1.1 | 2.139 | 1.4E-08 |
| *R05D7.3* | WBGene00011028 | protein_coding_gene | 1.8 | 3.5144 | 1.2E-04 | 1.5 | 2.845 | 4.0E-05 |
| *gtr-1* | WBGene00017779 | protein_coding_gene | 1.8 | 3.5135 | 4.6E-03 | 1.2 | 2.372 | 2.7E-02 |
| *F15A4.5* | WBGene00008840 | protein_coding_gene | 1.8 | 3.5108 | 1.5E-05 | 1.7 | 3.155 | 1.1E-06 |
| ***prkl-1*** | **WBGene00022727** | **protein_coding_gene** | **1.8** | **3.5066** | **9.5E-07** | **-1.0** | **0.490** | **1.4E-05** |
| *aex-3* | WBGene00000086 | protein_coding_gene | 1.8 | 3.5058 | 2.6E-07 | 1.2 | 2.270 | 2.2E-11 |
| *fipr-9* | WBGene00044175 | protein_coding_gene | 1.8 | 3.5052 | 5.9E-09 | 1.8 | 3.443 | 1.9E-06 |
| *R07E3.1* | WBGene00011102 | protein_coding_gene | 1.8 | 3.5040 | 3.1E-15 | 1.7 | 3.297 | 1.4E-20 |
| *F18A11.5* | WBGene00008932 | protein_coding_gene | 1.8 | 3.5038 | 2.3E-10 | 1.6 | 3.109 | 1.9E-08 |
| *C26D10.7* | WBGene00023420 | protein_coding_gene | 1.8 | 3.5023 | 3.4E-03 | 1.7 | 3.275 | 9.3E-03 |
| *T11F9.1* | WBGene00011709 | protein_coding_gene | 1.8 | 3.5008 | 1.3E-04 | 1.4 | 2.732 | 1.0E-02 |
| *F57A8.4* | WBGene00010179 | protein_coding_gene | 1.8 | 3.5005 | 2.0E-04 | 1.2 | 2.281 | 2.5E-02 |
| *ZK180.6* | WBGene00022680 | protein_coding_gene | 1.8 | 3.4995 | 3.7E-06 | 3.8 | 14.144 | 7.3E-05 |
| *T24C12.4* | WBGene00020766 | protein_coding_gene | 1.8 | 3.4960 | 1.7E-02 | 2.3 | 5.060 | 2.2E-04 |
| *Y116F11B.6* | WBGene00013822 | protein_coding_gene | 1.8 | 3.4960 | 2.6E-04 | 1.0 | 2.069 | 2.2E-02 |
| *dhp-1* | WBGene00000963 | protein_coding_gene | 1.8 | 3.4954 | 3.1E-09 | 1.3 | 2.483 | 2.2E-09 |
| *snf-11* | WBGene00004910 | protein_coding_gene | 1.8 | 3.4944 | 5.4E-07 | 1.0 | 2.050 | 2.5E-05 |
| *T05C1.2* | WBGene00020249 | protein_coding_gene | 1.8 | 3.4899 | 4.1E-03 | 2.7 | 6.365 | 2.6E-04 |
| *F41E6.7* | WBGene00018288 | protein_coding_gene | 1.8 | 3.4863 | 5.7E-04 | 1.4 | 2.702 | 1.7E-02 |
| *F28E10.1* | WBGene00017900 | protein_coding_gene | 1.8 | 3.4830 | 1.5E-08 | 1.4 | 2.640 | 1.9E-12 |
| *lat-2* | WBGene00002252 | protein_coding_gene | 1.8 | 3.4828 | 1.8E-10 | 2.2 | 4.750 | 9.5E-16 |
| *M162.5* | WBGene00010931 | protein_coding_gene | 1.8 | 3.4816 | 6.5E-07 | 1.4 | 2.704 | 1.4E-07 |
| *F14D12.1* | WBGene00017463 | protein_coding_gene | 1.8 | 3.4806 | 9.0E-09 | 1.7 | 3.243 | 6.0E-13 |
| *F14D7.8* | WBGene00008795 | protein_coding_gene | 1.8 | 3.4786 | 2.3E-02 | 2.1 | 4.214 | 1.1E-02 |
| *B0403.5* | WBGene00015169 | protein_coding_gene | 1.8 | 3.4767 | 1.3E-03 | 1.5 | 2.841 | 2.0E-04 |
| *mls-1* | WBGene00003376 | protein_coding_gene | 1.8 | 3.4744 | 1.1E-03 | 1.4 | 2.673 | 5.9E-03 |
| *F14B6.2* | WBGene00008783 | protein_coding_gene | 1.8 | 3.4730 | 1.5E-09 | 1.5 | 2.875 | 2.1E-07 |
| *rpl-24.2* | WBGene00004437 | protein_coding_gene | 1.8 | 3.4697 | 2.3E-33 | 1.6 | 3.123 | 1.0E-30 |
| *nhr-231* | WBGene00012449 | protein_coding_gene | 1.8 | 3.4670 | 1.7E-04 | 1.8 | 3.600 | 1.2E-05 |
| *eat-2* | WBGene00001133 | protein_coding_gene | 1.8 | 3.4635 | 8.0E-03 | 1.9 | 3.638 | 5.3E-04 |
| *ins-3* | WBGene00002086 | protein_coding_gene | 1.8 | 3.4619 | 5.0E-05 | 1.1 | 2.190 | 8.0E-03 |
| *F19C7.2* | WBGene00017592 | protein_coding_gene | 1.8 | 3.4617 | 4.1E-08 | 1.4 | 2.627 | 1.3E-13 |
| ***C01G6.9*** | **WBGene00007229** | **protein_coding_gene** | **1.8** | **3.4566** | **1.8E-04** | **-3.8** | **0.070** | **2.7E-10** |
| *myo-1* | WBGene00002348 | protein_coding_gene | 1.8 | 3.4558 | 7.7E-08 | 2.1 | 4.214 | 5.8E-32 |
| *slc-17.3* | WBGene00015333 | protein_coding_gene | 1.8 | 3.4528 | 3.2E-13 | 2.4 | 5.120 | 2.2E-19 |
| *nspc-10* | WBGene00009640 | protein_coding_gene | 1.8 | 3.4519 | 8.4E-12 | 1.2 | 2.252 | 4.8E-13 |
| *F49C5.11* | WBGene00044163 | protein_coding_gene | 1.8 | 3.4510 | 5.8E-03 | 4.5 | 22.609 | 9.1E-07 |
| *Y73F4A.2* | WBGene00013515 | protein_coding_gene | 1.8 | 3.4474 | 5.2E-09 | 1.1 | 2.210 | 1.7E-10 |
| *ift-74* | WBGene00016005 | protein_coding_gene | 1.8 | 3.4445 | 1.6E-06 | 1.0 | 2.037 | 4.2E-03 |
| *flp-8* | WBGene00001451 | protein_coding_gene | 1.8 | 3.4442 | 1.9E-10 | 1.4 | 2.564 | 2.3E-17 |
| *twk-25* | WBGene00006678 | protein_coding_gene | 1.8 | 3.4424 | 5.8E-08 | 1.7 | 3.155 | 3.1E-16 |
| *T23D8.3* | WBGene00011944 | protein_coding_gene | 1.8 | 3.4421 | 9.7E-39 | 1.2 | 2.367 | 9.0E-22 |
| *nhr-14* | WBGene00003613 | protein_coding_gene | 1.8 | 3.4421 | 6.5E-12 | 1.7 | 3.327 | 4.7E-28 |
| ***ZK287.1*** | **WBGene00013962** | **protein_coding_gene** | **1.8** | **3.4376** | **1.5E-05** | **-1.1** | **0.475** | **6.2E-04** |
| *mec-9* | WBGene00003173 | protein_coding_gene | 1.8 | 3.4344 | 6.3E-08 | 1.7 | 3.146 | 2.2E-05 |
| *F35B12.10* | WBGene00009389 | protein_coding_gene | 1.8 | 3.4335 | 2.9E-11 | 1.7 | 3.153 | 1.6E-17 |
| *zig-2* | WBGene00006979 | protein_coding_gene | 1.8 | 3.4304 | 8.0E-14 | 1.7 | 3.236 | 2.8E-09 |
| *crm-1* | WBGene00007103 | protein_coding_gene | 1.8 | 3.4261 | 1.1E-14 | 1.3 | 2.442 | 6.7E-18 |
| *maf-1* | WBGene00077521 | protein_coding_gene | 1.8 | 3.4257 | 1.6E-09 | 2.0 | 4.029 | 1.5E-09 |
| *W01A8.7* | WBGene00044764 | pseudogene | 1.8 | 3.4225 | 2.9E-02 | 1.7 | 3.332 | 4.2E-02 |
| *gcy-19* | WBGene00001544 | protein_coding_gene | 1.8 | 3.4224 | 2.7E-05 | 1.4 | 2.625 | 2.4E-03 |
| *pepm-1* | WBGene00009807 | protein_coding_gene | 1.8 | 3.4203 | 3.7E-04 | 2.5 | 5.610 | 1.8E-03 |
| *ckr-2* | WBGene00021439 | protein_coding_gene | 1.8 | 3.4134 | 1.2E-04 | 1.2 | 2.341 | 2.7E-04 |

| | | | | | | | | |
|---|---|---|---|---|---|---|---|---|
| *sem-4* | WBGene00004773 | protein_coding_gene | 1.8 | 3.4134 | 3.0E-09 | 1.3 | 2.461 | 1.1E-14 |
| *pqn-31* | WBGene00004119 | protein_coding_gene | 1.8 | 3.4120 | 6.3E-06 | 1.1 | 2.103 | 2.8E-10 |
| *T06D8.10* | WBGene00011530 | protein_coding_gene | 1.8 | 3.4083 | 6.2E-05 | 1.7 | 3.146 | 2.6E-02 |
| *nhr-31* | WBGene00003625 | protein_coding_gene | 1.8 | 3.4063 | 2.8E-07 | 2.3 | 4.973 | 2.0E-36 |
| *C23F12.9* | WBGene00304782 | protein_coding_gene | 1.8 | 3.4048 | 7.7E-03 | 1.5 | 2.912 | 1.4E-02 |
| *Y53F4B.1* | WBGene00013149 | protein_coding_gene | 1.8 | 3.4047 | 4.0E-10 | 1.3 | 2.521 | 9.1E-07 |
| *sma-3* | WBGene00004857 | protein_coding_gene | 1.8 | 3.4038 | 1.0E-18 | 1.4 | 2.720 | 1.6E-12 |
| *rga-6* | WBGene00015303 | protein_coding_gene | 1.8 | 3.4035 | 2.6E-06 | 1.2 | 2.295 | 2.8E-04 |
| *ric-4* | WBGene00004364 | protein_coding_gene | 1.8 | 3.3941 | 5.6E-19 | 1.1 | 2.219 | 2.2E-15 |
| *cdo-1* | WBGene00018986 | protein_coding_gene | 1.8 | 3.3937 | 5.9E-15 | 1.3 | 2.448 | 3.3E-18 |
| *rgs-6* | WBGene00004349 | protein_coding_gene | 1.8 | 3.3926 | 7.0E-05 | 1.6 | 3.048 | 1.5E-10 |
| *del-9* | WBGene00015957 | protein_coding_gene | 1.8 | 3.3906 | 8.6E-04 | 1.7 | 3.199 | 2.1E-05 |
| *clec-166* | WBGene00009518 | protein_coding_gene | 1.8 | 3.3845 | 2.0E-11 | 1.1 | 2.205 | 1.3E-12 |
| *lgc-37* | WBGene00013914 | protein_coding_gene | 1.8 | 3.3838 | 5.1E-03 | 2.4 | 5.316 | 2.0E-03 |
| *VM106R.1* | WBGene00014848 | protein_coding_gene | 1.8 | 3.3836 | 2.6E-07 | 1.9 | 3.852 | 4.0E-10 |
| *nlp-5* | WBGene00003743 | protein_coding_gene | 1.8 | 3.3835 | 1.0E-04 | 1.3 | 2.393 | 2.5E-04 |
| *cps-6* | WBGene00000787 | protein_coding_gene | 1.8 | 3.3817 | 1.8E-22 | 1.6 | 3.078 | 1.6E-33 |
| ***F59C6.16*** | **WBGene00077764** | **protein_coding_gene** | **1.8** | **3.3812** | **8.9E-04** | **-1.1** | **0.458** | **1.5E-07** |
| *F25E5.16* | WBGene00017792 | protein_coding_gene | 1.8 | 3.3812 | 3.4E-05 | 1.4 | 2.597 | 3.2E-03 |
| *Y37E11AL.1* | WBGene00021357 | protein_coding_gene | 1.8 | 3.3789 | 3.7E-03 | 2.0 | 3.891 | 7.0E-03 |
| ***arl-6*** | **WBGene00000193** | **protein_coding_gene** | **1.8** | **3.3704** | **3.7E-02** | **-1.4** | **0.389** | **2.9E-02** |
| *cutl-19* | WBGene00022533 | protein_coding_gene | 1.8 | 3.3698 | 2.3E-02 | 1.5 | 2.912 | 3.6E-02 |
| *M195.2* | WBGene00010948 | protein_coding_gene | 1.8 | 3.3678 | 3.7E-04 | 1.9 | 3.649 | 3.0E-02 |
| *col-14* | WBGene00000603 | protein_coding_gene | 1.7 | 3.3612 | 8.5E-05 | 2.3 | 4.812 | 1.8E-02 |
| *nhr-170* | WBGene00008309 | protein_coding_gene | 1.7 | 3.3573 | 1.2E-06 | 1.7 | 3.172 | 1.2E-06 |
| *T07D4.2* | WBGene00011579 | protein_coding_gene | 1.7 | 3.3557 | 3.6E-11 | 1.6 | 2.983 | 2.3E-13 |
| *gab-1* | WBGene00001512 | protein_coding_gene | 1.7 | 3.3536 | 2.5E-04 | 2.0 | 4.133 | 8.4E-05 |
| *casy-1* | WBGene00000403 | protein_coding_gene | 1.7 | 3.3519 | 1.5E-14 | 1.5 | 2.790 | 2.5E-24 |
| *unc-8* | WBGene00006748 | protein_coding_gene | 1.7 | 3.3509 | 1.4E-03 | 1.3 | 2.532 | 3.6E-02 |
| *pes-8* | WBGene00003981 | protein_coding_gene | 1.7 | 3.3480 | 1.8E-07 | 2.6 | 5.985 | 6.2E-04 |
| *pqn-26* | WBGene00004115 | protein_coding_gene | 1.7 | 3.3476 | 6.2E-04 | 1.3 | 2.398 | 1.9E-02 |
| *F14F9.3* | WBGene00017466 | protein_coding_gene | 1.7 | 3.3461 | 4.6E-02 | 1.2 | 2.316 | 1.7E-02 |
| *lon-8* | WBGene00013352 | protein_coding_gene | 1.7 | 3.3418 | 2.3E-10 | 1.8 | 3.418 | 2.3E-10 |
| *C34E7.3* | WBGene00007933 | protein_coding_gene | 1.7 | 3.3409 | 3.1E-03 | 2.1 | 4.268 | 2.7E-03 |
| *ugt-24* | WBGene00016762 | protein_coding_gene | 1.7 | 3.3405 | 6.9E-07 | 1.6 | 3.050 | 1.0E-12 |
| *clp-9* | WBGene00011705 | protein_coding_gene | 1.7 | 3.3401 | 4.1E-10 | 1.3 | 2.464 | 1.5E-07 |
| *T28F3.8* | WBGene00012134 | protein_coding_gene | 1.7 | 3.3397 | 1.4E-19 | 1.4 | 2.640 | 9.7E-18 |
| *cpna-1* | WBGene00006495 | protein_coding_gene | 1.7 | 3.3392 | 1.8E-07 | 1.1 | 2.218 | 8.5E-16 |
| *R03H10.1* | WBGene00019854 | protein_coding_gene | 1.7 | 3.3391 | 1.3E-08 | 1.3 | 2.526 | 2.7E-07 |
| *C49A9.5* | WBGene00016759 | protein_coding_gene | 1.7 | 3.3369 | 1.9E-05 | 2.3 | 4.981 | 2.1E-37 |
| *glb-6* | WBGene00015969 | protein_coding_gene | 1.7 | 3.3367 | 7.1E-07 | 2.0 | 3.886 | 2.4E-08 |
| *zmp-6* | WBGene00020646 | protein_coding_gene | 1.7 | 3.3359 | 2.2E-02 | 1.9 | 3.607 | 3.2E-02 |
| *nhr-146* | WBGene00045255 | protein_coding_gene | 1.7 | 3.3323 | 1.7E-11 | 1.9 | 3.674 | 7.2E-15 |
| *akt-2* | WBGene00000103 | protein_coding_gene | 1.7 | 3.3298 | 5.0E-09 | 1.1 | 2.125 | 6.2E-11 |
| *T06E4.8* | WBGene00011535 | protein_coding_gene | 1.7 | 3.3279 | 3.3E-02 | 1.8 | 3.584 | 4.5E-02 |
| *npr-2* | WBGene00003808 | protein_coding_gene | 1.7 | 3.3267 | 3.3E-05 | 1.4 | 2.576 | 6.4E-04 |
| *C18G1.6* | WBGene00015984 | protein_coding_gene | 1.7 | 3.3257 | 1.2E-03 | 1.2 | 2.268 | 4.7E-02 |
| *cpx-1* | WBGene00022271 | protein_coding_gene | 1.7 | 3.3252 | 6.4E-10 | 1.5 | 2.791 | 4.4E-14 |
| *C25G4.2* | WBGene00007730 | protein_coding_gene | 1.7 | 3.3242 | 1.7E-13 | 1.0 | 2.058 | 4.4E-11 |
| *src-2* | WBGene00005078 | protein_coding_gene | 1.7 | 3.3217 | 1.8E-12 | 1.5 | 2.804 | 4.7E-09 |
| *ifb-2* | WBGene00002054 | protein_coding_gene | 1.7 | 3.3181 | 5.5E-14 | 1.7 | 3.360 | 4.9E-35 |
| *cgt-1* | WBGene00011517 | protein_coding_gene | 1.7 | 3.3145 | 7.4E-08 | 1.6 | 2.987 | 6.6E-08 |
| *axl-1* | WBGene00010497 | protein_coding_gene | 1.7 | 3.3135 | 1.0E-07 | 1.2 | 2.232 | 3.0E-04 |
| *ZC196.5* | WBGene00022549 | protein_coding_gene | 1.7 | 3.3130 | 2.1E-11 | 1.1 | 2.165 | 2.6E-07 |
| *lntl-1* | WBGene00016289 | protein_coding_gene | 1.7 | 3.3109 | 4.0E-09 | 1.2 | 2.226 | 4.7E-12 |
| *C09F12.3* | WBGene00007483 | protein_coding_gene | 1.7 | 3.3086 | 2.4E-02 | 2.0 | 3.984 | 8.5E-03 |
| *cebp-1* | WBGene00016997 | protein_coding_gene | 1.7 | 3.3066 | 4.4E-07 | 1.2 | 2.333 | 8.8E-08 |
| *tom-1* | WBGene00006594 | protein_coding_gene | 1.7 | 3.3028 | 1.4E-20 | 1.5 | 2.915 | 2.6E-27 |
| *lge-1* | WBGene00010716 | protein_coding_gene | 1.7 | 3.3024 | 6.2E-03 | 1.7 | 3.362 | 7.3E-03 |
| *gpc-1* | WBGene00001681 | protein_coding_gene | 1.7 | 3.2992 | 5.4E-05 | 1.1 | 2.140 | 2.5E-02 |
| *F41E7.7* | WBGene00009623 | protein_coding_gene | 1.7 | 3.2956 | 6.9E-05 | 1.4 | 2.597 | 4.5E-06 |
| *tkr-1* | WBGene00006576 | protein_coding_gene | 1.7 | 3.2951 | 9.2E-03 | 1.1 | 2.193 | 2.5E-02 |
| *F52E1.9* | WBGene00018697 | protein_coding_gene | 1.7 | 3.2947 | 1.2E-08 | 1.3 | 2.469 | 3.8E-09 |
| *bcc-1* | WBGene00000246 | protein_coding_gene | 1.7 | 3.2865 | 5.4E-13 | 2.0 | 4.063 | 1.0E-12 |
| *nhr-213* | WBGene00011520 | protein_coding_gene | 1.7 | 3.2840 | 4.1E-07 | 1.1 | 2.117 | 2.4E-03 |
| *zig-6* | WBGene00006983 | protein_coding_gene | 1.7 | 3.2839 | 4.7E-11 | 1.4 | 2.705 | 1.2E-08 |
| *nhr-54* | WBGene00003644 | protein_coding_gene | 1.7 | 3.2826 | 2.4E-05 | 1.4 | 2.675 | 1.1E-08 |
| *col-34* | WBGene00000611 | protein_coding_gene | 1.7 | 3.2822 | 9.4E-09 | 1.8 | 3.601 | 2.1E-18 |
| *R01H2.7* | WBGene00023414 | protein_coding_gene | 1.7 | 3.2820 | 1.4E-12 | 1.1 | 2.086 | 1.9E-05 |
| *lin-1* | WBGene00002990 | protein_coding_gene | 1.7 | 3.2818 | 9.2E-09 | 1.1 | 2.084 | 1.3E-06 |
| *C08G5.6* | WBGene00015618 | protein_coding_gene | 1.7 | 3.2817 | 2.9E-02 | 2.1 | 4.238 | 7.0E-03 |
| *linc-125* | WBGene00219686 | lincRNA_gene | 1.7 | 3.2805 | 2.9E-02 | 2.4 | 5.193 | 5.7E-03 |
| *Y43C5A.2* | WBGene00012782 | protein_coding_gene | 1.7 | 3.2784 | 9.2E-19 | 1.0 | 2.011 | 9.5E-10 |
| *F09B9.4* | WBGene00008603 | protein_coding_gene | 1.7 | 3.2744 | 3.3E-06 | 1.4 | 2.595 | 1.4E-21 |
| *nhr-125* | WBGene00003715 | protein_coding_gene | 1.7 | 3.2717 | 1.1E-07 | 1.2 | 2.362 | 2.1E-04 |
| *ugt-1* | WBGene00007072 | protein_coding_gene | 1.7 | 3.2693 | 1.4E-05 | 1.4 | 2.696 | 3.3E-13 |
| *W05H12.1* | WBGene00012289 | protein_coding_gene | 1.7 | 3.2634 | 1.1E-12 | 1.6 | 2.938 | 3.5E-12 |
| *B0244.7* | WBGene00015082 | protein_coding_gene | 1.7 | 3.2633 | 6.1E-03 | 3.2 | 9.411 | 6.1E-06 |
| *glb-9* | WBGene00016184 | protein_coding_gene | 1.7 | 3.2607 | 3.7E-04 | 1.9 | 3.660 | 8.7E-05 |
| *twk-21* | WBGene00006674 | protein_coding_gene | 1.7 | 3.2605 | 2.1E-03 | 2.0 | 3.866 | 6.3E-04 |

| | | | | | | | | |
|---|---|---|---|---|---|---|---|---|
| *T06E4.12* | WBGene00044011 | protein_coding_gene | 1.7 | 3.2599 | 3.0E-02 | 2.1 | 4.405 | 2.2E-02 |
| *Y34B4A.10* | WBGene00021326 | protein_coding_gene | 1.7 | 3.2547 | 3.5E-06 | 1.6 | 3.135 | 9.7E-05 |
| *mab-9* | WBGene00003106 | protein_coding_gene | 1.7 | 3.2530 | 8.6E-04 | 1.5 | 2.863 | 3.8E-03 |
| *F59G1.4* | WBGene00019128 | protein_coding_gene | 1.7 | 3.2487 | 2.5E-06 | 1.1 | 2.082 | 4.3E-03 |
| *F14B6.4* | WBGene00008785 | protein_coding_gene | 1.7 | 3.2461 | 6.5E-03 | 1.3 | 2.480 | 2.9E-02 |
| *D2023.3* | WBGene00008407 | protein_coding_gene | 1.7 | 3.2449 | 1.8E-04 | 1.2 | 2.292 | 1.4E-02 |
| *T28F3.9* | WBGene00012135 | protein_coding_gene | 1.7 | 3.2420 | 6.5E-03 | 1.5 | 2.744 | 3.2E-02 |
| *nhr-201* | WBGene00019741 | protein_coding_gene | 1.7 | 3.2393 | 3.7E-09 | 1.2 | 2.234 | 1.3E-05 |
| *nlp-82* | WBGene00044006 | protein_coding_gene | 1.7 | 3.2378 | 1.4E-04 | 1.5 | 2.835 | 8.4E-04 |
| ***cnc-2*** | **WBGene00000556** | **protein_coding_gene** | **1.7** | **3.2360** | **3.0E-02** | **-1.4** | **0.383** | **4.3E-03** |
| *F42A9.9* | WBGene00018338 | protein_coding_gene | 1.7 | 3.2323 | 1.6E-02 | 2.3 | 4.782 | 2.2E-03 |
| *tre-3* | WBGene00006609 | protein_coding_gene | 1.7 | 3.2321 | 1.0E-13 | 1.6 | 2.938 | 1.0E-17 |
| ***M03B6.3*** | **WBGene00010835** | **protein_coding_gene** | **1.7** | **3.2318** | **3.4E-04** | **-1.3** | **0.395** | **3.3E-02** |
| *C33D3.3* | WBGene00007894 | protein_coding_gene | 1.7 | 3.2291 | 1.9E-04 | 3.6 | 12.529 | 2.5E-11 |
| *C28C12.4* | WBGene00016180 | protein_coding_gene | 1.7 | 3.2285 | 6.5E-07 | 1.6 | 2.982 | 2.1E-05 |
| *cng-2* | WBGene00022295 | protein_coding_gene | 1.7 | 3.2274 | 1.8E-02 | 1.6 | 3.108 | 2.0E-02 |
| *ugt-56* | WBGene00011453 | protein_coding_gene | 1.7 | 3.2265 | 2.1E-02 | 1.6 | 2.983 | 2.3E-02 |
| *F07H5.7* | WBGene00008558 | protein_coding_gene | 1.7 | 3.2110 | 1.1E-04 | 1.4 | 2.720 | 3.3E-03 |
| *ZK596.1* | WBGene00014006 | protein_coding_gene | 1.7 | 3.2096 | 9.3E-05 | 2.5 | 5.695 | 7.5E-53 |
| *nhr-121* | WBGene00003711 | protein_coding_gene | 1.7 | 3.2094 | 2.0E-08 | 1.1 | 2.181 | 1.1E-08 |
| *F49B2.6* | WBGene00009865 | protein_coding_gene | 1.7 | 3.2073 | 3.7E-08 | 1.8 | 3.534 | 1.4E-10 |
| *Y44A6D.3* | WBGene00012854 | protein_coding_gene | 1.7 | 3.2068 | 1.8E-09 | 1.2 | 2.231 | 2.4E-12 |
| *bli-6* | WBGene00000256 | protein_coding_gene | 1.7 | 3.2066 | 3.6E-04 | 3.1 | 8.292 | 2.5E-03 |
| *Y73F8A.35* | WBGene00013539 | protein_coding_gene | 1.7 | 3.2058 | 1.8E-05 | 1.3 | 2.517 | 3.5E-04 |
| *C28H8.5* | WBGene00016196 | protein_coding_gene | 1.7 | 3.2040 | 3.8E-02 | 2.4 | 5.191 | 1.4E-02 |
| *F14D7.6* | WBGene00008793 | protein_coding_gene | 1.7 | 3.2032 | 2.2E-08 | 2.5 | 5.798 | 9.2E-25 |
| *R07E4.1* | WBGene00019935 | protein_coding_gene | 1.7 | 3.2018 | 2.9E-14 | 2.3 | 4.845 | 3.4E-31 |
| *unc-96* | WBGene00006825 | protein_coding_gene | 1.7 | 3.2016 | 8.3E-08 | 1.0 | 2.012 | 4.1E-10 |
| *nspe-4* | WBGene00012589 | protein_coding_gene | 1.7 | 3.1988 | 2.4E-03 | 4.5 | 22.146 | 4.9E-17 |
| *igeg-1* | WBGene00017901 | protein_coding_gene | 1.7 | 3.1980 | 3.9E-11 | 1.2 | 2.254 | 6.8E-11 |
| *ZK54.3* | WBGene00022648 | protein_coding_gene | 1.7 | 3.1962 | 3.7E-03 | 1.5 | 2.913 | 2.6E-02 |
| *grl-10* | WBGene00001719 | protein_coding_gene | 1.7 | 3.1953 | 3.5E-05 | 1.2 | 2.334 | 8.0E-04 |
| *Y54G2A.49* | WBGene00044492 | protein_coding_gene | 1.7 | 3.1945 | 2.0E-05 | 1.3 | 2.428 | 3.4E-03 |
| *ifo-1* | WBGene00018350 | protein_coding_gene | 1.7 | 3.1937 | 3.6E-05 | 1.8 | 3.405 | 4.0E-42 |
| *F14F9.4* | WBGene00017467 | protein_coding_gene | 1.7 | 3.1933 | 5.0E-03 | 1.6 | 2.973 | 9.9E-14 |
| *dmsr-7* | WBGene00016428 | protein_coding_gene | 1.7 | 3.1930 | 7.8E-05 | 1.5 | 2.797 | 1.3E-04 |
| *nas-31* | WBGene00003549 | protein_coding_gene | 1.7 | 3.1891 | 8.9E-10 | 1.4 | 2.592 | 7.2E-06 |
| *cln-3.3* | WBGene00000541 | protein_coding_gene | 1.7 | 3.1883 | 1.5E-05 | 2.5 | 5.841 | 6.2E-22 |
| *best-9* | WBGene00016613 | protein_coding_gene | 1.7 | 3.1874 | 6.3E-08 | 1.1 | 2.115 | 2.3E-04 |
| *F19C6.3* | WBGene00008952 | protein_coding_gene | 1.7 | 3.1863 | 1.8E-05 | 1.4 | 2.722 | 8.4E-05 |
| *T19D12.4* | WBGene00020579 | protein_coding_gene | 1.7 | 3.1841 | 2.8E-05 | 1.9 | 3.798 | 8.3E-08 |
| *arrd-18* | WBGene00018060 | protein_coding_gene | 1.7 | 3.1808 | 1.2E-08 | 2.4 | 5.212 | 8.1E-19 |
| *Y102A11A.3* | WBGene00022413 | protein_coding_gene | 1.7 | 3.1787 | 8.7E-10 | 2.5 | 5.488 | 4.7E-54 |
| *Y54E2A.10* | WBGene00013195 | protein_coding_gene | 1.7 | 3.1784 | 1.5E-03 | 1.6 | 3.103 | 5.1E-04 |
| *C56G2.9* | WBGene00016982 | protein_coding_gene | 1.7 | 3.1769 | 1.6E-06 | 1.3 | 2.416 | 1.4E-03 |
| *F54F12.2* | WBGene00010073 | protein_coding_gene | 1.7 | 3.1731 | 7.5E-04 | 1.2 | 2.346 | 1.3E-02 |
| *ttr-28* | WBGene00011285 | protein_coding_gene | 1.7 | 3.1689 | 2.0E-04 | 1.3 | 2.401 | 1.9E-03 |
| *Y51H7C.13* | WBGene00021791 | protein_coding_gene | 1.7 | 3.1642 | 4.8E-05 | 3.8 | 14.248 | 4.6E-06 |
| ***F09F9.4*** | **WBGene00017309** | **protein_coding_gene** | **1.7** | **3.1634** | **1.3E-03** | **-1.2** | **0.431** | **5.5E-04** |
| *R102.6* | WBGene00011293 | protein_coding_gene | 1.7 | 3.1624 | 2.8E-04 | 2.0 | 4.129 | 2.6E-07 |
| *sorb-1* | WBGene00012891 | protein_coding_gene | 1.7 | 3.1591 | 3.7E-07 | 1.1 | 2.195 | 2.8E-07 |
| *cyp-34A5* | WBGene00015040 | protein_coding_gene | 1.7 | 3.1589 | 6.0E-04 | 1.6 | 2.982 | 1.9E-02 |
| *col-170* | WBGene00000743 | protein_coding_gene | 1.7 | 3.1574 | 7.9E-04 | 3.9 | 15.253 | 2.0E-11 |
| *nlp-50* | WBGene00018298 | protein_coding_gene | 1.7 | 3.1566 | 5.9E-08 | 1.0 | 2.042 | 1.6E-05 |
| *cri-1* | WBGene00010614 | protein_coding_gene | 1.7 | 3.1564 | 6.4E-04 | 2.6 | 6.002 | 7.8E-22 |
| *twk-48* | WBGene00010784 | protein_coding_gene | 1.7 | 3.1556 | 2.6E-03 | 1.9 | 3.703 | 3.0E-06 |
| *K07E8.6* | WBGene00019497 | protein_coding_gene | 1.7 | 3.1545 | 1.3E-03 | 1.6 | 2.982 | 3.0E-04 |
| *kgb-2* | WBGene00002188 | protein_coding_gene | 1.7 | 3.1539 | 6.0E-03 | 1.1 | 2.142 | 1.6E-05 |
| *cutl-23* | WBGene00021343 | protein_coding_gene | 1.7 | 3.1502 | 2.9E-06 | 1.5 | 2.848 | 9.0E-05 |
| *nhr-133* | WBGene00003723 | protein_coding_gene | 1.7 | 3.1472 | 6.7E-09 | 1.1 | 2.113 | 5.7E-06 |
| *K07E3.9* | WBGene00194670 | protein_coding_gene | 1.7 | 3.1463 | 1.2E-02 | 1.8 | 3.451 | 4.3E-02 |
| *glb-13* | WBGene00008957 | protein_coding_gene | 1.7 | 3.1461 | 6.4E-06 | 2.0 | 4.131 | 4.2E-06 |
| *F22E5.13* | WBGene00017711 | protein_coding_gene | 1.7 | 3.1418 | 1.7E-03 | 1.7 | 3.205 | 3.6E-03 |
| *lim-6* | WBGene00002988 | protein_coding_gene | 1.7 | 3.1417 | 1.1E-07 | 1.4 | 2.597 | 2.6E-08 |
| *C05D12.3* | WBGene00014666 | protein_coding_gene | 1.7 | 3.1411 | 1.2E-02 | 4.0 | 15.671 | 1.9E-104 |
| *T06G6.3* | WBGene00011545 | protein_coding_gene | 1.7 | 3.1404 | 5.7E-03 | 1.5 | 2.888 | 1.3E-02 |
| *Y53H1B.2* | WBGene00013181 | protein_coding_gene | 1.6 | 3.1305 | 3.6E-14 | 1.4 | 2.693 | 4.9E-19 |
| *lrp-1* | WBGene00003071 | protein_coding_gene | 1.6 | 3.1292 | 3.3E-08 | 1.6 | 3.109 | 2.1E-32 |
| *nhr-183* | WBGene00018266 | protein_coding_gene | 1.6 | 3.1284 | 5.4E-07 | 1.4 | 2.731 | 4.0E-02 |
| *far-3* | WBGene00001387 | protein_coding_gene | 1.6 | 3.1272 | 4.1E-05 | 2.2 | 4.466 | 3.1E-40 |
| *pqn-74* | WBGene00004156 | protein_coding_gene | 1.6 | 3.1271 | 2.8E-04 | 1.5 | 2.860 | 3.7E-02 |
| *F58E2.3* | WBGene00019047 | protein_coding_gene | 1.6 | 3.1263 | 1.7E-06 | 2.0 | 3.886 | 3.8E-07 |
| *nlp-41* | WBGene00007316 | protein_coding_gene | 1.6 | 3.1256 | 1.2E-04 | 1.4 | 2.728 | 7.8E-08 |
| *F25E5.8* | WBGene00017789 | protein_coding_gene | 1.6 | 3.1232 | 1.5E-13 | 1.6 | 2.953 | 1.9E-14 |
| *tnt-4* | WBGene00006589 | protein_coding_gene | 1.6 | 3.1165 | 7.6E-06 | 1.6 | 3.001 | 7.6E-27 |
| *ugt-23* | WBGene00007650 | protein_coding_gene | 1.6 | 3.1154 | 1.0E-25 | 1.1 | 2.167 | 5.8E-14 |
| *daf-38* | WBGene00013642 | protein_coding_gene | 1.6 | 3.1129 | 4.5E-05 | 1.6 | 3.131 | 2.1E-04 |
| *D1054.5* | WBGene00008372 | protein_coding_gene | 1.6 | 3.1114 | 2.9E-12 | 1.7 | 3.198 | 2.0E-13 |
| *lmp-2* | WBGene00015471 | protein_coding_gene | 1.6 | 3.1104 | 5.9E-15 | 2.2 | 4.599 | 1.5E-26 |
| *ins-29* | WBGene00002112 | protein_coding_gene | 1.6 | 3.1097 | 1.7E-02 | 1.9 | 3.735 | 2.9E-05 |

| | | | | | | | | |
|---|---|---|---|---|---|---|---|---|
| *sma-5* | WBGene00004859 | protein_coding_gene | 1.6 | 3.1093 | 4.6E-07 | 1.3 | 2.460 | 9.5E-20 |
| *glb-7* | WBGene00016024 | protein_coding_gene | 1.6 | 3.1073 | 1.4E-02 | 2.1 | 4.230 | 9.0E-04 |
| *myo-5* | WBGene00019064 | protein_coding_gene | 1.6 | 3.1054 | 1.5E-09 | 1.3 | 2.528 | 2.6E-17 |
| *otpl-3* | WBGene00017449 | protein_coding_gene | 1.6 | 3.1048 | 1.8E-05 | 2.0 | 4.079 | 7.5E-17 |
| *B0393.4* | WBGene00007169 | protein_coding_gene | 1.6 | 3.1033 | 2.6E-05 | 2.3 | 5.055 | 1.1E-21 |
| ***igcm-2*** | **WBGene00020130** | **protein_coding_gene** | **1.6** | **3.1031** | **1.2E-03** | **-1.0** | **0.486** | **5.1E-03** |
| *F13E6.2* | WBGene00008746 | protein_coding_gene | 1.6 | 3.1029 | 3.7E-04 | 1.2 | 2.316 | 1.6E-02 |
| *sup-9* | WBGene00006318 | protein_coding_gene | 1.6 | 3.1001 | 8.3E-04 | 1.0 | 2.049 | 1.7E-02 |
| *hot-9* | WBGene00044484 | protein_coding_gene | 1.6 | 3.0999 | 2.5E-04 | 1.1 | 2.096 | 1.5E-02 |
| *rgs-3* | WBGene00004346 | protein_coding_gene | 1.6 | 3.0998 | 1.2E-05 | 1.3 | 2.460 | 2.6E-04 |
| *K02D10.8* | WBGene00219329 | protein_coding_gene | 1.6 | 3.0969 | 3.1E-05 | 1.1 | 2.148 | 1.5E-02 |
| *prhg-1* | WBGene00022391 | protein_coding_gene | 1.6 | 3.0953 | 3.4E-06 | 1.6 | 3.113 | 2.8E-06 |
| *F43C1.5* | WBGene00009649 | protein_coding_gene | 1.6 | 3.0945 | 3.7E-06 | 2.0 | 3.933 | 3.0E-06 |
| *T23B12.5* | WBGene00020720 | protein_coding_gene | 1.6 | 3.0921 | 8.1E-03 | 2.3 | 4.810 | 1.9E-03 |
| *nhr-128* | WBGene00003718 | protein_coding_gene | 1.6 | 3.0921 | 7.2E-06 | 1.1 | 2.150 | 1.4E-04 |
| *byn-1* | WBGene00000276 | protein_coding_gene | 1.6 | 3.0920 | 3.1E-60 | 1.4 | 2.716 | 1.0E-33 |
| *C06E1.7* | WBGene00015522 | protein_coding_gene | 1.6 | 3.0887 | 3.9E-04 | 1.8 | 3.388 | 2.2E-05 |
| ***plg-1*** | **WBGene00004041** | **pseudogene** | **1.6** | **3.0825** | **8.9E-03** | **-5.3** | **0.025** | **3.2E-70** |
| *T11B7.5* | WBGene00011708 | protein_coding_gene | 1.6 | 3.0824 | 2.1E-09 | 1.6 | 2.965 | 8.3E-14 |
| *F29C6.1* | WBGene00009243 | protein_coding_gene | 1.6 | 3.0821 | 8.7E-11 | 1.9 | 3.651 | 1.1E-22 |
| *T19D2.2* | WBGene00004184 | protein_coding_gene | 1.6 | 3.0753 | 9.6E-06 | 1.2 | 2.227 | 2.5E-17 |
| *F16F9.3* | WBGene00017514 | protein_coding_gene | 1.6 | 3.0747 | 6.8E-05 | 1.6 | 2.985 | 4.7E-04 |
| *acs-12* | WBGene00009106 | protein_coding_gene | 1.6 | 3.0713 | 2.1E-16 | 1.3 | 2.395 | 4.1E-14 |
| *asp-8* | WBGene00019105 | protein_coding_gene | 1.6 | 3.0705 | 2.6E-10 | 2.8 | 6.865 | 8.3E-11 |
| *Y46G5A.15* | WBGene00012906 | protein_coding_gene | 1.6 | 3.0676 | 1.3E-10 | 1.3 | 2.500 | 4.4E-08 |
| *F38H4.3* | WBGene00009547 | protein_coding_gene | 1.6 | 3.0655 | 2.8E-10 | 1.6 | 3.013 | 1.2E-08 |
| *F19C7.8* | WBGene00017597 | protein_coding_gene | 1.6 | 3.0638 | 2.4E-05 | 1.6 | 2.973 | 1.2E-07 |
| *glb-19* | WBGene00010146 | protein_coding_gene | 1.6 | 3.0615 | 1.4E-05 | 1.6 | 3.069 | 2.4E-04 |
| *odr-2* | WBGene00003849 | protein_coding_gene | 1.6 | 3.0576 | 1.2E-05 | 1.1 | 2.095 | 6.2E-03 |
| *F22E12.3* | WBGene00009060 | protein_coding_gene | 1.6 | 3.0522 | 5.6E-04 | 1.2 | 2.220 | 3.7E-02 |
| *T10B5.7* | WBGene00020393 | protein_coding_gene | 1.6 | 3.0520 | 1.9E-16 | 1.4 | 2.582 | 1.2E-20 |
| *gst-40* | WBGene00001788 | protein_coding_gene | 1.6 | 3.0486 | 9.8E-03 | 1.1 | 2.171 | 3.9E-02 |
| *aexr-3* | WBGene00016748 | protein_coding_gene | 1.6 | 3.0482 | 1.2E-02 | 1.9 | 3.680 | 1.0E-02 |
| *F57B9.3* | WBGene00018997 | protein_coding_gene | 1.6 | 3.0437 | 1.0E-02 | 2.5 | 5.598 | 1.4E-04 |
| *Y37E11B.1* | WBGene00021374 | protein_coding_gene | 1.6 | 3.0427 | 2.0E-08 | 1.1 | 2.146 | 1.4E-04 |
| *F22B5.4* | WBGene00009042 | protein_coding_gene | 1.6 | 3.0424 | 9.2E-04 | 1.0 | 2.022 | 2.9E-04 |
| *rpl-11.2* | WBGene00004423 | protein_coding_gene | 1.6 | 3.0400 | 1.4E-16 | 2.0 | 3.905 | 4.8E-54 |
| *kcnl-2* | WBGene00008570 | protein_coding_gene | 1.6 | 3.0395 | 3.3E-07 | 1.5 | 2.913 | 5.9E-10 |
| *mec-1* | WBGene00003165 | protein_coding_gene | 1.6 | 3.0386 | 5.4E-06 | 1.6 | 3.097 | 2.9E-10 |
| *col-97* | WBGene00000672 | protein_coding_gene | 1.6 | 3.0385 | 5.6E-05 | 2.9 | 7.582 | 8.3E-04 |
| *W07G4.5* | WBGene00012339 | protein_coding_gene | 1.6 | 3.0373 | 1.4E-33 | 1.5 | 2.897 | 1.2E-16 |
| *Y47D3B.6* | WBGene00012942 | protein_coding_gene | 1.6 | 3.0364 | 6.2E-04 | 1.9 | 3.631 | 3.7E-02 |
| *dmd-6* | WBGene00007058 | protein_coding_gene | 1.6 | 3.0355 | 1.0E-08 | 1.2 | 2.300 | 1.5E-24 |
| *ngp-1* | WBGene00003596 | protein_coding_gene | 1.6 | 3.0341 | 1.5E-54 | 1.6 | 2.935 | 3.8E-30 |
| *D1086.18* | WBGene00045405 | protein_coding_gene | 1.6 | 3.0290 | 6.1E-04 | 1.1 | 2.121 | 7.4E-03 |
| *ram-2* | WBGene00004300 | protein_coding_gene | 1.6 | 3.0289 | 1.1E-04 | 3.4 | 10.671 | 1.5E-58 |
| *T23F2.2* | WBGene00020735 | protein_coding_gene | 1.6 | 3.0269 | 9.4E-07 | 1.7 | 3.143 | 1.7E-17 |
| *T20B5.2* | WBGene00020595 | protein_coding_gene | 1.6 | 3.0265 | 9.3E-08 | 1.4 | 2.670 | 1.5E-05 |
| *gst-23* | WBGene00001771 | protein_coding_gene | 1.6 | 3.0253 | 1.2E-02 | 1.3 | 2.402 | 2.1E-02 |
| *F47B10.5* | WBGene00009816 | protein_coding_gene | 1.6 | 3.0238 | 9.8E-10 | 1.4 | 2.580 | 4.2E-07 |
| *cat-2* | WBGene00000296 | protein_coding_gene | 1.6 | 3.0217 | 6.4E-03 | 1.4 | 2.620 | 1.7E-02 |
| *gst-35* | WBGene00001783 | protein_coding_gene | 1.6 | 3.0188 | 6.5E-06 | 1.9 | 3.674 | 1.7E-07 |
| *acdh-6* | WBGene00015335 | protein_coding_gene | 1.6 | 3.0179 | 2.9E-03 | 1.8 | 3.458 | 1.4E-02 |
| *str-176* | WBGene00006220 | protein_coding_gene | 1.6 | 3.0178 | 1.2E-17 | 1.8 | 3.515 | 6.9E-18 |
| *nhr-105* | WBGene00003695 | protein_coding_gene | 1.6 | 3.0162 | 1.9E-19 | 1.3 | 2.497 | 1.0E-14 |
| *gcy-21* | WBGene00001546 | protein_coding_gene | 1.6 | 3.0144 | 5.2E-06 | 1.6 | 3.082 | 1.2E-04 |
| *F14B8.6* | WBGene00017448 | protein_coding_gene | 1.6 | 3.0133 | 2.2E-11 | 1.7 | 3.230 | 1.1E-25 |
| *F18A1.1* | WBGene00017545 | protein_coding_gene | 1.6 | 3.0129 | 2.4E-03 | 1.6 | 3.015 | 2.6E-04 |
| *col-61* | WBGene00000637 | protein_coding_gene | 1.6 | 3.0121 | 1.4E-02 | 2.3 | 4.797 | 2.0E-02 |
| *otpl-7* | WBGene00018480 | protein_coding_gene | 1.6 | 3.0118 | 1.7E-02 | 1.6 | 3.012 | 4.4E-02 |
| *abhd-14* | WBGene00021927 | protein_coding_gene | 1.6 | 3.0058 | 7.1E-07 | 1.2 | 2.275 | 1.4E-07 |
| *sqst-5* | WBGene00269434 | protein_coding_gene | 1.6 | 3.0058 | 6.7E-06 | 2.2 | 4.642 | 3.2E-03 |
| *rei-2* | WBGene00019365 | protein_coding_gene | 1.6 | 3.0029 | 1.5E-10 | 1.0 | 2.035 | 3.0E-12 |
| *Y51A2D.21* | WBGene00044210 | protein_coding_gene | 1.6 | 3.0018 | 9.5E-10 | 1.1 | 2.149 | 3.1E-04 |
| *gmeb-3* | WBGene00008092 | protein_coding_gene | 1.6 | 3.0012 | 4.6E-03 | 2.2 | 4.601 | 1.8E-11 |
| *F58E2.4* | WBGene00019048 | protein_coding_gene | 1.6 | 2.9988 | 1.6E-06 | 1.9 | 3.750 | 3.5E-08 |
| *kmo-2* | WBGene00011088 | protein_coding_gene | 1.6 | 2.9975 | 1.4E-04 | 1.1 | 2.179 | 5.9E-05 |
| *D1007.19* | WBGene00045048 | protein_coding_gene | 1.6 | 2.9923 | 3.6E-03 | 2.1 | 4.145 | 9.4E-04 |
| *twk-49* | WBGene00019904 | protein_coding_gene | 1.6 | 2.9896 | 6.0E-14 | 1.9 | 3.717 | 2.2E-10 |
| *F27C1.11* | WBGene00017858 | protein_coding_gene | 1.6 | 2.9884 | 2.0E-05 | 1.4 | 2.726 | 1.7E-05 |
| *frpr-3* | WBGene00016149 | protein_coding_gene | 1.6 | 2.9830 | 4.1E-02 | 1.8 | 3.426 | 2.5E-02 |
| *best-23* | WBGene00014989 | protein_coding_gene | 1.6 | 2.9802 | 1.3E-04 | 1.0 | 2.048 | 1.5E-03 |
| *F38E9.6* | WBGene00018188 | protein_coding_gene | 1.6 | 2.9799 | 5.1E-06 | 1.3 | 2.530 | 1.4E-05 |
| *nhr-136* | WBGene00003726 | protein_coding_gene | 1.6 | 2.9738 | 1.5E-07 | 1.2 | 2.350 | 2.5E-09 |
| *B0511.6* | WBGene00015232 | protein_coding_gene | 1.6 | 2.9731 | 4.0E-35 | 1.2 | 2.319 | 8.0E-21 |
| *R10E11.5* | WBGene00011217 | protein_coding_gene | 1.6 | 2.9728 | 1.1E-07 | 1.6 | 3.048 | 2.0E-07 |
| *col-125* | WBGene00000699 | protein_coding_gene | 1.6 | 2.9727 | 1.6E-02 | 3.3 | 9.760 | 9.1E-13 |
| *F32B5.6* | WBGene00017979 | protein_coding_gene | 1.6 | 2.9720 | 3.6E-14 | 1.5 | 2.833 | 3.4E-38 |
| *B0034.1* | WBGene00015002 | protein_coding_gene | 1.6 | 2.9714 | 1.4E-05 | 2.7 | 6.514 | 2.0E-15 |
| *fbxa-209* | WBGene00012533 | protein_coding_gene | 1.6 | 2.9700 | 6.2E-06 | 1.2 | 2.359 | 3.1E-04 |

| | | | | | | | | |
|---|---|---|---|---|---|---|---|---|
| *fbxa-146* | WBGene00018807 | pseudogene | 1.6 | 2.9694 | 3.3E-10 | 1.4 | 2.671 | 1.1E-09 |
| *F40F12.8* | WBGene00014755 | pseudogene | 1.6 | 2.9692 | 1.8E-02 | 1.7 | 3.224 | 2.1E-06 |
| ***plpr-1*** | **WBGene00011524** | **protein_coding_gene** | **1.6** | **2.9691** | **1.1E-03** | **-1.2** | **0.439** | **3.5E-03** |
| *unc-47* | WBGene00006783 | protein_coding_gene | 1.6 | 2.9646 | 1.7E-04 | 1.7 | 3.297 | 2.3E-03 |
| *Y15E3A.4* | WBGene00012443 | protein_coding_gene | 1.6 | 2.9615 | 1.7E-20 | 1.7 | 3.359 | 1.2E-34 |
| *W05B5.1* | WBGene00012274 | protein_coding_gene | 1.6 | 2.9591 | 1.3E-05 | 1.1 | 2.139 | 2.9E-11 |
| *gbb-1* | WBGene00021528 | protein_coding_gene | 1.6 | 2.9554 | 7.5E-07 | 1.6 | 3.040 | 1.3E-11 |
| *unc-79* | WBGene00006811 | protein_coding_gene | 1.6 | 2.9522 | 2.9E-04 | 1.8 | 3.399 | 4.4E-17 |
| *PDB1.1* | WBGene00019803 | protein_coding_gene | 1.6 | 2.9513 | 5.3E-07 | 2.0 | 3.908 | 3.5E-22 |
| ***ins-23*** | **WBGene00002106** | **protein_coding_gene** | **1.6** | **2.9506** | **2.3E-02** | **-1.4** | **0.377** | **8.5E-03** |
| *kcnl-4* | WBGene00008265 | protein_coding_gene | 1.6 | 2.9484 | 1.9E-04 | 1.4 | 2.623 | 1.5E-04 |
| *pqn-8* | WBGene00004100 | protein_coding_gene | 1.6 | 2.9468 | 3.5E-07 | 1.3 | 2.523 | 1.2E-04 |
| *abu-11* | WBGene00000034 | protein_coding_gene | 1.6 | 2.9460 | 7.3E-03 | 2.3 | 4.945 | 5.9E-04 |
| *ssu-1* | WBGene00013748 | protein_coding_gene | 1.6 | 2.9453 | 4.6E-03 | 1.6 | 3.022 | 1.5E-02 |
| *lgc-53* | WBGene00020657 | protein_coding_gene | 1.6 | 2.9447 | 5.4E-04 | 1.8 | 3.379 | 9.5E-04 |
| *epac-1* | WBGene00004255 | protein_coding_gene | 1.6 | 2.9425 | 3.7E-03 | 1.6 | 3.136 | 4.6E-03 |
| *unc-31* | WBGene00006767 | protein_coding_gene | 1.6 | 2.9418 | 3.0E-09 | 1.8 | 3.401 | 3.0E-16 |
| *sto-2* | WBGene00006064 | protein_coding_gene | 1.6 | 2.9414 | 1.2E-03 | 1.7 | 3.300 | 3.6E-04 |
| *glb-27* | WBGene00012178 | protein_coding_gene | 1.6 | 2.9413 | 1.9E-03 | 1.4 | 2.722 | 6.7E-03 |
| *Y94H6A.5* | WBGene00022378 | protein_coding_gene | 1.6 | 2.9385 | 7.4E-11 | 1.2 | 2.223 | 6.1E-16 |
| *mec-10* | WBGene00003174 | protein_coding_gene | 1.6 | 2.9353 | 5.4E-04 | 1.3 | 2.505 | 4.1E-03 |
| *K04F10.1* | WBGene00019394 | protein_coding_gene | 1.6 | 2.9329 | 7.1E-14 | 1.5 | 2.855 | 3.4E-11 |
| *K04G2.7* | WBGene00010566 | protein_coding_gene | 1.6 | 2.9308 | 1.9E-03 | 2.8 | 6.865 | 1.2E-04 |
| *Y56A3A.33* | WBGene00013244 | protein_coding_gene | 1.6 | 2.9308 | 2.2E-05 | 2.1 | 4.154 | 1.8E-11 |
| *abt-4* | WBGene00000022 | protein_coding_gene | 1.6 | 2.9304 | 1.1E-06 | 1.5 | 2.891 | 2.5E-04 |
| *arrd-15* | WBGene00014033 | protein_coding_gene | 1.6 | 2.9293 | 6.9E-04 | 1.0 | 2.005 | 2.1E-02 |
| *ptr-8* | WBGene00004222 | protein_coding_gene | 1.5 | 2.9269 | 4.4E-05 | 1.1 | 2.144 | 4.5E-06 |
| *ZC247.1* | WBGene00013859 | protein_coding_gene | 1.5 | 2.9265 | 1.3E-06 | 2.0 | 3.983 | 1.5E-11 |
| *klo-2* | WBGene00017103 | protein_coding_gene | 1.5 | 2.9261 | 2.6E-12 | 1.5 | 2.809 | 6.9E-08 |
| *ndnf-1* | WBGene00021893 | protein_coding_gene | 1.5 | 2.9261 | 3.5E-07 | 2.0 | 3.971 | 1.1E-13 |
| *myo-6* | WBGene00009730 | protein_coding_gene | 1.5 | 2.9261 | 2.9E-11 | 1.1 | 2.156 | 9.7E-07 |
| *M04C9.1* | WBGene00010856 | protein_coding_gene | 1.5 | 2.9251 | 7.7E-07 | 1.4 | 2.591 | 2.7E-05 |
| *ncs-3* | WBGene00003565 | protein_coding_gene | 1.5 | 2.9250 | 4.2E-08 | 1.5 | 2.792 | 1.4E-06 |
| *itx-1* | WBGene00002174 | protein_coding_gene | 1.5 | 2.9178 | 4.3E-05 | 1.4 | 2.608 | 2.0E-28 |
| ***T20F5.4*** | **WBGene00020626** | **protein_coding_gene** | **1.5** | **2.9123** | **2.6E-03** | **-1.7** | **0.303** | **1.4E-06** |
| *Y34F4.4* | WBGene00021339 | protein_coding_gene | 1.5 | 2.9077 | 1.8E-04 | 1.5 | 2.921 | 7.7E-06 |
| *ztf-13* | WBGene00018704 | protein_coding_gene | 1.5 | 2.9001 | 7.8E-07 | 1.2 | 2.309 | 4.0E-11 |
| *ncx-2* | WBGene00003567 | protein_coding_gene | 1.5 | 2.8967 | 8.8E-06 | 1.6 | 3.050 | 6.0E-29 |
| *pept-3* | WBGene00003878 | protein_coding_gene | 1.5 | 2.8958 | 1.1E-03 | 1.2 | 2.342 | 2.2E-07 |
| *exp-1* | WBGene00001373 | protein_coding_gene | 1.5 | 2.8949 | 4.2E-03 | 1.4 | 2.671 | 1.6E-02 |
| *R11F4.2* | WBGene00020008 | protein_coding_gene | 1.5 | 2.8948 | 2.0E-02 | 1.6 | 3.037 | 3.5E-02 |
| *C15C7.6* | WBGene00015792 | protein_coding_gene | 1.5 | 2.8930 | 5.4E-03 | 2.7 | 6.476 | 1.4E-04 |
| *K11G9.5* | WBGene00019655 | protein_coding_gene | 1.5 | 2.8925 | 1.8E-04 | 1.7 | 3.358 | 1.0E-07 |
| *C13A10.1* | WBGene00015728 | protein_coding_gene | 1.5 | 2.8895 | 3.0E-10 | 1.1 | 2.099 | 2.2E-04 |
| *lgc-39* | WBGene00017314 | protein_coding_gene | 1.5 | 2.8890 | 3.4E-05 | 1.3 | 2.504 | 4.1E-06 |
| *F56A12.2* | WBGene00010143 | protein_coding_gene | 1.5 | 2.8856 | 8.7E-03 | 1.5 | 2.748 | 1.5E-02 |
| ***sqt-3*** | **WBGene00005018** | **protein_coding_gene** | **1.5** | **2.8851** | **8.2E-05** | **-1.2** | **0.439** | **4.7E-02** |
| *ZK84.1* | WBGene00022649 | protein_coding_gene | 1.5 | 2.8797 | 1.4E-06 | 3.8 | 13.538 | 1.3E-21 |
| *Y116A8A.6* | WBGene00013776 | protein_coding_gene | 1.5 | 2.8794 | 1.8E-02 | 1.9 | 3.694 | 3.8E-04 |
| *nhr-85* | WBGene00003675 | protein_coding_gene | 1.5 | 2.8744 | 4.0E-06 | 2.1 | 4.298 | 1.6E-08 |
| *klf-3* | WBGene00003480 | protein_coding_gene | 1.5 | 2.8716 | 6.0E-06 | 1.4 | 2.619 | 3.3E-09 |
| *bus-19* | WBGene00011590 | protein_coding_gene | 1.5 | 2.8699 | 6.7E-07 | 1.5 | 2.780 | 1.8E-07 |
| *Y38C1AA.9* | WBGene00021400 | protein_coding_gene | 1.5 | 2.8686 | 1.1E-05 | 1.2 | 2.365 | 2.9E-05 |
| *klp-13* | WBGene00002224 | protein_coding_gene | 1.5 | 2.8686 | 7.6E-04 | 2.0 | 4.049 | 2.1E-05 |
| *acy-2* | WBGene00000069 | protein_coding_gene | 1.5 | 2.8685 | 7.6E-04 | 1.3 | 2.434 | 2.1E-03 |
| *trhr-1* | WBGene00016265 | protein_coding_gene | 1.5 | 2.8669 | 2.5E-06 | 1.6 | 3.010 | 1.2E-08 |
| *ilcr-1* | WBGene00013415 | protein_coding_gene | 1.5 | 2.8656 | 1.1E-14 | 1.6 | 3.063 | 7.4E-15 |
| *T24B8.4* | WBGene00011978 | protein_coding_gene | 1.5 | 2.8650 | 2.4E-05 | 1.3 | 2.411 | 1.1E-19 |
| *F58D5.6* | WBGene00010245 | protein_coding_gene | 1.5 | 2.8623 | 3.2E-02 | 2.0 | 3.967 | 1.1E-07 |
| *C16B8.3* | WBGene00015823 | protein_coding_gene | 1.5 | 2.8617 | 2.6E-10 | 1.2 | 2.350 | 6.6E-13 |
| *eri-12* | WBGene00007091 | protein_coding_gene | 1.5 | 2.8607 | 2.7E-30 | 1.2 | 2.321 | 5.4E-25 |
| *fpn-1.2* | WBGene00019977 | protein_coding_gene | 1.5 | 2.8603 | 1.2E-06 | 1.7 | 3.188 | 1.9E-05 |
| *M03E7.1* | WBGene00019753 | protein_coding_gene | 1.5 | 2.8581 | 1.0E-03 | 1.2 | 2.335 | 2.5E-02 |
| *Y53G8AR.7* | WBGene00021814 | protein_coding_gene | 1.5 | 2.8556 | 4.6E-10 | 2.5 | 5.637 | 6.6E-35 |
| *cyp-33C5* | WBGene00018261 | protein_coding_gene | 1.5 | 2.8555 | 4.7E-04 | 1.7 | 3.190 | 7.6E-04 |
| *Y4C6B.3* | WBGene00021157 | protein_coding_gene | 1.5 | 2.8520 | 7.1E-07 | 1.1 | 2.150 | 1.1E-05 |
| *abts-3* | WBGene00019018 | protein_coding_gene | 1.5 | 2.8497 | 1.9E-05 | 1.5 | 2.839 | 7.4E-17 |
| *C54G4.5* | WBGene00008315 | protein_coding_gene | 1.5 | 2.8487 | 3.1E-03 | 1.3 | 2.435 | 1.4E-02 |
| ***try-5*** | **WBGene00006623** | **protein_coding_gene** | **1.5** | **2.8482** | **2.3E-02** | **-1.5** | **0.352** | **3.6E-05** |
| *fbxa-155* | WBGene00007323 | protein_coding_gene | 1.5 | 2.8474 | 1.7E-02 | 2.2 | 4.684 | 1.1E-10 |
| *fos-1* | WBGene00001345 | protein_coding_gene | 1.5 | 2.8464 | 1.7E-05 | 1.2 | 2.255 | 2.3E-15 |
| *K02E2.6* | WBGene00010507 | protein_coding_gene | 1.5 | 2.8450 | 1.0E-17 | 1.1 | 2.175 | 2.3E-07 |
| *mlt-2* | WBGene00007252 | protein_coding_gene | 1.5 | 2.8393 | 2.1E-07 | 1.6 | 2.990 | 1.8E-07 |
| *glb-14* | WBGene00008996 | protein_coding_gene | 1.5 | 2.8392 | 5.4E-03 | 1.4 | 2.639 | 6.6E-03 |
| *ser-2* | WBGene00004777 | protein_coding_gene | 1.5 | 2.8390 | 5.7E-04 | 1.5 | 2.847 | 1.7E-03 |
| *rgs-2* | WBGene00004345 | protein_coding_gene | 1.5 | 2.8368 | 4.8E-05 | 1.6 | 2.947 | 7.6E-12 |
| *linc-82* | WBGene00219570 | lincRNA_gene | 1.5 | 2.8344 | 2.4E-02 | 1.9 | 3.624 | 4.9E-04 |
| *W05F2.4* | WBGene00021036 | protein_coding_gene | 1.5 | 2.8338 | 2.3E-05 | 1.5 | 2.918 | 1.4E-27 |
| *glb-17* | WBGene00009887 | protein_coding_gene | 1.5 | 2.8296 | 6.2E-03 | 1.1 | 2.165 | 1.2E-02 |
| *acr-25* | WBGene00043066 | protein_coding_gene | 1.5 | 2.8295 | 8.0E-03 | 2.3 | 5.003 | 1.1E-05 |

| | | | | | | | | |
|---|---|---|---|---|---|---|---|---|
| *gnrr-3* | WBGene00013871 | protein_coding_gene | 1.5 | 2.8262 | 1.8E-02 | 1.4 | 2.592 | 4.8E-02 |
| ***F43D9.1*** | **WBGene00009653** | **protein_coding_gene** | **1.5** | **2.8236** | **5.5E-03** | **-1.7** | **0.301** | **8.8E-06** |
| *ifc-2* | WBGene00002056 | protein_coding_gene | 1.5 | 2.8223 | 4.0E-14 | 1.7 | 3.194 | 8.2E-10 |
| *F35E2.5* | WBGene00009412 | protein_coding_gene | 1.5 | 2.8218 | 1.2E-03 | 1.9 | 3.842 | 1.6E-09 |
| *srx-45* | WBGene00005936 | protein_coding_gene | 1.5 | 2.8199 | 1.1E-02 | 1.8 | 3.387 | 2.1E-03 |
| *F39G3.5* | WBGene00018210 | protein_coding_gene | 1.5 | 2.8182 | 3.3E-06 | 2.0 | 4.021 | 1.9E-16 |
| *pqn-11* | WBGene00004103 | pseudogene | 1.5 | 2.8153 | 1.6E-02 | 3.6 | 11.768 | 4.9E-04 |
| ***mtl-2*** | **WBGene00003474** | **protein_coding_gene** | **1.5** | **2.8111** | **1.1E-06** | **-1.5** | **0.361** | **6.0E-21** |
| *T08G2.2* | WBGene00020365 | protein_coding_gene | 1.5 | 2.8088 | 3.1E-03 | 1.3 | 2.524 | 1.4E-03 |
| *irg-3* | WBGene00018760 | protein_coding_gene | 1.5 | 2.8077 | 1.9E-06 | 1.2 | 2.332 | 1.8E-06 |
| *sta-1* | WBGene00013111 | protein_coding_gene | 1.5 | 2.8062 | 1.0E-08 | 1.1 | 2.194 | 4.8E-09 |
| *spp-16* | WBGene00005001 | protein_coding_gene | 1.5 | 2.8028 | 1.7E-12 | 1.2 | 2.317 | 5.4E-10 |
| *nhr-41* | WBGene00022423 | protein_coding_gene | 1.5 | 2.7988 | 6.5E-05 | 1.1 | 2.177 | 3.3E-07 |
| *stg-2* | WBGene00017400 | protein_coding_gene | 1.5 | 2.7980 | 2.4E-02 | 1.5 | 2.836 | 3.6E-02 |
| *F14D7.12* | WBGene00045331 | protein_coding_gene | 1.5 | 2.7972 | 9.8E-03 | 2.7 | 6.426 | 8.3E-06 |
| *twk-2* | WBGene00006657 | protein_coding_gene | 1.5 | 2.7961 | 1.7E-04 | 1.2 | 2.327 | 1.5E-05 |
| *K05F1.10* | WBGene00019411 | protein_coding_gene | 1.5 | 2.7912 | 1.2E-13 | 1.6 | 3.040 | 1.1E-16 |
| *W02A2.4* | WBGene00012189 | pseudogene | 1.5 | 2.7896 | 4.5E-02 | 1.4 | 2.560 | 2.1E-05 |
| *apt-9* | WBGene00000166 | protein_coding_gene | 1.5 | 2.7889 | 1.8E-03 | 1.6 | 2.930 | 5.4E-04 |
| *F09F7.5* | WBGene00017302 | protein_coding_gene | 1.5 | 2.7847 | 1.2E-09 | 1.7 | 3.208 | 1.3E-24 |
| *T04A8.18* | WBGene00119203 | protein_coding_gene | 1.5 | 2.7836 | 1.6E-05 | 2.0 | 3.874 | 5.2E-03 |
| *F07F6.2* | WBGene00017216 | protein_coding_gene | 1.5 | 2.7818 | 1.8E-04 | 1.4 | 2.676 | 5.3E-04 |
| *F36D4.4* | WBGene00018089 | protein_coding_gene | 1.5 | 2.7810 | 7.6E-03 | 1.6 | 2.934 | 5.3E-03 |
| *kin-1* | WBGene00002189 | protein_coding_gene | 1.5 | 2.7788 | 4.5E-06 | 1.3 | 2.546 | 2.8E-27 |
| *ins-26* | WBGene00002109 | protein_coding_gene | 1.5 | 2.7764 | 2.1E-06 | 1.0 | 2.039 | 1.0E-02 |
| ***vap-2*** | **WBGene00011462** | **protein_coding_gene** | **1.5** | **2.7764** | **5.3E-03** | **-1.2** | **0.436** | **2.8E-05** |
| *cyp-33A1* | WBGene00015709 | protein_coding_gene | 1.5 | 2.7756 | 3.9E-17 | 1.5 | 2.858 | 2.0E-13 |
| *ZK721.3* | WBGene00022810 | protein_coding_gene | 1.5 | 2.7746 | 7.3E-03 | 1.3 | 2.486 | 8.5E-03 |
| *pno-1* | WBGene00013144 | protein_coding_gene | 1.5 | 2.7701 | 5.6E-33 | 1.4 | 2.567 | 1.1E-22 |
| *npr-4* | WBGene00007635 | protein_coding_gene | 1.5 | 2.7680 | 1.6E-04 | 1.5 | 2.823 | 1.3E-04 |
| *F48B9.1* | WBGene00018587 | protein_coding_gene | 1.5 | 2.7672 | 3.9E-03 | 1.1 | 2.113 | 1.1E-02 |
| *slc-25A29* | WBGene00008320 | protein_coding_gene | 1.5 | 2.7624 | 8.2E-07 | 1.8 | 3.551 | 3.5E-15 |
| *clip-1* | WBGene00010796 | protein_coding_gene | 1.5 | 2.7602 | 2.3E-08 | 1.1 | 2.157 | 4.8E-12 |
| *F12E12.11* | WBGene00044811 | protein_coding_gene | 1.5 | 2.7600 | 5.4E-03 | 2.7 | 6.630 | 2.1E-18 |
| *F40A3.7* | WBGene00018222 | protein_coding_gene | 1.5 | 2.7560 | 1.1E-04 | 1.4 | 2.585 | 6.2E-04 |
| *F20C5.4* | WBGene00008969 | protein_coding_gene | 1.5 | 2.7551 | 1.1E-07 | 1.1 | 2.082 | 1.5E-07 |
| *ncx-4* | WBGene00003569 | protein_coding_gene | 1.5 | 2.7543 | 6.5E-06 | 1.8 | 3.537 | 2.7E-09 |
| *Y16B4A.2* | WBGene00012445 | protein_coding_gene | 1.5 | 2.7542 | 8.1E-05 | 3.0 | 7.747 | 1.8E-28 |
| *lgc-49* | WBGene00010741 | protein_coding_gene | 1.5 | 2.7519 | 2.2E-02 | 2.4 | 5.273 | 3.9E-04 |
| *ZK792.4* | WBGene00014077 | protein_coding_gene | 1.5 | 2.7489 | 5.0E-05 | 1.5 | 2.781 | 6.3E-07 |
| *nas-8* | WBGene00003527 | protein_coding_gene | 1.5 | 2.7488 | 4.0E-03 | 1.7 | 3.282 | 3.8E-04 |
| *dgk-1* | WBGene00000958 | protein_coding_gene | 1.5 | 2.7470 | 2.6E-05 | 1.1 | 2.110 | 5.4E-06 |
| *Y53G8B.3* | WBGene00021819 | pseudogene | 1.5 | 2.7467 | 5.9E-03 | 1.8 | 3.585 | 1.2E-04 |
| *Y54G2A.41* | WBGene00044485 | protein_coding_gene | 1.5 | 2.7457 | 2.7E-06 | 2.1 | 4.213 | 1.6E-13 |
| *C39H7.4* | WBGene00016542 | protein_coding_gene | 1.5 | 2.7453 | 1.9E-08 | 1.4 | 2.609 | 9.0E-22 |
| *unc-82* | WBGene00006814 | protein_coding_gene | 1.5 | 2.7431 | 2.4E-05 | 1.5 | 2.880 | 3.8E-20 |
| ***cutl-9*** | **WBGene00010893** | **protein_coding_gene** | **1.5** | **2.7406** | **2.0E-02** | **-2.8** | **0.148** | **2.7E-18** |
| *acr-2* | WBGene00000042 | protein_coding_gene | 1.5 | 2.7403 | 2.5E-03 | 1.5 | 2.791 | 9.4E-03 |
| *col-104* | WBGene00000678 | protein_coding_gene | 1.5 | 2.7399 | 2.8E-03 | 2.3 | 5.051 | 3.2E-04 |
| *nhr-212* | WBGene00020153 | protein_coding_gene | 1.5 | 2.7398 | 1.4E-04 | 1.5 | 2.735 | 4.1E-11 |
| *W09C5.1* | WBGene00012351 | protein_coding_gene | 1.5 | 2.7394 | 4.6E-27 | 1.3 | 2.424 | 2.1E-22 |
| *T25G12.3* | WBGene00020811 | protein_coding_gene | 1.5 | 2.7347 | 3.6E-05 | 2.2 | 4.714 | 2.5E-15 |
| *R05D8.7* | WBGene00019885 | protein_coding_gene | 1.5 | 2.7339 | 1.4E-04 | 1.3 | 2.538 | 1.1E-04 |
| ***E03A3.10*** | **WBGene00189944** | **pseudogene** | **1.5** | **2.7328** | **3.4E-02** | **-1.0** | **0.485** | **3.3E-02** |
| ***cpt-4*** | **WBGene00019644** | **protein_coding_gene** | **1.4** | **2.7320** | **2.2E-02** | **-1.8** | **0.287** | **1.0E-14** |
| *rab-37* | WBGene00004285 | protein_coding_gene | 1.4 | 2.7312 | 2.6E-04 | 1.2 | 2.311 | 8.0E-03 |
| *K02E10.7* | WBGene00019321 | protein_coding_gene | 1.4 | 2.7308 | 8.0E-05 | 2.2 | 4.756 | 1.0E-08 |
| *lron-4* | WBGene00008051 | protein_coding_gene | 1.4 | 2.7285 | 3.8E-05 | 1.5 | 2.839 | 2.0E-05 |
| *egl-47* | WBGene00001211 | protein_coding_gene | 1.4 | 2.7284 | 8.7E-03 | 1.5 | 2.819 | 2.8E-02 |
| *C55A6.6* | WBGene00008335 | protein_coding_gene | 1.4 | 2.7265 | 4.8E-10 | 1.2 | 2.240 | 6.7E-06 |
| *ZC21.10* | WBGene00044443 | protein_coding_gene | 1.4 | 2.7265 | 1.5E-02 | 1.3 | 2.432 | 1.1E-02 |
| *T19D12.1* | WBGene00020576 | protein_coding_gene | 1.4 | 2.7249 | 1.5E-04 | 1.9 | 3.691 | 1.1E-26 |
| *Y41E3.22* | WBGene00045146 | protein_coding_gene | 1.4 | 2.7236 | 4.6E-03 | 1.2 | 2.358 | 1.3E-02 |
| *C55A6.11* | WBGene00008340 | protein_coding_gene | 1.4 | 2.7206 | 9.1E-03 | 1.7 | 3.156 | 6.7E-05 |
| *nlp-52* | WBGene00018363 | protein_coding_gene | 1.4 | 2.7149 | 1.4E-02 | 1.2 | 2.267 | 1.7E-02 |
| *F16C3.3* | WBGene00023415 | protein_coding_gene | 1.4 | 2.7148 | 1.3E-06 | 1.1 | 2.159 | 1.4E-05 |
| *ZC155.2* | WBGene00022530 | protein_coding_gene | 1.4 | 2.7135 | 7.0E-03 | 2.1 | 4.163 | 3.8E-07 |
| *del-1* | WBGene00000952 | protein_coding_gene | 1.4 | 2.7120 | 6.7E-03 | 1.4 | 2.687 | 2.3E-02 |
| *D2085.7* | WBGene00008432 | protein_coding_gene | 1.4 | 2.7113 | 2.3E-04 | 1.4 | 2.705 | 1.1E-06 |
| *glb-10* | WBGene00016215 | protein_coding_gene | 1.4 | 2.7103 | 1.2E-05 | 1.5 | 2.733 | 2.4E-04 |
| *gap-1* | WBGene00001515 | protein_coding_gene | 1.4 | 2.7072 | 1.2E-07 | 1.2 | 2.356 | 2.3E-06 |
| *ptr-11* | WBGene00004225 | protein_coding_gene | 1.4 | 2.7055 | 5.2E-09 | 1.5 | 2.753 | 3.1E-10 |
| *H20J04.1* | WBGene00019216 | protein_coding_gene | 1.4 | 2.6995 | 1.1E-03 | 2.0 | 4.047 | 1.0E-16 |
| *cdd-1* | WBGene00000391 | protein_coding_gene | 1.4 | 2.6958 | 7.3E-10 | 1.4 | 2.688 | 2.5E-11 |
| *F55C12.4* | WBGene00018871 | protein_coding_gene | 1.4 | 2.6932 | 1.5E-04 | 1.4 | 2.731 | 7.5E-05 |
| *mtd-1* | WBGene00003471 | protein_coding_gene | 1.4 | 2.6925 | 2.7E-04 | 1.9 | 3.739 | 1.4E-05 |
| *Y105E8A.1* | WBGene00013666 | protein_coding_gene | 1.4 | 2.6909 | 5.2E-05 | 1.8 | 3.545 | 5.6E-15 |
| *osm-3* | WBGene00003884 | protein_coding_gene | 1.4 | 2.6870 | 3.8E-08 | 1.3 | 2.410 | 5.4E-03 |
| *nhr-109* | WBGene00003699 | protein_coding_gene | 1.4 | 2.6869 | 6.0E-07 | 1.0 | 2.020 | 1.4E-06 |
| *C49C8.2* | WBGene00016766 | protein_coding_gene | 1.4 | 2.6868 | 5.7E-04 | 1.2 | 2.327 | 1.2E-02 |

| | | | | | | | | |
|---|---|---|---|---|---|---|---|---|
| *spe-49* | WBGene00019667 | protein_coding_gene | 1.4 | 2.6823 | 8.8E-03 | 2.8 | 6.743 | 3.7E-20 |
| ***F07A5.4*** | **WBGene00008543** | **protein_coding_gene** | **1.4** | **2.6811** | **1.1E-02** | **-1.4** | **0.381** | **6.5E-03** |
| *ztf-27* | WBGene00011661 | protein_coding_gene | 1.4 | 2.6810 | 6.6E-11 | 1.2 | 2.305 | 2.1E-11 |
| *F55D10.4* | WBGene00018879 | protein_coding_gene | 1.4 | 2.6806 | 3.5E-03 | 2.0 | 4.036 | 4.9E-05 |
| *arrd-26* | WBGene00019873 | protein_coding_gene | 1.4 | 2.6791 | 3.3E-04 | 2.0 | 4.131 | 2.3E-06 |
| *col-77* | WBGene00000653 | protein_coding_gene | 1.4 | 2.6790 | 5.9E-03 | 2.4 | 5.457 | 5.6E-03 |
| *dpy-31* | WBGene00006592 | protein_coding_gene | 1.4 | 2.6778 | 5.7E-11 | 2.1 | 4.294 | 4.3E-49 |
| *acc-4* | WBGene00012099 | protein_coding_gene | 1.4 | 2.6775 | 1.2E-04 | 1.1 | 2.113 | 3.2E-03 |
| *Y102A11A.2* | WBGene00022412 | protein_coding_gene | 1.4 | 2.6736 | 1.4E-03 | 1.6 | 3.103 | 3.1E-11 |
| *npr-31* | WBGene00020319 | protein_coding_gene | 1.4 | 2.6683 | 1.5E-02 | 1.1 | 2.145 | 2.8E-02 |
| *ZK470.6* | WBGene00022747 | protein_coding_gene | 1.4 | 2.6683 | 4.3E-02 | 2.5 | 5.594 | 5.8E-03 |
| ***Y110A2AL.4*** | **WBGene00022441** | **protein_coding_gene** | **1.4** | **2.6682** | **6.1E-03** | **-2.4** | **0.196** | **2.0E-08** |
| *nas-30* | WBGene00003548 | protein_coding_gene | 1.4 | 2.6667 | 2.6E-03 | 2.2 | 4.604 | 2.6E-04 |
| *pry-1* | WBGene00004202 | protein_coding_gene | 1.4 | 2.6662 | 8.7E-10 | 1.1 | 2.136 | 4.5E-08 |
| *unc-36* | WBGene00006772 | protein_coding_gene | 1.4 | 2.6655 | 5.3E-11 | 1.1 | 2.108 | 5.9E-08 |
| *catp-5* | WBGene00019493 | protein_coding_gene | 1.4 | 2.6632 | 1.0E-05 | 1.8 | 3.493 | 1.5E-22 |
| *C18A11.3* | WBGene00015949 | protein_coding_gene | 1.4 | 2.6629 | 4.8E-08 | 1.9 | 3.632 | 3.6E-08 |
| *C31H5.6* | WBGene00007857 | protein_coding_gene | 1.4 | 2.6628 | 3.7E-06 | 1.1 | 2.130 | 2.6E-07 |
| *twk-32* | WBGene00006684 | protein_coding_gene | 1.4 | 2.6622 | 5.4E-03 | 1.1 | 2.163 | 4.1E-02 |
| *zzz-1* | WBGene00009333 | protein_coding_gene | 1.4 | 2.6608 | 7.3E-08 | 1.1 | 2.075 | 4.1E-05 |
| *daf-37* | WBGene00016246 | protein_coding_gene | 1.4 | 2.6605 | 8.8E-05 | 1.6 | 2.936 | 9.2E-06 |
| *acr-9* | WBGene00000048 | protein_coding_gene | 1.4 | 2.6601 | 1.9E-02 | 1.4 | 2.730 | 3.6E-02 |
| *T28H10.2* | WBGene00012143 | protein_coding_gene | 1.4 | 2.6598 | 3.1E-07 | 1.5 | 2.751 | 2.9E-08 |
| *add-2* | WBGene00000073 | protein_coding_gene | 1.4 | 2.6592 | 3.9E-14 | 1.6 | 2.976 | 1.2E-23 |
| *acp-5* | WBGene00017427 | protein_coding_gene | 1.4 | 2.6588 | 7.6E-09 | 1.1 | 2.133 | 4.3E-06 |
| *W03A5.1* | WBGene00020966 | protein_coding_gene | 1.4 | 2.6581 | 1.8E-04 | 2.3 | 4.800 | 3.7E-22 |
| *lact-7* | WBGene00008047 | protein_coding_gene | 1.4 | 2.6580 | 7.7E-03 | 1.6 | 3.041 | 3.5E-03 |
| *ZK622.4* | WBGene00022782 | protein_coding_gene | 1.4 | 2.6568 | 4.5E-07 | 1.0 | 2.003 | 3.1E-05 |
| *F47G3.1* | WBGene00018576 | protein_coding_gene | 1.4 | 2.6555 | 6.4E-09 | 2.2 | 4.548 | 2.0E-17 |
| ***clec-221*** | **WBGene00008916** | **protein_coding_gene** | **1.4** | **2.6553** | **3.8E-02** | **-2.5** | **0.182** | **2.7E-39** |
| *F56D2.5* | WBGene00018966 | protein_coding_gene | 1.4 | 2.6539 | 1.8E-03 | 1.1 | 2.075 | 9.8E-06 |
| *acy-3* | WBGene00000070 | protein_coding_gene | 1.4 | 2.6458 | 7.3E-09 | 1.0 | 2.023 | 2.4E-08 |
| *jun-1* | WBGene00012005 | protein_coding_gene | 1.4 | 2.6378 | 1.1E-07 | 1.2 | 2.276 | 5.2E-12 |
| *nlp-78* | WBGene00195143 | protein_coding_gene | 1.4 | 2.6347 | 2.2E-02 | 1.0 | 2.048 | 3.7E-02 |
| *C30H6.5* | WBGene00007823 | protein_coding_gene | 1.4 | 2.6347 | 5.7E-03 | 2.0 | 4.060 | 1.7E-02 |
| ***T02E9.5*** | **WBGene00011383** | **protein_coding_gene** | **1.4** | **2.6320** | **9.4E-04** | **-2.3** | **0.202** | **5.1E-34** |
| *kri-1* | WBGene00013955 | protein_coding_gene | 1.4 | 2.6297 | 8.4E-05 | 1.2 | 2.337 | 1.4E-03 |
| *Y53F4B.51* | WBGene00194905 | protein_coding_gene | 1.4 | 2.6294 | 8.2E-03 | 1.8 | 3.386 | 3.2E-03 |
| *unc-38* | WBGene00006774 | protein_coding_gene | 1.4 | 2.6293 | 1.7E-05 | 1.1 | 2.103 | 2.0E-03 |
| *T04C12.11* | WBGene00077693 | protein_coding_gene | 1.4 | 2.6272 | 1.8E-09 | 1.2 | 2.327 | 7.7E-11 |
| *twk-34* | WBGene00006686 | protein_coding_gene | 1.4 | 2.6245 | 3.0E-08 | 1.5 | 2.896 | 1.1E-10 |
| *R04A9.3* | WBGene00019863 | protein_coding_gene | 1.4 | 2.6168 | 8.8E-11 | 1.4 | 2.594 | 2.9E-10 |
| *T22F7.4* | WBGene00020703 | protein_coding_gene | 1.4 | 2.6155 | 1.1E-04 | 2.1 | 4.437 | 3.4E-08 |
| *col-130* | WBGene00000704 | protein_coding_gene | 1.4 | 2.6144 | 8.5E-03 | 2.0 | 4.118 | 2.3E-02 |
| *F26D2.3* | WBGene00009148 | protein_coding_gene | 1.4 | 2.6128 | 2.2E-02 | 2.9 | 7.280 | 1.3E-04 |
| *C02B8.12* | WBGene00219375 | protein_coding_gene | 1.4 | 2.6119 | 2.1E-04 | 1.4 | 2.554 | 2.3E-03 |
| *pes-7* | WBGene00003980 | protein_coding_gene | 1.4 | 2.6104 | 1.7E-07 | 1.3 | 2.477 | 8.5E-18 |
| *mec-18* | WBGene00003179 | protein_coding_gene | 1.4 | 2.6090 | 9.3E-05 | 1.8 | 3.466 | 8.7E-06 |
| *twk-39* | WBGene00006690 | protein_coding_gene | 1.4 | 2.6085 | 7.3E-04 | 1.2 | 2.333 | 2.1E-03 |
| *F23H11.7* | WBGene00017762 | protein_coding_gene | 1.4 | 2.6083 | 3.9E-05 | 1.8 | 3.414 | 1.4E-06 |
| *abcx-1* | WBGene00006522 | protein_coding_gene | 1.4 | 2.6074 | 4.0E-03 | 1.2 | 2.290 | 4.1E-02 |
| *cest-31* | WBGene00019653 | protein_coding_gene | 1.4 | 2.6061 | 6.5E-07 | 1.6 | 3.006 | 3.9E-09 |
| *gst-11* | WBGene00001759 | protein_coding_gene | 1.4 | 2.6012 | 1.3E-02 | 1.2 | 2.302 | 2.5E-02 |
| *C55A6.7* | WBGene00008336 | protein_coding_gene | 1.4 | 2.6009 | 7.4E-06 | 1.2 | 2.320 | 1.6E-05 |
| *C49F8.3* | WBGene00008215 | protein_coding_gene | 1.4 | 2.6008 | 2.9E-04 | 1.4 | 2.730 | 4.6E-05 |
| ***EEED8.2*** | **WBGene00017133** | **protein_coding_gene** | **1.4** | **2.6001** | **3.0E-05** | **-1.3** | **0.394** | **9.2E-08** |
| *oac-48* | WBGene00011960 | protein_coding_gene | 1.4 | 2.5967 | 5.0E-02 | 2.1 | 4.364 | 2.0E-04 |
| *plc-1* | WBGene00004036 | protein_coding_gene | 1.4 | 2.5965 | 7.9E-05 | 1.5 | 2.876 | 9.2E-23 |
| *R11F4.3* | WBGene00020009 | protein_coding_gene | 1.4 | 2.5957 | 1.7E-05 | 1.7 | 3.211 | 9.1E-09 |
| *nlp-79* | WBGene00008626 | protein_coding_gene | 1.4 | 2.5949 | 1.5E-04 | 1.2 | 2.352 | 2.8E-03 |
| *unc-1* | WBGene00006741 | protein_coding_gene | 1.4 | 2.5943 | 8.6E-06 | 1.0 | 2.059 | 6.5E-08 |
| *F13H8.5* | WBGene00017438 | protein_coding_gene | 1.4 | 2.5879 | 8.7E-04 | 2.0 | 3.933 | 1.0E-03 |
| *twk-40* | WBGene00006691 | protein_coding_gene | 1.4 | 2.5845 | 2.3E-03 | 1.7 | 3.236 | 2.2E-03 |
| *lgc-43* | WBGene00008076 | protein_coding_gene | 1.4 | 2.5818 | 2.9E-09 | 1.6 | 3.119 | 8.5E-11 |
| *daf-12* | WBGene00000908 | protein_coding_gene | 1.4 | 2.5811 | 5.0E-04 | 1.4 | 2.717 | 5.1E-18 |
| *C29F5.3* | WBGene00016212 | protein_coding_gene | 1.4 | 2.5789 | 1.8E-03 | 1.8 | 3.374 | 2.0E-08 |
| *let-756* | WBGene00002881 | protein_coding_gene | 1.4 | 2.5776 | 1.5E-04 | 1.2 | 2.240 | 1.2E-06 |
| *gar-2* | WBGene00001518 | protein_coding_gene | 1.4 | 2.5756 | 5.7E-07 | 1.7 | 3.292 | 5.5E-10 |
| *glt-3* | WBGene00001621 | protein_coding_gene | 1.4 | 2.5714 | 1.8E-06 | 1.1 | 2.073 | 4.6E-05 |
| *fbxa-54* | WBGene00020449 | protein_coding_gene | 1.4 | 2.5705 | 3.8E-07 | 1.6 | 3.018 | 2.5E-03 |
| *clh-4* | WBGene00000531 | protein_coding_gene | 1.4 | 2.5705 | 2.6E-03 | 1.6 | 3.138 | 1.4E-05 |
| *M01F1.3* | WBGene00010809 | protein_coding_gene | 1.4 | 2.5689 | 1.2E-22 | 1.3 | 2.429 | 4.0E-19 |
| *dmsr-1* | WBGene00010191 | protein_coding_gene | 1.4 | 2.5687 | 2.4E-08 | 1.3 | 2.443 | 8.6E-10 |
| *F08D12.7* | WBGene00017250 | protein_coding_gene | 1.4 | 2.5676 | 2.3E-07 | 2.1 | 4.311 | 1.7E-14 |
| *K10C9.9* | WBGene00044515 | pseudogene | 1.4 | 2.5658 | 2.8E-02 | 2.3 | 4.918 | 3.0E-03 |
| *col-118* | WBGene00000692 | protein_coding_gene | 1.4 | 2.5649 | 4.9E-05 | 2.0 | 3.986 | 7.9E-36 |
| *H43E16.1* | WBGene00019273 | protein_coding_gene | 1.4 | 2.5639 | 5.1E-03 | 2.0 | 4.042 | 1.5E-23 |
| *lido-17* | WBGene00007378 | protein_coding_gene | 1.4 | 2.5603 | 4.8E-03 | 1.0 | 2.008 | 7.1E-03 |
| *clec-205* | WBGene00021602 | protein_coding_gene | 1.4 | 2.5597 | 4.3E-06 | 2.2 | 4.568 | 9.3E-09 |
| *lido-16* | WBGene00020898 | protein_coding_gene | 1.4 | 2.5573 | 4.5E-03 | 2.6 | 6.226 | 3.1E-09 |

| | | | | | | | | |
|---|---|---|---|---|---|---|---|---|
| *nhr-3* | WBGene00003602 | protein_coding_gene | 1.4 | 2.5532 | 1.9E-05 | 1.5 | 2.839 | 3.1E-14 |
| *C14B9.3* | WBGene00015753 | protein_coding_gene | 1.4 | 2.5529 | 1.4E-06 | 1.8 | 3.378 | 3.3E-07 |
| *unc-49* | WBGene00006784 | protein_coding_gene | 1.4 | 2.5526 | 5.1E-05 | 1.2 | 2.357 | 4.4E-12 |
| *T05C12.11* | WBGene00011473 | protein_coding_gene | 1.4 | 2.5524 | 1.1E-02 | 1.3 | 2.527 | 3.2E-02 |
| *irk-1* | WBGene00002149 | protein_coding_gene | 1.4 | 2.5516 | 3.1E-02 | 1.9 | 3.675 | 1.2E-04 |
| *T19D2.3* | WBGene00020568 | protein_coding_gene | 1.4 | 2.5510 | 6.7E-04 | 1.2 | 2.353 | 9.9E-03 |
| ***F46H6.5*** | **WBGene00023460** | **protein_coding_gene** | **1.4** | **2.5498** | **2.6E-02** | **-1.3** | **0.404** | **2.3E-02** |
| *hum-8* | WBGene00002041 | protein_coding_gene | 1.3 | 2.5474 | 4.3E-12 | 1.1 | 2.123 | 9.4E-17 |
| *F28H7.3* | WBGene00009237 | protein_coding_gene | 1.3 | 2.5469 | 1.4E-05 | 2.2 | 4.712 | 3.6E-10 |
| *F25E5.5* | WBGene00017786 | protein_coding_gene | 1.3 | 2.5454 | 1.4E-05 | 1.1 | 2.131 | 2.3E-09 |
| *kin-16* | WBGene00002200 | protein_coding_gene | 1.3 | 2.5449 | 1.8E-08 | 1.4 | 2.730 | 3.3E-09 |
| *Y69A2AR.22* | WBGene00022093 | protein_coding_gene | 1.3 | 2.5420 | 2.6E-08 | 1.0 | 2.015 | 2.2E-08 |
| *Y71F9B.13* | WBGene00022131 | protein_coding_gene | 1.3 | 2.5343 | 1.5E-10 | 1.4 | 2.687 | 4.1E-16 |
| *arrd-3* | WBGene00006429 | protein_coding_gene | 1.3 | 2.5326 | 3.2E-02 | 1.7 | 3.308 | 4.5E-02 |
| *F22F1.3* | WBGene00017715 | protein_coding_gene | 1.3 | 2.5319 | 2.5E-02 | 1.9 | 3.717 | 8.8E-04 |
| *H10D12.2* | WBGene00019176 | pseudogene | 1.3 | 2.5280 | 1.9E-05 | 1.1 | 2.097 | 7.8E-05 |
| *Y37H2A.14* | WBGene00045416 | protein_coding_gene | 1.3 | 2.5266 | 4.1E-03 | 2.4 | 5.305 | 7.7E-12 |
| *trp-2* | WBGene00006615 | protein_coding_gene | 1.3 | 2.5241 | 2.0E-04 | 1.2 | 2.263 | 2.0E-02 |
| *F17H10.2* | WBGene00008926 | protein_coding_gene | 1.3 | 2.5234 | 1.5E-09 | 1.2 | 2.284 | 2.7E-13 |
| *gst-24* | WBGene00001772 | protein_coding_gene | 1.3 | 2.5209 | 1.3E-05 | 1.7 | 3.275 | 3.1E-05 |
| *unc-80* | WBGene00006812 | protein_coding_gene | 1.3 | 2.5189 | 3.9E-06 | 1.4 | 2.707 | 4.4E-15 |
| ***F13H10.5*** | **WBGene00008776** | **protein_coding_gene** | **1.3** | **2.5136** | **1.4E-02** | **-2.3** | **0.199** | **4.2E-09** |
| *kcnl-1* | WBGene00007176 | protein_coding_gene | 1.3 | 2.5125 | 5.5E-05 | 1.9 | 3.626 | 6.8E-10 |
| *B0507.2* | WBGene00015219 | protein_coding_gene | 1.3 | 2.5104 | 5.4E-20 | 1.0 | 2.011 | 1.4E-14 |
| *ZK1067.4* | WBGene00014210 | protein_coding_gene | 1.3 | 2.5101 | 7.1E-05 | 1.3 | 2.499 | 1.9E-04 |
| ***ccep-290*** | **WBGene00021643** | **protein_coding_gene** | **1.3** | **2.5099** | **7.2E-03** | **-1.1** | **0.465** | **2.5E-03** |
| *K03B8.6* | WBGene00010524 | protein_coding_gene | 1.3 | 2.5066 | 1.4E-06 | 1.3 | 2.490 | 9.4E-06 |
| *pan-1* | WBGene00003915 | protein_coding_gene | 1.3 | 2.5036 | 5.0E-11 | 1.5 | 2.784 | 1.9E-19 |
| *dhs-31* | WBGene00011424 | protein_coding_gene | 1.3 | 2.5033 | 5.7E-06 | 1.2 | 2.298 | 8.5E-08 |
| *F13D12.10* | WBGene00008744 | protein_coding_gene | 1.3 | 2.5014 | 6.4E-03 | 1.1 | 2.133 | 1.2E-02 |
| *K03H6.5* | WBGene00019370 | protein_coding_gene | 1.3 | 2.5003 | 6.5E-03 | 1.6 | 3.030 | 3.4E-03 |
| *C27D8.3* | WBGene00007779 | protein_coding_gene | 1.3 | 2.4993 | 4.1E-08 | 1.0 | 2.011 | 2.1E-11 |
| *pbo-5* | WBGene00012857 | protein_coding_gene | 1.3 | 2.4993 | 1.8E-06 | 1.2 | 2.308 | 1.7E-05 |
| *nep-22* | WBGene00020788 | protein_coding_gene | 1.3 | 2.4992 | 1.3E-03 | 2.9 | 7.235 | 2.0E-56 |
| *nlp-10* | WBGene00003748 | protein_coding_gene | 1.3 | 2.4952 | 5.7E-07 | 1.2 | 2.320 | 3.2E-11 |
| *cysl-2* | WBGene00010759 | protein_coding_gene | 1.3 | 2.4931 | 4.2E-09 | 1.2 | 2.250 | 5.7E-10 |
| *pck-1* | WBGene00021043 | protein_coding_gene | 1.3 | 2.4912 | 1.7E-08 | 1.5 | 2.914 | 3.1E-36 |
| *F29B9.12* | WBGene00044482 | protein_coding_gene | 1.3 | 2.4896 | 3.4E-02 | 1.7 | 3.332 | 1.1E-02 |
| *E02C12.6* | WBGene00017092 | protein_coding_gene | 1.3 | 2.4892 | 2.0E-02 | 2.8 | 6.743 | 2.9E-18 |
| *Y4C6B.4* | WBGene00021158 | protein_coding_gene | 1.3 | 2.4848 | 8.1E-06 | 1.2 | 2.229 | 2.8E-04 |
| *gur-4* | WBGene00001805 | protein_coding_gene | 1.3 | 2.4824 | 1.2E-02 | 2.3 | 4.852 | 1.4E-05 |
| *spp-14* | WBGene00004999 | protein_coding_gene | 1.3 | 2.4796 | 1.3E-09 | 1.8 | 3.569 | 1.5E-69 |
| *myrf-1* | WBGene00004134 | protein_coding_gene | 1.3 | 2.4760 | 6.4E-05 | 1.5 | 2.867 | 4.7E-13 |
| *ZK813.4* | WBGene00022823 | protein_coding_gene | 1.3 | 2.4726 | 1.5E-07 | 1.4 | 2.711 | 7.2E-08 |
| *nhr-211* | WBGene00020152 | protein_coding_gene | 1.3 | 2.4718 | 3.8E-03 | 1.2 | 2.317 | 2.5E-04 |
| *nlp-16* | WBGene00003754 | protein_coding_gene | 1.3 | 2.4711 | 3.7E-11 | 1.4 | 2.559 | 5.8E-16 |
| *E02H9.2* | WBGene00017100 | protein_coding_gene | 1.3 | 2.4668 | 6.9E-04 | 1.7 | 3.276 | 8.6E-07 |
| ***oac-51*** | **WBGene00020976** | **protein_coding_gene** | **1.3** | **2.4664** | **1.4E-02** | **-1.2** | **0.447** | **4.4E-03** |
| *saeg-1* | WBGene00010012 | protein_coding_gene | 1.3 | 2.4662 | 4.0E-06 | 1.4 | 2.620 | 1.5E-26 |
| *Y54G11A.4* | WBGene00013215 | protein_coding_gene | 1.3 | 2.4655 | 1.8E-04 | 1.0 | 2.019 | 2.0E-04 |
| *W08F4.5* | WBGene00021094 | protein_coding_gene | 1.3 | 2.4626 | 4.5E-03 | 1.9 | 3.789 | 1.6E-03 |
| *Y39B6A.25* | WBGene00012684 | protein_coding_gene | 1.3 | 2.4626 | 1.7E-02 | 2.0 | 3.865 | 1.9E-11 |
| *rsbp-1* | WBGene00011531 | protein_coding_gene | 1.3 | 2.4598 | 4.7E-06 | 1.0 | 2.044 | 1.3E-09 |
| *F08D12.3* | WBGene00017247 | protein_coding_gene | 1.3 | 2.4581 | 1.2E-03 | 1.0 | 2.052 | 9.5E-07 |
| *T20H4.2* | WBGene00020635 | protein_coding_gene | 1.3 | 2.4563 | 9.0E-03 | 1.8 | 3.445 | 7.9E-09 |
| *pcp-1* | WBGene00003956 | protein_coding_gene | 1.3 | 2.4532 | 6.6E-16 | 1.4 | 2.617 | 2.4E-36 |
| *Y73B6A.1* | WBGene00022228 | protein_coding_gene | 1.3 | 2.4504 | 1.5E-02 | 1.2 | 2.227 | 8.0E-04 |
| *nlp-47* | WBGene00018117 | protein_coding_gene | 1.3 | 2.4461 | 2.0E-05 | 1.1 | 2.196 | 1.5E-07 |
| *M05B5.3* | WBGene00010871 | protein_coding_gene | 1.3 | 2.4450 | 5.9E-04 | 1.8 | 3.456 | 1.5E-11 |
| *F16B4.5* | WBGene00017507 | protein_coding_gene | 1.3 | 2.4445 | 1.4E-05 | 1.7 | 3.228 | 1.8E-11 |
| *unc-17* | WBGene00006756 | protein_coding_gene | 1.3 | 2.4423 | 7.9E-05 | 1.7 | 3.354 | 1.5E-05 |
| *F22F7.2* | WBGene00017720 | protein_coding_gene | 1.3 | 2.4409 | 8.9E-06 | 1.5 | 2.777 | 3.0E-05 |
| *C17H12.10* | WBGene00015935 | protein_coding_gene | 1.3 | 2.4394 | 2.7E-03 | 1.4 | 2.693 | 2.1E-03 |
| *C44H4.8* | WBGene00008096 | protein_coding_gene | 1.3 | 2.4384 | 4.8E-02 | 1.6 | 3.005 | 9.8E-03 |
| *T28H11.8* | WBGene00020906 | protein_coding_gene | 1.3 | 2.4377 | 3.4E-05 | 1.2 | 2.240 | 1.8E-03 |
| *T21F4.1* | WBGene00020658 | protein_coding_gene | 1.3 | 2.4359 | 5.2E-08 | 1.2 | 2.224 | 1.9E-19 |
| *nipa-1* | WBGene00021820 | protein_coding_gene | 1.3 | 2.4342 | 6.0E-29 | 1.1 | 2.100 | 6.2E-18 |
| *tag-180* | WBGene00007041 | protein_coding_gene | 1.3 | 2.4336 | 1.3E-06 | 1.8 | 3.554 | 5.9E-12 |
| *C13G3.1* | WBGene00007553 | protein_coding_gene | 1.3 | 2.4320 | 2.9E-05 | 1.9 | 3.781 | 3.4E-13 |
| *F54G2.1* | WBGene00018837 | protein_coding_gene | 1.3 | 2.4308 | 6.5E-08 | 1.6 | 3.112 | 4.6E-17 |
| *F41B4.2* | WBGene00018258 | protein_coding_gene | 1.3 | 2.4304 | 1.6E-05 | 1.1 | 2.096 | 7.9E-08 |
| *ZK185.2* | WBGene00022682 | protein_coding_gene | 1.3 | 2.4296 | 8.7E-04 | 1.5 | 2.733 | 3.2E-04 |
| *T03D8.6* | WBGene00011393 | protein_coding_gene | 1.3 | 2.4280 | 3.8E-11 | 1.3 | 2.504 | 1.5E-11 |
| *C49C3.10* | WBGene00008200 | protein_coding_gene | 1.3 | 2.4269 | 2.4E-10 | 1.1 | 2.088 | 1.4E-07 |
| *R13H4.5* | WBGene00011263 | protein_coding_gene | 1.3 | 2.4242 | 5.4E-05 | 1.2 | 2.304 | 1.6E-07 |
| *F23D12.3* | WBGene00009087 | protein_coding_gene | 1.3 | 2.4237 | 4.5E-05 | 1.9 | 3.851 | 1.0E-02 |
| ***B0393.5*** | **WBGene00007170** | **protein_coding_gene** | **1.3** | **2.4233** | **1.5E-03** | **-1.6** | **0.340** | **3.0E-03** |
| *T02B11.8* | WBGene00044776 | protein_coding_gene | 1.3 | 2.4231 | 9.6E-05 | 1.0 | 2.009 | 1.2E-02 |
| *cest-19* | WBGene00020125 | protein_coding_gene | 1.3 | 2.4227 | 2.6E-02 | 1.2 | 2.289 | 1.9E-02 |
| *sipa-1* | WBGene00012100 | protein_coding_gene | 1.3 | 2.4224 | 3.0E-10 | 1.6 | 2.944 | 4.6E-20 |

| | | | | | | | | |
|---|---|---|---|---|---|---|---|---|
| *F49E2.2* | WBGene00009886 | protein_coding_gene | 1.3 | 2.4192 | 3.8E-12 | 1.6 | 2.963 | 1.7E-28 |
| *C50C3.2* | WBGene00016800 | protein_coding_gene | 1.3 | 2.4169 | 5.3E-03 | 2.0 | 4.095 | 2.4E-24 |
| *M163.8* | WBGene00010940 | protein_coding_gene | 1.3 | 2.4151 | 2.3E-05 | 2.0 | 3.986 | 1.6E-14 |
| *gtl-1* | WBGene00001795 | protein_coding_gene | 1.3 | 2.4127 | 1.5E-05 | 1.2 | 2.328 | 8.6E-19 |
| *tmc-1* | WBGene00020490 | protein_coding_gene | 1.3 | 2.4127 | 5.6E-05 | 1.3 | 2.546 | 1.3E-06 |
| *tre-4* | WBGene00006610 | protein_coding_gene | 1.3 | 2.4126 | 8.8E-07 | 1.7 | 3.243 | 1.3E-11 |
| *Y60A3A.24* | WBGene00023519 | protein_coding_gene | 1.3 | 2.4122 | 7.0E-04 | 1.2 | 2.229 | 6.1E-03 |
| *dhhc-2* | WBGene00012948 | protein_coding_gene | 1.3 | 2.4121 | 1.1E-06 | 1.1 | 2.158 | 2.2E-07 |
| *F32B5.7* | WBGene00017980 | protein_coding_gene | 1.3 | 2.4101 | 1.0E-05 | 1.3 | 2.483 | 1.1E-13 |
| *gst-41* | WBGene00001789 | protein_coding_gene | 1.3 | 2.4086 | 2.2E-03 | 3.1 | 8.309 | 7.6E-13 |
| *dat-1* | WBGene00000934 | protein_coding_gene | 1.3 | 2.4083 | 2.8E-03 | 2.2 | 4.513 | 7.7E-06 |
| *faah-6* | WBGene00012750 | protein_coding_gene | 1.3 | 2.4073 | 1.2E-04 | 1.3 | 2.457 | 1.5E-04 |
| *npr-22* | WBGene00022004 | protein_coding_gene | 1.3 | 2.4070 | 8.3E-06 | 1.4 | 2.608 | 5.5E-05 |
| *pcyt-2.1* | WBGene00021352 | protein_coding_gene | 1.3 | 2.4069 | 7.8E-22 | 1.0 | 2.038 | 9.2E-13 |
| *cnp-2* | WBGene00012901 | protein_coding_gene | 1.3 | 2.4066 | 1.1E-05 | 1.4 | 2.622 | 1.4E-02 |
| *sfxn-5* | WBGene00044049 | protein_coding_gene | 1.3 | 2.4055 | 1.7E-12 | 1.0 | 2.057 | 2.1E-09 |
| *scl-6* | WBGene00008028 | protein_coding_gene | 1.3 | 2.4054 | 5.3E-05 | 5.3 | 38.727 | 2.0E-21 |
| *sqrd-1* | WBGene00008538 | protein_coding_gene | 1.3 | 2.4011 | 5.4E-09 | 1.1 | 2.085 | 9.0E-08 |
| *sdz-3* | WBGene00016211 | pseudogene | 1.3 | 2.4005 | 4.7E-02 | 1.5 | 2.788 | 6.1E-03 |
| *pals-25* | WBGene00020540 | protein_coding_gene | 1.3 | 2.3976 | 1.6E-07 | 1.1 | 2.168 | 5.4E-03 |
| *mlk-1* | WBGene00003374 | protein_coding_gene | 1.3 | 2.3975 | 5.2E-05 | 1.1 | 2.212 | 6.5E-13 |
| *F54C1.1* | WBGene00018789 | protein_coding_gene | 1.3 | 2.3965 | 1.1E-04 | 2.5 | 5.587 | 1.1E-10 |
| *F33E2.4* | WBGene00009360 | protein_coding_gene | 1.3 | 2.3950 | 1.4E-03 | 2.0 | 4.127 | 1.5E-04 |
| *tep-1* | WBGene00013969 | protein_coding_gene | 1.3 | 2.3944 | 9.8E-05 | 1.1 | 2.085 | 1.1E-16 |
| *tra-1* | WBGene00006604 | protein_coding_gene | 1.3 | 2.3938 | 9.8E-05 | 1.4 | 2.639 | 2.2E-18 |
| *tag-151* | WBGene00006497 | protein_coding_gene | 1.3 | 2.3930 | 1.9E-21 | 1.3 | 2.533 | 1.8E-23 |
| *cutl-15* | WBGene00011888 | protein_coding_gene | 1.3 | 2.3925 | 2.5E-04 | 1.3 | 2.486 | 4.7E-06 |
| *F09A5.2* | WBGene00008599 | protein_coding_gene | 1.3 | 2.3887 | 1.2E-03 | 1.2 | 2.245 | 1.5E-04 |
| *arrd-17* | WBGene00011732 | protein_coding_gene | 1.3 | 2.3868 | 3.7E-04 | 1.2 | 2.285 | 4.8E-09 |
| *chil-12* | WBGene00010799 | protein_coding_gene | 1.3 | 2.3862 | 2.1E-07 | 1.6 | 3.132 | 1.2E-06 |
| *oig-5* | WBGene00023520 | protein_coding_gene | 1.3 | 2.3859 | 1.4E-02 | 2.1 | 4.411 | 5.4E-05 |
| *kcc-2* | WBGene00019205 | protein_coding_gene | 1.3 | 2.3855 | 5.0E-07 | 1.1 | 2.198 | 1.9E-22 |
| ***C26F1.1*** | **WBGene00016146** | **protein_coding_gene** | **1.3** | **2.3818** | **6.5E-03** | **-1.9** | **0.275** | **3.5E-08** |
| *lin-31* | WBGene00003017 | protein_coding_gene | 1.3 | 2.3814 | 6.5E-03 | 2.5 | 5.683 | 3.9E-06 |
| *che-7* | WBGene00000488 | protein_coding_gene | 1.2 | 2.3767 | 6.1E-03 | 1.1 | 2.214 | 1.1E-02 |
| *C50D2.2* | WBGene00016806 | protein_coding_gene | 1.2 | 2.3745 | 9.2E-05 | 1.4 | 2.641 | 1.0E-03 |
| *F56D6.17* | WBGene00219307 | pseudogene | 1.2 | 2.3741 | 4.9E-07 | 1.4 | 2.595 | 2.3E-06 |
| *slc-36.2* | WBGene00020837 | protein_coding_gene | 1.2 | 2.3741 | 5.1E-07 | 1.1 | 2.106 | 6.4E-07 |
| *ppw-2* | WBGene00004094 | protein_coding_gene | 1.2 | 2.3718 | 1.6E-10 | 2.4 | 5.423 | 1.4E-56 |
| *nekl-4* | WBGene00017033 | protein_coding_gene | 1.2 | 2.3709 | 8.1E-03 | 1.2 | 2.336 | 1.1E-02 |
| *kin-29* | WBGene00002210 | protein_coding_gene | 1.2 | 2.3698 | 1.5E-07 | 1.3 | 2.484 | 2.8E-24 |
| *wago-10* | WBGene00020707 | protein_coding_gene | 1.2 | 2.3684 | 1.4E-03 | 2.9 | 7.672 | 7.6E-41 |
| *ifc-1* | WBGene00002055 | protein_coding_gene | 1.2 | 2.3682 | 3.4E-05 | 1.5 | 2.776 | 2.3E-35 |
| *F28E10.5* | WBGene00044302 | pseudogene | 1.2 | 2.3679 | 7.7E-03 | 1.8 | 3.436 | 3.4E-09 |
| *F46C5.4* | WBGene00009780 | protein_coding_gene | 1.2 | 2.3628 | 4.6E-02 | 1.1 | 2.213 | 4.6E-02 |
| *amx-2* | WBGene00000138 | protein_coding_gene | 1.2 | 2.3627 | 4.3E-08 | 1.3 | 2.545 | 8.6E-15 |
| *asp-12* | WBGene00017678 | protein_coding_gene | 1.2 | 2.3626 | 1.3E-03 | 1.3 | 2.459 | 3.7E-07 |
| *srb-16* | WBGene00019025 | protein_coding_gene | 1.2 | 2.3599 | 3.0E-02 | 1.3 | 2.453 | 2.3E-02 |
| *C09G1.5* | WBGene00007487 | protein_coding_gene | 1.2 | 2.3597 | 3.9E-04 | 1.7 | 3.218 | 4.0E-06 |
| *oig-8* | WBGene00020582 | protein_coding_gene | 1.2 | 2.3593 | 3.4E-03 | 1.6 | 2.944 | 6.2E-04 |
| *T05H4.7* | WBGene00020270 | protein_coding_gene | 1.2 | 2.3577 | 4.3E-02 | 2.1 | 4.287 | 2.8E-05 |
| *D2089.3* | WBGene00008434 | protein_coding_gene | 1.2 | 2.3571 | 2.4E-02 | 1.2 | 2.356 | 4.9E-03 |
| *F48G7.4* | WBGene00018615 | protein_coding_gene | 1.2 | 2.3552 | 2.4E-04 | 1.1 | 2.086 | 2.0E-02 |
| *B0252.8* | WBGene00044615 | protein_coding_gene | 1.2 | 2.3526 | 1.2E-03 | 1.1 | 2.095 | 4.3E-03 |
| *nhl-1* | WBGene00003597 | protein_coding_gene | 1.2 | 2.3523 | 2.7E-03 | 1.6 | 2.949 | 1.2E-17 |
| *dac-1* | WBGene00000895 | protein_coding_gene | 1.2 | 2.3516 | 1.0E-04 | 1.3 | 2.509 | 3.3E-04 |
| *K10D2.8* | WBGene00189952 | protein_coding_gene | 1.2 | 2.3483 | 2.4E-03 | 1.9 | 3.675 | 2.3E-14 |
| *K01A11.1* | WBGene00010449 | protein_coding_gene | 1.2 | 2.3481 | 3.2E-07 | 1.4 | 2.604 | 1.2E-07 |
| *C08D8.7* | WBGene00235325 | protein_coding_gene | 1.2 | 2.3473 | 1.0E-02 | 1.1 | 2.203 | 3.6E-02 |
| *try-4* | WBGene00006622 | protein_coding_gene | 1.2 | 2.3437 | 2.6E-02 | 2.1 | 4.436 | 5.1E-03 |
| *R07B7.9* | WBGene00011093 | protein_coding_gene | 1.2 | 2.3400 | 1.1E-07 | 1.2 | 2.270 | 8.5E-05 |
| *T05A1.5* | WBGene00011456 | protein_coding_gene | 1.2 | 2.3367 | 6.2E-03 | 1.9 | 3.611 | 2.7E-04 |
| ***F20C5.6*** | **WBGene00008971** | **protein_coding_gene** | **1.2** | **2.3356** | **5.3E-05** | **-1.3** | **0.400** | **9.2E-08** |
| *C44B11.6* | WBGene00016635 | protein_coding_gene | 1.2 | 2.3341 | 4.4E-05 | 1.0 | 2.045 | 1.7E-03 |
| *B0250.4* | WBGene00007121 | pseudogene | 1.2 | 2.3336 | 3.6E-05 | 1.7 | 3.251 | 4.6E-10 |
| *Y56A3A.6* | WBGene00013227 | protein_coding_gene | 1.2 | 2.3321 | 7.6E-06 | 1.8 | 3.480 | 5.4E-20 |
| *lev-9* | WBGene00002976 | protein_coding_gene | 1.2 | 2.3286 | 1.8E-05 | 2.1 | 4.217 | 9.0E-10 |
| *crp-1* | WBGene00012532 | protein_coding_gene | 1.2 | 2.3222 | 6.0E-07 | 1.0 | 2.030 | 2.3E-05 |
| *ctsa-2* | WBGene00019617 | protein_coding_gene | 1.2 | 2.3217 | 1.2E-03 | 2.3 | 5.082 | 1.5E-08 |
| *kvs-1* | WBGene00002242 | protein_coding_gene | 1.2 | 2.3214 | 1.0E-02 | 1.3 | 2.428 | 2.0E-03 |
| *Y41C4A.12* | WBGene00012758 | protein_coding_gene | 1.2 | 2.3193 | 2.6E-06 | 1.5 | 2.917 | 9.2E-09 |
| *gpa-1* | WBGene00001663 | protein_coding_gene | 1.2 | 2.3179 | 5.1E-04 | 1.2 | 2.269 | 1.8E-02 |
| *pdfr-1* | WBGene00015735 | protein_coding_gene | 1.2 | 2.3142 | 1.1E-07 | 2.0 | 3.945 | 3.6E-25 |
| *T06A4.3* | WBGene00020283 | protein_coding_gene | 1.2 | 2.3124 | 1.1E-03 | 2.0 | 4.104 | 1.6E-06 |
| *aex-2* | WBGene00000085 | protein_coding_gene | 1.2 | 2.3121 | 1.6E-02 | 1.0 | 2.046 | 3.4E-02 |
| *gyg-1* | WBGene00006863 | protein_coding_gene | 1.2 | 2.3117 | 4.4E-07 | 1.7 | 3.348 | 1.0E-29 |
| *F27D4.6* | WBGene00009190 | protein_coding_gene | 1.2 | 2.3108 | 2.3E-08 | 1.0 | 2.006 | 5.5E-08 |
| *C05C12.6* | WBGene00007338 | protein_coding_gene | 1.2 | 2.3087 | 2.8E-02 | 1.2 | 2.250 | 3.3E-02 |
| *phat-4* | WBGene00020237 | protein_coding_gene | 1.2 | 2.3076 | 9.4E-03 | 1.7 | 3.328 | 5.5E-04 |
| ***igcm-1*** | **WBGene00018215** | **protein_coding_gene** | **1.2** | **2.3068** | **3.1E-04** | **-1.1** | **0.477** | **1.6E-04** |

| | | | | | | | | |
|---|---|---|---|---|---|---|---|---|
| *eef-1A.2* | WBGene00001169 | protein_coding_gene | 1.2 | 2.3065 | 2.4E-09 | 1.2 | 2.250 | 2.6E-26 |
| ***F15B9.8*** | **WBGene00008851** | **protein_coding_gene** | **1.2** | **2.3060** | **2.9E-02** | **-2.0** | **0.257** | **1.5E-02** |
| *egl-13* | WBGene00001182 | protein_coding_gene | 1.2 | 2.3052 | 1.7E-03 | 1.5 | 2.758 | 2.7E-15 |
| *flr-4* | WBGene00001468 | protein_coding_gene | 1.2 | 2.3048 | 2.5E-05 | 1.9 | 3.682 | 6.9E-09 |
| *dmd-7* | WBGene00019521 | protein_coding_gene | 1.2 | 2.3031 | 2.9E-07 | 1.4 | 2.688 | 1.4E-36 |
| *B0244.10* | WBGene00015085 | protein_coding_gene | 1.2 | 2.3024 | 7.0E-04 | 2.1 | 4.386 | 6.3E-10 |
| *W02C12.2* | WBGene00020929 | protein_coding_gene | 1.2 | 2.3017 | 4.5E-03 | 1.0 | 2.016 | 1.1E-02 |
| *clec-204* | WBGene00021603 | protein_coding_gene | 1.2 | 2.3014 | 4.9E-05 | 2.7 | 6.527 | 2.5E-14 |
| *K04F1.9* | WBGene00019388 | protein_coding_gene | 1.2 | 2.2995 | 8.3E-06 | 1.6 | 2.958 | 4.8E-10 |
| *F27E5.7* | WBGene00044035 | protein_coding_gene | 1.2 | 2.2985 | 7.6E-04 | 1.3 | 2.494 | 3.2E-03 |
| *ZK673.1* | WBGene00014057 | protein_coding_gene | 1.2 | 2.2978 | 2.7E-07 | 1.5 | 2.806 | 2.7E-16 |
| *nhr-7* | WBGene00003606 | protein_coding_gene | 1.2 | 2.2951 | 9.0E-04 | 1.8 | 3.435 | 8.8E-13 |
| *cyp-29A2* | WBGene00011830 | protein_coding_gene | 1.2 | 2.2933 | 2.0E-07 | 1.1 | 2.089 | 1.1E-10 |
| *ZK1055.2* | WBGene00022843 | protein_coding_gene | 1.2 | 2.2918 | 2.8E-02 | 1.9 | 3.804 | 9.7E-03 |
| *mpz-1* | WBGene00003404 | protein_coding_gene | 1.2 | 2.2915 | 3.2E-04 | 1.1 | 2.134 | 1.3E-12 |
| *R10H10.3* | WBGene00011222 | protein_coding_gene | 1.2 | 2.2891 | 7.4E-09 | 1.0 | 2.002 | 9.1E-17 |
| *nipi-4* | WBGene00077712 | protein_coding_gene | 1.2 | 2.2842 | 1.0E-04 | 1.2 | 2.246 | 3.0E-04 |
| *pde-4* | WBGene00020114 | protein_coding_gene | 1.2 | 2.2784 | 1.4E-05 | 1.0 | 2.007 | 1.9E-12 |
| *mfsd-6* | WBGene00020051 | protein_coding_gene | 1.2 | 2.2774 | 1.8E-08 | 1.2 | 2.313 | 1.8E-07 |
| *nlp-7* | WBGene00003745 | protein_coding_gene | 1.2 | 2.2749 | 2.2E-07 | 1.0 | 2.052 | 6.9E-05 |
| *Y57G11A.4* | WBGene00013292 | protein_coding_gene | 1.2 | 2.2718 | 2.7E-11 | 1.5 | 2.786 | 3.9E-14 |
| *ins-6* | WBGene00002089 | protein_coding_gene | 1.2 | 2.2714 | 8.8E-03 | 1.8 | 3.401 | 4.1E-08 |
| *Y40C5A.4* | WBGene00021497 | protein_coding_gene | 1.2 | 2.2700 | 1.1E-09 | 1.9 | 3.805 | 9.3E-07 |
| *oac-58* | WBGene00019580 | protein_coding_gene | 1.2 | 2.2674 | 2.5E-03 | 1.1 | 2.081 | 2.8E-03 |
| *ZK262.3* | WBGene00013950 | protein_coding_gene | 1.2 | 2.2671 | 1.1E-02 | 1.1 | 2.080 | 9.4E-03 |
| *grd-1* | WBGene00001690 | protein_coding_gene | 1.2 | 2.2630 | 2.1E-03 | 1.6 | 3.070 | 3.2E-03 |
| *pcp-2* | WBGene00003957 | protein_coding_gene | 1.2 | 2.2617 | 1.3E-02 | 2.4 | 5.346 | 1.2E-07 |
| *mab-10* | WBGene00003107 | protein_coding_gene | 1.2 | 2.2599 | 6.8E-04 | 2.2 | 4.452 | 5.1E-17 |
| *Y48G10A.6* | WBGene00013023 | protein_coding_gene | 1.2 | 2.2593 | 7.6E-03 | 1.1 | 2.143 | 1.1E-04 |
| *abcf-2* | WBGene00012097 | protein_coding_gene | 1.2 | 2.2576 | 2.7E-15 | 1.1 | 2.187 | 1.9E-24 |
| *cest-12* | WBGene00013540 | protein_coding_gene | 1.2 | 2.2557 | 1.3E-02 | 1.3 | 2.406 | 2.2E-08 |
| ***ddr-2*** | **WBGene00017381** | **protein_coding_gene** | **1.2** | **2.2548** | **5.2E-04** | **-1.1** | **0.455** | **5.2E-07** |
| *F36H12.14* | WBGene00018127 | pseudogene | 1.2 | 2.2542 | 2.5E-02 | 2.0 | 3.927 | 3.4E-10 |
| *T06F4.1* | WBGene00020294 | protein_coding_gene | 1.2 | 2.2542 | 1.9E-03 | 1.0 | 2.039 | 3.2E-05 |
| *ncs-6* | WBGene00021116 | protein_coding_gene | 1.2 | 2.2515 | 1.7E-02 | 1.3 | 2.522 | 6.8E-03 |
| *glr-5* | WBGene00001616 | protein_coding_gene | 1.2 | 2.2500 | 2.3E-04 | 1.4 | 2.685 | 3.0E-06 |
| *H01G02.3* | WBGene00010350 | protein_coding_gene | 1.2 | 2.2491 | 1.6E-02 | 1.0 | 2.005 | 3.2E-02 |
| *ctl-3* | WBGene00013220 | protein_coding_gene | 1.2 | 2.2472 | 5.9E-06 | 1.2 | 2.236 | 2.1E-06 |
| *Y71F9AL.2* | WBGene00022108 | protein_coding_gene | 1.2 | 2.2451 | 7.5E-03 | 1.8 | 3.450 | 2.6E-14 |
| *sqst-2* | WBGene00021495 | protein_coding_gene | 1.2 | 2.2445 | 1.1E-04 | 1.4 | 2.569 | 1.3E-11 |
| *bgal-2* | WBGene00019225 | protein_coding_gene | 1.2 | 2.2443 | 1.7E-06 | 1.4 | 2.629 | 6.8E-16 |
| *pde-1* | WBGene00011433 | protein_coding_gene | 1.2 | 2.2426 | 1.4E-05 | 1.4 | 2.589 | 2.2E-06 |
| *Y71G12B.17* | WBGene00022155 | protein_coding_gene | 1.2 | 2.2417 | 1.0E-03 | 1.2 | 2.245 | 4.4E-05 |
| ***F13E9.11*** | **WBGene00008760** | **protein_coding_gene** | **1.2** | **2.2402** | **3.7E-03** | **-2.0** | **0.259** | **3.4E-16** |
| *F21A3.4* | WBGene00008994 | protein_coding_gene | 1.2 | 2.2372 | 4.1E-02 | 1.5 | 2.878 | 3.9E-05 |
| *cest-15* | WBGene00012202 | pseudogene | 1.2 | 2.2363 | 2.2E-08 | 1.7 | 3.162 | 8.0E-10 |
| *klp-4* | WBGene00002217 | protein_coding_gene | 1.2 | 2.2356 | 8.3E-08 | 1.0 | 2.064 | 2.5E-17 |
| ***agr-1*** | **WBGene00018304** | **protein_coding_gene** | **1.2** | **2.2355** | **5.4E-06** | **-1.1** | **0.456** | **1.2E-05** |
| *zipt-16* | WBGene00006486 | protein_coding_gene | 1.2 | 2.2307 | 4.4E-03 | 1.6 | 2.961 | 8.1E-07 |
| *unc-2* | WBGene00006742 | protein_coding_gene | 1.2 | 2.2240 | 2.0E-03 | 1.3 | 2.392 | 1.0E-11 |
| *gst-42* | WBGene00001790 | protein_coding_gene | 1.2 | 2.2195 | 5.8E-04 | 1.2 | 2.334 | 7.0E-11 |
| ***T15B7.10*** | **WBGene00020522** | **pseudogene** | **1.1** | **2.2162** | **3.3E-03** | **-2.2** | **0.218** | **2.2E-04** |
| *F58F9.4* | WBGene00019058 | protein_coding_gene | 1.1 | 2.2122 | 4.9E-05 | 1.4 | 2.589 | 1.8E-06 |
| *avr-15* | WBGene00000233 | protein_coding_gene | 1.1 | 2.2113 | 4.1E-09 | 1.1 | 2.092 | 1.5E-10 |
| *frpr-17* | WBGene00011765 | protein_coding_gene | 1.1 | 2.2111 | 2.5E-03 | 1.4 | 2.588 | 2.2E-03 |
| *H03A11.2* | WBGene00010357 | protein_coding_gene | 1.1 | 2.2076 | 7.1E-07 | 2.1 | 4.197 | 2.8E-26 |
| *pll-1* | WBGene00004045 | protein_coding_gene | 1.1 | 2.2030 | 4.6E-03 | 1.5 | 2.735 | 1.2E-08 |
| *ugt-31* | WBGene00021464 | protein_coding_gene | 1.1 | 2.2020 | 8.8E-03 | 1.7 | 3.315 | 1.9E-04 |
| *T04F8.3* | WBGene00011442 | protein_coding_gene | 1.1 | 2.2012 | 3.0E-02 | 1.6 | 2.974 | 2.5E-02 |
| ***fkb-7*** | **WBGene00001432** | **protein_coding_gene** | **1.1** | **2.2009** | **2.2E-03** | **-1.5** | **0.344** | **1.6E-08** |
| *fmo-3* | WBGene00001478 | protein_coding_gene | 1.1 | 2.1987 | 1.9E-03 | 2.3 | 4.835 | 1.3E-24 |
| *Y106G6H.1* | WBGene00013716 | protein_coding_gene | 1.1 | 2.1981 | 1.4E-05 | 1.5 | 2.896 | 8.2E-13 |
| *F25B3.4* | WBGene00009101 | protein_coding_gene | 1.1 | 2.1980 | 1.5E-02 | 1.6 | 3.053 | 4.9E-09 |
| *qui-1* | WBGene00004265 | protein_coding_gene | 1.1 | 2.1952 | 1.4E-04 | 1.5 | 2.921 | 2.6E-12 |
| *W10G11.19* | WBGene00021148 | protein_coding_gene | 1.1 | 2.1918 | 1.0E-04 | 1.2 | 2.337 | 6.8E-08 |
| *hecw-1* | WBGene00009738 | protein_coding_gene | 1.1 | 2.1917 | 4.8E-03 | 1.2 | 2.235 | 1.2E-06 |
| *gsa-1* | WBGene00001745 | protein_coding_gene | 1.1 | 2.1899 | 2.1E-03 | 1.0 | 2.014 | 1.2E-09 |
| *ceh-60* | WBGene00017690 | protein_coding_gene | 1.1 | 2.1887 | 3.1E-06 | 1.2 | 2.228 | 1.7E-09 |
| *F10D7.11* | WBGene00303423 | protein_coding_gene | 1.1 | 2.1877 | 2.2E-02 | 1.2 | 2.267 | 2.6E-02 |
| *ept-1* | WBGene00018797 | protein_coding_gene | 1.1 | 2.1858 | 1.3E-23 | 1.3 | 2.412 | 4.4E-17 |
| *anc-1* | WBGene00000140 | protein_coding_gene | 1.1 | 2.1855 | 4.2E-06 | 1.9 | 3.771 | 3.9E-25 |
| *unc-104* | WBGene00006831 | protein_coding_gene | 1.1 | 2.1839 | 2.3E-07 | 1.2 | 2.374 | 1.5E-15 |
| *viro-2* | WBGene00015107 | protein_coding_gene | 1.1 | 2.1822 | 5.0E-04 | 1.1 | 2.208 | 2.5E-05 |
| *nhr-141* | WBGene00017787 | protein_coding_gene | 1.1 | 2.1819 | 9.9E-03 | 1.5 | 2.860 | 1.2E-04 |
| *clhm-1* | WBGene00016626 | protein_coding_gene | 1.1 | 2.1812 | 4.2E-03 | 1.2 | 2.236 | 4.7E-03 |
| *fbxa-74* | WBGene00021262 | protein_coding_gene | 1.1 | 2.1783 | 3.1E-03 | 1.1 | 2.071 | 1.6E-06 |
| *lys-5* | WBGene00003094 | protein_coding_gene | 1.1 | 2.1755 | 3.1E-05 | 1.5 | 2.822 | 3.6E-06 |
| *nekl-3* | WBGene00008956 | protein_coding_gene | 1.1 | 2.1639 | 4.1E-04 | 1.1 | 2.100 | 7.0E-07 |
| *kqt-2* | WBGene00002234 | protein_coding_gene | 1.1 | 2.1630 | 3.1E-04 | 1.1 | 2.148 | 2.1E-07 |
| ***ins-7*** | **WBGene00002090** | **protein_coding_gene** | **1.1** | **2.1625** | **6.5E-03** | **-1.3** | **0.409** | **1.7E-04** |

| | | | | | | | | |
|---|---|---|---|---|---|---|---|---|
| *lgc-10* | WBGene00022247 | protein_coding_gene | 1.1 | 2.1619 | 1.5E-02 | 1.2 | 2.225 | 7.4E-03 |
| *tyra-3* | WBGene00006475 | protein_coding_gene | 1.1 | 2.1612 | 3.2E-03 | 1.4 | 2.722 | 5.7E-04 |
| *ptr-12* | WBGene00004226 | protein_coding_gene | 1.1 | 2.1549 | 1.1E-04 | 1.3 | 2.535 | 2.1E-08 |
| *E04F6.6* | WBGene00017126 | protein_coding_gene | 1.1 | 2.1537 | 5.1E-07 | 1.3 | 2.424 | 3.4E-15 |
| ***zig-10*** | **WBGene00020800** | **protein_coding_gene** | **1.1** | **2.1532** | **5.0E-02** | **-1.9** | **0.269** | **2.0E-07** |
| *pals-16* | WBGene00022337 | protein_coding_gene | 1.1 | 2.1504 | 2.0E-03 | 1.1 | 2.202 | 3.8E-03 |
| *C17G10.7* | WBGene00015919 | protein_coding_gene | 1.1 | 2.1491 | 1.8E-03 | 1.5 | 2.759 | 1.1E-04 |
| *Y105C5B.11* | WBGene00013651 | protein_coding_gene | 1.1 | 2.1477 | 1.8E-02 | 1.3 | 2.477 | 4.8E-05 |
| *pcdr-1* | WBGene00010329 | protein_coding_gene | 1.1 | 2.1475 | 1.9E-03 | 1.1 | 2.092 | 6.7E-03 |
| *T13H5.1* | WBGene00011755 | protein_coding_gene | 1.1 | 2.1456 | 5.1E-05 | 1.1 | 2.083 | 1.3E-09 |
| *cnc-3* | WBGene00000557 | protein_coding_gene | 1.1 | 2.1444 | 5.1E-03 | 1.6 | 2.956 | 1.3E-07 |
| *acr-17* | WBGene00000056 | protein_coding_gene | 1.1 | 2.1434 | 2.4E-02 | 1.4 | 2.630 | 1.1E-02 |
| *F57C9.6* | WBGene00019012 | protein_coding_gene | 1.1 | 2.1430 | 5.0E-03 | 1.1 | 2.093 | 3.7E-02 |
| *F42G4.5* | WBGene00009642 | protein_coding_gene | 1.1 | 2.1423 | 5.0E-07 | 1.0 | 2.063 | 2.1E-06 |
| *wrt-3* | WBGene00006949 | protein_coding_gene | 1.1 | 2.1413 | 7.4E-04 | 1.8 | 3.574 | 1.3E-05 |
| *osm-7* | WBGene00003887 | protein_coding_gene | 1.1 | 2.1412 | 6.6E-05 | 1.7 | 3.144 | 1.5E-24 |
| *unc-103* | WBGene00006830 | protein_coding_gene | 1.1 | 2.1408 | 9.3E-05 | 1.3 | 2.418 | 5.0E-08 |
| *che-10* | WBGene00018051 | protein_coding_gene | 1.1 | 2.1371 | 3.3E-04 | 1.8 | 3.428 | 2.7E-28 |
| *rcs-1* | WBGene00016190 | protein_coding_gene | 1.1 | 2.1340 | 6.6E-04 | 1.3 | 2.506 | 7.1E-14 |
| *slo-2* | WBGene00004831 | protein_coding_gene | 1.1 | 2.1335 | 1.3E-07 | 1.2 | 2.291 | 1.3E-13 |
| *ptc-3* | WBGene00004210 | protein_coding_gene | 1.1 | 2.1329 | 1.0E-04 | 1.2 | 2.241 | 3.3E-14 |
| *lips-15* | WBGene00012585 | protein_coding_gene | 1.1 | 2.1327 | 5.4E-09 | 1.7 | 3.332 | 2.0E-18 |
| *mod-1* | WBGene00003386 | protein_coding_gene | 1.1 | 2.1327 | 2.2E-03 | 1.3 | 2.539 | 5.8E-04 |
| ***let-381*** | **WBGene00002601** | **protein_coding_gene** | **1.1** | **2.1309** | **4.0E-02** | **-1.4** | **0.379** | **1.1E-03** |
| *C14A11.6* | WBGene00015751 | protein_coding_gene | 1.1 | 2.1295 | 6.0E-04 | 1.7 | 3.200 | 3.3E-05 |
| *H14N18.4* | WBGene00019204 | protein_coding_gene | 1.1 | 2.1276 | 3.8E-06 | 1.5 | 2.894 | 1.1E-20 |
| *F17B5.8* | WBGene00045458 | protein_coding_gene | 1.1 | 2.1253 | 4.2E-02 | 1.3 | 2.380 | 3.4E-02 |
| *E02C12.8* | WBGene00017093 | protein_coding_gene | 1.1 | 2.1247 | 1.5E-02 | 1.8 | 3.497 | 2.1E-10 |
| *F39B2.8* | WBGene00009558 | protein_coding_gene | 1.1 | 2.1246 | 1.5E-04 | 1.3 | 2.438 | 1.2E-06 |
| *C08E8.10* | WBGene00077692 | protein_coding_gene | 1.1 | 2.1235 | 2.3E-02 | 1.0 | 2.050 | 4.1E-02 |
| *lyst-1* | WBGene00011361 | protein_coding_gene | 1.1 | 2.1232 | 7.4E-06 | 1.2 | 2.235 | 4.8E-15 |
| *aex-1* | WBGene00000084 | protein_coding_gene | 1.1 | 2.1219 | 7.1E-07 | 1.1 | 2.117 | 2.7E-09 |
| *F41C6.7* | WBGene00018280 | protein_coding_gene | 1.1 | 2.1198 | 3.2E-04 | 1.2 | 2.286 | 2.0E-04 |
| *rol-3* | WBGene00004395 | protein_coding_gene | 1.1 | 2.1195 | 1.2E-03 | 1.4 | 2.562 | 8.8E-10 |
| *C07F11.2* | WBGene00015570 | protein_coding_gene | 1.1 | 2.1191 | 3.6E-02 | 1.3 | 2.379 | 5.0E-02 |
| *fbxa-31* | WBGene00013838 | protein_coding_gene | 1.1 | 2.1188 | 1.1E-03 | 1.2 | 2.255 | 1.7E-04 |
| *cls-1* | WBGene00015580 | protein_coding_gene | 1.1 | 2.1180 | 3.7E-09 | 1.1 | 2.124 | 7.2E-15 |
| *R119.5* | WBGene00020090 | protein_coding_gene | 1.1 | 2.1165 | 4.6E-11 | 1.2 | 2.259 | 1.5E-08 |
| *ZC581.9* | WBGene00022635 | protein_coding_gene | 1.1 | 2.1112 | 6.8E-07 | 1.7 | 3.275 | 4.2E-11 |
| *C07A12.7* | WBGene00015561 | protein_coding_gene | 1.1 | 2.1109 | 1.9E-07 | 1.2 | 2.265 | 2.5E-15 |
| *F41H10.5* | WBGene00018318 | protein_coding_gene | 1.1 | 2.1100 | 5.4E-03 | 1.9 | 3.860 | 3.2E-12 |
| *nlp-20* | WBGene00003758 | protein_coding_gene | 1.1 | 2.1031 | 2.2E-06 | 1.0 | 2.004 | 1.5E-08 |
| *F57H12.5* | WBGene00019020 | protein_coding_gene | 1.1 | 2.1012 | 1.7E-02 | 1.8 | 3.537 | 1.8E-07 |
| *osm-11* | WBGene00003891 | protein_coding_gene | 1.1 | 2.1000 | 1.9E-05 | 1.7 | 3.237 | 1.1E-38 |
| *B0554.5* | WBGene00015258 | protein_coding_gene | 1.1 | 2.0896 | 2.1E-02 | 1.9 | 3.644 | 1.2E-05 |
| *rpl-21* | WBGene00004433 | protein_coding_gene | 1.1 | 2.0879 | 3.0E-19 | 1.5 | 2.767 | 6.9E-32 |
| *Y46G5A.29* | WBGene00012917 | protein_coding_gene | 1.1 | 2.0872 | 2.6E-02 | 3.3 | 9.945 | 2.1E-05 |
| *tag-68* | WBGene00006445 | protein_coding_gene | 1.1 | 2.0867 | 8.4E-04 | 1.2 | 2.239 | 1.9E-04 |
| *C02F5.14* | WBGene00044445 | protein_coding_gene | 1.1 | 2.0853 | 1.3E-04 | 1.1 | 2.205 | 9.6E-04 |
| *acox-1.6* | WBGene00010336 | protein_coding_gene | 1.1 | 2.0835 | 2.9E-04 | 1.0 | 2.031 | 5.6E-07 |
| *fbxa-34* | WBGene00013842 | pseudogene | 1.1 | 2.0815 | 4.0E-02 | 1.0 | 2.004 | 6.3E-03 |
| *cyp-33E2* | WBGene00018334 | protein_coding_gene | 1.1 | 2.0815 | 3.8E-05 | 1.2 | 2.259 | 1.9E-07 |
| *math-18* | WBGene00016544 | protein_coding_gene | 1.1 | 2.0775 | 7.3E-03 | 2.0 | 3.881 | 3.8E-06 |
| *ceh-18* | WBGene00000441 | protein_coding_gene | 1.1 | 2.0725 | 4.0E-06 | 1.4 | 2.684 | 5.8E-14 |
| *lec-8* | WBGene00002271 | protein_coding_gene | 1.1 | 2.0721 | 3.5E-04 | 2.0 | 3.979 | 6.7E-11 |
| *C18B2.5* | WBGene00015956 | protein_coding_gene | 1.0 | 2.0701 | 5.7E-07 | 1.4 | 2.731 | 2.6E-36 |
| *R02F2.8* | WBGene00019837 | protein_coding_gene | 1.0 | 2.0691 | 2.9E-04 | 1.1 | 2.103 | 1.2E-05 |
| *ZC416.2* | WBGene00022609 | protein_coding_gene | 1.0 | 2.0673 | 1.5E-02 | 1.2 | 2.307 | 1.2E-02 |
| *AC3.5* | WBGene00007071 | protein_coding_gene | 1.0 | 2.0662 | 8.0E-03 | 3.0 | 8.255 | 2.7E-81 |
| *Y32F6A.4* | WBGene00012529 | protein_coding_gene | 1.0 | 2.0648 | 1.7E-08 | 1.7 | 3.316 | 7.2E-34 |
| *sox-4* | WBGene00015716 | protein_coding_gene | 1.0 | 2.0592 | 3.4E-04 | 1.3 | 2.414 | 1.7E-05 |
| *tatn-1* | WBGene00009628 | protein_coding_gene | 1.0 | 2.0560 | 1.2E-02 | 1.4 | 2.574 | 5.8E-16 |
| *F35H10.10* | WBGene00018073 | protein_coding_gene | 1.0 | 2.0553 | 4.0E-05 | 1.8 | 3.411 | 1.5E-16 |
| *F13D2.1* | WBGene00008735 | protein_coding_gene | 1.0 | 2.0548 | 3.6E-05 | 1.1 | 2.163 | 1.2E-11 |
| *haf-9* | WBGene00001819 | protein_coding_gene | 1.0 | 2.0531 | 9.3E-06 | 1.6 | 2.976 | 1.4E-17 |
| *egl-8* | WBGene00001177 | protein_coding_gene | 1.0 | 2.0517 | 7.7E-05 | 1.3 | 2.380 | 1.2E-24 |
| *Y37F4.7* | WBGene00021388 | protein_coding_gene | 1.0 | 2.0483 | 7.8E-03 | 1.8 | 3.540 | 7.1E-06 |
| *Y106G6G.2* | WBGene00013711 | protein_coding_gene | 1.0 | 2.0476 | 6.3E-04 | 1.0 | 2.061 | 4.5E-04 |
| *twk-26* | WBGene00006679 | protein_coding_gene | 1.0 | 2.0463 | 6.2E-04 | 1.5 | 2.831 | 2.4E-08 |
| *pmk-2* | WBGene00004056 | protein_coding_gene | 1.0 | 2.0457 | 5.4E-09 | 1.0 | 2.022 | 2.7E-07 |
| *F35C11.6* | WBGene00009405 | protein_coding_gene | 1.0 | 2.0431 | 6.5E-04 | 1.0 | 2.007 | 4.6E-03 |
| *F55H12.3* | WBGene00010134 | protein_coding_gene | 1.0 | 2.0400 | 3.1E-04 | 1.4 | 2.597 | 7.2E-17 |
| *madf-4* | WBGene00011575 | protein_coding_gene | 1.0 | 2.0379 | 1.1E-03 | 1.2 | 2.300 | 1.4E-03 |
| *rnp-9* | WBGene00007396 | protein_coding_gene | 1.0 | 2.0330 | 2.6E-03 | 1.6 | 3.040 | 4.7E-09 |
| *ifd-2* | WBGene00002058 | protein_coding_gene | 1.0 | 2.0309 | 1.5E-02 | 1.9 | 3.621 | 5.5E-20 |
| *npr-6* | WBGene00009619 | protein_coding_gene | 1.0 | 2.0303 | 4.9E-02 | 1.1 | 2.176 | 2.3E-02 |
| *ZC449.7* | WBGene00045365 | protein_coding_gene | 1.0 | 2.0271 | 1.2E-02 | 1.8 | 3.394 | 1.2E-05 |
| *aak-2* | WBGene00020142 | protein_coding_gene | 1.0 | 2.0266 | 3.9E-08 | 1.2 | 2.282 | 5.2E-22 |
| *lec-5* | WBGene00002268 | protein_coding_gene | 1.0 | 2.0266 | 2.1E-06 | 1.1 | 2.155 | 7.9E-20 |
| *R08E3.1* | WBGene00019957 | protein_coding_gene | 1.0 | 2.0260 | 8.7E-04 | 1.1 | 2.207 | 1.1E-13 |

| | | | | | | | | |
|---|---|---|---|---|---|---|---|---|
| *cut-4* | WBGene00008482 | protein_coding_gene | 1.0 | 2.0260 | 1.9E-03 | 1.6 | 3.110 | 2.0E-03 |
| ***C03A3.1*** | **WBGene00007266** | **protein_coding_gene** | **1.0** | **2.0242** | **3.5E-02** | **-1.0** | **0.485** | **2.6E-03** |
| *R07B1.11* | WBGene00011082 | protein_coding_gene | 1.0 | 2.0237 | 3.9E-02 | 1.2 | 2.315 | 8.1E-03 |
| *swt-6* | WBGene00011190 | protein_coding_gene | 1.0 | 2.0236 | 1.0E-02 | 2.0 | 4.047 | 8.2E-11 |
| *avr-14* | WBGene00000232 | protein_coding_gene | 1.0 | 2.0221 | 3.0E-05 | 1.3 | 2.459 | 1.3E-07 |
| *F11D5.7* | WBGene00043980 | protein_coding_gene | 1.0 | 2.0199 | 4.7E-03 | 1.3 | 2.443 | 1.9E-04 |
| *twk-30* | WBGene00006682 | protein_coding_gene | 1.0 | 2.0196 | 5.7E-04 | 1.3 | 2.454 | 3.6E-06 |
| *gbb-2* | WBGene00022675 | protein_coding_gene | 1.0 | 2.0182 | 1.2E-04 | 1.7 | 3.184 | 1.9E-12 |
| *alh-12* | WBGene00000118 | protein_coding_gene | 1.0 | 2.0176 | 1.3E-05 | 1.1 | 2.082 | 5.3E-17 |
| *toh-1* | WBGene00006591 | protein_coding_gene | 1.0 | 2.0174 | 6.0E-06 | 1.6 | 3.110 | 5.6E-15 |
| ***T24B8.3*** | **WBGene00011977** | **protein_coding_gene** | **1.0** | **2.0150** | **4.1E-07** | **-1.2** | **0.447** | **6.3E-19** |
| *F57F5.1* | WBGene00010204 | protein_coding_gene | 1.0 | 2.0134 | 4.1E-10 | 1.4 | 2.690 | 1.5E-05 |
| *nhr-95* | WBGene00003685 | protein_coding_gene | 1.0 | 2.0124 | 2.9E-08 | 1.1 | 2.078 | 8.7E-06 |
| *R09B5.11* | WBGene00019979 | protein_coding_gene | 1.0 | 2.0109 | 3.9E-03 | 2.2 | 4.713 | 7.4E-21 |
| *T19C3.3* | WBGene00020561 | protein_coding_gene | 1.0 | 2.0079 | 3.9E-02 | 1.4 | 2.555 | 2.0E-03 |
| ***Y65A5A.1*** | **WBGene00013418** | **protein_coding_gene** | **1.0** | **2.0071** | **2.0E-03** | **-1.0** | **0.496** | **6.8E-06** |
| *D2013.6* | WBGene00008404 | protein_coding_gene | 1.0 | 2.0055 | 2.7E-08 | 1.3 | 2.443 | 8.2E-24 |
| *metl-17* | WBGene00009305 | protein_coding_gene | -1.0 | 0.4987 | 4.1E-12 | -1.0 | 0.499 | 1.4E-14 |
| *ddl-1* | WBGene00019125 | protein_coding_gene | -1.0 | 0.4982 | 7.4E-06 | -1.0 | 0.494 | 1.4E-09 |
| *leo-1* | WBGene00007110 | protein_coding_gene | -1.0 | 0.4978 | 1.3E-12 | -1.2 | 0.429 | 2.0E-33 |
| *B0304.2* | WBGene00015134 | protein_coding_gene | -1.0 | 0.4975 | 7.0E-08 | -1.1 | 0.468 | 2.9E-14 |
| *F16B4.4* | WBGene00017506 | protein_coding_gene | -1.0 | 0.4973 | 2.7E-03 | -1.4 | 0.379 | 3.7E-17 |
| *Y97E10AR.3* | WBGene00022398 | protein_coding_gene | -1.0 | 0.4965 | 1.0E-05 | -1.2 | 0.421 | 5.0E-22 |
| *bir-2* | WBGene00000250 | protein_coding_gene | -1.0 | 0.4959 | 2.7E-08 | -1.1 | 0.483 | 7.2E-13 |
| *BE0003N10.3* | WBGene00015268 | protein_coding_gene | -1.0 | 0.4959 | 1.9E-02 | -1.6 | 0.334 | 3.1E-06 |
| *E01A2.5* | WBGene00017087 | protein_coding_gene | -1.0 | 0.4957 | 1.3E-09 | -1.1 | 0.469 | 8.0E-10 |
| ***C55A6.4*** | **WBGene00008333** | **protein_coding_gene** | **-1.0** | **0.4957** | **2.9E-02** | **1.5** | **2.899** | **5.2E-12** |
| *B0361.2* | WBGene00015156 | protein_coding_gene | -1.0 | 0.4953 | 2.6E-08 | -1.2 | 0.444 | 3.1E-21 |
| ***rft-1*** | **WBGene00044637** | **protein_coding_gene** | **-1.0** | **0.4950** | **1.8E-02** | **1.1** | **2.076** | **1.8E-04** |
| *C41D11.9* | WBGene00016567 | protein_coding_gene | -1.0 | 0.4948 | 2.3E-05 | -1.6 | 0.330 | 1.1E-20 |
| *B0238.11* | WBGene00015075 | protein_coding_gene | -1.0 | 0.4943 | 2.0E-16 | -1.0 | 0.491 | 1.9E-19 |
| *pri-1* | WBGene00004180 | protein_coding_gene | -1.0 | 0.4943 | 5.7E-14 | -1.3 | 0.420 | 1.1E-19 |
| *taf-3* | WBGene00006384 | protein_coding_gene | -1.0 | 0.4938 | 6.4E-05 | -1.2 | 0.435 | 1.6E-18 |
| *C06A6.4* | WBGene00015509 | protein_coding_gene | -1.0 | 0.4937 | 2.6E-12 | -1.3 | 0.409 | 1.4E-24 |
| *C06G3.8* | WBGene00015554 | protein_coding_gene | -1.0 | 0.4936 | 1.9E-11 | -1.1 | 0.466 | 9.5E-15 |
| *B0285.3* | WBGene00007136 | protein_coding_gene | -1.0 | 0.4934 | 4.2E-07 | -1.6 | 0.338 | 8.1E-21 |
| *hda-2* | WBGene00001835 | protein_coding_gene | -1.0 | 0.4927 | 1.5E-09 | -1.3 | 0.407 | 4.6E-27 |
| *ZK1320.7* | WBGene00014256 | protein_coding_gene | -1.0 | 0.4924 | 3.8E-09 | -1.2 | 0.421 | 7.4E-17 |
| *F14B4.1* | WBGene00008779 | protein_coding_gene | -1.0 | 0.4921 | 8.1E-03 | -3.1 | 0.120 | 6.8E-28 |
| *dod-18* | WBGene00008316 | protein_coding_gene | -1.0 | 0.4920 | 7.4E-07 | -1.1 | 0.472 | 1.1E-08 |
| *B0416.5* | WBGene00015181 | protein_coding_gene | -1.0 | 0.4919 | 2.2E-06 | -1.2 | 0.450 | 2.8E-25 |
| *mdt-18* | WBGene00007018 | protein_coding_gene | -1.0 | 0.4913 | 7.0E-06 | -1.2 | 0.448 | 1.9E-15 |
| *C16C8.20* | WBGene00195070 | protein_coding_gene | -1.0 | 0.4910 | 4.0E-03 | -1.2 | 0.430 | 2.0E-05 |
| *dsb-3* | WBGene00016701 | protein_coding_gene | -1.0 | 0.4907 | 8.1E-11 | -1.0 | 0.494 | 1.1E-10 |
| *pot-1* | WBGene00015105 | protein_coding_gene | -1.0 | 0.4904 | 1.9E-10 | -1.0 | 0.498 | 1.6E-11 |
| *cdc-26* | WBGene00015235 | protein_coding_gene | -1.0 | 0.4901 | 8.5E-10 | -1.0 | 0.488 | 4.6E-19 |
| *Y67H2A.10* | WBGene00013465 | protein_coding_gene | -1.0 | 0.4898 | 1.2E-02 | -1.2 | 0.442 | 2.7E-19 |
| *moa-2* | WBGene00015205 | protein_coding_gene | -1.0 | 0.4897 | 3.6E-07 | -1.1 | 0.463 | 2.1E-12 |
| *unc-76* | WBGene00006808 | protein_coding_gene | -1.0 | 0.4896 | 1.3E-05 | -1.0 | 0.493 | 5.6E-20 |
| *F07H5.13* | WBGene00008563 | protein_coding_gene | -1.0 | 0.4894 | 1.5E-04 | -3.0 | 0.125 | 2.5E-33 |
| *F26A3.7* | WBGene00009145 | protein_coding_gene | -1.0 | 0.4894 | 6.3E-09 | -1.1 | 0.459 | 1.3E-15 |
| *rfp-1* | WBGene00007008 | protein_coding_gene | -1.0 | 0.4892 | 2.1E-08 | -1.2 | 0.436 | 2.7E-25 |
| *pot-2* | WBGene00010195 | protein_coding_gene | -1.0 | 0.4890 | 4.1E-06 | -1.5 | 0.351 | 1.7E-18 |
| *Y39B6A.13* | WBGene00012675 | protein_coding_gene | -1.0 | 0.4884 | 1.1E-12 | -1.5 | 0.361 | 1.6E-18 |
| *Y73F8A.13* | WBGene00013522 | protein_coding_gene | -1.0 | 0.4879 | 1.8E-05 | -1.5 | 0.353 | 3.4E-08 |
| *ZK1067.3* | WBGene00014209 | protein_coding_gene | -1.0 | 0.4879 | 6.7E-12 | -1.0 | 0.493 | 1.3E-16 |
| *lin-26* | WBGene00003012 | protein_coding_gene | -1.0 | 0.4874 | 1.0E-05 | -1.3 | 0.415 | 1.0E-26 |
| *C15F1.8* | WBGene00015797 | protein_coding_gene | -1.0 | 0.4871 | 4.0E-06 | -1.4 | 0.378 | 1.8E-08 |
| *rde-2* | WBGene00004324 | protein_coding_gene | -1.0 | 0.4869 | 1.9E-12 | -1.0 | 0.494 | 5.5E-18 |
| *C16C8.16* | WBGene00015853 | protein_coding_gene | -1.0 | 0.4869 | 4.7E-14 | -1.1 | 0.458 | 5.0E-16 |
| *M199.2* | WBGene00010951 | protein_coding_gene | -1.0 | 0.4865 | 2.5E-06 | -2.1 | 0.238 | 3.7E-21 |
| *T04B2.7* | WBGene00011425 | protein_coding_gene | -1.0 | 0.4865 | 3.8E-02 | -1.5 | 0.352 | 1.6E-05 |
| *cyn-9* | WBGene00000885 | protein_coding_gene | -1.0 | 0.4862 | 9.8E-15 | -1.1 | 0.472 | 1.0E-16 |
| *ssp-35* | WBGene00010091 | protein_coding_gene | -1.0 | 0.4861 | 2.7E-02 | -1.1 | 0.471 | 6.9E-07 |
| *clec-218* | WBGene00020938 | protein_coding_gene | -1.0 | 0.4861 | 2.1E-02 | -3.1 | 0.116 | 2.1E-67 |
| *rnp-2* | WBGene00004385 | protein_coding_gene | -1.0 | 0.4860 | 6.4E-15 | -1.3 | 0.414 | 8.9E-19 |
| *fbxc-52* | WBGene00016710 | protein_coding_gene | -1.0 | 0.4845 | 2.7E-05 | -1.1 | 0.460 | 2.9E-09 |
| *gadr-6* | WBGene00009823 | protein_coding_gene | -1.0 | 0.4838 | 1.8E-08 | -1.2 | 0.428 | 3.1E-11 |
| *C25A1.1* | WBGene00007705 | protein_coding_gene | -1.0 | 0.4831 | 7.8E-10 | -1.2 | 0.447 | 2.9E-21 |
| *K07A1.1* | WBGene00010608 | protein_coding_gene | -1.1 | 0.4828 | 7.0E-07 | -1.5 | 0.364 | 1.7E-33 |
| *Y95D11A.3* | WBGene00045434 | protein_coding_gene | -1.1 | 0.4823 | 1.9E-11 | -1.1 | 0.458 | 6.2E-19 |
| *F28C1.1* | WBGene00009200 | protein_coding_gene | -1.1 | 0.4821 | 1.2E-13 | -1.1 | 0.482 | 3.5E-20 |
| *mcrs-1* | WBGene00010420 | protein_coding_gene | -1.1 | 0.4816 | 1.3E-14 | -1.1 | 0.480 | 1.8E-15 |
| ***Y51A2D.18*** | **WBGene00013083** | **protein_coding_gene** | **-1.1** | **0.4814** | **3.1E-02** | **1.5** | **2.906** | **1.2E-12** |
| *orc-3* | WBGene00022488 | protein_coding_gene | -1.1 | 0.4809 | 5.7E-13 | -1.2 | 0.441 | 2.2E-16 |
| *T16H12.1* | WBGene00011811 | protein_coding_gene | -1.1 | 0.4806 | 1.0E-05 | -1.5 | 0.352 | 5.3E-25 |
| *amx-1* | WBGene00000137 | protein_coding_gene | -1.1 | 0.4803 | 1.5E-05 | -1.1 | 0.462 | 8.6E-10 |
| ***mua-6*** | **WBGene00003485** | **protein_coding_gene** | **-1.1** | **0.4796** | **2.9E-02** | **1.3** | **2.466** | **4.6E-09** |
| *nrde-3* | WBGene00019862 | protein_coding_gene | -1.1 | 0.4789 | 2.4E-05 | -1.7 | 0.310 | 3.6E-41 |
| *F25H8.1* | WBGene00009131 | protein_coding_gene | -1.1 | 0.4789 | 8.3E-14 | -1.2 | 0.424 | 2.9E-21 |

| | | | | | | | | |
|---|---|---|---|---|---|---|---|---|
| *set-17* | WBGene00011887 | protein_coding_gene | -1.1 | 0.4785 | 4.1E-07 | -1.0 | 0.494 | 9.8E-17 |
| *parp-2* | WBGene00004050 | protein_coding_gene | -1.1 | 0.4783 | 1.3E-09 | -1.1 | 0.464 | 1.4E-24 |
| *hmg-5* | WBGene00001975 | protein_coding_gene | -1.1 | 0.4783 | 2.0E-08 | -1.2 | 0.447 | 2.7E-27 |
| *nspd-6* | WBGene00043147 | protein_coding_gene | -1.1 | 0.4778 | 3.4E-02 | -1.1 | 0.471 | 6.9E-05 |
| *T23B12.6* | WBGene00020721 | protein_coding_gene | -1.1 | 0.4777 | 7.4E-15 | -1.1 | 0.457 | 1.7E-20 |
| *F40H3.6* | WBGene00018252 | protein_coding_gene | -1.1 | 0.4772 | 3.5E-04 | -1.5 | 0.349 | 1.7E-06 |
| *M05D6.2* | WBGene00010875 | protein_coding_gene | -1.1 | 0.4770 | 1.6E-09 | -1.1 | 0.464 | 5.3E-25 |
| *san-1* | WBGene00004721 | protein_coding_gene | -1.1 | 0.4765 | 3.7E-09 | -1.6 | 0.341 | 6.5E-36 |
| *smg-4* | WBGene00004882 | protein_coding_gene | -1.1 | 0.4761 | 3.6E-12 | -1.0 | 0.490 | 3.0E-17 |
| *rde-8* | WBGene00022620 | protein_coding_gene | -1.1 | 0.4760 | 9.4E-08 | -1.4 | 0.389 | 7.5E-26 |
| *R05D11.5* | WBGene00011033 | protein_coding_gene | -1.1 | 0.4748 | 2.4E-05 | -1.1 | 0.471 | 3.6E-15 |
| *gadr-5* | WBGene00045058 | protein_coding_gene | -1.1 | 0.4745 | 7.3E-05 | -1.9 | 0.259 | 3.2E-09 |
| *Y54F10BM.9* | WBGene00021863 | protein_coding_gene | -1.1 | 0.4738 | 2.4E-05 | -1.0 | 0.498 | 4.1E-08 |
| *C13F10.5* | WBGene00015744 | protein_coding_gene | -1.1 | 0.4735 | 1.4E-08 | -1.3 | 0.416 | 1.9E-18 |
| *C50F4.4* | WBGene00008232 | protein_coding_gene | -1.1 | 0.4734 | 3.8E-03 | -1.2 | 0.428 | 6.0E-04 |
| *szy-5* | WBGene00016154 | protein_coding_gene | -1.1 | 0.4732 | 3.4E-08 | -1.4 | 0.378 | 6.8E-38 |
| *Y5F2A.4* | WBGene00012385 | protein_coding_gene | -1.1 | 0.4729 | 5.1E-09 | -1.3 | 0.419 | 2.7E-22 |
| *F02E9.1* | WBGene00008528 | protein_coding_gene | -1.1 | 0.4727 | 2.0E-11 | -1.5 | 0.347 | 4.5E-20 |
| *sup-36* | WBGene00018164 | protein_coding_gene | -1.1 | 0.4725 | 1.8E-07 | -1.4 | 0.389 | 2.0E-20 |
| *bcas-2* | WBGene00020441 | protein_coding_gene | -1.1 | 0.4723 | 7.4E-12 | -1.0 | 0.484 | 5.4E-17 |
| *anat-1* | WBGene00015938 | protein_coding_gene | -1.1 | 0.4717 | 4.0E-09 | -1.3 | 0.404 | 1.4E-35 |
| *lin-28* | WBGene00003014 | protein_coding_gene | -1.1 | 0.4715 | 1.2E-13 | -1.0 | 0.496 | 2.1E-14 |
| *W01C8.5* | WBGene00020921 | protein_coding_gene | -1.1 | 0.4715 | 5.3E-06 | -1.1 | 0.474 | 3.0E-08 |
| *F40F11.3* | WBGene00009588 | protein_coding_gene | -1.1 | 0.4712 | 1.3E-10 | -1.4 | 0.380 | 7.0E-20 |
| *fbxc-33* | WBGene00019924 | protein_coding_gene | -1.1 | 0.4699 | 3.2E-04 | -1.7 | 0.308 | 4.4E-09 |
| *mdt-22* | WBGene00007022 | protein_coding_gene | -1.1 | 0.4698 | 3.4E-10 | -1.1 | 0.480 | 6.7E-16 |
| *F33H2.3* | WBGene00009367 | protein_coding_gene | -1.1 | 0.4696 | 2.7E-12 | -1.6 | 0.333 | 4.0E-47 |
| *bmk-1* | WBGene00000257 | protein_coding_gene | -1.1 | 0.4696 | 7.3E-10 | -1.2 | 0.447 | 1.3E-23 |
| *F36D3.1* | WBGene00009469 | protein_coding_gene | -1.1 | 0.4690 | 8.6E-12 | -1.6 | 0.331 | 1.3E-16 |
| *C43G2.3* | WBGene00016612 | protein_coding_gene | -1.1 | 0.4690 | 4.8E-02 | -1.3 | 0.420 | 6.9E-06 |
| *B0207.6* | WBGene00015029 | protein_coding_gene | -1.1 | 0.4690 | 3.1E-06 | -1.1 | 0.472 | 9.8E-17 |
| *C46C11.4* | WBGene00044567 | protein_coding_gene | -1.1 | 0.4688 | 4.8E-02 | -2.5 | 0.178 | 9.7E-08 |
| *D1054.3* | WBGene00008371 | protein_coding_gene | -1.1 | 0.4680 | 1.8E-06 | -1.1 | 0.468 | 2.5E-16 |
| ***grsp-4*** | **WBGene00017201** | **protein_coding_gene** | **-1.1** | **0.4676** | **1.1E-02** | **2.3** | **4.916** | **9.9E-09** |
| *tftc-5* | WBGene00011368 | protein_coding_gene | -1.1 | 0.4668 | 2.1E-13 | -1.2 | 0.445 | 1.2E-23 |
| *set-1* | WBGene00004781 | protein_coding_gene | -1.1 | 0.4668 | 2.1E-10 | -1.6 | 0.324 | 2.3E-49 |
| *Y34F4.2* | WBGene00021337 | protein_coding_gene | -1.1 | 0.4657 | 4.4E-03 | -1.0 | 0.491 | 1.7E-04 |
| ***C04G6.6*** | **WBGene00015456** | **protein_coding_gene** | **-1.1** | **0.4654** | **2.9E-08** | **1.6** | **2.974** | **1.5E-03** |
| *F43G9.13* | WBGene00009673 | protein_coding_gene | -1.1 | 0.4650 | 5.4E-05 | -1.2 | 0.429 | 3.2E-15 |
| *mdt-1.2* | WBGene00007011 | protein_coding_gene | -1.1 | 0.4646 | 2.6E-07 | -1.1 | 0.452 | 2.7E-08 |
| *heh-1* | WBGene00006452 | protein_coding_gene | -1.1 | 0.4644 | 1.6E-10 | -1.1 | 0.468 | 8.2E-14 |
| *T28A8.6* | WBGene00012112 | protein_coding_gene | -1.1 | 0.4643 | 6.2E-10 | -1.0 | 0.489 | 5.8E-15 |
| *pfkb-1.1* | WBGene00022456 | protein_coding_gene | -1.1 | 0.4642 | 1.9E-21 | -1.2 | 0.442 | 7.2E-30 |
| *R10H10.6* | WBGene00011224 | protein_coding_gene | -1.1 | 0.4641 | 1.2E-07 | -1.6 | 0.329 | 1.1E-15 |
| *F52E1.14* | WBGene00044696 | protein_coding_gene | -1.1 | 0.4641 | 1.6E-05 | -1.2 | 0.444 | 2.0E-10 |
| *apn-1* | WBGene00000151 | protein_coding_gene | -1.1 | 0.4640 | 1.0E-11 | -1.2 | 0.434 | 2.5E-26 |
| *Y66D12A.7* | WBGene00013433 | protein_coding_gene | -1.1 | 0.4639 | 4.1E-06 | -1.6 | 0.337 | 1.6E-13 |
| *F12A10.8* | WBGene00017397 | protein_coding_gene | -1.1 | 0.4639 | 3.8E-08 | -1.1 | 0.458 | 1.5E-11 |
| *try-10* | WBGene00008849 | protein_coding_gene | -1.1 | 0.4636 | 2.1E-03 | -2.1 | 0.233 | 3.9E-11 |
| *nspd-9* | WBGene00017808 | protein_coding_gene | -1.1 | 0.4635 | 1.9E-02 | -2.4 | 0.191 | 5.2E-24 |
| *ttb-1* | WBGene00006648 | protein_coding_gene | -1.1 | 0.4633 | 5.0E-09 | -1.2 | 0.441 | 9.0E-20 |
| *C16E9.2* | WBGene00015866 | protein_coding_gene | -1.1 | 0.4630 | 2.8E-10 | -1.2 | 0.438 | 3.7E-11 |
| *Y106G6H.8* | WBGene00013720 | protein_coding_gene | -1.1 | 0.4629 | 2.3E-07 | -1.1 | 0.469 | 5.4E-10 |
| *mis-12* | WBGene00021648 | protein_coding_gene | -1.1 | 0.4625 | 2.0E-10 | -1.1 | 0.452 | 1.3E-16 |
| *C31C9.2* | WBGene00007836 | protein_coding_gene | -1.1 | 0.4625 | 6.7E-07 | -1.2 | 0.424 | 1.3E-14 |
| *blos-1* | WBGene00011872 | protein_coding_gene | -1.1 | 0.4625 | 3.3E-10 | -1.4 | 0.369 | 4.6E-22 |
| *tim-1* | WBGene00006571 | protein_coding_gene | -1.1 | 0.4621 | 3.4E-20 | -1.1 | 0.451 | 3.2E-28 |
| *C41G11.1* | WBGene00016568 | protein_coding_gene | -1.1 | 0.4621 | 1.8E-04 | -2.9 | 0.132 | 9.8E-43 |
| *F56G4.4* | WBGene00010159 | protein_coding_gene | -1.1 | 0.4611 | 1.6E-12 | -1.3 | 0.408 | 1.5E-22 |
| *F52D10.2* | WBGene00009930 | protein_coding_gene | -1.1 | 0.4611 | 7.7E-11 | -1.2 | 0.440 | 2.7E-14 |
| *R09F10.3* | WBGene00019987 | protein_coding_gene | -1.1 | 0.4611 | 4.0E-06 | -1.3 | 0.416 | 1.7E-11 |
| *mog-2* | WBGene00003390 | protein_coding_gene | -1.1 | 0.4605 | 1.2E-15 | -1.2 | 0.438 | 1.9E-21 |
| *flh-2* | WBGene00016138 | protein_coding_gene | -1.1 | 0.4603 | 5.4E-06 | -1.0 | 0.486 | 1.8E-12 |
| *pafo-1* | WBGene00008338 | protein_coding_gene | -1.1 | 0.4599 | 4.2E-11 | -1.4 | 0.387 | 4.2E-30 |
| *unc-45* | WBGene00006781 | protein_coding_gene | -1.1 | 0.4595 | 1.0E-14 | -1.1 | 0.478 | 9.6E-21 |
| *elt-1* | WBGene00001249 | protein_coding_gene | -1.1 | 0.4595 | 4.3E-05 | -1.5 | 0.359 | 2.7E-23 |
| *vrk-1* | WBGene00017895 | protein_coding_gene | -1.1 | 0.4594 | 5.7E-10 | -1.4 | 0.388 | 2.3E-31 |
| *best-20* | WBGene00021368 | protein_coding_gene | -1.1 | 0.4593 | 7.2E-05 | -1.7 | 0.298 | 5.2E-18 |
| *C16C8.5* | WBGene00015843 | protein_coding_gene | -1.1 | 0.4587 | 4.5E-09 | -1.3 | 0.394 | 6.2E-12 |
| *T23F11.4* | WBGene00011956 | protein_coding_gene | -1.1 | 0.4586 | 1.7E-05 | -1.4 | 0.368 | 1.0E-14 |
| *M04F3.5* | WBGene00019771 | protein_coding_gene | -1.1 | 0.4585 | 5.0E-09 | -1.2 | 0.427 | 1.3E-25 |
| *Y105E8A.21* | WBGene00013679 | protein_coding_gene | -1.1 | 0.4582 | 1.0E-15 | -1.1 | 0.481 | 3.6E-18 |
| *C26F1.3* | WBGene00016148 | protein_coding_gene | -1.1 | 0.4579 | 1.1E-11 | -1.1 | 0.452 | 2.7E-19 |
| *F28H6.4* | WBGene00009233 | protein_coding_gene | -1.1 | 0.4575 | 1.3E-07 | -1.2 | 0.431 | 2.1E-13 |
| *T14B4.3* | WBGene00020500 | protein_coding_gene | -1.1 | 0.4572 | 1.2E-13 | -1.1 | 0.482 | 8.9E-17 |
| *unc-26* | WBGene00006763 | protein_coding_gene | -1.1 | 0.4569 | 2.6E-11 | -1.0 | 0.487 | 4.6E-14 |
| *B0272.3* | WBGene00007129 | protein_coding_gene | -1.1 | 0.4562 | 6.1E-09 | -1.3 | 0.415 | 2.6E-21 |
| *arp-6* | WBGene00007434 | protein_coding_gene | -1.1 | 0.4562 | 1.2E-11 | -1.1 | 0.478 | 1.0E-18 |
| *scbp-2* | WBGene00010569 | protein_coding_gene | -1.1 | 0.4561 | 5.8E-07 | -1.4 | 0.382 | 8.7E-12 |
| *M106.2* | WBGene00010910 | protein_coding_gene | -1.1 | 0.4549 | 1.2E-11 | -1.2 | 0.446 | 6.3E-21 |

| | | | | | | | | |
|---|---|---|---|---|---|---|---|---|
| *Y45F10C.4* | WBGene00012880 | protein_coding_gene | -1.1 | 0.4546 | 3.8E-03 | -2.1 | 0.232 | 1.6E-17 |
| *mrt-2* | WBGene00003417 | protein_coding_gene | -1.1 | 0.4542 | 8.4E-10 | -1.1 | 0.460 | 6.2E-11 |
| *mmab-1* | WBGene00016144 | protein_coding_gene | -1.1 | 0.4540 | 3.8E-06 | -1.2 | 0.443 | 2.2E-16 |
| *mutd-1* | WBGene00016404 | protein_coding_gene | -1.1 | 0.4539 | 2.1E-08 | -1.3 | 0.406 | 7.0E-19 |
| *C01F1.6* | WBGene00015301 | protein_coding_gene | -1.1 | 0.4537 | 1.1E-10 | -1.0 | 0.493 | 2.9E-20 |
| *lmtr-2* | WBGene00022402 | protein_coding_gene | -1.1 | 0.4530 | 1.2E-04 | -1.2 | 0.442 | 1.3E-14 |
| *hpl-2* | WBGene00001996 | protein_coding_gene | -1.1 | 0.4523 | 6.7E-16 | -1.2 | 0.438 | 2.6E-22 |
| *slrp-1* | WBGene00008224 | protein_coding_gene | -1.1 | 0.4509 | 6.4E-15 | -1.2 | 0.435 | 4.4E-23 |
| *C34B7.2* | WBGene00007912 | protein_coding_gene | -1.1 | 0.4509 | 6.0E-10 | -1.1 | 0.455 | 1.8E-18 |
| *K07B1.4* | WBGene00019464 | protein_coding_gene | -1.2 | 0.4502 | 1.1E-06 | -1.1 | 0.455 | 2.9E-16 |
| *gck-2* | WBGene00022603 | protein_coding_gene | -1.2 | 0.4501 | 3.5E-12 | -1.0 | 0.495 | 1.4E-20 |
| *mre-11* | WBGene00003405 | protein_coding_gene | -1.2 | 0.4497 | 1.4E-13 | -1.1 | 0.468 | 8.5E-23 |
| *nhr-66* | WBGene00003656 | protein_coding_gene | -1.2 | 0.4496 | 8.9E-15 | -1.3 | 0.404 | 7.6E-31 |
| *stl-1* | WBGene00006061 | protein_coding_gene | -1.2 | 0.4487 | 3.4E-24 | -1.1 | 0.482 | 2.7E-21 |
| *psf-3* | WBGene00022043 | protein_coding_gene | -1.2 | 0.4486 | 6.9E-09 | -1.2 | 0.444 | 3.5E-20 |
| *baf-1* | WBGene00000235 | protein_coding_gene | -1.2 | 0.4485 | 3.6E-14 | -1.1 | 0.479 | 2.0E-20 |
| *R10H10.7* | WBGene00011225 | protein_coding_gene | -1.2 | 0.4485 | 9.8E-16 | -1.2 | 0.429 | 4.2E-25 |
| *T05H10.1* | WBGene00011507 | protein_coding_gene | -1.2 | 0.4484 | 1.1E-11 | -1.0 | 0.499 | 1.7E-22 |
| *Y71A12B.12* | WBGene00013509 | protein_coding_gene | -1.2 | 0.4479 | 3.6E-04 | -2.2 | 0.220 | 2.0E-13 |
| *fecl-1* | WBGene00010645 | protein_coding_gene | -1.2 | 0.4477 | 2.1E-17 | -1.0 | 0.484 | 7.7E-20 |
| *gst-16* | WBGene00001764 | protein_coding_gene | -1.2 | 0.4474 | 3.5E-04 | -1.1 | 0.472 | 5.9E-03 |
| *nasp-1* | WBGene00007500 | protein_coding_gene | -1.2 | 0.4468 | 1.6E-21 | -1.1 | 0.472 | 2.5E-22 |
| *tlk-1* | WBGene00006579 | protein_coding_gene | -1.2 | 0.4465 | 1.5E-04 | -1.1 | 0.464 | 1.5E-29 |
| *mys-1* | WBGene00007029 | protein_coding_gene | -1.2 | 0.4461 | 5.0E-08 | -1.2 | 0.437 | 4.1E-25 |
| *nova-1* | WBGene00013347 | protein_coding_gene | -1.2 | 0.4461 | 2.1E-14 | -1.1 | 0.476 | 3.7E-22 |
| *C02F5.13* | WBGene00015353 | protein_coding_gene | -1.2 | 0.4457 | 1.8E-07 | -1.3 | 0.414 | 2.6E-16 |
| *kup-1* | WBGene00002241 | protein_coding_gene | -1.2 | 0.4455 | 9.1E-23 | -1.2 | 0.428 | 5.1E-31 |
| *flcn-1* | WBGene00017699 | protein_coding_gene | -1.2 | 0.4454 | 1.0E-06 | -1.2 | 0.421 | 4.7E-29 |
| *M117.6* | WBGene00044253 | protein_coding_gene | -1.2 | 0.4448 | 1.1E-03 | -1.2 | 0.444 | 1.3E-04 |
| *src-1* | WBGene00005077 | protein_coding_gene | -1.2 | 0.4447 | 3.0E-09 | -1.3 | 0.400 | 4.9E-37 |
| *egl-26* | WBGene00001193 | protein_coding_gene | -1.2 | 0.4442 | 4.4E-07 | -1.1 | 0.479 | 1.4E-10 |
| *F55G11.4* | WBGene00010124 | protein_coding_gene | -1.2 | 0.4442 | 4.7E-04 | -4.2 | 0.054 | 1.1E-15 |
| *pacs-1* | WBGene00044077 | protein_coding_gene | -1.2 | 0.4440 | 7.3E-10 | -1.0 | 0.499 | 1.9E-25 |
| *C35B1.4* | WBGene00016434 | protein_coding_gene | -1.2 | 0.4440 | 2.0E-03 | -1.6 | 0.333 | 1.5E-13 |
| *Y49E10.4* | WBGene00013030 | protein_coding_gene | -1.2 | 0.4440 | 5.4E-12 | -1.1 | 0.478 | 3.6E-16 |
| *smc-6* | WBGene00010056 | protein_coding_gene | -1.2 | 0.4437 | 3.8E-13 | -1.1 | 0.459 | 3.0E-31 |
| *pfd-4* | WBGene00007107 | protein_coding_gene | -1.2 | 0.4436 | 7.3E-17 | -1.2 | 0.420 | 4.0E-23 |
| *algn-1* | WBGene00020820 | protein_coding_gene | -1.2 | 0.4431 | 2.9E-17 | -1.2 | 0.444 | 6.2E-21 |
| *lsm-3* | WBGene00003077 | protein_coding_gene | -1.2 | 0.4429 | 1.1E-06 | -1.4 | 0.369 | 7.3E-24 |
| *syx-4* | WBGene00006374 | protein_coding_gene | -1.2 | 0.4428 | 1.9E-16 | -1.2 | 0.450 | 1.2E-12 |
| *snpc-1.1* | WBGene00015513 | protein_coding_gene | -1.2 | 0.4424 | 3.2E-21 | -1.2 | 0.433 | 3.4E-19 |
| *cwc-15* | WBGene00011687 | protein_coding_gene | -1.2 | 0.4423 | 3.5E-20 | -1.2 | 0.423 | 6.1E-31 |
| *fbxb-7* | WBGene00012492 | protein_coding_gene | -1.2 | 0.4421 | 1.4E-07 | -1.3 | 0.420 | 5.3E-08 |
| *Y53C12A.10* | WBGene00175030 | protein_coding_gene | -1.2 | 0.4420 | 5.5E-08 | -1.1 | 0.474 | 1.8E-14 |
| *zip-8* | WBGene00017755 | protein_coding_gene | -1.2 | 0.4413 | 6.0E-08 | -1.7 | 0.315 | 8.4E-21 |
| *C54G4.9* | WBGene00008318 | protein_coding_gene | -1.2 | 0.4412 | 9.3E-09 | -1.1 | 0.456 | 4.2E-18 |
| *bra-2* | WBGene00017757 | protein_coding_gene | -1.2 | 0.4411 | 3.2E-09 | -1.4 | 0.370 | 1.2E-41 |
| ***Y27F2A.6*** | **WBGene00021298** | **protein_coding_gene** | **-1.2** | **0.4406** | **4.5E-02** | **1.1** | **2.120** | **2.4E-02** |
| *ZK632.2* | WBGene00014011 | protein_coding_gene | -1.2 | 0.4402 | 6.6E-17 | -1.2 | 0.435 | 6.9E-28 |
| *ztf-20* | WBGene00012702 | protein_coding_gene | -1.2 | 0.4398 | 5.7E-07 | -1.1 | 0.452 | 1.1E-09 |
| *cpf-1* | WBGene00000773 | protein_coding_gene | -1.2 | 0.4396 | 2.2E-11 | -1.2 | 0.446 | 1.9E-21 |
| *F35D11.4* | WBGene00018045 | protein_coding_gene | -1.2 | 0.4391 | 1.1E-04 | -1.3 | 0.416 | 1.6E-13 |
| *C35D10.7* | WBGene00016444 | protein_coding_gene | -1.2 | 0.4391 | 1.9E-11 | -1.3 | 0.409 | 4.2E-22 |
| *C06A5.8* | WBGene00015505 | protein_coding_gene | -1.2 | 0.4390 | 1.4E-12 | -1.3 | 0.394 | 3.5E-19 |
| *T24D1.2* | WBGene00011985 | protein_coding_gene | -1.2 | 0.4388 | 2.6E-21 | -1.3 | 0.396 | 9.5E-32 |
| *tofu-5* | WBGene00012167 | protein_coding_gene | -1.2 | 0.4384 | 1.3E-12 | -1.2 | 0.441 | 8.1E-25 |
| *jhdm-1* | WBGene00020821 | protein_coding_gene | -1.2 | 0.4383 | 1.3E-06 | -1.1 | 0.455 | 2.1E-32 |
| *trpp-3* | WBGene00014222 | protein_coding_gene | -1.2 | 0.4382 | 1.0E-11 | -1.1 | 0.469 | 1.0E-12 |
| *lmd-1* | WBGene00009665 | protein_coding_gene | -1.2 | 0.4380 | 1.2E-12 | -1.2 | 0.437 | 9.9E-16 |
| *snrp-27* | WBGene00011035 | protein_coding_gene | -1.2 | 0.4379 | 6.6E-08 | -1.4 | 0.383 | 2.2E-27 |
| *rbm-17* | WBGene00010233 | protein_coding_gene | -1.2 | 0.4378 | 3.5E-13 | -1.2 | 0.440 | 1.2E-16 |
| *F10E9.5* | WBGene00017357 | protein_coding_gene | -1.2 | 0.4373 | 8.4E-10 | -1.4 | 0.370 | 1.3E-19 |
| *ZK1236.1* | WBGene00022862 | protein_coding_gene | -1.2 | 0.4365 | 7.0E-22 | -1.1 | 0.460 | 6.6E-15 |
| *rbx-1* | WBGene00004320 | protein_coding_gene | -1.2 | 0.4356 | 4.6E-16 | -1.2 | 0.431 | 5.8E-30 |
| *par-2* | WBGene00003917 | protein_coding_gene | -1.2 | 0.4354 | 6.1E-09 | -1.0 | 0.498 | 1.4E-12 |
| *tbcb-1* | WBGene00009987 | protein_coding_gene | -1.2 | 0.4350 | 9.1E-17 | -1.2 | 0.432 | 6.9E-26 |
| *F52A8.1* | WBGene00009915 | protein_coding_gene | -1.2 | 0.4349 | 8.2E-08 | -1.2 | 0.433 | 6.4E-25 |
| *mlc-6* | WBGene00010554 | protein_coding_gene | -1.2 | 0.4348 | 7.2E-03 | -1.3 | 0.405 | 2.6E-02 |
| *cdc-6* | WBGene00000382 | protein_coding_gene | -1.2 | 0.4344 | 6.3E-17 | -1.2 | 0.441 | 3.8E-29 |
| *C08G9.2* | WBGene00015620 | protein_coding_gene | -1.2 | 0.4342 | 7.6E-04 | -1.6 | 0.341 | 2.8E-04 |
| *C01G5.5* | WBGene00015307 | protein_coding_gene | -1.2 | 0.4331 | 1.3E-09 | -1.2 | 0.440 | 4.0E-17 |
| *Y55D9A.2* | WBGene00013224 | protein_coding_gene | -1.2 | 0.4331 | 4.5E-21 | -1.2 | 0.441 | 1.1E-21 |
| *damt-1* | WBGene00015939 | protein_coding_gene | -1.2 | 0.4324 | 1.6E-10 | -1.3 | 0.404 | 7.9E-17 |
| *W04D2.4* | WBGene00012243 | protein_coding_gene | -1.2 | 0.4324 | 1.7E-10 | -1.3 | 0.405 | 1.2E-29 |
| *ZK154.5* | WBGene00022667 | protein_coding_gene | -1.2 | 0.4314 | 2.2E-07 | -1.2 | 0.430 | 8.1E-16 |
| *F08F1.4* | WBGene00017258 | protein_coding_gene | -1.2 | 0.4312 | 5.8E-04 | -1.7 | 0.298 | 2.6E-09 |
| *wve-1* | WBGene00006958 | protein_coding_gene | -1.2 | 0.4312 | 5.1E-12 | -1.0 | 0.485 | 3.4E-17 |
| *H06H21.11* | WBGene00044483 | protein_coding_gene | -1.2 | 0.4311 | 6.7E-16 | -1.4 | 0.385 | 5.7E-21 |
| *T25G3.1* | WBGene00012029 | protein_coding_gene | -1.2 | 0.4310 | 5.1E-15 | -1.6 | 0.340 | 2.1E-27 |
| *toe-2* | WBGene00016971 | protein_coding_gene | -1.2 | 0.4309 | 3.0E-18 | -1.4 | 0.389 | 5.2E-31 |

| | | | | | | | | |
|---|---|---|---|---|---|---|---|---|
| *cku-80* | WBGene00000520 | protein_coding_gene | -1.2 | 0.4306 | 2.3E-18 | -1.0 | 0.491 | 4.2E-17 |
| *metl-6* | WBGene00014205 | protein_coding_gene | -1.2 | 0.4298 | 5.1E-09 | -1.1 | 0.472 | 2.5E-12 |
| *C16C8.4* | WBGene00015842 | protein_coding_gene | -1.2 | 0.4295 | 1.8E-15 | -1.6 | 0.324 | 3.5E-23 |
| *pigs-1* | WBGene00011776 | protein_coding_gene | -1.2 | 0.4295 | 9.3E-22 | -1.1 | 0.465 | 1.2E-20 |
| *ZK673.4* | WBGene00014060 | protein_coding_gene | -1.2 | 0.4291 | 6.5E-05 | -1.1 | 0.460 | 9.6E-04 |
| ***twk-22*** | **WBGene00006675** | **protein_coding_gene** | **-1.2** | **0.4291** | **8.8E-05** | **1.1** | **2.126** | **6.7E-05** |
| *spat-3* | WBGene00020496 | protein_coding_gene | -1.2 | 0.4290 | 6.0E-04 | -1.3 | 0.408 | 2.4E-20 |
| *T07F8.4* | WBGene00020320 | protein_coding_gene | -1.2 | 0.4290 | 6.1E-07 | -1.1 | 0.462 | 1.7E-09 |
| *dscc-1* | WBGene00019595 | protein_coding_gene | -1.2 | 0.4287 | 1.5E-13 | -1.3 | 0.392 | 1.6E-21 |
| *M106.3* | WBGene00010911 | protein_coding_gene | -1.2 | 0.4287 | 7.8E-12 | -1.1 | 0.461 | 2.8E-20 |
| *C45G9.5* | WBGene00016676 | protein_coding_gene | -1.2 | 0.4284 | 1.4E-15 | -1.2 | 0.440 | 2.8E-18 |
| *sorf-1* | WBGene00022769 | protein_coding_gene | -1.2 | 0.4280 | 7.0E-11 | -1.0 | 0.494 | 1.2E-08 |
| *F58H1.8* | WBGene00138720 | protein_coding_gene | -1.2 | 0.4275 | 5.8E-10 | -1.3 | 0.395 | 4.3E-14 |
| *top-1* | WBGene00006595 | protein_coding_gene | -1.2 | 0.4270 | 2.5E-06 | -1.4 | 0.379 | 5.7E-44 |
| *pkc-3* | WBGene00004034 | protein_coding_gene | -1.2 | 0.4270 | 1.5E-08 | -1.0 | 0.491 | 1.3E-20 |
| *let-504* | WBGene00002696 | protein_coding_gene | -1.2 | 0.4270 | 1.0E-13 | -1.1 | 0.468 | 2.9E-18 |
| *let-49* | WBGene00002324 | protein_coding_gene | -1.2 | 0.4267 | 3.5E-13 | -1.2 | 0.448 | 1.5E-14 |
| *rpb-10* | WBGene00021347 | protein_coding_gene | -1.2 | 0.4267 | 1.6E-12 | -1.1 | 0.477 | 2.1E-11 |
| *mage-1* | WBGene00018230 | protein_coding_gene | -1.2 | 0.4262 | 9.4E-11 | -1.3 | 0.411 | 2.9E-24 |
| *snf-5* | WBGene00004904 | protein_coding_gene | -1.2 | 0.4260 | 2.1E-02 | -1.1 | 0.457 | 1.0E-11 |
| *C55C3.4* | WBGene00016954 | protein_coding_gene | -1.2 | 0.4253 | 4.4E-02 | -1.1 | 0.462 | 7.5E-04 |
| *chup-1* | WBGene00006477 | protein_coding_gene | -1.2 | 0.4252 | 6.4E-09 | -1.4 | 0.378 | 1.4E-12 |
| *ZK945.4* | WBGene00014166 | protein_coding_gene | -1.2 | 0.4252 | 3.1E-06 | -1.2 | 0.434 | 8.8E-05 |
| *F55A11.8* | WBGene00010081 | protein_coding_gene | -1.2 | 0.4247 | 2.4E-14 | -1.1 | 0.460 | 3.0E-12 |
| *stam-1* | WBGene00004109 | protein_coding_gene | -1.2 | 0.4246 | 3.0E-12 | -1.0 | 0.483 | 1.5E-21 |
| *fem-2* | WBGene00001412 | protein_coding_gene | -1.2 | 0.4244 | 1.5E-17 | -1.1 | 0.453 | 1.4E-21 |
| *rpb-5* | WBGene00019246 | protein_coding_gene | -1.2 | 0.4243 | 3.7E-19 | -1.1 | 0.466 | 1.8E-24 |
| *tgn-38* | WBGene00011742 | protein_coding_gene | -1.2 | 0.4241 | 7.8E-10 | -1.2 | 0.443 | 1.5E-25 |
| *lido-8* | WBGene00019511 | protein_coding_gene | -1.2 | 0.4237 | 3.1E-11 | -1.2 | 0.428 | 1.7E-11 |
| ***nspc-1*** | **WBGene00023477** | **protein_coding_gene** | **-1.2** | **0.4225** | **2.8E-02** | **1.6** | **3.117** | **2.8E-04** |
| *smut-1* | WBGene00021391 | protein_coding_gene | -1.2 | 0.4223 | 5.8E-11 | -1.2 | 0.429 | 2.5E-24 |
| *C14B1.12* | WBGene00184990 | protein_coding_gene | -1.2 | 0.4221 | 4.7E-12 | -1.3 | 0.392 | 2.0E-20 |
| *mek-5* | WBGene00017098 | protein_coding_gene | -1.2 | 0.4219 | 2.5E-14 | -1.2 | 0.449 | 3.4E-29 |
| *tomm-22* | WBGene00021133 | protein_coding_gene | -1.2 | 0.4214 | 5.6E-06 | -1.0 | 0.486 | 1.4E-15 |
| *arv-1* | WBGene00011040 | protein_coding_gene | -1.2 | 0.4212 | 4.1E-10 | -1.3 | 0.400 | 2.1E-17 |
| *mics-1* | WBGene00011890 | protein_coding_gene | -1.2 | 0.4211 | 1.5E-12 | -1.1 | 0.465 | 1.8E-25 |
| *lsm-7* | WBGene00003081 | protein_coding_gene | -1.2 | 0.4210 | 8.2E-08 | -1.4 | 0.376 | 4.8E-28 |
| *Y40A1A.3* | WBGene00021485 | protein_coding_gene | -1.2 | 0.4208 | 1.4E-02 | -1.1 | 0.451 | 1.9E-02 |
| *pas-6* | WBGene00003927 | protein_coding_gene | -1.2 | 0.4208 | 2.6E-13 | -1.0 | 0.498 | 2.1E-16 |
| *faah-4* | WBGene00013232 | protein_coding_gene | -1.3 | 0.4203 | 3.8E-07 | -1.2 | 0.437 | 2.0E-18 |
| *com-1* | WBGene00008082 | protein_coding_gene | -1.3 | 0.4201 | 1.0E-10 | -1.2 | 0.424 | 8.8E-19 |
| *F02E9.10* | WBGene00008533 | protein_coding_gene | -1.3 | 0.4199 | 1.0E-16 | -1.1 | 0.460 | 1.4E-19 |
| *fipr-21* | WBGene00009621 | protein_coding_gene | -1.3 | 0.4198 | 1.1E-04 | -1.3 | 0.407 | 2.0E-14 |
| *fubl-4* | WBGene00019692 | protein_coding_gene | -1.3 | 0.4193 | 3.1E-12 | -1.2 | 0.424 | 2.3E-18 |
| *T22C1.5* | WBGene00011916 | protein_coding_gene | -1.3 | 0.4191 | 2.3E-13 | -1.3 | 0.407 | 4.6E-29 |
| *lin-61* | WBGene00003041 | protein_coding_gene | -1.3 | 0.4190 | 5.3E-10 | -1.1 | 0.476 | 5.0E-20 |
| *Y80D3A.9* | WBGene00013587 | protein_coding_gene | -1.3 | 0.4189 | 2.6E-06 | -1.1 | 0.466 | 1.8E-09 |
| *prcc-1* | WBGene00008034 | protein_coding_gene | -1.3 | 0.4189 | 1.4E-28 | -1.3 | 0.401 | 1.1E-28 |
| *exoc-8* | WBGene00013687 | protein_coding_gene | -1.3 | 0.4189 | 2.9E-26 | -1.1 | 0.473 | 6.9E-18 |
| *R12C12.5* | WBGene00020025 | protein_coding_gene | -1.3 | 0.4188 | 4.8E-10 | -1.4 | 0.375 | 6.2E-26 |
| *swsn-7* | WBGene00007433 | protein_coding_gene | -1.3 | 0.4187 | 5.1E-06 | -1.2 | 0.430 | 1.2E-29 |
| *clh-6* | WBGene00000533 | protein_coding_gene | -1.3 | 0.4186 | 2.0E-13 | -1.0 | 0.490 | 3.1E-14 |
| *smg-5* | WBGene00004883 | protein_coding_gene | -1.3 | 0.4186 | 1.4E-16 | -1.4 | 0.385 | 1.4E-24 |
| *lido-11* | WBGene00016574 | protein_coding_gene | -1.3 | 0.4185 | 5.1E-09 | -1.8 | 0.283 | 2.2E-15 |
| *ztf-8* | WBGene00022598 | protein_coding_gene | -1.3 | 0.4184 | 2.6E-08 | -1.1 | 0.473 | 1.5E-19 |
| *paxt-1* | WBGene00011034 | protein_coding_gene | -1.3 | 0.4181 | 1.4E-12 | -1.2 | 0.439 | 5.7E-22 |
| *ndc-80* | WBGene00003576 | protein_coding_gene | -1.3 | 0.4177 | 5.9E-19 | -1.3 | 0.394 | 4.7E-42 |
| ***drd-1*** | **WBGene00009902** | **protein_coding_gene** | **-1.3** | **0.4176** | **1.0E-03** | **1.0** | **2.034** | **1.9E-11** |
| *tbce-1* | WBGene00019503 | protein_coding_gene | -1.3 | 0.4173 | 6.5E-15 | -1.3 | 0.406 | 1.0E-23 |
| *F31A3.5* | WBGene00017939 | protein_coding_gene | -1.3 | 0.4169 | 1.3E-06 | -1.4 | 0.368 | 3.0E-16 |
| *fut-3* | WBGene00006402 | protein_coding_gene | -1.3 | 0.4168 | 2.5E-11 | -1.6 | 0.338 | 5.0E-39 |
| *zip-4* | WBGene00021552 | protein_coding_gene | -1.3 | 0.4166 | 3.6E-25 | -1.0 | 0.491 | 5.6E-14 |
| *F42G8.19* | WBGene00249817 | protein_coding_gene | -1.3 | 0.4165 | 1.2E-02 | -2.0 | 0.242 | 2.4E-04 |
| *unc-37* | WBGene00006773 | protein_coding_gene | -1.3 | 0.4158 | 3.7E-12 | -1.6 | 0.328 | 1.3E-42 |
| *Y50D4C.6* | WBGene00021750 | protein_coding_gene | -1.3 | 0.4157 | 6.5E-08 | -1.4 | 0.392 | 9.4E-12 |
| *lin-15B* | WBGene00023497 | protein_coding_gene | -1.3 | 0.4156 | 4.9E-06 | -1.2 | 0.422 | 1.7E-15 |
| *mrps-23* | WBGene00014224 | protein_coding_gene | -1.3 | 0.4153 | 2.6E-08 | -1.2 | 0.424 | 1.0E-15 |
| ***gst-26*** | **WBGene00001774** | **protein_coding_gene** | **-1.3** | **0.4153** | **1.8E-02** | **1.5** | **2.868** | **6.6E-12** |
| *C08B6.8* | WBGene00007429 | protein_coding_gene | -1.3 | 0.4152 | 5.8E-07 | -1.1 | 0.467 | 7.3E-17 |
| *trpp-9* | WBGene00007956 | protein_coding_gene | -1.3 | 0.4151 | 2.9E-12 | -1.1 | 0.471 | 2.2E-14 |
| *prp-38* | WBGene00008380 | protein_coding_gene | -1.3 | 0.4150 | 2.8E-13 | -1.3 | 0.408 | 4.3E-25 |
| *ugt-14* | WBGene00019233 | protein_coding_gene | -1.3 | 0.4146 | 2.5E-02 | -2.8 | 0.146 | 1.8E-09 |
| *csk-1* | WBGene00000812 | protein_coding_gene | -1.3 | 0.4145 | 1.1E-15 | -1.1 | 0.471 | 4.9E-20 |
| *hsb-1* | WBGene00002002 | protein_coding_gene | -1.3 | 0.4145 | 7.7E-11 | -1.3 | 0.409 | 2.0E-25 |
| *R11E3.2* | WBGene00020004 | protein_coding_gene | -1.3 | 0.4144 | 3.0E-06 | -1.0 | 0.492 | 2.7E-03 |
| *aph-2* | WBGene00000148 | protein_coding_gene | -1.3 | 0.4143 | 1.6E-22 | -1.2 | 0.434 | 1.6E-23 |
| *aap-1* | WBGene00000001 | protein_coding_gene | -1.3 | 0.4140 | 2.4E-09 | -1.0 | 0.487 | 4.7E-10 |
| *nth-1* | WBGene00011201 | protein_coding_gene | -1.3 | 0.4139 | 9.8E-10 | -1.3 | 0.396 | 3.8E-12 |
| *rib-1* | WBGene00004360 | protein_coding_gene | -1.3 | 0.4138 | 2.6E-10 | -1.1 | 0.453 | 2.2E-20 |
| *C50F2.2* | WBGene00016836 | protein_coding_gene | -1.3 | 0.4136 | 1.1E-09 | -1.3 | 0.411 | 9.7E-24 |

| | | | | | | | | |
|---|---|---|---|---|---|---|---|---|
| *F58E6.5* | WBGene00010254 | protein_coding_gene | -1.3 | 0.4135 | 9.4E-03 | -1.5 | 0.366 | 6.4E-11 |
| *set-14* | WBGene00011068 | protein_coding_gene | -1.3 | 0.4132 | 2.7E-16 | -1.0 | 0.495 | 5.4E-11 |
| *C36B1.14* | WBGene00044614 | protein_coding_gene | -1.3 | 0.4130 | 3.0E-06 | -1.8 | 0.296 | 6.3E-28 |
| *R05F9.9* | WBGene00019892 | protein_coding_gene | -1.3 | 0.4130 | 1.6E-12 | -1.3 | 0.417 | 2.6E-25 |
| *ced-2* | WBGene00000416 | protein_coding_gene | -1.3 | 0.4129 | 4.5E-22 | -1.1 | 0.477 | 2.3E-20 |
| *ssup-72* | WBGene00020480 | protein_coding_gene | -1.3 | 0.4126 | 7.0E-09 | -1.4 | 0.390 | 1.0E-19 |
| ***amt-1*** | **WBGene00000133** | **protein_coding_gene** | **-1.3** | **0.4124** | **2.3E-02** | **2.6** | **6.000** | **1.1E-07** |
| *K05C4.4* | WBGene00010581 | protein_coding_gene | -1.3 | 0.4123 | 1.5E-07 | -1.5 | 0.366 | 1.2E-08 |
| *sec-11* | WBGene00021844 | protein_coding_gene | -1.3 | 0.4121 | 7.8E-13 | -1.0 | 0.488 | 1.3E-19 |
| *pxf-1* | WBGene00004254 | protein_coding_gene | -1.3 | 0.4120 | 1.3E-06 | -1.0 | 0.487 | 2.1E-18 |
| *apm-3* | WBGene00000164 | protein_coding_gene | -1.3 | 0.4117 | 2.2E-09 | -1.5 | 0.354 | 3.5E-21 |
| *F42H10.2* | WBGene00018366 | protein_coding_gene | -1.3 | 0.4112 | 4.8E-13 | -1.6 | 0.328 | 3.6E-23 |
| *K09E9.3* | WBGene00010726 | protein_coding_gene | -1.3 | 0.4111 | 6.1E-09 | -1.4 | 0.369 | 7.8E-17 |
| *snrp-40.1* | WBGene00008586 | protein_coding_gene | -1.3 | 0.4110 | 2.1E-09 | -1.5 | 0.348 | 1.7E-25 |
| *Y62E10A.23* | WBGene00302964 | protein_coding_gene | -1.3 | 0.4109 | 2.1E-05 | -1.1 | 0.457 | 1.1E-07 |
| *pola-1* | WBGene00012936 | protein_coding_gene | -1.3 | 0.4108 | 4.1E-10 | -1.1 | 0.479 | 3.7E-25 |
| *trpp-10* | WBGene00022138 | protein_coding_gene | -1.3 | 0.4107 | 3.1E-15 | -1.2 | 0.442 | 4.5E-15 |
| *age-1* | WBGene00000090 | protein_coding_gene | -1.3 | 0.4106 | 8.1E-12 | -1.2 | 0.446 | 1.1E-22 |
| *cids-2* | WBGene00009451 | protein_coding_gene | -1.3 | 0.4105 | 4.1E-06 | -1.1 | 0.471 | 4.8E-27 |
| *stip-1* | WBGene00007412 | protein_coding_gene | -1.3 | 0.4102 | 5.5E-14 | -1.0 | 0.489 | 2.0E-19 |
| *ZK546.5* | WBGene00022762 | protein_coding_gene | -1.3 | 0.4099 | 2.4E-20 | -1.3 | 0.417 | 4.0E-18 |
| *cdk-8* | WBGene00000409 | protein_coding_gene | -1.3 | 0.4097 | 2.3E-07 | -1.3 | 0.405 | 1.0E-28 |
| *chaf-2* | WBGene00022141 | protein_coding_gene | -1.3 | 0.4097 | 8.1E-19 | -1.2 | 0.437 | 2.3E-18 |
| *C24D10.6* | WBGene00016057 | protein_coding_gene | -1.3 | 0.4090 | 4.6E-21 | -1.2 | 0.430 | 2.0E-23 |
| *taf-9* | WBGene00006391 | protein_coding_gene | -1.3 | 0.4085 | 8.4E-13 | -1.3 | 0.396 | 3.9E-24 |
| *C35D10.1* | WBGene00016439 | protein_coding_gene | -1.3 | 0.4085 | 9.0E-11 | -1.2 | 0.446 | 7.6E-23 |
| *R11H6.5* | WBGene00011253 | protein_coding_gene | -1.3 | 0.4081 | 3.1E-16 | -1.2 | 0.424 | 3.8E-27 |
| *M04B2.4* | WBGene00010847 | protein_coding_gene | -1.3 | 0.4075 | 8.5E-17 | -1.2 | 0.421 | 5.0E-21 |
| *ceh-74* | WBGene00013876 | protein_coding_gene | -1.3 | 0.4075 | 1.7E-05 | -1.7 | 0.307 | 6.0E-10 |
| *ent-7* | WBGene00009686 | protein_coding_gene | -1.3 | 0.4074 | 1.6E-03 | -1.0 | 0.496 | 7.2E-08 |
| *F02E9.7* | WBGene00008531 | protein_coding_gene | -1.3 | 0.4072 | 5.4E-17 | -1.3 | 0.401 | 2.4E-28 |
| *ceh-86* | WBGene00018355 | protein_coding_gene | -1.3 | 0.4069 | 1.3E-06 | -3.0 | 0.124 | 5.8E-39 |
| *B0464.9* | WBGene00007188 | protein_coding_gene | -1.3 | 0.4067 | 8.2E-20 | -1.0 | 0.486 | 3.7E-15 |
| *pcaf-1* | WBGene00021636 | protein_coding_gene | -1.3 | 0.4063 | 4.1E-15 | -1.1 | 0.466 | 2.0E-18 |
| *F55A11.4* | WBGene00010077 | protein_coding_gene | -1.3 | 0.4057 | 3.0E-16 | -1.1 | 0.458 | 9.5E-18 |
| *erh-1* | WBGene00011892 | protein_coding_gene | -1.3 | 0.4056 | 5.9E-11 | -1.5 | 0.353 | 9.5E-36 |
| *apc-2* | WBGene00000143 | protein_coding_gene | -1.3 | 0.4054 | 6.6E-23 | -1.1 | 0.475 | 8.3E-19 |
| *ceh-93* | WBGene00019864 | protein_coding_gene | -1.3 | 0.4053 | 6.0E-10 | -1.2 | 0.423 | 1.3E-14 |
| *M02D8.3* | WBGene00019729 | protein_coding_gene | -1.3 | 0.4051 | 6.4E-11 | -1.3 | 0.408 | 4.4E-12 |
| *F33H1.3* | WBGene00009364 | protein_coding_gene | -1.3 | 0.4050 | 3.3E-10 | -1.3 | 0.405 | 1.7E-31 |
| *eme-1* | WBGene00018923 | protein_coding_gene | -1.3 | 0.4048 | 5.4E-16 | -1.0 | 0.489 | 2.4E-14 |
| *T23G5.2* | WBGene00011962 | protein_coding_gene | -1.3 | 0.4040 | 2.3E-09 | -1.1 | 0.474 | 1.6E-30 |
| *C32E8.5* | WBGene00016323 | protein_coding_gene | -1.3 | 0.4040 | 3.2E-25 | -1.3 | 0.397 | 2.9E-32 |
| *EEED8.13* | WBGene00017141 | protein_coding_gene | -1.3 | 0.4037 | 7.2E-05 | -1.6 | 0.333 | 1.8E-07 |
| *T24H10.1* | WBGene00012000 | protein_coding_gene | -1.3 | 0.4031 | 4.5E-19 | -1.1 | 0.457 | 1.4E-23 |
| *T04A8.8* | WBGene00011410 | protein_coding_gene | -1.3 | 0.4027 | 1.3E-15 | -1.3 | 0.404 | 5.4E-28 |
| *natc-1* | WBGene00020719 | protein_coding_gene | -1.3 | 0.4027 | 1.1E-13 | -1.2 | 0.432 | 3.4E-30 |
| *F01F1.2* | WBGene00017159 | protein_coding_gene | -1.3 | 0.4018 | 3.7E-15 | -1.2 | 0.428 | 3.0E-23 |
| *bub-1* | WBGene00000275 | protein_coding_gene | -1.3 | 0.4018 | 3.3E-28 | -1.1 | 0.477 | 9.1E-24 |
| *pigo-1* | WBGene00016159 | protein_coding_gene | -1.3 | 0.4017 | 3.6E-20 | -1.0 | 0.486 | 6.4E-15 |
| *rog-1* | WBGene00018819 | protein_coding_gene | -1.3 | 0.4014 | 1.3E-06 | -1.1 | 0.472 | 4.1E-17 |
| *F55A11.1* | WBGene00010075 | protein_coding_gene | -1.3 | 0.4012 | 5.6E-10 | -1.1 | 0.458 | 6.9E-30 |
| *nip-1* | WBGene00018386 | protein_coding_gene | -1.3 | 0.4012 | 8.9E-11 | -1.8 | 0.294 | 7.3E-23 |
| *rars-2* | WBGene00004680 | protein_coding_gene | -1.3 | 0.4007 | 4.7E-21 | -1.4 | 0.382 | 9.8E-28 |
| *prmt-9* | WBGene00011939 | protein_coding_gene | -1.3 | 0.4006 | 6.0E-11 | -1.1 | 0.469 | 1.3E-23 |
| *spd-5* | WBGene00004955 | protein_coding_gene | -1.3 | 0.4006 | 5.2E-17 | -1.1 | 0.475 | 3.3E-28 |
| *vps-33.2* | WBGene00016960 | protein_coding_gene | -1.3 | 0.4005 | 4.4E-24 | -1.1 | 0.480 | 1.0E-16 |
| *T09B9.4* | WBGene00011645 | protein_coding_gene | -1.3 | 0.4003 | 3.3E-07 | -1.1 | 0.475 | 2.1E-09 |
| ***C50A2.3*** | **WBGene00016797** | **protein_coding_gene** | **-1.3** | **0.4003** | **6.2E-03** | **1.0** | **2.042** | **9.0E-04** |
| *vab-15* | WBGene00006881 | protein_coding_gene | -1.3 | 0.3999 | 2.9E-03 | -2.2 | 0.215 | 1.2E-07 |
| *F37A4.2* | WBGene00018132 | protein_coding_gene | -1.3 | 0.3999 | 1.2E-12 | -1.4 | 0.375 | 1.1E-10 |
| *him-18* | WBGene00011415 | protein_coding_gene | -1.3 | 0.3998 | 1.2E-15 | -1.3 | 0.409 | 7.4E-27 |
| *C50F4.6* | WBGene00001910 | protein_coding_gene | -1.3 | 0.3997 | 4.0E-10 | -2.2 | 0.212 | 1.0E-37 |
| *Y69A2AR.3* | WBGene00022075 | protein_coding_gene | -1.3 | 0.3996 | 7.0E-06 | -1.0 | 0.495 | 4.9E-12 |
| *F30F8.10* | WBGene00009277 | protein_coding_gene | -1.3 | 0.3995 | 8.9E-14 | -1.2 | 0.446 | 9.0E-25 |
| *far-2* | WBGene00001386 | protein_coding_gene | -1.3 | 0.3993 | 3.0E-08 | -1.2 | 0.434 | 4.8E-18 |
| *cec-3* | WBGene00011636 | protein_coding_gene | -1.3 | 0.3993 | 3.6E-11 | -1.3 | 0.402 | 7.5E-33 |
| *Y38A10A.7* | WBGene00021392 | protein_coding_gene | -1.3 | 0.3989 | 2.1E-17 | -1.5 | 0.362 | 8.3E-33 |
| *C49A9.10* | WBGene00016764 | protein_coding_gene | -1.3 | 0.3988 | 9.6E-17 | -1.2 | 0.431 | 8.8E-12 |
| ***nspd-7*** | **WBGene00012781** | **protein_coding_gene** | **-1.3** | **0.3988** | **1.5E-02** | **1.3** | **2.419** | **4.6E-02** |
| *ehbp-1* | WBGene00009098 | protein_coding_gene | -1.3 | 0.3987 | 1.6E-15 | -1.0 | 0.486 | 5.7E-21 |
| *hcp-6* | WBGene00001833 | protein_coding_gene | -1.3 | 0.3986 | 1.2E-12 | -1.4 | 0.378 | 5.9E-36 |
| *lbp-4* | WBGene00002256 | protein_coding_gene | -1.3 | 0.3983 | 7.1E-14 | -1.3 | 0.395 | 1.5E-26 |
| *clik-1* | WBGene00020808 | protein_coding_gene | -1.3 | 0.3979 | 9.1E-09 | -1.1 | 0.464 | 2.2E-14 |
| *unc-61* | WBGene00006795 | protein_coding_gene | -1.3 | 0.3979 | 1.5E-09 | -1.2 | 0.425 | 1.3E-29 |
| *gadr-3* | WBGene00018561 | protein_coding_gene | -1.3 | 0.3979 | 1.7E-05 | -2.1 | 0.241 | 8.0E-12 |
| *Y87G2A.13* | WBGene00013601 | protein_coding_gene | -1.3 | 0.3978 | 9.0E-14 | -1.0 | 0.490 | 2.5E-20 |
| *rpt-2* | WBGene00004502 | protein_coding_gene | -1.3 | 0.3974 | 2.3E-16 | -1.1 | 0.456 | 2.4E-23 |
| *cogc-5* | WBGene00016608 | protein_coding_gene | -1.3 | 0.3972 | 2.1E-18 | -1.2 | 0.433 | 2.4E-27 |
| *mes-4* | WBGene00003222 | protein_coding_gene | -1.3 | 0.3971 | 1.5E-10 | -1.0 | 0.487 | 1.8E-19 |

| | | | | | | | | |
|---|---|---|---|---|---|---|---|---|
| *rab-30* | WBGene00004282 | protein_coding_gene | -1.3 | 0.3970 | 3.1E-18 | -1.2 | 0.450 | 5.6E-10 |
| *ref-2* | WBGene00004335 | protein_coding_gene | -1.3 | 0.3959 | 8.1E-03 | -3.1 | 0.113 | 1.8E-15 |
| *F44G4.3* | WBGene00009713 | protein_coding_gene | -1.3 | 0.3959 | 4.1E-08 | -1.2 | 0.428 | 7.5E-12 |
| *R74.7* | WBGene00011281 | protein_coding_gene | -1.3 | 0.3958 | 1.7E-23 | -1.3 | 0.417 | 3.7E-24 |
| *wapl-1* | WBGene00019953 | protein_coding_gene | -1.3 | 0.3957 | 1.0E-18 | -1.2 | 0.442 | 3.3E-27 |
| *F48C1.5* | WBGene00018598 | protein_coding_gene | -1.3 | 0.3957 | 1.9E-14 | -1.5 | 0.357 | 5.1E-20 |
| *K04F10.3* | WBGene00019396 | protein_coding_gene | -1.3 | 0.3957 | 1.4E-13 | -1.2 | 0.437 | 1.6E-17 |
| *hmp-1* | WBGene00001978 | protein_coding_gene | -1.3 | 0.3951 | 3.2E-09 | -1.1 | 0.458 | 1.5E-30 |
| *rnp-4* | WBGene00004387 | protein_coding_gene | -1.3 | 0.3947 | 1.6E-20 | -1.2 | 0.427 | 2.0E-39 |
| *eya-1* | WBGene00001377 | protein_coding_gene | -1.3 | 0.3946 | 4.3E-04 | -3.1 | 0.116 | 1.7E-36 |
| *F10D11.2* | WBGene00008648 | protein_coding_gene | -1.3 | 0.3946 | 1.3E-11 | -1.4 | 0.383 | 1.6E-26 |
| *smn-1* | WBGene00004887 | protein_coding_gene | -1.3 | 0.3943 | 1.6E-25 | -1.4 | 0.367 | 2.7E-34 |
| *F55A12.5* | WBGene00018864 | protein_coding_gene | -1.3 | 0.3942 | 4.3E-13 | -1.3 | 0.403 | 6.0E-34 |
| *prx-14* | WBGene00004199 | protein_coding_gene | -1.3 | 0.3936 | 4.4E-17 | -1.5 | 0.344 | 1.2E-35 |
| *ced-11* | WBGene00000425 | protein_coding_gene | -1.3 | 0.3936 | 3.6E-06 | -2.7 | 0.158 | 8.6E-28 |
| *K08D8.6* | WBGene00010660 | protein_coding_gene | -1.3 | 0.3936 | 2.4E-03 | -1.2 | 0.438 | 1.7E-03 |
| *atg-10* | WBGene00008427 | protein_coding_gene | -1.3 | 0.3936 | 2.3E-05 | -1.2 | 0.426 | 1.0E-03 |
| *cdk-11.2* | WBGene00013917 | protein_coding_gene | -1.3 | 0.3934 | 2.1E-11 | -1.3 | 0.392 | 4.3E-15 |
| *Y40H7A.10* | WBGene00012747 | protein_coding_gene | -1.3 | 0.3933 | 2.6E-07 | -1.5 | 0.362 | 6.2E-26 |
| *M02B1.3* | WBGene00010828 | protein_coding_gene | -1.3 | 0.3933 | 2.0E-10 | -1.2 | 0.423 | 1.9E-29 |
| *R151.8* | WBGene00020111 | protein_coding_gene | -1.3 | 0.3931 | 1.7E-10 | -1.2 | 0.427 | 1.3E-21 |
| *C10C6.9* | WBGene00045175 | pseudogene | -1.3 | 0.3930 | 3.5E-02 | -1.5 | 0.358 | 2.5E-03 |
| *ZK1098.2* | WBGene00014219 | protein_coding_gene | -1.3 | 0.3928 | 2.2E-16 | -1.0 | 0.492 | 1.6E-18 |
| *vps-33.1* | WBGene00015130 | protein_coding_gene | -1.3 | 0.3928 | 3.8E-15 | -1.4 | 0.391 | 5.4E-25 |
| *lat-1* | WBGene00002251 | protein_coding_gene | -1.3 | 0.3928 | 1.9E-12 | -1.3 | 0.415 | 2.2E-40 |
| *cpn-1* | WBGene00000777 | protein_coding_gene | -1.4 | 0.3922 | 2.8E-07 | -1.1 | 0.475 | 9.1E-24 |
| *B0205.1* | WBGene00015017 | protein_coding_gene | -1.4 | 0.3921 | 2.0E-14 | -1.2 | 0.437 | 1.0E-29 |
| *thoc-7* | WBGene00007195 | protein_coding_gene | -1.4 | 0.3915 | 4.2E-15 | -1.5 | 0.347 | 8.4E-35 |
| *gtf-2H3* | WBGene00014230 | protein_coding_gene | -1.4 | 0.3912 | 1.0E-17 | -1.2 | 0.435 | 2.0E-19 |
| *mbk-2* | WBGene00003150 | protein_coding_gene | -1.4 | 0.3911 | 5.1E-12 | -1.0 | 0.491 | 4.8E-26 |
| *pgp-4* | WBGene00003998 | protein_coding_gene | -1.4 | 0.3910 | 3.5E-06 | -1.7 | 0.318 | 1.0E-16 |
| *nas-20* | WBGene00003539 | protein_coding_gene | -1.4 | 0.3909 | 3.9E-04 | -2.7 | 0.156 | 2.3E-27 |
| *iff-1* | WBGene00002064 | protein_coding_gene | -1.4 | 0.3909 | 1.9E-08 | -1.1 | 0.452 | 1.6E-17 |
| *sna-2* | WBGene00011747 | protein_coding_gene | -1.4 | 0.3907 | 7.4E-13 | -1.1 | 0.470 | 1.8E-28 |
| *C17H11.2* | WBGene00015922 | protein_coding_gene | -1.4 | 0.3907 | 9.0E-11 | -1.0 | 0.498 | 3.4E-08 |
| ***kin-15*** | **WBGene00002199** | **protein_coding_gene** | **-1.4** | **0.3906** | **1.3E-02** | **1.7** | **3.153** | **1.2E-07** |
| *rpt-5* | WBGene00004505 | protein_coding_gene | -1.4 | 0.3905 | 2.2E-16 | -1.1 | 0.464 | 2.3E-23 |
| *lsy-13* | WBGene00020287 | protein_coding_gene | -1.4 | 0.3904 | 2.3E-20 | -1.1 | 0.452 | 4.3E-19 |
| *tag-307* | WBGene00022787 | protein_coding_gene | -1.4 | 0.3903 | 8.0E-18 | -1.2 | 0.439 | 1.0E-23 |
| *hira-1* | WBGene00019627 | protein_coding_gene | -1.4 | 0.3900 | 2.0E-15 | -1.2 | 0.431 | 8.1E-25 |
| *ceh-41* | WBGene00000462 | protein_coding_gene | -1.4 | 0.3895 | 1.7E-11 | -1.4 | 0.378 | 3.3E-12 |
| *T28A8.3* | WBGene00012109 | protein_coding_gene | -1.4 | 0.3893 | 1.7E-16 | -1.4 | 0.382 | 3.4E-28 |
| *orc-4* | WBGene00012651 | protein_coding_gene | -1.4 | 0.3891 | 9.1E-18 | -1.5 | 0.346 | 3.3E-27 |
| *Y14H12B.1* | WBGene00021191 | protein_coding_gene | -1.4 | 0.3890 | 2.3E-18 | -1.1 | 0.453 | 8.2E-19 |
| *hmg-4* | WBGene00001974 | protein_coding_gene | -1.4 | 0.3889 | 2.5E-10 | -1.7 | 0.317 | 7.1E-55 |
| *C31E10.5* | WBGene00007846 | protein_coding_gene | -1.4 | 0.3887 | 7.5E-13 | -1.5 | 0.354 | 2.6E-22 |
| *F33H1.4* | WBGene00009365 | protein_coding_gene | -1.4 | 0.3885 | 7.7E-16 | -1.2 | 0.425 | 7.6E-37 |
| *Y69H2.7* | WBGene00013482 | protein_coding_gene | -1.4 | 0.3883 | 8.2E-21 | -1.1 | 0.483 | 9.7E-12 |
| *dpff-1* | WBGene00016200 | protein_coding_gene | -1.4 | 0.3881 | 1.5E-16 | -1.3 | 0.407 | 2.0E-29 |
| *B0334.4* | WBGene00007144 | protein_coding_gene | -1.4 | 0.3879 | 2.2E-10 | -1.6 | 0.331 | 1.9E-42 |
| *isy-1* | WBGene00009966 | protein_coding_gene | -1.4 | 0.3878 | 6.3E-26 | -1.1 | 0.458 | 4.2E-20 |
| *F25H2.4* | WBGene00009118 | protein_coding_gene | -1.4 | 0.3874 | 3.1E-20 | -1.1 | 0.466 | 1.1E-15 |
| *mdf-1* | WBGene00003160 | protein_coding_gene | -1.4 | 0.3871 | 4.5E-12 | -1.1 | 0.459 | 8.7E-27 |
| *nxf-2* | WBGene00003835 | protein_coding_gene | -1.4 | 0.3869 | 1.8E-13 | -1.0 | 0.485 | 7.2E-13 |
| *dnj-22* | WBGene00001040 | protein_coding_gene | -1.4 | 0.3867 | 6.8E-17 | -1.3 | 0.407 | 2.6E-27 |
| *vps-29* | WBGene00014234 | protein_coding_gene | -1.4 | 0.3862 | 4.9E-10 | -1.3 | 0.399 | 3.1E-24 |
| *ced-9* | WBGene00000423 | protein_coding_gene | -1.4 | 0.3861 | 1.2E-26 | -1.5 | 0.356 | 1.3E-28 |
| *zhp-3* | WBGene00006976 | protein_coding_gene | -1.4 | 0.3861 | 3.9E-09 | -1.3 | 0.401 | 4.7E-24 |
| *M106.8* | WBGene00077658 | protein_coding_gene | -1.4 | 0.3860 | 5.2E-10 | -1.3 | 0.417 | 1.2E-16 |
| *ZK1307.4* | WBGene00014247 | protein_coding_gene | -1.4 | 0.3859 | 1.7E-02 | -1.2 | 0.425 | 6.1E-07 |
| *mrpl-20* | WBGene00012992 | protein_coding_gene | -1.4 | 0.3855 | 4.1E-16 | -1.1 | 0.457 | 2.4E-11 |
| *uba-2* | WBGene00006700 | protein_coding_gene | -1.4 | 0.3853 | 2.6E-23 | -1.0 | 0.497 | 2.6E-24 |
| *lin-32* | WBGene00003018 | protein_coding_gene | -1.4 | 0.3851 | 4.9E-02 | -3.6 | 0.082 | 8.7E-12 |
| *T28A8.8* | WBGene00194852 | protein_coding_gene | -1.4 | 0.3851 | 4.4E-07 | -1.3 | 0.418 | 4.2E-07 |
| ***C38D9.13*** | **WBGene00235090** | **ncRNA_gene** | **-1.4** | **0.3851** | **1.8E-08** | **2.9** | **7.255** | **4.4E-03** |
| *F45F2.10* | WBGene00018482 | protein_coding_gene | -1.4 | 0.3844 | 3.3E-07 | -1.0 | 0.494 | 1.3E-19 |
| *C25H3.3* | WBGene00016112 | protein_coding_gene | -1.4 | 0.3843 | 3.9E-10 | -1.2 | 0.426 | 3.0E-18 |
| *set-22* | WBGene00012527 | protein_coding_gene | -1.4 | 0.3843 | 7.9E-09 | -1.2 | 0.439 | 1.1E-17 |
| *Y39A3CL.3* | WBGene00021441 | protein_coding_gene | -1.4 | 0.3838 | 1.5E-06 | -1.1 | 0.463 | 1.5E-13 |
| *thk-1* | WBGene00006568 | protein_coding_gene | -1.4 | 0.3838 | 4.7E-06 | -1.6 | 0.327 | 7.3E-36 |
| *spr-1* | WBGene00005006 | protein_coding_gene | -1.4 | 0.3837 | 1.0E-18 | -1.2 | 0.448 | 6.5E-21 |
| *hst-6* | WBGene00002031 | protein_coding_gene | -1.4 | 0.3834 | 1.2E-09 | -1.7 | 0.318 | 3.1E-16 |
| *rpb-4* | WBGene00018391 | protein_coding_gene | -1.4 | 0.3833 | 2.6E-16 | -1.4 | 0.391 | 2.8E-27 |
| *Y10G11A.1* | WBGene00012423 | protein_coding_gene | -1.4 | 0.3831 | 8.4E-16 | -1.2 | 0.438 | 6.2E-23 |
| *npp-9* | WBGene00003795 | protein_coding_gene | -1.4 | 0.3830 | 2.3E-09 | -1.0 | 0.490 | 2.7E-21 |
| *zip-7* | WBGene00013100 | protein_coding_gene | -1.4 | 0.3829 | 5.4E-03 | -3.2 | 0.110 | 9.7E-19 |
| *fbxa-169* | WBGene00045289 | protein_coding_gene | -1.4 | 0.3827 | 1.7E-02 | -1.8 | 0.293 | 2.8E-03 |
| *B0001.8* | WBGene00007093 | protein_coding_gene | -1.4 | 0.3826 | 4.9E-13 | -1.0 | 0.492 | 1.0E-18 |
| *eri-7* | WBGene00016566 | protein_coding_gene | -1.4 | 0.3825 | 1.7E-10 | -1.3 | 0.419 | 6.3E-17 |
| *sas-6* | WBGene00012888 | protein_coding_gene | -1.4 | 0.3825 | 1.6E-23 | -1.2 | 0.429 | 2.0E-26 |

| | | | | | | | | |
|---|---|---|---|---|---|---|---|---|
| *klp-18* | WBGene00002228 | protein_coding_gene | -1.4 | 0.3825 | 1.8E-11 | -1.1 | 0.482 | 9.9E-21 |
| *T03F1.12* | WBGene00020193 | protein_coding_gene | -1.4 | 0.3822 | 1.2E-09 | -1.5 | 0.351 | 7.2E-20 |
| *alfa-1* | WBGene00017547 | protein_coding_gene | -1.4 | 0.3820 | 2.5E-27 | -1.2 | 0.444 | 2.8E-18 |
| *mdt-9* | WBGene00014938 | protein_coding_gene | -1.4 | 0.3818 | 2.4E-05 | -1.3 | 0.411 | 2.0E-19 |
| *mdt-11* | WBGene00007015 | protein_coding_gene | -1.4 | 0.3816 | 2.6E-08 | -1.9 | 0.264 | 5.9E-50 |
| *R02F11.4* | WBGene00019842 | protein_coding_gene | -1.4 | 0.3810 | 2.2E-27 | -1.0 | 0.488 | 2.6E-15 |
| *bag-1* | WBGene00000236 | protein_coding_gene | -1.4 | 0.3808 | 1.1E-19 | -1.4 | 0.381 | 6.3E-27 |
| *vps-26* | WBGene00006931 | protein_coding_gene | -1.4 | 0.3807 | 9.1E-18 | -1.1 | 0.465 | 3.8E-21 |
| *C17G10.1* | WBGene00015915 | protein_coding_gene | -1.4 | 0.3807 | 1.1E-19 | -1.0 | 0.491 | 2.6E-19 |
| *Y106G6H.14* | WBGene00013724 | protein_coding_gene | -1.4 | 0.3807 | 3.1E-11 | -1.3 | 0.419 | 6.0E-14 |
| *lin-45* | WBGene00003030 | protein_coding_gene | -1.4 | 0.3806 | 6.9E-08 | -1.0 | 0.484 | 3.1E-26 |
| *C43E11.12* | WBGene00016609 | protein_coding_gene | -1.4 | 0.3806 | 1.3E-16 | -1.4 | 0.382 | 5.7E-21 |
| *hda-3* | WBGene00001836 | protein_coding_gene | -1.4 | 0.3806 | 5.0E-17 | -1.2 | 0.444 | 2.0E-22 |
| *R144.6* | WBGene00020096 | protein_coding_gene | -1.4 | 0.3805 | 3.1E-12 | -1.2 | 0.440 | 6.9E-13 |
| *clec-3* | WBGene00016577 | protein_coding_gene | -1.4 | 0.3805 | 4.0E-02 | -3.4 | 0.093 | 1.9E-14 |
| *C28C12.12* | WBGene00016182 | protein_coding_gene | -1.4 | 0.3804 | 5.7E-14 | -1.3 | 0.408 | 1.8E-28 |
| *F25G6.8* | WBGene00017799 | protein_coding_gene | -1.4 | 0.3804 | 9.5E-26 | -1.3 | 0.413 | 2.8E-23 |
| *R09H10.7* | WBGene00011184 | protein_coding_gene | -1.4 | 0.3803 | 2.5E-02 | -2.0 | 0.242 | 4.0E-05 |
| *C48D1.5* | WBGene00008178 | protein_coding_gene | -1.4 | 0.3799 | 6.7E-05 | -1.2 | 0.427 | 3.3E-03 |
| *vps-39* | WBGene00011625 | protein_coding_gene | -1.4 | 0.3798 | 3.2E-17 | -1.1 | 0.475 | 9.3E-24 |
| *K07A1.15* | WBGene00010619 | protein_coding_gene | -1.4 | 0.3796 | 1.4E-04 | -1.7 | 0.303 | 5.2E-19 |
| *mdf-2* | WBGene00003161 | protein_coding_gene | -1.4 | 0.3795 | 1.3E-15 | -1.9 | 0.265 | 2.8E-60 |
| *cyn-12* | WBGene00000888 | protein_coding_gene | -1.4 | 0.3793 | 1.2E-16 | -1.3 | 0.410 | 1.7E-20 |
| *T10B10.3* | WBGene00011680 | protein_coding_gene | -1.4 | 0.3790 | 2.0E-08 | -1.2 | 0.422 | 1.1E-17 |
| *row-1* | WBGene00009508 | protein_coding_gene | -1.4 | 0.3788 | 1.0E-19 | -1.5 | 0.360 | 7.1E-33 |
| *gon-4* | WBGene00001653 | protein_coding_gene | -1.4 | 0.3788 | 5.8E-11 | -1.5 | 0.352 | 1.6E-36 |
| *xpa-1* | WBGene00006963 | protein_coding_gene | -1.4 | 0.3787 | 7.8E-21 | -1.1 | 0.462 | 5.2E-17 |
| *K08E4.2* | WBGene00010666 | protein_coding_gene | -1.4 | 0.3785 | 1.1E-07 | -1.8 | 0.284 | 3.8E-18 |
| *T09A5.15* | WBGene00011641 | protein_coding_gene | -1.4 | 0.3778 | 7.1E-09 | -1.1 | 0.472 | 1.2E-07 |
| *F56D1.1* | WBGene00018959 | protein_coding_gene | -1.4 | 0.3776 | 2.2E-12 | -1.5 | 0.361 | 9.3E-33 |
| *T21B10.3* | WBGene00011885 | protein_coding_gene | -1.4 | 0.3773 | 3.3E-17 | -1.2 | 0.446 | 1.1E-26 |
| *C16C10.4* | WBGene00007625 | protein_coding_gene | -1.4 | 0.3773 | 3.1E-24 | -1.2 | 0.442 | 1.7E-25 |
| *C29E4.13* | WBGene00043051 | protein_coding_gene | -1.4 | 0.3771 | 1.0E-15 | -1.3 | 0.396 | 2.2E-17 |
| *Y37D8A.25* | WBGene00012560 | protein_coding_gene | -1.4 | 0.3771 | 5.9E-10 | -1.6 | 0.336 | 4.8E-12 |
| *rfc-1* | WBGene00004337 | protein_coding_gene | -1.4 | 0.3770 | 4.1E-16 | -1.3 | 0.398 | 3.5E-40 |
| *cisd-3.1* | WBGene00010909 | protein_coding_gene | -1.4 | 0.3770 | 1.2E-19 | -1.1 | 0.454 | 2.0E-18 |
| *sgo-1* | WBGene00016381 | protein_coding_gene | -1.4 | 0.3769 | 5.7E-11 | -1.6 | 0.339 | 1.6E-30 |
| *mcm-10* | WBGene00012935 | protein_coding_gene | -1.4 | 0.3765 | 7.3E-27 | -1.1 | 0.467 | 2.2E-16 |
| *knl-2* | WBGene00019432 | protein_coding_gene | -1.4 | 0.3764 | 2.0E-14 | -1.4 | 0.377 | 2.2E-37 |
| *fog-2* | WBGene00001482 | protein_coding_gene | -1.4 | 0.3763 | 3.1E-14 | -1.4 | 0.377 | 6.3E-24 |
| *drp-1* | WBGene00001093 | protein_coding_gene | -1.4 | 0.3761 | 1.3E-25 | -1.1 | 0.456 | 2.3E-26 |
| *cul-1* | WBGene00000836 | protein_coding_gene | -1.4 | 0.3760 | 9.2E-20 | -1.1 | 0.469 | 1.3E-22 |
| *mxl-1* | WBGene00003509 | protein_coding_gene | -1.4 | 0.3757 | 1.6E-13 | -1.3 | 0.407 | 1.4E-27 |
| *pat-3* | WBGene00003930 | protein_coding_gene | -1.4 | 0.3757 | 2.2E-13 | -1.5 | 0.356 | 9.0E-49 |
| *ubxn-6* | WBGene00019163 | protein_coding_gene | -1.4 | 0.3755 | 3.2E-24 | -1.2 | 0.448 | 6.4E-25 |
| *ari-1.2* | WBGene00016157 | protein_coding_gene | -1.4 | 0.3751 | 1.2E-25 | -1.4 | 0.391 | 1.9E-26 |
| *Y71F9B.6* | WBGene00022128 | protein_coding_gene | -1.4 | 0.3751 | 1.8E-24 | -1.0 | 0.500 | 1.0E-13 |
| *bir-1* | WBGene00000249 | protein_coding_gene | -1.4 | 0.3750 | 1.9E-17 | -1.3 | 0.399 | 8.3E-19 |
| *him-8* | WBGene00001867 | protein_coding_gene | -1.4 | 0.3750 | 7.2E-17 | -1.4 | 0.370 | 3.1E-32 |
| *vps-16* | WBGene00006516 | protein_coding_gene | -1.4 | 0.3749 | 3.4E-19 | -1.0 | 0.492 | 5.3E-15 |
| *C18G1.9* | WBGene00015987 | protein_coding_gene | -1.4 | 0.3748 | 3.4E-02 | -1.5 | 0.349 | 1.9E-05 |
| *Y54G2A.75* | WBGene00235102 | protein_coding_gene | -1.4 | 0.3747 | 8.9E-12 | -1.2 | 0.436 | 1.9E-17 |
| *lipl-4* | WBGene00019376 | protein_coding_gene | -1.4 | 0.3747 | 8.5E-03 | -2.3 | 0.206 | 1.3E-05 |
| *dnj-23* | WBGene00001041 | protein_coding_gene | -1.4 | 0.3739 | 1.4E-14 | -1.7 | 0.314 | 1.0E-42 |
| ***C45B2.2*** | **WBGene00016659** | **protein_coding_gene** | **-1.4** | **0.3738** | **1.1E-02** | **1.0** | **2.041** | **2.2E-02** |
| *pigc-1* | WBGene00011860 | protein_coding_gene | -1.4 | 0.3736 | 1.5E-14 | -1.2 | 0.433 | 2.6E-23 |
| *F59E12.1* | WBGene00019118 | protein_coding_gene | -1.4 | 0.3733 | 1.9E-19 | -1.2 | 0.421 | 1.7E-23 |
| *sna-1* | WBGene00020951 | protein_coding_gene | -1.4 | 0.3732 | 6.7E-22 | -1.0 | 0.486 | 3.7E-16 |
| *D2030.7* | WBGene00008417 | protein_coding_gene | -1.4 | 0.3731 | 4.5E-14 | -1.4 | 0.369 | 3.0E-35 |
| *nspd-1* | WBGene00022754 | protein_coding_gene | -1.4 | 0.3730 | 1.1E-02 | -1.5 | 0.351 | 6.8E-17 |
| *smcl-1* | WBGene00008085 | protein_coding_gene | -1.4 | 0.3725 | 1.0E-10 | -1.5 | 0.351 | 9.5E-12 |
| ***C40A11.8*** | **WBGene00016551** | **protein_coding_gene** | **-1.4** | **0.3723** | **4.5E-03** | **1.5** | **2.921** | **1.8E-03** |
| *lsm-6* | WBGene00003080 | protein_coding_gene | -1.4 | 0.3721 | 8.8E-17 | -1.4 | 0.392 | 1.6E-16 |
| *rfc-4* | WBGene00004340 | protein_coding_gene | -1.4 | 0.3717 | 3.2E-11 | -1.2 | 0.446 | 7.6E-28 |
| *sip-1* | WBGene00004798 | protein_coding_gene | -1.4 | 0.3715 | 2.9E-18 | -2.1 | 0.230 | 1.5E-64 |
| *Y87G2A.18* | WBGene00013604 | protein_coding_gene | -1.4 | 0.3714 | 8.1E-09 | -1.4 | 0.384 | 4.7E-10 |
| *rilp-1* | WBGene00007860 | protein_coding_gene | -1.4 | 0.3712 | 1.9E-17 | -1.2 | 0.428 | 1.0E-25 |
| *sel-12* | WBGene00004769 | protein_coding_gene | -1.4 | 0.3712 | 1.1E-10 | -1.4 | 0.377 | 7.2E-15 |
| *rnf-5* | WBGene00004381 | protein_coding_gene | -1.4 | 0.3711 | 5.2E-15 | -1.1 | 0.465 | 8.7E-20 |
| *nfx-1* | WBGene00007048 | protein_coding_gene | -1.4 | 0.3711 | 3.7E-17 | -1.1 | 0.469 | 4.3E-23 |
| *ZK418.5* | WBGene00022734 | protein_coding_gene | -1.4 | 0.3707 | 7.8E-10 | -1.2 | 0.431 | 1.2E-25 |
| *lin-52* | WBGene00003035 | protein_coding_gene | -1.4 | 0.3704 | 1.6E-18 | -1.6 | 0.323 | 7.5E-34 |
| *F12F6.7* | WBGene00008722 | protein_coding_gene | -1.4 | 0.3701 | 2.9E-19 | -1.4 | 0.384 | 6.5E-34 |
| *F22E5.9* | WBGene00017708 | protein_coding_gene | -1.4 | 0.3698 | 1.5E-13 | -1.1 | 0.470 | 2.4E-09 |
| *ppfr-2* | WBGene00017064 | protein_coding_gene | -1.4 | 0.3698 | 3.8E-21 | -1.0 | 0.494 | 2.1E-19 |
| *rfc-2* | WBGene00004338 | protein_coding_gene | -1.4 | 0.3697 | 8.6E-19 | -1.2 | 0.432 | 5.3E-20 |
| *hex-4* | WBGene00021766 | protein_coding_gene | -1.4 | 0.3694 | 6.9E-23 | -1.1 | 0.473 | 7.1E-17 |
| *mcm-4* | WBGene00003156 | protein_coding_gene | -1.4 | 0.3690 | 1.1E-12 | -2.0 | 0.255 | 1.8E-51 |
| *T04F8.2* | WBGene00011441 | protein_coding_gene | -1.4 | 0.3690 | 1.2E-14 | -1.0 | 0.488 | 8.4E-15 |
| *ssna-1* | WBGene00020300 | protein_coding_gene | -1.4 | 0.3689 | 6.5E-16 | -2.0 | 0.257 | 9.3E-39 |

| | | | | | | | | |
|---|---|---|---|---|---|---|---|---|
| *frh-1* | WBGene00001486 | protein_coding_gene | -1.4 | 0.3685 | 8.6E-11 | -1.3 | 0.395 | 7.6E-17 |
| *tbck-1* | WBGene00016352 | protein_coding_gene | -1.4 | 0.3684 | 1.9E-14 | -1.1 | 0.469 | 8.6E-09 |
| *erg-28* | WBGene00007589 | protein_coding_gene | -1.4 | 0.3681 | 1.0E-06 | -1.8 | 0.290 | 7.0E-16 |
| *dif-1* | WBGene00000996 | protein_coding_gene | -1.4 | 0.3681 | 1.9E-18 | -1.2 | 0.423 | 6.8E-23 |
| *eral-1* | WBGene00008456 | protein_coding_gene | -1.4 | 0.3679 | 6.2E-20 | -1.1 | 0.456 | 6.2E-14 |
| *gly-20* | WBGene00001645 | protein_coding_gene | -1.4 | 0.3679 | 6.3E-19 | -1.4 | 0.371 | 3.3E-44 |
| *hcf-1* | WBGene00001827 | protein_coding_gene | -1.4 | 0.3677 | 2.0E-16 | -1.2 | 0.447 | 1.0E-32 |
| *cat-4* | WBGene00000298 | protein_coding_gene | -1.4 | 0.3675 | 5.7E-06 | -1.5 | 0.366 | 4.3E-30 |
| *cyn-7* | WBGene00000883 | protein_coding_gene | -1.4 | 0.3675 | 2.9E-09 | -1.1 | 0.482 | 2.9E-18 |
| *tag-342* | WBGene00007187 | protein_coding_gene | -1.4 | 0.3673 | 2.4E-19 | -1.8 | 0.295 | 6.5E-42 |
| *nstp-2* | WBGene00017480 | protein_coding_gene | -1.4 | 0.3672 | 1.4E-22 | -1.2 | 0.449 | 1.3E-20 |
| *pigg-1* | WBGene00009204 | protein_coding_gene | -1.4 | 0.3672 | 3.5E-19 | -1.0 | 0.489 | 2.5E-14 |
| *brd-1* | WBGene00000265 | protein_coding_gene | -1.4 | 0.3669 | 1.5E-19 | -1.1 | 0.471 | 5.3E-15 |
| *rpn-13* | WBGene00016981 | protein_coding_gene | -1.4 | 0.3668 | 1.9E-16 | -1.0 | 0.485 | 4.7E-18 |
| *C28A5.1* | WBGene00007788 | protein_coding_gene | -1.4 | 0.3661 | 5.5E-12 | -1.5 | 0.364 | 8.6E-19 |
| *tdp-1* | WBGene00006514 | protein_coding_gene | -1.5 | 0.3657 | 4.1E-19 | -1.2 | 0.422 | 1.8E-32 |
| *ulp-1* | WBGene00006736 | protein_coding_gene | -1.5 | 0.3656 | 7.9E-14 | -1.2 | 0.423 | 1.2E-36 |
| *W05H9.4* | WBGene00021051 | protein_coding_gene | -1.5 | 0.3655 | 4.8E-16 | -1.5 | 0.355 | 1.7E-25 |
| ***col-149*** | **WBGene00000722** | **protein_coding_gene** | **-1.5** | **0.3653** | **1.7E-03** | **2.5** | **5.722** | **1.5E-07** |
| *lin-5* | WBGene00002994 | protein_coding_gene | -1.5 | 0.3652 | 9.5E-22 | -1.6 | 0.331 | 1.9E-41 |
| *R10D12.13* | WBGene00011194 | protein_coding_gene | -1.5 | 0.3648 | 2.5E-15 | -1.3 | 0.407 | 5.5E-17 |
| ***M03B6.1*** | **WBGene00010833** | **protein_coding_gene** | **-1.5** | **0.3639** | **3.1E-02** | **1.2** | **2.357** | **1.7E-03** |
| *msp-57* | WBGene00003450 | protein_coding_gene | -1.5 | 0.3639 | 6.6E-03 | -1.4 | 0.372 | 1.5E-15 |
| *B0511.2* | WBGene00015228 | protein_coding_gene | -1.5 | 0.3635 | 2.7E-04 | -1.8 | 0.284 | 3.4E-05 |
| *R12C12.6* | WBGene00020026 | protein_coding_gene | -1.5 | 0.3635 | 9.0E-14 | -1.5 | 0.364 | 7.5E-28 |
| *Y24F12A.1* | WBGene00012496 | protein_coding_gene | -1.5 | 0.3634 | 8.7E-28 | -1.2 | 0.448 | 5.1E-23 |
| *emb-27* | WBGene00001281 | protein_coding_gene | -1.5 | 0.3634 | 9.6E-16 | -1.3 | 0.402 | 1.1E-24 |
| *cuc-1* | WBGene00000835 | protein_coding_gene | -1.5 | 0.3634 | 2.5E-07 | -1.1 | 0.462 | 1.2E-16 |
| *knl-1* | WBGene00002231 | protein_coding_gene | -1.5 | 0.3633 | 1.8E-13 | -1.3 | 0.395 | 1.2E-32 |
| *H31G24.3* | WBGene00019254 | protein_coding_gene | -1.5 | 0.3633 | 3.3E-14 | -1.6 | 0.341 | 5.3E-13 |
| *C01G10.7* | WBGene00007234 | protein_coding_gene | -1.5 | 0.3626 | 6.3E-10 | -1.1 | 0.477 | 1.4E-13 |
| *lin-53* | WBGene00003036 | protein_coding_gene | -1.5 | 0.3625 | 2.1E-17 | -1.4 | 0.390 | 3.6E-34 |
| *Y47D3B.12* | WBGene00185055 | protein_coding_gene | -1.5 | 0.3624 | 6.9E-03 | -2.6 | 0.167 | 1.4E-07 |
| *D1014.2* | WBGene00017015 | protein_coding_gene | -1.5 | 0.3624 | 1.4E-04 | -1.6 | 0.338 | 5.9E-05 |
| *ubq-1* | WBGene00006727 | protein_coding_gene | -1.5 | 0.3622 | 2.7E-18 | -1.3 | 0.417 | 2.6E-23 |
| *W02D7.5* | WBGene00020941 | protein_coding_gene | -1.5 | 0.3621 | 3.9E-09 | -1.5 | 0.349 | 2.1E-08 |
| *rpt-1* | WBGene00004501 | protein_coding_gene | -1.5 | 0.3621 | 7.6E-17 | -1.1 | 0.474 | 3.1E-24 |
| *ocrl-1* | WBGene00007620 | protein_coding_gene | -1.5 | 0.3620 | 1.8E-27 | -1.2 | 0.430 | 4.4E-24 |
| *fbxa-179* | WBGene00045292 | protein_coding_gene | -1.5 | 0.3619 | 1.5E-02 | -2.9 | 0.136 | 3.4E-06 |
| *igcm-4* | WBGene00022418 | protein_coding_gene | -1.5 | 0.3618 | 9.7E-07 | -2.2 | 0.211 | 1.5E-29 |
| *rabn-5* | WBGene00017161 | protein_coding_gene | -1.5 | 0.3618 | 1.3E-17 | -1.1 | 0.460 | 4.6E-17 |
| *Y71F9AL.6* | WBGene00022111 | protein_coding_gene | -1.5 | 0.3618 | 5.1E-03 | -2.5 | 0.174 | 8.2E-07 |
| *T07F10.3* | WBGene00011589 | protein_coding_gene | -1.5 | 0.3615 | 2.3E-19 | -1.6 | 0.335 | 1.3E-36 |
| *ceh-58* | WBGene00007417 | protein_coding_gene | -1.5 | 0.3612 | 1.4E-11 | -2.0 | 0.257 | 2.0E-26 |
| *T10B11.8* | WBGene00020404 | protein_coding_gene | -1.5 | 0.3611 | 2.6E-12 | -1.1 | 0.451 | 6.7E-14 |
| *isw-1* | WBGene00002169 | protein_coding_gene | -1.5 | 0.3611 | 4.7E-14 | -1.0 | 0.496 | 2.0E-18 |
| *cdk-4* | WBGene00000406 | protein_coding_gene | -1.5 | 0.3610 | 3.4E-09 | -1.9 | 0.272 | 9.2E-17 |
| *polq-1* | WBGene00020964 | protein_coding_gene | -1.5 | 0.3609 | 1.2E-16 | -1.2 | 0.444 | 4.4E-23 |
| *C35D10.5* | WBGene00016442 | protein_coding_gene | -1.5 | 0.3608 | 3.0E-16 | -1.3 | 0.417 | 1.2E-16 |
| *F31B12.3* | WBGene00001495 | protein_coding_gene | -1.5 | 0.3606 | 2.5E-03 | -2.6 | 0.165 | 5.3E-08 |
| *nspd-3* | WBGene00016058 | protein_coding_gene | -1.5 | 0.3602 | 1.5E-03 | -1.5 | 0.351 | 1.2E-05 |
| *B0563.1* | WBGene00015261 | protein_coding_gene | -1.5 | 0.3601 | 4.7E-02 | -2.0 | 0.251 | 1.9E-02 |
| *dlc-1* | WBGene00001005 | protein_coding_gene | -1.5 | 0.3593 | 2.4E-17 | -1.5 | 0.365 | 6.2E-46 |
| *C53A3.4* | WBGene00303231 | protein_coding_gene | -1.5 | 0.3591 | 8.1E-03 | -3.2 | 0.108 | 7.3E-09 |
| *spcs-2* | WBGene00012550 | protein_coding_gene | -1.5 | 0.3590 | 1.8E-16 | -1.1 | 0.483 | 6.7E-18 |
| *taf-11.1* | WBGene00006393 | protein_coding_gene | -1.5 | 0.3590 | 2.6E-12 | -1.7 | 0.303 | 1.3E-15 |
| *kel-8* | WBGene00020952 | protein_coding_gene | -1.5 | 0.3589 | 3.5E-10 | -1.4 | 0.372 | 1.4E-15 |
| *hpr-17* | WBGene00001998 | protein_coding_gene | -1.5 | 0.3589 | 1.1E-19 | -1.6 | 0.335 | 9.4E-32 |
| *pas-7* | WBGene00003928 | protein_coding_gene | -1.5 | 0.3588 | 8.9E-23 | -1.1 | 0.481 | 7.3E-24 |
| *F13E9.1* | WBGene00008750 | protein_coding_gene | -1.5 | 0.3587 | 1.2E-20 | -1.1 | 0.453 | 2.0E-22 |
| *chaf-1* | WBGene00011532 | protein_coding_gene | -1.5 | 0.3586 | 7.2E-18 | -1.8 | 0.284 | 8.3E-66 |
| *bud-31* | WBGene00007400 | protein_coding_gene | -1.5 | 0.3584 | 2.9E-26 | -1.4 | 0.370 | 8.6E-30 |
| *C16A3.2* | WBGene00015807 | protein_coding_gene | -1.5 | 0.3584 | 3.2E-19 | -1.8 | 0.296 | 1.1E-25 |
| *swd-2.2* | WBGene00016374 | protein_coding_gene | -1.5 | 0.3583 | 1.6E-24 | -1.1 | 0.455 | 3.0E-23 |
| *C45G9.2* | WBGene00016674 | protein_coding_gene | -1.5 | 0.3582 | 2.6E-21 | -1.4 | 0.388 | 2.2E-17 |
| *ntp-1* | WBGene00016380 | protein_coding_gene | -1.5 | 0.3581 | 4.4E-14 | -1.6 | 0.338 | 8.0E-28 |
| *wht-2* | WBGene00007513 | protein_coding_gene | -1.5 | 0.3581 | 3.3E-17 | -1.3 | 0.410 | 4.0E-27 |
| *C15A11.4* | WBGene00007598 | protein_coding_gene | -1.5 | 0.3580 | 6.3E-03 | -2.1 | 0.237 | 9.5E-07 |
| *T26A5.8* | WBGene00020823 | protein_coding_gene | -1.5 | 0.3580 | 9.1E-08 | -1.9 | 0.277 | 8.8E-33 |
| *Y73F8A.27* | WBGene00013532 | protein_coding_gene | -1.5 | 0.3579 | 9.6E-24 | -1.0 | 0.491 | 4.9E-17 |
| *Y73B6BL.12* | WBGene00022236 | protein_coding_gene | -1.5 | 0.3570 | 3.1E-02 | -1.3 | 0.409 | 9.5E-03 |
| *pen-2* | WBGene00003975 | protein_coding_gene | -1.5 | 0.3569 | 3.7E-16 | -1.1 | 0.476 | 2.9E-09 |
| *lsy-22* | WBGene00009188 | protein_coding_gene | -1.5 | 0.3568 | 2.4E-09 | -1.5 | 0.359 | 4.9E-42 |
| ***F41E6.12*** | **WBGene00018293** | **protein_coding_gene** | **-1.5** | **0.3554** | **1.4E-02** | **1.5** | **2.861** | **6.0E-06** |
| *Y73B6BL.23* | WBGene00022244 | protein_coding_gene | -1.5 | 0.3553 | 9.6E-14 | -1.2 | 0.439 | 9.0E-25 |
| *rsp-3* | WBGene00004700 | protein_coding_gene | -1.5 | 0.3552 | 1.5E-19 | -1.1 | 0.459 | 3.4E-24 |
| *ife-3* | WBGene00002061 | protein_coding_gene | -1.5 | 0.3551 | 3.2E-19 | -1.1 | 0.463 | 5.1E-20 |
| *Y37E11B.6* | WBGene00021378 | protein_coding_gene | -1.5 | 0.3550 | 8.8E-16 | -1.6 | 0.324 | 3.0E-12 |
| *snx-3* | WBGene00006503 | protein_coding_gene | -1.5 | 0.3548 | 1.5E-26 | -1.2 | 0.446 | 3.5E-22 |
| *R04F11.3* | WBGene00011016 | protein_coding_gene | -1.5 | 0.3547 | 9.1E-14 | -1.6 | 0.322 | 4.2E-48 |

| | | | | | | | | |
|---|---|---|---|---|---|---|---|---|
| *R17.3* | WBGene00011269 | protein_coding_gene | -1.5 | 0.3546 | 5.5E-06 | -1.3 | 0.411 | 2.7E-04 |
| *gcc-2* | WBGene00011503 | protein_coding_gene | -1.5 | 0.3546 | 3.7E-22 | -1.2 | 0.444 | 5.5E-25 |
| *ssp-9* | WBGene00006038 | protein_coding_gene | -1.5 | 0.3546 | 2.5E-02 | -1.7 | 0.314 | 4.6E-06 |
| *ceh-38* | WBGene00000459 | protein_coding_gene | -1.5 | 0.3539 | 2.2E-11 | -1.3 | 0.414 | 8.8E-35 |
| *hda-1* | WBGene00001834 | protein_coding_gene | -1.5 | 0.3534 | 1.4E-12 | -1.3 | 0.395 | 3.1E-42 |
| *C17E4.20* | WBGene00219313 | protein_coding_gene | -1.5 | 0.3533 | 8.4E-09 | -1.8 | 0.287 | 2.7E-60 |
| *rpa-1* | WBGene00017546 | protein_coding_gene | -1.5 | 0.3533 | 2.0E-17 | -1.3 | 0.404 | 4.7E-40 |
| *hst-2* | WBGene00002029 | protein_coding_gene | -1.5 | 0.3532 | 6.3E-15 | -1.4 | 0.368 | 6.8E-13 |
| *lir-3* | WBGene00003046 | protein_coding_gene | -1.5 | 0.3532 | 1.6E-06 | -1.9 | 0.270 | 1.4E-09 |
| *hrpf-1* | WBGene00020936 | protein_coding_gene | -1.5 | 0.3531 | 3.8E-22 | -1.2 | 0.429 | 2.8E-18 |
| *F32D8.14* | WBGene00009336 | protein_coding_gene | -1.5 | 0.3530 | 5.9E-21 | -1.0 | 0.492 | 2.8E-15 |
| *F17A9.2* | WBGene00017534 | protein_coding_gene | -1.5 | 0.3526 | 1.0E-18 | -1.2 | 0.445 | 1.5E-20 |
| *H05L14.2* | WBGene00010367 | protein_coding_gene | -1.5 | 0.3525 | 1.1E-10 | -1.1 | 0.465 | 2.5E-16 |
| *exos-2* | WBGene00022232 | protein_coding_gene | -1.5 | 0.3524 | 2.6E-15 | -1.3 | 0.398 | 5.1E-24 |
| *dtmk-1* | WBGene00011272 | protein_coding_gene | -1.5 | 0.3524 | 7.3E-16 | -1.7 | 0.309 | 1.0E-36 |
| *T26C12.3* | WBGene00020833 | protein_coding_gene | -1.5 | 0.3523 | 4.0E-03 | -4.0 | 0.062 | 2.5E-24 |
| *fbxb-4* | WBGene00010921 | pseudogene | -1.5 | 0.3523 | 2.7E-02 | -1.7 | 0.300 | 2.0E-02 |
| *F21D5.6* | WBGene00009011 | protein_coding_gene | -1.5 | 0.3522 | 1.7E-14 | -1.5 | 0.349 | 1.5E-30 |
| *msh-2* | WBGene00003418 | protein_coding_gene | -1.5 | 0.3520 | 2.9E-19 | -1.1 | 0.467 | 4.4E-19 |
| ***ugt-19*** | **WBGene00019515** | **protein_coding_gene** | **-1.5** | **0.3520** | **3.0E-03** | **1.2** | **2.355** | **8.1E-08** |
| *blos-2* | WBGene00022251 | protein_coding_gene | -1.5 | 0.3520 | 8.3E-16 | -1.4 | 0.382 | 2.3E-27 |
| *mdt-8* | WBGene00007013 | protein_coding_gene | -1.5 | 0.3519 | 4.4E-09 | -1.9 | 0.260 | 6.9E-48 |
| *rnh-2* | WBGene00004383 | protein_coding_gene | -1.5 | 0.3517 | 1.3E-12 | -1.2 | 0.448 | 1.1E-20 |
| *F10E7.11* | WBGene00017352 | protein_coding_gene | -1.5 | 0.3517 | 4.0E-04 | -1.5 | 0.355 | 4.6E-03 |
| *rskd-1* | WBGene00010096 | protein_coding_gene | -1.5 | 0.3516 | 9.8E-26 | -1.3 | 0.397 | 8.8E-36 |
| *mob-4* | WBGene00016238 | protein_coding_gene | -1.5 | 0.3516 | 4.3E-07 | -1.4 | 0.387 | 6.3E-39 |
| *coa-5* | WBGene00012483 | protein_coding_gene | -1.5 | 0.3514 | 1.2E-16 | -1.1 | 0.457 | 6.8E-11 |
| *dnc-6* | WBGene00021827 | protein_coding_gene | -1.5 | 0.3510 | 4.7E-18 | -1.3 | 0.406 | 1.5E-21 |
| *R06F6.12* | WBGene00023418 | protein_coding_gene | -1.5 | 0.3510 | 7.3E-20 | -1.4 | 0.367 | 3.3E-29 |
| *die-1* | WBGene00000995 | protein_coding_gene | -1.5 | 0.3509 | 3.8E-07 | -2.5 | 0.182 | 3.6E-64 |
| *ttr-2* | WBGene00010539 | protein_coding_gene | -1.5 | 0.3507 | 8.1E-09 | -2.2 | 0.224 | 9.2E-40 |
| *dhfr-1* | WBGene00007974 | protein_coding_gene | -1.5 | 0.3507 | 1.8E-07 | -1.3 | 0.415 | 2.1E-25 |
| *lsm-4* | WBGene00003078 | protein_coding_gene | -1.5 | 0.3506 | 1.1E-14 | -1.5 | 0.358 | 3.4E-18 |
| *tofu-6* | WBGene00017132 | protein_coding_gene | -1.5 | 0.3504 | 2.0E-12 | -1.5 | 0.364 | 6.6E-29 |
| *usp-46* | WBGene00011216 | protein_coding_gene | -1.5 | 0.3503 | 2.1E-28 | -1.4 | 0.369 | 5.9E-31 |
| *efn-4* | WBGene00001165 | protein_coding_gene | -1.5 | 0.3502 | 3.2E-07 | -2.9 | 0.138 | 2.1E-46 |
| *num-1* | WBGene00003830 | protein_coding_gene | -1.5 | 0.3501 | 2.2E-19 | -1.0 | 0.491 | 1.8E-20 |
| *ubc-7* | WBGene00006704 | protein_coding_gene | -1.5 | 0.3500 | 6.6E-31 | -1.2 | 0.447 | 3.0E-26 |
| *C07E3.9* | WBGene00007419 | protein_coding_gene | -1.5 | 0.3499 | 6.5E-06 | -1.6 | 0.328 | 1.2E-17 |
| *spas-1* | WBGene00016045 | protein_coding_gene | -1.5 | 0.3497 | 1.6E-22 | -1.0 | 0.487 | 2.9E-16 |
| *K10C2.14* | WBGene00206388 | protein_coding_gene | -1.5 | 0.3491 | 7.0E-03 | -3.5 | 0.088 | 1.9E-29 |
| *T24D3.2* | WBGene00023492 | protein_coding_gene | -1.5 | 0.3490 | 1.4E-03 | -1.1 | 0.458 | 6.9E-03 |
| *C02F5.10* | WBGene00015351 | protein_coding_gene | -1.5 | 0.3490 | 1.6E-03 | -1.2 | 0.436 | 7.1E-04 |
| *C47G2.4* | WBGene00008165 | protein_coding_gene | -1.5 | 0.3489 | 4.9E-22 | -1.3 | 0.395 | 1.6E-30 |
| *K08F9.4* | WBGene00010687 | protein_coding_gene | -1.5 | 0.3483 | 1.4E-25 | -1.7 | 0.308 | 9.1E-34 |
| *such-1* | WBGene00013443 | protein_coding_gene | -1.5 | 0.3483 | 6.6E-24 | -1.0 | 0.500 | 2.7E-12 |
| *cdh-4* | WBGene00000396 | protein_coding_gene | -1.5 | 0.3482 | 2.7E-06 | -2.1 | 0.227 | 1.3E-55 |
| *CC8.2* | WBGene00016987 | protein_coding_gene | -1.5 | 0.3477 | 1.3E-14 | -1.4 | 0.373 | 6.4E-13 |
| *ssp-31* | WBGene00006048 | protein_coding_gene | -1.5 | 0.3475 | 2.3E-02 | -1.6 | 0.334 | 1.5E-07 |
| *madf-11* | WBGene00015468 | protein_coding_gene | -1.5 | 0.3474 | 1.7E-19 | -1.3 | 0.393 | 1.7E-26 |
| *rba-1* | WBGene00004312 | protein_coding_gene | -1.5 | 0.3473 | 3.2E-19 | -1.3 | 0.400 | 1.2E-37 |
| *cup-2* | WBGene00000843 | protein_coding_gene | -1.5 | 0.3473 | 3.7E-11 | -1.1 | 0.458 | 7.0E-22 |
| *brc-2* | WBGene00020316 | protein_coding_gene | -1.5 | 0.3471 | 2.6E-18 | -1.1 | 0.461 | 1.0E-16 |
| *hal-2* | WBGene00011819 | protein_coding_gene | -1.5 | 0.3470 | 3.8E-13 | -1.3 | 0.400 | 8.8E-23 |
| *mlst-8* | WBGene00015697 | protein_coding_gene | -1.5 | 0.3461 | 6.2E-15 | -1.2 | 0.421 | 4.8E-24 |
| *psr-1* | WBGene00004205 | protein_coding_gene | -1.5 | 0.3460 | 3.9E-21 | -1.2 | 0.431 | 4.4E-20 |
| *rpn-7* | WBGene00004463 | protein_coding_gene | -1.5 | 0.3454 | 4.3E-16 | -1.2 | 0.446 | 5.4E-25 |
| *hmp-2* | WBGene00001979 | protein_coding_gene | -1.5 | 0.3453 | 2.0E-17 | -1.2 | 0.444 | 3.5E-21 |
| *acaa-2* | WBGene00009952 | protein_coding_gene | -1.5 | 0.3452 | 3.1E-20 | -1.0 | 0.488 | 1.1E-15 |
| *Y110A7A.9* | WBGene00022459 | protein_coding_gene | -1.5 | 0.3449 | 3.6E-19 | -1.2 | 0.428 | 9.7E-31 |
| *F32D1.5* | WBGene00017984 | protein_coding_gene | -1.5 | 0.3448 | 6.2E-14 | -1.1 | 0.460 | 2.6E-19 |
| *F08B6.1* | WBGene00017239 | protein_coding_gene | -1.5 | 0.3448 | 8.9E-10 | -1.6 | 0.341 | 3.4E-10 |
| *lin-37* | WBGene00003022 | protein_coding_gene | -1.5 | 0.3448 | 1.7E-19 | -1.4 | 0.390 | 6.6E-26 |
| *ksr-2* | WBGene00002240 | protein_coding_gene | -1.5 | 0.3447 | 1.4E-15 | -1.1 | 0.453 | 2.8E-17 |
| *B0545.4* | WBGene00015247 | protein_coding_gene | -1.5 | 0.3441 | 2.9E-04 | -1.3 | 0.417 | 4.8E-04 |
| *F17H10.1* | WBGene00008925 | protein_coding_gene | -1.5 | 0.3440 | 4.4E-29 | -1.1 | 0.480 | 5.6E-10 |
| *mel-26* | WBGene00003209 | protein_coding_gene | -1.5 | 0.3439 | 2.8E-11 | -1.5 | 0.346 | 3.7E-46 |
| *madf-8* | WBGene00008118 | protein_coding_gene | -1.5 | 0.3435 | 1.5E-15 | -1.7 | 0.301 | 2.9E-28 |
| *bub-3* | WBGene00013209 | protein_coding_gene | -1.5 | 0.3433 | 1.7E-21 | -1.3 | 0.412 | 3.1E-27 |
| *T05H4.11* | WBGene00020274 | protein_coding_gene | -1.5 | 0.3432 | 2.0E-13 | -1.5 | 0.360 | 8.7E-33 |
| *T10E9.2* | WBGene00020412 | protein_coding_gene | -1.5 | 0.3432 | 1.0E-25 | -1.4 | 0.384 | 2.9E-32 |
| *hrpk-1* | WBGene00017816 | protein_coding_gene | -1.5 | 0.3431 | 4.6E-16 | -1.1 | 0.472 | 8.1E-26 |
| *mpk-1* | WBGene00003401 | protein_coding_gene | -1.5 | 0.3431 | 9.4E-14 | -1.1 | 0.480 | 3.3E-24 |
| *C56G2.15* | WBGene00016983 | protein_coding_gene | -1.5 | 0.3430 | 6.9E-10 | -1.5 | 0.347 | 1.4E-08 |
| *C48B6.9* | WBGene00016745 | protein_coding_gene | -1.5 | 0.3430 | 1.9E-07 | -2.4 | 0.186 | 4.1E-26 |
| *tag-175* | WBGene00006520 | protein_coding_gene | -1.5 | 0.3427 | 6.6E-14 | -1.1 | 0.469 | 9.5E-09 |
| *Y71H2AM.2* | WBGene00022167 | protein_coding_gene | -1.5 | 0.3426 | 7.0E-14 | -1.4 | 0.377 | 8.3E-26 |
| *strl-1* | WBGene00009940 | protein_coding_gene | -1.5 | 0.3422 | 1.1E-17 | -1.2 | 0.441 | 7.5E-21 |
| *ZC404.16* | WBGene00195226 | protein_coding_gene | -1.5 | 0.3422 | 2.6E-02 | -2.2 | 0.213 | 1.4E-02 |
| *K07C5.6* | WBGene00010629 | protein_coding_gene | -1.5 | 0.3421 | 2.2E-21 | -1.1 | 0.458 | 3.5E-24 |

| | | | | | | | | |
|---|---|---|---|---|---|---|---|---|
| *C17H12.2* | WBGene00015928 | protein_coding_gene | -1.5 | 0.3421 | 8.5E-19 | -1.0 | 0.491 | 1.5E-24 |
| *Y51F10.3* | WBGene00021764 | protein_coding_gene | -1.5 | 0.3421 | 1.2E-18 | -1.1 | 0.462 | 7.0E-22 |
| *htz-1* | WBGene00019947 | protein_coding_gene | -1.5 | 0.3419 | 4.5E-14 | -1.4 | 0.388 | 2.0E-32 |
| *farl-11* | WBGene00017349 | protein_coding_gene | -1.5 | 0.3419 | 4.3E-16 | -1.2 | 0.431 | 7.4E-26 |
| *vps-53* | WBGene00011502 | protein_coding_gene | -1.5 | 0.3418 | 4.2E-18 | -1.2 | 0.449 | 3.4E-20 |
| *gfi-3* | WBGene00001583 | protein_coding_gene | -1.5 | 0.3416 | 1.4E-12 | -1.1 | 0.451 | 2.3E-09 |
| *E01A2.1* | WBGene00017084 | protein_coding_gene | -1.5 | 0.3416 | 1.1E-15 | -1.7 | 0.315 | 1.9E-36 |
| *F43G9.4* | WBGene00009667 | protein_coding_gene | -1.6 | 0.3415 | 2.0E-18 | -1.4 | 0.376 | 3.5E-27 |
| *usp-14* | WBGene00006856 | protein_coding_gene | -1.6 | 0.3415 | 5.0E-14 | -1.1 | 0.452 | 2.5E-23 |
| *lin-7* | WBGene00002996 | protein_coding_gene | -1.6 | 0.3414 | 1.8E-24 | -1.2 | 0.433 | 1.0E-13 |
| *C04F12.1* | WBGene00007297 | protein_coding_gene | -1.6 | 0.3414 | 2.4E-12 | -1.4 | 0.371 | 1.2E-27 |
| *ZK1010.10* | WBGene00164985 | protein_coding_gene | -1.6 | 0.3412 | 2.6E-05 | -1.5 | 0.359 | 3.9E-05 |
| *grp-1* | WBGene00001743 | protein_coding_gene | -1.6 | 0.3407 | 1.0E-16 | -1.4 | 0.372 | 5.2E-38 |
| *swsn-4* | WBGene00004204 | protein_coding_gene | -1.6 | 0.3405 | 2.4E-12 | -1.5 | 0.359 | 1.1E-47 |
| *hmg-20* | WBGene00012209 | protein_coding_gene | -1.6 | 0.3402 | 6.6E-23 | -1.0 | 0.498 | 9.8E-10 |
| *F16B12.6* | WBGene00008882 | protein_coding_gene | -1.6 | 0.3401 | 2.1E-17 | -1.4 | 0.386 | 2.0E-22 |
| *F37D6.4* | WBGene00009510 | protein_coding_gene | -1.6 | 0.3399 | 4.9E-03 | -1.5 | 0.354 | 6.0E-03 |
| *ZK593.3* | WBGene00014003 | protein_coding_gene | -1.6 | 0.3399 | 1.3E-04 | -2.0 | 0.258 | 1.6E-08 |
| *csn-3* | WBGene00000815 | protein_coding_gene | -1.6 | 0.3397 | 4.1E-21 | -1.2 | 0.422 | 3.5E-31 |
| *zyg-12* | WBGene00006997 | protein_coding_gene | -1.6 | 0.3396 | 1.4E-17 | -1.2 | 0.432 | 2.3E-34 |
| *tra-2* | WBGene00006605 | protein_coding_gene | -1.6 | 0.3394 | 4.1E-10 | -1.1 | 0.474 | 7.2E-15 |
| *C05C10.2* | WBGene00007329 | protein_coding_gene | -1.6 | 0.3394 | 1.1E-21 | -1.4 | 0.385 | 2.2E-24 |
| *Y47D3A.1* | WBGene00012922 | protein_coding_gene | -1.6 | 0.3390 | 2.6E-06 | -2.6 | 0.164 | 6.1E-17 |
| *B0511.7* | WBGene00015233 | protein_coding_gene | -1.6 | 0.3390 | 7.6E-13 | -1.4 | 0.369 | 9.7E-26 |
| *clr-1* | WBGene00000548 | protein_coding_gene | -1.6 | 0.3389 | 5.3E-16 | -1.2 | 0.428 | 1.5E-27 |
| *lin-35* | WBGene00003020 | protein_coding_gene | -1.6 | 0.3385 | 3.5E-23 | -1.4 | 0.379 | 3.8E-37 |
| *emc-6* | WBGene00017999 | protein_coding_gene | -1.6 | 0.3384 | 1.6E-09 | -1.3 | 0.411 | 2.2E-17 |
| *R05F9.6* | WBGene00019890 | protein_coding_gene | -1.6 | 0.3384 | 9.1E-15 | -1.2 | 0.435 | 2.2E-22 |
| *F26H9.5* | WBGene00009177 | protein_coding_gene | -1.6 | 0.3382 | 1.9E-13 | -1.3 | 0.412 | 1.4E-22 |
| *dnsn-1* | WBGene00016074 | protein_coding_gene | -1.6 | 0.3382 | 6.9E-33 | -1.3 | 0.416 | 2.8E-17 |
| *pole-4* | WBGene00013150 | protein_coding_gene | -1.6 | 0.3381 | 1.6E-31 | -1.3 | 0.407 | 9.0E-16 |
| *vab-1* | WBGene00006868 | protein_coding_gene | -1.6 | 0.3380 | 3.0E-10 | -1.7 | 0.314 | 2.3E-48 |
| *ccnk-1* | WBGene00009650 | protein_coding_gene | -1.6 | 0.3377 | 8.2E-19 | -1.4 | 0.383 | 6.7E-27 |
| *flap-1* | WBGene00018998 | protein_coding_gene | -1.6 | 0.3375 | 6.2E-20 | -1.3 | 0.409 | 2.5E-23 |
| *kin-4* | WBGene00002192 | protein_coding_gene | -1.6 | 0.3370 | 1.5E-08 | -1.1 | 0.471 | 6.8E-23 |
| *mig-32* | WBGene00008684 | protein_coding_gene | -1.6 | 0.3364 | 2.4E-16 | -1.7 | 0.314 | 4.8E-30 |
| *bmy-1* | WBGene00017293 | protein_coding_gene | -1.6 | 0.3362 | 3.8E-24 | -1.8 | 0.293 | 2.2E-43 |
| *apd-3* | WBGene00000162 | protein_coding_gene | -1.6 | 0.3361 | 2.9E-18 | -1.1 | 0.464 | 1.9E-16 |
| *clh-5* | WBGene00000532 | protein_coding_gene | -1.6 | 0.3361 | 2.2E-24 | -1.0 | 0.489 | 2.0E-20 |
| *arl-8* | WBGene00000192 | protein_coding_gene | -1.6 | 0.3360 | 1.9E-14 | -1.2 | 0.446 | 5.7E-29 |
| *lsm-8* | WBGene00003082 | protein_coding_gene | -1.6 | 0.3359 | 2.6E-18 | -1.4 | 0.370 | 1.1E-28 |
| *D1044.6* | WBGene00017031 | protein_coding_gene | -1.6 | 0.3358 | 3.5E-12 | -1.2 | 0.444 | 1.5E-24 |
| *mek-1* | WBGene00003185 | protein_coding_gene | -1.6 | 0.3358 | 1.2E-12 | -1.2 | 0.424 | 1.1E-13 |
| *C06C3.7* | WBGene00007377 | protein_coding_gene | -1.6 | 0.3355 | 1.6E-02 | -1.9 | 0.271 | 2.7E-03 |
| *gsr-1* | WBGene00008117 | protein_coding_gene | -1.6 | 0.3353 | 4.9E-12 | -1.3 | 0.409 | 1.3E-26 |
| *C50F4.12* | WBGene00008236 | protein_coding_gene | -1.6 | 0.3351 | 5.0E-20 | -1.3 | 0.409 | 1.9E-20 |
| *C07D10.5* | WBGene00015569 | protein_coding_gene | -1.6 | 0.3350 | 8.7E-10 | -3.0 | 0.125 | 4.2E-62 |
| *gss-1* | WBGene00010941 | protein_coding_gene | -1.6 | 0.3348 | 6.4E-20 | -1.3 | 0.418 | 1.8E-25 |
| *brc-1* | WBGene00000264 | protein_coding_gene | -1.6 | 0.3348 | 4.3E-21 | -1.3 | 0.408 | 4.6E-20 |
| *R11A5.3* | WBGene00011231 | protein_coding_gene | -1.6 | 0.3347 | 1.0E-03 | -1.9 | 0.274 | 3.0E-07 |
| *H06A10.1* | WBGene00010368 | protein_coding_gene | -1.6 | 0.3346 | 3.9E-05 | -1.6 | 0.327 | 4.7E-04 |
| *lido-3* | WBGene00015588 | pseudogene | -1.6 | 0.3344 | 5.5E-03 | -2.1 | 0.230 | 6.5E-04 |
| *F31D5.6* | WBGene00017949 | protein_coding_gene | -1.6 | 0.3344 | 2.4E-02 | -3.6 | 0.082 | 1.7E-08 |
| *sel-5* | WBGene00004762 | protein_coding_gene | -1.6 | 0.3334 | 7.0E-14 | -1.3 | 0.411 | 2.1E-32 |
| *F28C6.10* | WBGene00009209 | protein_coding_gene | -1.6 | 0.3334 | 1.4E-02 | -6.1 | 0.014 | 8.1E-23 |
| *C26C6.6* | WBGene00007740 | protein_coding_gene | -1.6 | 0.3334 | 3.3E-02 | -1.1 | 0.453 | 4.4E-02 |
| *ceh-36* | WBGene00000457 | protein_coding_gene | -1.6 | 0.3329 | 2.9E-03 | -2.2 | 0.224 | 5.9E-08 |
| *C38D4.4* | WBGene00008005 | protein_coding_gene | -1.6 | 0.3328 | 7.2E-16 | -1.1 | 0.466 | 2.4E-17 |
| *W04A8.1* | WBGene00012236 | protein_coding_gene | -1.6 | 0.3328 | 2.8E-23 | -1.9 | 0.269 | 9.8E-30 |
| *lag-1* | WBGene00002245 | protein_coding_gene | -1.6 | 0.3324 | 5.8E-13 | -1.3 | 0.413 | 1.0E-30 |
| *bath-42* | WBGene00016803 | protein_coding_gene | -1.6 | 0.3323 | 7.2E-20 | -1.2 | 0.431 | 4.4E-30 |
| *rsa-2* | WBGene00012972 | protein_coding_gene | -1.6 | 0.3315 | 2.7E-26 | -1.3 | 0.402 | 1.6E-27 |
| *bre-5* | WBGene00000270 | protein_coding_gene | -1.6 | 0.3315 | 1.4E-15 | -1.2 | 0.449 | 7.3E-12 |
| *Y43E12A.3* | WBGene00012795 | protein_coding_gene | -1.6 | 0.3314 | 1.2E-15 | -1.3 | 0.420 | 2.8E-12 |
| *spo-11* | WBGene00004985 | protein_coding_gene | -1.6 | 0.3314 | 5.4E-18 | -1.7 | 0.310 | 3.8E-34 |
| *arl-5* | WBGene00000189 | protein_coding_gene | -1.6 | 0.3311 | 3.7E-17 | -1.6 | 0.334 | 3.9E-19 |
| *ncbp-1* | WBGene00018156 | protein_coding_gene | -1.6 | 0.3311 | 1.6E-21 | -1.2 | 0.437 | 3.7E-30 |
| *gfl-1* | WBGene00001585 | protein_coding_gene | -1.6 | 0.3310 | 1.3E-22 | -1.7 | 0.309 | 4.6E-33 |
| *hat-1* | WBGene00044069 | protein_coding_gene | -1.6 | 0.3305 | 1.8E-34 | -1.2 | 0.424 | 2.1E-28 |
| *algn-10* | WBGene00007043 | protein_coding_gene | -1.6 | 0.3304 | 4.5E-22 | -1.3 | 0.409 | 3.1E-21 |
| *F27D4.7* | WBGene00009191 | protein_coding_gene | -1.6 | 0.3303 | 8.7E-20 | -1.2 | 0.443 | 6.2E-17 |
| *rpn-5* | WBGene00004461 | protein_coding_gene | -1.6 | 0.3303 | 1.5E-20 | -1.2 | 0.447 | 9.5E-24 |
| *F30A10.13* | WBGene00044680 | protein_coding_gene | -1.6 | 0.3303 | 4.9E-04 | -3.5 | 0.091 | 1.5E-10 |
| *prp-31* | WBGene00022458 | protein_coding_gene | -1.6 | 0.3302 | 8.9E-37 | -1.4 | 0.366 | 1.8E-36 |
| *Y62E10A.13* | WBGene00013379 | protein_coding_gene | -1.6 | 0.3302 | 1.8E-09 | -1.2 | 0.449 | 5.6E-17 |
| *B0001.2* | WBGene00007087 | protein_coding_gene | -1.6 | 0.3300 | 2.1E-30 | -1.6 | 0.338 | 3.0E-48 |
| *thoc-5* | WBGene00021311 | protein_coding_gene | -1.6 | 0.3300 | 2.6E-36 | -1.3 | 0.411 | 3.8E-23 |
| *ooc-3* | WBGene00003868 | protein_coding_gene | -1.6 | 0.3296 | 3.5E-36 | -1.2 | 0.450 | 5.1E-18 |
| *aph-1* | WBGene00000147 | protein_coding_gene | -1.6 | 0.3295 | 1.6E-24 | -1.4 | 0.376 | 5.3E-24 |
| *cdt-1* | WBGene00000413 | protein_coding_gene | -1.6 | 0.3294 | 4.9E-14 | -1.5 | 0.357 | 8.6E-28 |

| | | | | | | | | |
|---|---|---|---|---|---|---|---|---|
| *rpn-12* | WBGene00004468 | protein_coding_gene | -1.6 | 0.3293 | 7.0E-16 | -1.2 | 0.423 | 5.7E-23 |
| *F01F1.11* | WBGene00017165 | protein_coding_gene | -1.6 | 0.3289 | 1.3E-26 | -1.6 | 0.326 | 4.7E-40 |
| *ntl-11* | WBGene00013530 | protein_coding_gene | -1.6 | 0.3285 | 9.0E-17 | -1.2 | 0.448 | 1.4E-23 |
| *ttc-17* | WBGene00020553 | protein_coding_gene | -1.6 | 0.3283 | 8.9E-19 | -1.1 | 0.458 | 7.0E-26 |
| *gst-7* | WBGene00001755 | protein_coding_gene | -1.6 | 0.3283 | 1.8E-07 | -2.2 | 0.219 | 2.5E-50 |
| *zwl-1* | WBGene00021460 | protein_coding_gene | -1.6 | 0.3282 | 1.2E-20 | -1.1 | 0.463 | 7.5E-18 |
| *F09G8.5* | WBGene00017320 | protein_coding_gene | -1.6 | 0.3282 | 4.7E-03 | -1.3 | 0.393 | 8.8E-05 |
| *snr-1* | WBGene00004914 | protein_coding_gene | -1.6 | 0.3281 | 7.2E-35 | -1.1 | 0.476 | 1.1E-21 |
| *npp-5* | WBGene00003791 | protein_coding_gene | -1.6 | 0.3279 | 3.5E-25 | -1.1 | 0.477 | 3.0E-21 |
| *C01G12.3* | WBGene00007244 | protein_coding_gene | -1.6 | 0.3279 | 7.8E-08 | -1.0 | 0.495 | 2.9E-04 |
| *cec-8* | WBGene00021913 | protein_coding_gene | -1.6 | 0.3278 | 1.1E-06 | -3.2 | 0.109 | 5.3E-47 |
| *slx-1* | WBGene00018909 | protein_coding_gene | -1.6 | 0.3277 | 2.0E-32 | -1.4 | 0.374 | 1.4E-30 |
| *rnh-1.2* | WBGene00014163 | protein_coding_gene | -1.6 | 0.3275 | 1.8E-14 | -1.3 | 0.400 | 8.3E-18 |
| *F35H10.5* | WBGene00018070 | protein_coding_gene | -1.6 | 0.3269 | 4.4E-11 | -1.4 | 0.373 | 1.4E-13 |
| *phf-30* | WBGene00020716 | protein_coding_gene | -1.6 | 0.3268 | 3.2E-19 | -1.5 | 0.346 | 1.8E-27 |
| *pas-4* | WBGene00003925 | protein_coding_gene | -1.6 | 0.3267 | 6.2E-21 | -1.1 | 0.460 | 4.5E-21 |
| ***C38D9.2*** | **WBGene00008010** | **protein_coding_gene** | **-1.6** | **0.3266** | **1.4E-22** | **3.4** | **10.795** | **1.6E-31** |
| *nspd-4* | WBGene00020715 | protein_coding_gene | -1.6 | 0.3261 | 2.4E-03 | -2.2 | 0.212 | 2.1E-06 |
| *mob-1* | WBGene00009551 | protein_coding_gene | -1.6 | 0.3256 | 1.0E-20 | -1.7 | 0.303 | 2.6E-32 |
| *nop-1* | WBGene00017774 | protein_coding_gene | -1.6 | 0.3255 | 1.2E-21 | -1.6 | 0.334 | 1.4E-50 |
| *rpa-2* | WBGene00019767 | protein_coding_gene | -1.6 | 0.3252 | 4.4E-12 | -1.2 | 0.426 | 2.0E-25 |
| *tbcd-1* | WBGene00008887 | protein_coding_gene | -1.6 | 0.3250 | 1.0E-31 | -1.5 | 0.361 | 2.6E-41 |
| *tag-115* | WBGene00006466 | protein_coding_gene | -1.6 | 0.3250 | 3.1E-17 | -1.6 | 0.328 | 1.2E-32 |
| *ubc-18* | WBGene00006713 | protein_coding_gene | -1.6 | 0.3245 | 7.2E-17 | -1.6 | 0.341 | 1.3E-42 |
| *nstp-10* | WBGene00008237 | protein_coding_gene | -1.6 | 0.3244 | 5.7E-16 | -1.0 | 0.488 | 1.8E-22 |
| *F57A10.4* | WBGene00010185 | protein_coding_gene | -1.6 | 0.3244 | 4.9E-13 | -2.2 | 0.223 | 1.2E-29 |
| *czw-1* | WBGene00017643 | protein_coding_gene | -1.6 | 0.3243 | 6.8E-21 | -1.4 | 0.367 | 2.4E-32 |
| *B0464.6* | WBGene00007186 | protein_coding_gene | -1.6 | 0.3240 | 1.2E-16 | -1.5 | 0.356 | 7.1E-45 |
| *eaf-1* | WBGene00017011 | protein_coding_gene | -1.6 | 0.3239 | 1.9E-16 | -1.5 | 0.343 | 1.5E-43 |
| *Y4C6B.1* | WBGene00021155 | protein_coding_gene | -1.6 | 0.3237 | 2.5E-19 | -1.5 | 0.344 | 3.2E-36 |
| *pst-2* | WBGene00018827 | protein_coding_gene | -1.6 | 0.3237 | 1.1E-12 | -1.1 | 0.478 | 5.7E-12 |
| *wdr-5.1* | WBGene00006474 | protein_coding_gene | -1.6 | 0.3233 | 1.8E-27 | -1.3 | 0.417 | 1.9E-24 |
| *cnep-1* | WBGene00018474 | protein_coding_gene | -1.6 | 0.3232 | 1.2E-17 | -1.1 | 0.464 | 3.3E-25 |
| *snr-5* | WBGene00004918 | protein_coding_gene | -1.6 | 0.3232 | 2.0E-12 | -1.3 | 0.398 | 7.4E-31 |
| *T24H7.3* | WBGene00020782 | protein_coding_gene | -1.6 | 0.3231 | 2.2E-08 | -1.6 | 0.336 | 4.5E-33 |
| *rsp-4* | WBGene00004701 | protein_coding_gene | -1.6 | 0.3230 | 3.0E-34 | -1.5 | 0.356 | 2.2E-44 |
| *tbc-2* | WBGene00022880 | protein_coding_gene | -1.6 | 0.3229 | 1.9E-13 | -1.3 | 0.416 | 2.1E-27 |
| *mop-25.3* | WBGene00020858 | protein_coding_gene | -1.6 | 0.3228 | 1.2E-28 | -1.6 | 0.338 | 4.2E-24 |
| *rpt-3* | WBGene00004503 | protein_coding_gene | -1.6 | 0.3227 | 1.4E-13 | -1.2 | 0.425 | 5.6E-28 |
| *pbs-3* | WBGene00003949 | protein_coding_gene | -1.6 | 0.3227 | 1.4E-12 | -1.3 | 0.411 | 6.9E-30 |
| *D2024.5* | WBGene00017053 | protein_coding_gene | -1.6 | 0.3227 | 5.2E-21 | -1.4 | 0.372 | 3.4E-26 |
| *asg-1* | WBGene00000209 | protein_coding_gene | -1.6 | 0.3225 | 3.8E-10 | -1.1 | 0.462 | 8.1E-16 |
| *lst-1* | WBGene00003083 | protein_coding_gene | -1.6 | 0.3223 | 1.4E-12 | -1.3 | 0.402 | 2.2E-26 |
| *taf-8* | WBGene00006390 | protein_coding_gene | -1.6 | 0.3219 | 3.5E-34 | -1.5 | 0.364 | 4.4E-35 |
| *cogc-2* | WBGene00000585 | protein_coding_gene | -1.6 | 0.3217 | 1.0E-25 | -1.0 | 0.499 | 1.2E-15 |
| *tre-1* | WBGene00006607 | protein_coding_gene | -1.6 | 0.3215 | 1.7E-35 | -1.2 | 0.422 | 8.4E-39 |
| *sao-1* | WBGene00011195 | protein_coding_gene | -1.6 | 0.3213 | 4.2E-09 | -1.0 | 0.484 | 5.8E-20 |
| *Y111B2A.2* | WBGene00013728 | protein_coding_gene | -1.6 | 0.3213 | 1.8E-31 | -1.4 | 0.387 | 6.4E-30 |
| *rde-4* | WBGene00004326 | protein_coding_gene | -1.6 | 0.3213 | 2.2E-18 | -1.6 | 0.336 | 5.3E-38 |
| *F57C9.4* | WBGene00019011 | protein_coding_gene | -1.6 | 0.3212 | 6.9E-27 | -1.5 | 0.352 | 5.3E-36 |
| *msp-40* | WBGene00003435 | protein_coding_gene | -1.6 | 0.3208 | 2.3E-03 | -1.1 | 0.479 | 4.1E-09 |
| *fbxb-80* | WBGene00017117 | pseudogene | -1.6 | 0.3204 | 7.0E-03 | -2.4 | 0.184 | 4.4E-06 |
| *cdk-5* | WBGene00000407 | protein_coding_gene | -1.6 | 0.3197 | 1.1E-18 | -1.4 | 0.378 | 1.5E-20 |
| *C03B8.2* | WBGene00015385 | protein_coding_gene | -1.6 | 0.3197 | 3.4E-04 | -1.7 | 0.315 | 8.1E-04 |
| *acl-8* | WBGene00020264 | protein_coding_gene | -1.6 | 0.3195 | 1.3E-29 | -1.1 | 0.476 | 1.4E-15 |
| *cyp-14A2* | WBGene00010706 | protein_coding_gene | -1.6 | 0.3192 | 1.6E-04 | -1.7 | 0.315 | 1.3E-04 |
| *K09E4.1* | WBGene00010719 | protein_coding_gene | -1.6 | 0.3192 | 6.4E-03 | -1.1 | 0.453 | 9.1E-07 |
| *metr-1* | WBGene00010988 | protein_coding_gene | -1.6 | 0.3187 | 1.5E-25 | -2.0 | 0.247 | 3.7E-43 |
| *pas-5* | WBGene00003926 | protein_coding_gene | -1.6 | 0.3187 | 1.1E-21 | -1.3 | 0.405 | 1.4E-32 |
| *ints-6* | WBGene00000994 | protein_coding_gene | -1.7 | 0.3184 | 5.7E-15 | -1.1 | 0.475 | 2.7E-16 |
| *rpb-3* | WBGene00007971 | protein_coding_gene | -1.7 | 0.3178 | 1.3E-21 | -1.3 | 0.407 | 3.6E-33 |
| *msp-3* | WBGene00003424 | protein_coding_gene | -1.7 | 0.3172 | 4.0E-04 | -2.0 | 0.251 | 6.0E-21 |
| *F30A10.14* | WBGene00045473 | protein_coding_gene | -1.7 | 0.3170 | 1.8E-07 | -2.9 | 0.132 | 1.8E-27 |
| *cyn-10* | WBGene00000886 | protein_coding_gene | -1.7 | 0.3169 | 8.3E-27 | -1.4 | 0.372 | 4.0E-22 |
| *sms-2* | WBGene00004893 | protein_coding_gene | -1.7 | 0.3167 | 8.5E-11 | -1.5 | 0.354 | 3.1E-13 |
| *sup-35* | WBGene00012976 | protein_coding_gene | -1.7 | 0.3166 | 1.7E-12 | -1.2 | 0.430 | 1.3E-09 |
| *C49C3.6* | WBGene00008196 | protein_coding_gene | -1.7 | 0.3164 | 7.8E-14 | -1.7 | 0.300 | 7.2E-37 |
| *K07A1.17* | WBGene00044987 | protein_coding_gene | -1.7 | 0.3163 | 1.1E-13 | -1.5 | 0.357 | 3.8E-38 |
| *T20F7.1* | WBGene00020630 | protein_coding_gene | -1.7 | 0.3162 | 5.5E-26 | -1.6 | 0.337 | 5.4E-25 |
| *bath-29* | WBGene00017452 | protein_coding_gene | -1.7 | 0.3161 | 2.1E-14 | -1.4 | 0.389 | 1.7E-08 |
| *duo-3* | WBGene00001112 | protein_coding_gene | -1.7 | 0.3158 | 2.3E-23 | -1.4 | 0.381 | 2.2E-35 |
| *snr-3* | WBGene00004916 | protein_coding_gene | -1.7 | 0.3158 | 6.3E-35 | -1.2 | 0.432 | 3.6E-25 |
| *D1081.6* | WBGene00008384 | protein_coding_gene | -1.7 | 0.3158 | 1.6E-07 | -1.3 | 0.401 | 1.5E-10 |
| *vacl-14* | WBGene00010565 | protein_coding_gene | -1.7 | 0.3156 | 2.4E-25 | -1.3 | 0.399 | 7.5E-25 |
| *C47E12.2* | WBGene00008147 | protein_coding_gene | -1.7 | 0.3154 | 1.1E-26 | -1.3 | 0.411 | 1.0E-28 |
| *swsn-2.2* | WBGene00015971 | protein_coding_gene | -1.7 | 0.3154 | 7.3E-18 | -1.2 | 0.425 | 2.3E-18 |
| *cnk-1* | WBGene00000564 | protein_coding_gene | -1.7 | 0.3151 | 2.9E-19 | -1.2 | 0.444 | 7.5E-26 |
| *F58A4.1* | WBGene00010226 | protein_coding_gene | -1.7 | 0.3150 | 3.1E-02 | -5.9 | 0.017 | 3.8E-21 |
| *pqbp-1.2* | WBGene00020295 | pseudogene | -1.7 | 0.3149 | 3.9E-10 | -1.1 | 0.461 | 3.4E-06 |
| *tag-266* | WBGene00044319 | protein_coding_gene | -1.7 | 0.3148 | 2.9E-13 | -1.1 | 0.460 | 7.4E-11 |

| | | | | | | | | |
|---|---|---|---|---|---|---|---|---|
| *him-10* | WBGene00001869 | protein_coding_gene | -1.7 | 0.3147 | 1.3E-17 | -1.5 | 0.354 | 2.1E-37 |
| *egl-6* | WBGene00001175 | protein_coding_gene | -1.7 | 0.3146 | 2.9E-13 | -1.1 | 0.463 | 1.9E-05 |
| *F26G1.1* | WBGene00017837 | protein_coding_gene | -1.7 | 0.3146 | 3.2E-21 | -1.4 | 0.387 | 7.4E-27 |
| *tres-1* | WBGene00022750 | protein_coding_gene | -1.7 | 0.3144 | 2.2E-18 | -1.4 | 0.389 | 2.7E-40 |
| *npp-1* | WBGene00003787 | protein_coding_gene | -1.7 | 0.3144 | 4.6E-23 | -1.5 | 0.365 | 1.2E-38 |
| ***aat-4*** | **WBGene00000005** | **protein_coding_gene** | **-1.7** | **0.3142** | **6.7E-06** | **1.5** | **2.782** | **2.1E-05** |
| *ZK1248.15* | WBGene00022883 | protein_coding_gene | -1.7 | 0.3140 | 1.6E-21 | -1.3 | 0.405 | 1.9E-22 |
| *cchl-1* | WBGene00011527 | protein_coding_gene | -1.7 | 0.3138 | 6.2E-31 | -1.1 | 0.469 | 2.1E-23 |
| *Y105E8A.14* | WBGene00013674 | protein_coding_gene | -1.7 | 0.3137 | 2.5E-16 | -1.2 | 0.432 | 1.6E-25 |
| *zyg-11* | WBGene00006996 | protein_coding_gene | -1.7 | 0.3136 | 2.0E-33 | -1.1 | 0.478 | 8.1E-19 |
| *F55G1.6* | WBGene00018902 | protein_coding_gene | -1.7 | 0.3136 | 8.1E-15 | -1.3 | 0.420 | 4.4E-22 |
| *swsn-9* | WBGene00007256 | protein_coding_gene | -1.7 | 0.3136 | 1.7E-25 | -1.5 | 0.357 | 6.4E-34 |
| *mans-1* | WBGene00008411 | protein_coding_gene | -1.7 | 0.3135 | 1.8E-31 | -1.2 | 0.425 | 1.3E-36 |
| *T19B4.5* | WBGene00020558 | protein_coding_gene | -1.7 | 0.3134 | 1.0E-17 | -1.2 | 0.437 | 1.5E-25 |
| *F23F12.12* | WBGene00017756 | protein_coding_gene | -1.7 | 0.3133 | 2.8E-03 | -1.1 | 0.478 | 1.8E-05 |
| *cids-1* | WBGene00015347 | protein_coding_gene | -1.7 | 0.3129 | 4.4E-18 | -1.4 | 0.380 | 1.6E-26 |
| *yop-1* | WBGene00022127 | protein_coding_gene | -1.7 | 0.3128 | 9.3E-23 | -1.3 | 0.416 | 4.6E-19 |
| *pigt-1* | WBGene00008918 | protein_coding_gene | -1.7 | 0.3128 | 1.7E-28 | -1.5 | 0.361 | 1.1E-38 |
| *aipl-1* | WBGene00010685 | protein_coding_gene | -1.7 | 0.3127 | 9.3E-28 | -1.3 | 0.398 | 1.5E-37 |
| *vps-54* | WBGene00006934 | protein_coding_gene | -1.7 | 0.3125 | 1.3E-21 | -1.1 | 0.469 | 1.2E-25 |
| *samp-1* | WBGene00011994 | protein_coding_gene | -1.7 | 0.3125 | 5.7E-14 | -1.4 | 0.373 | 1.6E-32 |
| *oef-1* | WBGene00018636 | protein_coding_gene | -1.7 | 0.3123 | 7.1E-27 | -1.4 | 0.368 | 3.5E-29 |
| *his-32* | WBGene00001906 | protein_coding_gene | -1.7 | 0.3120 | 7.9E-04 | -3.0 | 0.128 | 1.5E-07 |
| *K08E5.1* | WBGene00014801 | protein_coding_gene | -1.7 | 0.3118 | 2.6E-05 | -1.1 | 0.455 | 4.5E-07 |
| *slc-17.2* | WBGene00008000 | protein_coding_gene | -1.7 | 0.3111 | 2.1E-22 | -1.3 | 0.397 | 3.7E-32 |
| *R09E10.2* | WBGene00011172 | protein_coding_gene | -1.7 | 0.3107 | 1.4E-03 | -2.7 | 0.157 | 1.8E-09 |
| *M142.5* | WBGene00010922 | protein_coding_gene | -1.7 | 0.3106 | 2.1E-30 | -1.1 | 0.469 | 5.4E-14 |
| *pri-2* | WBGene00004181 | protein_coding_gene | -1.7 | 0.3106 | 7.0E-26 | -1.2 | 0.430 | 3.5E-18 |
| *acdh-1* | WBGene00016943 | protein_coding_gene | -1.7 | 0.3105 | 7.5E-15 | -2.3 | 0.210 | 1.3E-13 |
| *asp-13* | WBGene00017881 | protein_coding_gene | -1.7 | 0.3103 | 1.3E-20 | -2.6 | 0.170 | 3.7E-135 |
| *mrp-7* | WBGene00003413 | protein_coding_gene | -1.7 | 0.3102 | 3.8E-15 | -1.3 | 0.403 | 8.0E-21 |
| *gst-1* | WBGene00001749 | protein_coding_gene | -1.7 | 0.3102 | 1.5E-06 | -1.4 | 0.391 | 3.0E-27 |
| *C50C3.1* | WBGene00016799 | protein_coding_gene | -1.7 | 0.3099 | 9.2E-20 | -1.2 | 0.441 | 1.1E-22 |
| *vps-35* | WBGene00006933 | protein_coding_gene | -1.7 | 0.3097 | 4.5E-23 | -1.2 | 0.443 | 7.9E-25 |
| *lst-4* | WBGene00003086 | protein_coding_gene | -1.7 | 0.3096 | 1.8E-23 | -1.1 | 0.457 | 3.7E-23 |
| *ZK632.12* | WBGene00014019 | protein_coding_gene | -1.7 | 0.3094 | 2.2E-21 | -1.3 | 0.420 | 2.2E-23 |
| *E01A2.2* | WBGene00017085 | protein_coding_gene | -1.7 | 0.3094 | 1.2E-29 | -1.4 | 0.376 | 1.2E-48 |
| *msp-63* | WBGene00003456 | protein_coding_gene | -1.7 | 0.3092 | 1.8E-02 | -1.4 | 0.374 | 2.6E-02 |
| *txl-1* | WBGene00021826 | protein_coding_gene | -1.7 | 0.3090 | 6.5E-19 | -1.2 | 0.439 | 2.3E-22 |
| *kxd-1* | WBGene00015742 | protein_coding_gene | -1.7 | 0.3089 | 3.4E-10 | -2.1 | 0.238 | 1.4E-22 |
| *Y11D7A.7* | WBGene00012431 | protein_coding_gene | -1.7 | 0.3088 | 3.8E-19 | -2.0 | 0.247 | 1.4E-46 |
| *exos-3* | WBGene00010325 | protein_coding_gene | -1.7 | 0.3086 | 3.0E-19 | -1.3 | 0.408 | 1.6E-23 |
| *blos-8* | WBGene00010981 | protein_coding_gene | -1.7 | 0.3078 | 9.4E-08 | -1.7 | 0.304 | 1.2E-07 |
| *pid-3* | WBGene00021270 | protein_coding_gene | -1.7 | 0.3077 | 4.0E-30 | -1.2 | 0.435 | 4.9E-25 |
| *B0205.9* | WBGene00015023 | protein_coding_gene | -1.7 | 0.3077 | 1.1E-24 | -1.5 | 0.366 | 3.1E-32 |
| *pak-2* | WBGene00006443 | protein_coding_gene | -1.7 | 0.3076 | 1.9E-13 | -1.3 | 0.417 | 1.0E-27 |
| *Y39G8B.2* | WBGene00012723 | protein_coding_gene | -1.7 | 0.3076 | 1.0E-08 | -1.7 | 0.308 | 3.5E-12 |
| *swan-1* | WBGene00009977 | protein_coding_gene | -1.7 | 0.3068 | 9.3E-13 | -1.7 | 0.314 | 8.7E-34 |
| *div-1* | WBGene00001002 | protein_coding_gene | -1.7 | 0.3064 | 6.0E-23 | -1.5 | 0.360 | 6.3E-30 |
| *aly-2* | WBGene00000121 | protein_coding_gene | -1.7 | 0.3063 | 6.8E-32 | -1.6 | 0.326 | 2.9E-48 |
| *K11H3.3* | WBGene00010780 | protein_coding_gene | -1.7 | 0.3058 | 3.3E-25 | -1.5 | 0.349 | 4.8E-49 |
| *acdh-12* | WBGene00017125 | protein_coding_gene | -1.7 | 0.3057 | 4.1E-31 | -1.1 | 0.462 | 1.7E-26 |
| *F12F6.1* | WBGene00008720 | protein_coding_gene | -1.7 | 0.3057 | 2.3E-27 | -1.3 | 0.398 | 1.0E-35 |
| *nfyc-1* | WBGene00017742 | protein_coding_gene | -1.7 | 0.3054 | 4.5E-21 | -1.5 | 0.359 | 9.4E-27 |
| *mbd-2* | WBGene00023409 | protein_coding_gene | -1.7 | 0.3049 | 7.0E-18 | -1.6 | 0.320 | 4.6E-28 |
| *B0024.13* | WBGene00007102 | protein_coding_gene | -1.7 | 0.3046 | 3.5E-12 | -1.4 | 0.387 | 1.9E-25 |
| *msp-51* | WBGene00003444 | protein_coding_gene | -1.7 | 0.3043 | 1.4E-04 | -1.6 | 0.327 | 9.1E-16 |
| *Y39G10AR.32* | WBGene00219275 | protein_coding_gene | -1.7 | 0.3042 | 2.6E-21 | -2.0 | 0.253 | 9.4E-36 |
| *ztf-23* | WBGene00021846 | protein_coding_gene | -1.7 | 0.3039 | 3.0E-32 | -1.6 | 0.334 | 5.9E-25 |
| *usip-1* | WBGene00014124 | protein_coding_gene | -1.7 | 0.3033 | 1.7E-22 | -1.5 | 0.361 | 1.0E-46 |
| *Y48B6A.13* | WBGene00012984 | protein_coding_gene | -1.7 | 0.3032 | 3.1E-24 | -1.3 | 0.419 | 6.6E-15 |
| *K11H3.8* | WBGene00185002 | protein_coding_gene | -1.7 | 0.3032 | 6.9E-15 | -2.0 | 0.245 | 1.2E-48 |
| *zhp-2* | WBGene00008387 | protein_coding_gene | -1.7 | 0.3027 | 3.0E-21 | -1.1 | 0.472 | 1.2E-07 |
| *piga-1* | WBGene00008431 | protein_coding_gene | -1.7 | 0.3027 | 1.3E-16 | -1.4 | 0.368 | 8.4E-23 |
| *pes-5* | WBGene00003979 | protein_coding_gene | -1.7 | 0.3026 | 1.7E-13 | -1.9 | 0.263 | 6.9E-21 |
| ***F42A8.1*** | **WBGene00009626** | **protein_coding_gene** | **-1.7** | **0.3026** | **2.8E-02** | **2.5** | **5.717** | **5.9E-41** |
| *pqn-20* | WBGene00004110 | protein_coding_gene | -1.7 | 0.3024 | 9.9E-06 | -1.1 | 0.454 | 3.5E-21 |
| *ufd-3* | WBGene00007333 | protein_coding_gene | -1.7 | 0.3023 | 1.1E-27 | -1.1 | 0.459 | 1.5E-27 |
| *R07E5.11* | WBGene00011117 | protein_coding_gene | -1.7 | 0.3021 | 6.6E-10 | -1.7 | 0.316 | 1.1E-20 |
| *C15C6.4* | WBGene00007603 | protein_coding_gene | -1.7 | 0.3019 | 3.8E-13 | -1.4 | 0.379 | 2.0E-19 |
| *F52B5.7* | WBGene00009923 | protein_coding_gene | -1.7 | 0.3016 | 4.6E-15 | -2.0 | 0.245 | 3.6E-20 |
| *hsr-9* | WBGene00002027 | protein_coding_gene | -1.7 | 0.3016 | 6.6E-14 | -1.1 | 0.453 | 2.0E-34 |
| *hcp-3* | WBGene00001831 | protein_coding_gene | -1.7 | 0.3012 | 2.4E-24 | -1.2 | 0.422 | 1.6E-26 |
| *F10E9.3* | WBGene00017355 | protein_coding_gene | -1.7 | 0.3012 | 1.6E-25 | -1.4 | 0.369 | 1.5E-36 |
| *Y48G1C.12* | WBGene00044345 | protein_coding_gene | -1.7 | 0.3012 | 5.9E-26 | -1.1 | 0.480 | 4.9E-15 |
| *crn-1* | WBGene00000794 | protein_coding_gene | -1.7 | 0.3012 | 2.0E-20 | -1.7 | 0.317 | 1.6E-43 |
| *orc-2* | WBGene00003882 | protein_coding_gene | -1.7 | 0.3008 | 7.1E-19 | -1.6 | 0.334 | 2.1E-35 |
| *B0511.19* | WBGene00255714 | protein_coding_gene | -1.7 | 0.3008 | 5.1E-17 | -1.8 | 0.286 | 1.6E-39 |
| *C27F2.10* | WBGene00016171 | protein_coding_gene | -1.7 | 0.3008 | 9.0E-19 | -1.6 | 0.328 | 1.9E-39 |
| *pqn-82* | WBGene00004163 | protein_coding_gene | -1.7 | 0.3006 | 8.9E-12 | -1.4 | 0.390 | 4.3E-12 |

| | | | | | | | | |
|---|---|---|---|---|---|---|---|---|
| *R02D3.7* | WBGene00019824 | protein_coding_gene | -1.7 | 0.3005 | 1.6E-28 | -1.6 | 0.328 | 6.6E-38 |
| *C18E3.9* | WBGene00015976 | protein_coding_gene | -1.7 | 0.3004 | 3.9E-15 | -1.3 | 0.395 | 2.9E-22 |
| *mtm-9* | WBGene00003479 | protein_coding_gene | -1.7 | 0.3003 | 4.9E-28 | -1.4 | 0.382 | 2.2E-24 |
| *unc-40* | WBGene00006776 | protein_coding_gene | -1.7 | 0.3003 | 4.5E-11 | -1.1 | 0.474 | 4.3E-22 |
| *uri-1* | WBGene00016944 | protein_coding_gene | -1.7 | 0.2996 | 1.6E-25 | -1.6 | 0.319 | 2.3E-46 |
| *athp-1* | WBGene00008081 | protein_coding_gene | -1.7 | 0.2996 | 9.3E-24 | -1.2 | 0.439 | 4.3E-19 |
| *rsp-5* | WBGene00004702 | protein_coding_gene | -1.7 | 0.2994 | 7.0E-18 | -1.2 | 0.441 | 1.7E-17 |
| *efl-1* | WBGene00001161 | protein_coding_gene | -1.7 | 0.2992 | 1.0E-28 | -1.6 | 0.333 | 2.5E-41 |
| *wrm-1* | WBGene00006943 | protein_coding_gene | -1.7 | 0.2991 | 4.4E-18 | -1.4 | 0.376 | 9.5E-30 |
| *CD4.10* | WBGene00022970 | ncRNA_gene | -1.7 | 0.2990 | 2.7E-08 | -1.4 | 0.375 | 2.4E-08 |
| *nlp-39* | WBGene00008295 | protein_coding_gene | -1.7 | 0.2987 | 4.2E-02 | -6.4 | 0.012 | 2.7E-11 |
| *F39H2.3* | WBGene00009563 | protein_coding_gene | -1.7 | 0.2987 | 1.6E-27 | -1.2 | 0.439 | 2.2E-21 |
| *sand-1* | WBGene00018321 | protein_coding_gene | -1.7 | 0.2986 | 5.8E-26 | -1.8 | 0.286 | 5.9E-56 |
| *let-711* | WBGene00002845 | protein_coding_gene | -1.7 | 0.2986 | 7.3E-11 | -1.1 | 0.458 | 4.1E-26 |
| *rpt-4* | WBGene00004504 | protein_coding_gene | -1.7 | 0.2985 | 2.9E-15 | -1.3 | 0.402 | 3.1E-28 |
| *sumv-1* | WBGene00016409 | protein_coding_gene | -1.7 | 0.2984 | 2.2E-13 | -1.1 | 0.458 | 1.0E-20 |
| *Y43F4B.10* | WBGene00012807 | protein_coding_gene | -1.7 | 0.2983 | 1.8E-19 | -1.8 | 0.285 | 1.4E-37 |
| *epg-2* | WBGene00021468 | protein_coding_gene | -1.7 | 0.2983 | 4.0E-07 | -2.9 | 0.133 | 9.6E-101 |
| *tdpo-1* | WBGene00018678 | protein_coding_gene | -1.7 | 0.2982 | 1.7E-33 | -1.1 | 0.481 | 4.4E-11 |
| *mzt-1* | WBGene00044431 | protein_coding_gene | -1.7 | 0.2981 | 4.6E-14 | -1.8 | 0.297 | 9.1E-28 |
| *C29H12.2* | WBGene00016235 | protein_coding_gene | -1.7 | 0.2980 | 5.0E-28 | -1.2 | 0.439 | 7.3E-25 |
| *tpp-2* | WBGene00017686 | protein_coding_gene | -1.7 | 0.2980 | 1.9E-30 | -1.4 | 0.388 | 7.5E-37 |
| *baz-2* | WBGene00001470 | protein_coding_gene | -1.7 | 0.2979 | 2.7E-17 | -2.1 | 0.233 | 3.9E-57 |
| *ttr-53* | WBGene00011773 | protein_coding_gene | -1.7 | 0.2978 | 2.6E-13 | -1.4 | 0.389 | 4.7E-28 |
| *mom-4* | WBGene00003396 | protein_coding_gene | -1.7 | 0.2977 | 6.8E-31 | -1.4 | 0.378 | 1.4E-24 |
| *nspd-10* | WBGene00020860 | protein_coding_gene | -1.7 | 0.2975 | 2.9E-03 | -1.7 | 0.303 | 4.1E-10 |
| *rsp-6* | WBGene00004703 | protein_coding_gene | -1.7 | 0.2974 | 2.7E-27 | -1.3 | 0.405 | 1.7E-30 |
| *spr-2* | WBGene00005007 | protein_coding_gene | -1.7 | 0.2974 | 2.3E-26 | -1.5 | 0.364 | 3.7E-49 |
| *tag-63* | WBGene00006440 | protein_coding_gene | -1.8 | 0.2965 | 1.6E-28 | -1.3 | 0.406 | 6.3E-23 |
| *nprl-2* | WBGene00018635 | protein_coding_gene | -1.8 | 0.2960 | 2.6E-25 | -1.5 | 0.349 | 2.3E-28 |
| *T24H10.4* | WBGene00012002 | protein_coding_gene | -1.8 | 0.2960 | 2.3E-22 | -1.3 | 0.401 | 2.9E-19 |
| *swsn-6* | WBGene00022774 | protein_coding_gene | -1.8 | 0.2960 | 2.3E-15 | -1.6 | 0.329 | 1.9E-39 |
| *icp-1* | WBGene00002047 | protein_coding_gene | -1.8 | 0.2959 | 4.0E-27 | -1.2 | 0.427 | 4.6E-21 |
| *R10E4.11* | WBGene00011206 | protein_coding_gene | -1.8 | 0.2958 | 4.4E-18 | -1.4 | 0.383 | 9.9E-31 |
| *F09G2.8* | WBGene00017316 | protein_coding_gene | -1.8 | 0.2956 | 1.6E-23 | -1.8 | 0.296 | 1.0E-44 |
| *C47E12.3* | WBGene00008148 | protein_coding_gene | -1.8 | 0.2955 | 6.5E-23 | -1.1 | 0.458 | 4.1E-24 |
| *mtm-6* | WBGene00003478 | protein_coding_gene | -1.8 | 0.2955 | 6.3E-55 | -1.4 | 0.374 | 1.2E-31 |
| *ssp-33* | WBGene00011134 | protein_coding_gene | -1.8 | 0.2955 | 2.3E-03 | -1.5 | 0.359 | 9.2E-06 |
| *abf-1* | WBGene00000012 | protein_coding_gene | -1.8 | 0.2952 | 4.2E-02 | -4.8 | 0.036 | 2.2E-11 |
| *C13B4.1* | WBGene00007545 | protein_coding_gene | -1.8 | 0.2951 | 1.3E-24 | -1.1 | 0.456 | 8.6E-18 |
| *sax-7* | WBGene00004732 | protein_coding_gene | -1.8 | 0.2947 | 2.6E-13 | -1.3 | 0.402 | 1.4E-52 |
| *R151.2* | WBGene00020107 | protein_coding_gene | -1.8 | 0.2944 | 4.4E-20 | -1.4 | 0.385 | 3.9E-51 |
| *F41E6.15* | WBGene00044554 | protein_coding_gene | -1.8 | 0.2942 | 1.2E-13 | -1.5 | 0.345 | 4.6E-09 |
| *M110.3* | WBGene00010913 | protein_coding_gene | -1.8 | 0.2941 | 4.8E-13 | -1.4 | 0.382 | 5.2E-14 |
| *mmaa-1* | WBGene00020169 | protein_coding_gene | -1.8 | 0.2940 | 5.4E-31 | -1.9 | 0.260 | 8.6E-46 |
| *txdc-9* | WBGene00006515 | protein_coding_gene | -1.8 | 0.2940 | 7.3E-19 | -1.5 | 0.346 | 1.2E-37 |
| *pgap-2* | WBGene00007045 | protein_coding_gene | -1.8 | 0.2938 | 1.8E-18 | -1.3 | 0.417 | 5.4E-21 |
| *ucp-4* | WBGene00006729 | protein_coding_gene | -1.8 | 0.2938 | 1.1E-19 | -1.3 | 0.404 | 2.8E-14 |
| *F49C12.9* | WBGene00009878 | protein_coding_gene | -1.8 | 0.2935 | 1.2E-25 | -1.3 | 0.396 | 6.4E-26 |
| *F53H1.4* | WBGene00018778 | protein_coding_gene | -1.8 | 0.2934 | 2.7E-15 | -1.5 | 0.349 | 1.1E-43 |
| *lex-1* | WBGene00008682 | protein_coding_gene | -1.8 | 0.2934 | 8.5E-21 | -1.2 | 0.428 | 4.1E-28 |
| *C01B12.8* | WBGene00015291 | protein_coding_gene | -1.8 | 0.2932 | 1.5E-21 | -2.0 | 0.243 | 1.1E-32 |
| *hrde-2* | WBGene00011324 | protein_coding_gene | -1.8 | 0.2929 | 1.9E-27 | -1.1 | 0.471 | 5.7E-15 |
| *rpb-8* | WBGene00017830 | protein_coding_gene | -1.8 | 0.2927 | 2.7E-19 | -1.4 | 0.372 | 9.1E-34 |
| *kbp-3* | WBGene00009179 | protein_coding_gene | -1.8 | 0.2927 | 1.6E-31 | -1.8 | 0.286 | 2.1E-48 |
| *ufd-2* | WBGene00006734 | protein_coding_gene | -1.8 | 0.2926 | 4.6E-33 | -1.1 | 0.456 | 2.5E-32 |
| *F20G4.2* | WBGene00008991 | protein_coding_gene | -1.8 | 0.2923 | 8.9E-06 | -1.5 | 0.356 | 9.7E-13 |
| *sas-5* | WBGene00009385 | protein_coding_gene | -1.8 | 0.2923 | 2.0E-31 | -2.0 | 0.243 | 2.5E-62 |
| *Y48G10A.2* | WBGene00013019 | protein_coding_gene | -1.8 | 0.2921 | 1.5E-17 | -1.6 | 0.334 | 3.3E-28 |
| *F21H7.3* | WBGene00009029 | protein_coding_gene | -1.8 | 0.2920 | 3.5E-02 | -2.4 | 0.184 | 9.7E-03 |
| *ran-4* | WBGene00004305 | protein_coding_gene | -1.8 | 0.2918 | 3.4E-20 | -1.3 | 0.395 | 8.9E-34 |
| *wee-1.3* | WBGene00006940 | protein_coding_gene | -1.8 | 0.2917 | 1.2E-26 | -1.3 | 0.393 | 2.0E-30 |
| *pbs-7* | WBGene00003953 | protein_coding_gene | -1.8 | 0.2917 | 2.5E-15 | -1.3 | 0.401 | 5.7E-29 |
| *bath-15* | WBGene00015590 | protein_coding_gene | -1.8 | 0.2914 | 5.3E-16 | -2.0 | 0.251 | 1.2E-33 |
| *acl-3* | WBGene00006491 | protein_coding_gene | -1.8 | 0.2908 | 1.7E-23 | -1.5 | 0.361 | 3.5E-26 |
| *T01C3.2* | WBGene00011319 | protein_coding_gene | -1.8 | 0.2905 | 1.5E-36 | -1.9 | 0.263 | 2.3E-51 |
| *tag-321* | WBGene00044325 | protein_coding_gene | -1.8 | 0.2904 | 3.9E-23 | -1.6 | 0.320 | 1.0E-36 |
| *aho-3* | WBGene00045192 | protein_coding_gene | -1.8 | 0.2903 | 4.2E-32 | -1.5 | 0.350 | 1.5E-46 |
| *gut-2* | WBGene00001808 | protein_coding_gene | -1.8 | 0.2902 | 7.4E-18 | -1.6 | 0.323 | 7.6E-31 |
| *K08B4.2* | WBGene00019514 | protein_coding_gene | -1.8 | 0.2902 | 2.6E-02 | -5.4 | 0.023 | 5.8E-12 |
| *msp-76* | WBGene00003463 | protein_coding_gene | -1.8 | 0.2901 | 4.1E-04 | -1.0 | 0.490 | 7.9E-09 |
| *hil-4* | WBGene00001855 | protein_coding_gene | -1.8 | 0.2899 | 8.1E-29 | -1.2 | 0.435 | 1.5E-18 |
| *F32H2.10* | WBGene00009346 | protein_coding_gene | -1.8 | 0.2894 | 1.4E-16 | -1.9 | 0.275 | 1.5E-32 |
| *snr-6* | WBGene00004919 | protein_coding_gene | -1.8 | 0.2890 | 4.4E-44 | -1.7 | 0.309 | 1.5E-55 |
| *F13A7.14* | WBGene00045357 | protein_coding_gene | -1.8 | 0.2890 | 1.1E-32 | -1.6 | 0.339 | 4.2E-26 |
| *ins-2* | WBGene00002085 | protein_coding_gene | -1.8 | 0.2888 | 4.1E-02 | -4.4 | 0.049 | 2.5E-12 |
| *smo-1* | WBGene00004888 | protein_coding_gene | -1.8 | 0.2887 | 1.8E-33 | -1.6 | 0.324 | 3.1E-40 |
| *rod-1* | WBGene00018900 | protein_coding_gene | -1.8 | 0.2883 | 4.2E-20 | -1.2 | 0.427 | 2.2E-29 |
| *snr-4* | WBGene00004917 | protein_coding_gene | -1.8 | 0.2882 | 2.0E-32 | -1.5 | 0.363 | 1.1E-48 |
| *glh-2* | WBGene00001599 | protein_coding_gene | -1.8 | 0.2882 | 2.4E-33 | -1.7 | 0.299 | 1.6E-55 |

| | | | | | | | | |
|---|---|---|---|---|---|---|---|---|
| *W02D3.4* | WBGene00020933 | protein_coding_gene | -1.8 | 0.2878 | 2.5E-28 | -1.1 | 0.469 | 7.6E-21 |
| *B0035.3* | WBGene00007106 | protein_coding_gene | -1.8 | 0.2875 | 3.6E-33 | -1.2 | 0.430 | 2.3E-21 |
| *pfkb-1.2* | WBGene00019295 | protein_coding_gene | -1.8 | 0.2875 | 5.9E-22 | -1.4 | 0.383 | 2.0E-28 |
| *ent-6* | WBGene00009499 | protein_coding_gene | -1.8 | 0.2874 | 7.9E-21 | -1.4 | 0.386 | 5.6E-31 |
| ***T16G12.3*** | **WBGene00011804** | **protein_coding_gene** | **-1.8** | **0.2873** | **1.1E-15** | **1.5** | **2.882** | **4.4E-03** |
| *rpn-1* | WBGene00004458 | protein_coding_gene | -1.8 | 0.2871 | 4.7E-19 | -1.1 | 0.466 | 3.4E-19 |
| *Y105C5B.19* | WBGene00013658 | protein_coding_gene | -1.8 | 0.2865 | 8.3E-13 | -2.0 | 0.242 | 2.9E-24 |
| *T04A8.7* | WBGene00011409 | protein_coding_gene | -1.8 | 0.2864 | 3.5E-16 | -1.5 | 0.358 | 2.6E-35 |
| *ztf-15* | WBGene00011066 | protein_coding_gene | -1.8 | 0.2863 | 2.2E-29 | -1.8 | 0.285 | 3.4E-51 |
| *T05H10.3* | WBGene00011508 | protein_coding_gene | -1.8 | 0.2862 | 7.6E-03 | -4.3 | 0.050 | 3.1E-17 |
| *F59D12.5* | WBGene00010332 | protein_coding_gene | -1.8 | 0.2861 | 2.7E-07 | -1.8 | 0.281 | 9.3E-11 |
| *natb-1* | WBGene00022394 | protein_coding_gene | -1.8 | 0.2858 | 2.5E-18 | -1.2 | 0.429 | 1.2E-17 |
| *B0261.7* | WBGene00015095 | protein_coding_gene | -1.8 | 0.2855 | 3.5E-26 | -1.7 | 0.302 | 2.5E-61 |
| *glp-1* | WBGene00001609 | protein_coding_gene | -1.8 | 0.2853 | 1.6E-16 | -1.4 | 0.378 | 2.4E-32 |
| *sdc-1* | WBGene00004745 | protein_coding_gene | -1.8 | 0.2852 | 6.1E-27 | -1.3 | 0.403 | 9.3E-14 |
| *zhit-1* | WBGene00016992 | protein_coding_gene | -1.8 | 0.2850 | 2.5E-18 | -1.9 | 0.267 | 5.7E-29 |
| *pmp-3* | WBGene00004060 | protein_coding_gene | -1.8 | 0.2848 | 2.3E-20 | -1.6 | 0.339 | 3.0E-54 |
| *K01D12.6* | WBGene00010465 | protein_coding_gene | -1.8 | 0.2848 | 3.2E-33 | -1.1 | 0.455 | 8.3E-17 |
| *fbxb-101* | WBGene00013404 | protein_coding_gene | -1.8 | 0.2842 | 2.4E-02 | -3.5 | 0.087 | 8.3E-11 |
| *suds-3* | WBGene00044204 | protein_coding_gene | -1.8 | 0.2842 | 1.7E-26 | -1.4 | 0.369 | 1.2E-21 |
| *C16C8.12* | WBGene00015850 | protein_coding_gene | -1.8 | 0.2841 | 8.6E-12 | -2.0 | 0.249 | 1.4E-26 |
| *dpl-1* | WBGene00001061 | protein_coding_gene | -1.8 | 0.2839 | 1.2E-19 | -1.6 | 0.326 | 1.3E-43 |
| *C06E1.1* | WBGene00015518 | protein_coding_gene | -1.8 | 0.2838 | 6.5E-09 | -2.6 | 0.164 | 3.5E-36 |
| *rab-6.1* | WBGene00004269 | protein_coding_gene | -1.8 | 0.2838 | 4.5E-20 | -1.3 | 0.408 | 3.4E-29 |
| *clec-12* | WBGene00012103 | protein_coding_gene | -1.8 | 0.2836 | 2.1E-08 | -1.8 | 0.289 | 7.2E-08 |
| *F28C10.3* | WBGene00017898 | protein_coding_gene | -1.8 | 0.2835 | 2.1E-12 | -1.6 | 0.327 | 1.6E-17 |
| *utx-1* | WBGene00017046 | protein_coding_gene | -1.8 | 0.2834 | 6.9E-15 | -2.2 | 0.210 | 1.7E-42 |
| *slc-30A5* | WBGene00013668 | protein_coding_gene | -1.8 | 0.2832 | 1.6E-23 | -1.0 | 0.484 | 2.1E-20 |
| *hal-3* | WBGene00011788 | protein_coding_gene | -1.8 | 0.2830 | 2.1E-29 | -1.6 | 0.337 | 4.7E-22 |
| *K07A1.13* | WBGene00010617 | protein_coding_gene | -1.8 | 0.2829 | 3.1E-14 | -2.1 | 0.239 | 1.5E-45 |
| *ztf-4* | WBGene00020399 | protein_coding_gene | -1.8 | 0.2826 | 3.5E-28 | -1.3 | 0.412 | 1.9E-21 |
| *F35C8.8* | WBGene00018038 | protein_coding_gene | -1.8 | 0.2824 | 1.7E-02 | -2.2 | 0.218 | 1.2E-02 |
| *hphd-1* | WBGene00012608 | protein_coding_gene | -1.8 | 0.2822 | 5.3E-10 | -1.4 | 0.382 | 2.7E-04 |
| *rpl-41.1* | WBGene00023067 | protein_coding_gene | -1.8 | 0.2819 | 8.3E-11 | -1.5 | 0.350 | 1.9E-28 |
| *rbm-3.1* | WBGene00011155 | protein_coding_gene | -1.8 | 0.2813 | 1.2E-22 | -1.3 | 0.399 | 1.2E-31 |
| *eri-9* | WBGene00016143 | protein_coding_gene | -1.8 | 0.2811 | 4.9E-31 | -1.2 | 0.435 | 9.4E-20 |
| *gcna-1* | WBGene00022694 | protein_coding_gene | -1.8 | 0.2809 | 1.3E-22 | -1.6 | 0.332 | 4.9E-36 |
| *lid-1* | WBGene00007711 | protein_coding_gene | -1.8 | 0.2808 | 1.1E-30 | -1.5 | 0.345 | 1.8E-35 |
| *Y54G2A.12* | WBGene00021877 | protein_coding_gene | -1.8 | 0.2806 | 3.2E-23 | -2.0 | 0.249 | 3.7E-47 |
| *ragc-1* | WBGene00012497 | protein_coding_gene | -1.8 | 0.2805 | 1.9E-24 | -1.6 | 0.339 | 1.1E-52 |
| *wdr-4* | WBGene00022420 | protein_coding_gene | -1.8 | 0.2804 | 4.5E-19 | -1.8 | 0.293 | 8.2E-56 |
| *tyms-1* | WBGene00022455 | protein_coding_gene | -1.8 | 0.2802 | 4.7E-20 | -2.4 | 0.191 | 2.5E-69 |
| *pde-12* | WBGene00011268 | protein_coding_gene | -1.8 | 0.2802 | 1.1E-48 | -1.2 | 0.443 | 4.0E-19 |
| *cam-1* | WBGene00000289 | protein_coding_gene | -1.8 | 0.2801 | 3.0E-34 | -2.1 | 0.232 | 9.0E-77 |
| *C50E3.5* | WBGene00016816 | protein_coding_gene | -1.8 | 0.2800 | 3.2E-18 | -1.7 | 0.301 | 1.4E-33 |
| *C34D4.3* | WBGene00016399 | protein_coding_gene | -1.8 | 0.2797 | 1.1E-03 | -1.2 | 0.445 | 1.2E-06 |
| *cyp-44A1* | WBGene00000375 | protein_coding_gene | -1.8 | 0.2796 | 2.6E-25 | -1.8 | 0.283 | 2.7E-41 |
| *ugt-17* | WBGene00007422 | protein_coding_gene | -1.8 | 0.2795 | 4.1E-26 | -1.6 | 0.339 | 6.2E-15 |
| *ccr-4* | WBGene00000376 | protein_coding_gene | -1.8 | 0.2795 | 1.4E-13 | -1.7 | 0.317 | 1.5E-61 |
| *pas-2* | WBGene00003923 | protein_coding_gene | -1.8 | 0.2795 | 1.0E-15 | -1.5 | 0.362 | 9.4E-33 |
| *rnf-145* | WBGene00022471 | protein_coding_gene | -1.8 | 0.2794 | 1.3E-39 | -1.6 | 0.329 | 4.5E-17 |
| *asb-1* | WBGene00000206 | protein_coding_gene | -1.8 | 0.2794 | 5.5E-20 | -1.2 | 0.439 | 1.1E-25 |
| *npp-19* | WBGene00003805 | protein_coding_gene | -1.8 | 0.2792 | 3.6E-19 | -1.2 | 0.439 | 6.2E-23 |
| *asfl-1* | WBGene00007277 | protein_coding_gene | -1.8 | 0.2792 | 3.2E-13 | -1.7 | 0.303 | 4.7E-38 |
| *rad-23* | WBGene00013924 | protein_coding_gene | -1.8 | 0.2792 | 6.3E-28 | -1.0 | 0.488 | 8.7E-15 |
| *coq-5* | WBGene00000765 | protein_coding_gene | -1.8 | 0.2791 | 8.6E-21 | -1.3 | 0.395 | 2.1E-25 |
| *R12E2.8* | WBGene00020034 | protein_coding_gene | -1.8 | 0.2791 | 2.0E-02 | -2.2 | 0.218 | 1.1E-03 |
| *cdl-1* | WBGene00000411 | protein_coding_gene | -1.8 | 0.2785 | 1.9E-21 | -1.5 | 0.355 | 1.3E-36 |
| *C11D2.4* | WBGene00015702 | protein_coding_gene | -1.8 | 0.2784 | 1.5E-18 | -1.1 | 0.461 | 1.5E-14 |
| *erfa-3* | WBGene00010405 | protein_coding_gene | -1.8 | 0.2783 | 3.8E-24 | -1.3 | 0.396 | 5.7E-44 |
| *ctsa-1.1* | WBGene00018271 | protein_coding_gene | -1.8 | 0.2783 | 3.3E-14 | -1.3 | 0.400 | 5.4E-15 |
| *maco-1* | WBGene00017066 | protein_coding_gene | -1.8 | 0.2783 | 8.4E-14 | -1.5 | 0.361 | 2.4E-58 |
| *let-391* | WBGene00006492 | protein_coding_gene | -1.8 | 0.2781 | 1.4E-24 | -1.8 | 0.281 | 4.5E-50 |
| *R07E4.5* | WBGene00019938 | protein_coding_gene | -1.8 | 0.2780 | 2.8E-14 | -1.6 | 0.324 | 1.7E-22 |
| *ints-11* | WBGene00008642 | protein_coding_gene | -1.8 | 0.2779 | 3.0E-44 | -1.1 | 0.469 | 3.4E-19 |
| *B0001.7* | WBGene00007092 | protein_coding_gene | -1.8 | 0.2779 | 2.6E-21 | -1.3 | 0.394 | 1.2E-27 |
| *sur-2* | WBGene00006349 | protein_coding_gene | -1.8 | 0.2777 | 1.4E-07 | -1.4 | 0.374 | 1.3E-25 |
| *ulp-2* | WBGene00006737 | protein_coding_gene | -1.8 | 0.2775 | 6.7E-21 | -1.2 | 0.444 | 3.2E-21 |
| *set-12* | WBGene00019584 | protein_coding_gene | -1.8 | 0.2775 | 2.5E-09 | -3.6 | 0.084 | 1.6E-16 |
| *F10E9.7* | WBGene00017358 | protein_coding_gene | -1.8 | 0.2775 | 6.0E-16 | -1.7 | 0.308 | 9.3E-30 |
| *ect-2* | WBGene00002297 | protein_coding_gene | -1.9 | 0.2773 | 3.3E-14 | -1.1 | 0.462 | 1.6E-20 |
| *ran-1* | WBGene00004302 | protein_coding_gene | -1.9 | 0.2773 | 2.1E-18 | -1.6 | 0.324 | 2.5E-53 |
| *ZK973.1* | WBGene00022830 | protein_coding_gene | -1.9 | 0.2771 | 2.2E-43 | -1.8 | 0.290 | 3.0E-72 |
| *cpsf-4* | WBGene00044329 | protein_coding_gene | -1.9 | 0.2770 | 1.4E-37 | -1.6 | 0.338 | 7.9E-32 |
| *Y56A3A.16* | WBGene00013235 | protein_coding_gene | -1.9 | 0.2769 | 1.8E-25 | -1.6 | 0.321 | 3.5E-39 |
| *linc-4* | WBGene00219306 | lincRNA_gene | -1.9 | 0.2769 | 9.8E-07 | -1.8 | 0.282 | 1.5E-05 |
| *T15B7.1* | WBGene00020516 | protein_coding_gene | -1.9 | 0.2768 | 1.6E-10 | -1.4 | 0.389 | 8.6E-05 |
| *fan-1* | WBGene00015310 | protein_coding_gene | -1.9 | 0.2768 | 9.3E-38 | -1.7 | 0.317 | 5.9E-35 |
| *Y39B6A.42* | WBGene00012700 | protein_coding_gene | -1.9 | 0.2767 | 2.0E-22 | -1.6 | 0.332 | 3.6E-25 |
| *F42A9.8* | WBGene00018337 | protein_coding_gene | -1.9 | 0.2765 | 4.2E-11 | -1.8 | 0.278 | 4.8E-28 |

| | | | | | | | | |
|---|---|---|---|---|---|---|---|---|
| *mrg-1* | WBGene00003406 | protein_coding_gene | -1.9 | 0.2763 | 1.8E-27 | -1.5 | 0.354 | 1.9E-37 |
| *dro-1* | WBGene00001092 | protein_coding_gene | -1.9 | 0.2762 | 6.8E-56 | -1.5 | 0.344 | 1.5E-33 |
| *ari-1.3* | WBGene00016156 | protein_coding_gene | -1.9 | 0.2761 | 4.0E-29 | -1.5 | 0.350 | 2.6E-36 |
| *cup-5* | WBGene00000846 | protein_coding_gene | -1.9 | 0.2759 | 1.7E-27 | -1.6 | 0.326 | 4.7E-39 |
| *her-1* | WBGene00001842 | protein_coding_gene | -1.9 | 0.2754 | 1.1E-03 | -4.0 | 0.062 | 5.5E-14 |
| *hasp-1* | WBGene00007258 | protein_coding_gene | -1.9 | 0.2754 | 5.1E-29 | -1.5 | 0.351 | 1.8E-28 |
| *gon-14* | WBGene00002148 | protein_coding_gene | -1.9 | 0.2751 | 2.6E-18 | -1.6 | 0.339 | 8.4E-39 |
| *orc-5* | WBGene00013857 | protein_coding_gene | -1.9 | 0.2748 | 2.1E-30 | -1.5 | 0.342 | 2.1E-44 |
| *kbp-4* | WBGene00022357 | protein_coding_gene | -1.9 | 0.2747 | 5.5E-34 | -2.1 | 0.236 | 3.1E-43 |
| *trcs-2* | WBGene00011312 | protein_coding_gene | -1.9 | 0.2747 | 2.6E-20 | -1.1 | 0.470 | 5.5E-23 |
| *T19H12.2* | WBGene00020588 | protein_coding_gene | -1.9 | 0.2746 | 2.4E-30 | -1.6 | 0.323 | 3.9E-39 |
| *cec-4* | WBGene00017990 | protein_coding_gene | -1.9 | 0.2745 | 8.8E-26 | -1.7 | 0.300 | 8.2E-43 |
| *sun-1* | WBGene00006311 | protein_coding_gene | -1.9 | 0.2739 | 4.6E-30 | -1.4 | 0.378 | 1.0E-33 |
| *btb-12* | WBGene00022555 | protein_coding_gene | -1.9 | 0.2735 | 5.8E-04 | -5.4 | 0.023 | 1.3E-09 |
| *tag-340* | WBGene00017826 | protein_coding_gene | -1.9 | 0.2730 | 3.6E-50 | -1.5 | 0.344 | 1.2E-38 |
| *nlp-24* | WBGene00003762 | protein_coding_gene | -1.9 | 0.2730 | 4.7E-06 | -1.5 | 0.345 | 3.0E-03 |
| *nhr-12* | WBGene00003611 | protein_coding_gene | -1.9 | 0.2727 | 1.1E-12 | -1.4 | 0.385 | 3.7E-12 |
| *ska-3* | WBGene00018830 | protein_coding_gene | -1.9 | 0.2723 | 3.3E-30 | -1.9 | 0.265 | 1.3E-53 |
| *sqv-2* | WBGene00005020 | protein_coding_gene | -1.9 | 0.2718 | 9.2E-28 | -1.3 | 0.419 | 9.6E-17 |
| *F52D2.7* | WBGene00018689 | protein_coding_gene | -1.9 | 0.2718 | 1.4E-12 | -2.3 | 0.203 | 1.7E-27 |
| *dhhc-6* | WBGene00010892 | protein_coding_gene | -1.9 | 0.2717 | 1.2E-21 | -1.5 | 0.356 | 3.9E-37 |
| *kbp-2* | WBGene00017832 | protein_coding_gene | -1.9 | 0.2716 | 6.4E-20 | -1.9 | 0.259 | 9.3E-39 |
| *R11H6.2* | WBGene00011250 | protein_coding_gene | -1.9 | 0.2715 | 3.3E-19 | -1.6 | 0.321 | 6.5E-45 |
| *tcab-1* | WBGene00013669 | protein_coding_gene | -1.9 | 0.2714 | 1.3E-29 | -1.2 | 0.427 | 1.3E-21 |
| *B0416.11* | WBGene00206483 | protein_coding_gene | -1.9 | 0.2714 | 7.3E-03 | -2.3 | 0.210 | 7.6E-03 |
| *C48B4.9* | WBGene00008172 | protein_coding_gene | -1.9 | 0.2713 | 5.5E-29 | -1.0 | 0.499 | 2.8E-11 |
| *rpn-9* | WBGene00004465 | protein_coding_gene | -1.9 | 0.2712 | 2.3E-22 | -1.5 | 0.355 | 2.5E-33 |
| *gmn-1* | WBGene00013552 | protein_coding_gene | -1.9 | 0.2709 | 1.1E-45 | -2.0 | 0.252 | 1.0E-51 |
| *F23F1.5* | WBGene00017746 | protein_coding_gene | -1.9 | 0.2707 | 2.0E-27 | -1.8 | 0.281 | 3.0E-50 |
| *F33G12.3* | WBGene00018015 | protein_coding_gene | -1.9 | 0.2706 | 1.1E-28 | -1.8 | 0.279 | 1.3E-51 |
| *msp-78* | WBGene00003465 | protein_coding_gene | -1.9 | 0.2705 | 1.2E-03 | -1.5 | 0.347 | 5.5E-12 |
| *Y62F5A.12* | WBGene00077453 | protein_coding_gene | -1.9 | 0.2704 | 1.0E-35 | -1.5 | 0.353 | 2.7E-24 |
| *msp-36* | WBGene00003432 | protein_coding_gene | -1.9 | 0.2704 | 2.8E-04 | -1.5 | 0.349 | 4.9E-15 |
| *spr-3* | WBGene00005008 | protein_coding_gene | -1.9 | 0.2703 | 3.0E-16 | -1.8 | 0.292 | 2.5E-23 |
| *tba-1* | WBGene00006528 | protein_coding_gene | -1.9 | 0.2702 | 1.9E-21 | -1.5 | 0.343 | 1.1E-40 |
| *skpt-1* | WBGene00018613 | protein_coding_gene | -1.9 | 0.2701 | 1.0E-25 | -1.5 | 0.349 | 2.7E-34 |
| *acs-19* | WBGene00007969 | protein_coding_gene | -1.9 | 0.2699 | 1.3E-15 | -1.8 | 0.289 | 3.5E-40 |
| *Y47G6A.25* | WBGene00021649 | protein_coding_gene | -1.9 | 0.2699 | 1.0E-25 | -1.9 | 0.263 | 1.3E-33 |
| *gpaa-1* | WBGene00018006 | protein_coding_gene | -1.9 | 0.2697 | 1.9E-18 | -1.7 | 0.308 | 1.7E-47 |
| *ceh-5* | WBGene00000430 | protein_coding_gene | -1.9 | 0.2696 | 1.1E-02 | -5.1 | 0.030 | 6.8E-08 |
| *R13A5.7* | WBGene00020050 | protein_coding_gene | -1.9 | 0.2695 | 2.3E-16 | -1.2 | 0.450 | 1.5E-11 |
| *cosa-1* | WBGene00022172 | protein_coding_gene | -1.9 | 0.2690 | 8.0E-28 | -1.3 | 0.393 | 3.3E-14 |
| *abi-1* | WBGene00015146 | protein_coding_gene | -1.9 | 0.2690 | 7.5E-28 | -1.7 | 0.314 | 4.6E-56 |
| *algn-6* | WBGene00007435 | protein_coding_gene | -1.9 | 0.2689 | 8.4E-24 | -1.5 | 0.345 | 3.4E-29 |
| *trt-1* | WBGene00006618 | protein_coding_gene | -1.9 | 0.2689 | 1.5E-27 | -1.9 | 0.273 | 5.3E-42 |
| *R74.8* | WBGene00011282 | protein_coding_gene | -1.9 | 0.2683 | 2.6E-30 | -1.5 | 0.351 | 9.8E-43 |
| *spat-1* | WBGene00010198 | protein_coding_gene | -1.9 | 0.2681 | 1.6E-27 | -1.4 | 0.378 | 2.2E-26 |
| *cdc-48.1* | WBGene00007352 | protein_coding_gene | -1.9 | 0.2678 | 1.6E-26 | -1.2 | 0.435 | 8.2E-25 |
| *ctf-4* | WBGene00008921 | protein_coding_gene | -1.9 | 0.2676 | 4.9E-24 | -1.8 | 0.282 | 1.8E-63 |
| *mut-2* | WBGene00003499 | protein_coding_gene | -1.9 | 0.2675 | 6.9E-30 | -1.6 | 0.338 | 7.2E-39 |
| *anmt-2* | WBGene00018340 | protein_coding_gene | -1.9 | 0.2673 | 9.2E-34 | -1.2 | 0.421 | 8.3E-10 |
| *daf-41* | WBGene00022599 | protein_coding_gene | -1.9 | 0.2673 | 4.1E-23 | -1.5 | 0.364 | 3.8E-41 |
| *ZK686.1* | WBGene00022791 | protein_coding_gene | -1.9 | 0.2673 | 9.9E-24 | -1.8 | 0.284 | 4.4E-29 |
| *T12C9.7* | WBGene00020461 | protein_coding_gene | -1.9 | 0.2671 | 7.7E-23 | -1.3 | 0.401 | 5.7E-16 |
| *atx-3* | WBGene00006446 | protein_coding_gene | -1.9 | 0.2671 | 1.6E-18 | -1.6 | 0.339 | 5.3E-29 |
| *R107.5* | WBGene00011300 | protein_coding_gene | -1.9 | 0.2670 | 3.5E-20 | -1.5 | 0.345 | 5.0E-28 |
| *ccz-1* | WBGene00010283 | protein_coding_gene | -1.9 | 0.2667 | 4.2E-31 | -1.5 | 0.351 | 8.4E-32 |
| *F33E2.5* | WBGene00009361 | protein_coding_gene | -1.9 | 0.2666 | 2.6E-10 | -3.1 | 0.118 | 6.5E-53 |
| *acbp-6* | WBGene00012457 | protein_coding_gene | -1.9 | 0.2665 | 9.5E-05 | -1.6 | 0.339 | 1.4E-04 |
| *apr-1* | WBGene00000156 | protein_coding_gene | -1.9 | 0.2665 | 1.7E-18 | -2.0 | 0.253 | 6.3E-78 |
| *nud-1* | WBGene00003829 | protein_coding_gene | -1.9 | 0.2662 | 4.6E-26 | -1.7 | 0.317 | 1.1E-46 |
| *hpo-9* | WBGene00015463 | protein_coding_gene | -1.9 | 0.2655 | 2.4E-31 | -1.7 | 0.318 | 2.5E-53 |
| *T07D4.5* | WBGene00045407 | protein_coding_gene | -1.9 | 0.2654 | 4.0E-22 | -1.9 | 0.260 | 9.2E-41 |
| *rpb-11* | WBGene00012187 | protein_coding_gene | -1.9 | 0.2649 | 2.1E-29 | -1.5 | 0.356 | 1.4E-25 |
| *C06H2.7* | WBGene00007391 | protein_coding_gene | -1.9 | 0.2647 | 8.6E-18 | -2.6 | 0.166 | 6.5E-46 |
| *zfp-2* | WBGene00009448 | protein_coding_gene | -1.9 | 0.2644 | 1.3E-36 | -1.4 | 0.373 | 7.8E-31 |
| *K04G7.1* | WBGene00019400 | protein_coding_gene | -1.9 | 0.2643 | 2.0E-27 | -1.8 | 0.280 | 7.0E-54 |
| *edc-3* | WBGene00011036 | protein_coding_gene | -1.9 | 0.2639 | 7.2E-25 | -1.6 | 0.324 | 6.9E-50 |
| *alg-2* | WBGene00000106 | protein_coding_gene | -1.9 | 0.2636 | 1.1E-15 | -1.8 | 0.292 | 8.1E-40 |
| *wdr-20* | WBGene00007428 | protein_coding_gene | -1.9 | 0.2634 | 6.8E-22 | -1.1 | 0.453 | 3.5E-21 |
| *elli-1* | WBGene00008968 | protein_coding_gene | -1.9 | 0.2632 | 1.1E-16 | -1.6 | 0.334 | 9.3E-38 |
| *W03D8.7* | WBGene00020988 | protein_coding_gene | -1.9 | 0.2632 | 5.4E-03 | -2.9 | 0.131 | 1.4E-06 |
| *mom-5* | WBGene00003397 | protein_coding_gene | -1.9 | 0.2632 | 1.2E-19 | -1.4 | 0.372 | 4.8E-41 |
| *nra-2* | WBGene00011488 | protein_coding_gene | -1.9 | 0.2629 | 3.2E-34 | -1.2 | 0.422 | 9.9E-32 |
| *tsp-12* | WBGene00006638 | protein_coding_gene | -1.9 | 0.2629 | 9.4E-16 | -2.0 | 0.246 | 2.9E-64 |
| *Y69A2AR.21* | WBGene00022092 | protein_coding_gene | -1.9 | 0.2629 | 2.7E-55 | -1.8 | 0.284 | 1.7E-45 |
| *lin-54* | WBGene00003037 | protein_coding_gene | -1.9 | 0.2627 | 7.9E-22 | -1.8 | 0.283 | 1.9E-59 |
| *hse-5* | WBGene00002003 | protein_coding_gene | -1.9 | 0.2626 | 2.3E-38 | -1.9 | 0.262 | 7.8E-65 |
| *sygl-1* | WBGene00012104 | protein_coding_gene | -1.9 | 0.2623 | 5.4E-14 | -1.4 | 0.390 | 8.7E-21 |
| *nemp-1* | WBGene00017328 | protein_coding_gene | -1.9 | 0.2619 | 2.2E-52 | -1.6 | 0.327 | 4.3E-47 |

| | | | | | | | | |
|---|---|---|---|---|---|---|---|---|
| *mdh-2* | WBGene00003162 | protein_coding_gene | -1.9 | 0.2619 | 3.9E-25 | -1.3 | 0.415 | 3.8E-22 |
| *T27F6.7* | WBGene00012106 | protein_coding_gene | -1.9 | 0.2618 | 1.3E-19 | -1.4 | 0.370 | 1.3E-16 |
| *lbp-9* | WBGene00021486 | protein_coding_gene | -1.9 | 0.2614 | 7.1E-11 | -1.6 | 0.336 | 1.7E-37 |
| *fbxc-44* | WBGene00015819 | protein_coding_gene | -1.9 | 0.2614 | 2.8E-14 | -1.1 | 0.464 | 7.0E-10 |
| *mtp-18* | WBGene00077690 | protein_coding_gene | -1.9 | 0.2614 | 2.1E-08 | -1.0 | 0.494 | 1.8E-03 |
| *dnj-11* | WBGene00001029 | protein_coding_gene | -1.9 | 0.2613 | 2.7E-23 | -1.5 | 0.359 | 4.0E-49 |
| *C53H9.3* | WBGene00044746 | protein_coding_gene | -1.9 | 0.2613 | 9.3E-11 | -1.4 | 0.374 | 3.1E-13 |
| *ugt-64* | WBGene00015577 | protein_coding_gene | -1.9 | 0.2613 | 5.2E-22 | -1.5 | 0.349 | 2.4E-22 |
| *T02H6.5* | WBGene00020175 | protein_coding_gene | -1.9 | 0.2612 | 1.3E-03 | -2.4 | 0.192 | 1.4E-05 |
| *ttx-7* | WBGene00008765 | protein_coding_gene | -1.9 | 0.2611 | 3.3E-12 | -1.1 | 0.466 | 6.4E-19 |
| *fbxa-46* | WBGene00018690 | pseudogene | -1.9 | 0.2611 | 1.3E-03 | -3.0 | 0.124 | 3.3E-04 |
| *rfc-3* | WBGene00004339 | protein_coding_gene | -1.9 | 0.2609 | 2.0E-27 | -2.2 | 0.224 | 9.3E-65 |
| *F53A2.3* | WBGene00009951 | protein_coding_gene | -1.9 | 0.2609 | 8.5E-12 | -2.2 | 0.213 | 3.1E-16 |
| *B0336.13* | WBGene00015150 | protein_coding_gene | -1.9 | 0.2608 | 1.4E-19 | -1.7 | 0.300 | 1.7E-41 |
| *arle-14* | WBGene00015145 | protein_coding_gene | -1.9 | 0.2608 | 1.3E-36 | -1.6 | 0.319 | 1.5E-29 |
| *sys-1* | WBGene00006379 | protein_coding_gene | -1.9 | 0.2608 | 1.2E-21 | -2.3 | 0.203 | 3.2E-57 |
| *T07F12.2* | WBGene00020322 | protein_coding_gene | -1.9 | 0.2607 | 3.0E-21 | -1.4 | 0.384 | 7.1E-17 |
| *zen-4* | WBGene00006974 | protein_coding_gene | -1.9 | 0.2607 | 2.8E-24 | -1.8 | 0.297 | 1.1E-63 |
| *K05C4.7* | WBGene00010583 | protein_coding_gene | -1.9 | 0.2606 | 3.5E-36 | -1.6 | 0.324 | 8.8E-40 |
| *ung-1* | WBGene00013241 | protein_coding_gene | -1.9 | 0.2604 | 2.4E-33 | -1.5 | 0.347 | 1.4E-24 |
| *npp-23* | WBGene00007493 | protein_coding_gene | -1.9 | 0.2603 | 4.2E-28 | -1.6 | 0.332 | 1.0E-38 |
| *Y92H12BR.3* | WBGene00022369 | protein_coding_gene | -1.9 | 0.2596 | 2.2E-24 | -1.5 | 0.365 | 8.1E-40 |
| *xpo-2* | WBGene00002079 | protein_coding_gene | -1.9 | 0.2594 | 5.7E-17 | -1.4 | 0.388 | 7.7E-28 |
| *aka-1* | WBGene00000101 | protein_coding_gene | -1.9 | 0.2591 | 6.4E-17 | -1.1 | 0.458 | 1.4E-30 |
| *C15C8.4* | WBGene00007606 | protein_coding_gene | -1.9 | 0.2589 | 1.9E-39 | -1.6 | 0.322 | 2.2E-45 |
| *M03B6.4* | WBGene00010836 | protein_coding_gene | -1.9 | 0.2588 | 1.0E-15 | -2.4 | 0.196 | 3.2E-19 |
| *F01D4.5* | WBGene00008488 | protein_coding_gene | -2.0 | 0.2580 | 1.4E-33 | -1.8 | 0.280 | 2.3E-37 |
| *Y75B8A.16* | WBGene00013551 | protein_coding_gene | -2.0 | 0.2577 | 8.2E-42 | -1.5 | 0.354 | 3.4E-22 |
| *nfya-1* | WBGene00011614 | protein_coding_gene | -2.0 | 0.2576 | 9.4E-12 | -2.2 | 0.212 | 1.5E-28 |
| *apc-11* | WBGene00000145 | protein_coding_gene | -2.0 | 0.2572 | 1.2E-15 | -2.2 | 0.219 | 1.3E-45 |
| *sel-10* | WBGene00004767 | protein_coding_gene | -2.0 | 0.2571 | 6.7E-32 | -1.2 | 0.442 | 7.6E-33 |
| *psf-2* | WBGene00009287 | protein_coding_gene | -2.0 | 0.2569 | 9.8E-41 | -2.1 | 0.233 | 4.5E-51 |
| *C53D6.4* | WBGene00008281 | protein_coding_gene | -2.0 | 0.2568 | 5.6E-31 | -1.6 | 0.334 | 2.8E-29 |
| *F44B9.8* | WBGene00018409 | protein_coding_gene | -2.0 | 0.2566 | 1.3E-19 | -1.4 | 0.370 | 8.2E-35 |
| *tost-1* | WBGene00016449 | protein_coding_gene | -2.0 | 0.2562 | 1.0E-20 | -1.9 | 0.275 | 1.8E-52 |
| *toc-1* | WBGene00006590 | protein_coding_gene | -2.0 | 0.2557 | 2.4E-19 | -1.0 | 0.485 | 4.2E-18 |
| *mcm-6* | WBGene00003158 | protein_coding_gene | -2.0 | 0.2555 | 4.2E-20 | -1.9 | 0.264 | 1.3E-59 |
| *gld-4* | WBGene00014115 | protein_coding_gene | -2.0 | 0.2554 | 1.2E-08 | -1.7 | 0.311 | 2.1E-56 |
| *drd-10* | WBGene00016361 | protein_coding_gene | -2.0 | 0.2554 | 8.9E-05 | -2.1 | 0.233 | 5.1E-03 |
| *cdc-25.1* | WBGene00000386 | protein_coding_gene | -2.0 | 0.2551 | 1.3E-22 | -1.7 | 0.305 | 6.7E-52 |
| *laat-1* | WBGene00021546 | protein_coding_gene | -2.0 | 0.2549 | 2.3E-36 | -1.2 | 0.449 | 2.6E-18 |
| *rmh-2* | WBGene00011576 | protein_coding_gene | -2.0 | 0.2546 | 4.6E-22 | -1.8 | 0.285 | 3.8E-32 |
| *F33E11.3* | WBGene00018011 | protein_coding_gene | -2.0 | 0.2545 | 4.4E-45 | -1.8 | 0.290 | 1.6E-46 |
| *C27F2.7* | WBGene00016169 | protein_coding_gene | -2.0 | 0.2541 | 3.1E-28 | -1.4 | 0.371 | 4.9E-14 |
| *sucg-1* | WBGene00016844 | protein_coding_gene | -2.0 | 0.2540 | 2.2E-26 | -1.0 | 0.500 | 3.1E-18 |
| *nyn-2* | WBGene00013596 | protein_coding_gene | -2.0 | 0.2538 | 8.0E-33 | -1.4 | 0.382 | 1.6E-22 |
| *F25G6.9* | WBGene00017800 | protein_coding_gene | -2.0 | 0.2537 | 6.8E-32 | -1.5 | 0.349 | 2.0E-45 |
| *C53B7.2* | WBGene00016893 | protein_coding_gene | -2.0 | 0.2536 | 2.6E-08 | -2.4 | 0.186 | 5.6E-23 |
| *tram-1* | WBGene00007696 | protein_coding_gene | -2.0 | 0.2532 | 1.8E-19 | -1.1 | 0.478 | 6.3E-24 |
| *ZK688.9* | WBGene00022803 | protein_coding_gene | -2.0 | 0.2530 | 4.9E-49 | -1.7 | 0.316 | 2.4E-35 |
| *fbxa-83* | WBGene00013453 | protein_coding_gene | -2.0 | 0.2528 | 3.0E-11 | -2.6 | 0.170 | 1.9E-21 |
| *K08E3.5* | WBGene00010665 | protein_coding_gene | -2.0 | 0.2526 | 1.0E-39 | -1.6 | 0.332 | 2.0E-45 |
| *vang-1* | WBGene00015171 | protein_coding_gene | -2.0 | 0.2519 | 8.5E-23 | -2.4 | 0.187 | 4.8E-37 |
| *chdp-1* | WBGene00015687 | protein_coding_gene | -2.0 | 0.2518 | 1.5E-26 | -1.9 | 0.268 | 5.3E-80 |
| *skr-17* | WBGene00004823 | protein_coding_gene | -2.0 | 0.2517 | 3.7E-29 | -1.4 | 0.383 | 2.7E-26 |
| *F28F8.9* | WBGene00044188 | protein_coding_gene | -2.0 | 0.2516 | 6.8E-34 | -1.7 | 0.301 | 1.1E-30 |
| *ilcr-2* | WBGene00018960 | protein_coding_gene | -2.0 | 0.2513 | 2.0E-22 | -1.4 | 0.372 | 2.0E-27 |
| *F08B4.8* | WBGene00255375 | protein_coding_gene | -2.0 | 0.2512 | 3.2E-04 | -1.3 | 0.411 | 3.4E-08 |
| *algn-9* | WBGene00007556 | protein_coding_gene | -2.0 | 0.2510 | 8.6E-32 | -1.2 | 0.447 | 2.3E-26 |
| *ZK973.9* | WBGene00022835 | protein_coding_gene | -2.0 | 0.2508 | 9.3E-13 | -2.0 | 0.250 | 7.8E-40 |
| *cutl-25* | WBGene00021919 | protein_coding_gene | -2.0 | 0.2508 | 1.0E-03 | -3.3 | 0.100 | 1.8E-14 |
| *F25H9.6* | WBGene00009138 | protein_coding_gene | -2.0 | 0.2507 | 3.1E-33 | -1.5 | 0.345 | 4.7E-25 |
| *snpn-1* | WBGene00015327 | protein_coding_gene | -2.0 | 0.2507 | 3.0E-27 | -2.0 | 0.251 | 6.6E-41 |
| *nspd-2* | WBGene00010633 | protein_coding_gene | -2.0 | 0.2505 | 1.1E-04 | -1.5 | 0.342 | 1.8E-18 |
| *msp-31* | WBGene00003429 | protein_coding_gene | -2.0 | 0.2505 | 1.6E-04 | -1.1 | 0.456 | 4.1E-11 |
| *R06B9.5* | WBGene00011056 | protein_coding_gene | -2.0 | 0.2504 | 3.3E-17 | -1.8 | 0.283 | 8.7E-15 |
| *swd-2.1* | WBGene00016373 | protein_coding_gene | -2.0 | 0.2503 | 3.6E-39 | -1.6 | 0.330 | 6.2E-34 |
| *D1007.5* | WBGene00017003 | protein_coding_gene | -2.0 | 0.2503 | 6.6E-31 | -1.5 | 0.360 | 2.5E-29 |
| *F54D10.5* | WBGene00018806 | protein_coding_gene | -2.0 | 0.2500 | 2.3E-20 | -1.8 | 0.291 | 9.2E-25 |
| *dnj-28* | WBGene00001046 | protein_coding_gene | -2.0 | 0.2496 | 9.6E-33 | -1.1 | 0.473 | 6.8E-15 |
| *rnp-8* | WBGene00020091 | protein_coding_gene | -2.0 | 0.2490 | 4.8E-17 | -1.6 | 0.323 | 3.0E-35 |
| *mec-15* | WBGene00003177 | protein_coding_gene | -2.0 | 0.2488 | 1.3E-31 | -1.8 | 0.284 | 7.5E-57 |
| *F35G2.3* | WBGene00009437 | protein_coding_gene | -2.0 | 0.2485 | 3.0E-03 | -2.8 | 0.139 | 1.5E-05 |
| *C32B5.15* | WBGene00016308 | protein_coding_gene | -2.0 | 0.2479 | 1.7E-02 | -4.0 | 0.061 | 2.3E-05 |
| *F19H6.3* | WBGene00008958 | protein_coding_gene | -2.0 | 0.2477 | 5.0E-02 | -1.9 | 0.273 | 3.7E-02 |
| *F58A6.9* | WBGene00019026 | protein_coding_gene | -2.0 | 0.2476 | 1.4E-04 | -2.3 | 0.209 | 4.4E-28 |
| *sld-2* | WBGene00020466 | protein_coding_gene | -2.0 | 0.2476 | 3.8E-30 | -1.9 | 0.272 | 4.3E-37 |
| *C46C2.7* | WBGene00045078 | protein_coding_gene | -2.0 | 0.2472 | 9.1E-14 | -1.6 | 0.335 | 3.6E-09 |
| *ubc-9* | WBGene00006706 | protein_coding_gene | -2.0 | 0.2471 | 5.0E-26 | -1.7 | 0.304 | 3.2E-62 |
| *C31H2.4* | WBGene00016294 | protein_coding_gene | -2.0 | 0.2470 | 1.7E-07 | -1.5 | 0.349 | 4.3E-05 |

| | | | | | | | | |
|---|---|---|---|---|---|---|---|---|
| ***msp-64*** | **WBGene00003457** | **protein_coding_gene** | **-2.0** | **0.2470** | **1.7E-04** | **2.2** | **4.689** | **5.8E-04** |
| *nspd-5* | WBGene00017386 | protein_coding_gene | -2.0 | 0.2469 | 2.3E-04 | -1.8 | 0.287 | 1.7E-16 |
| *D2023.6* | WBGene00008410 | protein_coding_gene | -2.0 | 0.2467 | 6.3E-39 | -1.3 | 0.415 | 5.6E-26 |
| *srpr-1.2* | WBGene00045168 | ncRNA_gene | -2.0 | 0.2464 | 5.0E-02 | -2.9 | 0.135 | 1.1E-03 |
| *T05A7.1* | WBGene00020219 | protein_coding_gene | -2.0 | 0.2460 | 2.0E-21 | -1.7 | 0.303 | 1.3E-31 |
| *acly-2* | WBGene00007150 | protein_coding_gene | -2.0 | 0.2460 | 4.1E-25 | -1.8 | 0.288 | 1.1E-61 |
| *B0041.5* | WBGene00015009 | protein_coding_gene | -2.0 | 0.2459 | 1.0E-40 | -1.4 | 0.368 | 3.1E-36 |
| *flh-1* | WBGene00012435 | protein_coding_gene | -2.0 | 0.2458 | 2.4E-30 | -2.0 | 0.251 | 2.1E-56 |
| *T27F6.10* | WBGene00077508 | protein_coding_gene | -2.0 | 0.2458 | 3.4E-02 | -3.2 | 0.109 | 7.3E-04 |
| ***D1086.3*** | **WBGene00008390** | **protein_coding_gene** | **-2.0** | **0.2458** | **7.0E-04** | **1.3** | **2.466** | **1.9E-03** |
| *ssp-10* | WBGene00006039 | protein_coding_gene | -2.0 | 0.2457 | 1.2E-04 | -2.5 | 0.179 | 1.4E-31 |
| *B0212.3* | WBGene00015037 | protein_coding_gene | -2.0 | 0.2455 | 1.5E-26 | -1.8 | 0.296 | 5.4E-21 |
| *pcyt-1* | WBGene00017241 | protein_coding_gene | -2.0 | 0.2454 | 1.7E-12 | -1.0 | 0.492 | 1.2E-12 |
| *sdz-27* | WBGene00011124 | protein_coding_gene | -2.0 | 0.2452 | 2.9E-20 | -2.2 | 0.213 | 2.0E-56 |
| *F27D4.8* | WBGene00077543 | protein_coding_gene | -2.0 | 0.2451 | 5.1E-15 | -2.4 | 0.191 | 2.2E-26 |
| *C36A4.4* | WBGene00007965 | protein_coding_gene | -2.0 | 0.2449 | 3.9E-33 | -2.0 | 0.255 | 9.3E-65 |
| *impt-1* | WBGene00013122 | protein_coding_gene | -2.0 | 0.2449 | 1.7E-33 | -1.7 | 0.315 | 2.6E-27 |
| *imb-2* | WBGene00002076 | protein_coding_gene | -2.0 | 0.2447 | 1.3E-26 | -1.4 | 0.381 | 1.7E-32 |
| *evl-18* | WBGene00009372 | protein_coding_gene | -2.0 | 0.2447 | 4.7E-24 | -2.1 | 0.231 | 2.7E-68 |
| *F17E9.4* | WBGene00017541 | protein_coding_gene | -2.0 | 0.2440 | 4.4E-14 | -4.0 | 0.063 | 6.7E-110 |
| *C33F10.4* | WBGene00016353 | protein_coding_gene | -2.0 | 0.2440 | 7.6E-23 | -2.1 | 0.234 | 1.8E-40 |
| *Y71F9AL.7* | WBGene00022112 | protein_coding_gene | -2.0 | 0.2440 | 4.6E-06 | -3.3 | 0.099 | 4.7E-16 |
| *vha-7* | WBGene00006916 | protein_coding_gene | -2.0 | 0.2434 | 2.3E-22 | -1.8 | 0.286 | 2.7E-49 |
| *T05B9.1* | WBGene00011464 | protein_coding_gene | -2.0 | 0.2433 | 2.9E-33 | -1.9 | 0.274 | 1.3E-68 |
| *C42C1.12* | WBGene00016590 | protein_coding_gene | -2.0 | 0.2431 | 2.3E-06 | -1.9 | 0.268 | 1.3E-12 |
| *gls-1* | WBGene00007975 | protein_coding_gene | -2.0 | 0.2429 | 1.4E-12 | -1.2 | 0.427 | 5.2E-29 |
| *F30F8.1* | WBGene00009270 | protein_coding_gene | -2.0 | 0.2429 | 5.1E-23 | -1.5 | 0.348 | 1.0E-40 |
| *M01E11.1* | WBGene00019710 | protein_coding_gene | -2.0 | 0.2428 | 1.4E-17 | -1.6 | 0.327 | 6.5E-26 |
| *dsl-3* | WBGene00001105 | protein_coding_gene | -2.0 | 0.2425 | 3.6E-04 | -5.9 | 0.017 | 6.8E-23 |
| *C09G12.5* | WBGene00015655 | protein_coding_gene | -2.0 | 0.2423 | 4.6E-02 | -6.9 | 0.008 | 7.2E-13 |
| *ssq-2* | WBGene00006051 | protein_coding_gene | -2.0 | 0.2422 | 2.0E-03 | -1.2 | 0.429 | 3.7E-17 |
| *C35D10.10* | WBGene00016446 | protein_coding_gene | -2.0 | 0.2422 | 2.3E-31 | -1.6 | 0.341 | 2.8E-40 |
| *Y53C10A.6* | WBGene00013137 | protein_coding_gene | -2.0 | 0.2421 | 1.1E-21 | -1.4 | 0.367 | 1.2E-25 |
| *srgp-1* | WBGene00006406 | protein_coding_gene | -2.0 | 0.2420 | 2.8E-24 | -1.8 | 0.289 | 2.9E-74 |
| *ubxn-4* | WBGene00022703 | protein_coding_gene | -2.0 | 0.2416 | 4.2E-41 | -1.4 | 0.372 | 3.1E-41 |
| *mdt-6* | WBGene00003164 | protein_coding_gene | -2.0 | 0.2415 | 1.8E-31 | -2.0 | 0.255 | 2.6E-49 |
| *Y54E10BR.2* | WBGene00021841 | protein_coding_gene | -2.1 | 0.2414 | 5.4E-35 | -1.5 | 0.360 | 5.1E-25 |
| *btb-19* | WBGene00010194 | protein_coding_gene | -2.1 | 0.2409 | 2.1E-15 | -1.7 | 0.299 | 3.3E-31 |
| *T12F5.2* | WBGene00020467 | protein_coding_gene | -2.1 | 0.2401 | 8.7E-30 | -1.8 | 0.290 | 2.4E-22 |
| *Y47G6A.31* | WBGene00044436 | protein_coding_gene | -2.1 | 0.2401 | 1.4E-07 | -3.9 | 0.066 | 1.8E-18 |
| *cpt-6* | WBGene00020911 | protein_coding_gene | -2.1 | 0.2400 | 8.3E-15 | -1.3 | 0.396 | 6.3E-07 |
| *ptc-1* | WBGene00004208 | protein_coding_gene | -2.1 | 0.2397 | 6.3E-18 | -1.4 | 0.387 | 7.5E-38 |
| *K08F4.3* | WBGene00010678 | protein_coding_gene | -2.1 | 0.2396 | 5.9E-29 | -1.2 | 0.422 | 4.5E-23 |
| *hst-1* | WBGene00002028 | protein_coding_gene | -2.1 | 0.2396 | 9.4E-43 | -1.6 | 0.321 | 5.1E-37 |
| *phf-33* | WBGene00010708 | protein_coding_gene | -2.1 | 0.2394 | 2.8E-02 | -2.5 | 0.175 | 2.0E-03 |
| *ubc-12* | WBGene00006707 | protein_coding_gene | -2.1 | 0.2393 | 5.7E-18 | -1.6 | 0.327 | 4.4E-26 |
| *K11H12.6* | WBGene00019662 | protein_coding_gene | -2.1 | 0.2391 | 3.6E-02 | -2.3 | 0.208 | 4.9E-03 |
| *mbtr-1* | WBGene00021661 | protein_coding_gene | -2.1 | 0.2386 | 1.1E-40 | -1.4 | 0.377 | 4.6E-14 |
| *C41G7.3* | WBGene00008061 | protein_coding_gene | -2.1 | 0.2384 | 1.2E-47 | -1.4 | 0.367 | 2.9E-31 |
| *msrp-6* | WBGene00017058 | protein_coding_gene | -2.1 | 0.2383 | 5.3E-04 | -2.2 | 0.220 | 2.8E-15 |
| *exo-3* | WBGene00001372 | protein_coding_gene | -2.1 | 0.2382 | 1.3E-54 | -1.5 | 0.353 | 1.9E-30 |
| *C55A6.1* | WBGene00008330 | protein_coding_gene | -2.1 | 0.2371 | 7.4E-27 | -1.8 | 0.280 | 6.9E-45 |
| *D1046.2* | WBGene00008363 | protein_coding_gene | -2.1 | 0.2370 | 1.3E-40 | -1.7 | 0.305 | 1.5E-33 |
| *dnj-2* | WBGene00001020 | protein_coding_gene | -2.1 | 0.2369 | 6.0E-20 | -1.3 | 0.404 | 4.1E-22 |
| *F25E2.2* | WBGene00017780 | protein_coding_gene | -2.1 | 0.2369 | 6.9E-33 | -1.7 | 0.311 | 8.9E-13 |
| *T22B2.2* | WBGene00020666 | protein_coding_gene | -2.1 | 0.2367 | 4.6E-02 | -2.9 | 0.134 | 5.6E-03 |
| *F43C11.10* | WBGene00018387 | protein_coding_gene | -2.1 | 0.2356 | 4.0E-02 | -3.7 | 0.074 | 5.6E-05 |
| *F45F2.11* | WBGene00018483 | protein_coding_gene | -2.1 | 0.2354 | 1.1E-20 | -1.5 | 0.349 | 4.3E-31 |
| *F36A4.5* | WBGene00018084 | protein_coding_gene | -2.1 | 0.2350 | 2.1E-05 | -1.7 | 0.301 | 1.9E-03 |
| *fbxb-16* | WBGene00022560 | protein_coding_gene | -2.1 | 0.2349 | 9.0E-03 | -3.9 | 0.066 | 5.6E-05 |
| *T25D3.4* | WBGene00020798 | protein_coding_gene | -2.1 | 0.2347 | 4.1E-38 | -1.4 | 0.379 | 1.2E-35 |
| *snr-7* | WBGene00004920 | protein_coding_gene | -2.1 | 0.2345 | 9.8E-16 | -1.9 | 0.265 | 5.8E-38 |
| *K08C9.1* | WBGene00010650 | protein_coding_gene | -2.1 | 0.2342 | 1.9E-03 | -1.2 | 0.447 | 2.4E-04 |
| *F32D8.5* | WBGene00009330 | protein_coding_gene | -2.1 | 0.2342 | 2.0E-18 | -1.5 | 0.360 | 2.6E-17 |
| *ctf-8* | WBGene00011915 | protein_coding_gene | -2.1 | 0.2341 | 1.1E-20 | -2.5 | 0.175 | 2.8E-21 |
| *F26A1.1* | WBGene00017801 | protein_coding_gene | -2.1 | 0.2340 | 3.2E-37 | -1.8 | 0.283 | 1.7E-44 |
| *C44B12.3* | WBGene00016637 | protein_coding_gene | -2.1 | 0.2340 | 5.1E-03 | -4.4 | 0.048 | 1.5E-14 |
| *msrp-1* | WBGene00017978 | protein_coding_gene | -2.1 | 0.2339 | 1.3E-05 | -1.6 | 0.327 | 3.0E-18 |
| *npp-22* | WBGene00007118 | protein_coding_gene | -2.1 | 0.2336 | 2.7E-32 | -1.9 | 0.273 | 9.9E-60 |
| *F09E10.1* | WBGene00017294 | protein_coding_gene | -2.1 | 0.2333 | 2.1E-08 | -1.9 | 0.274 | 2.2E-14 |
| *F11G11.15* | WBGene00271822 | protein_coding_gene | -2.1 | 0.2332 | 3.7E-04 | -2.8 | 0.146 | 9.2E-04 |
| *air-2* | WBGene00000099 | protein_coding_gene | -2.1 | 0.2327 | 3.4E-37 | -2.0 | 0.250 | 8.9E-71 |
| *pch-2* | WBGene00008641 | protein_coding_gene | -2.1 | 0.2325 | 2.0E-23 | -1.9 | 0.264 | 1.9E-49 |
| *Y52E8A.2* | WBGene00021795 | protein_coding_gene | -2.1 | 0.2324 | 5.4E-18 | -1.7 | 0.311 | 2.5E-21 |
| *T09B4.2* | WBGene00020376 | protein_coding_gene | -2.1 | 0.2323 | 5.6E-22 | -2.0 | 0.255 | 2.6E-49 |
| *D1086.19* | WBGene00045406 | protein_coding_gene | -2.1 | 0.2323 | 3.6E-03 | -2.1 | 0.227 | 3.4E-03 |
| *pign-1* | WBGene00021840 | protein_coding_gene | -2.1 | 0.2321 | 7.7E-33 | -1.3 | 0.411 | 1.5E-18 |
| *C04G2.9* | WBGene00007308 | protein_coding_gene | -2.1 | 0.2321 | 3.3E-05 | -1.2 | 0.427 | 4.4E-15 |
| *Y76B12C.6* | WBGene00022300 | protein_coding_gene | -2.1 | 0.2319 | 6.0E-14 | -1.7 | 0.317 | 2.3E-27 |
| *ZK809.5* | WBGene00014087 | protein_coding_gene | -2.1 | 0.2318 | 1.1E-31 | -1.7 | 0.313 | 4.0E-40 |

| | | | | | | | | |
|---|---|---|---|---|---|---|---|---|
| *phf-31* | WBGene00008747 | protein_coding_gene | -2.1 | 0.2317 | 2.9E-12 | -2.0 | 0.247 | 2.0E-10 |
| *snpc-1.2* | WBGene00012715 | protein_coding_gene | -2.1 | 0.2312 | 6.5E-46 | -1.3 | 0.412 | 6.7E-27 |
| *memb-1* | WBGene00007128 | protein_coding_gene | -2.1 | 0.2311 | 8.6E-12 | -2.0 | 0.244 | 9.8E-17 |
| *mom-1* | WBGene00003394 | protein_coding_gene | -2.1 | 0.2309 | 4.0E-11 | -2.6 | 0.165 | 2.1E-24 |
| *C02F12.8* | WBGene00015357 | protein_coding_gene | -2.1 | 0.2306 | 3.0E-25 | -1.8 | 0.282 | 1.3E-17 |
| *F55G11.2* | WBGene00010123 | protein_coding_gene | -2.1 | 0.2305 | 2.3E-04 | -2.8 | 0.140 | 2.5E-06 |
| *D2096.13* | WBGene00044953 | protein_coding_gene | -2.1 | 0.2304 | 7.7E-04 | -4.8 | 0.037 | 5.0E-07 |
| *cya-1* | WBGene00000863 | protein_coding_gene | -2.1 | 0.2303 | 1.4E-26 | -2.1 | 0.232 | 2.3E-79 |
| *D2030.8* | WBGene00008418 | protein_coding_gene | -2.1 | 0.2301 | 6.0E-30 | -1.6 | 0.329 | 4.0E-34 |
| *scc-1* | WBGene00004737 | protein_coding_gene | -2.1 | 0.2296 | 5.0E-37 | -1.2 | 0.428 | 2.9E-21 |
| *T23F2.5* | WBGene00020738 | protein_coding_gene | -2.1 | 0.2294 | 5.3E-36 | -2.1 | 0.230 | 1.6E-46 |
| *dsl-6* | WBGene00001108 | protein_coding_gene | -2.1 | 0.2285 | 2.4E-02 | -6.7 | 0.010 | 1.9E-23 |
| *metl-9* | WBGene00020199 | protein_coding_gene | -2.1 | 0.2285 | 5.6E-23 | -2.0 | 0.243 | 3.5E-31 |
| ***F46F2.5*** | **WBGene00009789** | **protein_coding_gene** | **-2.1** | **0.2282** | **3.4E-04** | **1.8** | **3.382** | **4.9E-03** |
| *klp-19* | WBGene00002229 | protein_coding_gene | -2.1 | 0.2280 | 6.2E-29 | -1.5 | 0.352 | 1.8E-44 |
| *cest-2.3* | WBGene00013873 | protein_coding_gene | -2.1 | 0.2278 | 1.6E-03 | -4.0 | 0.061 | 5.0E-07 |
| *T09B4.5* | WBGene00020379 | protein_coding_gene | -2.1 | 0.2277 | 1.9E-09 | -4.3 | 0.051 | 1.4E-182 |
| *npp-26* | WBGene00021198 | protein_coding_gene | -2.1 | 0.2273 | 8.7E-04 | -3.1 | 0.119 | 8.0E-06 |
| *C13F10.6* | WBGene00015745 | protein_coding_gene | -2.1 | 0.2272 | 3.9E-28 | -2.1 | 0.233 | 1.9E-59 |
| *mans-4* | WBGene00013881 | protein_coding_gene | -2.1 | 0.2271 | 7.3E-30 | -1.7 | 0.306 | 2.0E-42 |
| *zipt-11* | WBGene00019077 | protein_coding_gene | -2.1 | 0.2268 | 6.9E-32 | -1.4 | 0.380 | 2.4E-32 |
| *set-4* | WBGene00016313 | protein_coding_gene | -2.1 | 0.2265 | 6.6E-32 | -1.7 | 0.304 | 2.7E-47 |
| *ugt-36* | WBGene00017315 | protein_coding_gene | -2.1 | 0.2263 | 2.6E-03 | -3.9 | 0.068 | 1.8E-05 |
| *F25C8.1* | WBGene00009104 | pseudogene | -2.1 | 0.2263 | 3.9E-02 | -3.3 | 0.100 | 1.9E-05 |
| *sqv-7* | WBGene00005025 | protein_coding_gene | -2.1 | 0.2262 | 1.1E-24 | -2.1 | 0.233 | 4.2E-78 |
| *Y53F4B.8* | WBGene00013155 | pseudogene | -2.1 | 0.2262 | 3.4E-03 | -2.8 | 0.139 | 5.3E-09 |
| *msd-4* | WBGene00016447 | protein_coding_gene | -2.1 | 0.2259 | 9.2E-04 | -1.6 | 0.331 | 2.2E-22 |
| *F59A3.12* | WBGene00185086 | protein_coding_gene | -2.1 | 0.2256 | 6.3E-32 | -1.6 | 0.319 | 3.3E-34 |
| *vha-18* | WBGene00018698 | protein_coding_gene | -2.2 | 0.2249 | 3.6E-42 | -1.6 | 0.338 | 1.6E-41 |
| *C29F7.3* | WBGene00007812 | protein_coding_gene | -2.2 | 0.2244 | 1.1E-18 | -1.5 | 0.347 | 4.7E-22 |
| *mogs-1* | WBGene00008775 | protein_coding_gene | -2.2 | 0.2243 | 2.5E-38 | -1.6 | 0.334 | 1.5E-49 |
| *F44E7.9* | WBGene00018431 | protein_coding_gene | -2.2 | 0.2243 | 2.7E-17 | -1.5 | 0.360 | 9.3E-33 |
| *npax-4* | WBGene00007496 | protein_coding_gene | -2.2 | 0.2233 | 1.8E-05 | -1.9 | 0.273 | 3.3E-03 |
| *ced-3* | WBGene00000417 | protein_coding_gene | -2.2 | 0.2227 | 3.9E-14 | -1.9 | 0.271 | 2.0E-71 |
| *T16G12.4* | WBGene00011805 | protein_coding_gene | -2.2 | 0.2225 | 7.9E-30 | -2.5 | 0.180 | 3.2E-59 |
| *dhhc-3* | WBGene00018009 | protein_coding_gene | -2.2 | 0.2220 | 1.8E-29 | -1.5 | 0.363 | 2.0E-33 |
| *F55A11.7* | WBGene00010080 | protein_coding_gene | -2.2 | 0.2219 | 7.7E-37 | -2.1 | 0.238 | 9.3E-45 |
| *Y65B4BL.4* | WBGene00022036 | protein_coding_gene | -2.2 | 0.2216 | 2.2E-23 | -1.2 | 0.447 | 1.8E-09 |
| *Y57G11B.8* | WBGene00045061 | protein_coding_gene | -2.2 | 0.2216 | 3.4E-02 | -2.6 | 0.165 | 9.8E-03 |
| *tbc-20* | WBGene00013196 | protein_coding_gene | -2.2 | 0.2213 | 7.6E-40 | -1.3 | 0.404 | 7.9E-24 |
| *alg-5* | WBGene00011945 | protein_coding_gene | -2.2 | 0.2212 | 9.9E-51 | -2.0 | 0.243 | 1.4E-62 |
| *F58G11.3* | WBGene00010281 | protein_coding_gene | -2.2 | 0.2212 | 4.9E-20 | -1.4 | 0.387 | 2.0E-26 |
| *F56D6.20* | WBGene00235297 | protein_coding_gene | -2.2 | 0.2210 | 6.2E-03 | -1.1 | 0.467 | 3.8E-02 |
| *F56C9.6* | WBGene00018950 | protein_coding_gene | -2.2 | 0.2209 | 1.4E-41 | -1.6 | 0.326 | 3.3E-39 |
| *idh-1* | WBGene00010317 | protein_coding_gene | -2.2 | 0.2208 | 3.6E-18 | -1.3 | 0.396 | 1.7E-19 |
| *M7.9* | WBGene00010885 | protein_coding_gene | -2.2 | 0.2208 | 3.8E-02 | -7.9 | 0.004 | 4.4E-31 |
| *dnj-5* | WBGene00001023 | protein_coding_gene | -2.2 | 0.2203 | 1.6E-26 | -1.7 | 0.308 | 6.6E-56 |
| *ari-1.4* | WBGene00013538 | protein_coding_gene | -2.2 | 0.2199 | 3.1E-56 | -1.9 | 0.261 | 1.3E-46 |
| *mcm-5* | WBGene00003157 | protein_coding_gene | -2.2 | 0.2194 | 8.6E-25 | -2.0 | 0.244 | 1.3E-70 |
| *F56C9.11* | WBGene00018954 | protein_coding_gene | -2.2 | 0.2193 | 1.3E-24 | -1.6 | 0.319 | 1.3E-26 |
| *C49H3.6* | WBGene00016792 | protein_coding_gene | -2.2 | 0.2192 | 1.4E-37 | -1.4 | 0.375 | 7.0E-27 |
| *ZK218.1* | WBGene00013935 | protein_coding_gene | -2.2 | 0.2191 | 3.6E-04 | -1.1 | 0.466 | 3.5E-02 |
| *polh-1* | WBGene00018721 | protein_coding_gene | -2.2 | 0.2189 | 4.7E-69 | -1.6 | 0.334 | 1.2E-35 |
| *hut-1* | WBGene00013740 | protein_coding_gene | -2.2 | 0.2188 | 4.2E-31 | -1.2 | 0.439 | 2.2E-15 |
| *bath-30* | WBGene00017460 | protein_coding_gene | -2.2 | 0.2185 | 1.1E-49 | -1.5 | 0.346 | 3.6E-07 |
| *cyb-1* | WBGene00000865 | protein_coding_gene | -2.2 | 0.2181 | 1.4E-47 | -1.5 | 0.350 | 3.8E-36 |
| *F53F1.2* | WBGene00009980 | protein_coding_gene | -2.2 | 0.2179 | 1.2E-09 | -1.1 | 0.479 | 3.2E-10 |
| *col-74* | WBGene00000650 | protein_coding_gene | -2.2 | 0.2178 | 1.3E-02 | -5.2 | 0.028 | 1.0E-26 |
| *B0041.8* | WBGene00015011 | protein_coding_gene | -2.2 | 0.2178 | 8.9E-26 | -2.0 | 0.253 | 2.3E-59 |
| *F11A10.5* | WBGene00008686 | protein_coding_gene | -2.2 | 0.2176 | 7.6E-41 | -1.8 | 0.288 | 2.1E-55 |
| *T05E12.8* | WBGene00045382 | protein_coding_gene | -2.2 | 0.2173 | 5.8E-04 | -1.9 | 0.260 | 1.3E-02 |
| *F19H6.5* | WBGene00008960 | protein_coding_gene | -2.2 | 0.2173 | 4.5E-02 | -4.2 | 0.056 | 3.5E-06 |
| *cept-2* | WBGene00013024 | protein_coding_gene | -2.2 | 0.2172 | 4.4E-27 | -1.5 | 0.359 | 3.8E-40 |
| *Y59A8B.21* | WBGene00013353 | protein_coding_gene | -2.2 | 0.2171 | 5.0E-26 | -1.1 | 0.464 | 1.6E-13 |
| *algn-12* | WBGene00022629 | protein_coding_gene | -2.2 | 0.2168 | 2.3E-34 | -1.7 | 0.317 | 7.5E-40 |
| *Y53C12A.3* | WBGene00013139 | protein_coding_gene | -2.2 | 0.2166 | 2.9E-24 | -1.5 | 0.360 | 1.2E-23 |
| *kbp-1* | WBGene00020064 | protein_coding_gene | -2.2 | 0.2164 | 6.3E-17 | -2.1 | 0.226 | 3.6E-36 |
| *pig-1* | WBGene00021012 | protein_coding_gene | -2.2 | 0.2163 | 5.8E-35 | -2.1 | 0.227 | 1.2E-55 |
| *Y53C12A.6* | WBGene00013141 | protein_coding_gene | -2.2 | 0.2161 | 2.4E-08 | -1.0 | 0.486 | 4.7E-03 |
| *C08F8.3* | WBGene00007445 | protein_coding_gene | -2.2 | 0.2158 | 1.5E-42 | -1.7 | 0.297 | 4.4E-41 |
| *har-2* | WBGene00011988 | protein_coding_gene | -2.2 | 0.2156 | 6.8E-44 | -2.5 | 0.173 | 9.0E-92 |
| *msd-3* | WBGene00009684 | protein_coding_gene | -2.2 | 0.2156 | 2.9E-02 | -2.5 | 0.174 | 1.9E-03 |
| *aptf-3* | WBGene00009203 | protein_coding_gene | -2.2 | 0.2156 | 1.7E-28 | -1.9 | 0.267 | 3.8E-31 |
| *unc-59* | WBGene00006793 | protein_coding_gene | -2.2 | 0.2156 | 2.3E-38 | -2.1 | 0.237 | 9.8E-78 |
| *Y20C6A.4* | WBGene00077495 | protein_coding_gene | -2.2 | 0.2155 | 5.0E-02 | -5.9 | 0.016 | 4.4E-11 |
| *T16A1.10* | WBGene00269431 | protein_coding_gene | -2.2 | 0.2153 | 2.4E-02 | -2.6 | 0.169 | 2.2E-04 |
| *F46F2.1* | WBGene00009786 | pseudogene | -2.2 | 0.2151 | 6.8E-03 | -2.0 | 0.256 | 1.7E-02 |
| *C32B5.18* | WBGene00235351 | protein_coding_gene | -2.2 | 0.2149 | 3.3E-02 | -4.4 | 0.047 | 3.0E-06 |
| *pdp-1* | WBGene00022832 | protein_coding_gene | -2.2 | 0.2149 | 4.4E-74 | -1.6 | 0.331 | 1.1E-30 |
| *F48A11.4* | WBGene00018585 | protein_coding_gene | -2.2 | 0.2144 | 4.0E-34 | -1.6 | 0.337 | 7.4E-28 |

| | | | | | | | | |
|---|---|---|---|---|---|---|---|---|
| *F55B12.11* | WBGene00077570 | pseudogene | -2.2 | 0.2143 | 3.0E-03 | -2.4 | 0.183 | 4.9E-07 |
| *plk-1* | WBGene00004042 | protein_coding_gene | -2.2 | 0.2136 | 8.3E-24 | -1.8 | 0.292 | 2.2E-53 |
| *arid-1* | WBGene00044689 | protein_coding_gene | -2.2 | 0.2135 | 3.7E-17 | -1.6 | 0.339 | 1.2E-47 |
| *csb-1* | WBGene00000803 | protein_coding_gene | -2.2 | 0.2135 | 4.5E-17 | -2.0 | 0.249 | 7.1E-24 |
| *cee-1* | WBGene00017825 | protein_coding_gene | -2.2 | 0.2133 | 2.1E-33 | -1.7 | 0.307 | 7.0E-42 |
| *C17E7.13* | WBGene00015906 | protein_coding_gene | -2.2 | 0.2131 | 2.3E-02 | -4.6 | 0.041 | 9.5E-07 |
| *ooc-5* | WBGene00003870 | protein_coding_gene | -2.2 | 0.2130 | 4.3E-33 | -1.9 | 0.268 | 2.0E-56 |
| *Y38E10A.34* | WBGene00304775 | protein_coding_gene | -2.2 | 0.2128 | 3.6E-02 | -2.1 | 0.225 | 4.1E-02 |
| *T18D3.1* | WBGene00011820 | protein_coding_gene | -2.2 | 0.2127 | 2.3E-19 | -2.4 | 0.195 | 1.3E-38 |
| *R04A9.9* | WBGene00195183 | protein_coding_gene | -2.2 | 0.2126 | 4.5E-06 | -1.2 | 0.429 | 1.2E-03 |
| *Y62E10A.2* | WBGene00013373 | protein_coding_gene | -2.2 | 0.2123 | 2.4E-11 | -1.6 | 0.328 | 3.9E-05 |
| *Y106G6H.6* | WBGene00013719 | protein_coding_gene | -2.2 | 0.2122 | 2.0E-29 | -1.9 | 0.270 | 7.5E-43 |
| *ceh-21* | WBGene00000444 | protein_coding_gene | -2.2 | 0.2121 | 1.2E-25 | -1.9 | 0.262 | 1.1E-20 |
| *fbxa-153* | WBGene00007162 | protein_coding_gene | -2.2 | 0.2118 | 2.0E-10 | -1.3 | 0.407 | 8.6E-08 |
| *plin-1* | WBGene00007024 | protein_coding_gene | -2.2 | 0.2116 | 1.8E-31 | -1.9 | 0.264 | 5.5E-52 |
| *sdz-23* | WBGene00019066 | protein_coding_gene | -2.2 | 0.2113 | 5.0E-02 | -2.5 | 0.183 | 2.2E-02 |
| *hop-1* | WBGene00001985 | protein_coding_gene | -2.2 | 0.2108 | 3.5E-51 | -1.9 | 0.268 | 4.8E-36 |
| *F28B3.5* | WBGene00017886 | protein_coding_gene | -2.2 | 0.2105 | 1.8E-31 | -1.3 | 0.406 | 9.7E-24 |
| *F25B4.4* | WBGene00017767 | protein_coding_gene | -2.3 | 0.2100 | 1.3E-06 | -1.7 | 0.315 | 2.5E-08 |
| *pqn-10* | WBGene00004102 | protein_coding_gene | -2.3 | 0.2099 | 2.9E-03 | -2.1 | 0.233 | 9.3E-03 |
| *cgt-3* | WBGene00019127 | protein_coding_gene | -2.3 | 0.2097 | 2.8E-76 | -1.5 | 0.354 | 1.7E-48 |
| *patr-1* | WBGene00009661 | protein_coding_gene | -2.3 | 0.2096 | 1.3E-24 | -1.7 | 0.312 | 3.8E-54 |
| *fbxa-104* | WBGene00011542 | protein_coding_gene | -2.3 | 0.2095 | 1.3E-02 | -2.4 | 0.184 | 6.7E-04 |
| *zipt-1* | WBGene00009286 | protein_coding_gene | -2.3 | 0.2095 | 1.6E-62 | -2.1 | 0.233 | 8.3E-95 |
| *T15D6.12* | WBGene00011786 | protein_coding_gene | -2.3 | 0.2091 | 3.0E-02 | -2.2 | 0.219 | 2.9E-02 |
| *msp-38* | WBGene00003434 | protein_coding_gene | -2.3 | 0.2089 | 1.4E-04 | -3.0 | 0.125 | 2.1E-34 |
| *K04H8.3* | WBGene00010578 | protein_coding_gene | -2.3 | 0.2089 | 4.0E-02 | -5.9 | 0.016 | 8.4E-11 |
| *his-68* | WBGene00001942 | protein_coding_gene | -2.3 | 0.2086 | 3.1E-14 | -1.5 | 0.344 | 3.6E-08 |
| *fbxb-66* | WBGene00044066 | protein_coding_gene | -2.3 | 0.2085 | 1.1E-05 | -3.9 | 0.067 | 6.3E-23 |
| *ech-6* | WBGene00001155 | protein_coding_gene | -2.3 | 0.2085 | 3.8E-22 | -1.5 | 0.363 | 4.1E-09 |
| *W06A7.4* | WBGene00012293 | protein_coding_gene | -2.3 | 0.2083 | 6.8E-11 | -1.2 | 0.445 | 1.3E-03 |
| *fbxc-15* | WBGene00016297 | protein_coding_gene | -2.3 | 0.2078 | 3.6E-02 | -2.8 | 0.139 | 5.0E-03 |
| *tpxl-1* | WBGene00021470 | protein_coding_gene | -2.3 | 0.2075 | 1.6E-15 | -1.4 | 0.370 | 6.5E-30 |
| *mtrr-1* | WBGene00006510 | protein_coding_gene | -2.3 | 0.2071 | 8.5E-49 | -1.6 | 0.324 | 4.1E-43 |
| *msp-45* | WBGene00003438 | protein_coding_gene | -2.3 | 0.2068 | 6.3E-06 | -1.8 | 0.286 | 9.7E-27 |
| *srpr-2.1* | WBGene00195016 | ncRNA_gene | -2.3 | 0.2067 | 7.7E-03 | -2.6 | 0.160 | 2.1E-03 |
| *R07G3.7* | WBGene00019943 | protein_coding_gene | -2.3 | 0.2066 | 9.2E-43 | -1.7 | 0.313 | 5.6E-49 |
| *tbb-1* | WBGene00006536 | protein_coding_gene | -2.3 | 0.2065 | 1.4E-33 | -1.6 | 0.333 | 4.1E-46 |
| *F52D2.6* | WBGene00018688 | protein_coding_gene | -2.3 | 0.2062 | 8.3E-03 | -1.6 | 0.336 | 2.8E-02 |
| *spe-39* | WBGene00004975 | protein_coding_gene | -2.3 | 0.2058 | 1.2E-34 | -1.3 | 0.404 | 1.6E-26 |
| *C56E6.2* | WBGene00016970 | protein_coding_gene | -2.3 | 0.2050 | 3.9E-25 | -2.5 | 0.180 | 7.2E-41 |
| *K03H1.7* | WBGene00010542 | protein_coding_gene | -2.3 | 0.2048 | 7.9E-16 | -2.3 | 0.210 | 1.8E-31 |
| *C52D10.10* | WBGene00023259 | pseudogene | -2.3 | 0.2047 | 1.3E-02 | -2.8 | 0.139 | 1.0E-02 |
| *clec-94* | WBGene00013928 | protein_coding_gene | -2.3 | 0.2046 | 2.5E-02 | -2.8 | 0.145 | 8.8E-03 |
| *hcp-1* | WBGene00001829 | protein_coding_gene | -2.3 | 0.2038 | 3.5E-31 | -1.6 | 0.335 | 9.1E-42 |
| *ima-1* | WBGene00002072 | protein_coding_gene | -2.3 | 0.2035 | 8.3E-44 | -1.8 | 0.290 | 4.9E-52 |
| *ZK666.2* | WBGene00014042 | protein_coding_gene | -2.3 | 0.2033 | 3.3E-02 | -2.0 | 0.247 | 3.5E-02 |
| *egl-18* | WBGene00001186 | protein_coding_gene | -2.3 | 0.2033 | 6.5E-14 | -2.7 | 0.159 | 6.1E-70 |
| *F54E12.2* | WBGene00010061 | protein_coding_gene | -2.3 | 0.2026 | 4.9E-41 | -2.1 | 0.239 | 1.7E-85 |
| *ifet-1* | WBGene00004132 | protein_coding_gene | -2.3 | 0.2025 | 1.3E-15 | -1.8 | 0.284 | 3.3E-74 |
| *tcc-1* | WBGene00013340 | protein_coding_gene | -2.3 | 0.2022 | 6.4E-28 | -1.7 | 0.318 | 7.0E-52 |
| *zipt-2.4* | WBGene00018897 | protein_coding_gene | -2.3 | 0.2021 | 7.5E-25 | -1.6 | 0.335 | 4.8E-35 |
| *bath-41* | WBGene00018137 | protein_coding_gene | -2.3 | 0.2019 | 2.2E-33 | -1.9 | 0.273 | 2.4E-57 |
| *his-63* | WBGene00001937 | protein_coding_gene | -2.3 | 0.2017 | 2.2E-02 | -3.5 | 0.087 | 1.9E-04 |
| *C14E2.11* | WBGene00206378 | protein_coding_gene | -2.3 | 0.2017 | 2.6E-02 | -3.5 | 0.091 | 7.0E-04 |
| *W04A8.6* | WBGene00012241 | protein_coding_gene | -2.3 | 0.2015 | 1.3E-52 | -2.0 | 0.248 | 9.8E-55 |
| *nos-1* | WBGene00003783 | protein_coding_gene | -2.3 | 0.2013 | 1.6E-29 | -1.9 | 0.271 | 5.7E-18 |
| *mel-32* | WBGene00003214 | protein_coding_gene | -2.3 | 0.2013 | 1.1E-21 | -1.4 | 0.373 | 7.4E-21 |
| *K07H8.7* | WBGene00019507 | protein_coding_gene | -2.3 | 0.2010 | 1.9E-04 | -2.2 | 0.212 | 2.3E-05 |
| *Y105C5A.1271* | WBGene00194667 | pseudogene | -2.3 | 0.2007 | 2.6E-02 | -2.2 | 0.223 | 2.4E-02 |
| *dod-24* | WBGene00007875 | protein_coding_gene | -2.3 | 0.2006 | 3.6E-03 | -4.3 | 0.049 | 1.1E-129 |
| *fbxb-85* | WBGene00011267 | protein_coding_gene | -2.3 | 0.2000 | 9.6E-05 | -2.1 | 0.237 | 1.1E-05 |
| *F33H2.2* | WBGene00009366 | protein_coding_gene | -2.3 | 0.1998 | 4.0E-56 | -1.9 | 0.263 | 9.5E-53 |
| *W09C3.7* | WBGene00021114 | protein_coding_gene | -2.3 | 0.1985 | 1.1E-13 | -1.3 | 0.402 | 6.6E-15 |
| *ckb-2* | WBGene00000512 | protein_coding_gene | -2.3 | 0.1984 | 3.6E-10 | -1.1 | 0.464 | 1.1E-07 |
| *blos-4* | WBGene00020783 | protein_coding_gene | -2.3 | 0.1983 | 5.5E-12 | -2.4 | 0.183 | 1.6E-33 |
| *pfn-1* | WBGene00003989 | protein_coding_gene | -2.3 | 0.1983 | 1.3E-39 | -1.6 | 0.341 | 4.0E-29 |
| *D2005.7* | WBGene00086551 | protein_coding_gene | -2.3 | 0.1966 | 2.5E-38 | -1.5 | 0.345 | 7.1E-21 |
| *polk-1* | WBGene00017696 | protein_coding_gene | -2.3 | 0.1965 | 1.1E-40 | -2.4 | 0.193 | 1.1E-69 |
| *cdd-2* | WBGene00000392 | protein_coding_gene | -2.3 | 0.1963 | 6.3E-31 | -2.0 | 0.245 | 7.9E-32 |
| *chin-1* | WBGene00015267 | protein_coding_gene | -2.3 | 0.1962 | 3.0E-51 | -1.8 | 0.288 | 1.2E-39 |
| *K07H8.2* | WBGene00019504 | protein_coding_gene | -2.4 | 0.1960 | 6.5E-38 | -1.7 | 0.308 | 3.7E-56 |
| *B0238.9* | WBGene00015073 | protein_coding_gene | -2.4 | 0.1959 | 2.0E-23 | -1.5 | 0.347 | 3.9E-37 |
| *mig-14* | WBGene00003246 | protein_coding_gene | -2.4 | 0.1953 | 2.1E-27 | -1.8 | 0.290 | 1.7E-35 |
| *ces-1* | WBGene00000468 | protein_coding_gene | -2.4 | 0.1952 | 7.9E-03 | -4.3 | 0.050 | 1.4E-08 |
| *Y75B12A.2* | WBGene00013566 | protein_coding_gene | -2.4 | 0.1951 | 3.9E-02 | -2.5 | 0.180 | 1.5E-02 |
| *Y71F9B.15* | WBGene00022133 | protein_coding_gene | -2.4 | 0.1951 | 1.5E-03 | -2.0 | 0.258 | 2.4E-03 |
| *F38A5.22* | WBGene00235391 | protein_coding_gene | -2.4 | 0.1950 | 3.6E-09 | -3.4 | 0.094 | 1.4E-20 |
| *T24C4.5* | WBGene00020761 | protein_coding_gene | -2.4 | 0.1949 | 1.2E-59 | -1.9 | 0.261 | 4.3E-49 |
| *Y106G6H.16* | WBGene00013726 | protein_coding_gene | -2.4 | 0.1949 | 1.9E-06 | -2.2 | 0.217 | 2.8E-06 |

| | | | | | | | | |
|---|---|---|---|---|---|---|---|---|
| *C08A9.10* | WBGene00044773 | protein_coding_gene | -2.4 | 0.1947 | 4.3E-09 | -5.2 | 0.028 | 2.4E-53 |
| *F15B10.3* | WBGene00017481 | protein_coding_gene | -2.4 | 0.1947 | 1.3E-09 | -2.4 | 0.186 | 2.1E-27 |
| *T13B5.9* | WBGene00020476 | protein_coding_gene | -2.4 | 0.1944 | 3.5E-06 | -2.9 | 0.131 | 2.7E-08 |
| *lrr-1* | WBGene00018016 | protein_coding_gene | -2.4 | 0.1939 | 3.3E-45 | -2.1 | 0.237 | 1.2E-62 |
| *Y11D7A.3* | WBGene00012428 | protein_coding_gene | -2.4 | 0.1938 | 1.4E-17 | -1.7 | 0.310 | 8.6E-25 |
| *lin-3* | WBGene00002992 | protein_coding_gene | -2.4 | 0.1935 | 3.1E-26 | -2.0 | 0.250 | 6.2E-46 |
| *ssq-1* | WBGene00006050 | protein_coding_gene | -2.4 | 0.1935 | 6.1E-06 | -1.3 | 0.394 | 2.5E-13 |
| *clec-207* | WBGene00018096 | protein_coding_gene | -2.4 | 0.1935 | 2.7E-02 | -7.8 | 0.004 | 2.2E-18 |
| *bcl-7* | WBGene00016192 | protein_coding_gene | -2.4 | 0.1934 | 1.8E-29 | -2.5 | 0.178 | 1.7E-62 |
| *cyp-13A9* | WBGene00014838 | pseudogene | -2.4 | 0.1934 | 2.4E-05 | -3.2 | 0.109 | 2.7E-06 |
| *snr-2* | WBGene00004915 | protein_coding_gene | -2.4 | 0.1932 | 9.3E-61 | -1.6 | 0.321 | 1.4E-55 |
| *rbg-3* | WBGene00007167 | protein_coding_gene | -2.4 | 0.1928 | 1.9E-32 | -1.9 | 0.263 | 4.1E-69 |
| *ztf-1* | WBGene00018833 | protein_coding_gene | -2.4 | 0.1920 | 6.8E-22 | -1.8 | 0.289 | 1.4E-21 |
| *C48B4.11* | WBGene00008174 | protein_coding_gene | -2.4 | 0.1915 | 1.1E-37 | -1.7 | 0.300 | 9.3E-40 |
| *ZK546.19* | WBGene00195246 | protein_coding_gene | -2.4 | 0.1912 | 2.8E-02 | -2.2 | 0.220 | 1.7E-02 |
| *H37A05.4* | WBGene00044200 | protein_coding_gene | -2.4 | 0.1911 | 1.4E-17 | -2.7 | 0.153 | 2.7E-24 |
| *wee-1.1* | WBGene00006938 | protein_coding_gene | -2.4 | 0.1907 | 2.0E-05 | -2.6 | 0.166 | 1.5E-05 |
| *ZC434.8* | WBGene00013894 | protein_coding_gene | -2.4 | 0.1905 | 2.3E-16 | -1.6 | 0.328 | 9.5E-41 |
| *cdc-7* | WBGene00016421 | protein_coding_gene | -2.4 | 0.1903 | 1.0E-29 | -2.2 | 0.223 | 1.7E-69 |
| *T13F2.6* | WBGene00011746 | protein_coding_gene | -2.4 | 0.1903 | 4.3E-34 | -1.9 | 0.276 | 5.4E-62 |
| *F31D5.7* | WBGene00206414 | protein_coding_gene | -2.4 | 0.1901 | 2.8E-03 | -3.5 | 0.090 | 6.7E-06 |
| *orc-1* | WBGene00012650 | protein_coding_gene | -2.4 | 0.1898 | 1.7E-56 | -2.0 | 0.249 | 3.6E-67 |
| *srd-14* | WBGene00005092 | pseudogene | -2.4 | 0.1898 | 5.9E-07 | -3.1 | 0.118 | 1.7E-06 |
| *C16C10.1* | WBGene00007622 | protein_coding_gene | -2.4 | 0.1890 | 7.6E-50 | -1.4 | 0.375 | 1.0E-23 |
| *C14B1.7* | WBGene00007578 | pseudogene | -2.4 | 0.1886 | 6.4E-40 | -1.4 | 0.376 | 7.8E-21 |
| *rrn-2.1* | WBGene00004567 | rRNA_gene | -2.4 | 0.1886 | 1.3E-02 | -4.1 | 0.059 | 4.0E-07 |
| *tsn-1* | WBGene00006626 | protein_coding_gene | -2.4 | 0.1882 | 4.2E-30 | -1.1 | 0.478 | 1.3E-16 |
| *H34C03.2* | WBGene00019259 | protein_coding_gene | -2.4 | 0.1879 | 1.8E-24 | -1.4 | 0.376 | 4.0E-34 |
| *K02C4.3* | WBGene00010502 | protein_coding_gene | -2.4 | 0.1874 | 8.4E-36 | -1.8 | 0.292 | 1.1E-64 |
| *col-95* | WBGene00000670 | protein_coding_gene | -2.4 | 0.1867 | 2.7E-06 | -2.0 | 0.244 | 3.9E-42 |
| *sqv-6* | WBGene00005024 | protein_coding_gene | -2.4 | 0.1865 | 2.0E-34 | -1.7 | 0.315 | 2.0E-32 |
| *set-13* | WBGene00019690 | protein_coding_gene | -2.4 | 0.1862 | 4.7E-05 | -1.5 | 0.362 | 3.1E-03 |
| *C04A11.2* | WBGene00007286 | protein_coding_gene | -2.4 | 0.1862 | 8.3E-16 | -4.0 | 0.065 | 2.6E-25 |
| *pgap-1* | WBGene00011835 | protein_coding_gene | -2.4 | 0.1860 | 8.6E-48 | -1.8 | 0.278 | 4.4E-48 |
| *fsn-1* | WBGene00001499 | protein_coding_gene | -2.4 | 0.1860 | 9.9E-34 | -1.9 | 0.267 | 1.1E-53 |
| *Y25C1A.14* | WBGene00194985 | protein_coding_gene | -2.4 | 0.1857 | 1.5E-11 | -2.3 | 0.200 | 1.5E-05 |
| *W10G11.2* | WBGene00021135 | protein_coding_gene | -2.4 | 0.1856 | 4.6E-04 | -2.3 | 0.205 | 5.7E-05 |
| *B0393.6* | WBGene00007171 | protein_coding_gene | -2.4 | 0.1856 | 1.3E-16 | -1.7 | 0.315 | 9.4E-27 |
| *act-2* | WBGene00000064 | protein_coding_gene | -2.4 | 0.1853 | 5.0E-23 | -1.7 | 0.308 | 6.5E-32 |
| *otub-3* | WBGene00009007 | protein_coding_gene | -2.4 | 0.1848 | 3.4E-57 | -2.1 | 0.234 | 1.8E-73 |
| *D1086.6* | WBGene00008393 | protein_coding_gene | -2.4 | 0.1848 | 4.7E-17 | -3.6 | 0.080 | 5.5E-107 |
| *lido-4* | WBGene00015589 | protein_coding_gene | -2.4 | 0.1848 | 8.0E-05 | -2.1 | 0.226 | 2.9E-04 |
| *perm-5* | WBGene00016955 | protein_coding_gene | -2.4 | 0.1845 | 1.9E-36 | -2.1 | 0.238 | 1.2E-71 |
| *fkh-2* | WBGene00001434 | protein_coding_gene | -2.4 | 0.1843 | 2.6E-06 | -3.6 | 0.082 | 1.3E-18 |
| *bath-27* | WBGene00017450 | protein_coding_gene | -2.4 | 0.1841 | 4.3E-16 | -2.0 | 0.255 | 2.1E-02 |
| *F43G6.3* | WBGene00009656 | pseudogene | -2.4 | 0.1835 | 2.4E-38 | -2.7 | 0.155 | 7.5E-43 |
| *W10D9.2* | WBGene00021130 | pseudogene | -2.4 | 0.1832 | 2.0E-04 | -3.7 | 0.077 | 4.5E-05 |
| *fitm-2* | WBGene00044094 | protein_coding_gene | -2.5 | 0.1829 | 9.2E-14 | -1.2 | 0.438 | 5.2E-21 |
| *pie-1* | WBGene00004027 | protein_coding_gene | -2.5 | 0.1827 | 6.7E-25 | -2.0 | 0.249 | 6.1E-36 |
| *sdz-19* | WBGene00009728 | protein_coding_gene | -2.5 | 0.1822 | 1.1E-02 | -2.4 | 0.191 | 1.6E-02 |
| *car-1* | WBGene00012484 | protein_coding_gene | -2.5 | 0.1817 | 2.5E-33 | -1.8 | 0.283 | 2.6E-61 |
| *cut-3* | WBGene00009041 | protein_coding_gene | -2.5 | 0.1809 | 6.1E-03 | -4.9 | 0.033 | 2.3E-23 |
| *kbp-5* | WBGene00016387 | protein_coding_gene | -2.5 | 0.1808 | 4.1E-38 | -2.2 | 0.212 | 1.2E-61 |
| *ZC155.4* | WBGene00022532 | protein_coding_gene | -2.5 | 0.1808 | 1.1E-29 | -2.5 | 0.182 | 2.7E-97 |
| *F23A7.8* | WBGene00044638 | protein_coding_gene | -2.5 | 0.1807 | 6.8E-17 | -2.7 | 0.156 | 1.0E-71 |
| *rsa-1* | WBGene00007710 | protein_coding_gene | -2.5 | 0.1806 | 4.2E-41 | -2.6 | 0.162 | 9.2E-107 |
| *ZK643.2* | WBGene00014034 | protein_coding_gene | -2.5 | 0.1804 | 9.5E-27 | -2.4 | 0.194 | 1.3E-43 |
| *cdk-1* | WBGene00000405 | protein_coding_gene | -2.5 | 0.1795 | 6.5E-31 | -2.2 | 0.219 | 1.2E-89 |
| *swah-1* | WBGene00007985 | protein_coding_gene | -2.5 | 0.1792 | 5.1E-34 | -2.1 | 0.230 | 1.1E-42 |
| *C42C1.8* | WBGene00016586 | protein_coding_gene | -2.5 | 0.1791 | 1.4E-32 | -2.0 | 0.252 | 1.7E-63 |
| *F07H5.10* | WBGene00008561 | protein_coding_gene | -2.5 | 0.1790 | 2.5E-26 | -1.7 | 0.298 | 5.1E-49 |
| *fbxc-40* | WBGene00016295 | protein_coding_gene | -2.5 | 0.1790 | 1.7E-07 | -2.3 | 0.205 | 5.4E-06 |
| *nas-17* | WBGene00003536 | protein_coding_gene | -2.5 | 0.1788 | 3.8E-02 | -7.4 | 0.006 | 1.4E-15 |
| *rrn-3.56* | WBGene00004677 | pseudogene | -2.5 | 0.1785 | 8.7E-03 | -3.1 | 0.118 | 2.0E-04 |
| *mcm-2* | WBGene00003154 | protein_coding_gene | -2.5 | 0.1782 | 1.9E-32 | -2.1 | 0.240 | 7.9E-60 |
| *gla-3* | WBGene00011376 | protein_coding_gene | -2.5 | 0.1778 | 1.7E-40 | -1.5 | 0.363 | 2.7E-41 |
| *R01H10.7* | WBGene00010975 | protein_coding_gene | -2.5 | 0.1775 | 3.3E-54 | -1.6 | 0.328 | 8.6E-31 |
| *R05H5.3* | WBGene00011038 | protein_coding_gene | -2.5 | 0.1772 | 2.3E-32 | -2.6 | 0.169 | 5.0E-104 |
| *sup-17* | WBGene00006324 | protein_coding_gene | -2.5 | 0.1769 | 1.4E-26 | -2.0 | 0.252 | 4.7E-69 |
| *K10C2.7* | WBGene00019622 | protein_coding_gene | -2.5 | 0.1767 | 3.2E-06 | -1.6 | 0.333 | 1.4E-02 |
| *F30A10.15* | WBGene00195244 | protein_coding_gene | -2.5 | 0.1766 | 2.9E-04 | -2.3 | 0.207 | 2.2E-02 |
| *cmt-1* | WBGene00016564 | protein_coding_gene | -2.5 | 0.1761 | 4.9E-42 | -2.5 | 0.180 | 3.4E-104 |
| *F23D12.11* | WBGene00194674 | protein_coding_gene | -2.5 | 0.1761 | 6.6E-08 | -1.1 | 0.451 | 2.9E-04 |
| *pup-2* | WBGene00019628 | protein_coding_gene | -2.5 | 0.1753 | 7.8E-17 | -2.0 | 0.250 | 1.1E-68 |
| *msp-50* | WBGene00003443 | protein_coding_gene | -2.5 | 0.1748 | 6.6E-08 | -1.5 | 0.349 | 1.8E-19 |
| *Y116A8C.44* | WBGene00013815 | protein_coding_gene | -2.5 | 0.1746 | 1.5E-02 | -5.7 | 0.019 | 4.1E-10 |
| *ifa-3* | WBGene00002051 | protein_coding_gene | -2.5 | 0.1746 | 5.1E-04 | -6.1 | 0.014 | 1.2E-24 |
| *R148.4* | WBGene00020103 | protein_coding_gene | -2.5 | 0.1743 | 3.4E-60 | -1.5 | 0.362 | 2.8E-20 |
| *zif-1* | WBGene00006977 | protein_coding_gene | -2.5 | 0.1734 | 1.4E-35 | -2.1 | 0.236 | 2.7E-50 |
| *ZC449.8* | WBGene00195189 | protein_coding_gene | -2.5 | 0.1731 | 3.6E-13 | -1.4 | 0.376 | 2.8E-03 |

| | | | | | | | | |
|---|---|---|---|---|---|---|---|---|
| *capg-2* | WBGene00010093 | protein_coding_gene | -2.5 | 0.1723 | 8.4E-37 | -2.0 | 0.256 | 1.7E-56 |
| *fbxb-65* | WBGene00008066 | protein_coding_gene | -2.5 | 0.1720 | 2.8E-04 | -4.9 | 0.034 | 3.7E-07 |
| *fbxb-99* | WBGene00021174 | protein_coding_gene | -2.5 | 0.1712 | 9.5E-03 | -3.8 | 0.072 | 5.9E-05 |
| *F45D3.1* | WBGene00009721 | protein_coding_gene | -2.6 | 0.1708 | 1.9E-04 | -4.2 | 0.054 | 1.6E-07 |
| *wdr-48* | WBGene00009441 | protein_coding_gene | -2.6 | 0.1705 | 3.5E-42 | -1.8 | 0.296 | 1.1E-39 |
| *F42A9.6* | WBGene00018335 | protein_coding_gene | -2.6 | 0.1697 | 3.6E-14 | -1.3 | 0.408 | 1.4E-11 |
| *T21B4.21* | WBGene00220076 | ncRNA_gene | -2.6 | 0.1697 | 3.2E-06 | -4.3 | 0.051 | 4.3E-21 |
| *C06E2.5* | WBGene00015529 | protein_coding_gene | -2.6 | 0.1693 | 1.1E-18 | -3.3 | 0.102 | 1.2E-31 |
| *F52B5.2* | WBGene00009921 | protein_coding_gene | -2.6 | 0.1682 | 1.0E-48 | -2.0 | 0.254 | 5.6E-69 |
| *C50B6.3* | WBGene00008219 | protein_coding_gene | -2.6 | 0.1682 | 4.5E-63 | -2.4 | 0.189 | 6.4E-96 |
| *K10C2.12* | WBGene00206386 | protein_coding_gene | -2.6 | 0.1681 | 4.1E-05 | -1.9 | 0.277 | 8.0E-03 |
| *C07D8.6* | WBGene00015565 | protein_coding_gene | -2.6 | 0.1679 | 2.7E-30 | -2.4 | 0.191 | 3.2E-60 |
| *Y22D7AL.10* | WBGene00021248 | protein_coding_gene | -2.6 | 0.1678 | 6.5E-29 | -1.6 | 0.334 | 2.6E-43 |
| *lsm-1* | WBGene00003076 | protein_coding_gene | -2.6 | 0.1677 | 6.1E-52 | -2.4 | 0.194 | 2.5E-76 |
| *col-184* | WBGene00000757 | protein_coding_gene | -2.6 | 0.1666 | 1.6E-23 | -3.3 | 0.102 | 2.2E-159 |
| *C10A4.1* | WBGene00015662 | protein_coding_gene | -2.6 | 0.1665 | 1.0E-03 | -3.1 | 0.114 | 7.0E-06 |
| *ugt-40* | WBGene00017331 | protein_coding_gene | -2.6 | 0.1661 | 1.5E-09 | -2.0 | 0.250 | 2.0E-05 |
| *D1086.11* | WBGene00044081 | protein_coding_gene | -2.6 | 0.1661 | 3.0E-28 | -4.1 | 0.058 | 6.9E-186 |
| *mig-5* | WBGene00003241 | protein_coding_gene | -2.6 | 0.1660 | 1.4E-38 | -2.0 | 0.245 | 4.3E-61 |
| *sqv-1* | WBGene00005019 | protein_coding_gene | -2.6 | 0.1658 | 9.3E-68 | -2.1 | 0.238 | 2.6E-93 |
| *F23D12.10* | WBGene00045291 | protein_coding_gene | -2.6 | 0.1657 | 1.4E-14 | -3.5 | 0.088 | 5.8E-17 |
| *hxk-1* | WBGene00008780 | protein_coding_gene | -2.6 | 0.1656 | 2.3E-47 | -2.2 | 0.224 | 4.3E-92 |
| *mcm-3* | WBGene00003155 | protein_coding_gene | -2.6 | 0.1655 | 1.3E-30 | -2.3 | 0.202 | 1.6E-81 |
| *gld-1* | WBGene00001595 | protein_coding_gene | -2.6 | 0.1653 | 2.4E-21 | -1.5 | 0.347 | 6.4E-58 |
| *F11E6.7* | WBGene00008710 | protein_coding_gene | -2.6 | 0.1652 | 9.4E-36 | -2.5 | 0.180 | 1.1E-83 |
| *adm-4* | WBGene00000075 | protein_coding_gene | -2.6 | 0.1651 | 2.2E-39 | -1.7 | 0.302 | 3.2E-20 |
| *C30F12.4* | WBGene00016263 | protein_coding_gene | -2.6 | 0.1650 | 1.5E-42 | -2.0 | 0.250 | 3.3E-52 |
| *B0563.8* | WBGene00015265 | protein_coding_gene | -2.6 | 0.1648 | 1.1E-02 | -3.5 | 0.089 | 5.2E-04 |
| *C05B5.11* | WBGene00045242 | protein_coding_gene | -2.6 | 0.1645 | 1.6E-02 | -7.1 | 0.007 | 2.8E-15 |
| *air-1* | WBGene00000098 | protein_coding_gene | -2.6 | 0.1645 | 4.1E-32 | -2.0 | 0.243 | 1.1E-78 |
| *cutc-1* | WBGene00022702 | protein_coding_gene | -2.6 | 0.1643 | 2.5E-35 | -2.0 | 0.257 | 3.0E-64 |
| *F53B1.2* | WBGene00018735 | protein_coding_gene | -2.6 | 0.1639 | 4.3E-22 | -2.1 | 0.237 | 4.2E-29 |
| *Y54G9A.5* | WBGene00013208 | protein_coding_gene | -2.6 | 0.1638 | 2.3E-48 | -2.3 | 0.210 | 1.8E-55 |
| *deps-1* | WBGene00022034 | protein_coding_gene | -2.6 | 0.1633 | 8.1E-55 | -1.5 | 0.348 | 2.0E-33 |
| *M117.7* | WBGene00044768 | pseudogene | -2.6 | 0.1632 | 6.2E-04 | -3.0 | 0.123 | 4.0E-05 |
| *cdc-25.2* | WBGene00000387 | protein_coding_gene | -2.6 | 0.1631 | 2.3E-31 | -3.0 | 0.128 | 5.1E-85 |
| *cup-15* | WBGene00009627 | protein_coding_gene | -2.6 | 0.1628 | 5.1E-47 | -2.3 | 0.203 | 2.1E-58 |
| *fbxc-39* | WBGene00018456 | protein_coding_gene | -2.6 | 0.1628 | 2.7E-05 | -1.6 | 0.340 | 2.0E-02 |
| *crn-4* | WBGene00000797 | protein_coding_gene | -2.6 | 0.1627 | 1.7E-28 | -2.1 | 0.239 | 3.7E-21 |
| *F56F11.4* | WBGene00018991 | protein_coding_gene | -2.6 | 0.1624 | 4.7E-47 | -2.5 | 0.172 | 3.2E-78 |
| *xol-1* | WBGene00006962 | protein_coding_gene | -2.6 | 0.1621 | 2.6E-09 | -2.9 | 0.134 | 5.4E-09 |
| *F08F8.4* | WBGene00017269 | protein_coding_gene | -2.6 | 0.1621 | 8.8E-60 | -2.4 | 0.196 | 1.9E-67 |
| *cyp-13A4* | WBGene00011671 | protein_coding_gene | -2.6 | 0.1620 | 1.4E-08 | -2.5 | 0.180 | 1.2E-08 |
| *col-106* | WBGene00000680 | protein_coding_gene | -2.6 | 0.1619 | 3.1E-19 | -3.2 | 0.111 | 1.9E-92 |
| *dut-1* | WBGene00010609 | protein_coding_gene | -2.6 | 0.1618 | 5.7E-43 | -1.9 | 0.270 | 4.3E-74 |
| *F14H12.6* | WBGene00017473 | protein_coding_gene | -2.6 | 0.1618 | 9.5E-04 | -3.7 | 0.075 | 5.8E-05 |
| ***abhd-3.2*** | **WBGene00016644** | **protein_coding_gene** | **-2.6** | **0.1617** | **2.6E-06** | **1.3** | **2.431** | **6.4E-06** |
| *sqv-3* | WBGene00005021 | protein_coding_gene | -2.6 | 0.1615 | 1.5E-55 | -1.9 | 0.277 | 4.6E-32 |
| *C06A5.6* | WBGene00015504 | protein_coding_gene | -2.6 | 0.1613 | 1.2E-31 | -1.4 | 0.372 | 6.0E-23 |
| *msp-152* | WBGene00003470 | protein_coding_gene | -2.6 | 0.1610 | 3.2E-08 | -2.4 | 0.183 | 1.2E-30 |
| *C25A11.2* | WBGene00016087 | protein_coding_gene | -2.6 | 0.1608 | 2.8E-33 | -2.9 | 0.136 | 8.5E-52 |
| *sdz-38* | WBGene00014131 | protein_coding_gene | -2.6 | 0.1606 | 1.1E-04 | -2.8 | 0.140 | 1.8E-05 |
| *plk-3* | WBGene00004044 | protein_coding_gene | -2.6 | 0.1604 | 4.6E-46 | -1.3 | 0.392 | 3.0E-42 |
| *R10E4.6* | WBGene00011202 | protein_coding_gene | -2.6 | 0.1601 | 1.3E-08 | -3.4 | 0.093 | 1.1E-13 |
| *C43D7.8* | WBGene00008070 | protein_coding_gene | -2.6 | 0.1601 | 1.2E-02 | -5.1 | 0.030 | 2.5E-08 |
| *maph-1.3* | WBGene00007966 | protein_coding_gene | -2.6 | 0.1600 | 1.2E-31 | -1.7 | 0.306 | 2.1E-22 |
| *cyp-31A3* | WBGene00021200 | protein_coding_gene | -2.6 | 0.1600 | 2.8E-49 | -2.1 | 0.230 | 1.0E-47 |
| *mltn-10* | WBGene00008965 | protein_coding_gene | -2.6 | 0.1597 | 1.5E-04 | -2.1 | 0.231 | 2.7E-05 |
| *cin-4* | WBGene00022854 | protein_coding_gene | -2.6 | 0.1597 | 5.9E-22 | -1.6 | 0.337 | 1.0E-21 |
| *ule-2* | WBGene00045038 | protein_coding_gene | -2.6 | 0.1596 | 1.6E-22 | -2.9 | 0.137 | 7.5E-51 |
| *hinf-1* | WBGene00009553 | protein_coding_gene | -2.7 | 0.1593 | 2.5E-28 | -2.4 | 0.191 | 7.3E-93 |
| *W08F4.3* | WBGene00021093 | protein_coding_gene | -2.7 | 0.1591 | 2.4E-47 | -1.8 | 0.288 | 2.2E-32 |
| *Y54F10BM.6* | WBGene00021861 | protein_coding_gene | -2.7 | 0.1591 | 8.2E-05 | -3.9 | 0.065 | 9.4E-07 |
| *Y57G11C.1144* | WBGene00219443 | protein_coding_gene | -2.7 | 0.1590 | 1.2E-02 | -2.7 | 0.154 | 8.5E-03 |
| *Y59A8B.12* | WBGene00013349 | protein_coding_gene | -2.7 | 0.1583 | 1.5E-68 | -2.5 | 0.183 | 7.1E-57 |
| *gei-14* | WBGene00001571 | protein_coding_gene | -2.7 | 0.1575 | 4.8E-52 | -2.2 | 0.221 | 3.5E-62 |
| *Y53C12A.11* | WBGene00194906 | protein_coding_gene | -2.7 | 0.1567 | 4.8E-18 | -2.1 | 0.238 | 7.5E-16 |
| *F31E9.6* | WBGene00009299 | protein_coding_gene | -2.7 | 0.1567 | 1.3E-12 | -1.4 | 0.381 | 3.1E-02 |
| *gld-3* | WBGene00001597 | protein_coding_gene | -2.7 | 0.1565 | 2.1E-31 | -2.3 | 0.207 | 1.8E-99 |
| *cpg-4* | WBGene00015677 | protein_coding_gene | -2.7 | 0.1565 | 8.1E-32 | -2.4 | 0.186 | 2.8E-77 |
| *C47D12.4* | WBGene00008138 | protein_coding_gene | -2.7 | 0.1564 | 6.1E-06 | -5.5 | 0.022 | 5.3E-13 |
| *fzy-1* | WBGene00001511 | protein_coding_gene | -2.7 | 0.1563 | 3.0E-46 | -2.1 | 0.241 | 4.8E-69 |
| *dpy-17* | WBGene00001076 | protein_coding_gene | -2.7 | 0.1556 | 4.9E-09 | -6.2 | 0.014 | 9.2E-83 |
| *dop-4* | WBGene00016872 | protein_coding_gene | -2.7 | 0.1554 | 9.0E-09 | -1.8 | 0.285 | 4.4E-05 |
| *ssp-37* | WBGene00012211 | protein_coding_gene | -2.7 | 0.1553 | 1.7E-07 | -4.4 | 0.049 | 5.1E-18 |
| *Y49E10.29* | WBGene00045239 | protein_coding_gene | -2.7 | 0.1553 | 4.8E-19 | -2.6 | 0.170 | 1.5E-10 |
| *T27E7.10* | WBGene00044268 | pseudogene | -2.7 | 0.1552 | 8.5E-04 | -1.7 | 0.308 | 6.4E-03 |
| *tbx-40* | WBGene00006559 | protein_coding_gene | -2.7 | 0.1549 | 6.9E-03 | -2.9 | 0.131 | 2.1E-03 |
| *fbxb-9* | WBGene00007447 | protein_coding_gene | -2.7 | 0.1543 | 5.1E-03 | -3.4 | 0.097 | 8.6E-04 |
| *rnr-1* | WBGene00004391 | protein_coding_gene | -2.7 | 0.1543 | 2.8E-32 | -2.2 | 0.223 | 9.6E-60 |

| | | | | | | | | |
|---|---|---|---|---|---|---|---|---|
| *picc-1* | WBGene00017931 | protein_coding_gene | -2.7 | 0.1540 | 9.7E-31 | -2.7 | 0.151 | 4.4E-94 |
| *F40G9.15* | WBGene00077531 | protein_coding_gene | -2.7 | 0.1532 | 1.6E-05 | -5.4 | 0.024 | 6.4E-09 |
| *Y53F4B.62* | WBGene00219491 | protein_coding_gene | -2.7 | 0.1532 | 4.4E-04 | -3.4 | 0.093 | 2.0E-07 |
| *F59A6.5* | WBGene00019087 | protein_coding_gene | -2.7 | 0.1532 | 2.2E-49 | -2.8 | 0.143 | 2.0E-93 |
| *col-122* | WBGene00000696 | protein_coding_gene | -2.7 | 0.1532 | 1.0E-22 | -3.1 | 0.114 | 6.0E-44 |
| *H24K24.2* | WBGene00019239 | protein_coding_gene | -2.7 | 0.1530 | 2.0E-04 | -5.0 | 0.032 | 2.0E-08 |
| *F08F1.21* | WBGene00304789 | ncRNA_gene | -2.7 | 0.1527 | 1.4E-05 | -3.2 | 0.112 | 4.2E-07 |
| *T22C8.4* | WBGene00011925 | protein_coding_gene | -2.7 | 0.1526 | 1.5E-07 | -2.8 | 0.139 | 2.9E-08 |
| *hach-1* | WBGene00017301 | protein_coding_gene | -2.7 | 0.1523 | 3.9E-79 | -2.1 | 0.237 | 1.8E-07 |
| *T27A1.2* | WBGene00020834 | protein_coding_gene | -2.7 | 0.1520 | 1.9E-09 | -2.6 | 0.164 | 2.4E-29 |
| *rga-3* | WBGene00019600 | protein_coding_gene | -2.7 | 0.1520 | 2.5E-28 | -1.8 | 0.279 | 1.9E-52 |
| *gras-1* | WBGene00009272 | protein_coding_gene | -2.7 | 0.1518 | 6.4E-59 | -2.1 | 0.232 | 6.0E-46 |
| *fbxb-102* | WBGene00008847 | protein_coding_gene | -2.7 | 0.1516 | 4.3E-03 | -4.3 | 0.051 | 7.2E-06 |
| *C11G6.2* | WBGene00007523 | protein_coding_gene | -2.7 | 0.1514 | 2.0E-02 | -3.4 | 0.093 | 3.8E-04 |
| ***T08A9.6*** | **WBGene00020338** | **pseudogene** | **-2.7** | **0.1514** | **1.0E-07** | **1.7** | **3.245** | **4.9E-05** |
| *wago-1* | WBGene00011061 | protein_coding_gene | -2.7 | 0.1512 | 6.6E-42 | -1.6 | 0.329 | 1.2E-46 |
| *T25D3.3* | WBGene00020797 | protein_coding_gene | -2.7 | 0.1511 | 8.3E-16 | -1.9 | 0.269 | 2.8E-06 |
| *cey-3* | WBGene00000474 | protein_coding_gene | -2.7 | 0.1505 | 1.9E-44 | -2.3 | 0.197 | 2.7E-104 |
| *inx-14* | WBGene00002136 | protein_coding_gene | -2.7 | 0.1503 | 2.4E-49 | -2.2 | 0.219 | 5.8E-75 |
| *F23A7.4* | WBGene00009069 | protein_coding_gene | -2.7 | 0.1501 | 1.7E-09 | -2.0 | 0.250 | 1.8E-33 |
| *C06A1.12* | WBGene00219803 | ncRNA_gene | -2.7 | 0.1499 | 3.5E-16 | -1.3 | 0.404 | 2.6E-06 |
| *F22G12.3* | WBGene00009063 | protein_coding_gene | -2.7 | 0.1497 | 3.9E-03 | -3.3 | 0.099 | 5.2E-04 |
| *mei-1* | WBGene00003183 | protein_coding_gene | -2.7 | 0.1497 | 2.4E-59 | -2.4 | 0.192 | 3.3E-62 |
| *T28C6.3* | WBGene00012119 | protein_coding_gene | -2.7 | 0.1496 | 3.2E-08 | -3.1 | 0.119 | 9.7E-12 |
| *ekl-5* | WBGene00012512 | protein_coding_gene | -2.7 | 0.1495 | 1.3E-27 | -2.4 | 0.187 | 6.1E-54 |
| *F35E2.2* | WBGene00009410 | protein_coding_gene | -2.7 | 0.1495 | 3.6E-03 | -3.4 | 0.096 | 4.4E-04 |
| *Y116F11A.4* | WBGene00014615 | pseudogene | -2.7 | 0.1491 | 9.8E-04 | -4.0 | 0.065 | 1.5E-05 |
| *hrde-1* | WBGene00007624 | protein_coding_gene | -2.7 | 0.1489 | 8.6E-43 | -1.6 | 0.341 | 3.4E-52 |
| *col-181* | WBGene00000754 | protein_coding_gene | -2.7 | 0.1487 | 2.8E-18 | -3.2 | 0.106 | 4.4E-137 |
| *mcm-7* | WBGene00003159 | protein_coding_gene | -2.8 | 0.1486 | 6.9E-37 | -2.6 | 0.162 | 2.1E-90 |
| *C44B7.12* | WBGene00016632 | protein_coding_gene | -2.8 | 0.1481 | 3.2E-65 | -2.1 | 0.227 | 1.0E-78 |
| *klc-1* | WBGene00002214 | protein_coding_gene | -2.8 | 0.1479 | 2.5E-51 | -2.4 | 0.183 | 3.4E-86 |
| *ztf-11* | WBGene00009939 | protein_coding_gene | -2.8 | 0.1477 | 6.3E-20 | -4.4 | 0.046 | 5.5E-103 |
| *C14C11.2* | WBGene00015766 | protein_coding_gene | -2.8 | 0.1476 | 2.5E-47 | -1.4 | 0.384 | 1.8E-12 |
| *C10A4.6* | WBGene00015667 | protein_coding_gene | -2.8 | 0.1476 | 3.0E-05 | -2.1 | 0.230 | 1.3E-02 |
| *col-80* | WBGene00000656 | protein_coding_gene | -2.8 | 0.1473 | 3.4E-13 | -2.2 | 0.211 | 1.4E-17 |
| *erh-2* | WBGene00009444 | protein_coding_gene | -2.8 | 0.1471 | 1.3E-24 | -2.6 | 0.168 | 1.3E-51 |
| *lsd-1* | WBGene00011615 | protein_coding_gene | -2.8 | 0.1468 | 3.7E-33 | -2.3 | 0.210 | 3.0E-29 |
| *acs-13* | WBGene00022037 | protein_coding_gene | -2.8 | 0.1466 | 1.0E-33 | -2.5 | 0.174 | 7.0E-104 |
| *B0393.3* | WBGene00007168 | protein_coding_gene | -2.8 | 0.1462 | 2.1E-37 | -2.1 | 0.226 | 4.6E-64 |
| *W08F4.20* | WBGene00268207 | pseudogene | -2.8 | 0.1461 | 1.5E-02 | -2.9 | 0.131 | 4.7E-03 |
| *C24H12.1* | WBGene00016071 | protein_coding_gene | -2.8 | 0.1461 | 2.4E-06 | -4.5 | 0.043 | 9.2E-17 |
| *T03F6.3* | WBGene00011399 | protein_coding_gene | -2.8 | 0.1459 | 1.0E-65 | -2.4 | 0.184 | 5.7E-59 |
| *Y17G9A.4* | WBGene00021197 | protein_coding_gene | -2.8 | 0.1458 | 8.0E-05 | -6.1 | 0.015 | 1.2E-35 |
| *end-1* | WBGene00001310 | protein_coding_gene | -2.8 | 0.1456 | 2.9E-05 | -4.8 | 0.036 | 1.1E-07 |
| *msrp-5* | WBGene00017059 | protein_coding_gene | -2.8 | 0.1454 | 1.4E-06 | -2.2 | 0.214 | 1.7E-18 |
| *D1081.7* | WBGene00008385 | protein_coding_gene | -2.8 | 0.1452 | 2.4E-70 | -1.7 | 0.313 | 7.7E-60 |
| *nhr-181* | WBGene00018189 | protein_coding_gene | -2.8 | 0.1451 | 2.8E-19 | -3.1 | 0.121 | 4.2E-18 |
| *C08F1.11* | WBGene00044417 | protein_coding_gene | -2.8 | 0.1450 | 2.4E-03 | -3.3 | 0.098 | 2.8E-04 |
| *W03C9.2* | WBGene00012220 | protein_coding_gene | -2.8 | 0.1450 | 1.5E-57 | -2.4 | 0.193 | 2.8E-70 |
| *dsh-2* | WBGene00001102 | protein_coding_gene | -2.8 | 0.1442 | 1.3E-48 | -2.5 | 0.178 | 2.1E-76 |
| *C09B8.5* | WBGene00015624 | protein_coding_gene | -2.8 | 0.1429 | 2.2E-20 | -3.6 | 0.082 | 1.2E-23 |
| *ZK662.5* | WBGene00014040 | protein_coding_gene | -2.8 | 0.1425 | 4.5E-03 | -2.5 | 0.177 | 7.5E-03 |
| *Y39G10AR.9* | WBGene00021467 | protein_coding_gene | -2.8 | 0.1420 | 2.8E-61 | -2.1 | 0.238 | 5.5E-54 |
| *rskn-1* | WBGene00011352 | protein_coding_gene | -2.8 | 0.1419 | 3.3E-36 | -2.3 | 0.202 | 1.2E-92 |
| *sex-1* | WBGene00004786 | protein_coding_gene | -2.8 | 0.1416 | 3.3E-24 | -3.8 | 0.070 | 1.3E-94 |
| *tric-1B.2* | WBGene00013268 | protein_coding_gene | -2.8 | 0.1415 | 6.2E-60 | -1.6 | 0.335 | 1.9E-24 |
| *F44F1.5* | WBGene00009697 | pseudogene | -2.8 | 0.1402 | 4.7E-11 | -4.3 | 0.051 | 1.0E-21 |
| *sdz-20* | WBGene00009950 | protein_coding_gene | -2.8 | 0.1400 | 1.4E-02 | -2.4 | 0.195 | 2.4E-02 |
| *cpna-3* | WBGene00007221 | protein_coding_gene | -2.8 | 0.1399 | 9.0E-45 | -2.3 | 0.209 | 6.1E-64 |
| *tbx-8* | WBGene00006545 | protein_coding_gene | -2.8 | 0.1397 | 8.9E-05 | -2.6 | 0.162 | 6.4E-09 |
| *Y23H5B.8* | WBGene00021279 | protein_coding_gene | -2.8 | 0.1392 | 2.0E-05 | -5.0 | 0.031 | 6.9E-20 |
| *T07E3.3* | WBGene00020314 | protein_coding_gene | -2.8 | 0.1387 | 5.0E-37 | -1.7 | 0.317 | 3.6E-44 |
| *fbxb-64* | WBGene00008071 | protein_coding_gene | -2.9 | 0.1387 | 1.4E-02 | -4.0 | 0.062 | 4.4E-06 |
| *F44E5.3* | WBGene00009690 | protein_coding_gene | -2.9 | 0.1386 | 3.7E-03 | -2.2 | 0.225 | 1.1E-06 |
| *let-99* | WBGene00002368 | protein_coding_gene | -2.9 | 0.1380 | 3.8E-41 | -2.3 | 0.209 | 1.3E-62 |
| *C37C3.9* | WBGene00016501 | protein_coding_gene | -2.9 | 0.1375 | 8.2E-52 | -2.2 | 0.213 | 7.9E-69 |
| *T05B11.4* | WBGene00020247 | protein_coding_gene | -2.9 | 0.1374 | 4.3E-24 | -2.2 | 0.223 | 4.8E-19 |
| *Y40A1A.6* | WBGene00304815 | ncRNA_gene | -2.9 | 0.1373 | 3.2E-23 | -2.7 | 0.157 | 4.0E-52 |
| *W08F4.7* | WBGene00021096 | protein_coding_gene | -2.9 | 0.1368 | 1.9E-04 | -3.6 | 0.082 | 7.5E-11 |
| *pcn-1* | WBGene00003955 | protein_coding_gene | -2.9 | 0.1362 | 1.9E-32 | -2.8 | 0.141 | 1.5E-114 |
| *F41G3.6* | WBGene00018302 | protein_coding_gene | -2.9 | 0.1358 | 6.0E-51 | -2.6 | 0.170 | 3.5E-80 |
| *Y17G9B.2* | WBGene00021199 | protein_coding_gene | -2.9 | 0.1357 | 1.8E-05 | -4.5 | 0.046 | 1.7E-09 |
| *T06D4.6* | WBGene00206385 | protein_coding_gene | -2.9 | 0.1354 | 5.8E-03 | -3.6 | 0.081 | 1.8E-04 |
| *Y37E11AL.9* | WBGene00021364 | protein_coding_gene | -2.9 | 0.1353 | 4.0E-03 | -2.5 | 0.173 | 1.0E-02 |
| *F44E7.5* | WBGene00018427 | protein_coding_gene | -2.9 | 0.1348 | 4.4E-20 | -1.8 | 0.293 | 9.7E-30 |
| *Y38H6A.4* | WBGene00014874 | pseudogene | -2.9 | 0.1345 | 5.0E-04 | -2.6 | 0.170 | 4.7E-04 |
| *pmt-2* | WBGene00018811 | protein_coding_gene | -2.9 | 0.1341 | 8.6E-27 | -1.4 | 0.387 | 3.9E-20 |
| *C45B2.1* | WBGene00016658 | protein_coding_gene | -2.9 | 0.1336 | 6.6E-10 | -3.9 | 0.068 | 8.2E-31 |
| *ify-1* | WBGene00002068 | protein_coding_gene | -2.9 | 0.1332 | 2.4E-53 | -2.1 | 0.231 | 1.2E-55 |

| | | | | | | | | |
|---|---|---|---|---|---|---|---|---|
| *F58D2.3* | WBGene00044978 | protein_coding_gene | -2.9 | 0.1331 | 5.8E-03 | -3.6 | 0.084 | 2.2E-04 |
| *C25A11.10* | WBGene00219374 | pseudogene | -2.9 | 0.1331 | 7.6E-19 | -4.0 | 0.063 | 2.0E-21 |
| *drsh-1* | WBGene00009163 | protein_coding_gene | -2.9 | 0.1327 | 2.1E-71 | -2.6 | 0.168 | 1.1E-91 |
| *Y73F8A.36* | WBGene00044259 | protein_coding_gene | -2.9 | 0.1326 | 7.4E-03 | -2.2 | 0.212 | 3.9E-02 |
| *zipt-7.1* | WBGene00044067 | protein_coding_gene | -2.9 | 0.1320 | 1.4E-61 | -1.4 | 0.369 | 1.9E-32 |
| *C27C12.1* | WBGene00007771 | protein_coding_gene | -2.9 | 0.1319 | 9.7E-50 | -2.1 | 0.227 | 4.4E-22 |
| *C36B1.11* | WBGene00007978 | protein_coding_gene | -2.9 | 0.1319 | 6.1E-44 | -2.5 | 0.181 | 2.5E-81 |
| *F41C6.4* | WBGene00018278 | protein_coding_gene | -2.9 | 0.1309 | 2.2E-04 | -3.1 | 0.115 | 1.0E-03 |
| *msp-65* | WBGene00003458 | protein_coding_gene | -2.9 | 0.1299 | 6.8E-06 | -2.6 | 0.164 | 3.4E-13 |
| *klp-16* | WBGene00002226 | protein_coding_gene | -2.9 | 0.1298 | 2.6E-46 | -2.3 | 0.209 | 4.1E-72 |
| *W09G12.3* | WBGene00023163 | pseudogene | -2.9 | 0.1295 | 3.4E-03 | -2.7 | 0.155 | 3.6E-03 |
| *F18E9.7* | WBGene00017573 | protein_coding_gene | -3.0 | 0.1293 | 4.4E-03 | -8.0 | 0.004 | 4.8E-16 |
| *C31H1.8* | WBGene00016291 | protein_coding_gene | -3.0 | 0.1290 | 3.1E-55 | -2.3 | 0.209 | 2.6E-70 |
| *msp-49* | WBGene00003442 | protein_coding_gene | -3.0 | 0.1286 | 7.7E-09 | -1.5 | 0.348 | 8.5E-12 |
| *cutl-2* | WBGene00013145 | protein_coding_gene | -3.0 | 0.1278 | 2.6E-04 | -6.3 | 0.013 | 3.4E-23 |
| *acly-1* | WBGene00016995 | protein_coding_gene | -3.0 | 0.1277 | 7.6E-37 | -1.7 | 0.303 | 2.0E-45 |
| *C29A12.2* | WBGene00007798 | pseudogene | -3.0 | 0.1274 | 1.5E-03 | -1.5 | 0.358 | 2.3E-02 |
| *ZK105.1* | WBGene00022653 | protein_coding_gene | -3.0 | 0.1264 | 2.0E-26 | -2.7 | 0.157 | 5.5E-60 |
| *F44E5.2* | WBGene00009689 | protein_coding_gene | -3.0 | 0.1261 | 3.0E-03 | -2.5 | 0.182 | 9.2E-04 |
| *F56C9.3* | WBGene00018948 | protein_coding_gene | -3.0 | 0.1259 | 1.8E-71 | -1.9 | 0.269 | 5.9E-64 |
| *F13H8.3* | WBGene00017436 | protein_coding_gene | -3.0 | 0.1258 | 3.6E-14 | -1.3 | 0.411 | 9.2E-09 |
| *F17C11.13* | WBGene00023503 | protein_coding_gene | -3.0 | 0.1255 | 3.0E-05 | -1.8 | 0.282 | 1.6E-02 |
| *Y110A2AR.1* | WBGene00022452 | protein_coding_gene | -3.0 | 0.1253 | 2.2E-55 | -2.2 | 0.211 | 6.5E-55 |
| *sqv-4* | WBGene00005022 | protein_coding_gene | -3.0 | 0.1253 | 3.9E-46 | -2.0 | 0.244 | 1.0E-76 |
| *Y42H9B.3* | WBGene00021541 | protein_coding_gene | -3.0 | 0.1251 | 8.2E-57 | -2.5 | 0.177 | 1.2E-95 |
| *Y46H3C.5* | WBGene00021605 | protein_coding_gene | -3.0 | 0.1250 | 1.0E-21 | -3.4 | 0.093 | 1.2E-18 |
| *Y116A8B.1* | WBGene00013779 | protein_coding_gene | -3.0 | 0.1250 | 1.8E-03 | -3.5 | 0.091 | 2.8E-05 |
| *R11G10.4* | WBGene00050968 | protein_coding_gene | -3.0 | 0.1242 | 2.2E-02 | -5.6 | 0.021 | 2.8E-09 |
| *col-19* | WBGene00000608 | protein_coding_gene | -3.0 | 0.1242 | 7.0E-22 | -3.6 | 0.081 | 3.7E-52 |
| *lin-41* | WBGene00003026 | protein_coding_gene | -3.0 | 0.1240 | 8.6E-23 | -2.9 | 0.135 | 1.6E-121 |
| *tbx-32* | WBGene00006551 | protein_coding_gene | -3.0 | 0.1239 | 4.9E-03 | -2.7 | 0.150 | 3.7E-03 |
| *ima-2* | WBGene00002073 | protein_coding_gene | -3.0 | 0.1228 | 9.8E-35 | -2.2 | 0.225 | 7.2E-69 |
| *ZK6.11* | WBGene00022645 | protein_coding_gene | -3.0 | 0.1228 | 1.4E-10 | -1.5 | 0.354 | 2.2E-03 |
| *K11D12.6* | WBGene00019646 | protein_coding_gene | -3.0 | 0.1227 | 3.1E-03 | -7.2 | 0.007 | 5.4E-21 |
| *ule-4* | WBGene00007458 | protein_coding_gene | -3.0 | 0.1227 | 1.4E-15 | -4.7 | 0.039 | 6.5E-76 |
| *C02F12.5* | WBGene00015355 | protein_coding_gene | -3.0 | 0.1225 | 5.8E-11 | -5.0 | 0.031 | 1.2E-17 |
| *lips-12* | WBGene00020473 | protein_coding_gene | -3.0 | 0.1220 | 2.9E-03 | -2.2 | 0.223 | 4.1E-02 |
| *fbxc-51* | WBGene00021576 | protein_coding_gene | -3.0 | 0.1217 | 3.0E-17 | -4.6 | 0.042 | 2.9E-84 |
| *kin-33* | WBGene00017083 | protein_coding_gene | -3.0 | 0.1210 | 3.0E-14 | -2.9 | 0.133 | 1.7E-15 |
| *M01D1.12* | WBGene00269373 | protein_coding_gene | -3.0 | 0.1208 | 6.2E-04 | -1.7 | 0.318 | 7.0E-04 |
| *ergo-1* | WBGene00019971 | protein_coding_gene | -3.1 | 0.1207 | 4.4E-40 | -2.8 | 0.144 | 3.2E-128 |
| *pygl-1* | WBGene00020696 | protein_coding_gene | -3.1 | 0.1200 | 1.2E-49 | -2.3 | 0.206 | 1.9E-89 |
| *F36H1.12* | WBGene00009497 | protein_coding_gene | -3.1 | 0.1200 | 3.3E-03 | -2.0 | 0.257 | 1.6E-03 |
| *ref-1* | WBGene00004334 | protein_coding_gene | -3.1 | 0.1195 | 3.2E-21 | -3.0 | 0.121 | 3.0E-20 |
| *R03H10.11* | WBGene00271824 | protein_coding_gene | -3.1 | 0.1194 | 5.8E-07 | -3.9 | 0.065 | 6.1E-10 |
| *ZK813.2* | WBGene00022821 | protein_coding_gene | -3.1 | 0.1191 | 1.3E-31 | -3.9 | 0.067 | 9.4E-93 |
| *K07F5.12* | WBGene00010637 | protein_coding_gene | -3.1 | 0.1186 | 5.9E-31 | -3.1 | 0.117 | 8.2E-55 |
| *col-140* | WBGene00000713 | protein_coding_gene | -3.1 | 0.1184 | 1.4E-21 | -3.4 | 0.092 | 4.2E-98 |
| *bath-33* | WBGene00015595 | protein_coding_gene | -3.1 | 0.1181 | 2.1E-03 | -2.8 | 0.146 | 4.4E-05 |
| *F55B11.2* | WBGene00010084 | protein_coding_gene | -3.1 | 0.1176 | 4.9E-30 | -3.9 | 0.069 | 5.7E-105 |
| *C04C11.25* | WBGene00284855 | protein_coding_gene | -3.1 | 0.1168 | 1.6E-04 | -5.4 | 0.023 | 9.4E-09 |
| *Y52B11A.8* | WBGene00013127 | protein_coding_gene | -3.1 | 0.1163 | 2.3E-43 | -2.4 | 0.184 | 9.7E-38 |
| *Y82E9BL.5* | WBGene00022322 | protein_coding_gene | -3.1 | 0.1162 | 1.1E-03 | -2.6 | 0.166 | 1.1E-02 |
| *C32B5.6* | WBGene00016299 | protein_coding_gene | -3.1 | 0.1162 | 1.3E-03 | -2.9 | 0.131 | 2.8E-03 |
| *C27A7.6* | WBGene00007757 | protein_coding_gene | -3.1 | 0.1161 | 7.1E-42 | -2.4 | 0.193 | 2.0E-46 |
| *hlh-2* | WBGene00001949 | protein_coding_gene | -3.1 | 0.1160 | 2.0E-36 | -3.8 | 0.072 | 4.7E-90 |
| *mes-1* | WBGene00003219 | protein_coding_gene | -3.1 | 0.1159 | 2.8E-19 | -2.5 | 0.173 | 2.7E-41 |
| *cgh-1* | WBGene00000479 | protein_coding_gene | -3.1 | 0.1156 | 1.7E-29 | -2.3 | 0.205 | 3.4E-121 |
| *apx-1* | WBGene00000168 | protein_coding_gene | -3.1 | 0.1152 | 4.0E-57 | -2.5 | 0.171 | 5.0E-74 |
| *F52D2.12* | WBGene00219451 | protein_coding_gene | -3.1 | 0.1151 | 8.7E-37 | -2.4 | 0.193 | 5.9E-28 |
| *F44E5.15* | WBGene00206377 | protein_coding_gene | -3.1 | 0.1149 | 2.6E-05 | -1.3 | 0.405 | 3.3E-02 |
| *F12F6.8* | WBGene00008723 | pseudogene | -3.1 | 0.1143 | 1.4E-07 | -4.1 | 0.060 | 9.5E-12 |
| *F22F4.5* | WBGene00017718 | protein_coding_gene | -3.1 | 0.1140 | 4.1E-11 | -3.5 | 0.088 | 2.2E-05 |
| *dvc-1* | WBGene00011834 | protein_coding_gene | -3.1 | 0.1140 | 1.5E-108 | -2.4 | 0.195 | 4.8E-54 |
| *F22H10.5* | WBGene00017729 | pseudogene | -3.1 | 0.1135 | 1.3E-03 | -2.8 | 0.148 | 1.2E-03 |
| *wdr-5.3* | WBGene00013862 | protein_coding_gene | -3.1 | 0.1132 | 1.8E-44 | -2.9 | 0.129 | 4.5E-86 |
| *F40G9.14* | WBGene00018248 | protein_coding_gene | -3.1 | 0.1132 | 2.0E-03 | -3.5 | 0.090 | 1.3E-04 |
| *lab-1* | WBGene00007278 | protein_coding_gene | -3.1 | 0.1128 | 1.8E-57 | -2.4 | 0.191 | 7.3E-53 |
| *W02B8.2* | WBGene00012198 | protein_coding_gene | -3.1 | 0.1128 | 1.1E-33 | -3.0 | 0.127 | 2.0E-35 |
| *fbxa-72* | WBGene00017962 | protein_coding_gene | -3.1 | 0.1127 | 2.0E-09 | -1.7 | 0.314 | 3.8E-05 |
| *inx-8* | WBGene00002130 | protein_coding_gene | -3.2 | 0.1126 | 3.4E-38 | -2.5 | 0.172 | 2.7E-106 |
| *C05D12.5* | WBGene00007342 | protein_coding_gene | -3.2 | 0.1124 | 3.5E-08 | -4.9 | 0.034 | 2.9E-08 |
| *Y71F9AL.25* | WBGene00304792 | protein_coding_gene | -3.2 | 0.1122 | 7.9E-06 | -5.1 | 0.029 | 6.4E-13 |
| *fbxb-6* | WBGene00017207 | protein_coding_gene | -3.2 | 0.1120 | 1.0E-03 | -3.0 | 0.123 | 3.3E-03 |
| *T10B5.8* | WBGene00020394 | protein_coding_gene | -3.2 | 0.1120 | 4.8E-19 | -2.5 | 0.175 | 2.3E-12 |
| *ced-8* | WBGene00000422 | protein_coding_gene | -3.2 | 0.1116 | 2.3E-13 | -4.8 | 0.037 | 8.6E-23 |
| *F47G3.3* | WBGene00018578 | protein_coding_gene | -3.2 | 0.1116 | 1.5E-31 | -2.5 | 0.171 | 1.3E-24 |
| *gpd-4* | WBGene00001686 | protein_coding_gene | -3.2 | 0.1115 | 6.3E-53 | -2.3 | 0.205 | 1.2E-42 |
| *fbxa-184* | WBGene00018439 | protein_coding_gene | -3.2 | 0.1113 | 9.8E-05 | -1.9 | 0.265 | 1.2E-03 |
| *F55C9.15* | WBGene00206500 | protein_coding_gene | -3.2 | 0.1113 | 1.4E-03 | -3.6 | 0.080 | 2.3E-04 |

| | | | | | | | | |
|---|---|---|---|---|---|---|---|---|
| *T05H10.4* | WBGene00011509 | protein_coding_gene | -3.2 | 0.1104 | 1.7E-37 | -3.6 | 0.084 | 1.7E-116 |
| *clec-157* | WBGene00017810 | protein_coding_gene | -3.2 | 0.1101 | 2.2E-03 | -7.2 | 0.007 | 6.3E-16 |
| *F53H4.2* | WBGene00010007 | protein_coding_gene | -3.2 | 0.1099 | 9.4E-14 | -4.5 | 0.044 | 9.4E-90 |
| *Y47G6A.33* | WBGene00045399 | protein_coding_gene | -3.2 | 0.1095 | 3.6E-22 | -2.1 | 0.241 | 9.0E-20 |
| *hlh-14* | WBGene00001958 | protein_coding_gene | -3.2 | 0.1095 | 1.6E-07 | -5.7 | 0.020 | 3.3E-22 |
| *ZK673.5* | WBGene00014061 | protein_coding_gene | -3.2 | 0.1094 | 2.5E-05 | -2.5 | 0.176 | 5.4E-04 |
| *fbxb-94* | WBGene00013410 | pseudogene | -3.2 | 0.1086 | 3.1E-05 | -2.0 | 0.245 | 2.3E-05 |
| *K03B8.14* | WBGene00194783 | protein_coding_gene | -3.2 | 0.1086 | 3.4E-03 | -5.9 | 0.017 | 1.5E-09 |
| *sra-14* | WBGene00005040 | protein_coding_gene | -3.2 | 0.1083 | 1.8E-05 | -1.8 | 0.295 | 6.9E-03 |
| *col-116* | WBGene00000690 | protein_coding_gene | -3.2 | 0.1082 | 3.1E-06 | -5.0 | 0.032 | 3.1E-08 |
| *fbxb-8* | WBGene00009861 | protein_coding_gene | -3.2 | 0.1070 | 1.2E-03 | -4.5 | 0.046 | 7.3E-07 |
| *Y37H2A.13* | WBGene00045417 | protein_coding_gene | -3.2 | 0.1062 | 5.3E-04 | -4.6 | 0.041 | 1.5E-07 |
| *fbxb-59* | WBGene00013759 | protein_coding_gene | -3.2 | 0.1061 | 5.3E-04 | -3.8 | 0.074 | 6.2E-05 |
| *K04A8.20* | WBGene00235332 | protein_coding_gene | -3.2 | 0.1059 | 1.5E-05 | -3.8 | 0.073 | 7.6E-13 |
| *ssp-19* | WBGene00006047 | protein_coding_gene | -3.2 | 0.1057 | 1.9E-06 | -2.9 | 0.136 | 1.8E-34 |
| *msra-1* | WBGene00018393 | protein_coding_gene | -3.2 | 0.1054 | 4.5E-14 | -2.8 | 0.148 | 8.1E-09 |
| *W08F4.12* | WBGene00021101 | protein_coding_gene | -3.3 | 0.1050 | 7.2E-04 | -3.4 | 0.093 | 2.4E-04 |
| *sir-2.3* | WBGene00004802 | protein_coding_gene | -3.3 | 0.1050 | 2.2E-17 | -1.6 | 0.326 | 2.2E-05 |
| *C04E7.5* | WBGene00045270 | protein_coding_gene | -3.3 | 0.1049 | 1.1E-03 | -2.2 | 0.215 | 1.5E-02 |
| *F40G9.12* | WBGene00018247 | protein_coding_gene | -3.3 | 0.1046 | 1.7E-03 | -3.4 | 0.096 | 8.5E-04 |
| *cest-10* | WBGene00016595 | protein_coding_gene | -3.3 | 0.1045 | 3.2E-20 | -2.4 | 0.196 | 5.2E-11 |
| *Y48A6C.10* | WBGene00220168 | snoRNA_gene | -3.3 | 0.1040 | 4.8E-03 | -2.4 | 0.184 | 2.3E-02 |
| *F26D2.16* | WBGene00009154 | protein_coding_gene | -3.3 | 0.1033 | 1.5E-03 | -8.7 | 0.002 | 3.7E-30 |
| *pash-1* | WBGene00011908 | protein_coding_gene | -3.3 | 0.1030 | 1.9E-71 | -2.5 | 0.178 | 2.2E-92 |
| *imp-1* | WBGene00007979 | protein_coding_gene | -3.3 | 0.1030 | 4.4E-52 | -2.8 | 0.141 | 2.8E-135 |
| *C29A12.1* | WBGene00007797 | protein_coding_gene | -3.3 | 0.1029 | 1.1E-88 | -2.4 | 0.190 | 3.4E-66 |
| *Y46H3C.7* | WBGene00021607 | protein_coding_gene | -3.3 | 0.1021 | 1.4E-13 | -3.1 | 0.117 | 6.4E-11 |
| *F55C5.10* | WBGene00010098 | protein_coding_gene | -3.3 | 0.1018 | 1.3E-04 | -4.1 | 0.058 | 2.9E-08 |
| *T08B6.1* | WBGene00020351 | protein_coding_gene | -3.3 | 0.1016 | 1.1E-03 | -3.2 | 0.107 | 1.4E-03 |
| *his-67* | WBGene00001941 | protein_coding_gene | -3.3 | 0.1015 | 1.1E-11 | -3.5 | 0.089 | 9.6E-09 |
| *pal-1* | WBGene00003912 | protein_coding_gene | -3.3 | 0.1011 | 9.0E-45 | -2.9 | 0.135 | 9.3E-138 |
| *ZK228.12* | WBGene00077652 | pseudogene | -3.3 | 0.1010 | 1.4E-03 | -3.5 | 0.086 | 8.9E-05 |
| *T05E12.6* | WBGene00011487 | protein_coding_gene | -3.3 | 0.1005 | 8.3E-18 | -3.0 | 0.121 | 4.6E-31 |
| *srh-237* | WBGene00005444 | protein_coding_gene | -3.3 | 0.1002 | 3.0E-13 | -2.7 | 0.152 | 1.7E-17 |
| *fbxc-14* | WBGene00016298 | protein_coding_gene | -3.3 | 0.1002 | 5.6E-05 | -4.9 | 0.033 | 2.3E-07 |
| *pup-3* | WBGene00019082 | protein_coding_gene | -3.3 | 0.1001 | 1.2E-20 | -3.3 | 0.101 | 1.3E-24 |
| *BE10.3* | WBGene00007211 | protein_coding_gene | -3.3 | 0.0998 | 4.4E-08 | -5.2 | 0.028 | 5.4E-08 |
| *col-20* | WBGene00000609 | protein_coding_gene | -3.3 | 0.0997 | 2.4E-22 | -3.3 | 0.105 | 1.4E-113 |
| *bath-9* | WBGene00021725 | protein_coding_gene | -3.3 | 0.0993 | 4.9E-15 | -4.6 | 0.042 | 1.6E-28 |
| *unc-71* | WBGene00006804 | protein_coding_gene | -3.3 | 0.0991 | 3.1E-97 | -2.8 | 0.143 | 1.2E-105 |
| *F53C3.11* | WBGene00018754 | protein_coding_gene | -3.3 | 0.0990 | 1.0E-05 | -3.6 | 0.083 | 9.1E-06 |
| *psf-1* | WBGene00011275 | protein_coding_gene | -3.3 | 0.0989 | 1.2E-38 | -3.1 | 0.116 | 2.1E-88 |
| *W04G3.13* | WBGene00194793 | protein_coding_gene | -3.3 | 0.0989 | 5.7E-04 | -3.4 | 0.093 | 1.8E-04 |
| *pgph-2* | WBGene00018424 | protein_coding_gene | -3.3 | 0.0984 | 4.1E-19 | -1.9 | 0.266 | 1.4E-14 |
| *T19H5.7* | WBGene00045414 | protein_coding_gene | -3.4 | 0.0979 | 8.3E-05 | -4.2 | 0.056 | 5.8E-06 |
| *ZK938.11* | WBGene00270290 | protein_coding_gene | -3.4 | 0.0974 | 2.5E-04 | -4.1 | 0.058 | 9.7E-06 |
| *clec-74* | WBGene00021585 | protein_coding_gene | -3.4 | 0.0973 | 5.2E-04 | -3.5 | 0.090 | 1.1E-12 |
| *Y48G1C.13* | WBGene00189949 | protein_coding_gene | -3.4 | 0.0967 | 4.0E-19 | -3.1 | 0.116 | 6.4E-22 |
| *F46C3.5* | WBGene00044703 | pseudogene | -3.4 | 0.0961 | 1.5E-03 | -2.7 | 0.152 | 9.0E-03 |
| *C14E2.12* | WBGene00219816 | snoRNA_gene | -3.4 | 0.0958 | 9.7E-10 | -1.1 | 0.464 | 2.0E-02 |
| *Y48G1A.2* | WBGene00021658 | protein_coding_gene | -3.4 | 0.0958 | 9.0E-80 | -3.6 | 0.081 | 4.3E-88 |
| *ces-2* | WBGene00000469 | protein_coding_gene | -3.4 | 0.0956 | 1.3E-06 | -7.0 | 0.008 | 2.4E-28 |
| *C24H12.11* | WBGene00016080 | pseudogene | -3.4 | 0.0956 | 2.3E-05 | -4.7 | 0.039 | 2.3E-09 |
| *perm-1* | WBGene00011350 | protein_coding_gene | -3.4 | 0.0954 | 3.4E-51 | -3.0 | 0.129 | 2.0E-117 |
| *col-124* | WBGene00000698 | protein_coding_gene | -3.4 | 0.0950 | 1.3E-28 | -4.0 | 0.062 | 1.5E-34 |
| *F07E5.13* | WBGene00304173 | protein_coding_gene | -3.4 | 0.0949 | 2.1E-04 | -4.2 | 0.054 | 4.1E-06 |
| *Y24F12A.3* | WBGene00012498 | pseudogene | -3.4 | 0.0947 | 1.4E-05 | -2.5 | 0.180 | 4.5E-05 |
| *F21D5.1* | WBGene00009006 | protein_coding_gene | -3.4 | 0.0940 | 1.6E-44 | -2.9 | 0.137 | 2.8E-115 |
| *fbxc-24* | WBGene00019927 | protein_coding_gene | -3.4 | 0.0937 | 2.0E-04 | -5.1 | 0.030 | 2.3E-08 |
| *sdz-7* | WBGene00016779 | protein_coding_gene | -3.4 | 0.0935 | 4.4E-04 | -3.3 | 0.102 | 4.4E-04 |
| *F27B3.6* | WBGene00017848 | protein_coding_gene | -3.4 | 0.0933 | 5.4E-08 | -4.2 | 0.054 | 2.2E-09 |
| *F54H12.5* | WBGene00018845 | protein_coding_gene | -3.4 | 0.0931 | 4.2E-29 | -2.8 | 0.142 | 7.7E-32 |
| *sqv-5* | WBGene00005023 | protein_coding_gene | -3.4 | 0.0926 | 1.9E-62 | -2.1 | 0.226 | 6.3E-78 |
| *W02D9.6* | WBGene00012212 | protein_coding_gene | -3.4 | 0.0926 | 2.9E-15 | -4.1 | 0.057 | 4.2E-69 |
| *F21C10.9* | WBGene00017658 | protein_coding_gene | -3.4 | 0.0924 | 1.4E-11 | -1.2 | 0.439 | 1.1E-19 |
| *K07A1.6* | WBGene00010613 | protein_coding_gene | -3.4 | 0.0922 | 4.3E-44 | -5.6 | 0.020 | 6.5E-195 |
| *maph-1.2* | WBGene00009113 | protein_coding_gene | -3.4 | 0.0917 | 1.3E-71 | -2.2 | 0.222 | 2.8E-48 |
| *R04A9.7* | WBGene00044281 | protein_coding_gene | -3.5 | 0.0910 | 1.4E-10 | -5.7 | 0.019 | 1.1E-10 |
| *T22D1.5* | WBGene00020684 | protein_coding_gene | -3.5 | 0.0910 | 7.1E-69 | -2.5 | 0.180 | 1.1E-89 |
| *col-178* | WBGene00000751 | protein_coding_gene | -3.5 | 0.0908 | 7.4E-25 | -3.8 | 0.071 | 2.3E-153 |
| *Y82E9BL.2* | WBGene00022319 | protein_coding_gene | -3.5 | 0.0908 | 6.5E-04 | -3.4 | 0.096 | 4.8E-04 |
| *gadr-4* | WBGene00021053 | protein_coding_gene | -3.5 | 0.0907 | 3.1E-08 | -4.1 | 0.057 | 4.6E-10 |
| *mex-3* | WBGene00003229 | protein_coding_gene | -3.5 | 0.0906 | 1.3E-36 | -3.0 | 0.125 | 7.0E-116 |
| *fbxb-84* | WBGene00020544 | protein_coding_gene | -3.5 | 0.0904 | 4.1E-12 | -4.7 | 0.038 | 1.4E-07 |
| *sdz-13* | WBGene00008755 | protein_coding_gene | -3.5 | 0.0901 | 3.3E-04 | -3.3 | 0.102 | 2.6E-04 |
| *fbxb-93* | WBGene00013408 | protein_coding_gene | -3.5 | 0.0900 | 7.1E-07 | -3.2 | 0.112 | 4.6E-12 |
| ***nhr-235*** | **WBGene00012597** | **protein_coding_gene** | **-3.5** | **0.0898** | **3.7E-03** | **3.0** | **8.004** | **1.1E-02** |
| *lido-12* | WBGene00016573 | protein_coding_gene | -3.5 | 0.0896 | 1.0E-40 | -3.4 | 0.097 | 1.3E-55 |
| *Y41D4B.26* | WBGene00021525 | protein_coding_gene | -3.5 | 0.0889 | 1.5E-04 | -6.8 | 0.009 | 2.0E-12 |
| *F54E4.2* | WBGene00010058 | protein_coding_gene | -3.5 | 0.0885 | 9.1E-04 | -2.7 | 0.159 | 1.1E-04 |

| | | | | | | | | |
|---|---|---|---|---|---|---|---|---|
| *T24E12.13* | WBGene00195077 | protein_coding_gene | -3.5 | 0.0868 | 3.8E-04 | -4.1 | 0.058 | 7.7E-06 |
| *tbx-39* | WBGene00006558 | protein_coding_gene | -3.5 | 0.0868 | 7.4E-04 | -2.2 | 0.214 | 3.5E-02 |
| *M151.10* | WBGene00303365 | protein_coding_gene | -3.5 | 0.0861 | 3.5E-04 | -7.9 | 0.004 | 1.1E-17 |
| *tbx-37* | WBGene00006556 | protein_coding_gene | -3.5 | 0.0858 | 2.8E-04 | -2.5 | 0.173 | 2.2E-03 |
| *W02D9.7* | WBGene00012213 | protein_coding_gene | -3.5 | 0.0857 | 2.6E-25 | -4.0 | 0.064 | 1.1E-117 |
| *C24H12.8* | WBGene00016077 | protein_coding_gene | -3.6 | 0.0850 | 3.9E-04 | -4.1 | 0.057 | 5.1E-06 |
| *fbxb-25* | WBGene00019034 | protein_coding_gene | -3.6 | 0.0849 | 1.3E-04 | -3.9 | 0.067 | 2.5E-05 |
| *ZK813.7* | WBGene00045394 | protein_coding_gene | -3.6 | 0.0847 | 2.9E-68 | -3.7 | 0.079 | 2.4E-128 |
| *fbxb-117* | WBGene00010099 | protein_coding_gene | -3.6 | 0.0847 | 2.3E-04 | -4.5 | 0.044 | 3.9E-07 |
| *fbxb-60* | WBGene00010104 | protein_coding_gene | -3.6 | 0.0842 | 1.4E-04 | -5.0 | 0.031 | 2.4E-08 |
| *pmp-5* | WBGene00004062 | protein_coding_gene | -3.6 | 0.0840 | 5.5E-22 | -2.6 | 0.169 | 2.7E-51 |
| *C49F5.6* | WBGene00008210 | protein_coding_gene | -3.6 | 0.0839 | 1.8E-46 | -4.1 | 0.058 | 5.5E-104 |
| *fbxb-86* | WBGene00012098 | pseudogene | -3.6 | 0.0835 | 2.8E-04 | -4.3 | 0.051 | 1.4E-06 |
| *klp-15* | WBGene00002225 | protein_coding_gene | -3.6 | 0.0834 | 1.2E-46 | -2.0 | 0.244 | 2.1E-59 |
| *Y105C5B.20* | WBGene00013659 | protein_coding_gene | -3.6 | 0.0831 | 4.2E-20 | -3.2 | 0.110 | 3.5E-33 |
| *C35D6.4* | WBGene00007961 | protein_coding_gene | -3.6 | 0.0830 | 2.2E-04 | -4.3 | 0.051 | 3.9E-06 |
| *F23A7.6* | WBGene00009071 | pseudogene | -3.6 | 0.0829 | 5.2E-17 | -4.2 | 0.054 | 5.1E-19 |
| *gpd-1* | WBGene00001683 | protein_coding_gene | -3.6 | 0.0826 | 5.9E-61 | -2.7 | 0.155 | 7.5E-59 |
| *R02F2.4* | WBGene00019833 | protein_coding_gene | -3.6 | 0.0826 | 4.9E-72 | -2.6 | 0.167 | 4.4E-57 |
| *chtl-1* | WBGene00018037 | protein_coding_gene | -3.6 | 0.0824 | 1.1E-43 | -3.0 | 0.124 | 4.6E-76 |
| *Y43D4A.6* | WBGene00012792 | protein_coding_gene | -3.6 | 0.0818 | 3.3E-04 | -3.2 | 0.110 | 7.2E-04 |
| *col-119* | WBGene00000693 | protein_coding_gene | -3.6 | 0.0817 | 2.9E-36 | -3.2 | 0.109 | 1.0E-143 |
| *C09G9.8* | WBGene00007497 | protein_coding_gene | -3.6 | 0.0812 | 4.3E-04 | -2.8 | 0.139 | 4.3E-03 |
| *gadr-1* | WBGene00010240 | pseudogene | -3.6 | 0.0812 | 2.6E-04 | -4.1 | 0.058 | 3.7E-07 |
| *C17G1.2* | WBGene00007649 | protein_coding_gene | -3.6 | 0.0809 | 1.9E-38 | -3.3 | 0.101 | 4.9E-66 |
| *R03G8.6* | WBGene00011000 | protein_coding_gene | -3.6 | 0.0803 | 3.1E-22 | -5.4 | 0.024 | 1.5E-39 |
| *ZK899.5* | WBGene00014143 | protein_coding_gene | -3.7 | 0.0788 | 9.5E-05 | -3.8 | 0.071 | 3.6E-05 |
| *sams-2* | WBGene00013652 | pseudogene | -3.7 | 0.0786 | 6.6E-09 | -2.4 | 0.190 | 2.8E-10 |
| *cey-2* | WBGene00000473 | protein_coding_gene | -3.7 | 0.0782 | 7.3E-81 | -2.9 | 0.137 | 2.5E-137 |
| *ngn-1* | WBGene00003595 | protein_coding_gene | -3.7 | 0.0781 | 4.8E-05 | -5.7 | 0.019 | 9.5E-11 |
| *Y60A9.2* | WBGene00013369 | pseudogene | -3.7 | 0.0779 | 2.7E-04 | -3.8 | 0.070 | 1.6E-05 |
| *T27A1.3* | WBGene00020835 | protein_coding_gene | -3.7 | 0.0778 | 9.1E-06 | -3.7 | 0.076 | 2.8E-09 |
| *F19C7.3* | WBGene00017593 | protein_coding_gene | -3.7 | 0.0771 | 4.9E-05 | -3.9 | 0.068 | 6.2E-10 |
| *C14B1.2* | WBGene00007573 | protein_coding_gene | -3.7 | 0.0770 | 7.6E-67 | -3.2 | 0.108 | 6.4E-114 |
| *sdz-32* | WBGene00021717 | protein_coding_gene | -3.7 | 0.0769 | 2.7E-04 | -2.4 | 0.188 | 2.2E-02 |
| *Y70G10A.1* | WBGene00014946 | pseudogene | -3.7 | 0.0767 | 1.0E-04 | -3.3 | 0.099 | 4.1E-05 |
| *B0334.9* | WBGene00007147 | pseudogene | -3.7 | 0.0765 | 1.0E-04 | -4.1 | 0.059 | 2.8E-06 |
| *trcs-1* | WBGene00009186 | protein_coding_gene | -3.7 | 0.0765 | 2.6E-81 | -3.5 | 0.087 | 1.3E-174 |
| *gfat-1* | WBGene00008546 | protein_coding_gene | -3.7 | 0.0765 | 1.9E-53 | -2.3 | 0.210 | 2.4E-61 |
| *fbxb-100* | WBGene00013403 | protein_coding_gene | -3.7 | 0.0764 | 1.1E-04 | -3.9 | 0.069 | 5.3E-09 |
| *col-121* | WBGene00000695 | protein_coding_gene | -3.7 | 0.0762 | 3.8E-05 | -7.6 | 0.005 | 1.2E-27 |
| *str-179* | WBGene00006223 | pseudogene | -3.7 | 0.0761 | 3.0E-04 | -2.7 | 0.149 | 9.8E-03 |
| *F54D5.5* | WBGene00010051 | protein_coding_gene | -3.7 | 0.0752 | 3.1E-59 | -4.5 | 0.043 | 8.8E-171 |
| *fbxb-24* | WBGene00013234 | protein_coding_gene | -3.7 | 0.0751 | 5.1E-12 | -4.4 | 0.046 | 3.0E-16 |
| *W04A8.9* | WBGene00195155 | protein_coding_gene | -3.7 | 0.0751 | 1.6E-04 | -2.8 | 0.141 | 6.3E-05 |
| *F21D9.5* | WBGene00009019 | pseudogene | -3.7 | 0.0749 | 1.1E-04 | -6.9 | 0.008 | 6.4E-15 |
| *F57C2.4* | WBGene00010196 | protein_coding_gene | -3.7 | 0.0746 | 4.7E-27 | -4.4 | 0.048 | 7.6E-92 |
| *ugt-41* | WBGene00017336 | protein_coding_gene | -3.7 | 0.0745 | 7.4E-33 | -2.4 | 0.192 | 5.9E-21 |
| *dpy-14* | WBGene00001075 | protein_coding_gene | -3.8 | 0.0742 | 1.4E-08 | -7.2 | 0.007 | 3.4E-63 |
| *lido-9* | WBGene00016576 | protein_coding_gene | -3.8 | 0.0740 | 5.9E-55 | -3.5 | 0.086 | 1.2E-53 |
| *sdz-37* | WBGene00014064 | protein_coding_gene | -3.8 | 0.0739 | 4.2E-06 | -3.8 | 0.072 | 3.6E-05 |
| *ceh-39* | WBGene00000460 | protein_coding_gene | -3.8 | 0.0737 | 1.0E-35 | -3.7 | 0.075 | 7.8E-61 |
| *mei-2* | WBGene00003184 | protein_coding_gene | -3.8 | 0.0737 | 1.3E-106 | -2.9 | 0.134 | 2.4E-91 |
| *F13A2.4* | WBGene00017413 | protein_coding_gene | -3.8 | 0.0737 | 4.6E-18 | -1.8 | 0.286 | 1.5E-06 |
| *fbxc-30* | WBGene00045302 | protein_coding_gene | -3.8 | 0.0732 | 1.3E-04 | -3.6 | 0.081 | 7.0E-05 |
| *fbxb-71* | WBGene00018236 | protein_coding_gene | -3.8 | 0.0730 | 1.1E-04 | -4.7 | 0.039 | 7.9E-08 |
| *Y82E9BL.6* | WBGene00022323 | protein_coding_gene | -3.8 | 0.0729 | 7.9E-05 | -4.4 | 0.046 | 3.2E-06 |
| *F32A11.5* | WBGene00009310 | pseudogene | -3.8 | 0.0729 | 3.7E-05 | -3.7 | 0.076 | 8.7E-06 |
| *clec-210* | WBGene00022261 | protein_coding_gene | -3.8 | 0.0729 | 1.1E-06 | -1.8 | 0.292 | 3.2E-05 |
| *C43D7.11* | WBGene00077664 | pseudogene | -3.8 | 0.0726 | 6.4E-05 | -5.8 | 0.018 | 7.0E-11 |
| *F21D9.11* | WBGene00219452 | protein_coding_gene | -3.8 | 0.0724 | 4.1E-05 | -5.1 | 0.029 | 1.0E-13 |
| *dod-23* | WBGene00009898 | protein_coding_gene | -3.8 | 0.0722 | 5.2E-30 | -4.8 | 0.035 | 3.7E-46 |
| *fem-3* | WBGene00001413 | protein_coding_gene | -3.8 | 0.0719 | 1.3E-145 | -3.2 | 0.108 | 1.1E-97 |
| *mom-2* | WBGene00003395 | protein_coding_gene | -3.8 | 0.0716 | 2.1E-61 | -2.9 | 0.132 | 1.8E-98 |
| *C09G5.14* | WBGene00271803 | protein_coding_gene | -3.8 | 0.0715 | 6.5E-05 | -3.1 | 0.113 | 8.1E-04 |
| *M151.9* | WBGene00219238 | pseudogene | -3.8 | 0.0713 | 1.2E-04 | -3.1 | 0.119 | 6.1E-04 |
| *fbxb-74* | WBGene00019952 | protein_coding_gene | -3.8 | 0.0711 | 6.9E-07 | -3.4 | 0.094 | 3.2E-09 |
| *F12E12.3* | WBGene00017404 | pseudogene | -3.8 | 0.0710 | 4.1E-05 | -3.2 | 0.105 | 4.5E-04 |
| *B0238.15* | WBGene00044537 | protein_coding_gene | -3.8 | 0.0708 | 2.4E-04 | -2.4 | 0.185 | 1.8E-02 |
| *F56A8.8* | WBGene00010142 | protein_coding_gene | -3.8 | 0.0705 | 8.2E-14 | -5.0 | 0.032 | 2.1E-24 |
| *F36F12.3* | WBGene00018094 | protein_coding_gene | -3.8 | 0.0702 | 5.4E-05 | -3.2 | 0.111 | 1.3E-04 |
| *hlh-3* | WBGene00001950 | protein_coding_gene | -3.8 | 0.0701 | 9.3E-12 | -5.6 | 0.020 | 2.8E-19 |
| *cyb-3* | WBGene00000868 | protein_coding_gene | -3.8 | 0.0695 | 2.9E-69 | -3.9 | 0.066 | 1.6E-217 |
| *W03D2.15* | WBGene00271791 | ncRNA_gene | -3.8 | 0.0695 | 6.7E-05 | -4.1 | 0.058 | 1.8E-05 |
| *inx-3* | WBGene00002125 | protein_coding_gene | -3.9 | 0.0690 | 1.9E-63 | -5.0 | 0.032 | 5.0E-246 |
| *C04H5.7* | WBGene00007315 | protein_coding_gene | -3.9 | 0.0686 | 3.6E-17 | -4.5 | 0.044 | 5.9E-46 |
| *linc-12* | WBGene00219687 | lincRNA_gene | -3.9 | 0.0685 | 3.6E-05 | -4.7 | 0.038 | 1.7E-07 |
| *taf-11.2* | WBGene00006394 | protein_coding_gene | -3.9 | 0.0684 | 1.5E-157 | -3.3 | 0.103 | 5.2E-99 |
| *col-179* | WBGene00000752 | protein_coding_gene | -3.9 | 0.0684 | 5.2E-32 | -3.0 | 0.121 | 3.4E-85 |
| *F27C1.4* | WBGene00017854 | protein_coding_gene | -3.9 | 0.0683 | 9.1E-17 | -3.4 | 0.097 | 1.8E-17 |

| | | | | | | | | |
|---|---|---|---|---|---|---|---|---|
| *Y9C9A.13* | WBGene00021180 | protein_coding_gene | -3.9 | 0.0672 | 6.2E-05 | -2.8 | 0.143 | 1.0E-05 |
| *C14B1.9* | WBGene00007580 | protein_coding_gene | -3.9 | 0.0671 | 4.5E-87 | -3.4 | 0.092 | 1.3E-166 |
| *daf-18* | WBGene00000913 | protein_coding_gene | -3.9 | 0.0670 | 3.0E-59 | -3.3 | 0.101 | 1.9E-162 |
| *fbxb-118* | WBGene00021054 | protein_coding_gene | -3.9 | 0.0668 | 4.3E-05 | -3.4 | 0.096 | 2.8E-04 |
| *F59A6.12* | WBGene00044665 | protein_coding_gene | -3.9 | 0.0668 | 4.7E-28 | -4.9 | 0.034 | 5.0E-31 |
| *sdz-26* | WBGene00011049 | protein_coding_gene | -3.9 | 0.0661 | 4.9E-10 | -4.2 | 0.056 | 5.5E-17 |
| *W06D11.1* | WBGene00012299 | protein_coding_gene | -3.9 | 0.0655 | 8.2E-27 | -3.8 | 0.071 | 8.6E-41 |
| *fbxb-35* | WBGene00017208 | protein_coding_gene | -3.9 | 0.0653 | 1.3E-06 | -2.9 | 0.132 | 3.8E-03 |
| *fkh-5* | WBGene00001437 | protein_coding_gene | -3.9 | 0.0650 | 2.0E-07 | -3.8 | 0.071 | 7.9E-10 |
| *ZK892.6* | WBGene00014130 | protein_coding_gene | -3.9 | 0.0648 | 4.6E-05 | -3.5 | 0.088 | 1.6E-04 |
| *neg-1* | WBGene00017985 | protein_coding_gene | -4.0 | 0.0647 | 4.8E-91 | -3.8 | 0.071 | 8.6E-105 |
| *nasp-2* | WBGene00008218 | protein_coding_gene | -4.0 | 0.0645 | 2.2E-79 | -3.7 | 0.076 | 4.7E-184 |
| *pph-4.2* | WBGene00004086 | protein_coding_gene | -4.0 | 0.0644 | 1.9E-94 | -2.9 | 0.130 | 4.9E-90 |
| *efn-3* | WBGene00001164 | protein_coding_gene | -4.0 | 0.0633 | 2.1E-16 | -3.6 | 0.085 | 1.1E-06 |
| *ZK666.4* | WBGene00014044 | protein_coding_gene | -4.0 | 0.0631 | 7.4E-05 | -4.0 | 0.061 | 3.0E-05 |
| *fbxc-37* | WBGene00017942 | pseudogene | -4.0 | 0.0630 | 3.1E-05 | -3.6 | 0.080 | 1.7E-04 |
| *T10C6.16* | WBGene00077607 | protein_coding_gene | -4.0 | 0.0630 | 5.2E-05 | -2.6 | 0.168 | 8.6E-03 |
| *fbxc-31* | WBGene00019925 | protein_coding_gene | -4.0 | 0.0626 | 4.5E-05 | -3.4 | 0.098 | 3.8E-04 |
| *wee-1.2* | WBGene00006939 | pseudogene | -4.0 | 0.0624 | 2.8E-09 | -3.1 | 0.116 | 3.7E-07 |
| *jmjd-3.2* | WBGene00009089 | protein_coding_gene | -4.0 | 0.0622 | 4.5E-12 | -6.0 | 0.015 | 7.2E-21 |
| *Y39G10AL.1* | WBGene00021458 | protein_coding_gene | -4.0 | 0.0621 | 1.3E-20 | -5.3 | 0.026 | 9.0E-40 |
| *scl-27* | WBGene00022638 | protein_coding_gene | -4.0 | 0.0619 | 4.4E-05 | -2.6 | 0.163 | 2.2E-02 |
| *C06A1.4* | WBGene00014667 | pseudogene | -4.0 | 0.0618 | 5.6E-49 | -1.9 | 0.270 | 3.3E-36 |
| *C49F5.9* | WBGene00045316 | pseudogene | -4.0 | 0.0618 | 1.2E-15 | -2.7 | 0.151 | 1.7E-09 |
| *M151.1* | WBGene00019793 | protein_coding_gene | -4.0 | 0.0615 | 8.6E-05 | -3.8 | 0.071 | 4.8E-05 |
| *C24H12.7* | WBGene00016076 | pseudogene | -4.0 | 0.0614 | 6.7E-06 | -4.7 | 0.039 | 7.0E-07 |
| *F35E2.8* | WBGene00009415 | protein_coding_gene | -4.0 | 0.0611 | 7.9E-05 | -4.7 | 0.037 | 1.1E-07 |
| *T24D5.5* | WBGene00011993 | pseudogene | -4.0 | 0.0611 | 3.6E-05 | -2.8 | 0.140 | 2.9E-03 |
| *cyp-31A5* | WBGene00013381 | protein_coding_gene | -4.0 | 0.0610 | 4.4E-50 | -3.2 | 0.110 | 1.4E-72 |
| *fbxb-30* | WBGene00018458 | protein_coding_gene | -4.0 | 0.0610 | 3.0E-05 | -4.8 | 0.035 | 9.2E-08 |
| *aptf-4* | WBGene00009202 | protein_coding_gene | -4.0 | 0.0609 | 3.3E-17 | -4.2 | 0.054 | 5.8E-26 |
| *C44B9.3* | WBGene00008080 | protein_coding_gene | -4.0 | 0.0606 | 3.9E-41 | -3.5 | 0.089 | 2.5E-61 |
| *fbxb-32* | WBGene00019798 | protein_coding_gene | -4.0 | 0.0605 | 1.3E-05 | -4.0 | 0.064 | 1.2E-05 |
| *F29G9.1* | WBGene00017930 | protein_coding_gene | -4.0 | 0.0604 | 2.6E-05 | -4.1 | 0.060 | 3.5E-08 |
| *fbxb-78* | WBGene00017116 | protein_coding_gene | -4.1 | 0.0602 | 7.0E-06 | -4.7 | 0.039 | 5.8E-09 |
| *Y27F2A.8* | WBGene00021299 | protein_coding_gene | -4.1 | 0.0601 | 1.9E-05 | -5.7 | 0.020 | 1.6E-10 |
| *B0205.13* | WBGene00044644 | protein_coding_gene | -4.1 | 0.0599 | 1.0E-16 | -1.1 | 0.453 | 1.3E-02 |
| *B0416.10* | WBGene00206482 | protein_coding_gene | -4.1 | 0.0598 | 8.6E-06 | -2.9 | 0.135 | 3.3E-05 |
| *Y22D7AR.10* | WBGene00021263 | protein_coding_gene | -4.1 | 0.0597 | 3.6E-15 | -5.5 | 0.022 | 1.8E-63 |
| *DY3.8* | WBGene00008440 | protein_coding_gene | -4.1 | 0.0595 | 4.4E-71 | -4.2 | 0.055 | 3.7E-155 |
| *F54F7.2* | WBGene00010064 | protein_coding_gene | -4.1 | 0.0594 | 3.9E-18 | -3.2 | 0.109 | 3.2E-41 |
| *ZK666.1* | WBGene00014041 | protein_coding_gene | -4.1 | 0.0592 | 2.7E-05 | -2.7 | 0.156 | 8.5E-03 |
| *ceh-51* | WBGene00013583 | protein_coding_gene | -4.1 | 0.0591 | 1.0E-05 | -3.2 | 0.111 | 3.0E-06 |
| *C27C7.7* | WBGene00007769 | protein_coding_gene | -4.1 | 0.0590 | 1.6E-05 | -2.4 | 0.188 | 1.3E-02 |
| *F36H5.13* | WBGene00206519 | protein_coding_gene | -4.1 | 0.0588 | 1.8E-05 | -4.9 | 0.033 | 4.0E-08 |
| *egg-4* | WBGene00020652 | protein_coding_gene | -4.1 | 0.0588 | 9.5E-84 | -3.5 | 0.089 | 6.9E-85 |
| *Y51H7C.3* | WBGene00021781 | protein_coding_gene | -4.1 | 0.0587 | 9.5E-91 | -4.1 | 0.059 | 2.9E-102 |
| *K02F6.7* | WBGene00019341 | protein_coding_gene | -4.1 | 0.0584 | 3.1E-67 | -6.0 | 0.015 | 7.2E-51 |
| *aptf-2* | WBGene00013383 | protein_coding_gene | -4.1 | 0.0582 | 8.2E-53 | -3.9 | 0.068 | 6.7E-67 |
| *F54D11.3* | WBGene00018813 | pseudogene | -4.1 | 0.0577 | 8.5E-33 | -3.2 | 0.112 | 1.6E-22 |
| *Y13C8A.1* | WBGene00021188 | protein_coding_gene | -4.1 | 0.0577 | 1.1E-06 | -3.8 | 0.070 | 5.5E-24 |
| *Y77E11A.23* | WBGene00235267 | protein_coding_gene | -4.1 | 0.0575 | 2.9E-07 | -5.8 | 0.017 | 8.4E-10 |
| *memi-1* | WBGene00003216 | protein_coding_gene | -4.1 | 0.0573 | 4.7E-67 | -2.5 | 0.171 | 1.2E-63 |
| *kca-1* | WBGene00015698 | protein_coding_gene | -4.1 | 0.0572 | 4.0E-89 | -2.9 | 0.132 | 1.8E-88 |
| *F35C11.5* | WBGene00009404 | protein_coding_gene | -4.1 | 0.0572 | 3.2E-67 | -3.1 | 0.113 | 3.2E-140 |
| *atz-1* | WBGene00009212 | protein_coding_gene | -4.2 | 0.0563 | 7.3E-26 | -5.9 | 0.016 | 1.6E-76 |
| *Y18D10A.11* | WBGene00012481 | protein_coding_gene | -4.2 | 0.0560 | 1.1E-76 | -3.4 | 0.098 | 3.4E-147 |
| *cpar-1* | WBGene00010036 | protein_coding_gene | -4.2 | 0.0558 | 1.7E-53 | -3.6 | 0.081 | 3.4E-103 |
| *fbxb-67* | WBGene00009862 | protein_coding_gene | -4.2 | 0.0556 | 2.1E-14 | -5.8 | 0.018 | 3.4E-11 |
| *sdz-33* | WBGene00013233 | protein_coding_gene | -4.2 | 0.0555 | 4.1E-13 | -4.2 | 0.054 | 3.0E-18 |
| *W08F4.13* | WBGene00044733 | protein_coding_gene | -4.2 | 0.0555 | 6.9E-06 | -4.5 | 0.043 | 9.7E-07 |
| *lido-2* | WBGene00015587 | protein_coding_gene | -4.2 | 0.0552 | 4.4E-13 | -4.9 | 0.033 | 2.6E-14 |
| *F55C9.14* | WBGene00077663 | protein_coding_gene | -4.2 | 0.0550 | 7.2E-06 | -4.7 | 0.038 | 1.1E-07 |
| *fbxa-215* | WBGene00012879 | protein_coding_gene | -4.2 | 0.0550 | 1.6E-128 | -4.2 | 0.054 | 5.5E-231 |
| *fbxb-5* | WBGene00018445 | protein_coding_gene | -4.2 | 0.0549 | 1.6E-05 | -3.5 | 0.087 | 1.8E-04 |
| *T04D3.5* | WBGene00011434 | protein_coding_gene | -4.2 | 0.0548 | 1.9E-72 | -5.0 | 0.032 | 6.1E-115 |
| *T04G9.7* | WBGene00020218 | protein_coding_gene | -4.2 | 0.0547 | 8.8E-29 | -5.0 | 0.030 | 3.1E-207 |
| *pigv-1* | WBGene00020375 | protein_coding_gene | -4.2 | 0.0544 | 1.4E-71 | -3.4 | 0.094 | 3.5E-169 |
| *F31F6.2* | WBGene00009301 | protein_coding_gene | -4.2 | 0.0538 | 1.8E-25 | -4.0 | 0.062 | 8.9E-30 |
| *fbxb-107* | WBGene00017912 | protein_coding_gene | -4.2 | 0.0537 | 4.5E-06 | -4.4 | 0.049 | 7.2E-08 |
| *fbxb-111* | WBGene00017254 | protein_coding_gene | -4.2 | 0.0535 | 1.7E-05 | -4.0 | 0.062 | 1.2E-05 |
| *clec-88* | WBGene00019606 | protein_coding_gene | -4.2 | 0.0533 | 9.6E-119 | -4.3 | 0.049 | 1.6E-292 |
| *fbxb-19* | WBGene00019040 | protein_coding_gene | -4.2 | 0.0532 | 4.8E-06 | -5.5 | 0.023 | 1.1E-09 |
| *rnr-2* | WBGene00004392 | protein_coding_gene | -4.2 | 0.0529 | 3.6E-50 | -4.0 | 0.061 | 6.5E-226 |
| *C24H12.9* | WBGene00016078 | pseudogene | -4.2 | 0.0529 | 6.6E-06 | -4.2 | 0.053 | 8.0E-06 |
| *R11H6.6* | WBGene00014822 | pseudogene | -4.2 | 0.0529 | 6.8E-18 | -3.3 | 0.102 | 1.1E-11 |
| *clec-196* | WBGene00009156 | protein_coding_gene | -4.2 | 0.0528 | 6.9E-19 | -6.1 | 0.014 | 1.0E-44 |
| *W06B4.1* | WBGene00021056 | protein_coding_gene | -4.2 | 0.0526 | 1.4E-116 | -3.7 | 0.077 | 2.9E-157 |
| *dsl-1* | WBGene00001103 | protein_coding_gene | -4.3 | 0.0525 | 3.0E-07 | -4.4 | 0.046 | 5.0E-11 |
| *ZC373.2* | WBGene00013867 | protein_coding_gene | -4.3 | 0.0524 | 5.4E-58 | -4.3 | 0.050 | 2.9E-177 |

| | | | | | | | | |
|---|---|---|---|---|---|---|---|---|
| *fbxb-34* | WBGene00021092 | protein_coding_gene | -4.3 | 0.0521 | 7.7E-06 | -4.6 | 0.042 | 4.1E-07 |
| *T11B7.1* | WBGene00011706 | protein_coding_gene | -4.3 | 0.0521 | 9.0E-27 | -4.7 | 0.040 | 1.6E-53 |
| *fbxb-1* | WBGene00012063 | protein_coding_gene | -4.3 | 0.0520 | 6.9E-06 | -3.8 | 0.069 | 1.4E-04 |
| *ZK813.1* | WBGene00022820 | protein_coding_gene | -4.3 | 0.0519 | 2.5E-61 | -4.3 | 0.052 | 5.6E-220 |
| *K09D9.12* | WBGene00019569 | pseudogene | -4.3 | 0.0519 | 1.5E-18 | -3.7 | 0.077 | 1.7E-15 |
| *fbxc-34* | WBGene00019921 | protein_coding_gene | -4.3 | 0.0519 | 8.0E-06 | -3.1 | 0.113 | 1.3E-03 |
| *inx-2* | WBGene00002124 | protein_coding_gene | -4.3 | 0.0518 | 2.7E-49 | -6.4 | 0.012 | 6.9E-152 |
| *rgs-10* | WBGene00004353 | protein_coding_gene | -4.3 | 0.0518 | 1.1E-07 | -3.6 | 0.085 | 2.1E-08 |
| *fbxb-2* | WBGene00012065 | protein_coding_gene | -4.3 | 0.0516 | 6.5E-06 | -5.1 | 0.029 | 6.8E-09 |
| *tbx-38* | WBGene00006557 | protein_coding_gene | -4.3 | 0.0513 | 2.3E-05 | -4.4 | 0.049 | 7.5E-07 |
| *fbxb-18* | WBGene00019039 | protein_coding_gene | -4.3 | 0.0508 | 3.3E-06 | -5.6 | 0.021 | 5.1E-10 |
| *bath-10* | WBGene00021726 | protein_coding_gene | -4.3 | 0.0508 | 2.3E-09 | -4.0 | 0.061 | 1.9E-42 |
| *spn-4* | WBGene00004984 | protein_coding_gene | -4.3 | 0.0508 | 6.2E-98 | -4.0 | 0.062 | 1.7E-201 |
| *Y37D8A.19* | WBGene00012557 | protein_coding_gene | -4.3 | 0.0506 | 2.9E-26 | -4.2 | 0.053 | 6.0E-126 |
| *H29C22.1* | WBGene00045261 | protein_coding_gene | -4.3 | 0.0505 | 5.0E-50 | -5.6 | 0.020 | 3.8E-108 |
| *F07E5.12* | WBGene00219315 | protein_coding_gene | -4.3 | 0.0504 | 8.1E-06 | -3.4 | 0.092 | 4.1E-04 |
| *C38D9.8* | WBGene00008016 | protein_coding_gene | -4.3 | 0.0504 | 2.7E-08 | -6.7 | 0.009 | 5.1E-14 |
| *F18A1.7* | WBGene00017548 | protein_coding_gene | -4.3 | 0.0503 | 5.1E-66 | -4.4 | 0.048 | 1.1E-189 |
| *ddo-3* | WBGene00017648 | protein_coding_gene | -4.3 | 0.0503 | 2.0E-36 | -4.5 | 0.045 | 4.9E-212 |
| *H04D03.4* | WBGene00010365 | protein_coding_gene | -4.3 | 0.0501 | 2.7E-12 | -2.7 | 0.150 | 1.4E-15 |
| *inx-22* | WBGene00002144 | protein_coding_gene | -4.3 | 0.0500 | 2.7E-84 | -3.3 | 0.103 | 1.1E-118 |
| *lido-5* | WBGene00019576 | protein_coding_gene | -4.3 | 0.0499 | 5.4E-18 | -3.9 | 0.067 | 7.2E-24 |
| *W04A8.3* | WBGene00012238 | protein_coding_gene | -4.3 | 0.0499 | 7.5E-06 | -4.6 | 0.041 | 3.7E-09 |
| *T16G12.8* | WBGene00011809 | protein_coding_gene | -4.3 | 0.0498 | 2.1E-39 | -4.9 | 0.033 | 9.8E-53 |
| *cyb-2.1* | WBGene00000866 | protein_coding_gene | -4.3 | 0.0495 | 3.7E-80 | -3.6 | 0.082 | 5.5E-160 |
| *Y82E9BR.1* | WBGene00022334 | protein_coding_gene | -4.3 | 0.0492 | 1.9E-18 | -6.9 | 0.009 | 1.1E-53 |
| *C44B7.5* | WBGene00016627 | protein_coding_gene | -4.4 | 0.0489 | 1.2E-47 | -4.8 | 0.035 | 8.0E-215 |
| *kin-34* | WBGene00019815 | protein_coding_gene | -4.4 | 0.0486 | 1.8E-07 | -5.6 | 0.020 | 2.4E-10 |
| *T24E12.11* | WBGene00020778 | protein_coding_gene | -4.4 | 0.0483 | 2.1E-17 | -3.3 | 0.104 | 1.7E-06 |
| *ceh-83* | WBGene00018446 | protein_coding_gene | -4.4 | 0.0480 | 5.0E-45 | -4.5 | 0.045 | 8.0E-49 |
| *F58E1.13* | WBGene00019044 | protein_coding_gene | -4.4 | 0.0478 | 2.3E-06 | -2.4 | 0.194 | 3.9E-02 |
| *Y75B12B.1* | WBGene00013567 | protein_coding_gene | -4.4 | 0.0478 | 5.8E-77 | -4.1 | 0.057 | 1.1E-210 |
| *tbx-43* | WBGene00044798 | protein_coding_gene | -4.4 | 0.0476 | 2.0E-06 | -2.0 | 0.252 | 5.2E-04 |
| *dgtr-1* | WBGene00020910 | protein_coding_gene | -4.4 | 0.0474 | 3.0E-51 | -3.5 | 0.091 | 7.6E-72 |
| *T28A11.22* | WBGene00044526 | protein_coding_gene | -4.4 | 0.0469 | 3.8E-06 | -5.4 | 0.023 | 8.8E-09 |
| *cyp-31A2* | WBGene00010354 | protein_coding_gene | -4.4 | 0.0467 | 7.0E-101 | -3.7 | 0.078 | 9.1E-184 |
| *ZK430.5* | WBGene00022741 | protein_coding_gene | -4.4 | 0.0466 | 4.7E-17 | -4.9 | 0.034 | 8.5E-22 |
| *sdz-12* | WBGene00017406 | protein_coding_gene | -4.4 | 0.0466 | 2.7E-06 | -2.2 | 0.217 | 2.1E-02 |
| *C35E7.3* | WBGene00016455 | protein_coding_gene | -4.4 | 0.0464 | 1.3E-12 | -4.9 | 0.034 | 4.4E-17 |
| *Y47H10A.2* | WBGene00012958 | protein_coding_gene | -4.4 | 0.0462 | 1.9E-11 | -3.6 | 0.080 | 3.2E-12 |
| *F23D12.4* | WBGene00009088 | protein_coding_gene | -4.4 | 0.0462 | 5.9E-06 | -5.1 | 0.029 | 6.0E-08 |
| *F55B11.3* | WBGene00010085 | protein_coding_gene | -4.4 | 0.0460 | 3.7E-45 | -4.6 | 0.042 | 1.4E-102 |
| *col-135* | WBGene00000708 | protein_coding_gene | -4.4 | 0.0459 | 7.3E-29 | -3.6 | 0.081 | 1.5E-70 |
| *fbxb-45* | WBGene00019705 | protein_coding_gene | -4.4 | 0.0459 | 1.2E-06 | -5.2 | 0.028 | 1.4E-08 |
| *gna-2* | WBGene00001647 | protein_coding_gene | -4.4 | 0.0458 | 3.6E-82 | -5.0 | 0.031 | 4.6E-217 |
| *T24D1.3* | WBGene00011986 | protein_coding_gene | -4.5 | 0.0456 | 2.1E-112 | -3.5 | 0.088 | 4.4E-88 |
| *fbxb-106* | WBGene00017911 | protein_coding_gene | -4.5 | 0.0456 | 1.7E-06 | -3.8 | 0.073 | 5.8E-05 |
| *fbxb-62* | WBGene00010105 | protein_coding_gene | -4.5 | 0.0456 | 1.4E-06 | -3.6 | 0.082 | 8.1E-05 |
| *Y49E10.30* | WBGene00194981 | protein_coding_gene | -4.5 | 0.0455 | 2.6E-06 | -2.9 | 0.134 | 5.7E-10 |
| *puf-10* | WBGene00004246 | pseudogene | -4.5 | 0.0453 | 3.3E-47 | -2.8 | 0.142 | 2.3E-43 |
| *H04M03.3* | WBGene00019153 | protein_coding_gene | -4.5 | 0.0453 | 5.7E-63 | -3.5 | 0.088 | 1.4E-57 |
| *ZK250.15* | WBGene00235355 | pseudogene | -4.5 | 0.0450 | 1.8E-06 | -4.7 | 0.039 | 1.1E-07 |
| *F19B10.2* | WBGene00017584 | protein_coding_gene | -4.5 | 0.0447 | 1.3E-38 | -2.5 | 0.182 | 1.4E-03 |
| *T05F1.2* | WBGene00011489 | protein_coding_gene | -4.5 | 0.0447 | 1.6E-147 | -4.5 | 0.043 | 8.7E-275 |
| *T15H9.6* | WBGene00011791 | protein_coding_gene | -4.5 | 0.0445 | 1.8E-09 | -5.7 | 0.019 | 1.3E-09 |
| *Y116A8C.20* | WBGene00013797 | protein_coding_gene | -4.5 | 0.0445 | 1.3E-06 | -4.7 | 0.039 | 2.1E-07 |
| *fbxc-22* | WBGene00019041 | protein_coding_gene | -4.5 | 0.0445 | 1.1E-06 | -5.0 | 0.031 | 2.1E-08 |
| *szy-4* | WBGene00077732 | protein_coding_gene | -4.5 | 0.0439 | 1.4E-170 | -3.7 | 0.078 | 5.7E-131 |
| *C27D9.1* | WBGene00016163 | protein_coding_gene | -4.5 | 0.0437 | 2.3E-111 | -3.7 | 0.078 | 1.8E-105 |
| *Y45G5AM.5* | WBGene00021557 | protein_coding_gene | -4.5 | 0.0436 | 2.2E-08 | -5.2 | 0.027 | 1.9E-16 |
| *Y51F10.2* | WBGene00021763 | protein_coding_gene | -4.5 | 0.0436 | 3.5E-108 | -4.4 | 0.048 | 5.6E-184 |
| *Y116F11A.1* | WBGene00013816 | protein_coding_gene | -4.5 | 0.0431 | 2.3E-06 | -2.4 | 0.195 | 8.4E-05 |
| *col-43* | WBGene00000620 | protein_coding_gene | -4.5 | 0.0429 | 4.1E-05 | -2.5 | 0.174 | 5.9E-03 |
| *bath-8* | WBGene00021723 | pseudogene | -4.6 | 0.0426 | 9.1E-07 | -5.0 | 0.031 | 8.4E-09 |
| *Y39A1A.16* | WBGene00012654 | protein_coding_gene | -4.6 | 0.0425 | 4.6E-06 | -3.3 | 0.105 | 1.0E-04 |
| *ZK1053.4* | WBGene00014199 | protein_coding_gene | -4.6 | 0.0419 | 1.2E-33 | -5.5 | 0.023 | 1.4E-78 |
| *fbxb-61* | WBGene00010106 | protein_coding_gene | -4.6 | 0.0419 | 1.0E-06 | -4.8 | 0.037 | 2.1E-07 |
| *W04A8.5* | WBGene00012240 | protein_coding_gene | -4.6 | 0.0418 | 9.3E-12 | -5.5 | 0.022 | 7.1E-09 |
| *T09B9.1* | WBGene00011642 | protein_coding_gene | -4.6 | 0.0418 | 2.1E-48 | -5.3 | 0.025 | 1.0E-50 |
| *D1054.10* | WBGene00008377 | protein_coding_gene | -4.6 | 0.0418 | 3.5E-52 | -4.5 | 0.045 | 1.7E-260 |
| *Y49F6A.11* | WBGene00303366 | protein_coding_gene | -4.6 | 0.0418 | 1.0E-06 | -3.6 | 0.081 | 4.1E-05 |
| *Y51F10.7* | WBGene00021768 | protein_coding_gene | -4.6 | 0.0416 | 5.9E-57 | -2.3 | 0.198 | 3.6E-37 |
| *B0513.4* | WBGene00007196 | protein_coding_gene | -4.6 | 0.0415 | 1.9E-46 | -3.6 | 0.082 | 8.3E-89 |
| *fbxb-3* | WBGene00009546 | protein_coding_gene | -4.6 | 0.0414 | 5.4E-07 | -5.5 | 0.021 | 4.3E-09 |
| *Y57A10C.11* | WBGene00044047 | protein_coding_gene | -4.6 | 0.0413 | 7.7E-07 | -4.8 | 0.037 | 1.1E-07 |
| *arrd-1* | WBGene00020205 | protein_coding_gene | -4.6 | 0.0413 | 1.9E-18 | -6.3 | 0.013 | 1.3E-30 |
| *fbxb-33* | WBGene00044314 | protein_coding_gene | -4.6 | 0.0408 | 1.2E-06 | -4.7 | 0.038 | 8.3E-07 |
| *egg-6* | WBGene00010621 | protein_coding_gene | -4.6 | 0.0408 | 1.3E-88 | -3.5 | 0.090 | 8.3E-175 |
| *fbxb-75* | WBGene00019956 | protein_coding_gene | -4.6 | 0.0408 | 8.8E-12 | -5.6 | 0.021 | 9.4E-16 |
| *F54F7.6* | WBGene00010066 | protein_coding_gene | -4.6 | 0.0407 | 6.3E-28 | -5.1 | 0.030 | 1.1E-50 |

| | | | | | | | | |
|---|---|---|---|---|---|---|---|---|
| *R02C2.6* | WBGene00019817 | protein_coding_gene | -4.6 | 0.0407 | 3.1E-06 | -3.4 | 0.096 | 3.0E-04 |
| *mex-5* | WBGene00003230 | protein_coding_gene | -4.6 | 0.0407 | 1.3E-71 | -3.5 | 0.085 | 1.1E-154 |
| *oma-1* | WBGene00003864 | protein_coding_gene | -4.6 | 0.0406 | 7.3E-103 | -3.7 | 0.076 | 4.1E-166 |
| *fbxb-47* | WBGene00019045 | protein_coding_gene | -4.6 | 0.0404 | 7.0E-07 | -4.7 | 0.037 | 9.7E-08 |
| *Y19D2B.2* | WBGene00012490 | protein_coding_gene | -4.6 | 0.0401 | 8.0E-07 | -1.3 | 0.414 | 2.4E-02 |
| *W04E12.2* | WBGene00012247 | protein_coding_gene | -4.6 | 0.0400 | 1.7E-10 | -4.2 | 0.055 | 2.9E-14 |
| *C43D7.10* | WBGene00077441 | pseudogene | -4.6 | 0.0399 | 8.6E-07 | -5.1 | 0.029 | 5.0E-09 |
| *ule-3* | WBGene00008378 | protein_coding_gene | -4.6 | 0.0398 | 8.9E-69 | -4.6 | 0.040 | 3.7E-210 |
| *T04B8.2* | WBGene00020204 | pseudogene | -4.7 | 0.0398 | 1.5E-06 | -5.1 | 0.029 | 1.3E-08 |
| *sdz-31* | WBGene00011704 | protein_coding_gene | -4.7 | 0.0396 | 7.1E-07 | -3.5 | 0.085 | 1.3E-04 |
| *C17E4.2* | WBGene00007642 | protein_coding_gene | -4.7 | 0.0396 | 4.1E-18 | -1.2 | 0.425 | 4.7E-13 |
| *EEED8.3* | WBGene00017134 | protein_coding_gene | -4.7 | 0.0391 | 1.9E-123 | -5.2 | 0.027 | 5.1E-237 |
| *mrp-4* | WBGene00003410 | protein_coding_gene | -4.7 | 0.0388 | 5.9E-66 | -3.7 | 0.077 | 5.3E-188 |
| *fbxb-10* | WBGene00020362 | protein_coding_gene | -4.7 | 0.0386 | 1.7E-07 | -2.3 | 0.209 | 1.6E-02 |
| *ptr-2* | WBGene00004217 | protein_coding_gene | -4.7 | 0.0386 | 2.9E-142 | -4.2 | 0.055 | 3.7E-157 |
| *Y57G7A.2* | WBGene00021966 | protein_coding_gene | -4.7 | 0.0380 | 9.1E-07 | -4.7 | 0.038 | 4.6E-07 |
| *F45D11.5* | WBGene00018452 | protein_coding_gene | -4.7 | 0.0379 | 1.9E-07 | -4.3 | 0.051 | 3.8E-06 |
| *clec-223* | WBGene00007282 | protein_coding_gene | -4.7 | 0.0378 | 8.8E-55 | -5.2 | 0.027 | 2.0E-69 |
| *K10C2.8* | WBGene00045265 | protein_coding_gene | -4.7 | 0.0377 | 7.0E-37 | -2.7 | 0.152 | 7.2E-14 |
| *fbxb-63* | WBGene00044197 | protein_coding_gene | -4.7 | 0.0374 | 8.1E-07 | -4.3 | 0.052 | 5.5E-06 |
| *fbxc-50* | WBGene00015912 | protein_coding_gene | -4.7 | 0.0373 | 1.9E-75 | -3.6 | 0.084 | 4.0E-25 |
| *fbxa-181* | WBGene00009427 | protein_coding_gene | -4.7 | 0.0372 | 1.3E-07 | -2.5 | 0.173 | 1.2E-03 |
| *T26E3.5* | WBGene00012038 | protein_coding_gene | -4.8 | 0.0371 | 1.5E-06 | -5.2 | 0.028 | 5.0E-08 |
| *Y62H9A.4* | WBGene00013392 | protein_coding_gene | -4.8 | 0.0370 | 5.1E-41 | -4.3 | 0.050 | 8.9E-82 |
| *W05F2.3* | WBGene00021035 | protein_coding_gene | -4.8 | 0.0369 | 7.2E-60 | -4.3 | 0.050 | 2.7E-199 |
| *tdc-1* | WBGene00006562 | protein_coding_gene | -4.8 | 0.0368 | 1.7E-102 | -4.0 | 0.062 | 6.0E-219 |
| *Y57A10C.1* | WBGene00013280 | protein_coding_gene | -4.8 | 0.0367 | 2.2E-07 | -5.4 | 0.023 | 1.3E-09 |
| *K11D12.13* | WBGene00044535 | protein_coding_gene | -4.8 | 0.0365 | 6.6E-53 | -5.5 | 0.022 | 4.1E-144 |
| *Y24F12A.4* | WBGene00012499 | pseudogene | -4.8 | 0.0364 | 6.1E-07 | -2.0 | 0.244 | 9.5E-03 |
| *T11F8.1* | WBGene00020433 | protein_coding_gene | -4.8 | 0.0364 | 1.8E-49 | -3.9 | 0.066 | 6.6E-94 |
| *fbxb-31* | WBGene00019797 | protein_coding_gene | -4.8 | 0.0364 | 7.7E-07 | -4.2 | 0.054 | 1.4E-05 |
| *fbxb-57* | WBGene00010101 | protein_coding_gene | -4.8 | 0.0361 | 3.8E-07 | -6.1 | 0.015 | 9.2E-12 |
| *F02D10.6* | WBGene00008526 | protein_coding_gene | -4.8 | 0.0359 | 1.1E-12 | -6.1 | 0.015 | 7.4E-11 |
| *col-8* | WBGene00000597 | protein_coding_gene | -4.8 | 0.0355 | 4.2E-50 | -5.0 | 0.030 | 1.7E-69 |
| *memi-3* | WBGene00010353 | protein_coding_gene | -4.8 | 0.0353 | 3.2E-122 | -2.9 | 0.131 | 4.7E-121 |
| *Y57A10B.6* | WBGene00013278 | protein_coding_gene | -4.8 | 0.0352 | 1.7E-11 | -6.2 | 0.014 | 1.5E-12 |
| *F40G9.18* | WBGene00189954 | protein_coding_gene | -4.8 | 0.0351 | 2.7E-07 | -5.0 | 0.031 | 2.4E-08 |
| *F36H5.4* | WBGene00018103 | protein_coding_gene | -4.8 | 0.0349 | 1.5E-07 | -5.6 | 0.020 | 6.0E-10 |
| *fbxc-26* | WBGene00019043 | protein_coding_gene | -4.8 | 0.0348 | 1.5E-07 | -5.2 | 0.028 | 8.3E-09 |
| *sdz-21* | WBGene00018828 | protein_coding_gene | -4.8 | 0.0348 | 9.9E-10 | -5.8 | 0.018 | 1.0E-09 |
| *fbxb-81* | WBGene00020541 | protein_coding_gene | -4.9 | 0.0341 | 3.1E-07 | -2.5 | 0.177 | 2.1E-02 |
| *marc-3* | WBGene00007643 | protein_coding_gene | -4.9 | 0.0341 | 1.0E-123 | -3.6 | 0.080 | 6.2E-164 |
| *ZK813.3* | WBGene00022822 | protein_coding_gene | -4.9 | 0.0340 | 3.7E-42 | -4.5 | 0.045 | 1.1E-164 |
| *C08F8.15* | WBGene00219217 | protein_coding_gene | -4.9 | 0.0334 | 5.5E-08 | -5.6 | 0.020 | 3.4E-10 |
| *Y49F6A.5* | WBGene00021713 | protein_coding_gene | -4.9 | 0.0334 | 1.8E-07 | -3.6 | 0.081 | 4.3E-05 |
| *sdz-10* | WBGene00017252 | protein_coding_gene | -4.9 | 0.0332 | 8.7E-08 | -5.5 | 0.023 | 8.2E-10 |
| *T21C9.13* | WBGene00011898 | protein_coding_gene | -4.9 | 0.0330 | 1.3E-78 | -4.5 | 0.043 | 1.4E-225 |
| *F22E5.20* | WBGene00017713 | protein_coding_gene | -4.9 | 0.0329 | 5.3E-08 | -5.7 | 0.019 | 1.3E-10 |
| *sprp-4* | WBGene00016189 | protein_coding_gene | -4.9 | 0.0325 | 4.1E-18 | -5.5 | 0.022 | 4.1E-21 |
| *dot-1.4* | WBGene00010120 | protein_coding_gene | -4.9 | 0.0325 | 3.5E-08 | -6.2 | 0.014 | 2.9E-12 |
| *egg-3* | WBGene00009701 | protein_coding_gene | -5.0 | 0.0323 | 1.9E-106 | -4.0 | 0.062 | 8.5E-178 |
| *F02H6.1* | WBGene00008534 | pseudogene | -5.0 | 0.0323 | 7.4E-13 | -4.7 | 0.039 | 3.2E-07 |
| *fbxc-21* | WBGene00019042 | protein_coding_gene | -5.0 | 0.0321 | 7.8E-08 | -4.8 | 0.036 | 5.0E-08 |
| *fbxc-25* | WBGene00019591 | protein_coding_gene | -5.0 | 0.0321 | 4.2E-08 | -2.5 | 0.174 | 5.0E-06 |
| *F12E12.6* | WBGene00017407 | protein_coding_gene | -5.0 | 0.0320 | 3.8E-08 | -5.7 | 0.019 | 3.7E-11 |
| *Y54F10AM.11* | WBGene00021853 | protein_coding_gene | -5.0 | 0.0319 | 6.6E-49 | -4.4 | 0.049 | 2.6E-54 |
| *W04A4.2* | WBGene00012231 | protein_coding_gene | -5.0 | 0.0319 | 8.5E-36 | -3.8 | 0.070 | 2.8E-50 |
| *hch-1* | WBGene00001828 | protein_coding_gene | -5.0 | 0.0319 | 2.7E-08 | -8.5 | 0.003 | 1.3E-20 |
| *fbxb-108* | WBGene00017249 | protein_coding_gene | -5.0 | 0.0317 | 1.5E-07 | -2.7 | 0.158 | 7.5E-03 |
| *F57G9.3* | WBGene00010219 | protein_coding_gene | -5.0 | 0.0317 | 4.1E-08 | -4.6 | 0.040 | 7.4E-07 |
| *T27E9.6* | WBGene00012096 | protein_coding_gene | -5.0 | 0.0315 | 6.1E-08 | -4.7 | 0.039 | 1.3E-07 |
| *tbh-1* | WBGene00006541 | protein_coding_gene | -5.0 | 0.0314 | 3.4E-56 | -4.6 | 0.040 | 2.7E-153 |
| *Y44A6C.2* | WBGene00012851 | protein_coding_gene | -5.0 | 0.0310 | 2.3E-16 | -4.6 | 0.042 | 4.0E-24 |
| *fbxb-11* | WBGene00018436 | protein_coding_gene | -5.0 | 0.0309 | 6.0E-08 | -5.2 | 0.026 | 3.2E-08 |
| *K09E3.7* | WBGene00019578 | protein_coding_gene | -5.0 | 0.0309 | 1.6E-36 | -4.3 | 0.051 | 6.1E-37 |
| *fbxb-92* | WBGene00044693 | protein_coding_gene | -5.0 | 0.0308 | 5.6E-08 | -3.5 | 0.090 | 1.5E-04 |
| *gln-6* | WBGene00001607 | protein_coding_gene | -5.0 | 0.0308 | 3.3E-156 | -4.8 | 0.037 | 3.3E-268 |
| *fbxb-82* | WBGene00020542 | protein_coding_gene | -5.0 | 0.0306 | 5.9E-08 | -4.9 | 0.033 | 6.6E-08 |
| *F45C12.17* | WBGene00269388 | protein_coding_gene | -5.0 | 0.0305 | 4.5E-08 | -5.9 | 0.017 | 5.9E-10 |
| *F40G9.9* | WBGene00018245 | protein_coding_gene | -5.0 | 0.0304 | 6.0E-08 | -4.6 | 0.041 | 1.2E-06 |
| *T05E12.10* | WBGene00077787 | ncRNA_gene | -5.0 | 0.0303 | 4.4E-08 | -4.2 | 0.053 | 1.4E-06 |
| *T24C4.2* | WBGene00020758 | protein_coding_gene | -5.0 | 0.0303 | 6.7E-32 | -5.3 | 0.026 | 6.4E-47 |
| *spp-17* | WBGene00005002 | protein_coding_gene | -5.1 | 0.0301 | 5.9E-54 | -3.6 | 0.084 | 6.9E-104 |
| *egg-5* | WBGene00020035 | protein_coding_gene | -5.1 | 0.0299 | 1.1E-88 | -3.6 | 0.081 | 6.1E-89 |
| *fbxb-83* | WBGene00020543 | protein_coding_gene | -5.1 | 0.0299 | 1.3E-07 | -3.6 | 0.081 | 7.2E-05 |
| *sdz-36* | WBGene00014242 | protein_coding_gene | -5.1 | 0.0298 | 6.1E-08 | -4.7 | 0.038 | 4.8E-07 |
| *F19F10.1* | WBGene00017598 | protein_coding_gene | -5.1 | 0.0297 | 2.2E-08 | -5.0 | 0.031 | 7.4E-09 |
| *acox-1.3* | WBGene00008566 | protein_coding_gene | -5.1 | 0.0297 | 1.0E-28 | -1.1 | 0.475 | 2.3E-03 |
| *clec-97* | WBGene00013931 | protein_coding_gene | -5.1 | 0.0296 | 9.0E-30 | -3.4 | 0.096 | 2.7E-25 |
| *F19B10.5* | WBGene00017587 | protein_coding_gene | -5.1 | 0.0295 | 3.0E-15 | -3.1 | 0.119 | 3.8E-10 |

| | | | | | | | | |
|---|---|---|---|---|---|---|---|---|
| *memi-2* | WBGene00013380 | protein_coding_gene | -5.1 | 0.0295 | 1.1E-82 | -3.7 | 0.076 | 1.9E-75 |
| *EEED8.12* | WBGene00017140 | protein_coding_gene | -5.1 | 0.0293 | 2.2E-08 | -5.3 | 0.025 | 4.2E-12 |
| *cht-3* | WBGene00016084 | protein_coding_gene | -5.1 | 0.0293 | 5.6E-55 | -5.8 | 0.018 | 1.1E-260 |
| *Y60A9.3* | WBGene00013370 | protein_coding_gene | -5.1 | 0.0292 | 1.2E-07 | -4.5 | 0.043 | 2.7E-06 |
| *C17F4.7* | WBGene00015913 | protein_coding_gene | -5.1 | 0.0292 | 4.4E-58 | -3.0 | 0.121 | 2.0E-18 |
| *F48E3.4* | WBGene00018605 | protein_coding_gene | -5.1 | 0.0291 | 9.1E-46 | -5.7 | 0.020 | 1.1E-180 |
| *C24D10.5* | WBGene00016056 | protein_coding_gene | -5.1 | 0.0290 | 4.9E-14 | -5.9 | 0.016 | 2.1E-15 |
| *fbxb-22* | WBGene00013230 | protein_coding_gene | -5.1 | 0.0284 | 4.1E-08 | -3.8 | 0.070 | 2.5E-10 |
| *C10G8.4* | WBGene00015683 | protein_coding_gene | -5.2 | 0.0280 | 2.5E-32 | -5.5 | 0.022 | 5.6E-155 |
| *nhr-2* | WBGene00003601 | protein_coding_gene | -5.2 | 0.0278 | 6.8E-47 | -4.8 | 0.036 | 2.0E-76 |
| *Y39E4B.5* | WBGene00012716 | protein_coding_gene | -5.2 | 0.0278 | 6.6E-65 | -3.8 | 0.071 | 1.7E-91 |
| *mex-1* | WBGene00003228 | protein_coding_gene | -5.2 | 0.0277 | 2.1E-111 | -4.8 | 0.035 | 1.2E-204 |
| *fbxb-90* | WBGene00044694 | protein_coding_gene | -5.2 | 0.0277 | 1.7E-08 | -3.1 | 0.115 | 2.7E-03 |
| *T11F8.2* | WBGene00020434 | pseudogene | -5.2 | 0.0276 | 1.1E-11 | -5.0 | 0.031 | 1.1E-15 |
| *mesp-1* | WBGene00016178 | protein_coding_gene | -5.2 | 0.0275 | 4.6E-116 | -3.9 | 0.066 | 1.2E-130 |
| *fbxb-48* | WBGene00018444 | protein_coding_gene | -5.2 | 0.0274 | 1.9E-08 | -5.1 | 0.029 | 6.4E-08 |
| *Y106G6D.1* | WBGene00013698 | protein_coding_gene | -5.2 | 0.0274 | 2.6E-14 | -5.8 | 0.019 | 2.2E-17 |
| *F44F1.6* | WBGene00009698 | protein_coding_gene | -5.2 | 0.0273 | 6.4E-09 | -7.3 | 0.007 | 4.5E-16 |
| *Y43B11AL.5* | WBGene00235272 | pseudogene | -5.2 | 0.0273 | 1.4E-08 | -4.1 | 0.058 | 6.7E-06 |
| *M151.2* | WBGene00019794 | protein_coding_gene | -5.2 | 0.0269 | 8.7E-09 | -4.9 | 0.035 | 3.0E-08 |
| *mex-6* | WBGene00003231 | protein_coding_gene | -5.2 | 0.0268 | 3.0E-141 | -4.2 | 0.053 | 5.5E-179 |
| *F10G2.4* | WBGene00017365 | protein_coding_gene | -5.2 | 0.0267 | 1.0E-08 | -2.9 | 0.134 | 2.5E-03 |
| *dct-13* | WBGene00013794 | protein_coding_gene | -5.2 | 0.0267 | 2.2E-08 | -4.8 | 0.036 | 8.9E-08 |
| *ceh-82* | WBGene00018433 | protein_coding_gene | -5.2 | 0.0265 | 1.2E-08 | -4.3 | 0.050 | 5.2E-07 |
| *fbxb-104* | WBGene00021719 | protein_coding_gene | -5.2 | 0.0263 | 6.9E-08 | -6.0 | 0.016 | 1.9E-11 |
| *W06D11.2* | WBGene00012300 | protein_coding_gene | -5.2 | 0.0263 | 3.3E-08 | -4.3 | 0.051 | 3.3E-06 |
| *Y43B11AL.1* | WBGene00021542 | protein_coding_gene | -5.2 | 0.0263 | 1.2E-08 | -5.4 | 0.023 | 2.0E-12 |
| *puf-7* | WBGene00004243 | protein_coding_gene | -5.3 | 0.0262 | 2.4E-73 | -4.2 | 0.056 | 4.5E-92 |
| *F48E3.6* | WBGene00018606 | protein_coding_gene | -5.3 | 0.0262 | 3.3E-11 | -6.8 | 0.009 | 1.1E-23 |
| *K08H2.3* | WBGene00010690 | protein_coding_gene | -5.3 | 0.0259 | 7.4E-34 | -3.2 | 0.108 | 3.5E-11 |
| *ceh-40* | WBGene00000461 | protein_coding_gene | -5.3 | 0.0259 | 6.7E-21 | -5.7 | 0.020 | 5.7E-28 |
| *F46C3.4* | WBGene00044702 | pseudogene | -5.3 | 0.0256 | 6.4E-12 | -2.8 | 0.147 | 1.3E-09 |
| *fbxb-119* | WBGene00007182 | protein_coding_gene | -5.3 | 0.0256 | 1.3E-08 | -4.7 | 0.039 | 3.3E-13 |
| *fbxa-170* | WBGene00016486 | protein_coding_gene | -5.3 | 0.0255 | 1.3E-27 | -4.4 | 0.048 | 1.8E-49 |
| *col-143* | WBGene00000716 | protein_coding_gene | -5.3 | 0.0254 | 3.3E-20 | -3.6 | 0.081 | 3.1E-29 |
| *sdz-5* | WBGene00016578 | protein_coding_gene | -5.3 | 0.0253 | 8.4E-09 | -6.2 | 0.014 | 4.6E-12 |
| *K08D9.9* | WBGene00235094 | protein_coding_gene | -5.3 | 0.0253 | 4.3E-10 | -6.1 | 0.014 | 5.7E-12 |
| *fbxc-29* | WBGene00019923 | protein_coding_gene | -5.3 | 0.0248 | 3.8E-09 | -5.2 | 0.026 | 3.6E-08 |
| *egg-2* | WBGene00019811 | protein_coding_gene | -5.4 | 0.0245 | 2.1E-128 | -5.3 | 0.025 | 1.6E-294 |
| *F38C2.7* | WBGene00009539 | protein_coding_gene | -5.4 | 0.0244 | 3.3E-08 | -5.2 | 0.026 | 6.7E-09 |
| *perm-4* | WBGene00016638 | protein_coding_gene | -5.4 | 0.0242 | 7.2E-70 | -4.0 | 0.061 | 7.0E-172 |
| *clec-190* | WBGene00010953 | protein_coding_gene | -5.4 | 0.0241 | 4.1E-07 | -5.9 | 0.016 | 1.5E-11 |
| *sdz-29* | WBGene00020081 | pseudogene | -5.4 | 0.0239 | 4.3E-09 | -3.1 | 0.118 | 4.8E-04 |
| *K02B12.2* | WBGene00010495 | protein_coding_gene | -5.4 | 0.0237 | 1.5E-18 | -5.8 | 0.018 | 6.6E-32 |
| *Y45F10C.2* | WBGene00012878 | protein_coding_gene | -5.4 | 0.0236 | 7.6E-08 | -6.0 | 0.016 | 1.5E-10 |
| *C05C10.5* | WBGene00007332 | protein_coding_gene | -5.4 | 0.0236 | 6.4E-174 | -4.9 | 0.034 | 1.2E-175 |
| *C39D10.7* | WBGene00016534 | protein_coding_gene | -5.4 | 0.0235 | 2.0E-57 | -4.7 | 0.038 | 1.2E-151 |
| *vet-1* | WBGene00020261 | protein_coding_gene | -5.4 | 0.0235 | 1.1E-29 | -5.5 | 0.022 | 1.5E-61 |
| *dot-1.5* | WBGene00022512 | protein_coding_gene | -5.4 | 0.0233 | 6.0E-09 | -4.2 | 0.054 | 1.1E-06 |
| *F46E10.2* | WBGene00018489 | protein_coding_gene | -5.4 | 0.0231 | 1.1E-08 | -7.8 | 0.004 | 4.1E-18 |
| *F34D6.1* | WBGene00018021 | protein_coding_gene | -5.4 | 0.0231 | 2.7E-09 | -6.1 | 0.015 | 5.0E-12 |
| *gadr-2* | WBGene00007887 | protein_coding_gene | -5.4 | 0.0231 | 2.0E-09 | -6.3 | 0.012 | 8.7E-13 |
| *fbxb-105* | WBGene00017251 | protein_coding_gene | -5.4 | 0.0230 | 4.9E-09 | -4.0 | 0.060 | 2.7E-05 |
| *ccch-2* | WBGene00009537 | protein_coding_gene | -5.4 | 0.0230 | 1.0E-08 | -5.7 | 0.019 | 1.3E-10 |
| *ule-5* | WBGene00013394 | protein_coding_gene | -5.4 | 0.0230 | 5.7E-108 | -5.4 | 0.023 | 6.9E-286 |
| *T10C6.10* | WBGene00011691 | protein_coding_gene | -5.5 | 0.0228 | 1.9E-36 | -3.0 | 0.125 | 5.0E-21 |
| *M04D5.1* | WBGene00014807 | protein_coding_gene | -5.5 | 0.0228 | 5.4E-09 | -4.9 | 0.034 | 4.3E-08 |
| *K08H2.2* | WBGene00010689 | protein_coding_gene | -5.5 | 0.0227 | 4.8E-09 | -4.9 | 0.033 | 2.1E-07 |
| *puf-6* | WBGene00004242 | protein_coding_gene | -5.5 | 0.0226 | 1.3E-64 | -3.9 | 0.065 | 2.1E-80 |
| *C46E10.8* | WBGene00016712 | protein_coding_gene | -5.5 | 0.0226 | 1.2E-09 | -4.9 | 0.033 | 1.1E-08 |
| *F31B9.3* | WBGene00009280 | protein_coding_gene | -5.5 | 0.0226 | 2.0E-27 | -4.4 | 0.047 | 2.9E-36 |
| *K12H6.7* | WBGene00019687 | protein_coding_gene | -5.5 | 0.0224 | 2.0E-09 | -3.8 | 0.074 | 1.7E-05 |
| *Y62H9A.3* | WBGene00013391 | protein_coding_gene | -5.5 | 0.0222 | 3.0E-58 | -4.8 | 0.036 | 6.6E-104 |
| *pos-1* | WBGene00004078 | protein_coding_gene | -5.5 | 0.0222 | 9.3E-124 | -5.2 | 0.027 | 0.0E+00 |
| *cyb-2.2* | WBGene00000867 | protein_coding_gene | -5.5 | 0.0221 | 2.8E-136 | -4.2 | 0.055 | 2.1E-197 |
| *perm-2* | WBGene00016636 | protein_coding_gene | -5.5 | 0.0219 | 5.0E-68 | -4.6 | 0.041 | 7.2E-196 |
| *F18A12.7* | WBGene00017556 | protein_coding_gene | -5.5 | 0.0219 | 1.5E-09 | -5.1 | 0.030 | 5.3E-08 |
| *cyp-13B2* | WBGene00010606 | protein_coding_gene | -5.5 | 0.0218 | 5.6E-13 | -4.3 | 0.049 | 4.1E-23 |
| *Y71A12B.11* | WBGene00013508 | protein_coding_gene | -5.5 | 0.0217 | 3.0E-24 | -7.2 | 0.007 | 3.5E-21 |
| *spsb-2* | WBGene00021596 | protein_coding_gene | -5.5 | 0.0217 | 7.2E-113 | -4.8 | 0.037 | 7.8E-114 |
| *R11.4* | WBGene00011229 | protein_coding_gene | -5.5 | 0.0214 | 1.1E-08 | -6.4 | 0.011 | 1.0E-11 |
| *Y5H2B.3* | WBGene00021165 | protein_coding_gene | -5.6 | 0.0213 | 2.8E-09 | -5.5 | 0.022 | 4.2E-10 |
| *M151.4* | WBGene00019796 | protein_coding_gene | -5.6 | 0.0213 | 6.9E-10 | -4.9 | 0.034 | 1.1E-08 |
| *skr-11* | WBGene00004817 | pseudogene | -5.6 | 0.0208 | 3.6E-10 | -7.1 | 0.007 | 9.8E-55 |
| *sgnh-1* | WBGene00020354 | protein_coding_gene | -5.6 | 0.0208 | 2.3E-09 | -5.7 | 0.019 | 9.6E-11 |
| *rmd-1* | WBGene00011501 | protein_coding_gene | -5.6 | 0.0207 | 1.5E-86 | -5.3 | 0.026 | 2.2E-301 |
| *btb-11* | WBGene00019700 | protein_coding_gene | -5.6 | 0.0207 | 1.2E-09 | -2.8 | 0.139 | 5.2E-15 |
| *era-1* | WBGene00020948 | protein_coding_gene | -5.6 | 0.0206 | 1.9E-110 | -4.9 | 0.033 | 5.9E-178 |
| *ttr-42* | WBGene00045268 | protein_coding_gene | -5.6 | 0.0204 | 2.8E-33 | -5.4 | 0.024 | 1.6E-43 |
| *fbxb-26* | WBGene00019035 | protein_coding_gene | -5.6 | 0.0204 | 1.4E-09 | -5.5 | 0.022 | 5.8E-09 |

| | | | | | | | | |
|---|---|---|---|---|---|---|---|---|
| *C46C2.5* | WBGene00008113 | protein_coding_gene | -5.6 | 0.0201 | 9.5E-24 | -6.2 | 0.013 | 1.1E-23 |
| *fbxa-200* | WBGene00020304 | pseudogene | -5.6 | 0.0201 | 3.3E-09 | -6.2 | 0.014 | 2.6E-12 |
| *fbxc-32* | WBGene00016303 | protein_coding_gene | -5.6 | 0.0200 | 1.2E-42 | -5.9 | 0.016 | 1.8E-51 |
| *Y62H9A.5* | WBGene00013393 | protein_coding_gene | -5.7 | 0.0199 | 3.5E-80 | -5.6 | 0.021 | 1.3E-261 |
| *C03D6.9* | WBGene00219796 | protein_coding_gene | -5.7 | 0.0198 | 3.3E-10 | -5.8 | 0.018 | 6.7E-10 |
| *Y38E10A.3* | WBGene00012581 | protein_coding_gene | -5.7 | 0.0197 | 1.1E-09 | -6.0 | 0.016 | 1.1E-11 |
| *ZK899.6* | WBGene00014144 | protein_coding_gene | -5.7 | 0.0197 | 1.4E-09 | -5.7 | 0.019 | 1.1E-10 |
| *clec-91* | WBGene00014117 | protein_coding_gene | -5.7 | 0.0196 | 2.0E-73 | -5.2 | 0.027 | 2.5E-215 |
| *arrd-2* | WBGene00013086 | protein_coding_gene | -5.7 | 0.0196 | 4.3E-12 | -6.6 | 0.010 | 9.8E-14 |
| *T12A2.3* | WBGene00020438 | protein_coding_gene | -5.7 | 0.0195 | 3.7E-10 | -5.2 | 0.028 | 4.5E-09 |
| *T27A8.2* | WBGene00012074 | protein_coding_gene | -5.7 | 0.0194 | 4.6E-13 | -6.0 | 0.015 | 2.5E-12 |
| *skpo-1* | WBGene00009897 | protein_coding_gene | -5.7 | 0.0193 | 2.0E-76 | -5.7 | 0.020 | 1.8E-216 |
| *T05D4.2* | WBGene00011475 | protein_coding_gene | -5.7 | 0.0192 | 4.3E-15 | -7.3 | 0.006 | 2.5E-16 |
| *F29G9.7* | WBGene00017933 | pseudogene | -5.7 | 0.0190 | 5.8E-14 | -4.3 | 0.051 | 1.4E-19 |
| *fbxb-40* | WBGene00019704 | protein_coding_gene | -5.7 | 0.0189 | 3.6E-10 | -5.3 | 0.025 | 2.0E-09 |
| *bath-31* | WBGene00017405 | protein_coding_gene | -5.7 | 0.0189 | 9.4E-10 | -5.9 | 0.016 | 1.5E-11 |
| *Y4C6A.3* | WBGene00021153 | protein_coding_gene | -5.7 | 0.0188 | 6.9E-80 | -4.5 | 0.046 | 1.1E-119 |
| *Y25C1A.6* | WBGene00021293 | protein_coding_gene | -5.7 | 0.0188 | 5.7E-10 | -5.3 | 0.026 | 2.7E-09 |
| *Y41C4A.32* | WBGene00270321 | protein_coding_gene | -5.7 | 0.0187 | 1.0E-20 | -3.9 | 0.066 | 7.6E-16 |
| *C08F11.12* | WBGene00007459 | protein_coding_gene | -5.7 | 0.0187 | 1.7E-09 | -6.0 | 0.016 | 2.6E-15 |
| *ins-34* | WBGene00002117 | protein_coding_gene | -5.8 | 0.0185 | 2.2E-10 | -5.8 | 0.018 | 5.5E-20 |
| *Y82E9BR.17* | WBGene00022349 | protein_coding_gene | -5.8 | 0.0185 | 4.6E-18 | -7.6 | 0.005 | 8.4E-35 |
| *hnd-1* | WBGene00001981 | protein_coding_gene | -5.8 | 0.0184 | 1.8E-10 | -6.6 | 0.011 | 1.3E-13 |
| *C52A10.3* | WBGene00016864 | protein_coding_gene | -5.8 | 0.0181 | 2.4E-10 | -4.6 | 0.043 | 6.5E-09 |
| *gyg-2* | WBGene00012020 | protein_coding_gene | -5.8 | 0.0179 | 1.2E-147 | -4.9 | 0.034 | 3.3E-274 |
| *fbxc-18* | WBGene00016307 | protein_coding_gene | -5.8 | 0.0179 | 2.7E-10 | -6.9 | 0.009 | 1.1E-14 |
| *clec-87* | WBGene00007709 | protein_coding_gene | -5.8 | 0.0179 | 1.0E-128 | -6.1 | 0.015 | 0.0E+00 |
| *fbxb-41* | WBGene00019703 | protein_coding_gene | -5.8 | 0.0178 | 1.4E-10 | -5.3 | 0.025 | 1.4E-09 |
| *C46E10.9* | WBGene00016713 | pseudogene | -5.8 | 0.0177 | 9.7E-11 | -4.8 | 0.035 | 2.8E-08 |
| *tbx-36* | WBGene00006555 | protein_coding_gene | -5.8 | 0.0177 | 3.4E-37 | -4.0 | 0.063 | 1.2E-45 |
| *cav-1* | WBGene00000301 | protein_coding_gene | -5.8 | 0.0176 | 2.9E-73 | -7.3 | 0.006 | 8.5E-252 |
| *ets-7* | WBGene00017601 | protein_coding_gene | -5.8 | 0.0175 | 2.8E-10 | -5.6 | 0.020 | 8.8E-11 |
| *sdz-28* | WBGene00020073 | protein_coding_gene | -5.8 | 0.0174 | 3.4E-18 | -3.4 | 0.097 | 2.0E-12 |
| *puf-11* | WBGene00022257 | protein_coding_gene | -5.9 | 0.0173 | 1.9E-101 | -4.9 | 0.034 | 4.3E-295 |
| *F15A4.2* | WBGene00008838 | protein_coding_gene | -5.9 | 0.0173 | 1.5E-09 | -5.4 | 0.023 | 6.6E-09 |
| *Y116A8C.19* | WBGene00013796 | protein_coding_gene | -5.9 | 0.0173 | 9.2E-10 | -5.9 | 0.017 | 4.7E-11 |
| *btb-7* | WBGene00018443 | protein_coding_gene | -5.9 | 0.0172 | 8.4E-10 | -6.8 | 0.009 | 9.9E-15 |
| *tbx-11* | WBGene00006547 | protein_coding_gene | -5.9 | 0.0172 | 3.1E-09 | -6.3 | 0.013 | 3.2E-11 |
| *pes-1* | WBGene00003976 | protein_coding_gene | -5.9 | 0.0170 | 1.4E-10 | -5.8 | 0.018 | 5.1E-11 |
| *M151.3* | WBGene00019795 | protein_coding_gene | -5.9 | 0.0168 | 7.8E-10 | -5.6 | 0.020 | 6.6E-11 |
| *tbx-33* | WBGene00006552 | protein_coding_gene | -5.9 | 0.0168 | 1.6E-17 | -6.0 | 0.015 | 5.3E-12 |
| *ugt-63* | WBGene00015449 | protein_coding_gene | -5.9 | 0.0168 | 9.6E-14 | -5.5 | 0.023 | 3.6E-42 |
| *F32D1.7* | WBGene00017986 | protein_coding_gene | -5.9 | 0.0168 | 6.0E-182 | -4.8 | 0.036 | 9.5E-161 |
| *sdz-16* | WBGene00009297 | protein_coding_gene | -5.9 | 0.0167 | 3.4E-10 | -3.6 | 0.082 | 6.4E-05 |
| *ZC53.1* | WBGene00022508 | protein_coding_gene | -5.9 | 0.0166 | 4.8E-25 | -5.4 | 0.023 | 4.8E-09 |
| *EEED8.4* | WBGene00017135 | protein_coding_gene | -5.9 | 0.0165 | 3.5E-09 | -6.1 | 0.015 | 1.1E-10 |
| *athp-3* | WBGene00013799 | protein_coding_gene | -5.9 | 0.0164 | 7.2E-11 | -5.8 | 0.018 | 2.6E-11 |
| *nip-2* | WBGene00011981 | protein_coding_gene | -5.9 | 0.0164 | 6.5E-11 | -5.8 | 0.018 | 2.4E-11 |
| *fbxb-17* | WBGene00019037 | protein_coding_gene | -5.9 | 0.0163 | 1.1E-10 | -6.8 | 0.009 | 2.6E-14 |
| *tag-276* | WBGene00013254 | protein_coding_gene | -6.0 | 0.0161 | 8.9E-10 | -5.6 | 0.020 | 1.5E-10 |
| *R02C2.1* | WBGene00019814 | protein_coding_gene | -6.0 | 0.0161 | 9.0E-11 | -6.0 | 0.016 | 3.3E-10 |
| *cpg-2* | WBGene00015102 | protein_coding_gene | -6.0 | 0.0161 | 1.2E-232 | -4.9 | 0.033 | 0.0E+00 |
| *skr-12* | WBGene00004818 | protein_coding_gene | -6.0 | 0.0159 | 7.5E-11 | -6.9 | 0.008 | 2.7E-25 |
| *sdz-4* | WBGene00016309 | protein_coding_gene | -6.0 | 0.0158 | 7.5E-11 | -6.2 | 0.013 | 4.8E-11 |
| *chs-1* | WBGene00000496 | protein_coding_gene | -6.0 | 0.0157 | 4.5E-144 | -5.2 | 0.027 | 5.5E-284 |
| *cpg-1* | WBGene00000465 | protein_coding_gene | -6.0 | 0.0157 | 1.6E-145 | -5.0 | 0.032 | 1.5E-294 |
| *F43G6.5* | WBGene00009658 | protein_coding_gene | -6.0 | 0.0156 | 3.0E-10 | -6.4 | 0.012 | 3.0E-13 |
| *vet-2* | WBGene00016453 | protein_coding_gene | -6.0 | 0.0156 | 4.1E-43 | -6.7 | 0.010 | 2.1E-74 |
| *C47F8.1* | WBGene00008157 | protein_coding_gene | -6.0 | 0.0155 | 3.7E-10 | -5.8 | 0.018 | 4.7E-10 |
| *clec-266* | WBGene00016088 | protein_coding_gene | -6.0 | 0.0154 | 4.6E-33 | -7.4 | 0.006 | 5.6E-89 |
| *F17A2.13* | WBGene00008904 | protein_coding_gene | -6.1 | 0.0150 | 7.5E-15 | -7.2 | 0.007 | 4.0E-16 |
| *Y73C8C.3* | WBGene00022262 | protein_coding_gene | -6.1 | 0.0150 | 6.1E-11 | -5.4 | 0.023 | 8.5E-10 |
| *C01G8.1* | WBGene00015312 | protein_coding_gene | -6.1 | 0.0147 | 2.7E-161 | -5.5 | 0.022 | 1.3E-192 |
| *gln-5* | WBGene00001606 | protein_coding_gene | -6.1 | 0.0143 | 2.5E-151 | -5.1 | 0.029 | 1.7E-295 |
| *puf-3* | WBGene00004239 | protein_coding_gene | -6.1 | 0.0142 | 7.4E-141 | -4.7 | 0.038 | 8.0E-290 |
| *try-1* | WBGene00006619 | protein_coding_gene | -6.2 | 0.0139 | 2.4E-63 | -6.2 | 0.014 | 2.6E-73 |
| *E02H9.7* | WBGene00017105 | protein_coding_gene | -6.2 | 0.0139 | 7.6E-21 | -5.9 | 0.017 | 2.5E-24 |
| *clec-222* | WBGene00007283 | protein_coding_gene | -6.2 | 0.0137 | 5.7E-43 | -5.9 | 0.017 | 9.3E-79 |
| *fbxb-103* | WBGene00018746 | protein_coding_gene | -6.2 | 0.0136 | 2.0E-11 | -6.8 | 0.009 | 2.8E-14 |
| *W06D11.3* | WBGene00012301 | protein_coding_gene | -6.2 | 0.0134 | 3.8E-38 | -6.8 | 0.009 | 7.0E-20 |
| *sdz-14* | WBGene00008782 | protein_coding_gene | -6.2 | 0.0133 | 1.2E-11 | -6.5 | 0.011 | 1.3E-13 |
| *cbd-1* | WBGene00010351 | protein_coding_gene | -6.2 | 0.0132 | 3.5E-107 | -5.1 | 0.029 | 0.0E+00 |
| *btb-8* | WBGene00018437 | protein_coding_gene | -6.3 | 0.0131 | 2.2E-11 | -6.4 | 0.012 | 1.2E-11 |
| *C29F7.10* | WBGene00206389 | protein_coding_gene | -6.3 | 0.0131 | 1.2E-10 | -5.0 | 0.032 | 1.2E-18 |
| *ccch-5* | WBGene00013319 | protein_coding_gene | -6.3 | 0.0131 | 9.7E-11 | -6.5 | 0.011 | 1.3E-13 |
| *M151.7* | WBGene00019799 | protein_coding_gene | -6.3 | 0.0131 | 8.2E-24 | -5.6 | 0.021 | 2.0E-22 |
| *EEED8.15* | WBGene00017143 | protein_coding_gene | -6.3 | 0.0131 | 6.1E-26 | -7.1 | 0.008 | 1.2E-61 |
| *fbxc-42* | WBGene00018455 | protein_coding_gene | -6.3 | 0.0131 | 3.5E-12 | -6.0 | 0.016 | 3.5E-23 |
| *C08F1.10* | WBGene00015613 | protein_coding_gene | -6.3 | 0.0130 | 2.8E-12 | -4.2 | 0.053 | 3.9E-16 |
| *sdz-9* | WBGene00017253 | protein_coding_gene | -6.3 | 0.0129 | 4.2E-11 | -6.5 | 0.011 | 3.3E-13 |

| | | | | | | | | |
|---|---|---|---|---|---|---|---|---|
| *Y49F6C.8* | WBGene00021730 | protein_coding_gene | -6.3 | 0.0129 | 5.9E-39 | -6.7 | 0.009 | 2.0E-14 |
| *T10C6.7* | WBGene00011689 | protein_coding_gene | -6.3 | 0.0129 | 1.2E-47 | -4.9 | 0.034 | 6.5E-40 |
| *fkh-4* | WBGene00001436 | protein_coding_gene | -6.3 | 0.0127 | 1.1E-11 | -6.5 | 0.011 | 1.8E-13 |
| *col-101* | WBGene00000675 | protein_coding_gene | -6.3 | 0.0126 | 1.7E-50 | -4.8 | 0.036 | 3.2E-34 |
| *T24E12.1* | WBGene00020771 | protein_coding_gene | -6.3 | 0.0126 | 1.9E-12 | -5.8 | 0.018 | 3.6E-10 |
| *meg-3* | WBGene00001569 | protein_coding_gene | -6.3 | 0.0125 | 3.9E-100 | -5.0 | 0.031 | 1.3E-131 |
| *F02H6.4* | WBGene00008537 | pseudogene | -6.3 | 0.0124 | 7.3E-51 | -6.4 | 0.012 | 7.2E-118 |
| *tbx-9* | WBGene00006546 | protein_coding_gene | -6.3 | 0.0124 | 2.0E-34 | -6.0 | 0.016 | 9.7E-36 |
| *T01C3.3* | WBGene00011320 | protein_coding_gene | -6.4 | 0.0122 | 2.4E-89 | -5.4 | 0.024 | 2.7E-232 |
| *C45H4.14* | WBGene00016695 | protein_coding_gene | -6.4 | 0.0121 | 1.8E-12 | -6.0 | 0.015 | 2.3E-12 |
| *C49F5.3* | WBGene00008207 | protein_coding_gene | -6.4 | 0.0120 | 2.7E-31 | -6.6 | 0.010 | 1.6E-27 |
| *F56F4.8* | WBGene00045498 | protein_coding_gene | -6.4 | 0.0119 | 1.4E-11 | -6.4 | 0.012 | 1.2E-11 |
| *Y46G5A.28* | WBGene00012916 | protein_coding_gene | -6.4 | 0.0119 | 1.4E-11 | -6.6 | 0.010 | 1.1E-13 |
| *C27C12.3* | WBGene00007773 | protein_coding_gene | -6.4 | 0.0117 | 7.5E-42 | -6.7 | 0.010 | 3.9E-32 |
| *cpg-3* | WBGene00011063 | protein_coding_gene | -6.4 | 0.0116 | 2.4E-189 | -6.3 | 0.012 | 0.0E+00 |
| *C08F1.6* | WBGene00015610 | protein_coding_gene | -6.4 | 0.0115 | 1.8E-12 | -5.1 | 0.029 | 1.7E-08 |
| *K01G12.3* | WBGene00010483 | protein_coding_gene | -6.5 | 0.0114 | 3.2E-11 | -5.0 | 0.032 | 4.7E-09 |
| *fkh-3* | WBGene00001435 | protein_coding_gene | -6.5 | 0.0112 | 3.5E-11 | -5.5 | 0.021 | 2.5E-09 |
| *fbxc-28* | WBGene00019922 | protein_coding_gene | -6.5 | 0.0112 | 3.6E-12 | -6.1 | 0.014 | 4.2E-19 |
| *E02H4.6* | WBGene00008464 | protein_coding_gene | -6.5 | 0.0111 | 3.7E-29 | -5.7 | 0.020 | 2.5E-31 |
| *gfat-2* | WBGene00009035 | protein_coding_gene | -6.5 | 0.0109 | 4.7E-152 | -4.8 | 0.037 | 2.5E-294 |
| *F31F6.1* | WBGene00009300 | protein_coding_gene | -6.5 | 0.0109 | 3.9E-11 | -6.0 | 0.016 | 1.7E-11 |
| *ZC308.4* | WBGene00022586 | protein_coding_gene | -6.5 | 0.0108 | 1.0E-121 | -6.1 | 0.015 | 1.2E-160 |
| *skr-13* | WBGene00004819 | protein_coding_gene | -6.5 | 0.0107 | 8.0E-13 | -5.2 | 0.026 | 2.0E-08 |
| *R04D3.4* | WBGene00011012 | protein_coding_gene | -6.5 | 0.0107 | 5.7E-21 | -7.3 | 0.006 | 9.4E-17 |
| *B0281.5* | WBGene00015112 | protein_coding_gene | -6.6 | 0.0106 | 1.6E-20 | -4.8 | 0.037 | 4.6E-07 |
| *F43D2.2* | WBGene00009651 | pseudogene | -6.6 | 0.0106 | 4.6E-56 | -6.1 | 0.015 | 1.2E-46 |
| *T01G5.7* | WBGene00011342 | protein_coding_gene | -6.6 | 0.0104 | 1.3E-17 | -6.4 | 0.012 | 1.6E-13 |
| *Y106G6D.2* | WBGene00013699 | protein_coding_gene | -6.6 | 0.0102 | 1.0E-12 | -6.8 | 0.009 | 1.9E-14 |
| *oma-2* | WBGene00003865 | protein_coding_gene | -6.6 | 0.0101 | 5.0E-172 | -6.1 | 0.015 | 0.0E+00 |
| *pes-10* | WBGene00003983 | protein_coding_gene | -6.6 | 0.0101 | 6.5E-12 | -6.5 | 0.011 | 2.9E-13 |
| *skr-9* | WBGene00004815 | protein_coding_gene | -6.6 | 0.0101 | 1.1E-12 | -7.3 | 0.006 | 2.0E-16 |
| *T12G3.6* | WBGene00011741 | protein_coding_gene | -6.6 | 0.0101 | 2.8E-135 | -6.1 | 0.015 | 1.0E-180 |
| *Y39B6A.10* | WBGene00012672 | protein_coding_gene | -6.7 | 0.0099 | 2.2E-82 | -5.0 | 0.031 | 7.1E-79 |
| *C31G12.1* | WBGene00007850 | protein_coding_gene | -6.7 | 0.0099 | 3.7E-13 | -6.7 | 0.010 | 3.5E-14 |
| *pzf-1* | WBGene00011505 | protein_coding_gene | -6.7 | 0.0098 | 3.0E-18 | -4.3 | 0.051 | 9.2E-16 |
| *rme-2* | WBGene00004374 | protein_coding_gene | -6.7 | 0.0097 | 1.3E-128 | -5.3 | 0.025 | 7.6E-296 |
| *R09A8.1* | WBGene00011152 | protein_coding_gene | -6.7 | 0.0096 | 9.8E-14 | -4.9 | 0.033 | 5.1E-23 |
| *Y75D11A.3* | WBGene00022292 | protein_coding_gene | -6.8 | 0.0092 | 3.6E-12 | -5.3 | 0.026 | 9.0E-18 |
| *Y73C8C.8* | WBGene00022265 | protein_coding_gene | -6.8 | 0.0092 | 5.7E-12 | -6.7 | 0.010 | 2.3E-12 |
| *T04D3.1* | WBGene00011431 | protein_coding_gene | -6.8 | 0.0091 | 7.9E-32 | -6.8 | 0.009 | 2.0E-36 |
| *rgs-9* | WBGene00004352 | protein_coding_gene | -6.8 | 0.0089 | 1.8E-27 | -5.2 | 0.027 | 2.3E-09 |
| *F56D5.5* | WBGene00010149 | protein_coding_gene | -6.8 | 0.0088 | 1.1E-13 | -6.4 | 0.012 | 3.2E-13 |
| *F14H3.5* | WBGene00008824 | protein_coding_gene | -6.9 | 0.0085 | 5.1E-19 | -6.7 | 0.010 | 2.2E-28 |
| *rgs-11* | WBGene00004354 | protein_coding_gene | -6.9 | 0.0085 | 1.4E-18 | -4.9 | 0.033 | 4.4E-13 |
| *F22E5.17* | WBGene00017712 | protein_coding_gene | -6.9 | 0.0084 | 5.8E-33 | -6.1 | 0.014 | 1.7E-43 |
| *R09F10.8* | WBGene00019989 | protein_coding_gene | -6.9 | 0.0082 | 9.6E-90 | -7.1 | 0.007 | 5.4E-125 |
| *flh-3* | WBGene00012436 | protein_coding_gene | -6.9 | 0.0081 | 1.9E-55 | -7.0 | 0.008 | 1.1E-67 |
| *F15A4.10* | WBGene00008844 | protein_coding_gene | -7.0 | 0.0080 | 1.8E-14 | -6.2 | 0.013 | 2.4E-11 |
| *ule-1* | WBGene00021005 | protein_coding_gene | -7.0 | 0.0078 | 5.7E-67 | -4.3 | 0.050 | 1.9E-86 |
| *F43G6.10* | WBGene00009662 | protein_coding_gene | -7.0 | 0.0078 | 1.3E-14 | -6.0 | 0.015 | 6.0E-11 |
| *egg-1* | WBGene00015083 | protein_coding_gene | -7.0 | 0.0078 | 3.0E-158 | -6.2 | 0.014 | 0.0E+00 |
| *Y57G11B.5* | WBGene00013297 | protein_coding_gene | -7.0 | 0.0078 | 1.6E-119 | -5.7 | 0.020 | 6.5E-210 |
| *puf-5* | WBGene00004241 | protein_coding_gene | -7.0 | 0.0077 | 1.0E-173 | -5.9 | 0.017 | 0.0E+00 |
| *F14B6.3* | WBGene00008784 | protein_coding_gene | -7.0 | 0.0077 | 2.2E-14 | -7.0 | 0.008 | 2.0E-15 |
| *T25E12.6* | WBGene00012021 | protein_coding_gene | -7.0 | 0.0076 | 2.1E-14 | -7.5 | 0.006 | 3.5E-17 |
| *T02G6.5* | WBGene00011388 | protein_coding_gene | -7.1 | 0.0075 | 5.9E-15 | -6.8 | 0.009 | 9.4E-20 |
| *F02H6.2* | WBGene00008535 | protein_coding_gene | -7.1 | 0.0075 | 7.1E-48 | -6.0 | 0.015 | 1.3E-64 |
| *skr-15* | WBGene00004821 | protein_coding_gene | -7.1 | 0.0074 | 1.9E-14 | -7.6 | 0.005 | 1.3E-17 |
| *F23D12.2* | WBGene00009086 | protein_coding_gene | -7.1 | 0.0074 | 2.8E-15 | -7.7 | 0.005 | 6.1E-18 |
| *F12E12.1* | WBGene00017402 | protein_coding_gene | -7.1 | 0.0073 | 1.5E-25 | -6.8 | 0.009 | 5.3E-20 |
| *ztf-25* | WBGene00012405 | protein_coding_gene | -7.1 | 0.0073 | 2.3E-13 | -6.3 | 0.013 | 1.2E-28 |
| *vet-6* | WBGene00006902 | protein_coding_gene | -7.1 | 0.0072 | 3.7E-20 | -8.6 | 0.003 | 4.0E-19 |
| *F53F8.3* | WBGene00010000 | protein_coding_gene | -7.1 | 0.0071 | 1.5E-61 | -6.4 | 0.012 | 1.1E-72 |
| *ent-5* | WBGene00017532 | protein_coding_gene | -7.2 | 0.0070 | 1.6E-24 | -6.6 | 0.010 | 3.7E-23 |
| *F08F3.6* | WBGene00017263 | protein_coding_gene | -7.2 | 0.0068 | 6.6E-74 | -6.4 | 0.012 | 3.9E-116 |
| *dsl-2* | WBGene00001104 | protein_coding_gene | -7.2 | 0.0068 | 3.5E-13 | -7.8 | 0.005 | 4.4E-16 |
| *ZK829.9* | WBGene00014097 | protein_coding_gene | -7.2 | 0.0067 | 1.5E-101 | -6.5 | 0.011 | 2.4E-151 |
| *C50E3.13* | WBGene00016824 | protein_coding_gene | -7.2 | 0.0067 | 1.7E-30 | -5.1 | 0.029 | 2.0E-08 |
| *B0416.4* | WBGene00015180 | protein_coding_gene | -7.2 | 0.0067 | 1.9E-15 | -6.6 | 0.011 | 1.4E-12 |
| *skr-10* | WBGene00004816 | protein_coding_gene | -7.3 | 0.0065 | 2.5E-13 | -8.1 | 0.004 | 1.7E-19 |
| *R04D3.2* | WBGene00011010 | protein_coding_gene | -7.3 | 0.0064 | 3.6E-57 | -7.2 | 0.007 | 3.6E-82 |
| *T22B2.1* | WBGene00020665 | protein_coding_gene | -7.3 | 0.0064 | 8.1E-15 | -7.0 | 0.008 | 4.2E-15 |
| *C35E7.5* | WBGene00016457 | protein_coding_gene | -7.3 | 0.0063 | 1.9E-53 | -7.8 | 0.004 | 5.5E-159 |
| *C01G6.3* | WBGene00007225 | protein_coding_gene | -7.3 | 0.0062 | 7.0E-108 | -8.5 | 0.003 | 9.5E-138 |
| *skr-8* | WBGene00004814 | protein_coding_gene | -7.4 | 0.0058 | 1.8E-15 | -7.8 | 0.005 | 3.3E-16 |
| *K04G2.10* | WBGene00010568 | protein_coding_gene | -7.5 | 0.0056 | 2.0E-14 | -7.2 | 0.007 | 3.6E-22 |
| *ttr-50* | WBGene00010234 | protein_coding_gene | -7.5 | 0.0055 | 5.1E-63 | -9.0 | 0.002 | 3.2E-137 |
| *F14H3.6* | WBGene00008825 | protein_coding_gene | -7.6 | 0.0052 | 1.6E-178 | -6.7 | 0.009 | 7.2E-227 |
| *C50E3.11* | WBGene00016822 | protein_coding_gene | -7.6 | 0.0052 | 2.6E-17 | -4.9 | 0.034 | 1.3E-18 |

| | | | | | | | | |
|---|---|---|---|---|---|---|---|---|
| *K03A11.1* | WBGene00010519 | protein_coding_gene | -7.6 | 0.0052 | 1.8E-16 | -7.0 | 0.008 | 1.4E-15 |
| *C17E7.12* | WBGene00015905 | protein_coding_gene | -7.6 | 0.0051 | 7.3E-15 | -6.4 | 0.011 | 7.1E-22 |
| *vit-6* | WBGene00006930 | protein_coding_gene | -7.7 | 0.0050 | 1.7E-102 | -6.5 | 0.011 | 6.9E-152 |
| *F02E8.4* | WBGene00017177 | protein_coding_gene | -7.7 | 0.0050 | 4.4E-97 | -6.8 | 0.009 | 1.8E-84 |
| *lsy-27* | WBGene00009834 | protein_coding_gene | -7.7 | 0.0050 | 1.8E-17 | -2.0 | 0.250 | 5.6E-11 |
| *fat-7* | WBGene00001399 | protein_coding_gene | -7.7 | 0.0049 | 4.3E-97 | -4.6 | 0.042 | 1.5E-38 |
| *C17E7.9* | WBGene00015903 | protein_coding_gene | -7.7 | 0.0049 | 2.1E-17 | -6.6 | 0.010 | 2.1E-27 |
| *meg-2* | WBGene00010493 | protein_coding_gene | -7.7 | 0.0048 | 7.0E-39 | -5.7 | 0.019 | 2.7E-67 |
| *sepa-1* | WBGene00010808 | protein_coding_gene | -7.7 | 0.0047 | 1.5E-14 | -8.3 | 0.003 | 4.8E-18 |
| *T06D4.1* | WBGene00020290 | protein_coding_gene | -7.7 | 0.0047 | 6.0E-68 | -7.7 | 0.005 | 1.3E-48 |
| *vit-2* | WBGene00006926 | protein_coding_gene | -7.7 | 0.0047 | 3.7E-136 | -6.9 | 0.009 | 0.0E+00 |
| *sdz-30* | WBGene00011432 | protein_coding_gene | -7.7 | 0.0046 | 3.7E-16 | -8.4 | 0.003 | 3.6E-18 |
| *F43G6.7* | WBGene00009659 | protein_coding_gene | -7.8 | 0.0046 | 4.9E-17 | -5.1 | 0.029 | 6.5E-24 |
| *meg-4* | WBGene00016485 | protein_coding_gene | -7.8 | 0.0045 | 1.8E-34 | -7.1 | 0.007 | 5.8E-51 |
| *F02H6.3* | WBGene00008536 | protein_coding_gene | -7.8 | 0.0044 | 8.9E-30 | -6.6 | 0.011 | 1.2E-60 |
| *btb-6* | WBGene00018438 | protein_coding_gene | -7.8 | 0.0044 | 3.7E-17 | -7.7 | 0.005 | 4.5E-16 |
| *F14D7.2* | WBGene00008789 | protein_coding_gene | -7.8 | 0.0044 | 9.7E-67 | -6.9 | 0.009 | 3.6E-81 |
| *ZK177.1* | WBGene00022669 | protein_coding_gene | -7.8 | 0.0044 | 4.6E-18 | -7.7 | 0.005 | 1.5E-18 |
| *R09A8.2* | WBGene00011153 | protein_coding_gene | -7.9 | 0.0043 | 6.3E-18 | -7.2 | 0.007 | 3.8E-22 |
| *C17E7.4* | WBGene00015899 | protein_coding_gene | -7.9 | 0.0043 | 4.8E-35 | -7.8 | 0.004 | 2.8E-19 |
| *R05G9.3* | WBGene00019905 | protein_coding_gene | -7.9 | 0.0042 | 1.4E-15 | -6.6 | 0.010 | 1.8E-31 |
| *F26G5.1* | WBGene00017843 | protein_coding_gene | -7.9 | 0.0042 | 9.3E-25 | -7.2 | 0.007 | 2.5E-42 |
| *skr-14* | WBGene00004820 | protein_coding_gene | -7.9 | 0.0041 | 1.5E-17 | -8.0 | 0.004 | 5.2E-17 |
| *C50E3.12* | WBGene00016823 | protein_coding_gene | -8.0 | 0.0040 | 8.7E-18 | -3.8 | 0.071 | 5.4E-07 |
| *sea-1* | WBGene00004750 | protein_coding_gene | -8.0 | 0.0040 | 4.1E-16 | -6.7 | 0.010 | 9.5E-13 |
| *skr-7* | WBGene00004813 | protein_coding_gene | -8.0 | 0.0039 | 3.1E-24 | -9.3 | 0.002 | 1.9E-24 |
| *F14H3.3* | WBGene00008822 | protein_coding_gene | -8.0 | 0.0039 | 3.0E-18 | -6.7 | 0.010 | 9.2E-20 |
| *ceh-49* | WBGene00017538 | protein_coding_gene | -8.0 | 0.0038 | 2.6E-16 | -7.4 | 0.006 | 2.2E-23 |
| *F19H6.4* | WBGene00008959 | protein_coding_gene | -8.1 | 0.0037 | 1.8E-16 | -6.9 | 0.008 | 2.2E-39 |
| *nos-2* | WBGene00003784 | protein_coding_gene | -8.2 | 0.0034 | 7.9E-40 | -6.4 | 0.012 | 7.8E-106 |
| *F40G12.11* | WBGene00009606 | protein_coding_gene | -8.4 | 0.0029 | 8.9E-28 | -7.2 | 0.007 | 2.3E-56 |
| *C04B4.2* | WBGene00007290 | protein_coding_gene | -8.6 | 0.0026 | 5.2E-35 | -8.0 | 0.004 | 4.9E-46 |
| *K04C1.5* | WBGene00010555 | protein_coding_gene | -8.9 | 0.0021 | 1.0E-22 | -8.5 | 0.003 | 4.7E-38 |
| *cdc-25.3* | WBGene00000388 | protein_coding_gene | -9.0 | 0.0020 | 1.1E-21 | -7.1 | 0.007 | 5.8E-36 |
| *meg-1* | WBGene00010492 | protein_coding_gene | -9.0 | 0.0019 | 3.7E-84 | -8.0 | 0.004 | 3.2E-110 |
| *F14H3.4* | WBGene00008823 | protein_coding_gene | -9.1 | 0.0018 | 1.4E-22 | -7.7 | 0.005 | 6.9E-19 |
| *R04D3.3* | WBGene00011011 | protein_coding_gene | -9.9 | 0.0011 | 1.2E-37 | -8.3 | 0.003 | 6.4E-83 |
| *vit-1* | WBGene00006925 | protein_coding_gene | -10.1 | 0.0009 | 9.8E-69 | -6.0 | 0.016 | 1.0E-18 |
| *vit-5* | WBGene00006929 | protein_coding_gene | -11.4 | 0.0004 | 3.0E-158 | -9.6 | 0.001 | 8.0E-298 |
| *vit-4* | WBGene00006928 | protein_coding_gene | -11.9 | 0.0003 | 6.6E-28 | -8.5 | 0.003 | 1.9E-24 |
| *vit-3* | WBGene00006927 | protein_coding_gene | -12.7 | 0.0001 | 5.5E-37 | -8.0 | 0.004 | 3.8E-22 |

Transcription Factors Regulated by *S. aureus* Infection

| Gene Name | WormBase Gene ID | Sequence Name | N2 $log_2$ FC | CB4856 $log_2$ FC |
|---|---|---|---|---|
| *aly-2* | WBGene00000121 | F23B2.6 | -1.7 | -1.6 |
| *bro-1* | WBGene00000272 | F56A3.5 | 2.2 | -1.2 |
| *ceh-5* | WBGene00000430 | C16C2.1 | -1.9 | -5.1 |
| *ceh-18* | WBGene00000441 | ZC64.3 | 1.1 | 1.4 |
| *ceh-21* | WBGene00000444 | T26C11.6 | -2.2 | -1.9 |
| *ceh-23* | WBGene00000446 | ZK652.5 | 3.4 | 2.9 |
| *ceh-36* | WBGene00000457 | C37E2.4 | -1.6 | -2.2 |
| *ceh-38* | WBGene00000459 | F22D3.1 | -1.5 | -1.3 |
| *ceh-39* | WBGene00000460 | T26C11.7 | -3.8 | -3.7 |
| *ceh-40* | WBGene00000461 | F17A2.5 | -5.3 | -5.7 |
| *ceh-41* | WBGene00000462 | T26C11.5 | -1.4 | -1.4 |
| *ces-1* | WBGene00000468 | F43G9.11 | -2.4 | -4.3 |
| *ces-2* | WBGene00000469 | ZK909.4 | -3.4 | -7.0 |
| *cey-2* | WBGene00000473 | F46F11.2 | -3.7 | -2.9 |
| *cey-3* | WBGene00000474 | M01E11.5 | -2.7 | -2.3 |
| *cky-1* | WBGene00000521 | C15C8.2 | 3.7 | -1.9 |
| *dac-1* | WBGene00000895 | B0412.1 | 1.2 | 1.3 |
| *daf-12* | WBGene00000908 | F11A1.3 | 1.4 | 1.4 |
| *dbl-1* | WBGene00000936 | T25F10.2 | 2.6 | 2.4 |
| *die-1* | WBGene00000995 | C18D1.1 | -1.5 | -2.5 |
| *dnj-11* | WBGene00001029 | F38A5.13 | -1.9 | -1.5 |
| *dpl-1* | WBGene00001061 | T23G7.1 | -1.8 | -1.6 |
| *dpy-20* | WBGene00001079 | T22B3.1 | 2.9 | 4.0 |
| *dro-1* | WBGene00001092 | F53A2.5 | -1.9 | -1.5 |
| *eff-1* | WBGene00001159 | C26D10.5 | 2.1 | 1.1 |
| *efl-1* | WBGene00001161 | Y102A5C.18 | -1.7 | -1.6 |
| *egl-13* | WBGene00001182 | T22B7.1 | 1.2 | 1.5 |
| *egl-18* | WBGene00001186 | F55A8.1 | -2.3 | -2.7 |
| *elt-1* | WBGene00001249 | W09C2.1 | -1.1 | -1.5 |
| *elt-2* | WBGene00001250 | C33D3.1 | 1.8 | 1.4 |
| *end-1* | WBGene00001310 | F58E10.2 | -2.8 | -4.8 |
| *fos-1* | WBGene00001345 | F29G9.4 | 1.5 | 1.2 |
| *fkh-2* | WBGene00001434 | T14G12.4 | -2.4 | -3.6 |
| *fkh-3* | WBGene00001435 | C29F7.4 | -6.5 | -5.5 |
| *fkh-4* | WBGene00001436 | C29F7.5 | -6.3 | -6.5 |
| *fkh-5* | WBGene00001437 | F26A1.2 | -3.9 | -3.8 |

| | | | | |
|---|---|---|---|---|
| *fkh-6* | WBGene00001438 | B0286.5 | 2.4 | 1.8 |
| *fkh-7* | WBGene00001439 | F26D12.1 | 2.3 | 1.0 |
| *fkh-9* | WBGene00001441 | K03C7.2 | 2.2 | 1.3 |
| *hcf-1* | WBGene00001827 | C46A5.9 | -1.4 | -1.2 |
| *him-8* | WBGene00001867 | T07G12.12 | -1.4 | -1.4 |
| *hlh-2* | WBGene00001949 | M05B5.5 | -3.1 | -3.8 |
| *hlh-3* | WBGene00001950 | T24B8.6 | -3.8 | -5.6 |
| *hlh-11* | WBGene00001955 | F58A4.7 | 1.9 | 2.0 |
| *hlh-14* | WBGene00001958 | C18A3.8 | -3.2 | -5.7 |
| *hmg-4* | WBGene00001974 | T20B12.8 | -1.4 | -1.7 |
| *hmg-5* | WBGene00001975 | F45E4.9 | -1.1 | -1.2 |
| *hnd-1* | WBGene00001981 | C44C10.8 | -5.8 | -6.6 |
| *hsb-1* | WBGene00002002 | K08E7.2 | -1.3 | -1.3 |
| *isw-1* | WBGene00002169 | F37A4.8 | -1.5 | -1.0 |
| *lag-1* | WBGene00002245 | K08B4.1 | -1.6 | -1.3 |
| *let-381* | WBGene00002601 | F26B1.7 | 1.1 | -1.4 |
| *lim-6* | WBGene00002988 | K03E6.1 | 1.7 | 1.4 |
| *lin-1* | WBGene00002990 | C37F5.1 | 1.7 | 1.1 |
| *lin-11* | WBGene00003000 | ZC247.3 | 2.8 | 2.1 |
| *lin-26* | WBGene00003012 | F18A1.2 | -1.0 | -1.3 |
| *lin-28* | WBGene00003014 | F02E9.2 | -1.1 | -1.0 |
| *lin-29* | WBGene00003015 | W03C9.4 | 2.0 | 2.6 |
| *lin-31* | WBGene00003017 | K10G6.1 | 1.3 | 2.5 |
| *lin-32* | WBGene00003018 | T14F9.5 | -1.4 | -3.6 |
| *lin-48* | WBGene00003033 | F34D10.5 | 2.6 | 1.3 |
| *lin-54* | WBGene00003037 | JC8.6 | -1.9 | -1.8 |
| *lir-3* | WBGene00003046 | F37H8.1 | -1.5 | -1.9 |
| *mab-9* | WBGene00003106 | T27A1.6 | 1.7 | 1.5 |
| *mab-10* | WBGene00003107 | R166.1 | 1.2 | 2.2 |
| *mdl-1* | WBGene00003163 | R03E9.1 | 2.4 | 1.3 |
| *mef-2* | WBGene00003182 | W10D5.1 | 2.1 | 1.5 |
| *mex-1* | WBGene00003228 | W03C9.7 | -5.2 | -4.8 |
| *mex-5* | WBGene00003230 | W02A2.7 | -4.6 | -3.5 |
| *mex-6* | WBGene00003231 | AH6.5 | -5.2 | -4.2 |
| *mig-5* | WBGene00003241 | T05C12.6 | -2.6 | -2.0 |
| *mls-1* | WBGene00003376 | H14A12.4 | 1.8 | 1.4 |
| *moe-3* | WBGene00003388 | F32A11.6 | 2.4 | 1.3 |
| *mrg-1* | WBGene00003406 | Y37D8A.9 | -1.9 | -1.5 |
| *klf-3* | WBGene00003480 | F54H5.4 | 1.5 | 1.4 |
| *mxl-1* | WBGene00003509 | T19B10.11 | -1.4 | -1.3 |
| *ngn-1* | WBGene00003595 | Y69A2AR.29 | -3.7 | -5.7 |
| *nhr-2* | WBGene00003601 | C32F10.6 | -5.2 | -4.8 |
| *nhr-3* | WBGene00003602 | H01A20.1 | 1.4 | 1.5 |
| *nhr-6* | WBGene00003605 | C48D5.1 | 3.2 | 2.1 |
| *nhr-7* | WBGene00003606 | F54D1.4 | 1.2 | 1.8 |
| *nhr-12* | WBGene00003611 | R04B5.4 | -1.9 | -1.4 |
| *nhr-14* | WBGene00003613 | T01B10.4 | 1.8 | 1.7 |
| *nhr-18* | WBGene00003617 | F44C8.3 | 3.2 | 1.5 |
| *nhr-19* | WBGene00003618 | E02H1.7 | 3.5 | 1.5 |
| *nhr-31* | WBGene00003625 | C26B2.3 | 1.8 | 2.3 |
| *nhr-35* | WBGene00003628 | C07A12.3 | 2.9 | 1.1 |
| *nhr-38* | WBGene00003629 | K01H12.3 | 3.0 | 3.3 |
| *nhr-44* | WBGene00003634 | T19A5.4 | 2.3 | 1.2 |
| *nhr-50* | WBGene00003640 | C06C6.5 | 1.8 | 1.2 |
| *nhr-54* | WBGene00003644 | F36D3.2 | 1.7 | 1.4 |
| *nhr-59* | WBGene00003649 | T27B7.1 | 2.0 | 1.2 |
| *nhr-63* | WBGene00003653 | C06C6.4 | 1.8 | 1.3 |
| *nhr-66* | WBGene00003656 | T09A12.4 | -1.2 | -1.3 |
| *nhr-85* | WBGene00003675 | W05B5.3 | 1.5 | 2.1 |
| *nhr-87* | WBGene00003677 | Y41D4B.7 | 1.9 | 1.4 |
| *nhr-88* | WBGene00003678 | K08A2.5 | 2.0 | 1.4 |
| *nhr-95* | WBGene00003685 | Y39B6A.17 | 1.0 | 1.1 |
| *nhr-99* | WBGene00003689 | M02H5.1 | 4.1 | 2.0 |
| *nhr-103* | WBGene00003693 | F44C8.4 | 2.1 | 1.1 |
| *nhr-105* | WBGene00003695 | C06G3.1 | 1.6 | 1.3 |
| *nhr-109* | WBGene00003699 | T12C9.5 | 1.4 | 1.0 |
| *nhr-117* | WBGene00003707 | F16B4.12 | 3.2 | 1.6 |
| *nhr-120* | WBGene00003710 | C25B8.6 | 2.4 | 1.5 |
| *nhr-121* | WBGene00003711 | E02H9.8 | 1.7 | 1.1 |
| *nhr-125* | WBGene00003715 | R02D1.1 | 1.7 | 1.2 |
| *nhr-126* | WBGene00003716 | F44C8.10 | 1.9 | 1.5 |
| *nhr-128* | WBGene00003718 | F44C8.5 | 1.6 | 1.1 |
| *nhr-129* | WBGene00003719 | C50B6.14 | 2.2 | 1.8 |
| *nhr-133* | WBGene00003723 | F44C8.8 | 1.7 | 1.1 |
| *nhr-136* | WBGene00003726 | C13C4.3 | 1.6 | 1.2 |
| *oma-1* | WBGene00003864 | C09G9.6 | -4.6 | -3.7 |
| *oma-2* | WBGene00003865 | ZC513.6 | -6.6 | -6.1 |
| *pal-1* | WBGene00003912 | C38D4.6 | -3.3 | -2.9 |
| *pax-2* | WBGene00003938 | K06B9.5 | 2.5 | 2.5 |
| *peb-1* | WBGene00003968 | T14F9.4 | 3.7 | 1.2 |

| | | | | |
|---|---|---|---|---|
| *pes-1* | WBGene00003976 | T28H11.4 | -5.9 | -5.8 |
| *php-3* | WBGene00004024 | Y75B8A.1 | 2.3 | 1.0 |
| *pie-1* | WBGene00004027 | Y49E10.14 | -2.5 | -2.0 |
| *pos-1* | WBGene00004078 | F52E1.1 | -5.5 | -5.2 |
| *pqm-1* | WBGene00004096 | F40F8.7 | 3.1 | 1.6 |
| *myrf-1* | WBGene00004134 | F59B10.1 | 1.3 | 1.5 |
| *swsn-4* | WBGene00004204 | F01G4.1 | -1.6 | -1.5 |
| *ref-1* | WBGene00004334 | T01E8.2 | -3.1 | -3.0 |
| *ref-2* | WBGene00004335 | C47C12.3 | -1.3 | -3.1 |
| *rnt-1* | WBGene00004393 | B0414.2 | 2.2 | 1.4 |
| *rsp-6* | WBGene00004703 | C33H5.12 | -1.7 | -1.3 |
| *sdc-1* | WBGene00004745 | F52E10.1 | -1.8 | -1.3 |
| *sea-1* | WBGene00004750 | F19B10.9 | -8.0 | -6.7 |
| *sem-4* | WBGene00004773 | F15C11.1 | 1.8 | 1.3 |
| *sex-1* | WBGene00004786 | F44A6.2 | -2.8 | -3.8 |
| *sma-2* | WBGene00004856 | ZK370.2 | 2.4 | 1.4 |
| *sma-3* | WBGene00004857 | R13F6.9 | 1.8 | 1.4 |
| *snf-12* | WBGene00004911 | T25B6.7 | 1.9 | 1.9 |
| *sox-3* | WBGene00004950 | F40E10.2 | 2.3 | 1.4 |
| *spr-1* | WBGene00005006 | D1014.8 | -1.4 | -1.2 |
| *spr-3* | WBGene00005008 | C07A12.5 | -1.9 | -1.8 |
| *sys-1* | WBGene00006379 | T23D8.9 | -1.9 | -2.3 |
| *taf-3* | WBGene00006384 | C11G6.1 | -1.0 | -1.2 |
| *taf-8* | WBGene00006390 | ZK1320.12 | -1.6 | -1.5 |
| *taf-9* | WBGene00006391 | T12D8.7 | -1.3 | -1.3 |
| *taf-11.1* | WBGene00006393 | F48D6.1 | -1.5 | -1.7 |
| *taf-11.2* | WBGene00006394 | K10D3.3 | -3.9 | -3.3 |
| *tag-68* | WBGene00006445 | F37D6.6 | 1.1 | 1.2 |
| *svh-5* | WBGene00006462 | C33A11.4 | 1.9 | 1.4 |
| *wdr-5.1* | WBGene00006474 | C14B1.4 | -1.6 | -1.3 |
| *let-391* | WBGene00006492 | C27A12.3 | -1.8 | -1.8 |
| *tbx-8* | WBGene00006545 | T07C4.2 | -2.8 | -2.6 |
| *tbx-9* | WBGene00006546 | T07C4.6 | -6.3 | -6.0 |
| *tbx-11* | WBGene00006547 | F40H6.4 | -5.9 | -6.3 |
| *tbx-32* | WBGene00006551 | ZK380.1 | -3.0 | -2.7 |
| *tbx-33* | WBGene00006552 | Y66A7A.8 | -5.9 | -6.0 |
| *tbx-36* | WBGene00006555 | ZK829.5 | -5.8 | -4.0 |
| *tbx-37* | WBGene00006556 | Y47D3A.12 | -3.5 | -2.5 |
| *tbx-38* | WBGene00006557 | C24H11.3 | -4.3 | -4.4 |
| *tbx-39* | WBGene00006558 | Y73F8A.16 | -3.5 | -2.2 |
| *tbx-40* | WBGene00006559 | Y73F8A.17 | -2.7 | -2.9 |
| *tra-1* | WBGene00006604 | Y47D3A.6 | 1.3 | 1.4 |
| *ttb-1* | WBGene00006648 | W03F9.5 | -1.1 | -1.2 |
| *ubc-9* | WBGene00006706 | F29B9.6 | -2.0 | -1.7 |
| *unc-4* | WBGene00006744 | F26C11.2 | 1.9 | 1.5 |
| *unc-37* | WBGene00006773 | W02D3.9 | -1.3 | -1.6 |
| *unc-55* | WBGene00006790 | F55D12.4 | 2.1 | 1.5 |
| *vab-15* | WBGene00006881 | R07B1.1 | -1.3 | -2.2 |
| *wrm-1* | WBGene00006943 | B0336.1 | -1.7 | -1.4 |
| *nfx-1* | WBGene00007048 | C16A3.7 | -1.4 | -1.1 |
| *dmd-6* | WBGene00007058 | F13G11.1 | 1.6 | 1.2 |
| *B0250.4* | WBGene00007121 | B0250.4 | 1.2 | 1.7 |
| *thoc-7* | WBGene00007195 | B0513.2 | -1.4 | -1.5 |
| *ceh-57* | WBGene00007416 | C07E3.5 | 2.7 | 1.6 |
| *ceh-58* | WBGene00007417 | C07E3.6 | -1.5 | -2.0 |
| *swsn-7* | WBGene00007433 | C08B11.3 | -1.3 | -1.2 |
| *npax-4* | WBGene00007496 | C09G9.7 | -2.2 | -1.9 |
| *C16C10.4* | WBGene00007625 | C16C10.4 | -1.4 | -1.2 |
| *nfki-1* | WBGene00007877 | C33A11.1 | 1.9 | 1.5 |
| *dmd-10* | WBGene00007929 | C34D1.1 | 3.9 | 2.0 |
| *C35D6.4* | WBGene00007961 | C35D6.4 | -3.6 | -4.3 |
| *athp-1* | WBGene00008081 | C44B9.4 | -1.7 | -1.2 |
| *gmeb-3* | WBGene00008092 | C44F1.2 | 1.6 | 2.2 |
| *madf-8* | WBGene00008118 | C46F11.3 | -1.5 | -1.7 |
| *nhr-167* | WBGene00008208 | C49F5.4 | 2.4 | 1.5 |
| *nhr-168* | WBGene00008221 | C50B6.8 | 2.7 | 2.7 |
| *nhr-170* | WBGene00008309 | C54E10.5 | 1.7 | 1.7 |
| *pafo-1* | WBGene00008338 | C55A6.9 | -1.1 | -1.4 |
| *D1046.2* | WBGene00008363 | D1046.2 | -2.1 | -1.7 |
| *D2030.7* | WBGene00008417 | D2030.7 | -1.4 | -1.4 |
| *F10B5.3* | WBGene00008640 | F10B5.3 | 7.1 | 7.9 |
| *ztf-2* | WBGene00008762 | F13G3.1 | 2.9 | 3.1 |
| *F16B12.6* | WBGene00008882 | F16B12.6 | -1.6 | -1.4 |
| *F23A7.6* | WBGene00009071 | F23A7.6 | -3.6 | -4.2 |
| *jmjd-3.2* | WBGene00009089 | F23D12.5 | -4.0 | -6.0 |
| *lsy-22* | WBGene00009188 | F27D4.2 | -1.5 | -1.5 |
| *F27D4.6* | WBGene00009190 | F27D4.6 | 1.2 | 1.0 |
| *aptf-4* | WBGene00009202 | F28C6.1 | -4.0 | -4.2 |
| *aptf-3* | WBGene00009203 | F28C6.2 | -2.2 | -1.9 |
| *F33H1.4* | WBGene00009365 | F33H1.4 | -1.4 | -1.2 |

| | | | | |
|---|---|---|---|---|
| *zfp-2* | WBGene00009448 | F35H8.3 | -1.9 | -1.4 |
| *row-1* | WBGene00009508 | F37D6.2 | -1.4 | -1.5 |
| *ccch-2* | WBGene00009537 | F38C2.5 | -5.4 | -5.7 |
| *F38C2.7* | WBGene00009539 | F38C2.7 | -5.4 | -5.2 |
| *hinf-1* | WBGene00009553 | F39B2.1 | -2.7 | -2.4 |
| *ccnk-1* | WBGene00009650 | F43D2.1 | -1.6 | -1.4 |
| *lsy-27* | WBGene00009834 | F47H4.1 | -7.7 | -2.0 |
| *F52B5.7* | WBGene00009923 | F52B5.7 | -1.7 | -2.0 |
| *ztf-11* | WBGene00009939 | F52F12.6 | -2.8 | -4.4 |
| *saeg-1* | WBGene00010012 | F53H10.2 | 1.3 | 1.4 |
| *fozi-1* | WBGene00010453 | K01B6.1 | 3.4 | 1.2 |
| *dhhc-6* | WBGene00010892 | M18.8 | -1.9 | -1.5 |
| *ztf-14* | WBGene00010936 | M163.2 | 2.3 | 2.1 |
| *ztf-15* | WBGene00011066 | R06C7.9 | -1.8 | -1.8 |
| *nhr-206* | WBGene00011097 | R07B7.13 | 3.0 | 1.4 |
| *nhr-207* | WBGene00011098 | R07B7.14 | 2.5 | 3.0 |
| *nhr-208* | WBGene00011099 | R07B7.15 | 2.9 | 1.9 |
| *zip-6* | WBGene00011130 | R07H5.10 | 3.8 | 2.6 |
| *R10E4.11* | WBGene00011206 | R10E4.11 | -1.8 | -1.4 |
| *T01C3.2* | WBGene00011319 | T01C3.2 | -1.8 | -1.9 |
| *tftc-5* | WBGene00011368 | T02C12.3 | -1.1 | -1.2 |
| *gla-3* | WBGene00011376 | T02E1.3 | -2.5 | -1.5 |
| *pzf-1* | WBGene00011505 | T05G11.1 | -6.7 | -4.3 |
| *nhr-213* | WBGene00011520 | T06C12.13 | 1.7 | 1.1 |
| *madf-4* | WBGene00011575 | T07C12.11 | 1.0 | 1.2 |
| *nfya-1* | WBGene00011614 | T08D10.1 | -2.0 | -2.2 |
| *lsd-1* | WBGene00011615 | T08D10.2 | -2.8 | -2.3 |
| *ztf-27* | WBGene00011661 | T09F3.1 | 1.4 | 1.2 |
| *T22C8.4* | WBGene00011925 | T22C8.4 | -2.7 | -2.8 |
| *sptf-2* | WBGene00011926 | T22C8.5 | 2.5 | 2.2 |
| *T23F11.4* | WBGene00011956 | T23F11.4 | -1.1 | -1.4 |
| *T24H10.1* | WBGene00012000 | T24H10.1 | -1.3 | -1.1 |
| *jun-1* | WBGene00012005 | T24H10.7 | 1.4 | 1.2 |
| *T27A8.2* | WBGene00012074 | T27A8.2 | -5.7 | -6.0 |
| *zip-10* | WBGene00012101 | T27F2.4 | 2.4 | 2.0 |
| *hmg-20* | WBGene00012209 | W02D9.3 | -1.6 | -1.0 |
| *W04D2.4* | WBGene00012243 | W04D2.4 | -1.2 | -1.3 |
| *Y5F2A.4* | WBGene00012385 | Y5F2A.4 | -1.1 | -1.3 |
| *ztf-25* | WBGene00012405 | Y6G8.3 | -7.1 | -6.3 |
| *flh-1* | WBGene00012435 | Y11D7A.12 | -2.0 | -2.0 |
| *flh-3* | WBGene00012436 | Y11D7A.13 | -6.9 | -7.0 |
| *nhr-231* | WBGene00012449 | Y17D7B.1 | 1.8 | 1.8 |
| *ztf-20* | WBGene00012702 | Y39B6A.46 | -1.2 | -1.1 |
| *snpc-1.2* | WBGene00012715 | Y39E4B.2 | -2.1 | -1.3 |
| *dhhc-2* | WBGene00012948 | Y47H9C.2 | 1.3 | 1.1 |
| *sup-35* | WBGene00012976 | Y48A6C.3 | -1.7 | -1.2 |
| *zip-7* | WBGene00013100 | Y51H4A.4 | -1.4 | -3.2 |
| *sta-1* | WBGene00013111 | Y51H4A.17 | 1.5 | 1.1 |
| *pole-4* | WBGene00013150 | Y53F4B.3 | -1.6 | -1.3 |
| *znf-593* | WBGene00013236 | Y56A3A.18 | 1.8 | 1.7 |
| *ccch-5* | WBGene00013319 | Y57G11C.25 | -6.3 | -6.5 |
| *Y60A9.3* | WBGene00013370 | Y60A9.3 | -5.1 | -4.5 |
| *aptf-2* | WBGene00013383 | Y62E10A.17 | -4.1 | -3.9 |
| *Y67H2A.10* | WBGene00013465 | Y67H2A.10 | -1.0 | -1.2 |
| *ceh-51* | WBGene00013583 | Y80D3A.3 | -4.1 | -3.2 |
| *dct-13* | WBGene00013794 | Y116A8C.17 | -5.2 | -4.8 |
| *Y116A8C.19* | WBGene00013796 | Y116A8C.19 | -5.9 | -5.9 |
| *Y116A8C.20* | WBGene00013797 | Y116A8C.20 | -4.5 | -4.7 |
| *athp-3* | WBGene00013799 | Y116A8C.22 | -5.9 | -5.8 |
| *ceh-74* | WBGene00013876 | ZC376.4 | -1.3 | -1.7 |
| *klu-1* | WBGene00013970 | ZK337.2 | 3.0 | 2.7 |
| *ZK673.4* | WBGene00014060 | ZK673.4 | -1.2 | -1.1 |
| *sdz-38* | WBGene00014131 | ZK892.7 | -2.6 | -2.8 |
| *nhr-247* | WBGene00014193 | ZK1037.5 | 3.6 | 1.9 |
| *gtf-2H3* | WBGene00014230 | ZK1128.4 | -1.4 | -1.2 |
| *B0336.13* | WBGene00015150 | B0336.13 | -1.9 | -1.7 |
| *madf-11* | WBGene00015468 | C05D2.6 | -1.5 | -1.3 |
| *C08G9.2* | WBGene00015620 | C08G9.2 | -1.2 | -1.6 |
| *sox-4* | WBGene00015716 | C12D12.5 | 1.0 | 1.3 |
| *swsn-2.2* | WBGene00015971 | C18E3.2 | -1.7 | -1.2 |
| *flh-2* | WBGene00016138 | C26E6.2 | -1.1 | -1.0 |
| *szy-5* | WBGene00016154 | C27A12.2 | -1.1 | -1.4 |
| *sprp-4* | WBGene00016189 | C28G1.4 | -4.9 | -5.5 |
| *dpff-1* | WBGene00016200 | C28H8.9 | -1.4 | -1.3 |
| *nhr-139* | WBGene00016365 | C33G8.8 | 1.9 | 1.2 |
| *nhr-162* | WBGene00016367 | C33G8.10 | 2.0 | 1.4 |
| *unk-1* | WBGene00016407 | C34D10.2 | 3.1 | 1.6 |
| *C46E10.8* | WBGene00016712 | C46E10.8 | -5.5 | -4.9 |
| *C46E10.9* | WBGene00016713 | C46E10.9 | -5.8 | -4.8 |
| *ets-10* | WBGene00016870 | C52B9.11 | 2.6 | 1.8 |

| | | | | |
|---|---|---|---|---|
| *cebp-1* | WBGene00016997 | D1005.3 | 1.7 | 1.2 |
| *eaf-1* | WBGene00017011 | D1007.16 | -1.6 | -1.5 |
| *F10E7.11* | WBGene00017352 | F10E7.11 | -1.5 | -1.5 |
| *sdz-12* | WBGene00017406 | F12E12.5 | -4.4 | -2.2 |
| *bcl-11* | WBGene00017430 | F13H6.1 | 2.0 | 1.8 |
| *nhr-179* | WBGene00017512 | F16B4.11 | 2.7 | 3.2 |
| *ceh-49* | WBGene00017538 | F17A9.6 | -8.0 | -7.4 |
| *F19F10.1* | WBGene00017598 | F19F10.1 | -5.1 | -5.0 |
| *ets-7* | WBGene00017601 | F19F10.5 | -5.8 | -5.6 |
| *ets-4* | WBGene00017687 | F22A3.1 | 2.1 | 2.2 |
| *ceh-60* | WBGene00017690 | F22A3.5 | 1.1 | 1.2 |
| *nfyc-1* | WBGene00017742 | F23F1.1 | -1.7 | -1.5 |
| *zip-8* | WBGene00017755 | F23F12.9 | -1.2 | -1.7 |
| *nhr-141* | WBGene00017787 | F25E5.6 | 1.1 | 1.5 |
| *nhr-180* | WBGene00017961 | F31F4.12 | 2.3 | 1.3 |
| *dhhc-3* | WBGene00018009 | F33D11.12 | -2.2 | -1.5 |
| *nhr-181* | WBGene00018189 | F38H12.3 | -2.8 | -3.1 |
| *F40G9.14* | WBGene00018248 | F40G9.14 | -3.1 | -3.5 |
| *nhr-183* | WBGene00018266 | F41B5.10 | 1.6 | 1.4 |
| *ceh-86* | WBGene00018355 | F42G2.6 | -1.3 | -3.0 |
| *rpb-4* | WBGene00018391 | F43E2.2 | -1.4 | -1.4 |
| *ceh-82* | WBGene00018433 | F45C12.2 | -5.2 | -4.3 |
| *ceh-83* | WBGene00018446 | F45C12.15 | -4.4 | -4.5 |
| *oef-1* | WBGene00018636 | F49E8.2 | -1.7 | -1.4 |
| *ztf-13* | WBGene00018704 | F52E4.8 | 1.5 | 1.2 |
| *ztf-1* | WBGene00018833 | F54F2.5 | -2.4 | -1.8 |
| *F56D1.1* | WBGene00018959 | F56D1.1 | -1.4 | -1.5 |
| *klf-1* | WBGene00018990 | F56F11.3 | 1.8 | 1.8 |
| *F57C9.4* | WBGene00019011 | F57C9.4 | -1.6 | -1.5 |
| *nhr-143* | WBGene00019116 | F59E11.11 | 2.1 | 1.2 |
| *zip-2* | WBGene00019327 | K02F3.4 | 1.8 | 1.1 |
| *dmd-7* | WBGene00019521 | K08B12.2 | 1.2 | 1.4 |
| *nhr-201* | WBGene00019741 | M02H5.3 | 1.7 | 1.2 |
| *R02D3.7* | WBGene00019824 | R02D3.7 | -1.7 | -1.6 |
| *ceh-93* | WBGene00019864 | R04A9.5 | -1.3 | -1.2 |
| *R151.8* | WBGene00020111 | R151.8 | -1.3 | -1.2 |
| *nhr-211* | WBGene00020152 | T01G6.5 | 1.3 | 1.2 |
| *nhr-212* | WBGene00020153 | T01G6.6 | 1.5 | 1.5 |
| *T07F8.4* | WBGene00020320 | T07F8.4 | -1.2 | -1.1 |
| *nhr-216* | WBGene00020385 | T09D3.4 | 2.3 | 3.0 |
| *ztf-4* | WBGene00020399 | T10B11.3 | -1.8 | -1.3 |
| *ssup-72* | WBGene00020480 | T13C2.4 | -1.3 | -1.4 |
| *T20F7.1* | WBGene00020630 | T20F7.1 | -1.7 | -1.6 |
| *ztf-31* | WBGene00020635 | T20H4.2 | 1.3 | 1.8 |
| *dmd-8* | WBGene00020708 | T22H9.4 | 2.5 | 2.9 |
| *nhr-221* | WBGene00020748 | T24A6.8 | 3.2 | 1.6 |
| *T26A5.8* | WBGene00020823 | T26A5.8 | -1.5 | -1.9 |
| *hlh-30* | WBGene00020930 | W02C12.3 | 2.2 | 1.1 |
| *nhr-275* | WBGene00021163 | Y5H2A.2 | 2.4 | 1.5 |
| *Y22D7AL.16* | WBGene00021254 | Y22D7AL.16 | 2.6 | 2.8 |
| *thoc-5* | WBGene00021311 | Y32H12A.2 | -1.6 | -1.3 |
| *Y37E11B.1* | WBGene00021374 | Y37E11B.1 | 1.6 | 1.1 |
| *nhr-236* | WBGene00021417 | Y38F2AL.5 | 2.3 | 1.5 |
| *hlh-33* | WBGene00021446 | Y39A3CR.6 | 2.3 | 1.6 |
| *zip-4* | WBGene00021552 | Y44E3B.1 | -1.3 | -1.0 |
| *pcaf-1* | WBGene00021636 | Y47G6A.6 | -1.3 | -1.1 |
| *ztf-23* | WBGene00021846 | Y54E10BR.8 | -1.7 | -1.6 |
| *znf-782* | WBGene00021931 | Y55F3AM.14 | 2.2 | 4.3 |
| *Y82E9BR.1* | WBGene00022334 | Y82E9BR.1 | -4.3 | -6.9 |
| *Y82E9BR.17* | WBGene00022349 | Y82E9BR.17 | -5.8 | -7.6 |
| *nhr-277* | WBGene00022374 | Y94H6A.1 | 2.8 | 2.5 |
| *nhr-41* | WBGene00022423 | Y104H12A.1 | 1.5 | 1.1 |
| *ztf-8* | WBGene00022598 | ZC395.8 | -1.3 | -1.1 |
| *madf-9* | WBGene00022608 | ZC416.1 | 2.3 | 2.4 |
| *nhr-250* | WBGene00022755 | ZK488.1 | 2.1 | 1.4 |
| *ZK546.5* | WBGene00022762 | ZK546.5 | -1.3 | -1.3 |
| *dve-1* | WBGene00022861 | ZK1193.5 | 2.1 | 1.1 |
| *lin-15B* | WBGene00023497 | ZK662.4 | -1.3 | -1.2 |
| *tag-243* | WBGene00044078 | T04A8.4 | 4.1 | 2.4 |
| *bnc-1* | WBGene00044791 | F55C5.11 | 3.2 | 2.0 |
| *tbx-43* | WBGene00044798 | Y46E12A.4 | -4.4 | -2.0 |
| *nhr-146* | WBGene00045255 | Y41D4B.27 | 1.7 | 1.9 |
| *maf-1* | WBGene00077521 | F45H11.6 | 1.8 | 2.0 |
| | | N2 Specific | | |
| *ahr-1* | WBGene00000096 | C41G7.5 | 1.6 | Not DE |
| *atf-6* | WBGene00000222 | F45E6.2 | 1.2 | Not DE |
| *ceh-2* | WBGene00000429 | C27A12.5 | 1.7 | Not DE |
| *ceh-6* | WBGene00000431 | K02B12.1 | 1.5 | Not DE |
| *ceh-9* | WBGene00000434 | Y65B4BR.9 | 2.8 | Not DE |
| *ceh-17* | WBGene00000440 | D1007.1 | 1.6 | Not DE |

| | | | | |
|---|---|---|---|---|
| *ceh-19* | WBGene00000442 | F20D12.6 | 1.4 | Not DE |
| *ceh-22* | WBGene00000445 | F29F11.5 | 2.0 | Not DE |
| *ceh-24* | WBGene00000447 | F55B12.1 | 2.0 | Not DE |
| *pros-1* | WBGene00000448 | K12H4.1 | 2.0 | Not DE |
| *ceh-27* | WBGene00000449 | F46F3.1 | 1.0 | Not DE |
| *ceh-31* | WBGene00000452 | C33D12.1 | 1.6 | Not DE |
| *ceh-33* | WBGene00000454 | C10G8.7 | 1.8 | Not DE |
| *ceh-34* | WBGene00000455 | C10G8.6 | 1.6 | Not DE |
| *ceh-37* | WBGene00000458 | C37E2.5 | 1.6 | Not DE |
| *cfi-1* | WBGene00000476 | T23D8.8 | 2.0 | Not DE |
| *cog-1* | WBGene00000584 | R03C1.3 | 1.6 | Not DE |
| *daf-5* | WBGene00000901 | W01G7.1 | 1.0 | Not DE |
| *daf-14* | WBGene00000910 | F01G10.8 | 2.1 | Not DE |
| *dnj-17* | WBGene00001035 | T03F6.2 | -1.0 | Not DE |
| *dsc-1* | WBGene00001096 | C18B12.3 | 2.2 | Not DE |
| *egl-5* | WBGene00001174 | C08C3.1 | 1.3 | Not DE |
| *egl-38* | WBGene00001204 | C04G2.7 | 1.4 | Not DE |
| *egl-43* | WBGene00001207 | R53.3 | 2.2 | Not DE |
| *egl-44* | WBGene00001208 | F28B12.2 | 2.1 | Not DE |
| *elt-3* | WBGene00001251 | K02B9.4 | 1.3 | Not DE |
| *elt-4* | WBGene00001252 | C39B10.6 | 2.7 | Not DE |
| *elt-6* | WBGene00001253 | F52C12.5 | 1.2 | Not DE |
| *eor-1* | WBGene00001324 | R11E3.6 | -1.1 | Not DE |
| *fax-1* | WBGene00001400 | F56E3.4 | 1.7 | Not DE |
| *fkh-8* | WBGene00001440 | F40H3.4 | 1.9 | Not DE |
| *fkh-10* | WBGene00001442 | C25A1.2 | 2.5 | Not DE |
| *frm-7* | WBGene00001493 | C51F7.1 | 1.2 | Not DE |
| *xnd-1* | WBGene00001514 | C05D2.5 | -1.6 | Not DE |
| *gei-13* | WBGene00001570 | F58A4.11 | 1.9 | Not DE |
| *grh-1* | WBGene00001707 | Y48G8AR.1 | 2.3 | Not DE |
| *ham-2* | WBGene00001821 | C07A12.1 | 1.6 | Not DE |
| *hlh-6* | WBGene00001952 | T15H9.3 | 2.1 | Not DE |
| *hlh-8* | WBGene00001953 | C02B8.4 | 2.0 | Not DE |
| *hlh-12* | WBGene00001956 | C28C12.8 | 1.3 | Not DE |
| *hlh-13* | WBGene00001957 | F48D6.3 | 3.6 | Not DE |
| *hlh-17* | WBGene00001961 | F38C2.2 | 1.9 | Not DE |
| *hlh-26* | WBGene00001965 | C17C3.8 | -3.2 | Not DE |
| *hmg-12* | WBGene00001977 | Y17G7A.1 | -1.5 | Not DE |
| *hsf-1* | WBGene00002004 | Y53C10A.12 | -1.0 | Not DE |
| *hsp-60* | WBGene00002025 | Y22D7AL.5 | -2.8 | Not DE |
| *lim-4* | WBGene00002987 | ZC64.4 | 3.5 | Not DE |
| *lin-22* | WBGene00003008 | Y54G2A.1 | 1.5 | Not DE |
| *lin-36* | WBGene00003021 | F44B9.6 | -1.2 | Not DE |
| *lin-39* | WBGene00003024 | C07H6.7 | 1.6 | Not DE |
| *mab-3* | WBGene00003100 | Y53C12B.5 | 2.2 | Not DE |
| *mab-5* | WBGene00003102 | C08C3.3 | 2.6 | Not DE |
| *mbf-1* | WBGene00003148 | H21P03.1 | -1.4 | Not DE |
| *mec-3* | WBGene00003167 | F01D4.6 | 2.7 | Not DE |
| *med-2* | WBGene00003181 | K04C2.6 | 3.1 | Not DE |
| *mel-28* | WBGene00003210 | C38D4.3 | -1.0 | Not DE |
| *mep-1* | WBGene00003218 | M04B2.1 | -1.1 | Not DE |
| *mxl-3* | WBGene00003511 | F46G10.6 | -2.0 | Not DE |
| *nhr-4* | WBGene00003603 | F32B6.1 | 1.4 | Not DE |
| *nhr-5* | WBGene00003604 | Y73F8A.21 | 1.1 | Not DE |
| *nhr-16* | WBGene00003615 | T12C9.6 | 1.1 | Not DE |
| *nhr-21* | WBGene00003620 | F21D12.1 | 2.8 | Not DE |
| *nhr-25* | WBGene00003623 | F11C1.6 | 1.3 | Not DE |
| *nhr-32* | WBGene00003626 | K08H2.8 | 2.2 | Not DE |
| *nhr-42* | WBGene00003632 | C33G8.6 | 2.2 | Not DE |
| *nhr-43* | WBGene00003633 | C29E6.5 | 1.0 | Not DE |
| *nhr-45* | WBGene00003635 | F16H11.5 | 1.5 | Not DE |
| *nhr-53* | WBGene00003643 | K06B4.11 | 1.9 | Not DE |
| *nhr-55* | WBGene00003645 | T01G6.7 | 1.7 | Not DE |
| *nhr-56* | WBGene00003646 | F44C8.6 | 1.2 | Not DE |
| *nhr-58* | WBGene00003648 | R11G11.2 | 2.0 | Not DE |
| *nhr-60* | WBGene00003650 | F57A10.5 | 1.6 | Not DE |
| *nhr-61* | WBGene00003651 | W01D2.2 | 1.1 | Not DE |
| *nhr-64* | WBGene00003654 | C45E1.1 | 1.4 | Not DE |
| *nhr-69* | WBGene00003659 | T23H4.2 | 1.6 | Not DE |
| *nhr-74* | WBGene00003664 | C27C7.3 | -1.5 | Not DE |
| *nhr-83* | WBGene00003673 | F48G7.3 | 1.1 | Not DE |
| *nhr-90* | WBGene00003680 | ZK488.2 | 1.5 | Not DE |
| *nhr-94* | WBGene00003684 | C12D5.8 | 2.2 | Not DE |
| *nhr-96* | WBGene00003686 | F44C8.11 | 1.6 | Not DE |
| *nhr-100* | WBGene00003690 | C28D4.1 | 1.1 | Not DE |
| *nhr-101* | WBGene00003691 | H12C20.6 | 1.2 | Not DE |
| *nhr-106* | WBGene00003696 | T01G6.4 | 1.6 | Not DE |
| *nhr-107* | WBGene00003697 | C33G8.11 | 2.1 | Not DE |
| *nhr-108* | WBGene00003698 | F35E8.12 | 1.0 | Not DE |
| *nhr-110* | WBGene00003700 | Y46H3D.5 | 1.4 | Not DE |

| | | | | |
|---|---|---|---|---|
| *nhr-112* | WBGene00003702 | Y70C5C.6 | 2.6 | Not DE |
| *nhr-113* | WBGene00003703 | ZK1025.9 | 4.9 | Not DE |
| *nhr-114* | WBGene00003704 | Y45G5AM.1 | -2.9 | Not DE |
| *nhr-116* | WBGene00003706 | F09C6.9 | 2.4 | Not DE |
| *nhr-119* | WBGene00003709 | K12H6.1 | 1.3 | Not DE |
| *nhr-122* | WBGene00003712 | Y41D4B.9 | 1.6 | Not DE |
| *nhr-123* | WBGene00003713 | M02H5.7 | 1.5 | Not DE |
| *nhr-127* | WBGene00003717 | T13F3.3 | 2.7 | Not DE |
| *nhr-131* | WBGene00003721 | T01G6.2 | 1.0 | Not DE |
| *nhr-134* | WBGene00003724 | F44C8.2 | 1.4 | Not DE |
| *nhr-135* | WBGene00003725 | VC5.5 | 1.1 | Not DE |
| *nob-1* | WBGene00003779 | Y75B8A.2 | 1.1 | Not DE |
| *blmp-1* | WBGene00003847 | F25D7.3 | 1.2 | Not DE |
| *pag-3* | WBGene00003909 | F45B8.4 | 1.8 | Not DE |
| *pat-9* | WBGene00003933 | T27B1.2 | 1.3 | Not DE |
| *pha-2* | WBGene00004011 | M6.3 | 1.7 | Not DE |
| *pha-4* | WBGene00004013 | F38A6.1 | 2.3 | Not DE |
| *plx-2* | WBGene00004048 | K04B12.1 | 1.5 | Not DE |
| *swsn-1* | WBGene00004203 | Y113G7B.23 | -1.2 | Not DE |
| *sma-4* | WBGene00004858 | R12B2.1 | 1.4 | Not DE |
| *sma-6* | WBGene00004860 | C32D5.2 | 1.0 | Not DE |
| *sma-9* | WBGene00004862 | T05A10.1 | 1.2 | Not DE |
| *sox-2* | WBGene00004949 | K08A8.2 | 1.9 | Not DE |
| *sup-37* | WBGene00006342 | C01B7.1 | 1.1 | Not DE |
| *taf-6.1* | WBGene00006387 | W09B6.2 | -1.2 | Not DE |
| *taf-10* | WBGene00006392 | K03B4.3 | -1.0 | Not DE |
| *ctbp-1* | WBGene00006424 | F49E10.5 | 2.2 | Not DE |
| *tbx-7* | WBGene00006544 | ZK328.8 | 1.8 | Not DE |
| *tbx-35* | WBGene00006554 | ZK177.10 | -2.6 | Not DE |
| *tlf-1* | WBGene00006577 | F39H11.2 | -1.0 | Not DE |
| *ttx-1* | WBGene00006652 | Y113G7A.6 | 1.9 | Not DE |
| *ttx-3* | WBGene00006654 | C40H5.5 | 1.9 | Not DE |
| *unc-3* | WBGene00006743 | Y16B4A.1 | 2.3 | Not DE |
| *unc-30* | WBGene00006766 | B0564.10 | 1.3 | Not DE |
| *unc-42* | WBGene00006778 | F58E6.10 | 3.4 | Not DE |
| *unc-86* | WBGene00006818 | C30A5.7 | 2.3 | Not DE |
| *unc-120* | WBGene00006844 | D1081.2 | 1.1 | Not DE |
| *vab-3* | WBGene00006870 | F14F3.1 | 1.9 | Not DE |
| *zag-1* | WBGene00006970 | F28F9.1 | 2.1 | Not DE |
| *zip-1* | WBGene00006986 | Y75B8A.35 | 1.5 | Not DE |
| *pbrm-1* | WBGene00007042 | C26C6.1 | -1.0 | Not DE |
| *nhr-150* | WBGene00007367 | C06B8.1 | 2.6 | Not DE |
| *jmjd-5* | WBGene00007387 | C06H2.3 | -1.2 | Not DE |
| *nhr-153* | WBGene00007546 | C13C4.1 | 1.2 | Not DE |
| *dhhc-7* | WBGene00007637 | C17D12.1 | -1.5 | Not DE |
| *C17E4.6* | WBGene00007645 | C17E4.6 | -1.2 | Not DE |
| *ceh-79* | WBGene00007749 | C26E1.3 | 1.5 | Not DE |
| *dmd-4* | WBGene00007776 | C27C12.6 | 1.7 | Not DE |
| *zip-5* | WBGene00007932 | C34D1.5 | 1.5 | Not DE |
| *yap-1* | WBGene00008748 | F13E6.4 | 1.9 | Not DE |
| *nhr-176* | WBGene00008830 | F14H3.11 | 1.0 | Not DE |
| *plp-2* | WBGene00008976 | F20D1.4 | 1.1 | Not DE |
| *myrf-2* | WBGene00008999 | F21A10.2 | 1.3 | Not DE |
| *F27D4.4* | WBGene00009189 | F27D4.4 | -1.0 | Not DE |
| *F36F2.1* | WBGene00009475 | F36F2.1 | 1.5 | Not DE |
| *ccch-1* | WBGene00009532 | F38B7.1 | 1.7 | Not DE |
| *psa-3* | WBGene00009560 | F39D8.2 | 2.2 | Not DE |
| *lsl-1* | WBGene00009937 | F52F12.4 | -1.7 | Not DE |
| *frpr-9* | WBGene00009965 | F53B7.2 | 1.4 | Not DE |
| *gmeb-2* | WBGene00010010 | F53H4.5 | 1.7 | Not DE |
| *nhr-193* | WBGene00010215 | F57G8.6 | 1.6 | Not DE |
| *sta-2* | WBGene00010251 | F58E6.1 | 1.8 | Not DE |
| *K11H3.4* | WBGene00010781 | K11H3.4 | -1.9 | Not DE |
| *somi-1* | WBGene00010868 | M04G12.4 | 1.7 | Not DE |
| *tceb-3* | WBGene00010990 | R03D7.4 | -1.1 | Not DE |
| *ceh-62* | WBGene00011069 | R06F6.6 | 1.3 | Not DE |
| *nhr-209* | WBGene00011100 | R07B7.16 | 1.5 | Not DE |
| *eea-1* | WBGene00011696 | T10G3.5 | -1.2 | Not DE |
| *rbm-22* | WBGene00011722 | T11G6.8 | -1.2 | Not DE |
| *tsct-1* | WBGene00011824 | T18D3.7 | 1.9 | Not DE |
| *T22C8.3* | WBGene00011924 | T22C8.3 | 1.2 | Not DE |
| *zip-3* | WBGene00012330 | W07G1.3 | -1.1 | Not DE |
| *Y17G7B.22* | WBGene00012473 | Y17G7B.22 | 2.1 | Not DE |
| *fezf-1* | WBGene00012639 | Y38H8A.5 | 2.6 | Not DE |
| *rabx-5* | WBGene00012644 | Y39A1A.5 | -1.1 | Not DE |
| *dmd-3* | WBGene00012832 | Y43F8C.10 | 6.1 | Not DE |
| *ztf-22* | WBGene00012988 | Y48C3A.4 | 1.4 | Not DE |
| *Y48C3A.12* | WBGene00012994 | Y48C3A.12 | -1.1 | Not DE |
| *mcd-1* | WBGene00013096 | Y51H1A.6 | -1.2 | Not DE |
| *hsf-2* | WBGene00013134 | Y53C10A.3 | -1.5 | Not DE |

| | | | | |
|---|---|---|---|---|
| *Y53H1A.2* | WBGene00013178 | Y53H1A.2 | -1.1 | Not DE |
| *Y56A3A.2* | WBGene00013225 | Y56A3A.2 | -1.1 | Not DE |
| *Y57A10A.31* | WBGene00013270 | Y57A10A.31 | -1.9 | Not DE |
| *slr-2* | WBGene00013350 | Y59A8B.13 | -1.1 | Not DE |
| *Y105C5A.15* | WBGene00013639 | Y105C5A.15 | 1.5 | Not DE |
| *madf-10* | WBGene00013717 | Y106G6H.4 | 1.6 | Not DE |
| *gtf-2A2* | WBGene00013736 | Y111B2A.13 | 2.0 | Not DE |
| *gmeb-1* | WBGene00015285 | C01B12.2 | 1.3 | Not DE |
| *nhr-147* | WBGene00015395 | C03G6.8 | 2.4 | Not DE |
| *C04F5.9* | WBGene00015451 | C04F5.9 | -1.1 | Not DE |
| *ztf-30* | WBGene00015523 | C06E1.8 | 2.6 | Not DE |
| *nhr-152* | WBGene00015705 | C12D5.2 | 1.5 | Not DE |
| *nhr-158* | WBGene00015901 | C17E7.6 | 1.5 | Not DE |
| *ceh-48* | WBGene00015934 | C17H12.9 | 1.3 | Not DE |
| *elt-7* | WBGene00015981 | C18G1.2 | 1.0 | Not DE |
| *nhr-258* | WBGene00016126 | C26B2.4 | 1.9 | Not DE |
| *nhr-290* | WBGene00016233 | C29G2.5 | 2.7 | Not DE |
| *nhr-140* | WBGene00016366 | C33G8.9 | 2.1 | Not DE |
| *nhr-163* | WBGene00016368 | C33G8.12 | 1.9 | Not DE |
| *ets-8* | WBGene00016798 | C50A2.4 | -2.7 | Not DE |
| *ets-9* | WBGene00016865 | C52B9.2 | 1.4 | Not DE |
| *znf-598* | WBGene00016888 | C52E12.1 | -1.3 | Not DE |
| *nhr-172* | WBGene00016927 | C54F6.9 | -2.0 | Not DE |
| *nhr-173* | WBGene00016975 | C56E10.1 | 1.9 | Not DE |
| *nhr-36* | WBGene00017198 | F07C3.10 | 1.6 | Not DE |
| *set-9* | WBGene00017482 | F15E6.1 | -1.0 | Not DE |
| *nhr-177* | WBGene00017503 | F16B4.1 | 1.1 | Not DE |
| *nhr-178* | WBGene00017510 | F16B4.9 | 3.4 | Not DE |
| *atf-8* | WBGene00017535 | F17A9.3 | 1.3 | Not DE |
| *F21A9.2* | WBGene00017651 | F21A9.2 | 1.2 | Not DE |
| *ubxn-1* | WBGene00017733 | F23C8.4 | -1.3 | Not DE |
| *F26A10.2* | WBGene00017814 | F26A10.2 | 2.5 | Not DE |
| *nhr-37* | WBGene00018412 | F44C4.2 | 3.9 | Not DE |
| *nhr-184* | WBGene00018415 | F44C8.9 | 1.3 | Not DE |
| *F44E2.7* | WBGene00018420 | F44E2.7 | -1.0 | Not DE |
| *mnat-1* | WBGene00018769 | F53G2.7 | -1.8 | Not DE |
| *hmbx-1* | WBGene00018786 | F54A5.1 | 1.1 | Not DE |
| *attf-3* | WBGene00018794 | F54C4.3 | -1.4 | Not DE |
| *nhr-195* | WBGene00019115 | F59E11.10 | 1.4 | Not DE |
| *snai-1* | WBGene00019299 | K02D7.2 | 2.0 | Not DE |
| *aptf-1* | WBGene00019424 | K06A1.1 | 2.1 | Not DE |
| *K09H9.7* | WBGene00019598 | K09H9.7 | -1.7 | Not DE |
| *K11D12.12* | WBGene00019651 | K11D12.12 | 1.5 | Not DE |
| *nhr-202* | WBGene00019742 | M02H5.4 | 1.0 | Not DE |
| *M03D4.4* | WBGene00019751 | M03D4.4 | 2.2 | Not DE |
| *nhr-204* | WBGene00019816 | R02C2.4 | 1.1 | Not DE |
| *ztf-16* | WBGene00019960 | R08E3.4 | 1.5 | Not DE |
| *nhr-270* | WBGene00020062 | R13D11.8 | 1.4 | Not DE |
| *camt-1* | WBGene00020251 | T05C1.4 | 1.4 | Not DE |
| *ast-1* | WBGene00020368 | T08H4.3 | 2.1 | Not DE |
| *T10D4.6* | WBGene00020408 | T10D4.6 | -1.2 | Not DE |
| *dhhc-11* | WBGene00020694 | T22E7.2 | 2.8 | Not DE |
| *ztf-18* | WBGene00020763 | T24C4.7 | -1.0 | Not DE |
| *nhr-225* | WBGene00020849 | T27B7.2 | 1.3 | Not DE |
| *nhr-227* | WBGene00020851 | T27B7.5 | 1.8 | Not DE |
| *W04B5.2* | WBGene00021019 | W04B5.2 | 2.1 | Not DE |
| *crtc-1* | WBGene00021237 | Y20F4.2 | 2.0 | Not DE |
| *taf-6.2* | WBGene00021363 | Y37E11AL.8 | -1.2 | Not DE |
| *cyh-1* | WBGene00021714 | Y49F6B.1 | -1.3 | Not DE |
| *rcor-1* | WBGene00022278 | Y74C9A.4 | -1.2 | Not DE |
| *zfh-2* | WBGene00022518 | ZC123.3 | 1.1 | Not DE |
| *ZK177.3* | WBGene00022671 | ZK177.3 | 2.6 | Not DE |
| *ceh-45* | WBGene00022837 | ZK993.1 | 1.5 | Not DE |
| *syd-9* | WBGene00044068 | ZK867.1 | 1.3 | Not DE |
| *ham-3* | WBGene00044072 | ZK1128.5 | -1.2 | Not DE |
| *alr-1* | WBGene00044330 | R08B4.2 | 2.0 | Not DE |
| | | CB4856 Specific | | |
| *atf-2* | WBGene00000220 | K08F8.2 | Not DE | -3.1 |
| *cdk-7* | WBGene00000408 | Y39G10AL.3 | Not DE | -1.0 |
| *ceh-8* | WBGene00000433 | ZK265.4 | Not DE | -2.4 |
| *ceh-16* | WBGene00000439 | C13G5.1 | Not DE | -2.2 |
| *ceh-30* | WBGene00000451 | C33D12.7 | Not DE | -2.8 |
| *ceh-43* | WBGene00000463 | C28A5.4 | Not DE | -2.4 |
| *cnd-1* | WBGene00000561 | C34E10.7 | Not DE | -3.7 |
| *egl-46* | WBGene00001210 | K11G9.4 | Not DE | -3.4 |
| *ehn-3* | WBGene00001223 | ZK616.10 | Not DE | -2.5 |
| *ham-1* | WBGene00001820 | F53B2.6 | Not DE | -1.5 |
| *hbl-1* | WBGene00001824 | F13D11.2 | Not DE | -2.9 |
| *hlh-1* | WBGene00001948 | B0304.1 | Not DE | -1.0 |
| *hlh-16* | WBGene00001960 | DY3.3 | Not DE | -3.9 |

| | | | | |
|---|---|---|---|---|
| *hmg-11* | WBGene00001976 | T05A7.4 | Not DE | -1.1 |
| *lir-2* | WBGene00003045 | F18A1.4 | Not DE | -1.0 |
| *mls-2* | WBGene00003377 | C39E6.4 | Not DE | -2.5 |
| *mnm-2* | WBGene00003380 | C10A4.8 | Not DE | -1.2 |
| *nhr-1* | WBGene00003600 | R09G11.2 | Not DE | 1.3 |
| *nhr-67* | WBGene00003657 | C08F8.8 | Not DE | -1.2 |
| *nhr-68* | WBGene00003658 | H12C20.3 | Not DE | -1.0 |
| *nhr-77* | WBGene00003667 | T15D6.6 | Not DE | -2.3 |
| *nhr-80* | WBGene00003670 | H10E21.3 | Not DE | 1.2 |
| *nhr-93* | WBGene00003683 | T24A6.9 | Not DE | 2.2 |
| *nhr-97* | WBGene00003687 | H27C11.1 | Not DE | 1.1 |
| *nhr-130* | WBGene00003720 | T01G6.8 | Not DE | 1.1 |
| *nhr-132* | WBGene00003722 | R11G11.1 | Not DE | 1.6 |
| *odd-1* | WBGene00003845 | B0280.4 | Not DE | -1.5 |
| *pax-1* | WBGene00003937 | K07C11.1 | Not DE | -3.5 |
| *pax-3* | WBGene00003939 | F27E5.2 | Not DE | -1.8 |
| *pop-1* | WBGene00004077 | W10C8.2 | Not DE | -1.4 |
| *pqn-75* | WBGene00004157 | W03D2.1 | Not DE | 1.8 |
| *sem-2* | WBGene00004771 | C32E12.5 | Not DE | -2.5 |
| *spe-10* | WBGene00004964 | AC3.10 | Not DE | 1.0 |
| *spt-4* | WBGene00005014 | F54C4.2 | Not DE | -1.2 |
| *tab-1* | WBGene00006380 | F31E8.3 | Not DE | -1.2 |
| *taf-7.1* | WBGene00006388 | F54F7.1 | Not DE | -2.0 |
| *taf-13* | WBGene00006397 | C14A4.10 | Not DE | -1.1 |
| *ubl-5* | WBGene00006726 | F46F11.4 | Not DE | -1.4 |
| *unc-39* | WBGene00006775 | F56A12.1 | Not DE | -2.2 |
| *C38D4.7* | WBGene00008007 | C38D4.7 | Not DE | 2.0 |
| *F21D5.9* | WBGene00009014 | F21D5.9 | Not DE | -2.8 |
| *F21G4.5* | WBGene00009026 | F21G4.5 | Not DE | 1.2 |
| *F28F8.7* | WBGene00009224 | F28F8.7 | Not DE | -1.4 |
| *hmg-6* | WBGene00009827 | F47G4.6 | Not DE | -2.3 |
| *efl-3* | WBGene00009899 | F49E12.6 | Not DE | -2.0 |
| *klf-2* | WBGene00009998 | F53F8.1 | Not DE | -4.0 |
| *dhhc-12* | WBGene00010323 | F59C6.2 | Not DE | 1.7 |
| *K04C1.3* | WBGene00010553 | K04C1.3 | Not DE | -1.0 |
| *nhr-198* | WBGene00010603 | K06B4.8 | Not DE | 2.2 |
| *K09A11.1* | WBGene00010704 | K09A11.1 | Not DE | -1.0 |
| *ccch-3* | WBGene00012277 | W05B10.2 | Not DE | -1.4 |
| *nhr-230* | WBGene00012446 | Y17D7A.1 | Not DE | 1.0 |
| *nhr-234* | WBGene00012596 | Y38E10A.18 | Not DE | 2.7 |
| *eyg-1* | WBGene00013147 | Y53C12C.1 | Not DE | -1.2 |
| *ztf-29* | WBGene00013438 | Y66D12A.12 | Not DE | -1.2 |
| *ekl-4* | WBGene00013676 | Y105E8A.17 | Not DE | -1.0 |
| *C06E2.1* | WBGene00015527 | C06E2.1 | Not DE | 1.5 |
| *ceh-53* | WBGene00015651 | C09G12.1 | Not DE | -1.9 |
| *nhr-155* | WBGene00015758 | C14C6.4 | Not DE | 4.1 |
| *nhr-157* | WBGene00015900 | C17E7.5 | Not DE | 2.4 |
| *nhr-30* | WBGene00016091 | C25E10.1 | Not DE | 1.8 |
| *nhr-260* | WBGene00016517 | C38C3.9 | Not DE | 1.7 |
| *scrt-1* | WBGene00016948 | C55C2.1 | Not DE | -2.1 |
| *dmd-5* | WBGene00017326 | F10C1.5 | Not DE | -3.1 |
| *F13C5.2* | WBGene00017423 | F13C5.2 | Not DE | -1.3 |
| *F37B4.10* | WBGene00018140 | F37B4.10 | Not DE | -1.2 |
| *cir-1* | WBGene00018892 | F55F8.4 | Not DE | -1.2 |
| *nhr-266* | WBGene00018993 | F56H1.2 | Not DE | 2.2 |
| *madf-3* | WBGene00019218 | H20J04.3 | Not DE | -1.2 |
| *dhhc-13* | WBGene00019257 | H32C10.3 | Not DE | 1.7 |
| *lect-2* | WBGene00019407 | K05F1.5 | Not DE | -1.3 |
| *dhhc-5* | WBGene00020066 | R13F6.5 | Not DE | 1.4 |
| *R144.3* | WBGene00020093 | R144.3 | Not DE | -1.1 |
| *ceh-54* | WBGene00020485 | T13C5.4 | Not DE | 2.4 |
| *nhr-219* | WBGene00020555 | T19A5.5 | Not DE | 3.3 |
| *W03F9.2* | WBGene00021000 | W03F9.2 | Not DE | -1.3 |
| *rabs-5* | WBGene00021538 | Y42H9AR.3 | Not DE | -1.2 |
| *duxl-1* | WBGene00022554 | ZC204.2 | Not DE | -1.8 |
| *klu-2* | WBGene00022592 | ZC328.2 | Not DE | 2.1 |
| *snu-23* | WBGene00022794 | ZK686.4 | Not DE | -1.1 |
| *ZK686.5* | WBGene00022795 | ZK686.5 | Not DE | 1.8 |
| *C27A2.7* | WBGene00044386 | C27A2.7 | Not DE | 2.3 |

**Supplemental Table 8**

**Lists of differentially expressed genes that are identified in either N2, CB4856, or both, after infection by *P. aeruginosa* (compared to *E. coli*)**

**Supplemental Table 8 - Differentially Expressed Genes in *P. aeruginosa* Infected Animals**

**N2**-exclusive DEGs in response to *P. aeruginosa* (vs. OP50)

| Gene Name | Wormbase Gene ID | Type | $log_2FC$ | FC | Adjusted P-value |
|---|---|---|---|---|---|
| *T27C5.8* | WBGene00012081 | protein_coding_gene | 5.1 | 33.134 | 7.1E-06 |
| *C25F9.11* | WBGene00045411 | protein_coding_gene | 5.0 | 30.976 | 6.6E-10 |
| *T26H5.9* | WBGene00044206 | protein_coding_gene | 4.8 | 28.647 | 2.1E-08 |
| *F53A9.8* | WBGene00018731 | protein_coding_gene | 4.6 | 24.314 | 3.0E-12 |
| *T26H5.14* | WBGene00235133 | protein_coding_gene | 4.4 | 21.608 | 7.3E-08 |
| *fbxa-161* | WBGene00015596 | pseudogene | 4.3 | 20.340 | 8.7E-05 |
| *T26H5.4* | WBGene00012069 | protein_coding_gene | 4.3 | 19.879 | 4.7E-06 |
| *W07G1.25* | WBGene00271643 | ncRNA_gene | 4.3 | 19.462 | 6.2E-04 |
| *C25F9.16* | WBGene00194803 | protein_coding_gene | 4.2 | 17.864 | 1.1E-04 |
| *fbxa-25* | WBGene00022321 | protein_coding_gene | 4.1 | 16.850 | 1.8E-04 |
| *fbxa-24* | WBGene00018803 | protein_coding_gene | 4.0 | 16.263 | 6.9E-10 |
| *B0250.8* | WBGene00007125 | protein_coding_gene | 4.0 | 16.125 | 5.3E-04 |
| *C35A5.11* | WBGene00044024 | protein_coding_gene | 3.9 | 14.839 | 4.8E-10 |
| *F21C10.11* | WBGene00017660 | protein_coding_gene | 3.8 | 14.408 | 1.6E-21 |
| *clec-46* | WBGene00017202 | pseudogene | 3.8 | 14.316 | 4.3E-02 |
| *W07G4.1* | WBGene00012335 | protein_coding_gene | 3.8 | 14.233 | 1.2E-03 |
| *Y37A1B.7* | WBGene00012540 | protein_coding_gene | 3.8 | 13.881 | 9.0E-10 |
| *T26H5.10* | WBGene00044783 | protein_coding_gene | 3.8 | 13.837 | 3.1E-05 |
| *C07A9.5* | WBGene00007401 | protein_coding_gene | 3.7 | 13.128 | 8.1E-04 |
| *C07G3.8* | WBGene00015576 | protein_coding_gene | 3.7 | 13.095 | 4.4E-04 |
| *F13A2.14* | WBGene00269436 | pseudogene | 3.7 | 12.896 | 1.2E-03 |
| *F18C5.9* | WBGene00017562 | protein_coding_gene | 3.7 | 12.774 | 8.4E-04 |
| *srw-86* | WBGene00005833 | protein_coding_gene | 3.6 | 11.833 | 7.1E-07 |
| *T04C12.1* | WBGene00011427 | protein_coding_gene | 3.5 | 11.586 | 5.3E-06 |
| *clec-42* | WBGene00008891 | protein_coding_gene | 3.5 | 11.432 | 1.3E-04 |
| *Y39A3A.3* | WBGene00021434 | protein_coding_gene | 3.5 | 11.387 | 1.2E-05 |
| *lipl-2* | WBGene00009773 | protein_coding_gene | 3.5 | 11.252 | 2.7E-05 |
| *lys-3* | WBGene00003092 | protein_coding_gene | 3.5 | 11.141 | 1.5E-06 |
| *F11C7.6* | WBGene00017378 | protein_coding_gene | 3.5 | 11.131 | 2.7E-08 |
| *Y74C9A.6* | WBGene00023193 | snoRNA_gene | 3.4 | 10.642 | 9.3E-03 |
| *linc-38* | WBGene00219415 | lincRNA_gene | 3.4 | 10.380 | 4.9E-03 |
| *sru-20* | WBGene00005683 | protein_coding_gene | 3.4 | 10.263 | 1.3E-03 |
| *ugt-18* | WBGene00013900 | protein_coding_gene | 3.3 | 10.177 | 2.8E-04 |
| *W01C9.2* | WBGene00012176 | protein_coding_gene | 3.3 | 10.027 | 5.4E-09 |
| *Y73B6BL.37* | WBGene00022256 | protein_coding_gene | 3.3 | 9.994 | 4.9E-04 |
| *kreg-1* | WBGene00018725 | protein_coding_gene | 3.3 | 9.901 | 5.7E-08 |
| *fezf-1* | WBGene00012639 | protein_coding_gene | 3.3 | 9.810 | 2.5E-03 |
| *ech-9* | WBGene00001158 | protein_coding_gene | 3.3 | 9.775 | 5.2E-07 |
| *ZK675.4* | WBGene00014067 | protein_coding_gene | 3.3 | 9.764 | 1.7E-04 |
| *C17B7.4* | WBGene00015878 | protein_coding_gene | 3.3 | 9.676 | 1.2E-06 |
| *C45B11.2* | WBGene00008105 | protein_coding_gene | 3.2 | 9.511 | 2.7E-06 |
| *grl-11* | WBGene00001720 | protein_coding_gene | 3.2 | 9.375 | 1.3E-02 |
| *Y57A10C.10* | WBGene00013288 | protein_coding_gene | 3.2 | 9.362 | 9.2E-03 |
| *F01D5.7* | WBGene00008497 | protein_coding_gene | 3.2 | 9.196 | 1.4E-03 |
| *Y43F8B.15* | WBGene00045415 | protein_coding_gene | 3.2 | 9.004 | 3.0E-06 |
| *T28A11.19* | WBGene00020881 | protein_coding_gene | 3.2 | 8.932 | 3.0E-07 |
| *B0250.11* | WBGene00050926 | pseudogene | 3.2 | 8.921 | 7.9E-03 |
| *Y54G2A.76* | WBGene00235268 | protein_coding_gene | 3.1 | 8.735 | 1.2E-05 |
| *phat-5* | WBGene00020242 | protein_coding_gene | 3.1 | 8.706 | 3.7E-06 |
| *fbxa-127* | WBGene00012027 | protein_coding_gene | 3.1 | 8.680 | 1.4E-05 |
| *C25F9.12* | WBGene00045412 | protein_coding_gene | 3.1 | 8.566 | 6.2E-04 |
| *Y71A12B.19* | WBGene00077443 | pseudogene | 3.1 | 8.365 | 8.9E-10 |
| *Y64H9A.2* | WBGene00022024 | protein_coding_gene | 3.1 | 8.294 | 1.7E-04 |
| *scl-14* | WBGene00008625 | protein_coding_gene | 3.1 | 8.289 | 2.0E-03 |
| *ZC395.5* | WBGene00022597 | protein_coding_gene | 3.0 | 8.276 | 2.4E-12 |
| *Y37E3.20* | WBGene00044925 | snoRNA_gene | 3.0 | 8.210 | 1.9E-02 |
| *C27A2.8* | WBGene00044387 | protein_coding_gene | 3.0 | 8.150 | 6.5E-03 |
| *F11A5.5* | WBGene00008673 | protein_coding_gene | 3.0 | 7.974 | 5.3E-03 |
| *ttr-23* | WBGene00011894 | protein_coding_gene | 3.0 | 7.854 | 3.4E-08 |
| *btb-21* | WBGene00018540 | pseudogene | 2.9 | 7.725 | 2.0E-13 |
| *C10F3.7* | WBGene00015679 | protein_coding_gene | 2.9 | 7.699 | 7.6E-05 |
| *C49C3.21* | WBGene00189993 | protein_coding_gene | 2.9 | 7.673 | 1.8E-02 |
| *T09E11.11* | WBGene00011660 | pseudogene | 2.9 | 7.659 | 6.9E-10 |
| *Y75B12B.12* | WBGene00045129 | snoRNA_gene | 2.9 | 7.622 | 2.9E-02 |
| *ZK813.6* | WBGene00044564 | protein_coding_gene | 2.9 | 7.480 | 1.4E-02 |
| *Y73B3A.16* | WBGene00022218 | pseudogene | 2.9 | 7.468 | 4.4E-02 |

| | | | | | |
|---|---|---|---|---|---|
| *T26E3.6* | WBGene00012039 | protein_coding_gene | 2.9 | 7.445 | 1.6E-02 |
| *C17B7.10* | WBGene00015884 | protein_coding_gene | 2.9 | 7.292 | 2.7E-03 |
| *fbxa-162* | WBGene00015597 | pseudogene | 2.8 | 7.192 | 9.6E-07 |
| *nhr-218* | WBGene00011750 | protein_coding_gene | 2.8 | 7.163 | 7.5E-03 |
| *K08B4.2* | WBGene00019514 | protein_coding_gene | 2.8 | 7.065 | 2.4E-06 |
| *T02G5.15* | WBGene00206521 | protein_coding_gene | 2.8 | 6.959 | 2.9E-06 |
| *cyp-32B1* | WBGene00021167 | protein_coding_gene | 2.8 | 6.926 | 1.3E-04 |
| *mir-238* | WBGene00003331 | miRNA_gene | 2.8 | 6.847 | 1.4E-02 |
| *Y39A3A.4* | WBGene00021435 | protein_coding_gene | 2.8 | 6.835 | 2.1E-06 |
| *R166.7* | WBGene00045074 | snoRNA_gene | 2.8 | 6.764 | 4.9E-02 |
| *hil-7* | WBGene00001858 | protein_coding_gene | 2.7 | 6.648 | 4.3E-04 |
| *C01G10.6* | WBGene00007233 | protein_coding_gene | 2.7 | 6.635 | 1.0E-03 |
| *srw-85* | WBGene00005832 | protein_coding_gene | 2.7 | 6.567 | 3.0E-15 |
| *Y102A5C.11* | WBGene00014956 | pseudogene | 2.7 | 6.535 | 2.0E-02 |
| *C50F4.9* | WBGene00008234 | protein_coding_gene | 2.7 | 6.476 | 3.8E-13 |
| *pax-1* | WBGene00003937 | protein_coding_gene | 2.7 | 6.469 | 1.3E-02 |
| *Y75B8A.33* | WBGene00013563 | protein_coding_gene | 2.7 | 6.388 | 1.1E-02 |
| *bath-23* | WBGene00021727 | protein_coding_gene | 2.7 | 6.369 | 3.8E-04 |
| *W02A2.9* | WBGene00044280 | protein_coding_gene | 2.7 | 6.323 | 1.7E-12 |
| *W05H7.2* | WBGene00021045 | pseudogene | 2.7 | 6.277 | 3.3E-02 |
| *Y49E10.7* | WBGene00013031 | pseudogene | 2.6 | 6.124 | 2.9E-06 |
| *M162.12* | WBGene00050946 | pseudogene | 2.6 | 6.095 | 8.9E-09 |
| *R08C7.4* | WBGene00019948 | protein_coding_gene | 2.6 | 6.039 | 3.8E-02 |
| *Y43F8B.23* | WBGene00194815 | protein_coding_gene | 2.6 | 5.995 | 4.9E-05 |
| *F35E12.13* | WBGene00235181 | ncRNA_gene | 2.6 | 5.988 | 2.7E-02 |
| *F31E3.12* | WBGene00219317 | protein_coding_gene | 2.6 | 5.973 | 2.3E-04 |
| *Y43F8B.14* | WBGene00012824 | protein_coding_gene | 2.6 | 5.957 | 3.8E-07 |
| *F10E9.12* | WBGene00017361 | protein_coding_gene | 2.6 | 5.893 | 4.8E-17 |
| *ces-1* | WBGene00000468 | protein_coding_gene | 2.6 | 5.878 | 9.6E-04 |
| *F13A7.11* | WBGene00008726 | protein_coding_gene | 2.5 | 5.852 | 6.6E-12 |
| *linc-47* | WBGene00219641 | lincRNA_gene | 2.5 | 5.826 | 1.6E-02 |
| *grl-32* | WBGene00044279 | pseudogene | 2.5 | 5.814 | 4.6E-02 |
| *pals-37* | WBGene00138721 | protein_coding_gene | 2.5 | 5.809 | 2.9E-03 |
| *F07E5.7* | WBGene00017212 | protein_coding_gene | 2.5 | 5.746 | 4.0E-02 |
| *pals-39* | WBGene00008301 | protein_coding_gene | 2.5 | 5.744 | 3.5E-04 |
| *hlh-32* | WBGene00013665 | protein_coding_gene | 2.5 | 5.723 | 3.8E-02 |
| *Y69A2AL.2* | WBGene00044468 | protein_coding_gene | 2.5 | 5.600 | 4.9E-12 |
| *Y41D4B.26* | WBGene00021525 | protein_coding_gene | 2.5 | 5.580 | 4.1E-06 |
| *sra-32* | WBGene00005058 | protein_coding_gene | 2.5 | 5.548 | 4.5E-02 |
| *F59A7.7* | WBGene00019094 | protein_coding_gene | 2.5 | 5.531 | 5.2E-05 |
| *Y82E9BR.6* | WBGene00022339 | protein_coding_gene | 2.5 | 5.486 | 1.6E-02 |
| *C30H6.12* | WBGene00044238 | protein_coding_gene | 2.5 | 5.467 | 3.0E-02 |
| *C36B1.13* | WBGene00044120 | protein_coding_gene | 2.4 | 5.438 | 3.5E-03 |
| *F46F11.12* | WBGene00045082 | snoRNA_gene | 2.4 | 5.433 | 2.8E-02 |
| *wrt-2* | WBGene00006948 | protein_coding_gene | 2.4 | 5.429 | 2.4E-05 |
| *F27E5.9* | WBGene00206354 | protein_coding_gene | 2.4 | 5.411 | 1.5E-07 |
| *Y55F3AM.21* | WBGene00219399 | protein_coding_gene | 2.4 | 5.397 | 8.4E-12 |
| *bath-11* | WBGene00018664 | pseudogene | 2.4 | 5.374 | 2.3E-18 |
| *F55C12.19* | WBGene00271817 | protein_coding_gene | 2.4 | 5.368 | 6.2E-06 |
| *Y110A2AL.4* | WBGene00022441 | protein_coding_gene | 2.4 | 5.365 | 1.7E-06 |
| *dao-2* | WBGene00000928 | protein_coding_gene | 2.4 | 5.364 | 5.7E-12 |
| *T05H10.3* | WBGene00011508 | protein_coding_gene | 2.4 | 5.352 | 1.3E-04 |
| *Y53F4B.45* | WBGene00044728 | ncRNA_gene | 2.4 | 5.329 | 5.9E-07 |
| *F19B10.13* | WBGene00194713 | protein_coding_gene | 2.4 | 5.319 | 7.3E-04 |
| *R12C12.10* | WBGene00044395 | protein_coding_gene | 2.4 | 5.294 | 5.6E-03 |
| *T19D12.3* | WBGene00020578 | protein_coding_gene | 2.4 | 5.292 | 1.9E-02 |
| *fbxc-20* | WBGene00018352 | protein_coding_gene | 2.4 | 5.275 | 4.5E-06 |
| *F14D2.14* | WBGene00017462 | protein_coding_gene | 2.4 | 5.242 | 4.1E-19 |
| *C30G12.11* | WBGene00284861 | ncRNA_gene | 2.4 | 5.215 | 1.6E-02 |
| *C08B6.4* | WBGene00007425 | protein_coding_gene | 2.4 | 5.214 | 9.7E-05 |
| *his-1* | WBGene00001875 | protein_coding_gene | 2.4 | 5.212 | 3.4E-02 |
| *F53A9.7* | WBGene00018730 | protein_coding_gene | 2.4 | 5.208 | 2.4E-09 |
| *ZK686.6* | WBGene00044799 | protein_coding_gene | 2.4 | 5.196 | 2.3E-03 |
| *fbxa-12* | WBGene00016525 | protein_coding_gene | 2.4 | 5.181 | 2.5E-02 |
| *H39E20.1* | WBGene00019267 | protein_coding_gene | 2.3 | 5.095 | 1.2E-04 |
| *ZC262.4* | WBGene00022581 | protein_coding_gene | 2.3 | 5.094 | 8.7E-03 |
| *C33D9.10* | WBGene00044239 | protein_coding_gene | 2.3 | 5.068 | 3.5E-02 |
| *Y5H2B.1* | WBGene00021164 | protein_coding_gene | 2.3 | 4.966 | 1.4E-02 |
| *fbxa-222* | WBGene00045295 | protein_coding_gene | 2.3 | 4.966 | 4.7E-02 |
| *fbxa-137* | WBGene00020853 | protein_coding_gene | 2.3 | 4.966 | 2.5E-02 |

| | | | | | |
|---|---|---|---|---|---|
| *Y54G2A.36* | WBGene00021898 | protein_coding_gene | 2.3 | 4.956 | 9.3E-08 |
| *R05H11.2* | WBGene00019908 | protein_coding_gene | 2.3 | 4.944 | 6.9E-03 |
| *cut-3* | WBGene00009041 | protein_coding_gene | 2.3 | 4.940 | 1.8E-02 |
| *col-74* | WBGene00000650 | protein_coding_gene | 2.3 | 4.936 | 1.4E-02 |
| *dsl-3* | WBGene00001105 | protein_coding_gene | 2.3 | 4.896 | 9.3E-08 |
| *oac-47* | WBGene00011933 | pseudogene | 2.3 | 4.829 | 3.8E-02 |
| *R12E2.8* | WBGene00020034 | protein_coding_gene | 2.3 | 4.826 | 3.2E-05 |
| *Y69A2AR.25* | WBGene00022096 | protein_coding_gene | 2.3 | 4.822 | 7.0E-04 |
| *M162.9* | WBGene00014811 | pseudogene | 2.3 | 4.806 | 3.2E-24 |
| *F47B7.6* | WBGene00018537 | pseudogene | 2.3 | 4.793 | 4.5E-02 |
| *fbxa-217* | WBGene00012954 | pseudogene | 2.3 | 4.778 | 3.2E-24 |
| *C46A5.4* | WBGene00016700 | protein_coding_gene | 2.3 | 4.777 | 1.1E-04 |
| *Y71G12B.42* | WBGene00220198 | ncRNA_gene | 2.2 | 4.738 | 5.2E-03 |
| *F08F1.9* | WBGene00017259 | protein_coding_gene | 2.2 | 4.699 | 6.4E-03 |
| *Y116A8C.25* | WBGene00013802 | protein_coding_gene | 2.2 | 4.692 | 4.1E-08 |
| *Y18D10A.30* | WBGene00271598 | pseudogene | 2.2 | 4.676 | 5.4E-06 |
| *C07E3.8* | WBGene00007418 | protein_coding_gene | 2.2 | 4.675 | 3.6E-02 |
| *C08E8.4* | WBGene00007440 | protein_coding_gene | 2.2 | 4.663 | 3.5E-03 |
| *Y54F10BM.12* | WBGene00021866 | protein_coding_gene | 2.2 | 4.661 | 9.8E-08 |
| *Y48G9A.7* | WBGene00021701 | protein_coding_gene | 2.2 | 4.644 | 3.6E-06 |
| *W06G6.20* | WBGene00219308 | protein_coding_gene | 2.2 | 4.627 | 2.8E-02 |
| *C24H10.3* | WBGene00016069 | protein_coding_gene | 2.2 | 4.624 | 5.6E-04 |
| *C50B8.4* | WBGene00008226 | protein_coding_gene | 2.2 | 4.620 | 2.0E-08 |
| *fbxa-30* | WBGene00013837 | protein_coding_gene | 2.2 | 4.602 | 7.4E-03 |
| *F57A10.2* | WBGene00010184 | protein_coding_gene | 2.2 | 4.601 | 2.4E-03 |
| *mboa-5* | WBGene00013995 | pseudogene | 2.2 | 4.599 | 2.3E-02 |
| *K09H11.16* | WBGene00269384 | ncRNA_gene | 2.2 | 4.588 | 4.6E-05 |
| *F35E12.9* | WBGene00009433 | protein_coding_gene | 2.2 | 4.583 | 3.0E-15 |
| *lipl-1* | WBGene00010062 | protein_coding_gene | 2.2 | 4.582 | 2.6E-03 |
| *his-59* | WBGene00001933 | protein_coding_gene | 2.2 | 4.578 | 3.7E-08 |
| *djr-1.2* | WBGene00016789 | protein_coding_gene | 2.2 | 4.572 | 9.0E-03 |
| *F14D2.11* | WBGene00017459 | protein_coding_gene | 2.2 | 4.538 | 9.3E-15 |
| *C16C4.18* | WBGene00194998 | protein_coding_gene | 2.2 | 4.518 | 6.3E-12 |
| *egl-1* | WBGene00001170 | protein_coding_gene | 2.2 | 4.508 | 4.4E-06 |
| *Y23H5B.8* | WBGene00021279 | protein_coding_gene | 2.2 | 4.501 | 1.5E-06 |
| *col-64* | WBGene00000640 | protein_coding_gene | 2.2 | 4.479 | 2.1E-02 |
| *T21D12.7* | WBGene00020648 | protein_coding_gene | 2.2 | 4.448 | 2.2E-02 |
| *C54D10.13* | WBGene00045400 | protein_coding_gene | 2.2 | 4.448 | 1.9E-06 |
| *C01G6.10* | WBGene00014663 | pseudogene | 2.2 | 4.446 | 5.2E-05 |
| *msp-113* | WBGene00003468 | protein_coding_gene | 2.2 | 4.444 | 3.7E-02 |
| *ZC190.10* | WBGene00022544 | protein_coding_gene | 2.1 | 4.427 | 3.8E-03 |
| *hlh-16* | WBGene00001960 | protein_coding_gene | 2.1 | 4.426 | 8.8E-04 |
| *fbxa-136* | WBGene00007672 | protein_coding_gene | 2.1 | 4.422 | 2.8E-02 |
| *fbxa-164* | WBGene00015599 | protein_coding_gene | 2.1 | 4.413 | 3.2E-02 |
| *skr-11* | WBGene00004817 | pseudogene | 2.1 | 4.409 | 1.7E-21 |
| *F58H1.2* | WBGene00010285 | protein_coding_gene | 2.1 | 4.401 | 6.8E-04 |
| *T10D4.15* | WBGene00077522 | protein_coding_gene | 2.1 | 4.396 | 1.7E-02 |
| *K11H12.11* | WBGene00044723 | protein_coding_gene | 2.1 | 4.386 | 8.5E-03 |
| *bath-24* | WBGene00015014 | protein_coding_gene | 2.1 | 4.384 | 3.2E-11 |
| *skr-5* | WBGene00004811 | protein_coding_gene | 2.1 | 4.380 | 1.5E-04 |
| *C07E3.3* | WBGene00007414 | protein_coding_gene | 2.1 | 4.364 | 6.0E-07 |
| *F13H8.1* | WBGene00017434 | protein_coding_gene | 2.1 | 4.354 | 1.1E-06 |
| *Y23H5B.12* | WBGene00194701 | protein_coding_gene | 2.1 | 4.352 | 3.7E-02 |
| *F49H12.7* | WBGene00044753 | protein_coding_gene | 2.1 | 4.349 | 5.3E-03 |
| *ttr-29* | WBGene00011286 | protein_coding_gene | 2.1 | 4.345 | 2.4E-06 |
| *M04C3.5* | WBGene00206418 | protein_coding_gene | 2.1 | 4.342 | 2.4E-08 |
| *ZK993.2* | WBGene00022838 | protein_coding_gene | 2.1 | 4.334 | 3.3E-02 |
| *fbxa-48* | WBGene00021862 | protein_coding_gene | 2.1 | 4.327 | 4.3E-03 |
| *Y6D1A.2* | WBGene00012396 | protein_coding_gene | 2.1 | 4.313 | 1.8E-06 |
| *linc-60* | WBGene00219618 | lincRNA_gene | 2.1 | 4.306 | 4.6E-02 |
| *clec-140* | WBGene00020220 | protein_coding_gene | 2.1 | 4.297 | 4.1E-02 |
| *bath-18* | WBGene00018224 | pseudogene | 2.1 | 4.297 | 1.9E-03 |
| *Y111B2A.21* | WBGene00013741 | protein_coding_gene | 2.1 | 4.272 | 6.4E-11 |
| *MTCE.33* | WBGene00014472 | rRNA_gene | 2.1 | 4.271 | 3.4E-17 |
| *F28E10.5* | WBGene00044302 | pseudogene | 2.1 | 4.264 | 3.1E-06 |
| *ZK666.11* | WBGene00014049 | protein_coding_gene | 2.1 | 4.263 | 2.8E-02 |
| *fbxa-144* | WBGene00009709 | protein_coding_gene | 2.1 | 4.219 | 2.8E-04 |
| *tag-234* | WBGene00018874 | protein_coding_gene | 2.1 | 4.194 | 2.8E-11 |
| *Y73F8A.37* | WBGene00044632 | pseudogene | 2.1 | 4.193 | 9.2E-04 |
| *K10D11.2* | WBGene00010746 | protein_coding_gene | 2.1 | 4.166 | 3.6E-02 |

| | | | | | |
|---|---|---|---|---|---|
| *F18C5.5* | WBGene00017560 | protein_coding_gene | 2.1 | 4.163 | 1.0E-03 |
| *T28H10.1* | WBGene00012142 | protein_coding_gene | 2.1 | 4.157 | 1.3E-19 |
| *C42D4.19* | WBGene00235295 | protein_coding_gene | 2.1 | 4.154 | 4.6E-02 |
| *C14A4.9* | WBGene00007560 | protein_coding_gene | 2.0 | 4.139 | 1.0E-03 |
| *F11E6.9* | WBGene00008712 | protein_coding_gene | 2.0 | 4.125 | 7.6E-03 |
| *C47G2.8* | WBGene00044161 | ncRNA_gene | 2.0 | 4.109 | 1.4E-02 |
| *fbxa-212* | WBGene00012568 | pseudogene | 2.0 | 4.106 | 1.1E-16 |
| *fbxa-21* | WBGene00022487 | protein_coding_gene | 2.0 | 4.083 | 3.9E-05 |
| *F52C6.4* | WBGene00018661 | protein_coding_gene | 2.0 | 4.077 | 2.6E-14 |
| *Y69A2AR.12* | WBGene00022083 | protein_coding_gene | 2.0 | 4.069 | 4.7E-06 |
| *gpx-4* | WBGene00022377 | protein_coding_gene | 2.0 | 4.068 | 6.0E-08 |
| *C14F11.6* | WBGene00015781 | protein_coding_gene | 2.0 | 4.054 | 1.5E-04 |
| *C52E2.4* | WBGene00016883 | protein_coding_gene | 2.0 | 4.027 | 8.9E-05 |
| *F13A7.12* | WBGene00008727 | protein_coding_gene | 2.0 | 4.023 | 7.6E-03 |
| *W08G11.6* | WBGene00023429 | snoRNA_gene | 2.0 | 4.009 | 2.9E-02 |
| *ins-11* | WBGene00002094 | protein_coding_gene | 2.0 | 3.970 | 3.0E-02 |
| *Y52B11A.11* | WBGene00014914 | protein_coding_gene | 2.0 | 3.955 | 6.0E-03 |
| *C35D6.5* | WBGene00007962 | pseudogene | 2.0 | 3.954 | 1.8E-02 |
| *T03G6.1* | WBGene00020194 | protein_coding_gene | 2.0 | 3.948 | 4.1E-04 |
| *Y61A9LA.4* | WBGene00022016 | protein_coding_gene | 2.0 | 3.944 | 5.0E-02 |
| *F30H5.4* | WBGene00023020 | snoRNA_gene | 2.0 | 3.932 | 3.3E-02 |
| *csp-1* | WBGene00000819 | protein_coding_gene | 2.0 | 3.916 | 1.3E-05 |
| *fbxa-140* | WBGene00008130 | protein_coding_gene | 2.0 | 3.912 | 5.9E-14 |
| *R09A8.9* | WBGene00219278 | protein_coding_gene | 2.0 | 3.911 | 1.4E-04 |
| *K08E7.10* | WBGene00044907 | pseudogene | 2.0 | 3.906 | 5.7E-03 |
| *F09F9.2* | WBGene00017307 | protein_coding_gene | 2.0 | 3.885 | 9.3E-04 |
| *R09B3.6* | WBGene00014820 | pseudogene | 2.0 | 3.882 | 9.6E-15 |
| *Y43B11AR.3* | WBGene00021544 | protein_coding_gene | 2.0 | 3.868 | 1.8E-12 |
| *F52C6.3* | WBGene00018660 | protein_coding_gene | 1.9 | 3.792 | 2.2E-14 |
| *bath-1* | WBGene00019139 | protein_coding_gene | 1.9 | 3.784 | 4.6E-12 |
| *Y37H2A.18* | WBGene00219210 | protein_coding_gene | 1.9 | 3.773 | 1.4E-13 |
| *C17E4.11* | WBGene00044236 | protein_coding_gene | 1.9 | 3.773 | 3.9E-20 |
| *Y50D4A.5* | WBGene00021739 | protein_coding_gene | 1.9 | 3.771 | 1.7E-14 |
| *Y17D7C.4* | WBGene00050903 | pseudogene | 1.9 | 3.763 | 2.0E-03 |
| *EEED8.13* | WBGene00017141 | protein_coding_gene | 1.9 | 3.754 | 5.2E-12 |
| *F23H12.10* | WBGene00044218 | protein_coding_gene | 1.9 | 3.751 | 2.5E-09 |
| *anr-8* | WBGene00219575 | antisense_lncRNA_gene | 1.9 | 3.735 | 1.0E-15 |
| *C03F11.2* | WBGene00015388 | protein_coding_gene | 1.9 | 3.731 | 2.3E-03 |
| *his-62* | WBGene00001936 | protein_coding_gene | 1.9 | 3.731 | 1.4E-03 |
| *T26F2.2* | WBGene00012058 | pseudogene | 1.9 | 3.720 | 1.9E-09 |
| *W08E12.6* | WBGene00021087 | protein_coding_gene | 1.9 | 3.711 | 5.0E-02 |
| *F14D2.8* | WBGene00017456 | protein_coding_gene | 1.9 | 3.704 | 1.2E-17 |
| *F53A9.6* | WBGene00018729 | protein_coding_gene | 1.9 | 3.689 | 1.0E-05 |
| *Y40H7A.7* | WBGene00012745 | pseudogene | 1.9 | 3.686 | 6.0E-03 |
| *Y71A12B.15* | WBGene00013511 | protein_coding_gene | 1.9 | 3.677 | 2.8E-02 |
| *F37C12.21* | WBGene00194742 | protein_coding_gene | 1.9 | 3.667 | 2.3E-03 |
| *C07H6.9* | WBGene00015582 | protein_coding_gene | 1.9 | 3.640 | 2.7E-04 |
| *spp-13* | WBGene00004998 | protein_coding_gene | 1.9 | 3.622 | 7.5E-04 |
| *C52E12.11* | WBGene00304189 | pseudogene | 1.9 | 3.613 | 6.4E-08 |
| *K08B12.1* | WBGene00019520 | protein_coding_gene | 1.9 | 3.608 | 1.5E-03 |
| *F36H1.11* | WBGene00009496 | protein_coding_gene | 1.8 | 3.593 | 4.0E-02 |
| *ZK678.3* | WBGene00014069 | pseudogene | 1.8 | 3.583 | 4.4E-02 |
| *pals-23* | WBGene00016218 | protein_coding_gene | 1.8 | 3.583 | 4.5E-06 |
| *Y53F4B.38* | WBGene00014920 | pseudogene | 1.8 | 3.581 | 1.4E-03 |
| *btb-3* | WBGene00015013 | pseudogene | 1.8 | 3.568 | 3.6E-10 |
| *C53A5.2* | WBGene00008263 | protein_coding_gene | 1.8 | 3.541 | 3.4E-03 |
| *srpr-2.1* | WBGene00195016 | ncRNA_gene | 1.8 | 3.531 | 4.0E-02 |
| *W01A11.7* | WBGene00020912 | protein_coding_gene | 1.8 | 3.521 | 1.5E-05 |
| *F54C4.4* | WBGene00018795 | protein_coding_gene | 1.8 | 3.519 | 2.2E-19 |
| *K08F9.3* | WBGene00010686 | protein_coding_gene | 1.8 | 3.503 | 3.1E-03 |
| *C33H5.13* | WBGene00016379 | protein_coding_gene | 1.8 | 3.500 | 2.9E-15 |
| *fbxa-211* | WBGene00012566 | protein_coding_gene | 1.8 | 3.497 | 2.2E-19 |
| *fbxa-207* | WBGene00013612 | pseudogene | 1.8 | 3.495 | 7.1E-12 |
| *lpr-4* | WBGene00012257 | protein_coding_gene | 1.8 | 3.491 | 2.2E-04 |
| *ins-2* | WBGene00002085 | protein_coding_gene | 1.8 | 3.483 | 1.1E-02 |
| *F37A4.3* | WBGene00018133 | protein_coding_gene | 1.8 | 3.472 | 2.9E-03 |
| *D2023.8* | WBGene00050970 | pseudogene | 1.8 | 3.470 | 4.9E-03 |
| *E03H4.8* | WBGene00008476 | protein_coding_gene | 1.8 | 3.465 | 8.4E-05 |
| *W03D2.9* | WBGene00020983 | protein_coding_gene | 1.8 | 3.464 | 3.4E-02 |
| *fbxa-3* | WBGene00020363 | protein_coding_gene | 1.8 | 3.454 | 5.1E-05 |

| | | | | | |
|---|---|---|---|---|---|
| *fbxa-193* | WBGene00010295 | pseudogene | 1.8 | 3.450 | 9.2E-17 |
| *Y81G3A.6* | WBGene00195150 | protein_coding_gene | 1.8 | 3.434 | 2.5E-02 |
| *EEED8.2* | WBGene00017133 | protein_coding_gene | 1.8 | 3.423 | 5.3E-08 |
| *ZK250.2* | WBGene00022685 | protein_coding_gene | 1.8 | 3.418 | 3.4E-03 |
| *bath-14* | WBGene00015016 | pseudogene | 1.8 | 3.411 | 1.8E-07 |
| *srr-2* | WBGene00005653 | protein_coding_gene | 1.8 | 3.411 | 2.4E-02 |
| *Y51H7C.12* | WBGene00021790 | protein_coding_gene | 1.8 | 3.405 | 1.1E-02 |
| *dmsr-3* | WBGene00012993 | protein_coding_gene | 1.8 | 3.403 | 5.8E-05 |
| *K02E7.5* | WBGene00019310 | pseudogene | 1.8 | 3.401 | 3.1E-03 |
| *bath-22* | WBGene00018658 | pseudogene | 1.8 | 3.398 | 3.3E-13 |
| *nhr-240* | WBGene00013363 | pseudogene | 1.8 | 3.397 | 4.7E-02 |
| *R13A1.10* | WBGene00044504 | protein_coding_gene | 1.8 | 3.370 | 1.2E-05 |
| *gst-22* | WBGene00001770 | protein_coding_gene | 1.8 | 3.369 | 4.0E-11 |
| *F59B2.8* | WBGene00010310 | protein_coding_gene | 1.8 | 3.367 | 9.6E-04 |
| *C06B3.6* | WBGene00007365 | protein_coding_gene | 1.8 | 3.365 | 6.8E-05 |
| *T24H10.5* | WBGene00012003 | protein_coding_gene | 1.8 | 3.364 | 2.6E-04 |
| *F54F11.1* | WBGene00010069 | protein_coding_gene | 1.7 | 3.360 | 4.8E-02 |
| *ces-2* | WBGene00000469 | protein_coding_gene | 1.7 | 3.352 | 2.1E-04 |
| *fbxa-59* | WBGene00020455 | protein_coding_gene | 1.7 | 3.350 | 4.8E-08 |
| *C24F3.2* | WBGene00007697 | protein_coding_gene | 1.7 | 3.346 | 3.0E-34 |
| *fbxa-157* | WBGene00007393 | protein_coding_gene | 1.7 | 3.344 | 2.3E-02 |
| *nspd-9* | WBGene00017808 | protein_coding_gene | 1.7 | 3.327 | 1.1E-04 |
| *C08F11.13* | WBGene00007460 | protein_coding_gene | 1.7 | 3.322 | 1.5E-03 |
| *F13H10.1* | WBGene00008773 | protein_coding_gene | 1.7 | 3.307 | 3.9E-02 |
| *bath-45* | WBGene00009247 | protein_coding_gene | 1.7 | 3.304 | 3.2E-05 |
| *pstk-1* | WBGene00013041 | protein_coding_gene | 1.7 | 3.294 | 2.9E-19 |
| *ift-43* | WBGene00016121 | protein_coding_gene | 1.7 | 3.281 | 3.5E-11 |
| *K08F9.1* | WBGene00010684 | protein_coding_gene | 1.7 | 3.277 | 8.3E-04 |
| *lea-1* | WBGene00002263 | protein_coding_gene | 1.7 | 3.272 | 3.2E-11 |
| *F46F3.3* | WBGene00009791 | protein_coding_gene | 1.7 | 3.272 | 2.4E-02 |
| *his-64* | WBGene00001938 | protein_coding_gene | 1.7 | 3.269 | 5.9E-05 |
| *F55G1.7* | WBGene00018903 | protein_coding_gene | 1.7 | 3.261 | 1.9E-10 |
| *C32H11.7* | WBGene00007870 | pseudogene | 1.7 | 3.258 | 2.4E-04 |
| *Y37H9A.5* | WBGene00014871 | protein_coding_gene | 1.7 | 3.257 | 6.4E-05 |
| *T09F3.4* | WBGene00011663 | protein_coding_gene | 1.7 | 3.249 | 8.7E-25 |
| *scbp-2* | WBGene00010569 | protein_coding_gene | 1.7 | 3.243 | 1.7E-16 |
| *T21G5.2* | WBGene00020660 | protein_coding_gene | 1.7 | 3.231 | 1.7E-02 |
| *F12E12.3* | WBGene00017404 | pseudogene | 1.7 | 3.230 | 9.3E-03 |
| *F25H2.12* | WBGene00009123 | protein_coding_gene | 1.7 | 3.229 | 1.4E-17 |
| *math-34* | WBGene00019699 | protein_coding_gene | 1.7 | 3.226 | 2.2E-03 |
| *bath-21* | WBGene00019138 | protein_coding_gene | 1.7 | 3.217 | 3.8E-11 |
| *bath-27* | WBGene00017450 | protein_coding_gene | 1.7 | 3.211 | 1.4E-10 |
| *spp-6* | WBGene00004991 | protein_coding_gene | 1.7 | 3.195 | 3.3E-02 |
| *oac-31* | WBGene00018707 | protein_coding_gene | 1.7 | 3.194 | 1.9E-03 |
| *pals-24* | WBGene00016219 | protein_coding_gene | 1.7 | 3.173 | 8.5E-06 |
| *ZK856.4* | WBGene00014105 | protein_coding_gene | 1.7 | 3.173 | 9.3E-03 |
| *ceh-16* | WBGene00000439 | protein_coding_gene | 1.7 | 3.169 | 3.3E-03 |
| *R06C7.6* | WBGene00011065 | protein_coding_gene | 1.7 | 3.161 | 8.2E-14 |
| *F53B2.8* | WBGene00009957 | protein_coding_gene | 1.7 | 3.160 | 3.4E-02 |
| *spp-2* | WBGene00004987 | protein_coding_gene | 1.7 | 3.154 | 1.0E-04 |
| *F13G3.6* | WBGene00008766 | protein_coding_gene | 1.7 | 3.147 | 1.3E-26 |
| *fbxa-117* | WBGene00014907 | pseudogene | 1.7 | 3.147 | 6.2E-07 |
| *btb-16* | WBGene00018195 | protein_coding_gene | 1.7 | 3.143 | 4.0E-08 |
| *T16G12.12* | WBGene00269433 | protein_coding_gene | 1.6 | 3.133 | 3.5E-04 |
| *ZC449.1* | WBGene00022611 | protein_coding_gene | 1.6 | 3.129 | 1.8E-02 |
| *fbxa-108* | WBGene00010293 | protein_coding_gene | 1.6 | 3.123 | 6.3E-16 |
| *ZK938.11* | WBGene00270290 | protein_coding_gene | 1.6 | 3.120 | 8.5E-04 |
| *mtcu-1* | WBGene00009557 | protein_coding_gene | 1.6 | 3.118 | 4.0E-30 |
| *Y71F9AL.8* | WBGene00022113 | protein_coding_gene | 1.6 | 3.108 | 1.3E-22 |
| *nmad-1* | WBGene00017304 | protein_coding_gene | 1.6 | 3.108 | 7.6E-13 |
| *his-48* | WBGene00001922 | protein_coding_gene | 1.6 | 3.108 | 5.9E-03 |
| *T04D3.8* | WBGene00011435 | protein_coding_gene | 1.6 | 3.100 | 8.1E-13 |
| *F53A9.1* | WBGene00018724 | protein_coding_gene | 1.6 | 3.098 | 1.4E-12 |
| *gbh-2* | WBGene00001523 | protein_coding_gene | 1.6 | 3.083 | 8.6E-05 |
| *K07A1.15* | WBGene00010619 | protein_coding_gene | 1.6 | 3.077 | 7.0E-07 |
| *spch-2* | WBGene00015689 | protein_coding_gene | 1.6 | 3.075 | 1.9E-02 |
| *F08B4.3* | WBGene00017235 | pseudogene | 1.6 | 3.072 | 2.2E-17 |
| *Y102A5C.2* | WBGene00013611 | protein_coding_gene | 1.6 | 3.069 | 5.0E-26 |
| *T11F9.10* | WBGene00011712 | protein_coding_gene | 1.6 | 3.064 | 3.7E-02 |
| *F40E3.7* | WBGene00077781 | pseudogene | 1.6 | 3.062 | 1.1E-02 |

| | | | | | |
|---|---|---|---|---|---|
| *twk-32* | WBGene00006684 | protein_coding_gene | 1.6 | 3.052 | 1.4E-03 |
| *faah-5* | WBGene00013226 | protein_coding_gene | 1.6 | 3.050 | 4.5E-02 |
| *F27D4.8* | WBGene00077543 | protein_coding_gene | 1.6 | 3.049 | 6.7E-13 |
| *atf-8* | WBGene00017535 | protein_coding_gene | 1.6 | 3.038 | 5.4E-03 |
| *F35G12.5* | WBGene00009442 | protein_coding_gene | 1.6 | 3.033 | 5.9E-16 |
| *fbxa-129* | WBGene00010294 | pseudogene | 1.6 | 3.022 | 3.7E-16 |
| *R10D12.18* | WBGene00194678 | ncRNA_gene | 1.6 | 3.019 | 2.8E-02 |
| *C08F11.6* | WBGene00014672 | pseudogene | 1.6 | 3.014 | 6.1E-11 |
| *fbxa-200* | WBGene00020304 | pseudogene | 1.6 | 3.001 | 2.0E-10 |
| *his-61* | WBGene00001935 | protein_coding_gene | 1.6 | 3.000 | 1.0E-06 |
| *C09B9.85* | WBGene00220250 | protein_coding_gene | 1.6 | 2.987 | 3.1E-02 |
| *math-21* | WBGene00016556 | protein_coding_gene | 1.6 | 2.986 | 4.0E-02 |
| *B0281.4* | WBGene00015111 | protein_coding_gene | 1.6 | 2.985 | 3.5E-04 |
| *fbxa-158* | WBGene00015602 | protein_coding_gene | 1.6 | 2.981 | 3.6E-02 |
| *C40A11.3* | WBGene00016546 | protein_coding_gene | 1.6 | 2.978 | 1.2E-03 |
| *W02D7.5* | WBGene00020941 | protein_coding_gene | 1.6 | 2.965 | 1.7E-14 |
| *C39B5.14* | WBGene00189950 | protein_coding_gene | 1.6 | 2.964 | 3.4E-02 |
| *Y75B12B.3* | WBGene00013568 | protein_coding_gene | 1.6 | 2.963 | 5.6E-06 |
| *JC8.4* | WBGene00010436 | protein_coding_gene | 1.6 | 2.961 | 7.2E-13 |
| *ZK1248.19* | WBGene00022885 | protein_coding_gene | 1.6 | 2.960 | 1.1E-18 |
| *K11D12.12* | WBGene00019651 | protein_coding_gene | 1.6 | 2.956 | 5.3E-14 |
| *T10C6.15* | WBGene00050899 | protein_coding_gene | 1.6 | 2.951 | 1.1E-02 |
| *fbxa-179* | WBGene00045292 | protein_coding_gene | 1.6 | 2.949 | 1.3E-03 |
| *Y7A5A.25* | WBGene00304824 | protein_coding_gene | 1.6 | 2.947 | 3.0E-02 |
| *ttr-52* | WBGene00008719 | protein_coding_gene | 1.6 | 2.942 | 1.7E-02 |
| *F29G9.7* | WBGene00017933 | pseudogene | 1.6 | 2.940 | 5.5E-14 |
| *F22D6.8* | WBGene00043279 | protein_coding_gene | 1.6 | 2.940 | 3.7E-11 |
| *D2030.11* | WBGene00008420 | protein_coding_gene | 1.6 | 2.938 | 3.4E-11 |
| *F31E3.6* | WBGene00017952 | protein_coding_gene | 1.6 | 2.929 | 6.9E-07 |
| *C25G4.2* | WBGene00007730 | protein_coding_gene | 1.5 | 2.920 | 7.8E-11 |
| *W03B1.3* | WBGene00020973 | protein_coding_gene | 1.5 | 2.914 | 1.7E-05 |
| *ZK1127.12* | WBGene00022857 | protein_coding_gene | 1.5 | 2.907 | 2.0E-13 |
| *Y42A5A.5* | WBGene00012780 | protein_coding_gene | 1.5 | 2.902 | 8.6E-15 |
| *W05G11.2* | WBGene00021040 | protein_coding_gene | 1.5 | 2.901 | 1.0E-05 |
| *Y55F3BL.4* | WBGene00185088 | protein_coding_gene | 1.5 | 2.889 | 1.1E-09 |
| *C30G12.4* | WBGene00016276 | protein_coding_gene | 1.5 | 2.882 | 4.9E-02 |
| *F11A5.18* | WBGene00045511 | protein_coding_gene | 1.5 | 2.882 | 1.1E-02 |
| *B0432.10* | WBGene00015191 | protein_coding_gene | 1.5 | 2.878 | 3.4E-19 |
| *F47D12.6* | WBGene00018562 | protein_coding_gene | 1.5 | 2.871 | 7.1E-05 |
| *C35D10.3* | WBGene00016441 | protein_coding_gene | 1.5 | 2.868 | 8.7E-03 |
| *his-60* | WBGene00001934 | protein_coding_gene | 1.5 | 2.868 | 4.0E-08 |
| *fbxc-58* | WBGene00007506 | protein_coding_gene | 1.5 | 2.867 | 1.7E-02 |
| *Y48G8AL.15* | WBGene00021692 | protein_coding_gene | 1.5 | 2.865 | 2.4E-16 |
| *F13G3.3* | WBGene00008763 | protein_coding_gene | 1.5 | 2.861 | 1.1E-02 |
| *ttr-26* | WBGene00013079 | protein_coding_gene | 1.5 | 2.833 | 2.7E-04 |
| *fbxc-45* | WBGene00021887 | protein_coding_gene | 1.5 | 2.822 | 8.2E-15 |
| *tsen-34* | WBGene00019535 | protein_coding_gene | 1.5 | 2.820 | 6.3E-13 |
| *tsen-2* | WBGene00013231 | protein_coding_gene | 1.5 | 2.815 | 1.9E-24 |
| *Y52E8A.2* | WBGene00021795 | protein_coding_gene | 1.5 | 2.814 | 7.2E-11 |
| *efsc-1* | WBGene00008133 | protein_coding_gene | 1.5 | 2.810 | 5.7E-18 |
| *bath-19* | WBGene00019132 | protein_coding_gene | 1.5 | 2.808 | 1.2E-11 |
| *F56D2.3* | WBGene00018965 | protein_coding_gene | 1.5 | 2.808 | 4.6E-03 |
| *C10B5.1* | WBGene00015670 | protein_coding_gene | 1.5 | 2.807 | 3.8E-21 |
| *Y75B8A.18* | WBGene00013553 | protein_coding_gene | 1.5 | 2.804 | 7.4E-23 |
| *K06B9.2* | WBGene00019439 | pseudogene | 1.5 | 2.801 | 2.9E-03 |
| *Y105E8B.6* | WBGene00013691 | protein_coding_gene | 1.5 | 2.800 | 9.1E-13 |
| *F46F11.7* | WBGene00018510 | protein_coding_gene | 1.5 | 2.798 | 1.9E-05 |
| *C46E10.9* | WBGene00016713 | pseudogene | 1.5 | 2.796 | 1.5E-07 |
| *B0035.18* | WBGene00044290 | protein_coding_gene | 1.5 | 2.796 | 2.4E-14 |
| *R05D3.3* | WBGene00019878 | protein_coding_gene | 1.5 | 2.782 | 5.9E-10 |
| *fbxb-102* | WBGene00008847 | protein_coding_gene | 1.5 | 2.781 | 3.4E-02 |
| *F42A9.8* | WBGene00018337 | protein_coding_gene | 1.5 | 2.779 | 4.9E-08 |
| *Y11D7A.8* | WBGene00012432 | protein_coding_gene | 1.5 | 2.774 | 3.2E-02 |
| *W07E6.5* | WBGene00021076 | protein_coding_gene | 1.5 | 2.771 | 5.9E-16 |
| *F22D6.9* | WBGene00009054 | protein_coding_gene | 1.5 | 2.770 | 5.5E-13 |
| *metl-5* | WBGene00008008 | protein_coding_gene | 1.5 | 2.763 | 1.4E-16 |
| *F59H6.3* | WBGene00019134 | pseudogene | 1.5 | 2.759 | 1.0E-02 |
| *T15B7.15* | WBGene00020527 | protein_coding_gene | 1.5 | 2.758 | 3.3E-02 |
| *osm-3* | WBGene00003884 | protein_coding_gene | 1.5 | 2.756 | 1.5E-08 |
| *T04F8.7* | WBGene00011445 | protein_coding_gene | 1.5 | 2.755 | 1.1E-04 |

| | | | | | |
|---|---|---|---|---|---|
| *T06D10.3* | WBGene00077558 | protein_coding_gene | 1.5 | 2.749 | 2.5E-02 |
| *fbxa-145* | WBGene00018595 | protein_coding_gene | 1.5 | 2.745 | 1.4E-07 |
| *T04H1.5* | WBGene00011451 | protein_coding_gene | 1.5 | 2.743 | 7.0E-18 |
| *F55B11.6* | WBGene00044249 | protein_coding_gene | 1.5 | 2.737 | 4.8E-08 |
| *Y11D7A.5* | WBGene00012429 | protein_coding_gene | 1.4 | 2.732 | 4.0E-02 |
| *rip-1* | WBGene00010973 | protein_coding_gene | 1.4 | 2.729 | 7.6E-05 |
| *Y57G7A.8* | WBGene00021971 | pseudogene | 1.4 | 2.727 | 9.7E-04 |
| *F52C6.12* | WBGene00018669 | protein_coding_gene | 1.4 | 2.726 | 3.7E-07 |
| *col-45* | WBGene00000622 | protein_coding_gene | 1.4 | 2.726 | 5.0E-02 |
| *dpy-10* | WBGene00001072 | protein_coding_gene | 1.4 | 2.723 | 7.8E-05 |
| *hus-1* | WBGene00002042 | protein_coding_gene | 1.4 | 2.722 | 1.9E-14 |
| *memb-2* | WBGene00019757 | protein_coding_gene | 1.4 | 2.721 | 9.5E-15 |
| *T23F11.6* | WBGene00011957 | protein_coding_gene | 1.4 | 2.720 | 1.6E-03 |
| *T16H12.1* | WBGene00011811 | protein_coding_gene | 1.4 | 2.719 | 5.5E-13 |
| *F58E1.13* | WBGene00019044 | protein_coding_gene | 1.4 | 2.713 | 1.4E-02 |
| *Y57G11C.8* | WBGene00013306 | protein_coding_gene | 1.4 | 2.712 | 7.4E-08 |
| *cdr-1* | WBGene00000412 | protein_coding_gene | 1.4 | 2.711 | 3.3E-02 |
| *sptf-2* | WBGene00011926 | protein_coding_gene | 1.4 | 2.706 | 2.5E-03 |
| *Y51H4A.1* | WBGene00013098 | protein_coding_gene | 1.4 | 2.705 | 2.4E-02 |
| *F08A8.5* | WBGene00008568 | protein_coding_gene | 1.4 | 2.697 | 2.1E-02 |
| *bbs-5* | WBGene00010974 | protein_coding_gene | 1.4 | 2.696 | 4.4E-04 |
| *btb-4* | WBGene00019142 | protein_coding_gene | 1.4 | 2.694 | 1.2E-09 |
| *W04A8.2* | WBGene00012237 | protein_coding_gene | 1.4 | 2.687 | 1.3E-13 |
| *C40A11.2* | WBGene00016545 | protein_coding_gene | 1.4 | 2.685 | 1.4E-03 |
| *C10A4.1* | WBGene00015662 | protein_coding_gene | 1.4 | 2.677 | 1.1E-02 |
| *rnh-1.2* | WBGene00014163 | protein_coding_gene | 1.4 | 2.674 | 1.0E-12 |
| *mdt-31* | WBGene00007026 | protein_coding_gene | 1.4 | 2.673 | 1.2E-10 |
| *F59B2.11* | WBGene00010313 | pseudogene | 1.4 | 2.673 | 2.7E-02 |
| *clec-66* | WBGene00009397 | protein_coding_gene | 1.4 | 2.671 | 4.9E-05 |
| *F38H4.6* | WBGene00009550 | protein_coding_gene | 1.4 | 2.667 | 3.2E-02 |
| *gst-30* | WBGene00001778 | protein_coding_gene | 1.4 | 2.666 | 1.5E-03 |
| *ZC204.12* | WBGene00022562 | protein_coding_gene | 1.4 | 2.660 | 7.6E-08 |
| *gmeb-3* | WBGene00008092 | protein_coding_gene | 1.4 | 2.658 | 1.4E-02 |
| *msp-65* | WBGene00003458 | protein_coding_gene | 1.4 | 2.651 | 1.4E-02 |
| *F52F12.9* | WBGene00009942 | protein_coding_gene | 1.4 | 2.650 | 3.9E-03 |
| *glod-4* | WBGene00006448 | protein_coding_gene | 1.4 | 2.647 | 1.4E-07 |
| *H20J04.9* | WBGene00044310 | protein_coding_gene | 1.4 | 2.647 | 5.0E-19 |
| *ftr-1* | WBGene00013756 | protein_coding_gene | 1.4 | 2.645 | 1.6E-10 |
| *lin-32* | WBGene00003018 | protein_coding_gene | 1.4 | 2.639 | 1.8E-02 |
| *Y55F3BR.10* | WBGene00044742 | protein_coding_gene | 1.4 | 2.635 | 1.3E-06 |
| *W09D6.1* | WBGene00012356 | protein_coding_gene | 1.4 | 2.631 | 1.6E-12 |
| *fbxa-87* | WBGene00044660 | pseudogene | 1.4 | 2.627 | 1.8E-02 |
| *F52C12.6* | WBGene00138717 | protein_coding_gene | 1.4 | 2.627 | 1.4E-12 |
| *Y61A9LA.12* | WBGene00044586 | protein_coding_gene | 1.4 | 2.626 | 2.9E-02 |
| *F14D2.5* | WBGene00017453 | protein_coding_gene | 1.4 | 2.622 | 3.5E-04 |
| *M18.3* | WBGene00010889 | protein_coding_gene | 1.4 | 2.617 | 2.0E-13 |
| *ZC404.16* | WBGene00195226 | protein_coding_gene | 1.4 | 2.616 | 1.7E-02 |
| *C16C8.11* | WBGene00015849 | protein_coding_gene | 1.4 | 2.615 | 3.6E-08 |
| *F45F2.9* | WBGene00018481 | protein_coding_gene | 1.4 | 2.615 | 3.0E-16 |
| *M04B2.2* | WBGene00010846 | protein_coding_gene | 1.4 | 2.615 | 7.3E-20 |
| *dhhc-13* | WBGene00019257 | protein_coding_gene | 1.4 | 2.610 | 9.8E-03 |
| *san-1* | WBGene00004721 | protein_coding_gene | 1.4 | 2.610 | 4.3E-15 |
| *tpk-1* | WBGene00014027 | protein_coding_gene | 1.4 | 2.610 | 1.3E-08 |
| *Y102A11A.5* | WBGene00022415 | protein_coding_gene | 1.4 | 2.606 | 4.9E-03 |
| *dohh-1* | WBGene00007555 | protein_coding_gene | 1.4 | 2.605 | 7.6E-11 |
| *fbxa-223* | WBGene00022630 | pseudogene | 1.4 | 2.603 | 4.8E-02 |
| *C06A5.12* | WBGene00044414 | protein_coding_gene | 1.4 | 2.600 | 1.1E-08 |
| *C27C7.1* | WBGene00007766 | protein_coding_gene | 1.4 | 2.599 | 1.1E-15 |
| *F41C3.7* | WBGene00018273 | protein_coding_gene | 1.4 | 2.598 | 2.1E-02 |
| *C34H4.5* | WBGene00016427 | protein_coding_gene | 1.4 | 2.593 | 2.5E-10 |
| *hyls-1* | WBGene00015466 | protein_coding_gene | 1.4 | 2.591 | 1.4E-13 |
| *josd-1* | WBGene00022190 | protein_coding_gene | 1.4 | 2.591 | 2.6E-12 |
| *fbxc-24* | WBGene00019927 | protein_coding_gene | 1.4 | 2.589 | 1.9E-02 |
| *F53A3.7* | WBGene00018723 | protein_coding_gene | 1.4 | 2.589 | 4.7E-10 |
| *icmt-1* | WBGene00017673 | protein_coding_gene | 1.4 | 2.581 | 1.7E-02 |
| *K08C9.8* | WBGene00045218 | pseudogene | 1.4 | 2.577 | 2.6E-03 |
| *C18A11.1* | WBGene00015947 | protein_coding_gene | 1.4 | 2.574 | 1.6E-03 |
| *ascc-1* | WBGene00016019 | protein_coding_gene | 1.4 | 2.570 | 9.3E-22 |
| *mks-6* | WBGene00010642 | protein_coding_gene | 1.4 | 2.567 | 4.2E-03 |
| *C09E7.5* | WBGene00015639 | protein_coding_gene | 1.4 | 2.564 | 4.9E-02 |

| | | | | | |
|---|---|---|---|---|---|
| *F32E10.5* | WBGene00017992 | protein_coding_gene | 1.4 | 2.563 | 2.0E-19 |
| *F59D6.1* | WBGene00019103 | protein_coding_gene | 1.4 | 2.560 | 4.1E-02 |
| *F40H6.6* | WBGene00044412 | pseudogene | 1.4 | 2.558 | 1.1E-08 |
| *nuaf-1* | WBGene00008225 | protein_coding_gene | 1.4 | 2.557 | 2.0E-15 |
| *F58F12.12* | WBGene00271812 | protein_coding_gene | 1.4 | 2.555 | 2.2E-02 |
| *metl-6* | WBGene00014205 | protein_coding_gene | 1.4 | 2.553 | 9.0E-12 |
| *F10G8.9* | WBGene00008667 | protein_coding_gene | 1.4 | 2.553 | 1.1E-15 |
| *Y38F2AL.12* | WBGene00219413 | protein_coding_gene | 1.4 | 2.550 | 6.1E-09 |
| *C30G12.1* | WBGene00016273 | protein_coding_gene | 1.3 | 2.547 | 9.2E-03 |
| *M03C11.6* | WBGene00010843 | protein_coding_gene | 1.3 | 2.545 | 1.2E-11 |
| *Y60A3A.8* | WBGene00013359 | protein_coding_gene | 1.3 | 2.545 | 2.4E-13 |
| *F08B6.1* | WBGene00017239 | protein_coding_gene | 1.3 | 2.544 | 1.5E-10 |
| *D1007.10* | WBGene00017007 | protein_coding_gene | 1.3 | 2.542 | 1.6E-08 |
| *evl-20* | WBGene00001358 | protein_coding_gene | 1.3 | 2.541 | 2.5E-17 |
| *EEED8.14* | WBGene00017142 | protein_coding_gene | 1.3 | 2.540 | 4.3E-20 |
| *trpp-6* | WBGene00010699 | protein_coding_gene | 1.3 | 2.535 | 1.5E-17 |
| *dod-18* | WBGene00008316 | protein_coding_gene | 1.3 | 2.535 | 9.5E-13 |
| *wrt-10* | WBGene00006956 | protein_coding_gene | 1.3 | 2.533 | 2.3E-04 |
| *fbxa-113* | WBGene00013757 | protein_coding_gene | 1.3 | 2.531 | 2.1E-02 |
| *ZK632.14* | WBGene00014020 | protein_coding_gene | 1.3 | 2.527 | 4.5E-22 |
| *T21C12.8* | WBGene00043998 | protein_coding_gene | 1.3 | 2.527 | 5.0E-05 |
| *Y47H10A.6* | WBGene00077537 | pseudogene | 1.3 | 2.523 | 1.4E-03 |
| *F46B6.4* | WBGene00009769 | protein_coding_gene | 1.3 | 2.521 | 1.5E-09 |
| *dyf-17* | WBGene00012673 | protein_coding_gene | 1.3 | 2.514 | 9.4E-07 |
| *elo-7* | WBGene00001245 | protein_coding_gene | 1.3 | 2.512 | 3.6E-02 |
| *BE10.1* | WBGene00007209 | protein_coding_gene | 1.3 | 2.509 | 1.2E-02 |
| *bath-20* | WBGene00015012 | protein_coding_gene | 1.3 | 2.508 | 1.2E-02 |
| *C16A3.2* | WBGene00015807 | protein_coding_gene | 1.3 | 2.506 | 1.2E-18 |
| *F12E12.6* | WBGene00017407 | protein_coding_gene | 1.3 | 2.506 | 4.9E-04 |
| *Y49A3A.3* | WBGene00013026 | protein_coding_gene | 1.3 | 2.506 | 2.0E-19 |
| *K07H8.7* | WBGene00019507 | protein_coding_gene | 1.3 | 2.505 | 2.5E-03 |
| *T10F2.5* | WBGene00044604 | protein_coding_gene | 1.3 | 2.501 | 5.3E-10 |
| *C53A5.16* | WBGene00194706 | protein_coding_gene | 1.3 | 2.499 | 2.2E-07 |
| *cps-6* | WBGene00000787 | protein_coding_gene | 1.3 | 2.498 | 4.9E-13 |
| *tth-1* | WBGene00006649 | protein_coding_gene | 1.3 | 2.496 | 9.0E-08 |
| *T07D10.8* | WBGene00045509 | pseudogene | 1.3 | 2.496 | 4.4E-09 |
| *T16H12.3* | WBGene00011813 | protein_coding_gene | 1.3 | 2.495 | 4.3E-20 |
| *E02H1.1* | WBGene00008455 | protein_coding_gene | 1.3 | 2.492 | 3.8E-21 |
| *F46A9.1* | WBGene00009754 | protein_coding_gene | 1.3 | 2.491 | 3.2E-02 |
| *erg-28* | WBGene00007589 | protein_coding_gene | 1.3 | 2.490 | 9.6E-07 |
| *ZK973.8* | WBGene00022834 | protein_coding_gene | 1.3 | 2.490 | 8.2E-06 |
| *dhs-11* | WBGene00000974 | protein_coding_gene | 1.3 | 2.489 | 7.6E-09 |
| *exos-8* | WBGene00018305 | protein_coding_gene | 1.3 | 2.489 | 4.0E-15 |
| *F36H12.14* | WBGene00018127 | pseudogene | 1.3 | 2.488 | 1.3E-02 |
| *C48D1.5* | WBGene00008178 | protein_coding_gene | 1.3 | 2.488 | 5.5E-05 |
| *glf-1* | WBGene00019154 | protein_coding_gene | 1.3 | 2.487 | 2.2E-02 |
| *C35D10.6* | WBGene00016443 | protein_coding_gene | 1.3 | 2.486 | 8.7E-23 |
| *T28C6.10* | WBGene00045183 | protein_coding_gene | 1.3 | 2.486 | 3.7E-02 |
| *F55A3.2* | WBGene00018848 | protein_coding_gene | 1.3 | 2.485 | 2.1E-22 |
| *ZC123.1* | WBGene00022517 | protein_coding_gene | 1.3 | 2.480 | 1.7E-02 |
| *blos-4* | WBGene00020783 | protein_coding_gene | 1.3 | 2.480 | 7.4E-05 |
| *prmt-7* | WBGene00012298 | protein_coding_gene | 1.3 | 2.480 | 1.0E-24 |
| *B0564.2* | WBGene00007202 | protein_coding_gene | 1.3 | 2.477 | 2.0E-12 |
| *fbxa-70* | WBGene00020458 | pseudogene | 1.3 | 2.477 | 3.1E-02 |
| *C56A3.10* | WBGene00077568 | pseudogene | 1.3 | 2.477 | 1.2E-02 |
| *K07H8.9* | WBGene00019509 | protein_coding_gene | 1.3 | 2.476 | 6.3E-17 |
| *clec-63* | WBGene00009394 | protein_coding_gene | 1.3 | 2.476 | 2.4E-05 |
| *C42C1.12* | WBGene00016590 | protein_coding_gene | 1.3 | 2.476 | 1.8E-03 |
| *B0462.5* | WBGene00194695 | protein_coding_gene | 1.3 | 2.475 | 7.5E-03 |
| *dph-3* | WBGene00010484 | protein_coding_gene | 1.3 | 2.472 | 1.3E-10 |
| *rpn-6.2* | WBGene00010309 | protein_coding_gene | 1.3 | 2.472 | 3.5E-12 |
| *Y1A5A.1* | WBGene00012379 | protein_coding_gene | 1.3 | 2.471 | 2.9E-02 |
| *Y57A10A.7* | WBGene00013252 | protein_coding_gene | 1.3 | 2.468 | 3.8E-09 |
| *ZK973.11* | WBGene00022836 | protein_coding_gene | 1.3 | 2.464 | 3.0E-09 |
| *C35D10.8* | WBGene00016445 | protein_coding_gene | 1.3 | 2.464 | 7.2E-06 |
| *ceh-45* | WBGene00022837 | protein_coding_gene | 1.3 | 2.463 | 4.5E-02 |
| *K12H4.2* | WBGene00019677 | protein_coding_gene | 1.3 | 2.459 | 1.4E-13 |
| *W10D9.6* | WBGene00185079 | protein_coding_gene | 1.3 | 2.455 | 7.4E-22 |
| *C54G4.9* | WBGene00008318 | protein_coding_gene | 1.3 | 2.453 | 1.0E-10 |
| *spi-1* | WBGene00023416 | protein_coding_gene | 1.3 | 2.453 | 2.9E-02 |

| | | | | | |
|---|---|---|---|---|---|
| *W04E12.4* | WBGene00012249 | protein_coding_gene | 1.3 | 2.452 | 7.5E-03 |
| *btb-7* | WBGene00018443 | protein_coding_gene | 1.3 | 2.448 | 1.1E-09 |
| *F25H9.7* | WBGene00009139 | protein_coding_gene | 1.3 | 2.448 | 7.8E-16 |
| *E02H1.6* | WBGene00008458 | protein_coding_gene | 1.3 | 2.445 | 1.1E-14 |
| *R10H10.6* | WBGene00011224 | protein_coding_gene | 1.3 | 2.444 | 3.8E-12 |
| *F40F11.3* | WBGene00009588 | protein_coding_gene | 1.3 | 2.443 | 1.7E-15 |
| *F45G2.7* | WBGene00009733 | protein_coding_gene | 1.3 | 2.439 | 2.1E-10 |
| *H04D03.6* | WBGene00194861 | protein_coding_gene | 1.3 | 2.438 | 2.7E-03 |
| *K06H7.7* | WBGene00019458 | protein_coding_gene | 1.3 | 2.437 | 2.2E-21 |
| *C36B1.14* | WBGene00044614 | protein_coding_gene | 1.3 | 2.436 | 1.6E-06 |
| *blos-7* | WBGene00021376 | protein_coding_gene | 1.3 | 2.434 | 8.9E-13 |
| *F54H5.5* | WBGene00018841 | protein_coding_gene | 1.3 | 2.432 | 2.8E-02 |
| *ZK1307.9* | WBGene00014250 | protein_coding_gene | 1.3 | 2.430 | 4.2E-20 |
| *C29G2.2* | WBGene00016230 | pseudogene | 1.3 | 2.425 | 4.6E-07 |
| *gtf-2E2* | WBGene00010054 | protein_coding_gene | 1.3 | 2.424 | 9.5E-18 |
| *ears-2* | WBGene00001338 | protein_coding_gene | 1.3 | 2.424 | 2.6E-13 |
| *C49A9.10* | WBGene00016764 | protein_coding_gene | 1.3 | 2.419 | 4.7E-19 |
| *pzf-1* | WBGene00011505 | protein_coding_gene | 1.3 | 2.418 | 3.6E-09 |
| *zip-7* | WBGene00013100 | protein_coding_gene | 1.3 | 2.418 | 4.4E-03 |
| *lgg-3* | WBGene00002982 | protein_coding_gene | 1.3 | 2.412 | 3.1E-16 |
| *B0019.2* | WBGene00007094 | protein_coding_gene | 1.3 | 2.410 | 2.0E-11 |
| *W01A8.2* | WBGene00012164 | protein_coding_gene | 1.3 | 2.409 | 5.6E-10 |
| *D1007.4* | WBGene00017002 | protein_coding_gene | 1.3 | 2.407 | 5.6E-10 |
| *elpc-4* | WBGene00016128 | protein_coding_gene | 1.3 | 2.406 | 3.7E-15 |
| *Y39B6A.49* | WBGene00077457 | pseudogene | 1.3 | 2.405 | 1.7E-08 |
| *Y54E5B.2* | WBGene00013204 | protein_coding_gene | 1.3 | 2.404 | 2.9E-15 |
| *C15C6.4* | WBGene00007603 | protein_coding_gene | 1.3 | 2.400 | 4.0E-08 |
| *ctl-3* | WBGene00013220 | protein_coding_gene | 1.3 | 2.398 | 7.6E-07 |
| *D2089.8* | WBGene00271706 | protein_coding_gene | 1.3 | 2.397 | 8.9E-08 |
| *wdr-60* | WBGene00016165 | protein_coding_gene | 1.3 | 2.394 | 1.8E-03 |
| *lir-2* | WBGene00003045 | protein_coding_gene | 1.3 | 2.392 | 2.6E-14 |
| *F15D4.2* | WBGene00008859 | protein_coding_gene | 1.3 | 2.390 | 2.1E-18 |
| *B0334.4* | WBGene00007144 | protein_coding_gene | 1.3 | 2.387 | 4.3E-09 |
| *ssp-11* | WBGene00006040 | protein_coding_gene | 1.3 | 2.387 | 4.7E-02 |
| *fbxa-101* | WBGene00008569 | protein_coding_gene | 1.3 | 2.387 | 2.1E-12 |
| *F10E7.2* | WBGene00017344 | protein_coding_gene | 1.3 | 2.387 | 1.7E-08 |
| *cht-2* | WBGene00011757 | protein_coding_gene | 1.3 | 2.383 | 3.0E-02 |
| *Y59E9AL.3* | WBGene00021993 | protein_coding_gene | 1.3 | 2.383 | 1.8E-02 |
| *W10D9.3* | WBGene00021131 | protein_coding_gene | 1.3 | 2.381 | 1.8E-07 |
| *grl-15* | WBGene00001724 | protein_coding_gene | 1.3 | 2.380 | 4.5E-02 |
| *H06I04.1* | WBGene00019167 | protein_coding_gene | 1.3 | 2.379 | 1.5E-20 |
| *C16C8.21* | WBGene00195071 | protein_coding_gene | 1.2 | 2.378 | 4.6E-02 |
| *K03B4.1* | WBGene00019353 | protein_coding_gene | 1.2 | 2.378 | 1.6E-14 |
| *tbce-1* | WBGene00019503 | protein_coding_gene | 1.2 | 2.377 | 1.5E-15 |
| *wht-6* | WBGene00007060 | protein_coding_gene | 1.2 | 2.377 | 6.0E-03 |
| *rde-8* | WBGene00022620 | protein_coding_gene | 1.2 | 2.377 | 2.5E-10 |
| *R07H5.11* | WBGene00011131 | protein_coding_gene | 1.2 | 2.374 | 4.6E-07 |
| *D1014.4* | WBGene00017017 | protein_coding_gene | 1.2 | 2.371 | 4.5E-11 |
| *pgs-1* | WBGene00021677 | protein_coding_gene | 1.2 | 2.370 | 2.2E-21 |
| *arl-5* | WBGene00000189 | protein_coding_gene | 1.2 | 2.368 | 5.8E-13 |
| *agt-1* | WBGene00000093 | protein_coding_gene | 1.2 | 2.367 | 7.5E-16 |
| *skr-21* | WBGene00004827 | protein_coding_gene | 1.2 | 2.365 | 3.3E-05 |
| *mdt-11* | WBGene00007015 | protein_coding_gene | 1.2 | 2.365 | 4.5E-07 |
| *F56G4.6* | WBGene00010161 | protein_coding_gene | 1.2 | 2.362 | 6.2E-08 |
| *C05C8.1* | WBGene00015460 | protein_coding_gene | 1.2 | 2.361 | 4.3E-08 |
| *C27B7.2* | WBGene00007760 | protein_coding_gene | 1.2 | 2.358 | 8.6E-10 |
| *F26F4.6* | WBGene00017828 | protein_coding_gene | 1.2 | 2.356 | 1.5E-09 |
| *C42C1.19* | WBGene00302978 | protein_coding_gene | 1.2 | 2.351 | 6.6E-12 |
| *F52F12.8* | WBGene00009941 | protein_coding_gene | 1.2 | 2.346 | 3.3E-02 |
| *F45G2.9* | WBGene00009735 | protein_coding_gene | 1.2 | 2.343 | 3.6E-10 |
| *him-8* | WBGene00001867 | protein_coding_gene | 1.2 | 2.341 | 8.0E-14 |
| *C02F5.13* | WBGene00015353 | protein_coding_gene | 1.2 | 2.341 | 1.3E-08 |
| *Y97E10AR.3* | WBGene00022398 | protein_coding_gene | 1.2 | 2.340 | 5.1E-08 |
| *C24D10.6* | WBGene00016057 | protein_coding_gene | 1.2 | 2.337 | 8.2E-21 |
| *K02A11.2* | WBGene00010487 | protein_coding_gene | 1.2 | 2.335 | 4.4E-02 |
| *F53F10.8* | WBGene00044423 | protein_coding_gene | 1.2 | 2.334 | 3.1E-09 |
| *dnc-5* | WBGene00022118 | protein_coding_gene | 1.2 | 2.334 | 7.5E-20 |
| *Y50D4A.10* | WBGene00235315 | protein_coding_gene | 1.2 | 2.332 | 2.7E-03 |
| *pelo-1* | WBGene00011280 | protein_coding_gene | 1.2 | 2.332 | 3.9E-18 |
| *B0304.2* | WBGene00015134 | protein_coding_gene | 1.2 | 2.330 | 1.4E-11 |

| | | | | | |
|---|---|---|---|---|---|
| *F10B5.2* | WBGene00008639 | protein_coding_gene | 1.2 | 2.328 | 4.0E-23 |
| *M02E1.3* | WBGene00023458 | protein_coding_gene | 1.2 | 2.328 | 3.6E-02 |
| *Y48G9A.11* | WBGene00021704 | protein_coding_gene | 1.2 | 2.325 | 1.9E-09 |
| *fbxl-1* | WBGene00015350 | protein_coding_gene | 1.2 | 2.325 | 1.7E-04 |
| *bath-28* | WBGene00017461 | protein_coding_gene | 1.2 | 2.324 | 5.5E-10 |
| *mut-2* | WBGene00003499 | protein_coding_gene | 1.2 | 2.324 | 9.6E-14 |
| *T12G3.4* | WBGene00011739 | protein_coding_gene | 1.2 | 2.319 | 5.5E-10 |
| *oac-23* | WBGene00018143 | protein_coding_gene | 1.2 | 2.319 | 4.1E-02 |
| *mrpl-14* | WBGene00018475 | protein_coding_gene | 1.2 | 2.316 | 4.4E-14 |
| *mafr-1* | WBGene00016622 | protein_coding_gene | 1.2 | 2.315 | 7.4E-13 |
| *F46B6.12* | WBGene00009776 | protein_coding_gene | 1.2 | 2.314 | 1.4E-06 |
| *btb-19* | WBGene00010194 | protein_coding_gene | 1.2 | 2.314 | 1.8E-06 |
| *K07A12.5* | WBGene00010623 | protein_coding_gene | 1.2 | 2.313 | 2.2E-04 |
| *R144.3* | WBGene00020093 | protein_coding_gene | 1.2 | 2.312 | 1.2E-09 |
| *trpp-5* | WBGene00013259 | protein_coding_gene | 1.2 | 2.309 | 2.5E-10 |
| *Y27F2A.8* | WBGene00021299 | protein_coding_gene | 1.2 | 2.309 | 5.5E-03 |
| *C42C1.9* | WBGene00016587 | protein_coding_gene | 1.2 | 2.309 | 9.1E-03 |
| *F55C5.2* | WBGene00010092 | pseudogene | 1.2 | 2.309 | 2.0E-02 |
| *Y49E10.4* | WBGene00013030 | protein_coding_gene | 1.2 | 2.308 | 2.6E-13 |
| *trip-4* | WBGene00000208 | protein_coding_gene | 1.2 | 2.308 | 4.0E-11 |
| *F49E8.7* | WBGene00018638 | protein_coding_gene | 1.2 | 2.307 | 3.1E-11 |
| *jmjd-4* | WBGene00011563 | protein_coding_gene | 1.2 | 2.306 | 6.3E-05 |
| *C18D4.8* | WBGene00007677 | pseudogene | 1.2 | 2.302 | 6.1E-03 |
| *dpy-3* | WBGene00001065 | protein_coding_gene | 1.2 | 2.301 | 3.1E-02 |
| *T04D1.2* | WBGene00020212 | protein_coding_gene | 1.2 | 2.300 | 4.6E-03 |
| *B0250.12* | WBGene00050927 | pseudogene | 1.2 | 2.299 | 5.6E-03 |
| *ZC204.14* | WBGene00022564 | protein_coding_gene | 1.2 | 2.298 | 3.5E-09 |
| *arp-11* | WBGene00016793 | protein_coding_gene | 1.2 | 2.296 | 6.5E-17 |
| *ubh-2* | WBGene00006722 | protein_coding_gene | 1.2 | 2.296 | 4.8E-11 |
| *sfxn-1.4* | WBGene00008137 | protein_coding_gene | 1.2 | 2.295 | 9.4E-04 |
| *Y43E12A.3* | WBGene00012795 | protein_coding_gene | 1.2 | 2.295 | 2.6E-11 |
| *arv-1* | WBGene00011040 | protein_coding_gene | 1.2 | 2.293 | 3.1E-10 |
| *ZK353.9* | WBGene00022704 | protein_coding_gene | 1.2 | 2.293 | 3.4E-06 |
| *F23A7.1* | WBGene00009067 | protein_coding_gene | 1.2 | 2.292 | 3.9E-02 |
| *C34C6.10* | WBGene00206355 | protein_coding_gene | 1.2 | 2.291 | 2.0E-02 |
| *secs-1* | WBGene00008379 | protein_coding_gene | 1.2 | 2.290 | 1.2E-16 |
| *B0393.4* | WBGene00007169 | protein_coding_gene | 1.2 | 2.290 | 3.0E-03 |
| *ZK1320.7* | WBGene00014256 | protein_coding_gene | 1.2 | 2.288 | 4.5E-13 |
| *F30A10.9* | WBGene00009266 | protein_coding_gene | 1.2 | 2.285 | 1.5E-11 |
| *R11G1.7* | WBGene00044725 | protein_coding_gene | 1.2 | 2.284 | 9.2E-04 |
| *F23D12.10* | WBGene00045291 | protein_coding_gene | 1.2 | 2.284 | 1.9E-06 |
| *F45G2.10* | WBGene00009736 | protein_coding_gene | 1.2 | 2.284 | 3.0E-20 |
| *Y50E8A.12* | WBGene00013055 | protein_coding_gene | 1.2 | 2.281 | 1.4E-02 |
| *F25B5.5* | WBGene00017776 | protein_coding_gene | 1.2 | 2.281 | 5.8E-15 |
| *F14H3.12* | WBGene00008831 | protein_coding_gene | 1.2 | 2.281 | 9.9E-04 |
| *algn-13* | WBGene00011193 | protein_coding_gene | 1.2 | 2.281 | 2.4E-14 |
| *mks-1* | WBGene00020100 | protein_coding_gene | 1.2 | 2.278 | 3.0E-02 |
| *zip-10* | WBGene00012101 | protein_coding_gene | 1.2 | 2.277 | 2.7E-02 |
| *F56C4.4* | WBGene00044977 | protein_coding_gene | 1.2 | 2.277 | 1.7E-02 |
| *ZK643.2* | WBGene00014034 | protein_coding_gene | 1.2 | 2.276 | 3.1E-08 |
| *Y45F10B.3* | WBGene00012871 | protein_coding_gene | 1.2 | 2.275 | 4.1E-02 |
| *Y46G5A.7* | WBGene00012899 | protein_coding_gene | 1.2 | 2.275 | 5.5E-05 |
| *nyn-1* | WBGene00011959 | protein_coding_gene | 1.2 | 2.271 | 5.5E-16 |
| *F59B2.9* | WBGene00010311 | protein_coding_gene | 1.2 | 2.271 | 3.8E-03 |
| *moc-3* | WBGene00018357 | protein_coding_gene | 1.2 | 2.270 | 2.7E-09 |
| *cpn-1* | WBGene00000777 | protein_coding_gene | 1.2 | 2.269 | 9.6E-06 |
| *Y102A5C.35* | WBGene00044220 | pseudogene | 1.2 | 2.267 | 4.3E-02 |
| *Y73B6A.1* | WBGene00022228 | protein_coding_gene | 1.2 | 2.267 | 3.2E-02 |
| *ZK632.11* | WBGene00014018 | protein_coding_gene | 1.2 | 2.267 | 9.0E-22 |
| *ercc-1* | WBGene00008665 | protein_coding_gene | 1.2 | 2.265 | 5.5E-13 |
| *gip-2* | WBGene00044008 | protein_coding_gene | 1.2 | 2.264 | 1.1E-11 |
| *C05C10.7* | WBGene00007334 | protein_coding_gene | 1.2 | 2.264 | 1.2E-10 |
| *clec-196* | WBGene00009156 | protein_coding_gene | 1.2 | 2.261 | 3.6E-04 |
| *W03F8.3* | WBGene00020993 | protein_coding_gene | 1.2 | 2.259 | 1.3E-26 |
| *B0207.6* | WBGene00015029 | protein_coding_gene | 1.2 | 2.258 | 3.9E-07 |
| *T23G7.3* | WBGene00011966 | protein_coding_gene | 1.2 | 2.257 | 6.5E-06 |
| *ZK616.2* | WBGene00022772 | protein_coding_gene | 1.2 | 2.255 | 9.5E-07 |
| *ttc-4* | WBGene00015916 | protein_coding_gene | 1.2 | 2.254 | 1.2E-19 |
| *B0495.9* | WBGene00015208 | protein_coding_gene | 1.2 | 2.253 | 2.5E-18 |
| *gtsf-1* | WBGene00020286 | protein_coding_gene | 1.2 | 2.253 | 1.0E-16 |

| | | | | | |
|---|---|---|---|---|---|
| *ZK1128.1* | WBGene00014227 | protein_coding_gene | 1.2 | 2.251 | 5.3E-21 |
| *rpap-2* | WBGene00011143 | protein_coding_gene | 1.2 | 2.251 | 9.1E-19 |
| *slc-25A18.2* | WBGene00018901 | protein_coding_gene | 1.2 | 2.251 | 7.5E-13 |
| *dnj-18* | WBGene00001036 | protein_coding_gene | 1.2 | 2.251 | 1.7E-10 |
| *K03H6.5* | WBGene00019370 | protein_coding_gene | 1.2 | 2.250 | 1.9E-02 |
| *Y17G9B.8* | WBGene00021205 | protein_coding_gene | 1.2 | 2.250 | 4.2E-11 |
| *R02D5.8* | WBGene00050937 | protein_coding_gene | 1.2 | 2.250 | 1.6E-08 |
| *Y57G11C.36* | WBGene00013325 | protein_coding_gene | 1.2 | 2.249 | 3.4E-07 |
| *F39B2.5* | WBGene00009556 | protein_coding_gene | 1.2 | 2.249 | 2.6E-18 |
| *tbck-1* | WBGene00016352 | protein_coding_gene | 1.2 | 2.248 | 2.3E-11 |
| *elf-1* | WBGene00016029 | protein_coding_gene | 1.2 | 2.248 | 5.4E-03 |
| *T26A5.8* | WBGene00020823 | protein_coding_gene | 1.2 | 2.247 | 2.6E-05 |
| *nog-1* | WBGene00020297 | protein_coding_gene | 1.2 | 2.245 | 7.9E-10 |
| *Y57A10A.6* | WBGene00013251 | pseudogene | 1.2 | 2.242 | 1.1E-04 |
| *pot-2* | WBGene00010195 | protein_coding_gene | 1.2 | 2.241 | 9.4E-08 |
| *clec-72* | WBGene00021583 | protein_coding_gene | 1.2 | 2.241 | 5.5E-05 |
| *F07H5.10* | WBGene00008561 | protein_coding_gene | 1.2 | 2.240 | 5.3E-07 |
| *W09G3.1* | WBGene00012365 | protein_coding_gene | 1.2 | 2.237 | 7.4E-13 |
| *F23C8.8* | WBGene00017737 | protein_coding_gene | 1.2 | 2.237 | 1.5E-02 |
| *Y102A5C.6* | WBGene00014955 | pseudogene | 1.2 | 2.236 | 1.3E-07 |
| *nhr-63* | WBGene00003653 | protein_coding_gene | 1.2 | 2.234 | 1.9E-02 |
| *fubl-4* | WBGene00019692 | protein_coding_gene | 1.2 | 2.232 | 3.4E-11 |
| *Y5F2A.4* | WBGene00012385 | protein_coding_gene | 1.2 | 2.232 | 1.7E-10 |
| *F11C3.1* | WBGene00008695 | protein_coding_gene | 1.2 | 2.232 | 1.1E-04 |
| *K12D12.4* | WBGene00010786 | protein_coding_gene | 1.2 | 2.231 | 3.9E-02 |
| *moa-2* | WBGene00015205 | protein_coding_gene | 1.2 | 2.231 | 1.6E-09 |
| *coq-2* | WBGene00000762 | protein_coding_gene | 1.2 | 2.229 | 2.3E-11 |
| *blos-9* | WBGene00010767 | protein_coding_gene | 1.2 | 2.228 | 3.0E-12 |
| *F32H2.10* | WBGene00009346 | protein_coding_gene | 1.2 | 2.228 | 2.5E-08 |
| *smg-7* | WBGene00004885 | protein_coding_gene | 1.2 | 2.225 | 9.3E-13 |
| *tag-115* | WBGene00006466 | protein_coding_gene | 1.2 | 2.224 | 5.9E-10 |
| *F22D6.14* | WBGene00009055 | pseudogene | 1.2 | 2.223 | 8.6E-06 |
| *cdc-48.3* | WBGene00010562 | protein_coding_gene | 1.2 | 2.222 | 4.4E-19 |
| *ssp-16* | WBGene00006044 | protein_coding_gene | 1.2 | 2.221 | 3.9E-02 |
| *dus-2* | WBGene00013201 | protein_coding_gene | 1.2 | 2.221 | 1.1E-11 |
| *E01A2.5* | WBGene00017087 | protein_coding_gene | 1.2 | 2.219 | 4.2E-14 |
| *apc-10* | WBGene00000144 | protein_coding_gene | 1.1 | 2.219 | 2.6E-11 |
| *Y71A12B.12* | WBGene00013509 | protein_coding_gene | 1.1 | 2.217 | 3.7E-05 |
| *sqt-3* | WBGene00005018 | protein_coding_gene | 1.1 | 2.217 | 4.5E-03 |
| *taf-13* | WBGene00006397 | protein_coding_gene | 1.1 | 2.216 | 1.5E-05 |
| *C01A2.4* | WBGene00007216 | protein_coding_gene | 1.1 | 2.216 | 1.1E-24 |
| *H20J04.1* | WBGene00019216 | protein_coding_gene | 1.1 | 2.216 | 1.2E-02 |
| *math-41* | WBGene00020360 | protein_coding_gene | 1.1 | 2.215 | 1.1E-04 |
| *T25G3.1* | WBGene00012029 | protein_coding_gene | 1.1 | 2.215 | 2.9E-15 |
| *T23G11.4* | WBGene00011969 | protein_coding_gene | 1.1 | 2.213 | 6.8E-18 |
| *C38C10.3* | WBGene00008001 | protein_coding_gene | 1.1 | 2.212 | 3.7E-02 |
| *Y37D8A.25* | WBGene00012560 | protein_coding_gene | 1.1 | 2.212 | 1.6E-08 |
| *E02H1.5* | WBGene00008457 | protein_coding_gene | 1.1 | 2.211 | 1.1E-14 |
| *C01G5.5* | WBGene00015307 | protein_coding_gene | 1.1 | 2.211 | 2.7E-09 |
| *C54C6.6* | WBGene00008288 | protein_coding_gene | 1.1 | 2.210 | 1.1E-10 |
| *C18E3.1* | WBGene00015970 | protein_coding_gene | 1.1 | 2.210 | 1.1E-02 |
| *F59A2.5* | WBGene00010305 | protein_coding_gene | 1.1 | 2.208 | 5.4E-13 |
| *F21D5.5* | WBGene00009010 | protein_coding_gene | 1.1 | 2.207 | 1.7E-16 |
| *ZK1127.3* | WBGene00022850 | protein_coding_gene | 1.1 | 2.207 | 9.4E-12 |
| *mdmh-35* | WBGene00011635 | protein_coding_gene | 1.1 | 2.206 | 5.8E-09 |
| *pdcd-2* | WBGene00011116 | protein_coding_gene | 1.1 | 2.206 | 1.5E-15 |
| *C13G5.2* | WBGene00015747 | protein_coding_gene | 1.1 | 2.205 | 3.8E-16 |
| *pir-1* | WBGene00011967 | protein_coding_gene | 1.1 | 2.205 | 4.1E-27 |
| *Y111B2A.10* | WBGene00013734 | protein_coding_gene | 1.1 | 2.205 | 4.0E-11 |
| *psmd-9* | WBGene00016623 | protein_coding_gene | 1.1 | 2.204 | 5.5E-08 |
| *Y54H5A.2* | WBGene00021900 | protein_coding_gene | 1.1 | 2.203 | 2.2E-12 |
| *C17F4.2* | WBGene00015910 | protein_coding_gene | 1.1 | 2.202 | 4.6E-02 |
| *F15B9.8* | WBGene00008851 | protein_coding_gene | 1.1 | 2.197 | 5.0E-02 |
| *T20H4.2* | WBGene00020635 | protein_coding_gene | 1.1 | 2.197 | 2.8E-02 |
| *Y53C12A.10* | WBGene00175030 | protein_coding_gene | 1.1 | 2.195 | 8.2E-08 |
| *fbxa-206* | WBGene00013610 | protein_coding_gene | 1.1 | 2.194 | 2.2E-11 |
| *K11H12.5* | WBGene00019661 | protein_coding_gene | 1.1 | 2.193 | 1.3E-03 |
| *rpoa-12* | WBGene00007616 | protein_coding_gene | 1.1 | 2.193 | 4.2E-08 |
| *Y38H8A.3* | WBGene00012637 | protein_coding_gene | 1.1 | 2.190 | 1.4E-02 |
| *kbp-2* | WBGene00017832 | protein_coding_gene | 1.1 | 2.188 | 1.1E-08 |

| | | | | | |
|---|---|---|---|---|---|
| *C41D11.9* | WBGene00016567 | protein_coding_gene | 1.1 | 2.188 | 9.5E-07 |
| *rde-11* | WBGene00023421 | protein_coding_gene | 1.1 | 2.187 | 1.3E-07 |
| *polk-1* | WBGene00017696 | protein_coding_gene | 1.1 | 2.185 | 7.6E-11 |
| *trpp-4* | WBGene00018088 | protein_coding_gene | 1.1 | 2.185 | 1.0E-13 |
| *gst-16* | WBGene00001764 | protein_coding_gene | 1.1 | 2.182 | 8.2E-05 |
| *T07C4.12* | WBGene00044891 | protein_coding_gene | 1.1 | 2.181 | 3.8E-15 |
| *Y39B6A.25* | WBGene00012684 | protein_coding_gene | 1.1 | 2.181 | 4.7E-02 |
| *T11F8.1* | WBGene00020433 | protein_coding_gene | 1.1 | 2.181 | 4.5E-07 |
| *mob-1* | WBGene00009551 | protein_coding_gene | 1.1 | 2.180 | 5.5E-12 |
| *mrps-30* | WBGene00044321 | protein_coding_gene | 1.1 | 2.180 | 4.4E-15 |
| *T09B4.2* | WBGene00020376 | protein_coding_gene | 1.1 | 2.180 | 1.9E-07 |
| *C01G12.12* | WBGene00044053 | pseudogene | 1.1 | 2.179 | 2.6E-05 |
| *K11H3.8* | WBGene00185002 | protein_coding_gene | 1.1 | 2.179 | 2.3E-07 |
| *Y43F4B.9* | WBGene00012806 | protein_coding_gene | 1.1 | 2.178 | 2.2E-15 |
| *taf-11.3* | WBGene00006395 | protein_coding_gene | 1.1 | 2.178 | 6.6E-05 |
| *ndx-6* | WBGene00003583 | protein_coding_gene | 1.1 | 2.177 | 1.9E-08 |
| *tut-2* | WBGene00009256 | protein_coding_gene | 1.1 | 2.176 | 4.7E-12 |
| *Y102A5C.36* | WBGene00044213 | protein_coding_gene | 1.1 | 2.175 | 1.2E-03 |
| *cnnm-4* | WBGene00007255 | protein_coding_gene | 1.1 | 2.174 | 2.1E-09 |
| *C25H3.7* | WBGene00016116 | protein_coding_gene | 1.1 | 2.174 | 2.8E-06 |
| *math-44* | WBGene00021193 | protein_coding_gene | 1.1 | 2.174 | 3.2E-04 |
| *C26D10.3* | WBGene00007744 | protein_coding_gene | 1.1 | 2.174 | 1.6E-08 |
| *F54E7.9* | WBGene00018831 | protein_coding_gene | 1.1 | 2.174 | 1.1E-11 |
| *F25H8.1* | WBGene00009131 | protein_coding_gene | 1.1 | 2.172 | 2.1E-16 |
| *Y54H5A.5* | WBGene00050920 | pseudogene | 1.1 | 2.171 | 2.5E-02 |
| *W02D9.4* | WBGene00012210 | protein_coding_gene | 1.1 | 2.171 | 1.1E-12 |
| *M01F1.9* | WBGene00010815 | protein_coding_gene | 1.1 | 2.171 | 7.6E-16 |
| *EEED8.3* | WBGene00017134 | protein_coding_gene | 1.1 | 2.170 | 9.1E-10 |
| *pqn-68* | WBGene00004151 | protein_coding_gene | 1.1 | 2.170 | 2.2E-04 |
| *T05H4.10* | WBGene00020273 | protein_coding_gene | 1.1 | 2.170 | 3.2E-26 |
| *xpg-1* | WBGene00019004 | protein_coding_gene | 1.1 | 2.169 | 2.1E-13 |
| *oac-20* | WBGene00009488 | protein_coding_gene | 1.1 | 2.169 | 4.8E-02 |
| *F10E9.11* | WBGene00017360 | protein_coding_gene | 1.1 | 2.166 | 1.6E-06 |
| *Y39B6A.3* | WBGene00012666 | protein_coding_gene | 1.1 | 2.165 | 2.5E-14 |
| *fbxb-97* | WBGene00016885 | protein_coding_gene | 1.1 | 2.165 | 2.3E-02 |
| *glrx-22* | WBGene00044608 | protein_coding_gene | 1.1 | 2.165 | 6.5E-14 |
| *phf-30* | WBGene00020716 | protein_coding_gene | 1.1 | 2.164 | 1.4E-10 |
| *Y37H2A.12* | WBGene00044758 | protein_coding_gene | 1.1 | 2.164 | 3.5E-08 |
| *cic-1* | WBGene00000506 | protein_coding_gene | 1.1 | 2.162 | 8.3E-11 |
| *D2005.3* | WBGene00008398 | protein_coding_gene | 1.1 | 2.160 | 1.0E-06 |
| *Y82E9BR.22* | WBGene00022354 | protein_coding_gene | 1.1 | 2.160 | 8.1E-07 |
| *Y50D7A.10* | WBGene00021758 | protein_coding_gene | 1.1 | 2.160 | 5.5E-13 |
| *F58A4.6* | WBGene00010229 | protein_coding_gene | 1.1 | 2.160 | 2.2E-09 |
| *gtf-2H1* | WBGene00019821 | protein_coding_gene | 1.1 | 2.159 | 3.8E-18 |
| *mrpl-4* | WBGene00020717 | protein_coding_gene | 1.1 | 2.158 | 2.5E-16 |
| *pigt-1* | WBGene00008918 | protein_coding_gene | 1.1 | 2.158 | 6.3E-14 |
| *C16C8.20* | WBGene00195070 | protein_coding_gene | 1.1 | 2.158 | 5.7E-04 |
| *ssup-72* | WBGene00020480 | protein_coding_gene | 1.1 | 2.156 | 2.6E-07 |
| *arp-6* | WBGene00007434 | protein_coding_gene | 1.1 | 2.156 | 1.5E-11 |
| *C08B6.8* | WBGene00007429 | protein_coding_gene | 1.1 | 2.154 | 1.3E-05 |
| *F10E9.7* | WBGene00017358 | protein_coding_gene | 1.1 | 2.154 | 1.0E-06 |
| *bath-3* | WBGene00019140 | protein_coding_gene | 1.1 | 2.153 | 8.9E-06 |
| *fbxb-67* | WBGene00009862 | protein_coding_gene | 1.1 | 2.152 | 7.0E-05 |
| *algn-11* | WBGene00015162 | protein_coding_gene | 1.1 | 2.148 | 1.2E-13 |
| *T07A9.15* | WBGene00045249 | protein_coding_gene | 1.1 | 2.147 | 9.4E-08 |
| *M110.3* | WBGene00010913 | protein_coding_gene | 1.1 | 2.147 | 2.9E-06 |
| *F26A1.1* | WBGene00017801 | protein_coding_gene | 1.1 | 2.143 | 6.1E-12 |
| *nol-9* | WBGene00010709 | protein_coding_gene | 1.1 | 2.143 | 1.1E-17 |
| *Y23H5A.2* | WBGene00021269 | protein_coding_gene | 1.1 | 2.141 | 8.6E-09 |
| *C31H1.1* | WBGene00016286 | protein_coding_gene | 1.1 | 2.141 | 2.3E-02 |
| *cup-15* | WBGene00009627 | protein_coding_gene | 1.1 | 2.139 | 1.6E-10 |
| *erl-1* | WBGene00016592 | protein_coding_gene | 1.1 | 2.138 | 1.2E-11 |
| *C37A5.7* | WBGene00007991 | protein_coding_gene | 1.1 | 2.137 | 2.8E-11 |
| *ZK632.5* | WBGene00014014 | protein_coding_gene | 1.1 | 2.137 | 5.9E-13 |
| *F58H1.3* | WBGene00010286 | protein_coding_gene | 1.1 | 2.136 | 2.6E-17 |
| *fbxa-13* | WBGene00021264 | protein_coding_gene | 1.1 | 2.133 | 1.9E-04 |
| *spe-44* | WBGene00007732 | protein_coding_gene | 1.1 | 2.133 | 1.1E-09 |
| *ntp-1* | WBGene00016380 | protein_coding_gene | 1.1 | 2.133 | 1.7E-08 |
| *rga-1* | WBGene00012203 | protein_coding_gene | 1.1 | 2.132 | 1.0E-11 |
| *tag-322* | WBGene00044326 | protein_coding_gene | 1.1 | 2.132 | 1.5E-09 |

| | | | | | |
|---|---|---|---|---|---|
| *F14B4.1* | WBGene00008779 | protein_coding_gene | 1.1 | 2.132 | 3.0E-03 |
| *mdt-20* | WBGene00007020 | protein_coding_gene | 1.1 | 2.131 | 6.2E-07 |
| *bath-10* | WBGene00021726 | protein_coding_gene | 1.1 | 2.131 | 4.6E-03 |
| *pms-2* | WBGene00004064 | protein_coding_gene | 1.1 | 2.131 | 2.6E-12 |
| *B0511.2* | WBGene00015228 | protein_coding_gene | 1.1 | 2.130 | 1.3E-03 |
| *rbx-2* | WBGene00019993 | protein_coding_gene | 1.1 | 2.129 | 1.7E-06 |
| *ulp-3* | WBGene00006738 | protein_coding_gene | 1.1 | 2.128 | 6.2E-11 |
| *T05G5.4* | WBGene00011499 | protein_coding_gene | 1.1 | 2.128 | 2.8E-03 |
| *C16D9.4* | WBGene00015859 | protein_coding_gene | 1.1 | 2.127 | 6.1E-03 |
| *Y61A9LA.11* | WBGene00022022 | protein_coding_gene | 1.1 | 2.126 | 5.0E-08 |
| *Y22D7AL.9* | WBGene00021247 | protein_coding_gene | 1.1 | 2.126 | 1.6E-10 |
| *ant-1.3* | WBGene00010485 | protein_coding_gene | 1.1 | 2.126 | 2.8E-02 |
| *cku-70* | WBGene00000519 | protein_coding_gene | 1.1 | 2.124 | 4.4E-16 |
| *C46C11.4* | WBGene00044567 | protein_coding_gene | 1.1 | 2.123 | 1.8E-02 |
| *F55G1.6* | WBGene00018902 | protein_coding_gene | 1.1 | 2.123 | 3.9E-07 |
| *trxr-2* | WBGene00014028 | protein_coding_gene | 1.1 | 2.121 | 1.2E-10 |
| *ceh-92* | WBGene00013431 | protein_coding_gene | 1.1 | 2.121 | 5.5E-15 |
| *F22E5.9* | WBGene00017708 | protein_coding_gene | 1.1 | 2.120 | 1.5E-09 |
| *C56A3.5* | WBGene00008344 | protein_coding_gene | 1.1 | 2.119 | 6.2E-10 |
| *M01F1.3* | WBGene00010809 | protein_coding_gene | 1.1 | 2.119 | 1.2E-14 |
| *mrps-11* | WBGene00012244 | protein_coding_gene | 1.1 | 2.116 | 4.6E-13 |
| *orc-4* | WBGene00012651 | protein_coding_gene | 1.1 | 2.116 | 1.4E-12 |
| *rabs-5* | WBGene00021538 | protein_coding_gene | 1.1 | 2.115 | 1.1E-16 |
| *T07F10.6* | WBGene00011592 | protein_coding_gene | 1.1 | 2.115 | 7.2E-03 |
| *C56C10.9* | WBGene00016965 | protein_coding_gene | 1.1 | 2.115 | 2.9E-17 |
| *eri-6* | WBGene00016561 | protein_coding_gene | 1.1 | 2.115 | 1.2E-06 |
| *C50B6.9* | WBGene00008222 | protein_coding_gene | 1.1 | 2.114 | 4.1E-03 |
| *smi-1* | WBGene00004886 | protein_coding_gene | 1.1 | 2.114 | 4.9E-15 |
| *pgap-3* | WBGene00019806 | protein_coding_gene | 1.1 | 2.113 | 1.1E-07 |
| *Y75B12B.1* | WBGene00013567 | protein_coding_gene | 1.1 | 2.113 | 4.7E-06 |
| *F36D1.9* | WBGene00009468 | protein_coding_gene | 1.1 | 2.111 | 7.1E-03 |
| *nxt-1* | WBGene00003836 | protein_coding_gene | 1.1 | 2.110 | 5.3E-07 |
| *frh-1* | WBGene00001486 | protein_coding_gene | 1.1 | 2.108 | 3.2E-07 |
| *F56A8.5* | WBGene00010141 | protein_coding_gene | 1.1 | 2.106 | 4.9E-12 |
| *D1054.1* | WBGene00008370 | protein_coding_gene | 1.1 | 2.103 | 9.4E-11 |
| *cdk-7* | WBGene00000408 | protein_coding_gene | 1.1 | 2.103 | 2.7E-13 |
| *mrps-21* | WBGene00017924 | protein_coding_gene | 1.1 | 2.102 | 1.0E-06 |
| *C50D2.10* | WBGene00044420 | protein_coding_gene | 1.1 | 2.101 | 3.1E-08 |
| *C30F12.3* | WBGene00016262 | protein_coding_gene | 1.1 | 2.101 | 1.9E-03 |
| *C24H11.5* | WBGene00007702 | protein_coding_gene | 1.1 | 2.101 | 5.0E-08 |
| *inx-7* | WBGene00002129 | protein_coding_gene | 1.1 | 2.100 | 1.6E-13 |
| *C56C10.7* | WBGene00016964 | protein_coding_gene | 1.1 | 2.099 | 8.9E-08 |
| *nekl-1* | WBGene00021461 | protein_coding_gene | 1.1 | 2.099 | 3.3E-02 |
| *hpo-9* | WBGene00015463 | protein_coding_gene | 1.1 | 2.099 | 6.4E-11 |
| *F01F1.11* | WBGene00017165 | protein_coding_gene | 1.1 | 2.098 | 3.9E-13 |
| *C13F10.5* | WBGene00015744 | protein_coding_gene | 1.1 | 2.098 | 6.2E-09 |
| *pho-9* | WBGene00022181 | protein_coding_gene | 1.1 | 2.098 | 1.2E-06 |
| *snpc-3.4* | WBGene00011367 | protein_coding_gene | 1.1 | 2.095 | 1.5E-16 |
| *acdh-4* | WBGene00020419 | protein_coding_gene | 1.1 | 2.095 | 1.0E-02 |
| *wht-5* | WBGene00008950 | protein_coding_gene | 1.1 | 2.095 | 3.9E-02 |
| *dph-2* | WBGene00007488 | protein_coding_gene | 1.1 | 2.095 | 8.9E-18 |
| *Y54F10AR.2* | WBGene00021855 | protein_coding_gene | 1.1 | 2.094 | 2.4E-14 |
| *psa-3* | WBGene00009560 | protein_coding_gene | 1.1 | 2.094 | 5.5E-03 |
| *mrpl-13* | WBGene00008769 | protein_coding_gene | 1.1 | 2.093 | 6.6E-14 |
| *Y38H6C.16* | WBGene00012628 | protein_coding_gene | 1.1 | 2.093 | 9.1E-03 |
| *exos-7* | WBGene00009289 | protein_coding_gene | 1.1 | 2.093 | 1.8E-13 |
| *ced-12* | WBGene00000426 | protein_coding_gene | 1.1 | 2.093 | 4.3E-20 |
| *Y40D12A.1* | WBGene00021502 | protein_coding_gene | 1.1 | 2.092 | 2.2E-08 |
| *Y71F9AL.6* | WBGene00022111 | protein_coding_gene | 1.1 | 2.091 | 1.2E-02 |
| *Y54G9A.9* | WBGene00013211 | protein_coding_gene | 1.1 | 2.091 | 9.2E-03 |
| *F56A8.4* | WBGene00010140 | protein_coding_gene | 1.1 | 2.091 | 1.5E-17 |
| *metl-18* | WBGene00010450 | protein_coding_gene | 1.1 | 2.089 | 1.6E-10 |
| *cki-1* | WBGene00000516 | protein_coding_gene | 1.1 | 2.089 | 4.1E-04 |
| *Y11D7A.7* | WBGene00012431 | protein_coding_gene | 1.1 | 2.087 | 2.3E-09 |
| *ZK688.11* | WBGene00044762 | protein_coding_gene | 1.1 | 2.087 | 1.1E-18 |
| *mrpl-18* | WBGene00017044 | protein_coding_gene | 1.1 | 2.086 | 2.3E-07 |
| *ztf-20* | WBGene00012702 | protein_coding_gene | 1.1 | 2.085 | 1.4E-06 |
| *R11H6.5* | WBGene00011253 | protein_coding_gene | 1.1 | 2.084 | 1.4E-11 |
| *T27A3.7* | WBGene00020843 | protein_coding_gene | 1.1 | 2.082 | 5.2E-12 |
| *lron-4* | WBGene00008051 | protein_coding_gene | 1.1 | 2.082 | 3.5E-03 |

| | | | | | |
|---|---|---|---|---|---|
| *B0464.9* | WBGene00007188 | protein_coding_gene | 1.1 | 2.082 | 3.1E-15 |
| *fbxa-115* | WBGene00013755 | protein_coding_gene | 1.1 | 2.082 | 3.4E-04 |
| *M142.8* | WBGene00043057 | protein_coding_gene | 1.1 | 2.081 | 6.2E-12 |
| *elpc-2* | WBGene00013738 | protein_coding_gene | 1.1 | 2.081 | 1.3E-11 |
| *vps-51* | WBGene00015176 | protein_coding_gene | 1.1 | 2.081 | 2.6E-14 |
| *C50F4.4* | WBGene00008232 | protein_coding_gene | 1.1 | 2.081 | 1.6E-03 |
| *rpap-3* | WBGene00016375 | protein_coding_gene | 1.1 | 2.079 | 9.4E-11 |
| *brap-2* | WBGene00017144 | protein_coding_gene | 1.1 | 2.078 | 7.6E-15 |
| *crn-5* | WBGene00000798 | protein_coding_gene | 1.1 | 2.077 | 2.0E-04 |
| *dnj-15* | WBGene00001033 | protein_coding_gene | 1.1 | 2.075 | 1.5E-18 |
| *C03B8.3* | WBGene00015386 | protein_coding_gene | 1.1 | 2.075 | 1.4E-02 |
| *R144.13* | WBGene00044648 | protein_coding_gene | 1.1 | 2.075 | 6.2E-07 |
| *F32B6.3* | WBGene00009320 | protein_coding_gene | 1.1 | 2.074 | 1.1E-05 |
| *T28A11.22* | WBGene00044526 | protein_coding_gene | 1.1 | 2.073 | 1.3E-02 |
| *hap-1* | WBGene00001823 | protein_coding_gene | 1.1 | 2.073 | 7.4E-07 |
| *C33F10.4* | WBGene00016353 | protein_coding_gene | 1.1 | 2.073 | 6.0E-08 |
| *dhps-1* | WBGene00012460 | protein_coding_gene | 1.1 | 2.072 | 2.0E-11 |
| *lpd-7* | WBGene00003063 | protein_coding_gene | 1.1 | 2.071 | 1.7E-13 |
| *lin-24* | WBGene00003010 | protein_coding_gene | 1.1 | 2.071 | 4.7E-09 |
| *lir-1* | WBGene00003044 | protein_coding_gene | 1.1 | 2.071 | 3.0E-08 |
| *C16C8.16* | WBGene00015853 | protein_coding_gene | 1.1 | 2.071 | 1.8E-15 |
| *puf-8* | WBGene00004244 | protein_coding_gene | 1.1 | 2.071 | 7.3E-11 |
| *T02E1.2* | WBGene00011375 | protein_coding_gene | 1.0 | 2.070 | 5.9E-14 |
| *mdt-9* | WBGene00014938 | protein_coding_gene | 1.0 | 2.069 | 1.7E-03 |
| *Y49F6B.2* | WBGene00021715 | protein_coding_gene | 1.0 | 2.069 | 5.8E-10 |
| *dhs-17* | WBGene00000980 | protein_coding_gene | 1.0 | 2.068 | 4.7E-08 |
| *Y106G6H.14* | WBGene00013724 | protein_coding_gene | 1.0 | 2.067 | 2.9E-07 |
| *T10G3.1* | WBGene00011692 | protein_coding_gene | 1.0 | 2.067 | 2.1E-02 |
| *fbxa-97* | WBGene00007667 | pseudogene | 1.0 | 2.067 | 3.1E-03 |
| *Y53G8B.2* | WBGene00021818 | protein_coding_gene | 1.0 | 2.066 | 1.0E-08 |
| *erh-1* | WBGene00011892 | protein_coding_gene | 1.0 | 2.064 | 1.3E-07 |
| *nyn-2* | WBGene00013596 | protein_coding_gene | 1.0 | 2.063 | 7.1E-11 |
| *C15F1.5* | WBGene00015796 | protein_coding_gene | 1.0 | 2.063 | 5.6E-03 |
| *phip-1* | WBGene00009455 | protein_coding_gene | 1.0 | 2.062 | 4.3E-13 |
| *Y37B11A.8* | WBGene00255596 | pseudogene | 1.0 | 2.062 | 1.8E-05 |
| *asps-1* | WBGene00007100 | protein_coding_gene | 1.0 | 2.061 | 1.4E-09 |
| *mett-10* | WBGene00014228 | protein_coding_gene | 1.0 | 2.061 | 3.4E-12 |
| *mrps-17* | WBGene00015487 | protein_coding_gene | 1.0 | 2.060 | 2.8E-18 |
| *fbxa-219* | WBGene00044403 | protein_coding_gene | 1.0 | 2.059 | 8.9E-03 |
| *gtf-2H3* | WBGene00014230 | protein_coding_gene | 1.0 | 2.058 | 1.1E-11 |
| *T24B8.2* | WBGene00011976 | protein_coding_gene | 1.0 | 2.056 | 6.9E-13 |
| *trx-2* | WBGene00007099 | protein_coding_gene | 1.0 | 2.056 | 6.9E-08 |
| *ZK856.11* | WBGene00014112 | protein_coding_gene | 1.0 | 2.054 | 2.2E-11 |
| *dcs-1* | WBGene00000940 | protein_coding_gene | 1.0 | 2.054 | 1.8E-10 |
| *F40G12.10* | WBGene00009605 | protein_coding_gene | 1.0 | 2.053 | 3.1E-02 |
| *F42G4.7* | WBGene00009644 | protein_coding_gene | 1.0 | 2.052 | 7.7E-03 |
| *T04A8.8* | WBGene00011410 | protein_coding_gene | 1.0 | 2.052 | 2.8E-10 |
| *Y39A1A.3* | WBGene00012643 | protein_coding_gene | 1.0 | 2.051 | 3.4E-11 |
| *cyp-44A1* | WBGene00000375 | protein_coding_gene | 1.0 | 2.051 | 1.4E-09 |
| *K08H2.4* | WBGene00010691 | protein_coding_gene | 1.0 | 2.051 | 1.5E-02 |
| *mes-6* | WBGene00003224 | protein_coding_gene | 1.0 | 2.050 | 1.7E-05 |
| *Y51H4A.15* | WBGene00013109 | protein_coding_gene | 1.0 | 2.050 | 2.7E-15 |
| *F59D12.5* | WBGene00010332 | protein_coding_gene | 1.0 | 2.049 | 1.5E-03 |
| *F02E9.1* | WBGene00008528 | protein_coding_gene | 1.0 | 2.049 | 1.2E-11 |
| *metl-17* | WBGene00009305 | protein_coding_gene | 1.0 | 2.049 | 2.5E-13 |
| *F10D11.2* | WBGene00008648 | protein_coding_gene | 1.0 | 2.048 | 1.5E-07 |
| *pch-2* | WBGene00008641 | protein_coding_gene | 1.0 | 2.048 | 9.9E-07 |
| *ZK1248.13* | WBGene00022882 | protein_coding_gene | 1.0 | 2.048 | 4.7E-11 |
| *npp-24* | WBGene00022672 | protein_coding_gene | 1.0 | 2.047 | 3.0E-15 |
| *cyp-31A5* | WBGene00013381 | protein_coding_gene | 1.0 | 2.046 | 7.6E-05 |
| *mrpl-23* | WBGene00020348 | protein_coding_gene | 1.0 | 2.045 | 4.4E-11 |
| *R10D12.8* | WBGene00011189 | protein_coding_gene | 1.0 | 2.045 | 1.1E-08 |
| *pgl-2* | WBGene00003993 | protein_coding_gene | 1.0 | 2.045 | 3.7E-14 |
| *arrd-13* | WBGene00011054 | protein_coding_gene | 1.0 | 2.044 | 3.1E-08 |
| *C50F4.16* | WBGene00008238 | protein_coding_gene | 1.0 | 2.044 | 6.3E-15 |
| *Y105C5B.20* | WBGene00013659 | protein_coding_gene | 1.0 | 2.044 | 3.7E-04 |
| *rpc-11* | WBGene00022309 | protein_coding_gene | 1.0 | 2.044 | 9.7E-05 |
| *smg-8* | WBGene00010551 | protein_coding_gene | 1.0 | 2.043 | 5.4E-13 |
| *Y80D4G.1* | WBGene00206459 | pseudogene | 1.0 | 2.043 | 1.7E-05 |
| *T10E9.14* | WBGene00271694 | protein_coding_gene | 1.0 | 2.042 | 2.1E-06 |

| | | | | | |
|---|---|---|---|---|---|
| *mrpl-10* | WBGene00010458 | protein_coding_gene | 1.0 | 2.042 | 6.6E-11 |
| *C29F5.3* | WBGene00016212 | protein_coding_gene | 1.0 | 2.042 | 2.5E-02 |
| *ndc-80* | WBGene00003576 | protein_coding_gene | 1.0 | 2.042 | 2.4E-13 |
| *C23H3.5* | WBGene00016021 | protein_coding_gene | 1.0 | 2.042 | 1.1E-14 |
| *jtr-1* | WBGene00002180 | protein_coding_gene | 1.0 | 2.042 | 1.0E-10 |
| *apc-11* | WBGene00000145 | protein_coding_gene | 1.0 | 2.042 | 2.2E-05 |
| *psf-1* | WBGene00011275 | protein_coding_gene | 1.0 | 2.041 | 4.0E-05 |
| *R01B10.6* | WBGene00019808 | protein_coding_gene | 1.0 | 2.041 | 3.9E-15 |
| *F59A7.8* | WBGene00019095 | pseudogene | 1.0 | 2.041 | 3.8E-06 |
| *zim-1* | WBGene00011597 | protein_coding_gene | 1.0 | 2.034 | 2.7E-08 |
| *F42A6.6* | WBGene00018328 | protein_coding_gene | 1.0 | 2.033 | 2.6E-07 |
| *fut-3* | WBGene00006402 | protein_coding_gene | 1.0 | 2.033 | 5.7E-08 |
| *clk-2* | WBGene00000537 | protein_coding_gene | 1.0 | 2.032 | 5.8E-08 |
| *vars-1* | WBGene00006935 | protein_coding_gene | 1.0 | 2.031 | 6.6E-13 |
| *mrpl-36* | WBGene00010783 | protein_coding_gene | 1.0 | 2.031 | 1.0E-06 |
| *M110.10* | WBGene00195142 | protein_coding_gene | 1.0 | 2.031 | 1.9E-03 |
| *T09A5.5* | WBGene00011634 | protein_coding_gene | 1.0 | 2.030 | 1.0E-20 |
| *mfap-1* | WBGene00009671 | protein_coding_gene | 1.0 | 2.028 | 4.4E-05 |
| *Y48A5A.1* | WBGene00021655 | protein_coding_gene | 1.0 | 2.028 | 3.9E-14 |
| *elpc-3* | WBGene00014123 | protein_coding_gene | 1.0 | 2.027 | 2.7E-11 |
| *Y62F5A.12* | WBGene00077453 | protein_coding_gene | 1.0 | 2.027 | 2.6E-13 |
| *Y39A1C.1* | WBGene00012662 | protein_coding_gene | 1.0 | 2.026 | 5.7E-11 |
| *mutd-1* | WBGene00016404 | protein_coding_gene | 1.0 | 2.025 | 3.3E-07 |
| *F14D2.19* | WBGene00206522 | protein_coding_gene | 1.0 | 2.024 | 9.8E-03 |
| *Y45G5AM.7* | WBGene00021559 | protein_coding_gene | 1.0 | 2.024 | 1.0E-15 |
| *gyg-3* | WBGene00013535 | pseudogene | 1.0 | 2.023 | 3.2E-02 |
| *asna-2* | WBGene00018577 | pseudogene | 1.0 | 2.023 | 1.7E-05 |
| *gna-2* | WBGene00001647 | protein_coding_gene | 1.0 | 2.023 | 8.1E-06 |
| *ZK896.5* | WBGene00014136 | protein_coding_gene | 1.0 | 2.022 | 5.4E-03 |
| *Y37D8A.16* | WBGene00012554 | protein_coding_gene | 1.0 | 2.022 | 2.3E-04 |
| *F46F11.8* | WBGene00018511 | protein_coding_gene | 1.0 | 2.021 | 2.2E-05 |
| *C23G10.7* | WBGene00016014 | protein_coding_gene | 1.0 | 2.021 | 5.2E-07 |
| *dhfr-1* | WBGene00007974 | protein_coding_gene | 1.0 | 2.020 | 6.6E-04 |
| *ctf-8* | WBGene00011915 | protein_coding_gene | 1.0 | 2.020 | 2.5E-09 |
| *dtmk-1* | WBGene00011272 | protein_coding_gene | 1.0 | 2.020 | 2.8E-08 |
| *tbcc-1* | WBGene00044609 | protein_coding_gene | 1.0 | 2.020 | 1.2E-14 |
| *ska-1* | WBGene00013725 | protein_coding_gene | 1.0 | 2.019 | 6.3E-10 |
| *his-67* | WBGene00001941 | protein_coding_gene | 1.0 | 2.019 | 4.9E-03 |
| *Y62F5A.10* | WBGene00013388 | protein_coding_gene | 1.0 | 2.018 | 2.8E-02 |
| *fbxa-33* | WBGene00013841 | pseudogene | 1.0 | 2.018 | 3.8E-02 |
| *tofu-5* | WBGene00012167 | protein_coding_gene | 1.0 | 2.016 | 1.2E-09 |
| *tppp-1* | WBGene00016321 | protein_coding_gene | 1.0 | 2.016 | 4.1E-14 |
| *cyn-10* | WBGene00000886 | protein_coding_gene | 1.0 | 2.016 | 4.6E-12 |
| *F02E9.7* | WBGene00008531 | protein_coding_gene | 1.0 | 2.015 | 1.6E-11 |
| *C34E10.10* | WBGene00016411 | protein_coding_gene | 1.0 | 2.015 | 2.3E-13 |
| *T07F10.3* | WBGene00011589 | protein_coding_gene | 1.0 | 2.015 | 2.4E-10 |
| *C01G10.7* | WBGene00007234 | protein_coding_gene | 1.0 | 2.013 | 2.0E-05 |
| *Y37H9A.1* | WBGene00012576 | protein_coding_gene | 1.0 | 2.013 | 1.1E-06 |
| *mrpl-49* | WBGene00011247 | protein_coding_gene | 1.0 | 2.012 | 5.2E-06 |
| *Y67D2.5* | WBGene00022055 | protein_coding_gene | 1.0 | 2.011 | 3.3E-07 |
| *mrps-10* | WBGene00012556 | protein_coding_gene | 1.0 | 2.011 | 3.0E-14 |
| *K04G2.10* | WBGene00010568 | protein_coding_gene | 1.0 | 2.011 | 5.9E-04 |
| *rpc-25* | WBGene00014111 | protein_coding_gene | 1.0 | 2.010 | 6.2E-11 |
| *B0361.2* | WBGene00015156 | protein_coding_gene | 1.0 | 2.009 | 2.3E-08 |
| *F44E2.9* | WBGene00018422 | protein_coding_gene | 1.0 | 2.009 | 6.3E-07 |
| *pot-3* | WBGene00007065 | protein_coding_gene | 1.0 | 2.008 | 7.6E-06 |
| *kel-3* | WBGene00002185 | protein_coding_gene | 1.0 | 2.007 | 4.8E-09 |
| *tads-1* | WBGene00007217 | protein_coding_gene | 1.0 | 2.006 | 1.7E-15 |
| *K08F4.3* | WBGene00010678 | protein_coding_gene | 1.0 | 2.006 | 5.1E-08 |
| *C03A7.13* | WBGene00015371 | protein_coding_gene | 1.0 | 2.006 | 3.2E-02 |
| *T26A5.2* | WBGene00020819 | protein_coding_gene | 1.0 | 2.005 | 3.4E-08 |
| *C05C8.5* | WBGene00015462 | protein_coding_gene | 1.0 | 2.005 | 6.4E-09 |
| *lcmt-1* | WBGene00007137 | protein_coding_gene | 1.0 | 2.005 | 1.3E-05 |
| *Y56A3A.33* | WBGene00013244 | protein_coding_gene | 1.0 | 2.005 | 9.0E-03 |
| *sft-1* | WBGene00004787 | protein_coding_gene | 1.0 | 2.003 | 1.0E-05 |
| *T28C6.5* | WBGene00012120 | protein_coding_gene | 1.0 | 2.003 | 3.2E-02 |
| *Y62F5A.9* | WBGene00013387 | protein_coding_gene | 1.0 | 2.003 | 2.5E-02 |
| *dph-1* | WBGene00007576 | protein_coding_gene | 1.0 | 2.003 | 2.1E-07 |
| *K06A4.7* | WBGene00010597 | protein_coding_gene | 1.0 | 2.002 | 2.7E-03 |
| *ZK688.7* | WBGene00022802 | protein_coding_gene | 1.0 | 2.002 | 1.7E-06 |

| | | | | | |
|---|---|---|---|---|---|
| *F22B5.5* | WBGene00009043 | protein_coding_gene | 1.0 | 2.002 | 3.9E-02 |
| *Y50D4C.2* | WBGene00021747 | pseudogene | 1.0 | 2.002 | 1.0E-03 |
| *C33A12.4* | WBGene00007883 | protein_coding_gene | 1.0 | 2.001 | 6.3E-03 |
| *sas-5* | WBGene00009385 | protein_coding_gene | 1.0 | 2.000 | 2.6E-11 |
| *act-1* | WBGene00000063 | protein_coding_gene | -1.0 | 0.499 | 3.1E-03 |
| *hsp-60* | WBGene00002025 | protein_coding_gene | -1.0 | 0.499 | 1.1E-04 |
| *arc-1* | WBGene00000180 | protein_coding_gene | -1.0 | 0.499 | 4.0E-10 |
| *klu-2* | WBGene00022592 | protein_coding_gene | -1.0 | 0.499 | 4.1E-05 |
| *mca-3* | WBGene00003153 | protein_coding_gene | -1.0 | 0.498 | 2.2E-05 |
| *F14B6.2* | WBGene00008783 | protein_coding_gene | -1.0 | 0.498 | 3.6E-03 |
| *C14H10.2* | WBGene00007592 | protein_coding_gene | -1.0 | 0.497 | 2.5E-04 |
| *clp-9* | WBGene00011705 | protein_coding_gene | -1.0 | 0.497 | 1.3E-03 |
| *H12D21.10* | WBGene00010386 | protein_coding_gene | -1.0 | 0.496 | 2.3E-06 |
| *lipl-7* | WBGene00008510 | protein_coding_gene | -1.0 | 0.496 | 8.2E-06 |
| *sgca-1* | WBGene00019227 | protein_coding_gene | -1.0 | 0.496 | 2.5E-05 |
| *B0403.3* | WBGene00015167 | protein_coding_gene | -1.0 | 0.496 | 3.5E-03 |
| *W07G4.5* | WBGene00012339 | protein_coding_gene | -1.0 | 0.496 | 8.5E-13 |
| *gcy-21* | WBGene00001546 | protein_coding_gene | -1.0 | 0.495 | 1.2E-02 |
| *nkat-3* | WBGene00010984 | protein_coding_gene | -1.0 | 0.495 | 5.9E-04 |
| *unc-78* | WBGene00006810 | protein_coding_gene | -1.0 | 0.495 | 1.4E-08 |
| *nhr-146* | WBGene00045255 | protein_coding_gene | -1.0 | 0.495 | 4.5E-04 |
| *F55C10.5* | WBGene00010109 | protein_coding_gene | -1.0 | 0.494 | 1.8E-04 |
| *inpp-1* | WBGene00012016 | protein_coding_gene | -1.0 | 0.494 | 4.9E-03 |
| *spin-4* | WBGene00013739 | protein_coding_gene | -1.0 | 0.494 | 4.4E-05 |
| *dpy-23* | WBGene00001082 | protein_coding_gene | -1.0 | 0.493 | 4.5E-07 |
| *H37A05.2* | WBGene00010426 | protein_coding_gene | -1.0 | 0.493 | 2.2E-02 |
| *K07C11.8* | WBGene00019480 | protein_coding_gene | -1.0 | 0.492 | 1.3E-03 |
| *nrx-1* | WBGene00007799 | protein_coding_gene | -1.0 | 0.492 | 1.8E-03 |
| *C25F9.5* | WBGene00007725 | protein_coding_gene | -1.0 | 0.492 | 8.0E-03 |
| *F58F9.4* | WBGene00019058 | protein_coding_gene | -1.0 | 0.492 | 9.4E-04 |
| *twf-2* | WBGene00018187 | protein_coding_gene | -1.0 | 0.491 | 1.1E-04 |
| *C39D10.6* | WBGene00016533 | protein_coding_gene | -1.0 | 0.491 | 2.0E-02 |
| *erd-2.1* | WBGene00001331 | protein_coding_gene | -1.0 | 0.491 | 1.5E-05 |
| *F47G9.4* | WBGene00009831 | protein_coding_gene | -1.0 | 0.491 | 2.4E-03 |
| *Y39G10AL.1* | WBGene00021458 | protein_coding_gene | -1.0 | 0.490 | 3.4E-03 |
| *hlh-11* | WBGene00001955 | protein_coding_gene | -1.0 | 0.490 | 1.3E-03 |
| *swt-7* | WBGene00019645 | protein_coding_gene | -1.0 | 0.490 | 3.8E-07 |
| *klo-2* | WBGene00017103 | protein_coding_gene | -1.0 | 0.489 | 1.6E-05 |
| *T19H12.3* | WBGene00020589 | protein_coding_gene | -1.0 | 0.489 | 3.4E-06 |
| *daf-7* | WBGene00000903 | protein_coding_gene | -1.0 | 0.489 | 1.6E-03 |
| *chil-25* | WBGene00011846 | protein_coding_gene | -1.0 | 0.489 | 2.7E-02 |
| *C49A9.6* | WBGene00016760 | protein_coding_gene | -1.0 | 0.488 | 9.1E-07 |
| *lin-14* | WBGene00003003 | protein_coding_gene | -1.0 | 0.488 | 6.6E-03 |
| *lgc-33* | WBGene00021941 | protein_coding_gene | -1.0 | 0.488 | 4.8E-02 |
| *rhy-1* | WBGene00012324 | protein_coding_gene | -1.0 | 0.488 | 6.4E-04 |
| *rfip-1* | WBGene00018869 | protein_coding_gene | -1.0 | 0.488 | 1.6E-05 |
| *cla-1* | WBGene00018468 | protein_coding_gene | -1.0 | 0.487 | 3.0E-02 |
| *kat-1* | WBGene00002183 | protein_coding_gene | -1.0 | 0.487 | 2.8E-10 |
| *F52G2.3* | WBGene00009943 | protein_coding_gene | -1.0 | 0.487 | 4.3E-06 |
| *zipt-7.2* | WBGene00006494 | protein_coding_gene | -1.0 | 0.486 | 3.1E-09 |
| *F52H2.7* | WBGene00018719 | protein_coding_gene | -1.0 | 0.484 | 4.8E-07 |
| *E02A10.3* | WBGene00008453 | protein_coding_gene | -1.0 | 0.484 | 1.4E-02 |
| *ifo-1* | WBGene00018350 | protein_coding_gene | -1.0 | 0.484 | 1.5E-02 |
| *idh-1* | WBGene00010317 | protein_coding_gene | -1.0 | 0.484 | 5.7E-05 |
| *C38D9.8* | WBGene00008016 | protein_coding_gene | -1.0 | 0.483 | 4.9E-02 |
| *T02G5.7* | WBGene00020166 | protein_coding_gene | -1.1 | 0.482 | 5.3E-08 |
| *K05F6.4* | WBGene00019415 | protein_coding_gene | -1.1 | 0.482 | 4.3E-02 |
| *ncl-1* | WBGene00003559 | protein_coding_gene | -1.1 | 0.482 | 2.5E-10 |
| *Y42G9A.2* | WBGene00021532 | protein_coding_gene | -1.1 | 0.482 | 1.6E-03 |
| *ZC449.7* | WBGene00045365 | protein_coding_gene | -1.1 | 0.482 | 1.6E-02 |
| *F38A1.9* | WBGene00009522 | protein_coding_gene | -1.1 | 0.481 | 2.8E-02 |
| *let-767* | WBGene00002891 | protein_coding_gene | -1.1 | 0.481 | 1.8E-06 |
| *fbp-1* | WBGene00001404 | protein_coding_gene | -1.1 | 0.480 | 1.5E-05 |
| *maf-1* | WBGene00077521 | protein_coding_gene | -1.1 | 0.480 | 1.7E-03 |
| *R07E3.1* | WBGene00011102 | protein_coding_gene | -1.1 | 0.480 | 9.7E-06 |
| *F54F7.3* | WBGene00010065 | protein_coding_gene | -1.1 | 0.480 | 3.6E-03 |
| *K09E9.4* | WBGene00044143 | protein_coding_gene | -1.1 | 0.480 | 1.9E-02 |
| *Y56A3A.30* | WBGene00013242 | protein_coding_gene | -1.1 | 0.480 | 4.2E-05 |
| *C01G10.9* | WBGene00007236 | protein_coding_gene | -1.1 | 0.480 | 4.6E-03 |
| *elo-1* | WBGene00001239 | protein_coding_gene | -1.1 | 0.480 | 1.5E-10 |

| | | | | | |
|---|---|---|---|---|---|
| *trap-4* | WBGene00013238 | protein_coding_gene | -1.1 | 0.480 | 1.7E-06 |
| *R09H10.5* | WBGene00011182 | protein_coding_gene | -1.1 | 0.479 | 2.0E-05 |
| *ftn-2* | WBGene00001501 | protein_coding_gene | -1.1 | 0.478 | 2.5E-12 |
| *flp-26* | WBGene00020126 | protein_coding_gene | -1.1 | 0.478 | 4.6E-03 |
| *sir-2.3* | WBGene00004802 | protein_coding_gene | -1.1 | 0.478 | 4.6E-04 |
| *lgc-31* | WBGene00008997 | protein_coding_gene | -1.1 | 0.478 | 2.3E-03 |
| *ZK6.11* | WBGene00022645 | protein_coding_gene | -1.1 | 0.477 | 3.8E-02 |
| *cpt-6* | WBGene00020911 | protein_coding_gene | -1.1 | 0.477 | 4.6E-05 |
| *Y46G5A.28* | WBGene00012916 | protein_coding_gene | -1.1 | 0.476 | 3.9E-02 |
| *ttr-34* | WBGene00045245 | protein_coding_gene | -1.1 | 0.476 | 1.1E-02 |
| *T14B4.5* | WBGene00020501 | protein_coding_gene | -1.1 | 0.476 | 1.5E-03 |
| *flp-14* | WBGene00001457 | protein_coding_gene | -1.1 | 0.476 | 2.7E-06 |
| *dhs-21* | WBGene00000984 | protein_coding_gene | -1.1 | 0.476 | 3.0E-04 |
| *hum-7* | WBGene00002040 | protein_coding_gene | -1.1 | 0.476 | 5.6E-05 |
| *E04F6.15* | WBGene00017131 | protein_coding_gene | -1.1 | 0.476 | 1.4E-05 |
| *Y66A7A.7* | WBGene00013426 | protein_coding_gene | -1.1 | 0.475 | 1.5E-04 |
| *cal-4* | WBGene00000288 | protein_coding_gene | -1.1 | 0.475 | 1.2E-03 |
| *pnk-4* | WBGene00006862 | protein_coding_gene | -1.1 | 0.475 | 4.4E-08 |
| *R102.4* | WBGene00011291 | protein_coding_gene | -1.1 | 0.474 | 1.0E-03 |
| *ncx-6* | WBGene00003571 | protein_coding_gene | -1.1 | 0.473 | 3.2E-03 |
| *ifd-1* | WBGene00002057 | protein_coding_gene | -1.1 | 0.473 | 7.2E-06 |
| *ZK185.2* | WBGene00022682 | protein_coding_gene | -1.1 | 0.473 | 1.3E-02 |
| *ets-4* | WBGene00017687 | protein_coding_gene | -1.1 | 0.473 | 1.9E-04 |
| *Y71G12B.6* | WBGene00022146 | protein_coding_gene | -1.1 | 0.473 | 1.6E-05 |
| *pab-2* | WBGene00003903 | protein_coding_gene | -1.1 | 0.473 | 1.6E-04 |
| *nhr-34* | WBGene00003627 | protein_coding_gene | -1.1 | 0.472 | 1.8E-08 |
| *ZC373.5* | WBGene00013870 | protein_coding_gene | -1.1 | 0.472 | 3.3E-08 |
| *T28F3.8* | WBGene00012134 | protein_coding_gene | -1.1 | 0.472 | 7.8E-08 |
| *R10E11.5* | WBGene00011217 | protein_coding_gene | -1.1 | 0.472 | 1.1E-03 |
| *ttr-38* | WBGene00044788 | protein_coding_gene | -1.1 | 0.472 | 4.6E-02 |
| *trp-2* | WBGene00006615 | protein_coding_gene | -1.1 | 0.471 | 5.3E-03 |
| *pkn-1* | WBGene00009793 | protein_coding_gene | -1.1 | 0.471 | 2.9E-05 |
| *set-30* | WBGene00022499 | protein_coding_gene | -1.1 | 0.471 | 4.8E-05 |
| *sdc-2* | WBGene00004746 | protein_coding_gene | -1.1 | 0.471 | 5.5E-06 |
| *pdf-1* | WBGene00020317 | protein_coding_gene | -1.1 | 0.471 | 1.2E-05 |
| *C15B12.4* | WBGene00015786 | protein_coding_gene | -1.1 | 0.470 | 3.5E-03 |
| *frpr-4* | WBGene00016909 | protein_coding_gene | -1.1 | 0.470 | 3.2E-02 |
| *trap-2* | WBGene00020216 | protein_coding_gene | -1.1 | 0.469 | 2.0E-05 |
| *H13N06.2* | WBGene00010396 | protein_coding_gene | -1.1 | 0.469 | 1.7E-02 |
| *F09A5.2* | WBGene00008599 | protein_coding_gene | -1.1 | 0.469 | 8.7E-03 |
| *fat-1* | WBGene00001393 | protein_coding_gene | -1.1 | 0.469 | 2.2E-08 |
| *C49A1.10* | WBGene00008191 | protein_coding_gene | -1.1 | 0.468 | 1.2E-03 |
| *lin-10* | WBGene00002999 | protein_coding_gene | -1.1 | 0.468 | 4.1E-08 |
| *gst-3* | WBGene00001751 | protein_coding_gene | -1.1 | 0.468 | 1.7E-02 |
| *vha-19* | WBGene00021952 | protein_coding_gene | -1.1 | 0.468 | 3.4E-08 |
| *sid-3* | WBGene00002207 | protein_coding_gene | -1.1 | 0.467 | 4.8E-05 |
| *B0244.10* | WBGene00015085 | protein_coding_gene | -1.1 | 0.467 | 4.2E-03 |
| *C05D10.4* | WBGene00015480 | protein_coding_gene | -1.1 | 0.467 | 2.7E-07 |
| *cal-3* | WBGene00000287 | protein_coding_gene | -1.1 | 0.467 | 2.5E-03 |
| *stau-1* | WBGene00018857 | protein_coding_gene | -1.1 | 0.467 | 1.7E-05 |
| *abu-14* | WBGene00004174 | protein_coding_gene | -1.1 | 0.466 | 2.1E-02 |
| *dmd-7* | WBGene00019521 | protein_coding_gene | -1.1 | 0.466 | 4.9E-06 |
| *ccb-1* | WBGene00000368 | protein_coding_gene | -1.1 | 0.466 | 2.3E-06 |
| *F15A4.6* | WBGene00008841 | protein_coding_gene | -1.1 | 0.466 | 7.8E-07 |
| *Y37D8A.2* | WBGene00012544 | protein_coding_gene | -1.1 | 0.466 | 2.3E-12 |
| *F58G1.7* | WBGene00010269 | protein_coding_gene | -1.1 | 0.465 | 2.3E-02 |
| *C07A12.2* | WBGene00015560 | protein_coding_gene | -1.1 | 0.465 | 3.6E-02 |
| *lido-8* | WBGene00019511 | protein_coding_gene | -1.1 | 0.465 | 2.3E-09 |
| *unc-112* | WBGene00006836 | protein_coding_gene | -1.1 | 0.465 | 7.5E-11 |
| *F47B7.2* | WBGene00018533 | protein_coding_gene | -1.1 | 0.465 | 1.1E-06 |
| *twk-40* | WBGene00006691 | protein_coding_gene | -1.1 | 0.464 | 3.4E-02 |
| *inx-16* | WBGene00002138 | protein_coding_gene | -1.1 | 0.464 | 3.7E-06 |
| *F43D2.7* | WBGene00045516 | protein_coding_gene | -1.1 | 0.464 | 1.2E-02 |
| *K08H2.10* | WBGene00164968 | protein_coding_gene | -1.1 | 0.464 | 7.5E-03 |
| *nhr-7* | WBGene00003606 | protein_coding_gene | -1.1 | 0.464 | 4.0E-03 |
| *ZK470.2* | WBGene00022745 | protein_coding_gene | -1.1 | 0.464 | 2.1E-04 |
| *F25E5.16* | WBGene00017792 | protein_coding_gene | -1.1 | 0.463 | 2.8E-02 |
| *linc-34* | WBGene00194814 | lincRNA_gene | -1.1 | 0.463 | 3.0E-02 |
| *suox-1* | WBGene00010397 | protein_coding_gene | -1.1 | 0.462 | 2.2E-07 |
| *npr-4* | WBGene00007635 | protein_coding_gene | -1.1 | 0.462 | 1.3E-02 |

| | | | | | |
|---|---|---|---|---|---|
| *F39G3.2* | WBGene00018207 | protein_coding_gene | -1.1 | 0.462 | 9.6E-05 |
| *acl-4* | WBGene00018657 | protein_coding_gene | -1.1 | 0.462 | 1.2E-11 |
| *Y39H10B.2* | WBGene00044527 | protein_coding_gene | -1.1 | 0.461 | 2.4E-10 |
| *Y11D7A.9* | WBGene00012433 | protein_coding_gene | -1.1 | 0.461 | 5.8E-03 |
| *ugt-50* | WBGene00011564 | protein_coding_gene | -1.1 | 0.461 | 6.7E-06 |
| *VM106R.1* | WBGene00014848 | protein_coding_gene | -1.1 | 0.460 | 3.7E-03 |
| *F28E10.1* | WBGene00017900 | protein_coding_gene | -1.1 | 0.460 | 1.1E-03 |
| *fbxa-71* | WBGene00020639 | protein_coding_gene | -1.1 | 0.459 | 8.7E-03 |
| *B0507.6* | WBGene00015223 | protein_coding_gene | -1.1 | 0.459 | 4.2E-02 |
| *R12E2.15* | WBGene00020040 | protein_coding_gene | -1.1 | 0.459 | 4.8E-02 |
| *lon-8* | WBGene00013352 | protein_coding_gene | -1.1 | 0.458 | 1.6E-04 |
| *slc-36.2* | WBGene00020837 | protein_coding_gene | -1.1 | 0.458 | 2.0E-05 |
| *ptrn-1* | WBGene00004121 | protein_coding_gene | -1.1 | 0.458 | 3.0E-05 |
| *slo-2* | WBGene00004831 | protein_coding_gene | -1.1 | 0.458 | 1.6E-07 |
| *lrch-1* | WBGene00015779 | protein_coding_gene | -1.1 | 0.458 | 3.0E-03 |
| *anr-35* | WBGene00219589 | antisense_lncRNA_gene | -1.1 | 0.457 | 8.6E-03 |
| *F57F5.1* | WBGene00010204 | protein_coding_gene | -1.1 | 0.457 | 3.4E-12 |
| *lrk-1* | WBGene00003068 | protein_coding_gene | -1.1 | 0.457 | 1.6E-06 |
| *F44E5.15* | WBGene00206377 | protein_coding_gene | -1.1 | 0.456 | 4.0E-02 |
| *dop-3* | WBGene00020506 | protein_coding_gene | -1.1 | 0.456 | 1.8E-02 |
| *F25H5.8* | WBGene00009130 | protein_coding_gene | -1.1 | 0.456 | 9.2E-06 |
| *flp-24* | WBGene00016028 | protein_coding_gene | -1.1 | 0.455 | 3.7E-09 |
| *unc-73* | WBGene00006805 | protein_coding_gene | -1.1 | 0.455 | 3.0E-06 |
| *linc-11* | WBGene00219758 | lincRNA_gene | -1.1 | 0.455 | 1.6E-02 |
| *cest-15* | WBGene00012202 | pseudogene | -1.1 | 0.454 | 5.0E-07 |
| *ncs-3* | WBGene00003565 | protein_coding_gene | -1.1 | 0.454 | 3.3E-04 |
| *fut-1* | WBGene00001505 | protein_coding_gene | -1.1 | 0.454 | 1.7E-10 |
| *ttm-5* | WBGene00013197 | protein_coding_gene | -1.1 | 0.453 | 1.3E-06 |
| *cyp-34A6* | WBGene00015041 | protein_coding_gene | -1.1 | 0.453 | 6.8E-03 |
| *M04C9.3* | WBGene00010858 | protein_coding_gene | -1.1 | 0.453 | 2.0E-04 |
| *C25E10.8* | WBGene00016097 | protein_coding_gene | -1.1 | 0.453 | 1.0E-03 |
| *T19D12.1* | WBGene00020576 | protein_coding_gene | -1.1 | 0.453 | 4.2E-03 |
| *Y57G11C.40* | WBGene00013329 | protein_coding_gene | -1.1 | 0.453 | 5.2E-03 |
| *F28C1.3* | WBGene00009201 | protein_coding_gene | -1.1 | 0.453 | 1.6E-05 |
| *C49C8.2* | WBGene00016766 | protein_coding_gene | -1.1 | 0.453 | 1.6E-02 |
| *lips-5* | WBGene00011725 | protein_coding_gene | -1.1 | 0.453 | 2.7E-03 |
| *T02D1.8* | WBGene00011374 | protein_coding_gene | -1.1 | 0.452 | 9.2E-07 |
| *K02E11.6* | WBGene00010515 | protein_coding_gene | -1.1 | 0.452 | 2.4E-04 |
| *C24A1.3* | WBGene00016030 | protein_coding_gene | -1.1 | 0.452 | 4.7E-06 |
| *F27D9.7* | WBGene00017865 | protein_coding_gene | -1.1 | 0.452 | 3.8E-03 |
| *H28G03.2* | WBGene00019250 | protein_coding_gene | -1.1 | 0.452 | 7.5E-06 |
| *K07A12.8* | WBGene00086557 | protein_coding_gene | -1.1 | 0.452 | 4.3E-03 |
| *ZK265.7* | WBGene00013959 | protein_coding_gene | -1.1 | 0.451 | 1.2E-03 |
| *wrt-6* | WBGene00006952 | protein_coding_gene | -1.2 | 0.450 | 3.4E-02 |
| *ftt-2* | WBGene00001502 | protein_coding_gene | -1.2 | 0.449 | 1.4E-09 |
| *eif-2alpha* | WBGene00021351 | protein_coding_gene | -1.2 | 0.449 | 3.2E-14 |
| *cdr-6* | WBGene00010471 | protein_coding_gene | -1.2 | 0.449 | 2.1E-06 |
| *F36H5.14* | WBGene00219271 | protein_coding_gene | -1.2 | 0.448 | 2.0E-02 |
| *cnp-2* | WBGene00012901 | protein_coding_gene | -1.2 | 0.448 | 1.2E-04 |
| *R05A10.3* | WBGene00011020 | protein_coding_gene | -1.2 | 0.448 | 2.7E-03 |
| *T04A8.18* | WBGene00119203 | protein_coding_gene | -1.2 | 0.448 | 3.5E-03 |
| *sec-61.A* | WBGene00013311 | protein_coding_gene | -1.2 | 0.446 | 6.5E-06 |
| *wee-1.1* | WBGene00006938 | protein_coding_gene | -1.2 | 0.446 | 2.7E-02 |
| *oct-2* | WBGene00003843 | protein_coding_gene | -1.2 | 0.446 | 8.3E-05 |
| *Y14H12A.1* | WBGene00021190 | protein_coding_gene | -1.2 | 0.446 | 1.6E-05 |
| *pmp-2* | WBGene00004059 | protein_coding_gene | -1.2 | 0.445 | 2.9E-13 |
| *frpr-7* | WBGene00018191 | protein_coding_gene | -1.2 | 0.445 | 5.0E-03 |
| *lonp-2* | WBGene00013541 | protein_coding_gene | -1.2 | 0.445 | 1.7E-06 |
| *mrp-5* | WBGene00003411 | protein_coding_gene | -1.2 | 0.445 | 1.3E-07 |
| *C07F11.2* | WBGene00015570 | protein_coding_gene | -1.2 | 0.444 | 4.9E-02 |
| *H01G02.3* | WBGene00010350 | protein_coding_gene | -1.2 | 0.444 | 3.1E-02 |
| *F53B1.2* | WBGene00018735 | protein_coding_gene | -1.2 | 0.444 | 8.6E-06 |
| *sek-1* | WBGene00004758 | protein_coding_gene | -1.2 | 0.443 | 3.9E-15 |
| *K09F6.6* | WBGene00019589 | protein_coding_gene | -1.2 | 0.443 | 2.2E-03 |
| *cyp-14A5* | WBGene00000373 | protein_coding_gene | -1.2 | 0.443 | 2.1E-06 |
| *R02F11.2* | WBGene00019840 | protein_coding_gene | -1.2 | 0.442 | 3.0E-02 |
| *C39D10.2* | WBGene00016530 | protein_coding_gene | -1.2 | 0.442 | 2.0E-02 |
| *nhr-153* | WBGene00007546 | protein_coding_gene | -1.2 | 0.442 | 1.4E-04 |
| *fipr-5* | WBGene00044174 | protein_coding_gene | -1.2 | 0.441 | 1.4E-03 |
| *fbxa-168* | WBGene00007838 | pseudogene | -1.2 | 0.441 | 3.0E-03 |

| | | | | | |
|---|---|---|---|---|---|
| *sdz-30* | WBGene00011432 | protein_coding_gene | -1.2 | 0.441 | 1.1E-02 |
| *dop-5* | WBGene00011382 | protein_coding_gene | -1.2 | 0.441 | 8.3E-03 |
| *F39C12.1* | WBGene00018193 | protein_coding_gene | -1.2 | 0.440 | 8.1E-04 |
| *dhhc-14* | WBGene00044071 | protein_coding_gene | -1.2 | 0.440 | 1.5E-09 |
| *T23C6.4* | WBGene00020726 | protein_coding_gene | -1.2 | 0.440 | 5.1E-05 |
| *twk-43* | WBGene00006693 | protein_coding_gene | -1.2 | 0.440 | 2.0E-04 |
| *elt-2* | WBGene00001250 | protein_coding_gene | -1.2 | 0.440 | 1.5E-05 |
| *amph-1* | WBGene00010272 | protein_coding_gene | -1.2 | 0.439 | 6.5E-24 |
| *acly-1* | WBGene00016995 | protein_coding_gene | -1.2 | 0.439 | 6.9E-07 |
| *gpa-2* | WBGene00001664 | protein_coding_gene | -1.2 | 0.439 | 3.8E-02 |
| *pqn-31* | WBGene00004119 | protein_coding_gene | -1.2 | 0.439 | 4.1E-03 |
| *nhr-162* | WBGene00016367 | protein_coding_gene | -1.2 | 0.438 | 2.3E-02 |
| *cytb-5.1* | WBGene00007848 | protein_coding_gene | -1.2 | 0.438 | 8.8E-07 |
| *F55G1.15* | WBGene00018907 | protein_coding_gene | -1.2 | 0.438 | 1.2E-07 |
| *seb-3* | WBGene00007664 | protein_coding_gene | -1.2 | 0.437 | 4.1E-03 |
| *Y49E10.30* | WBGene00194981 | protein_coding_gene | -1.2 | 0.437 | 2.0E-02 |
| *C06A8.8* | WBGene00015517 | protein_coding_gene | -1.2 | 0.437 | 9.2E-07 |
| *rgl-1* | WBGene00017891 | protein_coding_gene | -1.2 | 0.437 | 1.2E-12 |
| *dlc-2* | WBGene00010888 | protein_coding_gene | -1.2 | 0.437 | 1.7E-06 |
| *mrp-8* | WBGene00003414 | protein_coding_gene | -1.2 | 0.436 | 3.4E-09 |
| *ccdc-149* | WBGene00009258 | protein_coding_gene | -1.2 | 0.436 | 4.7E-03 |
| *B0222.1* | WBGene00015053 | protein_coding_gene | -1.2 | 0.436 | 4.2E-02 |
| *F09E10.15* | WBGene00219886 | ncRNA_gene | -1.2 | 0.436 | 2.6E-03 |
| *R07B1.11* | WBGene00011082 | protein_coding_gene | -1.2 | 0.435 | 2.5E-02 |
| *flp-33* | WBGene00011586 | protein_coding_gene | -1.2 | 0.434 | 7.5E-05 |
| *C15C7.4* | WBGene00015790 | protein_coding_gene | -1.2 | 0.434 | 1.8E-02 |
| *pcca-1* | WBGene00017864 | protein_coding_gene | -1.2 | 0.434 | 3.9E-12 |
| *mif-2* | WBGene00003235 | protein_coding_gene | -1.2 | 0.434 | 5.8E-07 |
| *glb-3* | WBGene00007389 | protein_coding_gene | -1.2 | 0.433 | 1.0E-02 |
| *flp-3* | WBGene00001446 | protein_coding_gene | -1.2 | 0.433 | 2.4E-06 |
| *Y73B6A.3* | WBGene00022230 | protein_coding_gene | -1.2 | 0.433 | 2.1E-02 |
| *epg-3* | WBGene00012559 | protein_coding_gene | -1.2 | 0.432 | 2.4E-16 |
| *F11C7.2* | WBGene00017376 | protein_coding_gene | -1.2 | 0.432 | 3.9E-03 |
| *unc-63* | WBGene00006797 | protein_coding_gene | -1.2 | 0.432 | 1.2E-04 |
| *ile-2* | WBGene00002071 | protein_coding_gene | -1.2 | 0.432 | 2.5E-12 |
| *linc-7* | WBGene00219609 | lincRNA_gene | -1.2 | 0.432 | 1.4E-06 |
| *tre-4* | WBGene00006610 | protein_coding_gene | -1.2 | 0.432 | 1.2E-05 |
| *hpo-26* | WBGene00018492 | protein_coding_gene | -1.2 | 0.432 | 1.5E-03 |
| *galt-1* | WBGene00019763 | protein_coding_gene | -1.2 | 0.431 | 8.6E-03 |
| *C08F11.1* | WBGene00007450 | protein_coding_gene | -1.2 | 0.431 | 3.2E-03 |
| *ptr-18* | WBGene00004232 | protein_coding_gene | -1.2 | 0.430 | 5.3E-03 |
| *T13G4.4* | WBGene00020491 | protein_coding_gene | -1.2 | 0.430 | 1.3E-07 |
| *T25C12.3* | WBGene00012018 | protein_coding_gene | -1.2 | 0.430 | 8.5E-05 |
| *ZK380.6* | WBGene00304812 | protein_coding_gene | -1.2 | 0.430 | 3.2E-04 |
| *arrd-22* | WBGene00020612 | protein_coding_gene | -1.2 | 0.429 | 9.6E-04 |
| *sipa-1* | WBGene00012100 | protein_coding_gene | -1.2 | 0.429 | 8.5E-09 |
| *F42A10.9* | WBGene00023489 | protein_coding_gene | -1.2 | 0.429 | 3.4E-04 |
| *lgx-1* | WBGene00002983 | protein_coding_gene | -1.2 | 0.429 | 1.2E-03 |
| *catp-5* | WBGene00019493 | protein_coding_gene | -1.2 | 0.428 | 2.4E-04 |
| *exp-2* | WBGene00001374 | protein_coding_gene | -1.2 | 0.428 | 5.5E-04 |
| *Y106G6G.2* | WBGene00013711 | protein_coding_gene | -1.2 | 0.428 | 1.8E-04 |
| *unc-49* | WBGene00006784 | protein_coding_gene | -1.2 | 0.427 | 4.4E-04 |
| *ZK380.t2* | WBGene00023401 | pseudogene | -1.2 | 0.427 | 4.0E-03 |
| *klu-1* | WBGene00013970 | protein_coding_gene | -1.2 | 0.427 | 1.0E-02 |
| *zig-9* | WBGene00022474 | protein_coding_gene | -1.2 | 0.427 | 1.0E-05 |
| *klf-3* | WBGene00003480 | protein_coding_gene | -1.2 | 0.426 | 5.2E-04 |
| *K09A9.8* | WBGene00194849 | protein_coding_gene | -1.2 | 0.426 | 3.7E-03 |
| *F52E10.4* | WBGene00009935 | protein_coding_gene | -1.2 | 0.426 | 3.2E-03 |
| *dnj-20* | WBGene00001038 | protein_coding_gene | -1.2 | 0.426 | 6.8E-07 |
| *plpp-1.3* | WBGene00008749 | protein_coding_gene | -1.2 | 0.425 | 2.7E-08 |
| *pghm-1* | WBGene00022144 | protein_coding_gene | -1.2 | 0.425 | 6.9E-07 |
| *hosl-1* | WBGene00016704 | protein_coding_gene | -1.2 | 0.425 | 4.4E-07 |
| *M153.1* | WBGene00010924 | protein_coding_gene | -1.2 | 0.425 | 3.6E-04 |
| *H10D18.5* | WBGene00019180 | protein_coding_gene | -1.2 | 0.425 | 9.4E-04 |
| *vet-6* | WBGene00006902 | protein_coding_gene | -1.2 | 0.425 | 1.9E-02 |
| *ZC239.6* | WBGene00022569 | protein_coding_gene | -1.2 | 0.425 | 1.7E-04 |
| *sng-1* | WBGene00004912 | protein_coding_gene | -1.2 | 0.425 | 5.1E-04 |
| *ins-12* | WBGene00002095 | protein_coding_gene | -1.2 | 0.424 | 4.0E-02 |
| *set-18* | WBGene00044070 | protein_coding_gene | -1.2 | 0.424 | 5.8E-08 |
| *C18B2.5* | WBGene00015956 | protein_coding_gene | -1.2 | 0.424 | 5.4E-09 |

| | | | | | |
|---|---|---|---|---|---|
| *Y38C1BA.1* | WBGene00021410 | protein_coding_gene | -1.2 | 0.424 | 1.8E-02 |
| *ttr-27* | WBGene00011284 | protein_coding_gene | -1.2 | 0.424 | 8.6E-03 |
| *C24B5.1* | WBGene00016044 | protein_coding_gene | -1.2 | 0.423 | 8.8E-03 |
| *K11C4.2* | WBGene00019643 | protein_coding_gene | -1.2 | 0.423 | 9.0E-05 |
| *axl-1* | WBGene00010497 | protein_coding_gene | -1.2 | 0.423 | 7.2E-04 |
| *F42G4.5* | WBGene00009642 | protein_coding_gene | -1.2 | 0.423 | 5.3E-08 |
| *ldp-1* | WBGene00017719 | protein_coding_gene | -1.2 | 0.423 | 4.6E-06 |
| *bus-19* | WBGene00011590 | protein_coding_gene | -1.2 | 0.422 | 1.5E-04 |
| *cest-9.2* | WBGene00007695 | protein_coding_gene | -1.2 | 0.422 | 1.3E-05 |
| *C04G2.10* | WBGene00007309 | protein_coding_gene | -1.2 | 0.422 | 4.4E-05 |
| *C31C9.2* | WBGene00007836 | protein_coding_gene | -1.2 | 0.422 | 2.7E-08 |
| *pyr-1* | WBGene00004259 | protein_coding_gene | -1.2 | 0.421 | 1.8E-08 |
| *hlb-1* | WBGene00011904 | protein_coding_gene | -1.2 | 0.421 | 1.8E-09 |
| *F47H4.2* | WBGene00009835 | protein_coding_gene | -1.3 | 0.420 | 2.2E-02 |
| *art-1* | WBGene00000198 | protein_coding_gene | -1.3 | 0.420 | 7.7E-09 |
| *svh-1* | WBGene00006620 | protein_coding_gene | -1.3 | 0.419 | 3.1E-03 |
| *nlp-6* | WBGene00003744 | protein_coding_gene | -1.3 | 0.419 | 9.4E-08 |
| *pdfr-1* | WBGene00015735 | protein_coding_gene | -1.3 | 0.419 | 1.7E-07 |
| *tni-1* | WBGene00006584 | protein_coding_gene | -1.3 | 0.419 | 6.6E-06 |
| *R07E3.7* | WBGene00011108 | protein_coding_gene | -1.3 | 0.419 | 5.3E-03 |
| *mel-32* | WBGene00003214 | protein_coding_gene | -1.3 | 0.419 | 4.3E-07 |
| *egl-36* | WBGene00001202 | protein_coding_gene | -1.3 | 0.419 | 9.0E-04 |
| *anc-1* | WBGene00000140 | protein_coding_gene | -1.3 | 0.418 | 4.3E-07 |
| *arrd-15* | WBGene00014033 | protein_coding_gene | -1.3 | 0.418 | 1.4E-02 |
| *trap-1* | WBGene00022122 | protein_coding_gene | -1.3 | 0.418 | 1.4E-08 |
| *srr-1* | WBGene00005652 | protein_coding_gene | -1.3 | 0.418 | 1.6E-02 |
| *slcf-1* | WBGene00010340 | protein_coding_gene | -1.3 | 0.417 | 9.6E-07 |
| *idh-2* | WBGene00007942 | protein_coding_gene | -1.3 | 0.417 | 3.8E-14 |
| *pfn-2* | WBGene00003990 | protein_coding_gene | -1.3 | 0.417 | 5.8E-14 |
| *C04A11.1* | WBGene00007285 | protein_coding_gene | -1.3 | 0.417 | 3.1E-02 |
| *pqn-25* | WBGene00004114 | protein_coding_gene | -1.3 | 0.416 | 1.4E-04 |
| *F22F7.7* | WBGene00017724 | protein_coding_gene | -1.3 | 0.416 | 1.4E-06 |
| *Y69A2AR.22* | WBGene00022093 | protein_coding_gene | -1.3 | 0.416 | 6.4E-07 |
| *F25E5.5* | WBGene00017786 | protein_coding_gene | -1.3 | 0.415 | 8.9E-05 |
| *C49A9.9* | WBGene00016763 | protein_coding_gene | -1.3 | 0.415 | 1.8E-09 |
| *Y47H10A.4* | WBGene00012960 | protein_coding_gene | -1.3 | 0.415 | 3.1E-03 |
| *acdh-6* | WBGene00015335 | protein_coding_gene | -1.3 | 0.415 | 3.8E-02 |
| *Y9C2UA.1* | WBGene00012421 | protein_coding_gene | -1.3 | 0.415 | 1.1E-06 |
| *flr-2* | WBGene00001466 | protein_coding_gene | -1.3 | 0.415 | 1.0E-04 |
| *clh-2* | WBGene00000529 | protein_coding_gene | -1.3 | 0.415 | 3.8E-03 |
| *nspc-18* | WBGene00008358 | protein_coding_gene | -1.3 | 0.415 | 3.8E-04 |
| *txdc-12.2* | WBGene00018656 | protein_coding_gene | -1.3 | 0.414 | 3.5E-07 |
| *CE7X_3.2* | WBGene00008350 | pseudogene | -1.3 | 0.414 | 2.8E-02 |
| *pitr-5* | WBGene00017312 | protein_coding_gene | -1.3 | 0.414 | 1.0E-04 |
| *Y106G6H.1* | WBGene00013716 | protein_coding_gene | -1.3 | 0.413 | 2.0E-06 |
| *nspc-7* | WBGene00015392 | protein_coding_gene | -1.3 | 0.413 | 1.5E-02 |
| *C06C3.10* | WBGene00044169 | protein_coding_gene | -1.3 | 0.413 | 3.6E-02 |
| *bcl-11* | WBGene00017430 | protein_coding_gene | -1.3 | 0.413 | 5.0E-04 |
| *lem-2* | WBGene00002275 | protein_coding_gene | -1.3 | 0.413 | 9.1E-11 |
| *clec-199* | WBGene00007820 | protein_coding_gene | -1.3 | 0.412 | 3.4E-02 |
| *apa-2* | WBGene00000161 | protein_coding_gene | -1.3 | 0.412 | 7.3E-08 |
| *acl-1* | WBGene00010339 | protein_coding_gene | -1.3 | 0.412 | 1.2E-10 |
| *F59C6.10* | WBGene00014789 | pseudogene | -1.3 | 0.411 | 2.4E-02 |
| *ghi-1* | WBGene00013923 | protein_coding_gene | -1.3 | 0.410 | 1.3E-02 |
| *unc-23* | WBGene00006760 | protein_coding_gene | -1.3 | 0.410 | 6.6E-04 |
| *C37H5.13* | WBGene00016511 | protein_coding_gene | -1.3 | 0.410 | 6.4E-13 |
| *flp-22* | WBGene00009562 | protein_coding_gene | -1.3 | 0.410 | 1.4E-07 |
| *ndnf-1* | WBGene00021893 | protein_coding_gene | -1.3 | 0.410 | 1.1E-04 |
| *T19D2.3* | WBGene00020568 | protein_coding_gene | -1.3 | 0.409 | 5.5E-03 |
| *C15H9.4* | WBGene00015800 | protein_coding_gene | -1.3 | 0.409 | 2.9E-14 |
| *atrn-1* | WBGene00006432 | protein_coding_gene | -1.3 | 0.409 | 1.1E-09 |
| *gst-28* | WBGene00001776 | protein_coding_gene | -1.3 | 0.409 | 2.0E-04 |
| *ctsa-3.2* | WBGene00007462 | protein_coding_gene | -1.3 | 0.408 | 3.1E-02 |
| *F36D4.4* | WBGene00018089 | protein_coding_gene | -1.3 | 0.408 | 4.8E-02 |
| *zfh-2* | WBGene00022518 | protein_coding_gene | -1.3 | 0.407 | 1.0E-04 |
| *Y57A10B.6* | WBGene00013278 | protein_coding_gene | -1.3 | 0.407 | 1.7E-04 |
| *tyra-2* | WBGene00017157 | protein_coding_gene | -1.3 | 0.407 | 6.2E-03 |
| *F58E2.4* | WBGene00019048 | protein_coding_gene | -1.3 | 0.406 | 6.0E-04 |
| *W02D9.10* | WBGene00012216 | protein_coding_gene | -1.3 | 0.406 | 1.5E-07 |
| *C49G7.3* | WBGene00016781 | protein_coding_gene | -1.3 | 0.406 | 9.3E-07 |

| | | | | | |
|---|---|---|---|---|---|
| *pcyt-1* | WBGene00017241 | protein_coding_gene | -1.3 | 0.405 | 8.3E-06 |
| *flp-4* | WBGene00001447 | protein_coding_gene | -1.3 | 0.404 | 2.0E-05 |
| *lgc-39* | WBGene00017314 | protein_coding_gene | -1.3 | 0.404 | 1.1E-03 |
| *ida-1* | WBGene00002048 | protein_coding_gene | -1.3 | 0.404 | 9.9E-05 |
| *npr-29* | WBGene00013848 | protein_coding_gene | -1.3 | 0.404 | 8.2E-03 |
| *ahcy-1* | WBGene00019322 | protein_coding_gene | -1.3 | 0.403 | 1.4E-06 |
| *asg-2* | WBGene00000210 | protein_coding_gene | -1.3 | 0.402 | 5.5E-06 |
| *C33G8.4* | WBGene00016362 | protein_coding_gene | -1.3 | 0.402 | 4.6E-07 |
| *F11E6.6* | WBGene00008709 | protein_coding_gene | -1.3 | 0.402 | 1.9E-02 |
| *nlp-7* | WBGene00003745 | protein_coding_gene | -1.3 | 0.402 | 2.5E-07 |
| *pgp-15* | WBGene00004009 | pseudogene | -1.3 | 0.402 | 1.2E-02 |
| *cdkl-1* | WBGene00012779 | protein_coding_gene | -1.3 | 0.402 | 4.3E-02 |
| *C14H10.3* | WBGene00007593 | protein_coding_gene | -1.3 | 0.402 | 1.4E-08 |
| *M04C9.4* | WBGene00010859 | protein_coding_gene | -1.3 | 0.402 | 6.8E-05 |
| *R03D7.3* | WBGene00014814 | pseudogene | -1.3 | 0.401 | 3.5E-02 |
| *unc-25* | WBGene00006762 | protein_coding_gene | -1.3 | 0.401 | 2.8E-05 |
| *fbxa-180* | WBGene00009295 | pseudogene | -1.3 | 0.401 | 4.3E-05 |
| *kvs-2* | WBGene00007862 | protein_coding_gene | -1.3 | 0.401 | 6.5E-03 |
| *nlp-55* | WBGene00008820 | protein_coding_gene | -1.3 | 0.400 | 1.0E-04 |
| *klp-13* | WBGene00002224 | protein_coding_gene | -1.3 | 0.400 | 1.2E-02 |
| *nhr-80* | WBGene00003670 | protein_coding_gene | -1.3 | 0.400 | 7.3E-09 |
| *acr-12* | WBGene00000051 | protein_coding_gene | -1.3 | 0.399 | 5.9E-04 |
| *F27B10.1* | WBGene00017850 | protein_coding_gene | -1.3 | 0.399 | 3.7E-03 |
| *Y34B4A.5* | WBGene00021321 | protein_coding_gene | -1.3 | 0.399 | 1.9E-05 |
| *Y57G7A.1* | WBGene00021965 | protein_coding_gene | -1.3 | 0.399 | 1.5E-03 |
| *T25B9.9* | WBGene00012015 | protein_coding_gene | -1.3 | 0.398 | 2.1E-07 |
| *egl-47* | WBGene00001211 | protein_coding_gene | -1.3 | 0.398 | 3.9E-02 |
| *aakg-2* | WBGene00020633 | protein_coding_gene | -1.3 | 0.398 | 8.3E-14 |
| *aat-5* | WBGene00000006 | protein_coding_gene | -1.3 | 0.397 | 7.5E-04 |
| *grk-2* | WBGene00001709 | protein_coding_gene | -1.3 | 0.397 | 6.6E-09 |
| *abf-6* | WBGene00000017 | protein_coding_gene | -1.3 | 0.397 | 1.9E-04 |
| *alp-1* | WBGene00001132 | protein_coding_gene | -1.3 | 0.397 | 1.2E-06 |
| *trpp-12* | WBGene00016614 | protein_coding_gene | -1.3 | 0.396 | 2.2E-08 |
| *aqp-9* | WBGene00000177 | protein_coding_gene | -1.3 | 0.396 | 3.3E-06 |
| *haf-4* | WBGene00001814 | protein_coding_gene | -1.3 | 0.395 | 6.3E-13 |
| *ugt-26* | WBGene00015695 | protein_coding_gene | -1.3 | 0.394 | 1.3E-05 |
| *acr-24* | WBGene00013534 | protein_coding_gene | -1.3 | 0.394 | 2.8E-02 |
| *Y102A11A.3* | WBGene00022413 | protein_coding_gene | -1.3 | 0.393 | 1.4E-06 |
| *srb-16* | WBGene00019025 | protein_coding_gene | -1.3 | 0.393 | 3.9E-02 |
| *Y55D5A.4* | WBGene00021918 | protein_coding_gene | -1.3 | 0.393 | 1.5E-02 |
| *F31B12.3* | WBGene00001495 | protein_coding_gene | -1.3 | 0.393 | 4.8E-03 |
| *C25G6.3* | WBGene00016108 | protein_coding_gene | -1.3 | 0.393 | 4.6E-02 |
| *cri-1* | WBGene00010614 | protein_coding_gene | -1.4 | 0.392 | 2.0E-02 |
| *F09C8.2* | WBGene00008622 | protein_coding_gene | -1.4 | 0.391 | 1.7E-06 |
| *nspc-9* | WBGene00009634 | protein_coding_gene | -1.4 | 0.391 | 1.0E-04 |
| *flp-5* | WBGene00001448 | protein_coding_gene | -1.4 | 0.391 | 2.1E-07 |
| *F29G9.1* | WBGene00017930 | protein_coding_gene | -1.4 | 0.391 | 2.3E-02 |
| *ttr-33* | WBGene00016505 | protein_coding_gene | -1.4 | 0.391 | 2.6E-06 |
| *R13A1.5* | WBGene00020043 | protein_coding_gene | -1.4 | 0.391 | 3.7E-03 |
| *F57A8.4* | WBGene00010179 | protein_coding_gene | -1.4 | 0.391 | 2.3E-02 |
| *gon-2* | WBGene00001651 | protein_coding_gene | -1.4 | 0.391 | 3.2E-09 |
| *glb-18* | WBGene00009917 | protein_coding_gene | -1.4 | 0.391 | 1.3E-05 |
| *R07E3.2* | WBGene00011103 | protein_coding_gene | -1.4 | 0.390 | 3.8E-02 |
| *gln-1* | WBGene00001602 | protein_coding_gene | -1.4 | 0.390 | 2.7E-16 |
| *C27A2.12* | WBGene00206373 | protein_coding_gene | -1.4 | 0.390 | 5.3E-04 |
| *asp-3* | WBGene00000216 | protein_coding_gene | -1.4 | 0.390 | 5.5E-15 |
| *mctp-1* | WBGene00017063 | protein_coding_gene | -1.4 | 0.390 | 4.6E-09 |
| *odr-3* | WBGene00003850 | protein_coding_gene | -1.4 | 0.389 | 1.6E-03 |
| *sid-2* | WBGene00004796 | protein_coding_gene | -1.4 | 0.389 | 1.1E-06 |
| *C10G8.8* | WBGene00015684 | protein_coding_gene | -1.4 | 0.389 | 1.2E-07 |
| *aat-6* | WBGene00000007 | protein_coding_gene | -1.4 | 0.388 | 1.6E-10 |
| *dct-6* | WBGene00017773 | protein_coding_gene | -1.4 | 0.388 | 7.3E-14 |
| *aex-1* | WBGene00000084 | protein_coding_gene | -1.4 | 0.388 | 2.1E-09 |
| *F57B7.2* | WBGene00010192 | protein_coding_gene | -1.4 | 0.388 | 6.5E-06 |
| *D1044.7* | WBGene00017032 | protein_coding_gene | -1.4 | 0.388 | 7.7E-03 |
| *DH11.5* | WBGene00008438 | protein_coding_gene | -1.4 | 0.388 | 7.8E-08 |
| *nlp-43* | WBGene00016684 | protein_coding_gene | -1.4 | 0.388 | 7.2E-06 |
| *unc-17* | WBGene00006756 | protein_coding_gene | -1.4 | 0.387 | 2.5E-04 |
| *pll-1* | WBGene00004045 | protein_coding_gene | -1.4 | 0.387 | 1.3E-03 |
| *H10E21.5* | WBGene00019185 | protein_coding_gene | -1.4 | 0.387 | 2.0E-04 |

| | | | | | |
|---|---|---|---|---|---|
| *twk-9* | WBGene00006664 | protein_coding_gene | -1.4 | 0.386 | 2.8E-02 |
| *C18B2.4* | WBGene00015955 | protein_coding_gene | -1.4 | 0.386 | 1.4E-12 |
| *F35F10.4* | WBGene00018054 | pseudogene | -1.4 | 0.386 | 1.3E-02 |
| *ZK652.8* | WBGene00022786 | protein_coding_gene | -1.4 | 0.386 | 3.5E-07 |
| *snt-4* | WBGene00004924 | protein_coding_gene | -1.4 | 0.386 | 4.6E-06 |
| *suca-1* | WBGene00009812 | protein_coding_gene | -1.4 | 0.385 | 1.4E-11 |
| *nhr-263* | WBGene00017367 | protein_coding_gene | -1.4 | 0.385 | 3.4E-02 |
| *rbc-1* | WBGene00004313 | protein_coding_gene | -1.4 | 0.385 | 9.2E-06 |
| *mec-2* | WBGene00003166 | protein_coding_gene | -1.4 | 0.385 | 9.3E-05 |
| *abu-12* | WBGene00004127 | protein_coding_gene | -1.4 | 0.385 | 1.3E-02 |
| *mab-9* | WBGene00003106 | protein_coding_gene | -1.4 | 0.384 | 2.5E-02 |
| *twk-48* | WBGene00010784 | protein_coding_gene | -1.4 | 0.384 | 2.3E-02 |
| *cest-26* | WBGene00017478 | protein_coding_gene | -1.4 | 0.384 | 2.9E-06 |
| *K02E11.5* | WBGene00010514 | protein_coding_gene | -1.4 | 0.383 | 5.0E-05 |
| *mocs-1* | WBGene00010983 | protein_coding_gene | -1.4 | 0.383 | 1.6E-09 |
| *gab-1* | WBGene00001512 | protein_coding_gene | -1.4 | 0.383 | 1.8E-02 |
| *M7.12* | WBGene00010887 | protein_coding_gene | -1.4 | 0.382 | 2.7E-07 |
| *F54G2.1* | WBGene00018837 | protein_coding_gene | -1.4 | 0.382 | 2.7E-08 |
| *srx-125* | WBGene00006016 | protein_coding_gene | -1.4 | 0.382 | 4.2E-02 |
| *C35B1.7* | WBGene00016436 | protein_coding_gene | -1.4 | 0.382 | 3.5E-02 |
| *C50D2.6* | WBGene00016809 | protein_coding_gene | -1.4 | 0.382 | 1.0E-04 |
| *T12G3.2* | WBGene00011738 | protein_coding_gene | -1.4 | 0.381 | 7.7E-13 |
| *Y32G9B.1* | WBGene00021309 | protein_coding_gene | -1.4 | 0.381 | 2.2E-03 |
| *ugt-16* | WBGene00013901 | protein_coding_gene | -1.4 | 0.381 | 2.1E-06 |
| *npr-30* | WBGene00019183 | protein_coding_gene | -1.4 | 0.380 | 3.8E-03 |
| *tmc-1* | WBGene00020490 | protein_coding_gene | -1.4 | 0.380 | 4.9E-05 |
| *dlk-1* | WBGene00001008 | protein_coding_gene | -1.4 | 0.380 | 5.5E-11 |
| *Y39B6A.41* | WBGene00012699 | protein_coding_gene | -1.4 | 0.380 | 1.5E-02 |
| *acr-25* | WBGene00043066 | protein_coding_gene | -1.4 | 0.380 | 2.8E-02 |
| *F38B2.6* | WBGene00194663 | protein_coding_gene | -1.4 | 0.380 | 9.5E-03 |
| *his-40* | WBGene00001914 | protein_coding_gene | -1.4 | 0.379 | 9.4E-03 |
| *pgrn-1* | WBGene00011936 | protein_coding_gene | -1.4 | 0.379 | 3.5E-13 |
| *unc-103* | WBGene00006830 | protein_coding_gene | -1.4 | 0.379 | 2.0E-06 |
| *glb-29* | WBGene00012461 | protein_coding_gene | -1.4 | 0.379 | 1.8E-02 |
| *ZK822.4* | WBGene00014091 | protein_coding_gene | -1.4 | 0.379 | 1.7E-14 |
| *nhr-147* | WBGene00015395 | protein_coding_gene | -1.4 | 0.379 | 3.4E-02 |
| *T19D2.2* | WBGene00004184 | protein_coding_gene | -1.4 | 0.378 | 2.2E-04 |
| *ZK622.4* | WBGene00022782 | protein_coding_gene | -1.4 | 0.378 | 5.0E-06 |
| *E02A10.4* | WBGene00008454 | protein_coding_gene | -1.4 | 0.378 | 2.3E-03 |
| *oac-59* | WBGene00016585 | protein_coding_gene | -1.4 | 0.378 | 1.2E-10 |
| *rdy-2* | WBGene00004328 | protein_coding_gene | -1.4 | 0.377 | 1.9E-02 |
| *dat-1* | WBGene00000934 | protein_coding_gene | -1.4 | 0.377 | 3.3E-03 |
| *F46C8.8* | WBGene00018487 | protein_coding_gene | -1.4 | 0.377 | 3.0E-04 |
| *ccm-2* | WBGene00019462 | protein_coding_gene | -1.4 | 0.377 | 5.3E-05 |
| *C55A6.6* | WBGene00008335 | protein_coding_gene | -1.4 | 0.377 | 1.1E-07 |
| *C31H5.5* | WBGene00007856 | protein_coding_gene | -1.4 | 0.377 | 7.4E-06 |
| *Y45G12C.1* | WBGene00021565 | protein_coding_gene | -1.4 | 0.377 | 1.2E-02 |
| *F10G2.1* | WBGene00017362 | protein_coding_gene | -1.4 | 0.377 | 1.5E-17 |
| *H18N23.2* | WBGene00019211 | protein_coding_gene | -1.4 | 0.377 | 2.9E-05 |
| *B0361.4* | WBGene00015158 | protein_coding_gene | -1.4 | 0.376 | 1.3E-02 |
| *T05E7.1* | WBGene00020258 | protein_coding_gene | -1.4 | 0.376 | 2.1E-11 |
| *nlg-1* | WBGene00006412 | protein_coding_gene | -1.4 | 0.376 | 6.2E-05 |
| *M03E7.1* | WBGene00019753 | protein_coding_gene | -1.4 | 0.375 | 8.1E-03 |
| *H41C03.3* | WBGene00019270 | protein_coding_gene | -1.4 | 0.375 | 7.3E-07 |
| *Y119D3B.21* | WBGene00022497 | protein_coding_gene | -1.4 | 0.375 | 7.0E-05 |
| *vha-13* | WBGene00013025 | protein_coding_gene | -1.4 | 0.375 | 9.6E-09 |
| *inx-10* | WBGene00002132 | protein_coding_gene | -1.4 | 0.375 | 9.7E-15 |
| *F54F12.2* | WBGene00010073 | protein_coding_gene | -1.4 | 0.374 | 1.3E-02 |
| *C44B7.7* | WBGene00016629 | protein_coding_gene | -1.4 | 0.373 | 1.6E-04 |
| *C02B8.12* | WBGene00219375 | protein_coding_gene | -1.4 | 0.373 | 1.3E-03 |
| *nlp-3* | WBGene00003741 | protein_coding_gene | -1.4 | 0.373 | 6.0E-07 |
| *ZK1240.11* | WBGene00206413 | protein_coding_gene | -1.4 | 0.372 | 9.6E-03 |
| *nlp-19* | WBGene00003757 | protein_coding_gene | -1.4 | 0.372 | 3.0E-02 |
| *arrd-17* | WBGene00011732 | protein_coding_gene | -1.4 | 0.371 | 1.0E-04 |
| *daf-11* | WBGene00000907 | protein_coding_gene | -1.4 | 0.371 | 3.2E-02 |
| *C36B7.5* | WBGene00016468 | protein_coding_gene | -1.4 | 0.371 | 6.0E-03 |
| *fbxb-105* | WBGene00017251 | protein_coding_gene | -1.4 | 0.371 | 1.8E-03 |
| *nspc-10* | WBGene00009640 | protein_coding_gene | -1.4 | 0.371 | 1.9E-07 |
| *cyp-25A1* | WBGene00007963 | protein_coding_gene | -1.4 | 0.371 | 3.0E-02 |
| *B0416.11* | WBGene00206483 | protein_coding_gene | -1.4 | 0.371 | 3.1E-02 |

| | | | | | |
|---|---|---|---|---|---|
| *frpr-17* | WBGene00011765 | protein_coding_gene | -1.4 | 0.370 | 1.2E-03 |
| *Y55D5A.3* | WBGene00021917 | protein_coding_gene | -1.4 | 0.370 | 4.5E-04 |
| *feh-1* | WBGene00001410 | protein_coding_gene | -1.4 | 0.370 | 6.6E-07 |
| *F36A4.5* | WBGene00018084 | protein_coding_gene | -1.4 | 0.370 | 1.4E-03 |
| *nhr-1* | WBGene00003600 | protein_coding_gene | -1.4 | 0.370 | 6.1E-10 |
| *D1044.1* | WBGene00017027 | protein_coding_gene | -1.4 | 0.369 | 4.7E-06 |
| *cyp-33C4* | WBGene00018413 | protein_coding_gene | -1.4 | 0.369 | 6.6E-03 |
| *C03H5.5* | WBGene00015407 | protein_coding_gene | -1.4 | 0.369 | 3.6E-10 |
| *Y75B7AR.1* | WBGene00022287 | protein_coding_gene | -1.4 | 0.369 | 4.6E-04 |
| *C44B11.6* | WBGene00016635 | protein_coding_gene | -1.4 | 0.369 | 9.2E-06 |
| *tnc-2* | WBGene00006583 | protein_coding_gene | -1.4 | 0.369 | 6.4E-07 |
| *Y26D4A.21* | WBGene00077684 | protein_coding_gene | -1.4 | 0.368 | 3.8E-02 |
| *gsto-1* | WBGene00016204 | protein_coding_gene | -1.4 | 0.368 | 3.1E-06 |
| *C09G9.5* | WBGene00007495 | protein_coding_gene | -1.4 | 0.367 | 5.5E-04 |
| *H10E21.4* | WBGene00019184 | protein_coding_gene | -1.4 | 0.367 | 5.7E-03 |
| *F17C11.2* | WBGene00008914 | protein_coding_gene | -1.4 | 0.367 | 9.3E-10 |
| *F15D4.4* | WBGene00008861 | protein_coding_gene | -1.5 | 0.366 | 1.5E-04 |
| *ace-1* | WBGene00000035 | protein_coding_gene | -1.5 | 0.365 | 1.3E-06 |
| *K08E7.8* | WBGene00010675 | protein_coding_gene | -1.5 | 0.365 | 2.1E-03 |
| *T09B4.8* | WBGene00020382 | protein_coding_gene | -1.5 | 0.365 | 1.4E-09 |
| *W10C8.5* | WBGene00021128 | protein_coding_gene | -1.5 | 0.364 | 6.5E-18 |
| *F31F7.2* | WBGene00017965 | protein_coding_gene | -1.5 | 0.363 | 3.7E-02 |
| *ZK1307.1* | WBGene00014244 | protein_coding_gene | -1.5 | 0.363 | 7.1E-06 |
| *ZK909.3* | WBGene00014148 | protein_coding_gene | -1.5 | 0.363 | 8.5E-13 |
| *lev-9* | WBGene00002976 | protein_coding_gene | -1.5 | 0.363 | 6.5E-06 |
| *ric-4* | WBGene00004364 | protein_coding_gene | -1.5 | 0.363 | 6.1E-12 |
| *arrd-18* | WBGene00018060 | protein_coding_gene | -1.5 | 0.362 | 6.3E-06 |
| *mai-1* | WBGene00003124 | protein_coding_gene | -1.5 | 0.362 | 1.9E-06 |
| *glr-1* | WBGene00001612 | protein_coding_gene | -1.5 | 0.362 | 7.2E-03 |
| *F13D12.8* | WBGene00008742 | protein_coding_gene | -1.5 | 0.361 | 9.7E-05 |
| *haf-9* | WBGene00001819 | protein_coding_gene | -1.5 | 0.361 | 4.9E-10 |
| *Y38C1AA.9* | WBGene00021400 | protein_coding_gene | -1.5 | 0.361 | 1.9E-04 |
| *nlp-13* | WBGene00003751 | protein_coding_gene | -1.5 | 0.361 | 9.8E-09 |
| *T02E1.8* | WBGene00011380 | protein_coding_gene | -1.5 | 0.361 | 2.8E-03 |
| *gnrr-1* | WBGene00018798 | protein_coding_gene | -1.5 | 0.361 | 3.0E-02 |
| *unc-38* | WBGene00006774 | protein_coding_gene | -1.5 | 0.360 | 8.6E-05 |
| *F52E1.2* | WBGene00018692 | protein_coding_gene | -1.5 | 0.360 | 2.8E-06 |
| *clhm-1* | WBGene00016626 | protein_coding_gene | -1.5 | 0.360 | 1.1E-03 |
| *npr-26* | WBGene00011372 | protein_coding_gene | -1.5 | 0.360 | 9.6E-03 |
| *atp-6* | WBGene00010960 | protein_coding_gene | -1.5 | 0.360 | 9.8E-07 |
| *gly-12* | WBGene00001637 | protein_coding_gene | -1.5 | 0.360 | 1.9E-11 |
| *F23F12.3* | WBGene00017751 | protein_coding_gene | -1.5 | 0.359 | 7.3E-04 |
| *Y119C1B.6* | WBGene00022472 | protein_coding_gene | -1.5 | 0.359 | 1.1E-03 |
| *slc-9B.2* | WBGene00010214 | protein_coding_gene | -1.5 | 0.359 | 1.2E-04 |
| *F55A4.7* | WBGene00018859 | protein_coding_gene | -1.5 | 0.358 | 1.8E-02 |
| *dyc-1* | WBGene00001116 | protein_coding_gene | -1.5 | 0.358 | 2.3E-05 |
| *C53D5.5* | WBGene00016906 | protein_coding_gene | -1.5 | 0.358 | 1.6E-13 |
| *T02B11.3* | WBGene00020155 | protein_coding_gene | -1.5 | 0.357 | 1.2E-13 |
| *vap-1* | WBGene00006886 | protein_coding_gene | -1.5 | 0.357 | 5.4E-08 |
| *xbx-5* | WBGene00011974 | protein_coding_gene | -1.5 | 0.357 | 7.6E-04 |
| *F57C7.4* | WBGene00010200 | protein_coding_gene | -1.5 | 0.357 | 1.2E-02 |
| *sto-5* | WBGene00006067 | protein_coding_gene | -1.5 | 0.357 | 1.8E-05 |
| *math-14* | WBGene00015828 | protein_coding_gene | -1.5 | 0.356 | 3.2E-08 |
| *R03C1.1* | WBGene00010987 | protein_coding_gene | -1.5 | 0.356 | 4.4E-02 |
| *R10E8.7* | WBGene00014821 | pseudogene | -1.5 | 0.356 | 4.8E-08 |
| *T23F2.2* | WBGene00020735 | protein_coding_gene | -1.5 | 0.356 | 1.6E-05 |
| *C49A9.3* | WBGene00016757 | protein_coding_gene | -1.5 | 0.355 | 2.4E-08 |
| *unc-97* | WBGene00006826 | protein_coding_gene | -1.5 | 0.355 | 3.5E-15 |
| *gnrr-3* | WBGene00013871 | protein_coding_gene | -1.5 | 0.355 | 4.7E-02 |
| *F58B6.1* | WBGene00019029 | protein_coding_gene | -1.5 | 0.355 | 8.6E-03 |
| *Y32F6A.4* | WBGene00012529 | protein_coding_gene | -1.5 | 0.355 | 2.0E-15 |
| *C40A11.8* | WBGene00016551 | protein_coding_gene | -1.5 | 0.354 | 3.3E-03 |
| *egl-10* | WBGene00001179 | protein_coding_gene | -1.5 | 0.354 | 5.3E-16 |
| *del-1* | WBGene00000952 | protein_coding_gene | -1.5 | 0.353 | 1.9E-02 |
| *F11A5.15* | WBGene00008680 | protein_coding_gene | -1.5 | 0.353 | 1.7E-02 |
| *dct-15* | WBGene00012340 | protein_coding_gene | -1.5 | 0.353 | 1.3E-02 |
| *pde-4* | WBGene00020114 | protein_coding_gene | -1.5 | 0.352 | 6.2E-08 |
| *glb-6* | WBGene00015969 | protein_coding_gene | -1.5 | 0.352 | 1.7E-04 |
| *F40E10.6* | WBGene00009574 | protein_coding_gene | -1.5 | 0.352 | 6.5E-07 |
| *trpa-2* | WBGene00010873 | protein_coding_gene | -1.5 | 0.352 | 8.0E-08 |

| | | | | | |
|---|---|---|---|---|---|
| *F36H9.5* | WBGene00018113 | protein_coding_gene | -1.5 | 0.352 | 2.9E-03 |
| *rgs-6* | WBGene00004349 | protein_coding_gene | -1.5 | 0.352 | 1.6E-03 |
| *pgal-1* | WBGene00017671 | protein_coding_gene | -1.5 | 0.352 | 6.8E-11 |
| *D1007.18* | WBGene00044421 | protein_coding_gene | -1.5 | 0.352 | 2.4E-03 |
| *sft-4* | WBGene00004788 | protein_coding_gene | -1.5 | 0.351 | 5.3E-14 |
| *cest-2.2* | WBGene00013874 | protein_coding_gene | -1.5 | 0.351 | 8.7E-11 |
| *fipr-2* | WBGene00009097 | protein_coding_gene | -1.5 | 0.351 | 2.0E-07 |
| *T20B6.3* | WBGene00020599 | protein_coding_gene | -1.5 | 0.350 | 1.1E-02 |
| *rund-1* | WBGene00020572 | protein_coding_gene | -1.5 | 0.350 | 7.1E-08 |
| *K08D10.14* | WBGene00219216 | protein_coding_gene | -1.5 | 0.350 | 3.9E-06 |
| *H40L08.3* | WBGene00010434 | protein_coding_gene | -1.5 | 0.350 | 1.4E-04 |
| *acl-12* | WBGene00015295 | protein_coding_gene | -1.5 | 0.349 | 1.2E-12 |
| *F22E12.3* | WBGene00009060 | protein_coding_gene | -1.5 | 0.349 | 5.6E-03 |
| *unc-10* | WBGene00006750 | protein_coding_gene | -1.5 | 0.349 | 6.7E-05 |
| *sma-5* | WBGene00004859 | protein_coding_gene | -1.5 | 0.348 | 5.3E-06 |
| *ZK669.2* | WBGene00014052 | protein_coding_gene | -1.5 | 0.348 | 6.3E-07 |
| *acd-3* | WBGene00007775 | protein_coding_gene | -1.5 | 0.348 | 3.4E-07 |
| *C09G1.4* | WBGene00007486 | protein_coding_gene | -1.5 | 0.348 | 4.5E-05 |
| *gpc-1* | WBGene00001681 | protein_coding_gene | -1.5 | 0.348 | 2.5E-03 |
| *M02B1.2* | WBGene00010827 | protein_coding_gene | -1.5 | 0.347 | 8.0E-10 |
| *R07B1.13* | WBGene00138711 | protein_coding_gene | -1.5 | 0.346 | 8.1E-03 |
| *H38K22.9* | WBGene00284844 | protein_coding_gene | -1.5 | 0.346 | 7.1E-03 |
| *lin-2* | WBGene00002991 | protein_coding_gene | -1.5 | 0.346 | 1.7E-12 |
| *ins-3* | WBGene00002086 | protein_coding_gene | -1.5 | 0.345 | 2.6E-03 |
| *R13H4.5* | WBGene00011263 | protein_coding_gene | -1.5 | 0.345 | 4.4E-06 |
| *M03A8.3* | WBGene00019749 | protein_coding_gene | -1.5 | 0.345 | 1.6E-08 |
| *M03F8.5* | WBGene00019764 | protein_coding_gene | -1.5 | 0.344 | 1.6E-02 |
| *F46C8.1* | WBGene00018484 | protein_coding_gene | -1.5 | 0.344 | 1.3E-02 |
| *F35E12.10* | WBGene00009434 | protein_coding_gene | -1.5 | 0.344 | 6.5E-05 |
| *frm-8* | WBGene00001494 | protein_coding_gene | -1.5 | 0.343 | 9.0E-10 |
| *kvs-3* | WBGene00018402 | protein_coding_gene | -1.5 | 0.343 | 6.7E-03 |
| *pudl-1* | WBGene00044489 | protein_coding_gene | -1.5 | 0.343 | 2.3E-05 |
| *T14G8.3* | WBGene00011771 | protein_coding_gene | -1.5 | 0.342 | 7.3E-07 |
| *lron-2* | WBGene00022789 | protein_coding_gene | -1.5 | 0.342 | 6.5E-03 |
| *ttr-36* | WBGene00044605 | protein_coding_gene | -1.5 | 0.342 | 3.2E-09 |
| *hum-5* | WBGene00002038 | protein_coding_gene | -1.6 | 0.341 | 6.1E-13 |
| *F56C9.7* | WBGene00018951 | protein_coding_gene | -1.6 | 0.341 | 2.2E-09 |
| *pfk-1.1* | WBGene00022199 | protein_coding_gene | -1.6 | 0.341 | 3.4E-19 |
| *cutl-15* | WBGene00011888 | protein_coding_gene | -1.6 | 0.341 | 3.1E-05 |
| *kin-9* | WBGene00002195 | protein_coding_gene | -1.6 | 0.341 | 1.3E-04 |
| *PDB1.1* | WBGene00019803 | protein_coding_gene | -1.6 | 0.341 | 1.2E-06 |
| *W05H12.1* | WBGene00012289 | protein_coding_gene | -1.6 | 0.341 | 6.4E-09 |
| *F15A4.5* | WBGene00008840 | protein_coding_gene | -1.6 | 0.340 | 1.1E-03 |
| *H23N18.5* | WBGene00019236 | protein_coding_gene | -1.6 | 0.340 | 1.6E-02 |
| *dpy-20* | WBGene00001079 | protein_coding_gene | -1.6 | 0.340 | 4.2E-02 |
| *C17H12.10* | WBGene00015935 | protein_coding_gene | -1.6 | 0.340 | 1.4E-03 |
| *cyp-33C5* | WBGene00018261 | protein_coding_gene | -1.6 | 0.340 | 1.5E-03 |
| *unc-31* | WBGene00006767 | protein_coding_gene | -1.6 | 0.339 | 7.7E-08 |
| *F14H12.3* | WBGene00017471 | protein_coding_gene | -1.6 | 0.338 | 5.6E-05 |
| *M05D6.9* | WBGene00010881 | protein_coding_gene | -1.6 | 0.338 | 3.2E-02 |
| *F10A3.4* | WBGene00008634 | protein_coding_gene | -1.6 | 0.338 | 2.0E-05 |
| *T22F3.7* | WBGene00020697 | protein_coding_gene | -1.6 | 0.338 | 8.4E-03 |
| *del-4* | WBGene00012116 | protein_coding_gene | -1.6 | 0.338 | 4.6E-02 |
| *fbxa-42* | WBGene00044676 | protein_coding_gene | -1.6 | 0.338 | 3.5E-03 |
| *F41F3.3* | WBGene00018297 | protein_coding_gene | -1.6 | 0.338 | 2.3E-03 |
| *C12D5.9* | WBGene00015710 | protein_coding_gene | -1.6 | 0.338 | 6.8E-09 |
| *twk-49* | WBGene00019904 | protein_coding_gene | -1.6 | 0.338 | 4.3E-10 |
| *Y102A11A.7* | WBGene00022417 | protein_coding_gene | -1.6 | 0.337 | 2.4E-02 |
| *clec-50* | WBGene00012253 | protein_coding_gene | -1.6 | 0.337 | 5.6E-07 |
| *scd-2* | WBGene00004740 | protein_coding_gene | -1.6 | 0.337 | 1.8E-04 |
| *T08E11.8* | WBGene00020364 | protein_coding_gene | -1.6 | 0.337 | 3.5E-02 |
| *T05H10.8* | WBGene00011512 | protein_coding_gene | -1.6 | 0.335 | 3.6E-04 |
| *eppl-1* | WBGene00020139 | protein_coding_gene | -1.6 | 0.335 | 2.3E-13 |
| *flp-11* | WBGene00001454 | protein_coding_gene | -1.6 | 0.334 | 8.6E-07 |
| *ZC239.13* | WBGene00022571 | protein_coding_gene | -1.6 | 0.334 | 1.5E-02 |
| *F13H8.5* | WBGene00017438 | protein_coding_gene | -1.6 | 0.334 | 2.6E-04 |
| *F49H12.4* | WBGene00018655 | protein_coding_gene | -1.6 | 0.334 | 4.0E-02 |
| *cgt-2* | WBGene00017625 | protein_coding_gene | -1.6 | 0.333 | 2.8E-09 |
| *C18E9.7* | WBGene00007687 | protein_coding_gene | -1.6 | 0.333 | 8.2E-04 |
| *Y41C4A.17* | WBGene00012760 | protein_coding_gene | -1.6 | 0.333 | 2.2E-02 |

| | | | | | |
|---|---|---|---|---|---|
| *Y57G11B.5* | WBGene00013297 | protein_coding_gene | -1.6 | 0.332 | 2.8E-10 |
| *F07H5.7* | WBGene00008558 | protein_coding_gene | -1.6 | 0.332 | 2.3E-03 |
| *nhr-172* | WBGene00016927 | protein_coding_gene | -1.6 | 0.332 | 4.7E-02 |
| *unc-8* | WBGene00006748 | protein_coding_gene | -1.6 | 0.332 | 1.7E-02 |
| *cbs-1* | WBGene00013866 | protein_coding_gene | -1.6 | 0.332 | 4.7E-20 |
| *arrd-26* | WBGene00019873 | protein_coding_gene | -1.6 | 0.331 | 6.9E-04 |
| *kin-29* | WBGene00002210 | protein_coding_gene | -1.6 | 0.331 | 5.1E-11 |
| *T11B7.5* | WBGene00011708 | protein_coding_gene | -1.6 | 0.331 | 3.2E-08 |
| *flp-18* | WBGene00001461 | protein_coding_gene | -1.6 | 0.331 | 1.3E-08 |
| *twk-24* | WBGene00006677 | protein_coding_gene | -1.6 | 0.331 | 2.5E-09 |
| *Y82E9BR.5* | WBGene00022338 | protein_coding_gene | -1.6 | 0.331 | 1.6E-05 |
| *pde-1* | WBGene00011433 | protein_coding_gene | -1.6 | 0.331 | 1.3E-07 |
| *C38C3.10* | WBGene00045390 | protein_coding_gene | -1.6 | 0.331 | 1.4E-03 |
| *srx-45* | WBGene00005936 | protein_coding_gene | -1.6 | 0.330 | 2.0E-02 |
| *nucb-1* | WBGene00009674 | protein_coding_gene | -1.6 | 0.330 | 1.7E-13 |
| *pqn-35* | WBGene00004122 | protein_coding_gene | -1.6 | 0.330 | 1.5E-08 |
| *F58E2.3* | WBGene00019047 | protein_coding_gene | -1.6 | 0.330 | 1.2E-04 |
| *D1086.7* | WBGene00008394 | protein_coding_gene | -1.6 | 0.330 | 8.2E-09 |
| *C03H5.9* | WBGene00206366 | protein_coding_gene | -1.6 | 0.329 | 4.7E-02 |
| *cest-13* | WBGene00010116 | protein_coding_gene | -1.6 | 0.329 | 4.9E-07 |
| *unc-80* | WBGene00006812 | protein_coding_gene | -1.6 | 0.329 | 7.3E-08 |
| *dop-1* | WBGene00001052 | protein_coding_gene | -1.6 | 0.329 | 6.7E-04 |
| *npr-22* | WBGene00022004 | protein_coding_gene | -1.6 | 0.329 | 1.8E-06 |
| *T05C12.11* | WBGene00011473 | protein_coding_gene | -1.6 | 0.329 | 8.6E-03 |
| *gbb-1* | WBGene00021528 | protein_coding_gene | -1.6 | 0.329 | 3.5E-06 |
| *F57E7.1* | WBGene00010202 | protein_coding_gene | -1.6 | 0.328 | 1.4E-03 |
| *flp-13* | WBGene00001456 | protein_coding_gene | -1.6 | 0.328 | 2.0E-08 |
| *chw-1* | WBGene00009059 | protein_coding_gene | -1.6 | 0.328 | 5.3E-08 |
| *ZK1067.4* | WBGene00014210 | protein_coding_gene | -1.6 | 0.327 | 2.7E-05 |
| *F15E6.6* | WBGene00017487 | protein_coding_gene | -1.6 | 0.327 | 1.1E-09 |
| *crn-6* | WBGene00000799 | protein_coding_gene | -1.6 | 0.327 | 2.1E-11 |
| *nlp-12* | WBGene00003750 | protein_coding_gene | -1.6 | 0.327 | 9.6E-06 |
| *vha-11* | WBGene00006920 | protein_coding_gene | -1.6 | 0.326 | 3.2E-10 |
| *T16A9.3* | WBGene00011793 | protein_coding_gene | -1.6 | 0.326 | 6.5E-05 |
| *F41G3.10* | WBGene00018303 | protein_coding_gene | -1.6 | 0.326 | 2.8E-10 |
| *vha-20* | WBGene00010993 | protein_coding_gene | -1.6 | 0.326 | 3.8E-14 |
| *Y38F1A.4* | WBGene00012607 | protein_coding_gene | -1.6 | 0.325 | 2.1E-02 |
| *nrf-6* | WBGene00003813 | protein_coding_gene | -1.6 | 0.325 | 1.4E-17 |
| *C08B6.11* | WBGene00007431 | protein_coding_gene | -1.6 | 0.325 | 1.2E-08 |
| *mab-10* | WBGene00003107 | protein_coding_gene | -1.6 | 0.324 | 4.1E-06 |
| *srp-6* | WBGene00005647 | protein_coding_gene | -1.6 | 0.324 | 5.9E-14 |
| *C15F1.2* | WBGene00015795 | protein_coding_gene | -1.6 | 0.323 | 3.1E-03 |
| *lgc-10* | WBGene00022247 | protein_coding_gene | -1.6 | 0.323 | 2.0E-03 |
| *idhb-1* | WBGene00007993 | protein_coding_gene | -1.6 | 0.323 | 1.1E-22 |
| *Y51H7BR.4* | WBGene00021776 | protein_coding_gene | -1.6 | 0.323 | 2.3E-02 |
| *M02D8.2* | WBGene00019728 | protein_coding_gene | -1.6 | 0.323 | 6.1E-05 |
| *C09F12.2* | WBGene00007482 | protein_coding_gene | -1.6 | 0.323 | 1.5E-02 |
| *hsp-25* | WBGene00002023 | protein_coding_gene | -1.6 | 0.323 | 3.3E-10 |
| *Y60A9.3* | WBGene00013370 | protein_coding_gene | -1.6 | 0.322 | 2.7E-03 |
| *T04F8.3* | WBGene00011442 | protein_coding_gene | -1.6 | 0.322 | 6.6E-03 |
| *R02E4.2* | WBGene00044021 | protein_coding_gene | -1.6 | 0.322 | 4.8E-03 |
| *ins-6* | WBGene00002089 | protein_coding_gene | -1.6 | 0.321 | 8.2E-04 |
| *sbt-1* | WBGene00011392 | protein_coding_gene | -1.6 | 0.321 | 2.2E-12 |
| *F35F10.5* | WBGene00018055 | protein_coding_gene | -1.6 | 0.321 | 4.5E-03 |
| *gst-23* | WBGene00001771 | protein_coding_gene | -1.6 | 0.320 | 3.1E-02 |
| *T19D7.7* | WBGene00020575 | protein_coding_gene | -1.6 | 0.320 | 1.3E-02 |
| *C49A9.4* | WBGene00016758 | protein_coding_gene | -1.6 | 0.320 | 7.0E-10 |
| *lys-8* | WBGene00003097 | protein_coding_gene | -1.6 | 0.320 | 1.2E-16 |
| *hot-8* | WBGene00020489 | protein_coding_gene | -1.6 | 0.319 | 7.0E-05 |
| *tbx-37* | WBGene00006556 | protein_coding_gene | -1.6 | 0.319 | 1.1E-02 |
| *immt-1* | WBGene00020511 | protein_coding_gene | -1.7 | 0.318 | 1.8E-22 |
| *ldh-1* | WBGene00002262 | protein_coding_gene | -1.7 | 0.318 | 3.2E-09 |
| *lgc-35* | WBGene00012915 | protein_coding_gene | -1.7 | 0.318 | 1.2E-02 |
| *F57B1.5* | WBGene00010188 | protein_coding_gene | -1.7 | 0.317 | 4.1E-08 |
| *rsbp-1* | WBGene00011531 | protein_coding_gene | -1.7 | 0.317 | 1.5E-08 |
| *C49C3.4* | WBGene00008194 | protein_coding_gene | -1.7 | 0.317 | 1.2E-07 |
| *nlp-14* | WBGene00003752 | protein_coding_gene | -1.7 | 0.317 | 1.6E-06 |
| *pdi-6* | WBGene00015168 | protein_coding_gene | -1.7 | 0.317 | 9.6E-07 |
| *gcy-23* | WBGene00001548 | protein_coding_gene | -1.7 | 0.316 | 5.6E-03 |
| *Y17D7B.10* | WBGene00194982 | protein_coding_gene | -1.7 | 0.316 | 2.8E-02 |

| | | | | | |
|---|---|---|---|---|---|
| *nas-8* | WBGene00003527 | protein_coding_gene | -1.7 | 0.316 | 2.7E-03 |
| *pks-1* | WBGene00016558 | protein_coding_gene | -1.7 | 0.316 | 7.7E-05 |
| *phat-3* | WBGene00016782 | protein_coding_gene | -1.7 | 0.315 | 2.1E-08 |
| *Y53G8AR.7* | WBGene00021814 | protein_coding_gene | -1.7 | 0.314 | 1.6E-10 |
| *F46G10.4* | WBGene00009798 | protein_coding_gene | -1.7 | 0.314 | 1.4E-06 |
| *Y102A11A.1* | WBGene00022411 | protein_coding_gene | -1.7 | 0.314 | 4.4E-03 |
| *T28C6.7* | WBGene00012121 | protein_coding_gene | -1.7 | 0.313 | 2.0E-09 |
| *dhcr-7* | WBGene00007126 | protein_coding_gene | -1.7 | 0.313 | 1.5E-03 |
| *T08B1.1* | WBGene00020340 | protein_coding_gene | -1.7 | 0.313 | 4.7E-12 |
| *hyl-2* | WBGene00002044 | protein_coding_gene | -1.7 | 0.313 | 2.4E-21 |
| *F36G3.1* | WBGene00009482 | protein_coding_gene | -1.7 | 0.312 | 6.1E-12 |
| *F52D10.4* | WBGene00009931 | protein_coding_gene | -1.7 | 0.312 | 2.4E-02 |
| *paqr-3* | WBGene00013457 | protein_coding_gene | -1.7 | 0.312 | 4.2E-04 |
| *smoc-1* | WBGene00011437 | protein_coding_gene | -1.7 | 0.312 | 2.7E-06 |
| *W05B5.1* | WBGene00012274 | protein_coding_gene | -1.7 | 0.310 | 4.6E-06 |
| *vha-12* | WBGene00006921 | protein_coding_gene | -1.7 | 0.310 | 7.2E-08 |
| *ncx-4* | WBGene00003569 | protein_coding_gene | -1.7 | 0.310 | 6.1E-06 |
| *nlp-10* | WBGene00003748 | protein_coding_gene | -1.7 | 0.309 | 1.8E-09 |
| *ugt-61* | WBGene00018206 | protein_coding_gene | -1.7 | 0.309 | 8.6E-11 |
| *nas-1* | WBGene00003520 | protein_coding_gene | -1.7 | 0.309 | 2.6E-02 |
| *sul-1* | WBGene00006308 | protein_coding_gene | -1.7 | 0.309 | 7.7E-07 |
| *F30A10.14* | WBGene00045473 | protein_coding_gene | -1.7 | 0.308 | 4.2E-08 |
| *fipr-3* | WBGene00009090 | protein_coding_gene | -1.7 | 0.308 | 3.9E-05 |
| *F13B6.2* | WBGene00017417 | protein_coding_gene | -1.7 | 0.307 | 2.6E-05 |
| *T07F12.1* | WBGene00020321 | protein_coding_gene | -1.7 | 0.307 | 4.4E-12 |
| *lec-3* | WBGene00002266 | protein_coding_gene | -1.7 | 0.307 | 3.7E-09 |
| *inx-1* | WBGene00002123 | protein_coding_gene | -1.7 | 0.307 | 1.3E-07 |
| *T11F9.12* | WBGene00011713 | protein_coding_gene | -1.7 | 0.306 | 7.8E-07 |
| *C29F3.7* | WBGene00007807 | protein_coding_gene | -1.7 | 0.306 | 5.4E-07 |
| *kcnl-4* | WBGene00008265 | protein_coding_gene | -1.7 | 0.306 | 3.7E-04 |
| *F15G9.6* | WBGene00008869 | protein_coding_gene | -1.7 | 0.305 | 3.7E-02 |
| *ZK813.4* | WBGene00022823 | protein_coding_gene | -1.7 | 0.305 | 2.1E-09 |
| *fbl-1* | WBGene00001403 | protein_coding_gene | -1.7 | 0.305 | 5.2E-10 |
| *gcy-32* | WBGene00001552 | protein_coding_gene | -1.7 | 0.305 | 1.4E-02 |
| *hsp-3* | WBGene00002007 | protein_coding_gene | -1.7 | 0.304 | 1.5E-10 |
| *C01G5.25* | WBGene00235291 | protein_coding_gene | -1.7 | 0.304 | 1.8E-03 |
| *M153.2* | WBGene00010925 | protein_coding_gene | -1.7 | 0.304 | 3.9E-06 |
| *lgc-53* | WBGene00020657 | protein_coding_gene | -1.7 | 0.304 | 2.0E-03 |
| *nspc-4* | WBGene00044339 | protein_coding_gene | -1.7 | 0.304 | 3.1E-06 |
| *C17C3.1* | WBGene00015887 | protein_coding_gene | -1.7 | 0.303 | 2.4E-21 |
| *Y38A10A.2* | WBGene00021389 | protein_coding_gene | -1.7 | 0.303 | 6.6E-04 |
| *Y54G11A.7* | WBGene00013216 | protein_coding_gene | -1.7 | 0.303 | 1.2E-12 |
| *R10H10.3* | WBGene00011222 | protein_coding_gene | -1.7 | 0.303 | 1.6E-16 |
| *F37H8.2* | WBGene00009511 | protein_coding_gene | -1.7 | 0.303 | 5.7E-08 |
| *D1046.4* | WBGene00008365 | protein_coding_gene | -1.7 | 0.302 | 1.7E-06 |
| *daf-37* | WBGene00016246 | protein_coding_gene | -1.7 | 0.302 | 5.5E-05 |
| *ptr-16* | WBGene00004230 | protein_coding_gene | -1.7 | 0.302 | 5.4E-04 |
| *fat-6* | WBGene00001398 | protein_coding_gene | -1.7 | 0.302 | 1.0E-14 |
| *Y37A1A.3* | WBGene00012536 | protein_coding_gene | -1.7 | 0.302 | 2.1E-09 |
| *B0457.6* | WBGene00007180 | protein_coding_gene | -1.7 | 0.301 | 2.1E-09 |
| *B0238.12* | WBGene00015076 | protein_coding_gene | -1.7 | 0.301 | 2.2E-03 |
| *F19C6.3* | WBGene00008952 | protein_coding_gene | -1.7 | 0.300 | 1.9E-04 |
| *egl-23* | WBGene00001190 | protein_coding_gene | -1.7 | 0.300 | 4.2E-08 |
| *acox-1.1* | WBGene00008564 | protein_coding_gene | -1.7 | 0.300 | 3.5E-19 |
| *F59B10.3* | WBGene00010319 | protein_coding_gene | -1.7 | 0.299 | 4.1E-03 |
| *F32D8.7* | WBGene00009331 | protein_coding_gene | -1.7 | 0.299 | 1.6E-03 |
| *C09G9.8* | WBGene00007497 | protein_coding_gene | -1.7 | 0.299 | 1.9E-02 |
| *dnj-7* | WBGene00001025 | protein_coding_gene | -1.7 | 0.298 | 3.8E-10 |
| *Y48G10A.6* | WBGene00013023 | protein_coding_gene | -1.7 | 0.298 | 1.8E-04 |
| *ubc-23* | WBGene00006718 | protein_coding_gene | -1.7 | 0.298 | 2.0E-05 |
| *EGAP4.1* | WBGene00017149 | protein_coding_gene | -1.7 | 0.298 | 5.6E-03 |
| *lgc-30* | WBGene00019069 | protein_coding_gene | -1.7 | 0.298 | 1.0E-03 |
| *cpn-4* | WBGene00000780 | protein_coding_gene | -1.7 | 0.298 | 9.8E-17 |
| *F16F9.1* | WBGene00017513 | protein_coding_gene | -1.7 | 0.298 | 7.3E-09 |
| *C23H5.11* | WBGene00044649 | protein_coding_gene | -1.7 | 0.298 | 2.8E-02 |
| *pcdr-1* | WBGene00010329 | protein_coding_gene | -1.7 | 0.297 | 2.0E-05 |
| *unc-9* | WBGene00006749 | protein_coding_gene | -1.7 | 0.297 | 2.6E-11 |
| *B0395.3* | WBGene00007175 | protein_coding_gene | -1.8 | 0.297 | 1.1E-19 |
| *fpn-1.2* | WBGene00019977 | protein_coding_gene | -1.8 | 0.297 | 1.1E-06 |
| *F26F12.8* | WBGene00044544 | protein_coding_gene | -1.8 | 0.297 | 4.8E-04 |

| | | | | | |
|---|---|---|---|---|---|
| *cht-4* | WBGene00021252 | protein_coding_gene | -1.8 | 0.297 | 8.5E-03 |
| *F46A8.11* | WBGene00014761 | protein_coding_gene | -1.8 | 0.296 | 3.1E-04 |
| *C45G9.8* | WBGene00016679 | protein_coding_gene | -1.8 | 0.296 | 1.9E-02 |
| *F46C5.6* | WBGene00009781 | protein_coding_gene | -1.8 | 0.296 | 2.3E-09 |
| *Y102A11A.9* | WBGene00022419 | protein_coding_gene | -1.8 | 0.295 | 5.5E-08 |
| *H09G03.1* | WBGene00019174 | protein_coding_gene | -1.8 | 0.295 | 5.6E-03 |
| *clec-223* | WBGene00007282 | protein_coding_gene | -1.8 | 0.295 | 3.7E-11 |
| *R05A10.2* | WBGene00011019 | protein_coding_gene | -1.8 | 0.295 | 1.5E-12 |
| *C04G2.14* | WBGene00255426 | ncRNA_gene | -1.8 | 0.295 | 9.6E-06 |
| *unc-105* | WBGene00006832 | protein_coding_gene | -1.8 | 0.295 | 1.1E-09 |
| *F53H4.3* | WBGene00010008 | protein_coding_gene | -1.8 | 0.295 | 9.1E-05 |
| *snet-1* | WBGene00015354 | protein_coding_gene | -1.8 | 0.295 | 1.9E-07 |
| *tre-3* | WBGene00006609 | protein_coding_gene | -1.8 | 0.293 | 6.0E-14 |
| *F46C5.2* | WBGene00009779 | protein_coding_gene | -1.8 | 0.293 | 4.0E-04 |
| *F27C1.11* | WBGene00017858 | protein_coding_gene | -1.8 | 0.293 | 2.7E-05 |
| *C26B9.5* | WBGene00016134 | protein_coding_gene | -1.8 | 0.293 | 1.1E-08 |
| *C30G7.2* | WBGene00044176 | protein_coding_gene | -1.8 | 0.293 | 2.1E-06 |
| *asp-9* | WBGene00015700 | protein_coding_gene | -1.8 | 0.292 | 3.4E-02 |
| *scl-3* | WBGene00009896 | protein_coding_gene | -1.8 | 0.292 | 5.0E-07 |
| *phf-33* | WBGene00010708 | protein_coding_gene | -1.8 | 0.292 | 3.7E-02 |
| *D1025.10* | WBGene00044134 | protein_coding_gene | -1.8 | 0.291 | 2.8E-06 |
| *F09E5.16* | WBGene00017292 | protein_coding_gene | -1.8 | 0.291 | 2.8E-07 |
| *spds-1* | WBGene00012909 | protein_coding_gene | -1.8 | 0.291 | 7.6E-10 |
| *dhc-4* | WBGene00012272 | protein_coding_gene | -1.8 | 0.291 | 7.0E-07 |
| *ncx-8* | WBGene00003573 | protein_coding_gene | -1.8 | 0.291 | 7.0E-09 |
| *odc-1* | WBGene00003844 | protein_coding_gene | -1.8 | 0.290 | 1.8E-28 |
| *tni-3* | WBGene00006585 | protein_coding_gene | -1.8 | 0.290 | 9.1E-11 |
| *clec-224* | WBGene00044790 | protein_coding_gene | -1.8 | 0.290 | 1.6E-02 |
| *pcp-4* | WBGene00003959 | protein_coding_gene | -1.8 | 0.290 | 2.0E-08 |
| *nspc-17* | WBGene00008359 | protein_coding_gene | -1.8 | 0.290 | 5.8E-07 |
| *egl-3* | WBGene00001172 | protein_coding_gene | -1.8 | 0.289 | 3.4E-08 |
| *rhr-1* | WBGene00004358 | protein_coding_gene | -1.8 | 0.289 | 7.6E-11 |
| *C26D10.6* | WBGene00007746 | protein_coding_gene | -1.8 | 0.289 | 1.9E-13 |
| *flp-16* | WBGene00001459 | protein_coding_gene | -1.8 | 0.289 | 1.0E-14 |
| *pgp-13* | WBGene00004007 | protein_coding_gene | -1.8 | 0.289 | 2.8E-11 |
| *T24C12.4* | WBGene00020766 | protein_coding_gene | -1.8 | 0.289 | 3.3E-02 |
| *nekl-4* | WBGene00017033 | protein_coding_gene | -1.8 | 0.288 | 9.4E-04 |
| *exp-1* | WBGene00001373 | protein_coding_gene | -1.8 | 0.288 | 4.8E-03 |
| *sto-2* | WBGene00006064 | protein_coding_gene | -1.8 | 0.288 | 2.2E-03 |
| *glb-22* | WBGene00011251 | protein_coding_gene | -1.8 | 0.288 | 8.6E-03 |
| *F49E12.8* | WBGene00009901 | protein_coding_gene | -1.8 | 0.288 | 1.3E-02 |
| *frpr-5* | WBGene00008342 | protein_coding_gene | -1.8 | 0.287 | 2.4E-05 |
| *prmt-6* | WBGene00016023 | protein_coding_gene | -1.8 | 0.286 | 9.1E-05 |
| *T18D3.9* | WBGene00011826 | protein_coding_gene | -1.8 | 0.285 | 7.0E-03 |
| *C11H1.5* | WBGene00007530 | protein_coding_gene | -1.8 | 0.285 | 2.9E-02 |
| *fkb-3* | WBGene00001428 | protein_coding_gene | -1.8 | 0.285 | 4.2E-07 |
| *fbxa-187* | WBGene00009838 | pseudogene | -1.8 | 0.285 | 5.9E-03 |
| *pdi-2* | WBGene00003963 | protein_coding_gene | -1.8 | 0.285 | 1.3E-12 |
| *F23H11.7* | WBGene00017762 | protein_coding_gene | -1.8 | 0.285 | 4.5E-06 |
| *pgp-4* | WBGene00003998 | protein_coding_gene | -1.8 | 0.285 | 3.0E-10 |
| *C36C5.5* | WBGene00016474 | protein_coding_gene | -1.8 | 0.284 | 6.6E-14 |
| *ilcr-1* | WBGene00013415 | protein_coding_gene | -1.8 | 0.284 | 4.0E-15 |
| *H28G03.1* | WBGene00019249 | protein_coding_gene | -1.8 | 0.284 | 1.0E-07 |
| *B0310.2* | WBGene00015138 | protein_coding_gene | -1.8 | 0.283 | 1.5E-08 |
| *F34H10.5* | WBGene00009382 | protein_coding_gene | -1.8 | 0.283 | 3.1E-02 |
| *dyla-1* | WBGene00018307 | protein_coding_gene | -1.8 | 0.283 | 6.4E-06 |
| *C06E7.2* | WBGene00015539 | protein_coding_gene | -1.8 | 0.282 | 6.5E-13 |
| *clec-118* | WBGene00021032 | protein_coding_gene | -1.8 | 0.282 | 3.4E-02 |
| *C02F5.14* | WBGene00044445 | protein_coding_gene | -1.8 | 0.282 | 2.1E-09 |
| *prx-1* | WBGene00004191 | protein_coding_gene | -1.8 | 0.281 | 1.7E-19 |
| *col-38* | WBGene00000615 | protein_coding_gene | -1.8 | 0.281 | 2.0E-08 |
| *fut-4* | WBGene00001507 | protein_coding_gene | -1.8 | 0.281 | 2.8E-04 |
| *R13A5.10* | WBGene00020052 | protein_coding_gene | -1.8 | 0.281 | 1.2E-05 |
| *clec-246* | WBGene00008892 | protein_coding_gene | -1.8 | 0.280 | 3.5E-02 |
| *pxmp-4* | WBGene00017574 | protein_coding_gene | -1.8 | 0.280 | 2.4E-18 |
| *calu-1* | WBGene00019760 | protein_coding_gene | -1.8 | 0.280 | 1.3E-12 |
| *twk-26* | WBGene00006679 | protein_coding_gene | -1.8 | 0.280 | 4.2E-09 |
| *grl-8* | WBGene00001717 | protein_coding_gene | -1.8 | 0.279 | 4.1E-03 |
| *npr-2* | WBGene00003808 | protein_coding_gene | -1.8 | 0.279 | 2.6E-04 |
| *T05A1.5* | WBGene00011456 | protein_coding_gene | -1.8 | 0.279 | 4.3E-04 |

| | | | | | |
|---|---|---|---|---|---|
| *C26C6.9* | WBGene00007743 | protein_coding_gene | -1.8 | 0.278 | 4.7E-09 |
| *ins-24* | WBGene00002107 | protein_coding_gene | -1.8 | 0.278 | 2.8E-04 |
| *K10C8.4* | WBGene00010737 | protein_coding_gene | -1.8 | 0.278 | 1.9E-02 |
| *gpx-3* | WBGene00007517 | protein_coding_gene | -1.8 | 0.277 | 3.1E-11 |
| *dbl-1* | WBGene00000936 | protein_coding_gene | -1.8 | 0.277 | 2.1E-08 |
| *F58F9.1* | WBGene00019055 | protein_coding_gene | -1.8 | 0.277 | 1.8E-02 |
| *F35A5.4* | WBGene00018026 | protein_coding_gene | -1.9 | 0.277 | 5.8E-03 |
| *daf-36* | WBGene00007536 | protein_coding_gene | -1.9 | 0.277 | 2.6E-05 |
| *myo-1* | WBGene00002348 | protein_coding_gene | -1.9 | 0.277 | 4.2E-08 |
| *drn-1* | WBGene00016911 | protein_coding_gene | -1.9 | 0.277 | 3.5E-08 |
| *F35H10.3* | WBGene00018069 | protein_coding_gene | -1.9 | 0.277 | 3.5E-05 |
| *nlp-49* | WBGene00019160 | protein_coding_gene | -1.9 | 0.277 | 2.2E-10 |
| *bli-5* | WBGene00000255 | protein_coding_gene | -1.9 | 0.277 | 7.5E-03 |
| *alh-9* | WBGene00000115 | protein_coding_gene | -1.9 | 0.276 | 1.4E-08 |
| *ent-4* | WBGene00008123 | protein_coding_gene | -1.9 | 0.276 | 9.1E-15 |
| *mrp-2* | WBGene00003408 | protein_coding_gene | -1.9 | 0.276 | 1.3E-21 |
| *T07G12.2* | WBGene00011593 | protein_coding_gene | -1.9 | 0.276 | 3.7E-02 |
| *F41E6.17* | WBGene00194909 | protein_coding_gene | -1.9 | 0.276 | 1.4E-02 |
| *lipl-5* | WBGene00022642 | protein_coding_gene | -1.9 | 0.275 | 5.1E-15 |
| *C18A11.3* | WBGene00015949 | protein_coding_gene | -1.9 | 0.275 | 1.2E-11 |
| *chil-1* | WBGene00007465 | protein_coding_gene | -1.9 | 0.275 | 1.0E-02 |
| *ZK105.1* | WBGene00022653 | protein_coding_gene | -1.9 | 0.274 | 3.1E-11 |
| *frpr-6* | WBGene00017661 | protein_coding_gene | -1.9 | 0.274 | 1.1E-02 |
| *F09B9.5* | WBGene00008604 | protein_coding_gene | -1.9 | 0.274 | 1.0E-03 |
| *ucr-2.2* | WBGene00011679 | protein_coding_gene | -1.9 | 0.273 | 9.7E-30 |
| *jud-4* | WBGene00007262 | protein_coding_gene | -1.9 | 0.273 | 3.7E-02 |
| *slc-25A18.1* | WBGene00008979 | protein_coding_gene | -1.9 | 0.273 | 1.2E-19 |
| *nhx-4* | WBGene00003732 | protein_coding_gene | -1.9 | 0.273 | 1.2E-10 |
| *F55G7.1* | WBGene00010119 | protein_coding_gene | -1.9 | 0.273 | 1.5E-03 |
| *otpl-7* | WBGene00018480 | protein_coding_gene | -1.9 | 0.272 | 2.4E-02 |
| *gcy-12* | WBGene00001538 | protein_coding_gene | -1.9 | 0.272 | 7.5E-12 |
| *nlp-80* | WBGene00011428 | protein_coding_gene | -1.9 | 0.272 | 1.2E-10 |
| *gar-2* | WBGene00001518 | protein_coding_gene | -1.9 | 0.272 | 1.9E-09 |
| *rft-1* | WBGene00044637 | protein_coding_gene | -1.9 | 0.272 | 8.3E-06 |
| *F26D2.3* | WBGene00009148 | protein_coding_gene | -1.9 | 0.272 | 7.7E-03 |
| *F53F4.17* | WBGene00045195 | protein_coding_gene | -1.9 | 0.272 | 6.1E-03 |
| *F11D5.7* | WBGene00043980 | protein_coding_gene | -1.9 | 0.271 | 1.6E-06 |
| *cpz-1* | WBGene00000788 | protein_coding_gene | -1.9 | 0.271 | 4.1E-18 |
| *C53C7.5* | WBGene00014294 | pseudogene | -1.9 | 0.271 | 4.5E-09 |
| *F54D5.17* | WBGene00195146 | protein_coding_gene | -1.9 | 0.270 | 5.1E-04 |
| *B0334.13* | WBGene00007149 | protein_coding_gene | -1.9 | 0.270 | 1.0E-02 |
| *F52G3.3* | WBGene00018711 | pseudogene | -1.9 | 0.270 | 4.5E-02 |
| *shk-1* | WBGene00014261 | protein_coding_gene | -1.9 | 0.270 | 6.2E-10 |
| *hot-9* | WBGene00044484 | protein_coding_gene | -1.9 | 0.270 | 5.0E-04 |
| *C04E6.8* | WBGene00015424 | protein_coding_gene | -1.9 | 0.270 | 2.5E-02 |
| *frpr-10* | WBGene00019019 | protein_coding_gene | -1.9 | 0.269 | 4.0E-02 |
| *E04F6.10* | WBGene00017129 | protein_coding_gene | -1.9 | 0.269 | 1.4E-03 |
| *F28H7.3* | WBGene00009237 | protein_coding_gene | -1.9 | 0.269 | 1.2E-09 |
| *T05C1.2* | WBGene00020249 | protein_coding_gene | -1.9 | 0.268 | 1.4E-02 |
| *T06F4.1* | WBGene00020294 | protein_coding_gene | -1.9 | 0.268 | 1.9E-06 |
| *tni-4* | WBGene00006586 | protein_coding_gene | -1.9 | 0.268 | 2.8E-15 |
| *myo-5* | WBGene00019064 | protein_coding_gene | -1.9 | 0.268 | 6.2E-12 |
| *linc-56* | WBGene00219650 | lincRNA_gene | -1.9 | 0.268 | 2.9E-02 |
| *hacd-1* | WBGene00019978 | protein_coding_gene | -1.9 | 0.267 | 9.4E-08 |
| *nep-12* | WBGene00017842 | protein_coding_gene | -1.9 | 0.267 | 6.4E-07 |
| *arrd-7* | WBGene00012464 | protein_coding_gene | -1.9 | 0.267 | 9.9E-11 |
| *C45E5.4* | WBGene00016667 | protein_coding_gene | -1.9 | 0.267 | 4.8E-09 |
| *gst-40* | WBGene00001788 | protein_coding_gene | -1.9 | 0.266 | 8.3E-03 |
| *avr-14* | WBGene00000232 | protein_coding_gene | -1.9 | 0.266 | 3.5E-12 |
| *hsd-2* | WBGene00022498 | protein_coding_gene | -1.9 | 0.266 | 1.4E-03 |
| *H11E01.3* | WBGene00019188 | protein_coding_gene | -1.9 | 0.266 | 1.5E-09 |
| *ZK896.1* | WBGene00014132 | protein_coding_gene | -1.9 | 0.265 | 1.8E-04 |
| *ZK721.3* | WBGene00022810 | protein_coding_gene | -1.9 | 0.265 | 1.6E-03 |
| *C18E9.5* | WBGene00007685 | protein_coding_gene | -1.9 | 0.265 | 2.5E-12 |
| *pmp-1* | WBGene00004058 | protein_coding_gene | -1.9 | 0.265 | 1.8E-12 |
| *T01B7.8* | WBGene00011313 | protein_coding_gene | -1.9 | 0.265 | 2.7E-02 |
| *Y77E11A.12* | WBGene00022313 | protein_coding_gene | -1.9 | 0.265 | 4.2E-08 |
| *T27A10.2* | WBGene00020844 | protein_coding_gene | -1.9 | 0.264 | 2.2E-03 |
| *gst-19* | WBGene00001767 | protein_coding_gene | -1.9 | 0.264 | 2.6E-02 |
| *T24C12.3* | WBGene00020765 | protein_coding_gene | -1.9 | 0.264 | 7.7E-15 |

| | | | | | |
|---|---|---|---|---|---|
| *srm-6* | WBGene00009810 | protein_coding_gene | -1.9 | 0.264 | 3.8E-02 |
| *ZK816.1* | WBGene00022825 | protein_coding_gene | -1.9 | 0.264 | 2.3E-05 |
| *vha-15* | WBGene00020507 | protein_coding_gene | -1.9 | 0.264 | 5.2E-16 |
| *C33E10.8* | WBGene00016349 | protein_coding_gene | -1.9 | 0.264 | 3.1E-04 |
| *cest-1.1* | WBGene00011362 | protein_coding_gene | -1.9 | 0.263 | 5.9E-04 |
| *Y113G7B.12* | WBGene00013762 | protein_coding_gene | -1.9 | 0.262 | 1.8E-12 |
| *hpo-34* | WBGene00009259 | protein_coding_gene | -1.9 | 0.262 | 1.6E-15 |
| *oig-8* | WBGene00020582 | protein_coding_gene | -1.9 | 0.262 | 9.6E-05 |
| *daf-12* | WBGene00000908 | protein_coding_gene | -1.9 | 0.262 | 1.2E-06 |
| *T03F7.7* | WBGene00011404 | protein_coding_gene | -1.9 | 0.261 | 4.2E-19 |
| *T10E10.3* | WBGene00020420 | protein_coding_gene | -1.9 | 0.261 | 4.7E-02 |
| *itr-1* | WBGene00002173 | protein_coding_gene | -1.9 | 0.261 | 1.0E-14 |
| *flr-4* | WBGene00001468 | protein_coding_gene | -1.9 | 0.260 | 3.6E-10 |
| *cubn-1* | WBGene00013855 | protein_coding_gene | -1.9 | 0.260 | 8.5E-08 |
| *AH9.6* | WBGene00044701 | protein_coding_gene | -1.9 | 0.260 | 1.2E-04 |
| *C18B2.1* | WBGene00015952 | protein_coding_gene | -1.9 | 0.259 | 3.5E-02 |
| *clec-80* | WBGene00043062 | protein_coding_gene | -2.0 | 0.259 | 9.6E-16 |
| *lgc-40* | WBGene00020767 | protein_coding_gene | -2.0 | 0.258 | 3.9E-07 |
| *pqn-8* | WBGene00004100 | protein_coding_gene | -2.0 | 0.258 | 1.3E-07 |
| *F44G4.6* | WBGene00009715 | protein_coding_gene | -2.0 | 0.258 | 8.9E-04 |
| *F56D6.16* | WBGene00219304 | pseudogene | -2.0 | 0.258 | 9.9E-07 |
| *tmem-135* | WBGene00019345 | protein_coding_gene | -2.0 | 0.258 | 4.3E-24 |
| *ttr-28* | WBGene00011285 | protein_coding_gene | -2.0 | 0.258 | 2.6E-04 |
| *F16H6.7* | WBGene00008897 | protein_coding_gene | -2.0 | 0.258 | 6.5E-04 |
| *lat-2* | WBGene00002252 | protein_coding_gene | -2.0 | 0.258 | 6.2E-09 |
| *K03B4.4* | WBGene00019355 | protein_coding_gene | -2.0 | 0.257 | 4.6E-06 |
| *C43F9.7* | WBGene00008075 | protein_coding_gene | -2.0 | 0.256 | 2.7E-02 |
| *F22F1.2* | WBGene00017714 | protein_coding_gene | -2.0 | 0.256 | 3.4E-03 |
| *C01H6.4* | WBGene00007254 | protein_coding_gene | -2.0 | 0.256 | 1.1E-08 |
| *C04E12.2* | WBGene00015433 | protein_coding_gene | -2.0 | 0.256 | 6.6E-10 |
| *nlp-67* | WBGene00011227 | protein_coding_gene | -2.0 | 0.255 | 2.4E-04 |
| *F48B9.3* | WBGene00018589 | protein_coding_gene | -2.0 | 0.255 | 1.1E-05 |
| *phy-2* | WBGene00004025 | protein_coding_gene | -2.0 | 0.255 | 6.1E-11 |
| *gnrr-7* | WBGene00008737 | protein_coding_gene | -2.0 | 0.255 | 1.1E-02 |
| *T20G5.8* | WBGene00011870 | protein_coding_gene | -2.0 | 0.255 | 1.2E-08 |
| *K09F6.13* | WBGene00235164 | protein_coding_gene | -2.0 | 0.255 | 2.1E-02 |
| *B0310.6* | WBGene00015142 | protein_coding_gene | -2.0 | 0.255 | 3.1E-02 |
| *C34E11.2* | WBGene00007935 | protein_coding_gene | -2.0 | 0.255 | 1.1E-04 |
| *unc-47* | WBGene00006783 | protein_coding_gene | -2.0 | 0.255 | 1.6E-04 |
| *ver-4* | WBGene00006897 | protein_coding_gene | -2.0 | 0.255 | 3.2E-06 |
| *spp-19* | WBGene00005004 | protein_coding_gene | -2.0 | 0.254 | 1.1E-02 |
| *nas-15* | WBGene00003534 | protein_coding_gene | -2.0 | 0.253 | 4.6E-03 |
| *Y39G8B.5* | WBGene00012726 | protein_coding_gene | -2.0 | 0.252 | 7.5E-03 |
| *Y39A3B.3* | WBGene00021438 | protein_coding_gene | -2.0 | 0.252 | 1.5E-02 |
| *K03B4.9* | WBGene00219236 | protein_coding_gene | -2.0 | 0.252 | 7.2E-04 |
| *F02C12.1* | WBGene00008515 | protein_coding_gene | -2.0 | 0.252 | 6.9E-14 |
| *C31C9.7* | WBGene00007841 | protein_coding_gene | -2.0 | 0.251 | 7.9E-07 |
| *ant-1.2* | WBGene00007057 | protein_coding_gene | -2.0 | 0.251 | 5.9E-03 |
| *T20B5.2* | WBGene00020595 | protein_coding_gene | -2.0 | 0.251 | 6.4E-08 |
| *mpst-7* | WBGene00011307 | protein_coding_gene | -2.0 | 0.251 | 2.2E-13 |
| *hen-1* | WBGene00001841 | protein_coding_gene | -2.0 | 0.250 | 1.2E-02 |
| *wrt-9* | WBGene00006955 | protein_coding_gene | -2.0 | 0.250 | 5.4E-03 |
| *ttr-35* | WBGene00017688 | protein_coding_gene | -2.0 | 0.250 | 3.8E-10 |
| *T24A6.20* | WBGene00020756 | protein_coding_gene | -2.0 | 0.250 | 1.1E-10 |
| *clec-187* | WBGene00014137 | protein_coding_gene | -2.0 | 0.250 | 4.5E-11 |
| *exc-1* | WBGene00001362 | protein_coding_gene | -2.0 | 0.249 | 6.0E-04 |
| *srr-3* | WBGene00005654 | protein_coding_gene | -2.0 | 0.249 | 2.2E-02 |
| *F19D8.2* | WBGene00008955 | protein_coding_gene | -2.0 | 0.249 | 3.1E-04 |
| *fipr-9* | WBGene00044175 | protein_coding_gene | -2.0 | 0.249 | 9.6E-08 |
| *F26G1.5* | WBGene00017841 | protein_coding_gene | -2.0 | 0.248 | 2.3E-05 |
| *C14B9.3* | WBGene00015753 | protein_coding_gene | -2.0 | 0.248 | 1.2E-09 |
| *C14B4.2* | WBGene00007583 | protein_coding_gene | -2.0 | 0.248 | 8.9E-10 |
| *srsx-20* | WBGene00019853 | pseudogene | -2.0 | 0.247 | 2.5E-02 |
| *C55A6.4* | WBGene00008333 | protein_coding_gene | -2.0 | 0.247 | 9.1E-06 |
| *C04E7.5* | WBGene00045270 | protein_coding_gene | -2.0 | 0.247 | 4.3E-03 |
| *fbxa-50* | WBGene00013844 | protein_coding_gene | -2.0 | 0.247 | 2.9E-03 |
| *E02C12.6* | WBGene00017092 | protein_coding_gene | -2.0 | 0.247 | 8.4E-04 |
| *lips-7* | WBGene00015645 | protein_coding_gene | -2.0 | 0.246 | 2.6E-03 |
| *vha-6* | WBGene00006915 | protein_coding_gene | -2.0 | 0.246 | 7.1E-17 |
| *gst-11* | WBGene00001759 | protein_coding_gene | -2.0 | 0.246 | 2.3E-03 |

| | | | | | |
|---|---|---|---|---|---|
| *dac-1* | WBGene00000895 | protein_coding_gene | -2.0 | 0.246 | 2.7E-08 |
| *R02D1.2* | WBGene00235310 | protein_coding_gene | -2.0 | 0.246 | 1.1E-05 |
| *Y43F8C.11* | WBGene00012833 | protein_coding_gene | -2.0 | 0.246 | 4.4E-03 |
| *C37C3.11* | WBGene00016503 | protein_coding_gene | -2.0 | 0.245 | 1.5E-02 |
| *F35H12.1* | WBGene00018074 | protein_coding_gene | -2.0 | 0.245 | 2.8E-04 |
| *C53D6.11* | WBGene00044241 | protein_coding_gene | -2.0 | 0.245 | 3.7E-07 |
| *C44H4.6* | WBGene00008095 | protein_coding_gene | -2.0 | 0.245 | 2.7E-03 |
| *acr-3* | WBGene00000043 | protein_coding_gene | -2.0 | 0.244 | 1.5E-02 |
| *ZC581.9* | WBGene00022635 | protein_coding_gene | -2.0 | 0.244 | 7.3E-16 |
| *F41G4.5* | WBGene00018309 | pseudogene | -2.0 | 0.244 | 1.0E-02 |
| *flp-6* | WBGene00001449 | protein_coding_gene | -2.0 | 0.244 | 1.3E-15 |
| *C05D12.3* | WBGene00014666 | protein_coding_gene | -2.0 | 0.243 | 2.1E-03 |
| *his-38* | WBGene00001912 | protein_coding_gene | -2.0 | 0.243 | 1.6E-03 |
| *F19C7.8* | WBGene00017597 | protein_coding_gene | -2.0 | 0.243 | 2.8E-06 |
| *F01E11.18* | WBGene00219373 | protein_coding_gene | -2.0 | 0.243 | 1.6E-04 |
| *arrd-28* | WBGene00009000 | protein_coding_gene | -2.0 | 0.243 | 4.8E-02 |
| *zmp-6* | WBGene00020646 | protein_coding_gene | -2.0 | 0.242 | 1.8E-02 |
| *daf-38* | WBGene00013642 | protein_coding_gene | -2.0 | 0.242 | 6.3E-05 |
| *C10A4.9* | WBGene00045456 | protein_coding_gene | -2.0 | 0.242 | 5.0E-04 |
| *ivns-1* | WBGene00006493 | protein_coding_gene | -2.0 | 0.242 | 5.8E-11 |
| *R11D1.13* | WBGene00045517 | protein_coding_gene | -2.0 | 0.242 | 1.3E-02 |
| *nex-3* | WBGene00003590 | protein_coding_gene | -2.0 | 0.242 | 1.0E-08 |
| *dhs-3* | WBGene00000967 | protein_coding_gene | -2.1 | 0.241 | 1.2E-09 |
| *C05D2.8* | WBGene00015469 | protein_coding_gene | -2.1 | 0.241 | 1.1E-20 |
| *unc-18* | WBGene00006757 | protein_coding_gene | -2.1 | 0.241 | 6.3E-26 |
| *cest-32* | WBGene00016862 | protein_coding_gene | -2.1 | 0.241 | 8.6E-15 |
| *unc-46* | WBGene00006782 | protein_coding_gene | -2.1 | 0.240 | 1.4E-03 |
| *gfi-1* | WBGene00001581 | protein_coding_gene | -2.1 | 0.240 | 7.1E-07 |
| *dhs-7* | WBGene00000971 | protein_coding_gene | -2.1 | 0.240 | 9.6E-11 |
| *acr-8* | WBGene00000047 | protein_coding_gene | -2.1 | 0.239 | 1.8E-05 |
| *C06H5.6* | WBGene00007394 | protein_coding_gene | -2.1 | 0.239 | 1.5E-10 |
| *T06G6.6* | WBGene00011548 | protein_coding_gene | -2.1 | 0.239 | 1.1E-02 |
| *slc-17.5* | WBGene00017681 | protein_coding_gene | -2.1 | 0.239 | 2.0E-09 |
| *ttn-1* | WBGene00006436 | protein_coding_gene | -2.1 | 0.239 | 6.7E-06 |
| *npr-19* | WBGene00015364 | protein_coding_gene | -2.1 | 0.238 | 4.8E-02 |
| *npr-7* | WBGene00018067 | protein_coding_gene | -2.1 | 0.238 | 2.0E-03 |
| *ZC116.1* | WBGene00013853 | protein_coding_gene | -2.1 | 0.238 | 4.9E-08 |
| *alh-13* | WBGene00011938 | protein_coding_gene | -2.1 | 0.238 | 3.8E-15 |
| *nmat-1* | WBGene00012295 | protein_coding_gene | -2.1 | 0.238 | 3.8E-09 |
| *E01G6.1* | WBGene00008449 | protein_coding_gene | -2.1 | 0.237 | 2.5E-04 |
| *otpl-8* | WBGene00010403 | protein_coding_gene | -2.1 | 0.237 | 1.9E-02 |
| *glb-10* | WBGene00016215 | protein_coding_gene | -2.1 | 0.237 | 2.1E-07 |
| *lev-1* | WBGene00002974 | protein_coding_gene | -2.1 | 0.237 | 5.3E-09 |
| *slc-36.4* | WBGene00010421 | protein_coding_gene | -2.1 | 0.236 | 8.5E-13 |
| *F31F4.1* | WBGene00017957 | protein_coding_gene | -2.1 | 0.236 | 6.0E-14 |
| *Y53F4B.51* | WBGene00194905 | protein_coding_gene | -2.1 | 0.236 | 1.9E-03 |
| *nspc-14* | WBGene00009638 | protein_coding_gene | -2.1 | 0.236 | 2.8E-08 |
| *nspc-5* | WBGene00044340 | protein_coding_gene | -2.1 | 0.236 | 1.8E-02 |
| *hlh-10* | WBGene00001954 | protein_coding_gene | -2.1 | 0.236 | 3.3E-02 |
| *fbxa-36* | WBGene00022479 | protein_coding_gene | -2.1 | 0.236 | 4.1E-02 |
| *otpl-1* | WBGene00015035 | protein_coding_gene | -2.1 | 0.235 | 4.1E-06 |
| *C01B9.1* | WBGene00007220 | protein_coding_gene | -2.1 | 0.235 | 8.0E-08 |
| *col-174* | WBGene00000747 | protein_coding_gene | -2.1 | 0.235 | 4.4E-03 |
| *fah-1* | WBGene00019620 | protein_coding_gene | -2.1 | 0.235 | 3.2E-16 |
| *C35B1.5* | WBGene00016435 | protein_coding_gene | -2.1 | 0.234 | 9.2E-14 |
| *T05B11.4* | WBGene00020247 | protein_coding_gene | -2.1 | 0.234 | 3.0E-16 |
| *nhx-2* | WBGene00003730 | protein_coding_gene | -2.1 | 0.234 | 5.7E-23 |
| *M02D8.5* | WBGene00019731 | protein_coding_gene | -2.1 | 0.234 | 4.8E-09 |
| *ctl-1* | WBGene00000830 | protein_coding_gene | -2.1 | 0.234 | 2.1E-03 |
| *add-1* | WBGene00000072 | protein_coding_gene | -2.1 | 0.234 | 1.1E-13 |
| *lev-10* | WBGene00002977 | protein_coding_gene | -2.1 | 0.234 | 1.9E-02 |
| *T10B5.10* | WBGene00020396 | protein_coding_gene | -2.1 | 0.234 | 1.6E-05 |
| *cyp-36A1* | WBGene00007913 | protein_coding_gene | -2.1 | 0.233 | 3.0E-07 |
| *M195.2* | WBGene00010948 | protein_coding_gene | -2.1 | 0.233 | 5.2E-04 |
| *H03E18.1* | WBGene00019148 | protein_coding_gene | -2.1 | 0.233 | 1.7E-07 |
| *tag-180* | WBGene00007041 | protein_coding_gene | -2.1 | 0.233 | 7.8E-12 |
| *C14E2.12* | WBGene00219816 | snoRNA_gene | -2.1 | 0.232 | 9.9E-06 |
| *W03G11.3* | WBGene00012225 | protein_coding_gene | -2.1 | 0.232 | 1.0E-15 |
| *marg-1* | WBGene00018538 | protein_coding_gene | -2.1 | 0.232 | 8.6E-08 |
| *xbx-4* | WBGene00016025 | protein_coding_gene | -2.1 | 0.231 | 2.8E-03 |

| | | | | | |
|---|---|---|---|---|---|
| *K07E3.9* | WBGene00194670 | protein_coding_gene | -2.1 | 0.231 | 5.7E-03 |
| *F54F7.8* | WBGene00010068 | protein_coding_gene | -2.1 | 0.230 | 3.8E-02 |
| *Y41D4B.1* | WBGene00021511 | protein_coding_gene | -2.1 | 0.230 | 5.6E-04 |
| *cnnm-1* | WBGene00016879 | protein_coding_gene | -2.1 | 0.230 | 4.1E-24 |
| *Y54F10AM.8* | WBGene00021852 | protein_coding_gene | -2.1 | 0.230 | 2.1E-15 |
| *K12B6.11* | WBGene00195164 | protein_coding_gene | -2.1 | 0.230 | 3.1E-03 |
| *mbl-1* | WBGene00019347 | protein_coding_gene | -2.1 | 0.229 | 8.4E-10 |
| *mans-2* | WBGene00008258 | protein_coding_gene | -2.1 | 0.229 | 8.5E-11 |
| *B0294.1* | WBGene00015117 | protein_coding_gene | -2.1 | 0.228 | 1.8E-04 |
| *pbo-5* | WBGene00012857 | protein_coding_gene | -2.1 | 0.228 | 2.7E-11 |
| *F16F9.3* | WBGene00017514 | protein_coding_gene | -2.1 | 0.228 | 2.4E-05 |
| *C25E10.7* | WBGene00016096 | protein_coding_gene | -2.1 | 0.228 | 3.0E-02 |
| *mxt-1* | WBGene00012478 | protein_coding_gene | -2.1 | 0.228 | 4.7E-16 |
| *T18D3.5* | WBGene00011822 | protein_coding_gene | -2.1 | 0.227 | 2.4E-03 |
| *dsc-4* | WBGene00001099 | protein_coding_gene | -2.1 | 0.227 | 8.5E-11 |
| *Y26E6A.3* | WBGene00012514 | protein_coding_gene | -2.1 | 0.227 | 1.3E-18 |
| *R13D7.2* | WBGene00020054 | protein_coding_gene | -2.1 | 0.227 | 2.1E-10 |
| *pqn-88* | WBGene00004168 | protein_coding_gene | -2.1 | 0.227 | 1.3E-10 |
| *F35F10.6* | WBGene00018056 | protein_coding_gene | -2.1 | 0.227 | 2.2E-05 |
| *pepm-1* | WBGene00009807 | protein_coding_gene | -2.1 | 0.227 | 1.2E-03 |
| *ugt-7* | WBGene00015739 | protein_coding_gene | -2.1 | 0.226 | 1.3E-10 |
| *C37A5.11* | WBGene00077520 | protein_coding_gene | -2.1 | 0.226 | 2.2E-02 |
| *ser-6* | WBGene00021897 | protein_coding_gene | -2.1 | 0.226 | 1.9E-04 |
| *D1007.13* | WBGene00017009 | protein_coding_gene | -2.2 | 0.225 | 3.6E-03 |
| *dgat-2* | WBGene00010296 | protein_coding_gene | -2.2 | 0.225 | 1.8E-08 |
| *F41E6.15* | WBGene00044554 | protein_coding_gene | -2.2 | 0.225 | 3.4E-20 |
| *mtd-1* | WBGene00003471 | protein_coding_gene | -2.2 | 0.224 | 1.1E-05 |
| *trpa-1* | WBGene00007801 | protein_coding_gene | -2.2 | 0.223 | 5.1E-16 |
| *T22F7.4* | WBGene00020703 | protein_coding_gene | -2.2 | 0.223 | 1.4E-07 |
| *gst-36* | WBGene00001784 | protein_coding_gene | -2.2 | 0.223 | 2.1E-08 |
| *nlf-1* | WBGene00018854 | protein_coding_gene | -2.2 | 0.222 | 9.6E-16 |
| *F42C5.6* | WBGene00018348 | protein_coding_gene | -2.2 | 0.222 | 1.5E-02 |
| *T06G6.3* | WBGene00011545 | protein_coding_gene | -2.2 | 0.222 | 2.7E-03 |
| *F56A12.2* | WBGene00010143 | protein_coding_gene | -2.2 | 0.222 | 2.7E-03 |
| *nlp-42* | WBGene00013588 | protein_coding_gene | -2.2 | 0.221 | 1.6E-07 |
| *R07B1.8* | WBGene00001608 | protein_coding_gene | -2.2 | 0.221 | 2.9E-04 |
| *ser-2* | WBGene00004777 | protein_coding_gene | -2.2 | 0.220 | 7.5E-05 |
| *F48G7.8* | WBGene00018619 | protein_coding_gene | -2.2 | 0.219 | 2.9E-02 |
| *nlp-70* | WBGene00044513 | protein_coding_gene | -2.2 | 0.219 | 3.6E-05 |
| *T04B2.3* | WBGene00011422 | protein_coding_gene | -2.2 | 0.219 | 6.7E-12 |
| *M6.4* | WBGene00019776 | protein_coding_gene | -2.2 | 0.219 | 6.5E-03 |
| *F58H7.1* | WBGene00019067 | protein_coding_gene | -2.2 | 0.219 | 1.9E-03 |
| *C11E4.8* | WBGene00007522 | protein_coding_gene | -2.2 | 0.218 | 5.7E-03 |
| *T28B8.3* | WBGene00012114 | protein_coding_gene | -2.2 | 0.218 | 1.0E-03 |
| *gcp-2.1* | WBGene00020082 | protein_coding_gene | -2.2 | 0.218 | 2.2E-23 |
| *npr-6* | WBGene00009619 | protein_coding_gene | -2.2 | 0.217 | 2.3E-04 |
| *ZK105.12* | WBGene00235113 | protein_coding_gene | -2.2 | 0.217 | 1.2E-02 |
| *gst-27* | WBGene00001775 | protein_coding_gene | -2.2 | 0.217 | 1.4E-11 |
| *C49F8.15* | WBGene00304787 | protein_coding_gene | -2.2 | 0.217 | 4.5E-02 |
| *K04C2.5* | WBGene00019382 | protein_coding_gene | -2.2 | 0.216 | 7.2E-03 |
| *C28H8.5* | WBGene00016196 | protein_coding_gene | -2.2 | 0.216 | 2.3E-02 |
| *irk-1* | WBGene00002149 | protein_coding_gene | -2.2 | 0.216 | 1.7E-03 |
| *rig-6* | WBGene00016354 | protein_coding_gene | -2.2 | 0.215 | 1.6E-17 |
| *oig-5* | WBGene00023520 | protein_coding_gene | -2.2 | 0.215 | 2.3E-04 |
| *C44H4.8* | WBGene00008096 | protein_coding_gene | -2.2 | 0.215 | 3.3E-03 |
| *K03B8.6* | WBGene00010524 | protein_coding_gene | -2.2 | 0.215 | 1.7E-13 |
| *clec-205* | WBGene00021602 | protein_coding_gene | -2.2 | 0.215 | 4.1E-11 |
| *eat-18* | WBGene00001147 | protein_coding_gene | -2.2 | 0.215 | 4.4E-02 |
| *xbx-9* | WBGene00007604 | protein_coding_gene | -2.2 | 0.214 | 2.6E-06 |
| *K04G2.7* | WBGene00010566 | protein_coding_gene | -2.2 | 0.214 | 2.8E-04 |
| *R102.11* | WBGene00077714 | protein_coding_gene | -2.2 | 0.214 | 3.0E-02 |
| *Y45F10D.14* | WBGene00012892 | protein_coding_gene | -2.2 | 0.214 | 1.1E-02 |
| *glr-4* | WBGene00001615 | protein_coding_gene | -2.2 | 0.213 | 4.0E-10 |
| *myo-2* | WBGene00003514 | protein_coding_gene | -2.2 | 0.213 | 8.3E-11 |
| *unc-58* | WBGene00006792 | protein_coding_gene | -2.2 | 0.213 | 7.4E-18 |
| *K09F6.10* | WBGene00019593 | protein_coding_gene | -2.2 | 0.213 | 4.3E-05 |
| *nlp-63* | WBGene00044028 | protein_coding_gene | -2.2 | 0.212 | 2.6E-03 |
| *R05C11.4* | WBGene00044467 | protein_coding_gene | -2.2 | 0.212 | 1.6E-12 |
| *shl-1* | WBGene00022240 | protein_coding_gene | -2.2 | 0.212 | 1.6E-10 |
| *C50E3.6* | WBGene00016817 | protein_coding_gene | -2.2 | 0.212 | 6.4E-08 |

| | | | | | |
|---|---|---|---|---|---|
| *C54D2.2* | WBGene00016916 | protein_coding_gene | -2.2 | 0.212 | 4.6E-04 |
| *clik-2* | WBGene00016898 | protein_coding_gene | -2.2 | 0.211 | 3.4E-14 |
| *Y52E8A.6* | WBGene00021798 | protein_coding_gene | -2.2 | 0.210 | 4.8E-02 |
| *C03B1.5* | WBGene00015376 | protein_coding_gene | -2.2 | 0.210 | 6.7E-03 |
| *T25E4.1* | WBGene00020803 | protein_coding_gene | -2.3 | 0.210 | 1.6E-04 |
| *K02E2.1* | WBGene00010506 | protein_coding_gene | -2.3 | 0.210 | 3.7E-02 |
| *ZK470.6* | WBGene00022747 | protein_coding_gene | -2.3 | 0.209 | 6.2E-03 |
| *chil-3* | WBGene00007467 | protein_coding_gene | -2.3 | 0.209 | 8.2E-04 |
| *nas-23* | WBGene00003542 | protein_coding_gene | -2.3 | 0.209 | 1.1E-02 |
| *srd-5* | WBGene00005083 | protein_coding_gene | -2.3 | 0.209 | 5.0E-02 |
| *cyp-33C8* | WBGene00019967 | protein_coding_gene | -2.3 | 0.209 | 1.7E-07 |
| *Y19D10B.6* | WBGene00021235 | protein_coding_gene | -2.3 | 0.209 | 2.6E-09 |
| *kcnl-1* | WBGene00007176 | protein_coding_gene | -2.3 | 0.209 | 2.7E-09 |
| *nhr-157* | WBGene00015900 | protein_coding_gene | -2.3 | 0.208 | 3.6E-02 |
| *comt-3* | WBGene00021487 | protein_coding_gene | -2.3 | 0.208 | 9.7E-10 |
| *C03B1.6* | WBGene00015377 | protein_coding_gene | -2.3 | 0.208 | 4.5E-02 |
| *Y43F8B.18* | WBGene00050911 | protein_coding_gene | -2.3 | 0.208 | 4.3E-02 |
| *ins-30* | WBGene00002113 | protein_coding_gene | -2.3 | 0.208 | 7.1E-06 |
| *slc-17.3* | WBGene00015333 | protein_coding_gene | -2.3 | 0.208 | 5.0E-15 |
| *math-18* | WBGene00016544 | protein_coding_gene | -2.3 | 0.208 | 1.6E-07 |
| *F14D7.12* | WBGene00045331 | protein_coding_gene | -2.3 | 0.207 | 1.6E-03 |
| *F49C5.3* | WBGene00009866 | protein_coding_gene | -2.3 | 0.207 | 4.9E-02 |
| *pde-6* | WBGene00022389 | protein_coding_gene | -2.3 | 0.207 | 3.0E-14 |
| *pals-1* | WBGene00008858 | protein_coding_gene | -2.3 | 0.206 | 5.3E-04 |
| *F10D7.11* | WBGene00303423 | protein_coding_gene | -2.3 | 0.206 | 1.4E-04 |
| *D1005.10* | WBGene00206384 | protein_coding_gene | -2.3 | 0.206 | 3.4E-02 |
| *Y70D2A.1* | WBGene00013496 | protein_coding_gene | -2.3 | 0.206 | 1.2E-03 |
| *T14G8.6* | WBGene00304822 | protein_coding_gene | -2.3 | 0.206 | 1.1E-02 |
| *Y54G2A.48* | WBGene00044491 | protein_coding_gene | -2.3 | 0.206 | 1.4E-05 |
| *ckc-1* | WBGene00000515 | protein_coding_gene | -2.3 | 0.206 | 2.0E-11 |
| *lact-1* | WBGene00018523 | protein_coding_gene | -2.3 | 0.205 | 2.0E-05 |
| *W03G11.2* | WBGene00012224 | protein_coding_gene | -2.3 | 0.205 | 1.9E-05 |
| *aco-1* | WBGene00000040 | protein_coding_gene | -2.3 | 0.205 | 1.3E-26 |
| *col-65* | WBGene00000641 | protein_coding_gene | -2.3 | 0.205 | 1.2E-05 |
| *Y73E7A.3* | WBGene00022270 | protein_coding_gene | -2.3 | 0.205 | 2.9E-11 |
| *comt-1* | WBGene00018741 | protein_coding_gene | -2.3 | 0.204 | 3.1E-02 |
| *C24A3.1* | WBGene00016032 | protein_coding_gene | -2.3 | 0.204 | 2.8E-02 |
| *acbp-6* | WBGene00012457 | protein_coding_gene | -2.3 | 0.204 | 1.9E-06 |
| *npr-16* | WBGene00006864 | protein_coding_gene | -2.3 | 0.204 | 3.4E-06 |
| *T24F1.4* | WBGene00011996 | protein_coding_gene | -2.3 | 0.203 | 4.9E-03 |
| *C31H2.4* | WBGene00016294 | protein_coding_gene | -2.3 | 0.203 | 1.5E-09 |
| *F20D1.22* | WBGene00304908 | protein_coding_gene | -2.3 | 0.203 | 1.8E-10 |
| *scav-4* | WBGene00008692 | protein_coding_gene | -2.3 | 0.203 | 1.2E-11 |
| *linc-45* | WBGene00219661 | lincRNA_gene | -2.3 | 0.202 | 1.7E-02 |
| *nrfl-1* | WBGene00006438 | protein_coding_gene | -2.3 | 0.202 | 4.2E-14 |
| *ctsa-3.1* | WBGene00017969 | protein_coding_gene | -2.3 | 0.202 | 7.9E-15 |
| *clec-53* | WBGene00020191 | protein_coding_gene | -2.3 | 0.202 | 4.7E-09 |
| *T06G6.4* | WBGene00011546 | protein_coding_gene | -2.3 | 0.202 | 2.5E-05 |
| *nac-3* | WBGene00003519 | protein_coding_gene | -2.3 | 0.202 | 5.2E-16 |
| *K02F2.5* | WBGene00019324 | protein_coding_gene | -2.3 | 0.201 | 2.2E-16 |
| *cyp-35D1* | WBGene00008829 | protein_coding_gene | -2.3 | 0.201 | 5.6E-06 |
| *C10G8.3* | WBGene00015682 | protein_coding_gene | -2.3 | 0.200 | 3.5E-02 |
| *Y53C10A.5* | WBGene00013136 | protein_coding_gene | -2.3 | 0.200 | 5.6E-16 |
| *ctsa-2* | WBGene00019617 | protein_coding_gene | -2.3 | 0.200 | 2.9E-10 |
| *slc-17.8* | WBGene00017798 | protein_coding_gene | -2.3 | 0.199 | 5.5E-04 |
| *folt-3* | WBGene00044738 | protein_coding_gene | -2.3 | 0.199 | 4.0E-02 |
| *nhr-30* | WBGene00016091 | protein_coding_gene | -2.3 | 0.199 | 4.4E-02 |
| *inx-18* | WBGene00002140 | protein_coding_gene | -2.3 | 0.199 | 8.1E-06 |
| *ceh-54* | WBGene00020485 | protein_coding_gene | -2.3 | 0.199 | 3.4E-02 |
| *Y37F4.7* | WBGene00021388 | protein_coding_gene | -2.3 | 0.199 | 1.8E-07 |
| *ceh-14* | WBGene00000438 | protein_coding_gene | -2.3 | 0.198 | 4.6E-10 |
| *F02E11.2* | WBGene00017180 | protein_coding_gene | -2.3 | 0.198 | 1.3E-04 |
| *piit-1* | WBGene00009384 | protein_coding_gene | -2.3 | 0.198 | 5.2E-06 |
| *F52F10.2* | WBGene00018706 | protein_coding_gene | -2.3 | 0.197 | 5.3E-04 |
| *srx-58* | WBGene00005949 | protein_coding_gene | -2.3 | 0.197 | 1.8E-02 |
| *C29F9.6* | WBGene00016221 | protein_coding_gene | -2.3 | 0.196 | 9.9E-03 |
| *C02G6.1* | WBGene00015359 | pseudogene | -2.3 | 0.196 | 2.3E-11 |
| *pgp-12* | WBGene00004006 | protein_coding_gene | -2.4 | 0.196 | 4.3E-12 |
| *sri-22* | WBGene00005534 | pseudogene | -2.4 | 0.196 | 4.5E-02 |
| *C10E2.1* | WBGene00015672 | protein_coding_gene | -2.4 | 0.196 | 8.2E-04 |

| | | | | | |
|---|---|---|---|---|---|
| *nhr-148* | WBGene00015396 | protein_coding_gene | -2.4 | 0.196 | 1.4E-03 |
| *clec-229* | WBGene00010355 | protein_coding_gene | -2.4 | 0.195 | 3.7E-02 |
| *act-5* | WBGene00000067 | protein_coding_gene | -2.4 | 0.194 | 9.8E-09 |
| *ugt-48* | WBGene00015965 | protein_coding_gene | -2.4 | 0.194 | 1.1E-12 |
| *C33C12.11* | WBGene00044360 | protein_coding_gene | -2.4 | 0.194 | 1.4E-02 |
| *nlp-59* | WBGene00008522 | protein_coding_gene | -2.4 | 0.194 | 9.5E-13 |
| *ugt-47* | WBGene00011006 | protein_coding_gene | -2.4 | 0.194 | 7.2E-09 |
| *gst-42* | WBGene00001790 | protein_coding_gene | -2.4 | 0.193 | 1.7E-12 |
| *F55C10.4* | WBGene00010108 | protein_coding_gene | -2.4 | 0.192 | 3.5E-02 |
| *cest-7* | WBGene00007693 | protein_coding_gene | -2.4 | 0.192 | 1.6E-04 |
| *Y47D3B.6* | WBGene00012942 | protein_coding_gene | -2.4 | 0.192 | 1.2E-06 |
| *D1005.4* | WBGene00016998 | protein_coding_gene | -2.4 | 0.192 | 1.4E-09 |
| *glb-13* | WBGene00008957 | protein_coding_gene | -2.4 | 0.192 | 1.8E-07 |
| *F48C5.2* | WBGene00009843 | protein_coding_gene | -2.4 | 0.191 | 1.2E-10 |
| *pho-8* | WBGene00011262 | protein_coding_gene | -2.4 | 0.191 | 1.8E-04 |
| *C54D2.1* | WBGene00016915 | protein_coding_gene | -2.4 | 0.191 | 1.3E-05 |
| *F41C6.4* | WBGene00018278 | protein_coding_gene | -2.4 | 0.191 | 7.7E-04 |
| *C33D3.4* | WBGene00007895 | protein_coding_gene | -2.4 | 0.190 | 3.7E-02 |
| *nep-17* | WBGene00010070 | protein_coding_gene | -2.4 | 0.190 | 4.7E-16 |
| *btb-22* | WBGene00269387 | protein_coding_gene | -2.4 | 0.190 | 3.3E-02 |
| *Y43F8C.13* | WBGene00012834 | protein_coding_gene | -2.4 | 0.189 | 3.0E-09 |
| *ctsa-1.2* | WBGene00008741 | protein_coding_gene | -2.4 | 0.189 | 2.9E-16 |
| *srv-1* | WBGene00005712 | protein_coding_gene | -2.4 | 0.189 | 3.1E-29 |
| *B0244.17* | WBGene00271715 | protein_coding_gene | -2.4 | 0.189 | 2.6E-04 |
| *ZK328.6* | WBGene00022695 | protein_coding_gene | -2.4 | 0.189 | 7.3E-13 |
| *F41E7.6* | WBGene00009622 | protein_coding_gene | -2.4 | 0.189 | 3.4E-02 |
| *K01D12.8* | WBGene00010467 | protein_coding_gene | -2.4 | 0.188 | 1.6E-02 |
| *K08C7.7* | WBGene00010649 | protein_coding_gene | -2.4 | 0.188 | 6.3E-07 |
| *C35C5.10* | WBGene00007959 | protein_coding_gene | -2.4 | 0.187 | 1.6E-27 |
| *kvs-5* | WBGene00021948 | protein_coding_gene | -2.4 | 0.187 | 1.2E-08 |
| *T28F4.8* | WBGene00271567 | protein_coding_gene | -2.4 | 0.187 | 4.3E-02 |
| *gem-1* | WBGene00008214 | protein_coding_gene | -2.4 | 0.186 | 2.3E-17 |
| *Y5H2A.4* | WBGene00044521 | protein_coding_gene | -2.4 | 0.186 | 4.7E-02 |
| *F14H8.4* | WBGene00008834 | protein_coding_gene | -2.4 | 0.186 | 2.0E-02 |
| *ZK185.3* | WBGene00022683 | protein_coding_gene | -2.4 | 0.186 | 1.6E-10 |
| *dpy-9* | WBGene00001071 | protein_coding_gene | -2.4 | 0.186 | 1.4E-07 |
| *F40G12.2* | WBGene00009597 | protein_coding_gene | -2.4 | 0.186 | 4.7E-03 |
| *srb-14* | WBGene00016338 | pseudogene | -2.4 | 0.185 | 1.2E-02 |
| *srx-123* | WBGene00006014 | pseudogene | -2.4 | 0.185 | 4.3E-02 |
| *Y11D7A.1* | WBGene00012426 | pseudogene | -2.4 | 0.185 | 2.8E-03 |
| *R10E8.10* | WBGene00195374 | ncRNA_gene | -2.4 | 0.184 | 4.1E-02 |
| *B0563.6* | WBGene00015263 | protein_coding_gene | -2.4 | 0.184 | 7.4E-11 |
| *W06D12.8* | WBGene00198082 | ncRNA_gene | -2.4 | 0.184 | 2.7E-02 |
| *glb-16* | WBGene00018486 | protein_coding_gene | -2.4 | 0.184 | 2.7E-03 |
| *ZC334.13* | WBGene00086553 | protein_coding_gene | -2.4 | 0.184 | 2.6E-02 |
| *mab-23* | WBGene00003114 | protein_coding_gene | -2.4 | 0.184 | 3.8E-02 |
| *F21F8.6* | WBGene00017680 | pseudogene | -2.4 | 0.183 | 8.8E-05 |
| *btb-14* | WBGene00013275 | protein_coding_gene | -2.5 | 0.183 | 1.1E-20 |
| *K02H11.4* | WBGene00019348 | protein_coding_gene | -2.5 | 0.183 | 3.5E-02 |
| *T03G11.10* | WBGene00044732 | protein_coding_gene | -2.5 | 0.183 | 9.2E-03 |
| *bca-1* | WBGene00000245 | protein_coding_gene | -2.5 | 0.182 | 1.1E-15 |
| *lgc-12* | WBGene00020048 | protein_coding_gene | -2.5 | 0.182 | 2.5E-04 |
| *C04G6.2* | WBGene00015453 | protein_coding_gene | -2.5 | 0.182 | 1.3E-04 |
| *gst-21* | WBGene00001769 | protein_coding_gene | -2.5 | 0.181 | 2.1E-08 |
| *F17C11.12* | WBGene00008923 | protein_coding_gene | -2.5 | 0.181 | 2.6E-17 |
| *F35E2.7* | WBGene00009414 | pseudogene | -2.5 | 0.181 | 2.5E-02 |
| *clec-204* | WBGene00021603 | protein_coding_gene | -2.5 | 0.180 | 8.1E-14 |
| *cyp-34A5* | WBGene00015040 | protein_coding_gene | -2.5 | 0.180 | 1.7E-05 |
| *Y60A3A.25* | WBGene00044211 | protein_coding_gene | -2.5 | 0.180 | 1.1E-10 |
| *Y110A7A.23* | WBGene00201165 | ncRNA_gene | -2.5 | 0.179 | 4.8E-02 |
| *col-182* | WBGene00000755 | pseudogene | -2.5 | 0.179 | 1.8E-05 |
| *R09H10.6* | WBGene00011183 | protein_coding_gene | -2.5 | 0.179 | 3.6E-02 |
| *gst-10* | WBGene00001758 | protein_coding_gene | -2.5 | 0.179 | 1.9E-08 |
| *C05D9.12* | WBGene00219800 | ncRNA_gene | -2.5 | 0.178 | 4.4E-06 |
| *nep-22* | WBGene00020788 | protein_coding_gene | -2.5 | 0.178 | 8.1E-10 |
| *aakg-3* | WBGene00021527 | protein_coding_gene | -2.5 | 0.177 | 1.1E-08 |
| *C33D3.5* | WBGene00007896 | protein_coding_gene | -2.5 | 0.176 | 3.1E-02 |
| *rgs-2* | WBGene00004345 | protein_coding_gene | -2.5 | 0.176 | 3.4E-10 |
| *ZK1025.4* | WBGene00014184 | protein_coding_gene | -2.5 | 0.176 | 3.3E-02 |
| *C02B8.3* | WBGene00015322 | protein_coding_gene | -2.5 | 0.175 | 3.5E-08 |

| | | | | | |
|---|---|---|---|---|---|
| *F22H10.6* | WBGene00017730 | protein_coding_gene | -2.5 | 0.175 | 7.4E-07 |
| *K02E10.7* | WBGene00019321 | protein_coding_gene | -2.5 | 0.175 | 5.2E-08 |
| *dmsr-16* | WBGene00194821 | protein_coding_gene | -2.5 | 0.174 | 1.4E-02 |
| *pudl-2* | WBGene00044490 | protein_coding_gene | -2.5 | 0.174 | 1.8E-06 |
| *W04G3.11* | WBGene00044153 | protein_coding_gene | -2.5 | 0.173 | 4.0E-03 |
| *vha-3* | WBGene00006912 | protein_coding_gene | -2.5 | 0.173 | 2.7E-06 |
| *gst-9* | WBGene00001757 | protein_coding_gene | -2.5 | 0.173 | 3.7E-08 |
| *sptl-3* | WBGene00011932 | protein_coding_gene | -2.5 | 0.172 | 4.7E-23 |
| *nlp-16* | WBGene00003754 | protein_coding_gene | -2.5 | 0.172 | 8.7E-29 |
| *Y52B11B.1* | WBGene00013130 | protein_coding_gene | -2.5 | 0.171 | 3.2E-02 |
| *ZK105.6* | WBGene00022657 | protein_coding_gene | -2.5 | 0.171 | 9.4E-05 |
| *pqn-75* | WBGene00004157 | protein_coding_gene | -2.6 | 0.171 | 3.3E-18 |
| *twk-17* | WBGene00006671 | protein_coding_gene | -2.6 | 0.170 | 5.1E-10 |
| *sri-8* | WBGene00005520 | pseudogene | -2.6 | 0.170 | 1.1E-02 |
| *cah-3* | WBGene00000281 | protein_coding_gene | -2.6 | 0.170 | 3.3E-31 |
| *C44C10.11* | WBGene00043985 | protein_coding_gene | -2.6 | 0.170 | 4.7E-17 |
| *K02B12.9* | WBGene00044679 | protein_coding_gene | -2.6 | 0.170 | 7.6E-03 |
| *Y48G8AL.16* | WBGene00195068 | protein_coding_gene | -2.6 | 0.169 | 4.5E-02 |
| *gst-2* | WBGene00001750 | protein_coding_gene | -2.6 | 0.169 | 3.9E-05 |
| *glb-30* | WBGene00021259 | protein_coding_gene | -2.6 | 0.169 | 1.5E-02 |
| *F59F5.5* | WBGene00010344 | protein_coding_gene | -2.6 | 0.169 | 5.1E-05 |
| *aat-4* | WBGene00000005 | protein_coding_gene | -2.6 | 0.169 | 4.5E-12 |
| *gpa-10* | WBGene00001672 | protein_coding_gene | -2.6 | 0.168 | 2.6E-03 |
| *F32E10.8* | WBGene00017995 | protein_coding_gene | -2.6 | 0.168 | 8.6E-04 |
| *K09H11.6* | WBGene00019603 | protein_coding_gene | -2.6 | 0.168 | 5.0E-03 |
| *K05F1.14* | WBGene00271818 | protein_coding_gene | -2.6 | 0.168 | 3.5E-04 |
| *grl-9* | WBGene00001718 | protein_coding_gene | -2.6 | 0.168 | 2.8E-02 |
| *C34F6.1* | WBGene00007938 | protein_coding_gene | -2.6 | 0.168 | 3.3E-05 |
| *ZK742.3* | WBGene00022813 | protein_coding_gene | -2.6 | 0.168 | 7.9E-17 |
| *F52H2.6* | WBGene00018718 | protein_coding_gene | -2.6 | 0.168 | 2.5E-10 |
| *ZK822.10* | WBGene00219456 | protein_coding_gene | -2.6 | 0.167 | 1.7E-02 |
| *F07H5.8* | WBGene00008559 | protein_coding_gene | -2.6 | 0.167 | 1.3E-05 |
| *F47F2.1* | WBGene00018569 | protein_coding_gene | -2.6 | 0.166 | 1.2E-19 |
| *Y16B4A.2* | WBGene00012445 | protein_coding_gene | -2.6 | 0.166 | 2.0E-12 |
| *cyp-13A2* | WBGene00011676 | protein_coding_gene | -2.6 | 0.166 | 5.6E-11 |
| *scl-16* | WBGene00009425 | pseudogene | -2.6 | 0.166 | 3.6E-02 |
| *C36A4.14* | WBGene00194650 | pseudogene | -2.6 | 0.166 | 2.1E-02 |
| *Y67H2A.9* | WBGene00014941 | pseudogene | -2.6 | 0.166 | 4.1E-13 |
| *fbxa-38* | WBGene00016522 | protein_coding_gene | -2.6 | 0.165 | 2.0E-09 |
| *wrt-4* | WBGene00006950 | protein_coding_gene | -2.6 | 0.165 | 5.2E-09 |
| *lgc-21* | WBGene00007903 | protein_coding_gene | -2.6 | 0.164 | 1.8E-07 |
| *nas-7* | WBGene00003526 | protein_coding_gene | -2.6 | 0.164 | 3.1E-10 |
| *T28C12.2* | WBGene00020889 | protein_coding_gene | -2.6 | 0.164 | 1.1E-06 |
| *T13C5.9* | WBGene00195168 | protein_coding_gene | -2.6 | 0.164 | 2.5E-02 |
| *R02E12.5* | WBGene00019829 | protein_coding_gene | -2.6 | 0.163 | 4.7E-12 |
| *F55D10.4* | WBGene00018879 | protein_coding_gene | -2.6 | 0.163 | 4.4E-05 |
| *aexr-3* | WBGene00016748 | protein_coding_gene | -2.6 | 0.163 | 2.2E-03 |
| *ZK662.2* | WBGene00014039 | protein_coding_gene | -2.6 | 0.162 | 2.2E-07 |
| *fipr-7* | WBGene00007543 | protein_coding_gene | -2.6 | 0.162 | 6.0E-04 |
| *K08H2.7* | WBGene00010693 | protein_coding_gene | -2.6 | 0.162 | 2.6E-04 |
| *nas-30* | WBGene00003548 | protein_coding_gene | -2.6 | 0.161 | 7.0E-07 |
| *che-6* | WBGene00000487 | protein_coding_gene | -2.6 | 0.161 | 1.2E-02 |
| *clec-168* | WBGene00009520 | protein_coding_gene | -2.6 | 0.161 | 1.8E-02 |
| *gba-4* | WBGene00021160 | protein_coding_gene | -2.6 | 0.161 | 1.4E-08 |
| *col-58* | WBGene00000634 | protein_coding_gene | -2.6 | 0.161 | 3.1E-06 |
| *ifa-1* | WBGene00002050 | protein_coding_gene | -2.6 | 0.161 | 1.2E-23 |
| *B0511.17* | WBGene00219234 | protein_coding_gene | -2.6 | 0.161 | 2.6E-02 |
| *Y45F10B.13* | WBGene00012875 | protein_coding_gene | -2.6 | 0.160 | 5.5E-33 |
| *C07G1.7* | WBGene00015573 | protein_coding_gene | -2.6 | 0.160 | 6.4E-06 |
| *T25G12.6* | WBGene00020813 | protein_coding_gene | -2.6 | 0.160 | 1.1E-14 |
| *F31D5.1* | WBGene00017944 | protein_coding_gene | -2.6 | 0.160 | 2.2E-05 |
| *ttr-14* | WBGene00011460 | protein_coding_gene | -2.6 | 0.160 | 2.6E-14 |
| *C35A11.4* | WBGene00016431 | protein_coding_gene | -2.6 | 0.160 | 7.0E-10 |
| *21ur-8714* | WBGene00174936 | piRNA_gene | -2.6 | 0.160 | 9.9E-03 |
| *apt-9* | WBGene00000166 | protein_coding_gene | -2.7 | 0.159 | 4.3E-06 |
| *ppat-1* | WBGene00011407 | protein_coding_gene | -2.7 | 0.159 | 3.9E-10 |
| *21ur-14029* | WBGene00173683 | piRNA_gene | -2.7 | 0.158 | 2.0E-02 |
| *F45B8.5* | WBGene00194794 | protein_coding_gene | -2.7 | 0.157 | 5.9E-04 |
| *F35A5.1* | WBGene00018024 | protein_coding_gene | -2.7 | 0.156 | 9.6E-15 |
| *Y66C5A.1* | WBGene00044156 | protein_coding_gene | -2.7 | 0.156 | 6.2E-08 |

| | | | | | |
|---|---|---|---|---|---|
| *F19C6.4* | WBGene00008953 | protein_coding_gene | -2.7 | 0.156 | 8.7E-20 |
| *F17C11.13* | WBGene00023503 | protein_coding_gene | -2.7 | 0.155 | 6.3E-05 |
| *col-117* | WBGene00000691 | protein_coding_gene | -2.7 | 0.155 | 4.2E-03 |
| *gst-39* | WBGene00001787 | protein_coding_gene | -2.7 | 0.155 | 5.7E-14 |
| *gldc-1* | WBGene00020022 | protein_coding_gene | -2.7 | 0.154 | 4.6E-15 |
| *try-4* | WBGene00006622 | protein_coding_gene | -2.7 | 0.154 | 3.8E-04 |
| *T22B2.6* | WBGene00020670 | protein_coding_gene | -2.7 | 0.154 | 2.2E-02 |
| *gst-26* | WBGene00001774 | protein_coding_gene | -2.7 | 0.154 | 2.5E-07 |
| *ZK1037.6* | WBGene00014194 | protein_coding_gene | -2.7 | 0.154 | 4.1E-12 |
| *Y70C5A.2* | WBGene00013490 | protein_coding_gene | -2.7 | 0.154 | 1.3E-02 |
| *bas-1* | WBGene00000239 | protein_coding_gene | -2.7 | 0.153 | 2.1E-09 |
| *arrd-9* | WBGene00008843 | protein_coding_gene | -2.7 | 0.153 | 1.6E-04 |
| *elo-9* | WBGene00001247 | protein_coding_gene | -2.7 | 0.152 | 1.6E-09 |
| *K01D12.5* | WBGene00010464 | protein_coding_gene | -2.7 | 0.152 | 4.1E-04 |
| *lin-29* | WBGene00003015 | protein_coding_gene | -2.7 | 0.152 | 1.2E-12 |
| *ZC302.3* | WBGene00013863 | protein_coding_gene | -2.7 | 0.151 | 1.3E-06 |
| *nkb-3* | WBGene00010117 | protein_coding_gene | -2.7 | 0.151 | 2.5E-27 |
| *Y38E10A.8* | WBGene00012586 | protein_coding_gene | -2.7 | 0.151 | 3.1E-02 |
| *sra-37* | WBGene00005063 | protein_coding_gene | -2.7 | 0.150 | 2.9E-03 |
| *col-108* | WBGene00000682 | protein_coding_gene | -2.7 | 0.150 | 2.1E-02 |
| *srm-5* | WBGene00044112 | protein_coding_gene | -2.7 | 0.150 | 1.6E-02 |
| *chil-26* | WBGene00011847 | protein_coding_gene | -2.7 | 0.150 | 4.4E-03 |
| *col-107* | WBGene00000681 | protein_coding_gene | -2.7 | 0.149 | 4.9E-10 |
| *glc-4* | WBGene00001594 | protein_coding_gene | -2.8 | 0.148 | 1.9E-10 |
| *nhr-31* | WBGene00003625 | protein_coding_gene | -2.8 | 0.148 | 2.0E-15 |
| *linc-125* | WBGene00219686 | lincRNA_gene | -2.8 | 0.148 | 7.8E-03 |
| *ins-9* | WBGene00002092 | protein_coding_gene | -2.8 | 0.147 | 1.0E-02 |
| *C33E10.10* | WBGene00016350 | protein_coding_gene | -2.8 | 0.147 | 1.4E-02 |
| *sptl-2* | WBGene00018398 | protein_coding_gene | -2.8 | 0.147 | 2.3E-11 |
| *smd-1* | WBGene00004875 | protein_coding_gene | -2.8 | 0.147 | 1.1E-12 |
| *T11B7.2* | WBGene00011707 | protein_coding_gene | -2.8 | 0.146 | 1.0E-06 |
| *ceh-18* | WBGene00000441 | protein_coding_gene | -2.8 | 0.146 | 1.2E-28 |
| *F14D7.5* | WBGene00008792 | protein_coding_gene | -2.8 | 0.146 | 1.7E-02 |
| *ZK180.5* | WBGene00022679 | protein_coding_gene | -2.8 | 0.146 | 1.1E-09 |
| *T02G6.1* | WBGene00011384 | protein_coding_gene | -2.8 | 0.146 | 5.4E-03 |
| *pqn-74* | WBGene00004156 | protein_coding_gene | -2.8 | 0.146 | 1.7E-09 |
| *cdh-7* | WBGene00000399 | protein_coding_gene | -2.8 | 0.145 | 5.3E-05 |
| *Y50D4B.7* | WBGene00021746 | pseudogene | -2.8 | 0.145 | 2.0E-02 |
| *F57B1.10* | WBGene00196471 | ncRNA_gene | -2.8 | 0.144 | 3.7E-02 |
| *tbb-6* | WBGene00006539 | protein_coding_gene | -2.8 | 0.144 | 6.6E-08 |
| *ZK177.9* | WBGene00022674 | protein_coding_gene | -2.8 | 0.144 | 2.3E-02 |
| *C33D3.3* | WBGene00007894 | protein_coding_gene | -2.8 | 0.143 | 1.7E-07 |
| *bli-1* | WBGene00000251 | protein_coding_gene | -2.8 | 0.142 | 5.7E-04 |
| *F43D2.3* | WBGene00009652 | protein_coding_gene | -2.8 | 0.142 | 1.1E-03 |
| *F53B1.3* | WBGene00018736 | protein_coding_gene | -2.8 | 0.141 | 2.4E-03 |
| *cutl-18* | WBGene00009830 | protein_coding_gene | -2.8 | 0.140 | 2.0E-07 |
| *C03F11.4* | WBGene00015390 | protein_coding_gene | -2.8 | 0.140 | 2.5E-20 |
| *pcp-3* | WBGene00003958 | protein_coding_gene | -2.8 | 0.140 | 3.7E-12 |
| *ZK1086.2* | WBGene00014216 | protein_coding_gene | -2.8 | 0.140 | 9.2E-04 |
| *F23F1.2* | WBGene00017743 | protein_coding_gene | -2.8 | 0.140 | 5.8E-03 |
| *cdh-10* | WBGene00000402 | protein_coding_gene | -2.8 | 0.139 | 2.8E-06 |
| *C01B10.3* | WBGene00015278 | protein_coding_gene | -2.8 | 0.139 | 4.6E-19 |
| *C08H9.18* | WBGene00206395 | protein_coding_gene | -2.8 | 0.139 | 2.6E-02 |
| *B0222.11* | WBGene00044582 | protein_coding_gene | -2.8 | 0.139 | 7.8E-07 |
| *nhr-115* | WBGene00003705 | protein_coding_gene | -2.9 | 0.138 | 2.9E-14 |
| *gcy-9* | WBGene00001536 | protein_coding_gene | -2.9 | 0.138 | 4.0E-08 |
| *nhr-72* | WBGene00003662 | protein_coding_gene | -2.9 | 0.137 | 1.2E-02 |
| *acr-2* | WBGene00000042 | protein_coding_gene | -2.9 | 0.137 | 2.0E-05 |
| *B0280.7* | WBGene00015103 | protein_coding_gene | -2.9 | 0.137 | 6.5E-04 |
| *F26F2.10* | WBGene00138719 | protein_coding_gene | -2.9 | 0.137 | 6.7E-03 |
| *B0244.7* | WBGene00015082 | protein_coding_gene | -2.9 | 0.136 | 1.2E-04 |
| *lon-3* | WBGene00003057 | protein_coding_gene | -2.9 | 0.136 | 1.3E-07 |
| *nspb-6* | WBGene00019152 | protein_coding_gene | -2.9 | 0.135 | 1.1E-03 |
| *linc-21* | WBGene00219688 | lincRNA_gene | -2.9 | 0.134 | 3.1E-05 |
| *F56D5.3* | WBGene00010148 | protein_coding_gene | -2.9 | 0.132 | 3.0E-17 |
| *C15H9.9* | WBGene00015803 | protein_coding_gene | -2.9 | 0.132 | 4.2E-50 |
| *Y60A3A.23* | WBGene00013367 | protein_coding_gene | -2.9 | 0.132 | 2.1E-02 |
| *pals-4* | WBGene00007658 | protein_coding_gene | -2.9 | 0.132 | 8.7E-05 |
| *elo-6* | WBGene00001244 | protein_coding_gene | -2.9 | 0.132 | 3.2E-06 |
| *srt-49* | WBGene00012772 | protein_coding_gene | -2.9 | 0.132 | 1.7E-02 |

| | | | | | |
|---|---|---|---|---|---|
| *clec-22* | WBGene00009855 | protein_coding_gene | -2.9 | 0.131 | 8.9E-03 |
| *nhr-235* | WBGene00012597 | protein_coding_gene | -2.9 | 0.130 | 2.1E-02 |
| *gur-4* | WBGene00001805 | protein_coding_gene | -2.9 | 0.130 | 5.4E-06 |
| *linc-41* | WBGene00219689 | lincRNA_gene | -2.9 | 0.130 | 8.2E-03 |
| *AC3.13* | WBGene00077504 | pseudogene | -2.9 | 0.130 | 1.5E-02 |
| *gon-1* | WBGene00001650 | protein_coding_gene | -3.0 | 0.129 | 7.3E-25 |
| *F11G11.13* | WBGene00017388 | protein_coding_gene | -3.0 | 0.129 | 8.0E-03 |
| *cdr-4* | WBGene00010470 | protein_coding_gene | -3.0 | 0.129 | 1.6E-14 |
| *srz-27* | WBGene00021181 | pseudogene | -3.0 | 0.128 | 1.8E-02 |
| *srv-15* | WBGene00005726 | protein_coding_gene | -3.0 | 0.128 | 3.6E-05 |
| *sra-36* | WBGene00005062 | protein_coding_gene | -3.0 | 0.128 | 4.5E-03 |
| *dmsr-11* | WBGene00022605 | protein_coding_gene | -3.0 | 0.127 | 2.6E-03 |
| *ZK822.8* | WBGene00194665 | antisense_lncRNA_gene | -3.0 | 0.127 | 4.4E-04 |
| *T20D4.7* | WBGene00020613 | protein_coding_gene | -3.0 | 0.127 | 1.1E-03 |
| *K08D9.6* | WBGene00019527 | protein_coding_gene | -3.0 | 0.127 | 1.8E-02 |
| *T02G6.2* | WBGene00011385 | protein_coding_gene | -3.0 | 0.126 | 2.6E-03 |
| *C37H5.14* | WBGene00045308 | protein_coding_gene | -3.0 | 0.126 | 1.6E-13 |
| *R09H10.1* | WBGene00011179 | protein_coding_gene | -3.0 | 0.126 | 1.6E-03 |
| *acox-1.4* | WBGene00008567 | protein_coding_gene | -3.0 | 0.126 | 1.2E-09 |
| *pept-1* | WBGene00003877 | protein_coding_gene | -3.0 | 0.124 | 1.6E-21 |
| *acbp-7* | WBGene00044603 | protein_coding_gene | -3.0 | 0.123 | 8.3E-03 |
| *C14F11.4* | WBGene00015780 | protein_coding_gene | -3.0 | 0.123 | 1.2E-29 |
| *ram-5* | WBGene00004301 | protein_coding_gene | -3.0 | 0.123 | 1.1E-03 |
| *C18A11.4* | WBGene00015950 | protein_coding_gene | -3.0 | 0.122 | 2.5E-06 |
| *C04C3.14* | WBGene00219410 | protein_coding_gene | -3.0 | 0.122 | 2.0E-02 |
| *paic-1* | WBGene00015116 | protein_coding_gene | -3.1 | 0.120 | 5.1E-12 |
| *F49D11.3* | WBGene00018627 | protein_coding_gene | -3.1 | 0.120 | 1.1E-03 |
| *srd-55* | WBGene00005133 | protein_coding_gene | -3.1 | 0.119 | 4.5E-03 |
| *Y43D4A.1* | WBGene00012787 | protein_coding_gene | -3.1 | 0.119 | 7.4E-03 |
| *fbxa-45* | WBGene00008618 | protein_coding_gene | -3.1 | 0.119 | 4.7E-03 |
| *twk-42* | WBGene00006692 | protein_coding_gene | -3.1 | 0.119 | 1.7E-18 |
| *clec-56* | WBGene00008595 | protein_coding_gene | -3.1 | 0.117 | 1.1E-23 |
| *F56C3.4* | WBGene00018941 | protein_coding_gene | -3.1 | 0.116 | 8.6E-03 |
| *egl-17* | WBGene00001185 | protein_coding_gene | -3.1 | 0.116 | 9.6E-04 |
| *W01A8.8* | WBGene00044988 | protein_coding_gene | -3.1 | 0.116 | 5.0E-10 |
| *C17B7.5* | WBGene00015879 | protein_coding_gene | -3.1 | 0.114 | 4.8E-06 |
| *col-94* | WBGene00000669 | protein_coding_gene | -3.1 | 0.114 | 6.8E-03 |
| *M01H9.2* | WBGene00019718 | protein_coding_gene | -3.1 | 0.114 | 3.7E-04 |
| *R07E3.4* | WBGene00011105 | protein_coding_gene | -3.1 | 0.114 | 1.9E-14 |
| *T06E4.14* | WBGene00077691 | protein_coding_gene | -3.1 | 0.114 | 4.8E-06 |
| *fbxa-96* | WBGene00010360 | protein_coding_gene | -3.1 | 0.114 | 5.1E-03 |
| *dmd-8* | WBGene00020708 | protein_coding_gene | -3.1 | 0.114 | 1.0E-03 |
| *tor-2* | WBGene00006598 | protein_coding_gene | -3.1 | 0.113 | 4.2E-03 |
| *abu-11* | WBGene00000034 | protein_coding_gene | -3.1 | 0.113 | 1.9E-07 |
| *Y70C5C.1* | WBGene00013492 | protein_coding_gene | -3.1 | 0.113 | 9.0E-11 |
| *DC2.5* | WBGene00017082 | protein_coding_gene | -3.1 | 0.113 | 8.8E-18 |
| *F31D4.8* | WBGene00009293 | protein_coding_gene | -3.2 | 0.113 | 5.2E-12 |
| *R05G9.5* | WBGene00206514 | protein_coding_gene | -3.2 | 0.112 | 2.9E-03 |
| *ncs-4* | WBGene00008998 | protein_coding_gene | -3.2 | 0.112 | 1.7E-09 |
| *F11F1.8* | WBGene00044399 | protein_coding_gene | -3.2 | 0.112 | 3.2E-03 |
| *F38B6.4* | WBGene00018174 | protein_coding_gene | -3.2 | 0.112 | 4.6E-11 |
| *mec-19* | WBGene00008216 | protein_coding_gene | -3.2 | 0.110 | 4.5E-07 |
| *C06E4.3* | WBGene00015532 | protein_coding_gene | -3.2 | 0.110 | 4.1E-04 |
| *tbc-19* | WBGene00007849 | protein_coding_gene | -3.2 | 0.110 | 1.4E-17 |
| *srm-3* | WBGene00010273 | protein_coding_gene | -3.2 | 0.110 | 3.4E-03 |
| *gst-34* | WBGene00001782 | protein_coding_gene | -3.2 | 0.110 | 1.2E-03 |
| *ugt-65* | WBGene00008583 | protein_coding_gene | -3.2 | 0.109 | 2.1E-08 |
| *F22F1.3* | WBGene00017715 | protein_coding_gene | -3.2 | 0.109 | 9.9E-05 |
| *T20D4.5* | WBGene00020611 | protein_coding_gene | -3.2 | 0.108 | 7.1E-09 |
| *stdh-2* | WBGene00008678 | protein_coding_gene | -3.2 | 0.108 | 2.2E-07 |
| *F16H6.4* | WBGene00008894 | protein_coding_gene | -3.2 | 0.107 | 9.2E-06 |
| *B0462.4* | WBGene00007183 | protein_coding_gene | -3.2 | 0.107 | 2.5E-03 |
| *T23B12.5* | WBGene00020720 | protein_coding_gene | -3.2 | 0.106 | 1.1E-03 |
| *col-48* | WBGene00000625 | protein_coding_gene | -3.2 | 0.106 | 7.5E-13 |
| *col-115* | WBGene00000689 | protein_coding_gene | -3.2 | 0.106 | 2.3E-07 |
| *F10D11.6* | WBGene00008652 | protein_coding_gene | -3.2 | 0.105 | 3.8E-15 |
| *F28A10.5* | WBGene00017873 | protein_coding_gene | -3.2 | 0.105 | 7.7E-03 |
| *C31B8.1* | WBGene00016279 | protein_coding_gene | -3.2 | 0.105 | 3.1E-03 |
| *ZK228.3* | WBGene00013946 | protein_coding_gene | -3.2 | 0.105 | 3.8E-21 |
| *phat-4* | WBGene00020237 | protein_coding_gene | -3.3 | 0.105 | 1.1E-11 |

| | | | | | |
|---|---|---|---|---|---|
| *Y49G5A.1* | WBGene00021731 | protein_coding_gene | -3.3 | 0.105 | 4.1E-03 |
| *ech-7* | WBGene00001156 | protein_coding_gene | -3.3 | 0.104 | 4.5E-16 |
| *him-4* | WBGene00001863 | protein_coding_gene | -3.3 | 0.104 | 7.4E-26 |
| *C53B7.6* | WBGene00016895 | protein_coding_gene | -3.3 | 0.104 | 7.2E-03 |
| *cnnm-3* | WBGene00016343 | protein_coding_gene | -3.3 | 0.104 | 1.3E-26 |
| *F07G6.10* | WBGene00045271 | protein_coding_gene | -3.3 | 0.103 | 2.8E-15 |
| *npax-3* | WBGene00011257 | protein_coding_gene | -3.3 | 0.103 | 1.2E-03 |
| *col-109* | WBGene00000683 | protein_coding_gene | -3.3 | 0.103 | 2.1E-13 |
| *R12E2.7* | WBGene00020033 | protein_coding_gene | -3.3 | 0.102 | 1.3E-07 |
| *F34H10.4* | WBGene00009381 | protein_coding_gene | -3.3 | 0.102 | 7.2E-04 |
| *F41E6.11* | WBGene00018292 | protein_coding_gene | -3.3 | 0.101 | 3.2E-06 |
| *cln-3.1* | WBGene00000539 | protein_coding_gene | -3.3 | 0.100 | 1.2E-19 |
| *acs-1* | WBGene00018488 | protein_coding_gene | -3.3 | 0.099 | 5.0E-15 |
| *F18E9.4* | WBGene00017570 | protein_coding_gene | -3.3 | 0.099 | 9.1E-05 |
| *nas-27* | WBGene00003545 | protein_coding_gene | -3.3 | 0.098 | 2.4E-04 |
| *gst-4* | WBGene00001752 | protein_coding_gene | -3.4 | 0.097 | 1.3E-10 |
| *acox-1.2* | WBGene00008565 | protein_coding_gene | -3.4 | 0.097 | 7.7E-22 |
| *nspb-7* | WBGene00007243 | protein_coding_gene | -3.4 | 0.097 | 9.0E-04 |
| *C05C8.8* | WBGene00015465 | protein_coding_gene | -3.4 | 0.097 | 1.6E-07 |
| *spp-23* | WBGene00021335 | protein_coding_gene | -3.4 | 0.096 | 1.7E-11 |
| *C30G12.10* | WBGene00284859 | ncRNA_gene | -3.4 | 0.096 | 1.3E-03 |
| *M03B6.1* | WBGene00010833 | protein_coding_gene | -3.4 | 0.096 | 4.2E-07 |
| *D1086.20* | WBGene00077448 | pseudogene | -3.4 | 0.096 | 2.0E-03 |
| *fpn-1.3* | WBGene00019969 | pseudogene | -3.4 | 0.096 | 2.7E-03 |
| *dct-5* | WBGene00017218 | protein_coding_gene | -3.4 | 0.095 | 3.6E-05 |
| *C54C6.5* | WBGene00008287 | protein_coding_gene | -3.4 | 0.095 | 2.1E-05 |
| *K09E9.7* | WBGene00194664 | protein_coding_gene | -3.4 | 0.095 | 2.8E-04 |
| *dct-16* | WBGene00012615 | protein_coding_gene | -3.4 | 0.095 | 1.4E-37 |
| *ncx-7* | WBGene00003572 | protein_coding_gene | -3.4 | 0.094 | 4.2E-16 |
| *abu-4* | WBGene00000027 | protein_coding_gene | -3.4 | 0.094 | 4.9E-03 |
| *C04F1.1* | WBGene00015442 | protein_coding_gene | -3.4 | 0.094 | 5.1E-06 |
| *ora-1* | WBGene00003879 | protein_coding_gene | -3.4 | 0.092 | 2.1E-06 |
| *Y17D7B.7* | WBGene00014562 | pseudogene | -3.5 | 0.091 | 8.1E-10 |
| *ins-27* | WBGene00002110 | protein_coding_gene | -3.5 | 0.090 | 4.5E-16 |
| *F57B1.9* | WBGene00045488 | protein_coding_gene | -3.5 | 0.090 | 1.0E-08 |
| *col-113* | WBGene00000687 | protein_coding_gene | -3.5 | 0.090 | 1.4E-11 |
| *T19D12.10* | WBGene00020584 | protein_coding_gene | -3.5 | 0.089 | 4.2E-04 |
| *T06E4.10* | WBGene00011537 | protein_coding_gene | -3.5 | 0.089 | 7.9E-05 |
| *sru-25* | WBGene00005688 | protein_coding_gene | -3.5 | 0.089 | 2.0E-03 |
| *Y32B12C.3* | WBGene00012526 | pseudogene | -3.5 | 0.089 | 6.8E-04 |
| *col-110* | WBGene00000684 | protein_coding_gene | -3.5 | 0.088 | 3.0E-08 |
| *C33H5.1* | WBGene00016370 | protein_coding_gene | -3.5 | 0.087 | 5.0E-03 |
| *Y95B8A.2* | WBGene00022383 | protein_coding_gene | -3.5 | 0.087 | 8.0E-12 |
| *W02H3.1* | WBGene00012217 | protein_coding_gene | -3.5 | 0.086 | 3.2E-18 |
| *C52D10.3* | WBGene00016877 | protein_coding_gene | -3.5 | 0.086 | 4.8E-08 |
| *comt-5* | WBGene00021492 | protein_coding_gene | -3.6 | 0.085 | 2.8E-06 |
| *nhr-93* | WBGene00003683 | protein_coding_gene | -3.6 | 0.084 | 4.6E-07 |
| *nspb-9* | WBGene00007251 | protein_coding_gene | -3.6 | 0.084 | 8.8E-04 |
| *cdr-2* | WBGene00008296 | protein_coding_gene | -3.6 | 0.083 | 2.2E-19 |
| *col-79* | WBGene00000655 | protein_coding_gene | -3.6 | 0.082 | 1.2E-07 |
| *F22F7.8* | WBGene00194908 | protein_coding_gene | -3.6 | 0.082 | 1.5E-30 |
| *col-104* | WBGene00000678 | protein_coding_gene | -3.6 | 0.082 | 2.4E-13 |
| *col-71* | WBGene00000647 | protein_coding_gene | -3.6 | 0.082 | 9.9E-16 |
| *C05E7.2* | WBGene00007344 | protein_coding_gene | -3.6 | 0.081 | 7.7E-09 |
| *grl-16* | WBGene00001725 | protein_coding_gene | -3.6 | 0.081 | 8.3E-14 |
| *abu-8* | WBGene00000031 | protein_coding_gene | -3.6 | 0.080 | 1.8E-09 |
| *col-152* | WBGene00000725 | protein_coding_gene | -3.6 | 0.080 | 9.2E-05 |
| *C04F6.7* | WBGene00195081 | protein_coding_gene | -3.7 | 0.079 | 6.3E-04 |
| *ZK6.6* | WBGene00022641 | protein_coding_gene | -3.7 | 0.078 | 4.2E-11 |
| *C30H6.5* | WBGene00007823 | protein_coding_gene | -3.7 | 0.078 | 7.0E-09 |
| *pqn-63* | WBGene00004146 | protein_coding_gene | -3.7 | 0.077 | 2.3E-07 |
| *C49F5.8* | WBGene00008212 | protein_coding_gene | -3.7 | 0.077 | 2.8E-04 |
| *bli-2* | WBGene00000252 | protein_coding_gene | -3.7 | 0.076 | 4.2E-08 |
| *nkat-1* | WBGene00009232 | protein_coding_gene | -3.7 | 0.076 | 1.2E-03 |
| *abhd-3.2* | WBGene00016644 | protein_coding_gene | -3.7 | 0.075 | 2.1E-11 |
| *aat-7* | WBGene00000008 | protein_coding_gene | -3.7 | 0.075 | 1.4E-11 |
| *clec-245* | WBGene00007851 | protein_coding_gene | -3.7 | 0.075 | 4.9E-06 |
| *pes-8* | WBGene00003981 | protein_coding_gene | -3.8 | 0.074 | 1.3E-20 |
| *ets-5* | WBGene00016600 | protein_coding_gene | -3.8 | 0.073 | 1.9E-05 |
| *K09H11.4* | WBGene00019601 | protein_coding_gene | -3.8 | 0.073 | 3.1E-04 |

| | | | | | |
|---|---|---|---|---|---|
| *T12B5.14* | WBGene00050913 | protein_coding_gene | -3.8 | 0.073 | 2.0E-07 |
| *chil-5* | WBGene00007470 | protein_coding_gene | -3.8 | 0.072 | 7.2E-05 |
| *grl-21* | WBGene00001730 | protein_coding_gene | -3.8 | 0.072 | 1.5E-11 |
| *asns-1* | WBGene00003816 | protein_coding_gene | -3.8 | 0.071 | 1.3E-12 |
| *T10E9.3* | WBGene00020413 | protein_coding_gene | -3.8 | 0.070 | 5.6E-19 |
| *W07A12.8* | WBGene00012325 | protein_coding_gene | -3.8 | 0.069 | 3.1E-14 |
| *T17H7.1* | WBGene00020550 | protein_coding_gene | -3.9 | 0.069 | 7.3E-20 |
| *dao-4* | WBGene00000930 | protein_coding_gene | -3.9 | 0.068 | 2.0E-06 |
| *T10G3.3* | WBGene00011694 | protein_coding_gene | -3.9 | 0.068 | 2.4E-06 |
| *linc-130* | WBGene00219651 | lincRNA_gene | -3.9 | 0.067 | 1.1E-03 |
| *Y47D7A.13* | WBGene00021625 | protein_coding_gene | -3.9 | 0.067 | 6.3E-13 |
| *linc-72* | WBGene00219590 | lincRNA_gene | -3.9 | 0.067 | 3.5E-04 |
| *H17B01.2* | WBGene00000646 | protein_coding_gene | -3.9 | 0.067 | 2.7E-03 |
| *21ur-8696* | WBGene00173920 | piRNA_gene | -3.9 | 0.066 | 8.8E-04 |
| *T12B5.15* | WBGene00050914 | protein_coding_gene | -3.9 | 0.066 | 8.5E-15 |
| *ZC250.4* | WBGene00045305 | protein_coding_gene | -3.9 | 0.065 | 4.2E-14 |
| *C46F2.1* | WBGene00016714 | protein_coding_gene | -4.0 | 0.065 | 3.0E-10 |
| *F49C5.11* | WBGene00044163 | protein_coding_gene | -4.0 | 0.064 | 1.2E-04 |
| *cut-4* | WBGene00008482 | protein_coding_gene | -4.0 | 0.064 | 4.8E-17 |
| *R12E2.6* | WBGene00020032 | protein_coding_gene | -4.0 | 0.063 | 7.5E-04 |
| *oac-37* | WBGene00010391 | protein_coding_gene | -4.0 | 0.062 | 1.5E-04 |
| *nas-38* | WBGene00003554 | protein_coding_gene | -4.0 | 0.061 | 4.0E-09 |
| *nspb-11* | WBGene00009398 | protein_coding_gene | -4.0 | 0.061 | 8.2E-05 |
| *abu-10* | WBGene00000033 | protein_coding_gene | -4.0 | 0.060 | 8.7E-11 |
| *nspb-10* | WBGene00007246 | protein_coding_gene | -4.1 | 0.059 | 5.6E-05 |
| *Y57G11C.42* | WBGene00013331 | protein_coding_gene | -4.1 | 0.059 | 5.6E-11 |
| *Y50D4B.6* | WBGene00021745 | protein_coding_gene | -4.1 | 0.058 | 4.0E-12 |
| *nas-22* | WBGene00003541 | protein_coding_gene | -4.1 | 0.057 | 9.0E-07 |
| *C04E12.4* | WBGene00015434 | protein_coding_gene | -4.1 | 0.057 | 3.2E-38 |
| *T06E4.9* | WBGene00011536 | protein_coding_gene | -4.1 | 0.057 | 2.4E-06 |
| *math-10* | WBGene00015839 | protein_coding_gene | -4.2 | 0.056 | 1.1E-05 |
| *clec-206* | WBGene00019089 | protein_coding_gene | -4.2 | 0.056 | 1.3E-04 |
| *F53B3.6* | WBGene00018744 | protein_coding_gene | -4.2 | 0.055 | 1.3E-09 |
| *ugt-25* | WBGene00015692 | protein_coding_gene | -4.2 | 0.055 | 1.8E-41 |
| *F59B1.4* | WBGene00019098 | protein_coding_gene | -4.2 | 0.054 | 4.7E-05 |
| *col-97* | WBGene00000672 | protein_coding_gene | -4.2 | 0.054 | 3.2E-23 |
| *ZK84.1* | WBGene00022649 | protein_coding_gene | -4.2 | 0.054 | 9.3E-28 |
| *H25K10.1* | WBGene00010411 | protein_coding_gene | -4.2 | 0.054 | 7.3E-08 |
| *E04D5.5* | WBGene00008484 | protein_coding_gene | -4.2 | 0.054 | 1.9E-16 |
| *nspb-12* | WBGene00017305 | protein_coding_gene | -4.2 | 0.053 | 1.3E-05 |
| *lbp-8* | WBGene00002260 | protein_coding_gene | -4.2 | 0.053 | 1.0E-04 |
| *sprr-2* | WBGene00009629 | protein_coding_gene | -4.2 | 0.053 | 1.4E-14 |
| *clec-210* | WBGene00022261 | protein_coding_gene | -4.2 | 0.053 | 1.7E-08 |
| *col-168* | WBGene00000741 | protein_coding_gene | -4.3 | 0.052 | 1.2E-12 |
| *D1086.3* | WBGene00008390 | protein_coding_gene | -4.3 | 0.051 | 1.2E-12 |
| *K10H10.4* | WBGene00010760 | protein_coding_gene | -4.3 | 0.051 | 5.5E-08 |
| *Y46G5A.29* | WBGene00012917 | protein_coding_gene | -4.3 | 0.051 | 1.2E-10 |
| *sri-40* | WBGene00005552 | protein_coding_gene | -4.3 | 0.050 | 1.1E-85 |
| *lips-14* | WBGene00019208 | pseudogene | -4.4 | 0.049 | 7.5E-05 |
| *col-157* | WBGene00000730 | protein_coding_gene | -4.4 | 0.049 | 6.8E-11 |
| *dpy-8* | WBGene00001070 | protein_coding_gene | -4.4 | 0.048 | 1.9E-08 |
| *Y47D7A.6* | WBGene00021619 | protein_coding_gene | -4.4 | 0.048 | 5.4E-07 |
| *T28D6.3* | WBGene00012123 | protein_coding_gene | -4.4 | 0.048 | 3.1E-23 |
| *Y43C5A.7* | WBGene00044922 | protein_coding_gene | -4.4 | 0.047 | 3.2E-12 |
| *pqn-2* | WBGene00004098 | protein_coding_gene | -4.4 | 0.047 | 2.1E-05 |
| *C04E12.5* | WBGene00015435 | protein_coding_gene | -4.4 | 0.047 | 2.4E-05 |
| *dpy-6* | WBGene00001068 | protein_coding_gene | -4.4 | 0.047 | 1.3E-11 |
| *col-61* | WBGene00000637 | protein_coding_gene | -4.4 | 0.046 | 4.5E-06 |
| *pqn-54* | WBGene00004139 | protein_coding_gene | -4.4 | 0.046 | 7.0E-11 |
| *twk-10* | WBGene00006665 | protein_coding_gene | -4.5 | 0.045 | 3.1E-06 |
| *rol-6* | WBGene00004397 | protein_coding_gene | -4.5 | 0.044 | 2.4E-16 |
| *F46F2.5* | WBGene00009789 | protein_coding_gene | -4.5 | 0.044 | 3.0E-12 |
| *col-14* | WBGene00000603 | protein_coding_gene | -4.5 | 0.044 | 6.1E-23 |
| *Y38H6C.23* | WBGene00012634 | protein_coding_gene | -4.5 | 0.043 | 4.3E-31 |
| *acp-6* | WBGene00022245 | protein_coding_gene | -4.6 | 0.043 | 2.4E-23 |
| *col-173* | WBGene00000746 | protein_coding_gene | -4.6 | 0.042 | 1.2E-09 |
| *abu-1* | WBGene00000024 | protein_coding_gene | -4.6 | 0.042 | 8.4E-06 |
| *R07E5.4* | WBGene00011112 | protein_coding_gene | -4.6 | 0.041 | 5.0E-07 |
| *F53F1.4* | WBGene00009982 | protein_coding_gene | -4.6 | 0.041 | 7.4E-13 |
| *K02F2.11* | WBGene00220000 | ncRNA_gene | -4.6 | 0.041 | 7.0E-05 |

| | | | | | |
|---|---|---|---|---|---|
| *nspb-8* | WBGene00007250 | protein_coding_gene | -4.6 | 0.040 | 1.5E-06 |
| *F58G6.3* | WBGene00010274 | protein_coding_gene | -4.6 | 0.040 | 1.5E-11 |
| *grd-4* | WBGene00001693 | protein_coding_gene | -4.7 | 0.039 | 7.1E-08 |
| *F42A8.1* | WBGene00009626 | protein_coding_gene | -4.7 | 0.039 | 4.0E-10 |
| *ZC239.5* | WBGene00022568 | protein_coding_gene | -4.7 | 0.038 | 1.1E-06 |
| *col-167* | WBGene00000740 | protein_coding_gene | -4.7 | 0.038 | 2.5E-20 |
| *plep-1* | WBGene00021797 | protein_coding_gene | -4.7 | 0.037 | 1.7E-09 |
| *T20D4.3* | WBGene00020609 | protein_coding_gene | -4.8 | 0.037 | 1.9E-06 |
| *T06E4.8* | WBGene00011535 | protein_coding_gene | -4.8 | 0.036 | 4.7E-06 |
| *col-41* | WBGene00000618 | protein_coding_gene | -5.0 | 0.032 | 1.5E-06 |
| *R08E5.1* | WBGene00019961 | protein_coding_gene | -5.0 | 0.032 | 6.9E-19 |
| *C04E12.10* | WBGene00015439 | pseudogene | -5.0 | 0.032 | 4.2E-06 |
| *abu-7* | WBGene00000030 | protein_coding_gene | -5.0 | 0.030 | 7.3E-07 |
| *ugt-19* | WBGene00019515 | protein_coding_gene | -5.1 | 0.030 | 5.9E-23 |
| *sqt-2* | WBGene00005017 | protein_coding_gene | -5.1 | 0.029 | 1.5E-18 |
| *grd-14* | WBGene00001703 | protein_coding_gene | -5.1 | 0.029 | 1.4E-11 |
| *E01G4.6* | WBGene00008448 | protein_coding_gene | -5.1 | 0.028 | 2.7E-21 |
| *sqt-1* | WBGene00005016 | protein_coding_gene | -5.2 | 0.028 | 2.1E-15 |
| *col-130* | WBGene00000704 | protein_coding_gene | -5.2 | 0.026 | 2.4E-21 |
| *ZK180.6* | WBGene00022680 | protein_coding_gene | -5.2 | 0.026 | 3.9E-24 |
| *col-90* | WBGene00000665 | protein_coding_gene | -5.3 | 0.026 | 4.1E-11 |
| *grd-6* | WBGene00001695 | protein_coding_gene | -5.3 | 0.025 | 2.9E-15 |
| *Y38E10A.14* | WBGene00012592 | protein_coding_gene | -5.3 | 0.025 | 1.0E-30 |
| *abu-15* | WBGene00004099 | protein_coding_gene | -5.3 | 0.025 | 1.8E-07 |
| *abu-6* | WBGene00000029 | protein_coding_gene | -5.4 | 0.024 | 1.6E-07 |
| *col-120* | WBGene00000694 | protein_coding_gene | -5.4 | 0.024 | 4.2E-19 |
| *col-60* | WBGene00000636 | protein_coding_gene | -5.4 | 0.024 | 3.6E-15 |
| *clec-10* | WBGene00015403 | protein_coding_gene | -5.4 | 0.024 | 9.8E-56 |
| *R03G8.4* | WBGene00010999 | pseudogene | -5.4 | 0.023 | 2.0E-07 |
| *Y48G8AL.12* | WBGene00021690 | protein_coding_gene | -5.5 | 0.022 | 2.5E-10 |
| *str-146* | WBGene00006193 | protein_coding_gene | -5.5 | 0.022 | 1.7E-07 |
| *T06E4.12* | WBGene00044011 | protein_coding_gene | -5.7 | 0.019 | 1.2E-08 |
| *col-73* | WBGene00000649 | protein_coding_gene | -5.8 | 0.018 | 1.3E-08 |
| *cest-28* | WBGene00019654 | protein_coding_gene | -5.8 | 0.018 | 7.0E-11 |
| *rol-8* | WBGene00004398 | protein_coding_gene | -5.8 | 0.018 | 4.4E-10 |
| *Y48E1B.8* | WBGene00013007 | protein_coding_gene | -5.9 | 0.017 | 3.5E-08 |
| *Y51H7C.1* | WBGene00021779 | protein_coding_gene | -6.0 | 0.016 | 2.1E-31 |
| *nhr-74* | WBGene00003664 | protein_coding_gene | -6.0 | 0.015 | 4.9E-09 |
| *col-162* | WBGene00000735 | protein_coding_gene | -6.1 | 0.014 | 1.3E-15 |
| *col-77* | WBGene00000653 | protein_coding_gene | -6.2 | 0.013 | 4.0E-32 |
| *dhs-26* | WBGene00000989 | protein_coding_gene | -6.3 | 0.013 | 6.5E-14 |
| *K02E11.10* | WBGene00044109 | protein_coding_gene | -6.7 | 0.010 | 1.3E-11 |
| *col-63* | WBGene00000639 | protein_coding_gene | -6.7 | 0.010 | 2.1E-25 |
| *col-17* | WBGene00000606 | protein_coding_gene | -6.7 | 0.009 | 1.5E-11 |
| *col-175* | WBGene00000748 | protein_coding_gene | -6.9 | 0.008 | 1.9E-24 |
| *dpy-4* | WBGene00001066 | protein_coding_gene | -7.1 | 0.007 | 8.9E-30 |
| *col-49* | WBGene00000626 | protein_coding_gene | -7.3 | 0.006 | 2.7E-18 |
| *bli-6* | WBGene00000256 | protein_coding_gene | -7.4 | 0.006 | 8.3E-35 |
| *dpy-5* | WBGene00001067 | protein_coding_gene | -7.4 | 0.006 | 2.0E-28 |
| *rol-1* | WBGene00004394 | protein_coding_gene | -7.6 | 0.005 | 1.5E-23 |
| *dpy-13* | WBGene00001074 | protein_coding_gene | -7.6 | 0.005 | 3.6E-23 |
| *col-138* | WBGene00000711 | protein_coding_gene | -7.7 | 0.005 | 6.8E-28 |
| *folt-2* | WBGene00018138 | protein_coding_gene | -8.4 | 0.003 | 3.8E-98 |
| *Y47D7A.15* | WBGene00021627 | protein_coding_gene | -8.5 | 0.003 | 2.1E-16 |

**CB4856**-exclusive DEGs in response to *P. aeruginosa* (vs. OP50)

| Gene Name | Wormbase Gene ID | Type | $log_2FC$ | FC | Adjusted P-value |
|---|---|---|---|---|---|
| *T24B8.5* | WBGene00011979 | protein_coding_gene | 5.9 | 61.463 | 1.9E-73 |
| *sri-74* | WBGene00005586 | protein_coding_gene | 5.4 | 41.481 | 1.6E-07 |
| *fbxa-20* | WBGene00022332 | protein_coding_gene | 5.1 | 33.924 | 8.9E-06 |
| *pqn-98* | WBGene00004177 | protein_coding_gene | 4.8 | 27.361 | 1.4E-05 |
| *C30E1.6* | WBGene00016255 | pseudogene | 4.6 | 23.535 | 7.0E-06 |
| *Y6G8.5* | WBGene00044207 | protein_coding_gene | 4.4 | 20.838 | 4.5E-05 |
| *rab-11.2* | WBGene00004275 | protein_coding_gene | 4.3 | 19.983 | 5.5E-05 |
| *F01D5.2* | WBGene00008493 | protein_coding_gene | 4.2 | 18.999 | 4.2E-05 |
| *F35E12.2* | WBGene00009426 | protein_coding_gene | 4.0 | 16.440 | 4.9E-04 |
| *Y37H2A.11* | WBGene00012571 | protein_coding_gene | 4.0 | 15.515 | 1.5E-03 |
| *srz-100* | WBGene00012026 | protein_coding_gene | 3.8 | 13.749 | 1.5E-08 |
| *sri-38* | WBGene00005550 | protein_coding_gene | 3.7 | 12.662 | 1.5E-03 |
| *C55B6.5* | WBGene00016941 | protein_coding_gene | 3.6 | 11.936 | 5.4E-04 |

| | | | | | |
|---|---|---|---|---|---|
| *pud-2.1* | WBGene00017490 | protein_coding_gene | 3.4 | 10.337 | 4.1E-03 |
| *drd-50* | WBGene00018643 | protein_coding_gene | 3.4 | 10.231 | 7.2E-25 |
| *Y53F4B.16* | WBGene00013162 | protein_coding_gene | 3.3 | 10.000 | 9.3E-07 |
| *C15H11.16* | WBGene00206494 | antisense_lncRNA_gene | 3.3 | 9.910 | 1.1E-03 |
| *F01D5.1* | WBGene00008492 | protein_coding_gene | 3.3 | 9.883 | 1.7E-21 |
| *F14H3.15* | WBGene00219414 | protein_coding_gene | 3.3 | 9.868 | 6.3E-03 |
| *F11D11.22* | WBGene00271804 | protein_coding_gene | 3.2 | 9.153 | 9.0E-03 |
| *F09C6.12* | WBGene00044182 | protein_coding_gene | 3.1 | 8.868 | 1.1E-02 |
| *F20G2.9* | WBGene00195065 | antisense_lncRNA_gene | 3.1 | 8.785 | 7.4E-03 |
| *ZK250.14* | WBGene00235354 | protein_coding_gene | 3.1 | 8.747 | 5.1E-07 |
| *math-47* | WBGene00022688 | pseudogene | 3.1 | 8.702 | 1.1E-24 |
| *T20B6.1* | WBGene00020597 | protein_coding_gene | 3.1 | 8.364 | 3.9E-05 |
| *arrd-3* | WBGene00006429 | protein_coding_gene | 3.0 | 8.120 | 4.5E-04 |
| *C35A5.10* | WBGene00044016 | protein_coding_gene | 3.0 | 7.879 | 5.7E-05 |
| *F55G11.2* | WBGene00010123 | protein_coding_gene | 3.0 | 7.863 | 1.5E-07 |
| *Y39G8B.9* | WBGene00012728 | protein_coding_gene | 3.0 | 7.784 | 3.7E-07 |
| *hsp-16.41* | WBGene00002018 | protein_coding_gene | 2.9 | 7.691 | 2.9E-04 |
| *hsp-17* | WBGene00002021 | protein_coding_gene | 2.9 | 7.551 | 1.9E-47 |
| *F09C6.13* | WBGene00044183 | protein_coding_gene | 2.9 | 7.405 | 6.9E-06 |
| *F22E5.6* | WBGene00017705 | protein_coding_gene | 2.9 | 7.317 | 6.3E-07 |
| *Y54G2A.45* | WBGene00044488 | protein_coding_gene | 2.9 | 7.303 | 2.0E-08 |
| *T01H10.4* | WBGene00011357 | pseudogene | 2.8 | 7.148 | 7.5E-03 |
| *Y106G6E.1* | WBGene00013705 | protein_coding_gene | 2.8 | 7.027 | 1.5E-03 |
| *M28.2* | WBGene00010894 | protein_coding_gene | 2.8 | 6.979 | 8.3E-03 |
| *C04G6.6* | WBGene00015456 | protein_coding_gene | 2.8 | 6.977 | 4.0E-09 |
| *K11H12.3* | WBGene00019659 | protein_coding_gene | 2.8 | 6.972 | 5.1E-05 |
| *Y9C9A.8* | WBGene00021178 | protein_coding_gene | 2.8 | 6.871 | 3.1E-11 |
| *Y53F4B.17* | WBGene00013163 | protein_coding_gene | 2.8 | 6.867 | 7.8E-04 |
| *F26D2.10* | WBGene00009149 | protein_coding_gene | 2.7 | 6.654 | 1.5E-02 |
| *oac-34* | WBGene00010157 | protein_coding_gene | 2.7 | 6.648 | 3.3E-13 |
| *str-131* | WBGene00006180 | protein_coding_gene | 2.7 | 6.632 | 4.1E-03 |
| *C27D9.2* | WBGene00016164 | protein_coding_gene | 2.7 | 6.618 | 5.1E-03 |
| *F28B1.10* | WBGene00050958 | pseudogene | 2.7 | 6.535 | 1.8E-02 |
| *srg-62* | WBGene00005219 | protein_coding_gene | 2.7 | 6.517 | 1.9E-02 |
| *abf-4* | WBGene00000015 | protein_coding_gene | 2.7 | 6.471 | 1.2E-04 |
| *F58G1.9* | WBGene00010271 | pseudogene | 2.7 | 6.334 | 2.6E-04 |
| *K09D9.11* | WBGene00019568 | pseudogene | 2.7 | 6.282 | 2.3E-05 |
| *Y57G11C.51* | WBGene00044258 | protein_coding_gene | 2.6 | 6.273 | 5.2E-34 |
| *clp-6* | WBGene00000546 | protein_coding_gene | 2.6 | 6.229 | 1.1E-06 |
| *arl-7* | WBGene00000191 | pseudogene | 2.6 | 6.045 | 1.2E-07 |
| *ZC15.3* | WBGene00013832 | protein_coding_gene | 2.6 | 6.019 | 4.1E-03 |
| *B0303.16* | WBGene00044678 | protein_coding_gene | 2.6 | 6.003 | 1.3E-02 |
| *Y53C10A.1* | WBGene00014915 | pseudogene | 2.6 | 5.941 | 1.7E-19 |
| *C17H12.8* | WBGene00015933 | protein_coding_gene | 2.5 | 5.841 | 7.5E-43 |
| *ferl-1* | WBGene00020262 | protein_coding_gene | 2.5 | 5.840 | 8.0E-04 |
| *K08D10.5* | WBGene00019529 | pseudogene | 2.5 | 5.761 | 1.1E-05 |
| *T08D2.7* | WBGene00011612 | protein_coding_gene | 2.5 | 5.733 | 1.4E-09 |
| *F28A10.9* | WBGene00017877 | pseudogene | 2.5 | 5.658 | 8.4E-03 |
| *F17C11.14* | WBGene00045151 | snoRNA_gene | 2.5 | 5.596 | 1.7E-02 |
| *K08H2.5* | WBGene00010692 | protein_coding_gene | 2.5 | 5.574 | 1.4E-02 |
| *clec-230* | WBGene00007806 | protein_coding_gene | 2.5 | 5.570 | 3.9E-03 |
| *Y82E9BL.12* | WBGene00022328 | protein_coding_gene | 2.5 | 5.538 | 4.1E-05 |
| *fbxa-178* | WBGene00017533 | pseudogene | 2.5 | 5.518 | 9.9E-07 |
| *nduo-2* | WBGene00010961 | protein_coding_gene | 2.4 | 5.401 | 1.8E-12 |
| *clpr-2* | WBGene00023410 | protein_coding_gene | 2.4 | 5.344 | 4.3E-04 |
| *dod-20* | WBGene00015259 | protein_coding_gene | 2.4 | 5.327 | 8.6E-04 |
| *hrg-4* | WBGene00009493 | protein_coding_gene | 2.4 | 5.326 | 9.3E-07 |
| *Y43F4B.12* | WBGene00045113 | snoRNA_gene | 2.4 | 5.303 | 3.0E-02 |
| *K09F6.3* | WBGene00019586 | protein_coding_gene | 2.4 | 5.209 | 3.4E-02 |
| *C32H11.6* | WBGene00007869 | protein_coding_gene | 2.4 | 5.166 | 4.1E-02 |
| *F21D9.2* | WBGene00009016 | protein_coding_gene | 2.4 | 5.139 | 2.2E-04 |
| *sdz-35* | WBGene00022570 | protein_coding_gene | 2.3 | 5.094 | 2.8E-02 |
| *F16B12.7* | WBGene00008883 | protein_coding_gene | 2.3 | 5.087 | 3.6E-02 |
| *F14H3.8* | WBGene00008827 | pseudogene | 2.3 | 5.080 | 3.3E-02 |
| *F17B5.8* | WBGene00045458 | protein_coding_gene | 2.3 | 5.078 | 2.7E-05 |
| *fbxa-100* | WBGene00008826 | protein_coding_gene | 2.3 | 5.056 | 1.2E-11 |
| *C14A6.5* | WBGene00007569 | protein_coding_gene | 2.3 | 5.015 | 5.2E-03 |
| *C28C12.4* | WBGene00016180 | protein_coding_gene | 2.3 | 5.001 | 9.2E-11 |
| *otpl-5* | WBGene00018478 | protein_coding_gene | 2.3 | 4.953 | 2.4E-02 |
| *srw-96* | WBGene00005843 | pseudogene | 2.3 | 4.941 | 3.7E-02 |

| | | | | | |
|---|---|---|---|---|---|
| *C08G5.3* | WBGene00015616 | pseudogene | 2.3 | 4.929 | 2.7E-03 |
| *cyp-35D2* | WBGene00014735 | pseudogene | 2.3 | 4.923 | 1.4E-02 |
| *hsp-16.2* | WBGene00002016 | protein_coding_gene | 2.3 | 4.911 | 5.5E-04 |
| *pqn-97* | WBGene00004176 | protein_coding_gene | 2.3 | 4.910 | 1.7E-02 |
| *Y37H9A.2* | WBGene00012577 | protein_coding_gene | 2.3 | 4.902 | 3.5E-02 |
| *H11L12.1* | WBGene00019189 | protein_coding_gene | 2.3 | 4.853 | 3.5E-05 |
| *F57G4.11* | WBGene00195165 | protein_coding_gene | 2.3 | 4.810 | 4.4E-02 |
| *F53C11.1* | WBGene00009971 | protein_coding_gene | 2.3 | 4.764 | 1.9E-22 |
| *F14F9.3* | WBGene00017466 | protein_coding_gene | 2.2 | 4.756 | 2.7E-06 |
| *Y44A6B.3* | WBGene00012848 | protein_coding_gene | 2.2 | 4.747 | 4.9E-03 |
| *Y26D4A.22* | WBGene00077789 | pseudogene | 2.2 | 4.742 | 4.9E-02 |
| *B0457.3* | WBGene00014660 | pseudogene | 2.2 | 4.733 | 2.9E-04 |
| *F54B3.4* | WBGene00175034 | protein_coding_gene | 2.2 | 4.708 | 7.4E-03 |
| *R193.6* | WBGene00304820 | protein_coding_gene | 2.2 | 4.658 | 4.4E-02 |
| *nhr-216* | WBGene00020385 | protein_coding_gene | 2.2 | 4.656 | 2.7E-03 |
| *T23E1.1* | WBGene00020728 | protein_coding_gene | 2.2 | 4.629 | 3.9E-03 |
| *Y43B11AR.7* | WBGene00045086 | snoRNA_gene | 2.2 | 4.617 | 7.9E-07 |
| *B0250.7* | WBGene00007124 | protein_coding_gene | 2.2 | 4.617 | 5.7E-23 |
| *C50H2.7* | WBGene00008243 | protein_coding_gene | 2.2 | 4.601 | 1.6E-03 |
| *fbxa-89* | WBGene00013555 | protein_coding_gene | 2.2 | 4.588 | 3.9E-06 |
| *enu-3.5* | WBGene00012552 | protein_coding_gene | 2.2 | 4.540 | 9.8E-06 |
| *pqn-29* | WBGene00004118 | protein_coding_gene | 2.2 | 4.534 | 5.1E-03 |
| *fbxa-74* | WBGene00021262 | protein_coding_gene | 2.2 | 4.518 | 3.2E-25 |
| *F13B6.1* | WBGene00017416 | protein_coding_gene | 2.2 | 4.515 | 2.3E-09 |
| *F27C1.18* | WBGene00219918 | ncRNA_gene | 2.2 | 4.496 | 3.1E-05 |
| *H25P06.5* | WBGene00219319 | protein_coding_gene | 2.2 | 4.492 | 7.2E-04 |
| *T28A11.3* | WBGene00020870 | protein_coding_gene | 2.2 | 4.482 | 1.7E-02 |
| *tyr-3* | WBGene00009001 | protein_coding_gene | 2.2 | 4.449 | 2.1E-02 |
| *K09E2.1* | WBGene00019570 | protein_coding_gene | 2.1 | 4.429 | 3.7E-02 |
| *oac-5* | WBGene00007829 | protein_coding_gene | 2.1 | 4.421 | 2.8E-04 |
| *chk-2* | WBGene00000499 | protein_coding_gene | 2.1 | 4.397 | 2.6E-25 |
| *ZC373.3* | WBGene00013868 | protein_coding_gene | 2.1 | 4.397 | 4.0E-03 |
| *T23B12.14* | WBGene00220084 | ncRNA_gene | 2.1 | 4.357 | 3.9E-03 |
| *W03F9.3* | WBGene00021001 | pseudogene | 2.1 | 4.345 | 4.9E-04 |
| *F12A10.2* | WBGene00017391 | pseudogene | 2.1 | 4.344 | 3.2E-02 |
| *ZC15.4* | WBGene00014977 | pseudogene | 2.1 | 4.341 | 4.3E-03 |
| *F14F8.8* | WBGene00008816 | protein_coding_gene | 2.1 | 4.307 | 2.4E-02 |
| *nhr-38* | WBGene00003629 | protein_coding_gene | 2.1 | 4.256 | 4.6E-02 |
| *nep-15* | WBGene00018227 | protein_coding_gene | 2.1 | 4.246 | 3.9E-02 |
| *B0041.1* | WBGene00015006 | protein_coding_gene | 2.1 | 4.233 | 9.8E-03 |
| *Y14H12A.4* | WBGene00220133 | ncRNA_gene | 2.1 | 4.223 | 1.7E-02 |
| *ZK856.14* | WBGene00044020 | protein_coding_gene | 2.1 | 4.189 | 4.4E-02 |
| *Y24F12A.4* | WBGene00012499 | pseudogene | 2.1 | 4.168 | 4.2E-04 |
| *F23B2.10* | WBGene00009077 | protein_coding_gene | 2.1 | 4.165 | 1.8E-02 |
| *F20D6.2* | WBGene00017634 | protein_coding_gene | 2.0 | 4.121 | 3.9E-02 |
| *ZK795.6* | WBGene00194644 | pseudogene | 2.0 | 4.107 | 5.3E-03 |
| *basl-2* | WBGene00017392 | pseudogene | 2.0 | 4.039 | 4.3E-06 |
| *wago-2* | WBGene00018862 | pseudogene | 2.0 | 4.034 | 8.1E-30 |
| *twk-28* | WBGene00006680 | protein_coding_gene | 2.0 | 4.009 | 1.1E-04 |
| *fbxa-84* | WBGene00011670 | protein_coding_gene | 2.0 | 3.999 | 6.7E-10 |
| *C07D8.2* | WBGene00015562 | pseudogene | 2.0 | 3.990 | 4.7E-02 |
| *T25B9.2* | WBGene00012008 | protein_coding_gene | 2.0 | 3.986 | 1.2E-02 |
| *F35F10.13* | WBGene00018062 | protein_coding_gene | 2.0 | 3.975 | 1.7E-11 |
| *F59E12.3* | WBGene00019119 | protein_coding_gene | 2.0 | 3.933 | 3.7E-03 |
| *R11D1.2* | WBGene00011243 | protein_coding_gene | 2.0 | 3.930 | 4.5E-02 |
| *oac-58* | WBGene00019580 | protein_coding_gene | 2.0 | 3.902 | 3.3E-09 |
| *irld-57* | WBGene00012401 | protein_coding_gene | 2.0 | 3.886 | 2.2E-02 |
| *C53D6.5* | WBGene00008282 | pseudogene | 2.0 | 3.878 | 1.6E-02 |
| *T20H9.6* | WBGene00020642 | protein_coding_gene | 2.0 | 3.873 | 4.7E-03 |
| *zipt-2.3* | WBGene00013207 | protein_coding_gene | 2.0 | 3.868 | 4.8E-04 |
| *C50E3.12* | WBGene00016823 | protein_coding_gene | 1.9 | 3.852 | 2.3E-07 |
| *Y48G1BM.6* | WBGene00021671 | pseudogene | 1.9 | 3.840 | 7.6E-12 |
| *F41G4.7* | WBGene00018310 | protein_coding_gene | 1.9 | 3.819 | 1.1E-02 |
| *gcy-7* | WBGene00001534 | protein_coding_gene | 1.9 | 3.813 | 4.0E-02 |
| *T08D2.4* | WBGene00011609 | protein_coding_gene | 1.9 | 3.806 | 4.9E-02 |
| *H20E11.3* | WBGene00019215 | protein_coding_gene | 1.9 | 3.797 | 3.1E-13 |
| *C06A1.12* | WBGene00219803 | ncRNA_gene | 1.9 | 3.795 | 2.1E-16 |
| *F40E12.2* | WBGene00018235 | protein_coding_gene | 1.9 | 3.778 | 2.6E-08 |
| *clec-265* | WBGene00019738 | protein_coding_gene | 1.9 | 3.756 | 8.1E-28 |
| *C47E8.10* | WBGene00044216 | pseudogene | 1.9 | 3.750 | 7.3E-05 |

| | | | | | |
|---|---|---|---|---|---|
| *gcy-29* | WBGene00007314 | protein_coding_gene | 1.9 | 3.738 | 2.4E-02 |
| *mpst-4* | WBGene00017387 | protein_coding_gene | 1.9 | 3.722 | 4.7E-06 |
| *T16H12.13* | WBGene00194725 | protein_coding_gene | 1.9 | 3.720 | 3.3E-02 |
| *dod-19* | WBGene00022644 | protein_coding_gene | 1.9 | 3.708 | 2.0E-06 |
| *F19B6.3* | WBGene00008949 | protein_coding_gene | 1.9 | 3.695 | 2.1E-05 |
| *hrg-3* | WBGene00010256 | protein_coding_gene | 1.9 | 3.692 | 5.6E-04 |
| *T13A10.1* | WBGene00020468 | protein_coding_gene | 1.9 | 3.688 | 6.6E-03 |
| *R02C2.8* | WBGene00255728 | pseudogene | 1.9 | 3.685 | 6.6E-04 |
| *C50E3.13* | WBGene00016824 | protein_coding_gene | 1.9 | 3.677 | 8.0E-10 |
| *M01E5.1* | WBGene00010804 | pseudogene | 1.9 | 3.673 | 1.2E-02 |
| *H43E16.1* | WBGene00019273 | protein_coding_gene | 1.9 | 3.667 | 1.4E-20 |
| *clec-152* | WBGene00008658 | pseudogene | 1.9 | 3.655 | 2.7E-07 |
| *F22B3.5* | WBGene00009036 | protein_coding_gene | 1.9 | 3.639 | 4.4E-14 |
| *F26A1.4* | WBGene00017803 | protein_coding_gene | 1.9 | 3.638 | 1.1E-05 |
| *D1081.10* | WBGene00045508 | protein_coding_gene | 1.9 | 3.623 | 2.3E-03 |
| *T05F1.5* | WBGene00011491 | protein_coding_gene | 1.8 | 3.598 | 7.2E-05 |
| *F58A4.12* | WBGene00014777 | pseudogene | 1.8 | 3.584 | 4.0E-02 |
| *ctc-3* | WBGene00010962 | protein_coding_gene | 1.8 | 3.562 | 3.8E-13 |
| *Y73B3A.18* | WBGene00022219 | pseudogene | 1.8 | 3.553 | 4.9E-11 |
| *F23C8.1* | WBGene00017731 | pseudogene | 1.8 | 3.552 | 8.7E-03 |
| *Y41D4A.1* | WBGene00021504 | protein_coding_gene | 1.8 | 3.539 | 1.2E-02 |
| *T20D3.2* | WBGene00011856 | protein_coding_gene | 1.8 | 3.536 | 4.5E-13 |
| *B0001.5* | WBGene00007090 | protein_coding_gene | 1.8 | 3.525 | 3.3E-47 |
| *Y34B4A.6* | WBGene00021322 | protein_coding_gene | 1.8 | 3.521 | 9.8E-10 |
| *R07C3.13* | WBGene00019929 | protein_coding_gene | 1.8 | 3.517 | 9.4E-06 |
| *C38C6.5* | WBGene00007998 | protein_coding_gene | 1.8 | 3.506 | 6.5E-04 |
| *math-2* | WBGene00015606 | protein_coding_gene | 1.8 | 3.498 | 1.1E-05 |
| *K01A6.5* | WBGene00010447 | protein_coding_gene | 1.8 | 3.492 | 1.9E-02 |
| *Y51A2B.5* | WBGene00013069 | protein_coding_gene | 1.8 | 3.487 | 2.6E-02 |
| *K08D8.12* | WBGene00194926 | protein_coding_gene | 1.8 | 3.481 | 5.7E-06 |
| *fbxa-173* | WBGene00044960 | pseudogene | 1.8 | 3.465 | 4.5E-02 |
| *Y102A5C.38* | WBGene00050883 | pseudogene | 1.8 | 3.450 | 1.6E-02 |
| *ift-74* | WBGene00016005 | protein_coding_gene | 1.8 | 3.447 | 1.2E-07 |
| *miro-3* | WBGene00021651 | pseudogene | 1.8 | 3.429 | 2.5E-03 |
| *oac-27* | WBGene00009609 | protein_coding_gene | 1.8 | 3.420 | 4.1E-03 |
| *wht-9* | WBGene00013032 | pseudogene | 1.8 | 3.420 | 2.9E-02 |
| *srd-64* | WBGene00005141 | protein_coding_gene | 1.8 | 3.408 | 1.1E-02 |
| *Y62E10A.3* | WBGene00013374 | protein_coding_gene | 1.8 | 3.407 | 7.7E-03 |
| *C38D9.5* | WBGene00008013 | pseudogene | 1.8 | 3.401 | 1.6E-05 |
| *fbxa-195* | WBGene00019976 | pseudogene | 1.8 | 3.387 | 3.7E-02 |
| *B0524.4* | WBGene00015243 | protein_coding_gene | 1.8 | 3.376 | 3.9E-02 |
| *C26E6.1* | WBGene00016137 | protein_coding_gene | 1.8 | 3.375 | 1.9E-02 |
| *F09C6.1* | WBGene00008615 | protein_coding_gene | 1.8 | 3.372 | 1.1E-03 |
| *nhr-234* | WBGene00012596 | protein_coding_gene | 1.8 | 3.368 | 4.5E-02 |
| *ZK218.8* | WBGene00013942 | pseudogene | 1.7 | 3.355 | 2.7E-05 |
| *Y54G11A.19* | WBGene00235256 | protein_coding_gene | 1.7 | 3.351 | 1.2E-05 |
| *ctc-1* | WBGene00010964 | protein_coding_gene | 1.7 | 3.347 | 5.6E-28 |
| *F10B5.3* | WBGene00008640 | protein_coding_gene | 1.7 | 3.344 | 1.0E-02 |
| *eat-2* | WBGene00001133 | protein_coding_gene | 1.7 | 3.341 | 1.2E-03 |
| *R08F11.7* | WBGene00019970 | protein_coding_gene | 1.7 | 3.337 | 9.7E-03 |
| *mth-1* | WBGene00017948 | protein_coding_gene | 1.7 | 3.333 | 5.5E-09 |
| *T25D10.1* | WBGene00020799 | protein_coding_gene | 1.7 | 3.332 | 8.8E-25 |
| *F22E5.1* | WBGene00017703 | protein_coding_gene | 1.7 | 3.326 | 1.0E-05 |
| *ppw-2* | WBGene00004094 | protein_coding_gene | 1.7 | 3.324 | 7.9E-29 |
| *C49C8.8* | WBGene00235271 | protein_coding_gene | 1.7 | 3.292 | 1.6E-02 |
| *K01G12.3* | WBGene00010483 | protein_coding_gene | 1.7 | 3.285 | 4.1E-06 |
| *snf-4* | WBGene00004903 | pseudogene | 1.7 | 3.280 | 5.6E-05 |
| *tag-244* | WBGene00044073 | protein_coding_gene | 1.7 | 3.279 | 1.9E-17 |
| *Y105E8A.13* | WBGene00013673 | protein_coding_gene | 1.7 | 3.263 | 4.8E-03 |
| *T23F6.1* | WBGene00011949 | protein_coding_gene | 1.7 | 3.243 | 3.6E-02 |
| *AC8.1* | WBGene00004054 | pseudogene | 1.7 | 3.239 | 2.2E-02 |
| *fbxa-8* | WBGene00016526 | pseudogene | 1.7 | 3.228 | 3.5E-02 |
| *C40H5.8* | WBGene00044585 | pseudogene | 1.7 | 3.224 | 8.4E-03 |
| *Y105C5B.15* | WBGene00013654 | protein_coding_gene | 1.7 | 3.222 | 2.6E-05 |
| *Y60A3A.11* | WBGene00014933 | pseudogene | 1.7 | 3.216 | 3.6E-02 |
| *T27E7.10* | WBGene00044268 | pseudogene | 1.7 | 3.216 | 8.3E-04 |
| *H01G02.4* | WBGene00045108 | protein_coding_gene | 1.7 | 3.214 | 1.4E-02 |
| *C01G10.1* | WBGene00007230 | protein_coding_gene | 1.7 | 3.208 | 7.2E-05 |
| *Y47H9C.12* | WBGene00012955 | protein_coding_gene | 1.7 | 3.190 | 1.4E-04 |
| *Y55B1AR.3* | WBGene00021907 | protein_coding_gene | 1.7 | 3.186 | 4.4E-02 |

| | | | | | |
|---|---|---|---|---|---|
| *linc-76* | WBGene00219619 | lincRNA_gene | 1.7 | 3.177 | 2.2E-03 |
| *gcsh-1* | WBGene00008354 | protein_coding_gene | 1.7 | 3.173 | 2.2E-09 |
| *Y67D8C.23* | WBGene00235353 | pseudogene | 1.7 | 3.172 | 1.6E-05 |
| *C18H2.3* | WBGene00015990 | pseudogene | 1.7 | 3.171 | 2.4E-04 |
| *fbxc-47* | WBGene00019723 | protein_coding_gene | 1.7 | 3.165 | 2.8E-03 |
| *C55C3.8* | WBGene00044501 | protein_coding_gene | 1.7 | 3.164 | 2.1E-02 |
| *dod-17* | WBGene00010745 | protein_coding_gene | 1.7 | 3.145 | 8.5E-03 |
| *timm-17B.2* | WBGene00017069 | protein_coding_gene | 1.6 | 3.136 | 1.0E-10 |
| *R155.4* | WBGene00020118 | protein_coding_gene | 1.6 | 3.135 | 1.3E-05 |
| *ZK783.6* | WBGene00044448 | protein_coding_gene | 1.6 | 3.128 | 3.0E-04 |
| *Y59E9AL.8* | WBGene00044346 | protein_coding_gene | 1.6 | 3.128 | 1.8E-02 |
| *Y39G8C.2* | WBGene00012731 | protein_coding_gene | 1.6 | 3.127 | 8.1E-06 |
| *Y111B2A.1* | WBGene00013727 | protein_coding_gene | 1.6 | 3.118 | 7.9E-13 |
| *F10C1.8* | WBGene00017327 | protein_coding_gene | 1.6 | 3.081 | 6.0E-03 |
| *glb-12* | WBGene00008256 | protein_coding_gene | 1.6 | 3.068 | 8.0E-03 |
| *ctb-1* | WBGene00000829 | protein_coding_gene | 1.6 | 3.063 | 1.1E-21 |
| *C49C8.5* | WBGene00016769 | protein_coding_gene | 1.6 | 3.058 | 4.2E-19 |
| *ptp-5.3* | WBGene00021207 | protein_coding_gene | 1.6 | 3.047 | 2.5E-03 |
| *Y69A2AR.14* | WBGene00022085 | pseudogene | 1.6 | 3.024 | 2.6E-05 |
| *T20B6.2* | WBGene00020598 | protein_coding_gene | 1.6 | 3.004 | 1.5E-03 |
| *W05F2.8* | WBGene00195045 | protein_coding_gene | 1.6 | 3.004 | 1.5E-04 |
| *Y65B4BL.4* | WBGene00022036 | protein_coding_gene | 1.6 | 3.001 | 4.7E-19 |
| *F44B9.9* | WBGene00018410 | protein_coding_gene | 1.6 | 2.983 | 1.1E-02 |
| *nbs-1* | WBGene00007504 | protein_coding_gene | 1.6 | 2.982 | 6.7E-28 |
| *C09B9.1* | WBGene00015626 | protein_coding_gene | 1.6 | 2.982 | 2.1E-02 |
| *Y73B3A.3* | WBGene00022205 | pseudogene | 1.6 | 2.975 | 1.1E-08 |
| *ugt-29* | WBGene00021709 | protein_coding_gene | 1.6 | 2.972 | 3.7E-08 |
| *F47H4.12* | WBGene00043053 | pseudogene | 1.6 | 2.971 | 3.5E-05 |
| *dot-1.3* | WBGene00012302 | protein_coding_gene | 1.6 | 2.968 | 1.8E-02 |
| *nduo-4* | WBGene00010963 | protein_coding_gene | 1.6 | 2.963 | 3.6E-22 |
| *T25E12.16* | WBGene00194715 | protein_coding_gene | 1.6 | 2.945 | 1.1E-02 |
| *lec-9* | WBGene00002272 | protein_coding_gene | 1.6 | 2.945 | 7.4E-17 |
| *acs-23* | WBGene00011174 | pseudogene | 1.6 | 2.939 | 4.1E-03 |
| *Y49F6A.5* | WBGene00021713 | protein_coding_gene | 1.6 | 2.935 | 1.5E-02 |
| *taf-11.2* | WBGene00006394 | protein_coding_gene | 1.6 | 2.929 | 1.5E-34 |
| *F19C7.1* | WBGene00017591 | protein_coding_gene | 1.5 | 2.925 | 4.6E-08 |
| *nep-13* | WBGene00018196 | protein_coding_gene | 1.5 | 2.914 | 2.0E-03 |
| *msh-4* | WBGene00003420 | pseudogene | 1.5 | 2.911 | 4.3E-07 |
| *C14B1.7* | WBGene00007578 | pseudogene | 1.5 | 2.909 | 4.4E-27 |
| *math-46* | WBGene00021175 | protein_coding_gene | 1.5 | 2.906 | 4.1E-02 |
| *K09H9.7* | WBGene00019598 | protein_coding_gene | 1.5 | 2.890 | 8.2E-04 |
| *Y32G9A.3* | WBGene00021302 | protein_coding_gene | 1.5 | 2.882 | 1.6E-02 |
| *M05D6.1* | WBGene00010874 | protein_coding_gene | 1.5 | 2.868 | 5.0E-03 |
| *Y59E9AL.5* | WBGene00021995 | pseudogene | 1.5 | 2.865 | 2.4E-03 |
| *best-12* | WBGene00008821 | protein_coding_gene | 1.5 | 2.851 | 1.6E-03 |
| *F27B3.7* | WBGene00017849 | pseudogene | 1.5 | 2.842 | 1.0E-10 |
| *nspd-7* | WBGene00012781 | protein_coding_gene | 1.5 | 2.837 | 1.6E-02 |
| *C24H12.2* | WBGene00016072 | protein_coding_gene | 1.5 | 2.835 | 1.1E-03 |
| *sdz-3* | WBGene00016211 | pseudogene | 1.5 | 2.835 | 4.9E-03 |
| *F19C7.4* | WBGene00017594 | protein_coding_gene | 1.5 | 2.835 | 2.0E-04 |
| *R155.3* | WBGene00020117 | protein_coding_gene | 1.5 | 2.829 | 4.1E-07 |
| *ZK1236.8* | WBGene00023223 | pseudogene | 1.5 | 2.829 | 3.6E-02 |
| *C30F2.2* | WBGene00007816 | protein_coding_gene | 1.5 | 2.811 | 4.8E-02 |
| *ttr-57* | WBGene00050892 | protein_coding_gene | 1.5 | 2.811 | 2.0E-02 |
| *Y48B6A.10* | WBGene00012981 | protein_coding_gene | 1.5 | 2.803 | 4.4E-02 |
| *C45G9.6* | WBGene00016677 | protein_coding_gene | 1.5 | 2.802 | 2.7E-16 |
| *T07A9.14* | WBGene00044789 | protein_coding_gene | 1.5 | 2.802 | 3.9E-28 |
| *nep-7* | WBGene00017552 | protein_coding_gene | 1.5 | 2.799 | 1.0E-02 |
| *T03F6.8* | WBGene00044229 | pseudogene | 1.5 | 2.798 | 3.3E-04 |
| *T04A8.13* | WBGene00011414 | protein_coding_gene | 1.5 | 2.796 | 2.4E-05 |
| *R144.10* | WBGene00020098 | protein_coding_gene | 1.5 | 2.794 | 1.9E-25 |
| *oac-19* | WBGene00009413 | protein_coding_gene | 1.5 | 2.791 | 2.0E-03 |
| *fbxa-196* | WBGene00020265 | protein_coding_gene | 1.5 | 2.791 | 1.2E-05 |
| *D1065.3* | WBGene00017035 | pseudogene | 1.5 | 2.781 | 1.9E-04 |
| *R11H6.7* | WBGene00077579 | pseudogene | 1.5 | 2.780 | 8.0E-03 |
| *nas-6* | WBGene00003525 | protein_coding_gene | 1.5 | 2.776 | 9.4E-03 |
| *C34B4.2* | WBGene00007907 | protein_coding_gene | 1.5 | 2.771 | 7.1E-20 |
| *fbxa-192* | WBGene00010212 | protein_coding_gene | 1.5 | 2.765 | 1.8E-08 |
| *T02C12.4* | WBGene00011369 | pseudogene | 1.5 | 2.764 | 2.1E-02 |
| *F14D7.6* | WBGene00008793 | protein_coding_gene | 1.5 | 2.754 | 7.4E-09 |

| | | | | | |
|---|---|---|---|---|---|
| *fbxa-31* | WBGene00013838 | protein_coding_gene | 1.5 | 2.750 | 1.4E-06 |
| *Y22D7AR.7* | WBGene00021261 | protein_coding_gene | 1.5 | 2.747 | 8.4E-04 |
| *tipn-1* | WBGene00017738 | protein_coding_gene | 1.5 | 2.744 | 6.0E-27 |
| *W01G7.2* | WBGene00014851 | pseudogene | 1.5 | 2.742 | 3.0E-03 |
| *C06E1.7* | WBGene00015522 | protein_coding_gene | 1.5 | 2.741 | 4.5E-04 |
| *F21H12.7* | WBGene00044287 | protein_coding_gene | 1.5 | 2.740 | 2.5E-02 |
| *F08H9.2* | WBGene00008590 | protein_coding_gene | 1.5 | 2.738 | 1.6E-11 |
| *tag-51* | WBGene00006430 | protein_coding_gene | 1.5 | 2.738 | 2.7E-19 |
| *cye-1* | WBGene00000871 | protein_coding_gene | 1.4 | 2.728 | 5.1E-33 |
| *fbxa-201* | WBGene00020342 | protein_coding_gene | 1.4 | 2.728 | 3.4E-02 |
| *C49G7.13* | WBGene00269425 | protein_coding_gene | 1.4 | 2.716 | 3.2E-04 |
| *Y54G2A.21* | WBGene00021886 | pseudogene | 1.4 | 2.713 | 5.7E-08 |
| *F25E5.2* | WBGene00017783 | protein_coding_gene | 1.4 | 2.709 | 1.7E-02 |
| *T05C3.2* | WBGene00020254 | protein_coding_gene | 1.4 | 2.709 | 4.7E-03 |
| *fbxa-190* | WBGene00018939 | protein_coding_gene | 1.4 | 2.704 | 7.3E-04 |
| *klp-20* | WBGene00002230 | protein_coding_gene | 1.4 | 2.703 | 3.3E-12 |
| *F01D4.3* | WBGene00008487 | pseudogene | 1.4 | 2.689 | 3.8E-04 |
| *nduo-5* | WBGene00010967 | protein_coding_gene | 1.4 | 2.688 | 6.2E-08 |
| *C25G4.7* | WBGene00007734 | protein_coding_gene | 1.4 | 2.687 | 1.4E-04 |
| *cdr-7* | WBGene00010472 | protein_coding_gene | 1.4 | 2.686 | 3.4E-08 |
| *F36G9.13* | WBGene00009489 | protein_coding_gene | 1.4 | 2.682 | 6.8E-05 |
| *hpr-9* | WBGene00001997 | protein_coding_gene | 1.4 | 2.677 | 9.3E-24 |
| *T09F5.10* | WBGene00011669 | protein_coding_gene | 1.4 | 2.677 | 3.1E-02 |
| *scrm-4* | WBGene00008681 | protein_coding_gene | 1.4 | 2.675 | 2.7E-06 |
| *Y50D4B.2* | WBGene00021741 | protein_coding_gene | 1.4 | 2.670 | 2.4E-04 |
| *K09B11.4* | WBGene00010711 | protein_coding_gene | 1.4 | 2.670 | 1.7E-02 |
| *zip-11* | WBGene00021082 | protein_coding_gene | 1.4 | 2.664 | 3.5E-10 |
| *C33H5.16* | WBGene00016382 | protein_coding_gene | 1.4 | 2.664 | 1.1E-03 |
| *C18D11.3* | WBGene00007681 | protein_coding_gene | 1.4 | 2.662 | 7.5E-33 |
| *B0334.5* | WBGene00007145 | protein_coding_gene | 1.4 | 2.662 | 6.6E-29 |
| *fbxa-79* | WBGene00022329 | protein_coding_gene | 1.4 | 2.658 | 2.3E-04 |
| *F18A11.6* | WBGene00008933 | pseudogene | 1.4 | 2.656 | 1.0E-04 |
| *K09C6.2* | WBGene00019556 | protein_coding_gene | 1.4 | 2.655 | 2.1E-05 |
| *set-24* | WBGene00012845 | protein_coding_gene | 1.4 | 2.647 | 2.2E-24 |
| *prde-1* | WBGene00008995 | protein_coding_gene | 1.4 | 2.645 | 2.9E-17 |
| *Y105C5B.14* | WBGene00013653 | protein_coding_gene | 1.4 | 2.638 | 4.5E-02 |
| *csp-2* | WBGene00000820 | protein_coding_gene | 1.4 | 2.638 | 2.1E-03 |
| *gpx-8* | WBGene00018850 | protein_coding_gene | 1.4 | 2.625 | 4.8E-02 |
| *Y73B3A.20* | WBGene00022221 | protein_coding_gene | 1.4 | 2.617 | 5.8E-12 |
| *E02H9.3* | WBGene00017101 | protein_coding_gene | 1.4 | 2.605 | 3.4E-19 |
| *C28D4.5* | WBGene00007793 | protein_coding_gene | 1.4 | 2.605 | 1.6E-02 |
| *F19F10.11* | WBGene00017607 | protein_coding_gene | 1.4 | 2.602 | 9.0E-31 |
| *nhr-168* | WBGene00008221 | protein_coding_gene | 1.4 | 2.596 | 2.9E-05 |
| *snpc-3.3* | WBGene00021662 | protein_coding_gene | 1.4 | 2.591 | 9.6E-07 |
| *C09E7.6* | WBGene00015640 | pseudogene | 1.4 | 2.588 | 5.7E-16 |
| *snpc-3.1* | WBGene00015098 | protein_coding_gene | 1.4 | 2.586 | 2.2E-05 |
| *enu-3.2* | WBGene00008003 | protein_coding_gene | 1.4 | 2.584 | 3.3E-11 |
| *T05E12.8* | WBGene00045382 | protein_coding_gene | 1.4 | 2.583 | 3.2E-02 |
| *rbmx-2* | WBGene00016245 | protein_coding_gene | 1.4 | 2.578 | 1.7E-32 |
| *gpx-1* | WBGene00009165 | protein_coding_gene | 1.4 | 2.577 | 1.7E-10 |
| *C09G4.4* | WBGene00044001 | protein_coding_gene | 1.4 | 2.575 | 2.0E-05 |
| *C06A5.6* | WBGene00015504 | protein_coding_gene | 1.4 | 2.575 | 4.1E-23 |
| *D1086.18* | WBGene00045405 | protein_coding_gene | 1.4 | 2.574 | 4.9E-04 |
| *clp-2* | WBGene00000543 | protein_coding_gene | 1.4 | 2.566 | 8.0E-26 |
| *F07E5.8* | WBGene00017213 | protein_coding_gene | 1.4 | 2.563 | 1.0E-02 |
| *ife-5* | WBGene00002063 | protein_coding_gene | 1.4 | 2.562 | 4.7E-20 |
| *F55D12.5* | WBGene00010113 | protein_coding_gene | 1.4 | 2.553 | 4.9E-33 |
| *pgph-2* | WBGene00018424 | protein_coding_gene | 1.3 | 2.543 | 2.3E-10 |
| *Y71A12B.13* | WBGene00013510 | pseudogene | 1.3 | 2.542 | 2.2E-02 |
| *clec-155* | WBGene00011405 | pseudogene | 1.3 | 2.541 | 1.2E-02 |
| *C05B5.2* | WBGene00007320 | protein_coding_gene | 1.3 | 2.535 | 3.6E-12 |
| *H04J21.1* | WBGene00019150 | protein_coding_gene | 1.3 | 2.534 | 3.1E-04 |
| *nduo-3* | WBGene00010966 | protein_coding_gene | 1.3 | 2.530 | 2.7E-02 |
| *wago-4* | WBGene00010263 | protein_coding_gene | 1.3 | 2.518 | 3.5E-24 |
| *syp-5* | WBGene00021832 | protein_coding_gene | 1.3 | 2.515 | 1.3E-25 |
| *ZK809.1* | WBGene00014085 | protein_coding_gene | 1.3 | 2.513 | 4.3E-02 |
| *F43G6.10* | WBGene00009662 | protein_coding_gene | 1.3 | 2.507 | 5.3E-10 |
| *ppw-1* | WBGene00004093 | protein_coding_gene | 1.3 | 2.506 | 1.4E-12 |
| *T08D2.9* | WBGene00044700 | pseudogene | 1.3 | 2.504 | 5.0E-03 |
| *R05D3.12* | WBGene00019884 | protein_coding_gene | 1.3 | 2.502 | 1.5E-22 |

| | | | | | |
|---|---|---|---|---|---|
| *D2005.7* | WBGene00086551 | protein_coding_gene | 1.3 | 2.502 | 4.9E-19 |
| *F10C2.5* | WBGene00008646 | protein_coding_gene | 1.3 | 2.501 | 2.0E-27 |
| *C25H3.1* | WBGene00016111 | protein_coding_gene | 1.3 | 2.495 | 2.0E-02 |
| *lsl-1* | WBGene00009937 | protein_coding_gene | 1.3 | 2.495 | 1.3E-21 |
| *tofu-7* | WBGene00022737 | protein_coding_gene | 1.3 | 2.494 | 1.4E-24 |
| *Y24F12A.3* | WBGene00012498 | pseudogene | 1.3 | 2.490 | 6.4E-03 |
| *C17C3.11* | WBGene00015893 | protein_coding_gene | 1.3 | 2.485 | 2.6E-02 |
| *nas-25* | WBGene00003544 | protein_coding_gene | 1.3 | 2.484 | 3.4E-03 |
| *F54H5.3* | WBGene00018840 | protein_coding_gene | 1.3 | 2.480 | 6.2E-04 |
| *cpb-1* | WBGene00000770 | protein_coding_gene | 1.3 | 2.478 | 1.4E-21 |
| *R04F11.5* | WBGene00011017 | protein_coding_gene | 1.3 | 2.475 | 4.3E-19 |
| *rpl-31* | WBGene00004445 | protein_coding_gene | 1.3 | 2.472 | 2.5E-24 |
| *M02B7.2* | WBGene00019724 | protein_coding_gene | 1.3 | 2.469 | 7.7E-11 |
| *nep-5* | WBGene00017393 | protein_coding_gene | 1.3 | 2.466 | 2.2E-02 |
| *cln-3.3* | WBGene00000541 | protein_coding_gene | 1.3 | 2.465 | 1.9E-06 |
| *T08D2.3* | WBGene00011608 | pseudogene | 1.3 | 2.463 | 1.1E-05 |
| *fbxa-183* | WBGene00018428 | protein_coding_gene | 1.3 | 2.463 | 1.7E-02 |
| *F14H3.3* | WBGene00008822 | protein_coding_gene | 1.3 | 2.459 | 8.9E-15 |
| *T10B11.8* | WBGene00020404 | protein_coding_gene | 1.3 | 2.456 | 1.0E-20 |
| *set-5* | WBGene00008145 | protein_coding_gene | 1.3 | 2.449 | 1.5E-18 |
| *nhr-266* | WBGene00018993 | pseudogene | 1.3 | 2.449 | 2.7E-02 |
| *C14C10.1* | WBGene00007584 | protein_coding_gene | 1.3 | 2.439 | 1.5E-11 |
| *scrm-7* | WBGene00013052 | protein_coding_gene | 1.3 | 2.434 | 2.8E-03 |
| *D2024.1* | WBGene00017050 | protein_coding_gene | 1.3 | 2.431 | 1.5E-02 |
| *Y53C10A.6* | WBGene00013137 | protein_coding_gene | 1.3 | 2.430 | 1.3E-24 |
| *F54A3.6* | WBGene00018785 | protein_coding_gene | 1.3 | 2.426 | 3.5E-16 |
| *C33F10.1* | WBGene00016351 | protein_coding_gene | 1.3 | 2.423 | 3.5E-10 |
| *C14C11.1* | WBGene00015765 | protein_coding_gene | 1.3 | 2.422 | 1.1E-05 |
| *ZK354.3* | WBGene00022706 | protein_coding_gene | 1.3 | 2.420 | 7.6E-06 |
| *Y52E8A.3* | WBGene00021796 | protein_coding_gene | 1.3 | 2.419 | 3.0E-03 |
| *Y73B6BL.12* | WBGene00022236 | protein_coding_gene | 1.3 | 2.416 | 2.8E-03 |
| *F22E5.7* | WBGene00017706 | protein_coding_gene | 1.3 | 2.412 | 4.3E-04 |
| *yars-2* | WBGene00006968 | protein_coding_gene | 1.3 | 2.409 | 1.1E-28 |
| *H25P19.1* | WBGene00019244 | protein_coding_gene | 1.3 | 2.409 | 1.7E-13 |
| *math-48* | WBGene00022689 | protein_coding_gene | 1.3 | 2.408 | 5.7E-04 |
| *CD4.8* | WBGene00016993 | protein_coding_gene | 1.3 | 2.408 | 7.3E-12 |
| *clec-143* | WBGene00014063 | protein_coding_gene | 1.3 | 2.406 | 2.8E-02 |
| *F36D1.6* | WBGene00009465 | protein_coding_gene | 1.3 | 2.403 | 4.6E-02 |
| *C08F11.14* | WBGene00007461 | pseudogene | 1.3 | 2.400 | 4.8E-05 |
| *linc-82* | WBGene00219570 | lincRNA_gene | 1.3 | 2.400 | 2.1E-02 |
| *hrg-7* | WBGene00007605 | protein_coding_gene | 1.3 | 2.393 | 9.9E-04 |
| *B0035.15* | WBGene00007113 | protein_coding_gene | 1.3 | 2.388 | 1.9E-25 |
| *C04E6.4* | WBGene00015421 | protein_coding_gene | 1.3 | 2.387 | 5.4E-03 |
| *set-32* | WBGene00008062 | protein_coding_gene | 1.3 | 2.385 | 3.5E-13 |
| *ubl-1* | WBGene00006725 | protein_coding_gene | 1.3 | 2.383 | 1.5E-21 |
| *D2096.11* | WBGene00017078 | protein_coding_gene | 1.3 | 2.383 | 1.3E-02 |
| *abt-1* | WBGene00000019 | protein_coding_gene | 1.3 | 2.380 | 2.1E-04 |
| *Y66D12A.3* | WBGene00013429 | protein_coding_gene | 1.3 | 2.380 | 2.3E-04 |
| *dnj-17* | WBGene00001035 | protein_coding_gene | 1.3 | 2.379 | 5.0E-23 |
| *Y94H6A.3* | WBGene00022376 | protein_coding_gene | 1.2 | 2.378 | 3.7E-15 |
| *fbxa-218* | WBGene00013036 | protein_coding_gene | 1.2 | 2.375 | 1.2E-02 |
| *T05F1.7* | WBGene00011492 | protein_coding_gene | 1.2 | 2.372 | 1.6E-02 |
| *Y57G11C.23* | WBGene00013318 | protein_coding_gene | 1.2 | 2.370 | 2.8E-02 |
| *elo-5* | WBGene00001243 | protein_coding_gene | 1.2 | 2.368 | 2.3E-15 |
| *Y97E10AR.8* | WBGene00044552 | protein_coding_gene | 1.2 | 2.367 | 3.2E-02 |
| *lip-1* | WBGene00003043 | protein_coding_gene | 1.2 | 2.366 | 3.2E-25 |
| *Y54G2A.73* | WBGene00219425 | protein_coding_gene | 1.2 | 2.365 | 6.4E-10 |
| *pigg-1* | WBGene00009204 | protein_coding_gene | 1.2 | 2.365 | 2.2E-23 |
| *Y75B12A.1* | WBGene00014949 | pseudogene | 1.2 | 2.360 | 8.3E-04 |
| *C34H4.2* | WBGene00016425 | protein_coding_gene | 1.2 | 2.359 | 1.8E-05 |
| *hrg-1* | WBGene00019830 | protein_coding_gene | 1.2 | 2.358 | 5.2E-09 |
| *K12C11.5* | WBGene00044896 | protein_coding_gene | 1.2 | 2.357 | 2.3E-02 |
| *mog-3* | WBGene00003391 | protein_coding_gene | 1.2 | 2.354 | 2.2E-20 |
| *M05D6.3* | WBGene00010876 | protein_coding_gene | 1.2 | 2.351 | 1.4E-02 |
| *aars-1* | WBGene00000196 | protein_coding_gene | 1.2 | 2.344 | 1.5E-18 |
| *clec-186* | WBGene00014138 | protein_coding_gene | 1.2 | 2.342 | 5.9E-14 |
| *puf-7* | WBGene00004243 | protein_coding_gene | 1.2 | 2.335 | 8.4E-16 |
| *pid-5* | WBGene00021555 | protein_coding_gene | 1.2 | 2.335 | 4.9E-22 |
| *fbxa-210* | WBGene00012565 | protein_coding_gene | 1.2 | 2.333 | 2.0E-12 |
| *Y73B3A.21* | WBGene00022222 | pseudogene | 1.2 | 2.327 | 1.4E-03 |

| | | | | | |
|---|---|---|---|---|---|
| *hil-5* | WBGene00001856 | protein_coding_gene | 1.2 | 2.326 | 5.3E-24 |
| *ZK938.1* | WBGene00014158 | protein_coding_gene | 1.2 | 2.324 | 1.7E-12 |
| *Y39B6A.18* | WBGene00012679 | pseudogene | 1.2 | 2.323 | 4.9E-03 |
| *Y48A6C.1* | WBGene00012974 | pseudogene | 1.2 | 2.320 | 8.2E-05 |
| *memi-2* | WBGene00013380 | protein_coding_gene | 1.2 | 2.312 | 5.7E-15 |
| *R08D7.1* | WBGene00011142 | protein_coding_gene | 1.2 | 2.312 | 4.0E-27 |
| *hil-4* | WBGene00001855 | protein_coding_gene | 1.2 | 2.305 | 3.6E-19 |
| *swp-1* | WBGene00006359 | protein_coding_gene | 1.2 | 2.304 | 5.1E-22 |
| *C06C6.6* | WBGene00007380 | pseudogene | 1.2 | 2.302 | 3.5E-02 |
| *T14G10.8* | WBGene00011777 | protein_coding_gene | 1.2 | 2.302 | 5.8E-10 |
| *Y39B6A.43* | WBGene00012701 | protein_coding_gene | 1.2 | 2.296 | 1.1E-14 |
| *trpl-5* | WBGene00004149 | protein_coding_gene | 1.2 | 2.295 | 1.1E-03 |
| *C17H12.12* | WBGene00015937 | protein_coding_gene | 1.2 | 2.289 | 2.3E-08 |
| *F54A3.2* | WBGene00018781 | protein_coding_gene | 1.2 | 2.288 | 3.1E-07 |
| *kup-1* | WBGene00002241 | protein_coding_gene | 1.2 | 2.288 | 1.5E-32 |
| *F53B2.5* | WBGene00009956 | protein_coding_gene | 1.2 | 2.284 | 3.0E-07 |
| *F56H1.3* | WBGene00018994 | protein_coding_gene | 1.2 | 2.284 | 4.4E-03 |
| *coa-1* | WBGene00195063 | protein_coding_gene | 1.2 | 2.284 | 1.0E-11 |
| *F56G4.4* | WBGene00010159 | protein_coding_gene | 1.2 | 2.281 | 1.9E-21 |
| *T28A11.25* | WBGene00269355 | protein_coding_gene | 1.2 | 2.280 | 4.9E-02 |
| *W03H9.2* | WBGene00012228 | protein_coding_gene | 1.2 | 2.279 | 2.1E-02 |
| *asp-14* | WBGene00019619 | protein_coding_gene | 1.2 | 2.279 | 8.6E-19 |
| *Y14H12B.2* | WBGene00021192 | protein_coding_gene | 1.2 | 2.279 | 5.4E-14 |
| *syx-16* | WBGene00022534 | protein_coding_gene | 1.2 | 2.278 | 8.9E-22 |
| *trul-1* | WBGene00015194 | protein_coding_gene | 1.2 | 2.278 | 1.0E-16 |
| *wip-1* | WBGene00020094 | protein_coding_gene | 1.2 | 2.278 | 1.6E-14 |
| *ulp-5* | WBGene00006740 | protein_coding_gene | 1.2 | 2.277 | 1.8E-16 |
| *F56C9.6* | WBGene00018950 | protein_coding_gene | 1.2 | 2.275 | 3.5E-23 |
| *Y22D7AR.12* | WBGene00021265 | pseudogene | 1.2 | 2.274 | 3.1E-04 |
| *Y67A10A.3* | WBGene00013452 | protein_coding_gene | 1.2 | 2.274 | 1.7E-04 |
| *K05F1.1* | WBGene00019405 | protein_coding_gene | 1.2 | 2.273 | 2.0E-02 |
| *F54D5.2* | WBGene00010048 | protein_coding_gene | 1.2 | 2.273 | 2.3E-17 |
| *W03F8.2* | WBGene00020992 | pseudogene | 1.2 | 2.271 | 3.4E-04 |
| *lido-18* | WBGene00015988 | protein_coding_gene | 1.2 | 2.266 | 5.7E-13 |
| *R144.11* | WBGene00020099 | protein_coding_gene | 1.2 | 2.264 | 2.2E-17 |
| *K03H9.3* | WBGene00019372 | protein_coding_gene | 1.2 | 2.264 | 4.6E-02 |
| *VH15N14R.1* | WBGene00012151 | protein_coding_gene | 1.2 | 2.262 | 1.2E-05 |
| *T13F3.8* | WBGene00044205 | protein_coding_gene | 1.2 | 2.262 | 1.6E-04 |
| *B0207.7* | WBGene00015030 | protein_coding_gene | 1.2 | 2.260 | 2.3E-04 |
| *rpl-28* | WBGene00004442 | protein_coding_gene | 1.2 | 2.259 | 5.1E-16 |
| *T28A8.5* | WBGene00012111 | protein_coding_gene | 1.2 | 2.255 | 2.2E-15 |
| *bgnt-1.4* | WBGene00015982 | protein_coding_gene | 1.2 | 2.255 | 2.0E-02 |
| *F01G4.4* | WBGene00008503 | protein_coding_gene | 1.2 | 2.254 | 2.0E-23 |
| *ZK809.5* | WBGene00014087 | protein_coding_gene | 1.2 | 2.253 | 3.5E-22 |
| *mbtr-1* | WBGene00021661 | protein_coding_gene | 1.2 | 2.250 | 1.8E-11 |
| *ints-11* | WBGene00008642 | protein_coding_gene | 1.2 | 2.249 | 2.2E-23 |
| *syp-6* | WBGene00019002 | protein_coding_gene | 1.2 | 2.249 | 2.3E-23 |
| *rps-22* | WBGene00004491 | protein_coding_gene | 1.2 | 2.247 | 9.8E-15 |
| *rps-0* | WBGene00004469 | protein_coding_gene | 1.2 | 2.247 | 1.5E-16 |
| *ZK1127.6* | WBGene00022853 | protein_coding_gene | 1.2 | 2.243 | 3.3E-12 |
| *him-6* | WBGene00001865 | protein_coding_gene | 1.2 | 2.241 | 4.7E-21 |
| *K12D12.5* | WBGene00010787 | protein_coding_gene | 1.2 | 2.238 | 1.6E-18 |
| *F10G7.9* | WBGene00017372 | protein_coding_gene | 1.2 | 2.238 | 3.4E-22 |
| *F54F2.9* | WBGene00018836 | protein_coding_gene | 1.2 | 2.237 | 8.7E-20 |
| *wago-11* | WBGene00021711 | protein_coding_gene | 1.2 | 2.237 | 1.9E-03 |
| *C48B4.11* | WBGene00008174 | protein_coding_gene | 1.2 | 2.235 | 5.0E-22 |
| *R06C1.4* | WBGene00011059 | protein_coding_gene | 1.2 | 2.234 | 2.3E-27 |
| *rnf-1* | WBGene00004380 | protein_coding_gene | 1.2 | 2.226 | 2.2E-20 |
| *zhp-4* | WBGene00012678 | protein_coding_gene | 1.2 | 2.223 | 2.2E-08 |
| *hrde-2* | WBGene00011324 | protein_coding_gene | 1.2 | 2.222 | 2.2E-18 |
| *tmed-12* | WBGene00011606 | protein_coding_gene | 1.2 | 2.221 | 1.7E-02 |
| *K07A1.17* | WBGene00044987 | protein_coding_gene | 1.2 | 2.221 | 2.5E-26 |
| *C49C3.2* | WBGene00008192 | pseudogene | 1.2 | 2.220 | 1.2E-02 |
| *rpl-17* | WBGene00004429 | protein_coding_gene | 1.1 | 2.218 | 4.8E-20 |
| *let-504* | WBGene00002696 | protein_coding_gene | 1.1 | 2.218 | 7.5E-21 |
| *Y106G6H.16* | WBGene00013726 | protein_coding_gene | 1.1 | 2.216 | 1.7E-03 |
| *K08H10.5* | WBGene00014805 | pseudogene | 1.1 | 2.216 | 5.0E-02 |
| *ubxn-5* | WBGene00011336 | protein_coding_gene | 1.1 | 2.214 | 9.9E-07 |
| *T07A5.7* | WBGene00045475 | protein_coding_gene | 1.1 | 2.213 | 1.8E-02 |
| *Y54G2A.15* | WBGene00021880 | protein_coding_gene | 1.1 | 2.211 | 6.1E-06 |

| | | | | | |
|---|---|---|---|---|---|
| *gly-10* | WBGene00001635 | protein_coding_gene | 1.1 | 2.210 | 6.9E-24 |
| *htp-1* | WBGene00002032 | protein_coding_gene | 1.1 | 2.209 | 1.8E-18 |
| *Y48G1A.1* | WBGene00021657 | protein_coding_gene | 1.1 | 2.205 | 1.6E-12 |
| *fbxc-50* | WBGene00015912 | protein_coding_gene | 1.1 | 2.205 | 1.6E-07 |
| *such-1* | WBGene00013443 | protein_coding_gene | 1.1 | 2.203 | 2.1E-16 |
| *T12E12.3* | WBGene00020464 | protein_coding_gene | 1.1 | 2.199 | 3.7E-19 |
| *coa-6* | WBGene00013893 | protein_coding_gene | 1.1 | 2.199 | 9.5E-21 |
| *F18C12.3* | WBGene00008934 | protein_coding_gene | 1.1 | 2.198 | 1.9E-22 |
| *Y75B8A.44* | WBGene00194855 | protein_coding_gene | 1.1 | 2.198 | 4.6E-03 |
| *T08D2.2* | WBGene00011607 | pseudogene | 1.1 | 2.196 | 2.2E-03 |
| *H14A12.3* | WBGene00019196 | protein_coding_gene | 1.1 | 2.192 | 1.4E-19 |
| *F43G6.3* | WBGene00009656 | pseudogene | 1.1 | 2.191 | 4.6E-12 |
| *R04B5.11* | WBGene00011008 | protein_coding_gene | 1.1 | 2.189 | 4.6E-02 |
| *gei-14* | WBGene00001571 | protein_coding_gene | 1.1 | 2.189 | 8.2E-21 |
| *fbf-2* | WBGene00001402 | protein_coding_gene | 1.1 | 2.188 | 5.0E-11 |
| *ZK1251.5* | WBGene00014996 | pseudogene | 1.1 | 2.188 | 4.6E-03 |
| *emr-1* | WBGene00001309 | protein_coding_gene | 1.1 | 2.187 | 1.1E-18 |
| *rrp-1* | WBGene00008151 | protein_coding_gene | 1.1 | 2.187 | 3.8E-18 |
| *ZK287.9* | WBGene00013968 | protein_coding_gene | 1.1 | 2.187 | 3.7E-07 |
| *slr-2* | WBGene00013350 | protein_coding_gene | 1.1 | 2.187 | 3.2E-11 |
| *C33F10.11* | WBGene00016357 | protein_coding_gene | 1.1 | 2.186 | 2.9E-06 |
| *znfx-1* | WBGene00014208 | protein_coding_gene | 1.1 | 2.185 | 1.5E-19 |
| *R148.4* | WBGene00020103 | protein_coding_gene | 1.1 | 2.185 | 6.6E-15 |
| *math-43* | WBGene00022432 | pseudogene | 1.1 | 2.184 | 3.2E-02 |
| *lido-16* | WBGene00020898 | protein_coding_gene | 1.1 | 2.183 | 1.7E-02 |
| *F47B3.4* | WBGene00018528 | protein_coding_gene | 1.1 | 2.183 | 2.7E-02 |
| *kca-1* | WBGene00015698 | protein_coding_gene | 1.1 | 2.182 | 3.9E-19 |
| *rnh-1.3* | WBGene00007303 | protein_coding_gene | 1.1 | 2.180 | 6.5E-06 |
| *duo-1* | WBGene00001110 | protein_coding_gene | 1.1 | 2.180 | 5.2E-21 |
| *gfl-1* | WBGene00001585 | protein_coding_gene | 1.1 | 2.180 | 2.9E-18 |
| *cus-2* | WBGene00022025 | protein_coding_gene | 1.1 | 2.179 | 3.4E-18 |
| *C06G4.1* | WBGene00015557 | protein_coding_gene | 1.1 | 2.178 | 5.5E-22 |
| *rpl-15* | WBGene00004427 | protein_coding_gene | 1.1 | 2.178 | 3.0E-17 |
| *ZK550.5* | WBGene00013999 | protein_coding_gene | 1.1 | 2.178 | 2.2E-05 |
| *T28A8.6* | WBGene00012112 | protein_coding_gene | 1.1 | 2.177 | 9.9E-20 |
| *klp-10* | WBGene00002221 | pseudogene | 1.1 | 2.175 | 5.9E-14 |
| *Y116F11B.8* | WBGene00013824 | protein_coding_gene | 1.1 | 2.171 | 2.8E-02 |
| *dpf-7* | WBGene00001060 | protein_coding_gene | 1.1 | 2.170 | 1.4E-18 |
| *vps-36* | WBGene00008919 | protein_coding_gene | 1.1 | 2.169 | 1.4E-22 |
| *F59A6.2* | WBGene00019084 | protein_coding_gene | 1.1 | 2.168 | 7.6E-04 |
| *plp-2* | WBGene00008976 | protein_coding_gene | 1.1 | 2.167 | 4.5E-10 |
| *ZK1290.14* | WBGene00044396 | protein_coding_gene | 1.1 | 2.165 | 6.7E-07 |
| *rps-29* | WBGene00004498 | protein_coding_gene | 1.1 | 2.163 | 5.0E-15 |
| *rps-4* | WBGene00004473 | protein_coding_gene | 1.1 | 2.163 | 2.6E-21 |
| *ZK1098.1* | WBGene00014218 | protein_coding_gene | 1.1 | 2.160 | 1.0E-29 |
| *igeg-2* | WBGene00009842 | protein_coding_gene | 1.1 | 2.160 | 5.0E-03 |
| *lurp-4* | WBGene00077698 | protein_coding_gene | 1.1 | 2.159 | 3.4E-02 |
| *C08B11.9* | WBGene00007436 | protein_coding_gene | 1.1 | 2.158 | 4.1E-15 |
| *aptf-3* | WBGene00009203 | protein_coding_gene | 1.1 | 2.158 | 1.9E-13 |
| *nud-1* | WBGene00003829 | protein_coding_gene | 1.1 | 2.157 | 2.4E-23 |
| *fbxa-51* | WBGene00020448 | protein_coding_gene | 1.1 | 2.156 | 1.2E-02 |
| *F55H2.7* | WBGene00010132 | protein_coding_gene | 1.1 | 2.153 | 3.3E-22 |
| *glh-1* | WBGene00001598 | protein_coding_gene | 1.1 | 2.151 | 3.8E-25 |
| *rtel-1* | WBGene00009124 | protein_coding_gene | 1.1 | 2.150 | 3.2E-19 |
| *R09E10.6* | WBGene00011176 | protein_coding_gene | 1.1 | 2.149 | 1.9E-11 |
| *Y44F5A.1* | WBGene00012858 | protein_coding_gene | 1.1 | 2.147 | 3.5E-15 |
| *T03D3.5* | WBGene00020183 | protein_coding_gene | 1.1 | 2.146 | 7.6E-09 |
| *Y73B3A.9* | WBGene00022211 | pseudogene | 1.1 | 2.145 | 1.1E-13 |
| *him-18* | WBGene00011415 | protein_coding_gene | 1.1 | 2.144 | 1.3E-21 |
| *clec-142* | WBGene00010460 | protein_coding_gene | 1.1 | 2.142 | 1.4E-02 |
| *mppe-1* | WBGene00015238 | protein_coding_gene | 1.1 | 2.141 | 5.2E-12 |
| *F56D6.12* | WBGene00044474 | protein_coding_gene | 1.1 | 2.139 | 4.3E-02 |
| *C36B1.11* | WBGene00007978 | protein_coding_gene | 1.1 | 2.136 | 1.3E-18 |
| *C05C10.5* | WBGene00007332 | protein_coding_gene | 1.1 | 2.130 | 2.3E-13 |
| *ekl-7* | WBGene00021465 | protein_coding_gene | 1.1 | 2.128 | 8.7E-14 |
| *F58B3.4* | WBGene00010231 | protein_coding_gene | 1.1 | 2.127 | 5.2E-23 |
| *C24G6.8* | WBGene00016062 | protein_coding_gene | 1.1 | 2.127 | 2.8E-16 |
| *ZK896.3* | WBGene00014134 | pseudogene | 1.1 | 2.126 | 6.3E-03 |
| *F08F8.6* | WBGene00017271 | protein_coding_gene | 1.1 | 2.126 | 1.4E-03 |
| *F35C11.2* | WBGene00009401 | protein_coding_gene | 1.1 | 2.126 | 2.8E-03 |

| | | | | | |
|---|---|---|---|---|---|
| *Y32B12B.2* | WBGene00012522 | protein_coding_gene | 1.1 | 2.125 | 4.0E-17 |
| *F46F5.1* | WBGene00018493 | protein_coding_gene | 1.1 | 2.125 | 3.1E-03 |
| *C45E1.4* | WBGene00016663 | protein_coding_gene | 1.1 | 2.124 | 3.4E-05 |
| *F38B2.2* | WBGene00009529 | protein_coding_gene | 1.1 | 2.123 | 1.2E-02 |
| *K11H3.4* | WBGene00010781 | protein_coding_gene | 1.1 | 2.121 | 1.9E-13 |
| *C50E10.1* | WBGene00016825 | protein_coding_gene | 1.1 | 2.121 | 1.2E-05 |
| *T08G3.4* | WBGene00011617 | pseudogene | 1.1 | 2.118 | 4.4E-04 |
| *R07E5.1* | WBGene00011109 | protein_coding_gene | 1.1 | 2.118 | 9.0E-20 |
| *C17E4.19* | WBGene00219310 | protein_coding_gene | 1.1 | 2.117 | 4.8E-11 |
| *ZC155.2* | WBGene00022530 | protein_coding_gene | 1.1 | 2.117 | 1.1E-02 |
| *mrpl-11* | WBGene00015133 | protein_coding_gene | 1.1 | 2.117 | 2.9E-17 |
| *cdk-11.1* | WBGene00015203 | protein_coding_gene | 1.1 | 2.116 | 3.3E-22 |
| *T06E6.1* | WBGene00011538 | protein_coding_gene | 1.1 | 2.115 | 1.6E-13 |
| *Y111B2A.3* | WBGene00013729 | protein_coding_gene | 1.1 | 2.114 | 1.6E-24 |
| *eif-2D* | WBGene00016113 | protein_coding_gene | 1.1 | 2.112 | 2.4E-17 |
| *eea-1* | WBGene00011696 | protein_coding_gene | 1.1 | 2.112 | 1.9E-17 |
| *gtf-2F2* | WBGene00012694 | protein_coding_gene | 1.1 | 2.111 | 3.1E-19 |
| *F25E5.1* | WBGene00017782 | protein_coding_gene | 1.1 | 2.110 | 3.8E-13 |
| *Y20F4.8* | WBGene00044435 | protein_coding_gene | 1.1 | 2.110 | 4.4E-02 |
| *spd-2* | WBGene00004953 | protein_coding_gene | 1.1 | 2.109 | 1.3E-19 |
| *Y4C6A.3* | WBGene00021153 | protein_coding_gene | 1.1 | 2.106 | 1.2E-16 |
| *syp-4* | WBGene00019247 | protein_coding_gene | 1.1 | 2.105 | 3.2E-27 |
| *Y110A7A.15* | WBGene00022462 | protein_coding_gene | 1.1 | 2.105 | 7.0E-18 |
| *daz-1* | WBGene00000935 | protein_coding_gene | 1.1 | 2.102 | 3.8E-27 |
| *pst-2* | WBGene00018827 | protein_coding_gene | 1.1 | 2.101 | 9.9E-14 |
| *zhp-2* | WBGene00008387 | protein_coding_gene | 1.1 | 2.101 | 8.3E-09 |
| *Y80D3A.8* | WBGene00013586 | protein_coding_gene | 1.1 | 2.101 | 7.7E-03 |
| *lin-9* | WBGene00002998 | protein_coding_gene | 1.1 | 2.098 | 1.7E-18 |
| *Y4C6A.1* | WBGene00021151 | pseudogene | 1.1 | 2.098 | 6.9E-03 |
| *F14H3.4* | WBGene00008823 | protein_coding_gene | 1.1 | 2.098 | 4.0E-11 |
| *F28A10.4* | WBGene00017872 | protein_coding_gene | 1.1 | 2.097 | 3.0E-02 |
| *pid-2* | WBGene00021676 | protein_coding_gene | 1.1 | 2.096 | 4.4E-10 |
| *rpl-36* | WBGene00004450 | protein_coding_gene | 1.1 | 2.095 | 1.5E-21 |
| *Y46G5A.43* | WBGene00077636 | pseudogene | 1.1 | 2.095 | 1.1E-02 |
| *hmg-20* | WBGene00012209 | protein_coding_gene | 1.1 | 2.094 | 1.2E-12 |
| *F43C1.5* | WBGene00009649 | protein_coding_gene | 1.1 | 2.093 | 1.6E-02 |
| *hal-3* | WBGene00011788 | protein_coding_gene | 1.1 | 2.093 | 7.7E-13 |
| *C18E3.9* | WBGene00015976 | protein_coding_gene | 1.1 | 2.093 | 7.7E-17 |
| *Y23H5B.6* | WBGene00021277 | protein_coding_gene | 1.1 | 2.093 | 4.8E-18 |
| *cogc-2* | WBGene00000585 | protein_coding_gene | 1.1 | 2.092 | 1.8E-18 |
| *Y48G1BL.5* | WBGene00021664 | pseudogene | 1.1 | 2.091 | 2.9E-05 |
| *F44E7.5* | WBGene00018427 | protein_coding_gene | 1.1 | 2.091 | 9.2E-14 |
| *set-9* | WBGene00017482 | protein_coding_gene | 1.1 | 2.091 | 1.9E-17 |
| *snpc-1.1* | WBGene00015513 | protein_coding_gene | 1.1 | 2.091 | 2.1E-16 |
| *him-10* | WBGene00001869 | protein_coding_gene | 1.1 | 2.091 | 2.7E-21 |
| *ztf-8* | WBGene00022598 | protein_coding_gene | 1.1 | 2.090 | 1.9E-20 |
| *Y47G6A.9* | WBGene00021638 | protein_coding_gene | 1.1 | 2.089 | 8.9E-17 |
| *C30G7.3* | WBGene00044177 | protein_coding_gene | 1.1 | 2.089 | 6.2E-07 |
| *cyp-31A3* | WBGene00021200 | protein_coding_gene | 1.1 | 2.088 | 2.8E-15 |
| *T02E1.6* | WBGene00011378 | protein_coding_gene | 1.1 | 2.087 | 1.5E-05 |
| *cebp-2* | WBGene00016754 | protein_coding_gene | 1.1 | 2.086 | 8.3E-11 |
| *lpd-2* | WBGene00003059 | protein_coding_gene | 1.1 | 2.086 | 4.4E-18 |
| *dnsn-1* | WBGene00016074 | protein_coding_gene | 1.1 | 2.085 | 8.6E-14 |
| *C15C8.4* | WBGene00007606 | protein_coding_gene | 1.1 | 2.085 | 6.0E-23 |
| *T08B2.12* | WBGene00020350 | protein_coding_gene | 1.1 | 2.081 | 2.3E-03 |
| *R07B7.2* | WBGene00011087 | protein_coding_gene | 1.1 | 2.081 | 5.4E-15 |
| *rbd-1* | WBGene00004315 | protein_coding_gene | 1.1 | 2.080 | 3.4E-20 |
| *fbxa-63* | WBGene00022485 | protein_coding_gene | 1.1 | 2.080 | 1.5E-03 |
| *K07C5.6* | WBGene00010629 | protein_coding_gene | 1.1 | 2.079 | 2.7E-22 |
| *mrps-14* | WBGene00011334 | protein_coding_gene | 1.1 | 2.078 | 7.6E-15 |
| *thoc-7* | WBGene00007195 | protein_coding_gene | 1.1 | 2.078 | 3.2E-19 |
| *K08F9.4* | WBGene00010687 | protein_coding_gene | 1.1 | 2.077 | 4.2E-15 |
| *F15D4.6* | WBGene00008863 | protein_coding_gene | 1.1 | 2.076 | 1.1E-02 |
| *F55A11.7* | WBGene00010080 | protein_coding_gene | 1.1 | 2.076 | 2.4E-14 |
| *F02H6.2* | WBGene00008535 | protein_coding_gene | 1.1 | 2.076 | 3.8E-14 |
| *aqp-4* | WBGene00000172 | protein_coding_gene | 1.1 | 2.076 | 4.0E-07 |
| *tofu-4* | WBGene00019013 | protein_coding_gene | 1.1 | 2.075 | 1.6E-11 |
| *T01G5.7* | WBGene00011342 | protein_coding_gene | 1.0 | 2.070 | 8.4E-08 |
| *T10C6.7* | WBGene00011689 | protein_coding_gene | 1.0 | 2.068 | 5.4E-08 |
| *fbxa-82* | WBGene00007716 | protein_coding_gene | 1.0 | 2.068 | 2.4E-03 |

| | | | | | |
|---|---|---|---|---|---|
| *F30F8.9* | WBGene00009276 | protein_coding_gene | 1.0 | 2.067 | 4.4E-16 |
| *Y48G1C.5* | WBGene00021678 | protein_coding_gene | 1.0 | 2.067 | 4.4E-03 |
| *nep-9* | WBGene00017554 | protein_coding_gene | 1.0 | 2.065 | 1.3E-02 |
| *ZK616.5* | WBGene00022775 | protein_coding_gene | 1.0 | 2.065 | 2.3E-15 |
| *Y71F9AL.2* | WBGene00022108 | protein_coding_gene | 1.0 | 2.063 | 1.4E-05 |
| *ZK632.2* | WBGene00014011 | protein_coding_gene | 1.0 | 2.063 | 6.3E-23 |
| *C16A11.4* | WBGene00015817 | protein_coding_gene | 1.0 | 2.062 | 5.4E-17 |
| *chaf-2* | WBGene00022141 | protein_coding_gene | 1.0 | 2.062 | 3.1E-15 |
| *polh-1* | WBGene00018721 | protein_coding_gene | 1.0 | 2.062 | 4.8E-18 |
| *cars-2* | WBGene00000801 | pseudogene | 1.0 | 2.062 | 9.1E-06 |
| *pole-4* | WBGene00013150 | protein_coding_gene | 1.0 | 2.062 | 4.3E-12 |
| *F48F5.2* | WBGene00009846 | pseudogene | 1.0 | 2.061 | 9.2E-05 |
| *B0001.3* | WBGene00007088 | protein_coding_gene | 1.0 | 2.060 | 2.1E-15 |
| *F09E5.8* | WBGene00017286 | protein_coding_gene | 1.0 | 2.059 | 2.4E-14 |
| *B0207.1* | WBGene00015026 | protein_coding_gene | 1.0 | 2.058 | 2.0E-03 |
| *F36A2.10* | WBGene00009457 | protein_coding_gene | 1.0 | 2.058 | 5.7E-07 |
| *F10G8.1* | WBGene00008661 | protein_coding_gene | 1.0 | 2.057 | 2.1E-04 |
| *coh-4* | WBGene00021560 | protein_coding_gene | 1.0 | 2.057 | 2.3E-11 |
| *gld-1* | WBGene00001595 | protein_coding_gene | 1.0 | 2.056 | 9.8E-28 |
| *ZK688.12* | WBGene00044763 | protein_coding_gene | 1.0 | 2.055 | 1.0E-02 |
| *ptps-1* | WBGene00015010 | protein_coding_gene | 1.0 | 2.054 | 2.7E-05 |
| *ceh-99* | WBGene00044032 | protein_coding_gene | 1.0 | 2.054 | 1.0E-05 |
| *T28A11.16* | WBGene00020878 | protein_coding_gene | 1.0 | 2.052 | 4.8E-02 |
| *cec-6* | WBGene00020463 | protein_coding_gene | 1.0 | 2.051 | 2.5E-17 |
| *Y47D9A.5* | WBGene00021632 | protein_coding_gene | 1.0 | 2.051 | 3.4E-02 |
| *cin-4* | WBGene00022854 | protein_coding_gene | 1.0 | 2.050 | 1.2E-11 |
| *K07F5.14* | WBGene00010638 | protein_coding_gene | 1.0 | 2.049 | 3.4E-18 |
| *lec-8* | WBGene00002271 | protein_coding_gene | 1.0 | 2.049 | 1.0E-03 |
| *hrde-4* | WBGene00020403 | protein_coding_gene | 1.0 | 2.048 | 1.6E-19 |
| *parp-1* | WBGene00004049 | protein_coding_gene | 1.0 | 2.047 | 8.3E-16 |
| *vrp-1* | WBGene00021869 | protein_coding_gene | 1.0 | 2.047 | 2.8E-21 |
| *ZK792.5* | WBGene00014078 | protein_coding_gene | 1.0 | 2.046 | 7.4E-22 |
| *R05H5.4* | WBGene00011039 | protein_coding_gene | 1.0 | 2.045 | 3.1E-11 |
| *prp-38* | WBGene00008380 | protein_coding_gene | 1.0 | 2.044 | 1.1E-17 |
| *Y47H9C.8* | WBGene00012951 | protein_coding_gene | 1.0 | 2.044 | 1.4E-14 |
| *C15C6.2* | WBGene00007601 | protein_coding_gene | 1.0 | 2.043 | 7.1E-08 |
| *snpc-1.3* | WBGene00017265 | protein_coding_gene | 1.0 | 2.043 | 2.9E-04 |
| *ZK418.7* | WBGene00022736 | protein_coding_gene | 1.0 | 2.042 | 9.9E-05 |
| *K06A9.1* | WBGene00019435 | protein_coding_gene | 1.0 | 2.042 | 3.8E-05 |
| *C08F8.3* | WBGene00007445 | protein_coding_gene | 1.0 | 2.042 | 1.2E-16 |
| *F54C9.9* | WBGene00010044 | protein_coding_gene | 1.0 | 2.041 | 2.6E-27 |
| *smgl-2* | WBGene00021365 | protein_coding_gene | 1.0 | 2.041 | 1.3E-17 |
| *pqbp-1.2* | WBGene00020295 | pseudogene | 1.0 | 2.040 | 2.6E-06 |
| *C15H11.1* | WBGene00007613 | protein_coding_gene | 1.0 | 2.040 | 5.5E-05 |
| *rpl-19* | WBGene00004431 | protein_coding_gene | 1.0 | 2.039 | 1.7E-15 |
| *ufm-1* | WBGene00044324 | protein_coding_gene | 1.0 | 2.037 | 4.9E-12 |
| *M151.3* | WBGene00019795 | protein_coding_gene | 1.0 | 2.037 | 6.2E-04 |
| *F26H11.4* | WBGene00009182 | protein_coding_gene | 1.0 | 2.037 | 3.5E-03 |
| *bre-5* | WBGene00000270 | protein_coding_gene | 1.0 | 2.036 | 3.8E-11 |
| *utp-20* | WBGene00007050 | protein_coding_gene | 1.0 | 2.035 | 2.7E-13 |
| *B0001.2* | WBGene00007087 | protein_coding_gene | 1.0 | 2.035 | 7.8E-23 |
| *F29D10.1* | WBGene00009250 | protein_coding_gene | 1.0 | 2.034 | 3.4E-02 |
| *spd-3* | WBGene00004954 | protein_coding_gene | 1.0 | 2.034 | 7.0E-12 |
| *puf-6* | WBGene00004242 | protein_coding_gene | 1.0 | 2.033 | 1.7E-11 |
| *T10B5.2* | WBGene00020388 | pseudogene | 1.0 | 2.032 | 1.9E-03 |
| *W03C9.2* | WBGene00012220 | protein_coding_gene | 1.0 | 2.030 | 1.9E-17 |
| *kri-1* | WBGene00013955 | protein_coding_gene | 1.0 | 2.030 | 7.6E-03 |
| *sumv-1* | WBGene00016409 | protein_coding_gene | 1.0 | 2.029 | 1.5E-18 |
| *abhd-11.2* | WBGene00011029 | protein_coding_gene | 1.0 | 2.028 | 3.4E-15 |
| *pgl-1* | WBGene00003992 | protein_coding_gene | 1.0 | 2.028 | 7.0E-21 |
| *xnd-1* | WBGene00001514 | protein_coding_gene | 1.0 | 2.028 | 4.8E-14 |
| *pie-1* | WBGene00004027 | protein_coding_gene | 1.0 | 2.026 | 2.3E-11 |
| *K10C3.5* | WBGene00010732 | protein_coding_gene | 1.0 | 2.026 | 8.1E-15 |
| *clec-236* | WBGene00013495 | pseudogene | 1.0 | 2.025 | 1.0E-02 |
| *nifk-1* | WBGene00011408 | protein_coding_gene | 1.0 | 2.025 | 3.8E-18 |
| *F44E2.6* | WBGene00018419 | protein_coding_gene | 1.0 | 2.025 | 6.8E-14 |
| *ZK1248.15* | WBGene00022883 | protein_coding_gene | 1.0 | 2.024 | 1.3E-15 |
| *F57H12.5* | WBGene00019020 | protein_coding_gene | 1.0 | 2.024 | 5.0E-03 |
| *Y17G7B.21* | WBGene00012472 | protein_coding_gene | 1.0 | 2.024 | 4.9E-20 |
| *cux-7* | WBGene00000854 | protein_coding_gene | 1.0 | 2.024 | 1.1E-14 |

| | | | | | |
|---|---|---|---|---|---|
| *plk-3* | WBGene00004044 | protein_coding_gene | 1.0 | 2.024 | 2.1E-25 |
| *cfim-1* | WBGene00009668 | protein_coding_gene | 1.0 | 2.023 | 4.5E-12 |
| *dvc-1* | WBGene00011834 | protein_coding_gene | 1.0 | 2.023 | 5.8E-12 |
| *C14C11.7* | WBGene00015769 | protein_coding_gene | 1.0 | 2.020 | 2.2E-07 |
| *C09G1.5* | WBGene00007487 | protein_coding_gene | 1.0 | 2.019 | 7.2E-03 |
| *ztf-28* | WBGene00018099 | protein_coding_gene | 1.0 | 2.019 | 2.1E-07 |
| *ZC477.3* | WBGene00022618 | protein_coding_gene | 1.0 | 2.018 | 4.6E-19 |
| *F54D5.9* | WBGene00010053 | protein_coding_gene | 1.0 | 2.018 | 4.6E-13 |
| *akir-1* | WBGene00017088 | protein_coding_gene | 1.0 | 2.018 | 6.6E-21 |
| *H06I04.5* | WBGene00019169 | protein_coding_gene | 1.0 | 2.018 | 5.2E-04 |
| *hex-4* | WBGene00021766 | protein_coding_gene | 1.0 | 2.018 | 4.3E-17 |
| *F09C8.1* | WBGene00008621 | protein_coding_gene | 1.0 | 2.017 | 1.6E-02 |
| *pfd-2* | WBGene00019220 | protein_coding_gene | 1.0 | 2.017 | 6.8E-16 |
| *ZK546.14* | WBGene00022765 | protein_coding_gene | 1.0 | 2.016 | 1.8E-21 |
| *rps-10* | WBGene00004479 | protein_coding_gene | 1.0 | 2.015 | 3.0E-15 |
| *T16H12.2* | WBGene00011812 | protein_coding_gene | 1.0 | 2.014 | 1.9E-02 |
| *spe-48* | WBGene00021771 | protein_coding_gene | 1.0 | 2.013 | 1.6E-14 |
| *mcm-10* | WBGene00012935 | protein_coding_gene | 1.0 | 2.013 | 2.1E-15 |
| *pfn-1* | WBGene00003989 | protein_coding_gene | 1.0 | 2.013 | 3.4E-13 |
| *C09D4.4* | WBGene00015635 | protein_coding_gene | 1.0 | 2.012 | 3.6E-28 |
| *C26E6.12* | WBGene00016145 | protein_coding_gene | 1.0 | 2.011 | 5.2E-16 |
| *glh-4* | WBGene00001601 | protein_coding_gene | 1.0 | 2.011 | 4.3E-21 |
| *H08M01.1* | WBGene00010373 | protein_coding_gene | 1.0 | 2.011 | 1.7E-03 |
| *C47A4.1* | WBGene00008122 | protein_coding_gene | 1.0 | 2.010 | 5.2E-03 |
| *mtx-1* | WBGene00009559 | protein_coding_gene | 1.0 | 2.009 | 2.3E-14 |
| *pges-2* | WBGene00011239 | protein_coding_gene | 1.0 | 2.008 | 1.7E-25 |
| *F25G6.1* | WBGene00017796 | protein_coding_gene | 1.0 | 2.007 | 3.9E-04 |
| *hpr-17* | WBGene00001998 | protein_coding_gene | 1.0 | 2.006 | 1.2E-15 |
| *eif-3.J* | WBGene00012738 | protein_coding_gene | 1.0 | 2.006 | 3.3E-14 |
| *F26B1.5* | WBGene00017817 | protein_coding_gene | 1.0 | 2.006 | 2.7E-02 |
| *K06G5.1* | WBGene00010605 | protein_coding_gene | 1.0 | 2.006 | 1.4E-09 |
| *msh-5* | WBGene00003421 | protein_coding_gene | 1.0 | 2.005 | 1.7E-16 |
| *F25B4.4* | WBGene00017767 | protein_coding_gene | 1.0 | 2.005 | 9.9E-05 |
| *ubc-6* | WBGene00006703 | protein_coding_gene | 1.0 | 2.005 | 9.0E-18 |
| *ccm-3* | WBGene00007561 | protein_coding_gene | 1.0 | 2.005 | 1.5E-16 |
| *C09B9.7* | WBGene00015630 | protein_coding_gene | 1.0 | 2.005 | 7.6E-04 |
| *mom-2* | WBGene00003395 | protein_coding_gene | 1.0 | 2.004 | 5.8E-15 |
| *T19D12.5* | WBGene00020580 | protein_coding_gene | 1.0 | 2.004 | 2.2E-04 |
| *inft-2* | WBGene00008848 | protein_coding_gene | 1.0 | 2.003 | 3.6E-18 |
| *ent-1* | WBGene00001320 | protein_coding_gene | 1.0 | 2.002 | 5.2E-17 |
| *lec-10* | WBGene00002273 | protein_coding_gene | 1.0 | 2.002 | 2.6E-02 |
| *F27C1.1* | WBGene00017851 | protein_coding_gene | 1.0 | 2.002 | 4.6E-08 |
| *sel-13* | WBGene00011411 | protein_coding_gene | 1.0 | 2.002 | 1.4E-14 |
| *com-1* | WBGene00008082 | protein_coding_gene | 1.0 | 2.001 | 5.3E-14 |
| *B0035.3* | WBGene00007106 | protein_coding_gene | 1.0 | 2.001 | 2.2E-16 |
| *szy-5* | WBGene00016154 | protein_coding_gene | 1.0 | 2.000 | 6.6E-22 |
| *F44A6.4* | WBGene00009676 | protein_coding_gene | -1.0 | 0.500 | 8.1E-08 |
| *egl-38* | WBGene00001204 | protein_coding_gene | -1.0 | 0.500 | 3.8E-02 |
| *nhr-79* | WBGene00003669 | protein_coding_gene | -1.0 | 0.499 | 4.9E-06 |
| *anoh-2* | WBGene00013326 | protein_coding_gene | -1.0 | 0.499 | 1.8E-04 |
| *F59A6.12* | WBGene00044665 | protein_coding_gene | -1.0 | 0.499 | 3.2E-04 |
| *hpo-15* | WBGene00016061 | protein_coding_gene | -1.0 | 0.498 | 2.5E-10 |
| *2L52.1* | WBGene00007063 | protein_coding_gene | -1.0 | 0.498 | 2.0E-08 |
| *crb-1* | WBGene00000792 | protein_coding_gene | -1.0 | 0.498 | 1.2E-08 |
| *C23G10.14* | WBGene00271647 | ncRNA_gene | -1.0 | 0.497 | 3.8E-02 |
| *K08B5.2* | WBGene00019519 | protein_coding_gene | -1.0 | 0.497 | 8.4E-05 |
| *C18A11.2* | WBGene00015948 | protein_coding_gene | -1.0 | 0.496 | 1.6E-02 |
| *aagr-1* | WBGene00017071 | protein_coding_gene | -1.0 | 0.495 | 4.7E-09 |
| *F57A8.1* | WBGene00010177 | protein_coding_gene | -1.0 | 0.495 | 7.4E-12 |
| *fkh-2* | WBGene00001434 | protein_coding_gene | -1.0 | 0.495 | 8.2E-04 |
| *nhr-64* | WBGene00003654 | protein_coding_gene | -1.0 | 0.495 | 9.3E-09 |
| *nlp-57* | WBGene00008589 | protein_coding_gene | -1.0 | 0.495 | 6.8E-03 |
| *msp-152* | WBGene00003470 | protein_coding_gene | -1.0 | 0.494 | 4.5E-07 |
| *dhs-29* | WBGene00000992 | protein_coding_gene | -1.0 | 0.494 | 5.4E-06 |
| *C33A11.2* | WBGene00007878 | protein_coding_gene | -1.0 | 0.494 | 2.4E-05 |
| *C08A9.10* | WBGene00044773 | protein_coding_gene | -1.0 | 0.493 | 2.1E-08 |
| *C05G5.3* | WBGene00007349 | protein_coding_gene | -1.0 | 0.493 | 3.3E-04 |
| *dgn-1* | WBGene00000961 | protein_coding_gene | -1.0 | 0.493 | 6.0E-14 |
| *nep-24* | WBGene00013786 | protein_coding_gene | -1.0 | 0.492 | 3.5E-02 |
| *slcf-2* | WBGene00006865 | protein_coding_gene | -1.0 | 0.492 | 1.9E-03 |

| | | | | | |
|---|---|---|---|---|---|
| *D2045.8* | WBGene00008425 | protein_coding_gene | -1.0 | 0.491 | 1.7E-05 |
| *T04F8.9* | WBGene00011447 | protein_coding_gene | -1.0 | 0.490 | 3.1E-02 |
| *ncr-1* | WBGene00003561 | protein_coding_gene | -1.0 | 0.490 | 2.7E-10 |
| *T19C3.4* | WBGene00020562 | protein_coding_gene | -1.0 | 0.490 | 2.0E-16 |
| *K10D6.4* | WBGene00010744 | protein_coding_gene | -1.0 | 0.490 | 1.3E-08 |
| *T14A8.2* | WBGene00020497 | protein_coding_gene | -1.0 | 0.490 | 2.1E-03 |
| *F26F2.8* | WBGene00023504 | protein_coding_gene | -1.0 | 0.489 | 4.1E-02 |
| *iglr-1* | WBGene00008977 | protein_coding_gene | -1.0 | 0.489 | 1.1E-02 |
| *agr-1* | WBGene00018304 | protein_coding_gene | -1.0 | 0.489 | 5.0E-05 |
| *pqn-89* | WBGene00004169 | protein_coding_gene | -1.0 | 0.488 | 1.7E-11 |
| *T23F2.3* | WBGene00020736 | protein_coding_gene | -1.0 | 0.488 | 2.1E-05 |
| *dmd-5* | WBGene00017326 | protein_coding_gene | -1.0 | 0.488 | 8.9E-03 |
| *C35B8.3* | WBGene00016438 | protein_coding_gene | -1.0 | 0.488 | 1.2E-06 |
| *rcan-1* | WBGene00004321 | protein_coding_gene | -1.0 | 0.488 | 4.6E-10 |
| *F59C12.3* | WBGene00019102 | protein_coding_gene | -1.0 | 0.488 | 1.3E-12 |
| *cutl-25* | WBGene00021919 | protein_coding_gene | -1.0 | 0.488 | 5.4E-03 |
| *F09G2.1* | WBGene00017310 | protein_coding_gene | -1.0 | 0.487 | 1.9E-08 |
| *ags-3* | WBGene00000092 | protein_coding_gene | -1.0 | 0.487 | 8.7E-11 |
| *Y20C6A.1* | WBGene00012491 | protein_coding_gene | -1.0 | 0.487 | 5.6E-08 |
| *D1086.5* | WBGene00008392 | protein_coding_gene | -1.0 | 0.486 | 4.5E-06 |
| *C16H3.3* | WBGene00015868 | protein_coding_gene | -1.0 | 0.486 | 6.4E-10 |
| *lin-33* | WBGene00003019 | protein_coding_gene | -1.0 | 0.486 | 2.6E-07 |
| *F13C5.1* | WBGene00017422 | protein_coding_gene | -1.0 | 0.486 | 4.9E-06 |
| *sol-1* | WBGene00004944 | protein_coding_gene | -1.0 | 0.486 | 1.5E-02 |
| *F13D12.10* | WBGene00008744 | protein_coding_gene | -1.0 | 0.486 | 2.5E-02 |
| *K10C9.1* | WBGene00019623 | protein_coding_gene | -1.0 | 0.486 | 3.4E-02 |
| *Y41G9A.5* | WBGene00021529 | protein_coding_gene | -1.0 | 0.485 | 2.2E-02 |
| *somi-1* | WBGene00010868 | protein_coding_gene | -1.0 | 0.485 | 1.0E-09 |
| *clec-83* | WBGene00021879 | protein_coding_gene | -1.0 | 0.485 | 1.4E-10 |
| *rig-5* | WBGene00004372 | protein_coding_gene | -1.0 | 0.485 | 7.1E-04 |
| *nfm-1* | WBGene00003593 | protein_coding_gene | -1.0 | 0.484 | 6.9E-16 |
| *cpna-4* | WBGene00006441 | protein_coding_gene | -1.0 | 0.484 | 1.4E-02 |
| *dct-11* | WBGene00021060 | protein_coding_gene | -1.0 | 0.484 | 1.3E-11 |
| *rrf-2* | WBGene00004509 | protein_coding_gene | -1.0 | 0.483 | 2.8E-16 |
| *ugt-28* | WBGene00015693 | protein_coding_gene | -1.0 | 0.483 | 1.3E-02 |
| *F53C11.9* | WBGene00044921 | protein_coding_gene | -1.1 | 0.483 | 7.3E-04 |
| *R03E9.2* | WBGene00019843 | protein_coding_gene | -1.1 | 0.483 | 7.8E-04 |
| *C36A4.11* | WBGene00044167 | protein_coding_gene | -1.1 | 0.483 | 9.6E-03 |
| *H12D21.9* | WBGene00010385 | protein_coding_gene | -1.1 | 0.483 | 9.7E-03 |
| *Y75B8A.19* | WBGene00013554 | protein_coding_gene | -1.1 | 0.482 | 7.2E-06 |
| *R11H6.4* | WBGene00011252 | protein_coding_gene | -1.1 | 0.482 | 9.9E-08 |
| *epg-9* | WBGene00022078 | protein_coding_gene | -1.1 | 0.482 | 1.3E-11 |
| *vem-1* | WBGene00006890 | protein_coding_gene | -1.1 | 0.481 | 1.3E-14 |
| *ttbk-7* | WBGene00011283 | protein_coding_gene | -1.1 | 0.481 | 1.4E-08 |
| *ceh-31* | WBGene00000452 | protein_coding_gene | -1.1 | 0.481 | 4.0E-02 |
| *hst-3.2* | WBGene00044303 | protein_coding_gene | -1.1 | 0.481 | 1.0E-03 |
| *K10G6.9* | WBGene00206374 | protein_coding_gene | -1.1 | 0.480 | 3.1E-02 |
| *C36C5.15* | WBGene00016484 | protein_coding_gene | -1.1 | 0.480 | 3.9E-02 |
| *ceh-62* | WBGene00011069 | protein_coding_gene | -1.1 | 0.480 | 4.9E-02 |
| *ssr-2* | WBGene00006055 | protein_coding_gene | -1.1 | 0.480 | 2.9E-02 |
| *nphp-4* | WBGene00011261 | protein_coding_gene | -1.1 | 0.480 | 4.7E-03 |
| *ser-4* | WBGene00004779 | protein_coding_gene | -1.1 | 0.480 | 5.8E-04 |
| *ddr-1* | WBGene00016104 | protein_coding_gene | -1.1 | 0.480 | 1.5E-05 |
| *dhs-27* | WBGene00000990 | protein_coding_gene | -1.1 | 0.479 | 7.6E-08 |
| *cutl-21* | WBGene00018861 | protein_coding_gene | -1.1 | 0.478 | 4.1E-03 |
| *mthf-1* | WBGene00015512 | protein_coding_gene | -1.1 | 0.478 | 2.2E-34 |
| *bre-2* | WBGene00000267 | protein_coding_gene | -1.1 | 0.478 | 4.2E-05 |
| *F59F5.7* | WBGene00010345 | protein_coding_gene | -1.1 | 0.477 | 2.3E-02 |
| *ZK512.11* | WBGene00013989 | protein_coding_gene | -1.1 | 0.477 | 1.3E-03 |
| *cwn-2* | WBGene00000858 | protein_coding_gene | -1.1 | 0.477 | 1.1E-06 |
| *ins-17* | WBGene00002100 | protein_coding_gene | -1.1 | 0.477 | 1.7E-07 |
| *mnm-2* | WBGene00003380 | protein_coding_gene | -1.1 | 0.475 | 1.4E-02 |
| *ceh-6* | WBGene00000431 | protein_coding_gene | -1.1 | 0.475 | 1.6E-03 |
| *C05E11.3* | WBGene00015494 | protein_coding_gene | -1.1 | 0.473 | 1.7E-07 |
| *pho-6* | WBGene00018696 | protein_coding_gene | -1.1 | 0.473 | 3.3E-05 |
| *igcm-3* | WBGene00020160 | protein_coding_gene | -1.1 | 0.473 | 1.8E-13 |
| *kel-1* | WBGene00002184 | protein_coding_gene | -1.1 | 0.472 | 1.4E-03 |
| *zmp-2* | WBGene00019212 | protein_coding_gene | -1.1 | 0.472 | 1.7E-03 |
| *ceh-37* | WBGene00000458 | protein_coding_gene | -1.1 | 0.472 | 2.0E-08 |
| *ttr-10* | WBGene00017600 | protein_coding_gene | -1.1 | 0.472 | 1.0E-04 |

| | | | | | |
|---|---|---|---|---|---|
| *lad-2* | WBGene00002243 | protein_coding_gene | -1.1 | 0.471 | 6.8E-04 |
| *B0228.1* | WBGene00015059 | protein_coding_gene | -1.1 | 0.471 | 2.8E-07 |
| *fbxa-69* | WBGene00010018 | protein_coding_gene | -1.1 | 0.470 | 6.0E-09 |
| *K02E10.1* | WBGene00019317 | protein_coding_gene | -1.1 | 0.470 | 1.2E-07 |
| *nmtn-1* | WBGene00019434 | protein_coding_gene | -1.1 | 0.470 | 2.2E-03 |
| *wht-1* | WBGene00015479 | protein_coding_gene | -1.1 | 0.470 | 1.9E-04 |
| *C18C4.7* | WBGene00015968 | protein_coding_gene | -1.1 | 0.470 | 7.8E-16 |
| *Y65B4A.2* | WBGene00022026 | protein_coding_gene | -1.1 | 0.469 | 1.0E-05 |
| *lron-15* | WBGene00016974 | protein_coding_gene | -1.1 | 0.469 | 2.9E-04 |
| *rnst-2* | WBGene00019624 | protein_coding_gene | -1.1 | 0.468 | 3.7E-13 |
| *unc-120* | WBGene00006844 | protein_coding_gene | -1.1 | 0.468 | 2.0E-08 |
| *C38C3.4* | WBGene00016513 | protein_coding_gene | -1.1 | 0.468 | 5.0E-08 |
| *F56H9.2* | WBGene00010174 | protein_coding_gene | -1.1 | 0.468 | 2.7E-05 |
| *Y57A10A.26* | WBGene00013266 | protein_coding_gene | -1.1 | 0.466 | 1.2E-11 |
| *C25F6.7* | WBGene00016106 | protein_coding_gene | -1.1 | 0.466 | 6.6E-03 |
| *mig-18* | WBGene00003249 | protein_coding_gene | -1.1 | 0.466 | 8.3E-07 |
| *ZK262.3* | WBGene00013950 | protein_coding_gene | -1.1 | 0.465 | 1.2E-02 |
| *fkb-7* | WBGene00001432 | protein_coding_gene | -1.1 | 0.465 | 3.8E-05 |
| *F07A5.4* | WBGene00008543 | protein_coding_gene | -1.1 | 0.465 | 2.8E-02 |
| *linc-15* | WBGene00219571 | lincRNA_gene | -1.1 | 0.465 | 2.5E-05 |
| *tub-2* | WBGene00022139 | protein_coding_gene | -1.1 | 0.464 | 1.7E-17 |
| *mks-3* | WBGene00018042 | protein_coding_gene | -1.1 | 0.464 | 4.1E-02 |
| *M01H9.4* | WBGene00019720 | protein_coding_gene | -1.1 | 0.464 | 2.9E-06 |
| *fkh-8* | WBGene00001440 | protein_coding_gene | -1.1 | 0.463 | 6.2E-03 |
| *ttx-1* | WBGene00006652 | protein_coding_gene | -1.1 | 0.463 | 3.5E-05 |
| *aakb-1* | WBGene00010115 | protein_coding_gene | -1.1 | 0.463 | 6.4E-13 |
| *F56C3.9* | WBGene00018946 | protein_coding_gene | -1.1 | 0.462 | 2.5E-02 |
| *tag-304* | WBGene00018757 | protein_coding_gene | -1.1 | 0.462 | 3.5E-16 |
| *sma-1* | WBGene00004855 | protein_coding_gene | -1.1 | 0.461 | 2.8E-22 |
| *C32E12.4* | WBGene00016330 | protein_coding_gene | -1.1 | 0.461 | 1.5E-13 |
| *lron-11* | WBGene00022129 | protein_coding_gene | -1.1 | 0.461 | 2.0E-09 |
| *eya-1* | WBGene00001377 | protein_coding_gene | -1.1 | 0.461 | 7.4E-08 |
| *F14B6.6* | WBGene00008787 | protein_coding_gene | -1.1 | 0.461 | 2.6E-04 |
| *F20C5.6* | WBGene00008971 | protein_coding_gene | -1.1 | 0.460 | 5.5E-06 |
| *clc-2* | WBGene00000523 | protein_coding_gene | -1.1 | 0.460 | 4.2E-04 |
| *ham-1* | WBGene00001820 | protein_coding_gene | -1.1 | 0.460 | 5.1E-09 |
| *K02C4.5* | WBGene00010503 | protein_coding_gene | -1.1 | 0.458 | 1.2E-12 |
| *aff-1* | WBGene00016625 | protein_coding_gene | -1.1 | 0.458 | 5.7E-03 |
| *ZC123.4* | WBGene00022519 | protein_coding_gene | -1.1 | 0.458 | 7.2E-06 |
| *F59F5.3* | WBGene00010342 | protein_coding_gene | -1.1 | 0.457 | 8.0E-04 |
| *pals-34* | WBGene00017823 | protein_coding_gene | -1.1 | 0.457 | 1.7E-03 |
| *ugt-39* | WBGene00017329 | protein_coding_gene | -1.1 | 0.457 | 1.8E-04 |
| *let-381* | WBGene00002601 | protein_coding_gene | -1.1 | 0.456 | 6.4E-03 |
| *F46H6.5* | WBGene00023460 | protein_coding_gene | -1.1 | 0.456 | 4.1E-02 |
| *W08E3.4* | WBGene00012345 | protein_coding_gene | -1.1 | 0.456 | 1.6E-02 |
| *marc-4* | WBGene00016903 | protein_coding_gene | -1.1 | 0.456 | 1.6E-04 |
| *dlg-1* | WBGene00001006 | protein_coding_gene | -1.1 | 0.455 | 1.4E-18 |
| *pals-14* | WBGene00009061 | protein_coding_gene | -1.1 | 0.455 | 4.3E-02 |
| *C37E2.2* | WBGene00007994 | protein_coding_gene | -1.1 | 0.455 | 3.2E-02 |
| *jkk-1* | WBGene00002177 | protein_coding_gene | -1.1 | 0.455 | 3.1E-02 |
| *mab-5* | WBGene00003102 | protein_coding_gene | -1.1 | 0.455 | 1.2E-03 |
| *F07H5.4* | WBGene00014308 | pseudogene | -1.1 | 0.455 | 2.0E-13 |
| *mec-6* | WBGene00003170 | protein_coding_gene | -1.1 | 0.454 | 8.3E-03 |
| *egas-4* | WBGene00018906 | protein_coding_gene | -1.1 | 0.453 | 1.0E-02 |
| *aqp-2* | WBGene00000170 | protein_coding_gene | -1.1 | 0.452 | 9.8E-10 |
| *shn-1* | WBGene00006444 | protein_coding_gene | -1.1 | 0.452 | 1.7E-11 |
| *phlp-2* | WBGene00009647 | protein_coding_gene | -1.1 | 0.452 | 1.4E-02 |
| *snx-27* | WBGene00009116 | protein_coding_gene | -1.1 | 0.452 | 1.5E-13 |
| *atg-2* | WBGene00019748 | protein_coding_gene | -1.1 | 0.451 | 3.5E-16 |
| *hum-6* | WBGene00002039 | protein_coding_gene | -1.2 | 0.451 | 2.5E-07 |
| *nhr-110* | WBGene00003700 | protein_coding_gene | -1.2 | 0.451 | 3.2E-09 |
| *daao-1* | WBGene00022076 | protein_coding_gene | -1.2 | 0.450 | 5.6E-05 |
| *frm-7* | WBGene00001493 | protein_coding_gene | -1.2 | 0.450 | 4.2E-20 |
| *Y38F1A.8* | WBGene00012610 | protein_coding_gene | -1.2 | 0.450 | 2.0E-05 |
| *Y49A3A.4* | WBGene00013027 | protein_coding_gene | -1.2 | 0.450 | 1.5E-02 |
| *cdh-9* | WBGene00000401 | protein_coding_gene | -1.2 | 0.449 | 3.0E-03 |
| *Y62H9A.16* | WBGene00304791 | protein_coding_gene | -1.2 | 0.449 | 9.7E-03 |
| *trhr-1* | WBGene00016265 | protein_coding_gene | -1.2 | 0.448 | 1.8E-04 |
| *unc-5* | WBGene00006745 | protein_coding_gene | -1.2 | 0.448 | 4.3E-15 |
| *C05A9.2* | WBGene00007317 | protein_coding_gene | -1.2 | 0.447 | 8.2E-08 |

| | | | | | |
|---|---|---|---|---|---|
| *K12B6.9* | WBGene00023432 | protein_coding_gene | -1.2 | 0.447 | 1.2E-10 |
| *K02E10.6* | WBGene00019320 | protein_coding_gene | -1.2 | 0.447 | 7.4E-04 |
| *T05H4.4* | WBGene00020267 | protein_coding_gene | -1.2 | 0.447 | 1.9E-02 |
| *memo-1* | WBGene00016500 | protein_coding_gene | -1.2 | 0.447 | 5.4E-18 |
| *ugt-57* | WBGene00017154 | protein_coding_gene | -1.2 | 0.446 | 2.8E-02 |
| *T12A7.6* | WBGene00011727 | protein_coding_gene | -1.2 | 0.446 | 9.4E-09 |
| *F59C6.8* | WBGene00010327 | protein_coding_gene | -1.2 | 0.446 | 7.6E-03 |
| *grdn-1* | WBGene00013082 | protein_coding_gene | -1.2 | 0.445 | 1.3E-12 |
| *F10G8.8* | WBGene00008666 | protein_coding_gene | -1.2 | 0.445 | 7.8E-12 |
| *F53A2.1* | WBGene00009949 | protein_coding_gene | -1.2 | 0.445 | 5.4E-03 |
| *lim-4* | WBGene00002987 | protein_coding_gene | -1.2 | 0.445 | 1.5E-02 |
| *ncx-3* | WBGene00003568 | protein_coding_gene | -1.2 | 0.444 | 2.6E-09 |
| *dhs-18* | WBGene00000981 | protein_coding_gene | -1.2 | 0.444 | 9.6E-18 |
| *hhat-2* | WBGene00013312 | protein_coding_gene | -1.2 | 0.444 | 1.1E-03 |
| *T09B9.3* | WBGene00011644 | protein_coding_gene | -1.2 | 0.444 | 4.5E-02 |
| *F58H1.7* | WBGene00010290 | protein_coding_gene | -1.2 | 0.444 | 2.7E-07 |
| *H29C22.1* | WBGene00045261 | protein_coding_gene | -1.2 | 0.444 | 1.5E-11 |
| *lag-2* | WBGene00002246 | protein_coding_gene | -1.2 | 0.444 | 4.6E-02 |
| *nhr-134* | WBGene00003724 | protein_coding_gene | -1.2 | 0.444 | 2.7E-08 |
| *vab-15* | WBGene00006881 | protein_coding_gene | -1.2 | 0.443 | 2.5E-03 |
| *F59B10.5* | WBGene00010321 | protein_coding_gene | -1.2 | 0.443 | 4.1E-03 |
| *clec-264* | WBGene00009291 | protein_coding_gene | -1.2 | 0.443 | 1.2E-06 |
| *acs-21* | WBGene00016849 | protein_coding_gene | -1.2 | 0.442 | 1.3E-03 |
| *cest-35.2* | WBGene00016863 | protein_coding_gene | -1.2 | 0.442 | 1.8E-03 |
| *ssp-19* | WBGene00006047 | protein_coding_gene | -1.2 | 0.442 | 4.8E-08 |
| *K01F9.2* | WBGene00010475 | protein_coding_gene | -1.2 | 0.442 | 3.4E-06 |
| *nhr-103* | WBGene00003693 | protein_coding_gene | -1.2 | 0.442 | 1.7E-06 |
| *bro-1* | WBGene00000272 | protein_coding_gene | -1.2 | 0.442 | 1.6E-03 |
| *ins-4* | WBGene00002087 | protein_coding_gene | -1.2 | 0.441 | 4.3E-04 |
| *lbp-1* | WBGene00002253 | protein_coding_gene | -1.2 | 0.441 | 2.9E-13 |
| *F53F10.1* | WBGene00018763 | protein_coding_gene | -1.2 | 0.441 | 4.4E-07 |
| *msp-10* | WBGene00003425 | protein_coding_gene | -1.2 | 0.440 | 2.5E-03 |
| *F57C12.2* | WBGene00019014 | protein_coding_gene | -1.2 | 0.440 | 5.2E-05 |
| *ZC168.2* | WBGene00013856 | protein_coding_gene | -1.2 | 0.440 | 1.6E-02 |
| *sym-2* | WBGene00006367 | protein_coding_gene | -1.2 | 0.440 | 1.4E-07 |
| *cest-34* | WBGene00019652 | protein_coding_gene | -1.2 | 0.440 | 1.5E-02 |
| *unc-6* | WBGene00006746 | protein_coding_gene | -1.2 | 0.440 | 3.0E-08 |
| *F28H7.2* | WBGene00009236 | protein_coding_gene | -1.2 | 0.440 | 4.2E-02 |
| *crn-7* | WBGene00007056 | protein_coding_gene | -1.2 | 0.439 | 6.3E-07 |
| *clec-1* | WBGene00017772 | protein_coding_gene | -1.2 | 0.439 | 1.2E-09 |
| *Y34D9A.8* | WBGene00021333 | protein_coding_gene | -1.2 | 0.439 | 3.2E-06 |
| *acdh-10* | WBGene00020366 | protein_coding_gene | -1.2 | 0.439 | 2.9E-22 |
| *madd-2* | WBGene00016539 | protein_coding_gene | -1.2 | 0.438 | 1.1E-11 |
| *cyp-33C3* | WBGene00018262 | protein_coding_gene | -1.2 | 0.438 | 6.3E-06 |
| *ccep-290* | WBGene00021643 | protein_coding_gene | -1.2 | 0.438 | 6.9E-04 |
| *unc-3* | WBGene00006743 | protein_coding_gene | -1.2 | 0.438 | 9.5E-04 |
| *dpy-1* | WBGene00001063 | protein_coding_gene | -1.2 | 0.437 | 9.1E-03 |
| *snt-7* | WBGene00011682 | protein_coding_gene | -1.2 | 0.437 | 2.3E-02 |
| *fkb-4* | WBGene00001429 | protein_coding_gene | -1.2 | 0.437 | 1.3E-03 |
| *F53C11.3* | WBGene00009973 | protein_coding_gene | -1.2 | 0.436 | 5.2E-26 |
| *subs-4* | WBGene00012939 | protein_coding_gene | -1.2 | 0.436 | 1.3E-02 |
| *Y67A10A.7* | WBGene00013456 | protein_coding_gene | -1.2 | 0.436 | 1.8E-02 |
| *cnd-1* | WBGene00000561 | protein_coding_gene | -1.2 | 0.435 | 2.8E-04 |
| *Y55D5A.1* | WBGene00021915 | protein_coding_gene | -1.2 | 0.435 | 1.4E-06 |
| *bus-12* | WBGene00010442 | protein_coding_gene | -1.2 | 0.435 | 2.0E-02 |
| *R06C1.6* | WBGene00011060 | protein_coding_gene | -1.2 | 0.434 | 5.7E-07 |
| *igdb-3* | WBGene00020691 | protein_coding_gene | -1.2 | 0.433 | 2.6E-03 |
| *Y54E5A.8* | WBGene00013203 | protein_coding_gene | -1.2 | 0.432 | 1.9E-06 |
| *ppm-1.H* | WBGene00016580 | protein_coding_gene | -1.2 | 0.432 | 3.0E-02 |
| *tba-4* | WBGene00006530 | protein_coding_gene | -1.2 | 0.432 | 2.0E-10 |
| *T13H5.6* | WBGene00011760 | protein_coding_gene | -1.2 | 0.432 | 1.5E-06 |
| *ltd-1* | WBGene00003089 | protein_coding_gene | -1.2 | 0.432 | 4.8E-06 |
| *ceh-32* | WBGene00000453 | protein_coding_gene | -1.2 | 0.432 | 4.4E-11 |
| *cpn-2* | WBGene00000778 | protein_coding_gene | -1.2 | 0.432 | 2.3E-02 |
| *R05G9R.1* | WBGene00019906 | protein_coding_gene | -1.2 | 0.432 | 1.2E-02 |
| *prkl-1* | WBGene00022727 | protein_coding_gene | -1.2 | 0.431 | 1.6E-07 |
| *unc-129* | WBGene00006852 | protein_coding_gene | -1.2 | 0.430 | 7.7E-07 |
| *F47G4.4* | WBGene00009825 | protein_coding_gene | -1.2 | 0.430 | 6.5E-12 |
| *ceh-30* | WBGene00000451 | protein_coding_gene | -1.2 | 0.429 | 1.9E-02 |
| *T22F3.10* | WBGene00020699 | protein_coding_gene | -1.2 | 0.429 | 3.9E-02 |

| | | | | | |
|---|---|---|---|---|---|
| *ttr-20* | WBGene00018085 | protein_coding_gene | -1.2 | 0.429 | 3.7E-04 |
| *rsf-1* | WBGene00011995 | protein_coding_gene | -1.2 | 0.429 | 5.6E-13 |
| *mig-13* | WBGene00003245 | protein_coding_gene | -1.2 | 0.428 | 7.6E-08 |
| *Y57A10A.14* | WBGene00013257 | protein_coding_gene | -1.2 | 0.428 | 1.1E-05 |
| *K12G11.6* | WBGene00010792 | protein_coding_gene | -1.2 | 0.428 | 3.0E-02 |
| *BE0003N10.3* | WBGene00015268 | protein_coding_gene | -1.2 | 0.428 | 1.9E-04 |
| *Y48A6B.9* | WBGene00012970 | protein_coding_gene | -1.2 | 0.427 | 3.1E-10 |
| *twk-11* | WBGene00006666 | protein_coding_gene | -1.2 | 0.426 | 1.7E-03 |
| *magu-3* | WBGene00016841 | protein_coding_gene | -1.2 | 0.426 | 4.2E-04 |
| *T27E4.7* | WBGene00020865 | protein_coding_gene | -1.2 | 0.426 | 4.2E-11 |
| *let-653* | WBGene00002827 | protein_coding_gene | -1.2 | 0.426 | 5.8E-03 |
| *tag-243* | WBGene00044078 | protein_coding_gene | -1.2 | 0.426 | 5.9E-07 |
| *C28H8.2* | WBGene00016193 | protein_coding_gene | -1.2 | 0.425 | 7.9E-03 |
| *oig-3* | WBGene00003861 | protein_coding_gene | -1.2 | 0.425 | 5.4E-08 |
| *R03H10.1* | WBGene00019854 | protein_coding_gene | -1.2 | 0.425 | 1.1E-05 |
| *syd-9* | WBGene00044068 | protein_coding_gene | -1.2 | 0.425 | 1.1E-08 |
| *F21D9.4* | WBGene00009018 | protein_coding_gene | -1.2 | 0.425 | 7.4E-03 |
| *dyf-5* | WBGene00001121 | protein_coding_gene | -1.2 | 0.424 | 6.9E-03 |
| *R10H1.1* | WBGene00019998 | protein_coding_gene | -1.2 | 0.424 | 1.6E-11 |
| *Y54F10BM.3* | WBGene00021858 | protein_coding_gene | -1.2 | 0.424 | 1.5E-07 |
| *C08F11.12* | WBGene00007459 | protein_coding_gene | -1.2 | 0.424 | 4.8E-06 |
| *T01D1.8* | WBGene00044641 | protein_coding_gene | -1.2 | 0.423 | 2.5E-02 |
| *unc-30* | WBGene00006766 | protein_coding_gene | -1.2 | 0.423 | 2.0E-03 |
| *Y67D8C.9* | WBGene00022073 | protein_coding_gene | -1.2 | 0.423 | 6.5E-10 |
| *K10B3.5* | WBGene00019611 | protein_coding_gene | -1.2 | 0.422 | 1.0E-10 |
| *C02D5.4* | WBGene00043097 | protein_coding_gene | -1.2 | 0.422 | 3.4E-21 |
| *B0334.10* | WBGene00007148 | protein_coding_gene | -1.2 | 0.422 | 4.5E-02 |
| *H03G16.7* | WBGene00304793 | protein_coding_gene | -1.2 | 0.421 | 4.6E-02 |
| *cpna-5* | WBGene00007044 | protein_coding_gene | -1.2 | 0.421 | 2.6E-12 |
| *Y22D7AL.6* | WBGene00021245 | protein_coding_gene | -1.2 | 0.421 | 2.1E-02 |
| *plpr-1* | WBGene00011524 | protein_coding_gene | -1.2 | 0.421 | 1.7E-03 |
| *C17G1.5* | WBGene00007652 | protein_coding_gene | -1.2 | 0.421 | 7.8E-07 |
| *C26B9.6* | WBGene00016135 | protein_coding_gene | -1.2 | 0.421 | 4.4E-07 |
| *Y79H2A.2* | WBGene00013579 | protein_coding_gene | -1.2 | 0.421 | 5.5E-08 |
| *K02E7.6* | WBGene00019311 | pseudogene | -1.3 | 0.420 | 3.5E-02 |
| *ctn-1* | WBGene00000832 | protein_coding_gene | -1.3 | 0.419 | 3.3E-12 |
| *cuti-1* | WBGene00022591 | protein_coding_gene | -1.3 | 0.418 | 2.2E-03 |
| *F10E7.9* | WBGene00017350 | protein_coding_gene | -1.3 | 0.417 | 6.6E-07 |
| *F36F12.1* | WBGene00018092 | protein_coding_gene | -1.3 | 0.417 | 4.6E-02 |
| *kcc-3* | WBGene00019289 | protein_coding_gene | -1.3 | 0.417 | 7.5E-06 |
| *hlh-33* | WBGene00021446 | protein_coding_gene | -1.3 | 0.416 | 7.7E-06 |
| *C06B8.7* | WBGene00007372 | protein_coding_gene | -1.3 | 0.416 | 1.5E-16 |
| *nhr-88* | WBGene00003678 | protein_coding_gene | -1.3 | 0.416 | 3.3E-12 |
| *cyp-13A3* | WBGene00011675 | protein_coding_gene | -1.3 | 0.415 | 7.5E-03 |
| *K01A2.4* | WBGene00019279 | protein_coding_gene | -1.3 | 0.415 | 1.1E-06 |
| *trx-3* | WBGene00019717 | protein_coding_gene | -1.3 | 0.415 | 1.3E-07 |
| *slt-1* | WBGene00004854 | protein_coding_gene | -1.3 | 0.414 | 3.6E-07 |
| *xdh-1* | WBGene00010083 | protein_coding_gene | -1.3 | 0.414 | 7.9E-17 |
| *cyp-13A6* | WBGene00011673 | protein_coding_gene | -1.3 | 0.414 | 2.6E-03 |
| *F58G11.4* | WBGene00010282 | protein_coding_gene | -1.3 | 0.414 | 1.2E-02 |
| *pes-4* | WBGene00003978 | protein_coding_gene | -1.3 | 0.414 | 1.2E-14 |
| *R08A2.7* | WBGene00011137 | protein_coding_gene | -1.3 | 0.414 | 2.4E-02 |
| *Y48A6B.7* | WBGene00012968 | protein_coding_gene | -1.3 | 0.414 | 2.2E-19 |
| *tmed-13* | WBGene00022255 | protein_coding_gene | -1.3 | 0.414 | 7.4E-04 |
| *W07E11.1* | WBGene00012326 | protein_coding_gene | -1.3 | 0.414 | 8.1E-28 |
| *gsto-3* | WBGene00019636 | protein_coding_gene | -1.3 | 0.413 | 7.1E-09 |
| *Y116F11A.6* | WBGene00013819 | protein_coding_gene | -1.3 | 0.413 | 1.4E-02 |
| *clec-25* | WBGene00044331 | protein_coding_gene | -1.3 | 0.412 | 4.2E-03 |
| *C15B12.8* | WBGene00015788 | protein_coding_gene | -1.3 | 0.412 | 2.9E-03 |
| *4R79.2* | WBGene00007067 | protein_coding_gene | -1.3 | 0.412 | 2.6E-03 |
| *bus-17* | WBGene00044630 | protein_coding_gene | -1.3 | 0.412 | 8.2E-03 |
| *gly-19* | WBGene00001644 | protein_coding_gene | -1.3 | 0.411 | 6.3E-07 |
| *vab-3* | WBGene00006870 | protein_coding_gene | -1.3 | 0.411 | 1.6E-09 |
| *kal-1* | WBGene00002181 | protein_coding_gene | -1.3 | 0.411 | 4.7E-10 |
| *D2030.2* | WBGene00008412 | protein_coding_gene | -1.3 | 0.411 | 2.8E-39 |
| *F42G2.7* | WBGene00044362 | protein_coding_gene | -1.3 | 0.410 | 5.9E-03 |
| *npr-31* | WBGene00020319 | protein_coding_gene | -1.3 | 0.410 | 1.8E-02 |
| *F57C9.6* | WBGene00019012 | protein_coding_gene | -1.3 | 0.409 | 1.9E-02 |
| *chil-28* | WBGene00008842 | pseudogene | -1.3 | 0.409 | 6.3E-03 |
| *nlr-1* | WBGene00003772 | protein_coding_gene | -1.3 | 0.409 | 1.9E-09 |

| | | | | | |
|---|---|---|---|---|---|
| *mam-5* | WBGene00011572 | protein_coding_gene | -1.3 | 0.409 | 3.4E-02 |
| *lgc-55* | WBGene00013746 | protein_coding_gene | -1.3 | 0.409 | 2.5E-03 |
| *unc-24* | WBGene00006761 | protein_coding_gene | -1.3 | 0.408 | 1.0E-03 |
| *nhr-18* | WBGene00003617 | protein_coding_gene | -1.3 | 0.407 | 1.4E-08 |
| *lin-22* | WBGene00003008 | protein_coding_gene | -1.3 | 0.407 | 1.7E-02 |
| *F33G12.6* | WBGene00018018 | protein_coding_gene | -1.3 | 0.407 | 2.8E-17 |
| *D2005.6* | WBGene00008401 | protein_coding_gene | -1.3 | 0.407 | 1.1E-04 |
| *ceh-22* | WBGene00000445 | protein_coding_gene | -1.3 | 0.406 | 1.3E-03 |
| *F54C8.6* | WBGene00010039 | protein_coding_gene | -1.3 | 0.406 | 1.1E-05 |
| *ncam-1* | WBGene00017184 | protein_coding_gene | -1.3 | 0.406 | 3.5E-12 |
| *F38B6.6* | WBGene00018175 | protein_coding_gene | -1.3 | 0.406 | 3.1E-05 |
| *cutl-17* | WBGene00009541 | protein_coding_gene | -1.3 | 0.406 | 3.3E-02 |
| *C26E1.2* | WBGene00007748 | protein_coding_gene | -1.3 | 0.406 | 8.2E-15 |
| *F02A9.7* | WBGene00044406 | protein_coding_gene | -1.3 | 0.406 | 3.6E-02 |
| *sox-2* | WBGene00004949 | protein_coding_gene | -1.3 | 0.406 | 1.3E-15 |
| *lipl-6* | WBGene00021963 | protein_coding_gene | -1.3 | 0.405 | 7.8E-03 |
| *cht-1* | WBGene00000503 | protein_coding_gene | -1.3 | 0.405 | 1.4E-20 |
| *zig-4* | WBGene00006981 | protein_coding_gene | -1.3 | 0.405 | 1.3E-05 |
| *K09E4.4* | WBGene00010722 | protein_coding_gene | -1.3 | 0.404 | 5.8E-05 |
| *C14A11.2* | WBGene00015748 | protein_coding_gene | -1.3 | 0.404 | 5.6E-03 |
| *nlp-52* | WBGene00018363 | protein_coding_gene | -1.3 | 0.404 | 1.6E-02 |
| *F32D8.3* | WBGene00009328 | protein_coding_gene | -1.3 | 0.404 | 4.5E-02 |
| *B0478.3* | WBGene00015202 | protein_coding_gene | -1.3 | 0.403 | 5.4E-03 |
| *Y79H2A.12* | WBGene00043997 | protein_coding_gene | -1.3 | 0.403 | 4.1E-02 |
| *sulp-8* | WBGene00013963 | protein_coding_gene | -1.3 | 0.403 | 5.6E-09 |
| *F35D2.3* | WBGene00018041 | protein_coding_gene | -1.3 | 0.402 | 6.0E-09 |
| *ZK809.9* | WBGene00194934 | protein_coding_gene | -1.3 | 0.400 | 4.5E-06 |
| *F13B12.4* | WBGene00008732 | protein_coding_gene | -1.3 | 0.400 | 3.1E-02 |
| *unc-86* | WBGene00006818 | protein_coding_gene | -1.3 | 0.399 | 6.7E-05 |
| *M03D4.4* | WBGene00019751 | protein_coding_gene | -1.3 | 0.399 | 7.5E-08 |
| *T21B4.15* | WBGene00043054 | protein_coding_gene | -1.3 | 0.399 | 3.8E-02 |
| *C27B7.7* | WBGene00007764 | protein_coding_gene | -1.3 | 0.399 | 5.8E-11 |
| *B0365.9* | WBGene00219208 | protein_coding_gene | -1.3 | 0.398 | 6.4E-03 |
| *cutl-13* | WBGene00021025 | protein_coding_gene | -1.3 | 0.398 | 1.1E-02 |
| *F14F9.5* | WBGene00017468 | protein_coding_gene | -1.3 | 0.398 | 2.7E-05 |
| *C35E7.4* | WBGene00016456 | protein_coding_gene | -1.3 | 0.398 | 8.5E-07 |
| *dhs-16* | WBGene00000979 | protein_coding_gene | -1.3 | 0.397 | 2.2E-02 |
| *F26G1.11* | WBGene00045455 | protein_coding_gene | -1.3 | 0.397 | 2.6E-02 |
| *lin-17* | WBGene00003006 | protein_coding_gene | -1.3 | 0.397 | 1.2E-09 |
| *best-18* | WBGene00011868 | protein_coding_gene | -1.3 | 0.397 | 3.2E-08 |
| *C18F10.2* | WBGene00015977 | protein_coding_gene | -1.3 | 0.396 | 4.5E-03 |
| *F55B12.10* | WBGene00077439 | protein_coding_gene | -1.3 | 0.396 | 1.4E-02 |
| *T14B1.1* | WBGene00011763 | protein_coding_gene | -1.3 | 0.395 | 3.6E-09 |
| *sma-2* | WBGene00004856 | protein_coding_gene | -1.3 | 0.394 | 1.0E-09 |
| *C10G8.4* | WBGene00015683 | protein_coding_gene | -1.3 | 0.394 | 2.3E-20 |
| *mop-25.1* | WBGene00019827 | protein_coding_gene | -1.3 | 0.393 | 2.1E-33 |
| *ZC196.3* | WBGene00022547 | protein_coding_gene | -1.4 | 0.391 | 2.0E-02 |
| *ZK822.1* | WBGene00014089 | protein_coding_gene | -1.4 | 0.391 | 3.1E-05 |
| *nhr-32* | WBGene00003626 | protein_coding_gene | -1.4 | 0.391 | 3.5E-17 |
| *Y58A7A.2* | WBGene00021976 | protein_coding_gene | -1.4 | 0.391 | 3.5E-02 |
| *K07A1.6* | WBGene00010613 | protein_coding_gene | -1.4 | 0.391 | 9.6E-20 |
| *Y51H4A.5* | WBGene00013101 | protein_coding_gene | -1.4 | 0.390 | 1.3E-04 |
| *tric-1B.1* | WBGene00013255 | protein_coding_gene | -1.4 | 0.389 | 1.4E-22 |
| *try-3* | WBGene00006621 | protein_coding_gene | -1.4 | 0.389 | 2.6E-05 |
| *ZC262.13* | WBGene00284846 | protein_coding_gene | -1.4 | 0.388 | 4.6E-03 |
| *Y47G6A.5* | WBGene00021635 | protein_coding_gene | -1.4 | 0.388 | 1.6E-19 |
| *cbp-2* | WBGene00010545 | protein_coding_gene | -1.4 | 0.388 | 1.5E-12 |
| *ttr-43* | WBGene00007590 | protein_coding_gene | -1.4 | 0.387 | 2.4E-04 |
| *C17C3.3* | WBGene00015889 | protein_coding_gene | -1.4 | 0.387 | 2.1E-02 |
| *tba-5* | WBGene00006531 | protein_coding_gene | -1.4 | 0.387 | 5.5E-03 |
| *nmur-3* | WBGene00017176 | protein_coding_gene | -1.4 | 0.387 | 1.8E-02 |
| *R12G8.1* | WBGene00011254 | protein_coding_gene | -1.4 | 0.387 | 3.3E-02 |
| *Y50E8A.5* | WBGene00013048 | protein_coding_gene | -1.4 | 0.387 | 3.9E-06 |
| *R11F4.1* | WBGene00020007 | protein_coding_gene | -1.4 | 0.387 | 1.3E-13 |
| *jmjd-3.3* | WBGene00007813 | protein_coding_gene | -1.4 | 0.386 | 2.8E-04 |
| *lgc-1* | WBGene00020236 | protein_coding_gene | -1.4 | 0.386 | 2.5E-02 |
| *Y48G8AL.13* | WBGene00021691 | protein_coding_gene | -1.4 | 0.386 | 3.5E-21 |
| *drr-1* | WBGene00009741 | protein_coding_gene | -1.4 | 0.386 | 4.4E-14 |
| *mab-21* | WBGene00003112 | protein_coding_gene | -1.4 | 0.385 | 1.2E-06 |
| *M01G12.9* | WBGene00010822 | protein_coding_gene | -1.4 | 0.385 | 3.3E-02 |

| | | | | | |
|---|---|---|---|---|---|
| *C45E5.3* | WBGene00016666 | pseudogene | -1.4 | 0.385 | 2.5E-04 |
| *T05E11.9* | WBGene00014505 | ncRNA_gene | -1.4 | 0.384 | 2.8E-20 |
| *F49C12.6* | WBGene00009876 | protein_coding_gene | -1.4 | 0.384 | 8.3E-05 |
| *unc-122* | WBGene00006845 | protein_coding_gene | -1.4 | 0.384 | 1.2E-02 |
| *Y39B6A.7* | WBGene00012669 | protein_coding_gene | -1.4 | 0.384 | 3.4E-02 |
| *egl-43* | WBGene00001207 | protein_coding_gene | -1.4 | 0.383 | 1.3E-08 |
| *nhr-171* | WBGene00016926 | protein_coding_gene | -1.4 | 0.383 | 1.2E-02 |
| *fipr-4* | WBGene00007541 | protein_coding_gene | -1.4 | 0.383 | 4.2E-02 |
| *lam-3* | WBGene00002248 | protein_coding_gene | -1.4 | 0.383 | 1.4E-14 |
| *Y6B3B.7* | WBGene00012391 | protein_coding_gene | -1.4 | 0.383 | 1.5E-02 |
| *R57.2* | WBGene00020083 | protein_coding_gene | -1.4 | 0.383 | 9.8E-03 |
| *ztf-11* | WBGene00009939 | protein_coding_gene | -1.4 | 0.382 | 9.7E-19 |
| *D1069.1* | WBGene00017036 | protein_coding_gene | -1.4 | 0.382 | 1.6E-02 |
| *T27D12.6* | WBGene00194848 | protein_coding_gene | -1.4 | 0.382 | 4.2E-02 |
| *F55A4.4* | WBGene00018856 | pseudogene | -1.4 | 0.382 | 1.8E-02 |
| *fkh-7* | WBGene00001439 | protein_coding_gene | -1.4 | 0.382 | 4.8E-20 |
| *linc-84* | WBGene00219592 | lincRNA_gene | -1.4 | 0.380 | 4.0E-02 |
| *Y43F8B.2* | WBGene00012813 | protein_coding_gene | -1.4 | 0.380 | 1.5E-26 |
| *F56C9.8* | WBGene00018952 | protein_coding_gene | -1.4 | 0.380 | 1.1E-13 |
| *cest-16* | WBGene00020330 | protein_coding_gene | -1.4 | 0.380 | 4.5E-04 |
| *C07B5.4* | WBGene00007409 | protein_coding_gene | -1.4 | 0.379 | 7.8E-09 |
| *tub-1* | WBGene00006655 | protein_coding_gene | -1.4 | 0.379 | 1.6E-03 |
| *metr-1* | WBGene00010988 | protein_coding_gene | -1.4 | 0.379 | 2.8E-21 |
| *cest-17* | WBGene00019477 | protein_coding_gene | -1.4 | 0.378 | 2.8E-11 |
| *T16G1.2* | WBGene00011796 | protein_coding_gene | -1.4 | 0.378 | 7.4E-03 |
| *ZK1073.1* | WBGene00014213 | protein_coding_gene | -1.4 | 0.378 | 9.6E-21 |
| *glb-23* | WBGene00020045 | protein_coding_gene | -1.4 | 0.377 | 2.2E-02 |
| *lgc-20* | WBGene00007191 | protein_coding_gene | -1.4 | 0.377 | 1.1E-02 |
| *ZC239.16* | WBGene00022574 | protein_coding_gene | -1.4 | 0.376 | 3.4E-02 |
| *T19A6.4* | WBGene00011829 | protein_coding_gene | -1.4 | 0.375 | 1.4E-04 |
| *ham-2* | WBGene00001821 | protein_coding_gene | -1.4 | 0.375 | 4.7E-12 |
| *Y87G2A.25* | WBGene00219288 | protein_coding_gene | -1.4 | 0.375 | 1.6E-03 |
| *C06G3.5* | WBGene00015551 | protein_coding_gene | -1.4 | 0.374 | 4.4E-16 |
| *col-53* | WBGene00000630 | protein_coding_gene | -1.4 | 0.374 | 2.3E-03 |
| *cut-6* | WBGene00000853 | protein_coding_gene | -1.4 | 0.373 | 5.6E-04 |
| *tsp-16* | WBGene00006642 | protein_coding_gene | -1.4 | 0.372 | 2.9E-06 |
| *F40G9.5* | WBGene00018241 | protein_coding_gene | -1.4 | 0.372 | 3.5E-07 |
| *cyp-23A1* | WBGene00015135 | protein_coding_gene | -1.4 | 0.372 | 1.8E-02 |
| *mam-1* | WBGene00022504 | protein_coding_gene | -1.4 | 0.372 | 1.1E-02 |
| *Y71H2AM.15* | WBGene00022180 | protein_coding_gene | -1.4 | 0.372 | 1.0E-05 |
| *nhr-120* | WBGene00003710 | protein_coding_gene | -1.4 | 0.371 | 4.7E-10 |
| *K01D12.1* | WBGene00010461 | protein_coding_gene | -1.4 | 0.371 | 1.4E-02 |
| *F08G2.8* | WBGene00008579 | protein_coding_gene | -1.4 | 0.371 | 1.3E-02 |
| *eva-1* | WBGene00009304 | protein_coding_gene | -1.4 | 0.371 | 8.0E-10 |
| *F09F9.1* | WBGene00017306 | protein_coding_gene | -1.4 | 0.370 | 1.9E-08 |
| *skr-19* | WBGene00004825 | protein_coding_gene | -1.4 | 0.370 | 1.4E-17 |
| *F58E6.11* | WBGene00010258 | protein_coding_gene | -1.4 | 0.370 | 4.4E-03 |
| *dep-1* | WBGene00009717 | protein_coding_gene | -1.4 | 0.369 | 1.2E-13 |
| *F28G4.2* | WBGene00009227 | protein_coding_gene | -1.4 | 0.369 | 2.1E-02 |
| *R148.5* | WBGene00020104 | protein_coding_gene | -1.4 | 0.368 | 1.5E-21 |
| *F01D5.6* | WBGene00008496 | protein_coding_gene | -1.4 | 0.368 | 4.5E-03 |
| *fozi-1* | WBGene00010453 | protein_coding_gene | -1.4 | 0.368 | 6.6E-15 |
| *cdh-12* | WBGene00022103 | protein_coding_gene | -1.4 | 0.367 | 2.1E-02 |
| *C26B9.3* | WBGene00016133 | protein_coding_gene | -1.4 | 0.367 | 2.5E-02 |
| *ace-3* | WBGene00000037 | protein_coding_gene | -1.4 | 0.366 | 1.8E-05 |
| *K01A12.3* | WBGene00019287 | protein_coding_gene | -1.5 | 0.366 | 4.9E-02 |
| *ZK287.1* | WBGene00013962 | protein_coding_gene | -1.5 | 0.366 | 2.4E-06 |
| *pals-33* | WBGene00021081 | protein_coding_gene | -1.5 | 0.365 | 9.0E-04 |
| *F08F1.21* | WBGene00304789 | ncRNA_gene | -1.5 | 0.365 | 4.0E-03 |
| *cest-27* | WBGene00018958 | protein_coding_gene | -1.5 | 0.365 | 3.1E-16 |
| *hbl-1* | WBGene00001824 | protein_coding_gene | -1.5 | 0.364 | 3.6E-07 |
| *efn-4* | WBGene00001165 | protein_coding_gene | -1.5 | 0.364 | 5.2E-18 |
| *Y75B8A.6* | WBGene00013543 | protein_coding_gene | -1.5 | 0.364 | 3.8E-15 |
| *dma-1* | WBGene00011345 | protein_coding_gene | -1.5 | 0.363 | 1.3E-10 |
| *C24A8.5* | WBGene00016039 | protein_coding_gene | -1.5 | 0.363 | 2.0E-02 |
| *klf-1* | WBGene00018990 | protein_coding_gene | -1.5 | 0.362 | 1.8E-17 |
| *H10D18.1* | WBGene00019177 | protein_coding_gene | -1.5 | 0.362 | 9.8E-03 |
| *F55B11.5* | WBGene00010087 | protein_coding_gene | -1.5 | 0.362 | 4.1E-05 |
| *K12C11.7* | WBGene00045053 | protein_coding_gene | -1.5 | 0.362 | 3.8E-07 |
| *aex-3* | WBGene00000086 | protein_coding_gene | -1.5 | 0.361 | 1.1E-14 |

| | | | | | |
|---|---|---|---|---|---|
| *K10D3.4* | WBGene00010738 | protein_coding_gene | -1.5 | 0.361 | 7.6E-03 |
| *C29A12.6* | WBGene00007800 | protein_coding_gene | -1.5 | 0.361 | 3.1E-03 |
| *F17C8.3* | WBGene00008910 | protein_coding_gene | -1.5 | 0.361 | 7.2E-08 |
| *nhr-25* | WBGene00003623 | protein_coding_gene | -1.5 | 0.361 | 1.8E-12 |
| *glb-5* | WBGene00015964 | protein_coding_gene | -1.5 | 0.361 | 2.4E-03 |
| *C36E8.4* | WBGene00007983 | protein_coding_gene | -1.5 | 0.360 | 2.7E-10 |
| *Y82E9BL.19* | WBGene00194735 | protein_coding_gene | -1.5 | 0.360 | 2.1E-02 |
| *siah-1* | WBGene00021369 | protein_coding_gene | -1.5 | 0.360 | 1.2E-26 |
| *F23A7.5* | WBGene00009070 | protein_coding_gene | -1.5 | 0.360 | 7.6E-04 |
| *zmp-5* | WBGene00010423 | protein_coding_gene | -1.5 | 0.359 | 1.2E-02 |
| *Y6E2A.5* | WBGene00012399 | protein_coding_gene | -1.5 | 0.359 | 9.1E-06 |
| *Y69A2AR.31* | WBGene00022100 | protein_coding_gene | -1.5 | 0.359 | 3.3E-08 |
| *ptr-17* | WBGene00004231 | protein_coding_gene | -1.5 | 0.359 | 8.2E-03 |
| *cwp-4* | WBGene00000862 | protein_coding_gene | -1.5 | 0.359 | 4.0E-03 |
| *F46A8.1* | WBGene00014759 | pseudogene | -1.5 | 0.359 | 2.9E-05 |
| *F55H12.2* | WBGene00010133 | protein_coding_gene | -1.5 | 0.358 | 5.8E-16 |
| *C24H10.1* | WBGene00016067 | protein_coding_gene | -1.5 | 0.358 | 5.7E-05 |
| *K09F5.6* | WBGene00044698 | protein_coding_gene | -1.5 | 0.358 | 6.3E-15 |
| *T17H7.7* | WBGene00020551 | protein_coding_gene | -1.5 | 0.358 | 1.3E-03 |
| *R05G6.1* | WBGene00019896 | protein_coding_gene | -1.5 | 0.357 | 1.5E-03 |
| *F07C3.9* | WBGene00017197 | protein_coding_gene | -1.5 | 0.357 | 4.0E-07 |
| *C29H12.6* | WBGene00016237 | protein_coding_gene | -1.5 | 0.356 | 1.8E-07 |
| *ZC487.1* | WBGene00022623 | protein_coding_gene | -1.5 | 0.356 | 2.3E-04 |
| *osm-6* | WBGene00003886 | protein_coding_gene | -1.5 | 0.356 | 1.7E-03 |
| *hlh-30* | WBGene00020930 | protein_coding_gene | -1.5 | 0.355 | 4.2E-35 |
| *dhrs-4* | WBGene00010063 | protein_coding_gene | -1.5 | 0.355 | 1.4E-22 |
| *E02H4.4* | WBGene00008462 | protein_coding_gene | -1.5 | 0.355 | 1.8E-20 |
| *W05B2.2* | WBGene00012271 | protein_coding_gene | -1.5 | 0.355 | 1.9E-02 |
| *clec-260* | WBGene00013357 | protein_coding_gene | -1.5 | 0.354 | 5.6E-05 |
| *T13C2.3* | WBGene00020479 | protein_coding_gene | -1.5 | 0.354 | 1.2E-06 |
| *C13E3.1* | WBGene00015740 | protein_coding_gene | -1.5 | 0.353 | 3.0E-09 |
| *F40G9.20* | WBGene00195034 | ncRNA_gene | -1.5 | 0.353 | 6.3E-03 |
| *ttr-17* | WBGene00012383 | protein_coding_gene | -1.5 | 0.353 | 3.7E-20 |
| *rei-2* | WBGene00019365 | protein_coding_gene | -1.5 | 0.353 | 3.7E-22 |
| *gtr-1* | WBGene00017779 | protein_coding_gene | -1.5 | 0.353 | 1.7E-02 |
| *C27A12.4* | WBGene00016155 | protein_coding_gene | -1.5 | 0.352 | 3.9E-02 |
| *unc-29* | WBGene00006765 | protein_coding_gene | -1.5 | 0.352 | 6.6E-06 |
| *igcm-2* | WBGene00020130 | protein_coding_gene | -1.5 | 0.352 | 3.0E-05 |
| *F32D1.11* | WBGene00044530 | protein_coding_gene | -1.5 | 0.352 | 1.2E-03 |
| *C06C3.4* | WBGene00007375 | protein_coding_gene | -1.5 | 0.351 | 9.2E-03 |
| *F38E9.1* | WBGene00018184 | protein_coding_gene | -1.5 | 0.351 | 8.1E-24 |
| *ugt-51* | WBGene00015369 | protein_coding_gene | -1.5 | 0.351 | 5.8E-08 |
| *ZC449.2* | WBGene00022612 | protein_coding_gene | -1.5 | 0.351 | 7.0E-04 |
| *F49B2.4* | WBGene00009864 | pseudogene | -1.5 | 0.350 | 2.8E-02 |
| *F15E6.9* | WBGene00017489 | protein_coding_gene | -1.5 | 0.350 | 1.0E-04 |
| *bkip-1* | WBGene00014569 | protein_coding_gene | -1.5 | 0.349 | 2.0E-03 |
| *T03G11.9* | WBGene00044584 | protein_coding_gene | -1.5 | 0.348 | 4.2E-02 |
| *F46G11.6* | WBGene00045269 | protein_coding_gene | -1.5 | 0.348 | 3.5E-02 |
| *igcm-1* | WBGene00018215 | protein_coding_gene | -1.5 | 0.347 | 4.2E-08 |
| *F48E3.8* | WBGene00018607 | protein_coding_gene | -1.5 | 0.347 | 3.8E-11 |
| *Y51A2D.13* | WBGene00013080 | protein_coding_gene | -1.5 | 0.347 | 9.9E-30 |
| *pals-19* | WBGene00044761 | protein_coding_gene | -1.5 | 0.347 | 2.1E-03 |
| *F52D1.2* | WBGene00018683 | protein_coding_gene | -1.5 | 0.347 | 1.6E-06 |
| *T05F1.11* | WBGene00011495 | protein_coding_gene | -1.5 | 0.347 | 1.6E-09 |
| *C54F6.18* | WBGene00235093 | protein_coding_gene | -1.5 | 0.347 | 2.5E-02 |
| *ZK1128.7* | WBGene00014233 | protein_coding_gene | -1.5 | 0.346 | 9.3E-21 |
| *F08F1.3* | WBGene00017257 | protein_coding_gene | -1.5 | 0.346 | 3.0E-07 |
| *nsy-4* | WBGene00021415 | protein_coding_gene | -1.5 | 0.346 | 3.7E-10 |
| *cest-19* | WBGene00020125 | protein_coding_gene | -1.5 | 0.344 | 7.0E-03 |
| *cdf-2* | WBGene00011821 | protein_coding_gene | -1.5 | 0.344 | 1.5E-12 |
| *cpt-1* | WBGene00012907 | protein_coding_gene | -1.5 | 0.344 | 7.9E-24 |
| *ceh-48* | WBGene00015934 | protein_coding_gene | -1.5 | 0.343 | 5.9E-07 |
| *lgc-27* | WBGene00022283 | protein_coding_gene | -1.5 | 0.343 | 9.8E-05 |
| *F43G6.17* | WBGene00219378 | protein_coding_gene | -1.5 | 0.343 | 5.3E-04 |
| *tag-138* | WBGene00006484 | protein_coding_gene | -1.5 | 0.343 | 1.6E-09 |
| *F32D1.3* | WBGene00017983 | protein_coding_gene | -1.5 | 0.342 | 4.4E-10 |
| *zig-10* | WBGene00020800 | protein_coding_gene | -1.5 | 0.342 | 9.8E-06 |
| *mks-5* | WBGene00007490 | protein_coding_gene | -1.5 | 0.342 | 1.7E-03 |
| *gei-13* | WBGene00001570 | protein_coding_gene | -1.5 | 0.342 | 2.1E-05 |
| *osm-5* | WBGene00003885 | protein_coding_gene | -1.6 | 0.341 | 1.1E-03 |

| | | | | | |
|---|---|---|---|---|---|
| *T28F4.5* | WBGene00012140 | protein_coding_gene | -1.6 | 0.340 | 1.9E-18 |
| *F30H5.3* | WBGene00017937 | protein_coding_gene | -1.6 | 0.340 | 1.3E-02 |
| *dex-1* | WBGene00017028 | protein_coding_gene | -1.6 | 0.340 | 4.7E-15 |
| *cgt-1* | WBGene00011517 | protein_coding_gene | -1.6 | 0.340 | 1.8E-06 |
| *lido-15* | WBGene00018804 | protein_coding_gene | -1.6 | 0.339 | 3.2E-08 |
| *T04A6.2* | WBGene00020201 | protein_coding_gene | -1.6 | 0.339 | 4.7E-03 |
| *tpra-1* | WBGene00008366 | protein_coding_gene | -1.6 | 0.339 | 2.8E-24 |
| *F53E10.1* | WBGene00018758 | protein_coding_gene | -1.6 | 0.338 | 9.4E-23 |
| *F20D6.10* | WBGene00017639 | protein_coding_gene | -1.6 | 0.338 | 3.6E-08 |
| *ddo-1* | WBGene00008127 | protein_coding_gene | -1.6 | 0.338 | 1.2E-11 |
| *dct-1* | WBGene00015776 | protein_coding_gene | -1.6 | 0.337 | 1.5E-24 |
| *F10E7.3* | WBGene00017345 | pseudogene | -1.6 | 0.337 | 3.1E-02 |
| *C32D5.12* | WBGene00016319 | protein_coding_gene | -1.6 | 0.337 | 2.4E-06 |
| *ZK721.4* | WBGene00022811 | protein_coding_gene | -1.6 | 0.337 | 4.2E-02 |
| *F45E1.5* | WBGene00018466 | protein_coding_gene | -1.6 | 0.336 | 7.0E-03 |
| *nhr-91* | WBGene00003681 | protein_coding_gene | -1.6 | 0.336 | 1.4E-06 |
| *C03A3.1* | WBGene00007266 | protein_coding_gene | -1.6 | 0.336 | 3.5E-06 |
| *ZK836.3* | WBGene00014099 | protein_coding_gene | -1.6 | 0.335 | 1.3E-06 |
| *spp-8* | WBGene00004993 | protein_coding_gene | -1.6 | 0.335 | 1.1E-10 |
| *cpr-2* | WBGene00000782 | protein_coding_gene | -1.6 | 0.335 | 1.9E-02 |
| *Y65B4BL.1* | WBGene00022033 | protein_coding_gene | -1.6 | 0.334 | 5.8E-04 |
| *F56F11.1* | WBGene00018988 | protein_coding_gene | -1.6 | 0.333 | 2.5E-02 |
| *mksr-1* | WBGene00019364 | protein_coding_gene | -1.6 | 0.332 | 6.0E-03 |
| *noah-1* | WBGene00016422 | protein_coding_gene | -1.6 | 0.332 | 8.4E-03 |
| *comt-4* | WBGene00021491 | protein_coding_gene | -1.6 | 0.330 | 3.7E-02 |
| *Y105C5A.15* | WBGene00013639 | protein_coding_gene | -1.6 | 0.330 | 7.3E-16 |
| *tmem-231* | WBGene00020825 | protein_coding_gene | -1.6 | 0.329 | 2.3E-03 |
| *R02E12.12* | WBGene00303425 | protein_coding_gene | -1.6 | 0.329 | 2.6E-02 |
| *F09F9.4* | WBGene00017309 | protein_coding_gene | -1.6 | 0.329 | 3.0E-06 |
| *ZK381.2* | WBGene00022725 | protein_coding_gene | -1.6 | 0.329 | 4.3E-02 |
| *cutl-16* | WBGene00019426 | protein_coding_gene | -1.6 | 0.329 | 8.4E-09 |
| *Y97E10AR.1* | WBGene00022396 | protein_coding_gene | -1.6 | 0.328 | 1.8E-11 |
| *arrd-8* | WBGene00012467 | protein_coding_gene | -1.6 | 0.328 | 1.1E-02 |
| *C07A4.2* | WBGene00007397 | protein_coding_gene | -1.6 | 0.328 | 5.8E-03 |
| *fmil-1* | WBGene00021362 | protein_coding_gene | -1.6 | 0.328 | 1.1E-07 |
| *unk-1* | WBGene00016407 | protein_coding_gene | -1.6 | 0.328 | 5.0E-29 |
| *wrt-1* | WBGene00006947 | protein_coding_gene | -1.6 | 0.328 | 5.0E-02 |
| *noah-2* | WBGene00009926 | protein_coding_gene | -1.6 | 0.327 | 1.5E-02 |
| *C44H9.5* | WBGene00008100 | protein_coding_gene | -1.6 | 0.327 | 1.3E-18 |
| *F25C8.1* | WBGene00009104 | pseudogene | -1.6 | 0.326 | 1.3E-03 |
| *ssp-37* | WBGene00012211 | protein_coding_gene | -1.6 | 0.326 | 9.3E-08 |
| *ttr-54* | WBGene00011774 | protein_coding_gene | -1.6 | 0.326 | 1.7E-11 |
| *K03H1.5* | WBGene00010540 | protein_coding_gene | -1.6 | 0.326 | 1.8E-23 |
| *C48B4.3* | WBGene00008168 | protein_coding_gene | -1.6 | 0.326 | 7.8E-05 |
| *C07A4.3* | WBGene00007398 | protein_coding_gene | -1.6 | 0.325 | 1.9E-05 |
| *pef-1* | WBGene00003969 | protein_coding_gene | -1.6 | 0.325 | 1.7E-10 |
| *sups-1* | WBGene00017716 | protein_coding_gene | -1.6 | 0.324 | 1.4E-02 |
| *T28H10.3* | WBGene00012144 | protein_coding_gene | -1.6 | 0.324 | 2.6E-47 |
| *T01D3.3* | WBGene00011328 | protein_coding_gene | -1.6 | 0.324 | 8.7E-08 |
| *F17C11.4* | WBGene00008915 | protein_coding_gene | -1.6 | 0.324 | 5.2E-20 |
| *ZC84.1* | WBGene00013846 | protein_coding_gene | -1.6 | 0.323 | 2.1E-03 |
| *W10C8.4* | WBGene00021127 | protein_coding_gene | -1.6 | 0.323 | 4.4E-04 |
| *F23H12.5* | WBGene00009094 | protein_coding_gene | -1.6 | 0.323 | 1.0E-02 |
| *gcy-5* | WBGene00001532 | protein_coding_gene | -1.6 | 0.322 | 3.2E-02 |
| *T19C4.1* | WBGene00011838 | protein_coding_gene | -1.6 | 0.322 | 1.1E-06 |
| *W03G1.5* | WBGene00021011 | protein_coding_gene | -1.6 | 0.322 | 1.8E-06 |
| *osm-9* | WBGene00003889 | protein_coding_gene | -1.6 | 0.322 | 7.8E-03 |
| *acs-16* | WBGene00018579 | protein_coding_gene | -1.6 | 0.322 | 2.0E-06 |
| *R05A10.4* | WBGene00011021 | protein_coding_gene | -1.6 | 0.321 | 3.4E-03 |
| *mlt-11* | WBGene00012186 | protein_coding_gene | -1.6 | 0.321 | 4.1E-02 |
| *F43G6.8* | WBGene00009660 | protein_coding_gene | -1.6 | 0.320 | 3.2E-23 |
| *exc-5* | WBGene00001366 | protein_coding_gene | -1.6 | 0.320 | 2.2E-33 |
| *R06B10.2* | WBGene00019913 | protein_coding_gene | -1.6 | 0.320 | 2.5E-02 |
| *Y46C8AM.1* | WBGene00271795 | protein_coding_gene | -1.6 | 0.320 | 2.6E-03 |
| *ptr-15* | WBGene00004229 | protein_coding_gene | -1.6 | 0.320 | 1.2E-04 |
| *F22B7.9* | WBGene00017698 | protein_coding_gene | -1.6 | 0.320 | 4.6E-17 |
| *bath-25* | WBGene00020859 | protein_coding_gene | -1.6 | 0.319 | 5.3E-03 |
| *Y43F8C.3* | WBGene00012825 | protein_coding_gene | -1.6 | 0.319 | 2.9E-25 |
| *ttm-2* | WBGene00017840 | protein_coding_gene | -1.7 | 0.318 | 1.5E-10 |
| *T14B4.19* | WBGene00302993 | protein_coding_gene | -1.7 | 0.318 | 2.4E-02 |

| | | | | | |
|---|---|---|---|---|---|
| *max-1* | WBGene00003143 | protein_coding_gene | -1.7 | 0.318 | 3.3E-12 |
| *ceh-5* | WBGene00000430 | protein_coding_gene | -1.7 | 0.318 | 4.0E-04 |
| *M03F4.6* | WBGene00019759 | protein_coding_gene | -1.7 | 0.317 | 4.6E-05 |
| *T14E8.4* | WBGene00043981 | protein_coding_gene | -1.7 | 0.317 | 8.3E-04 |
| *zag-1* | WBGene00006970 | protein_coding_gene | -1.7 | 0.316 | 3.9E-12 |
| *ZK593.2* | WBGene00014002 | protein_coding_gene | -1.7 | 0.316 | 3.6E-08 |
| *linc-103* | WBGene00219743 | lincRNA_gene | -1.7 | 0.315 | 2.4E-02 |
| *W08G11.1* | WBGene00012346 | protein_coding_gene | -1.7 | 0.314 | 1.9E-02 |
| *F58F12.4* | WBGene00044756 | protein_coding_gene | -1.7 | 0.314 | 9.5E-05 |
| *glb-8* | WBGene00007741 | protein_coding_gene | -1.7 | 0.314 | 1.5E-04 |
| *F53A2.9* | WBGene00009953 | protein_coding_gene | -1.7 | 0.313 | 3.3E-13 |
| *C07E3.9* | WBGene00007419 | protein_coding_gene | -1.7 | 0.312 | 5.9E-20 |
| *Y57G11C.22* | WBGene00013317 | protein_coding_gene | -1.7 | 0.312 | 2.1E-09 |
| *tig-2* | WBGene00006570 | protein_coding_gene | -1.7 | 0.312 | 6.2E-11 |
| *ugt-5* | WBGene00013906 | protein_coding_gene | -1.7 | 0.312 | 2.4E-20 |
| *plx-2* | WBGene00004048 | protein_coding_gene | -1.7 | 0.312 | 4.6E-14 |
| *pros-1* | WBGene00000448 | protein_coding_gene | -1.7 | 0.312 | 8.3E-11 |
| *B0393.5* | WBGene00007170 | protein_coding_gene | -1.7 | 0.311 | 1.2E-03 |
| *gst-15* | WBGene00001763 | protein_coding_gene | -1.7 | 0.311 | 2.6E-09 |
| *K04A8.21* | WBGene00271815 | protein_coding_gene | -1.7 | 0.311 | 9.1E-03 |
| *nphp-1* | WBGene00010898 | protein_coding_gene | -1.7 | 0.311 | 6.3E-05 |
| *lpr-5* | WBGene00012256 | protein_coding_gene | -1.7 | 0.310 | 3.9E-02 |
| *srp-5* | WBGene00005646 | pseudogene | -1.7 | 0.310 | 2.5E-06 |
| *F15H9.7* | WBGene00077535 | pseudogene | -1.7 | 0.309 | 1.9E-02 |
| *K07G5.5* | WBGene00010644 | protein_coding_gene | -1.7 | 0.308 | 1.6E-10 |
| *nhr-69* | WBGene00003659 | protein_coding_gene | -1.7 | 0.308 | 3.7E-23 |
| *kpc-1* | WBGene00002232 | protein_coding_gene | -1.7 | 0.308 | 5.4E-39 |
| *clc-5* | WBGene00000526 | protein_coding_gene | -1.7 | 0.307 | 2.0E-08 |
| *R07E3.6* | WBGene00011107 | protein_coding_gene | -1.7 | 0.307 | 4.9E-02 |
| *rhgf-2* | WBGene00006476 | protein_coding_gene | -1.7 | 0.306 | 3.7E-14 |
| *nab-1* | WBGene00003516 | protein_coding_gene | -1.7 | 0.306 | 1.5E-13 |
| *K02D7.1* | WBGene00019298 | protein_coding_gene | -1.7 | 0.306 | 6.8E-32 |
| *F31D5.6* | WBGene00017949 | protein_coding_gene | -1.7 | 0.305 | 5.8E-04 |
| *M7.8* | WBGene00010884 | protein_coding_gene | -1.7 | 0.304 | 3.0E-13 |
| *K08E7.6* | WBGene00010674 | protein_coding_gene | -1.7 | 0.303 | 2.8E-04 |
| *C13C12.2* | WBGene00007552 | protein_coding_gene | -1.7 | 0.303 | 7.6E-04 |
| *F42C5.4* | WBGene00018346 | protein_coding_gene | -1.7 | 0.302 | 5.1E-14 |
| *ddr-2* | WBGene00017381 | protein_coding_gene | -1.7 | 0.302 | 1.4E-14 |
| *C29F7.2* | WBGene00007811 | protein_coding_gene | -1.7 | 0.302 | 3.8E-38 |
| *R03H10.6* | WBGene00019858 | protein_coding_gene | -1.7 | 0.301 | 8.2E-12 |
| *clec-68* | WBGene00018970 | protein_coding_gene | -1.7 | 0.301 | 4.0E-02 |
| *glc-3* | WBGene00001593 | protein_coding_gene | -1.7 | 0.301 | 7.7E-03 |
| *F16G10.15* | WBGene00017529 | pseudogene | -1.7 | 0.301 | 3.4E-10 |
| *ncx-5* | WBGene00003570 | protein_coding_gene | -1.7 | 0.300 | 3.7E-02 |
| *bgnt-1.3* | WBGene00009032 | protein_coding_gene | -1.7 | 0.299 | 2.7E-02 |
| *nhr-180* | WBGene00017961 | protein_coding_gene | -1.7 | 0.299 | 4.9E-09 |
| *F36D1.7* | WBGene00009466 | protein_coding_gene | -1.7 | 0.299 | 8.5E-10 |
| *W05B10.4* | WBGene00012279 | protein_coding_gene | -1.7 | 0.299 | 5.1E-07 |
| *W03F9.4* | WBGene00021002 | protein_coding_gene | -1.7 | 0.299 | 2.0E-05 |
| *F33H2.8* | WBGene00009371 | protein_coding_gene | -1.7 | 0.297 | 1.3E-04 |
| *dpy-2* | WBGene00001064 | protein_coding_gene | -1.8 | 0.297 | 2.0E-02 |
| *D1086.2* | WBGene00008389 | protein_coding_gene | -1.8 | 0.296 | 1.7E-06 |
| *acs-3* | WBGene00020343 | protein_coding_gene | -1.8 | 0.295 | 9.8E-16 |
| *cyp-33C12* | WBGene00021168 | protein_coding_gene | -1.8 | 0.295 | 1.5E-02 |
| *F48F7.3* | WBGene00009849 | protein_coding_gene | -1.8 | 0.294 | 8.8E-03 |
| *tmem-218* | WBGene00044771 | protein_coding_gene | -1.8 | 0.294 | 4.2E-02 |
| *F36F2.2* | WBGene00009476 | protein_coding_gene | -1.8 | 0.293 | 6.7E-16 |
| *ZC84.6* | WBGene00013849 | protein_coding_gene | -1.8 | 0.292 | 7.0E-04 |
| *K11H12.6* | WBGene00019662 | protein_coding_gene | -1.8 | 0.292 | 1.7E-02 |
| *ZK180.8* | WBGene00195011 | protein_coding_gene | -1.8 | 0.291 | 2.9E-02 |
| *F33E2.6* | WBGene00009362 | protein_coding_gene | -1.8 | 0.290 | 2.9E-02 |
| *F13D11.3* | WBGene00017428 | protein_coding_gene | -1.8 | 0.289 | 7.6E-19 |
| *C54F6.3* | WBGene00016921 | protein_coding_gene | -1.8 | 0.289 | 4.6E-02 |
| *F22F7.3* | WBGene00017721 | protein_coding_gene | -1.8 | 0.289 | 1.4E-03 |
| *M04C9.2* | WBGene00010857 | protein_coding_gene | -1.8 | 0.288 | 6.4E-03 |
| *cyp-33C2* | WBGene00016697 | protein_coding_gene | -1.8 | 0.288 | 4.5E-03 |
| *snb-5* | WBGene00004899 | protein_coding_gene | -1.8 | 0.288 | 9.1E-03 |
| *chpf-2* | WBGene00019108 | protein_coding_gene | -1.8 | 0.288 | 2.4E-11 |
| *F11A5.9* | WBGene00008677 | protein_coding_gene | -1.8 | 0.288 | 7.1E-16 |
| *afmd-2* | WBGene00012866 | protein_coding_gene | -1.8 | 0.288 | 4.5E-12 |

| | | | | | |
|---|---|---|---|---|---|
| *F21A3.11* | WBGene00219316 | protein_coding_gene | -1.8 | 0.288 | 3.5E-18 |
| *tbb-4* | WBGene00006538 | protein_coding_gene | -1.8 | 0.288 | 5.9E-11 |
| *cpr-4* | WBGene00000784 | protein_coding_gene | -1.8 | 0.288 | 3.9E-11 |
| *C17G10.10* | WBGene00195182 | protein_coding_gene | -1.8 | 0.287 | 5.1E-03 |
| *acn-1* | WBGene00000039 | protein_coding_gene | -1.8 | 0.287 | 1.7E-02 |
| *C09F9.2* | WBGene00007479 | protein_coding_gene | -1.8 | 0.287 | 2.4E-05 |
| *K02G10.15* | WBGene00206361 | protein_coding_gene | -1.8 | 0.286 | 1.9E-02 |
| *ZK470.14* | WBGene00219362 | protein_coding_gene | -1.8 | 0.286 | 2.0E-03 |
| *ptr-10* | WBGene00004224 | protein_coding_gene | -1.8 | 0.286 | 8.5E-06 |
| *col-121* | WBGene00000695 | protein_coding_gene | -1.8 | 0.286 | 7.6E-10 |
| *Y116A8C.40* | WBGene00013812 | protein_coding_gene | -1.8 | 0.286 | 1.9E-02 |
| *swip-10* | WBGene00018738 | protein_coding_gene | -1.8 | 0.285 | 4.1E-05 |
| *tor-1* | WBGene00006597 | protein_coding_gene | -1.8 | 0.285 | 2.5E-02 |
| *ZC239.14* | WBGene00022572 | protein_coding_gene | -1.8 | 0.285 | 7.3E-04 |
| *R11D1.3* | WBGene00011244 | protein_coding_gene | -1.8 | 0.285 | 1.6E-07 |
| *F38B2.3* | WBGene00009530 | protein_coding_gene | -1.8 | 0.284 | 4.8E-02 |
| *F18C5.10* | WBGene00017563 | protein_coding_gene | -1.8 | 0.284 | 4.5E-53 |
| *K10D6.2* | WBGene00010742 | protein_coding_gene | -1.8 | 0.284 | 4.3E-13 |
| *C55A1.6* | WBGene00008323 | protein_coding_gene | -1.8 | 0.283 | 2.8E-03 |
| *Y48G9A.6* | WBGene00021700 | protein_coding_gene | -1.8 | 0.283 | 3.9E-02 |
| *hot-2* | WBGene00001987 | protein_coding_gene | -1.8 | 0.283 | 2.2E-03 |
| *C38H2.3* | WBGene00008020 | protein_coding_gene | -1.8 | 0.282 | 2.6E-07 |
| *glr-3* | WBGene00001614 | protein_coding_gene | -1.8 | 0.282 | 4.8E-02 |
| *T10B10.8* | WBGene00011684 | protein_coding_gene | -1.8 | 0.282 | 1.2E-11 |
| *C55C3.7* | WBGene00043065 | protein_coding_gene | -1.8 | 0.281 | 3.9E-02 |
| *anr-20* | WBGene00219581 | antisense_lncRNA_gene | -1.8 | 0.281 | 2.7E-02 |
| *pxd-1* | WBGene00007982 | protein_coding_gene | -1.8 | 0.281 | 2.0E-35 |
| *W02D7.8* | WBGene00020943 | protein_coding_gene | -1.8 | 0.281 | 8.1E-08 |
| *C01C4.2* | WBGene00015292 | protein_coding_gene | -1.8 | 0.281 | 3.6E-02 |
| *F13E9.11* | WBGene00008760 | protein_coding_gene | -1.8 | 0.281 | 1.2E-14 |
| *cut-5* | WBGene00011104 | protein_coding_gene | -1.8 | 0.279 | 1.8E-04 |
| *moe-3* | WBGene00003388 | protein_coding_gene | -1.8 | 0.279 | 3.8E-21 |
| *F53H2.1* | WBGene00010004 | protein_coding_gene | -1.8 | 0.279 | 5.5E-04 |
| *gcy-8* | WBGene00001535 | protein_coding_gene | -1.8 | 0.278 | 2.1E-03 |
| *F43D9.1* | WBGene00009653 | protein_coding_gene | -1.8 | 0.277 | 1.8E-06 |
| *ZK105.13* | WBGene00235114 | protein_coding_gene | -1.8 | 0.277 | 5.5E-05 |
| *T04C12.8* | WBGene00011430 | protein_coding_gene | -1.9 | 0.277 | 3.4E-08 |
| *bbs-1* | WBGene00000241 | protein_coding_gene | -1.9 | 0.277 | 2.0E-03 |
| *F21H7.3* | WBGene00009029 | protein_coding_gene | -1.9 | 0.276 | 2.9E-02 |
| *C02F4.4* | WBGene00007264 | protein_coding_gene | -1.9 | 0.276 | 1.5E-03 |
| *insc-1* | WBGene00018392 | protein_coding_gene | -1.9 | 0.275 | 2.3E-16 |
| *R07G3.8* | WBGene00019944 | protein_coding_gene | -1.9 | 0.275 | 4.5E-19 |
| *F58G6.9* | WBGene00044107 | protein_coding_gene | -1.9 | 0.275 | 1.5E-03 |
| *Y77E11A.14* | WBGene00022314 | protein_coding_gene | -1.9 | 0.274 | 8.3E-08 |
| *fax-1* | WBGene00001400 | protein_coding_gene | -1.9 | 0.273 | 2.6E-03 |
| *rga-2* | WBGene00013135 | protein_coding_gene | -1.9 | 0.273 | 1.8E-12 |
| *F19H6.6* | WBGene00008961 | protein_coding_gene | -1.9 | 0.273 | 4.1E-03 |
| *Y53F4B.20* | WBGene00013166 | protein_coding_gene | -1.9 | 0.273 | 6.7E-04 |
| *tbc-6* | WBGene00015410 | protein_coding_gene | -1.9 | 0.272 | 4.7E-23 |
| *Y6G8.2* | WBGene00012404 | protein_coding_gene | -1.9 | 0.271 | 2.1E-08 |
| *ZK550.2* | WBGene00013996 | protein_coding_gene | -1.9 | 0.271 | 1.4E-12 |
| *grh-1* | WBGene00001707 | protein_coding_gene | -1.9 | 0.271 | 1.6E-12 |
| *F40F8.12* | WBGene00044917 | protein_coding_gene | -1.9 | 0.270 | 1.1E-02 |
| *C16C10.9* | WBGene00007629 | protein_coding_gene | -1.9 | 0.270 | 9.9E-03 |
| *F16B12.1* | WBGene00008879 | protein_coding_gene | -1.9 | 0.269 | 2.1E-02 |
| *cfz-2* | WBGene00000478 | protein_coding_gene | -1.9 | 0.268 | 7.2E-11 |
| *F58A6.1* | WBGene00019022 | protein_coding_gene | -1.9 | 0.268 | 2.4E-20 |
| *hpo-36* | WBGene00020855 | protein_coding_gene | -1.9 | 0.268 | 2.9E-04 |
| *C31H5.7* | WBGene00007858 | protein_coding_gene | -1.9 | 0.267 | 9.9E-04 |
| *pals-31* | WBGene00018614 | protein_coding_gene | -1.9 | 0.267 | 2.5E-03 |
| *T24C2.5* | WBGene00011984 | protein_coding_gene | -1.9 | 0.267 | 4.4E-15 |
| *T01C3.11* | WBGene00011325 | protein_coding_gene | -1.9 | 0.266 | 3.4E-02 |
| *Y52B11A.4* | WBGene00013124 | protein_coding_gene | -1.9 | 0.266 | 2.4E-10 |
| *F32D8.15* | WBGene00009325 | protein_coding_gene | -1.9 | 0.265 | 2.3E-04 |
| *C17H12.37* | WBGene00236786 | protein_coding_gene | -1.9 | 0.265 | 7.8E-03 |
| *T01B10.5* | WBGene00020137 | protein_coding_gene | -1.9 | 0.265 | 7.7E-06 |
| *crtc-1* | WBGene00021237 | protein_coding_gene | -1.9 | 0.265 | 4.0E-26 |
| *C06G3.3* | WBGene00015549 | protein_coding_gene | -1.9 | 0.265 | 1.0E-06 |
| *npr-20* | WBGene00011578 | protein_coding_gene | -1.9 | 0.265 | 5.5E-10 |
| *F10D7.4* | WBGene00017341 | protein_coding_gene | -1.9 | 0.264 | 6.3E-03 |

| | | | | | |
|---|---|---|---|---|---|
| *trx-1* | WBGene00015062 | protein_coding_gene | -1.9 | 0.263 | 2.8E-02 |
| *K12C11.6* | WBGene00045052 | protein_coding_gene | -1.9 | 0.263 | 2.6E-15 |
| *hda-5* | WBGene00009663 | protein_coding_gene | -1.9 | 0.263 | 2.0E-14 |
| *F34H10.3* | WBGene00009380 | protein_coding_gene | -1.9 | 0.262 | 1.8E-37 |
| *unc-75* | WBGene00006807 | protein_coding_gene | -1.9 | 0.262 | 6.0E-05 |
| *Y57G11B.2* | WBGene00013295 | protein_coding_gene | -1.9 | 0.262 | 1.6E-15 |
| *ptr-9* | WBGene00004223 | protein_coding_gene | -1.9 | 0.261 | 2.4E-02 |
| *cutl-9* | WBGene00010893 | protein_coding_gene | -1.9 | 0.261 | 4.2E-11 |
| *fmi-1* | WBGene00001475 | protein_coding_gene | -1.9 | 0.261 | 3.1E-15 |
| *M151.1* | WBGene00019793 | protein_coding_gene | -1.9 | 0.261 | 1.4E-02 |
| *Y71G12B.31* | WBGene00044348 | protein_coding_gene | -1.9 | 0.261 | 5.2E-06 |
| *mab-3* | WBGene00003100 | protein_coding_gene | -1.9 | 0.261 | 1.4E-09 |
| *her-1* | WBGene00001842 | protein_coding_gene | -1.9 | 0.260 | 5.0E-07 |
| *F25D7.5* | WBGene00009114 | protein_coding_gene | -1.9 | 0.260 | 1.4E-03 |
| *F57B1.6* | WBGene00010189 | protein_coding_gene | -2.0 | 0.257 | 7.5E-07 |
| *ubc-8* | WBGene00006705 | protein_coding_gene | -2.0 | 0.257 | 1.7E-56 |
| *R08E5.4* | WBGene00019964 | protein_coding_gene | -2.0 | 0.257 | 1.4E-02 |
| *arl-6* | WBGene00000193 | protein_coding_gene | -2.0 | 0.257 | 1.3E-03 |
| *K01A6.8* | WBGene00194869 | protein_coding_gene | -2.0 | 0.256 | 4.1E-03 |
| *M01G12.10* | WBGene00010823 | protein_coding_gene | -2.0 | 0.255 | 2.5E-05 |
| *F08F1.4* | WBGene00017258 | protein_coding_gene | -2.0 | 0.255 | 5.4E-12 |
| *aqp-3* | WBGene00000171 | protein_coding_gene | -2.0 | 0.255 | 1.2E-04 |
| *F31C3.6* | WBGene00009288 | protein_coding_gene | -2.0 | 0.254 | 8.5E-19 |
| *Y46H3A.5* | WBGene00021601 | protein_coding_gene | -2.0 | 0.254 | 4.7E-03 |
| *akt-2* | WBGene00000103 | protein_coding_gene | -2.0 | 0.254 | 1.7E-28 |
| *clec-55* | WBGene00008597 | protein_coding_gene | -2.0 | 0.254 | 4.1E-09 |
| *ifa-3* | WBGene00002051 | protein_coding_gene | -2.0 | 0.253 | 8.1E-06 |
| *T20B3.1* | WBGene00011850 | protein_coding_gene | -2.0 | 0.253 | 3.0E-20 |
| *nas-19* | WBGene00003538 | protein_coding_gene | -2.0 | 0.253 | 2.9E-02 |
| *cpt-4* | WBGene00019644 | protein_coding_gene | -2.0 | 0.252 | 3.4E-18 |
| *nlp-45* | WBGene00020136 | protein_coding_gene | -2.0 | 0.252 | 1.0E-07 |
| *F02D10.3* | WBGene00008524 | protein_coding_gene | -2.0 | 0.251 | 4.9E-02 |
| *T18D3.7* | WBGene00011824 | protein_coding_gene | -2.0 | 0.250 | 4.7E-26 |
| *nep-14* | WBGene00018225 | pseudogene | -2.0 | 0.249 | 3.1E-11 |
| *nlp-46* | WBGene00012145 | protein_coding_gene | -2.0 | 0.249 | 1.7E-03 |
| *C36B1.6* | WBGene00007973 | protein_coding_gene | -2.0 | 0.249 | 2.2E-18 |
| *T20F5.8* | WBGene00077599 | protein_coding_gene | -2.0 | 0.249 | 4.1E-02 |
| *rocf-1* | WBGene00011948 | protein_coding_gene | -2.0 | 0.248 | 1.1E-05 |
| *F59A1.2* | WBGene00014779 | pseudogene | -2.0 | 0.248 | 4.9E-02 |
| *T19E7.6* | WBGene00044500 | protein_coding_gene | -2.0 | 0.248 | 1.8E-19 |
| *folr-1* | WBGene00007648 | protein_coding_gene | -2.0 | 0.247 | 1.1E-02 |
| *Y39H10B.3* | WBGene00194902 | protein_coding_gene | -2.0 | 0.247 | 2.2E-02 |
| *F17B5.1* | WBGene00008905 | protein_coding_gene | -2.0 | 0.246 | 2.3E-03 |
| *C09F5.1* | WBGene00015647 | protein_coding_gene | -2.0 | 0.246 | 3.1E-20 |
| *F32H2.11* | WBGene00043049 | protein_coding_gene | -2.0 | 0.245 | 1.6E-02 |
| *klp-11* | WBGene00002222 | protein_coding_gene | -2.0 | 0.245 | 6.0E-14 |
| *linc-116* | WBGene00219726 | lincRNA_gene | -2.0 | 0.245 | 2.4E-02 |
| *clec-78* | WBGene00018547 | protein_coding_gene | -2.0 | 0.245 | 1.5E-14 |
| *clec-44* | WBGene00019928 | protein_coding_gene | -2.0 | 0.244 | 1.7E-03 |
| *Y46E12BR.1* | WBGene00021598 | protein_coding_gene | -2.0 | 0.242 | 3.3E-04 |
| *W05B10.3* | WBGene00012278 | protein_coding_gene | -2.0 | 0.242 | 1.0E-02 |
| *F01G12.18* | WBGene00206382 | protein_coding_gene | -2.0 | 0.242 | 2.5E-02 |
| *ZK180.7* | WBGene00045248 | protein_coding_gene | -2.0 | 0.242 | 2.7E-03 |
| *srr-6* | WBGene00005657 | protein_coding_gene | -2.1 | 0.240 | 2.5E-09 |
| *ztf-30* | WBGene00015523 | protein_coding_gene | -2.1 | 0.240 | 2.8E-06 |
| *lgc-18* | WBGene00011360 | protein_coding_gene | -2.1 | 0.240 | 1.8E-02 |
| *AC3.9* | WBGene00044022 | pseudogene | -2.1 | 0.239 | 2.1E-02 |
| *F09E10.6* | WBGene00017296 | protein_coding_gene | -2.1 | 0.239 | 5.7E-04 |
| *srt-63* | WBGene00020877 | protein_coding_gene | -2.1 | 0.238 | 7.5E-03 |
| *lgc-23* | WBGene00007594 | protein_coding_gene | -2.1 | 0.238 | 2.1E-02 |
| *F11C1.9* | WBGene00195213 | protein_coding_gene | -2.1 | 0.238 | 4.9E-02 |
| *Y47D7A.18* | WBGene00235165 | protein_coding_gene | -2.1 | 0.238 | 4.4E-02 |
| *clec-23* | WBGene00011851 | pseudogene | -2.1 | 0.237 | 2.5E-04 |
| *fbn-1* | WBGene00022816 | protein_coding_gene | -2.1 | 0.237 | 1.6E-06 |
| *Y6E2A.1* | WBGene00012397 | protein_coding_gene | -2.1 | 0.236 | 2.3E-02 |
| *R160.4* | WBGene00020121 | protein_coding_gene | -2.1 | 0.233 | 2.3E-03 |
| *F52C9.5* | WBGene00018675 | protein_coding_gene | -2.1 | 0.232 | 6.8E-04 |
| *B0310.3* | WBGene00015139 | protein_coding_gene | -2.1 | 0.232 | 2.2E-10 |
| *clec-40* | WBGene00011853 | protein_coding_gene | -2.1 | 0.232 | 1.2E-04 |
| *K09C8.7* | WBGene00010717 | protein_coding_gene | -2.1 | 0.232 | 5.8E-04 |

| | | | | | |
|---|---|---|---|---|---|
| *C39D10.5* | WBGene00016532 | protein_coding_gene | -2.1 | 0.232 | 4.0E-03 |
| *Y37F4.8* | WBGene00044289 | protein_coding_gene | -2.1 | 0.231 | 6.5E-07 |
| *H08J19.1* | WBGene00010372 | protein_coding_gene | -2.1 | 0.230 | 4.7E-02 |
| *F34H10.9* | WBGene00219417 | pseudogene | -2.1 | 0.230 | 4.1E-02 |
| *M01A8.1* | WBGene00010795 | protein_coding_gene | -2.1 | 0.230 | 2.1E-15 |
| *unc-51* | WBGene00006786 | protein_coding_gene | -2.1 | 0.228 | 4.7E-52 |
| *F53F4.2* | WBGene00009986 | protein_coding_gene | -2.1 | 0.227 | 4.4E-02 |
| *dlc-5* | WBGene00043067 | protein_coding_gene | -2.1 | 0.225 | 1.5E-05 |
| *R10F2.5* | WBGene00019996 | protein_coding_gene | -2.1 | 0.225 | 3.5E-02 |
| *C43H6.6* | WBGene00016619 | protein_coding_gene | -2.2 | 0.225 | 1.9E-03 |
| *F26F12.3* | WBGene00017834 | protein_coding_gene | -2.2 | 0.225 | 6.0E-91 |
| *K11D5.2* | WBGene00235318 | ncRNA_gene | -2.2 | 0.224 | 4.5E-02 |
| *F21C10.10* | WBGene00017659 | protein_coding_gene | -2.2 | 0.224 | 1.4E-06 |
| *H23N18.4* | WBGene00019235 | protein_coding_gene | -2.2 | 0.223 | 2.2E-03 |
| *F44E2.4* | WBGene00018418 | protein_coding_gene | -2.2 | 0.223 | 1.6E-04 |
| *Y2H9A.6* | WBGene00086568 | protein_coding_gene | -2.2 | 0.223 | 5.5E-06 |
| *T04C12.32* | WBGene00235086 | ncRNA_gene | -2.2 | 0.223 | 4.4E-03 |
| *ugt-54* | WBGene00012013 | protein_coding_gene | -2.2 | 0.222 | 5.2E-26 |
| *F10G2.2* | WBGene00017363 | protein_coding_gene | -2.2 | 0.222 | 3.7E-02 |
| *B0462.1* | WBGene00007181 | pseudogene | -2.2 | 0.222 | 4.3E-08 |
| *C54F6.6* | WBGene00016924 | protein_coding_gene | -2.2 | 0.222 | 8.8E-03 |
| *dhp-1* | WBGene00000963 | protein_coding_gene | -2.2 | 0.221 | 5.5E-21 |
| *gstk-2* | WBGene00017054 | protein_coding_gene | -2.2 | 0.221 | 1.4E-13 |
| *aqp-5* | WBGene00000173 | protein_coding_gene | -2.2 | 0.221 | 2.6E-03 |
| *chil-16* | WBGene00011158 | protein_coding_gene | -2.2 | 0.220 | 2.3E-02 |
| *C32E8.13* | WBGene00284853 | protein_coding_gene | -2.2 | 0.220 | 4.3E-02 |
| *R03H10.5* | WBGene00019857 | protein_coding_gene | -2.2 | 0.220 | 4.7E-02 |
| *B0222.15* | WBGene00235316 | protein_coding_gene | -2.2 | 0.219 | 3.7E-02 |
| *K06A4.10* | WBGene00077497 | protein_coding_gene | -2.2 | 0.219 | 9.7E-03 |
| *F59A1.18* | WBGene00206506 | protein_coding_gene | -2.2 | 0.218 | 3.0E-02 |
| *mltn-8* | WBGene00021451 | protein_coding_gene | -2.2 | 0.218 | 3.2E-02 |
| *hsp-12.6* | WBGene00002013 | protein_coding_gene | -2.2 | 0.217 | 5.5E-11 |
| *F23F12.8* | WBGene00017754 | protein_coding_gene | -2.2 | 0.217 | 1.9E-04 |
| *ugt-52* | WBGene00018931 | protein_coding_gene | -2.2 | 0.216 | 3.2E-16 |
| *clec-2* | WBGene00015198 | protein_coding_gene | -2.2 | 0.216 | 3.4E-02 |
| *F58E6.4* | WBGene00010253 | protein_coding_gene | -2.2 | 0.215 | 2.6E-02 |
| *pitr-2* | WBGene00015054 | protein_coding_gene | -2.2 | 0.215 | 3.3E-02 |
| *ptr-4* | WBGene00004219 | protein_coding_gene | -2.2 | 0.214 | 1.7E-04 |
| *gcy-34* | WBGene00001554 | protein_coding_gene | -2.2 | 0.214 | 4.3E-04 |
| *srab-3* | WBGene00015448 | protein_coding_gene | -2.2 | 0.213 | 3.2E-02 |
| *C23G10.12* | WBGene00219827 | snoRNA_gene | -2.2 | 0.213 | 3.1E-02 |
| *T10H4.13* | WBGene00045339 | protein_coding_gene | -2.2 | 0.211 | 1.1E-03 |
| *mboa-1* | WBGene00007174 | protein_coding_gene | -2.2 | 0.210 | 4.7E-11 |
| *T21B4.21* | WBGene00220076 | ncRNA_gene | -2.2 | 0.210 | 2.2E-13 |
| *C39E6.2* | WBGene00016537 | protein_coding_gene | -2.3 | 0.210 | 2.8E-02 |
| *F41C6.6* | WBGene00018279 | protein_coding_gene | -2.3 | 0.210 | 7.7E-03 |
| *C14C6.13* | WBGene00015764 | protein_coding_gene | -2.3 | 0.209 | 3.8E-02 |
| *gst-29* | WBGene00001777 | protein_coding_gene | -2.3 | 0.209 | 2.4E-05 |
| *M199.9* | WBGene00044998 | protein_coding_gene | -2.3 | 0.208 | 1.9E-02 |
| *ptr-19* | WBGene00004233 | protein_coding_gene | -2.3 | 0.208 | 1.7E-04 |
| *nlp-23* | WBGene00003761 | protein_coding_gene | -2.3 | 0.207 | 3.0E-02 |
| *C03A7.2* | WBGene00015366 | protein_coding_gene | -2.3 | 0.206 | 2.2E-04 |
| *R10D12.1* | WBGene00011185 | protein_coding_gene | -2.3 | 0.206 | 2.2E-10 |
| *F57C12.6* | WBGene00194687 | protein_coding_gene | -2.3 | 0.206 | 1.6E-07 |
| *tig-3* | WBGene00021594 | protein_coding_gene | -2.3 | 0.206 | 2.2E-06 |
| *T06H11.11* | WBGene00197930 | ncRNA_gene | -2.3 | 0.205 | 2.9E-02 |
| *srbc-20* | WBGene00016693 | protein_coding_gene | -2.3 | 0.205 | 3.2E-02 |
| *T24E12.3* | WBGene00020773 | protein_coding_gene | -2.3 | 0.204 | 2.1E-02 |
| *E04A4.3* | WBGene00017118 | pseudogene | -2.3 | 0.204 | 1.9E-03 |
| *F16H6.8* | WBGene00008898 | protein_coding_gene | -2.3 | 0.203 | 3.0E-02 |
| *Y42A5A.3* | WBGene00012778 | protein_coding_gene | -2.3 | 0.203 | 3.4E-07 |
| *ZC443.3* | WBGene00013898 | protein_coding_gene | -2.3 | 0.202 | 2.3E-53 |
| *W04A4.3* | WBGene00012232 | protein_coding_gene | -2.3 | 0.201 | 3.1E-02 |
| *K10H10.10* | WBGene00010765 | protein_coding_gene | -2.3 | 0.200 | 3.6E-04 |
| *C25B8.8* | WBGene00044297 | protein_coding_gene | -2.3 | 0.200 | 4.5E-04 |
| *nas-3* | WBGene00003522 | protein_coding_gene | -2.3 | 0.200 | 9.2E-03 |
| *clec-11* | WBGene00008294 | protein_coding_gene | -2.3 | 0.199 | 4.5E-02 |
| *clec-242* | WBGene00013471 | protein_coding_gene | -2.3 | 0.199 | 1.3E-03 |
| *Y6E2A.4* | WBGene00012398 | protein_coding_gene | -2.3 | 0.199 | 2.8E-19 |
| *F47G9.6* | WBGene00009833 | protein_coding_gene | -2.3 | 0.199 | 3.0E-02 |

| | | | | | |
|---|---|---|---|---|---|
| *F20A1.8* | WBGene00017619 | protein_coding_gene | -2.3 | 0.199 | 1.4E-03 |
| *R10E8.8* | WBGene00011213 | protein_coding_gene | -2.3 | 0.198 | 7.8E-19 |
| *R05A10.5* | WBGene00011022 | pseudogene | -2.3 | 0.198 | 2.4E-02 |
| *ZK1307.2* | WBGene00014245 | protein_coding_gene | -2.3 | 0.197 | 3.0E-07 |
| *K08E3.2* | WBGene00010662 | protein_coding_gene | -2.3 | 0.197 | 2.0E-04 |
| *pag-3* | WBGene00003909 | protein_coding_gene | -2.3 | 0.196 | 1.1E-05 |
| *Y53C12B.7* | WBGene00013146 | protein_coding_gene | -2.3 | 0.196 | 3.9E-19 |
| *C54F6.5* | WBGene00016923 | protein_coding_gene | -2.4 | 0.196 | 1.6E-16 |
| *hmit-1.3* | WBGene00010811 | protein_coding_gene | -2.4 | 0.195 | 1.3E-54 |
| *W05H9.3* | WBGene00021050 | ncRNA_gene | -2.4 | 0.195 | 1.0E-42 |
| *chil-11* | WBGene00016665 | protein_coding_gene | -2.4 | 0.194 | 6.2E-04 |
| *trf-1* | WBGene00006612 | protein_coding_gene | -2.4 | 0.194 | 5.0E-04 |
| *dod-23* | WBGene00009898 | protein_coding_gene | -2.4 | 0.194 | 3.3E-12 |
| *M03A1.8* | WBGene00077490 | protein_coding_gene | -2.4 | 0.194 | 1.6E-07 |
| *W06D12.6* | WBGene00012305 | protein_coding_gene | -2.4 | 0.194 | 2.6E-02 |
| *F19H6.5* | WBGene00008960 | protein_coding_gene | -2.4 | 0.193 | 3.7E-04 |
| *dos-3* | WBGene00019331 | protein_coding_gene | -2.4 | 0.192 | 1.7E-02 |
| *cyp-35A1* | WBGene00015399 | protein_coding_gene | -2.4 | 0.192 | 4.4E-03 |
| *coel-1* | WBGene00016866 | protein_coding_gene | -2.4 | 0.191 | 5.8E-49 |
| *pals-7* | WBGene00007661 | protein_coding_gene | -2.4 | 0.191 | 2.2E-03 |
| *F11D5.5* | WBGene00017382 | protein_coding_gene | -2.4 | 0.190 | 1.2E-10 |
| *C54C8.4* | WBGene00008292 | protein_coding_gene | -2.4 | 0.190 | 2.6E-03 |
| *F16H6.10* | WBGene00008900 | protein_coding_gene | -2.4 | 0.189 | 1.9E-03 |
| *T26H5.8* | WBGene00012070 | protein_coding_gene | -2.4 | 0.188 | 1.9E-24 |
| *ZK402.5* | WBGene00022731 | protein_coding_gene | -2.4 | 0.187 | 1.9E-04 |
| *C38C5.1* | WBGene00016518 | protein_coding_gene | -2.4 | 0.186 | 1.4E-02 |
| *T03E6.8* | WBGene00011397 | protein_coding_gene | -2.4 | 0.185 | 1.6E-06 |
| *R09E10.5* | WBGene00011175 | protein_coding_gene | -2.4 | 0.185 | 1.5E-05 |
| *ZC15.5* | WBGene00013833 | protein_coding_gene | -2.4 | 0.184 | 1.9E-02 |
| *T09D3.8* | WBGene00044538 | protein_coding_gene | -2.4 | 0.183 | 5.5E-03 |
| *C29F9.5* | WBGene00016220 | protein_coding_gene | -2.4 | 0.183 | 1.4E-02 |
| *nmr-1* | WBGene00003774 | protein_coding_gene | -2.4 | 0.183 | 1.8E-03 |
| *linc-23* | WBGene00219692 | lincRNA_gene | -2.5 | 0.181 | 1.6E-02 |
| *C04H5.9* | WBGene00044054 | pseudogene | -2.5 | 0.179 | 1.4E-02 |
| *clec-109* | WBGene00008907 | protein_coding_gene | -2.5 | 0.177 | 1.3E-02 |
| *F38A1.11* | WBGene00009524 | protein_coding_gene | -2.5 | 0.177 | 1.3E-02 |
| *F02H6.7* | WBGene00008540 | protein_coding_gene | -2.5 | 0.176 | 2.0E-02 |
| *F17B5.6* | WBGene00044058 | protein_coding_gene | -2.5 | 0.176 | 4.0E-03 |
| *K09C4.4* | WBGene00019549 | protein_coding_gene | -2.5 | 0.176 | 3.5E-03 |
| *chil-9* | WBGene00014162 | protein_coding_gene | -2.5 | 0.176 | 1.5E-02 |
| *F14D7.11* | WBGene00008798 | protein_coding_gene | -2.5 | 0.175 | 1.7E-02 |
| *ins-19* | WBGene00002102 | protein_coding_gene | -2.5 | 0.175 | 9.8E-06 |
| *F17E9.15* | WBGene00194833 | protein_coding_gene | -2.5 | 0.175 | 1.6E-02 |
| *clec-216* | WBGene00017822 | protein_coding_gene | -2.5 | 0.175 | 2.3E-02 |
| *C34C12.7* | WBGene00007926 | protein_coding_gene | -2.5 | 0.174 | 1.1E-21 |
| *F13H6.5* | WBGene00017433 | protein_coding_gene | -2.5 | 0.174 | 3.2E-14 |
| *F42C5.3* | WBGene00018345 | protein_coding_gene | -2.5 | 0.173 | 1.5E-08 |
| *F18G5.5* | WBGene00017581 | pseudogene | -2.5 | 0.171 | 1.7E-02 |
| *K09C4.1* | WBGene00019547 | protein_coding_gene | -2.5 | 0.171 | 1.1E-19 |
| *K01A2.10* | WBGene00019284 | protein_coding_gene | -2.6 | 0.171 | 7.3E-34 |
| *nlp-48* | WBGene00008002 | protein_coding_gene | -2.6 | 0.170 | 6.3E-06 |
| *cutl-11* | WBGene00011443 | protein_coding_gene | -2.6 | 0.170 | 7.6E-06 |
| *Y19D10B.1* | WBGene00021230 | pseudogene | -2.6 | 0.168 | 2.8E-03 |
| *T09A12.8* | WBGene00235275 | protein_coding_gene | -2.6 | 0.168 | 5.7E-05 |
| *E02C12.12* | WBGene00017097 | protein_coding_gene | -2.6 | 0.168 | 1.4E-02 |
| *rom-2* | WBGene00004401 | protein_coding_gene | -2.6 | 0.167 | 1.3E-08 |
| *Y71F9B.9* | WBGene00022130 | protein_coding_gene | -2.6 | 0.166 | 1.2E-30 |
| *Y51H4A.937* | WBGene00194927 | protein_coding_gene | -2.6 | 0.166 | 1.7E-02 |
| *slc-17.9* | WBGene00011643 | protein_coding_gene | -2.6 | 0.166 | 1.6E-08 |
| *T16G1.4* | WBGene00011798 | protein_coding_gene | -2.6 | 0.166 | 5.2E-13 |
| *C44C10.15* | WBGene00219301 | protein_coding_gene | -2.6 | 0.165 | 1.8E-02 |
| *T19B10.12* | WBGene00050898 | protein_coding_gene | -2.6 | 0.165 | 1.6E-02 |
| *pals-32* | WBGene00016281 | protein_coding_gene | -2.6 | 0.165 | 3.8E-19 |
| *Y7A5A.5* | WBGene00012411 | protein_coding_gene | -2.6 | 0.164 | 2.1E-02 |
| *F52E1.12* | WBGene00018699 | pseudogene | -2.6 | 0.164 | 7.3E-03 |
| *F19C7.2* | WBGene00017592 | protein_coding_gene | -2.6 | 0.164 | 2.1E-35 |
| *F45D3.15* | WBGene00201625 | ncRNA_gene | -2.6 | 0.164 | 1.7E-02 |
| *Y54G2A.11* | WBGene00021876 | protein_coding_gene | -2.6 | 0.164 | 1.4E-49 |
| *Y47D7A.12* | WBGene00021624 | protein_coding_gene | -2.6 | 0.163 | 2.1E-06 |
| *C05C9.1* | WBGene00007325 | protein_coding_gene | -2.6 | 0.163 | 8.0E-04 |

| | | | | | |
|---|---|---|---|---|---|
| *Y50E8A.1* | WBGene00013046 | protein_coding_gene | -2.6 | 0.163 | 1.7E-14 |
| *ZK546.19* | WBGene00195246 | protein_coding_gene | -2.6 | 0.162 | 3.5E-03 |
| *fbxb-1* | WBGene00012063 | protein_coding_gene | -2.6 | 0.162 | 4.8E-03 |
| *xtr-2* | WBGene00006966 | protein_coding_gene | -2.6 | 0.162 | 4.4E-03 |
| *Y47H10A.5* | WBGene00012961 | protein_coding_gene | -2.6 | 0.161 | 1.3E-16 |
| *M01B2.13* | WBGene00045338 | protein_coding_gene | -2.6 | 0.161 | 1.5E-11 |
| *R09D1.12* | WBGene00011168 | protein_coding_gene | -2.6 | 0.160 | 1.5E-10 |
| *F33E2.7* | WBGene00009363 | protein_coding_gene | -2.7 | 0.159 | 9.9E-03 |
| *pals-5* | WBGene00007659 | protein_coding_gene | -2.7 | 0.159 | 5.9E-03 |
| *F40H6.2* | WBGene00018254 | protein_coding_gene | -2.7 | 0.159 | 1.5E-02 |
| *C08D8.1* | WBGene00015592 | protein_coding_gene | -2.7 | 0.156 | 2.6E-12 |
| *Y70C5A.3* | WBGene00050904 | protein_coding_gene | -2.7 | 0.155 | 1.1E-20 |
| *C49C8.6* | WBGene00016770 | protein_coding_gene | -2.7 | 0.155 | 3.3E-03 |
| *btb-15* | WBGene00015594 | pseudogene | -2.7 | 0.154 | 4.7E-04 |
| *B0228.8* | WBGene00015065 | protein_coding_gene | -2.7 | 0.154 | 2.4E-03 |
| *C25H3.10* | WBGene00016119 | protein_coding_gene | -2.7 | 0.154 | 3.5E-39 |
| *F54F3.2* | WBGene00014770 | pseudogene | -2.7 | 0.152 | 1.1E-02 |
| *clec-69* | WBGene00044642 | pseudogene | -2.7 | 0.151 | 1.4E-04 |
| *F17A2.16* | WBGene00304215 | protein_coding_gene | -2.7 | 0.151 | 5.5E-03 |
| *T11F9.22* | WBGene00194784 | protein_coding_gene | -2.7 | 0.150 | 9.1E-06 |
| *F53B7.7* | WBGene00009969 | protein_coding_gene | -2.8 | 0.148 | 1.9E-04 |
| *nspe-6* | WBGene00012603 | protein_coding_gene | -2.8 | 0.147 | 1.2E-04 |
| *cyp-13A11* | WBGene00008809 | protein_coding_gene | -2.8 | 0.145 | 4.9E-03 |
| *K06A9.2* | WBGene00019436 | protein_coding_gene | -2.8 | 0.145 | 5.3E-30 |
| *C54F6.12* | WBGene00016929 | protein_coding_gene | -2.8 | 0.144 | 6.3E-08 |
| *Y57E12B.10* | WBGene00202501 | protein_coding_gene | -2.8 | 0.144 | 5.1E-04 |
| *F16G10.2* | WBGene00017517 | protein_coding_gene | -2.8 | 0.143 | 5.7E-03 |
| *F41C3.2* | WBGene00018268 | protein_coding_gene | -2.8 | 0.143 | 5.4E-28 |
| *clec-77* | WBGene00044343 | pseudogene | -2.8 | 0.142 | 7.3E-73 |
| *K07E8.1* | WBGene00019494 | pseudogene | -2.8 | 0.142 | 5.3E-03 |
| *K04F1.8* | WBGene00019387 | protein_coding_gene | -2.8 | 0.141 | 5.7E-03 |
| *C35C5.9* | WBGene00007958 | protein_coding_gene | -2.8 | 0.141 | 2.5E-36 |
| *C44B12.9* | WBGene00044462 | protein_coding_gene | -2.8 | 0.140 | 8.1E-13 |
| *R160.10* | WBGene00219369 | protein_coding_gene | -2.8 | 0.139 | 3.0E-06 |
| *C04A11.5* | WBGene00007288 | protein_coding_gene | -2.9 | 0.138 | 6.2E-11 |
| *ugt-4* | WBGene00013905 | protein_coding_gene | -2.9 | 0.138 | 2.5E-32 |
| *scl-18* | WBGene00011724 | protein_coding_gene | -2.9 | 0.138 | 2.7E-04 |
| *C42D4.1* | WBGene00016594 | protein_coding_gene | -2.9 | 0.138 | 8.0E-79 |
| *F29C4.4* | WBGene00017927 | protein_coding_gene | -2.9 | 0.138 | 9.2E-03 |
| *R09E10.13* | WBGene00194658 | protein_coding_gene | -2.9 | 0.138 | 2.3E-07 |
| *cyp-13A5* | WBGene00011672 | protein_coding_gene | -2.9 | 0.138 | 2.1E-65 |
| *Y32B12C.1* | WBGene00012525 | protein_coding_gene | -2.9 | 0.137 | 2.3E-03 |
| *C27A7.8* | WBGene00007758 | protein_coding_gene | -2.9 | 0.137 | 1.5E-27 |
| *C39B5.10* | WBGene00016528 | protein_coding_gene | -2.9 | 0.136 | 7.1E-44 |
| *cyp-33C7* | WBGene00018260 | protein_coding_gene | -2.9 | 0.135 | 2.3E-41 |
| *cyp-35B1* | WBGene00019472 | protein_coding_gene | -2.9 | 0.135 | 8.4E-11 |
| *faah-2* | WBGene00015048 | protein_coding_gene | -2.9 | 0.133 | 1.3E-20 |
| *F49C5.12* | WBGene00184956 | protein_coding_gene | -2.9 | 0.133 | 1.5E-03 |
| *cyp-13B1* | WBGene00008519 | protein_coding_gene | -2.9 | 0.132 | 3.9E-19 |
| *Y70G10A.2* | WBGene00013498 | protein_coding_gene | -2.9 | 0.130 | 1.1E-03 |
| *W05H9.1* | WBGene00021048 | protein_coding_gene | -2.9 | 0.130 | 5.5E-100 |
| *W08F4.7* | WBGene00021096 | protein_coding_gene | -2.9 | 0.130 | 1.6E-10 |
| *Y54G2A.37* | WBGene00023484 | protein_coding_gene | -3.0 | 0.129 | 5.7E-10 |
| *F49C5.10* | WBGene00044162 | protein_coding_gene | -3.0 | 0.128 | 2.4E-03 |
| *tyr-5* | WBGene00021553 | protein_coding_gene | -3.0 | 0.128 | 1.7E-23 |
| *C54E10.10* | WBGene00206523 | protein_coding_gene | -3.0 | 0.127 | 4.0E-03 |
| *Y71A12B.24* | WBGene00195228 | protein_coding_gene | -3.0 | 0.126 | 8.3E-06 |
| *Y32B12C.5* | WBGene00194769 | pseudogene | -3.0 | 0.124 | 2.3E-13 |
| *Y50E8A.8* | WBGene00013051 | pseudogene | -3.0 | 0.124 | 1.2E-03 |
| *F10D7.3* | WBGene00017340 | protein_coding_gene | -3.0 | 0.123 | 1.4E-45 |
| *F25E5.4* | WBGene00017785 | protein_coding_gene | -3.0 | 0.123 | 4.7E-70 |
| *F56H9.9* | WBGene00206460 | protein_coding_gene | -3.0 | 0.122 | 3.5E-04 |
| *clec-52* | WBGene00015052 | protein_coding_gene | -3.0 | 0.122 | 1.3E-15 |
| *ugt-64* | WBGene00015577 | protein_coding_gene | -3.0 | 0.122 | 9.2E-76 |
| *C29F7.11* | WBGene00206531 | ncRNA_gene | -3.0 | 0.121 | 5.3E-04 |
| *F55G11.10* | WBGene00010129 | protein_coding_gene | -3.1 | 0.121 | 5.1E-03 |
| *oac-29* | WBGene00018295 | protein_coding_gene | -3.1 | 0.120 | 9.5E-08 |
| *clec-93* | WBGene00013929 | protein_coding_gene | -3.1 | 0.118 | 1.6E-03 |
| *str-163* | WBGene00006208 | protein_coding_gene | -3.1 | 0.118 | 2.1E-07 |
| *nspe-5* | WBGene00012594 | protein_coding_gene | -3.1 | 0.118 | 4.9E-04 |

| | | | | | |
|---|---|---|---|---|---|
| *clec-33* | WBGene00009859 | protein_coding_gene | -3.1 | 0.117 | 8.6E-07 |
| *sodh-1* | WBGene00010790 | protein_coding_gene | -3.1 | 0.116 | 3.9E-11 |
| *asp-11* | WBGene00017677 | pseudogene | -3.1 | 0.116 | 4.9E-08 |
| *C35A11.3* | WBGene00016430 | protein_coding_gene | -3.1 | 0.115 | 1.7E-03 |
| *Y39H10A.1* | WBGene00021478 | protein_coding_gene | -3.1 | 0.115 | 4.8E-04 |
| *Y69H2.10* | WBGene00013485 | protein_coding_gene | -3.1 | 0.113 | 3.8E-08 |
| *srbc-82* | WBGene00007832 | protein_coding_gene | -3.2 | 0.112 | 1.6E-03 |
| *clec-27* | WBGene00013607 | protein_coding_gene | -3.2 | 0.112 | 1.6E-03 |
| *chil-18* | WBGene00011161 | protein_coding_gene | -3.2 | 0.112 | 9.8E-06 |
| *Y18H1A.8* | WBGene00021212 | protein_coding_gene | -3.2 | 0.111 | 5.5E-14 |
| *aat-8* | WBGene00000009 | protein_coding_gene | -3.2 | 0.111 | 7.1E-12 |
| *F55C9.12* | WBGene00044196 | pseudogene | -3.2 | 0.111 | 1.5E-03 |
| *Y62H9A.8* | WBGene00013396 | protein_coding_gene | -3.2 | 0.110 | 1.8E-03 |
| *bigr-1* | WBGene00009512 | protein_coding_gene | -3.2 | 0.110 | 2.0E-06 |
| *K08C9.10* | WBGene00086564 | protein_coding_gene | -3.2 | 0.109 | 1.1E-03 |
| *T19H5.6* | WBGene00044807 | protein_coding_gene | -3.2 | 0.108 | 6.4E-24 |
| *ZK512.7* | WBGene00013986 | protein_coding_gene | -3.2 | 0.107 | 4.7E-18 |
| *Y71F9AM.9* | WBGene00271781 | lincRNA_gene | -3.2 | 0.107 | 1.7E-06 |
| *C23G10.11* | WBGene00016017 | protein_coding_gene | -3.2 | 0.106 | 2.6E-04 |
| *T08G5.1* | WBGene00011622 | protein_coding_gene | -3.2 | 0.106 | 2.1E-05 |
| *ZC204.17* | WBGene00194714 | protein_coding_gene | -3.2 | 0.105 | 1.8E-03 |
| *fil-1* | WBGene00011321 | protein_coding_gene | -3.2 | 0.105 | 4.4E-41 |
| *Y39A1A.27* | WBGene00194731 | protein_coding_gene | -3.3 | 0.105 | 4.1E-14 |
| *F25B3.2* | WBGene00009099 | protein_coding_gene | -3.3 | 0.104 | 1.8E-07 |
| *lov-1* | WBGene00003058 | protein_coding_gene | -3.3 | 0.104 | 6.1E-05 |
| *Y26G10.3* | WBGene00050973 | pseudogene | -3.3 | 0.104 | 1.2E-03 |
| *W10G11.1* | WBGene00021134 | protein_coding_gene | -3.3 | 0.103 | 7.4E-05 |
| *C07G3.10* | WBGene00015578 | protein_coding_gene | -3.3 | 0.103 | 7.6E-04 |
| *swt-1* | WBGene00019300 | protein_coding_gene | -3.3 | 0.102 | 3.4E-40 |
| *F13E9.4* | WBGene00008753 | protein_coding_gene | -3.3 | 0.102 | 1.6E-04 |
| *C05B5.1* | WBGene00007319 | protein_coding_gene | -3.3 | 0.102 | 1.1E-03 |
| *clec-75* | WBGene00021586 | protein_coding_gene | -3.3 | 0.100 | 5.3E-07 |
| *F09F3.8* | WBGene00014732 | pseudogene | -3.3 | 0.100 | 7.4E-07 |
| *Y19D10B.4* | WBGene00021233 | pseudogene | -3.3 | 0.099 | 1.2E-04 |
| *D2096.5* | WBGene00017072 | protein_coding_gene | -3.3 | 0.098 | 5.1E-23 |
| *sri-23* | WBGene00005535 | pseudogene | -3.4 | 0.098 | 5.0E-04 |
| *F28B1.3* | WBGene00009197 | pseudogene | -3.4 | 0.098 | 8.7E-04 |
| *E02H9.1* | WBGene00017099 | protein_coding_gene | -3.4 | 0.097 | 7.4E-19 |
| *Y57G11C.1132* | WBGene00194666 | antisense_lncRNA_gene | -3.4 | 0.097 | 2.7E-04 |
| *ftn-1* | WBGene00001500 | protein_coding_gene | -3.4 | 0.097 | 9.4E-15 |
| *Y43C5A.3* | WBGene00012783 | protein_coding_gene | -3.4 | 0.097 | 5.4E-82 |
| *K01A6.7* | WBGene00045096 | protein_coding_gene | -3.4 | 0.096 | 9.5E-06 |
| *grsp-3* | WBGene00016403 | protein_coding_gene | -3.4 | 0.096 | 7.9E-21 |
| *K01A6.4* | WBGene00010446 | protein_coding_gene | -3.4 | 0.095 | 3.1E-08 |
| *C45B2.3* | WBGene00016660 | protein_coding_gene | -3.4 | 0.095 | 6.9E-04 |
| *F11E6.11* | WBGene00023501 | protein_coding_gene | -3.4 | 0.095 | 1.1E-41 |
| *cwp-5* | WBGene00009844 | protein_coding_gene | -3.4 | 0.095 | 1.7E-04 |
| *ZK666.14* | WBGene00194952 | protein_coding_gene | -3.4 | 0.094 | 5.0E-05 |
| *C14E2.4* | WBGene00015773 | protein_coding_gene | -3.4 | 0.092 | 2.4E-04 |
| *clec-32* | WBGene00009860 | protein_coding_gene | -3.4 | 0.092 | 1.5E-12 |
| *F28B1.2* | WBGene00009196 | protein_coding_gene | -3.5 | 0.091 | 5.1E-04 |
| *abf-1* | WBGene00000012 | protein_coding_gene | -3.5 | 0.091 | 1.4E-14 |
| *dod-3* | WBGene00016052 | protein_coding_gene | -3.5 | 0.090 | 2.8E-14 |
| *F29A7.3* | WBGene00017913 | protein_coding_gene | -3.5 | 0.089 | 1.5E-04 |
| *Y17G7B.23* | WBGene00044030 | protein_coding_gene | -3.5 | 0.089 | 8.5E-09 |
| *C41G7.15* | WBGene00219284 | protein_coding_gene | -3.5 | 0.088 | 7.4E-05 |
| *tre-5* | WBGene00006611 | protein_coding_gene | -3.5 | 0.086 | 5.4E-43 |
| *C54E10.11* | WBGene00206524 | protein_coding_gene | -3.5 | 0.086 | 5.3E-04 |
| *Y75B12B.13* | WBGene00045402 | protein_coding_gene | -3.6 | 0.085 | 3.4E-04 |
| *F21H7.12* | WBGene00009033 | protein_coding_gene | -3.6 | 0.085 | 1.0E-05 |
| *C37A5.6* | WBGene00014700 | protein_coding_gene | -3.6 | 0.085 | 5.7E-04 |
| *hil-1* | WBGene00001852 | protein_coding_gene | -3.6 | 0.084 | 2.3E-30 |
| *fipr-24* | WBGene00007992 | protein_coding_gene | -3.6 | 0.083 | 3.1E-08 |
| *T10D4.7* | WBGene00020409 | protein_coding_gene | -3.6 | 0.083 | 5.6E-04 |
| *F28C6.10* | WBGene00009209 | protein_coding_gene | -3.6 | 0.083 | 2.5E-40 |
| *clec-94* | WBGene00013928 | protein_coding_gene | -3.6 | 0.082 | 2.4E-04 |
| *pitr-4* | WBGene00007141 | protein_coding_gene | -3.6 | 0.082 | 2.6E-06 |
| *C27D6.15* | WBGene00303362 | protein_coding_gene | -3.6 | 0.081 | 3.3E-04 |
| *clec-162* | WBGene00007399 | protein_coding_gene | -3.6 | 0.081 | 2.1E-08 |
| *far-7* | WBGene00001391 | protein_coding_gene | -3.6 | 0.081 | 8.0E-56 |

| | | | | | |
|---|---|---|---|---|---|
| *F26C11.3* | WBGene00009147 | protein_coding_gene | -3.6 | 0.080 | 5.0E-05 |
| *T12A2.6* | WBGene00020440 | protein_coding_gene | -3.7 | 0.079 | 3.1E-04 |
| *Y43F8C.23* | WBGene00045344 | protein_coding_gene | -3.7 | 0.078 | 2.0E-04 |
| *F59C6.16* | WBGene00077764 | protein_coding_gene | -3.7 | 0.077 | 3.9E-66 |
| *F02H6.6* | WBGene00008539 | protein_coding_gene | -3.7 | 0.077 | 1.8E-16 |
| *H32K16.2* | WBGene00010422 | protein_coding_gene | -3.7 | 0.077 | 9.9E-20 |
| *F16B4.7* | WBGene00017509 | protein_coding_gene | -3.7 | 0.077 | 3.2E-11 |
| *F46B3.14* | WBGene00009765 | protein_coding_gene | -3.7 | 0.076 | 1.5E-04 |
| *F46C5.1* | WBGene00009778 | protein_coding_gene | -3.7 | 0.076 | 8.1E-11 |
| *W07G1.7* | WBGene00012334 | protein_coding_gene | -3.7 | 0.075 | 1.3E-06 |
| *E02C12.10* | WBGene00017095 | protein_coding_gene | -3.7 | 0.075 | 1.9E-04 |
| *M02E1.4* | WBGene00284848 | protein_coding_gene | -3.7 | 0.075 | 6.3E-12 |
| *F13E9.9* | WBGene00008758 | protein_coding_gene | -3.8 | 0.074 | 5.2E-10 |
| *C18D11.6* | WBGene00007682 | protein_coding_gene | -3.8 | 0.072 | 1.0E-08 |
| *fipr-26* | WBGene00009963 | protein_coding_gene | -3.8 | 0.072 | 1.8E-05 |
| *try-5* | WBGene00006623 | protein_coding_gene | -3.8 | 0.072 | 3.0E-18 |
| *Y73F8A.10* | WBGene00013519 | protein_coding_gene | -3.8 | 0.071 | 2.7E-08 |
| *Y64G10A.10* | WBGene00045144 | protein_coding_gene | -3.8 | 0.070 | 6.9E-05 |
| *C16C8.17* | WBGene00015854 | protein_coding_gene | -3.9 | 0.069 | 7.6E-07 |
| *T02E9.6* | WBGene00044203 | protein_coding_gene | -3.9 | 0.069 | 6.2E-06 |
| *clec-232* | WBGene00009487 | protein_coding_gene | -3.9 | 0.069 | 1.4E-33 |
| *clec-24* | WBGene00009856 | protein_coding_gene | -3.9 | 0.068 | 5.6E-36 |
| *C35B1.8* | WBGene00016437 | protein_coding_gene | -3.9 | 0.068 | 8.2E-13 |
| *C37A5.3* | WBGene00014698 | protein_coding_gene | -3.9 | 0.067 | 4.1E-05 |
| *cnc-11* | WBGene00044900 | protein_coding_gene | -3.9 | 0.067 | 1.4E-14 |
| *F59A6.3* | WBGene00019085 | protein_coding_gene | -3.9 | 0.066 | 1.9E-05 |
| *clec-183* | WBGene00011855 | protein_coding_gene | -3.9 | 0.065 | 2.5E-05 |
| *F49F1.2* | WBGene00235363 | protein_coding_gene | -4.0 | 0.064 | 4.7E-05 |
| *cwp-2* | WBGene00000860 | protein_coding_gene | -4.0 | 0.063 | 4.6E-05 |
| *T16G12.10* | WBGene00044222 | protein_coding_gene | -4.0 | 0.062 | 1.0E-68 |
| *C05C10.8* | WBGene00086567 | protein_coding_gene | -4.0 | 0.062 | 2.8E-05 |
| *C06A12.8* | WBGene00007359 | protein_coding_gene | -4.0 | 0.061 | 6.2E-05 |
| *F46A8.4* | WBGene00009747 | protein_coding_gene | -4.1 | 0.059 | 3.7E-07 |
| *LLC1.121* | WBGene00235121 | pseudogene | -4.1 | 0.057 | 3.5E-05 |
| *T07G12.5* | WBGene00011596 | protein_coding_gene | -4.1 | 0.057 | 1.5E-123 |
| *F56D6.9* | WBGene00044471 | protein_coding_gene | -4.2 | 0.056 | 2.8E-05 |
| *C17G10.6* | WBGene00015918 | protein_coding_gene | -4.2 | 0.055 | 6.5E-06 |
| *B0207.5* | WBGene00015028 | protein_coding_gene | -4.2 | 0.055 | 4.2E-88 |
| *Y46D2A.3* | WBGene00021591 | protein_coding_gene | -4.2 | 0.055 | 1.2E-11 |
| *clec-98* | WBGene00013932 | protein_coding_gene | -4.2 | 0.054 | 3.1E-05 |
| *scl-26* | WBGene00016821 | protein_coding_gene | -4.2 | 0.054 | 5.4E-10 |
| *Y37F4.3* | WBGene00021384 | protein_coding_gene | -4.2 | 0.054 | 1.1E-05 |
| *F58F9.9* | WBGene00044307 | protein_coding_gene | -4.2 | 0.053 | 3.6E-06 |
| *F46B3.15* | WBGene00009766 | protein_coding_gene | -4.2 | 0.053 | 5.2E-06 |
| *ugt-34* | WBGene00009255 | protein_coding_gene | -4.3 | 0.052 | 4.2E-05 |
| *T28C12.15* | WBGene00271821 | protein_coding_gene | -4.3 | 0.051 | 1.3E-05 |
| *C08E3.14* | WBGene00044415 | protein_coding_gene | -4.3 | 0.050 | 2.0E-05 |
| *scl-2* | WBGene00009895 | protein_coding_gene | -4.3 | 0.049 | 1.1E-147 |
| *F57A8.7* | WBGene00010182 | protein_coding_gene | -4.4 | 0.049 | 2.1E-05 |
| *Y47D7A.11* | WBGene00021623 | protein_coding_gene | -4.4 | 0.048 | 2.0E-06 |
| *C10G8.2* | WBGene00015681 | protein_coding_gene | -4.4 | 0.048 | 5.4E-06 |
| *ZC204.6* | WBGene00022556 | protein_coding_gene | -4.4 | 0.048 | 1.6E-05 |
| *ZC328.5* | WBGene00022594 | protein_coding_gene | -4.4 | 0.046 | 9.1E-31 |
| *fipr-23* | WBGene00007990 | protein_coding_gene | -4.4 | 0.046 | 2.4E-07 |
| *F56D2.8* | WBGene00018968 | protein_coding_gene | -4.5 | 0.045 | 9.9E-16 |
| *thn-1* | WBGene00009213 | protein_coding_gene | -4.5 | 0.044 | 4.9E-16 |
| *R11E3.19* | WBGene00235374 | protein_coding_gene | -4.5 | 0.043 | 7.1E-08 |
| *F17E9.19* | WBGene00249825 | pseudogene | -4.5 | 0.043 | 1.1E-06 |
| *nspe-1* | WBGene00012591 | protein_coding_gene | -4.5 | 0.043 | 4.3E-10 |
| *T23E7.6* | WBGene00044772 | protein_coding_gene | -4.6 | 0.043 | 2.8E-42 |
| *Y116A8C.44* | WBGene00013815 | protein_coding_gene | -4.6 | 0.042 | 9.0E-13 |
| *F13H6.4* | WBGene00017432 | pseudogene | -4.6 | 0.041 | 1.0E-17 |
| *Y43F8C.16* | WBGene00012837 | protein_coding_gene | -4.6 | 0.040 | 1.9E-06 |
| *clec-129* | WBGene00021145 | protein_coding_gene | -4.6 | 0.040 | 1.9E-06 |
| *F56D6.8* | WBGene00044292 | protein_coding_gene | -4.6 | 0.040 | 2.4E-09 |
| *F47C12.8* | WBGene00018551 | protein_coding_gene | -4.7 | 0.040 | 1.8E-26 |
| *T02D1.7* | WBGene00011373 | protein_coding_gene | -4.7 | 0.039 | 1.1E-06 |
| *Y57E12AL.4* | WBGene00021958 | protein_coding_gene | -4.7 | 0.039 | 2.6E-06 |
| *F48C1.11* | WBGene00195179 | protein_coding_gene | -4.7 | 0.039 | 2.4E-06 |
| *C41D11.10* | WBGene00195180 | protein_coding_gene | -4.7 | 0.038 | 1.1E-11 |

| | | | | | |
|---|---|---|---|---|---|
| *Y116F11A.3* | WBGene00013817 | protein_coding_gene | -4.7 | 0.038 | 2.3E-07 |
| *F26H11.8* | WBGene00195149 | protein_coding_gene | -4.7 | 0.038 | 2.4E-25 |
| *F17E9.18* | WBGene00235282 | protein_coding_gene | -4.7 | 0.038 | 8.5E-07 |
| *C16C8.8* | WBGene00015846 | protein_coding_gene | -4.7 | 0.037 | 9.4E-07 |
| *flp-23* | WBGene00017693 | protein_coding_gene | -4.8 | 0.037 | 2.9E-07 |
| *fip-7* | WBGene00013105 | protein_coding_gene | -4.8 | 0.037 | 3.4E-07 |
| *C16C8.10* | WBGene00015848 | protein_coding_gene | -4.8 | 0.036 | 4.8E-07 |
| *clec-221* | WBGene00008916 | protein_coding_gene | -4.8 | 0.036 | 5.4E-93 |
| *F58F9.8* | WBGene00044306 | protein_coding_gene | -4.8 | 0.036 | 3.6E-14 |
| *K09C8.2* | WBGene00010715 | protein_coding_gene | -4.8 | 0.035 | 2.9E-15 |
| *C06E2.9* | WBGene00044579 | protein_coding_gene | -4.8 | 0.035 | 7.7E-16 |
| *F58A4.1* | WBGene00010226 | protein_coding_gene | -4.8 | 0.035 | 2.1E-40 |
| *R05G6.3* | WBGene00043466 | protein_coding_gene | -4.9 | 0.034 | 2.7E-07 |
| *F59A6.11* | WBGene00044664 | protein_coding_gene | -4.9 | 0.034 | 1.1E-07 |
| *Y6G8.16* | WBGene00194801 | protein_coding_gene | -4.9 | 0.033 | 6.8E-07 |
| *pkd-2* | WBGene00004035 | protein_coding_gene | -4.9 | 0.033 | 8.3E-08 |
| *T22C1.12* | WBGene00011921 | protein_coding_gene | -5.0 | 0.032 | 2.4E-13 |
| *C14F11.7* | WBGene00015782 | protein_coding_gene | -5.0 | 0.032 | 4.0E-50 |
| *ZC178.2* | WBGene00022536 | protein_coding_gene | -5.0 | 0.032 | 6.2E-08 |
| *F59A1.16* | WBGene00010301 | protein_coding_gene | -5.0 | 0.032 | 7.5E-08 |
| *Y82E9BR.8* | WBGene00022341 | protein_coding_gene | -5.0 | 0.031 | 1.1E-07 |
| *nspf-1* | WBGene00044364 | protein_coding_gene | -5.0 | 0.031 | 1.7E-07 |
| *F56C3.8* | WBGene00018945 | protein_coding_gene | -5.0 | 0.031 | 3.0E-24 |
| *T04C12.7* | WBGene00011429 | protein_coding_gene | -5.0 | 0.031 | 8.5E-19 |
| *F59A1.6* | WBGene00010292 | protein_coding_gene | -5.0 | 0.031 | 4.5E-07 |
| *fipr-19* | WBGene00044451 | protein_coding_gene | -5.0 | 0.031 | 1.2E-07 |
| *Y110A2AL.6* | WBGene00022443 | protein_coding_gene | -5.0 | 0.031 | 4.2E-14 |
| *R11G10.4* | WBGene00050968 | protein_coding_gene | -5.0 | 0.031 | 1.3E-09 |
| *C45G9.10* | WBGene00016681 | protein_coding_gene | -5.0 | 0.031 | 3.0E-33 |
| *R09E10.8* | WBGene00011177 | protein_coding_gene | -5.0 | 0.030 | 1.0E-29 |
| *nas-24* | WBGene00003543 | protein_coding_gene | -5.1 | 0.030 | 1.3E-62 |
| *W09G12.6* | WBGene00021120 | protein_coding_gene | -5.1 | 0.030 | 7.6E-08 |
| *glc-1* | WBGene00001591 | protein_coding_gene | -5.1 | 0.029 | 4.7E-122 |
| *oac-4* | WBGene00015873 | protein_coding_gene | -5.1 | 0.029 | 2.0E-08 |
| *Y64G10A.2* | WBGene00013412 | protein_coding_gene | -5.1 | 0.029 | 2.9E-08 |
| *F47C12.6* | WBGene00018549 | protein_coding_gene | -5.1 | 0.029 | 4.5E-08 |
| *fipr-16* | WBGene00015521 | protein_coding_gene | -5.1 | 0.028 | 3.1E-35 |
| *F47C12.12* | WBGene00219377 | pseudogene | -5.2 | 0.028 | 1.2E-24 |
| *linc-1* | WBGene00194659 | lincRNA_gene | -5.2 | 0.028 | 2.9E-45 |
| *clec-100* | WBGene00013179 | protein_coding_gene | -5.2 | 0.028 | 1.6E-08 |
| *clec-108* | WBGene00012504 | protein_coding_gene | -5.2 | 0.027 | 1.7E-08 |
| *T14G12.11* | WBGene00219411 | protein_coding_gene | -5.2 | 0.027 | 2.4E-19 |
| *F30A10.11* | WBGene00009268 | protein_coding_gene | -5.2 | 0.026 | 2.3E-36 |
| *Y17G9A.4* | WBGene00021197 | protein_coding_gene | -5.3 | 0.026 | 5.0E-47 |
| *F40G9.7* | WBGene00018243 | protein_coding_gene | -5.3 | 0.026 | 9.5E-09 |
| *B0222.10* | WBGene00015058 | ncRNA_gene | -5.3 | 0.026 | 3.1E-62 |
| *C49C3.11* | WBGene00008201 | protein_coding_gene | -5.3 | 0.026 | 7.6E-113 |
| *T12A2.5* | WBGene00020439 | protein_coding_gene | -5.3 | 0.026 | 5.0E-30 |
| *F13E9.19* | WBGene00194922 | protein_coding_gene | -5.3 | 0.026 | 1.8E-11 |
| *nspf-2* | WBGene00044365 | protein_coding_gene | -5.3 | 0.025 | 1.2E-08 |
| *fip-3* | WBGene00015520 | protein_coding_gene | -5.3 | 0.025 | 6.8E-08 |
| *T12A7.10* | WBGene00194933 | protein_coding_gene | -5.4 | 0.024 | 6.1E-09 |
| *scl-8* | WBGene00008030 | protein_coding_gene | -5.4 | 0.023 | 3.7E-20 |
| *ins-8* | WBGene00002091 | protein_coding_gene | -5.4 | 0.023 | 3.7E-07 |
| *F40G9.15* | WBGene00077531 | protein_coding_gene | -5.5 | 0.023 | 3.5E-08 |
| *F35C5.3* | WBGene00009391 | protein_coding_gene | -5.5 | 0.022 | 1.8E-08 |
| *F13E9.10* | WBGene00008759 | protein_coding_gene | -5.5 | 0.022 | 2.7E-29 |
| *F16H6.9* | WBGene00008899 | protein_coding_gene | -5.5 | 0.022 | 1.9E-12 |
| *Y51A2D.1* | WBGene00013072 | protein_coding_gene | -5.6 | 0.021 | 7.3E-09 |
| *clec-130* | WBGene00021144 | protein_coding_gene | -5.6 | 0.021 | 1.0E-11 |
| *B0513.90* | WBGene00235301 | protein_coding_gene | -5.6 | 0.021 | 8.4E-09 |
| *W02B12.16* | WBGene00195148 | protein_coding_gene | -5.6 | 0.021 | 3.5E-17 |
| *scl-9* | WBGene00009890 | protein_coding_gene | -5.6 | 0.020 | 8.8E-33 |
| *clec-126* | WBGene00021117 | protein_coding_gene | -5.6 | 0.020 | 1.5E-35 |
| *Y43F8C.24* | WBGene00045345 | pseudogene | -5.7 | 0.019 | 1.2E-09 |
| *clec-157* | WBGene00017810 | protein_coding_gene | -5.7 | 0.019 | 1.9E-57 |
| *K08C9.6* | WBGene00010653 | protein_coding_gene | -5.8 | 0.019 | 5.8E-10 |
| *R53.8* | WBGene00011277 | protein_coding_gene | -5.8 | 0.018 | 1.7E-38 |
| *F07G6.8* | WBGene00017227 | protein_coding_gene | -5.8 | 0.018 | 2.0E-21 |
| *T22F3.11* | WBGene00020700 | protein_coding_gene | -5.8 | 0.018 | 7.0E-13 |

| | | | | | |
|---|---|---|---|---|---|
| *cwp-1* | WBGene00000859 | protein_coding_gene | -5.8 | 0.018 | 2.2E-09 |
| *clec-136* | WBGene00018047 | protein_coding_gene | -5.9 | 0.017 | 8.0E-14 |
| *K03B8.14* | WBGene00194783 | protein_coding_gene | -5.9 | 0.017 | 2.1E-13 |
| *K12H6.5* | WBGene00019685 | protein_coding_gene | -5.9 | 0.017 | 7.8E-11 |
| *F46A8.3* | WBGene00009746 | protein_coding_gene | -5.9 | 0.017 | 3.4E-19 |
| *F17E9.3* | WBGene00017540 | protein_coding_gene | -5.9 | 0.017 | 1.4E-13 |
| *Y49F6B.13* | WBGene00044377 | protein_coding_gene | -5.9 | 0.016 | 3.5E-10 |
| *clec-181* | WBGene00044511 | protein_coding_gene | -5.9 | 0.016 | 1.4E-09 |
| *F25E5.3* | WBGene00017784 | protein_coding_gene | -5.9 | 0.016 | 4.6E-44 |
| *Y43F8B.19* | WBGene00077688 | protein_coding_gene | -5.9 | 0.016 | 1.2E-10 |
| *Y18D10A.2* | WBGene00012475 | protein_coding_gene | -6.0 | 0.016 | 1.6E-14 |
| *T26A8.3* | WBGene00020826 | protein_coding_gene | -6.0 | 0.016 | 2.3E-31 |
| *F38E1.10* | WBGene00018182 | protein_coding_gene | -6.0 | 0.016 | 7.9E-10 |
| *clec-105* | WBGene00012488 | protein_coding_gene | -6.0 | 0.016 | 9.3E-11 |
| *D1022.2* | WBGene00017021 | protein_coding_gene | -6.0 | 0.016 | 2.2E-17 |
| *C35E7.7* | WBGene00016459 | protein_coding_gene | -6.0 | 0.016 | 9.4E-14 |
| *R05H10.7* | WBGene00011046 | protein_coding_gene | -6.0 | 0.016 | 2.8E-33 |
| *ZK1225.10* | WBGene00044714 | pseudogene | -6.0 | 0.016 | 9.7E-11 |
| *H25K10.4* | WBGene00010413 | protein_coding_gene | -6.0 | 0.015 | 7.6E-19 |
| *clec-135* | WBGene00015821 | protein_coding_gene | -6.0 | 0.015 | 5.9E-21 |
| *F26C11.4* | WBGene00044159 | protein_coding_gene | -6.0 | 0.015 | 9.8E-18 |
| *clec-132* | WBGene00021143 | protein_coding_gene | -6.1 | 0.015 | 2.0E-17 |
| *W02D7.12* | WBGene00044356 | protein_coding_gene | -6.1 | 0.015 | 3.1E-11 |
| *K03B8.11* | WBGene00010526 | protein_coding_gene | -6.1 | 0.015 | 4.0E-10 |
| *K12H6.8* | WBGene00019688 | protein_coding_gene | -6.1 | 0.014 | 1.3E-11 |
| *F45D3.4* | WBGene00009724 | protein_coding_gene | -6.1 | 0.014 | 6.9E-95 |
| *F46A8.8* | WBGene00009751 | protein_coding_gene | -6.2 | 0.014 | 2.9E-10 |
| *clec-141* | WBGene00017139 | protein_coding_gene | -6.2 | 0.013 | 2.0E-32 |
| *Y43F8C.15* | WBGene00012836 | protein_coding_gene | -6.2 | 0.013 | 1.3E-11 |
| *K11D12.6* | WBGene00019646 | protein_coding_gene | -6.3 | 0.013 | 1.4E-54 |
| *fipr-17* | WBGene00012965 | protein_coding_gene | -6.3 | 0.013 | 6.9E-66 |
| *Y6G8.15* | WBGene00189937 | protein_coding_gene | -6.3 | 0.013 | 3.0E-22 |
| *clec-261* | WBGene00013834 | protein_coding_gene | -6.3 | 0.013 | 1.4E-10 |
| *F28B1.9* | WBGene00045381 | protein_coding_gene | -6.3 | 0.013 | 1.0E-11 |
| *Y6G8.14* | WBGene00138715 | protein_coding_gene | -6.3 | 0.013 | 4.8E-12 |
| *dct-7* | WBGene00017488 | protein_coding_gene | -6.3 | 0.013 | 2.0E-09 |
| *F11A6.16* | WBGene00271780 | protein_coding_gene | -6.3 | 0.012 | 9.3E-11 |
| *C47A10.12* | WBGene00008129 | protein_coding_gene | -6.3 | 0.012 | 1.6E-16 |
| *Y49F6B.6* | WBGene00021718 | protein_coding_gene | -6.4 | 0.012 | 2.3E-12 |
| *F16G10.10* | WBGene00017525 | protein_coding_gene | -6.4 | 0.012 | 5.1E-16 |
| *F48C1.9* | WBGene00018601 | protein_coding_gene | -6.4 | 0.012 | 1.0E-27 |
| *clec-263* | WBGene00009171 | protein_coding_gene | -6.4 | 0.012 | 4.4E-12 |
| *Y74C10AR.2* | WBGene00022282 | protein_coding_gene | -6.4 | 0.012 | 2.5E-75 |
| *C09G12.5* | WBGene00015655 | protein_coding_gene | -6.4 | 0.012 | 3.6E-28 |
| *nspf-3* | WBGene00044366 | protein_coding_gene | -6.5 | 0.011 | 1.6E-12 |
| *C16C8.19* | WBGene00015856 | protein_coding_gene | -6.5 | 0.011 | 1.3E-16 |
| *Y49F6B.20* | WBGene00303369 | protein_coding_gene | -6.5 | 0.011 | 2.4E-12 |
| *M7.10* | WBGene00010886 | protein_coding_gene | -6.6 | 0.010 | 1.0E-26 |
| *tag-329* | WBGene00008231 | protein_coding_gene | -6.6 | 0.010 | 6.6E-67 |
| *try-8* | WBGene00017791 | protein_coding_gene | -6.6 | 0.010 | 1.2E-51 |
| *M151.10* | WBGene00303365 | protein_coding_gene | -6.6 | 0.010 | 2.2E-42 |
| *F43C11.2* | WBGene00018379 | protein_coding_gene | -6.6 | 0.010 | 1.1E-21 |
| *C16C8.7* | WBGene00015845 | protein_coding_gene | -6.6 | 0.010 | 1.9E-17 |
| *clec-133* | WBGene00021142 | protein_coding_gene | -6.6 | 0.010 | 5.3E-17 |
| *F59B2.12* | WBGene00010314 | protein_coding_gene | -6.7 | 0.010 | 2.2E-17 |
| *clec-92* | WBGene00013125 | protein_coding_gene | -6.7 | 0.010 | 7.3E-18 |
| *C05B5.11* | WBGene00045242 | protein_coding_gene | -6.7 | 0.009 | 1.5E-34 |
| *ZK1290.11* | WBGene00022892 | protein_coding_gene | -6.8 | 0.009 | 2.6E-26 |
| *clec-138* | WBGene00018049 | protein_coding_gene | -6.8 | 0.009 | 4.0E-14 |
| *M176.10* | WBGene00010946 | protein_coding_gene | -6.8 | 0.009 | 6.6E-35 |
| *fmo-2* | WBGene00001477 | protein_coding_gene | -6.8 | 0.009 | 4.0E-19 |
| *clec-161* | WBGene00010228 | protein_coding_gene | -6.8 | 0.009 | 1.5E-31 |
| *F42A6.2* | WBGene00018324 | protein_coding_gene | -6.8 | 0.009 | 1.6E-13 |
| *clec-106* | WBGene00012482 | protein_coding_gene | -6.9 | 0.008 | 4.3E-14 |
| *F25E5.7* | WBGene00017788 | protein_coding_gene | -6.9 | 0.008 | 2.3E-49 |
| *T23G4.5* | WBGene00011961 | protein_coding_gene | -6.9 | 0.008 | 4.2E-41 |
| *hpo-2* | WBGene00012500 | protein_coding_gene | -6.9 | 0.008 | 3.0E-54 |
| *Y25C1A.2* | WBGene00021289 | protein_coding_gene | -7.0 | 0.008 | 1.8E-19 |
| *F46A8.5* | WBGene00009748 | protein_coding_gene | -7.0 | 0.008 | 4.6E-13 |
| *cpt-3* | WBGene00021703 | protein_coding_gene | -7.0 | 0.008 | 6.7E-25 |

| | | | | | |
|---|---|---|---|---|---|
| *K04H8.3* | WBGene00010578 | protein_coding_gene | -7.1 | 0.008 | 2.2E-14 |
| *C33G3.5* | WBGene00007905 | protein_coding_gene | -7.1 | 0.008 | 1.4E-24 |
| *C16C8.9* | WBGene00015847 | protein_coding_gene | -7.2 | 0.007 | 3.7E-43 |
| *dct-8* | WBGene00044472 | protein_coding_gene | -7.2 | 0.007 | 5.2E-12 |
| *ins-31* | WBGene00002114 | protein_coding_gene | -7.2 | 0.007 | 1.6E-60 |
| *clec-208* | WBGene00018097 | protein_coding_gene | -7.2 | 0.007 | 8.4E-82 |
| *fipr-28* | WBGene00013120 | protein_coding_gene | -7.3 | 0.007 | 6.2E-26 |
| *F26D10.13* | WBGene00044244 | protein_coding_gene | -7.3 | 0.007 | 1.3E-45 |
| *clec-217* | WBGene00017824 | protein_coding_gene | -7.3 | 0.006 | 7.7E-16 |
| *clec-110* | WBGene00008909 | protein_coding_gene | -7.3 | 0.006 | 5.6E-36 |
| *clec-207* | WBGene00018096 | protein_coding_gene | -7.3 | 0.006 | 1.5E-50 |
| *ZK1290.1* | WBGene00022886 | protein_coding_gene | -7.3 | 0.006 | 9.9E-22 |
| *clec-99* | WBGene00013933 | protein_coding_gene | -7.3 | 0.006 | 3.3E-41 |
| *clec-96* | WBGene00013930 | protein_coding_gene | -7.4 | 0.006 | 4.0E-14 |
| *T10B10.9* | WBGene00011685 | protein_coding_gene | -7.4 | 0.006 | 4.5E-14 |
| *Y43F8C.17* | WBGene00012838 | protein_coding_gene | -7.5 | 0.006 | 2.1E-14 |
| *clec-124* | WBGene00015909 | protein_coding_gene | -7.5 | 0.006 | 1.3E-22 |
| *F26D2.16* | WBGene00009154 | protein_coding_gene | -7.6 | 0.005 | 3.5E-98 |
| *fipr-20* | WBGene00044449 | protein_coding_gene | -7.6 | 0.005 | 3.0E-23 |
| *fipr-18* | WBGene00044450 | protein_coding_gene | -7.7 | 0.005 | 1.4E-39 |
| *Y110A2AL.7* | WBGene00022444 | protein_coding_gene | -7.7 | 0.005 | 2.4E-17 |
| *C48B4.12* | WBGene00008175 | protein_coding_gene | -7.7 | 0.005 | 9.7E-35 |
| *clec-137* | WBGene00018048 | protein_coding_gene | -7.7 | 0.005 | 3.3E-46 |
| *F54D12.9* | WBGene00018822 | protein_coding_gene | -7.8 | 0.005 | 7.9E-18 |
| *T22G5.1* | WBGene00011929 | protein_coding_gene | -7.8 | 0.004 | 1.8E-15 |
| *nas-17* | WBGene00003536 | protein_coding_gene | -7.8 | 0.004 | 2.8E-17 |
| *clec-95* | WBGene00013927 | protein_coding_gene | -7.8 | 0.004 | 6.7E-42 |
| *K12H6.4* | WBGene00019684 | protein_coding_gene | -7.9 | 0.004 | 2.3E-30 |
| *clec-197* | WBGene00008202 | protein_coding_gene | -8.0 | 0.004 | 1.1E-97 |
| *pqn-84* | WBGene00004165 | protein_coding_gene | -8.1 | 0.004 | 2.3E-26 |
| *F18E9.7* | WBGene00017573 | protein_coding_gene | -8.2 | 0.003 | 1.1E-19 |
| *K12H6.9* | WBGene00019689 | protein_coding_gene | -8.3 | 0.003 | 3.4E-27 |
| *plg-1* | WBGene00004041 | pseudogene | -8.4 | 0.003 | 4.2E-21 |
| *M7.9* | WBGene00010885 | protein_coding_gene | -8.4 | 0.003 | 1.9E-35 |
| *clec-219* | WBGene00020945 | protein_coding_gene | -9.3 | 0.002 | 3.8E-43 |
| *ilys-3* | WBGene00016670 | protein_coding_gene | -9.5 | 0.001 | 2.6E-35 |

Shared (N2&CB4856) DEGs in response to *P. aeruginosa* (vs. OP50)

| | | | N2 | | | CB4856 | | |
|---|---|---|---|---|---|---|---|---|
| Gene Name | Wormbase Gene ID | Type | $log_2$FC | FC | Adjusted P-value | $log_2$FC | FC | Adjusted P-value |
| *dod-21* | WBGene00007873 | protein_coding_gene | 9.1 | 559.6949 | 1.4E-15 | 3.7 | 12.7068 | 8.0E-03 |
| *clec-45* | WBGene00017199 | protein_coding_gene | 8.5 | 350.2918 | 2.9E-13 | 4.2 | 17.7972 | 2.7E-04 |
| *Y53G8AM.5* | WBGene00021805 | protein_coding_gene | 6.2 | 75.0094 | 1.1E-08 | 6.9 | 123.2343 | 1.2E-12 |
| *F49H6.13* | WBGene00009914 | protein_coding_gene | 6.2 | 71.8589 | 7.3E-08 | 4.7 | 26.8964 | 1.6E-05 |
| *Y26D4A.3* | WBGene00012501 | protein_coding_gene | 5.9 | 60.8088 | 7.4E-09 | 3.5 | 11.3835 | 1.7E-03 |
| *sri-36* | WBGene00005548 | protein_coding_gene | 5.6 | 48.7611 | 7.2E-07 | 4.7 | 25.8998 | 1.5E-15 |
| *B0024.4* | WBGene00007097 | protein_coding_gene | 5.6 | 47.3158 | 3.4E-26 | 4.1 | 17.5952 | 2.4E-25 |
| *Y65B4BR.1* | WBGene00022040 | protein_coding_gene | 5.4 | 42.1401 | 3.3E-39 | 3.6 | 11.9993 | 3.5E-13 |
| *F08G2.5* | WBGene00008577 | protein_coding_gene | 5.3 | 38.6876 | 4.9E-46 | 2.9 | 7.2677 | 1.7E-11 |
| *F44G3.10* | WBGene00009710 | protein_coding_gene | 5.2 | 36.2579 | 5.1E-24 | 4.0 | 15.6717 | 1.7E-31 |
| *C49G7.12* | WBGene00077593 | protein_coding_gene | 5.2 | 35.9355 | 2.3E-12 | 5.6 | 49.8879 | 2.0E-97 |
| *clec-17* | WBGene00008477 | protein_coding_gene | 5.2 | 35.5994 | 7.7E-10 | 2.7 | 6.3797 | 1.1E-05 |
| *T24C4.4* | WBGene00020760 | protein_coding_gene | 5.1 | 34.0092 | 5.7E-33 | 3.0 | 8.2199 | 1.7E-27 |
| *dod-22* | WBGene00010125 | protein_coding_gene | 4.9 | 30.3545 | 9.0E-63 | 3.2 | 9.2674 | 6.4E-61 |
| *srj-29* | WBGene00005616 | protein_coding_gene | 4.9 | 30.1617 | 6.0E-07 | 4.1 | 16.7911 | 7.8E-12 |
| *F55G11.6* | WBGene00010126 | protein_coding_gene | 4.9 | 29.5689 | 2.1E-05 | 5.8 | 56.5495 | 2.0E-07 |
| *tsp-1* | WBGene00006627 | protein_coding_gene | 4.9 | 29.0994 | 5.2E-23 | 1.7 | 3.3296 | 1.1E-03 |
| *srz-99* | WBGene00012028 | protein_coding_gene | 4.6 | 24.8204 | 8.8E-06 | 2.7 | 6.3325 | 4.0E-08 |
| *linc-37* | WBGene00219709 | lincRNA_gene | 4.5 | 22.9330 | 8.9E-07 | 2.3 | 4.8831 | 1.5E-05 |
| ***nlp-39*** | **WBGene00008295** | **protein_coding_gene** | **4.5** | **21.9995** | **4.6E-08** | **-2.4** | **0.1915** | **1.0E-05** |
| *F55G11.7* | WBGene00010127 | protein_coding_gene | 4.4 | 21.7414 | 1.7E-05 | 4.1 | 16.7561 | 7.8E-12 |
| *H02F09.2* | WBGene00019145 | protein_coding_gene | 4.4 | 21.7323 | 2.6E-05 | 4.1 | 17.4140 | 6.3E-05 |
| *Y17D7B.3* | WBGene00012451 | protein_coding_gene | 4.3 | 20.1340 | 3.9E-05 | 3.7 | 12.8848 | 1.4E-07 |
| *tsp-2* | WBGene00006628 | protein_coding_gene | 4.3 | 19.2325 | 8.1E-20 | 1.9 | 3.7041 | 4.1E-11 |
| *irg-6* | WBGene00007864 | protein_coding_gene | 4.2 | 18.4142 | 4.4E-07 | 1.7 | 3.2813 | 2.4E-07 |
| *irg-5* | WBGene00009429 | protein_coding_gene | 4.2 | 18.3050 | 4.7E-22 | 7.4 | 173.2678 | 2.1E-102 |
| *ZK1290.10* | WBGene00022891 | protein_coding_gene | 4.1 | 16.9597 | 4.5E-22 | 2.8 | 7.1351 | 6.5E-16 |
| ***cyp-37B1*** | **WBGene00009226** | **protein_coding_gene** | **4.1** | **16.8781** | **4.0E-07** | **-2.1** | **0.2330** | **6.0E-11** |
| *irg-1* | WBGene00015574 | protein_coding_gene | 4.1 | 16.7257 | 1.2E-10 | 4.4 | 21.2921 | 2.2E-28 |
| *F35E12.4* | WBGene00009428 | pseudogene | 4.1 | 16.5717 | 7.6E-08 | 2.7 | 6.6381 | 1.0E-19 |
| *Y51B9A.8* | WBGene00013090 | protein_coding_gene | 4.0 | 16.1533 | 5.6E-04 | 5.8 | 56.0187 | 3.9E-08 |

| | | | | | | | | |
|---|---|---|---|---|---|---|---|---|
| *F11D11.3* | WBGene00008698 | protein_coding_gene | 4.0 | 16.0696 | 1.2E-02 | 5.0 | 32.9922 | 5.5E-10 |
| *H02F09.3* | WBGene00019146 | protein_coding_gene | 4.0 | 15.9989 | 6.9E-21 | 6.7 | 105.5755 | 1.5E-30 |
| *fbxa-88* | WBGene00008632 | protein_coding_gene | 4.0 | 15.8853 | 4.3E-07 | 4.1 | 17.3574 | 1.1E-13 |
| *F10A3.1* | WBGene00008631 | protein_coding_gene | 4.0 | 15.6472 | 3.3E-02 | 3.2 | 9.3058 | 1.3E-09 |
| ***C35B1.3*** | **WBGene00016433** | **protein_coding_gene** | **4.0** | **15.6260** | **1.5E-03** | **-4.9** | **0.0327** | **7.0E-16** |
| ***C50F7.5*** | **WBGene00016845** | **protein_coding_gene** | **3.9** | **15.3149** | **7.0E-06** | **-3.8** | **0.0702** | **4.6E-127** |
| *C01G10.5* | WBGene00007232 | protein_coding_gene | 3.8 | 13.9320 | 1.4E-03 | 5.1 | 34.3685 | 5.4E-17 |
| *C04G6.5* | WBGene00015455 | protein_coding_gene | 3.7 | 13.2724 | 6.2E-09 | 1.6 | 3.0109 | 3.8E-05 |
| ***F49F1.15*** | **WBGene00235365** | **protein_coding_gene** | **3.7** | **13.0111** | **5.4E-04** | **-6.0** | **0.0161** | **4.7E-26** |
| ***clec-158*** | **WBGene00020067** | **protein_coding_gene** | **3.6** | **12.5502** | **7.7E-05** | **-5.7** | **0.0193** | **2.9E-23** |
| ***F15E6.10*** | **WBGene00044479** | **protein_coding_gene** | **3.6** | **12.2289** | **9.4E-04** | **-1.8** | **0.2778** | **2.1E-03** |
| ***fbxa-163*** | **WBGene00015598** | **protein_coding_gene** | **3.6** | **12.1261** | **6.2E-09** | **-1.4** | **0.3855** | **1.9E-05** |
| *gst-38* | WBGene00001786 | protein_coding_gene | 3.6 | 12.0414 | 2.5E-16 | 7.8 | 222.9372 | 7.1E-14 |
| *dod-24* | WBGene00007875 | protein_coding_gene | 3.6 | 11.9628 | 1.0E-05 | 3.0 | 7.7920 | 1.7E-69 |
| ***C06E7.88*** | **WBGene00189995** | **protein_coding_gene** | **3.6** | **11.9074** | **1.9E-06** | **-1.1** | **0.4819** | **2.3E-02** |
| *C18C4.17* | WBGene00219824 | ncRNA_gene | 3.6 | 11.7828 | 6.1E-03 | 3.5 | 11.3854 | 3.4E-03 |
| *irg-3* | WBGene00018760 | protein_coding_gene | 3.5 | 11.2574 | 5.7E-31 | 3.9 | 14.4804 | 1.0E-57 |
| *lys-2* | WBGene00003091 | protein_coding_gene | 3.5 | 11.1075 | 1.5E-27 | 3.3 | 9.5529 | 3.2E-13 |
| *C17H12.6* | WBGene00015932 | protein_coding_gene | 3.4 | 10.8892 | 2.8E-18 | 6.6 | 100.1148 | 6.5E-15 |
| *ZC47.11* | WBGene00014979 | pseudogene | 3.4 | 10.8319 | 4.5E-03 | 2.5 | 5.7891 | 2.7E-02 |
| *F33H12.7* | WBGene00045457 | protein_coding_gene | 3.4 | 10.6929 | 9.0E-05 | 2.6 | 6.0687 | 3.0E-08 |
| *clec-174* | WBGene00021580 | protein_coding_gene | 3.4 | 10.5723 | 5.4E-08 | 3.8 | 14.1352 | 8.9E-13 |
| *T24A6.7* | WBGene00020747 | protein_coding_gene | 3.4 | 10.5177 | 3.0E-05 | 3.4 | 10.5719 | 1.3E-04 |
| *C06B3.7* | WBGene00007366 | protein_coding_gene | 3.4 | 10.4956 | 1.6E-04 | 3.2 | 8.9872 | 6.6E-12 |
| ***tts-1*** | **WBGene00006650** | **ncRNA_gene** | **3.3** | **10.1468** | **2.1E-06** | **-1.9** | **0.2725** | **1.2E-12** |
| *Y41D4B.15* | WBGene00021517 | protein_coding_gene | 3.3 | 10.0746 | 1.1E-03 | 2.8 | 6.8875 | 2.7E-03 |
| *C01G10.17* | WBGene00044658 | protein_coding_gene | 3.3 | 9.9635 | 1.1E-03 | 3.1 | 8.5677 | 5.2E-11 |
| *Y17D7B.2* | WBGene00012450 | protein_coding_gene | 3.2 | 9.4479 | 7.0E-04 | 1.6 | 3.1156 | 1.2E-03 |
| *clec-67* | WBGene00018971 | protein_coding_gene | 3.2 | 9.2921 | 1.8E-11 | 3.4 | 10.3364 | 1.9E-14 |
| ***F46A8.13*** | **WBGene00219950** | **ncRNA_gene** | **3.2** | **9.0095** | **8.2E-04** | **-1.5** | **0.3457** | **4.7E-05** |
| *F15B9.6* | WBGene00008850 | protein_coding_gene | 3.2 | 8.9490 | 9.4E-10 | 1.8 | 3.5175 | 1.3E-03 |
| ***F47B8.4*** | **WBGene00009805** | **protein_coding_gene** | **3.1** | **8.8697** | **1.5E-03** | **-2.2** | **0.2228** | **5.9E-11** |
| *dct-17* | WBGene00009431 | protein_coding_gene | 3.1 | 8.7497 | 1.6E-09 | 3.1 | 8.7431 | 1.1E-07 |
| *Y46H3C.6* | WBGene00021606 | pseudogene | 3.1 | 8.7484 | 5.3E-07 | 3.6 | 11.9808 | 9.1E-07 |
| ***scl-7*** | **WBGene00008029** | **protein_coding_gene** | **3.1** | **8.6499** | **6.6E-03** | **-6.4** | **0.0120** | **4.5E-30** |
| *fbxa-128* | WBGene00012923 | protein_coding_gene | 3.1 | 8.4709 | 2.7E-21 | 2.6 | 6.2611 | 1.6E-18 |
| *C32H11.4* | WBGene00007867 | protein_coding_gene | 3.1 | 8.4450 | 2.5E-33 | 3.8 | 13.9616 | 9.5E-30 |
| *valv-1* | WBGene00015216 | protein_coding_gene | 3.1 | 8.2885 | 2.5E-20 | 1.7 | 3.3310 | 1.4E-15 |
| ***F47B8.10*** | **WBGene00009811** | **protein_coding_gene** | **3.0** | **8.2785** | **7.5E-04** | **-1.9** | **0.2713** | **1.8E-05** |
| *F20G2.5* | WBGene00008988 | protein_coding_gene | 3.0 | 8.2451 | 1.9E-08 | 2.9 | 7.5771 | 4.4E-10 |
| *Y51H4A.25* | WBGene00013119 | protein_coding_gene | 3.0 | 7.8105 | 8.7E-29 | 2.7 | 6.3174 | 2.2E-31 |
| *comt-2* | WBGene00012518 | protein_coding_gene | 3.0 | 7.7357 | 5.3E-07 | 3.1 | 8.3642 | 1.0E-05 |
| *C01G10.4* | WBGene00007231 | protein_coding_gene | 2.9 | 7.7200 | 2.0E-04 | 5.1 | 35.0394 | 8.7E-22 |
| *C56G3.2* | WBGene00016985 | protein_coding_gene | 2.9 | 7.6137 | 8.0E-03 | 3.1 | 8.3954 | 7.3E-04 |
| *pgp-5* | WBGene00003999 | protein_coding_gene | 2.9 | 7.5214 | 1.5E-05 | 3.5 | 11.3407 | 4.2E-24 |
| *Y22D7AL.15* | WBGene00021253 | protein_coding_gene | 2.9 | 7.4634 | 3.2E-24 | 1.3 | 2.4514 | 2.3E-10 |
| *T24C4.8* | WBGene00044681 | protein_coding_gene | 2.9 | 7.3864 | 2.8E-07 | 2.3 | 4.9983 | 1.9E-06 |
| *Y47H9C.1* | WBGene00012947 | protein_coding_gene | 2.8 | 7.1352 | 1.4E-06 | 3.4 | 10.3553 | 3.1E-10 |
| *F54B8.4* | WBGene00010019 | protein_coding_gene | 2.8 | 7.0521 | 5.4E-23 | 1.1 | 2.1435 | 2.6E-05 |
| *best-5* | WBGene00007404 | protein_coding_gene | 2.8 | 6.8990 | 1.8E-08 | 1.2 | 2.2474 | 5.7E-07 |
| ***Y7A5A.6*** | **WBGene00012412** | **protein_coding_gene** | **2.8** | **6.8808** | **1.5E-03** | **-1.1** | **0.4736** | **2.1E-02** |
| ***C54C8.12*** | **WBGene00044242** | **protein_coding_gene** | **2.8** | **6.8574** | **1.5E-02** | **-2.5** | **0.1782** | **1.6E-02** |
| *W03D8.7* | WBGene00020988 | protein_coding_gene | 2.8 | 6.8366 | 2.3E-09 | 1.9 | 3.8130 | 3.9E-07 |
| ***F54E2.2*** | **WBGene00018824** | **protein_coding_gene** | **2.8** | **6.8292** | **1.1E-02** | **-1.5** | **0.3638** | **1.7E-02** |
| *irg-2* | WBGene00016783 | protein_coding_gene | 2.8 | 6.7730 | 2.1E-03 | 1.2 | 2.3322 | 3.7E-02 |
| ***Y41D4B.6*** | **WBGene00021513** | **protein_coding_gene** | **2.7** | **6.4915** | **6.2E-04** | **-1.5** | **0.3462** | **4.0E-05** |
| *oac-18* | WBGene00009230 | pseudogene | 2.7 | 6.4291 | 4.6E-03 | 1.6 | 3.1306 | 2.9E-02 |
| ***F46B6.13*** | **WBGene00044192** | **protein_coding_gene** | **2.7** | **6.3898** | **1.9E-02** | **-7.3** | **0.0061** | **7.1E-16** |
| ***F49H6.5*** | **WBGene00009908** | **protein_coding_gene** | **2.7** | **6.3865** | **1.2E-06** | **-2.0** | **0.2564** | **3.6E-12** |
| *F53F4.4* | WBGene00009988 | protein_coding_gene | 2.7 | 6.3328 | 3.0E-12 | 2.5 | 5.7895 | 6.1E-07 |
| *E02C12.8* | WBGene00017093 | protein_coding_gene | 2.7 | 6.2839 | 7.1E-10 | 2.6 | 6.2398 | 2.1E-21 |
| *fbxc-43* | WBGene00045297 | protein_coding_gene | 2.6 | 6.2554 | 6.9E-21 | 2.2 | 4.4916 | 7.2E-07 |
| *F40D4.13* | WBGene00009572 | protein_coding_gene | 2.6 | 6.2414 | 3.9E-04 | 2.1 | 4.3417 | 2.0E-06 |
| *W09H1.4* | WBGene00012374 | protein_coding_gene | 2.6 | 6.2202 | 4.9E-09 | 2.1 | 4.3438 | 7.8E-05 |
| *fbxa-66* | WBGene00021865 | protein_coding_gene | 2.6 | 6.1873 | 5.0E-03 | 1.4 | 2.7043 | 1.3E-02 |
| *linc-40* | WBGene00219770 | lincRNA_gene | 2.6 | 6.1003 | 1.2E-09 | 2.5 | 5.8307 | 2.6E-07 |
| *cpr-3* | WBGene00000783 | protein_coding_gene | 2.6 | 6.0867 | 1.0E-14 | 2.5 | 5.7379 | 7.2E-36 |
| *C34H4.1* | WBGene00016424 | protein_coding_gene | 2.6 | 6.0568 | 5.4E-08 | 3.3 | 9.8157 | 5.5E-26 |
| *Y58A7A.4* | WBGene00021978 | protein_coding_gene | 2.6 | 6.0551 | 2.9E-03 | 1.9 | 3.8314 | 1.2E-03 |
| *F43C1.7* | WBGene00023423 | protein_coding_gene | 2.6 | 6.0277 | 8.4E-04 | 3.1 | 8.7470 | 4.5E-07 |

| | | | | | | | | |
|---|---|---|---|---|---|---|---|---|
| ***nhr-127*** | **WBGene00003717** | **protein_coding_gene** | **2.6** | **5.9585** | **2.8E-03** | **-1.1** | **0.4621** | **7.2E-03** |
| *Y41D4B.17* | WBGene00021519 | protein_coding_gene | 2.5 | 5.8518 | 3.9E-12 | 3.5 | 11.6403 | 1.3E-17 |
| *Y52B11A.12* | WBGene00077539 | protein_coding_gene | 2.5 | 5.7458 | 1.8E-07 | 1.3 | 2.3803 | 5.4E-05 |
| *ZK930.5* | WBGene00014155 | protein_coding_gene | 2.5 | 5.7392 | 1.2E-05 | 1.3 | 2.5111 | 4.4E-05 |
| ***W09G12.7*** | **WBGene00021121** | **protein_coding_gene** | **2.5** | **5.6729** | **2.5E-06** | **-1.0** | **0.4904** | **1.3E-05** |
| *B0334.9* | WBGene00007147 | pseudogene | 2.5 | 5.6023 | 9.7E-08 | 1.4 | 2.6127 | 1.2E-02 |
| ***W02D7.11*** | **WBGene00020946** | **protein_coding_gene** | **2.5** | **5.5832** | **6.1E-03** | **-3.1** | **0.1163** | **3.2E-15** |
| *gly-14* | WBGene00001639 | protein_coding_gene | 2.5 | 5.5823 | 3.8E-14 | 1.6 | 3.0018 | 7.8E-07 |
| *T05F1.9* | WBGene00011494 | protein_coding_gene | 2.5 | 5.5138 | 2.5E-07 | 3.2 | 9.5016 | 1.8E-11 |
| ***ins-20*** | **WBGene00002103** | **protein_coding_gene** | **2.5** | **5.4882** | **1.1E-02** | **-2.3** | **0.2029** | **8.9E-10** |
| ***clec-60*** | **WBGene00014046** | **protein_coding_gene** | **2.4** | **5.4417** | **1.6E-03** | **-2.1** | **0.2386** | **6.6E-07** |
| ***Y105C5B.7*** | **WBGene00013648** | **pseudogene** | **2.4** | **5.4236** | **1.3E-02** | **-1.1** | **0.4661** | **4.7E-02** |
| ***T05A7.11*** | **WBGene00044383** | **protein_coding_gene** | **2.4** | **5.3843** | **6.1E-03** | **-1.8** | **0.2848** | **9.5E-08** |
| *Y71G12B.32* | WBGene00044349 | protein_coding_gene | 2.4 | 5.3583 | 2.6E-06 | 2.7 | 6.5624 | 7.6E-08 |
| *srv-7* | WBGene00005718 | protein_coding_gene | 2.4 | 5.3581 | 4.7E-08 | 1.6 | 3.0491 | 1.7E-04 |
| *C02B8.1* | WBGene00015320 | protein_coding_gene | 2.4 | 5.3311 | 2.8E-08 | 1.2 | 2.2471 | 1.6E-02 |
| *irg-4* | WBGene00008584 | protein_coding_gene | 2.4 | 5.2929 | 7.2E-09 | 5.5 | 44.8972 | 2.0E-106 |
| ***dpy-14*** | **WBGene00001075** | **protein_coding_gene** | **2.4** | **5.2302** | **3.6E-04** | **-1.1** | **0.4679** | **1.2E-02** |
| *C06E1.11* | WBGene00015526 | protein_coding_gene | 2.4 | 5.1550 | 2.9E-13 | 1.3 | 2.4029 | 2.9E-12 |
| *C15H7.2* | WBGene00014678 | pseudogene | 2.3 | 5.0635 | 1.0E-02 | 1.7 | 3.2174 | 1.8E-03 |
| *Y48B6A.16* | WBGene00050948 | pseudogene | 2.3 | 5.0528 | 1.6E-12 | 1.5 | 2.8250 | 3.6E-06 |
| *hic-1* | WBGene00020887 | protein_coding_gene | 2.3 | 4.9732 | 2.0E-10 | 2.8 | 7.0521 | 2.1E-31 |
| *F25A2.1* | WBGene00017764 | protein_coding_gene | 2.3 | 4.9243 | 1.2E-04 | 2.7 | 6.3051 | 8.7E-09 |
| ***dsl-6*** | **WBGene00001108** | **protein_coding_gene** | **2.3** | **4.9152** | **4.0E-03** | **-1.6** | **0.3267** | **1.7E-03** |
| *fbxa-198* | WBGene00011514 | pseudogene | 2.3 | 4.8878 | 3.1E-19 | 1.1 | 2.1135 | 1.0E-06 |
| *F21C10.18* | WBGene00235330 | protein_coding_gene | 2.3 | 4.8698 | 7.0E-06 | 2.6 | 6.0493 | 6.7E-09 |
| *bath-35* | WBGene00012196 | protein_coding_gene | 2.3 | 4.8223 | 6.3E-18 | 1.4 | 2.6577 | 3.6E-04 |
| *Y32G9A.13* | WBGene00044517 | pseudogene | 2.3 | 4.8186 | 3.7E-12 | 1.2 | 2.2439 | 1.5E-03 |
| *fbxa-184* | WBGene00018439 | protein_coding_gene | 2.3 | 4.8047 | 4.3E-07 | 1.7 | 3.1649 | 2.9E-04 |
| *T11F8.2* | WBGene00020434 | pseudogene | 2.3 | 4.7803 | 4.8E-21 | 1.2 | 2.2884 | 4.0E-06 |
| ***C44H9.7*** | **WBGene00008102** | **protein_coding_gene** | **2.3** | **4.7658** | **7.2E-04** | **-1.4** | **0.3707** | **1.1E-04** |
| ***C03C10.5*** | **WBGene00007271** | **protein_coding_gene** | **2.2** | **4.7325** | **1.5E-03** | **-1.1** | **0.4693** | **5.3E-03** |
| *Y67D8C.6* | WBGene00022070 | protein_coding_gene | 2.2 | 4.6932 | 1.7E-02 | 1.8 | 3.3965 | 7.8E-04 |
| *fbxa-181* | WBGene00009427 | protein_coding_gene | 2.2 | 4.6931 | 2.8E-07 | 2.0 | 4.0884 | 2.8E-04 |
| ***K10G6.4*** | **WBGene00019641** | **protein_coding_gene** | **2.2** | **4.6765** | **1.5E-02** | **-1.3** | **0.4195** | **1.1E-02** |
| *Y71H2B.11* | WBGene00022198 | protein_coding_gene | 2.2 | 4.6331 | 2.5E-30 | 2.2 | 4.6968 | 4.5E-27 |
| *BE0003N10.6* | WBGene00255598 | protein_coding_gene | 2.2 | 4.6031 | 2.2E-07 | 1.3 | 2.4317 | 2.0E-03 |
| *Y54G11A.14* | WBGene00013221 | protein_coding_gene | 2.2 | 4.5790 | 2.3E-10 | 3.2 | 9.0175 | 3.3E-32 |
| *F32D8.13* | WBGene00009335 | protein_coding_gene | 2.2 | 4.5400 | 3.1E-11 | 1.0 | 2.0541 | 2.9E-04 |
| *F41B5.1* | WBGene00023284 | protein_coding_gene | 2.2 | 4.5131 | 2.8E-22 | 1.9 | 3.8167 | 5.6E-11 |
| ***cky-1*** | **WBGene00000521** | **protein_coding_gene** | **2.2** | **4.5081** | **1.9E-02** | **-1.9** | **0.2623** | **5.1E-03** |
| *Y71G12B.2* | WBGene00022142 | protein_coding_gene | 2.2 | 4.4715 | 2.4E-05 | 3.1 | 8.5515 | 2.1E-08 |
| *F48G7.13* | WBGene00018624 | protein_coding_gene | 2.2 | 4.4660 | 1.5E-02 | 3.5 | 11.1811 | 2.3E-05 |
| *fbxc-46* | WBGene00012569 | pseudogene | 2.2 | 4.4453 | 8.1E-25 | 1.4 | 2.5714 | 1.1E-08 |
| *parg-2* | WBGene00004052 | protein_coding_gene | 2.1 | 4.3772 | 1.5E-03 | 2.4 | 5.1672 | 3.3E-07 |
| *F40D4.17* | WBGene00194682 | protein_coding_gene | 2.1 | 4.3654 | 7.3E-16 | 1.3 | 2.4392 | 4.8E-07 |
| *fbxa-94* | WBGene00009225 | protein_coding_gene | 2.1 | 4.3591 | 2.2E-05 | 1.7 | 3.2547 | 2.4E-02 |
| *fbxa-93* | WBGene00012368 | pseudogene | 2.1 | 4.3487 | 1.0E-16 | 1.3 | 2.5059 | 3.0E-05 |
| *B0348.2* | WBGene00015152 | protein_coding_gene | 2.1 | 4.3431 | 2.3E-03 | 1.7 | 3.2887 | 1.8E-06 |
| *C49G7.7* | WBGene00016785 | protein_coding_gene | 2.1 | 4.3377 | 9.0E-04 | 1.9 | 3.7687 | 7.2E-06 |
| *linc-20* | WBGene00219736 | lincRNA_gene | 2.1 | 4.2845 | 4.0E-07 | 1.3 | 2.5418 | 9.3E-04 |
| *E04D5.4* | WBGene00008483 | protein_coding_gene | 2.1 | 4.2322 | 1.5E-04 | 1.7 | 3.2795 | 1.3E-03 |
| *T26C12.2* | WBGene00020832 | protein_coding_gene | 2.1 | 4.2213 | 4.6E-03 | 1.4 | 2.7315 | 7.1E-03 |
| *Y55F3AM.11* | WBGene00021928 | protein_coding_gene | 2.1 | 4.2193 | 6.4E-20 | 1.0 | 2.0644 | 3.6E-06 |
| ***C08F1.11*** | **WBGene00044417** | **protein_coding_gene** | **2.1** | **4.2184** | **1.0E-03** | **-1.5** | **0.3499** | **4.4E-02** |
| *C49C8.3* | WBGene00016767 | protein_coding_gene | 2.1 | 4.2154 | 1.1E-24 | 1.4 | 2.5780 | 1.6E-10 |
| *T24A6.1* | WBGene00020744 | protein_coding_gene | 2.1 | 4.1951 | 1.0E-05 | 1.6 | 3.0942 | 4.1E-03 |
| *his-63* | WBGene00001937 | protein_coding_gene | 2.1 | 4.1876 | 5.1E-03 | 1.5 | 2.7699 | 4.0E-02 |
| *hpo-38* | WBGene00021755 | protein_coding_gene | 2.1 | 4.1851 | 2.4E-07 | 2.0 | 3.9735 | 7.6E-08 |
| *inx-20* | WBGene00002142 | protein_coding_gene | 2.1 | 4.1833 | 1.6E-04 | 1.0 | 2.0271 | 3.2E-02 |
| *F54D10.4* | WBGene00018805 | protein_coding_gene | 2.1 | 4.1828 | 2.4E-10 | 1.7 | 3.2197 | 6.1E-08 |
| ***ZC443.2*** | **WBGene00013897** | **protein_coding_gene** | **2.0** | **4.1142** | **1.6E-02** | **-1.6** | **0.3195** | **2.0E-04** |
| *D2096.10* | WBGene00017077 | protein_coding_gene | 2.0 | 4.1098 | 1.1E-17 | 1.6 | 3.0720 | 7.1E-14 |
| *C07D10.6* | WBGene00044673 | protein_coding_gene | 2.0 | 4.1031 | 2.6E-02 | 1.5 | 2.8708 | 4.7E-02 |
| *Y17G9B.4* | WBGene00021201 | protein_coding_gene | 2.0 | 4.0965 | 3.7E-21 | 1.6 | 2.9895 | 2.3E-16 |
| *fbxa-203* | WBGene00021037 | pseudogene | 2.0 | 4.0963 | 2.9E-18 | 2.1 | 4.3947 | 5.0E-22 |
| *hpo-6* | WBGene00021518 | protein_coding_gene | 2.0 | 4.0574 | 3.1E-12 | 3.0 | 8.2243 | 1.2E-11 |
| *bath-7* | WBGene00018667 | protein_coding_gene | 2.0 | 4.0414 | 1.9E-14 | 2.6 | 6.1113 | 1.7E-02 |
| *dyf-3* | WBGene00001119 | protein_coding_gene | 2.0 | 4.0366 | 2.8E-07 | 1.3 | 2.4529 | 9.9E-12 |
| *C18H7.11* | WBGene00044457 | protein_coding_gene | 2.0 | 4.0087 | 4.5E-05 | 1.1 | 2.1769 | 7.6E-03 |

| | | | | | | | | |
|---|---|---|---|---|---|---|---|---|
| *M02H5.8* | WBGene00019744 | protein_coding_gene | 2.0 | 3.9787 | 2.3E-15 | 1.0 | 2.0554 | 7.4E-03 |
| *C37A2.6* | WBGene00016492 | protein_coding_gene | 2.0 | 3.9445 | 6.7E-14 | 1.2 | 2.3485 | 5.1E-05 |
| *linc-63* | WBGene00219725 | lincRNA_gene | 2.0 | 3.9224 | 6.9E-04 | 2.2 | 4.6416 | 3.0E-03 |
| *Y34F4.3* | WBGene00021338 | protein_coding_gene | 2.0 | 3.9194 | 2.4E-04 | 1.2 | 2.3151 | 4.8E-04 |
| *MTCE.7* | WBGene00014454 | rRNA_gene | 2.0 | 3.9189 | 3.2E-09 | 1.7 | 3.2551 | 4.5E-02 |
| *selt-1.2* | WBGene00009238 | protein_coding_gene | 2.0 | 3.9019 | 9.5E-06 | 1.6 | 2.9902 | 5.3E-05 |
| *hsf-2* | WBGene00013134 | pseudogene | 2.0 | 3.8842 | 4.2E-08 | 1.5 | 2.7845 | 4.2E-03 |
| *linc-61* | WBGene00195170 | lincRNA_gene | 2.0 | 3.8653 | 7.5E-03 | 3.2 | 9.2460 | 9.2E-07 |
| *Y37H2A.14* | WBGene00045416 | protein_coding_gene | 2.0 | 3.8642 | 2.6E-05 | 4.3 | 19.6254 | 2.9E-35 |
| *asm-3* | WBGene00000213 | protein_coding_gene | 1.9 | 3.8610 | 8.3E-03 | 2.1 | 4.2460 | 4.0E-08 |
| *clc-1* | WBGene00000522 | protein_coding_gene | 1.9 | 3.8520 | 1.1E-07 | 1.5 | 2.8556 | 2.2E-05 |
| *crn-2* | WBGene00000795 | protein_coding_gene | 1.9 | 3.8501 | 7.0E-32 | 1.5 | 2.8166 | 6.0E-17 |
| *btb-1* | WBGene00001963 | pseudogene | 1.9 | 3.8075 | 2.6E-07 | 1.3 | 2.4487 | 5.3E-03 |
| *fbxa-18* | WBGene00008633 | protein_coding_gene | 1.9 | 3.8051 | 1.9E-02 | 4.1 | 17.6176 | 1.1E-08 |
| *T28A8.8* | WBGene00194852 | protein_coding_gene | 1.9 | 3.8042 | 6.3E-18 | 1.1 | 2.2054 | 1.3E-07 |
| *K11D2.1* | WBGene00010768 | protein_coding_gene | 1.9 | 3.7944 | 1.3E-13 | 1.1 | 2.1982 | 7.7E-05 |
| *fbxa-105* | WBGene00015362 | protein_coding_gene | 1.9 | 3.7854 | 4.3E-09 | 1.7 | 3.2467 | 2.6E-15 |
| ***abf-2*** | **WBGene00000013** | **protein_coding_gene** | **1.9** | **3.7675** | **1.7E-03** | **-1.9** | **0.2757** | **3.8E-08** |
| ***T20F5.4*** | **WBGene00020626** | **protein_coding_gene** | **1.9** | **3.7567** | **1.9E-04** | **-1.3** | **0.4149** | **3.3E-04** |
| ***dpy-17*** | **WBGene00001076** | **protein_coding_gene** | **1.9** | **3.7565** | **3.2E-05** | **-1.2** | **0.4208** | **9.2E-05** |
| ***B0205.4*** | **WBGene00015019** | **protein_coding_gene** | **1.9** | **3.7545** | **2.5E-03** | **-1.0** | **0.4949** | **1.1E-02** |
| *linc-29* | WBGene00189951 | lincRNA_gene | 1.9 | 3.7383 | 9.2E-09 | 2.5 | 5.7384 | 1.6E-22 |
| *F57A10.4* | WBGene00010185 | protein_coding_gene | 1.9 | 3.7246 | 5.9E-21 | 1.1 | 2.1645 | 9.4E-12 |
| *F14F9.4* | WBGene00017467 | protein_coding_gene | 1.9 | 3.7116 | 1.8E-03 | 1.1 | 2.2028 | 9.7E-08 |
| *mul-1* | WBGene00018646 | protein_coding_gene | 1.9 | 3.7107 | 8.1E-05 | 1.6 | 3.0482 | 2.8E-06 |
| *C15H9.11* | WBGene00015805 | protein_coding_gene | 1.9 | 3.7003 | 6.1E-20 | 2.5 | 5.8129 | 3.0E-50 |
| *Y58A7A.3* | WBGene00021977 | protein_coding_gene | 1.9 | 3.6850 | 4.2E-05 | 1.3 | 2.3863 | 2.5E-11 |
| *F02H6.1* | WBGene00008534 | pseudogene | 1.9 | 3.6714 | 1.8E-15 | 1.9 | 3.7271 | 8.1E-08 |
| *nduo-6* | WBGene00010957 | protein_coding_gene | 1.9 | 3.6641 | 2.5E-11 | 1.6 | 3.0639 | 7.7E-11 |
| *Y66D12A.6* | WBGene00013432 | protein_coding_gene | 1.9 | 3.6557 | 9.9E-21 | 1.9 | 3.8329 | 6.5E-19 |
| *anr-32* | WBGene00219683 | antisense_lncRNA_gene | 1.9 | 3.6432 | 3.8E-02 | 2.6 | 5.9812 | 3.1E-04 |
| *T16G1.5* | WBGene00011799 | protein_coding_gene | 1.9 | 3.6411 | 2.0E-04 | 2.0 | 3.9583 | 2.3E-10 |
| *bath-6* | WBGene00018666 | pseudogene | 1.9 | 3.6297 | 2.3E-11 | 2.0 | 3.8951 | 6.4E-03 |
| *bath-34* | WBGene00012195 | protein_coding_gene | 1.9 | 3.6132 | 3.7E-18 | 1.5 | 2.8256 | 1.4E-08 |
| *enu-3.6* | WBGene00010477 | protein_coding_gene | 1.9 | 3.6120 | 4.0E-05 | 2.1 | 4.3505 | 6.5E-08 |
| ***cutl-28*** | **WBGene00018256** | **protein_coding_gene** | **1.8** | **3.5948** | **1.5E-03** | **-1.4** | **0.3725** | **2.1E-02** |
| *C48D1.9* | WBGene00194643 | protein_coding_gene | 1.8 | 3.5945 | 2.1E-04 | 1.2 | 2.3361 | 7.1E-04 |
| *oac-57* | WBGene00019579 | protein_coding_gene | 1.8 | 3.5841 | 9.2E-07 | 2.1 | 4.3221 | 1.5E-13 |
| *Y48G1BM.8* | WBGene00021673 | pseudogene | 1.8 | 3.5682 | 5.2E-05 | 2.3 | 4.7696 | 1.3E-06 |
| ***C18H7.1*** | **WBGene00015993** | **protein_coding_gene** | **1.8** | **3.5626** | **2.2E-02** | **-5.8** | **0.0179** | **3.2E-31** |
| *acd-4* | WBGene00017879 | protein_coding_gene | 1.8 | 3.5611 | 2.9E-02 | 2.4 | 5.2974 | 8.4E-04 |
| *fbxa-124* | WBGene00014916 | pseudogene | 1.8 | 3.5508 | 3.4E-21 | 1.0 | 2.0248 | 1.3E-07 |
| *F40F12.8* | WBGene00014755 | pseudogene | 1.8 | 3.5418 | 6.4E-03 | 1.5 | 2.8212 | 2.6E-05 |
| ***F55H12.7*** | **WBGene00219283** | **protein_coding_gene** | **1.8** | **3.5307** | **3.5E-02** | **-1.6** | **0.3307** | **4.2E-03** |
| *sws-1* | WBGene00012698 | protein_coding_gene | 1.8 | 3.5301 | 3.4E-19 | 1.3 | 2.3810 | 2.5E-13 |
| ***unc-42*** | **WBGene00006778** | **protein_coding_gene** | **1.8** | **3.5141** | **3.1E-03** | **-1.1** | **0.4641** | **5.6E-03** |
| ***E02H9.9*** | **WBGene00017106** | **protein_coding_gene** | **1.8** | **3.5063** | **1.5E-03** | **-3.7** | **0.0792** | **1.3E-31** |
| *Y68A4A.13* | WBGene00044212 | protein_coding_gene | 1.8 | 3.4874 | 1.4E-02 | 2.4 | 5.1507 | 4.6E-02 |
| *ugt-31* | WBGene00021464 | protein_coding_gene | 1.8 | 3.4731 | 2.6E-05 | 2.5 | 5.8328 | 1.3E-08 |
| *C15F1.8* | WBGene00015797 | protein_coding_gene | 1.8 | 3.4509 | 2.1E-20 | 1.7 | 3.2179 | 3.8E-15 |
| *F55G11.8* | WBGene00010128 | protein_coding_gene | 1.8 | 3.4459 | 4.1E-05 | 2.3 | 4.8482 | 4.8E-18 |
| *fbxa-191* | WBGene00010209 | pseudogene | 1.8 | 3.4430 | 3.3E-10 | 1.2 | 2.3245 | 3.2E-08 |
| *spe-5* | WBGene00004959 | protein_coding_gene | 1.8 | 3.4184 | 4.1E-20 | 1.0 | 2.0203 | 5.5E-10 |
| *clec-74* | WBGene00021585 | protein_coding_gene | 1.8 | 3.4183 | 4.0E-03 | 1.9 | 3.7036 | 7.5E-09 |
| *Y17G7B.8* | WBGene00012462 | protein_coding_gene | 1.8 | 3.4164 | 9.1E-12 | 2.5 | 5.5663 | 1.0E-36 |
| *C10C6.9* | WBGene00045175 | pseudogene | 1.8 | 3.4062 | 7.6E-04 | 1.1 | 2.1230 | 1.2E-02 |
| *T08B2.4* | WBGene00020345 | protein_coding_gene | 1.8 | 3.3983 | 4.1E-08 | 1.2 | 2.2305 | 9.1E-04 |
| *F53G12.4* | WBGene00018772 | protein_coding_gene | 1.8 | 3.3929 | 5.0E-04 | 1.1 | 2.1038 | 4.3E-02 |
| ***F54B11.10*** | **WBGene00010033** | **protein_coding_gene** | **1.8** | **3.3921** | **2.0E-02** | **-1.5** | **0.3441** | **1.9E-02** |
| ***mtl-1*** | **WBGene00003473** | **protein_coding_gene** | **1.8** | **3.3886** | **4.1E-02** | **-5.3** | **0.0248** | **9.0E-44** |
| *trp-4* | WBGene00006616 | protein_coding_gene | 1.7 | 3.3605 | 2.3E-06 | 2.8 | 7.2015 | 5.0E-09 |
| *C38D4.7* | WBGene00008007 | protein_coding_gene | 1.7 | 3.3570 | 2.4E-09 | 1.9 | 3.6793 | 3.2E-07 |
| ***cutl-2*** | **WBGene00013145** | **protein_coding_gene** | **1.7** | **3.3456** | **9.7E-04** | **-1.2** | **0.4370** | **8.9E-06** |
| *linc-9* | WBGene00219678 | lincRNA_gene | 1.7 | 3.3366 | 1.3E-06 | 1.3 | 2.4814 | 3.8E-06 |
| *Y53G8B.3* | WBGene00021819 | pseudogene | 1.7 | 3.3180 | 1.1E-03 | 1.7 | 3.3557 | 2.4E-04 |
| *E02D9.3* | WBGene00304811 | protein_coding_gene | 1.7 | 3.3174 | 1.2E-13 | 1.1 | 2.1745 | 3.8E-09 |
| *B0304.4* | WBGene00015136 | protein_coding_gene | 1.7 | 3.3123 | 2.0E-27 | 1.3 | 2.5471 | 1.2E-17 |
| *fbxa-95* | WBGene00009222 | protein_coding_gene | 1.7 | 3.2986 | 3.9E-09 | 1.1 | 2.2093 | 4.3E-03 |
| *F58D5.3* | WBGene00010243 | protein_coding_gene | 1.7 | 3.2936 | 6.0E-03 | 1.3 | 2.4799 | 3.5E-02 |
| *Y71A12B.14* | WBGene00014947 | pseudogene | 1.7 | 3.2889 | 1.0E-08 | 1.4 | 2.7021 | 1.0E-06 |

| | | | | | | | | |
|---|---|---|---|---|---|---|---|---|
| *B0416.10* | WBGene00206482 | protein_coding_gene | 1.7 | 3.2888 | 1.2E-05 | 1.4 | 2.5625 | 3.8E-03 |
| *F53G12.9* | WBGene00018774 | protein_coding_gene | 1.7 | 3.2877 | 2.8E-05 | 1.3 | 2.4882 | 2.0E-03 |
| *Y38F2AL.6* | WBGene00021418 | pseudogene | 1.7 | 3.2780 | 1.5E-03 | 1.6 | 2.9596 | 2.3E-03 |
| *kbp-1* | WBGene00020064 | protein_coding_gene | 1.7 | 3.2732 | 3.0E-12 | 1.1 | 2.1450 | 5.5E-14 |
| *bath-13* | WBGene00018223 | protein_coding_gene | 1.7 | 3.2720 | 1.0E-03 | 2.3 | 4.8897 | 4.1E-04 |
| *fbxa-123* | WBGene00014918 | pseudogene | 1.7 | 3.2707 | 4.5E-18 | 1.2 | 2.3598 | 1.6E-10 |
| *T07F10.5* | WBGene00011591 | protein_coding_gene | 1.7 | 3.2634 | 2.1E-08 | 1.6 | 3.0811 | 5.1E-09 |
| *E_BE45912.2* | WBGene00017153 | protein_coding_gene | 1.7 | 3.2614 | 2.3E-14 | 1.0 | 2.0401 | 3.8E-07 |
| *Y37H9A.4* | WBGene00014870 | pseudogene | 1.7 | 3.2548 | 1.8E-05 | 1.5 | 2.9026 | 1.0E-03 |
| ***oac-14*** | **WBGene00008602** | **protein_coding_gene** | **1.7** | **3.2469** | **3.8E-02** | **-2.2** | **0.2151** | **1.1E-15** |
| *fbxa-122* | WBGene00014917 | pseudogene | 1.7 | 3.2204 | 1.9E-11 | 1.1 | 2.0921 | 9.1E-06 |
| *F07F6.7* | WBGene00017219 | protein_coding_gene | 1.7 | 3.2200 | 2.1E-08 | 1.8 | 3.4111 | 5.6E-10 |
| *T19D12.4* | WBGene00020579 | protein_coding_gene | 1.7 | 3.2075 | 3.3E-05 | 2.2 | 4.7432 | 3.7E-10 |
| *fbxa-15* | WBGene00022327 | protein_coding_gene | 1.7 | 3.1986 | 8.3E-04 | 5.0 | 31.6273 | 1.7E-07 |
| *K10D2.8* | WBGene00189952 | protein_coding_gene | 1.7 | 3.1897 | 2.8E-05 | 1.0 | 2.0346 | 5.3E-05 |
| *F26F4.8* | WBGene00005011 | protein_coding_gene | 1.7 | 3.1893 | 2.2E-20 | 1.0 | 2.0700 | 6.5E-13 |
| *F54H5.2* | WBGene00018839 | protein_coding_gene | 1.7 | 3.1883 | 9.0E-03 | 1.7 | 3.1695 | 8.7E-06 |
| *T08D2.5* | WBGene00011610 | protein_coding_gene | 1.7 | 3.1867 | 1.1E-04 | 2.0 | 3.8809 | 1.1E-07 |
| ***F31D4.5*** | **WBGene00009292** | **protein_coding_gene** | **1.7** | **3.1841** | **1.2E-03** | **-1.7** | **0.3127** | **2.1E-03** |
| *nhr-112* | WBGene00003702 | protein_coding_gene | 1.7 | 3.1719 | 4.4E-06 | 1.0 | 2.0403 | 1.1E-04 |
| *K09D9.12* | WBGene00019569 | pseudogene | 1.7 | 3.1704 | 6.6E-14 | 1.5 | 2.8116 | 1.6E-07 |
| *fbxa-121* | WBGene00012485 | pseudogene | 1.7 | 3.1641 | 1.5E-07 | 1.1 | 2.0832 | 3.7E-04 |
| *Y71F9AL.4* | WBGene00022110 | protein_coding_gene | 1.7 | 3.1622 | 3.3E-13 | 1.2 | 2.2651 | 5.2E-10 |
| *H04D03.3* | WBGene00010364 | protein_coding_gene | 1.7 | 3.1604 | 3.2E-04 | 2.2 | 4.5801 | 6.2E-04 |
| ***C06E4.8*** | **WBGene00015537** | **protein_coding_gene** | **1.7** | **3.1450** | **1.1E-04** | **-1.4** | **0.3673** | **2.8E-09** |
| *ndfl-4* | WBGene00010958 | protein_coding_gene | 1.6 | 3.1347 | 1.6E-03 | 1.6 | 3.1248 | 1.3E-03 |
| *C29A12.2* | WBGene00007798 | pseudogene | 1.6 | 3.1267 | 6.1E-03 | 1.2 | 2.3217 | 3.0E-02 |
| *fbxc-44* | WBGene00015819 | protein_coding_gene | 1.6 | 3.1118 | 4.7E-11 | 1.7 | 3.2493 | 1.6E-25 |
| *Y48G8AL.7* | WBGene00021687 | protein_coding_gene | 1.6 | 3.1036 | 5.8E-13 | 1.2 | 2.3732 | 1.6E-11 |
| *C46E10.8* | WBGene00016712 | protein_coding_gene | 1.6 | 3.0977 | 3.2E-08 | 1.4 | 2.6126 | 5.3E-04 |
| *C18H2.2* | WBGene00015989 | pseudogene | 1.6 | 3.0973 | 3.0E-09 | 2.0 | 4.1159 | 8.8E-19 |
| *bir-2* | WBGene00000250 | protein_coding_gene | 1.6 | 3.0962 | 4.7E-21 | 1.7 | 3.3537 | 1.9E-39 |
| *fbxa-106* | WBGene00013348 | protein_coding_gene | 1.6 | 3.0892 | 1.6E-11 | 1.1 | 2.0815 | 1.1E-08 |
| *T08B2.11* | WBGene00020349 | protein_coding_gene | 1.6 | 3.0873 | 2.6E-21 | 1.7 | 3.3064 | 6.0E-23 |
| *Y37H2C.1* | WBGene00012573 | pseudogene | 1.6 | 3.0838 | 9.6E-04 | 2.8 | 7.0748 | 3.3E-12 |
| *Y116A8A.10* | WBGene00014975 | pseudogene | 1.6 | 3.0492 | 3.3E-04 | 1.0 | 2.0432 | 3.8E-04 |
| *nhr-236* | WBGene00021417 | protein_coding_gene | 1.6 | 3.0479 | 6.6E-03 | 2.0 | 3.9589 | 9.0E-03 |
| *C09G5.7* | WBGene00007489 | protein_coding_gene | 1.6 | 3.0424 | 2.9E-12 | 1.0 | 2.0110 | 1.7E-03 |
| ***pqn-73*** | **WBGene00004155** | **protein_coding_gene** | **1.6** | **3.0370** | **4.2E-02** | **-1.6** | **0.3413** | **2.2E-03** |
| *ZC47.8* | WBGene00014978 | pseudogene | 1.6 | 3.0275 | 3.7E-04 | 1.7 | 3.1806 | 9.4E-06 |
| *bath-12* | WBGene00019131 | pseudogene | 1.6 | 3.0265 | 5.1E-12 | 2.2 | 4.5183 | 7.7E-23 |
| *W03F9.2* | WBGene00021000 | protein_coding_gene | 1.6 | 3.0237 | 2.1E-13 | 1.4 | 2.5744 | 3.4E-09 |
| *T25B9.8* | WBGene00012014 | protein_coding_gene | 1.6 | 3.0231 | 1.0E-22 | 1.2 | 2.2443 | 6.5E-14 |
| *chst-1* | WBGene00008050 | protein_coding_gene | 1.6 | 3.0091 | 1.4E-04 | 1.4 | 2.6054 | 2.9E-02 |
| *F01G10.6* | WBGene00008509 | protein_coding_gene | 1.6 | 2.9977 | 2.2E-03 | 2.5 | 5.4706 | 6.7E-10 |
| *C14C10.2* | WBGene00007585 | protein_coding_gene | 1.6 | 2.9916 | 5.8E-19 | 1.0 | 2.0019 | 3.7E-09 |
| *cld-9* | WBGene00009432 | protein_coding_gene | 1.6 | 2.9836 | 4.2E-06 | 3.7 | 12.7242 | 1.0E-49 |
| ***Y65B4BM.3*** | **WBGene00302974** | **protein_coding_gene** | **1.6** | **2.9806** | **1.1E-04** | **-1.0** | **0.4915** | **7.7E-03** |
| *C16C8.14* | WBGene00015852 | protein_coding_gene | 1.6 | 2.9793 | 1.4E-21 | 1.3 | 2.4651 | 2.2E-11 |
| *F32A11.5* | WBGene00009310 | pseudogene | 1.6 | 2.9770 | 1.8E-04 | 2.2 | 4.5706 | 9.7E-06 |
| *spp-18* | WBGene00005003 | protein_coding_gene | 1.6 | 2.9710 | 7.4E-09 | 1.2 | 2.2400 | 7.0E-17 |
| *R119.1* | WBGene00020087 | pseudogene | 1.6 | 2.9708 | 2.2E-19 | 1.3 | 2.4893 | 3.2E-17 |
| *damt-1* | WBGene00015939 | protein_coding_gene | 1.6 | 2.9694 | 2.5E-19 | 1.1 | 2.1991 | 1.3E-15 |
| ***peb-1*** | **WBGene00003968** | **protein_coding_gene** | **1.6** | **2.9685** | **3.7E-03** | **-1.0** | **0.4855** | **3.9E-06** |
| *C30H6.8* | WBGene00007825 | protein_coding_gene | 1.6 | 2.9559 | 1.9E-16 | 1.0 | 2.0688 | 2.8E-16 |
| *fbxa-114* | WBGene00013758 | protein_coding_gene | 1.6 | 2.9526 | 3.9E-26 | 1.5 | 2.7798 | 2.3E-14 |
| *F28F8.10* | WBGene00044219 | pseudogene | 1.6 | 2.9517 | 9.7E-11 | 1.6 | 3.0055 | 7.3E-11 |
| *F56D2.5* | WBGene00018966 | protein_coding_gene | 1.6 | 2.9517 | 6.4E-04 | 1.4 | 2.7120 | 5.2E-10 |
| ***W01C9.1*** | **WBGene00012175** | **protein_coding_gene** | **1.6** | **2.9516** | **3.6E-02** | **-1.4** | **0.3760** | **1.9E-02** |
| *C16A11.5* | WBGene00015818 | protein_coding_gene | 1.6 | 2.9516 | 9.2E-28 | 1.5 | 2.9122 | 4.4E-22 |
| ***C28C12.3*** | **WBGene00016179** | **protein_coding_gene** | **1.6** | **2.9510** | **4.0E-02** | **-5.7** | **0.0195** | **3.1E-62** |
| *pot-1* | WBGene00015105 | protein_coding_gene | 1.5 | 2.9216 | 2.2E-25 | 1.2 | 2.3217 | 4.5E-19 |
| *C46C2.3* | WBGene00008111 | protein_coding_gene | 1.5 | 2.9199 | 6.0E-03 | 2.2 | 4.6715 | 1.6E-06 |
| *clec-65* | WBGene00009396 | protein_coding_gene | 1.5 | 2.9147 | 7.6E-25 | 1.8 | 3.3786 | 2.8E-30 |
| ***oac-51*** | **WBGene00020976** | **protein_coding_gene** | **1.5** | **2.9092** | **4.0E-03** | **-1.2** | **0.4307** | **2.4E-03** |
| ***M01D1.12*** | **WBGene00269373** | **protein_coding_gene** | **1.5** | **2.9037** | **1.8E-02** | **-1.1** | **0.4778** | **2.7E-02** |
| *F20D6.1* | WBGene00017633 | protein_coding_gene | 1.5 | 2.9033 | 2.6E-02 | 1.2 | 2.3509 | 1.6E-02 |
| *lido-4* | WBGene00015589 | protein_coding_gene | 1.5 | 2.8945 | 3.7E-04 | 1.3 | 2.4072 | 1.0E-02 |
| *F19B2.12* | WBGene00194812 | pseudogene | 1.5 | 2.8896 | 4.7E-12 | 1.1 | 2.1661 | 4.3E-06 |
| *Y67D2.4* | WBGene00022054 | protein_coding_gene | 1.5 | 2.8808 | 6.3E-19 | 1.3 | 2.3951 | 3.8E-16 |

| | | | | | | | | |
|---|---|---|---|---|---|---|---|---|
| ***C44B12.3*** | **WBGene00016637** | **protein_coding_gene** | **1.5** | **2.8760** | **1.1E-02** | **-1.3** | **0.3977** | **1.9E-05** |
| ***T02H6.9*** | **WBGene00020179** | **protein_coding_gene** | **1.5** | **2.8708** | **5.0E-02** | **-2.3** | **0.2092** | **1.5E-12** |
| *Y37H2A.7* | WBGene00012567 | protein_coding_gene | 1.5 | 2.8687 | 4.0E-07 | 1.2 | 2.2563 | 1.6E-08 |
| *ccch-3* | WBGene00012277 | protein_coding_gene | 1.5 | 2.8632 | 2.1E-25 | 1.0 | 2.0246 | 7.2E-21 |
| ***cup-16*** | **WBGene00007919** | **protein_coding_gene** | **1.5** | **2.8625** | **2.6E-03** | **-1.9** | **0.2598** | **3.3E-20** |
| *C16C8.13* | WBGene00015851 | protein_coding_gene | 1.5 | 2.8489 | 3.0E-09 | 1.2 | 2.2477 | 3.5E-05 |
| *prg-2* | WBGene00004179 | pseudogene | 1.5 | 2.8476 | 8.6E-12 | 2.4 | 5.1852 | 1.3E-58 |
| *ave-1* | WBGene00044341 | protein_coding_gene | 1.5 | 2.8432 | 2.3E-13 | 1.1 | 2.1420 | 1.7E-08 |
| *C03E10.3* | WBGene00007281 | protein_coding_gene | 1.5 | 2.8331 | 2.0E-04 | 1.3 | 2.4577 | 2.7E-03 |
| *C16C8.5* | WBGene00015843 | protein_coding_gene | 1.5 | 2.8312 | 4.6E-18 | 1.4 | 2.6696 | 3.2E-17 |
| *Y54E5A.2* | WBGene00013198 | protein_coding_gene | 1.5 | 2.8251 | 6.0E-06 | 1.1 | 2.0773 | 3.8E-06 |
| *F14D2.15* | WBGene00023427 | protein_coding_gene | 1.5 | 2.8229 | 2.9E-16 | 1.5 | 2.7479 | 1.8E-02 |
| *kbp-4* | WBGene00022357 | protein_coding_gene | 1.5 | 2.8202 | 8.4E-27 | 1.0 | 2.0311 | 1.3E-13 |
| *T08B6.2* | WBGene00020352 | pseudogene | 1.5 | 2.8103 | 6.0E-03 | 1.8 | 3.5385 | 3.9E-04 |
| *K07D4.1* | WBGene00023318 | pseudogene | 1.5 | 2.8082 | 2.3E-08 | 1.4 | 2.6584 | 2.1E-07 |
| *C18H2.4* | WBGene00015991 | pseudogene | 1.5 | 2.8081 | 3.3E-05 | 1.6 | 3.0502 | 2.2E-05 |
| *rpl-25.2* | WBGene00004439 | protein_coding_gene | 1.5 | 2.8055 | 2.2E-17 | 1.5 | 2.8226 | 6.5E-35 |
| ***F37A8.5*** | **WBGene00009503** | **protein_coding_gene** | **1.5** | **2.8044** | **1.3E-02** | **-1.9** | **0.2622** | **3.7E-32** |
| *Y39B6A.37* | WBGene00012695 | protein_coding_gene | 1.5 | 2.8009 | 1.1E-09 | 1.1 | 2.1686 | 4.8E-12 |
| *C48B4.7* | WBGene00008170 | protein_coding_gene | 1.5 | 2.8004 | 1.5E-10 | 1.6 | 3.0407 | 1.0E-20 |
| *C46A5.6* | WBGene00016702 | protein_coding_gene | 1.5 | 2.7987 | 5.5E-28 | 1.3 | 2.4174 | 2.6E-22 |
| *mttu-1* | WBGene00007114 | protein_coding_gene | 1.5 | 2.7985 | 2.9E-12 | 1.1 | 2.0927 | 1.0E-07 |
| *ZK616.3* | WBGene00022773 | protein_coding_gene | 1.5 | 2.7973 | 3.0E-17 | 1.1 | 2.1596 | 7.1E-13 |
| *nos-1* | WBGene00003783 | protein_coding_gene | 1.5 | 2.7969 | 4.4E-17 | 1.7 | 3.2132 | 1.8E-19 |
| *C08F11.7* | WBGene00007454 | protein_coding_gene | 1.5 | 2.7957 | 1.2E-14 | 2.0 | 4.0128 | 1.6E-02 |
| *sip-1* | WBGene00004798 | protein_coding_gene | 1.5 | 2.7937 | 2.0E-19 | 1.4 | 2.6853 | 4.7E-30 |
| *blos-1* | WBGene00011872 | protein_coding_gene | 1.5 | 2.7936 | 1.4E-19 | 1.0 | 2.0222 | 5.8E-14 |
| *W06A11.4* | WBGene00021055 | protein_coding_gene | 1.5 | 2.7921 | 2.8E-14 | 1.7 | 3.3351 | 3.9E-15 |
| *rpl-21* | WBGene00004433 | protein_coding_gene | 1.5 | 2.7879 | 4.5E-36 | 1.6 | 3.1027 | 5.1E-39 |
| *parn-1* | WBGene00010734 | protein_coding_gene | 1.5 | 2.7855 | 4.2E-12 | 1.9 | 3.6901 | 2.3E-26 |
| *Y57A10A.3* | WBGene00013248 | protein_coding_gene | 1.5 | 2.7851 | 2.0E-13 | 1.9 | 3.8146 | 1.1E-16 |
| *C56A3.8* | WBGene00008346 | protein_coding_gene | 1.5 | 2.7839 | 2.1E-21 | 1.1 | 2.1180 | 7.1E-09 |
| *hil-8* | WBGene00001859 | protein_coding_gene | 1.5 | 2.7833 | 4.0E-03 | 1.9 | 3.6481 | 1.4E-03 |
| *T07D4.5* | WBGene00045407 | protein_coding_gene | 1.5 | 2.7782 | 3.4E-15 | 1.1 | 2.1676 | 4.2E-18 |
| *F15D4.5* | WBGene00008862 | protein_coding_gene | 1.5 | 2.7764 | 7.3E-04 | 2.2 | 4.4889 | 4.5E-13 |
| *F55A3.7* | WBGene00018852 | pseudogene | 1.5 | 2.7758 | 2.2E-09 | 3.6 | 12.0302 | 1.2E-17 |
| *Y94H6A.2* | WBGene00022375 | protein_coding_gene | 1.5 | 2.7738 | 1.6E-02 | 1.1 | 2.0770 | 2.8E-02 |
| *Y39B6A.13* | WBGene00012675 | protein_coding_gene | 1.5 | 2.7714 | 1.9E-29 | 1.2 | 2.3017 | 3.8E-15 |
| *nsun-5* | WBGene00013151 | protein_coding_gene | 1.5 | 2.7693 | 1.1E-13 | 1.2 | 2.3596 | 8.6E-14 |
| ***C32F10.4*** | **WBGene00016332** | **protein_coding_gene** | **1.5** | **2.7673** | **3.1E-06** | **-1.0** | **0.4898** | **6.0E-04** |
| *Y37B11A.3* | WBGene00021345 | protein_coding_gene | 1.5 | 2.7661 | 4.0E-05 | 1.8 | 3.4610 | 1.9E-04 |
| *nth-1* | WBGene00011201 | protein_coding_gene | 1.5 | 2.7638 | 5.7E-14 | 1.2 | 2.3659 | 3.2E-12 |
| *sir-2.4* | WBGene00004803 | protein_coding_gene | 1.5 | 2.7627 | 4.5E-17 | 1.1 | 2.0948 | 8.1E-15 |
| *ogr-2* | WBGene00009946 | protein_coding_gene | 1.5 | 2.7567 | 3.4E-20 | 1.0 | 2.0577 | 7.3E-13 |
| *C34B7.1* | WBGene00007911 | protein_coding_gene | 1.5 | 2.7552 | 7.4E-06 | 2.0 | 3.9796 | 5.2E-12 |
| *ZK1225.7* | WBGene00014994 | pseudogene | 1.5 | 2.7523 | 1.3E-05 | 1.7 | 3.1601 | 3.9E-04 |
| *C16C8.12* | WBGene00015850 | protein_coding_gene | 1.5 | 2.7523 | 1.8E-09 | 1.1 | 2.0809 | 2.6E-11 |
| *trt-1* | WBGene00006618 | protein_coding_gene | 1.5 | 2.7483 | 1.7E-18 | 1.3 | 2.5251 | 2.6E-25 |
| *Y54G2A.50* | WBGene00044493 | protein_coding_gene | 1.5 | 2.7424 | 8.6E-20 | 1.3 | 2.4169 | 1.0E-13 |
| *C14B1.12* | WBGene00184990 | protein_coding_gene | 1.5 | 2.7409 | 2.6E-18 | 1.2 | 2.2875 | 1.1E-19 |
| *F59C6.12* | WBGene00044251 | protein_coding_gene | 1.5 | 2.7330 | 2.4E-12 | 1.0 | 2.0539 | 3.3E-06 |
| *C49H3.4* | WBGene00016791 | protein_coding_gene | 1.4 | 2.7312 | 1.2E-14 | 1.1 | 2.0794 | 3.2E-10 |
| *F55A11.8* | WBGene00010081 | protein_coding_gene | 1.4 | 2.7260 | 1.1E-21 | 1.2 | 2.2726 | 2.0E-15 |
| *T08A9.6* | WBGene00020338 | pseudogene | 1.4 | 2.7250 | 2.3E-05 | 1.6 | 3.1128 | 7.8E-05 |
| *C34D4.10* | WBGene00016402 | protein_coding_gene | 1.4 | 2.7247 | 2.2E-17 | 1.5 | 2.7974 | 3.2E-25 |
| *R06C7.2* | WBGene00011062 | protein_coding_gene | 1.4 | 2.7240 | 4.9E-14 | 1.5 | 2.7706 | 1.8E-17 |
| *him-5* | WBGene00001864 | protein_coding_gene | 1.4 | 2.7223 | 6.1E-15 | 1.3 | 2.4490 | 5.1E-17 |
| *K03H6.2* | WBGene00019368 | protein_coding_gene | 1.4 | 2.7193 | 3.8E-04 | 2.2 | 4.5909 | 1.1E-08 |
| *R02D3.8* | WBGene00019825 | protein_coding_gene | 1.4 | 2.7075 | 1.6E-42 | 1.2 | 2.3345 | 3.8E-17 |
| ***R03H10.2*** | **WBGene00019855** | **protein_coding_gene** | **1.4** | **2.7007** | **1.5E-03** | **-1.1** | **0.4714** | **2.6E-05** |
| *Y57A10A.2* | WBGene00013247 | protein_coding_gene | 1.4 | 2.6986 | 5.7E-15 | 1.5 | 2.7382 | 1.7E-23 |
| *R11D1.1* | WBGene00011242 | protein_coding_gene | 1.4 | 2.6982 | 2.2E-17 | 1.1 | 2.0963 | 2.5E-19 |
| *Y57A10A.4* | WBGene00013249 | protein_coding_gene | 1.4 | 2.6974 | 9.6E-10 | 1.7 | 3.2370 | 9.3E-10 |
| *madf-8* | WBGene00008118 | protein_coding_gene | 1.4 | 2.6949 | 3.6E-15 | 1.2 | 2.2503 | 4.9E-17 |
| *C27H6.9* | WBGene00077500 | protein_coding_gene | 1.4 | 2.6899 | 3.2E-15 | 1.2 | 2.2653 | 9.6E-20 |
| *C46G7.5* | WBGene00044502 | protein_coding_gene | 1.4 | 2.6873 | 1.6E-04 | 1.7 | 3.2397 | 4.5E-03 |
| *madf-6* | WBGene00016930 | protein_coding_gene | 1.4 | 2.6851 | 1.2E-24 | 1.3 | 2.4536 | 6.1E-14 |
| *C28A5.1* | WBGene00007788 | protein_coding_gene | 1.4 | 2.6836 | 5.5E-13 | 1.8 | 3.5008 | 2.1E-34 |
| *R05F9.11* | WBGene00019894 | protein_coding_gene | 1.4 | 2.6747 | 1.2E-19 | 1.7 | 3.3427 | 1.3E-16 |
| *F01D5.3* | WBGene00008494 | protein_coding_gene | 1.4 | 2.6728 | 2.8E-02 | 5.7 | 51.3393 | 9.9E-26 |

| | | | | | | | | |
|---|---|---|---|---|---|---|---|---|
| *Y38H6C.9* | WBGene00012622 | protein_coding_gene | 1.4 | 2.6720 | 8.6E-04 | 1.9 | 3.8042 | 2.0E-03 |
| *agt-2* | WBGene00000094 | protein_coding_gene | 1.4 | 2.6670 | 6.5E-10 | 1.5 | 2.7706 | 1.0E-20 |
| *ndx-7* | WBGene00003584 | protein_coding_gene | 1.4 | 2.6665 | 3.2E-19 | 1.2 | 2.3644 | 5.4E-17 |
| *zhp-1* | WBGene00018867 | protein_coding_gene | 1.4 | 2.6604 | 1.1E-20 | 1.2 | 2.2356 | 6.6E-14 |
| *Y105E8A.28* | WBGene00013685 | protein_coding_gene | 1.4 | 2.6583 | 2.8E-03 | 1.7 | 3.1969 | 4.8E-04 |
| *clec-4* | WBGene00012583 | protein_coding_gene | 1.4 | 2.6573 | 1.1E-03 | 1.0 | 2.0071 | 2.2E-02 |
| ***T12D8.5*** | **WBGene00011733** | **protein_coding_gene** | **1.4** | **2.6564** | **6.4E-03** | **-2.6** | **0.1637** | **1.7E-17** |
| *Y66D12A.7* | WBGene00013433 | protein_coding_gene | 1.4 | 2.6554 | 2.8E-10 | 1.0 | 2.0039 | 2.5E-07 |
| *C16C8.4* | WBGene00015842 | protein_coding_gene | 1.4 | 2.6509 | 1.6E-23 | 1.1 | 2.1101 | 4.1E-13 |
| *D2096.7* | WBGene00017074 | protein_coding_gene | 1.4 | 2.6502 | 7.1E-35 | 1.2 | 2.2311 | 4.8E-22 |
| *tgt-1* | WBGene00006566 | protein_coding_gene | 1.4 | 2.6499 | 8.8E-32 | 1.1 | 2.1622 | 4.0E-18 |
| *kxd-1* | WBGene00015742 | protein_coding_gene | 1.4 | 2.6498 | 2.9E-08 | 1.2 | 2.3400 | 7.5E-12 |
| *rev-3* | WBGene00021344 | protein_coding_gene | 1.4 | 2.6493 | 1.4E-13 | 1.0 | 2.0263 | 2.9E-10 |
| *F46C3.4* | WBGene00044702 | pseudogene | 1.4 | 2.6469 | 9.8E-09 | 1.5 | 2.8959 | 1.3E-06 |
| *C49C3.9* | WBGene00008199 | protein_coding_gene | 1.4 | 2.6423 | 2.2E-03 | 1.2 | 2.3088 | 4.3E-03 |
| *F57B9.3* | WBGene00018997 | protein_coding_gene | 1.4 | 2.6264 | 3.2E-02 | 2.4 | 5.4103 | 1.7E-04 |
| *C49G7.10* | WBGene00016788 | protein_coding_gene | 1.4 | 2.6197 | 3.3E-03 | 1.5 | 2.8108 | 3.8E-04 |
| ***Y43F8B.9*** | **WBGene00012819** | **protein_coding_gene** | **1.4** | **2.6166** | **5.9E-03** | **-2.7** | **0.1527** | **2.7E-05** |
| *fbxa-125* | WBGene00015229 | protein_coding_gene | 1.4 | 2.6115 | 1.4E-02 | 1.7 | 3.2915 | 4.1E-05 |
| *W02D7.6* | WBGene00020942 | protein_coding_gene | 1.4 | 2.6099 | 1.4E-14 | 1.1 | 2.0933 | 1.6E-14 |
| *Y54G2A.49* | WBGene00044492 | protein_coding_gene | 1.4 | 2.6065 | 5.6E-04 | 1.2 | 2.2346 | 7.7E-03 |
| *let-49* | WBGene00002324 | protein_coding_gene | 1.4 | 2.6050 | 3.7E-18 | 1.4 | 2.6483 | 3.0E-24 |
| *C18D4.6* | WBGene00007675 | protein_coding_gene | 1.4 | 2.6049 | 1.3E-09 | 1.1 | 2.1861 | 1.3E-12 |
| *dsb-2* | WBGene00194892 | protein_coding_gene | 1.4 | 2.6016 | 5.2E-15 | 1.2 | 2.2394 | 2.4E-17 |
| *F17C11.11* | WBGene00008922 | protein_coding_gene | 1.4 | 2.5989 | 1.4E-06 | 1.3 | 2.5227 | 1.1E-04 |
| *C37A2.8* | WBGene00016494 | protein_coding_gene | 1.4 | 2.5984 | 5.6E-11 | 1.3 | 2.5440 | 1.8E-30 |
| *F26F4.9* | WBGene00017829 | protein_coding_gene | 1.4 | 2.5940 | 9.0E-19 | 1.3 | 2.4260 | 4.0E-21 |
| ***hch-1*** | **WBGene00001828** | **protein_coding_gene** | **1.4** | **2.5938** | **3.1E-02** | **-1.1** | **0.4579** | **1.2E-06** |
| *F59G1.8* | WBGene00019129 | protein_coding_gene | 1.4 | 2.5855 | 5.6E-14 | 1.0 | 2.0417 | 8.7E-16 |
| *Y32H12A.6* | WBGene00021314 | protein_coding_gene | 1.4 | 2.5845 | 2.9E-09 | 1.1 | 2.2107 | 7.9E-11 |
| *T02G5.4* | WBGene00020164 | pseudogene | 1.4 | 2.5821 | 1.5E-18 | 2.6 | 5.8702 | 2.0E-26 |
| ***F18G5.6*** | **WBGene00017582** | **protein_coding_gene** | **1.4** | **2.5778** | **2.1E-03** | **-1.0** | **0.4887** | **9.1E-13** |
| *Y50D4A.6* | WBGene00195169 | protein_coding_gene | 1.4 | 2.5758 | 7.5E-03 | 1.5 | 2.7617 | 4.1E-04 |
| *Y34B4A.9* | WBGene00021325 | protein_coding_gene | 1.4 | 2.5739 | 1.3E-07 | 2.2 | 4.4414 | 2.2E-18 |
| *spg-20* | WBGene00019006 | protein_coding_gene | 1.4 | 2.5704 | 1.9E-19 | 1.1 | 2.1430 | 6.3E-17 |
| *C42C1.13* | WBGene00016591 | protein_coding_gene | 1.4 | 2.5639 | 1.2E-15 | 1.1 | 2.1109 | 1.9E-11 |
| *set-23* | WBGene00021515 | protein_coding_gene | 1.4 | 2.5636 | 1.6E-16 | 1.2 | 2.3641 | 1.9E-16 |
| *zhit-2* | WBGene00009648 | protein_coding_gene | 1.4 | 2.5626 | 2.7E-15 | 1.1 | 2.1159 | 6.9E-12 |
| ***ins-7*** | **WBGene00002090** | **protein_coding_gene** | **1.4** | **2.5541** | **8.2E-04** | **-1.7** | **0.3074** | **3.8E-07** |
| *R12C12.7* | WBGene00020027 | protein_coding_gene | 1.4 | 2.5520 | 5.9E-16 | 1.3 | 2.5056 | 3.6E-17 |
| *B0391.10* | WBGene00007165 | pseudogene | 1.4 | 2.5504 | 2.6E-07 | 2.9 | 7.6778 | 3.2E-32 |
| *F33G12.2* | WBGene00018014 | protein_coding_gene | 1.4 | 2.5500 | 5.7E-21 | 1.1 | 2.0850 | 1.7E-17 |
| *rde-4* | WBGene00004326 | protein_coding_gene | 1.4 | 2.5495 | 3.5E-13 | 1.5 | 2.7460 | 2.1E-36 |
| *C49C3.6* | WBGene00008196 | protein_coding_gene | 1.4 | 2.5492 | 5.8E-10 | 1.1 | 2.1567 | 1.4E-18 |
| *C27F2.7* | WBGene00016169 | protein_coding_gene | 1.3 | 2.5478 | 1.3E-18 | 1.2 | 2.3446 | 3.7E-13 |
| *slc-25A32* | WBGene00010459 | protein_coding_gene | 1.3 | 2.5458 | 1.8E-14 | 1.0 | 2.0341 | 3.6E-10 |
| *Y57G11C.1130* | WBGene00194649 | protein_coding_gene | 1.3 | 2.5435 | 1.1E-03 | 1.2 | 2.2448 | 1.9E-03 |
| *K11H12.4* | WBGene00019660 | protein_coding_gene | 1.3 | 2.5421 | 4.2E-04 | 1.9 | 3.7360 | 1.7E-03 |
| *Y37E11B.6* | WBGene00021378 | protein_coding_gene | 1.3 | 2.5420 | 2.0E-17 | 1.4 | 2.5789 | 2.2E-11 |
| *C06A5.8* | WBGene00015505 | protein_coding_gene | 1.3 | 2.5410 | 2.0E-17 | 1.3 | 2.4679 | 2.3E-21 |
| *C30H6.9* | WBGene00007826 | protein_coding_gene | 1.3 | 2.5362 | 4.0E-16 | 1.3 | 2.5193 | 2.2E-17 |
| ***tsp-17*** | **WBGene00006643** | **protein_coding_gene** | **1.3** | **2.5284** | **3.1E-02** | **-1.5** | **0.3647** | **2.0E-04** |
| *rbm-17* | WBGene00010233 | protein_coding_gene | 1.3 | 2.5279 | 7.5E-18 | 1.3 | 2.4497 | 2.0E-22 |
| *enu-3.4* | WBGene00021034 | protein_coding_gene | 1.3 | 2.5271 | 9.4E-04 | 2.1 | 4.3062 | 2.0E-06 |
| *R10H10.7* | WBGene00011225 | protein_coding_gene | 1.3 | 2.5251 | 3.9E-22 | 1.1 | 2.1045 | 5.6E-22 |
| *ZK945.4* | WBGene00014166 | protein_coding_gene | 1.3 | 2.5196 | 1.5E-08 | 1.5 | 2.8155 | 1.2E-08 |
| *ZK858.5* | WBGene00014118 | protein_coding_gene | 1.3 | 2.5159 | 1.3E-13 | 1.1 | 2.1851 | 1.1E-15 |
| *K08D8.4* | WBGene00010658 | protein_coding_gene | 1.3 | 2.5052 | 2.6E-03 | 2.4 | 5.1339 | 6.5E-06 |
| *B0432.8* | WBGene00015189 | protein_coding_gene | 1.3 | 2.5027 | 4.1E-15 | 1.2 | 2.2979 | 6.9E-14 |
| *C16A11.3* | WBGene00015816 | protein_coding_gene | 1.3 | 2.5006 | 1.4E-14 | 1.1 | 2.0973 | 3.2E-09 |
| *H31G24.3* | WBGene00019254 | protein_coding_gene | 1.3 | 2.4999 | 2.2E-15 | 1.1 | 2.1480 | 5.7E-09 |
| *tofu-1* | WBGene00007785 | protein_coding_gene | 1.3 | 2.4972 | 1.1E-14 | 1.1 | 2.0911 | 8.3E-19 |
| *R05G6.4* | WBGene00019898 | protein_coding_gene | 1.3 | 2.4946 | 7.3E-17 | 1.0 | 2.0680 | 1.3E-11 |
| *K05C4.4* | WBGene00010581 | protein_coding_gene | 1.3 | 2.4944 | 1.9E-09 | 1.1 | 2.1864 | 3.2E-07 |
| *M18.6* | WBGene00010891 | protein_coding_gene | 1.3 | 2.4889 | 5.3E-19 | 1.0 | 2.0301 | 4.1E-13 |
| ***C40H1.2*** | **WBGene00008038** | **protein_coding_gene** | **1.3** | **2.4842** | **5.5E-03** | **-1.2** | **0.4372** | **1.5E-04** |
| *F43D2.2* | WBGene00009651 | pseudogene | 1.3 | 2.4815 | 1.5E-11 | 1.2 | 2.2497 | 7.3E-13 |
| *rpl-26* | WBGene00004440 | protein_coding_gene | 1.3 | 2.4810 | 7.3E-09 | 1.5 | 2.8817 | 9.8E-32 |
| *K07B1.7* | WBGene00019467 | protein_coding_gene | 1.3 | 2.4770 | 6.8E-09 | 1.3 | 2.4983 | 7.8E-17 |
| *pro-1* | WBGene00004185 | protein_coding_gene | 1.3 | 2.4763 | 1.2E-17 | 1.0 | 2.0251 | 1.5E-16 |

| | | | | | | | | |
|---|---|---|---|---|---|---|---|---|
| *bath-29* | WBGene00017452 | protein_coding_gene | 1.3 | 2.4755 | 2.7E-11 | 1.2 | 2.2771 | 2.5E-08 |
| *Y66D12A.16* | WBGene00013442 | protein_coding_gene | 1.3 | 2.4748 | 6.6E-17 | 1.6 | 2.9633 | 4.8E-19 |
| *rec-1* | WBGene00004332 | protein_coding_gene | 1.3 | 2.4748 | 6.4E-25 | 1.6 | 2.9310 | 5.4E-29 |
| *rdl-1* | WBGene00016703 | protein_coding_gene | 1.3 | 2.4728 | 3.2E-07 | 1.3 | 2.4311 | 7.8E-04 |
| *T26E3.7* | WBGene00012040 | protein_coding_gene | 1.3 | 2.4719 | 2.8E-02 | 1.6 | 2.9623 | 4.2E-03 |
| *K12B6.8* | WBGene00019672 | pseudogene | 1.3 | 2.4642 | 9.3E-12 | 1.3 | 2.4049 | 7.4E-08 |
| *T20G5.9* | WBGene00011871 | protein_coding_gene | 1.3 | 2.4589 | 2.5E-14 | 1.3 | 2.4405 | 1.6E-19 |
| *T23E1.2* | WBGene00020729 | pseudogene | 1.3 | 2.4553 | 3.6E-08 | 1.1 | 2.1581 | 4.2E-08 |
| *fbxa-197* | WBGene00011513 | protein_coding_gene | 1.3 | 2.4522 | 2.9E-10 | 1.1 | 2.2125 | 8.2E-07 |
| *C53A5.17* | WBGene00194707 | protein_coding_gene | 1.3 | 2.4519 | 3.6E-22 | 1.0 | 2.0074 | 2.9E-11 |
| *pri-1* | WBGene00004180 | protein_coding_gene | 1.3 | 2.4488 | 6.7E-24 | 1.0 | 2.0375 | 6.7E-15 |
| *fbxa-34* | WBGene00013842 | pseudogene | 1.3 | 2.4487 | 1.2E-02 | 1.5 | 2.8601 | 1.4E-05 |
| *T07H8.11* | WBGene00304201 | protein_coding_gene | 1.3 | 2.4451 | 3.9E-02 | 1.9 | 3.6734 | 2.2E-08 |
| *lin-56* | WBGene00003038 | protein_coding_gene | 1.3 | 2.4449 | 1.7E-18 | 1.2 | 2.3563 | 1.0E-17 |
| *C29E4.12* | WBGene00016209 | protein_coding_gene | 1.3 | 2.4425 | 1.8E-24 | 1.1 | 2.0743 | 5.0E-16 |
| *bath-15* | WBGene00015590 | protein_coding_gene | 1.3 | 2.4416 | 9.6E-10 | 1.1 | 2.1876 | 3.3E-15 |
| *fbxa-53* | WBGene00017226 | protein_coding_gene | 1.3 | 2.4373 | 1.6E-02 | 1.5 | 2.8259 | 1.9E-05 |
| *dnj-3* | WBGene00001021 | protein_coding_gene | 1.3 | 2.4358 | 2.2E-11 | 1.0 | 2.0245 | 1.7E-08 |
| *oxy-4* | WBGene00021902 | protein_coding_gene | 1.3 | 2.4352 | 2.3E-18 | 1.0 | 2.0257 | 2.7E-14 |
| *gro-1* | WBGene00001740 | protein_coding_gene | 1.3 | 2.4328 | 2.2E-16 | 1.1 | 2.0891 | 1.4E-14 |
| *C45H4.14* | WBGene00016695 | protein_coding_gene | 1.3 | 2.4265 | 2.6E-07 | 1.5 | 2.8743 | 5.1E-10 |
| *cng-1* | WBGene00000562 | protein_coding_gene | 1.3 | 2.4260 | 6.7E-05 | 2.0 | 3.9536 | 6.0E-07 |
| ***F13H10.5*** | **WBGene00008776** | **protein_coding_gene** | **1.3** | **2.4259** | **2.1E-02** | **-1.8** | **0.2805** | **1.3E-06** |
| *syx-18* | WBGene00020425 | protein_coding_gene | 1.3 | 2.4248 | 6.2E-14 | 1.2 | 2.2739 | 3.8E-25 |
| *Y110A2AL.3* | WBGene00022440 | protein_coding_gene | 1.3 | 2.4234 | 3.6E-03 | 1.1 | 2.1089 | 4.3E-02 |
| *tbca-1* | WBGene00021475 | protein_coding_gene | 1.3 | 2.4225 | 5.2E-06 | 1.2 | 2.2676 | 1.3E-15 |
| *uev-3* | WBGene00006732 | protein_coding_gene | 1.3 | 2.4208 | 7.3E-18 | 1.1 | 2.1218 | 4.6E-18 |
| *R11D1.10* | WBGene00011248 | protein_coding_gene | 1.3 | 2.4198 | 4.0E-15 | 1.4 | 2.6063 | 4.4E-25 |
| *F10E9.5* | WBGene00017357 | protein_coding_gene | 1.3 | 2.4196 | 1.2E-11 | 1.1 | 2.2174 | 9.2E-15 |
| ***lpr-6*** | **WBGene00012255** | **protein_coding_gene** | **1.3** | **2.4195** | **2.6E-02** | **-1.6** | **0.3322** | **3.6E-02** |
| *F25H2.4* | WBGene00009118 | protein_coding_gene | 1.3 | 2.4183 | 4.3E-19 | 1.0 | 2.0084 | 2.5E-14 |
| *nmat-2* | WBGene00009176 | protein_coding_gene | 1.3 | 2.4182 | 3.3E-12 | 1.0 | 2.0583 | 4.9E-10 |
| *use-1* | WBGene00022460 | protein_coding_gene | 1.3 | 2.4148 | 7.0E-22 | 1.1 | 2.1254 | 9.5E-14 |
| *F28F8.9* | WBGene00044188 | protein_coding_gene | 1.3 | 2.4106 | 9.6E-18 | 1.3 | 2.5460 | 2.4E-25 |
| *R08D7.5* | WBGene00011145 | protein_coding_gene | 1.3 | 2.4097 | 2.0E-10 | 1.1 | 2.0728 | 3.8E-16 |
| *lab-1* | WBGene00007278 | protein_coding_gene | 1.3 | 2.4089 | 2.2E-13 | 1.2 | 2.3765 | 7.9E-21 |
| *spo-11* | WBGene00004985 | protein_coding_gene | 1.3 | 2.4065 | 2.0E-12 | 1.4 | 2.5570 | 4.1E-26 |
| *F26D11.1* | WBGene00017818 | protein_coding_gene | 1.3 | 2.4025 | 7.8E-07 | 1.0 | 2.0537 | 1.4E-05 |
| *heri-1* | WBGene00016236 | protein_coding_gene | 1.3 | 2.3984 | 5.9E-21 | 1.4 | 2.6444 | 9.5E-28 |
| *ZK1058.3* | WBGene00014203 | protein_coding_gene | 1.3 | 2.3944 | 1.4E-13 | 1.0 | 2.0205 | 7.3E-09 |
| *syp-3* | WBGene00006377 | protein_coding_gene | 1.3 | 2.3926 | 1.5E-22 | 1.2 | 2.2309 | 5.3E-18 |
| *dom-3* | WBGene00001050 | protein_coding_gene | 1.3 | 2.3839 | 6.9E-11 | 1.3 | 2.4140 | 9.4E-23 |
| *ddl-1* | WBGene00019125 | protein_coding_gene | 1.3 | 2.3800 | 5.3E-09 | 1.1 | 2.2122 | 8.6E-14 |
| *piga-1* | WBGene00008431 | protein_coding_gene | 1.2 | 2.3775 | 4.8E-10 | 1.1 | 2.1877 | 1.6E-17 |
| *ZC317.7* | WBGene00022590 | protein_coding_gene | 1.2 | 2.3773 | 4.4E-04 | 1.3 | 2.4144 | 1.1E-10 |
| ***lpr-1*** | **WBGene00002393** | **protein_coding_gene** | **1.2** | **2.3772** | **2.5E-02** | **-1.4** | **0.3891** | **7.1E-03** |
| *C32H11.3* | WBGene00007866 | protein_coding_gene | 1.2 | 2.3721 | 1.1E-02 | 1.2 | 2.2519 | 1.1E-02 |
| *F12A10.8* | WBGene00017397 | protein_coding_gene | 1.2 | 2.3708 | 1.2E-10 | 1.2 | 2.2593 | 2.3E-14 |
| *Y82E9BR.2* | WBGene00022335 | protein_coding_gene | 1.2 | 2.3666 | 8.4E-15 | 1.6 | 2.9467 | 1.8E-17 |
| *T16G12.3* | WBGene00011804 | protein_coding_gene | 1.2 | 2.3648 | 6.1E-09 | 3.4 | 10.9263 | 1.4E-11 |
| *F08F3.8* | WBGene00017264 | protein_coding_gene | 1.2 | 2.3645 | 2.3E-02 | 1.6 | 2.9633 | 1.5E-02 |
| ***F43C11.7*** | **WBGene00018384** | **protein_coding_gene** | **1.2** | **2.3601** | **6.4E-03** | **-2.0** | **0.2456** | **2.3E-21** |
| *zim-2* | WBGene00011600 | protein_coding_gene | 1.2 | 2.3562 | 2.9E-18 | 1.3 | 2.4649 | 6.7E-30 |
| *Y87G2A.18* | WBGene00013604 | protein_coding_gene | 1.2 | 2.3539 | 1.2E-07 | 1.1 | 2.1213 | 3.3E-08 |
| ***hsp-12.1*** | **WBGene00011906** | **protein_coding_gene** | **1.2** | **2.3539** | **9.4E-03** | **-1.2** | **0.4206** | **5.8E-04** |
| ***dpy-7*** | **WBGene00001069** | **protein_coding_gene** | **1.2** | **2.3535** | **1.4E-02** | **-1.6** | **0.3403** | **4.6E-02** |
| *set-17* | WBGene00011887 | protein_coding_gene | 1.2 | 2.3521 | 2.5E-09 | 1.2 | 2.2425 | 9.0E-24 |
| *nfya-2* | WBGene00043056 | protein_coding_gene | 1.2 | 2.3512 | 2.6E-07 | 1.4 | 2.6452 | 2.8E-16 |
| *K08E4.2* | WBGene00010666 | protein_coding_gene | 1.2 | 2.3504 | 1.3E-06 | 1.0 | 2.0480 | 1.2E-08 |
| *Y73B3B.5* | WBGene00022227 | pseudogene | 1.2 | 2.3499 | 8.5E-09 | 1.5 | 2.7775 | 1.1E-13 |
| *C08H9.16* | WBGene00086566 | protein_coding_gene | 1.2 | 2.3495 | 1.2E-12 | 1.5 | 2.9145 | 9.0E-41 |
| *R10D12.13* | WBGene00011194 | protein_coding_gene | 1.2 | 2.3492 | 1.8E-12 | 1.3 | 2.4695 | 3.5E-20 |
| *T23B5.4* | WBGene00011941 | protein_coding_gene | 1.2 | 2.3444 | 5.8E-09 | 1.3 | 2.4987 | 8.1E-19 |
| *ivph-3* | WBGene00022067 | protein_coding_gene | 1.2 | 2.3433 | 2.3E-16 | 1.3 | 2.4071 | 6.0E-12 |
| *H21P03.2* | WBGene00010409 | protein_coding_gene | 1.2 | 2.3413 | 9.2E-19 | 1.0 | 2.0304 | 8.8E-19 |
| *W02F12.1* | WBGene00023355 | pseudogene | 1.2 | 2.3404 | 9.7E-06 | 1.4 | 2.5712 | 1.9E-04 |
| *C17G10.13* | WBGene00270322 | protein_coding_gene | 1.2 | 2.3401 | 3.4E-27 | 1.2 | 2.3117 | 8.4E-20 |
| *snpc-3.2* | WBGene00021667 | protein_coding_gene | 1.2 | 2.3388 | 6.6E-04 | 1.7 | 3.1431 | 4.3E-04 |
| *C35B1.2* | WBGene00016432 | protein_coding_gene | 1.2 | 2.3364 | 6.9E-22 | 1.3 | 2.3859 | 2.3E-24 |
| *chs-2* | WBGene00000497 | protein_coding_gene | 1.2 | 2.3314 | 4.4E-06 | 1.4 | 2.6051 | 2.3E-03 |

| | | | | | | | | |
|---|---|---|---|---|---|---|---|---|
| ***F09F7.6*** | **WBGene00017303** | **protein_coding_gene** | **1.2** | **2.3310** | **7.5E-03** | **-2.0** | **0.2521** | **1.7E-24** |
| *eri-5* | WBGene00021419 | protein_coding_gene | 1.2 | 2.3302 | 4.3E-15 | 1.1 | 2.1060 | 1.9E-11 |
| *cas-2* | WBGene00015975 | protein_coding_gene | 1.2 | 2.3284 | 6.3E-12 | 1.2 | 2.2481 | 3.0E-23 |
| *H06H21.11* | WBGene00044483 | protein_coding_gene | 1.2 | 2.3283 | 6.0E-18 | 1.0 | 2.0183 | 1.3E-13 |
| ***W04A8.4*** | **WBGene00012239** | **protein_coding_gene** | **1.2** | **2.3265** | **5.1E-04** | **-1.3** | **0.4143** | **3.8E-10** |
| *W03C9.1* | WBGene00012219 | protein_coding_gene | 1.2 | 2.3244 | 1.3E-02 | 1.2 | 2.3491 | 6.2E-04 |
| *F46F11.10* | WBGene00018513 | protein_coding_gene | 1.2 | 2.3238 | 5.8E-10 | 1.2 | 2.2204 | 6.0E-10 |
| *K08F11.6* | WBGene00023323 | pseudogene | 1.2 | 2.3186 | 2.9E-06 | 1.5 | 2.7336 | 3.5E-08 |
| *F10E7.11* | WBGene00017352 | protein_coding_gene | 1.2 | 2.3169 | 1.3E-03 | 1.1 | 2.1028 | 3.0E-02 |
| *mdt-22* | WBGene00007022 | protein_coding_gene | 1.2 | 2.3154 | 4.9E-13 | 1.1 | 2.0911 | 2.4E-18 |
| *K06B9.4* | WBGene00019441 | pseudogene | 1.2 | 2.3137 | 7.0E-05 | 2.0 | 4.1227 | 7.6E-14 |
| *set-22* | WBGene00012527 | protein_coding_gene | 1.2 | 2.3131 | 3.5E-07 | 1.0 | 2.0326 | 8.7E-15 |
| *Y27F2A.6* | WBGene00021298 | protein_coding_gene | 1.2 | 2.3117 | 2.1E-02 | 1.0 | 2.0242 | 3.4E-02 |
| *F18A1.1* | WBGene00017545 | protein_coding_gene | 1.2 | 2.3110 | 2.7E-02 | 1.6 | 2.9467 | 3.1E-04 |
| *mrt-1* | WBGene00045237 | protein_coding_gene | 1.2 | 2.3106 | 1.2E-24 | 1.3 | 2.3851 | 3.6E-26 |
| *Y69A2AR.32* | WBGene00022101 | protein_coding_gene | 1.2 | 2.3094 | 9.4E-14 | 1.1 | 2.1958 | 2.8E-13 |
| *F32E10.9* | WBGene00043068 | protein_coding_gene | 1.2 | 2.3093 | 9.7E-11 | 1.3 | 2.3957 | 1.8E-09 |
| *T24D1.3* | WBGene00011986 | protein_coding_gene | 1.2 | 2.3083 | 4.8E-12 | 1.2 | 2.3336 | 9.3E-15 |
| *syp-2* | WBGene00006376 | protein_coding_gene | 1.2 | 2.3050 | 2.2E-10 | 1.2 | 2.2203 | 1.2E-20 |
| *T14B4.1* | WBGene00020498 | protein_coding_gene | 1.2 | 2.3031 | 4.3E-23 | 1.1 | 2.1849 | 1.0E-17 |
| *enu-3.3* | WBGene00021015 | protein_coding_gene | 1.2 | 2.3026 | 9.5E-04 | 1.9 | 3.6986 | 3.6E-12 |
| *C44B12.6* | WBGene00016639 | protein_coding_gene | 1.2 | 2.2995 | 2.9E-02 | 1.6 | 3.0186 | 2.5E-02 |
| *F27D4.3* | WBGene00014746 | pseudogene | 1.2 | 2.2980 | 1.6E-04 | 1.6 | 3.0419 | 3.1E-06 |
| *W09C3.4* | WBGene00021112 | protein_coding_gene | 1.2 | 2.2971 | 7.9E-14 | 1.0 | 2.0335 | 8.3E-14 |
| *C01B12.8* | WBGene00015291 | protein_coding_gene | 1.2 | 2.2935 | 6.1E-12 | 1.3 | 2.3877 | 4.0E-16 |
| *Y50D4C.6* | WBGene00021750 | protein_coding_gene | 1.2 | 2.2927 | 5.7E-08 | 1.5 | 2.8842 | 2.6E-18 |
| *Y53G8AR.5* | WBGene00021812 | protein_coding_gene | 1.2 | 2.2925 | 2.0E-19 | 1.1 | 2.1115 | 5.7E-12 |
| *zim-3* | WBGene00011601 | protein_coding_gene | 1.2 | 2.2896 | 3.5E-17 | 1.1 | 2.0975 | 1.4E-20 |
| *Y17G7B.12* | WBGene00012465 | protein_coding_gene | 1.2 | 2.2889 | 2.8E-18 | 1.1 | 2.0928 | 6.6E-12 |
| *Y22D7AL.7* | WBGene00021246 | protein_coding_gene | 1.2 | 2.2880 | 9.6E-08 | 1.1 | 2.0945 | 7.2E-07 |
| *Y17G9B.5* | WBGene00021202 | protein_coding_gene | 1.2 | 2.2878 | 1.6E-19 | 1.2 | 2.2669 | 2.1E-21 |
| *pnn-1* | WBGene00011308 | protein_coding_gene | 1.2 | 2.2828 | 2.6E-11 | 1.2 | 2.2629 | 3.4E-24 |
| *fncm-1* | WBGene00018908 | protein_coding_gene | 1.2 | 2.2818 | 2.8E-12 | 1.2 | 2.2600 | 1.0E-18 |
| *Y54E2A.8* | WBGene00013193 | protein_coding_gene | 1.2 | 2.2813 | 8.6E-08 | 1.4 | 2.6101 | 3.3E-21 |
| *C03B8.2* | WBGene00015385 | protein_coding_gene | 1.2 | 2.2802 | 2.7E-03 | 1.5 | 2.8419 | 2.3E-04 |
| *bath-46* | WBGene00020325 | protein_coding_gene | 1.2 | 2.2792 | 3.4E-02 | 2.8 | 7.1010 | 1.2E-11 |
| *F33E11.2* | WBGene00018010 | protein_coding_gene | 1.2 | 2.2767 | 6.8E-09 | 1.1 | 2.1609 | 7.9E-14 |
| ***M03B6.3*** | **WBGene00010835** | **protein_coding_gene** | **1.2** | **2.2762** | **1.6E-02** | **-1.9** | **0.2600** | **1.5E-03** |
| *K06A5.1* | WBGene00019429 | protein_coding_gene | 1.2 | 2.2728 | 6.8E-20 | 1.4 | 2.7183 | 4.5E-35 |
| *R07H5.9* | WBGene00011129 | protein_coding_gene | 1.2 | 2.2699 | 1.1E-02 | 1.2 | 2.3076 | 1.4E-04 |
| *mrrf-1* | WBGene00020625 | protein_coding_gene | 1.2 | 2.2684 | 8.0E-14 | 1.2 | 2.3327 | 1.1E-15 |
| *ZK892.6* | WBGene00014130 | protein_coding_gene | 1.2 | 2.2668 | 1.4E-02 | 1.5 | 2.9059 | 8.7E-03 |
| ***hhat-1*** | **WBGene00013852** | **protein_coding_gene** | **1.2** | **2.2648** | **2.7E-02** | **-1.8** | **0.2836** | **1.7E-03** |
| *zgpa-1* | WBGene00016383 | protein_coding_gene | 1.2 | 2.2640 | 2.4E-14 | 1.1 | 2.1760 | 2.7E-15 |
| *coh-3* | WBGene00000592 | protein_coding_gene | 1.2 | 2.2629 | 5.2E-16 | 1.2 | 2.3174 | 5.2E-24 |
| *syp-1* | WBGene00006375 | protein_coding_gene | 1.2 | 2.2611 | 3.0E-12 | 1.1 | 2.1770 | 1.8E-19 |
| *lys-1* | WBGene00003090 | protein_coding_gene | 1.2 | 2.2571 | 2.9E-05 | 1.8 | 3.5604 | 2.4E-10 |
| *C30F2.4* | WBGene00007818 | protein_coding_gene | 1.2 | 2.2564 | 4.5E-02 | 2.8 | 6.8266 | 1.8E-10 |
| *Y95D11A.2* | WBGene00044262 | pseudogene | 1.2 | 2.2537 | 6.6E-04 | 1.3 | 2.3889 | 4.1E-04 |
| ***T22B7.3*** | **WBGene00020672** | **protein_coding_gene** | **1.2** | **2.2531** | **5.8E-03** | **-1.2** | **0.4298** | **1.8E-04** |
| *gst-44* | WBGene00001792 | protein_coding_gene | 1.2 | 2.2528 | 7.0E-03 | 1.1 | 2.1162 | 1.7E-02 |
| *B0391.8* | WBGene00014643 | pseudogene | 1.2 | 2.2524 | 5.7E-07 | 1.2 | 2.3686 | 1.7E-10 |
| *F59A3.2* | WBGene00019075 | protein_coding_gene | 1.2 | 2.2520 | 2.7E-12 | 1.5 | 2.7419 | 5.6E-22 |
| *C39B5.6* | WBGene00016524 | protein_coding_gene | 1.2 | 2.2496 | 5.7E-13 | 1.1 | 2.1523 | 4.0E-15 |
| ***egl-46*** | **WBGene00001210** | **protein_coding_gene** | **1.2** | **2.2474** | **2.3E-02** | **-1.3** | **0.4092** | **1.1E-06** |
| *dnj-4* | WBGene00001022 | protein_coding_gene | 1.2 | 2.2443 | 1.5E-13 | 1.2 | 2.3276 | 5.4E-21 |
| *CD4.10* | WBGene00022970 | ncRNA_gene | 1.2 | 2.2438 | 1.0E-04 | 1.6 | 2.9549 | 6.2E-13 |
| *M151.4* | WBGene00019796 | protein_coding_gene | 1.2 | 2.2434 | 1.5E-04 | 1.4 | 2.6977 | 1.4E-04 |
| *F58H1.8* | WBGene00138720 | protein_coding_gene | 1.2 | 2.2434 | 3.8E-10 | 1.1 | 2.1608 | 6.1E-12 |
| *Y39A1A.20* | WBGene00012657 | protein_coding_gene | 1.2 | 2.2410 | 5.0E-05 | 1.0 | 2.0259 | 2.3E-05 |
| *snu-23* | WBGene00022794 | protein_coding_gene | 1.2 | 2.2393 | 2.0E-07 | 1.3 | 2.4205 | 1.4E-33 |
| *prx-10* | WBGene00006946 | protein_coding_gene | 1.2 | 2.2368 | 5.8E-12 | 1.0 | 2.0187 | 5.3E-15 |
| *dnj-26* | WBGene00001044 | protein_coding_gene | 1.2 | 2.2364 | 3.2E-09 | 1.0 | 2.0456 | 1.9E-12 |
| ***F49F1.5*** | **WBGene00018645** | **protein_coding_gene** | **1.2** | **2.2363** | **9.1E-03** | **-2.2** | **0.2192** | **4.4E-17** |
| *K09D9.1* | WBGene00019564 | protein_coding_gene | 1.2 | 2.2357 | 2.9E-02 | 1.4 | 2.5665 | 6.7E-03 |
| *bath-4* | WBGene00018665 | protein_coding_gene | 1.2 | 2.2327 | 1.4E-05 | 1.6 | 3.0131 | 3.3E-06 |
| *mdt-10* | WBGene00007014 | protein_coding_gene | 1.2 | 2.2310 | 5.9E-13 | 1.2 | 2.2338 | 9.6E-18 |
| *Y57A10A.1* | WBGene00013246 | protein_coding_gene | 1.2 | 2.2296 | 3.7E-09 | 1.1 | 2.1893 | 4.1E-21 |
| *E01B7.2* | WBGene00008442 | protein_coding_gene | 1.2 | 2.2293 | 1.8E-12 | 1.0 | 2.0546 | 3.1E-11 |
| *H35B03.2* | WBGene00019264 | protein_coding_gene | 1.2 | 2.2289 | 1.3E-21 | 1.0 | 2.0694 | 1.8E-15 |

| | | | | | | | | |
|---|---|---|---|---|---|---|---|---|
| *spin-3* | WBGene00008598 | protein_coding_gene | 1.2 | 2.2280 | 4.8E-05 | 2.0 | 3.9836 | 5.0E-09 |
| *ztf-23* | WBGene00021846 | protein_coding_gene | 1.2 | 2.2253 | 3.7E-18 | 1.1 | 2.1614 | 5.0E-15 |
| *lrg-1* | WBGene00003067 | pseudogene | 1.2 | 2.2252 | 1.7E-15 | 2.8 | 6.9965 | 1.0E-38 |
| *polq-1* | WBGene00020964 | protein_coding_gene | 1.2 | 2.2225 | 6.1E-11 | 1.1 | 2.2186 | 2.3E-24 |
| *C45G9.2* | WBGene00016674 | protein_coding_gene | 1.1 | 2.2180 | 1.7E-15 | 1.0 | 2.0327 | 5.4E-12 |
| ***clec-125*** | **WBGene00021118** | **protein_coding_gene** | **1.1** | **2.2139** | **3.4E-02** | **-4.5** | **0.0428** | **5.0E-82** |
| *B0024.13* | WBGene00007102 | protein_coding_gene | 1.1 | 2.2134 | 3.4E-06 | 1.1 | 2.1901 | 8.5E-21 |
| *ZK384.7* | WBGene00269383 | ncRNA_gene | 1.1 | 2.2099 | 6.7E-03 | 1.1 | 2.1111 | 1.5E-05 |
| *idi-1* | WBGene00019460 | protein_coding_gene | 1.1 | 2.2078 | 1.2E-11 | 1.3 | 2.4451 | 1.1E-30 |
| *tbx-36* | WBGene00006555 | protein_coding_gene | 1.1 | 2.2078 | 1.3E-06 | 1.1 | 2.1259 | 6.3E-11 |
| *clec-84* | WBGene00021895 | protein_coding_gene | 1.1 | 2.2053 | 3.1E-07 | 2.4 | 5.2566 | 3.4E-51 |
| *Y82E9BL.18* | WBGene00044402 | protein_coding_gene | 1.1 | 2.2013 | 8.4E-03 | 2.6 | 6.2638 | 2.6E-03 |
| *R11A8.1* | WBGene00011236 | protein_coding_gene | 1.1 | 2.2010 | 5.3E-09 | 2.5 | 5.5374 | 1.9E-41 |
| *tsen-54* | WBGene00018680 | protein_coding_gene | 1.1 | 2.1991 | 3.2E-10 | 1.4 | 2.5978 | 1.1E-13 |
| *C48B4.6* | WBGene00008169 | protein_coding_gene | 1.1 | 2.1991 | 3.8E-08 | 1.1 | 2.0742 | 3.7E-09 |
| *fbxa-14* | WBGene00022326 | protein_coding_gene | 1.1 | 2.1986 | 6.8E-03 | 3.1 | 8.8508 | 3.2E-17 |
| *cec-7* | WBGene00012551 | protein_coding_gene | 1.1 | 2.1979 | 5.6E-22 | 1.0 | 2.0272 | 7.0E-16 |
| *F49C12.9* | WBGene00009878 | protein_coding_gene | 1.1 | 2.1971 | 7.1E-12 | 1.2 | 2.3271 | 9.4E-25 |
| *gbas-1* | WBGene00019130 | protein_coding_gene | 1.1 | 2.1961 | 1.9E-16 | 1.0 | 2.0183 | 2.0E-13 |
| *tag-72* | WBGene00006447 | protein_coding_gene | 1.1 | 2.1949 | 2.4E-12 | 1.1 | 2.1446 | 1.1E-17 |
| *C29E4.13* | WBGene00043051 | protein_coding_gene | 1.1 | 2.1919 | 1.4E-11 | 1.2 | 2.2632 | 5.3E-16 |
| *B0285.3* | WBGene00007136 | protein_coding_gene | 1.1 | 2.1902 | 6.5E-09 | 1.2 | 2.2843 | 5.4E-15 |
| *C25G4.3* | WBGene00007731 | protein_coding_gene | 1.1 | 2.1900 | 7.4E-15 | 1.0 | 2.0498 | 1.1E-10 |
| *T01C3.2* | WBGene00011319 | protein_coding_gene | 1.1 | 2.1886 | 3.8E-17 | 1.0 | 2.0494 | 8.1E-19 |
| *T24H10.4* | WBGene00012002 | protein_coding_gene | 1.1 | 2.1885 | 1.6E-10 | 1.4 | 2.5864 | 1.4E-23 |
| *vps-33.2* | WBGene00016960 | protein_coding_gene | 1.1 | 2.1874 | 1.4E-19 | 1.1 | 2.1764 | 3.2E-20 |
| *rpl-16* | WBGene00004428 | protein_coding_gene | 1.1 | 2.1831 | 1.5E-16 | 1.3 | 2.4624 | 4.7E-25 |
| *brc-1* | WBGene00000264 | protein_coding_gene | 1.1 | 2.1827 | 1.6E-12 | 1.2 | 2.2260 | 5.0E-19 |
| *Y53G8AR.8* | WBGene00021815 | protein_coding_gene | 1.1 | 2.1824 | 1.8E-14 | 1.1 | 2.1887 | 2.1E-16 |
| *sid-1* | WBGene00004795 | protein_coding_gene | 1.1 | 2.1822 | 2.7E-06 | 2.5 | 5.8157 | 7.5E-49 |
| *T10E9.1* | WBGene00020411 | protein_coding_gene | 1.1 | 2.1816 | 9.4E-08 | 1.0 | 2.0085 | 1.3E-15 |
| *rps-21* | WBGene00004490 | protein_coding_gene | 1.1 | 2.1769 | 2.9E-08 | 1.3 | 2.4515 | 5.7E-21 |
| ***T02E9.5*** | **WBGene00011383** | **protein_coding_gene** | **1.1** | **2.1722** | **1.1E-02** | **-1.2** | **0.4351** | **1.8E-10** |
| *ekl-4* | WBGene00013676 | protein_coding_gene | 1.1 | 2.1711 | 5.4E-13 | 1.0 | 2.0408 | 5.2E-19 |
| *Y73F8A.1173* | WBGene00206491 | pseudogene | 1.1 | 2.1696 | 2.8E-04 | 1.1 | 2.1430 | 1.1E-02 |
| *fan-1* | WBGene00015310 | protein_coding_gene | 1.1 | 2.1665 | 7.1E-16 | 1.2 | 2.3251 | 3.2E-22 |
| *W05F2.6* | WBGene00021038 | protein_coding_gene | 1.1 | 2.1660 | 3.2E-20 | 1.3 | 2.4262 | 1.3E-21 |
| *C10C6.7* | WBGene00007515 | protein_coding_gene | 1.1 | 2.1657 | 1.2E-03 | 1.1 | 2.1625 | 1.8E-02 |
| *C14B1.8* | WBGene00007579 | protein_coding_gene | 1.1 | 2.1655 | 6.5E-15 | 1.2 | 2.2786 | 2.7E-20 |
| *T10C6.10* | WBGene00011691 | protein_coding_gene | 1.1 | 2.1647 | 6.4E-08 | 1.4 | 2.5544 | 1.8E-08 |
| *ztf-1* | WBGene00018833 | protein_coding_gene | 1.1 | 2.1643 | 1.7E-06 | 1.4 | 2.6577 | 2.1E-18 |
| *Y57G11C.43* | WBGene00013332 | protein_coding_gene | 1.1 | 2.1643 | 5.7E-13 | 1.2 | 2.2422 | 4.5E-20 |
| *Y71H2AM.14* | WBGene00022179 | protein_coding_gene | 1.1 | 2.1621 | 2.4E-02 | 2.1 | 4.3835 | 5.8E-06 |
| *C03H5.3* | WBGene00015405 | protein_coding_gene | 1.1 | 2.1620 | 2.6E-09 | 1.2 | 2.2568 | 1.2E-16 |
| *F36H12.2* | WBGene00018118 | protein_coding_gene | 1.1 | 2.1617 | 9.0E-11 | 1.4 | 2.6430 | 5.5E-17 |
| *rnp-3* | WBGene00004386 | protein_coding_gene | 1.1 | 2.1609 | 6.1E-20 | 1.0 | 2.0089 | 4.0E-16 |
| *Y58A7A.5* | WBGene00021979 | protein_coding_gene | 1.1 | 2.1608 | 4.7E-02 | 3.0 | 7.7314 | 2.2E-14 |
| *mrpl-19* | WBGene00022470 | protein_coding_gene | 1.1 | 2.1607 | 2.1E-12 | 1.0 | 2.0575 | 8.4E-15 |
| *Y71A12B.10* | WBGene00013507 | protein_coding_gene | 1.1 | 2.1604 | 3.2E-02 | 1.8 | 3.5627 | 1.7E-03 |
| *swsn-2.2* | WBGene00015971 | protein_coding_gene | 1.1 | 2.1598 | 7.4E-09 | 1.0 | 2.0175 | 1.6E-13 |
| *orc-5* | WBGene00013857 | protein_coding_gene | 1.1 | 2.1572 | 4.0E-12 | 1.1 | 2.1809 | 2.6E-27 |
| *K08E4.7* | WBGene00045036 | protein_coding_gene | 1.1 | 2.1531 | 4.1E-02 | 1.5 | 2.8076 | 1.6E-05 |
| *fbxa-107* | WBGene00012564 | protein_coding_gene | 1.1 | 2.1497 | 5.1E-07 | 1.0 | 2.0278 | 2.1E-06 |
| *C49C3.7* | WBGene00008197 | protein_coding_gene | 1.1 | 2.1489 | 1.8E-12 | 1.1 | 2.1143 | 5.5E-24 |
| *C30G12.6* | WBGene00016277 | protein_coding_gene | 1.1 | 2.1489 | 6.3E-19 | 1.3 | 2.4239 | 2.6E-29 |
| *Y47G6A.25* | WBGene00021649 | protein_coding_gene | 1.1 | 2.1472 | 9.0E-11 | 1.1 | 2.1840 | 3.7E-15 |
| *T02G5.11* | WBGene00020167 | pseudogene | 1.1 | 2.1460 | 9.9E-03 | 1.1 | 2.0738 | 3.6E-02 |
| *smut-1* | WBGene00021391 | protein_coding_gene | 1.1 | 2.1448 | 5.0E-09 | 1.0 | 2.0186 | 7.1E-19 |
| *T28D9.4* | WBGene00020896 | protein_coding_gene | 1.1 | 2.1424 | 4.9E-04 | 1.4 | 2.5586 | 2.5E-23 |
| *rmh-1* | WBGene00019712 | protein_coding_gene | 1.1 | 2.1385 | 2.7E-14 | 1.5 | 2.8298 | 5.6E-28 |
| *xpa-1* | WBGene00006963 | protein_coding_gene | 1.1 | 2.1365 | 1.4E-14 | 1.2 | 2.2271 | 4.0E-21 |
| *btb-20* | WBGene00010193 | protein_coding_gene | 1.1 | 2.1363 | 1.2E-11 | 1.1 | 2.1655 | 2.5E-12 |
| *F25H8.2* | WBGene00009132 | protein_coding_gene | 1.1 | 2.1349 | 6.7E-12 | 1.2 | 2.3504 | 4.1E-26 |
| *him-3* | WBGene00001862 | protein_coding_gene | 1.1 | 2.1325 | 3.0E-14 | 1.2 | 2.2289 | 1.5E-21 |
| *R11H6.6* | WBGene00014822 | pseudogene | 1.1 | 2.1324 | 5.0E-06 | 1.5 | 2.7787 | 5.3E-06 |
| *Y65B4BL.3* | WBGene00022035 | protein_coding_gene | 1.1 | 2.1315 | 2.3E-12 | 1.1 | 2.1339 | 1.8E-09 |
| *F44G4.3* | WBGene00009713 | protein_coding_gene | 1.1 | 2.1294 | 4.3E-06 | 1.2 | 2.2627 | 3.8E-13 |
| *Y105E8A.20* | WBGene00013678 | protein_coding_gene | 1.1 | 2.1279 | 9.3E-09 | 1.0 | 2.0423 | 1.2E-15 |
| *rmh-2* | WBGene00011576 | protein_coding_gene | 1.1 | 2.1276 | 4.8E-08 | 1.1 | 2.1503 | 4.9E-15 |
| *C50E3.11* | WBGene00016822 | protein_coding_gene | 1.1 | 2.1244 | 2.5E-08 | 1.0 | 2.0038 | 1.2E-04 |

| | | | | | | | | |
|---|---|---|---|---|---|---|---|---|
| *ets-6* | WBGene00017606 | protein_coding_gene | 1.1 | 2.1242 | 1.6E-11 | 1.0 | 2.0107 | 4.4E-12 |
| *C04F5.9* | WBGene00015451 | protein_coding_gene | 1.1 | 2.1215 | 2.6E-11 | 1.3 | 2.3833 | 3.1E-17 |
| ***C01G6.9*** | **WBGene00007229** | **protein_coding_gene** | **1.1** | **2.1194** | **3.2E-02** | **-1.4** | **0.3860** | **5.4E-03** |
| *nduo-1* | WBGene00010959 | protein_coding_gene | 1.1 | 2.1188 | 2.1E-07 | 1.7 | 3.3408 | 3.5E-40 |
| *hpo-40* | WBGene00014113 | protein_coding_gene | 1.1 | 2.1183 | 8.1E-08 | 1.3 | 2.4553 | 1.8E-27 |
| *rps-11* | WBGene00004480 | protein_coding_gene | 1.1 | 2.1164 | 2.4E-05 | 1.2 | 2.2504 | 1.9E-21 |
| *apn-1* | WBGene00000151 | protein_coding_gene | 1.1 | 2.1161 | 1.8E-11 | 1.2 | 2.3658 | 1.0E-30 |
| *mans-4* | WBGene00013881 | protein_coding_gene | 1.1 | 2.1161 | 4.5E-09 | 1.1 | 2.0901 | 1.6E-20 |
| *F41H10.4* | WBGene00018317 | protein_coding_gene | 1.1 | 2.1160 | 1.4E-15 | 1.0 | 2.0104 | 5.4E-16 |
| *dnj-22* | WBGene00001040 | protein_coding_gene | 1.1 | 2.1133 | 3.4E-11 | 1.3 | 2.4693 | 2.4E-30 |
| *Y39E4B.14* | WBGene00014884 | pseudogene | 1.1 | 2.1124 | 4.0E-02 | 1.6 | 3.0974 | 4.4E-03 |
| *bath-30* | WBGene00017460 | protein_coding_gene | 1.1 | 2.1106 | 2.8E-16 | 1.1 | 2.1011 | 5.8E-05 |
| *algn-1* | WBGene00020820 | protein_coding_gene | 1.1 | 2.1105 | 1.5E-15 | 1.2 | 2.2388 | 1.3E-22 |
| *gras-1* | WBGene00009272 | protein_coding_gene | 1.1 | 2.1103 | 2.6E-11 | 1.1 | 2.1206 | 1.6E-14 |
| *zhit-1* | WBGene00016992 | protein_coding_gene | 1.1 | 2.1094 | 3.5E-08 | 1.0 | 2.0017 | 3.3E-11 |
| *chaf-1* | WBGene00011532 | protein_coding_gene | 1.1 | 2.1082 | 3.8E-10 | 1.1 | 2.1850 | 2.6E-29 |
| *coa-7* | WBGene00013925 | protein_coding_gene | 1.1 | 2.1073 | 8.7E-12 | 1.2 | 2.3436 | 3.3E-19 |
| *F28F8.7* | WBGene00009224 | protein_coding_gene | 1.1 | 2.1062 | 7.7E-05 | 1.3 | 2.3816 | 2.2E-15 |
| *F57B10.8* | WBGene00019005 | protein_coding_gene | 1.1 | 2.1055 | 4.0E-11 | 1.1 | 2.1658 | 7.8E-13 |
| *swah-1* | WBGene00007985 | protein_coding_gene | 1.1 | 2.1039 | 2.9E-08 | 1.2 | 2.2330 | 1.9E-18 |
| *C50E3.5* | WBGene00016816 | protein_coding_gene | 1.1 | 2.1015 | 3.4E-07 | 1.0 | 2.0514 | 1.7E-15 |
| *F57C9.4* | WBGene00019011 | protein_coding_gene | 1.1 | 2.0956 | 1.6E-12 | 1.1 | 2.0807 | 8.4E-20 |
| *F43E2.1* | WBGene00018390 | protein_coding_gene | 1.1 | 2.0948 | 3.1E-10 | 1.5 | 2.8719 | 3.2E-24 |
| *ZK637.6* | WBGene00014987 | pseudogene | 1.1 | 2.0928 | 1.5E-10 | 1.9 | 3.7953 | 7.3E-18 |
| *K10D2.5* | WBGene00019631 | protein_coding_gene | 1.1 | 2.0900 | 5.9E-10 | 1.1 | 2.1938 | 5.2E-15 |
| *rec-8* | WBGene00004333 | protein_coding_gene | 1.1 | 2.0892 | 2.5E-11 | 1.1 | 2.1666 | 3.3E-20 |
| *tag-124* | WBGene00006473 | protein_coding_gene | 1.1 | 2.0882 | 4.2E-13 | 1.1 | 2.1091 | 9.8E-14 |
| *F56C9.11* | WBGene00018954 | protein_coding_gene | 1.1 | 2.0882 | 1.8E-07 | 1.1 | 2.2021 | 6.2E-16 |
| *Y50D4A.1* | WBGene00021735 | protein_coding_gene | 1.1 | 2.0879 | 1.5E-22 | 1.3 | 2.4029 | 6.1E-19 |
| *dsb-3* | WBGene00016701 | protein_coding_gene | 1.1 | 2.0877 | 5.3E-13 | 1.1 | 2.1089 | 2.2E-13 |
| *mei-2* | WBGene00003184 | protein_coding_gene | 1.1 | 2.0870 | 7.8E-11 | 1.3 | 2.4123 | 6.5E-22 |
| *F52C6.2* | WBGene00018659 | protein_coding_gene | 1.1 | 2.0821 | 2.0E-07 | 2.0 | 3.8685 | 2.1E-14 |
| *gst-6* | WBGene00001754 | protein_coding_gene | 1.1 | 2.0805 | 2.3E-04 | 2.5 | 5.7991 | 3.0E-34 |
| *C09E7.8* | WBGene00015642 | protein_coding_gene | 1.1 | 2.0775 | 1.1E-09 | 1.6 | 3.0546 | 7.8E-33 |
| *Y116F11B.10* | WBGene00013826 | pseudogene | 1.1 | 2.0774 | 5.6E-04 | 2.1 | 4.2753 | 1.7E-09 |
| *C25A11.10* | WBGene00219374 | pseudogene | 1.1 | 2.0762 | 1.6E-04 | 1.2 | 2.3092 | 6.1E-06 |
| *Y54G2A.12* | WBGene00021877 | protein_coding_gene | 1.1 | 2.0761 | 1.3E-09 | 1.2 | 2.2408 | 3.5E-21 |
| *zip-4* | WBGene00021552 | protein_coding_gene | 1.1 | 2.0759 | 2.6E-20 | 1.0 | 2.0530 | 1.3E-15 |
| *C55A6.1* | WBGene00008330 | protein_coding_gene | 1.1 | 2.0731 | 5.4E-08 | 1.0 | 2.0110 | 7.2E-16 |
| *cyn-9* | WBGene00000885 | protein_coding_gene | 1.1 | 2.0713 | 4.2E-16 | 1.1 | 2.0848 | 1.7E-17 |
| *C28A5.2* | WBGene00007789 | protein_coding_gene | 1.1 | 2.0712 | 2.4E-09 | 1.5 | 2.8079 | 8.8E-19 |
| *gcc-2* | WBGene00011503 | protein_coding_gene | 1.1 | 2.0709 | 7.4E-12 | 1.2 | 2.2493 | 3.0E-27 |
| *bcs-1* | WBGene00010042 | protein_coding_gene | 1.0 | 2.0678 | 2.4E-13 | 1.1 | 2.1252 | 1.8E-20 |
| *bath-36* | WBGene00013569 | protein_coding_gene | 1.0 | 2.0665 | 1.6E-05 | 1.0 | 2.0054 | 4.5E-06 |
| *enu-3.1* | WBGene00010362 | protein_coding_gene | 1.0 | 2.0647 | 2.5E-07 | 1.5 | 2.8854 | 2.8E-17 |
| *ZC262.2* | WBGene00022579 | protein_coding_gene | 1.0 | 2.0640 | 7.8E-23 | 1.1 | 2.1108 | 2.4E-20 |
| *eri-3* | WBGene00021103 | protein_coding_gene | 1.0 | 2.0634 | 1.2E-18 | 1.2 | 2.3301 | 1.1E-19 |
| *sygl-1* | WBGene00012104 | protein_coding_gene | 1.0 | 2.0615 | 6.7E-05 | 1.1 | 2.1983 | 4.6E-16 |
| *clp-3* | WBGene00000544 | protein_coding_gene | 1.0 | 2.0588 | 2.4E-03 | 1.2 | 2.3764 | 1.5E-02 |
| *nars-2* | WBGene00013447 | protein_coding_gene | 1.0 | 2.0578 | 3.2E-06 | 1.2 | 2.2619 | 2.5E-14 |
| *F56D2.2* | WBGene00018964 | protein_coding_gene | 1.0 | 2.0577 | 6.4E-15 | 1.1 | 2.0820 | 7.1E-16 |
| *K12H6.2* | WBGene00019683 | protein_coding_gene | 1.0 | 2.0539 | 1.6E-03 | 1.2 | 2.3052 | 4.7E-06 |
| *gkow-1* | WBGene00011237 | protein_coding_gene | 1.0 | 2.0527 | 8.4E-11 | 1.1 | 2.2133 | 1.7E-25 |
| *F56D5.4* | WBGene00014772 | pseudogene | 1.0 | 2.0521 | 5.6E-04 | 1.4 | 2.5829 | 5.2E-20 |
| *C17E4.4* | WBGene00007644 | protein_coding_gene | 1.0 | 2.0510 | 8.8E-07 | 1.1 | 2.1154 | 4.3E-13 |
| *Y43F11A.6* | WBGene00014576 | pseudogene | 1.0 | 2.0509 | 3.7E-07 | 2.0 | 4.1201 | 2.1E-23 |
| *F14H3.5* | WBGene00008824 | protein_coding_gene | 1.0 | 2.0507 | 1.7E-05 | 1.2 | 2.3603 | 3.3E-13 |
| *M04F3.2* | WBGene00019768 | protein_coding_gene | 1.0 | 2.0505 | 5.8E-05 | 1.5 | 2.8263 | 5.9E-09 |
| *R03D7.2* | WBGene00010989 | protein_coding_gene | 1.0 | 2.0495 | 1.1E-06 | 2.1 | 4.3511 | 9.9E-11 |
| *suds-3* | WBGene00044204 | protein_coding_gene | 1.0 | 2.0491 | 3.3E-10 | 1.2 | 2.3743 | 6.6E-19 |
| *C24D10.4* | WBGene00016055 | protein_coding_gene | 1.0 | 2.0454 | 4.2E-16 | 1.2 | 2.3603 | 1.3E-21 |
| *bpl-1* | WBGene00000259 | protein_coding_gene | 1.0 | 2.0450 | 2.8E-19 | 1.1 | 2.1247 | 2.4E-18 |
| *hal-2* | WBGene00011819 | protein_coding_gene | 1.0 | 2.0438 | 5.1E-07 | 1.2 | 2.3447 | 3.1E-24 |
| ***C08F1.10*** | **WBGene00015613** | **protein_coding_gene** | **1.0** | **2.0437** | **2.7E-03** | **-1.1** | **0.4562** | **1.5E-03** |
| *rpl-30* | WBGene00004444 | protein_coding_gene | 1.0 | 2.0436 | 1.1E-10 | 1.0 | 2.0030 | 3.0E-17 |
| *sas-4* | WBGene00004726 | protein_coding_gene | 1.0 | 2.0415 | 1.9E-06 | 1.2 | 2.2193 | 2.1E-21 |
| *B0416.4* | WBGene00015180 | protein_coding_gene | 1.0 | 2.0393 | 3.6E-07 | 1.2 | 2.2806 | 1.3E-10 |
| *F26A3.7* | WBGene00009145 | protein_coding_gene | 1.0 | 2.0384 | 2.5E-09 | 1.1 | 2.1168 | 1.5E-16 |
| *npl-4.1* | WBGene00019120 | protein_coding_gene | 1.0 | 2.0333 | 2.1E-05 | 1.0 | 2.0622 | 1.3E-03 |
| *fbf-1* | WBGene00001401 | protein_coding_gene | 1.0 | 2.0327 | 3.5E-10 | 1.4 | 2.5936 | 6.3E-14 |

| | | | | | | | | |
|---|---|---|---|---|---|---|---|---|
| *mage-1* | WBGene00018230 | protein_coding_gene | 1.0 | 2.0320 | 4.8E-08 | 1.0 | 2.0167 | 1.7E-17 |
| *hop-1* | WBGene00001985 | protein_coding_gene | 1.0 | 2.0307 | 2.9E-14 | 1.0 | 2.0108 | 1.4E-13 |
| *rps-5* | WBGene00004474 | protein_coding_gene | 1.0 | 2.0270 | 7.2E-08 | 1.4 | 2.6490 | 5.6E-24 |
| *mtcu-2* | WBGene00009944 | protein_coding_gene | 1.0 | 2.0262 | 1.7E-17 | 1.3 | 2.4221 | 3.6E-24 |
| *C29A12.1* | WBGene00007797 | protein_coding_gene | 1.0 | 2.0260 | 4.6E-12 | 1.1 | 2.1531 | 3.2E-18 |
| *F09E5.10* | WBGene00017288 | protein_coding_gene | 1.0 | 2.0258 | 8.3E-03 | 1.1 | 2.1060 | 3.2E-03 |
| *Y53C12A.6* | WBGene00013141 | protein_coding_gene | 1.0 | 2.0239 | 3.5E-03 | 1.6 | 3.0923 | 1.0E-07 |
| *C53H9.2* | WBGene00016907 | protein_coding_gene | 1.0 | 2.0226 | 1.2E-09 | 1.1 | 2.1935 | 7.5E-19 |
| *Y51F10.2* | WBGene00021763 | protein_coding_gene | 1.0 | 2.0222 | 1.9E-07 | 1.1 | 2.0741 | 3.2E-17 |
| *micu-1* | WBGene00013462 | protein_coding_gene | 1.0 | 2.0220 | 2.8E-11 | 1.1 | 2.1792 | 6.4E-25 |
| *unc-69* | WBGene00006802 | protein_coding_gene | 1.0 | 2.0214 | 3.5E-10 | 1.1 | 2.0991 | 5.1E-14 |
| *ani-3* | WBGene00012835 | protein_coding_gene | 1.0 | 2.0180 | 1.0E-09 | 1.4 | 2.7311 | 5.8E-24 |
| *B0244.6* | WBGene00015081 | protein_coding_gene | 1.0 | 2.0177 | 5.5E-03 | 1.5 | 2.7480 | 4.6E-04 |
| *T12C9.7* | WBGene00020461 | protein_coding_gene | 1.0 | 2.0147 | 3.7E-08 | 1.2 | 2.2806 | 2.8E-15 |
| *F08F8.4* | WBGene00017269 | protein_coding_gene | 1.0 | 2.0091 | 7.6E-11 | 1.1 | 2.2098 | 2.8E-20 |
| *Y71G12B.10* | WBGene00022150 | protein_coding_gene | 1.0 | 2.0081 | 1.6E-07 | 1.2 | 2.2617 | 1.1E-24 |
| *uvs-1* | WBGene00022812 | protein_coding_gene | 1.0 | 2.0081 | 9.3E-11 | 1.3 | 2.3993 | 5.9E-19 |
| *B0491.6* | WBGene00007193 | protein_coding_gene | 1.0 | 2.0072 | 5.1E-17 | 1.1 | 2.1197 | 2.1E-15 |
| *eri-9* | WBGene00016143 | protein_coding_gene | 1.0 | 2.0066 | 1.2E-10 | 1.4 | 2.5887 | 2.0E-28 |
| *F39G3.5* | WBGene00018210 | protein_coding_gene | 1.0 | 2.0064 | 2.7E-03 | 1.9 | 3.8509 | 1.2E-15 |
| *rps-27* | WBGene00004496 | protein_coding_gene | 1.0 | 2.0021 | 1.9E-06 | 1.0 | 2.0622 | 2.2E-13 |
| *B0025.4* | WBGene00015001 | protein_coding_gene | 1.0 | 2.0016 | 3.9E-04 | 1.1 | 2.1814 | 6.0E-17 |
| *cep-1* | WBGene00000467 | protein_coding_gene | 1.0 | 2.0004 | 1.8E-08 | 1.4 | 2.5617 | 1.9E-29 |
| *che-14* | WBGene00000493 | protein_coding_gene | -1.0 | 0.4997 | 9.3E-03 | -1.4 | 0.3718 | 1.9E-04 |
| *lfe-2* | WBGene00002979 | protein_coding_gene | -1.0 | 0.4994 | 1.8E-07 | -1.5 | 0.3593 | 5.0E-29 |
| *ZK563.7* | WBGene00044355 | protein_coding_gene | -1.0 | 0.4993 | 5.5E-03 | -1.1 | 0.4663 | 1.1E-02 |
| *F16C3.2* | WBGene00008886 | protein_coding_gene | -1.0 | 0.4986 | 2.3E-02 | -3.9 | 0.0655 | 1.7E-72 |
| *anmt-1* | WBGene00015124 | protein_coding_gene | -1.0 | 0.4985 | 1.8E-03 | -1.3 | 0.4125 | 1.8E-04 |
| *Y34F4.1* | WBGene00021336 | protein_coding_gene | -1.0 | 0.4982 | 4.0E-03 | -2.1 | 0.2339 | 8.4E-23 |
| *Y54G2A.7* | WBGene00043702 | protein_coding_gene | -1.0 | 0.4978 | 2.7E-03 | -2.2 | 0.2163 | 1.1E-19 |
| *dyf-7* | WBGene00001123 | protein_coding_gene | -1.0 | 0.4973 | 4.6E-02 | -1.8 | 0.2901 | 1.1E-03 |
| *C07H4.1* | WBGene00007421 | protein_coding_gene | -1.0 | 0.4973 | 1.0E-02 | -1.5 | 0.3577 | 3.1E-10 |
| *Y62H9A.5* | WBGene00013393 | protein_coding_gene | -1.0 | 0.4964 | 1.1E-03 | -1.5 | 0.3546 | 4.6E-26 |
| *syd-1* | WBGene00006363 | protein_coding_gene | -1.0 | 0.4961 | 3.5E-02 | -1.4 | 0.3848 | 2.0E-05 |
| *magu-4* | WBGene00009678 | protein_coding_gene | -1.0 | 0.4953 | 2.1E-07 | -1.1 | 0.4726 | 4.5E-13 |
| *nhr-106* | WBGene00003696 | protein_coding_gene | -1.0 | 0.4952 | 1.4E-02 | -1.1 | 0.4765 | 1.8E-04 |
| *F16B3.3* | WBGene00044774 | protein_coding_gene | -1.0 | 0.4949 | 2.9E-02 | -1.1 | 0.4813 | 4.0E-04 |
| *W10C8.6* | WBGene00044433 | protein_coding_gene | -1.0 | 0.4945 | 4.2E-02 | -1.9 | 0.2685 | 3.0E-08 |
| *svh-2* | WBGene00020504 | protein_coding_gene | -1.0 | 0.4935 | 4.4E-04 | -1.1 | 0.4669 | 3.3E-20 |
| *lin-1* | WBGene00002990 | protein_coding_gene | -1.0 | 0.4933 | 1.7E-03 | -1.2 | 0.4289 | 1.5E-07 |
| *C27D8.3* | WBGene00007779 | protein_coding_gene | -1.0 | 0.4931 | 5.0E-05 | -1.3 | 0.3965 | 1.7E-17 |
| *ttr-5* | WBGene00008040 | protein_coding_gene | -1.0 | 0.4927 | 1.8E-06 | -2.1 | 0.2379 | 2.2E-24 |
| ***Y39D8A.1*** | **WBGene00021448** | **protein_coding_gene** | **-1.0** | **0.4927** | **3.7E-02** | **1.7** | **3.1896** | **7.2E-11** |
| *droe-4* | WBGene00010262 | protein_coding_gene | -1.0 | 0.4927 | 3.5E-03 | -2.4 | 0.1858 | 6.7E-45 |
| *his-71* | WBGene00001945 | protein_coding_gene | -1.0 | 0.4926 | 1.1E-04 | -1.6 | 0.3373 | 1.6E-23 |
| *glo-3* | WBGene00010341 | protein_coding_gene | -1.0 | 0.4922 | 1.3E-04 | -1.6 | 0.3343 | 9.8E-09 |
| *F41E7.2* | WBGene00009618 | protein_coding_gene | -1.0 | 0.4909 | 3.7E-03 | -1.8 | 0.2949 | 4.0E-08 |
| *B0272.4* | WBGene00007130 | protein_coding_gene | -1.0 | 0.4904 | 6.3E-03 | -1.6 | 0.3277 | 2.0E-12 |
| *C10B5.3* | WBGene00015671 | protein_coding_gene | -1.0 | 0.4890 | 2.3E-05 | -1.0 | 0.4974 | 2.4E-08 |
| *twk-7* | WBGene00006662 | protein_coding_gene | -1.0 | 0.4889 | 3.0E-02 | -2.0 | 0.2459 | 7.8E-19 |
| *glb-25* | WBGene00020279 | protein_coding_gene | -1.0 | 0.4880 | 1.3E-02 | -1.2 | 0.4281 | 3.2E-04 |
| *osg-1* | WBGene00019832 | protein_coding_gene | -1.0 | 0.4880 | 8.4E-03 | -1.6 | 0.3227 | 4.4E-09 |
| *ztf-29* | WBGene00013438 | protein_coding_gene | -1.0 | 0.4874 | 9.1E-07 | -1.1 | 0.4652 | 1.9E-09 |
| *nhr-135* | WBGene00003725 | protein_coding_gene | -1.0 | 0.4873 | 4.1E-04 | -1.3 | 0.4153 | 5.7E-08 |
| *F45D3.2* | WBGene00009722 | protein_coding_gene | -1.0 | 0.4858 | 1.1E-02 | -1.0 | 0.4846 | 9.4E-04 |
| *sma-6* | WBGene00004860 | protein_coding_gene | -1.0 | 0.4848 | 5.4E-07 | -1.2 | 0.4470 | 1.3E-11 |
| *dhp-2* | WBGene00000964 | protein_coding_gene | -1.0 | 0.4847 | 1.1E-03 | -1.3 | 0.3971 | 4.3E-10 |
| *nlp-50* | WBGene00018298 | protein_coding_gene | -1.0 | 0.4845 | 2.1E-03 | -1.1 | 0.4631 | 9.3E-06 |
| *C17H11.6* | WBGene00015926 | protein_coding_gene | -1.0 | 0.4843 | 3.9E-07 | -1.4 | 0.3880 | 5.6E-22 |
| *lips-15* | WBGene00012585 | protein_coding_gene | -1.0 | 0.4838 | 6.8E-08 | -1.6 | 0.3294 | 2.9E-14 |
| *syd-2* | WBGene00006364 | protein_coding_gene | -1.0 | 0.4837 | 6.1E-05 | -1.2 | 0.4271 | 2.9E-14 |
| *ipmk-1* | WBGene00014081 | protein_coding_gene | -1.0 | 0.4833 | 5.5E-06 | -1.0 | 0.4945 | 2.3E-05 |
| *crm-1* | WBGene00007103 | protein_coding_gene | -1.0 | 0.4831 | 1.4E-05 | -1.5 | 0.3491 | 3.6E-22 |
| *dre-1* | WBGene00001089 | protein_coding_gene | -1.0 | 0.4831 | 4.1E-07 | -1.1 | 0.4682 | 1.0E-09 |
| *ins-18* | WBGene00002101 | protein_coding_gene | -1.1 | 0.4826 | 2.5E-02 | -1.3 | 0.4081 | 3.5E-04 |
| *jnk-1* | WBGene00002178 | protein_coding_gene | -1.1 | 0.4817 | 9.5E-04 | -1.1 | 0.4813 | 1.6E-04 |
| *ztf-13* | WBGene00018704 | protein_coding_gene | -1.1 | 0.4812 | 1.4E-03 | -1.2 | 0.4275 | 1.5E-10 |
| *C16B8.4* | WBGene00015824 | protein_coding_gene | -1.1 | 0.4812 | 9.9E-03 | -1.3 | 0.4063 | 1.8E-07 |
| *spp-15* | WBGene00005000 | protein_coding_gene | -1.1 | 0.4809 | 2.7E-04 | -1.1 | 0.4685 | 1.3E-06 |
| *cyp-33A1* | WBGene00015709 | protein_coding_gene | -1.1 | 0.4808 | 1.4E-08 | -1.6 | 0.3371 | 6.4E-13 |

| | | | | | | | | |
|---|---|---|---|---|---|---|---|---|
| *daf-8* | WBGene00000904 | protein_coding_gene | -1.1 | 0.4807 | 8.5E-07 | -1.0 | 0.4945 | 6.2E-08 |
| *T04A6.1* | WBGene00020200 | protein_coding_gene | -1.1 | 0.4796 | 7.0E-05 | -1.9 | 0.2718 | 9.6E-27 |
| *cyp-25A3* | WBGene00014697 | protein_coding_gene | -1.1 | 0.4791 | 5.9E-03 | -1.8 | 0.2837 | 2.1E-17 |
| *ttr-51* | WBGene00010440 | protein_coding_gene | -1.1 | 0.4777 | 7.3E-06 | -2.0 | 0.2476 | 1.4E-38 |
| *cex-2* | WBGene00023408 | protein_coding_gene | -1.1 | 0.4775 | 8.2E-03 | -1.5 | 0.3555 | 4.0E-17 |
| *frpr-11* | WBGene00019444 | protein_coding_gene | -1.1 | 0.4772 | 8.7E-03 | -1.7 | 0.3106 | 5.4E-09 |
| *sma-4* | WBGene00004858 | protein_coding_gene | -1.1 | 0.4741 | 1.8E-05 | -2.2 | 0.2174 | 2.6E-49 |
| *W02C12.2* | WBGene00020929 | protein_coding_gene | -1.1 | 0.4739 | 2.1E-02 | -1.7 | 0.3142 | 7.1E-05 |
| *F56H11.2* | WBGene00010175 | protein_coding_gene | -1.1 | 0.4738 | 2.0E-02 | -2.3 | 0.2008 | 1.5E-13 |
| *C34C12.6* | WBGene00007925 | protein_coding_gene | -1.1 | 0.4735 | 2.9E-03 | -1.5 | 0.3574 | 1.0E-04 |
| *rga-6* | WBGene00015303 | protein_coding_gene | -1.1 | 0.4733 | 1.4E-02 | -1.2 | 0.4256 | 6.6E-04 |
| *swm-1* | WBGene00023417 | protein_coding_gene | -1.1 | 0.4733 | 5.9E-07 | -1.3 | 0.4146 | 6.3E-16 |
| *Y51B9A.9* | WBGene00013091 | protein_coding_gene | -1.1 | 0.4724 | 1.9E-02 | -1.9 | 0.2764 | 1.5E-12 |
| *C50F2.7* | WBGene00016840 | protein_coding_gene | -1.1 | 0.4723 | 1.7E-02 | -1.4 | 0.3868 | 3.2E-04 |
| *F42C5.9* | WBGene00018349 | protein_coding_gene | -1.1 | 0.4722 | 1.5E-11 | -1.4 | 0.3699 | 8.1E-20 |
| *F35F10.1* | WBGene00018052 | protein_coding_gene | -1.1 | 0.4716 | 4.5E-02 | -1.3 | 0.4087 | 4.8E-04 |
| *tag-196* | WBGene00007055 | protein_coding_gene | -1.1 | 0.4715 | 2.7E-04 | -1.4 | 0.3664 | 9.6E-27 |
| *nnt-1* | WBGene00003778 | protein_coding_gene | -1.1 | 0.4714 | 3.6E-02 | -3.3 | 0.1022 | 2.2E-74 |
| *fipr-1* | WBGene00009096 | protein_coding_gene | -1.1 | 0.4707 | 1.3E-06 | -1.4 | 0.3695 | 2.3E-15 |
| *acox-3* | WBGene00019060 | protein_coding_gene | -1.1 | 0.4696 | 1.2E-02 | -1.5 | 0.3600 | 2.9E-11 |
| *nex-2* | WBGene00003589 | protein_coding_gene | -1.1 | 0.4685 | 9.4E-07 | -1.6 | 0.3269 | 7.4E-46 |
| *nhr-43* | WBGene00003633 | protein_coding_gene | -1.1 | 0.4682 | 3.8E-03 | -1.4 | 0.3921 | 8.3E-06 |
| *hid-1* | WBGene00001844 | protein_coding_gene | -1.1 | 0.4680 | 2.9E-07 | -1.2 | 0.4262 | 6.6E-18 |
| *F43G6.4* | WBGene00009657 | protein_coding_gene | -1.1 | 0.4679 | 2.9E-02 | -1.2 | 0.4310 | 6.9E-07 |
| *Y55B1BL.1* | WBGene00021909 | protein_coding_gene | -1.1 | 0.4673 | 4.2E-08 | -2.3 | 0.2025 | 8.9E-46 |
| ***F57F4.4*** | **WBGene00019017** | **protein_coding_gene** | **-1.1** | **0.4669** | **4.4E-04** | **1.7** | **3.2093** | **4.1E-03** |
| *M02D8.6* | WBGene00019732 | protein_coding_gene | -1.1 | 0.4663 | 1.2E-03 | -1.7 | 0.3176 | 3.2E-14 |
| *frpr-18* | WBGene00020586 | protein_coding_gene | -1.1 | 0.4659 | 2.4E-02 | -1.2 | 0.4440 | 7.6E-03 |
| ***pcp-2*** | **WBGene00003957** | **protein_coding_gene** | **-1.1** | **0.4659** | **2.7E-02** | **2.8** | **6.9212** | **9.4E-10** |
| *chil-10* | WBGene00016642 | protein_coding_gene | -1.1 | 0.4657 | 4.9E-10 | -1.0 | 0.4900 | 2.0E-11 |
| *nhr-62* | WBGene00003652 | protein_coding_gene | -1.1 | 0.4649 | 1.5E-08 | -1.0 | 0.4861 | 1.6E-05 |
| *let-413* | WBGene00002632 | protein_coding_gene | -1.1 | 0.4647 | 8.8E-07 | -1.1 | 0.4692 | 6.5E-17 |
| *sma-10* | WBGene00020649 | protein_coding_gene | -1.1 | 0.4647 | 1.4E-03 | -2.1 | 0.2284 | 2.1E-39 |
| ***C41H7.1*** | **WBGene00016571** | **protein_coding_gene** | **-1.1** | **0.4645** | **5.7E-03** | **1.4** | **2.7256** | **8.7E-03** |
| *F41G4.8* | WBGene00018311 | protein_coding_gene | -1.1 | 0.4643 | 3.6E-03 | -1.5 | 0.3634 | 1.5E-08 |
| *twk-31* | WBGene00006683 | protein_coding_gene | -1.1 | 0.4638 | 9.4E-03 | -1.3 | 0.3945 | 1.5E-07 |
| *ttr-2* | WBGene00010539 | protein_coding_gene | -1.1 | 0.4633 | 3.5E-05 | -1.7 | 0.3028 | 2.9E-26 |
| *Y64G10A.7* | WBGene00013416 | protein_coding_gene | -1.1 | 0.4628 | 1.4E-03 | -1.2 | 0.4353 | 2.1E-05 |
| *flp-12* | WBGene00001455 | protein_coding_gene | -1.1 | 0.4625 | 5.9E-04 | -1.2 | 0.4431 | 2.5E-11 |
| *snt-6* | WBGene00004926 | protein_coding_gene | -1.1 | 0.4624 | 8.1E-03 | -1.0 | 0.4941 | 5.7E-04 |
| *rab-28* | WBGene00004281 | protein_coding_gene | -1.1 | 0.4622 | 4.4E-02 | -1.4 | 0.3678 | 3.4E-03 |
| *jac-1* | WBGene00002175 | protein_coding_gene | -1.1 | 0.4619 | 1.4E-07 | -1.3 | 0.4141 | 7.5E-23 |
| *F26A1.19* | WBGene00270319 | protein_coding_gene | -1.1 | 0.4615 | 3.3E-02 | -1.1 | 0.4706 | 6.8E-03 |
| *sul-2* | WBGene00006309 | protein_coding_gene | -1.1 | 0.4614 | 2.0E-02 | -1.4 | 0.3724 | 3.7E-07 |
| *nhr-121* | WBGene00003711 | protein_coding_gene | -1.1 | 0.4605 | 5.2E-04 | -1.1 | 0.4631 | 1.3E-07 |
| *exc-7* | WBGene00001368 | protein_coding_gene | -1.1 | 0.4593 | 8.3E-05 | -1.4 | 0.3670 | 1.5E-12 |
| *C03B1.13* | WBGene00015381 | protein_coding_gene | -1.1 | 0.4586 | 8.8E-06 | -1.5 | 0.3567 | 1.8E-10 |
| *chd-3* | WBGene00000482 | protein_coding_gene | -1.1 | 0.4582 | 2.5E-04 | -1.1 | 0.4779 | 1.5E-14 |
| ***sth-1*** | **WBGene00006060** | **protein_coding_gene** | **-1.1** | **0.4581** | **5.8E-03** | **1.2** | **2.3201** | **3.0E-04** |
| *Y57G11C.38* | WBGene00013327 | protein_coding_gene | -1.1 | 0.4573 | 6.7E-04 | -1.5 | 0.3564 | 4.2E-07 |
| *K01A2.5* | WBGene00019280 | protein_coding_gene | -1.1 | 0.4568 | 4.3E-04 | -1.5 | 0.3512 | 4.0E-14 |
| *C32D5.1* | WBGene00016310 | protein_coding_gene | -1.1 | 0.4564 | 1.6E-03 | -1.2 | 0.4230 | 6.8E-05 |
| *T07D1.2* | WBGene00020301 | protein_coding_gene | -1.1 | 0.4563 | 6.7E-05 | -1.2 | 0.4302 | 1.7E-14 |
| *ule-5* | WBGene00013394 | protein_coding_gene | -1.1 | 0.4557 | 6.9E-06 | -1.4 | 0.3913 | 4.4E-23 |
| *F20G2.7* | WBGene00044920 | pseudogene | -1.1 | 0.4546 | 5.9E-04 | -1.0 | 0.4903 | 1.6E-03 |
| *M60.4* | WBGene00019780 | protein_coding_gene | -1.1 | 0.4543 | 4.6E-03 | -3.2 | 0.1104 | 1.3E-108 |
| *mod-5* | WBGene00003387 | protein_coding_gene | -1.1 | 0.4537 | 1.2E-02 | -2.3 | 0.1963 | 9.0E-12 |
| *ttr-37* | WBGene00017667 | protein_coding_gene | -1.1 | 0.4531 | 1.9E-03 | -2.8 | 0.1446 | 1.1E-31 |
| *pho-4* | WBGene00004023 | protein_coding_gene | -1.1 | 0.4526 | 1.6E-09 | -1.4 | 0.3713 | 6.1E-15 |
| *qua-1* | WBGene00004264 | protein_coding_gene | -1.1 | 0.4525 | 8.2E-03 | -1.9 | 0.2721 | 1.1E-03 |
| *Y48G9A.9* | WBGene00021702 | protein_coding_gene | -1.1 | 0.4524 | 6.9E-05 | -1.3 | 0.4014 | 1.3E-18 |
| *Y25C1A.14* | WBGene00194985 | protein_coding_gene | -1.1 | 0.4523 | 7.6E-05 | -1.7 | 0.2978 | 3.3E-04 |
| *fbxa-32* | WBGene00013839 | protein_coding_gene | -1.1 | 0.4520 | 4.4E-03 | -1.0 | 0.4845 | 1.1E-02 |
| *rig-1* | WBGene00004369 | protein_coding_gene | -1.1 | 0.4515 | 3.1E-05 | -1.3 | 0.3932 | 4.8E-20 |
| *F45E10.2* | WBGene00009729 | protein_coding_gene | -1.1 | 0.4512 | 1.8E-03 | -2.2 | 0.2244 | 9.2E-18 |
| *F45E4.5* | WBGene00018470 | protein_coding_gene | -1.1 | 0.4508 | 2.7E-03 | -1.9 | 0.2697 | 2.2E-08 |
| *lite-1* | WBGene00001803 | protein_coding_gene | -1.1 | 0.4507 | 5.6E-03 | -1.3 | 0.4035 | 8.5E-05 |
| *fat-3* | WBGene00001395 | protein_coding_gene | -1.2 | 0.4501 | 4.9E-05 | -2.2 | 0.2127 | 4.1E-14 |
| *flp-27* | WBGene00016114 | protein_coding_gene | -1.2 | 0.4497 | 3.6E-03 | -1.4 | 0.3787 | 4.2E-09 |
| *T24B8.4* | WBGene00011978 | protein_coding_gene | -1.2 | 0.4497 | 2.2E-03 | -1.3 | 0.4008 | 7.6E-19 |

| | | | | | | | | |
|---|---|---|---|---|---|---|---|---|
| *kgb-1* | WBGene00002187 | protein_coding_gene | -1.2 | 0.4491 | 1.1E-09 | -1.0 | 0.4932 | 6.3E-18 |
| *T16H12.9* | WBGene00011817 | protein_coding_gene | -1.2 | 0.4491 | 2.9E-02 | -2.2 | 0.2202 | 1.6E-12 |
| *C29F7.1* | WBGene00007810 | protein_coding_gene | -1.2 | 0.4490 | 3.2E-05 | -1.2 | 0.4467 | 5.8E-06 |
| *rnf-145* | WBGene00022471 | protein_coding_gene | -1.2 | 0.4485 | 6.1E-18 | -1.3 | 0.4129 | 1.6E-11 |
| *nhr-173* | WBGene00016975 | protein_coding_gene | -1.2 | 0.4484 | 2.9E-03 | -1.1 | 0.4557 | 3.3E-07 |
| *klo-1* | WBGene00016848 | protein_coding_gene | -1.2 | 0.4481 | 3.2E-13 | -1.5 | 0.3631 | 4.9E-16 |
| *spc-1* | WBGene00004951 | protein_coding_gene | -1.2 | 0.4477 | 6.9E-06 | -1.2 | 0.4293 | 9.5E-19 |
| *K02D10.8* | WBGene00219329 | protein_coding_gene | -1.2 | 0.4473 | 1.2E-02 | -1.8 | 0.2924 | 3.6E-04 |
| *F59E11.7* | WBGene00019113 | protein_coding_gene | -1.2 | 0.4472 | 4.1E-02 | -2.7 | 0.1568 | 4.1E-14 |
| *twk-2* | WBGene00006657 | protein_coding_gene | -1.2 | 0.4472 | 8.1E-03 | -1.3 | 0.4096 | 3.2E-05 |
| *odd-1* | WBGene00003845 | protein_coding_gene | -1.2 | 0.4467 | 5.3E-03 | -1.3 | 0.3994 | 2.3E-03 |
| *lron-12* | WBGene00013542 | protein_coding_gene | -1.2 | 0.4461 | 3.7E-03 | -1.5 | 0.3577 | 2.8E-05 |
| *tsp-20* | WBGene00006646 | protein_coding_gene | -1.2 | 0.4458 | 1.4E-04 | -1.4 | 0.3918 | 3.5E-07 |
| *cutl-12* | WBGene00011928 | protein_coding_gene | -1.2 | 0.4455 | 3.5E-02 | -1.3 | 0.3944 | 2.3E-02 |
| *C30G7.4* | WBGene00044178 | protein_coding_gene | -1.2 | 0.4451 | 2.8E-03 | -1.5 | 0.3424 | 9.1E-07 |
| *bam-2* | WBGene00000237 | protein_coding_gene | -1.2 | 0.4445 | 2.6E-05 | -1.7 | 0.3163 | 1.2E-21 |
| *dhs-5* | WBGene00000969 | protein_coding_gene | -1.2 | 0.4443 | 2.5E-07 | -1.2 | 0.4429 | 2.6E-06 |
| *syg-1* | WBGene00006365 | protein_coding_gene | -1.2 | 0.4438 | 2.4E-03 | -1.3 | 0.4077 | 1.2E-07 |
| *ZC581.3* | WBGene00022633 | protein_coding_gene | -1.2 | 0.4434 | 4.7E-03 | -1.7 | 0.3091 | 7.9E-26 |
| *nhr-131* | WBGene00003721 | protein_coding_gene | -1.2 | 0.4431 | 5.6E-03 | -1.5 | 0.3558 | 1.1E-09 |
| *fkh-9* | WBGene00001441 | protein_coding_gene | -1.2 | 0.4429 | 6.4E-03 | -1.3 | 0.3932 | 1.6E-22 |
| *C53B7.3* | WBGene00016894 | protein_coding_gene | -1.2 | 0.4429 | 1.9E-06 | -1.4 | 0.3841 | 2.9E-15 |
| *dyf-1* | WBGene00001117 | protein_coding_gene | -1.2 | 0.4427 | 4.0E-03 | -1.6 | 0.3250 | 5.2E-05 |
| *F35C5.12* | WBGene00009400 | protein_coding_gene | -1.2 | 0.4424 | 4.5E-02 | -1.1 | 0.4673 | 5.7E-03 |
| *nhr-36* | WBGene00017198 | protein_coding_gene | -1.2 | 0.4422 | 1.1E-03 | -1.6 | 0.3197 | 3.4E-08 |
| *octr-1* | WBGene00006411 | protein_coding_gene | -1.2 | 0.4421 | 1.5E-02 | -1.4 | 0.3753 | 7.6E-03 |
| *ZC190.4* | WBGene00022538 | protein_coding_gene | -1.2 | 0.4421 | 1.4E-04 | -1.1 | 0.4815 | 8.1E-11 |
| *tag-273* | WBGene00013289 | protein_coding_gene | -1.2 | 0.4416 | 7.0E-05 | -1.3 | 0.4057 | 4.6E-10 |
| *C46C2.2* | WBGene00008110 | protein_coding_gene | -1.2 | 0.4412 | 1.9E-13 | -1.1 | 0.4517 | 3.9E-16 |
| *abhd-5.1* | WBGene00016506 | protein_coding_gene | -1.2 | 0.4401 | 8.5E-03 | -2.1 | 0.2386 | 9.2E-13 |
| *catp-7* | WBGene00022010 | protein_coding_gene | -1.2 | 0.4400 | 8.1E-04 | -2.1 | 0.2385 | 1.2E-51 |
| *dve-1* | WBGene00022861 | protein_coding_gene | -1.2 | 0.4398 | 8.2E-04 | -1.1 | 0.4698 | 7.0E-11 |
| *cfap-36* | WBGene00022435 | protein_coding_gene | -1.2 | 0.4393 | 2.1E-03 | -1.1 | 0.4753 | 4.2E-03 |
| *ssu-2* | WBGene00013455 | protein_coding_gene | -1.2 | 0.4391 | 2.1E-02 | -1.9 | 0.2629 | 3.3E-05 |
| *ule-2* | WBGene00045038 | protein_coding_gene | -1.2 | 0.4390 | 1.8E-05 | -1.1 | 0.4700 | 3.6E-10 |
| *cnnm-2* | WBGene00019869 | protein_coding_gene | -1.2 | 0.4389 | 2.2E-09 | -1.3 | 0.4113 | 3.5E-11 |
| *T28D6.4* | WBGene00012124 | protein_coding_gene | -1.2 | 0.4383 | 8.6E-04 | -1.3 | 0.4113 | 1.6E-05 |
| *ajm-1* | WBGene00000100 | protein_coding_gene | -1.2 | 0.4382 | 1.6E-04 | -1.2 | 0.4238 | 1.2E-23 |
| *tax-6* | WBGene00006527 | protein_coding_gene | -1.2 | 0.4379 | 3.2E-04 | -1.7 | 0.3079 | 3.0E-43 |
| ***Y55F3AM.14*** | **WBGene00021931** | **protein_coding_gene** | **-1.2** | **0.4378** | **4.3E-02** | **2.5** | **5.6184** | **2.8E-03** |
| *glt-5* | WBGene00001623 | protein_coding_gene | -1.2 | 0.4376 | 2.3E-02 | -1.9 | 0.2704 | 1.8E-11 |
| *F23F12.13* | WBGene00044455 | protein_coding_gene | -1.2 | 0.4376 | 5.2E-03 | -1.6 | 0.3343 | 8.8E-06 |
| *tsp-6* | WBGene00006632 | protein_coding_gene | -1.2 | 0.4368 | 4.9E-02 | -3.2 | 0.1105 | 2.0E-09 |
| *F37C4.4* | WBGene00018144 | protein_coding_gene | -1.2 | 0.4368 | 3.4E-04 | -1.3 | 0.4072 | 9.6E-06 |
| *lron-7* | WBGene00019351 | protein_coding_gene | -1.2 | 0.4365 | 3.0E-10 | -1.2 | 0.4249 | 4.4E-27 |
| *anp-1* | WBGene00011587 | protein_coding_gene | -1.2 | 0.4362 | 1.3E-06 | -1.2 | 0.4496 | 3.3E-17 |
| *gcl-1* | WBGene00013382 | protein_coding_gene | -1.2 | 0.4358 | 6.1E-04 | -2.3 | 0.1996 | 4.1E-32 |
| ***let-383*** | **WBGene00044898** | **protein_coding_gene** | **-1.2** | **0.4358** | **4.7E-04** | **1.1** | **2.0837** | **6.3E-03** |
| *nsy-1* | WBGene00003822 | protein_coding_gene | -1.2 | 0.4354 | 1.0E-05 | -1.2 | 0.4293 | 7.4E-23 |
| *Y42G9A.3* | WBGene00021533 | protein_coding_gene | -1.2 | 0.4352 | 1.6E-03 | -1.7 | 0.3126 | 2.1E-06 |
| *nlp-15* | WBGene00003753 | protein_coding_gene | -1.2 | 0.4351 | 6.1E-04 | -1.1 | 0.4695 | 2.0E-08 |
| *atgp-1* | WBGene00000224 | protein_coding_gene | -1.2 | 0.4349 | 1.6E-07 | -1.7 | 0.3060 | 4.5E-40 |
| *cutl-3* | WBGene00008978 | protein_coding_gene | -1.2 | 0.4348 | 7.2E-03 | -2.0 | 0.2544 | 3.0E-06 |
| *chd-7* | WBGene00007053 | protein_coding_gene | -1.2 | 0.4348 | 1.8E-03 | -1.3 | 0.4020 | 1.4E-24 |
| *ztf-16* | WBGene00019960 | protein_coding_gene | -1.2 | 0.4346 | 4.1E-04 | -1.2 | 0.4469 | 2.5E-09 |
| *dhhc-2* | WBGene00012948 | protein_coding_gene | -1.2 | 0.4345 | 1.9E-05 | -1.1 | 0.4651 | 2.1E-06 |
| *E02H9.7* | WBGene00017105 | protein_coding_gene | -1.2 | 0.4340 | 1.6E-04 | -1.1 | 0.4576 | 1.3E-06 |
| *ric-3* | WBGene00004363 | protein_coding_gene | -1.2 | 0.4337 | 2.4E-04 | -1.6 | 0.3198 | 1.9E-10 |
| *zig-7* | WBGene00006984 | protein_coding_gene | -1.2 | 0.4335 | 2.3E-04 | -1.7 | 0.3018 | 7.2E-35 |
| *H06H21.8* | WBGene00019164 | protein_coding_gene | -1.2 | 0.4334 | 9.4E-07 | -1.5 | 0.3442 | 1.6E-25 |
| *C44E12.1* | WBGene00016657 | protein_coding_gene | -1.2 | 0.4329 | 7.9E-06 | -2.5 | 0.1727 | 3.9E-25 |
| *olrn-1* | WBGene00007260 | protein_coding_gene | -1.2 | 0.4328 | 2.8E-03 | -1.1 | 0.4815 | 1.7E-05 |
| *clec-222* | WBGene00007283 | protein_coding_gene | -1.2 | 0.4328 | 4.4E-08 | -1.2 | 0.4488 | 5.7E-11 |
| *C54D10.10* | WBGene00008304 | protein_coding_gene | -1.2 | 0.4322 | 3.8E-03 | -1.5 | 0.3542 | 3.6E-03 |
| *cest-1.2* | WBGene00011364 | protein_coding_gene | -1.2 | 0.4322 | 3.2E-02 | -4.1 | 0.0567 | 1.0E-76 |
| *sma-9* | WBGene00004862 | protein_coding_gene | -1.2 | 0.4320 | 5.3E-04 | -1.4 | 0.3829 | 1.9E-25 |
| *let-23* | WBGene00002299 | protein_coding_gene | -1.2 | 0.4316 | 1.1E-04 | -1.4 | 0.3785 | 1.8E-23 |
| *msrp-7* | WBGene00011070 | protein_coding_gene | -1.2 | 0.4316 | 1.1E-04 | -1.2 | 0.4457 | 4.2E-07 |
| *F32B4.5* | WBGene00009315 | protein_coding_gene | -1.2 | 0.4315 | 2.3E-02 | -2.5 | 0.1822 | 5.8E-09 |
| *Y87G2A.16* | WBGene00013602 | protein_coding_gene | -1.2 | 0.4312 | 7.3E-03 | -2.3 | 0.2051 | 6.1E-05 |

| | | | | | | | | |
|---|---|---|---|---|---|---|---|---|
| *F54C9.11* | WBGene00010045 | protein_coding_gene | -1.2 | 0.4310 | 6.5E-05 | -1.0 | 0.5000 | 8.2E-05 |
| *M60.6* | WBGene00019781 | protein_coding_gene | -1.2 | 0.4310 | 7.6E-09 | -1.2 | 0.4404 | 1.8E-07 |
| *unc-79* | WBGene00006811 | protein_coding_gene | -1.2 | 0.4300 | 8.6E-03 | -1.5 | 0.3618 | 1.4E-09 |
| *Y71H2B.8* | WBGene00022197 | protein_coding_gene | -1.2 | 0.4282 | 6.2E-03 | -3.0 | 0.1248 | 5.2E-17 |
| *K09C4.10* | WBGene00019554 | protein_coding_gene | -1.2 | 0.4282 | 1.8E-07 | -1.4 | 0.3920 | 2.6E-10 |
| *clp-4* | WBGene00000545 | protein_coding_gene | -1.2 | 0.4282 | 2.9E-06 | -1.7 | 0.2993 | 1.2E-31 |
| *F22B7.1* | WBGene00017692 | protein_coding_gene | -1.2 | 0.4281 | 4.0E-03 | -1.7 | 0.3054 | 4.7E-09 |
| *C07A12.7* | WBGene00015561 | protein_coding_gene | -1.2 | 0.4278 | 8.5E-09 | -1.1 | 0.4642 | 1.4E-12 |
| *C10A4.13* | WBGene00206485 | protein_coding_gene | -1.2 | 0.4272 | 9.2E-03 | -1.1 | 0.4680 | 1.1E-02 |
| *W02B8.1* | WBGene00012197 | protein_coding_gene | -1.2 | 0.4270 | 7.3E-03 | -1.3 | 0.3990 | 3.9E-09 |
| *best-1* | WBGene00007203 | protein_coding_gene | -1.2 | 0.4267 | 2.5E-03 | -1.9 | 0.2702 | 5.0E-04 |
| *C54E4.4* | WBGene00016919 | protein_coding_gene | -1.2 | 0.4265 | 2.3E-04 | -1.2 | 0.4252 | 1.4E-03 |
| *ampd-1* | WBGene00016415 | protein_coding_gene | -1.2 | 0.4265 | 3.4E-12 | -1.1 | 0.4548 | 1.1E-23 |
| *C01A2.2* | WBGene00007214 | protein_coding_gene | -1.2 | 0.4258 | 1.9E-03 | -1.9 | 0.2739 | 4.5E-09 |
| *bicd-1* | WBGene00016611 | protein_coding_gene | -1.2 | 0.4250 | 3.4E-07 | -1.2 | 0.4364 | 1.1E-17 |
| *cof-2* | WBGene00000583 | protein_coding_gene | -1.2 | 0.4247 | 2.1E-03 | -1.1 | 0.4521 | 8.8E-03 |
| *ech-8* | WBGene00001157 | protein_coding_gene | -1.2 | 0.4242 | 4.9E-05 | -1.5 | 0.3424 | 2.2E-22 |
| *unc-53* | WBGene00006788 | protein_coding_gene | -1.2 | 0.4240 | 1.3E-03 | -1.5 | 0.3557 | 6.9E-20 |
| *E01G6.2* | WBGene00008450 | protein_coding_gene | -1.2 | 0.4237 | 4.5E-05 | -1.3 | 0.3964 | 3.0E-07 |
| *F40F8.5* | WBGene00009578 | protein_coding_gene | -1.2 | 0.4236 | 6.5E-05 | -1.7 | 0.3141 | 4.6E-30 |
| *F02A9.1* | WBGene00008513 | protein_coding_gene | -1.2 | 0.4233 | 4.5E-02 | -1.6 | 0.3413 | 3.5E-04 |
| ***nmy-3*** | **WBGene00012437** | **protein_coding_gene** | **-1.2** | **0.4233** | **3.3E-03** | **1.2** | **2.2193** | **3.0E-05** |
| *sulp-3* | WBGene00018283 | protein_coding_gene | -1.2 | 0.4226 | 2.5E-02 | -1.7 | 0.3149 | 1.6E-05 |
| *C56G2.4* | WBGene00016979 | protein_coding_gene | -1.2 | 0.4224 | 4.7E-04 | -1.8 | 0.2960 | 2.1E-07 |
| *T15B7.10* | WBGene00020522 | pseudogene | -1.2 | 0.4222 | 3.9E-03 | -3.1 | 0.1164 | 3.4E-07 |
| *vav-1* | WBGene00006887 | protein_coding_gene | -1.2 | 0.4218 | 2.5E-09 | -1.0 | 0.4968 | 2.1E-13 |
| *scd-1* | WBGene00004739 | protein_coding_gene | -1.2 | 0.4217 | 5.5E-05 | -1.1 | 0.4523 | 3.0E-20 |
| *R06F6.14* | WBGene00269421 | protein_coding_gene | -1.3 | 0.4202 | 1.1E-05 | -1.6 | 0.3205 | 1.5E-10 |
| *endu-1* | WBGene00010488 | protein_coding_gene | -1.3 | 0.4200 | 6.2E-08 | -1.7 | 0.2974 | 3.3E-33 |
| *C34C6.3* | WBGene00007916 | protein_coding_gene | -1.3 | 0.4184 | 3.1E-03 | -1.6 | 0.3345 | 2.4E-05 |
| *smf-3* | WBGene00004878 | protein_coding_gene | -1.3 | 0.4184 | 1.1E-02 | -4.0 | 0.0636 | 5.9E-29 |
| *W03G11.4* | WBGene00012226 | protein_coding_gene | -1.3 | 0.4181 | 1.5E-04 | -1.5 | 0.3484 | 1.7E-15 |
| *sta-1* | WBGene00013111 | protein_coding_gene | -1.3 | 0.4181 | 6.7E-06 | -1.3 | 0.4120 | 9.7E-10 |
| *C05B5.8* | WBGene00007324 | protein_coding_gene | -1.3 | 0.4181 | 9.4E-03 | -2.3 | 0.2047 | 8.6E-21 |
| *ccch-1* | WBGene00009532 | protein_coding_gene | -1.3 | 0.4179 | 2.7E-03 | -1.3 | 0.4026 | 2.7E-23 |
| *Y45F10C.2* | WBGene00012878 | protein_coding_gene | -1.3 | 0.4177 | 2.8E-02 | -1.3 | 0.3967 | 6.1E-06 |
| *cfi-1* | WBGene00000476 | protein_coding_gene | -1.3 | 0.4177 | 5.1E-04 | -1.8 | 0.2907 | 3.7E-11 |
| *sek-6* | WBGene00012162 | protein_coding_gene | -1.3 | 0.4174 | 1.5E-03 | -1.3 | 0.3948 | 1.4E-04 |
| *Y40B10A.9* | WBGene00021493 | protein_coding_gene | -1.3 | 0.4166 | 3.3E-06 | -2.1 | 0.2305 | 2.3E-23 |
| *F23A7.3* | WBGene00009068 | protein_coding_gene | -1.3 | 0.4156 | 5.0E-03 | -1.8 | 0.2907 | 3.3E-08 |
| *ptr-3* | WBGene00004218 | protein_coding_gene | -1.3 | 0.4156 | 8.9E-03 | -2.9 | 0.1310 | 4.9E-20 |
| *lntl-1* | WBGene00016289 | protein_coding_gene | -1.3 | 0.4147 | 4.4E-05 | -1.1 | 0.4540 | 1.4E-10 |
| *F52G3.1* | WBGene00018710 | protein_coding_gene | -1.3 | 0.4130 | 6.1E-05 | -1.2 | 0.4455 | 4.2E-25 |
| *cex-1* | WBGene00023407 | protein_coding_gene | -1.3 | 0.4127 | 9.5E-04 | -2.2 | 0.2120 | 6.4E-31 |
| *gst-13* | WBGene00001761 | protein_coding_gene | -1.3 | 0.4126 | 2.2E-06 | -1.0 | 0.4936 | 1.6E-14 |
| *vab-19* | WBGene00006882 | protein_coding_gene | -1.3 | 0.4125 | 8.4E-06 | -1.5 | 0.3609 | 1.7E-14 |
| *rgs-4* | WBGene00004347 | protein_coding_gene | -1.3 | 0.4115 | 5.7E-07 | -1.8 | 0.2862 | 2.4E-11 |
| *ctbp-1* | WBGene00006424 | protein_coding_gene | -1.3 | 0.4113 | 1.8E-03 | -1.4 | 0.3700 | 9.0E-08 |
| *C08G9.1* | WBGene00015619 | protein_coding_gene | -1.3 | 0.4112 | 8.1E-05 | -1.7 | 0.3019 | 1.1E-18 |
| *nhr-202* | WBGene00019742 | protein_coding_gene | -1.3 | 0.4111 | 1.3E-04 | -1.6 | 0.3215 | 1.6E-11 |
| *B0393.9* | WBGene00044636 | protein_coding_gene | -1.3 | 0.4105 | 1.2E-08 | -1.2 | 0.4220 | 8.7E-10 |
| *daf-6* | WBGene00000902 | protein_coding_gene | -1.3 | 0.4098 | 2.2E-02 | -1.3 | 0.4133 | 3.5E-02 |
| *Y41E3.22* | WBGene00045146 | protein_coding_gene | -1.3 | 0.4097 | 2.5E-02 | -2.6 | 0.1665 | 1.7E-05 |
| *ceeh-2* | WBGene00010628 | protein_coding_gene | -1.3 | 0.4096 | 1.7E-07 | -2.1 | 0.2329 | 3.4E-24 |
| ***ZK228.4*** | **WBGene00013947** | **protein_coding_gene** | **-1.3** | **0.4095** | **1.5E-06** | **2.7** | **6.5929** | **8.0E-38** |
| *T01E8.1* | WBGene00011331 | protein_coding_gene | -1.3 | 0.4091 | 3.7E-04 | -2.0 | 0.2458 | 2.3E-29 |
| *F40A3.1* | WBGene00018216 | protein_coding_gene | -1.3 | 0.4090 | 6.0E-12 | -1.0 | 0.4926 | 6.9E-10 |
| *dlc-3* | WBGene00012424 | protein_coding_gene | -1.3 | 0.4085 | 1.6E-03 | -1.8 | 0.2935 | 3.6E-08 |
| *gtl-1* | WBGene00001795 | protein_coding_gene | -1.3 | 0.4082 | 1.5E-05 | -1.1 | 0.4577 | 1.6E-15 |
| *dgk-2* | WBGene00000959 | protein_coding_gene | -1.3 | 0.4080 | 7.4E-12 | -1.7 | 0.3125 | 1.4E-35 |
| *acox-1.6* | WBGene00010336 | protein_coding_gene | -1.3 | 0.4078 | 1.7E-05 | -1.4 | 0.3679 | 2.4E-11 |
| *Y51A2D.14* | WBGene00013081 | protein_coding_gene | -1.3 | 0.4076 | 1.2E-04 | -1.4 | 0.3797 | 6.7E-17 |
| *T25B2.2* | WBGene00020786 | protein_coding_gene | -1.3 | 0.4075 | 4.2E-05 | -1.3 | 0.4196 | 1.8E-11 |
| *tsp-9* | WBGene00006635 | protein_coding_gene | -1.3 | 0.4073 | 5.7E-08 | -1.2 | 0.4452 | 1.6E-11 |
| *T05E7.4* | WBGene00020260 | protein_coding_gene | -1.3 | 0.4072 | 2.3E-02 | -2.1 | 0.2358 | 3.3E-05 |
| *twk-29* | WBGene00006681 | protein_coding_gene | -1.3 | 0.4072 | 1.4E-02 | -1.4 | 0.3818 | 8.0E-04 |
| *uggt-1* | WBGene00018604 | protein_coding_gene | -1.3 | 0.4070 | 4.7E-08 | -1.1 | 0.4659 | 5.3E-21 |
| *ZK355.2* | WBGene00022712 | protein_coding_gene | -1.3 | 0.4069 | 5.6E-13 | -1.2 | 0.4469 | 2.8E-12 |
| *F52E4.5* | WBGene00018702 | protein_coding_gene | -1.3 | 0.4067 | 5.0E-04 | -1.3 | 0.4111 | 4.4E-07 |
| *R07D5.2* | WBGene00011101 | protein_coding_gene | -1.3 | 0.4064 | 6.5E-03 | -1.4 | 0.3886 | 1.4E-03 |

| | | | | | | | | |
|---|---|---|---|---|---|---|---|---|
| *asp-12* | WBGene00017678 | protein_coding_gene | -1.3 | 0.4054 | 1.0E-03 | -1.6 | 0.3301 | 3.0E-09 |
| *nhr-92* | WBGene00003682 | protein_coding_gene | -1.3 | 0.4052 | 1.6E-03 | -2.0 | 0.2498 | 2.1E-08 |
| *nhr-142* | WBGene00018430 | protein_coding_gene | -1.3 | 0.4051 | 7.4E-04 | -1.0 | 0.4887 | 1.6E-02 |
| *sod-4* | WBGene00004933 | protein_coding_gene | -1.3 | 0.4050 | 1.1E-06 | -2.3 | 0.2102 | 5.0E-32 |
| *gad-2* | WBGene00015666 | protein_coding_gene | -1.3 | 0.4047 | 2.0E-02 | -1.4 | 0.3743 | 1.1E-04 |
| *col-99* | WBGene00000674 | protein_coding_gene | -1.3 | 0.4041 | 1.7E-03 | -1.8 | 0.2849 | 1.7E-14 |
| *ZC373.2* | WBGene00013867 | protein_coding_gene | -1.3 | 0.4038 | 1.0E-06 | -1.3 | 0.4173 | 1.3E-20 |
| *Y87G2A.19* | WBGene00044260 | protein_coding_gene | -1.3 | 0.4036 | 3.5E-04 | -1.4 | 0.3753 | 2.5E-15 |
| *din-1* | WBGene00008549 | protein_coding_gene | -1.3 | 0.4033 | 1.7E-04 | -1.1 | 0.4515 | 9.4E-21 |
| *K02D10.4* | WBGene00019304 | protein_coding_gene | -1.3 | 0.4024 | 5.0E-04 | -1.8 | 0.2918 | 7.7E-11 |
| *T10H9.8* | WBGene00044779 | protein_coding_gene | -1.3 | 0.4020 | 1.2E-03 | -3.0 | 0.1251 | 1.7E-32 |
| *pqn-18* | WBGene00004108 | protein_coding_gene | -1.3 | 0.4019 | 3.9E-04 | -1.1 | 0.4735 | 2.8E-10 |
| *hlh-8* | WBGene00001953 | protein_coding_gene | -1.3 | 0.4018 | 3.6E-02 | -1.6 | 0.3401 | 1.0E-02 |
| *C08G9.2* | WBGene00015620 | protein_coding_gene | -1.3 | 0.4018 | 1.3E-04 | -1.2 | 0.4490 | 5.6E-03 |
| *F58B4.6* | WBGene00010239 | protein_coding_gene | -1.3 | 0.4015 | 9.6E-03 | -1.0 | 0.4892 | 2.2E-03 |
| *aak-2* | WBGene00020142 | protein_coding_gene | -1.3 | 0.4015 | 3.2E-12 | -1.5 | 0.3657 | 2.7E-29 |
| *acs-22* | WBGene00017012 | protein_coding_gene | -1.3 | 0.4012 | 2.9E-05 | -2.3 | 0.2079 | 2.6E-74 |
| *C10C5.7* | WBGene00007511 | protein_coding_gene | -1.3 | 0.4009 | 6.0E-03 | -1.5 | 0.3547 | 4.1E-05 |
| *T05A8.3* | WBGene00020229 | protein_coding_gene | -1.3 | 0.4002 | 2.0E-04 | -1.3 | 0.4043 | 2.5E-14 |
| *egl-8* | WBGene00001177 | protein_coding_gene | -1.3 | 0.4001 | 5.7E-07 | -1.3 | 0.4065 | 6.7E-25 |
| *D1054.10* | WBGene00008377 | protein_coding_gene | -1.3 | 0.4000 | 2.3E-05 | -1.4 | 0.3801 | 4.0E-30 |
| *F25F8.1* | WBGene00017795 | protein_coding_gene | -1.3 | 0.3998 | 7.1E-06 | -1.4 | 0.3684 | 2.7E-08 |
| *Y48A6B.8* | WBGene00012969 | protein_coding_gene | -1.3 | 0.3997 | 1.9E-02 | -2.3 | 0.2092 | 1.5E-07 |
| *tag-241* | WBGene00044079 | protein_coding_gene | -1.3 | 0.3997 | 5.5E-04 | -1.4 | 0.3691 | 5.7E-13 |
| *C27H5.4* | WBGene00016174 | protein_coding_gene | -1.3 | 0.3975 | 7.6E-08 | -1.1 | 0.4753 | 9.5E-08 |
| *mlc-7* | WBGene00023451 | protein_coding_gene | -1.3 | 0.3973 | 4.0E-04 | -1.1 | 0.4802 | 8.9E-10 |
| *T28C6.8* | WBGene00012122 | protein_coding_gene | -1.3 | 0.3968 | 7.6E-12 | -1.5 | 0.3655 | 2.7E-12 |
| *nhr-45* | WBGene00003635 | protein_coding_gene | -1.3 | 0.3965 | 1.9E-06 | -1.2 | 0.4304 | 2.2E-16 |
| *cyp-33E2* | WBGene00018334 | protein_coding_gene | -1.3 | 0.3962 | 4.6E-07 | -1.0 | 0.4849 | 1.7E-05 |
| *nhr-238* | WBGene00021611 | protein_coding_gene | -1.3 | 0.3962 | 7.4E-04 | -2.9 | 0.1315 | 2.7E-07 |
| ***bcmo-2*** | **WBGene00018755** | **protein_coding_gene** | **-1.3** | **0.3956** | **6.9E-11** | **1.3** | **2.3861** | **1.1E-06** |
| *ckr-2* | WBGene00021439 | protein_coding_gene | -1.3 | 0.3954 | 9.8E-03 | -1.3 | 0.4150 | 4.6E-04 |
| *F53B3.3* | WBGene00018742 | protein_coding_gene | -1.3 | 0.3944 | 7.4E-03 | -1.3 | 0.4202 | 2.9E-03 |
| *rrc-1* | WBGene00009800 | protein_coding_gene | -1.3 | 0.3943 | 7.5E-05 | -1.6 | 0.3247 | 2.3E-09 |
| *R12C12.9* | WBGene00020029 | protein_coding_gene | -1.3 | 0.3940 | 1.5E-04 | -1.8 | 0.2949 | 5.6E-14 |
| *Y97E10AR.2* | WBGene00022397 | protein_coding_gene | -1.3 | 0.3938 | 6.4E-03 | -1.1 | 0.4642 | 1.0E-02 |
| *sem-4* | WBGene00004773 | protein_coding_gene | -1.3 | 0.3936 | 2.1E-05 | -1.1 | 0.4621 | 8.8E-10 |
| *F42H10.3* | WBGene00018367 | protein_coding_gene | -1.3 | 0.3932 | 3.5E-08 | -1.4 | 0.3886 | 1.1E-09 |
| *dpy-18* | WBGene00001077 | protein_coding_gene | -1.3 | 0.3931 | 1.6E-07 | -1.1 | 0.4582 | 1.9E-08 |
| *Y39G10AR.11* | WBGene00021469 | protein_coding_gene | -1.3 | 0.3929 | 1.7E-05 | -2.7 | 0.1560 | 1.0E-76 |
| *F32A5.8* | WBGene00017971 | protein_coding_gene | -1.3 | 0.3928 | 5.0E-08 | -1.0 | 0.4936 | 1.0E-08 |
| *ZC455.1* | WBGene00013902 | protein_coding_gene | -1.4 | 0.3920 | 2.7E-04 | -1.4 | 0.3906 | 8.4E-11 |
| *dpy-31* | WBGene00006592 | protein_coding_gene | -1.4 | 0.3919 | 1.7E-09 | -2.1 | 0.2371 | 3.1E-38 |
| *sdpn-1* | WBGene00018467 | protein_coding_gene | -1.4 | 0.3918 | 4.1E-12 | -1.1 | 0.4693 | 2.0E-13 |
| ***F28B4.3*** | **WBGene00017892** | **protein_coding_gene** | **-1.4** | **0.3913** | **6.1E-05** | **1.1** | **2.1636** | **4.8E-02** |
| *M04B2.6* | WBGene00010848 | protein_coding_gene | -1.4 | 0.3913 | 2.3E-03 | -1.3 | 0.4073 | 5.5E-04 |
| *dgk-5* | WBGene00019428 | protein_coding_gene | -1.4 | 0.3911 | 3.4E-06 | -1.4 | 0.3774 | 2.0E-13 |
| ***ctsa-4.2*** | **WBGene00021503** | **protein_coding_gene** | **-1.4** | **0.3906** | **4.2E-05** | **1.4** | **2.6219** | **2.3E-05** |
| *Y71G12B.26* | WBGene00022161 | protein_coding_gene | -1.4 | 0.3903 | 2.0E-02 | -1.7 | 0.3094 | 1.8E-04 |
| *C10C5.5* | WBGene00007509 | protein_coding_gene | -1.4 | 0.3902 | 4.7E-04 | -1.4 | 0.3888 | 4.4E-04 |
| *F38E9.6* | WBGene00018188 | protein_coding_gene | -1.4 | 0.3900 | 7.2E-04 | -1.8 | 0.2828 | 1.8E-07 |
| *nlp-82* | WBGene00044006 | protein_coding_gene | -1.4 | 0.3882 | 9.8E-03 | -1.7 | 0.2983 | 8.6E-04 |
| *ensh-1* | WBGene00017013 | protein_coding_gene | -1.4 | 0.3880 | 5.2E-03 | -2.0 | 0.2429 | 1.8E-06 |
| *C48E7.6* | WBGene00016751 | protein_coding_gene | -1.4 | 0.3879 | 2.0E-04 | -1.3 | 0.4116 | 7.4E-10 |
| *itx-1* | WBGene00002174 | protein_coding_gene | -1.4 | 0.3875 | 4.5E-04 | -1.6 | 0.3301 | 2.0E-33 |
| *ceh-79* | WBGene00007749 | protein_coding_gene | -1.4 | 0.3873 | 2.5E-03 | -2.1 | 0.2328 | 2.3E-17 |
| *che-3* | WBGene00000485 | protein_coding_gene | -1.4 | 0.3868 | 2.1E-03 | -1.5 | 0.3494 | 1.6E-08 |
| *tpa-1* | WBGene00006599 | protein_coding_gene | -1.4 | 0.3864 | 8.5E-08 | -1.5 | 0.3606 | 6.9E-25 |
| *nuc-1* | WBGene00003828 | protein_coding_gene | -1.4 | 0.3863 | 5.2E-09 | -1.3 | 0.4073 | 2.5E-26 |
| *F49H12.3* | WBGene00018654 | protein_coding_gene | -1.4 | 0.3861 | 2.6E-02 | -1.5 | 0.3637 | 8.0E-03 |
| *drl-1* | WBGene00017578 | protein_coding_gene | -1.4 | 0.3860 | 1.6E-08 | -1.3 | 0.4001 | 9.8E-12 |
| *T05A7.7* | WBGene00020224 | protein_coding_gene | -1.4 | 0.3853 | 7.9E-03 | -1.6 | 0.3391 | 2.6E-04 |
| *hum-4* | WBGene00002037 | protein_coding_gene | -1.4 | 0.3831 | 1.7E-06 | -1.1 | 0.4812 | 1.1E-09 |
| *T19A5.3* | WBGene00020554 | protein_coding_gene | -1.4 | 0.3827 | 2.6E-04 | -2.0 | 0.2506 | 2.8E-03 |
| *sgk-1* | WBGene00004789 | protein_coding_gene | -1.4 | 0.3826 | 2.6E-09 | -1.7 | 0.3054 | 2.7E-36 |
| *vpat-1* | WBGene00018860 | protein_coding_gene | -1.4 | 0.3816 | 1.5E-11 | -1.3 | 0.3936 | 4.1E-15 |
| *T02C5.1* | WBGene00020158 | protein_coding_gene | -1.4 | 0.3805 | 1.8E-05 | -1.9 | 0.2657 | 2.6E-53 |
| *C39B10.1* | WBGene00008021 | protein_coding_gene | -1.4 | 0.3805 | 2.0E-03 | -1.1 | 0.4557 | 4.8E-03 |
| *ndx-1* | WBGene00003578 | protein_coding_gene | -1.4 | 0.3804 | 2.0E-05 | -1.2 | 0.4304 | 7.3E-07 |
| *K10D3.6* | WBGene00010740 | protein_coding_gene | -1.4 | 0.3802 | 7.0E-06 | -2.4 | 0.1888 | 5.8E-19 |

| | | | | | | | | |
|---|---|---|---|---|---|---|---|---|
| *Y106G6D.8* | WBGene00013704 | protein_coding_gene | -1.4 | 0.3802 | 1.1E-05 | -1.3 | 0.4032 | 6.8E-10 |
| *F58A3.3* | WBGene00010223 | protein_coding_gene | -1.4 | 0.3798 | 5.9E-04 | -2.3 | 0.2044 | 8.4E-09 |
| *W06H8.6* | WBGene00021068 | protein_coding_gene | -1.4 | 0.3794 | 4.6E-03 | -1.7 | 0.2990 | 2.5E-35 |
| *H04M03.12* | WBGene00019156 | protein_coding_gene | -1.4 | 0.3790 | 3.3E-03 | -1.1 | 0.4805 | 7.7E-04 |
| *ZK550.6* | WBGene00014000 | protein_coding_gene | -1.4 | 0.3782 | 1.3E-03 | -2.7 | 0.1537 | 2.3E-43 |
| ***arf-1.1*** | **WBGene00000190** | **protein_coding_gene** | **-1.4** | **0.3779** | **2.0E-03** | **1.7** | **3.1680** | **5.8E-05** |
| *W03F8.6* | WBGene00020995 | protein_coding_gene | -1.4 | 0.3777 | 2.5E-03 | -2.2 | 0.2212 | 3.2E-14 |
| *VF13D12L.3* | WBGene00012149 | protein_coding_gene | -1.4 | 0.3776 | 3.6E-18 | -1.0 | 0.4933 | 5.7E-09 |
| *C32D5.6* | WBGene00016314 | protein_coding_gene | -1.4 | 0.3772 | 2.5E-06 | -1.3 | 0.4155 | 2.5E-07 |
| *D1086.1* | WBGene00008388 | protein_coding_gene | -1.4 | 0.3771 | 6.8E-05 | -1.5 | 0.3487 | 1.1E-08 |
| *capa-1* | WBGene00021268 | protein_coding_gene | -1.4 | 0.3765 | 5.1E-04 | -1.6 | 0.3372 | 1.2E-09 |
| *D2096.13* | WBGene00044953 | protein_coding_gene | -1.4 | 0.3764 | 1.3E-02 | -2.1 | 0.2357 | 1.6E-05 |
| *mab-7* | WBGene00003104 | protein_coding_gene | -1.4 | 0.3761 | 3.7E-03 | -1.9 | 0.2643 | 1.1E-05 |
| *Y116F11A.1* | WBGene00013816 | protein_coding_gene | -1.4 | 0.3757 | 3.7E-02 | -1.1 | 0.4556 | 3.5E-02 |
| *F32B5.6* | WBGene00017979 | protein_coding_gene | -1.4 | 0.3756 | 4.9E-11 | -1.4 | 0.3816 | 3.1E-29 |
| *ZK185.5* | WBGene00044481 | protein_coding_gene | -1.4 | 0.3756 | 1.2E-10 | -1.5 | 0.3598 | 3.1E-11 |
| *E04F6.9* | WBGene00017128 | protein_coding_gene | -1.4 | 0.3746 | 1.0E-10 | -2.0 | 0.2524 | 3.6E-42 |
| *zig-1* | WBGene00006978 | protein_coding_gene | -1.4 | 0.3744 | 8.5E-08 | -1.3 | 0.3941 | 3.2E-11 |
| *W05F2.4* | WBGene00021036 | protein_coding_gene | -1.4 | 0.3743 | 1.0E-04 | -1.3 | 0.3933 | 5.2E-19 |
| *unc-36* | WBGene00006772 | protein_coding_gene | -1.4 | 0.3739 | 7.9E-10 | -1.0 | 0.4831 | 4.8E-07 |
| *lin-46* | WBGene00003031 | protein_coding_gene | -1.4 | 0.3736 | 1.0E-03 | -1.3 | 0.4060 | 8.7E-03 |
| *sbp-1* | WBGene00004735 | protein_coding_gene | -1.4 | 0.3734 | 5.5E-06 | -1.2 | 0.4371 | 4.4E-17 |
| *mig-17* | WBGene00003248 | protein_coding_gene | -1.4 | 0.3718 | 3.9E-06 | -1.2 | 0.4310 | 4.5E-06 |
| *nas-39* | WBGene00003555 | protein_coding_gene | -1.4 | 0.3718 | 3.5E-05 | -1.1 | 0.4785 | 2.6E-05 |
| *gpx-6* | WBGene00020373 | protein_coding_gene | -1.4 | 0.3717 | 3.1E-05 | -1.5 | 0.3509 | 2.4E-04 |
| *W07A12.4* | WBGene00012322 | protein_coding_gene | -1.4 | 0.3711 | 5.8E-03 | -1.8 | 0.2776 | 5.4E-07 |
| *F36F2.1* | WBGene00009475 | protein_coding_gene | -1.4 | 0.3709 | 1.2E-06 | -1.2 | 0.4446 | 4.1E-11 |
| *T11F9.1* | WBGene00011709 | protein_coding_gene | -1.4 | 0.3705 | 1.6E-02 | -1.3 | 0.4011 | 3.4E-02 |
| *ZK899.2* | WBGene00014141 | protein_coding_gene | -1.4 | 0.3705 | 1.7E-10 | -1.0 | 0.4989 | 2.7E-10 |
| *cyp-37A1* | WBGene00008499 | protein_coding_gene | -1.4 | 0.3703 | 6.6E-04 | -3.1 | 0.1163 | 4.3E-57 |
| *C18G1.1* | WBGene00015980 | protein_coding_gene | -1.4 | 0.3699 | 6.9E-03 | -1.9 | 0.2678 | 1.3E-04 |
| ***pqn-36*** | **WBGene00004123** | **protein_coding_gene** | **-1.4** | **0.3698** | **5.5E-03** | **1.5** | **2.8100** | **1.6E-05** |
| *C14A4.7* | WBGene00007558 | protein_coding_gene | -1.4 | 0.3698 | 1.6E-04 | -1.3 | 0.3988 | 3.9E-06 |
| *ephx-1* | WBGene00019487 | protein_coding_gene | -1.4 | 0.3697 | 6.4E-16 | -1.6 | 0.3275 | 1.6E-31 |
| *W02D7.3* | WBGene00020939 | protein_coding_gene | -1.4 | 0.3695 | 1.8E-02 | -1.1 | 0.4736 | 4.7E-02 |
| *K12C11.3* | WBGene00019674 | protein_coding_gene | -1.4 | 0.3694 | 8.8E-03 | -1.2 | 0.4405 | 4.3E-03 |
| *F55B11.2* | WBGene00010084 | protein_coding_gene | -1.4 | 0.3693 | 1.6E-07 | -1.4 | 0.3805 | 7.3E-18 |
| *ZK813.3* | WBGene00022822 | protein_coding_gene | -1.4 | 0.3689 | 8.2E-05 | -1.3 | 0.4016 | 8.9E-22 |
| ***D2096.6*** | **WBGene00017073** | **protein_coding_gene** | **-1.4** | **0.3688** | **4.7E-03** | **1.2** | **2.2823** | **2.3E-02** |
| *abts-4* | WBGene00019844 | protein_coding_gene | -1.4 | 0.3677 | 7.7E-10 | -1.7 | 0.3144 | 1.7E-33 |
| *ctl-2* | WBGene00000831 | protein_coding_gene | -1.4 | 0.3673 | 3.8E-08 | -2.2 | 0.2129 | 4.0E-72 |
| *twk-8* | WBGene00006663 | protein_coding_gene | -1.4 | 0.3672 | 1.4E-06 | -1.3 | 0.3960 | 2.2E-11 |
| *dhs-15* | WBGene00000978 | protein_coding_gene | -1.4 | 0.3669 | 2.3E-02 | -2.2 | 0.2114 | 2.4E-09 |
| *arrd-6* | WBGene00009579 | protein_coding_gene | -1.4 | 0.3668 | 8.9E-03 | -1.3 | 0.3973 | 2.5E-03 |
| *F47G3.4* | WBGene00194641 | protein_coding_gene | -1.4 | 0.3661 | 6.3E-07 | -3.3 | 0.1013 | 1.6E-14 |
| *slc-28.2* | WBGene00017868 | protein_coding_gene | -1.5 | 0.3658 | 6.4E-03 | -2.8 | 0.1388 | 3.8E-11 |
| *lgc-50* | WBGene00020605 | protein_coding_gene | -1.5 | 0.3652 | 5.1E-04 | -1.5 | 0.3630 | 1.0E-06 |
| *cdh-8* | WBGene00017576 | protein_coding_gene | -1.5 | 0.3651 | 7.2E-05 | -1.3 | 0.4140 | 3.9E-05 |
| *nca-2* | WBGene00003558 | protein_coding_gene | -1.5 | 0.3650 | 5.8E-05 | -1.3 | 0.4139 | 5.2E-04 |
| *F23F12.12* | WBGene00017756 | protein_coding_gene | -1.5 | 0.3649 | 1.2E-02 | -1.8 | 0.2954 | 7.4E-13 |
| *ubc-17* | WBGene00006712 | protein_coding_gene | -1.5 | 0.3648 | 1.1E-03 | -1.0 | 0.4856 | 5.7E-03 |
| *F52H2.4* | WBGene00018716 | protein_coding_gene | -1.5 | 0.3644 | 5.2E-05 | -1.5 | 0.3589 | 9.7E-06 |
| *acy-2* | WBGene00000069 | protein_coding_gene | -1.5 | 0.3641 | 5.1E-03 | -1.1 | 0.4574 | 1.5E-02 |
| *Y62H9A.3* | WBGene00013391 | protein_coding_gene | -1.5 | 0.3639 | 7.5E-06 | -1.1 | 0.4693 | 6.8E-11 |
| *T19B10.2* | WBGene00011831 | protein_coding_gene | -1.5 | 0.3639 | 1.7E-05 | -1.6 | 0.3310 | 5.5E-04 |
| *ugt-20* | WBGene00019516 | protein_coding_gene | -1.5 | 0.3639 | 2.2E-06 | -1.7 | 0.3135 | 3.0E-06 |
| *K08D9.9* | WBGene00235094 | protein_coding_gene | -1.5 | 0.3638 | 6.4E-03 | -2.2 | 0.2214 | 3.1E-13 |
| *acp-5* | WBGene00017427 | protein_coding_gene | -1.5 | 0.3637 | 2.3E-08 | -2.1 | 0.2283 | 5.1E-17 |
| *Y43F4A.1* | WBGene00012796 | protein_coding_gene | -1.5 | 0.3636 | 1.2E-08 | -1.5 | 0.3660 | 1.1E-07 |
| *F07C6.3* | WBGene00008554 | protein_coding_gene | -1.5 | 0.3636 | 1.2E-07 | -1.2 | 0.4358 | 2.3E-09 |
| *gar-1* | WBGene00001517 | protein_coding_gene | -1.5 | 0.3636 | 1.6E-03 | -1.2 | 0.4403 | 4.4E-04 |
| *T08G11.3* | WBGene00011630 | protein_coding_gene | -1.5 | 0.3633 | 3.1E-02 | -1.7 | 0.3146 | 1.8E-02 |
| *npr-18* | WBGene00008065 | protein_coding_gene | -1.5 | 0.3628 | 2.5E-02 | -1.3 | 0.4185 | 4.2E-02 |
| *gana-1* | WBGene00011095 | protein_coding_gene | -1.5 | 0.3627 | 2.8E-06 | -1.3 | 0.3980 | 1.9E-14 |
| *gpa-7* | WBGene00001669 | protein_coding_gene | -1.5 | 0.3624 | 4.6E-08 | -1.5 | 0.3547 | 1.8E-09 |
| *prx-5* | WBGene00004194 | protein_coding_gene | -1.5 | 0.3621 | 8.5E-15 | -1.3 | 0.3949 | 9.1E-17 |
| *pqn-94* | WBGene00004173 | protein_coding_gene | -1.5 | 0.3621 | 2.1E-05 | -1.2 | 0.4358 | 3.3E-09 |
| *rme-4* | WBGene00004375 | protein_coding_gene | -1.5 | 0.3618 | 8.9E-15 | -1.1 | 0.4752 | 5.4E-15 |
| *bgal-1* | WBGene00011832 | protein_coding_gene | -1.5 | 0.3611 | 4.6E-13 | -1.2 | 0.4337 | 1.4E-10 |
| *W04G3.5* | WBGene00012258 | protein_coding_gene | -1.5 | 0.3609 | 2.0E-11 | -1.1 | 0.4802 | 1.2E-12 |

| | | | | | | | | |
|---|---|---|---|---|---|---|---|---|
| *maoc-1* | WBGene00017123 | protein_coding_gene | -1.5 | 0.3606 | 2.7E-11 | -1.7 | 0.3112 | 2.0E-52 |
| *daam-1* | WBGene00018976 | protein_coding_gene | -1.5 | 0.3604 | 2.5E-04 | -2.2 | 0.2183 | 2.0E-11 |
| *nlp-21* | WBGene00003759 | protein_coding_gene | -1.5 | 0.3604 | 5.6E-05 | -1.1 | 0.4662 | 1.3E-09 |
| *pqn-48* | WBGene00004135 | protein_coding_gene | -1.5 | 0.3603 | 4.4E-07 | -1.1 | 0.4766 | 9.0E-07 |
| *rncs-1* | WBGene00023069 | ncRNA_gene | -1.5 | 0.3596 | 6.2E-04 | -3.9 | 0.0669 | 1.7E-32 |
| *mtl-2* | WBGene00003474 | protein_coding_gene | -1.5 | 0.3591 | 2.3E-06 | -4.0 | 0.0638 | 5.3E-125 |
| *flp-20* | WBGene00001463 | protein_coding_gene | -1.5 | 0.3590 | 2.7E-04 | -1.5 | 0.3641 | 3.3E-05 |
| *F25B3.5* | WBGene00009102 | protein_coding_gene | -1.5 | 0.3589 | 2.1E-03 | -2.0 | 0.2424 | 7.7E-11 |
| *ZK813.2* | WBGene00022821 | protein_coding_gene | -1.5 | 0.3586 | 9.1E-09 | -1.6 | 0.3293 | 1.0E-26 |
| ***dct-18*** | **WBGene00010266** | **protein_coding_gene** | **-1.5** | **0.3586** | **2.8E-04** | **1.7** | **3.2493** | **1.4E-46** |
| *lpr-2* | WBGene00011726 | protein_coding_gene | -1.5 | 0.3586 | 1.7E-03 | -2.1 | 0.2402 | 3.8E-13 |
| *cyp-33D1* | WBGene00010589 | protein_coding_gene | -1.5 | 0.3583 | 2.7E-02 | -1.8 | 0.2873 | 3.4E-02 |
| *C35A5.6* | WBGene00007950 | protein_coding_gene | -1.5 | 0.3577 | 2.3E-05 | -2.5 | 0.1785 | 2.7E-30 |
| *Y43F8A.1* | WBGene00012808 | protein_coding_gene | -1.5 | 0.3577 | 2.6E-02 | -2.2 | 0.2144 | 5.3E-05 |
| *shw-3* | WBGene00004793 | protein_coding_gene | -1.5 | 0.3568 | 4.7E-04 | -1.8 | 0.2781 | 6.8E-05 |
| *twk-46* | WBGene00006696 | protein_coding_gene | -1.5 | 0.3568 | 3.8E-05 | -1.2 | 0.4371 | 5.1E-06 |
| *lep-5* | WBGene00010424 | lincRNA_gene | -1.5 | 0.3566 | 9.6E-21 | -1.1 | 0.4703 | 4.3E-18 |
| *ist-1* | WBGene00002163 | protein_coding_gene | -1.5 | 0.3565 | 4.9E-05 | -2.0 | 0.2424 | 1.0E-22 |
| *W02D9.6* | WBGene00012212 | protein_coding_gene | -1.5 | 0.3559 | 4.6E-04 | -2.2 | 0.2104 | 1.9E-34 |
| *casy-1* | WBGene00000403 | protein_coding_gene | -1.5 | 0.3558 | 2.1E-10 | -1.0 | 0.4905 | 1.1E-11 |
| *sgn-1* | WBGene00004790 | protein_coding_gene | -1.5 | 0.3557 | 5.8E-05 | -2.0 | 0.2539 | 3.2E-11 |
| *T15B12.1* | WBGene00020530 | protein_coding_gene | -1.5 | 0.3556 | 2.6E-09 | -1.4 | 0.3769 | 2.3E-25 |
| *T01G1.2* | WBGene00011337 | protein_coding_gene | -1.5 | 0.3554 | 1.0E-06 | -1.4 | 0.3681 | 8.5E-08 |
| *flp-25* | WBGene00010575 | protein_coding_gene | -1.5 | 0.3552 | 9.9E-04 | -1.5 | 0.3605 | 1.1E-04 |
| *C35E7.6* | WBGene00016458 | protein_coding_gene | -1.5 | 0.3547 | 1.1E-09 | -2.1 | 0.2263 | 5.7E-53 |
| *klp-4* | WBGene00002217 | protein_coding_gene | -1.5 | 0.3546 | 1.2E-11 | -1.2 | 0.4466 | 1.4E-19 |
| *sms-5* | WBGene00021075 | protein_coding_gene | -1.5 | 0.3545 | 1.3E-04 | -1.1 | 0.4516 | 1.4E-03 |
| *ugt-49* | WBGene00007070 | protein_coding_gene | -1.5 | 0.3544 | 3.6E-15 | -1.7 | 0.3145 | 4.3E-27 |
| *math-38* | WBGene00020326 | protein_coding_gene | -1.5 | 0.3539 | 1.2E-02 | -2.0 | 0.2416 | 2.0E-25 |
| *T27A1.3* | WBGene00020835 | protein_coding_gene | -1.5 | 0.3537 | 4.5E-03 | -1.5 | 0.3510 | 6.8E-04 |
| *clec-150* | WBGene00019914 | protein_coding_gene | -1.5 | 0.3536 | 1.9E-09 | -1.2 | 0.4221 | 1.8E-24 |
| *nhr-130* | WBGene00003720 | protein_coding_gene | -1.5 | 0.3533 | 9.2E-09 | -1.1 | 0.4689 | 2.3E-05 |
| *B0252.8* | WBGene00044615 | protein_coding_gene | -1.5 | 0.3532 | 5.5E-04 | -1.8 | 0.2817 | 1.1E-05 |
| *Y116A8C.3* | WBGene00013784 | protein_coding_gene | -1.5 | 0.3531 | 2.1E-04 | -1.8 | 0.2926 | 9.8E-06 |
| *F14H12.6* | WBGene00017473 | protein_coding_gene | -1.5 | 0.3528 | 3.5E-02 | -1.3 | 0.4194 | 3.6E-02 |
| *ZC374.2* | WBGene00013872 | protein_coding_gene | -1.5 | 0.3524 | 4.8E-04 | -1.4 | 0.3805 | 2.5E-05 |
| *F09E10.1* | WBGene00017294 | protein_coding_gene | -1.5 | 0.3523 | 2.4E-05 | -2.4 | 0.1907 | 6.5E-23 |
| *K07H8.11* | WBGene00044510 | protein_coding_gene | -1.5 | 0.3522 | 3.9E-02 | -1.1 | 0.4710 | 2.7E-02 |
| *F23F1.4* | WBGene00017745 | protein_coding_gene | -1.5 | 0.3515 | 5.8E-19 | -1.1 | 0.4558 | 2.2E-10 |
| *nlp-51* | WBGene00011997 | protein_coding_gene | -1.5 | 0.3513 | 2.4E-06 | -1.0 | 0.4980 | 3.2E-08 |
| *F53F4.13* | WBGene00009995 | protein_coding_gene | -1.5 | 0.3509 | 1.4E-09 | -1.6 | 0.3317 | 2.1E-31 |
| *fshr-1* | WBGene00008239 | protein_coding_gene | -1.5 | 0.3498 | 9.1E-10 | -1.4 | 0.3715 | 3.8E-24 |
| *wdfy-3* | WBGene00007752 | protein_coding_gene | -1.5 | 0.3490 | 1.3E-06 | -1.4 | 0.3799 | 1.5E-26 |
| *R06A10.4* | WBGene00019911 | protein_coding_gene | -1.5 | 0.3487 | 1.3E-06 | -1.2 | 0.4228 | 1.3E-06 |
| *C01B10.11* | WBGene00044294 | protein_coding_gene | -1.5 | 0.3487 | 5.6E-07 | -1.9 | 0.2716 | 2.5E-18 |
| *nspc-1* | WBGene00023477 | protein_coding_gene | -1.5 | 0.3487 | 7.6E-03 | -1.5 | 0.3474 | 2.8E-03 |
| *F10D7.10* | WBGene00271701 | protein_coding_gene | -1.5 | 0.3487 | 1.9E-02 | -2.0 | 0.2490 | 2.1E-05 |
| *gcy-15* | WBGene00001541 | protein_coding_gene | -1.5 | 0.3486 | 1.6E-02 | -1.9 | 0.2721 | 1.4E-02 |
| *plr-1* | WBGene00023404 | protein_coding_gene | -1.5 | 0.3485 | 7.6E-06 | -1.3 | 0.4189 | 2.2E-07 |
| *T05A10.6* | WBGene00011463 | protein_coding_gene | -1.5 | 0.3484 | 5.4E-03 | -2.0 | 0.2574 | 1.5E-04 |
| *flr-1* | WBGene00001465 | protein_coding_gene | -1.5 | 0.3482 | 1.4E-07 | -1.3 | 0.4151 | 4.4E-07 |
| *nlp-79* | WBGene00008626 | protein_coding_gene | -1.5 | 0.3481 | 4.0E-04 | -1.3 | 0.4107 | 5.1E-03 |
| *npr-24* | WBGene00020086 | protein_coding_gene | -1.5 | 0.3480 | 5.0E-04 | -1.9 | 0.2658 | 3.8E-05 |
| *F47D12.3* | WBGene00018560 | protein_coding_gene | -1.5 | 0.3478 | 4.8E-03 | -1.3 | 0.3933 | 1.9E-03 |
| *frpr-16* | WBGene00020023 | protein_coding_gene | -1.5 | 0.3471 | 4.4E-03 | -1.6 | 0.3284 | 3.1E-03 |
| *clec-145* | WBGene00013013 | protein_coding_gene | -1.5 | 0.3470 | 4.1E-02 | -2.7 | 0.1578 | 1.3E-04 |
| *F13H10.6* | WBGene00008777 | protein_coding_gene | -1.5 | 0.3462 | 3.0E-07 | -1.3 | 0.4182 | 2.6E-11 |
| *zipt-2.1* | WBGene00014669 | protein_coding_gene | -1.5 | 0.3458 | 1.3E-05 | -2.5 | 0.1783 | 3.4E-54 |
| *flp-32* | WBGene00010982 | protein_coding_gene | -1.5 | 0.3454 | 3.0E-05 | -1.3 | 0.4139 | 8.2E-08 |
| *pat-4* | WBGene00003931 | protein_coding_gene | -1.5 | 0.3453 | 8.4E-09 | -1.4 | 0.3886 | 3.6E-14 |
| *best-4* | WBGene00015288 | protein_coding_gene | -1.5 | 0.3450 | 5.4E-03 | -1.8 | 0.2853 | 4.1E-03 |
| *F23B12.4* | WBGene00009081 | protein_coding_gene | -1.5 | 0.3441 | 1.1E-12 | -1.2 | 0.4419 | 6.7E-13 |
| *sago-1* | WBGene00019666 | protein_coding_gene | -1.5 | 0.3441 | 1.2E-13 | -1.6 | 0.3238 | 1.4E-45 |
| *T05E12.3* | WBGene00011486 | protein_coding_gene | -1.5 | 0.3439 | 3.6E-06 | -3.7 | 0.0760 | 2.6E-116 |
| *acs-12* | WBGene00009106 | protein_coding_gene | -1.5 | 0.3435 | 1.2E-12 | -2.0 | 0.2541 | 1.5E-25 |
| *hlh-12* | WBGene00001956 | protein_coding_gene | -1.5 | 0.3429 | 3.7E-02 | -2.1 | 0.2334 | 5.5E-03 |
| *clec-184* | WBGene00023502 | protein_coding_gene | -1.5 | 0.3428 | 2.8E-03 | -1.6 | 0.3233 | 1.8E-03 |
| *pxl-1* | WBGene00016197 | protein_coding_gene | -1.5 | 0.3423 | 1.5E-11 | -1.3 | 0.4102 | 1.9E-22 |
| *ncx-10* | WBGene00003575 | protein_coding_gene | -1.5 | 0.3419 | 3.2E-03 | -1.9 | 0.2655 | 9.6E-06 |
| *dhs-28* | WBGene00000991 | protein_coding_gene | -1.6 | 0.3415 | 3.9E-11 | -1.2 | 0.4438 | 1.3E-13 |

| | | | | | | | | |
|---|---|---|---|---|---|---|---|---|
| ***C26B2.8*** | **WBGene00016130** | **protein_coding_gene** | **-1.6** | **0.3414** | **7.8E-03** | **1.9** | **3.7206** | **1.5E-02** |
| *C34E7.4* | WBGene00007934 | protein_coding_gene | -1.6 | 0.3414 | 7.5E-06 | -1.2 | 0.4489 | 4.4E-12 |
| *stn-2* | WBGene00017866 | protein_coding_gene | -1.6 | 0.3413 | 3.4E-06 | -1.2 | 0.4292 | 8.1E-05 |
| *bar-1* | WBGene00000238 | protein_coding_gene | -1.6 | 0.3409 | 2.0E-10 | -1.4 | 0.3678 | 1.4E-15 |
| *nspc-3* | WBGene00044338 | protein_coding_gene | -1.6 | 0.3408 | 7.3E-03 | -3.2 | 0.1113 | 1.8E-10 |
| *twk-39* | WBGene00006690 | protein_coding_gene | -1.6 | 0.3408 | 9.5E-04 | -1.8 | 0.2934 | 8.6E-05 |
| *B0207.2* | WBGene00015027 | protein_coding_gene | -1.6 | 0.3404 | 8.8E-03 | -1.4 | 0.3862 | 1.7E-02 |
| *Y42A5A.1* | WBGene00012776 | protein_coding_gene | -1.6 | 0.3400 | 5.8E-05 | -1.7 | 0.3101 | 5.4E-09 |
| *pho-11* | WBGene00007331 | protein_coding_gene | -1.6 | 0.3399 | 9.5E-06 | -1.5 | 0.3543 | 2.1E-08 |
| *pdf-2* | WBGene00018590 | protein_coding_gene | -1.6 | 0.3397 | 4.5E-05 | -1.6 | 0.3189 | 3.9E-16 |
| *asd-2* | WBGene00006423 | protein_coding_gene | -1.6 | 0.3395 | 8.6E-06 | -1.1 | 0.4527 | 8.1E-11 |
| *Y9C12A.1* | WBGene00021185 | protein_coding_gene | -1.6 | 0.3389 | 1.5E-05 | -1.6 | 0.3267 | 4.9E-06 |
| *Y57E12AR.1* | WBGene00044612 | protein_coding_gene | -1.6 | 0.3389 | 1.3E-04 | -1.6 | 0.3415 | 8.4E-06 |
| *F07A11.4* | WBGene00008547 | protein_coding_gene | -1.6 | 0.3387 | 2.9E-08 | -1.2 | 0.4430 | 8.8E-13 |
| *F53H4.2* | WBGene00010007 | protein_coding_gene | -1.6 | 0.3383 | 4.2E-04 | -1.4 | 0.3896 | 8.9E-13 |
| *lim-7* | WBGene00002989 | protein_coding_gene | -1.6 | 0.3378 | 2.3E-09 | -1.3 | 0.4189 | 3.8E-07 |
| *C04E7.3* | WBGene00015430 | protein_coding_gene | -1.6 | 0.3373 | 1.2E-05 | -1.4 | 0.3821 | 3.3E-13 |
| *nlp-1* | WBGene00003739 | protein_coding_gene | -1.6 | 0.3370 | 2.7E-07 | -1.0 | 0.4844 | 2.5E-05 |
| *viro-2* | WBGene00015107 | protein_coding_gene | -1.6 | 0.3369 | 5.3E-06 | -1.5 | 0.3465 | 1.0E-07 |
| *Y43F8B.11* | WBGene00012821 | protein_coding_gene | -1.6 | 0.3363 | 1.1E-03 | -1.8 | 0.2821 | 2.9E-03 |
| *B0198.3* | WBGene00007116 | protein_coding_gene | -1.6 | 0.3363 | 1.7E-05 | -1.4 | 0.3860 | 1.6E-06 |
| *C05E7.1* | WBGene00007343 | protein_coding_gene | -1.6 | 0.3362 | 1.6E-04 | -1.8 | 0.2828 | 8.7E-07 |
| *lev-8* | WBGene00002975 | protein_coding_gene | -1.6 | 0.3352 | 1.6E-03 | -1.7 | 0.3143 | 7.9E-06 |
| *arrd-24* | WBGene00009852 | protein_coding_gene | -1.6 | 0.3347 | 1.3E-02 | -1.6 | 0.3273 | 5.8E-03 |
| *kcc-2* | WBGene00019205 | protein_coding_gene | -1.6 | 0.3344 | 4.3E-10 | -1.3 | 0.4202 | 3.5E-25 |
| *phat-6* | WBGene00011683 | protein_coding_gene | -1.6 | 0.3344 | 6.1E-06 | -1.7 | 0.3177 | 9.3E-13 |
| *hot-6* | WBGene00001991 | protein_coding_gene | -1.6 | 0.3343 | 7.6E-06 | -1.7 | 0.3001 | 2.5E-09 |
| *F20A1.6* | WBGene00017618 | protein_coding_gene | -1.6 | 0.3341 | 5.5E-06 | -2.8 | 0.1392 | 4.5E-35 |
| *pes-7* | WBGene00003980 | protein_coding_gene | -1.6 | 0.3341 | 5.1E-09 | -1.1 | 0.4794 | 2.7E-11 |
| *Y8A9A.2* | WBGene00021171 | protein_coding_gene | -1.6 | 0.3336 | 6.9E-03 | -1.7 | 0.2986 | 3.0E-10 |
| ***F20G2.1*** | **WBGene00008985** | **protein_coding_gene** | **-1.6** | **0.3336** | **2.1E-03** | **1.4** | **2.7112** | **5.9E-04** |
| *dao-3* | WBGene00000929 | protein_coding_gene | -1.6 | 0.3335 | 4.4E-08 | -1.3 | 0.4028 | 5.3E-13 |
| *ule-1* | WBGene00021005 | protein_coding_gene | -1.6 | 0.3334 | 1.0E-06 | -2.3 | 0.2101 | 5.6E-35 |
| *dys-1* | WBGene00001131 | protein_coding_gene | -1.6 | 0.3332 | 1.1E-08 | -1.5 | 0.3508 | 6.9E-27 |
| *tsp-11* | WBGene00006637 | protein_coding_gene | -1.6 | 0.3331 | 5.4E-05 | -1.2 | 0.4418 | 4.4E-04 |
| *ugt-37* | WBGene00017332 | protein_coding_gene | -1.6 | 0.3329 | 1.3E-03 | -1.2 | 0.4367 | 2.7E-02 |
| *T28F3.5* | WBGene00012131 | protein_coding_gene | -1.6 | 0.3327 | 1.7E-04 | -1.2 | 0.4469 | 3.0E-03 |
| *F01D4.8* | WBGene00008490 | protein_coding_gene | -1.6 | 0.3326 | 5.1E-04 | -1.1 | 0.4686 | 1.5E-02 |
| *glb-33* | WBGene00022284 | protein_coding_gene | -1.6 | 0.3319 | 1.2E-09 | -1.1 | 0.4801 | 3.8E-05 |
| *F15E6.3* | WBGene00017484 | protein_coding_gene | -1.6 | 0.3314 | 5.6E-03 | -6.3 | 0.0128 | 3.1E-82 |
| *C17H12.11* | WBGene00015936 | protein_coding_gene | -1.6 | 0.3313 | 2.1E-05 | -2.4 | 0.1833 | 5.7E-18 |
| *lgc-25* | WBGene00010994 | protein_coding_gene | -1.6 | 0.3313 | 8.4E-04 | -2.1 | 0.2351 | 5.5E-07 |
| *Y34B4A.4* | WBGene00021320 | protein_coding_gene | -1.6 | 0.3306 | 6.0E-08 | -1.1 | 0.4818 | 3.4E-05 |
| *dpy-22* | WBGene00001081 | protein_coding_gene | -1.6 | 0.3306 | 2.2E-04 | -1.2 | 0.4426 | 5.1E-19 |
| *lim-6* | WBGene00002988 | protein_coding_gene | -1.6 | 0.3305 | 4.6E-06 | -1.1 | 0.4609 | 4.5E-05 |
| *unc-77* | WBGene00006809 | protein_coding_gene | -1.6 | 0.3305 | 5.2E-08 | -1.1 | 0.4762 | 3.3E-04 |
| *M176.11* | WBGene00010947 | protein_coding_gene | -1.6 | 0.3299 | 3.2E-10 | -1.2 | 0.4214 | 7.5E-10 |
| *K07H8.12* | WBGene00044509 | protein_coding_gene | -1.6 | 0.3299 | 6.4E-05 | -1.7 | 0.3027 | 1.0E-06 |
| *gba-1* | WBGene00016335 | protein_coding_gene | -1.6 | 0.3296 | 7.7E-05 | -2.0 | 0.2415 | 5.4E-20 |
| *elp-1* | WBGene00001248 | protein_coding_gene | -1.6 | 0.3296 | 3.4E-11 | -1.7 | 0.3167 | 3.0E-14 |
| *nas-32* | WBGene00003550 | protein_coding_gene | -1.6 | 0.3294 | 7.6E-05 | -1.8 | 0.2802 | 3.9E-06 |
| *Y66H1A.5* | WBGene00022047 | protein_coding_gene | -1.6 | 0.3293 | 6.5E-11 | -1.1 | 0.4779 | 7.2E-12 |
| *cyp-34A2* | WBGene00011699 | protein_coding_gene | -1.6 | 0.3292 | 1.7E-02 | -3.8 | 0.0732 | 2.0E-17 |
| *acd-5* | WBGene00020903 | protein_coding_gene | -1.6 | 0.3288 | 6.0E-08 | -1.1 | 0.4684 | 5.7E-06 |
| *tmbi-4* | WBGene00006479 | protein_coding_gene | -1.6 | 0.3286 | 2.0E-20 | -1.4 | 0.3910 | 5.0E-25 |
| *F41E6.5* | WBGene00018286 | protein_coding_gene | -1.6 | 0.3281 | 1.1E-04 | -3.1 | 0.1182 | 1.7E-41 |
| *unc-104* | WBGene00006831 | protein_coding_gene | -1.6 | 0.3280 | 1.1E-12 | -1.1 | 0.4635 | 3.5E-11 |
| *poml-4* | WBGene00044697 | protein_coding_gene | -1.6 | 0.3278 | 9.1E-03 | -2.5 | 0.1819 | 5.7E-07 |
| *H01G02.1* | WBGene00010348 | protein_coding_gene | -1.6 | 0.3277 | 5.3E-05 | -1.4 | 0.3826 | 8.3E-11 |
| *Y73F8A.5* | WBGene00013518 | protein_coding_gene | -1.6 | 0.3277 | 1.5E-04 | -1.1 | 0.4725 | 7.4E-04 |
| *K08C7.6* | WBGene00010648 | protein_coding_gene | -1.6 | 0.3276 | 1.3E-08 | -1.3 | 0.4113 | 6.5E-11 |
| *slc-36.3* | WBGene00012629 | protein_coding_gene | -1.6 | 0.3274 | 1.2E-02 | -3.0 | 0.1253 | 2.6E-14 |
| *C44B11.4* | WBGene00016634 | protein_coding_gene | -1.6 | 0.3273 | 5.2E-06 | -1.2 | 0.4420 | 2.5E-04 |
| *ZK1073.2* | WBGene00014214 | protein_coding_gene | -1.6 | 0.3272 | 2.0E-07 | -1.3 | 0.4162 | 2.1E-05 |
| *Y66D12A.13* | WBGene00013439 | protein_coding_gene | -1.6 | 0.3272 | 6.9E-06 | -2.4 | 0.1953 | 2.7E-21 |
| *nra-4* | WBGene00015344 | protein_coding_gene | -1.6 | 0.3271 | 2.7E-10 | -1.0 | 0.4930 | 5.0E-21 |
| *F19C6.5* | WBGene00008954 | protein_coding_gene | -1.6 | 0.3269 | 3.7E-06 | -1.4 | 0.3872 | 4.3E-04 |
| *Y59C2A.1* | WBGene00021984 | protein_coding_gene | -1.6 | 0.3268 | 2.5E-07 | -1.6 | 0.3386 | 9.4E-11 |
| *cest-33* | WBGene00017431 | protein_coding_gene | -1.6 | 0.3267 | 3.9E-11 | -1.9 | 0.2769 | 3.5E-16 |
| *K09H9.8* | WBGene00045387 | protein_coding_gene | -1.6 | 0.3267 | 5.9E-09 | -2.6 | 0.1596 | 1.3E-22 |

| | | | | | | | | |
|---|---|---|---|---|---|---|---|---|
| *T27A10.6* | WBGene00020846 | protein_coding_gene | -1.6 | 0.3264 | 1.1E-05 | -1.7 | 0.3026 | 8.1E-05 |
| *tep-1* | WBGene00013969 | protein_coding_gene | -1.6 | 0.3261 | 7.5E-07 | -2.0 | 0.2573 | 4.9E-49 |
| *slc-17.6* | WBGene00017398 | protein_coding_gene | -1.6 | 0.3261 | 5.2E-10 | -1.2 | 0.4251 | 2.5E-10 |
| *rsef-1* | WBGene00016344 | protein_coding_gene | -1.6 | 0.3258 | 1.4E-03 | -2.2 | 0.2156 | 1.4E-08 |
| *R09D1.9* | WBGene00011165 | pseudogene | -1.6 | 0.3256 | 5.0E-03 | -1.9 | 0.2599 | 9.9E-14 |
| *F14H12.8* | WBGene00017475 | protein_coding_gene | -1.6 | 0.3255 | 3.3E-03 | -1.8 | 0.2855 | 3.8E-04 |
| *C44H9.4* | WBGene00008099 | protein_coding_gene | -1.6 | 0.3248 | 8.6E-09 | -1.4 | 0.3689 | 5.1E-17 |
| *snf-11* | WBGene00004910 | protein_coding_gene | -1.6 | 0.3248 | 2.8E-05 | -1.6 | 0.3357 | 1.8E-09 |
| *rege-1* | WBGene00016260 | protein_coding_gene | -1.6 | 0.3243 | 1.8E-07 | -1.6 | 0.3276 | 8.8E-15 |
| *cyp-42A1* | WBGene00013585 | protein_coding_gene | -1.6 | 0.3242 | 1.1E-04 | -2.2 | 0.2109 | 4.1E-16 |
| *rbf-1* | WBGene00004316 | protein_coding_gene | -1.6 | 0.3241 | 4.1E-06 | -1.5 | 0.3651 | 4.1E-12 |
| *pod-2* | WBGene00004076 | protein_coding_gene | -1.6 | 0.3240 | 6.7E-11 | -1.1 | 0.4594 | 2.8E-23 |
| *Y45F3A.4* | WBGene00012861 | protein_coding_gene | -1.6 | 0.3240 | 7.9E-07 | -1.1 | 0.4784 | 5.4E-05 |
| *F07C3.2* | WBGene00017193 | protein_coding_gene | -1.6 | 0.3239 | 4.0E-06 | -1.7 | 0.3090 | 1.1E-06 |
| *cab-1* | WBGene00000277 | protein_coding_gene | -1.6 | 0.3234 | 7.3E-09 | -1.0 | 0.4867 | 1.3E-18 |
| *F07A11.5* | WBGene00008548 | protein_coding_gene | -1.6 | 0.3233 | 1.8E-12 | -1.2 | 0.4253 | 2.2E-10 |
| *nhr-41* | WBGene00022423 | protein_coding_gene | -1.6 | 0.3230 | 3.4E-05 | -1.6 | 0.3410 | 7.5E-11 |
| *faah-3* | WBGene00019068 | protein_coding_gene | -1.6 | 0.3230 | 1.2E-15 | -1.4 | 0.3880 | 5.4E-15 |
| *acc-3* | WBGene00018880 | protein_coding_gene | -1.6 | 0.3228 | 3.6E-02 | -1.5 | 0.3520 | 1.4E-03 |
| *ZK563.2* | WBGene00022767 | protein_coding_gene | -1.6 | 0.3226 | 1.0E-03 | -1.7 | 0.3055 | 3.3E-04 |
| *glt-7* | WBGene00001625 | protein_coding_gene | -1.6 | 0.3225 | 2.9E-03 | -1.4 | 0.3891 | 4.4E-02 |
| *far-8* | WBGene00001392 | protein_coding_gene | -1.6 | 0.3223 | 1.7E-13 | -1.4 | 0.3904 | 1.4E-21 |
| *lys-7* | WBGene00003096 | protein_coding_gene | -1.6 | 0.3222 | 5.2E-06 | -1.5 | 0.3510 | 2.0E-07 |
| *Y71H2B.4* | WBGene00022194 | protein_coding_gene | -1.6 | 0.3221 | 3.2E-08 | -1.3 | 0.4151 | 1.0E-10 |
| *clec-82* | WBGene00021873 | protein_coding_gene | -1.6 | 0.3221 | 1.0E-02 | -4.1 | 0.0601 | 4.2E-13 |
| *K11D12.13* | WBGene00044535 | protein_coding_gene | -1.6 | 0.3215 | 4.2E-08 | -1.2 | 0.4460 | 1.2E-12 |
| *ras-2* | WBGene00004311 | protein_coding_gene | -1.6 | 0.3210 | 5.2E-18 | -1.4 | 0.3921 | 5.6E-21 |
| *sup-9* | WBGene00006318 | protein_coding_gene | -1.6 | 0.3207 | 4.7E-03 | -1.3 | 0.3989 | 3.8E-03 |
| *C46C11.3* | WBGene00016706 | protein_coding_gene | -1.6 | 0.3206 | 8.9E-08 | -1.1 | 0.4716 | 1.8E-05 |
| *tag-209* | WBGene00011072 | protein_coding_gene | -1.6 | 0.3205 | 1.0E-05 | -1.6 | 0.3412 | 1.5E-17 |
| *Y43F8A.5* | WBGene00012811 | protein_coding_gene | -1.6 | 0.3205 | 1.2E-04 | -1.0 | 0.4906 | 3.9E-03 |
| *sfxn-5* | WBGene00044049 | protein_coding_gene | -1.6 | 0.3201 | 2.4E-17 | -1.3 | 0.4145 | 3.9E-12 |
| *F55D12.1* | WBGene00010110 | protein_coding_gene | -1.6 | 0.3201 | 3.4E-05 | -1.4 | 0.3889 | 1.4E-04 |
| *glt-3* | WBGene00001621 | protein_coding_gene | -1.6 | 0.3198 | 1.7E-07 | -2.1 | 0.2298 | 2.8E-13 |
| *Y39H10A.6* | WBGene00021482 | protein_coding_gene | -1.6 | 0.3197 | 3.9E-11 | -1.9 | 0.2685 | 2.0E-39 |
| *msi-1* | WBGene00003423 | protein_coding_gene | -1.6 | 0.3193 | 8.2E-07 | -2.0 | 0.2564 | 2.7E-38 |
| *pcp-5* | WBGene00022801 | protein_coding_gene | -1.6 | 0.3191 | 8.8E-15 | -1.1 | 0.4525 | 3.3E-15 |
| *Y53H1B.2* | WBGene00013181 | protein_coding_gene | -1.7 | 0.3185 | 2.5E-13 | -2.3 | 0.2009 | 4.0E-43 |
| *M04G7.1* | WBGene00019772 | protein_coding_gene | -1.7 | 0.3185 | 3.7E-07 | -1.3 | 0.4143 | 2.1E-10 |
| *dhs-14* | WBGene00000977 | protein_coding_gene | -1.7 | 0.3182 | 5.7E-09 | -1.4 | 0.3881 | 6.4E-15 |
| *F19G12.10* | WBGene00195186 | lincRNA_gene | -1.7 | 0.3181 | 2.9E-06 | -1.2 | 0.4342 | 1.2E-04 |
| *gly-18* | WBGene00001643 | protein_coding_gene | -1.7 | 0.3178 | 1.4E-02 | -1.4 | 0.3713 | 2.6E-02 |
| *spp-10* | WBGene00004995 | protein_coding_gene | -1.7 | 0.3177 | 6.3E-22 | -1.3 | 0.3997 | 7.1E-43 |
| *mgl-1* | WBGene00003232 | protein_coding_gene | -1.7 | 0.3177 | 1.8E-03 | -1.3 | 0.4101 | 1.3E-02 |
| *Y47D3A.32* | WBGene00012938 | protein_coding_gene | -1.7 | 0.3177 | 4.8E-09 | -1.5 | 0.3416 | 3.6E-08 |
| *nhr-85* | WBGene00003675 | protein_coding_gene | -1.7 | 0.3176 | 3.0E-05 | -1.1 | 0.4715 | 1.2E-02 |
| *prx-3* | WBGene00004193 | protein_coding_gene | -1.7 | 0.3175 | 3.0E-11 | -1.4 | 0.3661 | 9.5E-14 |
| *R08F11.4* | WBGene00019968 | protein_coding_gene | -1.7 | 0.3172 | 3.2E-04 | -1.1 | 0.4693 | 1.4E-02 |
| *exc-6* | WBGene00019030 | protein_coding_gene | -1.7 | 0.3171 | 2.3E-07 | -1.3 | 0.4200 | 3.4E-06 |
| *C24A3.4* | WBGene00016034 | protein_coding_gene | -1.7 | 0.3171 | 6.7E-12 | -1.2 | 0.4387 | 1.2E-05 |
| *C10A4.10* | WBGene00050957 | protein_coding_gene | -1.7 | 0.3158 | 1.4E-06 | -1.2 | 0.4296 | 5.1E-06 |
| *gmeb-2* | WBGene00010010 | protein_coding_gene | -1.7 | 0.3154 | 4.3E-03 | -1.1 | 0.4638 | 1.1E-02 |
| *ilys-2* | WBGene00016669 | protein_coding_gene | -1.7 | 0.3153 | 3.7E-02 | -5.6 | 0.0208 | 5.4E-10 |
| *R11G1.6* | WBGene00020012 | protein_coding_gene | -1.7 | 0.3152 | 4.1E-09 | -1.8 | 0.2919 | 1.3E-24 |
| *D1053.4* | WBGene00008369 | protein_coding_gene | -1.7 | 0.3144 | 3.2E-06 | -1.1 | 0.4699 | 3.4E-03 |
| *R09B5.11* | WBGene00019979 | protein_coding_gene | -1.7 | 0.3143 | 5.1E-06 | -2.4 | 0.1948 | 1.9E-14 |
| *dmsr-7* | WBGene00016428 | protein_coding_gene | -1.7 | 0.3143 | 5.8E-04 | -1.3 | 0.3990 | 2.2E-03 |
| *oac-8* | WBGene00008077 | protein_coding_gene | -1.7 | 0.3138 | 5.2E-03 | -2.3 | 0.2064 | 6.6E-05 |
| *nfki-1* | WBGene00007877 | protein_coding_gene | -1.7 | 0.3137 | 2.0E-08 | -1.3 | 0.3945 | 9.2E-20 |
| *Y47G6A.36* | WBGene00220165 | ncRNA_gene | -1.7 | 0.3137 | 1.6E-04 | -1.8 | 0.2784 | 1.7E-07 |
| *C33D9.3* | WBGene00007898 | protein_coding_gene | -1.7 | 0.3134 | 2.2E-07 | -1.4 | 0.3681 | 9.1E-14 |
| *mlk-1* | WBGene00003374 | protein_coding_gene | -1.7 | 0.3130 | 1.3E-07 | -1.3 | 0.4063 | 1.3E-14 |
| *C06H5.8* | WBGene00044795 | protein_coding_gene | -1.7 | 0.3130 | 1.8E-07 | -1.8 | 0.2821 | 1.5E-06 |
| *C23H3.9* | WBGene00016022 | protein_coding_gene | -1.7 | 0.3129 | 2.8E-07 | -1.4 | 0.3867 | 3.0E-07 |
| *C27D6.12* | WBGene00044389 | protein_coding_gene | -1.7 | 0.3118 | 9.0E-09 | -1.6 | 0.3266 | 3.7E-09 |
| *F49E10.4* | WBGene00018641 | protein_coding_gene | -1.7 | 0.3116 | 2.7E-03 | -1.1 | 0.4720 | 2.7E-02 |
| *ttr-25* | WBGene00013078 | protein_coding_gene | -1.7 | 0.3099 | 1.4E-10 | -1.7 | 0.2980 | 2.8E-27 |
| *C14A4.6* | WBGene00007557 | protein_coding_gene | -1.7 | 0.3097 | 1.6E-05 | -1.7 | 0.3041 | 9.2E-14 |
| *cpd-2* | WBGene00012073 | protein_coding_gene | -1.7 | 0.3097 | 4.1E-09 | -1.3 | 0.3963 | 1.7E-07 |
| *hrg-2* | WBGene00010473 | protein_coding_gene | -1.7 | 0.3096 | 4.4E-03 | -2.5 | 0.1781 | 3.8E-03 |

| | | | | | | | | |
|---|---|---|---|---|---|---|---|---|
| *tsp-15* | WBGene00006641 | protein_coding_gene | -1.7 | 0.3093 | 5.0E-07 | -1.4 | 0.3736 | 1.0E-12 |
| *pals-26* | WBGene00007131 | protein_coding_gene | -1.7 | 0.3093 | 1.8E-02 | -2.2 | 0.2244 | 5.4E-06 |
| *cyp-13A10* | WBGene00014254 | protein_coding_gene | -1.7 | 0.3091 | 1.9E-03 | -1.2 | 0.4422 | 1.2E-03 |
| *K08C7.4* | WBGene00010647 | protein_coding_gene | -1.7 | 0.3085 | 1.2E-02 | -1.0 | 0.4961 | 9.5E-03 |
| *F26C11.1* | WBGene00009146 | protein_coding_gene | -1.7 | 0.3079 | 2.4E-04 | -1.2 | 0.4495 | 3.4E-02 |
| *K09C6.9* | WBGene00019563 | protein_coding_gene | -1.7 | 0.3079 | 4.4E-06 | -1.3 | 0.3995 | 8.9E-09 |
| *pitr-3* | WBGene00015055 | protein_coding_gene | -1.7 | 0.3067 | 2.5E-05 | -2.1 | 0.2355 | 2.5E-17 |
| *C45G7.13* | WBGene00235340 | protein_coding_gene | -1.7 | 0.3066 | 2.6E-05 | -1.1 | 0.4706 | 9.8E-04 |
| *ggr-3* | WBGene00001588 | protein_coding_gene | -1.7 | 0.3066 | 4.3E-04 | -1.3 | 0.4106 | 1.1E-02 |
| *F55A11.6* | WBGene00010079 | protein_coding_gene | -1.7 | 0.3061 | 1.5E-10 | -1.3 | 0.3973 | 4.8E-14 |
| *F40F8.4* | WBGene00009577 | protein_coding_gene | -1.7 | 0.3059 | 1.5E-11 | -1.2 | 0.4485 | 2.4E-11 |
| *F08D12.3* | WBGene00017247 | protein_coding_gene | -1.7 | 0.3057 | 3.4E-05 | -1.2 | 0.4346 | 5.9E-08 |
| *spv-1* | WBGene00014051 | protein_coding_gene | -1.7 | 0.3057 | 1.4E-09 | -1.7 | 0.3082 | 1.3E-19 |
| *acds-10* | WBGene00019599 | protein_coding_gene | -1.7 | 0.3056 | 1.4E-12 | -1.2 | 0.4286 | 1.6E-17 |
| *Y4C6B.2* | WBGene00021156 | protein_coding_gene | -1.7 | 0.3055 | 4.5E-04 | -2.9 | 0.1365 | 1.5E-16 |
| *tmc-2* | WBGene00015177 | protein_coding_gene | -1.7 | 0.3055 | 4.1E-07 | -1.1 | 0.4592 | 3.4E-05 |
| *T21C9.6* | WBGene00011893 | protein_coding_gene | -1.7 | 0.3051 | 6.0E-11 | -1.6 | 0.3413 | 2.4E-16 |
| *M04B2.7* | WBGene00010849 | protein_coding_gene | -1.7 | 0.3050 | 8.3E-03 | -1.5 | 0.3578 | 2.3E-02 |
| *gei-1* | WBGene00001559 | protein_coding_gene | -1.7 | 0.3043 | 2.0E-09 | -1.8 | 0.2954 | 9.6E-29 |
| *F27D9.2* | WBGene00017861 | protein_coding_gene | -1.7 | 0.3043 | 9.9E-12 | -1.3 | 0.4203 | 4.4E-08 |
| *C54F6.20* | WBGene00271796 | protein_coding_gene | -1.7 | 0.3042 | 1.1E-02 | -1.9 | 0.2717 | 8.2E-03 |
| *vab-10* | WBGene00006876 | protein_coding_gene | -1.7 | 0.3039 | 2.9E-09 | -1.1 | 0.4825 | 8.9E-17 |
| *kqt-3* | WBGene00002235 | protein_coding_gene | -1.7 | 0.3036 | 5.9E-09 | -1.3 | 0.4095 | 4.5E-06 |
| *lec-2* | WBGene00002265 | protein_coding_gene | -1.7 | 0.3035 | 4.1E-05 | -2.5 | 0.1720 | 2.3E-18 |
| *R10E4.6* | WBGene00011202 | protein_coding_gene | -1.7 | 0.3034 | 7.8E-06 | -3.3 | 0.1023 | 2.5E-15 |
| *F42G8.19* | WBGene00249817 | protein_coding_gene | -1.7 | 0.3028 | 5.3E-04 | -2.3 | 0.2041 | 2.1E-05 |
| *srab-4* | WBGene00016363 | protein_coding_gene | -1.7 | 0.3024 | 4.8E-03 | -1.9 | 0.2742 | 7.7E-04 |
| *rimb-1* | WBGene00006513 | protein_coding_gene | -1.7 | 0.3019 | 7.0E-10 | -1.0 | 0.4856 | 1.9E-04 |
| *F28F9.3* | WBGene00017907 | protein_coding_gene | -1.7 | 0.3016 | 7.9E-03 | -1.5 | 0.3522 | 1.4E-03 |
| *clip-1* | WBGene00010796 | protein_coding_gene | -1.7 | 0.3015 | 1.8E-10 | -1.4 | 0.3905 | 7.7E-16 |
| *gsa-1* | WBGene00001745 | protein_coding_gene | -1.7 | 0.3014 | 3.2E-06 | -1.0 | 0.4860 | 3.2E-09 |
| *R02C2.7* | WBGene00019818 | protein_coding_gene | -1.7 | 0.3013 | 9.7E-12 | -1.0 | 0.4943 | 1.2E-10 |
| *Y48C3A.3* | WBGene00012987 | protein_coding_gene | -1.7 | 0.3011 | 8.4E-21 | -1.2 | 0.4274 | 2.3E-09 |
| *flp-10* | WBGene00001453 | protein_coding_gene | -1.7 | 0.3010 | 1.5E-04 | -2.3 | 0.2081 | 1.5E-34 |
| *W08A12.2* | WBGene00021079 | protein_coding_gene | -1.7 | 0.3007 | 1.7E-04 | -3.5 | 0.0894 | 5.9E-58 |
| *T27A3.8* | WBGene00044611 | protein_coding_gene | -1.7 | 0.3006 | 3.5E-02 | -2.1 | 0.2323 | 3.4E-04 |
| *ptr-22* | WBGene00004236 | protein_coding_gene | -1.7 | 0.3006 | 1.7E-02 | -5.2 | 0.0276 | 1.1E-97 |
| *madd-4* | WBGene00009958 | protein_coding_gene | -1.7 | 0.3004 | 5.6E-09 | -1.7 | 0.3107 | 6.9E-11 |
| *F20D6.8* | WBGene00017637 | protein_coding_gene | -1.7 | 0.3002 | 2.4E-02 | -2.3 | 0.1984 | 1.5E-06 |
| *best-8* | WBGene00007988 | protein_coding_gene | -1.7 | 0.2996 | 7.0E-05 | -1.9 | 0.2671 | 3.1E-06 |
| *C01C4.3* | WBGene00015293 | protein_coding_gene | -1.7 | 0.2993 | 4.4E-05 | -1.0 | 0.4955 | 1.5E-06 |
| *C27A7.3* | WBGene00007755 | protein_coding_gene | -1.7 | 0.2988 | 9.4E-07 | -1.4 | 0.3827 | 1.3E-04 |
| *slc-36.5* | WBGene00016628 | protein_coding_gene | -1.7 | 0.2987 | 3.6E-04 | -4.7 | 0.0396 | 3.6E-47 |
| *Y47A7.2* | WBGene00021614 | protein_coding_gene | -1.7 | 0.2987 | 4.0E-08 | -1.6 | 0.3401 | 1.5E-15 |
| *epi-1* | WBGene00001328 | protein_coding_gene | -1.7 | 0.2987 | 3.7E-09 | -1.0 | 0.4868 | 2.7E-13 |
| *clec-28* | WBGene00009857 | protein_coding_gene | -1.7 | 0.2986 | 3.0E-02 | -3.6 | 0.0804 | 9.3E-12 |
| *irk-2* | WBGene00002150 | protein_coding_gene | -1.7 | 0.2985 | 1.5E-04 | -1.0 | 0.4922 | 4.3E-02 |
| *C36C9.10* | WBGene00219849 | ncRNA_gene | -1.7 | 0.2982 | 5.5E-05 | -1.4 | 0.3814 | 7.4E-04 |
| *sup-1* | WBGene00012759 | protein_coding_gene | -1.7 | 0.2976 | 6.6E-12 | -1.5 | 0.3460 | 2.4E-16 |
| *ZK856.7* | WBGene00014108 | protein_coding_gene | -1.7 | 0.2974 | 2.6E-19 | -1.0 | 0.4846 | 8.9E-19 |
| *arrd-23* | WBGene00020323 | protein_coding_gene | -1.8 | 0.2971 | 7.0E-03 | -1.4 | 0.3689 | 4.7E-02 |
| *heh-1* | WBGene00006452 | protein_coding_gene | -1.8 | 0.2966 | 1.6E-24 | -1.2 | 0.4267 | 2.4E-17 |
| *F11E6.8* | WBGene00008711 | protein_coding_gene | -1.8 | 0.2962 | 1.3E-07 | -1.6 | 0.3357 | 4.7E-10 |
| *C01H6.8* | WBGene00007257 | protein_coding_gene | -1.8 | 0.2961 | 3.9E-08 | -1.4 | 0.3764 | 3.7E-06 |
| *lin-44* | WBGene00003029 | protein_coding_gene | -1.8 | 0.2952 | 3.6E-04 | -1.4 | 0.3846 | 2.3E-04 |
| *C49A9.2* | WBGene00016756 | protein_coding_gene | -1.8 | 0.2949 | 1.3E-15 | -1.1 | 0.4560 | 6.1E-11 |
| *nlp-41* | WBGene00007316 | protein_coding_gene | -1.8 | 0.2945 | 1.7E-04 | -1.5 | 0.3610 | 1.6E-06 |
| *myo-3* | WBGene00003515 | protein_coding_gene | -1.8 | 0.2939 | 1.1E-11 | -1.8 | 0.2819 | 4.9E-36 |
| *B0361.9* | WBGene00015163 | protein_coding_gene | -1.8 | 0.2930 | 1.6E-21 | -2.8 | 0.1395 | 2.1E-65 |
| *apl-1* | WBGene00000149 | protein_coding_gene | -1.8 | 0.2926 | 2.7E-08 | -1.7 | 0.3128 | 2.5E-47 |
| *D1005.2* | WBGene00016996 | protein_coding_gene | -1.8 | 0.2924 | 2.3E-05 | -1.6 | 0.3295 | 7.1E-08 |
| *cyp-25A2* | WBGene00007964 | protein_coding_gene | -1.8 | 0.2924 | 5.6E-08 | -1.1 | 0.4626 | 1.2E-06 |
| *cutl-26* | WBGene00021950 | protein_coding_gene | -1.8 | 0.2923 | 1.1E-16 | -1.9 | 0.2624 | 3.3E-11 |
| *F58A3.4* | WBGene00010224 | protein_coding_gene | -1.8 | 0.2921 | 6.1E-03 | -1.4 | 0.3908 | 1.7E-02 |
| *F20A1.1* | WBGene00017613 | protein_coding_gene | -1.8 | 0.2919 | 1.6E-11 | -1.5 | 0.3420 | 2.8E-29 |
| *R01H10.4* | WBGene00010972 | protein_coding_gene | -1.8 | 0.2914 | 7.6E-07 | -2.9 | 0.1363 | 2.6E-36 |
| *Y37D8A.19* | WBGene00012557 | protein_coding_gene | -1.8 | 0.2909 | 1.9E-05 | -1.9 | 0.2724 | 6.2E-31 |
| *blmp-1* | WBGene00003847 | protein_coding_gene | -1.8 | 0.2906 | 2.6E-06 | -1.7 | 0.2993 | 3.6E-05 |
| *egl-21* | WBGene00001189 | protein_coding_gene | -1.8 | 0.2905 | 8.3E-18 | -1.3 | 0.4073 | 7.7E-23 |
| *gnrr-4* | WBGene00016570 | protein_coding_gene | -1.8 | 0.2905 | 2.1E-05 | -1.6 | 0.3220 | 3.2E-05 |

| | | | | | | | | |
|---|---|---|---|---|---|---|---|---|
| *W05G11.8* | WBGene00271636 | ncRNA_gene | -1.8 | 0.2903 | 5.1E-03 | -1.2 | 0.4294 | 1.0E-03 |
| *col-177* | WBGene00000750 | protein_coding_gene | -1.8 | 0.2903 | 2.1E-03 | -1.7 | 0.2991 | 5.7E-03 |
| *C15B12.1* | WBGene00015783 | protein_coding_gene | -1.8 | 0.2902 | 1.2E-10 | -1.5 | 0.3479 | 2.4E-09 |
| *cpna-1* | WBGene00006495 | protein_coding_gene | -1.8 | 0.2898 | 1.6E-07 | -2.1 | 0.2414 | 5.7E-43 |
| *aat-1* | WBGene00000002 | protein_coding_gene | -1.8 | 0.2897 | 6.3E-12 | -1.6 | 0.3201 | 2.4E-13 |
| *upb-1* | WBGene00017440 | protein_coding_gene | -1.8 | 0.2895 | 3.1E-32 | -1.6 | 0.3247 | 3.4E-32 |
| *F46C3.2* | WBGene00009777 | protein_coding_gene | -1.8 | 0.2891 | 9.5E-08 | -1.1 | 0.4626 | 1.5E-03 |
| *T07E3.4* | WBGene00020315 | protein_coding_gene | -1.8 | 0.2890 | 1.9E-11 | -1.3 | 0.4064 | 1.7E-13 |
| *grl-14* | WBGene00001723 | protein_coding_gene | -1.8 | 0.2888 | 3.7E-03 | -1.6 | 0.3348 | 7.6E-03 |
| *F52E1.5* | WBGene00018694 | protein_coding_gene | -1.8 | 0.2888 | 8.1E-04 | -2.7 | 0.1532 | 1.3E-17 |
| *prx-12* | WBGene00004197 | protein_coding_gene | -1.8 | 0.2884 | 8.4E-12 | -1.3 | 0.3997 | 3.4E-08 |
| *tbc-16* | WBGene00018639 | protein_coding_gene | -1.8 | 0.2884 | 3.1E-04 | -2.1 | 0.2357 | 2.5E-05 |
| *snt-5* | WBGene00004925 | protein_coding_gene | -1.8 | 0.2880 | 5.5E-17 | -1.2 | 0.4214 | 7.3E-09 |
| ***T15B7.14*** | **WBGene00020526** | **protein_coding_gene** | **-1.8** | **0.2880** | **1.5E-02** | **1.3** | **2.4584** | **1.0E-02** |
| *nlp-9* | WBGene00003747 | protein_coding_gene | -1.8 | 0.2879 | 1.7E-06 | -1.3 | 0.3928 | 2.5E-11 |
| *svh-5* | WBGene00006462 | protein_coding_gene | -1.8 | 0.2875 | 5.3E-10 | -1.5 | 0.3519 | 1.8E-09 |
| *cft-1* | WBGene00000477 | protein_coding_gene | -1.8 | 0.2873 | 6.5E-08 | -2.1 | 0.2413 | 3.8E-08 |
| *ttr-15* | WBGene00011561 | protein_coding_gene | -1.8 | 0.2870 | 6.6E-23 | -2.2 | 0.2125 | 1.1E-80 |
| *D1086.12* | WBGene00044082 | protein_coding_gene | -1.8 | 0.2869 | 1.7E-20 | -1.0 | 0.4985 | 4.4E-07 |
| *Y18H1A.9* | WBGene00021213 | protein_coding_gene | -1.8 | 0.2866 | 4.3E-04 | -1.3 | 0.4188 | 6.9E-03 |
| *pamn-1* | WBGene00020556 | protein_coding_gene | -1.8 | 0.2866 | 1.9E-22 | -1.9 | 0.2760 | 1.6E-28 |
| *unc-43* | WBGene00006779 | protein_coding_gene | -1.8 | 0.2866 | 1.2E-11 | -1.4 | 0.3859 | 3.7E-23 |
| *C01F1.3* | WBGene00015298 | protein_coding_gene | -1.8 | 0.2864 | 5.9E-06 | -1.2 | 0.4308 | 5.3E-06 |
| *gpn-1* | WBGene00001687 | protein_coding_gene | -1.8 | 0.2863 | 1.3E-06 | -1.3 | 0.3954 | 5.8E-04 |
| *tag-260* | WBGene00010867 | protein_coding_gene | -1.8 | 0.2863 | 2.4E-09 | -1.5 | 0.3617 | 1.4E-15 |
| *C04C11.1* | WBGene00007295 | protein_coding_gene | -1.8 | 0.2861 | 5.0E-06 | -1.2 | 0.4403 | 1.1E-02 |
| *C49C3.10* | WBGene00008200 | protein_coding_gene | -1.8 | 0.2860 | 2.3E-15 | -1.6 | 0.3255 | 3.4E-13 |
| *K05F1.6* | WBGene00019408 | protein_coding_gene | -1.8 | 0.2859 | 1.4E-15 | -1.0 | 0.4924 | 1.6E-07 |
| *C24A3.2* | WBGene00016033 | protein_coding_gene | -1.8 | 0.2858 | 5.6E-14 | -1.2 | 0.4295 | 3.3E-13 |
| *B0252.1* | WBGene00015087 | protein_coding_gene | -1.8 | 0.2857 | 1.8E-08 | -1.1 | 0.4580 | 3.5E-04 |
| *Y39G8B.1* | WBGene00012722 | protein_coding_gene | -1.8 | 0.2857 | 3.6E-20 | -1.1 | 0.4689 | 6.7E-15 |
| *mec-1* | WBGene00003165 | protein_coding_gene | -1.8 | 0.2856 | 2.6E-06 | -1.9 | 0.2768 | 2.3E-09 |
| *Y48G1C.13* | WBGene00189949 | protein_coding_gene | -1.8 | 0.2855 | 3.8E-09 | -1.9 | 0.2718 | 2.9E-12 |
| *F09E5.12* | WBGene00017290 | protein_coding_gene | -1.8 | 0.2855 | 4.6E-03 | -1.3 | 0.4200 | 4.9E-02 |
| *C49H3.12* | WBGene00016795 | protein_coding_gene | -1.8 | 0.2854 | 1.3E-03 | -3.9 | 0.0650 | 1.2E-11 |
| *K01A11.1* | WBGene00010449 | protein_coding_gene | -1.8 | 0.2854 | 1.0E-11 | -1.6 | 0.3370 | 6.4E-08 |
| *plc-3* | WBGene00004038 | protein_coding_gene | -1.8 | 0.2853 | 2.9E-13 | -1.7 | 0.3096 | 1.5E-25 |
| *Y43B11AR.1* | WBGene00021543 | protein_coding_gene | -1.8 | 0.2851 | 2.2E-22 | -2.4 | 0.1882 | 4.9E-49 |
| *marc-5* | WBGene00013273 | protein_coding_gene | -1.8 | 0.2848 | 3.9E-06 | -1.6 | 0.3297 | 2.5E-10 |
| *F17A9.4* | WBGene00017536 | protein_coding_gene | -1.8 | 0.2845 | 5.7E-31 | -1.5 | 0.3629 | 1.8E-19 |
| *F19F10.3* | WBGene00017599 | protein_coding_gene | -1.8 | 0.2843 | 2.0E-05 | -1.5 | 0.3503 | 5.3E-06 |
| *T03D8.6* | WBGene00011393 | protein_coding_gene | -1.8 | 0.2843 | 4.3E-19 | -1.4 | 0.3809 | 1.0E-11 |
| *K04G2.9* | WBGene00010567 | protein_coding_gene | -1.8 | 0.2843 | 7.2E-09 | -1.1 | 0.4734 | 2.3E-06 |
| *C48E7.1* | WBGene00016749 | protein_coding_gene | -1.8 | 0.2839 | 1.5E-13 | -1.9 | 0.2710 | 2.0E-26 |
| *F29C6.1* | WBGene00009243 | protein_coding_gene | -1.8 | 0.2838 | 1.6E-12 | -1.2 | 0.4251 | 1.3E-09 |
| *arrd-25* | WBGene00007296 | protein_coding_gene | -1.8 | 0.2838 | 6.5E-20 | -1.4 | 0.3662 | 4.4E-23 |
| *F53F1.6* | WBGene00009984 | protein_coding_gene | -1.8 | 0.2837 | 7.8E-04 | -2.2 | 0.2203 | 1.4E-21 |
| *R09F10.5* | WBGene00019988 | protein_coding_gene | -1.8 | 0.2837 | 4.4E-10 | -1.7 | 0.3091 | 3.2E-07 |
| *C16D9.1* | WBGene00015857 | protein_coding_gene | -1.8 | 0.2836 | 1.2E-05 | -2.8 | 0.1456 | 5.4E-07 |
| *gba-3* | WBGene00008706 | protein_coding_gene | -1.8 | 0.2836 | 1.2E-14 | -1.3 | 0.4030 | 4.6E-13 |
| *mec-9* | WBGene00003173 | protein_coding_gene | -1.8 | 0.2835 | 6.5E-06 | -1.0 | 0.4898 | 2.1E-02 |
| *ZK6.8* | WBGene00022643 | protein_coding_gene | -1.8 | 0.2835 | 7.5E-04 | -4.6 | 0.0423 | 6.8E-29 |
| *Y71H2B.1* | WBGene00022191 | protein_coding_gene | -1.8 | 0.2833 | 1.5E-12 | -1.5 | 0.3447 | 1.4E-08 |
| *K09H9.5* | WBGene00019597 | protein_coding_gene | -1.8 | 0.2832 | 7.7E-09 | -1.3 | 0.4031 | 5.0E-08 |
| *T16G1.6* | WBGene00011800 | protein_coding_gene | -1.8 | 0.2829 | 3.7E-04 | -3.6 | 0.0845 | 2.5E-55 |
| *F43C9.1* | WBGene00018375 | protein_coding_gene | -1.8 | 0.2828 | 2.1E-03 | -1.2 | 0.4361 | 1.4E-03 |
| *gas-1* | WBGene00001520 | protein_coding_gene | -1.8 | 0.2823 | 3.5E-49 | -1.1 | 0.4533 | 5.3E-26 |
| *pals-38* | WBGene00008302 | protein_coding_gene | -1.8 | 0.2822 | 2.0E-02 | -3.1 | 0.1192 | 2.0E-05 |
| *F43C9.2* | WBGene00018376 | protein_coding_gene | -1.8 | 0.2821 | 6.6E-09 | -1.3 | 0.4147 | 1.2E-15 |
| *alh-8* | WBGene00000114 | protein_coding_gene | -1.8 | 0.2820 | 6.8E-29 | -1.4 | 0.3787 | 4.6E-28 |
| *C52B9.8* | WBGene00016868 | protein_coding_gene | -1.8 | 0.2819 | 1.9E-02 | -1.4 | 0.3676 | 1.6E-02 |
| *F36H12.17* | WBGene00018130 | protein_coding_gene | -1.8 | 0.2817 | 4.8E-03 | -2.0 | 0.2574 | 1.4E-03 |
| *F35A5.2* | WBGene00018025 | protein_coding_gene | -1.8 | 0.2814 | 5.9E-07 | -1.5 | 0.3643 | 2.7E-07 |
| *gob-1* | WBGene00001649 | protein_coding_gene | -1.8 | 0.2813 | 5.7E-34 | -1.1 | 0.4727 | 2.6E-18 |
| *glb-24* | WBGene00011287 | protein_coding_gene | -1.8 | 0.2813 | 8.3E-05 | -1.2 | 0.4240 | 1.2E-02 |
| *dnj-24* | WBGene00001042 | protein_coding_gene | -1.8 | 0.2811 | 1.9E-07 | -1.2 | 0.4282 | 6.8E-09 |
| *M03A1.3* | WBGene00019746 | protein_coding_gene | -1.8 | 0.2810 | 7.3E-07 | -2.0 | 0.2521 | 9.3E-06 |
| *zzz-1* | WBGene00009333 | protein_coding_gene | -1.8 | 0.2808 | 8.3E-10 | -1.2 | 0.4418 | 2.4E-05 |
| *T02B11.6* | WBGene00020157 | protein_coding_gene | -1.8 | 0.2808 | 5.2E-03 | -2.6 | 0.1672 | 1.0E-06 |
| *Y71G12B.25* | WBGene00022160 | protein_coding_gene | -1.8 | 0.2806 | 2.9E-05 | -1.4 | 0.3694 | 1.4E-05 |

| | | | | | | | | |
|---|---|---|---|---|---|---|---|---|
| *F41G3.3* | WBGene00018300 | protein_coding_gene | -1.8 | 0.2803 | 1.3E-02 | -1.4 | 0.3727 | 4.1E-02 |
| *F59F4.3* | WBGene00010338 | protein_coding_gene | -1.8 | 0.2799 | 6.7E-28 | -1.7 | 0.3101 | 2.5E-16 |
| *T28H11.8* | WBGene00020906 | protein_coding_gene | -1.8 | 0.2797 | 4.9E-07 | -2.4 | 0.1935 | 3.3E-08 |
| *R05H10.1* | WBGene00011042 | protein_coding_gene | -1.8 | 0.2795 | 9.8E-04 | -4.2 | 0.0546 | 2.5E-59 |
| *rab-3* | WBGene00004267 | protein_coding_gene | -1.8 | 0.2794 | 5.8E-14 | -2.0 | 0.2431 | 2.2E-28 |
| *F16F9.4* | WBGene00017515 | protein_coding_gene | -1.8 | 0.2794 | 8.6E-09 | -1.3 | 0.4055 | 7.8E-08 |
| *slo-1* | WBGene00004830 | protein_coding_gene | -1.8 | 0.2794 | 2.8E-07 | -1.3 | 0.4004 | 1.5E-08 |
| *C40H1.7* | WBGene00008042 | protein_coding_gene | -1.8 | 0.2792 | 2.2E-11 | -1.4 | 0.3830 | 6.3E-16 |
| *F02D8.4* | WBGene00008521 | protein_coding_gene | -1.8 | 0.2791 | 2.4E-14 | -1.1 | 0.4731 | 3.2E-15 |
| *cnc-7* | WBGene00010005 | protein_coding_gene | -1.8 | 0.2789 | 3.3E-03 | -6.0 | 0.0154 | 4.8E-14 |
| *T19B10.5* | WBGene00011833 | protein_coding_gene | -1.8 | 0.2786 | 1.9E-05 | -1.4 | 0.3741 | 4.8E-06 |
| *ptr-5* | WBGene00004220 | protein_coding_gene | -1.8 | 0.2781 | 2.2E-06 | -1.9 | 0.2622 | 2.7E-21 |
| *C05E11.6* | WBGene00015495 | protein_coding_gene | -1.8 | 0.2780 | 7.5E-04 | -1.1 | 0.4705 | 1.7E-02 |
| *unc-96* | WBGene00006825 | protein_coding_gene | -1.8 | 0.2779 | 1.6E-08 | -1.2 | 0.4315 | 9.8E-13 |
| *acs-14* | WBGene00008669 | protein_coding_gene | -1.8 | 0.2778 | 2.2E-16 | -1.9 | 0.2633 | 5.7E-39 |
| *mec-18* | WBGene00003179 | protein_coding_gene | -1.8 | 0.2776 | 1.1E-05 | -1.1 | 0.4531 | 1.6E-02 |
| *Y45F10A.7* | WBGene00012869 | protein_coding_gene | -1.8 | 0.2776 | 2.1E-08 | -1.4 | 0.3745 | 1.7E-10 |
| *M163.8* | WBGene00010940 | protein_coding_gene | -1.9 | 0.2772 | 6.3E-08 | -1.5 | 0.3419 | 2.0E-07 |
| *R186.1* | WBGene00011305 | protein_coding_gene | -1.9 | 0.2767 | 9.7E-10 | -1.6 | 0.3384 | 1.6E-05 |
| *egl-13* | WBGene00001182 | protein_coding_gene | -1.9 | 0.2765 | 1.7E-06 | -1.0 | 0.4949 | 4.0E-07 |
| *C14A4.12* | WBGene00007562 | protein_coding_gene | -1.9 | 0.2764 | 2.9E-16 | -1.4 | 0.3854 | 2.1E-10 |
| *irk-3* | WBGene00002151 | protein_coding_gene | -1.9 | 0.2761 | 4.6E-08 | -1.3 | 0.4107 | 3.9E-05 |
| *F48A9.2* | WBGene00018582 | protein_coding_gene | -1.9 | 0.2761 | 1.3E-09 | -1.5 | 0.3567 | 1.6E-12 |
| *mak-1* | WBGene00010681 | protein_coding_gene | -1.9 | 0.2759 | 4.9E-26 | -1.5 | 0.3576 | 1.7E-32 |
| *sad-1* | WBGene00004719 | protein_coding_gene | -1.9 | 0.2758 | 2.5E-08 | -1.5 | 0.3494 | 2.7E-08 |
| *mec-17* | WBGene00003178 | protein_coding_gene | -1.9 | 0.2756 | 3.1E-09 | -2.0 | 0.2528 | 6.1E-19 |
| *pqn-62* | WBGene00004145 | protein_coding_gene | -1.9 | 0.2755 | 8.4E-09 | -1.0 | 0.4945 | 2.6E-05 |
| *R04B3.3* | WBGene00019868 | protein_coding_gene | -1.9 | 0.2755 | 7.9E-04 | -1.5 | 0.3519 | 8.5E-05 |
| *npr-12* | WBGene00020689 | protein_coding_gene | -1.9 | 0.2750 | 8.8E-06 | -1.1 | 0.4683 | 7.3E-03 |
| *nlp-31* | WBGene00003769 | protein_coding_gene | -1.9 | 0.2750 | 2.7E-07 | -3.1 | 0.1148 | 7.4E-60 |
| *ttr-41* | WBGene00017374 | protein_coding_gene | -1.9 | 0.2750 | 1.0E-13 | -1.6 | 0.3248 | 1.6E-24 |
| *C29F5.1* | WBGene00016210 | protein_coding_gene | -1.9 | 0.2747 | 2.3E-13 | -1.4 | 0.3906 | 1.6E-18 |
| *spon-1* | WBGene00006893 | protein_coding_gene | -1.9 | 0.2746 | 2.3E-15 | -1.2 | 0.4428 | 1.6E-12 |
| *C06A1.2* | WBGene00007353 | protein_coding_gene | -1.9 | 0.2743 | 3.7E-07 | -1.2 | 0.4236 | 3.3E-05 |
| *ttr-24* | WBGene00013077 | protein_coding_gene | -1.9 | 0.2743 | 8.0E-14 | -1.6 | 0.3266 | 3.0E-33 |
| ***ZK662.6*** | **WBGene00045494** | **protein_coding_gene** | **-1.9** | **0.2741** | **1.6E-02** | **2.3** | **4.8354** | **4.7E-03** |
| *Y73F4A.2* | WBGene00013515 | protein_coding_gene | -1.9 | 0.2740 | 2.7E-09 | -1.5 | 0.3496 | 1.6E-16 |
| *nep-21* | WBGene00011794 | protein_coding_gene | -1.9 | 0.2740 | 4.8E-09 | -2.4 | 0.1871 | 5.8E-28 |
| *srr-4* | WBGene00005655 | protein_coding_gene | -1.9 | 0.2740 | 4.6E-09 | -2.2 | 0.2139 | 5.2E-41 |
| *sucl-1* | WBGene00007350 | protein_coding_gene | -1.9 | 0.2739 | 2.9E-50 | -1.1 | 0.4764 | 2.0E-18 |
| *ttr-45* | WBGene00044737 | protein_coding_gene | -1.9 | 0.2738 | 2.7E-13 | -2.3 | 0.2032 | 2.9E-38 |
| *famk-1* | WBGene00010356 | protein_coding_gene | -1.9 | 0.2737 | 6.0E-28 | -1.1 | 0.4680 | 8.4E-14 |
| *gei-18* | WBGene00001575 | protein_coding_gene | -1.9 | 0.2737 | 9.0E-04 | -1.6 | 0.3274 | 1.6E-02 |
| *C53B7.2* | WBGene00016893 | protein_coding_gene | -1.9 | 0.2736 | 1.2E-07 | -1.5 | 0.3491 | 5.5E-12 |
| *dmsr-5* | WBGene00012962 | protein_coding_gene | -1.9 | 0.2732 | 1.5E-03 | -1.6 | 0.3207 | 5.4E-03 |
| *nlp-38* | WBGene00007219 | protein_coding_gene | -1.9 | 0.2731 | 1.2E-07 | -2.0 | 0.2473 | 6.7E-12 |
| *F21C10.4* | WBGene00017654 | protein_coding_gene | -1.9 | 0.2725 | 3.2E-04 | -1.4 | 0.3823 | 6.2E-03 |
| *R04A9.6* | WBGene00019865 | protein_coding_gene | -1.9 | 0.2722 | 1.8E-06 | -1.9 | 0.2757 | 4.9E-04 |
| *nlp-30* | WBGene00003768 | protein_coding_gene | -1.9 | 0.2721 | 1.2E-03 | -3.6 | 0.0803 | 2.9E-24 |
| *C44C1.1* | WBGene00016641 | protein_coding_gene | -1.9 | 0.2719 | 1.1E-10 | -1.1 | 0.4540 | 1.5E-05 |
| *ZK370.6* | WBGene00022720 | protein_coding_gene | -1.9 | 0.2716 | 3.0E-04 | -1.4 | 0.3715 | 8.5E-03 |
| *nhr-119* | WBGene00003709 | protein_coding_gene | -1.9 | 0.2715 | 6.4E-09 | -2.6 | 0.1597 | 1.1E-40 |
| *C41G11.1* | WBGene00016568 | protein_coding_gene | -1.9 | 0.2713 | 1.7E-10 | -3.4 | 0.0977 | 3.3E-57 |
| *nas-31* | WBGene00003549 | protein_coding_gene | -1.9 | 0.2711 | 1.4E-08 | -1.7 | 0.2986 | 4.8E-07 |
| *che-7* | WBGene00000488 | protein_coding_gene | -1.9 | 0.2707 | 2.7E-04 | -1.0 | 0.4845 | 3.4E-02 |
| *mec-10* | WBGene00003174 | protein_coding_gene | -1.9 | 0.2704 | 3.9E-04 | -1.6 | 0.3397 | 2.4E-03 |
| *ZC266.1* | WBGene00022584 | protein_coding_gene | -1.9 | 0.2702 | 1.4E-10 | -1.1 | 0.4528 | 2.4E-02 |
| *clc-3* | WBGene00000524 | protein_coding_gene | -1.9 | 0.2702 | 9.7E-08 | -1.5 | 0.3658 | 3.2E-04 |
| *C05D9.9* | WBGene00015476 | protein_coding_gene | -1.9 | 0.2698 | 2.9E-04 | -4.3 | 0.0500 | 6.9E-81 |
| *B0403.6* | WBGene00043982 | protein_coding_gene | -1.9 | 0.2698 | 2.1E-05 | -2.4 | 0.1928 | 1.8E-12 |
| *igdb-2* | WBGene00007736 | protein_coding_gene | -1.9 | 0.2698 | 2.5E-14 | -1.4 | 0.3747 | 1.1E-18 |
| *asp-2* | WBGene00000215 | protein_coding_gene | -1.9 | 0.2697 | 2.2E-21 | -1.0 | 0.4921 | 2.3E-10 |
| ***elo-2*** | **WBGene00001240** | **protein_coding_gene** | **-1.9** | **0.2695** | **4.6E-05** | **1.8** | **3.4660** | **8.2E-12** |
| *C53A5.13* | WBGene00008270 | protein_coding_gene | -1.9 | 0.2693 | 1.2E-24 | -2.3 | 0.2080 | 3.8E-20 |
| *gap-1* | WBGene00001515 | protein_coding_gene | -1.9 | 0.2690 | 1.8E-09 | -2.2 | 0.2142 | 2.9E-13 |
| *F13H8.11* | WBGene00017443 | protein_coding_gene | -1.9 | 0.2685 | 1.4E-11 | -1.8 | 0.2780 | 1.6E-09 |
| *lin-18* | WBGene00003007 | protein_coding_gene | -1.9 | 0.2685 | 5.0E-15 | -1.0 | 0.4886 | 3.5E-06 |
| *F44E7.7* | WBGene00018429 | protein_coding_gene | -1.9 | 0.2680 | 4.9E-03 | -1.5 | 0.3433 | 4.7E-02 |
| *T14G8.4* | WBGene00011772 | protein_coding_gene | -1.9 | 0.2680 | 3.5E-03 | -1.4 | 0.3781 | 1.2E-02 |
| *ndx-8* | WBGene00003585 | protein_coding_gene | -1.9 | 0.2677 | 1.4E-09 | -1.3 | 0.3926 | 4.0E-05 |

| | | | | | | | | |
|---|---|---|---|---|---|---|---|---|
| *T19C3.5* | WBGene00020563 | protein_coding_gene | -1.9 | 0.2676 | 2.5E-14 | -1.5 | 0.3496 | 5.3E-10 |
| *F46H5.7* | WBGene00018522 | protein_coding_gene | -1.9 | 0.2674 | 3.5E-19 | -1.5 | 0.3524 | 8.5E-42 |
| *Y44A6D.2* | WBGene00012853 | protein_coding_gene | -1.9 | 0.2672 | 1.5E-07 | -1.1 | 0.4724 | 4.1E-05 |
| *pmt-1* | WBGene00022781 | protein_coding_gene | -1.9 | 0.2669 | 4.3E-16 | -2.1 | 0.2352 | 4.3E-75 |
| *F11C1.4* | WBGene00008693 | protein_coding_gene | -1.9 | 0.2667 | 4.6E-03 | -1.3 | 0.4029 | 2.9E-02 |
| *unc-2* | WBGene00006742 | protein_coding_gene | -1.9 | 0.2666 | 4.7E-07 | -1.1 | 0.4811 | 6.3E-08 |
| *nep-25* | WBGene00022874 | protein_coding_gene | -1.9 | 0.2665 | 1.3E-02 | -4.6 | 0.0416 | 3.3E-25 |
| *nrf-5* | WBGene00003812 | protein_coding_gene | -1.9 | 0.2664 | 1.8E-16 | -1.7 | 0.3149 | 1.0E-27 |
| *ttc-36* | WBGene00009947 | protein_coding_gene | -1.9 | 0.2664 | 1.5E-11 | -2.2 | 0.2181 | 7.7E-30 |
| *C39E9.7* | WBGene00008031 | protein_coding_gene | -1.9 | 0.2662 | 1.2E-06 | -3.2 | 0.1080 | 8.4E-16 |
| *Y40H7A.10* | WBGene00012747 | protein_coding_gene | -1.9 | 0.2660 | 2.1E-13 | -1.1 | 0.4540 | 3.2E-17 |
| *C35C5.8* | WBGene00007957 | protein_coding_gene | -1.9 | 0.2660 | 1.9E-05 | -4.0 | 0.0610 | 1.5E-30 |
| *F59B8.1* | WBGene00010316 | protein_coding_gene | -1.9 | 0.2659 | 4.0E-04 | -1.5 | 0.3502 | 2.4E-03 |
| *rgef-1* | WBGene00009100 | protein_coding_gene | -1.9 | 0.2656 | 1.3E-06 | -1.7 | 0.2981 | 1.4E-07 |
| *F44A2.5* | WBGene00018405 | protein_coding_gene | -1.9 | 0.2656 | 1.5E-09 | -1.7 | 0.3101 | 5.3E-18 |
| *cgr-1* | WBGene00020847 | protein_coding_gene | -1.9 | 0.2655 | 2.2E-17 | -1.2 | 0.4225 | 2.4E-04 |
| *F21D5.3* | WBGene00009008 | protein_coding_gene | -1.9 | 0.2655 | 1.6E-10 | -1.1 | 0.4719 | 8.8E-04 |
| *F26A1.9* | WBGene00017807 | protein_coding_gene | -1.9 | 0.2653 | 5.3E-04 | -2.8 | 0.1477 | 1.4E-07 |
| *ucr-2.1* | WBGene00012158 | protein_coding_gene | -1.9 | 0.2652 | 1.7E-31 | -1.2 | 0.4284 | 8.7E-28 |
| *acox-1.5* | WBGene00008167 | protein_coding_gene | -1.9 | 0.2652 | 3.1E-12 | -1.9 | 0.2662 | 2.9E-11 |
| *B0285.6* | WBGene00007138 | protein_coding_gene | -1.9 | 0.2652 | 4.9E-05 | -1.3 | 0.4004 | 5.9E-03 |
| *M176.4* | WBGene00010943 | protein_coding_gene | -1.9 | 0.2648 | 2.4E-46 | -2.5 | 0.1774 | 3.8E-84 |
| *F45E1.1* | WBGene00018462 | protein_coding_gene | -1.9 | 0.2645 | 4.3E-11 | -1.3 | 0.4001 | 1.0E-04 |
| *F52E1.9* | WBGene00018697 | protein_coding_gene | -1.9 | 0.2643 | 6.5E-09 | -1.2 | 0.4293 | 5.7E-07 |
| *lam-1* | WBGene00002247 | protein_coding_gene | -1.9 | 0.2642 | 3.5E-16 | -1.3 | 0.4085 | 5.1E-21 |
| *pph-1* | WBGene00004083 | protein_coding_gene | -1.9 | 0.2641 | 6.1E-11 | -1.6 | 0.3310 | 1.5E-16 |
| *ugt-58* | WBGene00009450 | protein_coding_gene | -1.9 | 0.2640 | 1.4E-29 | -1.7 | 0.3102 | 4.5E-23 |
| *maph-1.1* | WBGene00009306 | protein_coding_gene | -1.9 | 0.2638 | 6.7E-20 | -1.2 | 0.4281 | 1.9E-19 |
| *asm-1* | WBGene00000211 | protein_coding_gene | -1.9 | 0.2637 | 7.1E-12 | -1.9 | 0.2709 | 6.4E-23 |
| *ace-4* | WBGene00000038 | protein_coding_gene | -1.9 | 0.2637 | 1.0E-02 | -1.5 | 0.3572 | 1.8E-02 |
| *F57G12.1* | WBGene00010221 | protein_coding_gene | -1.9 | 0.2634 | 1.3E-07 | -1.5 | 0.3543 | 5.9E-24 |
| *F41B5.11* | WBGene00023039 | pseudogene | -1.9 | 0.2632 | 4.7E-03 | -2.7 | 0.1539 | 2.2E-08 |
| *F37C4.5* | WBGene00018145 | protein_coding_gene | -1.9 | 0.2631 | 6.1E-17 | -1.5 | 0.3509 | 1.5E-16 |
| *Y9C9A.16* | WBGene00021183 | protein_coding_gene | -1.9 | 0.2630 | 2.1E-02 | -1.3 | 0.3992 | 2.9E-02 |
| *msa-1* | WBGene00013882 | protein_coding_gene | -1.9 | 0.2630 | 2.3E-13 | -2.5 | 0.1806 | 5.5E-52 |
| *R08D7.7* | WBGene00011147 | protein_coding_gene | -1.9 | 0.2629 | 2.0E-11 | -1.6 | 0.3204 | 1.7E-09 |
| *E04F6.4* | WBGene00017124 | protein_coding_gene | -1.9 | 0.2626 | 3.6E-12 | -2.4 | 0.1952 | 1.6E-20 |
| *H03E18.2* | WBGene00019149 | protein_coding_gene | -1.9 | 0.2625 | 3.7E-02 | -1.7 | 0.3140 | 2.4E-02 |
| *slc-17.1* | WBGene00022647 | protein_coding_gene | -1.9 | 0.2625 | 9.5E-14 | -1.1 | 0.4513 | 6.1E-07 |
| *F46C3.6* | WBGene00045253 | protein_coding_gene | -1.9 | 0.2621 | 7.0E-07 | -2.4 | 0.1917 | 1.9E-10 |
| *Y32F6B.1* | WBGene00012531 | protein_coding_gene | -1.9 | 0.2619 | 4.9E-07 | -3.1 | 0.1158 | 5.7E-93 |
| *pck-2* | WBGene00011232 | protein_coding_gene | -1.9 | 0.2619 | 4.7E-22 | -2.0 | 0.2464 | 3.4E-77 |
| *C15A11.4* | WBGene00007598 | protein_coding_gene | -1.9 | 0.2614 | 3.4E-04 | -1.9 | 0.2727 | 1.6E-06 |
| *srh-2* | WBGene00005228 | protein_coding_gene | -1.9 | 0.2612 | 1.9E-02 | -3.9 | 0.0661 | 6.1E-08 |
| *klp-3* | WBGene00002216 | protein_coding_gene | -1.9 | 0.2611 | 8.4E-30 | -1.3 | 0.4017 | 1.3E-10 |
| *cest-2.1* | WBGene00013875 | protein_coding_gene | -1.9 | 0.2606 | 7.7E-15 | -1.4 | 0.3770 | 6.9E-09 |
| *clec-54* | WBGene00008596 | protein_coding_gene | -1.9 | 0.2599 | 1.3E-12 | -1.9 | 0.2674 | 1.3E-36 |
| *des-2* | WBGene00000955 | protein_coding_gene | -1.9 | 0.2594 | 5.6E-05 | -1.9 | 0.2770 | 1.5E-07 |
| *cyp-33E1* | WBGene00016768 | protein_coding_gene | -1.9 | 0.2594 | 2.5E-12 | -2.6 | 0.1667 | 1.3E-32 |
| *ZC196.4* | WBGene00022548 | protein_coding_gene | -1.9 | 0.2592 | 1.2E-05 | -1.9 | 0.2592 | 1.4E-06 |
| *F13D11.4* | WBGene00017429 | protein_coding_gene | -1.9 | 0.2592 | 2.0E-08 | -2.3 | 0.2073 | 2.5E-30 |
| *tln-1* | WBGene00006771 | protein_coding_gene | -1.9 | 0.2591 | 5.2E-12 | -1.7 | 0.3161 | 2.4E-43 |
| *flp-2* | WBGene00001445 | protein_coding_gene | -1.9 | 0.2591 | 2.4E-08 | -2.1 | 0.2317 | 9.2E-10 |
| *acs-17* | WBGene00016716 | protein_coding_gene | -2.0 | 0.2587 | 1.6E-05 | -2.5 | 0.1778 | 6.7E-75 |
| *C18H9.5* | WBGene00016003 | protein_coding_gene | -2.0 | 0.2582 | 6.4E-04 | -4.2 | 0.0550 | 5.7E-41 |
| *acy-1* | WBGene00000068 | protein_coding_gene | -2.0 | 0.2582 | 2.0E-09 | -1.4 | 0.3908 | 5.0E-10 |
| *hsp-12.2* | WBGene00002011 | protein_coding_gene | -2.0 | 0.2577 | 5.3E-12 | -1.8 | 0.2912 | 1.7E-32 |
| *B0198.2* | WBGene00007115 | protein_coding_gene | -2.0 | 0.2576 | 9.7E-06 | -1.5 | 0.3564 | 8.1E-04 |
| *tps-2* | WBGene00006603 | protein_coding_gene | -2.0 | 0.2574 | 3.7E-31 | -2.1 | 0.2358 | 7.9E-43 |
| *sulp-4* | WBGene00010788 | protein_coding_gene | -2.0 | 0.2569 | 1.3E-07 | -2.0 | 0.2463 | 6.9E-11 |
| *pho-14* | WBGene00020471 | protein_coding_gene | -2.0 | 0.2569 | 2.0E-08 | -1.6 | 0.3379 | 1.5E-26 |
| *hir-1* | WBGene00016059 | protein_coding_gene | -2.0 | 0.2569 | 9.0E-35 | -1.2 | 0.4226 | 1.2E-13 |
| *kin-2* | WBGene00002190 | protein_coding_gene | -2.0 | 0.2566 | 2.7E-09 | -1.1 | 0.4809 | 5.4E-17 |
| *F20A1.10* | WBGene00017621 | protein_coding_gene | -2.0 | 0.2566 | 2.4E-13 | -2.0 | 0.2458 | 1.3E-26 |
| *dop-4* | WBGene00016872 | protein_coding_gene | -2.0 | 0.2565 | 8.1E-06 | -3.1 | 0.1169 | 1.3E-11 |
| *T24C12.1* | WBGene00020764 | protein_coding_gene | -2.0 | 0.2561 | 2.0E-03 | -2.0 | 0.2567 | 7.9E-04 |
| *pmp-4* | WBGene00004061 | protein_coding_gene | -2.0 | 0.2561 | 1.5E-14 | -1.6 | 0.3393 | 3.2E-13 |
| *W10G11.2* | WBGene00021135 | protein_coding_gene | -2.0 | 0.2560 | 1.4E-03 | -4.9 | 0.0330 | 3.8E-11 |
| *W02F12.2* | WBGene00020947 | protein_coding_gene | -2.0 | 0.2558 | 1.1E-10 | -2.3 | 0.1975 | 3.1E-15 |
| *F36H2.3* | WBGene00009500 | protein_coding_gene | -2.0 | 0.2557 | 8.7E-07 | -1.6 | 0.3241 | 7.7E-08 |

| | | | | | | | | |
|---|---|---|---|---|---|---|---|---|
| *hif-1* | WBGene00001851 | protein_coding_gene | -2.0 | 0.2557 | 2.8E-16 | -1.5 | 0.3541 | 7.6E-44 |
| *F13B12.2* | WBGene00008730 | protein_coding_gene | -2.0 | 0.2556 | 2.0E-05 | -2.3 | 0.1978 | 2.6E-19 |
| *unc-7* | WBGene00006747 | protein_coding_gene | -2.0 | 0.2555 | 6.0E-06 | -1.0 | 0.4970 | 1.5E-04 |
| *Y67A10A.10* | WBGene00013459 | protein_coding_gene | -2.0 | 0.2554 | 9.0E-08 | -1.2 | 0.4364 | 1.2E-04 |
| *bli-3* | WBGene00000253 | protein_coding_gene | -2.0 | 0.2552 | 2.5E-26 | -1.5 | 0.3483 | 2.9E-20 |
| *unc-44* | WBGene00006780 | protein_coding_gene | -2.0 | 0.2551 | 7.3E-12 | -1.6 | 0.3392 | 1.1E-28 |
| *F25E2.3* | WBGene00017781 | protein_coding_gene | -2.0 | 0.2550 | 3.3E-06 | -3.1 | 0.1183 | 4.8E-27 |
| *pxn-2* | WBGene00004257 | protein_coding_gene | -2.0 | 0.2547 | 3.2E-07 | -1.0 | 0.4961 | 1.6E-05 |
| *Y15E3A.4* | WBGene00012443 | protein_coding_gene | -2.0 | 0.2544 | 1.4E-27 | -1.4 | 0.3897 | 6.3E-19 |
| *F53B1.8* | WBGene00018739 | protein_coding_gene | -2.0 | 0.2544 | 1.5E-23 | -1.8 | 0.2941 | 4.2E-20 |
| *prk-2* | WBGene00004183 | protein_coding_gene | -2.0 | 0.2543 | 1.8E-14 | -2.4 | 0.1880 | 1.0E-23 |
| *B0545.4* | WBGene00015247 | protein_coding_gene | -2.0 | 0.2541 | 2.1E-06 | -2.1 | 0.2403 | 1.1E-08 |
| *M03D4.3* | WBGene00019750 | protein_coding_gene | -2.0 | 0.2540 | 1.2E-03 | -2.0 | 0.2489 | 5.7E-06 |
| *clec-36* | WBGene00044332 | protein_coding_gene | -2.0 | 0.2540 | 3.4E-05 | -2.5 | 0.1742 | 2.5E-14 |
| *F41C3.11* | WBGene00018275 | protein_coding_gene | -2.0 | 0.2535 | 7.6E-13 | -1.8 | 0.2941 | 2.4E-08 |
| *clh-4* | WBGene00000531 | protein_coding_gene | -2.0 | 0.2530 | 9.1E-05 | -1.7 | 0.3103 | 1.2E-04 |
| *atgp-2* | WBGene00000225 | protein_coding_gene | -2.0 | 0.2530 | 1.1E-19 | -1.4 | 0.3841 | 3.1E-14 |
| *tps-1* | WBGene00006602 | protein_coding_gene | -2.0 | 0.2527 | 6.3E-13 | -2.1 | 0.2301 | 1.0E-34 |
| *flp-19* | WBGene00001462 | protein_coding_gene | -2.0 | 0.2526 | 1.1E-13 | -1.9 | 0.2616 | 2.6E-16 |
| *abt-4* | WBGene00000022 | protein_coding_gene | -2.0 | 0.2524 | 5.9E-10 | -1.2 | 0.4297 | 4.3E-03 |
| *B0416.3* | WBGene00015179 | protein_coding_gene | -2.0 | 0.2522 | 1.0E-06 | -1.2 | 0.4290 | 2.6E-04 |
| *zipt-16* | WBGene00006486 | protein_coding_gene | -2.0 | 0.2520 | 1.0E-05 | -1.1 | 0.4567 | 1.8E-03 |
| *ptr-11* | WBGene00004225 | protein_coding_gene | -2.0 | 0.2518 | 7.7E-13 | -1.5 | 0.3645 | 9.8E-09 |
| *F28D1.6* | WBGene00009216 | protein_coding_gene | -2.0 | 0.2518 | 2.7E-06 | -1.4 | 0.3814 | 2.4E-04 |
| *F32B5.7* | WBGene00017980 | protein_coding_gene | -2.0 | 0.2517 | 6.0E-11 | -1.0 | 0.4968 | 1.4E-07 |
| *C34D1.4* | WBGene00007931 | protein_coding_gene | -2.0 | 0.2517 | 7.6E-09 | -2.7 | 0.1516 | 8.8E-23 |
| *M79.2* | WBGene00010902 | protein_coding_gene | -2.0 | 0.2515 | 1.0E-19 | -1.9 | 0.2643 | 4.5E-32 |
| *pgp-3* | WBGene00003997 | protein_coding_gene | -2.0 | 0.2515 | 2.1E-25 | -2.0 | 0.2537 | 7.1E-25 |
| *F54E4.3* | WBGene00010059 | protein_coding_gene | -2.0 | 0.2512 | 4.8E-13 | -1.4 | 0.3832 | 3.1E-09 |
| *dhs-2* | WBGene00000966 | protein_coding_gene | -2.0 | 0.2511 | 8.1E-07 | -3.4 | 0.0942 | 2.3E-42 |
| *lbp-3* | WBGene00002255 | protein_coding_gene | -2.0 | 0.2505 | 1.1E-26 | -1.2 | 0.4259 | 1.2E-12 |
| *glb-9* | WBGene00016184 | protein_coding_gene | -2.0 | 0.2505 | 4.6E-04 | -1.4 | 0.3693 | 1.0E-02 |
| *aexr-1* | WBGene00016089 | protein_coding_gene | -2.0 | 0.2504 | 7.8E-03 | -1.8 | 0.2907 | 8.5E-03 |
| *Y45G5AM.6* | WBGene00021558 | protein_coding_gene | -2.0 | 0.2502 | 2.1E-16 | -1.3 | 0.4071 | 1.0E-07 |
| *K09C8.8* | WBGene00010718 | protein_coding_gene | -2.0 | 0.2500 | 4.5E-02 | -1.7 | 0.3110 | 1.5E-02 |
| *clec-49* | WBGene00012251 | protein_coding_gene | -2.0 | 0.2499 | 6.6E-13 | -2.4 | 0.1889 | 1.2E-24 |
| *C46C2.5* | WBGene00008113 | protein_coding_gene | -2.0 | 0.2497 | 2.2E-08 | -2.3 | 0.2078 | 2.7E-21 |
| *W01B11.6* | WBGene00020917 | protein_coding_gene | -2.0 | 0.2497 | 6.3E-10 | -3.1 | 0.1151 | 7.8E-54 |
| *zip-9* | WBGene00077761 | protein_coding_gene | -2.0 | 0.2492 | 1.9E-15 | -1.6 | 0.3244 | 2.3E-38 |
| *inx-13* | WBGene00002135 | protein_coding_gene | -2.0 | 0.2491 | 5.4E-20 | -1.4 | 0.3877 | 2.2E-15 |
| *F22G12.8* | WBGene00044712 | pseudogene | -2.0 | 0.2489 | 1.2E-05 | -1.1 | 0.4640 | 1.3E-02 |
| *flp-1* | WBGene00001444 | protein_coding_gene | -2.0 | 0.2488 | 2.4E-09 | -1.6 | 0.3411 | 2.5E-18 |
| *F47B10.5* | WBGene00009816 | protein_coding_gene | -2.0 | 0.2488 | 6.3E-11 | -1.7 | 0.3005 | 2.4E-08 |
| *saeg-1* | WBGene00010012 | protein_coding_gene | -2.0 | 0.2486 | 1.4E-12 | -1.6 | 0.3203 | 4.8E-33 |
| *fard-1* | WBGene00022200 | protein_coding_gene | -2.0 | 0.2481 | 3.0E-16 | -1.6 | 0.3327 | 1.3E-17 |
| *F14D12.1* | WBGene00017463 | protein_coding_gene | -2.0 | 0.2479 | 7.6E-09 | -1.6 | 0.3367 | 7.9E-09 |
| *mlt-2* | WBGene00007252 | protein_coding_gene | -2.0 | 0.2477 | 1.9E-09 | -1.6 | 0.3207 | 2.7E-06 |
| *VC5.2* | WBGene00020908 | protein_coding_gene | -2.0 | 0.2476 | 1.4E-14 | -2.2 | 0.2193 | 1.6E-36 |
| *nep-11* | WBGene00017557 | protein_coding_gene | -2.0 | 0.2475 | 6.5E-08 | -1.8 | 0.2821 | 7.4E-18 |
| *nhx-9* | WBGene00003736 | protein_coding_gene | -2.0 | 0.2474 | 1.4E-07 | -2.8 | 0.1482 | 4.5E-27 |
| *dapk-1* | WBGene00003400 | protein_coding_gene | -2.0 | 0.2471 | 2.9E-28 | -1.4 | 0.3873 | 4.4E-19 |
| *flp-7* | WBGene00001450 | protein_coding_gene | -2.0 | 0.2468 | 6.4E-12 | -1.1 | 0.4611 | 4.5E-06 |
| *lin-11* | WBGene00003000 | protein_coding_gene | -2.0 | 0.2467 | 6.2E-06 | -1.5 | 0.3610 | 7.5E-06 |
| *ttr-48* | WBGene00022515 | protein_coding_gene | -2.0 | 0.2462 | 2.5E-34 | -2.1 | 0.2350 | 1.4E-50 |
| *nhr-156* | WBGene00015897 | protein_coding_gene | -2.0 | 0.2460 | 2.0E-07 | -2.3 | 0.2026 | 1.8E-13 |
| *gst-35* | WBGene00001783 | protein_coding_gene | -2.0 | 0.2458 | 8.9E-07 | -1.4 | 0.3720 | 9.7E-04 |
| *pgp-11* | WBGene00004005 | protein_coding_gene | -2.0 | 0.2457 | 3.9E-18 | -3.5 | 0.0884 | 4.0E-56 |
| *Y47D3B.12* | WBGene00185055 | protein_coding_gene | -2.0 | 0.2456 | 1.5E-04 | -2.0 | 0.2574 | 4.0E-06 |
| *tmd-2* | WBGene00006582 | protein_coding_gene | -2.0 | 0.2454 | 6.3E-07 | -1.3 | 0.4105 | 2.9E-14 |
| *M03F8.6* | WBGene00019765 | protein_coding_gene | -2.0 | 0.2451 | 2.7E-04 | -1.6 | 0.3285 | 1.3E-03 |
| *frpr-8* | WBGene00018728 | protein_coding_gene | -2.0 | 0.2449 | 9.0E-04 | -1.7 | 0.3004 | 3.3E-03 |
| *F36A2.3* | WBGene00009453 | protein_coding_gene | -2.0 | 0.2448 | 1.5E-18 | -2.3 | 0.2099 | 3.8E-49 |
| *lim-9* | WBGene00006407 | protein_coding_gene | -2.0 | 0.2444 | 6.2E-15 | -1.1 | 0.4557 | 1.0E-21 |
| *nhr-97* | WBGene00003687 | protein_coding_gene | -2.0 | 0.2443 | 5.3E-35 | -1.2 | 0.4215 | 1.5E-26 |
| *C25E10.10* | WBGene00016099 | protein_coding_gene | -2.0 | 0.2440 | 4.0E-10 | -1.2 | 0.4472 | 2.5E-05 |
| *nekl-2* | WBGene00022631 | protein_coding_gene | -2.0 | 0.2440 | 8.5E-09 | -2.7 | 0.1559 | 8.2E-26 |
| *Y41G9A.10* | WBGene00044901 | protein_coding_gene | -2.0 | 0.2434 | 1.2E-09 | -4.7 | 0.0380 | 9.8E-30 |
| *dlhd-1* | WBGene00020741 | protein_coding_gene | -2.0 | 0.2431 | 2.6E-07 | -2.8 | 0.1465 | 1.1E-06 |
| *cyp-29A2* | WBGene00011830 | protein_coding_gene | -2.0 | 0.2425 | 2.4E-17 | -2.1 | 0.2390 | 3.5E-32 |
| *T03F6.4* | WBGene00011400 | protein_coding_gene | -2.0 | 0.2425 | 7.4E-06 | -1.3 | 0.4131 | 4.3E-04 |

| | | | | | | | | |
|---|---|---|---|---|---|---|---|---|
| *ZK970.7* | WBGene00014173 | protein_coding_gene | -2.0 | 0.2424 | 2.1E-03 | -6.2 | 0.0134 | 2.0E-17 |
| *mpz-1* | WBGene00003404 | protein_coding_gene | -2.0 | 0.2423 | 1.6E-09 | -1.3 | 0.4160 | 1.1E-14 |
| *dur-1* | WBGene00001113 | protein_coding_gene | -2.0 | 0.2423 | 5.4E-15 | -1.4 | 0.3793 | 2.5E-13 |
| *Y113G7A.16* | WBGene00013753 | protein_coding_gene | -2.0 | 0.2419 | 3.9E-02 | -1.8 | 0.2792 | 2.9E-02 |
| *aagr-2* | WBGene00019895 | protein_coding_gene | -2.0 | 0.2415 | 7.5E-23 | -1.1 | 0.4737 | 2.3E-05 |
| *C24B5.4* | WBGene00016046 | protein_coding_gene | -2.1 | 0.2415 | 1.3E-15 | -1.7 | 0.2989 | 2.1E-11 |
| *lips-16* | WBGene00012588 | pseudogene | -2.1 | 0.2414 | 1.2E-05 | -2.4 | 0.1864 | 4.1E-22 |
| *K11D12.7* | WBGene00019647 | protein_coding_gene | -2.1 | 0.2413 | 5.8E-28 | -1.5 | 0.3426 | 3.7E-18 |
| *ttr-1* | WBGene00010541 | protein_coding_gene | -2.1 | 0.2411 | 1.5E-13 | -2.4 | 0.1933 | 2.6E-10 |
| *alh-5* | WBGene00000111 | protein_coding_gene | -2.1 | 0.2411 | 1.1E-05 | -5.0 | 0.0319 | 4.5E-102 |
| *aakg-4* | WBGene00020135 | protein_coding_gene | -2.1 | 0.2398 | 8.6E-07 | -3.5 | 0.0884 | 4.5E-62 |
| *T04B8.5* | WBGene00020207 | protein_coding_gene | -2.1 | 0.2395 | 1.2E-17 | -1.0 | 0.4879 | 8.7E-15 |
| *C43F9.5* | WBGene00008073 | protein_coding_gene | -2.1 | 0.2393 | 3.1E-05 | -1.6 | 0.3317 | 7.4E-04 |
| *aagr-4* | WBGene00018682 | protein_coding_gene | -2.1 | 0.2392 | 9.0E-11 | -1.2 | 0.4261 | 5.0E-04 |
| *kvs-1* | WBGene00002242 | protein_coding_gene | -2.1 | 0.2388 | 6.0E-05 | -1.6 | 0.3278 | 4.9E-04 |
| *let-268* | WBGene00002497 | protein_coding_gene | -2.1 | 0.2388 | 7.0E-18 | -1.4 | 0.3758 | 3.3E-13 |
| *ugt-53* | WBGene00020182 | protein_coding_gene | -2.1 | 0.2387 | 8.4E-11 | -1.5 | 0.3553 | 3.2E-05 |
| *cpx-1* | WBGene00022271 | protein_coding_gene | -2.1 | 0.2385 | 2.7E-11 | -1.3 | 0.4068 | 4.1E-09 |
| *K04F10.9* | WBGene00220006 | ncRNA_gene | -2.1 | 0.2385 | 4.7E-02 | -2.6 | 0.1676 | 6.8E-04 |
| *lgc-42* | WBGene00021437 | protein_coding_gene | -2.1 | 0.2384 | 3.2E-03 | -1.4 | 0.3810 | 3.7E-02 |
| *F30A10.13* | WBGene00044680 | protein_coding_gene | -2.1 | 0.2379 | 4.4E-06 | -1.2 | 0.4214 | 3.3E-03 |
| *T16G1.7* | WBGene00011801 | protein_coding_gene | -2.1 | 0.2374 | 4.3E-03 | -4.9 | 0.0345 | 7.0E-164 |
| *F39H12.2* | WBGene00018213 | protein_coding_gene | -2.1 | 0.2374 | 5.6E-03 | -1.1 | 0.4730 | 4.6E-02 |
| *F49E2.5* | WBGene00009888 | protein_coding_gene | -2.1 | 0.2370 | 1.3E-07 | -1.6 | 0.3373 | 1.8E-29 |
| *D1086.8* | WBGene00008395 | protein_coding_gene | -2.1 | 0.2370 | 9.5E-19 | -1.1 | 0.4538 | 3.0E-06 |
| *sesn-1* | WBGene00022279 | protein_coding_gene | -2.1 | 0.2367 | 1.1E-24 | -1.4 | 0.3814 | 9.7E-19 |
| *F41E7.7* | WBGene00009623 | protein_coding_gene | -2.1 | 0.2366 | 2.8E-05 | -1.6 | 0.3326 | 1.9E-06 |
| *ptr-1* | WBGene00004216 | protein_coding_gene | -2.1 | 0.2364 | 4.4E-06 | -1.9 | 0.2682 | 6.8E-04 |
| *sdha-1* | WBGene00015391 | protein_coding_gene | -2.1 | 0.2363 | 1.2E-22 | -1.3 | 0.3975 | 1.9E-21 |
| *lgc-28* | WBGene00044301 | protein_coding_gene | -2.1 | 0.2362 | 1.8E-05 | -2.3 | 0.2037 | 3.0E-08 |
| *inx-12* | WBGene00002134 | protein_coding_gene | -2.1 | 0.2360 | 9.5E-23 | -1.6 | 0.3281 | 1.3E-27 |
| *K08B5.1* | WBGene00019518 | protein_coding_gene | -2.1 | 0.2358 | 2.5E-04 | -2.4 | 0.1959 | 3.6E-08 |
| *ins-22* | WBGene00002105 | protein_coding_gene | -2.1 | 0.2358 | 1.1E-04 | -1.3 | 0.4029 | 4.4E-03 |
| *nlp-69* | WBGene00013752 | protein_coding_gene | -2.1 | 0.2354 | 6.3E-04 | -1.9 | 0.2665 | 7.4E-04 |
| *acsd-1* | WBGene00022104 | protein_coding_gene | -2.1 | 0.2344 | 2.6E-09 | -1.1 | 0.4804 | 2.5E-04 |
| *nep-2* | WBGene00020230 | protein_coding_gene | -2.1 | 0.2344 | 1.4E-08 | -1.4 | 0.3839 | 9.2E-16 |
| *ins-26* | WBGene00002109 | protein_coding_gene | -2.1 | 0.2343 | 2.2E-08 | -1.2 | 0.4477 | 6.7E-03 |
| *Y50D4B.1* | WBGene00021740 | protein_coding_gene | -2.1 | 0.2337 | 1.9E-11 | -1.1 | 0.4785 | 3.4E-05 |
| *ent-2* | WBGene00010701 | protein_coding_gene | -2.1 | 0.2337 | 4.0E-20 | -1.0 | 0.4934 | 1.7E-14 |
| *prdh-1* | WBGene00007197 | protein_coding_gene | -2.1 | 0.2337 | 2.1E-16 | -1.3 | 0.4011 | 1.9E-19 |
| *C34C12.9* | WBGene00007928 | protein_coding_gene | -2.1 | 0.2336 | 1.1E-10 | -1.3 | 0.4095 | 9.1E-05 |
| *Y95B8A.6* | WBGene00022386 | protein_coding_gene | -2.1 | 0.2333 | 1.4E-06 | -2.9 | 0.1363 | 9.2E-36 |
| *unc-82* | WBGene00006814 | protein_coding_gene | -2.1 | 0.2331 | 1.8E-09 | -1.3 | 0.4154 | 4.3E-13 |
| *C50F4.8* | WBGene00008233 | protein_coding_gene | -2.1 | 0.2330 | 1.1E-09 | -3.5 | 0.0911 | 5.1E-65 |
| *T13H5.1* | WBGene00011755 | protein_coding_gene | -2.1 | 0.2330 | 8.1E-14 | -1.2 | 0.4468 | 2.8E-10 |
| *ZK622.5* | WBGene00044667 | protein_coding_gene | -2.1 | 0.2330 | 2.0E-12 | -1.2 | 0.4417 | 1.3E-04 |
| *F14H12.7* | WBGene00017474 | protein_coding_gene | -2.1 | 0.2327 | 3.9E-03 | -2.2 | 0.2161 | 8.6E-06 |
| *mod-1* | WBGene00003386 | protein_coding_gene | -2.1 | 0.2325 | 8.3E-07 | -1.6 | 0.3344 | 3.5E-04 |
| *pes-23* | WBGene00003987 | protein_coding_gene | -2.1 | 0.2323 | 5.3E-10 | -1.4 | 0.3743 | 3.0E-07 |
| *bcmo-1* | WBGene00012914 | protein_coding_gene | -2.1 | 0.2323 | 6.5E-06 | -1.5 | 0.3613 | 4.3E-03 |
| *ncs-5* | WBGene00008307 | protein_coding_gene | -2.1 | 0.2322 | 1.7E-03 | -1.3 | 0.4005 | 1.6E-02 |
| *linc-67* | WBGene00219648 | lincRNA_gene | -2.1 | 0.2312 | 2.9E-02 | -3.2 | 0.1093 | 7.8E-04 |
| *C10C5.3* | WBGene00007507 | protein_coding_gene | -2.1 | 0.2311 | 1.1E-06 | -3.1 | 0.1141 | 2.8E-11 |
| *asp-4* | WBGene00000217 | protein_coding_gene | -2.1 | 0.2310 | 1.7E-17 | -1.8 | 0.2942 | 1.7E-64 |
| *zig-6* | WBGene00006983 | protein_coding_gene | -2.1 | 0.2309 | 8.0E-12 | -2.0 | 0.2508 | 1.8E-13 |
| *ech-6* | WBGene00001155 | protein_coding_gene | -2.1 | 0.2308 | 2.5E-19 | -1.1 | 0.4725 | 1.8E-05 |
| *kin-1* | WBGene00002189 | protein_coding_gene | -2.1 | 0.2306 | 6.2E-11 | -1.0 | 0.4852 | 1.7E-15 |
| *Y38F2AR.12* | WBGene00021430 | protein_coding_gene | -2.1 | 0.2303 | 6.8E-25 | -1.7 | 0.3127 | 8.0E-20 |
| *ZC247.2* | WBGene00013860 | protein_coding_gene | -2.1 | 0.2302 | 9.2E-11 | -1.3 | 0.4000 | 2.5E-08 |
| *Y54G2A.41* | WBGene00044485 | protein_coding_gene | -2.1 | 0.2301 | 3.7E-09 | -1.3 | 0.4197 | 1.3E-04 |
| *gar-3* | WBGene00001519 | protein_coding_gene | -2.1 | 0.2299 | 4.6E-08 | -1.5 | 0.3605 | 1.8E-07 |
| *F54H5.14* | WBGene00255600 | protein_coding_gene | -2.1 | 0.2299 | 3.4E-07 | -1.1 | 0.4807 | 1.3E-02 |
| *ZC513.7* | WBGene00023209 | pseudogene | -2.1 | 0.2298 | 2.2E-17 | -2.3 | 0.2040 | 1.1E-64 |
| *R04B5.6* | WBGene00011004 | protein_coding_gene | -2.1 | 0.2295 | 9.1E-06 | -2.3 | 0.1968 | 1.2E-03 |
| *T22B11.4* | WBGene00020678 | protein_coding_gene | -2.1 | 0.2293 | 1.4E-14 | -1.0 | 0.4871 | 3.5E-12 |
| *oac-46* | WBGene00011768 | protein_coding_gene | -2.1 | 0.2292 | 1.5E-06 | -2.1 | 0.2395 | 3.1E-30 |
| *ule-3* | WBGene00008378 | protein_coding_gene | -2.1 | 0.2292 | 1.9E-15 | -2.1 | 0.2290 | 7.8E-52 |
| *unc-70* | WBGene00006803 | protein_coding_gene | -2.1 | 0.2290 | 8.9E-13 | -1.7 | 0.3141 | 2.0E-36 |
| *R09H10.2* | WBGene00011180 | protein_coding_gene | -2.1 | 0.2289 | 2.0E-07 | -2.5 | 0.1812 | 9.3E-09 |
| *F23D12.7* | WBGene00009091 | protein_coding_gene | -2.1 | 0.2288 | 8.6E-12 | -1.5 | 0.3452 | 2.2E-09 |

| | | | | | | | | |
|---|---|---|---|---|---|---|---|---|
| *lact-8* | WBGene00012777 | protein_coding_gene | -2.1 | 0.2284 | 9.4E-13 | -1.3 | 0.4028 | 4.2E-06 |
| *anmt-3* | WBGene00011573 | protein_coding_gene | -2.1 | 0.2284 | 3.4E-19 | -1.4 | 0.3684 | 5.5E-20 |
| *ZC449.5* | WBGene00022615 | protein_coding_gene | -2.1 | 0.2284 | 3.8E-10 | -1.2 | 0.4422 | 1.9E-12 |
| *mrp-1* | WBGene00003407 | protein_coding_gene | -2.1 | 0.2283 | 1.2E-18 | -1.2 | 0.4490 | 2.9E-23 |
| *clec-165* | WBGene00009523 | protein_coding_gene | -2.1 | 0.2278 | 2.4E-06 | -2.1 | 0.2372 | 8.5E-07 |
| *unc-1* | WBGene00006741 | protein_coding_gene | -2.1 | 0.2278 | 7.0E-11 | -1.3 | 0.3972 | 1.6E-10 |
| *gur-5* | WBGene00001806 | protein_coding_gene | -2.1 | 0.2277 | 2.8E-03 | -2.5 | 0.1750 | 2.3E-03 |
| *F55B11.3* | WBGene00010085 | protein_coding_gene | -2.1 | 0.2275 | 6.4E-13 | -1.3 | 0.3948 | 2.5E-21 |
| *snt-1* | WBGene00004921 | protein_coding_gene | -2.1 | 0.2269 | 2.6E-12 | -2.2 | 0.2188 | 7.3E-29 |
| *nipi-4* | WBGene00077712 | protein_coding_gene | -2.1 | 0.2269 | 8.7E-10 | -2.2 | 0.2189 | 3.2E-09 |
| *mlp-1* | WBGene00003375 | protein_coding_gene | -2.1 | 0.2266 | 9.0E-16 | -1.3 | 0.3964 | 1.2E-14 |
| *M28.10* | WBGene00010901 | protein_coding_gene | -2.1 | 0.2261 | 1.9E-17 | -1.0 | 0.4845 | 1.6E-11 |
| ***R193.2*** | **WBGene00020128** | **protein_coding_gene** | **-2.1** | **0.2257** | **2.1E-03** | **2.4** | **5.2710** | **2.9E-20** |
| *mboa-3* | WBGene00016934 | protein_coding_gene | -2.2 | 0.2253 | 9.4E-24 | -1.0 | 0.4911 | 6.9E-10 |
| *ZC412.3* | WBGene00013884 | protein_coding_gene | -2.2 | 0.2251 | 2.0E-11 | -1.6 | 0.3371 | 1.5E-20 |
| *F35B12.9* | WBGene00009388 | protein_coding_gene | -2.2 | 0.2244 | 2.2E-09 | -2.2 | 0.2215 | 4.1E-20 |
| *gei-3* | WBGene00001560 | protein_coding_gene | -2.2 | 0.2243 | 5.4E-09 | -1.2 | 0.4224 | 3.2E-17 |
| *F36G3.2* | WBGene00009483 | protein_coding_gene | -2.2 | 0.2242 | 1.1E-20 | -1.3 | 0.3984 | 1.2E-08 |
| *K09E2.2* | WBGene00019571 | protein_coding_gene | -2.2 | 0.2239 | 2.6E-02 | -2.4 | 0.1913 | 1.9E-03 |
| *D1086.6* | WBGene00008393 | protein_coding_gene | -2.2 | 0.2238 | 1.1E-13 | -1.5 | 0.3571 | 5.1E-24 |
| *ptb-1* | WBGene00004207 | protein_coding_gene | -2.2 | 0.2237 | 1.5E-14 | -1.3 | 0.4001 | 1.1E-10 |
| *Y69H2.15* | WBGene00050884 | protein_coding_gene | -2.2 | 0.2236 | 2.5E-05 | -1.9 | 0.2660 | 1.7E-03 |
| *K10H10.12* | WBGene00045403 | protein_coding_gene | -2.2 | 0.2230 | 1.6E-03 | -1.5 | 0.3650 | 7.1E-03 |
| *C53B4.3* | WBGene00008273 | protein_coding_gene | -2.2 | 0.2229 | 1.2E-20 | -1.9 | 0.2663 | 1.3E-24 |
| *mlck-1* | WBGene00013869 | protein_coding_gene | -2.2 | 0.2229 | 7.5E-15 | -1.2 | 0.4288 | 4.9E-13 |
| *clec-8* | WBGene00012582 | protein_coding_gene | -2.2 | 0.2228 | 2.1E-09 | -4.0 | 0.0642 | 2.0E-19 |
| *mig-39* | WBGene00018369 | protein_coding_gene | -2.2 | 0.2227 | 4.1E-10 | -1.2 | 0.4500 | 7.4E-06 |
| ***F09B12.3*** | **WBGene00008607** | **protein_coding_gene** | **-2.2** | **0.2226** | **3.4E-07** | **1.2** | **2.2902** | **1.5E-11** |
| *C05D11.5* | WBGene00015483 | protein_coding_gene | -2.2 | 0.2224 | 6.4E-11 | -1.9 | 0.2648 | 8.2E-15 |
| *seb-2* | WBGene00014035 | protein_coding_gene | -2.2 | 0.2223 | 4.9E-09 | -2.5 | 0.1761 | 1.1E-21 |
| *asb-2* | WBGene00000207 | protein_coding_gene | -2.2 | 0.2221 | 9.5E-51 | -1.0 | 0.4888 | 3.2E-19 |
| *C05D9.3* | WBGene00015472 | protein_coding_gene | -2.2 | 0.2221 | 5.7E-09 | -1.4 | 0.3793 | 6.5E-14 |
| *R07B1.9* | WBGene00011081 | protein_coding_gene | -2.2 | 0.2220 | 2.7E-07 | -1.4 | 0.3665 | 4.4E-04 |
| *Y39B6A.21* | WBGene00012680 | pseudogene | -2.2 | 0.2220 | 8.5E-04 | -2.7 | 0.1503 | 8.4E-08 |
| *K09E2.3* | WBGene00019572 | protein_coding_gene | -2.2 | 0.2218 | 2.4E-18 | -2.4 | 0.1893 | 1.8E-54 |
| *B0554.5* | WBGene00015258 | protein_coding_gene | -2.2 | 0.2216 | 3.2E-05 | -1.3 | 0.3975 | 7.2E-03 |
| *ace-2* | WBGene00000036 | protein_coding_gene | -2.2 | 0.2214 | 2.3E-14 | -1.5 | 0.3520 | 3.1E-19 |
| *F13G3.12* | WBGene00008770 | protein_coding_gene | -2.2 | 0.2210 | 8.6E-09 | -1.4 | 0.3736 | 1.4E-04 |
| *nhl-1* | WBGene00003597 | protein_coding_gene | -2.2 | 0.2209 | 1.7E-07 | -1.6 | 0.3379 | 4.7E-15 |
| *F41B4.2* | WBGene00018258 | protein_coding_gene | -2.2 | 0.2208 | 1.9E-11 | -1.3 | 0.4136 | 1.6E-09 |
| *unc-68* | WBGene00006801 | protein_coding_gene | -2.2 | 0.2208 | 1.9E-10 | -1.1 | 0.4540 | 7.8E-16 |
| *atn-1* | WBGene00000228 | protein_coding_gene | -2.2 | 0.2207 | 2.7E-24 | -1.4 | 0.3716 | 2.0E-19 |
| *C03G6.17* | WBGene00015402 | protein_coding_gene | -2.2 | 0.2205 | 1.8E-09 | -2.7 | 0.1593 | 3.1E-25 |
| *plc-1* | WBGene00004036 | protein_coding_gene | -2.2 | 0.2202 | 7.3E-10 | -1.0 | 0.4865 | 4.4E-10 |
| *glb-15* | WBGene00009387 | protein_coding_gene | -2.2 | 0.2202 | 4.7E-04 | -2.5 | 0.1741 | 2.9E-07 |
| *F53C3.8* | WBGene00018752 | protein_coding_gene | -2.2 | 0.2200 | 2.7E-02 | -1.5 | 0.3600 | 2.6E-03 |
| *acr-16* | WBGene00000055 | protein_coding_gene | -2.2 | 0.2200 | 4.6E-05 | -1.0 | 0.4857 | 2.0E-02 |
| *F28A10.13* | WBGene00284851 | protein_coding_gene | -2.2 | 0.2198 | 5.8E-03 | -1.3 | 0.4062 | 4.1E-02 |
| *dach-1* | WBGene00020386 | protein_coding_gene | -2.2 | 0.2197 | 7.1E-07 | -3.5 | 0.0858 | 4.7E-20 |
| *cyp-14A4* | WBGene00011009 | protein_coding_gene | -2.2 | 0.2196 | 5.9E-03 | -2.5 | 0.1759 | 2.6E-04 |
| *gly-2* | WBGene00001627 | protein_coding_gene | -2.2 | 0.2196 | 1.7E-16 | -1.4 | 0.3908 | 2.9E-10 |
| *fut-8* | WBGene00001508 | protein_coding_gene | -2.2 | 0.2194 | 2.9E-12 | -1.0 | 0.4868 | 5.9E-07 |
| *gpx-7* | WBGene00019846 | protein_coding_gene | -2.2 | 0.2191 | 5.4E-20 | -1.5 | 0.3569 | 3.4E-11 |
| *Y51A2D.8* | WBGene00013076 | protein_coding_gene | -2.2 | 0.2187 | 1.6E-02 | -7.3 | 0.0065 | 1.9E-64 |
| *vglu-2* | WBGene00010755 | protein_coding_gene | -2.2 | 0.2185 | 2.3E-16 | -2.5 | 0.1795 | 7.4E-46 |
| *egl-15* | WBGene00001184 | protein_coding_gene | -2.2 | 0.2181 | 4.3E-07 | -3.2 | 0.1100 | 8.4E-69 |
| *qui-1* | WBGene00004265 | protein_coding_gene | -2.2 | 0.2181 | 2.8E-10 | -1.5 | 0.3501 | 2.5E-09 |
| ***lact-7*** | **WBGene00008047** | **protein_coding_gene** | **-2.2** | **0.2179** | **3.9E-04** | **1.1** | **2.1712** | **4.7E-02** |
| *unc-60* | WBGene00006794 | protein_coding_gene | -2.2 | 0.2179 | 1.6E-23 | -1.5 | 0.3507 | 2.5E-27 |
| *ZC449.4* | WBGene00022614 | protein_coding_gene | -2.2 | 0.2178 | 3.7E-08 | -1.8 | 0.2877 | 2.7E-09 |
| *tag-297* | WBGene00007999 | protein_coding_gene | -2.2 | 0.2177 | 9.8E-05 | -1.9 | 0.2607 | 2.2E-02 |
| *C47D12.5* | WBGene00008139 | protein_coding_gene | -2.2 | 0.2176 | 7.0E-14 | -3.0 | 0.1245 | 7.8E-37 |
| *wrk-1* | WBGene00006942 | protein_coding_gene | -2.2 | 0.2172 | 2.6E-14 | -1.4 | 0.3779 | 7.0E-27 |
| *F45E1.4* | WBGene00018465 | protein_coding_gene | -2.2 | 0.2171 | 6.7E-08 | -1.8 | 0.2880 | 5.2E-11 |
| *F48C1.8* | WBGene00018600 | protein_coding_gene | -2.2 | 0.2171 | 4.3E-02 | -3.0 | 0.1290 | 6.8E-04 |
| *rig-3* | WBGene00004370 | protein_coding_gene | -2.2 | 0.2170 | 1.3E-13 | -2.0 | 0.2573 | 5.8E-25 |
| *C23H4.8* | WBGene00044132 | protein_coding_gene | -2.2 | 0.2170 | 7.8E-05 | -3.3 | 0.1042 | 5.8E-10 |
| *acl-7* | WBGene00012911 | protein_coding_gene | -2.2 | 0.2168 | 1.9E-26 | -1.5 | 0.3625 | 2.0E-18 |
| *pqn-24* | WBGene00004113 | protein_coding_gene | -2.2 | 0.2166 | 2.8E-21 | -1.2 | 0.4263 | 2.6E-13 |
| *glb-28* | WBGene00012441 | protein_coding_gene | -2.2 | 0.2165 | 1.7E-02 | -1.9 | 0.2671 | 1.6E-02 |

| | | | | | | | | |
|---|---|---|---|---|---|---|---|---|
| *T04G9.7* | WBGene00020218 | protein_coding_gene | -2.2 | 0.2165 | 4.6E-09 | -2.9 | 0.1311 | 2.8E-106 |
| *F26G1.2* | WBGene00017838 | protein_coding_gene | -2.2 | 0.2164 | 3.7E-11 | -3.7 | 0.0796 | 5.2E-26 |
| *F08D12.7* | WBGene00017250 | protein_coding_gene | -2.2 | 0.2160 | 1.5E-13 | -1.1 | 0.4773 | 6.9E-04 |
| *eff-1* | WBGene00001159 | protein_coding_gene | -2.2 | 0.2158 | 3.7E-10 | -1.8 | 0.2945 | 1.2E-06 |
| *srd-23* | WBGene00005101 | protein_coding_gene | -2.2 | 0.2153 | 3.5E-02 | -2.7 | 0.1567 | 5.8E-03 |
| *C04E6.7* | WBGene00015423 | protein_coding_gene | -2.2 | 0.2152 | 1.4E-12 | -1.3 | 0.4032 | 2.6E-09 |
| *F32H5.4* | WBGene00009350 | protein_coding_gene | -2.2 | 0.2152 | 4.4E-08 | -1.8 | 0.2851 | 3.2E-06 |
| *Y62H9A.4* | WBGene00013392 | protein_coding_gene | -2.2 | 0.2151 | 1.6E-10 | -1.2 | 0.4364 | 3.4E-11 |
| *ule-4* | WBGene00007458 | protein_coding_gene | -2.2 | 0.2149 | 8.4E-09 | -2.3 | 0.2000 | 1.2E-20 |
| *C10C5.4* | WBGene00007508 | protein_coding_gene | -2.2 | 0.2149 | 6.0E-06 | -2.9 | 0.1335 | 4.9E-61 |
| *abcx-1* | WBGene00006522 | protein_coding_gene | -2.2 | 0.2148 | 2.4E-04 | -1.2 | 0.4213 | 4.6E-02 |
| *cri-2* | WBGene00019478 | protein_coding_gene | -2.2 | 0.2147 | 1.4E-13 | -2.0 | 0.2539 | 1.6E-26 |
| *Y105E8A.1* | WBGene00013666 | protein_coding_gene | -2.2 | 0.2144 | 5.1E-09 | -1.3 | 0.3934 | 5.1E-07 |
| *got-1.2* | WBGene00020146 | protein_coding_gene | -2.2 | 0.2142 | 3.1E-24 | -1.7 | 0.3062 | 1.0E-40 |
| *F21G4.5* | WBGene00009026 | protein_coding_gene | -2.2 | 0.2140 | 1.1E-06 | -2.3 | 0.2019 | 3.3E-06 |
| *Y2H9A.4* | WBGene00012381 | protein_coding_gene | -2.2 | 0.2138 | 4.4E-07 | -2.7 | 0.1589 | 5.9E-16 |
| *F47B8.8* | WBGene00009809 | protein_coding_gene | -2.2 | 0.2131 | 1.4E-12 | -2.6 | 0.1597 | 6.0E-26 |
| *srp-2* | WBGene00005643 | protein_coding_gene | -2.2 | 0.2130 | 3.4E-26 | -2.8 | 0.1413 | 2.5E-41 |
| *aex-2* | WBGene00000085 | protein_coding_gene | -2.2 | 0.2130 | 2.3E-04 | -1.8 | 0.2789 | 5.9E-04 |
| *K06A4.2* | WBGene00010592 | protein_coding_gene | -2.2 | 0.2129 | 1.1E-03 | -2.6 | 0.1705 | 2.4E-05 |
| *cpx-2* | WBGene00010520 | protein_coding_gene | -2.2 | 0.2128 | 1.2E-08 | -1.8 | 0.2862 | 4.9E-08 |
| *C44C1.6* | WBGene00016645 | protein_coding_gene | -2.2 | 0.2127 | 6.7E-05 | -4.6 | 0.0405 | 1.2E-33 |
| *nhr-76* | WBGene00015497 | protein_coding_gene | -2.2 | 0.2126 | 1.1E-21 | -1.3 | 0.4078 | 1.6E-08 |
| *clc-4* | WBGene00000525 | protein_coding_gene | -2.2 | 0.2122 | 4.4E-02 | -5.6 | 0.0201 | 6.4E-16 |
| *dgk-1* | WBGene00000958 | protein_coding_gene | -2.2 | 0.2121 | 2.1E-09 | -1.8 | 0.2832 | 1.1E-12 |
| *nlp-58* | WBGene00077775 | protein_coding_gene | -2.2 | 0.2120 | 2.7E-09 | -1.4 | 0.3722 | 1.0E-06 |
| *T28B4.1* | WBGene00020884 | protein_coding_gene | -2.2 | 0.2120 | 4.3E-14 | -1.2 | 0.4391 | 4.0E-09 |
| *Y57G11C.31* | WBGene00013321 | protein_coding_gene | -2.2 | 0.2119 | 6.8E-30 | -1.9 | 0.2655 | 5.1E-29 |
| *slc-25A10* | WBGene00019656 | protein_coding_gene | -2.2 | 0.2118 | 6.2E-34 | -1.4 | 0.3851 | 2.4E-30 |
| *Y22D7AR.10* | WBGene00021263 | protein_coding_gene | -2.2 | 0.2118 | 9.9E-06 | -2.5 | 0.1781 | 1.3E-44 |
| *M04G7.3* | WBGene00019774 | protein_coding_gene | -2.2 | 0.2113 | 5.9E-10 | -1.1 | 0.4519 | 1.2E-04 |
| *R05D8.7* | WBGene00019885 | protein_coding_gene | -2.2 | 0.2112 | 1.5E-07 | -1.5 | 0.3622 | 1.3E-04 |
| *npr-23* | WBGene00021328 | protein_coding_gene | -2.2 | 0.2111 | 2.3E-06 | -1.7 | 0.3133 | 2.5E-04 |
| *pkc-2* | WBGene00004033 | protein_coding_gene | -2.2 | 0.2111 | 1.6E-26 | -1.4 | 0.3760 | 6.5E-26 |
| *deg-3* | WBGene00000951 | protein_coding_gene | -2.2 | 0.2109 | 2.3E-07 | -1.2 | 0.4383 | 6.6E-04 |
| *mltn-1* | WBGene00009311 | protein_coding_gene | -2.2 | 0.2103 | 9.3E-06 | -2.4 | 0.1931 | 7.8E-05 |
| *pssy-1* | WBGene00013920 | protein_coding_gene | -2.3 | 0.2101 | 7.3E-37 | -1.3 | 0.4030 | 2.8E-20 |
| *gei-15* | WBGene00001572 | protein_coding_gene | -2.3 | 0.2101 | 2.0E-15 | -1.4 | 0.3724 | 3.7E-13 |
| *F45D3.3* | WBGene00009723 | protein_coding_gene | -2.3 | 0.2100 | 1.5E-16 | -3.5 | 0.0900 | 2.6E-25 |
| *C24B9.3* | WBGene00016048 | protein_coding_gene | -2.3 | 0.2098 | 9.1E-25 | -1.9 | 0.2753 | 1.4E-11 |
| *clec-61* | WBGene00014047 | protein_coding_gene | -2.3 | 0.2098 | 1.1E-02 | -6.3 | 0.0127 | 2.0E-20 |
| *sod-3* | WBGene00004932 | protein_coding_gene | -2.3 | 0.2098 | 3.1E-10 | -1.3 | 0.4032 | 6.4E-05 |
| *acr-15* | WBGene00000054 | protein_coding_gene | -2.3 | 0.2097 | 1.4E-08 | -1.3 | 0.4188 | 3.9E-04 |
| *R04B5.5* | WBGene00011003 | protein_coding_gene | -2.3 | 0.2094 | 1.4E-09 | -3.4 | 0.0980 | 1.6E-64 |
| *M04C9.1* | WBGene00010856 | protein_coding_gene | -2.3 | 0.2092 | 1.9E-09 | -1.8 | 0.2805 | 1.3E-06 |
| *F49E12.10* | WBGene00009903 | protein_coding_gene | -2.3 | 0.2090 | 1.1E-09 | -3.8 | 0.0739 | 1.5E-21 |
| *unc-13* | WBGene00006752 | protein_coding_gene | -2.3 | 0.2090 | 8.0E-11 | -2.3 | 0.2033 | 3.3E-43 |
| *C34F6.10* | WBGene00007944 | protein_coding_gene | -2.3 | 0.2089 | 9.0E-17 | -1.3 | 0.4181 | 1.7E-12 |
| *mec-7* | WBGene00003171 | protein_coding_gene | -2.3 | 0.2087 | 1.2E-21 | -2.3 | 0.2046 | 2.0E-24 |
| *dig-1* | WBGene00000998 | protein_coding_gene | -2.3 | 0.2085 | 3.1E-09 | -1.1 | 0.4730 | 7.7E-08 |
| *cpi-1* | WBGene00000535 | protein_coding_gene | -2.3 | 0.2084 | 1.0E-11 | -1.9 | 0.2669 | 1.6E-38 |
| *Y72A10A.1* | WBGene00022202 | protein_coding_gene | -2.3 | 0.2082 | 6.4E-08 | -1.3 | 0.3961 | 3.3E-04 |
| *nlp-34* | WBGene00015046 | protein_coding_gene | -2.3 | 0.2082 | 3.8E-04 | -4.9 | 0.0337 | 2.3E-21 |
| *daf-22* | WBGene00013284 | protein_coding_gene | -2.3 | 0.2080 | 7.1E-22 | -2.1 | 0.2357 | 5.1E-41 |
| *Y18D10A.3* | WBGene00012476 | protein_coding_gene | -2.3 | 0.2078 | 3.9E-15 | -1.8 | 0.2884 | 1.2E-10 |
| *cysl-1* | WBGene00007653 | protein_coding_gene | -2.3 | 0.2078 | 7.1E-21 | -1.5 | 0.3557 | 1.4E-21 |
| *C15A11.7* | WBGene00007599 | protein_coding_gene | -2.3 | 0.2078 | 4.5E-16 | -3.4 | 0.0956 | 5.9E-46 |
| *ccg-1* | WBGene00000370 | protein_coding_gene | -2.3 | 0.2076 | 3.6E-32 | -1.7 | 0.3079 | 2.5E-53 |
| *F59B10.4* | WBGene00010320 | protein_coding_gene | -2.3 | 0.2076 | 8.2E-07 | -3.3 | 0.0986 | 6.2E-26 |
| *nhr-5* | WBGene00003604 | protein_coding_gene | -2.3 | 0.2073 | 3.0E-19 | -1.8 | 0.2873 | 3.6E-19 |
| *K02F3.9* | WBGene00019332 | protein_coding_gene | -2.3 | 0.2072 | 8.2E-21 | -1.8 | 0.2872 | 1.0E-24 |
| *C46C11.2* | WBGene00016705 | protein_coding_gene | -2.3 | 0.2070 | 2.1E-25 | -1.4 | 0.3762 | 7.6E-10 |
| *F13C5.5* | WBGene00017426 | protein_coding_gene | -2.3 | 0.2069 | 1.1E-09 | -1.8 | 0.2968 | 8.0E-20 |
| *glb-19* | WBGene00010146 | protein_coding_gene | -2.3 | 0.2069 | 1.5E-06 | -1.4 | 0.3853 | 5.7E-03 |
| *W04B5.1* | WBGene00021018 | pseudogene | -2.3 | 0.2066 | 6.3E-16 | -1.7 | 0.3021 | 5.9E-08 |
| *D2089.3* | WBGene00008434 | protein_coding_gene | -2.3 | 0.2064 | 3.8E-04 | -1.2 | 0.4429 | 1.7E-02 |
| *arrd-14* | WBGene00011055 | protein_coding_gene | -2.3 | 0.2063 | 2.6E-29 | -2.7 | 0.1522 | 1.2E-36 |
| *C06G1.1* | WBGene00015545 | protein_coding_gene | -2.3 | 0.2060 | 1.7E-06 | -1.8 | 0.2818 | 8.6E-08 |
| *rhgf-1* | WBGene00006468 | protein_coding_gene | -2.3 | 0.2060 | 6.3E-11 | -1.5 | 0.3632 | 1.1E-09 |
| *cka-2* | WBGene00000510 | protein_coding_gene | -2.3 | 0.2059 | 2.8E-13 | -1.6 | 0.3234 | 3.7E-08 |

| | | | | | | | | |
|---|---|---|---|---|---|---|---|---|
| *atf-4* | WBGene00000221 | protein_coding_gene | -2.3 | 0.2054 | 3.6E-21 | -1.4 | 0.3870 | 5.5E-31 |
| *K07E3.4* | WBGene00019492 | protein_coding_gene | -2.3 | 0.2049 | 1.8E-25 | -2.2 | 0.2187 | 8.9E-66 |
| *C07A9.12* | WBGene00043994 | protein_coding_gene | -2.3 | 0.2048 | 4.1E-07 | -2.3 | 0.2062 | 2.0E-10 |
| *pgp-2* | WBGene00003996 | protein_coding_gene | -2.3 | 0.2048 | 2.5E-32 | -1.7 | 0.3106 | 1.8E-23 |
| *pgp-10* | WBGene00004004 | protein_coding_gene | -2.3 | 0.2047 | 2.0E-09 | -3.6 | 0.0824 | 1.0E-36 |
| *C29G2.6* | WBGene00016234 | protein_coding_gene | -2.3 | 0.2045 | 9.3E-13 | -1.4 | 0.3753 | 6.6E-12 |
| *unc-98* | WBGene00006827 | protein_coding_gene | -2.3 | 0.2044 | 1.7E-13 | -1.4 | 0.3793 | 1.9E-11 |
| *flp-8* | WBGene00001451 | protein_coding_gene | -2.3 | 0.2044 | 1.0E-12 | -2.4 | 0.1934 | 4.2E-36 |
| *Y69E1A.5* | WBGene00013477 | protein_coding_gene | -2.3 | 0.2043 | 2.4E-14 | -2.6 | 0.1606 | 3.5E-47 |
| *R09E12.9* | WBGene00044514 | protein_coding_gene | -2.3 | 0.2040 | 9.0E-08 | -2.3 | 0.1993 | 3.5E-09 |
| *Y67A10A.9* | WBGene00013458 | protein_coding_gene | -2.3 | 0.2036 | 1.3E-07 | -2.1 | 0.2409 | 4.2E-11 |
| *che-10* | WBGene00018051 | protein_coding_gene | -2.3 | 0.2033 | 6.9E-14 | -1.2 | 0.4460 | 2.2E-11 |
| *sta-2* | WBGene00010251 | protein_coding_gene | -2.3 | 0.2032 | 4.3E-12 | -3.4 | 0.0955 | 5.1E-48 |
| *F37H8.5* | WBGene00009514 | protein_coding_gene | -2.3 | 0.2030 | 1.6E-33 | -1.5 | 0.3518 | 3.6E-17 |
| *F35C12.3* | WBGene00009408 | protein_coding_gene | -2.3 | 0.2030 | 5.5E-06 | -1.3 | 0.4121 | 1.9E-03 |
| *cyp-34A9* | WBGene00015044 | protein_coding_gene | -2.3 | 0.2029 | 2.0E-03 | -5.8 | 0.0178 | 1.2E-284 |
| *deb-1* | WBGene00000942 | protein_coding_gene | -2.3 | 0.2027 | 1.2E-29 | -1.6 | 0.3204 | 1.5E-40 |
| *lron-9* | WBGene00011971 | protein_coding_gene | -2.3 | 0.2026 | 1.7E-15 | -2.0 | 0.2456 | 4.1E-13 |
| *plpp-1.2* | WBGene00020895 | protein_coding_gene | -2.3 | 0.2026 | 3.2E-41 | -2.0 | 0.2455 | 3.4E-26 |
| *ttr-3* | WBGene00010538 | protein_coding_gene | -2.3 | 0.2023 | 4.6E-04 | -2.0 | 0.2420 | 1.3E-02 |
| *T19C3.3* | WBGene00020561 | protein_coding_gene | -2.3 | 0.2022 | 2.9E-05 | -1.5 | 0.3532 | 2.4E-03 |
| *ttr-47* | WBGene00019203 | protein_coding_gene | -2.3 | 0.2020 | 7.1E-19 | -2.1 | 0.2365 | 3.5E-62 |
| *unc-115* | WBGene00006839 | protein_coding_gene | -2.3 | 0.2019 | 5.6E-20 | -1.5 | 0.3425 | 1.2E-27 |
| *F59B2.13* | WBGene00010315 | protein_coding_gene | -2.3 | 0.2018 | 2.8E-11 | -1.9 | 0.2763 | 3.5E-12 |
| *C31B8.12* | WBGene00016285 | protein_coding_gene | -2.3 | 0.2016 | 3.9E-02 | -5.2 | 0.0274 | 6.0E-08 |
| *apy-1* | WBGene00017244 | protein_coding_gene | -2.3 | 0.2014 | 1.6E-37 | -1.1 | 0.4604 | 2.4E-18 |
| *acr-18* | WBGene00000057 | protein_coding_gene | -2.3 | 0.2013 | 1.3E-09 | -1.8 | 0.2815 | 1.1E-06 |
| *acy-4* | WBGene00000071 | protein_coding_gene | -2.3 | 0.2013 | 3.6E-33 | -1.4 | 0.3797 | 2.3E-11 |
| *rap-3* | WBGene00004309 | protein_coding_gene | -2.3 | 0.2012 | 5.6E-06 | -1.5 | 0.3434 | 3.1E-03 |
| *ugt-1* | WBGene00007072 | protein_coding_gene | -2.3 | 0.2010 | 2.5E-08 | -2.8 | 0.1427 | 5.2E-33 |
| *T01E8.8* | WBGene00011335 | protein_coding_gene | -2.3 | 0.2010 | 1.5E-11 | -1.3 | 0.4161 | 2.9E-08 |
| *aqp-11* | WBGene00000179 | protein_coding_gene | -2.3 | 0.2004 | 8.0E-17 | -1.1 | 0.4740 | 2.8E-15 |
| *nhr-3* | WBGene00003602 | protein_coding_gene | -2.3 | 0.2001 | 1.2E-12 | -2.2 | 0.2248 | 4.7E-24 |
| *F55D12.2* | WBGene00010111 | protein_coding_gene | -2.3 | 0.1999 | 4.3E-15 | -2.2 | 0.2226 | 2.1E-23 |
| *nlp-20* | WBGene00003758 | protein_coding_gene | -2.3 | 0.1998 | 3.0E-21 | -1.9 | 0.2660 | 2.5E-23 |
| *glt-4* | WBGene00001622 | protein_coding_gene | -2.3 | 0.1996 | 7.4E-11 | -2.2 | 0.2131 | 5.9E-29 |
| *inx-5* | WBGene00002127 | protein_coding_gene | -2.3 | 0.1996 | 1.1E-18 | -1.5 | 0.3529 | 1.6E-15 |
| *pcyt-2.2* | WBGene00016531 | protein_coding_gene | -2.3 | 0.1995 | 4.2E-14 | -2.1 | 0.2392 | 8.2E-23 |
| *thn-2* | WBGene00009215 | protein_coding_gene | -2.3 | 0.1994 | 4.0E-06 | -4.7 | 0.0377 | 2.9E-95 |
| *W01C8.5* | WBGene00020921 | protein_coding_gene | -2.3 | 0.1993 | 6.9E-23 | -1.3 | 0.4053 | 1.4E-11 |
| *ceeh-1* | WBGene00019329 | protein_coding_gene | -2.3 | 0.1992 | 1.5E-27 | -1.1 | 0.4663 | 1.3E-09 |
| *cest-3* | WBGene00008451 | protein_coding_gene | -2.3 | 0.1991 | 1.2E-18 | -1.8 | 0.2809 | 3.2E-10 |
| *twk-5* | WBGene00006660 | protein_coding_gene | -2.3 | 0.1987 | 1.5E-19 | -2.0 | 0.2428 | 4.8E-21 |
| *ztf-26* | WBGene00010137 | protein_coding_gene | -2.3 | 0.1984 | 6.0E-16 | -1.7 | 0.3101 | 3.5E-11 |
| *fgt-1* | WBGene00019207 | protein_coding_gene | -2.3 | 0.1982 | 1.9E-44 | -1.5 | 0.3494 | 1.4E-32 |
| *lipl-3* | WBGene00020016 | protein_coding_gene | -2.3 | 0.1979 | 3.1E-03 | -7.8 | 0.0045 | 3.1E-34 |
| *ttr-6* | WBGene00020886 | protein_coding_gene | -2.3 | 0.1979 | 7.2E-16 | -1.8 | 0.2775 | 2.0E-30 |
| *eat-4* | WBGene00001135 | protein_coding_gene | -2.3 | 0.1978 | 2.2E-10 | -1.7 | 0.2985 | 3.2E-05 |
| *del-9* | WBGene00015957 | protein_coding_gene | -2.3 | 0.1978 | 1.4E-04 | -2.1 | 0.2281 | 6.6E-06 |
| *F47B7.1* | WBGene00018532 | protein_coding_gene | -2.3 | 0.1977 | 3.6E-24 | -2.0 | 0.2575 | 3.2E-45 |
| *nmy-1* | WBGene00003776 | protein_coding_gene | -2.3 | 0.1973 | 7.4E-21 | -1.5 | 0.3627 | 1.2E-20 |
| *F21C10.17* | WBGene00235309 | pseudogene | -2.3 | 0.1971 | 3.1E-02 | -3.3 | 0.0993 | 3.4E-04 |
| *C49F5.11* | WBGene00219859 | ncRNA_gene | -2.3 | 0.1967 | 1.9E-05 | -2.5 | 0.1722 | 7.2E-07 |
| *lys-5* | WBGene00003094 | protein_coding_gene | -2.3 | 0.1967 | 2.2E-12 | -3.4 | 0.0959 | 3.9E-14 |
| *hot-1* | WBGene00001986 | protein_coding_gene | -2.3 | 0.1965 | 5.8E-12 | -1.3 | 0.3980 | 5.0E-05 |
| *T13B5.9* | WBGene00020476 | protein_coding_gene | -2.3 | 0.1965 | 8.5E-07 | -1.7 | 0.3065 | 1.5E-04 |
| *E02H4.5* | WBGene00008463 | protein_coding_gene | -2.3 | 0.1965 | 2.0E-02 | -1.6 | 0.3341 | 6.4E-03 |
| *F22F7.2* | WBGene00017720 | protein_coding_gene | -2.4 | 0.1960 | 2.2E-10 | -1.6 | 0.3263 | 6.2E-05 |
| *ugt-2* | WBGene00007073 | protein_coding_gene | -2.4 | 0.1960 | 4.7E-09 | -1.7 | 0.3068 | 3.1E-23 |
| *chil-27* | WBGene00011848 | protein_coding_gene | -2.4 | 0.1959 | 3.9E-02 | -2.4 | 0.1911 | 1.4E-02 |
| *Y53G8B.1* | WBGene00021817 | protein_coding_gene | -2.4 | 0.1958 | 8.3E-14 | -2.0 | 0.2464 | 4.5E-15 |
| *C13G3.1* | WBGene00007553 | protein_coding_gene | -2.4 | 0.1958 | 1.4E-11 | -1.2 | 0.4281 | 4.6E-05 |
| *F48E3.2* | WBGene00018603 | protein_coding_gene | -2.4 | 0.1957 | 3.9E-06 | -1.8 | 0.2970 | 9.8E-05 |
| *F38A6.5* | WBGene00086569 | protein_coding_gene | -2.4 | 0.1956 | 4.9E-18 | -1.1 | 0.4819 | 3.0E-04 |
| *R11F4.3* | WBGene00020009 | protein_coding_gene | -2.4 | 0.1956 | 6.3E-10 | -2.3 | 0.2005 | 4.4E-11 |
| *W05E10.1* | WBGene00012280 | protein_coding_gene | -2.4 | 0.1956 | 5.5E-07 | -1.7 | 0.3051 | 5.2E-08 |
| *egl-2* | WBGene00001171 | protein_coding_gene | -2.4 | 0.1954 | 4.3E-09 | -2.0 | 0.2519 | 7.8E-08 |
| *sup-10* | WBGene00006319 | protein_coding_gene | -2.4 | 0.1954 | 3.8E-07 | -1.0 | 0.4850 | 1.3E-02 |
| *Y45F10D.2* | WBGene00012883 | protein_coding_gene | -2.4 | 0.1954 | 1.4E-04 | -2.1 | 0.2269 | 9.7E-03 |
| *T06E6.10* | WBGene00011540 | protein_coding_gene | -2.4 | 0.1951 | 3.0E-18 | -1.9 | 0.2656 | 2.1E-16 |

| | | | | | | | | |
|---|---|---|---|---|---|---|---|---|
| *tyra-3* | WBGene00006475 | protein_coding_gene | -2.4 | 0.1949 | 3.7E-07 | -1.3 | 0.3933 | 5.0E-03 |
| *act-4* | WBGene00000066 | protein_coding_gene | -2.4 | 0.1944 | 1.1E-22 | -1.3 | 0.4102 | 1.7E-29 |
| *lgc-44* | WBGene00009790 | protein_coding_gene | -2.4 | 0.1944 | 5.7E-19 | -2.4 | 0.1850 | 8.9E-16 |
| *mrp-3* | WBGene00003409 | protein_coding_gene | -2.4 | 0.1943 | 1.7E-29 | -2.2 | 0.2105 | 1.0E-32 |
| *F25E5.8* | WBGene00017789 | protein_coding_gene | -2.4 | 0.1943 | 5.3E-24 | -2.3 | 0.2033 | 1.5E-26 |
| *npax-4* | WBGene00007496 | protein_coding_gene | -2.4 | 0.1936 | 5.9E-07 | -2.3 | 0.1987 | 1.8E-04 |
| *cdh-1* | WBGene00019994 | protein_coding_gene | -2.4 | 0.1935 | 2.0E-08 | -2.4 | 0.1917 | 7.3E-27 |
| *lgc-22* | WBGene00017483 | protein_coding_gene | -2.4 | 0.1933 | 2.6E-11 | -3.4 | 0.0944 | 4.8E-45 |
| *slc-25A29* | WBGene00008320 | protein_coding_gene | -2.4 | 0.1931 | 3.1E-13 | -2.3 | 0.1982 | 6.3E-18 |
| *mlc-6* | WBGene00010554 | protein_coding_gene | -2.4 | 0.1930 | 2.5E-07 | -1.4 | 0.3717 | 1.1E-02 |
| *spin-1* | WBGene00007549 | protein_coding_gene | -2.4 | 0.1928 | 9.9E-28 | -1.3 | 0.4003 | 9.5E-21 |
| *C02B10.3* | WBGene00015328 | protein_coding_gene | -2.4 | 0.1928 | 4.7E-26 | -2.0 | 0.2420 | 3.1E-20 |
| *zyx-1* | WBGene00006999 | protein_coding_gene | -2.4 | 0.1928 | 1.1E-20 | -1.1 | 0.4632 | 1.8E-21 |
| *F15G9.1* | WBGene00008865 | protein_coding_gene | -2.4 | 0.1927 | 3.1E-14 | -2.0 | 0.2448 | 2.6E-39 |
| *Y62H9A.13* | WBGene00013401 | protein_coding_gene | -2.4 | 0.1927 | 3.8E-02 | -2.2 | 0.2253 | 1.9E-03 |
| *old-1* | WBGene00003862 | protein_coding_gene | -2.4 | 0.1923 | 1.3E-05 | -2.5 | 0.1713 | 9.4E-04 |
| *F12A10.1* | WBGene00017390 | protein_coding_gene | -2.4 | 0.1922 | 1.4E-06 | -4.8 | 0.0359 | 4.7E-46 |
| *ifa-4* | WBGene00002052 | protein_coding_gene | -2.4 | 0.1922 | 5.0E-23 | -1.2 | 0.4310 | 6.9E-14 |
| *Y51A2D.21* | WBGene00044210 | protein_coding_gene | -2.4 | 0.1920 | 1.4E-12 | -2.2 | 0.2118 | 1.3E-10 |
| *T04G9.4* | WBGene00020215 | protein_coding_gene | -2.4 | 0.1919 | 9.5E-16 | -1.5 | 0.3618 | 5.5E-12 |
| *cat-1* | WBGene00000295 | protein_coding_gene | -2.4 | 0.1917 | 4.3E-06 | -2.3 | 0.2015 | 5.9E-06 |
| *dmsr-1* | WBGene00010191 | protein_coding_gene | -2.4 | 0.1916 | 7.4E-17 | -1.4 | 0.3814 | 3.1E-09 |
| *pitp-1* | WBGene00010813 | protein_coding_gene | -2.4 | 0.1914 | 2.1E-19 | -1.5 | 0.3569 | 3.4E-16 |
| *sdz-27* | WBGene00011124 | protein_coding_gene | -2.4 | 0.1912 | 7.0E-28 | -2.4 | 0.1912 | 1.7E-68 |
| *flp-17* | WBGene00001460 | protein_coding_gene | -2.4 | 0.1912 | 5.2E-11 | -1.8 | 0.2926 | 1.3E-18 |
| *lbp-2* | WBGene00002254 | protein_coding_gene | -2.4 | 0.1911 | 9.9E-20 | -2.0 | 0.2447 | 1.6E-31 |
| *abts-3* | WBGene00019018 | protein_coding_gene | -2.4 | 0.1910 | 7.9E-11 | -1.4 | 0.3779 | 1.5E-12 |
| *asns-2* | WBGene00019730 | protein_coding_gene | -2.4 | 0.1909 | 1.4E-06 | -2.9 | 0.1300 | 9.9E-28 |
| *F56F10.2* | WBGene00018985 | protein_coding_gene | -2.4 | 0.1907 | 1.1E-11 | -1.7 | 0.3084 | 1.6E-06 |
| *col-116* | WBGene00000690 | protein_coding_gene | -2.4 | 0.1904 | 2.7E-05 | -2.7 | 0.1541 | 1.5E-10 |
| *lron-14* | WBGene00011454 | protein_coding_gene | -2.4 | 0.1902 | 3.7E-09 | -1.9 | 0.2732 | 6.2E-06 |
| *nhr-14* | WBGene00003613 | protein_coding_gene | -2.4 | 0.1900 | 3.7E-18 | -1.4 | 0.3739 | 5.7E-16 |
| *sulp-7* | WBGene00012259 | protein_coding_gene | -2.4 | 0.1900 | 7.6E-25 | -2.0 | 0.2450 | 4.1E-27 |
| *cht-3* | WBGene00016084 | protein_coding_gene | -2.4 | 0.1899 | 1.1E-13 | -2.6 | 0.1637 | 8.0E-83 |
| *twk-18* | WBGene00006672 | protein_coding_gene | -2.4 | 0.1897 | 2.8E-12 | -1.9 | 0.2607 | 5.4E-31 |
| *lrp-1* | WBGene00003071 | protein_coding_gene | -2.4 | 0.1896 | 1.2E-15 | -2.3 | 0.2064 | 1.4E-58 |
| *mig-6* | WBGene00003242 | protein_coding_gene | -2.4 | 0.1896 | 5.6E-14 | -1.9 | 0.2755 | 3.6E-54 |
| *R11E3.2* | WBGene00020004 | protein_coding_gene | -2.4 | 0.1894 | 7.9E-18 | -1.6 | 0.3316 | 1.7E-06 |
| *F26E4.3* | WBGene00009158 | protein_coding_gene | -2.4 | 0.1890 | 3.4E-14 | -1.8 | 0.2925 | 3.7E-13 |
| *pfas-1* | WBGene00008654 | protein_coding_gene | -2.4 | 0.1889 | 3.0E-27 | -1.2 | 0.4356 | 1.0E-13 |
| *idhg-2* | WBGene00016266 | protein_coding_gene | -2.4 | 0.1889 | 4.5E-32 | -1.8 | 0.2778 | 1.4E-29 |
| *srd-53* | WBGene00005131 | protein_coding_gene | -2.4 | 0.1884 | 4.1E-04 | -1.4 | 0.3682 | 2.3E-02 |
| *F54E4.4* | WBGene00010060 | protein_coding_gene | -2.4 | 0.1881 | 2.5E-03 | -1.8 | 0.2862 | 4.5E-02 |
| *wrt-3* | WBGene00006949 | protein_coding_gene | -2.4 | 0.1880 | 4.4E-10 | -1.5 | 0.3627 | 2.5E-03 |
| *cest-4* | WBGene00015930 | protein_coding_gene | -2.4 | 0.1878 | 1.6E-22 | -1.7 | 0.3017 | 6.0E-23 |
| *rab-37* | WBGene00004285 | protein_coding_gene | -2.4 | 0.1878 | 2.0E-06 | -1.0 | 0.4848 | 4.0E-02 |
| *F16C3.1* | WBGene00008885 | protein_coding_gene | -2.4 | 0.1878 | 1.2E-07 | -1.7 | 0.3162 | 1.8E-04 |
| *jph-1* | WBGene00002179 | protein_coding_gene | -2.4 | 0.1877 | 1.6E-17 | -1.4 | 0.3772 | 9.3E-20 |
| *best-3* | WBGene00015286 | protein_coding_gene | -2.4 | 0.1876 | 1.6E-14 | -2.8 | 0.1457 | 1.1E-30 |
| *rskn-2* | WBGene00008311 | protein_coding_gene | -2.4 | 0.1875 | 2.0E-24 | -2.5 | 0.1788 | 7.9E-84 |
| *npr-5* | WBGene00021983 | protein_coding_gene | -2.4 | 0.1871 | 3.0E-20 | -1.4 | 0.3681 | 3.6E-08 |
| *T28D6.5* | WBGene00012125 | protein_coding_gene | -2.4 | 0.1871 | 1.6E-13 | -1.9 | 0.2640 | 4.0E-22 |
| *T07D4.2* | WBGene00011579 | protein_coding_gene | -2.4 | 0.1869 | 1.2E-15 | -1.6 | 0.3262 | 4.0E-11 |
| *chil-4* | WBGene00007469 | protein_coding_gene | -2.4 | 0.1868 | 2.0E-02 | -2.0 | 0.2430 | 4.5E-02 |
| *gst-33* | WBGene00001781 | protein_coding_gene | -2.4 | 0.1865 | 1.6E-04 | -3.4 | 0.0936 | 1.7E-12 |
| *hsp-43* | WBGene00002024 | protein_coding_gene | -2.4 | 0.1865 | 4.7E-27 | -1.2 | 0.4245 | 7.0E-21 |
| *C04C11.25* | WBGene00284855 | protein_coding_gene | -2.4 | 0.1860 | 3.2E-04 | -5.3 | 0.0257 | 2.0E-13 |
| *Y41C4A.6* | WBGene00012753 | protein_coding_gene | -2.4 | 0.1855 | 3.4E-07 | -1.7 | 0.3101 | 4.6E-05 |
| *cyp-33C9* | WBGene00016860 | protein_coding_gene | -2.4 | 0.1853 | 2.6E-25 | -1.9 | 0.2595 | 6.5E-20 |
| *F41C3.1* | WBGene00018267 | protein_coding_gene | -2.4 | 0.1851 | 1.7E-02 | -4.0 | 0.0607 | 1.3E-06 |
| *F17H10.1* | WBGene00008925 | protein_coding_gene | -2.4 | 0.1850 | 3.8E-70 | -2.2 | 0.2155 | 7.8E-39 |
| *F55C12.4* | WBGene00018871 | protein_coding_gene | -2.4 | 0.1849 | 5.3E-08 | -1.1 | 0.4636 | 7.0E-03 |
| *C55A6.12* | WBGene00044019 | protein_coding_gene | -2.4 | 0.1845 | 5.3E-25 | -1.8 | 0.2781 | 6.3E-20 |
| *egl-19* | WBGene00001187 | protein_coding_gene | -2.4 | 0.1845 | 2.6E-12 | -1.8 | 0.2907 | 3.4E-21 |
| *srh-46* | WBGene00005269 | protein_coding_gene | -2.4 | 0.1844 | 2.6E-02 | -2.3 | 0.2074 | 1.2E-02 |
| *gyg-1* | WBGene00006863 | protein_coding_gene | -2.4 | 0.1843 | 1.2E-23 | -1.3 | 0.4066 | 1.7E-15 |
| *K09E9.1* | WBGene00010724 | protein_coding_gene | -2.4 | 0.1841 | 4.5E-29 | -1.4 | 0.3694 | 5.7E-17 |
| *F44B9.10* | WBGene00018411 | protein_coding_gene | -2.4 | 0.1834 | 2.7E-03 | -1.5 | 0.3647 | 1.3E-02 |
| *T21D12.14* | WBGene00045177 | protein_coding_gene | -2.4 | 0.1834 | 7.6E-04 | -3.5 | 0.0910 | 5.2E-10 |
| *K01B6.3* | WBGene00010454 | protein_coding_gene | -2.4 | 0.1834 | 1.1E-11 | -1.5 | 0.3657 | 1.2E-04 |

| | | | | | | | | |
|---|---|---|---|---|---|---|---|---|
| *C51E3.6* | WBGene00008252 | protein_coding_gene | -2.4 | 0.1834 | 9.4E-21 | -1.9 | 0.2619 | 1.3E-14 |
| *clec-51* | WBGene00015050 | protein_coding_gene | -2.4 | 0.1834 | 1.5E-25 | -1.6 | 0.3226 | 7.1E-17 |
| *Y116F11B.6* | WBGene00013822 | protein_coding_gene | -2.4 | 0.1831 | 3.6E-05 | -1.9 | 0.2651 | 1.0E-04 |
| *F09B9.4* | WBGene00008603 | protein_coding_gene | -2.4 | 0.1830 | 9.1E-11 | -1.9 | 0.2736 | 1.7E-31 |
| *ads-1* | WBGene00000081 | protein_coding_gene | -2.5 | 0.1830 | 3.3E-31 | -2.2 | 0.2196 | 1.3E-77 |
| *kin-30* | WBGene00002211 | protein_coding_gene | -2.5 | 0.1822 | 3.0E-07 | -2.3 | 0.2066 | 3.0E-06 |
| ***T16G12.1*** | **WBGene00011803** | **protein_coding_gene** | **-2.5** | **0.1821** | **1.4E-10** | **1.6** | **3.0430** | **8.7E-07** |
| *tph-1* | WBGene00006600 | protein_coding_gene | -2.5 | 0.1820 | 1.5E-03 | -1.7 | 0.3038 | 3.1E-02 |
| *lec-4* | WBGene00002267 | protein_coding_gene | -2.5 | 0.1816 | 1.8E-19 | -2.1 | 0.2382 | 4.1E-58 |
| *lec-5* | WBGene00002268 | protein_coding_gene | -2.5 | 0.1815 | 8.8E-31 | -1.8 | 0.2812 | 1.1E-48 |
| *nid-1* | WBGene00003738 | protein_coding_gene | -2.5 | 0.1813 | 1.0E-16 | -1.1 | 0.4797 | 9.2E-11 |
| *Y70G10A.3* | WBGene00013499 | protein_coding_gene | -2.5 | 0.1812 | 3.0E-09 | -3.5 | 0.0896 | 5.5E-60 |
| *mbr-1* | WBGene00011315 | protein_coding_gene | -2.5 | 0.1811 | 8.1E-11 | -2.2 | 0.2190 | 1.1E-06 |
| *F09E10.7* | WBGene00017297 | protein_coding_gene | -2.5 | 0.1810 | 9.2E-06 | -1.1 | 0.4736 | 8.9E-03 |
| *ptc-3* | WBGene00004210 | protein_coding_gene | -2.5 | 0.1810 | 5.1E-18 | -2.6 | 0.1627 | 6.3E-56 |
| *F23D12.3* | WBGene00009087 | protein_coding_gene | -2.5 | 0.1805 | 9.4E-11 | -2.2 | 0.2151 | 1.4E-02 |
| *B0228.10* | WBGene00219780 | snoRNA_gene | -2.5 | 0.1804 | 1.3E-03 | -1.8 | 0.2842 | 1.5E-02 |
| *C54D1.7* | WBGene00016914 | protein_coding_gene | -2.5 | 0.1799 | 3.8E-08 | -3.1 | 0.1159 | 1.0E-08 |
| *rnp-9* | WBGene00007396 | protein_coding_gene | -2.5 | 0.1792 | 3.0E-11 | -1.2 | 0.4366 | 1.2E-04 |
| *C26D10.7* | WBGene00023420 | protein_coding_gene | -2.5 | 0.1791 | 1.6E-03 | -1.9 | 0.2730 | 1.6E-02 |
| ***F22E5.8*** | **WBGene00017707** | **protein_coding_gene** | **-2.5** | **0.1790** | **4.1E-05** | **1.5** | **2.7497** | **3.3E-04** |
| *far-6* | WBGene00001390 | protein_coding_gene | -2.5 | 0.1790 | 2.7E-16 | -2.6 | 0.1664 | 1.7E-57 |
| *lrp-2* | WBGene00003072 | protein_coding_gene | -2.5 | 0.1788 | 3.5E-13 | -1.9 | 0.2703 | 1.3E-46 |
| *T18D3.6* | WBGene00011823 | protein_coding_gene | -2.5 | 0.1787 | 3.0E-24 | -1.4 | 0.3665 | 5.9E-09 |
| *W06A11.1* | WBGene00021052 | protein_coding_gene | -2.5 | 0.1782 | 2.3E-03 | -3.4 | 0.0980 | 5.9E-05 |
| *K11D9.3* | WBGene00010772 | protein_coding_gene | -2.5 | 0.1780 | 1.1E-30 | -1.7 | 0.2997 | 8.1E-16 |
| *C40H1.8* | WBGene00008043 | protein_coding_gene | -2.5 | 0.1780 | 2.6E-06 | -2.5 | 0.1721 | 9.2E-08 |
| *F28A12.3* | WBGene00017880 | protein_coding_gene | -2.5 | 0.1779 | 5.7E-08 | -1.6 | 0.3202 | 2.2E-06 |
| *Y71H10B.1* | WBGene00022201 | protein_coding_gene | -2.5 | 0.1779 | 3.0E-26 | -1.2 | 0.4351 | 2.0E-13 |
| *glna-1* | WBGene00007480 | protein_coding_gene | -2.5 | 0.1778 | 5.6E-10 | -2.1 | 0.2257 | 4.2E-13 |
| *M117.1* | WBGene00010918 | protein_coding_gene | -2.5 | 0.1778 | 6.3E-28 | -2.2 | 0.2142 | 2.8E-32 |
| ***K10C9.9*** | **WBGene00044515** | **pseudogene** | **-2.5** | **0.1776** | **1.4E-03** | **2.2** | **4.5902** | **4.2E-03** |
| *F31D5.2* | WBGene00017945 | protein_coding_gene | -2.5 | 0.1769 | 4.9E-16 | -1.7 | 0.3115 | 4.8E-10 |
| *mlc-2* | WBGene00003370 | protein_coding_gene | -2.5 | 0.1768 | 2.5E-36 | -1.6 | 0.3269 | 8.1E-45 |
| *F21C10.7* | WBGene00017657 | protein_coding_gene | -2.5 | 0.1768 | 5.7E-18 | -1.5 | 0.3599 | 3.1E-23 |
| *W04E12.7* | WBGene00012252 | protein_coding_gene | -2.5 | 0.1764 | 8.1E-20 | -1.4 | 0.3813 | 3.9E-05 |
| *B0513.4* | WBGene00007196 | protein_coding_gene | -2.5 | 0.1763 | 1.0E-15 | -2.6 | 0.1635 | 1.3E-53 |
| *F55G7.4* | WBGene00045493 | protein_coding_gene | -2.5 | 0.1760 | 1.6E-02 | -3.6 | 0.0814 | 3.0E-04 |
| *hach-1* | WBGene00017301 | protein_coding_gene | -2.5 | 0.1760 | 5.6E-68 | -1.6 | 0.3308 | 7.4E-05 |
| *elt-3* | WBGene00001251 | protein_coding_gene | -2.5 | 0.1759 | 2.6E-15 | -1.5 | 0.3453 | 8.2E-21 |
| *F14E5.1* | WBGene00008799 | protein_coding_gene | -2.5 | 0.1755 | 1.1E-10 | -2.5 | 0.1740 | 4.9E-09 |
| *pat-12* | WBGene00003936 | protein_coding_gene | -2.5 | 0.1754 | 6.8E-23 | -1.3 | 0.4150 | 1.0E-18 |
| *ptr-12* | WBGene00004226 | protein_coding_gene | -2.5 | 0.1752 | 2.4E-14 | -1.9 | 0.2738 | 6.1E-11 |
| *lam-2* | WBGene00016913 | protein_coding_gene | -2.5 | 0.1749 | 1.2E-22 | -2.0 | 0.2552 | 7.3E-41 |
| *R02F11.3* | WBGene00019841 | protein_coding_gene | -2.5 | 0.1748 | 3.0E-21 | -1.2 | 0.4461 | 1.6E-08 |
| *F08D12.2* | WBGene00017246 | protein_coding_gene | -2.5 | 0.1746 | 1.7E-15 | -2.2 | 0.2189 | 8.4E-06 |
| *vap-2* | WBGene00011462 | protein_coding_gene | -2.5 | 0.1745 | 7.7E-06 | -2.3 | 0.2089 | 5.6E-15 |
| *lron-3* | WBGene00007261 | protein_coding_gene | -2.5 | 0.1743 | 3.9E-39 | -1.4 | 0.3753 | 2.5E-14 |
| *C06A12.3* | WBGene00007357 | protein_coding_gene | -2.5 | 0.1742 | 1.4E-18 | -2.3 | 0.2068 | 4.3E-24 |
| *pcp-1* | WBGene00003956 | protein_coding_gene | -2.5 | 0.1741 | 5.9E-53 | -1.4 | 0.3890 | 6.1E-32 |
| *F57G9.6* | WBGene00044041 | pseudogene | -2.5 | 0.1740 | 1.1E-04 | -2.4 | 0.1942 | 2.0E-13 |
| *ugt-12* | WBGene00020592 | protein_coding_gene | -2.5 | 0.1739 | 1.3E-21 | -1.9 | 0.2608 | 1.7E-28 |
| *glb-17* | WBGene00009887 | protein_coding_gene | -2.5 | 0.1738 | 1.1E-04 | -1.4 | 0.3707 | 3.4E-03 |
| *F55A12.6* | WBGene00018865 | protein_coding_gene | -2.5 | 0.1735 | 7.9E-04 | -4.1 | 0.0572 | 1.6E-09 |
| *spp-1* | WBGene00004986 | protein_coding_gene | -2.5 | 0.1734 | 5.5E-14 | -3.5 | 0.0906 | 3.9E-33 |
| *F43B10.1* | WBGene00018373 | protein_coding_gene | -2.5 | 0.1733 | 3.6E-15 | -1.3 | 0.3925 | 1.7E-13 |
| *molo-1* | WBGene00017299 | protein_coding_gene | -2.5 | 0.1731 | 2.4E-05 | -2.0 | 0.2522 | 2.6E-04 |
| *F46C5.10* | WBGene00009785 | protein_coding_gene | -2.5 | 0.1729 | 4.7E-10 | -2.8 | 0.1436 | 3.9E-25 |
| *C31H5.6* | WBGene00007857 | protein_coding_gene | -2.5 | 0.1727 | 3.6E-16 | -2.7 | 0.1550 | 1.3E-33 |
| *T22E5.1* | WBGene00020690 | protein_coding_gene | -2.5 | 0.1722 | 6.8E-09 | -1.1 | 0.4553 | 1.8E-02 |
| *Y102E9.5* | WBGene00044452 | protein_coding_gene | -2.5 | 0.1721 | 1.2E-15 | -1.1 | 0.4671 | 2.6E-03 |
| *C53A3.4* | WBGene00303231 | protein_coding_gene | -2.5 | 0.1714 | 1.0E-05 | -1.7 | 0.3075 | 1.6E-04 |
| *F32H5.1* | WBGene00009347 | protein_coding_gene | -2.5 | 0.1714 | 5.3E-32 | -1.3 | 0.4076 | 5.1E-15 |
| *timp-1* | WBGene00019476 | protein_coding_gene | -2.5 | 0.1711 | 2.6E-21 | -2.4 | 0.1885 | 1.2E-23 |
| *T27F6.8* | WBGene00012107 | protein_coding_gene | -2.5 | 0.1711 | 3.6E-10 | -3.1 | 0.1188 | 1.0E-21 |
| *ptr-6* | WBGene00004221 | protein_coding_gene | -2.5 | 0.1710 | 1.1E-14 | -2.5 | 0.1763 | 6.5E-18 |
| *frm-5.1* | WBGene00001492 | protein_coding_gene | -2.6 | 0.1707 | 2.2E-10 | -1.9 | 0.2651 | 2.1E-08 |
| *tkr-3* | WBGene00006428 | protein_coding_gene | -2.6 | 0.1707 | 4.6E-06 | -1.1 | 0.4512 | 3.9E-02 |
| *cyp-43A1* | WBGene00017108 | protein_coding_gene | -2.6 | 0.1706 | 1.2E-20 | -2.2 | 0.2248 | 1.0E-22 |
| *Y62H9A.12* | WBGene00013400 | protein_coding_gene | -2.6 | 0.1706 | 5.3E-04 | -2.1 | 0.2297 | 1.3E-03 |

| | | | | | | | | |
|---|---|---|---|---|---|---|---|---|
| *F09C12.6* | WBGene00017278 | protein_coding_gene | -2.6 | 0.1703 | 1.1E-02 | -2.0 | 0.2451 | 3.3E-02 |
| *C51E3.9* | WBGene00008254 | protein_coding_gene | -2.6 | 0.1703 | 3.3E-16 | -1.2 | 0.4454 | 3.5E-09 |
| *B0252.3* | WBGene00015088 | protein_coding_gene | -2.6 | 0.1702 | 1.0E-22 | -1.7 | 0.3169 | 2.4E-19 |
| *kcnl-2* | WBGene00008570 | protein_coding_gene | -2.6 | 0.1701 | 2.1E-12 | -1.6 | 0.3357 | 2.6E-08 |
| *fil-2* | WBGene00019668 | protein_coding_gene | -2.6 | 0.1701 | 6.3E-03 | -2.0 | 0.2544 | 1.6E-03 |
| *F23F1.6* | WBGene00017747 | protein_coding_gene | -2.6 | 0.1698 | 1.7E-10 | -2.0 | 0.2506 | 2.0E-11 |
| *best-20* | WBGene00021368 | protein_coding_gene | -2.6 | 0.1697 | 1.3E-18 | -3.0 | 0.1246 | 1.5E-44 |
| *cor-1* | WBGene00000768 | protein_coding_gene | -2.6 | 0.1696 | 3.2E-24 | -1.9 | 0.2626 | 6.1E-16 |
| *C52G5.2* | WBGene00008260 | protein_coding_gene | -2.6 | 0.1692 | 2.3E-10 | -1.4 | 0.3688 | 1.3E-05 |
| *Y106G6D.6* | WBGene00013702 | protein_coding_gene | -2.6 | 0.1690 | 3.4E-08 | -3.0 | 0.1251 | 3.4E-13 |
| *C45G9.12* | WBGene00016683 | protein_coding_gene | -2.6 | 0.1689 | 1.8E-04 | -2.1 | 0.2293 | 3.7E-04 |
| *R13D11.4* | WBGene00020058 | protein_coding_gene | -2.6 | 0.1689 | 2.8E-13 | -2.1 | 0.2374 | 1.5E-13 |
| *ins-23* | WBGene00002106 | protein_coding_gene | -2.6 | 0.1685 | 2.9E-03 | -3.1 | 0.1151 | 4.1E-08 |
| *dmsr-12* | WBGene00019262 | protein_coding_gene | -2.6 | 0.1684 | 7.8E-04 | -4.4 | 0.0462 | 3.6E-20 |
| *gbb-2* | WBGene00022675 | protein_coding_gene | -2.6 | 0.1684 | 7.4E-18 | -1.1 | 0.4557 | 2.7E-05 |
| *clik-3* | WBGene00019361 | protein_coding_gene | -2.6 | 0.1682 | 1.1E-25 | -1.5 | 0.3648 | 3.5E-25 |
| *del-10* | WBGene00020897 | protein_coding_gene | -2.6 | 0.1680 | 2.3E-18 | -1.3 | 0.4053 | 1.6E-05 |
| *pfn-3* | WBGene00003991 | protein_coding_gene | -2.6 | 0.1680 | 1.3E-16 | -1.2 | 0.4484 | 3.7E-07 |
| *tnt-4* | WBGene00006589 | protein_coding_gene | -2.6 | 0.1679 | 3.7E-12 | -1.0 | 0.4969 | 1.3E-10 |
| *F23G4.1* | WBGene00044574 | protein_coding_gene | -2.6 | 0.1676 | 9.4E-05 | -3.4 | 0.0956 | 1.5E-07 |
| *mltn-9* | WBGene00008964 | protein_coding_gene | -2.6 | 0.1676 | 1.3E-11 | -2.2 | 0.2186 | 1.9E-08 |
| *catp-1* | WBGene00013672 | protein_coding_gene | -2.6 | 0.1674 | 6.7E-23 | -3.2 | 0.1076 | 3.7E-69 |
| *ugt-32* | WBGene00018543 | protein_coding_gene | -2.6 | 0.1673 | 1.6E-03 | -2.8 | 0.1483 | 2.4E-05 |
| *M162.5* | WBGene00010931 | protein_coding_gene | -2.6 | 0.1669 | 3.2E-09 | -4.0 | 0.0624 | 1.6E-27 |
| *scrm-2* | WBGene00014200 | protein_coding_gene | -2.6 | 0.1669 | 5.2E-04 | -1.6 | 0.3382 | 8.7E-03 |
| *nhr-137* | WBGene00003727 | protein_coding_gene | -2.6 | 0.1669 | 4.6E-13 | -1.1 | 0.4736 | 1.4E-04 |
| *C49G9.2* | WBGene00045474 | protein_coding_gene | -2.6 | 0.1669 | 1.1E-05 | -2.1 | 0.2265 | 1.9E-03 |
| *ptr-21* | WBGene00004235 | protein_coding_gene | -2.6 | 0.1668 | 3.6E-27 | -1.8 | 0.2808 | 3.2E-36 |
| *ttr-16* | WBGene00012382 | protein_coding_gene | -2.6 | 0.1667 | 1.4E-28 | -1.6 | 0.3195 | 2.2E-46 |
| *F37C4.6* | WBGene00018146 | protein_coding_gene | -2.6 | 0.1666 | 9.5E-19 | -1.1 | 0.4790 | 5.7E-09 |
| *ctsa-4.1* | WBGene00019605 | protein_coding_gene | -2.6 | 0.1666 | 1.6E-10 | -1.1 | 0.4576 | 2.6E-07 |
| *cal-5* | WBGene00006861 | protein_coding_gene | -2.6 | 0.1663 | 9.6E-31 | -1.0 | 0.4899 | 3.3E-13 |
| *K07C11.7* | WBGene00019479 | protein_coding_gene | -2.6 | 0.1663 | 6.6E-25 | -2.0 | 0.2450 | 5.0E-39 |
| *smp-2* | WBGene00004890 | protein_coding_gene | -2.6 | 0.1662 | 6.5E-22 | -2.0 | 0.2432 | 8.3E-48 |
| *rhr-2* | WBGene00004359 | protein_coding_gene | -2.6 | 0.1657 | 6.1E-12 | -2.4 | 0.1950 | 2.4E-10 |
| *cca-1* | WBGene00000367 | protein_coding_gene | -2.6 | 0.1654 | 3.3E-20 | -1.2 | 0.4303 | 5.3E-09 |
| *hex-1* | WBGene00020509 | protein_coding_gene | -2.6 | 0.1651 | 2.1E-24 | -2.8 | 0.1420 | 1.2E-47 |
| *C18D11.1* | WBGene00007679 | protein_coding_gene | -2.6 | 0.1649 | 7.8E-14 | -1.6 | 0.3381 | 7.9E-11 |
| *cest-25* | WBGene00022178 | protein_coding_gene | -2.6 | 0.1648 | 8.6E-27 | -1.8 | 0.2819 | 4.3E-25 |
| *trk-1* | WBGene00017039 | protein_coding_gene | -2.6 | 0.1647 | 8.2E-09 | -1.8 | 0.2970 | 1.5E-05 |
| *avr-15* | WBGene00000233 | protein_coding_gene | -2.6 | 0.1646 | 4.7E-32 | -1.4 | 0.3912 | 3.1E-14 |
| *cpg-7* | WBGene00010723 | protein_coding_gene | -2.6 | 0.1644 | 6.0E-04 | -4.2 | 0.0557 | 1.1E-62 |
| *B0310.1* | WBGene00015137 | protein_coding_gene | -2.6 | 0.1643 | 1.5E-15 | -1.4 | 0.3708 | 8.5E-10 |
| *F36H1.12* | WBGene00009497 | protein_coding_gene | -2.6 | 0.1641 | 2.4E-03 | -2.3 | 0.2043 | 1.4E-04 |
| *srx-21* | WBGene00005912 | protein_coding_gene | -2.6 | 0.1640 | 3.1E-02 | -3.1 | 0.1135 | 1.2E-03 |
| *lgc-26* | WBGene00021981 | protein_coding_gene | -2.6 | 0.1640 | 3.7E-11 | -1.7 | 0.3069 | 2.0E-06 |
| *cnc-6* | WBGene00000560 | protein_coding_gene | -2.6 | 0.1639 | 1.1E-02 | -4.7 | 0.0396 | 1.2E-06 |
| *fbxa-39* | WBGene00021859 | protein_coding_gene | -2.6 | 0.1636 | 8.9E-05 | -1.2 | 0.4232 | 2.2E-02 |
| *fln-2* | WBGene00016006 | protein_coding_gene | -2.6 | 0.1634 | 2.5E-13 | -2.0 | 0.2429 | 2.4E-46 |
| *T10H9.1* | WBGene00020424 | protein_coding_gene | -2.6 | 0.1629 | 1.3E-07 | -1.4 | 0.3692 | 1.4E-02 |
| *ser-1* | WBGene00004776 | protein_coding_gene | -2.6 | 0.1626 | 6.6E-10 | -1.0 | 0.4860 | 1.2E-02 |
| *snf-12* | WBGene00004911 | protein_coding_gene | -2.6 | 0.1625 | 3.0E-12 | -3.1 | 0.1173 | 4.5E-14 |
| *pyk-2* | WBGene00014001 | protein_coding_gene | -2.6 | 0.1615 | 2.0E-29 | -1.7 | 0.3020 | 1.2E-38 |
| *F20G2.3* | WBGene00008987 | protein_coding_gene | -2.6 | 0.1615 | 1.0E-18 | -1.8 | 0.2858 | 1.1E-09 |
| *chil-6* | WBGene00007471 | protein_coding_gene | -2.6 | 0.1612 | 1.8E-03 | -2.8 | 0.1461 | 2.6E-03 |
| *ddn-1* | WBGene00015227 | protein_coding_gene | -2.6 | 0.1612 | 1.4E-07 | -1.7 | 0.3166 | 1.6E-03 |
| *nlp-25* | WBGene00003763 | protein_coding_gene | -2.6 | 0.1610 | 3.2E-04 | -4.8 | 0.0371 | 7.0E-11 |
| *F09A5.3* | WBGene00008600 | protein_coding_gene | -2.6 | 0.1608 | 8.8E-03 | -1.8 | 0.2807 | 3.3E-02 |
| *T20F7.5* | WBGene00020632 | pseudogene | -2.6 | 0.1608 | 4.8E-15 | -1.5 | 0.3480 | 8.2E-06 |
| *T07D10.3* | WBGene00011583 | protein_coding_gene | -2.6 | 0.1607 | 1.4E-07 | -3.5 | 0.0898 | 6.9E-08 |
| *M01G12.14* | WBGene00010825 | protein_coding_gene | -2.6 | 0.1606 | 6.8E-05 | -2.8 | 0.1428 | 3.2E-08 |
| *glt-6* | WBGene00001624 | protein_coding_gene | -2.6 | 0.1605 | 5.1E-11 | -3.0 | 0.1254 | 5.8E-14 |
| *lips-17* | WBGene00019939 | protein_coding_gene | -2.6 | 0.1603 | 6.0E-13 | -1.9 | 0.2764 | 4.6E-09 |
| *B0379.1* | WBGene00007155 | protein_coding_gene | -2.6 | 0.1603 | 8.0E-27 | -2.3 | 0.2012 | 5.7E-33 |
| *T22E5.6* | WBGene00020692 | protein_coding_gene | -2.6 | 0.1600 | 1.1E-10 | -1.5 | 0.3471 | 1.0E-04 |
| *F17H10.2* | WBGene00008926 | protein_coding_gene | -2.6 | 0.1599 | 1.5E-28 | -3.4 | 0.0960 | 3.5E-70 |
| ***scl-6*** | **WBGene00008028** | **protein_coding_gene** | **-2.6** | **0.1597** | **6.1E-12** | **2.9** | **7.5343** | **8.6E-08** |
| *F16B4.4* | WBGene00017506 | protein_coding_gene | -2.6 | 0.1597 | 7.6E-16 | -3.0 | 0.1230 | 1.5E-71 |
| *tbc-7* | WBGene00016292 | protein_coding_gene | -2.6 | 0.1597 | 4.9E-25 | -1.9 | 0.2692 | 7.2E-27 |
| *nlp-29* | WBGene00003767 | protein_coding_gene | -2.6 | 0.1597 | 8.4E-08 | -5.8 | 0.0176 | 4.3E-125 |

| | | | | | | | | |
|---|---|---|---|---|---|---|---|---|
| *prmn-1* | WBGene00008571 | protein_coding_gene | -2.7 | 0.1589 | 2.5E-24 | -2.1 | 0.2396 | 3.5E-21 |
| *T22C8.6* | WBGene00011927 | protein_coding_gene | -2.7 | 0.1588 | 1.7E-07 | -3.2 | 0.1070 | 4.4E-17 |
| *gad-3* | WBGene00015057 | protein_coding_gene | -2.7 | 0.1586 | 1.2E-25 | -2.8 | 0.1441 | 1.2E-110 |
| *nmur-1* | WBGene00016747 | protein_coding_gene | -2.7 | 0.1584 | 2.8E-03 | -1.5 | 0.3630 | 2.2E-02 |
| *npr-28* | WBGene00018886 | protein_coding_gene | -2.7 | 0.1584 | 2.1E-14 | -1.6 | 0.3210 | 4.1E-09 |
| *C11G10.1* | WBGene00007525 | protein_coding_gene | -2.7 | 0.1584 | 4.3E-07 | -4.4 | 0.0486 | 3.7E-23 |
| *C12D5.3* | WBGene00015706 | protein_coding_gene | -2.7 | 0.1583 | 5.0E-03 | -1.9 | 0.2719 | 1.0E-04 |
| *lips-10* | WBGene00008803 | protein_coding_gene | -2.7 | 0.1582 | 4.2E-14 | -4.3 | 0.0493 | 1.2E-55 |
| *F47G3.1* | WBGene00018576 | protein_coding_gene | -2.7 | 0.1579 | 2.9E-19 | -2.0 | 0.2559 | 7.3E-10 |
| *C08F11.3* | WBGene00007452 | protein_coding_gene | -2.7 | 0.1577 | 3.2E-04 | -3.9 | 0.0650 | 1.6E-10 |
| *F53F8.4* | WBGene00010001 | protein_coding_gene | -2.7 | 0.1577 | 1.0E-15 | -4.0 | 0.0638 | 1.3E-58 |
| *chhy-1* | WBGene00011923 | protein_coding_gene | -2.7 | 0.1577 | 4.8E-28 | -1.8 | 0.2807 | 2.9E-14 |
| *gly-6* | WBGene00001631 | protein_coding_gene | -2.7 | 0.1576 | 8.5E-20 | -2.4 | 0.1881 | 1.4E-60 |
| *mgl-2* | WBGene00003233 | protein_coding_gene | -2.7 | 0.1576 | 2.0E-10 | -2.0 | 0.2483 | 1.2E-10 |
| *mec-12* | WBGene00003175 | protein_coding_gene | -2.7 | 0.1572 | 2.0E-30 | -1.5 | 0.3555 | 4.1E-16 |
| *chil-13* | WBGene00010945 | protein_coding_gene | -2.7 | 0.1567 | 1.7E-27 | -3.1 | 0.1150 | 2.0E-31 |
| *oac-33* | WBGene00009972 | pseudogene | -2.7 | 0.1561 | 5.0E-04 | -2.0 | 0.2463 | 9.7E-04 |
| *C08B6.2* | WBGene00007423 | protein_coding_gene | -2.7 | 0.1560 | 2.5E-03 | -2.8 | 0.1443 | 7.4E-06 |
| *R102.2* | WBGene00011289 | protein_coding_gene | -2.7 | 0.1560 | 8.5E-25 | -1.2 | 0.4239 | 2.1E-14 |
| *pat-2* | WBGene00003929 | protein_coding_gene | -2.7 | 0.1559 | 2.8E-22 | -1.9 | 0.2711 | 7.7E-41 |
| *unc-95* | WBGene00006824 | protein_coding_gene | -2.7 | 0.1558 | 5.3E-25 | -1.1 | 0.4541 | 1.9E-05 |
| *mlc-3* | WBGene00003371 | protein_coding_gene | -2.7 | 0.1557 | 4.2E-30 | -1.8 | 0.2966 | 2.2E-52 |
| *F49E2.1* | WBGene00009885 | protein_coding_gene | -2.7 | 0.1556 | 1.4E-31 | -1.3 | 0.3983 | 1.7E-09 |
| *Y37E11B.7* | WBGene00021379 | protein_coding_gene | -2.7 | 0.1556 | 5.9E-06 | -2.5 | 0.1822 | 2.9E-07 |
| *lin-31* | WBGene00003017 | protein_coding_gene | -2.7 | 0.1555 | 9.7E-07 | -2.6 | 0.1652 | 1.2E-03 |
| *cpin-1* | WBGene00000775 | protein_coding_gene | -2.7 | 0.1554 | 9.4E-27 | -2.3 | 0.2059 | 6.8E-17 |
| *C49F5.7* | WBGene00008211 | protein_coding_gene | -2.7 | 0.1548 | 1.6E-18 | -4.3 | 0.0500 | 7.2E-116 |
| *cyp-13A7* | WBGene00000372 | protein_coding_gene | -2.7 | 0.1547 | 3.2E-03 | -5.2 | 0.0269 | 7.8E-25 |
| *F38B2.4* | WBGene00009531 | protein_coding_gene | -2.7 | 0.1546 | 6.9E-29 | -1.5 | 0.3502 | 1.7E-19 |
| *C29F7.3* | WBGene00007812 | protein_coding_gene | -2.7 | 0.1545 | 2.0E-28 | -1.7 | 0.3042 | 3.3E-28 |
| *K07A1.13* | WBGene00010617 | protein_coding_gene | -2.7 | 0.1542 | 8.6E-29 | -2.7 | 0.1584 | 1.1E-74 |
| *drd-1* | WBGene00009902 | protein_coding_gene | -2.7 | 0.1541 | 7.8E-13 | -1.8 | 0.2807 | 5.8E-26 |
| *acp-1* | WBGene00022770 | protein_coding_gene | -2.7 | 0.1540 | 7.6E-19 | -1.1 | 0.4640 | 2.7E-04 |
| *K09A9.6* | WBGene00010703 | protein_coding_gene | -2.7 | 0.1534 | 3.3E-17 | -1.3 | 0.4145 | 8.3E-10 |
| *K02E7.11* | WBGene00019315 | protein_coding_gene | -2.7 | 0.1533 | 2.0E-03 | -1.8 | 0.2795 | 8.2E-03 |
| *lron-8* | WBGene00020693 | protein_coding_gene | -2.7 | 0.1531 | 1.5E-27 | -1.5 | 0.3418 | 9.1E-11 |
| *mct-6* | WBGene00015676 | protein_coding_gene | -2.7 | 0.1528 | 5.4E-36 | -1.5 | 0.3585 | 2.5E-32 |
| *F59A7.2* | WBGene00019090 | protein_coding_gene | -2.7 | 0.1524 | 3.1E-05 | -3.4 | 0.0969 | 9.5E-28 |
| *nlp-18* | WBGene00003756 | protein_coding_gene | -2.7 | 0.1521 | 1.3E-18 | -1.4 | 0.3891 | 2.7E-11 |
| *Y7A5A.1* | WBGene00012407 | protein_coding_gene | -2.7 | 0.1519 | 2.9E-27 | -1.1 | 0.4762 | 1.3E-08 |
| *nlp-47* | WBGene00018117 | protein_coding_gene | -2.7 | 0.1518 | 1.3E-16 | -1.4 | 0.3887 | 3.0E-09 |
| *K02E11.7* | WBGene00010516 | protein_coding_gene | -2.7 | 0.1518 | 3.6E-15 | -1.6 | 0.3247 | 2.6E-05 |
| *F13D2.1* | WBGene00008735 | protein_coding_gene | -2.7 | 0.1517 | 5.9E-24 | -1.7 | 0.3124 | 3.4E-20 |
| *R01H2.7* | WBGene00023414 | protein_coding_gene | -2.7 | 0.1514 | 6.7E-18 | -2.3 | 0.2072 | 4.7E-16 |
| *W08F4.5* | WBGene00021094 | protein_coding_gene | -2.7 | 0.1513 | 6.8E-06 | -2.4 | 0.1954 | 2.2E-03 |
| *kmo-1* | WBGene00011089 | protein_coding_gene | -2.7 | 0.1513 | 2.6E-25 | -1.7 | 0.3184 | 1.4E-18 |
| *T15B7.1* | WBGene00020516 | protein_coding_gene | -2.7 | 0.1512 | 3.7E-21 | -1.8 | 0.2897 | 1.9E-07 |
| *Y32F6A.5* | WBGene00012530 | protein_coding_gene | -2.7 | 0.1512 | 7.5E-43 | -1.3 | 0.4091 | 9.1E-26 |
| *F26F12.5* | WBGene00017836 | protein_coding_gene | -2.7 | 0.1510 | 3.7E-06 | -2.8 | 0.1469 | 1.3E-06 |
| *ser-7* | WBGene00004780 | protein_coding_gene | -2.7 | 0.1508 | 4.7E-09 | -2.3 | 0.2081 | 2.7E-08 |
| *B0403.5* | WBGene00015169 | protein_coding_gene | -2.7 | 0.1508 | 4.1E-06 | -1.6 | 0.3253 | 3.6E-04 |
| *nekl-3* | WBGene00008956 | protein_coding_gene | -2.7 | 0.1507 | 5.3E-16 | -2.7 | 0.1544 | 2.3E-25 |
| *T19D12.9* | WBGene00020583 | protein_coding_gene | -2.7 | 0.1507 | 5.1E-05 | -2.0 | 0.2573 | 1.5E-02 |
| *glb-20* | WBGene00010970 | protein_coding_gene | -2.7 | 0.1506 | 1.4E-09 | -2.0 | 0.2416 | 2.2E-07 |
| *C14A11.6* | WBGene00015751 | protein_coding_gene | -2.7 | 0.1506 | 6.1E-12 | -1.7 | 0.3010 | 3.1E-04 |
| *ZC449.8* | WBGene00195189 | protein_coding_gene | -2.7 | 0.1505 | 1.8E-16 | -1.6 | 0.3395 | 5.4E-04 |
| *T23B7.3* | WBGene00044767 | protein_coding_gene | -2.7 | 0.1502 | 1.1E-07 | -1.0 | 0.4875 | 1.9E-02 |
| *srg-34* | WBGene00005191 | protein_coding_gene | -2.7 | 0.1499 | 8.8E-05 | -3.8 | 0.0695 | 6.4E-10 |
| *pbo-6* | WBGene00017375 | protein_coding_gene | -2.7 | 0.1496 | 1.1E-08 | -2.0 | 0.2508 | 4.1E-06 |
| *mltn-10* | WBGene00008965 | protein_coding_gene | -2.7 | 0.1493 | 4.3E-05 | -2.9 | 0.1366 | 1.7E-08 |
| *T11G6.3* | WBGene00011718 | protein_coding_gene | -2.7 | 0.1493 | 2.7E-11 | -2.7 | 0.1565 | 4.5E-20 |
| *vit-6* | WBGene00006930 | protein_coding_gene | -2.7 | 0.1491 | 6.5E-14 | -2.4 | 0.1862 | 5.2E-22 |
| *cpn-3* | WBGene00000779 | protein_coding_gene | -2.7 | 0.1490 | 5.3E-32 | -1.8 | 0.2799 | 4.4E-43 |
| *C09D4.1* | WBGene00015632 | protein_coding_gene | -2.7 | 0.1490 | 2.9E-33 | -2.6 | 0.1669 | 3.5E-61 |
| *cth-2* | WBGene00022856 | protein_coding_gene | -2.7 | 0.1490 | 1.6E-18 | -1.2 | 0.4265 | 5.7E-37 |
| *dpyd-1* | WBGene00016103 | protein_coding_gene | -2.7 | 0.1489 | 8.9E-25 | -1.4 | 0.3766 | 2.4E-33 |
| *ttr-46* | WBGene00011571 | protein_coding_gene | -2.7 | 0.1489 | 8.2E-35 | -1.9 | 0.2675 | 2.0E-65 |
| *mlt-3* | WBGene00012704 | protein_coding_gene | -2.7 | 0.1488 | 9.7E-19 | -2.8 | 0.1426 | 1.1E-18 |
| *lev-11* | WBGene00002978 | protein_coding_gene | -2.7 | 0.1487 | 2.2E-34 | -2.0 | 0.2450 | 1.9E-73 |
| *pkg-2* | WBGene00015650 | protein_coding_gene | -2.8 | 0.1486 | 6.5E-33 | -1.6 | 0.3387 | 2.1E-31 |

| | | | | | | | | |
|---|---|---|---|---|---|---|---|---|
| *fip-6* | WBGene00009964 | protein_coding_gene | -2.8 | 0.1483 | 2.2E-09 | -3.0 | 0.1259 | 3.2E-08 |
| *C10A4.4* | WBGene00015665 | protein_coding_gene | -2.8 | 0.1482 | 1.6E-08 | -1.5 | 0.3557 | 1.1E-02 |
| *C33A12.19* | WBGene00007890 | protein_coding_gene | -2.8 | 0.1477 | 3.3E-27 | -3.6 | 0.0806 | 4.1E-43 |
| *F22F4.4* | WBGene00017717 | protein_coding_gene | -2.8 | 0.1477 | 3.3E-55 | -1.4 | 0.3708 | 5.6E-25 |
| *ins-29* | WBGene00002112 | protein_coding_gene | -2.8 | 0.1475 | 4.9E-04 | -2.5 | 0.1753 | 1.8E-05 |
| *F17C11.6* | WBGene00008917 | protein_coding_gene | -2.8 | 0.1475 | 1.5E-19 | -1.8 | 0.2899 | 6.0E-12 |
| *Y45F10D.6* | WBGene00012886 | protein_coding_gene | -2.8 | 0.1473 | 2.9E-06 | -5.1 | 0.0292 | 1.3E-21 |
| *tag-290* | WBGene00009386 | protein_coding_gene | -2.8 | 0.1473 | 4.2E-19 | -1.5 | 0.3467 | 6.7E-14 |
| *vwa-8* | WBGene00008694 | protein_coding_gene | -2.8 | 0.1468 | 1.1E-69 | -1.5 | 0.3498 | 2.1E-21 |
| *ugt-11* | WBGene00020593 | protein_coding_gene | -2.8 | 0.1466 | 2.9E-18 | -2.2 | 0.2204 | 5.0E-15 |
| *T26C5.4* | WBGene00012034 | protein_coding_gene | -2.8 | 0.1465 | 1.1E-09 | -2.4 | 0.1929 | 2.4E-11 |
| *F54C1.1* | WBGene00018789 | protein_coding_gene | -2.8 | 0.1464 | 1.3E-10 | -1.1 | 0.4683 | 2.2E-02 |
| *ncx-2* | WBGene00003567 | protein_coding_gene | -2.8 | 0.1462 | 1.1E-15 | -1.8 | 0.2834 | 1.3E-33 |
| *T21B6.3* | WBGene00011880 | protein_coding_gene | -2.8 | 0.1457 | 1.2E-17 | -1.6 | 0.3263 | 3.2E-20 |
| *W04G3.13* | WBGene00194793 | protein_coding_gene | -2.8 | 0.1456 | 1.2E-04 | -4.2 | 0.0562 | 2.2E-05 |
| *faah-6* | WBGene00012750 | protein_coding_gene | -2.8 | 0.1456 | 1.2E-11 | -3.7 | 0.0751 | 4.4E-14 |
| *C05G5.1* | WBGene00007347 | protein_coding_gene | -2.8 | 0.1456 | 5.0E-30 | -1.9 | 0.2691 | 2.0E-19 |
| *ttr-30* | WBGene00020335 | protein_coding_gene | -2.8 | 0.1455 | 2.7E-19 | -4.1 | 0.0602 | 1.2E-29 |
| *Y4C6B.3* | WBGene00021157 | protein_coding_gene | -2.8 | 0.1454 | 6.9E-15 | -2.2 | 0.2179 | 3.9E-15 |
| *swt-3* | WBGene00007384 | protein_coding_gene | -2.8 | 0.1452 | 2.1E-14 | -2.6 | 0.1668 | 6.1E-31 |
| *F49C12.14* | WBGene00009883 | protein_coding_gene | -2.8 | 0.1451 | 3.6E-18 | -1.7 | 0.3165 | 1.8E-07 |
| *oig-2* | WBGene00003860 | protein_coding_gene | -2.8 | 0.1451 | 3.3E-15 | -1.8 | 0.2777 | 2.8E-18 |
| *npr-10* | WBGene00008278 | protein_coding_gene | -2.8 | 0.1450 | 1.2E-08 | -1.3 | 0.3971 | 1.0E-03 |
| *F13E9.14* | WBGene00045207 | protein_coding_gene | -2.8 | 0.1449 | 7.4E-03 | -2.1 | 0.2357 | 2.8E-03 |
| *ZC21.3* | WBGene00022505 | protein_coding_gene | -2.8 | 0.1446 | 6.1E-09 | -1.4 | 0.3922 | 9.4E-20 |
| *nlp-28* | WBGene00003766 | protein_coding_gene | -2.8 | 0.1446 | 1.2E-12 | -5.0 | 0.0309 | 1.2E-135 |
| *gtl-2* | WBGene00001796 | protein_coding_gene | -2.8 | 0.1444 | 2.0E-20 | -2.3 | 0.1976 | 3.1E-15 |
| *T01D1.4* | WBGene00020149 | protein_coding_gene | -2.8 | 0.1441 | 7.5E-15 | -1.7 | 0.3145 | 2.4E-12 |
| *W06A7.4* | WBGene00012293 | protein_coding_gene | -2.8 | 0.1441 | 4.1E-16 | -1.9 | 0.2659 | 1.1E-07 |
| *best-7* | WBGene00007808 | protein_coding_gene | -2.8 | 0.1440 | 1.4E-52 | -2.1 | 0.2303 | 4.0E-19 |
| *ipp-5* | WBGene00002146 | protein_coding_gene | -2.8 | 0.1439 | 8.5E-24 | -2.4 | 0.1833 | 6.0E-27 |
| *dim-1* | WBGene00001000 | protein_coding_gene | -2.8 | 0.1439 | 2.2E-42 | -1.2 | 0.4228 | 1.3E-25 |
| *ncx-1* | WBGene00003566 | protein_coding_gene | -2.8 | 0.1436 | 1.4E-43 | -2.3 | 0.2051 | 9.1E-61 |
| *ndg-4* | WBGene00003577 | protein_coding_gene | -2.8 | 0.1436 | 1.3E-21 | -2.1 | 0.2268 | 1.1E-48 |
| *T10B10.4* | WBGene00011681 | protein_coding_gene | -2.8 | 0.1436 | 1.3E-34 | -1.1 | 0.4799 | 1.6E-07 |
| *C08B6.10* | WBGene00007430 | protein_coding_gene | -2.8 | 0.1429 | 1.2E-12 | -2.1 | 0.2376 | 2.7E-08 |
| *cpr-6* | WBGene00000786 | protein_coding_gene | -2.8 | 0.1426 | 1.4E-39 | -2.1 | 0.2382 | 1.9E-60 |
| *F11E6.4* | WBGene00008708 | protein_coding_gene | -2.8 | 0.1425 | 1.2E-05 | -2.9 | 0.1330 | 1.6E-05 |
| *aat-2* | WBGene00000003 | protein_coding_gene | -2.8 | 0.1424 | 3.4E-18 | -1.8 | 0.2856 | 6.1E-13 |
| *M04D8.7* | WBGene00010865 | protein_coding_gene | -2.8 | 0.1422 | 2.1E-06 | -1.2 | 0.4493 | 3.9E-03 |
| *lfi-1* | WBGene00022500 | protein_coding_gene | -2.8 | 0.1418 | 6.7E-24 | -1.8 | 0.2786 | 7.7E-39 |
| *vit-2* | WBGene00006926 | protein_coding_gene | -2.8 | 0.1417 | 1.0E-18 | -2.9 | 0.1318 | 1.7E-78 |
| *srp-3* | WBGene00005644 | protein_coding_gene | -2.8 | 0.1414 | 5.2E-09 | -1.2 | 0.4225 | 1.3E-05 |
| *gln-3* | WBGene00001604 | protein_coding_gene | -2.8 | 0.1414 | 3.5E-16 | -2.6 | 0.1671 | 3.9E-73 |
| *clec-146* | WBGene00013008 | protein_coding_gene | -2.8 | 0.1413 | 4.3E-21 | -3.7 | 0.0766 | 1.3E-52 |
| *T19H12.15* | WBGene00271827 | protein_coding_gene | -2.8 | 0.1413 | 3.6E-10 | -1.8 | 0.2909 | 1.6E-07 |
| *F41C6.7* | WBGene00018280 | protein_coding_gene | -2.8 | 0.1411 | 3.0E-13 | -2.0 | 0.2509 | 4.8E-08 |
| *dop-6* | WBGene00016037 | protein_coding_gene | -2.8 | 0.1410 | 9.7E-04 | -2.4 | 0.1923 | 4.1E-04 |
| *H35B03.1* | WBGene00019263 | protein_coding_gene | -2.8 | 0.1409 | 1.1E-35 | -2.8 | 0.1450 | 3.0E-33 |
| *unc-15* | WBGene00006754 | protein_coding_gene | -2.8 | 0.1408 | 4.4E-19 | -2.1 | 0.2265 | 4.0E-70 |
| *C55C3.1* | WBGene00016951 | protein_coding_gene | -2.8 | 0.1406 | 5.1E-11 | -1.9 | 0.2763 | 5.6E-04 |
| *C30F12.5* | WBGene00016264 | protein_coding_gene | -2.8 | 0.1404 | 2.1E-34 | -2.5 | 0.1733 | 5.1E-28 |
| *ptr-23* | WBGene00004237 | protein_coding_gene | -2.8 | 0.1398 | 2.8E-30 | -2.2 | 0.2230 | 2.0E-48 |
| *F37B12.1* | WBGene00009504 | protein_coding_gene | -2.8 | 0.1396 | 1.8E-09 | -1.3 | 0.4118 | 1.2E-03 |
| *F44B9.2* | WBGene00018407 | protein_coding_gene | -2.8 | 0.1396 | 1.9E-17 | -1.0 | 0.4997 | 1.4E-06 |
| *K10C2.12* | WBGene00206386 | protein_coding_gene | -2.8 | 0.1394 | 2.9E-06 | -2.0 | 0.2519 | 2.9E-03 |
| *R13H4.2* | WBGene00014826 | protein_coding_gene | -2.8 | 0.1394 | 7.8E-33 | -2.6 | 0.1629 | 7.0E-46 |
| *pgp-14* | WBGene00004008 | protein_coding_gene | -2.8 | 0.1392 | 1.4E-12 | -1.5 | 0.3633 | 1.1E-15 |
| *F16C3.3* | WBGene00023415 | protein_coding_gene | -2.8 | 0.1389 | 2.4E-14 | -2.7 | 0.1518 | 1.7E-17 |
| *grd-1* | WBGene00001690 | protein_coding_gene | -2.8 | 0.1389 | 5.1E-11 | -1.5 | 0.3592 | 1.1E-02 |
| *osm-7* | WBGene00003887 | protein_coding_gene | -2.8 | 0.1388 | 1.2E-22 | -2.7 | 0.1530 | 2.4E-40 |
| *Y54E10A.17* | WBGene00021835 | protein_coding_gene | -2.8 | 0.1388 | 6.7E-16 | -1.4 | 0.3902 | 8.2E-12 |
| *F22F4.5* | WBGene00017718 | protein_coding_gene | -2.9 | 0.1385 | 6.2E-12 | -1.5 | 0.3433 | 7.8E-03 |
| *aman-1* | WBGene00018877 | protein_coding_gene | -2.9 | 0.1383 | 1.0E-19 | -2.1 | 0.2281 | 7.1E-18 |
| *F40A3.6* | WBGene00018221 | protein_coding_gene | -2.9 | 0.1380 | 3.1E-22 | -1.4 | 0.3909 | 5.7E-10 |
| *ssu-1* | WBGene00013748 | protein_coding_gene | -2.9 | 0.1379 | 1.8E-04 | -1.6 | 0.3239 | 2.5E-02 |
| *F44D12.2* | WBGene00009679 | protein_coding_gene | -2.9 | 0.1379 | 2.5E-31 | -2.9 | 0.1325 | 2.9E-29 |
| *unc-87* | WBGene00006819 | protein_coding_gene | -2.9 | 0.1378 | 1.1E-47 | -1.9 | 0.2724 | 2.5E-55 |
| *hgo-1* | WBGene00001843 | protein_coding_gene | -2.9 | 0.1377 | 1.7E-32 | -1.7 | 0.3122 | 1.5E-23 |
| *unc-89* | WBGene00006820 | protein_coding_gene | -2.9 | 0.1370 | 3.1E-14 | -1.3 | 0.3972 | 8.1E-07 |

| | | | | | | | | |
|---|---|---|---|---|---|---|---|---|
| ***pho-1*** | **WBGene00004020** | **protein_coding_gene** | **-2.9** | **0.1365** | **1.4E-15** | **1.5** | **2.8306** | **2.4E-04** |
| *Y73B6BL.44* | WBGene00044350 | protein_coding_gene | -2.9 | 0.1361 | 2.5E-15 | -3.5 | 0.0891 | 7.1E-36 |
| *W01A8.6* | WBGene00012168 | protein_coding_gene | -2.9 | 0.1360 | 1.2E-14 | -1.9 | 0.2706 | 9.5E-08 |
| *srap-1* | WBGene00011522 | protein_coding_gene | -2.9 | 0.1360 | 8.1E-10 | -3.9 | 0.0648 | 5.7E-101 |
| *ZK1058.9* | WBGene00014207 | protein_coding_gene | -2.9 | 0.1359 | 3.0E-32 | -2.0 | 0.2491 | 1.7E-49 |
| *T27D12.1* | WBGene00012085 | protein_coding_gene | -2.9 | 0.1358 | 8.5E-45 | -2.6 | 0.1691 | 1.2E-47 |
| *col-135* | WBGene00000708 | protein_coding_gene | -2.9 | 0.1358 | 5.7E-13 | -1.1 | 0.4686 | 2.5E-09 |
| *acs-5* | WBGene00013575 | protein_coding_gene | -2.9 | 0.1356 | 6.2E-70 | -1.6 | 0.3228 | 2.0E-31 |
| *W06A7.2* | WBGene00012291 | protein_coding_gene | -2.9 | 0.1355 | 1.4E-18 | -2.1 | 0.2399 | 3.1E-18 |
| *mct-4* | WBGene00003986 | protein_coding_gene | -2.9 | 0.1353 | 9.2E-19 | -1.2 | 0.4360 | 5.9E-15 |
| *clec-48* | WBGene00007565 | protein_coding_gene | -2.9 | 0.1352 | 8.1E-47 | -2.6 | 0.1642 | 4.4E-82 |
| *twk-25* | WBGene00006678 | protein_coding_gene | -2.9 | 0.1348 | 2.2E-15 | -2.3 | 0.2080 | 1.3E-19 |
| *F38A6.4* | WBGene00009528 | protein_coding_gene | -2.9 | 0.1346 | 5.6E-06 | -2.0 | 0.2422 | 5.7E-03 |
| *cle-1* | WBGene00000527 | protein_coding_gene | -2.9 | 0.1344 | 1.8E-61 | -1.9 | 0.2689 | 6.7E-36 |
| *rpy-1* | WBGene00004507 | protein_coding_gene | -2.9 | 0.1343 | 1.6E-25 | -2.3 | 0.2025 | 3.5E-18 |
| *F43D2.6* | WBGene00044191 | protein_coding_gene | -2.9 | 0.1341 | 2.5E-38 | -1.3 | 0.4020 | 9.5E-11 |
| *tag-18* | WBGene00006408 | protein_coding_gene | -2.9 | 0.1340 | 4.3E-46 | -1.6 | 0.3354 | 5.6E-25 |
| *twk-13* | WBGene00006668 | protein_coding_gene | -2.9 | 0.1339 | 2.2E-13 | -1.0 | 0.4854 | 2.8E-03 |
| *cup-4* | WBGene00000845 | protein_coding_gene | -2.9 | 0.1335 | 2.1E-13 | -2.0 | 0.2578 | 8.9E-10 |
| *C37C3.7* | WBGene00016499 | protein_coding_gene | -2.9 | 0.1334 | 1.1E-13 | -2.8 | 0.1388 | 3.1E-14 |
| *srp-1* | WBGene00005642 | protein_coding_gene | -2.9 | 0.1333 | 8.3E-30 | -2.5 | 0.1715 | 2.5E-24 |
| *R05D3.9* | WBGene00019882 | protein_coding_gene | -2.9 | 0.1330 | 1.7E-54 | -2.1 | 0.2369 | 3.1E-29 |
| *ZK809.8* | WBGene00014088 | protein_coding_gene | -2.9 | 0.1329 | 6.9E-30 | -2.1 | 0.2413 | 4.4E-51 |
| *C04F12.8* | WBGene00007302 | protein_coding_gene | -2.9 | 0.1328 | 1.5E-14 | -2.3 | 0.2085 | 1.5E-19 |
| *dcar-1* | WBGene00007395 | protein_coding_gene | -2.9 | 0.1325 | 4.7E-16 | -3.5 | 0.0876 | 1.3E-100 |
| *afmd-1* | WBGene00017051 | protein_coding_gene | -2.9 | 0.1325 | 5.7E-18 | -2.2 | 0.2160 | 1.5E-06 |
| *K02G10.5* | WBGene00019346 | protein_coding_gene | -2.9 | 0.1324 | 8.4E-10 | -2.0 | 0.2525 | 1.1E-06 |
| *cyp-14A3* | WBGene00010707 | protein_coding_gene | -2.9 | 0.1318 | 4.6E-03 | -4.5 | 0.0435 | 2.0E-11 |
| *suro-1* | WBGene00011235 | protein_coding_gene | -2.9 | 0.1318 | 7.5E-10 | -1.2 | 0.4396 | 1.4E-02 |
| *abts-2* | WBGene00009929 | protein_coding_gene | -2.9 | 0.1316 | 6.6E-15 | -2.7 | 0.1494 | 1.6E-13 |
| *test-1* | WBGene00016918 | protein_coding_gene | -2.9 | 0.1314 | 3.6E-27 | -1.5 | 0.3583 | 4.1E-32 |
| *F39H12.3* | WBGene00018214 | protein_coding_gene | -2.9 | 0.1312 | 3.8E-30 | -2.1 | 0.2276 | 7.9E-34 |
| *twk-36* | WBGene00006688 | protein_coding_gene | -2.9 | 0.1310 | 5.6E-04 | -2.8 | 0.1429 | 4.5E-03 |
| *nhx-1* | WBGene00003729 | protein_coding_gene | -2.9 | 0.1308 | 7.7E-10 | -1.6 | 0.3382 | 1.7E-05 |
| *adt-1* | WBGene00000082 | protein_coding_gene | -2.9 | 0.1307 | 8.8E-08 | -1.3 | 0.3967 | 3.6E-03 |
| *Y71F9B.1* | WBGene00022125 | protein_coding_gene | -2.9 | 0.1306 | 2.9E-17 | -2.3 | 0.1970 | 7.1E-18 |
| *cyp-13A8* | WBGene00011674 | protein_coding_gene | -2.9 | 0.1306 | 1.9E-03 | -3.1 | 0.1193 | 1.7E-03 |
| *C46G7.2* | WBGene00016722 | protein_coding_gene | -2.9 | 0.1304 | 1.4E-66 | -2.2 | 0.2149 | 4.6E-61 |
| *C08E8.3* | WBGene00007439 | protein_coding_gene | -2.9 | 0.1300 | 5.2E-03 | -4.7 | 0.0379 | 4.9E-10 |
| *F01G10.9* | WBGene00008511 | protein_coding_gene | -2.9 | 0.1299 | 3.3E-12 | -1.7 | 0.3060 | 1.8E-03 |
| *dpf-1* | WBGene00001054 | protein_coding_gene | -2.9 | 0.1295 | 1.6E-19 | -1.9 | 0.2649 | 1.6E-12 |
| *far-2* | WBGene00001386 | protein_coding_gene | -3.0 | 0.1294 | 4.4E-36 | -2.3 | 0.2010 | 1.9E-63 |
| *nhr-53* | WBGene00003643 | protein_coding_gene | -3.0 | 0.1293 | 8.3E-13 | -2.0 | 0.2476 | 2.5E-11 |
| *Y71F9B.13* | WBGene00022131 | protein_coding_gene | -3.0 | 0.1289 | 2.7E-35 | -3.6 | 0.0805 | 9.1E-53 |
| *K11H12.7* | WBGene00019663 | protein_coding_gene | -3.0 | 0.1288 | 5.8E-31 | -1.7 | 0.3037 | 3.5E-23 |
| *npr-13* | WBGene00013883 | protein_coding_gene | -3.0 | 0.1288 | 7.3E-06 | -2.5 | 0.1796 | 3.4E-03 |
| *C17G10.7* | WBGene00015919 | protein_coding_gene | -3.0 | 0.1287 | 3.8E-11 | -2.1 | 0.2393 | 5.1E-06 |
| *T01B4.3* | WBGene00011310 | protein_coding_gene | -3.0 | 0.1287 | 3.7E-20 | -1.7 | 0.3150 | 5.9E-10 |
| *cest-31* | WBGene00019653 | protein_coding_gene | -3.0 | 0.1287 | 4.6E-17 | -1.8 | 0.2895 | 8.4E-09 |
| *amt-4* | WBGene00000136 | protein_coding_gene | -3.0 | 0.1285 | 4.5E-18 | -1.7 | 0.3029 | 1.1E-18 |
| *T23B12.11* | WBGene00044553 | protein_coding_gene | -3.0 | 0.1284 | 2.1E-20 | -2.4 | 0.1852 | 6.4E-30 |
| *F17C11.26* | WBGene00303011 | protein_coding_gene | -3.0 | 0.1282 | 4.5E-33 | -2.6 | 0.1689 | 8.2E-39 |
| *cest-9.1* | WBGene00007692 | protein_coding_gene | -3.0 | 0.1282 | 5.1E-23 | -1.2 | 0.4422 | 1.1E-05 |
| *T03G6.3* | WBGene00020195 | protein_coding_gene | -3.0 | 0.1279 | 2.6E-35 | -2.4 | 0.1832 | 6.3E-53 |
| *lgc-4* | WBGene00017580 | protein_coding_gene | -3.0 | 0.1279 | 8.3E-13 | -2.3 | 0.1986 | 2.1E-05 |
| *dpf-2* | WBGene00001055 | protein_coding_gene | -3.0 | 0.1279 | 1.0E-30 | -2.6 | 0.1634 | 1.4E-54 |
| *T01C8.2* | WBGene00020143 | protein_coding_gene | -3.0 | 0.1276 | 2.2E-17 | -2.0 | 0.2523 | 1.6E-20 |
| *chil-7* | WBGene00007472 | protein_coding_gene | -3.0 | 0.1276 | 2.0E-04 | -3.0 | 0.1224 | 3.6E-05 |
| *toh-1* | WBGene00006591 | protein_coding_gene | -3.0 | 0.1275 | 1.8E-33 | -1.9 | 0.2723 | 1.5E-14 |
| *hsp-12.3* | WBGene00002012 | protein_coding_gene | -3.0 | 0.1272 | 6.6E-18 | -2.5 | 0.1763 | 2.0E-20 |
| *twk-12* | WBGene00006667 | protein_coding_gene | -3.0 | 0.1272 | 1.4E-14 | -1.3 | 0.4067 | 2.2E-03 |
| *far-1* | WBGene00001385 | protein_coding_gene | -3.0 | 0.1271 | 1.2E-42 | -2.2 | 0.2213 | 3.4E-66 |
| *fbxa-27* | WBGene00022333 | protein_coding_gene | -3.0 | 0.1271 | 9.9E-12 | -2.2 | 0.2199 | 3.8E-04 |
| *csq-1* | WBGene00000822 | protein_coding_gene | -3.0 | 0.1271 | 1.1E-47 | -2.5 | 0.1745 | 1.1E-94 |
| *F23F1.7* | WBGene00017748 | protein_coding_gene | -3.0 | 0.1263 | 7.3E-24 | -1.9 | 0.2656 | 7.2E-09 |
| *dpf-6* | WBGene00001059 | protein_coding_gene | -3.0 | 0.1260 | 1.4E-20 | -1.3 | 0.4119 | 2.9E-04 |
| *F58B4.5* | WBGene00010238 | protein_coding_gene | -3.0 | 0.1259 | 8.0E-31 | -1.6 | 0.3310 | 2.1E-16 |
| *mrp-6* | WBGene00003412 | protein_coding_gene | -3.0 | 0.1258 | 1.7E-25 | -2.4 | 0.1849 | 9.9E-24 |
| *B0034.1* | WBGene00015002 | protein_coding_gene | -3.0 | 0.1256 | 2.6E-11 | -1.4 | 0.3745 | 8.4E-04 |
| *R11A5.6* | WBGene00011234 | protein_coding_gene | -3.0 | 0.1256 | 6.9E-05 | -1.3 | 0.4125 | 4.9E-02 |

| | | | | | | | | |
|---|---|---|---|---|---|---|---|---|
| *fasn-1* | WBGene00009342 | protein_coding_gene | -3.0 | 0.1255 | 1.4E-23 | -2.2 | 0.2183 | 4.9E-16 |
| *F48G7.10* | WBGene00018621 | protein_coding_gene | -3.0 | 0.1254 | 2.0E-03 | -1.6 | 0.3258 | 4.8E-02 |
| *cnc-2* | WBGene00000556 | protein_coding_gene | -3.0 | 0.1253 | 3.7E-03 | -5.6 | 0.0199 | 8.5E-10 |
| *ost-1* | WBGene00003893 | protein_coding_gene | -3.0 | 0.1251 | 4.9E-30 | -1.7 | 0.3169 | 1.0E-49 |
| *pck-1* | WBGene00021043 | protein_coding_gene | -3.0 | 0.1250 | 2.3E-37 | -2.6 | 0.1685 | 4.5E-82 |
| *clec-190* | WBGene00010953 | protein_coding_gene | -3.0 | 0.1247 | 1.7E-05 | -2.4 | 0.1929 | 7.9E-17 |
| *zmp-4* | WBGene00012364 | protein_coding_gene | -3.0 | 0.1245 | 5.2E-52 | -2.8 | 0.1418 | 2.2E-87 |
| *amx-2* | WBGene00000138 | protein_coding_gene | -3.0 | 0.1244 | 2.6E-32 | -1.9 | 0.2675 | 3.7E-22 |
| *Y42H9AR.5* | WBGene00021540 | protein_coding_gene | -3.0 | 0.1242 | 5.0E-09 | -3.3 | 0.1037 | 6.0E-06 |
| *ZK909.6* | WBGene00014150 | protein_coding_gene | -3.0 | 0.1242 | 1.5E-21 | -1.7 | 0.3006 | 3.6E-09 |
| *T19D12.6* | WBGene00020581 | protein_coding_gene | -3.0 | 0.1241 | 5.4E-50 | -2.0 | 0.2522 | 2.2E-39 |
| *Y69H2.3* | WBGene00013481 | protein_coding_gene | -3.0 | 0.1240 | 8.6E-42 | -2.1 | 0.2393 | 1.6E-46 |
| *try-9* | WBGene00023425 | protein_coding_gene | -3.0 | 0.1236 | 1.8E-03 | -6.2 | 0.0141 | 1.6E-10 |
| *ugt-66* | WBGene00016013 | protein_coding_gene | -3.0 | 0.1235 | 9.8E-08 | -5.5 | 0.0218 | 3.1E-29 |
| *tsp-8* | WBGene00006634 | protein_coding_gene | -3.0 | 0.1234 | 4.6E-23 | -1.8 | 0.2805 | 1.6E-15 |
| *Y7A9A.1* | WBGene00012416 | protein_coding_gene | -3.0 | 0.1234 | 7.4E-19 | -2.1 | 0.2261 | 6.7E-11 |
| *C52B11.5* | WBGene00016874 | protein_coding_gene | -3.0 | 0.1230 | 3.0E-29 | -1.2 | 0.4443 | 2.0E-09 |
| *col-95* | WBGene00000670 | protein_coding_gene | -3.0 | 0.1229 | 4.7E-09 | -5.0 | 0.0320 | 1.3E-177 |
| *cpna-2* | WBGene00015061 | protein_coding_gene | -3.0 | 0.1229 | 2.9E-21 | -1.6 | 0.3263 | 7.5E-31 |
| *F40A3.2* | WBGene00018217 | protein_coding_gene | -3.0 | 0.1228 | 5.4E-33 | -2.4 | 0.1944 | 5.9E-32 |
| *T12A2.1* | WBGene00020436 | protein_coding_gene | -3.0 | 0.1227 | 6.7E-30 | -2.2 | 0.2214 | 9.1E-15 |
| ***daf-28*** | **WBGene00000920** | **protein_coding_gene** | **-3.0** | **0.1226** | **1.5E-11** | **1.4** | **2.5685** | **1.1E-04** |
| *T03F6.10* | WBGene00185005 | protein_coding_gene | -3.0 | 0.1226 | 1.3E-07 | -1.8 | 0.2900 | 8.1E-05 |
| *B0507.8* | WBGene00015225 | protein_coding_gene | -3.0 | 0.1225 | 1.1E-05 | -2.0 | 0.2503 | 2.8E-03 |
| *C17G1.2* | WBGene00007649 | protein_coding_gene | -3.0 | 0.1225 | 1.3E-30 | -3.1 | 0.1202 | 6.4E-66 |
| *F35H10.10* | WBGene00018073 | protein_coding_gene | -3.0 | 0.1225 | 1.7E-32 | -1.6 | 0.3342 | 1.8E-12 |
| *ttr-44* | WBGene00008341 | protein_coding_gene | -3.0 | 0.1224 | 2.9E-23 | -4.0 | 0.0631 | 2.5E-31 |
| *K11D12.8* | WBGene00019648 | protein_coding_gene | -3.0 | 0.1222 | 5.7E-05 | -3.8 | 0.0738 | 5.3E-07 |
| *F28H1.4* | WBGene00017909 | protein_coding_gene | -3.0 | 0.1217 | 2.9E-06 | -2.0 | 0.2460 | 1.1E-03 |
| *pept-2* | WBGene00003876 | protein_coding_gene | -3.0 | 0.1215 | 4.3E-29 | -2.4 | 0.1900 | 1.5E-28 |
| *F48B9.1* | WBGene00018587 | protein_coding_gene | -3.0 | 0.1213 | 1.2E-06 | -3.3 | 0.0995 | 3.1E-09 |
| *C35B1.4* | WBGene00016434 | protein_coding_gene | -3.0 | 0.1213 | 5.1E-16 | -1.5 | 0.3640 | 1.9E-12 |
| *Y119D3B.13* | WBGene00022490 | protein_coding_gene | -3.0 | 0.1212 | 1.9E-48 | -1.2 | 0.4288 | 2.1E-09 |
| *F38G1.3* | WBGene00044556 | protein_coding_gene | -3.1 | 0.1206 | 2.5E-10 | -2.3 | 0.2073 | 1.1E-08 |
| *C31H2.14* | WBGene00206484 | protein_coding_gene | -3.1 | 0.1205 | 4.3E-10 | -1.9 | 0.2691 | 6.2E-05 |
| *C09B8.8* | WBGene00015625 | protein_coding_gene | -3.1 | 0.1202 | 3.6E-05 | -2.5 | 0.1733 | 4.2E-04 |
| *mca-2* | WBGene00019875 | protein_coding_gene | -3.1 | 0.1201 | 3.0E-40 | -2.7 | 0.1493 | 1.2E-74 |
| *alh-10* | WBGene00000116 | protein_coding_gene | -3.1 | 0.1200 | 3.2E-38 | -1.8 | 0.2783 | 5.9E-23 |
| *F41D9.2* | WBGene00018282 | protein_coding_gene | -3.1 | 0.1198 | 9.5E-10 | -1.4 | 0.3840 | 1.7E-05 |
| *ttr-32* | WBGene00020047 | protein_coding_gene | -3.1 | 0.1196 | 2.6E-20 | -3.3 | 0.1050 | 4.3E-27 |
| ***C36E6.8*** | **WBGene00044683** | **protein_coding_gene** | **-3.1** | **0.1192** | **1.5E-04** | **2.1** | **4.4349** | **2.0E-03** |
| *abch-1* | WBGene00016973 | protein_coding_gene | -3.1 | 0.1192 | 2.1E-03 | -1.7 | 0.3035 | 2.1E-02 |
| *lon-1* | WBGene00003055 | protein_coding_gene | -3.1 | 0.1192 | 7.9E-15 | -1.2 | 0.4330 | 1.5E-03 |
| *D2023.1* | WBGene00014300 | protein_coding_gene | -3.1 | 0.1191 | 1.1E-35 | -1.8 | 0.2908 | 2.5E-10 |
| *C45B2.8* | WBGene00016662 | protein_coding_gene | -3.1 | 0.1181 | 6.4E-03 | -2.8 | 0.1405 | 5.9E-03 |
| *F20D6.5* | WBGene00017635 | protein_coding_gene | -3.1 | 0.1180 | 1.2E-14 | -1.4 | 0.3828 | 1.9E-02 |
| *mua-3* | WBGene00003482 | protein_coding_gene | -3.1 | 0.1180 | 1.0E-16 | -1.3 | 0.4067 | 5.5E-04 |
| *ugt-9* | WBGene00020587 | protein_coding_gene | -3.1 | 0.1180 | 3.1E-17 | -5.0 | 0.0317 | 2.3E-42 |
| *cyp-29A3* | WBGene00021412 | protein_coding_gene | -3.1 | 0.1177 | 1.4E-12 | -3.0 | 0.1249 | 1.3E-13 |
| *zip-3* | WBGene00012330 | protein_coding_gene | -3.1 | 0.1177 | 1.0E-17 | -1.2 | 0.4235 | 1.0E-11 |
| *C44B7.5* | WBGene00016627 | protein_coding_gene | -3.1 | 0.1175 | 5.7E-25 | -3.5 | 0.0908 | 5.8E-132 |
| *W10G11.4* | WBGene00021137 | protein_coding_gene | -3.1 | 0.1174 | 6.2E-03 | -5.1 | 0.0286 | 1.2E-06 |
| *pah-1* | WBGene00000240 | protein_coding_gene | -3.1 | 0.1172 | 5.0E-61 | -2.1 | 0.2391 | 2.8E-58 |
| *daf-9* | WBGene00000905 | protein_coding_gene | -3.1 | 0.1171 | 1.6E-09 | -1.4 | 0.3880 | 1.3E-02 |
| *F46H5.3* | WBGene00018519 | protein_coding_gene | -3.1 | 0.1171 | 2.6E-64 | -2.5 | 0.1796 | 2.1E-83 |
| *chil-8* | WBGene00007473 | protein_coding_gene | -3.1 | 0.1169 | 9.6E-06 | -3.7 | 0.0783 | 4.4E-08 |
| *R07E4.1* | WBGene00019935 | protein_coding_gene | -3.1 | 0.1168 | 3.0E-35 | -1.2 | 0.4301 | 2.8E-08 |
| *ZC443.1* | WBGene00013896 | protein_coding_gene | -3.1 | 0.1168 | 2.7E-25 | -1.7 | 0.3049 | 5.0E-13 |
| *lact-6* | WBGene00007358 | protein_coding_gene | -3.1 | 0.1168 | 1.1E-03 | -2.9 | 0.1340 | 1.6E-04 |
| *K08B4.7* | WBGene00023486 | protein_coding_gene | -3.1 | 0.1167 | 6.9E-11 | -2.1 | 0.2272 | 6.7E-07 |
| *asic-1* | WBGene00022815 | protein_coding_gene | -3.1 | 0.1165 | 5.5E-06 | -1.3 | 0.3950 | 1.7E-02 |
| *F15G9.5* | WBGene00008868 | protein_coding_gene | -3.1 | 0.1165 | 2.3E-15 | -1.4 | 0.3706 | 5.8E-05 |
| *ttr-8* | WBGene00020049 | protein_coding_gene | -3.1 | 0.1163 | 1.3E-18 | -3.1 | 0.1145 | 1.2E-25 |
| *C25E10.12* | WBGene00016101 | protein_coding_gene | -3.1 | 0.1162 | 1.1E-41 | -2.6 | 0.1644 | 7.7E-37 |
| *col-166* | WBGene00000739 | protein_coding_gene | -3.1 | 0.1158 | 1.0E-19 | -2.8 | 0.1473 | 5.3E-23 |
| *Y47D7A.7* | WBGene00021620 | protein_coding_gene | -3.1 | 0.1157 | 4.4E-03 | -3.6 | 0.0819 | 2.5E-07 |
| *nhr-205* | WBGene00011002 | protein_coding_gene | -3.1 | 0.1156 | 1.2E-10 | -2.5 | 0.1752 | 1.6E-07 |
| *best-15* | WBGene00011258 | protein_coding_gene | -3.1 | 0.1153 | 1.3E-18 | -1.5 | 0.3597 | 4.7E-07 |
| *aldo-1* | WBGene00011474 | protein_coding_gene | -3.1 | 0.1149 | 2.0E-32 | -2.3 | 0.2031 | 3.1E-50 |
| *col-144* | WBGene00000717 | protein_coding_gene | -3.1 | 0.1148 | 4.6E-08 | -1.9 | 0.2615 | 5.5E-03 |

| | | | | | | | | |
|---|---|---|---|---|---|---|---|---|
| *zig-2* | WBGene00006979 | protein_coding_gene | -3.1 | 0.1145 | 2.1E-20 | -2.2 | 0.2110 | 5.7E-11 |
| *Y7A9A.79* | WBGene00185076 | ncRNA_gene | -3.1 | 0.1144 | 3.3E-06 | -2.1 | 0.2261 | 2.0E-03 |
| *folt-1* | WBGene00007388 | protein_coding_gene | -3.1 | 0.1143 | 2.7E-23 | -1.2 | 0.4446 | 1.0E-05 |
| *lbp-6* | WBGene00002258 | protein_coding_gene | -3.1 | 0.1142 | 5.5E-16 | -1.4 | 0.3666 | 2.6E-29 |
| *cat-4* | WBGene00000298 | protein_coding_gene | -3.1 | 0.1142 | 3.0E-23 | -1.3 | 0.3989 | 3.1E-26 |
| *ltah-1.2* | WBGene00022610 | protein_coding_gene | -3.1 | 0.1141 | 2.4E-26 | -1.3 | 0.4133 | 8.2E-19 |
| *zipt-2.2* | WBGene00017936 | protein_coding_gene | -3.1 | 0.1139 | 1.2E-20 | -1.6 | 0.3291 | 3.6E-08 |
| *F53F4.8* | WBGene00009991 | pseudogene | -3.1 | 0.1137 | 1.9E-04 | -4.1 | 0.0593 | 8.5E-18 |
| *W02H5.8* | WBGene00020962 | protein_coding_gene | -3.1 | 0.1136 | 1.2E-33 | -2.1 | 0.2272 | 5.3E-38 |
| *ges-1* | WBGene00001578 | protein_coding_gene | -3.1 | 0.1136 | 8.8E-34 | -2.8 | 0.1439 | 5.5E-78 |
| *W01F3.2* | WBGene00012185 | protein_coding_gene | -3.1 | 0.1134 | 5.0E-44 | -3.0 | 0.1276 | 2.0E-74 |
| *flp-34* | WBGene00019975 | protein_coding_gene | -3.1 | 0.1132 | 2.2E-09 | -4.3 | 0.0514 | 7.5E-28 |
| *clec-139* | WBGene00018050 | protein_coding_gene | -3.1 | 0.1132 | 8.8E-06 | -1.4 | 0.3899 | 2.1E-02 |
| *clec-7* | WBGene00017364 | protein_coding_gene | -3.1 | 0.1132 | 5.2E-10 | -5.1 | 0.0295 | 9.5E-67 |
| *gsnl-1* | WBGene00010593 | protein_coding_gene | -3.1 | 0.1130 | 5.4E-34 | -2.5 | 0.1738 | 3.0E-48 |
| *cth-1* | WBGene00009048 | protein_coding_gene | -3.1 | 0.1127 | 2.1E-09 | -1.3 | 0.4059 | 8.7E-07 |
| *twk-1* | WBGene00006656 | protein_coding_gene | -3.2 | 0.1125 | 1.6E-39 | -2.4 | 0.1868 | 6.2E-25 |
| *Y18D10A.23* | WBGene00012487 | protein_coding_gene | -3.2 | 0.1122 | 3.5E-40 | -2.5 | 0.1796 | 1.9E-27 |
| *ddo-3* | WBGene00017648 | protein_coding_gene | -3.2 | 0.1122 | 1.1E-20 | -3.6 | 0.0839 | 8.2E-170 |
| *H41C03.1* | WBGene00019268 | protein_coding_gene | -3.2 | 0.1121 | 3.1E-24 | -2.2 | 0.2163 | 5.1E-17 |
| *inx-9* | WBGene00002131 | protein_coding_gene | -3.2 | 0.1119 | 1.2E-33 | -3.5 | 0.0903 | 7.9E-81 |
| *F33C8.4* | WBGene00009358 | protein_coding_gene | -3.2 | 0.1116 | 1.1E-28 | -1.2 | 0.4310 | 4.2E-07 |
| *Y45F10C.4* | WBGene00012880 | protein_coding_gene | -3.2 | 0.1114 | 3.9E-16 | -1.6 | 0.3209 | 2.7E-12 |
| *let-805* | WBGene00002915 | protein_coding_gene | -3.2 | 0.1113 | 1.6E-15 | -1.6 | 0.3393 | 6.9E-21 |
| *D2092.4* | WBGene00017065 | protein_coding_gene | -3.2 | 0.1110 | 1.2E-30 | -1.9 | 0.2672 | 1.5E-34 |
| *nas-33* | WBGene00003551 | protein_coding_gene | -3.2 | 0.1110 | 3.0E-06 | -3.6 | 0.0812 | 3.0E-13 |
| *hex-5* | WBGene00013497 | protein_coding_gene | -3.2 | 0.1107 | 8.5E-19 | -2.1 | 0.2306 | 9.9E-13 |
| *oct-1* | WBGene00003842 | protein_coding_gene | -3.2 | 0.1097 | 4.0E-20 | -1.9 | 0.2758 | 3.8E-10 |
| *F47E1.2* | WBGene00018566 | protein_coding_gene | -3.2 | 0.1097 | 2.0E-17 | -2.9 | 0.1376 | 2.2E-14 |
| *pat-10* | WBGene00003934 | protein_coding_gene | -3.2 | 0.1095 | 4.2E-30 | -2.6 | 0.1628 | 2.9E-125 |
| *ttr-59* | WBGene00020339 | protein_coding_gene | -3.2 | 0.1093 | 5.4E-34 | -2.8 | 0.1448 | 1.0E-35 |
| *Y6B3B.1* | WBGene00012387 | protein_coding_gene | -3.2 | 0.1088 | 8.3E-14 | -2.0 | 0.2505 | 1.1E-05 |
| *gta-1* | WBGene00001794 | protein_coding_gene | -3.2 | 0.1085 | 7.0E-42 | -2.5 | 0.1782 | 3.5E-97 |
| *C41A3.2* | WBGene00016559 | protein_coding_gene | -3.2 | 0.1082 | 3.5E-30 | -1.4 | 0.3892 | 7.2E-09 |
| *F22E12.1* | WBGene00009058 | protein_coding_gene | -3.2 | 0.1082 | 1.3E-26 | -2.0 | 0.2517 | 3.3E-22 |
| *npa-1* | WBGene00003786 | protein_coding_gene | -3.2 | 0.1081 | 3.3E-41 | -3.0 | 0.1288 | 1.8E-129 |
| *sra-14* | WBGene00005040 | protein_coding_gene | -3.2 | 0.1080 | 3.1E-06 | -3.2 | 0.1106 | 6.5E-06 |
| *C14C6.2* | WBGene00015756 | protein_coding_gene | -3.2 | 0.1077 | 1.4E-39 | -5.3 | 0.0258 | 7.2E-37 |
| *cyp-32A1* | WBGene00016147 | protein_coding_gene | -3.2 | 0.1076 | 9.4E-18 | -4.7 | 0.0382 | 1.2E-43 |
| *ZK1321.4* | WBGene00014262 | protein_coding_gene | -3.2 | 0.1073 | 2.3E-45 | -1.6 | 0.3309 | 3.3E-22 |
| *C25E10.4* | WBGene00016094 | protein_coding_gene | -3.2 | 0.1072 | 3.3E-16 | -1.2 | 0.4363 | 1.3E-03 |
| *gnrr-6* | WBGene00008736 | protein_coding_gene | -3.2 | 0.1072 | 1.2E-04 | -1.4 | 0.3722 | 2.4E-02 |
| *argk-1* | WBGene00009706 | protein_coding_gene | -3.2 | 0.1071 | 1.0E-03 | -2.4 | 0.1871 | 1.5E-04 |
| *aqp-1* | WBGene00000169 | protein_coding_gene | -3.2 | 0.1071 | 9.7E-25 | -2.3 | 0.2016 | 1.6E-09 |
| *C27A7.1* | WBGene00007753 | protein_coding_gene | -3.2 | 0.1069 | 2.4E-31 | -3.3 | 0.1016 | 1.0E-78 |
| *F09G8.5* | WBGene00017320 | protein_coding_gene | -3.2 | 0.1067 | 1.9E-08 | -3.2 | 0.1069 | 2.1E-17 |
| *ZK673.1* | WBGene00014057 | protein_coding_gene | -3.2 | 0.1067 | 2.2E-34 | -2.0 | 0.2420 | 3.7E-22 |
| *F25E2.2* | WBGene00017780 | protein_coding_gene | -3.2 | 0.1066 | 2.7E-75 | -1.6 | 0.3257 | 4.4E-12 |
| *D1054.6* | WBGene00008373 | pseudogene | -3.2 | 0.1063 | 2.1E-03 | -1.9 | 0.2639 | 4.4E-02 |
| *ZK596.3* | WBGene00014008 | protein_coding_gene | -3.2 | 0.1063 | 2.1E-08 | -1.8 | 0.2924 | 1.1E-03 |
| *inx-8* | WBGene00002130 | protein_coding_gene | -3.2 | 0.1062 | 5.7E-41 | -3.5 | 0.0878 | 4.7E-192 |
| *ugt-6* | WBGene00013904 | protein_coding_gene | -3.2 | 0.1056 | 1.4E-13 | -1.4 | 0.3758 | 3.9E-07 |
| *F41B4.1* | WBGene00018257 | protein_coding_gene | -3.2 | 0.1054 | 4.9E-17 | -2.2 | 0.2163 | 3.7E-16 |
| *spp-4* | WBGene00004989 | protein_coding_gene | -3.2 | 0.1052 | 2.0E-19 | -2.1 | 0.2358 | 7.8E-05 |
| *F32D8.12* | WBGene00009334 | protein_coding_gene | -3.3 | 0.1050 | 2.1E-62 | -3.2 | 0.1080 | 2.8E-60 |
| *cyp-35A5* | WBGene00019473 | protein_coding_gene | -3.3 | 0.1049 | 6.4E-18 | -2.1 | 0.2368 | 2.9E-14 |
| *gpd-3* | WBGene00001685 | protein_coding_gene | -3.3 | 0.1047 | 8.1E-51 | -2.7 | 0.1586 | 6.4E-109 |
| *C53D6.7* | WBGene00008284 | protein_coding_gene | -3.3 | 0.1046 | 7.0E-36 | -3.5 | 0.0876 | 2.2E-31 |
| *cpl-1* | WBGene00000776 | protein_coding_gene | -3.3 | 0.1043 | 2.1E-34 | -2.3 | 0.1964 | 1.5E-48 |
| *ugt-13* | WBGene00019232 | protein_coding_gene | -3.3 | 0.1043 | 9.0E-25 | -3.4 | 0.0976 | 5.0E-79 |
| *F19H8.2* | WBGene00008963 | protein_coding_gene | -3.3 | 0.1041 | 2.6E-23 | -2.0 | 0.2527 | 2.5E-09 |
| *acdh-9* | WBGene00017874 | protein_coding_gene | -3.3 | 0.1040 | 2.2E-63 | -1.7 | 0.3064 | 4.1E-26 |
| *R04A9.9* | WBGene00195183 | protein_coding_gene | -3.3 | 0.1038 | 3.3E-11 | -1.9 | 0.2710 | 3.9E-07 |
| *clec-178* | WBGene00020585 | protein_coding_gene | -3.3 | 0.1036 | 2.7E-10 | -1.2 | 0.4313 | 6.8E-03 |
| *ugt-33* | WBGene00007946 | protein_coding_gene | -3.3 | 0.1035 | 2.1E-06 | -4.0 | 0.0646 | 9.0E-14 |
| *ZK669.3* | WBGene00014053 | protein_coding_gene | -3.3 | 0.1031 | 3.7E-16 | -4.7 | 0.0385 | 1.9E-133 |
| *C05D12.2* | WBGene00007340 | protein_coding_gene | -3.3 | 0.1030 | 6.5E-25 | -3.5 | 0.0866 | 3.9E-30 |
| *tba-9* | WBGene00006535 | protein_coding_gene | -3.3 | 0.1030 | 5.2E-23 | -1.1 | 0.4676 | 2.4E-04 |
| *F28H7.8* | WBGene00009241 | protein_coding_gene | -3.3 | 0.1030 | 1.5E-15 | -2.3 | 0.2095 | 1.8E-09 |
| *T04C12.11* | WBGene00077693 | protein_coding_gene | -3.3 | 0.1030 | 5.5E-34 | -3.0 | 0.1239 | 2.0E-39 |

| | | | | | | | | |
|---|---|---|---|---|---|---|---|---|
| *R02F2.8* | WBGene00019837 | protein_coding_gene | -3.3 | 0.1028 | 3.2E-23 | -2.6 | 0.1623 | 1.9E-18 |
| *best-22* | WBGene00013921 | protein_coding_gene | -3.3 | 0.1028 | 1.2E-39 | -2.3 | 0.2076 | 8.3E-25 |
| *Y51H4A.7* | WBGene00013103 | protein_coding_gene | -3.3 | 0.1027 | 1.2E-68 | -3.4 | 0.0949 | 1.1E-84 |
| *F36G9.3* | WBGene00009485 | protein_coding_gene | -3.3 | 0.1026 | 1.8E-13 | -4.6 | 0.0405 | 2.1E-07 |
| *ZK1193.2* | WBGene00022858 | protein_coding_gene | -3.3 | 0.1026 | 4.9E-45 | -1.5 | 0.3423 | 1.4E-11 |
| *cpt-5* | WBGene00008629 | protein_coding_gene | -3.3 | 0.1024 | 2.4E-47 | -1.7 | 0.3027 | 1.7E-14 |
| *F38B7.2* | WBGene00009533 | protein_coding_gene | -3.3 | 0.1023 | 6.6E-10 | -2.8 | 0.1433 | 1.4E-10 |
| *Y17G9A.3* | WBGene00021196 | protein_coding_gene | -3.3 | 0.1023 | 3.5E-05 | -2.8 | 0.1409 | 2.2E-03 |
| *C01C10.2* | WBGene00015294 | protein_coding_gene | -3.3 | 0.1022 | 4.4E-34 | -2.2 | 0.2106 | 1.8E-07 |
| *B0416.7* | WBGene00015182 | protein_coding_gene | -3.3 | 0.1021 | 6.5E-18 | -2.9 | 0.1306 | 2.7E-11 |
| *C18B2.3* | WBGene00015954 | protein_coding_gene | -3.3 | 0.1019 | 4.0E-20 | -1.4 | 0.3682 | 1.4E-08 |
| *B0410.3* | WBGene00015172 | protein_coding_gene | -3.3 | 0.1018 | 5.8E-19 | -1.8 | 0.2811 | 6.7E-21 |
| *C11H1.9* | WBGene00023500 | protein_coding_gene | -3.3 | 0.1016 | 1.6E-15 | -1.8 | 0.2899 | 4.4E-12 |
| *alh-12* | WBGene00000118 | protein_coding_gene | -3.3 | 0.1016 | 2.2E-45 | -1.7 | 0.3154 | 1.8E-36 |
| *zig-12* | WBGene00019727 | protein_coding_gene | -3.3 | 0.1014 | 3.5E-52 | -1.4 | 0.3921 | 1.1E-16 |
| *T23E7.2* | WBGene00020732 | protein_coding_gene | -3.3 | 0.1013 | 4.6E-23 | -1.6 | 0.3257 | 6.6E-34 |
| *aqp-8* | WBGene00000176 | protein_coding_gene | -3.3 | 0.1012 | 3.3E-27 | -2.9 | 0.1322 | 6.8E-87 |
| *ptr-24* | WBGene00004238 | protein_coding_gene | -3.3 | 0.1011 | 7.8E-36 | -1.6 | 0.3226 | 3.0E-25 |
| *C05D12.1* | WBGene00007339 | protein_coding_gene | -3.3 | 0.1010 | 3.7E-15 | -2.1 | 0.2354 | 6.8E-08 |
| *ZK616.1* | WBGene00022771 | protein_coding_gene | -3.3 | 0.1009 | 1.1E-06 | -1.9 | 0.2735 | 3.1E-04 |
| *lgc-43* | WBGene00008076 | protein_coding_gene | -3.3 | 0.1007 | 7.4E-28 | -2.6 | 0.1704 | 3.0E-16 |
| *Y105C5A.8* | WBGene00013633 | protein_coding_gene | -3.3 | 0.1006 | 6.3E-43 | -4.1 | 0.0602 | 4.1E-72 |
| *icl-1* | WBGene00001564 | protein_coding_gene | -3.3 | 0.1004 | 1.0E-15 | -4.2 | 0.0546 | 4.3E-131 |
| *T04F3.1* | WBGene00011436 | protein_coding_gene | -3.3 | 0.1003 | 3.7E-39 | -1.8 | 0.2955 | 6.5E-40 |
| *cah-4* | WBGene00000282 | protein_coding_gene | -3.3 | 0.1003 | 8.3E-37 | -3.5 | 0.0883 | 1.7E-113 |
| *cyp-35A2* | WBGene00015400 | protein_coding_gene | -3.3 | 0.1000 | 1.9E-16 | -3.6 | 0.0804 | 1.3E-30 |
| *Y43F8B.1* | WBGene00012812 | protein_coding_gene | -3.3 | 0.1000 | 3.0E-50 | -2.0 | 0.2552 | 1.4E-38 |
| *col-34* | WBGene00000611 | protein_coding_gene | -3.3 | 0.0998 | 2.3E-21 | -1.6 | 0.3297 | 5.9E-11 |
| *far-4* | WBGene00001388 | protein_coding_gene | -3.3 | 0.0998 | 6.3E-04 | -1.9 | 0.2770 | 4.4E-02 |
| *F59A1.13* | WBGene00010299 | protein_coding_gene | -3.3 | 0.0998 | 6.2E-05 | -2.5 | 0.1758 | 7.9E-10 |
| *col-89* | WBGene00000664 | protein_coding_gene | -3.3 | 0.0997 | 1.1E-08 | -1.7 | 0.3114 | 4.5E-04 |
| *T12D8.9* | WBGene00011736 | protein_coding_gene | -3.3 | 0.0997 | 5.6E-33 | -2.2 | 0.2142 | 5.4E-45 |
| *col-186* | WBGene00000759 | protein_coding_gene | -3.3 | 0.0996 | 3.1E-08 | -2.7 | 0.1564 | 4.4E-06 |
| *ketn-1* | WBGene00004130 | protein_coding_gene | -3.3 | 0.0993 | 6.5E-22 | -1.2 | 0.4397 | 9.4E-13 |
| *sul-3* | WBGene00006310 | protein_coding_gene | -3.3 | 0.0991 | 1.8E-19 | -2.2 | 0.2214 | 1.1E-08 |
| *R03A10.5* | WBGene00010985 | protein_coding_gene | -3.3 | 0.0989 | 9.9E-43 | -2.9 | 0.1315 | 1.2E-40 |
| *R03G8.6* | WBGene00011000 | protein_coding_gene | -3.3 | 0.0989 | 8.1E-20 | -2.0 | 0.2541 | 7.1E-13 |
| *F27E5.7* | WBGene00044035 | protein_coding_gene | -3.3 | 0.0988 | 8.5E-13 | -2.2 | 0.2222 | 3.4E-05 |
| *T21F4.1* | WBGene00020658 | protein_coding_gene | -3.3 | 0.0987 | 2.8E-43 | -3.5 | 0.0866 | 1.3E-127 |
| *T25F10.3* | WBGene00020806 | protein_coding_gene | -3.3 | 0.0986 | 3.7E-24 | -2.6 | 0.1602 | 6.1E-32 |
| *far-5* | WBGene00001389 | protein_coding_gene | -3.3 | 0.0986 | 7.5E-15 | -1.6 | 0.3370 | 2.2E-06 |
| *nlp-77* | WBGene00015514 | protein_coding_gene | -3.3 | 0.0984 | 1.7E-26 | -1.3 | 0.4081 | 1.3E-28 |
| *C34F11.8* | WBGene00016417 | protein_coding_gene | -3.3 | 0.0981 | 2.4E-16 | -4.1 | 0.0583 | 2.1E-57 |
| *tdo-2* | WBGene00016201 | protein_coding_gene | -3.3 | 0.0981 | 6.9E-23 | -2.4 | 0.1939 | 5.6E-27 |
| *cyp-34A7* | WBGene00015042 | protein_coding_gene | -3.4 | 0.0974 | 1.2E-06 | -1.1 | 0.4602 | 4.4E-02 |
| *F35H8.2* | WBGene00009447 | protein_coding_gene | -3.4 | 0.0973 | 2.2E-25 | -3.3 | 0.1020 | 7.2E-26 |
| *pqn-72* | WBGene00004154 | protein_coding_gene | -3.4 | 0.0972 | 1.3E-15 | -2.3 | 0.2013 | 7.0E-08 |
| ***ins-32*** | **WBGene00002115** | **protein_coding_gene** | **-3.4** | **0.0972** | **2.6E-03** | **2.0** | **4.1093** | **1.9E-02** |
| *C08A9.11* | WBGene00045273 | pseudogene | -3.4 | 0.0968 | 7.7E-09 | -3.3 | 0.1027 | 1.1E-08 |
| *B0454.6* | WBGene00015197 | protein_coding_gene | -3.4 | 0.0968 | 1.2E-24 | -4.6 | 0.0407 | 1.6E-60 |
| *unc-22* | WBGene00006759 | protein_coding_gene | -3.4 | 0.0968 | 3.3E-17 | -1.6 | 0.3320 | 4.9E-10 |
| *pgp-9* | WBGene00004003 | protein_coding_gene | -3.4 | 0.0965 | 1.4E-14 | -1.1 | 0.4784 | 3.3E-05 |
| *rol-3* | WBGene00004395 | protein_coding_gene | -3.4 | 0.0959 | 5.0E-21 | -1.8 | 0.2958 | 5.3E-13 |
| *clec-26* | WBGene00011852 | protein_coding_gene | -3.4 | 0.0959 | 7.2E-13 | -3.2 | 0.1060 | 5.7E-17 |
| *best-21* | WBGene00013520 | protein_coding_gene | -3.4 | 0.0959 | 2.6E-14 | -4.0 | 0.0630 | 1.7E-18 |
| *unc-52* | WBGene00006787 | protein_coding_gene | -3.4 | 0.0958 | 7.5E-31 | -1.5 | 0.3540 | 4.0E-27 |
| *F18E9.3* | WBGene00017569 | protein_coding_gene | -3.4 | 0.0957 | 1.5E-18 | -3.7 | 0.0764 | 1.3E-37 |
| *T25G12.3* | WBGene00020811 | protein_coding_gene | -3.4 | 0.0956 | 1.2E-16 | -2.3 | 0.2020 | 1.2E-10 |
| *F08F3.4* | WBGene00017262 | protein_coding_gene | -3.4 | 0.0954 | 1.2E-15 | -1.9 | 0.2614 | 3.8E-44 |
| *K11G9.5* | WBGene00019655 | protein_coding_gene | -3.4 | 0.0953 | 4.9E-11 | -2.9 | 0.1356 | 2.6E-11 |
| *cpr-8* | WBGene00021070 | protein_coding_gene | -3.4 | 0.0952 | 1.3E-18 | -3.7 | 0.0781 | 4.0E-16 |
| *ram-2* | WBGene00004300 | protein_coding_gene | -3.4 | 0.0951 | 3.0E-16 | -1.3 | 0.4160 | 1.3E-07 |
| *K10C2.14* | WBGene00206388 | protein_coding_gene | -3.4 | 0.0949 | 4.2E-07 | -1.8 | 0.2880 | 1.5E-15 |
| *dhs-31* | WBGene00011424 | protein_coding_gene | -3.4 | 0.0948 | 3.4E-15 | -2.9 | 0.1348 | 3.3E-26 |
| *str-176* | WBGene00006220 | protein_coding_gene | -3.4 | 0.0948 | 4.9E-53 | -3.2 | 0.1126 | 9.6E-28 |
| *atic-1* | WBGene00016957 | protein_coding_gene | -3.4 | 0.0946 | 2.3E-12 | -1.0 | 0.4877 | 9.8E-08 |
| *sgcb-1* | WBGene00019277 | protein_coding_gene | -3.4 | 0.0946 | 8.8E-19 | -1.7 | 0.3125 | 2.2E-06 |
| *W07B8.4* | WBGene00021072 | protein_coding_gene | -3.4 | 0.0945 | 1.8E-06 | -3.9 | 0.0683 | 5.2E-20 |
| *emb-9* | WBGene00001263 | protein_coding_gene | -3.4 | 0.0945 | 6.0E-28 | -2.4 | 0.1904 | 1.9E-78 |
| *best-16* | WBGene00020046 | protein_coding_gene | -3.4 | 0.0943 | 1.2E-05 | -3.0 | 0.1292 | 7.3E-08 |

| | | | | | | | | |
|---|---|---|---|---|---|---|---|---|
| *bcc-1* | WBGene00000246 | protein_coding_gene | -3.4 | 0.0940 | 7.0E-22 | -3.2 | 0.1100 | 3.7E-15 |
| *unc-54* | WBGene00006789 | protein_coding_gene | -3.4 | 0.0938 | 1.9E-28 | -2.9 | 0.1315 | 1.3E-80 |
| *sulp-2* | WBGene00017464 | protein_coding_gene | -3.4 | 0.0937 | 1.4E-31 | -4.9 | 0.0337 | 4.1E-47 |
| *W03D8.8* | WBGene00020989 | protein_coding_gene | -3.4 | 0.0935 | 9.7E-21 | -3.1 | 0.1200 | 1.9E-17 |
| *K04F1.9* | WBGene00019388 | protein_coding_gene | -3.4 | 0.0933 | 1.1E-21 | -2.9 | 0.1381 | 2.2E-19 |
| *glna-2* | WBGene00008435 | protein_coding_gene | -3.4 | 0.0932 | 2.8E-64 | -3.0 | 0.1213 | 1.7E-58 |
| *bcat-1* | WBGene00001149 | protein_coding_gene | -3.4 | 0.0932 | 5.8E-47 | -1.8 | 0.2859 | 3.7E-29 |
| *zig-3* | WBGene00006980 | protein_coding_gene | -3.4 | 0.0932 | 7.9E-16 | -2.4 | 0.1930 | 3.4E-13 |
| *snf-3* | WBGene00004902 | protein_coding_gene | -3.4 | 0.0931 | 1.4E-50 | -2.6 | 0.1637 | 2.8E-34 |
| *T06A4.3* | WBGene00020283 | protein_coding_gene | -3.4 | 0.0929 | 3.7E-15 | -1.2 | 0.4318 | 1.2E-02 |
| *K08C7.1* | WBGene00010646 | protein_coding_gene | -3.4 | 0.0927 | 4.2E-48 | -2.0 | 0.2454 | 2.5E-16 |
| *K09G1.1* | WBGene00010727 | protein_coding_gene | -3.4 | 0.0925 | 2.0E-40 | -1.8 | 0.2878 | 1.1E-37 |
| *unc-27* | WBGene00006764 | protein_coding_gene | -3.4 | 0.0924 | 8.8E-68 | -2.0 | 0.2567 | 1.4E-67 |
| *mlc-1* | WBGene00003369 | protein_coding_gene | -3.4 | 0.0922 | 7.1E-36 | -2.2 | 0.2187 | 2.0E-85 |
| *haao-1* | WBGene00010595 | protein_coding_gene | -3.4 | 0.0922 | 1.9E-14 | -1.4 | 0.3718 | 2.3E-09 |
| *fig-1* | WBGene00009968 | protein_coding_gene | -3.4 | 0.0921 | 1.3E-19 | -2.4 | 0.1953 | 4.1E-34 |
| *C08E8.10* | WBGene00077692 | protein_coding_gene | -3.4 | 0.0921 | 2.1E-07 | -2.7 | 0.1541 | 6.0E-06 |
| *ddo-2* | WBGene00017565 | protein_coding_gene | -3.4 | 0.0917 | 3.2E-40 | -2.4 | 0.1895 | 7.7E-33 |
| *far-3* | WBGene00001387 | protein_coding_gene | -3.4 | 0.0915 | 3.6E-16 | -3.7 | 0.0747 | 1.6E-65 |
| *nlp-33* | WBGene00003771 | protein_coding_gene | -3.5 | 0.0915 | 1.8E-24 | -2.9 | 0.1307 | 2.8E-12 |
| *gpd-2* | WBGene00001684 | protein_coding_gene | -3.5 | 0.0914 | 7.0E-64 | -1.9 | 0.2679 | 5.8E-47 |
| *cdo-1* | WBGene00018986 | protein_coding_gene | -3.5 | 0.0912 | 2.8E-40 | -4.1 | 0.0602 | 1.6E-86 |
| *R102.6* | WBGene00011293 | protein_coding_gene | -3.5 | 0.0910 | 3.1E-08 | -3.6 | 0.0854 | 2.2E-08 |
| *cima-1* | WBGene00018473 | protein_coding_gene | -3.5 | 0.0910 | 4.0E-17 | -1.8 | 0.2913 | 2.9E-06 |
| *ugt-60* | WBGene00007402 | protein_coding_gene | -3.5 | 0.0909 | 4.3E-12 | -1.4 | 0.3818 | 4.1E-06 |
| *F31A3.3* | WBGene00017938 | protein_coding_gene | -3.5 | 0.0906 | 7.1E-12 | -1.3 | 0.4159 | 4.2E-02 |
| *T01D1.3* | WBGene00020148 | protein_coding_gene | -3.5 | 0.0901 | 2.3E-10 | -2.4 | 0.1948 | 1.3E-12 |
| *sulp-5* | WBGene00010789 | protein_coding_gene | -3.5 | 0.0901 | 9.0E-10 | -3.0 | 0.1216 | 1.6E-07 |
| *F17C8.9* | WBGene00185014 | protein_coding_gene | -3.5 | 0.0900 | 9.0E-19 | -2.3 | 0.2089 | 2.8E-21 |
| *chil-12* | WBGene00010799 | protein_coding_gene | -3.5 | 0.0898 | 6.7E-22 | -2.1 | 0.2383 | 2.6E-07 |
| *oac-53* | WBGene00012308 | protein_coding_gene | -3.5 | 0.0897 | 3.0E-04 | -2.8 | 0.1453 | 6.5E-04 |
| *srsx-18* | WBGene00015419 | protein_coding_gene | -3.5 | 0.0895 | 2.1E-04 | -2.9 | 0.1324 | 3.5E-03 |
| *ins-28* | WBGene00002111 | protein_coding_gene | -3.5 | 0.0895 | 2.6E-10 | -2.3 | 0.1973 | 3.3E-06 |
| *F55C12.6* | WBGene00018873 | protein_coding_gene | -3.5 | 0.0893 | 2.9E-05 | -3.6 | 0.0852 | 2.8E-06 |
| *T25B9.1* | WBGene00012007 | protein_coding_gene | -3.5 | 0.0893 | 1.2E-17 | -1.2 | 0.4238 | 5.7E-07 |
| *F48E3.4* | WBGene00018605 | protein_coding_gene | -3.5 | 0.0886 | 3.9E-23 | -3.9 | 0.0656 | 2.8E-129 |
| *fkh-6* | WBGene00001438 | protein_coding_gene | -3.5 | 0.0886 | 2.5E-10 | -2.1 | 0.2305 | 1.4E-07 |
| *F46G11.2* | WBGene00018515 | protein_coding_gene | -3.5 | 0.0879 | 3.0E-09 | -1.9 | 0.2739 | 2.7E-03 |
| *pept-3* | WBGene00003878 | protein_coding_gene | -3.5 | 0.0876 | 1.1E-12 | -4.0 | 0.0646 | 3.1E-43 |
| *C25E10.5* | WBGene00016095 | protein_coding_gene | -3.5 | 0.0875 | 3.8E-24 | -2.2 | 0.2138 | 2.1E-14 |
| *C06E2.1* | WBGene00015527 | protein_coding_gene | -3.5 | 0.0873 | 5.2E-10 | -1.4 | 0.3767 | 6.9E-03 |
| *D1054.8* | WBGene00008375 | protein_coding_gene | -3.5 | 0.0867 | 1.2E-11 | -1.3 | 0.3946 | 4.7E-13 |
| *grd-13* | WBGene00001702 | protein_coding_gene | -3.5 | 0.0864 | 2.5E-10 | -2.0 | 0.2567 | 7.8E-04 |
| *R08F11.1* | WBGene00019965 | protein_coding_gene | -3.5 | 0.0859 | 3.2E-47 | -2.1 | 0.2327 | 1.2E-42 |
| *mlt-4* | WBGene00013835 | protein_coding_gene | -3.5 | 0.0858 | 3.4E-11 | -2.0 | 0.2453 | 3.3E-05 |
| *brp-1* | WBGene00000273 | protein_coding_gene | -3.5 | 0.0857 | 5.3E-44 | -3.7 | 0.0796 | 2.3E-170 |
| *C54E4.5* | WBGene00016920 | protein_coding_gene | -3.5 | 0.0856 | 2.2E-19 | -1.4 | 0.3775 | 4.9E-11 |
| *cnc-3* | WBGene00000557 | protein_coding_gene | -3.5 | 0.0856 | 1.2E-15 | -3.7 | 0.0751 | 3.3E-22 |
| *nspe-7* | WBGene00012593 | protein_coding_gene | -3.5 | 0.0854 | 5.1E-04 | -4.5 | 0.0442 | 6.2E-09 |
| *F42G10.1* | WBGene00009645 | protein_coding_gene | -3.6 | 0.0854 | 4.7E-32 | -1.1 | 0.4724 | 2.5E-05 |
| *best-24* | WBGene00022797 | protein_coding_gene | -3.6 | 0.0848 | 1.0E-21 | -4.4 | 0.0490 | 4.2E-110 |
| *T28H10.2* | WBGene00012143 | protein_coding_gene | -3.6 | 0.0848 | 1.4E-25 | -2.0 | 0.2482 | 6.3E-11 |
| *F59E11.5* | WBGene00019111 | protein_coding_gene | -3.6 | 0.0845 | 1.3E-26 | -3.7 | 0.0757 | 3.6E-60 |
| *cyp-35A4* | WBGene00016786 | protein_coding_gene | -3.6 | 0.0844 | 4.9E-08 | -1.9 | 0.2665 | 2.6E-02 |
| *acp-7* | WBGene00022246 | protein_coding_gene | -3.6 | 0.0844 | 8.2E-28 | -2.6 | 0.1614 | 2.7E-20 |
| *nep-4* | WBGene00016896 | protein_coding_gene | -3.6 | 0.0844 | 4.4E-22 | -1.9 | 0.2617 | 3.0E-08 |
| *aass-1* | WBGene00019819 | protein_coding_gene | -3.6 | 0.0843 | 9.8E-46 | -1.6 | 0.3272 | 5.2E-36 |
| *mup-2* | WBGene00003495 | protein_coding_gene | -3.6 | 0.0838 | 2.3E-59 | -2.1 | 0.2258 | 4.1E-60 |
| *lys-6* | WBGene00003095 | protein_coding_gene | -3.6 | 0.0838 | 8.8E-12 | -4.1 | 0.0573 | 3.2E-29 |
| *lgc-45* | WBGene00021146 | protein_coding_gene | -3.6 | 0.0836 | 5.9E-28 | -1.9 | 0.2705 | 1.8E-14 |
| *C44C10.9* | WBGene00008089 | protein_coding_gene | -3.6 | 0.0830 | 6.6E-49 | -2.1 | 0.2371 | 1.2E-26 |
| *F33E2.4* | WBGene00009360 | protein_coding_gene | -3.6 | 0.0828 | 8.9E-11 | -2.3 | 0.2018 | 1.0E-03 |
| *gpx-5* | WBGene00007516 | protein_coding_gene | -3.6 | 0.0826 | 4.2E-29 | -1.8 | 0.2939 | 3.6E-19 |
| *anmt-2* | WBGene00018340 | protein_coding_gene | -3.6 | 0.0826 | 1.5E-97 | -1.5 | 0.3469 | 2.3E-14 |
| *F41E7.20* | WBGene00219546 | ncRNA_gene | -3.6 | 0.0824 | 2.0E-04 | -2.7 | 0.1587 | 3.5E-03 |
| *F57B1.1* | WBGene00010187 | protein_coding_gene | -3.6 | 0.0820 | 5.5E-05 | -2.6 | 0.1613 | 2.3E-03 |
| *let-2* | WBGene00002280 | protein_coding_gene | -3.6 | 0.0819 | 2.2E-33 | -2.4 | 0.1863 | 5.8E-87 |
| *B0457.2* | WBGene00007178 | protein_coding_gene | -3.6 | 0.0818 | 1.1E-51 | -3.9 | 0.0686 | 2.1E-53 |
| *F54B11.11* | WBGene00010034 | protein_coding_gene | -3.6 | 0.0816 | 7.7E-21 | -3.7 | 0.0769 | 5.4E-27 |
| *K08D8.3* | WBGene00010657 | protein_coding_gene | -3.6 | 0.0814 | 1.0E-33 | -1.0 | 0.4923 | 3.3E-05 |

| | | | | | | | | |
|---|---|---|---|---|---|---|---|---|
| *Y105C5B.5* | WBGene00013647 | protein_coding_gene | -3.6 | 0.0814 | 3.4E-54 | -2.8 | 0.1475 | 2.1E-98 |
| *Y48G1BR.1* | WBGene00021675 | protein_coding_gene | -3.6 | 0.0812 | 4.5E-26 | -2.7 | 0.1573 | 2.8E-22 |
| *hpd-1* | WBGene00001993 | protein_coding_gene | -3.6 | 0.0811 | 2.2E-16 | -1.4 | 0.3804 | 1.1E-17 |
| *C50A2.3* | WBGene00016797 | protein_coding_gene | -3.6 | 0.0807 | 1.9E-13 | -1.6 | 0.3273 | 3.6E-06 |
| *exc-15* | WBGene00020369 | protein_coding_gene | -3.6 | 0.0807 | 1.3E-73 | -2.4 | 0.1887 | 5.2E-72 |
| *gpa-12* | WBGene00001674 | protein_coding_gene | -3.6 | 0.0806 | 2.7E-21 | -2.3 | 0.2024 | 4.7E-14 |
| *frpr-1* | WBGene00015323 | protein_coding_gene | -3.6 | 0.0805 | 1.2E-03 | -2.4 | 0.1869 | 2.0E-03 |
| *col-147* | WBGene00000720 | protein_coding_gene | -3.6 | 0.0804 | 4.1E-08 | -2.6 | 0.1660 | 8.9E-14 |
| *ifb-1* | WBGene00002053 | protein_coding_gene | -3.6 | 0.0803 | 4.2E-65 | -1.6 | 0.3359 | 2.2E-26 |
| *Y34F4.6* | WBGene00189956 | protein_coding_gene | -3.6 | 0.0797 | 5.6E-09 | -2.5 | 0.1746 | 1.2E-02 |
| *cest-24* | WBGene00020891 | protein_coding_gene | -3.7 | 0.0795 | 1.9E-40 | -4.1 | 0.0590 | 3.5E-32 |
| *C08H9.15* | WBGene00007474 | protein_coding_gene | -3.7 | 0.0792 | 9.8E-23 | -3.2 | 0.1121 | 5.4E-31 |
| *F11F1.1* | WBGene00008714 | protein_coding_gene | -3.7 | 0.0792 | 1.3E-28 | -2.1 | 0.2331 | 5.6E-11 |
| *lgc-34* | WBGene00020836 | protein_coding_gene | -3.7 | 0.0791 | 2.2E-36 | -2.4 | 0.1833 | 1.5E-24 |
| *nas-37* | WBGene00003553 | protein_coding_gene | -3.7 | 0.0789 | 6.6E-21 | -4.5 | 0.0447 | 2.8E-20 |
| *R05D8.9* | WBGene00019886 | protein_coding_gene | -3.7 | 0.0786 | 1.7E-03 | -4.3 | 0.0515 | 7.5E-06 |
| *C23H5.15* | WBGene00219423 | protein_coding_gene | -3.7 | 0.0786 | 8.2E-17 | -5.1 | 0.0287 | 1.2E-21 |
| *lys-4* | WBGene00003093 | protein_coding_gene | -3.7 | 0.0785 | 5.4E-66 | -5.4 | 0.0244 | 1.5E-183 |
| *asp-8* | WBGene00019105 | protein_coding_gene | -3.7 | 0.0782 | 1.3E-35 | -1.1 | 0.4517 | 1.3E-02 |
| *R160.3* | WBGene00020120 | protein_coding_gene | -3.7 | 0.0779 | 3.2E-10 | -2.1 | 0.2314 | 2.2E-05 |
| *ins-33* | WBGene00002116 | protein_coding_gene | -3.7 | 0.0778 | 4.9E-11 | -2.9 | 0.1341 | 1.6E-06 |
| *H14N18.4* | WBGene00019204 | protein_coding_gene | -3.7 | 0.0772 | 1.5E-41 | -2.7 | 0.1554 | 2.6E-39 |
| *fbxa-62* | WBGene00016521 | protein_coding_gene | -3.7 | 0.0770 | 1.4E-04 | -2.8 | 0.1468 | 2.1E-03 |
| *C26G2.2* | WBGene00007751 | protein_coding_gene | -3.7 | 0.0769 | 1.5E-42 | -3.4 | 0.0978 | 3.1E-50 |
| *C23H3.2* | WBGene00016018 | protein_coding_gene | -3.7 | 0.0769 | 1.2E-51 | -1.9 | 0.2628 | 1.6E-27 |
| *R17.3* | WBGene00011269 | protein_coding_gene | -3.7 | 0.0768 | 1.2E-24 | -2.4 | 0.1836 | 1.8E-11 |
| *perm-2* | WBGene00016636 | protein_coding_gene | -3.7 | 0.0767 | 6.7E-32 | -3.7 | 0.0770 | 6.9E-139 |
| *best-14* | WBGene00009340 | protein_coding_gene | -3.7 | 0.0767 | 1.5E-36 | -2.4 | 0.1832 | 1.2E-45 |
| *T10H10.2* | WBGene00020426 | protein_coding_gene | -3.7 | 0.0766 | 3.2E-19 | -3.5 | 0.0897 | 2.3E-29 |
| *ugt-3* | WBGene00013903 | protein_coding_gene | -3.7 | 0.0765 | 3.3E-04 | -2.1 | 0.2379 | 4.9E-03 |
| *Y59H11AR.4* | WBGene00022012 | protein_coding_gene | -3.7 | 0.0763 | 3.6E-20 | -3.2 | 0.1061 | 4.2E-30 |
| *srsx-34* | WBGene00016093 | protein_coding_gene | -3.7 | 0.0759 | 3.7E-42 | -4.0 | 0.0636 | 4.6E-50 |
| *col-153* | WBGene00000726 | protein_coding_gene | -3.7 | 0.0759 | 1.9E-21 | -2.9 | 0.1323 | 3.3E-30 |
| *lron-5* | WBGene00008656 | protein_coding_gene | -3.7 | 0.0758 | 4.6E-15 | -3.4 | 0.0949 | 4.1E-13 |
| *fip-5* | WBGene00009620 | protein_coding_gene | -3.7 | 0.0757 | 4.9E-16 | -4.0 | 0.0645 | 5.5E-11 |
| *C50F2.4* | WBGene00016838 | protein_coding_gene | -3.7 | 0.0754 | 4.8E-34 | -3.0 | 0.1259 | 7.6E-46 |
| *F47E1.4* | WBGene00018568 | protein_coding_gene | -3.7 | 0.0752 | 3.4E-16 | -3.8 | 0.0739 | 9.9E-30 |
| *tnt-2* | WBGene00006587 | protein_coding_gene | -3.7 | 0.0752 | 5.1E-52 | -2.3 | 0.1974 | 3.3E-52 |
| *Y22D7AL.11* | WBGene00021249 | protein_coding_gene | -3.7 | 0.0748 | 2.9E-19 | -1.6 | 0.3284 | 4.6E-04 |
| *skpo-1* | WBGene00009897 | protein_coding_gene | -3.7 | 0.0747 | 8.4E-36 | -3.8 | 0.0720 | 6.2E-127 |
| *C15F1.1* | WBGene00015794 | protein_coding_gene | -3.7 | 0.0745 | 3.5E-25 | -2.4 | 0.1863 | 2.6E-16 |
| *fipr-11* | WBGene00008244 | protein_coding_gene | -3.7 | 0.0743 | 2.3E-08 | -1.5 | 0.3557 | 5.7E-03 |
| *T24D3.2* | WBGene00023492 | protein_coding_gene | -3.8 | 0.0743 | 3.0E-13 | -3.3 | 0.0989 | 1.6E-12 |
| *D1086.10* | WBGene00044080 | protein_coding_gene | -3.8 | 0.0743 | 8.3E-54 | -2.4 | 0.1928 | 1.5E-17 |
| *cnc-4* | WBGene00000558 | protein_coding_gene | -3.8 | 0.0743 | 1.1E-07 | -6.7 | 0.0095 | 7.0E-22 |
| *grsp-1* | WBGene00012840 | protein_coding_gene | -3.8 | 0.0742 | 6.4E-12 | -1.7 | 0.3154 | 1.3E-04 |
| *drd-5* | WBGene00018885 | protein_coding_gene | -3.8 | 0.0742 | 1.0E-44 | -1.9 | 0.2741 | 1.1E-10 |
| *arrd-10* | WBGene00010268 | protein_coding_gene | -3.8 | 0.0741 | 2.4E-14 | -1.8 | 0.2846 | 1.2E-05 |
| *T01G5.1* | WBGene00011339 | protein_coding_gene | -3.8 | 0.0740 | 1.1E-20 | -2.0 | 0.2566 | 7.7E-13 |
| *C45B2.2* | WBGene00016659 | protein_coding_gene | -3.8 | 0.0738 | 4.8E-11 | -2.5 | 0.1734 | 3.6E-06 |
| *osm-11* | WBGene00003891 | protein_coding_gene | -3.8 | 0.0736 | 4.5E-53 | -3.2 | 0.1083 | 1.3E-122 |
| *ugt-46* | WBGene00015141 | protein_coding_gene | -3.8 | 0.0735 | 2.9E-23 | -1.3 | 0.4060 | 9.3E-11 |
| *ugt-36* | WBGene00017315 | protein_coding_gene | -3.8 | 0.0735 | 2.0E-05 | -4.3 | 0.0495 | 1.0E-06 |
| *nas-9* | WBGene00003528 | protein_coding_gene | -3.8 | 0.0733 | 2.9E-29 | -4.4 | 0.0482 | 3.8E-61 |
| *best-17* | WBGene00020559 | protein_coding_gene | -3.8 | 0.0731 | 7.8E-27 | -2.5 | 0.1811 | 2.3E-11 |
| *grd-10* | WBGene00001699 | protein_coding_gene | -3.8 | 0.0728 | 2.5E-16 | -2.0 | 0.2467 | 1.1E-07 |
| *Y73B6BL.31* | WBGene00022252 | protein_coding_gene | -3.8 | 0.0727 | 7.7E-46 | -1.5 | 0.3546 | 1.5E-16 |
| *nac-1* | WBGene00003517 | protein_coding_gene | -3.8 | 0.0725 | 8.0E-41 | -2.6 | 0.1618 | 2.1E-25 |
| *oat-1* | WBGene00003837 | protein_coding_gene | -3.8 | 0.0722 | 9.1E-22 | -2.6 | 0.1698 | 1.7E-12 |
| *B0303.11* | WBGene00015131 | protein_coding_gene | -3.8 | 0.0722 | 3.1E-59 | -2.7 | 0.1498 | 6.5E-40 |
| *ttr-31* | WBGene00010225 | protein_coding_gene | -3.8 | 0.0715 | 3.8E-15 | -2.3 | 0.1999 | 2.8E-14 |
| *ctsa-1.1* | WBGene00018271 | protein_coding_gene | -3.8 | 0.0714 | 3.9E-56 | -3.2 | 0.1118 | 2.9E-79 |
| *ttr-7* | WBGene00020566 | protein_coding_gene | -3.8 | 0.0712 | 5.9E-18 | -3.2 | 0.1058 | 3.8E-18 |
| *F43H9.4* | WBGene00018400 | protein_coding_gene | -3.8 | 0.0711 | 7.3E-20 | -3.8 | 0.0722 | 7.7E-74 |
| *ZK822.5* | WBGene00014092 | protein_coding_gene | -3.8 | 0.0706 | 7.3E-73 | -2.3 | 0.2019 | 3.5E-35 |
| *F23D12.11* | WBGene00194674 | protein_coding_gene | -3.8 | 0.0705 | 1.4E-16 | -2.6 | 0.1658 | 8.3E-17 |
| *cyp-35C1* | WBGene00007362 | protein_coding_gene | -3.8 | 0.0705 | 1.1E-31 | -2.4 | 0.1875 | 1.4E-07 |
| *Y57G11A.4* | WBGene00013292 | protein_coding_gene | -3.8 | 0.0702 | 5.5E-47 | -2.4 | 0.1919 | 9.4E-23 |
| *C27B7.9* | WBGene00007765 | protein_coding_gene | -3.8 | 0.0702 | 5.8E-23 | -6.8 | 0.0091 | 2.2E-223 |
| *clec-170* | WBGene00009515 | protein_coding_gene | -3.8 | 0.0701 | 1.5E-27 | -3.6 | 0.0842 | 2.6E-56 |

| | | | | | | | | |
|---|---|---|---|---|---|---|---|---|
| *glt-1* | WBGene00001620 | protein_coding_gene | -3.8 | 0.0696 | 1.1E-48 | -2.6 | 0.1688 | 6.8E-87 |
| *dhs-23* | WBGene00000986 | protein_coding_gene | -3.8 | 0.0693 | 2.6E-15 | -4.9 | 0.0330 | 7.9E-34 |
| *Y17G9A.2* | WBGene00021195 | protein_coding_gene | -3.9 | 0.0689 | 3.8E-06 | -2.2 | 0.2235 | 1.0E-03 |
| *poml-3* | WBGene00077701 | protein_coding_gene | -3.9 | 0.0687 | 2.7E-56 | -3.0 | 0.1255 | 5.9E-33 |
| *clec-31* | WBGene00009854 | protein_coding_gene | -3.9 | 0.0686 | 5.9E-10 | -5.4 | 0.0239 | 9.4E-15 |
| *F14B8.4* | WBGene00017446 | protein_coding_gene | -3.9 | 0.0685 | 1.3E-17 | -2.9 | 0.1373 | 2.7E-13 |
| *gly-8* | WBGene00001633 | protein_coding_gene | -3.9 | 0.0683 | 2.0E-17 | -1.7 | 0.3058 | 1.1E-10 |
| *twk-33* | WBGene00006685 | protein_coding_gene | -3.9 | 0.0681 | 2.2E-30 | -3.0 | 0.1223 | 4.1E-15 |
| *clec-43* | WBGene00019917 | protein_coding_gene | -3.9 | 0.0681 | 6.0E-12 | -3.3 | 0.1039 | 6.2E-08 |
| *chil-22* | WBGene00011166 | protein_coding_gene | -3.9 | 0.0679 | 8.4E-29 | -3.3 | 0.1037 | 1.6E-30 |
| *lmd-4* | WBGene00017233 | protein_coding_gene | -3.9 | 0.0679 | 1.3E-05 | -1.7 | 0.3107 | 2.3E-02 |
| *tnt-3* | WBGene00006588 | protein_coding_gene | -3.9 | 0.0677 | 1.7E-41 | -2.1 | 0.2395 | 9.2E-41 |
| *Y37D8A.3* | WBGene00012545 | protein_coding_gene | -3.9 | 0.0671 | 6.9E-56 | -2.2 | 0.2199 | 3.3E-21 |
| *W10G11.3* | WBGene00021136 | protein_coding_gene | -3.9 | 0.0670 | 1.8E-15 | -3.0 | 0.1239 | 1.1E-04 |
| *grsp-2* | WBGene00012118 | protein_coding_gene | -3.9 | 0.0669 | 7.3E-30 | -3.3 | 0.1031 | 1.3E-40 |
| *Y4C6B.7* | WBGene00021161 | protein_coding_gene | -3.9 | 0.0669 | 2.4E-57 | -3.4 | 0.0975 | 3.8E-49 |
| *slc-9B.1* | WBGene00009617 | protein_coding_gene | -3.9 | 0.0669 | 5.7E-21 | -3.1 | 0.1181 | 1.5E-22 |
| *nhr-68* | WBGene00003658 | protein_coding_gene | -3.9 | 0.0668 | 2.4E-79 | -3.2 | 0.1098 | 2.5E-119 |
| *cyp-34A10* | WBGene00015045 | protein_coding_gene | -3.9 | 0.0667 | 5.9E-05 | -6.8 | 0.0092 | 5.9E-22 |
| *D1022.3* | WBGene00017022 | protein_coding_gene | -3.9 | 0.0661 | 5.3E-09 | -2.4 | 0.1955 | 6.0E-04 |
| *Y11D7A.3* | WBGene00012428 | protein_coding_gene | -3.9 | 0.0660 | 1.5E-44 | -2.7 | 0.1504 | 3.2E-60 |
| *haly-1* | WBGene00009813 | protein_coding_gene | -3.9 | 0.0660 | 2.6E-47 | -3.5 | 0.0873 | 1.9E-121 |
| *mec-5* | WBGene00003169 | protein_coding_gene | -3.9 | 0.0660 | 2.8E-51 | -1.3 | 0.4143 | 7.0E-06 |
| *oac-54* | WBGene00012323 | protein_coding_gene | -3.9 | 0.0659 | 1.2E-11 | -1.7 | 0.2995 | 1.2E-03 |
| *vha-5* | WBGene00006914 | protein_coding_gene | -3.9 | 0.0658 | 1.9E-38 | -2.8 | 0.1465 | 7.5E-86 |
| *F29B9.8* | WBGene00017923 | protein_coding_gene | -3.9 | 0.0658 | 5.7E-69 | -2.7 | 0.1540 | 1.2E-50 |
| *T12B3.2* | WBGene00020445 | protein_coding_gene | -3.9 | 0.0658 | 6.7E-23 | -2.8 | 0.1410 | 4.9E-34 |
| *pqn-22* | WBGene00004112 | protein_coding_gene | -3.9 | 0.0658 | 7.6E-39 | -1.2 | 0.4325 | 9.6E-11 |
| *C05C8.7* | WBGene00015464 | protein_coding_gene | -3.9 | 0.0651 | 8.7E-29 | -2.2 | 0.2144 | 5.8E-26 |
| *tyr-6* | WBGene00022231 | protein_coding_gene | -3.9 | 0.0649 | 9.5E-44 | -3.4 | 0.0969 | 2.8E-39 |
| *clec-70* | WBGene00021581 | protein_coding_gene | -3.9 | 0.0649 | 4.0E-11 | -2.4 | 0.1913 | 1.3E-10 |
| *Y39F10A.1* | WBGene00021452 | protein_coding_gene | -3.9 | 0.0649 | 4.7E-10 | -1.8 | 0.2932 | 8.8E-03 |
| *K03B8.8* | WBGene00010525 | protein_coding_gene | -3.9 | 0.0648 | 1.2E-28 | -2.7 | 0.1508 | 4.5E-12 |
| *clik-1* | WBGene00020808 | protein_coding_gene | -4.0 | 0.0644 | 1.6E-67 | -2.9 | 0.1348 | 5.5E-90 |
| *cpr-1* | WBGene00000781 | protein_coding_gene | -4.0 | 0.0642 | 2.7E-35 | -4.8 | 0.0364 | 2.2E-246 |
| *tbh-1* | WBGene00006541 | protein_coding_gene | -4.0 | 0.0638 | 3.5E-37 | -3.6 | 0.0835 | 1.5E-106 |
| *acd-2* | WBGene00016066 | protein_coding_gene | -4.0 | 0.0635 | 1.1E-30 | -3.5 | 0.0908 | 2.2E-26 |
| *Y47G6A.33* | WBGene00045399 | protein_coding_gene | -4.0 | 0.0633 | 1.6E-33 | -2.2 | 0.2159 | 3.4E-24 |
| *Y39B6A.27* | WBGene00012686 | protein_coding_gene | -4.0 | 0.0630 | 3.1E-18 | -2.0 | 0.2421 | 4.6E-04 |
| *pxn-1* | WBGene00004256 | protein_coding_gene | -4.0 | 0.0626 | 1.5E-55 | -1.7 | 0.2996 | 5.2E-22 |
| *col-118* | WBGene00000692 | protein_coding_gene | -4.0 | 0.0619 | 6.1E-31 | -2.5 | 0.1768 | 9.3E-43 |
| *lipl-4* | WBGene00019376 | protein_coding_gene | -4.0 | 0.0618 | 2.8E-08 | -4.4 | 0.0472 | 3.0E-10 |
| *F48D6.4* | WBGene00018602 | protein_coding_gene | -4.0 | 0.0616 | 1.6E-106 | -2.9 | 0.1385 | 5.3E-83 |
| *R09H10.7* | WBGene00011184 | protein_coding_gene | -4.0 | 0.0615 | 9.9E-10 | -3.9 | 0.0650 | 2.3E-12 |
| *mnr-1* | WBGene00012184 | protein_coding_gene | -4.0 | 0.0611 | 6.2E-66 | -2.7 | 0.1561 | 1.6E-42 |
| *Y49E10.29* | WBGene00045239 | protein_coding_gene | -4.0 | 0.0608 | 2.1E-34 | -1.9 | 0.2641 | 1.9E-08 |
| *cnc-8* | WBGene00009624 | protein_coding_gene | -4.0 | 0.0606 | 1.1E-25 | -3.2 | 0.1096 | 1.3E-55 |
| *ugt-21* | WBGene00007885 | protein_coding_gene | -4.0 | 0.0605 | 2.4E-25 | -1.6 | 0.3290 | 2.1E-06 |
| *lgc-54* | WBGene00020528 | protein_coding_gene | -4.1 | 0.0603 | 1.9E-13 | -2.0 | 0.2581 | 3.4E-06 |
| *C10A4.6* | WBGene00015667 | protein_coding_gene | -4.1 | 0.0603 | 7.9E-08 | -2.8 | 0.1394 | 1.1E-03 |
| *perm-4* | WBGene00016638 | protein_coding_gene | -4.1 | 0.0598 | 7.0E-41 | -4.1 | 0.0581 | 1.1E-181 |
| *gst-41* | WBGene00001789 | protein_coding_gene | -4.1 | 0.0597 | 2.9E-15 | -2.3 | 0.2047 | 6.5E-05 |
| *C05C9.3* | WBGene00007327 | protein_coding_gene | -4.1 | 0.0596 | 1.2E-15 | -1.1 | 0.4708 | 4.2E-03 |
| *kynu-1* | WBGene00015802 | protein_coding_gene | -4.1 | 0.0596 | 1.5E-21 | -1.6 | 0.3241 | 9.2E-23 |
| *ugt-8* | WBGene00019234 | protein_coding_gene | -4.1 | 0.0595 | 3.8E-31 | -4.7 | 0.0383 | 5.0E-58 |
| *VB0395L.1* | WBGene00044150 | protein_coding_gene | -4.1 | 0.0592 | 6.4E-16 | -2.0 | 0.2486 | 6.6E-05 |
| *zipt-13* | WBGene00007591 | protein_coding_gene | -4.1 | 0.0589 | 1.2E-42 | -2.6 | 0.1696 | 3.7E-33 |
| *clec-227* | WBGene00008593 | protein_coding_gene | -4.1 | 0.0587 | 9.2E-14 | -3.6 | 0.0825 | 7.4E-12 |
| *T21D12.12* | WBGene00020651 | protein_coding_gene | -4.1 | 0.0587 | 3.4E-53 | -2.6 | 0.1707 | 2.5E-26 |
| *vit-5* | WBGene00006929 | protein_coding_gene | -4.1 | 0.0586 | 6.7E-24 | -3.4 | 0.0929 | 1.2E-44 |
| *K10C2.7* | WBGene00019622 | protein_coding_gene | -4.1 | 0.0585 | 1.3E-11 | -1.4 | 0.3792 | 2.2E-02 |
| *grl-22* | WBGene00001731 | protein_coding_gene | -4.1 | 0.0583 | 9.7E-06 | -2.4 | 0.1845 | 6.3E-03 |
| *fipr-10* | WBGene00008245 | protein_coding_gene | -4.1 | 0.0582 | 5.4E-11 | -1.7 | 0.3157 | 1.1E-04 |
| *F54F7.2* | WBGene00010064 | protein_coding_gene | -4.1 | 0.0579 | 7.7E-19 | -4.4 | 0.0470 | 2.3E-63 |
| *nas-20* | WBGene00003539 | protein_coding_gene | -4.1 | 0.0579 | 5.6E-26 | -3.0 | 0.1229 | 1.0E-36 |
| *F13H8.3* | WBGene00017436 | protein_coding_gene | -4.1 | 0.0577 | 6.5E-25 | -1.9 | 0.2666 | 8.1E-18 |
| *cah-5* | WBGene00000283 | protein_coding_gene | -4.1 | 0.0576 | 5.0E-37 | -2.2 | 0.2187 | 2.4E-22 |
| *Y58A7A.1* | WBGene00021975 | protein_coding_gene | -4.1 | 0.0574 | 5.5E-15 | -2.1 | 0.2395 | 2.3E-12 |
| *Y73F4A.1* | WBGene00013514 | protein_coding_gene | -4.1 | 0.0568 | 3.5E-44 | -3.7 | 0.0744 | 8.1E-46 |
| *W04A4.2* | WBGene00012231 | protein_coding_gene | -4.1 | 0.0564 | 9.9E-33 | -3.5 | 0.0881 | 6.7E-54 |

| | | | | | | | | |
|---|---|---|---|---|---|---|---|---|
| *F57H12.6* | WBGene00019021 | protein_coding_gene | -4.2 | 0.0559 | 7.6E-07 | -3.4 | 0.0969 | 8.3E-07 |
| *ZK829.1* | WBGene00014093 | protein_coding_gene | -4.2 | 0.0558 | 7.2E-05 | -4.7 | 0.0397 | 1.1E-06 |
| *cept-1* | WBGene00009057 | protein_coding_gene | -4.2 | 0.0558 | 3.2E-26 | -1.5 | 0.3567 | 5.5E-18 |
| *clec-160* | WBGene00017322 | protein_coding_gene | -4.2 | 0.0556 | 1.7E-54 | -2.7 | 0.1555 | 1.3E-33 |
| *C39D10.8* | WBGene00016535 | protein_coding_gene | -4.2 | 0.0555 | 2.8E-41 | -1.8 | 0.2910 | 3.0E-13 |
| *ptr-20* | WBGene00004234 | protein_coding_gene | -4.2 | 0.0553 | 4.0E-32 | -3.9 | 0.0648 | 8.9E-55 |
| *adt-3* | WBGene00020567 | protein_coding_gene | -4.2 | 0.0553 | 2.9E-25 | -1.9 | 0.2642 | 2.9E-07 |
| *DH11.2* | WBGene00008436 | protein_coding_gene | -4.2 | 0.0553 | 3.8E-39 | -4.9 | 0.0344 | 1.0E-221 |
| *clec-169* | WBGene00009526 | protein_coding_gene | -4.2 | 0.0551 | 4.1E-11 | -4.4 | 0.0458 | 2.6E-10 |
| *mct-3* | WBGene00010834 | protein_coding_gene | -4.2 | 0.0549 | 1.2E-54 | -2.2 | 0.2108 | 2.4E-62 |
| *twk-16* | WBGene00006670 | protein_coding_gene | -4.2 | 0.0547 | 9.2E-12 | -1.7 | 0.3063 | 3.4E-03 |
| *clh-1* | WBGene00000528 | protein_coding_gene | -4.2 | 0.0545 | 4.3E-45 | -3.6 | 0.0823 | 6.5E-117 |
| *F42A10.7* | WBGene00018343 | protein_coding_gene | -4.2 | 0.0543 | 6.3E-59 | -2.8 | 0.1457 | 2.6E-13 |
| *del-6* | WBGene00011891 | protein_coding_gene | -4.2 | 0.0543 | 5.1E-73 | -2.9 | 0.1314 | 3.2E-73 |
| *pmt-2* | WBGene00018811 | protein_coding_gene | -4.2 | 0.0540 | 5.6E-55 | -2.8 | 0.1474 | 1.8E-77 |
| *nspe-2* | WBGene00012604 | protein_coding_gene | -4.2 | 0.0539 | 3.5E-05 | -4.5 | 0.0450 | 4.1E-12 |
| *papl-1* | WBGene00008936 | protein_coding_gene | -4.2 | 0.0534 | 2.7E-25 | -2.5 | 0.1737 | 2.1E-57 |
| *doxa-1* | WBGene00015519 | protein_coding_gene | -4.2 | 0.0528 | 9.7E-26 | -1.5 | 0.3445 | 4.1E-06 |
| *F11G11.15* | WBGene00271822 | protein_coding_gene | -4.3 | 0.0524 | 7.3E-08 | -1.9 | 0.2594 | 3.4E-03 |
| *Y12A6A.1* | WBGene00012439 | protein_coding_gene | -4.3 | 0.0523 | 3.7E-53 | -3.2 | 0.1105 | 9.5E-28 |
| *Y51B9A.6* | WBGene00013088 | protein_coding_gene | -4.3 | 0.0521 | 1.3E-05 | -2.7 | 0.1558 | 1.8E-03 |
| *fmo-4* | WBGene00001479 | protein_coding_gene | -4.3 | 0.0520 | 7.5E-47 | -3.8 | 0.0734 | 2.2E-127 |
| *spl-2* | WBGene00006418 | protein_coding_gene | -4.3 | 0.0519 | 1.3E-14 | -4.5 | 0.0456 | 1.2E-30 |
| *Y73C8C.10* | WBGene00022267 | protein_coding_gene | -4.3 | 0.0518 | 4.8E-05 | -2.8 | 0.1444 | 4.7E-05 |
| *ifc-1* | WBGene00002055 | protein_coding_gene | -4.3 | 0.0513 | 4.6E-41 | -3.7 | 0.0753 | 9.1E-105 |
| *pho-7* | WBGene00011879 | protein_coding_gene | -4.3 | 0.0513 | 7.3E-18 | -1.7 | 0.3075 | 2.3E-05 |
| *C16E9.1* | WBGene00015865 | protein_coding_gene | -4.3 | 0.0512 | 2.4E-19 | -3.6 | 0.0800 | 7.5E-10 |
| *lys-10* | WBGene00003099 | protein_coding_gene | -4.3 | 0.0512 | 1.1E-06 | -5.7 | 0.0189 | 2.6E-09 |
| *ptr-8* | WBGene00004222 | protein_coding_gene | -4.3 | 0.0508 | 7.1E-23 | -5.0 | 0.0305 | 2.7E-59 |
| *F12E12.11* | WBGene00044811 | protein_coding_gene | -4.3 | 0.0504 | 2.6E-16 | -1.8 | 0.2907 | 7.5E-07 |
| *F59B1.2* | WBGene00019097 | protein_coding_gene | -4.3 | 0.0503 | 1.3E-63 | -3.1 | 0.1188 | 3.7E-118 |
| *Y41C4A.32* | WBGene00270321 | protein_coding_gene | -4.3 | 0.0501 | 9.1E-14 | -1.1 | 0.4622 | 1.9E-03 |
| *pqn-37* | WBGene00004124 | protein_coding_gene | -4.3 | 0.0500 | 4.9E-22 | -2.7 | 0.1516 | 3.8E-07 |
| *adt-2* | WBGene00000083 | protein_coding_gene | -4.3 | 0.0499 | 8.5E-25 | -1.7 | 0.3058 | 5.8E-22 |
| ***Y47G6A.15*** | **WBGene00021641** | **protein_coding_gene** | **-4.3** | **0.0494** | **2.3E-26** | **2.4** | **5.2497** | **1.8E-04** |
| *ttr-18* | WBGene00007142 | protein_coding_gene | -4.3 | 0.0491 | 3.4E-45 | -3.2 | 0.1075 | 4.8E-57 |
| *nlp-24* | WBGene00003762 | protein_coding_gene | -4.4 | 0.0482 | 8.0E-27 | -3.1 | 0.1188 | 1.3E-09 |
| *ugt-22* | WBGene00007455 | protein_coding_gene | -4.4 | 0.0482 | 3.9E-32 | -1.6 | 0.3291 | 7.0E-06 |
| *tag-10* | WBGene00006404 | protein_coding_gene | -4.4 | 0.0482 | 3.4E-47 | -2.0 | 0.2431 | 3.5E-26 |
| *nas-11* | WBGene00003530 | protein_coding_gene | -4.4 | 0.0481 | 3.6E-67 | -3.6 | 0.0808 | 5.2E-116 |
| *tyr-1* | WBGene00015332 | protein_coding_gene | -4.4 | 0.0480 | 1.0E-33 | -5.4 | 0.0234 | 4.8E-84 |
| *F25D1.5* | WBGene00009110 | protein_coding_gene | -4.4 | 0.0479 | 4.7E-37 | -3.1 | 0.1146 | 1.8E-18 |
| *twk-22* | WBGene00006675 | protein_coding_gene | -4.4 | 0.0478 | 1.1E-29 | -1.8 | 0.2972 | 4.4E-08 |
| *cbl-1* | WBGene00007533 | protein_coding_gene | -4.4 | 0.0478 | 2.0E-27 | -1.3 | 0.3935 | 3.8E-13 |
| *W02B3.4* | WBGene00020924 | protein_coding_gene | -4.4 | 0.0477 | 3.4E-21 | -3.4 | 0.0957 | 1.7E-12 |
| *dos-1* | WBGene00013980 | protein_coding_gene | -4.4 | 0.0475 | 1.6E-33 | -4.8 | 0.0351 | 1.1E-42 |
| *H11E01.2* | WBGene00019187 | protein_coding_gene | -4.4 | 0.0474 | 1.1E-25 | -3.4 | 0.0951 | 2.4E-17 |
| *Y4C6B.4* | WBGene00021158 | protein_coding_gene | -4.4 | 0.0473 | 1.5E-16 | -2.4 | 0.1913 | 2.9E-10 |
| *C42D4.3* | WBGene00016596 | protein_coding_gene | -4.4 | 0.0472 | 1.0E-12 | -1.2 | 0.4294 | 4.4E-02 |
| *K04A8.20* | WBGene00235332 | protein_coding_gene | -4.4 | 0.0471 | 1.5E-07 | -6.1 | 0.0149 | 8.2E-14 |
| *pgph-3* | WBGene00016892 | protein_coding_gene | -4.4 | 0.0469 | 5.7E-13 | -1.8 | 0.2802 | 4.0E-06 |
| *mxl-3* | WBGene00003511 | protein_coding_gene | -4.4 | 0.0469 | 2.4E-16 | -2.0 | 0.2521 | 8.1E-05 |
| *C39D10.7* | WBGene00016534 | protein_coding_gene | -4.4 | 0.0467 | 4.5E-39 | -4.2 | 0.0558 | 7.2E-125 |
| *slc-17.4* | WBGene00012130 | protein_coding_gene | -4.4 | 0.0467 | 3.0E-49 | -2.1 | 0.2281 | 3.7E-19 |
| *fmo-3* | WBGene00001478 | protein_coding_gene | -4.4 | 0.0466 | 5.9E-27 | -2.3 | 0.1995 | 1.1E-15 |
| *ugt-30* | WBGene00011340 | protein_coding_gene | -4.4 | 0.0464 | 1.6E-24 | -2.4 | 0.1876 | 1.7E-04 |
| *grl-4* | WBGene00001713 | protein_coding_gene | -4.4 | 0.0461 | 1.3E-18 | -2.2 | 0.2231 | 1.2E-11 |
| *str-180* | WBGene00006224 | protein_coding_gene | -4.4 | 0.0460 | 9.2E-06 | -2.7 | 0.1496 | 1.6E-04 |
| *cyp-35A3* | WBGene00019565 | protein_coding_gene | -4.4 | 0.0460 | 2.2E-09 | -4.3 | 0.0509 | 5.2E-09 |
| *acs-2* | WBGene00009221 | protein_coding_gene | -4.4 | 0.0459 | 1.2E-28 | -3.0 | 0.1252 | 3.3E-14 |
| *ZK455.5* | WBGene00013975 | protein_coding_gene | -4.5 | 0.0453 | 1.3E-09 | -2.4 | 0.1833 | 2.1E-05 |
| *ptr-13* | WBGene00004227 | protein_coding_gene | -4.5 | 0.0452 | 9.4E-53 | -4.1 | 0.0582 | 8.0E-92 |
| *F13D12.3* | WBGene00008739 | protein_coding_gene | -4.5 | 0.0450 | 6.2E-50 | -4.2 | 0.0553 | 1.6E-42 |
| *F58G6.7* | WBGene00010277 | protein_coding_gene | -4.5 | 0.0450 | 2.0E-15 | -2.0 | 0.2495 | 2.1E-05 |
| *AC3.5* | WBGene00007071 | protein_coding_gene | -4.5 | 0.0449 | 2.9E-31 | -1.6 | 0.3304 | 3.0E-19 |
| *ugt-15* | WBGene00008097 | protein_coding_gene | -4.5 | 0.0448 | 1.0E-06 | -4.0 | 0.0609 | 7.0E-18 |
| *mlt-7* | WBGene00022743 | protein_coding_gene | -4.5 | 0.0446 | 1.0E-45 | -4.1 | 0.0596 | 6.3E-78 |
| *spp-17* | WBGene00005002 | protein_coding_gene | -4.5 | 0.0446 | 6.0E-44 | -3.7 | 0.0777 | 1.0E-114 |
| *glc-2* | WBGene00001592 | protein_coding_gene | -4.5 | 0.0442 | 4.7E-23 | -3.5 | 0.0869 | 4.5E-32 |
| *nhx-3* | WBGene00003731 | protein_coding_gene | -4.5 | 0.0440 | 5.8E-32 | -2.7 | 0.1587 | 1.5E-14 |

| | | | | | | | | |
|---|---|---|---|---|---|---|---|---|
| *kel-8* | WBGene00020952 | protein_coding_gene | -4.5 | 0.0438 | 5.5E-73 | -3.5 | 0.0867 | 2.8E-76 |
| *kin-16* | WBGene00002200 | protein_coding_gene | -4.5 | 0.0438 | 2.0E-33 | -3.1 | 0.1168 | 1.2E-22 |
| *ugt-14* | WBGene00019233 | protein_coding_gene | -4.5 | 0.0437 | 1.9E-07 | -4.3 | 0.0495 | 3.5E-16 |
| *grsp-4* | WBGene00017201 | protein_coding_gene | -4.5 | 0.0437 | 2.9E-25 | -2.1 | 0.2307 | 4.2E-05 |
| *cyp-14A2* | WBGene00010706 | protein_coding_gene | -4.5 | 0.0436 | 1.2E-14 | -4.6 | 0.0408 | 5.0E-13 |
| *nlp-27* | WBGene00003765 | protein_coding_gene | -4.5 | 0.0435 | 6.3E-35 | -4.0 | 0.0615 | 7.2E-101 |
| *poml-2* | WBGene00017089 | protein_coding_gene | -4.5 | 0.0429 | 5.9E-32 | -2.2 | 0.2172 | 1.4E-10 |
| *C06E8.5* | WBGene00015544 | protein_coding_gene | -4.5 | 0.0428 | 3.5E-14 | -2.6 | 0.1700 | 7.7E-06 |
| *ugt-17* | WBGene00007422 | protein_coding_gene | -4.6 | 0.0426 | 9.1E-107 | -5.2 | 0.0270 | 1.2E-113 |
| *mup-4* | WBGene00003497 | protein_coding_gene | -4.6 | 0.0420 | 7.6E-42 | -2.4 | 0.1902 | 3.1E-52 |
| *cyp-34A8* | WBGene00015043 | protein_coding_gene | -4.6 | 0.0420 | 9.6E-70 | -3.0 | 0.1221 | 5.6E-55 |
| *E04F6.6* | WBGene00017126 | protein_coding_gene | -4.6 | 0.0418 | 3.5E-55 | -4.2 | 0.0547 | 1.1E-58 |
| *F41E6.12* | WBGene00018293 | protein_coding_gene | -4.6 | 0.0416 | 3.9E-14 | -1.3 | 0.4122 | 1.1E-03 |
| *Y37A1A.2* | WBGene00012535 | protein_coding_gene | -4.6 | 0.0414 | 6.4E-27 | -3.7 | 0.0752 | 3.7E-20 |
| *ptr-14* | WBGene00004228 | protein_coding_gene | -4.6 | 0.0408 | 4.3E-26 | -3.5 | 0.0868 | 3.3E-43 |
| *ent-7* | WBGene00009686 | protein_coding_gene | -4.6 | 0.0406 | 1.3E-27 | -2.7 | 0.1502 | 7.1E-42 |
| *Y37A1B.5* | WBGene00012538 | protein_coding_gene | -4.6 | 0.0405 | 1.1E-93 | -3.5 | 0.0858 | 1.7E-70 |
| *Y51A2D.18* | WBGene00013083 | protein_coding_gene | -4.6 | 0.0405 | 1.1E-20 | -2.4 | 0.1838 | 6.3E-18 |
| *Y50D4B.4* | WBGene00021743 | protein_coding_gene | -4.6 | 0.0404 | 3.6E-32 | -2.7 | 0.1488 | 9.1E-25 |
| *F21A3.3* | WBGene00008993 | protein_coding_gene | -4.6 | 0.0403 | 2.1E-26 | -1.9 | 0.2647 | 9.1E-07 |
| *col-170* | WBGene00000743 | protein_coding_gene | -4.6 | 0.0403 | 6.7E-08 | -1.7 | 0.3019 | 2.5E-02 |
| *K01C8.1* | WBGene00010456 | protein_coding_gene | -4.7 | 0.0397 | 6.9E-54 | -5.3 | 0.0249 | 1.8E-179 |
| *cyp-13A9* | WBGene00014838 | pseudogene | -4.7 | 0.0397 | 3.2E-10 | -4.4 | 0.0468 | 2.0E-08 |
| *cpg-9* | WBGene00022072 | protein_coding_gene | -4.7 | 0.0389 | 8.9E-36 | -1.2 | 0.4281 | 3.3E-07 |
| *K07E1.1* | WBGene00019489 | protein_coding_gene | -4.7 | 0.0385 | 2.9E-47 | -4.4 | 0.0460 | 1.1E-83 |
| *C02F12.5* | WBGene00015355 | protein_coding_gene | -4.7 | 0.0384 | 7.3E-21 | -3.6 | 0.0845 | 1.9E-14 |
| *T10B5.8* | WBGene00020394 | protein_coding_gene | -4.7 | 0.0381 | 1.8E-30 | -3.8 | 0.0718 | 2.7E-23 |
| *vit-4* | WBGene00006928 | protein_coding_gene | -4.7 | 0.0380 | 1.4E-10 | -3.6 | 0.0832 | 9.0E-07 |
| *pho-13* | WBGene00008560 | protein_coding_gene | -4.7 | 0.0380 | 6.4E-19 | -1.5 | 0.3453 | 7.2E-03 |
| *C44C10.3* | WBGene00008084 | protein_coding_gene | -4.7 | 0.0380 | 1.0E-23 | -2.6 | 0.1689 | 4.0E-12 |
| *F17A9.5* | WBGene00017537 | protein_coding_gene | -4.7 | 0.0378 | 9.9E-106 | -3.5 | 0.0867 | 1.8E-42 |
| *C39E9.8* | WBGene00008032 | protein_coding_gene | -4.7 | 0.0377 | 2.5E-98 | -5.1 | 0.0301 | 8.4E-159 |
| *col-150* | WBGene00000723 | protein_coding_gene | -4.7 | 0.0375 | 1.7E-27 | -1.6 | 0.3297 | 7.2E-07 |
| *F35B3.4* | WBGene00018031 | protein_coding_gene | -4.7 | 0.0374 | 6.4E-08 | -2.3 | 0.2091 | 1.7E-03 |
| *C45E5.1* | WBGene00016664 | protein_coding_gene | -4.7 | 0.0374 | 1.2E-17 | -1.9 | 0.2615 | 1.3E-11 |
| *ttll-15* | WBGene00010630 | protein_coding_gene | -4.7 | 0.0372 | 1.4E-32 | -2.9 | 0.1336 | 4.1E-15 |
| *cest-10* | WBGene00016595 | protein_coding_gene | -4.8 | 0.0367 | 1.0E-34 | -1.6 | 0.3254 | 3.8E-07 |
| *catp-3* | WBGene00015660 | protein_coding_gene | -4.8 | 0.0365 | 1.0E-25 | -4.2 | 0.0537 | 6.9E-30 |
| *kin-15* | WBGene00002199 | protein_coding_gene | -4.8 | 0.0362 | 6.5E-17 | -2.5 | 0.1808 | 1.7E-08 |
| *F44A2.3* | WBGene00018403 | protein_coding_gene | -4.8 | 0.0361 | 1.1E-53 | -4.2 | 0.0541 | 9.1E-52 |
| *ZC434.9* | WBGene00013895 | protein_coding_gene | -4.8 | 0.0359 | 3.5E-47 | -2.7 | 0.1553 | 2.0E-32 |
| *ZC196.2* | WBGene00022546 | protein_coding_gene | -4.8 | 0.0356 | 2.8E-10 | -2.3 | 0.2091 | 5.4E-05 |
| *col-39* | WBGene00000616 | protein_coding_gene | -4.8 | 0.0353 | 8.8E-23 | -3.1 | 0.1196 | 9.2E-41 |
| *grd-5* | WBGene00001694 | protein_coding_gene | -4.8 | 0.0353 | 2.7E-31 | -3.1 | 0.1204 | 2.0E-12 |
| *twk-34* | WBGene00006686 | protein_coding_gene | -4.8 | 0.0352 | 2.6E-45 | -3.4 | 0.0916 | 5.9E-23 |
| *nlp-26* | WBGene00003764 | protein_coding_gene | -4.9 | 0.0345 | 2.4E-25 | -4.1 | 0.0576 | 6.4E-17 |
| *col-169* | WBGene00000742 | protein_coding_gene | -4.9 | 0.0339 | 3.1E-09 | -1.9 | 0.2770 | 6.6E-03 |
| *clec-218* | WBGene00020938 | protein_coding_gene | -4.9 | 0.0338 | 7.2E-24 | -5.5 | 0.0225 | 4.4E-144 |
| *F36H9.4* | WBGene00018112 | protein_coding_gene | -4.9 | 0.0334 | 4.2E-32 | -4.2 | 0.0553 | 2.7E-47 |
| *decr-1.1* | WBGene00018380 | protein_coding_gene | -4.9 | 0.0330 | 3.8E-17 | -3.2 | 0.1054 | 4.9E-18 |
| *fipr-21* | WBGene00009621 | protein_coding_gene | -4.9 | 0.0330 | 9.5E-52 | -3.8 | 0.0724 | 4.1E-100 |
| *aqp-7* | WBGene00000175 | protein_coding_gene | -4.9 | 0.0329 | 1.9E-118 | -2.8 | 0.1486 | 7.7E-59 |
| *Y59E9AL.4* | WBGene00021994 | protein_coding_gene | -4.9 | 0.0326 | 5.1E-24 | -3.1 | 0.1141 | 7.0E-10 |
| *ugt-40* | WBGene00017331 | protein_coding_gene | -4.9 | 0.0326 | 1.1E-18 | -4.2 | 0.0542 | 2.9E-12 |
| *F54E4.2* | WBGene00010058 | protein_coding_gene | -4.9 | 0.0324 | 1.6E-06 | -4.7 | 0.0388 | 8.9E-07 |
| *clec-97* | WBGene00013931 | protein_coding_gene | -5.0 | 0.0323 | 8.5E-38 | -3.2 | 0.1081 | 4.0E-29 |
| *K09C4.5* | WBGene00019550 | protein_coding_gene | -5.0 | 0.0322 | 4.6E-08 | -2.0 | 0.2445 | 5.9E-05 |
| *T21C9.9* | WBGene00011895 | protein_coding_gene | -5.0 | 0.0320 | 3.3E-12 | -1.6 | 0.3279 | 1.2E-02 |
| *ugt-42* | WBGene00017959 | protein_coding_gene | -5.0 | 0.0310 | 2.3E-07 | -3.6 | 0.0809 | 4.9E-06 |
| *K10C2.8* | WBGene00045265 | protein_coding_gene | -5.1 | 0.0295 | 6.8E-48 | -2.2 | 0.2239 | 3.6E-12 |
| *msra-1* | WBGene00018393 | protein_coding_gene | -5.1 | 0.0295 | 2.0E-32 | -4.6 | 0.0399 | 9.6E-23 |
| *F55H12.4* | WBGene00010135 | protein_coding_gene | -5.1 | 0.0293 | 7.3E-89 | -4.3 | 0.0520 | 9.1E-97 |
| *Y47G6A.19* | WBGene00021645 | protein_coding_gene | -5.1 | 0.0292 | 4.2E-40 | -2.3 | 0.2029 | 1.0E-19 |
| *tatn-1* | WBGene00009628 | protein_coding_gene | -5.1 | 0.0287 | 1.3E-37 | -3.2 | 0.1102 | 3.8E-73 |
| *tyr-4* | WBGene00016419 | protein_coding_gene | -5.1 | 0.0284 | 8.6E-140 | -4.4 | 0.0489 | 5.4E-176 |
| *lgc-32* | WBGene00020569 | protein_coding_gene | -5.2 | 0.0281 | 3.8E-25 | -4.2 | 0.0539 | 1.3E-18 |
| *Y53F4B.23* | WBGene00013169 | protein_coding_gene | -5.2 | 0.0277 | 2.2E-08 | -2.5 | 0.1751 | 6.4E-06 |
| *C23H5.8* | WBGene00016027 | protein_coding_gene | -5.2 | 0.0275 | 7.7E-50 | -1.1 | 0.4821 | 1.4E-02 |
| *H06A10.1* | WBGene00010368 | protein_coding_gene | -5.2 | 0.0267 | 2.1E-22 | -4.1 | 0.0598 | 5.7E-09 |
| *C39B5.5* | WBGene00016523 | protein_coding_gene | -5.2 | 0.0266 | 2.0E-22 | -1.9 | 0.2676 | 3.7E-03 |

| | | | | | | | | |
|---|---|---|---|---|---|---|---|---|
| *vglu-3* | WBGene00011556 | protein_coding_gene | -5.2 | 0.0265 | 2.9E-36 | -5.0 | 0.0311 | 1.8E-24 |
| *col-141* | WBGene00000714 | protein_coding_gene | -5.2 | 0.0264 | 1.8E-10 | -2.2 | 0.2151 | 1.4E-02 |
| *F44A6.5* | WBGene00009677 | protein_coding_gene | -5.2 | 0.0263 | 2.1E-40 | -5.0 | 0.0305 | 1.1E-73 |
| *linc-107* | WBGene00219655 | lincRNA_gene | -5.2 | 0.0263 | 3.6E-07 | -2.8 | 0.1460 | 2.9E-03 |
| *acox-1.3* | WBGene00008566 | protein_coding_gene | -5.3 | 0.0262 | 3.2E-37 | -2.2 | 0.2229 | 1.1E-09 |
| *col-125* | WBGene00000699 | protein_coding_gene | -5.3 | 0.0262 | 1.9E-16 | -1.3 | 0.4118 | 1.0E-02 |
| *drd-10* | WBGene00016361 | protein_coding_gene | -5.3 | 0.0261 | 3.5E-14 | -4.1 | 0.0563 | 1.8E-05 |
| *cyp-13A12* | WBGene00008810 | protein_coding_gene | -5.3 | 0.0258 | 3.2E-07 | -2.4 | 0.1895 | 3.3E-03 |
| *T25G12.13* | WBGene00219274 | protein_coding_gene | -5.3 | 0.0256 | 7.1E-08 | -2.3 | 0.1977 | 2.2E-02 |
| *fat-7* | WBGene00001399 | protein_coding_gene | -5.3 | 0.0253 | 4.4E-59 | -4.2 | 0.0534 | 1.5E-34 |
| *efn-3* | WBGene00001164 | protein_coding_gene | -5.3 | 0.0253 | 1.3E-21 | -2.8 | 0.1395 | 2.1E-06 |
| *skpo-3* | WBGene00017968 | protein_coding_gene | -5.3 | 0.0247 | 2.2E-65 | -2.8 | 0.1445 | 2.0E-64 |
| *sams-1* | WBGene00008205 | protein_coding_gene | -5.3 | 0.0247 | 2.9E-43 | -2.7 | 0.1514 | 1.7E-100 |
| *cyp-14A1* | WBGene00010705 | protein_coding_gene | -5.4 | 0.0244 | 7.3E-17 | -3.4 | 0.0972 | 1.2E-08 |
| *srd-14* | WBGene00005092 | pseudogene | -5.4 | 0.0244 | 5.9E-14 | -5.5 | 0.0226 | 2.0E-09 |
| *F18E3.13* | WBGene00045276 | protein_coding_gene | -5.4 | 0.0243 | 1.8E-13 | -2.1 | 0.2363 | 3.9E-03 |
| *pgp-1* | WBGene00003995 | protein_coding_gene | -5.4 | 0.0243 | 9.2E-46 | -3.7 | 0.0794 | 2.1E-12 |
| *dhs-25* | WBGene00000988 | protein_coding_gene | -5.4 | 0.0243 | 1.7E-49 | -1.4 | 0.3821 | 4.6E-04 |
| *F40H3.2* | WBGene00018250 | protein_coding_gene | -5.4 | 0.0235 | 1.2E-15 | -1.5 | 0.3534 | 5.7E-04 |
| *cpr-5* | WBGene00000785 | protein_coding_gene | -5.4 | 0.0234 | 3.1E-213 | -4.8 | 0.0356 | 1.3E-40 |
| *col-106* | WBGene00000680 | protein_coding_gene | -5.4 | 0.0234 | 1.0E-77 | -3.9 | 0.0678 | 8.0E-138 |
| *clec-57* | WBGene00008594 | protein_coding_gene | -5.4 | 0.0233 | 1.8E-28 | -3.2 | 0.1077 | 3.5E-17 |
| *ugt-62* | WBGene00010904 | protein_coding_gene | -5.4 | 0.0233 | 1.0E-50 | -2.9 | 0.1380 | 3.8E-19 |
| *mua-6* | WBGene00003485 | protein_coding_gene | -5.4 | 0.0232 | 3.8E-30 | -2.8 | 0.1441 | 3.3E-26 |
| *col-184* | WBGene00000757 | protein_coding_gene | -5.4 | 0.0232 | 3.4E-99 | -4.5 | 0.0432 | 2.7E-295 |
| *B0222.5* | WBGene00015056 | protein_coding_gene | -5.4 | 0.0230 | 1.5E-46 | -3.6 | 0.0845 | 9.1E-46 |
| *E02H4.7* | WBGene00077757 | protein_coding_gene | -5.4 | 0.0229 | 5.7E-30 | -2.8 | 0.1440 | 2.3E-34 |
| *col-155* | WBGene00000728 | protein_coding_gene | -5.4 | 0.0229 | 3.7E-36 | -3.3 | 0.1025 | 5.8E-23 |
| *fipr-13* | WBGene00010183 | protein_coding_gene | -5.5 | 0.0228 | 1.7E-19 | -4.0 | 0.0605 | 3.9E-30 |
| *C11E4.7* | WBGene00007521 | protein_coding_gene | -5.5 | 0.0228 | 5.7E-27 | -3.2 | 0.1093 | 4.4E-22 |
| *C49F5.9* | WBGene00045316 | pseudogene | -5.5 | 0.0228 | 4.0E-18 | -2.9 | 0.1342 | 1.1E-11 |
| *B0205.13* | WBGene00044644 | protein_coding_gene | -5.5 | 0.0226 | 8.6E-28 | -2.1 | 0.2316 | 2.9E-06 |
| *col-96* | WBGene00000671 | protein_coding_gene | -5.5 | 0.0222 | 3.6E-35 | -4.1 | 0.0572 | 9.8E-27 |
| *col-176* | WBGene00000749 | protein_coding_gene | -5.5 | 0.0221 | 1.1E-09 | -2.1 | 0.2342 | 1.9E-02 |
| *col-119* | WBGene00000693 | protein_coding_gene | -5.5 | 0.0221 | 2.2E-82 | -4.1 | 0.0595 | 3.6E-230 |
| *T04F8.8* | WBGene00011446 | protein_coding_gene | -5.5 | 0.0220 | 1.4E-121 | -4.9 | 0.0326 | 2.1E-214 |
| *col-43* | WBGene00000620 | protein_coding_gene | -5.5 | 0.0218 | 1.6E-07 | -2.5 | 0.1817 | 1.3E-02 |
| *ugt-41* | WBGene00017336 | protein_coding_gene | -5.5 | 0.0217 | 6.4E-55 | -5.1 | 0.0294 | 5.2E-56 |
| *T07A5.1* | WBGene00011554 | protein_coding_gene | -5.5 | 0.0217 | 1.3E-13 | -3.2 | 0.1057 | 9.6E-06 |
| *K08H10.6* | WBGene00010698 | protein_coding_gene | -5.5 | 0.0216 | 1.8E-08 | -3.1 | 0.1165 | 2.6E-04 |
| *C09B8.4* | WBGene00015623 | protein_coding_gene | -5.6 | 0.0213 | 2.8E-53 | -3.1 | 0.1172 | 1.0E-20 |
| *F32H5.3* | WBGene00009349 | protein_coding_gene | -5.6 | 0.0213 | 8.9E-36 | -5.4 | 0.0237 | 2.7E-20 |
| *clec-166* | WBGene00009518 | protein_coding_gene | -5.6 | 0.0210 | 3.4E-32 | -6.2 | 0.0134 | 1.6E-45 |
| *ZK593.3* | WBGene00014003 | protein_coding_gene | -5.7 | 0.0198 | 2.3E-38 | -5.6 | 0.0209 | 1.5E-51 |
| *col-62* | WBGene00000638 | protein_coding_gene | -5.7 | 0.0195 | 2.5E-11 | -3.9 | 0.0687 | 1.8E-06 |
| *gpdh-1* | WBGene00009824 | protein_coding_gene | -5.7 | 0.0194 | 1.0E-11 | -2.0 | 0.2494 | 4.6E-07 |
| *hphd-1* | WBGene00012608 | protein_coding_gene | -5.7 | 0.0193 | 2.6E-85 | -4.8 | 0.0359 | 2.2E-38 |
| ***Y51H7C.13*** | **WBGene00021791** | **protein_coding_gene** | **-5.7** | **0.0191** | **4.0E-25** | **1.9** | **3.7583** | **2.5E-02** |
| *zmp-3* | WBGene00016283 | protein_coding_gene | -5.7 | 0.0190 | 1.5E-48 | -5.1 | 0.0284 | 4.0E-23 |
| *col-154* | WBGene00000727 | protein_coding_gene | -5.7 | 0.0187 | 8.6E-13 | -1.9 | 0.2673 | 3.0E-05 |
| *F18E3.11* | WBGene00017567 | protein_coding_gene | -5.8 | 0.0186 | 7.9E-11 | -2.6 | 0.1647 | 1.0E-03 |
| *F46F2.3* | WBGene00009787 | protein_coding_gene | -5.8 | 0.0185 | 1.0E-10 | -2.0 | 0.2509 | 9.2E-06 |
| *B0454.5* | WBGene00015196 | protein_coding_gene | -5.8 | 0.0181 | 2.8E-24 | -4.7 | 0.0382 | 2.0E-13 |
| *clec-5* | WBGene00016450 | protein_coding_gene | -5.8 | 0.0181 | 2.2E-63 | -6.1 | 0.0147 | 1.7E-57 |
| *amt-1* | WBGene00000133 | protein_coding_gene | -5.8 | 0.0178 | 5.1E-11 | -3.8 | 0.0715 | 3.2E-06 |
| *clec-47* | WBGene00011668 | protein_coding_gene | -5.8 | 0.0178 | 3.6E-62 | -5.6 | 0.0200 | 2.6E-50 |
| *tyr-2* | WBGene00010661 | protein_coding_gene | -5.9 | 0.0171 | 2.3E-43 | -2.5 | 0.1709 | 4.6E-16 |
| *Y38H6C.21* | WBGene00012633 | protein_coding_gene | -5.9 | 0.0162 | 1.2E-09 | -6.0 | 0.0158 | 1.2E-11 |
| *col-122* | WBGene00000696 | protein_coding_gene | -6.0 | 0.0159 | 1.0E-105 | -4.7 | 0.0376 | 2.2E-98 |
| *vit-3* | WBGene00006927 | protein_coding_gene | -6.0 | 0.0159 | 4.2E-25 | -4.8 | 0.0354 | 2.2E-10 |
| *vit-1* | WBGene00006925 | protein_coding_gene | -6.0 | 0.0157 | 5.5E-26 | -3.1 | 0.1193 | 1.3E-05 |
| *Y38H6C.19* | WBGene00012631 | protein_coding_gene | -6.0 | 0.0151 | 1.4E-11 | -3.1 | 0.1199 | 5.7E-04 |
| *col-178* | WBGene00000751 | protein_coding_gene | -6.1 | 0.0149 | 7.4E-73 | -4.7 | 0.0390 | 2.9E-222 |
| *srh-237* | WBGene00005444 | protein_coding_gene | -6.1 | 0.0141 | 1.4E-28 | -3.9 | 0.0647 | 2.2E-31 |
| *col-181* | WBGene00000754 | protein_coding_gene | -6.2 | 0.0141 | 1.6E-86 | -5.3 | 0.0261 | 0.0E+00 |
| *hmit-1.2* | WBGene00013074 | protein_coding_gene | -6.2 | 0.0138 | 7.3E-29 | -4.2 | 0.0556 | 1.7E-19 |
| *col-143* | WBGene00000716 | protein_coding_gene | -6.2 | 0.0136 | 4.4E-27 | -4.2 | 0.0532 | 3.0E-39 |
| *ZK218.1* | WBGene00013935 | protein_coding_gene | -6.2 | 0.0135 | 3.5E-09 | -3.8 | 0.0715 | 1.7E-07 |
| *col-19* | WBGene00000608 | protein_coding_gene | -6.2 | 0.0134 | 6.3E-89 | -4.7 | 0.0386 | 3.3E-86 |
| *grd-3* | WBGene00001692 | protein_coding_gene | -6.2 | 0.0132 | 9.8E-67 | -4.2 | 0.0550 | 2.0E-21 |

| | | | | | | | | |
|---|---|---|---|---|---|---|---|---|
| *col-7* | WBGene00000596 | protein_coding_gene | -6.3 | 0.0131 | 1.0E-10 | -4.4 | 0.0482 | 7.2E-06 |
| *col-145* | WBGene00000718 | protein_coding_gene | -6.3 | 0.0129 | 5.8E-14 | -2.3 | 0.2031 | 2.3E-02 |
| *col-179* | WBGene00000752 | protein_coding_gene | -6.3 | 0.0129 | 3.0E-79 | -4.1 | 0.0578 | 2.6E-151 |
| *col-140* | WBGene00000713 | protein_coding_gene | -6.3 | 0.0128 | 6.5E-86 | -4.9 | 0.0326 | 5.2E-198 |
| *col-20* | WBGene00000609 | protein_coding_gene | -6.3 | 0.0128 | 2.0E-76 | -4.9 | 0.0341 | 3.6E-248 |
| *ugt-63* | WBGene00015449 | protein_coding_gene | -6.3 | 0.0127 | 1.0E-15 | -6.2 | 0.0138 | 1.2E-51 |
| *K08D12.6* | WBGene00019540 | protein_coding_gene | -6.3 | 0.0125 | 5.6E-53 | -3.6 | 0.0825 | 3.3E-26 |
| *C45B2.1* | WBGene00016658 | protein_coding_gene | -6.3 | 0.0124 | 7.8E-40 | -3.7 | 0.0789 | 3.3E-28 |
| *T05E12.6* | WBGene00011487 | protein_coding_gene | -6.3 | 0.0124 | 1.1E-58 | -4.1 | 0.0597 | 1.4E-53 |
| *col-156* | WBGene00000729 | protein_coding_gene | -6.4 | 0.0117 | 9.8E-13 | -2.7 | 0.1548 | 4.7E-04 |
| *col-160* | WBGene00000733 | protein_coding_gene | -6.5 | 0.0111 | 7.1E-73 | -4.6 | 0.0399 | 2.5E-215 |
| *R08E5.3* | WBGene00019963 | protein_coding_gene | -6.5 | 0.0110 | 1.2E-42 | -2.1 | 0.2398 | 1.4E-23 |
| *col-137* | WBGene00000710 | protein_coding_gene | -6.5 | 0.0109 | 1.2E-09 | -3.1 | 0.1146 | 1.8E-03 |
| *col-91* | WBGene00000666 | protein_coding_gene | -6.6 | 0.0107 | 1.3E-20 | -3.7 | 0.0756 | 5.8E-09 |
| *F54D8.10* | WBGene00284847 | protein_coding_gene | -6.6 | 0.0102 | 8.5E-10 | -3.1 | 0.1133 | 1.7E-04 |
| *F11E6.3* | WBGene00008707 | protein_coding_gene | -6.6 | 0.0101 | 2.4E-86 | -3.9 | 0.0671 | 6.7E-129 |
| *col-101* | WBGene00000675 | protein_coding_gene | -6.6 | 0.0101 | 9.0E-56 | -4.9 | 0.0338 | 1.1E-35 |
| *nhr-114* | WBGene00003704 | protein_coding_gene | -6.7 | 0.0098 | 1.1E-56 | -3.6 | 0.0824 | 3.4E-103 |
| *col-180* | WBGene00000753 | protein_coding_gene | -6.7 | 0.0096 | 7.0E-19 | -2.4 | 0.1859 | 8.0E-04 |
| *col-8* | WBGene00000597 | protein_coding_gene | -6.7 | 0.0096 | 6.7E-95 | -4.3 | 0.0521 | 5.4E-51 |
| *fbxa-72* | WBGene00017962 | protein_coding_gene | -6.8 | 0.0090 | 1.7E-32 | -4.4 | 0.0471 | 5.8E-23 |
| *Y34F4.2* | WBGene00021337 | protein_coding_gene | -6.8 | 0.0090 | 2.4E-23 | -6.6 | 0.0102 | 1.6E-25 |
| *F18E3.12* | WBGene00045275 | protein_coding_gene | -6.8 | 0.0089 | 3.2E-12 | -3.3 | 0.1031 | 1.6E-04 |
| *asp-13* | WBGene00017881 | protein_coding_gene | -6.9 | 0.0086 | 1.8E-237 | -5.2 | 0.0271 | 0.0E+00 |
| *cyp-13A4* | WBGene00011671 | protein_coding_gene | -6.9 | 0.0083 | 4.8E-13 | -6.3 | 0.0130 | 6.9E-15 |
| *col-12* | WBGene00000601 | protein_coding_gene | -7.1 | 0.0072 | 3.6E-30 | -2.6 | 0.1653 | 1.5E-06 |
| *fat-5* | WBGene00001397 | protein_coding_gene | -7.2 | 0.0069 | 3.7E-104 | -4.1 | 0.0584 | 1.8E-19 |
| *ilys-5* | WBGene00017691 | protein_coding_gene | -7.2 | 0.0067 | 3.1E-96 | -2.7 | 0.1582 | 4.0E-09 |
| *col-124* | WBGene00000698 | protein_coding_gene | -7.2 | 0.0066 | 1.8E-124 | -5.2 | 0.0269 | 2.9E-57 |
| *F21C10.9* | WBGene00017658 | protein_coding_gene | -7.3 | 0.0064 | 5.4E-34 | -5.8 | 0.0176 | 3.3E-106 |
| *col-146* | WBGene00000719 | protein_coding_gene | -7.4 | 0.0059 | 3.5E-12 | -3.1 | 0.1131 | 9.1E-07 |
| *col-142* | WBGene00000715 | protein_coding_gene | -7.5 | 0.0057 | 1.8E-92 | -5.4 | 0.0234 | 1.7E-94 |
| *col-42* | WBGene00013489 | protein_coding_gene | -7.5 | 0.0056 | 1.3E-182 | -4.4 | 0.0490 | 2.4E-135 |
| *C30G12.2* | WBGene00016274 | protein_coding_gene | -7.6 | 0.0053 | 2.7E-71 | -5.0 | 0.0307 | 1.4E-28 |
| *col-103* | WBGene00000677 | protein_coding_gene | -7.6 | 0.0051 | 2.5E-113 | -4.3 | 0.0509 | 8.9E-170 |
| *F15E6.4* | WBGene00017485 | protein_coding_gene | -7.6 | 0.0051 | 2.6E-19 | -7.8 | 0.0044 | 1.3E-14 |
| *col-80* | WBGene00000656 | protein_coding_gene | -7.6 | 0.0050 | 2.2E-90 | -5.7 | 0.0195 | 1.1E-101 |
| *T22B7.7* | WBGene00020674 | protein_coding_gene | -7.6 | 0.0050 | 2.5E-39 | -2.3 | 0.1986 | 1.1E-05 |
| *C35A5.3* | WBGene00007947 | protein_coding_gene | -7.7 | 0.0047 | 4.0E-66 | -5.0 | 0.0320 | 2.0E-17 |
| *pmp-5* | WBGene00004062 | protein_coding_gene | -7.7 | 0.0047 | 1.5E-85 | -6.1 | 0.0147 | 7.2E-176 |
| *col-98* | WBGene00000673 | protein_coding_gene | -7.8 | 0.0046 | 9.8E-106 | -4.3 | 0.0502 | 5.9E-168 |
| *col-88* | WBGene00000663 | protein_coding_gene | -7.9 | 0.0041 | 1.6E-14 | -2.4 | 0.1952 | 5.5E-03 |
| *col-161* | WBGene00000734 | protein_coding_gene | -8.3 | 0.0032 | 9.0E-16 | -1.8 | 0.2779 | 3.1E-02 |
| *col-139* | WBGene00000712 | protein_coding_gene | -8.3 | 0.0031 | 5.8E-39 | -4.4 | 0.0480 | 8.9E-30 |
| *col-149* | WBGene00000722 | protein_coding_gene | -8.4 | 0.0030 | 8.9E-16 | -2.0 | 0.2462 | 4.7E-04 |
| *col-159* | WBGene00000732 | protein_coding_gene | -8.5 | 0.0028 | 2.8E-54 | -4.8 | 0.0364 | 2.7E-40 |
| *col-133* | WBGene00000707 | protein_coding_gene | -8.6 | 0.0026 | 5.8E-17 | -2.7 | 0.1501 | 4.4E-04 |
| *col-81* | WBGene00000657 | protein_coding_gene | -9.0 | 0.0020 | 2.3E-40 | -5.1 | 0.0299 | 4.9E-32 |
| *acdh-2* | WBGene00015894 | protein_coding_gene | -9.2 | 0.0017 | 8.7E-54 | -6.6 | 0.0106 | 1.2E-27 |
| *col-129* | WBGene00000703 | protein_coding_gene | -9.3 | 0.0016 | 2.8E-70 | -5.4 | 0.0244 | 2.3E-57 |
| *acdh-1* | WBGene00016943 | protein_coding_gene | -13.9 | 0.0001 | 5.4E-71 | -10.7 | 0.0006 | 5.7E-195 |

Differentially Expressed Transcription Factors in *P. aeuruginosa* Infected Animals

| Gene Name | WormBase Gene ID | Sequence Name | N2 $log_2$FC | CB4856 $log_2$FC |
|---|---|---|---|---|
| *atf-4* | WBGene00000221 | T04C10.4 | -2.3 | -1.4 |
| *cep-1* | WBGene00000467 | F52B5.5 | 1.0 | 1.4 |
| *cfi-1* | WBGene00000476 | T23D8.8 | -1.3 | -1.8 |
| *cky-1* | WBGene00000521 | C15C8.2 | 2.2 | -1.9 |
| *daf-8* | WBGene00000904 | R05D11.1 | -1.1 | -1.0 |
| *dpy-22* | WBGene00001081 | F47A4.2 | -1.6 | -1.2 |
| *eff-1* | WBGene00001159 | C26D10.5 | -2.2 | -1.8 |
| *egl-13* | WBGene00001182 | T22B7.1 | -1.9 | -1.0 |
| *egl-46* | WBGene00001210 | K11G9.4 | 1.2 | -1.3 |
| *elt-3* | WBGene00001251 | K02B9.4 | -2.5 | -1.5 |
| *fkh-6* | WBGene00001438 | B0286.5 | -3.5 | -2.1 |
| *fkh-9* | WBGene00001441 | K03C7.2 | -1.2 | -1.3 |
| *gei-3* | WBGene00001560 | T22H6.6 | -2.2 | -1.2 |
| *hif-1* | WBGene00001851 | F38A6.3 | -2.0 | -1.5 |
| *hlh-8* | WBGene00001953 | C02B8.4 | -1.3 | -1.6 |
| *hlh-12* | WBGene00001956 | C28C12.8 | -1.5 | -2.1 |
| *lim-6* | WBGene00002988 | K03E6.1 | -1.6 | -1.1 |

| | | | | |
|---|---|---|---|---|
| *lim-7* | WBGene00002989 | C04F1.3 | -1.6 | -1.3 |
| *lin-1* | WBGene00002990 | C37F5.1 | -1.0 | -1.2 |
| *lin-11* | WBGene00003000 | ZC247.3 | -2.0 | -1.5 |
| *lin-31* | WBGene00003017 | K10G6.1 | -2.7 | -2.6 |
| *mgl-2* | WBGene00003233 | F45H11.4 | -2.7 | -2.0 |
| *mxl-3* | WBGene00003511 | F46G10.6 | -4.4 | -2.0 |
| *nhr-3* | WBGene00003602 | H01A20.1 | -2.3 | -2.2 |
| *nhr-5* | WBGene00003604 | Y73F8A.21 | -2.3 | -1.8 |
| *nhr-14* | WBGene00003613 | T01B10.4 | -2.4 | -1.4 |
| *nhr-43* | WBGene00003633 | C29E6.5 | -1.1 | -1.4 |
| *nhr-45* | WBGene00003635 | F16H11.5 | -1.3 | -1.2 |
| *nhr-53* | WBGene00003643 | K06B4.11 | -3.0 | -2.0 |
| *nhr-62* | WBGene00003652 | Y67A6A.2 | -1.1 | -1.0 |
| *nhr-68* | WBGene00003658 | H12C20.3 | -3.9 | -3.2 |
| *nhr-85* | WBGene00003675 | W05B5.3 | -1.7 | -1.1 |
| *nhr-92* | WBGene00003682 | Y41D4B.8 | -1.3 | -2.0 |
| *nhr-97* | WBGene00003687 | H27C11.1 | -2.0 | -1.2 |
| *nhr-106* | WBGene00003696 | T01G6.4 | -1.0 | -1.1 |
| *nhr-112* | WBGene00003702 | Y70C5C.6 | 1.7 | 1.0 |
| *nhr-114* | WBGene00003704 | Y45G5AM.1 | -6.7 | -3.6 |
| *nhr-119* | WBGene00003709 | K12H6.1 | -1.9 | -2.6 |
| *nhr-121* | WBGene00003711 | E02H9.8 | -1.1 | -1.1 |
| *nhr-127* | WBGene00003717 | T13F3.3 | 2.6 | -1.1 |
| *nhr-130* | WBGene00003720 | T01G6.8 | -1.5 | -1.1 |
| *nhr-131* | WBGene00003721 | T01G6.2 | -1.2 | -1.5 |
| *nhr-135* | WBGene00003725 | VC5.5 | -1.0 | -1.3 |
| *nhr-137* | WBGene00003727 | C56E10.4 | -2.6 | -1.1 |
| *odd-1* | WBGene00003845 | B0280.4 | -1.2 | -1.3 |
| *blmp-1* | WBGene00003847 | F25D7.3 | -1.8 | -1.7 |
| *peb-1* | WBGene00003968 | T14F9.4 | 1.6 | -1.0 |
| *sbp-1* | WBGene00004735 | Y47D3B.7 | -1.4 | -1.2 |
| *sem-4* | WBGene00004773 | F15C11.1 | -1.3 | -1.1 |
| *sir-2.4* | WBGene00004803 | C06A5.11 | 1.5 | 1.1 |
| *sma-4* | WBGene00004858 | R12B2.1 | -1.1 | -2.2 |
| *sma-6* | WBGene00004860 | C32D5.2 | -1.0 | -1.2 |
| *sma-9* | WBGene00004862 | T05A10.1 | -1.2 | -1.4 |
| *snf-12* | WBGene00004911 | T25B6.7 | -2.6 | -3.1 |
| *F26F4.8* | WBGene00005011 | F26F4.8 | 1.7 | 1.0 |
| *ctbp-1* | WBGene00006424 | F49E10.5 | -1.3 | -1.4 |
| *svh-5* | WBGene00006462 | C33A11.4 | -1.8 | -1.5 |
| *tbx-36* | WBGene00006555 | ZK829.5 | 1.1 | 1.1 |
| *unc-42* | WBGene00006778 | F58E6.10 | 1.8 | -1.1 |
| *unc-98* | WBGene00006827 | F08C6.7 | -2.3 | -1.4 |
| *chd-7* | WBGene00007053 | T04D1.4 | -1.2 | -1.3 |
| *npax-4* | WBGene00007496 | C09G9.7 | -2.4 | -2.3 |
| *ceh-79* | WBGene00007749 | C26E1.3 | -1.4 | -2.1 |
| *nfki-1* | WBGene00007877 | C33A11.1 | -1.7 | -1.3 |
| *C38D4.7* | WBGene00008007 | C38D4.7 | 1.7 | 1.9 |
| *madf-8* | WBGene00008118 | C46F11.3 | 1.4 | 1.2 |
| *F21G4.5* | WBGene00009026 | F21G4.5 | -2.2 | -2.3 |
| *F28F8.7* | WBGene00009224 | F28F8.7 | 1.1 | 1.3 |
| *F36F2.1* | WBGene00009475 | F36F2.1 | -1.4 | -1.2 |
| *ccch-1* | WBGene00009532 | F38B7.1 | -1.3 | -1.3 |
| *gmeb-2* | WBGene00010010 | F53H4.5 | -1.7 | -1.1 |
| *saeg-1* | WBGene00010012 | F53H10.2 | -2.0 | -1.6 |
| *ztf-26* | WBGene00010137 | F55H12.6 | -2.3 | -1.7 |
| *sta-2* | WBGene00010251 | F58E6.1 | -2.3 | -3.4 |
| *nhr-205* | WBGene00011002 | R04B5.3 | -3.1 | -2.5 |
| *mbr-1* | WBGene00011315 | T01C1.2 | -2.5 | -2.2 |
| *T01C3.2* | WBGene00011319 | T01C3.2 | 1.1 | 1.0 |
| *T07D10.3* | WBGene00011583 | T07D10.3 | -2.6 | -3.5 |
| *zim-2* | WBGene00011600 | T07G12.10 | 1.2 | 1.3 |
| *zim-3* | WBGene00011601 | T07G12.11 | 1.2 | 1.1 |
| *ccch-3* | WBGene00012277 | W05B10.2 | 1.5 | 1.0 |
| *zip-3* | WBGene00012330 | W07G1.3 | -3.1 | -1.2 |
| *dhhc-2* | WBGene00012948 | Y47H9C.2 | -1.2 | -1.1 |
| *sta-1* | WBGene00013111 | Y51H4A.17 | -1.3 | -1.3 |
| *hsf-2* | WBGene00013134 | Y53C10A.3 | 2.0 | 1.5 |
| *ztf-29* | WBGene00013438 | Y66D12A.12 | -1.0 | -1.1 |
| *ekl-4* | WBGene00013676 | Y105E8A.17 | 1.1 | 1.0 |

| | | | | |
|---|---|---|---|---|
| *C04F5.9* | WBGene00015451 | C04F5.9 | 1.1 | 1.3 |
| *nhr-76* | WBGene00015497 | C05G6.1 | -2.2 | -1.3 |
| *C06E2.1* | WBGene00015527 | C06E2.1 | -3.5 | -1.4 |
| *C08G9.2* | WBGene00015620 | C08G9.2 | -1.3 | -1.2 |
| *nhr-156* | WBGene00015897 | C17E7.1 | -2.0 | -2.3 |
| *swsn-2.2* | WBGene00015971 | C18E3.2 | 1.1 | 1.0 |
| *zgpa-1* | WBGene00016383 | C33H5.17 | 1.2 | 1.1 |
| *C46E10.8* | WBGene00016712 | C46E10.8 | 1.6 | 1.4 |
| *madf-6* | WBGene00016930 | C54G6.1 | 1.4 | 1.3 |
| *nhr-173* | WBGene00016975 | C56E10.1 | -1.2 | -1.1 |
| *nhr-36* | WBGene00017198 | F07C3.10 | -1.2 | -1.6 |
| *F10E7.11* | WBGene00017352 | F10E7.11 | 1.2 | 1.1 |
| *ets-6* | WBGene00017606 | F19F10.10 | 1.1 | 1.0 |
| *nhr-142* | WBGene00018430 | F44E7.8 | -1.3 | -1.0 |
| *ztf-13* | WBGene00018704 | F52E4.8 | -1.1 | -1.2 |
| *ztf-1* | WBGene00018833 | F54F2.5 | 1.1 | 1.4 |
| *F57C9.4* | WBGene00019011 | F57C9.4 | 1.1 | 1.1 |
| *nhr-202* | WBGene00019742 | M02H5.4 | -1.3 | -1.6 |
| *ztf-16* | WBGene00019960 | R08E3.4 | -1.2 | -1.2 |
| *W02D7.6* | WBGene00020942 | W02D7.6 | 1.4 | 1.1 |
| *W03F9.2* | WBGene00021000 | W03F9.2 | 1.6 | 1.4 |
| *nhr-236* | WBGene00021417 | Y38F2AL.5 | 1.6 | 2.0 |
| *zip-4* | WBGene00021552 | Y44E3B.1 | 1.1 | 1.0 |
| *nhr-238* | WBGene00021611 | Y46H3D.7 | -1.3 | -2.9 |
| *Y53G8AR.5* | WBGene00021812 | Y53G8AR.5 | 1.2 | 1.1 |
| *ztf-23* | WBGene00021846 | Y54E10BR.8 | 1.2 | 1.1 |
| *znf-782* | WBGene00021931 | Y55F3AM.14 | -1.2 | 2.5 |
| *nhr-41* | WBGene00022423 | Y104H12A.1 | -1.6 | -1.6 |
| *lfi-1* | WBGene00022500 | ZC8.4 | -2.8 | -1.8 |
| *snu-23* | WBGene00022794 | ZK686.4 | 1.2 | 1.3 |
| *dve-1* | WBGene00022861 | ZK1193.5 | -1.2 | -1.1 |
| *nfya-2* | WBGene00043056 | Y53H1A.5 | 1.2 | 1.4 |
| *zip-9* | WBGene00077761 | F17C11.17 | -2.0 | -1.6 |

**Supplemental Table 9**

**Lists of differentially expressed genes that are identified in either N2, CB4856, or both, after infection by *P. rettgeri* (compared to *E. coli*)**

**Supplemental Table 9 - Differntially Expressed Genes in *P. rettgeri* Condition**

N2-exclusive DEGs in response to *P. rettgeri* (vs. OP50)

| Gene Name | Wormbase Gene ID | Type | $log_2FC$ | FC | Adj P-value |
|---|---|---|---|---|---|
| *fmo-2* | WBGene00001477 | protein_coding_gene | 6.5 | 87.59 | 8.1E-10 |
| *fipr-22* | WBGene00007989 | protein_coding_gene | 4.5 | 22.76 | 2.0E-05 |
| *F35E12.4* | WBGene00009428 | pseudogene | 3.7 | 12.93 | 4.0E-05 |
| *K05B2.4* | WBGene00019404 | protein_coding_gene | 3.7 | 12.70 | 2.5E-05 |
| *pgp-5* | WBGene00003999 | protein_coding_gene | 3.1 | 8.49 | 1.2E-04 |
| *C25F9.11* | WBGene00045411 | protein_coding_gene | 3.0 | 7.79 | 2.8E-03 |
| *W01C8.1* | WBGene00020918 | protein_coding_gene | 2.8 | 6.98 | 3.1E-02 |
| *Y47H10A.3* | WBGene00012959 | protein_coding_gene | 2.7 | 6.50 | 1.8E-02 |
| *C49G7.12* | WBGene00077593 | protein_coding_gene | 2.6 | 5.95 | 6.0E-03 |
| *sqst-3* | WBGene00012067 | protein_coding_gene | 2.5 | 5.52 | 4.9E-02 |
| *C25F9.12* | WBGene00045412 | protein_coding_gene | 2.4 | 5.19 | 4.0E-02 |
| *T10D4.15* | WBGene00077522 | protein_coding_gene | 2.4 | 5.13 | 4.2E-02 |
| *F14F9.3* | WBGene00017466 | protein_coding_gene | 2.2 | 4.72 | 4.9E-02 |
| *pgp-6* | WBGene00004000 | protein_coding_gene | 2.2 | 4.59 | 1.2E-04 |
| *clec-76* | WBGene00021587 | protein_coding_gene | 2.2 | 4.53 | 2.8E-02 |
| *cyp-13B1* | WBGene00008519 | protein_coding_gene | 2.2 | 4.50 | 1.4E-03 |
| *srw-86* | WBGene00005833 | protein_coding_gene | 2.0 | 4.06 | 3.2E-02 |
| *F11A5.15* | WBGene00008680 | protein_coding_gene | 2.0 | 4.04 | 2.6E-03 |
| *F11C7.6* | WBGene00017378 | protein_coding_gene | 2.0 | 4.00 | 1.4E-02 |
| *T27A3.8* | WBGene00044611 | protein_coding_gene | 1.9 | 3.77 | 2.2E-02 |
| *acs-7* | WBGene00007228 | protein_coding_gene | 1.9 | 3.71 | 6.3E-04 |
| *E03H4.8* | WBGene00008476 | protein_coding_gene | 1.9 | 3.64 | 7.2E-04 |
| *ZC239.22* | WBGene00194717 | protein_coding_gene | 1.9 | 3.63 | 2.5E-02 |
| *ech-1.1* | WBGene00001150 | protein_coding_gene | 1.8 | 3.51 | 4.5E-05 |
| *ceh-45* | WBGene00022837 | protein_coding_gene | 1.8 | 3.51 | 2.5E-02 |
| *C34H4.1* | WBGene00016424 | protein_coding_gene | 1.8 | 3.37 | 3.6E-03 |
| *nhr-58* | WBGene00003648 | protein_coding_gene | 1.8 | 3.37 | 5.6E-03 |
| *Y43F8B.23* | WBGene00194815 | protein_coding_gene | 1.8 | 3.37 | 3.9E-02 |
| *acox-3* | WBGene00019060 | protein_coding_gene | 1.8 | 3.37 | 4.3E-04 |
| *E02H4.4* | WBGene00008462 | protein_coding_gene | 1.7 | 3.36 | 1.1E-02 |
| *E02C12.8* | WBGene00017093 | protein_coding_gene | 1.7 | 3.33 | 1.3E-03 |
| *C10C5.5* | WBGene00007509 | protein_coding_gene | 1.7 | 3.27 | 7.2E-05 |
| *C52E2.4* | WBGene00016883 | protein_coding_gene | 1.7 | 3.18 | 1.1E-02 |
| *F12A10.1* | WBGene00017390 | protein_coding_gene | 1.6 | 3.09 | 6.5E-03 |
| *F07E5.13* | WBGene00304173 | protein_coding_gene | 1.6 | 3.07 | 7.2E-03 |
| *cest-34* | WBGene00019652 | protein_coding_gene | 1.6 | 3.07 | 9.9E-03 |
| *Y51A2D.13* | WBGene00013080 | protein_coding_gene | 1.6 | 2.98 | 9.3E-03 |
| *C46C11.4* | WBGene00044567 | protein_coding_gene | 1.6 | 2.96 | 4.7E-03 |
| *nhr-171* | WBGene00016926 | protein_coding_gene | 1.6 | 2.94 | 4.9E-02 |
| *K08B5.1* | WBGene00019518 | protein_coding_gene | 1.5 | 2.89 | 6.7E-03 |
| *ctl-1* | WBGene00000830 | protein_coding_gene | 1.5 | 2.73 | 4.9E-02 |
| *asns-2* | WBGene00019730 | protein_coding_gene | 1.4 | 2.72 | 2.5E-02 |
| *fbxb-87* | WBGene00043996 | protein_coding_gene | 1.4 | 2.72 | 4.6E-02 |
| *B0272.4* | WBGene00007130 | protein_coding_gene | 1.4 | 2.68 | 1.1E-03 |
| *C40A11.4* | WBGene00016547 | protein_coding_gene | 1.4 | 2.67 | 3.3E-02 |
| *nhr-116* | WBGene00003706 | protein_coding_gene | 1.4 | 2.66 | 2.1E-03 |
| *fbxb-44* | WBGene00019416 | protein_coding_gene | 1.4 | 2.63 | 2.4E-02 |
| *ugt-31* | WBGene00021464 | protein_coding_gene | 1.4 | 2.61 | 1.2E-02 |
| *B0250.12* | WBGene00050927 | pseudogene | 1.4 | 2.61 | 1.2E-02 |
| *Y58A7A.5* | WBGene00021979 | protein_coding_gene | 1.4 | 2.60 | 4.9E-02 |
| *K08B5.2* | WBGene00019519 | protein_coding_gene | 1.4 | 2.59 | 2.0E-03 |
| *T05F1.9* | WBGene00011494 | protein_coding_gene | 1.4 | 2.58 | 3.8E-02 |
| *cpr-4* | WBGene00000784 | protein_coding_gene | 1.4 | 2.56 | 5.9E-04 |
| *fbxa-60* | WBGene00020456 | protein_coding_gene | 1.3 | 2.54 | 2.0E-02 |

| | | | | | |
|---|---|---|---|---|---|
| *F45E10.2* | WBGene00009729 | protein_coding_gene | 1.3 | 2.54 | 1.5E-03 |
| *swt-6* | WBGene00011190 | protein_coding_gene | 1.3 | 2.51 | 6.8E-03 |
| *ref-2* | WBGene00004335 | protein_coding_gene | 1.3 | 2.49 | 2.7E-02 |
| *C25D7.5* | WBGene00007717 | protein_coding_gene | 1.3 | 2.48 | 3.7E-02 |
| *zeel-1* | WBGene00021463 | protein_coding_gene | 1.3 | 2.48 | 1.7E-02 |
| *F31F7.1* | WBGene00017964 | protein_coding_gene | 1.3 | 2.48 | 7.8E-05 |
| *cyp-34A5* | WBGene00015040 | protein_coding_gene | 1.3 | 2.47 | 4.4E-02 |
| *pept-3* | WBGene00003878 | protein_coding_gene | 1.3 | 2.44 | 3.9E-02 |
| *pals-23* | WBGene00016218 | protein_coding_gene | 1.3 | 2.42 | 1.4E-02 |
| *zip-8* | WBGene00017755 | protein_coding_gene | 1.2 | 2.37 | 8.8E-08 |
| *Y111B2A.21* | WBGene00013741 | protein_coding_gene | 1.2 | 2.35 | 3.0E-03 |
| *zig-4* | WBGene00006981 | protein_coding_gene | 1.2 | 2.32 | 4.0E-02 |
| *R11F4.1* | WBGene00020007 | protein_coding_gene | 1.2 | 2.31 | 5.3E-04 |
| *F45D3.3* | WBGene00009723 | protein_coding_gene | 1.2 | 2.31 | 1.4E-04 |
| *F58A6.1* | WBGene00019022 | protein_coding_gene | 1.2 | 2.31 | 6.5E-03 |
| *fbxc-29* | WBGene00019923 | protein_coding_gene | 1.2 | 2.29 | 2.3E-02 |
| *nhr-21* | WBGene00003620 | protein_coding_gene | 1.2 | 2.29 | 3.4E-03 |
| *T04F8.7* | WBGene00011445 | protein_coding_gene | 1.2 | 2.28 | 1.4E-02 |
| *F44F1.6* | WBGene00009698 | protein_coding_gene | 1.2 | 2.26 | 4.2E-03 |
| *ZK757.2* | WBGene00014074 | protein_coding_gene | 1.2 | 2.26 | 4.4E-02 |
| *F35B3.7* | WBGene00018033 | protein_coding_gene | 1.2 | 2.26 | 2.4E-02 |
| *prx-11* | WBGene00004196 | protein_coding_gene | 1.2 | 2.25 | 2.5E-03 |
| *cpt-1* | WBGene00012907 | protein_coding_gene | 1.2 | 2.24 | 2.6E-05 |
| *F21D12.3* | WBGene00017663 | protein_coding_gene | 1.2 | 2.24 | 1.8E-02 |
| *D2063.1* | WBGene00017060 | protein_coding_gene | 1.2 | 2.23 | 2.2E-02 |
| *pals-24* | WBGene00016219 | protein_coding_gene | 1.2 | 2.23 | 1.7E-02 |
| *endu-2* | WBGene00019779 | protein_coding_gene | 1.2 | 2.23 | 1.3E-03 |
| *fbxb-63* | WBGene00044197 | protein_coding_gene | 1.1 | 2.21 | 2.6E-02 |
| *jmjd-3.3* | WBGene00007813 | protein_coding_gene | 1.1 | 2.20 | 4.3E-02 |
| *sdz-28* | WBGene00020073 | protein_coding_gene | 1.1 | 2.18 | 1.7E-02 |
| *M162.5* | WBGene00010931 | protein_coding_gene | 1.1 | 2.18 | 2.0E-02 |
| *F32D8.13* | WBGene00009335 | protein_coding_gene | 1.1 | 2.16 | 1.3E-02 |
| *C18H9.6* | WBGene00016004 | protein_coding_gene | 1.1 | 2.16 | 5.6E-03 |
| *Y41D4A.7* | WBGene00021509 | protein_coding_gene | 1.1 | 2.16 | 1.8E-02 |
| *dod-22* | WBGene00010125 | protein_coding_gene | 1.1 | 2.14 | 9.4E-03 |
| *R03H10.11* | WBGene00271824 | protein_coding_gene | 1.1 | 2.13 | 3.4E-02 |
| *Y70G10A.3* | WBGene00013499 | protein_coding_gene | 1.1 | 2.12 | 4.6E-02 |
| *btb-16* | WBGene00018195 | protein_coding_gene | 1.1 | 2.11 | 6.4E-03 |
| *B0403.6* | WBGene00043982 | protein_coding_gene | 1.1 | 2.09 | 4.6E-02 |
| *clec-52* | WBGene00015052 | protein_coding_gene | 1.1 | 2.09 | 7.1E-08 |
| *fbxc-40* | WBGene00016295 | protein_coding_gene | 1.1 | 2.08 | 3.2E-02 |
| *T05D4.2* | WBGene00011475 | protein_coding_gene | 1.0 | 2.06 | 3.7E-02 |
| *Y71A12B.11* | WBGene00013508 | protein_coding_gene | 1.0 | 2.06 | 4.1E-02 |
| *cng-1* | WBGene00000562 | protein_coding_gene | 1.0 | 2.06 | 1.4E-02 |
| *Y46G5A.7* | WBGene00012899 | protein_coding_gene | 1.0 | 2.04 | 6.1E-03 |
| *fmo-5* | WBGene00001480 | protein_coding_gene | 1.0 | 2.04 | 1.7E-02 |
| *T12A7.2* | WBGene00011723 | protein_coding_gene | 1.0 | 2.03 | 1.8E-02 |
| *C16C4.18* | WBGene00194998 | protein_coding_gene | 1.0 | 2.02 | 1.8E-02 |
| *fbxb-41* | WBGene00019703 | protein_coding_gene | 1.0 | 2.01 | 3.1E-02 |
| *pgp-10* | WBGene00004004 | protein_coding_gene | 1.0 | 2.01 | 3.3E-02 |
| *F39F10.4* | WBGene00018204 | protein_coding_gene | 1.0 | 2.00 | 2.8E-02 |
| *tdo-2* | WBGene00016201 | protein_coding_gene | -1.0 | 0.50 | 2.7E-02 |
| *F25D1.5* | WBGene00009110 | protein_coding_gene | -1.0 | 0.50 | 1.1E-02 |
| *clec-51* | WBGene00015050 | protein_coding_gene | -1.0 | 0.50 | 4.3E-04 |
| *H03A11.2* | WBGene00010357 | protein_coding_gene | -1.0 | 0.50 | 7.3E-04 |
| *T01H3.5* | WBGene00011351 | protein_coding_gene | -1.0 | 0.50 | 4.2E-02 |
| *pmt-2* | WBGene00018811 | protein_coding_gene | -1.0 | 0.49 | 3.6E-03 |
| *F41B4.3* | WBGene00018259 | protein_coding_gene | -1.0 | 0.49 | 2.2E-02 |

| | | | | | |
|---|---|---|---|---|---|
| *F37H8.2* | WBGene00009511 | protein_coding_gene | -1.0 | 0.49 | 1.5E-02 |
| *spp-4* | WBGene00004989 | protein_coding_gene | -1.0 | 0.49 | 3.2E-02 |
| *aat-2* | WBGene00000003 | protein_coding_gene | -1.0 | 0.49 | 1.2E-02 |
| *K09F6.4* | WBGene00019587 | protein_coding_gene | -1.0 | 0.49 | 8.1E-03 |
| *bcmo-2* | WBGene00018755 | protein_coding_gene | -1.0 | 0.49 | 8.4E-05 |
| *F17C8.9* | WBGene00185014 | protein_coding_gene | -1.0 | 0.49 | 4.4E-02 |
| *F58B4.5* | WBGene00010238 | protein_coding_gene | -1.0 | 0.48 | 1.3E-03 |
| *C25F9.4* | WBGene00007724 | protein_coding_gene | -1.1 | 0.48 | 6.6E-03 |
| *D2023.1* | WBGene00014300 | protein_coding_gene | -1.1 | 0.48 | 5.3E-05 |
| *Y38E10A.14* | WBGene00012592 | protein_coding_gene | -1.1 | 0.48 | 4.0E-02 |
| *lron-8* | WBGene00020693 | protein_coding_gene | -1.1 | 0.48 | 3.1E-04 |
| *pmp-5* | WBGene00004062 | protein_coding_gene | -1.1 | 0.48 | 3.4E-02 |
| *shl-1* | WBGene00022240 | protein_coding_gene | -1.1 | 0.48 | 1.6E-02 |
| *C31H2.4* | WBGene00016294 | protein_coding_gene | -1.1 | 0.48 | 3.3E-02 |
| *arrd-18* | WBGene00018060 | protein_coding_gene | -1.1 | 0.48 | 1.3E-02 |
| *K03B8.6* | WBGene00010524 | protein_coding_gene | -1.1 | 0.47 | 4.3E-03 |
| *mct-3* | WBGene00010834 | protein_coding_gene | -1.1 | 0.47 | 1.2E-03 |
| *C08H9.15* | WBGene00007474 | protein_coding_gene | -1.1 | 0.47 | 2.0E-02 |
| *ndx-2* | WBGene00003579 | protein_coding_gene | -1.1 | 0.47 | 1.1E-02 |
| *ZC239.6* | WBGene00022569 | protein_coding_gene | -1.1 | 0.47 | 1.2E-02 |
| *hacd-1* | WBGene00019978 | protein_coding_gene | -1.1 | 0.47 | 1.9E-02 |
| *fkb-3* | WBGene00001428 | protein_coding_gene | -1.1 | 0.47 | 1.8E-02 |
| *tbc-19* | WBGene00007849 | protein_coding_gene | -1.1 | 0.47 | 2.1E-02 |
| *cnp-2* | WBGene00012901 | protein_coding_gene | -1.1 | 0.46 | 3.5E-03 |
| *cest-10* | WBGene00016595 | protein_coding_gene | -1.1 | 0.46 | 6.7E-03 |
| *K10D11.5* | WBGene00010749 | protein_coding_gene | -1.1 | 0.46 | 2.5E-02 |
| *F27E5.7* | WBGene00044035 | protein_coding_gene | -1.1 | 0.46 | 3.3E-02 |
| *R13D11.4* | WBGene00020058 | protein_coding_gene | -1.1 | 0.46 | 9.7E-03 |
| *T04B2.3* | WBGene00011422 | protein_coding_gene | -1.1 | 0.46 | 4.7E-03 |
| *pfn-3* | WBGene00003991 | protein_coding_gene | -1.1 | 0.45 | 3.1E-03 |
| *F34D10.9* | WBGene00045484 | protein_coding_gene | -1.1 | 0.45 | 2.3E-02 |
| *Y34F4.2* | WBGene00021337 | protein_coding_gene | -1.1 | 0.45 | 2.2E-02 |
| *C15C7.5* | WBGene00015791 | protein_coding_gene | -1.1 | 0.45 | 1.9E-03 |
| *T20D4.11* | WBGene00020617 | protein_coding_gene | -1.2 | 0.45 | 3.5E-02 |
| *wago-10* | WBGene00020707 | protein_coding_gene | -1.2 | 0.45 | 4.0E-02 |
| *nas-20* | WBGene00003539 | protein_coding_gene | -1.2 | 0.45 | 1.9E-02 |
| *bath-26* | WBGene00020329 | protein_coding_gene | -1.2 | 0.44 | 3.1E-02 |
| *T22E5.1* | WBGene00020690 | protein_coding_gene | -1.2 | 0.44 | 3.2E-02 |
| *R11E3.1* | WBGene00020003 | protein_coding_gene | -1.2 | 0.44 | 4.4E-02 |
| *msp-51* | WBGene00003444 | protein_coding_gene | -1.2 | 0.44 | 5.0E-02 |
| *spe-41* | WBGene00004976 | protein_coding_gene | -1.2 | 0.44 | 2.9E-02 |
| *tba-9* | WBGene00006535 | protein_coding_gene | -1.2 | 0.44 | 1.8E-03 |
| *alg-4* | WBGene00006449 | protein_coding_gene | -1.2 | 0.44 | 4.7E-02 |
| *Y43C5B.3* | WBGene00012786 | protein_coding_gene | -1.2 | 0.44 | 3.3E-02 |
| *Y57G11C.40* | WBGene00013329 | protein_coding_gene | -1.2 | 0.44 | 3.4E-02 |
| *let-805* | WBGene00002915 | protein_coding_gene | -1.2 | 0.43 | 2.3E-02 |
| *T28F4.3* | WBGene00012138 | protein_coding_gene | -1.2 | 0.43 | 4.1E-02 |
| *ttr-14* | WBGene00011460 | protein_coding_gene | -1.2 | 0.43 | 6.7E-03 |
| *dac-1* | WBGene00000895 | protein_coding_gene | -1.2 | 0.43 | 1.0E-02 |
| *F46C8.8* | WBGene00018487 | protein_coding_gene | -1.2 | 0.43 | 2.1E-02 |
| *dop-4* | WBGene00016872 | protein_coding_gene | -1.2 | 0.43 | 3.7E-02 |
| *W06A7.4* | WBGene00012293 | protein_coding_gene | -1.2 | 0.43 | 4.2E-03 |
| *C55A6.6* | WBGene00008335 | protein_coding_gene | -1.2 | 0.43 | 4.6E-04 |
| *K03B8.8* | WBGene00010525 | protein_coding_gene | -1.2 | 0.43 | 6.4E-04 |
| *B0252.5* | WBGene00015089 | protein_coding_gene | -1.2 | 0.43 | 4.1E-02 |
| *ttr-7* | WBGene00020566 | protein_coding_gene | -1.2 | 0.42 | 5.9E-03 |
| *C11H1.9* | WBGene00023500 | protein_coding_gene | -1.2 | 0.42 | 1.9E-02 |
| *C50D2.6* | WBGene00016809 | protein_coding_gene | -1.2 | 0.42 | 6.9E-03 |

| | | | | | |
|---|---|---|---|---|---|
| *F54C1.1* | WBGene00018789 | protein_coding_gene | -1.2 | 0.42 | 1.4E-02 |
| *C09D4.3* | WBGene00015634 | protein_coding_gene | -1.3 | 0.42 | 3.4E-02 |
| *K09A9.8* | WBGene00194849 | protein_coding_gene | -1.3 | 0.42 | 2.9E-02 |
| *abu-14* | WBGene00004174 | protein_coding_gene | -1.3 | 0.42 | 3.8E-02 |
| *R07E5.15* | WBGene00011120 | protein_coding_gene | -1.3 | 0.42 | 4.8E-02 |
| *irld-6* | WBGene00009239 | protein_coding_gene | -1.3 | 0.42 | 4.2E-02 |
| *F33E2.4* | WBGene00009360 | protein_coding_gene | -1.3 | 0.42 | 3.4E-02 |
| *C33F10.8* | WBGene00016356 | protein_coding_gene | -1.3 | 0.42 | 4.8E-02 |
| *K01B6.3* | WBGene00010454 | protein_coding_gene | -1.3 | 0.41 | 4.1E-03 |
| *lon-8* | WBGene00013352 | protein_coding_gene | -1.3 | 0.41 | 7.3E-04 |
| *fpn-1.2* | WBGene00019977 | protein_coding_gene | -1.3 | 0.41 | 6.6E-03 |
| *best-17* | WBGene00020559 | protein_coding_gene | -1.3 | 0.41 | 3.9E-04 |
| *pgp-14* | WBGene00004008 | protein_coding_gene | -1.3 | 0.41 | 1.4E-02 |
| *col-17* | WBGene00000606 | protein_coding_gene | -1.3 | 0.41 | 3.2E-02 |
| *C47E12.11* | WBGene00008154 | protein_coding_gene | -1.3 | 0.40 | 3.6E-02 |
| *pgp-9* | WBGene00004003 | protein_coding_gene | -1.3 | 0.40 | 2.3E-02 |
| *F07A5.2* | WBGene00008541 | protein_coding_gene | -1.3 | 0.40 | 5.0E-02 |
| *clec-57* | WBGene00008594 | protein_coding_gene | -1.3 | 0.40 | 1.0E-02 |
| *C09H10.9* | WBGene00007503 | protein_coding_gene | -1.3 | 0.40 | 2.0E-02 |
| *spe-4* | WBGene00004958 | protein_coding_gene | -1.3 | 0.40 | 3.3E-02 |
| *C18E9.8* | WBGene00007688 | protein_coding_gene | -1.3 | 0.40 | 4.8E-02 |
| *ZK418.2* | WBGene00022732 | protein_coding_gene | -1.3 | 0.40 | 3.1E-02 |
| *cpr-5* | WBGene00000785 | protein_coding_gene | -1.3 | 0.39 | 7.4E-13 |
| *F40G12.10* | WBGene00009605 | protein_coding_gene | -1.3 | 0.39 | 3.3E-02 |
| *T13F2.9* | WBGene00011748 | pseudogene | -1.4 | 0.39 | 4.8E-02 |
| *W01A8.6* | WBGene00012168 | protein_coding_gene | -1.4 | 0.39 | 2.5E-03 |
| *ugt-21* | WBGene00007885 | protein_coding_gene | -1.4 | 0.39 | 2.2E-03 |
| *msp-56* | WBGene00003449 | protein_coding_gene | -1.4 | 0.39 | 3.3E-02 |
| *F54C9.14* | WBGene00175035 | protein_coding_gene | -1.4 | 0.39 | 2.3E-02 |
| *ilys-5* | WBGene00017691 | protein_coding_gene | -1.4 | 0.39 | 2.0E-03 |
| *M176.9* | WBGene00014813 | protein_coding_gene | -1.4 | 0.39 | 3.6E-02 |
| *H05L14.1* | WBGene00010366 | protein_coding_gene | -1.4 | 0.39 | 4.6E-02 |
| *spch-1* | WBGene00007307 | protein_coding_gene | -1.4 | 0.39 | 5.0E-02 |
| *F58H1.6* | WBGene00010289 | protein_coding_gene | -1.4 | 0.39 | 2.2E-02 |
| *T20D4.5* | WBGene00020611 | protein_coding_gene | -1.4 | 0.39 | 2.9E-02 |
| *ugt-41* | WBGene00017336 | protein_coding_gene | -1.4 | 0.39 | 9.4E-05 |
| *F45E4.6* | WBGene00018471 | protein_coding_gene | -1.4 | 0.39 | 4.5E-02 |
| *gst-39* | WBGene00001787 | protein_coding_gene | -1.4 | 0.39 | 1.9E-03 |
| *suro-1* | WBGene00011235 | protein_coding_gene | -1.4 | 0.38 | 2.5E-02 |
| *F36A4.4* | WBGene00018083 | protein_coding_gene | -1.4 | 0.38 | 3.1E-02 |
| *ZK6.6* | WBGene00022641 | protein_coding_gene | -1.4 | 0.38 | 4.4E-02 |
| *math-22* | WBGene00016717 | protein_coding_gene | -1.4 | 0.38 | 4.7E-04 |
| *Y55B1AR.4* | WBGene00021908 | protein_coding_gene | -1.4 | 0.38 | 2.1E-02 |
| *F55C5.2* | WBGene00010092 | pseudogene | -1.4 | 0.38 | 4.6E-02 |
| *lgc-21* | WBGene00007903 | protein_coding_gene | -1.4 | 0.38 | 3.1E-02 |
| *F31A3.3* | WBGene00017938 | protein_coding_gene | -1.4 | 0.38 | 1.3E-02 |
| *ZK858.8* | WBGene00014121 | protein_coding_gene | -1.4 | 0.38 | 2.5E-02 |
| *F19B2.5* | WBGene00008944 | protein_coding_gene | -1.4 | 0.38 | 2.1E-02 |
| *F41F3.3* | WBGene00018297 | protein_coding_gene | -1.4 | 0.38 | 3.2E-02 |
| *ent-3* | WBGene00010510 | protein_coding_gene | -1.4 | 0.38 | 2.7E-02 |
| *ZK455.5* | WBGene00013975 | protein_coding_gene | -1.4 | 0.37 | 4.6E-02 |
| *F09C12.8* | WBGene00017279 | protein_coding_gene | -1.4 | 0.37 | 2.9E-02 |
| *rnh-1.1* | WBGene00022888 | protein_coding_gene | -1.4 | 0.37 | 2.6E-02 |
| *msra-1* | WBGene00018393 | protein_coding_gene | -1.4 | 0.37 | 1.2E-02 |
| *T03F6.6* | WBGene00011401 | protein_coding_gene | -1.4 | 0.37 | 3.5E-02 |
| *F34D10.8* | WBGene00009377 | protein_coding_gene | -1.4 | 0.37 | 2.0E-02 |
| *spch-3* | WBGene00020840 | protein_coding_gene | -1.4 | 0.37 | 4.3E-02 |
| *Y69E1A.8* | WBGene00013478 | protein_coding_gene | -1.4 | 0.37 | 2.2E-02 |

| | | | | | |
|---|---|---|---|---|---|
| *Y38H8A.4* | WBGene00012638 | protein_coding_gene | -1.4 | 0.37 | 3.8E-02 |
| *ptr-14* | WBGene00004228 | protein_coding_gene | -1.4 | 0.37 | 5.7E-03 |
| *ZC581.2* | WBGene00022632 | protein_coding_gene | -1.4 | 0.37 | 4.1E-02 |
| *T28B8.4* | WBGene00012115 | protein_coding_gene | -1.5 | 0.36 | 4.0E-02 |
| *nhx-1* | WBGene00003729 | protein_coding_gene | -1.5 | 0.36 | 1.6E-02 |
| *F56D5.3* | WBGene00010148 | protein_coding_gene | -1.5 | 0.36 | 3.7E-04 |
| *clec-43* | WBGene00019917 | protein_coding_gene | -1.5 | 0.36 | 1.0E-02 |
| *ctsa-4.1* | WBGene00019605 | protein_coding_gene | -1.5 | 0.36 | 4.0E-03 |
| *acr-8* | WBGene00000047 | protein_coding_gene | -1.5 | 0.36 | 2.1E-02 |
| *F55G11.4* | WBGene00010124 | protein_coding_gene | -1.5 | 0.36 | 3.4E-04 |
| *C33C12.4* | WBGene00016336 | protein_coding_gene | -1.5 | 0.36 | 3.1E-02 |
| *T19D12.5* | WBGene00020580 | protein_coding_gene | -1.5 | 0.36 | 3.8E-02 |
| *DC2.5* | WBGene00017082 | protein_coding_gene | -1.5 | 0.36 | 1.0E-03 |
| *C09B9.4* | WBGene00015629 | protein_coding_gene | -1.5 | 0.36 | 2.2E-02 |
| *adt-3* | WBGene00020567 | protein_coding_gene | -1.5 | 0.36 | 1.5E-04 |
| *C08F11.14* | WBGene00007461 | pseudogene | -1.5 | 0.36 | 4.9E-02 |
| *Y47G6A.15* | WBGene00021641 | protein_coding_gene | -1.5 | 0.35 | 2.0E-03 |
| *col-39* | WBGene00000616 | protein_coding_gene | -1.5 | 0.35 | 1.9E-02 |
| *C54D2.1* | WBGene00016915 | protein_coding_gene | -1.5 | 0.35 | 3.7E-02 |
| *clec-170* | WBGene00009515 | protein_coding_gene | -1.5 | 0.35 | 6.7E-06 |
| *C54E4.5* | WBGene00016920 | protein_coding_gene | -1.5 | 0.35 | 1.9E-03 |
| *ttbk-4* | WBGene00012169 | protein_coding_gene | -1.5 | 0.35 | 3.0E-02 |
| *cest-1.1* | WBGene00011362 | protein_coding_gene | -1.5 | 0.35 | 4.9E-02 |
| *F54H12.7* | WBGene00044633 | protein_coding_gene | -1.5 | 0.35 | 3.3E-02 |
| *C15C6.2* | WBGene00007601 | protein_coding_gene | -1.5 | 0.35 | 4.5E-02 |
| *zipt-17* | WBGene00006487 | protein_coding_gene | -1.5 | 0.35 | 3.6E-02 |
| *R06B10.1* | WBGene00019912 | pseudogene | -1.5 | 0.35 | 1.8E-02 |
| *C27D8.1* | WBGene00007777 | protein_coding_gene | -1.5 | 0.35 | 2.6E-02 |
| *grl-21* | WBGene00001730 | protein_coding_gene | -1.5 | 0.34 | 2.4E-02 |
| *B0034.1* | WBGene00015002 | protein_coding_gene | -1.5 | 0.34 | 3.6E-03 |
| *nas-38* | WBGene00003554 | protein_coding_gene | -1.5 | 0.34 | 4.4E-02 |
| *cbs-2* | WBGene00018783 | protein_coding_gene | -1.6 | 0.34 | 1.0E-02 |
| *Y66C5A.1* | WBGene00044156 | protein_coding_gene | -1.6 | 0.34 | 1.3E-02 |
| *K08H10.6* | WBGene00010698 | protein_coding_gene | -1.6 | 0.34 | 4.1E-02 |
| *T10B5.8* | WBGene00020394 | protein_coding_gene | -1.6 | 0.34 | 2.6E-05 |
| *H20J04.1* | WBGene00019216 | protein_coding_gene | -1.6 | 0.34 | 2.0E-02 |
| *sod-3* | WBGene00004932 | protein_coding_gene | -1.6 | 0.34 | 5.2E-04 |
| *old-1* | WBGene00003862 | protein_coding_gene | -1.6 | 0.34 | 2.9E-02 |
| *F36D3.5* | WBGene00009471 | protein_coding_gene | -1.6 | 0.34 | 4.5E-02 |
| *Y34F4.6* | WBGene00189956 | protein_coding_gene | -1.6 | 0.34 | 1.3E-02 |
| *F59A1.15* | WBGene00010300 | protein_coding_gene | -1.6 | 0.33 | 1.8E-02 |
| *col-38* | WBGene00000615 | protein_coding_gene | -1.6 | 0.33 | 6.2E-05 |
| *C33E10.8* | WBGene00016349 | protein_coding_gene | -1.6 | 0.33 | 2.9E-02 |
| *col-115* | WBGene00000689 | protein_coding_gene | -1.6 | 0.33 | 4.5E-02 |
| *F35F10.6* | WBGene00018056 | protein_coding_gene | -1.6 | 0.33 | 1.7E-02 |
| *F13H8.5* | WBGene00017438 | protein_coding_gene | -1.6 | 0.33 | 3.6E-03 |
| *R13A1.3* | WBGene00020042 | protein_coding_gene | -1.6 | 0.33 | 3.3E-02 |
| *C16C8.18* | WBGene00015855 | protein_coding_gene | -1.6 | 0.33 | 1.8E-02 |
| *cyp-33C8* | WBGene00019967 | protein_coding_gene | -1.6 | 0.33 | 2.9E-03 |
| *T02E1.7* | WBGene00011379 | protein_coding_gene | -1.6 | 0.33 | 2.0E-02 |
| *ugt-13* | WBGene00019232 | protein_coding_gene | -1.6 | 0.33 | 2.1E-05 |
| *acdh-5* | WBGene00016491 | protein_coding_gene | -1.6 | 0.32 | 1.2E-02 |
| *gska-3* | WBGene00007977 | protein_coding_gene | -1.6 | 0.32 | 3.0E-02 |
| *T02B11.9* | WBGene00044777 | protein_coding_gene | -1.6 | 0.32 | 2.8E-02 |
| *F27E5.3* | WBGene00009193 | protein_coding_gene | -1.6 | 0.32 | 2.6E-02 |
| *Y6E2A.10* | WBGene00050915 | protein_coding_gene | -1.6 | 0.32 | 3.6E-02 |
| *msp-65* | WBGene00003458 | protein_coding_gene | -1.6 | 0.32 | 3.8E-02 |
| *ugt-30* | WBGene00011340 | protein_coding_gene | -1.6 | 0.32 | 1.2E-05 |

| | | | | | |
|---|---|---|---|---|---|
| *F11E6.6* | WBGene00008709 | protein_coding_gene | -1.7 | 0.31 | 3.8E-02 |
| *ant-1.4* | WBGene00020140 | protein_coding_gene | -1.7 | 0.31 | 1.3E-02 |
| *F10D11.6* | WBGene00008652 | protein_coding_gene | -1.7 | 0.31 | 1.0E-03 |
| *col-109* | WBGene00000683 | protein_coding_gene | -1.7 | 0.31 | 2.2E-03 |
| *C35A5.4* | WBGene00007948 | protein_coding_gene | -1.7 | 0.31 | 2.2E-02 |
| *Y39B6A.31* | WBGene00012690 | pseudogene | -1.7 | 0.31 | 4.4E-02 |
| *T28C12.2* | WBGene00020889 | protein_coding_gene | -1.7 | 0.31 | 1.3E-02 |
| *abu-10* | WBGene00000033 | protein_coding_gene | -1.7 | 0.31 | 1.7E-02 |
| *Y47D3B.6* | WBGene00012942 | protein_coding_gene | -1.7 | 0.30 | 6.2E-03 |
| *Y67A10A.3* | WBGene00013452 | protein_coding_gene | -1.7 | 0.30 | 2.1E-02 |
| *aat-7* | WBGene00000008 | protein_coding_gene | -1.7 | 0.30 | 1.6E-03 |
| *F16H6.4* | WBGene00008894 | protein_coding_gene | -1.7 | 0.30 | 5.0E-02 |
| *T06E4.14* | WBGene00077691 | protein_coding_gene | -1.7 | 0.30 | 5.0E-02 |
| *C49F8.3* | WBGene00008215 | protein_coding_gene | -1.7 | 0.30 | 7.2E-04 |
| *D2062.4* | WBGene00017056 | protein_coding_gene | -1.8 | 0.30 | 4.8E-02 |
| *C17B7.5* | WBGene00015879 | protein_coding_gene | -1.8 | 0.30 | 2.7E-02 |
| *ZK783.6* | WBGene00044448 | protein_coding_gene | -1.8 | 0.29 | 4.4E-02 |
| *spe-12* | WBGene00004966 | protein_coding_gene | -1.8 | 0.29 | 1.8E-02 |
| *fis-1* | WBGene00001424 | protein_coding_gene | -1.8 | 0.29 | 3.4E-02 |
| *bath-46* | WBGene00020325 | protein_coding_gene | -1.8 | 0.29 | 4.7E-02 |
| *dpy-4* | WBGene00001066 | protein_coding_gene | -1.8 | 0.29 | 3.4E-04 |
| *col-169* | WBGene00000742 | protein_coding_gene | -1.8 | 0.29 | 4.9E-03 |
| *oac-19* | WBGene00009413 | protein_coding_gene | -1.8 | 0.29 | 1.7E-02 |
| *T04A8.13* | WBGene00011414 | protein_coding_gene | -1.8 | 0.29 | 3.6E-02 |
| *C30H6.5* | WBGene00007823 | protein_coding_gene | -1.8 | 0.29 | 1.3E-02 |
| *dpy-5* | WBGene00001067 | protein_coding_gene | -1.8 | 0.29 | 1.1E-03 |
| *T19C9.8* | WBGene00011844 | protein_coding_gene | -1.8 | 0.29 | 3.0E-02 |
| *R155.3* | WBGene00020117 | protein_coding_gene | -1.8 | 0.29 | 1.6E-02 |
| *H10E21.4* | WBGene00019184 | protein_coding_gene | -1.8 | 0.28 | 5.5E-03 |
| *ssp-9* | WBGene00006038 | protein_coding_gene | -1.8 | 0.28 | 4.2E-02 |
| *K03H1.9* | WBGene00010544 | protein_coding_gene | -1.8 | 0.28 | 3.8E-02 |
| *C06C6.7* | WBGene00007381 | protein_coding_gene | -1.8 | 0.28 | 1.9E-02 |
| *abu-11* | WBGene00000034 | protein_coding_gene | -1.8 | 0.28 | 1.9E-02 |
| *ugt-40* | WBGene00017331 | protein_coding_gene | -1.8 | 0.28 | 2.5E-04 |
| *ZK180.5* | WBGene00022679 | protein_coding_gene | -1.8 | 0.28 | 1.2E-03 |
| *mpst-4* | WBGene00017387 | protein_coding_gene | -1.8 | 0.28 | 1.2E-02 |
| *T03F6.8* | WBGene00044229 | pseudogene | -1.8 | 0.28 | 3.9E-02 |
| *hrg-2* | WBGene00010473 | protein_coding_gene | -1.9 | 0.28 | 2.4E-02 |
| *T10E9.3* | WBGene00020413 | protein_coding_gene | -1.9 | 0.28 | 2.1E-04 |
| *pqn-63* | WBGene00004146 | protein_coding_gene | -1.9 | 0.28 | 1.9E-02 |
| *col-90* | WBGene00000665 | protein_coding_gene | -1.9 | 0.27 | 1.1E-02 |
| *wrt-4* | WBGene00006950 | protein_coding_gene | -1.9 | 0.27 | 7.4E-04 |
| *abu-8* | WBGene00000031 | protein_coding_gene | -1.9 | 0.27 | 1.2E-02 |
| *T20D4.3* | WBGene00020609 | protein_coding_gene | -1.9 | 0.27 | 1.4E-02 |
| *tsp-19* | WBGene00006645 | protein_coding_gene | -1.9 | 0.27 | 9.7E-03 |
| *col-71* | WBGene00000647 | protein_coding_gene | -1.9 | 0.26 | 4.7E-04 |
| *nhr-115* | WBGene00003705 | protein_coding_gene | -1.9 | 0.26 | 1.9E-05 |
| *col-12* | WBGene00000601 | protein_coding_gene | -1.9 | 0.26 | 4.3E-03 |
| *F10D11.4* | WBGene00008650 | protein_coding_gene | -1.9 | 0.26 | 1.2E-02 |
| *T19D12.9* | WBGene00020583 | protein_coding_gene | -1.9 | 0.26 | 2.8E-02 |
| *col-113* | WBGene00000687 | protein_coding_gene | -1.9 | 0.26 | 1.8E-03 |
| *nas-30* | WBGene00003548 | protein_coding_gene | -2.0 | 0.26 | 4.0E-03 |
| *clec-236* | WBGene00013495 | pseudogene | -2.0 | 0.26 | 2.1E-02 |
| *Y39G10AR.15* | WBGene00021471 | protein_coding_gene | -2.0 | 0.26 | 3.6E-02 |
| *fipr-10* | WBGene00008245 | protein_coding_gene | -2.0 | 0.26 | 2.0E-03 |
| *col-14* | WBGene00000603 | protein_coding_gene | -2.0 | 0.26 | 3.4E-04 |
| *F26G1.5* | WBGene00017841 | protein_coding_gene | -2.0 | 0.25 | 1.0E-03 |
| *F17B5.8* | WBGene00045458 | protein_coding_gene | -2.0 | 0.25 | 1.9E-02 |

| | | | | | |
|---|---|---|---|---|---|
| *pqn-54* | WBGene00004139 | protein_coding_gene | -2.0 | 0.25 | 1.1E-02 |
| *amt-1* | WBGene00000133 | protein_coding_gene | -2.0 | 0.25 | 5.8E-03 |
| *col-73* | WBGene00000649 | protein_coding_gene | -2.0 | 0.25 | 5.6E-03 |
| *T25G12.13* | WBGene00219274 | protein_coding_gene | -2.0 | 0.25 | 2.3E-02 |
| *cyp-14A1* | WBGene00010705 | protein_coding_gene | -2.0 | 0.25 | 3.4E-03 |
| *rol-1* | WBGene00004394 | protein_coding_gene | -2.0 | 0.25 | 2.5E-04 |
| *K11H12.3* | WBGene00019659 | protein_coding_gene | -2.0 | 0.25 | 1.4E-02 |
| *col-65* | WBGene00000641 | protein_coding_gene | -2.0 | 0.25 | 2.0E-03 |
| *ZK180.6* | WBGene00022680 | protein_coding_gene | -2.0 | 0.25 | 5.6E-05 |
| *F11G11.15* | WBGene00271822 | protein_coding_gene | -2.0 | 0.25 | 7.7E-03 |
| *F29G6.1* | WBGene00009257 | protein_coding_gene | -2.0 | 0.25 | 7.3E-04 |
| *H23N18.5* | WBGene00019236 | protein_coding_gene | -2.0 | 0.24 | 1.3E-02 |
| *C45E5.4* | WBGene00016667 | protein_coding_gene | -2.0 | 0.24 | 4.7E-07 |
| *col-175* | WBGene00000748 | protein_coding_gene | -2.0 | 0.24 | 3.3E-04 |
| *F10A3.4* | WBGene00008634 | protein_coding_gene | -2.0 | 0.24 | 1.8E-05 |
| *col-58* | WBGene00000634 | protein_coding_gene | -2.0 | 0.24 | 4.7E-03 |
| *col-138* | WBGene00000711 | protein_coding_gene | -2.0 | 0.24 | 1.3E-04 |
| *asp-8* | WBGene00019105 | protein_coding_gene | -2.0 | 0.24 | 1.3E-10 |
| *B0244.4* | WBGene00015079 | protein_coding_gene | -2.0 | 0.24 | 3.1E-02 |
| *ZK662.2* | WBGene00014039 | protein_coding_gene | -2.1 | 0.24 | 1.3E-03 |
| *C09B9.7* | WBGene00015630 | protein_coding_gene | -2.1 | 0.24 | 9.4E-03 |
| *K10H10.4* | WBGene00010760 | protein_coding_gene | -2.1 | 0.24 | 1.6E-02 |
| *R02E4.3* | WBGene00044724 | protein_coding_gene | -2.1 | 0.24 | 1.7E-03 |
| *col-161* | WBGene00000734 | protein_coding_gene | -2.1 | 0.24 | 3.5E-02 |
| *K02E11.10* | WBGene00044109 | protein_coding_gene | -2.1 | 0.24 | 1.1E-02 |
| *col-104* | WBGene00000678 | protein_coding_gene | -2.1 | 0.24 | 5.1E-04 |
| *pqn-74* | WBGene00004156 | protein_coding_gene | -2.1 | 0.23 | 2.1E-04 |
| *fat-7* | WBGene00001399 | protein_coding_gene | -2.1 | 0.23 | 4.3E-08 |
| *T01G5.1* | WBGene00011339 | protein_coding_gene | -2.1 | 0.23 | 7.6E-06 |
| *ram-2* | WBGene00004300 | protein_coding_gene | -2.1 | 0.23 | 2.6E-05 |
| *col-97* | WBGene00000672 | protein_coding_gene | -2.1 | 0.23 | 1.9E-05 |
| *arf-1.1* | WBGene00000190 | protein_coding_gene | -2.1 | 0.23 | 1.5E-04 |
| *dpy-13* | WBGene00001074 | protein_coding_gene | -2.1 | 0.23 | 2.7E-04 |
| *col-168* | WBGene00000741 | protein_coding_gene | -2.1 | 0.23 | 3.9E-04 |
| *col-63* | WBGene00000639 | protein_coding_gene | -2.1 | 0.23 | 7.2E-05 |
| *C42D4.3* | WBGene00016596 | protein_coding_gene | -2.1 | 0.23 | 3.7E-03 |
| *R12E2.15* | WBGene00020040 | protein_coding_gene | -2.1 | 0.23 | 1.8E-03 |
| *C06E8.5* | WBGene00015544 | protein_coding_gene | -2.2 | 0.23 | 5.4E-04 |
| *col-170* | WBGene00000743 | protein_coding_gene | -2.2 | 0.22 | 4.5E-03 |
| *Y51H7C.13* | WBGene00021791 | protein_coding_gene | -2.2 | 0.22 | 2.5E-05 |
| *Y46G5A.29* | WBGene00012917 | protein_coding_gene | -2.2 | 0.22 | 1.3E-03 |
| *C15H7.4* | WBGene00007611 | protein_coding_gene | -2.2 | 0.22 | 1.8E-02 |
| *nduo-2* | WBGene00010961 | protein_coding_gene | -2.2 | 0.22 | 3.3E-02 |
| *T06E4.12* | WBGene00044011 | protein_coding_gene | -2.2 | 0.22 | 3.8E-02 |
| *asns-1* | WBGene00003816 | protein_coding_gene | -2.2 | 0.22 | 4.0E-04 |
| *abu-15* | WBGene00004099 | protein_coding_gene | -2.2 | 0.21 | 7.1E-03 |
| *T06E4.8* | WBGene00011535 | protein_coding_gene | -2.2 | 0.21 | 4.5E-02 |
| *pqn-2* | WBGene00004098 | protein_coding_gene | -2.2 | 0.21 | 3.1E-02 |
| *col-125* | WBGene00000699 | protein_coding_gene | -2.3 | 0.21 | 5.2E-03 |
| *Y95B8A.2* | WBGene00022383 | protein_coding_gene | -2.3 | 0.21 | 2.3E-04 |
| *col-107* | WBGene00000681 | protein_coding_gene | -2.3 | 0.21 | 1.9E-05 |
| *T21C9.9* | WBGene00011895 | protein_coding_gene | -2.3 | 0.21 | 9.4E-04 |
| *F35B3.4* | WBGene00018031 | protein_coding_gene | -2.3 | 0.21 | 4.1E-02 |
| *C04E12.5* | WBGene00015435 | protein_coding_gene | -2.3 | 0.21 | 3.3E-02 |
| *tbb-6* | WBGene00006539 | protein_coding_gene | -2.3 | 0.20 | 3.1E-04 |
| *bli-6* | WBGene00000256 | protein_coding_gene | -2.3 | 0.20 | 5.0E-05 |
| *F13A7.7* | WBGene00008725 | protein_coding_gene | -2.3 | 0.20 | 6.5E-03 |
| *col-144* | WBGene00000717 | protein_coding_gene | -2.3 | 0.20 | 1.1E-03 |

| | | | | | |
|---|---|---|---|---|---|
| *col-157* | WBGene00000730 | protein_coding_gene | -2.3 | 0.20 | 4.3E-03 |
| *col-167* | WBGene00000740 | protein_coding_gene | -2.3 | 0.20 | 5.0E-05 |
| *gln-4* | WBGene00001605 | pseudogene | -2.3 | 0.20 | 1.0E-02 |
| *C34F11.1* | WBGene00016413 | protein_coding_gene | -2.4 | 0.19 | 3.9E-02 |
| *wrt-8* | WBGene00006954 | protein_coding_gene | -2.4 | 0.19 | 3.5E-03 |
| *cdr-4* | WBGene00010470 | protein_coding_gene | -2.4 | 0.19 | 8.8E-08 |
| *T06E4.9* | WBGene00011536 | protein_coding_gene | -2.4 | 0.19 | 2.1E-02 |
| *C04G6.2* | WBGene00015453 | protein_coding_gene | -2.4 | 0.19 | 7.4E-03 |
| *gmeb-3* | WBGene00008092 | protein_coding_gene | -2.4 | 0.19 | 2.8E-02 |
| *Y47D7A.15* | WBGene00021627 | protein_coding_gene | -2.4 | 0.19 | 1.0E-02 |
| *col-49* | WBGene00000626 | protein_coding_gene | -2.4 | 0.18 | 3.2E-05 |
| *cyp-13A12* | WBGene00008810 | protein_coding_gene | -2.4 | 0.18 | 9.9E-03 |
| *col-162* | WBGene00000735 | protein_coding_gene | -2.5 | 0.18 | 9.9E-03 |
| *dct-5* | WBGene00017218 | protein_coding_gene | -2.5 | 0.18 | 1.9E-02 |
| *Y47D7A.13* | WBGene00021625 | protein_coding_gene | -2.5 | 0.18 | 1.8E-04 |
| *T20D4.10* | WBGene00020616 | protein_coding_gene | -2.5 | 0.18 | 3.7E-02 |
| *grl-16* | WBGene00001725 | protein_coding_gene | -2.5 | 0.17 | 1.6E-05 |
| *C02E7.7* | WBGene00015340 | protein_coding_gene | -2.5 | 0.17 | 1.1E-02 |
| *abu-6* | WBGene00000029 | protein_coding_gene | -2.6 | 0.17 | 2.4E-02 |
| *oac-1* | WBGene00007177 | protein_coding_gene | -2.6 | 0.17 | 3.8E-02 |
| *col-60* | WBGene00000636 | protein_coding_gene | -2.6 | 0.17 | 2.0E-03 |
| *col-152* | WBGene00000725 | protein_coding_gene | -2.6 | 0.17 | 2.6E-02 |
| *col-130* | WBGene00000704 | protein_coding_gene | -2.6 | 0.17 | 4.0E-05 |
| *C30G12.2* | WBGene00016274 | protein_coding_gene | -2.6 | 0.16 | 2.1E-08 |
| *C05E7.2* | WBGene00007344 | protein_coding_gene | -2.7 | 0.16 | 5.4E-04 |
| *col-77* | WBGene00000653 | protein_coding_gene | -2.7 | 0.16 | 1.2E-05 |
| *clec-169* | WBGene00009526 | protein_coding_gene | -2.7 | 0.16 | 2.4E-04 |
| *C04E12.10* | WBGene00015439 | pseudogene | -2.7 | 0.16 | 1.1E-02 |
| *col-180* | WBGene00000753 | protein_coding_gene | -2.7 | 0.16 | 3.0E-05 |
| *dhs-26* | WBGene00000989 | protein_coding_gene | -2.7 | 0.15 | 4.1E-07 |
| *col-120* | WBGene00000694 | protein_coding_gene | -2.8 | 0.15 | 7.2E-05 |
| *R12E2.7* | WBGene00020033 | protein_coding_gene | -2.8 | 0.15 | 2.9E-04 |
| *zmp-6* | WBGene00020646 | protein_coding_gene | -2.8 | 0.15 | 4.1E-02 |
| *grd-6* | WBGene00001695 | protein_coding_gene | -2.8 | 0.14 | 1.7E-04 |
| *col-145* | WBGene00000718 | protein_coding_gene | -2.8 | 0.14 | 8.7E-04 |
| *ZK662.6* | WBGene00045494 | protein_coding_gene | -2.8 | 0.14 | 2.5E-02 |
| *T22B7.7* | WBGene00020674 | protein_coding_gene | -2.9 | 0.13 | 5.6E-08 |
| *F35F10.5* | WBGene00018055 | protein_coding_gene | -3.0 | 0.13 | 1.2E-02 |
| *col-156* | WBGene00000729 | protein_coding_gene | -3.0 | 0.13 | 6.4E-05 |
| *Y48G8AL.12* | WBGene00021690 | protein_coding_gene | -3.0 | 0.13 | 5.0E-04 |
| *col-88* | WBGene00000663 | protein_coding_gene | -3.0 | 0.12 | 1.2E-03 |
| *col-137* | WBGene00000710 | protein_coding_gene | -3.1 | 0.12 | 4.4E-03 |
| *nhr-74* | WBGene00003664 | protein_coding_gene | -3.1 | 0.12 | 9.9E-04 |
| *C07G1.7* | WBGene00015573 | protein_coding_gene | -3.5 | 0.09 | 1.8E-05 |
| *Y41C4A.32* | WBGene00270321 | protein_coding_gene | -3.8 | 0.07 | 2.4E-08 |
| *pgp-1* | WBGene00003995 | protein_coding_gene | -4.6 | 0.04 | 1.3E-30 |

CB4856-exclusive DEGs in response to *P. rettgeri* (vs. OP50)

| Gene Name | Wormbase Gene ID | Type | $log_2FC$ | FC | Adj P-value |
|---|---|---|---|---|---|
| F46F2.3 | WBGene00009787 | protein_coding_gene | 6.1 | 68.33 | 1.0E-51 |
| Y48E1B.8 | WBGene00013007 | protein_coding_gene | 5.9 | 59.84 | 2.0E-26 |
| Y19D10B.1 | WBGene00021230 | pseudogene | 5.2 | 37.79 | 9.0E-13 |
| argk-1 | WBGene00009706 | protein_coding_gene | 5.0 | 32.45 | 7.0E-36 |
| Y19D10B.4 | WBGene00021233 | pseudogene | 5.0 | 31.59 | 1.7E-17 |
| F42A8.1 | WBGene00009626 | protein_coding_gene | 4.6 | 24.35 | 4.2E-138 |
| H25K10.1 | WBGene00010411 | protein_coding_gene | 4.0 | 15.63 | 7.2E-11 |
| hsd-2 | WBGene00022498 | protein_coding_gene | 3.9 | 14.84 | 7.7E-04 |
| nas-3 | WBGene00003522 | protein_coding_gene | 3.8 | 14.28 | 8.7E-10 |

| | | | | | |
|---|---|---|---|---|---|
| K02F2.11 | WBGene00220000 | ncRNA_gene | 3.8 | 13.52 | 3.2E-03 |
| H20E11.2 | WBGene00019214 | protein_coding_gene | 3.7 | 13.36 | 1.0E-03 |
| F18E3.12 | WBGene00045275 | protein_coding_gene | 3.7 | 13.02 | 2.4E-07 |
| F18E3.11 | WBGene00017567 | protein_coding_gene | 3.6 | 12.12 | 7.0E-07 |
| linc-130 | WBGene00219651 | lincRNA_gene | 3.6 | 12.05 | 6.1E-03 |
| K09C4.5 | WBGene00019550 | protein_coding_gene | 3.6 | 11.92 | 1.8E-20 |
| F01D5.3 | WBGene00008494 | protein_coding_gene | 3.6 | 11.88 | 3.8E-10 |
| F58G6.3 | WBGene00010274 | protein_coding_gene | 3.5 | 11.65 | 3.1E-22 |
| drd-50 | WBGene00018643 | protein_coding_gene | 3.5 | 11.54 | 7.1E-26 |
| R07E5.4 | WBGene00011112 | protein_coding_gene | 3.5 | 11.28 | 9.7E-05 |
| gpdh-1 | WBGene00009824 | protein_coding_gene | 3.5 | 11.28 | 9.2E-27 |
| D1086.3 | WBGene00008390 | protein_coding_gene | 3.4 | 10.73 | 3.8E-17 |
| zipt-2.3 | WBGene00013207 | protein_coding_gene | 3.4 | 10.36 | 8.1E-09 |
| ins-32 | WBGene00002115 | protein_coding_gene | 3.4 | 10.28 | 1.4E-03 |
| C50E3.9 | WBGene00016820 | protein_coding_gene | 3.3 | 9.71 | 8.9E-10 |
| col-158 | WBGene00000731 | protein_coding_gene | 3.3 | 9.70 | 1.7E-07 |
| col-94 | WBGene00000669 | protein_coding_gene | 3.3 | 9.59 | 1.0E-02 |
| clec-245 | WBGene00007851 | protein_coding_gene | 3.3 | 9.55 | 4.5E-04 |
| H02F09.3 | WBGene00019146 | protein_coding_gene | 3.2 | 9.45 | 7.2E-08 |
| C49G7.13 | WBGene00269425 | protein_coding_gene | 3.2 | 9.37 | 4.5E-16 |
| Y39B6A.21 | WBGene00012680 | pseudogene | 3.2 | 9.35 | 1.0E-09 |
| sur-5 | WBGene00006351 | protein_coding_gene | 3.2 | 9.28 | 5.6E-106 |
| fbxa-15 | WBGene00022327 | protein_coding_gene | 3.2 | 9.19 | 1.2E-03 |
| T24B8.5 | WBGene00011979 | protein_coding_gene | 3.2 | 9.15 | 1.2E-20 |
| lbp-8 | WBGene00002260 | protein_coding_gene | 3.2 | 9.09 | 4.4E-05 |
| pcp-2 | WBGene00003957 | protein_coding_gene | 3.2 | 9.01 | 1.6E-11 |
| F17C11.14 | WBGene00045151 | snoRNA_gene | 3.1 | 8.63 | 7.2E-03 |
| elo-5 | WBGene00001243 | protein_coding_gene | 3.1 | 8.54 | 1.2E-89 |
| gst-19 | WBGene00001767 | protein_coding_gene | 3.1 | 8.43 | 4.9E-04 |
| T10G3.3 | WBGene00011694 | protein_coding_gene | 3.1 | 8.41 | 1.4E-28 |
| F18E3.13 | WBGene00045276 | protein_coding_gene | 3.1 | 8.32 | 2.1E-05 |
| F54D10.8 | WBGene00018809 | protein_coding_gene | 3.0 | 8.18 | 1.5E-02 |
| vit-1 | WBGene00006925 | protein_coding_gene | 3.0 | 8.06 | 9.2E-05 |
| lips-14 | WBGene00019208 | pseudogene | 3.0 | 8.01 | 1.8E-03 |
| lgc-54 | WBGene00020528 | protein_coding_gene | 3.0 | 8.01 | 1.6E-19 |
| srx-12 | WBGene00005903 | protein_coding_gene | 3.0 | 7.94 | 1.1E-05 |
| F22B8.4 | WBGene00009047 | pseudogene | 2.9 | 7.59 | 2.3E-02 |
| gst-26 | WBGene00001774 | protein_coding_gene | 2.9 | 7.57 | 1.8E-40 |
| F01D5.2 | WBGene00008493 | protein_coding_gene | 2.9 | 7.47 | 9.3E-03 |
| acp-6 | WBGene00022245 | protein_coding_gene | 2.9 | 7.44 | 7.1E-79 |
| irg-4 | WBGene00008584 | protein_coding_gene | 2.9 | 7.39 | 4.0E-27 |
| pqn-98 | WBGene00004177 | protein_coding_gene | 2.9 | 7.37 | 1.0E-02 |
| folt-3 | WBGene00044738 | protein_coding_gene | 2.9 | 7.25 | 6.6E-05 |
| M03B6.1 | WBGene00010833 | protein_coding_gene | 2.8 | 7.15 | 2.1E-13 |
| gcsh-1 | WBGene00008354 | protein_coding_gene | 2.8 | 7.12 | 9.5E-24 |
| F57F4.4 | WBGene00019017 | protein_coding_gene | 2.8 | 7.09 | 2.5E-06 |
| clec-134 | WBGene00021141 | protein_coding_gene | 2.8 | 6.98 | 3.6E-02 |
| F54E7.11 | WBGene00044942 | snoRNA_gene | 2.8 | 6.84 | 1.2E-02 |
| F07C6.6 | WBGene00045067 | protein_coding_gene | 2.8 | 6.81 | 3.4E-02 |
| rrn-4.2 | WBGene00077465 | rRNA_gene | 2.8 | 6.81 | 1.6E-02 |
| F22H10.6 | WBGene00017730 | protein_coding_gene | 2.7 | 6.71 | 1.5E-11 |
| F55G11.2 | WBGene00010123 | protein_coding_gene | 2.7 | 6.70 | 4.8E-06 |
| Y32B12C.3 | WBGene00012526 | pseudogene | 2.7 | 6.66 | 5.4E-03 |
| F49D11.3 | WBGene00018627 | protein_coding_gene | 2.7 | 6.64 | 3.2E-02 |
| F54D8.10 | WBGene00284847 | protein_coding_gene | 2.7 | 6.56 | 5.3E-06 |
| acdh-2 | WBGene00015894 | protein_coding_gene | 2.7 | 6.49 | 2.1E-07 |
| C49F5.8 | WBGene00008212 | protein_coding_gene | 2.7 | 6.46 | 1.5E-02 |
| Y75B12B.12 | WBGene00045129 | snoRNA_gene | 2.7 | 6.33 | 4.2E-02 |

| | | | | | |
|---|---|---|---|---|---|
| Y58A7A.1 | WBGene00021975 | protein_coding_gene | 2.6 | 6.18 | 4.8E-30 |
| hsp-17 | WBGene00002021 | protein_coding_gene | 2.6 | 6.16 | 3.7E-36 |
| B0462.4 | WBGene00007183 | protein_coding_gene | 2.6 | 6.05 | 6.2E-05 |
| col-43 | WBGene00000620 | protein_coding_gene | 2.6 | 6.05 | 3.5E-03 |
| sdz-35 | WBGene00022570 | protein_coding_gene | 2.6 | 6.00 | 2.9E-02 |
| ins-19 | WBGene00002102 | protein_coding_gene | 2.6 | 5.97 | 6.3E-07 |
| rrn-3.56 | WBGene00004677 | pseudogene | 2.6 | 5.96 | 1.3E-02 |
| ZK185.9 | WBGene00220238 | snoRNA_gene | 2.6 | 5.90 | 2.1E-02 |
| col-147 | WBGene00000720 | protein_coding_gene | 2.6 | 5.89 | 4.9E-24 |
| Y43F4B.12 | WBGene00045113 | snoRNA_gene | 2.6 | 5.88 | 3.5E-02 |
| far-4 | WBGene00001388 | protein_coding_gene | 2.6 | 5.87 | 3.1E-03 |
| F11A10.9 | WBGene00045153 | snoRNA_gene | 2.6 | 5.87 | 4.0E-02 |
| Y39D8A.1 | WBGene00021448 | protein_coding_gene | 2.5 | 5.85 | 1.1E-22 |
| F52D4.1 | WBGene00045391 | ncRNA_gene | 2.5 | 5.80 | 3.4E-02 |
| gst-4 | WBGene00001752 | protein_coding_gene | 2.5 | 5.80 | 8.0E-25 |
| C05D12.3 | WBGene00014666 | protein_coding_gene | 2.5 | 5.79 | 5.2E-41 |
| Y26D4A.21 | WBGene00077684 | protein_coding_gene | 2.5 | 5.72 | 1.0E-02 |
| drd-1 | WBGene00009902 | protein_coding_gene | 2.5 | 5.70 | 4.5E-64 |
| ZK228.4 | WBGene00013947 | protein_coding_gene | 2.5 | 5.68 | 1.0E-30 |
| ZK218.1 | WBGene00013935 | protein_coding_gene | 2.5 | 5.67 | 3.9E-08 |
| ncx-7 | WBGene00003572 | protein_coding_gene | 2.5 | 5.55 | 3.8E-34 |
| atic-1 | WBGene00016957 | protein_coding_gene | 2.5 | 5.52 | 4.7E-41 |
| F26D10.24 | WBGene00219917 | snoRNA_gene | 2.5 | 5.51 | 2.5E-02 |
| pqn-11 | WBGene00004103 | pseudogene | 2.5 | 5.49 | 2.3E-02 |
| K11H12.12 | WBGene00044948 | snoRNA_gene | 2.5 | 5.49 | 2.3E-02 |
| clec-118 | WBGene00021032 | protein_coding_gene | 2.4 | 5.42 | 5.9E-03 |
| F25C8.1 | WBGene00009104 | pseudogene | 2.4 | 5.39 | 4.8E-08 |
| cyp-36A1 | WBGene00007913 | protein_coding_gene | 2.4 | 5.33 | 9.3E-23 |
| rrn-2.1 | WBGene00004567 | rRNA_gene | 2.4 | 5.33 | 2.6E-02 |
| Y45F10B.15 | WBGene00012876 | pseudogene | 2.4 | 5.30 | 4.9E-02 |
| grl-4 | WBGene00001713 | protein_coding_gene | 2.4 | 5.26 | 2.9E-20 |
| srv-32 | WBGene00005743 | protein_coding_gene | 2.4 | 5.24 | 4.3E-02 |
| clec-1 | WBGene00017772 | protein_coding_gene | 2.4 | 5.22 | 3.1E-36 |
| twk-10 | WBGene00006665 | protein_coding_gene | 2.4 | 5.17 | 2.9E-04 |
| R08E5.3 | WBGene00019963 | protein_coding_gene | 2.4 | 5.13 | 7.3E-35 |
| H06I04.9 | WBGene00045092 | snoRNA_gene | 2.4 | 5.12 | 9.6E-03 |
| C18B2.1 | WBGene00015952 | protein_coding_gene | 2.3 | 5.09 | 4.0E-02 |
| cysl-3 | WBGene00019962 | protein_coding_gene | 2.3 | 5.08 | 2.9E-22 |
| C05C8.8 | WBGene00015465 | protein_coding_gene | 2.3 | 5.07 | 5.6E-03 |
| dod-19 | WBGene00022644 | protein_coding_gene | 2.3 | 5.04 | 1.5E-08 |
| R166.7 | WBGene00045074 | snoRNA_gene | 2.3 | 5.00 | 2.7E-02 |
| linc-49 | WBGene00219731 | lincRNA_gene | 2.3 | 4.98 | 2.5E-02 |
| ctsa-2 | WBGene00019617 | protein_coding_gene | 2.3 | 4.97 | 1.0E-07 |
| Y105C5B.15 | WBGene00013654 | protein_coding_gene | 2.3 | 4.86 | 3.7E-08 |
| comt-5 | WBGene00021492 | protein_coding_gene | 2.3 | 4.85 | 2.7E-04 |
| srpr-2.1 | WBGene00195016 | ncRNA_gene | 2.3 | 4.85 | 3.5E-02 |
| lon-1 | WBGene00003055 | protein_coding_gene | 2.3 | 4.84 | 1.7E-09 |
| Y24D9A.9 | WBGene00044952 | snoRNA_gene | 2.3 | 4.82 | 3.7E-02 |
| Y51A2D.18 | WBGene00013083 | protein_coding_gene | 2.3 | 4.77 | 1.2E-24 |
| F56C9.12 | WBGene00023071 | snoRNA_gene | 2.2 | 4.76 | 2.8E-02 |
| clec-258 | WBGene00010928 | protein_coding_gene | 2.2 | 4.73 | 8.5E-03 |
| ppat-1 | WBGene00011407 | protein_coding_gene | 2.2 | 4.72 | 4.5E-21 |
| F20G2.1 | WBGene00008985 | protein_coding_gene | 2.2 | 4.71 | 4.8E-07 |
| Y50D4B.7 | WBGene00021746 | pseudogene | 2.2 | 4.71 | 3.3E-03 |
| gst-27 | WBGene00001775 | protein_coding_gene | 2.2 | 4.65 | 2.4E-03 |
| Y82E9BL.18 | WBGene00044402 | protein_coding_gene | 2.2 | 4.58 | 2.7E-02 |
| mth-1 | WBGene00017948 | protein_coding_gene | 2.2 | 4.58 | 2.6E-12 |
| smd-1 | WBGene00004875 | protein_coding_gene | 2.2 | 4.57 | 2.7E-15 |

| | | | | | |
|---|---|---|---|---|---|
| K10C9.9 | WBGene00044515 | pseudogene | 2.2 | 4.50 | 1.3E-02 |
| acox-1.2 | WBGene00008565 | protein_coding_gene | 2.2 | 4.48 | 2.5E-13 |
| col-110 | WBGene00000684 | protein_coding_gene | 2.2 | 4.45 | 3.8E-02 |
| C01G10.17 | WBGene00044658 | protein_coding_gene | 2.2 | 4.45 | 4.8E-05 |
| F09C8.1 | WBGene00008621 | protein_coding_gene | 2.2 | 4.44 | 3.2E-07 |
| Y37H2A.13 | WBGene00045417 | protein_coding_gene | 2.1 | 4.42 | 7.0E-07 |
| Y17D7B.7 | WBGene00014562 | pseudogene | 2.1 | 4.41 | 2.3E-04 |
| AC3.5 | WBGene00007071 | protein_coding_gene | 2.1 | 4.35 | 1.2E-37 |
| Y37E3.22 | WBGene00044932 | snoRNA_gene | 2.1 | 4.34 | 3.7E-02 |
| ets-5 | WBGene00016600 | protein_coding_gene | 2.1 | 4.31 | 2.0E-02 |
| hsp-16.41 | WBGene00002018 | protein_coding_gene | 2.1 | 4.29 | 2.2E-02 |
| Y49G5A.1 | WBGene00021731 | protein_coding_gene | 2.1 | 4.27 | 2.3E-02 |
| F58G6.9 | WBGene00044107 | protein_coding_gene | 2.1 | 4.23 | 7.7E-04 |
| gfi-1 | WBGene00001581 | protein_coding_gene | 2.1 | 4.22 | 5.2E-04 |
| col-143 | WBGene00000716 | protein_coding_gene | 2.1 | 4.17 | 1.0E-09 |
| daf-36 | WBGene00007536 | protein_coding_gene | 2.1 | 4.15 | 6.7E-20 |
| his-40 | WBGene00001914 | protein_coding_gene | 2.0 | 4.11 | 1.2E-02 |
| C06E1.7 | WBGene00015522 | protein_coding_gene | 2.0 | 4.06 | 6.0E-06 |
| fbxa-84 | WBGene00011670 | protein_coding_gene | 2.0 | 4.06 | 7.9E-09 |
| C05C9.3 | WBGene00007327 | protein_coding_gene | 2.0 | 4.05 | 3.9E-09 |
| twk-16 | WBGene00006670 | protein_coding_gene | 2.0 | 4.05 | 1.2E-04 |
| cest-28 | WBGene00019654 | protein_coding_gene | 2.0 | 4.04 | 1.8E-02 |
| F14H12.3 | WBGene00017471 | protein_coding_gene | 2.0 | 4.01 | 8.3E-13 |
| let-383 | WBGene00044898 | protein_coding_gene | 2.0 | 3.98 | 6.2E-07 |
| Y38F1A.4 | WBGene00012607 | protein_coding_gene | 2.0 | 3.98 | 4.3E-02 |
| F54E4.2 | WBGene00010058 | protein_coding_gene | 2.0 | 3.97 | 1.5E-03 |
| frpr-1 | WBGene00015323 | protein_coding_gene | 2.0 | 3.97 | 4.1E-03 |
| vit-4 | WBGene00006928 | protein_coding_gene | 2.0 | 3.96 | 2.0E-02 |
| mxl-3 | WBGene00003511 | protein_coding_gene | 2.0 | 3.96 | 1.9E-04 |
| F23F12.3 | WBGene00017751 | protein_coding_gene | 2.0 | 3.95 | 5.3E-06 |
| C09B8.4 | WBGene00015623 | protein_coding_gene | 2.0 | 3.93 | 8.6E-11 |
| pho-7 | WBGene00011879 | protein_coding_gene | 2.0 | 3.91 | 2.2E-08 |
| acox-1.4 | WBGene00008567 | protein_coding_gene | 2.0 | 3.90 | 5.8E-08 |
| clec-265 | WBGene00019738 | protein_coding_gene | 1.9 | 3.86 | 4.6E-27 |
| dct-18 | WBGene00010266 | protein_coding_gene | 1.9 | 3.86 | 3.3E-59 |
| his-38 | WBGene00001912 | protein_coding_gene | 1.9 | 3.85 | 9.8E-03 |
| cah-5 | WBGene00000283 | protein_coding_gene | 1.9 | 3.82 | 2.5E-21 |
| ctsa-3.1 | WBGene00017969 | protein_coding_gene | 1.9 | 3.81 | 3.1E-04 |
| F48E3.2 | WBGene00018603 | protein_coding_gene | 1.9 | 3.80 | 3.1E-06 |
| F35E2.7 | WBGene00009414 | pseudogene | 1.9 | 3.80 | 3.9E-02 |
| F57H12.6 | WBGene00019021 | protein_coding_gene | 1.9 | 3.79 | 7.3E-05 |
| col-96 | WBGene00000671 | protein_coding_gene | 1.9 | 3.78 | 7.1E-15 |
| C23H5.8 | WBGene00016027 | protein_coding_gene | 1.9 | 3.77 | 9.8E-06 |
| ugt-47 | WBGene00011006 | protein_coding_gene | 1.9 | 3.75 | 8.2E-47 |
| H43E16.1 | WBGene00019273 | protein_coding_gene | 1.9 | 3.72 | 3.4E-19 |
| F34H10.4 | WBGene00009381 | protein_coding_gene | 1.9 | 3.68 | 4.3E-02 |
| H06A10.1 | WBGene00010368 | protein_coding_gene | 1.9 | 3.68 | 1.7E-06 |
| T15B7.14 | WBGene00020526 | protein_coding_gene | 1.9 | 3.67 | 6.6E-04 |
| T17H7.1 | WBGene00020550 | protein_coding_gene | 1.9 | 3.65 | 2.8E-02 |
| pes-8 | WBGene00003981 | protein_coding_gene | 1.9 | 3.65 | 2.8E-02 |
| cyp-25A1 | WBGene00007963 | protein_coding_gene | 1.9 | 3.62 | 5.2E-08 |
| C34E7.3 | WBGene00007933 | protein_coding_gene | 1.8 | 3.58 | 2.1E-02 |
| nlp-24 | WBGene00003762 | protein_coding_gene | 1.8 | 3.56 | 8.0E-04 |
| F14D7.6 | WBGene00008793 | protein_coding_gene | 1.8 | 3.56 | 7.8E-12 |
| ins-30 | WBGene00002113 | protein_coding_gene | 1.8 | 3.56 | 2.5E-02 |
| Y24F12A.4 | WBGene00012499 | pseudogene | 1.8 | 3.55 | 5.5E-03 |
| K04G2.7 | WBGene00010566 | protein_coding_gene | 1.8 | 3.55 | 3.0E-02 |
| W01C9.2 | WBGene00012176 | protein_coding_gene | 1.8 | 3.54 | 7.2E-03 |

| | | | | | |
|---|---|---|---|---|---|
| oac-58 | WBGene00019580 | protein_coding_gene | 1.8 | 3.51 | 7.0E-07 |
| Y54G2A.45 | WBGene00044488 | protein_coding_gene | 1.8 | 3.49 | 2.5E-03 |
| T06A1.5 | WBGene00020280 | protein_coding_gene | 1.8 | 3.48 | 4.6E-20 |
| F41G4.5 | WBGene00018309 | pseudogene | 1.8 | 3.47 | 3.6E-02 |
| C35A5.3 | WBGene00007947 | protein_coding_gene | 1.8 | 3.47 | 1.8E-03 |
| F25H2.14 | WBGene00045138 | snoRNA_gene | 1.8 | 3.47 | 1.5E-02 |
| linc-61 | WBGene00195170 | lincRNA_gene | 1.8 | 3.46 | 2.0E-02 |
| col-176 | WBGene00000749 | protein_coding_gene | 1.8 | 3.44 | 2.5E-02 |
| his-63 | WBGene00001937 | protein_coding_gene | 1.8 | 3.43 | 2.2E-02 |
| clec-174 | WBGene00021580 | protein_coding_gene | 1.8 | 3.43 | 4.7E-03 |
| C14E2.12 | WBGene00219816 | snoRNA_gene | 1.8 | 3.40 | 6.1E-05 |
| ctsa-4.2 | WBGene00021503 | protein_coding_gene | 1.8 | 3.39 | 6.2E-07 |
| D1054.8 | WBGene00008375 | protein_coding_gene | 1.8 | 3.38 | 5.3E-24 |
| hmit-1.2 | WBGene00013074 | protein_coding_gene | 1.8 | 3.38 | 3.2E-09 |
| ugt-65 | WBGene00008583 | protein_coding_gene | 1.8 | 3.37 | 2.4E-04 |
| grsp-1 | WBGene00012840 | protein_coding_gene | 1.8 | 3.37 | 4.5E-05 |
| pgp-8 | WBGene00004002 | protein_coding_gene | 1.7 | 3.35 | 9.1E-03 |
| F28H7.3 | WBGene00009237 | protein_coding_gene | 1.7 | 3.35 | 4.8E-06 |
| T01C8.3 | WBGene00020144 | protein_coding_gene | 1.7 | 3.35 | 5.3E-04 |
| lurp-4 | WBGene00077698 | protein_coding_gene | 1.7 | 3.33 | 2.1E-03 |
| E04D5.5 | WBGene00008484 | protein_coding_gene | 1.7 | 3.32 | 1.2E-07 |
| Y47G6A.33 | WBGene00045399 | protein_coding_gene | 1.7 | 3.31 | 1.8E-17 |
| C45B2.1 | WBGene00016658 | protein_coding_gene | 1.7 | 3.30 | 7.4E-07 |
| his-62 | WBGene00001936 | protein_coding_gene | 1.7 | 3.30 | 4.2E-02 |
| Y45F10C.4 | WBGene00012880 | protein_coding_gene | 1.7 | 3.30 | 5.3E-14 |
| oac-48 | WBGene00011960 | protein_coding_gene | 1.7 | 3.29 | 9.2E-03 |
| R13A5.10 | WBGene00020052 | protein_coding_gene | 1.7 | 3.28 | 3.4E-07 |
| F58G1.9 | WBGene00010271 | pseudogene | 1.7 | 3.28 | 4.6E-02 |
| T25C12.3 | WBGene00012018 | protein_coding_gene | 1.7 | 3.27 | 5.3E-05 |
| C16E9.1 | WBGene00015865 | protein_coding_gene | 1.7 | 3.25 | 7.7E-07 |
| cysl-2 | WBGene00010759 | protein_coding_gene | 1.7 | 3.25 | 8.6E-19 |
| gur-4 | WBGene00001805 | protein_coding_gene | 1.7 | 3.24 | 4.8E-03 |
| T15B7.1 | WBGene00020516 | protein_coding_gene | 1.7 | 3.22 | 3.0E-06 |
| hpd-1 | WBGene00001993 | protein_coding_gene | 1.7 | 3.22 | 3.2E-25 |
| F46F2.5 | WBGene00009789 | protein_coding_gene | 1.7 | 3.21 | 1.8E-02 |
| T10B5.10 | WBGene00020396 | protein_coding_gene | 1.7 | 3.15 | 1.4E-02 |
| Y73C8C.10 | WBGene00022267 | protein_coding_gene | 1.6 | 3.14 | 4.3E-03 |
| C41H7.1 | WBGene00016571 | protein_coding_gene | 1.6 | 3.13 | 8.3E-03 |
| col-101 | WBGene00000675 | protein_coding_gene | 1.6 | 3.12 | 1.3E-04 |
| K08H2.7 | WBGene00010693 | protein_coding_gene | 1.6 | 3.12 | 2.8E-02 |
| clec-186 | WBGene00014138 | protein_coding_gene | 1.6 | 3.11 | 9.5E-23 |
| nlp-77 | WBGene00015514 | protein_coding_gene | 1.6 | 3.10 | 2.8E-45 |
| gst-9 | WBGene00001757 | protein_coding_gene | 1.6 | 3.10 | 1.0E-03 |
| F55G11.8 | WBGene00010128 | protein_coding_gene | 1.6 | 3.09 | 7.4E-09 |
| K08D12.6 | WBGene00019540 | protein_coding_gene | 1.6 | 3.08 | 6.1E-06 |
| F28B4.3 | WBGene00017892 | protein_coding_gene | 1.6 | 3.08 | 6.3E-03 |
| W08A12.3 | WBGene00021080 | protein_coding_gene | 1.6 | 3.06 | 6.4E-07 |
| Y39B6A.41 | WBGene00012699 | protein_coding_gene | 1.6 | 3.05 | 1.3E-04 |
| C12D12.1 | WBGene00015713 | protein_coding_gene | 1.6 | 3.05 | 3.3E-07 |
| pmp-1 | WBGene00004058 | protein_coding_gene | 1.6 | 3.04 | 1.4E-21 |
| lin-31 | WBGene00003017 | protein_coding_gene | 1.6 | 3.03 | 1.2E-02 |
| str-180 | WBGene00006224 | protein_coding_gene | 1.6 | 3.03 | 1.3E-02 |
| C18B2.3 | WBGene00015954 | protein_coding_gene | 1.6 | 3.02 | 5.2E-11 |
| rft-1 | WBGene00044637 | protein_coding_gene | 1.6 | 3.02 | 5.2E-08 |
| hsp-70 | WBGene00002026 | protein_coding_gene | 1.6 | 2.99 | 2.0E-04 |
| Y81B9A.5 | WBGene00304906 | ncRNA_gene | 1.6 | 2.99 | 3.0E-02 |
| pks-1 | WBGene00016558 | protein_coding_gene | 1.6 | 2.99 | 4.3E-11 |
| oac-34 | WBGene00010157 | protein_coding_gene | 1.6 | 2.98 | 2.6E-04 |

| | | | | | |
|---|---|---|---|---|---|
| ugt-32 | WBGene00018543 | protein_coding_gene | 1.6 | 2.97 | 9.2E-03 |
| lgc-4 | WBGene00017580 | protein_coding_gene | 1.6 | 2.96 | 1.4E-03 |
| linc-7 | WBGene00219609 | lincRNA_gene | 1.6 | 2.95 | 6.0E-18 |
| ZC449.8 | WBGene00195189 | protein_coding_gene | 1.6 | 2.94 | 3.9E-04 |
| rhr-1 | WBGene00004358 | protein_coding_gene | 1.6 | 2.94 | 1.8E-40 |
| C46F2.1 | WBGene00016714 | protein_coding_gene | 1.6 | 2.94 | 1.3E-06 |
| srj-29 | WBGene00005616 | protein_coding_gene | 1.6 | 2.94 | 3.2E-02 |
| him-4 | WBGene00001863 | protein_coding_gene | 1.5 | 2.93 | 1.6E-16 |
| C06A1.12 | WBGene00219803 | ncRNA_gene | 1.5 | 2.92 | 9.4E-10 |
| amt-4 | WBGene00000136 | protein_coding_gene | 1.5 | 2.88 | 1.4E-15 |
| F10C1.8 | WBGene00017327 | protein_coding_gene | 1.5 | 2.87 | 2.8E-02 |
| slc-17.4 | WBGene00012130 | protein_coding_gene | 1.5 | 2.87 | 6.8E-13 |
| Y53F4B.23 | WBGene00013169 | protein_coding_gene | 1.5 | 2.87 | 3.1E-03 |
| sqrd-1 | WBGene00008538 | protein_coding_gene | 1.5 | 2.85 | 1.9E-13 |
| T05E12.6 | WBGene00011487 | protein_coding_gene | 1.5 | 2.84 | 4.0E-08 |
| F21C10.9 | WBGene00017658 | protein_coding_gene | 1.5 | 2.84 | 5.4E-35 |
| dct-16 | WBGene00012615 | protein_coding_gene | 1.5 | 2.84 | 1.0E-04 |
| ZK105.3 | WBGene00022654 | protein_coding_gene | 1.5 | 2.83 | 1.2E-02 |
| gpx-5 | WBGene00007516 | protein_coding_gene | 1.5 | 2.81 | 1.2E-14 |
| ent-4 | WBGene00008123 | protein_coding_gene | 1.5 | 2.80 | 6.9E-09 |
| R02D1.2 | WBGene00235310 | protein_coding_gene | 1.5 | 2.78 | 3.8E-04 |
| col-8 | WBGene00000597 | protein_coding_gene | 1.5 | 2.78 | 1.3E-06 |
| F20D1.22 | WBGene00304908 | protein_coding_gene | 1.5 | 2.77 | 1.8E-02 |
| inx-18 | WBGene00002140 | protein_coding_gene | 1.5 | 2.77 | 1.1E-02 |
| Y49E10.29 | WBGene00045239 | protein_coding_gene | 1.5 | 2.76 | 6.0E-06 |
| dod-17 | WBGene00010745 | protein_coding_gene | 1.5 | 2.76 | 3.7E-02 |
| F43C1.5 | WBGene00009649 | protein_coding_gene | 1.5 | 2.75 | 2.9E-03 |
| T22F7.4 | WBGene00020703 | protein_coding_gene | 1.5 | 2.74 | 1.3E-03 |
| ZK6.11 | WBGene00022645 | protein_coding_gene | 1.5 | 2.74 | 6.5E-03 |
| ZK228.3 | WBGene00013946 | protein_coding_gene | 1.5 | 2.74 | 6.4E-09 |
| Y41D4B.17 | WBGene00021519 | protein_coding_gene | 1.4 | 2.73 | 4.1E-03 |
| B0416.2 | WBGene00015178 | protein_coding_gene | 1.4 | 2.73 | 1.6E-02 |
| clec-56 | WBGene00008595 | protein_coding_gene | 1.4 | 2.73 | 4.1E-16 |
| nit-1 | WBGene00014206 | protein_coding_gene | 1.4 | 2.72 | 9.3E-08 |
| T04B2.1 | WBGene00011421 | pseudogene | 1.4 | 2.72 | 3.6E-02 |
| H09G03.1 | WBGene00019174 | protein_coding_gene | 1.4 | 2.72 | 1.1E-02 |
| dsc-4 | WBGene00001099 | protein_coding_gene | 1.4 | 2.71 | 4.9E-22 |
| fbxa-88 | WBGene00008632 | protein_coding_gene | 1.4 | 2.71 | 3.5E-02 |
| enu-3.5 | WBGene00012552 | protein_coding_gene | 1.4 | 2.70 | 1.4E-02 |
| acr-16 | WBGene00000055 | protein_coding_gene | 1.4 | 2.70 | 1.0E-03 |
| C45E5.1 | WBGene00016664 | protein_coding_gene | 1.4 | 2.69 | 3.7E-08 |
| Y119D3B.21 | WBGene00022497 | protein_coding_gene | 1.4 | 2.69 | 1.2E-19 |
| hrg-6 | WBGene00009495 | protein_coding_gene | 1.4 | 2.69 | 6.3E-03 |
| ncs-4 | WBGene00008998 | protein_coding_gene | 1.4 | 2.69 | 7.7E-03 |
| K08C7.1 | WBGene00010646 | protein_coding_gene | 1.4 | 2.69 | 2.2E-10 |
| spds-1 | WBGene00012909 | protein_coding_gene | 1.4 | 2.68 | 2.3E-13 |
| his-67 | WBGene00001941 | protein_coding_gene | 1.4 | 2.67 | 8.5E-03 |
| C18E9.5 | WBGene00007685 | protein_coding_gene | 1.4 | 2.66 | 9.0E-16 |
| Y34B4A.6 | WBGene00021322 | protein_coding_gene | 1.4 | 2.66 | 8.9E-06 |
| T07A5.1 | WBGene00011554 | protein_coding_gene | 1.4 | 2.66 | 1.1E-02 |
| scav-5 | WBGene00011076 | protein_coding_gene | 1.4 | 2.65 | 4.1E-06 |
| lips-5 | WBGene00011725 | protein_coding_gene | 1.4 | 2.64 | 1.1E-03 |
| F01D5.1 | WBGene00008492 | protein_coding_gene | 1.4 | 2.64 | 6.8E-04 |
| pqn-22 | WBGene00004112 | protein_coding_gene | 1.4 | 2.63 | 1.3E-13 |
| T23B7.3 | WBGene00044767 | protein_coding_gene | 1.4 | 2.62 | 1.4E-03 |
| Y7A9A.79 | WBGene00185076 | ncRNA_gene | 1.4 | 2.62 | 3.6E-02 |
| C35B1.4 | WBGene00016434 | protein_coding_gene | 1.4 | 2.61 | 1.3E-11 |
| Y102E9.5 | WBGene00044452 | protein_coding_gene | 1.4 | 2.60 | 1.5E-04 |

| | | | | | |
|---|---|---|---|---|---|
| F19C7.1 | WBGene00017591 | protein_coding_gene | 1.4 | 2.60 | 4.7E-06 |
| T08A9.6 | WBGene00020338 | pseudogene | 1.4 | 2.59 | 4.1E-03 |
| F54F7.2 | WBGene00010064 | protein_coding_gene | 1.4 | 2.59 | 1.5E-12 |
| twk-26 | WBGene00006679 | protein_coding_gene | 1.4 | 2.59 | 3.3E-06 |
| lec-9 | WBGene00002272 | protein_coding_gene | 1.4 | 2.59 | 1.7E-12 |
| C45G9.12 | WBGene00016683 | protein_coding_gene | 1.4 | 2.59 | 1.5E-02 |
| kri-1 | WBGene00013955 | protein_coding_gene | 1.4 | 2.58 | 1.3E-03 |
| ptps-1 | WBGene00015010 | protein_coding_gene | 1.4 | 2.58 | 2.8E-07 |
| F20B6.7 | WBGene00017631 | pseudogene | 1.4 | 2.57 | 3.9E-02 |
| ctsa-1.2 | WBGene00008741 | protein_coding_gene | 1.4 | 2.57 | 3.2E-12 |
| acdh-9 | WBGene00017874 | protein_coding_gene | 1.4 | 2.57 | 6.5E-17 |
| haao-1 | WBGene00010595 | protein_coding_gene | 1.4 | 2.56 | 1.0E-08 |
| oac-3 | WBGene00007361 | protein_coding_gene | 1.4 | 2.56 | 4.6E-02 |
| ZK512.7 | WBGene00013986 | protein_coding_gene | 1.4 | 2.56 | 7.8E-04 |
| ncx-8 | WBGene00003573 | protein_coding_gene | 1.4 | 2.56 | 6.8E-03 |
| exp-1 | WBGene00001373 | protein_coding_gene | 1.4 | 2.56 | 4.7E-02 |
| Y43B11AR.7 | WBGene00045086 | snoRNA_gene | 1.4 | 2.55 | 1.1E-02 |
| fbxa-184 | WBGene00018439 | protein_coding_gene | 1.3 | 2.54 | 1.1E-02 |
| ugt-62 | WBGene00010904 | protein_coding_gene | 1.3 | 2.54 | 6.7E-05 |
| T05C3.2 | WBGene00020254 | protein_coding_gene | 1.3 | 2.54 | 2.3E-02 |
| T08B1.1 | WBGene00020340 | protein_coding_gene | 1.3 | 2.54 | 6.7E-04 |
| gpd-2 | WBGene00001684 | protein_coding_gene | 1.3 | 2.54 | 2.4E-24 |
| B0391.10 | WBGene00007165 | pseudogene | 1.3 | 2.54 | 2.3E-06 |
| B0294.1 | WBGene00015117 | protein_coding_gene | 1.3 | 2.52 | 3.8E-02 |
| hpo-6 | WBGene00021518 | protein_coding_gene | 1.3 | 2.51 | 9.6E-03 |
| cpt-5 | WBGene00008629 | protein_coding_gene | 1.3 | 2.51 | 1.4E-09 |
| B0410.3 | WBGene00015172 | protein_coding_gene | 1.3 | 2.50 | 5.7E-13 |
| cbl-1 | WBGene00007533 | protein_coding_gene | 1.3 | 2.50 | 2.6E-13 |
| aqp-8 | WBGene00000176 | protein_coding_gene | 1.3 | 2.50 | 3.5E-23 |
| Y54E10A.17 | WBGene00021835 | protein_coding_gene | 1.3 | 2.50 | 1.2E-11 |
| prg-2 | WBGene00004179 | pseudogene | 1.3 | 2.50 | 5.2E-18 |
| cln-3.3 | WBGene00000541 | protein_coding_gene | 1.3 | 2.50 | 1.3E-05 |
| ldh-1 | WBGene00002262 | protein_coding_gene | 1.3 | 2.49 | 1.4E-21 |
| oig-5 | WBGene00023520 | protein_coding_gene | 1.3 | 2.49 | 4.1E-02 |
| Y105C5B.7 | WBGene00013648 | pseudogene | 1.3 | 2.49 | 2.1E-02 |
| PDB1.1 | WBGene00019803 | protein_coding_gene | 1.3 | 2.49 | 2.7E-09 |
| K08D8.3 | WBGene00010657 | protein_coding_gene | 1.3 | 2.49 | 1.4E-07 |
| ugt-6 | WBGene00013904 | protein_coding_gene | 1.3 | 2.49 | 6.2E-07 |
| papl-1 | WBGene00008936 | protein_coding_gene | 1.3 | 2.49 | 2.0E-17 |
| tba-4 | WBGene00006530 | protein_coding_gene | 1.3 | 2.48 | 5.8E-12 |
| vha-12 | WBGene00006921 | protein_coding_gene | 1.3 | 2.47 | 3.9E-17 |
| Y42H9AR.5 | WBGene00021540 | protein_coding_gene | 1.3 | 2.47 | 1.4E-02 |
| hphd-1 | WBGene00012608 | protein_coding_gene | 1.3 | 2.47 | 1.7E-03 |
| C17C3.15 | WBGene00015896 | protein_coding_gene | 1.3 | 2.45 | 2.0E-05 |
| kvs-5 | WBGene00021948 | protein_coding_gene | 1.3 | 2.45 | 5.4E-08 |
| ugt-16 | WBGene00013901 | protein_coding_gene | 1.3 | 2.44 | 1.7E-06 |
| C11E4.7 | WBGene00007521 | protein_coding_gene | 1.3 | 2.44 | 7.0E-07 |
| wago-2 | WBGene00018862 | pseudogene | 1.3 | 2.43 | 8.0E-12 |
| folt-1 | WBGene00007388 | protein_coding_gene | 1.3 | 2.42 | 1.1E-06 |
| chk-2 | WBGene00000499 | protein_coding_gene | 1.3 | 2.41 | 4.6E-08 |
| cdr-6 | WBGene00010471 | protein_coding_gene | 1.3 | 2.41 | 1.5E-21 |
| T19D12.1 | WBGene00020576 | protein_coding_gene | 1.3 | 2.40 | 2.5E-11 |
| ZK185.3 | WBGene00022683 | protein_coding_gene | 1.3 | 2.39 | 4.2E-06 |
| unc-23 | WBGene00006760 | protein_coding_gene | 1.3 | 2.39 | 3.2E-09 |
| gst-36 | WBGene00001784 | protein_coding_gene | 1.3 | 2.39 | 4.4E-11 |
| F41G3.21 | WBGene00044316 | protein_coding_gene | 1.3 | 2.38 | 4.4E-03 |
| sptl-3 | WBGene00011932 | protein_coding_gene | 1.2 | 2.38 | 2.0E-25 |
| nhr-31 | WBGene00003625 | protein_coding_gene | 1.2 | 2.38 | 2.6E-10 |

| | | | | | |
|---|---|---|---|---|---|
| Y43F8C.13 | WBGene00012834 | protein_coding_gene | 1.2 | 2.36 | 9.9E-12 |
| ZK742.3 | WBGene00022813 | protein_coding_gene | 1.2 | 2.36 | 1.2E-04 |
| F56F10.1 | WBGene00018984 | protein_coding_gene | 1.2 | 2.36 | 5.2E-08 |
| F13H8.3 | WBGene00017436 | protein_coding_gene | 1.2 | 2.35 | 7.3E-09 |
| bas-1 | WBGene00000239 | protein_coding_gene | 1.2 | 2.35 | 1.0E-07 |
| K01G12.3 | WBGene00010483 | protein_coding_gene | 1.2 | 2.35 | 4.0E-03 |
| R09H10.5 | WBGene00011182 | protein_coding_gene | 1.2 | 2.35 | 3.2E-04 |
| bca-1 | WBGene00000245 | protein_coding_gene | 1.2 | 2.35 | 3.2E-09 |
| hach-1 | WBGene00017301 | protein_coding_gene | 1.2 | 2.35 | 5.6E-03 |
| T10H9.1 | WBGene00020424 | protein_coding_gene | 1.2 | 2.34 | 4.1E-02 |
| C01B10.3 | WBGene00015278 | protein_coding_gene | 1.2 | 2.34 | 2.2E-12 |
| C05C10.3 | WBGene00007330 | protein_coding_gene | 1.2 | 2.34 | 3.5E-18 |
| eppl-1 | WBGene00020139 | protein_coding_gene | 1.2 | 2.34 | 8.4E-14 |
| rhy-1 | WBGene00012324 | protein_coding_gene | 1.2 | 2.33 | 1.2E-09 |
| cyp-34A8 | WBGene00015043 | protein_coding_gene | 1.2 | 2.33 | 3.0E-11 |
| trp-4 | WBGene00006616 | protein_coding_gene | 1.2 | 2.32 | 4.6E-02 |
| pgph-2 | WBGene00018424 | protein_coding_gene | 1.2 | 2.32 | 1.3E-07 |
| Y54G11A.14 | WBGene00013221 | protein_coding_gene | 1.2 | 2.31 | 1.6E-04 |
| C15H9.9 | WBGene00015803 | protein_coding_gene | 1.2 | 2.31 | 6.1E-26 |
| C06A1.4 | WBGene00014667 | pseudogene | 1.2 | 2.31 | 4.7E-16 |
| F54E2.1 | WBGene00018823 | protein_coding_gene | 1.2 | 2.31 | 1.0E-02 |
| F19B6.3 | WBGene00008949 | protein_coding_gene | 1.2 | 2.30 | 2.7E-02 |
| F52H2.6 | WBGene00018718 | protein_coding_gene | 1.2 | 2.30 | 6.4E-09 |
| C34H4.2 | WBGene00016425 | protein_coding_gene | 1.2 | 2.30 | 1.5E-04 |
| spin-3 | WBGene00008598 | protein_coding_gene | 1.2 | 2.30 | 2.8E-03 |
| Y43C5A.7 | WBGene00044922 | protein_coding_gene | 1.2 | 2.30 | 1.8E-02 |
| ugt-63 | WBGene00015449 | protein_coding_gene | 1.2 | 2.29 | 9.0E-04 |
| ech-6 | WBGene00001155 | protein_coding_gene | 1.2 | 2.29 | 6.2E-06 |
| ltah-1.2 | WBGene00022610 | protein_coding_gene | 1.2 | 2.29 | 1.4E-16 |
| pck-1 | WBGene00021043 | protein_coding_gene | 1.2 | 2.29 | 1.2E-20 |
| C15B12.8 | WBGene00015788 | protein_coding_gene | 1.2 | 2.29 | 5.1E-03 |
| cth-2 | WBGene00022856 | protein_coding_gene | 1.2 | 2.28 | 3.4E-35 |
| C44B7.7 | WBGene00016629 | protein_coding_gene | 1.2 | 2.28 | 1.0E-05 |
| clec-187 | WBGene00014137 | protein_coding_gene | 1.2 | 2.28 | 3.5E-06 |
| aqp-7 | WBGene00000175 | protein_coding_gene | 1.2 | 2.28 | 4.7E-12 |
| trap-1 | WBGene00022122 | protein_coding_gene | 1.2 | 2.28 | 3.7E-19 |
| C37H5.14 | WBGene00045308 | protein_coding_gene | 1.2 | 2.27 | 2.3E-02 |
| F57F5.1 | WBGene00010204 | protein_coding_gene | 1.2 | 2.27 | 9.7E-04 |
| pes-9 | WBGene00003982 | protein_coding_gene | 1.2 | 2.27 | 1.6E-23 |
| irg-6 | WBGene00007864 | protein_coding_gene | 1.2 | 2.26 | 2.0E-03 |
| aass-1 | WBGene00019819 | protein_coding_gene | 1.2 | 2.26 | 7.0E-20 |
| F44B9.2 | WBGene00018407 | protein_coding_gene | 1.2 | 2.26 | 1.3E-08 |
| Y13C8A.4 | WBGene00219303 | pseudogene | 1.2 | 2.26 | 7.1E-05 |
| F09E10.1 | WBGene00017294 | protein_coding_gene | 1.2 | 2.26 | 5.0E-07 |
| unc-105 | WBGene00006832 | protein_coding_gene | 1.2 | 2.26 | 4.3E-05 |
| C49A9.4 | WBGene00016758 | protein_coding_gene | 1.2 | 2.25 | 3.8E-04 |
| ugt-26 | WBGene00015695 | protein_coding_gene | 1.2 | 2.25 | 3.4E-05 |
| Y43F11A.6 | WBGene00014576 | pseudogene | 1.2 | 2.24 | 2.9E-07 |
| Y6B3B.1 | WBGene00012387 | protein_coding_gene | 1.2 | 2.24 | 9.6E-03 |
| T07D3.9 | WBGene00020311 | protein_coding_gene | 1.2 | 2.24 | 1.6E-06 |
| cyp-25A2 | WBGene00007964 | protein_coding_gene | 1.2 | 2.24 | 5.6E-07 |
| pho-8 | WBGene00011262 | protein_coding_gene | 1.2 | 2.23 | 9.5E-03 |
| Y38H6C.21 | WBGene00012633 | protein_coding_gene | 1.2 | 2.23 | 6.9E-04 |
| K10C2.8 | WBGene00045265 | protein_coding_gene | 1.2 | 2.23 | 1.0E-04 |
| sec-61.B | WBGene00021427 | protein_coding_gene | 1.2 | 2.23 | 2.1E-13 |
| vha-11 | WBGene00006920 | protein_coding_gene | 1.2 | 2.23 | 2.9E-14 |
| C09F12.2 | WBGene00007482 | protein_coding_gene | 1.2 | 2.23 | 4.3E-02 |
| Y51F10.7 | WBGene00021768 | protein_coding_gene | 1.2 | 2.23 | 9.7E-10 |

| | | | | | |
|---|---|---|---|---|---|
| F42G10.1 | WBGene00009645 | protein_coding_gene | 1.2 | 2.22 | 1.3E-05 |
| B0244.17 | WBGene00271715 | protein_coding_gene | 1.2 | 2.22 | 3.6E-02 |
| pgp-12 | WBGene00004006 | protein_coding_gene | 1.2 | 2.22 | 2.4E-02 |
| Y77E11A.12 | WBGene00022313 | protein_coding_gene | 1.1 | 2.22 | 2.3E-02 |
| klu-2 | WBGene00022592 | protein_coding_gene | 1.1 | 2.22 | 1.7E-06 |
| set-9 | WBGene00017482 | protein_coding_gene | 1.1 | 2.22 | 1.2E-18 |
| C26B9.5 | WBGene00016134 | protein_coding_gene | 1.1 | 2.21 | 1.2E-11 |
| hmit-1.1 | WBGene00013073 | protein_coding_gene | 1.1 | 2.21 | 9.7E-04 |
| R03G8.3 | WBGene00010998 | protein_coding_gene | 1.1 | 2.20 | 6.9E-06 |
| Y22D7AR.10 | WBGene00021263 | protein_coding_gene | 1.1 | 2.20 | 3.7E-11 |
| twk-17 | WBGene00006671 | protein_coding_gene | 1.1 | 2.20 | 9.1E-03 |
| F40E12.2 | WBGene00018235 | protein_coding_gene | 1.1 | 2.20 | 6.6E-03 |
| slc-9B.2 | WBGene00010214 | protein_coding_gene | 1.1 | 2.19 | 1.7E-02 |
| puf-7 | WBGene00004243 | protein_coding_gene | 1.1 | 2.19 | 9.9E-13 |
| glr-4 | WBGene00001615 | protein_coding_gene | 1.1 | 2.18 | 3.4E-04 |
| D1005.t1 | WBGene00023263 | pseudogene | 1.1 | 2.18 | 2.2E-02 |
| pals-16 | WBGene00022337 | protein_coding_gene | 1.1 | 2.17 | 1.2E-02 |
| idh-1 | WBGene00010317 | protein_coding_gene | 1.1 | 2.17 | 2.2E-13 |
| pgph-3 | WBGene00016892 | protein_coding_gene | 1.1 | 2.17 | 8.7E-03 |
| pat-12 | WBGene00003936 | protein_coding_gene | 1.1 | 2.17 | 1.0E-14 |
| T08D2.3 | WBGene00011608 | pseudogene | 1.1 | 2.17 | 9.9E-04 |
| mans-2 | WBGene00008258 | protein_coding_gene | 1.1 | 2.16 | 8.7E-18 |
| cth-1 | WBGene00009048 | protein_coding_gene | 1.1 | 2.16 | 4.0E-05 |
| Y53C10A.5 | WBGene00013136 | protein_coding_gene | 1.1 | 2.16 | 5.4E-11 |
| rig-6 | WBGene00016354 | protein_coding_gene | 1.1 | 2.16 | 6.1E-10 |
| vha-5 | WBGene00006914 | protein_coding_gene | 1.1 | 2.16 | 9.8E-16 |
| C17C3.1 | WBGene00015887 | protein_coding_gene | 1.1 | 2.16 | 1.5E-07 |
| B0250.7 | WBGene00007124 | protein_coding_gene | 1.1 | 2.15 | 1.4E-05 |
| ketn-1 | WBGene00004130 | protein_coding_gene | 1.1 | 2.15 | 1.1E-10 |
| clec-173 | WBGene00044469 | protein_coding_gene | 1.1 | 2.15 | 7.6E-06 |
| sec-61.A | WBGene00013311 | protein_coding_gene | 1.1 | 2.14 | 3.6E-19 |
| R06C1.4 | WBGene00011059 | protein_coding_gene | 1.1 | 2.14 | 2.6E-23 |
| lev-9 | WBGene00002976 | protein_coding_gene | 1.1 | 2.14 | 7.3E-03 |
| lrg-1 | WBGene00003067 | pseudogene | 1.1 | 2.14 | 1.4E-05 |
| C02F12.5 | WBGene00015355 | protein_coding_gene | 1.1 | 2.14 | 2.9E-02 |
| memi-2 | WBGene00013380 | protein_coding_gene | 1.1 | 2.14 | 1.5E-11 |
| M02D8.5 | WBGene00019731 | protein_coding_gene | 1.1 | 2.14 | 1.6E-04 |
| glb-3 | WBGene00007389 | protein_coding_gene | 1.1 | 2.14 | 1.4E-02 |
| F55H12.4 | WBGene00010135 | protein_coding_gene | 1.1 | 2.14 | 1.3E-13 |
| anc-1 | WBGene00000140 | protein_coding_gene | 1.1 | 2.13 | 3.5E-08 |
| lev-1 | WBGene00002974 | protein_coding_gene | 1.1 | 2.13 | 7.2E-04 |
| Y57G11C.51 | WBGene00044258 | protein_coding_gene | 1.1 | 2.13 | 1.4E-05 |
| fbxa-190 | WBGene00018939 | protein_coding_gene | 1.1 | 2.12 | 3.5E-02 |
| K08D8.12 | WBGene00194926 | protein_coding_gene | 1.1 | 2.12 | 2.5E-02 |
| vha-20 | WBGene00010993 | protein_coding_gene | 1.1 | 2.12 | 1.1E-18 |
| Y45G12C.1 | WBGene00021565 | protein_coding_gene | 1.1 | 2.12 | 1.8E-02 |
| C23H3.2 | WBGene00016018 | protein_coding_gene | 1.1 | 2.12 | 6.9E-10 |
| twk-12 | WBGene00006667 | protein_coding_gene | 1.1 | 2.12 | 1.4E-02 |
| hpo-26 | WBGene00018492 | protein_coding_gene | 1.1 | 2.12 | 1.6E-03 |
| Y73B3A.1 | WBGene00022203 | protein_coding_gene | 1.1 | 2.11 | 1.4E-11 |
| F09E10.7 | WBGene00017297 | protein_coding_gene | 1.1 | 2.11 | 1.1E-02 |
| Y73B6BL.31 | WBGene00022252 | protein_coding_gene | 1.1 | 2.10 | 2.7E-09 |
| vha-13 | WBGene00013025 | protein_coding_gene | 1.1 | 2.10 | 1.9E-19 |
| C27D9.1 | WBGene00016163 | protein_coding_gene | 1.1 | 2.10 | 1.9E-14 |
| col-20 | WBGene00000609 | protein_coding_gene | 1.1 | 2.10 | 1.1E-12 |
| col-119 | WBGene00000693 | protein_coding_gene | 1.1 | 2.09 | 2.3E-16 |
| Y69H2.3 | WBGene00013481 | protein_coding_gene | 1.1 | 2.09 | 3.9E-13 |
| cnnm-3 | WBGene00016343 | protein_coding_gene | 1.1 | 2.09 | 1.0E-08 |

| | | | | | |
|---|---|---|---|---|---|
| C34B4.2 | WBGene00007907 | protein_coding_gene | 1.1 | 2.09 | 1.0E-09 |
| T07A9.14 | WBGene00044789 | protein_coding_gene | 1.1 | 2.09 | 2.2E-13 |
| dmsr-2 | WBGene00021275 | protein_coding_gene | 1.1 | 2.09 | 1.5E-02 |
| zig-12 | WBGene00019727 | protein_coding_gene | 1.1 | 2.08 | 1.6E-10 |
| C50E3.12 | WBGene00016823 | protein_coding_gene | 1.1 | 2.07 | 2.0E-02 |
| Y53C10A.1 | WBGene00014915 | pseudogene | 1.1 | 2.07 | 2.2E-03 |
| dim-1 | WBGene00001000 | protein_coding_gene | 1.0 | 2.07 | 3.5E-18 |
| T05B11.4 | WBGene00020247 | protein_coding_gene | 1.0 | 2.06 | 1.5E-06 |
| dhs-3 | WBGene00000967 | protein_coding_gene | 1.0 | 2.06 | 2.2E-09 |
| vha-19 | WBGene00021952 | protein_coding_gene | 1.0 | 2.06 | 4.0E-17 |
| F25E2.2 | WBGene00017780 | protein_coding_gene | 1.0 | 2.06 | 2.5E-05 |
| cin-4 | WBGene00022854 | protein_coding_gene | 1.0 | 2.06 | 9.2E-11 |
| hprt-1 | WBGene00013690 | protein_coding_gene | 1.0 | 2.06 | 2.8E-15 |
| col-122 | WBGene00000696 | protein_coding_gene | 1.0 | 2.06 | 2.3E-05 |
| E01G4.3 | WBGene00008446 | protein_coding_gene | 1.0 | 2.06 | 1.8E-09 |
| C29G2.6 | WBGene00016234 | protein_coding_gene | 1.0 | 2.06 | 4.7E-07 |
| vha-8 | WBGene00006917 | protein_coding_gene | 1.0 | 2.05 | 2.4E-18 |
| F53C11.1 | WBGene00009971 | protein_coding_gene | 1.0 | 2.05 | 7.9E-05 |
| lact-1 | WBGene00018523 | protein_coding_gene | 1.0 | 2.04 | 1.4E-02 |
| R03D7.2 | WBGene00010989 | protein_coding_gene | 1.0 | 2.04 | 8.9E-03 |
| C28A5.1 | WBGene00007788 | protein_coding_gene | 1.0 | 2.04 | 1.3E-10 |
| prmt-6 | WBGene00016023 | protein_coding_gene | 1.0 | 2.04 | 4.8E-03 |
| sid-1 | WBGene00004795 | protein_coding_gene | 1.0 | 2.03 | 1.1E-07 |
| enu-3.6 | WBGene00010477 | protein_coding_gene | 1.0 | 2.03 | 3.9E-02 |
| nkb-3 | WBGene00010117 | protein_coding_gene | 1.0 | 2.03 | 1.2E-15 |
| ins-27 | WBGene00002110 | protein_coding_gene | 1.0 | 2.03 | 1.8E-02 |
| mai-1 | WBGene00003124 | protein_coding_gene | 1.0 | 2.02 | 4.1E-04 |
| C25A11.10 | WBGene00219374 | pseudogene | 1.0 | 2.02 | 6.5E-04 |
| dod-24 | WBGene00007875 | protein_coding_gene | 1.0 | 2.02 | 1.8E-08 |
| ttn-1 | WBGene00006436 | protein_coding_gene | 1.0 | 2.02 | 1.2E-02 |
| lec-8 | WBGene00002271 | protein_coding_gene | 1.0 | 2.02 | 3.2E-03 |
| Y37F4.7 | WBGene00021388 | protein_coding_gene | 1.0 | 2.01 | 4.2E-02 |
| pid-5 | WBGene00021555 | protein_coding_gene | 1.0 | 2.01 | 4.8E-14 |
| glc-4 | WBGene00001594 | protein_coding_gene | 1.0 | 2.01 | 2.2E-03 |
| K10C2.14 | WBGene00206388 | protein_coding_gene | 1.0 | 2.01 | 6.1E-06 |
| Y39E4B.5 | WBGene00012716 | protein_coding_gene | 1.0 | 2.01 | 5.2E-10 |
| hrde-1 | WBGene00007624 | protein_coding_gene | 1.0 | 2.01 | 3.4E-22 |
| C14B1.7 | WBGene00007578 | pseudogene | 1.0 | 2.01 | 3.1E-11 |
| tnt-3 | WBGene00006588 | protein_coding_gene | 1.0 | 2.01 | 2.5E-10 |
| pxn-1 | WBGene00004256 | protein_coding_gene | 1.0 | 2.01 | 1.2E-08 |
| F09E10.15 | WBGene00219886 | ncRNA_gene | 1.0 | 2.01 | 8.6E-03 |
| col-140 | WBGene00000713 | protein_coding_gene | 1.0 | 2.01 | 7.9E-09 |
| cyp-35A5 | WBGene00019473 | protein_coding_gene | 1.0 | 2.01 | 5.1E-04 |
| Y14H12A.1 | WBGene00021190 | protein_coding_gene | 1.0 | 2.00 | 4.5E-03 |
| zif-1 | WBGene00006977 | protein_coding_gene | 1.0 | 2.00 | 7.7E-13 |
| T20D3.2 | WBGene00011856 | protein_coding_gene | 1.0 | 2.00 | 2.9E-04 |
| C32E12.4 | WBGene00016330 | protein_coding_gene | -1.0 | 0.50 | 4.1E-09 |
| slc-28.2 | WBGene00017868 | protein_coding_gene | -1.0 | 0.50 | 3.4E-02 |
| C50F4.10 | WBGene00008235 | protein_coding_gene | -1.0 | 0.50 | 4.8E-03 |
| ZK673.2 | WBGene00014058 | protein_coding_gene | -1.0 | 0.50 | 1.8E-19 |
| pmt-1 | WBGene00022781 | protein_coding_gene | -1.0 | 0.50 | 3.4E-17 |
| K03H1.13 | WBGene00044903 | protein_coding_gene | -1.0 | 0.50 | 1.9E-02 |
| cec-1 | WBGene00000414 | protein_coding_gene | -1.0 | 0.50 | 3.6E-13 |
| vab-2 | WBGene00006869 | protein_coding_gene | -1.0 | 0.50 | 1.1E-03 |
| nhr-92 | WBGene00003682 | protein_coding_gene | -1.0 | 0.50 | 1.8E-02 |
| hda-5 | WBGene00009663 | protein_coding_gene | -1.0 | 0.50 | 4.7E-04 |
| ptp-1 | WBGene00004213 | protein_coding_gene | -1.0 | 0.50 | 6.4E-07 |
| M79.2 | WBGene00010902 | protein_coding_gene | -1.0 | 0.50 | 1.4E-08 |

| | | | | | |
|---|---|---|---|---|---|
| ins-4 | WBGene00002087 | protein_coding_gene | -1.0 | 0.50 | 1.3E-02 |
| C01A2.2 | WBGene00007214 | protein_coding_gene | -1.0 | 0.50 | 6.9E-03 |
| K01A2.5 | WBGene00019280 | protein_coding_gene | -1.0 | 0.50 | 5.7E-06 |
| Y43E12A.2 | WBGene00012794 | protein_coding_gene | -1.0 | 0.50 | 1.9E-07 |
| ceh-79 | WBGene00007749 | protein_coding_gene | -1.0 | 0.50 | 3.2E-04 |
| R10D12.1 | WBGene00011185 | protein_coding_gene | -1.0 | 0.50 | 1.5E-02 |
| W10C8.6 | WBGene00044433 | protein_coding_gene | -1.0 | 0.50 | 1.3E-02 |
| Y62F5A.10 | WBGene00013388 | protein_coding_gene | -1.0 | 0.50 | 6.3E-03 |
| B0454.6 | WBGene00015197 | protein_coding_gene | -1.0 | 0.50 | 4.2E-06 |
| M02H5.8 | WBGene00019744 | protein_coding_gene | -1.0 | 0.50 | 2.0E-02 |
| pes-4 | WBGene00003978 | protein_coding_gene | -1.0 | 0.50 | 2.7E-08 |
| nhr-55 | WBGene00003645 | protein_coding_gene | -1.0 | 0.50 | 5.2E-06 |
| ncx-3 | WBGene00003568 | protein_coding_gene | -1.0 | 0.50 | 2.6E-06 |
| nhr-107 | WBGene00003697 | protein_coding_gene | -1.0 | 0.50 | 2.9E-05 |
| F49E12.10 | WBGene00009903 | protein_coding_gene | -1.0 | 0.49 | 1.5E-02 |
| unc-51 | WBGene00006786 | protein_coding_gene | -1.0 | 0.49 | 1.4E-11 |
| nex-2 | WBGene00003589 | protein_coding_gene | -1.0 | 0.49 | 7.9E-17 |
| daf-14 | WBGene00000910 | protein_coding_gene | -1.0 | 0.49 | 1.0E-06 |
| Y57A10A.14 | WBGene00013257 | protein_coding_gene | -1.0 | 0.49 | 2.1E-03 |
| Y71F9AL.25 | WBGene00304792 | protein_coding_gene | -1.0 | 0.49 | 1.9E-02 |
| F13B12.2 | WBGene00008730 | protein_coding_gene | -1.0 | 0.49 | 4.3E-04 |
| ttr-17 | WBGene00012383 | protein_coding_gene | -1.0 | 0.49 | 5.1E-09 |
| R03H10.1 | WBGene00019854 | protein_coding_gene | -1.0 | 0.49 | 2.9E-03 |
| math-37 | WBGene00020080 | protein_coding_gene | -1.0 | 0.49 | 1.2E-02 |
| dhs-15 | WBGene00000978 | protein_coding_gene | -1.0 | 0.49 | 2.0E-02 |
| W02D7.4 | WBGene00020940 | protein_coding_gene | -1.0 | 0.49 | 1.8E-06 |
| K07E3.2 | WBGene00019491 | protein_coding_gene | -1.0 | 0.49 | 3.5E-05 |
| lact-5 | WBGene00016355 | protein_coding_gene | -1.0 | 0.49 | 1.5E-04 |
| lron-15 | WBGene00016974 | protein_coding_gene | -1.0 | 0.49 | 5.7E-03 |
| F57C12.6 | WBGene00194687 | protein_coding_gene | -1.0 | 0.49 | 4.7E-02 |
| mrp-3 | WBGene00003409 | protein_coding_gene | -1.0 | 0.49 | 6.0E-07 |
| tag-275 | WBGene00016423 | protein_coding_gene | -1.0 | 0.49 | 6.3E-03 |
| sma-2 | WBGene00004856 | protein_coding_gene | -1.0 | 0.49 | 7.7E-05 |
| Y43B11AR.3 | WBGene00021544 | protein_coding_gene | -1.0 | 0.49 | 4.1E-07 |
| ptp-4 | WBGene00010558 | protein_coding_gene | -1.0 | 0.49 | 3.7E-03 |
| Y73F4A.2 | WBGene00013515 | protein_coding_gene | -1.0 | 0.49 | 3.3E-07 |
| D1044.1 | WBGene00017027 | protein_coding_gene | -1.0 | 0.49 | 8.5E-03 |
| nhx-9 | WBGene00003736 | protein_coding_gene | -1.0 | 0.49 | 1.7E-04 |
| elt-7 | WBGene00015981 | protein_coding_gene | -1.0 | 0.49 | 3.4E-06 |
| hpo-15 | WBGene00016061 | protein_coding_gene | -1.0 | 0.49 | 7.7E-10 |
| F55H12.5 | WBGene00010136 | protein_coding_gene | -1.0 | 0.49 | 8.7E-05 |
| Y64G10A.7 | WBGene00013416 | protein_coding_gene | -1.0 | 0.49 | 1.4E-03 |
| che-14 | WBGene00000493 | protein_coding_gene | -1.0 | 0.49 | 2.4E-02 |
| T16G1.5 | WBGene00011799 | protein_coding_gene | -1.0 | 0.49 | 2.8E-02 |
| F19C7.2 | WBGene00017592 | protein_coding_gene | -1.0 | 0.49 | 4.2E-06 |
| sulp-8 | WBGene00013963 | protein_coding_gene | -1.0 | 0.49 | 5.3E-05 |
| F39E9.28 | WBGene00271716 | pseudogene | -1.0 | 0.49 | 1.6E-03 |
| crb-1 | WBGene00000792 | protein_coding_gene | -1.0 | 0.49 | 1.3E-07 |
| W04A8.5 | WBGene00012240 | protein_coding_gene | -1.0 | 0.49 | 2.6E-02 |
| F33E2.5 | WBGene00009361 | protein_coding_gene | -1.0 | 0.49 | 9.8E-08 |
| M04D8.7 | WBGene00010865 | protein_coding_gene | -1.0 | 0.49 | 4.1E-02 |
| ttx-1 | WBGene00006652 | protein_coding_gene | -1.0 | 0.49 | 1.3E-03 |
| nhr-195 | WBGene00019115 | protein_coding_gene | -1.0 | 0.49 | 6.7E-03 |
| nhr-41 | WBGene00022423 | protein_coding_gene | -1.0 | 0.49 | 2.1E-04 |
| F54C9.3 | WBGene00010041 | protein_coding_gene | -1.0 | 0.49 | 1.6E-12 |
| kap-1 | WBGene00002182 | protein_coding_gene | -1.0 | 0.49 | 2.9E-03 |
| F44E5.3 | WBGene00009690 | protein_coding_gene | -1.0 | 0.49 | 3.5E-02 |
| lec-11 | WBGene00002274 | protein_coding_gene | -1.0 | 0.48 | 1.8E-04 |

| | | | | | |
|---|---|---|---|---|---|
| B0546.4 | WBGene00015251 | protein_coding_gene | -1.0 | 0.48 | 3.1E-09 |
| nhr-90 | WBGene00003680 | protein_coding_gene | -1.0 | 0.48 | 1.1E-05 |
| Y20C6A.4 | WBGene00077495 | protein_coding_gene | -1.0 | 0.48 | 2.8E-02 |
| F08F1.4 | WBGene00017258 | protein_coding_gene | -1.0 | 0.48 | 1.4E-03 |
| K02A11.4 | WBGene00044607 | protein_coding_gene | -1.0 | 0.48 | 4.3E-04 |
| cec-8 | WBGene00021913 | protein_coding_gene | -1.1 | 0.48 | 6.0E-07 |
| grh-1 | WBGene00001707 | protein_coding_gene | -1.1 | 0.48 | 5.1E-04 |
| Y71A12B.18 | WBGene00045077 | pseudogene | -1.1 | 0.48 | 2.6E-02 |
| kal-1 | WBGene00002181 | protein_coding_gene | -1.1 | 0.48 | 5.7E-06 |
| K12C11.6 | WBGene00045052 | protein_coding_gene | -1.1 | 0.48 | 1.5E-04 |
| gst-15 | WBGene00001763 | protein_coding_gene | -1.1 | 0.48 | 2.2E-03 |
| dcar-1 | WBGene00007395 | protein_coding_gene | -1.1 | 0.48 | 8.6E-11 |
| C05E11.3 | WBGene00015494 | protein_coding_gene | -1.1 | 0.48 | 6.9E-06 |
| jmjd-4 | WBGene00011563 | protein_coding_gene | -1.1 | 0.48 | 6.7E-04 |
| cal-2 | WBGene00000286 | protein_coding_gene | -1.1 | 0.48 | 4.4E-06 |
| ZK673.1 | WBGene00014057 | protein_coding_gene | -1.1 | 0.48 | 8.1E-06 |
| daf-10 | WBGene00000906 | protein_coding_gene | -1.1 | 0.48 | 3.0E-02 |
| K02C4.5 | WBGene00010503 | protein_coding_gene | -1.1 | 0.48 | 2.3E-09 |
| F09F9.1 | WBGene00017306 | protein_coding_gene | -1.1 | 0.48 | 4.8E-04 |
| C34B4.3 | WBGene00007908 | protein_coding_gene | -1.1 | 0.48 | 2.4E-02 |
| hnd-1 | WBGene00001981 | protein_coding_gene | -1.1 | 0.48 | 1.2E-03 |
| sams-5 | WBGene00006416 | protein_coding_gene | -1.1 | 0.48 | 2.2E-06 |
| F20C5.11 | WBGene00303015 | protein_coding_gene | -1.1 | 0.48 | 3.8E-04 |
| F08C6.5 | WBGene00017243 | protein_coding_gene | -1.1 | 0.48 | 2.3E-02 |
| odr-3 | WBGene00003850 | protein_coding_gene | -1.1 | 0.48 | 3.8E-02 |
| B0507.6 | WBGene00015223 | protein_coding_gene | -1.1 | 0.48 | 1.0E-02 |
| clc-5 | WBGene00000526 | protein_coding_gene | -1.1 | 0.48 | 2.8E-03 |
| F25B3.5 | WBGene00009102 | protein_coding_gene | -1.1 | 0.48 | 3.2E-03 |
| pqn-89 | WBGene00004169 | protein_coding_gene | -1.1 | 0.48 | 7.8E-10 |
| ric-3 | WBGene00004363 | protein_coding_gene | -1.1 | 0.48 | 2.5E-04 |
| ilys-4 | WBGene00016958 | protein_coding_gene | -1.1 | 0.48 | 1.3E-04 |
| C33G3.4 | WBGene00007904 | protein_coding_gene | -1.1 | 0.48 | 1.5E-09 |
| T25D10.4 | WBGene00020801 | protein_coding_gene | -1.1 | 0.47 | 3.8E-06 |
| T19H12.15 | WBGene00271827 | protein_coding_gene | -1.1 | 0.47 | 1.0E-02 |
| F59C6.11 | WBGene00010328 | protein_coding_gene | -1.1 | 0.47 | 5.0E-03 |
| Y71A12B.19 | WBGene00077443 | pseudogene | -1.1 | 0.47 | 2.8E-05 |
| ZK688.7 | WBGene00022802 | protein_coding_gene | -1.1 | 0.47 | 2.4E-05 |
| cest-35.2 | WBGene00016863 | protein_coding_gene | -1.1 | 0.47 | 2.1E-02 |
| tsp-20 | WBGene00006646 | protein_coding_gene | -1.1 | 0.47 | 5.8E-04 |
| ptr-8 | WBGene00004222 | protein_coding_gene | -1.1 | 0.47 | 6.8E-05 |
| ttc-36 | WBGene00009947 | protein_coding_gene | -1.1 | 0.47 | 3.4E-07 |
| B0286.1 | WBGene00015115 | protein_coding_gene | -1.1 | 0.47 | 2.5E-04 |
| bath-38 | WBGene00009945 | protein_coding_gene | -1.1 | 0.47 | 1.4E-08 |
| ZC190.4 | WBGene00022538 | protein_coding_gene | -1.1 | 0.47 | 4.1E-09 |
| nfm-1 | WBGene00003593 | protein_coding_gene | -1.1 | 0.47 | 7.9E-14 |
| T19H5.4 | WBGene00011849 | protein_coding_gene | -1.1 | 0.47 | 7.5E-12 |
| F52B10.3 | WBGene00044304 | protein_coding_gene | -1.1 | 0.47 | 1.2E-02 |
| cex-1 | WBGene00023407 | protein_coding_gene | -1.1 | 0.47 | 2.5E-07 |
| B0507.3 | WBGene00015220 | protein_coding_gene | -1.1 | 0.47 | 1.3E-02 |
| Y49A10A.2 | WBGene00044949 | protein_coding_gene | -1.1 | 0.47 | 7.2E-06 |
| ceh-20 | WBGene00000443 | protein_coding_gene | -1.1 | 0.47 | 2.6E-10 |
| B0284.3 | WBGene00007133 | protein_coding_gene | -1.1 | 0.47 | 3.2E-03 |
| pdl-1 | WBGene00003966 | protein_coding_gene | -1.1 | 0.47 | 2.0E-02 |
| T23B3.2 | WBGene00020710 | protein_coding_gene | -1.1 | 0.47 | 2.4E-09 |
| ZK593.2 | WBGene00014002 | protein_coding_gene | -1.1 | 0.47 | 1.8E-03 |
| C30G7.4 | WBGene00044178 | protein_coding_gene | -1.1 | 0.47 | 2.8E-03 |
| BE0003N10.3 | WBGene00015268 | protein_coding_gene | -1.1 | 0.47 | 4.5E-03 |
| Y57G11B.2 | WBGene00013295 | protein_coding_gene | -1.1 | 0.47 | 6.8E-05 |

| | | | | | |
|---|---|---|---|---|---|
| rhgf-2 | WBGene00006476 | protein_coding_gene | -1.1 | 0.47 | 1.1E-05 |
| frm-2 | WBGene00001489 | protein_coding_gene | -1.1 | 0.47 | 1.7E-10 |
| asm-1 | WBGene00000211 | protein_coding_gene | -1.1 | 0.47 | 2.0E-07 |
| mdl-1 | WBGene00003163 | protein_coding_gene | -1.1 | 0.47 | 1.5E-09 |
| fbxa-167 | WBGene00007837 | protein_coding_gene | -1.1 | 0.47 | 2.0E-02 |
| tli-1 | WBGene00006578 | protein_coding_gene | -1.1 | 0.46 | 3.2E-08 |
| nhr-94 | WBGene00003684 | protein_coding_gene | -1.1 | 0.46 | 1.8E-03 |
| T19D7.6 | WBGene00020574 | protein_coding_gene | -1.1 | 0.46 | 2.7E-03 |
| snb-2 | WBGene00004898 | protein_coding_gene | -1.1 | 0.46 | 5.2E-11 |
| grdn-1 | WBGene00013082 | protein_coding_gene | -1.1 | 0.46 | 7.0E-10 |
| nhr-140 | WBGene00016366 | protein_coding_gene | -1.1 | 0.46 | 4.1E-03 |
| tbx-2 | WBGene00006543 | protein_coding_gene | -1.1 | 0.46 | 1.1E-09 |
| K09C8.2 | WBGene00010715 | protein_coding_gene | -1.1 | 0.46 | 2.0E-02 |
| Y52B11A.4 | WBGene00013124 | protein_coding_gene | -1.1 | 0.46 | 1.8E-03 |
| F46C3.6 | WBGene00045253 | protein_coding_gene | -1.1 | 0.46 | 9.7E-03 |
| C10C5.7 | WBGene00007511 | protein_coding_gene | -1.1 | 0.46 | 1.4E-02 |
| ugt-29 | WBGene00021709 | protein_coding_gene | -1.1 | 0.46 | 2.7E-03 |
| K02A6.3 | WBGene00019294 | protein_coding_gene | -1.1 | 0.46 | 3.1E-07 |
| dlc-3 | WBGene00012424 | protein_coding_gene | -1.1 | 0.46 | 3.7E-03 |
| egas-4 | WBGene00018906 | protein_coding_gene | -1.1 | 0.46 | 4.1E-02 |
| ZK930.2 | WBGene00014152 | protein_coding_gene | -1.1 | 0.46 | 2.2E-06 |
| C05C12.4 | WBGene00007336 | protein_coding_gene | -1.1 | 0.46 | 8.3E-17 |
| Y22D7AL.15 | WBGene00021253 | protein_coding_gene | -1.1 | 0.46 | 1.0E-06 |
| F57A8.1 | WBGene00010177 | protein_coding_gene | -1.1 | 0.46 | 1.2E-11 |
| nhr-210 | WBGene00020015 | protein_coding_gene | -1.1 | 0.46 | 9.2E-03 |
| C40H1.9 | WBGene00008044 | protein_coding_gene | -1.1 | 0.46 | 4.8E-02 |
| tsp-16 | WBGene00006642 | protein_coding_gene | -1.1 | 0.46 | 2.5E-03 |
| Y55F3BR.11 | WBGene00044757 | protein_coding_gene | -1.1 | 0.46 | 1.2E-03 |
| tub-1 | WBGene00006655 | protein_coding_gene | -1.1 | 0.46 | 3.9E-02 |
| C33H5.13 | WBGene00016379 | protein_coding_gene | -1.1 | 0.46 | 2.9E-08 |
| Y95B8A.6 | WBGene00022386 | protein_coding_gene | -1.1 | 0.46 | 2.8E-06 |
| T10B11.5 | WBGene00020401 | protein_coding_gene | -1.1 | 0.46 | 2.9E-04 |
| hot-6 | WBGene00001991 | protein_coding_gene | -1.1 | 0.46 | 9.0E-04 |
| C02D5.4 | WBGene00043097 | protein_coding_gene | -1.1 | 0.46 | 1.6E-15 |
| R11D1.3 | WBGene00011244 | protein_coding_gene | -1.1 | 0.46 | 7.2E-03 |
| T22C8.6 | WBGene00011927 | protein_coding_gene | -1.1 | 0.46 | 3.5E-03 |
| irx-1 | WBGene00007984 | protein_coding_gene | -1.1 | 0.46 | 1.4E-08 |
| T19B10.5 | WBGene00011833 | protein_coding_gene | -1.1 | 0.46 | 1.1E-03 |
| ZK512.11 | WBGene00013989 | protein_coding_gene | -1.1 | 0.45 | 4.9E-03 |
| pho-4 | WBGene00004023 | protein_coding_gene | -1.1 | 0.45 | 1.6E-08 |
| fbxb-100 | WBGene00013403 | protein_coding_gene | -1.1 | 0.45 | 4.7E-02 |
| syg-2 | WBGene00007750 | protein_coding_gene | -1.1 | 0.45 | 2.5E-06 |
| gpa-9 | WBGene00001671 | protein_coding_gene | -1.1 | 0.45 | 5.6E-04 |
| rig-5 | WBGene00004372 | protein_coding_gene | -1.1 | 0.45 | 1.8E-03 |
| osm-12 | WBGene00003892 | protein_coding_gene | -1.1 | 0.45 | 4.8E-02 |
| cutl-24 | WBGene00021396 | protein_coding_gene | -1.1 | 0.45 | 4.6E-03 |
| E02C12.6 | WBGene00017092 | protein_coding_gene | -1.1 | 0.45 | 1.8E-02 |
| T21B4.21 | WBGene00220076 | ncRNA_gene | -1.1 | 0.45 | 1.1E-03 |
| acl-12 | WBGene00015295 | protein_coding_gene | -1.1 | 0.45 | 3.6E-11 |
| gln-3 | WBGene00001604 | protein_coding_gene | -1.1 | 0.45 | 2.1E-14 |
| C33D9.5 | WBGene00007899 | pseudogene | -1.1 | 0.45 | 1.1E-04 |
| cyp-14A3 | WBGene00010707 | protein_coding_gene | -1.1 | 0.45 | 3.2E-02 |
| W08F4.7 | WBGene00021096 | protein_coding_gene | -1.1 | 0.45 | 2.6E-02 |
| C50B6.9 | WBGene00008222 | protein_coding_gene | -1.1 | 0.45 | 1.1E-02 |
| Y2H9A.4 | WBGene00012381 | protein_coding_gene | -1.1 | 0.45 | 1.3E-03 |
| ztf-11 | WBGene00009939 | protein_coding_gene | -1.1 | 0.45 | 1.4E-11 |
| C08F8.15 | WBGene00219217 | protein_coding_gene | -1.2 | 0.45 | 3.0E-02 |
| strm-1 | WBGene00019198 | protein_coding_gene | -1.2 | 0.45 | 9.4E-03 |

| | | | | | |
|---|---|---|---|---|---|
| W03G11.4 | WBGene00012226 | protein_coding_gene | -1.2 | 0.45 | 3.2E-08 |
| nhr-110 | WBGene00003700 | protein_coding_gene | -1.2 | 0.45 | 3.4E-07 |
| ztf-13 | WBGene00018704 | protein_coding_gene | -1.2 | 0.45 | 1.1E-07 |
| F47B8.2 | WBGene00009803 | protein_coding_gene | -1.2 | 0.45 | 6.0E-08 |
| kgb-2 | WBGene00002188 | protein_coding_gene | -1.2 | 0.45 | 2.7E-04 |
| nhr-35 | WBGene00003628 | protein_coding_gene | -1.2 | 0.45 | 1.5E-14 |
| exc-5 | WBGene00001366 | protein_coding_gene | -1.2 | 0.45 | 2.3E-14 |
| C33D9.13 | WBGene00194651 | protein_coding_gene | -1.2 | 0.45 | 2.9E-05 |
| cest-27 | WBGene00018958 | protein_coding_gene | -1.2 | 0.45 | 1.4E-08 |
| T05E11.8 | WBGene00011484 | protein_coding_gene | -1.2 | 0.45 | 2.9E-03 |
| F49C12.6 | WBGene00009876 | protein_coding_gene | -1.2 | 0.45 | 7.2E-03 |
| nhr-131 | WBGene00003721 | protein_coding_gene | -1.2 | 0.45 | 5.7E-05 |
| afmd-2 | WBGene00012866 | protein_coding_gene | -1.2 | 0.45 | 1.2E-04 |
| Y77E11A.23 | WBGene00235267 | protein_coding_gene | -1.2 | 0.45 | 5.6E-03 |
| K10D6.4 | WBGene00010744 | protein_coding_gene | -1.2 | 0.45 | 1.7E-08 |
| F11E6.8 | WBGene00008711 | protein_coding_gene | -1.2 | 0.45 | 8.1E-05 |
| Y105E8B.9 | WBGene00013693 | protein_coding_gene | -1.2 | 0.45 | 3.7E-05 |
| C08G5.7 | WBGene00044418 | protein_coding_gene | -1.2 | 0.45 | 3.5E-04 |
| F58A3.3 | WBGene00010223 | protein_coding_gene | -1.2 | 0.45 | 1.2E-02 |
| Y48A6B.8 | WBGene00012969 | protein_coding_gene | -1.2 | 0.45 | 2.2E-02 |
| Y42A5A.3 | WBGene00012778 | protein_coding_gene | -1.2 | 0.45 | 2.8E-02 |
| nlp-2 | WBGene00003740 | protein_coding_gene | -1.2 | 0.44 | 1.1E-02 |
| tag-278 | WBGene00015356 | protein_coding_gene | -1.2 | 0.44 | 1.2E-07 |
| cyp-33E1 | WBGene00016768 | protein_coding_gene | -1.2 | 0.44 | 4.7E-07 |
| Y47H10A.5 | WBGene00012961 | protein_coding_gene | -1.2 | 0.44 | 8.2E-04 |
| K08D8.7 | WBGene00185092 | protein_coding_gene | -1.2 | 0.44 | 7.5E-04 |
| ttr-51 | WBGene00010440 | protein_coding_gene | -1.2 | 0.44 | 5.2E-13 |
| clh-4 | WBGene00000531 | protein_coding_gene | -1.2 | 0.44 | 3.3E-02 |
| F13E9.15 | WBGene00045208 | protein_coding_gene | -1.2 | 0.44 | 3.0E-07 |
| ZK1290.13 | WBGene00022893 | protein_coding_gene | -1.2 | 0.44 | 1.8E-08 |
| hum-6 | WBGene00002039 | protein_coding_gene | -1.2 | 0.44 | 1.4E-06 |
| igcm-3 | WBGene00020160 | protein_coding_gene | -1.2 | 0.44 | 1.6E-14 |
| F56C9.8 | WBGene00018952 | protein_coding_gene | -1.2 | 0.44 | 1.5E-08 |
| rrc-1 | WBGene00009800 | protein_coding_gene | -1.2 | 0.44 | 1.8E-04 |
| tag-273 | WBGene00013289 | protein_coding_gene | -1.2 | 0.44 | 1.2E-07 |
| dyf-1 | WBGene00001117 | protein_coding_gene | -1.2 | 0.44 | 1.4E-02 |
| frpr-18 | WBGene00020586 | protein_coding_gene | -1.2 | 0.44 | 3.3E-02 |
| W03B1.3 | WBGene00020973 | protein_coding_gene | -1.2 | 0.44 | 6.9E-04 |
| T24B8.3 | WBGene00011977 | protein_coding_gene | -1.2 | 0.44 | 2.4E-18 |
| nlr-1 | WBGene00003772 | protein_coding_gene | -1.2 | 0.44 | 9.9E-07 |
| F54D1.6 | WBGene00010047 | protein_coding_gene | -1.2 | 0.44 | 7.9E-21 |
| F53C11.9 | WBGene00044921 | protein_coding_gene | -1.2 | 0.44 | 1.0E-03 |
| zip-10 | WBGene00012101 | protein_coding_gene | -1.2 | 0.44 | 6.0E-03 |
| ceeh-1 | WBGene00019329 | protein_coding_gene | -1.2 | 0.44 | 5.7E-09 |
| C45G7.13 | WBGene00235340 | protein_coding_gene | -1.2 | 0.44 | 3.7E-03 |
| F10G8.8 | WBGene00008666 | protein_coding_gene | -1.2 | 0.44 | 1.4E-10 |
| lect-2 | WBGene00019407 | protein_coding_gene | -1.2 | 0.44 | 1.1E-07 |
| Y58A7A.3 | WBGene00021977 | protein_coding_gene | -1.2 | 0.43 | 1.4E-08 |
| sym-2 | WBGene00006367 | protein_coding_gene | -1.2 | 0.43 | 1.4E-06 |
| catp-7 | WBGene00022010 | protein_coding_gene | -1.2 | 0.43 | 7.1E-17 |
| C50D2.1 | WBGene00016805 | protein_coding_gene | -1.2 | 0.43 | 2.6E-02 |
| lea-1 | WBGene00002263 | protein_coding_gene | -1.2 | 0.43 | 1.3E-17 |
| C24H10.1 | WBGene00016067 | protein_coding_gene | -1.2 | 0.43 | 5.9E-03 |
| hmg-11 | WBGene00001976 | protein_coding_gene | -1.2 | 0.43 | 1.3E-09 |
| Y41G9A.5 | WBGene00021529 | protein_coding_gene | -1.2 | 0.43 | 3.2E-02 |
| glb-8 | WBGene00007741 | protein_coding_gene | -1.2 | 0.43 | 2.6E-02 |
| T13C2.3 | WBGene00020479 | protein_coding_gene | -1.2 | 0.43 | 7.3E-04 |
| C56G2.4 | WBGene00016979 | protein_coding_gene | -1.2 | 0.43 | 2.3E-03 |

| | | | | | |
|---|---|---|---|---|---|
| tag-243 | WBGene00044078 | protein_coding_gene | -1.2 | 0.43 | 3.8E-05 |
| C41G7.8 | WBGene00077525 | protein_coding_gene | -1.2 | 0.43 | 6.3E-04 |
| K01A2.4 | WBGene00019279 | protein_coding_gene | -1.2 | 0.43 | 4.2E-05 |
| C56E10.3 | WBGene00016976 | protein_coding_gene | -1.2 | 0.43 | 8.5E-03 |
| cest-31 | WBGene00019653 | protein_coding_gene | -1.2 | 0.43 | 1.1E-03 |
| rpi-2 | WBGene00019536 | protein_coding_gene | -1.2 | 0.43 | 4.6E-02 |
| nhr-120 | WBGene00003710 | protein_coding_gene | -1.2 | 0.43 | 6.8E-06 |
| Y52B11A.12 | WBGene00077539 | protein_coding_gene | -1.2 | 0.43 | 4.1E-03 |
| akt-2 | WBGene00000103 | protein_coding_gene | -1.2 | 0.43 | 5.6E-10 |
| R03E9.2 | WBGene00019843 | protein_coding_gene | -1.2 | 0.43 | 6.7E-04 |
| F20C5.5 | WBGene00008970 | protein_coding_gene | -1.2 | 0.43 | 3.2E-03 |
| lips-10 | WBGene00008803 | protein_coding_gene | -1.2 | 0.43 | 1.2E-05 |
| gcl-1 | WBGene00013382 | protein_coding_gene | -1.2 | 0.43 | 9.5E-09 |
| lido-17 | WBGene00007378 | protein_coding_gene | -1.2 | 0.43 | 1.2E-02 |
| pho-6 | WBGene00018696 | protein_coding_gene | -1.2 | 0.43 | 1.5E-04 |
| F13B6.1 | WBGene00017416 | protein_coding_gene | -1.2 | 0.43 | 4.6E-03 |
| R05D8.7 | WBGene00019885 | protein_coding_gene | -1.2 | 0.43 | 1.0E-02 |
| F18A11.2 | WBGene00008929 | protein_coding_gene | -1.2 | 0.43 | 2.4E-02 |
| F48E3.9 | WBGene00018608 | protein_coding_gene | -1.2 | 0.43 | 1.8E-02 |
| F38B6.6 | WBGene00018175 | protein_coding_gene | -1.2 | 0.42 | 9.1E-04 |
| bath-47 | WBGene00006464 | protein_coding_gene | -1.2 | 0.42 | 1.8E-04 |
| C35E7.6 | WBGene00016458 | protein_coding_gene | -1.2 | 0.42 | 2.0E-16 |
| K08E7.6 | WBGene00010674 | protein_coding_gene | -1.2 | 0.42 | 3.6E-02 |
| nhr-69 | WBGene00003659 | protein_coding_gene | -1.2 | 0.42 | 5.2E-11 |
| K03H1.5 | WBGene00010540 | protein_coding_gene | -1.2 | 0.42 | 4.2E-12 |
| jmjd-3.1 | WBGene00017571 | protein_coding_gene | -1.2 | 0.42 | 2.5E-10 |
| ckr-2 | WBGene00021439 | protein_coding_gene | -1.2 | 0.42 | 5.5E-03 |
| osg-1 | WBGene00019832 | protein_coding_gene | -1.2 | 0.42 | 9.2E-05 |
| B0478.3 | WBGene00015202 | protein_coding_gene | -1.2 | 0.42 | 4.2E-02 |
| fbxa-79 | WBGene00022329 | protein_coding_gene | -1.2 | 0.42 | 2.5E-02 |
| Y65B4BL.7 | WBGene00022039 | protein_coding_gene | -1.2 | 0.42 | 1.3E-02 |
| nep-14 | WBGene00018225 | pseudogene | -1.2 | 0.42 | 2.9E-04 |
| T01D3.6 | WBGene00011330 | protein_coding_gene | -1.2 | 0.42 | 3.0E-06 |
| W01F3.2 | WBGene00012185 | protein_coding_gene | -1.2 | 0.42 | 1.3E-13 |
| daam-1 | WBGene00018976 | protein_coding_gene | -1.2 | 0.42 | 9.6E-04 |
| K10B4.3 | WBGene00019615 | protein_coding_gene | -1.2 | 0.42 | 6.6E-11 |
| cyp-25A3 | WBGene00014697 | protein_coding_gene | -1.2 | 0.42 | 2.3E-07 |
| twk-11 | WBGene00006666 | protein_coding_gene | -1.2 | 0.42 | 7.4E-03 |
| nhr-117 | WBGene00003707 | protein_coding_gene | -1.2 | 0.42 | 1.4E-05 |
| cfi-1 | WBGene00000476 | protein_coding_gene | -1.2 | 0.42 | 6.9E-05 |
| ceh-36 | WBGene00000457 | protein_coding_gene | -1.2 | 0.42 | 4.5E-03 |
| col-99 | WBGene00000674 | protein_coding_gene | -1.2 | 0.42 | 2.3E-06 |
| madf-10 | WBGene00013717 | protein_coding_gene | -1.2 | 0.42 | 7.3E-03 |
| T02C12.5 | WBGene00194692 | protein_coding_gene | -1.3 | 0.42 | 3.2E-02 |
| W04B5.2 | WBGene00021019 | protein_coding_gene | -1.3 | 0.42 | 2.2E-04 |
| C17H12.11 | WBGene00015936 | protein_coding_gene | -1.3 | 0.42 | 6.5E-05 |
| F08B12.4 | WBGene00008572 | protein_coding_gene | -1.3 | 0.42 | 2.6E-11 |
| clec-47 | WBGene00011668 | protein_coding_gene | -1.3 | 0.42 | 3.1E-05 |
| chpf-2 | WBGene00019108 | protein_coding_gene | -1.3 | 0.42 | 5.1E-05 |
| vap-2 | WBGene00011462 | protein_coding_gene | -1.3 | 0.42 | 1.0E-04 |
| ZC487.1 | WBGene00022623 | protein_coding_gene | -1.3 | 0.42 | 1.3E-02 |
| R10E4.6 | WBGene00011202 | protein_coding_gene | -1.3 | 0.42 | 4.0E-03 |
| C35A11.2 | WBGene00016429 | protein_coding_gene | -1.3 | 0.42 | 2.0E-02 |
| Y37D8A.16 | WBGene00012554 | protein_coding_gene | -1.3 | 0.42 | 3.2E-12 |
| C54E10.6 | WBGene00008310 | protein_coding_gene | -1.3 | 0.42 | 3.6E-04 |
| inx-6 | WBGene00002128 | protein_coding_gene | -1.3 | 0.41 | 5.3E-05 |
| Y75B8A.6 | WBGene00013543 | protein_coding_gene | -1.3 | 0.41 | 4.9E-10 |
| F13G3.3 | WBGene00008763 | protein_coding_gene | -1.3 | 0.41 | 2.4E-02 |

| | | | | | |
|---|---|---|---|---|---|
| sma-10 | WBGene00020649 | protein_coding_gene | -1.3 | 0.41 | 1.7E-13 |
| F53A2.9 | WBGene00009953 | protein_coding_gene | -1.3 | 0.41 | 1.3E-06 |
| fbxb-75 | WBGene00019956 | protein_coding_gene | -1.3 | 0.41 | 2.6E-04 |
| Y6G8.2 | WBGene00012404 | protein_coding_gene | -1.3 | 0.41 | 1.7E-03 |
| Y57A10A.26 | WBGene00013266 | protein_coding_gene | -1.3 | 0.41 | 3.6E-13 |
| R04B5.5 | WBGene00011003 | protein_coding_gene | -1.3 | 0.41 | 3.4E-11 |
| ift-81 | WBGene00017973 | protein_coding_gene | -1.3 | 0.41 | 4.7E-02 |
| bath-10 | WBGene00021726 | protein_coding_gene | -1.3 | 0.41 | 2.1E-07 |
| zip-5 | WBGene00007932 | protein_coding_gene | -1.3 | 0.41 | 6.8E-07 |
| C32H11.3 | WBGene00007866 | protein_coding_gene | -1.3 | 0.41 | 4.5E-02 |
| Y97E10AL.1 | WBGene00022392 | protein_coding_gene | -1.3 | 0.41 | 1.7E-02 |
| sox-2 | WBGene00004949 | protein_coding_gene | -1.3 | 0.41 | 1.4E-12 |
| F31D5.6 | WBGene00017949 | protein_coding_gene | -1.3 | 0.41 | 3.4E-02 |
| dep-1 | WBGene00009717 | protein_coding_gene | -1.3 | 0.41 | 9.7E-10 |
| ttr-44 | WBGene00008341 | protein_coding_gene | -1.3 | 0.41 | 4.4E-04 |
| eol-1 | WBGene00045401 | protein_coding_gene | -1.3 | 0.41 | 1.1E-02 |
| cfz-2 | WBGene00000478 | protein_coding_gene | -1.3 | 0.41 | 1.1E-04 |
| W04C9.8 | WBGene00235387 | ncRNA_gene | -1.3 | 0.41 | 3.5E-02 |
| F58H10.1 | WBGene00010291 | protein_coding_gene | -1.3 | 0.41 | 5.0E-03 |
| Y71F9AL.6 | WBGene00022111 | protein_coding_gene | -1.3 | 0.41 | 2.1E-02 |
| R148.5 | WBGene00020104 | protein_coding_gene | -1.3 | 0.41 | 5.9E-15 |
| Y39C12A.9 | WBGene00012710 | protein_coding_gene | -1.3 | 0.41 | 4.6E-02 |
| T14B1.1 | WBGene00011763 | protein_coding_gene | -1.3 | 0.41 | 3.6E-07 |
| cutl-21 | WBGene00018861 | protein_coding_gene | -1.3 | 0.41 | 4.3E-03 |
| C07A9.12 | WBGene00043994 | protein_coding_gene | -1.3 | 0.41 | 1.7E-03 |
| cest-16 | WBGene00020330 | protein_coding_gene | -1.3 | 0.41 | 7.2E-03 |
| ugt-1 | WBGene00007072 | protein_coding_gene | -1.3 | 0.41 | 1.3E-07 |
| elt-6 | WBGene00001253 | protein_coding_gene | -1.3 | 0.40 | 1.9E-04 |
| C51E3.10 | WBGene00008255 | protein_coding_gene | -1.3 | 0.40 | 2.6E-03 |
| Y54F10BM.3 | WBGene00021858 | protein_coding_gene | -1.3 | 0.40 | 2.1E-06 |
| best-3 | WBGene00015286 | protein_coding_gene | -1.3 | 0.40 | 2.7E-07 |
| pat-9 | WBGene00003933 | protein_coding_gene | -1.3 | 0.40 | 7.8E-09 |
| D1054.18 | WBGene00077783 | protein_coding_gene | -1.3 | 0.40 | 3.1E-02 |
| F56H9.2 | WBGene00010174 | protein_coding_gene | -1.3 | 0.40 | 1.5E-06 |
| T04F8.9 | WBGene00011447 | protein_coding_gene | -1.3 | 0.40 | 2.6E-02 |
| col-76 | WBGene00000652 | protein_coding_gene | -1.3 | 0.40 | 6.4E-08 |
| F13D11.3 | WBGene00017428 | protein_coding_gene | -1.3 | 0.40 | 1.2E-08 |
| ZK863.8 | WBGene00014125 | protein_coding_gene | -1.3 | 0.40 | 1.9E-09 |
| F56H11.2 | WBGene00010175 | protein_coding_gene | -1.3 | 0.40 | 2.1E-04 |
| F53B3.3 | WBGene00018742 | protein_coding_gene | -1.3 | 0.40 | 1.1E-02 |
| unc-120 | WBGene00006844 | protein_coding_gene | -1.3 | 0.40 | 1.0E-09 |
| ddn-1 | WBGene00015227 | protein_coding_gene | -1.3 | 0.40 | 3.8E-02 |
| ZK105.13 | WBGene00235114 | protein_coding_gene | -1.3 | 0.40 | 1.9E-02 |
| C34E11.2 | WBGene00007935 | protein_coding_gene | -1.3 | 0.40 | 1.5E-02 |
| R10H1.1 | WBGene00019998 | protein_coding_gene | -1.3 | 0.40 | 3.7E-10 |
| clec-264 | WBGene00009291 | protein_coding_gene | -1.3 | 0.40 | 2.1E-06 |
| tab-1 | WBGene00006380 | protein_coding_gene | -1.3 | 0.40 | 4.4E-02 |
| cpn-2 | WBGene00000778 | protein_coding_gene | -1.3 | 0.39 | 2.5E-02 |
| C31H5.4 | WBGene00007855 | protein_coding_gene | -1.3 | 0.39 | 7.3E-18 |
| clec-196 | WBGene00009156 | protein_coding_gene | -1.3 | 0.39 | 9.1E-10 |
| cyp-42A1 | WBGene00013585 | protein_coding_gene | -1.3 | 0.39 | 1.2E-05 |
| ugt-2 | WBGene00007073 | protein_coding_gene | -1.3 | 0.39 | 3.5E-12 |
| insc-1 | WBGene00018392 | protein_coding_gene | -1.3 | 0.39 | 1.6E-07 |
| K12B6.9 | WBGene00023432 | protein_coding_gene | -1.3 | 0.39 | 5.9E-11 |
| nhr-59 | WBGene00003649 | protein_coding_gene | -1.3 | 0.39 | 6.3E-04 |
| ceh-48 | WBGene00015934 | protein_coding_gene | -1.4 | 0.39 | 1.5E-04 |
| famh-161 | WBGene00012626 | protein_coding_gene | -1.4 | 0.39 | 1.9E-02 |
| glb-15 | WBGene00009387 | protein_coding_gene | -1.4 | 0.39 | 1.8E-02 |

| | | | | | |
|---|---|---|---|---|---|
| cgt-1 | WBGene00011517 | protein_coding_gene | -1.4 | 0.39 | 6.9E-04 |
| pef-1 | WBGene00003969 | protein_coding_gene | -1.4 | 0.39 | 8.2E-06 |
| T27A10.6 | WBGene00020846 | protein_coding_gene | -1.4 | 0.39 | 5.9E-03 |
| C05A9.2 | WBGene00007317 | protein_coding_gene | -1.4 | 0.39 | 1.4E-08 |
| C06E4.2 | WBGene00015531 | protein_coding_gene | -1.4 | 0.39 | 1.1E-02 |
| C05G5.5 | WBGene00007351 | protein_coding_gene | -1.4 | 0.39 | 1.9E-02 |
| nhr-103 | WBGene00003693 | protein_coding_gene | -1.4 | 0.39 | 4.4E-06 |
| nhr-25 | WBGene00003623 | protein_coding_gene | -1.4 | 0.39 | 7.8E-09 |
| R10E8.6 | WBGene00011212 | protein_coding_gene | -1.4 | 0.39 | 2.0E-11 |
| F26F2.8 | WBGene00023504 | protein_coding_gene | -1.4 | 0.39 | 2.8E-02 |
| T04A6.1 | WBGene00020200 | protein_coding_gene | -1.4 | 0.39 | 2.3E-12 |
| tmed-13 | WBGene00022255 | protein_coding_gene | -1.4 | 0.39 | 3.9E-03 |
| Y47G6A.5 | WBGene00021635 | protein_coding_gene | -1.4 | 0.39 | 3.0E-16 |
| F53H2.1 | WBGene00010004 | protein_coding_gene | -1.4 | 0.39 | 4.1E-02 |
| asp-12 | WBGene00017678 | protein_coding_gene | -1.4 | 0.39 | 1.1E-05 |
| chil-22 | WBGene00011166 | protein_coding_gene | -1.4 | 0.39 | 3.1E-06 |
| F08G2.5 | WBGene00008577 | protein_coding_gene | -1.4 | 0.39 | 9.5E-03 |
| C26F1.1 | WBGene00016146 | protein_coding_gene | -1.4 | 0.39 | 2.8E-04 |
| let-653 | WBGene00002827 | protein_coding_gene | -1.4 | 0.39 | 5.3E-03 |
| C34D1.4 | WBGene00007931 | protein_coding_gene | -1.4 | 0.39 | 4.3E-06 |
| C14C6.5 | WBGene00015759 | protein_coding_gene | -1.4 | 0.39 | 2.7E-03 |
| dyf-19 | WBGene00012826 | protein_coding_gene | -1.4 | 0.39 | 4.7E-02 |
| B0365.9 | WBGene00219208 | protein_coding_gene | -1.4 | 0.38 | 1.4E-02 |
| F47E1.4 | WBGene00018568 | protein_coding_gene | -1.4 | 0.38 | 1.7E-05 |
| T19A6.4 | WBGene00011829 | protein_coding_gene | -1.4 | 0.38 | 2.6E-03 |
| Y43F8A.1 | WBGene00012808 | protein_coding_gene | -1.4 | 0.38 | 3.8E-02 |
| hst-3.2 | WBGene00044303 | protein_coding_gene | -1.4 | 0.38 | 3.6E-04 |
| eya-1 | WBGene00001377 | protein_coding_gene | -1.4 | 0.38 | 5.0E-09 |
| lron-2 | WBGene00022789 | protein_coding_gene | -1.4 | 0.38 | 1.6E-02 |
| F56C3.9 | WBGene00018946 | protein_coding_gene | -1.4 | 0.38 | 3.1E-02 |
| osm-5 | WBGene00003885 | protein_coding_gene | -1.4 | 0.38 | 1.9E-02 |
| Y71H2B.8 | WBGene00022197 | protein_coding_gene | -1.4 | 0.38 | 2.8E-04 |
| dlg-1 | WBGene00001006 | protein_coding_gene | -1.4 | 0.38 | 8.3E-25 |
| tag-196 | WBGene00007055 | protein_coding_gene | -1.4 | 0.38 | 2.8E-22 |
| Y94H6A.2 | WBGene00022375 | protein_coding_gene | -1.4 | 0.38 | 3.6E-02 |
| slc-36.3 | WBGene00012629 | protein_coding_gene | -1.4 | 0.38 | 8.6E-04 |
| Y6E2A.5 | WBGene00012399 | protein_coding_gene | -1.4 | 0.38 | 8.6E-04 |
| F12E12.11 | WBGene00044811 | protein_coding_gene | -1.4 | 0.38 | 1.3E-03 |
| C49H3.12 | WBGene00016795 | protein_coding_gene | -1.4 | 0.38 | 1.0E-02 |
| ceh-31 | WBGene00000452 | protein_coding_gene | -1.4 | 0.38 | 2.8E-02 |
| kel-1 | WBGene00002184 | protein_coding_gene | -1.4 | 0.38 | 4.4E-04 |
| C06G3.6 | WBGene00015552 | protein_coding_gene | -1.4 | 0.38 | 3.8E-15 |
| Y65A5A.1 | WBGene00013418 | protein_coding_gene | -1.4 | 0.38 | 3.1E-08 |
| fbxb-66 | WBGene00044066 | protein_coding_gene | -1.4 | 0.38 | 2.1E-05 |
| ceh-32 | WBGene00000453 | protein_coding_gene | -1.4 | 0.38 | 6.8E-12 |
| C54D10.13 | WBGene00045400 | protein_coding_gene | -1.4 | 0.38 | 7.9E-04 |
| ZK970.7 | WBGene00014173 | protein_coding_gene | -1.4 | 0.38 | 2.7E-03 |
| F23F12.8 | WBGene00017754 | protein_coding_gene | -1.4 | 0.38 | 4.6E-02 |
| R05H10.1 | WBGene00011042 | protein_coding_gene | -1.4 | 0.38 | 3.0E-09 |
| dlc-5 | WBGene00043067 | protein_coding_gene | -1.4 | 0.38 | 1.8E-02 |
| nhr-56 | WBGene00003646 | protein_coding_gene | -1.4 | 0.38 | 1.5E-05 |
| C13C4.7 | WBGene00007551 | protein_coding_gene | -1.4 | 0.38 | 1.4E-04 |
| ncam-1 | WBGene00017184 | protein_coding_gene | -1.4 | 0.38 | 1.8E-12 |
| ceh-6 | WBGene00000431 | protein_coding_gene | -1.4 | 0.38 | 6.8E-04 |
| pha-4 | WBGene00004013 | protein_coding_gene | -1.4 | 0.38 | 7.2E-17 |
| gba-1 | WBGene00016335 | protein_coding_gene | -1.4 | 0.38 | 9.3E-09 |
| F59B10.4 | WBGene00010320 | protein_coding_gene | -1.4 | 0.38 | 1.2E-05 |
| B0462.5 | WBGene00194695 | protein_coding_gene | -1.4 | 0.38 | 2.9E-04 |

| | | | | | |
|---|---|---|---|---|---|
| sqst-1 | WBGene00011737 | protein_coding_gene | -1.4 | 0.38 | 3.0E-37 |
| nhr-246 | WBGene00014192 | protein_coding_gene | -1.4 | 0.37 | 3.6E-02 |
| sta-2 | WBGene00010251 | protein_coding_gene | -1.4 | 0.37 | 1.2E-09 |
| unc-130 | WBGene00006853 | protein_coding_gene | -1.4 | 0.37 | 1.5E-06 |
| lipl-6 | WBGene00021963 | protein_coding_gene | -1.4 | 0.37 | 1.6E-02 |
| nhr-134 | WBGene00003724 | protein_coding_gene | -1.4 | 0.37 | 1.0E-08 |
| T01D1.8 | WBGene00044641 | protein_coding_gene | -1.4 | 0.37 | 3.3E-02 |
| pax-3 | WBGene00003939 | protein_coding_gene | -1.4 | 0.37 | 3.5E-04 |
| sodh-2 | WBGene00010791 | protein_coding_gene | -1.4 | 0.37 | 3.4E-02 |
| zmp-2 | WBGene00019212 | protein_coding_gene | -1.4 | 0.37 | 7.9E-04 |
| tag-343 | WBGene00018374 | protein_coding_gene | -1.4 | 0.37 | 4.4E-11 |
| sulp-3 | WBGene00018283 | protein_coding_gene | -1.4 | 0.37 | 3.2E-03 |
| ist-1 | WBGene00002163 | protein_coding_gene | -1.4 | 0.37 | 7.5E-10 |
| F57C12.2 | WBGene00019014 | protein_coding_gene | -1.4 | 0.37 | 5.8E-05 |
| R11E3.19 | WBGene00235374 | protein_coding_gene | -1.4 | 0.37 | 4.9E-02 |
| let-522 | WBGene00010870 | protein_coding_gene | -1.4 | 0.37 | 1.7E-10 |
| shn-1 | WBGene00006444 | protein_coding_gene | -1.4 | 0.37 | 1.3E-13 |
| aagr-1 | WBGene00017071 | protein_coding_gene | -1.4 | 0.37 | 2.8E-15 |
| F59F5.7 | WBGene00010345 | protein_coding_gene | -1.4 | 0.37 | 1.0E-02 |
| nlp-34 | WBGene00015046 | protein_coding_gene | -1.4 | 0.37 | 1.6E-03 |
| pha-2 | WBGene00004011 | protein_coding_gene | -1.4 | 0.37 | 1.9E-02 |
| gasr-8 | WBGene00008144 | protein_coding_gene | -1.4 | 0.37 | 3.0E-02 |
| nhr-152 | WBGene00015705 | protein_coding_gene | -1.5 | 0.37 | 6.7E-05 |
| lys-10 | WBGene00003099 | protein_coding_gene | -1.5 | 0.37 | 2.5E-03 |
| C53C11.5 | WBGene00044770 | protein_coding_gene | -1.5 | 0.37 | 2.8E-10 |
| vab-23 | WBGene00014153 | protein_coding_gene | -1.5 | 0.37 | 6.9E-06 |
| Y41E3.18 | WBGene00044003 | protein_coding_gene | -1.5 | 0.37 | 1.4E-02 |
| cest-17 | WBGene00019477 | protein_coding_gene | -1.5 | 0.36 | 2.2E-10 |
| math-42 | WBGene00020532 | protein_coding_gene | -1.5 | 0.36 | 1.4E-02 |
| C08G9.2 | WBGene00015620 | protein_coding_gene | -1.5 | 0.36 | 3.0E-03 |
| T25B9.3 | WBGene00012009 | protein_coding_gene | -1.5 | 0.36 | 2.4E-02 |
| W05H9.3 | WBGene00021050 | ncRNA_gene | -1.5 | 0.36 | 4.1E-16 |
| spi-1 | WBGene00023416 | protein_coding_gene | -1.5 | 0.36 | 8.4E-03 |
| marc-4 | WBGene00016903 | protein_coding_gene | -1.5 | 0.36 | 9.0E-05 |
| spp-1 | WBGene00004986 | protein_coding_gene | -1.5 | 0.36 | 1.9E-06 |
| che-12 | WBGene00000491 | protein_coding_gene | -1.5 | 0.36 | 7.1E-04 |
| C27B7.7 | WBGene00007764 | protein_coding_gene | -1.5 | 0.36 | 2.6E-10 |
| C34E11.4 | WBGene00007937 | protein_coding_gene | -1.5 | 0.36 | 2.6E-03 |
| F48E3.6 | WBGene00018606 | protein_coding_gene | -1.5 | 0.36 | 5.7E-10 |
| ZK669.3 | WBGene00014053 | protein_coding_gene | -1.5 | 0.36 | 3.0E-21 |
| R07G3.8 | WBGene00019944 | protein_coding_gene | -1.5 | 0.36 | 9.7E-10 |
| ceh-2 | WBGene00000429 | protein_coding_gene | -1.5 | 0.36 | 4.5E-02 |
| F34H10.3 | WBGene00009380 | protein_coding_gene | -1.5 | 0.36 | 1.8E-19 |
| C06G3.5 | WBGene00015551 | protein_coding_gene | -1.5 | 0.36 | 4.3E-16 |
| clc-2 | WBGene00000523 | protein_coding_gene | -1.5 | 0.36 | 8.7E-05 |
| phlp-2 | WBGene00009647 | protein_coding_gene | -1.5 | 0.36 | 9.9E-03 |
| F18C5.5 | WBGene00017560 | protein_coding_gene | -1.5 | 0.36 | 4.9E-02 |
| clec-172 | WBGene00044719 | protein_coding_gene | -1.5 | 0.36 | 5.2E-04 |
| vab-7 | WBGene00006873 | protein_coding_gene | -1.5 | 0.36 | 4.3E-03 |
| mod-5 | WBGene00003387 | protein_coding_gene | -1.5 | 0.35 | 1.7E-04 |
| fbxa-146 | WBGene00018807 | pseudogene | -1.5 | 0.35 | 1.8E-06 |
| tpra-1 | WBGene00008366 | protein_coding_gene | -1.5 | 0.35 | 7.9E-19 |
| fbxa-172 | WBGene00008014 | protein_coding_gene | -1.5 | 0.35 | 9.3E-04 |
| C45B11.6 | WBGene00045379 | protein_coding_gene | -1.5 | 0.35 | 1.1E-03 |
| M60.4 | WBGene00019780 | protein_coding_gene | -1.5 | 0.35 | 1.6E-24 |
| ugt-66 | WBGene00016013 | protein_coding_gene | -1.5 | 0.35 | 1.6E-06 |
| Y51B9A.9 | WBGene00013091 | protein_coding_gene | -1.5 | 0.35 | 1.8E-06 |
| C10F3.7 | WBGene00015679 | protein_coding_gene | -1.5 | 0.35 | 4.1E-02 |

| | | | | | |
|---|---|---|---|---|---|
| bus-17 | WBGene00044630 | protein_coding_gene | -1.5 | 0.35 | 8.6E-03 |
| K08F9.1 | WBGene00010684 | protein_coding_gene | -1.5 | 0.35 | 6.4E-04 |
| Y32F6B.1 | WBGene00012531 | protein_coding_gene | -1.5 | 0.35 | 2.5E-22 |
| C39B5.14 | WBGene00189950 | protein_coding_gene | -1.5 | 0.35 | 3.7E-02 |
| C36E8.4 | WBGene00007983 | protein_coding_gene | -1.5 | 0.35 | 1.6E-08 |
| srg-34 | WBGene00005191 | protein_coding_gene | -1.5 | 0.35 | 8.5E-03 |
| Y87G2A.16 | WBGene00013602 | protein_coding_gene | -1.5 | 0.35 | 2.1E-02 |
| Y44E3A.1 | WBGene00021547 | protein_coding_gene | -1.5 | 0.35 | 4.1E-06 |
| sulp-2 | WBGene00017464 | protein_coding_gene | -1.5 | 0.35 | 1.0E-06 |
| T02G5.14 | WBGene00020170 | protein_coding_gene | -1.5 | 0.35 | 5.6E-04 |
| ZC455.1 | WBGene00013902 | protein_coding_gene | -1.5 | 0.35 | 1.3E-09 |
| sel-7 | WBGene00004764 | protein_coding_gene | -1.5 | 0.35 | 6.3E-03 |
| W05B10.4 | WBGene00012279 | protein_coding_gene | -1.5 | 0.35 | 3.3E-04 |
| xtr-1 | WBGene00006965 | protein_coding_gene | -1.5 | 0.35 | 5.9E-03 |
| F52E1.5 | WBGene00018694 | protein_coding_gene | -1.5 | 0.35 | 1.2E-05 |
| ttr-50 | WBGene00010234 | protein_coding_gene | -1.5 | 0.35 | 6.7E-27 |
| cyp-13A1 | WBGene00011677 | protein_coding_gene | -1.5 | 0.34 | 3.7E-02 |
| ceh-37 | WBGene00000458 | protein_coding_gene | -1.5 | 0.34 | 1.8E-11 |
| ceh-10 | WBGene00000435 | protein_coding_gene | -1.5 | 0.34 | 9.7E-03 |
| nhr-232 | WBGene00012494 | protein_coding_gene | -1.5 | 0.34 | 1.6E-09 |
| nlp-30 | WBGene00003768 | protein_coding_gene | -1.5 | 0.34 | 1.6E-05 |
| Y42G9A.1 | WBGene00021531 | protein_coding_gene | -1.5 | 0.34 | 1.4E-07 |
| T02B11.6 | WBGene00020157 | protein_coding_gene | -1.5 | 0.34 | 1.3E-02 |
| ptr-19 | WBGene00004233 | protein_coding_gene | -1.5 | 0.34 | 3.1E-02 |
| mltn-12 | WBGene00008277 | protein_coding_gene | -1.5 | 0.34 | 4.6E-02 |
| R02D5.3 | WBGene00010977 | protein_coding_gene | -1.5 | 0.34 | 9.8E-08 |
| magu-3 | WBGene00016841 | protein_coding_gene | -1.5 | 0.34 | 1.8E-04 |
| lgc-1 | WBGene00020236 | protein_coding_gene | -1.5 | 0.34 | 4.1E-02 |
| C54F6.12 | WBGene00016929 | protein_coding_gene | -1.5 | 0.34 | 7.9E-03 |
| plx-2 | WBGene00004048 | protein_coding_gene | -1.5 | 0.34 | 3.9E-10 |
| acs-3 | WBGene00020343 | protein_coding_gene | -1.6 | 0.34 | 3.8E-10 |
| cutl-11 | WBGene00011443 | protein_coding_gene | -1.6 | 0.34 | 2.2E-02 |
| nmad-1 | WBGene00017304 | protein_coding_gene | -1.6 | 0.34 | 5.9E-16 |
| EEED8.15 | WBGene00017143 | protein_coding_gene | -1.6 | 0.34 | 6.7E-19 |
| vab-3 | WBGene00006870 | protein_coding_gene | -1.6 | 0.34 | 2.3E-10 |
| clec-40 | WBGene00011853 | protein_coding_gene | -1.6 | 0.34 | 2.0E-02 |
| F20D6.10 | WBGene00017639 | protein_coding_gene | -1.6 | 0.34 | 6.0E-06 |
| vab-15 | WBGene00006881 | protein_coding_gene | -1.6 | 0.34 | 8.1E-04 |
| F21H7.2 | WBGene00009028 | protein_coding_gene | -1.6 | 0.34 | 1.2E-03 |
| ZC196.3 | WBGene00022547 | protein_coding_gene | -1.6 | 0.34 | 3.3E-02 |
| M01H9.4 | WBGene00019720 | protein_coding_gene | -1.6 | 0.34 | 1.5E-08 |
| C06E7.4 | WBGene00015541 | protein_coding_gene | -1.6 | 0.34 | 2.8E-03 |
| C35A5.6 | WBGene00007950 | protein_coding_gene | -1.6 | 0.34 | 3.5E-11 |
| F56D3.1 | WBGene00018969 | protein_coding_gene | -1.6 | 0.34 | 8.4E-03 |
| tsp-6 | WBGene00006632 | protein_coding_gene | -1.6 | 0.34 | 5.9E-03 |
| F58E6.11 | WBGene00010258 | protein_coding_gene | -1.6 | 0.34 | 1.0E-02 |
| C33D9.6 | WBGene00007900 | protein_coding_gene | -1.6 | 0.34 | 7.8E-11 |
| tbx-7 | WBGene00006544 | protein_coding_gene | -1.6 | 0.34 | 9.7E-03 |
| twk-7 | WBGene00006662 | protein_coding_gene | -1.6 | 0.34 | 1.3E-09 |
| skr-19 | WBGene00004825 | protein_coding_gene | -1.6 | 0.34 | 1.1E-18 |
| M03F4.6 | WBGene00019759 | protein_coding_gene | -1.6 | 0.34 | 1.0E-03 |
| F36D1.7 | WBGene00009466 | protein_coding_gene | -1.6 | 0.34 | 1.2E-06 |
| R02C2.7 | WBGene00019818 | protein_coding_gene | -1.6 | 0.33 | 6.3E-18 |
| pag-3 | WBGene00003909 | protein_coding_gene | -1.6 | 0.33 | 1.3E-02 |
| lbp-7 | WBGene00002259 | protein_coding_gene | -1.6 | 0.33 | 1.5E-16 |
| C26B9.6 | WBGene00016135 | protein_coding_gene | -1.6 | 0.33 | 1.7E-08 |
| pals-33 | WBGene00021081 | protein_coding_gene | -1.6 | 0.33 | 4.9E-03 |
| ccep-290 | WBGene00021643 | protein_coding_gene | -1.6 | 0.33 | 2.9E-04 |

| | | | | | |
|---|---|---|---|---|---|
| Y75B8A.28 | WBGene00013559 | protein_coding_gene | -1.6 | 0.33 | 1.7E-07 |
| C18D11.6 | WBGene00007682 | protein_coding_gene | -1.6 | 0.33 | 1.1E-02 |
| F54B8.4 | WBGene00010019 | protein_coding_gene | -1.6 | 0.33 | 9.1E-07 |
| dpy-1 | WBGene00001063 | protein_coding_gene | -1.6 | 0.33 | 3.4E-03 |
| C36A4.11 | WBGene00044167 | protein_coding_gene | -1.6 | 0.33 | 1.5E-03 |
| F17C8.3 | WBGene00008910 | protein_coding_gene | -1.6 | 0.33 | 1.8E-06 |
| C37E2.2 | WBGene00007994 | protein_coding_gene | -1.6 | 0.33 | 1.3E-02 |
| F36H5.4 | WBGene00018103 | protein_coding_gene | -1.6 | 0.33 | 3.8E-03 |
| C16H3.3 | WBGene00015868 | protein_coding_gene | -1.6 | 0.33 | 1.7E-17 |
| unc-129 | WBGene00006852 | protein_coding_gene | -1.6 | 0.33 | 6.4E-09 |
| F13C5.1 | WBGene00017422 | protein_coding_gene | -1.6 | 0.33 | 3.8E-09 |
| pals-38 | WBGene00008302 | protein_coding_gene | -1.6 | 0.33 | 3.9E-02 |
| ins-7 | WBGene00002090 | protein_coding_gene | -1.6 | 0.33 | 4.5E-05 |
| ceh-22 | WBGene00000445 | protein_coding_gene | -1.6 | 0.33 | 1.0E-03 |
| ZK822.1 | WBGene00014089 | protein_coding_gene | -1.6 | 0.33 | 1.6E-05 |
| cex-2 | WBGene00023408 | protein_coding_gene | -1.6 | 0.33 | 8.3E-15 |
| mam-3 | WBGene00010714 | protein_coding_gene | -1.6 | 0.33 | 1.2E-02 |
| Y71H2AM.15 | WBGene00022180 | protein_coding_gene | -1.6 | 0.33 | 1.5E-05 |
| aff-1 | WBGene00016625 | protein_coding_gene | -1.6 | 0.33 | 1.2E-03 |
| zag-1 | WBGene00006970 | protein_coding_gene | -1.6 | 0.32 | 3.9E-09 |
| F46H6.5 | WBGene00023460 | protein_coding_gene | -1.6 | 0.32 | 2.0E-02 |
| Y17G9B.2 | WBGene00021199 | protein_coding_gene | -1.6 | 0.32 | 2.9E-03 |
| D2096.6 | WBGene00017073 | protein_coding_gene | -1.6 | 0.32 | 2.1E-02 |
| C07D10.5 | WBGene00015569 | protein_coding_gene | -1.6 | 0.32 | 5.7E-20 |
| fbxa-135 | WBGene00011539 | protein_coding_gene | -1.6 | 0.32 | 1.5E-02 |
| del-2 | WBGene00000953 | protein_coding_gene | -1.6 | 0.32 | 1.3E-02 |
| cht-4 | WBGene00021252 | protein_coding_gene | -1.6 | 0.32 | 6.9E-03 |
| T16A1.10 | WBGene00269431 | protein_coding_gene | -1.6 | 0.32 | 3.5E-02 |
| nhr-150 | WBGene00007367 | protein_coding_gene | -1.6 | 0.32 | 7.4E-03 |
| F41C3.2 | WBGene00018268 | protein_coding_gene | -1.6 | 0.32 | 2.2E-09 |
| F43G6.17 | WBGene00219378 | protein_coding_gene | -1.6 | 0.32 | 4.3E-03 |
| Y69H2.10 | WBGene00013485 | protein_coding_gene | -1.7 | 0.32 | 7.8E-03 |
| ddr-2 | WBGene00017381 | protein_coding_gene | -1.7 | 0.32 | 4.0E-11 |
| fbxa-136 | WBGene00007672 | protein_coding_gene | -1.7 | 0.32 | 4.3E-03 |
| lin-32 | WBGene00003018 | protein_coding_gene | -1.7 | 0.32 | 4.5E-04 |
| dyf-11 | WBGene00001127 | protein_coding_gene | -1.7 | 0.32 | 2.5E-03 |
| spp-22 | WBGene00044283 | protein_coding_gene | -1.7 | 0.32 | 1.8E-06 |
| Y6D1A.2 | WBGene00012396 | protein_coding_gene | -1.7 | 0.32 | 1.7E-03 |
| Y54G2A.10 | WBGene00021875 | protein_coding_gene | -1.7 | 0.32 | 2.9E-05 |
| ZK512.1 | WBGene00013982 | protein_coding_gene | -1.7 | 0.32 | 1.7E-07 |
| cnd-1 | WBGene00000561 | protein_coding_gene | -1.7 | 0.32 | 3.5E-05 |
| ZC239.14 | WBGene00022572 | protein_coding_gene | -1.7 | 0.31 | 1.5E-02 |
| T08G5.3 | WBGene00011624 | protein_coding_gene | -1.7 | 0.31 | 3.7E-02 |
| F43H9.4 | WBGene00018400 | protein_coding_gene | -1.7 | 0.31 | 4.6E-17 |
| Y37F4.8 | WBGene00044289 | protein_coding_gene | -1.7 | 0.31 | 1.0E-03 |
| Y46D2A.1 | WBGene00021589 | protein_coding_gene | -1.7 | 0.31 | 2.4E-02 |
| cyp-13A4 | WBGene00011671 | protein_coding_gene | -1.7 | 0.31 | 4.2E-04 |
| lips-3 | WBGene00008655 | protein_coding_gene | -1.7 | 0.31 | 1.5E-02 |
| B0563.5 | WBGene00015262 | protein_coding_gene | -1.7 | 0.31 | 2.6E-03 |
| C14C6.2 | WBGene00015756 | protein_coding_gene | -1.7 | 0.31 | 1.3E-08 |
| ceh-34 | WBGene00000455 | protein_coding_gene | -1.7 | 0.31 | 4.9E-07 |
| hsp-12.6 | WBGene00002013 | protein_coding_gene | -1.7 | 0.31 | 4.4E-05 |
| F47B8.3 | WBGene00009804 | protein_coding_gene | -1.7 | 0.31 | 6.1E-10 |
| Y51H7C.10 | WBGene00021788 | protein_coding_gene | -1.7 | 0.31 | 1.5E-02 |
| egl-38 | WBGene00001204 | protein_coding_gene | -1.7 | 0.31 | 4.1E-03 |
| fat-3 | WBGene00001395 | protein_coding_gene | -1.7 | 0.31 | 5.5E-08 |
| efl-3 | WBGene00009899 | protein_coding_gene | -1.7 | 0.31 | 8.5E-14 |
| abhd-5.1 | WBGene00016506 | protein_coding_gene | -1.7 | 0.31 | 5.3E-07 |

| | | | | | |
|---|---|---|---|---|---|
| F22B7.1 | WBGene00017692 | protein_coding_gene | -1.7 | 0.31 | 1.7E-06 |
| Y2H9A.6 | WBGene00086568 | protein_coding_gene | -1.7 | 0.31 | 3.8E-03 |
| fbxa-137 | WBGene00020853 | protein_coding_gene | -1.7 | 0.31 | 4.1E-02 |
| R11G1.2 | WBGene00020011 | protein_coding_gene | -1.7 | 0.31 | 2.3E-04 |
| lam-3 | WBGene00002248 | protein_coding_gene | -1.7 | 0.31 | 1.5E-19 |
| Y54G2A.36 | WBGene00021898 | protein_coding_gene | -1.7 | 0.31 | 1.1E-20 |
| mec-6 | WBGene00003170 | protein_coding_gene | -1.7 | 0.31 | 7.5E-04 |
| npr-20 | WBGene00011578 | protein_coding_gene | -1.7 | 0.31 | 4.4E-06 |
| Y71G12B.31 | WBGene00044348 | protein_coding_gene | -1.7 | 0.31 | 1.4E-03 |
| oac-51 | WBGene00020976 | protein_coding_gene | -1.7 | 0.31 | 2.2E-04 |
| F13B12.4 | WBGene00008732 | protein_coding_gene | -1.7 | 0.31 | 1.8E-02 |
| lec-2 | WBGene00002265 | protein_coding_gene | -1.7 | 0.31 | 2.7E-08 |
| nhr-44 | WBGene00003634 | protein_coding_gene | -1.7 | 0.31 | 3.1E-06 |
| bkip-1 | WBGene00014569 | protein_coding_gene | -1.7 | 0.31 | 6.0E-03 |
| Y50E8A.5 | WBGene00013048 | protein_coding_gene | -1.7 | 0.31 | 1.2E-06 |
| C09F5.1 | WBGene00015647 | protein_coding_gene | -1.7 | 0.31 | 3.7E-12 |
| pals-31 | WBGene00018614 | protein_coding_gene | -1.7 | 0.31 | 3.1E-02 |
| clec-69 | WBGene00044642 | pseudogene | -1.7 | 0.30 | 3.9E-02 |
| T18D3.7 | WBGene00011824 | protein_coding_gene | -1.7 | 0.30 | 7.1E-16 |
| F59B2.9 | WBGene00010311 | protein_coding_gene | -1.7 | 0.30 | 6.7E-04 |
| ltd-1 | WBGene00003089 | protein_coding_gene | -1.7 | 0.30 | 7.5E-09 |
| eva-1 | WBGene00009304 | protein_coding_gene | -1.7 | 0.30 | 1.4E-10 |
| T01D3.3 | WBGene00011328 | protein_coding_gene | -1.7 | 0.30 | 1.2E-06 |
| phg-1 | WBGene00004017 | protein_coding_gene | -1.7 | 0.30 | 1.4E-02 |
| fbxa-175 | WBGene00045293 | pseudogene | -1.7 | 0.30 | 7.7E-06 |
| fbxb-88 | WBGene00016359 | protein_coding_gene | -1.7 | 0.30 | 9.7E-09 |
| ZK666.15 | WBGene00195145 | protein_coding_gene | -1.7 | 0.30 | 4.8E-02 |
| Y58A7A.2 | WBGene00021976 | protein_coding_gene | -1.7 | 0.30 | 3.1E-02 |
| F41G4.8 | WBGene00018311 | protein_coding_gene | -1.7 | 0.30 | 3.7E-08 |
| fbxb-77 | WBGene00021204 | protein_coding_gene | -1.7 | 0.30 | 1.6E-02 |
| hhat-2 | WBGene00013312 | protein_coding_gene | -1.7 | 0.30 | 9.3E-05 |
| fbxc-58 | WBGene00007506 | protein_coding_gene | -1.7 | 0.30 | 7.3E-04 |
| E03A3.10 | WBGene00189944 | pseudogene | -1.7 | 0.30 | 4.2E-03 |
| F57F5.3 | WBGene00010205 | protein_coding_gene | -1.7 | 0.30 | 4.5E-02 |
| igdb-3 | WBGene00020691 | protein_coding_gene | -1.7 | 0.30 | 7.2E-04 |
| T08A9.13 | WBGene00044442 | protein_coding_gene | -1.7 | 0.30 | 8.8E-03 |
| F55B12.10 | WBGene00077439 | protein_coding_gene | -1.7 | 0.30 | 2.6E-02 |
| cbp-3 | WBGene00009595 | protein_coding_gene | -1.7 | 0.30 | 3.8E-16 |
| prkl-1 | WBGene00022727 | protein_coding_gene | -1.7 | 0.30 | 7.0E-11 |
| nhr-42 | WBGene00003632 | protein_coding_gene | -1.7 | 0.30 | 2.8E-15 |
| gstk-2 | WBGene00017054 | protein_coding_gene | -1.7 | 0.30 | 7.2E-07 |
| C07A4.3 | WBGene00007398 | protein_coding_gene | -1.7 | 0.30 | 1.3E-04 |
| fbxb-61 | WBGene00010106 | protein_coding_gene | -1.7 | 0.30 | 2.3E-02 |
| Y53F4B.20 | WBGene00013166 | protein_coding_gene | -1.8 | 0.30 | 1.1E-02 |
| ZK180.7 | WBGene00045248 | protein_coding_gene | -1.8 | 0.30 | 4.4E-02 |
| F58H1.2 | WBGene00010285 | protein_coding_gene | -1.8 | 0.30 | 3.9E-02 |
| C35C5.9 | WBGene00007958 | protein_coding_gene | -1.8 | 0.30 | 3.6E-13 |
| B0507.8 | WBGene00015225 | protein_coding_gene | -1.8 | 0.30 | 3.4E-02 |
| nhr-67 | WBGene00003657 | protein_coding_gene | -1.8 | 0.30 | 1.7E-04 |
| faah-2 | WBGene00015048 | protein_coding_gene | -1.8 | 0.29 | 1.2E-07 |
| lron-1 | WBGene00008093 | protein_coding_gene | -1.8 | 0.29 | 8.7E-03 |
| T23F11.6 | WBGene00011957 | protein_coding_gene | -1.8 | 0.29 | 1.4E-03 |
| Y105E8A.34 | WBGene00077689 | pseudogene | -1.8 | 0.29 | 3.4E-02 |
| nhr-180 | WBGene00017961 | protein_coding_gene | -1.8 | 0.29 | 1.2E-06 |
| lido-15 | WBGene00018804 | protein_coding_gene | -1.8 | 0.29 | 3.4E-07 |
| C44C1.6 | WBGene00016645 | protein_coding_gene | -1.8 | 0.29 | 4.5E-08 |
| T09A12.8 | WBGene00235275 | protein_coding_gene | -1.8 | 0.29 | 2.1E-02 |
| F42C5.4 | WBGene00018346 | protein_coding_gene | -1.8 | 0.29 | 4.4E-11 |

| | | | | | |
|---|---|---|---|---|---|
| C08D8.1 | WBGene00015592 | protein_coding_gene | -1.8 | 0.29 | 4.7E-05 |
| osm-6 | WBGene00003886 | protein_coding_gene | -1.8 | 0.29 | 3.2E-03 |
| F44A6.5 | WBGene00009677 | protein_coding_gene | -1.8 | 0.29 | 2.1E-14 |
| T03E6.8 | WBGene00011397 | protein_coding_gene | -1.8 | 0.29 | 4.2E-03 |
| C33A12.4 | WBGene00007883 | protein_coding_gene | -1.8 | 0.29 | 8.9E-10 |
| F08A7.1 | WBGene00045013 | protein_coding_gene | -1.8 | 0.29 | 1.1E-08 |
| clec-180 | WBGene00017991 | protein_coding_gene | -1.8 | 0.29 | 4.9E-02 |
| F22F7.3 | WBGene00017721 | protein_coding_gene | -1.8 | 0.29 | 9.1E-03 |
| cyp-23A1 | WBGene00015135 | protein_coding_gene | -1.8 | 0.29 | 1.5E-02 |
| lgc-27 | WBGene00022283 | protein_coding_gene | -1.8 | 0.29 | 6.6E-05 |
| F25E2.3 | WBGene00017781 | protein_coding_gene | -1.8 | 0.29 | 3.9E-09 |
| hsp-12.1 | WBGene00011906 | protein_coding_gene | -1.8 | 0.29 | 1.6E-05 |
| dmd-4 | WBGene00007776 | protein_coding_gene | -1.8 | 0.29 | 1.5E-02 |
| F52D1.2 | WBGene00018683 | protein_coding_gene | -1.8 | 0.29 | 5.1E-06 |
| F41C3.1 | WBGene00018267 | protein_coding_gene | -1.8 | 0.29 | 1.8E-02 |
| nphp-4 | WBGene00011261 | protein_coding_gene | -1.8 | 0.29 | 1.4E-04 |
| F23H12.5 | WBGene00009094 | protein_coding_gene | -1.8 | 0.29 | 1.1E-02 |
| F13D11.4 | WBGene00017429 | protein_coding_gene | -1.8 | 0.29 | 3.9E-15 |
| W03F8.6 | WBGene00020995 | protein_coding_gene | -1.8 | 0.29 | 4.8E-08 |
| fkh-8 | WBGene00001440 | protein_coding_gene | -1.8 | 0.29 | 3.4E-04 |
| fkb-4 | WBGene00001429 | protein_coding_gene | -1.8 | 0.29 | 4.3E-05 |
| W05B2.2 | WBGene00012271 | protein_coding_gene | -1.8 | 0.29 | 2.4E-02 |
| dma-1 | WBGene00011345 | protein_coding_gene | -1.8 | 0.29 | 4.0E-11 |
| F53F10.1 | WBGene00018763 | protein_coding_gene | -1.8 | 0.29 | 1.6E-10 |
| F53F1.6 | WBGene00009984 | protein_coding_gene | -1.8 | 0.28 | 7.9E-12 |
| glo-3 | WBGene00010341 | protein_coding_gene | -1.8 | 0.28 | 1.4E-08 |
| Y110A2AL.3 | WBGene00022440 | protein_coding_gene | -1.8 | 0.28 | 2.7E-02 |
| syd-9 | WBGene00044068 | protein_coding_gene | -1.8 | 0.28 | 7.9E-15 |
| C26B9.3 | WBGene00016133 | protein_coding_gene | -1.8 | 0.28 | 9.9E-03 |
| F37A4.3 | WBGene00018133 | protein_coding_gene | -1.8 | 0.28 | 2.3E-02 |
| C38C3.4 | WBGene00016513 | protein_coding_gene | -1.8 | 0.28 | 2.8E-14 |
| F20D6.8 | WBGene00017637 | protein_coding_gene | -1.8 | 0.28 | 1.6E-03 |
| K01A2.3 | WBGene00019278 | pseudogene | -1.8 | 0.28 | 7.3E-06 |
| lpr-7 | WBGene00020571 | protein_coding_gene | -1.8 | 0.28 | 6.2E-03 |
| Y82E9BR.1 | WBGene00022334 | protein_coding_gene | -1.8 | 0.28 | 2.5E-21 |
| F58E6.13 | WBGene00077697 | protein_coding_gene | -1.8 | 0.28 | 2.0E-02 |
| klp-11 | WBGene00002222 | protein_coding_gene | -1.8 | 0.28 | 2.1E-09 |
| C36B1.13 | WBGene00044120 | protein_coding_gene | -1.8 | 0.28 | 4.8E-03 |
| maph-9 | WBGene00016397 | protein_coding_gene | -1.8 | 0.28 | 6.3E-03 |
| swip-10 | WBGene00018738 | protein_coding_gene | -1.8 | 0.28 | 4.3E-04 |
| pals-26 | WBGene00007131 | protein_coding_gene | -1.8 | 0.28 | 2.6E-03 |
| ZK353.2 | WBGene00022698 | protein_coding_gene | -1.8 | 0.28 | 3.0E-02 |
| F57G12.2 | WBGene00010222 | protein_coding_gene | -1.8 | 0.28 | 3.7E-02 |
| C40H1.2 | WBGene00008038 | protein_coding_gene | -1.8 | 0.28 | 2.3E-06 |
| btb-15 | WBGene00015594 | pseudogene | -1.8 | 0.28 | 4.9E-02 |
| clec-100 | WBGene00013179 | protein_coding_gene | -1.9 | 0.28 | 8.3E-03 |
| cdh-5 | WBGene00000397 | protein_coding_gene | -1.9 | 0.28 | 8.9E-03 |
| C54C6.7 | WBGene00044166 | protein_coding_gene | -1.9 | 0.28 | 2.7E-04 |
| dyf-5 | WBGene00001121 | protein_coding_gene | -1.9 | 0.28 | 1.3E-03 |
| mam-1 | WBGene00022504 | protein_coding_gene | -1.9 | 0.28 | 6.3E-03 |
| C46A5.4 | WBGene00016700 | protein_coding_gene | -1.9 | 0.27 | 3.8E-04 |
| C28G1.6 | WBGene00016191 | protein_coding_gene | -1.9 | 0.27 | 3.9E-10 |
| Y54G2A.52 | WBGene00045397 | protein_coding_gene | -1.9 | 0.27 | 1.6E-16 |
| M01D1.12 | WBGene00269373 | protein_coding_gene | -1.9 | 0.27 | 9.4E-04 |
| hot-5 | WBGene00001990 | protein_coding_gene | -1.9 | 0.27 | 8.5E-03 |
| agr-1 | WBGene00018304 | protein_coding_gene | -1.9 | 0.27 | 2.1E-10 |
| bbs-9 | WBGene00016744 | protein_coding_gene | -1.9 | 0.27 | 2.1E-03 |
| F49B2.4 | WBGene00009864 | pseudogene | -1.9 | 0.27 | 3.1E-02 |

| | | | | | |
|---|---|---|---|---|---|
| clec-17 | WBGene00008477 | protein_coding_gene | -1.9 | 0.27 | 1.2E-02 |
| C54F6.17 | WBGene00235092 | protein_coding_gene | -1.9 | 0.27 | 3.6E-02 |
| cpr-2 | WBGene00000782 | protein_coding_gene | -1.9 | 0.27 | 4.4E-02 |
| unc-30 | WBGene00006766 | protein_coding_gene | -1.9 | 0.27 | 2.4E-04 |
| gst-33 | WBGene00001781 | protein_coding_gene | -1.9 | 0.27 | 2.1E-04 |
| E04A4.3 | WBGene00017118 | pseudogene | -1.9 | 0.27 | 4.2E-02 |
| ZK154.1 | WBGene00022665 | protein_coding_gene | -1.9 | 0.27 | 2.3E-02 |
| Y32B12C.5 | WBGene00194769 | pseudogene | -1.9 | 0.27 | 3.2E-05 |
| C50B6.7 | WBGene00008220 | protein_coding_gene | -1.9 | 0.27 | 1.8E-15 |
| ugt-36 | WBGene00017315 | protein_coding_gene | -1.9 | 0.27 | 2.3E-02 |
| Y58A7A.4 | WBGene00021978 | protein_coding_gene | -1.9 | 0.27 | 4.0E-02 |
| ttm-2 | WBGene00017840 | protein_coding_gene | -1.9 | 0.27 | 2.3E-10 |
| B0228.6 | WBGene00015063 | protein_coding_gene | -1.9 | 0.27 | 8.0E-12 |
| lim-4 | WBGene00002987 | protein_coding_gene | -1.9 | 0.27 | 1.8E-03 |
| abu-13 | WBGene00004133 | protein_coding_gene | -1.9 | 0.27 | 2.8E-02 |
| slc-28.1 | WBGene00017867 | protein_coding_gene | -1.9 | 0.27 | 2.1E-07 |
| cpg-7 | WBGene00010723 | protein_coding_gene | -1.9 | 0.27 | 1.0E-14 |
| Y11D7A.5 | WBGene00012429 | protein_coding_gene | -1.9 | 0.27 | 3.1E-02 |
| C18H7.11 | WBGene00044457 | protein_coding_gene | -1.9 | 0.27 | 1.4E-04 |
| tag-234 | WBGene00018874 | protein_coding_gene | -1.9 | 0.27 | 1.4E-21 |
| nsy-4 | WBGene00021415 | protein_coding_gene | -1.9 | 0.26 | 3.3E-12 |
| gad-3 | WBGene00015057 | protein_coding_gene | -1.9 | 0.26 | 1.5E-48 |
| W04E12.9 | WBGene00012254 | protein_coding_gene | -1.9 | 0.26 | 2.4E-03 |
| dyf-7 | WBGene00001123 | protein_coding_gene | -1.9 | 0.26 | 1.4E-03 |
| hlh-6 | WBGene00001952 | protein_coding_gene | -1.9 | 0.26 | 3.1E-03 |
| Y54G2A.37 | WBGene00023484 | protein_coding_gene | -1.9 | 0.26 | 3.9E-04 |
| Y77E11A.14 | WBGene00022314 | protein_coding_gene | -1.9 | 0.26 | 1.8E-05 |
| ZK971.1 | WBGene00014175 | protein_coding_gene | -1.9 | 0.26 | 3.9E-07 |
| K09D9.1 | WBGene00019564 | protein_coding_gene | -1.9 | 0.26 | 5.9E-04 |
| cpr-1 | WBGene00000781 | protein_coding_gene | -1.9 | 0.26 | 1.5E-40 |
| fbxa-157 | WBGene00007393 | protein_coding_gene | -1.9 | 0.26 | 2.0E-04 |
| T26C5.2 | WBGene00012032 | protein_coding_gene | -1.9 | 0.26 | 4.3E-02 |
| T07G12.3 | WBGene00011594 | protein_coding_gene | -1.9 | 0.26 | 2.3E-02 |
| F15H10.8 | WBGene00008875 | protein_coding_gene | -1.9 | 0.26 | 2.6E-02 |
| plpr-1 | WBGene00011524 | protein_coding_gene | -1.9 | 0.26 | 4.2E-05 |
| F26A10.2 | WBGene00017814 | protein_coding_gene | -1.9 | 0.26 | 1.6E-08 |
| F53C3.6 | WBGene00018750 | protein_coding_gene | -1.9 | 0.26 | 5.1E-03 |
| C17B7.2 | WBGene00015876 | pseudogene | -1.9 | 0.26 | 1.2E-02 |
| mab-7 | WBGene00003104 | protein_coding_gene | -1.9 | 0.26 | 1.3E-04 |
| cwn-2 | WBGene00000858 | protein_coding_gene | -1.9 | 0.26 | 9.2E-14 |
| fbxa-30 | WBGene00013837 | protein_coding_gene | -1.9 | 0.26 | 6.6E-07 |
| R05G9R.1 | WBGene00019906 | protein_coding_gene | -1.9 | 0.26 | 7.0E-04 |
| ZC123.1 | WBGene00022517 | protein_coding_gene | -1.9 | 0.26 | 2.0E-02 |
| F13A7.11 | WBGene00008726 | protein_coding_gene | -1.9 | 0.26 | 8.8E-08 |
| F49E10.2 | WBGene00018640 | protein_coding_gene | -2.0 | 0.26 | 4.2E-02 |
| C08E8.4 | WBGene00007440 | protein_coding_gene | -2.0 | 0.26 | 4.0E-12 |
| mec-3 | WBGene00003167 | protein_coding_gene | -2.0 | 0.26 | 3.4E-04 |
| che-13 | WBGene00000492 | protein_coding_gene | -2.0 | 0.26 | 6.4E-04 |
| let-381 | WBGene00002601 | protein_coding_gene | -2.0 | 0.26 | 1.8E-04 |
| T01B6.1 | WBGene00020134 | protein_coding_gene | -2.0 | 0.26 | 1.6E-05 |
| F32H2.6 | WBGene00009343 | protein_coding_gene | -2.0 | 0.26 | 8.7E-04 |
| R160.4 | WBGene00020121 | protein_coding_gene | -2.0 | 0.26 | 2.7E-02 |
| C25F6.7 | WBGene00016106 | protein_coding_gene | -2.0 | 0.26 | 2.8E-05 |
| F41C3.7 | WBGene00018273 | protein_coding_gene | -2.0 | 0.26 | 3.0E-03 |
| T05E7.4 | WBGene00020260 | protein_coding_gene | -2.0 | 0.26 | 1.4E-03 |
| C08B6.4 | WBGene00007425 | protein_coding_gene | -2.0 | 0.25 | 1.1E-03 |
| F07H5.13 | WBGene00008563 | protein_coding_gene | -2.0 | 0.25 | 1.5E-14 |
| cut-6 | WBGene00000853 | protein_coding_gene | -2.0 | 0.25 | 8.2E-05 |

| | | | | | |
|---|---|---|---|---|---|
| W10G11.2 | WBGene00021135 | protein_coding_gene | -2.0 | 0.25 | 2.4E-03 |
| R13A5.15 | WBGene00194726 | protein_coding_gene | -2.0 | 0.25 | 1.5E-03 |
| bbs-1 | WBGene00000241 | protein_coding_gene | -2.0 | 0.25 | 1.1E-02 |
| R06B10.2 | WBGene00019913 | protein_coding_gene | -2.0 | 0.25 | 3.0E-02 |
| spp-8 | WBGene00004993 | protein_coding_gene | -2.0 | 0.25 | 3.7E-15 |
| fax-1 | WBGene00001400 | protein_coding_gene | -2.0 | 0.25 | 9.9E-03 |
| rga-2 | WBGene00013135 | protein_coding_gene | -2.0 | 0.25 | 1.6E-11 |
| chil-23 | WBGene00011167 | protein_coding_gene | -2.0 | 0.25 | 5.8E-08 |
| ugt-9 | WBGene00020587 | protein_coding_gene | -2.0 | 0.25 | 3.0E-11 |
| nlp-25 | WBGene00003763 | protein_coding_gene | -2.0 | 0.25 | 1.5E-03 |
| gei-13 | WBGene00001570 | protein_coding_gene | -2.0 | 0.25 | 2.8E-07 |
| srr-4 | WBGene00005655 | protein_coding_gene | -2.0 | 0.25 | 1.1E-25 |
| ZK550.2 | WBGene00013996 | protein_coding_gene | -2.0 | 0.25 | 2.4E-11 |
| fbxb-78 | WBGene00017116 | protein_coding_gene | -2.0 | 0.25 | 3.1E-04 |
| phat-5 | WBGene00020242 | protein_coding_gene | -2.0 | 0.25 | 9.4E-03 |
| ZK287.1 | WBGene00013962 | protein_coding_gene | -2.0 | 0.25 | 9.0E-09 |
| Y57G11C.39 | WBGene00013328 | protein_coding_gene | -2.0 | 0.25 | 4.6E-02 |
| fbxa-127 | WBGene00012027 | protein_coding_gene | -2.0 | 0.25 | 9.0E-03 |
| nhr-218 | WBGene00011750 | protein_coding_gene | -2.0 | 0.25 | 4.4E-02 |
| bus-4 | WBGene00044620 | protein_coding_gene | -2.0 | 0.25 | 1.5E-02 |
| F59D6.1 | WBGene00019103 | protein_coding_gene | -2.0 | 0.25 | 1.6E-02 |
| F52C9.5 | WBGene00018675 | protein_coding_gene | -2.0 | 0.25 | 9.8E-03 |
| JC8.16 | WBGene00194915 | protein_coding_gene | -2.0 | 0.25 | 3.0E-02 |
| igcm-2 | WBGene00020130 | protein_coding_gene | -2.0 | 0.25 | 4.8E-06 |
| Y73E7A.8 | WBGene00022274 | protein_coding_gene | -2.0 | 0.25 | 1.2E-02 |
| pax-1 | WBGene00003937 | protein_coding_gene | -2.0 | 0.25 | 4.9E-02 |
| clec-218 | WBGene00020938 | protein_coding_gene | -2.0 | 0.24 | 4.1E-29 |
| T01B10.5 | WBGene00020137 | protein_coding_gene | -2.0 | 0.24 | 9.0E-05 |
| R57.2 | WBGene00020083 | protein_coding_gene | -2.0 | 0.24 | 2.0E-03 |
| R07E3.6 | WBGene00011107 | protein_coding_gene | -2.0 | 0.24 | 3.3E-02 |
| M03A1.8 | WBGene00077490 | protein_coding_gene | -2.0 | 0.24 | 2.7E-04 |
| F46G11.6 | WBGene00045269 | protein_coding_gene | -2.0 | 0.24 | 1.7E-02 |
| ast-1 | WBGene00020368 | protein_coding_gene | -2.0 | 0.24 | 3.1E-07 |
| T19H5.6 | WBGene00044807 | protein_coding_gene | -2.0 | 0.24 | 2.4E-09 |
| C10A4.3 | WBGene00015664 | protein_coding_gene | -2.0 | 0.24 | 2.7E-03 |
| R01H2.8 | WBGene00045190 | protein_coding_gene | -2.0 | 0.24 | 3.8E-02 |
| zig-10 | WBGene00020800 | protein_coding_gene | -2.0 | 0.24 | 9.1E-07 |
| lpr-1 | WBGene00002393 | protein_coding_gene | -2.1 | 0.24 | 4.0E-04 |
| aat-8 | WBGene00000009 | protein_coding_gene | -2.1 | 0.24 | 6.6E-05 |
| F20G2.5 | WBGene00008988 | protein_coding_gene | -2.1 | 0.24 | 1.2E-02 |
| fbxa-161 | WBGene00015596 | pseudogene | -2.1 | 0.24 | 2.6E-04 |
| lips-9 | WBGene00010267 | protein_coding_gene | -2.1 | 0.24 | 8.3E-03 |
| mks-5 | WBGene00007490 | protein_coding_gene | -2.1 | 0.24 | 6.5E-04 |
| C14A11.2 | WBGene00015748 | protein_coding_gene | -2.1 | 0.24 | 4.4E-04 |
| Y53F4B.25 | WBGene00013171 | protein_coding_gene | -2.1 | 0.24 | 2.3E-02 |
| T04C12.8 | WBGene00011430 | protein_coding_gene | -2.1 | 0.24 | 3.5E-07 |
| Y5H2B.1 | WBGene00021164 | protein_coding_gene | -2.1 | 0.24 | 1.6E-02 |
| jbts-14 | WBGene00018727 | protein_coding_gene | -2.1 | 0.24 | 1.2E-02 |
| zmp-5 | WBGene00010423 | protein_coding_gene | -2.1 | 0.24 | 7.5E-03 |
| ZC84.6 | WBGene00013849 | protein_coding_gene | -2.1 | 0.24 | 7.8E-04 |
| C04E6.12 | WBGene00015426 | protein_coding_gene | -2.1 | 0.24 | 2.9E-02 |
| F07C3.9 | WBGene00017197 | protein_coding_gene | -2.1 | 0.24 | 2.7E-08 |
| unc-86 | WBGene00006818 | protein_coding_gene | -2.1 | 0.24 | 3.6E-07 |
| F13H6.5 | WBGene00017433 | protein_coding_gene | -2.1 | 0.24 | 1.5E-07 |
| Y71F9B.9 | WBGene00022130 | protein_coding_gene | -2.1 | 0.24 | 7.2E-17 |
| clec-8 | WBGene00012582 | protein_coding_gene | -2.1 | 0.24 | 1.0E-06 |
| C35E7.4 | WBGene00016456 | protein_coding_gene | -2.1 | 0.23 | 4.3E-11 |
| Y22D7AL.6 | WBGene00021245 | protein_coding_gene | -2.1 | 0.23 | 3.3E-03 |

| | | | | | |
|---|---|---|---|---|---|
| srh-234 | WBGene00005441 | protein_coding_gene | -2.1 | 0.23 | 4.6E-02 |
| F32D8.1 | WBGene00009326 | protein_coding_gene | -2.1 | 0.23 | 3.9E-02 |
| F55H12.7 | WBGene00219283 | protein_coding_gene | -2.1 | 0.23 | 4.6E-03 |
| ifta-2 | WBGene00012132 | protein_coding_gene | -2.1 | 0.23 | 1.5E-02 |
| T09E11.11 | WBGene00011660 | pseudogene | -2.1 | 0.23 | 2.6E-10 |
| F30H5.3 | WBGene00017937 | protein_coding_gene | -2.1 | 0.23 | 1.8E-03 |
| T19E7.6 | WBGene00044500 | protein_coding_gene | -2.1 | 0.23 | 5.7E-16 |
| skr-21 | WBGene00004827 | protein_coding_gene | -2.1 | 0.23 | 1.2E-27 |
| C03A3.1 | WBGene00007266 | protein_coding_gene | -2.1 | 0.23 | 1.3E-07 |
| asah-1 | WBGene00010769 | protein_coding_gene | -2.1 | 0.23 | 2.2E-11 |
| R02F11.1 | WBGene00019839 | protein_coding_gene | -2.1 | 0.23 | 3.4E-02 |
| tth-1 | WBGene00006649 | protein_coding_gene | -2.1 | 0.23 | 2.1E-43 |
| F30A10.2 | WBGene00009261 | protein_coding_gene | -2.1 | 0.23 | 2.6E-02 |
| F41G3.3 | WBGene00018300 | protein_coding_gene | -2.1 | 0.23 | 1.9E-02 |
| acn-1 | WBGene00000039 | protein_coding_gene | -2.1 | 0.23 | 9.1E-03 |
| D1086.2 | WBGene00008389 | protein_coding_gene | -2.1 | 0.23 | 9.7E-07 |
| nab-1 | WBGene00003516 | protein_coding_gene | -2.1 | 0.23 | 4.4E-18 |
| T21D12.7 | WBGene00020648 | protein_coding_gene | -2.1 | 0.23 | 2.5E-03 |
| ceh-8 | WBGene00000433 | protein_coding_gene | -2.1 | 0.23 | 5.7E-03 |
| dhs-16 | WBGene00000979 | protein_coding_gene | -2.1 | 0.23 | 3.3E-03 |
| linc-103 | WBGene00219743 | lincRNA_gene | -2.1 | 0.23 | 3.0E-02 |
| arl-6 | WBGene00000193 | protein_coding_gene | -2.1 | 0.23 | 5.8E-03 |
| R07E5.17 | WBGene00011121 | protein_coding_gene | -2.1 | 0.23 | 7.3E-03 |
| ptr-3 | WBGene00004218 | protein_coding_gene | -2.1 | 0.23 | 2.6E-09 |
| ZK836.3 | WBGene00014099 | protein_coding_gene | -2.1 | 0.23 | 3.1E-07 |
| T17H7.7 | WBGene00020551 | protein_coding_gene | -2.1 | 0.23 | 9.7E-05 |
| aakg-4 | WBGene00020135 | protein_coding_gene | -2.1 | 0.23 | 6.1E-23 |
| fbxa-42 | WBGene00044676 | protein_coding_gene | -2.1 | 0.23 | 1.3E-04 |
| phat-2 | WBGene00016733 | protein_coding_gene | -2.1 | 0.23 | 1.8E-02 |
| C26B9.7 | WBGene00016136 | protein_coding_gene | -2.1 | 0.23 | 3.6E-02 |
| bath-25 | WBGene00020859 | protein_coding_gene | -2.1 | 0.23 | 9.1E-03 |
| Y87G2A.25 | WBGene00219288 | protein_coding_gene | -2.1 | 0.23 | 1.3E-04 |
| Y50E8A.1 | WBGene00013046 | protein_coding_gene | -2.2 | 0.22 | 1.2E-07 |
| cyp-13A9 | WBGene00014838 | pseudogene | -2.2 | 0.22 | 2.7E-03 |
| nhr-91 | WBGene00003681 | protein_coding_gene | -2.2 | 0.22 | 6.6E-09 |
| nhr-208 | WBGene00011099 | protein_coding_gene | -2.2 | 0.22 | 8.4E-04 |
| F46F3.3 | WBGene00009791 | protein_coding_gene | -2.2 | 0.22 | 1.1E-02 |
| C17E7.13 | WBGene00015906 | protein_coding_gene | -2.2 | 0.22 | 1.4E-02 |
| F09F9.2 | WBGene00017307 | protein_coding_gene | -2.2 | 0.22 | 2.7E-02 |
| osm-8 | WBGene00003888 | protein_coding_gene | -2.2 | 0.22 | 7.4E-03 |
| C53A5.2 | WBGene00008263 | protein_coding_gene | -2.2 | 0.22 | 1.1E-04 |
| C32D5.12 | WBGene00016319 | protein_coding_gene | -2.2 | 0.22 | 4.2E-08 |
| cutl-16 | WBGene00019426 | protein_coding_gene | -2.2 | 0.22 | 1.1E-11 |
| T05C1.3 | WBGene00020250 | protein_coding_gene | -2.2 | 0.22 | 1.7E-02 |
| K10D3.6 | WBGene00010740 | protein_coding_gene | -2.2 | 0.22 | 3.6E-12 |
| B0391.14 | WBGene00206462 | protein_coding_gene | -2.2 | 0.22 | 1.4E-02 |
| rom-2 | WBGene00004401 | protein_coding_gene | -2.2 | 0.22 | 5.9E-05 |
| rgba-1 | WBGene00077766 | protein_coding_gene | -2.2 | 0.22 | 4.3E-02 |
| W10G11.4 | WBGene00021137 | protein_coding_gene | -2.2 | 0.22 | 1.8E-02 |
| K10H10.10 | WBGene00010765 | protein_coding_gene | -2.2 | 0.22 | 1.2E-02 |
| T21D12.14 | WBGene00045177 | protein_coding_gene | -2.2 | 0.22 | 3.1E-04 |
| M04C9.2 | WBGene00010857 | protein_coding_gene | -2.2 | 0.22 | 2.4E-02 |
| K02E10.4 | WBGene00019318 | protein_coding_gene | -2.2 | 0.22 | 1.8E-03 |
| cky-1 | WBGene00000521 | protein_coding_gene | -2.2 | 0.22 | 9.6E-03 |
| pros-1 | WBGene00000448 | protein_coding_gene | -2.2 | 0.22 | 2.1E-13 |
| ZC449.1 | WBGene00022611 | protein_coding_gene | -2.2 | 0.22 | 2.8E-02 |
| C01G6.9 | WBGene00007229 | protein_coding_gene | -2.2 | 0.22 | 1.7E-04 |
| C17G10.6 | WBGene00015918 | protein_coding_gene | -2.2 | 0.22 | 3.6E-02 |

| | | | | | |
|---|---|---|---|---|---|
| Y37A1B.7 | WBGene00012540 | protein_coding_gene | -2.2 | 0.22 | 1.9E-02 |
| Y47G6A.32 | WBGene00044437 | protein_coding_gene | -2.2 | 0.22 | 2.6E-02 |
| C24H10.3 | WBGene00016069 | protein_coding_gene | -2.2 | 0.21 | 4.0E-04 |
| C11D2.3 | WBGene00015701 | protein_coding_gene | -2.2 | 0.21 | 4.5E-04 |
| F33H2.8 | WBGene00009371 | protein_coding_gene | -2.2 | 0.21 | 8.1E-05 |
| F57B1.6 | WBGene00010189 | protein_coding_gene | -2.2 | 0.21 | 2.4E-06 |
| R12A1.3 | WBGene00020017 | protein_coding_gene | -2.2 | 0.21 | 2.6E-02 |
| F40G9.20 | WBGene00195034 | ncRNA_gene | -2.2 | 0.21 | 1.4E-03 |
| F26F12.4 | WBGene00017835 | protein_coding_gene | -2.2 | 0.21 | 1.2E-04 |
| Y39C12A.2 | WBGene00014883 | pseudogene | -2.2 | 0.21 | 3.6E-02 |
| F38B6.7 | WBGene00018176 | protein_coding_gene | -2.3 | 0.21 | 3.7E-02 |
| F59E11.7 | WBGene00019113 | protein_coding_gene | -2.3 | 0.21 | 3.7E-08 |
| srbc-20 | WBGene00016693 | protein_coding_gene | -2.3 | 0.21 | 4.5E-02 |
| fbxa-99 | WBGene00009490 | protein_coding_gene | -2.3 | 0.21 | 3.6E-03 |
| R09D1.12 | WBGene00011168 | protein_coding_gene | -2.3 | 0.21 | 6.4E-06 |
| C18A11.1 | WBGene00015947 | protein_coding_gene | -2.3 | 0.21 | 7.9E-21 |
| T16G1.2 | WBGene00011796 | protein_coding_gene | -2.3 | 0.21 | 7.7E-04 |
| K10D6.2 | WBGene00010742 | protein_coding_gene | -2.3 | 0.21 | 1.4E-16 |
| mtl-2 | WBGene00003474 | protein_coding_gene | -2.3 | 0.20 | 2.2E-42 |
| ptr-15 | WBGene00004229 | protein_coding_gene | -2.3 | 0.20 | 6.5E-06 |
| C43H6.6 | WBGene00016619 | protein_coding_gene | -2.3 | 0.20 | 1.0E-02 |
| cut-2 | WBGene00009983 | protein_coding_gene | -2.3 | 0.20 | 5.0E-03 |
| M02G9.2 | WBGene00010831 | protein_coding_gene | -2.3 | 0.20 | 8.4E-03 |
| clec-24 | WBGene00009856 | protein_coding_gene | -2.3 | 0.20 | 2.0E-13 |
| pho-11 | WBGene00007331 | protein_coding_gene | -2.3 | 0.20 | 2.6E-16 |
| F28H7.2 | WBGene00009236 | protein_coding_gene | -2.3 | 0.20 | 8.3E-03 |
| bus-8 | WBGene00044623 | protein_coding_gene | -2.3 | 0.20 | 6.7E-03 |
| Y38H8A.1 | WBGene00012635 | protein_coding_gene | -2.3 | 0.20 | 1.5E-03 |
| F59F5.3 | WBGene00010342 | protein_coding_gene | -2.3 | 0.20 | 1.2E-07 |
| K11H12.11 | WBGene00044723 | protein_coding_gene | -2.3 | 0.20 | 5.5E-03 |
| F56H9.9 | WBGene00206460 | protein_coding_gene | -2.3 | 0.20 | 2.5E-02 |
| F57A8.7 | WBGene00010182 | protein_coding_gene | -2.3 | 0.20 | 2.1E-02 |
| clec-31 | WBGene00009854 | protein_coding_gene | -2.3 | 0.20 | 1.5E-07 |
| W03D2.9 | WBGene00020983 | protein_coding_gene | -2.3 | 0.20 | 5.8E-03 |
| T19H12.6 | WBGene00020590 | protein_coding_gene | -2.3 | 0.20 | 1.5E-02 |
| Y97E10AR.1 | WBGene00022396 | protein_coding_gene | -2.3 | 0.20 | 1.7E-16 |
| dhhc-11 | WBGene00020694 | protein_coding_gene | -2.3 | 0.20 | 2.4E-02 |
| R04B5.6 | WBGene00011004 | protein_coding_gene | -2.3 | 0.20 | 1.1E-02 |
| slc-17.9 | WBGene00011643 | protein_coding_gene | -2.4 | 0.20 | 2.7E-05 |
| W06G6.20 | WBGene00219308 | protein_coding_gene | -2.4 | 0.20 | 2.5E-02 |
| bbs-2 | WBGene00000242 | protein_coding_gene | -2.4 | 0.20 | 1.2E-03 |
| W06G6.11 | WBGene00012313 | protein_coding_gene | -2.4 | 0.19 | 1.0E-02 |
| B0393.5 | WBGene00007170 | protein_coding_gene | -2.4 | 0.19 | 3.1E-05 |
| W08A12.2 | WBGene00021079 | protein_coding_gene | -2.4 | 0.19 | 4.7E-25 |
| pals-19 | WBGene00044761 | protein_coding_gene | -2.4 | 0.19 | 3.6E-04 |
| fbxc-53 | WBGene00016711 | protein_coding_gene | -2.4 | 0.19 | 7.0E-03 |
| C14A4.9 | WBGene00007560 | protein_coding_gene | -2.4 | 0.19 | 1.6E-02 |
| subs-4 | WBGene00012939 | protein_coding_gene | -2.4 | 0.19 | 2.8E-05 |
| T09F5.1 | WBGene00011665 | protein_coding_gene | -2.4 | 0.19 | 1.7E-02 |
| T24C4.4 | WBGene00020760 | protein_coding_gene | -2.4 | 0.19 | 3.7E-08 |
| nhr-172 | WBGene00016927 | protein_coding_gene | -2.4 | 0.19 | 4.7E-02 |
| C13C12.2 | WBGene00007552 | protein_coding_gene | -2.4 | 0.19 | 7.2E-05 |
| fbxa-68 | WBGene00020641 | pseudogene | -2.4 | 0.19 | 1.6E-02 |
| Y70C5A.3 | WBGene00050904 | protein_coding_gene | -2.4 | 0.19 | 1.2E-12 |
| tbb-4 | WBGene00006538 | protein_coding_gene | -2.4 | 0.19 | 4.2E-13 |
| mlt-8 | WBGene00021095 | protein_coding_gene | -2.4 | 0.19 | 7.9E-04 |
| M03B6.5 | WBGene00010837 | protein_coding_gene | -2.4 | 0.19 | 3.6E-03 |
| mks-2 | WBGene00194710 | protein_coding_gene | -2.4 | 0.19 | 4.0E-02 |

| | | | | | |
|---|---|---|---|---|---|
| K10G4.13 | WBGene00077629 | pseudogene | -2.4 | 0.19 | 2.7E-02 |
| K07D4.6 | WBGene00019486 | protein_coding_gene | -2.4 | 0.19 | 2.4E-02 |
| F01D5.6 | WBGene00008496 | protein_coding_gene | -2.4 | 0.19 | 5.6E-05 |
| ZC449.2 | WBGene00022612 | protein_coding_gene | -2.4 | 0.19 | 1.9E-06 |
| F44E2.4 | WBGene00018418 | protein_coding_gene | -2.4 | 0.19 | 7.8E-05 |
| F48C1.11 | WBGene00195179 | protein_coding_gene | -2.4 | 0.19 | 2.7E-02 |
| ZK678.3 | WBGene00014069 | pseudogene | -2.4 | 0.19 | 3.7E-02 |
| C18H9.5 | WBGene00016003 | protein_coding_gene | -2.4 | 0.19 | 1.1E-14 |
| K04H4.2 | WBGene00010573 | protein_coding_gene | -2.4 | 0.19 | 1.3E-04 |
| F54F11.1 | WBGene00010069 | protein_coding_gene | -2.4 | 0.18 | 3.1E-02 |
| oac-29 | WBGene00018295 | protein_coding_gene | -2.4 | 0.18 | 4.7E-04 |
| srv-13 | WBGene00005724 | protein_coding_gene | -2.4 | 0.18 | 4.2E-02 |
| F41G3.2 | WBGene00018299 | protein_coding_gene | -2.5 | 0.18 | 2.8E-02 |
| ins-23 | WBGene00002106 | protein_coding_gene | -2.5 | 0.18 | 2.7E-04 |
| cnc-6 | WBGene00000560 | protein_coding_gene | -2.5 | 0.18 | 3.1E-02 |
| F48G7.5 | WBGene00018616 | protein_coding_gene | -2.5 | 0.18 | 2.1E-02 |
| col-74 | WBGene00000650 | protein_coding_gene | -2.5 | 0.18 | 6.3E-13 |
| mks-3 | WBGene00018042 | protein_coding_gene | -2.5 | 0.18 | 6.7E-04 |
| F08H9.3 | WBGene00008591 | protein_coding_gene | -2.5 | 0.18 | 3.3E-05 |
| R04B3.1 | WBGene00019866 | protein_coding_gene | -2.5 | 0.18 | 1.3E-02 |
| tmem-231 | WBGene00020825 | protein_coding_gene | -2.5 | 0.18 | 3.7E-04 |
| C16D9.3 | WBGene00015858 | protein_coding_gene | -2.5 | 0.18 | 1.7E-02 |
| lhfp-4 | WBGene00009155 | protein_coding_gene | -2.5 | 0.18 | 4.6E-03 |
| bgnt-1.3 | WBGene00009032 | protein_coding_gene | -2.5 | 0.18 | 1.8E-02 |
| thn-1 | WBGene00009213 | protein_coding_gene | -2.5 | 0.18 | 1.4E-06 |
| cyp-34A7 | WBGene00015042 | protein_coding_gene | -2.5 | 0.18 | 2.6E-03 |
| nep-16 | WBGene00018472 | protein_coding_gene | -2.5 | 0.18 | 7.9E-04 |
| F54B11.10 | WBGene00010033 | protein_coding_gene | -2.5 | 0.18 | 2.9E-03 |
| M01G12.14 | WBGene00010825 | protein_coding_gene | -2.5 | 0.18 | 9.3E-05 |
| nhr-165 | WBGene00008158 | protein_coding_gene | -2.5 | 0.18 | 3.1E-02 |
| math-38 | WBGene00020326 | protein_coding_gene | -2.5 | 0.18 | 4.7E-23 |
| acr-14 | WBGene00000053 | protein_coding_gene | -2.5 | 0.18 | 1.5E-02 |
| nspe-7 | WBGene00012593 | protein_coding_gene | -2.5 | 0.18 | 3.0E-03 |
| pqn-16 | WBGene00004107 | protein_coding_gene | -2.5 | 0.18 | 2.0E-02 |
| chil-18 | WBGene00011161 | protein_coding_gene | -2.5 | 0.18 | 4.8E-03 |
| atf-8 | WBGene00017535 | protein_coding_gene | -2.5 | 0.17 | 1.3E-02 |
| hmg-6 | WBGene00009827 | protein_coding_gene | -2.5 | 0.17 | 2.2E-02 |
| spp-6 | WBGene00004991 | protein_coding_gene | -2.5 | 0.17 | 2.7E-03 |
| ttr-52 | WBGene00008719 | protein_coding_gene | -2.5 | 0.17 | 7.3E-05 |
| bus-18 | WBGene00044631 | protein_coding_gene | -2.5 | 0.17 | 6.1E-04 |
| F36F12.1 | WBGene00018092 | protein_coding_gene | -2.5 | 0.17 | 2.4E-03 |
| tag-297 | WBGene00007999 | protein_coding_gene | -2.5 | 0.17 | 5.3E-03 |
| ceh-5 | WBGene00000430 | protein_coding_gene | -2.5 | 0.17 | 2.8E-05 |
| trx-3 | WBGene00019717 | protein_coding_gene | -2.6 | 0.17 | 2.6E-16 |
| C17C3.3 | WBGene00015889 | protein_coding_gene | -2.6 | 0.17 | 2.9E-03 |
| K01A6.4 | WBGene00010446 | protein_coding_gene | -2.6 | 0.17 | 3.4E-04 |
| C35A5.5 | WBGene00007949 | protein_coding_gene | -2.6 | 0.17 | 2.2E-03 |
| F15B9.8 | WBGene00008851 | protein_coding_gene | -2.6 | 0.17 | 3.0E-03 |
| cutl-8 | WBGene00013960 | protein_coding_gene | -2.6 | 0.17 | 7.6E-03 |
| C02E7.6 | WBGene00015339 | protein_coding_gene | -2.6 | 0.17 | 8.8E-03 |
| smf-3 | WBGene00004878 | protein_coding_gene | -2.6 | 0.17 | 2.8E-12 |
| pqn-13 | WBGene00004104 | protein_coding_gene | -2.6 | 0.17 | 1.3E-03 |
| ins-2 | WBGene00002085 | protein_coding_gene | -2.6 | 0.17 | 1.3E-06 |
| cbn-1 | WBGene00019285 | protein_coding_gene | -2.6 | 0.17 | 2.9E-08 |
| grl-2 | WBGene00001711 | protein_coding_gene | -2.6 | 0.17 | 3.0E-02 |
| ces-1 | WBGene00000468 | protein_coding_gene | -2.6 | 0.17 | 7.3E-05 |
| W02D7.8 | WBGene00020943 | protein_coding_gene | -2.6 | 0.17 | 8.7E-08 |
| F21C10.10 | WBGene00017659 | protein_coding_gene | -2.6 | 0.16 | 2.4E-08 |

| | | | | | |
|---|---|---|---|---|---|
| H32K16.2 | WBGene00010422 | protein_coding_gene | -2.6 | 0.16 | 3.5E-09 |
| F13H6.4 | WBGene00017432 | pseudogene | -2.6 | 0.16 | 9.0E-08 |
| pals-7 | WBGene00007661 | protein_coding_gene | -2.6 | 0.16 | 1.5E-02 |
| srbc-82 | WBGene00007832 | protein_coding_gene | -2.6 | 0.16 | 3.2E-02 |
| F49D11.6 | WBGene00018630 | protein_coding_gene | -2.6 | 0.16 | 5.2E-03 |
| W07G4.1 | WBGene00012335 | protein_coding_gene | -2.6 | 0.16 | 1.5E-02 |
| tts-2 | WBGene00006651 | ncRNA_gene | -2.6 | 0.16 | 1.3E-05 |
| dhs-23 | WBGene00000986 | protein_coding_gene | -2.6 | 0.16 | 1.1E-12 |
| dlhd-1 | WBGene00020741 | protein_coding_gene | -2.6 | 0.16 | 7.0E-04 |
| wrt-1 | WBGene00006947 | protein_coding_gene | -2.6 | 0.16 | 2.1E-03 |
| linc-37 | WBGene00219709 | lincRNA_gene | -2.6 | 0.16 | 5.5E-03 |
| clec-82 | WBGene00021873 | protein_coding_gene | -2.6 | 0.16 | 9.9E-06 |
| C45E5.3 | WBGene00016666 | pseudogene | -2.6 | 0.16 | 4.2E-08 |
| C06E4.8 | WBGene00015537 | protein_coding_gene | -2.7 | 0.16 | 2.1E-15 |
| Y54G2A.76 | WBGene00235268 | protein_coding_gene | -2.7 | 0.16 | 5.6E-03 |
| spp-13 | WBGene00004998 | protein_coding_gene | -2.7 | 0.16 | 7.1E-05 |
| nhr-81 | WBGene00003671 | protein_coding_gene | -2.7 | 0.16 | 7.8E-03 |
| spp-7 | WBGene00004992 | protein_coding_gene | -2.7 | 0.16 | 1.1E-02 |
| F16B3.2 | WBGene00017502 | protein_coding_gene | -2.7 | 0.15 | 2.1E-04 |
| T16G1.6 | WBGene00011800 | protein_coding_gene | -2.7 | 0.15 | 2.8E-29 |
| F35G2.5 | WBGene00009438 | protein_coding_gene | -2.7 | 0.15 | 1.3E-02 |
| C05D9.4 | WBGene00015473 | protein_coding_gene | -2.7 | 0.15 | 9.3E-03 |
| F07A5.4 | WBGene00008543 | protein_coding_gene | -2.7 | 0.15 | 1.9E-05 |
| cyp-13A3 | WBGene00011675 | protein_coding_gene | -2.7 | 0.15 | 2.0E-05 |
| H39E23.2 | WBGene00010432 | protein_coding_gene | -2.7 | 0.15 | 1.6E-02 |
| F16B4.7 | WBGene00017509 | protein_coding_gene | -2.7 | 0.15 | 1.3E-05 |
| K01D12.1 | WBGene00010461 | protein_coding_gene | -2.7 | 0.15 | 1.8E-04 |
| ZK813.6 | WBGene00044564 | protein_coding_gene | -2.7 | 0.15 | 1.3E-02 |
| C28G1.2 | WBGene00016187 | protein_coding_gene | -2.7 | 0.15 | 3.8E-03 |
| T14E8.4 | WBGene00043981 | protein_coding_gene | -2.7 | 0.15 | 9.0E-05 |
| Y23H5B.8 | WBGene00021279 | protein_coding_gene | -2.8 | 0.15 | 1.9E-10 |
| hlh-13 | WBGene00001957 | protein_coding_gene | -2.8 | 0.15 | 9.1E-03 |
| nhr-221 | WBGene00020748 | protein_coding_gene | -2.8 | 0.15 | 1.1E-02 |
| ztf-30 | WBGene00015523 | protein_coding_gene | -2.8 | 0.15 | 3.5E-06 |
| nphp-1 | WBGene00010898 | protein_coding_gene | -2.8 | 0.15 | 6.3E-07 |
| F42C5.3 | WBGene00018345 | protein_coding_gene | -2.8 | 0.15 | 1.5E-05 |
| fipr-26 | WBGene00009963 | protein_coding_gene | -2.8 | 0.15 | 3.2E-03 |
| C54F6.18 | WBGene00235093 | protein_coding_gene | -2.8 | 0.15 | 1.4E-03 |
| fut-2 | WBGene00001506 | protein_coding_gene | -2.8 | 0.15 | 7.4E-06 |
| cuti-1 | WBGene00022591 | protein_coding_gene | -2.8 | 0.14 | 2.8E-09 |
| chil-28 | WBGene00008842 | pseudogene | -2.8 | 0.14 | 6.4E-04 |
| gcy-8 | WBGene00001535 | protein_coding_gene | -2.8 | 0.14 | 1.6E-03 |
| pitr-6 | WBGene00012285 | protein_coding_gene | -2.8 | 0.14 | 1.5E-02 |
| W02A2.9 | WBGene00044280 | protein_coding_gene | -2.8 | 0.14 | 2.5E-22 |
| lys-3 | WBGene00003092 | protein_coding_gene | -2.8 | 0.14 | 2.2E-08 |
| K09C4.4 | WBGene00019549 | protein_coding_gene | -2.8 | 0.14 | 1.3E-02 |
| cutl-13 | WBGene00021025 | protein_coding_gene | -2.8 | 0.14 | 1.0E-04 |
| B0284.5 | WBGene00045188 | pseudogene | -2.8 | 0.14 | 4.5E-03 |
| C26E1.2 | WBGene00007748 | protein_coding_gene | -2.9 | 0.14 | 2.6E-35 |
| fbxb-101 | WBGene00013404 | protein_coding_gene | -2.9 | 0.14 | 7.3E-07 |
| sru-40 | WBGene00005703 | protein_coding_gene | -2.9 | 0.14 | 2.0E-03 |
| K10H10.13 | WBGene00194888 | protein_coding_gene | -2.9 | 0.14 | 9.8E-03 |
| K01A6.7 | WBGene00045096 | protein_coding_gene | -2.9 | 0.14 | 3.3E-03 |
| Y46H3A.5 | WBGene00021601 | protein_coding_gene | -2.9 | 0.14 | 2.2E-03 |
| Y51H7C.12 | WBGene00021790 | protein_coding_gene | -2.9 | 0.14 | 3.1E-03 |
| T21E8.6 | WBGene00164971 | protein_coding_gene | -2.9 | 0.13 | 1.4E-02 |
| F11D5.5 | WBGene00017382 | protein_coding_gene | -2.9 | 0.13 | 1.5E-09 |
| C03F11.2 | WBGene00015388 | protein_coding_gene | -2.9 | 0.13 | 1.6E-04 |

| | | | | | |
|---|---|---|---|---|---|
| cut-3 | WBGene00009041 | protein_coding_gene | -2.9 | 0.13 | 2.4E-10 |
| rocf-1 | WBGene00011948 | protein_coding_gene | -2.9 | 0.13 | 1.1E-06 |
| Y45F10D.6 | WBGene00012886 | protein_coding_gene | -2.9 | 0.13 | 3.2E-10 |
| Y39H10A.1 | WBGene00021478 | protein_coding_gene | -2.9 | 0.13 | 1.6E-03 |
| T05H10.3 | WBGene00011508 | protein_coding_gene | -2.9 | 0.13 | 1.8E-08 |
| F17B5.6 | WBGene00044058 | protein_coding_gene | -2.9 | 0.13 | 1.1E-02 |
| K08B4.2 | WBGene00019514 | protein_coding_gene | -3.0 | 0.13 | 3.0E-09 |
| E02C12.10 | WBGene00017095 | protein_coding_gene | -3.0 | 0.13 | 1.0E-02 |
| bro-1 | WBGene00000272 | protein_coding_gene | -3.0 | 0.13 | 1.4E-08 |
| C54C8.4 | WBGene00008292 | protein_coding_gene | -3.0 | 0.13 | 3.6E-03 |
| fbxa-162 | WBGene00015597 | pseudogene | -3.0 | 0.13 | 1.2E-09 |
| T28A11.25 | WBGene00269355 | protein_coding_gene | -3.0 | 0.13 | 1.4E-03 |
| R160.10 | WBGene00219369 | protein_coding_gene | -3.0 | 0.13 | 2.6E-04 |
| dao-6 | WBGene00000932 | protein_coding_gene | -3.0 | 0.13 | 2.2E-04 |
| droe-8 | WBGene00022550 | protein_coding_gene | -3.0 | 0.13 | 3.1E-03 |
| K08F9.3 | WBGene00010686 | protein_coding_gene | -3.0 | 0.13 | 5.6E-06 |
| F09E10.6 | WBGene00017296 | protein_coding_gene | -3.0 | 0.12 | 1.4E-04 |
| C30F8.3 | WBGene00016259 | protein_coding_gene | -3.0 | 0.12 | 2.7E-03 |
| T09D3.3 | WBGene00020384 | protein_coding_gene | -3.0 | 0.12 | 6.5E-03 |
| R09E10.13 | WBGene00194658 | protein_coding_gene | -3.0 | 0.12 | 4.0E-04 |
| fbxa-163 | WBGene00015598 | protein_coding_gene | -3.0 | 0.12 | 4.1E-11 |
| mboa-1 | WBGene00007174 | protein_coding_gene | -3.0 | 0.12 | 1.5E-10 |
| H25K10.4 | WBGene00010413 | protein_coding_gene | -3.0 | 0.12 | 2.1E-10 |
| oac-43 | WBGene00011653 | protein_coding_gene | -3.0 | 0.12 | 4.1E-03 |
| W08G11.1 | WBGene00012346 | protein_coding_gene | -3.1 | 0.12 | 6.8E-04 |
| fbxb-114 | WBGene00016254 | protein_coding_gene | -3.1 | 0.12 | 7.5E-03 |
| F13H10.5 | WBGene00008776 | protein_coding_gene | -3.1 | 0.12 | 1.1E-10 |
| B0205.4 | WBGene00015019 | protein_coding_gene | -3.1 | 0.12 | 1.1E-06 |
| W04H10.5 | WBGene00206416 | protein_coding_gene | -3.1 | 0.12 | 3.2E-03 |
| C14F11.6 | WBGene00015781 | protein_coding_gene | -3.1 | 0.12 | 9.3E-07 |
| T19A5.3 | WBGene00020554 | protein_coding_gene | -3.1 | 0.12 | 1.7E-05 |
| F54E2.2 | WBGene00018824 | protein_coding_gene | -3.1 | 0.11 | 2.2E-04 |
| F33H12.7 | WBGene00045457 | protein_coding_gene | -3.1 | 0.11 | 1.5E-03 |
| bus-12 | WBGene00010442 | protein_coding_gene | -3.2 | 0.11 | 4.0E-06 |
| C37A5.6 | WBGene00014700 | protein_coding_gene | -3.2 | 0.11 | 3.7E-03 |
| C31H5.1 | WBGene00007854 | protein_coding_gene | -3.2 | 0.11 | 1.3E-03 |
| clec-32 | WBGene00009860 | protein_coding_gene | -3.2 | 0.11 | 1.8E-07 |
| F49F1.5 | WBGene00018645 | protein_coding_gene | -3.2 | 0.11 | 5.6E-23 |
| C03C10.5 | WBGene00007271 | protein_coding_gene | -3.2 | 0.11 | 8.9E-11 |
| glf-1 | WBGene00019154 | protein_coding_gene | -3.2 | 0.11 | 7.0E-04 |
| tsp-1 | WBGene00006627 | protein_coding_gene | -3.2 | 0.11 | 2.0E-07 |
| F53B2.8 | WBGene00009957 | protein_coding_gene | -3.2 | 0.11 | 2.4E-16 |
| T04C12.32 | WBGene00235086 | ncRNA_gene | -3.2 | 0.11 | 4.6E-03 |
| F18C5.9 | WBGene00017562 | protein_coding_gene | -3.2 | 0.11 | 4.2E-03 |
| F48F7.3 | WBGene00009849 | protein_coding_gene | -3.2 | 0.11 | 1.0E-03 |
| col-165 | WBGene00000738 | protein_coding_gene | -3.3 | 0.10 | 5.2E-18 |
| C08F11.3 | WBGene00007452 | protein_coding_gene | -3.3 | 0.10 | 1.0E-05 |
| K04A8.21 | WBGene00271815 | protein_coding_gene | -3.3 | 0.10 | 6.2E-04 |
| ZK896.5 | WBGene00014136 | protein_coding_gene | -3.3 | 0.10 | 3.3E-07 |
| clec-77 | WBGene00044343 | pseudogene | -3.3 | 0.10 | 4.3E-69 |
| ugt-33 | WBGene00007946 | protein_coding_gene | -3.3 | 0.10 | 8.6E-08 |
| D1014.5 | WBGene00017018 | protein_coding_gene | -3.3 | 0.10 | 1.5E-04 |
| K09C8.7 | WBGene00010717 | protein_coding_gene | -3.3 | 0.10 | 1.4E-03 |
| C29A12.6 | WBGene00007800 | protein_coding_gene | -3.4 | 0.10 | 5.4E-07 |
| nstp-3 | WBGene00022577 | protein_coding_gene | -3.4 | 0.10 | 2.9E-03 |
| K08C9.5 | WBGene00010652 | protein_coding_gene | -3.4 | 0.09 | 5.8E-04 |
| F15E6.10 | WBGene00044479 | protein_coding_gene | -3.4 | 0.09 | 4.4E-04 |
| W09G12.7 | WBGene00021121 | protein_coding_gene | -3.4 | 0.09 | 1.3E-29 |

| | | | | | |
|---|---|---|---|---|---|
| T26H5.8 | WBGene00012070 | protein_coding_gene | -3.4 | 0.09 | 2.2E-20 |
| T04C12.1 | WBGene00011427 | protein_coding_gene | -3.5 | 0.09 | 1.1E-05 |
| C23G10.11 | WBGene00016017 | protein_coding_gene | -3.5 | 0.09 | 4.5E-04 |
| B0462.1 | WBGene00007181 | pseudogene | -3.5 | 0.09 | 3.0E-08 |
| alh-5 | WBGene00000111 | protein_coding_gene | -3.5 | 0.09 | 3.4E-52 |
| C54C8.12 | WBGene00044242 | protein_coding_gene | -3.5 | 0.09 | 3.9E-03 |
| col-121 | WBGene00000695 | protein_coding_gene | -3.5 | 0.09 | 6.2E-26 |
| clec-60 | WBGene00014046 | protein_coding_gene | -3.5 | 0.09 | 2.4E-11 |
| ZK1307.2 | WBGene00014245 | protein_coding_gene | -3.5 | 0.09 | 1.3E-06 |
| T22F3.8 | WBGene00020698 | protein_coding_gene | -3.6 | 0.08 | 2.6E-04 |
| pitr-4 | WBGene00007141 | protein_coding_gene | -3.6 | 0.08 | 5.7E-04 |
| tsp-2 | WBGene00006628 | protein_coding_gene | -3.6 | 0.08 | 3.3E-14 |
| W10G11.1 | WBGene00021134 | protein_coding_gene | -3.6 | 0.08 | 2.0E-03 |
| C08E8.3 | WBGene00007439 | protein_coding_gene | -3.7 | 0.08 | 1.1E-05 |
| nspe-6 | WBGene00012603 | protein_coding_gene | -3.7 | 0.08 | 7.8E-04 |
| clec-23 | WBGene00011851 | pseudogene | -3.7 | 0.08 | 4.6E-05 |
| ZC84.1 | WBGene00013846 | protein_coding_gene | -3.7 | 0.07 | 4.9E-06 |
| cut-5 | WBGene00011104 | protein_coding_gene | -3.7 | 0.07 | 4.0E-07 |
| pals-32 | WBGene00016281 | protein_coding_gene | -3.8 | 0.07 | 1.8E-14 |
| ugt-34 | WBGene00009255 | protein_coding_gene | -3.8 | 0.07 | 2.0E-03 |
| T10C6.15 | WBGene00050899 | protein_coding_gene | -3.8 | 0.07 | 1.3E-11 |
| F17B5.1 | WBGene00008905 | protein_coding_gene | -3.9 | 0.07 | 1.3E-06 |
| fbxa-164 | WBGene00015599 | protein_coding_gene | -3.9 | 0.07 | 4.8E-05 |
| F21H7.12 | WBGene00009033 | protein_coding_gene | -4.0 | 0.06 | 1.0E-04 |
| W07B8.4 | WBGene00021072 | protein_coding_gene | -4.0 | 0.06 | 9.8E-09 |
| C06E7.88 | WBGene00189995 | protein_coding_gene | -4.0 | 0.06 | 1.2E-09 |
| str-163 | WBGene00006208 | protein_coding_gene | -4.0 | 0.06 | 4.8E-05 |
| F49H6.5 | WBGene00009908 | protein_coding_gene | -4.1 | 0.06 | 4.7E-20 |
| dct-7 | WBGene00017488 | protein_coding_gene | -4.1 | 0.06 | 1.1E-06 |
| F56D6.9 | WBGene00044471 | protein_coding_gene | -4.1 | 0.06 | 1.5E-04 |
| cyp-13A7 | WBGene00000372 | protein_coding_gene | -4.1 | 0.06 | 3.4E-14 |
| ins-8 | WBGene00002091 | protein_coding_gene | -4.7 | 0.04 | 1.1E-04 |
| T16G1.7 | WBGene00011801 | protein_coding_gene | -4.7 | 0.04 | 2.8E-97 |
| F56D6.8 | WBGene00044292 | protein_coding_gene | -4.8 | 0.04 | 3.1E-05 |
| clec-61 | WBGene00014047 | protein_coding_gene | -4.8 | 0.04 | 1.6E-15 |
| M02E1.4 | WBGene00284848 | protein_coding_gene | -4.8 | 0.04 | 3.2E-07 |
| W02D7.11 | WBGene00020946 | protein_coding_gene | -5.1 | 0.03 | 4.2E-07 |
| cyp-13A5 | WBGene00011672 | protein_coding_gene | -5.1 | 0.03 | 2.1E-102 |
| dct-8 | WBGene00044472 | protein_coding_gene | -5.8 | 0.02 | 3.2E-09 |

Shared (N2&CB4856) DEGs in response to *P. rettgeri* (vs. OP50)

| | | | | N2 | | | CB4856 | |
|---|---|---|---|---|---|---|---|---|
| Gene Name | Wormbase Gene ID | Type | $log_2FC$ | FC | Adj P-value | $log_2FC$ | FC | Adj P-value |
| *F46C5.1* | WBGene00009778 | protein_coding_gene | 5.4 | 41.77 | 4.3E-08 | -4.1 | 0.06 | 4.5E-11 |
| *mtl-1* | WBGene00003473 | protein_coding_gene | 4.9 | 30.50 | 6.4E-07 | -6.6 | 0.01 | 4.9E-44 |
| *F48C1.9* | WBGene00018601 | protein_coding_gene | 4.9 | 29.92 | 8.6E-05 | -2.6 | 0.16 | 8.5E-09 |
| *F45D3.4* | WBGene00009724 | protein_coding_gene | 4.7 | 25.65 | 3.7E-09 | -4.5 | 0.04 | 4.7E-51 |
| *T07G12.5* | WBGene00011596 | protein_coding_gene | 4.6 | 23.44 | 1.9E-05 | -5.6 | 0.02 | 4.2E-70 |
| *fipr-24* | WBGene00007992 | protein_coding_gene | 4.4 | 21.06 | 9.9E-04 | -2.4 | 0.18 | 6.6E-04 |
| *ugt-18* | WBGene00013900 | protein_coding_gene | 4.2 | 17.94 | 1.9E-04 | -4.9 | 0.03 | 3.9E-21 |
| *C37A5.3* | WBGene00014698 | protein_coding_gene | 4.2 | 17.92 | 2.0E-06 | -2.3 | 0.20 | 3.4E-02 |
| *cest-29* | WBGene00017205 | pseudogene | 4.1 | 17.70 | 1.2E-03 | -4.3 | 0.05 | 5.6E-17 |
| *tts-1* | WBGene00006650 | ncRNA_gene | 4.1 | 16.61 | 6.2E-07 | -4.8 | 0.04 | 2.2E-74 |
| *C18H7.1* | WBGene00015993 | protein_coding_gene | 4.0 | 16.50 | 2.8E-05 | -5.9 | 0.02 | 7.7E-13 |
| *F47B8.4* | WBGene00009805 | protein_coding_gene | 4.0 | 16.18 | 1.2E-03 | -5.6 | 0.02 | 3.0E-10 |
| *T26H5.9* | WBGene00044206 | protein_coding_gene | 4.0 | 15.67 | 1.1E-04 | -4.7 | 0.04 | 1.7E-65 |
| *C06B3.7* | WBGene00007366 | protein_coding_gene | 3.9 | 14.81 | 3.1E-04 | -2.0 | 0.25 | 4.1E-04 |
| *cpt-3* | WBGene00021703 | protein_coding_gene | 3.7 | 12.82 | 5.0E-03 | -3.6 | 0.08 | 8.0E-15 |
| *C50F7.5* | WBGene00016845 | protein_coding_gene | 3.6 | 12.44 | 5.6E-04 | -5.6 | 0.02 | 1.3E-109 |

| | | | | | | | | |
|---|---|---|---|---|---|---|---|---|
| *srh-2* | WBGene00005228 | protein_coding_gene | 3.6 | 12.02 | 4.7E-06 | -2.4 | 0.19 | 1.6E-03 |
| *cyp-37B1* | WBGene00009226 | protein_coding_gene | 3.6 | 11.96 | 2.1E-04 | -5.5 | 0.02 | 2.1E-52 |
| *F46A8.13* | WBGene00219950 | ncRNA_gene | 3.5 | 11.65 | 2.7E-03 | -2.5 | 0.18 | 2.1E-07 |
| *scl-2* | WBGene00009895 | protein_coding_gene | 3.5 | 11.34 | 2.3E-04 | -5.0 | 0.03 | 6.7E-157 |
| *T26H5.4* | WBGene00012069 | protein_coding_gene | 3.5 | 10.97 | 1.8E-03 | -3.3 | 0.10 | 1.4E-05 |
| *fipr-23* | WBGene00007990 | protein_coding_gene | 3.4 | 10.83 | 4.0E-03 | -2.8 | 0.14 | 1.6E-03 |
| *lipl-2* | WBGene00009773 | protein_coding_gene | 3.4 | 10.51 | 7.6E-04 | -4.6 | 0.04 | 2.9E-32 |
| *cyp-32B1* | WBGene00021167 | protein_coding_gene | 3.3 | 9.81 | 1.5E-04 | -3.5 | 0.09 | 1.1E-13 |
| *ilys-3* | WBGene00016670 | protein_coding_gene | 3.2 | 9.49 | 2.7E-03 | -7.2 | 0.01 | 5.8E-55 |
| *T20B3.1* | WBGene00011850 | protein_coding_gene | 3.2 | 9.47 | 3.9E-08 | -2.8 | 0.15 | 4.7E-31 |
| *irg-2* | WBGene00016783 | protein_coding_gene | 3.2 | 9.24 | 3.2E-03 | -2.6 | 0.17 | 1.1E-04 |
| *F53A9.8* | WBGene00018731 | protein_coding_gene | 3.2 | 8.93 | 7.2E-05 | -4.0 | 0.06 | 3.2E-42 |
| *cnc-11* | WBGene00044900 | protein_coding_gene | 3.1 | 8.82 | 1.7E-02 | -4.5 | 0.04 | 7.8E-06 |
| *fbxa-24* | WBGene00018803 | protein_coding_gene | 3.1 | 8.70 | 7.1E-05 | -4.2 | 0.05 | 4.5E-33 |
| *T23E7.6* | WBGene00044772 | protein_coding_gene | 3.1 | 8.70 | 1.6E-04 | -3.7 | 0.08 | 1.1E-22 |
| *tre-5* | WBGene00006611 | protein_coding_gene | 3.1 | 8.62 | 4.4E-06 | -3.4 | 0.09 | 4.1E-27 |
| *cnc-4* | WBGene00000558 | protein_coding_gene | 3.0 | 8.11 | 1.2E-07 | -2.4 | 0.19 | 1.6E-04 |
| *F47B8.10* | WBGene00009811 | protein_coding_gene | 3.0 | 8.08 | 8.1E-03 | -2.7 | 0.15 | 1.5E-05 |
| *ech-9* | WBGene00001158 | protein_coding_gene | 3.0 | 8.04 | 1.4E-04 | -2.4 | 0.18 | 6.9E-04 |
| *ptr-22* | WBGene00004236 | protein_coding_gene | 3.0 | 7.88 | 1.7E-04 | -1.8 | 0.29 | 2.4E-18 |
| *F15E6.3* | WBGene00017484 | protein_coding_gene | 3.0 | 7.87 | 7.2E-07 | -4.1 | 0.06 | 2.7E-41 |
| *nhr-127* | WBGene00003717 | protein_coding_gene | 3.0 | 7.81 | 6.0E-03 | -2.2 | 0.22 | 4.2E-05 |
| *cyp-34A9* | WBGene00015044 | protein_coding_gene | 3.0 | 7.81 | 1.0E-03 | -3.5 | 0.09 | 4.1E-129 |
| *ZC443.3* | WBGene00013898 | protein_coding_gene | 3.0 | 7.78 | 1.1E-06 | -3.8 | 0.07 | 6.0E-91 |
| *T26H5.14* | WBGene00235133 | protein_coding_gene | 2.9 | 7.68 | 3.1E-03 | -3.2 | 0.11 | 1.7E-07 |
| *swt-1* | WBGene00019300 | protein_coding_gene | 2.9 | 7.67 | 1.5E-07 | -2.0 | 0.24 | 5.6E-15 |
| *irg-1* | WBGene00015574 | protein_coding_gene | 2.9 | 7.60 | 1.4E-04 | -1.8 | 0.30 | 1.1E-02 |
| *coel-1* | WBGene00016866 | protein_coding_gene | 2.9 | 7.59 | 1.2E-05 | -3.1 | 0.11 | 2.4E-56 |
| *K10D11.2* | WBGene00010746 | protein_coding_gene | 2.9 | 7.51 | 2.7E-02 | -2.9 | 0.13 | 3.7E-03 |
| *K10G6.4* | WBGene00019641 | protein_coding_gene | 2.9 | 7.38 | 1.3E-02 | -2.9 | 0.13 | 8.7E-06 |
| *Y43F8B.9* | WBGene00012819 | protein_coding_gene | 2.9 | 7.32 | 2.5E-07 | -1.7 | 0.30 | 2.4E-02 |
| *F11E6.11* | WBGene00023501 | protein_coding_gene | 2.9 | 7.30 | 2.5E-05 | -4.9 | 0.03 | 4.5E-33 |
| *Y41D4B.6* | WBGene00021513 | protein_coding_gene | 2.8 | 7.20 | 3.4E-03 | -3.7 | 0.08 | 7.3E-14 |
| *F09F7.6* | WBGene00017303 | protein_coding_gene | 2.8 | 6.96 | 2.1E-08 | -2.6 | 0.16 | 1.9E-33 |
| *F10D7.3* | WBGene00017340 | protein_coding_gene | 2.8 | 6.84 | 7.8E-06 | -2.7 | 0.15 | 1.1E-30 |
| *C25H3.10* | WBGene00016119 | protein_coding_gene | 2.8 | 6.83 | 1.0E-06 | -3.2 | 0.11 | 6.9E-43 |
| *cpt-4* | WBGene00019644 | protein_coding_gene | 2.8 | 6.79 | 2.1E-04 | -2.7 | 0.16 | 8.5E-23 |
| *ilys-2* | WBGene00016669 | protein_coding_gene | 2.8 | 6.75 | 1.8E-03 | -3.0 | 0.13 | 4.0E-05 |
| *oac-14* | WBGene00008602 | protein_coding_gene | 2.7 | 6.64 | 4.8E-03 | -5.5 | 0.02 | 4.2E-39 |
| *EGAP1.1* | WBGene00017146 | protein_coding_gene | 2.7 | 6.62 | 3.4E-03 | -2.2 | 0.22 | 4.3E-03 |
| *lipl-3* | WBGene00020016 | protein_coding_gene | 2.7 | 6.57 | 1.8E-03 | -7.4 | 0.01 | 7.7E-26 |
| *cyp-34A2* | WBGene00011699 | protein_coding_gene | 2.6 | 6.19 | 4.3E-04 | -3.0 | 0.13 | 2.8E-09 |
| *sodh-1* | WBGene00010790 | protein_coding_gene | 2.6 | 6.17 | 2.5E-07 | -2.8 | 0.14 | 1.1E-08 |
| *nnt-1* | WBGene00003778 | protein_coding_gene | 2.6 | 6.14 | 1.3E-06 | -3.1 | 0.12 | 1.8E-46 |
| *ins-20* | WBGene00002103 | protein_coding_gene | 2.6 | 6.12 | 3.7E-02 | -3.6 | 0.08 | 2.4E-11 |
| *F37A8.5* | WBGene00009503 | protein_coding_gene | 2.6 | 5.95 | 1.9E-04 | -3.4 | 0.10 | 8.2E-49 |
| *T12D8.5* | WBGene00011733 | protein_coding_gene | 2.6 | 5.95 | 1.2E-05 | -1.9 | 0.27 | 7.4E-09 |
| *F09F3.8* | WBGene00014732 | pseudogene | 2.6 | 5.94 | 1.2E-02 | -1.5 | 0.35 | 3.9E-02 |
| *Y65B4BL.1* | WBGene00022033 | protein_coding_gene | 2.6 | 5.93 | 3.7E-02 | -2.7 | 0.15 | 4.4E-05 |
| *srr-6* | WBGene00005657 | protein_coding_gene | 2.6 | 5.89 | 1.2E-02 | -3.8 | 0.07 | 5.6E-11 |
| *fbxa-48* | WBGene00021862 | protein_coding_gene | 2.6 | 5.89 | 5.6E-03 | -3.2 | 0.11 | 7.7E-09 |
| *wrt-2* | WBGene00006948 | protein_coding_gene | 2.5 | 5.82 | 3.0E-04 | -2.6 | 0.16 | 9.2E-03 |
| *T04C12.7* | WBGene00011429 | protein_coding_gene | 2.5 | 5.80 | 3.7E-04 | -2.8 | 0.15 | 2.6E-07 |
| *fil-1* | WBGene00011321 | protein_coding_gene | 2.5 | 5.79 | 1.2E-03 | -5.3 | 0.03 | 4.2E-39 |
| *fmo-1* | WBGene00001476 | protein_coding_gene | 2.5 | 5.72 | 3.0E-03 | -1.1 | 0.45 | 1.1E-10 |
| *T05A7.13* | WBGene00271415 | protein_coding_gene | 2.5 | 5.57 | 9.4E-03 | -2.2 | 0.22 | 8.3E-06 |
| *W03F9.4* | WBGene00021002 | protein_coding_gene | 2.5 | 5.47 | 2.1E-08 | -1.2 | 0.42 | 7.6E-03 |
| *C08F11.13* | WBGene00007460 | protein_coding_gene | 2.4 | 5.45 | 1.2E-04 | -3.4 | 0.10 | 7.6E-59 |

| | | | | | | | | |
|---|---|---|---|---|---|---|---|---|
| *unc-42* | WBGene00006778 | protein_coding_gene | 2.4 | 5.44 | 1.1E-03 | -2.0 | 0.25 | 4.2E-05 |
| *T22F3.11* | WBGene00020700 | protein_coding_gene | 2.4 | 5.43 | 2.3E-02 | -5.3 | 0.02 | 1.9E-09 |
| *C06B3.6* | WBGene00007365 | protein_coding_gene | 2.4 | 5.42 | 1.3E-06 | -2.1 | 0.24 | 4.7E-23 |
| *ZK675.4* | WBGene00014067 | protein_coding_gene | 2.4 | 5.41 | 3.1E-02 | -3.2 | 0.11 | 6.7E-05 |
| *T05A7.11* | WBGene00044383 | protein_coding_gene | 2.4 | 5.32 | 3.5E-02 | -3.9 | 0.07 | 1.0E-14 |
| *T08G5.1* | WBGene00011622 | protein_coding_gene | 2.4 | 5.27 | 3.0E-02 | -1.9 | 0.26 | 2.8E-02 |
| *Y47H9C.1* | WBGene00012947 | protein_coding_gene | 2.4 | 5.25 | 9.8E-04 | 4.4 | 20.89 | 2.9E-15 |
| *ZC443.2* | WBGene00013897 | protein_coding_gene | 2.4 | 5.23 | 3.0E-02 | -3.1 | 0.12 | 2.2E-06 |
| *Y7A5A.6* | WBGene00012412 | protein_coding_gene | 2.4 | 5.20 | 3.5E-02 | -2.4 | 0.19 | 1.5E-04 |
| *cyp-34A10* | WBGene00015045 | protein_coding_gene | 2.4 | 5.19 | 1.3E-02 | -4.6 | 0.04 | 1.3E-19 |
| *nlp-39* | WBGene00008295 | protein_coding_gene | 2.4 | 5.19 | 2.5E-02 | -3.4 | 0.10 | 5.3E-07 |
| *cup-16* | WBGene00007919 | protein_coding_gene | 2.3 | 4.99 | 7.2E-05 | -3.0 | 0.12 | 4.8E-32 |
| *ttr-23* | WBGene00011894 | protein_coding_gene | 2.3 | 4.99 | 4.7E-04 | -1.6 | 0.33 | 6.0E-09 |
| *T16G1.4* | WBGene00011798 | protein_coding_gene | 2.3 | 4.95 | 2.4E-02 | -4.7 | 0.04 | 2.3E-25 |
| *parg-2* | WBGene00004052 | protein_coding_gene | 2.3 | 4.94 | 5.1E-03 | -2.0 | 0.26 | 3.4E-03 |
| *F59C6.16* | WBGene00077764 | protein_coding_gene | 2.3 | 4.90 | 3.3E-04 | -2.9 | 0.14 | 1.2E-37 |
| *hch-1* | WBGene00001828 | protein_coding_gene | 2.3 | 4.90 | 1.9E-03 | -2.5 | 0.17 | 6.9E-23 |
| *Y46E12BR.1* | WBGene00021598 | protein_coding_gene | 2.3 | 4.87 | 3.6E-02 | -3.1 | 0.11 | 4.7E-04 |
| *K01F9.2* | WBGene00010475 | protein_coding_gene | 2.3 | 4.81 | 1.0E-04 | -2.1 | 0.23 | 1.7E-12 |
| *Y53F4B.45* | WBGene00044728 | ncRNA_gene | 2.3 | 4.81 | 9.3E-05 | -4.1 | 0.06 | 6.1E-62 |
| *Y53C12B.7* | WBGene00013146 | protein_coding_gene | 2.3 | 4.78 | 1.4E-03 | -2.1 | 0.23 | 7.2E-12 |
| *nhr-99* | WBGene00003689 | protein_coding_gene | 2.2 | 4.64 | 1.9E-02 | -3.2 | 0.11 | 2.9E-21 |
| *ftn-1* | WBGene00001500 | protein_coding_gene | 2.2 | 4.64 | 2.0E-02 | -3.7 | 0.08 | 4.5E-15 |
| *cutl-9* | WBGene00010893 | protein_coding_gene | 2.2 | 4.62 | 4.0E-03 | -2.8 | 0.15 | 4.6E-15 |
| *Y17G9A.4* | WBGene00021197 | protein_coding_gene | 2.2 | 4.58 | 3.7E-04 | -3.2 | 0.11 | 8.7E-21 |
| *cutl-28* | WBGene00018256 | protein_coding_gene | 2.2 | 4.56 | 2.0E-03 | -2.5 | 0.17 | 1.3E-04 |
| *far-7* | WBGene00001391 | protein_coding_gene | 2.2 | 4.53 | 4.7E-04 | -3.8 | 0.07 | 2.3E-44 |
| *C08F1.11* | WBGene00044417 | protein_coding_gene | 2.2 | 4.52 | 5.6E-03 | -2.0 | 0.26 | 3.9E-02 |
| *spp-2* | WBGene00004987 | protein_coding_gene | 2.2 | 4.51 | 1.4E-05 | -3.1 | 0.12 | 2.7E-61 |
| *dsl-6* | WBGene00001108 | protein_coding_gene | 2.2 | 4.51 | 3.4E-02 | -3.7 | 0.08 | 8.6E-12 |
| *C44H9.7* | WBGene00008102 | protein_coding_gene | 2.2 | 4.49 | 1.0E-02 | -3.2 | 0.11 | 1.3E-10 |
| *dod-3* | WBGene00016052 | protein_coding_gene | 2.2 | 4.47 | 4.5E-02 | -4.3 | 0.05 | 2.9E-12 |
| *asp-11* | WBGene00017677 | pseudogene | 2.2 | 4.46 | 2.3E-02 | -3.4 | 0.09 | 7.0E-05 |
| *dsl-3* | WBGene00001105 | protein_coding_gene | 2.2 | 4.45 | 3.1E-05 | -1.8 | 0.28 | 5.5E-08 |
| *F43C11.7* | WBGene00018384 | protein_coding_gene | 2.1 | 4.39 | 3.9E-05 | -3.7 | 0.08 | 1.6E-49 |
| *F58F12.12* | WBGene00271812 | protein_coding_gene | 2.1 | 4.34 | 2.9E-03 | -2.3 | 0.20 | 4.6E-06 |
| *C54F6.5* | WBGene00016923 | protein_coding_gene | 2.1 | 4.31 | 1.1E-02 | -2.7 | 0.16 | 3.2E-12 |
| *C35C5.8* | WBGene00007957 | protein_coding_gene | 2.1 | 4.31 | 2.0E-05 | -2.7 | 0.16 | 2.8E-13 |
| *peb-1* | WBGene00003968 | protein_coding_gene | 2.1 | 4.28 | 1.2E-03 | -2.7 | 0.16 | 1.8E-19 |
| *best-5* | WBGene00007404 | protein_coding_gene | 2.1 | 4.26 | 7.0E-04 | -1.3 | 0.40 | 2.9E-05 |
| *clec-25* | WBGene00044331 | protein_coding_gene | 2.1 | 4.25 | 3.6E-02 | -1.4 | 0.38 | 1.0E-02 |
| *tsp-17* | WBGene00006643 | protein_coding_gene | 2.1 | 4.22 | 5.6E-03 | -2.2 | 0.22 | 1.0E-05 |
| *ubc-8* | WBGene00006705 | protein_coding_gene | 2.1 | 4.19 | 2.0E-05 | -2.2 | 0.22 | 3.5E-54 |
| *clec-78* | WBGene00018547 | protein_coding_gene | 2.1 | 4.15 | 4.6E-03 | -2.5 | 0.17 | 1.2E-18 |
| *dpy-14* | WBGene00001075 | protein_coding_gene | 2.1 | 4.15 | 1.6E-02 | -3.0 | 0.13 | 4.5E-12 |
| *H42K12.3* | WBGene00019272 | protein_coding_gene | 2.1 | 4.15 | 4.4E-04 | -3.0 | 0.13 | 4.0E-04 |
| *Y54G2A.11* | WBGene00021876 | protein_coding_gene | 2.0 | 4.12 | 2.5E-04 | -2.0 | 0.25 | 1.6E-26 |
| *egl-46* | WBGene00001210 | protein_coding_gene | 2.0 | 4.11 | 5.6E-04 | -2.7 | 0.16 | 6.1E-20 |
| *mlt-11* | WBGene00012186 | protein_coding_gene | 2.0 | 4.09 | 7.8E-04 | -2.4 | 0.19 | 5.0E-03 |
| *daao-1* | WBGene00022076 | protein_coding_gene | 2.0 | 4.07 | 5.3E-05 | -1.1 | 0.48 | 1.8E-03 |
| *F11E6.9* | WBGene00008712 | protein_coding_gene | 2.0 | 4.07 | 4.2E-02 | -3.7 | 0.08 | 1.1E-05 |
| *M01B2.13* | WBGene00045338 | protein_coding_gene | 2.0 | 4.03 | 4.0E-02 | -3.1 | 0.12 | 4.7E-09 |
| *cdf-2* | WBGene00011821 | protein_coding_gene | 2.0 | 4.02 | 5.0E-04 | -2.6 | 0.17 | 2.9E-22 |
| *hil-7* | WBGene00001858 | protein_coding_gene | 2.0 | 4.02 | 4.8E-02 | -3.3 | 0.10 | 2.2E-19 |
| *gbh-2* | WBGene00001523 | protein_coding_gene | 2.0 | 3.98 | 4.3E-05 | -2.0 | 0.26 | 6.3E-19 |
| *ZK6.8* | WBGene00022643 | protein_coding_gene | 2.0 | 3.98 | 1.5E-03 | -2.4 | 0.19 | 1.6E-09 |
| *oac-6* | WBGene00007833 | protein_coding_gene | 2.0 | 3.96 | 1.5E-04 | -2.0 | 0.25 | 1.2E-05 |
| *R09E10.5* | WBGene00011175 | protein_coding_gene | 2.0 | 3.94 | 1.6E-02 | -3.1 | 0.12 | 2.6E-07 |

| | | | | | | | | |
|---|---|---|---|---|---|---|---|---|
| *C35A5.11* | WBGene00044024 | protein_coding_gene | 2.0 | 3.93 | 1.7E-02 | -3.7 | 0.08 | 1.2E-11 |
| *F31D4.5* | WBGene00009292 | protein_coding_gene | 2.0 | 3.90 | 1.9E-03 | -3.4 | 0.10 | 1.8E-08 |
| *C05D9.9* | WBGene00015476 | protein_coding_gene | 2.0 | 3.89 | 1.7E-03 | -2.2 | 0.22 | 1.1E-23 |
| *lipl-1* | WBGene00010062 | protein_coding_gene | 2.0 | 3.88 | 3.7E-02 | -5.0 | 0.03 | 2.0E-16 |
| *R10E8.8* | WBGene00011213 | protein_coding_gene | 2.0 | 3.88 | 1.9E-03 | -2.1 | 0.24 | 2.5E-11 |
| *M60.7* | WBGene00019782 | protein_coding_gene | 2.0 | 3.87 | 8.1E-03 | -1.9 | 0.27 | 1.3E-05 |
| *pqn-73* | WBGene00004155 | protein_coding_gene | 1.9 | 3.86 | 4.9E-02 | -2.9 | 0.13 | 1.0E-07 |
| *K01A2.10* | WBGene00019284 | protein_coding_gene | 1.9 | 3.85 | 7.2E-04 | -2.4 | 0.20 | 5.1E-21 |
| *F43D9.1* | WBGene00009653 | protein_coding_gene | 1.9 | 3.84 | 3.6E-03 | -3.0 | 0.12 | 2.6E-13 |
| *K06A9.2* | WBGene00019436 | protein_coding_gene | 1.9 | 3.83 | 3.6E-03 | -2.7 | 0.15 | 2.1E-20 |
| *F31E3.12* | WBGene00219317 | protein_coding_gene | 1.9 | 3.81 | 3.8E-02 | -2.1 | 0.23 | 7.1E-03 |
| *fbxa-21* | WBGene00022487 | protein_coding_gene | 1.9 | 3.73 | 1.9E-03 | -2.9 | 0.14 | 8.9E-21 |
| *K08B12.1* | WBGene00019520 | protein_coding_gene | 1.9 | 3.72 | 9.9E-03 | -2.7 | 0.15 | 2.1E-03 |
| *nhr-19* | WBGene00003618 | protein_coding_gene | 1.9 | 3.72 | 1.6E-04 | -1.8 | 0.29 | 5.7E-18 |
| *dpy-17* | WBGene00001076 | protein_coding_gene | 1.9 | 3.71 | 6.7E-04 | -2.7 | 0.15 | 3.0E-17 |
| *icl-1* | WBGene00001564 | protein_coding_gene | 1.9 | 3.71 | 1.9E-04 | -1.7 | 0.30 | 6.7E-23 |
| *fbxc-24* | WBGene00019927 | protein_coding_gene | 1.9 | 3.71 | 7.1E-03 | -1.7 | 0.31 | 7.7E-04 |
| *cyp-35D1* | WBGene00008829 | protein_coding_gene | 1.9 | 3.68 | 2.7E-04 | 2.0 | 3.95 | 6.0E-05 |
| *dpy-3* | WBGene00001065 | protein_coding_gene | 1.9 | 3.67 | 4.5E-03 | -2.2 | 0.22 | 2.1E-02 |
| *nlp-28* | WBGene00003766 | protein_coding_gene | 1.9 | 3.64 | 5.5E-05 | -1.3 | 0.39 | 3.9E-12 |
| *siah-1* | WBGene00021369 | protein_coding_gene | 1.9 | 3.62 | 4.7E-04 | -1.1 | 0.46 | 7.8E-13 |
| *lpr-4* | WBGene00012257 | protein_coding_gene | 1.8 | 3.60 | 2.0E-03 | -3.0 | 0.12 | 8.8E-04 |
| *C27A7.8* | WBGene00007758 | protein_coding_gene | 1.8 | 3.59 | 1.9E-05 | -1.4 | 0.37 | 3.4E-07 |
| *F59B2.8* | WBGene00010310 | protein_coding_gene | 1.8 | 3.59 | 5.7E-03 | -1.8 | 0.29 | 1.0E-04 |
| *nlp-29* | WBGene00003767 | protein_coding_gene | 1.8 | 3.55 | 2.5E-03 | -1.5 | 0.35 | 2.0E-14 |
| *F14F9.4* | WBGene00017467 | protein_coding_gene | 1.8 | 3.54 | 1.7E-02 | -1.1 | 0.47 | 6.2E-05 |
| *F53A9.7* | WBGene00018730 | protein_coding_gene | 1.8 | 3.51 | 2.2E-04 | -2.5 | 0.18 | 4.0E-27 |
| *ifa-3* | WBGene00002051 | protein_coding_gene | 1.8 | 3.49 | 2.6E-02 | -3.1 | 0.12 | 2.0E-10 |
| *oac-31* | WBGene00018707 | protein_coding_gene | 1.8 | 3.48 | 7.8E-03 | -1.2 | 0.44 | 1.8E-02 |
| *lpr-3* | WBGene00012261 | protein_coding_gene | 1.8 | 3.47 | 6.4E-04 | -2.4 | 0.20 | 4.3E-03 |
| *dpy-7* | WBGene00001069 | protein_coding_gene | 1.8 | 3.45 | 2.8E-03 | -2.5 | 0.18 | 2.1E-03 |
| *unk-1* | WBGene00016407 | protein_coding_gene | 1.8 | 3.43 | 5.5E-03 | -2.0 | 0.25 | 1.2E-39 |
| *glc-1* | WBGene00001591 | protein_coding_gene | 1.8 | 3.42 | 1.4E-04 | -3.8 | 0.07 | 5.2E-66 |
| *T28H10.3* | WBGene00012144 | protein_coding_gene | 1.8 | 3.41 | 2.0E-03 | -2.8 | 0.15 | 2.2E-122 |
| *T14B4.19* | WBGene00302993 | protein_coding_gene | 1.8 | 3.39 | 6.3E-03 | -2.4 | 0.19 | 2.1E-03 |
| *F53F4.8* | WBGene00009991 | pseudogene | 1.8 | 3.39 | 4.0E-02 | -3.1 | 0.12 | 5.5E-09 |
| *ceh-30* | WBGene00000451 | protein_coding_gene | 1.8 | 3.39 | 2.2E-02 | -1.3 | 0.41 | 4.7E-02 |
| *C04A11.5* | WBGene00007288 | protein_coding_gene | 1.8 | 3.39 | 4.3E-02 | -2.3 | 0.20 | 9.7E-06 |
| *Y51H4A.5* | WBGene00013101 | protein_coding_gene | 1.8 | 3.39 | 1.4E-02 | -2.7 | 0.15 | 6.5E-09 |
| *Y44A6C.1* | WBGene00012850 | protein_coding_gene | 1.8 | 3.37 | 1.0E-05 | -1.7 | 0.31 | 7.2E-24 |
| *F13E9.11* | WBGene00008760 | protein_coding_gene | 1.7 | 3.36 | 2.6E-04 | -2.5 | 0.18 | 6.6E-22 |
| *kreg-1* | WBGene00018725 | protein_coding_gene | 1.7 | 3.36 | 3.3E-02 | -3.0 | 0.13 | 1.0E-05 |
| *D2045.8* | WBGene00008425 | protein_coding_gene | 1.7 | 3.35 | 3.7E-04 | -1.4 | 0.39 | 1.6E-06 |
| *T12A7.6* | WBGene00011727 | protein_coding_gene | 1.7 | 3.33 | 5.5E-03 | -2.4 | 0.20 | 1.1E-19 |
| *cutl-2* | WBGene00013145 | protein_coding_gene | 1.7 | 3.33 | 9.3E-03 | -2.6 | 0.16 | 4.5E-16 |
| *T28F4.5* | WBGene00012140 | protein_coding_gene | 1.7 | 3.32 | 1.4E-04 | -1.7 | 0.31 | 3.4E-20 |
| *ugt-52* | WBGene00018931 | protein_coding_gene | 1.7 | 3.32 | 1.1E-02 | -1.7 | 0.30 | 3.9E-08 |
| *R03H10.2* | WBGene00019855 | protein_coding_gene | 1.7 | 3.32 | 1.6E-03 | -2.0 | 0.24 | 7.9E-12 |
| *lpr-5* | WBGene00012256 | protein_coding_gene | 1.7 | 3.32 | 2.3E-03 | -2.6 | 0.16 | 2.1E-03 |
| *C50F4.9* | WBGene00008234 | protein_coding_gene | 1.7 | 3.31 | 2.6E-04 | -1.4 | 0.39 | 1.0E-03 |
| *dhs-18* | WBGene00000981 | protein_coding_gene | 1.7 | 3.30 | 4.0E-04 | -1.4 | 0.38 | 7.7E-20 |
| *F56D2.3* | WBGene00018965 | protein_coding_gene | 1.7 | 3.29 | 9.5E-03 | -2.2 | 0.22 | 1.9E-02 |
| *ceh-16* | WBGene00000439 | protein_coding_gene | 1.7 | 3.27 | 1.9E-02 | -1.5 | 0.36 | 1.6E-03 |
| *zip-7* | WBGene00013100 | protein_coding_gene | 1.7 | 3.27 | 1.4E-03 | -1.6 | 0.33 | 3.6E-06 |
| *noah-1* | WBGene00016422 | protein_coding_gene | 1.7 | 3.27 | 1.6E-04 | -2.3 | 0.21 | 2.9E-04 |
| *hlh-16* | WBGene00001960 | protein_coding_gene | 1.7 | 3.25 | 4.8E-02 | -2.0 | 0.25 | 4.7E-05 |
| *noah-2* | WBGene00009926 | protein_coding_gene | 1.7 | 3.24 | 7.0E-04 | -2.4 | 0.20 | 6.9E-04 |
| *T02G5.15* | WBGene00206521 | protein_coding_gene | 1.7 | 3.21 | 3.7E-02 | -1.5 | 0.34 | 9.3E-04 |

| | | | | | | | | |
|---|---|---|---|---|---|---|---|---|
| *ugt-54* | WBGene00012013 | protein_coding_gene | 1.7 | 3.21 | 4.1E-03 | -3.0 | 0.13 | 7.9E-35 |
| *dpy-2* | WBGene00001064 | protein_coding_gene | 1.7 | 3.20 | 4.4E-03 | -2.6 | 0.16 | 7.9E-04 |
| *slc-36.5* | WBGene00016628 | protein_coding_gene | 1.7 | 3.19 | 2.2E-03 | -2.4 | 0.19 | 2.5E-16 |
| *ces-2* | WBGene00000469 | protein_coding_gene | 1.7 | 3.19 | 4.4E-03 | -2.4 | 0.19 | 9.2E-18 |
| *C09F9.2* | WBGene00007479 | protein_coding_gene | 1.7 | 3.19 | 1.4E-03 | -2.3 | 0.20 | 3.4E-07 |
| *cutl-29* | WBGene00017421 | protein_coding_gene | 1.7 | 3.17 | 4.8E-02 | -1.3 | 0.39 | 4.6E-02 |
| *F41E6.5* | WBGene00018286 | protein_coding_gene | 1.7 | 3.16 | 5.9E-04 | -1.6 | 0.32 | 7.2E-12 |
| *hbl-1* | WBGene00001824 | protein_coding_gene | 1.7 | 3.15 | 1.4E-02 | -2.4 | 0.19 | 6.6E-16 |
| *ugt-57* | WBGene00017154 | protein_coding_gene | 1.7 | 3.15 | 2.3E-02 | -2.0 | 0.25 | 4.2E-04 |
| *F18G5.6* | WBGene00017582 | protein_coding_gene | 1.7 | 3.15 | 2.0E-03 | -2.5 | 0.18 | 4.8E-54 |
| *ikb-1* | WBGene00002069 | protein_coding_gene | 1.6 | 3.13 | 9.0E-03 | -2.1 | 0.23 | 9.1E-24 |
| *B0310.3* | WBGene00015139 | protein_coding_gene | 1.6 | 3.13 | 8.2E-03 | -1.8 | 0.29 | 5.2E-06 |
| *F16C3.2* | WBGene00008886 | protein_coding_gene | 1.6 | 3.12 | 1.0E-03 | -2.7 | 0.15 | 9.9E-33 |
| *math-34* | WBGene00019699 | protein_coding_gene | 1.6 | 3.11 | 2.2E-02 | -2.5 | 0.18 | 1.0E-08 |
| *sym-1* | WBGene00006366 | protein_coding_gene | 1.6 | 3.08 | 1.9E-04 | -2.0 | 0.24 | 2.2E-03 |
| *C29F7.2* | WBGene00007811 | protein_coding_gene | 1.6 | 3.07 | 5.1E-04 | -1.6 | 0.32 | 2.7E-30 |
| *cyp-33C7* | WBGene00018260 | protein_coding_gene | 1.6 | 3.06 | 3.3E-03 | -1.3 | 0.41 | 7.8E-09 |
| *btb-21* | WBGene00018540 | pseudogene | 1.6 | 3.05 | 2.0E-03 | -2.0 | 0.25 | 7.9E-12 |
| *ptr-10* | WBGene00004224 | protein_coding_gene | 1.6 | 3.05 | 1.9E-02 | -2.8 | 0.15 | 4.9E-08 |
| *W05H9.1* | WBGene00021048 | protein_coding_gene | 1.6 | 3.04 | 3.3E-03 | -2.5 | 0.17 | 1.8E-68 |
| *C06G3.3* | WBGene00015549 | protein_coding_gene | 1.6 | 3.04 | 1.8E-02 | -2.0 | 0.26 | 1.4E-04 |
| *hil-1* | WBGene00001852 | protein_coding_gene | 1.6 | 3.03 | 3.9E-03 | -2.7 | 0.16 | 5.9E-16 |
| *K09C4.1* | WBGene00019547 | protein_coding_gene | 1.6 | 3.03 | 1.0E-02 | -2.3 | 0.20 | 3.4E-14 |
| *Y6E2A.4* | WBGene00012398 | protein_coding_gene | 1.6 | 3.02 | 3.1E-02 | -2.6 | 0.16 | 5.1E-15 |
| *T05F1.11* | WBGene00011495 | protein_coding_gene | 1.6 | 3.02 | 3.3E-03 | -2.8 | 0.14 | 1.7E-17 |
| *dpy-10* | WBGene00001072 | protein_coding_gene | 1.6 | 3.01 | 3.1E-04 | -2.3 | 0.20 | 1.8E-03 |
| *F21A3.11* | WBGene00219316 | protein_coding_gene | 1.6 | 3.01 | 8.7E-05 | -2.2 | 0.21 | 1.4E-23 |
| *dmd-5* | WBGene00017326 | protein_coding_gene | 1.6 | 3.00 | 2.8E-02 | -1.8 | 0.30 | 2.6E-04 |
| *K10D3.4* | WBGene00010738 | protein_coding_gene | 1.6 | 3.00 | 2.4E-03 | -2.3 | 0.21 | 7.7E-05 |
| *C11G10.1* | WBGene00007525 | protein_coding_gene | 1.6 | 3.00 | 3.5E-03 | -2.3 | 0.20 | 1.2E-08 |
| *clec-7* | WBGene00017364 | protein_coding_gene | 1.6 | 3.00 | 1.0E-02 | -2.0 | 0.25 | 2.1E-16 |
| *ZK550.6* | WBGene00014000 | protein_coding_gene | 1.6 | 3.00 | 2.7E-03 | -1.7 | 0.31 | 1.5E-16 |
| *Y110A2AL.4* | WBGene00022441 | protein_coding_gene | 1.6 | 2.99 | 1.7E-02 | -2.2 | 0.22 | 3.1E-06 |
| *T03G6.1* | WBGene00020194 | protein_coding_gene | 1.6 | 2.98 | 3.3E-02 | -4.3 | 0.05 | 3.1E-07 |
| *hhat-1* | WBGene00013852 | protein_coding_gene | 1.6 | 2.95 | 1.8E-02 | -2.8 | 0.14 | 9.8E-06 |
| *T26C12.3* | WBGene00020833 | protein_coding_gene | 1.6 | 2.95 | 4.9E-03 | -1.1 | 0.46 | 7.7E-04 |
| *F26F12.3* | WBGene00017834 | protein_coding_gene | 1.6 | 2.95 | 1.2E-03 | -2.4 | 0.19 | 1.5E-105 |
| *Y41D4B.26* | WBGene00021525 | protein_coding_gene | 1.6 | 2.94 | 3.1E-02 | -2.6 | 0.16 | 9.4E-12 |
| *rncs-1* | WBGene00023069 | ncRNA_gene | 1.6 | 2.93 | 2.8E-03 | -1.6 | 0.32 | 3.7E-06 |
| *C48B4.3* | WBGene00008168 | protein_coding_gene | 1.6 | 2.93 | 1.8E-02 | -2.7 | 0.16 | 1.8E-06 |
| *K08D9.4* | WBGene00019525 | protein_coding_gene | 1.5 | 2.92 | 2.2E-02 | -1.4 | 0.39 | 7.3E-04 |
| *gst-35* | WBGene00001783 | protein_coding_gene | 1.5 | 2.92 | 3.8E-04 | 1.9 | 3.72 | 1.0E-06 |
| *M01A8.1* | WBGene00010795 | protein_coding_gene | 1.5 | 2.91 | 3.0E-02 | -2.4 | 0.18 | 1.3E-14 |
| *Y7A9D.1* | WBGene00012420 | protein_coding_gene | 1.5 | 2.90 | 1.4E-03 | -1.7 | 0.30 | 1.3E-13 |
| *cutl-25* | WBGene00021919 | protein_coding_gene | 1.5 | 2.89 | 1.0E-02 | -1.6 | 0.33 | 2.0E-04 |
| *wrt-10* | WBGene00006956 | protein_coding_gene | 1.5 | 2.89 | 4.6E-04 | -2.3 | 0.20 | 3.7E-03 |
| *Y65B4BM.3* | WBGene00302974 | protein_coding_gene | 1.5 | 2.88 | 2.7E-03 | -2.3 | 0.20 | 1.6E-08 |
| *D1086.5* | WBGene00008392 | protein_coding_gene | 1.5 | 2.88 | 3.7E-03 | -1.8 | 0.28 | 8.5E-12 |
| *lpr-6* | WBGene00012255 | protein_coding_gene | 1.5 | 2.88 | 3.4E-02 | -3.0 | 0.13 | 1.5E-04 |
| *D2030.2* | WBGene00008412 | protein_coding_gene | 1.5 | 2.87 | 1.6E-05 | -1.5 | 0.36 | 3.3E-48 |
| *ugt-4* | WBGene00013905 | protein_coding_gene | 1.5 | 2.85 | 7.9E-03 | -2.9 | 0.13 | 2.6E-23 |
| *T19C4.1* | WBGene00011838 | protein_coding_gene | 1.5 | 2.85 | 2.9E-02 | -2.6 | 0.17 | 2.1E-11 |
| *F18C5.10* | WBGene00017563 | protein_coding_gene | 1.5 | 2.83 | 3.4E-04 | -2.0 | 0.25 | 2.7E-58 |
| *tyr-1* | WBGene00015332 | protein_coding_gene | 1.5 | 2.83 | 5.3E-05 | -1.1 | 0.46 | 1.7E-06 |
| *F46E10.2* | WBGene00018489 | protein_coding_gene | 1.5 | 2.81 | 1.1E-03 | -1.5 | 0.35 | 3.1E-10 |
| *C10C5.3* | WBGene00007507 | protein_coding_gene | 1.5 | 2.81 | 5.6E-03 | -1.6 | 0.32 | 1.7E-03 |
| *R05G6.10* | WBGene00019902 | protein_coding_gene | 1.5 | 2.80 | 2.3E-03 | -1.9 | 0.27 | 6.6E-22 |
| *F21C10.11* | WBGene00017660 | protein_coding_gene | 1.5 | 2.80 | 6.3E-03 | -3.0 | 0.13 | 2.1E-07 |

| | | | | | | | | |
|---|---|---|---|---|---|---|---|---|
| *clec-55* | WBGene00008597 | protein_coding_gene | 1.5 | 2.79 | 3.3E-02 | -2.3 | 0.20 | 1.8E-07 |
| *hlh-14* | WBGene00001958 | protein_coding_gene | 1.5 | 2.79 | 1.9E-03 | -1.5 | 0.35 | 1.2E-07 |
| *glt-5* | WBGene00001623 | protein_coding_gene | 1.5 | 2.79 | 1.6E-02 | -1.9 | 0.26 | 3.9E-09 |
| *ceh-27* | WBGene00000449 | protein_coding_gene | 1.5 | 2.77 | 3.3E-03 | -1.6 | 0.32 | 3.2E-08 |
| *irg-5* | WBGene00009429 | protein_coding_gene | 1.5 | 2.77 | 1.0E-02 | 2.6 | 6.26 | 1.2E-12 |
| *M03B6.3* | WBGene00010835 | protein_coding_gene | 1.5 | 2.77 | 1.8E-02 | -2.7 | 0.15 | 4.5E-05 |
| *T19C4.5* | WBGene00011839 | protein_coding_gene | 1.5 | 2.76 | 2.0E-03 | -1.6 | 0.34 | 6.8E-09 |
| *let-4* | WBGene00002282 | protein_coding_gene | 1.5 | 2.75 | 9.1E-03 | -2.0 | 0.25 | 3.7E-03 |
| *Y43C5A.3* | WBGene00012783 | protein_coding_gene | 1.5 | 2.74 | 1.3E-04 | -1.5 | 0.37 | 2.2E-16 |
| *atf-2* | WBGene00000220 | protein_coding_gene | 1.4 | 2.72 | 3.0E-02 | -2.3 | 0.20 | 4.5E-04 |
| *T02E9.5* | WBGene00011383 | protein_coding_gene | 1.4 | 2.71 | 6.9E-03 | -2.3 | 0.21 | 2.9E-30 |
| *bigr-1* | WBGene00009512 | protein_coding_gene | 1.4 | 2.71 | 5.4E-04 | -2.6 | 0.17 | 4.6E-04 |
| *C36B1.6* | WBGene00007973 | protein_coding_gene | 1.4 | 2.69 | 7.2E-03 | -2.5 | 0.18 | 1.5E-18 |
| *ngn-1* | WBGene00003595 | protein_coding_gene | 1.4 | 2.68 | 1.8E-02 | -1.3 | 0.42 | 6.9E-03 |
| *C10C5.4* | WBGene00007508 | protein_coding_gene | 1.4 | 2.68 | 2.5E-02 | -1.9 | 0.27 | 1.9E-24 |
| *C42D4.1* | WBGene00016594 | protein_coding_gene | 1.4 | 2.67 | 9.4E-04 | -1.6 | 0.32 | 9.0E-26 |
| *fbn-1* | WBGene00022816 | protein_coding_gene | 1.4 | 2.67 | 1.3E-02 | -2.0 | 0.25 | 1.3E-05 |
| *acs-17* | WBGene00016716 | protein_coding_gene | 1.4 | 2.66 | 1.4E-02 | -2.0 | 0.25 | 6.1E-44 |
| *ZC395.5* | WBGene00022597 | protein_coding_gene | 1.4 | 2.66 | 1.5E-02 | -2.5 | 0.18 | 1.2E-22 |
| *acs-22* | WBGene00017012 | protein_coding_gene | 1.4 | 2.66 | 1.6E-04 | -1.5 | 0.36 | 5.8E-30 |
| *T20F5.4* | WBGene00020626 | protein_coding_gene | 1.4 | 2.66 | 3.7E-02 | -2.8 | 0.14 | 3.0E-12 |
| *hmit-1.3* | WBGene00010811 | protein_coding_gene | 1.4 | 2.65 | 7.6E-03 | -1.6 | 0.32 | 5.0E-24 |
| *M03D4.4* | WBGene00019751 | protein_coding_gene | 1.4 | 2.63 | 6.8E-03 | -2.2 | 0.22 | 3.3E-14 |
| *F53A9.6* | WBGene00018729 | protein_coding_gene | 1.4 | 2.63 | 1.1E-02 | -1.6 | 0.33 | 5.8E-17 |
| *mfb-1* | WBGene00008439 | protein_coding_gene | 1.4 | 2.62 | 1.7E-03 | -2.1 | 0.24 | 5.2E-20 |
| *Y55D5A.1* | WBGene00021915 | protein_coding_gene | 1.4 | 2.62 | 7.8E-03 | -1.1 | 0.46 | 1.2E-04 |
| *cest-1.2* | WBGene00011364 | protein_coding_gene | 1.4 | 2.61 | 4.9E-02 | -3.6 | 0.08 | 2.7E-46 |
| *unc-3* | WBGene00006743 | protein_coding_gene | 1.4 | 2.60 | 3.5E-02 | -1.2 | 0.44 | 6.3E-03 |
| *F56C4.4* | WBGene00044977 | protein_coding_gene | 1.4 | 2.60 | 3.1E-02 | -1.3 | 0.39 | 5.5E-03 |
| *F09F9.4* | WBGene00017309 | protein_coding_gene | 1.4 | 2.58 | 5.0E-02 | -1.9 | 0.27 | 4.4E-06 |
| *thn-2* | WBGene00009215 | protein_coding_gene | 1.4 | 2.57 | 3.7E-02 | -2.6 | 0.17 | 1.9E-34 |
| *Y48A6B.7* | WBGene00012968 | protein_coding_gene | 1.4 | 2.56 | 6.0E-05 | -1.7 | 0.31 | 4.1E-27 |
| *F43G6.8* | WBGene00009660 | protein_coding_gene | 1.4 | 2.56 | 9.3E-03 | -2.0 | 0.26 | 2.6E-24 |
| *T10G3.1* | WBGene00011692 | protein_coding_gene | 1.4 | 2.56 | 1.6E-02 | -1.7 | 0.31 | 5.1E-10 |
| *M163.1* | WBGene00010935 | protein_coding_gene | 1.4 | 2.56 | 4.3E-03 | -1.9 | 0.26 | 2.7E-23 |
| *W04A8.4* | WBGene00012239 | protein_coding_gene | 1.4 | 2.56 | 1.4E-03 | -2.4 | 0.18 | 4.7E-26 |
| *C18D4.8* | WBGene00007677 | pseudogene | 1.4 | 2.55 | 1.4E-02 | -1.6 | 0.32 | 2.7E-09 |
| *C40A11.3* | WBGene00016546 | protein_coding_gene | 1.3 | 2.54 | 3.6E-02 | -1.5 | 0.35 | 5.0E-06 |
| *T22B7.3* | WBGene00020672 | protein_coding_gene | 1.3 | 2.53 | 1.2E-02 | -2.6 | 0.17 | 2.6E-10 |
| *mls-2* | WBGene00003377 | protein_coding_gene | 1.3 | 2.53 | 3.1E-02 | -1.1 | 0.48 | 2.8E-02 |
| *dao-2* | WBGene00000928 | protein_coding_gene | 1.3 | 2.51 | 3.1E-03 | -2.5 | 0.17 | 1.6E-03 |
| *his-24* | WBGene00001898 | protein_coding_gene | 1.3 | 2.50 | 1.4E-03 | -1.3 | 0.40 | 1.0E-20 |
| *igcm-1* | WBGene00018215 | protein_coding_gene | 1.3 | 2.49 | 1.5E-03 | -1.9 | 0.28 | 1.9E-09 |
| *fbxc-51* | WBGene00021576 | protein_coding_gene | 1.3 | 2.49 | 9.0E-04 | -1.5 | 0.35 | 3.3E-14 |
| *fkb-7* | WBGene00001432 | protein_coding_gene | 1.3 | 2.47 | 5.1E-03 | -2.5 | 0.18 | 2.1E-15 |
| *Y39G10AR.11* | WBGene00021469 | protein_coding_gene | 1.3 | 2.47 | 5.2E-04 | -2.0 | 0.24 | 2.2E-41 |
| *M7.8* | WBGene00010884 | protein_coding_gene | 1.3 | 2.46 | 1.5E-02 | -1.7 | 0.31 | 8.1E-10 |
| *C50B8.4* | WBGene00008226 | protein_coding_gene | 1.3 | 2.45 | 1.6E-02 | -2.4 | 0.19 | 5.2E-09 |
| *fozi-1* | WBGene00010453 | protein_coding_gene | 1.3 | 2.45 | 3.9E-02 | -1.9 | 0.27 | 3.5E-17 |
| *R04A9.7* | WBGene00044281 | protein_coding_gene | 1.3 | 2.44 | 3.2E-03 | -1.1 | 0.47 | 3.2E-03 |
| *cyp-35B1* | WBGene00019472 | protein_coding_gene | 1.3 | 2.43 | 9.4E-03 | -1.2 | 0.44 | 1.3E-02 |
| *ptr-4* | WBGene00004219 | protein_coding_gene | 1.3 | 2.43 | 4.8E-02 | -3.0 | 0.12 | 1.7E-06 |
| *nlp-31* | WBGene00003769 | protein_coding_gene | 1.3 | 2.42 | 4.1E-03 | -1.3 | 0.41 | 1.3E-10 |
| *clec-266* | WBGene00016088 | protein_coding_gene | 1.3 | 2.42 | 9.2E-03 | -1.5 | 0.36 | 3.2E-18 |
| *F54C8.6* | WBGene00010039 | protein_coding_gene | 1.3 | 2.40 | 1.9E-02 | -1.8 | 0.28 | 1.8E-07 |
| *C08F1.10* | WBGene00015613 | protein_coding_gene | 1.3 | 2.40 | 2.3E-03 | -1.8 | 0.28 | 1.6E-05 |
| *F48E3.8* | WBGene00018607 | protein_coding_gene | 1.3 | 2.40 | 1.3E-02 | -1.7 | 0.30 | 1.9E-10 |
| *dhrs-4* | WBGene00010063 | protein_coding_gene | 1.3 | 2.40 | 3.8E-03 | -1.6 | 0.34 | 4.7E-20 |

| | | | | | | | | |
|---|---|---|---|---|---|---|---|---|
| *F14B4.1* | WBGene00008779 | protein_coding_gene | 1.3 | 2.39 | 5.3E-03 | -2.4 | 0.19 | 8.2E-17 |
| *T09B4.5* | WBGene00020379 | protein_coding_gene | 1.3 | 2.39 | 5.0E-03 | -1.3 | 0.41 | 4.5E-22 |
| *F13H8.1* | WBGene00017434 | protein_coding_gene | 1.3 | 2.39 | 3.3E-02 | -1.9 | 0.27 | 2.7E-09 |
| *dct-1* | WBGene00015776 | protein_coding_gene | 1.3 | 2.39 | 5.3E-05 | -1.9 | 0.27 | 1.3E-30 |
| *unc-39* | WBGene00006775 | protein_coding_gene | 1.3 | 2.38 | 3.2E-02 | -1.2 | 0.44 | 1.2E-02 |
| *egl-15* | WBGene00001184 | protein_coding_gene | 1.2 | 2.37 | 2.4E-02 | -1.8 | 0.28 | 2.6E-22 |
| *ham-2* | WBGene00001821 | protein_coding_gene | 1.2 | 2.37 | 8.9E-03 | -1.5 | 0.35 | 2.8E-10 |
| *Y71F9AL.7* | WBGene00022112 | protein_coding_gene | 1.2 | 2.37 | 2.3E-03 | -1.0 | 0.50 | 6.8E-03 |
| *lgc-22* | WBGene00017483 | protein_coding_gene | 1.2 | 2.37 | 4.1E-03 | -1.2 | 0.44 | 2.5E-06 |
| *Y82E9BR.17* | WBGene00022349 | protein_coding_gene | 1.2 | 2.37 | 4.9E-03 | -1.9 | 0.27 | 1.1E-18 |
| *sptf-1* | WBGene00009743 | protein_coding_gene | 1.2 | 2.36 | 1.9E-03 | -1.2 | 0.44 | 5.7E-06 |
| *nhr-206* | WBGene00011097 | protein_coding_gene | 1.2 | 2.36 | 4.5E-02 | -2.3 | 0.21 | 6.3E-07 |
| *sqt-3* | WBGene00005018 | protein_coding_gene | 1.2 | 2.35 | 1.4E-02 | -2.5 | 0.18 | 5.1E-05 |
| *mnp-1* | WBGene00007139 | protein_coding_gene | 1.2 | 2.35 | 1.6E-04 | -1.4 | 0.39 | 7.1E-13 |
| *F53B3.5* | WBGene00018743 | protein_coding_gene | 1.2 | 2.35 | 5.5E-03 | -2.2 | 0.22 | 2.7E-22 |
| *arrd-4* | WBGene00014159 | protein_coding_gene | 1.2 | 2.34 | 1.7E-02 | -1.3 | 0.41 | 1.3E-05 |
| *best-1* | WBGene00007203 | protein_coding_gene | 1.2 | 2.34 | 5.0E-03 | -1.5 | 0.35 | 1.5E-02 |
| *moe-3* | WBGene00003388 | protein_coding_gene | 1.2 | 2.33 | 2.0E-02 | -1.9 | 0.26 | 9.1E-18 |
| *fkh-7* | WBGene00001439 | protein_coding_gene | 1.2 | 2.33 | 4.9E-03 | -1.7 | 0.31 | 2.2E-24 |
| *cht-1* | WBGene00000503 | protein_coding_gene | 1.2 | 2.33 | 4.3E-03 | -1.3 | 0.40 | 3.3E-19 |
| *fmi-1* | WBGene00001475 | protein_coding_gene | 1.2 | 2.32 | 3.8E-02 | -2.0 | 0.26 | 1.7E-13 |
| *F17C11.4* | WBGene00008915 | protein_coding_gene | 1.2 | 2.32 | 1.2E-02 | -1.9 | 0.26 | 1.1E-23 |
| *ham-1* | WBGene00001820 | protein_coding_gene | 1.2 | 2.32 | 2.1E-03 | -1.2 | 0.44 | 2.2E-08 |
| *C24H12.1* | WBGene00016071 | protein_coding_gene | 1.2 | 2.30 | 2.9E-02 | -1.4 | 0.37 | 5.8E-04 |
| *aqp-2* | WBGene00000170 | protein_coding_gene | 1.2 | 2.28 | 7.5E-03 | -1.9 | 0.27 | 5.9E-23 |
| *C27B7.9* | WBGene00007765 | protein_coding_gene | 1.2 | 2.28 | 1.6E-02 | -1.6 | 0.33 | 3.7E-31 |
| *fbxl-1* | WBGene00015350 | protein_coding_gene | 1.2 | 2.27 | 2.9E-03 | -1.1 | 0.46 | 1.1E-19 |
| *dach-1* | WBGene00020386 | protein_coding_gene | 1.2 | 2.27 | 3.8E-02 | -2.9 | 0.13 | 1.1E-10 |
| *T24C2.5* | WBGene00011984 | protein_coding_gene | 1.2 | 2.27 | 5.6E-03 | -2.4 | 0.19 | 1.2E-16 |
| *C07E3.3* | WBGene00007414 | protein_coding_gene | 1.2 | 2.27 | 4.0E-02 | -1.6 | 0.33 | 2.2E-13 |
| *EEED8.2* | WBGene00017133 | protein_coding_gene | 1.2 | 2.26 | 5.5E-03 | -2.4 | 0.18 | 2.2E-18 |
| *F40F12.9* | WBGene00043995 | protein_coding_gene | 1.2 | 2.25 | 2.0E-02 | -1.1 | 0.47 | 1.8E-02 |
| *T28B8.1* | WBGene00012113 | protein_coding_gene | 1.2 | 2.24 | 1.8E-03 | -1.7 | 0.31 | 4.5E-27 |
| *cyp-37A1* | WBGene00008499 | protein_coding_gene | 1.2 | 2.24 | 3.0E-02 | -1.5 | 0.35 | 3.9E-14 |
| *ttr-37* | WBGene00017667 | protein_coding_gene | 1.2 | 2.24 | 9.4E-03 | -1.4 | 0.37 | 2.0E-08 |
| *dhs-8* | WBGene00000972 | protein_coding_gene | 1.2 | 2.23 | 4.8E-02 | -1.2 | 0.44 | 4.1E-02 |
| *arrd-1* | WBGene00020205 | protein_coding_gene | 1.2 | 2.23 | 2.9E-02 | -1.1 | 0.46 | 6.9E-06 |
| *hil-3* | WBGene00001854 | protein_coding_gene | 1.2 | 2.23 | 8.8E-04 | -1.3 | 0.41 | 1.0E-21 |
| *nhr-32* | WBGene00003626 | protein_coding_gene | 1.2 | 2.22 | 7.8E-04 | -1.2 | 0.44 | 1.6E-10 |
| *qua-1* | WBGene00004264 | protein_coding_gene | 1.1 | 2.20 | 3.1E-02 | -1.8 | 0.28 | 4.1E-03 |
| *F20C5.6* | WBGene00008971 | protein_coding_gene | 1.1 | 2.20 | 2.8E-03 | -1.8 | 0.28 | 3.9E-12 |
| *ceh-43* | WBGene00000463 | protein_coding_gene | 1.1 | 2.19 | 7.2E-03 | -1.5 | 0.36 | 2.0E-11 |
| *memo-1* | WBGene00016500 | protein_coding_gene | 1.1 | 2.19 | 5.1E-03 | -1.5 | 0.35 | 1.3E-24 |
| *jmjd-3.2* | WBGene00009089 | protein_coding_gene | 1.1 | 2.19 | 1.6E-02 | -1.2 | 0.44 | 8.2E-04 |
| *dhs-2* | WBGene00000966 | protein_coding_gene | 1.1 | 2.19 | 2.5E-02 | -1.4 | 0.38 | 1.4E-08 |
| *Y43F8C.3* | WBGene00012825 | protein_coding_gene | 1.1 | 2.18 | 2.4E-02 | -1.6 | 0.34 | 1.4E-18 |
| *pxd-1* | WBGene00007982 | protein_coding_gene | 1.1 | 2.18 | 7.8E-05 | -1.8 | 0.29 | 8.0E-28 |
| *nhr-18* | WBGene00003617 | protein_coding_gene | 1.1 | 2.18 | 1.4E-02 | -2.0 | 0.25 | 1.5E-13 |
| *F31C3.6* | WBGene00009288 | protein_coding_gene | 1.1 | 2.18 | 1.0E-02 | -1.4 | 0.39 | 4.8E-08 |
| *C13C4.6* | WBGene00007550 | protein_coding_gene | 1.1 | 2.17 | 3.7E-02 | -1.1 | 0.47 | 1.0E-04 |
| *C07B5.4* | WBGene00007409 | protein_coding_gene | 1.1 | 2.17 | 2.6E-02 | -1.2 | 0.44 | 1.9E-05 |
| *Y15E3A.5* | WBGene00012444 | protein_coding_gene | 1.1 | 2.17 | 2.8E-02 | -1.5 | 0.36 | 7.9E-11 |
| *frpr-11* | WBGene00019444 | protein_coding_gene | 1.1 | 2.17 | 1.9E-02 | -1.3 | 0.41 | 1.6E-04 |
| *cbp-2* | WBGene00010545 | protein_coding_gene | 1.1 | 2.16 | 1.5E-02 | -1.7 | 0.30 | 2.5E-16 |
| *egl-43* | WBGene00001207 | protein_coding_gene | 1.1 | 2.16 | 1.6E-02 | -1.4 | 0.39 | 1.5E-06 |
| *F35D2.3* | WBGene00018041 | protein_coding_gene | 1.1 | 2.15 | 5.8E-03 | -1.8 | 0.29 | 8.5E-13 |
| *C44H9.5* | WBGene00008100 | protein_coding_gene | 1.1 | 2.14 | 1.9E-03 | -1.6 | 0.32 | 1.5E-15 |
| *F55H12.2* | WBGene00010133 | protein_coding_gene | 1.1 | 2.14 | 2.3E-02 | -2.1 | 0.23 | 1.8E-26 |

| | | | | | | | | |
|---|---|---|---|---|---|---|---|---|
| *droe-4* | WBGene00010262 | protein_coding_gene | 1.1 | 2.13 | 8.7E-03 | -1.0 | 0.48 | 1.2E-08 |
| *D1053.3* | WBGene00008368 | protein_coding_gene | 1.1 | 2.13 | 4.2E-02 | -1.1 | 0.48 | 4.0E-04 |
| *epg-2* | WBGene00021468 | protein_coding_gene | 1.1 | 2.13 | 1.5E-02 | -1.5 | 0.35 | 1.5E-27 |
| *Y69A2AL.2* | WBGene00044468 | protein_coding_gene | 1.1 | 2.11 | 2.6E-02 | -1.1 | 0.48 | 1.4E-06 |
| *F10E9.12* | WBGene00017361 | protein_coding_gene | 1.1 | 2.11 | 8.9E-03 | -1.8 | 0.28 | 5.9E-17 |
| *kpc-1* | WBGene00002232 | protein_coding_gene | 1.1 | 2.10 | 1.6E-02 | -1.6 | 0.34 | 1.8E-28 |
| *asp-14* | WBGene00019619 | protein_coding_gene | 1.1 | 2.09 | 1.9E-02 | 1.3 | 2.43 | 1.1E-20 |
| *C30G4.4* | WBGene00016269 | protein_coding_gene | 1.1 | 2.09 | 1.9E-02 | -1.3 | 0.40 | 4.2E-12 |
| *Y73C8B.1* | WBGene00022258 | protein_coding_gene | 1.1 | 2.09 | 4.7E-02 | -1.3 | 0.39 | 1.3E-03 |
| *C32F10.4* | WBGene00016332 | protein_coding_gene | 1.1 | 2.08 | 8.6E-03 | -2.0 | 0.25 | 9.9E-11 |
| *ins-34* | WBGene00002117 | protein_coding_gene | 1.1 | 2.08 | 4.9E-02 | -1.1 | 0.46 | 2.6E-04 |
| *epg-9* | WBGene00022078 | protein_coding_gene | 1.1 | 2.08 | 4.6E-03 | -1.4 | 0.39 | 3.7E-16 |
| *ZC443.4* | WBGene00013899 | protein_coding_gene | 1.1 | 2.07 | 2.9E-02 | -1.8 | 0.28 | 5.1E-22 |
| *T10H9.8* | WBGene00044779 | protein_coding_gene | 1.1 | 2.07 | 3.7E-02 | -1.8 | 0.29 | 3.2E-11 |
| *crtc-1* | WBGene00021237 | protein_coding_gene | 1.0 | 2.07 | 2.4E-02 | -1.2 | 0.44 | 2.2E-09 |
| *cav-1* | WBGene00000301 | protein_coding_gene | 1.0 | 2.07 | 4.0E-03 | -1.3 | 0.40 | 9.0E-21 |
| *sup-26* | WBGene00006331 | protein_coding_gene | 1.0 | 2.07 | 2.5E-02 | -1.1 | 0.46 | 2.1E-21 |
| *ZK1128.7* | WBGene00014233 | protein_coding_gene | 1.0 | 2.07 | 1.7E-02 | -1.8 | 0.29 | 1.8E-21 |
| *fkh-2* | WBGene00001434 | protein_coding_gene | 1.0 | 2.05 | 4.1E-02 | -1.6 | 0.33 | 1.5E-05 |
| *sto-1* | WBGene00006063 | protein_coding_gene | 1.0 | 2.05 | 3.9E-03 | -1.1 | 0.45 | 6.9E-13 |
| *T01E8.1* | WBGene00011331 | protein_coding_gene | 1.0 | 2.05 | 2.3E-02 | -1.2 | 0.44 | 1.2E-09 |
| *dex-1* | WBGene00017028 | protein_coding_gene | 1.0 | 2.04 | 4.4E-02 | -1.6 | 0.33 | 1.9E-13 |
| *atz-1* | WBGene00009212 | protein_coding_gene | 1.0 | 2.04 | 1.7E-02 | -1.2 | 0.43 | 9.2E-13 |
| *T05E12.3* | WBGene00011486 | protein_coding_gene | 1.0 | 2.04 | 1.1E-02 | -2.4 | 0.19 | 2.6E-46 |
| *lgg-1* | WBGene00002980 | protein_coding_gene | 1.0 | 2.03 | 7.2E-05 | -1.4 | 0.39 | 2.2E-45 |
| *ugt-51* | WBGene00015369 | protein_coding_gene | 1.0 | 2.03 | 4.5E-02 | -2.2 | 0.22 | 9.2E-10 |
| *C08F1.6* | WBGene00015610 | protein_coding_gene | 1.0 | 2.02 | 3.6E-02 | -1.6 | 0.34 | 1.5E-02 |
| *gem-4* | WBGene00001577 | protein_coding_gene | 1.0 | 2.02 | 2.9E-03 | -1.7 | 0.30 | 1.0E-14 |
| *cki-1* | WBGene00000516 | protein_coding_gene | 1.0 | 2.02 | 7.7E-03 | -1.8 | 0.29 | 8.1E-18 |
| *F36F2.2* | WBGene00009476 | protein_coding_gene | 1.0 | 2.02 | 8.7E-03 | -2.1 | 0.23 | 2.9E-19 |
| *C35E7.5* | WBGene00016457 | protein_coding_gene | 1.0 | 2.01 | 2.2E-02 | -1.0 | 0.49 | 7.1E-13 |
| *F13C5.2* | WBGene00017423 | protein_coding_gene | 1.0 | 2.01 | 1.9E-02 | -1.2 | 0.44 | 4.3E-12 |
| *lgg-2* | WBGene00002981 | protein_coding_gene | 1.0 | 2.01 | 2.0E-03 | -1.1 | 0.48 | 3.0E-20 |
| *F38E9.1* | WBGene00018184 | protein_coding_gene | 1.0 | 2.01 | 5.7E-03 | -1.3 | 0.42 | 5.8E-15 |
| *C28G1.10* | WBGene00219376 | protein_coding_gene | 1.0 | 2.00 | 6.3E-03 | -1.1 | 0.47 | 4.1E-04 |
| *nlp-27* | WBGene00003765 | protein_coding_gene | -1.0 | 0.50 | 4.0E-02 | 1.2 | 2.37 | 1.0E-14 |
| *C18A11.3* | WBGene00015949 | protein_coding_gene | -1.0 | 0.50 | 3.9E-03 | 1.1 | 2.07 | 7.2E-03 |
| *aat-4* | WBGene00000005 | protein_coding_gene | -1.0 | 0.49 | 3.9E-02 | 2.1 | 4.24 | 7.8E-09 |
| *mtp-18* | WBGene00077690 | protein_coding_gene | -1.0 | 0.49 | 2.2E-02 | 1.3 | 2.54 | 1.2E-05 |
| *C14B4.2* | WBGene00007583 | protein_coding_gene | -1.0 | 0.49 | 1.8E-02 | 1.1 | 2.11 | 8.0E-10 |
| *oat-1* | WBGene00003837 | protein_coding_gene | -1.0 | 0.49 | 3.1E-02 | 1.3 | 2.47 | 2.1E-05 |
| *ugt-8* | WBGene00019234 | protein_coding_gene | -1.0 | 0.49 | 6.3E-03 | -1.9 | 0.27 | 8.7E-13 |
| *slc-17.5* | WBGene00017681 | protein_coding_gene | -1.0 | 0.49 | 2.4E-02 | 1.6 | 2.98 | 1.7E-10 |
| *Y53G8AR.7* | WBGene00021814 | protein_coding_gene | -1.0 | 0.49 | 2.0E-03 | 1.2 | 2.32 | 1.5E-07 |
| *poml-2* | WBGene00017089 | protein_coding_gene | -1.0 | 0.49 | 9.7E-03 | 1.3 | 2.45 | 3.9E-05 |
| *ldp-1* | WBGene00017719 | protein_coding_gene | -1.0 | 0.49 | 2.0E-03 | 1.6 | 3.07 | 1.4E-54 |
| *lgc-45* | WBGene00021146 | protein_coding_gene | -1.0 | 0.49 | 9.8E-03 | 1.7 | 3.16 | 2.0E-14 |
| *twk-22* | WBGene00006675 | protein_coding_gene | -1.0 | 0.49 | 9.3E-03 | 1.8 | 3.40 | 2.9E-10 |
| *F55A3.7* | WBGene00018852 | pseudogene | -1.1 | 0.48 | 3.4E-03 | 1.3 | 2.46 | 1.2E-02 |
| *ugt-48* | WBGene00015965 | protein_coding_gene | -1.1 | 0.48 | 1.5E-02 | 2.0 | 4.03 | 1.2E-19 |
| *col-42* | WBGene00013489 | protein_coding_gene | -1.1 | 0.48 | 6.7E-04 | 2.0 | 4.03 | 2.3E-33 |
| *F11E6.3* | WBGene00008707 | protein_coding_gene | -1.1 | 0.48 | 1.6E-02 | 1.1 | 2.18 | 6.3E-13 |
| *Y50D4B.2* | WBGene00021741 | protein_coding_gene | -1.1 | 0.48 | 2.6E-02 | 1.3 | 2.53 | 2.2E-03 |
| *scav-4* | WBGene00008692 | protein_coding_gene | -1.1 | 0.48 | 1.7E-02 | 2.6 | 6.01 | 3.8E-20 |
| *mua-3* | WBGene00003482 | protein_coding_gene | -1.1 | 0.48 | 3.3E-02 | 1.0 | 2.06 | 1.2E-02 |
| *F23C8.7* | WBGene00017736 | protein_coding_gene | -1.1 | 0.48 | 2.2E-02 | 1.1 | 2.09 | 1.2E-04 |
| *Y73E7A.3* | WBGene00022270 | protein_coding_gene | -1.1 | 0.48 | 1.7E-02 | 1.4 | 2.68 | 9.1E-07 |
| *C06H5.6* | WBGene00007394 | protein_coding_gene | -1.1 | 0.47 | 9.9E-03 | 1.1 | 2.11 | 2.0E-06 |

| | | | | | | | | |
|---|---|---|---|---|---|---|---|---|
| *C39D10.8* | WBGene00016535 | protein_coding_gene | -1.1 | 0.47 | 1.2E-03 | 1.7 | 3.30 | 3.8E-16 |
| *col-166* | WBGene00000739 | protein_coding_gene | -1.1 | 0.47 | 1.3E-02 | 1.2 | 2.22 | 7.2E-07 |
| *cld-9* | WBGene00009432 | protein_coding_gene | -1.1 | 0.47 | 2.0E-02 | 1.2 | 2.34 | 2.6E-05 |
| *C52G5.2* | WBGene00008260 | protein_coding_gene | -1.1 | 0.47 | 4.4E-02 | 1.0 | 2.02 | 4.2E-03 |
| *zip-3* | WBGene00012330 | protein_coding_gene | -1.1 | 0.47 | 2.1E-02 | 1.6 | 2.97 | 3.5E-19 |
| *Y67H2A.9* | WBGene00014941 | pseudogene | -1.1 | 0.47 | 1.9E-02 | 1.9 | 3.78 | 2.5E-23 |
| *col-179* | WBGene00000752 | protein_coding_gene | -1.1 | 0.46 | 1.1E-02 | 1.3 | 2.49 | 7.7E-17 |
| *nlp-26* | WBGene00003764 | protein_coding_gene | -1.1 | 0.46 | 3.0E-02 | 1.4 | 2.59 | 3.0E-05 |
| *ifc-1* | WBGene00002055 | protein_coding_gene | -1.1 | 0.46 | 3.7E-03 | 1.2 | 2.28 | 2.2E-21 |
| *pqn-60* | WBGene00004144 | protein_coding_gene | -1.1 | 0.46 | 1.2E-02 | 1.1 | 2.12 | 1.7E-03 |
| *ugt-22* | WBGene00007455 | protein_coding_gene | -1.1 | 0.46 | 2.5E-02 | 1.6 | 3.01 | 2.4E-05 |
| *C02G6.1* | WBGene00015359 | pseudogene | -1.1 | 0.46 | 1.1E-02 | 3.4 | 10.29 | 1.9E-04 |
| *comt-3* | WBGene00021487 | protein_coding_gene | -1.1 | 0.46 | 2.1E-02 | 1.4 | 2.70 | 8.2E-11 |
| *T16G12.1* | WBGene00011803 | protein_coding_gene | -1.1 | 0.45 | 2.5E-02 | 2.5 | 5.60 | 7.0E-14 |
| *Y16B4A.2* | WBGene00012445 | protein_coding_gene | -1.1 | 0.45 | 1.9E-02 | 2.3 | 5.05 | 2.6E-17 |
| *cima-1* | WBGene00018473 | protein_coding_gene | -1.1 | 0.45 | 2.3E-02 | 1.6 | 3.03 | 4.2E-06 |
| *twk-42* | WBGene00006692 | protein_coding_gene | -1.1 | 0.45 | 5.5E-03 | 1.9 | 3.75 | 2.7E-06 |
| *kynu-1* | WBGene00015802 | protein_coding_gene | -1.1 | 0.45 | 4.9E-02 | 1.4 | 2.63 | 6.5E-19 |
| *clec-205* | WBGene00021602 | protein_coding_gene | -1.1 | 0.45 | 6.8E-03 | 1.5 | 2.91 | 3.7E-04 |
| *F21A3.3* | WBGene00008993 | protein_coding_gene | -1.1 | 0.45 | 1.9E-02 | 1.8 | 3.38 | 5.5E-07 |
| *gon-1* | WBGene00001650 | protein_coding_gene | -1.1 | 0.45 | 1.2E-03 | 1.2 | 2.31 | 6.5E-11 |
| *Y59E9AL.4* | WBGene00021994 | protein_coding_gene | -1.2 | 0.45 | 4.7E-03 | 1.5 | 2.89 | 3.9E-05 |
| *T28D6.3* | WBGene00012123 | protein_coding_gene | -1.2 | 0.45 | 4.9E-02 | 3.0 | 7.93 | 3.0E-42 |
| *B0244.10* | WBGene00015085 | protein_coding_gene | -1.2 | 0.45 | 2.2E-02 | 1.1 | 2.19 | 5.4E-03 |
| *C35A11.4* | WBGene00016431 | protein_coding_gene | -1.2 | 0.44 | 4.0E-02 | 3.2 | 9.46 | 1.1E-37 |
| *F19C7.4* | WBGene00017594 | protein_coding_gene | -1.2 | 0.44 | 1.9E-02 | 1.1 | 2.15 | 2.2E-02 |
| *Y73B3A.18* | WBGene00022219 | pseudogene | -1.2 | 0.44 | 5.1E-03 | 1.5 | 2.88 | 9.6E-07 |
| *aqp-4* | WBGene00000172 | protein_coding_gene | -1.2 | 0.44 | 4.3E-04 | 1.2 | 2.27 | 1.9E-07 |
| *col-141* | WBGene00000714 | protein_coding_gene | -1.2 | 0.44 | 4.9E-02 | 2.0 | 4.08 | 1.2E-02 |
| *W02H3.1* | WBGene00012217 | protein_coding_gene | -1.2 | 0.43 | 1.8E-02 | 1.6 | 3.02 | 2.2E-05 |
| *grd-5* | WBGene00001694 | protein_coding_gene | -1.2 | 0.43 | 3.0E-02 | 1.9 | 3.79 | 2.9E-05 |
| *ZC449.7* | WBGene00045365 | protein_coding_gene | -1.2 | 0.43 | 3.8E-02 | 1.0 | 2.02 | 4.2E-02 |
| *pept-1* | WBGene00003877 | protein_coding_gene | -1.2 | 0.43 | 2.7E-03 | 1.7 | 3.18 | 2.8E-22 |
| *ttr-18* | WBGene00007142 | protein_coding_gene | -1.2 | 0.43 | 1.2E-03 | 1.4 | 2.64 | 3.7E-17 |
| *F54B11.11* | WBGene00010034 | protein_coding_gene | -1.2 | 0.43 | 1.1E-02 | -1.9 | 0.27 | 4.5E-08 |
| *unc-52* | WBGene00006787 | protein_coding_gene | -1.2 | 0.42 | 7.7E-04 | 1.5 | 2.80 | 7.2E-27 |
| *F42A10.7* | WBGene00018343 | protein_coding_gene | -1.2 | 0.42 | 3.5E-06 | 1.7 | 3.21 | 7.1E-06 |
| *clec-97* | WBGene00013931 | protein_coding_gene | -1.2 | 0.42 | 1.5E-03 | 1.0 | 2.00 | 8.5E-05 |
| *D1005.4* | WBGene00016998 | protein_coding_gene | -1.2 | 0.42 | 1.3E-02 | 1.0 | 2.07 | 1.4E-02 |
| *B0454.5* | WBGene00015196 | protein_coding_gene | -1.2 | 0.42 | 1.0E-02 | 1.2 | 2.34 | 5.4E-04 |
| *nmy-3* | WBGene00012437 | protein_coding_gene | -1.3 | 0.42 | 2.4E-02 | 1.2 | 2.28 | 1.7E-04 |
| *Y39F10A.1* | WBGene00021452 | protein_coding_gene | -1.3 | 0.42 | 4.6E-02 | 1.5 | 2.90 | 1.8E-02 |
| *C01B9.1* | WBGene00007220 | protein_coding_gene | -1.3 | 0.42 | 1.3E-02 | 1.4 | 2.62 | 1.2E-05 |
| *C01H6.4* | WBGene00007254 | protein_coding_gene | -1.3 | 0.42 | 3.9E-03 | 1.2 | 2.32 | 1.8E-06 |
| *lys-5* | WBGene00003094 | protein_coding_gene | -1.3 | 0.42 | 2.0E-03 | -1.3 | 0.41 | 3.9E-03 |
| *C50A2.3* | WBGene00016797 | protein_coding_gene | -1.3 | 0.42 | 4.9E-02 | 1.8 | 3.36 | 3.7E-08 |
| *daf-28* | WBGene00000920 | protein_coding_gene | -1.3 | 0.42 | 1.9E-02 | 1.6 | 2.98 | 5.7E-05 |
| *ttr-31* | WBGene00010225 | protein_coding_gene | -1.3 | 0.41 | 4.6E-02 | 1.9 | 3.66 | 8.4E-13 |
| *pqn-37* | WBGene00004124 | protein_coding_gene | -1.3 | 0.41 | 7.9E-03 | 1.8 | 3.40 | 2.2E-05 |
| *grd-3* | WBGene00001692 | protein_coding_gene | -1.3 | 0.41 | 2.9E-03 | 2.0 | 4.05 | 1.1E-05 |
| *gldc-1* | WBGene00020022 | protein_coding_gene | -1.3 | 0.41 | 3.4E-03 | 1.2 | 2.24 | 1.7E-25 |
| *sams-1* | WBGene00008205 | protein_coding_gene | -1.3 | 0.41 | 1.2E-02 | 1.4 | 2.64 | 1.4E-28 |
| *skpo-3* | WBGene00017968 | protein_coding_gene | -1.3 | 0.41 | 3.1E-04 | 1.2 | 2.23 | 1.6E-17 |
| *cept-1* | WBGene00009057 | protein_coding_gene | -1.3 | 0.41 | 1.1E-02 | 1.7 | 3.34 | 1.3E-26 |
| *F11F1.1* | WBGene00008714 | protein_coding_gene | -1.3 | 0.41 | 6.7E-04 | 1.3 | 2.45 | 7.9E-06 |
| *Y50D4B.4* | WBGene00021743 | protein_coding_gene | -1.3 | 0.41 | 9.9E-03 | 2.3 | 5.06 | 3.7E-26 |
| *pho-1* | WBGene00004020 | protein_coding_gene | -1.3 | 0.40 | 4.4E-03 | 3.2 | 9.46 | 1.7E-15 |
| *adt-2* | WBGene00000083 | protein_coding_gene | -1.3 | 0.40 | 1.7E-02 | 2.0 | 4.06 | 2.1E-36 |

| | | | | | | | | |
|---|---|---|---|---|---|---|---|---|
| *col-98* | WBGene00000673 | protein_coding_gene | -1.3 | 0.40 | 3.2E-03 | 2.5 | 5.78 | 3.6E-78 |
| *F09B12.3* | WBGene00008607 | protein_coding_gene | -1.3 | 0.40 | 1.8E-02 | 2.3 | 4.91 | 4.3E-38 |
| *acs-1* | WBGene00018488 | protein_coding_gene | -1.3 | 0.40 | 2.0E-02 | 2.4 | 5.42 | 2.5E-51 |
| *col-142* | WBGene00000715 | protein_coding_gene | -1.3 | 0.40 | 8.9E-04 | 1.2 | 2.32 | 3.3E-06 |
| *ugt-46* | WBGene00015141 | protein_coding_gene | -1.3 | 0.40 | 6.0E-03 | 1.1 | 2.21 | 1.7E-08 |
| *T19C3.2* | WBGene00020560 | protein_coding_gene | -1.3 | 0.39 | 4.1E-02 | -2.7 | 0.16 | 6.7E-03 |
| *Y22D7AL.11* | WBGene00021249 | protein_coding_gene | -1.3 | 0.39 | 1.9E-03 | 2.1 | 4.18 | 5.9E-07 |
| *doxa-1* | WBGene00015519 | protein_coding_gene | -1.4 | 0.39 | 2.0E-03 | 2.0 | 4.05 | 5.1E-12 |
| *col-103* | WBGene00000677 | protein_coding_gene | -1.4 | 0.39 | 9.7E-04 | 2.4 | 5.32 | 9.3E-74 |
| *Y37H2A.14* | WBGene00045416 | protein_coding_gene | -1.4 | 0.39 | 2.9E-02 | 1.6 | 3.03 | 3.5E-05 |
| *col-80* | WBGene00000656 | protein_coding_gene | -1.4 | 0.39 | 5.2E-03 | 1.5 | 2.87 | 4.2E-08 |
| *mup-4* | WBGene00003497 | protein_coding_gene | -1.4 | 0.39 | 1.2E-03 | 1.7 | 3.19 | 8.5E-29 |
| *ugt-60* | WBGene00007402 | protein_coding_gene | -1.4 | 0.39 | 3.6E-02 | 1.3 | 2.43 | 2.3E-05 |
| *cut-4* | WBGene00008482 | protein_coding_gene | -1.4 | 0.39 | 3.0E-03 | 1.4 | 2.60 | 2.3E-02 |
| *arrd-10* | WBGene00010268 | protein_coding_gene | -1.4 | 0.38 | 1.8E-02 | 1.4 | 2.58 | 4.8E-04 |
| *nep-22* | WBGene00020788 | protein_coding_gene | -1.4 | 0.38 | 8.1E-03 | 1.8 | 3.54 | 6.2E-22 |
| *ZC250.4* | WBGene00045305 | protein_coding_gene | -1.4 | 0.38 | 4.4E-02 | 2.9 | 7.44 | 1.7E-32 |
| *T25G12.6* | WBGene00020813 | protein_coding_gene | -1.4 | 0.38 | 7.6E-04 | 1.4 | 2.67 | 7.8E-05 |
| *nas-7* | WBGene00003526 | protein_coding_gene | -1.4 | 0.38 | 7.0E-03 | 1.3 | 2.49 | 4.7E-03 |
| *paic-1* | WBGene00015116 | protein_coding_gene | -1.4 | 0.38 | 1.5E-02 | 2.5 | 5.85 | 5.8E-74 |
| *fmo-3* | WBGene00001478 | protein_coding_gene | -1.4 | 0.38 | 2.5E-03 | 2.0 | 3.87 | 1.1E-16 |
| *cln-3.1* | WBGene00000539 | protein_coding_gene | -1.4 | 0.37 | 1.5E-03 | 1.8 | 3.54 | 2.4E-09 |
| *Y51H7C.1* | WBGene00021779 | protein_coding_gene | -1.4 | 0.37 | 2.1E-03 | 2.6 | 5.88 | 3.9E-03 |
| *F41D9.2* | WBGene00018282 | protein_coding_gene | -1.4 | 0.37 | 3.0E-02 | 1.0 | 2.05 | 1.9E-03 |
| *T15B7.10* | WBGene00020522 | pseudogene | -1.4 | 0.37 | 1.5E-02 | -1.8 | 0.29 | 9.7E-03 |
| *ugt-19* | WBGene00019515 | protein_coding_gene | -1.4 | 0.37 | 3.1E-02 | 2.0 | 4.13 | 1.5E-18 |
| *phat-4* | WBGene00020237 | protein_coding_gene | -1.4 | 0.37 | 1.9E-02 | 1.8 | 3.47 | 9.9E-04 |
| *F37B12.1* | WBGene00009504 | protein_coding_gene | -1.4 | 0.37 | 1.8E-02 | 1.7 | 3.21 | 1.3E-05 |
| *R07E3.4* | WBGene00011105 | protein_coding_gene | -1.4 | 0.37 | 5.5E-03 | 1.3 | 2.40 | 1.5E-06 |
| *T06A4.3* | WBGene00020283 | protein_coding_gene | -1.4 | 0.37 | 4.1E-03 | 1.0 | 2.03 | 4.9E-02 |
| *elo-9* | WBGene00001247 | protein_coding_gene | -1.5 | 0.36 | 1.1E-02 | 1.2 | 2.32 | 1.8E-05 |
| *act-5* | WBGene00000067 | protein_coding_gene | -1.5 | 0.36 | 4.8E-03 | 2.1 | 4.30 | 4.4E-06 |
| *fat-5* | WBGene00001397 | protein_coding_gene | -1.5 | 0.35 | 9.1E-06 | 1.7 | 3.15 | 4.5E-05 |
| *pho-13* | WBGene00008560 | protein_coding_gene | -1.5 | 0.35 | 3.1E-02 | 2.2 | 4.67 | 1.7E-04 |
| *elo-2* | WBGene00001240 | protein_coding_gene | -1.5 | 0.35 | 9.4E-03 | 3.0 | 8.22 | 4.7E-31 |
| *col-160* | WBGene00000733 | protein_coding_gene | -1.5 | 0.35 | 7.2E-04 | 1.9 | 3.76 | 2.8E-46 |
| *fbxa-72* | WBGene00017962 | protein_coding_gene | -1.5 | 0.35 | 2.8E-02 | 1.7 | 3.28 | 4.0E-05 |
| *ZC434.9* | WBGene00013895 | protein_coding_gene | -1.5 | 0.34 | 2.6E-05 | 1.8 | 3.58 | 1.1E-25 |
| *tag-10* | WBGene00006404 | protein_coding_gene | -1.5 | 0.34 | 2.0E-05 | 1.6 | 3.11 | 3.5E-20 |
| *cpg-9* | WBGene00022072 | protein_coding_gene | -1.5 | 0.34 | 6.5E-04 | 2.3 | 5.09 | 1.4E-24 |
| *decr-1.1* | WBGene00018380 | protein_coding_gene | -1.6 | 0.34 | 3.6E-03 | 1.2 | 2.36 | 8.3E-06 |
| *T01D1.3* | WBGene00020148 | protein_coding_gene | -1.6 | 0.34 | 2.6E-02 | 1.6 | 3.04 | 2.5E-09 |
| *tyr-2* | WBGene00010661 | protein_coding_gene | -1.6 | 0.33 | 4.6E-04 | 2.0 | 4.08 | 5.6E-13 |
| *T25B9.1* | WBGene00012007 | protein_coding_gene | -1.6 | 0.33 | 1.4E-03 | 1.3 | 2.40 | 2.2E-07 |
| *kin-15* | WBGene00002199 | protein_coding_gene | -1.6 | 0.33 | 2.2E-02 | 3.5 | 10.96 | 1.2E-29 |
| *VB0395L.1* | WBGene00044150 | protein_coding_gene | -1.6 | 0.33 | 3.5E-03 | 1.7 | 3.14 | 3.7E-04 |
| *Y47G6A.19* | WBGene00021645 | protein_coding_gene | -1.6 | 0.33 | 3.4E-04 | 2.8 | 6.79 | 2.4E-41 |
| *spp-23* | WBGene00021335 | protein_coding_gene | -1.6 | 0.33 | 1.2E-02 | 4.0 | 15.85 | 6.1E-14 |
| *gsto-1* | WBGene00016204 | protein_coding_gene | -1.7 | 0.32 | 1.6E-05 | 1.0 | 2.03 | 1.1E-03 |
| *dhs-25* | WBGene00000988 | protein_coding_gene | -1.7 | 0.32 | 2.1E-04 | 3.2 | 8.89 | 6.4E-16 |
| *mua-6* | WBGene00003485 | protein_coding_gene | -1.7 | 0.32 | 5.3E-03 | 2.6 | 6.12 | 1.8E-32 |
| *dpy-6* | WBGene00001068 | protein_coding_gene | -1.7 | 0.32 | 2.9E-02 | 3.1 | 8.33 | 3.9E-06 |
| *nhr-114* | WBGene00003704 | protein_coding_gene | -1.7 | 0.31 | 1.4E-03 | 2.2 | 4.45 | 2.7E-48 |
| *abhd-3.2* | WBGene00016644 | protein_coding_gene | -1.7 | 0.31 | 2.3E-02 | 3.3 | 10.14 | 1.4E-34 |
| *F22E5.8* | WBGene00017707 | protein_coding_gene | -1.7 | 0.31 | 3.4E-02 | 1.3 | 2.42 | 7.2E-03 |
| *C52D10.3* | WBGene00016877 | protein_coding_gene | -1.7 | 0.31 | 4.8E-02 | 1.0 | 2.06 | 2.1E-02 |
| *sptl-2* | WBGene00018398 | protein_coding_gene | -1.7 | 0.31 | 9.4E-04 | 2.1 | 4.34 | 7.2E-27 |
| *ZK84.1* | WBGene00022649 | protein_coding_gene | -1.7 | 0.31 | 2.9E-05 | 1.1 | 2.15 | 2.3E-02 |

| | | | | | | | | |
|---|---|---|---|---|---|---|---|---|
| *C15F1.1* | WBGene00015794 | protein_coding_gene | -1.7 | 0.30 | 4.3E-05 | 1.9 | 3.63 | 1.6E-14 |
| *elo-6* | WBGene00001244 | protein_coding_gene | -1.7 | 0.30 | 3.9E-02 | 2.5 | 5.60 | 6.6E-12 |
| *C55A6.4* | WBGene00008333 | protein_coding_gene | -1.7 | 0.30 | 2.2E-03 | 1.0 | 2.04 | 5.4E-05 |
| *F41E6.12* | WBGene00018293 | protein_coding_gene | -1.7 | 0.30 | 2.7E-02 | 3.5 | 11.23 | 1.1E-26 |
| *F58G6.7* | WBGene00010277 | protein_coding_gene | -1.7 | 0.30 | 1.7E-02 | 2.3 | 4.90 | 1.2E-06 |
| *F53B3.6* | WBGene00018744 | protein_coding_gene | -1.7 | 0.30 | 4.8E-02 | 3.5 | 11.38 | 2.6E-15 |
| *mpst-7* | WBGene00011307 | protein_coding_gene | -1.8 | 0.30 | 3.0E-07 | 2.1 | 4.36 | 6.2E-06 |
| *col-159* | WBGene00000732 | protein_coding_gene | -1.8 | 0.29 | 4.3E-04 | 2.0 | 4.03 | 3.0E-15 |
| *Y50D4B.6* | WBGene00021745 | protein_coding_gene | -1.8 | 0.29 | 4.2E-03 | 3.0 | 7.77 | 1.9E-06 |
| *gst-10* | WBGene00001758 | protein_coding_gene | -1.8 | 0.28 | 8.8E-04 | 2.7 | 6.33 | 2.7E-24 |
| *vha-3* | WBGene00006912 | protein_coding_gene | -1.8 | 0.28 | 7.9E-03 | 1.2 | 2.28 | 6.4E-04 |
| *F40H3.2* | WBGene00018250 | protein_coding_gene | -1.8 | 0.28 | 2.4E-02 | 3.0 | 8.27 | 1.9E-17 |
| *C39B5.5* | WBGene00016523 | protein_coding_gene | -1.9 | 0.28 | 2.0E-03 | 1.6 | 2.98 | 2.9E-02 |
| *C45B2.2* | WBGene00016659 | protein_coding_gene | -1.9 | 0.27 | 8.6E-03 | 2.7 | 6.57 | 5.5E-10 |
| *pcp-3* | WBGene00003958 | protein_coding_gene | -1.9 | 0.27 | 1.7E-04 | 1.2 | 2.25 | 8.3E-12 |
| *cdr-2* | WBGene00008296 | protein_coding_gene | -1.9 | 0.27 | 9.2E-05 | 1.6 | 3.04 | 8.0E-12 |
| *ech-7* | WBGene00001156 | protein_coding_gene | -1.9 | 0.27 | 9.9E-05 | 2.0 | 4.11 | 1.1E-30 |
| *F38B6.4* | WBGene00018174 | protein_coding_gene | -1.9 | 0.27 | 1.4E-03 | 2.4 | 5.14 | 1.9E-26 |
| *col-149* | WBGene00000722 | protein_coding_gene | -1.9 | 0.26 | 6.9E-04 | 3.3 | 9.61 | 4.0E-11 |
| *clec-204* | WBGene00021603 | protein_coding_gene | -1.9 | 0.26 | 1.7E-06 | 2.0 | 3.90 | 4.6E-07 |
| *ugt-15* | WBGene00008097 | protein_coding_gene | -2.0 | 0.25 | 5.6E-03 | -1.9 | 0.27 | 1.4E-05 |
| *W07A12.8* | WBGene00012325 | protein_coding_gene | -2.0 | 0.25 | 2.1E-04 | 2.6 | 6.03 | 2.6E-03 |
| *K01D12.9* | WBGene00010468 | protein_coding_gene | -2.0 | 0.24 | 8.9E-03 | -2.0 | 0.25 | 3.7E-02 |
| *hrg-7* | WBGene00007605 | protein_coding_gene | -2.1 | 0.24 | 2.6E-04 | 1.2 | 2.29 | 4.3E-03 |
| *Y47D7A.6* | WBGene00021619 | protein_coding_gene | -2.1 | 0.24 | 1.4E-02 | 1.8 | 3.43 | 7.0E-05 |
| *C40H1.8* | WBGene00008043 | protein_coding_gene | -2.1 | 0.23 | 2.2E-03 | -3.1 | 0.12 | 3.8E-07 |
| *grd-10* | WBGene00001699 | protein_coding_gene | -2.1 | 0.23 | 9.9E-05 | 2.6 | 6.16 | 1.0E-15 |
| *F57B1.9* | WBGene00045488 | protein_coding_gene | -2.2 | 0.22 | 3.6E-03 | 2.1 | 4.19 | 6.7E-05 |
| *ins-33* | WBGene00002116 | protein_coding_gene | -2.2 | 0.22 | 2.9E-04 | 2.1 | 4.43 | 4.9E-07 |
| *C33D3.3* | WBGene00007894 | protein_coding_gene | -2.2 | 0.22 | 1.2E-03 | 1.8 | 3.51 | 4.7E-03 |
| *R193.2* | WBGene00020128 | protein_coding_gene | -2.2 | 0.21 | 9.7E-03 | 4.2 | 18.80 | 3.9E-60 |
| *clec-71* | WBGene00021582 | protein_coding_gene | -2.3 | 0.21 | 5.8E-03 | -2.3 | 0.21 | 6.5E-13 |
| *scl-6* | WBGene00008028 | protein_coding_gene | -2.3 | 0.21 | 4.4E-06 | 3.8 | 14.34 | 3.0E-11 |
| *T28A11.6* | WBGene00020873 | pseudogene | -2.3 | 0.21 | 4.4E-02 | 2.3 | 4.92 | 3.1E-02 |
| *F53F1.4* | WBGene00009982 | protein_coding_gene | -2.3 | 0.20 | 4.3E-03 | 2.7 | 6.51 | 3.7E-05 |
| *R12E2.14* | WBGene00020039 | protein_coding_gene | -2.4 | 0.19 | 3.4E-03 | -2.1 | 0.24 | 4.1E-02 |
| *grd-13* | WBGene00001702 | protein_coding_gene | -2.4 | 0.19 | 3.6E-04 | 2.8 | 6.97 | 2.6E-09 |
| *col-91* | WBGene00000666 | protein_coding_gene | -2.5 | 0.17 | 5.0E-04 | -1.3 | 0.41 | 3.8E-02 |
| *grd-14* | WBGene00001703 | protein_coding_gene | -2.5 | 0.17 | 7.9E-03 | 3.3 | 9.92 | 2.8E-04 |
| *hrg-4* | WBGene00009493 | protein_coding_gene | -2.7 | 0.15 | 4.7E-04 | 3.2 | 9.51 | 4.3E-10 |
| *clec-70* | WBGene00021581 | protein_coding_gene | -2.9 | 0.14 | 1.9E-05 | -2.4 | 0.19 | 5.6E-10 |
| *col-154* | WBGene00000727 | protein_coding_gene | -3.0 | 0.12 | 1.9E-05 | 1.2 | 2.35 | 4.4E-03 |
| *B0205.13* | WBGene00044644 | protein_coding_gene | -4.1 | 0.06 | 1.3E-13 | 1.1 | 2.18 | 2.2E-02 |

**Transcription Factors Enriched in *P. rettgeri* Infected Animals**

Shared (N2&CB4856) DEGs in response to P. rettgeri (vs. OP50)

| Gene Name | WormBase Gene ID | Sequence Name | N2 $log_2$FC | CB4856 $log_2$FC |
|---|---|---|---|---|
| *atf-2* | WBGene00000220 | K08F8.2 | 1.4 | -2.3 |
| *ceh-16* | WBGene00000439 | C13G5.1 | 1.7 | -1.5 |
| *ceh-27* | WBGene00000449 | F46F3.1 | 1.5 | -1.6 |
| *ceh-30* | WBGene00000451 | C33D12.7 | 1.8 | -1.3 |
| *ceh-43* | WBGene00000463 | C28A5.4 | 1.1 | -1.5 |
| *ces-2* | WBGene00000469 | ZK909.4 | 1.7 | -2.4 |
| *egl-43* | WBGene00001207 | R53.3 | 1.1 | -1.4 |
| *egl-46* | WBGene00001210 | K11G9.4 | 2.0 | -2.7 |
| *fkh-2* | WBGene00001434 | T14G12.4 | 1.0 | -1.6 |
| *fkh-7* | WBGene00001439 | F26D12.1 | 1.2 | -1.7 |
| *ham-1* | WBGene00001820 | F53B2.6 | 1.2 | -1.2 |

| | | | | |
|---|---|---|---|---|
| *ham-2* | WBGene00001821 | C07A12.1 | 1.2 | -1.5 |
| *hbl-1* | WBGene00001824 | F13D11.2 | 1.7 | -2.4 |
| *hlh-14* | WBGene00001958 | C18A3.8 | 1.5 | -1.5 |
| *hlh-16* | WBGene00001960 | DY3.3 | 1.7 | -2.0 |
| *mls-2* | WBGene00003377 | C39E6.4 | 1.3 | -1.1 |
| *moe-3* | WBGene00003388 | F32A11.6 | 1.2 | -1.9 |
| *ngn-1* | WBGene00003595 | Y69A2AR.29 | 1.4 | -1.3 |
| *nhr-18* | WBGene00003617 | F44C8.3 | 1.1 | -2.0 |
| *nhr-19* | WBGene00003618 | E02H1.7 | 1.9 | -1.8 |
| *nhr-32* | WBGene00003626 | K08H2.8 | 1.2 | -1.2 |
| *nhr-99* | WBGene00003689 | M02H5.1 | 2.2 | -3.2 |
| *nhr-114* | WBGene00003704 | Y45G5AM.1 | -1.7 | 2.2 |
| *nhr-127* | WBGene00003717 | T13F3.3 | 3.0 | -2.2 |
| *peb-1* | WBGene00003968 | T14F9.4 | 2.1 | -2.7 |
| *unc-3* | WBGene00006743 | Y16B4A.1 | 1.4 | -1.2 |
| *unc-39* | WBGene00006775 | F56A12.1 | 1.3 | -1.2 |
| *unc-42* | WBGene00006778 | F58E6.10 | 2.4 | -2.0 |
| *jmjd-3.2* | WBGene00009089 | F23D12.5 | 1.1 | -1.2 |
| *sptf-1* | WBGene00009743 | F45H11.1 | 1.2 | -1.2 |
| *fozi-1* | WBGene00010453 | K01B6.1 | 1.3 | -1.9 |
| *nhr-206* | WBGene00011097 | R07B7.13 | 1.2 | -2.3 |
| *zip-3* | WBGene00012330 | W07G1.3 | -1.1 | 1.6 |
| *zip-7* | WBGene00013100 | Y51H4A.4 | 1.7 | -1.6 |
| *unk-1* | WBGene00016407 | C34D10.2 | 1.8 | -2.0 |
| *dmd-5* | WBGene00017326 | F10C1.5 | 1.6 | -1.8 |
| *F13C5.2* | WBGene00017423 | F13C5.2 | 1.0 | -1.2 |
| *M03D4.4* | WBGene00019751 | M03D4.4 | 1.4 | -2.2 |
| *crtc-1* | WBGene00021237 | Y20F4.2 | 1.0 | -1.2 |
| *Y82E9BR.17* | WBGene00022349 | Y82E9BR.17 | 1.2 | -1.9 |
| | | N2-Specific | | |
| *dac-1* | WBGene00000895 | B0412.1 | -1.2 | Not DE |
| *nhr-21* | WBGene00003620 | F21D12.1 | 1.2 | Not DE |
| *nhr-58* | WBGene00003648 | R11G11.2 | 1.8 | Not DE |
| *nhr-115* | WBGene00003705 | T27B7.4 | -1.9 | Not DE |
| *nhr-116* | WBGene00003706 | F09C6.9 | 1.4 | Not DE |
| *ref-2* | WBGene00004335 | C47C12.3 | 1.3 | Not DE |
| *jmjd-3.3* | WBGene00007813 | C29F7.6 | 1.1 | Not DE |
| *gmeb-3* | WBGene00008092 | C44F1.2 | -2.4 | Not DE |
| *nhr-171* | WBGene00016926 | C54F6.8 | 1.6 | Not DE |
| *zip-8* | WBGene00017755 | F23F12.9 | 1.2 | Not DE |
| *ceh-45* | WBGene00022837 | ZK993.1 | 1.8 | Not DE |
| *nhr-74* | WBGene00003664 | C27C7.3 | -3.1 | Not DE |
| | | CB4856-Specific | | |
| *bro-1* | WBGene00000272 | F56A3.5 | Not DE | -3.0 |
| *ceh-2* | WBGene00000429 | C27A12.5 | Not DE | -1.5 |
| *ceh-5* | WBGene00000430 | C16C2.1 | Not DE | -2.5 |
| *ceh-6* | WBGene00000431 | K02B12.1 | Not DE | -1.4 |
| *ceh-8* | WBGene00000433 | ZK265.4 | Not DE | -2.1 |
| *ceh-10* | WBGene00000435 | W03A3.1 | Not DE | -1.5 |
| *ceh-20* | WBGene00000443 | F31E3.1 | Not DE | -1.1 |
| *ceh-22* | WBGene00000445 | F29F11.5 | Not DE | -1.6 |
| *pros-1* | WBGene00000448 | K12H4.1 | Not DE | -2.2 |
| *ceh-31* | WBGene00000452 | C33D12.1 | Not DE | -1.4 |
| *ceh-32* | WBGene00000453 | W05E10.3 | Not DE | -1.4 |
| *ceh-34* | WBGene00000455 | C10G8.6 | Not DE | -1.7 |
| *ceh-36* | WBGene00000457 | C37E2.4 | Not DE | -1.2 |
| *ceh-37* | WBGene00000458 | C37E2.5 | Not DE | -1.5 |
| *ces-1* | WBGene00000468 | F43G9.11 | Not DE | -2.6 |

| | | | | |
|---|---|---|---|---|
| *cfi-1* | WBGene00000476 | T23D8.8 | Not DE | -1.2 |
| *cky-1* | WBGene00000521 | C15C8.2 | Not DE | -2.2 |
| *cnd-1* | WBGene00000561 | C34E10.7 | Not DE | -1.7 |
| *daf-14* | WBGene00000910 | F01G10.8 | Not DE | -1.0 |
| *egl-38* | WBGene00001204 | C04G2.7 | Not DE | -1.7 |
| *elt-6* | WBGene00001253 | F52C12.5 | Not DE | -1.3 |
| *fax-1* | WBGene00001400 | F56E3.4 | Not DE | -2.0 |
| *fkh-8* | WBGene00001440 | F40H3.4 | Not DE | -1.8 |
| *gei-13* | WBGene00001570 | F58A4.11 | Not DE | -2.0 |
| *grh-1* | WBGene00001707 | Y48G8AR.1 | Not DE | -1.1 |
| *hlh-6* | WBGene00001952 | T15H9.3 | Not DE | -1.9 |
| *hlh-13* | WBGene00001957 | F48D6.3 | Not DE | -2.8 |
| *hmg-11* | WBGene00001976 | T05A7.4 | Not DE | -1.2 |
| *hnd-1* | WBGene00001981 | C44C10.8 | Not DE | -1.1 |
| *let-381* | WBGene00002601 | F26B1.7 | Not DE | -2.0 |
| *lim-4* | WBGene00002987 | ZC64.4 | Not DE | -1.9 |
| *lin-31* | WBGene00003017 | K10G6.1 | Not DE | 1.6 |
| *lin-32* | WBGene00003018 | T14F9.5 | Not DE | -1.7 |
| *mdl-1* | WBGene00003163 | R03E9.1 | Not DE | -1.1 |
| *mec-3* | WBGene00003167 | F01D4.6 | Not DE | -2.0 |
| *mxl-3* | WBGene00003511 | F46G10.6 | Not DE | 2.0 |
| *nhr-25* | WBGene00003623 | F11C1.6 | Not DE | -1.4 |
| *nhr-31* | WBGene00003625 | C26B2.3 | Not DE | 1.2 |
| *nhr-35* | WBGene00003628 | C07A12.3 | Not DE | -1.2 |
| *nhr-42* | WBGene00003632 | C33G8.6 | Not DE | -1.7 |
| *nhr-44* | WBGene00003634 | T19A5.4 | Not DE | -1.7 |
| *nhr-55* | WBGene00003645 | T01G6.7 | Not DE | -1.0 |
| *nhr-56* | WBGene00003646 | F44C8.6 | Not DE | -1.4 |
| *nhr-59* | WBGene00003649 | T27B7.1 | Not DE | -1.3 |
| *nhr-67* | WBGene00003657 | C08F8.8 | Not DE | -1.8 |
| *nhr-69* | WBGene00003659 | T23H4.2 | Not DE | -1.2 |
| *nhr-81* | WBGene00003671 | C47F8.8 | Not DE | -2.7 |
| *nhr-90* | WBGene00003680 | ZK488.2 | Not DE | -1.0 |
| *nhr-91* | WBGene00003681 | Y15E3A.1 | Not DE | -2.2 |
| *nhr-92* | WBGene00003682 | Y41D4B.8 | Not DE | -1.0 |
| *nhr-94* | WBGene00003684 | C12D5.8 | Not DE | -1.1 |
| *nhr-103* | WBGene00003693 | F44C8.4 | Not DE | -1.4 |
| *nhr-107* | WBGene00003697 | C33G8.11 | Not DE | -1.0 |
| *nhr-110* | WBGene00003700 | Y46H3D.5 | Not DE | -1.2 |
| *nhr-117* | WBGene00003707 | F16B4.12 | Not DE | -1.2 |
| *nhr-120* | WBGene00003710 | C25B8.6 | Not DE | -1.2 |
| *nhr-131* | WBGene00003721 | T01G6.2 | Not DE | -1.2 |
| *nhr-134* | WBGene00003724 | F44C8.2 | Not DE | -1.4 |
| *pag-3* | WBGene00003909 | F45B8.4 | Not DE | -1.6 |
| *pat-9* | WBGene00003933 | T27B1.2 | Not DE | -1.3 |
| *pax-1* | WBGene00003937 | K07C11.1 | Not DE | -2.0 |
| *pax-3* | WBGene00003939 | F27E5.2 | Not DE | -1.4 |
| *pha-2* | WBGene00004011 | M6.3 | Not DE | -1.4 |
| *pha-4* | WBGene00004013 | F38A6.1 | Not DE | -1.4 |
| *plx-2* | WBGene00004048 | K04B12.1 | Not DE | -1.5 |
| *sma-2* | WBGene00004856 | ZK370.2 | Not DE | -1.0 |
| *sox-2* | WBGene00004949 | K08A8.2 | Not DE | -1.3 |
| *tab-1* | WBGene00006380 | F31E8.3 | Not DE | -1.3 |
| *rhgf-2* | WBGene00006476 | T08H4.1 | Not DE | -1.1 |
| *tbx-2* | WBGene00006543 | F21H11.3 | Not DE | -1.1 |
| *tbx-7* | WBGene00006544 | ZK328.8 | Not DE | -1.6 |
| *ttx-1* | WBGene00006652 | Y113G7A.6 | Not DE | -1.0 |
| *unc-30* | WBGene00006766 | B0564.10 | Not DE | -1.9 |

| | | | | |
|---|---|---|---|---|
| *unc-86* | WBGene00006818 | C30A5.7 | Not DE | -2.1 |
| *unc-120* | WBGene00006844 | D1081.2 | Not DE | -1.3 |
| *unc-130* | WBGene00006853 | C47G2.2 | Not DE | -1.4 |
| *vab-3* | WBGene00006870 | F14F3.1 | Not DE | -1.6 |
| *vab-7* | WBGene00006873 | M142.4 | Not DE | -1.5 |
| *vab-15* | WBGene00006881 | R07B1.1 | Not DE | -1.6 |
| *zag-1* | WBGene00006970 | F28F9.1 | Not DE | -1.6 |
| *nhr-150* | WBGene00007367 | C06B8.1 | Not DE | -1.6 |
| *ceh-79* | WBGene00007749 | C26E1.3 | Not DE | -1.0 |
| *dmd-4* | WBGene00007776 | C27C12.6 | Not DE | -1.8 |
| *C34B4.2* | WBGene00007907 | C34B4.2 | Not DE | 1.1 |
| *zip-5* | WBGene00007932 | C34D1.5 | Not DE | -1.3 |
| *irx-1* | WBGene00007984 | C36F7.1 | Not DE | -1.1 |
| *nhr-165* | WBGene00008158 | C47F8.2 | Not DE | -2.5 |
| *hmg-6* | WBGene00009827 | F47G4.6 | Not DE | -2.5 |
| *efl-3* | WBGene00009899 | F49E12.6 | Not DE | -1.7 |
| *ztf-11* | WBGene00009939 | F52F12.6 | Not DE | -1.1 |
| *sta-2* | WBGene00010251 | F58E6.1 | Not DE | -1.4 |
| *nhr-208* | WBGene00011099 | R07B7.15 | Not DE | -2.2 |
| *jmjd-4* | WBGene00011563 | T07C4.11 | Not DE | -1.1 |
| *nhr-218* | WBGene00011750 | T13F3.2 | Not DE | -2.0 |
| *tsct-1* | WBGene00011824 | T18D3.7 | Not DE | -1.7 |
| *zip-10* | WBGene00012101 | T27F2.4 | Not DE | -1.2 |
| *nhr-232* | WBGene00012494 | Y22F5A.1 | Not DE | -1.5 |
| *madf-10* | WBGene00013717 | Y106G6H.4 | Not DE | -1.2 |
| *ztf-30* | WBGene00015523 | C06E1.8 | Not DE | -2.8 |
| *C08G9.2* | WBGene00015620 | C08G9.2 | Not DE | -1.5 |
| *nhr-152* | WBGene00015705 | C12D5.2 | Not DE | -1.5 |
| *ceh-48* | WBGene00015934 | C17H12.9 | Not DE | -1.4 |
| *elt-7* | WBGene00015981 | C18G1.2 | Not DE | -1.0 |
| *nhr-140* | WBGene00016366 | C33G8.9 | Not DE | -1.1 |
| *ets-5* | WBGene00016600 | C42D8.4 | Not DE | 2.1 |
| *nhr-172* | WBGene00016927 | C54F6.9 | Not DE | -2.4 |
| *set-9* | WBGene00017482 | F15E6.1 | Not DE | 1.1 |
| *atf-8* | WBGene00017535 | F17A9.3 | Not DE | -2.5 |
| *F26A10.2* | WBGene00017814 | F26A10.2 | Not DE | -1.9 |
| *nhr-180* | WBGene00017961 | F31F4.12 | Not DE | -1.8 |
| *ztf-13* | WBGene00018704 | F52E4.8 | Not DE | -1.2 |
| *nhr-195* | WBGene00019115 | F59E11.10 | Not DE | -1.0 |
| *lect-2* | WBGene00019407 | K05F1.5 | Not DE | -1.2 |
| *nhr-210* | WBGene00020015 | R11G11.12 | Not DE | -1.1 |
| *ast-1* | WBGene00020368 | T08H4.3 | Not DE | -2.0 |
| *dhhc-11* | WBGene00020694 | T22E7.2 | Not DE | -2.3 |
| *nhr-221* | WBGene00020748 | T24A6.8 | Not DE | -2.8 |
| *W04B5.2* | WBGene00021019 | W04B5.2 | Not DE | -1.3 |
| *Y82E9BR.1* | WBGene00022334 | Y82E9BR.1 | Not DE | -1.8 |
| *nhr-41* | WBGene00022423 | Y104H12A.1 | Not DE | -1.0 |
| *klu-2* | WBGene00022592 | ZC328.2 | Not DE | 1.1 |
| *syd-9* | WBGene00044068 | ZK867.1 | Not DE | -1.8 |
| *tag-243* | WBGene00044078 | T04A8.4 | Not DE | -1.2 |

**Supplemental Table 10**

**Lists of differentially expressed genes that are identified in either N2, CB4856, or both, after infection by *S. epidermidis* (compared to *S. aureus*)**

**Supplemental Table 10 - Differentially Expressed Genes in the *S. epidermidis* Condition**

**N2**-specific genes exclusively expressed following *S. epidermidis* infection (vs. OP50) or DE in opposing directions following infection with *S. epidermidis* or *S. auerus* (vs. OP50)

| Gene Name | Wormbase Gene ID | Type | $log_2FC$ | FC | Adj P-value |
|---|---|---|---|---|---|
| *C14C6.8* | WBGene00015762 | protein_coding_gene | 4.6 | 25.07 | 1.2E-03 |
| *cyp-35B3* | WBGene00019470 | protein_coding_gene | 4.5 | 23.02 | 2.0E-03 |
| *clec-2* | WBGene00015198 | protein_coding_gene | 4.3 | 19.19 | 2.8E-04 |
| *Y41D4B.6* | WBGene00021513 | protein_coding_gene | 4.0 | 16.47 | 1.3E-06 |
| *M7.10* | WBGene00010886 | protein_coding_gene | 4.0 | 15.85 | 1.4E-03 |
| *F54E2.2* | WBGene00018824 | protein_coding_gene | 3.8 | 13.93 | 1.8E-03 |
| *T05A7.11* | WBGene00044383 | protein_coding_gene | 3.8 | 13.86 | 8.5E-05 |
| *C35A5.11* | WBGene00044024 | protein_coding_gene | 3.5 | 11.05 | 2.4E-07 |
| *K10G6.4* | WBGene00019641 | protein_coding_gene | 3.4 | 10.78 | 6.2E-04 |
| *T05A7.13* | WBGene00271415 | protein_coding_gene | 3.4 | 10.64 | 2.3E-05 |
| *F18C5.9* | WBGene00017562 | protein_coding_gene | 3.4 | 10.37 | 4.3E-03 |
| *ZK675.4* | WBGene00014067 | protein_coding_gene | 3.4 | 10.22 | 3.8E-04 |
| *dsl-6* | WBGene00001108 | protein_coding_gene | 3.3 | 10.04 | 7.6E-05 |
| *aat-7* | WBGene00000008 | protein_coding_gene | 3.3 | 9.82 | 3.3E-23 |
| *pqn-73* | WBGene00004155 | protein_coding_gene | 3.2 | 9.31 | 6.4E-05 |
| *pax-1* | WBGene00003937 | protein_coding_gene | 3.2 | 9.16 | 7.5E-03 |
| *hil-7* | WBGene00001858 | protein_coding_gene | 3.2 | 9.03 | 1.3E-04 |
| *F47B7.6* | WBGene00018537 | pseudogene | 3.1 | 8.84 | 1.2E-02 |
| *clec-78* | WBGene00018547 | protein_coding_gene | 3.1 | 8.81 | 2.7E-07 |
| *nhr-218* | WBGene00011750 | protein_coding_gene | 3.1 | 8.79 | 7.2E-03 |
| *dpy-14* | WBGene00001075 | protein_coding_gene | 3.1 | 8.57 | 1.1E-05 |
| *F21C10.10* | WBGene00017659 | protein_coding_gene | 3.1 | 8.46 | 2.0E-21 |
| *C06E7.88* | WBGene00189995 | protein_coding_gene | 3.0 | 8.28 | 1.9E-04 |
| *ZK381.2* | WBGene00022725 | protein_coding_gene | 3.0 | 8.19 | 1.9E-02 |
| *ZK813.6* | WBGene00044564 | protein_coding_gene | 3.0 | 7.95 | 1.9E-02 |
| *T02G6.7* | WBGene00011390 | protein_coding_gene | 3.0 | 7.94 | 1.8E-02 |
| *R09E10.5* | WBGene00011175 | protein_coding_gene | 3.0 | 7.83 | 1.3E-05 |
| *F17B5.6* | WBGene00044058 | protein_coding_gene | 3.0 | 7.81 | 2.1E-02 |
| *lips-6* | WBGene00014009 | protein_coding_gene | 2.9 | 7.64 | 8.5E-04 |
| *nlp-39* | WBGene00008295 | protein_coding_gene | 2.9 | 7.57 | 1.1E-03 |
| *scl-15* | WBGene00017183 | protein_coding_gene | 2.9 | 7.50 | 2.0E-02 |
| *col-121* | WBGene00000695 | protein_coding_gene | 2.9 | 7.41 | 4.9E-03 |
| *F46B6.13* | WBGene00044192 | protein_coding_gene | 2.9 | 7.37 | 1.9E-02 |
| *egl-46* | WBGene00001210 | protein_coding_gene | 2.9 | 7.31 | 1.8E-08 |
| *clec-263* | WBGene00009171 | protein_coding_gene | 2.9 | 7.29 | 3.6E-02 |
| *dpy-17* | WBGene00001076 | protein_coding_gene | 2.9 | 7.25 | 2.1E-09 |
| *M176.10* | WBGene00010946 | protein_coding_gene | 2.8 | 6.95 | 1.9E-02 |
| *bgnt-1.3* | WBGene00009032 | protein_coding_gene | 2.8 | 6.84 | 3.5E-02 |
| *ifa-3* | WBGene00002051 | protein_coding_gene | 2.8 | 6.84 | 2.8E-05 |
| *T04C12.1* | WBGene00011427 | protein_coding_gene | 2.8 | 6.83 | 1.1E-03 |
| *hlh-32* | WBGene00013665 | protein_coding_gene | 2.8 | 6.75 | 3.6E-02 |
| *C01C4.2* | WBGene00015292 | protein_coding_gene | 2.8 | 6.75 | 2.4E-02 |
| *T09D3.3* | WBGene00020384 | protein_coding_gene | 2.7 | 6.60 | 3.8E-02 |
| *math-10* | WBGene00015839 | protein_coding_gene | 2.7 | 6.58 | 1.3E-05 |
| *hch-1* | WBGene00001828 | protein_coding_gene | 2.7 | 6.47 | 2.8E-05 |
| *scl-8* | WBGene00008030 | protein_coding_gene | 2.7 | 6.43 | 1.8E-02 |
| *W07G4.1* | WBGene00012335 | protein_coding_gene | 2.7 | 6.28 | 3.3E-02 |

| | | | | | |
|---|---|---|---|---|---|
| *C03C10.5* | WBGene00007271 | protein_coding_gene | 2.6 | 6.18 | 5.2E-04 |
| *R160.4* | WBGene00020121 | protein_coding_gene | 2.6 | 6.05 | 1.2E-02 |
| *cutl-13* | WBGene00021025 | protein_coding_gene | 2.6 | 6.02 | 4.3E-03 |
| *C27A2.8* | WBGene00044387 | protein_coding_gene | 2.6 | 5.92 | 3.0E-02 |
| *atf-2* | WBGene00000220 | protein_coding_gene | 2.6 | 5.87 | 1.9E-06 |
| *F31D4.5* | WBGene00009292 | protein_coding_gene | 2.5 | 5.85 | 2.3E-06 |
| *hbl-1* | WBGene00001824 | protein_coding_gene | 2.5 | 5.80 | 5.6E-06 |
| *str-173* | WBGene00006217 | protein_coding_gene | 2.5 | 5.70 | 3.2E-02 |
| *clec-208* | WBGene00018097 | protein_coding_gene | 2.5 | 5.66 | 1.3E-02 |
| *E02H9.6* | WBGene00017104 | protein_coding_gene | 2.4 | 5.46 | 9.0E-03 |
| *dpy-10* | WBGene00001072 | protein_coding_gene | 2.4 | 5.32 | 1.4E-10 |
| *T15D6.12* | WBGene00011786 | protein_coding_gene | 2.4 | 5.25 | 2.3E-02 |
| *C29A12.6* | WBGene00007800 | protein_coding_gene | 2.4 | 5.25 | 2.8E-03 |
| *ces-2* | WBGene00000469 | protein_coding_gene | 2.4 | 5.23 | 1.4E-06 |
| *C08F1.11* | WBGene00044417 | protein_coding_gene | 2.4 | 5.20 | 4.6E-04 |
| *K11H12.11* | WBGene00044723 | protein_coding_gene | 2.4 | 5.19 | 6.5E-03 |
| *C07D10.6* | WBGene00044673 | protein_coding_gene | 2.4 | 5.17 | 1.6E-02 |
| *ZC84.6* | WBGene00013849 | protein_coding_gene | 2.4 | 5.14 | 2.0E-03 |
| *rocf-1* | WBGene00011948 | protein_coding_gene | 2.4 | 5.14 | 1.2E-02 |
| *dao-2* | WBGene00000928 | protein_coding_gene | 2.3 | 5.05 | 3.7E-10 |
| *F55B11.5* | WBGene00010087 | protein_coding_gene | 2.3 | 4.99 | 4.2E-03 |
| *tmem-231* | WBGene00020825 | protein_coding_gene | 2.3 | 4.97 | 2.7E-03 |
| *cut-5* | WBGene00011104 | protein_coding_gene | 2.3 | 4.97 | 1.6E-03 |
| *fipr-17* | WBGene00012965 | protein_coding_gene | 2.3 | 4.88 | 2.4E-02 |
| *dsl-3* | WBGene00001105 | protein_coding_gene | 2.3 | 4.83 | 9.0E-07 |
| *ces-1* | WBGene00000468 | protein_coding_gene | 2.3 | 4.82 | 7.5E-03 |
| *icl-1* | WBGene00001564 | protein_coding_gene | 2.3 | 4.80 | 4.0E-07 |
| *F23F12.8* | WBGene00017754 | protein_coding_gene | 2.2 | 4.73 | 9.6E-04 |
| *ins-2* | WBGene00002085 | protein_coding_gene | 2.2 | 4.68 | 3.0E-03 |
| *C14C6.5* | WBGene00015759 | protein_coding_gene | 2.2 | 4.64 | 3.1E-03 |
| *cutl-2* | WBGene00013145 | protein_coding_gene | 2.2 | 4.63 | 8.1E-05 |
| *noah-2* | WBGene00009926 | protein_coding_gene | 2.2 | 4.62 | 3.6E-07 |
| *K08B4.2* | WBGene00019514 | protein_coding_gene | 2.2 | 4.60 | 9.1E-04 |
| *K11H12.6* | WBGene00019662 | protein_coding_gene | 2.2 | 4.59 | 1.2E-02 |
| *F31E3.12* | WBGene00219317 | protein_coding_gene | 2.2 | 4.57 | 4.5E-03 |
| *clec-232* | WBGene00009487 | protein_coding_gene | 2.2 | 4.52 | 5.0E-02 |
| *C27A7.8* | WBGene00007758 | protein_coding_gene | 2.2 | 4.47 | 1.0E-08 |
| *Y41D4B.26* | WBGene00021525 | protein_coding_gene | 2.2 | 4.45 | 2.7E-04 |
| *fmi-1* | WBGene00001475 | protein_coding_gene | 2.2 | 4.44 | 4.8E-06 |
| *T14B4.19* | WBGene00302993 | protein_coding_gene | 2.1 | 4.34 | 1.2E-04 |
| *clec-161* | WBGene00010228 | protein_coding_gene | 2.1 | 4.30 | 4.6E-02 |
| *nphp-1* | WBGene00010898 | protein_coding_gene | 2.1 | 4.29 | 3.3E-04 |
| *C03A7.2* | WBGene00015366 | protein_coding_gene | 2.1 | 4.26 | 2.7E-02 |
| *F57B1.6* | WBGene00010189 | protein_coding_gene | 2.1 | 4.23 | 1.7E-03 |
| *noah-1* | WBGene00016422 | protein_coding_gene | 2.1 | 4.21 | 1.9E-07 |
| *F53C3.3* | WBGene00018747 | protein_coding_gene | 2.1 | 4.19 | 7.7E-03 |
| *cut-6* | WBGene00000853 | protein_coding_gene | 2.1 | 4.15 | 2.0E-03 |
| *hlh-16* | WBGene00001960 | protein_coding_gene | 2.0 | 4.08 | 4.1E-03 |
| *hpo-2* | WBGene00012500 | protein_coding_gene | 2.0 | 4.01 | 4.3E-02 |
| *T05H10.3* | WBGene00011508 | protein_coding_gene | 2.0 | 3.99 | 4.0E-03 |
| *wrt-10* | WBGene00006956 | protein_coding_gene | 2.0 | 3.95 | 1.7E-07 |
| *math-14* | WBGene00015828 | protein_coding_gene | 2.0 | 3.90 | 7.7E-15 |

| | | | | | |
|---|---|---|---|---|---|
| *F41C3.1* | WBGene00018267 | protein_coding_gene | 1.9 | 3.82 | 2.0E-02 |
| *mam-5* | WBGene00011572 | protein_coding_gene | 1.9 | 3.80 | 4.7E-02 |
| *R04B3.1* | WBGene00019866 | protein_coding_gene | 1.9 | 3.77 | 4.6E-02 |
| *ceh-8* | WBGene00000433 | protein_coding_gene | 1.9 | 3.74 | 2.3E-02 |
| *K01D12.1* | WBGene00010461 | protein_coding_gene | 1.9 | 3.67 | 4.6E-02 |
| *F22F7.3* | WBGene00017721 | protein_coding_gene | 1.9 | 3.66 | 7.9E-03 |
| *F54B11.10* | WBGene00010033 | protein_coding_gene | 1.9 | 3.64 | 2.3E-02 |
| *ZC84.1* | WBGene00013846 | protein_coding_gene | 1.9 | 3.61 | 3.4E-03 |
| *del-3* | WBGene00009144 | protein_coding_gene | 1.9 | 3.61 | 4.4E-02 |
| *R160.10* | WBGene00219369 | protein_coding_gene | 1.8 | 3.59 | 1.9E-02 |
| *Y43F8C.18* | WBGene00012839 | ncRNA_gene | 1.8 | 3.59 | 4.1E-02 |
| *hrg-1* | WBGene00019830 | protein_coding_gene | 1.8 | 3.59 | 7.0E-06 |
| *C40H1.2* | WBGene00008038 | protein_coding_gene | 1.8 | 3.57 | 2.2E-04 |
| *let-4* | WBGene00002282 | protein_coding_gene | 1.8 | 3.55 | 1.1E-04 |
| *Y23H5B.8* | WBGene00021279 | protein_coding_gene | 1.8 | 3.54 | 2.4E-04 |
| *M01D1.12* | WBGene00269373 | protein_coding_gene | 1.8 | 3.53 | 8.9E-03 |
| *ptr-19* | WBGene00004233 | protein_coding_gene | 1.8 | 3.53 | 1.8E-03 |
| *sym-1* | WBGene00006366 | protein_coding_gene | 1.8 | 3.52 | 2.4E-06 |
| *C35A5.6* | WBGene00007950 | protein_coding_gene | 1.8 | 3.52 | 2.8E-07 |
| *ceh-16* | WBGene00000439 | protein_coding_gene | 1.8 | 3.46 | 3.6E-03 |
| *gsto-1* | WBGene00016204 | protein_coding_gene | 1.8 | 3.45 | 1.3E-08 |
| *C26C6.4* | WBGene00007739 | protein_coding_gene | 1.8 | 3.44 | 2.1E-02 |
| *M01H9.2* | WBGene00019718 | protein_coding_gene | 1.8 | 3.42 | 6.5E-03 |
| *cdh-5* | WBGene00000397 | protein_coding_gene | 1.8 | 3.41 | 1.6E-04 |
| *W01C9.1* | WBGene00012175 | protein_coding_gene | 1.8 | 3.39 | 2.7E-02 |
| *F38E9.1* | WBGene00018184 | protein_coding_gene | 1.7 | 3.36 | 6.8E-09 |
| *ZK822.1* | WBGene00014089 | protein_coding_gene | 1.7 | 3.34 | 1.2E-03 |
| *Y44E3A.1* | WBGene00021547 | protein_coding_gene | 1.7 | 3.30 | 4.9E-08 |
| *ZK593.9* | WBGene00014005 | pseudogene | 1.7 | 3.30 | 6.4E-03 |
| *T05E12.3* | WBGene00011486 | protein_coding_gene | 1.7 | 3.29 | 2.2E-07 |
| *Y82E9BL.3* | WBGene00022320 | protein_coding_gene | 1.7 | 3.29 | 4.6E-02 |
| *bus-18* | WBGene00044631 | protein_coding_gene | 1.7 | 3.24 | 2.2E-03 |
| *C03A7.13* | WBGene00015371 | protein_coding_gene | 1.7 | 3.20 | 4.9E-04 |
| *rig-5* | WBGene00004372 | protein_coding_gene | 1.7 | 3.18 | 1.5E-03 |
| *Y46H3C.6* | WBGene00021606 | pseudogene | 1.7 | 3.14 | 2.1E-02 |
| *ZC449.2* | WBGene00022612 | protein_coding_gene | 1.6 | 3.12 | 7.4E-03 |
| *F14B4.1* | WBGene00008779 | protein_coding_gene | 1.6 | 3.09 | 2.3E-05 |
| *C28C12.3* | WBGene00016179 | protein_coding_gene | 1.6 | 3.05 | 5.0E-02 |
| *Y51H7C.12* | WBGene00021790 | protein_coding_gene | 1.6 | 3.04 | 3.6E-02 |
| *clec-147* | WBGene00007313 | protein_coding_gene | 1.6 | 3.04 | 2.9E-11 |
| *nab-1* | WBGene00003516 | protein_coding_gene | 1.6 | 3.03 | 7.2E-05 |
| *tbb-4* | WBGene00006538 | protein_coding_gene | 1.6 | 3.02 | 6.4E-04 |
| *fbn-1* | WBGene00022816 | protein_coding_gene | 1.6 | 2.98 | 1.2E-03 |
| *cutl-25* | WBGene00021919 | protein_coding_gene | 1.6 | 2.97 | 2.2E-03 |
| *nhr-171* | WBGene00016926 | protein_coding_gene | 1.6 | 2.94 | 1.9E-02 |
| *fbxb-101* | WBGene00013404 | protein_coding_gene | 1.5 | 2.92 | 3.5E-02 |
| *E01G4.7* | WBGene00045410 | pseudogene | 1.5 | 2.92 | 1.2E-09 |
| *ceh-30* | WBGene00000451 | protein_coding_gene | 1.5 | 2.90 | 2.1E-02 |
| *hsp-12.1* | WBGene00011906 | protein_coding_gene | 1.5 | 2.88 | 2.6E-03 |
| *dyf-7* | WBGene00001123 | protein_coding_gene | 1.5 | 2.88 | 2.3E-03 |
| *dpy-1* | WBGene00001063 | protein_coding_gene | 1.5 | 2.87 | 2.6E-02 |
| *C17E4.19* | WBGene00219310 | protein_coding_gene | 1.5 | 2.84 | 4.8E-02 |

| | | | | | |
|---|---|---|---|---|---|
| *srx-125* | WBGene00006016 | protein_coding_gene | 1.5 | 2.80 | 2.1E-02 |
| *F55G1.7* | WBGene00018903 | protein_coding_gene | 1.5 | 2.78 | 4.8E-07 |
| *Y51H4A.1* | WBGene00013098 | protein_coding_gene | 1.5 | 2.76 | 3.6E-02 |
| *F35D2.3* | WBGene00018041 | protein_coding_gene | 1.5 | 2.74 | 1.8E-05 |
| *T19C9.8* | WBGene00011844 | protein_coding_gene | 1.4 | 2.72 | 3.3E-02 |
| *dmd-5* | WBGene00017326 | protein_coding_gene | 1.4 | 2.72 | 1.9E-02 |
| *zip-7* | WBGene00013100 | protein_coding_gene | 1.4 | 2.66 | 3.3E-03 |
| *Y7A9D.1* | WBGene00012420 | protein_coding_gene | 1.4 | 2.65 | 1.0E-03 |
| *T26C12.3* | WBGene00020833 | protein_coding_gene | 1.4 | 2.65 | 4.0E-03 |
| *F53B3.5* | WBGene00018743 | protein_coding_gene | 1.4 | 2.64 | 2.6E-04 |
| *ddn-1* | WBGene00015227 | protein_coding_gene | 1.4 | 2.63 | 2.9E-03 |
| *F19C7.4* | WBGene00017594 | protein_coding_gene | 1.4 | 2.63 | 1.0E-04 |
| *C35D10.3* | WBGene00016441 | protein_coding_gene | 1.4 | 2.62 | 3.1E-02 |
| *dex-1* | WBGene00017028 | protein_coding_gene | 1.4 | 2.62 | 9.6E-04 |
| *K07E3.4* | WBGene00019492 | protein_coding_gene | 1.4 | 2.61 | 2.8E-09 |
| *best-16* | WBGene00020046 | protein_coding_gene | 1.4 | 2.60 | 1.5E-02 |
| *lgc-22* | WBGene00017483 | protein_coding_gene | 1.4 | 2.55 | 3.3E-04 |
| *F46E10.2* | WBGene00018489 | protein_coding_gene | 1.3 | 2.50 | 1.2E-03 |
| *Y38H6C.21* | WBGene00012633 | protein_coding_gene | 1.3 | 2.50 | 1.0E-03 |
| *nas-33* | WBGene00003551 | protein_coding_gene | 1.3 | 2.48 | 2.7E-02 |
| *flp-34* | WBGene00019975 | protein_coding_gene | 1.3 | 2.48 | 7.6E-03 |
| *rga-2* | WBGene00013135 | protein_coding_gene | 1.3 | 2.47 | 6.0E-03 |
| *C46C11.4* | WBGene00044567 | protein_coding_gene | 1.3 | 2.46 | 8.4E-03 |
| *F59B2.9* | WBGene00010311 | protein_coding_gene | 1.3 | 2.45 | 3.7E-03 |
| *hrg-7* | WBGene00007605 | protein_coding_gene | 1.3 | 2.45 | 1.4E-02 |
| *R07B7.6* | WBGene00011090 | protein_coding_gene | 1.3 | 2.43 | 9.0E-06 |
| *clec-66* | WBGene00009397 | protein_coding_gene | 1.3 | 2.42 | 8.2E-04 |
| *Y57A10A.26* | WBGene00013266 | protein_coding_gene | 1.3 | 2.41 | 1.3E-05 |
| *fat-1* | WBGene00001393 | protein_coding_gene | 1.3 | 2.41 | 6.7E-10 |
| *D2005.6* | WBGene00008401 | protein_coding_gene | 1.2 | 2.38 | 2.6E-02 |
| *C30G7.4* | WBGene00044178 | protein_coding_gene | 1.2 | 2.37 | 1.9E-03 |
| *K02E10.4* | WBGene00019318 | protein_coding_gene | 1.2 | 2.35 | 1.5E-02 |
| *lad-2* | WBGene00002243 | protein_coding_gene | 1.2 | 2.35 | 2.8E-03 |
| *acox-3* | WBGene00019060 | protein_coding_gene | 1.2 | 2.34 | 7.3E-03 |
| *nas-37* | WBGene00003553 | protein_coding_gene | 1.2 | 2.34 | 3.6E-03 |
| *B0454.6* | WBGene00015197 | protein_coding_gene | 1.2 | 2.33 | 1.1E-04 |
| *ptp-4* | WBGene00010558 | protein_coding_gene | 1.2 | 2.32 | 7.7E-03 |
| *ZK896.1* | WBGene00014132 | protein_coding_gene | 1.2 | 2.31 | 1.0E-02 |
| *T02G5.11* | WBGene00020167 | pseudogene | 1.2 | 2.29 | 9.6E-03 |
| *cdh-8* | WBGene00017576 | protein_coding_gene | 1.2 | 2.29 | 1.2E-03 |
| *C41G11.1* | WBGene00016568 | protein_coding_gene | 1.2 | 2.29 | 1.6E-04 |
| *C44E12.1* | WBGene00016657 | protein_coding_gene | 1.2 | 2.28 | 1.3E-05 |
| *jmjd-3.3* | WBGene00007813 | protein_coding_gene | 1.2 | 2.28 | 1.2E-02 |
| *T25B2.2* | WBGene00020786 | protein_coding_gene | 1.2 | 2.28 | 4.0E-04 |
| *ZK287.4* | WBGene00013965 | protein_coding_gene | 1.2 | 2.27 | 3.4E-02 |
| *hex-1* | WBGene00020509 | protein_coding_gene | 1.2 | 2.25 | 2.1E-05 |
| *K01A2.3* | WBGene00019278 | pseudogene | 1.2 | 2.25 | 1.9E-02 |
| *B0228.6* | WBGene00015063 | protein_coding_gene | 1.2 | 2.24 | 4.3E-05 |
| *ceh-32* | WBGene00000453 | protein_coding_gene | 1.2 | 2.23 | 1.3E-03 |
| *Y59H11AR.4* | WBGene00022012 | protein_coding_gene | 1.2 | 2.23 | 1.2E-03 |
| *ech-1.1* | WBGene00001150 | protein_coding_gene | 1.1 | 2.22 | 6.1E-03 |
| *C02B8.1* | WBGene00015320 | protein_coding_gene | 1.1 | 2.21 | 2.8E-02 |

| | | | | | |
|---|---|---|---|---|---|
| *F47E1.4* | WBGene00018568 | protein_coding_gene | 1.1 | 2.20 | 4.9E-03 |
| *C40A11.3* | WBGene00016546 | protein_coding_gene | 1.1 | 2.20 | 4.0E-02 |
| *pudl-2* | WBGene00044490 | protein_coding_gene | 1.1 | 2.20 | 2.2E-02 |
| *mks-5* | WBGene00007490 | protein_coding_gene | 1.1 | 2.20 | 3.0E-02 |
| *col-99* | WBGene00000674 | protein_coding_gene | 1.1 | 2.19 | 1.1E-02 |
| *cest-16* | WBGene00020330 | protein_coding_gene | 1.1 | 2.18 | 1.1E-02 |
| *spin-2* | WBGene00008033 | protein_coding_gene | 1.1 | 2.18 | 3.5E-03 |
| *lipl-4* | WBGene00019376 | protein_coding_gene | 1.1 | 2.18 | 3.7E-02 |
| *bam-2* | WBGene00000237 | protein_coding_gene | 1.1 | 2.17 | 1.4E-04 |
| *W02D7.4* | WBGene00020940 | protein_coding_gene | 1.1 | 2.17 | 1.1E-06 |
| *arrd-4* | WBGene00014159 | protein_coding_gene | 1.1 | 2.16 | 1.2E-02 |
| *mig-13* | WBGene00003245 | protein_coding_gene | 1.1 | 2.16 | 5.7E-04 |
| *Y62F5A.9* | WBGene00013387 | protein_coding_gene | 1.1 | 2.15 | 2.3E-02 |
| *hlh-14* | WBGene00001958 | protein_coding_gene | 1.1 | 2.15 | 1.1E-02 |
| *nhr-91* | WBGene00003681 | protein_coding_gene | 1.1 | 2.15 | 7.3E-03 |
| *cng-1* | WBGene00000562 | protein_coding_gene | 1.1 | 2.14 | 2.2E-03 |
| *fbxa-175* | WBGene00045293 | pseudogene | 1.1 | 2.14 | 2.5E-02 |
| *C08F1.10* | WBGene00015613 | protein_coding_gene | 1.1 | 2.13 | 3.2E-03 |
| *F23F1.6* | WBGene00017747 | protein_coding_gene | 1.1 | 2.13 | 9.1E-03 |
| *clec-54* | WBGene00008596 | protein_coding_gene | 1.1 | 2.12 | 9.3E-05 |
| *C23G10.14* | WBGene00271647 | ncRNA_gene | 1.1 | 2.12 | 4.3E-02 |
| *skr-21* | WBGene00004827 | protein_coding_gene | 1.1 | 2.11 | 1.1E-03 |
| *C11H1.7* | WBGene00007531 | protein_coding_gene | 1.1 | 2.10 | 4.1E-02 |
| *nlr-1* | WBGene00003772 | protein_coding_gene | 1.1 | 2.09 | 8.4E-05 |
| *nlp-73* | WBGene00013964 | protein_coding_gene | 1.1 | 2.09 | 1.8E-03 |
| *syg-1* | WBGene00006365 | protein_coding_gene | 1.1 | 2.09 | 8.3E-03 |
| *F13C5.5* | WBGene00017426 | protein_coding_gene | 1.1 | 2.09 | 9.0E-03 |
| *D1044.7* | WBGene00017032 | protein_coding_gene | 1.1 | 2.09 | 3.4E-02 |
| *W03G11.4* | WBGene00012226 | protein_coding_gene | 1.1 | 2.07 | 3.2E-03 |
| *B0403.6* | WBGene00043982 | protein_coding_gene | 1.0 | 2.06 | 2.0E-02 |
| *glt-6* | WBGene00001624 | protein_coding_gene | 1.0 | 2.06 | 3.8E-03 |
| *fat-4* | WBGene00001396 | protein_coding_gene | 1.0 | 2.05 | 2.4E-07 |
| *syg-2* | WBGene00007750 | protein_coding_gene | 1.0 | 2.04 | 5.9E-03 |
| *R09A1.3* | WBGene00019973 | protein_coding_gene | 1.0 | 2.04 | 1.2E-04 |
| *abts-4* | WBGene00019844 | protein_coding_gene | 1.0 | 2.04 | 4.1E-05 |
| *T09B4.5* | WBGene00020379 | protein_coding_gene | 1.0 | 2.03 | 1.0E-02 |
| *hsp-12.2* | WBGene00002011 | protein_coding_gene | 1.0 | 2.03 | 1.0E-03 |
| *kal-1* | WBGene00002181 | protein_coding_gene | 1.0 | 2.03 | 4.7E-04 |
| *catp-7* | WBGene00022010 | protein_coding_gene | 1.0 | 2.02 | 8.3E-03 |
| *DH11.2* | WBGene00008436 | protein_coding_gene | 1.0 | 2.01 | 3.8E-03 |
| *sto-1* | WBGene00006063 | protein_coding_gene | 1.0 | 2.01 | 1.4E-03 |
| *ddo-2* | WBGene00017565 | protein_coding_gene | 1.0 | 2.00 | 3.2E-04 |
| *lam-3* | WBGene00002248 | protein_coding_gene | 1.0 | 2.00 | 5.2E-03 |
| *mnp-1* | WBGene00007139 | protein_coding_gene | 1.0 | 2.00 | 8.1E-04 |
| *gst-39* | WBGene00001787 | protein_coding_gene | -1.0 | 0.50 | 1.1E-02 |
| *tyr-2* | WBGene00010661 | protein_coding_gene | -1.0 | 0.50 | 1.6E-02 |
| *F49D11.10* | WBGene00045433 | protein_coding_gene | -1.0 | 0.50 | 3.0E-05 |
| *oat-1* | WBGene00003837 | protein_coding_gene | -1.0 | 0.50 | 1.2E-02 |
| *cct-5* | WBGene00000380 | protein_coding_gene | -1.0 | 0.50 | 2.7E-05 |
| *cct-3* | WBGene00018782 | protein_coding_gene | -1.0 | 0.49 | 5.6E-08 |
| *nit-1* | WBGene00014206 | protein_coding_gene | -1.0 | 0.49 | 1.2E-03 |
| *cest-4* | WBGene00015930 | protein_coding_gene | -1.0 | 0.49 | 1.8E-04 |

| | | | | | |
|---|---|---|---|---|---|
| *aspm-1* | WBGene00008107 | protein_coding_gene | -1.0 | 0.49 | 2.4E-09 |
| *Y116F11B.10* | WBGene00013826 | pseudogene | -1.0 | 0.49 | 7.0E-03 |
| *W02H5.8* | WBGene00020962 | protein_coding_gene | -1.0 | 0.49 | 3.0E-04 |
| *col-109* | WBGene00000683 | protein_coding_gene | -1.0 | 0.49 | 3.9E-02 |
| *fah-1* | WBGene00019620 | protein_coding_gene | -1.0 | 0.49 | 3.0E-04 |
| *F31A3.3* | WBGene00017938 | protein_coding_gene | -1.0 | 0.49 | 3.0E-02 |
| *duo-1* | WBGene00001110 | protein_coding_gene | -1.0 | 0.49 | 7.5E-07 |
| *acl-1* | WBGene00010339 | protein_coding_gene | -1.0 | 0.49 | 1.7E-06 |
| *cln-3.1* | WBGene00000539 | protein_coding_gene | -1.0 | 0.49 | 9.6E-03 |
| *elpc-1* | WBGene00022463 | protein_coding_gene | -1.0 | 0.49 | 3.5E-17 |
| *twk-42* | WBGene00006692 | protein_coding_gene | -1.0 | 0.49 | 3.4E-03 |
| *atfs-1* | WBGene00013878 | protein_coding_gene | -1.1 | 0.48 | 2.5E-13 |
| *haly-1* | WBGene00009813 | protein_coding_gene | -1.1 | 0.48 | 3.2E-04 |
| *cima-1* | WBGene00018473 | protein_coding_gene | -1.1 | 0.48 | 1.3E-02 |
| *F21A3.3* | WBGene00008993 | protein_coding_gene | -1.1 | 0.48 | 1.1E-02 |
| *mrpl-24* | WBGene00019076 | protein_coding_gene | -1.1 | 0.48 | 6.3E-06 |
| *gpa-4* | WBGene00001666 | protein_coding_gene | -1.1 | 0.48 | 4.9E-02 |
| *cra-1* | WBGene00020068 | protein_coding_gene | -1.1 | 0.48 | 2.0E-07 |
| *C10C6.7* | WBGene00007515 | protein_coding_gene | -1.1 | 0.47 | 1.9E-02 |
| *gst-20* | WBGene00001768 | protein_coding_gene | -1.1 | 0.47 | 2.0E-03 |
| *T28A11.2* | WBGene00020869 | protein_coding_gene | -1.1 | 0.47 | 2.0E-05 |
| *tofu-4* | WBGene00019013 | protein_coding_gene | -1.1 | 0.47 | 2.3E-07 |
| *Y54E10A.17* | WBGene00021835 | protein_coding_gene | -1.1 | 0.47 | 6.0E-03 |
| *B0391.10* | WBGene00007165 | pseudogene | -1.1 | 0.47 | 2.6E-04 |
| *heri-1* | WBGene00016236 | protein_coding_gene | -1.1 | 0.47 | 1.1E-12 |
| *F28H7.8* | WBGene00009241 | protein_coding_gene | -1.1 | 0.47 | 1.0E-02 |
| *K12H4.7* | WBGene00019682 | protein_coding_gene | -1.1 | 0.47 | 1.9E-06 |
| *oig-2* | WBGene00003860 | protein_coding_gene | -1.1 | 0.46 | 4.2E-03 |
| *K01C8.1* | WBGene00010456 | protein_coding_gene | -1.1 | 0.46 | 6.5E-04 |
| *pqn-37* | WBGene00004124 | protein_coding_gene | -1.1 | 0.46 | 6.8E-03 |
| *col-38* | WBGene00000615 | protein_coding_gene | -1.1 | 0.46 | 2.0E-03 |
| *clec-265* | WBGene00019738 | protein_coding_gene | -1.1 | 0.46 | 2.1E-02 |
| *cth-2* | WBGene00022856 | protein_coding_gene | -1.1 | 0.46 | 1.4E-03 |
| *bath-14* | WBGene00015016 | pseudogene | -1.1 | 0.46 | 6.0E-03 |
| *D1044.1* | WBGene00017027 | protein_coding_gene | -1.1 | 0.46 | 1.4E-03 |
| *Y50D4B.6* | WBGene00021745 | protein_coding_gene | -1.1 | 0.46 | 3.6E-02 |
| *C54E4.5* | WBGene00016920 | protein_coding_gene | -1.1 | 0.46 | 8.9E-03 |
| *ugt-4* | WBGene00013905 | protein_coding_gene | -1.1 | 0.45 | 3.2E-02 |
| *grd-10* | WBGene00001699 | protein_coding_gene | -1.1 | 0.45 | 2.2E-02 |
| *eif-2gamma* | WBGene00021466 | protein_coding_gene | -1.2 | 0.45 | 2.3E-11 |
| *grl-21* | WBGene00001730 | protein_coding_gene | -1.2 | 0.45 | 5.0E-02 |
| *mua-3* | WBGene00003482 | protein_coding_gene | -1.2 | 0.45 | 5.7E-03 |
| *sprr-2* | WBGene00009629 | protein_coding_gene | -1.2 | 0.45 | 1.6E-02 |
| *cct-2* | WBGene00000378 | protein_coding_gene | -1.2 | 0.44 | 7.1E-10 |
| *cdr-2* | WBGene00008296 | protein_coding_gene | -1.2 | 0.44 | 8.4E-03 |
| *R160.3* | WBGene00020120 | protein_coding_gene | -1.2 | 0.44 | 3.4E-02 |
| *C44F1.1* | WBGene00008091 | protein_coding_gene | -1.2 | 0.44 | 2.4E-03 |
| *nas-36* | WBGene00003552 | protein_coding_gene | -1.2 | 0.43 | 2.1E-02 |
| *Y47D3B.6* | WBGene00012942 | protein_coding_gene | -1.2 | 0.43 | 2.8E-02 |
| *col-17* | WBGene00000606 | protein_coding_gene | -1.2 | 0.43 | 1.8E-02 |
| *col-12* | WBGene00000601 | protein_coding_gene | -1.2 | 0.43 | 4.8E-02 |
| *nhr-115* | WBGene00003705 | protein_coding_gene | -1.2 | 0.43 | 2.2E-03 |

| | | | | | |
|---|---|---|---|---|---|
| *C39E9.8* | WBGene00008032 | protein_coding_gene | -1.2 | 0.43 | 6.6E-08 |
| *pcp-2* | WBGene00003957 | protein_coding_gene | -1.2 | 0.43 | 2.1E-02 |
| *col-176* | WBGene00000749 | protein_coding_gene | -1.2 | 0.43 | 1.2E-02 |
| *nmat-1* | WBGene00012295 | protein_coding_gene | -1.2 | 0.43 | 1.5E-03 |
| *T28C12.2* | WBGene00020889 | protein_coding_gene | -1.2 | 0.42 | 3.8E-02 |
| *col-107* | WBGene00000681 | protein_coding_gene | -1.2 | 0.42 | 1.3E-02 |
| *H10E21.4* | WBGene00019184 | protein_coding_gene | -1.2 | 0.42 | 3.2E-02 |
| *col-63* | WBGene00000639 | protein_coding_gene | -1.2 | 0.42 | 1.2E-02 |
| *ugt-42* | WBGene00017959 | protein_coding_gene | -1.2 | 0.42 | 4.4E-02 |
| *cld-9* | WBGene00009432 | protein_coding_gene | -1.2 | 0.42 | 1.6E-03 |
| *C49C3.9* | WBGene00008199 | protein_coding_gene | -1.3 | 0.42 | 1.3E-02 |
| *B0554.5* | WBGene00015258 | protein_coding_gene | -1.3 | 0.42 | 2.8E-02 |
| *R02E4.3* | WBGene00044724 | protein_coding_gene | -1.3 | 0.41 | 2.2E-02 |
| *cutl-18* | WBGene00009830 | protein_coding_gene | -1.3 | 0.41 | 3.4E-02 |
| *gst-24* | WBGene00001772 | protein_coding_gene | -1.3 | 0.41 | 6.0E-04 |
| *mig-21* | WBGene00003252 | protein_coding_gene | -1.3 | 0.40 | 5.3E-03 |
| *sdz-8* | WBGene00008334 | protein_coding_gene | -1.3 | 0.40 | 5.3E-04 |
| *pho-1* | WBGene00004020 | protein_coding_gene | -1.3 | 0.40 | 1.1E-03 |
| *col-110* | WBGene00000684 | protein_coding_gene | -1.3 | 0.40 | 2.5E-02 |
| *col-115* | WBGene00000689 | protein_coding_gene | -1.3 | 0.40 | 4.6E-02 |
| *mpst-7* | WBGene00011307 | protein_coding_gene | -1.3 | 0.40 | 7.0E-06 |
| *cyp-35C1* | WBGene00007362 | protein_coding_gene | -1.3 | 0.40 | 2.1E-04 |
| *suro-1* | WBGene00011235 | protein_coding_gene | -1.3 | 0.39 | 1.1E-02 |
| *Y55F3AM.14* | WBGene00021931 | protein_coding_gene | -1.3 | 0.39 | 4.5E-02 |
| *col-14* | WBGene00000603 | protein_coding_gene | -1.4 | 0.39 | 7.0E-03 |
| *F19B2.5* | WBGene00008944 | protein_coding_gene | -1.4 | 0.39 | 1.0E-02 |
| *C25E10.10* | WBGene00016099 | protein_coding_gene | -1.4 | 0.39 | 1.6E-04 |
| *clec-186* | WBGene00014138 | protein_coding_gene | -1.4 | 0.39 | 3.3E-04 |
| *ZK180.6* | WBGene00022680 | protein_coding_gene | -1.4 | 0.39 | 2.6E-03 |
| *F41B4.3* | WBGene00018259 | protein_coding_gene | -1.4 | 0.39 | 2.8E-04 |
| *col-7* | WBGene00000596 | protein_coding_gene | -1.4 | 0.38 | 3.5E-02 |
| *C33D3.3* | WBGene00007894 | protein_coding_gene | -1.4 | 0.38 | 1.3E-02 |
| *Y47G6A.15* | WBGene00021641 | protein_coding_gene | -1.4 | 0.38 | 9.4E-04 |
| *K01D12.9* | WBGene00010468 | protein_coding_gene | -1.4 | 0.37 | 3.7E-02 |
| *miro-3* | WBGene00021651 | pseudogene | -1.4 | 0.37 | 8.8E-03 |
| *C36C5.5* | WBGene00016474 | protein_coding_gene | -1.4 | 0.37 | 9.4E-08 |
| *dpy-13* | WBGene00001074 | protein_coding_gene | -1.4 | 0.37 | 7.5E-03 |
| *cdr-4* | WBGene00010470 | protein_coding_gene | -1.4 | 0.37 | 8.8E-04 |
| *hpd-1* | WBGene00001993 | protein_coding_gene | -1.4 | 0.37 | 4.3E-03 |
| *gst-12* | WBGene00001760 | protein_coding_gene | -1.4 | 0.37 | 1.4E-03 |
| *pho-12* | WBGene00016152 | protein_coding_gene | -1.4 | 0.37 | 1.9E-02 |
| *arl-7* | WBGene00000191 | pseudogene | -1.4 | 0.37 | 1.3E-02 |
| *Y47D7A.13* | WBGene00021625 | protein_coding_gene | -1.4 | 0.37 | 1.9E-02 |
| *Y119D3B.21* | WBGene00022497 | protein_coding_gene | -1.4 | 0.37 | 1.8E-04 |
| *dct-18* | WBGene00010266 | protein_coding_gene | -1.4 | 0.37 | 1.0E-03 |
| *F09B12.3* | WBGene00008607 | protein_coding_gene | -1.4 | 0.37 | 2.2E-03 |
| *dpy-5* | WBGene00001067 | protein_coding_gene | -1.5 | 0.36 | 3.3E-03 |
| *C17C3.1* | WBGene00015887 | protein_coding_gene | -1.5 | 0.36 | 3.0E-14 |
| *T21C9.9* | WBGene00011895 | protein_coding_gene | -1.5 | 0.36 | 1.5E-02 |
| *col-97* | WBGene00000672 | protein_coding_gene | -1.5 | 0.36 | 1.1E-03 |
| *C07A9.9* | WBGene00007405 | protein_coding_gene | -1.5 | 0.36 | 5.2E-06 |
| *his-40* | WBGene00001914 | protein_coding_gene | -1.5 | 0.36 | 1.7E-02 |

| | | | | | |
|---|---|---|---|---|---|
| *col-144* | WBGene00000717 | protein_coding_gene | -1.5 | 0.36 | 9.6E-03 |
| *Y58A7A.1* | WBGene00021975 | protein_coding_gene | -1.5 | 0.36 | 1.1E-02 |
| *cyp-35A4* | WBGene00016786 | protein_coding_gene | -1.5 | 0.36 | 4.4E-02 |
| *T01D1.3* | WBGene00020148 | protein_coding_gene | -1.5 | 0.36 | 1.4E-02 |
| *T10E9.3* | WBGene00020413 | protein_coding_gene | -1.5 | 0.36 | 8.4E-04 |
| *gly-8* | WBGene00001633 | protein_coding_gene | -1.5 | 0.35 | 2.9E-03 |
| *C32H11.4* | WBGene00007867 | protein_coding_gene | -1.5 | 0.35 | 1.4E-05 |
| *C17H12.6* | WBGene00015932 | protein_coding_gene | -1.5 | 0.35 | 1.6E-02 |
| *C23H5.8* | WBGene00016027 | protein_coding_gene | -1.5 | 0.35 | 8.7E-05 |
| *F54D5.4* | WBGene00010050 | protein_coding_gene | -1.5 | 0.35 | 1.1E-10 |
| *C33E10.8* | WBGene00016349 | protein_coding_gene | -1.5 | 0.35 | 1.2E-02 |
| *col-71* | WBGene00000647 | protein_coding_gene | -1.5 | 0.35 | 2.3E-03 |
| *C30H6.5* | WBGene00007823 | protein_coding_gene | -1.5 | 0.35 | 1.3E-02 |
| *Y51B9A.6* | WBGene00013088 | protein_coding_gene | -1.5 | 0.35 | 3.4E-02 |
| *F45E1.4* | WBGene00018465 | protein_coding_gene | -1.5 | 0.35 | 6.7E-04 |
| *C26B2.8* | WBGene00016130 | protein_coding_gene | -1.5 | 0.35 | 2.0E-02 |
| *R02D1.2* | WBGene00235310 | protein_coding_gene | -1.5 | 0.34 | 2.8E-03 |
| *C35A11.4* | WBGene00016431 | protein_coding_gene | -1.5 | 0.34 | 1.1E-03 |
| *F08F3.4* | WBGene00017262 | protein_coding_gene | -1.6 | 0.34 | 9.4E-04 |
| *pcp-4* | WBGene00003959 | protein_coding_gene | -1.6 | 0.34 | 8.0E-06 |
| *K08D8.12* | WBGene00194926 | protein_coding_gene | -1.6 | 0.34 | 2.4E-03 |
| *col-169* | WBGene00000742 | protein_coding_gene | -1.6 | 0.34 | 3.3E-03 |
| *R12E2.7* | WBGene00020033 | protein_coding_gene | -1.6 | 0.33 | 2.2E-02 |
| *dpy-4* | WBGene00001066 | protein_coding_gene | -1.6 | 0.33 | 3.6E-04 |
| *Y43C5A.7* | WBGene00044922 | protein_coding_gene | -1.6 | 0.33 | 1.1E-02 |
| *col-104* | WBGene00000678 | protein_coding_gene | -1.6 | 0.33 | 3.2E-03 |
| *col-175* | WBGene00000748 | protein_coding_gene | -1.6 | 0.33 | 1.8E-03 |
| *pgp-8* | WBGene00004002 | protein_coding_gene | -1.6 | 0.33 | 4.1E-04 |
| *col-130* | WBGene00000704 | protein_coding_gene | -1.6 | 0.33 | 5.9E-03 |
| *ugt-37* | WBGene00017332 | protein_coding_gene | -1.6 | 0.33 | 4.9E-03 |
| *col-125* | WBGene00000699 | protein_coding_gene | -1.6 | 0.32 | 2.4E-02 |
| *Y47G6A.19* | WBGene00021645 | protein_coding_gene | -1.6 | 0.32 | 4.5E-05 |
| *cyp-33C8* | WBGene00019967 | protein_coding_gene | -1.7 | 0.32 | 5.0E-04 |
| *Y39B6A.21* | WBGene00012680 | pseudogene | -1.7 | 0.32 | 2.1E-02 |
| *grl-16* | WBGene00001725 | protein_coding_gene | -1.7 | 0.32 | 2.1E-03 |
| *col-73* | WBGene00000649 | protein_coding_gene | -1.7 | 0.31 | 6.4E-03 |
| *col-90* | WBGene00000665 | protein_coding_gene | -1.7 | 0.31 | 7.4E-03 |
| *ttr-31* | WBGene00010225 | protein_coding_gene | -1.7 | 0.31 | 1.4E-03 |
| *H23N18.5* | WBGene00019236 | protein_coding_gene | -1.7 | 0.31 | 1.6E-02 |
| *col-129* | WBGene00000703 | protein_coding_gene | -1.7 | 0.31 | 1.6E-03 |
| *grsp-1* | WBGene00012840 | protein_coding_gene | -1.7 | 0.31 | 3.4E-03 |
| *col-139* | WBGene00000712 | protein_coding_gene | -1.7 | 0.31 | 1.8E-02 |
| *ram-2* | WBGene00004300 | protein_coding_gene | -1.7 | 0.31 | 1.8E-04 |
| *dpy-6* | WBGene00001068 | protein_coding_gene | -1.7 | 0.30 | 7.5E-03 |
| *col-49* | WBGene00000626 | protein_coding_gene | -1.7 | 0.30 | 1.2E-03 |
| *F41G3.21* | WBGene00044316 | protein_coding_gene | -1.7 | 0.30 | 3.9E-04 |
| *rol-1* | WBGene00004394 | protein_coding_gene | -1.7 | 0.30 | 4.1E-04 |
| *Y51H7C.1* | WBGene00021779 | protein_coding_gene | -1.7 | 0.30 | 1.6E-05 |
| *T20D4.3* | WBGene00020609 | protein_coding_gene | -1.8 | 0.30 | 5.7E-03 |
| *col-138* | WBGene00000711 | protein_coding_gene | -1.8 | 0.30 | 2.8E-04 |
| *mec-19* | WBGene00008216 | protein_coding_gene | -1.8 | 0.30 | 6.8E-03 |
| *ugt-46* | WBGene00015141 | protein_coding_gene | -1.8 | 0.29 | 1.8E-05 |

| | | | | | |
|---|---|---|---|---|---|
| *C42D4.3* | WBGene00016596 | protein_coding_gene | -1.8 | 0.29 | 5.8E-03 |
| *lact-1* | WBGene00018523 | protein_coding_gene | -1.8 | 0.29 | 2.5E-03 |
| *T06E4.9* | WBGene00011536 | protein_coding_gene | -1.8 | 0.29 | 3.9E-02 |
| *Y94H6A.10* | WBGene00022382 | protein_coding_gene | -1.8 | 0.29 | 1.8E-06 |
| *col-77* | WBGene00000653 | protein_coding_gene | -1.8 | 0.28 | 1.0E-03 |
| *dod-20* | WBGene00015259 | protein_coding_gene | -1.8 | 0.28 | 4.7E-02 |
| *T25G12.13* | WBGene00219274 | protein_coding_gene | -1.9 | 0.27 | 9.5E-03 |
| *Y46G5A.29* | WBGene00012917 | protein_coding_gene | -1.9 | 0.27 | 1.2E-03 |
| *bli-6* | WBGene00000256 | protein_coding_gene | -1.9 | 0.27 | 3.2E-04 |
| *F16H6.4* | WBGene00008894 | protein_coding_gene | -1.9 | 0.27 | 1.0E-02 |
| *wago-11* | WBGene00021711 | protein_coding_gene | -1.9 | 0.26 | 6.9E-05 |
| *Y51H7C.13* | WBGene00021791 | protein_coding_gene | -1.9 | 0.26 | 2.4E-05 |
| *R04B5.6* | WBGene00011004 | protein_coding_gene | -2.0 | 0.26 | 3.2E-04 |
| *ugt-9* | WBGene00020587 | protein_coding_gene | -2.0 | 0.26 | 1.2E-07 |
| *oac-20* | WBGene00009488 | protein_coding_gene | -2.0 | 0.26 | 2.1E-03 |
| *dod-17* | WBGene00010745 | protein_coding_gene | -2.0 | 0.26 | 8.3E-05 |
| *col-60* | WBGene00000636 | protein_coding_gene | -2.0 | 0.25 | 8.6E-03 |
| *col-81* | WBGene00000657 | protein_coding_gene | -2.0 | 0.25 | 5.9E-03 |
| *col-161* | WBGene00000734 | protein_coding_gene | -2.0 | 0.25 | 1.8E-02 |
| *swt-6* | WBGene00011190 | protein_coding_gene | -2.0 | 0.25 | 3.1E-05 |
| *Y39B6A.27* | WBGene00012686 | protein_coding_gene | -2.0 | 0.25 | 5.2E-06 |
| *col-152* | WBGene00000725 | protein_coding_gene | -2.0 | 0.25 | 3.2E-02 |
| *R03H10.12* | WBGene00303363 | pseudogene | -2.0 | 0.25 | 2.6E-02 |
| *C04E12.5* | WBGene00015435 | protein_coding_gene | -2.0 | 0.25 | 2.1E-02 |
| *gst-10* | WBGene00001758 | protein_coding_gene | -2.0 | 0.24 | 2.3E-05 |
| *R08E5.3* | WBGene00019963 | protein_coding_gene | -2.0 | 0.24 | 1.3E-05 |
| *clec-53* | WBGene00020191 | protein_coding_gene | -2.0 | 0.24 | 3.2E-06 |
| *col-120* | WBGene00000694 | protein_coding_gene | -2.1 | 0.24 | 1.0E-03 |
| *cyp-29A3* | WBGene00021412 | protein_coding_gene | -2.1 | 0.23 | 5.0E-06 |
| *col-167* | WBGene00000740 | protein_coding_gene | -2.1 | 0.23 | 3.3E-05 |
| *F52C6.14* | WBGene00018671 | protein_coding_gene | -2.1 | 0.23 | 3.3E-02 |
| *cyp-13A12* | WBGene00008810 | protein_coding_gene | -2.2 | 0.22 | 7.1E-03 |
| *col-168* | WBGene00000741 | protein_coding_gene | -2.2 | 0.21 | 2.5E-05 |
| *col-133* | WBGene00000707 | protein_coding_gene | -2.2 | 0.21 | 9.0E-03 |
| *col-170* | WBGene00000743 | protein_coding_gene | -2.2 | 0.21 | 6.0E-04 |
| *F57B9.3* | WBGene00018997 | protein_coding_gene | -2.2 | 0.21 | 1.4E-02 |
| *gba-4* | WBGene00021160 | protein_coding_gene | -2.3 | 0.21 | 5.9E-06 |
| *col-91* | WBGene00000666 | protein_coding_gene | -2.3 | 0.21 | 4.5E-04 |
| *pgp-12* | WBGene00004006 | protein_coding_gene | -2.3 | 0.20 | 2.6E-09 |
| *F35F10.13* | WBGene00018062 | protein_coding_gene | -2.3 | 0.20 | 3.1E-02 |
| *mth-1* | WBGene00017948 | protein_coding_gene | -2.3 | 0.20 | 8.9E-08 |
| *F52G3.3* | WBGene00018711 | pseudogene | -2.4 | 0.20 | 3.8E-02 |
| *ugt-30* | WBGene00011340 | protein_coding_gene | -2.4 | 0.19 | 1.4E-11 |
| *srm-6* | WBGene00009810 | protein_coding_gene | -2.4 | 0.19 | 3.3E-02 |
| *ugt-16* | WBGene00013901 | protein_coding_gene | -2.4 | 0.19 | 1.9E-14 |
| *col-162* | WBGene00000735 | protein_coding_gene | -2.4 | 0.19 | 3.6E-03 |
| *T20D4.7* | WBGene00020613 | protein_coding_gene | -2.4 | 0.19 | 1.7E-02 |
| *Y45G12C.1* | WBGene00021565 | protein_coding_gene | -2.5 | 0.18 | 1.0E-04 |
| *R07E5.4* | WBGene00011112 | protein_coding_gene | -2.5 | 0.18 | 1.9E-02 |
| *far-3* | WBGene00001387 | protein_coding_gene | -2.5 | 0.18 | 3.2E-08 |
| *K11H12.3* | WBGene00019659 | protein_coding_gene | -2.5 | 0.17 | 5.5E-04 |
| *C55C3.7* | WBGene00043065 | protein_coding_gene | -2.5 | 0.17 | 3.7E-02 |

| | | | | | |
|---|---|---|---|---|---|
| *col-157* | WBGene00000730 | protein_coding_gene | -2.6 | 0.17 | 2.6E-04 |
| *B0205.14* | WBGene00044646 | protein_coding_gene | -2.7 | 0.16 | 6.6E-03 |
| *grd-14* | WBGene00001703 | protein_coding_gene | -2.7 | 0.15 | 1.0E-03 |
| *Y47D7A.15* | WBGene00021627 | protein_coding_gene | -2.7 | 0.15 | 8.0E-04 |
| *grd-13* | WBGene00001702 | protein_coding_gene | -2.8 | 0.15 | 4.6E-06 |
| *Y48G8AL.12* | WBGene00021690 | protein_coding_gene | -2.8 | 0.15 | 2.5E-04 |
| *Y75B8A.39* | WBGene00023483 | protein_coding_gene | -2.8 | 0.15 | 1.4E-02 |
| *col-159* | WBGene00000732 | protein_coding_gene | -2.8 | 0.15 | 2.3E-10 |
| *C04E12.10* | WBGene00015439 | pseudogene | -2.8 | 0.15 | 1.9E-03 |
| *col-88* | WBGene00000663 | protein_coding_gene | -2.8 | 0.14 | 6.2E-04 |
| *zipt-2.3* | WBGene00013207 | protein_coding_gene | -2.9 | 0.14 | 8.6E-04 |
| *col-154* | WBGene00000727 | protein_coding_gene | -2.9 | 0.13 | 2.9E-06 |
| *C49G7.13* | WBGene00269425 | protein_coding_gene | -2.9 | 0.13 | 1.3E-02 |
| *F53F1.4* | WBGene00009982 | protein_coding_gene | -3.0 | 0.13 | 2.1E-05 |
| *pgp-1* | WBGene00003995 | protein_coding_gene | -3.0 | 0.13 | 6.6E-15 |
| *col-145* | WBGene00000718 | protein_coding_gene | -3.1 | 0.12 | 4.0E-05 |
| *col-156* | WBGene00000729 | protein_coding_gene | -3.1 | 0.12 | 3.6E-06 |
| *col-180* | WBGene00000753 | protein_coding_gene | -3.1 | 0.12 | 6.9E-08 |
| *F35E12.2* | WBGene00009426 | protein_coding_gene | -3.1 | 0.12 | 9.5E-03 |
| *gpdh-1* | WBGene00009824 | protein_coding_gene | -3.3 | 0.10 | 2.0E-04 |

**CB4856**-specific genes exclusively expressed following *S. epidermidis* infection (vs. OP50) or DE

| Gene Name | Wormbase Gene ID | Type | $log_2FC$ | FC | Adj P-value |
|---|---|---|---|---|---|
| *F56H6.7* | WBGene00010168 | protein_coding_gene | 4.3 | 19.76 | 1.3E-06 |
| *oac-35* | WBGene00010171 | protein_coding_gene | 3.6 | 12.36 | 7.2E-04 |
| *F56H6.9* | WBGene00010170 | protein_coding_gene | 3.3 | 9.92 | 3.3E-03 |
| *C47F8.6* | WBGene00008162 | protein_coding_gene | 3.3 | 9.88 | 2.6E-03 |
| *hsd-2* | WBGene00022498 | protein_coding_gene | 3.2 | 9.40 | 3.1E-03 |
| *col-94* | WBGene00000669 | protein_coding_gene | 3.2 | 9.18 | 8.9E-03 |
| *H25K10.1* | WBGene00010411 | protein_coding_gene | 3.0 | 8.16 | 9.3E-07 |
| *Y106G6E.1* | WBGene00013705 | protein_coding_gene | 3.0 | 8.11 | 1.2E-03 |
| *elo-6* | WBGene00001244 | protein_coding_gene | 2.9 | 7.56 | 6.1E-16 |
| *C49F5.8* | WBGene00008212 | protein_coding_gene | 2.9 | 7.29 | 6.4E-03 |
| *elo-5* | WBGene00001243 | protein_coding_gene | 2.8 | 7.12 | 6.8E-75 |
| *elo-2* | WBGene00001240 | protein_coding_gene | 2.8 | 6.98 | 4.2E-26 |
| *thn-2* | WBGene00009215 | protein_coding_gene | 2.7 | 6.31 | 1.3E-53 |
| *T28D6.3* | WBGene00012123 | protein_coding_gene | 2.7 | 6.28 | 2.7E-33 |
| *clec-174* | WBGene00021580 | protein_coding_gene | 2.6 | 6.07 | 5.1E-06 |
| *oac-45* | WBGene00011656 | protein_coding_gene | 2.5 | 5.71 | 7.5E-03 |
| *tag-10* | WBGene00006404 | protein_coding_gene | 2.5 | 5.69 | 1.2E-48 |
| *H06A10.1* | WBGene00010368 | protein_coding_gene | 2.5 | 5.53 | 4.2E-11 |
| *clec-100* | WBGene00013179 | protein_coding_gene | 2.5 | 5.52 | 6.8E-08 |
| *acox-1.4* | WBGene00008567 | protein_coding_gene | 2.5 | 5.49 | 2.1E-12 |
| *F11A5.17* | WBGene00045510 | protein_coding_gene | 2.5 | 5.48 | 8.9E-03 |
| *F49D11.3* | WBGene00018627 | protein_coding_gene | 2.4 | 5.32 | 4.4E-02 |
| *T10H10.2* | WBGene00020426 | protein_coding_gene | 2.4 | 5.18 | 3.4E-22 |
| *C54D2.1* | WBGene00016915 | protein_coding_gene | 2.4 | 5.15 | 1.6E-03 |
| *C54F6.12* | WBGene00016929 | protein_coding_gene | 2.3 | 5.10 | 4.8E-09 |
| *lys-7* | WBGene00003096 | protein_coding_gene | 2.3 | 5.01 | 3.2E-15 |
| *Y54G2A.45* | WBGene00044488 | protein_coding_gene | 2.3 | 5.00 | 2.2E-05 |
| *T22F3.11* | WBGene00020700 | protein_coding_gene | 2.3 | 4.99 | 1.1E-02 |
| *F46B3.14* | WBGene00009765 | protein_coding_gene | 2.3 | 4.81 | 6.8E-04 |
| *F58F9.9* | WBGene00044307 | protein_coding_gene | 2.3 | 4.79 | 3.2E-04 |

| | | | | | |
|---|---|---|---|---|---|
| *Y53F4B.23* | WBGene00013169 | protein_coding_gene | 2.2 | 4.74 | 2.0E-06 |
| *abu-1* | WBGene00000024 | protein_coding_gene | 2.2 | 4.72 | 4.0E-02 |
| *Y45F10D.2* | WBGene00012883 | protein_coding_gene | 2.2 | 4.65 | 1.4E-03 |
| *oac-44* | WBGene00011654 | protein_coding_gene | 2.2 | 4.64 | 2.4E-02 |
| *C28F5.4* | WBGene00016185 | protein_coding_gene | 2.2 | 4.59 | 1.7E-02 |
| *decr-1.1* | WBGene00018380 | protein_coding_gene | 2.1 | 4.43 | 2.2E-17 |
| *Y50D4B.4* | WBGene00021743 | protein_coding_gene | 2.1 | 4.43 | 3.5E-22 |
| *F29A7.3* | WBGene00017913 | protein_coding_gene | 2.1 | 4.41 | 7.1E-03 |
| *pqn-71* | WBGene00004153 | protein_coding_gene | 2.1 | 4.41 | 3.0E-02 |
| *F11A5.9* | WBGene00008677 | protein_coding_gene | 2.1 | 4.39 | 3.2E-23 |
| *tor-2* | WBGene00006598 | protein_coding_gene | 2.1 | 4.39 | 1.5E-02 |
| *str-146* | WBGene00006193 | protein_coding_gene | 2.1 | 4.39 | 2.9E-02 |
| *F46B3.15* | WBGene00009766 | protein_coding_gene | 2.1 | 4.34 | 2.0E-03 |
| *W03F11.5* | WBGene00021008 | protein_coding_gene | 2.1 | 4.34 | 1.4E-04 |
| *F11C7.6* | WBGene00017378 | protein_coding_gene | 2.1 | 4.32 | 2.6E-13 |
| *Y38H6C.19* | WBGene00012631 | protein_coding_gene | 2.1 | 4.29 | 1.1E-02 |
| *F59A6.3* | WBGene00019085 | protein_coding_gene | 2.1 | 4.15 | 3.8E-03 |
| *nspb-12* | WBGene00017305 | protein_coding_gene | 2.0 | 4.13 | 4.9E-02 |
| *F40H6.5* | WBGene00018255 | protein_coding_gene | 2.0 | 4.12 | 2.5E-02 |
| *ets-5* | WBGene00016600 | protein_coding_gene | 2.0 | 4.09 | 1.7E-02 |
| *T25E4.1* | WBGene00020803 | protein_coding_gene | 2.0 | 3.99 | 1.2E-02 |
| *col-58* | WBGene00000634 | protein_coding_gene | 2.0 | 3.99 | 9.6E-03 |
| *cwp-5* | WBGene00009844 | protein_coding_gene | 2.0 | 3.97 | 6.8E-04 |
| *F59B1.4* | WBGene00019098 | protein_coding_gene | 2.0 | 3.96 | 3.2E-02 |
| *F37A4.6* | WBGene00018136 | protein_coding_gene | 2.0 | 3.96 | 1.3E-02 |
| *R03C1.1* | WBGene00010987 | protein_coding_gene | 2.0 | 3.96 | 2.2E-02 |
| *clec-1* | WBGene00017772 | protein_coding_gene | 2.0 | 3.94 | 1.0E-24 |
| *K02D3.2* | WBGene00010505 | protein_coding_gene | 2.0 | 3.92 | 2.9E-03 |
| *clec-245* | WBGene00007851 | protein_coding_gene | 2.0 | 3.88 | 3.1E-02 |
| *Y47D7A.2* | WBGene00021616 | protein_coding_gene | 2.0 | 3.88 | 4.5E-02 |
| *F46F2.3* | WBGene00009787 | protein_coding_gene | 2.0 | 3.87 | 6.6E-06 |
| *abu-8* | WBGene00000031 | protein_coding_gene | 2.0 | 3.87 | 1.3E-02 |
| *ech-7* | WBGene00001156 | protein_coding_gene | 1.9 | 3.85 | 4.0E-28 |
| *fipr-6* | WBGene00007544 | protein_coding_gene | 1.9 | 3.81 | 3.9E-02 |
| *drd-5* | WBGene00018885 | protein_coding_gene | 1.9 | 3.77 | 3.4E-12 |
| *fbxa-73* | WBGene00020640 | protein_coding_gene | 1.9 | 3.77 | 2.3E-03 |
| *Y39F10A.1* | WBGene00021452 | protein_coding_gene | 1.9 | 3.74 | 1.3E-03 |
| *D1054.6* | WBGene00008373 | pseudogene | 1.9 | 3.72 | 3.3E-02 |
| *W01C8.1* | WBGene00020918 | protein_coding_gene | 1.9 | 3.70 | 1.3E-02 |
| *F22H10.6* | WBGene00017730 | protein_coding_gene | 1.9 | 3.67 | 6.8E-06 |
| *C50E3.9* | WBGene00016820 | protein_coding_gene | 1.9 | 3.65 | 8.0E-04 |
| *F01D5.7* | WBGene00008497 | protein_coding_gene | 1.9 | 3.63 | 6.5E-03 |
| *Y82E9BL.18* | WBGene00044402 | protein_coding_gene | 1.9 | 3.63 | 4.5E-02 |
| *Y38F1A.4* | WBGene00012607 | protein_coding_gene | 1.8 | 3.60 | 4.2E-02 |
| *M03E7.4* | WBGene00019756 | protein_coding_gene | 1.8 | 3.59 | 4.1E-02 |
| *F25C8.1* | WBGene00009104 | pseudogene | 1.8 | 3.59 | 4.3E-05 |
| *mup-4* | WBGene00003497 | protein_coding_gene | 1.8 | 3.58 | 9.8E-35 |
| *Y67D8C.7* | WBGene00022071 | protein_coding_gene | 1.8 | 3.56 | 3.7E-02 |
| *C10F3.7* | WBGene00015679 | protein_coding_gene | 1.8 | 3.54 | 7.7E-04 |
| *C08G5.3* | WBGene00015616 | pseudogene | 1.8 | 3.53 | 2.6E-02 |
| *inx-18* | WBGene00002140 | protein_coding_gene | 1.8 | 3.49 | 6.3E-04 |
| *abu-10* | WBGene00000033 | protein_coding_gene | 1.8 | 3.48 | 4.8E-02 |

| | | | | | |
|---|---|---|---|---|---|
| *daao-1* | WBGene00022076 | protein_coding_gene | 1.8 | 3.46 | 6.4E-11 |
| *ugt-21* | WBGene00007885 | protein_coding_gene | 1.8 | 3.45 | 7.8E-09 |
| *F33E2.6* | WBGene00009362 | protein_coding_gene | 1.8 | 3.45 | 1.5E-02 |
| *C35A5.10* | WBGene00044016 | protein_coding_gene | 1.8 | 3.41 | 2.7E-02 |
| *lgc-54* | WBGene00020528 | protein_coding_gene | 1.8 | 3.40 | 4.5E-07 |
| *C18F10.2* | WBGene00015977 | protein_coding_gene | 1.8 | 3.40 | 4.1E-05 |
| *ncx-7* | WBGene00003572 | protein_coding_gene | 1.7 | 3.35 | 5.9E-17 |
| *Y57G11C.42* | WBGene00013331 | protein_coding_gene | 1.7 | 3.34 | 4.9E-03 |
| *F57B1.9* | WBGene00045488 | protein_coding_gene | 1.7 | 3.31 | 6.9E-04 |
| *C30A5.10* | WBGene00016242 | protein_coding_gene | 1.7 | 3.30 | 2.9E-02 |
| *F54E4.4* | WBGene00010060 | protein_coding_gene | 1.7 | 3.29 | 3.8E-02 |
| *srh-254* | WBGene00005460 | pseudogene | 1.7 | 3.28 | 1.3E-09 |
| *cah-5* | WBGene00000283 | protein_coding_gene | 1.7 | 3.27 | 9.6E-17 |
| *F19H6.6* | WBGene00008961 | protein_coding_gene | 1.7 | 3.26 | 4.1E-03 |
| *ZK228.4* | WBGene00013947 | protein_coding_gene | 1.7 | 3.25 | 3.7E-14 |
| *pho-13* | WBGene00008560 | protein_coding_gene | 1.7 | 3.24 | 3.9E-03 |
| *atic-1* | WBGene00016957 | protein_coding_gene | 1.7 | 3.23 | 2.9E-19 |
| *asns-1* | WBGene00003816 | protein_coding_gene | 1.7 | 3.23 | 1.5E-03 |
| *mct-6* | WBGene00015676 | protein_coding_gene | 1.7 | 3.22 | 8.6E-44 |
| *Y73C8C.10* | WBGene00022267 | protein_coding_gene | 1.7 | 3.22 | 2.0E-03 |
| *R05H10.7* | WBGene00011046 | protein_coding_gene | 1.7 | 3.20 | 2.7E-15 |
| *F19H6.5* | WBGene00008960 | protein_coding_gene | 1.7 | 3.18 | 5.2E-03 |
| *F07H5.8* | WBGene00008559 | protein_coding_gene | 1.7 | 3.18 | 1.9E-02 |
| *daf-36* | WBGene00007536 | protein_coding_gene | 1.7 | 3.17 | 2.7E-13 |
| *smd-1* | WBGene00004875 | protein_coding_gene | 1.7 | 3.17 | 5.1E-09 |
| *ldp-1* | WBGene00017719 | protein_coding_gene | 1.7 | 3.16 | 6.9E-58 |
| *haao-1* | WBGene00010595 | protein_coding_gene | 1.7 | 3.16 | 5.8E-13 |
| *T07G12.3* | WBGene00011594 | protein_coding_gene | 1.7 | 3.15 | 1.5E-02 |
| *T10B11.4* | WBGene00020400 | protein_coding_gene | 1.6 | 3.13 | 1.7E-02 |
| *K08D8.3* | WBGene00010657 | protein_coding_gene | 1.6 | 3.13 | 1.3E-11 |
| *twk-16* | WBGene00006670 | protein_coding_gene | 1.6 | 3.12 | 1.4E-03 |
| *F56C3.9* | WBGene00018946 | protein_coding_gene | 1.6 | 3.12 | 3.3E-04 |
| *Y7A5A.11* | WBGene00044155 | pseudogene | 1.6 | 3.11 | 1.7E-03 |
| *F26G1.5* | WBGene00017841 | protein_coding_gene | 1.6 | 3.10 | 3.4E-02 |
| *cpt-5* | WBGene00008629 | protein_coding_gene | 1.6 | 3.10 | 2.0E-14 |
| *C01G10.17* | WBGene00044658 | protein_coding_gene | 1.6 | 3.09 | 1.8E-03 |
| *Y23H5B.12* | WBGene00194701 | protein_coding_gene | 1.6 | 3.06 | 3.5E-02 |
| *Y55D5A.4* | WBGene00021918 | protein_coding_gene | 1.6 | 3.05 | 1.0E-03 |
| *F56D6.12* | WBGene00044474 | protein_coding_gene | 1.6 | 3.05 | 3.4E-03 |
| *mgl-2* | WBGene00003233 | protein_coding_gene | 1.6 | 3.05 | 3.7E-09 |
| *Y41D4B.1* | WBGene00021511 | protein_coding_gene | 1.6 | 3.03 | 2.5E-02 |
| *lon-1* | WBGene00003055 | protein_coding_gene | 1.6 | 3.03 | 3.4E-05 |
| *acp-6* | WBGene00022245 | protein_coding_gene | 1.6 | 3.03 | 9.9E-24 |
| *C29F4.3* | WBGene00007809 | protein_coding_gene | 1.6 | 3.02 | 4.6E-02 |
| *W02H3.1* | WBGene00012217 | protein_coding_gene | 1.6 | 3.02 | 1.2E-05 |
| *C15F1.1* | WBGene00015794 | protein_coding_gene | 1.6 | 3.02 | 6.3E-11 |
| *pkd-2* | WBGene00004035 | protein_coding_gene | 1.6 | 3.02 | 1.4E-04 |
| *F37C12.21* | WBGene00194742 | protein_coding_gene | 1.6 | 3.01 | 1.7E-03 |
| *tph-1* | WBGene00006600 | protein_coding_gene | 1.6 | 3.01 | 3.0E-02 |
| *C39B5.5* | WBGene00016523 | protein_coding_gene | 1.6 | 3.00 | 1.9E-02 |
| *Y105C5B.7* | WBGene00013648 | pseudogene | 1.6 | 2.98 | 2.3E-03 |
| *F55A3.7* | WBGene00018852 | pseudogene | 1.6 | 2.98 | 7.4E-04 |

| | | | | | |
|---|---|---|---|---|---|
| *ppat-1* | WBGene00011407 | protein_coding_gene | 1.6 | 2.96 | 1.5E-10 |
| *col-165* | WBGene00000738 | protein_coding_gene | 1.6 | 2.96 | 7.4E-07 |
| *C11E4.7* | WBGene00007521 | protein_coding_gene | 1.6 | 2.95 | 3.0E-10 |
| *col-143* | WBGene00000716 | protein_coding_gene | 1.6 | 2.94 | 5.8E-06 |
| *col-160* | WBGene00000733 | protein_coding_gene | 1.6 | 2.94 | 2.6E-30 |
| *Y43F8C.16* | WBGene00012837 | protein_coding_gene | 1.6 | 2.93 | 4.0E-03 |
| *cyp-36A1* | WBGene00007913 | protein_coding_gene | 1.5 | 2.91 | 1.7E-09 |
| *C46F2.1* | WBGene00016714 | protein_coding_gene | 1.5 | 2.88 | 1.0E-06 |
| *T02D1.7* | WBGene00011373 | protein_coding_gene | 1.5 | 2.86 | 2.8E-02 |
| *lov-1* | WBGene00003058 | protein_coding_gene | 1.5 | 2.86 | 2.1E-02 |
| *T22F3.7* | WBGene00020697 | protein_coding_gene | 1.5 | 2.83 | 3.0E-02 |
| *inpp-1* | WBGene00012016 | protein_coding_gene | 1.5 | 2.83 | 4.4E-08 |
| *F54H5.3* | WBGene00018840 | protein_coding_gene | 1.5 | 2.82 | 2.0E-04 |
| *R13A5.10* | WBGene00020052 | protein_coding_gene | 1.5 | 2.82 | 6.5E-06 |
| *K12H6.5* | WBGene00019685 | protein_coding_gene | 1.5 | 2.82 | 8.7E-05 |
| *Y47D7A.7* | WBGene00021620 | protein_coding_gene | 1.5 | 2.79 | 2.4E-03 |
| *Y6B3B.1* | WBGene00012387 | protein_coding_gene | 1.5 | 2.78 | 3.2E-04 |
| *lgc-21* | WBGene00007903 | protein_coding_gene | 1.5 | 2.76 | 2.0E-02 |
| *C47A10.12* | WBGene00008129 | protein_coding_gene | 1.5 | 2.76 | 2.6E-08 |
| *nas-22* | WBGene00003541 | protein_coding_gene | 1.5 | 2.76 | 1.9E-02 |
| *acr-8* | WBGene00000047 | protein_coding_gene | 1.5 | 2.76 | 4.1E-04 |
| *pqn-22* | WBGene00004112 | protein_coding_gene | 1.5 | 2.76 | 8.8E-15 |
| *T04A8.13* | WBGene00011414 | protein_coding_gene | 1.5 | 2.75 | 9.8E-05 |
| *clec-155* | WBGene00011405 | pseudogene | 1.5 | 2.75 | 9.6E-03 |
| *F13H8.6* | WBGene00017439 | protein_coding_gene | 1.5 | 2.74 | 3.3E-02 |
| *T01C8.3* | WBGene00020144 | protein_coding_gene | 1.5 | 2.74 | 2.7E-03 |
| *ugt-32* | WBGene00018543 | protein_coding_gene | 1.5 | 2.74 | 1.1E-02 |
| *tor-1* | WBGene00006597 | protein_coding_gene | 1.5 | 2.74 | 4.9E-02 |
| *W06H8.2* | WBGene00021066 | protein_coding_gene | 1.5 | 2.74 | 2.3E-04 |
| *Y40H7A.11* | WBGene00012748 | protein_coding_gene | 1.4 | 2.72 | 4.9E-02 |
| *clec-132* | WBGene00021143 | protein_coding_gene | 1.4 | 2.71 | 1.2E-05 |
| *T12A2.5* | WBGene00020439 | protein_coding_gene | 1.4 | 2.71 | 3.5E-07 |
| *F25D7.5* | WBGene00009114 | protein_coding_gene | 1.4 | 2.70 | 1.1E-02 |
| *ugt-3* | WBGene00013903 | protein_coding_gene | 1.4 | 2.70 | 3.2E-02 |
| *F15H9.7* | WBGene00077535 | pseudogene | 1.4 | 2.68 | 3.7E-02 |
| *unc-52* | WBGene00006787 | protein_coding_gene | 1.4 | 2.68 | 1.9E-24 |
| *ctsa-3.1* | WBGene00017969 | protein_coding_gene | 1.4 | 2.67 | 7.8E-03 |
| *pde-5* | WBGene00016328 | protein_coding_gene | 1.4 | 2.66 | 3.6E-02 |
| *col-98* | WBGene00000673 | protein_coding_gene | 1.4 | 2.66 | 5.3E-24 |
| *clec-105* | WBGene00012488 | protein_coding_gene | 1.4 | 2.64 | 6.3E-04 |
| *T10H9.1* | WBGene00020424 | protein_coding_gene | 1.4 | 2.63 | 1.1E-02 |
| *ZK402.5* | WBGene00022731 | protein_coding_gene | 1.4 | 2.63 | 2.1E-02 |
| *C31B8.12* | WBGene00016285 | protein_coding_gene | 1.4 | 2.62 | 3.0E-03 |
| *C17H12.37* | WBGene00236786 | protein_coding_gene | 1.4 | 2.62 | 4.4E-02 |
| *Y51A2D.1* | WBGene00013072 | protein_coding_gene | 1.4 | 2.61 | 3.8E-04 |
| *clec-108* | WBGene00012504 | protein_coding_gene | 1.4 | 2.60 | 4.2E-03 |
| *cyp-35D1* | WBGene00008829 | protein_coding_gene | 1.4 | 2.59 | 5.4E-03 |
| *ctsa-3.2* | WBGene00007462 | protein_coding_gene | 1.4 | 2.58 | 5.9E-03 |
| *T02G5.7* | WBGene00020166 | protein_coding_gene | 1.4 | 2.58 | 6.8E-23 |
| *col-96* | WBGene00000671 | protein_coding_gene | 1.4 | 2.57 | 6.8E-08 |
| *W10C8.5* | WBGene00021128 | protein_coding_gene | 1.4 | 2.57 | 4.4E-16 |
| *T26H5.10* | WBGene00044783 | protein_coding_gene | 1.4 | 2.56 | 1.3E-03 |

| | | | | | |
|---|---|---|---|---|---|
| *ilys-2* | WBGene00016669 | protein_coding_gene | 1.4 | 2.56 | 3.7E-03 |
| *Y43F8C.17* | WBGene00012838 | protein_coding_gene | 1.4 | 2.55 | 1.0E-10 |
| *vha-5* | WBGene00006914 | protein_coding_gene | 1.4 | 2.55 | 3.6E-23 |
| *clec-165* | WBGene00009523 | protein_coding_gene | 1.4 | 2.55 | 6.7E-04 |
| *unc-41* | WBGene00006777 | protein_coding_gene | 1.4 | 2.55 | 7.2E-07 |
| *C34F11.1* | WBGene00016413 | protein_coding_gene | 1.4 | 2.55 | 8.9E-03 |
| *cah-4* | WBGene00000282 | protein_coding_gene | 1.3 | 2.54 | 3.8E-18 |
| *F09E5.16* | WBGene00017292 | protein_coding_gene | 1.3 | 2.53 | 4.0E-07 |
| *cwp-1* | WBGene00000859 | protein_coding_gene | 1.3 | 2.53 | 2.3E-04 |
| *B0416.2* | WBGene00015178 | protein_coding_gene | 1.3 | 2.52 | 1.8E-02 |
| *cwp-2* | WBGene00000860 | protein_coding_gene | 1.3 | 2.52 | 4.7E-02 |
| *pat-12* | WBGene00003936 | protein_coding_gene | 1.3 | 2.52 | 5.4E-21 |
| *bgnt-1.6* | WBGene00010167 | protein_coding_gene | 1.3 | 2.52 | 2.6E-02 |
| *nlp-26* | WBGene00003764 | protein_coding_gene | 1.3 | 2.51 | 3.0E-05 |
| *pqn-72* | WBGene00004154 | protein_coding_gene | 1.3 | 2.50 | 7.3E-04 |
| *C35B1.4* | WBGene00016434 | protein_coding_gene | 1.3 | 2.49 | 1.1E-10 |
| *VB0395L.1* | WBGene00044150 | protein_coding_gene | 1.3 | 2.48 | 3.8E-03 |
| *glr-1* | WBGene00001612 | protein_coding_gene | 1.3 | 2.48 | 2.2E-03 |
| *amt-4* | WBGene00000136 | protein_coding_gene | 1.3 | 2.48 | 1.3E-11 |
| *cyp-25A1* | WBGene00007963 | protein_coding_gene | 1.3 | 2.48 | 1.5E-04 |
| *Y116F11A.3* | WBGene00013817 | protein_coding_gene | 1.3 | 2.47 | 2.7E-02 |
| *mab-3* | WBGene00003100 | protein_coding_gene | 1.3 | 2.47 | 5.9E-06 |
| *Y51H4A.935* | WBGene00194647 | protein_coding_gene | 1.3 | 2.47 | 2.4E-02 |
| *dmsr-3* | WBGene00012993 | protein_coding_gene | 1.3 | 2.47 | 1.5E-02 |
| *C28D4.8* | WBGene00007795 | protein_coding_gene | 1.3 | 2.47 | 1.3E-02 |
| *C18B2.3* | WBGene00015954 | protein_coding_gene | 1.3 | 2.47 | 1.1E-07 |
| *F43C11.2* | WBGene00018379 | protein_coding_gene | 1.3 | 2.46 | 9.4E-08 |
| *F46A9.2* | WBGene00009755 | protein_coding_gene | 1.3 | 2.46 | 4.4E-06 |
| *Y53C10A.15* | WBGene00044805 | protein_coding_gene | 1.3 | 2.45 | 2.4E-02 |
| *clec-135* | WBGene00015821 | protein_coding_gene | 1.3 | 2.45 | 1.7E-06 |
| *kcc-3* | WBGene00019289 | protein_coding_gene | 1.3 | 2.45 | 2.1E-06 |
| *sur-5* | WBGene00006351 | protein_coding_gene | 1.3 | 2.44 | 1.3E-16 |
| *scl-26* | WBGene00016821 | protein_coding_gene | 1.3 | 2.44 | 8.5E-03 |
| *aqp-8* | WBGene00000176 | protein_coding_gene | 1.3 | 2.43 | 7.2E-22 |
| *fbxa-172* | WBGene00008014 | protein_coding_gene | 1.3 | 2.42 | 1.4E-04 |
| *Y66C5A.1* | WBGene00044156 | protein_coding_gene | 1.3 | 2.42 | 3.4E-02 |
| *F59A6.11* | WBGene00044664 | protein_coding_gene | 1.3 | 2.42 | 9.2E-03 |
| *F57A8.6* | WBGene00010181 | protein_coding_gene | 1.3 | 2.40 | 2.6E-03 |
| *ugt-43* | WBGene00008485 | protein_coding_gene | 1.3 | 2.40 | 1.4E-05 |
| *F52H2.6* | WBGene00018718 | protein_coding_gene | 1.3 | 2.40 | 6.7E-10 |
| *ZK1193.2* | WBGene00022858 | protein_coding_gene | 1.3 | 2.39 | 6.0E-08 |
| *sams-1* | WBGene00008205 | protein_coding_gene | 1.3 | 2.39 | 5.5E-23 |
| *frpr-6* | WBGene00017661 | protein_coding_gene | 1.3 | 2.39 | 2.8E-02 |
| *aqp-7* | WBGene00000175 | protein_coding_gene | 1.3 | 2.39 | 2.5E-13 |
| *best-24* | WBGene00022797 | protein_coding_gene | 1.3 | 2.39 | 8.8E-19 |
| *R07E3.4* | WBGene00011105 | protein_coding_gene | 1.3 | 2.39 | 7.6E-07 |
| *F40F9.3* | WBGene00009581 | protein_coding_gene | 1.3 | 2.38 | 2.4E-05 |
| *T02E9.6* | WBGene00044203 | protein_coding_gene | 1.2 | 2.38 | 4.1E-02 |
| *F38A1.9* | WBGene00009522 | protein_coding_gene | 1.2 | 2.37 | 4.4E-04 |
| *F58B4.5* | WBGene00010238 | protein_coding_gene | 1.2 | 2.36 | 8.2E-11 |
| *dpf-2* | WBGene00001055 | protein_coding_gene | 1.2 | 2.36 | 2.8E-15 |
| *cpn-4* | WBGene00000780 | protein_coding_gene | 1.2 | 2.35 | 2.6E-17 |

| | | | | | |
|---|---|---|---|---|---|
| *Y42G9A.3* | WBGene00021533 | protein_coding_gene | 1.2 | 2.35 | 8.4E-04 |
| *unc-23* | WBGene00006760 | protein_coding_gene | 1.2 | 2.34 | 6.0E-09 |
| *clec-92* | WBGene00013125 | protein_coding_gene | 1.2 | 2.33 | 9.0E-06 |
| *poml-3* | WBGene00077701 | protein_coding_gene | 1.2 | 2.33 | 9.7E-08 |
| *W03D8.5* | WBGene00020987 | protein_coding_gene | 1.2 | 2.33 | 5.4E-04 |
| *Y87G2A.16* | WBGene00013602 | protein_coding_gene | 1.2 | 2.33 | 4.6E-02 |
| *C01B10.44* | WBGene00194839 | protein_coding_gene | 1.2 | 2.33 | 3.5E-02 |
| *Y65B4A.7* | WBGene00022030 | protein_coding_gene | 1.2 | 2.32 | 2.5E-02 |
| *aco-1* | WBGene00000040 | protein_coding_gene | 1.2 | 2.32 | 1.9E-17 |
| *unc-46* | WBGene00006782 | protein_coding_gene | 1.2 | 2.32 | 5.0E-02 |
| *cat-1* | WBGene00000295 | protein_coding_gene | 1.2 | 2.32 | 6.8E-03 |
| *hex-5* | WBGene00013497 | protein_coding_gene | 1.2 | 2.31 | 2.8E-05 |
| *C01B10.3* | WBGene00015278 | protein_coding_gene | 1.2 | 2.31 | 2.0E-12 |
| *F37C4.6* | WBGene00018146 | protein_coding_gene | 1.2 | 2.31 | 1.5E-11 |
| *Y110A2AL.6* | WBGene00022443 | protein_coding_gene | 1.2 | 2.30 | 5.9E-04 |
| *F33C8.4* | WBGene00009358 | protein_coding_gene | 1.2 | 2.30 | 2.1E-07 |
| *cal-6* | WBGene00022214 | pseudogene | 1.2 | 2.30 | 4.7E-02 |
| *clec-99* | WBGene00013933 | protein_coding_gene | 1.2 | 2.29 | 2.3E-09 |
| *unc-25* | WBGene00006762 | protein_coding_gene | 1.2 | 2.29 | 5.6E-09 |
| *idh-1* | WBGene00010317 | protein_coding_gene | 1.2 | 2.29 | 4.6E-15 |
| *C12D5.9* | WBGene00015710 | protein_coding_gene | 1.2 | 2.29 | 4.6E-05 |
| *C35C5.8* | WBGene00007957 | protein_coding_gene | 1.2 | 2.29 | 7.9E-04 |
| *bca-1* | WBGene00000245 | protein_coding_gene | 1.2 | 2.29 | 7.8E-09 |
| *M02D8.5* | WBGene00019731 | protein_coding_gene | 1.2 | 2.28 | 1.5E-05 |
| *Y39B6A.41* | WBGene00012699 | protein_coding_gene | 1.2 | 2.28 | 4.2E-03 |
| *flp-23* | WBGene00017693 | protein_coding_gene | 1.2 | 2.27 | 2.2E-02 |
| *clec-130* | WBGene00021144 | protein_coding_gene | 1.2 | 2.27 | 1.0E-02 |
| *F58B6.1* | WBGene00019029 | protein_coding_gene | 1.2 | 2.27 | 1.4E-02 |
| *Y46D2A.3* | WBGene00021591 | protein_coding_gene | 1.2 | 2.27 | 4.8E-03 |
| *T09B4.3* | WBGene00020377 | protein_coding_gene | 1.2 | 2.26 | 1.1E-02 |
| *clec-56* | WBGene00008595 | protein_coding_gene | 1.2 | 2.26 | 6.5E-11 |
| *Y6G8.16* | WBGene00194801 | protein_coding_gene | 1.2 | 2.26 | 3.3E-02 |
| *F40H3.2* | WBGene00018250 | protein_coding_gene | 1.2 | 2.26 | 2.8E-03 |
| *F44G3.10* | WBGene00009710 | protein_coding_gene | 1.2 | 2.26 | 2.9E-03 |
| *mct-3* | WBGene00010834 | protein_coding_gene | 1.2 | 2.26 | 3.7E-20 |
| *sng-1* | WBGene00004912 | protein_coding_gene | 1.2 | 2.25 | 2.4E-03 |
| *Y6G8.15* | WBGene00189937 | protein_coding_gene | 1.2 | 2.25 | 5.2E-05 |
| *M02D8.6* | WBGene00019732 | protein_coding_gene | 1.2 | 2.25 | 2.7E-08 |
| *Y71H2AM.12* | WBGene00022177 | protein_coding_gene | 1.2 | 2.25 | 1.3E-04 |
| *F46A9.1* | WBGene00009754 | protein_coding_gene | 1.2 | 2.25 | 8.8E-07 |
| *F38E1.10* | WBGene00018182 | protein_coding_gene | 1.2 | 2.25 | 8.8E-04 |
| *F42G10.1* | WBGene00009645 | protein_coding_gene | 1.2 | 2.24 | 6.7E-06 |
| *Y43F8B.19* | WBGene00077688 | protein_coding_gene | 1.2 | 2.24 | 1.1E-02 |
| *inx-1* | WBGene00002123 | protein_coding_gene | 1.2 | 2.24 | 9.4E-06 |
| *W01G7.2* | WBGene00014851 | pseudogene | 1.2 | 2.24 | 2.9E-02 |
| *K01B6.3* | WBGene00010454 | protein_coding_gene | 1.2 | 2.23 | 1.2E-03 |
| *asp-11* | WBGene00017677 | pseudogene | 1.2 | 2.23 | 2.0E-02 |
| *gon-1* | WBGene00001650 | protein_coding_gene | 1.2 | 2.23 | 2.6E-10 |
| *Y64G10A.2* | WBGene00013412 | protein_coding_gene | 1.2 | 2.23 | 3.6E-02 |
| *mxt-1* | WBGene00012478 | protein_coding_gene | 1.2 | 2.23 | 1.6E-08 |
| *clec-138* | WBGene00018049 | protein_coding_gene | 1.2 | 2.23 | 3.7E-06 |
| *Y53C10A.5* | WBGene00013136 | protein_coding_gene | 1.1 | 2.22 | 6.5E-12 |

| | | | | | |
|---|---|---|---|---|---|
| *clec-110* | WBGene00008909 | protein_coding_gene | 1.1 | 2.21 | 1.3E-08 |
| *F46F5.1* | WBGene00018493 | protein_coding_gene | 1.1 | 2.21 | 3.2E-03 |
| *F37B12.1* | WBGene00009504 | protein_coding_gene | 1.1 | 2.21 | 3.2E-03 |
| *F31D5.2* | WBGene00017945 | protein_coding_gene | 1.1 | 2.21 | 6.2E-06 |
| *C08G5.1* | WBGene00015614 | pseudogene | 1.1 | 2.21 | 5.8E-03 |
| *T26H5.8* | WBGene00012070 | protein_coding_gene | 1.1 | 2.21 | 7.4E-08 |
| *ugt-48* | WBGene00015965 | protein_coding_gene | 1.1 | 2.20 | 8.9E-07 |
| *F58F9.8* | WBGene00044306 | protein_coding_gene | 1.1 | 2.20 | 2.6E-03 |
| *C36B7.5* | WBGene00016468 | protein_coding_gene | 1.1 | 2.20 | 4.0E-02 |
| *W03G11.2* | WBGene00012224 | protein_coding_gene | 1.1 | 2.20 | 1.3E-02 |
| *frpr-5* | WBGene00008342 | protein_coding_gene | 1.1 | 2.20 | 1.8E-03 |
| *M02D8.2* | WBGene00019728 | protein_coding_gene | 1.1 | 2.20 | 3.9E-02 |
| *F10C1.9* | WBGene00044296 | protein_coding_gene | 1.1 | 2.20 | 6.5E-11 |
| *argk-1* | WBGene00009706 | protein_coding_gene | 1.1 | 2.19 | 1.6E-02 |
| *aldo-1* | WBGene00011474 | protein_coding_gene | 1.1 | 2.19 | 1.3E-13 |
| *madd-4* | WBGene00009958 | protein_coding_gene | 1.1 | 2.19 | 6.3E-06 |
| *fipr-29* | WBGene00019171 | protein_coding_gene | 1.1 | 2.19 | 3.6E-02 |
| *Y51F10.7* | WBGene00021768 | protein_coding_gene | 1.1 | 2.18 | 2.8E-09 |
| *plg-1* | WBGene00004041 | pseudogene | 1.1 | 2.18 | 3.2E-11 |
| *mec-12* | WBGene00003175 | protein_coding_gene | 1.1 | 2.18 | 7.2E-11 |
| *F16H6.9* | WBGene00008899 | protein_coding_gene | 1.1 | 2.18 | 6.5E-04 |
| *sdha-1* | WBGene00015391 | protein_coding_gene | 1.1 | 2.17 | 3.1E-15 |
| *F58A4.1* | WBGene00010226 | protein_coding_gene | 1.1 | 2.17 | 2.0E-06 |
| *K05F1.9* | WBGene00019410 | protein_coding_gene | 1.1 | 2.17 | 8.9E-08 |
| *T14A8.2* | WBGene00020497 | protein_coding_gene | 1.1 | 2.17 | 6.4E-04 |
| *pept-2* | WBGene00003876 | protein_coding_gene | 1.1 | 2.17 | 4.4E-08 |
| *Y34B4A.4* | WBGene00021320 | protein_coding_gene | 1.1 | 2.17 | 5.6E-06 |
| *spin-1* | WBGene00007549 | protein_coding_gene | 1.1 | 2.16 | 5.4E-16 |
| *nas-20* | WBGene00003539 | protein_coding_gene | 1.1 | 2.16 | 1.1E-07 |
| *B0513.90* | WBGene00235301 | protein_coding_gene | 1.1 | 2.16 | 8.6E-03 |
| *F08H9.2* | WBGene00008590 | protein_coding_gene | 1.1 | 2.16 | 1.3E-06 |
| *clec-95* | WBGene00013927 | protein_coding_gene | 1.1 | 2.15 | 1.2E-08 |
| *fipr-5* | WBGene00044174 | protein_coding_gene | 1.1 | 2.15 | 3.0E-05 |
| *Y67H2A.9* | WBGene00014941 | pseudogene | 1.1 | 2.15 | 4.1E-08 |
| *unc-9* | WBGene00006749 | protein_coding_gene | 1.1 | 2.15 | 9.5E-07 |
| *C14C11.1* | WBGene00015765 | protein_coding_gene | 1.1 | 2.14 | 4.4E-04 |
| *F02H6.6* | WBGene00008539 | protein_coding_gene | 1.1 | 2.14 | 2.0E-03 |
| *C33F10.11* | WBGene00016357 | protein_coding_gene | 1.1 | 2.14 | 1.7E-05 |
| *K09A9.6* | WBGene00010703 | protein_coding_gene | 1.1 | 2.14 | 4.6E-08 |
| *Y45F10C.4* | WBGene00012880 | protein_coding_gene | 1.1 | 2.14 | 3.5E-06 |
| *D2045.5* | WBGene00008423 | protein_coding_gene | 1.1 | 2.14 | 3.3E-03 |
| *cyp-35A2* | WBGene00015400 | protein_coding_gene | 1.1 | 2.13 | 1.2E-03 |
| *col-113* | WBGene00000687 | protein_coding_gene | 1.1 | 2.13 | 1.3E-02 |
| *Y2H9A.6* | WBGene00086568 | protein_coding_gene | 1.1 | 2.13 | 1.5E-02 |
| *tbc-19* | WBGene00007849 | protein_coding_gene | 1.1 | 2.13 | 1.1E-05 |
| *F40G9.7* | WBGene00018243 | protein_coding_gene | 1.1 | 2.13 | 1.1E-02 |
| *C03F11.4* | WBGene00015390 | protein_coding_gene | 1.1 | 2.13 | 4.1E-06 |
| *aat-1* | WBGene00000002 | protein_coding_gene | 1.1 | 2.13 | 1.0E-06 |
| *F38B2.4* | WBGene00009531 | protein_coding_gene | 1.1 | 2.13 | 4.6E-11 |
| *K05F1.1* | WBGene00019405 | protein_coding_gene | 1.1 | 2.12 | 4.5E-02 |
| *F36G3.1* | WBGene00009482 | protein_coding_gene | 1.1 | 2.12 | 1.8E-08 |
| *C17H12.12* | WBGene00015937 | protein_coding_gene | 1.1 | 2.12 | 1.7E-06 |

| | | | | | |
|---|---|---|---|---|---|
| *pod-2* | WBGene00004076 | protein_coding_gene | 1.1 | 2.12 | 5.1E-21 |
| *prmt-6* | WBGene00016023 | protein_coding_gene | 1.1 | 2.11 | 1.4E-03 |
| *F22G12.8* | WBGene00044712 | pseudogene | 1.1 | 2.11 | 1.6E-02 |
| *ifb-1* | WBGene00002053 | protein_coding_gene | 1.1 | 2.11 | 1.1E-12 |
| *ivns-1* | WBGene00006493 | protein_coding_gene | 1.1 | 2.11 | 3.0E-17 |
| *Y49F6B.8* | WBGene00021720 | protein_coding_gene | 1.1 | 2.11 | 1.9E-02 |
| *C17F3.3* | WBGene00015908 | protein_coding_gene | 1.1 | 2.11 | 4.6E-04 |
| *C33G3.5* | WBGene00007905 | protein_coding_gene | 1.1 | 2.11 | 1.5E-06 |
| *B0222.10* | WBGene00015058 | ncRNA_gene | 1.1 | 2.10 | 3.8E-07 |
| *Y71G12B.5* | WBGene00022145 | protein_coding_gene | 1.1 | 2.10 | 1.9E-02 |
| *scav-6* | WBGene00008542 | protein_coding_gene | 1.1 | 2.10 | 5.4E-06 |
| *col-186* | WBGene00000759 | protein_coding_gene | 1.1 | 2.10 | 3.9E-02 |
| *T14G12.11* | WBGene00219411 | protein_coding_gene | 1.1 | 2.10 | 7.8E-04 |
| *Y116A8C.44* | WBGene00013815 | protein_coding_gene | 1.1 | 2.09 | 2.1E-02 |
| *F59E11.2* | WBGene00019109 | protein_coding_gene | 1.1 | 2.09 | 8.6E-03 |
| *C45E5.1* | WBGene00016664 | protein_coding_gene | 1.1 | 2.09 | 4.1E-05 |
| *T08B2.12* | WBGene00020350 | protein_coding_gene | 1.1 | 2.09 | 3.7E-03 |
| *tba-5* | WBGene00006531 | protein_coding_gene | 1.1 | 2.09 | 2.8E-02 |
| *cpr-4* | WBGene00000784 | protein_coding_gene | 1.1 | 2.09 | 2.6E-04 |
| *C49C3.11* | WBGene00008201 | protein_coding_gene | 1.1 | 2.09 | 1.5E-09 |
| *mrp-5* | WBGene00003411 | protein_coding_gene | 1.1 | 2.08 | 7.2E-22 |
| *T11F9.12* | WBGene00011713 | protein_coding_gene | 1.1 | 2.08 | 2.0E-12 |
| *Y7A5A.1* | WBGene00012407 | protein_coding_gene | 1.1 | 2.08 | 4.8E-08 |
| *cpr-8* | WBGene00021070 | protein_coding_gene | 1.1 | 2.08 | 1.1E-03 |
| *T13F3.8* | WBGene00044205 | protein_coding_gene | 1.1 | 2.08 | 1.7E-03 |
| *F46A8.5* | WBGene00009748 | protein_coding_gene | 1.1 | 2.07 | 1.0E-04 |
| *F32D8.12* | WBGene00009334 | protein_coding_gene | 1.0 | 2.07 | 2.0E-07 |
| *afmd-1* | WBGene00017051 | protein_coding_gene | 1.0 | 2.07 | 1.2E-02 |
| *rig-6* | WBGene00016354 | protein_coding_gene | 1.0 | 2.06 | 7.0E-09 |
| *rab-3* | WBGene00004267 | protein_coding_gene | 1.0 | 2.06 | 2.6E-09 |
| *acs-18* | WBGene00011173 | protein_coding_gene | 1.0 | 2.06 | 3.3E-07 |
| *F25E5.3* | WBGene00017784 | protein_coding_gene | 1.0 | 2.06 | 3.9E-07 |
| *ugt-47* | WBGene00011006 | protein_coding_gene | 1.0 | 2.06 | 5.9E-14 |
| *T01B4.3* | WBGene00011310 | protein_coding_gene | 1.0 | 2.05 | 7.3E-05 |
| *Y43F8C.15* | WBGene00012836 | protein_coding_gene | 1.0 | 2.05 | 1.6E-02 |
| *W03F9.4* | WBGene00021002 | protein_coding_gene | 1.0 | 2.05 | 1.7E-02 |
| *try-8* | WBGene00017791 | protein_coding_gene | 1.0 | 2.05 | 3.8E-03 |
| *R102.3* | WBGene00011290 | protein_coding_gene | 1.0 | 2.05 | 2.2E-02 |
| *wht-4* | WBGene00017179 | protein_coding_gene | 1.0 | 2.05 | 1.9E-03 |
| *Y18D10A.2* | WBGene00012475 | protein_coding_gene | 1.0 | 2.04 | 5.5E-04 |
| *del-10* | WBGene00020897 | protein_coding_gene | 1.0 | 2.04 | 4.2E-04 |
| *R05G6.1* | WBGene00019896 | protein_coding_gene | 1.0 | 2.04 | 2.6E-02 |
| *C41A3.2* | WBGene00016559 | protein_coding_gene | 1.0 | 2.04 | 5.8E-06 |
| *F10D7.10* | WBGene00271701 | protein_coding_gene | 1.0 | 2.04 | 2.2E-02 |
| *F11E6.3* | WBGene00008707 | protein_coding_gene | 1.0 | 2.04 | 6.4E-11 |
| *clec-106* | WBGene00012482 | protein_coding_gene | 1.0 | 2.04 | 9.1E-04 |
| *C43F9.5* | WBGene00008073 | protein_coding_gene | 1.0 | 2.04 | 2.3E-02 |
| *ver-4* | WBGene00006897 | protein_coding_gene | 1.0 | 2.04 | 2.1E-02 |
| *T21B6.3* | WBGene00011880 | protein_coding_gene | 1.0 | 2.04 | 2.2E-09 |
| *Y6G8.14* | WBGene00138715 | protein_coding_gene | 1.0 | 2.04 | 2.5E-03 |
| *F07C3.2* | WBGene00017193 | protein_coding_gene | 1.0 | 2.04 | 2.0E-03 |
| *cal-5* | WBGene00006861 | protein_coding_gene | 1.0 | 2.04 | 2.6E-13 |

| | | | | | |
|---|---|---|---|---|---|
| *Y43F8C.13* | WBGene00012834 | protein_coding_gene | 1.0 | 2.03 | 2.1E-08 |
| *C35B1.8* | WBGene00016437 | protein_coding_gene | 1.0 | 2.03 | 1.7E-02 |
| *poml-2* | WBGene00017089 | protein_coding_gene | 1.0 | 2.03 | 8.9E-04 |
| *F35C5.3* | WBGene00009391 | protein_coding_gene | 1.0 | 2.03 | 2.6E-02 |
| *C35B1.3* | WBGene00016433 | protein_coding_gene | 1.0 | 2.03 | 1.2E-02 |
| *ZK616.61* | WBGene00194816 | protein_coding_gene | 1.0 | 2.03 | 5.0E-04 |
| *C49C3.4* | WBGene00008194 | protein_coding_gene | 1.0 | 2.03 | 1.7E-08 |
| *F18E9.7* | WBGene00017573 | protein_coding_gene | 1.0 | 2.03 | 2.2E-05 |
| *C04C11.1* | WBGene00007295 | protein_coding_gene | 1.0 | 2.03 | 2.6E-02 |
| *ugt-22* | WBGene00007455 | protein_coding_gene | 1.0 | 2.02 | 7.7E-03 |
| *Y51A2D.8* | WBGene00013076 | protein_coding_gene | 1.0 | 2.02 | 9.7E-07 |
| *spe-12* | WBGene00004966 | protein_coding_gene | 1.0 | 2.02 | 9.2E-03 |
| *Y39B6A.18* | WBGene00012679 | pseudogene | 1.0 | 2.02 | 3.0E-02 |
| *acr-18* | WBGene00000057 | protein_coding_gene | 1.0 | 2.02 | 4.2E-03 |
| *crn-6* | WBGene00000799 | protein_coding_gene | 1.0 | 2.02 | 5.8E-07 |
| *C25D7.15* | WBGene00007722 | protein_coding_gene | 1.0 | 2.01 | 3.6E-02 |
| *F49E11.7* | WBGene00009893 | protein_coding_gene | 1.0 | 2.01 | 6.5E-03 |
| *oac-59* | WBGene00016585 | protein_coding_gene | 1.0 | 2.01 | 1.3E-06 |
| *ins-31* | WBGene00002114 | protein_coding_gene | 1.0 | 2.01 | 3.2E-03 |
| *F36A2.10* | WBGene00009457 | protein_coding_gene | 1.0 | 2.01 | 5.1E-06 |
| *clec-124* | WBGene00015909 | protein_coding_gene | 1.0 | 2.01 | 1.4E-06 |
| *F59B2.11* | WBGene00010313 | pseudogene | 1.0 | 2.01 | 3.0E-02 |
| *C48B4.12* | WBGene00008175 | protein_coding_gene | 1.0 | 2.01 | 4.9E-07 |
| *C45G9.10* | WBGene00016681 | protein_coding_gene | 1.0 | 2.01 | 2.4E-04 |
| *rnh-1.1* | WBGene00022888 | protein_coding_gene | 1.0 | 2.00 | 7.2E-05 |
| *ent-7* | WBGene00009686 | protein_coding_gene | 1.0 | 2.00 | 4.9E-08 |
| *exc-12* | WBGene00011476 | protein_coding_gene | 1.0 | 2.00 | 3.3E-02 |
| *fbp-1* | WBGene00001404 | protein_coding_gene | 1.0 | 2.00 | 1.0E-15 |
| *dhc-3* | WBGene00007154 | protein_coding_gene | 1.0 | 2.00 | 4.4E-02 |
| *ZK1290.1* | WBGene00022886 | protein_coding_gene | 1.0 | 2.00 | 1.3E-05 |
| *Y37E11AL.4* | WBGene00021360 | pseudogene | 1.0 | 2.00 | 5.0E-05 |
| *clec-162* | WBGene00007399 | protein_coding_gene | 1.0 | 2.00 | 4.4E-02 |
| *pxd-1* | WBGene00007982 | protein_coding_gene | -1.0 | 0.50 | 1.1E-10 |
| *glc-1* | WBGene00001591 | protein_coding_gene | -1.0 | 0.50 | 2.1E-09 |
| *F44E2.9* | WBGene00018422 | protein_coding_gene | -1.0 | 0.50 | 3.3E-08 |
| *F47B8.10* | WBGene00009811 | protein_coding_gene | -1.0 | 0.50 | 3.2E-02 |
| *mrps-17* | WBGene00015487 | protein_coding_gene | -1.0 | 0.50 | 2.2E-10 |
| *cpg-8* | WBGene00019357 | protein_coding_gene | -1.0 | 0.50 | 1.6E-13 |
| *glo-3* | WBGene00010341 | protein_coding_gene | -1.0 | 0.50 | 6.7E-04 |
| *F53F10.1* | WBGene00018763 | protein_coding_gene | -1.0 | 0.50 | 7.3E-05 |
| *C44C1.6* | WBGene00016645 | protein_coding_gene | -1.0 | 0.50 | 6.8E-04 |
| *eif-2alpha* | WBGene00021351 | protein_coding_gene | -1.0 | 0.50 | 3.6E-18 |
| *C06G3.5* | WBGene00015551 | protein_coding_gene | -1.0 | 0.50 | 4.1E-08 |
| *anr-8* | WBGene00219575 | antisense_lncRNA_gene | -1.0 | 0.50 | 1.2E-03 |
| *tsen-34* | WBGene00019535 | protein_coding_gene | -1.0 | 0.50 | 4.2E-05 |
| *ZK380.6* | WBGene00304812 | protein_coding_gene | -1.0 | 0.49 | 6.6E-13 |
| *mfb-1* | WBGene00008439 | protein_coding_gene | -1.0 | 0.49 | 2.2E-06 |
| *acdh-13* | WBGene00001566 | protein_coding_gene | -1.0 | 0.49 | 1.1E-14 |
| *bath-35* | WBGene00012196 | protein_coding_gene | -1.0 | 0.49 | 3.1E-02 |
| *F33H2.8* | WBGene00009371 | protein_coding_gene | -1.0 | 0.49 | 3.7E-02 |
| *C36B1.6* | WBGene00007973 | protein_coding_gene | -1.0 | 0.49 | 1.9E-05 |
| *M162.5* | WBGene00010931 | protein_coding_gene | -1.0 | 0.49 | 9.3E-04 |

| | | | | | |
|---|---|---|---|---|---|
| *R05D3.3* | WBGene00019878 | protein_coding_gene | -1.0 | 0.49 | 2.9E-07 |
| *fbxb-116* | WBGene00012066 | protein_coding_gene | -1.0 | 0.48 | 2.2E-02 |
| *aakg-4* | WBGene00020135 | protein_coding_gene | -1.1 | 0.48 | 3.2E-07 |
| *K12C11.6* | WBGene00045052 | protein_coding_gene | -1.1 | 0.48 | 4.2E-05 |
| *eif-3.H* | WBGene00001231 | protein_coding_gene | -1.1 | 0.48 | 4.1E-17 |
| *F52C12.2* | WBGene00018679 | protein_coding_gene | -1.1 | 0.48 | 1.5E-09 |
| *F56G4.6* | WBGene00010161 | protein_coding_gene | -1.1 | 0.48 | 2.8E-10 |
| *Y111B2A.21* | WBGene00013741 | protein_coding_gene | -1.1 | 0.48 | 3.7E-02 |
| *dyf-17* | WBGene00012673 | protein_coding_gene | -1.1 | 0.48 | 3.1E-03 |
| *F46C5.10* | WBGene00009785 | protein_coding_gene | -1.1 | 0.48 | 6.4E-05 |
| *ins-20* | WBGene00002103 | protein_coding_gene | -1.1 | 0.48 | 7.1E-03 |
| *T08A9.13* | WBGene00044442 | protein_coding_gene | -1.1 | 0.48 | 4.6E-02 |
| *fbxa-157* | WBGene00007393 | protein_coding_gene | -1.1 | 0.48 | 1.7E-02 |
| *zip-11* | WBGene00021082 | protein_coding_gene | -1.1 | 0.48 | 9.4E-05 |
| *tyr-1* | WBGene00015332 | protein_coding_gene | -1.1 | 0.48 | 2.2E-06 |
| *fbxc-45* | WBGene00021887 | protein_coding_gene | -1.1 | 0.48 | 1.2E-06 |
| *M01B2.13* | WBGene00045338 | protein_coding_gene | -1.1 | 0.48 | 6.9E-03 |
| *K06H7.7* | WBGene00019458 | protein_coding_gene | -1.1 | 0.47 | 4.6E-11 |
| *F13B12.2* | WBGene00008730 | protein_coding_gene | -1.1 | 0.47 | 4.4E-05 |
| *R06C7.2* | WBGene00011062 | protein_coding_gene | -1.1 | 0.47 | 1.4E-07 |
| *ZK1127.12* | WBGene00022857 | protein_coding_gene | -1.1 | 0.47 | 7.8E-08 |
| *K11D12.12* | WBGene00019651 | protein_coding_gene | -1.1 | 0.47 | 1.6E-10 |
| *F46A8.13* | WBGene00219950 | ncRNA_gene | -1.1 | 0.47 | 7.3E-03 |
| *M01G12.14* | WBGene00010825 | protein_coding_gene | -1.1 | 0.47 | 3.0E-02 |
| *csa-1* | WBGene00008403 | protein_coding_gene | -1.1 | 0.47 | 1.4E-04 |
| *ZK185.5* | WBGene00044481 | protein_coding_gene | -1.1 | 0.47 | 4.0E-06 |
| *dnj-9* | WBGene00001027 | protein_coding_gene | -1.1 | 0.47 | 1.8E-17 |
| *F59B10.4* | WBGene00010320 | protein_coding_gene | -1.1 | 0.46 | 1.9E-04 |
| *ttr-23* | WBGene00011894 | protein_coding_gene | -1.1 | 0.46 | 4.4E-06 |
| *C08F11.13* | WBGene00007460 | protein_coding_gene | -1.1 | 0.46 | 3.8E-10 |
| *abf-2* | WBGene00000013 | protein_coding_gene | -1.1 | 0.46 | 2.0E-03 |
| *slc-17.9* | WBGene00011643 | protein_coding_gene | -1.1 | 0.46 | 1.7E-02 |
| *B0478.3* | WBGene00015202 | protein_coding_gene | -1.1 | 0.46 | 3.2E-02 |
| *clec-146* | WBGene00013008 | protein_coding_gene | -1.1 | 0.46 | 2.2E-07 |
| *F59B1.8* | WBGene00019100 | protein_coding_gene | -1.1 | 0.46 | 1.7E-08 |
| *Y49A3A.3* | WBGene00013026 | protein_coding_gene | -1.1 | 0.46 | 7.4E-13 |
| *F40G9.17* | WBGene00185078 | protein_coding_gene | -1.1 | 0.46 | 1.3E-11 |
| *him-14* | WBGene00001872 | protein_coding_gene | -1.1 | 0.45 | 2.7E-13 |
| *F59C12.4* | WBGene00045272 | protein_coding_gene | -1.1 | 0.45 | 2.3E-08 |
| *dlc-5* | WBGene00043067 | protein_coding_gene | -1.1 | 0.45 | 3.1E-02 |
| *fbxa-212* | WBGene00012568 | pseudogene | -1.1 | 0.45 | 1.3E-05 |
| *F35E12.9* | WBGene00009433 | protein_coding_gene | -1.1 | 0.45 | 4.6E-04 |
| *R10E8.8* | WBGene00011213 | protein_coding_gene | -1.1 | 0.45 | 2.3E-05 |
| *fbxa-145* | WBGene00018595 | protein_coding_gene | -1.1 | 0.45 | 6.8E-04 |
| *atad-3* | WBGene00010015 | protein_coding_gene | -1.1 | 0.45 | 1.7E-18 |
| *cyp-34A9* | WBGene00015044 | protein_coding_gene | -1.2 | 0.45 | 3.8E-18 |
| *Y48A6B.7* | WBGene00012968 | protein_coding_gene | -1.2 | 0.45 | 1.0E-14 |
| *W04A8.2* | WBGene00012237 | protein_coding_gene | -1.2 | 0.45 | 9.5E-06 |
| *F52D1.2* | WBGene00018683 | protein_coding_gene | -1.2 | 0.44 | 7.3E-04 |
| *tsp-2* | WBGene00006628 | protein_coding_gene | -1.2 | 0.44 | 4.2E-04 |
| *fbxa-198* | WBGene00011514 | pseudogene | -1.2 | 0.44 | 6.7E-06 |
| *Y57G11B.2* | WBGene00013295 | protein_coding_gene | -1.2 | 0.44 | 4.3E-06 |

| | | | | | |
|---|---|---|---|---|---|
| *C05C10.7* | WBGene00007334 | protein_coding_gene | -1.2 | 0.44 | 2.4E-10 |
| *M03A1.8* | WBGene00077490 | protein_coding_gene | -1.2 | 0.44 | 1.1E-02 |
| *D2007.1* | WBGene00017041 | protein_coding_gene | -1.2 | 0.44 | 4.3E-05 |
| *ZK688.7* | WBGene00022802 | protein_coding_gene | -1.2 | 0.44 | 6.3E-07 |
| *T07G12.8* | WBGene00011599 | pseudogene | -1.2 | 0.44 | 8.4E-06 |
| *K07C5.13* | WBGene00050947 | protein_coding_gene | -1.2 | 0.43 | 2.5E-02 |
| *D2089.8* | WBGene00271706 | protein_coding_gene | -1.2 | 0.43 | 1.3E-06 |
| *T12D8.5* | WBGene00011733 | protein_coding_gene | -1.2 | 0.43 | 2.3E-04 |
| *C05D9.9* | WBGene00015476 | protein_coding_gene | -1.2 | 0.43 | 6.3E-09 |
| *F42H10.6* | WBGene00018370 | protein_coding_gene | -1.2 | 0.43 | 4.3E-11 |
| *cyp-13A3* | WBGene00011675 | protein_coding_gene | -1.2 | 0.43 | 1.7E-02 |
| *F21C10.11* | WBGene00017660 | protein_coding_gene | -1.2 | 0.43 | 2.2E-02 |
| *F13C5.1* | WBGene00017422 | protein_coding_gene | -1.2 | 0.43 | 8.6E-07 |
| *K02A11.4* | WBGene00044607 | protein_coding_gene | -1.2 | 0.43 | 9.5E-06 |
| *F41C3.2* | WBGene00018268 | protein_coding_gene | -1.2 | 0.43 | 2.0E-06 |
| *K01A6.4* | WBGene00010446 | protein_coding_gene | -1.2 | 0.42 | 3.4E-02 |
| *Y73B3A.21* | WBGene00022222 | pseudogene | -1.2 | 0.42 | 7.3E-03 |
| *spd-3* | WBGene00004954 | protein_coding_gene | -1.2 | 0.42 | 3.2E-13 |
| *cest-29* | WBGene00017205 | pseudogene | -1.2 | 0.42 | 3.2E-05 |
| *T28F4.5* | WBGene00012140 | protein_coding_gene | -1.2 | 0.42 | 2.1E-11 |
| *F36F2.2* | WBGene00009476 | protein_coding_gene | -1.2 | 0.42 | 7.8E-08 |
| *faah-2* | WBGene00015048 | protein_coding_gene | -1.2 | 0.42 | 2.0E-04 |
| *F08B4.3* | WBGene00017235 | pseudogene | -1.2 | 0.42 | 2.2E-09 |
| *T06A1.1* | WBGene00020277 | protein_coding_gene | -1.3 | 0.42 | 1.2E-02 |
| *C18A11.1* | WBGene00015947 | protein_coding_gene | -1.3 | 0.42 | 2.8E-09 |
| *C11D2.3* | WBGene00015701 | protein_coding_gene | -1.3 | 0.42 | 1.8E-02 |
| *fbxa-122* | WBGene00014917 | pseudogene | -1.3 | 0.42 | 5.8E-06 |
| *F18C5.10* | WBGene00017563 | protein_coding_gene | -1.3 | 0.42 | 9.3E-25 |
| *amt-1* | WBGene00000133 | protein_coding_gene | -1.3 | 0.42 | 4.2E-02 |
| *F01D5.5* | WBGene00008495 | protein_coding_gene | -1.3 | 0.42 | 7.1E-04 |
| *alh-5* | WBGene00000111 | protein_coding_gene | -1.3 | 0.42 | 2.3E-12 |
| *fbxa-95* | WBGene00009222 | protein_coding_gene | -1.3 | 0.41 | 1.1E-02 |
| *ZK550.2* | WBGene00013996 | protein_coding_gene | -1.3 | 0.41 | 6.6E-06 |
| *ZK896.4* | WBGene00014135 | protein_coding_gene | -1.3 | 0.41 | 1.1E-03 |
| *K06A9.2* | WBGene00019436 | protein_coding_gene | -1.3 | 0.41 | 2.1E-07 |
| *C54F6.5* | WBGene00016923 | protein_coding_gene | -1.3 | 0.41 | 9.9E-06 |
| *jmjd-4* | WBGene00011563 | protein_coding_gene | -1.3 | 0.40 | 5.4E-06 |
| *K10G6.9* | WBGene00206374 | protein_coding_gene | -1.3 | 0.40 | 1.6E-02 |
| *cyp-13A5* | WBGene00011672 | protein_coding_gene | -1.3 | 0.40 | 3.7E-14 |
| *F59A7.2* | WBGene00019090 | protein_coding_gene | -1.3 | 0.40 | 1.7E-06 |
| *W05H9.3* | WBGene00021050 | ncRNA_gene | -1.3 | 0.40 | 2.6E-13 |
| *D2045.8* | WBGene00008425 | protein_coding_gene | -1.3 | 0.40 | 5.0E-07 |
| *Y58A7A.3* | WBGene00021977 | protein_coding_gene | -1.3 | 0.40 | 1.0E-10 |
| *Y34F4.3* | WBGene00021338 | protein_coding_gene | -1.3 | 0.40 | 1.6E-03 |
| *C08F11.3* | WBGene00007452 | protein_coding_gene | -1.3 | 0.40 | 1.3E-02 |
| *pot-3* | WBGene00007065 | protein_coding_gene | -1.3 | 0.40 | 3.1E-11 |
| *W01F3.2* | WBGene00012185 | protein_coding_gene | -1.3 | 0.40 | 3.0E-16 |
| *C08E8.10* | WBGene00077692 | protein_coding_gene | -1.3 | 0.39 | 2.4E-02 |
| *F45E10.2* | WBGene00009729 | protein_coding_gene | -1.3 | 0.39 | 3.8E-07 |
| *C18H9.5* | WBGene00016003 | protein_coding_gene | -1.3 | 0.39 | 6.6E-06 |
| *dct-17* | WBGene00009431 | protein_coding_gene | -1.3 | 0.39 | 3.7E-02 |
| *dach-1* | WBGene00020386 | protein_coding_gene | -1.4 | 0.39 | 1.2E-04 |

| | | | | | |
|---|---|---|---|---|---|
| *spp-22* | WBGene00044283 | protein_coding_gene | -1.4 | 0.39 | 1.9E-05 |
| *ftn-1* | WBGene00001500 | protein_coding_gene | -1.4 | 0.39 | 3.2E-03 |
| *F09F3.8* | WBGene00014732 | pseudogene | -1.4 | 0.39 | 3.0E-02 |
| *frpr-11* | WBGene00019444 | protein_coding_gene | -1.4 | 0.39 | 1.1E-05 |
| *fbxa-140* | WBGene00008130 | protein_coding_gene | -1.4 | 0.39 | 8.0E-09 |
| *C36C9.10* | WBGene00219849 | ncRNA_gene | -1.4 | 0.38 | 2.2E-03 |
| *W01B11.6* | WBGene00020917 | protein_coding_gene | -1.4 | 0.38 | 4.8E-11 |
| *fbxa-162* | WBGene00015597 | pseudogene | -1.4 | 0.38 | 1.3E-04 |
| *C51E3.10* | WBGene00008255 | protein_coding_gene | -1.4 | 0.38 | 4.1E-04 |
| *F48D6.4* | WBGene00018602 | protein_coding_gene | -1.4 | 0.38 | 7.3E-20 |
| *R09B3.6* | WBGene00014820 | pseudogene | -1.4 | 0.38 | 6.8E-08 |
| *ZK380.t2* | WBGene00023401 | pseudogene | -1.4 | 0.37 | 1.7E-05 |
| *M03B6.5* | WBGene00010837 | protein_coding_gene | -1.4 | 0.37 | 2.7E-02 |
| *ugt-29* | WBGene00021709 | protein_coding_gene | -1.4 | 0.37 | 2.6E-05 |
| *fbxa-99* | WBGene00009490 | protein_coding_gene | -1.4 | 0.37 | 2.1E-02 |
| *fbxa-163* | WBGene00015598 | protein_coding_gene | -1.5 | 0.37 | 3.8E-05 |
| *Y37A1A.2* | WBGene00012535 | protein_coding_gene | -1.5 | 0.36 | 6.8E-05 |
| *C49G7.12* | WBGene00077593 | protein_coding_gene | -1.5 | 0.36 | 4.5E-04 |
| *F15E6.10* | WBGene00044479 | protein_coding_gene | -1.5 | 0.36 | 2.4E-02 |
| *lips-10* | WBGene00008803 | protein_coding_gene | -1.5 | 0.36 | 5.1E-08 |
| *F13A7.11* | WBGene00008726 | protein_coding_gene | -1.5 | 0.36 | 1.3E-05 |
| *M163.1* | WBGene00010935 | protein_coding_gene | -1.5 | 0.36 | 4.6E-15 |
| *F08H9.3* | WBGene00008591 | protein_coding_gene | -1.5 | 0.36 | 7.5E-04 |
| *M60.7* | WBGene00019782 | protein_coding_gene | -1.5 | 0.35 | 8.4E-05 |
| *T22B7.3* | WBGene00020672 | protein_coding_gene | -1.5 | 0.35 | 2.2E-05 |
| *F21H7.12* | WBGene00009033 | protein_coding_gene | -1.5 | 0.35 | 3.8E-02 |
| *tbc-6* | WBGene00015410 | protein_coding_gene | -1.5 | 0.35 | 4.0E-14 |
| *clec-40* | WBGene00011853 | protein_coding_gene | -1.5 | 0.34 | 9.2E-03 |
| *Y47G6A.14* | WBGene00021640 | protein_coding_gene | -1.5 | 0.34 | 2.4E-06 |
| *ugt-18* | WBGene00013900 | protein_coding_gene | -1.6 | 0.34 | 7.8E-04 |
| *cnc-7* | WBGene00010005 | protein_coding_gene | -1.6 | 0.34 | 7.1E-04 |
| *mtl-1* | WBGene00003473 | protein_coding_gene | -1.6 | 0.34 | 1.1E-04 |
| *fipr-26* | WBGene00009963 | protein_coding_gene | -1.6 | 0.34 | 8.3E-03 |
| *F40G9.20* | WBGene00195034 | ncRNA_gene | -1.6 | 0.34 | 9.3E-03 |
| *C06B3.6* | WBGene00007365 | protein_coding_gene | -1.6 | 0.34 | 4.6E-14 |
| *ugt-33* | WBGene00007946 | protein_coding_gene | -1.6 | 0.34 | 1.5E-03 |
| *catp-3* | WBGene00015660 | protein_coding_gene | -1.6 | 0.33 | 3.4E-06 |
| *swt-1* | WBGene00019300 | protein_coding_gene | -1.6 | 0.33 | 4.4E-10 |
| *T23F11.6* | WBGene00011957 | protein_coding_gene | -1.6 | 0.33 | 1.3E-03 |
| *T04C12.8* | WBGene00011430 | protein_coding_gene | -1.6 | 0.33 | 1.0E-05 |
| *C06B3.7* | WBGene00007366 | protein_coding_gene | -1.6 | 0.33 | 2.0E-03 |
| *F45D3.4* | WBGene00009724 | protein_coding_gene | -1.6 | 0.32 | 1.5E-07 |
| *Y53F4B.38* | WBGene00014920 | pseudogene | -1.6 | 0.32 | 4.2E-02 |
| *D2024.10* | WBGene00044505 | pseudogene | -1.6 | 0.32 | 3.2E-02 |
| *F37A8.5* | WBGene00009503 | protein_coding_gene | -1.7 | 0.31 | 5.7E-22 |
| *F11D5.5* | WBGene00017382 | protein_coding_gene | -1.7 | 0.31 | 2.2E-05 |
| *Y37H2A.12* | WBGene00044758 | protein_coding_gene | -1.7 | 0.31 | 2.7E-05 |
| *acdh-2* | WBGene00015894 | protein_coding_gene | -1.7 | 0.31 | 1.5E-03 |
| *Y8A9A.2* | WBGene00021171 | protein_coding_gene | -1.7 | 0.31 | 1.5E-08 |
| *dao-6* | WBGene00000932 | protein_coding_gene | -1.7 | 0.30 | 9.0E-03 |
| *tre-5* | WBGene00006611 | protein_coding_gene | -1.7 | 0.30 | 2.6E-12 |
| *clec-23* | WBGene00011851 | pseudogene | -1.7 | 0.30 | 5.1E-03 |

| | | | | | |
|---|---|---|---|---|---|
| *T10C6.15* | WBGene00050899 | protein_coding_gene | -1.7 | 0.30 | 3.5E-07 |
| *Y37F4.8* | WBGene00044289 | protein_coding_gene | -1.7 | 0.30 | 1.7E-04 |
| *H32K16.2* | WBGene00010422 | protein_coding_gene | -1.8 | 0.30 | 6.4E-06 |
| *pqn-16* | WBGene00004107 | protein_coding_gene | -1.8 | 0.29 | 4.0E-02 |
| *C18H7.1* | WBGene00015993 | protein_coding_gene | -1.8 | 0.29 | 3.8E-08 |
| *T04C12.7* | WBGene00011429 | protein_coding_gene | -1.8 | 0.29 | 2.5E-04 |
| *irg-5* | WBGene00009429 | protein_coding_gene | -1.8 | 0.29 | 1.9E-04 |
| *W05H9.1* | WBGene00021048 | protein_coding_gene | -1.8 | 0.29 | 2.7E-37 |
| *oac-54* | WBGene00012323 | protein_coding_gene | -1.8 | 0.28 | 2.5E-03 |
| *K10D11.2* | WBGene00010746 | protein_coding_gene | -1.8 | 0.28 | 4.3E-02 |
| *F10D7.3* | WBGene00017340 | protein_coding_gene | -1.8 | 0.28 | 5.6E-17 |
| *clec-145* | WBGene00013013 | protein_coding_gene | -1.8 | 0.28 | 1.3E-02 |
| *pho-9* | WBGene00022181 | protein_coding_gene | -1.8 | 0.28 | 8.0E-11 |
| *ttr-37* | WBGene00017667 | protein_coding_gene | -1.8 | 0.28 | 9.2E-14 |
| *C09B8.4* | WBGene00015623 | protein_coding_gene | -1.9 | 0.27 | 3.2E-08 |
| *F16B4.7* | WBGene00017509 | protein_coding_gene | -1.9 | 0.27 | 3.3E-04 |
| *K12B6.11* | WBGene00195164 | protein_coding_gene | -1.9 | 0.27 | 4.2E-02 |
| *nlp-31* | WBGene00003769 | protein_coding_gene | -1.9 | 0.26 | 1.9E-23 |
| *F53B2.8* | WBGene00009957 | protein_coding_gene | -1.9 | 0.26 | 1.1E-06 |
| *R09E10.13* | WBGene00194658 | protein_coding_gene | -2.0 | 0.26 | 4.9E-04 |
| *K01A6.7* | WBGene00045096 | protein_coding_gene | -2.0 | 0.26 | 4.4E-03 |
| *irg-2* | WBGene00016783 | protein_coding_gene | -2.0 | 0.26 | 1.4E-03 |
| *nspe-6* | WBGene00012603 | protein_coding_gene | -2.0 | 0.25 | 9.2E-03 |
| *ugt-54* | WBGene00012013 | protein_coding_gene | -2.0 | 0.25 | 4.0E-20 |
| *srd-28* | WBGene00005106 | protein_coding_gene | -2.0 | 0.24 | 4.6E-02 |
| *K09D9.1* | WBGene00019564 | protein_coding_gene | -2.1 | 0.24 | 1.1E-04 |
| *ZK1307.2* | WBGene00014245 | protein_coding_gene | -2.1 | 0.24 | 3.4E-05 |
| *C54F6.18* | WBGene00235093 | protein_coding_gene | -2.1 | 0.23 | 4.4E-03 |
| *ZK6.8* | WBGene00022643 | protein_coding_gene | -2.1 | 0.23 | 2.5E-08 |
| *nlp-34* | WBGene00015046 | protein_coding_gene | -2.1 | 0.23 | 1.0E-06 |
| *sri-36* | WBGene00005548 | protein_coding_gene | -2.1 | 0.23 | 1.9E-02 |
| *M04C9.2* | WBGene00010857 | protein_coding_gene | -2.1 | 0.23 | 8.6E-03 |
| *Y45F10D.6* | WBGene00012886 | protein_coding_gene | -2.1 | 0.23 | 6.8E-08 |
| *C31E10.11* | WBGene00304783 | protein_coding_gene | -2.2 | 0.22 | 8.0E-03 |
| *btb-15* | WBGene00015594 | pseudogene | -2.2 | 0.22 | 8.8E-03 |
| *ugt-34* | WBGene00009255 | protein_coding_gene | -2.2 | 0.22 | 1.5E-02 |
| *nlp-30* | WBGene00003768 | protein_coding_gene | -2.2 | 0.22 | 1.8E-10 |
| *F47B8.4* | WBGene00009805 | protein_coding_gene | -2.2 | 0.22 | 1.1E-09 |
| *ptr-22* | WBGene00004236 | protein_coding_gene | -2.2 | 0.22 | 7.6E-28 |
| *clec-24* | WBGene00009856 | protein_coding_gene | -2.2 | 0.21 | 2.0E-14 |
| *T04C12.32* | WBGene00235086 | ncRNA_gene | -2.3 | 0.21 | 8.9E-03 |
| *F54D8.10* | WBGene00284847 | protein_coding_gene | -2.3 | 0.20 | 5.0E-03 |
| *Y24D9A.9* | WBGene00044952 | snoRNA_gene | -2.3 | 0.20 | 1.3E-02 |
| *clec-32* | WBGene00009860 | protein_coding_gene | -2.3 | 0.20 | 1.9E-06 |
| *Y46H3A.5* | WBGene00021601 | protein_coding_gene | -2.4 | 0.19 | 2.5E-03 |
| *W08A12.2* | WBGene00021079 | protein_coding_gene | -2.4 | 0.19 | 7.6E-28 |
| *Y71F9AM.9* | WBGene00271781 | lincRNA_gene | -2.4 | 0.19 | 7.3E-04 |
| *dod-22* | WBGene00010125 | protein_coding_gene | -2.4 | 0.19 | 7.6E-24 |
| *F11E6.11* | WBGene00023501 | protein_coding_gene | -2.4 | 0.18 | 7.2E-22 |
| *pitr-4* | WBGene00007141 | protein_coding_gene | -2.5 | 0.18 | 1.4E-03 |
| *cnc-6* | WBGene00000560 | protein_coding_gene | -2.5 | 0.18 | 1.6E-02 |
| *fipr-23* | WBGene00007990 | protein_coding_gene | -2.5 | 0.17 | 1.6E-03 |

| | | | | | |
|---|---|---|---|---|---|
| *F54E2.1* | WBGene00018823 | protein_coding_gene | -2.5 | 0.17 | 6.7E-07 |
| *W10G11.1* | WBGene00021134 | protein_coding_gene | -2.6 | 0.17 | 3.3E-03 |
| *C49G7.7* | WBGene00016785 | protein_coding_gene | -2.6 | 0.17 | 4.8E-06 |
| *fil-1* | WBGene00011321 | protein_coding_gene | -2.6 | 0.17 | 5.7E-25 |
| *C37A5.3* | WBGene00014698 | protein_coding_gene | -2.6 | 0.16 | 8.4E-03 |
| *T23E7.6* | WBGene00044772 | protein_coding_gene | -2.7 | 0.16 | 8.6E-21 |
| *srr-6* | WBGene00005657 | protein_coding_gene | -2.7 | 0.15 | 5.4E-11 |
| *C50F7.5* | WBGene00016845 | protein_coding_gene | -2.8 | 0.15 | 6.3E-68 |
| *fipr-24* | WBGene00007992 | protein_coding_gene | -3.0 | 0.13 | 1.2E-05 |
| *C18D11.6* | WBGene00007682 | protein_coding_gene | -3.0 | 0.12 | 6.2E-06 |
| *C08E8.3* | WBGene00007439 | protein_coding_gene | -3.1 | 0.12 | 1.1E-05 |
| *C37A5.6* | WBGene00014700 | protein_coding_gene | -3.1 | 0.12 | 4.2E-03 |
| *ZK970.7* | WBGene00014173 | protein_coding_gene | -3.2 | 0.11 | 1.3E-11 |
| *cnc-11* | WBGene00044900 | protein_coding_gene | -3.3 | 0.10 | 7.1E-10 |
| *nlp-29* | WBGene00003767 | protein_coding_gene | -3.3 | 0.10 | 3.3E-62 |
| *cpt-3* | WBGene00021703 | protein_coding_gene | -3.3 | 0.10 | 8.6E-15 |
| *F56D6.9* | WBGene00044471 | protein_coding_gene | -3.4 | 0.10 | 5.3E-04 |
| *F48C1.11* | WBGene00195179 | protein_coding_gene | -3.8 | 0.07 | 7.3E-04 |
| *W02D7.11* | WBGene00020946 | protein_coding_gene | -3.9 | 0.07 | 4.9E-13 |
| *fmo-2* | WBGene00001477 | protein_coding_gene | -3.9 | 0.07 | 2.0E-07 |
| *F56D6.8* | WBGene00044292 | protein_coding_gene | -4.9 | 0.03 | 4.1E-06 |
| *dct-7* | WBGene00017488 | protein_coding_gene | -5.2 | 0.03 | 4.0E-08 |

N2 & CB4856 share genes exclusively expressed following *S. epidermidis* infection (vs. OP50) or DE in opposing directions

| | | | N2 | | | CB4856 | | |
|---|---|---|---|---|---|---|---|---|
| Gene Name | Wormbase Gene ID | Type | $log_2FC$ | FC | Adj P-value | $log_2FC$ | FC | Adj P-value |
| *lys-10* | WBGene00003099 | protein_coding_gene | 6.5 | 93.17 | 3.8E-31 | 2.8 | 6.91 | 1.1E-15 |
| *aqp-1* | WBGene00000169 | protein_coding_gene | 4.0 | 15.48 | 5.7E-39 | 3.6 | 12.10 | 5.0E-21 |
| *acs-2* | WBGene00009221 | protein_coding_gene | 3.8 | 14.03 | 5.0E-21 | 2.5 | 5.77 | 7.6E-10 |
| *cyp-13A7* | WBGene00000372 | protein_coding_gene | 3.0 | 8.17 | 4.4E-03 | 2.0 | 4.05 | 3.4E-12 |
| *clec-158* | WBGene00020067 | protein_coding_gene | 2.9 | 7.35 | 4.1E-03 | 1.4 | 2.63 | 8.1E-03 |
| *C35D6.5* | WBGene00007962 | pseudogene | 2.8 | 7.18 | 1.4E-03 | 1.3 | 2.49 | 3.9E-03 |
| ***mir-238*** | **WBGene00003331** | **miRNA_gene** | **2.7** | **6.58** | **2.6E-02** | **-2.3** | **0.21** | **2.6E-02** |
| *F22F7.7* | WBGene00017724 | protein_coding_gene | 2.5 | 5.65 | 2.4E-24 | 2.0 | 4.05 | 1.8E-19 |
| *hacd-1* | WBGene00019978 | protein_coding_gene | 2.5 | 5.63 | 1.4E-11 | 3.7 | 12.98 | 3.0E-13 |
| *scl-7* | WBGene00008029 | protein_coding_gene | 2.5 | 5.57 | 4.6E-02 | 1.2 | 2.26 | 2.8E-06 |
| *F49F1.15* | WBGene00235365 | protein_coding_gene | 2.5 | 5.49 | 3.3E-02 | 1.1 | 2.11 | 4.2E-05 |
| *C46A5.4* | WBGene00016700 | protein_coding_gene | 2.4 | 5.16 | 1.8E-04 | 1.1 | 2.15 | 8.9E-03 |
| *cut-3* | WBGene00009041 | protein_coding_gene | 2.3 | 5.08 | 2.4E-02 | 1.4 | 2.67 | 1.1E-03 |
| *F43C9.1* | WBGene00018375 | protein_coding_gene | 2.3 | 5.07 | 4.7E-05 | 1.0 | 2.02 | 8.2E-03 |
| *col-74* | WBGene00000650 | protein_coding_gene | 2.3 | 5.07 | 2.0E-02 | 1.8 | 3.55 | 9.3E-11 |
| *dmsr-16* | WBGene00194821 | protein_coding_gene | 2.3 | 4.90 | 5.4E-03 | 1.5 | 2.87 | 2.7E-02 |
| *fbxa-50* | WBGene00013844 | protein_coding_gene | 2.2 | 4.57 | 3.2E-04 | 1.5 | 2.82 | 3.2E-03 |
| *C08B6.4* | WBGene00007425 | protein_coding_gene | 2.1 | 4.39 | 1.4E-03 | 1.4 | 2.67 | 6.4E-03 |
| *lgc-13* | WBGene00011354 | protein_coding_gene | 2.1 | 4.32 | 4.4E-03 | 2.1 | 4.37 | 1.2E-03 |
| *pqn-84* | WBGene00004165 | protein_coding_gene | 1.8 | 3.38 | 1.1E-02 | 1.2 | 2.33 | 1.9E-10 |
| *Y47D7A.6* | WBGene00021619 | protein_coding_gene | 1.7 | 3.23 | 6.7E-03 | 2.2 | 4.63 | 1.7E-07 |
| *aat-2* | WBGene00000003 | protein_coding_gene | 1.7 | 3.23 | 1.5E-07 | 1.7 | 3.24 | 2.7E-12 |
| *gst-34* | WBGene00001782 | protein_coding_gene | 1.6 | 2.99 | 2.9E-02 | 3.0 | 7.75 | 1.1E-02 |
| *fat-2* | WBGene00001394 | protein_coding_gene | 1.5 | 2.89 | 8.0E-15 | 1.2 | 2.35 | 1.5E-21 |
| *pho-8* | WBGene00011262 | protein_coding_gene | 1.5 | 2.78 | 2.4E-02 | 1.9 | 3.71 | 1.9E-06 |
| *acer-1* | WBGene00016630 | protein_coding_gene | 1.4 | 2.69 | 3.1E-10 | 1.1 | 2.15 | 5.3E-14 |
| *pitr-5* | WBGene00017312 | protein_coding_gene | 1.4 | 2.66 | 1.5E-05 | 2.0 | 3.99 | 1.1E-09 |

| | | | | | | | | |
|---|---|---|---|---|---|---|---|---|
| *Y17D7B.7* | WBGene00014562 | pseudogene | 1.4 | 2.63 | 1.2E-02 | 1.7 | 3.22 | 3.1E-03 |
| *acs-19* | WBGene00007969 | protein_coding_gene | 1.4 | 2.58 | 8.9E-08 | 1.2 | 2.26 | 3.0E-17 |
| *Y37A1B.5* | WBGene00012538 | protein_coding_gene | 1.3 | 2.42 | 5.2E-08 | 1.1 | 2.21 | 8.6E-09 |
| *lys-4* | WBGene00003093 | protein_coding_gene | 1.2 | 2.35 | 1.4E-07 | 1.3 | 2.50 | 2.4E-12 |
| *aman-3* | WBGene00018594 | protein_coding_gene | 1.1 | 2.19 | 1.1E-04 | 1.8 | 3.40 | 1.5E-20 |
| *tag-290* | WBGene00009386 | protein_coding_gene | 1.1 | 2.17 | 2.3E-04 | 1.3 | 2.45 | 1.3E-11 |
| *F41E6.15* | WBGene00044554 | protein_coding_gene | 1.1 | 2.11 | 8.2E-06 | 1.4 | 2.70 | 2.7E-08 |
| *F41G3.10* | WBGene00018303 | protein_coding_gene | 1.1 | 2.11 | 9.6E-05 | 1.2 | 2.30 | 1.0E-16 |
| *pgp-13* | WBGene00004007 | protein_coding_gene | 1.1 | 2.08 | 7.3E-05 | 1.6 | 3.00 | 4.5E-13 |
| *fat-5* | WBGene00001397 | protein_coding_gene | 1.0 | 2.01 | 1.4E-03 | 3.5 | 11.71 | 9.0E-21 |
| *gstk-1* | WBGene00014251 | protein_coding_gene | -1.0 | 0.49 | 2.7E-07 | -1.4 | 0.38 | 3.4E-14 |
| *trd-1* | WBGene00020600 | protein_coding_gene | -1.0 | 0.49 | 1.7E-13 | -1.1 | 0.48 | 8.9E-14 |
| *F21F3.4* | WBGene00017674 | protein_coding_gene | -1.0 | 0.49 | 8.9E-05 | -1.2 | 0.43 | 1.6E-10 |
| *clec-150* | WBGene00019914 | protein_coding_gene | -1.1 | 0.48 | 1.0E-04 | -1.3 | 0.41 | 7.0E-24 |
| *mrps-22* | WBGene00007564 | protein_coding_gene | -1.1 | 0.47 | 4.4E-18 | -1.1 | 0.47 | 7.7E-18 |
| *C29F3.7* | WBGene00007807 | protein_coding_gene | -1.1 | 0.47 | 3.8E-03 | -1.6 | 0.33 | 3.3E-12 |
| *ZK1307.1* | WBGene00014244 | protein_coding_gene | -1.1 | 0.47 | 2.2E-03 | -1.1 | 0.46 | 5.1E-08 |
| *F12A10.1* | WBGene00017390 | protein_coding_gene | -1.1 | 0.46 | 4.6E-02 | -2.6 | 0.17 | 2.2E-19 |
| *eel-1* | WBGene00022069 | protein_coding_gene | -1.1 | 0.46 | 1.4E-03 | -1.0 | 0.49 | 1.3E-13 |
| *D2063.1* | WBGene00017060 | protein_coding_gene | -1.1 | 0.46 | 1.6E-02 | -1.2 | 0.45 | 2.5E-05 |
| *irg-4* | WBGene00008584 | protein_coding_gene | -1.2 | 0.45 | 1.7E-02 | -1.5 | 0.35 | 1.0E-04 |
| *F55G11.8* | WBGene00010128 | protein_coding_gene | -1.3 | 0.40 | 1.2E-02 | -1.2 | 0.43 | 7.0E-05 |
| *F49E12.10* | WBGene00009903 | protein_coding_gene | -1.4 | 0.38 | 5.8E-04 | -1.3 | 0.41 | 7.6E-04 |
| *fbxc-58* | WBGene00007506 | protein_coding_gene | -1.4 | 0.38 | 4.6E-02 | -1.1 | 0.45 | 1.0E-02 |
| *clec-86* | WBGene00016912 | protein_coding_gene | -1.4 | 0.38 | 1.0E-05 | -1.9 | 0.26 | 1.4E-16 |
| *C40H1.8* | WBGene00008043 | protein_coding_gene | -1.4 | 0.37 | 1.3E-02 | -1.1 | 0.47 | 3.2E-02 |
| *C18H7.11* | WBGene00044457 | protein_coding_gene | -1.4 | 0.37 | 1.3E-02 | -1.4 | 0.37 | 1.8E-03 |
| *F07G6.11* | WBGene00304813 | ncRNA_gene | -1.5 | 0.37 | 2.2E-02 | -1.8 | 0.29 | 2.0E-04 |
| *F21D5.3* | WBGene00009008 | protein_coding_gene | -1.5 | 0.35 | 2.4E-06 | -1.0 | 0.49 | 3.0E-03 |
| ***T23F6.1*** | **WBGene00011949** | **protein_coding_gene** | **-1.6** | **0.32** | **3.0E-02** | **1.7** | **3.17** | **4.9E-02** |
| *nlp-33* | WBGene00003771 | protein_coding_gene | -1.6 | 0.32 | 8.0E-06 | -1.3 | 0.40 | 3.8E-03 |
| *C49G7.10* | WBGene00016788 | protein_coding_gene | -1.7 | 0.31 | 1.3E-03 | -1.5 | 0.36 | 9.4E-04 |
| *cnc-4* | WBGene00000558 | protein_coding_gene | -1.7 | 0.30 | 6.7E-03 | -5.7 | 0.02 | 5.9E-17 |
| ***C05E7.2*** | **WBGene00007344** | **protein_coding_gene** | **-2.1** | **0.23** | **2.0E-03** | **1.8** | **3.39** | **5.3E-03** |
| ***grd-6*** | **WBGene00001695** | **protein_coding_gene** | **-2.1** | **0.23** | **1.6E-03** | **1.9** | **3.84** | **1.3E-02** |
| ***B0334.13*** | **WBGene00007149** | **protein_coding_gene** | **-2.2** | **0.22** | **8.4E-03** | **2.1** | **4.18** | **2.7E-02** |
| *asp-12* | WBGene00017678 | protein_coding_gene | -2.3 | 0.20 | 2.2E-08 | -2.2 | 0.21 | 3.0E-13 |
| *C10C5.3* | WBGene00007507 | protein_coding_gene | -2.4 | 0.19 | 3.7E-07 | -2.2 | 0.21 | 6.7E-06 |
| *F23F12.3* | WBGene00017751 | protein_coding_gene | -2.5 | 0.18 | 9.5E-07 | -1.3 | 0.40 | 1.2E-02 |
| *C17H12.8* | WBGene00015933 | protein_coding_gene | -2.8 | 0.14 | 5.2E-13 | -2.0 | 0.25 | 5.6E-21 |
| *nlp-25* | WBGene00003763 | protein_coding_gene | -2.8 | 0.14 | 1.1E-03 | -3.6 | 0.08 | 1.1E-07 |
| *F33H12.7* | WBGene00045457 | protein_coding_gene | -3.2 | 0.11 | 7.5E-04 | -2.6 | 0.16 | 3.6E-04 |
| *F15E6.4* | WBGene00017485 | protein_coding_gene | -3.2 | 0.11 | 2.2E-09 | -4.4 | 0.05 | 3.8E-11 |
| *cnc-3* | WBGene00000557 | protein_coding_gene | -3.3 | 0.10 | 1.3E-11 | -2.3 | 0.20 | 2.4E-10 |
| *K11H12.4* | WBGene00019660 | protein_coding_gene | -3.3 | 0.10 | 1.3E-11 | -2.1 | 0.24 | 1.4E-03 |
| *fip-6* | WBGene00009964 | protein_coding_gene | -3.8 | 0.07 | 4.9E-08 | -2.1 | 0.24 | 2.0E-04 |

**Supplemental Table 11**

**GO terms enriched in either N2, CB4856, or both, after infection by *S. epidermidis* (compared to *S. aureus*)**

**Supplemental Table 11 - GO Terms Enriched in *S. epidermidis* infected N2 or CB4856 animals**

N2 Specific Categories

| Category | Number of DEGs | Genes in Geneset | Downregulated Genes in Geneset | Upregulated Genes in Geneset | p-value | FDR-adjusted p-value | Differentially Expressed Genes within a Given Geneset |
|---|---|---|---|---|---|---|---|
| Extracellular material: collagen | 55 | 185 | 50 | 5 | 5.3E-46 | 4.5E-44 | WBGene00000256,WBGene00000596,WBGene00000601,WBGene00000603,WBGene00000606,WBGene00000615,WBGene00000626,WBGene00000636,WBGene00000639,WBGene00000647,WBGene00000649,WBGene00000653,WBGene00000657,WBGene00000663,WBGene00000665,WBGene00000666,WBGene00000672,WBGene00000674,WBGene00000678,WBGene00000681,WBGene00000683,WBGene00000684,WBGene00000689,WBGene00000694,WBGene00000695,WBGene00000699,WBGene00000703,WBGene00000704,WBGene00000707,WBGene00000711,WBGene00000712,WBGene00000717,WBGene00000718,WBGene00000725,WBGene00000727,WBGene00000729,WBGene00000730,WBGene00000732,WBGene00000734,WBGene00000735,WBGene00000740,WBGene00000741,WBGene00000742,WBGene00000743,WBGene00000748,WBGene00000749,WBGene00000753,WBGene00001066,WBGene00001067,WBGene00001072,WBGene00001074,WBGene00001075,WBGene00001076,WBGene00004300,WBGene00004394 |
| Extracellular material: cuticlin | 7 | 35 | 1 | 6 | 1.8E-06 | 1.5E-04 | WBGene00000853,WBGene00001063,WBGene00009830,WBGene00011104,WBGene00013145,WBGene00021025,WBGene00021919 |
| Stress response: detoxification | 13 | 151 | 11 | 1 | 3.2E-06 | 2.8E-04 | WBGene00007362,WBGene00008810,WBGene00011340,WBGene00013901,WBGene00013905,WBGene00015141,WBGene00016786,WBGene00017332,WBGene00017959,WBGene00019470,WBGene00019967,WBGene00020587,WBGene00021412 |

CB4856 Specific Categories

| Category | Number of DEGs | Genes in Geneset | Downregulated Genes in Geneset | Upregulated Genes in Geneset | p-value | FDR-adjusted p-value | Differentially Expressed Genes within a Given Geneset |
|---|---|---|---|---|---|---|---|
| Unknown | 293 | 8160 | 106 | 187 | 1.7E-12 | 1.7E-10 | WBGene00000860,WBGene00000932,WBGene00004107,WBGene00004112,WBGene00004153,WBGene00004154,WBGene00005657,WBGene00006493,WBGene00007295,WBGene00007334,WBGene00007366,WBGene00007439,WBGene00007452,WBGene00007460,WBGene00007521,WBGene00007544,WBGene00007682,WBGene00007722,WBGene00007795,WBGene00007809,WBGene00007849,WBGene00007905,WBGene00007957,WBGene00007973,WBGene00007982,WBGene00007990,WBGene00007992,WBGene00008073,WBGene00008129,WBGene00008175,WBGene00008194,WBGene00008212,WBGene00008255,WBGene00008403,WBGene00008497,WBGene00008539,WBGene00008559,WBGene00008590,WBGene00008677,WBGene00008707,WBGene00008899,WBGene00008960,WBGene00008961,WBGene00009033,WBGene00009114,WBGene00009358,WBGene00009362,WBGene00009371,WBGene00009433,WBGene00009457,WBGene00009476,WBGene00009482,WBGene00009503,WBGene00009504,WBGene00009522,WBGene00009581,WBGene00009724,WBGene00009754,WBGene00009755,WBGene00009765,WBGene00009766,WBGene00009785,WBGene00009787,WBGene00009957,WBGene00009963,WBGene00010060,WBGene00010161,WBGene00010168,WBGene00010170,WBGene00010181,WBGene00010226,WBGene00010238,WBGene00010320,WBGene00010368,WBGene00010422,WBGene00010446,WBGene00010454,WBGene00010825,WBGene00010837,WBGene00010935,WBGene00010987,WBGene00011046,WBGene00011062,WBGene00011213,WBGene00011290,WBGene00011310,WBGene00011373,WBGene00011414,WBGene00011429,WBGene00011430,WBGene00011476,WBGene00011594,WBGene00011713,WBGene00011880,WBGene00011894,WBGene00011957,WBGene00012123,WBGene00012140,WBGene00012185,WBGene00012196,WBGene00012217,WBGene00012237,WBGene00012387,WBGene00012475,WBGene00012603,WBGene00012607,WBGene00012748,WBGene00012834,WBGene00012836,WBGene00012837,WBGene00012838,WBGene00012880,WBGene00012883,WBGene00012886,WBGene00013026,WBGene00013169,WBGene00013295,WBGene00013331,WBGene00013412,WBGene00013741,WBGene00013815,WBGene00013817,WBGene00014173,WBGene00014245,WBGene00014698,WBGene00014700,WBGene00015058,WBGene00015178,WBGene00015390,WBGene00015410,WBGene00015476,WBGene00015679,WBGene00015701,WBGene00015710,WBGene00015765,WBGene00015794,WBGene00015908,WBGene00015937,WBGene00015947,WBGene00015954,WBGene00015977,WBGene00015993,WBGene00016185,WBGene00016285,WBGene00016413,WBGene00016433,WBGene00016437,WBGene00016468,WBGene00016559,WBGene00016645,WBGene00016681,WBGene00016714,WBGene00016785,WBGene00016820,WBGene00016845,WBGene00016915,WBGene00016923,WBGene00016929,WBGene00017193,WBGene00017205,WBGene00017292,WBGene00017305,WBGene00017378,WBGene00017439,WBGene00017488,WBGene00017509,WBGene00017563,WBGene00017573,WBGene00017660,WBGene00017730,WBGene00017784,WBGene00017841,WBGene00017913,WBGene00018136,WBGene00018182,WBGene00018243,WBGene00018250,WBGene00018379,WBGene00018380,WBGene00018422,WBGene00018493,WBGene00018602,WBGene00018683,WBGene00018763,WBGene00018823,WBGene00018946,WBGene00019029,WBGene00019085,WBGene00019109,WBGene00019171,WBGene00019405,WBGene00019410,WBGene00019458,WBGene00019564,WBGene00019685,WBGene00019728,WBGene00019731,WBGene00019732,WBGene00019878,WBGene00019896,WBGene00020144,WBGene00020277,WBGene00020350,WBGene00020377,WBGene00020400,WBGene00020439,WBGene00020697,WBGene00020700,WBGene00020803,WBGene00020918,WBGene00020946,WBGene00020987,WBGene00021008,WBGene00021048,WBGene00021050,WBGene00021066,WBGene00021079,WBGene00021134,WBGene00021452,WBGene00021511,WBGene00021533,WBGene00021591,WBGene00021601,WBGene00021616,WBGene00021620,WBGene00021640,WBGene00021720,WBGene00021743,WBGene00021768,WBGene00021918,WBGene00021977,WBGene00022030,WBGene00022071,WBGene00022145,WBGene00022267,WBGene00022443,WBGene00022731,WBGene00022802,WBGene00022857,WBGene00022858,WBGene00022886,WBGene00023501,WBGene00044016,WBGene00044156,WBGene00044174,WBGene00044203,WBGene00044205,WBGene00044283,WBGene00044289,WBGene00044292,WBGene00044296,WBGene00044306,WBGene00044307,WBGene00044402,WBGene00044442,WBGene00044471,WBGene00044474,WBGene00044479,WBGene00044488,WBGene00044607 WBGene00044658,WBGene00044664,WBGene00044772,WBGene00044783,WBGene00045096,WBGene00045272,WBGene00045338,WBGene00045488,WBGene00045510,WBGene00050899,WBGene00050947,WBGene00077593,WBGene00077692,WBGene00086568,WBGene00138715,WBGene00189937,WBGene00194647,WBGene00194658,WBGene00194701,WBGene00194742,WBGene00194801,WBGene00194816,WBGene00194839,WBGene00195164,WBGene00195179,WBGene00206374,WBGene00219411,WBGene00235093,WBGene00235301,WBGene00236786,WBGene00271701,WBGene00271706,WBGene00284847 |
| Stress response: pathogen | 22 | 177 | 17 | 5 | 4.5E-10 | 4.6E-08 | WBGene00000013,WBGene00000560,WBGene00003096,WBGene00003768,WBGene00003769,WBGene00008495,WBGene00008726,WBGene00009215,WBGene00009429,WBGene00009431,WBGene00009805,WBGene00010005,WBGene00010125,WBGene00010657,WBGene00010746,WBGene00013947,WBGene00014135,WBGene00016669,WBGene00016783,WBGene00017041,WBGene00019090,WBGene00044900 |
| Stress response: C-type Lectin | 26 | 257 | 5 | 21 | 7.5E-10 | 7.8E-08 | WBGene00007399,WBGene00007851,WBGene00008201,WBGene00008595,WBGene00008909,WBGene00009523,WBGene00009844,WBGene00009856,WBGene00009860,WBGene00011853,WBGene00012482,WBGene00012488,WBGene00012504,WBGene00013008,WBGene00013013,WBGene00013125,WBGene00013179,WBGene00013927,WBGene00013933,WBGene00015821,WBGene00015909,WBGene00017772,WBGene00018049,WBGene00021143,WBGene00021144,WBGene00021580 |
| Stress response: detoxification | 20 | 151 | 9 | 11 | 9.9E-10 | 1.0E-07 | WBGene00007455,WBGene00007885,WBGene00007913,WBGene00007946,WBGene00007963,WBGene00008485,WBGene00008829,WBGene00009255,WBGene00011006,WBGene00011672,WBGene00011675,WBGene00012013,WBGene00013900,WBGene00013903,WBGene00015044,WBGene00015400,WBGene00015965,WBGene00018543,WBGene00020386,WBGene00021709 |
| Metabolism: lipid | 30 | 525 | 8 | 22 | 5.0E-06 | 5.2E-04 | WBGene00001156,WBGene00001240,WBGene00001243,WBGene00001244,WBGene00004076,WBGene00006351,WBGene00007365,WBGene00007536,WBGene00007549,WBGene00008567,WBGene00008629,WBGene00008803,WBGene00010171,WBGene00010505,WBGene00010595,WBGene00011173,WBGene00011321,WBGene00011654,WBGene00011656,WBGene00012323,WBGene00012407,WBGene00015048,WBGene00015894,WBGene00016585,WBGene00018370,WBGene00018718,WBGene00020166,WBGene00021002,WBGene00021703,WBGene00022498 |

N2 and CB4856 Shared

| Category | Number of DEGs | Genes in Geneset | Downregulated Genes in Geneset | Upregulated Genes in Geneset | p-value | FDR-adjusted p-value | Differentially Expressed Genes within a Given Geneset |
|---|---|---|---|---|---|---|---|
| Stress response: pathogen | 7 | 177 | 7 (N2); 7 (CB4856) | 0 (N2); 0 (CB4856) | 4.36E-07 | 1.4E-05 | WBGene00000557,WBGene00000558,WBGene00003771,WBGene00007506,WBGene00007807,WBGene00008584,WBGene00010128 |

**Supplemental Table 12**

**Lists of differentially expressed genes in CB4856 fed *E. coli* (compared to N2)**

**Supplemental Table 12 - Genes expressed in CB4856 (vs. N2) fed *E. coli* OP50**

| Gene Name | Wormbase Gene ID | Type | $log_2FC$ | FC | Adj P-value |
|---|---|---|---|---|---|
| *nspa-1* | WBGene00010377 | protein_coding_gene | 8.1 | 271.084 | 8.9E-20 |
| *clec-77* | WBGene00044343 | pseudogene | 7.8 | 219.595 | 2.3E-25 |
| *cyp-37B1* | WBGene00009226 | protein_coding_gene | 7.7 | 206.507 | 4.8E-60 |
| *C50F7.5* | WBGene00016845 | protein_coding_gene | 7.3 | 156.348 | 2.2E-70 |
| *pud-2.2* | WBGene00017500 | protein_coding_gene | 7.3 | 153.739 | 1.0E-16 |
| *K02F6.4* | WBGene00019338 | protein_coding_gene | 7.2 | 151.450 | 1.6E-16 |
| *clec-110* | WBGene00008909 | protein_coding_gene | 7.1 | 141.223 | 3.8E-16 |
| *clec-208* | WBGene00018097 | protein_coding_gene | 7.1 | 140.507 | 3.7E-36 |
| *fmo-2* | WBGene00001477 | protein_coding_gene | 7.0 | 129.383 | 1.8E-23 |
| *fipr-18* | WBGene00044450 | protein_coding_gene | 6.9 | 117.843 | 2.3E-20 |
| *C39B5.10* | WBGene00016528 | protein_coding_gene | 6.8 | 107.731 | 2.2E-34 |
| *T07G12.5* | WBGene00011596 | protein_coding_gene | 6.7 | 106.385 | 2.0E-51 |
| *T23G4.5* | WBGene00011961 | protein_coding_gene | 6.6 | 96.062 | 4.0E-14 |
| *clec-124* | WBGene00015909 | protein_coding_gene | 6.4 | 85.625 | 2.2E-13 |
| *M7.10* | WBGene00010886 | protein_coding_gene | 6.4 | 83.926 | 2.1E-13 |
| *fipr-17* | WBGene00012965 | protein_coding_gene | 6.3 | 80.244 | 1.8E-31 |
| *T10D4.1* | WBGene00020406 | protein_coding_gene | 6.3 | 77.877 | 2.0E-11 |
| *ZK1290.1* | WBGene00022886 | protein_coding_gene | 6.2 | 74.969 | 1.2E-12 |
| *lipl-2* | WBGene00009773 | protein_coding_gene | 6.2 | 72.425 | 3.9E-45 |
| *clec-158* | WBGene00020067 | protein_coding_gene | 6.2 | 71.598 | 4.3E-16 |
| *M176.10* | WBGene00010946 | protein_coding_gene | 6.2 | 71.438 | 2.0E-16 |
| *Y110A2AL.7* | WBGene00022444 | protein_coding_gene | 6.1 | 70.435 | 2.2E-16 |
| *fipr-20* | WBGene00044449 | protein_coding_gene | 6.1 | 70.085 | 1.2E-19 |
| *Y43F8C.17* | WBGene00012838 | protein_coding_gene | 6.1 | 67.164 | 8.4E-11 |
| *T10B10.9* | WBGene00011685 | protein_coding_gene | 6.0 | 66.234 | 9.5E-11 |
| *mtl-1* | WBGene00003473 | protein_coding_gene | 6.0 | 64.429 | 1.3E-26 |
| *F46B6.13* | WBGene00044192 | protein_coding_gene | 6.0 | 64.373 | 7.3E-12 |
| *R03H10.6* | WBGene00019858 | protein_coding_gene | 6.0 | 63.165 | 2.7E-57 |
| *scl-7* | WBGene00008029 | protein_coding_gene | 6.0 | 62.814 | 1.1E-11 |
| *T26H5.9* | WBGene00044206 | protein_coding_gene | 6.0 | 62.767 | 3.4E-47 |
| *clec-141* | WBGene00017139 | protein_coding_gene | 5.9 | 61.679 | 1.0E-11 |
| *hpo-2* | WBGene00012500 | protein_coding_gene | 5.9 | 60.038 | 7.3E-32 |
| *Y25C1A.2* | WBGene00021289 | protein_coding_gene | 5.9 | 59.343 | 1.9E-11 |
| *ZK1053.3* | WBGene00014198 | protein_coding_gene | 5.9 | 59.049 | 2.9E-46 |
| *ilys-3* | WBGene00016670 | protein_coding_gene | 5.9 | 58.694 | 9.4E-25 |
| *E02H4.4* | WBGene00008462 | protein_coding_gene | 5.8 | 55.944 | 1.1E-51 |
| *F48C1.9* | WBGene00018601 | protein_coding_gene | 5.8 | 55.279 | 1.7E-13 |
| *cest-29* | WBGene00017205 | pseudogene | 5.8 | 55.114 | 5.6E-10 |
| *clec-232* | WBGene00009487 | protein_coding_gene | 5.8 | 53.938 | 3.9E-14 |
| *clec-161* | WBGene00010228 | protein_coding_gene | 5.7 | 53.636 | 5.1E-20 |
| *clec-133* | WBGene00021142 | protein_coding_gene | 5.7 | 53.234 | 7.3E-10 |
| *C49C3.11* | WBGene00008201 | protein_coding_gene | 5.7 | 52.668 | 2.4E-62 |
| *B0250.3* | WBGene00007120 | pseudogene | 5.7 | 52.173 | 6.8E-104 |
| *fipr-28* | WBGene00013120 | protein_coding_gene | 5.7 | 51.238 | 2.1E-22 |
| *F46A8.5* | WBGene00009748 | protein_coding_gene | 5.7 | 50.381 | 1.2E-09 |
| *clec-132* | WBGene00021143 | protein_coding_gene | 5.6 | 49.716 | 6.6E-11 |
| *lipl-3* | WBGene00020016 | protein_coding_gene | 5.6 | 49.010 | 1.4E-17 |
| *scl-9* | WBGene00009890 | protein_coding_gene | 5.6 | 47.603 | 1.7E-13 |
| *fil-1* | WBGene00011321 | protein_coding_gene | 5.5 | 46.597 | 3.1E-39 |
| *F46C5.1* | WBGene00009778 | protein_coding_gene | 5.5 | 46.219 | 4.5E-19 |
| *tts-1* | WBGene00006650 | ncRNA_gene | 5.5 | 46.014 | 1.6E-26 |
| *clec-24* | WBGene00009856 | protein_coding_gene | 5.5 | 45.709 | 1.9E-18 |
| *Y6G8.15* | WBGene00189937 | protein_coding_gene | 5.5 | 45.586 | 4.4E-10 |
| *F49F1.15* | WBGene00235365 | protein_coding_gene | 5.5 | 44.924 | 3.2E-10 |
| *C17E4.19* | WBGene00219310 | protein_coding_gene | 5.5 | 44.778 | 3.3E-32 |
| *F45D3.4* | WBGene00009724 | protein_coding_gene | 5.4 | 43.195 | 5.0E-99 |
| *F07G6.8* | WBGene00017227 | protein_coding_gene | 5.4 | 42.889 | 5.3E-09 |

| | | | | | |
|---|---|---|---|---|---|
| *pqn-84* | WBGene00004165 | protein_coding_gene | 5.4 | 42.343 | 2.7E-42 |
| *Y59H11AM.1* | WBGene00022005 | protein_coding_gene | 5.4 | 42.329 | 1.6E-12 |
| *R53.8* | WBGene00011277 | protein_coding_gene | 5.4 | 42.279 | 7.2E-20 |
| *ZK1290.11* | WBGene00022892 | protein_coding_gene | 5.4 | 41.770 | 1.6E-19 |
| *Y82E9BL.9* | WBGene00023395 | pseudogene | 5.4 | 41.268 | 1.0E-18 |
| *tag-329* | WBGene00008231 | protein_coding_gene | 5.4 | 40.930 | 1.6E-37 |
| *nspc-12* | WBGene00023493 | protein_coding_gene | 5.3 | 40.560 | 5.3E-14 |
| *R09E10.8* | WBGene00011177 | protein_coding_gene | 5.3 | 40.200 | 3.2E-12 |
| *F43C11.2* | WBGene00018379 | protein_coding_gene | 5.3 | 39.272 | 1.4E-14 |
| *F42A6.2* | WBGene00018324 | protein_coding_gene | 5.3 | 39.242 | 7.3E-10 |
| *D1022.2* | WBGene00017021 | protein_coding_gene | 5.3 | 39.237 | 1.2E-08 |
| *clec-138* | WBGene00018049 | protein_coding_gene | 5.3 | 38.891 | 4.1E-12 |
| *F16G10.10* | WBGene00017525 | protein_coding_gene | 5.2 | 37.650 | 2.1E-09 |
| *C45G9.10* | WBGene00016681 | protein_coding_gene | 5.2 | 37.248 | 3.3E-14 |
| *F47C12.12* | WBGene00219377 | pseudogene | 5.2 | 37.076 | 3.1E-09 |
| *scl-2* | WBGene00009895 | protein_coding_gene | 5.2 | 36.375 | 1.2E-103 |
| *fbxa-24* | WBGene00018803 | protein_coding_gene | 5.2 | 35.683 | 3.0E-43 |
| *C18H7.1* | WBGene00015993 | protein_coding_gene | 5.2 | 35.522 | 9.9E-15 |
| *F11A6.16* | WBGene00271780 | protein_coding_gene | 5.1 | 35.397 | 2.1E-09 |
| *Y6G8.14* | WBGene00138715 | protein_coding_gene | 5.1 | 35.388 | 2.9E-08 |
| *clec-125* | WBGene00021118 | protein_coding_gene | 5.1 | 35.338 | 6.1E-51 |
| *T05A7.11* | WBGene00044383 | protein_coding_gene | 5.1 | 35.157 | 3.5E-11 |
| *B0222.10* | WBGene00015058 | ncRNA_gene | 5.1 | 35.104 | 7.3E-28 |
| *T26A8.3* | WBGene00020826 | protein_coding_gene | 5.1 | 34.850 | 2.2E-17 |
| *plg-1* | WBGene00004041 | pseudogene | 5.1 | 34.769 | 3.9E-51 |
| *F47B8.4* | WBGene00009805 | protein_coding_gene | 5.1 | 34.310 | 8.6E-11 |
| *F13E9.10* | WBGene00008759 | protein_coding_gene | 5.1 | 34.109 | 2.0E-13 |
| *C28C12.3* | WBGene00016179 | protein_coding_gene | 5.1 | 34.032 | 1.2E-25 |
| *nas-24* | WBGene00003543 | protein_coding_gene | 5.1 | 33.855 | 1.8E-26 |
| *C35B1.3* | WBGene00016433 | protein_coding_gene | 5.1 | 33.628 | 3.5E-09 |
| *B0513.7* | WBGene00007199 | pseudogene | 5.0 | 33.051 | 4.0E-09 |
| *ins-20* | WBGene00002103 | protein_coding_gene | 5.0 | 32.770 | 6.1E-08 |
| *C47A10.12* | WBGene00008129 | protein_coding_gene | 5.0 | 32.703 | 9.4E-11 |
| *C16C8.7* | WBGene00015845 | protein_coding_gene | 5.0 | 32.556 | 5.2E-13 |
| *T16G12.10* | WBGene00044222 | protein_coding_gene | 5.0 | 31.742 | 2.0E-30 |
| *F22B3.5* | WBGene00009036 | protein_coding_gene | 5.0 | 31.698 | 6.1E-09 |
| *tag-138* | WBGene00006484 | protein_coding_gene | 5.0 | 31.628 | 9.4E-11 |
| *clec-130* | WBGene00021144 | protein_coding_gene | 5.0 | 31.389 | 6.8E-09 |
| *F47C12.8* | WBGene00018551 | protein_coding_gene | 5.0 | 31.263 | 1.0E-10 |
| *H25K10.4* | WBGene00010413 | protein_coding_gene | 4.9 | 30.684 | 2.0E-12 |
| *clec-32* | WBGene00009860 | protein_coding_gene | 4.9 | 30.573 | 8.6E-09 |
| *dct-8* | WBGene00044472 | protein_coding_gene | 4.9 | 30.465 | 2.0E-10 |
| *R05H10.7* | WBGene00011046 | protein_coding_gene | 4.9 | 30.025 | 4.3E-17 |
| *F26C11.4* | WBGene00044159 | protein_coding_gene | 4.9 | 29.900 | 1.8E-10 |
| *W02D7.12* | WBGene00044356 | protein_coding_gene | 4.9 | 29.092 | 1.5E-07 |
| *try-8* | WBGene00017791 | protein_coding_gene | 4.9 | 29.007 | 2.5E-36 |
| *Y40H7A.147* | WBGene00077755 | pseudogene | 4.8 | 28.812 | 1.5E-08 |
| *F01D5.4* | WBGene00014730 | pseudogene | 4.8 | 28.475 | 2.0E-07 |
| *Y110A2AL.6* | WBGene00022443 | protein_coding_gene | 4.8 | 28.010 | 2.0E-08 |
| *F38E1.10* | WBGene00018182 | protein_coding_gene | 4.8 | 27.880 | 2.1E-08 |
| *T14G12.11* | WBGene00219411 | protein_coding_gene | 4.8 | 27.640 | 4.7E-08 |
| *C14F11.7* | WBGene00015782 | protein_coding_gene | 4.8 | 27.543 | 4.2E-22 |
| *B0207.5* | WBGene00015028 | protein_coding_gene | 4.8 | 27.449 | 7.4E-32 |
| *F53A9.8* | WBGene00018731 | protein_coding_gene | 4.8 | 27.394 | 1.6E-52 |
| *C09G12.5* | WBGene00015655 | protein_coding_gene | 4.8 | 27.190 | 7.1E-17 |
| *T26H5.8* | WBGene00012070 | protein_coding_gene | 4.8 | 27.104 | 1.7E-16 |
| *fipr-16* | WBGene00015521 | protein_coding_gene | 4.7 | 26.796 | 7.7E-18 |
| *clec-181* | WBGene00044511 | protein_coding_gene | 4.7 | 26.754 | 3.1E-08 |
| *Y43F8C.15* | WBGene00012836 | protein_coding_gene | 4.7 | 26.675 | 3.2E-08 |
| *F26H11.8* | WBGene00195149 | protein_coding_gene | 4.7 | 26.576 | 2.9E-11 |

| | | | | | |
|---|---|---|---|---|---|
| *F02H6.6* | WBGene00008539 | protein_coding_gene | 4.7 | 26.017 | 4.0E-07 |
| *F46A8.3* | WBGene00009746 | protein_coding_gene | 4.7 | 25.860 | 5.7E-11 |
| *M151.10* | WBGene00303365 | protein_coding_gene | 4.7 | 25.680 | 5.0E-27 |
| *K09C8.2* | WBGene00010715 | protein_coding_gene | 4.7 | 25.670 | 4.6E-07 |
| *Y39A1A.27* | WBGene00194731 | protein_coding_gene | 4.7 | 25.454 | 4.8E-07 |
| *K03B8.11* | WBGene00010526 | protein_coding_gene | 4.7 | 25.194 | 5.3E-07 |
| *clec-157* | WBGene00017810 | protein_coding_gene | 4.6 | 24.948 | 6.2E-30 |
| *E02H9.1* | WBGene00017099 | protein_coding_gene | 4.6 | 24.853 | 2.2E-12 |
| *ftn-1* | WBGene00001500 | protein_coding_gene | 4.6 | 24.799 | 3.5E-21 |
| *F15E6.3* | WBGene00017484 | protein_coding_gene | 4.6 | 24.669 | 2.8E-36 |
| *clec-92* | WBGene00013125 | protein_coding_gene | 4.6 | 24.592 | 1.6E-14 |
| *W02B12.16* | WBGene00195148 | protein_coding_gene | 4.6 | 24.496 | 3.8E-09 |
| *F46A8.13* | WBGene00219950 | ncRNA_gene | 4.6 | 24.360 | 2.1E-07 |
| *srh-254* | WBGene00005460 | pseudogene | 4.6 | 23.879 | 8.1E-07 |
| *Y41D4B.6* | WBGene00021513 | protein_coding_gene | 4.6 | 23.878 | 1.1E-12 |
| *clec-61* | WBGene00014047 | protein_coding_gene | 4.6 | 23.826 | 5.2E-13 |
| *clec-263* | WBGene00009171 | protein_coding_gene | 4.6 | 23.745 | 8.5E-07 |
| *F49H6.5* | WBGene00009908 | protein_coding_gene | 4.6 | 23.736 | 1.4E-15 |
| *linc-1* | WBGene00194659 | lincRNA_gene | 4.5 | 23.277 | 8.0E-25 |
| *T12A2.5* | WBGene00020439 | protein_coding_gene | 4.5 | 23.272 | 1.5E-13 |
| *scl-8* | WBGene00008030 | protein_coding_gene | 4.5 | 23.004 | 2.4E-10 |
| *ptr-22* | WBGene00004236 | protein_coding_gene | 4.5 | 22.995 | 1.3E-44 |
| *C16C8.19* | WBGene00015856 | protein_coding_gene | 4.5 | 22.862 | 1.1E-12 |
| *F17E9.3* | WBGene00017540 | protein_coding_gene | 4.5 | 22.826 | 6.9E-09 |
| *clec-219* | WBGene00020945 | protein_coding_gene | 4.5 | 22.305 | 3.7E-10 |
| *fbxa-58* | WBGene00016225 | pseudogene | 4.5 | 22.251 | 1.6E-08 |
| *F28B1.9* | WBGene00045381 | protein_coding_gene | 4.5 | 22.211 | 1.3E-06 |
| *nspf-3* | WBGene00044366 | protein_coding_gene | 4.5 | 22.196 | 1.2E-11 |
| *Y53F4B.45* | WBGene00044728 | ncRNA_gene | 4.4 | 21.778 | 6.1E-61 |
| *F46A8.8* | WBGene00009751 | protein_coding_gene | 4.4 | 21.564 | 6.1E-07 |
| *Y51A2D.1* | WBGene00013072 | protein_coding_gene | 4.4 | 21.509 | 2.3E-07 |
| *Y49F6B.6* | WBGene00021718 | protein_coding_gene | 4.4 | 21.347 | 8.4E-10 |
| *Y43F8B.19* | WBGene00077688 | protein_coding_gene | 4.4 | 20.939 | 2.9E-07 |
| *C06E2.9* | WBGene00044579 | protein_coding_gene | 4.4 | 20.926 | 6.2E-07 |
| *ZK1225.10* | WBGene00044714 | pseudogene | 4.4 | 20.864 | 3.0E-07 |
| *Y18D10A.2* | WBGene00012475 | protein_coding_gene | 4.4 | 20.779 | 9.4E-10 |
| *F49F1.5* | WBGene00018645 | protein_coding_gene | 4.4 | 20.765 | 2.1E-21 |
| *F25E5.3* | WBGene00017784 | protein_coding_gene | 4.3 | 20.371 | 2.9E-24 |
| *clec-126* | WBGene00021117 | protein_coding_gene | 4.3 | 20.127 | 7.6E-22 |
| *hpo-7* | WBGene00016548 | protein_coding_gene | 4.3 | 20.062 | 4.2E-07 |
| *cwp-1* | WBGene00000859 | protein_coding_gene | 4.3 | 19.967 | 2.8E-06 |
| *F30A10.11* | WBGene00009268 | protein_coding_gene | 4.3 | 19.870 | 7.2E-20 |
| *clec-105* | WBGene00012488 | protein_coding_gene | 4.3 | 19.844 | 4.7E-07 |
| *F56D6.8* | WBGene00044292 | protein_coding_gene | 4.3 | 19.811 | 4.7E-07 |
| *cnc-11* | WBGene00044900 | protein_coding_gene | 4.3 | 19.469 | 5.5E-07 |
| *clec-135* | WBGene00015821 | protein_coding_gene | 4.3 | 19.381 | 4.9E-12 |
| *R13.5* | WBGene00014825 | pseudogene | 4.3 | 19.337 | 5.8E-07 |
| *clec-261* | WBGene00013834 | protein_coding_gene | 4.3 | 19.313 | 4.6E-08 |
| *dsl-6* | WBGene00001108 | protein_coding_gene | 4.3 | 19.195 | 3.5E-09 |
| *F11E6.11* | WBGene00023501 | protein_coding_gene | 4.3 | 19.187 | 1.9E-35 |
| *clec-70* | WBGene00021581 | protein_coding_gene | 4.3 | 19.083 | 1.1E-25 |
| *W02D7.11* | WBGene00020946 | protein_coding_gene | 4.2 | 18.898 | 1.6E-06 |
| *T22C1.12* | WBGene00011921 | protein_coding_gene | 4.2 | 18.854 | 1.9E-06 |
| *F26D2.16* | WBGene00009154 | protein_coding_gene | 4.2 | 18.835 | 1.6E-09 |
| *nlp-39* | WBGene00008295 | protein_coding_gene | 4.2 | 18.666 | 2.4E-08 |
| *F47B8.10* | WBGene00009811 | protein_coding_gene | 4.2 | 18.662 | 7.8E-07 |
| *T12A7.10* | WBGene00194933 | protein_coding_gene | 4.2 | 18.470 | 5.0E-06 |
| *C41D11.10* | WBGene00195180 | protein_coding_gene | 4.2 | 18.465 | 5.0E-06 |
| *msp-79* | WBGene00003466 | protein_coding_gene | 4.2 | 18.312 | 1.4E-07 |
| *cutl-9* | WBGene00010893 | protein_coding_gene | 4.2 | 18.264 | 1.1E-16 |

| | | | | | |
|---|---|---|---|---|---|
| *F13E9.19* | WBGene00194922 | protein_coding_gene | 4.2 | 17.909 | 2.8E-06 |
| *nspa-4* | WBGene00044272 | protein_coding_gene | 4.1 | 17.738 | 6.8E-10 |
| *clec-106* | WBGene00012482 | protein_coding_gene | 4.1 | 17.721 | 1.1E-10 |
| *srr-6* | WBGene00005657 | protein_coding_gene | 4.1 | 17.661 | 2.9E-09 |
| *B0513.90* | WBGene00235301 | protein_coding_gene | 4.1 | 17.356 | 7.8E-06 |
| *F13E9.9* | WBGene00008758 | protein_coding_gene | 4.1 | 17.195 | 1.6E-06 |
| *far-7* | WBGene00001391 | protein_coding_gene | 4.1 | 17.130 | 3.2E-35 |
| *K08C9.6* | WBGene00010653 | protein_coding_gene | 4.1 | 17.035 | 1.7E-06 |
| *T23E7.6* | WBGene00044772 | protein_coding_gene | 4.1 | 16.801 | 7.4E-20 |
| *fbxa-161* | WBGene00015596 | pseudogene | 4.0 | 16.560 | 1.3E-05 |
| *fbxc-20* | WBGene00018352 | protein_coding_gene | 4.0 | 16.520 | 3.4E-14 |
| *K01A6.4* | WBGene00010446 | protein_coding_gene | 4.0 | 16.480 | 2.2E-06 |
| *F56D6.11* | WBGene00044473 | protein_coding_gene | 4.0 | 16.443 | 1.3E-05 |
| *F59A6.11* | WBGene00044664 | protein_coding_gene | 4.0 | 16.434 | 2.3E-06 |
| *srp-5* | WBGene00005646 | pseudogene | 4.0 | 16.285 | 6.4E-06 |
| *ZC328.5* | WBGene00022594 | protein_coding_gene | 4.0 | 16.165 | 8.2E-16 |
| *cpt-3* | WBGene00021703 | protein_coding_gene | 4.0 | 16.151 | 3.0E-17 |
| *Y43F8C.24* | WBGene00045345 | pseudogene | 4.0 | 16.142 | 2.6E-06 |
| *F40G9.7* | WBGene00018243 | protein_coding_gene | 4.0 | 16.020 | 1.4E-05 |
| *K12H6.5* | WBGene00019685 | protein_coding_gene | 4.0 | 15.883 | 1.5E-05 |
| *dct-7* | WBGene00017488 | protein_coding_gene | 4.0 | 15.826 | 6.7E-06 |
| *C17E4.12* | WBGene00044769 | pseudogene | 4.0 | 15.700 | 5.6E-09 |
| *fbxa-163* | WBGene00015598 | protein_coding_gene | 4.0 | 15.636 | 1.6E-10 |
| *K10G6.4* | WBGene00019641 | protein_coding_gene | 4.0 | 15.579 | 9.9E-07 |
| *nspc-17* | WBGene00008359 | protein_coding_gene | 4.0 | 15.470 | 3.3E-50 |
| *E02H9.9* | WBGene00017106 | protein_coding_gene | 3.9 | 15.378 | 1.5E-18 |
| *K04H8.3* | WBGene00010578 | protein_coding_gene | 3.9 | 15.327 | 1.3E-11 |
| *T26H5.14* | WBGene00235133 | protein_coding_gene | 3.9 | 15.327 | 1.0E-06 |
| *Y71A12B.19* | WBGene00077443 | pseudogene | 3.9 | 15.269 | 5.8E-22 |
| *T26H5.4* | WBGene00012069 | protein_coding_gene | 3.9 | 15.202 | 4.0E-06 |
| *Y17G9A.4* | WBGene00021197 | protein_coding_gene | 3.9 | 15.114 | 1.9E-24 |
| *fipr-23* | WBGene00007990 | protein_coding_gene | 3.9 | 15.078 | 3.8E-06 |
| *F47C12.6* | WBGene00018549 | protein_coding_gene | 3.9 | 14.951 | 2.2E-05 |
| *flp-23* | WBGene00017693 | protein_coding_gene | 3.9 | 14.863 | 5.2E-06 |
| *T05E12.3* | WBGene00011486 | protein_coding_gene | 3.9 | 14.821 | 3.6E-75 |
| *F37A8.5* | WBGene00009503 | protein_coding_gene | 3.9 | 14.690 | 5.6E-45 |
| *C35A5.11* | WBGene00044024 | protein_coding_gene | 3.9 | 14.659 | 3.1E-09 |
| *msp-71* | WBGene00003460 | pseudogene | 3.9 | 14.643 | 7.1E-27 |
| *clec-108* | WBGene00012504 | protein_coding_gene | 3.9 | 14.523 | 1.9E-05 |
| *Y20C6A.4* | WBGene00077495 | protein_coding_gene | 3.9 | 14.502 | 3.4E-08 |
| *plp-2* | WBGene00008976 | protein_coding_gene | 3.9 | 14.464 | 1.8E-20 |
| *M01G12.10* | WBGene00010823 | protein_coding_gene | 3.9 | 14.433 | 6.5E-06 |
| *F56D2.8* | WBGene00018968 | protein_coding_gene | 3.8 | 14.384 | 2.0E-07 |
| *nspd-8* | WBGene00021280 | protein_coding_gene | 3.8 | 14.369 | 6.8E-06 |
| *cutl-28* | WBGene00018256 | protein_coding_gene | 3.8 | 14.339 | 8.2E-14 |
| *C06E7.88* | WBGene00189995 | protein_coding_gene | 3.8 | 14.274 | 6.0E-08 |
| *F58F9.8* | WBGene00044306 | protein_coding_gene | 3.8 | 14.096 | 3.2E-07 |
| *F28C6.10* | WBGene00009209 | protein_coding_gene | 3.8 | 14.085 | 1.4E-22 |
| *T03G6.1* | WBGene00020194 | protein_coding_gene | 3.8 | 13.996 | 6.1E-13 |
| *3R5.2* | WBGene00007066 | pseudogene | 3.8 | 13.968 | 8.4E-06 |
| *K12H6.9* | WBGene00019689 | protein_coding_gene | 3.8 | 13.786 | 4.0E-07 |
| *egl-46* | WBGene00001210 | protein_coding_gene | 3.8 | 13.752 | 4.7E-17 |
| *nhr-127* | WBGene00003717 | protein_coding_gene | 3.8 | 13.726 | 3.7E-06 |
| *C16C8.9* | WBGene00015847 | protein_coding_gene | 3.8 | 13.720 | 5.3E-07 |
| *C44H9.7* | WBGene00008102 | protein_coding_gene | 3.8 | 13.535 | 7.9E-10 |
| *wrt-2* | WBGene00006948 | protein_coding_gene | 3.7 | 13.389 | 1.9E-11 |
| *clec-137* | WBGene00018048 | protein_coding_gene | 3.7 | 13.367 | 5.3E-07 |
| *Y49F6B.13* | WBGene00044377 | protein_coding_gene | 3.7 | 13.362 | 3.1E-05 |
| *ugt-18* | WBGene00013900 | protein_coding_gene | 3.7 | 13.356 | 6.8E-07 |
| *F56C3.8* | WBGene00018945 | protein_coding_gene | 3.7 | 13.346 | 3.8E-13 |

| | | | | | |
|---|---|---|---|---|---|
| *Y46D2A.3* | WBGene00021591 | protein_coding_gene | 3.7 | 13.336 | 3.2E-05 |
| *swt-1* | WBGene00019300 | protein_coding_gene | 3.7 | 13.326 | 3.6E-27 |
| *F35C5.3* | WBGene00009391 | protein_coding_gene | 3.7 | 13.321 | 3.1E-05 |
| *cutl-2* | WBGene00013145 | protein_coding_gene | 3.7 | 13.240 | 9.0E-20 |
| *dpy-14* | WBGene00001075 | protein_coding_gene | 3.7 | 13.195 | 6.2E-10 |
| *F16B4.7* | WBGene00017509 | protein_coding_gene | 3.7 | 13.085 | 6.7E-06 |
| *C35E7.7* | WBGene00016459 | protein_coding_gene | 3.7 | 13.040 | 5.0E-10 |
| *dod-3* | WBGene00016052 | protein_coding_gene | 3.7 | 13.016 | 3.5E-09 |
| *F59A1.6* | WBGene00010292 | protein_coding_gene | 3.7 | 12.959 | 1.5E-05 |
| *C49G7.13* | WBGene00269425 | protein_coding_gene | 3.7 | 12.900 | 1.7E-06 |
| *ZC443.2* | WBGene00013897 | protein_coding_gene | 3.7 | 12.766 | 7.8E-06 |
| *clec-221* | WBGene00008916 | protein_coding_gene | 3.7 | 12.764 | 2.1E-07 |
| *dpy-7* | WBGene00001069 | protein_coding_gene | 3.7 | 12.759 | 8.3E-20 |
| *H42K12.3* | WBGene00019272 | protein_coding_gene | 3.7 | 12.708 | 5.1E-18 |
| *F58A4.1* | WBGene00010226 | protein_coding_gene | 3.7 | 12.705 | 5.5E-16 |
| *D2096.5* | WBGene00017072 | protein_coding_gene | 3.7 | 12.694 | 9.5E-11 |
| *C25H3.10* | WBGene00016119 | protein_coding_gene | 3.7 | 12.614 | 2.5E-24 |
| *M142.3* | WBGene00014809 | pseudogene | 3.6 | 12.508 | 8.8E-10 |
| *F54E2.2* | WBGene00018824 | protein_coding_gene | 3.6 | 12.494 | 7.0E-05 |
| *C36C5.14* | WBGene00016483 | protein_coding_gene | 3.6 | 12.367 | 1.6E-06 |
| *W07G1.7* | WBGene00012334 | protein_coding_gene | 3.6 | 12.340 | 2.2E-05 |
| *fbxa-48* | WBGene00021862 | protein_coding_gene | 3.6 | 12.311 | 4.5E-08 |
| *T05A7.13* | WBGene00271415 | protein_coding_gene | 3.6 | 12.236 | 3.9E-08 |
| *scl-26* | WBGene00016821 | protein_coding_gene | 3.6 | 12.118 | 6.4E-05 |
| *T20B3.1* | WBGene00011850 | protein_coding_gene | 3.6 | 12.111 | 1.3E-26 |
| *math-38* | WBGene00020326 | protein_coding_gene | 3.6 | 12.070 | 7.3E-26 |
| *B0462.1* | WBGene00007181 | pseudogene | 3.6 | 11.855 | 4.5E-06 |
| *C01G5.7* | WBGene00015309 | protein_coding_gene | 3.6 | 11.842 | 1.2E-07 |
| *arrd-11* | WBGene00011052 | protein_coding_gene | 3.6 | 11.839 | 1.6E-07 |
| *fip-3* | WBGene00015520 | protein_coding_gene | 3.6 | 11.776 | 8.0E-05 |
| *lpr-5* | WBGene00012256 | protein_coding_gene | 3.6 | 11.775 | 3.3E-18 |
| *lpr-4* | WBGene00012257 | protein_coding_gene | 3.6 | 11.769 | 1.8E-13 |
| *Y54G2A.37* | WBGene00023484 | protein_coding_gene | 3.6 | 11.757 | 1.1E-04 |
| *ces-2* | WBGene00000469 | protein_coding_gene | 3.6 | 11.747 | 5.0E-22 |
| *F25E5.7* | WBGene00017788 | protein_coding_gene | 3.6 | 11.738 | 1.3E-06 |
| *F45D11.1* | WBGene00018448 | pseudogene | 3.6 | 11.731 | 1.6E-07 |
| *cyp-34A9* | WBGene00015044 | protein_coding_gene | 3.5 | 11.711 | 1.2E-127 |
| *Y37E11AL.4* | WBGene00021360 | pseudogene | 3.5 | 11.708 | 1.1E-13 |
| *Y6E2A.4* | WBGene00012398 | protein_coding_gene | 3.5 | 11.684 | 2.0E-13 |
| *F43D9.1* | WBGene00009653 | protein_coding_gene | 3.5 | 11.681 | 5.5E-16 |
| *F16G10.15* | WBGene00017529 | pseudogene | 3.5 | 11.550 | 5.6E-15 |
| *T12D8.5* | WBGene00011733 | protein_coding_gene | 3.5 | 11.549 | 2.3E-36 |
| *clc-4* | WBGene00000525 | protein_coding_gene | 3.5 | 11.545 | 1.4E-10 |
| *T02H6.9* | WBGene00020179 | protein_coding_gene | 3.5 | 11.516 | 5.6E-10 |
| *sodh-1* | WBGene00010790 | protein_coding_gene | 3.5 | 11.469 | 1.8E-08 |
| *hch-1* | WBGene00001828 | protein_coding_gene | 3.5 | 11.465 | 1.0E-16 |
| *oac-14* | WBGene00008602 | protein_coding_gene | 3.5 | 11.453 | 2.1E-18 |
| *fbxa-100* | WBGene00008826 | protein_coding_gene | 3.5 | 11.449 | 3.9E-05 |
| *dpy-2* | WBGene00001064 | protein_coding_gene | 3.5 | 11.434 | 9.9E-17 |
| *glf-1* | WBGene00019154 | protein_coding_gene | 3.5 | 11.429 | 2.1E-12 |
| *cnc-4* | WBGene00000558 | protein_coding_gene | 3.5 | 11.398 | 9.8E-13 |
| *F59C6.16* | WBGene00077764 | protein_coding_gene | 3.5 | 11.295 | 6.4E-33 |
| *mlt-11* | WBGene00012186 | protein_coding_gene | 3.5 | 11.214 | 1.5E-31 |
| *ZC395.5* | WBGene00022597 | protein_coding_gene | 3.5 | 11.177 | 1.4E-23 |
| *F55H12.7* | WBGene00219283 | protein_coding_gene | 3.5 | 11.157 | 4.6E-05 |
| *pals-32* | WBGene00016281 | protein_coding_gene | 3.5 | 11.156 | 4.2E-11 |
| *T19H5.6* | WBGene00044807 | protein_coding_gene | 3.5 | 11.144 | 2.1E-10 |
| *Y73F8A.10* | WBGene00013519 | protein_coding_gene | 3.5 | 11.128 | 1.6E-04 |
| *F17B5.1* | WBGene00008905 | protein_coding_gene | 3.5 | 11.106 | 1.4E-06 |
| *dpy-3* | WBGene00001065 | protein_coding_gene | 3.5 | 11.070 | 2.0E-15 |

| | | | | | |
|---|---|---|---|---|---|
| *bro-1* | WBGene00000272 | protein_coding_gene | 3.5 | 11.010 | 2.4E-08 |
| *Y65B4BL.1* | WBGene00022033 | protein_coding_gene | 3.5 | 10.958 | 1.1E-04 |
| *C03C10.5* | WBGene00007271 | protein_coding_gene | 3.5 | 10.938 | 9.2E-09 |
| *dpy-10* | WBGene00001072 | protein_coding_gene | 3.4 | 10.877 | 4.8E-24 |
| *Y102A5C.4* | WBGene00013613 | protein_coding_gene | 3.4 | 10.868 | 8.6E-10 |
| *dao-2* | WBGene00000928 | protein_coding_gene | 3.4 | 10.846 | 2.6E-25 |
| *F41C3.2* | WBGene00018268 | protein_coding_gene | 3.4 | 10.828 | 2.4E-16 |
| *ZC443.3* | WBGene00013898 | protein_coding_gene | 3.4 | 10.782 | 1.7E-61 |
| *Y75B12A.1* | WBGene00014949 | pseudogene | 3.4 | 10.749 | 1.8E-04 |
| *Y64G10A.2* | WBGene00013412 | protein_coding_gene | 3.4 | 10.719 | 6.3E-05 |
| *clec-136* | WBGene00018047 | protein_coding_gene | 3.4 | 10.672 | 1.4E-09 |
| *dod-22* | WBGene00010125 | protein_coding_gene | 3.4 | 10.627 | 4.2E-20 |
| *Y41D4B.26* | WBGene00021525 | protein_coding_gene | 3.4 | 10.606 | 4.8E-16 |
| *W09G12.6* | WBGene00021120 | protein_coding_gene | 3.4 | 10.595 | 6.9E-05 |
| *W09G12.7* | WBGene00021121 | protein_coding_gene | 3.4 | 10.584 | 2.2E-18 |
| *lpr-3* | WBGene00012261 | protein_coding_gene | 3.4 | 10.583 | 2.0E-21 |
| *K08B12.1* | WBGene00019520 | protein_coding_gene | 3.4 | 10.573 | 8.7E-11 |
| *lpr-6* | WBGene00012255 | protein_coding_gene | 3.4 | 10.514 | 2.9E-12 |
| *F46A8.4* | WBGene00009747 | protein_coding_gene | 3.4 | 10.473 | 2.2E-04 |
| *K03B8.14* | WBGene00194783 | protein_coding_gene | 3.4 | 10.417 | 3.8E-09 |
| *clec-60* | WBGene00014046 | protein_coding_gene | 3.4 | 10.395 | 4.0E-10 |
| *fipr-24* | WBGene00007992 | protein_coding_gene | 3.4 | 10.359 | 7.8E-05 |
| *ZC178.2* | WBGene00022536 | protein_coding_gene | 3.4 | 10.320 | 8.3E-05 |
| *Y116A8C.44* | WBGene00013815 | protein_coding_gene | 3.4 | 10.275 | 2.8E-07 |
| *Y54G2A.76* | WBGene00235268 | protein_coding_gene | 3.4 | 10.214 | 2.3E-06 |
| *fbxa-30* | WBGene00013837 | protein_coding_gene | 3.4 | 10.214 | 1.0E-08 |
| *abf-2* | WBGene00000013 | protein_coding_gene | 3.3 | 10.137 | 1.0E-08 |
| *Y71A12B.24* | WBGene00195228 | protein_coding_gene | 3.3 | 10.131 | 9.3E-05 |
| *Y57E12AL.4* | WBGene00021958 | protein_coding_gene | 3.3 | 10.122 | 9.2E-05 |
| *noah-2* | WBGene00009926 | protein_coding_gene | 3.3 | 10.099 | 3.7E-32 |
| *F15E6.10* | WBGene00044479 | protein_coding_gene | 3.3 | 10.076 | 3.2E-04 |
| *cky-1* | WBGene00000521 | protein_coding_gene | 3.3 | 10.074 | 8.0E-05 |
| *dpy-17* | WBGene00001076 | protein_coding_gene | 3.3 | 10.064 | 5.4E-09 |
| *F09F7.6* | WBGene00017303 | protein_coding_gene | 3.3 | 10.064 | 5.3E-31 |
| *F31D4.5* | WBGene00009292 | protein_coding_gene | 3.3 | 10.031 | 9.1E-10 |
| *C14F11.6* | WBGene00015781 | protein_coding_gene | 3.3 | 10.030 | 2.4E-13 |
| *C06E4.8* | WBGene00015537 | protein_coding_gene | 3.3 | 10.023 | 1.2E-11 |
| *F17E9.19* | WBGene00249825 | pseudogene | 3.3 | 9.954 | 3.5E-04 |
| *Y82E9BR.8* | WBGene00022341 | protein_coding_gene | 3.3 | 9.940 | 1.1E-04 |
| *pkd-2* | WBGene00004035 | protein_coding_gene | 3.3 | 9.934 | 2.2E-04 |
| *F43G6.17* | WBGene00219378 | protein_coding_gene | 3.3 | 9.930 | 2.7E-04 |
| *oac-4* | WBGene00015873 | protein_coding_gene | 3.3 | 9.917 | 2.2E-05 |
| *K08B4.2* | WBGene00019514 | protein_coding_gene | 3.3 | 9.766 | 5.8E-10 |
| *chil-28* | WBGene00008842 | pseudogene | 3.3 | 9.713 | 1.3E-04 |
| *lim-4* | WBGene00002987 | protein_coding_gene | 3.3 | 9.708 | 1.0E-04 |
| *T04C12.1* | WBGene00011427 | protein_coding_gene | 3.3 | 9.647 | 1.1E-05 |
| *T26H5.10* | WBGene00044783 | protein_coding_gene | 3.3 | 9.622 | 3.8E-04 |
| *Y7A5A.6* | WBGene00012412 | protein_coding_gene | 3.3 | 9.571 | 1.1E-04 |
| *fbxa-136* | WBGene00007672 | protein_coding_gene | 3.3 | 9.571 | 3.8E-04 |
| *ZK105.13* | WBGene00235114 | protein_coding_gene | 3.3 | 9.563 | 3.3E-04 |
| *ugt-57* | WBGene00017154 | protein_coding_gene | 3.3 | 9.538 | 7.1E-12 |
| *ZK970.7* | WBGene00014173 | protein_coding_gene | 3.2 | 9.479 | 3.4E-10 |
| *F21D9.5* | WBGene00009019 | pseudogene | 3.2 | 9.463 | 7.0E-16 |
| *F53A9.7* | WBGene00018730 | protein_coding_gene | 3.2 | 9.462 | 3.2E-25 |
| *F22E5.7* | WBGene00017706 | protein_coding_gene | 3.2 | 9.442 | 4.5E-04 |
| *K10D3.4* | WBGene00010738 | protein_coding_gene | 3.2 | 9.434 | 8.6E-22 |
| *coel-1* | WBGene00016866 | protein_coding_gene | 3.2 | 9.423 | 3.7E-52 |
| *F56D2.3* | WBGene00018965 | protein_coding_gene | 3.2 | 9.387 | 3.5E-10 |
| *atf-2* | WBGene00000220 | protein_coding_gene | 3.2 | 9.340 | 4.1E-16 |
| *R03H10.2* | WBGene00019855 | protein_coding_gene | 3.2 | 9.316 | 1.2E-15 |

| | | | | | |
|---|---|---|---|---|---|
| *unc-42* | WBGene00006778 | protein_coding_gene | 3.2 | 9.305 | 2.0E-06 |
| *R05G6.3* | WBGene00043466 | protein_coding_gene | 3.2 | 9.293 | 1.7E-04 |
| *T02E9.5* | WBGene00011383 | protein_coding_gene | 3.2 | 9.282 | 6.8E-28 |
| *nspf-1* | WBGene00044364 | protein_coding_gene | 3.2 | 9.240 | 1.8E-04 |
| *W04E12.9* | WBGene00012254 | protein_coding_gene | 3.2 | 9.236 | 3.6E-04 |
| *Y74C10AR.2* | WBGene00022282 | protein_coding_gene | 3.2 | 9.222 | 1.2E-05 |
| *F11C7.6* | WBGene00017378 | protein_coding_gene | 3.2 | 9.210 | 5.0E-06 |
| *Y49F6B.20* | WBGene00303369 | protein_coding_gene | 3.2 | 9.203 | 1.4E-05 |
| *fbxa-21* | WBGene00022487 | protein_coding_gene | 3.2 | 9.175 | 7.1E-18 |
| *dsl-3* | WBGene00001105 | protein_coding_gene | 3.2 | 9.113 | 5.0E-17 |
| *try-5* | WBGene00006623 | protein_coding_gene | 3.2 | 9.108 | 8.4E-09 |
| *hhat-1* | WBGene00013852 | protein_coding_gene | 3.2 | 9.102 | 3.8E-11 |
| *F13E9.11* | WBGene00008760 | protein_coding_gene | 3.2 | 9.093 | 2.3E-22 |
| *spp-2* | WBGene00004987 | protein_coding_gene | 3.2 | 9.093 | 4.6E-41 |
| *atf-8* | WBGene00017535 | protein_coding_gene | 3.2 | 9.084 | 5.3E-08 |
| *F13B6.1* | WBGene00017416 | protein_coding_gene | 3.2 | 9.065 | 1.2E-08 |
| *wrt-10* | WBGene00006956 | protein_coding_gene | 3.2 | 9.052 | 8.8E-27 |
| *fbxa-103* | WBGene00008612 | protein_coding_gene | 3.2 | 9.015 | 3.4E-14 |
| *M03B6.3* | WBGene00010835 | protein_coding_gene | 3.2 | 9.004 | 2.7E-10 |
| *hil-1* | WBGene00001852 | protein_coding_gene | 3.2 | 8.994 | 1.2E-20 |
| *Y37F4.3* | WBGene00021384 | protein_coding_gene | 3.2 | 8.953 | 2.1E-04 |
| *C45E5.3* | WBGene00016666 | pseudogene | 3.2 | 8.941 | 5.7E-09 |
| *hlh-16* | WBGene00001960 | protein_coding_gene | 3.2 | 8.924 | 1.6E-08 |
| *F49B2.4* | WBGene00009864 | pseudogene | 3.2 | 8.914 | 6.7E-04 |
| *Y18H1A.8* | WBGene00021212 | protein_coding_gene | 3.2 | 8.911 | 9.2E-08 |
| *T10C6.15* | WBGene00050899 | protein_coding_gene | 3.1 | 8.867 | 4.2E-08 |
| *C17E4.20* | WBGene00219313 | protein_coding_gene | 3.1 | 8.835 | 1.8E-114 |
| *ptr-10* | WBGene00004224 | protein_coding_gene | 3.1 | 8.797 | 4.6E-08 |
| *ZK218.8* | WBGene00013942 | pseudogene | 3.1 | 8.772 | 2.5E-04 |
| *T02E9.6* | WBGene00044203 | protein_coding_gene | 3.1 | 8.767 | 2.5E-04 |
| *ins-2* | WBGene00002085 | protein_coding_gene | 3.1 | 8.738 | 7.1E-08 |
| *peb-1* | WBGene00003968 | protein_coding_gene | 3.1 | 8.714 | 2.0E-18 |
| *C25A6.1* | WBGene00016081 | protein_coding_gene | 3.1 | 8.692 | 1.1E-07 |
| *Y43F8C.16* | WBGene00012837 | protein_coding_gene | 3.1 | 8.677 | 8.0E-04 |
| *Y37A1B.7* | WBGene00012540 | protein_coding_gene | 3.1 | 8.668 | 1.3E-06 |
| *C36C5.15* | WBGene00016484 | protein_coding_gene | 3.1 | 8.668 | 7.7E-04 |
| *nep-25* | WBGene00022874 | protein_coding_gene | 3.1 | 8.649 | 1.1E-11 |
| *K12H6.4* | WBGene00019684 | protein_coding_gene | 3.1 | 8.645 | 3.1E-05 |
| *F48F7.3* | WBGene00009849 | protein_coding_gene | 3.1 | 8.606 | 6.7E-04 |
| *Y65B4BM.3* | WBGene00302974 | protein_coding_gene | 3.1 | 8.576 | 1.5E-14 |
| *thn-1* | WBGene00009213 | protein_coding_gene | 3.1 | 8.549 | 3.5E-05 |
| *ces-1* | WBGene00000468 | protein_coding_gene | 3.1 | 8.548 | 7.0E-06 |
| *clec-25* | WBGene00044331 | protein_coding_gene | 3.1 | 8.545 | 4.1E-06 |
| *T21D12.7* | WBGene00020648 | protein_coding_gene | 3.1 | 8.494 | 6.0E-04 |
| *nnt-1* | WBGene00003778 | protein_coding_gene | 3.1 | 8.487 | 3.3E-33 |
| *ptr-4* | WBGene00004219 | protein_coding_gene | 3.1 | 8.471 | 8.4E-13 |
| *noah-1* | WBGene00016422 | protein_coding_gene | 3.1 | 8.440 | 1.1E-33 |
| *fipr-19* | WBGene00044451 | protein_coding_gene | 3.1 | 8.434 | 9.3E-04 |
| *dod-24* | WBGene00007875 | protein_coding_gene | 3.1 | 8.422 | 6.0E-07 |
| *C08E8.3* | WBGene00007439 | protein_coding_gene | 3.1 | 8.346 | 7.1E-05 |
| *clec-99* | WBGene00013933 | protein_coding_gene | 3.1 | 8.315 | 7.6E-05 |
| *T11F9.22* | WBGene00194784 | protein_coding_gene | 3.1 | 8.302 | 9.9E-04 |
| *K06A9.2* | WBGene00019436 | protein_coding_gene | 3.1 | 8.294 | 4.6E-24 |
| *ZC204.6* | WBGene00022556 | protein_coding_gene | 3.1 | 8.291 | 3.6E-04 |
| *oac-29* | WBGene00018295 | protein_coding_gene | 3.0 | 8.234 | 4.0E-04 |
| *ttr-29* | WBGene00011286 | protein_coding_gene | 3.0 | 8.202 | 3.8E-10 |
| *R09E10.13* | WBGene00194658 | protein_coding_gene | 3.0 | 8.199 | 3.3E-04 |
| *skr-11* | WBGene00004817 | pseudogene | 3.0 | 8.196 | 1.1E-36 |
| *clec-162* | WBGene00007399 | protein_coding_gene | 3.0 | 8.172 | 3.6E-04 |
| *cup-16* | WBGene00007919 | protein_coding_gene | 3.0 | 8.170 | 1.5E-21 |

| | | | | | |
|---|---|---|---|---|---|
| *clec-78* | WBGene00018547 | protein_coding_gene | 3.0 | 8.166 | 6.0E-06 |
| *clec-23* | WBGene00011851 | pseudogene | 3.0 | 8.123 | 1.2E-04 |
| *btb-15* | WBGene00015594 | pseudogene | 3.0 | 8.040 | 4.3E-04 |
| *Y116F11A.3* | WBGene00013817 | protein_coding_gene | 3.0 | 8.011 | 4.6E-04 |
| *C08E3.14* | WBGene00044415 | protein_coding_gene | 3.0 | 8.010 | 4.6E-04 |
| *ins-8* | WBGene00002091 | protein_coding_gene | 3.0 | 7.964 | 1.1E-03 |
| *C16C8.8* | WBGene00015846 | protein_coding_gene | 3.0 | 7.956 | 3.6E-04 |
| *T16H12.1* | WBGene00011811 | protein_coding_gene | 3.0 | 7.949 | 4.8E-57 |
| *ugt-54* | WBGene00012013 | protein_coding_gene | 3.0 | 7.949 | 1.1E-36 |
| *2RSSE.3* | WBGene00044951 | pseudogene | 3.0 | 7.932 | 4.8E-04 |
| *Y105C5B.7* | WBGene00013648 | pseudogene | 3.0 | 7.884 | 4.7E-04 |
| *clec-17* | WBGene00008477 | protein_coding_gene | 3.0 | 7.875 | 7.2E-04 |
| *W02A2.9* | WBGene00044280 | protein_coding_gene | 3.0 | 7.859 | 1.7E-18 |
| *Y59E9AR.1* | WBGene00021997 | protein_coding_gene | 3.0 | 7.839 | 5.0E-04 |
| *Y39A3A.4* | WBGene00021435 | protein_coding_gene | 3.0 | 7.836 | 3.9E-07 |
| *dct-10* | WBGene00012618 | pseudogene | 3.0 | 7.818 | 2.7E-04 |
| *F17E9.18* | WBGene00235282 | protein_coding_gene | 3.0 | 7.806 | 5.2E-04 |
| *fbxa-157* | WBGene00007393 | protein_coding_gene | 3.0 | 7.797 | 4.0E-06 |
| *Y66H1A.8* | WBGene00235263 | protein_coding_gene | 3.0 | 7.785 | 1.5E-05 |
| *hbl-1* | WBGene00001824 | protein_coding_gene | 3.0 | 7.759 | 1.4E-15 |
| *btb-21* | WBGene00018540 | pseudogene | 3.0 | 7.754 | 1.6E-12 |
| *W06G6.11* | WBGene00012313 | protein_coding_gene | 3.0 | 7.737 | 5.4E-04 |
| *nspf-2* | WBGene00044365 | protein_coding_gene | 2.9 | 7.716 | 1.1E-03 |
| *C09F9.2* | WBGene00007479 | protein_coding_gene | 2.9 | 7.714 | 4.2E-39 |
| *nlp-45* | WBGene00020136 | protein_coding_gene | 2.9 | 7.706 | 7.0E-06 |
| *W01G7.2* | WBGene00014851 | pseudogene | 2.9 | 7.703 | 5.9E-04 |
| *ifa-3* | WBGene00002051 | protein_coding_gene | 2.9 | 7.701 | 5.3E-05 |
| *Y17G7B.23* | WBGene00044030 | protein_coding_gene | 2.9 | 7.689 | 2.8E-04 |
| *R09E10.5* | WBGene00011175 | protein_coding_gene | 2.9 | 7.665 | 1.2E-04 |
| *M01B2.13* | WBGene00045338 | protein_coding_gene | 2.9 | 7.654 | 2.1E-07 |
| *K07A1.6* | WBGene00010613 | protein_coding_gene | 2.9 | 7.631 | 1.5E-82 |
| *ttr-23* | WBGene00011894 | protein_coding_gene | 2.9 | 7.631 | 6.6E-10 |
| *try-9* | WBGene00023425 | protein_coding_gene | 2.9 | 7.631 | 4.4E-08 |
| *tre-5* | WBGene00006611 | protein_coding_gene | 2.9 | 7.610 | 3.5E-21 |
| *lipl-1* | WBGene00010062 | protein_coding_gene | 2.9 | 7.609 | 4.3E-10 |
| *lys-3* | WBGene00003092 | protein_coding_gene | 2.9 | 7.609 | 7.4E-06 |
| *F46F5.1* | WBGene00018493 | protein_coding_gene | 2.9 | 7.594 | 4.3E-05 |
| *T20F5.4* | WBGene00020626 | protein_coding_gene | 2.9 | 7.589 | 1.7E-07 |
| *clec-207* | WBGene00018096 | protein_coding_gene | 2.9 | 7.582 | 7.0E-05 |
| *F26C11.3* | WBGene00009147 | protein_coding_gene | 2.9 | 7.569 | 6.3E-04 |
| *K12H6.8* | WBGene00019688 | protein_coding_gene | 2.9 | 7.567 | 5.5E-05 |
| *cyp-34A10* | WBGene00015045 | protein_coding_gene | 2.9 | 7.559 | 7.7E-12 |
| *M01D1.12* | WBGene00269373 | protein_coding_gene | 2.9 | 7.553 | 3.3E-06 |
| *F57B1.6* | WBGene00010189 | protein_coding_gene | 2.9 | 7.537 | 7.2E-08 |
| *F32B5.3* | WBGene00017977 | protein_coding_gene | 2.9 | 7.536 | 7.1E-05 |
| *clec-71* | WBGene00021582 | protein_coding_gene | 2.9 | 7.520 | 1.6E-08 |
| *F13H10.5* | WBGene00008776 | protein_coding_gene | 2.9 | 7.501 | 6.3E-07 |
| *ech-9* | WBGene00001158 | protein_coding_gene | 2.9 | 7.454 | 5.4E-05 |
| *Y74C9A.6* | WBGene00023193 | snoRNA_gene | 2.9 | 7.444 | 1.4E-03 |
| *C06B3.6* | WBGene00007365 | protein_coding_gene | 2.9 | 7.441 | 1.6E-23 |
| *fbxa-137* | WBGene00020853 | protein_coding_gene | 2.9 | 7.434 | 1.7E-03 |
| *zmp-5* | WBGene00010423 | protein_coding_gene | 2.9 | 7.403 | 1.7E-03 |
| *clec-217* | WBGene00017824 | protein_coding_gene | 2.9 | 7.399 | 2.6E-04 |
| *F49F1.2* | WBGene00235363 | protein_coding_gene | 2.9 | 7.397 | 6.7E-04 |
| *Y81B9A.5* | WBGene00304906 | ncRNA_gene | 2.9 | 7.392 | 1.7E-03 |
| *tsp-17* | WBGene00006643 | protein_coding_gene | 2.9 | 7.382 | 3.3E-06 |
| *F21C10.11* | WBGene00017660 | protein_coding_gene | 2.9 | 7.368 | 1.7E-06 |
| *M01D1.11* | WBGene00220026 | snoRNA_gene | 2.9 | 7.363 | 2.0E-03 |
| *C18D11.6* | WBGene00007682 | protein_coding_gene | 2.9 | 7.362 | 7.7E-04 |
| *math-43* | WBGene00022432 | pseudogene | 2.9 | 7.355 | 7.8E-04 |

| | | | | | |
|---|---|---|---|---|---|
| *C27A7.8* | WBGene00007758 | protein_coding_gene | 2.9 | 7.353 | 2.0E-17 |
| *srab-25* | WBGene00022807 | protein_coding_gene | 2.9 | 7.340 | 8.0E-04 |
| *prkl-1* | WBGene00022727 | protein_coding_gene | 2.9 | 7.340 | 2.1E-18 |
| *C41G7.15* | WBGene00219284 | protein_coding_gene | 2.9 | 7.329 | 8.0E-04 |
| *zig-10* | WBGene00020800 | protein_coding_gene | 2.9 | 7.326 | 2.9E-08 |
| *Y51H4A.5* | WBGene00013101 | protein_coding_gene | 2.9 | 7.324 | 7.6E-07 |
| *C54F6.5* | WBGene00016923 | protein_coding_gene | 2.9 | 7.304 | 2.3E-08 |
| *T19C4.1* | WBGene00011838 | protein_coding_gene | 2.9 | 7.292 | 9.6E-10 |
| *ZC449.2* | WBGene00022612 | protein_coding_gene | 2.9 | 7.291 | 5.7E-08 |
| *str-163* | WBGene00006208 | protein_coding_gene | 2.9 | 7.262 | 1.4E-04 |
| *C54C8.4* | WBGene00008292 | protein_coding_gene | 2.9 | 7.243 | 1.8E-03 |
| *F56D6.9* | WBGene00044471 | protein_coding_gene | 2.9 | 7.239 | 7.5E-04 |
| *Y51A2D.8* | WBGene00013076 | protein_coding_gene | 2.9 | 7.236 | 2.6E-04 |
| *F55B11.5* | WBGene00010087 | protein_coding_gene | 2.9 | 7.223 | 5.6E-07 |
| *T09A12.8* | WBGene00235275 | protein_coding_gene | 2.9 | 7.215 | 7.8E-04 |
| *C29A12.6* | WBGene00007800 | protein_coding_gene | 2.9 | 7.213 | 7.0E-05 |
| *R09D1.12* | WBGene00011168 | protein_coding_gene | 2.8 | 7.188 | 8.9E-07 |
| *F42G2.7* | WBGene00044362 | protein_coding_gene | 2.8 | 7.170 | 4.4E-04 |
| *sqt-3* | WBGene00005018 | protein_coding_gene | 2.8 | 7.162 | 1.4E-20 |
| *F21D9.11* | WBGene00219452 | protein_coding_gene | 2.8 | 7.157 | 3.9E-09 |
| *M02E1.4* | WBGene00284848 | protein_coding_gene | 2.8 | 7.125 | 2.9E-05 |
| *F25E5.4* | WBGene00017785 | protein_coding_gene | 2.8 | 7.123 | 2.2E-31 |
| *T14B4.19* | WBGene00302993 | protein_coding_gene | 2.8 | 7.088 | 6.6E-09 |
| *tsp-1* | WBGene00006627 | protein_coding_gene | 2.8 | 7.027 | 2.7E-06 |
| *fip-7* | WBGene00013105 | protein_coding_gene | 2.8 | 6.997 | 2.2E-03 |
| *rncs-1* | WBGene00023069 | ncRNA_gene | 2.8 | 6.994 | 1.1E-19 |
| *F17B5.6* | WBGene00044058 | protein_coding_gene | 2.8 | 6.941 | 2.1E-03 |
| *F11E6.9* | WBGene00008712 | protein_coding_gene | 2.8 | 6.929 | 5.5E-04 |
| *C24H10.3* | WBGene00016069 | protein_coding_gene | 2.8 | 6.922 | 2.1E-05 |
| *C04F12.6* | WBGene00007300 | pseudogene | 2.8 | 6.883 | 2.6E-06 |
| *F36F12.1* | WBGene00018092 | protein_coding_gene | 2.8 | 6.881 | 1.3E-03 |
| *ZK381.2* | WBGene00022725 | protein_coding_gene | 2.8 | 6.877 | 2.6E-03 |
| *clec-96* | WBGene00013930 | protein_coding_gene | 2.8 | 6.862 | 3.1E-04 |
| *cpt-4* | WBGene00019644 | protein_coding_gene | 2.8 | 6.814 | 3.0E-07 |
| *fbn-1* | WBGene00022816 | protein_coding_gene | 2.8 | 6.801 | 4.5E-47 |
| *T02D1.7* | WBGene00011373 | protein_coding_gene | 2.8 | 6.800 | 1.2E-03 |
| *F25D7.5* | WBGene00009114 | protein_coding_gene | 2.8 | 6.792 | 2.9E-03 |
| *M01G12.7* | WBGene00010821 | protein_coding_gene | 2.8 | 6.768 | 1.2E-03 |
| *abf-1* | WBGene00000012 | protein_coding_gene | 2.8 | 6.743 | 5.5E-07 |
| *C16C8.10* | WBGene00015848 | protein_coding_gene | 2.8 | 6.738 | 2.6E-03 |
| *EEED8.2* | WBGene00017133 | protein_coding_gene | 2.8 | 6.735 | 5.1E-13 |
| *irg-6* | WBGene00007864 | protein_coding_gene | 2.8 | 6.730 | 2.7E-04 |
| *fbxa-135* | WBGene00011539 | protein_coding_gene | 2.7 | 6.726 | 1.6E-03 |
| *kreg-1* | WBGene00018725 | protein_coding_gene | 2.7 | 6.721 | 3.8E-05 |
| *clec-260* | WBGene00013357 | protein_coding_gene | 2.7 | 6.715 | 5.6E-06 |
| *ZK675.4* | WBGene00014067 | protein_coding_gene | 2.7 | 6.672 | 8.0E-04 |
| *F28A10.5* | WBGene00017873 | protein_coding_gene | 2.7 | 6.647 | 5.8E-05 |
| *best-5* | WBGene00007404 | protein_coding_gene | 2.7 | 6.634 | 1.0E-08 |
| *T22F3.11* | WBGene00020700 | protein_coding_gene | 2.7 | 6.615 | 4.7E-05 |
| *B0310.3* | WBGene00015139 | protein_coding_gene | 2.7 | 6.595 | 7.0E-09 |
| *fbxa-68* | WBGene00020641 | pseudogene | 2.7 | 6.587 | 1.4E-03 |
| *sym-1* | WBGene00006366 | protein_coding_gene | 2.7 | 6.565 | 2.1E-31 |
| *fmi-1* | WBGene00001475 | protein_coding_gene | 2.7 | 6.544 | 1.2E-17 |
| *C17E4.1* | WBGene00007641 | protein_coding_gene | 2.7 | 6.542 | 3.6E-10 |
| *clpr-1* | WBGene00012233 | protein_coding_gene | 2.7 | 6.525 | 3.9E-03 |
| *fbxa-164* | WBGene00015599 | protein_coding_gene | 2.7 | 6.524 | 4.0E-03 |
| *hil-7* | WBGene00001858 | protein_coding_gene | 2.7 | 6.524 | 6.5E-04 |
| *Y46H3A.5* | WBGene00021601 | protein_coding_gene | 2.7 | 6.512 | 2.8E-03 |
| *fbxa-217* | WBGene00012954 | pseudogene | 2.7 | 6.511 | 1.2E-31 |
| *slc-36.5* | WBGene00016628 | protein_coding_gene | 2.7 | 6.500 | 1.1E-15 |

| | | | | | |
|---|---|---|---|---|---|
| *C35C5.8* | WBGene00007957 | protein_coding_gene | 2.7 | 6.497 | 1.6E-17 |
| *clec-42* | WBGene00008891 | protein_coding_gene | 2.7 | 6.486 | 2.3E-03 |
| *bgnt-1.3* | WBGene00009032 | protein_coding_gene | 2.7 | 6.486 | 3.0E-03 |
| *F16H6.9* | WBGene00008899 | protein_coding_gene | 2.7 | 6.444 | 1.6E-06 |
| *cwp-2* | WBGene00000860 | protein_coding_gene | 2.7 | 6.441 | 1.7E-03 |
| *R09D1.9* | WBGene00011165 | pseudogene | 2.7 | 6.428 | 8.5E-12 |
| *K08F9.3* | WBGene00010686 | protein_coding_gene | 2.7 | 6.426 | 8.9E-05 |
| *cnc-2* | WBGene00000556 | protein_coding_gene | 2.7 | 6.413 | 1.5E-06 |
| *K10C2.14* | WBGene00206388 | protein_coding_gene | 2.7 | 6.401 | 1.0E-12 |
| *cut-5* | WBGene00011104 | protein_coding_gene | 2.7 | 6.399 | 9.8E-05 |
| *nphp-1* | WBGene00010898 | protein_coding_gene | 2.7 | 6.397 | 4.7E-06 |
| *clec-129* | WBGene00021145 | protein_coding_gene | 2.7 | 6.392 | 4.0E-03 |
| *T04C12.7* | WBGene00011429 | protein_coding_gene | 2.7 | 6.381 | 1.5E-05 |
| *Y41C4A.21* | WBGene00194733 | protein_coding_gene | 2.7 | 6.367 | 1.3E-03 |
| *pax-1* | WBGene00003937 | protein_coding_gene | 2.7 | 6.325 | 1.9E-03 |
| *T02G5.15* | WBGene00206521 | protein_coding_gene | 2.7 | 6.312 | 5.5E-05 |
| *clec-100* | WBGene00013179 | protein_coding_gene | 2.7 | 6.289 | 2.1E-03 |
| *F36D1.7* | WBGene00009466 | protein_coding_gene | 2.6 | 6.244 | 1.5E-09 |
| *sru-20* | WBGene00005683 | protein_coding_gene | 2.6 | 6.228 | 1.8E-03 |
| *dmsr-12* | WBGene00019262 | protein_coding_gene | 2.6 | 6.218 | 1.6E-06 |
| *ins-19* | WBGene00002102 | protein_coding_gene | 2.6 | 6.216 | 2.7E-03 |
| *gei-13* | WBGene00001570 | protein_coding_gene | 2.6 | 6.211 | 5.7E-12 |
| *nspa-6* | WBGene00044273 | protein_coding_gene | 2.6 | 6.206 | 1.8E-03 |
| *F18C5.9* | WBGene00017562 | protein_coding_gene | 2.6 | 6.191 | 4.5E-03 |
| *col-121* | WBGene00000695 | protein_coding_gene | 2.6 | 6.174 | 9.8E-04 |
| *F55H12.2* | WBGene00010133 | protein_coding_gene | 2.6 | 6.155 | 1.7E-22 |
| *C35B1.8* | WBGene00016437 | protein_coding_gene | 2.6 | 6.149 | 1.5E-05 |
| *F10D2.8* | WBGene00017334 | protein_coding_gene | 2.6 | 6.149 | 3.8E-06 |
| *Y57G11C.1132* | WBGene00194666 | antisense_lncRNA_gene | 2.6 | 6.122 | 2.2E-03 |
| *pitr-4* | WBGene00007141 | protein_coding_gene | 2.6 | 6.115 | 1.0E-03 |
| *let-4* | WBGene00002282 | protein_coding_gene | 2.6 | 6.115 | 3.3E-15 |
| *C10G8.2* | WBGene00015681 | protein_coding_gene | 2.6 | 6.101 | 2.4E-03 |
| *F10D7.4* | WBGene00017341 | protein_coding_gene | 2.6 | 6.089 | 5.1E-03 |
| *C31B8.12* | WBGene00016285 | protein_coding_gene | 2.6 | 6.086 | 6.1E-04 |
| *T04C12.32* | WBGene00235086 | ncRNA_gene | 2.6 | 6.074 | 5.2E-03 |
| *C40A11.3* | WBGene00016546 | protein_coding_gene | 2.6 | 6.062 | 2.2E-08 |
| *T05F1.11* | WBGene00011495 | protein_coding_gene | 2.6 | 6.061 | 3.2E-11 |
| *mboa-1* | WBGene00007174 | protein_coding_gene | 2.6 | 6.060 | 1.7E-06 |
| *F59B2.8* | WBGene00010310 | protein_coding_gene | 2.6 | 6.054 | 1.3E-05 |
| *Y82E9BR.1* | WBGene00022334 | protein_coding_gene | 2.6 | 6.049 | 3.5E-31 |
| *C16C8.17* | WBGene00015854 | protein_coding_gene | 2.6 | 6.047 | 2.6E-03 |
| *ttr-22* | WBGene00011450 | protein_coding_gene | 2.6 | 6.046 | 4.3E-03 |
| *fbxa-162* | WBGene00015597 | pseudogene | 2.6 | 6.039 | 4.9E-06 |
| *F13E9.4* | WBGene00008753 | protein_coding_gene | 2.6 | 6.039 | 2.5E-03 |
| *ZK287.1* | WBGene00013962 | protein_coding_gene | 2.6 | 6.023 | 5.7E-09 |
| *nlp-34* | WBGene00015046 | protein_coding_gene | 2.6 | 6.020 | 2.1E-05 |
| *C17B7.2* | WBGene00015876 | pseudogene | 2.6 | 6.017 | 5.3E-03 |
| *Y54G2A.24* | WBGene00021889 | pseudogene | 2.6 | 6.014 | 7.3E-09 |
| *ugt-52* | WBGene00018931 | protein_coding_gene | 2.6 | 5.996 | 2.5E-11 |
| *tmem-231* | WBGene00020825 | protein_coding_gene | 2.6 | 5.984 | 3.3E-04 |
| *Y7A9D.1* | WBGene00012420 | protein_coding_gene | 2.6 | 5.983 | 1.2E-19 |
| *bus-12* | WBGene00010442 | protein_coding_gene | 2.6 | 5.973 | 1.0E-04 |
| *K11D12.6* | WBGene00019646 | protein_coding_gene | 2.6 | 5.972 | 6.8E-04 |
| *ZC84.6* | WBGene00013849 | protein_coding_gene | 2.6 | 5.957 | 6.9E-05 |
| *B0205.4* | WBGene00015019 | protein_coding_gene | 2.6 | 5.941 | 4.9E-04 |
| *M03D4.4* | WBGene00019751 | protein_coding_gene | 2.6 | 5.937 | 1.8E-10 |
| *rig-5* | WBGene00004372 | protein_coding_gene | 2.6 | 5.928 | 5.9E-08 |
| *ppt-1* | WBGene00004092 | protein_coding_gene | 2.6 | 5.918 | 7.2E-27 |
| *scl-18* | WBGene00011724 | protein_coding_gene | 2.6 | 5.910 | 2.7E-03 |
| *F11D5.5* | WBGene00017382 | protein_coding_gene | 2.6 | 5.908 | 1.6E-07 |

| | | | | | |
|---|---|---|---|---|---|
| *T01D3.3* | WBGene00011328 | protein_coding_gene | 2.6 | 5.896 | 3.4E-08 |
| *F40G9.20* | WBGene00195034 | ncRNA_gene | 2.6 | 5.887 | 3.0E-04 |
| *K01A6.8* | WBGene00194869 | protein_coding_gene | 2.6 | 5.868 | 6.0E-03 |
| *F59A1.16* | WBGene00010301 | protein_coding_gene | 2.6 | 5.866 | 5.1E-03 |
| *C06E1.11* | WBGene00015526 | protein_coding_gene | 2.6 | 5.863 | 1.1E-13 |
| *ceh-16* | WBGene00000439 | protein_coding_gene | 2.5 | 5.847 | 3.5E-06 |
| *ZC123.1* | WBGene00022517 | protein_coding_gene | 2.5 | 5.846 | 1.5E-07 |
| *F10D7.3* | WBGene00017340 | protein_coding_gene | 2.5 | 5.846 | 3.4E-18 |
| *math-47* | WBGene00022688 | pseudogene | 2.5 | 5.840 | 1.2E-03 |
| *ceh-8* | WBGene00000433 | protein_coding_gene | 2.5 | 5.839 | 1.0E-03 |
| *lov-1* | WBGene00003058 | protein_coding_gene | 2.5 | 5.836 | 5.3E-03 |
| *sodh-2* | WBGene00010791 | protein_coding_gene | 2.5 | 5.835 | 1.0E-03 |
| *fbxc-49* | WBGene00020534 | protein_coding_gene | 2.5 | 5.820 | 2.6E-04 |
| *R09A8.9* | WBGene00219278 | protein_coding_gene | 2.5 | 5.818 | 2.3E-06 |
| *C14A11.2* | WBGene00015748 | protein_coding_gene | 2.5 | 5.817 | 8.7E-04 |
| *srt-63* | WBGene00020877 | protein_coding_gene | 2.5 | 5.816 | 4.0E-03 |
| *linc-140* | WBGene00219596 | lincRNA_gene | 2.5 | 5.815 | 6.1E-03 |
| *spp-7* | WBGene00004992 | protein_coding_gene | 2.5 | 5.812 | 6.8E-03 |
| *subs-4* | WBGene00012939 | protein_coding_gene | 2.5 | 5.809 | 1.1E-05 |
| *F58H1.2* | WBGene00010285 | protein_coding_gene | 2.5 | 5.805 | 1.6E-04 |
| *C48B4.3* | WBGene00008168 | protein_coding_gene | 2.5 | 5.792 | 1.3E-05 |
| *lpr-1* | WBGene00002393 | protein_coding_gene | 2.5 | 5.781 | 1.1E-05 |
| *F09F9.4* | WBGene00017309 | protein_coding_gene | 2.5 | 5.777 | 5.8E-07 |
| *C36B1.6* | WBGene00007973 | protein_coding_gene | 2.5 | 5.777 | 4.2E-18 |
| *T26C12.3* | WBGene00020833 | protein_coding_gene | 2.5 | 5.776 | 1.9E-11 |
| *nspe-6* | WBGene00012603 | protein_coding_gene | 2.5 | 5.766 | 5.2E-03 |
| *ubc-8* | WBGene00006705 | protein_coding_gene | 2.5 | 5.761 | 2.2E-49 |
| *Y47D7A.12* | WBGene00021624 | protein_coding_gene | 2.5 | 5.743 | 3.7E-03 |
| *R05H10.1* | WBGene00011042 | protein_coding_gene | 2.5 | 5.733 | 1.9E-13 |
| *F25B3.2* | WBGene00009099 | protein_coding_gene | 2.5 | 5.728 | 8.1E-04 |
| *tyr-5* | WBGene00021553 | protein_coding_gene | 2.5 | 5.726 | 1.9E-10 |
| *ZC449.1* | WBGene00022611 | protein_coding_gene | 2.5 | 5.723 | 7.6E-04 |
| *spp-6* | WBGene00004991 | protein_coding_gene | 2.5 | 5.723 | 2.5E-03 |
| *nspe-7* | WBGene00012593 | protein_coding_gene | 2.5 | 5.716 | 2.7E-04 |
| *C04F12.12* | WBGene00077685 | protein_coding_gene | 2.5 | 5.691 | 6.5E-11 |
| *ZK783.3* | WBGene00022818 | pseudogene | 2.5 | 5.683 | 2.0E-03 |
| *F16C3.2* | WBGene00008886 | protein_coding_gene | 2.5 | 5.681 | 3.8E-26 |
| *ttr-52* | WBGene00008719 | protein_coding_gene | 2.5 | 5.680 | 1.3E-04 |
| *C18D4.8* | WBGene00007677 | pseudogene | 2.5 | 5.668 | 3.0E-17 |
| *fbxb-101* | WBGene00013404 | protein_coding_gene | 2.5 | 5.665 | 3.9E-06 |
| *gbh-2* | WBGene00001523 | protein_coding_gene | 2.5 | 5.662 | 5.3E-17 |
| *cyp-32B1* | WBGene00021167 | protein_coding_gene | 2.5 | 5.656 | 1.5E-08 |
| *wrt-1* | WBGene00006947 | protein_coding_gene | 2.5 | 5.631 | 1.7E-05 |
| *F44E2.4* | WBGene00018418 | protein_coding_gene | 2.5 | 5.630 | 4.3E-11 |
| *cutl-16* | WBGene00019426 | protein_coding_gene | 2.5 | 5.630 | 5.5E-10 |
| *F18G5.6* | WBGene00017582 | protein_coding_gene | 2.5 | 5.615 | 4.2E-56 |
| *R10D12.18* | WBGene00194678 | ncRNA_gene | 2.5 | 5.610 | 2.0E-04 |
| *W03D8.10* | WBGene00020991 | pseudogene | 2.5 | 5.604 | 2.2E-08 |
| *spp-13* | WBGene00004998 | protein_coding_gene | 2.5 | 5.604 | 2.1E-04 |
| *F09E10.6* | WBGene00017296 | protein_coding_gene | 2.5 | 5.596 | 3.6E-04 |
| *F15B9.8* | WBGene00008851 | protein_coding_gene | 2.5 | 5.596 | 4.3E-06 |
| *igcm-2* | WBGene00020130 | protein_coding_gene | 2.5 | 5.595 | 2.7E-06 |
| *T26C5.2* | WBGene00012032 | protein_coding_gene | 2.5 | 5.593 | 6.9E-06 |
| *bath-10* | WBGene00021726 | protein_coding_gene | 2.5 | 5.593 | 1.1E-19 |
| *fbxb-39* | WBGene00019702 | protein_coding_gene | 2.5 | 5.588 | 7.8E-03 |
| *Y47D7A.11* | WBGene00021623 | protein_coding_gene | 2.5 | 5.586 | 7.8E-03 |
| *fbxa-113* | WBGene00013757 | protein_coding_gene | 2.5 | 5.569 | 1.1E-05 |
| *T12A7.6* | WBGene00011727 | protein_coding_gene | 2.5 | 5.563 | 7.3E-17 |
| *Y23H5B.8* | WBGene00021279 | protein_coding_gene | 2.5 | 5.548 | 8.3E-07 |
| *cutl-25* | WBGene00021919 | protein_coding_gene | 2.5 | 5.542 | 5.4E-09 |

| | | | | | |
|---|---|---|---|---|---|
| *Y6G8.16* | WBGene00194801 | protein_coding_gene | 2.5 | 5.536 | 3.0E-03 |
| *aqp-2* | WBGene00000170 | protein_coding_gene | 2.5 | 5.523 | 2.4E-23 |
| *F54C8.6* | WBGene00010039 | protein_coding_gene | 2.5 | 5.512 | 5.9E-08 |
| *Y97E10AL.1* | WBGene00022392 | protein_coding_gene | 2.5 | 5.512 | 7.1E-04 |
| *dmd-5* | WBGene00017326 | protein_coding_gene | 2.5 | 5.508 | 1.0E-06 |
| *math-34* | WBGene00019699 | protein_coding_gene | 2.5 | 5.507 | 5.0E-05 |
| *tsp-2* | WBGene00006628 | protein_coding_gene | 2.5 | 5.502 | 2.7E-06 |
| *nhr-218* | WBGene00011750 | protein_coding_gene | 2.5 | 5.498 | 4.3E-03 |
| *F54D12.9* | WBGene00018822 | protein_coding_gene | 2.5 | 5.496 | 4.9E-04 |
| *F58F12.12* | WBGene00271812 | protein_coding_gene | 2.5 | 5.490 | 1.2E-04 |
| *W03F9.4* | WBGene00021002 | protein_coding_gene | 2.5 | 5.479 | 4.5E-16 |
| *C04A11.5* | WBGene00007288 | protein_coding_gene | 2.5 | 5.471 | 1.5E-05 |
| *C44B12.9* | WBGene00044462 | protein_coding_gene | 2.4 | 5.463 | 7.8E-07 |
| *K01A2.3* | WBGene00019278 | pseudogene | 2.4 | 5.455 | 7.5E-07 |
| *nep-16* | WBGene00018472 | protein_coding_gene | 2.4 | 5.452 | 1.3E-03 |
| *fbxa-172* | WBGene00008014 | protein_coding_gene | 2.4 | 5.448 | 1.3E-04 |
| *C05B5.11* | WBGene00045242 | protein_coding_gene | 2.4 | 5.433 | 1.7E-03 |
| *K08E3.2* | WBGene00010662 | protein_coding_gene | 2.4 | 5.431 | 1.8E-03 |
| *T28H10.3* | WBGene00012144 | protein_coding_gene | 2.4 | 5.424 | 2.0E-29 |
| *dma-1* | WBGene00011345 | protein_coding_gene | 2.4 | 5.400 | 9.1E-14 |
| *sssh-1* | WBGene00044138 | protein_coding_gene | 2.4 | 5.398 | 9.0E-03 |
| *C06A12.8* | WBGene00007359 | protein_coding_gene | 2.4 | 5.393 | 7.8E-03 |
| *fmo-1* | WBGene00001476 | protein_coding_gene | 2.4 | 5.392 | 2.7E-22 |
| *chil-23* | WBGene00011167 | protein_coding_gene | 2.4 | 5.391 | 1.8E-07 |
| *F13A7.11* | WBGene00008726 | protein_coding_gene | 2.4 | 5.383 | 2.7E-07 |
| *C43D7.8* | WBGene00008070 | protein_coding_gene | 2.4 | 5.375 | 4.3E-04 |
| *M7.9* | WBGene00010885 | protein_coding_gene | 2.4 | 5.372 | 1.5E-03 |
| *K08F9.1* | WBGene00010684 | protein_coding_gene | 2.4 | 5.372 | 7.5E-05 |
| *zag-1* | WBGene00006970 | protein_coding_gene | 2.4 | 5.356 | 2.3E-13 |
| *K01A6.7* | WBGene00045096 | protein_coding_gene | 2.4 | 5.354 | 8.7E-03 |
| *F46B3.15* | WBGene00009766 | protein_coding_gene | 2.4 | 5.354 | 4.8E-03 |
| *acn-1* | WBGene00000039 | protein_coding_gene | 2.4 | 5.327 | 1.4E-11 |
| *F58F9.9* | WBGene00044307 | protein_coding_gene | 2.4 | 5.321 | 5.0E-03 |
| *W10G11.4* | WBGene00021137 | protein_coding_gene | 2.4 | 5.317 | 1.5E-03 |
| *F13H6.5* | WBGene00017433 | protein_coding_gene | 2.4 | 5.301 | 5.4E-08 |
| *str-173* | WBGene00006217 | protein_coding_gene | 2.4 | 5.299 | 9.2E-03 |
| *T16G1.4* | WBGene00011798 | protein_coding_gene | 2.4 | 5.296 | 9.7E-09 |
| *tbb-4* | WBGene00006538 | protein_coding_gene | 2.4 | 5.286 | 1.0E-08 |
| *C32F10.4* | WBGene00016332 | protein_coding_gene | 2.4 | 5.285 | 1.4E-15 |
| *Y71F9B.9* | WBGene00022130 | protein_coding_gene | 2.4 | 5.280 | 7.6E-12 |
| *clec-75* | WBGene00021586 | protein_coding_gene | 2.4 | 5.260 | 1.0E-03 |
| *F14B4.1* | WBGene00008779 | protein_coding_gene | 2.4 | 5.260 | 3.5E-13 |
| *Y37F4.8* | WBGene00044289 | protein_coding_gene | 2.4 | 5.251 | 4.1E-05 |
| *clec-242* | WBGene00013471 | protein_coding_gene | 2.4 | 5.249 | 1.0E-02 |
| *E03A3.10* | WBGene00189944 | pseudogene | 2.4 | 5.239 | 5.4E-04 |
| *cki-1* | WBGene00000516 | protein_coding_gene | 2.4 | 5.235 | 6.4E-19 |
| *mlt-8* | WBGene00021095 | protein_coding_gene | 2.4 | 5.225 | 1.2E-12 |
| *nas-17* | WBGene00003536 | protein_coding_gene | 2.4 | 5.225 | 2.7E-03 |
| *K08C9.5* | WBGene00010652 | protein_coding_gene | 2.4 | 5.224 | 1.0E-02 |
| *C33G3.5* | WBGene00007905 | protein_coding_gene | 2.4 | 5.219 | 9.9E-04 |
| *C43D7.11* | WBGene00077664 | pseudogene | 2.4 | 5.217 | 1.0E-05 |
| *F35D2.3* | WBGene00018041 | protein_coding_gene | 2.4 | 5.197 | 4.8E-14 |
| *T26F2.2* | WBGene00012058 | pseudogene | 2.4 | 5.197 | 5.6E-13 |
| *F59E11.7* | WBGene00019113 | protein_coding_gene | 2.4 | 5.193 | 1.1E-06 |
| *asp-11* | WBGene00017677 | pseudogene | 2.4 | 5.177 | 1.8E-03 |
| *F23F12.8* | WBGene00017754 | protein_coding_gene | 2.4 | 5.164 | 1.7E-04 |
| *sknr-1* | WBGene00020961 | protein_coding_gene | 2.4 | 5.161 | 1.1E-02 |
| *K01A2.4* | WBGene00019279 | protein_coding_gene | 2.4 | 5.160 | 1.9E-11 |
| *C01G6.9* | WBGene00007229 | protein_coding_gene | 2.4 | 5.154 | 4.6E-04 |
| *F46B3.14* | WBGene00009765 | protein_coding_gene | 2.4 | 5.142 | 6.0E-03 |

| | | | | | |
|---|---|---|---|---|---|
| *pros-1* | WBGene00000448 | protein_coding_gene | 2.4 | 5.125 | 1.9E-10 |
| *Y54G2A.36* | WBGene00021898 | protein_coding_gene | 2.4 | 5.124 | 2.0E-24 |
| *ZK512.1* | WBGene00013982 | protein_coding_gene | 2.4 | 5.117 | 3.9E-10 |
| *C13C12.2* | WBGene00007552 | protein_coding_gene | 2.4 | 5.115 | 3.2E-05 |
| *trpl-5* | WBGene00004149 | protein_coding_gene | 2.4 | 5.109 | 7.6E-04 |
| *cnc-6* | WBGene00000560 | protein_coding_gene | 2.4 | 5.104 | 6.2E-03 |
| *clec-82* | WBGene00021873 | protein_coding_gene | 2.4 | 5.102 | 4.6E-03 |
| *C23G10.11* | WBGene00016017 | protein_coding_gene | 2.4 | 5.102 | 1.2E-03 |
| *W07G4.1* | WBGene00012335 | protein_coding_gene | 2.3 | 5.097 | 9.7E-03 |
| *Y70C5A.3* | WBGene00050904 | protein_coding_gene | 2.3 | 5.094 | 2.6E-08 |
| *C54F6.18* | WBGene00235093 | protein_coding_gene | 2.3 | 5.078 | 3.7E-03 |
| *F56H9.9* | WBGene00206460 | protein_coding_gene | 2.3 | 5.066 | 1.2E-02 |
| *ast-1* | WBGene00020368 | protein_coding_gene | 2.3 | 5.065 | 2.2E-06 |
| *syd-9* | WBGene00044068 | protein_coding_gene | 2.3 | 5.061 | 2.3E-17 |
| *pqn-16* | WBGene00004107 | protein_coding_gene | 2.3 | 5.059 | 1.2E-02 |
| *nstp-3* | WBGene00022577 | protein_coding_gene | 2.3 | 5.056 | 3.7E-03 |
| *C44B12.3* | WBGene00016637 | protein_coding_gene | 2.3 | 5.054 | 5.9E-06 |
| *C14A4.9* | WBGene00007560 | protein_coding_gene | 2.3 | 5.050 | 3.3E-04 |
| *F53B3.5* | WBGene00018743 | protein_coding_gene | 2.3 | 5.049 | 3.1E-17 |
| *hlh-14* | WBGene00001958 | protein_coding_gene | 2.3 | 5.024 | 1.1E-12 |
| *lin-32* | WBGene00003018 | protein_coding_gene | 2.3 | 5.023 | 6.4E-06 |
| *F31E3.12* | WBGene00219317 | protein_coding_gene | 2.3 | 5.021 | 3.6E-03 |
| *ZK1307.2* | WBGene00014245 | protein_coding_gene | 2.3 | 5.021 | 1.7E-04 |
| *K08D9.4* | WBGene00019525 | protein_coding_gene | 2.3 | 5.017 | 5.6E-06 |
| *F10E9.12* | WBGene00017361 | protein_coding_gene | 2.3 | 5.016 | 7.9E-18 |
| *F57A8.7* | WBGene00010182 | protein_coding_gene | 2.3 | 5.005 | 9.5E-03 |
| *C37A5.3* | WBGene00014698 | protein_coding_gene | 2.3 | 4.997 | 4.7E-03 |
| *fkb-7* | WBGene00001432 | protein_coding_gene | 2.3 | 4.989 | 3.8E-11 |
| *gcy-34* | WBGene00001554 | protein_coding_gene | 2.3 | 4.986 | 8.8E-03 |
| *T22B7.3* | WBGene00020672 | protein_coding_gene | 2.3 | 4.984 | 4.8E-06 |
| *K10G4.5* | WBGene00010754 | protein_coding_gene | 2.3 | 4.984 | 1.5E-02 |
| *F18E9.7* | WBGene00017573 | protein_coding_gene | 2.3 | 4.979 | 1.7E-03 |
| *ssp-19* | WBGene00006047 | protein_coding_gene | 2.3 | 4.975 | 2.2E-06 |
| *Y44A6C.1* | WBGene00012850 | protein_coding_gene | 2.3 | 4.974 | 3.8E-24 |
| *K10D6.2* | WBGene00010742 | protein_coding_gene | 2.3 | 4.966 | 7.7E-11 |
| *Y53C12B.7* | WBGene00013146 | protein_coding_gene | 2.3 | 4.963 | 2.9E-09 |
| *hsp-12.1* | WBGene00011906 | protein_coding_gene | 2.3 | 4.960 | 8.3E-06 |
| *ZK813.6* | WBGene00044564 | protein_coding_gene | 2.3 | 4.960 | 1.0E-02 |
| *scl-5* | WBGene00008027 | protein_coding_gene | 2.3 | 4.954 | 5.6E-09 |
| *droe-4* | WBGene00010262 | protein_coding_gene | 2.3 | 4.946 | 1.2E-22 |
| *cutl-13* | WBGene00021025 | protein_coding_gene | 2.3 | 4.941 | 4.1E-03 |
| *tag-234* | WBGene00018874 | protein_coding_gene | 2.3 | 4.940 | 4.6E-19 |
| *F30A10.2* | WBGene00009261 | protein_coding_gene | 2.3 | 4.940 | 3.0E-04 |
| *Y6E2A.1* | WBGene00012397 | protein_coding_gene | 2.3 | 4.934 | 7.0E-03 |
| *C35D6.5* | WBGene00007962 | pseudogene | 2.3 | 4.926 | 3.0E-03 |
| *F30H5.3* | WBGene00017937 | protein_coding_gene | 2.3 | 4.925 | 1.9E-07 |
| *Y46D2A.1* | WBGene00021589 | protein_coding_gene | 2.3 | 4.921 | 8.2E-04 |
| *F46F3.3* | WBGene00009791 | protein_coding_gene | 2.3 | 4.920 | 2.4E-03 |
| *C08F11.13* | WBGene00007460 | protein_coding_gene | 2.3 | 4.913 | 3.1E-20 |
| *C29F7.11* | WBGene00206531 | ncRNA_gene | 2.3 | 4.911 | 1.1E-02 |
| *F23H12.5* | WBGene00009094 | protein_coding_gene | 2.3 | 4.910 | 2.9E-05 |
| *unc-3* | WBGene00006743 | protein_coding_gene | 2.3 | 4.901 | 1.6E-05 |
| *nlp-28* | WBGene00003766 | protein_coding_gene | 2.3 | 4.894 | 3.0E-18 |
| *C08D8.1* | WBGene00015592 | protein_coding_gene | 2.3 | 4.886 | 3.8E-06 |
| *nhr-246* | WBGene00014192 | protein_coding_gene | 2.3 | 4.876 | 2.4E-03 |
| *F46E10.2* | WBGene00018489 | protein_coding_gene | 2.3 | 4.870 | 2.7E-16 |
| *srz-56* | WBGene00016519 | protein_coding_gene | 2.3 | 4.864 | 8.0E-03 |
| *ddr-2* | WBGene00017381 | protein_coding_gene | 2.3 | 4.862 | 1.4E-19 |
| *F07H5.13* | WBGene00008563 | protein_coding_gene | 2.3 | 4.861 | 2.0E-18 |
| *C50F2.5* | WBGene00016839 | protein_coding_gene | 2.3 | 4.861 | 3.5E-05 |

| | | | | | |
|---|---|---|---|---|---|
| *T05C1.3* | WBGene00020250 | protein_coding_gene | 2.3 | 4.837 | 7.6E-03 |
| *Y82E9BL.19* | WBGene00194735 | protein_coding_gene | 2.3 | 4.834 | 1.1E-02 |
| *F46G11.6* | WBGene00045269 | protein_coding_gene | 2.3 | 4.821 | 2.1E-03 |
| *W09C3.1* | WBGene00021109 | protein_coding_gene | 2.3 | 4.820 | 1.1E-02 |
| *C53A5.2* | WBGene00008263 | protein_coding_gene | 2.3 | 4.819 | 5.7E-04 |
| *skr-21* | WBGene00004827 | protein_coding_gene | 2.3 | 4.813 | 1.7E-22 |
| *ugt-43* | WBGene00008485 | protein_coding_gene | 2.3 | 4.812 | 9.6E-07 |
| *fbxa-105* | WBGene00015362 | protein_coding_gene | 2.3 | 4.807 | 1.5E-09 |
| *igcm-1* | WBGene00018215 | protein_coding_gene | 2.3 | 4.800 | 4.5E-14 |
| *cuti-1* | WBGene00022591 | protein_coding_gene | 2.3 | 4.797 | 1.1E-06 |
| *B0393.5* | WBGene00007170 | protein_coding_gene | 2.3 | 4.783 | 5.6E-08 |
| *K09C4.1* | WBGene00019547 | protein_coding_gene | 2.3 | 4.782 | 1.8E-07 |
| *ZK666.14* | WBGene00194952 | protein_coding_gene | 2.3 | 4.777 | 1.6E-02 |
| *nab-1* | WBGene00003516 | protein_coding_gene | 2.3 | 4.771 | 7.1E-12 |
| *R08C7.4* | WBGene00019948 | protein_coding_gene | 2.3 | 4.763 | 8.9E-03 |
| *clec-95* | WBGene00013927 | protein_coding_gene | 2.3 | 4.762 | 3.1E-04 |
| *F53A9.6* | WBGene00018729 | protein_coding_gene | 2.3 | 4.759 | 1.1E-23 |
| *F53F4.8* | WBGene00009991 | pseudogene | 2.2 | 4.756 | 1.6E-05 |
| *ZK971.1* | WBGene00014175 | protein_coding_gene | 2.2 | 4.752 | 1.5E-05 |
| *F48C1.11* | WBGene00195179 | protein_coding_gene | 2.2 | 4.746 | 1.3E-02 |
| *his-1* | WBGene00001875 | protein_coding_gene | 2.2 | 4.738 | 1.1E-02 |
| *Y43C5A.3* | WBGene00012783 | protein_coding_gene | 2.2 | 4.733 | 1.9E-30 |
| *dyf-7* | WBGene00001123 | protein_coding_gene | 2.2 | 4.728 | 2.2E-09 |
| *ZC443.4* | WBGene00013899 | protein_coding_gene | 2.2 | 4.711 | 2.1E-18 |
| *ttr-26* | WBGene00013079 | protein_coding_gene | 2.2 | 4.703 | 7.2E-10 |
| *oac-6* | WBGene00007833 | protein_coding_gene | 2.2 | 4.703 | 1.1E-04 |
| *ins-31* | WBGene00002114 | protein_coding_gene | 2.2 | 4.701 | 7.9E-04 |
| *Y87G2A.16* | WBGene00013602 | protein_coding_gene | 2.2 | 4.701 | 4.8E-05 |
| *ptr-9* | WBGene00004223 | protein_coding_gene | 2.2 | 4.700 | 1.6E-02 |
| *math-42* | WBGene00020532 | protein_coding_gene | 2.2 | 4.692 | 4.0E-03 |
| *Y39B6A.21* | WBGene00012680 | pseudogene | 2.2 | 4.689 | 2.1E-03 |
| *F26F12.3* | WBGene00017834 | protein_coding_gene | 2.2 | 4.687 | 8.1E-27 |
| *C50B8.4* | WBGene00008226 | protein_coding_gene | 2.2 | 4.681 | 2.0E-06 |
| *cnc-7* | WBGene00010005 | protein_coding_gene | 2.2 | 4.680 | 5.4E-05 |
| *F53B7.7* | WBGene00009969 | protein_coding_gene | 2.2 | 4.678 | 3.3E-03 |
| *W02A2.4* | WBGene00012189 | pseudogene | 2.2 | 4.670 | 4.8E-04 |
| *clec-66* | WBGene00009397 | protein_coding_gene | 2.2 | 4.662 | 8.0E-12 |
| *srbc-82* | WBGene00007832 | protein_coding_gene | 2.2 | 4.660 | 1.6E-02 |
| *R11E3.19* | WBGene00235374 | protein_coding_gene | 2.2 | 4.659 | 8.4E-04 |
| *nsy-4* | WBGene00021415 | protein_coding_gene | 2.2 | 4.655 | 9.2E-10 |
| *ncx-9* | WBGene00003574 | protein_coding_gene | 2.2 | 4.650 | 2.1E-03 |
| *F20B6.1* | WBGene00017627 | pseudogene | 2.2 | 4.642 | 1.0E-02 |
| *B0348.2* | WBGene00015152 | protein_coding_gene | 2.2 | 4.641 | 6.2E-03 |
| *grsp-3* | WBGene00016403 | protein_coding_gene | 2.2 | 4.636 | 6.6E-08 |
| *gasr-8* | WBGene00008144 | protein_coding_gene | 2.2 | 4.636 | 4.8E-03 |
| *ikb-1* | WBGene00002069 | protein_coding_gene | 2.2 | 4.632 | 3.3E-18 |
| *F53H2.1* | WBGene00010004 | protein_coding_gene | 2.2 | 4.630 | 1.1E-02 |
| *C33A12.4* | WBGene00007883 | protein_coding_gene | 2.2 | 4.628 | 3.4E-09 |
| *mir-238* | WBGene00003331 | miRNA_gene | 2.2 | 4.625 | 7.9E-03 |
| *R160.10* | WBGene00219369 | protein_coding_gene | 2.2 | 4.623 | 1.4E-03 |
| *acd-1* | WBGene00016064 | protein_coding_gene | 2.2 | 4.617 | 3.1E-03 |
| *K01A2.10* | WBGene00019284 | protein_coding_gene | 2.2 | 4.615 | 1.4E-11 |
| *pals-7* | WBGene00007661 | protein_coding_gene | 2.2 | 4.613 | 1.0E-02 |
| *D1014.5* | WBGene00017018 | protein_coding_gene | 2.2 | 4.610 | 5.1E-04 |
| *C35C5.9* | WBGene00007958 | protein_coding_gene | 2.2 | 4.598 | 2.2E-17 |
| *F26F12.4* | WBGene00017835 | protein_coding_gene | 2.2 | 4.596 | 1.0E-03 |
| *nhr-99* | WBGene00003689 | protein_coding_gene | 2.2 | 4.586 | 2.7E-08 |
| *zig-4* | WBGene00006981 | protein_coding_gene | 2.2 | 4.583 | 1.2E-05 |
| *F59A7.2* | WBGene00019090 | protein_coding_gene | 2.2 | 4.575 | 2.0E-07 |
| *oac-51* | WBGene00020976 | protein_coding_gene | 2.2 | 4.558 | 4.1E-04 |

| | | | | | |
|---|---|---|---|---|---|
| *Y48A6B.7* | WBGene00012968 | protein_coding_gene | 2.2 | 4.550 | 2.5E-31 |
| *C54F6.6* | WBGene00016924 | protein_coding_gene | 2.2 | 4.549 | 9.9E-03 |
| *cdf-2* | WBGene00011821 | protein_coding_gene | 2.2 | 4.540 | 4.3E-15 |
| *Y38H8A.1* | WBGene00012635 | protein_coding_gene | 2.2 | 4.539 | 6.3E-03 |
| *W05F2.8* | WBGene00195045 | protein_coding_gene | 2.2 | 4.536 | 7.5E-03 |
| *B0462.5* | WBGene00194695 | protein_coding_gene | 2.2 | 4.535 | 3.7E-05 |
| *let-381* | WBGene00002601 | protein_coding_gene | 2.2 | 4.535 | 2.2E-04 |
| *cpr-4* | WBGene00000784 | protein_coding_gene | 2.2 | 4.534 | 1.3E-13 |
| *bath-25* | WBGene00020859 | protein_coding_gene | 2.2 | 4.531 | 1.5E-02 |
| *W03D2.9* | WBGene00020983 | protein_coding_gene | 2.2 | 4.530 | 9.5E-03 |
| *R10E8.6* | WBGene00011212 | protein_coding_gene | 2.2 | 4.529 | 5.2E-15 |
| *eva-1* | WBGene00009304 | protein_coding_gene | 2.2 | 4.525 | 9.5E-10 |
| *clec-170* | WBGene00009515 | protein_coding_gene | 2.2 | 4.522 | 9.0E-12 |
| *cbn-1* | WBGene00019285 | protein_coding_gene | 2.2 | 4.520 | 3.4E-04 |
| *ztf-30* | WBGene00015523 | protein_coding_gene | 2.2 | 4.513 | 8.0E-04 |
| *cnd-1* | WBGene00000561 | protein_coding_gene | 2.2 | 4.509 | 4.5E-07 |
| *K07A1.3* | WBGene00010610 | protein_coding_gene | 2.2 | 4.509 | 4.5E-16 |
| *glb-5* | WBGene00015964 | protein_coding_gene | 2.2 | 4.503 | 6.2E-03 |
| *W10G11.1* | WBGene00021134 | protein_coding_gene | 2.2 | 4.500 | 1.1E-02 |
| *cyp-14A3* | WBGene00010707 | protein_coding_gene | 2.2 | 4.500 | 9.0E-04 |
| *arl-6* | WBGene00000193 | protein_coding_gene | 2.2 | 4.494 | 8.6E-03 |
| *linc-103* | WBGene00219743 | lincRNA_gene | 2.2 | 4.493 | 2.3E-02 |
| *Y73B6BL.28* | WBGene00022249 | pseudogene | 2.2 | 4.492 | 2.0E-02 |
| *F38B6.3* | WBGene00018173 | protein_coding_gene | 2.2 | 4.492 | 2.0E-02 |
| *fbxa-176* | WBGene00008017 | protein_coding_gene | 2.2 | 4.480 | 1.1E-03 |
| *T09E11.11* | WBGene00011660 | pseudogene | 2.2 | 4.473 | 6.0E-05 |
| *F32H2.6* | WBGene00009343 | protein_coding_gene | 2.2 | 4.447 | 4.2E-04 |
| *dhs-16* | WBGene00000979 | protein_coding_gene | 2.2 | 4.439 | 4.9E-03 |
| *F54B11.10* | WBGene00010033 | protein_coding_gene | 2.1 | 4.435 | 8.6E-03 |
| *C44C1.6* | WBGene00016645 | protein_coding_gene | 2.1 | 4.430 | 5.0E-09 |
| *C39D10.5* | WBGene00016532 | protein_coding_gene | 2.1 | 4.418 | 1.3E-02 |
| *plx-2* | WBGene00004048 | protein_coding_gene | 2.1 | 4.410 | 8.4E-10 |
| *osm-8* | WBGene00003888 | protein_coding_gene | 2.1 | 4.404 | 3.7E-03 |
| *C17H12.37* | WBGene00236786 | protein_coding_gene | 2.1 | 4.391 | 2.4E-02 |
| *B0284.5* | WBGene00045188 | pseudogene | 2.1 | 4.390 | 2.4E-02 |
| *bigr-1* | WBGene00009512 | protein_coding_gene | 2.1 | 4.389 | 9.8E-03 |
| *fbxb-114* | WBGene00016254 | protein_coding_gene | 2.1 | 4.389 | 1.3E-02 |
| *T01B10.5* | WBGene00020137 | protein_coding_gene | 2.1 | 4.387 | 1.9E-04 |
| *clec-98* | WBGene00013932 | protein_coding_gene | 2.1 | 4.386 | 1.7E-02 |
| *fbxa-87* | WBGene00044660 | pseudogene | 2.1 | 4.377 | 5.9E-07 |
| *W03G1.5* | WBGene00021011 | protein_coding_gene | 2.1 | 4.369 | 6.3E-04 |
| *F10E7.3* | WBGene00017345 | pseudogene | 2.1 | 4.367 | 2.4E-02 |
| *ZK836.3* | WBGene00014099 | protein_coding_gene | 2.1 | 4.366 | 8.9E-06 |
| *spi-1* | WBGene00023416 | protein_coding_gene | 2.1 | 4.364 | 8.0E-04 |
| *eol-1* | WBGene00045401 | protein_coding_gene | 2.1 | 4.362 | 2.0E-03 |
| *F22F7.3* | WBGene00017721 | protein_coding_gene | 2.1 | 4.360 | 2.8E-03 |
| *clec-69* | WBGene00044642 | pseudogene | 2.1 | 4.360 | 1.2E-02 |
| *slc-17.9* | WBGene00011643 | protein_coding_gene | 2.1 | 4.350 | 6.4E-04 |
| *R155.4* | WBGene00020118 | protein_coding_gene | 2.1 | 4.347 | 9.8E-03 |
| *W02D7.8* | WBGene00020943 | protein_coding_gene | 2.1 | 4.346 | 2.7E-05 |
| *clec-28* | WBGene00009857 | protein_coding_gene | 2.1 | 4.344 | 2.5E-03 |
| *R160.4* | WBGene00020121 | protein_coding_gene | 2.1 | 4.343 | 1.6E-02 |
| *cut-6* | WBGene00000853 | protein_coding_gene | 2.1 | 4.342 | 2.5E-03 |
| *T10D4.7* | WBGene00020409 | protein_coding_gene | 2.1 | 4.342 | 2.0E-02 |
| *best-1* | WBGene00007203 | protein_coding_gene | 2.1 | 4.338 | 6.2E-05 |
| *C17E4.14* | WBGene00195884 | ncRNA_gene | 2.1 | 4.336 | 2.4E-02 |
| *F07A5.4* | WBGene00008543 | protein_coding_gene | 2.1 | 4.333 | 1.9E-03 |
| *F58A6.9* | WBGene00019026 | protein_coding_gene | 2.1 | 4.330 | 7.1E-08 |
| *fbxa-160* | WBGene00015604 | protein_coding_gene | 2.1 | 4.323 | 1.4E-02 |
| *W08G11.1* | WBGene00012346 | protein_coding_gene | 2.1 | 4.319 | 7.0E-03 |

| | | | | | |
|---|---|---|---|---|---|
| *pph-2* | WBGene00004084 | protein_coding_gene | 2.1 | 4.318 | 1.4E-03 |
| *Y34F4.3* | WBGene00021338 | protein_coding_gene | 2.1 | 4.314 | 8.0E-04 |
| *ZC84.1* | WBGene00013846 | protein_coding_gene | 2.1 | 4.310 | 4.9E-04 |
| *F21A3.11* | WBGene00219316 | protein_coding_gene | 2.1 | 4.305 | 2.0E-13 |
| *fipr-26* | WBGene00009963 | protein_coding_gene | 2.1 | 4.304 | 1.9E-02 |
| *lgc-27* | WBGene00022283 | protein_coding_gene | 2.1 | 4.299 | 9.0E-07 |
| *ham-2* | WBGene00001821 | protein_coding_gene | 2.1 | 4.297 | 5.1E-14 |
| *F36F2.2* | WBGene00009476 | protein_coding_gene | 2.1 | 4.292 | 8.4E-13 |
| *nhr-19* | WBGene00003618 | protein_coding_gene | 2.1 | 4.292 | 3.4E-15 |
| *F57G9.6* | WBGene00044041 | pseudogene | 2.1 | 4.291 | 1.7E-05 |
| *F53C3.8* | WBGene00018752 | protein_coding_gene | 2.1 | 4.291 | 1.9E-03 |
| *F49E10.2* | WBGene00018640 | protein_coding_gene | 2.1 | 4.289 | 6.3E-04 |
| *F20C5.6* | WBGene00008971 | protein_coding_gene | 2.1 | 4.288 | 1.1E-10 |
| *H32K16.2* | WBGene00010422 | protein_coding_gene | 2.1 | 4.283 | 1.2E-05 |
| *F22B7.3* | WBGene00017694 | protein_coding_gene | 2.1 | 4.278 | 1.5E-02 |
| *F59A6.3* | WBGene00019085 | protein_coding_gene | 2.1 | 4.278 | 1.5E-02 |
| *ceh-48* | WBGene00015934 | protein_coding_gene | 2.1 | 4.272 | 3.4E-06 |
| *Y17D7B.2* | WBGene00012450 | protein_coding_gene | 2.1 | 4.271 | 2.2E-02 |
| *T10G3.1* | WBGene00011692 | protein_coding_gene | 2.1 | 4.266 | 1.1E-09 |
| *F44E5.3* | WBGene00009690 | protein_coding_gene | 2.1 | 4.265 | 4.4E-04 |
| *Y97E10AR.1* | WBGene00022396 | protein_coding_gene | 2.1 | 4.264 | 2.6E-10 |
| *mks-3* | WBGene00018042 | protein_coding_gene | 2.1 | 4.264 | 4.3E-03 |
| *Y55D5A.1* | WBGene00021915 | protein_coding_gene | 2.1 | 4.261 | 6.0E-07 |
| *C02F4.4* | WBGene00007264 | protein_coding_gene | 2.1 | 4.260 | 4.4E-03 |
| *F32D1.3* | WBGene00017983 | protein_coding_gene | 2.1 | 4.256 | 3.1E-09 |
| *rocf-1* | WBGene00011948 | protein_coding_gene | 2.1 | 4.248 | 9.3E-03 |
| *sups-1* | WBGene00017716 | protein_coding_gene | 2.1 | 4.247 | 5.9E-05 |
| *T28C12.15* | WBGene00271821 | protein_coding_gene | 2.1 | 4.242 | 3.0E-02 |
| *T04B2.1* | WBGene00011421 | pseudogene | 2.1 | 4.241 | 2.3E-02 |
| *C54C8.12* | WBGene00044242 | protein_coding_gene | 2.1 | 4.234 | 2.7E-02 |
| *Y47D7A.7* | WBGene00021620 | protein_coding_gene | 2.1 | 4.234 | 6.3E-03 |
| *mks-2* | WBGene00194710 | protein_coding_gene | 2.1 | 4.230 | 2.5E-02 |
| *pag-3* | WBGene00003909 | protein_coding_gene | 2.1 | 4.227 | 2.0E-03 |
| *ceh-45* | WBGene00022837 | protein_coding_gene | 2.1 | 4.222 | 2.2E-03 |
| *C45B11.6* | WBGene00045379 | protein_coding_gene | 2.1 | 4.221 | 2.1E-04 |
| *clec-172* | WBGene00044719 | protein_coding_gene | 2.1 | 4.219 | 7.3E-04 |
| *C46A5.4* | WBGene00016700 | protein_coding_gene | 2.1 | 4.209 | 3.5E-03 |
| *K04H4.2* | WBGene00010573 | protein_coding_gene | 2.1 | 4.208 | 1.0E-06 |
| *Y65A5A.1* | WBGene00013418 | protein_coding_gene | 2.1 | 4.205 | 1.4E-13 |
| *Y57G7A.8* | WBGene00021971 | pseudogene | 2.1 | 4.201 | 6.5E-06 |
| *W05B2.2* | WBGene00012271 | protein_coding_gene | 2.1 | 4.197 | 8.4E-03 |
| *Y46E12BR.1* | WBGene00021598 | protein_coding_gene | 2.1 | 4.188 | 7.8E-03 |
| *F57A10.2* | WBGene00010184 | protein_coding_gene | 2.1 | 4.186 | 3.1E-03 |
| *D2024.10* | WBGene00044505 | pseudogene | 2.1 | 4.166 | 3.0E-02 |
| *T05H10.3* | WBGene00011508 | protein_coding_gene | 2.1 | 4.162 | 1.4E-02 |
| *F09F9.2* | WBGene00017307 | protein_coding_gene | 2.1 | 4.149 | 2.0E-03 |
| *C32D5.12* | WBGene00016319 | protein_coding_gene | 2.1 | 4.145 | 2.4E-06 |
| *C49C8.6* | WBGene00016770 | protein_coding_gene | 2.1 | 4.143 | 1.7E-02 |
| *C43H6.6* | WBGene00016619 | protein_coding_gene | 2.1 | 4.143 | 1.3E-02 |
| *E02C12.10* | WBGene00017095 | protein_coding_gene | 2.1 | 4.142 | 1.8E-02 |
| *clec-183* | WBGene00011855 | protein_coding_gene | 2.0 | 4.140 | 2.2E-02 |
| *clec-197* | WBGene00008202 | protein_coding_gene | 2.0 | 4.138 | 3.9E-03 |
| *cal-6* | WBGene00022214 | pseudogene | 2.0 | 4.134 | 2.9E-02 |
| *R11F4.1* | WBGene00020007 | protein_coding_gene | 2.0 | 4.134 | 1.8E-13 |
| *dct-1* | WBGene00015776 | protein_coding_gene | 2.0 | 4.133 | 6.7E-24 |
| *C32B5.18* | WBGene00235351 | protein_coding_gene | 2.0 | 4.126 | 1.3E-02 |
| *math-2* | WBGene00015606 | protein_coding_gene | 2.0 | 4.122 | 2.3E-02 |
| *B0228.8* | WBGene00015065 | protein_coding_gene | 2.0 | 4.122 | 2.9E-02 |
| *F20A1.8* | WBGene00017619 | protein_coding_gene | 2.0 | 4.118 | 2.8E-02 |
| *F21C10.10* | WBGene00017659 | protein_coding_gene | 2.0 | 4.118 | 1.9E-03 |

| | | | | | |
|---|---|---|---|---|---|
| *F17C11.4* | WBGene00008915 | protein_coding_gene | 2.0 | 4.113 | 2.5E-16 |
| *EGAP1.1* | WBGene00017146 | protein_coding_gene | 2.0 | 4.113 | 1.3E-02 |
| *Y52B11A.12* | WBGene00077539 | protein_coding_gene | 2.0 | 4.105 | 5.5E-04 |
| *mam-5* | WBGene00011572 | protein_coding_gene | 2.0 | 4.101 | 1.4E-02 |
| *unc-86* | WBGene00006818 | protein_coding_gene | 2.0 | 4.096 | 2.2E-04 |
| *fbxa-175* | WBGene00045293 | pseudogene | 2.0 | 4.089 | 6.5E-06 |
| *ZC168.2* | WBGene00013856 | protein_coding_gene | 2.0 | 4.085 | 5.6E-03 |
| *K02E10.4* | WBGene00019318 | protein_coding_gene | 2.0 | 4.084 | 6.2E-06 |
| *F21D9.1* | WBGene00009015 | protein_coding_gene | 2.0 | 4.078 | 9.9E-04 |
| *T28F4.5* | WBGene00012140 | protein_coding_gene | 2.0 | 4.078 | 2.7E-20 |
| *gpa-9* | WBGene00001671 | protein_coding_gene | 2.0 | 4.077 | 5.5E-06 |
| *ceh-34* | WBGene00000455 | protein_coding_gene | 2.0 | 4.072 | 3.1E-07 |
| *M163.1* | WBGene00010935 | protein_coding_gene | 2.0 | 4.068 | 1.6E-42 |
| *ins-7* | WBGene00002090 | protein_coding_gene | 2.0 | 4.068 | 9.4E-06 |
| *aakg-4* | WBGene00020135 | protein_coding_gene | 2.0 | 4.066 | 2.3E-23 |
| *lips-16* | WBGene00012588 | pseudogene | 2.0 | 4.060 | 1.2E-10 |
| *C27B7.9* | WBGene00007765 | protein_coding_gene | 2.0 | 4.058 | 1.1E-38 |
| *tbx-7* | WBGene00006544 | protein_coding_gene | 2.0 | 4.055 | 2.9E-03 |
| *R05G6.10* | WBGene00019902 | protein_coding_gene | 2.0 | 4.055 | 4.8E-22 |
| *alh-5* | WBGene00000111 | protein_coding_gene | 2.0 | 4.053 | 2.3E-19 |
| *M7.8* | WBGene00010884 | protein_coding_gene | 2.0 | 4.049 | 4.1E-10 |
| *C54F6.12* | WBGene00016929 | protein_coding_gene | 2.0 | 4.040 | 4.1E-03 |
| *F19G12.3* | WBGene00017611 | protein_coding_gene | 2.0 | 4.040 | 3.2E-02 |
| *ceh-30* | WBGene00000451 | protein_coding_gene | 2.0 | 4.038 | 1.1E-03 |
| *T16G1.7* | WBGene00011801 | protein_coding_gene | 2.0 | 4.033 | 5.8E-08 |
| *nlp-29* | WBGene00003767 | protein_coding_gene | 2.0 | 4.027 | 1.3E-09 |
| *fbxc-7* | WBGene00016302 | protein_coding_gene | 2.0 | 4.024 | 2.4E-02 |
| *H10D18.1* | WBGene00019177 | protein_coding_gene | 2.0 | 4.024 | 2.0E-02 |
| *K11H12.5* | WBGene00019661 | protein_coding_gene | 2.0 | 4.024 | 3.2E-06 |
| *C06G3.3* | WBGene00015549 | protein_coding_gene | 2.0 | 4.021 | 2.4E-03 |
| *C03A3.1* | WBGene00007266 | protein_coding_gene | 2.0 | 4.021 | 2.1E-05 |
| *F59F4.6* | WBGene00303371 | ncRNA_gene | 2.0 | 4.016 | 1.9E-02 |
| *arrd-4* | WBGene00014159 | protein_coding_gene | 2.0 | 4.016 | 9.9E-11 |
| *F26G1.15* | WBGene00270318 | protein_coding_gene | 2.0 | 4.016 | 3.7E-02 |
| *mec-3* | WBGene00003167 | protein_coding_gene | 2.0 | 4.011 | 2.0E-03 |
| *fax-1* | WBGene00001400 | protein_coding_gene | 2.0 | 4.011 | 1.1E-02 |
| *Y40H7A.7* | WBGene00012745 | pseudogene | 2.0 | 4.010 | 3.7E-03 |
| *F28G4.2* | WBGene00009227 | protein_coding_gene | 2.0 | 4.007 | 1.5E-02 |
| *F43C11.7* | WBGene00018384 | protein_coding_gene | 2.0 | 4.005 | 1.2E-13 |
| *his-64* | WBGene00001938 | protein_coding_gene | 2.0 | 4.004 | 6.1E-06 |
| *mec-8* | WBGene00003172 | protein_coding_gene | 2.0 | 4.002 | 3.9E-41 |
| *C31B8.9* | WBGene00016284 | protein_coding_gene | 2.0 | 4.002 | 3.2E-02 |
| *F28B1.2* | WBGene00009196 | protein_coding_gene | 2.0 | 3.999 | 1.8E-02 |
| *Y62H9A.16* | WBGene00304791 | protein_coding_gene | 2.0 | 3.999 | 4.3E-03 |
| *Y50E8A.5* | WBGene00013048 | protein_coding_gene | 2.0 | 3.997 | 7.8E-06 |
| *pef-1* | WBGene00003969 | protein_coding_gene | 2.0 | 3.997 | 4.7E-07 |
| *F59B2.12* | WBGene00010314 | protein_coding_gene | 2.0 | 3.992 | 2.7E-03 |
| *ceh-5* | WBGene00000430 | protein_coding_gene | 2.0 | 3.990 | 2.7E-04 |
| *C54F6.17* | WBGene00235092 | protein_coding_gene | 2.0 | 3.990 | 3.3E-02 |
| *F26F2.11* | WBGene00195158 | protein_coding_gene | 2.0 | 3.986 | 3.4E-02 |
| *C35A5.5* | WBGene00007949 | protein_coding_gene | 2.0 | 3.986 | 7.4E-03 |
| *F28B1.3* | WBGene00009197 | pseudogene | 2.0 | 3.980 | 2.1E-02 |
| *F49C5.10* | WBGene00044162 | protein_coding_gene | 2.0 | 3.978 | 2.0E-02 |
| *F59B10.4* | WBGene00010320 | protein_coding_gene | 2.0 | 3.974 | 4.8E-08 |
| *osm-12* | WBGene00003892 | protein_coding_gene | 2.0 | 3.967 | 3.3E-03 |
| *D1086.2* | WBGene00008389 | protein_coding_gene | 2.0 | 3.966 | 3.5E-04 |
| *Y57E12B.10* | WBGene00202501 | protein_coding_gene | 2.0 | 3.962 | 2.6E-02 |
| *cyp-23A1* | WBGene00015135 | protein_coding_gene | 2.0 | 3.954 | 1.5E-02 |
| *Y5H2B.1* | WBGene00021164 | protein_coding_gene | 2.0 | 3.948 | 2.8E-02 |
| *R13A1.1* | WBGene00020041 | protein_coding_gene | 2.0 | 3.948 | 2.2E-02 |

| | | | | | |
|---|---|---|---|---|---|
| *Y20C6A.1* | WBGene00012491 | protein_coding_gene | 2.0 | 3.948 | 2.4E-11 |
| *Y39B6A.31* | WBGene00012690 | pseudogene | 2.0 | 3.942 | 4.3E-04 |
| *C05D9.9* | WBGene00015476 | protein_coding_gene | 2.0 | 3.938 | 1.6E-13 |
| *C06A12.19* | WBGene00255702 | protein_coding_gene | 2.0 | 3.936 | 3.0E-02 |
| *rpr-1* | WBGene00045238 | ncRNA_gene | 2.0 | 3.933 | 1.6E-02 |
| *set-28* | WBGene00022224 | protein_coding_gene | 2.0 | 3.931 | 2.6E-02 |
| *F18C5.10* | WBGene00017563 | protein_coding_gene | 2.0 | 3.929 | 8.7E-40 |
| *ZK402.5* | WBGene00022731 | protein_coding_gene | 2.0 | 3.924 | 2.7E-02 |
| *rrn-1.2* | WBGene00004513 | rRNA_gene | 2.0 | 3.917 | 2.2E-02 |
| *nhr-150* | WBGene00007367 | protein_coding_gene | 2.0 | 3.917 | 8.9E-03 |
| *C36B1.13* | WBGene00044120 | protein_coding_gene | 2.0 | 3.908 | 2.1E-02 |
| *K10H10.13* | WBGene00194888 | protein_coding_gene | 2.0 | 3.907 | 3.9E-02 |
| *C27A2.8* | WBGene00044387 | protein_coding_gene | 2.0 | 3.906 | 3.1E-02 |
| *smf-3* | WBGene00004878 | protein_coding_gene | 2.0 | 3.904 | 9.7E-06 |
| *Y50E8A.1* | WBGene00013046 | protein_coding_gene | 2.0 | 3.903 | 1.1E-05 |
| *K08C9.10* | WBGene00086564 | protein_coding_gene | 2.0 | 3.891 | 3.6E-02 |
| *hst-3.2* | WBGene00044303 | protein_coding_gene | 2.0 | 3.886 | 2.0E-05 |
| *B0228.6* | WBGene00015063 | protein_coding_gene | 2.0 | 3.872 | 3.0E-08 |
| *fbxc-51* | WBGene00021576 | protein_coding_gene | 2.0 | 3.869 | 1.3E-22 |
| *nlp-66* | WBGene00011141 | protein_coding_gene | 2.0 | 3.868 | 5.5E-03 |
| *B0334.10* | WBGene00007148 | protein_coding_gene | 1.9 | 3.863 | 2.9E-02 |
| *C07E3.3* | WBGene00007414 | protein_coding_gene | 1.9 | 3.861 | 3.8E-11 |
| *zip-7* | WBGene00013100 | protein_coding_gene | 1.9 | 3.858 | 1.0E-06 |
| *R102.1* | WBGene00011288 | protein_coding_gene | 1.9 | 3.856 | 3.2E-02 |
| *pqn-73* | WBGene00004155 | protein_coding_gene | 1.9 | 3.852 | 2.2E-02 |
| *F59B2.11* | WBGene00010313 | pseudogene | 1.9 | 3.851 | 5.9E-03 |
| *C17E4.11* | WBGene00044236 | protein_coding_gene | 1.9 | 3.851 | 4.2E-17 |
| *Y38H6C.16* | WBGene00012628 | protein_coding_gene | 1.9 | 3.843 | 1.2E-04 |
| *M195.4* | WBGene00010949 | protein_coding_gene | 1.9 | 3.842 | 2.5E-02 |
| *T24C2.5* | WBGene00011984 | protein_coding_gene | 1.9 | 3.841 | 1.1E-08 |
| *cutl-21* | WBGene00018861 | protein_coding_gene | 1.9 | 3.837 | 2.1E-04 |
| *F21D5.9* | WBGene00009014 | protein_coding_gene | 1.9 | 3.832 | 5.6E-03 |
| *R11G10.4* | WBGene00050968 | protein_coding_gene | 1.9 | 3.832 | 1.5E-02 |
| *F31D5.6* | WBGene00017949 | protein_coding_gene | 1.9 | 3.826 | 8.1E-04 |
| *tig-3* | WBGene00021594 | protein_coding_gene | 1.9 | 3.824 | 1.3E-03 |
| *egas-4* | WBGene00018906 | protein_coding_gene | 1.9 | 3.823 | 3.0E-03 |
| *rga-2* | WBGene00013135 | protein_coding_gene | 1.9 | 3.817 | 3.5E-07 |
| *C39B10.7* | WBGene00164973 | protein_coding_gene | 1.9 | 3.815 | 3.7E-03 |
| *nlp-2* | WBGene00003740 | protein_coding_gene | 1.9 | 3.813 | 2.6E-03 |
| *M01A8.1* | WBGene00010795 | protein_coding_gene | 1.9 | 3.812 | 4.1E-06 |
| *K08E5.1* | WBGene00014801 | protein_coding_gene | 1.9 | 3.809 | 1.0E-07 |
| *igdb-3* | WBGene00020691 | protein_coding_gene | 1.9 | 3.804 | 1.2E-03 |
| *chpf-2* | WBGene00019108 | protein_coding_gene | 1.9 | 3.800 | 1.3E-07 |
| *C10A4.3* | WBGene00015664 | protein_coding_gene | 1.9 | 3.799 | 6.5E-03 |
| *pgp-11* | WBGene00004005 | protein_coding_gene | 1.9 | 3.788 | 1.8E-10 |
| *Y45F10B.59* | WBGene00194917 | protein_coding_gene | 1.9 | 3.785 | 3.9E-02 |
| *aagr-1* | WBGene00017071 | protein_coding_gene | 1.9 | 3.782 | 8.8E-10 |
| *nspe-2* | WBGene00012604 | protein_coding_gene | 1.9 | 3.781 | 2.3E-03 |
| *C36A4.11* | WBGene00044167 | protein_coding_gene | 1.9 | 3.778 | 2.5E-03 |
| *linc-37* | WBGene00219709 | lincRNA_gene | 1.9 | 3.774 | 3.0E-02 |
| *K11H12.11* | WBGene00044723 | protein_coding_gene | 1.9 | 3.768 | 2.4E-02 |
| *K01F9.2* | WBGene00010475 | protein_coding_gene | 1.9 | 3.766 | 1.1E-06 |
| *klp-11* | WBGene00002222 | protein_coding_gene | 1.9 | 3.764 | 5.0E-07 |
| *C18H9.5* | WBGene00016003 | protein_coding_gene | 1.9 | 3.758 | 1.8E-07 |
| *K01D12.1* | WBGene00010461 | protein_coding_gene | 1.9 | 3.754 | 2.0E-02 |
| *cyp-13B1* | WBGene00008519 | protein_coding_gene | 1.9 | 3.754 | 3.8E-06 |
| *nhr-165* | WBGene00008158 | protein_coding_gene | 1.9 | 3.753 | 4.1E-02 |
| *dlg-1* | WBGene00001006 | protein_coding_gene | 1.9 | 3.752 | 1.0E-24 |
| *F15H9.8* | WBGene00194652 | pseudogene | 1.9 | 3.752 | 5.7E-03 |
| *oac-43* | WBGene00011653 | protein_coding_gene | 1.9 | 3.752 | 4.2E-02 |

| | | | | | |
|---|---|---|---|---|---|
| *clec-218* | WBGene00020938 | protein_coding_gene | 1.9 | 3.751 | 1.3E-03 |
| *C17E7.13* | WBGene00015906 | protein_coding_gene | 1.9 | 3.750 | 1.4E-02 |
| *W08A12.2* | WBGene00021079 | protein_coding_gene | 1.9 | 3.748 | 8.9E-08 |
| *B0238.13* | WBGene00015077 | pseudogene | 1.9 | 3.746 | 4.1E-02 |
| *W03B1.3* | WBGene00020973 | protein_coding_gene | 1.9 | 3.744 | 3.4E-05 |
| *T12A2.6* | WBGene00020440 | protein_coding_gene | 1.9 | 3.739 | 2.8E-02 |
| *fbxc-41* | WBGene00016304 | pseudogene | 1.9 | 3.736 | 2.9E-02 |
| *srz-99* | WBGene00012028 | protein_coding_gene | 1.9 | 3.736 | 4.0E-02 |
| *clec-40* | WBGene00011853 | protein_coding_gene | 1.9 | 3.735 | 6.3E-03 |
| *tts-2* | WBGene00006651 | ncRNA_gene | 1.9 | 3.724 | 1.3E-05 |
| *T22G5.1* | WBGene00011929 | protein_coding_gene | 1.9 | 3.723 | 6.2E-03 |
| *F49D11.6* | WBGene00018630 | protein_coding_gene | 1.9 | 3.718 | 1.6E-02 |
| *C35E7.12* | WBGene00271779 | protein_coding_gene | 1.9 | 3.717 | 4.1E-02 |
| *Y39G10AR.11* | WBGene00021469 | protein_coding_gene | 1.9 | 3.714 | 4.2E-37 |
| *C08G5.1* | WBGene00015614 | pseudogene | 1.9 | 3.714 | 7.2E-03 |
| *Y69H2.10* | WBGene00013485 | protein_coding_gene | 1.9 | 3.714 | 7.4E-03 |
| *Y82E9BR.17* | WBGene00022349 | protein_coding_gene | 1.9 | 3.710 | 4.6E-25 |
| *egl-43* | WBGene00001207 | protein_coding_gene | 1.9 | 3.704 | 2.1E-07 |
| *C47E12.13* | WBGene00008156 | protein_coding_gene | 1.9 | 3.703 | 4.9E-02 |
| *F36H1.11* | WBGene00009496 | protein_coding_gene | 1.9 | 3.695 | 2.6E-02 |
| *Y47H9C.12* | WBGene00012955 | protein_coding_gene | 1.9 | 3.694 | 2.8E-02 |
| *cutl-8* | WBGene00013960 | protein_coding_gene | 1.9 | 3.690 | 2.4E-02 |
| *trx-3* | WBGene00019717 | protein_coding_gene | 1.9 | 3.689 | 1.3E-06 |
| *dex-1* | WBGene00017028 | protein_coding_gene | 1.9 | 3.681 | 2.3E-12 |
| *pho-6* | WBGene00018696 | protein_coding_gene | 1.9 | 3.681 | 2.6E-05 |
| *linc-43* | WBGene00219579 | lincRNA_gene | 1.9 | 3.680 | 3.0E-02 |
| *F42G4.6* | WBGene00009643 | protein_coding_gene | 1.9 | 3.675 | 9.7E-04 |
| *rom-2* | WBGene00004401 | protein_coding_gene | 1.9 | 3.675 | 1.1E-03 |
| *comt-4* | WBGene00021491 | protein_coding_gene | 1.9 | 3.674 | 4.2E-02 |
| *cest-1.2* | WBGene00011364 | protein_coding_gene | 1.9 | 3.674 | 1.3E-13 |
| *F58E6.11* | WBGene00010258 | protein_coding_gene | 1.9 | 3.672 | 7.6E-03 |
| *F22B7.1* | WBGene00017692 | protein_coding_gene | 1.9 | 3.671 | 5.5E-06 |
| *T27A1.2* | WBGene00020834 | protein_coding_gene | 1.9 | 3.670 | 1.3E-07 |
| *ins-23* | WBGene00002106 | protein_coding_gene | 1.9 | 3.668 | 7.0E-03 |
| *T21E8.6* | WBGene00164971 | protein_coding_gene | 1.9 | 3.668 | 3.9E-02 |
| *F21H7.12* | WBGene00009033 | protein_coding_gene | 1.9 | 3.665 | 2.0E-02 |
| *C26E1.2* | WBGene00007748 | protein_coding_gene | 1.9 | 3.665 | 2.0E-14 |
| *W08G11.6* | WBGene00023429 | snoRNA_gene | 1.9 | 3.665 | 1.3E-02 |
| *Y71G12B.5* | WBGene00022145 | protein_coding_gene | 1.9 | 3.665 | 8.0E-03 |
| *ceh-31* | WBGene00000452 | protein_coding_gene | 1.9 | 3.664 | 6.4E-03 |
| *F16G10.9* | WBGene00017524 | protein_coding_gene | 1.9 | 3.663 | 4.7E-02 |
| *ceh-2* | WBGene00000429 | protein_coding_gene | 1.9 | 3.661 | 1.8E-02 |
| *F53F10.1* | WBGene00018763 | protein_coding_gene | 1.9 | 3.661 | 8.6E-08 |
| *chil-18* | WBGene00011161 | protein_coding_gene | 1.9 | 3.659 | 2.9E-02 |
| *osr-1* | WBGene00016329 | protein_coding_gene | 1.9 | 3.659 | 9.3E-03 |
| *C37A5.6* | WBGene00014700 | protein_coding_gene | 1.9 | 3.657 | 4.0E-02 |
| *tth-1* | WBGene00006649 | protein_coding_gene | 1.9 | 3.656 | 1.1E-18 |
| *pha-2* | WBGene00004011 | protein_coding_gene | 1.9 | 3.656 | 7.0E-03 |
| *grdn-1* | WBGene00013082 | protein_coding_gene | 1.9 | 3.655 | 1.5E-18 |
| *clec-3* | WBGene00016577 | protein_coding_gene | 1.9 | 3.654 | 4.7E-04 |
| *D2030.2* | WBGene00008412 | protein_coding_gene | 1.9 | 3.647 | 5.2E-45 |
| *F23A7.1* | WBGene00009067 | protein_coding_gene | 1.9 | 3.647 | 3.2E-03 |
| *R10D12.10* | WBGene00011191 | protein_coding_gene | 1.9 | 3.645 | 1.9E-05 |
| *cdh-5* | WBGene00000397 | protein_coding_gene | 1.9 | 3.645 | 9.5E-05 |
| *slc-28.1* | WBGene00017867 | protein_coding_gene | 1.9 | 3.640 | 1.6E-05 |
| *clec-33* | WBGene00009859 | protein_coding_gene | 1.9 | 3.637 | 2.2E-02 |
| *ceh-27* | WBGene00000449 | protein_coding_gene | 1.9 | 3.635 | 2.2E-09 |
| *hlh-32* | WBGene00013665 | protein_coding_gene | 1.9 | 3.634 | 4.6E-02 |
| *F20A1.6* | WBGene00017618 | protein_coding_gene | 1.9 | 3.633 | 3.1E-09 |
| *Y15E3A.5* | WBGene00012444 | protein_coding_gene | 1.9 | 3.625 | 3.7E-12 |

| | | | | | |
|---|---|---|---|---|---|
| *F46F11.12* | WBGene00045082 | snoRNA_gene | 1.9 | 3.615 | 3.7E-02 |
| *magu-3* | WBGene00016841 | protein_coding_gene | 1.9 | 3.612 | 3.1E-04 |
| *T21E12.5* | WBGene00020656 | protein_coding_gene | 1.9 | 3.611 | 1.5E-02 |
| *fbxb-88* | WBGene00016359 | protein_coding_gene | 1.9 | 3.608 | 2.6E-08 |
| *dod-23* | WBGene00009898 | protein_coding_gene | 1.9 | 3.606 | 2.1E-13 |
| *thn-2* | WBGene00009215 | protein_coding_gene | 1.8 | 3.605 | 4.5E-11 |
| *fbxa-182* | WBGene00018353 | protein_coding_gene | 1.8 | 3.603 | 2.9E-07 |
| *Y75B12B.13* | WBGene00045402 | protein_coding_gene | 1.8 | 3.603 | 3.3E-02 |
| *maph-9* | WBGene00016397 | protein_coding_gene | 1.8 | 3.600 | 9.7E-03 |
| *mfb-1* | WBGene00008439 | protein_coding_gene | 1.8 | 3.599 | 4.7E-13 |
| *F22B7.9* | WBGene00017698 | protein_coding_gene | 1.8 | 3.599 | 1.2E-11 |
| *F46F5.4* | WBGene00018495 | pseudogene | 1.8 | 3.598 | 7.6E-03 |
| *F19H6.5* | WBGene00008960 | protein_coding_gene | 1.8 | 3.597 | 3.2E-02 |
| *C07A4.3* | WBGene00007398 | protein_coding_gene | 1.8 | 3.591 | 4.5E-05 |
| *cyp-13A5* | WBGene00011672 | protein_coding_gene | 1.8 | 3.591 | 7.5E-11 |
| *Y52D5A.2* | WBGene00021793 | protein_coding_gene | 1.8 | 3.586 | 1.1E-02 |
| *B0024.4* | WBGene00007097 | protein_coding_gene | 1.8 | 3.581 | 1.2E-04 |
| *W05H9.1* | WBGene00021048 | protein_coding_gene | 1.8 | 3.579 | 1.1E-45 |
| *ZK822.1* | WBGene00014089 | protein_coding_gene | 1.8 | 3.579 | 8.1E-05 |
| *lhfp-4* | WBGene00009155 | protein_coding_gene | 1.8 | 3.578 | 1.3E-02 |
| *spp-8* | WBGene00004993 | protein_coding_gene | 1.8 | 3.576 | 3.3E-12 |
| *ugt-66* | WBGene00016013 | protein_coding_gene | 1.8 | 3.576 | 1.6E-06 |
| *xtr-2* | WBGene00006966 | protein_coding_gene | 1.8 | 3.566 | 3.0E-02 |
| *marc-4* | WBGene00016903 | protein_coding_gene | 1.8 | 3.563 | 4.0E-04 |
| *K07A1.15* | WBGene00010619 | protein_coding_gene | 1.8 | 3.563 | 1.8E-09 |
| *msrp-1* | WBGene00017978 | protein_coding_gene | 1.8 | 3.557 | 4.8E-06 |
| *Y53F4B.8* | WBGene00013155 | pseudogene | 1.8 | 3.556 | 1.5E-03 |
| *B0478.3* | WBGene00015202 | protein_coding_gene | 1.8 | 3.555 | 1.1E-02 |
| *folr-1* | WBGene00007648 | protein_coding_gene | 1.8 | 3.555 | 4.8E-02 |
| *ccep-290* | WBGene00021643 | protein_coding_gene | 1.8 | 3.555 | 1.8E-04 |
| *F48E3.6* | WBGene00018606 | protein_coding_gene | 1.8 | 3.551 | 5.5E-12 |
| *F35G2.5* | WBGene00009438 | protein_coding_gene | 1.8 | 3.543 | 4.0E-02 |
| *C29F7.2* | WBGene00007811 | protein_coding_gene | 1.8 | 3.535 | 7.1E-25 |
| *fbxb-78* | WBGene00017116 | protein_coding_gene | 1.8 | 3.534 | 5.2E-04 |
| *F13G3.3* | WBGene00008763 | protein_coding_gene | 1.8 | 3.532 | 7.6E-03 |
| *ttr-17* | WBGene00012383 | protein_coding_gene | 1.8 | 3.531 | 1.6E-22 |
| *F21A9.2* | WBGene00017651 | protein_coding_gene | 1.8 | 3.531 | 5.0E-06 |
| *W03F8.6* | WBGene00020995 | protein_coding_gene | 1.8 | 3.531 | 2.1E-06 |
| *K07A1.13* | WBGene00010617 | protein_coding_gene | 1.8 | 3.528 | 4.7E-26 |
| *clc-2* | WBGene00000523 | protein_coding_gene | 1.8 | 3.524 | 1.2E-05 |
| *clec-74* | WBGene00021585 | protein_coding_gene | 1.8 | 3.522 | 3.3E-03 |
| *oac-31* | WBGene00018707 | protein_coding_gene | 1.8 | 3.520 | 4.8E-03 |
| *F35E12.9* | WBGene00009433 | protein_coding_gene | 1.8 | 3.519 | 1.2E-06 |
| *fbxa-168* | WBGene00007838 | pseudogene | 1.8 | 3.519 | 1.5E-07 |
| *nep-24* | WBGene00013786 | protein_coding_gene | 1.8 | 3.518 | 7.6E-03 |
| *C53A3.1* | WBGene00016891 | protein_coding_gene | 1.8 | 3.515 | 4.2E-02 |
| *fbxa-127* | WBGene00012027 | protein_coding_gene | 1.8 | 3.507 | 3.0E-02 |
| *bbs-1* | WBGene00000241 | protein_coding_gene | 1.8 | 3.498 | 2.2E-02 |
| *srh-149* | WBGene00005365 | protein_coding_gene | 1.8 | 3.495 | 3.8E-02 |
| *C38C3.4* | WBGene00016513 | protein_coding_gene | 1.8 | 3.492 | 1.4E-11 |
| *W09G10.9* | WBGene00271830 | protein_coding_gene | 1.8 | 3.485 | 3.0E-02 |
| *mrpr-1* | WBGene00045095 | ncRNA_gene | 1.8 | 3.482 | 1.8E-02 |
| *C24A8.5* | WBGene00016039 | protein_coding_gene | 1.8 | 3.482 | 2.6E-02 |
| *rrf-2* | WBGene00004509 | protein_coding_gene | 1.8 | 3.475 | 1.3E-17 |
| *Y95B8A.6* | WBGene00022386 | protein_coding_gene | 1.8 | 3.472 | 9.5E-12 |
| *C08B6.4* | WBGene00007425 | protein_coding_gene | 1.8 | 3.472 | 2.4E-02 |
| *M60.4* | WBGene00019780 | protein_coding_gene | 1.8 | 3.470 | 7.5E-20 |
| *T19C4.5* | WBGene00011839 | protein_coding_gene | 1.8 | 3.469 | 1.3E-07 |
| *ugt-4* | WBGene00013905 | protein_coding_gene | 1.8 | 3.467 | 7.8E-07 |
| *T14E8.4* | WBGene00043981 | protein_coding_gene | 1.8 | 3.465 | 6.7E-03 |

| | | | | | |
|---|---|---|---|---|---|
| *ceh-43* | WBGene00000463 | protein_coding_gene | 1.8 | 3.458 | 1.2E-18 |
| *F13C5.1* | WBGene00017422 | protein_coding_gene | 1.8 | 3.456 | 6.0E-09 |
| *Y6E2A.5* | WBGene00012399 | protein_coding_gene | 1.8 | 3.451 | 1.2E-03 |
| *Y45F10D.6* | WBGene00012886 | protein_coding_gene | 1.8 | 3.451 | 6.9E-05 |
| *clec-54* | WBGene00008596 | protein_coding_gene | 1.8 | 3.449 | 1.6E-16 |
| *F26A10.2* | WBGene00017814 | protein_coding_gene | 1.8 | 3.447 | 3.9E-07 |
| *F13A7.12* | WBGene00008727 | protein_coding_gene | 1.8 | 3.447 | 3.5E-02 |
| *Y64G10A.10* | WBGene00045144 | protein_coding_gene | 1.8 | 3.444 | 5.0E-02 |
| *F42C5.3* | WBGene00018345 | protein_coding_gene | 1.8 | 3.444 | 1.2E-02 |
| *glt-5* | WBGene00001623 | protein_coding_gene | 1.8 | 3.444 | 3.8E-08 |
| *math-37* | WBGene00020080 | protein_coding_gene | 1.8 | 3.440 | 9.1E-04 |
| *R04B3.1* | WBGene00019866 | protein_coding_gene | 1.8 | 3.438 | 4.2E-02 |
| *C05A9.2* | WBGene00007317 | protein_coding_gene | 1.8 | 3.435 | 2.7E-10 |
| *F41C3.1* | WBGene00018267 | protein_coding_gene | 1.8 | 3.434 | 2.1E-02 |
| *Y87G2A.25* | WBGene00219288 | protein_coding_gene | 1.8 | 3.433 | 1.5E-03 |
| *F48E3.8* | WBGene00018607 | protein_coding_gene | 1.8 | 3.432 | 3.8E-07 |
| *twk-7* | WBGene00006662 | protein_coding_gene | 1.8 | 3.431 | 7.7E-09 |
| *F10A3.1* | WBGene00008631 | protein_coding_gene | 1.8 | 3.428 | 3.4E-02 |
| *ptr-15* | WBGene00004229 | protein_coding_gene | 1.8 | 3.426 | 1.3E-03 |
| *R11D1.12* | WBGene00045336 | protein_coding_gene | 1.8 | 3.426 | 2.5E-02 |
| *W07B8.4* | WBGene00021072 | protein_coding_gene | 1.8 | 3.416 | 1.5E-03 |
| *C02B10.6* | WBGene00015331 | protein_coding_gene | 1.8 | 3.416 | 4.1E-02 |
| *Y17D7C.4* | WBGene00050903 | pseudogene | 1.8 | 3.413 | 1.9E-02 |
| *C50F4.9* | WBGene00008234 | protein_coding_gene | 1.8 | 3.412 | 2.5E-04 |
| *Y102A11A.5* | WBGene00022415 | protein_coding_gene | 1.8 | 3.411 | 2.7E-03 |
| *C24H12.1* | WBGene00016071 | protein_coding_gene | 1.8 | 3.407 | 6.7E-06 |
| *R05G9R.1* | WBGene00019906 | protein_coding_gene | 1.8 | 3.407 | 4.6E-03 |
| *F20D6.8* | WBGene00017637 | protein_coding_gene | 1.8 | 3.405 | 2.6E-03 |
| *C36F7.5* | WBGene00007986 | protein_coding_gene | 1.8 | 3.399 | 1.3E-04 |
| *Y51H4A.1* | WBGene00013098 | protein_coding_gene | 1.8 | 3.398 | 1.1E-02 |
| *nhr-42* | WBGene00003632 | protein_coding_gene | 1.8 | 3.396 | 1.0E-11 |
| *bus-8* | WBGene00044623 | protein_coding_gene | 1.8 | 3.395 | 2.2E-03 |
| *C04C11.25* | WBGene00284855 | protein_coding_gene | 1.8 | 3.387 | 7.3E-04 |
| *ham-1* | WBGene00001820 | protein_coding_gene | 1.8 | 3.386 | 2.4E-16 |
| *cutl-11* | WBGene00011443 | protein_coding_gene | 1.8 | 3.385 | 8.6E-03 |
| *unc-129* | WBGene00006852 | protein_coding_gene | 1.8 | 3.381 | 4.6E-08 |
| *glc-1* | WBGene00001591 | protein_coding_gene | 1.8 | 3.381 | 8.2E-16 |
| *T10H9.8* | WBGene00044779 | protein_coding_gene | 1.8 | 3.380 | 2.5E-09 |
| *F53C3.6* | WBGene00018750 | protein_coding_gene | 1.8 | 3.376 | 2.6E-02 |
| *ttr-50* | WBGene00010234 | protein_coding_gene | 1.8 | 3.374 | 2.3E-40 |
| *Y37H2A.7* | WBGene00012567 | protein_coding_gene | 1.8 | 3.373 | 2.1E-07 |
| *Y43F8C.3* | WBGene00012825 | protein_coding_gene | 1.8 | 3.372 | 2.5E-16 |
| *lam-3* | WBGene00002248 | protein_coding_gene | 1.8 | 3.371 | 3.1E-16 |
| *gad-3* | WBGene00015057 | protein_coding_gene | 1.8 | 3.371 | 5.7E-22 |
| *mnp-1* | WBGene00007139 | protein_coding_gene | 1.7 | 3.363 | 8.2E-25 |
| *clec-44* | WBGene00019928 | protein_coding_gene | 1.7 | 3.361 | 2.6E-02 |
| *C38C3.8* | WBGene00016516 | pseudogene | 1.7 | 3.361 | 6.3E-03 |
| *Y71H2AM.15* | WBGene00022180 | protein_coding_gene | 1.7 | 3.360 | 1.4E-04 |
| *T04F8.9* | WBGene00011447 | protein_coding_gene | 1.7 | 3.355 | 5.0E-03 |
| *ZK593.2* | WBGene00014002 | protein_coding_gene | 1.7 | 3.353 | 2.4E-05 |
| *C48B4.12* | WBGene00008175 | protein_coding_gene | 1.7 | 3.348 | 5.6E-03 |
| *Y13C8A.1* | WBGene00021188 | protein_coding_gene | 1.7 | 3.345 | 2.1E-04 |
| *Y71F9AL.25* | WBGene00304792 | protein_coding_gene | 1.7 | 3.344 | 3.6E-04 |
| *F13E9.12* | WBGene00008761 | protein_coding_gene | 1.7 | 3.338 | 3.4E-02 |
| *F33H2.8* | WBGene00009371 | protein_coding_gene | 1.7 | 3.337 | 3.5E-03 |
| *K10G6.9* | WBGene00206374 | protein_coding_gene | 1.7 | 3.325 | 1.3E-02 |
| *srh-2* | WBGene00005228 | protein_coding_gene | 1.7 | 3.323 | 1.8E-02 |
| *bus-18* | WBGene00044631 | protein_coding_gene | 1.7 | 3.323 | 1.1E-02 |
| *C12D5.3* | WBGene00015706 | protein_coding_gene | 1.7 | 3.320 | 1.5E-02 |
| *T25B9.3* | WBGene00012009 | protein_coding_gene | 1.7 | 3.320 | 1.2E-02 |

| | | | | | |
|---|---|---|---|---|---|
| *C25F6.7* | WBGene00016106 | protein_coding_gene | 1.7 | 3.319 | 2.6E-04 |
| *jbts-14* | WBGene00018727 | protein_coding_gene | 1.7 | 3.319 | 2.2E-02 |
| *clec-107* | WBGene00012502 | protein_coding_gene | 1.7 | 3.317 | 4.5E-02 |
| *AC3.9* | WBGene00044022 | pseudogene | 1.7 | 3.315 | 4.3E-02 |
| *cpr-1* | WBGene00000781 | protein_coding_gene | 1.7 | 3.311 | 8.4E-27 |
| *Y73E7A.8* | WBGene00022274 | protein_coding_gene | 1.7 | 3.309 | 4.3E-03 |
| *C07A9.5* | WBGene00007401 | protein_coding_gene | 1.7 | 3.305 | 4.7E-02 |
| *B0284.3* | WBGene00007133 | protein_coding_gene | 1.7 | 3.305 | 8.6E-05 |
| *fbxa-69* | WBGene00010018 | protein_coding_gene | 1.7 | 3.304 | 2.9E-12 |
| *nlp-48* | WBGene00008002 | protein_coding_gene | 1.7 | 3.304 | 1.4E-02 |
| *Y43F8B.2* | WBGene00012813 | protein_coding_gene | 1.7 | 3.304 | 9.8E-27 |
| *pitr-3* | WBGene00015055 | protein_coding_gene | 1.7 | 3.302 | 5.2E-06 |
| *fkh-2* | WBGene00001434 | protein_coding_gene | 1.7 | 3.300 | 1.8E-05 |
| *fbxa-165* | WBGene00015600 | protein_coding_gene | 1.7 | 3.297 | 4.8E-02 |
| *C40H1.9* | WBGene00008044 | protein_coding_gene | 1.7 | 3.296 | 1.4E-02 |
| *Y43C5B.2* | WBGene00012785 | protein_coding_gene | 1.7 | 3.294 | 5.1E-03 |
| *Y75B8A.28* | WBGene00013559 | protein_coding_gene | 1.7 | 3.293 | 1.8E-05 |
| *W01C9.1* | WBGene00012175 | protein_coding_gene | 1.7 | 3.292 | 2.3E-02 |
| *trf-1* | WBGene00006612 | protein_coding_gene | 1.7 | 3.290 | 4.7E-02 |
| *cutl-29* | WBGene00017421 | protein_coding_gene | 1.7 | 3.290 | 1.8E-02 |
| *C47D12.4* | WBGene00008138 | protein_coding_gene | 1.7 | 3.288 | 1.0E-05 |
| *F13E9.13* | WBGene00044797 | protein_coding_gene | 1.7 | 3.287 | 5.2E-06 |
| *clec-196* | WBGene00009156 | protein_coding_gene | 1.7 | 3.287 | 2.1E-15 |
| *C14E2.4* | WBGene00015773 | protein_coding_gene | 1.7 | 3.285 | 4.9E-02 |
| *F18C5.5* | WBGene00017560 | protein_coding_gene | 1.7 | 3.285 | 1.0E-02 |
| *kal-1* | WBGene00002181 | protein_coding_gene | 1.7 | 3.283 | 1.0E-10 |
| *W01B11.6* | WBGene00020917 | protein_coding_gene | 1.7 | 3.283 | 1.3E-07 |
| *Y62F5A.10* | WBGene00013388 | protein_coding_gene | 1.7 | 3.282 | 1.6E-04 |
| *osm-6* | WBGene00003886 | protein_coding_gene | 1.7 | 3.281 | 1.1E-02 |
| *fbxc-53* | WBGene00016711 | protein_coding_gene | 1.7 | 3.281 | 3.7E-02 |
| *her-1* | WBGene00001842 | protein_coding_gene | 1.7 | 3.280 | 2.1E-04 |
| *F57C12.6* | WBGene00194687 | protein_coding_gene | 1.7 | 3.279 | 1.9E-03 |
| *F52D1.2* | WBGene00018683 | protein_coding_gene | 1.7 | 3.278 | 7.1E-04 |
| *C26B9.6* | WBGene00016135 | protein_coding_gene | 1.7 | 3.274 | 5.6E-06 |
| *B0391.14* | WBGene00206462 | protein_coding_gene | 1.7 | 3.274 | 3.1E-02 |
| *T08A9.13* | WBGene00044442 | protein_coding_gene | 1.7 | 3.268 | 3.0E-02 |
| *Y1A5A.1* | WBGene00012379 | protein_coding_gene | 1.7 | 3.265 | 1.3E-03 |
| *ZK546.7* | WBGene00022763 | pseudogene | 1.7 | 3.263 | 9.3E-03 |
| *D1054.18* | WBGene00077783 | protein_coding_gene | 1.7 | 3.261 | 3.8E-02 |
| *Y53H1B.4* | WBGene00013182 | protein_coding_gene | 1.7 | 3.260 | 3.6E-02 |
| *F07H5.4* | WBGene00014308 | pseudogene | 1.7 | 3.255 | 6.6E-21 |
| *tab-1* | WBGene00006380 | protein_coding_gene | 1.7 | 3.255 | 1.5E-02 |
| *ceh-32* | WBGene00000453 | protein_coding_gene | 1.7 | 3.252 | 3.2E-16 |
| *sptf-1* | WBGene00009743 | protein_coding_gene | 1.7 | 3.250 | 8.9E-08 |
| *Y51H7C.12* | WBGene00021790 | protein_coding_gene | 1.7 | 3.248 | 2.6E-02 |
| *C26B9.3* | WBGene00016133 | protein_coding_gene | 1.7 | 3.247 | 2.4E-04 |
| *C11G10.1* | WBGene00007525 | protein_coding_gene | 1.7 | 3.246 | 9.3E-05 |
| *fkh-7* | WBGene00001439 | protein_coding_gene | 1.7 | 3.243 | 1.2E-20 |
| *Y71G12B.31* | WBGene00044348 | protein_coding_gene | 1.7 | 3.241 | 3.6E-03 |
| *C13C4.6* | WBGene00007550 | protein_coding_gene | 1.7 | 3.231 | 3.0E-05 |
| *F17C8.3* | WBGene00008910 | protein_coding_gene | 1.7 | 3.229 | 2.2E-05 |
| *Y53F4B.20* | WBGene00013166 | protein_coding_gene | 1.7 | 3.226 | 1.9E-02 |
| *F13H6.4* | WBGene00017432 | pseudogene | 1.7 | 3.225 | 2.1E-03 |
| *unk-1* | WBGene00016407 | protein_coding_gene | 1.7 | 3.222 | 3.5E-21 |
| *T18D3.7* | WBGene00011824 | protein_coding_gene | 1.7 | 3.215 | 6.0E-11 |
| *F13H8.1* | WBGene00017434 | protein_coding_gene | 1.7 | 3.213 | 1.1E-04 |
| *srpr-1.2* | WBGene00045168 | ncRNA_gene | 1.7 | 3.208 | 2.7E-02 |
| *F54B8.4* | WBGene00010019 | protein_coding_gene | 1.7 | 3.208 | 7.5E-06 |
| *Y77E11A.14* | WBGene00022314 | protein_coding_gene | 1.7 | 3.202 | 4.4E-03 |
| *F33D4.6* | WBGene00017998 | protein_coding_gene | 1.7 | 3.201 | 3.9E-04 |

| | | | | | |
|---|---|---|---|---|---|
| *slc-36.3* | WBGene00012629 | protein_coding_gene | 1.7 | 3.200 | 6.4E-04 |
| *frpr-11* | WBGene00019444 | protein_coding_gene | 1.7 | 3.194 | 1.1E-06 |
| *F08G2.5* | WBGene00008577 | protein_coding_gene | 1.7 | 3.194 | 8.4E-03 |
| *ZK930.5* | WBGene00014155 | protein_coding_gene | 1.7 | 3.193 | 1.0E-02 |
| *K06A4.8* | WBGene00044015 | pseudogene | 1.7 | 3.191 | 3.8E-03 |
| *Y71F9AM.9* | WBGene00271781 | lincRNA_gene | 1.7 | 3.190 | 3.1E-02 |
| *tpra-1* | WBGene00008366 | protein_coding_gene | 1.7 | 3.188 | 2.5E-14 |
| *C53C11.5* | WBGene00044770 | protein_coding_gene | 1.7 | 3.185 | 1.4E-08 |
| *F16B3.2* | WBGene00017502 | protein_coding_gene | 1.7 | 3.185 | 3.6E-02 |
| *kpc-1* | WBGene00002232 | protein_coding_gene | 1.7 | 3.183 | 9.7E-30 |
| *fbxa-219* | WBGene00044403 | protein_coding_gene | 1.7 | 3.182 | 2.5E-05 |
| *asp-14* | WBGene00019619 | protein_coding_gene | 1.7 | 3.182 | 7.4E-14 |
| *F44F1.6* | WBGene00009698 | protein_coding_gene | 1.7 | 3.177 | 3.0E-10 |
| *F41E6.5* | WBGene00018286 | protein_coding_gene | 1.7 | 3.176 | 4.8E-09 |
| *nhr-18* | WBGene00003617 | protein_coding_gene | 1.7 | 3.172 | 8.1E-09 |
| *F45D3.3* | WBGene00009723 | protein_coding_gene | 1.7 | 3.171 | 1.1E-08 |
| *fut-2* | WBGene00001506 | protein_coding_gene | 1.7 | 3.166 | 1.3E-02 |
| *nphp-4* | WBGene00011261 | protein_coding_gene | 1.7 | 3.155 | 2.6E-03 |
| *clec-55* | WBGene00008597 | protein_coding_gene | 1.7 | 3.154 | 9.6E-04 |
| *lipl-6* | WBGene00021963 | protein_coding_gene | 1.7 | 3.150 | 6.5E-03 |
| *pha-4* | WBGene00004013 | protein_coding_gene | 1.7 | 3.146 | 1.6E-14 |
| *qua-1* | WBGene00004264 | protein_coding_gene | 1.7 | 3.146 | 2.3E-04 |
| *T09B4.5* | WBGene00020379 | protein_coding_gene | 1.7 | 3.144 | 4.5E-35 |
| *C31H2.3* | WBGene00016293 | protein_coding_gene | 1.7 | 3.144 | 3.9E-02 |
| *ins-4* | WBGene00002087 | protein_coding_gene | 1.7 | 3.142 | 1.3E-03 |
| *jmjd-3.2* | WBGene00009089 | protein_coding_gene | 1.7 | 3.140 | 3.5E-08 |
| *F46B6.9* | WBGene00009774 | protein_coding_gene | 1.6 | 3.129 | 7.6E-03 |
| *Y105E8A.34* | WBGene00077689 | pseudogene | 1.6 | 3.124 | 4.5E-02 |
| *Y54G2A.52* | WBGene00045397 | protein_coding_gene | 1.6 | 3.122 | 8.2E-10 |
| *C06B3.7* | WBGene00007366 | protein_coding_gene | 1.6 | 3.115 | 6.4E-03 |
| *daf-10* | WBGene00000906 | protein_coding_gene | 1.6 | 3.115 | 5.0E-03 |
| *fkh-8* | WBGene00001440 | protein_coding_gene | 1.6 | 3.114 | 1.9E-03 |
| *nep-14* | WBGene00018225 | pseudogene | 1.6 | 3.113 | 6.8E-06 |
| *T06G6.11* | WBGene00011550 | pseudogene | 1.6 | 3.110 | 3.0E-02 |
| *ZK669.3* | WBGene00014053 | protein_coding_gene | 1.6 | 3.106 | 1.8E-19 |
| *W08F4.7* | WBGene00021096 | protein_coding_gene | 1.6 | 3.106 | 1.3E-03 |
| *K07H8.11* | WBGene00044510 | protein_coding_gene | 1.6 | 3.103 | 1.6E-02 |
| *efl-3* | WBGene00009899 | protein_coding_gene | 1.6 | 3.103 | 1.1E-18 |
| *egl-15* | WBGene00001184 | protein_coding_gene | 1.6 | 3.099 | 1.8E-14 |
| *lec-2* | WBGene00002265 | protein_coding_gene | 1.6 | 3.097 | 3.0E-08 |
| *unc-39* | WBGene00006775 | protein_coding_gene | 1.6 | 3.096 | 2.0E-03 |
| *F20D6.10* | WBGene00017639 | protein_coding_gene | 1.6 | 3.090 | 3.2E-04 |
| *C50D2.1* | WBGene00016805 | protein_coding_gene | 1.6 | 3.089 | 1.9E-02 |
| *Y8A9A.2* | WBGene00021171 | protein_coding_gene | 1.6 | 3.089 | 9.3E-08 |
| *T28B8.1* | WBGene00012113 | protein_coding_gene | 1.6 | 3.088 | 1.5E-22 |
| *dao-6* | WBGene00000932 | protein_coding_gene | 1.6 | 3.082 | 4.0E-02 |
| *C17C3.3* | WBGene00015889 | protein_coding_gene | 1.6 | 3.081 | 3.5E-02 |
| *D2005.6* | WBGene00008401 | protein_coding_gene | 1.6 | 3.080 | 1.5E-04 |
| *C08E8.4* | WBGene00007440 | protein_coding_gene | 1.6 | 3.078 | 7.8E-03 |
| *Y6D1A.2* | WBGene00012396 | protein_coding_gene | 1.6 | 3.074 | 1.5E-02 |
| *clec-7* | WBGene00017364 | protein_coding_gene | 1.6 | 3.074 | 3.0E-10 |
| *F38B6.6* | WBGene00018175 | protein_coding_gene | 1.6 | 3.071 | 4.8E-04 |
| *B0393.4* | WBGene00007169 | protein_coding_gene | 1.6 | 3.068 | 4.1E-04 |
| *C13C4.7* | WBGene00007551 | protein_coding_gene | 1.6 | 3.067 | 5.9E-04 |
| *C05E11.3* | WBGene00015494 | protein_coding_gene | 1.6 | 3.067 | 1.4E-06 |
| *agr-1* | WBGene00018304 | protein_coding_gene | 1.6 | 3.067 | 7.1E-06 |
| *T16A1.10* | WBGene00269431 | protein_coding_gene | 1.6 | 3.065 | 2.9E-02 |
| *F52E1.5* | WBGene00018694 | protein_coding_gene | 1.6 | 3.064 | 3.2E-05 |
| *C16H3.3* | WBGene00015868 | protein_coding_gene | 1.6 | 3.059 | 4.3E-19 |
| *T01B6.1* | WBGene00020134 | protein_coding_gene | 1.6 | 3.057 | 1.3E-03 |

| | | | | | |
|---|---|---|---|---|---|
| *flp-32* | WBGene00010982 | protein_coding_gene | 1.6 | 3.056 | 2.8E-05 |
| *dyf-5* | WBGene00001121 | protein_coding_gene | 1.6 | 3.054 | 1.6E-02 |
| *Y116A8C.40* | WBGene00013812 | protein_coding_gene | 1.6 | 3.053 | 4.5E-02 |
| *T24C4.4* | WBGene00020760 | protein_coding_gene | 1.6 | 3.052 | 5.8E-03 |
| *grh-1* | WBGene00001707 | protein_coding_gene | 1.6 | 3.051 | 1.3E-05 |
| *C33D9.5* | WBGene00007899 | pseudogene | 1.6 | 3.050 | 9.1E-06 |
| *K09F5.6* | WBGene00044698 | protein_coding_gene | 1.6 | 3.050 | 1.3E-07 |
| *nhr-178* | WBGene00017510 | protein_coding_gene | 1.6 | 3.050 | 3.5E-02 |
| *nhr-91* | WBGene00003681 | protein_coding_gene | 1.6 | 3.049 | 2.8E-06 |
| *hmit-1.3* | WBGene00010811 | protein_coding_gene | 1.6 | 3.048 | 1.7E-13 |
| *C28G1.6* | WBGene00016191 | protein_coding_gene | 1.6 | 3.042 | 2.3E-05 |
| *cri-1* | WBGene00010614 | protein_coding_gene | 1.6 | 3.037 | 6.3E-03 |
| *C55A1.6* | WBGene00008323 | protein_coding_gene | 1.6 | 3.036 | 4.1E-02 |
| *math-48* | WBGene00022689 | protein_coding_gene | 1.6 | 3.035 | 2.2E-02 |
| *fbxc-24* | WBGene00019927 | protein_coding_gene | 1.6 | 3.032 | 1.3E-03 |
| *ZK896.4* | WBGene00014135 | protein_coding_gene | 1.6 | 3.031 | 3.2E-03 |
| *eya-1* | WBGene00001377 | protein_coding_gene | 1.6 | 3.030 | 3.5E-11 |
| *M01H9.10* | WBGene00271788 | protein_coding_gene | 1.6 | 3.030 | 1.4E-02 |
| *nspc-10* | WBGene00009640 | protein_coding_gene | 1.6 | 3.021 | 1.1E-11 |
| *ZK6.8* | WBGene00022643 | protein_coding_gene | 1.6 | 3.018 | 6.6E-04 |
| *cbp-2* | WBGene00010545 | protein_coding_gene | 1.6 | 3.014 | 3.5E-09 |
| *skr-19* | WBGene00004825 | protein_coding_gene | 1.6 | 3.014 | 1.8E-11 |
| *eyg-1* | WBGene00013147 | protein_coding_gene | 1.6 | 3.013 | 2.7E-02 |
| *F01D5.6* | WBGene00008496 | protein_coding_gene | 1.6 | 3.011 | 1.1E-02 |
| *fbxb-66* | WBGene00044066 | protein_coding_gene | 1.6 | 3.011 | 2.4E-06 |
| *npr-20* | WBGene00011578 | protein_coding_gene | 1.6 | 3.010 | 5.1E-05 |
| *mls-2* | WBGene00003377 | protein_coding_gene | 1.6 | 3.007 | 2.1E-03 |
| *cdh-9* | WBGene00000401 | protein_coding_gene | 1.6 | 3.005 | 4.2E-03 |
| *K08B5.2* | WBGene00019519 | protein_coding_gene | 1.6 | 3.002 | 1.0E-05 |
| *sym-2* | WBGene00006367 | protein_coding_gene | 1.6 | 2.999 | 2.1E-07 |
| *fbxb-60* | WBGene00010104 | protein_coding_gene | 1.6 | 2.996 | 2.9E-02 |
| *pat-9* | WBGene00003933 | protein_coding_gene | 1.6 | 2.993 | 1.1E-09 |
| *hhat-2* | WBGene00013312 | protein_coding_gene | 1.6 | 2.992 | 1.9E-03 |
| *fbxb-69* | WBGene00008613 | protein_coding_gene | 1.6 | 2.991 | 2.4E-02 |
| *lys-10* | WBGene00003099 | protein_coding_gene | 1.6 | 2.989 | 2.1E-02 |
| *C31H5.4* | WBGene00007855 | protein_coding_gene | 1.6 | 2.988 | 3.1E-17 |
| *cyp-13A3* | WBGene00011675 | protein_coding_gene | 1.6 | 2.984 | 7.3E-03 |
| *Y40A1A.6* | WBGene00304815 | ncRNA_gene | 1.6 | 2.982 | 4.7E-15 |
| *F53A2.1* | WBGene00009949 | protein_coding_gene | 1.6 | 2.981 | 4.3E-03 |
| *math-45* | WBGene00021772 | protein_coding_gene | 1.6 | 2.979 | 1.4E-03 |
| *F25B3.5* | WBGene00009102 | protein_coding_gene | 1.6 | 2.978 | 7.6E-04 |
| *lron-1* | WBGene00008093 | protein_coding_gene | 1.6 | 2.977 | 3.5E-02 |
| *Y41G9A.10* | WBGene00044901 | protein_coding_gene | 1.6 | 2.976 | 1.3E-04 |
| *ttr-37* | WBGene00017667 | protein_coding_gene | 1.6 | 2.975 | 7.3E-11 |
| *hlh-6* | WBGene00001952 | protein_coding_gene | 1.6 | 2.971 | 2.7E-02 |
| *unc-75* | WBGene00006807 | protein_coding_gene | 1.6 | 2.970 | 5.6E-03 |
| *nlp-31* | WBGene00003769 | protein_coding_gene | 1.6 | 2.967 | 2.1E-07 |
| *cex-1* | WBGene00023407 | protein_coding_gene | 1.6 | 2.966 | 8.3E-11 |
| *Y75B8A.6* | WBGene00013543 | protein_coding_gene | 1.6 | 2.964 | 2.9E-15 |
| *mlt-9* | WBGene00008605 | protein_coding_gene | 1.6 | 2.963 | 2.0E-03 |
| *F02A9.1* | WBGene00008513 | protein_coding_gene | 1.6 | 2.962 | 1.3E-02 |
| *exc-13* | WBGene00015393 | protein_coding_gene | 1.6 | 2.962 | 3.6E-03 |
| *F40F9.10* | WBGene00009586 | protein_coding_gene | 1.6 | 2.960 | 2.1E-02 |
| *Y42A5A.3* | WBGene00012778 | protein_coding_gene | 1.6 | 2.958 | 1.7E-02 |
| *skr-12* | WBGene00004818 | protein_coding_gene | 1.6 | 2.958 | 1.3E-08 |
| *C11H1.7* | WBGene00007531 | protein_coding_gene | 1.6 | 2.957 | 3.7E-03 |
| *Y54G2A.11* | WBGene00021876 | protein_coding_gene | 1.6 | 2.954 | 9.0E-10 |
| *C06E7.4* | WBGene00015541 | protein_coding_gene | 1.6 | 2.947 | 3.5E-08 |
| *F56C4.4* | WBGene00044977 | protein_coding_gene | 1.6 | 2.946 | 4.9E-03 |
| *egl-38* | WBGene00001204 | protein_coding_gene | 1.6 | 2.945 | 1.2E-02 |

| | | | | | |
|---|---|---|---|---|---|
| *C46C11.4* | WBGene00044567 | protein_coding_gene | 1.6 | 2.941 | 8.2E-03 |
| *C17E4.10* | WBGene00007647 | protein_coding_gene | 1.6 | 2.940 | 1.8E-21 |
| *T19E7.6* | WBGene00044500 | protein_coding_gene | 1.6 | 2.940 | 1.3E-06 |
| *his-24* | WBGene00001898 | protein_coding_gene | 1.6 | 2.939 | 1.2E-39 |
| *igcm-3* | WBGene00020160 | protein_coding_gene | 1.6 | 2.935 | 7.4E-32 |
| *F52C9.5* | WBGene00018675 | protein_coding_gene | 1.6 | 2.932 | 5.0E-02 |
| *C42D4.1* | WBGene00016594 | protein_coding_gene | 1.6 | 2.932 | 2.0E-13 |
| *F59A1.13* | WBGene00010299 | protein_coding_gene | 1.6 | 2.929 | 5.1E-03 |
| *bkip-1* | WBGene00014569 | protein_coding_gene | 1.5 | 2.927 | 2.4E-02 |
| *che-2* | WBGene00000484 | protein_coding_gene | 1.5 | 2.923 | 4.0E-02 |
| *ugt-9* | WBGene00020587 | protein_coding_gene | 1.5 | 2.919 | 7.7E-09 |
| *dyf-11* | WBGene00001127 | protein_coding_gene | 1.5 | 2.919 | 1.1E-02 |
| *sox-2* | WBGene00004949 | protein_coding_gene | 1.5 | 2.917 | 3.7E-12 |
| *clec-36* | WBGene00044332 | protein_coding_gene | 1.5 | 2.913 | 1.2E-03 |
| *col-76* | WBGene00000652 | protein_coding_gene | 1.5 | 2.911 | 1.5E-06 |
| *Y64G10A.7* | WBGene00013416 | protein_coding_gene | 1.5 | 2.911 | 2.3E-05 |
| *ugt-5* | WBGene00013906 | protein_coding_gene | 1.5 | 2.904 | 2.1E-09 |
| *fbxb-57* | WBGene00010101 | protein_coding_gene | 1.5 | 2.902 | 2.6E-03 |
| *K12B6.9* | WBGene00023432 | protein_coding_gene | 1.5 | 2.901 | 2.7E-06 |
| *T22F3.8* | WBGene00020698 | protein_coding_gene | 1.5 | 2.900 | 4.8E-02 |
| *gcy-22* | WBGene00001547 | protein_coding_gene | 1.5 | 2.897 | 2.1E-02 |
| *col-99* | WBGene00000674 | protein_coding_gene | 1.5 | 2.897 | 1.2E-06 |
| *moe-3* | WBGene00003388 | protein_coding_gene | 1.5 | 2.897 | 2.0E-08 |
| *msp-55* | WBGene00003448 | protein_coding_gene | 1.5 | 2.896 | 2.2E-02 |
| *ncam-1* | WBGene00017184 | protein_coding_gene | 1.5 | 2.896 | 9.3E-12 |
| *F08H9.3* | WBGene00008591 | protein_coding_gene | 1.5 | 2.894 | 1.8E-02 |
| *C17E4.2* | WBGene00007642 | protein_coding_gene | 1.5 | 2.894 | 5.7E-10 |
| *T16G1.6* | WBGene00011800 | protein_coding_gene | 1.5 | 2.892 | 2.9E-07 |
| *W04C9.8* | WBGene00235387 | ncRNA_gene | 1.5 | 2.892 | 2.1E-02 |
| *Y11D7A.19* | WBGene00077756 | protein_coding_gene | 1.5 | 2.892 | 4.3E-02 |
| *pdf-2* | WBGene00018590 | protein_coding_gene | 1.5 | 2.892 | 6.3E-07 |
| *acc-3* | WBGene00018880 | protein_coding_gene | 1.5 | 2.890 | 3.4E-02 |
| *C07B5.4* | WBGene00007409 | protein_coding_gene | 1.5 | 2.889 | 6.6E-05 |
| *msp-57* | WBGene00003450 | protein_coding_gene | 1.5 | 2.889 | 3.6E-04 |
| *EEED8.15* | WBGene00017143 | protein_coding_gene | 1.5 | 2.889 | 8.7E-15 |
| *C40H1.2* | WBGene00008038 | protein_coding_gene | 1.5 | 2.888 | 1.1E-04 |
| *F13D11.3* | WBGene00017428 | protein_coding_gene | 1.5 | 2.888 | 3.1E-07 |
| *C49H3.12* | WBGene00016795 | protein_coding_gene | 1.5 | 2.888 | 1.4E-02 |
| *cpn-2* | WBGene00000778 | protein_coding_gene | 1.5 | 2.888 | 7.3E-05 |
| *nhr-25* | WBGene00003623 | protein_coding_gene | 1.5 | 2.884 | 1.2E-10 |
| *fbxc-48* | WBGene00015615 | protein_coding_gene | 1.5 | 2.883 | 2.9E-02 |
| *gstk-2* | WBGene00017054 | protein_coding_gene | 1.5 | 2.883 | 4.4E-04 |
| *C35D10.3* | WBGene00016441 | protein_coding_gene | 1.5 | 2.879 | 3.9E-02 |
| *cest-17* | WBGene00019477 | protein_coding_gene | 1.5 | 2.876 | 2.0E-08 |
| *F59B2.9* | WBGene00010311 | protein_coding_gene | 1.5 | 2.874 | 1.0E-02 |
| *B0391.8* | WBGene00014643 | pseudogene | 1.5 | 2.874 | 1.2E-06 |
| *F46H6.5* | WBGene00023460 | protein_coding_gene | 1.5 | 2.870 | 3.5E-02 |
| *F08A7.1* | WBGene00045013 | protein_coding_gene | 1.5 | 2.869 | 1.3E-05 |
| *sto-1* | WBGene00006063 | protein_coding_gene | 1.5 | 2.867 | 1.4E-15 |
| *M03A1.8* | WBGene00077490 | protein_coding_gene | 1.5 | 2.863 | 1.0E-02 |
| *che-11* | WBGene00000490 | protein_coding_gene | 1.5 | 2.862 | 1.1E-02 |
| *Y47G6A.26* | WBGene00021650 | protein_coding_gene | 1.5 | 2.862 | 1.3E-03 |
| *ins-37* | WBGene00002120 | protein_coding_gene | 1.5 | 2.861 | 2.3E-02 |
| *D1086.5* | WBGene00008392 | protein_coding_gene | 1.5 | 2.857 | 1.3E-07 |
| *nhr-112* | WBGene00003702 | protein_coding_gene | 1.5 | 2.857 | 9.6E-04 |
| *lys-6* | WBGene00003095 | protein_coding_gene | 1.5 | 2.857 | 8.4E-04 |
| *nlp-30* | WBGene00003768 | protein_coding_gene | 1.5 | 2.857 | 6.8E-04 |
| *C32H11.4* | WBGene00007867 | protein_coding_gene | 1.5 | 2.857 | 4.0E-04 |
| *zip-8* | WBGene00017755 | protein_coding_gene | 1.5 | 2.855 | 2.7E-14 |
| *unc-130* | WBGene00006853 | protein_coding_gene | 1.5 | 2.853 | 1.7E-06 |

| | | | | | |
|---|---|---|---|---|---|
| *C08B6.2* | WBGene00007423 | protein_coding_gene | 1.5 | 2.849 | 4.0E-02 |
| *M02D8.6* | WBGene00019732 | protein_coding_gene | 1.5 | 2.849 | 7.8E-07 |
| *F42C5.4* | WBGene00018346 | protein_coding_gene | 1.5 | 2.849 | 5.9E-08 |
| *C16B8.4* | WBGene00015824 | protein_coding_gene | 1.5 | 2.848 | 2.9E-06 |
| *linc-46* | WBGene00219748 | lincRNA_gene | 1.5 | 2.846 | 1.2E-02 |
| *F43E2.6* | WBGene00018394 | protein_coding_gene | 1.5 | 2.845 | 1.9E-03 |
| *vab-3* | WBGene00006870 | protein_coding_gene | 1.5 | 2.842 | 3.2E-07 |
| *nspd-9* | WBGene00017808 | protein_coding_gene | 1.5 | 2.840 | 2.0E-03 |
| *fbxb-18* | WBGene00019039 | protein_coding_gene | 1.5 | 2.840 | 7.5E-03 |
| *swip-10* | WBGene00018738 | protein_coding_gene | 1.5 | 2.840 | 4.2E-03 |
| *F31F7.1* | WBGene00017964 | protein_coding_gene | 1.5 | 2.837 | 8.9E-38 |
| *C35E7.4* | WBGene00016456 | protein_coding_gene | 1.5 | 2.833 | 6.8E-04 |
| *C09B9.4* | WBGene00015629 | protein_coding_gene | 1.5 | 2.826 | 1.7E-03 |
| *ZK757.2* | WBGene00014074 | protein_coding_gene | 1.5 | 2.826 | 1.5E-02 |
| *icl-1* | WBGene00001564 | protein_coding_gene | 1.5 | 2.823 | 2.1E-03 |
| *unc-120* | WBGene00006844 | protein_coding_gene | 1.5 | 2.823 | 2.0E-07 |
| *K12G11.6* | WBGene00010792 | protein_coding_gene | 1.5 | 2.822 | 4.5E-02 |
| *phlp-2* | WBGene00009647 | protein_coding_gene | 1.5 | 2.820 | 1.4E-02 |
| *gem-4* | WBGene00001577 | protein_coding_gene | 1.5 | 2.819 | 1.6E-11 |
| *pxd-1* | WBGene00007982 | protein_coding_gene | 1.5 | 2.817 | 2.2E-18 |
| *ceh-22* | WBGene00000445 | protein_coding_gene | 1.5 | 2.817 | 8.4E-03 |
| *Y67D8C.9* | WBGene00022073 | protein_coding_gene | 1.5 | 2.817 | 1.1E-08 |
| *F17C8.7* | WBGene00008912 | protein_coding_gene | 1.5 | 2.813 | 8.8E-04 |
| *hil-3* | WBGene00001854 | protein_coding_gene | 1.5 | 2.810 | 8.5E-27 |
| *igcm-4* | WBGene00022418 | protein_coding_gene | 1.5 | 2.810 | 2.2E-12 |
| *F45C12.17* | WBGene00269388 | protein_coding_gene | 1.5 | 2.809 | 1.9E-04 |
| *fbxb-19* | WBGene00019040 | protein_coding_gene | 1.5 | 2.809 | 1.0E-02 |
| *K10D3.6* | WBGene00010740 | protein_coding_gene | 1.5 | 2.808 | 1.5E-06 |
| *M01H9.4* | WBGene00019720 | protein_coding_gene | 1.5 | 2.808 | 6.5E-08 |
| *K04A8.20* | WBGene00235332 | protein_coding_gene | 1.5 | 2.808 | 1.0E-02 |
| *pqn-42* | WBGene00004129 | protein_coding_gene | 1.5 | 2.806 | 2.4E-03 |
| *Y71G12B.35* | WBGene00185098 | protein_coding_gene | 1.5 | 2.804 | 1.8E-04 |
| *cyp-42A1* | WBGene00013585 | protein_coding_gene | 1.5 | 2.804 | 7.9E-08 |
| *ngn-1* | WBGene00003595 | protein_coding_gene | 1.5 | 2.803 | 1.5E-03 |
| *pals-33* | WBGene00021081 | protein_coding_gene | 1.5 | 2.797 | 4.5E-02 |
| *fbxb-77* | WBGene00021204 | protein_coding_gene | 1.5 | 2.797 | 4.5E-02 |
| *W10G11.2* | WBGene00021135 | protein_coding_gene | 1.5 | 2.796 | 5.8E-03 |
| *F09F3.8* | WBGene00014732 | pseudogene | 1.5 | 2.790 | 3.7E-02 |
| *osm-5* | WBGene00003885 | protein_coding_gene | 1.5 | 2.790 | 1.9E-02 |
| *F46C8.3* | WBGene00018485 | protein_coding_gene | 1.5 | 2.790 | 2.3E-06 |
| *D1053.3* | WBGene00008368 | protein_coding_gene | 1.5 | 2.788 | 7.7E-04 |
| *C36C5.12* | WBGene00016481 | protein_coding_gene | 1.5 | 2.787 | 3.7E-03 |
| *T01D3.6* | WBGene00011330 | protein_coding_gene | 1.5 | 2.785 | 1.1E-09 |
| *ref-2* | WBGene00004335 | protein_coding_gene | 1.5 | 2.783 | 4.2E-04 |
| *F33E2.5* | WBGene00009361 | protein_coding_gene | 1.5 | 2.782 | 3.3E-13 |
| *plpr-1* | WBGene00011524 | protein_coding_gene | 1.5 | 2.781 | 4.9E-03 |
| *dep-1* | WBGene00009717 | protein_coding_gene | 1.5 | 2.781 | 1.9E-08 |
| *clec-266* | WBGene00016088 | protein_coding_gene | 1.5 | 2.780 | 6.0E-15 |
| *hmg-11* | WBGene00001976 | protein_coding_gene | 1.5 | 2.780 | 5.0E-12 |
| *C33D9.6* | WBGene00007900 | protein_coding_gene | 1.5 | 2.778 | 4.8E-06 |
| *dlc-5* | WBGene00043067 | protein_coding_gene | 1.5 | 2.778 | 1.4E-02 |
| *F57C12.2* | WBGene00019014 | protein_coding_gene | 1.5 | 2.777 | 4.2E-04 |
| *lgg-1* | WBGene00002980 | protein_coding_gene | 1.5 | 2.777 | 3.5E-19 |
| *cyp-34A2* | WBGene00011699 | protein_coding_gene | 1.5 | 2.775 | 1.8E-03 |
| *nep-19* | WBGene00020292 | protein_coding_gene | 1.5 | 2.774 | 4.0E-02 |
| *F55A12.6* | WBGene00018865 | protein_coding_gene | 1.5 | 2.772 | 2.0E-02 |
| *H37A05.2* | WBGene00010426 | protein_coding_gene | 1.5 | 2.772 | 3.5E-05 |
| *ptr-19* | WBGene00004233 | protein_coding_gene | 1.5 | 2.769 | 2.7E-02 |
| *F20C5.5* | WBGene00008970 | protein_coding_gene | 1.5 | 2.768 | 3.3E-03 |
| *K12C11.6* | WBGene00045052 | protein_coding_gene | 1.5 | 2.764 | 6.1E-06 |

| | | | | | |
|---|---|---|---|---|---|
| *W03G11.4* | WBGene00012226 | protein_coding_gene | 1.5 | 2.763 | 8.9E-09 |
| *C38D9.8* | WBGene00008016 | protein_coding_gene | 1.5 | 2.761 | 2.4E-03 |
| *F11C7.2* | WBGene00017376 | protein_coding_gene | 1.5 | 2.760 | 1.7E-04 |
| *ace-3* | WBGene00000037 | protein_coding_gene | 1.5 | 2.758 | 1.7E-03 |
| *Y54G2A.28* | WBGene00021892 | protein_coding_gene | 1.5 | 2.758 | 3.6E-02 |
| *W02B8.1* | WBGene00012197 | protein_coding_gene | 1.5 | 2.756 | 1.4E-05 |
| *T16G1.2* | WBGene00011796 | protein_coding_gene | 1.5 | 2.755 | 2.0E-02 |
| *bath-47* | WBGene00006464 | protein_coding_gene | 1.5 | 2.752 | 9.6E-03 |
| *nhr-67* | WBGene00003657 | protein_coding_gene | 1.5 | 2.751 | 5.0E-03 |
| *mks-6* | WBGene00010642 | protein_coding_gene | 1.5 | 2.751 | 2.9E-02 |
| *lips-10* | WBGene00008803 | protein_coding_gene | 1.5 | 2.749 | 3.0E-05 |
| *W05G11.8* | WBGene00271636 | ncRNA_gene | 1.5 | 2.747 | 3.9E-02 |
| *ugt-33* | WBGene00007946 | protein_coding_gene | 1.5 | 2.746 | 2.2E-03 |
| *W05H9.3* | WBGene00021050 | ncRNA_gene | 1.5 | 2.746 | 2.9E-12 |
| *K02D7.1* | WBGene00019298 | protein_coding_gene | 1.5 | 2.745 | 1.3E-10 |
| *ltd-1* | WBGene00003089 | protein_coding_gene | 1.5 | 2.744 | 3.4E-05 |
| *F45E10.2* | WBGene00009729 | protein_coding_gene | 1.5 | 2.744 | 8.7E-06 |
| *srpr-2.1* | WBGene00195016 | ncRNA_gene | 1.5 | 2.744 | 3.5E-02 |
| *sulp-8* | WBGene00013963 | protein_coding_gene | 1.5 | 2.739 | 1.1E-05 |
| *ZK1128.7* | WBGene00014233 | protein_coding_gene | 1.5 | 2.739 | 4.9E-12 |
| *Y58A7A.3* | WBGene00021977 | protein_coding_gene | 1.5 | 2.738 | 2.1E-05 |
| *T10E9.6* | WBGene00020416 | protein_coding_gene | 1.5 | 2.737 | 1.9E-02 |
| *daao-1* | WBGene00022076 | protein_coding_gene | 1.5 | 2.736 | 4.6E-03 |
| *linc-91* | WBGene00219746 | lincRNA_gene | 1.5 | 2.735 | 5.0E-02 |
| *F38E9.1* | WBGene00018184 | protein_coding_gene | 1.5 | 2.734 | 2.7E-13 |
| *F35E2.10* | WBGene00044270 | pseudogene | 1.5 | 2.734 | 3.6E-02 |
| *Y51H4A.8* | WBGene00013104 | protein_coding_gene | 1.5 | 2.734 | 7.8E-06 |
| *T02B11.4* | WBGene00020156 | protein_coding_gene | 1.4 | 2.732 | 3.1E-03 |
| *fbxb-100* | WBGene00013403 | protein_coding_gene | 1.4 | 2.730 | 7.1E-03 |
| *C36E8.4* | WBGene00007983 | protein_coding_gene | 1.4 | 2.727 | 1.9E-05 |
| *Y116A8A.10* | WBGene00014975 | pseudogene | 1.4 | 2.726 | 5.4E-03 |
| *Y38F1A.8* | WBGene00012610 | protein_coding_gene | 1.4 | 2.724 | 1.2E-03 |
| *cht-1* | WBGene00000503 | protein_coding_gene | 1.4 | 2.719 | 1.6E-17 |
| *F58H1.5* | WBGene00010288 | protein_coding_gene | 1.4 | 2.719 | 1.0E-07 |
| *ZK896.5* | WBGene00014136 | protein_coding_gene | 1.4 | 2.718 | 1.7E-02 |
| *dhs-15* | WBGene00000978 | protein_coding_gene | 1.4 | 2.718 | 1.5E-02 |
| *T21D12.14* | WBGene00045177 | protein_coding_gene | 1.4 | 2.715 | 4.9E-02 |
| *nhr-32* | WBGene00003626 | protein_coding_gene | 1.4 | 2.713 | 8.2E-12 |
| *F21D12.3* | WBGene00017663 | protein_coding_gene | 1.4 | 2.707 | 2.3E-08 |
| *capa-1* | WBGene00021268 | protein_coding_gene | 1.4 | 2.703 | 2.9E-04 |
| *Y27F2A.8* | WBGene00021299 | protein_coding_gene | 1.4 | 2.701 | 3.1E-03 |
| *linc-6* | WBGene00219700 | lincRNA_gene | 1.4 | 2.701 | 5.4E-04 |
| *T04C12.8* | WBGene00011430 | protein_coding_gene | 1.4 | 2.700 | 2.4E-03 |
| *sst-20* | WBGene00006058 | protein_coding_gene | 1.4 | 2.700 | 9.0E-03 |
| *T24B8.3* | WBGene00011977 | protein_coding_gene | 1.4 | 2.699 | 3.3E-24 |
| *fbxl-1* | WBGene00015350 | protein_coding_gene | 1.4 | 2.699 | 1.1E-16 |
| *dhs-31* | WBGene00011424 | protein_coding_gene | 1.4 | 2.698 | 6.5E-06 |
| *F53F1.6* | WBGene00009984 | protein_coding_gene | 1.4 | 2.697 | 9.8E-04 |
| *vab-7* | WBGene00006873 | protein_coding_gene | 1.4 | 2.695 | 7.7E-03 |
| *Y22D7AL.15* | WBGene00021253 | protein_coding_gene | 1.4 | 2.692 | 1.1E-07 |
| *nhr-54* | WBGene00003644 | protein_coding_gene | 1.4 | 2.690 | 6.3E-03 |
| *sta-2* | WBGene00010251 | protein_coding_gene | 1.4 | 2.688 | 6.9E-08 |
| *C54D10.13* | WBGene00045400 | protein_coding_gene | 1.4 | 2.686 | 4.8E-03 |
| *ttx-1* | WBGene00006652 | protein_coding_gene | 1.4 | 2.685 | 1.4E-03 |
| *hsp-12.6* | WBGene00002013 | protein_coding_gene | 1.4 | 2.683 | 6.1E-03 |
| *crb-1* | WBGene00000792 | protein_coding_gene | 1.4 | 2.682 | 2.5E-08 |
| *faah-2* | WBGene00015048 | protein_coding_gene | 1.4 | 2.682 | 7.1E-05 |
| *aff-1* | WBGene00016625 | protein_coding_gene | 1.4 | 2.679 | 1.6E-02 |
| *ugt-14* | WBGene00019233 | protein_coding_gene | 1.4 | 2.678 | 4.3E-03 |
| *osg-1* | WBGene00019832 | protein_coding_gene | 1.4 | 2.676 | 2.3E-04 |

| | | | | | |
|---|---|---|---|---|---|
| *cbp-3* | WBGene00009595 | protein_coding_gene | 1.4 | 2.673 | 1.1E-04 |
| *K08C7.4* | WBGene00010647 | protein_coding_gene | 1.4 | 2.673 | 2.6E-02 |
| *poml-4* | WBGene00044697 | protein_coding_gene | 1.4 | 2.673 | 4.0E-02 |
| *cpg-7* | WBGene00010723 | protein_coding_gene | 1.4 | 2.671 | 1.5E-03 |
| *C30G4.4* | WBGene00016269 | protein_coding_gene | 1.4 | 2.671 | 2.4E-08 |
| *Y55F3AM.21* | WBGene00219399 | protein_coding_gene | 1.4 | 2.670 | 3.3E-03 |
| *Y57G11B.2* | WBGene00013295 | protein_coding_gene | 1.4 | 2.669 | 6.0E-05 |
| *ttr-21* | WBGene00008628 | protein_coding_gene | 1.4 | 2.669 | 7.7E-08 |
| *ins-34* | WBGene00002117 | protein_coding_gene | 1.4 | 2.668 | 1.1E-04 |
| *xtr-1* | WBGene00006965 | protein_coding_gene | 1.4 | 2.667 | 4.1E-02 |
| *F43C9.1* | WBGene00018375 | protein_coding_gene | 1.4 | 2.666 | 4.4E-03 |
| *F21D9.4* | WBGene00009018 | protein_coding_gene | 1.4 | 2.666 | 2.9E-02 |
| *pax-3* | WBGene00003939 | protein_coding_gene | 1.4 | 2.666 | 1.2E-03 |
| *F13E9.15* | WBGene00045208 | protein_coding_gene | 1.4 | 2.666 | 9.4E-07 |
| *T16H12.9* | WBGene00011817 | protein_coding_gene | 1.4 | 2.665 | 5.5E-03 |
| *T23F11.6* | WBGene00011957 | protein_coding_gene | 1.4 | 2.664 | 1.6E-02 |
| *dpy-1* | WBGene00001063 | protein_coding_gene | 1.4 | 2.661 | 1.6E-02 |
| *K03H1.5* | WBGene00010540 | protein_coding_gene | 1.4 | 2.661 | 9.2E-10 |
| *psa-3* | WBGene00009560 | protein_coding_gene | 1.4 | 2.660 | 2.2E-03 |
| *nhr-21* | WBGene00003620 | protein_coding_gene | 1.4 | 2.659 | 8.8E-08 |
| *lgc-22* | WBGene00017483 | protein_coding_gene | 1.4 | 2.654 | 6.2E-14 |
| *C47E12.9* | WBGene00008152 | protein_coding_gene | 1.4 | 2.649 | 1.3E-02 |
| *T01D1.8* | WBGene00044641 | protein_coding_gene | 1.4 | 2.649 | 3.6E-02 |
| *M162.5* | WBGene00010931 | protein_coding_gene | 1.4 | 2.644 | 4.2E-04 |
| *F26D10.13* | WBGene00044244 | protein_coding_gene | 1.4 | 2.641 | 2.7E-02 |
| *F47B8.2* | WBGene00009803 | protein_coding_gene | 1.4 | 2.641 | 3.8E-06 |
| *F02H6.4* | WBGene00008537 | pseudogene | 1.4 | 2.638 | 9.6E-12 |
| *F32D1.11* | WBGene00044530 | protein_coding_gene | 1.4 | 2.636 | 1.1E-02 |
| *W08F4.13* | WBGene00044733 | protein_coding_gene | 1.4 | 2.630 | 2.2E-02 |
| *aqp-3* | WBGene00000171 | protein_coding_gene | 1.4 | 2.628 | 4.0E-02 |
| *cex-2* | WBGene00023408 | protein_coding_gene | 1.4 | 2.627 | 1.3E-07 |
| *dhrs-4* | WBGene00010063 | protein_coding_gene | 1.4 | 2.625 | 8.2E-14 |
| *T05B11.7* | WBGene00044342 | protein_coding_gene | 1.4 | 2.620 | 2.9E-02 |
| *ncr-2* | WBGene00003562 | protein_coding_gene | 1.4 | 2.617 | 7.5E-04 |
| *Y57A10A.26* | WBGene00013266 | protein_coding_gene | 1.4 | 2.617 | 2.6E-12 |
| *K08E7.6* | WBGene00010674 | protein_coding_gene | 1.4 | 2.617 | 4.5E-02 |
| *C05G5.5* | WBGene00007351 | protein_coding_gene | 1.4 | 2.615 | 2.4E-02 |
| *Y92H12BR.4* | WBGene00022370 | pseudogene | 1.4 | 2.614 | 4.0E-03 |
| *msp-42* | WBGene00003437 | pseudogene | 1.4 | 2.612 | 4.0E-02 |
| *madd-2* | WBGene00016539 | protein_coding_gene | 1.4 | 2.612 | 5.5E-10 |
| *F18A11.2* | WBGene00008929 | protein_coding_gene | 1.4 | 2.612 | 3.3E-02 |
| *C18H9.6* | WBGene00016004 | protein_coding_gene | 1.4 | 2.610 | 1.3E-05 |
| *dhs-2* | WBGene00000966 | protein_coding_gene | 1.4 | 2.607 | 4.1E-07 |
| *C35A11.2* | WBGene00016429 | protein_coding_gene | 1.4 | 2.604 | 4.1E-02 |
| *memo-1* | WBGene00016500 | protein_coding_gene | 1.4 | 2.601 | 1.7E-14 |
| *tos-1* | WBGene00019466 | protein_coding_gene | 1.4 | 2.601 | 2.0E-46 |
| *tag-243* | WBGene00044078 | protein_coding_gene | 1.4 | 2.596 | 3.5E-04 |
| *best-3* | WBGene00015286 | protein_coding_gene | 1.4 | 2.592 | 2.0E-05 |
| *T14B1.1* | WBGene00011763 | protein_coding_gene | 1.4 | 2.591 | 6.5E-05 |
| *F44A6.5* | WBGene00009677 | protein_coding_gene | 1.4 | 2.590 | 3.6E-07 |
| *Y50D4A.10* | WBGene00235315 | protein_coding_gene | 1.4 | 2.589 | 6.6E-03 |
| *F56H11.2* | WBGene00010175 | protein_coding_gene | 1.4 | 2.589 | 5.7E-04 |
| *K10D6.4* | WBGene00010744 | protein_coding_gene | 1.4 | 2.585 | 2.0E-09 |
| *F47B8.3* | WBGene00009804 | protein_coding_gene | 1.4 | 2.583 | 1.1E-05 |
| *C08G5.7* | WBGene00044418 | protein_coding_gene | 1.4 | 2.581 | 3.0E-03 |
| *vap-2* | WBGene00011462 | protein_coding_gene | 1.4 | 2.579 | 7.8E-04 |
| *C24H12.11* | WBGene00016080 | pseudogene | 1.4 | 2.575 | 5.0E-03 |
| *cav-1* | WBGene00000301 | protein_coding_gene | 1.4 | 2.573 | 1.6E-20 |
| *Y39F10A.3* | WBGene00021454 | protein_coding_gene | 1.4 | 2.573 | 4.4E-02 |
| *sma-10* | WBGene00020649 | protein_coding_gene | 1.4 | 2.571 | 4.6E-14 |

| | | | | | |
|---|---|---|---|---|---|
| *Y110A2AL.4* | WBGene00022441 | protein_coding_gene | 1.4 | 2.567 | 4.2E-02 |
| *nhr-117* | WBGene00003707 | protein_coding_gene | 1.4 | 2.566 | 2.2E-04 |
| *Y55F3BR.11* | WBGene00044757 | protein_coding_gene | 1.4 | 2.562 | 4.3E-03 |
| *ugt-15* | WBGene00008097 | protein_coding_gene | 1.4 | 2.562 | 3.1E-02 |
| *mod-5* | WBGene00003387 | protein_coding_gene | 1.4 | 2.561 | 2.0E-03 |
| *tag-89* | WBGene00006459 | protein_coding_gene | 1.4 | 2.559 | 7.4E-03 |
| *Y52B11A.4* | WBGene00013124 | protein_coding_gene | 1.4 | 2.557 | 3.3E-03 |
| *cfi-1* | WBGene00000476 | protein_coding_gene | 1.3 | 2.548 | 1.0E-03 |
| *Y50D4C.2* | WBGene00021747 | pseudogene | 1.3 | 2.547 | 1.0E-08 |
| *F52F12.9* | WBGene00009942 | protein_coding_gene | 1.3 | 2.546 | 2.3E-02 |
| *ZK809.9* | WBGene00194934 | protein_coding_gene | 1.3 | 2.545 | 1.6E-03 |
| *Y71F9AL.7* | WBGene00022112 | protein_coding_gene | 1.3 | 2.543 | 7.8E-04 |
| *ceh-10* | WBGene00000435 | protein_coding_gene | 1.3 | 2.543 | 3.3E-02 |
| *cwn-1* | WBGene00000857 | protein_coding_gene | 1.3 | 2.542 | 5.1E-07 |
| *ncx-3* | WBGene00003568 | protein_coding_gene | 1.3 | 2.542 | 1.6E-06 |
| *ZK384.7* | WBGene00269383 | ncRNA_gene | 1.3 | 2.540 | 1.4E-03 |
| *M60.7* | WBGene00019782 | protein_coding_gene | 1.3 | 2.540 | 5.8E-03 |
| *insc-1* | WBGene00018392 | protein_coding_gene | 1.3 | 2.540 | 1.6E-05 |
| *F41G4.8* | WBGene00018311 | protein_coding_gene | 1.3 | 2.539 | 6.4E-04 |
| *F59F5.3* | WBGene00010342 | protein_coding_gene | 1.3 | 2.536 | 8.5E-03 |
| *nlp-71* | WBGene00022254 | protein_coding_gene | 1.3 | 2.536 | 9.9E-04 |
| *Y17G9B.8* | WBGene00021205 | protein_coding_gene | 1.3 | 2.535 | 2.1E-12 |
| *slt-1* | WBGene00004854 | protein_coding_gene | 1.3 | 2.534 | 4.6E-04 |
| *F54D1.6* | WBGene00010047 | protein_coding_gene | 1.3 | 2.533 | 1.1E-16 |
| *Y73F4A.3* | WBGene00013516 | protein_coding_gene | 1.3 | 2.533 | 9.0E-04 |
| *mltn-12* | WBGene00008277 | protein_coding_gene | 1.3 | 2.529 | 4.7E-02 |
| *nmad-1* | WBGene00017304 | protein_coding_gene | 1.3 | 2.529 | 2.0E-06 |
| *cest-27* | WBGene00018958 | protein_coding_gene | 1.3 | 2.526 | 2.3E-07 |
| *spp-1* | WBGene00004986 | protein_coding_gene | 1.3 | 2.526 | 2.2E-02 |
| *inx-2* | WBGene00002124 | protein_coding_gene | 1.3 | 2.523 | 3.1E-13 |
| *ceh-20* | WBGene00000443 | protein_coding_gene | 1.3 | 2.518 | 2.7E-12 |
| *Y65B4BL.7* | WBGene00022039 | protein_coding_gene | 1.3 | 2.518 | 1.4E-02 |
| *C03A7.13* | WBGene00015371 | protein_coding_gene | 1.3 | 2.516 | 2.2E-03 |
| *T13C2.3* | WBGene00020479 | protein_coding_gene | 1.3 | 2.516 | 2.6E-03 |
| *R148.5* | WBGene00020104 | protein_coding_gene | 1.3 | 2.515 | 7.3E-11 |
| *ser-4* | WBGene00004779 | protein_coding_gene | 1.3 | 2.514 | 2.6E-03 |
| *F54H5.5* | WBGene00018841 | protein_coding_gene | 1.3 | 2.513 | 2.7E-02 |
| *F10G8.8* | WBGene00008666 | protein_coding_gene | 1.3 | 2.511 | 1.6E-07 |
| *unc-5* | WBGene00006745 | protein_coding_gene | 1.3 | 2.511 | 2.1E-10 |
| *K04F10.2* | WBGene00019395 | protein_coding_gene | 1.3 | 2.510 | 3.8E-02 |
| *C29H12.6* | WBGene00016237 | protein_coding_gene | 1.3 | 2.510 | 5.2E-04 |
| *F36H5.13* | WBGene00206519 | protein_coding_gene | 1.3 | 2.509 | 1.2E-02 |
| *mdl-1* | WBGene00003163 | protein_coding_gene | 1.3 | 2.507 | 1.8E-05 |
| *lad-2* | WBGene00002243 | protein_coding_gene | 1.3 | 2.506 | 4.5E-03 |
| *C52E2.4* | WBGene00016883 | protein_coding_gene | 1.3 | 2.505 | 1.7E-02 |
| *flp-8* | WBGene00001451 | protein_coding_gene | 1.3 | 2.505 | 3.0E-07 |
| *cpna-5* | WBGene00007044 | protein_coding_gene | 1.3 | 2.503 | 4.5E-08 |
| *C18A11.2* | WBGene00015948 | protein_coding_gene | 1.3 | 2.501 | 4.5E-02 |
| *F43G6.8* | WBGene00009660 | protein_coding_gene | 1.3 | 2.500 | 3.0E-12 |
| *acs-17* | WBGene00016716 | protein_coding_gene | 1.3 | 2.500 | 1.4E-09 |
| *C31H1.1* | WBGene00016286 | protein_coding_gene | 1.3 | 2.500 | 1.1E-02 |
| *mig-13* | WBGene00003245 | protein_coding_gene | 1.3 | 2.500 | 2.5E-06 |
| *T25D10.4* | WBGene00020801 | protein_coding_gene | 1.3 | 2.499 | 4.0E-07 |
| *C44H9.5* | WBGene00008100 | protein_coding_gene | 1.3 | 2.499 | 8.5E-09 |
| *fkb-4* | WBGene00001429 | protein_coding_gene | 1.3 | 2.498 | 2.2E-02 |
| *Y37D8A.16* | WBGene00012554 | protein_coding_gene | 1.3 | 2.497 | 5.5E-06 |
| *W04A8.4* | WBGene00012239 | protein_coding_gene | 1.3 | 2.496 | 1.9E-05 |
| *nhr-206* | WBGene00011097 | protein_coding_gene | 1.3 | 2.496 | 9.5E-03 |
| *F32B4.5* | WBGene00009315 | protein_coding_gene | 1.3 | 2.495 | 2.2E-02 |
| *F55C5.2* | WBGene00010092 | pseudogene | 1.3 | 2.494 | 1.3E-03 |

| | | | | | |
|---|---|---|---|---|---|
| *T26H2.10* | WBGene00044976 | pseudogene | 1.3 | 2.493 | 1.5E-02 |
| *C54E10.6* | WBGene00008310 | protein_coding_gene | 1.3 | 2.490 | 1.7E-03 |
| *R05G6.1* | WBGene00019896 | protein_coding_gene | 1.3 | 2.488 | 3.6E-02 |
| *T04A6.1* | WBGene00020200 | protein_coding_gene | 1.3 | 2.488 | 8.3E-09 |
| *fat-3* | WBGene00001395 | protein_coding_gene | 1.3 | 2.484 | 1.1E-02 |
| *clec-79* | WBGene00018548 | protein_coding_gene | 1.3 | 2.482 | 7.8E-03 |
| *C06G3.5* | WBGene00015551 | protein_coding_gene | 1.3 | 2.480 | 1.2E-07 |
| *slc-28.2* | WBGene00017868 | protein_coding_gene | 1.3 | 2.479 | 1.8E-02 |
| *tsp-6* | WBGene00006632 | protein_coding_gene | 1.3 | 2.475 | 2.8E-02 |
| *Y75B7AL.2* | WBGene00022285 | protein_coding_gene | 1.3 | 2.475 | 2.1E-02 |
| *T08B6.9* | WBGene00020356 | protein_coding_gene | 1.3 | 2.474 | 3.2E-02 |
| *BE0003N10.3* | WBGene00015268 | protein_coding_gene | 1.3 | 2.466 | 2.9E-04 |
| *ZC455.1* | WBGene00013902 | protein_coding_gene | 1.3 | 2.465 | 2.5E-05 |
| *C34B4.3* | WBGene00007908 | protein_coding_gene | 1.3 | 2.465 | 3.0E-02 |
| *C24H12.7* | WBGene00016076 | pseudogene | 1.3 | 2.461 | 2.7E-02 |
| *cest-24* | WBGene00020891 | protein_coding_gene | 1.3 | 2.461 | 3.5E-04 |
| *epg-2* | WBGene00021468 | protein_coding_gene | 1.3 | 2.459 | 2.9E-12 |
| *Y71F9AL.6* | WBGene00022111 | protein_coding_gene | 1.3 | 2.457 | 1.2E-02 |
| *B0507.6* | WBGene00015223 | protein_coding_gene | 1.3 | 2.456 | 1.8E-02 |
| *gst-33* | WBGene00001781 | protein_coding_gene | 1.3 | 2.455 | 2.5E-02 |
| *F12A10.1* | WBGene00017390 | protein_coding_gene | 1.3 | 2.455 | 2.6E-05 |
| *F48E3.9* | WBGene00018608 | protein_coding_gene | 1.3 | 2.449 | 3.4E-02 |
| *M01G12.14* | WBGene00010825 | protein_coding_gene | 1.3 | 2.447 | 3.7E-02 |
| *glb-15* | WBGene00009387 | protein_coding_gene | 1.3 | 2.446 | 3.6E-02 |
| *R08A2.1* | WBGene00011132 | protein_coding_gene | 1.3 | 2.446 | 2.4E-03 |
| *fbxc-18* | WBGene00016307 | protein_coding_gene | 1.3 | 2.442 | 2.7E-04 |
| *vab-2* | WBGene00006869 | protein_coding_gene | 1.3 | 2.442 | 6.9E-04 |
| *C06G3.6* | WBGene00015552 | protein_coding_gene | 1.3 | 2.440 | 1.8E-07 |
| *vab-15* | WBGene00006881 | protein_coding_gene | 1.3 | 2.440 | 5.3E-03 |
| *ZC487.1* | WBGene00022623 | protein_coding_gene | 1.3 | 2.439 | 2.9E-02 |
| *F39E9.28* | WBGene00271716 | pseudogene | 1.3 | 2.439 | 8.5E-04 |
| *ttr-44* | WBGene00008341 | protein_coding_gene | 1.3 | 2.438 | 6.0E-04 |
| *fbxb-107* | WBGene00017912 | protein_coding_gene | 1.3 | 2.436 | 2.3E-02 |
| *T28H10.1* | WBGene00012142 | protein_coding_gene | 1.3 | 2.436 | 6.2E-05 |
| *C35A5.6* | WBGene00007950 | protein_coding_gene | 1.3 | 2.435 | 7.7E-06 |
| *zmp-2* | WBGene00019212 | protein_coding_gene | 1.3 | 2.435 | 3.1E-02 |
| *flp-21* | WBGene00001464 | protein_coding_gene | 1.3 | 2.433 | 9.9E-04 |
| *MTCE.7* | WBGene00014454 | rRNA_gene | 1.3 | 2.433 | 3.2E-04 |
| *Y47G6A.5* | WBGene00021635 | protein_coding_gene | 1.3 | 2.432 | 2.7E-09 |
| *flp-12* | WBGene00001455 | protein_coding_gene | 1.3 | 2.431 | 7.7E-06 |
| *F19C7.2* | WBGene00017592 | protein_coding_gene | 1.3 | 2.430 | 1.1E-03 |
| *cyn-6* | WBGene00000882 | protein_coding_gene | 1.3 | 2.430 | 7.5E-10 |
| *T19A5.3* | WBGene00020554 | protein_coding_gene | 1.3 | 2.429 | 1.7E-02 |
| *Y51A2D.13* | WBGene00013080 | protein_coding_gene | 1.3 | 2.427 | 4.6E-14 |
| *ZK1290.13* | WBGene00022893 | protein_coding_gene | 1.3 | 2.425 | 2.4E-05 |
| *lact-5* | WBGene00016355 | protein_coding_gene | 1.3 | 2.424 | 3.0E-06 |
| *ttm-2* | WBGene00017840 | protein_coding_gene | 1.3 | 2.421 | 4.7E-04 |
| *atz-1* | WBGene00009212 | protein_coding_gene | 1.3 | 2.420 | 4.6E-11 |
| *F36A4.2* | WBGene00018081 | protein_coding_gene | 1.3 | 2.420 | 2.0E-02 |
| *sqst-1* | WBGene00011737 | protein_coding_gene | 1.3 | 2.419 | 1.3E-07 |
| *W02D7.4* | WBGene00020940 | protein_coding_gene | 1.3 | 2.419 | 2.6E-07 |
| *R13A5.15* | WBGene00194726 | protein_coding_gene | 1.3 | 2.416 | 2.7E-02 |
| *R11G1.2* | WBGene00020011 | protein_coding_gene | 1.3 | 2.416 | 2.4E-02 |
| *C18A11.1* | WBGene00015947 | protein_coding_gene | 1.3 | 2.414 | 2.1E-04 |
| *nspc-16* | WBGene00008357 | protein_coding_gene | 1.3 | 2.414 | 3.3E-02 |
| *msp-3* | WBGene00003424 | protein_coding_gene | 1.3 | 2.413 | 1.2E-03 |
| *syg-2* | WBGene00007750 | protein_coding_gene | 1.3 | 2.412 | 4.4E-05 |
| *tsp-16* | WBGene00006642 | protein_coding_gene | 1.3 | 2.412 | 9.4E-03 |
| *Y75B12B.3* | WBGene00013568 | protein_coding_gene | 1.3 | 2.410 | 2.5E-02 |
| *mab-7* | WBGene00003104 | protein_coding_gene | 1.3 | 2.410 | 3.7E-02 |

| | | | | | |
|---|---|---|---|---|---|
| *let-522* | WBGene00010870 | protein_coding_gene | 1.3 | 2.409 | 1.1E-06 |
| *clec-86* | WBGene00016912 | protein_coding_gene | 1.3 | 2.409 | 8.2E-05 |
| *C08F11.12* | WBGene00007459 | protein_coding_gene | 1.3 | 2.407 | 2.8E-04 |
| *kel-1* | WBGene00002184 | protein_coding_gene | 1.3 | 2.406 | 7.3E-03 |
| *C07D10.5* | WBGene00015569 | protein_coding_gene | 1.3 | 2.404 | 3.6E-11 |
| *vap-1* | WBGene00006886 | protein_coding_gene | 1.3 | 2.399 | 6.2E-07 |
| *hlh-1* | WBGene00001948 | protein_coding_gene | 1.3 | 2.397 | 2.9E-04 |
| *sulp-3* | WBGene00018283 | protein_coding_gene | 1.3 | 2.388 | 3.0E-02 |
| *R03E9.2* | WBGene00019843 | protein_coding_gene | 1.3 | 2.388 | 3.4E-04 |
| *pho-4* | WBGene00004023 | protein_coding_gene | 1.3 | 2.386 | 2.9E-06 |
| *nhr-43* | WBGene00003633 | protein_coding_gene | 1.3 | 2.386 | 8.0E-04 |
| *let-653* | WBGene00002827 | protein_coding_gene | 1.3 | 2.386 | 3.2E-06 |
| *F59F5.7* | WBGene00010345 | protein_coding_gene | 1.3 | 2.386 | 2.4E-02 |
| *Y77E11A.23* | WBGene00235267 | protein_coding_gene | 1.3 | 2.385 | 4.0E-03 |
| *tyr-1* | WBGene00015332 | protein_coding_gene | 1.3 | 2.381 | 4.4E-05 |
| *F26G1.2* | WBGene00017838 | protein_coding_gene | 1.3 | 2.380 | 3.5E-04 |
| *aqp-9* | WBGene00000177 | protein_coding_gene | 1.3 | 2.379 | 2.3E-07 |
| *lect-2* | WBGene00019407 | protein_coding_gene | 1.2 | 2.376 | 3.2E-07 |
| *ztf-11* | WBGene00009939 | protein_coding_gene | 1.2 | 2.375 | 1.2E-10 |
| *fbxb-80* | WBGene00017117 | pseudogene | 1.2 | 2.374 | 3.9E-02 |
| *T05E11.8* | WBGene00011484 | protein_coding_gene | 1.2 | 2.373 | 3.3E-03 |
| *srm-1* | WBGene00005638 | protein_coding_gene | 1.2 | 2.372 | 4.9E-02 |
| *Y62F5A.9* | WBGene00013387 | protein_coding_gene | 1.2 | 2.372 | 4.7E-02 |
| *madf-10* | WBGene00013717 | protein_coding_gene | 1.2 | 2.369 | 7.1E-03 |
| *Y53F4B.11* | WBGene00013158 | protein_coding_gene | 1.2 | 2.368 | 1.2E-02 |
| *K12C11.7* | WBGene00045053 | protein_coding_gene | 1.2 | 2.368 | 9.5E-04 |
| *T23F11.2* | WBGene00011954 | protein_coding_gene | 1.2 | 2.368 | 6.6E-03 |
| *C35E7.5* | WBGene00016457 | protein_coding_gene | 1.2 | 2.368 | 2.0E-08 |
| *zig-8* | WBGene00006985 | protein_coding_gene | 1.2 | 2.367 | 4.9E-02 |
| *B0507.1* | WBGene00015218 | protein_coding_gene | 1.2 | 2.364 | 3.1E-03 |
| *F20C5.11* | WBGene00303015 | protein_coding_gene | 1.2 | 2.360 | 2.9E-03 |
| *Y70G10A.3* | WBGene00013499 | protein_coding_gene | 1.2 | 2.360 | 1.1E-05 |
| *Y73B6A.1* | WBGene00022228 | protein_coding_gene | 1.2 | 2.360 | 3.0E-02 |
| *K02E10.1* | WBGene00019317 | protein_coding_gene | 1.2 | 2.359 | 1.2E-05 |
| *C34F11.8* | WBGene00016417 | protein_coding_gene | 1.2 | 2.356 | 2.8E-05 |
| *ugt-8* | WBGene00019234 | protein_coding_gene | 1.2 | 2.356 | 1.6E-09 |
| *F35B3.7* | WBGene00018033 | protein_coding_gene | 1.2 | 2.356 | 4.8E-03 |
| *Y41C4A.8* | WBGene00012755 | protein_coding_gene | 1.2 | 2.354 | 5.1E-05 |
| *pqn-13* | WBGene00004104 | protein_coding_gene | 1.2 | 2.352 | 1.9E-02 |
| *daam-1* | WBGene00018976 | protein_coding_gene | 1.2 | 2.350 | 5.6E-03 |
| *Y38C1AA.6* | WBGene00021397 | protein_coding_gene | 1.2 | 2.346 | 2.1E-02 |
| *nlr-1* | WBGene00003772 | protein_coding_gene | 1.2 | 2.346 | 7.6E-05 |
| *C05B5.5* | WBGene00007322 | protein_coding_gene | 1.2 | 2.344 | 2.6E-02 |
| *fozi-1* | WBGene00010453 | protein_coding_gene | 1.2 | 2.343 | 2.9E-04 |
| *bus-17* | WBGene00044630 | protein_coding_gene | 1.2 | 2.341 | 4.2E-02 |
| *sup-26* | WBGene00006331 | protein_coding_gene | 1.2 | 2.340 | 6.9E-22 |
| *R11G1.7* | WBGene00044725 | protein_coding_gene | 1.2 | 2.340 | 2.5E-03 |
| *dhhc-13* | WBGene00019257 | protein_coding_gene | 1.2 | 2.337 | 3.6E-02 |
| *T09A12.1* | WBGene00020372 | protein_coding_gene | 1.2 | 2.337 | 2.9E-02 |
| *epg-9* | WBGene00022078 | protein_coding_gene | 1.2 | 2.335 | 1.1E-08 |
| *cwn-2* | WBGene00000858 | protein_coding_gene | 1.2 | 2.332 | 8.0E-05 |
| *C01A2.1* | WBGene00007213 | protein_coding_gene | 1.2 | 2.332 | 4.1E-02 |
| *C06B8.2* | WBGene00007368 | protein_coding_gene | 1.2 | 2.331 | 2.1E-04 |
| *Y42G9A.1* | WBGene00021531 | protein_coding_gene | 1.2 | 2.327 | 7.0E-04 |
| *fbxb-116* | WBGene00012066 | protein_coding_gene | 1.2 | 2.327 | 1.9E-02 |
| *cal-2* | WBGene00000286 | protein_coding_gene | 1.2 | 2.325 | 3.5E-04 |
| *Y50D4A.5* | WBGene00021739 | protein_coding_gene | 1.2 | 2.324 | 1.1E-04 |
| *T27A10.6* | WBGene00020846 | protein_coding_gene | 1.2 | 2.324 | 3.0E-04 |
| *ssp-16* | WBGene00006044 | protein_coding_gene | 1.2 | 2.324 | 2.1E-02 |
| *lgg-2* | WBGene00002981 | protein_coding_gene | 1.2 | 2.319 | 1.7E-15 |

| | | | | | |
|---|---|---|---|---|---|
| *nhr-55* | WBGene00003645 | protein_coding_gene | 1.2 | 2.319 | 2.9E-05 |
| *sfrp-1* | WBGene00022242 | protein_coding_gene | 1.2 | 2.319 | 3.5E-03 |
| *ptr-3* | WBGene00004218 | protein_coding_gene | 1.2 | 2.319 | 2.4E-02 |
| *Y17G9B.2* | WBGene00021199 | protein_coding_gene | 1.2 | 2.318 | 1.4E-02 |
| *Y44E3A.1* | WBGene00021547 | protein_coding_gene | 1.2 | 2.317 | 3.4E-03 |
| *Y50E8A.12* | WBGene00013055 | protein_coding_gene | 1.2 | 2.315 | 2.3E-02 |
| *grl-7* | WBGene00001716 | protein_coding_gene | 1.2 | 2.314 | 3.6E-02 |
| *clec-180* | WBGene00017991 | protein_coding_gene | 1.2 | 2.312 | 4.7E-02 |
| *cest-16* | WBGene00020330 | protein_coding_gene | 1.2 | 2.308 | 2.9E-02 |
| *ptp-4* | WBGene00010558 | protein_coding_gene | 1.2 | 2.308 | 4.9E-03 |
| *trh-1* | WBGene00007827 | protein_coding_gene | 1.2 | 2.307 | 2.6E-02 |
| *ZK550.2* | WBGene00013996 | protein_coding_gene | 1.2 | 2.307 | 1.6E-03 |
| *nhr-110* | WBGene00003700 | protein_coding_gene | 1.2 | 2.304 | 2.0E-06 |
| *strm-1* | WBGene00019198 | protein_coding_gene | 1.2 | 2.303 | 1.7E-02 |
| *C47A4.1* | WBGene00008122 | protein_coding_gene | 1.2 | 2.303 | 4.2E-02 |
| *F37A8.1* | WBGene00009501 | protein_coding_gene | 1.2 | 2.299 | 2.9E-02 |
| *F55B12.11* | WBGene00077570 | pseudogene | 1.2 | 2.299 | 4.0E-02 |
| *M7.7* | WBGene00010883 | protein_coding_gene | 1.2 | 2.298 | 4.8E-02 |
| *C05C12.4* | WBGene00007336 | protein_coding_gene | 1.2 | 2.297 | 3.8E-12 |
| *T04F8.7* | WBGene00011445 | protein_coding_gene | 1.2 | 2.296 | 1.4E-05 |
| *C34D1.4* | WBGene00007931 | protein_coding_gene | 1.2 | 2.296 | 1.3E-03 |
| *fbxb-93* | WBGene00013408 | protein_coding_gene | 1.2 | 2.296 | 1.7E-02 |
| *F13C5.2* | WBGene00017423 | protein_coding_gene | 1.2 | 2.295 | 5.0E-15 |
| *C13E3.1* | WBGene00015740 | protein_coding_gene | 1.2 | 2.294 | 5.3E-03 |
| *T19H5.4* | WBGene00011849 | protein_coding_gene | 1.2 | 2.294 | 1.7E-13 |
| *tbx-8* | WBGene00006545 | protein_coding_gene | 1.2 | 2.293 | 1.8E-02 |
| *Y41D4A.3* | WBGene00021505 | protein_coding_gene | 1.2 | 2.293 | 2.2E-03 |
| *C49F5.7* | WBGene00008211 | protein_coding_gene | 1.2 | 2.292 | 6.0E-07 |
| *ZK863.8* | WBGene00014125 | protein_coding_gene | 1.2 | 2.291 | 4.8E-05 |
| *syg-1* | WBGene00006365 | protein_coding_gene | 1.2 | 2.291 | 4.6E-04 |
| *R07G3.8* | WBGene00019944 | protein_coding_gene | 1.2 | 2.291 | 9.1E-05 |
| *zig-6* | WBGene00006983 | protein_coding_gene | 1.2 | 2.289 | 3.6E-03 |
| *T28A11.22* | WBGene00044526 | protein_coding_gene | 1.2 | 2.286 | 1.3E-02 |
| *F08C6.5* | WBGene00017243 | protein_coding_gene | 1.2 | 2.282 | 3.7E-02 |
| *BE10.1* | WBGene00007209 | protein_coding_gene | 1.2 | 2.281 | 1.9E-02 |
| *nhr-152* | WBGene00015705 | protein_coding_gene | 1.2 | 2.278 | 3.5E-03 |
| *C50C3.2* | WBGene00016800 | protein_coding_gene | 1.2 | 2.278 | 2.2E-02 |
| *inx-6* | WBGene00002128 | protein_coding_gene | 1.2 | 2.275 | 5.9E-03 |
| *C18E9.8* | WBGene00007688 | protein_coding_gene | 1.2 | 2.275 | 2.2E-02 |
| *mig-18* | WBGene00003249 | protein_coding_gene | 1.2 | 2.274 | 3.2E-05 |
| *ptp-1* | WBGene00004213 | protein_coding_gene | 1.2 | 2.271 | 2.0E-05 |
| *F58H1.7* | WBGene00010290 | protein_coding_gene | 1.2 | 2.271 | 9.6E-04 |
| *Y13C8A.4* | WBGene00219303 | pseudogene | 1.2 | 2.270 | 7.7E-03 |
| *ins-18* | WBGene00002101 | protein_coding_gene | 1.2 | 2.266 | 5.0E-02 |
| *E02H4.7* | WBGene00077757 | protein_coding_gene | 1.2 | 2.266 | 1.0E-07 |
| *tag-275* | WBGene00016423 | protein_coding_gene | 1.2 | 2.262 | 2.4E-03 |
| *fbxa-211* | WBGene00012566 | protein_coding_gene | 1.2 | 2.261 | 2.1E-07 |
| *sdz-14* | WBGene00008782 | protein_coding_gene | 1.2 | 2.261 | 4.7E-04 |
| *F58A3.3* | WBGene00010223 | protein_coding_gene | 1.2 | 2.258 | 1.6E-02 |
| *fbxc-36* | WBGene00019918 | protein_coding_gene | 1.2 | 2.257 | 6.1E-07 |
| *elt-6* | WBGene00001253 | protein_coding_gene | 1.2 | 2.257 | 5.9E-03 |
| *R10E9.3* | WBGene00011215 | protein_coding_gene | 1.2 | 2.255 | 4.5E-03 |
| *Y41C4A.18* | WBGene00012761 | protein_coding_gene | 1.2 | 2.253 | 3.4E-02 |
| *vab-23* | WBGene00014153 | protein_coding_gene | 1.2 | 2.252 | 3.7E-03 |
| *ist-1* | WBGene00002163 | protein_coding_gene | 1.2 | 2.246 | 1.6E-05 |
| *DH11.2* | WBGene00008436 | protein_coding_gene | 1.2 | 2.246 | 7.3E-11 |
| *ceh-6* | WBGene00000431 | protein_coding_gene | 1.2 | 2.244 | 3.3E-02 |
| *C06A8.6* | WBGene00015516 | protein_coding_gene | 1.2 | 2.244 | 3.4E-02 |
| *C27B7.7* | WBGene00007764 | protein_coding_gene | 1.2 | 2.244 | 3.8E-06 |
| *cdh-8* | WBGene00017576 | protein_coding_gene | 1.2 | 2.243 | 4.6E-03 |

| | | | | | |
|---|---|---|---|---|---|
| *K11D12.12* | WBGene00019651 | protein_coding_gene | 1.2 | 2.242 | 7.4E-07 |
| *ZK154.1* | WBGene00022665 | protein_coding_gene | 1.2 | 2.237 | 4.8E-02 |
| *tag-273* | WBGene00013289 | protein_coding_gene | 1.2 | 2.236 | 2.9E-04 |
| *gad-2* | WBGene00015666 | protein_coding_gene | 1.2 | 2.236 | 4.8E-02 |
| *Y43B11AR.3* | WBGene00021544 | protein_coding_gene | 1.2 | 2.235 | 1.8E-05 |
| *clec-264* | WBGene00009291 | protein_coding_gene | 1.2 | 2.234 | 2.0E-03 |
| *spp-22* | WBGene00044283 | protein_coding_gene | 1.2 | 2.234 | 4.3E-04 |
| *nspc-9* | WBGene00009634 | protein_coding_gene | 1.2 | 2.234 | 2.2E-07 |
| *nspc-3* | WBGene00044338 | protein_coding_gene | 1.2 | 2.232 | 4.4E-02 |
| *tmed-13* | WBGene00022255 | protein_coding_gene | 1.2 | 2.229 | 3.7E-02 |
| *R04B5.5* | WBGene00011003 | protein_coding_gene | 1.2 | 2.229 | 3.2E-06 |
| *ceh-37* | WBGene00000458 | protein_coding_gene | 1.2 | 2.229 | 4.7E-06 |
| *acs-3* | WBGene00020343 | protein_coding_gene | 1.2 | 2.228 | 6.4E-06 |
| *pudl-1* | WBGene00044489 | protein_coding_gene | 1.2 | 2.227 | 4.1E-04 |
| *C18D4.6* | WBGene00007675 | protein_coding_gene | 1.2 | 2.227 | 1.5E-07 |
| *jmjd-3.1* | WBGene00017571 | protein_coding_gene | 1.2 | 2.226 | 9.7E-07 |
| *T01E8.1* | WBGene00011331 | protein_coding_gene | 1.2 | 2.224 | 3.0E-05 |
| *mec-6* | WBGene00003170 | protein_coding_gene | 1.2 | 2.224 | 4.6E-02 |
| *nhr-94* | WBGene00003684 | protein_coding_gene | 1.2 | 2.221 | 7.5E-03 |
| *efn-4* | WBGene00001165 | protein_coding_gene | 1.2 | 2.220 | 1.6E-08 |
| *Y49A10A.2* | WBGene00044949 | protein_coding_gene | 1.1 | 2.217 | 6.0E-07 |
| *flp-34* | WBGene00019975 | protein_coding_gene | 1.1 | 2.216 | 2.8E-03 |
| *F56C9.8* | WBGene00018952 | protein_coding_gene | 1.1 | 2.216 | 1.3E-05 |
| *F55G11.8* | WBGene00010128 | protein_coding_gene | 1.1 | 2.214 | 3.2E-02 |
| *F19C7.3* | WBGene00017593 | protein_coding_gene | 1.1 | 2.214 | 3.0E-02 |
| *F58B4.6* | WBGene00010239 | protein_coding_gene | 1.1 | 2.214 | 2.3E-02 |
| *C10C5.7* | WBGene00007511 | protein_coding_gene | 1.1 | 2.212 | 4.8E-02 |
| *F12E12.6* | WBGene00017407 | protein_coding_gene | 1.1 | 2.211 | 1.9E-02 |
| *max-1* | WBGene00003143 | protein_coding_gene | 1.1 | 2.210 | 6.5E-04 |
| *rhgf-2* | WBGene00006476 | protein_coding_gene | 1.1 | 2.208 | 2.5E-04 |
| *nhr-35* | WBGene00003628 | protein_coding_gene | 1.1 | 2.208 | 6.3E-07 |
| *cdh-12* | WBGene00022103 | protein_coding_gene | 1.1 | 2.207 | 2.7E-03 |
| *ZK418.2* | WBGene00022732 | protein_coding_gene | 1.1 | 2.204 | 3.3E-02 |
| *lgc-47* | WBGene00009799 | protein_coding_gene | 1.1 | 2.204 | 3.0E-02 |
| *F57A8.1* | WBGene00010177 | protein_coding_gene | 1.1 | 2.202 | 9.6E-07 |
| *Y57A10B.6* | WBGene00013278 | protein_coding_gene | 1.1 | 2.201 | 8.1E-04 |
| *lron-15* | WBGene00016974 | protein_coding_gene | 1.1 | 2.200 | 1.1E-02 |
| *T03F1.11* | WBGene00020192 | protein_coding_gene | 1.1 | 2.197 | 1.3E-04 |
| *F13B12.2* | WBGene00008730 | protein_coding_gene | 1.1 | 2.196 | 2.2E-03 |
| *clec-31* | WBGene00009854 | protein_coding_gene | 1.1 | 2.196 | 9.1E-03 |
| *srp-3* | WBGene00005644 | protein_coding_gene | 1.1 | 2.196 | 2.2E-02 |
| *ZK250.13* | WBGene00235352 | protein_coding_gene | 1.1 | 2.194 | 1.4E-02 |
| *ant-1.4* | WBGene00020140 | protein_coding_gene | 1.1 | 2.193 | 1.7E-02 |
| *lev-8* | WBGene00002975 | protein_coding_gene | 1.1 | 2.193 | 4.6E-02 |
| *F54C9.3* | WBGene00010041 | protein_coding_gene | 1.1 | 2.191 | 1.2E-12 |
| *C35E7.6* | WBGene00016458 | protein_coding_gene | 1.1 | 2.189 | 2.2E-11 |
| *msp-65* | WBGene00003458 | protein_coding_gene | 1.1 | 2.185 | 4.0E-02 |
| *linc-15* | WBGene00219571 | lincRNA_gene | 1.1 | 2.184 | 1.0E-03 |
| *C44E12.1* | WBGene00016657 | protein_coding_gene | 1.1 | 2.183 | 9.8E-06 |
| *magu-2* | WBGene00006467 | protein_coding_gene | 1.1 | 2.179 | 6.6E-06 |
| *C33G3.4* | WBGene00007904 | protein_coding_gene | 1.1 | 2.179 | 3.1E-11 |
| *che-12* | WBGene00000491 | protein_coding_gene | 1.1 | 2.179 | 3.5E-02 |
| *chil-22* | WBGene00011166 | protein_coding_gene | 1.1 | 2.176 | 1.9E-03 |
| *ptr-8* | WBGene00004222 | protein_coding_gene | 1.1 | 2.176 | 3.4E-03 |
| *nep-21* | WBGene00011794 | protein_coding_gene | 1.1 | 2.176 | 8.7E-04 |
| *T05E11.2* | WBGene00011479 | protein_coding_gene | 1.1 | 2.171 | 3.4E-02 |
| *R01H10.4* | WBGene00010972 | protein_coding_gene | 1.1 | 2.170 | 5.3E-05 |
| *jmjd-3.3* | WBGene00007813 | protein_coding_gene | 1.1 | 2.170 | 2.9E-02 |
| *C06B8.7* | WBGene00007372 | protein_coding_gene | 1.1 | 2.168 | 6.5E-12 |
| *D2063.1* | WBGene00017060 | protein_coding_gene | 1.1 | 2.168 | 2.2E-03 |

| | | | | | |
|---|---|---|---|---|---|
| *nspc-18* | WBGene00008358 | protein_coding_gene | 1.1 | 2.167 | 2.8E-03 |
| *ccch-2* | WBGene00009537 | protein_coding_gene | 1.1 | 2.166 | 3.4E-02 |
| *dcar-1* | WBGene00007395 | protein_coding_gene | 1.1 | 2.164 | 3.3E-08 |
| *glb-11* | WBGene00007981 | protein_coding_gene | 1.1 | 2.163 | 1.4E-02 |
| *T28C6.5* | WBGene00012120 | protein_coding_gene | 1.1 | 2.163 | 2.5E-02 |
| *unc-30* | WBGene00006766 | protein_coding_gene | 1.1 | 2.159 | 3.0E-02 |
| *Y71A12B.12* | WBGene00013509 | protein_coding_gene | 1.1 | 2.156 | 1.9E-03 |
| *clc-5* | WBGene00000526 | protein_coding_gene | 1.1 | 2.155 | 1.6E-02 |
| *C34D4.10* | WBGene00016402 | protein_coding_gene | 1.1 | 2.154 | 7.5E-08 |
| *Y73F8A.5* | WBGene00013518 | protein_coding_gene | 1.1 | 2.154 | 2.4E-02 |
| *exc-5* | WBGene00001366 | protein_coding_gene | 1.1 | 2.154 | 1.5E-08 |
| *siah-1* | WBGene00021369 | protein_coding_gene | 1.1 | 2.153 | 1.1E-10 |
| *W04B5.2* | WBGene00021019 | protein_coding_gene | 1.1 | 2.150 | 6.9E-03 |
| *Y45G5AM.5* | WBGene00021557 | protein_coding_gene | 1.1 | 2.146 | 2.4E-02 |
| *nhr-103* | WBGene00003693 | protein_coding_gene | 1.1 | 2.146 | 3.5E-03 |
| *F49C12.7* | WBGene00009877 | protein_coding_gene | 1.1 | 2.145 | 1.4E-03 |
| *R11.4* | WBGene00011229 | protein_coding_gene | 1.1 | 2.145 | 6.1E-03 |
| *flp-27* | WBGene00016114 | protein_coding_gene | 1.1 | 2.144 | 7.0E-03 |
| *cyp-25A3* | WBGene00014697 | protein_coding_gene | 1.1 | 2.143 | 5.0E-04 |
| *nhr-120* | WBGene00003710 | protein_coding_gene | 1.1 | 2.143 | 1.3E-03 |
| *dhp-2* | WBGene00000964 | protein_coding_gene | 1.1 | 2.141 | 4.2E-04 |
| *R10E8.8* | WBGene00011213 | protein_coding_gene | 1.1 | 2.135 | 1.1E-03 |
| *F08B4.3* | WBGene00017235 | pseudogene | 1.1 | 2.134 | 9.2E-06 |
| *EGAP9.3* | WBGene00017151 | protein_coding_gene | 1.1 | 2.131 | 3.0E-02 |
| *C06E1.1* | WBGene00015518 | protein_coding_gene | 1.1 | 2.130 | 3.6E-06 |
| *nlp-42* | WBGene00013588 | protein_coding_gene | 1.1 | 2.130 | 6.2E-03 |
| *alr-1* | WBGene00044330 | protein_coding_gene | 1.1 | 2.129 | 9.6E-04 |
| *H04M03.12* | WBGene00019156 | protein_coding_gene | 1.1 | 2.128 | 1.9E-02 |
| *atg-2* | WBGene00019748 | protein_coding_gene | 1.1 | 2.126 | 3.7E-10 |
| *cutl-20* | WBGene00018787 | protein_coding_gene | 1.1 | 2.126 | 2.0E-02 |
| *hlh-3* | WBGene00001950 | protein_coding_gene | 1.1 | 2.125 | 6.2E-03 |
| *M03F4.6* | WBGene00019759 | protein_coding_gene | 1.1 | 2.124 | 2.1E-02 |
| *fipr-1* | WBGene00009096 | protein_coding_gene | 1.1 | 2.121 | 6.7E-05 |
| *fbxa-33* | WBGene00013841 | pseudogene | 1.1 | 2.121 | 4.2E-02 |
| *nfm-1* | WBGene00003593 | protein_coding_gene | 1.1 | 2.121 | 5.6E-07 |
| *ZK550.6* | WBGene00014000 | protein_coding_gene | 1.1 | 2.119 | 5.8E-10 |
| *nhr-88* | WBGene00003678 | protein_coding_gene | 1.1 | 2.119 | 2.4E-04 |
| *nhr-136* | WBGene00003726 | protein_coding_gene | 1.1 | 2.119 | 1.2E-03 |
| *fbxb-17* | WBGene00019037 | protein_coding_gene | 1.1 | 2.118 | 1.6E-02 |
| *elt-7* | WBGene00015981 | protein_coding_gene | 1.1 | 2.117 | 4.8E-05 |
| *R02D5.1* | WBGene00010976 | protein_coding_gene | 1.1 | 2.117 | 2.4E-04 |
| *lido-15* | WBGene00018804 | protein_coding_gene | 1.1 | 2.115 | 1.0E-02 |
| *ilys-4* | WBGene00016958 | protein_coding_gene | 1.1 | 2.114 | 2.7E-03 |
| *F22E5.20* | WBGene00017713 | protein_coding_gene | 1.1 | 2.113 | 4.2E-02 |
| *R07D5.2* | WBGene00011101 | protein_coding_gene | 1.1 | 2.113 | 4.3E-02 |
| *aat-1* | WBGene00000002 | protein_coding_gene | 1.1 | 2.106 | 1.1E-03 |
| *pqm-1* | WBGene00004096 | protein_coding_gene | 1.1 | 2.106 | 9.1E-07 |
| *ZK84.2* | WBGene00022650 | protein_coding_gene | 1.1 | 2.106 | 1.2E-02 |
| *T05E11.9* | WBGene00014505 | ncRNA_gene | 1.1 | 2.105 | 3.7E-07 |
| *shn-1* | WBGene00006444 | protein_coding_gene | 1.1 | 2.103 | 2.7E-05 |
| *irx-1* | WBGene00007984 | protein_coding_gene | 1.1 | 2.101 | 1.3E-05 |
| *H29C22.1* | WBGene00045261 | protein_coding_gene | 1.1 | 2.099 | 1.7E-06 |
| *ZK1010.8* | WBGene00014181 | protein_coding_gene | 1.1 | 2.098 | 3.2E-02 |
| *sdz-5* | WBGene00016578 | protein_coding_gene | 1.1 | 2.097 | 2.8E-02 |
| *Y69A2AL.2* | WBGene00044468 | protein_coding_gene | 1.1 | 2.097 | 4.5E-03 |
| *C28G1.10* | WBGene00219376 | protein_coding_gene | 1.1 | 2.096 | 3.8E-03 |
| *myo-3* | WBGene00003515 | protein_coding_gene | 1.1 | 2.095 | 1.4E-08 |
| *lin-12* | WBGene00003001 | protein_coding_gene | 1.1 | 2.093 | 1.9E-06 |
| *hot-6* | WBGene00001991 | protein_coding_gene | 1.1 | 2.091 | 4.9E-03 |
| *F59E11.5* | WBGene00019111 | protein_coding_gene | 1.1 | 2.091 | 5.8E-05 |

| | | | | | |
|---|---|---|---|---|---|
| *C32E12.4* | WBGene00016330 | protein_coding_gene | 1.1 | 2.087 | 6.1E-08 |
| *T23B3.2* | WBGene00020710 | protein_coding_gene | 1.1 | 2.087 | 1.4E-06 |
| *tag-278* | WBGene00015356 | protein_coding_gene | 1.1 | 2.086 | 3.9E-04 |
| *F42G2.2* | WBGene00018351 | protein_coding_gene | 1.1 | 2.086 | 1.2E-05 |
| *C33D9.13* | WBGene00194651 | protein_coding_gene | 1.1 | 2.080 | 7.1E-04 |
| *crtc-1* | WBGene00021237 | protein_coding_gene | 1.1 | 2.079 | 2.5E-06 |
| *M02H5.8* | WBGene00019744 | protein_coding_gene | 1.1 | 2.078 | 2.3E-02 |
| *akt-2* | WBGene00000103 | protein_coding_gene | 1.1 | 2.077 | 4.1E-06 |
| *tcl-2* | WBGene00006561 | protein_coding_gene | 1.1 | 2.074 | 4.5E-03 |
| *eri-6* | WBGene00016561 | protein_coding_gene | 1.1 | 2.074 | 3.5E-06 |
| *fbxa-212* | WBGene00012568 | pseudogene | 1.1 | 2.074 | 4.8E-04 |
| *nac-2* | WBGene00003518 | protein_coding_gene | 1.0 | 2.070 | 1.1E-03 |
| *lin-17* | WBGene00003006 | protein_coding_gene | 1.0 | 2.070 | 4.6E-04 |
| *nex-2* | WBGene00003589 | protein_coding_gene | 1.0 | 2.068 | 1.4E-10 |
| *C40H1.7* | WBGene00008042 | protein_coding_gene | 1.0 | 2.068 | 6.1E-05 |
| *twk-31* | WBGene00006683 | protein_coding_gene | 1.0 | 2.065 | 2.8E-03 |
| *endu-2* | WBGene00019779 | protein_coding_gene | 1.0 | 2.064 | 1.5E-05 |
| *unc-6* | WBGene00006746 | protein_coding_gene | 1.0 | 2.061 | 1.9E-04 |
| *sdz-30* | WBGene00011432 | protein_coding_gene | 1.0 | 2.061 | 7.3E-04 |
| *Y34D9A.8* | WBGene00021333 | protein_coding_gene | 1.0 | 2.060 | 6.3E-03 |
| *MTCE.33* | WBGene00014472 | rRNA_gene | 1.0 | 2.059 | 9.4E-04 |
| *cdh-3* | WBGene00000395 | protein_coding_gene | 1.0 | 2.059 | 9.3E-05 |
| *ate-1* | WBGene00010615 | protein_coding_gene | 1.0 | 2.057 | 5.9E-14 |
| *T22C8.6* | WBGene00011927 | protein_coding_gene | 1.0 | 2.056 | 2.7E-02 |
| *F09C3.2* | WBGene00008610 | protein_coding_gene | 1.0 | 2.055 | 1.9E-03 |
| *C34C12.7* | WBGene00007926 | protein_coding_gene | 1.0 | 2.054 | 4.7E-03 |
| *Y54G2A.32* | WBGene00021894 | protein_coding_gene | 1.0 | 2.054 | 2.7E-03 |
| *nhr-44* | WBGene00003634 | protein_coding_gene | 1.0 | 2.053 | 2.6E-02 |
| *R10D12.1* | WBGene00011185 | protein_coding_gene | 1.0 | 2.052 | 4.5E-02 |
| *ugt-51* | WBGene00015369 | protein_coding_gene | 1.0 | 2.051 | 8.7E-03 |
| *Y73B6BL.44* | WBGene00044350 | protein_coding_gene | 1.0 | 2.051 | 2.3E-03 |
| *dhs-27* | WBGene00000990 | protein_coding_gene | 1.0 | 2.050 | 1.4E-04 |
| *R10H1.1* | WBGene00019998 | protein_coding_gene | 1.0 | 2.050 | 6.1E-04 |
| *oig-3* | WBGene00003861 | protein_coding_gene | 1.0 | 2.049 | 1.9E-04 |
| *mtl-2* | WBGene00003474 | protein_coding_gene | 1.0 | 2.049 | 3.7E-05 |
| *D2045.8* | WBGene00008425 | protein_coding_gene | 1.0 | 2.047 | 7.0E-04 |
| *F14F9.5* | WBGene00017468 | protein_coding_gene | 1.0 | 2.046 | 1.1E-02 |
| *T21B4.21* | WBGene00220076 | ncRNA_gene | 1.0 | 2.045 | 8.8E-03 |
| *valv-1* | WBGene00015216 | protein_coding_gene | 1.0 | 2.045 | 1.3E-02 |
| *C13C4.4* | WBGene00007548 | protein_coding_gene | 1.0 | 2.045 | 7.6E-04 |
| *Y51B9A.9* | WBGene00013091 | protein_coding_gene | 1.0 | 2.044 | 5.1E-03 |
| *K02A6.3* | WBGene00019294 | protein_coding_gene | 1.0 | 2.042 | 1.6E-04 |
| *exc-9* | WBGene00017644 | protein_coding_gene | 1.0 | 2.041 | 7.1E-04 |
| *F52B10.3* | WBGene00044304 | protein_coding_gene | 1.0 | 2.041 | 5.0E-02 |
| *C33H5.13* | WBGene00016379 | protein_coding_gene | 1.0 | 2.040 | 3.0E-04 |
| *C15H7.3* | WBGene00007610 | protein_coding_gene | 1.0 | 2.040 | 5.0E-02 |
| *Y32F6B.1* | WBGene00012531 | protein_coding_gene | 1.0 | 2.039 | 2.0E-07 |
| *srj-14* | WBGene00005602 | protein_coding_gene | 1.0 | 2.038 | 8.1E-03 |
| *mig-10* | WBGene00003243 | protein_coding_gene | 1.0 | 2.038 | 1.2E-07 |
| *ZK1053.4* | WBGene00014199 | protein_coding_gene | 1.0 | 2.037 | 1.8E-09 |
| *clec-151* | WBGene00008659 | protein_coding_gene | 1.0 | 2.036 | 4.6E-02 |
| *Y105E8B.9* | WBGene00013693 | protein_coding_gene | 1.0 | 2.035 | 1.4E-03 |
| *ins-17* | WBGene00002100 | protein_coding_gene | 1.0 | 2.034 | 2.3E-03 |
| *sulp-2* | WBGene00017464 | protein_coding_gene | 1.0 | 2.034 | 1.2E-02 |
| *Y57A10A.14* | WBGene00013257 | protein_coding_gene | 1.0 | 2.033 | 6.7E-03 |
| *hnd-1* | WBGene00001981 | protein_coding_gene | 1.0 | 2.033 | 3.8E-03 |
| *nspd-6* | WBGene00043147 | protein_coding_gene | 1.0 | 2.033 | 2.6E-02 |
| *nhr-119* | WBGene00003709 | protein_coding_gene | 1.0 | 2.030 | 3.7E-05 |
| *C41G7.8* | WBGene00077525 | protein_coding_gene | 1.0 | 2.028 | 1.7E-02 |
| *F56H9.2* | WBGene00010174 | protein_coding_gene | 1.0 | 2.027 | 3.5E-04 |

| | | | | | |
|---|---|---|---|---|---|
| *sul-2* | WBGene00006309 | protein_coding_gene | 1.0 | 2.025 | 1.6E-02 |
| *Y2H9A.4* | WBGene00012381 | protein_coding_gene | 1.0 | 2.022 | 1.3E-02 |
| *ric-3* | WBGene00004363 | protein_coding_gene | 1.0 | 2.022 | 1.1E-02 |
| *nhr-107* | WBGene00003697 | protein_coding_gene | 1.0 | 2.021 | 2.9E-03 |
| *T05G5.4* | WBGene00011499 | protein_coding_gene | 1.0 | 2.019 | 2.9E-02 |
| *F42G4.2* | WBGene00009641 | protein_coding_gene | 1.0 | 2.017 | 4.2E-02 |
| *F46G10.2* | WBGene00009797 | protein_coding_gene | 1.0 | 2.016 | 3.7E-05 |
| *clec-190* | WBGene00010953 | protein_coding_gene | 1.0 | 2.014 | 2.9E-02 |
| *nob-1* | WBGene00003779 | protein_coding_gene | 1.0 | 2.014 | 2.0E-02 |
| *C30G7.4* | WBGene00044178 | protein_coding_gene | 1.0 | 2.014 | 3.4E-02 |
| *zig-7* | WBGene00006984 | protein_coding_gene | 1.0 | 2.012 | 2.9E-07 |
| *R02D5.3* | WBGene00010977 | protein_coding_gene | 1.0 | 2.012 | 4.5E-03 |
| *C34E11.4* | WBGene00007937 | protein_coding_gene | 1.0 | 2.011 | 4.9E-02 |
| *mab-21* | WBGene00003112 | protein_coding_gene | 1.0 | 2.009 | 8.8E-03 |
| *cpz-2* | WBGene00000789 | protein_coding_gene | 1.0 | 2.008 | 2.0E-03 |
| *srap-1* | WBGene00011522 | protein_coding_gene | 1.0 | 2.008 | 2.3E-06 |
| *asah-2* | WBGene00009192 | protein_coding_gene | 1.0 | 2.008 | 6.3E-04 |
| *C35D10.8* | WBGene00016445 | protein_coding_gene | 1.0 | 2.007 | 4.8E-03 |
| *F53F8.4* | WBGene00010001 | protein_coding_gene | 1.0 | 2.006 | 4.6E-03 |
| *glo-3* | WBGene00010341 | protein_coding_gene | 1.0 | 2.004 | 1.7E-02 |
| *T19D7.6* | WBGene00020574 | protein_coding_gene | 1.0 | 2.004 | 2.8E-02 |
| *T05H4.15* | WBGene00020276 | protein_coding_gene | 1.0 | 2.004 | 3.5E-02 |
| *skr-20* | WBGene00004826 | protein_coding_gene | 1.0 | 2.002 | 1.9E-06 |
| *tsp-20* | WBGene00006646 | protein_coding_gene | 1.0 | 2.001 | 6.6E-03 |
| *lgc-32* | WBGene00020569 | protein_coding_gene | -1.0 | 0.500 | 4.6E-03 |
| *trap-3* | WBGene00021420 | protein_coding_gene | -1.0 | 0.500 | 1.6E-10 |
| *ifd-1* | WBGene00002057 | protein_coding_gene | -1.0 | 0.499 | 4.3E-03 |
| *M01G5.1* | WBGene00019715 | protein_coding_gene | -1.0 | 0.499 | 3.3E-14 |
| *cdr-7* | WBGene00010472 | protein_coding_gene | -1.0 | 0.499 | 2.0E-03 |
| *Y102A11A.3* | WBGene00022413 | protein_coding_gene | -1.0 | 0.498 | 1.7E-04 |
| *pid-5* | WBGene00021555 | protein_coding_gene | -1.0 | 0.498 | 8.2E-12 |
| *let-805* | WBGene00002915 | protein_coding_gene | -1.0 | 0.498 | 4.3E-06 |
| *col-150* | WBGene00000723 | protein_coding_gene | -1.0 | 0.497 | 5.8E-03 |
| *cyp-33E2* | WBGene00018334 | protein_coding_gene | -1.0 | 0.497 | 4.9E-03 |
| *cpn-4* | WBGene00000780 | protein_coding_gene | -1.0 | 0.496 | 3.7E-06 |
| *F54D5.12* | WBGene00010055 | protein_coding_gene | -1.0 | 0.496 | 1.3E-04 |
| *Y75B8A.24* | WBGene00013557 | protein_coding_gene | -1.0 | 0.496 | 2.5E-10 |
| *set-9* | WBGene00017482 | protein_coding_gene | -1.0 | 0.496 | 2.2E-13 |
| *tkt-1* | WBGene00008506 | protein_coding_gene | -1.0 | 0.495 | 6.4E-14 |
| *lsy-27* | WBGene00009834 | protein_coding_gene | -1.0 | 0.494 | 2.0E-04 |
| *C06A5.12* | WBGene00044414 | protein_coding_gene | -1.0 | 0.493 | 3.1E-02 |
| *rhy-1* | WBGene00012324 | protein_coding_gene | -1.0 | 0.493 | 6.6E-05 |
| *Y119D3B.21* | WBGene00022497 | protein_coding_gene | -1.0 | 0.491 | 6.3E-05 |
| *F53C11.1* | WBGene00009971 | protein_coding_gene | -1.0 | 0.491 | 3.6E-03 |
| *elo-1* | WBGene00001239 | protein_coding_gene | -1.0 | 0.490 | 6.2E-11 |
| *F55H12.4* | WBGene00010135 | protein_coding_gene | -1.0 | 0.490 | 1.3E-04 |
| *F35H10.10* | WBGene00018073 | protein_coding_gene | -1.0 | 0.490 | 3.6E-05 |
| *pfas-1* | WBGene00008654 | protein_coding_gene | -1.0 | 0.490 | 5.4E-06 |
| *mua-3* | WBGene00003482 | protein_coding_gene | -1.0 | 0.489 | 1.1E-05 |
| *memi-2* | WBGene00013380 | protein_coding_gene | -1.0 | 0.489 | 1.1E-05 |
| *C05D2.8* | WBGene00015469 | protein_coding_gene | -1.0 | 0.488 | 9.0E-06 |
| *C17C3.1* | WBGene00015887 | protein_coding_gene | -1.0 | 0.487 | 7.6E-04 |
| *vha-8* | WBGene00006917 | protein_coding_gene | -1.0 | 0.487 | 2.4E-11 |
| *C17H12.8* | WBGene00015933 | protein_coding_gene | -1.0 | 0.487 | 1.8E-02 |
| *tfg-1* | WBGene00006565 | protein_coding_gene | -1.0 | 0.486 | 1.4E-09 |
| *vha-6* | WBGene00006915 | protein_coding_gene | -1.0 | 0.486 | 1.2E-05 |
| *bckd-1B* | WBGene00006518 | protein_coding_gene | -1.0 | 0.486 | 9.3E-13 |
| *vha-19* | WBGene00021952 | protein_coding_gene | -1.0 | 0.486 | 5.2E-11 |
| *Y102E9.5* | WBGene00044452 | protein_coding_gene | -1.0 | 0.485 | 2.0E-02 |
| *T13C5.6* | WBGene00020486 | protein_coding_gene | -1.0 | 0.485 | 1.4E-04 |

| | | | | | |
|---|---|---|---|---|---|
| *ZK909.6* | WBGene00014150 | protein_coding_gene | -1.0 | 0.485 | 3.4E-04 |
| *acs-21* | WBGene00016849 | protein_coding_gene | -1.0 | 0.485 | 1.3E-02 |
| *papl-1* | WBGene00008936 | protein_coding_gene | -1.0 | 0.485 | 7.3E-07 |
| *glt-1* | WBGene00001620 | protein_coding_gene | -1.0 | 0.485 | 1.8E-10 |
| *T16H12.2* | WBGene00011812 | protein_coding_gene | -1.0 | 0.484 | 3.1E-02 |
| *oatr-1* | WBGene00015814 | protein_coding_gene | -1.0 | 0.483 | 2.3E-15 |
| *R06C1.4* | WBGene00011059 | protein_coding_gene | -1.0 | 0.483 | 4.8E-06 |
| *M02B7.2* | WBGene00019724 | protein_coding_gene | -1.0 | 0.483 | 5.6E-06 |
| *C11E4.7* | WBGene00007521 | protein_coding_gene | -1.0 | 0.483 | 1.1E-03 |
| *F54F7.2* | WBGene00010064 | protein_coding_gene | -1.1 | 0.482 | 4.1E-04 |
| *B0511.11* | WBGene00015236 | protein_coding_gene | -1.1 | 0.481 | 8.0E-03 |
| *C06A1.4* | WBGene00014667 | pseudogene | -1.1 | 0.481 | 4.7E-07 |
| *C52G5.2* | WBGene00008260 | protein_coding_gene | -1.1 | 0.480 | 2.1E-03 |
| *ccm-2* | WBGene00019462 | protein_coding_gene | -1.1 | 0.480 | 2.2E-02 |
| *F14H12.3* | WBGene00017471 | protein_coding_gene | -1.1 | 0.480 | 1.3E-02 |
| *F56F11.2* | WBGene00018989 | protein_coding_gene | -1.1 | 0.480 | 2.5E-02 |
| *ctsa-1.2* | WBGene00008741 | protein_coding_gene | -1.1 | 0.479 | 6.3E-05 |
| *C14B1.7* | WBGene00007578 | pseudogene | -1.1 | 0.479 | 4.4E-11 |
| *C14C11.2* | WBGene00015766 | protein_coding_gene | -1.1 | 0.479 | 1.7E-07 |
| *cal-5* | WBGene00006861 | protein_coding_gene | -1.1 | 0.479 | 3.5E-05 |
| *egg-5* | WBGene00020035 | protein_coding_gene | -1.1 | 0.478 | 1.5E-08 |
| *C14A11.6* | WBGene00015751 | protein_coding_gene | -1.1 | 0.478 | 4.2E-02 |
| *F17C8.9* | WBGene00185014 | protein_coding_gene | -1.1 | 0.478 | 2.4E-04 |
| *F47F2.1* | WBGene00018569 | protein_coding_gene | -1.1 | 0.477 | 7.3E-04 |
| *C10G8.8* | WBGene00015684 | protein_coding_gene | -1.1 | 0.477 | 1.4E-11 |
| *nas-20* | WBGene00003539 | protein_coding_gene | -1.1 | 0.476 | 4.7E-03 |
| *C55A6.12* | WBGene00044019 | protein_coding_gene | -1.1 | 0.476 | 4.2E-06 |
| *B0303.11* | WBGene00015131 | protein_coding_gene | -1.1 | 0.476 | 7.0E-10 |
| *del-6* | WBGene00011891 | protein_coding_gene | -1.1 | 0.475 | 2.4E-07 |
| *odc-1* | WBGene00003844 | protein_coding_gene | -1.1 | 0.475 | 2.4E-09 |
| *ltah-1.2* | WBGene00022610 | protein_coding_gene | -1.1 | 0.474 | 1.8E-11 |
| *T04F3.1* | WBGene00011436 | protein_coding_gene | -1.1 | 0.474 | 1.0E-06 |
| *C01G10.9* | WBGene00007236 | protein_coding_gene | -1.1 | 0.474 | 4.6E-05 |
| *Y53C10A.5* | WBGene00013136 | protein_coding_gene | -1.1 | 0.472 | 3.9E-06 |
| *H10E21.1* | WBGene00019182 | protein_coding_gene | -1.1 | 0.472 | 3.6E-02 |
| *ZK652.8* | WBGene00022786 | protein_coding_gene | -1.1 | 0.471 | 4.4E-03 |
| *tbc-10* | WBGene00019915 | protein_coding_gene | -1.1 | 0.471 | 3.9E-04 |
| *nceh-1* | WBGene00012810 | protein_coding_gene | -1.1 | 0.471 | 3.6E-02 |
| *vrp-1* | WBGene00021869 | protein_coding_gene | -1.1 | 0.470 | 9.2E-18 |
| *C48B4.7* | WBGene00008170 | protein_coding_gene | -1.1 | 0.469 | 8.3E-06 |
| *pmp-4* | WBGene00004061 | protein_coding_gene | -1.1 | 0.468 | 6.6E-06 |
| *srd-14* | WBGene00005092 | pseudogene | -1.1 | 0.468 | 1.6E-02 |
| *ZK105.1* | WBGene00022653 | protein_coding_gene | -1.1 | 0.467 | 6.5E-06 |
| *cest-9.1* | WBGene00007692 | protein_coding_gene | -1.1 | 0.467 | 2.1E-04 |
| *C26B9.5* | WBGene00016134 | protein_coding_gene | -1.1 | 0.467 | 8.2E-09 |
| *nduo-6* | WBGene00010957 | protein_coding_gene | -1.1 | 0.466 | 1.5E-03 |
| *Y111B2A.1* | WBGene00013727 | protein_coding_gene | -1.1 | 0.466 | 5.4E-04 |
| *ldh-1* | WBGene00002262 | protein_coding_gene | -1.1 | 0.465 | 9.4E-11 |
| *pgph-3* | WBGene00016892 | protein_coding_gene | -1.1 | 0.465 | 9.9E-03 |
| *K08C7.1* | WBGene00010646 | protein_coding_gene | -1.1 | 0.465 | 6.1E-05 |
| *cyp-25A2* | WBGene00007964 | protein_coding_gene | -1.1 | 0.465 | 1.1E-05 |
| *twk-17* | WBGene00006671 | protein_coding_gene | -1.1 | 0.465 | 2.4E-02 |
| *trpp-12* | WBGene00016614 | protein_coding_gene | -1.1 | 0.464 | 2.1E-06 |
| *slc-17.3* | WBGene00015333 | protein_coding_gene | -1.1 | 0.463 | 4.8E-03 |
| *coq-1* | WBGene00000761 | protein_coding_gene | -1.1 | 0.462 | 4.3E-11 |
| *B0222.5* | WBGene00015056 | protein_coding_gene | -1.1 | 0.461 | 8.2E-08 |
| *mct-4* | WBGene00003986 | protein_coding_gene | -1.1 | 0.461 | 7.9E-08 |
| *acs-5* | WBGene00013575 | protein_coding_gene | -1.1 | 0.461 | 1.2E-10 |
| *F35F10.1* | WBGene00018052 | protein_coding_gene | -1.1 | 0.461 | 3.0E-02 |
| *nhr-132* | WBGene00003722 | protein_coding_gene | -1.1 | 0.461 | 9.7E-03 |

| | | | | | |
|---|---|---|---|---|---|
| *ptr-16* | WBGene00004230 | protein_coding_gene | -1.1 | 0.461 | 2.2E-02 |
| *T01C8.2* | WBGene00020143 | protein_coding_gene | -1.1 | 0.461 | 4.4E-07 |
| *col-106* | WBGene00000680 | protein_coding_gene | -1.1 | 0.461 | 1.9E-11 |
| *W04B5.3* | WBGene00021020 | protein_coding_gene | -1.1 | 0.460 | 5.4E-04 |
| *mpc-1* | WBGene00011119 | protein_coding_gene | -1.1 | 0.460 | 3.3E-14 |
| *fah-1* | WBGene00019620 | protein_coding_gene | -1.1 | 0.459 | 7.7E-13 |
| *F58D5.2* | WBGene00010242 | protein_coding_gene | -1.1 | 0.459 | 2.9E-02 |
| *cest-13* | WBGene00010116 | protein_coding_gene | -1.1 | 0.458 | 9.8E-04 |
| *C31H2.4* | WBGene00016294 | protein_coding_gene | -1.1 | 0.458 | 3.4E-04 |
| *fbxc-44* | WBGene00015819 | protein_coding_gene | -1.1 | 0.458 | 4.9E-06 |
| *F31D4.8* | WBGene00009293 | protein_coding_gene | -1.1 | 0.457 | 2.6E-03 |
| *ugt-26* | WBGene00015695 | protein_coding_gene | -1.1 | 0.457 | 7.5E-04 |
| *C16E9.1* | WBGene00015865 | protein_coding_gene | -1.1 | 0.457 | 3.2E-02 |
| *Y43F11A.6* | WBGene00014576 | pseudogene | -1.1 | 0.457 | 2.7E-06 |
| *Y46H3C.7* | WBGene00021607 | protein_coding_gene | -1.1 | 0.456 | 1.3E-02 |
| *col-65* | WBGene00000641 | protein_coding_gene | -1.1 | 0.456 | 4.4E-02 |
| *tatn-1* | WBGene00009628 | protein_coding_gene | -1.1 | 0.455 | 1.0E-05 |
| *F13H8.8* | WBGene00017441 | protein_coding_gene | -1.1 | 0.455 | 1.1E-02 |
| *Y54E10A.17* | WBGene00021835 | protein_coding_gene | -1.1 | 0.454 | 7.7E-05 |
| *pat-12* | WBGene00003936 | protein_coding_gene | -1.1 | 0.454 | 2.9E-08 |
| *tald-1* | WBGene00021286 | protein_coding_gene | -1.1 | 0.453 | 4.0E-13 |
| *K10C2.12* | WBGene00206386 | protein_coding_gene | -1.1 | 0.452 | 5.0E-02 |
| *spds-1* | WBGene00012909 | protein_coding_gene | -1.1 | 0.452 | 3.6E-07 |
| *C23H3.2* | WBGene00016018 | protein_coding_gene | -1.1 | 0.452 | 2.6E-08 |
| *mpc-2* | WBGene00018765 | protein_coding_gene | -1.1 | 0.452 | 1.1E-08 |
| *R102.4* | WBGene00011291 | protein_coding_gene | -1.1 | 0.451 | 1.8E-11 |
| *him-4* | WBGene00001863 | protein_coding_gene | -1.1 | 0.451 | 7.4E-06 |
| *H43E16.1* | WBGene00019273 | protein_coding_gene | -1.1 | 0.451 | 3.2E-04 |
| *lbp-6* | WBGene00002258 | protein_coding_gene | -1.1 | 0.451 | 1.2E-12 |
| *Y14H12B.2* | WBGene00021192 | protein_coding_gene | -1.1 | 0.451 | 9.0E-10 |
| *F21H7.5* | WBGene00009031 | protein_coding_gene | -1.1 | 0.451 | 4.0E-02 |
| *gly-8* | WBGene00001633 | protein_coding_gene | -1.1 | 0.451 | 7.0E-08 |
| *DC2.5* | WBGene00017082 | protein_coding_gene | -1.2 | 0.450 | 2.1E-04 |
| *dhs-14* | WBGene00000977 | protein_coding_gene | -1.2 | 0.449 | 4.3E-09 |
| *Y46H3C.5* | WBGene00021605 | protein_coding_gene | -1.2 | 0.448 | 3.5E-03 |
| *C26C6.9* | WBGene00007743 | protein_coding_gene | -1.2 | 0.448 | 6.9E-07 |
| *R17.3* | WBGene00011269 | protein_coding_gene | -1.2 | 0.447 | 3.6E-03 |
| *clec-41* | WBGene00007153 | protein_coding_gene | -1.2 | 0.446 | 4.0E-07 |
| *C06A1.12* | WBGene00219803 | ncRNA_gene | -1.2 | 0.446 | 1.1E-03 |
| *T11F9.12* | WBGene00011713 | protein_coding_gene | -1.2 | 0.446 | 4.5E-08 |
| *scav-5* | WBGene00011076 | protein_coding_gene | -1.2 | 0.446 | 5.2E-03 |
| *H03A11.2* | WBGene00010357 | protein_coding_gene | -1.2 | 0.445 | 5.2E-05 |
| *acdh-9* | WBGene00017874 | protein_coding_gene | -1.2 | 0.444 | 1.2E-06 |
| *gale-1* | WBGene00008132 | protein_coding_gene | -1.2 | 0.443 | 5.0E-18 |
| *Y62H9A.4* | WBGene00013392 | protein_coding_gene | -1.2 | 0.443 | 2.6E-07 |
| *sec-61.A* | WBGene00013311 | protein_coding_gene | -1.2 | 0.443 | 3.1E-20 |
| *trap-1* | WBGene00022122 | protein_coding_gene | -1.2 | 0.441 | 1.7E-15 |
| *T22F7.4* | WBGene00020703 | protein_coding_gene | -1.2 | 0.440 | 3.3E-02 |
| *nhr-80* | WBGene00003670 | protein_coding_gene | -1.2 | 0.440 | 5.7E-05 |
| *cysl-3* | WBGene00019962 | protein_coding_gene | -1.2 | 0.439 | 2.3E-04 |
| *vha-13* | WBGene00013025 | protein_coding_gene | -1.2 | 0.438 | 6.3E-17 |
| *pqn-72* | WBGene00004154 | protein_coding_gene | -1.2 | 0.438 | 2.0E-02 |
| *lec-8* | WBGene00002271 | protein_coding_gene | -1.2 | 0.438 | 8.8E-05 |
| *R06B10.2* | WBGene00019913 | protein_coding_gene | -1.2 | 0.438 | 3.6E-02 |
| *gon-1* | WBGene00001650 | protein_coding_gene | -1.2 | 0.437 | 7.9E-12 |
| *sea-1* | WBGene00004750 | protein_coding_gene | -1.2 | 0.437 | 9.4E-05 |
| *cytb-5.1* | WBGene00007848 | protein_coding_gene | -1.2 | 0.436 | 1.9E-09 |
| *clec-52* | WBGene00015052 | protein_coding_gene | -1.2 | 0.436 | 1.2E-03 |
| *vha-5* | WBGene00006914 | protein_coding_gene | -1.2 | 0.436 | 1.2E-11 |
| *drd-5* | WBGene00018885 | protein_coding_gene | -1.2 | 0.436 | 7.2E-04 |

| | | | | | |
|---|---|---|---|---|---|
| *F39B2.5* | WBGene00009556 | protein_coding_gene | -1.2 | 0.433 | 8.7E-08 |
| *ZK1193.2* | WBGene00022858 | protein_coding_gene | -1.2 | 0.433 | 4.8E-05 |
| *F19C7.1* | WBGene00017591 | protein_coding_gene | -1.2 | 0.433 | 4.6E-04 |
| *C48D1.5* | WBGene00008178 | protein_coding_gene | -1.2 | 0.433 | 8.0E-03 |
| *mct-3* | WBGene00010834 | protein_coding_gene | -1.2 | 0.432 | 1.4E-09 |
| *nduo-1* | WBGene00010959 | protein_coding_gene | -1.2 | 0.431 | 5.4E-08 |
| *Y26E6A.3* | WBGene00012514 | protein_coding_gene | -1.2 | 0.431 | 2.2E-05 |
| *K03B8.8* | WBGene00010525 | protein_coding_gene | -1.2 | 0.431 | 4.3E-03 |
| *C52B11.5* | WBGene00016874 | protein_coding_gene | -1.2 | 0.429 | 1.7E-07 |
| *irg-5* | WBGene00009429 | protein_coding_gene | -1.2 | 0.429 | 2.1E-02 |
| *idh-1* | WBGene00010317 | protein_coding_gene | -1.2 | 0.429 | 3.5E-21 |
| *cnnm-3* | WBGene00016343 | protein_coding_gene | -1.2 | 0.428 | 8.6E-07 |
| *F20G2.2* | WBGene00008986 | protein_coding_gene | -1.2 | 0.427 | 3.5E-10 |
| *R09H10.5* | WBGene00011182 | protein_coding_gene | -1.2 | 0.427 | 9.0E-05 |
| *eri-7* | WBGene00016566 | protein_coding_gene | -1.2 | 0.426 | 1.1E-10 |
| *lsl-1* | WBGene00009937 | protein_coding_gene | -1.2 | 0.426 | 3.6E-18 |
| *C05C10.3* | WBGene00007330 | protein_coding_gene | -1.2 | 0.426 | 1.1E-10 |
| *gly-13* | WBGene00001638 | protein_coding_gene | -1.2 | 0.424 | 5.8E-12 |
| *C18B2.3* | WBGene00015954 | protein_coding_gene | -1.2 | 0.424 | 1.6E-03 |
| *pcyt-1* | WBGene00017241 | protein_coding_gene | -1.2 | 0.423 | 6.0E-15 |
| *F36G9.3* | WBGene00009485 | protein_coding_gene | -1.2 | 0.423 | 9.1E-03 |
| *fbp-1* | WBGene00001404 | protein_coding_gene | -1.2 | 0.423 | 2.0E-21 |
| *fbxb-84* | WBGene00020544 | protein_coding_gene | -1.2 | 0.423 | 1.1E-02 |
| *gba-4* | WBGene00021160 | protein_coding_gene | -1.2 | 0.422 | 5.1E-04 |
| *cyp-36A1* | WBGene00007913 | protein_coding_gene | -1.2 | 0.422 | 4.1E-05 |
| *suro-1* | WBGene00011235 | protein_coding_gene | -1.2 | 0.422 | 1.7E-02 |
| *toh-1* | WBGene00006591 | protein_coding_gene | -1.2 | 0.422 | 7.6E-05 |
| *add-1* | WBGene00000072 | protein_coding_gene | -1.2 | 0.421 | 1.3E-06 |
| *pxn-1* | WBGene00004256 | protein_coding_gene | -1.2 | 0.421 | 7.3E-08 |
| *Y73B3A.18* | WBGene00022219 | pseudogene | -1.2 | 0.421 | 3.1E-03 |
| *ugt-47* | WBGene00011006 | protein_coding_gene | -1.2 | 0.421 | 8.2E-14 |
| *adt-3* | WBGene00020567 | protein_coding_gene | -1.3 | 0.419 | 3.1E-03 |
| *asna-2* | WBGene00018577 | pseudogene | -1.3 | 0.419 | 4.1E-03 |
| *F56F10.1* | WBGene00018984 | protein_coding_gene | -1.3 | 0.418 | 1.3E-04 |
| *D2096.11* | WBGene00017078 | protein_coding_gene | -1.3 | 0.418 | 3.2E-02 |
| *col-96* | WBGene00000671 | protein_coding_gene | -1.3 | 0.418 | 1.2E-03 |
| *nkb-3* | WBGene00010117 | protein_coding_gene | -1.3 | 0.417 | 1.9E-11 |
| *kat-1* | WBGene00002183 | protein_coding_gene | -1.3 | 0.417 | 5.9E-08 |
| *ech-6* | WBGene00001155 | protein_coding_gene | -1.3 | 0.417 | 1.2E-05 |
| *let-767* | WBGene00002891 | protein_coding_gene | -1.3 | 0.416 | 3.1E-15 |
| *pmt-2* | WBGene00018811 | protein_coding_gene | -1.3 | 0.416 | 3.6E-09 |
| *D1054.1* | WBGene00008370 | protein_coding_gene | -1.3 | 0.416 | 5.6E-08 |
| *zmp-3* | WBGene00016283 | protein_coding_gene | -1.3 | 0.415 | 3.0E-05 |
| *C14E2.12* | WBGene00219816 | snoRNA_gene | -1.3 | 0.415 | 2.0E-02 |
| *asp-12* | WBGene00017678 | protein_coding_gene | -1.3 | 0.415 | 3.3E-03 |
| *D1054.8* | WBGene00008375 | protein_coding_gene | -1.3 | 0.415 | 8.8E-11 |
| *T01C8.3* | WBGene00020144 | protein_coding_gene | -1.3 | 0.415 | 3.3E-02 |
| *B0403.3* | WBGene00015167 | protein_coding_gene | -1.3 | 0.415 | 2.2E-03 |
| *F28H7.8* | WBGene00009241 | protein_coding_gene | -1.3 | 0.414 | 8.9E-04 |
| *cld-9* | WBGene00009432 | protein_coding_gene | -1.3 | 0.414 | 8.0E-03 |
| *R04A9.9* | WBGene00195183 | protein_coding_gene | -1.3 | 0.414 | 1.2E-04 |
| *sth-1* | WBGene00006060 | protein_coding_gene | -1.3 | 0.414 | 5.5E-03 |
| *ser-5* | WBGene00008890 | protein_coding_gene | -1.3 | 0.413 | 2.8E-02 |
| *dpf-6* | WBGene00001059 | protein_coding_gene | -1.3 | 0.413 | 6.4E-03 |
| *C06E2.1* | WBGene00015527 | protein_coding_gene | -1.3 | 0.412 | 2.0E-02 |
| *T08B1.1* | WBGene00020340 | protein_coding_gene | -1.3 | 0.412 | 1.1E-03 |
| *T08D2.7* | WBGene00011612 | protein_coding_gene | -1.3 | 0.411 | 4.6E-02 |
| *C44B7.7* | WBGene00016629 | protein_coding_gene | -1.3 | 0.411 | 8.2E-05 |
| *cth-2* | WBGene00022856 | protein_coding_gene | -1.3 | 0.411 | 7.5E-26 |
| *hprt-1* | WBGene00013690 | protein_coding_gene | -1.3 | 0.409 | 9.0E-20 |

| | | | | | |
|---|---|---|---|---|---|
| *his-38* | WBGene00001912 | protein_coding_gene | -1.3 | 0.409 | 4.0E-02 |
| *mans-2* | WBGene00008258 | protein_coding_gene | -1.3 | 0.409 | 4.8E-14 |
| *C18E9.5* | WBGene00007685 | protein_coding_gene | -1.3 | 0.408 | 4.1E-10 |
| *bcmo-2* | WBGene00018755 | protein_coding_gene | -1.3 | 0.408 | 6.9E-05 |
| *pmp-1* | WBGene00004058 | protein_coding_gene | -1.3 | 0.408 | 4.1E-11 |
| *afmd-1* | WBGene00017051 | protein_coding_gene | -1.3 | 0.407 | 1.7E-03 |
| *sdz-28* | WBGene00020073 | protein_coding_gene | -1.3 | 0.405 | 8.7E-04 |
| *D2096.6* | WBGene00017073 | protein_coding_gene | -1.3 | 0.405 | 3.5E-02 |
| *ctb-1* | WBGene00000829 | protein_coding_gene | -1.3 | 0.405 | 8.6E-12 |
| *ckc-1* | WBGene00000515 | protein_coding_gene | -1.3 | 0.405 | 6.5E-06 |
| *gst-36* | WBGene00001784 | protein_coding_gene | -1.3 | 0.404 | 4.5E-07 |
| *klu-2* | WBGene00022592 | protein_coding_gene | -1.3 | 0.404 | 5.4E-05 |
| *bas-1* | WBGene00000239 | protein_coding_gene | -1.3 | 0.403 | 1.6E-07 |
| *C18A11.3* | WBGene00015949 | protein_coding_gene | -1.3 | 0.403 | 1.2E-02 |
| *Y37H2A.14* | WBGene00045416 | protein_coding_gene | -1.3 | 0.402 | 4.3E-02 |
| *grsp-1* | WBGene00012840 | protein_coding_gene | -1.3 | 0.402 | 5.4E-04 |
| *ZC449.7* | WBGene00045365 | protein_coding_gene | -1.3 | 0.402 | 1.5E-02 |
| *K08D8.3* | WBGene00010657 | protein_coding_gene | -1.3 | 0.402 | 8.4E-06 |
| *R08F11.4* | WBGene00019968 | protein_coding_gene | -1.3 | 0.402 | 4.3E-02 |
| *col-113* | WBGene00000687 | protein_coding_gene | -1.3 | 0.401 | 1.6E-02 |
| *C01B10.3* | WBGene00015278 | protein_coding_gene | -1.3 | 0.401 | 4.2E-08 |
| *ttll-15* | WBGene00010630 | protein_coding_gene | -1.3 | 0.400 | 3.4E-04 |
| *ZK512.7* | WBGene00013986 | protein_coding_gene | -1.3 | 0.400 | 2.0E-02 |
| *Y49F6C.8* | WBGene00021730 | protein_coding_gene | -1.3 | 0.400 | 3.7E-06 |
| *slc-17.5* | WBGene00017681 | protein_coding_gene | -1.3 | 0.399 | 1.7E-05 |
| *F25D1.5* | WBGene00009110 | protein_coding_gene | -1.3 | 0.399 | 1.5E-04 |
| *ptr-24* | WBGene00004238 | protein_coding_gene | -1.3 | 0.399 | 1.3E-12 |
| *F42A9.6* | WBGene00018335 | protein_coding_gene | -1.3 | 0.398 | 9.2E-11 |
| *nspc-1* | WBGene00023477 | protein_coding_gene | -1.3 | 0.397 | 1.9E-02 |
| *nas-7* | WBGene00003526 | protein_coding_gene | -1.3 | 0.397 | 2.4E-03 |
| *ahcy-1* | WBGene00019322 | protein_coding_gene | -1.3 | 0.397 | 3.0E-17 |
| *aass-1* | WBGene00019819 | protein_coding_gene | -1.3 | 0.396 | 7.5E-12 |
| *ule-2* | WBGene00045038 | protein_coding_gene | -1.3 | 0.396 | 1.0E-13 |
| *qdpr-1* | WBGene00011398 | protein_coding_gene | -1.3 | 0.395 | 7.3E-21 |
| *daf-28* | WBGene00000920 | protein_coding_gene | -1.3 | 0.395 | 7.0E-03 |
| *tmem-135* | WBGene00019345 | protein_coding_gene | -1.3 | 0.395 | 3.7E-07 |
| *col-62* | WBGene00000638 | protein_coding_gene | -1.3 | 0.395 | 1.5E-02 |
| *Y49E10.29* | WBGene00045239 | protein_coding_gene | -1.3 | 0.395 | 1.8E-04 |
| *F23D12.11* | WBGene00194674 | protein_coding_gene | -1.3 | 0.395 | 5.6E-05 |
| *cest-2.2* | WBGene00013874 | protein_coding_gene | -1.3 | 0.394 | 2.8E-10 |
| *sptl-1* | WBGene00016020 | protein_coding_gene | -1.3 | 0.394 | 8.2E-17 |
| *gpx-5* | WBGene00007516 | protein_coding_gene | -1.4 | 0.392 | 2.0E-10 |
| *linc-7* | WBGene00219609 | lincRNA_gene | -1.4 | 0.391 | 2.5E-05 |
| *pgph-2* | WBGene00018424 | protein_coding_gene | -1.4 | 0.391 | 2.3E-06 |
| *F41D9.2* | WBGene00018282 | protein_coding_gene | -1.4 | 0.390 | 5.2E-04 |
| *T06A4.3* | WBGene00020283 | protein_coding_gene | -1.4 | 0.388 | 2.1E-02 |
| *K02F6.3* | WBGene00019337 | protein_coding_gene | -1.4 | 0.387 | 4.5E-03 |
| *F22F4.5* | WBGene00017718 | protein_coding_gene | -1.4 | 0.387 | 4.1E-03 |
| *dhc-4* | WBGene00012272 | protein_coding_gene | -1.4 | 0.387 | 5.2E-03 |
| *PDB1.1* | WBGene00019803 | protein_coding_gene | -1.4 | 0.387 | 4.9E-09 |
| *unc-52* | WBGene00006787 | protein_coding_gene | -1.4 | 0.386 | 4.0E-09 |
| *Y82E9BR.22* | WBGene00022354 | protein_coding_gene | -1.4 | 0.386 | 2.6E-07 |
| *lgc-4* | WBGene00017580 | protein_coding_gene | -1.4 | 0.386 | 1.2E-02 |
| *twk-26* | WBGene00006679 | protein_coding_gene | -1.4 | 0.386 | 1.1E-03 |
| *F56D6.16* | WBGene00219304 | pseudogene | -1.4 | 0.385 | 2.8E-02 |
| *pqn-22* | WBGene00004112 | protein_coding_gene | -1.4 | 0.385 | 2.6E-07 |
| *phat-4* | WBGene00020237 | protein_coding_gene | -1.4 | 0.385 | 2.2E-02 |
| *Y82E9BR.14* | WBGene00022347 | protein_coding_gene | -1.4 | 0.384 | 2.4E-19 |
| *gpd-2* | WBGene00001684 | protein_coding_gene | -1.4 | 0.384 | 3.2E-12 |
| *Y14H12A.1* | WBGene00021190 | protein_coding_gene | -1.4 | 0.384 | 2.1E-05 |

| | | | | | |
|---|---|---|---|---|---|
| *F26C11.1* | WBGene00009146 | protein_coding_gene | -1.4 | 0.384 | 1.3E-02 |
| *col-48* | WBGene00000625 | protein_coding_gene | -1.4 | 0.384 | 2.8E-04 |
| *nhr-31* | WBGene00003625 | protein_coding_gene | -1.4 | 0.383 | 4.0E-08 |
| *Y116A8C.33* | WBGene00013809 | protein_coding_gene | -1.4 | 0.383 | 1.4E-02 |
| *C50D2.6* | WBGene00016809 | protein_coding_gene | -1.4 | 0.383 | 7.8E-05 |
| *pmp-5* | WBGene00004062 | protein_coding_gene | -1.4 | 0.382 | 3.7E-09 |
| *tbc-19* | WBGene00007849 | protein_coding_gene | -1.4 | 0.382 | 9.2E-08 |
| *C17F4.7* | WBGene00015913 | protein_coding_gene | -1.4 | 0.382 | 6.0E-07 |
| *pqn-36* | WBGene00004123 | protein_coding_gene | -1.4 | 0.382 | 3.6E-03 |
| *C06H5.6* | WBGene00007394 | protein_coding_gene | -1.4 | 0.382 | 2.5E-10 |
| *aqp-4* | WBGene00000172 | protein_coding_gene | -1.4 | 0.381 | 1.3E-07 |
| *F42A10.7* | WBGene00018343 | protein_coding_gene | -1.4 | 0.381 | 8.3E-05 |
| *fbxa-14* | WBGene00022326 | protein_coding_gene | -1.4 | 0.381 | 8.5E-03 |
| *ZC239.16* | WBGene00022574 | protein_coding_gene | -1.4 | 0.381 | 4.2E-02 |
| *chk-2* | WBGene00000499 | protein_coding_gene | -1.4 | 0.380 | 1.5E-07 |
| *C32E8.9* | WBGene00016325 | protein_coding_gene | -1.4 | 0.380 | 2.5E-09 |
| *R06B10.1* | WBGene00019912 | pseudogene | -1.4 | 0.379 | 2.8E-02 |
| *sec-61.B* | WBGene00021427 | protein_coding_gene | -1.4 | 0.379 | 5.9E-21 |
| *math-18* | WBGene00016544 | protein_coding_gene | -1.4 | 0.379 | 6.2E-03 |
| *Y39F10A.1* | WBGene00021452 | protein_coding_gene | -1.4 | 0.378 | 4.7E-02 |
| *fbxb-108* | WBGene00017249 | protein_coding_gene | -1.4 | 0.378 | 3.3E-02 |
| *cest-10* | WBGene00016595 | protein_coding_gene | -1.4 | 0.378 | 1.9E-08 |
| *W04E12.7* | WBGene00012252 | protein_coding_gene | -1.4 | 0.378 | 1.2E-03 |
| *C09B8.4* | WBGene00015623 | protein_coding_gene | -1.4 | 0.377 | 7.5E-05 |
| *cpt-5* | WBGene00008629 | protein_coding_gene | -1.4 | 0.376 | 1.7E-06 |
| *aqp-7* | WBGene00000175 | protein_coding_gene | -1.4 | 0.376 | 2.6E-16 |
| *C14F11.4* | WBGene00015780 | protein_coding_gene | -1.4 | 0.376 | 8.5E-08 |
| *H10E21.4* | WBGene00019184 | protein_coding_gene | -1.4 | 0.376 | 1.5E-02 |
| *F43C1.5* | WBGene00009649 | protein_coding_gene | -1.4 | 0.376 | 1.7E-02 |
| *C06E8.5* | WBGene00015544 | protein_coding_gene | -1.4 | 0.375 | 2.7E-02 |
| *Y73E7A.3* | WBGene00022270 | protein_coding_gene | -1.4 | 0.375 | 3.5E-05 |
| *T03D3.5* | WBGene00020183 | protein_coding_gene | -1.4 | 0.375 | 1.2E-10 |
| *dod-19* | WBGene00022644 | protein_coding_gene | -1.4 | 0.374 | 2.0E-02 |
| *clec-205* | WBGene00021602 | protein_coding_gene | -1.4 | 0.374 | 8.3E-03 |
| *cah-5* | WBGene00000283 | protein_coding_gene | -1.4 | 0.374 | 2.5E-08 |
| *flr-4* | WBGene00001468 | protein_coding_gene | -1.4 | 0.374 | 5.2E-03 |
| *plep-1* | WBGene00021797 | protein_coding_gene | -1.4 | 0.373 | 1.7E-02 |
| *F31F4.1* | WBGene00017957 | protein_coding_gene | -1.4 | 0.373 | 1.9E-05 |
| *col-101* | WBGene00000675 | protein_coding_gene | -1.4 | 0.373 | 1.2E-02 |
| *Y32B12C.1* | WBGene00012525 | protein_coding_gene | -1.4 | 0.372 | 4.1E-02 |
| *art-1* | WBGene00000198 | protein_coding_gene | -1.4 | 0.372 | 5.3E-17 |
| *hmit-1.2* | WBGene00013074 | protein_coding_gene | -1.4 | 0.372 | 1.6E-04 |
| *pes-9* | WBGene00003982 | protein_coding_gene | -1.4 | 0.372 | 2.2E-26 |
| *K10C2.7* | WBGene00019622 | protein_coding_gene | -1.4 | 0.371 | 4.5E-03 |
| *ptps-1* | WBGene00015010 | protein_coding_gene | -1.4 | 0.371 | 3.4E-07 |
| *his-60* | WBGene00001934 | protein_coding_gene | -1.4 | 0.370 | 6.8E-03 |
| *wrt-4* | WBGene00006950 | protein_coding_gene | -1.4 | 0.370 | 1.2E-02 |
| *arrd-10* | WBGene00010268 | protein_coding_gene | -1.4 | 0.370 | 1.2E-03 |
| *F08F3.4* | WBGene00017262 | protein_coding_gene | -1.4 | 0.370 | 1.6E-18 |
| *nlp-26* | WBGene00003764 | protein_coding_gene | -1.4 | 0.369 | 1.2E-04 |
| *F23F12.3* | WBGene00017751 | protein_coding_gene | -1.4 | 0.369 | 5.6E-03 |
| *col-80* | WBGene00000656 | protein_coding_gene | -1.4 | 0.368 | 4.2E-06 |
| *C15H9.9* | WBGene00015803 | protein_coding_gene | -1.4 | 0.368 | 7.1E-22 |
| *acl-1* | WBGene00010339 | protein_coding_gene | -1.4 | 0.368 | 5.5E-21 |
| *bca-1* | WBGene00000245 | protein_coding_gene | -1.4 | 0.368 | 5.8E-08 |
| *gst-24* | WBGene00001772 | protein_coding_gene | -1.4 | 0.367 | 2.8E-03 |
| *C08F1.6* | WBGene00015610 | protein_coding_gene | -1.4 | 0.367 | 7.9E-04 |
| *F22E5.1* | WBGene00017703 | protein_coding_gene | -1.4 | 0.366 | 1.1E-02 |
| *F52H2.6* | WBGene00018718 | protein_coding_gene | -1.4 | 0.366 | 1.4E-17 |
| *ZK218.1* | WBGene00013935 | protein_coding_gene | -1.5 | 0.366 | 1.6E-03 |

| | | | | | |
|---|---|---|---|---|---|
| *R08E5.3* | WBGene00019963 | protein_coding_gene | -1.5 | 0.366 | 1.3E-07 |
| *fbxb-35* | WBGene00017208 | protein_coding_gene | -1.5 | 0.366 | 3.2E-02 |
| *Y57G11C.51* | WBGene00044258 | protein_coding_gene | -1.5 | 0.366 | 7.1E-08 |
| *T03E6.8* | WBGene00011397 | protein_coding_gene | -1.5 | 0.365 | 1.7E-03 |
| *vha-12* | WBGene00006921 | protein_coding_gene | -1.5 | 0.364 | 9.9E-23 |
| *R13A5.10* | WBGene00020052 | protein_coding_gene | -1.5 | 0.364 | 9.1E-04 |
| *W06A7.4* | WBGene00012293 | protein_coding_gene | -1.5 | 0.363 | 4.3E-05 |
| *piit-1* | WBGene00009384 | protein_coding_gene | -1.5 | 0.362 | 6.1E-04 |
| *C52D10.3* | WBGene00016877 | protein_coding_gene | -1.5 | 0.361 | 3.2E-05 |
| *mtp-18* | WBGene00077690 | protein_coding_gene | -1.5 | 0.361 | 6.0E-06 |
| *cnp-2* | WBGene00012901 | protein_coding_gene | -1.5 | 0.361 | 9.0E-04 |
| *ZK185.3* | WBGene00022683 | protein_coding_gene | -1.5 | 0.361 | 9.5E-07 |
| *F37B12.1* | WBGene00009504 | protein_coding_gene | -1.5 | 0.361 | 2.3E-03 |
| *gpx-1* | WBGene00009165 | protein_coding_gene | -1.5 | 0.360 | 9.5E-11 |
| *C04G2.14* | WBGene00255426 | ncRNA_gene | -1.5 | 0.360 | 9.1E-03 |
| *col-166* | WBGene00000739 | protein_coding_gene | -1.5 | 0.359 | 5.3E-05 |
| *ZK180.5* | WBGene00022679 | protein_coding_gene | -1.5 | 0.359 | 3.0E-03 |
| *clec-187* | WBGene00014137 | protein_coding_gene | -1.5 | 0.359 | 6.3E-09 |
| *folt-1* | WBGene00007388 | protein_coding_gene | -1.5 | 0.358 | 1.6E-07 |
| *pals-18* | WBGene00022353 | protein_coding_gene | -1.5 | 0.358 | 1.5E-02 |
| *col-144* | WBGene00000717 | protein_coding_gene | -1.5 | 0.358 | 4.0E-02 |
| *F42G10.1* | WBGene00009645 | protein_coding_gene | -1.5 | 0.358 | 5.5E-05 |
| *cyp-33C8* | WBGene00019967 | protein_coding_gene | -1.5 | 0.358 | 4.6E-03 |
| *clec-57* | WBGene00008594 | protein_coding_gene | -1.5 | 0.356 | 1.2E-04 |
| *T25B9.9* | WBGene00012015 | protein_coding_gene | -1.5 | 0.356 | 3.1E-20 |
| *alh-9* | WBGene00000115 | protein_coding_gene | -1.5 | 0.356 | 5.6E-21 |
| *str-176* | WBGene00006220 | protein_coding_gene | -1.5 | 0.355 | 2.4E-10 |
| *T25C12.3* | WBGene00012018 | protein_coding_gene | -1.5 | 0.354 | 8.0E-03 |
| *K01D12.9* | WBGene00010468 | protein_coding_gene | -1.5 | 0.354 | 1.6E-02 |
| *B0454.5* | WBGene00015196 | protein_coding_gene | -1.5 | 0.354 | 1.6E-05 |
| *C12D12.1* | WBGene00015713 | protein_coding_gene | -1.5 | 0.353 | 3.7E-06 |
| *ZC302.3* | WBGene00013863 | protein_coding_gene | -1.5 | 0.353 | 2.8E-02 |
| *ugt-48* | WBGene00015965 | protein_coding_gene | -1.5 | 0.352 | 4.8E-07 |
| *ncx-6* | WBGene00003571 | protein_coding_gene | -1.5 | 0.352 | 1.8E-04 |
| *gldc-1* | WBGene00020022 | protein_coding_gene | -1.5 | 0.351 | 2.2E-25 |
| *Y43F8C.13* | WBGene00012834 | protein_coding_gene | -1.5 | 0.351 | 4.0E-17 |
| *pzf-1* | WBGene00011505 | protein_coding_gene | -1.5 | 0.351 | 5.3E-08 |
| *slc-17.4* | WBGene00012130 | protein_coding_gene | -1.5 | 0.351 | 8.0E-09 |
| *dct-16* | WBGene00012615 | protein_coding_gene | -1.5 | 0.350 | 2.5E-03 |
| *K12H6.7* | WBGene00019687 | protein_coding_gene | -1.5 | 0.349 | 1.3E-02 |
| *oat-1* | WBGene00003837 | protein_coding_gene | -1.5 | 0.349 | 4.1E-06 |
| *dsc-4* | WBGene00001099 | protein_coding_gene | -1.5 | 0.349 | 6.2E-17 |
| *sprr-2* | WBGene00009629 | protein_coding_gene | -1.5 | 0.348 | 3.0E-03 |
| *acox-1.3* | WBGene00008566 | protein_coding_gene | -1.5 | 0.348 | 6.0E-04 |
| *pks-1* | WBGene00016558 | protein_coding_gene | -1.5 | 0.347 | 3.2E-07 |
| *Y65B4A.9* | WBGene00022032 | protein_coding_gene | -1.5 | 0.347 | 2.0E-02 |
| *dhs-3* | WBGene00000967 | protein_coding_gene | -1.5 | 0.347 | 1.2E-19 |
| *fut-4* | WBGene00001507 | protein_coding_gene | -1.5 | 0.346 | 2.1E-02 |
| *bath-4* | WBGene00018665 | protein_coding_gene | -1.5 | 0.346 | 1.6E-05 |
| *Y67H2A.9* | WBGene00014941 | pseudogene | -1.5 | 0.346 | 1.7E-13 |
| *ugt-60* | WBGene00007402 | protein_coding_gene | -1.5 | 0.346 | 4.6E-05 |
| *C39D10.8* | WBGene00016535 | protein_coding_gene | -1.5 | 0.346 | 1.8E-07 |
| *R12E2.15* | WBGene00020040 | protein_coding_gene | -1.5 | 0.345 | 5.1E-03 |
| *acr-16* | WBGene00000055 | protein_coding_gene | -1.5 | 0.344 | 7.1E-04 |
| *Y110A2AL.9* | WBGene00022445 | protein_coding_gene | -1.5 | 0.344 | 1.4E-02 |
| *clec-56* | WBGene00008595 | protein_coding_gene | -1.5 | 0.343 | 1.5E-13 |
| *F20D6.5* | WBGene00017635 | protein_coding_gene | -1.5 | 0.342 | 2.6E-03 |
| *lgc-54* | WBGene00020528 | protein_coding_gene | -1.6 | 0.341 | 1.8E-04 |
| *mec-19* | WBGene00008216 | protein_coding_gene | -1.6 | 0.341 | 1.6E-02 |
| *twk-34* | WBGene00006686 | protein_coding_gene | -1.6 | 0.340 | 5.7E-09 |

| | | | | | |
|---|---|---|---|---|---|
| *cln-3.1* | WBGene00000539 | protein_coding_gene | -1.6 | 0.340 | 8.3E-04 |
| *K08H10.6* | WBGene00010698 | protein_coding_gene | -1.6 | 0.339 | 3.8E-02 |
| *poml-2* | WBGene00017089 | protein_coding_gene | -1.6 | 0.338 | 2.6E-05 |
| *wrt-8* | WBGene00006954 | protein_coding_gene | -1.6 | 0.337 | 4.2E-02 |
| *hacd-1* | WBGene00019978 | protein_coding_gene | -1.6 | 0.337 | 3.5E-07 |
| *col-110* | WBGene00000684 | protein_coding_gene | -1.6 | 0.337 | 2.0E-02 |
| *C01H6.4* | WBGene00007254 | protein_coding_gene | -1.6 | 0.337 | 2.3E-05 |
| *C45E5.1* | WBGene00016664 | protein_coding_gene | -1.6 | 0.337 | 8.0E-09 |
| *F40D4.13* | WBGene00009572 | protein_coding_gene | -1.6 | 0.335 | 2.3E-02 |
| *K11H12.3* | WBGene00019659 | protein_coding_gene | -1.6 | 0.335 | 4.7E-02 |
| *K01G12.3* | WBGene00010483 | protein_coding_gene | -1.6 | 0.335 | 7.3E-04 |
| *cyp-13A12* | WBGene00008810 | protein_coding_gene | -1.6 | 0.334 | 3.0E-02 |
| *F29G6.1* | WBGene00009257 | protein_coding_gene | -1.6 | 0.334 | 3.0E-02 |
| *ttr-18* | WBGene00007142 | protein_coding_gene | -1.6 | 0.334 | 9.0E-12 |
| *T24E12.11* | WBGene00020778 | protein_coding_gene | -1.6 | 0.334 | 3.4E-04 |
| *ctsa-4.1* | WBGene00019605 | protein_coding_gene | -1.6 | 0.333 | 4.3E-08 |
| *Y22D7AL.11* | WBGene00021249 | protein_coding_gene | -1.6 | 0.333 | 5.7E-04 |
| *F13H8.3* | WBGene00017436 | protein_coding_gene | -1.6 | 0.332 | 1.1E-13 |
| *twk-22* | WBGene00006675 | protein_coding_gene | -1.6 | 0.332 | 2.4E-06 |
| *nmat-1* | WBGene00012295 | protein_coding_gene | -1.6 | 0.332 | 1.1E-04 |
| *ttr-14* | WBGene00011460 | protein_coding_gene | -1.6 | 0.332 | 9.9E-06 |
| *zip-3* | WBGene00012330 | protein_coding_gene | -1.6 | 0.331 | 1.5E-13 |
| *lgc-45* | WBGene00021146 | protein_coding_gene | -1.6 | 0.331 | 5.2E-11 |
| *efn-3* | WBGene00001164 | protein_coding_gene | -1.6 | 0.330 | 4.3E-04 |
| *ctsa-2* | WBGene00019617 | protein_coding_gene | -1.6 | 0.330 | 7.5E-03 |
| *F33E2.4* | WBGene00009360 | protein_coding_gene | -1.6 | 0.329 | 1.0E-02 |
| *nlp-77* | WBGene00015514 | protein_coding_gene | -1.6 | 0.328 | 4.6E-27 |
| *fmo-3* | WBGene00001478 | protein_coding_gene | -1.6 | 0.327 | 7.6E-06 |
| *Y54G11A.19* | WBGene00235256 | protein_coding_gene | -1.6 | 0.327 | 4.8E-04 |
| *col-135* | WBGene00000708 | protein_coding_gene | -1.6 | 0.326 | 5.0E-22 |
| *Y66C5A.1* | WBGene00044156 | protein_coding_gene | -1.6 | 0.325 | 6.2E-03 |
| *pgp-1* | WBGene00003995 | protein_coding_gene | -1.6 | 0.325 | 1.7E-02 |
| *H23N18.5* | WBGene00019236 | protein_coding_gene | -1.6 | 0.323 | 3.9E-03 |
| *sptl-3* | WBGene00011932 | protein_coding_gene | -1.6 | 0.322 | 9.5E-29 |
| *pals-17* | WBGene00022346 | protein_coding_gene | -1.6 | 0.322 | 1.3E-04 |
| *T25E4.1* | WBGene00020803 | protein_coding_gene | -1.6 | 0.321 | 1.7E-02 |
| *F26G1.5* | WBGene00017841 | protein_coding_gene | -1.6 | 0.321 | 1.6E-04 |
| *let-383* | WBGene00044898 | protein_coding_gene | -1.6 | 0.321 | 8.8E-05 |
| *col-155* | WBGene00000728 | protein_coding_gene | -1.6 | 0.320 | 1.1E-05 |
| *F31A3.3* | WBGene00017938 | protein_coding_gene | -1.6 | 0.320 | 6.7E-03 |
| *col-154* | WBGene00000727 | protein_coding_gene | -1.6 | 0.319 | 9.5E-03 |
| *ZK742.3* | WBGene00022813 | protein_coding_gene | -1.6 | 0.319 | 1.2E-05 |
| *fip-5* | WBGene00009620 | protein_coding_gene | -1.6 | 0.319 | 7.7E-07 |
| *Y24F12A.4* | WBGene00012499 | pseudogene | -1.6 | 0.319 | 7.6E-03 |
| *ZC53.1* | WBGene00022508 | protein_coding_gene | -1.6 | 0.319 | 3.9E-06 |
| *F10A3.4* | WBGene00008634 | protein_coding_gene | -1.7 | 0.318 | 1.2E-03 |
| *F10D11.6* | WBGene00008652 | protein_coding_gene | -1.7 | 0.318 | 8.3E-05 |
| *hpd-1* | WBGene00001993 | protein_coding_gene | -1.7 | 0.318 | 1.1E-12 |
| *clp-3* | WBGene00000544 | protein_coding_gene | -1.7 | 0.317 | 5.1E-03 |
| *ZK228.3* | WBGene00013946 | protein_coding_gene | -1.7 | 0.317 | 1.4E-09 |
| *ctsa-3.1* | WBGene00017969 | protein_coding_gene | -1.7 | 0.317 | 1.9E-02 |
| *nhr-65* | WBGene00003655 | protein_coding_gene | -1.7 | 0.316 | 1.1E-05 |
| *C26B2.8* | WBGene00016130 | protein_coding_gene | -1.7 | 0.316 | 2.2E-02 |
| *T25B9.1* | WBGene00012007 | protein_coding_gene | -1.7 | 0.316 | 1.1E-08 |
| *col-39* | WBGene00000616 | protein_coding_gene | -1.7 | 0.315 | 1.3E-02 |
| *lbp-8* | WBGene00002260 | protein_coding_gene | -1.7 | 0.315 | 4.9E-02 |
| *sdz-12* | WBGene00017406 | protein_coding_gene | -1.7 | 0.314 | 4.1E-02 |
| *linc-24* | WBGene00219771 | lincRNA_gene | -1.7 | 0.314 | 4.5E-02 |
| *B0244.7* | WBGene00015082 | protein_coding_gene | -1.7 | 0.314 | 4.5E-02 |
| *vha-11* | WBGene00006920 | protein_coding_gene | -1.7 | 0.313 | 5.3E-21 |

| | | | | | |
|---|---|---|---|---|---|
| *comt-3* | WBGene00021487 | protein_coding_gene | -1.7 | 0.313 | 3.8E-06 |
| *Y55F3BR.10* | WBGene00044742 | protein_coding_gene | -1.7 | 0.312 | 3.8E-04 |
| *T10B5.10* | WBGene00020396 | protein_coding_gene | -1.7 | 0.311 | 2.1E-02 |
| *nlp-24* | WBGene00003762 | protein_coding_gene | -1.7 | 0.311 | 1.7E-02 |
| *Y60A3A.11* | WBGene00014933 | pseudogene | -1.7 | 0.310 | 3.9E-02 |
| *col-115* | WBGene00000689 | protein_coding_gene | -1.7 | 0.309 | 2.3E-02 |
| *ets-5* | WBGene00016600 | protein_coding_gene | -1.7 | 0.308 | 4.6E-02 |
| *F20D1.22* | WBGene00304908 | protein_coding_gene | -1.7 | 0.308 | 1.9E-03 |
| *Y39B6A.27* | WBGene00012686 | protein_coding_gene | -1.7 | 0.308 | 8.7E-05 |
| *Y34F4.6* | WBGene00189956 | protein_coding_gene | -1.7 | 0.307 | 1.6E-02 |
| *nhr-93* | WBGene00003683 | protein_coding_gene | -1.7 | 0.307 | 3.5E-02 |
| *Y102A11A.7* | WBGene00022417 | protein_coding_gene | -1.7 | 0.307 | 4.4E-02 |
| *ZK1240.9* | WBGene00022873 | protein_coding_gene | -1.7 | 0.306 | 1.6E-04 |
| *F07G6.10* | WBGene00045271 | protein_coding_gene | -1.7 | 0.306 | 3.4E-03 |
| *T01D1.3* | WBGene00020148 | protein_coding_gene | -1.7 | 0.306 | 3.5E-08 |
| *elo-9* | WBGene00001247 | protein_coding_gene | -1.7 | 0.306 | 3.1E-09 |
| *fbxa-138* | WBGene00022345 | protein_coding_gene | -1.7 | 0.305 | 4.7E-02 |
| *srb-14* | WBGene00016338 | pseudogene | -1.7 | 0.305 | 4.4E-02 |
| *mir-8199* | WBGene00016553 | miRNA_gene | -1.7 | 0.305 | 4.9E-02 |
| *fbxb-83* | WBGene00020543 | protein_coding_gene | -1.7 | 0.305 | 3.4E-03 |
| *cima-1* | WBGene00018473 | protein_coding_gene | -1.7 | 0.305 | 2.3E-06 |
| *skr-5* | WBGene00004811 | protein_coding_gene | -1.7 | 0.304 | 4.5E-02 |
| *lgc-40* | WBGene00020767 | protein_coding_gene | -1.7 | 0.304 | 3.2E-03 |
| *Y59E9AL.4* | WBGene00021994 | protein_coding_gene | -1.7 | 0.304 | 4.7E-06 |
| *T20D4.19* | WBGene00020624 | protein_coding_gene | -1.7 | 0.304 | 4.6E-02 |
| *col-8* | WBGene00000597 | protein_coding_gene | -1.7 | 0.304 | 1.3E-10 |
| *fbxb-90* | WBGene00044694 | protein_coding_gene | -1.7 | 0.302 | 1.7E-02 |
| *VB0395L.1* | WBGene00044150 | protein_coding_gene | -1.7 | 0.302 | 3.4E-04 |
| *col-178* | WBGene00000751 | protein_coding_gene | -1.7 | 0.302 | 4.1E-37 |
| *skpo-3* | WBGene00017968 | protein_coding_gene | -1.7 | 0.302 | 1.5E-09 |
| *C27A2.5* | WBGene00016153 | protein_coding_gene | -1.7 | 0.302 | 4.7E-02 |
| *mup-4* | WBGene00003497 | protein_coding_gene | -1.7 | 0.301 | 1.1E-20 |
| *drd-10* | WBGene00016361 | protein_coding_gene | -1.7 | 0.301 | 1.0E-04 |
| *fat-7* | WBGene00001399 | protein_coding_gene | -1.7 | 0.300 | 2.1E-03 |
| *K06G5.1* | WBGene00010605 | protein_coding_gene | -1.7 | 0.299 | 4.4E-14 |
| *haao-1* | WBGene00010595 | protein_coding_gene | -1.7 | 0.299 | 8.9E-20 |
| *amt-1* | WBGene00000133 | protein_coding_gene | -1.7 | 0.298 | 1.1E-03 |
| *dao-4* | WBGene00000930 | protein_coding_gene | -1.7 | 0.298 | 4.2E-02 |
| *grd-5* | WBGene00001694 | protein_coding_gene | -1.7 | 0.298 | 2.3E-06 |
| *ugt-46* | WBGene00015141 | protein_coding_gene | -1.7 | 0.298 | 3.4E-13 |
| *EGAP4.1* | WBGene00017149 | protein_coding_gene | -1.7 | 0.297 | 3.2E-02 |
| *nas-38* | WBGene00003554 | protein_coding_gene | -1.8 | 0.297 | 1.5E-02 |
| *E04D5.5* | WBGene00008484 | protein_coding_gene | -1.8 | 0.297 | 1.1E-04 |
| *str-148* | WBGene00006195 | protein_coding_gene | -1.8 | 0.296 | 4.4E-02 |
| *nduo-3* | WBGene00010966 | protein_coding_gene | -1.8 | 0.296 | 6.5E-04 |
| *Y48D7A.1* | WBGene00021656 | protein_coding_gene | -1.8 | 0.295 | 4.1E-02 |
| *ugt-21* | WBGene00007885 | protein_coding_gene | -1.8 | 0.295 | 3.8E-05 |
| *pho-13* | WBGene00008560 | protein_coding_gene | -1.8 | 0.295 | 2.0E-02 |
| *nep-10* | WBGene00017555 | protein_coding_gene | -1.8 | 0.294 | 2.5E-02 |
| *Y17D7C.3* | WBGene00045366 | pseudogene | -1.8 | 0.294 | 1.3E-06 |
| *fbxb-21* | WBGene00022558 | protein_coding_gene | -1.8 | 0.294 | 4.1E-02 |
| *Y43F8B.23* | WBGene00194815 | protein_coding_gene | -1.8 | 0.294 | 4.3E-02 |
| *Y38H6C.23* | WBGene00012634 | protein_coding_gene | -1.8 | 0.294 | 1.0E-05 |
| *T04A8.18* | WBGene00119203 | protein_coding_gene | -1.8 | 0.294 | 1.5E-02 |
| *F26H11.4* | WBGene00009182 | protein_coding_gene | -1.8 | 0.293 | 3.2E-06 |
| *ttr-31* | WBGene00010225 | protein_coding_gene | -1.8 | 0.293 | 8.0E-08 |
| *rgs-11* | WBGene00004354 | protein_coding_gene | -1.8 | 0.292 | 3.2E-07 |
| *linc-130* | WBGene00219651 | lincRNA_gene | -1.8 | 0.292 | 4.1E-02 |
| *nas-36* | WBGene00003552 | protein_coding_gene | -1.8 | 0.292 | 5.8E-03 |
| *nas-1* | WBGene00003520 | protein_coding_gene | -1.8 | 0.292 | 4.3E-02 |

| | | | | | |
|---|---|---|---|---|---|
| *H16D19.5* | WBGene00195253 | pseudogene | -1.8 | 0.291 | 1.8E-02 |
| *C04G6.2* | WBGene00015453 | protein_coding_gene | -1.8 | 0.291 | 3.5E-02 |
| *K06B9.4* | WBGene00019441 | pseudogene | -1.8 | 0.291 | 3.7E-08 |
| *col-81* | WBGene00000657 | protein_coding_gene | -1.8 | 0.290 | 1.1E-02 |
| *Y35H6.3* | WBGene00021342 | pseudogene | -1.8 | 0.290 | 9.0E-07 |
| *cbl-1* | WBGene00007533 | protein_coding_gene | -1.8 | 0.290 | 9.1E-13 |
| *F49F1.12* | WBGene00018652 | protein_coding_gene | -1.8 | 0.290 | 2.4E-02 |
| *clec-204* | WBGene00021603 | protein_coding_gene | -1.8 | 0.290 | 4.6E-04 |
| *col-58* | WBGene00000634 | protein_coding_gene | -1.8 | 0.290 | 4.8E-03 |
| *F01D5.1* | WBGene00008492 | protein_coding_gene | -1.8 | 0.288 | 1.9E-04 |
| *F58B4.5* | WBGene00010238 | protein_coding_gene | -1.8 | 0.288 | 2.2E-10 |
| *col-169* | WBGene00000742 | protein_coding_gene | -1.8 | 0.288 | 1.4E-02 |
| *C46F2.1* | WBGene00016714 | protein_coding_gene | -1.8 | 0.287 | 1.2E-07 |
| *pqn-2* | WBGene00004098 | protein_coding_gene | -1.8 | 0.287 | 4.9E-02 |
| *oac-37* | WBGene00010391 | protein_coding_gene | -1.8 | 0.286 | 3.9E-02 |
| *twk-10* | WBGene00006665 | protein_coding_gene | -1.8 | 0.286 | 1.1E-02 |
| *F28H7.3* | WBGene00009237 | protein_coding_gene | -1.8 | 0.286 | 4.0E-07 |
| *C50E3.11* | WBGene00016822 | protein_coding_gene | -1.8 | 0.286 | 3.1E-13 |
| *Y57G11C.23* | WBGene00013318 | protein_coding_gene | -1.8 | 0.286 | 1.9E-02 |
| *C55A6.4* | WBGene00008333 | protein_coding_gene | -1.8 | 0.285 | 8.4E-10 |
| *ugt-31* | WBGene00021464 | protein_coding_gene | -1.8 | 0.285 | 1.4E-03 |
| *cpr-5* | WBGene00000785 | protein_coding_gene | -1.8 | 0.285 | 1.4E-07 |
| *K10C2.8* | WBGene00045265 | protein_coding_gene | -1.8 | 0.284 | 7.6E-10 |
| *pcp-3* | WBGene00003958 | protein_coding_gene | -1.8 | 0.284 | 3.3E-10 |
| *wago-2* | WBGene00018862 | pseudogene | -1.8 | 0.284 | 1.3E-18 |
| *math-10* | WBGene00015839 | protein_coding_gene | -1.8 | 0.283 | 3.2E-02 |
| *pept-1* | WBGene00003877 | protein_coding_gene | -1.8 | 0.283 | 7.5E-15 |
| *T02G6.11* | WBGene00086562 | protein_coding_gene | -1.8 | 0.283 | 3.5E-02 |
| *str-227* | WBGene00006258 | protein_coding_gene | -1.8 | 0.283 | 3.5E-02 |
| *F11E6.3* | WBGene00008707 | protein_coding_gene | -1.8 | 0.282 | 6.2E-11 |
| *col-170* | WBGene00000743 | protein_coding_gene | -1.8 | 0.282 | 2.0E-02 |
| *clec-80* | WBGene00043062 | protein_coding_gene | -1.8 | 0.281 | 1.5E-06 |
| *ZK662.2* | WBGene00014039 | protein_coding_gene | -1.8 | 0.281 | 3.8E-03 |
| *C18A11.4* | WBGene00015950 | protein_coding_gene | -1.8 | 0.281 | 1.2E-02 |
| *mth-1* | WBGene00017948 | protein_coding_gene | -1.8 | 0.280 | 5.6E-07 |
| *vit-4* | WBGene00006928 | protein_coding_gene | -1.8 | 0.280 | 2.9E-02 |
| *C15F1.1* | WBGene00015794 | protein_coding_gene | -1.8 | 0.280 | 1.7E-10 |
| *F57F4.4* | WBGene00019017 | protein_coding_gene | -1.8 | 0.280 | 1.5E-02 |
| *ZK228.4* | WBGene00013947 | protein_coding_gene | -1.8 | 0.279 | 1.5E-13 |
| *F22F1.2* | WBGene00017714 | protein_coding_gene | -1.8 | 0.279 | 3.7E-02 |
| *mpst-4* | WBGene00017387 | protein_coding_gene | -1.8 | 0.279 | 3.9E-03 |
| *sams-1* | WBGene00008205 | protein_coding_gene | -1.8 | 0.278 | 2.4E-45 |
| *cut-4* | WBGene00008482 | protein_coding_gene | -1.8 | 0.278 | 1.0E-03 |
| *R12E2.6* | WBGene00020032 | protein_coding_gene | -1.9 | 0.277 | 3.3E-02 |
| *ldp-1* | WBGene00017719 | protein_coding_gene | -1.9 | 0.276 | 3.5E-39 |
| *C05C9.3* | WBGene00007327 | protein_coding_gene | -1.9 | 0.276 | 9.0E-07 |
| *lrg-1* | WBGene00003067 | pseudogene | -1.9 | 0.276 | 9.4E-12 |
| *Y48B6A.10* | WBGene00012981 | protein_coding_gene | -1.9 | 0.276 | 5.1E-03 |
| *pqn-11* | WBGene00004103 | pseudogene | -1.9 | 0.276 | 3.2E-02 |
| *C41H7.1* | WBGene00016571 | protein_coding_gene | -1.9 | 0.275 | 1.6E-03 |
| *ZC449.8* | WBGene00195189 | protein_coding_gene | -1.9 | 0.275 | 7.0E-06 |
| *C17H12.6* | WBGene00015932 | protein_coding_gene | -1.9 | 0.275 | 4.4E-02 |
| *doxa-1* | WBGene00015519 | protein_coding_gene | -1.9 | 0.275 | 4.8E-06 |
| *pqn-54* | WBGene00004139 | protein_coding_gene | -1.9 | 0.275 | 1.1E-02 |
| *C50A2.3* | WBGene00016797 | protein_coding_gene | -1.9 | 0.274 | 3.9E-09 |
| *C01C10.2* | WBGene00015294 | protein_coding_gene | -1.9 | 0.274 | 1.9E-08 |
| *ZK218.5* | WBGene00013939 | protein_coding_gene | -1.9 | 0.274 | 3.5E-02 |
| *C53B7.6* | WBGene00016895 | protein_coding_gene | -1.9 | 0.274 | 2.8E-02 |
| *pgp-9* | WBGene00004003 | protein_coding_gene | -1.9 | 0.274 | 5.2E-03 |
| *K02E7.10* | WBGene00019314 | protein_coding_gene | -1.9 | 0.274 | 3.0E-02 |

| | | | | | |
|---|---|---|---|---|---|
| *sdz-29* | WBGene00020081 | pseudogene | -1.9 | 0.273 | 8.1E-05 |
| *T02G5.4* | WBGene00020164 | pseudogene | -1.9 | 0.272 | 1.3E-11 |
| *F56D5.3* | WBGene00010148 | protein_coding_gene | -1.9 | 0.272 | 1.0E-06 |
| *F35B3.4* | WBGene00018031 | protein_coding_gene | -1.9 | 0.272 | 7.0E-03 |
| *clec-206* | WBGene00019089 | protein_coding_gene | -1.9 | 0.272 | 3.2E-02 |
| *fbxa-199* | WBGene00005931 | protein_coding_gene | -1.9 | 0.271 | 4.0E-02 |
| *F35E12.2* | WBGene00009426 | protein_coding_gene | -1.9 | 0.271 | 3.8E-02 |
| *F44E5.16* | WBGene00219949 | ncRNA_gene | -1.9 | 0.271 | 2.9E-02 |
| *rmd-6* | WBGene00020070 | protein_coding_gene | -1.9 | 0.271 | 1.7E-02 |
| *ZC434.9* | WBGene00013895 | protein_coding_gene | -1.9 | 0.271 | 5.4E-09 |
| *abu-7* | WBGene00000030 | protein_coding_gene | -1.9 | 0.271 | 3.7E-02 |
| *Y47G6A.33* | WBGene00045399 | protein_coding_gene | -1.9 | 0.271 | 8.2E-14 |
| *F35F10.6* | WBGene00018056 | protein_coding_gene | -1.9 | 0.271 | 8.0E-03 |
| *ncx-7* | WBGene00003572 | protein_coding_gene | -1.9 | 0.271 | 2.9E-19 |
| *C02E7.7* | WBGene00015340 | protein_coding_gene | -1.9 | 0.270 | 4.7E-04 |
| *Y32B12C.3* | WBGene00012526 | pseudogene | -1.9 | 0.270 | 2.6E-02 |
| *nas-30* | WBGene00003548 | protein_coding_gene | -1.9 | 0.269 | 1.1E-02 |
| *ZK6.11* | WBGene00022645 | protein_coding_gene | -1.9 | 0.269 | 6.5E-03 |
| *drh-2* | WBGene00001091 | pseudogene | -1.9 | 0.268 | 3.3E-27 |
| *grd-10* | WBGene00001699 | protein_coding_gene | -1.9 | 0.268 | 1.2E-03 |
| *K10G4.10* | WBGene00044202 | protein_coding_gene | -1.9 | 0.267 | 2.9E-02 |
| *ppw-1* | WBGene00004093 | protein_coding_gene | -1.9 | 0.267 | 1.7E-18 |
| *clec-118* | WBGene00021032 | protein_coding_gene | -1.9 | 0.267 | 2.7E-02 |
| *dct-18* | WBGene00010266 | protein_coding_gene | -1.9 | 0.267 | 1.0E-28 |
| *C45B11.2* | WBGene00008105 | protein_coding_gene | -1.9 | 0.267 | 2.7E-02 |
| *F10C1.8* | WBGene00017327 | protein_coding_gene | -1.9 | 0.266 | 5.2E-03 |
| *col-143* | WBGene00000716 | protein_coding_gene | -1.9 | 0.266 | 8.3E-11 |
| *ptp-5.3* | WBGene00021207 | protein_coding_gene | -1.9 | 0.266 | 3.7E-03 |
| *Y37H2A.12* | WBGene00044758 | protein_coding_gene | -1.9 | 0.266 | 1.8E-10 |
| *C35A5.3* | WBGene00007947 | protein_coding_gene | -1.9 | 0.266 | 6.1E-03 |
| *osm-3* | WBGene00003884 | protein_coding_gene | -1.9 | 0.266 | 8.9E-05 |
| *R03H10.12* | WBGene00303363 | pseudogene | -1.9 | 0.265 | 2.4E-02 |
| *T25G12.6* | WBGene00020813 | protein_coding_gene | -1.9 | 0.265 | 7.6E-08 |
| *fbxa-55* | WBGene00020453 | protein_coding_gene | -1.9 | 0.264 | 7.3E-06 |
| *gsto-1* | WBGene00016204 | protein_coding_gene | -1.9 | 0.264 | 2.5E-07 |
| *R11A8.1* | WBGene00011236 | protein_coding_gene | -1.9 | 0.263 | 2.3E-20 |
| *R12C12.7* | WBGene00020027 | protein_coding_gene | -1.9 | 0.262 | 7.0E-25 |
| *egas-3* | WBGene00013480 | protein_coding_gene | -1.9 | 0.261 | 3.6E-02 |
| *F52C6.14* | WBGene00018671 | protein_coding_gene | -1.9 | 0.260 | 2.4E-02 |
| *K06B4.15* | WBGene00077450 | protein_coding_gene | -1.9 | 0.260 | 2.8E-02 |
| *pgp-8* | WBGene00004002 | protein_coding_gene | -1.9 | 0.259 | 1.8E-03 |
| *C49F5.8* | WBGene00008212 | protein_coding_gene | -1.9 | 0.259 | 4.2E-02 |
| *nhr-226* | WBGene00020850 | protein_coding_gene | -1.9 | 0.259 | 4.6E-03 |
| *Y57G11C.42* | WBGene00013331 | protein_coding_gene | -2.0 | 0.258 | 9.4E-04 |
| *T10E9.3* | WBGene00020413 | protein_coding_gene | -2.0 | 0.257 | 3.3E-04 |
| *F11F1.1* | WBGene00008714 | protein_coding_gene | -2.0 | 0.256 | 2.1E-09 |
| *asns-1* | WBGene00003816 | protein_coding_gene | -2.0 | 0.256 | 5.9E-04 |
| *H06A10.1* | WBGene00010368 | protein_coding_gene | -2.0 | 0.256 | 3.5E-06 |
| *T23F6.1* | WBGene00011949 | protein_coding_gene | -2.0 | 0.256 | 1.1E-02 |
| *grl-16* | WBGene00001725 | protein_coding_gene | -2.0 | 0.255 | 3.5E-03 |
| *ncx-8* | WBGene00003573 | protein_coding_gene | -2.0 | 0.255 | 2.3E-04 |
| *act-5* | WBGene00000067 | protein_coding_gene | -2.0 | 0.254 | 1.4E-03 |
| *W02H3.1* | WBGene00012217 | protein_coding_gene | -2.0 | 0.254 | 2.6E-05 |
| *fbxa-59* | WBGene00020455 | protein_coding_gene | -2.0 | 0.254 | 3.9E-06 |
| *Y37H2B.1* | WBGene00012572 | pseudogene | -2.0 | 0.254 | 2.3E-02 |
| *C49A9.4* | WBGene00016758 | protein_coding_gene | -2.0 | 0.254 | 1.2E-07 |
| *T07A5.1* | WBGene00011554 | protein_coding_gene | -2.0 | 0.254 | 5.0E-04 |
| *T06E4.8* | WBGene00011535 | protein_coding_gene | -2.0 | 0.252 | 2.2E-02 |
| *K05F6.12* | WBGene00045486 | protein_coding_gene | -2.0 | 0.252 | 3.1E-02 |
| *Y43C5A.7* | WBGene00044922 | protein_coding_gene | -2.0 | 0.252 | 1.1E-05 |

| | | | | | |
|---|---|---|---|---|---|
| *C16C4.17* | WBGene00044359 | protein_coding_gene | -2.0 | 0.251 | 2.1E-02 |
| *F58G6.9* | WBGene00044107 | protein_coding_gene | -2.0 | 0.251 | 9.1E-03 |
| *F41E6.11* | WBGene00018292 | protein_coding_gene | -2.0 | 0.251 | 1.1E-02 |
| *cpg-9* | WBGene00022072 | protein_coding_gene | -2.0 | 0.250 | 1.6E-07 |
| *comt-5* | WBGene00021492 | protein_coding_gene | -2.0 | 0.250 | 2.3E-03 |
| *abu-6* | WBGene00000029 | protein_coding_gene | -2.0 | 0.249 | 2.5E-02 |
| *fbxc-43* | WBGene00045297 | protein_coding_gene | -2.0 | 0.249 | 5.9E-05 |
| *C39B5.5* | WBGene00016523 | protein_coding_gene | -2.0 | 0.249 | 9.4E-05 |
| *ugt-29* | WBGene00021709 | protein_coding_gene | -2.0 | 0.249 | 2.6E-19 |
| *lips-14* | WBGene00019208 | pseudogene | -2.0 | 0.248 | 1.7E-02 |
| *F10G2.4* | WBGene00017365 | protein_coding_gene | -2.0 | 0.248 | 3.2E-03 |
| *col-160* | WBGene00000733 | protein_coding_gene | -2.0 | 0.248 | 3.7E-16 |
| *fbxa-72* | WBGene00017962 | protein_coding_gene | -2.0 | 0.248 | 1.9E-07 |
| *sid-1* | WBGene00004795 | protein_coding_gene | -2.0 | 0.248 | 9.9E-18 |
| *nep-8* | WBGene00017553 | protein_coding_gene | -2.0 | 0.247 | 1.7E-03 |
| *gst-41* | WBGene00001789 | protein_coding_gene | -2.0 | 0.247 | 4.2E-04 |
| *Y47H9C.14* | WBGene00012956 | pseudogene | -2.0 | 0.247 | 2.0E-02 |
| *fbxb-81* | WBGene00020541 | protein_coding_gene | -2.0 | 0.247 | 9.5E-03 |
| *timm-17B.2* | WBGene00017069 | protein_coding_gene | -2.0 | 0.247 | 5.2E-15 |
| *fbxa-3* | WBGene00020363 | protein_coding_gene | -2.0 | 0.247 | 3.5E-02 |
| *C05D12.3* | WBGene00014666 | protein_coding_gene | -2.0 | 0.245 | 1.1E-23 |
| *Y105C5B.15* | WBGene00013654 | protein_coding_gene | -2.0 | 0.245 | 8.2E-04 |
| *fbxb-36* | WBGene00017211 | protein_coding_gene | -2.0 | 0.244 | 1.2E-02 |
| *T21C9.9* | WBGene00011895 | protein_coding_gene | -2.0 | 0.244 | 1.9E-03 |
| *W10G11.3* | WBGene00021136 | protein_coding_gene | -2.0 | 0.243 | 5.0E-05 |
| *C04C3.14* | WBGene00219410 | protein_coding_gene | -2.0 | 0.242 | 2.4E-02 |
| *K08D12.6* | WBGene00019540 | protein_coding_gene | -2.0 | 0.242 | 1.4E-04 |
| *T15B7.14* | WBGene00020526 | protein_coding_gene | -2.1 | 0.241 | 4.1E-04 |
| *clec-233* | WBGene00009030 | protein_coding_gene | -2.1 | 0.241 | 1.6E-02 |
| *adt-2* | WBGene00000083 | protein_coding_gene | -2.1 | 0.241 | 2.7E-21 |
| *lact-1* | WBGene00018523 | protein_coding_gene | -2.1 | 0.241 | 3.0E-08 |
| *C28D4.5* | WBGene00007793 | protein_coding_gene | -2.1 | 0.241 | 5.2E-03 |
| *K02E7.6* | WBGene00019311 | pseudogene | -2.1 | 0.240 | 2.4E-05 |
| *Y73B6BL.31* | WBGene00022252 | protein_coding_gene | -2.1 | 0.239 | 4.0E-18 |
| *F54A3.2* | WBGene00018781 | protein_coding_gene | -2.1 | 0.239 | 4.9E-24 |
| *nhr-183* | WBGene00018266 | protein_coding_gene | -2.1 | 0.238 | 6.3E-04 |
| *arf-1.1* | WBGene00000190 | protein_coding_gene | -2.1 | 0.238 | 7.0E-06 |
| *F01D5.2* | WBGene00008493 | protein_coding_gene | -2.1 | 0.238 | 3.2E-02 |
| *fbxa-210* | WBGene00012565 | protein_coding_gene | -2.1 | 0.238 | 2.0E-34 |
| *Y55F3AM.14* | WBGene00021931 | protein_coding_gene | -2.1 | 0.238 | 1.2E-02 |
| *cyp-14A1* | WBGene00010705 | protein_coding_gene | -2.1 | 0.237 | 2.9E-06 |
| *C16C4.18* | WBGene00194998 | protein_coding_gene | -2.1 | 0.237 | 1.5E-04 |
| *cept-1* | WBGene00009057 | protein_coding_gene | -2.1 | 0.237 | 7.1E-45 |
| *tba-4* | WBGene00006530 | protein_coding_gene | -2.1 | 0.236 | 3.9E-19 |
| *C45B2.2* | WBGene00016659 | protein_coding_gene | -2.1 | 0.236 | 1.4E-03 |
| *col-42* | WBGene00013489 | protein_coding_gene | -2.1 | 0.235 | 9.4E-21 |
| *nspb-12* | WBGene00017305 | protein_coding_gene | -2.1 | 0.235 | 1.8E-02 |
| *fbxa-51* | WBGene00020448 | protein_coding_gene | -2.1 | 0.235 | 2.2E-06 |
| *grl-21* | WBGene00001730 | protein_coding_gene | -2.1 | 0.234 | 2.3E-03 |
| *T06E4.14* | WBGene00077691 | protein_coding_gene | -2.1 | 0.233 | 5.2E-03 |
| *nhr-263* | WBGene00017367 | protein_coding_gene | -2.1 | 0.233 | 2.7E-02 |
| *nas-27* | WBGene00003545 | protein_coding_gene | -2.1 | 0.233 | 1.1E-02 |
| *R03H10.7* | WBGene00019859 | protein_coding_gene | -2.1 | 0.233 | 1.5E-02 |
| *rol-8* | WBGene00004398 | protein_coding_gene | -2.1 | 0.233 | 1.7E-04 |
| *twk-42* | WBGene00006692 | protein_coding_gene | -2.1 | 0.232 | 8.6E-08 |
| *col-43* | WBGene00000620 | protein_coding_gene | -2.1 | 0.232 | 4.3E-03 |
| *mxl-3* | WBGene00003511 | protein_coding_gene | -2.1 | 0.232 | 3.7E-21 |
| *col-79* | WBGene00000655 | protein_coding_gene | -2.1 | 0.232 | 1.1E-02 |
| *F59H6.3* | WBGene00019134 | pseudogene | -2.1 | 0.231 | 1.4E-02 |
| *col-90* | WBGene00000665 | protein_coding_gene | -2.1 | 0.230 | 6.1E-04 |

| | | | | | |
|---|---|---|---|---|---|
| *vit-1* | WBGene00006925 | protein_coding_gene | -2.1 | 0.230 | 9.6E-03 |
| *Y47D3B.6* | WBGene00012942 | protein_coding_gene | -2.1 | 0.230 | 7.4E-04 |
| *trp-2* | WBGene00006615 | protein_coding_gene | -2.1 | 0.230 | 1.9E-05 |
| *pgp-12* | WBGene00004006 | protein_coding_gene | -2.1 | 0.229 | 5.9E-07 |
| *clec-47* | WBGene00011668 | protein_coding_gene | -2.1 | 0.229 | 2.0E-20 |
| *clec-1* | WBGene00017772 | protein_coding_gene | -2.1 | 0.228 | 4.4E-16 |
| *C30G12.10* | WBGene00284859 | ncRNA_gene | -2.1 | 0.228 | 1.4E-02 |
| *ZK637.6* | WBGene00014987 | pseudogene | -2.1 | 0.228 | 2.3E-20 |
| *C33D3.3* | WBGene00007894 | protein_coding_gene | -2.1 | 0.227 | 1.6E-03 |
| *col-104* | WBGene00000678 | protein_coding_gene | -2.1 | 0.227 | 8.1E-08 |
| *Y51A2D.18* | WBGene00013083 | protein_coding_gene | -2.1 | 0.227 | 1.0E-18 |
| *T05F1.5* | WBGene00011491 | protein_coding_gene | -2.1 | 0.227 | 4.1E-03 |
| *daf-36* | WBGene00007536 | protein_coding_gene | -2.1 | 0.226 | 7.3E-19 |
| *linc-21* | WBGene00219688 | lincRNA_gene | -2.1 | 0.226 | 1.5E-02 |
| *pcp-2* | WBGene00003957 | protein_coding_gene | -2.1 | 0.225 | 2.0E-03 |
| *fbxa-95* | WBGene00009222 | protein_coding_gene | -2.2 | 0.225 | 7.2E-09 |
| *M162.12* | WBGene00050946 | pseudogene | -2.2 | 0.225 | 1.9E-02 |
| *tyr-2* | WBGene00010661 | protein_coding_gene | -2.2 | 0.225 | 1.3E-09 |
| *pes-8* | WBGene00003981 | protein_coding_gene | -2.2 | 0.225 | 3.5E-06 |
| *F21A3.3* | WBGene00008993 | protein_coding_gene | -2.2 | 0.224 | 5.1E-08 |
| *nspc-4* | WBGene00044339 | protein_coding_gene | -2.2 | 0.223 | 6.7E-17 |
| *ctl-1* | WBGene00000830 | protein_coding_gene | -2.2 | 0.223 | 1.9E-02 |
| *H04D03.3* | WBGene00010364 | protein_coding_gene | -2.2 | 0.222 | 1.9E-03 |
| *aat-7* | WBGene00000008 | protein_coding_gene | -2.2 | 0.221 | 1.0E-03 |
| *C30H6.5* | WBGene00007823 | protein_coding_gene | -2.2 | 0.220 | 6.2E-03 |
| *col-107* | WBGene00000681 | protein_coding_gene | -2.2 | 0.220 | 6.2E-07 |
| *CE7X_3.1* | WBGene00008349 | pseudogene | -2.2 | 0.220 | 3.8E-04 |
| *cya-2* | WBGene00000864 | protein_coding_gene | -2.2 | 0.219 | 1.7E-02 |
| *scl-24* | WBGene00008575 | protein_coding_gene | -2.2 | 0.219 | 1.6E-02 |
| *irg-4* | WBGene00008584 | protein_coding_gene | -2.2 | 0.218 | 1.6E-08 |
| *rgs-9* | WBGene00004352 | protein_coding_gene | -2.2 | 0.217 | 1.7E-15 |
| *sqt-2* | WBGene00005017 | protein_coding_gene | -2.2 | 0.217 | 1.1E-05 |
| *sdz-16* | WBGene00009297 | protein_coding_gene | -2.2 | 0.217 | 5.8E-05 |
| *oac-32* | WBGene00018708 | protein_coding_gene | -2.2 | 0.216 | 1.6E-02 |
| *pqn-74* | WBGene00004156 | protein_coding_gene | -2.2 | 0.216 | 5.8E-06 |
| *cpr-8* | WBGene00021070 | protein_coding_gene | -2.2 | 0.216 | 2.9E-07 |
| *enu-3.5* | WBGene00012552 | protein_coding_gene | -2.2 | 0.216 | 2.7E-05 |
| *fbxa-187* | WBGene00009838 | pseudogene | -2.2 | 0.215 | 2.0E-02 |
| *ugt-65* | WBGene00008583 | protein_coding_gene | -2.2 | 0.215 | 6.8E-07 |
| *kynu-1* | WBGene00015802 | protein_coding_gene | -2.2 | 0.214 | 2.8E-41 |
| *F01D5.3* | WBGene00008494 | protein_coding_gene | -2.2 | 0.214 | 2.1E-03 |
| *Y16B4A.2* | WBGene00012445 | protein_coding_gene | -2.2 | 0.214 | 1.0E-19 |
| *F31E9.6* | WBGene00009299 | protein_coding_gene | -2.2 | 0.214 | 4.8E-06 |
| *W04E12.5* | WBGene00012250 | protein_coding_gene | -2.2 | 0.213 | 1.4E-02 |
| *fkb-8* | WBGene00001433 | protein_coding_gene | -2.2 | 0.212 | 7.3E-06 |
| *cdr-2* | WBGene00008296 | protein_coding_gene | -2.2 | 0.212 | 1.6E-06 |
| *ram-2* | WBGene00004300 | protein_coding_gene | -2.2 | 0.211 | 6.0E-08 |
| *C29F9.2* | WBGene00016217 | protein_coding_gene | -2.2 | 0.211 | 3.9E-03 |
| *abu-15* | WBGene00004099 | protein_coding_gene | -2.2 | 0.211 | 5.1E-03 |
| *fbxa-15* | WBGene00022327 | protein_coding_gene | -2.2 | 0.211 | 1.9E-02 |
| *Y17D7B.7* | WBGene00014562 | pseudogene | -2.2 | 0.211 | 6.3E-05 |
| *atic-1* | WBGene00016957 | protein_coding_gene | -2.2 | 0.211 | 4.3E-22 |
| *gfi-1* | WBGene00001581 | protein_coding_gene | -2.2 | 0.211 | 1.7E-03 |
| *T08E11.8* | WBGene00020364 | protein_coding_gene | -2.2 | 0.211 | 7.3E-03 |
| *T17H7.1* | WBGene00020550 | protein_coding_gene | -2.3 | 0.210 | 7.0E-05 |
| *F45B8.5* | WBGene00194794 | protein_coding_gene | -2.3 | 0.209 | 1.6E-02 |
| *acdh-2* | WBGene00015894 | protein_coding_gene | -2.3 | 0.209 | 1.8E-10 |
| *rft-1* | WBGene00044637 | protein_coding_gene | -2.3 | 0.209 | 1.8E-10 |
| *nep-22* | WBGene00020788 | protein_coding_gene | -2.3 | 0.209 | 4.2E-34 |
| *F58G6.7* | WBGene00010277 | protein_coding_gene | -2.3 | 0.208 | 6.5E-08 |

| | | | | | |
|---|---|---|---|---|---|
| *sqt-1* | WBGene00005016 | protein_coding_gene | -2.3 | 0.207 | 1.4E-04 |
| *C13G5.2* | WBGene00015747 | protein_coding_gene | -2.3 | 0.207 | 7.5E-40 |
| *F31D5.1* | WBGene00017944 | protein_coding_gene | -2.3 | 0.207 | 1.9E-03 |
| *pho-7* | WBGene00011879 | protein_coding_gene | -2.3 | 0.207 | 6.8E-09 |
| *sqst-5* | WBGene00269434 | protein_coding_gene | -2.3 | 0.206 | 1.4E-03 |
| *ppat-1* | WBGene00011407 | protein_coding_gene | -2.3 | 0.206 | 9.7E-23 |
| *B0391.10* | WBGene00007165 | pseudogene | -2.3 | 0.206 | 1.9E-17 |
| *ugt-62* | WBGene00010904 | protein_coding_gene | -2.3 | 0.204 | 1.2E-04 |
| *Y58A7A.1* | WBGene00021975 | protein_coding_gene | -2.3 | 0.204 | 9.7E-19 |
| *Y41C4A.32* | WBGene00270321 | protein_coding_gene | -2.3 | 0.204 | 1.1E-03 |
| *vit-3* | WBGene00006927 | protein_coding_gene | -2.3 | 0.204 | 4.5E-03 |
| *F11A5.18* | WBGene00045511 | protein_coding_gene | -2.3 | 0.202 | 6.5E-03 |
| *hrg-7* | WBGene00007605 | protein_coding_gene | -2.3 | 0.202 | 2.7E-04 |
| *C05D9.12* | WBGene00219800 | ncRNA_gene | -2.3 | 0.202 | 1.4E-04 |
| *ugt-22* | WBGene00007455 | protein_coding_gene | -2.3 | 0.202 | 1.4E-05 |
| *nhr-227* | WBGene00020851 | protein_coding_gene | -2.3 | 0.201 | 7.4E-03 |
| *pqn-37* | WBGene00004124 | protein_coding_gene | -2.3 | 0.201 | 1.3E-07 |
| *nspb-10* | WBGene00007246 | protein_coding_gene | -2.3 | 0.201 | 1.4E-02 |
| *F07C6.6* | WBGene00045067 | protein_coding_gene | -2.3 | 0.201 | 1.2E-02 |
| *abu-8* | WBGene00000031 | protein_coding_gene | -2.3 | 0.200 | 2.0E-04 |
| *ZC266.1* | WBGene00022584 | protein_coding_gene | -2.3 | 0.200 | 7.6E-10 |
| *oac-52* | WBGene00020977 | protein_coding_gene | -2.3 | 0.200 | 5.2E-03 |
| *hsp-17* | WBGene00002021 | protein_coding_gene | -2.3 | 0.200 | 1.9E-17 |
| *smd-1* | WBGene00004875 | protein_coding_gene | -2.3 | 0.198 | 2.0E-21 |
| *clec-73* | WBGene00021579 | protein_coding_gene | -2.3 | 0.198 | 2.8E-05 |
| *F54D8.10* | WBGene00284847 | protein_coding_gene | -2.3 | 0.197 | 1.4E-05 |
| *mec-5* | WBGene00003169 | protein_coding_gene | -2.3 | 0.197 | 9.3E-21 |
| *K08H2.3* | WBGene00010690 | protein_coding_gene | -2.3 | 0.197 | 1.6E-13 |
| *Y38H8A.8* | WBGene00045062 | pseudogene | -2.3 | 0.196 | 1.2E-02 |
| *col-73* | WBGene00000649 | protein_coding_gene | -2.4 | 0.196 | 8.0E-04 |
| *ins-30* | WBGene00002113 | protein_coding_gene | -2.4 | 0.196 | 3.2E-03 |
| *col-176* | WBGene00000749 | protein_coding_gene | -2.4 | 0.196 | 3.3E-03 |
| *col-109* | WBGene00000683 | protein_coding_gene | -2.4 | 0.195 | 1.3E-08 |
| *F08D12.2* | WBGene00017246 | protein_coding_gene | -2.4 | 0.195 | 2.8E-12 |
| *C30G12.2* | WBGene00016274 | protein_coding_gene | -2.4 | 0.195 | 1.6E-04 |
| *rol-6* | WBGene00004397 | protein_coding_gene | -2.4 | 0.195 | 2.4E-05 |
| *paic-1* | WBGene00015116 | protein_coding_gene | -2.4 | 0.194 | 9.1E-32 |
| *K02F2.11* | WBGene00220000 | ncRNA_gene | -2.4 | 0.194 | 6.0E-03 |
| *fbxa-64* | WBGene00020883 | protein_coding_gene | -2.4 | 0.194 | 1.6E-10 |
| *E01G4.7* | WBGene00045410 | pseudogene | -2.4 | 0.192 | 3.6E-12 |
| *Y46G5A.29* | WBGene00012917 | protein_coding_gene | -2.4 | 0.192 | 7.1E-04 |
| *Y95B8A.2* | WBGene00022383 | protein_coding_gene | -2.4 | 0.192 | 2.2E-05 |
| *ugt-30* | WBGene00011340 | protein_coding_gene | -2.4 | 0.192 | 7.0E-08 |
| *str-146* | WBGene00006193 | protein_coding_gene | -2.4 | 0.192 | 2.7E-03 |
| *K09F6.5* | WBGene00019588 | protein_coding_gene | -2.4 | 0.191 | 1.2E-02 |
| *peel-1* | WBGene00077563 | protein_coding_gene | -2.4 | 0.191 | 4.9E-03 |
| *nhr-114* | WBGene00003704 | protein_coding_gene | -2.4 | 0.191 | 3.0E-30 |
| *fbxa-192* | WBGene00010212 | protein_coding_gene | -2.4 | 0.191 | 2.3E-23 |
| *bath-13* | WBGene00018223 | protein_coding_gene | -2.4 | 0.190 | 5.6E-04 |
| *B0205.13* | WBGene00044644 | protein_coding_gene | -2.4 | 0.190 | 1.3E-03 |
| *C23H5.8* | WBGene00016027 | protein_coding_gene | -2.4 | 0.190 | 8.8E-06 |
| *C56A3.10* | WBGene00077568 | pseudogene | -2.4 | 0.190 | 5.2E-03 |
| *H11L12.1* | WBGene00019189 | protein_coding_gene | -2.4 | 0.190 | 1.4E-05 |
| *abu-10* | WBGene00000033 | protein_coding_gene | -2.4 | 0.189 | 1.3E-03 |
| *R10E8.3* | WBGene00011209 | protein_coding_gene | -2.4 | 0.189 | 4.0E-03 |
| *C25F9.6* | WBGene00007726 | protein_coding_gene | -2.4 | 0.189 | 5.1E-03 |
| *hil-6* | WBGene00001857 | protein_coding_gene | -2.4 | 0.188 | 5.1E-03 |
| *col-147* | WBGene00000720 | protein_coding_gene | -2.4 | 0.188 | 1.5E-03 |
| *col-168* | WBGene00000741 | protein_coding_gene | -2.4 | 0.188 | 2.6E-04 |
| *H25K10.1* | WBGene00010411 | protein_coding_gene | -2.4 | 0.186 | 3.2E-04 |

| | | | | | |
|---|---|---|---|---|---|
| *F35F10.5* | WBGene00018055 | protein_coding_gene | -2.4 | 0.186 | 4.8E-03 |
| *B0334.13* | WBGene00007149 | protein_coding_gene | -2.4 | 0.186 | 2.8E-03 |
| *grl-4* | WBGene00001713 | protein_coding_gene | -2.4 | 0.186 | 6.6E-18 |
| *C46G7.5* | WBGene00044502 | protein_coding_gene | -2.4 | 0.186 | 4.2E-05 |
| *fbxc-50* | WBGene00015912 | protein_coding_gene | -2.4 | 0.185 | 4.2E-27 |
| *ugt-19* | WBGene00019515 | protein_coding_gene | -2.4 | 0.185 | 3.6E-04 |
| *col-173* | WBGene00000746 | protein_coding_gene | -2.4 | 0.185 | 8.8E-04 |
| *AC3.5* | WBGene00007071 | protein_coding_gene | -2.4 | 0.184 | 3.3E-27 |
| *his-48* | WBGene00001922 | protein_coding_gene | -2.4 | 0.184 | 7.4E-03 |
| *scav-4* | WBGene00008692 | protein_coding_gene | -2.4 | 0.184 | 2.8E-10 |
| *col-17* | WBGene00000606 | protein_coding_gene | -2.4 | 0.184 | 1.6E-05 |
| *lon-1* | WBGene00003055 | protein_coding_gene | -2.4 | 0.183 | 6.5E-44 |
| *col-103* | WBGene00000677 | protein_coding_gene | -2.4 | 0.183 | 4.6E-49 |
| *bath-21* | WBGene00019138 | protein_coding_gene | -2.5 | 0.183 | 8.5E-06 |
| *ZC239.13* | WBGene00022571 | protein_coding_gene | -2.5 | 0.183 | 3.2E-03 |
| *ZK6.6* | WBGene00022641 | protein_coding_gene | -2.5 | 0.183 | 3.2E-05 |
| *ZK402.3* | WBGene00022730 | protein_coding_gene | -2.5 | 0.182 | 4.2E-03 |
| *F57B1.9* | WBGene00045488 | protein_coding_gene | -2.5 | 0.181 | 7.5E-07 |
| *clec-147* | WBGene00007313 | protein_coding_gene | -2.5 | 0.181 | 1.4E-07 |
| *fbxa-54* | WBGene00020449 | protein_coding_gene | -2.5 | 0.180 | 1.6E-06 |
| *col-156* | WBGene00000729 | protein_coding_gene | -2.5 | 0.180 | 5.3E-05 |
| *Y47G6A.14* | WBGene00021640 | protein_coding_gene | -2.5 | 0.180 | 1.9E-27 |
| *F09B12.3* | WBGene00008607 | protein_coding_gene | -2.5 | 0.179 | 5.6E-30 |
| *C42D4.3* | WBGene00016596 | protein_coding_gene | -2.5 | 0.179 | 2.0E-05 |
| *math-49* | WBGene00022690 | protein_coding_gene | -2.5 | 0.179 | 8.3E-03 |
| *col-137* | WBGene00000710 | protein_coding_gene | -2.5 | 0.178 | 8.2E-04 |
| *K10H10.4* | WBGene00010760 | protein_coding_gene | -2.5 | 0.178 | 1.6E-03 |
| *F55A3.7* | WBGene00018852 | pseudogene | -2.5 | 0.178 | 5.0E-07 |
| *gst-10* | WBGene00001758 | protein_coding_gene | -2.5 | 0.178 | 9.8E-18 |
| *F23D12.3* | WBGene00009087 | protein_coding_gene | -2.5 | 0.178 | 2.2E-04 |
| *T28B8.4* | WBGene00012115 | protein_coding_gene | -2.5 | 0.177 | 1.6E-04 |
| *F52C6.13* | WBGene00018670 | protein_coding_gene | -2.5 | 0.177 | 3.5E-03 |
| *cdr-4* | WBGene00010470 | protein_coding_gene | -2.5 | 0.176 | 2.0E-05 |
| *R07E3.4* | WBGene00011105 | protein_coding_gene | -2.5 | 0.175 | 1.2E-15 |
| *C35A11.4* | WBGene00016431 | protein_coding_gene | -2.5 | 0.174 | 1.5E-19 |
| *cest-7* | WBGene00007693 | protein_coding_gene | -2.5 | 0.174 | 1.5E-03 |
| *Y47D7A.13* | WBGene00021625 | protein_coding_gene | -2.5 | 0.174 | 1.4E-04 |
| *T06E4.9* | WBGene00011536 | protein_coding_gene | -2.5 | 0.173 | 3.4E-03 |
| *linc-72* | WBGene00219590 | lincRNA_gene | -2.5 | 0.173 | 3.1E-03 |
| *Y82E9BL.18* | WBGene00044402 | protein_coding_gene | -2.5 | 0.172 | 3.0E-03 |
| *F52C6.2* | WBGene00018659 | protein_coding_gene | -2.5 | 0.171 | 6.1E-30 |
| *col-60* | WBGene00000636 | protein_coding_gene | -2.5 | 0.171 | 4.8E-07 |
| *srw-85* | WBGene00005832 | protein_coding_gene | -2.6 | 0.171 | 3.0E-03 |
| *T16G12.1* | WBGene00011803 | protein_coding_gene | -2.6 | 0.170 | 8.8E-21 |
| *T06E4.12* | WBGene00044011 | protein_coding_gene | -2.6 | 0.170 | 2.3E-03 |
| *Y47H9C.9* | WBGene00012952 | protein_coding_gene | -2.6 | 0.170 | 2.8E-03 |
| *E01G4.6* | WBGene00008448 | protein_coding_gene | -2.6 | 0.170 | 5.9E-08 |
| *C32H11.7* | WBGene00007870 | pseudogene | -2.6 | 0.169 | 2.7E-03 |
| *cyp-35A4* | WBGene00016786 | protein_coding_gene | -2.6 | 0.169 | 1.6E-03 |
| *fbxb-53* | WBGene00018104 | protein_coding_gene | -2.6 | 0.168 | 9.1E-04 |
| *fbxb-12* | WBGene00018109 | pseudogene | -2.6 | 0.168 | 2.7E-03 |
| *twk-16* | WBGene00006670 | protein_coding_gene | -2.6 | 0.167 | 2.8E-08 |
| *col-125* | WBGene00000699 | protein_coding_gene | -2.6 | 0.167 | 1.4E-04 |
| *ZK84.1* | WBGene00022649 | protein_coding_gene | -2.6 | 0.166 | 4.1E-09 |
| *pqn-63* | WBGene00004146 | protein_coding_gene | -2.6 | 0.166 | 3.3E-03 |
| *K05F6.15* | WBGene00269419 | pseudogene | -2.6 | 0.166 | 3.1E-04 |
| *bath-20* | WBGene00015012 | protein_coding_gene | -2.6 | 0.164 | 2.3E-03 |
| *B0281.5* | WBGene00015112 | protein_coding_gene | -2.6 | 0.164 | 1.5E-18 |
| *nhr-220* | WBGene00020591 | pseudogene | -2.6 | 0.164 | 1.4E-03 |
| *nspc-5* | WBGene00044340 | protein_coding_gene | -2.6 | 0.163 | 5.1E-03 |

| | | | | | |
|---|---|---|---|---|---|
| *col-14* | WBGene00000603 | protein_coding_gene | -2.6 | 0.163 | 1.6E-10 |
| *R12E2.7* | WBGene00020033 | protein_coding_gene | -2.6 | 0.163 | 9.8E-07 |
| *fbxb-51* | WBGene00019414 | protein_coding_gene | -2.6 | 0.162 | 2.2E-03 |
| *F19G12.t1* | WBGene00023271 | pseudogene | -2.6 | 0.162 | 2.2E-03 |
| *ZC239.15* | WBGene00022573 | protein_coding_gene | -2.6 | 0.161 | 3.8E-04 |
| *Y23H5A.8* | WBGene00023172 | protein_coding_gene | -2.6 | 0.161 | 1.0E-03 |
| *gst-26* | WBGene00001774 | protein_coding_gene | -2.6 | 0.160 | 4.1E-38 |
| *tag-10* | WBGene00006404 | protein_coding_gene | -2.6 | 0.160 | 2.2E-25 |
| *clec-245* | WBGene00007851 | protein_coding_gene | -2.6 | 0.160 | 1.2E-03 |
| *col-158* | WBGene00000731 | protein_coding_gene | -2.7 | 0.159 | 1.6E-05 |
| *Y53C10A.1* | WBGene00014915 | pseudogene | -2.7 | 0.158 | 1.1E-23 |
| *F59E12.15* | WBGene00235269 | protein_coding_gene | -2.7 | 0.158 | 1.9E-03 |
| *R03D7.2* | WBGene00010989 | protein_coding_gene | -2.7 | 0.157 | 9.6E-22 |
| *gst-4* | WBGene00001752 | protein_coding_gene | -2.7 | 0.157 | 2.8E-21 |
| *Y40C5A.4* | WBGene00021497 | protein_coding_gene | -2.7 | 0.156 | 7.3E-11 |
| *nspc-19* | WBGene00008360 | protein_coding_gene | -2.7 | 0.154 | 1.3E-04 |
| *Y69A2AR.12* | WBGene00022083 | protein_coding_gene | -2.7 | 0.154 | 3.8E-03 |
| *col-71* | WBGene00000647 | protein_coding_gene | -2.7 | 0.154 | 9.9E-10 |
| *F21D9.2* | WBGene00009016 | protein_coding_gene | -2.7 | 0.154 | 2.1E-04 |
| *K09F6.13* | WBGene00235164 | protein_coding_gene | -2.7 | 0.153 | 1.6E-03 |
| *Y47G6A.19* | WBGene00021645 | protein_coding_gene | -2.7 | 0.153 | 4.9E-18 |
| *mpst-7* | WBGene00011307 | protein_coding_gene | -2.7 | 0.153 | 3.8E-09 |
| *col-159* | WBGene00000732 | protein_coding_gene | -2.7 | 0.152 | 1.2E-12 |
| *T26E3.11* | WBGene00044905 | pseudogene | -2.7 | 0.152 | 1.5E-03 |
| *F38B6.4* | WBGene00018174 | protein_coding_gene | -2.7 | 0.151 | 3.2E-25 |
| *Y50D4B.6* | WBGene00021745 | protein_coding_gene | -2.7 | 0.151 | 2.3E-05 |
| *npl-4.1* | WBGene00019120 | protein_coding_gene | -2.7 | 0.151 | 1.1E-22 |
| *scrm-3* | WBGene00015437 | protein_coding_gene | -2.7 | 0.151 | 1.2E-03 |
| *gpdh-1* | WBGene00009824 | protein_coding_gene | -2.7 | 0.151 | 1.1E-07 |
| *acox-1.2* | WBGene00008565 | protein_coding_gene | -2.7 | 0.150 | 5.1E-16 |
| *F36H5.17* | WBGene00270293 | protein_coding_gene | -2.7 | 0.150 | 1.4E-03 |
| *M03B6.1* | WBGene00010833 | protein_coding_gene | -2.7 | 0.150 | 6.5E-11 |
| *bath-30* | WBGene00017460 | protein_coding_gene | -2.7 | 0.150 | 5.7E-37 |
| *col-98* | WBGene00000673 | protein_coding_gene | -2.7 | 0.149 | 3.1E-51 |
| *F11A5.15* | WBGene00008680 | protein_coding_gene | -2.8 | 0.149 | 1.3E-03 |
| *F12E12.11* | WBGene00044811 | protein_coding_gene | -2.8 | 0.149 | 2.6E-09 |
| *Y102A5C.35* | WBGene00044220 | pseudogene | -2.8 | 0.148 | 1.3E-03 |
| *acp-6* | WBGene00022245 | protein_coding_gene | -2.8 | 0.147 | 3.9E-42 |
| *Y38H6C.19* | WBGene00012631 | protein_coding_gene | -2.8 | 0.146 | 6.7E-06 |
| *acs-1* | WBGene00018488 | protein_coding_gene | -2.8 | 0.145 | 1.7E-92 |
| *sdz-18* | WBGene00018442 | protein_coding_gene | -2.8 | 0.145 | 1.1E-03 |
| *bath-6* | WBGene00018666 | pseudogene | -2.8 | 0.143 | 1.5E-06 |
| *F22H10.6* | WBGene00017730 | protein_coding_gene | -2.8 | 0.143 | 5.8E-11 |
| *col-120* | WBGene00000694 | protein_coding_gene | -2.8 | 0.143 | 2.8E-08 |
| *kin-15* | WBGene00002199 | protein_coding_gene | -2.8 | 0.142 | 2.0E-15 |
| *fbxb-43* | WBGene00021774 | protein_coding_gene | -2.8 | 0.142 | 4.9E-04 |
| *arrd-20* | WBGene00015440 | pseudogene | -2.8 | 0.142 | 1.0E-03 |
| *C25F9.9* | WBGene00007728 | protein_coding_gene | -2.8 | 0.141 | 9.8E-04 |
| *F16H6.7* | WBGene00008897 | protein_coding_gene | -2.8 | 0.141 | 9.1E-04 |
| *cest-28* | WBGene00019654 | protein_coding_gene | -2.8 | 0.141 | 1.4E-05 |
| *CE7X_3.2* | WBGene00008350 | pseudogene | -2.8 | 0.140 | 2.3E-03 |
| *C05E7.2* | WBGene00007344 | protein_coding_gene | -2.8 | 0.140 | 1.2E-06 |
| *col-133* | WBGene00000707 | protein_coding_gene | -2.8 | 0.140 | 5.1E-05 |
| *ZC239.20* | WBGene00044371 | protein_coding_gene | -2.8 | 0.140 | 1.1E-04 |
| *ZC250.4* | WBGene00045305 | protein_coding_gene | -2.8 | 0.140 | 5.9E-25 |
| *grd-13* | WBGene00001702 | protein_coding_gene | -2.8 | 0.139 | 8.3E-07 |
| *C05C8.8* | WBGene00015465 | protein_coding_gene | -2.8 | 0.139 | 6.9E-05 |
| *mua-6* | WBGene00003485 | protein_coding_gene | -2.8 | 0.139 | 1.5E-31 |
| *col-161* | WBGene00000734 | protein_coding_gene | -2.8 | 0.139 | 1.1E-04 |
| *F13B6.3* | WBGene00017418 | protein_coding_gene | -2.9 | 0.139 | 8.8E-12 |

| | | | | | |
|---|---|---|---|---|---|
| *nduo-2* | WBGene00010961 | protein_coding_gene | -2.9 | 0.138 | 1.9E-07 |
| *ugt-28* | WBGene00015693 | protein_coding_gene | -2.9 | 0.138 | 3.0E-21 |
| *C17B7.5* | WBGene00015879 | protein_coding_gene | -2.9 | 0.138 | 8.2E-04 |
| *col-63* | WBGene00000639 | protein_coding_gene | -2.9 | 0.137 | 1.9E-11 |
| *spp-23* | WBGene00021335 | protein_coding_gene | -2.9 | 0.136 | 3.0E-08 |
| *D1022.17* | WBGene00271809 | ncRNA_gene | -2.9 | 0.136 | 2.7E-05 |
| *scrm-4* | WBGene00008681 | protein_coding_gene | -2.9 | 0.136 | 2.2E-22 |
| *folt-2* | WBGene00018138 | protein_coding_gene | -2.9 | 0.135 | 2.3E-04 |
| *fbxa-57* | WBGene00016224 | pseudogene | -2.9 | 0.135 | 7.7E-04 |
| *col-130* | WBGene00000704 | protein_coding_gene | -2.9 | 0.135 | 1.1E-08 |
| *C40A11.4* | WBGene00016547 | protein_coding_gene | -2.9 | 0.135 | 1.7E-03 |
| *fbxb-52* | WBGene00019417 | protein_coding_gene | -2.9 | 0.135 | 7.3E-04 |
| *col-91* | WBGene00000666 | protein_coding_gene | -2.9 | 0.134 | 2.1E-07 |
| *F57E7.1* | WBGene00010202 | protein_coding_gene | -2.9 | 0.134 | 7.1E-04 |
| *abu-11* | WBGene00000034 | protein_coding_gene | -2.9 | 0.134 | 4.5E-05 |
| *Y51H7C.1* | WBGene00021779 | protein_coding_gene | -2.9 | 0.134 | 7.0E-10 |
| *fbxc-19* | WBGene00020361 | protein_coding_gene | -2.9 | 0.133 | 6.5E-04 |
| *fbxb-13* | WBGene00015607 | protein_coding_gene | -2.9 | 0.133 | 2.9E-04 |
| *zipt-2.3* | WBGene00013207 | protein_coding_gene | -2.9 | 0.133 | 1.4E-06 |
| *col-77* | WBGene00000653 | protein_coding_gene | -2.9 | 0.133 | 2.0E-13 |
| *T28D6.3* | WBGene00012123 | protein_coding_gene | -2.9 | 0.132 | 4.0E-27 |
| *Y50D4B.4* | WBGene00021743 | protein_coding_gene | -2.9 | 0.132 | 1.2E-32 |
| *Y43F8B.14* | WBGene00012824 | protein_coding_gene | -2.9 | 0.131 | 6.2E-04 |
| *col-146* | WBGene00000719 | protein_coding_gene | -2.9 | 0.131 | 1.0E-07 |
| *fbxb-44* | WBGene00019416 | protein_coding_gene | -2.9 | 0.130 | 5.9E-04 |
| *bli-6* | WBGene00000256 | protein_coding_gene | -2.9 | 0.130 | 6.0E-12 |
| *clec-119* | WBGene00018575 | protein_coding_gene | -2.9 | 0.130 | 9.4E-06 |
| *Y45F10C.6* | WBGene00044256 | protein_coding_gene | -3.0 | 0.129 | 5.4E-04 |
| *drd-1* | WBGene00009902 | protein_coding_gene | -3.0 | 0.128 | 3.6E-58 |
| *col-141* | WBGene00000714 | protein_coding_gene | -3.0 | 0.128 | 3.3E-05 |
| *F40G9.6* | WBGene00018242 | protein_coding_gene | -3.0 | 0.126 | 4.7E-04 |
| *aat-4* | WBGene00000005 | protein_coding_gene | -3.0 | 0.125 | 7.8E-19 |
| *fbxb-97* | WBGene00016885 | protein_coding_gene | -3.0 | 0.125 | 4.5E-04 |
| *dhs-25* | WBGene00000988 | protein_coding_gene | -3.0 | 0.125 | 8.3E-08 |
| *F21D9.6* | WBGene00009020 | pseudogene | -3.0 | 0.125 | 4.6E-04 |
| *R10E8.1* | WBGene00011207 | protein_coding_gene | -3.0 | 0.124 | 2.7E-14 |
| *C50E3.13* | WBGene00016824 | protein_coding_gene | -3.0 | 0.123 | 8.6E-30 |
| *msp-53* | WBGene00003446 | protein_coding_gene | -3.0 | 0.123 | 2.2E-04 |
| *dpy-13* | WBGene00001074 | protein_coding_gene | -3.0 | 0.123 | 7.0E-12 |
| *ZK1240.11* | WBGene00206413 | protein_coding_gene | -3.0 | 0.123 | 4.2E-04 |
| *M01G12.9* | WBGene00010822 | protein_coding_gene | -3.0 | 0.122 | 4.0E-09 |
| *M04C3.5* | WBGene00206418 | protein_coding_gene | -3.0 | 0.122 | 3.9E-04 |
| *C40A11.8* | WBGene00016551 | protein_coding_gene | -3.0 | 0.121 | 9.3E-09 |
| *T20D4.3* | WBGene00020609 | protein_coding_gene | -3.1 | 0.120 | 3.4E-04 |
| *hsd-2* | WBGene00022498 | protein_coding_gene | -3.1 | 0.120 | 8.8E-04 |
| *Y39G10AR.16* | WBGene00021472 | protein_coding_gene | -3.1 | 0.120 | 1.6E-05 |
| *elo-6* | WBGene00001244 | protein_coding_gene | -3.1 | 0.120 | 1.1E-28 |
| *F18E3.13* | WBGene00045276 | protein_coding_gene | -3.1 | 0.119 | 9.4E-11 |
| *K08D10.9* | WBGene00019532 | pseudogene | -3.1 | 0.118 | 3.0E-04 |
| *pho-1* | WBGene00004020 | protein_coding_gene | -3.1 | 0.116 | 1.6E-07 |
| *sptl-2* | WBGene00018398 | protein_coding_gene | -3.1 | 0.116 | 2.1E-43 |
| *elo-2* | WBGene00001240 | protein_coding_gene | -3.1 | 0.115 | 7.0E-79 |
| *linc-11* | WBGene00219758 | lincRNA_gene | -3.1 | 0.115 | 7.6E-05 |
| *T10G3.3* | WBGene00011694 | protein_coding_gene | -3.1 | 0.114 | 1.9E-30 |
| *C33C12.7* | WBGene00016339 | protein_coding_gene | -3.1 | 0.113 | 2.3E-04 |
| *ZC239.22* | WBGene00194717 | protein_coding_gene | -3.1 | 0.113 | 6.9E-04 |
| *clp-6* | WBGene00000546 | protein_coding_gene | -3.1 | 0.113 | 2.4E-06 |
| *col-12* | WBGene00000601 | protein_coding_gene | -3.2 | 0.113 | 1.5E-06 |
| *F16H6.4* | WBGene00008894 | protein_coding_gene | -3.2 | 0.112 | 2.3E-04 |
| *F40H3.2* | WBGene00018250 | protein_coding_gene | -3.2 | 0.112 | 8.5E-16 |

| | | | | | |
|---|---|---|---|---|---|
| *fbxa-156* | WBGene00007392 | protein_coding_gene | -3.2 | 0.112 | 5.9E-05 |
| *T20D4.11* | WBGene00020617 | protein_coding_gene | -3.2 | 0.111 | 6.5E-04 |
| *scl-6* | WBGene00008028 | protein_coding_gene | -3.2 | 0.111 | 1.8E-06 |
| *grd-6* | WBGene00001695 | protein_coding_gene | -3.2 | 0.110 | 2.5E-07 |
| *fbxb-98* | WBGene00015830 | protein_coding_gene | -3.2 | 0.110 | 1.9E-04 |
| *T24E12.5* | WBGene00020774 | protein_coding_gene | -3.2 | 0.110 | 1.7E-04 |
| *F47H4.2* | WBGene00009835 | protein_coding_gene | -3.2 | 0.109 | 5.6E-04 |
| *F53B3.6* | WBGene00018744 | protein_coding_gene | -3.2 | 0.109 | 8.3E-11 |
| *col-97* | WBGene00000672 | protein_coding_gene | -3.2 | 0.109 | 1.3E-15 |
| *C50E3.12* | WBGene00016823 | protein_coding_gene | -3.2 | 0.108 | 2.1E-19 |
| *F09C6.12* | WBGene00044182 | protein_coding_gene | -3.2 | 0.108 | 5.1E-04 |
| *Y113G7A.13* | WBGene00013750 | protein_coding_gene | -3.2 | 0.108 | 1.7E-04 |
| *ZK1025.3* | WBGene00014183 | protein_coding_gene | -3.2 | 0.108 | 1.5E-04 |
| *C52E2.5* | WBGene00016884 | protein_coding_gene | -3.2 | 0.108 | 4.7E-04 |
| *ech-7* | WBGene00001156 | protein_coding_gene | -3.2 | 0.107 | 6.9E-33 |
| *nspa-5* | WBGene00013887 | protein_coding_gene | -3.2 | 0.106 | 1.1E-04 |
| *gst-38* | WBGene00001786 | protein_coding_gene | -3.2 | 0.106 | 1.5E-04 |
| *W02A11.6* | WBGene00014852 | pseudogene | -3.2 | 0.106 | 1.5E-04 |
| *B0281.4* | WBGene00015111 | protein_coding_gene | -3.2 | 0.106 | 1.4E-04 |
| *duxl-1* | WBGene00022554 | protein_coding_gene | -3.3 | 0.105 | 2.6E-26 |
| *grd-14* | WBGene00001703 | protein_coding_gene | -3.3 | 0.105 | 2.6E-13 |
| *dhs-26* | WBGene00000989 | protein_coding_gene | -3.3 | 0.104 | 1.8E-10 |
| *W05G11.2* | WBGene00021040 | protein_coding_gene | -3.3 | 0.104 | 9.0E-05 |
| *Y46B2A.4* | WBGene00284850 | protein_coding_gene | -3.3 | 0.104 | 1.1E-04 |
| *elo-5* | WBGene00001243 | protein_coding_gene | -3.3 | 0.104 | 1.9E-108 |
| *col-38* | WBGene00000615 | protein_coding_gene | -3.3 | 0.103 | 8.0E-25 |
| *C08F11.6* | WBGene00014672 | pseudogene | -3.3 | 0.103 | 2.7E-06 |
| *col-162* | WBGene00000735 | protein_coding_gene | -3.3 | 0.103 | 7.2E-06 |
| *kin-21* | WBGene00002204 | protein_coding_gene | -3.3 | 0.103 | 1.2E-04 |
| *F59H6.15* | WBGene00269394 | protein_coding_gene | -3.3 | 0.102 | 1.1E-04 |
| *acox-1.4* | WBGene00008567 | protein_coding_gene | -3.3 | 0.101 | 2.3E-28 |
| *F53F1.4* | WBGene00009982 | protein_coding_gene | -3.3 | 0.100 | 9.3E-24 |
| *col-175* | WBGene00000748 | protein_coding_gene | -3.3 | 0.100 | 4.5E-15 |
| *Y47H10A.6* | WBGene00077537 | pseudogene | -3.3 | 0.100 | 9.9E-05 |
| *fbxb-50* | WBGene00019413 | protein_coding_gene | -3.3 | 0.100 | 9.9E-05 |
| *dpy-4* | WBGene00001066 | protein_coding_gene | -3.3 | 0.099 | 5.5E-14 |
| *C40A11.6* | WBGene00016549 | protein_coding_gene | -3.3 | 0.099 | 9.0E-05 |
| *Y19D10B.6* | WBGene00021235 | protein_coding_gene | -3.3 | 0.099 | 3.9E-06 |
| *Y49E10.37* | WBGene00304219 | ncRNA_gene | -3.3 | 0.098 | 9.7E-16 |
| *T02G6.10* | WBGene00077772 | pseudogene | -3.3 | 0.098 | 1.1E-05 |
| *D1086.3* | WBGene00008390 | protein_coding_gene | -3.4 | 0.097 | 7.1E-15 |
| *F16H6.10* | WBGene00008900 | protein_coding_gene | -3.4 | 0.097 | 9.5E-15 |
| *Y47G7B.2* | WBGene00021654 | protein_coding_gene | -3.4 | 0.096 | 4.4E-05 |
| *arrd-22* | WBGene00020612 | protein_coding_gene | -3.4 | 0.095 | 1.2E-05 |
| *C04E12.5* | WBGene00015435 | protein_coding_gene | -3.4 | 0.095 | 7.1E-05 |
| *fbxb-42* | WBGene00021773 | protein_coding_gene | -3.4 | 0.095 | 2.2E-04 |
| *fbxb-15* | WBGene00022557 | protein_coding_gene | -3.4 | 0.095 | 5.3E-06 |
| *ZK180.6* | WBGene00022680 | protein_coding_gene | -3.4 | 0.095 | 1.6E-12 |
| *npr-30* | WBGene00019183 | protein_coding_gene | -3.4 | 0.095 | 2.5E-04 |
| *sdz-11* | WBGene00017408 | protein_coding_gene | -3.4 | 0.094 | 6.4E-05 |
| *btb-17* | WBGene00018200 | protein_coding_gene | -3.4 | 0.094 | 2.4E-06 |
| *W07A12.8* | WBGene00012325 | protein_coding_gene | -3.4 | 0.093 | 1.4E-06 |
| *stdh-2* | WBGene00008678 | protein_coding_gene | -3.4 | 0.093 | 5.7E-05 |
| *C04E12.10* | WBGene00015439 | pseudogene | -3.4 | 0.093 | 6.1E-05 |
| *Y47G6A.15* | WBGene00021641 | protein_coding_gene | -3.4 | 0.093 | 1.3E-14 |
| *Y60A3A.25* | WBGene00044211 | protein_coding_gene | -3.4 | 0.092 | 3.2E-05 |
| *dpy-6* | WBGene00001068 | protein_coding_gene | -3.5 | 0.091 | 1.8E-09 |
| *Y43F8B.24* | WBGene00195176 | protein_coding_gene | -3.5 | 0.091 | 4.9E-05 |
| *phat-3* | WBGene00016782 | protein_coding_gene | -3.5 | 0.090 | 6.6E-17 |
| *C17F4.8* | WBGene00015914 | protein_coding_gene | -3.5 | 0.089 | 4.6E-05 |

| | | | | | |
|---|---|---|---|---|---|
| *F26A3.5* | WBGene00009143 | protein_coding_gene | -3.5 | 0.089 | 1.4E-06 |
| *F14D2.5* | WBGene00017453 | protein_coding_gene | -3.5 | 0.089 | 4.6E-05 |
| *F36H5.14* | WBGene00219271 | protein_coding_gene | -3.5 | 0.089 | 4.4E-05 |
| *sdz-24* | WBGene00019495 | protein_coding_gene | -3.5 | 0.089 | 2.2E-05 |
| *F19B2.6* | WBGene00008945 | protein_coding_gene | -3.5 | 0.089 | 8.2E-07 |
| *col-138* | WBGene00000711 | protein_coding_gene | -3.5 | 0.088 | 2.5E-15 |
| *col-157* | WBGene00000730 | protein_coding_gene | -3.5 | 0.088 | 1.6E-07 |
| *fbxb-96* | WBGene00016886 | protein_coding_gene | -3.5 | 0.088 | 3.9E-05 |
| *col-88* | WBGene00000663 | protein_coding_gene | -3.5 | 0.087 | 4.2E-13 |
| *col-49* | WBGene00000626 | protein_coding_gene | -3.5 | 0.087 | 3.1E-11 |
| *F41E6.12* | WBGene00018293 | protein_coding_gene | -3.5 | 0.087 | 1.6E-23 |
| *bath-3* | WBGene00019140 | protein_coding_gene | -3.5 | 0.086 | 2.8E-13 |
| *F46F2.5* | WBGene00009789 | protein_coding_gene | -3.5 | 0.086 | 2.2E-12 |
| *asp-17* | WBGene00012683 | protein_coding_gene | -3.6 | 0.085 | 4.0E-06 |
| *F39F10.3* | WBGene00018203 | protein_coding_gene | -3.6 | 0.085 | 7.7E-10 |
| *fbxa-188* | WBGene00009839 | protein_coding_gene | -3.6 | 0.085 | 3.0E-05 |
| *dpy-5* | WBGene00001067 | protein_coding_gene | -3.6 | 0.083 | 6.9E-17 |
| *Y73B6BL.14* | WBGene00022237 | protein_coding_gene | -3.6 | 0.083 | 1.1E-04 |
| *ZC204.12* | WBGene00022562 | protein_coding_gene | -3.6 | 0.082 | 1.7E-07 |
| *ZK1240.5* | WBGene00022870 | protein_coding_gene | -3.6 | 0.081 | 2.2E-05 |
| *scrm-8* | WBGene00019530 | protein_coding_gene | -3.6 | 0.081 | 1.9E-05 |
| *T20D4.5* | WBGene00020611 | protein_coding_gene | -3.6 | 0.080 | 2.0E-05 |
| *Y51H7C.13* | WBGene00021791 | protein_coding_gene | -3.6 | 0.080 | 7.5E-13 |
| *nhr-74* | WBGene00003664 | protein_coding_gene | -3.6 | 0.080 | 7.4E-05 |
| *sur-5* | WBGene00006351 | protein_coding_gene | -3.6 | 0.080 | 6.7E-203 |
| *C04G6.6* | WBGene00015456 | protein_coding_gene | -3.7 | 0.079 | 1.7E-18 |
| *F58G6.3* | WBGene00010274 | protein_coding_gene | -3.7 | 0.079 | 7.0E-26 |
| *F54E2.1* | WBGene00018823 | protein_coding_gene | -3.7 | 0.076 | 5.1E-08 |
| *ZC239.21* | WBGene00044372 | protein_coding_gene | -3.7 | 0.076 | 1.3E-05 |
| *col-180* | WBGene00000753 | protein_coding_gene | -3.7 | 0.076 | 2.7E-11 |
| *C07G1.7* | WBGene00015573 | protein_coding_gene | -3.7 | 0.076 | 1.1E-06 |
| *F14D2.15* | WBGene00023427 | protein_coding_gene | -3.7 | 0.075 | 1.0E-20 |
| *msp-64* | WBGene00003457 | protein_coding_gene | -3.7 | 0.075 | 7.6E-07 |
| *hrg-4* | WBGene00009493 | protein_coding_gene | -3.8 | 0.074 | 2.3E-10 |
| *bath-27* | WBGene00017450 | protein_coding_gene | -3.8 | 0.074 | 4.6E-16 |
| *btb-3* | WBGene00015013 | pseudogene | -3.8 | 0.073 | 1.0E-05 |
| *K02E11.10* | WBGene00044109 | protein_coding_gene | -3.8 | 0.072 | 6.1E-08 |
| *F19B10.2* | WBGene00017584 | protein_coding_gene | -3.8 | 0.072 | 4.5E-22 |
| *F59H6.2* | WBGene00019133 | pseudogene | -3.8 | 0.072 | 8.7E-06 |
| *math-20* | WBGene00016555 | protein_coding_gene | -3.8 | 0.072 | 2.3E-10 |
| *fbxa-117* | WBGene00014907 | pseudogene | -3.8 | 0.072 | 8.3E-06 |
| *scrm-5* | WBGene00019531 | protein_coding_gene | -3.8 | 0.071 | 6.4E-06 |
| *T19C9.8* | WBGene00011844 | protein_coding_gene | -3.8 | 0.071 | 6.3E-06 |
| *Y48G8AL.12* | WBGene00021690 | protein_coding_gene | -3.8 | 0.071 | 2.9E-07 |
| *fbxa-10* | WBGene00020450 | protein_coding_gene | -3.8 | 0.070 | 7.1E-06 |
| *col-167* | WBGene00000740 | protein_coding_gene | -3.8 | 0.070 | 2.4E-11 |
| *fbxb-49* | WBGene00019412 | protein_coding_gene | -3.8 | 0.070 | 6.5E-06 |
| *nhr-115* | WBGene00003705 | protein_coding_gene | -3.8 | 0.070 | 4.4E-10 |
| *R03G8.4* | WBGene00010999 | pseudogene | -3.8 | 0.069 | 3.0E-05 |
| *col-149* | WBGene00000722 | protein_coding_gene | -3.9 | 0.069 | 4.8E-13 |
| *gcsh-1* | WBGene00008354 | protein_coding_gene | -3.9 | 0.069 | 7.2E-37 |
| *F18E3.11* | WBGene00017567 | protein_coding_gene | -3.9 | 0.069 | 1.5E-20 |
| *F52C6.4* | WBGene00018661 | protein_coding_gene | -3.9 | 0.068 | 2.7E-05 |
| *F10E7.1* | WBGene00017343 | protein_coding_gene | -3.9 | 0.068 | 5.6E-06 |
| *K09H9.7* | WBGene00019598 | protein_coding_gene | -3.9 | 0.068 | 6.2E-28 |
| *rol-1* | WBGene00004394 | protein_coding_gene | -3.9 | 0.067 | 9.9E-17 |
| *fat-5* | WBGene00001397 | protein_coding_gene | -3.9 | 0.067 | 3.0E-36 |
| *T12B5.14* | WBGene00050913 | protein_coding_gene | -3.9 | 0.066 | 4.4E-06 |
| *T26E3.8* | WBGene00012041 | protein_coding_gene | -3.9 | 0.066 | 4.4E-06 |
| *K09C4.5* | WBGene00019550 | protein_coding_gene | -4.0 | 0.064 | 3.5E-32 |

| | | | | | |
|---|---|---|---|---|---|
| *col-145* | WBGene00000718 | protein_coding_gene | -4.0 | 0.064 | 1.6E-09 |
| *nspd-7* | WBGene00012781 | protein_coding_gene | -4.0 | 0.063 | 8.5E-08 |
| *fbxa-11* | WBGene00020451 | protein_coding_gene | -4.0 | 0.063 | 2.9E-06 |
| *Y47D7A.15* | WBGene00021627 | protein_coding_gene | -4.0 | 0.063 | 1.6E-10 |
| *F42A8.1* | WBGene00009626 | protein_coding_gene | -4.0 | 0.062 | 1.9E-43 |
| *math-40* | WBGene00020359 | protein_coding_gene | -4.0 | 0.062 | 4.0E-08 |
| *F14D2.19* | WBGene00206522 | protein_coding_gene | -4.0 | 0.060 | 2.1E-06 |
| *fbxa-6* | WBGene00008352 | protein_coding_gene | -4.0 | 0.060 | 2.1E-06 |
| *Y57G11C.40* | WBGene00013329 | protein_coding_gene | -4.1 | 0.060 | 2.1E-06 |
| *Y9D1A.1* | WBGene00021186 | protein_coding_gene | -4.1 | 0.060 | 7.8E-16 |
| *pud-3* | WBGene00017501 | protein_coding_gene | -4.1 | 0.059 | 1.2E-06 |
| *K05F6.4* | WBGene00019415 | protein_coding_gene | -4.1 | 0.059 | 5.8E-09 |
| *F42G2.5* | WBGene00018354 | protein_coding_gene | -4.1 | 0.058 | 1.6E-06 |
| *grd-4* | WBGene00001693 | protein_coding_gene | -4.1 | 0.058 | 4.5E-06 |
| *C02G6.1* | WBGene00015359 | pseudogene | -4.1 | 0.057 | 2.1E-07 |
| *F31E9.11* | WBGene00195233 | ncRNA_gene | -4.1 | 0.057 | 7.1E-06 |
| *math-24* | WBGene00016719 | protein_coding_gene | -4.2 | 0.056 | 1.9E-09 |
| *F26D2.10* | WBGene00009149 | protein_coding_gene | -4.2 | 0.056 | 9.9E-07 |
| *fbxb-46* | WBGene00019420 | protein_coding_gene | -4.2 | 0.056 | 1.1E-06 |
| *ugt-25* | WBGene00015692 | protein_coding_gene | -4.2 | 0.055 | 2.0E-25 |
| *clec-72* | WBGene00021583 | protein_coding_gene | -4.2 | 0.055 | 9.8E-07 |
| *fbxb-20* | WBGene00022559 | protein_coding_gene | -4.2 | 0.054 | 8.6E-07 |
| *fbxa-38* | WBGene00016522 | protein_coding_gene | -4.2 | 0.054 | 8.5E-07 |
| *D1065.6* | WBGene00268212 | ncRNA_gene | -4.2 | 0.053 | 6.9E-08 |
| *fbxb-95* | WBGene00016880 | protein_coding_gene | -4.3 | 0.053 | 6.3E-07 |
| *nas-3* | WBGene00003522 | protein_coding_gene | -4.3 | 0.052 | 3.2E-16 |
| *F52C6.12* | WBGene00018669 | protein_coding_gene | -4.3 | 0.052 | 3.8E-06 |
| *Y75B8A.32* | WBGene00013562 | protein_coding_gene | -4.3 | 0.051 | 5.0E-07 |
| *F19B10.1* | WBGene00017583 | protein_coding_gene | -4.3 | 0.051 | 3.6E-14 |
| *abhd-3.2* | WBGene00016644 | protein_coding_gene | -4.3 | 0.051 | 1.9E-73 |
| *fbxa-60* | WBGene00020456 | protein_coding_gene | -4.3 | 0.050 | 1.4E-28 |
| *fbxb-54* | WBGene00019418 | protein_coding_gene | -4.4 | 0.048 | 2.9E-07 |
| *btb-14* | WBGene00013275 | protein_coding_gene | -4.4 | 0.048 | 6.0E-25 |
| *F52C6.3* | WBGene00018660 | protein_coding_gene | -4.4 | 0.047 | 2.6E-07 |
| *grsp-4* | WBGene00017201 | protein_coding_gene | -4.4 | 0.047 | 3.4E-47 |
| *M04C3.2* | WBGene00010851 | protein_coding_gene | -4.4 | 0.047 | 2.4E-07 |
| *ctsa-4.2* | WBGene00021503 | protein_coding_gene | -4.4 | 0.047 | 7.5E-44 |
| *K08D10.14* | WBGene00219216 | protein_coding_gene | -4.5 | 0.045 | 1.6E-07 |
| *R07E5.4* | WBGene00011112 | protein_coding_gene | -4.5 | 0.045 | 1.4E-27 |
| *Y73B6BL.288* | WBGene00219218 | protein_coding_gene | -4.5 | 0.045 | 3.1E-15 |
| *zeel-1* | WBGene00021463 | protein_coding_gene | -4.5 | 0.045 | 1.5E-07 |
| *C04E12.2* | WBGene00015433 | protein_coding_gene | -4.5 | 0.044 | 1.1E-06 |
| *fbxa-180* | WBGene00009295 | pseudogene | -4.5 | 0.043 | 9.0E-07 |
| *F18E3.12* | WBGene00045275 | protein_coding_gene | -4.5 | 0.043 | 6.8E-19 |
| *R07E5.15* | WBGene00011120 | protein_coding_gene | -4.6 | 0.042 | 3.5E-07 |
| *F49F1.7* | WBGene00018647 | protein_coding_gene | -4.6 | 0.041 | 7.1E-07 |
| *R193.2* | WBGene00020128 | protein_coding_gene | -4.6 | 0.040 | 1.4E-44 |
| *math-39* | WBGene00020358 | protein_coding_gene | -4.6 | 0.040 | 5.0E-07 |
| *F14D2.14* | WBGene00017462 | protein_coding_gene | -4.7 | 0.039 | 4.3E-08 |
| *bath-11* | WBGene00018664 | pseudogene | -4.7 | 0.039 | 4.0E-08 |
| *Y54G2A.45* | WBGene00044488 | protein_coding_gene | -4.7 | 0.038 | 3.9E-27 |
| *math-41* | WBGene00020360 | protein_coding_gene | -4.7 | 0.038 | 2.4E-11 |
| *T22B7.7* | WBGene00020674 | protein_coding_gene | -4.7 | 0.038 | 1.9E-30 |
| *T28A11.2* | WBGene00020869 | protein_coding_gene | -4.7 | 0.038 | 8.6E-15 |
| *F14D2.11* | WBGene00017459 | protein_coding_gene | -4.7 | 0.037 | 3.0E-08 |
| *fbxb-91* | WBGene00044692 | protein_coding_gene | -4.8 | 0.037 | 2.5E-08 |
| *Y82E9BR.5* | WBGene00022338 | protein_coding_gene | -4.8 | 0.037 | 2.4E-08 |
| *F46F2.3* | WBGene00009787 | protein_coding_gene | -4.8 | 0.036 | 1.6E-17 |
| *Y46G5A.7* | WBGene00012899 | protein_coding_gene | -4.8 | 0.035 | 1.7E-08 |
| *nspc-14* | WBGene00009638 | protein_coding_gene | -4.8 | 0.035 | 6.1E-31 |

| | | | | | |
|---|---|---|---|---|---|
| *fbxa-216* | WBGene00012953 | protein_coding_gene | -4.9 | 0.034 | 4.1E-10 |
| *T16G12.3* | WBGene00011804 | protein_coding_gene | -4.9 | 0.034 | 1.4E-40 |
| *Y7A9C.1* | WBGene00012418 | pseudogene | -4.9 | 0.033 | 9.2E-09 |
| *C25F9.10* | WBGene00044291 | protein_coding_gene | -4.9 | 0.033 | 9.1E-09 |
| *fbxa-65* | WBGene00015885 | protein_coding_gene | -5.0 | 0.032 | 1.3E-12 |
| *C25F9.14* | WBGene00045461 | protein_coding_gene | -5.0 | 0.032 | 6.5E-09 |
| *M162.9* | WBGene00014811 | pseudogene | -5.0 | 0.031 | 5.2E-09 |
| *btb-10* | WBGene00019313 | protein_coding_gene | -5.0 | 0.030 | 4.2E-09 |
| *C08F11.7* | WBGene00007454 | protein_coding_gene | -5.0 | 0.030 | 3.6E-17 |
| *R10E8.7* | WBGene00014821 | pseudogene | -5.1 | 0.030 | 6.6E-11 |
| *F14D2.8* | WBGene00017456 | protein_coding_gene | -5.1 | 0.030 | 3.6E-09 |
| *K09F6.15* | WBGene00271744 | protein_coding_gene | -5.1 | 0.030 | 3.6E-09 |
| *bath-7* | WBGene00018667 | protein_coding_gene | -5.1 | 0.030 | 7.4E-09 |
| *bath-22* | WBGene00018658 | pseudogene | -5.1 | 0.029 | 2.9E-09 |
| *bath-14* | WBGene00015016 | pseudogene | -5.1 | 0.029 | 2.7E-09 |
| *Y46G5A.8* | WBGene00012900 | protein_coding_gene | -5.2 | 0.028 | 1.7E-09 |
| *bath-24* | WBGene00015014 | protein_coding_gene | -5.2 | 0.028 | 1.7E-09 |
| *K09F6.3* | WBGene00019586 | protein_coding_gene | -5.2 | 0.028 | 9.9E-09 |
| *fbxa-189* | WBGene00009840 | protein_coding_gene | -5.2 | 0.027 | 1.6E-09 |
| *ZC239.6* | WBGene00022569 | protein_coding_gene | -5.2 | 0.027 | 1.4E-09 |
| *R08E5.1* | WBGene00019961 | protein_coding_gene | -5.2 | 0.027 | 1.3E-09 |
| *K09F6.7* | WBGene00019590 | protein_coding_gene | -5.3 | 0.026 | 9.0E-10 |
| *K09F6.6* | WBGene00019589 | protein_coding_gene | -5.3 | 0.026 | 7.1E-10 |
| *C25F9.4* | WBGene00007724 | protein_coding_gene | -5.3 | 0.025 | 6.4E-10 |
| *math-26* | WBGene00018100 | protein_coding_gene | -5.3 | 0.025 | 1.2E-21 |
| *K05C4.9* | WBGene00010585 | protein_coding_gene | -5.3 | 0.025 | 5.5E-10 |
| *K09F6.4* | WBGene00019587 | protein_coding_gene | -5.3 | 0.025 | 8.3E-09 |
| *C38D9.13* | WBGene00235090 | ncRNA_gene | -5.4 | 0.024 | 2.1E-12 |
| *skr-4* | WBGene00004810 | protein_coding_gene | -5.4 | 0.024 | 1.2E-23 |
| *Y46D2A.2* | WBGene00021590 | protein_coding_gene | -5.4 | 0.024 | 3.8E-10 |
| *K09F6.10* | WBGene00019593 | protein_coding_gene | -5.4 | 0.023 | 2.9E-10 |
| *msp-19* | WBGene00003426 | protein_coding_gene | -5.4 | 0.023 | 7.5E-10 |
| *fbxa-207* | WBGene00013612 | pseudogene | -5.5 | 0.022 | 1.8E-10 |
| *math-14* | WBGene00015828 | protein_coding_gene | -5.5 | 0.022 | 1.5E-10 |
| *nspa-8* | WBGene00012294 | protein_coding_gene | -5.5 | 0.022 | 1.7E-10 |
| *math-27* | WBGene00018101 | pseudogene | -5.5 | 0.022 | 1.7E-18 |
| *math-3* | WBGene00015608 | protein_coding_gene | -5.5 | 0.022 | 5.3E-26 |
| *F59E12.6* | WBGene00019122 | protein_coding_gene | -5.6 | 0.021 | 9.5E-11 |
| *K09F6.9* | WBGene00019592 | protein_coding_gene | -5.6 | 0.021 | 2.0E-09 |
| *gst-22* | WBGene00001770 | protein_coding_gene | -5.6 | 0.021 | 8.2E-11 |
| *E01G4.5* | WBGene00008447 | protein_coding_gene | -5.6 | 0.021 | 1.8E-10 |
| *fbxa-129* | WBGene00010294 | pseudogene | -5.6 | 0.020 | 7.0E-11 |
| *C29G2.2* | WBGene00016230 | pseudogene | -5.6 | 0.020 | 6.8E-11 |
| *fbxa-67* | WBGene00020457 | protein_coding_gene | -5.7 | 0.020 | 5.2E-11 |
| *T07D3.4* | WBGene00020307 | protein_coding_gene | -5.7 | 0.019 | 9.5E-10 |
| *Y102A5C.36* | WBGene00044213 | protein_coding_gene | -5.8 | 0.018 | 2.4E-11 |
| *pals-23* | WBGene00016218 | protein_coding_gene | -5.9 | 0.017 | 1.1E-11 |
| *Y48E1B.8* | WBGene00013007 | protein_coding_gene | -5.9 | 0.017 | 5.7E-31 |
| *bath-1* | WBGene00019139 | protein_coding_gene | -5.9 | 0.017 | 8.5E-12 |
| *ZK1240.2* | WBGene00022867 | protein_coding_gene | -5.9 | 0.016 | 7.0E-12 |
| *bath-19* | WBGene00019132 | protein_coding_gene | -5.9 | 0.016 | 6.7E-12 |
| *ZC204.14* | WBGene00022564 | protein_coding_gene | -5.9 | 0.016 | 6.5E-12 |
| *C25F9.5* | WBGene00007725 | protein_coding_gene | -6.0 | 0.016 | 4.6E-12 |
| *msp-33* | WBGene00003431 | protein_coding_gene | -6.1 | 0.015 | 7.8E-17 |
| *btb-4* | WBGene00019142 | protein_coding_gene | -6.1 | 0.015 | 2.2E-12 |
| *B0281.3* | WBGene00015110 | protein_coding_gene | -6.1 | 0.015 | 1.7E-12 |
| *pals-24* | WBGene00016219 | protein_coding_gene | -6.2 | 0.014 | 1.3E-12 |
| *Y113G7B.12* | WBGene00013762 | protein_coding_gene | -6.2 | 0.014 | 1.1E-12 |
| *fbxa-193* | WBGene00010295 | pseudogene | -6.2 | 0.014 | 8.5E-13 |
| *F10E7.2* | WBGene00017344 | protein_coding_gene | -6.2 | 0.014 | 8.5E-13 |

| | | | | | |
|---|---|---|---|---|---|
| *Y38E10A.14* | WBGene00012592 | protein_coding_gene | -6.2 | 0.013 | 1.2E-12 |
| *R07C12.2* | WBGene00019932 | pseudogene | -6.3 | 0.013 | 4.1E-13 |
| *C04E12.4* | WBGene00015434 | protein_coding_gene | -6.3 | 0.013 | 3.3E-13 |
| *Y113G7B.14* | WBGene00013763 | pseudogene | -6.4 | 0.012 | 2.6E-13 |
| *bath-2* | WBGene00018668 | protein_coding_gene | -6.4 | 0.011 | 9.3E-18 |
| *Y102A5C.2* | WBGene00013611 | protein_coding_gene | -6.5 | 0.011 | 7.1E-14 |
| *math-22* | WBGene00016717 | protein_coding_gene | -6.6 | 0.011 | 4.2E-14 |
| *C38D9.2* | WBGene00008010 | protein_coding_gene | -6.6 | 0.010 | 1.9E-182 |
| *bath-28* | WBGene00017461 | protein_coding_gene | -6.9 | 0.009 | 4.1E-15 |
| *math-4* | WBGene00015609 | protein_coding_gene | -6.9 | 0.008 | 2.0E-15 |
| *gst-27* | WBGene00001775 | protein_coding_gene | -7.1 | 0.007 | 2.8E-45 |
| *ZK1240.3* | WBGene00022868 | protein_coding_gene | -7.1 | 0.007 | 4.7E-16 |
| *F54D10.7* | WBGene00018808 | protein_coding_gene | -7.1 | 0.007 | 3.7E-16 |
| *clec-10* | WBGene00015403 | protein_coding_gene | -7.1 | 0.007 | 3.6E-16 |
| *F59A7.8* | WBGene00019095 | pseudogene | -7.2 | 0.007 | 2.1E-16 |
| *Y102A5C.5* | WBGene00014954 | pseudogene | -7.2 | 0.007 | 1.9E-16 |
| *F55B11.6* | WBGene00044249 | protein_coding_gene | -7.3 | 0.006 | 7.1E-17 |
| *bath-5* | WBGene00019141 | protein_coding_gene | -7.4 | 0.006 | 4.6E-17 |
| *T12B5.15* | WBGene00050914 | protein_coding_gene | -7.5 | 0.006 | 2.0E-17 |
| *F19B10.10* | WBGene00017589 | protein_coding_gene | -7.5 | 0.005 | 1.6E-15 |
| *fbxa-206* | WBGene00013610 | protein_coding_gene | -7.6 | 0.005 | 5.8E-18 |
| *C27C7.1* | WBGene00007766 | protein_coding_gene | -7.7 | 0.005 | 1.3E-18 |
| *M04C3.1* | WBGene00010850 | protein_coding_gene | -8.0 | 0.004 | 2.4E-19 |
| *F19B2.5* | WBGene00008944 | protein_coding_gene | -8.1 | 0.004 | 5.6E-20 |
| *sri-40* | WBGene00005552 | protein_coding_gene | -8.2 | 0.003 | 3.9E-18 |
| *Y102A5C.6* | WBGene00014955 | pseudogene | -8.3 | 0.003 | 6.9E-21 |
| *Y94H6A.10* | WBGene00022382 | protein_coding_gene | -8.6 | 0.002 | 3.6E-22 |

**Supplemental Table 13**

**Lists of differentially expressed genes that were conditionally different between N2 and CB4856 fed *E. coli* upon infection by *S. epidermidis***

**Supplemental Table 13 - Genes expressed in CB4856 (vs. N2) fed *S. epidermidis***

| Gene Name | Wormbase Gene ID | Type | $log_2FC$ | FC | Adj P-value |
|---|---|---|---|---|---|
| *clec-110* | WBGene00008909 | protein_coding_gene | 6.5 | 89.28 | 5.6E-37 |
| *nspa-1* | WBGene00010377 | protein_coding_gene | 6.4 | 85.47 | 5.9E-18 |
| *R03H10.6* | WBGene00019858 | protein_coding_gene | 6.3 | 81.21 | 6.3E-160 |
| *clec-124* | WBGene00015909 | protein_coding_gene | 6.2 | 73.80 | 2.2E-29 |
| *K02F6.4* | WBGene00019338 | protein_coding_gene | 6.0 | 64.68 | 7.4E-28 |
| *B0250.3* | WBGene00007120 | pseudogene | 5.8 | 55.68 | 9.7E-111 |
| *Y43F8C.17* | WBGene00012838 | protein_coding_gene | 5.8 | 55.60 | 5.2E-38 |
| *F26D10.13* | WBGene00044244 | protein_coding_gene | 5.7 | 51.99 | 7.0E-90 |
| *clec-219* | WBGene00020945 | protein_coding_gene | 5.7 | 50.62 | 7.2E-185 |
| *plg-1* | WBGene00004041 | pseudogene | 5.6 | 49.38 | 1.3E-108 |
| *Y6G8.15* | WBGene00189937 | protein_coding_gene | 5.6 | 47.60 | 9.3E-16 |
| *T23G4.5* | WBGene00011961 | protein_coding_gene | 5.5 | 45.69 | 2.5E-69 |
| *clec-208* | WBGene00018097 | protein_coding_gene | 5.5 | 44.46 | 1.6E-124 |
| *clec-77* | WBGene00044343 | pseudogene | 5.5 | 43.91 | 5.5E-136 |
| *F26D2.16* | WBGene00009154 | protein_coding_gene | 5.4 | 43.22 | 8.8E-158 |
| *Y59H11AM.1* | WBGene00022005 | protein_coding_gene | 5.4 | 41.59 | 1.9E-13 |
| *clec-132* | WBGene00021143 | protein_coding_gene | 5.3 | 40.17 | 5.7E-15 |
| *clec-125* | WBGene00021118 | protein_coding_gene | 5.3 | 40.12 | 4.0E-117 |
| *R05H10.7* | WBGene00011046 | protein_coding_gene | 5.3 | 38.52 | 1.2E-68 |
| *clec-95* | WBGene00013927 | protein_coding_gene | 5.2 | 37.63 | 1.3E-114 |
| *M7.9* | WBGene00010885 | protein_coding_gene | 5.2 | 36.79 | 9.9E-134 |
| *nas-17* | WBGene00003536 | protein_coding_gene | 5.2 | 36.67 | 1.7E-33 |
| *clec-141* | WBGene00017139 | protein_coding_gene | 5.2 | 36.11 | 1.2E-45 |
| *C33G3.5* | WBGene00007905 | protein_coding_gene | 5.2 | 35.98 | 2.0E-63 |
| *clec-137* | WBGene00018048 | protein_coding_gene | 5.2 | 35.66 | 1.1E-113 |
| *F47C12.8* | WBGene00018551 | protein_coding_gene | 5.2 | 35.53 | 1.5E-12 |
| *fipr-18* | WBGene00044450 | protein_coding_gene | 5.1 | 35.15 | 5.4E-79 |
| *Y74C10AR.2* | WBGene00022282 | protein_coding_gene | 5.1 | 34.43 | 1.0E-118 |
| *ins-31* | WBGene00002114 | protein_coding_gene | 5.1 | 34.29 | 2.8E-194 |
| *C49C3.11* | WBGene00008201 | protein_coding_gene | 5.1 | 34.28 | 6.5E-151 |
| *F18E9.7* | WBGene00017573 | protein_coding_gene | 5.1 | 33.84 | 3.5E-92 |
| *M151.10* | WBGene00303365 | protein_coding_gene | 5.1 | 33.83 | 2.7E-71 |
| *C47A10.12* | WBGene00008129 | protein_coding_gene | 5.1 | 33.62 | 2.0E-41 |
| *F59B2.12* | WBGene00010314 | protein_coding_gene | 5.1 | 33.41 | 0.0E+00 |
| *F54D12.9* | WBGene00018822 | protein_coding_gene | 5.1 | 33.24 | 7.9E-76 |
| *fbxa-58* | WBGene00016225 | pseudogene | 5.0 | 32.96 | 3.4E-12 |
| *srh-254* | WBGene00005460 | pseudogene | 5.0 | 32.87 | 2.4E-13 |
| *clec-96* | WBGene00013930 | protein_coding_gene | 5.0 | 32.27 | 8.2E-38 |
| *F26H11.8* | WBGene00195149 | protein_coding_gene | 5.0 | 32.27 | 3.1E-13 |
| *clec-99* | WBGene00013933 | protein_coding_gene | 5.0 | 31.48 | 7.6E-104 |
| *clec-158* | WBGene00020067 | protein_coding_gene | 5.0 | 31.13 | 1.5E-70 |
| *T14G12.11* | WBGene00219411 | protein_coding_gene | 5.0 | 31.05 | 5.3E-12 |
| *clec-157* | WBGene00017810 | protein_coding_gene | 4.9 | 30.72 | 5.5E-82 |
| *try-8* | WBGene00017791 | protein_coding_gene | 4.9 | 30.29 | 1.5E-155 |
| *scl-7* | WBGene00008029 | protein_coding_gene | 4.9 | 30.18 | 2.2E-52 |
| *B0207.5* | WBGene00015028 | protein_coding_gene | 4.9 | 29.44 | 3.0E-118 |
| *B0222.10* | WBGene00015058 | ncRNA_gene | 4.9 | 29.03 | 5.3E-90 |
| *T22G5.1* | WBGene00011929 | protein_coding_gene | 4.9 | 28.99 | 3.2E-57 |
| *F13E9.10* | WBGene00008759 | protein_coding_gene | 4.8 | 28.57 | 2.8E-37 |
| *clec-197* | WBGene00008202 | protein_coding_gene | 4.8 | 28.56 | 6.3E-169 |
| *F43C11.2* | WBGene00018379 | protein_coding_gene | 4.8 | 28.46 | 7.1E-56 |
| *pqn-84* | WBGene00004165 | protein_coding_gene | 4.8 | 28.30 | 6.7E-131 |
| *K11D12.6* | WBGene00019646 | protein_coding_gene | 4.8 | 28.18 | 1.4E-85 |
| *fipr-17* | WBGene00012965 | protein_coding_gene | 4.8 | 27.59 | 2.5E-78 |
| *clec-207* | WBGene00018096 | protein_coding_gene | 4.8 | 27.57 | 2.5E-101 |
| *fipr-20* | WBGene00044449 | protein_coding_gene | 4.8 | 27.55 | 2.0E-79 |
| *ZK1290.1* | WBGene00022886 | protein_coding_gene | 4.8 | 27.53 | 6.5E-70 |

| | | | | | |
|---|---|---|---|---|---|
| *F47C12.12* | WBGene00219377 | pseudogene | 4.8 | 27.37 | 5.3E-16 |
| *F25E5.3* | WBGene00017784 | protein_coding_gene | 4.8 | 27.35 | 8.5E-71 |
| *C14F11.7* | WBGene00015782 | protein_coding_gene | 4.8 | 27.30 | 1.3E-55 |
| *C16C8.9* | WBGene00015847 | protein_coding_gene | 4.8 | 27.29 | 2.3E-97 |
| *Y51A2D.8* | WBGene00013076 | protein_coding_gene | 4.8 | 27.08 | 4.3E-156 |
| *ZK1053.3* | WBGene00014198 | protein_coding_gene | 4.7 | 26.78 | 2.7E-60 |
| *Y25C1A.2* | WBGene00021289 | protein_coding_gene | 4.7 | 26.55 | 9.9E-36 |
| *F46A8.5* | WBGene00009748 | protein_coding_gene | 4.7 | 26.46 | 7.0E-39 |
| *T12A2.5* | WBGene00020439 | protein_coding_gene | 4.7 | 26.30 | 5.8E-55 |
| *T16G12.10* | WBGene00044222 | protein_coding_gene | 4.7 | 25.85 | 1.6E-88 |
| *F16G10.10* | WBGene00017525 | protein_coding_gene | 4.7 | 25.54 | 4.7E-25 |
| *R09E10.8* | WBGene00011177 | protein_coding_gene | 4.7 | 25.20 | 1.6E-34 |
| *Y82E9BL.9* | WBGene00023395 | pseudogene | 4.7 | 25.14 | 5.3E-25 |
| *C48B4.12* | WBGene00008175 | protein_coding_gene | 4.6 | 25.07 | 4.0E-93 |
| *clec-217* | WBGene00017824 | protein_coding_gene | 4.6 | 24.98 | 3.7E-51 |
| *fipr-28* | WBGene00013120 | protein_coding_gene | 4.6 | 24.97 | 1.2E-62 |
| *T10B10.9* | WBGene00011685 | protein_coding_gene | 4.6 | 24.85 | 6.1E-37 |
| *C05B5.11* | WBGene00045242 | protein_coding_gene | 4.6 | 24.62 | 2.5E-69 |
| *R53.8* | WBGene00011277 | protein_coding_gene | 4.6 | 24.46 | 1.7E-56 |
| *F26C11.4* | WBGene00044159 | protein_coding_gene | 4.6 | 24.28 | 8.2E-18 |
| *clec-106* | WBGene00012482 | protein_coding_gene | 4.6 | 24.21 | 1.9E-24 |
| *clec-100* | WBGene00013179 | protein_coding_gene | 4.6 | 23.95 | 5.3E-13 |
| *clec-161* | WBGene00010228 | protein_coding_gene | 4.6 | 23.83 | 5.4E-64 |
| *hpo-2* | WBGene00012500 | protein_coding_gene | 4.6 | 23.71 | 7.5E-81 |
| *nspc-12* | WBGene00023493 | protein_coding_gene | 4.5 | 22.97 | 4.5E-25 |
| *C16C8.7* | WBGene00015845 | protein_coding_gene | 4.5 | 22.88 | 4.3E-34 |
| *Y37E11AL.4* | WBGene00021360 | pseudogene | 4.5 | 22.68 | 5.6E-32 |
| *K12H6.9* | WBGene00019689 | protein_coding_gene | 4.5 | 22.53 | 1.1E-87 |
| *M176.10* | WBGene00010946 | protein_coding_gene | 4.5 | 22.37 | 1.4E-75 |
| *clec-133* | WBGene00021142 | protein_coding_gene | 4.5 | 22.14 | 3.4E-34 |
| *tag-329* | WBGene00008231 | protein_coding_gene | 4.5 | 21.87 | 2.1E-97 |
| *clec-232* | WBGene00009487 | protein_coding_gene | 4.4 | 21.80 | 5.4E-52 |
| *T10D4.1* | WBGene00020406 | protein_coding_gene | 4.4 | 21.61 | 1.6E-12 |
| *F49F1.15* | WBGene00235365 | protein_coding_gene | 4.4 | 21.33 | 2.1E-44 |
| *K12H6.4* | WBGene00019684 | protein_coding_gene | 4.4 | 21.32 | 1.9E-85 |
| *nas-24* | WBGene00003543 | protein_coding_gene | 4.4 | 21.11 | 5.3E-80 |
| *ZK1290.11* | WBGene00022892 | protein_coding_gene | 4.4 | 20.89 | 1.4E-49 |
| *clec-105* | WBGene00012488 | protein_coding_gene | 4.4 | 20.78 | 3.4E-10 |
| *msp-71* | WBGene00003460 | pseudogene | 4.4 | 20.75 | 5.1E-64 |
| *clec-135* | WBGene00015821 | protein_coding_gene | 4.3 | 20.26 | 3.0E-41 |
| *T26A8.3* | WBGene00020826 | protein_coding_gene | 4.3 | 20.19 | 1.2E-49 |
| *Y43F8C.15* | WBGene00012836 | protein_coding_gene | 4.3 | 19.98 | 2.2E-12 |
| *fipr-16* | WBGene00015521 | protein_coding_gene | 4.3 | 19.97 | 2.2E-34 |
| *D1022.2* | WBGene00017021 | protein_coding_gene | 4.3 | 19.97 | 1.1E-27 |
| *C45G9.10* | WBGene00016681 | protein_coding_gene | 4.3 | 19.73 | 2.3E-46 |
| *clec-138* | WBGene00018049 | protein_coding_gene | 4.3 | 19.72 | 1.1E-44 |
| *C39B5.10* | WBGene00016528 | protein_coding_gene | 4.3 | 19.22 | 7.1E-10 |
| *Y110A2AL.7* | WBGene00022444 | protein_coding_gene | 4.3 | 19.12 | 1.5E-61 |
| *K04H8.3* | WBGene00010578 | protein_coding_gene | 4.2 | 18.83 | 1.7E-26 |
| *clec-221* | WBGene00008916 | protein_coding_gene | 4.2 | 18.76 | 8.4E-127 |
| *clec-92* | WBGene00013125 | protein_coding_gene | 4.2 | 18.74 | 1.2E-52 |
| *scl-9* | WBGene00009890 | protein_coding_gene | 4.2 | 18.59 | 4.3E-39 |
| *C28C12.3* | WBGene00016179 | protein_coding_gene | 4.2 | 18.29 | 6.4E-73 |
| *Y6G8.14* | WBGene00138715 | protein_coding_gene | 4.2 | 17.99 | 3.8E-26 |
| *C16C8.19* | WBGene00015856 | protein_coding_gene | 4.2 | 17.91 | 5.9E-33 |
| *Y43F8B.19* | WBGene00077688 | protein_coding_gene | 4.2 | 17.81 | 1.3E-09 |
| *F46B6.13* | WBGene00044192 | protein_coding_gene | 4.1 | 17.69 | 6.9E-45 |
| *Y51A2D.1* | WBGene00013072 | protein_coding_gene | 4.1 | 17.59 | 7.4E-15 |
| *F07G6.8* | WBGene00017227 | protein_coding_gene | 4.1 | 17.51 | 1.8E-21 |
| *F13E9.19* | WBGene00194922 | protein_coding_gene | 4.1 | 17.44 | 4.4E-10 |

| | | | | | |
|---|---|---|---|---|---|
| *C35B1.3* | WBGene00016433 | protein_coding_gene | 4.1 | 17.37 | 3.1E-18 |
| *F11A6.16* | WBGene00271780 | protein_coding_gene | 4.1 | 17.30 | 1.2E-19 |
| *Y49F6B.6* | WBGene00021718 | protein_coding_gene | 4.1 | 17.29 | 6.6E-24 |
| *ZC328.5* | WBGene00022594 | protein_coding_gene | 4.1 | 17.25 | 2.4E-39 |
| *F02H6.6* | WBGene00008539 | protein_coding_gene | 4.1 | 17.06 | 5.9E-21 |
| *T22C1.12* | WBGene00011921 | protein_coding_gene | 4.1 | 17.03 | 1.8E-09 |
| *F28A10.5* | WBGene00017873 | protein_coding_gene | 4.1 | 16.81 | 2.1E-11 |
| *F46A8.3* | WBGene00009746 | protein_coding_gene | 4.1 | 16.72 | 2.5E-29 |
| *C09G12.5* | WBGene00015655 | protein_coding_gene | 4.1 | 16.71 | 2.9E-55 |
| *nep-25* | WBGene00022874 | protein_coding_gene | 4.1 | 16.61 | 2.1E-26 |
| *F42A6.2* | WBGene00018324 | protein_coding_gene | 4.0 | 16.55 | 4.3E-19 |
| *Y102A5C.4* | WBGene00013613 | protein_coding_gene | 4.0 | 16.50 | 1.3E-20 |
| *Y40H7A.147* | WBGene00077755 | pseudogene | 4.0 | 16.43 | 2.4E-09 |
| *M7.10* | WBGene00010886 | protein_coding_gene | 4.0 | 16.42 | 6.1E-41 |
| *ZK1225.10* | WBGene00044714 | pseudogene | 4.0 | 16.42 | 3.1E-11 |
| *K03B8.14* | WBGene00194783 | protein_coding_gene | 4.0 | 16.32 | 3.3E-14 |
| *F58A4.1* | WBGene00010226 | protein_coding_gene | 4.0 | 16.32 | 1.1E-59 |
| *Y110A2AL.6* | WBGene00022443 | protein_coding_gene | 4.0 | 15.87 | 2.6E-21 |
| *F58F9.8* | WBGene00044306 | protein_coding_gene | 4.0 | 15.79 | 5.6E-20 |
| *scl-26* | WBGene00016821 | protein_coding_gene | 4.0 | 15.67 | 2.7E-09 |
| *K03B8.11* | WBGene00010526 | protein_coding_gene | 4.0 | 15.64 | 7.9E-14 |
| *linc-1* | WBGene00194659 | lincRNA_gene | 4.0 | 15.47 | 8.5E-45 |
| *F22B3.5* | WBGene00009036 | protein_coding_gene | 4.0 | 15.47 | 1.1E-09 |
| *Y46D2A.3* | WBGene00021591 | protein_coding_gene | 3.9 | 15.23 | 1.0E-11 |
| *clec-181* | WBGene00044511 | protein_coding_gene | 3.9 | 15.15 | 7.4E-11 |
| *E02H9.9* | WBGene00017106 | protein_coding_gene | 3.9 | 14.81 | 1.3E-40 |
| *Y18D10A.2* | WBGene00012475 | protein_coding_gene | 3.9 | 14.75 | 3.6E-30 |
| *F25E5.7* | WBGene00017788 | protein_coding_gene | 3.9 | 14.64 | 1.6E-82 |
| *K09C8.2* | WBGene00010715 | protein_coding_gene | 3.9 | 14.61 | 3.0E-16 |
| *F56D6.11* | WBGene00044473 | protein_coding_gene | 3.9 | 14.55 | 9.4E-11 |
| *R13.5* | WBGene00014825 | pseudogene | 3.8 | 14.18 | 1.1E-08 |
| *cwp-1* | WBGene00000859 | protein_coding_gene | 3.8 | 13.87 | 3.8E-20 |
| *clec-261* | WBGene00013834 | protein_coding_gene | 3.8 | 13.82 | 5.3E-18 |
| *F38E1.10* | WBGene00018182 | protein_coding_gene | 3.8 | 13.49 | 1.2E-21 |
| *Y18H1A.8* | WBGene00021212 | protein_coding_gene | 3.7 | 13.41 | 1.4E-17 |
| *D2096.5* | WBGene00017072 | protein_coding_gene | 3.7 | 13.38 | 1.6E-33 |
| *clec-126* | WBGene00021117 | protein_coding_gene | 3.7 | 13.22 | 7.2E-41 |
| *F28B1.9* | WBGene00045381 | protein_coding_gene | 3.7 | 13.13 | 8.7E-14 |
| *pkd-2* | WBGene00004035 | protein_coding_gene | 3.7 | 13.11 | 2.7E-16 |
| *Y116A8C.44* | WBGene00013815 | protein_coding_gene | 3.7 | 12.81 | 1.1E-16 |
| *F56D2.8* | WBGene00018968 | protein_coding_gene | 3.7 | 12.81 | 5.9E-17 |
| *F28C6.10* | WBGene00009209 | protein_coding_gene | 3.7 | 12.72 | 2.9E-55 |
| *F40G9.7* | WBGene00018243 | protein_coding_gene | 3.7 | 12.67 | 1.2E-14 |
| *E02H9.1* | WBGene00017099 | protein_coding_gene | 3.7 | 12.60 | 2.9E-23 |
| *srp-5* | WBGene00005646 | pseudogene | 3.7 | 12.56 | 5.0E-10 |
| *Y39A1A.27* | WBGene00194731 | protein_coding_gene | 3.6 | 12.42 | 2.5E-18 |
| *F17E9.3* | WBGene00017540 | protein_coding_gene | 3.6 | 12.40 | 2.6E-21 |
| *C17E4.19* | WBGene00219310 | protein_coding_gene | 3.6 | 12.21 | 4.1E-46 |
| *F59A6.11* | WBGene00044664 | protein_coding_gene | 3.6 | 12.17 | 4.1E-13 |
| *K08C9.6* | WBGene00010653 | protein_coding_gene | 3.6 | 12.11 | 2.2E-08 |
| *W02D7.12* | WBGene00044356 | protein_coding_gene | 3.6 | 12.00 | 2.4E-19 |
| *plp-2* | WBGene00008976 | protein_coding_gene | 3.6 | 11.99 | 1.5E-26 |
| *W02B12.16* | WBGene00195148 | protein_coding_gene | 3.6 | 11.89 | 8.3E-19 |
| *clec-190* | WBGene00010953 | protein_coding_gene | 3.6 | 11.87 | 1.1E-24 |
| *C44B12.9* | WBGene00044462 | protein_coding_gene | 3.6 | 11.83 | 8.1E-20 |
| *msp-79* | WBGene00003466 | protein_coding_gene | 3.6 | 11.81 | 5.2E-15 |
| *T02H6.9* | WBGene00020179 | protein_coding_gene | 3.5 | 11.66 | 1.1E-23 |
| *clec-108* | WBGene00012504 | protein_coding_gene | 3.5 | 11.55 | 2.2E-15 |
| *Y49F6B.20* | WBGene00303369 | protein_coding_gene | 3.5 | 11.55 | 2.5E-20 |
| *T12A7.10* | WBGene00194933 | protein_coding_gene | 3.5 | 11.45 | 2.4E-08 |

| | | | | | |
|---|---|---|---|---|---|
| *clc-4* | WBGene00000525 | protein_coding_gene | 3.5 | 11.45 | 9.8E-22 |
| *F46A8.8* | WBGene00009751 | protein_coding_gene | 3.5 | 11.41 | 5.0E-17 |
| *M01G12.10* | WBGene00010823 | protein_coding_gene | 3.5 | 11.31 | 1.4E-09 |
| *K12H6.5* | WBGene00019685 | protein_coding_gene | 3.5 | 11.20 | 6.8E-21 |
| *F58F9.9* | WBGene00044307 | protein_coding_gene | 3.5 | 11.18 | 1.4E-09 |
| *Y43F8C.24* | WBGene00045345 | pseudogene | 3.5 | 11.13 | 1.4E-09 |
| *C04F12.6* | WBGene00007300 | pseudogene | 3.5 | 11.04 | 6.2E-19 |
| *nspa-4* | WBGene00044272 | protein_coding_gene | 3.5 | 10.94 | 3.8E-17 |
| *Y39B6A.21* | WBGene00012680 | pseudogene | 3.4 | 10.85 | 3.9E-82 |
| *C35E7.7* | WBGene00016459 | protein_coding_gene | 3.4 | 10.64 | 4.7E-22 |
| *F17E9.19* | WBGene00249825 | pseudogene | 3.4 | 10.62 | 2.6E-09 |
| *C49G7.13* | WBGene00269425 | protein_coding_gene | 3.4 | 10.57 | 2.6E-09 |
| *C31B8.12* | WBGene00016285 | protein_coding_gene | 3.4 | 10.51 | 1.1E-16 |
| *ssp-19* | WBGene00006047 | protein_coding_gene | 3.4 | 10.50 | 1.7E-42 |
| *Y39B6A.31* | WBGene00012690 | pseudogene | 3.4 | 10.44 | 3.1E-30 |
| *M142.3* | WBGene00014809 | pseudogene | 3.4 | 10.30 | 6.0E-23 |
| *C36C5.14* | WBGene00016483 | protein_coding_gene | 3.4 | 10.21 | 3.7E-09 |
| *F35C5.3* | WBGene00009391 | protein_coding_gene | 3.3 | 10.12 | 8.3E-14 |
| *nspd-8* | WBGene00021280 | protein_coding_gene | 3.3 | 10.04 | 4.4E-09 |
| *Y82E9BR.8* | WBGene00022341 | protein_coding_gene | 3.3 | 10.01 | 3.7E-09 |
| *tyr-5* | WBGene00021553 | protein_coding_gene | 3.3 | 9.99 | 1.4E-39 |
| *F25C8.1* | WBGene00009104 | pseudogene | 3.3 | 9.96 | 3.9E-13 |
| *C08G5.1* | WBGene00015614 | pseudogene | 3.3 | 9.96 | 2.2E-10 |
| *F16H6.9* | WBGene00008899 | protein_coding_gene | 3.3 | 9.85 | 7.5E-23 |
| *B0513.90* | WBGene00235301 | protein_coding_gene | 3.3 | 9.65 | 1.9E-16 |
| *C41D11.10* | WBGene00195180 | protein_coding_gene | 3.3 | 9.62 | 4.9E-11 |
| *flp-23* | WBGene00017693 | protein_coding_gene | 3.3 | 9.59 | 7.2E-12 |
| *ZK402.5* | WBGene00022731 | protein_coding_gene | 3.2 | 9.30 | 2.8E-07 |
| *clec-170* | WBGene00009515 | protein_coding_gene | 3.2 | 9.16 | 1.8E-69 |
| *nspf-3* | WBGene00044366 | protein_coding_gene | 3.2 | 9.14 | 8.7E-22 |
| *skr-11* | WBGene00004817 | pseudogene | 3.2 | 9.10 | 4.1E-63 |
| *Y49F6B.13* | WBGene00044377 | protein_coding_gene | 3.2 | 9.08 | 5.2E-07 |
| *C06E2.9* | WBGene00044579 | protein_coding_gene | 3.2 | 9.00 | 3.1E-13 |
| *Y64G10A.2* | WBGene00013412 | protein_coding_gene | 3.2 | 9.00 | 3.8E-07 |
| *Y73F8A.10* | WBGene00013519 | protein_coding_gene | 3.2 | 8.92 | 1.5E-09 |
| *clec-263* | WBGene00009171 | protein_coding_gene | 3.1 | 8.75 | 1.8E-19 |
| *Y43F8C.16* | WBGene00012837 | protein_coding_gene | 3.1 | 8.68 | 6.7E-11 |
| *C17E4.12* | WBGene00044769 | pseudogene | 3.1 | 8.55 | 6.0E-07 |
| *ZC178.2* | WBGene00022536 | protein_coding_gene | 3.1 | 8.51 | 5.3E-07 |
| *C16C8.8* | WBGene00015846 | protein_coding_gene | 3.1 | 8.51 | 3.1E-08 |
| *hpo-7* | WBGene00016548 | protein_coding_gene | 3.1 | 8.47 | 2.3E-08 |
| *Y54G2A.24* | WBGene00021889 | pseudogene | 3.1 | 8.46 | 4.4E-20 |
| *F47C12.6* | WBGene00018549 | protein_coding_gene | 3.1 | 8.43 | 2.5E-08 |
| *W03D8.10* | WBGene00020991 | pseudogene | 3.1 | 8.37 | 1.5E-27 |
| *Y71A12B.24* | WBGene00195228 | protein_coding_gene | 3.0 | 8.21 | 2.7E-08 |
| *F13B6.1* | WBGene00017416 | protein_coding_gene | 3.0 | 8.21 | 3.0E-31 |
| *oac-4* | WBGene00015873 | protein_coding_gene | 3.0 | 8.19 | 1.2E-06 |
| *abf-1* | WBGene00000012 | protein_coding_gene | 3.0 | 8.08 | 3.4E-18 |
| *gmd-2* | WBGene00010166 | protein_coding_gene | 3.0 | 8.02 | 3.7E-16 |
| *math-2* | WBGene00015606 | protein_coding_gene | 3.0 | 7.92 | 1.3E-06 |
| *nspc-17* | WBGene00008359 | protein_coding_gene | 3.0 | 7.92 | 9.3E-74 |
| *K12H6.8* | WBGene00019688 | protein_coding_gene | 3.0 | 7.87 | 1.5E-17 |
| *trpl-5* | WBGene00004149 | protein_coding_gene | 3.0 | 7.73 | 1.1E-07 |
| *col-182* | WBGene00000755 | pseudogene | 2.9 | 7.71 | 2.9E-23 |
| *Y116F11A.3* | WBGene00013817 | protein_coding_gene | 2.9 | 7.58 | 1.7E-06 |
| *lov-1* | WBGene00003058 | protein_coding_gene | 2.9 | 7.55 | 1.4E-07 |
| *B0513.7* | WBGene00007199 | pseudogene | 2.9 | 7.52 | 2.1E-06 |
| *W01G7.2* | WBGene00014851 | pseudogene | 2.9 | 7.51 | 6.1E-08 |
| *Y37F4.3* | WBGene00021384 | protein_coding_gene | 2.9 | 7.47 | 2.0E-06 |
| *F56C3.8* | WBGene00018945 | protein_coding_gene | 2.9 | 7.46 | 1.3E-19 |

| | | | | | |
|---|---|---|---|---|---|
| *F46B3.15* | WBGene00009766 | protein_coding_gene | 2.9 | 7.30 | 2.4E-08 |
| *T02D1.7* | WBGene00011373 | protein_coding_gene | 2.9 | 7.22 | 4.3E-08 |
| *try-5* | WBGene00006623 | protein_coding_gene | 2.9 | 7.21 | 2.8E-17 |
| *Y17G7B.23* | WBGene00044030 | protein_coding_gene | 2.8 | 7.10 | 3.7E-10 |
| *F46B3.14* | WBGene00009765 | protein_coding_gene | 2.8 | 7.07 | 8.6E-09 |
| *F46A8.4* | WBGene00009747 | protein_coding_gene | 2.8 | 7.00 | 1.4E-07 |
| *F32B5.3* | WBGene00017977 | protein_coding_gene | 2.8 | 6.99 | 3.8E-09 |
| *scl-8* | WBGene00008030 | protein_coding_gene | 2.8 | 6.96 | 2.8E-21 |
| *F59A6.3* | WBGene00019085 | protein_coding_gene | 2.8 | 6.89 | 4.8E-07 |
| *clec-260* | WBGene00013357 | protein_coding_gene | 2.8 | 6.88 | 2.3E-14 |
| *F01D5.4* | WBGene00014730 | pseudogene | 2.8 | 6.86 | 3.8E-07 |
| *F59A1.6* | WBGene00010292 | protein_coding_gene | 2.8 | 6.79 | 3.9E-07 |
| *C17H12.37* | WBGene00236786 | protein_coding_gene | 2.8 | 6.76 | 4.6E-06 |
| *T16H12.1* | WBGene00011811 | protein_coding_gene | 2.8 | 6.76 | 7.2E-46 |
| *T02E9.6* | WBGene00044203 | protein_coding_gene | 2.8 | 6.74 | 3.1E-08 |
| *E02H4.4* | WBGene00008462 | protein_coding_gene | 2.7 | 6.71 | 5.1E-109 |
| *arrd-11* | WBGene00011052 | protein_coding_gene | 2.7 | 6.71 | 1.9E-10 |
| *C50E3.9* | WBGene00016820 | protein_coding_gene | 2.7 | 6.71 | 5.8E-06 |
| *math-47* | WBGene00022688 | pseudogene | 2.7 | 6.62 | 1.7E-09 |
| *F10D2.8* | WBGene00017334 | protein_coding_gene | 2.7 | 6.62 | 2.2E-27 |
| *F56H6.7* | WBGene00010168 | protein_coding_gene | 2.7 | 6.52 | 3.8E-10 |
| *clec-136* | WBGene00018047 | protein_coding_gene | 2.7 | 6.49 | 1.6E-18 |
| *C04F12.12* | WBGene00077685 | protein_coding_gene | 2.7 | 6.37 | 5.0E-21 |
| *C35B1.8* | WBGene00016437 | protein_coding_gene | 2.6 | 6.26 | 5.9E-16 |
| *F42G2.7* | WBGene00044362 | protein_coding_gene | 2.6 | 6.17 | 3.7E-08 |
| *C44B12.3* | WBGene00016637 | protein_coding_gene | 2.6 | 6.17 | 1.0E-14 |
| *R10D12.18* | WBGene00194678 | ncRNA_gene | 2.6 | 6.11 | 2.6E-11 |
| *F46F5.4* | WBGene00018495 | pseudogene | 2.6 | 6.09 | 2.3E-08 |
| *F29A7.3* | WBGene00017913 | protein_coding_gene | 2.6 | 6.03 | 3.9E-07 |
| *F43G6.17* | WBGene00219378 | protein_coding_gene | 2.6 | 6.02 | 1.3E-05 |
| *clec-169* | WBGene00009526 | protein_coding_gene | 2.6 | 5.96 | 1.6E-23 |
| *fip-7* | WBGene00013105 | protein_coding_gene | 2.6 | 5.95 | 9.0E-06 |
| *cwp-2* | WBGene00000860 | protein_coding_gene | 2.6 | 5.94 | 2.3E-06 |
| *F59A1.16* | WBGene00010301 | protein_coding_gene | 2.6 | 5.93 | 1.3E-05 |
| *fbxa-136* | WBGene00007672 | protein_coding_gene | 2.6 | 5.87 | 3.7E-11 |
| *clec-162* | WBGene00007399 | protein_coding_gene | 2.6 | 5.87 | 7.2E-10 |
| *cyp-37B1* | WBGene00009226 | protein_coding_gene | 2.6 | 5.86 | 3.6E-14 |
| *fip-3* | WBGene00015520 | protein_coding_gene | 2.5 | 5.80 | 1.2E-07 |
| *R09D1.9* | WBGene00011165 | pseudogene | 2.5 | 5.80 | 1.4E-13 |
| *clpr-1* | WBGene00012233 | protein_coding_gene | 2.5 | 5.79 | 1.8E-05 |
| *F22E5.7* | WBGene00017706 | protein_coding_gene | 2.5 | 5.78 | 1.0E-06 |
| *W08F4.7* | WBGene00021096 | protein_coding_gene | 2.5 | 5.77 | 1.3E-10 |
| *F13E9.9* | WBGene00008758 | protein_coding_gene | 2.5 | 5.76 | 2.9E-08 |
| *K10C2.14* | WBGene00206388 | protein_coding_gene | 2.5 | 5.72 | 4.8E-26 |
| *mab-3* | WBGene00003100 | protein_coding_gene | 2.5 | 5.72 | 1.3E-18 |
| *R05G6.3* | WBGene00043466 | protein_coding_gene | 2.5 | 5.67 | 2.3E-05 |
| *F49F1.2* | WBGene00235363 | protein_coding_gene | 2.5 | 5.64 | 8.4E-07 |
| *Y71G12B.5* | WBGene00022145 | protein_coding_gene | 2.5 | 5.62 | 7.8E-11 |
| *F13E9.4* | WBGene00008753 | protein_coding_gene | 2.5 | 5.59 | 2.6E-05 |
| *clec-130* | WBGene00021144 | protein_coding_gene | 2.5 | 5.58 | 1.9E-09 |
| *F49F1.5* | WBGene00018645 | protein_coding_gene | 2.5 | 5.51 | 2.9E-28 |
| *C36C5.12* | WBGene00016481 | protein_coding_gene | 2.5 | 5.47 | 1.0E-13 |
| *T26F2.2* | WBGene00012058 | pseudogene | 2.4 | 5.44 | 3.9E-27 |
| *M01G12.7* | WBGene00010821 | protein_coding_gene | 2.4 | 5.44 | 1.4E-06 |
| *F19H6.5* | WBGene00008960 | protein_coding_gene | 2.4 | 5.41 | 3.2E-09 |
| *F26C11.3* | WBGene00009147 | protein_coding_gene | 2.4 | 5.40 | 1.7E-07 |
| *Y57G11C.1132* | WBGene00194666 | antisense_lncRNA_gene | 2.4 | 5.40 | 1.5E-06 |
| *ins-20* | WBGene00002103 | protein_coding_gene | 2.4 | 5.32 | 8.8E-08 |
| *clec-98* | WBGene00013932 | protein_coding_gene | 2.4 | 5.31 | 1.5E-06 |
| *C41G7.15* | WBGene00219284 | protein_coding_gene | 2.4 | 5.30 | 4.5E-05 |

| | | | | | |
|---|---|---|---|---|---|
| *C16C8.17* | WBGene00015854 | protein_coding_gene | 2.4 | 5.28 | 7.1E-07 |
| *F19H6.6* | WBGene00008961 | protein_coding_gene | 2.4 | 5.27 | 1.3E-07 |
| *F25E5.4* | WBGene00017785 | protein_coding_gene | 2.4 | 5.26 | 2.2E-35 |
| *fbxa-182* | WBGene00018353 | protein_coding_gene | 2.4 | 5.24 | 1.7E-33 |
| *far-3* | WBGene00001387 | protein_coding_gene | 2.4 | 5.22 | 1.4E-24 |
| *irg-6* | WBGene00007864 | protein_coding_gene | 2.4 | 5.15 | 9.0E-07 |
| *F25D7.5* | WBGene00009114 | protein_coding_gene | 2.4 | 5.15 | 1.7E-08 |
| *B0393.4* | WBGene00007169 | protein_coding_gene | 2.4 | 5.14 | 1.4E-18 |
| *oac-35* | WBGene00010171 | protein_coding_gene | 2.4 | 5.10 | 5.7E-05 |
| *Y57E12AL.4* | WBGene00021958 | protein_coding_gene | 2.3 | 5.08 | 1.1E-06 |
| *Y64G10A.10* | WBGene00045144 | protein_coding_gene | 2.3 | 5.06 | 2.6E-06 |
| *nspf-2* | WBGene00044365 | protein_coding_gene | 2.3 | 5.05 | 1.2E-06 |
| *F02H6.4* | WBGene00008537 | pseudogene | 2.3 | 5.04 | 3.9E-50 |
| *R11G10.4* | WBGene00050968 | protein_coding_gene | 2.3 | 5.04 | 2.6E-06 |
| *B0334.10* | WBGene00007148 | protein_coding_gene | 2.3 | 5.02 | 1.5E-05 |
| *F30A10.11* | WBGene00009268 | protein_coding_gene | 2.3 | 5.01 | 1.9E-08 |
| *bgnt-1.6* | WBGene00010167 | protein_coding_gene | 2.3 | 4.99 | 6.5E-06 |
| *F45E1.4* | WBGene00018465 | protein_coding_gene | 2.3 | 4.91 | 1.5E-24 |
| *clec-24* | WBGene00009856 | protein_coding_gene | 2.3 | 4.90 | 1.6E-07 |
| *clec-242* | WBGene00013471 | protein_coding_gene | 2.3 | 4.89 | 3.7E-06 |
| *ZK418.2* | WBGene00022732 | protein_coding_gene | 2.3 | 4.86 | 7.6E-27 |
| *C17E4.20* | WBGene00219313 | protein_coding_gene | 2.3 | 4.83 | 2.6E-34 |
| *K08C9.10* | WBGene00086564 | protein_coding_gene | 2.3 | 4.83 | 9.3E-05 |
| *ZC204.6* | WBGene00022556 | protein_coding_gene | 2.3 | 4.78 | 1.0E-04 |
| *W07G1.7* | WBGene00012334 | protein_coding_gene | 2.3 | 4.76 | 3.2E-06 |
| *Y75B12A.1* | WBGene00014949 | pseudogene | 2.2 | 4.75 | 4.8E-06 |
| *C17E4.1* | WBGene00007641 | protein_coding_gene | 2.2 | 4.74 | 5.7E-20 |
| *M195.4* | WBGene00010949 | protein_coding_gene | 2.2 | 4.73 | 5.3E-05 |
| *fbxc-49* | WBGene00020534 | protein_coding_gene | 2.2 | 4.69 | 2.1E-09 |
| *ZK666.14* | WBGene00194952 | protein_coding_gene | 2.2 | 4.63 | 4.8E-06 |
| *B0462.1* | WBGene00007181 | pseudogene | 2.2 | 4.62 | 4.1E-08 |
| *Y39E4A.1* | WBGene00012711 | protein_coding_gene | 2.2 | 4.59 | 1.3E-07 |
| *ZK218.8* | WBGene00013942 | pseudogene | 2.2 | 4.58 | 6.5E-06 |
| *Y47D7A.7* | WBGene00021620 | protein_coding_gene | 2.2 | 4.57 | 6.3E-09 |
| *F57A10.2* | WBGene00010184 | protein_coding_gene | 2.2 | 4.55 | 1.1E-06 |
| *Y55F3AM.21* | WBGene00219399 | protein_coding_gene | 2.2 | 4.54 | 5.1E-10 |
| *nlp-45* | WBGene00020136 | protein_coding_gene | 2.2 | 4.53 | 7.7E-11 |
| *F17E9.18* | WBGene00235282 | protein_coding_gene | 2.2 | 4.53 | 1.4E-04 |
| *H10D18.6* | WBGene00019181 | pseudogene | 2.2 | 4.52 | 3.3E-07 |
| *cal-6* | WBGene00022214 | pseudogene | 2.2 | 4.50 | 5.0E-05 |
| *fbxa-163* | WBGene00015598 | protein_coding_gene | 2.2 | 4.49 | 2.9E-06 |
| *Y41C4A.21* | WBGene00194733 | protein_coding_gene | 2.2 | 4.48 | 2.1E-05 |
| *3R5.2* | WBGene00007066 | pseudogene | 2.2 | 4.46 | 8.2E-06 |
| *Y105C5B.7* | WBGene00013648 | pseudogene | 2.2 | 4.45 | 2.0E-08 |
| *W02A2.4* | WBGene00012189 | pseudogene | 2.2 | 4.44 | 1.5E-07 |
| *ugt-66* | WBGene00016013 | protein_coding_gene | 2.1 | 4.43 | 5.4E-15 |
| *cyp-35A5* | WBGene00019473 | protein_coding_gene | 2.1 | 4.42 | 2.1E-20 |
| *C06A12.8* | WBGene00007359 | protein_coding_gene | 2.1 | 4.41 | 1.2E-04 |
| *C47F8.6* | WBGene00008162 | protein_coding_gene | 2.1 | 4.41 | 1.2E-04 |
| *Y59E9AR.1* | WBGene00021997 | protein_coding_gene | 2.1 | 4.41 | 4.0E-06 |
| *Y1A5A.1* | WBGene00012379 | protein_coding_gene | 2.1 | 4.35 | 3.7E-12 |
| *Y53F4B.11* | WBGene00013158 | protein_coding_gene | 2.1 | 4.30 | 1.8E-13 |
| *F58A6.9* | WBGene00019026 | protein_coding_gene | 2.1 | 4.28 | 4.4E-09 |
| *C08E3.14* | WBGene00044415 | protein_coding_gene | 2.1 | 4.27 | 2.4E-04 |
| *scl-18* | WBGene00011724 | protein_coding_gene | 2.1 | 4.27 | 2.5E-04 |
| *F35F10.13* | WBGene00018062 | protein_coding_gene | 2.1 | 4.26 | 2.4E-04 |
| *Y47D7A.11* | WBGene00021623 | protein_coding_gene | 2.1 | 4.26 | 1.0E-05 |
| *fbxa-217* | WBGene00012954 | pseudogene | 2.1 | 4.26 | 2.4E-26 |
| *Y20C6A.4* | WBGene00077495 | protein_coding_gene | 2.1 | 4.25 | 8.6E-05 |
| *F40F8.12* | WBGene00044917 | protein_coding_gene | 2.1 | 4.25 | 5.0E-05 |

| | | | | | |
|---|---|---|---|---|---|
| *K08E5.1* | WBGene00014801 | protein_coding_gene | 2.1 | 4.19 | 8.9E-13 |
| *B0228.8* | WBGene00015065 | protein_coding_gene | 2.1 | 4.17 | 2.9E-04 |
| *C09B9.4* | WBGene00015629 | protein_coding_gene | 2.1 | 4.17 | 2.7E-16 |
| *ZK849.1* | WBGene00014100 | protein_coding_gene | 2.1 | 4.15 | 3.3E-11 |
| *clec-93* | WBGene00013929 | protein_coding_gene | 2.1 | 4.14 | 9.9E-06 |
| *Y46D2A.1* | WBGene00021589 | protein_coding_gene | 2.0 | 4.08 | 4.9E-07 |
| *Y43F8C.23* | WBGene00045344 | protein_coding_gene | 2.0 | 4.07 | 1.9E-05 |
| *B0348.2* | WBGene00015152 | protein_coding_gene | 2.0 | 4.07 | 6.3E-07 |
| *rrf-2* | WBGene00004509 | protein_coding_gene | 2.0 | 4.07 | 4.0E-41 |
| *nspf-1* | WBGene00044364 | protein_coding_gene | 2.0 | 4.06 | 4.8E-05 |
| *F21D9.11* | WBGene00219452 | protein_coding_gene | 2.0 | 4.06 | 1.3E-07 |
| *F46F5.1* | WBGene00018493 | protein_coding_gene | 2.0 | 4.05 | 4.8E-08 |
| *R155.4* | WBGene00020118 | protein_coding_gene | 2.0 | 4.05 | 8.0E-07 |
| *F44G3.10* | WBGene00009710 | protein_coding_gene | 2.0 | 4.04 | 2.1E-06 |
| *R09A8.9* | WBGene00219278 | protein_coding_gene | 2.0 | 4.03 | 1.2E-08 |
| *F27E5.3* | WBGene00009193 | protein_coding_gene | 2.0 | 4.01 | 3.0E-10 |
| *fbxa-103* | WBGene00008612 | protein_coding_gene | 2.0 | 4.01 | 1.2E-09 |
| *Y73C8C.10* | WBGene00022267 | protein_coding_gene | 2.0 | 4.00 | 6.2E-07 |
| *C01G5.7* | WBGene00015309 | protein_coding_gene | 2.0 | 3.98 | 1.9E-07 |
| *fbxa-168* | WBGene00007838 | pseudogene | 2.0 | 3.97 | 4.3E-12 |
| *ZK783.3* | WBGene00022818 | pseudogene | 2.0 | 3.97 | 2.8E-06 |
| *math-48* | WBGene00022689 | protein_coding_gene | 2.0 | 3.96 | 7.9E-07 |
| *C10F3.7* | WBGene00015679 | protein_coding_gene | 2.0 | 3.95 | 5.0E-09 |
| *C36C5.15* | WBGene00016484 | protein_coding_gene | 2.0 | 3.95 | 2.5E-05 |
| *C09B9.85* | WBGene00220250 | protein_coding_gene | 2.0 | 3.95 | 1.0E-05 |
| *ppt-1* | WBGene00004092 | protein_coding_gene | 2.0 | 3.93 | 6.1E-23 |
| *C05B5.1* | WBGene00007319 | protein_coding_gene | 2.0 | 3.93 | 2.7E-04 |
| *fbxa-113* | WBGene00013757 | protein_coding_gene | 2.0 | 3.92 | 1.1E-07 |
| *F25B3.2* | WBGene00009099 | protein_coding_gene | 2.0 | 3.92 | 5.4E-05 |
| *R10D12.10* | WBGene00011191 | protein_coding_gene | 2.0 | 3.90 | 1.0E-14 |
| *Y6G8.16* | WBGene00194801 | protein_coding_gene | 2.0 | 3.90 | 1.9E-07 |
| *fipr-19* | WBGene00044451 | protein_coding_gene | 1.9 | 3.85 | 2.3E-05 |
| *ZK896.5* | WBGene00014136 | protein_coding_gene | 1.9 | 3.83 | 1.6E-12 |
| *Y43C5B.2* | WBGene00012785 | protein_coding_gene | 1.9 | 3.82 | 2.0E-09 |
| *ant-1.3* | WBGene00010485 | protein_coding_gene | 1.9 | 3.82 | 5.9E-18 |
| *R05G6.1* | WBGene00019896 | protein_coding_gene | 1.9 | 3.82 | 4.4E-07 |
| *Y39A3A.4* | WBGene00021435 | protein_coding_gene | 1.9 | 3.81 | 1.1E-07 |
| *F49H6.5* | WBGene00009908 | protein_coding_gene | 1.9 | 3.80 | 8.9E-11 |
| *F17E9.15* | WBGene00194833 | protein_coding_gene | 1.9 | 3.79 | 4.8E-04 |
| *T11F9.22* | WBGene00194784 | protein_coding_gene | 1.9 | 3.79 | 1.8E-05 |
| *C10G8.2* | WBGene00015681 | protein_coding_gene | 1.9 | 3.77 | 3.8E-04 |
| *col-154* | WBGene00000727 | protein_coding_gene | 1.9 | 3.75 | 3.3E-07 |
| *Y71G12B.35* | WBGene00185098 | protein_coding_gene | 1.9 | 3.75 | 5.8E-14 |
| *ZK546.7* | WBGene00022763 | pseudogene | 1.9 | 3.75 | 1.1E-06 |
| *msrp-1* | WBGene00017978 | protein_coding_gene | 1.9 | 3.72 | 3.9E-07 |
| *C34D4.2* | WBGene00016398 | protein_coding_gene | 1.9 | 3.70 | 3.2E-11 |
| *F20A1.8* | WBGene00017619 | protein_coding_gene | 1.9 | 3.70 | 3.7E-04 |
| *C06E1.11* | WBGene00015526 | protein_coding_gene | 1.9 | 3.70 | 6.2E-11 |
| *oac-23* | WBGene00018143 | protein_coding_gene | 1.9 | 3.70 | 3.4E-07 |
| *C07G3.10* | WBGene00015578 | protein_coding_gene | 1.9 | 3.68 | 1.1E-04 |
| *F56H6.9* | WBGene00010170 | protein_coding_gene | 1.9 | 3.68 | 2.3E-04 |
| *F46F5.11* | WBGene00018502 | protein_coding_gene | 1.9 | 3.67 | 6.5E-08 |
| *Y47D7A.12* | WBGene00021624 | protein_coding_gene | 1.9 | 3.67 | 9.3E-07 |
| *T10B11.4* | WBGene00020400 | protein_coding_gene | 1.9 | 3.66 | 9.4E-05 |
| *gmeb-3* | WBGene00008092 | protein_coding_gene | 1.9 | 3.65 | 3.8E-07 |
| *F32B5.1* | WBGene00017975 | protein_coding_gene | 1.9 | 3.65 | 1.5E-11 |
| *T07H8.11* | WBGene00304201 | protein_coding_gene | 1.9 | 3.63 | 2.2E-19 |
| *K09H11.4* | WBGene00019601 | protein_coding_gene | 1.9 | 3.63 | 4.7E-05 |
| *C17G10.10* | WBGene00195182 | protein_coding_gene | 1.9 | 3.63 | 4.4E-05 |
| *Y37H2A.7* | WBGene00012567 | protein_coding_gene | 1.9 | 3.62 | 1.7E-10 |

| | | | | | |
|---|---|---|---|---|---|
| *F28A10.1* | WBGene00017869 | protein_coding_gene | 1.9 | 3.61 | 2.4E-08 |
| *clec-210* | WBGene00022261 | protein_coding_gene | 1.8 | 3.57 | 2.4E-04 |
| *C14E2.4* | WBGene00015773 | protein_coding_gene | 1.8 | 3.56 | 9.3E-04 |
| *ZK1290.7* | WBGene00022889 | protein_coding_gene | 1.8 | 3.56 | 1.9E-04 |
| *Y47G6A.26* | WBGene00021650 | protein_coding_gene | 1.8 | 3.56 | 4.5E-10 |
| *F11C7.6* | WBGene00017378 | protein_coding_gene | 1.8 | 3.56 | 4.3E-16 |
| *ptr-22* | WBGene00004236 | protein_coding_gene | 1.8 | 3.56 | 1.5E-12 |
| *nep-19* | WBGene00020292 | protein_coding_gene | 1.8 | 3.55 | 7.3E-06 |
| *C01G10.17* | WBGene00044658 | protein_coding_gene | 1.8 | 3.55 | 8.0E-05 |
| *F46F5.14* | WBGene00018505 | protein_coding_gene | 1.8 | 3.54 | 1.7E-06 |
| *C49C8.6* | WBGene00016770 | protein_coding_gene | 1.8 | 3.54 | 1.2E-04 |
| *H25K10.4* | WBGene00010413 | protein_coding_gene | 1.8 | 3.54 | 2.4E-05 |
| *Y82E9BR.1* | WBGene00022334 | protein_coding_gene | 1.8 | 3.54 | 2.0E-22 |
| *F17C8.7* | WBGene00008912 | protein_coding_gene | 1.8 | 3.53 | 4.8E-10 |
| *math-23* | WBGene00016718 | protein_coding_gene | 1.8 | 3.53 | 2.5E-04 |
| *pph-2* | WBGene00004084 | protein_coding_gene | 1.8 | 3.52 | 8.5E-08 |
| *K06A4.8* | WBGene00044015 | pseudogene | 1.8 | 3.52 | 2.3E-09 |
| *Y66H1A.8* | WBGene00235263 | protein_coding_gene | 1.8 | 3.51 | 2.8E-04 |
| *ugt-32* | WBGene00018543 | protein_coding_gene | 1.8 | 3.49 | 1.5E-06 |
| *F52F12.9* | WBGene00009942 | protein_coding_gene | 1.8 | 3.49 | 6.1E-07 |
| *K07A1.6* | WBGene00010613 | protein_coding_gene | 1.8 | 3.49 | 5.3E-54 |
| *fbxc-48* | WBGene00015615 | protein_coding_gene | 1.8 | 3.48 | 1.5E-05 |
| *fbxa-161* | WBGene00015596 | pseudogene | 1.8 | 3.45 | 1.8E-04 |
| *nep-24* | WBGene00013786 | protein_coding_gene | 1.8 | 3.45 | 2.3E-06 |
| *sknr-1* | WBGene00020961 | protein_coding_gene | 1.8 | 3.45 | 5.0E-04 |
| *Y50D4B.7* | WBGene00021746 | pseudogene | 1.8 | 3.39 | 3.1E-04 |
| *trf-1* | WBGene00006612 | protein_coding_gene | 1.8 | 3.38 | 1.1E-04 |
| *fbxa-100* | WBGene00008826 | protein_coding_gene | 1.8 | 3.38 | 1.3E-03 |
| *msp-3* | WBGene00003424 | protein_coding_gene | 1.8 | 3.37 | 3.6E-13 |
| *fbxa-73* | WBGene00020640 | protein_coding_gene | 1.8 | 3.36 | 6.7E-05 |
| *fer-1* | WBGene00001414 | protein_coding_gene | 1.7 | 3.36 | 3.8E-13 |
| *dct-10* | WBGene00012618 | pseudogene | 1.7 | 3.36 | 8.4E-04 |
| *F28B1.2* | WBGene00009196 | protein_coding_gene | 1.7 | 3.36 | 1.7E-04 |
| *K07E8.1* | WBGene00019494 | pseudogene | 1.7 | 3.35 | 1.5E-03 |
| *F33E2.6* | WBGene00009362 | protein_coding_gene | 1.7 | 3.34 | 4.5E-04 |
| *F56D5.2* | WBGene00010147 | protein_coding_gene | 1.7 | 3.33 | 1.9E-05 |
| *F21D9.5* | WBGene00009019 | pseudogene | 1.7 | 3.33 | 5.0E-07 |
| *ZC513.3* | WBGene00022628 | protein_coding_gene | 1.7 | 3.32 | 8.0E-07 |
| *pcp-4* | WBGene00003959 | protein_coding_gene | 1.7 | 3.30 | 6.8E-15 |
| *F59A1.13* | WBGene00010299 | protein_coding_gene | 1.7 | 3.30 | 8.6E-08 |
| *F10E7.3* | WBGene00017345 | pseudogene | 1.7 | 3.30 | 9.4E-04 |
| *C50C3.2* | WBGene00016800 | protein_coding_gene | 1.7 | 3.30 | 1.4E-13 |
| *folr-1* | WBGene00007648 | protein_coding_gene | 1.7 | 3.29 | 7.0E-04 |
| *C31B8.9* | WBGene00016284 | protein_coding_gene | 1.7 | 3.29 | 1.8E-03 |
| *C16C8.10* | WBGene00015848 | protein_coding_gene | 1.7 | 3.27 | 6.8E-04 |
| *T04B2.1* | WBGene00011421 | pseudogene | 1.7 | 3.27 | 1.2E-03 |
| *Y62H9A.8* | WBGene00013396 | protein_coding_gene | 1.7 | 3.27 | 1.2E-04 |
| *C31H1.1* | WBGene00016286 | protein_coding_gene | 1.7 | 3.27 | 5.1E-09 |
| *R05H10.1* | WBGene00011042 | protein_coding_gene | 1.7 | 3.24 | 7.8E-11 |
| *Y116A8A.10* | WBGene00014975 | pseudogene | 1.7 | 3.23 | 1.7E-05 |
| *F07D3.3* | WBGene00008556 | protein_coding_gene | 1.7 | 3.23 | 8.1E-05 |
| *W06G6.11* | WBGene00012313 | protein_coding_gene | 1.7 | 3.22 | 1.4E-04 |
| *H20J04.1* | WBGene00019216 | protein_coding_gene | 1.7 | 3.22 | 3.0E-09 |
| *nkb-1* | WBGene00007646 | protein_coding_gene | 1.7 | 3.22 | 1.6E-79 |
| *linc-140* | WBGene00219596 | lincRNA_gene | 1.7 | 3.21 | 3.4E-04 |
| *F45D11.1* | WBGene00018448 | pseudogene | 1.7 | 3.20 | 1.5E-07 |
| *W05F2.8* | WBGene00195045 | protein_coding_gene | 1.7 | 3.19 | 7.8E-04 |
| *F55G11.1* | WBGene00010122 | protein_coding_gene | 1.7 | 3.19 | 7.1E-05 |
| *R13A1.1* | WBGene00020041 | protein_coding_gene | 1.7 | 3.18 | 1.0E-03 |
| *her-1* | WBGene00001842 | protein_coding_gene | 1.7 | 3.18 | 1.0E-05 |

| | | | | | |
|---|---|---|---|---|---|
| *C29F9.5* | WBGene00016220 | protein_coding_gene | 1.7 | 3.17 | 1.2E-03 |
| *Y39F10A.3* | WBGene00021454 | protein_coding_gene | 1.7 | 3.17 | 3.4E-05 |
| *eol-1* | WBGene00045401 | protein_coding_gene | 1.7 | 3.16 | 3.3E-07 |
| *C18E9.8* | WBGene00007688 | protein_coding_gene | 1.7 | 3.15 | 2.0E-09 |
| *clec-165* | WBGene00009523 | protein_coding_gene | 1.7 | 3.15 | 1.8E-08 |
| *F55C9.12* | WBGene00044196 | pseudogene | 1.7 | 3.14 | 1.0E-03 |
| *F55C5.2* | WBGene00010092 | pseudogene | 1.7 | 3.14 | 2.6E-08 |
| *clec-183* | WBGene00011855 | protein_coding_gene | 1.6 | 3.14 | 1.9E-03 |
| *Y69E1A.8* | WBGene00013478 | protein_coding_gene | 1.6 | 3.13 | 3.0E-08 |
| *C43D7.11* | WBGene00077664 | pseudogene | 1.6 | 3.13 | 1.9E-04 |
| *rpl-42* | WBGene00004455 | protein_coding_gene | 1.6 | 3.12 | 7.9E-06 |
| *F57G9.6* | WBGene00044041 | pseudogene | 1.6 | 3.11 | 2.2E-07 |
| *W09G12.6* | WBGene00021120 | protein_coding_gene | 1.6 | 3.09 | 7.2E-04 |
| *ant-1.4* | WBGene00020140 | protein_coding_gene | 1.6 | 3.09 | 1.0E-04 |
| *C08F11.14* | WBGene00007461 | pseudogene | 1.6 | 3.09 | 5.1E-06 |
| *F58A6.5* | WBGene00019024 | protein_coding_gene | 1.6 | 3.09 | 1.5E-09 |
| *Y69A2AR.19* | WBGene00022090 | protein_coding_gene | 1.6 | 3.09 | 5.4E-19 |
| *lips-6* | WBGene00014009 | protein_coding_gene | 1.6 | 3.09 | 3.8E-09 |
| *W05B2.3* | WBGene00014854 | pseudogene | 1.6 | 3.08 | 7.4E-04 |
| *Y41C4A.8* | WBGene00012755 | protein_coding_gene | 1.6 | 3.08 | 1.3E-12 |
| *R12E2.14* | WBGene00020039 | protein_coding_gene | 1.6 | 3.08 | 1.2E-08 |
| *clec-79* | WBGene00018548 | protein_coding_gene | 1.6 | 3.08 | 3.9E-13 |
| *msp-50* | WBGene00003443 | protein_coding_gene | 1.6 | 3.06 | 6.6E-17 |
| *ZC412.5* | WBGene00013886 | protein_coding_gene | 1.6 | 3.05 | 5.7E-07 |
| *tag-138* | WBGene00006484 | protein_coding_gene | 1.6 | 3.05 | 1.6E-06 |
| *C44B9.2* | WBGene00008079 | protein_coding_gene | 1.6 | 3.05 | 1.5E-10 |
| *cyc-2.2* | WBGene00013854 | protein_coding_gene | 1.6 | 3.05 | 1.2E-12 |
| *Y67D8C.7* | WBGene00022071 | protein_coding_gene | 1.6 | 3.05 | 1.4E-03 |
| *Y82E9BL.12* | WBGene00022328 | protein_coding_gene | 1.6 | 3.05 | 1.6E-03 |
| *R11E3.19* | WBGene00235374 | protein_coding_gene | 1.6 | 3.04 | 3.0E-04 |
| *cri-1* | WBGene00010614 | protein_coding_gene | 1.6 | 3.04 | 1.6E-06 |
| *srp-3* | WBGene00005644 | protein_coding_gene | 1.6 | 3.04 | 2.9E-08 |
| *Y7A5A.9* | WBGene00012415 | protein_coding_gene | 1.6 | 3.03 | 2.0E-03 |
| *try-9* | WBGene00023425 | protein_coding_gene | 1.6 | 3.03 | 4.7E-09 |
| *wago-10* | WBGene00020707 | protein_coding_gene | 1.6 | 3.03 | 4.8E-12 |
| *clec-42* | WBGene00008891 | protein_coding_gene | 1.6 | 3.02 | 4.8E-05 |
| *lips-16* | WBGene00012588 | pseudogene | 1.6 | 3.02 | 5.7E-12 |
| *haf-8* | WBGene00001818 | protein_coding_gene | 1.6 | 3.00 | 2.7E-04 |
| *F18F11.4* | WBGene00017577 | protein_coding_gene | 1.6 | 2.99 | 6.5E-04 |
| *cest-35.2* | WBGene00016863 | protein_coding_gene | 1.6 | 2.99 | 9.5E-08 |
| *C02B10.6* | WBGene00015331 | protein_coding_gene | 1.6 | 2.98 | 9.3E-04 |
| *gln-4* | WBGene00001605 | pseudogene | 1.6 | 2.98 | 4.7E-05 |
| *clec-195* | WBGene00013798 | protein_coding_gene | 1.6 | 2.98 | 3.0E-04 |
| *clec-129* | WBGene00021145 | protein_coding_gene | 1.6 | 2.97 | 1.3E-03 |
| *aqp-9* | WBGene00000177 | protein_coding_gene | 1.6 | 2.96 | 3.2E-21 |
| *F49C5.12* | WBGene00184956 | protein_coding_gene | 1.6 | 2.96 | 9.4E-04 |
| *F37A8.1* | WBGene00009501 | protein_coding_gene | 1.6 | 2.96 | 1.7E-07 |
| *nhr-219* | WBGene00020555 | protein_coding_gene | 1.6 | 2.96 | 2.4E-03 |
| *C02G6.2* | WBGene00015360 | protein_coding_gene | 1.6 | 2.95 | 2.4E-04 |
| *T09E11.12* | WBGene00045103 | protein_coding_gene | 1.6 | 2.95 | 3.1E-04 |
| *F28B1.3* | WBGene00009197 | pseudogene | 1.6 | 2.95 | 2.4E-03 |
| *C38D9.8* | WBGene00008016 | protein_coding_gene | 1.6 | 2.95 | 3.7E-05 |
| *K08C7.4* | WBGene00010647 | protein_coding_gene | 1.6 | 2.94 | 3.0E-05 |
| *fut-5* | WBGene00043986 | protein_coding_gene | 1.6 | 2.94 | 8.5E-04 |
| *W02B3.7* | WBGene00020927 | protein_coding_gene | 1.6 | 2.94 | 2.6E-03 |
| *cut-3* | WBGene00009041 | protein_coding_gene | 1.6 | 2.94 | 5.1E-13 |
| *nep-7* | WBGene00017552 | protein_coding_gene | 1.6 | 2.94 | 1.8E-03 |
| *fbxc-20* | WBGene00018352 | protein_coding_gene | 1.6 | 2.94 | 2.4E-08 |
| *Y13C8A.1* | WBGene00021188 | protein_coding_gene | 1.6 | 2.94 | 5.4E-06 |
| *R155.5* | WBGene00189953 | protein_coding_gene | 1.6 | 2.93 | 2.1E-03 |

| | | | | | |
|---|---|---|---|---|---|
| *Y50E8A.12* | WBGene00013055 | protein_coding_gene | 1.6 | 2.93 | 1.5E-06 |
| *W03F8.2* | WBGene00020992 | pseudogene | 1.5 | 2.93 | 1.9E-05 |
| *pax-2* | WBGene00003938 | protein_coding_gene | 1.5 | 2.93 | 1.2E-03 |
| *K07A1.13* | WBGene00010617 | protein_coding_gene | 1.5 | 2.92 | 9.1E-32 |
| *C50F2.5* | WBGene00016839 | protein_coding_gene | 1.5 | 2.92 | 3.1E-06 |
| *oac-45* | WBGene00011656 | protein_coding_gene | 1.5 | 2.92 | 8.2E-04 |
| *fbxa-172* | WBGene00008014 | protein_coding_gene | 1.5 | 2.91 | 1.1E-07 |
| *skr-12* | WBGene00004818 | protein_coding_gene | 1.5 | 2.90 | 2.5E-11 |
| *Y39E4B.11* | WBGene00012720 | protein_coding_gene | 1.5 | 2.90 | 5.4E-06 |
| *C53B4.2* | WBGene00008272 | protein_coding_gene | 1.5 | 2.89 | 6.1E-05 |
| *C08B6.4* | WBGene00007425 | protein_coding_gene | 1.5 | 2.89 | 1.2E-09 |
| *M05B5.3* | WBGene00010871 | protein_coding_gene | 1.5 | 2.89 | 4.9E-08 |
| *nspd-9* | WBGene00017808 | protein_coding_gene | 1.5 | 2.88 | 5.8E-11 |
| *K07E12.2* | WBGene00019501 | protein_coding_gene | 1.5 | 2.88 | 4.9E-07 |
| *Y53F4B.16* | WBGene00013162 | protein_coding_gene | 1.5 | 2.88 | 3.4E-04 |
| *F55F10.3* | WBGene00050916 | protein_coding_gene | 1.5 | 2.88 | 1.1E-04 |
| *C16C10.9* | WBGene00007629 | protein_coding_gene | 1.5 | 2.87 | 2.0E-03 |
| *T06D8.10* | WBGene00011530 | protein_coding_gene | 1.5 | 2.87 | 1.2E-07 |
| *F20A1.6* | WBGene00017618 | protein_coding_gene | 1.5 | 2.87 | 2.8E-15 |
| *F35E2.10* | WBGene00044270 | pseudogene | 1.5 | 2.87 | 1.2E-04 |
| *F22B3.10* | WBGene00045122 | protein_coding_gene | 1.5 | 2.86 | 4.1E-04 |
| *K11H12.5* | WBGene00019661 | protein_coding_gene | 1.5 | 2.86 | 5.7E-09 |
| *srab-25* | WBGene00022807 | protein_coding_gene | 1.5 | 2.86 | 4.3E-03 |
| *T05E12.3* | WBGene00011486 | protein_coding_gene | 1.5 | 2.85 | 1.9E-30 |
| *C06A6.7* | WBGene00015511 | protein_coding_gene | 1.5 | 2.85 | 7.1E-07 |
| *pals-6* | WBGene00007660 | protein_coding_gene | 1.5 | 2.85 | 2.3E-05 |
| *T25D1.2* | WBGene00020794 | protein_coding_gene | 1.5 | 2.84 | 1.9E-03 |
| *F54H5.5* | WBGene00018841 | protein_coding_gene | 1.5 | 2.84 | 1.6E-07 |
| *fbxa-30* | WBGene00013837 | protein_coding_gene | 1.5 | 2.84 | 1.1E-04 |
| *C47F8.5* | WBGene00008161 | protein_coding_gene | 1.5 | 2.84 | 3.1E-03 |
| *sre-6* | WBGene00009242 | protein_coding_gene | 1.5 | 2.84 | 5.5E-04 |
| *wht-5* | WBGene00008950 | protein_coding_gene | 1.5 | 2.83 | 2.2E-08 |
| *decr-1.3* | WBGene00011467 | protein_coding_gene | 1.5 | 2.83 | 5.3E-08 |
| *gad-2* | WBGene00015666 | protein_coding_gene | 1.5 | 2.83 | 1.9E-05 |
| *D1081.5* | WBGene00008383 | protein_coding_gene | 1.5 | 2.83 | 4.9E-07 |
| *F58D5.6* | WBGene00010245 | protein_coding_gene | 1.5 | 2.83 | 1.9E-04 |
| *clec-28* | WBGene00009857 | protein_coding_gene | 1.5 | 2.83 | 2.6E-04 |
| *T09D3.8* | WBGene00044538 | protein_coding_gene | 1.5 | 2.82 | 3.9E-03 |
| *msp-51* | WBGene00003444 | protein_coding_gene | 1.5 | 2.82 | 8.4E-11 |
| *K07A1.4* | WBGene00010611 | protein_coding_gene | 1.5 | 2.81 | 7.4E-10 |
| *F02H6.7* | WBGene00008540 | protein_coding_gene | 1.5 | 2.81 | 4.5E-03 |
| *col-74* | WBGene00000650 | protein_coding_gene | 1.5 | 2.81 | 3.6E-10 |
| *C17E4.2* | WBGene00007642 | protein_coding_gene | 1.5 | 2.81 | 3.1E-12 |
| *R102.3* | WBGene00011290 | protein_coding_gene | 1.5 | 2.81 | 1.7E-04 |
| *ZK1251.3* | WBGene00014241 | protein_coding_gene | 1.5 | 2.81 | 2.3E-04 |
| *F44E5.3* | WBGene00009690 | protein_coding_gene | 1.5 | 2.80 | 3.4E-04 |
| *C35E7.10* | WBGene00016462 | protein_coding_gene | 1.5 | 2.79 | 1.5E-11 |
| *clec-32* | WBGene00009860 | protein_coding_gene | 1.5 | 2.78 | 3.2E-03 |
| *F49C5.10* | WBGene00044162 | protein_coding_gene | 1.5 | 2.78 | 3.3E-03 |
| *C09H10.9* | WBGene00007503 | protein_coding_gene | 1.5 | 2.78 | 3.0E-06 |
| *C34D4.10* | WBGene00016402 | protein_coding_gene | 1.5 | 2.78 | 6.6E-21 |
| *K07F5.6* | WBGene00010634 | protein_coding_gene | 1.5 | 2.77 | 7.5E-07 |
| *Y18D10A.26* | WBGene00023459 | protein_coding_gene | 1.5 | 2.77 | 5.3E-04 |
| *fbxa-176* | WBGene00008017 | protein_coding_gene | 1.5 | 2.76 | 4.7E-05 |
| *dsl-6* | WBGene00001108 | protein_coding_gene | 1.5 | 2.76 | 1.3E-09 |
| *T25B9.6* | WBGene00012012 | protein_coding_gene | 1.5 | 2.75 | 1.7E-09 |
| *Y50D4C.2* | WBGene00021747 | pseudogene | 1.5 | 2.75 | 3.7E-11 |
| *Y57A10A.1* | WBGene00013246 | protein_coding_gene | 1.5 | 2.75 | 3.3E-22 |
| *C06A8.6* | WBGene00015516 | protein_coding_gene | 1.5 | 2.75 | 9.1E-09 |
| *msp-45* | WBGene00003438 | protein_coding_gene | 1.5 | 2.74 | 2.2E-12 |

| | | | | | |
|---|---|---|---|---|---|
| *acc-3* | WBGene00018880 | protein_coding_gene | 1.5 | 2.74 | 1.8E-04 |
| *inpp-1* | WBGene00012016 | protein_coding_gene | 1.5 | 2.74 | 5.6E-11 |
| *F08H9.4* | WBGene00008592 | protein_coding_gene | 1.4 | 2.73 | 3.0E-03 |
| *ttr-22* | WBGene00011450 | protein_coding_gene | 1.4 | 2.73 | 4.6E-03 |
| *acdh-8* | WBGene00019406 | protein_coding_gene | 1.4 | 2.73 | 1.4E-11 |
| *Y47H9C.12* | WBGene00012955 | protein_coding_gene | 1.4 | 2.73 | 4.3E-03 |
| *T07G12.3* | WBGene00011594 | protein_coding_gene | 1.4 | 2.72 | 4.7E-05 |
| *C09D4.3* | WBGene00015634 | protein_coding_gene | 1.4 | 2.71 | 9.5E-09 |
| *his-70* | WBGene00001944 | protein_coding_gene | 1.4 | 2.71 | 8.0E-06 |
| *Y40A1A.6* | WBGene00304815 | ncRNA_gene | 1.4 | 2.71 | 4.1E-14 |
| *F40H6.5* | WBGene00018255 | protein_coding_gene | 1.4 | 2.70 | 4.3E-03 |
| *T11F8.4* | WBGene00020435 | protein_coding_gene | 1.4 | 2.70 | 1.6E-07 |
| *col-65* | WBGene00000641 | protein_coding_gene | 1.4 | 2.70 | 2.7E-16 |
| *C25H3.16* | WBGene00044392 | protein_coding_gene | 1.4 | 2.69 | 2.1E-03 |
| *dhhc-12* | WBGene00010323 | protein_coding_gene | 1.4 | 2.69 | 2.0E-06 |
| *ZK484.7* | WBGene00022753 | protein_coding_gene | 1.4 | 2.69 | 1.8E-06 |
| *oac-44* | WBGene00011654 | protein_coding_gene | 1.4 | 2.69 | 3.2E-03 |
| *F44F1.1* | WBGene00009693 | pseudogene | 1.4 | 2.69 | 2.5E-03 |
| *ugt-9* | WBGene00020587 | protein_coding_gene | 1.4 | 2.68 | 6.3E-05 |
| *col-165* | WBGene00000738 | protein_coding_gene | 1.4 | 2.68 | 9.7E-12 |
| *T21E12.5* | WBGene00020656 | protein_coding_gene | 1.4 | 2.68 | 1.6E-03 |
| *F21F3.2* | WBGene00017672 | protein_coding_gene | 1.4 | 2.68 | 1.7E-05 |
| *C25A6.1* | WBGene00016081 | protein_coding_gene | 1.4 | 2.67 | 4.5E-05 |
| *msp-55* | WBGene00003448 | protein_coding_gene | 1.4 | 2.67 | 2.0E-04 |
| *F36H12.9* | WBGene00018123 | protein_coding_gene | 1.4 | 2.67 | 3.6E-03 |
| *nhr-54* | WBGene00003644 | protein_coding_gene | 1.4 | 2.67 | 6.4E-06 |
| *clec-67* | WBGene00018971 | protein_coding_gene | 1.4 | 2.66 | 3.4E-09 |
| *F09E10.1* | WBGene00017294 | protein_coding_gene | 1.4 | 2.66 | 4.7E-13 |
| *Y38F1A.7* | WBGene00012609 | protein_coding_gene | 1.4 | 2.66 | 1.1E-03 |
| *Y47D7A.2* | WBGene00021616 | protein_coding_gene | 1.4 | 2.66 | 4.1E-03 |
| *C35A11.3* | WBGene00016430 | protein_coding_gene | 1.4 | 2.66 | 4.1E-03 |
| *trf-2* | WBGene00022454 | protein_coding_gene | 1.4 | 2.66 | 1.2E-03 |
| *F01G10.5* | WBGene00008508 | protein_coding_gene | 1.4 | 2.65 | 2.2E-09 |
| *ZK896.4* | WBGene00014135 | protein_coding_gene | 1.4 | 2.65 | 1.8E-04 |
| *spe-47* | WBGene00021271 | protein_coding_gene | 1.4 | 2.65 | 2.3E-05 |
| *F36H1.3* | WBGene00009492 | protein_coding_gene | 1.4 | 2.64 | 1.0E-06 |
| *str-7* | WBGene00006075 | protein_coding_gene | 1.4 | 2.64 | 5.1E-03 |
| *kin-24* | WBGene00002206 | protein_coding_gene | 1.4 | 2.64 | 5.6E-06 |
| *C17E4.10* | WBGene00007647 | protein_coding_gene | 1.4 | 2.64 | 4.0E-50 |
| *Y2H9A.6* | WBGene00086568 | protein_coding_gene | 1.4 | 2.64 | 1.9E-05 |
| *F55B12.11* | WBGene00077570 | pseudogene | 1.4 | 2.64 | 1.3E-03 |
| *K01A6.4* | WBGene00010446 | protein_coding_gene | 1.4 | 2.64 | 5.3E-03 |
| *W09C3.1* | WBGene00021109 | protein_coding_gene | 1.4 | 2.64 | 4.1E-03 |
| *F49E11.7* | WBGene00009893 | protein_coding_gene | 1.4 | 2.63 | 2.7E-05 |
| *Y48B6A.5* | WBGene00012979 | protein_coding_gene | 1.4 | 2.63 | 2.0E-07 |
| *F54C8.1* | WBGene00010035 | protein_coding_gene | 1.4 | 2.63 | 6.1E-08 |
| *acdh-5* | WBGene00016491 | protein_coding_gene | 1.4 | 2.63 | 5.5E-08 |
| *K07A1.3* | WBGene00010610 | protein_coding_gene | 1.4 | 2.63 | 2.0E-14 |
| *T27F6.1* | WBGene00012102 | protein_coding_gene | 1.4 | 2.63 | 2.8E-04 |
| *T19H12.15* | WBGene00271827 | protein_coding_gene | 1.4 | 2.62 | 4.7E-18 |
| *clec-134* | WBGene00021141 | protein_coding_gene | 1.4 | 2.62 | 8.0E-03 |
| *spe-42* | WBGene00007117 | protein_coding_gene | 1.4 | 2.62 | 2.0E-04 |
| *T15D6.10* | WBGene00011784 | protein_coding_gene | 1.4 | 2.61 | 5.6E-03 |
| *F36H12.14* | WBGene00018127 | pseudogene | 1.4 | 2.61 | 4.6E-05 |
| *tsp-19* | WBGene00006645 | protein_coding_gene | 1.4 | 2.61 | 1.8E-04 |
| *W03G1.5* | WBGene00021011 | protein_coding_gene | 1.4 | 2.61 | 1.8E-05 |
| *T04A6.3* | WBGene00020202 | protein_coding_gene | 1.4 | 2.60 | 1.6E-03 |
| *T24C4.4* | WBGene00020760 | protein_coding_gene | 1.4 | 2.60 | 1.3E-04 |
| *T25B9.5* | WBGene00012011 | protein_coding_gene | 1.4 | 2.60 | 7.1E-05 |
| *C55C3.7* | WBGene00043065 | protein_coding_gene | 1.4 | 2.60 | 4.4E-03 |

| | | | | | |
|---|---|---|---|---|---|
| *F21A3.4* | WBGene00008994 | protein_coding_gene | 1.4 | 2.60 | 3.4E-04 |
| *clec-36* | WBGene00044332 | protein_coding_gene | 1.4 | 2.60 | 3.0E-06 |
| *K08D8.5* | WBGene00010659 | protein_coding_gene | 1.4 | 2.60 | 7.7E-07 |
| *T10D4.7* | WBGene00020409 | protein_coding_gene | 1.4 | 2.59 | 6.0E-03 |
| *C38C3.8* | WBGene00016516 | pseudogene | 1.4 | 2.59 | 6.3E-04 |
| *M05D6.1* | WBGene00010874 | protein_coding_gene | 1.4 | 2.59 | 2.7E-03 |
| *Y47D7A.18* | WBGene00235165 | protein_coding_gene | 1.4 | 2.59 | 9.8E-04 |
| *K07A1.10* | WBGene00010616 | protein_coding_gene | 1.4 | 2.59 | 5.3E-36 |
| *F38A1.9* | WBGene00009522 | protein_coding_gene | 1.4 | 2.59 | 1.2E-06 |
| *C16C8.18* | WBGene00015855 | protein_coding_gene | 1.4 | 2.58 | 5.7E-07 |
| *T28C6.5* | WBGene00012120 | protein_coding_gene | 1.4 | 2.58 | 3.4E-05 |
| *nhr-81* | WBGene00003671 | protein_coding_gene | 1.4 | 2.58 | 4.6E-03 |
| *tig-3* | WBGene00021594 | protein_coding_gene | 1.4 | 2.58 | 3.1E-03 |
| *Y66A7A.4* | WBGene00013424 | protein_coding_gene | 1.4 | 2.58 | 2.4E-04 |
| *ZK1010.6* | WBGene00014180 | pseudogene | 1.4 | 2.58 | 9.4E-03 |
| *F15D4.5* | WBGene00008862 | protein_coding_gene | 1.4 | 2.58 | 3.0E-05 |
| *ZK809.9* | WBGene00194934 | protein_coding_gene | 1.4 | 2.57 | 1.0E-04 |
| *F23B12.1* | WBGene00009079 | protein_coding_gene | 1.4 | 2.57 | 2.2E-05 |
| *F10C1.3* | WBGene00017325 | protein_coding_gene | 1.4 | 2.57 | 2.9E-06 |
| *T12A2.6* | WBGene00020440 | protein_coding_gene | 1.4 | 2.57 | 9.1E-03 |
| *B0207.7* | WBGene00015030 | protein_coding_gene | 1.4 | 2.57 | 3.8E-05 |
| *dhhc-13* | WBGene00019257 | protein_coding_gene | 1.4 | 2.57 | 1.4E-04 |
| *C47F8.3* | WBGene00008159 | protein_coding_gene | 1.4 | 2.57 | 8.9E-03 |
| *C39H7.1* | WBGene00016541 | protein_coding_gene | 1.4 | 2.56 | 2.1E-11 |
| *ZK892.4* | WBGene00014128 | protein_coding_gene | 1.4 | 2.56 | 3.3E-03 |
| *W03A5.1* | WBGene00020966 | protein_coding_gene | 1.4 | 2.56 | 4.0E-06 |
| *sst-20* | WBGene00006058 | protein_coding_gene | 1.4 | 2.56 | 5.9E-06 |
| *T20B6.2* | WBGene00020598 | protein_coding_gene | 1.4 | 2.56 | 2.4E-03 |
| *C27D6.15* | WBGene00303362 | protein_coding_gene | 1.4 | 2.56 | 9.7E-04 |
| *T06G6.11* | WBGene00011550 | pseudogene | 1.4 | 2.56 | 6.9E-03 |
| *2RSSE.3* | WBGene00044951 | pseudogene | 1.4 | 2.56 | 1.1E-03 |
| *Y39B6A.25* | WBGene00012684 | protein_coding_gene | 1.4 | 2.56 | 1.1E-04 |
| *mboa-4* | WBGene00007446 | protein_coding_gene | 1.4 | 2.56 | 7.5E-07 |
| *ugt-63* | WBGene00015449 | protein_coding_gene | 1.4 | 2.56 | 3.3E-14 |
| *T05H4.7* | WBGene00020270 | protein_coding_gene | 1.3 | 2.55 | 1.4E-03 |
| *tag-293* | WBGene00015398 | protein_coding_gene | 1.3 | 2.55 | 8.9E-03 |
| *Y82E9BL.19* | WBGene00194735 | protein_coding_gene | 1.3 | 2.54 | 1.0E-02 |
| *ZK354.2* | WBGene00022705 | protein_coding_gene | 1.3 | 2.54 | 6.5E-11 |
| *msp-57* | WBGene00003450 | protein_coding_gene | 1.3 | 2.54 | 1.5E-03 |
| *F36A2.14* | WBGene00023490 | protein_coding_gene | 1.3 | 2.54 | 6.4E-09 |
| *spch-2* | WBGene00015689 | protein_coding_gene | 1.3 | 2.54 | 6.5E-05 |
| *Y50D4B.9* | WBGene00268189 | pseudogene | 1.3 | 2.54 | 6.7E-03 |
| *Y38A10A.2* | WBGene00021389 | protein_coding_gene | 1.3 | 2.54 | 3.3E-06 |
| *F16C3.4* | WBGene00044243 | protein_coding_gene | 1.3 | 2.54 | 3.7E-04 |
| *irld-14* | WBGene00011322 | protein_coding_gene | 1.3 | 2.53 | 5.2E-05 |
| *Y53F4B.27* | WBGene00013173 | protein_coding_gene | 1.3 | 2.53 | 4.5E-05 |
| *LLC1.121* | WBGene00235121 | pseudogene | 1.3 | 2.53 | 7.4E-03 |
| *bath-10* | WBGene00021726 | protein_coding_gene | 1.3 | 2.53 | 7.2E-07 |
| *oac-48* | WBGene00011960 | protein_coding_gene | 1.3 | 2.53 | 2.2E-03 |
| *4R79.2* | WBGene00007067 | protein_coding_gene | 1.3 | 2.52 | 1.4E-04 |
| *xtr-2* | WBGene00006966 | protein_coding_gene | 1.3 | 2.52 | 8.5E-03 |
| *F15H9.7* | WBGene00077535 | pseudogene | 1.3 | 2.51 | 3.6E-03 |
| *T27A8.7* | WBGene00206527 | protein_coding_gene | 1.3 | 2.51 | 9.4E-03 |
| *F16G10.2* | WBGene00017517 | protein_coding_gene | 1.3 | 2.51 | 1.3E-03 |
| *msp-65* | WBGene00003458 | protein_coding_gene | 1.3 | 2.50 | 7.5E-05 |
| *C36H8.1* | WBGene00007987 | protein_coding_gene | 1.3 | 2.50 | 6.0E-08 |
| *M199.9* | WBGene00044998 | protein_coding_gene | 1.3 | 2.50 | 1.1E-03 |
| *C49A1.5* | WBGene00008187 | protein_coding_gene | 1.3 | 2.50 | 8.6E-03 |
| *C17H12.3* | WBGene00015929 | protein_coding_gene | 1.3 | 2.50 | 1.2E-07 |
| *spe-45* | WBGene00009217 | protein_coding_gene | 1.3 | 2.50 | 7.0E-04 |

| | | | | | |
|---|---|---|---|---|---|
| *msrp-6* | WBGene00017058 | protein_coding_gene | 1.3 | 2.50 | 1.4E-06 |
| *snb-5* | WBGene00004899 | protein_coding_gene | 1.3 | 2.50 | 6.9E-03 |
| *F54H12.8* | WBGene00045247 | protein_coding_gene | 1.3 | 2.49 | 1.0E-04 |
| *C54E10.10* | WBGene00206523 | protein_coding_gene | 1.3 | 2.49 | 1.3E-03 |
| *irld-6* | WBGene00009239 | protein_coding_gene | 1.3 | 2.49 | 6.0E-05 |
| *ugt-16* | WBGene00013901 | protein_coding_gene | 1.3 | 2.49 | 6.4E-11 |
| *K03H1.12* | WBGene00010547 | protein_coding_gene | 1.3 | 2.49 | 2.3E-05 |
| *spp-17* | WBGene00005002 | protein_coding_gene | 1.3 | 2.49 | 6.1E-04 |
| *R02F11.10* | WBGene00202497 | protein_coding_gene | 1.3 | 2.49 | 1.4E-03 |
| *nspd-5* | WBGene00017386 | protein_coding_gene | 1.3 | 2.49 | 7.2E-04 |
| *T23F11.2* | WBGene00011954 | protein_coding_gene | 1.3 | 2.49 | 3.8E-03 |
| *cyp-25A6* | WBGene00019438 | protein_coding_gene | 1.3 | 2.48 | 8.4E-03 |
| *T27A1.2* | WBGene00020834 | protein_coding_gene | 1.3 | 2.48 | 1.4E-09 |
| *glf-1* | WBGene00019154 | protein_coding_gene | 1.3 | 2.47 | 2.5E-08 |
| *Y71A12B.19* | WBGene00077443 | pseudogene | 1.3 | 2.47 | 4.1E-04 |
| *C05D12.3* | WBGene00014666 | protein_coding_gene | 1.3 | 2.47 | 2.4E-21 |
| *gasr-8* | WBGene00008144 | protein_coding_gene | 1.3 | 2.47 | 4.8E-03 |
| *zig-6* | WBGene00006983 | protein_coding_gene | 1.3 | 2.47 | 3.5E-06 |
| *F43E2.6* | WBGene00018394 | protein_coding_gene | 1.3 | 2.47 | 6.7E-05 |
| *oac-36* | WBGene00010172 | protein_coding_gene | 1.3 | 2.46 | 8.8E-03 |
| *C16D9.5* | WBGene00015860 | protein_coding_gene | 1.3 | 2.46 | 7.3E-04 |
| *T15D6.8* | WBGene00011782 | protein_coding_gene | 1.3 | 2.46 | 8.2E-03 |
| *F26F2.11* | WBGene00195158 | protein_coding_gene | 1.3 | 2.46 | 7.8E-03 |
| *F32H2.7* | WBGene00009344 | protein_coding_gene | 1.3 | 2.46 | 3.3E-12 |
| *uggt-2* | WBGene00009178 | protein_coding_gene | 1.3 | 2.46 | 2.8E-06 |
| *F46A9.2* | WBGene00009755 | protein_coding_gene | 1.3 | 2.46 | 1.7E-06 |
| *scl-6* | WBGene00008028 | protein_coding_gene | 1.3 | 2.46 | 2.5E-08 |
| *K08E7.4* | WBGene00010672 | protein_coding_gene | 1.3 | 2.46 | 7.5E-03 |
| *pud-2.2* | WBGene00017500 | protein_coding_gene | 1.3 | 2.45 | 1.4E-09 |
| *F53C3.8* | WBGene00018752 | protein_coding_gene | 1.3 | 2.45 | 1.7E-03 |
| *nlp-16* | WBGene00003754 | protein_coding_gene | 1.3 | 2.45 | 4.2E-23 |
| *oac-34* | WBGene00010157 | protein_coding_gene | 1.3 | 2.45 | 8.0E-03 |
| *col-170* | WBGene00000743 | protein_coding_gene | 1.3 | 2.45 | 5.7E-03 |
| *F37A4.5* | WBGene00018135 | protein_coding_gene | 1.3 | 2.45 | 2.0E-07 |
| *C14A4.8* | WBGene00007559 | protein_coding_gene | 1.3 | 2.44 | 1.3E-03 |
| *Y40B10A.4* | WBGene00021489 | pseudogene | 1.3 | 2.44 | 4.4E-03 |
| *T08G3.13* | WBGene00011621 | protein_coding_gene | 1.3 | 2.44 | 1.2E-02 |
| *C48D1.9* | WBGene00194643 | protein_coding_gene | 1.3 | 2.44 | 4.7E-03 |
| *gst-39* | WBGene00001787 | protein_coding_gene | 1.3 | 2.44 | 1.3E-08 |
| *F39B3.7* | WBGene00304237 | protein_coding_gene | 1.3 | 2.44 | 9.7E-03 |
| *Y62H9A.16* | WBGene00304791 | protein_coding_gene | 1.3 | 2.44 | 3.6E-03 |
| *rsef-1* | WBGene00016344 | protein_coding_gene | 1.3 | 2.43 | 2.6E-05 |
| *C33F10.8* | WBGene00016356 | protein_coding_gene | 1.3 | 2.43 | 5.2E-06 |
| *scrm-6* | WBGene00009753 | protein_coding_gene | 1.3 | 2.43 | 4.7E-05 |
| *F16G10.15* | WBGene00017529 | pseudogene | 1.3 | 2.43 | 2.7E-04 |
| *M28.9* | WBGene00010900 | protein_coding_gene | 1.3 | 2.43 | 1.6E-09 |
| *elo-7* | WBGene00001245 | protein_coding_gene | 1.3 | 2.43 | 3.8E-04 |
| *nas-19* | WBGene00003538 | protein_coding_gene | 1.3 | 2.42 | 9.3E-03 |
| *C33F10.12* | WBGene00016358 | protein_coding_gene | 1.3 | 2.42 | 6.9E-08 |
| *Y57A10B.6* | WBGene00013278 | protein_coding_gene | 1.3 | 2.42 | 2.0E-05 |
| *F13H6.4* | WBGene00017432 | pseudogene | 1.3 | 2.42 | 6.6E-05 |
| *C08F11.1* | WBGene00007450 | protein_coding_gene | 1.3 | 2.42 | 5.9E-04 |
| *fbxa-160* | WBGene00015604 | protein_coding_gene | 1.3 | 2.42 | 1.8E-03 |
| *C31B8.7* | WBGene00016282 | protein_coding_gene | 1.3 | 2.41 | 5.8E-06 |
| *Y47D3B.12* | WBGene00185055 | protein_coding_gene | 1.3 | 2.41 | 3.2E-04 |
| *F13D12.9* | WBGene00008743 | protein_coding_gene | 1.3 | 2.41 | 2.0E-07 |
| *F21D9.1* | WBGene00009015 | protein_coding_gene | 1.3 | 2.41 | 6.4E-03 |
| *qui-1* | WBGene00004265 | protein_coding_gene | 1.3 | 2.41 | 1.0E-08 |
| *C29E4.14* | WBGene00044634 | protein_coding_gene | 1.3 | 2.41 | 1.2E-03 |
| *clec-164* | WBGene00021949 | protein_coding_gene | 1.3 | 2.41 | 1.4E-02 |

| | | | | | |
|---|---|---|---|---|---|
| *F36D3.8* | WBGene00009473 | protein_coding_gene | 1.3 | 2.40 | 1.9E-03 |
| *dmsr-12* | WBGene00019262 | protein_coding_gene | 1.3 | 2.40 | 4.3E-05 |
| *F58E6.5* | WBGene00010254 | protein_coding_gene | 1.3 | 2.40 | 5.1E-09 |
| *clec-193* | WBGene00013774 | protein_coding_gene | 1.3 | 2.40 | 1.2E-02 |
| *T05H10.3* | WBGene00011508 | protein_coding_gene | 1.3 | 2.40 | 2.5E-10 |
| *E03H4.11* | WBGene00008478 | protein_coding_gene | 1.3 | 2.40 | 7.4E-03 |
| *B0507.6* | WBGene00015223 | protein_coding_gene | 1.3 | 2.40 | 6.4E-05 |
| *AC7.3* | WBGene00014997 | protein_coding_gene | 1.3 | 2.39 | 1.8E-03 |
| *arrd-8* | WBGene00012467 | protein_coding_gene | 1.3 | 2.39 | 6.3E-03 |
| *C01G5.3* | WBGene00015305 | protein_coding_gene | 1.3 | 2.39 | 6.3E-04 |
| *W06H8.4* | WBGene00021067 | pseudogene | 1.3 | 2.39 | 1.0E-02 |
| *W04E12.9* | WBGene00012254 | protein_coding_gene | 1.3 | 2.39 | 6.5E-03 |
| *Y26G10.3* | WBGene00050973 | pseudogene | 1.3 | 2.39 | 8.1E-03 |
| *R02D1.2* | WBGene00235310 | protein_coding_gene | 1.3 | 2.39 | 1.7E-03 |
| *ipla-5* | WBGene00019229 | protein_coding_gene | 1.3 | 2.39 | 2.3E-03 |
| *ram-2* | WBGene00004300 | protein_coding_gene | 1.3 | 2.38 | 1.9E-15 |
| *K07A3.3* | WBGene00019461 | protein_coding_gene | 1.3 | 2.38 | 1.4E-07 |
| *F10G7.12* | WBGene00044744 | protein_coding_gene | 1.3 | 2.38 | 1.9E-03 |
| *lips-9* | WBGene00010267 | protein_coding_gene | 1.3 | 2.38 | 1.7E-03 |
| *nhr-169* | WBGene00008289 | protein_coding_gene | 1.3 | 2.38 | 8.6E-03 |
| *M7.7* | WBGene00010883 | protein_coding_gene | 1.3 | 2.38 | 1.2E-05 |
| *csp-1* | WBGene00000819 | protein_coding_gene | 1.2 | 2.37 | 2.4E-03 |
| *F20B6.1* | WBGene00017627 | pseudogene | 1.2 | 2.37 | 7.4E-03 |
| *F57G8.7* | WBGene00010216 | protein_coding_gene | 1.2 | 2.37 | 1.5E-02 |
| *C35A5.4* | WBGene00007948 | protein_coding_gene | 1.2 | 2.36 | 1.2E-04 |
| *ptp-5.1* | WBGene00016053 | protein_coding_gene | 1.2 | 2.36 | 1.1E-06 |
| *K01D12.9* | WBGene00010468 | protein_coding_gene | 1.2 | 2.36 | 2.4E-04 |
| *msp-31* | WBGene00003429 | protein_coding_gene | 1.2 | 2.36 | 6.2E-11 |
| *ZK616.61* | WBGene00194816 | protein_coding_gene | 1.2 | 2.36 | 1.8E-06 |
| *C40A11.3* | WBGene00016546 | protein_coding_gene | 1.2 | 2.36 | 6.2E-05 |
| *spch-1* | WBGene00007307 | protein_coding_gene | 1.2 | 2.36 | 2.2E-03 |
| *C47E8.1* | WBGene00008141 | protein_coding_gene | 1.2 | 2.36 | 1.1E-03 |
| *irld-7* | WBGene00018168 | protein_coding_gene | 1.2 | 2.36 | 3.6E-03 |
| *C06A1.3* | WBGene00007354 | protein_coding_gene | 1.2 | 2.36 | 3.9E-07 |
| *mec-8* | WBGene00003172 | protein_coding_gene | 1.2 | 2.36 | 6.9E-16 |
| *T05A7.11* | WBGene00044383 | protein_coding_gene | 1.2 | 2.36 | 2.4E-04 |
| *Y38H8A.4* | WBGene00012638 | protein_coding_gene | 1.2 | 2.36 | 3.6E-06 |
| *F35E2.5* | WBGene00009412 | protein_coding_gene | 1.2 | 2.36 | 3.7E-04 |
| *ZK1248.5* | WBGene00022876 | protein_coding_gene | 1.2 | 2.36 | 5.7E-05 |
| *Y52D5A.2* | WBGene00021793 | protein_coding_gene | 1.2 | 2.36 | 3.8E-03 |
| *C54E4.5* | WBGene00016920 | protein_coding_gene | 1.2 | 2.35 | 1.2E-13 |
| *lys-6* | WBGene00003095 | protein_coding_gene | 1.2 | 2.35 | 8.9E-06 |
| *T10H4.13* | WBGene00045339 | protein_coding_gene | 1.2 | 2.35 | 6.1E-03 |
| *best-6* | WBGene00015628 | protein_coding_gene | 1.2 | 2.35 | 1.2E-02 |
| *C47A4.1* | WBGene00008122 | protein_coding_gene | 1.2 | 2.35 | 4.3E-03 |
| *best-10* | WBGene00008185 | protein_coding_gene | 1.2 | 2.35 | 3.7E-03 |
| *scl-5* | WBGene00008027 | protein_coding_gene | 1.2 | 2.34 | 2.2E-12 |
| *T05F1.8* | WBGene00011493 | protein_coding_gene | 1.2 | 2.34 | 8.0E-07 |
| *C09B9.7* | WBGene00015630 | protein_coding_gene | 1.2 | 2.34 | 6.7E-04 |
| *fbxb-64* | WBGene00008071 | protein_coding_gene | 1.2 | 2.34 | 1.3E-02 |
| *F58G1.3* | WBGene00010265 | protein_coding_gene | 1.2 | 2.34 | 8.5E-06 |
| *Y75B12B.13* | WBGene00045402 | protein_coding_gene | 1.2 | 2.34 | 9.2E-03 |
| *bgnt-1.8* | WBGene00008290 | protein_coding_gene | 1.2 | 2.34 | 1.5E-02 |
| *ssp-10* | WBGene00006039 | protein_coding_gene | 1.2 | 2.33 | 1.1E-03 |
| *ent-3* | WBGene00010510 | protein_coding_gene | 1.2 | 2.33 | 2.0E-06 |
| *paf-1* | WBGene00003906 | protein_coding_gene | 1.2 | 2.33 | 1.2E-03 |
| *T14A8.2* | WBGene00020497 | protein_coding_gene | 1.2 | 2.33 | 3.5E-06 |
| *W03D8.1* | WBGene00020984 | protein_coding_gene | 1.2 | 2.33 | 5.0E-03 |
| *K10G4.5* | WBGene00010754 | protein_coding_gene | 1.2 | 2.32 | 1.3E-02 |
| *ssq-2* | WBGene00006051 | protein_coding_gene | 1.2 | 2.32 | 1.5E-17 |

| | | | | | |
|---|---|---|---|---|---|
| *K07A1.15* | WBGene00010619 | protein_coding_gene | 1.2 | 2.32 | 4.0E-07 |
| *C30B5.6* | WBGene00016247 | protein_coding_gene | 1.2 | 2.32 | 1.2E-03 |
| *Y50E8A.6* | WBGene00013049 | protein_coding_gene | 1.2 | 2.32 | 1.1E-02 |
| *msp-81* | WBGene00003467 | protein_coding_gene | 1.2 | 2.32 | 4.2E-09 |
| *F44G4.5* | WBGene00009714 | protein_coding_gene | 1.2 | 2.32 | 5.1E-04 |
| *T27A3.5* | WBGene00020841 | protein_coding_gene | 1.2 | 2.32 | 3.3E-06 |
| *C17E4.14* | WBGene00195884 | ncRNA_gene | 1.2 | 2.31 | 1.0E-02 |
| *fbxb-17* | WBGene00019037 | protein_coding_gene | 1.2 | 2.31 | 1.9E-04 |
| *F56F3.4* | WBGene00010155 | protein_coding_gene | 1.2 | 2.31 | 2.0E-08 |
| *M163.22* | WBGene00304195 | protein_coding_gene | 1.2 | 2.31 | 2.4E-03 |
| *B0310.3* | WBGene00015139 | protein_coding_gene | 1.2 | 2.31 | 7.2E-04 |
| *ZK616.1* | WBGene00022771 | protein_coding_gene | 1.2 | 2.31 | 3.9E-03 |
| *nspc-3* | WBGene00044338 | protein_coding_gene | 1.2 | 2.30 | 1.5E-03 |
| *Y49G5A.1* | WBGene00021731 | protein_coding_gene | 1.2 | 2.30 | 1.5E-02 |
| *F35C5.4* | WBGene00009392 | protein_coding_gene | 1.2 | 2.30 | 2.3E-03 |
| *cdc-25.4* | WBGene00000389 | protein_coding_gene | 1.2 | 2.30 | 1.3E-03 |
| *F42G4.2* | WBGene00009641 | protein_coding_gene | 1.2 | 2.30 | 3.5E-05 |
| *Y32G9A.3* | WBGene00021302 | protein_coding_gene | 1.2 | 2.30 | 1.5E-02 |
| *nas-22* | WBGene00003541 | protein_coding_gene | 1.2 | 2.30 | 4.3E-03 |
| *wht-4* | WBGene00017179 | protein_coding_gene | 1.2 | 2.30 | 4.0E-04 |
| *nhr-246* | WBGene00014192 | protein_coding_gene | 1.2 | 2.30 | 1.7E-02 |
| *B0024.4* | WBGene00007097 | protein_coding_gene | 1.2 | 2.30 | 1.2E-02 |
| *K06A1.2* | WBGene00019425 | protein_coding_gene | 1.2 | 2.29 | 2.0E-03 |
| *try-7* | WBGene00006625 | protein_coding_gene | 1.2 | 2.29 | 4.2E-05 |
| *str-11* | WBGene00006079 | pseudogene | 1.2 | 2.29 | 1.8E-02 |
| *Y67A6A.1* | WBGene00013449 | protein_coding_gene | 1.2 | 2.29 | 5.9E-05 |
| *gly-1* | WBGene00001626 | protein_coding_gene | 1.2 | 2.29 | 1.8E-02 |
| *H37A05.2* | WBGene00010426 | protein_coding_gene | 1.2 | 2.29 | 1.2E-04 |
| *F58D5.7* | WBGene00010246 | protein_coding_gene | 1.2 | 2.29 | 5.4E-07 |
| *B0244.9* | WBGene00015084 | protein_coding_gene | 1.2 | 2.29 | 3.1E-04 |
| *F36G3.3* | WBGene00009484 | protein_coding_gene | 1.2 | 2.29 | 1.6E-02 |
| *grl-15* | WBGene00001724 | protein_coding_gene | 1.2 | 2.29 | 1.6E-04 |
| *Y17G9A.4* | WBGene00021197 | protein_coding_gene | 1.2 | 2.29 | 6.7E-05 |
| *grl-16* | WBGene00001725 | protein_coding_gene | 1.2 | 2.28 | 1.1E-10 |
| *F38C2.4* | WBGene00009536 | protein_coding_gene | 1.2 | 2.28 | 1.5E-02 |
| *K08F4.12* | WBGene00010680 | protein_coding_gene | 1.2 | 2.28 | 1.4E-02 |
| *F36D3.5* | WBGene00009471 | protein_coding_gene | 1.2 | 2.28 | 8.5E-05 |
| *F13H8.6* | WBGene00017439 | protein_coding_gene | 1.2 | 2.28 | 1.3E-02 |
| *dhhc-5* | WBGene00020066 | protein_coding_gene | 1.2 | 2.28 | 9.4E-03 |
| *R13H9.5* | WBGene00020071 | protein_coding_gene | 1.2 | 2.28 | 1.2E-03 |
| *T22B7.3* | WBGene00020672 | protein_coding_gene | 1.2 | 2.28 | 3.1E-03 |
| *T02B11.4* | WBGene00020156 | protein_coding_gene | 1.2 | 2.28 | 8.7E-05 |
| *B0391.8* | WBGene00014643 | pseudogene | 1.2 | 2.28 | 2.9E-06 |
| *C29F5.3* | WBGene00016212 | protein_coding_gene | 1.2 | 2.27 | 6.6E-04 |
| *col-115* | WBGene00000689 | protein_coding_gene | 1.2 | 2.27 | 3.3E-03 |
| *C36C9.4* | WBGene00016487 | protein_coding_gene | 1.2 | 2.27 | 1.7E-02 |
| *C32B5.13* | WBGene00016306 | protein_coding_gene | 1.2 | 2.27 | 1.9E-02 |
| *msp-142* | WBGene00003469 | protein_coding_gene | 1.2 | 2.27 | 2.5E-08 |
| *ZC262.4* | WBGene00022581 | protein_coding_gene | 1.2 | 2.27 | 1.8E-02 |
| *T21G5.1* | WBGene00020659 | protein_coding_gene | 1.2 | 2.27 | 1.5E-05 |
| *Y45F10B.3* | WBGene00012871 | protein_coding_gene | 1.2 | 2.27 | 9.2E-04 |
| *Y39B6A.9* | WBGene00012671 | protein_coding_gene | 1.2 | 2.27 | 8.3E-03 |
| *Y57E12B.10* | WBGene00202501 | protein_coding_gene | 1.2 | 2.26 | 2.0E-02 |
| *ZC581.2* | WBGene00022632 | protein_coding_gene | 1.2 | 2.26 | 1.3E-04 |
| *Y116F11B.11* | WBGene00013827 | pseudogene | 1.2 | 2.26 | 1.1E-02 |
| *Y37H2A.13* | WBGene00045417 | protein_coding_gene | 1.2 | 2.26 | 4.4E-04 |
| *glct-4* | WBGene00008160 | protein_coding_gene | 1.2 | 2.26 | 1.8E-02 |
| *C47E12.12* | WBGene00008155 | protein_coding_gene | 1.2 | 2.26 | 5.3E-03 |
| *msp-40* | WBGene00003435 | protein_coding_gene | 1.2 | 2.26 | 5.1E-03 |
| *fbxa-87* | WBGene00044660 | pseudogene | 1.2 | 2.26 | 8.4E-05 |

| | | | | | |
|---|---|---|---|---|---|
| *Y23H5B.12* | WBGene00194701 | protein_coding_gene | 1.2 | 2.26 | 1.8E-02 |
| *F56F4.4* | WBGene00018981 | protein_coding_gene | 1.2 | 2.26 | 1.4E-02 |
| *F28E10.4* | WBGene00017902 | protein_coding_gene | 1.2 | 2.26 | 1.7E-03 |
| *C34C6.3* | WBGene00007916 | protein_coding_gene | 1.2 | 2.26 | 1.3E-06 |
| *F35C11.2* | WBGene00009401 | protein_coding_gene | 1.2 | 2.26 | 4.2E-03 |
| *B0496.1* | WBGene00015210 | protein_coding_gene | 1.2 | 2.26 | 4.4E-03 |
| *C15H7.3* | WBGene00007610 | protein_coding_gene | 1.2 | 2.26 | 1.7E-05 |
| *bgnt-1.4* | WBGene00015982 | protein_coding_gene | 1.2 | 2.25 | 1.3E-02 |
| *C27A2.8* | WBGene00044387 | protein_coding_gene | 1.2 | 2.25 | 1.3E-02 |
| *gst-40* | WBGene00001788 | protein_coding_gene | 1.2 | 2.25 | 1.1E-02 |
| *Y45F10B.15* | WBGene00012876 | pseudogene | 1.2 | 2.25 | 2.0E-02 |
| *exc-13* | WBGene00015393 | protein_coding_gene | 1.2 | 2.25 | 1.3E-04 |
| *T10E9.6* | WBGene00020416 | protein_coding_gene | 1.2 | 2.25 | 3.2E-03 |
| *F10G2.11* | WBGene00271802 | pseudogene | 1.2 | 2.25 | 2.0E-02 |
| *smz-2* | WBGene00020661 | protein_coding_gene | 1.2 | 2.25 | 1.5E-07 |
| *wht-8* | WBGene00012925 | protein_coding_gene | 1.2 | 2.25 | 5.0E-06 |
| *D1081.12* | WBGene00086554 | protein_coding_gene | 1.2 | 2.25 | 1.4E-05 |
| *C06C6.6* | WBGene00007380 | pseudogene | 1.2 | 2.25 | 9.6E-03 |
| *F44B9.10* | WBGene00018411 | protein_coding_gene | 1.2 | 2.25 | 1.2E-02 |
| *C07B5.3* | WBGene00007408 | protein_coding_gene | 1.2 | 2.24 | 6.6E-03 |
| *C15B12.8* | WBGene00015788 | protein_coding_gene | 1.2 | 2.24 | 3.4E-05 |
| *F38H4.6* | WBGene00009550 | protein_coding_gene | 1.2 | 2.24 | 1.9E-03 |
| *K08E7.5* | WBGene00010673 | protein_coding_gene | 1.2 | 2.24 | 2.8E-03 |
| *F41G3.5* | WBGene00018301 | protein_coding_gene | 1.2 | 2.24 | 7.2E-05 |
| *C28H8.7* | WBGene00016198 | protein_coding_gene | 1.2 | 2.24 | 2.2E-02 |
| *frk-1* | WBGene00001487 | protein_coding_gene | 1.2 | 2.24 | 1.9E-06 |
| *pals-33* | WBGene00021081 | protein_coding_gene | 1.2 | 2.24 | 2.2E-04 |
| *F59C6.3* | WBGene00010324 | protein_coding_gene | 1.2 | 2.24 | 7.5E-04 |
| *F36G9.17* | WBGene00077605 | pseudogene | 1.2 | 2.23 | 1.2E-02 |
| *F33D11.1* | WBGene00018000 | protein_coding_gene | 1.2 | 2.23 | 2.1E-02 |
| *fbxb-57* | WBGene00010101 | protein_coding_gene | 1.2 | 2.23 | 8.5E-03 |
| *T23E7.6* | WBGene00044772 | protein_coding_gene | 1.2 | 2.23 | 9.6E-03 |
| *T23G11.10* | WBGene00011973 | protein_coding_gene | 1.2 | 2.23 | 1.7E-02 |
| *tag-191* | WBGene00007049 | protein_coding_gene | 1.2 | 2.23 | 5.3E-04 |
| *C08F11.12* | WBGene00007459 | protein_coding_gene | 1.2 | 2.23 | 1.3E-05 |
| *F46F5.16* | WBGene00018507 | protein_coding_gene | 1.2 | 2.23 | 2.1E-02 |
| *F55G11.8* | WBGene00010128 | protein_coding_gene | 1.2 | 2.23 | 1.7E-03 |
| *T02E1.7* | WBGene00011379 | protein_coding_gene | 1.2 | 2.23 | 5.9E-05 |
| *F53B2.5* | WBGene00009956 | protein_coding_gene | 1.2 | 2.22 | 8.0E-06 |
| *oac-43* | WBGene00011653 | protein_coding_gene | 1.2 | 2.22 | 1.4E-02 |
| *gsp-3* | WBGene00021113 | protein_coding_gene | 1.2 | 2.22 | 1.1E-07 |
| *C03C10.2* | WBGene00007269 | protein_coding_gene | 1.2 | 2.22 | 1.2E-03 |
| *Y105C5B.11* | WBGene00013651 | protein_coding_gene | 1.2 | 2.22 | 6.8E-04 |
| *F17C11.11* | WBGene00008922 | protein_coding_gene | 1.2 | 2.22 | 2.8E-15 |
| *F22G12.8* | WBGene00044712 | pseudogene | 1.2 | 2.22 | 7.4E-06 |
| *F32A11.3* | WBGene00009308 | protein_coding_gene | 1.2 | 2.22 | 3.8E-10 |
| *dod-23* | WBGene00009898 | protein_coding_gene | 1.2 | 2.22 | 5.0E-30 |
| *C46H11.6* | WBGene00016730 | protein_coding_gene | 1.1 | 2.22 | 2.0E-03 |
| *F18G5.5* | WBGene00017581 | pseudogene | 1.1 | 2.21 | 2.4E-02 |
| *F32B4.2* | WBGene00009313 | protein_coding_gene | 1.1 | 2.21 | 2.7E-06 |
| *M7.12* | WBGene00010887 | protein_coding_gene | 1.1 | 2.21 | 4.1E-07 |
| *W03D8.3* | WBGene00020986 | protein_coding_gene | 1.1 | 2.21 | 8.3E-03 |
| *R08C7.5* | WBGene00019949 | protein_coding_gene | 1.1 | 2.21 | 9.5E-05 |
| *R03D7.5* | WBGene00010991 | protein_coding_gene | 1.1 | 2.21 | 1.5E-03 |
| *F28A10.4* | WBGene00017872 | protein_coding_gene | 1.1 | 2.21 | 1.6E-02 |
| *ZK930.7* | WBGene00014157 | protein_coding_gene | 1.1 | 2.21 | 1.9E-03 |
| *ptp-5.2* | WBGene00013304 | protein_coding_gene | 1.1 | 2.20 | 4.0E-05 |
| *F09C12.8* | WBGene00017279 | protein_coding_gene | 1.1 | 2.20 | 3.8E-06 |
| *R05C11.2* | WBGene00019874 | protein_coding_gene | 1.1 | 2.20 | 2.4E-02 |
| *F19B6.3* | WBGene00008949 | protein_coding_gene | 1.1 | 2.20 | 1.8E-02 |

| | | | | | |
|---|---|---|---|---|---|
| *grd-16* | WBGene00001705 | protein_coding_gene | 1.1 | 2.20 | 1.3E-02 |
| *F31E9.8* | WBGene00045408 | pseudogene | 1.1 | 2.20 | 3.7E-03 |
| *C46A5.4* | WBGene00016700 | protein_coding_gene | 1.1 | 2.20 | 2.2E-04 |
| *F26F12.2* | WBGene00017833 | protein_coding_gene | 1.1 | 2.20 | 2.1E-02 |
| *F25B3.4* | WBGene00009101 | protein_coding_gene | 1.1 | 2.20 | 1.6E-03 |
| *F44D12.8* | WBGene00009685 | protein_coding_gene | 1.1 | 2.20 | 2.6E-04 |
| *T09F5.12* | WBGene00014836 | lincRNA_gene | 1.1 | 2.20 | 1.1E-05 |
| *msrp-4* | WBGene00018500 | protein_coding_gene | 1.1 | 2.20 | 1.3E-02 |
| *comp-1* | WBGene00018158 | protein_coding_gene | 1.1 | 2.19 | 3.4E-05 |
| *K07A1.20* | WBGene00219285 | protein_coding_gene | 1.1 | 2.19 | 2.4E-02 |
| *Y53F4B.8* | WBGene00013155 | pseudogene | 1.1 | 2.19 | 7.3E-03 |
| *R01H2.4* | WBGene00019812 | protein_coding_gene | 1.1 | 2.19 | 2.9E-03 |
| *F59A7.2* | WBGene00019090 | protein_coding_gene | 1.1 | 2.19 | 4.0E-03 |
| *clec-156* | WBGene00017809 | protein_coding_gene | 1.1 | 2.18 | 3.8E-03 |
| *C06A5.2* | WBGene00015500 | protein_coding_gene | 1.1 | 2.18 | 9.6E-04 |
| *Y62H9A.13* | WBGene00013401 | protein_coding_gene | 1.1 | 2.18 | 2.5E-02 |
| *clec-68* | WBGene00018970 | protein_coding_gene | 1.1 | 2.18 | 2.5E-02 |
| *F16B3.2* | WBGene00017502 | protein_coding_gene | 1.1 | 2.18 | 1.0E-02 |
| *ugt-41* | WBGene00017336 | protein_coding_gene | 1.1 | 2.18 | 1.3E-07 |
| *cut-2* | WBGene00009983 | protein_coding_gene | 1.1 | 2.18 | 4.4E-07 |
| *sdz-30* | WBGene00011432 | protein_coding_gene | 1.1 | 2.18 | 5.6E-05 |
| *Y57G7A.6* | WBGene00021969 | protein_coding_gene | 1.1 | 2.17 | 4.8E-06 |
| *tbx-34* | WBGene00006553 | protein_coding_gene | 1.1 | 2.17 | 2.4E-04 |
| *lgc-47* | WBGene00009799 | protein_coding_gene | 1.1 | 2.17 | 2.1E-05 |
| *alg-4* | WBGene00006449 | protein_coding_gene | 1.1 | 2.17 | 1.9E-03 |
| *C25F9.8* | WBGene00007727 | protein_coding_gene | 1.1 | 2.17 | 4.3E-03 |
| *ssp-9* | WBGene00006038 | protein_coding_gene | 1.1 | 2.17 | 3.5E-03 |
| *C10C6.3* | WBGene00007512 | protein_coding_gene | 1.1 | 2.17 | 3.3E-03 |
| *F37A8.2* | WBGene00009502 | protein_coding_gene | 1.1 | 2.17 | 1.2E-03 |
| *ZC477.2* | WBGene00022617 | protein_coding_gene | 1.1 | 2.17 | 2.3E-03 |
| *C27D6.11* | WBGene00044388 | protein_coding_gene | 1.1 | 2.17 | 3.9E-05 |
| *K10D2.8* | WBGene00189952 | protein_coding_gene | 1.1 | 2.17 | 6.5E-04 |
| *B0205.13* | WBGene00044644 | protein_coding_gene | 1.1 | 2.17 | 1.7E-02 |
| *Y7A5A.11* | WBGene00044155 | pseudogene | 1.1 | 2.17 | 3.7E-03 |
| *H32C10.1* | WBGene00019255 | protein_coding_gene | 1.1 | 2.16 | 7.9E-06 |
| *Y57G11A.2* | WBGene00013290 | protein_coding_gene | 1.1 | 2.16 | 9.0E-04 |
| *F12E12.6* | WBGene00017407 | protein_coding_gene | 1.1 | 2.16 | 1.0E-02 |
| *cwp-3* | WBGene00000861 | protein_coding_gene | 1.1 | 2.16 | 1.5E-02 |
| *F42G8.8* | WBGene00018359 | protein_coding_gene | 1.1 | 2.16 | 4.5E-06 |
| *ttbk-4* | WBGene00012169 | protein_coding_gene | 1.1 | 2.16 | 2.5E-06 |
| *ZK1248.20* | WBGene00044393 | protein_coding_gene | 1.1 | 2.16 | 6.5E-03 |
| *C27D8.1* | WBGene00007777 | protein_coding_gene | 1.1 | 2.16 | 2.9E-07 |
| *R08A2.1* | WBGene00011132 | protein_coding_gene | 1.1 | 2.16 | 2.1E-05 |
| *Y37D8A.8* | WBGene00012549 | protein_coding_gene | 1.1 | 2.16 | 5.3E-04 |
| *C17C3.9* | WBGene00015892 | protein_coding_gene | 1.1 | 2.16 | 2.3E-02 |
| *B0280.2* | WBGene00015100 | protein_coding_gene | 1.1 | 2.16 | 2.9E-03 |
| *ZC581.10* | WBGene00195178 | protein_coding_gene | 1.1 | 2.16 | 1.5E-03 |
| *pgp-12* | WBGene00004006 | protein_coding_gene | 1.1 | 2.16 | 4.4E-03 |
| *nspc-10* | WBGene00009640 | protein_coding_gene | 1.1 | 2.15 | 1.2E-12 |
| *T16G12.7* | WBGene00011808 | protein_coding_gene | 1.1 | 2.15 | 6.9E-04 |
| *glct-3* | WBGene00011781 | protein_coding_gene | 1.1 | 2.15 | 2.7E-02 |
| *C47E12.11* | WBGene00008154 | protein_coding_gene | 1.1 | 2.15 | 1.4E-04 |
| *T05G5.4* | WBGene00011499 | protein_coding_gene | 1.1 | 2.15 | 3.3E-03 |
| *msp-152* | WBGene00003470 | protein_coding_gene | 1.1 | 2.15 | 5.6E-03 |
| *skpo-2* | WBGene00015841 | protein_coding_gene | 1.1 | 2.15 | 4.1E-04 |
| *gsp-4* | WBGene00020187 | protein_coding_gene | 1.1 | 2.15 | 6.1E-09 |
| *ZK1128.3* | WBGene00014229 | protein_coding_gene | 1.1 | 2.15 | 1.9E-04 |
| *ZK84.2* | WBGene00022650 | protein_coding_gene | 1.1 | 2.15 | 1.4E-04 |
| *ssp-11* | WBGene00006040 | protein_coding_gene | 1.1 | 2.15 | 8.2E-03 |
| *C35D6.5* | WBGene00007962 | pseudogene | 1.1 | 2.15 | 1.1E-03 |

| | | | | | |
|---|---|---|---|---|---|
| *C30A5.10* | WBGene00016242 | protein_coding_gene | 1.1 | 2.15 | 2.0E-02 |
| *clec-123* | WBGene00021288 | protein_coding_gene | 1.1 | 2.15 | 2.5E-06 |
| *argk-1* | WBGene00009706 | protein_coding_gene | 1.1 | 2.15 | 1.9E-02 |
| *K08F8.5* | WBGene00010682 | protein_coding_gene | 1.1 | 2.15 | 4.5E-04 |
| *C14A6.10* | WBGene00045348 | pseudogene | 1.1 | 2.14 | 2.5E-02 |
| *msp-78* | WBGene00003465 | protein_coding_gene | 1.1 | 2.14 | 3.7E-05 |
| *Y71G12B.33* | WBGene00044358 | protein_coding_gene | 1.1 | 2.14 | 3.1E-02 |
| *C18F10.2* | WBGene00015977 | protein_coding_gene | 1.1 | 2.14 | 4.1E-04 |
| *F46F5.7* | WBGene00018498 | protein_coding_gene | 1.1 | 2.14 | 5.5E-03 |
| *col-168* | WBGene00000741 | protein_coding_gene | 1.1 | 2.14 | 1.5E-02 |
| *F38E1.3* | WBGene00018178 | protein_coding_gene | 1.1 | 2.14 | 1.1E-03 |
| *srr-3* | WBGene00005654 | protein_coding_gene | 1.1 | 2.14 | 2.8E-02 |
| *ssp-16* | WBGene00006044 | protein_coding_gene | 1.1 | 2.14 | 1.6E-02 |
| *C17H12.5* | WBGene00015931 | protein_coding_gene | 1.1 | 2.14 | 7.5E-06 |
| *ZK105.14* | WBGene00271707 | protein_coding_gene | 1.1 | 2.13 | 4.8E-03 |
| *F13A7.1* | WBGene00008724 | protein_coding_gene | 1.1 | 2.13 | 5.7E-06 |
| *W05B10.3* | WBGene00012278 | protein_coding_gene | 1.1 | 2.13 | 2.9E-02 |
| *C15H7.2* | WBGene00014678 | pseudogene | 1.1 | 2.13 | 2.7E-02 |
| *alg-3* | WBGene00011910 | protein_coding_gene | 1.1 | 2.13 | 2.1E-03 |
| *ZK180.5* | WBGene00022679 | protein_coding_gene | 1.1 | 2.13 | 3.8E-10 |
| *msd-4* | WBGene00016447 | protein_coding_gene | 1.1 | 2.13 | 8.8E-04 |
| *C53D6.5* | WBGene00008282 | pseudogene | 1.1 | 2.13 | 3.2E-02 |
| *F27C8.5* | WBGene00009185 | protein_coding_gene | 1.1 | 2.13 | 1.4E-04 |
| *dmsr-3* | WBGene00012993 | protein_coding_gene | 1.1 | 2.13 | 9.2E-03 |
| *cnnm-5* | WBGene00011260 | protein_coding_gene | 1.1 | 2.13 | 5.5E-03 |
| *cwp-5* | WBGene00009844 | protein_coding_gene | 1.1 | 2.13 | 3.4E-03 |
| *Y65B4A.7* | WBGene00022030 | protein_coding_gene | 1.1 | 2.13 | 1.3E-02 |
| *C55B7.10* | WBGene00016946 | protein_coding_gene | 1.1 | 2.13 | 2.0E-03 |
| *wrt-1* | WBGene00006947 | protein_coding_gene | 1.1 | 2.13 | 9.7E-07 |
| *F57F4.1* | WBGene00019015 | protein_coding_gene | 1.1 | 2.13 | 6.0E-06 |
| *ttbk-3* | WBGene00009324 | protein_coding_gene | 1.1 | 2.12 | 2.0E-04 |
| *Y6E2A.4* | WBGene00012398 | protein_coding_gene | 1.1 | 2.12 | 2.2E-03 |
| *F22B7.3* | WBGene00017694 | protein_coding_gene | 1.1 | 2.12 | 5.1E-03 |
| *B0218.5* | WBGene00015049 | protein_coding_gene | 1.1 | 2.12 | 8.1E-04 |
| *C09H5.7* | WBGene00015661 | protein_coding_gene | 1.1 | 2.12 | 4.9E-03 |
| *Y57G11B.3* | WBGene00013296 | protein_coding_gene | 1.1 | 2.12 | 5.0E-03 |
| *srb-3* | WBGene00005068 | protein_coding_gene | 1.1 | 2.12 | 2.9E-02 |
| *K11C4.1* | WBGene00019642 | protein_coding_gene | 1.1 | 2.12 | 1.9E-04 |
| *clec-236* | WBGene00013495 | pseudogene | 1.1 | 2.12 | 1.3E-02 |
| *abf-2* | WBGene00000013 | protein_coding_gene | 1.1 | 2.12 | 3.1E-03 |
| *Y23H5B.2* | WBGene00021273 | pseudogene | 1.1 | 2.12 | 2.0E-02 |
| *spe-11* | WBGene00004965 | protein_coding_gene | 1.1 | 2.12 | 1.2E-06 |
| *R07H5.9* | WBGene00011129 | protein_coding_gene | 1.1 | 2.12 | 3.5E-03 |
| *K05F1.1* | WBGene00019405 | protein_coding_gene | 1.1 | 2.12 | 1.3E-02 |
| *nhr-175* | WBGene00008630 | protein_coding_gene | 1.1 | 2.12 | 3.3E-02 |
| *M01D1.11* | WBGene00220026 | snoRNA_gene | 1.1 | 2.11 | 3.3E-02 |
| *Y6E2A.5* | WBGene00012399 | protein_coding_gene | 1.1 | 2.11 | 6.7E-03 |
| *pck-3* | WBGene00019151 | protein_coding_gene | 1.1 | 2.11 | 1.4E-05 |
| *cpb-2* | WBGene00000771 | protein_coding_gene | 1.1 | 2.11 | 5.5E-05 |
| *pals-7* | WBGene00007661 | protein_coding_gene | 1.1 | 2.11 | 3.3E-02 |
| *F58H1.6* | WBGene00010289 | protein_coding_gene | 1.1 | 2.11 | 8.4E-06 |
| *F07F6.1* | WBGene00017215 | protein_coding_gene | 1.1 | 2.11 | 1.0E-03 |
| *msp-59* | WBGene00003452 | protein_coding_gene | 1.1 | 2.11 | 1.3E-06 |
| *C55C3.4* | WBGene00016954 | protein_coding_gene | 1.1 | 2.11 | 2.6E-04 |
| *linc-23* | WBGene00219692 | lincRNA_gene | 1.1 | 2.11 | 1.8E-02 |
| *col-144* | WBGene00000717 | protein_coding_gene | 1.1 | 2.11 | 1.6E-02 |
| *ZC581.7* | WBGene00022634 | protein_coding_gene | 1.1 | 2.11 | 1.6E-03 |
| *C38C10.3* | WBGene00008001 | protein_coding_gene | 1.1 | 2.11 | 9.1E-04 |
| *swt-5* | WBGene00010594 | protein_coding_gene | 1.1 | 2.11 | 2.6E-02 |
| *nkb-2* | WBGene00008074 | protein_coding_gene | 1.1 | 2.11 | 1.2E-05 |

| | | | | | |
|---|---|---|---|---|---|
| *msp-76* | WBGene00003463 | protein_coding_gene | 1.1 | 2.11 | 1.5E-07 |
| *W07A12.4* | WBGene00012322 | protein_coding_gene | 1.1 | 2.11 | 7.7E-04 |
| *hdl-1* | WBGene00001839 | protein_coding_gene | 1.1 | 2.10 | 9.6E-05 |
| *C15A11.2* | WBGene00007597 | protein_coding_gene | 1.1 | 2.10 | 1.2E-03 |
| *acs-6* | WBGene00022849 | protein_coding_gene | 1.1 | 2.10 | 2.1E-05 |
| *rab-3* | WBGene00004267 | protein_coding_gene | 1.1 | 2.10 | 3.7E-13 |
| *C17G10.3* | WBGene00015917 | protein_coding_gene | 1.1 | 2.10 | 1.1E-03 |
| *snf-2* | WBGene00004901 | protein_coding_gene | 1.1 | 2.10 | 1.4E-04 |
| *Y53F4B.24* | WBGene00013170 | protein_coding_gene | 1.1 | 2.10 | 6.7E-03 |
| *ZC443.2* | WBGene00013897 | protein_coding_gene | 1.1 | 2.10 | 1.9E-02 |
| *F23C8.8* | WBGene00017737 | protein_coding_gene | 1.1 | 2.10 | 2.8E-03 |
| *F23F12.12* | WBGene00017756 | protein_coding_gene | 1.1 | 2.10 | 2.5E-02 |
| *T22B3.3* | WBGene00011911 | protein_coding_gene | 1.1 | 2.10 | 1.9E-07 |
| *R03D7.8* | WBGene00010992 | protein_coding_gene | 1.1 | 2.10 | 1.4E-03 |
| *F31E8.5* | WBGene00017955 | protein_coding_gene | 1.1 | 2.10 | 2.9E-05 |
| *Y7A5A.21* | WBGene00219352 | pseudogene | 1.1 | 2.10 | 3.6E-02 |
| *clp-10* | WBGene00021041 | protein_coding_gene | 1.1 | 2.10 | 3.3E-02 |
| *fbxb-68* | WBGene00008611 | protein_coding_gene | 1.1 | 2.10 | 1.9E-02 |
| *F21C10.9* | WBGene00017658 | protein_coding_gene | 1.1 | 2.09 | 9.5E-12 |
| *phf-33* | WBGene00010708 | protein_coding_gene | 1.1 | 2.09 | 2.9E-02 |
| *Y71F9B.9* | WBGene00022130 | protein_coding_gene | 1.1 | 2.09 | 2.3E-06 |
| *K04F1.8* | WBGene00019387 | protein_coding_gene | 1.1 | 2.09 | 5.8E-03 |
| *C45G9.4* | WBGene00016675 | protein_coding_gene | 1.1 | 2.09 | 2.1E-03 |
| *F13E9.5* | WBGene00008754 | protein_coding_gene | 1.1 | 2.09 | 2.8E-04 |
| *C32D5.4* | WBGene00016312 | protein_coding_gene | 1.1 | 2.09 | 1.2E-03 |
| *mam-3* | WBGene00010714 | protein_coding_gene | 1.1 | 2.09 | 6.9E-03 |
| *oac-9* | WBGene00016753 | protein_coding_gene | 1.1 | 2.09 | 2.3E-05 |
| *K03H1.9* | WBGene00010544 | protein_coding_gene | 1.1 | 2.09 | 6.9E-03 |
| *C18E3.1* | WBGene00015970 | protein_coding_gene | 1.1 | 2.09 | 1.9E-03 |
| *str-163* | WBGene00006208 | protein_coding_gene | 1.1 | 2.09 | 3.2E-02 |
| *Y38E10A.3* | WBGene00012581 | protein_coding_gene | 1.1 | 2.08 | 8.0E-03 |
| *ttr-56* | WBGene00050896 | protein_coding_gene | 1.1 | 2.08 | 2.0E-02 |
| *H10E21.4* | WBGene00019184 | protein_coding_gene | 1.1 | 2.08 | 7.5E-04 |
| *msrp-5* | WBGene00017059 | protein_coding_gene | 1.1 | 2.08 | 1.0E-04 |
| *acs-18* | WBGene00011173 | protein_coding_gene | 1.1 | 2.08 | 4.1E-06 |
| *Y4C6A.1* | WBGene00021151 | pseudogene | 1.1 | 2.08 | 1.4E-02 |
| *T04F3.3* | WBGene00011438 | protein_coding_gene | 1.1 | 2.08 | 4.0E-04 |
| *Y41C4A.18* | WBGene00012761 | protein_coding_gene | 1.1 | 2.08 | 3.0E-03 |
| *kin-5* | WBGene00002193 | protein_coding_gene | 1.1 | 2.08 | 1.8E-04 |
| *spe-49* | WBGene00019667 | protein_coding_gene | 1.1 | 2.08 | 5.4E-03 |
| *W08G11.1* | WBGene00012346 | protein_coding_gene | 1.1 | 2.08 | 1.3E-02 |
| *kvs-5* | WBGene00021948 | protein_coding_gene | 1.1 | 2.08 | 1.7E-06 |
| *C49C8.1* | WBGene00016765 | protein_coding_gene | 1.1 | 2.08 | 4.2E-03 |
| *R04B5.11* | WBGene00011008 | protein_coding_gene | 1.1 | 2.08 | 2.8E-02 |
| *twk-6* | WBGene00006661 | protein_coding_gene | 1.1 | 2.07 | 2.0E-02 |
| *F45C12.9* | WBGene00018440 | protein_coding_gene | 1.1 | 2.07 | 2.1E-02 |
| *M163.8* | WBGene00010940 | protein_coding_gene | 1.1 | 2.07 | 1.8E-06 |
| *T05A8.7* | WBGene00020233 | protein_coding_gene | 1.1 | 2.07 | 3.8E-02 |
| *klp-17* | WBGene00002227 | protein_coding_gene | 1.1 | 2.07 | 2.4E-03 |
| *R12E2.15* | WBGene00020040 | protein_coding_gene | 1.1 | 2.07 | 2.2E-05 |
| *F30A10.2* | WBGene00009261 | protein_coding_gene | 1.1 | 2.07 | 2.2E-03 |
| *best-11* | WBGene00008186 | protein_coding_gene | 1.1 | 2.07 | 1.4E-02 |
| *sfxn-1.2* | WBGene00009513 | protein_coding_gene | 1.1 | 2.07 | 3.9E-03 |
| *Y106G6A.4* | WBGene00013696 | protein_coding_gene | 1.1 | 2.07 | 2.3E-06 |
| *F41H10.5* | WBGene00018318 | protein_coding_gene | 1.0 | 2.07 | 3.3E-03 |
| *Y59E9AL.3* | WBGene00021993 | protein_coding_gene | 1.0 | 2.07 | 7.3E-04 |
| *F42C5.5* | WBGene00018347 | protein_coding_gene | 1.0 | 2.07 | 3.3E-04 |
| *Y40H4A.2* | WBGene00012741 | protein_coding_gene | 1.0 | 2.07 | 1.4E-03 |
| *W06D12.6* | WBGene00012305 | protein_coding_gene | 1.0 | 2.07 | 6.5E-03 |
| *Y73F8A.20* | WBGene00013526 | protein_coding_gene | 1.0 | 2.07 | 4.6E-04 |

| | | | | | |
|---|---|---|---|---|---|
| *ttr-9* | WBGene00007888 | protein_coding_gene | 1.0 | 2.07 | 1.3E-02 |
| *math-37* | WBGene00020080 | protein_coding_gene | 1.0 | 2.06 | 1.3E-04 |
| *ugt-42* | WBGene00017959 | protein_coding_gene | 1.0 | 2.06 | 1.8E-02 |
| *K09E4.1* | WBGene00010719 | protein_coding_gene | 1.0 | 2.06 | 5.4E-04 |
| *Y24D9B.1* | WBGene00021287 | protein_coding_gene | 1.0 | 2.06 | 2.8E-03 |
| *F33C8.4* | WBGene00009358 | protein_coding_gene | 1.0 | 2.06 | 2.3E-08 |
| *C26C6.6* | WBGene00007740 | protein_coding_gene | 1.0 | 2.06 | 1.7E-02 |
| *T09B4.3* | WBGene00020377 | protein_coding_gene | 1.0 | 2.06 | 3.5E-03 |
| *wht-6* | WBGene00007060 | protein_coding_gene | 1.0 | 2.06 | 4.7E-05 |
| *F58D5.8* | WBGene00010247 | protein_coding_gene | 1.0 | 2.06 | 1.3E-02 |
| *phat-4* | WBGene00020237 | protein_coding_gene | 1.0 | 2.06 | 7.6E-06 |
| *clec-97* | WBGene00013931 | protein_coding_gene | 1.0 | 2.06 | 7.6E-07 |
| *C28A5.6* | WBGene00007791 | protein_coding_gene | 1.0 | 2.06 | 3.4E-08 |
| *F57A8.7* | WBGene00010182 | protein_coding_gene | 1.0 | 2.06 | 2.1E-02 |
| *C24H12.2* | WBGene00016072 | protein_coding_gene | 1.0 | 2.06 | 3.8E-02 |
| *mboa-1* | WBGene00007174 | protein_coding_gene | 1.0 | 2.06 | 5.4E-03 |
| *rnh-1.1* | WBGene00022888 | protein_coding_gene | 1.0 | 2.06 | 3.2E-04 |
| *ZK970.8* | WBGene00014174 | protein_coding_gene | 1.0 | 2.06 | 1.4E-03 |
| *Y37E11AL.2* | WBGene00021358 | protein_coding_gene | 1.0 | 2.06 | 3.9E-05 |
| *clec-74* | WBGene00021585 | protein_coding_gene | 1.0 | 2.06 | 4.2E-02 |
| *F59A1.2* | WBGene00014779 | pseudogene | 1.0 | 2.06 | 6.1E-03 |
| *spe-10* | WBGene00004964 | protein_coding_gene | 1.0 | 2.06 | 2.1E-03 |
| *C36F7.5* | WBGene00007986 | protein_coding_gene | 1.0 | 2.06 | 4.1E-04 |
| *M05B5.1* | WBGene00010869 | protein_coding_gene | 1.0 | 2.05 | 3.1E-04 |
| *spe-9* | WBGene00004963 | protein_coding_gene | 1.0 | 2.05 | 6.5E-03 |
| *Y80D3A.8* | WBGene00013586 | protein_coding_gene | 1.0 | 2.05 | 1.6E-02 |
| *ZK616.65* | WBGene00235383 | protein_coding_gene | 1.0 | 2.05 | 3.6E-03 |
| *ttr-29* | WBGene00011286 | protein_coding_gene | 1.0 | 2.05 | 1.7E-03 |
| *ssp-33* | WBGene00011134 | protein_coding_gene | 1.0 | 2.05 | 4.2E-03 |
| *Y73F8A.15* | WBGene00013524 | protein_coding_gene | 1.0 | 2.05 | 9.0E-03 |
| *nhr-198* | WBGene00010603 | protein_coding_gene | 1.0 | 2.05 | 2.2E-02 |
| *R05G9R.1* | WBGene00019906 | protein_coding_gene | 1.0 | 2.05 | 5.0E-04 |
| *fbxa-42* | WBGene00044676 | protein_coding_gene | 1.0 | 2.05 | 1.0E-02 |
| *C05C12.1* | WBGene00007335 | protein_coding_gene | 1.0 | 2.05 | 3.7E-04 |
| *fbxa-88* | WBGene00008632 | protein_coding_gene | 1.0 | 2.05 | 3.5E-02 |
| *H05L14.1* | WBGene00010366 | protein_coding_gene | 1.0 | 2.05 | 3.1E-07 |
| *Y71G12B.3* | WBGene00022143 | protein_coding_gene | 1.0 | 2.05 | 1.3E-03 |
| *C28D4.8* | WBGene00007795 | protein_coding_gene | 1.0 | 2.05 | 1.2E-02 |
| *msp-38* | WBGene00003434 | protein_coding_gene | 1.0 | 2.04 | 1.8E-04 |
| *F41C3.2* | WBGene00018268 | protein_coding_gene | 1.0 | 2.04 | 1.4E-03 |
| *vap-1* | WBGene00006886 | protein_coding_gene | 1.0 | 2.04 | 2.3E-11 |
| *C32E12.1* | WBGene00016327 | protein_coding_gene | 1.0 | 2.04 | 1.1E-05 |
| *tsp-18* | WBGene00006644 | protein_coding_gene | 1.0 | 2.04 | 2.0E-02 |
| *Y53C12A.8* | WBGene00044158 | protein_coding_gene | 1.0 | 2.04 | 2.1E-02 |
| *ZK795.2* | WBGene00014082 | protein_coding_gene | 1.0 | 2.04 | 1.9E-05 |
| *C18A3.7* | WBGene00015944 | protein_coding_gene | 1.0 | 2.04 | 2.4E-03 |
| *fbxa-166* | WBGene00015601 | protein_coding_gene | 1.0 | 2.04 | 3.9E-02 |
| *F40H6.1* | WBGene00018253 | protein_coding_gene | 1.0 | 2.04 | 5.0E-05 |
| *Y6E2A.1* | WBGene00012397 | protein_coding_gene | 1.0 | 2.04 | 7.2E-03 |
| *F10A3.1* | WBGene00008631 | protein_coding_gene | 1.0 | 2.04 | 7.2E-03 |
| *nhr-251* | WBGene00022756 | protein_coding_gene | 1.0 | 2.04 | 2.2E-02 |
| *C35A11.2* | WBGene00016429 | protein_coding_gene | 1.0 | 2.04 | 1.9E-02 |
| *oac-56* | WBGene00013450 | protein_coding_gene | 1.0 | 2.04 | 4.2E-02 |
| *T19H5.6* | WBGene00044807 | protein_coding_gene | 1.0 | 2.04 | 2.5E-03 |
| *R02E4.3* | WBGene00044724 | protein_coding_gene | 1.0 | 2.04 | 1.0E-02 |
| *kin-26* | WBGene00002208 | protein_coding_gene | 1.0 | 2.03 | 1.4E-03 |
| *Y20C6A.1* | WBGene00012491 | protein_coding_gene | 1.0 | 2.03 | 9.2E-08 |
| *Y53F4B.17* | WBGene00013163 | protein_coding_gene | 1.0 | 2.03 | 6.7E-03 |
| *F57A8.6* | WBGene00010181 | protein_coding_gene | 1.0 | 2.03 | 3.7E-03 |
| *T15H9.4* | WBGene00011789 | protein_coding_gene | 1.0 | 2.03 | 2.4E-02 |

| | | | | | |
|---|---|---|---|---|---|
| *F55A12.6* | WBGene00018865 | protein_coding_gene | 1.0 | 2.03 | 9.0E-03 |
| *F59A6.4* | WBGene00019086 | protein_coding_gene | 1.0 | 2.03 | 8.7E-05 |
| *Y102A5C.38* | WBGene00050883 | pseudogene | 1.0 | 2.03 | 7.6E-03 |
| *T16A1.2* | WBGene00020533 | protein_coding_gene | 1.0 | 2.03 | 1.5E-02 |
| *T16A9.5* | WBGene00011795 | protein_coding_gene | 1.0 | 2.03 | 1.6E-05 |
| *ZK945.6* | WBGene00014168 | protein_coding_gene | 1.0 | 2.03 | 1.6E-03 |
| *gst-32* | WBGene00001780 | protein_coding_gene | 1.0 | 2.03 | 4.6E-02 |
| *clec-109* | WBGene00008907 | protein_coding_gene | 1.0 | 2.03 | 4.5E-02 |
| *AC3.9* | WBGene00044022 | pseudogene | 1.0 | 2.03 | 4.7E-02 |
| *nep-23* | WBGene00013785 | protein_coding_gene | 1.0 | 2.03 | 2.2E-04 |
| *lipl-2* | WBGene00009773 | protein_coding_gene | 1.0 | 2.03 | 1.5E-02 |
| *Y41E3.461* | WBGene00219444 | ncRNA_gene | 1.0 | 2.03 | 4.6E-02 |
| *bus-8* | WBGene00044623 | protein_coding_gene | 1.0 | 2.03 | 6.8E-05 |
| *gst-33* | WBGene00001781 | protein_coding_gene | 1.0 | 2.03 | 2.5E-03 |
| *cbs-2* | WBGene00018783 | protein_coding_gene | 1.0 | 2.02 | 2.0E-03 |
| *M04G7.2* | WBGene00019773 | protein_coding_gene | 1.0 | 2.02 | 2.3E-02 |
| *ate-1* | WBGene00010615 | protein_coding_gene | 1.0 | 2.02 | 1.2E-25 |
| *snf-5* | WBGene00004904 | protein_coding_gene | 1.0 | 2.02 | 6.3E-03 |
| *Y119C1B.1* | WBGene00022467 | protein_coding_gene | 1.0 | 2.02 | 3.0E-03 |
| *Y39A1A.2* | WBGene00012642 | protein_coding_gene | 1.0 | 2.02 | 8.2E-03 |
| *glna-1* | WBGene00007480 | protein_coding_gene | 1.0 | 2.02 | 4.9E-04 |
| *C09D4.9* | WBGene00206357 | protein_coding_gene | 1.0 | 2.02 | 1.6E-02 |
| *Y7A5A.20* | WBGene00219351 | protein_coding_gene | 1.0 | 2.02 | 4.1E-02 |
| *Y67D8B.5* | WBGene00022064 | protein_coding_gene | 1.0 | 2.02 | 4.5E-02 |
| *F11C7.2* | WBGene00017376 | protein_coding_gene | 1.0 | 2.02 | 3.7E-07 |
| *snf-4* | WBGene00004903 | pseudogene | 1.0 | 2.01 | 1.5E-02 |
| *F55B11.5* | WBGene00010087 | protein_coding_gene | 1.0 | 2.01 | 9.9E-03 |
| *clec-6* | WBGene00016451 | protein_coding_gene | 1.0 | 2.01 | 4.4E-02 |
| *T03F6.6* | WBGene00011401 | protein_coding_gene | 1.0 | 2.01 | 4.1E-05 |
| *BE10.1* | WBGene00007209 | protein_coding_gene | 1.0 | 2.01 | 3.2E-03 |
| *R13H4.8* | WBGene00011266 | protein_coding_gene | 1.0 | 2.01 | 4.7E-02 |
| *oac-20* | WBGene00009488 | protein_coding_gene | 1.0 | 2.01 | 3.1E-02 |
| *tor-2* | WBGene00006598 | protein_coding_gene | 1.0 | 2.01 | 4.5E-02 |
| *F13A2.2* | WBGene00017411 | protein_coding_gene | 1.0 | 2.01 | 4.5E-02 |
| *oig-2* | WBGene00003860 | protein_coding_gene | 1.0 | 2.01 | 5.5E-05 |
| *Y71G12B.18* | WBGene00022156 | protein_coding_gene | 1.0 | 2.01 | 4.3E-02 |
| *F13H8.5* | WBGene00017438 | protein_coding_gene | 1.0 | 2.01 | 4.3E-04 |
| *T16G1.13* | WBGene00194787 | protein_coding_gene | 1.0 | 2.01 | 1.3E-02 |
| *Y54G2A.20* | WBGene00021885 | pseudogene | 1.0 | 2.01 | 4.4E-02 |
| *nhr-113* | WBGene00003703 | protein_coding_gene | 1.0 | 2.01 | 4.4E-02 |
| *F35E2.9* | WBGene00009416 | protein_coding_gene | 1.0 | 2.01 | 5.7E-05 |
| *abu-12* | WBGene00004127 | protein_coding_gene | 1.0 | 2.01 | 1.5E-03 |
| *Y116A8C.24* | WBGene00013801 | protein_coding_gene | 1.0 | 2.00 | 2.6E-02 |
| *B0238.13* | WBGene00015077 | pseudogene | 1.0 | 2.00 | 4.6E-02 |
| *T28B8.6* | WBGene00044122 | protein_coding_gene | 1.0 | 2.00 | 2.3E-03 |
| *C17B7.2* | WBGene00015876 | pseudogene | 1.0 | 2.00 | 4.1E-02 |
| *F36A2.11* | WBGene00009458 | protein_coding_gene | 1.0 | 2.00 | 4.1E-03 |
| *F58E6.13* | WBGene00077697 | protein_coding_gene | 1.0 | 2.00 | 4.8E-04 |
| *K10G4.10* | WBGene00044202 | protein_coding_gene | -1.0 | 0.50 | 8.8E-03 |
| *T08E11.8* | WBGene00020364 | protein_coding_gene | -1.0 | 0.50 | 7.8E-03 |
| *fbxa-4* | WBGene00044672 | protein_coding_gene | -1.0 | 0.49 | 2.7E-03 |
| *fbxb-84* | WBGene00020544 | protein_coding_gene | -1.0 | 0.49 | 1.8E-02 |
| *skr-13* | WBGene00004819 | protein_coding_gene | -1.0 | 0.49 | 5.8E-04 |
| *Y49F6A.5* | WBGene00021713 | protein_coding_gene | -1.0 | 0.49 | 4.8E-02 |
| *acs-2* | WBGene00009221 | protein_coding_gene | -1.0 | 0.49 | 3.1E-03 |
| *Y69A2AR.8* | WBGene00022079 | pseudogene | -1.0 | 0.49 | 1.7E-02 |
| *fbxa-93* | WBGene00012368 | pseudogene | -1.0 | 0.49 | 6.0E-04 |
| *ncx-6* | WBGene00003571 | protein_coding_gene | -1.0 | 0.49 | 7.5E-06 |
| *his-48* | WBGene00001922 | protein_coding_gene | -1.0 | 0.49 | 7.5E-03 |
| *asp-12* | WBGene00017678 | protein_coding_gene | -1.0 | 0.49 | 1.9E-03 |

| | | | | | |
|---|---|---|---|---|---|
| *trm-2A* | WBGene00019241 | protein_coding_gene | -1.0 | 0.49 | 1.8E-12 |
| *C33C12.11* | WBGene00044360 | protein_coding_gene | -1.0 | 0.49 | 2.3E-02 |
| *C02G6.1* | WBGene00015359 | pseudogene | -1.0 | 0.49 | 5.3E-03 |
| *pals-17* | WBGene00022346 | protein_coding_gene | -1.0 | 0.49 | 9.9E-05 |
| *fbxa-185* | WBGene00009836 | protein_coding_gene | -1.0 | 0.49 | 7.7E-03 |
| *Y57G11C.51* | WBGene00044258 | protein_coding_gene | -1.0 | 0.49 | 8.4E-06 |
| *fip-5* | WBGene00009620 | protein_coding_gene | -1.0 | 0.49 | 1.9E-03 |
| *irld-27* | WBGene00077751 | protein_coding_gene | -1.0 | 0.49 | 7.6E-03 |
| *glc-1* | WBGene00001591 | protein_coding_gene | -1.0 | 0.49 | 3.1E-12 |
| *R11A8.1* | WBGene00011236 | protein_coding_gene | -1.0 | 0.49 | 5.4E-13 |
| *nhr-65* | WBGene00003655 | protein_coding_gene | -1.0 | 0.49 | 1.1E-03 |
| *T08E11.1* | WBGene00020357 | protein_coding_gene | -1.0 | 0.48 | 3.7E-02 |
| *nlp-31* | WBGene00003769 | protein_coding_gene | -1.0 | 0.48 | 6.8E-10 |
| *csn-4* | WBGene00000816 | protein_coding_gene | -1.0 | 0.48 | 2.9E-16 |
| *fbxb-108* | WBGene00017249 | protein_coding_gene | -1.1 | 0.48 | 3.7E-02 |
| *F41B5.1* | WBGene00023284 | protein_coding_gene | -1.1 | 0.48 | 2.4E-04 |
| *Y37A1B.5* | WBGene00012538 | protein_coding_gene | -1.1 | 0.48 | 5.6E-18 |
| *hil-2* | WBGene00001853 | protein_coding_gene | -1.1 | 0.48 | 2.3E-09 |
| *T28B8.4* | WBGene00012115 | protein_coding_gene | -1.1 | 0.48 | 3.7E-03 |
| *fbxa-29* | WBGene00013836 | protein_coding_gene | -1.1 | 0.48 | 1.9E-03 |
| *F57F4.2* | WBGene00019016 | protein_coding_gene | -1.1 | 0.48 | 3.4E-02 |
| *Y48D7A.1* | WBGene00021656 | protein_coding_gene | -1.1 | 0.48 | 6.5E-03 |
| *linc-72* | WBGene00219590 | lincRNA_gene | -1.1 | 0.48 | 3.2E-02 |
| *Y73B3A.1* | WBGene00022203 | protein_coding_gene | -1.1 | 0.48 | 5.7E-11 |
| *eri-5* | WBGene00021419 | protein_coding_gene | -1.1 | 0.48 | 5.0E-10 |
| *clec-75* | WBGene00021586 | protein_coding_gene | -1.1 | 0.48 | 1.1E-02 |
| *K02F6.3* | WBGene00019337 | protein_coding_gene | -1.1 | 0.48 | 7.8E-05 |
| *T09F5.10* | WBGene00011669 | protein_coding_gene | -1.1 | 0.48 | 2.0E-02 |
| *fbxa-108* | WBGene00010293 | protein_coding_gene | -1.1 | 0.48 | 2.7E-08 |
| *acs-21* | WBGene00016849 | protein_coding_gene | -1.1 | 0.48 | 1.0E-04 |
| *cdr-4* | WBGene00010470 | protein_coding_gene | -1.1 | 0.47 | 1.9E-12 |
| *thn-2* | WBGene00009215 | protein_coding_gene | -1.1 | 0.47 | 3.9E-17 |
| *nhr-123* | WBGene00003713 | protein_coding_gene | -1.1 | 0.47 | 2.8E-02 |
| *cest-1.1* | WBGene00011362 | protein_coding_gene | -1.1 | 0.47 | 3.4E-10 |
| *F49F1.7* | WBGene00018647 | protein_coding_gene | -1.1 | 0.47 | 2.0E-02 |
| *ctc-1* | WBGene00010964 | protein_coding_gene | -1.1 | 0.47 | 9.4E-21 |
| *D1044.7* | WBGene00017032 | protein_coding_gene | -1.1 | 0.47 | 9.9E-03 |
| *clec-225* | WBGene00007152 | protein_coding_gene | -1.1 | 0.47 | 2.2E-02 |
| *Y18D10A.22* | WBGene00014864 | protein_coding_gene | -1.1 | 0.47 | 1.8E-03 |
| *Y54G11A.19* | WBGene00235256 | protein_coding_gene | -1.1 | 0.47 | 6.9E-03 |
| *F10C1.8* | WBGene00017327 | protein_coding_gene | -1.1 | 0.47 | 1.8E-03 |
| *Y105C5B.25* | WBGene00013662 | protein_coding_gene | -1.1 | 0.47 | 3.1E-02 |
| *oac-54* | WBGene00012323 | protein_coding_gene | -1.1 | 0.47 | 2.0E-02 |
| *F59H6.3* | WBGene00019134 | pseudogene | -1.1 | 0.47 | 4.6E-03 |
| *C29F3.7* | WBGene00007807 | protein_coding_gene | -1.1 | 0.47 | 5.7E-11 |
| *B0250.12* | WBGene00050927 | pseudogene | -1.1 | 0.47 | 3.5E-03 |
| *Y8A9A.2* | WBGene00021171 | protein_coding_gene | -1.1 | 0.46 | 6.1E-04 |
| *pals-22* | WBGene00016216 | protein_coding_gene | -1.1 | 0.46 | 2.5E-03 |
| *zipt-1* | WBGene00009286 | protein_coding_gene | -1.1 | 0.46 | 1.5E-34 |
| *M162.12* | WBGene00050946 | pseudogene | -1.1 | 0.46 | 4.7E-03 |
| *nape-1* | WBGene00021371 | protein_coding_gene | -1.1 | 0.46 | 7.9E-07 |
| *K03D3.2* | WBGene00010528 | protein_coding_gene | -1.1 | 0.46 | 4.1E-03 |
| *fbxa-199* | WBGene00005931 | protein_coding_gene | -1.1 | 0.46 | 4.1E-03 |
| *lsl-1* | WBGene00009937 | protein_coding_gene | -1.1 | 0.46 | 1.5E-16 |
| *twk-34* | WBGene00006686 | protein_coding_gene | -1.1 | 0.46 | 4.1E-08 |
| *fbxc-40* | WBGene00016295 | protein_coding_gene | -1.1 | 0.46 | 1.4E-03 |
| *Y69A2AR.47* | WBGene00235294 | protein_coding_gene | -1.1 | 0.46 | 2.3E-02 |
| *bath-35* | WBGene00012196 | protein_coding_gene | -1.1 | 0.46 | 4.7E-03 |
| *C50F7.5* | WBGene00016845 | protein_coding_gene | -1.1 | 0.46 | 6.5E-07 |
| *C04G2.14* | WBGene00255426 | ncRNA_gene | -1.1 | 0.45 | 1.4E-03 |

| | | | | | |
|---|---|---|---|---|---|
| *fbxb-83* | WBGene00020543 | protein_coding_gene | -1.1 | 0.45 | 9.4E-03 |
| *K08H2.3* | WBGene00010690 | protein_coding_gene | -1.1 | 0.45 | 7.0E-05 |
| *Y56A3A.19* | WBGene00013237 | protein_coding_gene | -1.1 | 0.45 | 1.6E-25 |
| *nhr-132* | WBGene00003722 | protein_coding_gene | -1.2 | 0.45 | 6.1E-05 |
| *K12H6.7* | WBGene00019687 | protein_coding_gene | -1.2 | 0.45 | 9.5E-03 |
| *fbxc-39* | WBGene00018456 | protein_coding_gene | -1.2 | 0.45 | 1.2E-02 |
| *Y53C10A.1* | WBGene00014915 | pseudogene | -1.2 | 0.45 | 9.8E-07 |
| *oac-18* | WBGene00009230 | pseudogene | -1.2 | 0.45 | 1.5E-02 |
| *acox-1.2* | WBGene00008565 | protein_coding_gene | -1.2 | 0.45 | 4.8E-05 |
| *C49C8.2* | WBGene00016766 | protein_coding_gene | -1.2 | 0.45 | 6.1E-03 |
| *oac-14* | WBGene00008602 | protein_coding_gene | -1.2 | 0.45 | 1.5E-04 |
| *F58E1.13* | WBGene00019044 | protein_coding_gene | -1.2 | 0.45 | 3.8E-03 |
| *ver-1* | WBGene00006894 | protein_coding_gene | -1.2 | 0.45 | 2.0E-02 |
| *Y24F12A.4* | WBGene00012499 | pseudogene | -1.2 | 0.45 | 1.3E-02 |
| *cdr-1* | WBGene00000412 | protein_coding_gene | -1.2 | 0.45 | 1.4E-02 |
| *clec-80* | WBGene00043062 | protein_coding_gene | -1.2 | 0.44 | 5.1E-23 |
| *T20D4.13* | WBGene00020619 | pseudogene | -1.2 | 0.44 | 2.0E-02 |
| *C25F9.6* | WBGene00007726 | protein_coding_gene | -1.2 | 0.44 | 1.2E-02 |
| *Y73B6BL.14* | WBGene00022237 | protein_coding_gene | -1.2 | 0.44 | 3.4E-03 |
| *fbxb-45* | WBGene00019705 | protein_coding_gene | -1.2 | 0.44 | 5.9E-03 |
| *Y47G7B.2* | WBGene00021654 | protein_coding_gene | -1.2 | 0.44 | 2.0E-02 |
| *M02B7.2* | WBGene00019724 | protein_coding_gene | -1.2 | 0.44 | 5.3E-08 |
| *math-38* | WBGene00020326 | protein_coding_gene | -1.2 | 0.44 | 7.0E-13 |
| *F35F10.6* | WBGene00018056 | protein_coding_gene | -1.2 | 0.44 | 9.6E-03 |
| *btb-2* | WBGene00020802 | protein_coding_gene | -1.2 | 0.44 | 4.1E-03 |
| *C33G8.2* | WBGene00016360 | protein_coding_gene | -1.2 | 0.44 | 6.4E-15 |
| *nhr-80* | WBGene00003670 | protein_coding_gene | -1.2 | 0.44 | 2.8E-13 |
| *W04B5.1* | WBGene00021018 | pseudogene | -1.2 | 0.44 | 1.6E-07 |
| *Y47H9C.1* | WBGene00012947 | protein_coding_gene | -1.2 | 0.44 | 4.6E-03 |
| *F47G3.4* | WBGene00194641 | protein_coding_gene | -1.2 | 0.44 | 4.2E-06 |
| *T07D3.6* | WBGene00020309 | pseudogene | -1.2 | 0.44 | 1.7E-02 |
| *his-72* | WBGene00001946 | protein_coding_gene | -1.2 | 0.44 | 1.8E-28 |
| *T07D3.9* | WBGene00020311 | protein_coding_gene | -1.2 | 0.44 | 1.4E-11 |
| *Y57G11C.1143* | WBGene00219342 | pseudogene | -1.2 | 0.44 | 3.0E-03 |
| *str-176* | WBGene00006220 | protein_coding_gene | -1.2 | 0.44 | 4.4E-13 |
| *ugt-31* | WBGene00021464 | protein_coding_gene | -1.2 | 0.43 | 3.2E-04 |
| *T24A6.1* | WBGene00020744 | protein_coding_gene | -1.2 | 0.43 | 9.4E-03 |
| *F21H7.2* | WBGene00009028 | protein_coding_gene | -1.2 | 0.43 | 5.7E-06 |
| *nhr-72* | WBGene00003662 | protein_coding_gene | -1.2 | 0.43 | 1.8E-02 |
| *T04A8.18* | WBGene00119203 | protein_coding_gene | -1.2 | 0.43 | 1.5E-03 |
| *ttr-12* | WBGene00009759 | protein_coding_gene | -1.2 | 0.43 | 2.7E-03 |
| *Y43F8B.13* | WBGene00012823 | protein_coding_gene | -1.2 | 0.43 | 2.4E-03 |
| *Y43F8B.15* | WBGene00045415 | protein_coding_gene | -1.2 | 0.43 | 2.6E-03 |
| *ubc-19* | WBGene00006714 | protein_coding_gene | -1.2 | 0.43 | 1.6E-04 |
| *M116.2* | WBGene00019789 | pseudogene | -1.2 | 0.43 | 1.3E-10 |
| *C28C12.4* | WBGene00016180 | protein_coding_gene | -1.2 | 0.43 | 3.0E-03 |
| *mxl-2* | WBGene00003510 | protein_coding_gene | -1.2 | 0.43 | 7.7E-15 |
| *C02B8.6* | WBGene00015324 | protein_coding_gene | -1.2 | 0.43 | 1.8E-10 |
| *fbxa-107* | WBGene00012564 | protein_coding_gene | -1.2 | 0.43 | 4.4E-11 |
| *D1054.1* | WBGene00008370 | protein_coding_gene | -1.2 | 0.43 | 7.0E-10 |
| *Y82E9BR.14* | WBGene00022347 | protein_coding_gene | -1.2 | 0.42 | 1.5E-19 |
| *H16D19.6* | WBGene00195254 | pseudogene | -1.2 | 0.42 | 1.5E-02 |
| *W06D12.8* | WBGene00198082 | ncRNA_gene | -1.3 | 0.42 | 1.2E-02 |
| *fbxa-51* | WBGene00020448 | protein_coding_gene | -1.3 | 0.42 | 8.5E-05 |
| *Y6G8.9* | WBGene00050974 | pseudogene | -1.3 | 0.42 | 1.2E-02 |
| *T02G5.11* | WBGene00020167 | pseudogene | -1.3 | 0.42 | 6.2E-04 |
| *srh-2* | WBGene00005228 | protein_coding_gene | -1.3 | 0.42 | 4.0E-04 |
| *F47B10.3* | WBGene00009814 | protein_coding_gene | -1.3 | 0.42 | 8.2E-03 |
| *fat-7* | WBGene00001399 | protein_coding_gene | -1.3 | 0.42 | 2.5E-15 |
| *ZC239.5* | WBGene00022568 | protein_coding_gene | -1.3 | 0.41 | 1.2E-02 |

| | | | | | |
|---|---|---|---|---|---|
| *F09C6.3* | WBGene00008617 | protein_coding_gene | -1.3 | 0.41 | 1.3E-02 |
| *H04D03.3* | WBGene00010364 | protein_coding_gene | -1.3 | 0.41 | 3.0E-03 |
| *F52C6.13* | WBGene00018670 | protein_coding_gene | -1.3 | 0.41 | 6.4E-03 |
| *Y116F11B.7* | WBGene00013823 | protein_coding_gene | -1.3 | 0.41 | 1.1E-02 |
| *F42A9.6* | WBGene00018335 | protein_coding_gene | -1.3 | 0.41 | 2.1E-19 |
| *sid-1* | WBGene00004795 | protein_coding_gene | -1.3 | 0.41 | 1.5E-23 |
| *F55A3.7* | WBGene00018852 | pseudogene | -1.3 | 0.41 | 3.3E-06 |
| *W04A8.4* | WBGene00012239 | protein_coding_gene | -1.3 | 0.41 | 1.1E-13 |
| *F25D1.5* | WBGene00009110 | protein_coding_gene | -1.3 | 0.41 | 7.3E-08 |
| *nhr-148* | WBGene00015396 | protein_coding_gene | -1.3 | 0.41 | 1.1E-02 |
| *C31C9.7* | WBGene00007841 | protein_coding_gene | -1.3 | 0.41 | 1.2E-05 |
| *fbxb-81* | WBGene00020541 | protein_coding_gene | -1.3 | 0.41 | 9.0E-03 |
| *F31F4.1* | WBGene00017957 | protein_coding_gene | -1.3 | 0.41 | 2.5E-08 |
| *ZC239.13* | WBGene00022571 | protein_coding_gene | -1.3 | 0.41 | 1.7E-03 |
| *Y111B2A.21* | WBGene00013741 | protein_coding_gene | -1.3 | 0.40 | 5.3E-04 |
| *F49F1.12* | WBGene00018652 | protein_coding_gene | -1.3 | 0.40 | 1.1E-03 |
| *W07A12.8* | WBGene00012325 | protein_coding_gene | -1.3 | 0.40 | 1.5E-03 |
| *tbc-10* | WBGene00019915 | protein_coding_gene | -1.3 | 0.40 | 7.1E-11 |
| *C06A5.12* | WBGene00044414 | protein_coding_gene | -1.3 | 0.40 | 2.5E-04 |
| *sdz-28* | WBGene00020073 | protein_coding_gene | -1.3 | 0.40 | 1.1E-05 |
| *F14D2.5* | WBGene00017453 | protein_coding_gene | -1.3 | 0.40 | 1.4E-03 |
| *F01D5.5* | WBGene00008495 | protein_coding_gene | -1.3 | 0.40 | 3.8E-05 |
| *F58D5.3* | WBGene00010243 | protein_coding_gene | -1.3 | 0.39 | 1.1E-03 |
| *ZK637.6* | WBGene00014987 | pseudogene | -1.3 | 0.39 | 3.2E-11 |
| *M01H9.2* | WBGene00019718 | protein_coding_gene | -1.3 | 0.39 | 2.0E-03 |
| *osm-3* | WBGene00003884 | protein_coding_gene | -1.3 | 0.39 | 5.6E-05 |
| *C30G12.2* | WBGene00016274 | protein_coding_gene | -1.3 | 0.39 | 6.2E-11 |
| *fbxb-52* | WBGene00019417 | protein_coding_gene | -1.4 | 0.39 | 1.1E-03 |
| *fbxb-97* | WBGene00016885 | protein_coding_gene | -1.4 | 0.39 | 1.1E-03 |
| *Y113G7A.14* | WBGene00013751 | protein_coding_gene | -1.4 | 0.39 | 6.1E-03 |
| *Y51A2B.5* | WBGene00013069 | protein_coding_gene | -1.4 | 0.39 | 2.4E-03 |
| *fbxc-34* | WBGene00019921 | protein_coding_gene | -1.4 | 0.39 | 4.6E-03 |
| *F20D1.22* | WBGene00304908 | protein_coding_gene | -1.4 | 0.39 | 4.7E-04 |
| *clp-3* | WBGene00000544 | protein_coding_gene | -1.4 | 0.39 | 3.8E-05 |
| *bath-4* | WBGene00018665 | protein_coding_gene | -1.4 | 0.39 | 1.6E-05 |
| *F11A5.18* | WBGene00045511 | protein_coding_gene | -1.4 | 0.39 | 1.1E-03 |
| *nduo-1* | WBGene00010959 | protein_coding_gene | -1.4 | 0.38 | 6.1E-19 |
| *C14A6.16* | WBGene00206536 | pseudogene | -1.4 | 0.38 | 3.8E-03 |
| *fat-5* | WBGene00001397 | protein_coding_gene | -1.4 | 0.38 | 2.6E-23 |
| *ZK250.2* | WBGene00022685 | protein_coding_gene | -1.4 | 0.38 | 9.5E-04 |
| *enu-3.5* | WBGene00012552 | protein_coding_gene | -1.4 | 0.38 | 6.7E-04 |
| *fbxb-10* | WBGene00020362 | protein_coding_gene | -1.4 | 0.38 | 2.1E-03 |
| *gst-38* | WBGene00001786 | protein_coding_gene | -1.4 | 0.38 | 4.6E-03 |
| *F11A5.17* | WBGene00045510 | protein_coding_gene | -1.4 | 0.38 | 7.9E-05 |
| *nep-8* | WBGene00017553 | protein_coding_gene | -1.4 | 0.38 | 5.2E-05 |
| *Y110A2AL.4* | WBGene00022441 | protein_coding_gene | -1.4 | 0.38 | 2.2E-09 |
| *F10G2.4* | WBGene00017365 | protein_coding_gene | -1.4 | 0.38 | 4.1E-03 |
| *K08D12.6* | WBGene00019540 | protein_coding_gene | -1.4 | 0.38 | 6.9E-20 |
| *D2096.11* | WBGene00017078 | protein_coding_gene | -1.4 | 0.38 | 2.5E-04 |
| *Y80D4G.1* | WBGene00206459 | pseudogene | -1.4 | 0.38 | 1.5E-11 |
| *timm-17B.2* | WBGene00017069 | protein_coding_gene | -1.4 | 0.38 | 1.5E-09 |
| *Y51A2B.6* | WBGene00013070 | protein_coding_gene | -1.4 | 0.38 | 2.5E-03 |
| *Y43F8B.14* | WBGene00012824 | protein_coding_gene | -1.4 | 0.38 | 8.8E-04 |
| *ZC239.17* | WBGene00022575 | protein_coding_gene | -1.4 | 0.38 | 8.6E-04 |
| *Y116A8C.33* | WBGene00013809 | protein_coding_gene | -1.4 | 0.38 | 2.6E-06 |
| *T15B7.10* | WBGene00020522 | pseudogene | -1.4 | 0.37 | 1.2E-05 |
| *pzf-1* | WBGene00011505 | protein_coding_gene | -1.4 | 0.37 | 3.0E-08 |
| *F26H11.4* | WBGene00009182 | protein_coding_gene | -1.4 | 0.37 | 8.1E-07 |
| *C25F9.11* | WBGene00045411 | protein_coding_gene | -1.4 | 0.37 | 5.5E-04 |
| *F01D5.1* | WBGene00008492 | protein_coding_gene | -1.4 | 0.37 | 8.3E-04 |

| | | | | | |
|---|---|---|---|---|---|
| *C29F9.2* | WBGene00016217 | protein_coding_gene | -1.4 | 0.37 | 2.1E-03 |
| *W02A11.6* | WBGene00014852 | pseudogene | -1.4 | 0.37 | 7.0E-04 |
| *linc-16* | WBGene00219728 | lincRNA_gene | -1.4 | 0.37 | 5.4E-04 |
| *C25F9.15* | WBGene00050967 | protein_coding_gene | -1.4 | 0.37 | 6.3E-04 |
| *dhs-23* | WBGene00000986 | protein_coding_gene | -1.4 | 0.37 | 8.5E-10 |
| *gst-41* | WBGene00001789 | protein_coding_gene | -1.4 | 0.37 | 1.3E-08 |
| *Y102A5C.35* | WBGene00044220 | pseudogene | -1.5 | 0.36 | 3.8E-03 |
| *C40A11.2* | WBGene00016545 | protein_coding_gene | -1.5 | 0.36 | 2.1E-03 |
| *drh-2* | WBGene00001091 | pseudogene | -1.5 | 0.36 | 4.8E-05 |
| *Y110A2AL.3* | WBGene00022440 | protein_coding_gene | -1.5 | 0.36 | 2.0E-05 |
| *R07C12.1* | WBGene00019931 | protein_coding_gene | -1.5 | 0.36 | 2.9E-04 |
| *ZC266.1* | WBGene00022584 | protein_coding_gene | -1.5 | 0.36 | 6.8E-06 |
| *peel-1* | WBGene00077563 | protein_coding_gene | -1.5 | 0.36 | 3.8E-04 |
| *K10G4.3* | WBGene00010752 | protein_coding_gene | -1.5 | 0.36 | 2.4E-03 |
| *fbxa-26* | WBGene00022324 | protein_coding_gene | -1.5 | 0.36 | 1.0E-06 |
| *M02E1.3* | WBGene00023458 | protein_coding_gene | -1.5 | 0.36 | 3.3E-03 |
| *Y49F6C.8* | WBGene00021730 | protein_coding_gene | -1.5 | 0.36 | 7.3E-10 |
| *T03E6.9* | WBGene00045356 | protein_coding_gene | -1.5 | 0.36 | 5.3E-03 |
| *Y57G11C.52* | WBGene00044741 | protein_coding_gene | -1.5 | 0.36 | 3.2E-03 |
| *nhr-263* | WBGene00017367 | protein_coding_gene | -1.5 | 0.36 | 2.6E-03 |
| *bath-6* | WBGene00018666 | pseudogene | -1.5 | 0.36 | 1.2E-03 |
| *clec-7* | WBGene00017364 | protein_coding_gene | -1.5 | 0.36 | 3.7E-25 |
| *fbxb-12* | WBGene00018109 | pseudogene | -1.5 | 0.36 | 5.3E-04 |
| *B0281.4* | WBGene00015111 | protein_coding_gene | -1.5 | 0.36 | 4.9E-04 |
| *clec-2* | WBGene00015198 | protein_coding_gene | -1.5 | 0.36 | 1.2E-03 |
| *F59E12.15* | WBGene00235269 | protein_coding_gene | -1.5 | 0.35 | 4.5E-04 |
| *sea-1* | WBGene00004750 | protein_coding_gene | -1.5 | 0.35 | 8.4E-18 |
| *gpx-1* | WBGene00009165 | protein_coding_gene | -1.5 | 0.35 | 7.7E-16 |
| *Y35H6.3* | WBGene00021342 | pseudogene | -1.5 | 0.35 | 8.7E-06 |
| *F39B2.5* | WBGene00009556 | protein_coding_gene | -1.5 | 0.35 | 3.6E-13 |
| *F36H5.17* | WBGene00270293 | protein_coding_gene | -1.5 | 0.35 | 4.5E-04 |
| *F55C12.19* | WBGene00271817 | protein_coding_gene | -1.5 | 0.35 | 1.4E-03 |
| *fbxb-90* | WBGene00044694 | protein_coding_gene | -1.5 | 0.35 | 2.4E-03 |
| *ZK686.1* | WBGene00022791 | protein_coding_gene | -1.5 | 0.35 | 3.2E-29 |
| *F49F1.18* | WBGene00235368 | protein_coding_gene | -1.5 | 0.35 | 3.9E-04 |
| *ppw-1* | WBGene00004093 | protein_coding_gene | -1.5 | 0.35 | 5.5E-26 |
| *F59A7.7* | WBGene00019094 | protein_coding_gene | -1.5 | 0.35 | 3.5E-03 |
| *Y73B6A.6* | WBGene00044503 | pseudogene | -1.5 | 0.34 | 2.4E-03 |
| *F09C6.14* | WBGene00044278 | protein_coding_gene | -1.5 | 0.34 | 2.3E-03 |
| *cpr-2* | WBGene00000782 | protein_coding_gene | -1.5 | 0.34 | 1.2E-03 |
| *C17B7.4* | WBGene00015878 | protein_coding_gene | -1.5 | 0.34 | 3.6E-04 |
| *C14C6.2* | WBGene00015756 | protein_coding_gene | -1.5 | 0.34 | 1.8E-32 |
| *C23H5.8* | WBGene00016027 | protein_coding_gene | -1.5 | 0.34 | 8.4E-24 |
| *fbxb-49* | WBGene00019412 | protein_coding_gene | -1.6 | 0.34 | 2.9E-03 |
| *fbxb-98* | WBGene00015830 | protein_coding_gene | -1.6 | 0.34 | 3.2E-04 |
| *F31E9.3* | WBGene00009296 | protein_coding_gene | -1.6 | 0.34 | 5.8E-04 |
| *R06B10.2* | WBGene00019913 | protein_coding_gene | -1.6 | 0.34 | 3.6E-04 |
| *F59H6.2* | WBGene00019133 | pseudogene | -1.6 | 0.34 | 3.0E-04 |
| *W04E12.5* | WBGene00012250 | protein_coding_gene | -1.6 | 0.34 | 2.6E-03 |
| *srh-70* | WBGene00005291 | protein_coding_gene | -1.6 | 0.34 | 3.2E-03 |
| *fbxa-114* | WBGene00013758 | protein_coding_gene | -1.6 | 0.34 | 7.2E-16 |
| *F07E5.5* | WBGene00017210 | protein_coding_gene | -1.6 | 0.34 | 1.1E-25 |
| *egas-3* | WBGene00013480 | protein_coding_gene | -1.6 | 0.33 | 1.6E-03 |
| *ZK1240.1* | WBGene00022866 | protein_coding_gene | -1.6 | 0.33 | 1.0E-03 |
| *fbxa-85* | WBGene00044654 | protein_coding_gene | -1.6 | 0.33 | 1.9E-03 |
| *F26F2.1* | WBGene00009166 | protein_coding_gene | -1.6 | 0.33 | 2.8E-04 |
| *msp-33* | WBGene00003431 | protein_coding_gene | -1.6 | 0.33 | 9.2E-07 |
| *ZC239.21* | WBGene00044372 | protein_coding_gene | -1.6 | 0.33 | 2.3E-04 |
| *sdz-11* | WBGene00017408 | protein_coding_gene | -1.6 | 0.33 | 2.4E-04 |
| *H16D19.5* | WBGene00195253 | pseudogene | -1.6 | 0.33 | 6.3E-04 |

| | | | | | |
|---|---|---|---|---|---|
| *R03D7.2* | WBGene00010989 | protein_coding_gene | -1.6 | 0.33 | 1.5E-07 |
| *arrd-21* | WBGene00015441 | pseudogene | -1.6 | 0.33 | 8.2E-04 |
| *T05F1.5* | WBGene00011491 | protein_coding_gene | -1.6 | 0.33 | 1.0E-06 |
| *sdz-18* | WBGene00018442 | protein_coding_gene | -1.6 | 0.32 | 2.5E-04 |
| *cest-29* | WBGene00017205 | pseudogene | -1.6 | 0.32 | 2.6E-10 |
| *Y71H2AM.14* | WBGene00022179 | protein_coding_gene | -1.6 | 0.32 | 7.9E-05 |
| *pals-37* | WBGene00138721 | protein_coding_gene | -1.6 | 0.32 | 5.0E-04 |
| *F53C3.3* | WBGene00018747 | protein_coding_gene | -1.6 | 0.32 | 1.2E-03 |
| *F15B9.6* | WBGene00008850 | protein_coding_gene | -1.6 | 0.32 | 1.5E-05 |
| *Y47H10A.3* | WBGene00012959 | protein_coding_gene | -1.6 | 0.32 | 4.9E-04 |
| *lys-10* | WBGene00003099 | protein_coding_gene | -1.6 | 0.32 | 7.2E-06 |
| *C08F1.11* | WBGene00044417 | protein_coding_gene | -1.6 | 0.32 | 9.3E-05 |
| *col-12* | WBGene00000601 | protein_coding_gene | -1.6 | 0.32 | 6.2E-16 |
| *F52C6.2* | WBGene00018659 | protein_coding_gene | -1.6 | 0.32 | 6.1E-14 |
| *F41D3.9* | WBGene00009614 | protein_coding_gene | -1.7 | 0.32 | 2.0E-04 |
| *mir-8199* | WBGene00016553 | miRNA_gene | -1.7 | 0.32 | 1.8E-04 |
| *nhr-58* | WBGene00003648 | protein_coding_gene | -1.7 | 0.32 | 3.0E-06 |
| *F09C6.10* | WBGene00008620 | protein_coding_gene | -1.7 | 0.32 | 1.6E-04 |
| *bath-21* | WBGene00019138 | protein_coding_gene | -1.7 | 0.32 | 3.3E-05 |
| *col-141* | WBGene00000714 | protein_coding_gene | -1.7 | 0.32 | 1.2E-04 |
| *col-135* | WBGene00000708 | protein_coding_gene | -1.7 | 0.31 | 6.5E-26 |
| *linc-24* | WBGene00219771 | lincRNA_gene | -1.7 | 0.31 | 1.7E-04 |
| *fbxa-14* | WBGene00022326 | protein_coding_gene | -1.7 | 0.31 | 1.1E-05 |
| *Y32B12C.5* | WBGene00194769 | pseudogene | -1.7 | 0.31 | 5.1E-11 |
| *Y48E1B.8* | WBGene00013007 | protein_coding_gene | -1.7 | 0.31 | 4.3E-04 |
| *F16H6.7* | WBGene00008897 | protein_coding_gene | -1.7 | 0.31 | 1.3E-04 |
| *clec-52* | WBGene00015052 | protein_coding_gene | -1.7 | 0.31 | 6.8E-23 |
| *irg-5* | WBGene00009429 | protein_coding_gene | -1.7 | 0.31 | 8.2E-05 |
| *R12C12.7* | WBGene00020027 | protein_coding_gene | -1.7 | 0.31 | 2.4E-45 |
| *C01C10.2* | WBGene00015294 | protein_coding_gene | -1.7 | 0.31 | 6.3E-11 |
| *F54A3.2* | WBGene00018781 | protein_coding_gene | -1.7 | 0.31 | 1.6E-17 |
| *Y48G9A.7* | WBGene00021701 | protein_coding_gene | -1.7 | 0.31 | 1.1E-03 |
| *fbxb-35* | WBGene00017208 | protein_coding_gene | -1.7 | 0.31 | 1.7E-04 |
| *F07C4.6* | WBGene00017200 | protein_coding_gene | -1.7 | 0.31 | 4.1E-06 |
| *F58D5.2* | WBGene00010242 | protein_coding_gene | -1.7 | 0.30 | 2.1E-07 |
| *Y69A2AR.12* | WBGene00022083 | protein_coding_gene | -1.7 | 0.30 | 5.7E-04 |
| *sdz-29* | WBGene00020081 | pseudogene | -1.7 | 0.30 | 1.2E-04 |
| *ZK1240.11* | WBGene00206413 | protein_coding_gene | -1.7 | 0.30 | 1.1E-04 |
| *H11L12.1* | WBGene00019189 | protein_coding_gene | -1.7 | 0.30 | 1.9E-07 |
| *fbxc-43* | WBGene00045297 | protein_coding_gene | -1.7 | 0.30 | 3.7E-07 |
| *F31E9.6* | WBGene00009299 | protein_coding_gene | -1.7 | 0.30 | 2.7E-06 |
| *nhr-225* | WBGene00020849 | protein_coding_gene | -1.7 | 0.30 | 9.9E-04 |
| *C40A11.8* | WBGene00016551 | protein_coding_gene | -1.7 | 0.30 | 2.5E-07 |
| *fbxb-51* | WBGene00019414 | protein_coding_gene | -1.7 | 0.30 | 1.2E-04 |
| *Y46D2A.2* | WBGene00021590 | protein_coding_gene | -1.7 | 0.30 | 1.2E-04 |
| *fbxb-53* | WBGene00018104 | protein_coding_gene | -1.7 | 0.30 | 9.8E-04 |
| *fbxb-36* | WBGene00017211 | protein_coding_gene | -1.7 | 0.30 | 3.1E-04 |
| *Y45F10C.6* | WBGene00044256 | protein_coding_gene | -1.8 | 0.30 | 8.6E-05 |
| *C50E3.12* | WBGene00016823 | protein_coding_gene | -1.8 | 0.30 | 1.2E-08 |
| *K06G5.1* | WBGene00010605 | protein_coding_gene | -1.8 | 0.29 | 6.3E-93 |
| *rmd-6* | WBGene00020070 | protein_coding_gene | -1.8 | 0.29 | 1.0E-04 |
| *C32H11.7* | WBGene00007870 | pseudogene | -1.8 | 0.29 | 1.0E-04 |
| *Y37H2B.1* | WBGene00012572 | pseudogene | -1.8 | 0.29 | 7.8E-04 |
| *Y46C8AM.1* | WBGene00271795 | protein_coding_gene | -1.8 | 0.29 | 2.1E-07 |
| *sqst-5* | WBGene00269434 | protein_coding_gene | -1.8 | 0.29 | 1.4E-06 |
| *H16D19.4* | WBGene00010402 | pseudogene | -1.8 | 0.29 | 1.3E-03 |
| *Y6G8.2* | WBGene00012404 | protein_coding_gene | -1.8 | 0.29 | 2.7E-15 |
| *Y47H10A.6* | WBGene00077537 | pseudogene | -1.8 | 0.29 | 7.5E-05 |
| *dach-1* | WBGene00020386 | protein_coding_gene | -1.8 | 0.29 | 8.8E-13 |
| *nhr-226* | WBGene00020850 | protein_coding_gene | -1.8 | 0.29 | 1.5E-06 |

| | | | | | |
|---|---|---|---|---|---|
| *ZK218.5* | WBGene00013939 | protein_coding_gene | -1.8 | 0.28 | 1.8E-04 |
| *clec-37* | WBGene00013609 | protein_coding_gene | -1.8 | 0.28 | 1.2E-03 |
| *fbxb-43* | WBGene00021774 | protein_coding_gene | -1.8 | 0.28 | 5.7E-04 |
| *drd-10* | WBGene00016361 | protein_coding_gene | -1.8 | 0.28 | 2.6E-09 |
| *Y82E9BL.3* | WBGene00022320 | protein_coding_gene | -1.8 | 0.28 | 2.8E-04 |
| *math-49* | WBGene00022690 | protein_coding_gene | -1.8 | 0.28 | 7.8E-04 |
| *pals-19* | WBGene00044761 | protein_coding_gene | -1.9 | 0.28 | 4.7E-08 |
| *ZK1025.3* | WBGene00014183 | protein_coding_gene | -1.9 | 0.28 | 5.1E-04 |
| *T02G5.4* | WBGene00020164 | pseudogene | -1.9 | 0.28 | 1.3E-25 |
| *fbxb-50* | WBGene00019413 | protein_coding_gene | -1.9 | 0.28 | 4.3E-05 |
| *cyp-35B3* | WBGene00019470 | protein_coding_gene | -1.9 | 0.27 | 5.6E-04 |
| *folt-2* | WBGene00018138 | protein_coding_gene | -1.9 | 0.27 | 3.1E-35 |
| *F57E7.2* | WBGene00010203 | protein_coding_gene | -1.9 | 0.27 | 5.0E-05 |
| *fbxa-55* | WBGene00020453 | protein_coding_gene | -1.9 | 0.27 | 6.9E-11 |
| *CE7X_3.1* | WBGene00008349 | pseudogene | -1.9 | 0.27 | 1.6E-04 |
| *nspe-4* | WBGene00012589 | protein_coding_gene | -1.9 | 0.27 | 6.9E-07 |
| *trp-2* | WBGene00006615 | protein_coding_gene | -1.9 | 0.26 | 9.1E-10 |
| *rgs-11* | WBGene00004354 | protein_coding_gene | -1.9 | 0.26 | 3.7E-16 |
| *D1022.17* | WBGene00271809 | ncRNA_gene | -1.9 | 0.26 | 1.4E-04 |
| *scrm-3* | WBGene00015437 | protein_coding_gene | -1.9 | 0.26 | 3.0E-05 |
| *F23D12.3* | WBGene00009087 | protein_coding_gene | -1.9 | 0.26 | 2.8E-08 |
| *Y40C5A.4* | WBGene00021497 | protein_coding_gene | -1.9 | 0.26 | 3.3E-17 |
| *Y34F4.6* | WBGene00189956 | protein_coding_gene | -1.9 | 0.26 | 1.2E-06 |
| *gcy-15* | WBGene00001541 | protein_coding_gene | -1.9 | 0.26 | 1.3E-06 |
| *fbxb-46* | WBGene00019420 | protein_coding_gene | -1.9 | 0.26 | 3.2E-05 |
| *nhr-220* | WBGene00020591 | pseudogene | -1.9 | 0.26 | 2.5E-04 |
| *bath-13* | WBGene00018223 | protein_coding_gene | -1.9 | 0.26 | 1.3E-07 |
| *Y17D7B.7* | WBGene00014562 | pseudogene | -2.0 | 0.26 | 2.0E-12 |
| *fbxb-20* | WBGene00022559 | protein_coding_gene | -2.0 | 0.26 | 1.9E-04 |
| *K03D3.5* | WBGene00010531 | protein_coding_gene | -2.0 | 0.26 | 3.4E-05 |
| *R03H10.7* | WBGene00019859 | protein_coding_gene | -2.0 | 0.26 | 3.1E-05 |
| *fbxb-44* | WBGene00019416 | protein_coding_gene | -2.0 | 0.26 | 3.3E-05 |
| *C28D4.5* | WBGene00007793 | protein_coding_gene | -2.0 | 0.26 | 5.6E-10 |
| *CE7X_3.2* | WBGene00008350 | pseudogene | -2.0 | 0.26 | 2.1E-05 |
| *F21D9.2* | WBGene00009016 | protein_coding_gene | -2.0 | 0.26 | 4.8E-07 |
| *fbxa-54* | WBGene00020449 | protein_coding_gene | -2.0 | 0.26 | 5.7E-12 |
| *fbxb-13* | WBGene00015607 | protein_coding_gene | -2.0 | 0.25 | 1.8E-04 |
| *ptp-5.3* | WBGene00021207 | protein_coding_gene | -2.0 | 0.25 | 4.2E-09 |
| *C08F1.10* | WBGene00015613 | protein_coding_gene | -2.0 | 0.25 | 1.2E-13 |
| *K02E7.5* | WBGene00019310 | pseudogene | -2.0 | 0.25 | 4.9E-05 |
| *clec-233* | WBGene00009030 | protein_coding_gene | -2.0 | 0.25 | 2.2E-05 |
| *C50E3.11* | WBGene00016822 | protein_coding_gene | -2.0 | 0.25 | 2.1E-25 |
| *ZK1240.9* | WBGene00022873 | protein_coding_gene | -2.0 | 0.25 | 5.7E-12 |
| *ugt-28* | WBGene00015693 | protein_coding_gene | -2.0 | 0.25 | 3.3E-13 |
| *nhr-115* | WBGene00003705 | protein_coding_gene | -2.0 | 0.25 | 5.4E-07 |
| *F12E12.11* | WBGene00044811 | protein_coding_gene | -2.0 | 0.25 | 3.8E-09 |
| *npr-30* | WBGene00019183 | protein_coding_gene | -2.0 | 0.25 | 1.7E-06 |
| *skr-5* | WBGene00004811 | protein_coding_gene | -2.0 | 0.25 | 1.6E-05 |
| *bath-20* | WBGene00015012 | protein_coding_gene | -2.0 | 0.25 | 2.1E-05 |
| *arrd-22* | WBGene00020612 | protein_coding_gene | -2.0 | 0.25 | 1.3E-06 |
| *his-60* | WBGene00001934 | protein_coding_gene | -2.0 | 0.24 | 3.1E-08 |
| *scrm-4* | WBGene00008681 | protein_coding_gene | -2.0 | 0.24 | 2.7E-26 |
| *C14C6.8* | WBGene00015762 | protein_coding_gene | -2.0 | 0.24 | 1.3E-04 |
| *Y60A3A.25* | WBGene00044211 | protein_coding_gene | -2.0 | 0.24 | 3.7E-07 |
| *fbxa-59* | WBGene00020455 | protein_coding_gene | -2.0 | 0.24 | 2.9E-22 |
| *K06B4.15* | WBGene00077450 | protein_coding_gene | -2.0 | 0.24 | 7.7E-05 |
| *T20D4.11* | WBGene00020617 | protein_coding_gene | -2.1 | 0.24 | 2.5E-04 |
| *K09F6.5* | WBGene00019588 | protein_coding_gene | -2.1 | 0.24 | 1.8E-04 |
| *nhr-227* | WBGene00020851 | protein_coding_gene | -2.1 | 0.24 | 1.6E-05 |
| *irg-4* | WBGene00008584 | protein_coding_gene | -2.1 | 0.24 | 2.3E-08 |

| | | | | | |
|---|---|---|---|---|---|
| *Y23H5A.8* | WBGene00023172 | protein_coding_gene | -2.1 | 0.24 | 2.0E-04 |
| *F21D9.6* | WBGene00009020 | pseudogene | -2.1 | 0.23 | 2.5E-04 |
| *K08D10.9* | WBGene00019532 | pseudogene | -2.1 | 0.23 | 1.2E-05 |
| *fbxc-19* | WBGene00020361 | protein_coding_gene | -2.1 | 0.23 | 1.7E-04 |
| *Y46B2A.4* | WBGene00284850 | protein_coding_gene | -2.1 | 0.23 | 1.9E-04 |
| *Y113G7A.13* | WBGene00013750 | protein_coding_gene | -2.1 | 0.23 | 1.7E-04 |
| *Y49E10.37* | WBGene00304219 | ncRNA_gene | -2.1 | 0.23 | 2.9E-11 |
| *ZC239.20* | WBGene00044371 | protein_coding_gene | -2.1 | 0.23 | 9.2E-06 |
| *fbxa-187* | WBGene00009838 | pseudogene | -2.1 | 0.23 | 1.3E-04 |
| *C08F11.3* | WBGene00007452 | protein_coding_gene | -2.1 | 0.23 | 6.2E-11 |
| *C04G6.6* | WBGene00015456 | protein_coding_gene | -2.1 | 0.23 | 4.8E-15 |
| *F19B2.6* | WBGene00008945 | protein_coding_gene | -2.1 | 0.23 | 1.8E-08 |
| *fbxa-64* | WBGene00020883 | protein_coding_gene | -2.2 | 0.23 | 2.4E-14 |
| *R10E8.3* | WBGene00011209 | protein_coding_gene | -2.2 | 0.23 | 1.1E-04 |
| *C17B7.5* | WBGene00015879 | protein_coding_gene | -2.2 | 0.22 | 9.9E-06 |
| *grsp-4* | WBGene00017201 | protein_coding_gene | -2.2 | 0.22 | 1.3E-20 |
| *fbxb-96* | WBGene00016886 | protein_coding_gene | -2.2 | 0.22 | 7.5E-06 |
| *K02E7.6* | WBGene00019311 | pseudogene | -2.2 | 0.22 | 5.7E-13 |
| *Y17D7C.3* | WBGene00045366 | pseudogene | -2.2 | 0.22 | 2.8E-19 |
| *K05F6.10* | WBGene00019421 | protein_coding_gene | -2.2 | 0.22 | 5.8E-11 |
| *T22B7.7* | WBGene00020674 | protein_coding_gene | -2.2 | 0.22 | 7.2E-20 |
| *F40G9.6* | WBGene00018242 | protein_coding_gene | -2.2 | 0.22 | 1.1E-04 |
| *arrd-20* | WBGene00015440 | pseudogene | -2.2 | 0.22 | 1.3E-04 |
| *nhr-116* | WBGene00003706 | protein_coding_gene | -2.2 | 0.22 | 8.3E-09 |
| *F10E7.1* | WBGene00017343 | protein_coding_gene | -2.2 | 0.22 | 6.8E-06 |
| *rgs-9* | WBGene00004352 | protein_coding_gene | -2.2 | 0.22 | 4.2E-20 |
| *fbxa-117* | WBGene00014907 | pseudogene | -2.2 | 0.22 | 6.8E-06 |
| *C08F1.6* | WBGene00015610 | protein_coding_gene | -2.2 | 0.22 | 7.1E-12 |
| *Y73B6BL.288* | WBGene00219218 | protein_coding_gene | -2.2 | 0.21 | 7.6E-06 |
| *C13G5.2* | WBGene00015747 | protein_coding_gene | -2.2 | 0.21 | 3.2E-49 |
| *F11A5.15* | WBGene00008680 | protein_coding_gene | -2.2 | 0.21 | 6.5E-06 |
| *fkb-8* | WBGene00001433 | protein_coding_gene | -2.2 | 0.21 | 7.6E-12 |
| *fbxa-192* | WBGene00010212 | protein_coding_gene | -2.2 | 0.21 | 2.0E-23 |
| *Y82E9BR.22* | WBGene00022354 | protein_coding_gene | -2.2 | 0.21 | 1.3E-32 |
| *fbxa-210* | WBGene00012565 | protein_coding_gene | -2.2 | 0.21 | 1.8E-46 |
| *ZC239.16* | WBGene00022574 | protein_coding_gene | -2.2 | 0.21 | 2.6E-10 |
| *T26E3.8* | WBGene00012041 | protein_coding_gene | -2.2 | 0.21 | 8.7E-05 |
| *C17F4.8* | WBGene00015914 | protein_coding_gene | -2.2 | 0.21 | 5.9E-06 |
| *clec-47* | WBGene00011668 | protein_coding_gene | -2.2 | 0.21 | 3.6E-08 |
| *btb-3* | WBGene00015013 | pseudogene | -2.3 | 0.21 | 8.7E-05 |
| *K05F6.15* | WBGene00269419 | pseudogene | -2.3 | 0.21 | 2.3E-05 |
| *Y57G11C.40* | WBGene00013329 | protein_coding_gene | -2.3 | 0.21 | 4.3E-06 |
| *Y19D10B.6* | WBGene00021235 | protein_coding_gene | -2.3 | 0.21 | 1.1E-06 |
| *fbxc-50* | WBGene00015912 | protein_coding_gene | -2.3 | 0.21 | 3.0E-42 |
| *C46G7.5* | WBGene00044502 | protein_coding_gene | -2.3 | 0.20 | 5.4E-11 |
| *clp-6* | WBGene00000546 | protein_coding_gene | -2.3 | 0.20 | 1.7E-14 |
| *Y32B12C.1* | WBGene00012525 | protein_coding_gene | -2.3 | 0.20 | 7.8E-10 |
| *C52E12.11* | WBGene00304189 | pseudogene | -2.3 | 0.20 | 2.0E-08 |
| *fbxa-3* | WBGene00020363 | protein_coding_gene | -2.3 | 0.20 | 8.9E-05 |
| *nhr-183* | WBGene00018266 | protein_coding_gene | -2.3 | 0.20 | 1.5E-08 |
| *C16C4.18* | WBGene00194998 | protein_coding_gene | -2.3 | 0.20 | 6.9E-09 |
| *F40D4.13* | WBGene00009572 | protein_coding_gene | -2.3 | 0.20 | 5.0E-24 |
| *C40A11.4* | WBGene00016547 | protein_coding_gene | -2.3 | 0.20 | 1.4E-05 |
| *fbxb-42* | WBGene00021773 | protein_coding_gene | -2.3 | 0.20 | 2.7E-06 |
| *T20D4.5* | WBGene00020611 | protein_coding_gene | -2.3 | 0.20 | 3.1E-06 |
| *F53C3.4* | WBGene00018748 | protein_coding_gene | -2.4 | 0.19 | 7.4E-07 |
| *fbxa-57* | WBGene00016224 | pseudogene | -2.4 | 0.19 | 5.2E-05 |
| *F09C6.13* | WBGene00044183 | protein_coding_gene | -2.4 | 0.19 | 5.6E-11 |
| *ctl-1* | WBGene00000830 | protein_coding_gene | -2.4 | 0.19 | 2.5E-06 |
| *bath-14* | WBGene00015016 | pseudogene | -2.4 | 0.19 | 2.6E-05 |

| | | | | | |
|---|---|---|---|---|---|
| *F55G11.4* | WBGene00010124 | protein_coding_gene | -2.4 | 0.19 | 3.7E-13 |
| *stdh-2* | WBGene00008678 | protein_coding_gene | -2.4 | 0.19 | 2.4E-06 |
| *F13B6.3* | WBGene00017418 | protein_coding_gene | -2.4 | 0.19 | 4.4E-26 |
| *Y37H2A.12* | WBGene00044758 | protein_coding_gene | -2.4 | 0.19 | 3.1E-15 |
| *srw-85* | WBGene00005832 | protein_coding_gene | -2.4 | 0.19 | 4.5E-05 |
| *Y43F8B.23* | WBGene00194815 | protein_coding_gene | -2.4 | 0.19 | 3.4E-05 |
| *C33C12.7* | WBGene00016339 | protein_coding_gene | -2.4 | 0.19 | 2.1E-05 |
| *pals-18* | WBGene00022353 | protein_coding_gene | -2.4 | 0.18 | 1.4E-14 |
| *clec-73* | WBGene00021579 | protein_coding_gene | -2.4 | 0.18 | 2.8E-20 |
| *F09C6.11* | WBGene00044181 | protein_coding_gene | -2.5 | 0.18 | 1.4E-06 |
| *fbxb-95* | WBGene00016880 | protein_coding_gene | -2.5 | 0.18 | 1.4E-06 |
| *Y110A2AL.9* | WBGene00022445 | protein_coding_gene | -2.5 | 0.18 | 2.5E-16 |
| *F57E7.4* | WBGene00077616 | protein_coding_gene | -2.5 | 0.18 | 9.9E-07 |
| *clec-119* | WBGene00018575 | protein_coding_gene | -2.5 | 0.18 | 1.0E-09 |
| *bath-3* | WBGene00019140 | protein_coding_gene | -2.5 | 0.18 | 6.8E-10 |
| *Y54G2A.45* | WBGene00044488 | protein_coding_gene | -2.5 | 0.17 | 1.3E-30 |
| *npl-4.1* | WBGene00019120 | protein_coding_gene | -2.5 | 0.17 | 2.3E-25 |
| *T02G6.10* | WBGene00077772 | pseudogene | -2.5 | 0.17 | 1.8E-05 |
| *F59H6.15* | WBGene00269394 | protein_coding_gene | -2.6 | 0.17 | 1.3E-05 |
| *Y38H8A.8* | WBGene00045062 | pseudogene | -2.6 | 0.17 | 1.4E-05 |
| *ugt-29* | WBGene00021709 | protein_coding_gene | -2.6 | 0.17 | 1.1E-22 |
| *fbxb-15* | WBGene00022557 | protein_coding_gene | -2.6 | 0.17 | 1.8E-05 |
| *W05G11.2* | WBGene00021040 | protein_coding_gene | -2.6 | 0.17 | 1.5E-05 |
| *F52C6.4* | WBGene00018661 | protein_coding_gene | -2.6 | 0.17 | 2.5E-06 |
| *B0281.5* | WBGene00015112 | protein_coding_gene | -2.6 | 0.17 | 7.3E-17 |
| *msp-53* | WBGene00003446 | protein_coding_gene | -2.6 | 0.16 | 8.0E-07 |
| *bath-11* | WBGene00018664 | pseudogene | -2.6 | 0.16 | 1.3E-07 |
| *kin-21* | WBGene00002204 | protein_coding_gene | -2.6 | 0.16 | 9.2E-06 |
| *scrm-8* | WBGene00019530 | protein_coding_gene | -2.6 | 0.16 | 9.1E-06 |
| *fbxa-138* | WBGene00022345 | protein_coding_gene | -2.6 | 0.16 | 4.9E-07 |
| *Y43F8B.9* | WBGene00012819 | protein_coding_gene | -2.7 | 0.16 | 3.3E-17 |
| *ilys-2* | WBGene00016669 | protein_coding_gene | -2.7 | 0.16 | 9.7E-12 |
| *F08D12.2* | WBGene00017246 | protein_coding_gene | -2.7 | 0.15 | 2.6E-31 |
| *C40A11.6* | WBGene00016549 | protein_coding_gene | -2.7 | 0.15 | 2.3E-07 |
| *bath-30* | WBGene00017460 | protein_coding_gene | -2.7 | 0.15 | 4.5E-49 |
| *K05F6.4* | WBGene00019415 | protein_coding_gene | -2.8 | 0.15 | 3.9E-08 |
| *Y46G5A.8* | WBGene00012900 | protein_coding_gene | -2.8 | 0.15 | 3.2E-06 |
| *F14D7.10* | WBGene00008797 | protein_coding_gene | -2.8 | 0.15 | 9.8E-08 |
| *pals-39* | WBGene00008301 | protein_coding_gene | -2.8 | 0.14 | 4.2E-09 |
| *D1065.6* | WBGene00268212 | ncRNA_gene | -2.8 | 0.14 | 5.0E-06 |
| *fbxb-54* | WBGene00019418 | protein_coding_gene | -2.8 | 0.14 | 1.3E-07 |
| *ZK1240.5* | WBGene00022870 | protein_coding_gene | -2.8 | 0.14 | 1.6E-07 |
| *bath-22* | WBGene00018658 | pseudogene | -2.8 | 0.14 | 3.4E-06 |
| *C08F11.6* | WBGene00014672 | pseudogene | -2.8 | 0.14 | 1.4E-09 |
| *fbxb-91* | WBGene00044692 | protein_coding_gene | -2.8 | 0.14 | 2.0E-06 |
| *F26D2.10* | WBGene00009149 | protein_coding_gene | -2.9 | 0.14 | 1.1E-09 |
| *fbxa-156* | WBGene00007392 | protein_coding_gene | -2.9 | 0.14 | 8.2E-10 |
| *fbxa-95* | WBGene00009222 | protein_coding_gene | -2.9 | 0.14 | 1.2E-17 |
| *Y9D1A.1* | WBGene00021186 | protein_coding_gene | -2.9 | 0.14 | 3.7E-15 |
| *K02E7.10* | WBGene00019314 | protein_coding_gene | -2.9 | 0.13 | 1.6E-06 |
| *K09H9.7* | WBGene00019598 | protein_coding_gene | -2.9 | 0.13 | 3.8E-27 |
| *M01G12.9* | WBGene00010822 | protein_coding_gene | -2.9 | 0.13 | 1.3E-18 |
| *C14C6.5* | WBGene00015759 | protein_coding_gene | -2.9 | 0.13 | 3.0E-16 |
| *F39F10.3* | WBGene00018203 | protein_coding_gene | -2.9 | 0.13 | 5.0E-11 |
| *T28A11.2* | WBGene00020869 | protein_coding_gene | -2.9 | 0.13 | 5.0E-11 |
| *F57E7.1* | WBGene00010202 | protein_coding_gene | -3.0 | 0.13 | 6.5E-08 |
| *C50E3.13* | WBGene00016824 | protein_coding_gene | -3.0 | 0.13 | 4.7E-39 |
| *asp-17* | WBGene00012683 | protein_coding_gene | -3.0 | 0.13 | 9.0E-17 |
| *fbxa-11* | WBGene00020451 | protein_coding_gene | -3.0 | 0.13 | 5.5E-08 |
| *T12B5.14* | WBGene00050913 | protein_coding_gene | -3.0 | 0.12 | 3.0E-08 |

| | | | | | |
|---|---|---|---|---|---|
| *btb-14* | WBGene00013275 | protein_coding_gene | -3.1 | 0.12 | 2.1E-29 |
| *F36H5.14* | WBGene00219271 | protein_coding_gene | -3.1 | 0.12 | 2.1E-08 |
| *clec-72* | WBGene00021583 | protein_coding_gene | -3.1 | 0.12 | 2.6E-08 |
| *F14D2.14* | WBGene00017462 | protein_coding_gene | -3.1 | 0.12 | 2.5E-08 |
| *thn-1* | WBGene00009213 | protein_coding_gene | -3.1 | 0.12 | 2.3E-65 |
| *K09F6.13* | WBGene00235164 | protein_coding_gene | -3.1 | 0.12 | 1.8E-08 |
| *T03E6.8* | WBGene00011397 | protein_coding_gene | -3.1 | 0.12 | 1.7E-29 |
| *Y68A4A.13* | WBGene00044212 | protein_coding_gene | -3.1 | 0.12 | 1.7E-13 |
| *nspc-19* | WBGene00008360 | protein_coding_gene | -3.1 | 0.11 | 3.6E-21 |
| *Y39G10AR.16* | WBGene00021472 | protein_coding_gene | -3.1 | 0.11 | 5.0E-20 |
| *fbxa-188* | WBGene00009839 | protein_coding_gene | -3.2 | 0.11 | 1.4E-08 |
| *R10E8.1* | WBGene00011207 | protein_coding_gene | -3.2 | 0.11 | 1.5E-27 |
| *F14D2.8* | WBGene00017456 | protein_coding_gene | -3.2 | 0.11 | 6.6E-07 |
| *F52C6.3* | WBGene00018660 | protein_coding_gene | -3.2 | 0.11 | 3.4E-07 |
| *C52E2.5* | WBGene00016884 | protein_coding_gene | -3.2 | 0.11 | 5.4E-07 |
| *fbxa-10* | WBGene00020450 | protein_coding_gene | -3.2 | 0.11 | 4.7E-07 |
| *C38D9.13* | WBGene00235090 | ncRNA_gene | -3.2 | 0.11 | 5.2E-20 |
| *clec-147* | WBGene00007313 | protein_coding_gene | -3.2 | 0.11 | 5.4E-37 |
| *Y47G6A.14* | WBGene00021640 | protein_coding_gene | -3.2 | 0.11 | 2.1E-45 |
| *ctsa-4.2* | WBGene00021503 | protein_coding_gene | -3.2 | 0.11 | 1.8E-84 |
| *M162.9* | WBGene00014811 | pseudogene | -3.2 | 0.11 | 8.0E-09 |
| *Y43F8B.24* | WBGene00195176 | protein_coding_gene | -3.2 | 0.11 | 3.8E-07 |
| *Y38H6C.19* | WBGene00012631 | protein_coding_gene | -3.2 | 0.11 | 7.2E-35 |
| *ZC239.6* | WBGene00022569 | protein_coding_gene | -3.2 | 0.11 | 7.9E-09 |
| *bath-7* | WBGene00018667 | protein_coding_gene | -3.2 | 0.11 | 5.6E-11 |
| *C25F9.9* | WBGene00007728 | protein_coding_gene | -3.3 | 0.10 | 9.5E-09 |
| *math-41* | WBGene00020360 | protein_coding_gene | -3.3 | 0.10 | 1.6E-12 |
| *fbxa-216* | WBGene00012953 | protein_coding_gene | -3.3 | 0.10 | 6.0E-09 |
| *Y38H6C.23* | WBGene00012634 | protein_coding_gene | -3.3 | 0.10 | 6.5E-29 |
| *F14D2.11* | WBGene00017459 | protein_coding_gene | -3.3 | 0.10 | 1.6E-07 |
| *T16G12.3* | WBGene00011804 | protein_coding_gene | -3.4 | 0.10 | 1.4E-38 |
| *K09F6.9* | WBGene00019592 | protein_coding_gene | -3.4 | 0.10 | 1.7E-15 |
| *zeel-1* | WBGene00021463 | protein_coding_gene | -3.4 | 0.10 | 1.3E-09 |
| *btb-16* | WBGene00018195 | protein_coding_gene | -3.4 | 0.10 | 1.5E-59 |
| *math-40* | WBGene00020359 | protein_coding_gene | -3.4 | 0.10 | 7.3E-13 |
| *R08E5.1* | WBGene00019961 | protein_coding_gene | -3.4 | 0.09 | 6.3E-08 |
| *F52C6.12* | WBGene00018669 | protein_coding_gene | -3.4 | 0.09 | 3.3E-09 |
| *F26A3.5* | WBGene00009143 | protein_coding_gene | -3.4 | 0.09 | 8.9E-16 |
| *linc-11* | WBGene00219758 | lincRNA_gene | -3.4 | 0.09 | 1.5E-07 |
| *fbxa-60* | WBGene00020456 | protein_coding_gene | -3.4 | 0.09 | 2.0E-65 |
| *math-39* | WBGene00020358 | protein_coding_gene | -3.5 | 0.09 | 6.8E-15 |
| *F16H6.10* | WBGene00008900 | protein_coding_gene | -3.5 | 0.09 | 3.0E-22 |
| *ZC239.22* | WBGene00194717 | protein_coding_gene | -3.5 | 0.09 | 9.7E-10 |
| *bath-27* | WBGene00017450 | protein_coding_gene | -3.5 | 0.09 | 3.9E-17 |
| *btb-10* | WBGene00019313 | protein_coding_gene | -3.5 | 0.09 | 6.9E-08 |
| *F54E2.1* | WBGene00018823 | protein_coding_gene | -3.5 | 0.09 | 1.2E-26 |
| *fbxa-129* | WBGene00010294 | pseudogene | -3.5 | 0.09 | 1.8E-09 |
| *scrm-5* | WBGene00019531 | protein_coding_gene | -3.5 | 0.09 | 1.4E-09 |
| *R07E5.15* | WBGene00011120 | protein_coding_gene | -3.5 | 0.09 | 1.9E-14 |
| *fbxa-38* | WBGene00016522 | protein_coding_gene | -3.5 | 0.09 | 1.6E-09 |
| *pals-23* | WBGene00016218 | protein_coding_gene | -3.5 | 0.09 | 7.8E-16 |
| *fbxa-207* | WBGene00013612 | pseudogene | -3.5 | 0.09 | 5.7E-10 |
| *F19B10.1* | WBGene00017583 | protein_coding_gene | -3.5 | 0.09 | 5.8E-19 |
| *C08F11.7* | WBGene00007454 | protein_coding_gene | -3.5 | 0.09 | 4.9E-19 |
| *math-10* | WBGene00015839 | protein_coding_gene | -3.5 | 0.09 | 1.3E-09 |
| *math-20* | WBGene00016555 | protein_coding_gene | -3.5 | 0.09 | 7.2E-24 |
| *F31E9.11* | WBGene00195233 | ncRNA_gene | -3.6 | 0.09 | 1.1E-09 |
| *fbxa-6* | WBGene00008352 | protein_coding_gene | -3.6 | 0.08 | 2.3E-08 |
| *K09F6.4* | WBGene00019587 | protein_coding_gene | -3.6 | 0.08 | 2.7E-10 |
| *F59E12.6* | WBGene00019122 | protein_coding_gene | -3.6 | 0.08 | 7.5E-10 |

| | | | | | |
|---|---|---|---|---|---|
| *fbxa-65* | WBGene00015885 | protein_coding_gene | -3.6 | 0.08 | 2.2E-16 |
| *C29G2.2* | WBGene00016230 | pseudogene | -3.6 | 0.08 | 3.8E-08 |
| *F14D2.15* | WBGene00023427 | protein_coding_gene | -3.6 | 0.08 | 9.7E-24 |
| *nspa-5* | WBGene00013887 | protein_coding_gene | -3.7 | 0.08 | 2.3E-08 |
| *F19B10.2* | WBGene00017584 | protein_coding_gene | -3.7 | 0.08 | 2.4E-30 |
| *Y75B8A.32* | WBGene00013562 | protein_coding_gene | -3.7 | 0.08 | 4.1E-10 |
| *bath-24* | WBGene00015014 | protein_coding_gene | -3.7 | 0.07 | 5.8E-09 |
| *Y82E9BR.5* | WBGene00022338 | protein_coding_gene | -3.8 | 0.07 | 1.5E-08 |
| *fbxa-193* | WBGene00010295 | pseudogene | -3.8 | 0.07 | 5.0E-14 |
| *nspd-7* | WBGene00012781 | protein_coding_gene | -3.8 | 0.07 | 1.7E-40 |
| *duxl-1* | WBGene00022554 | protein_coding_gene | -3.8 | 0.07 | 4.8E-48 |
| *msp-64* | WBGene00003457 | protein_coding_gene | -3.8 | 0.07 | 1.3E-34 |
| *F09C6.12* | WBGene00044182 | protein_coding_gene | -3.8 | 0.07 | 1.9E-08 |
| *math-27* | WBGene00018101 | pseudogene | -3.8 | 0.07 | 2.8E-20 |
| *fbxa-180* | WBGene00009295 | pseudogene | -3.9 | 0.07 | 2.0E-09 |
| *F14D2.19* | WBGene00206522 | protein_coding_gene | -3.9 | 0.07 | 9.7E-11 |
| *Y46G5A.7* | WBGene00012899 | protein_coding_gene | -3.9 | 0.07 | 6.0E-09 |
| *C25F9.10* | WBGene00044291 | protein_coding_gene | -4.0 | 0.06 | 7.3E-11 |
| *K05C4.9* | WBGene00010585 | protein_coding_gene | -4.0 | 0.06 | 3.4E-09 |
| *K08D10.14* | WBGene00219216 | protein_coding_gene | -4.0 | 0.06 | 1.4E-09 |
| *C04E12.2* | WBGene00015433 | protein_coding_gene | -4.0 | 0.06 | 6.5E-11 |
| *math-22* | WBGene00016717 | protein_coding_gene | -4.0 | 0.06 | 3.5E-20 |
| *bath-19* | WBGene00019132 | protein_coding_gene | -4.0 | 0.06 | 5.6E-11 |
| *phat-3* | WBGene00016782 | protein_coding_gene | -4.0 | 0.06 | 2.6E-60 |
| *R10E8.7* | WBGene00014821 | pseudogene | -4.0 | 0.06 | 2.7E-12 |
| *ugt-25* | WBGene00015692 | protein_coding_gene | -4.0 | 0.06 | 6.5E-37 |
| *M04C3.5* | WBGene00206418 | protein_coding_gene | -4.0 | 0.06 | 3.9E-11 |
| *C25F9.14* | WBGene00045461 | protein_coding_gene | -4.0 | 0.06 | 4.4E-11 |
| *K09F6.15* | WBGene00271744 | protein_coding_gene | -4.1 | 0.06 | 3.0E-10 |
| *gst-22* | WBGene00001770 | protein_coding_gene | -4.2 | 0.05 | 1.3E-11 |
| *math-24* | WBGene00016719 | protein_coding_gene | -4.2 | 0.05 | 6.7E-22 |
| *E01G4.7* | WBGene00045410 | pseudogene | -4.2 | 0.05 | 2.7E-50 |
| *fbxa-189* | WBGene00009840 | protein_coding_gene | -4.2 | 0.05 | 9.5E-12 |
| *T07D3.4* | WBGene00020307 | protein_coding_gene | -4.3 | 0.05 | 1.1E-10 |
| *T24E12.5* | WBGene00020774 | protein_coding_gene | -4.3 | 0.05 | 4.1E-10 |
| *ZC204.14* | WBGene00022564 | protein_coding_gene | -4.3 | 0.05 | 6.8E-10 |
| *K09F6.6* | WBGene00019589 | protein_coding_gene | -4.3 | 0.05 | 4.0E-14 |
| *math-26* | WBGene00018100 | protein_coding_gene | -4.3 | 0.05 | 7.4E-42 |
| *math-3* | WBGene00015608 | protein_coding_gene | -4.3 | 0.05 | 6.4E-28 |
| *K09F6.7* | WBGene00019590 | protein_coding_gene | -4.3 | 0.05 | 4.7E-14 |
| *K09F6.3* | WBGene00019586 | protein_coding_gene | -4.3 | 0.05 | 2.0E-27 |
| *bath-2* | WBGene00018668 | protein_coding_gene | -4.3 | 0.05 | 7.8E-33 |
| *pud-3* | WBGene00017501 | protein_coding_gene | -4.4 | 0.05 | 1.6E-11 |
| *nspa-8* | WBGene00012294 | protein_coding_gene | -4.4 | 0.05 | 2.4E-10 |
| *M04C3.2* | WBGene00010851 | protein_coding_gene | -4.4 | 0.05 | 5.9E-11 |
| *Y102A5C.36* | WBGene00044213 | protein_coding_gene | -4.4 | 0.05 | 2.3E-12 |
| *Y102A5C.2* | WBGene00013611 | protein_coding_gene | -4.4 | 0.05 | 1.3E-16 |
| *btb-4* | WBGene00019142 | protein_coding_gene | -4.4 | 0.05 | 4.5E-14 |
| *F42G2.5* | WBGene00018354 | protein_coding_gene | -4.4 | 0.05 | 3.8E-11 |
| *btb-17* | WBGene00018200 | protein_coding_gene | -4.4 | 0.05 | 1.3E-16 |
| *T19C9.8* | WBGene00011844 | protein_coding_gene | -4.4 | 0.05 | 2.5E-10 |
| *E01G4.5* | WBGene00008447 | protein_coding_gene | -4.5 | 0.04 | 2.4E-17 |
| *bath-1* | WBGene00019139 | protein_coding_gene | -4.5 | 0.04 | 1.5E-10 |
| *C38D9.2* | WBGene00008010 | protein_coding_gene | -4.6 | 0.04 | 2.1E-193 |
| *fbxa-67* | WBGene00020457 | protein_coding_gene | -4.6 | 0.04 | 7.9E-11 |
| *msp-19* | WBGene00003426 | protein_coding_gene | -4.6 | 0.04 | 1.3E-20 |
| *Y7A9C.1* | WBGene00012418 | pseudogene | -4.7 | 0.04 | 5.3E-12 |
| *gst-27* | WBGene00001775 | protein_coding_gene | -4.7 | 0.04 | 1.7E-112 |
| *Y113G7B.12* | WBGene00013762 | protein_coding_gene | -4.7 | 0.04 | 3.6E-11 |
| *K09F6.10* | WBGene00019593 | protein_coding_gene | -4.8 | 0.04 | 1.3E-11 |

| | | | | | |
|---|---|---|---|---|---|
| *ZK1240.2* | WBGene00022867 | protein_coding_gene | -4.8 | 0.04 | 2.1E-12 |
| *C25F9.4* | WBGene00007724 | protein_coding_gene | -4.8 | 0.03 | 3.7E-16 |
| *F47H4.2* | WBGene00009835 | protein_coding_gene | -4.9 | 0.03 | 2.6E-14 |
| *R07C12.2* | WBGene00019932 | pseudogene | -4.9 | 0.03 | 2.0E-22 |
| *Y38E10A.14* | WBGene00012592 | protein_coding_gene | -4.9 | 0.03 | 1.4E-22 |
| *fbxa-206* | WBGene00013610 | protein_coding_gene | -4.9 | 0.03 | 1.0E-11 |
| *F19B10.10* | WBGene00017589 | protein_coding_gene | -4.9 | 0.03 | 1.3E-48 |
| *skr-4* | WBGene00004810 | protein_coding_gene | -4.9 | 0.03 | 8.5E-43 |
| *Y113G7B.14* | WBGene00013763 | pseudogene | -4.9 | 0.03 | 7.3E-12 |
| *F10E7.2* | WBGene00017344 | protein_coding_gene | -4.9 | 0.03 | 1.8E-25 |
| *bath-28* | WBGene00017461 | protein_coding_gene | -5.0 | 0.03 | 2.9E-28 |
| *clec-10* | WBGene00015403 | protein_coding_gene | -5.1 | 0.03 | 1.9E-12 |
| *C25F9.5* | WBGene00007725 | protein_coding_gene | -5.1 | 0.03 | 1.1E-14 |
| *F54D10.7* | WBGene00018808 | protein_coding_gene | -5.1 | 0.03 | 2.3E-12 |
| *B0281.3* | WBGene00015110 | protein_coding_gene | -5.2 | 0.03 | 6.1E-14 |
| *nspc-14* | WBGene00009638 | protein_coding_gene | -5.2 | 0.03 | 1.0E-105 |
| bath-5 | WBGene00019141 | protein_coding_gene | -5.3 | 0.03 | 7.0E-26 |
| *pals-24* | WBGene00016219 | protein_coding_gene | -5.3 | 0.02 | 1.1E-22 |
| *sri-40* | WBGene00005552 | protein_coding_gene | -5.3 | 0.02 | 2.9E-29 |
| *Y94H6A.10* | WBGene00022382 | protein_coding_gene | -5.4 | 0.02 | 3.6E-23 |
| *clec-76* | WBGene00021587 | protein_coding_gene | -5.4 | 0.02 | 4.0E-24 |
| *ZK1240.3* | WBGene00022868 | protein_coding_gene | -5.5 | 0.02 | 3.1E-31 |
| *F59A7.8* | WBGene00019095 | pseudogene | -5.5 | 0.02 | 9.7E-20 |
| *F19B2.5* | WBGene00008944 | protein_coding_gene | -5.6 | 0.02 | 2.6E-24 |
| *C04E12.4* | WBGene00015434 | protein_coding_gene | -5.6 | 0.02 | 2.4E-35 |
| *ZC204.12* | WBGene00022562 | protein_coding_gene | -5.6 | 0.02 | 3.1E-42 |
| *sdz-24* | WBGene00019495 | protein_coding_gene | -5.7 | 0.02 | 2.2E-101 |
| *T12B5.15* | WBGene00050914 | protein_coding_gene | -5.7 | 0.02 | 3.5E-39 |
| *math-4* | WBGene00015609 | protein_coding_gene | -5.7 | 0.02 | 1.5E-36 |
| *math-14* | WBGene00015828 | protein_coding_gene | -5.7 | 0.02 | 2.3E-25 |
| *Y102A5C.5* | WBGene00014954 | pseudogene | -5.7 | 0.02 | 5.0E-16 |
| *F55B11.6* | WBGene00044249 | protein_coding_gene | -5.9 | 0.02 | 1.0E-39 |
| *C27C7.1* | WBGene00007766 | protein_coding_gene | -6.0 | 0.02 | 8.3E-60 |
| *Y102A5C.6* | WBGene00014955 | pseudogene | -6.4 | 0.01 | 9.4E-55 |
| *M04C3.1* | WBGene00010850 | protein_coding_gene | -6.4 | 0.01 | 1.9E-55 |

**Supplemental Table 14**

**Lists of differentially expressed genes, by direction, that change after infection by *S. epidermidis* (compared to *E. coli*). Genes were specifically those that are different between N2 and CB4856 on *E. coli* prior to infection**

**Supplemental Table 14 - Differentially Expressed Genes Comparing the *E. coli* and *S. epidermidis* Conditions**

Genes exclusively expressed in CB4856 (vs. N2) fed *E. coli* OP50

| Gene Name | Wormbase Gene ID | Type | $log_2FC$ | FC | Adj P-value |
|---|---|---|---|---|---|
| *fmo-2* | WBGene00001477 | protein_coding_gene | 7.0 | 129.38 | 1.8E-23 |
| *T07G12.5* | WBGene00011596 | protein_coding_gene | 6.7 | 106.39 | 2.0E-51 |
| *mtl-1* | WBGene00003473 | protein_coding_gene | 6.0 | 64.43 | 1.3E-26 |
| *T26H5.9* | WBGene00044206 | protein_coding_gene | 6.0 | 62.77 | 3.4E-47 |
| *ilys-3* | WBGene00016670 | protein_coding_gene | 5.9 | 58.69 | 9.4E-25 |
| *F48C1.9* | WBGene00018601 | protein_coding_gene | 5.8 | 55.28 | 1.7E-13 |
| *lipl-3* | WBGene00020016 | protein_coding_gene | 5.6 | 49.01 | 1.4E-17 |
| *fil-1* | WBGene00011321 | protein_coding_gene | 5.5 | 46.60 | 3.1E-39 |
| *F46C5.1* | WBGene00009778 | protein_coding_gene | 5.5 | 46.22 | 4.5E-19 |
| *tts-1* | WBGene00006650 | ncRNA_gene | 5.5 | 46.01 | 1.6E-26 |
| *F45D3.4* | WBGene00009724 | protein_coding_gene | 5.4 | 43.19 | 5.0E-99 |
| *scl-2* | WBGene00009895 | protein_coding_gene | 5.2 | 36.38 | 1.2E-103 |
| *fbxa-24* | WBGene00018803 | protein_coding_gene | 5.2 | 35.68 | 3.0E-43 |
| *C18H7.1* | WBGene00015993 | protein_coding_gene | 5.2 | 35.52 | 9.9E-15 |
| *F47B8.4* | WBGene00009805 | protein_coding_gene | 5.1 | 34.31 | 8.6E-11 |
| *dct-8* | WBGene00044472 | protein_coding_gene | 4.9 | 30.46 | 2.0E-10 |
| *F53A9.8* | WBGene00018731 | protein_coding_gene | 4.8 | 27.39 | 1.6E-52 |
| *T26H5.8* | WBGene00012070 | protein_coding_gene | 4.8 | 27.10 | 1.7E-16 |
| *ftn-1* | WBGene00001500 | protein_coding_gene | 4.6 | 24.80 | 3.5E-21 |
| *F15E6.3* | WBGene00017484 | protein_coding_gene | 4.6 | 24.67 | 2.8E-36 |
| *F46A8.13* | WBGene00219950 | ncRNA_gene | 4.6 | 24.36 | 2.1E-07 |
| *Y41D4B.6* | WBGene00021513 | protein_coding_gene | 4.6 | 23.88 | 1.1E-12 |
| *clec-61* | WBGene00014047 | protein_coding_gene | 4.6 | 23.83 | 5.2E-13 |
| *Y53F4B.45* | WBGene00044728 | ncRNA_gene | 4.4 | 21.78 | 6.1E-61 |
| *F56D6.8* | WBGene00044292 | protein_coding_gene | 4.3 | 19.81 | 4.7E-07 |
| *cnc-11* | WBGene00044900 | protein_coding_gene | 4.3 | 19.47 | 5.5E-07 |
| *F11E6.11* | WBGene00023501 | protein_coding_gene | 4.3 | 19.19 | 1.9E-35 |
| *clec-70* | WBGene00021581 | protein_coding_gene | 4.3 | 19.08 | 1.1E-25 |
| *W02D7.11* | WBGene00020946 | protein_coding_gene | 4.2 | 18.90 | 1.6E-06 |
| *nlp-39* | WBGene00008295 | protein_coding_gene | 4.2 | 18.67 | 2.4E-08 |
| *F47B8.10* | WBGene00009811 | protein_coding_gene | 4.2 | 18.66 | 7.8E-07 |
| *cutl-9* | WBGene00010893 | protein_coding_gene | 4.2 | 18.26 | 1.1E-16 |
| *srr-6* | WBGene00005657 | protein_coding_gene | 4.1 | 17.66 | 2.9E-09 |
| *far-7* | WBGene00001391 | protein_coding_gene | 4.1 | 17.13 | 3.2E-35 |
| *cpt-3* | WBGene00021703 | protein_coding_gene | 4.0 | 16.15 | 3.0E-17 |
| *dct-7* | WBGene00017488 | protein_coding_gene | 4.0 | 15.83 | 6.7E-06 |
| *K10G6.4* | WBGene00019641 | protein_coding_gene | 4.0 | 15.58 | 9.9E-07 |
| *T26H5.14* | WBGene00235133 | protein_coding_gene | 3.9 | 15.33 | 1.0E-06 |
| *T26H5.4* | WBGene00012069 | protein_coding_gene | 3.9 | 15.20 | 4.0E-06 |
| *fipr-23* | WBGene00007990 | protein_coding_gene | 3.9 | 15.08 | 3.8E-06 |
| *F37A8.5* | WBGene00009503 | protein_coding_gene | 3.9 | 14.69 | 5.6E-45 |
| *C35A5.11* | WBGene00044024 | protein_coding_gene | 3.9 | 14.66 | 3.1E-09 |
| *cutl-28* | WBGene00018256 | protein_coding_gene | 3.8 | 14.34 | 8.2E-14 |
| *C06E7.88* | WBGene00189995 | protein_coding_gene | 3.8 | 14.27 | 6.0E-08 |
| *T03G6.1* | WBGene00020194 | protein_coding_gene | 3.8 | 14.00 | 6.1E-13 |
| *egl-46* | WBGene00001210 | protein_coding_gene | 3.8 | 13.75 | 4.7E-17 |
| *nhr-127* | WBGene00003717 | protein_coding_gene | 3.8 | 13.73 | 3.7E-06 |
| *C44H9.7* | WBGene00008102 | protein_coding_gene | 3.8 | 13.54 | 7.9E-10 |
| *wrt-2* | WBGene00006948 | protein_coding_gene | 3.7 | 13.39 | 1.9E-11 |
| *ugt-18* | WBGene00013900 | protein_coding_gene | 3.7 | 13.36 | 6.8E-07 |
| *swt-1* | WBGene00019300 | protein_coding_gene | 3.7 | 13.33 | 3.6E-27 |
| *cutl-2* | WBGene00013145 | protein_coding_gene | 3.7 | 13.24 | 9.0E-20 |
| *dpy-14* | WBGene00001075 | protein_coding_gene | 3.7 | 13.20 | 6.2E-10 |
| *F16B4.7* | WBGene00017509 | protein_coding_gene | 3.7 | 13.09 | 6.7E-06 |
| *dod-3* | WBGene00016052 | protein_coding_gene | 3.7 | 13.02 | 3.5E-09 |
| *dpy-7* | WBGene00001069 | protein_coding_gene | 3.7 | 12.76 | 8.3E-20 |
| *H42K12.3* | WBGene00019272 | protein_coding_gene | 3.7 | 12.71 | 5.1E-18 |
| *C25H3.10* | WBGene00016119 | protein_coding_gene | 3.7 | 12.61 | 2.5E-24 |
| *F54E2.2* | WBGene00018824 | protein_coding_gene | 3.6 | 12.49 | 7.0E-05 |
| *fbxa-48* | WBGene00021862 | protein_coding_gene | 3.6 | 12.31 | 4.5E-08 |
| *T05A7.13* | WBGene00271415 | protein_coding_gene | 3.6 | 12.24 | 3.9E-08 |

| | | | | | |
|---|---|---|---|---|---|
| *T20B3.1* | WBGene00011850 | protein_coding_gene | 3.6 | 12.11 | 1.3E-26 |
| *lpr-5* | WBGene00012256 | protein_coding_gene | 3.6 | 11.77 | 3.3E-18 |
| *lpr-4* | WBGene00012257 | protein_coding_gene | 3.6 | 11.77 | 1.8E-13 |
| *Y54G2A.37* | WBGene00023484 | protein_coding_gene | 3.6 | 11.76 | 1.1E-04 |
| *ces-2* | WBGene00000469 | protein_coding_gene | 3.6 | 11.75 | 5.0E-22 |
| *cyp-34A9* | WBGene00015044 | protein_coding_gene | 3.5 | 11.71 | 1.2E-127 |
| *F43D9.1* | WBGene00009653 | protein_coding_gene | 3.5 | 11.68 | 5.5E-16 |
| *T12D8.5* | WBGene00011733 | protein_coding_gene | 3.5 | 11.55 | 2.3E-36 |
| *sodh-1* | WBGene00010790 | protein_coding_gene | 3.5 | 11.47 | 1.8E-08 |
| *hch-1* | WBGene00001828 | protein_coding_gene | 3.5 | 11.47 | 1.0E-16 |
| *dpy-2* | WBGene00001064 | protein_coding_gene | 3.5 | 11.43 | 9.9E-17 |
| *cnc-4* | WBGene00000558 | protein_coding_gene | 3.5 | 11.40 | 9.8E-13 |
| *F59C6.16* | WBGene00077764 | protein_coding_gene | 3.5 | 11.29 | 6.4E-33 |
| *mlt-11* | WBGene00012186 | protein_coding_gene | 3.5 | 11.21 | 1.5E-31 |
| *ZC395.5* | WBGene00022597 | protein_coding_gene | 3.5 | 11.18 | 1.4E-23 |
| *F55H12.7* | WBGene00219283 | protein_coding_gene | 3.5 | 11.16 | 4.6E-05 |
| *pals-32* | WBGene00016281 | protein_coding_gene | 3.5 | 11.16 | 4.2E-11 |
| *F17B5.1* | WBGene00008905 | protein_coding_gene | 3.5 | 11.11 | 1.4E-06 |
| *dpy-3* | WBGene00001065 | protein_coding_gene | 3.5 | 11.07 | 2.0E-15 |
| *bro-1* | WBGene00000272 | protein_coding_gene | 3.5 | 11.01 | 2.4E-08 |
| *Y65B4BL.1* | WBGene00022033 | protein_coding_gene | 3.5 | 10.96 | 1.1E-04 |
| *C03C10.5* | WBGene00007271 | protein_coding_gene | 3.5 | 10.94 | 9.2E-09 |
| *dpy-10* | WBGene00001072 | protein_coding_gene | 3.4 | 10.88 | 4.8E-24 |
| *dao-2* | WBGene00000928 | protein_coding_gene | 3.4 | 10.85 | 2.6E-25 |
| *ZC443.3* | WBGene00013898 | protein_coding_gene | 3.4 | 10.78 | 1.7E-61 |
| *dod-22* | WBGene00010125 | protein_coding_gene | 3.4 | 10.63 | 4.2E-20 |
| *Y41D4B.26* | WBGene00021525 | protein_coding_gene | 3.4 | 10.61 | 4.8E-16 |
| *W09G12.7* | WBGene00021121 | protein_coding_gene | 3.4 | 10.58 | 2.2E-18 |
| *lpr-3* | WBGene00012261 | protein_coding_gene | 3.4 | 10.58 | 2.0E-21 |
| *K08B12.1* | WBGene00019520 | protein_coding_gene | 3.4 | 10.57 | 8.7E-11 |
| *lpr-6* | WBGene00012255 | protein_coding_gene | 3.4 | 10.51 | 2.9E-12 |
| *clec-60* | WBGene00014046 | protein_coding_gene | 3.4 | 10.39 | 4.0E-10 |
| *fipr-24* | WBGene00007992 | protein_coding_gene | 3.4 | 10.36 | 7.8E-05 |
| *Y54G2A.76* | WBGene00235268 | protein_coding_gene | 3.4 | 10.21 | 2.3E-06 |
| *noah-2* | WBGene00009926 | protein_coding_gene | 3.3 | 10.10 | 3.7E-32 |
| *F15E6.10* | WBGene00044479 | protein_coding_gene | 3.3 | 10.08 | 3.2E-04 |
| *cky-1* | WBGene00000521 | protein_coding_gene | 3.3 | 10.07 | 8.0E-05 |
| *dpy-17* | WBGene00001076 | protein_coding_gene | 3.3 | 10.06 | 5.4E-09 |
| *F09F7.6* | WBGene00017303 | protein_coding_gene | 3.3 | 10.06 | 5.3E-31 |
| *F31D4.5* | WBGene00009292 | protein_coding_gene | 3.3 | 10.03 | 9.1E-10 |
| *C14F11.6* | WBGene00015781 | protein_coding_gene | 3.3 | 10.03 | 2.4E-13 |
| *C06E4.8* | WBGene00015537 | protein_coding_gene | 3.3 | 10.02 | 1.2E-11 |
| *K08B4.2* | WBGene00019514 | protein_coding_gene | 3.3 | 9.77 | 5.8E-10 |
| *chil-28* | WBGene00008842 | pseudogene | 3.3 | 9.71 | 1.3E-04 |
| *lim-4* | WBGene00002987 | protein_coding_gene | 3.3 | 9.71 | 1.0E-04 |
| *T04C12.1* | WBGene00011427 | protein_coding_gene | 3.3 | 9.65 | 1.1E-05 |
| *T26H5.10* | WBGene00044783 | protein_coding_gene | 3.3 | 9.62 | 3.8E-04 |
| *Y7A5A.6* | WBGene00012412 | protein_coding_gene | 3.3 | 9.57 | 1.1E-04 |
| *ZK105.13* | WBGene00235114 | protein_coding_gene | 3.3 | 9.56 | 3.3E-04 |
| *ugt-57* | WBGene00017154 | protein_coding_gene | 3.3 | 9.54 | 7.1E-12 |
| *ZK970.7* | WBGene00014173 | protein_coding_gene | 3.2 | 9.48 | 3.4E-10 |
| *F53A9.7* | WBGene00018730 | protein_coding_gene | 3.2 | 9.46 | 3.2E-25 |
| *K10D3.4* | WBGene00010738 | protein_coding_gene | 3.2 | 9.43 | 8.6E-22 |
| *coel-1* | WBGene00016866 | protein_coding_gene | 3.2 | 9.42 | 3.7E-52 |
| *F56D2.3* | WBGene00018965 | protein_coding_gene | 3.2 | 9.39 | 3.5E-10 |
| *atf-2* | WBGene00000220 | protein_coding_gene | 3.2 | 9.34 | 4.1E-16 |
| *R03H10.2* | WBGene00019855 | protein_coding_gene | 3.2 | 9.32 | 1.2E-15 |
| *unc-42* | WBGene00006778 | protein_coding_gene | 3.2 | 9.30 | 2.0E-06 |
| *T02E9.5* | WBGene00011383 | protein_coding_gene | 3.2 | 9.28 | 6.8E-28 |
| *fbxa-21* | WBGene00022487 | protein_coding_gene | 3.2 | 9.17 | 7.1E-18 |
| *dsl-3* | WBGene00001105 | protein_coding_gene | 3.2 | 9.11 | 5.0E-17 |
| *hhat-1* | WBGene00013852 | protein_coding_gene | 3.2 | 9.10 | 3.8E-11 |
| *F13E9.11* | WBGene00008760 | protein_coding_gene | 3.2 | 9.09 | 2.3E-22 |
| *spp-2* | WBGene00004987 | protein_coding_gene | 3.2 | 9.09 | 4.6E-41 |
| *atf-8* | WBGene00017535 | protein_coding_gene | 3.2 | 9.08 | 5.3E-08 |

| | | | | | |
|---|---|---|---|---|---|
| *wrt-10* | WBGene00006956 | protein_coding_gene | 3.2 | 9.05 | 8.8E-27 |
| *M03B6.3* | WBGene00010835 | protein_coding_gene | 3.2 | 9.00 | 2.7E-10 |
| *hil-1* | WBGene00001852 | protein_coding_gene | 3.2 | 8.99 | 1.2E-20 |
| *C45E5.3* | WBGene00016666 | pseudogene | 3.2 | 8.94 | 5.7E-09 |
| *hlh-16* | WBGene00001960 | protein_coding_gene | 3.2 | 8.92 | 1.6E-08 |
| *F49B2.4* | WBGene00009864 | pseudogene | 3.2 | 8.91 | 6.7E-04 |
| *T10C6.15* | WBGene00050899 | protein_coding_gene | 3.1 | 8.87 | 4.2E-08 |
| *ptr-10* | WBGene00004224 | protein_coding_gene | 3.1 | 8.80 | 4.6E-08 |
| *ins-2* | WBGene00002085 | protein_coding_gene | 3.1 | 8.74 | 7.1E-08 |
| *peb-1* | WBGene00003968 | protein_coding_gene | 3.1 | 8.71 | 2.0E-18 |
| *Y37A1B.7* | WBGene00012540 | protein_coding_gene | 3.1 | 8.67 | 1.3E-06 |
| *F48F7.3* | WBGene00009849 | protein_coding_gene | 3.1 | 8.61 | 6.7E-04 |
| *Y65B4BM.3* | WBGene00302974 | protein_coding_gene | 3.1 | 8.58 | 1.5E-14 |
| *ces-1* | WBGene00000468 | protein_coding_gene | 3.1 | 8.55 | 7.0E-06 |
| *clec-25* | WBGene00044331 | protein_coding_gene | 3.1 | 8.54 | 4.1E-06 |
| *T21D12.7* | WBGene00020648 | protein_coding_gene | 3.1 | 8.49 | 6.0E-04 |
| *nnt-1* | WBGene00003778 | protein_coding_gene | 3.1 | 8.49 | 3.3E-33 |
| *ptr-4* | WBGene00004219 | protein_coding_gene | 3.1 | 8.47 | 8.4E-13 |
| *noah-1* | WBGene00016422 | protein_coding_gene | 3.1 | 8.44 | 1.1E-33 |
| *dod-24* | WBGene00007875 | protein_coding_gene | 3.1 | 8.42 | 6.0E-07 |
| *C08E8.3* | WBGene00007439 | protein_coding_gene | 3.1 | 8.35 | 7.1E-05 |
| *K06A9.2* | WBGene00019436 | protein_coding_gene | 3.1 | 8.29 | 4.6E-24 |
| *oac-29* | WBGene00018295 | protein_coding_gene | 3.0 | 8.23 | 4.0E-04 |
| *R09E10.13* | WBGene00194658 | protein_coding_gene | 3.0 | 8.20 | 3.3E-04 |
| *cup-16* | WBGene00007919 | protein_coding_gene | 3.0 | 8.17 | 1.5E-21 |
| *clec-78* | WBGene00018547 | protein_coding_gene | 3.0 | 8.17 | 6.0E-06 |
| *clec-23* | WBGene00011851 | pseudogene | 3.0 | 8.12 | 1.2E-04 |
| *btb-15* | WBGene00015594 | pseudogene | 3.0 | 8.04 | 4.3E-04 |
| *ins-8* | WBGene00002091 | protein_coding_gene | 3.0 | 7.96 | 1.1E-03 |
| *ugt-54* | WBGene00012013 | protein_coding_gene | 3.0 | 7.95 | 1.1E-36 |
| *clec-17* | WBGene00008477 | protein_coding_gene | 3.0 | 7.87 | 7.2E-04 |
| *W02A2.9* | WBGene00044280 | protein_coding_gene | 3.0 | 7.86 | 1.7E-18 |
| *fbxa-157* | WBGene00007393 | protein_coding_gene | 3.0 | 7.80 | 4.0E-06 |
| *hbl-1* | WBGene00001824 | protein_coding_gene | 3.0 | 7.76 | 1.4E-15 |
| *btb-21* | WBGene00018540 | pseudogene | 3.0 | 7.75 | 1.6E-12 |
| *C09F9.2* | WBGene00007479 | protein_coding_gene | 2.9 | 7.71 | 4.2E-39 |
| *ifa-3* | WBGene00002051 | protein_coding_gene | 2.9 | 7.70 | 5.3E-05 |
| *R09E10.5* | WBGene00011175 | protein_coding_gene | 2.9 | 7.67 | 1.2E-04 |
| *M01B2.13* | WBGene00045338 | protein_coding_gene | 2.9 | 7.65 | 2.1E-07 |
| *ttr-23* | WBGene00011894 | protein_coding_gene | 2.9 | 7.63 | 6.6E-10 |
| *tre-5* | WBGene00006611 | protein_coding_gene | 2.9 | 7.61 | 3.5E-21 |
| *lipl-1* | WBGene00010062 | protein_coding_gene | 2.9 | 7.61 | 4.3E-10 |
| *lys-3* | WBGene00003092 | protein_coding_gene | 2.9 | 7.61 | 7.4E-06 |
| *T20F5.4* | WBGene00020626 | protein_coding_gene | 2.9 | 7.59 | 1.7E-07 |
| *cyp-34A10* | WBGene00015045 | protein_coding_gene | 2.9 | 7.56 | 7.7E-12 |
| *M01D1.12* | WBGene00269373 | protein_coding_gene | 2.9 | 7.55 | 3.3E-06 |
| *F57B1.6* | WBGene00010189 | protein_coding_gene | 2.9 | 7.54 | 7.2E-08 |
| *clec-71* | WBGene00021582 | protein_coding_gene | 2.9 | 7.52 | 1.6E-08 |
| *F13H10.5* | WBGene00008776 | protein_coding_gene | 2.9 | 7.50 | 6.3E-07 |
| *ech-9* | WBGene00001158 | protein_coding_gene | 2.9 | 7.45 | 5.4E-05 |
| *Y74C9A.6* | WBGene00023193 | snoRNA_gene | 2.9 | 7.44 | 1.4E-03 |
| *C06B3.6* | WBGene00007365 | protein_coding_gene | 2.9 | 7.44 | 1.6E-23 |
| *fbxa-137* | WBGene00020853 | protein_coding_gene | 2.9 | 7.43 | 1.7E-03 |
| *zmp-5* | WBGene00010423 | protein_coding_gene | 2.9 | 7.40 | 1.7E-03 |
| *Y81B9A.5* | WBGene00304906 | ncRNA_gene | 2.9 | 7.39 | 1.7E-03 |
| *tsp-17* | WBGene00006643 | protein_coding_gene | 2.9 | 7.38 | 3.3E-06 |
| *F21C10.11* | WBGene00017660 | protein_coding_gene | 2.9 | 7.37 | 1.7E-06 |
| *C18D11.6* | WBGene00007682 | protein_coding_gene | 2.9 | 7.36 | 7.7E-04 |
| *math-43* | WBGene00022432 | pseudogene | 2.9 | 7.36 | 7.8E-04 |
| *C27A7.8* | WBGene00007758 | protein_coding_gene | 2.9 | 7.35 | 2.0E-17 |
| *prkl-1* | WBGene00022727 | protein_coding_gene | 2.9 | 7.34 | 2.1E-18 |
| *zig-10* | WBGene00020800 | protein_coding_gene | 2.9 | 7.33 | 2.9E-08 |
| *Y51H4A.5* | WBGene00013101 | protein_coding_gene | 2.9 | 7.32 | 7.6E-07 |
| *C54F6.5* | WBGene00016923 | protein_coding_gene | 2.9 | 7.30 | 2.3E-08 |
| *T19C4.1* | WBGene00011838 | protein_coding_gene | 2.9 | 7.29 | 9.6E-10 |

| | | | | | |
|---|---|---|---|---|---|
| *ZC449.2* | WBGene00022612 | protein_coding_gene | 2.9 | 7.29 | 5.7E-08 |
| *C54C8.4* | WBGene00008292 | protein_coding_gene | 2.9 | 7.24 | 1.8E-03 |
| *F56D6.9* | WBGene00044471 | protein_coding_gene | 2.9 | 7.24 | 7.5E-04 |
| *T09A12.8* | WBGene00235275 | protein_coding_gene | 2.9 | 7.21 | 7.8E-04 |
| *C29A12.6* | WBGene00007800 | protein_coding_gene | 2.9 | 7.21 | 7.0E-05 |
| *R09D1.12* | WBGene00011168 | protein_coding_gene | 2.8 | 7.19 | 8.9E-07 |
| *sqt-3* | WBGene00005018 | protein_coding_gene | 2.8 | 7.16 | 1.4E-20 |
| *M02E1.4* | WBGene00284848 | protein_coding_gene | 2.8 | 7.12 | 2.9E-05 |
| *T14B4.19* | WBGene00302993 | protein_coding_gene | 2.8 | 7.09 | 6.6E-09 |
| *tsp-1* | WBGene00006627 | protein_coding_gene | 2.8 | 7.03 | 2.7E-06 |
| *rncs-1* | WBGene00023069 | ncRNA_gene | 2.8 | 6.99 | 1.1E-19 |
| *F17B5.6* | WBGene00044058 | protein_coding_gene | 2.8 | 6.94 | 2.1E-03 |
| *F11E6.9* | WBGene00008712 | protein_coding_gene | 2.8 | 6.93 | 5.5E-04 |
| *C24H10.3* | WBGene00016069 | protein_coding_gene | 2.8 | 6.92 | 2.1E-05 |
| *F36F12.1* | WBGene00018092 | protein_coding_gene | 2.8 | 6.88 | 1.3E-03 |
| *ZK381.2* | WBGene00022725 | protein_coding_gene | 2.8 | 6.88 | 2.6E-03 |
| *cpt-4* | WBGene00019644 | protein_coding_gene | 2.8 | 6.81 | 3.0E-07 |
| *fbn-1* | WBGene00022816 | protein_coding_gene | 2.8 | 6.80 | 4.5E-47 |
| *EEED8.2* | WBGene00017133 | protein_coding_gene | 2.8 | 6.73 | 5.1E-13 |
| *fbxa-135* | WBGene00011539 | protein_coding_gene | 2.7 | 6.73 | 1.6E-03 |
| *kreg-1* | WBGene00018725 | protein_coding_gene | 2.7 | 6.72 | 3.8E-05 |
| *ZK675.4* | WBGene00014067 | protein_coding_gene | 2.7 | 6.67 | 8.0E-04 |
| *best-5* | WBGene00007404 | protein_coding_gene | 2.7 | 6.63 | 1.0E-08 |
| *T22F3.11* | WBGene00020700 | protein_coding_gene | 2.7 | 6.61 | 4.7E-05 |
| *fbxa-68* | WBGene00020641 | pseudogene | 2.7 | 6.59 | 1.4E-03 |
| *sym-1* | WBGene00006366 | protein_coding_gene | 2.7 | 6.56 | 2.1E-31 |
| *fmi-1* | WBGene00001475 | protein_coding_gene | 2.7 | 6.54 | 1.2E-17 |
| *fbxa-164* | WBGene00015599 | protein_coding_gene | 2.7 | 6.52 | 4.0E-03 |
| *hil-7* | WBGene00001858 | protein_coding_gene | 2.7 | 6.52 | 6.5E-04 |
| *Y46H3A.5* | WBGene00021601 | protein_coding_gene | 2.7 | 6.51 | 2.8E-03 |
| *slc-36.5* | WBGene00016628 | protein_coding_gene | 2.7 | 6.50 | 1.1E-15 |
| *C35C5.8* | WBGene00007957 | protein_coding_gene | 2.7 | 6.50 | 1.6E-17 |
| *bgnt-1.3* | WBGene00009032 | protein_coding_gene | 2.7 | 6.49 | 3.0E-03 |
| *K08F9.3* | WBGene00010686 | protein_coding_gene | 2.7 | 6.43 | 8.9E-05 |
| *cnc-2* | WBGene00000556 | protein_coding_gene | 2.7 | 6.41 | 1.5E-06 |
| *cut-5* | WBGene00011104 | protein_coding_gene | 2.7 | 6.40 | 9.8E-05 |
| *nphp-1* | WBGene00010898 | protein_coding_gene | 2.7 | 6.40 | 4.7E-06 |
| *T04C12.7* | WBGene00011429 | protein_coding_gene | 2.7 | 6.38 | 1.5E-05 |
| *pax-1* | WBGene00003937 | protein_coding_gene | 2.7 | 6.32 | 1.9E-03 |
| *T02G5.15* | WBGene00206521 | protein_coding_gene | 2.7 | 6.31 | 5.5E-05 |
| *F36D1.7* | WBGene00009466 | protein_coding_gene | 2.6 | 6.24 | 1.5E-09 |
| *sru-20* | WBGene00005683 | protein_coding_gene | 2.6 | 6.23 | 1.8E-03 |
| *ins-19* | WBGene00002102 | protein_coding_gene | 2.6 | 6.22 | 2.7E-03 |
| *gei-13* | WBGene00001570 | protein_coding_gene | 2.6 | 6.21 | 5.7E-12 |
| *nspa-6* | WBGene00044273 | protein_coding_gene | 2.6 | 6.21 | 1.8E-03 |
| *F18C5.9* | WBGene00017562 | protein_coding_gene | 2.6 | 6.19 | 4.5E-03 |
| *col-121* | WBGene00000695 | protein_coding_gene | 2.6 | 6.17 | 9.8E-04 |
| *F55H12.2* | WBGene00010133 | protein_coding_gene | 2.6 | 6.15 | 1.7E-22 |
| *pitr-4* | WBGene00007141 | protein_coding_gene | 2.6 | 6.12 | 1.0E-03 |
| *let-4* | WBGene00002282 | protein_coding_gene | 2.6 | 6.11 | 3.3E-15 |
| *F10D7.4* | WBGene00017341 | protein_coding_gene | 2.6 | 6.09 | 5.1E-03 |
| *T04C12.32* | WBGene00235086 | ncRNA_gene | 2.6 | 6.07 | 5.2E-03 |
| *T05F1.11* | WBGene00011495 | protein_coding_gene | 2.6 | 6.06 | 3.2E-11 |
| *F59B2.8* | WBGene00010310 | protein_coding_gene | 2.6 | 6.05 | 1.3E-05 |
| *fbxa-162* | WBGene00015597 | pseudogene | 2.6 | 6.04 | 4.9E-06 |
| *ZK287.1* | WBGene00013962 | protein_coding_gene | 2.6 | 6.02 | 5.7E-09 |
| *nlp-34* | WBGene00015046 | protein_coding_gene | 2.6 | 6.02 | 2.1E-05 |
| *ugt-52* | WBGene00018931 | protein_coding_gene | 2.6 | 6.00 | 2.5E-11 |
| *tmem-231* | WBGene00020825 | protein_coding_gene | 2.6 | 5.98 | 3.3E-04 |
| *Y7A9D.1* | WBGene00012420 | protein_coding_gene | 2.6 | 5.98 | 1.2E-19 |
| *bus-12* | WBGene00010442 | protein_coding_gene | 2.6 | 5.97 | 1.0E-04 |
| *ZC84.6* | WBGene00013849 | protein_coding_gene | 2.6 | 5.96 | 6.9E-05 |
| *B0205.4* | WBGene00015019 | protein_coding_gene | 2.6 | 5.94 | 4.9E-04 |
| *M03D4.4* | WBGene00019751 | protein_coding_gene | 2.6 | 5.94 | 1.8E-10 |
| *rig-5* | WBGene00004372 | protein_coding_gene | 2.6 | 5.93 | 5.9E-08 |

| | | | | | |
|---|---|---|---|---|---|
| *F11D5.5* | WBGene00017382 | protein_coding_gene | 2.6 | 5.91 | 1.6E-07 |
| *T01D3.3* | WBGene00011328 | protein_coding_gene | 2.6 | 5.90 | 3.4E-08 |
| *F40G9.20* | WBGene00195034 | ncRNA_gene | 2.6 | 5.89 | 3.0E-04 |
| *K01A6.8* | WBGene00194869 | protein_coding_gene | 2.6 | 5.87 | 6.0E-03 |
| *ceh-16* | WBGene00000439 | protein_coding_gene | 2.5 | 5.85 | 3.5E-06 |
| *ZC123.1* | WBGene00022517 | protein_coding_gene | 2.5 | 5.85 | 1.5E-07 |
| *F10D7.3* | WBGene00017340 | protein_coding_gene | 2.5 | 5.85 | 3.4E-18 |
| *ceh-8* | WBGene00000433 | protein_coding_gene | 2.5 | 5.84 | 1.0E-03 |
| *sodh-2* | WBGene00010791 | protein_coding_gene | 2.5 | 5.83 | 1.0E-03 |
| *C14A11.2* | WBGene00015748 | protein_coding_gene | 2.5 | 5.82 | 8.7E-04 |
| *srt-63* | WBGene00020877 | protein_coding_gene | 2.5 | 5.82 | 4.0E-03 |
| *spp-7* | WBGene00004992 | protein_coding_gene | 2.5 | 5.81 | 6.8E-03 |
| *subs-4* | WBGene00012939 | protein_coding_gene | 2.5 | 5.81 | 1.1E-05 |
| *F58H1.2* | WBGene00010285 | protein_coding_gene | 2.5 | 5.80 | 1.6E-04 |
| *C48B4.3* | WBGene00008168 | protein_coding_gene | 2.5 | 5.79 | 1.3E-05 |
| *lpr-1* | WBGene00002393 | protein_coding_gene | 2.5 | 5.78 | 1.1E-05 |
| *F09F9.4* | WBGene00017309 | protein_coding_gene | 2.5 | 5.78 | 5.8E-07 |
| *C36B1.6* | WBGene00007973 | protein_coding_gene | 2.5 | 5.78 | 4.2E-18 |
| *T26C12.3* | WBGene00020833 | protein_coding_gene | 2.5 | 5.78 | 1.9E-11 |
| *nspe-6* | WBGene00012603 | protein_coding_gene | 2.5 | 5.77 | 5.2E-03 |
| *ubc-8* | WBGene00006705 | protein_coding_gene | 2.5 | 5.76 | 2.2E-49 |
| *ZC449.1* | WBGene00022611 | protein_coding_gene | 2.5 | 5.72 | 7.6E-04 |
| *spp-6* | WBGene00004991 | protein_coding_gene | 2.5 | 5.72 | 2.5E-03 |
| *nspe-7* | WBGene00012593 | protein_coding_gene | 2.5 | 5.72 | 2.7E-04 |
| *F16C3.2* | WBGene00008886 | protein_coding_gene | 2.5 | 5.68 | 3.8E-26 |
| *ttr-52* | WBGene00008719 | protein_coding_gene | 2.5 | 5.68 | 1.3E-04 |
| *C18D4.8* | WBGene00007677 | pseudogene | 2.5 | 5.67 | 3.0E-17 |
| *fbxb-101* | WBGene00013404 | protein_coding_gene | 2.5 | 5.67 | 3.9E-06 |
| *gbh-2* | WBGene00001523 | protein_coding_gene | 2.5 | 5.66 | 5.3E-17 |
| *cyp-32B1* | WBGene00021167 | protein_coding_gene | 2.5 | 5.66 | 1.5E-08 |
| *F44E2.4* | WBGene00018418 | protein_coding_gene | 2.5 | 5.63 | 4.3E-11 |
| *cutl-16* | WBGene00019426 | protein_coding_gene | 2.5 | 5.63 | 5.5E-10 |
| *F18G5.6* | WBGene00017582 | protein_coding_gene | 2.5 | 5.61 | 4.2E-56 |
| *spp-13* | WBGene00004998 | protein_coding_gene | 2.5 | 5.60 | 2.1E-04 |
| *F09E10.6* | WBGene00017296 | protein_coding_gene | 2.5 | 5.60 | 3.6E-04 |
| *F15B9.8* | WBGene00008851 | protein_coding_gene | 2.5 | 5.60 | 4.3E-06 |
| *igcm-2* | WBGene00020130 | protein_coding_gene | 2.5 | 5.59 | 2.7E-06 |
| *T26C5.2* | WBGene00012032 | protein_coding_gene | 2.5 | 5.59 | 6.9E-06 |
| *fbxb-39* | WBGene00019702 | protein_coding_gene | 2.5 | 5.59 | 7.8E-03 |
| *T12A7.6* | WBGene00011727 | protein_coding_gene | 2.5 | 5.56 | 7.3E-17 |
| *Y23H5B.8* | WBGene00021279 | protein_coding_gene | 2.5 | 5.55 | 8.3E-07 |
| *cutl-25* | WBGene00021919 | protein_coding_gene | 2.5 | 5.54 | 5.4E-09 |
| *aqp-2* | WBGene00000170 | protein_coding_gene | 2.5 | 5.52 | 2.4E-23 |
| *F54C8.6* | WBGene00010039 | protein_coding_gene | 2.5 | 5.51 | 5.9E-08 |
| *Y97E10AL.1* | WBGene00022392 | protein_coding_gene | 2.5 | 5.51 | 7.1E-04 |
| *dmd-5* | WBGene00017326 | protein_coding_gene | 2.5 | 5.51 | 1.0E-06 |
| *math-34* | WBGene00019699 | protein_coding_gene | 2.5 | 5.51 | 5.0E-05 |
| *tsp-2* | WBGene00006628 | protein_coding_gene | 2.5 | 5.50 | 2.7E-06 |
| *nhr-218* | WBGene00011750 | protein_coding_gene | 2.5 | 5.50 | 4.3E-03 |
| *F58F12.12* | WBGene00271812 | protein_coding_gene | 2.5 | 5.49 | 1.2E-04 |
| *W03F9.4* | WBGene00021002 | protein_coding_gene | 2.5 | 5.48 | 4.5E-16 |
| *C04A11.5* | WBGene00007288 | protein_coding_gene | 2.5 | 5.47 | 1.5E-05 |
| *K01A2.3* | WBGene00019278 | pseudogene | 2.4 | 5.45 | 7.5E-07 |
| *nep-16* | WBGene00018472 | protein_coding_gene | 2.4 | 5.45 | 1.3E-03 |
| *K08E3.2* | WBGene00010662 | protein_coding_gene | 2.4 | 5.43 | 1.8E-03 |
| *T28H10.3* | WBGene00012144 | protein_coding_gene | 2.4 | 5.42 | 2.0E-29 |
| *dma-1* | WBGene00011345 | protein_coding_gene | 2.4 | 5.40 | 9.1E-14 |
| *sssh-1* | WBGene00044138 | protein_coding_gene | 2.4 | 5.40 | 9.0E-03 |
| *fmo-1* | WBGene00001476 | protein_coding_gene | 2.4 | 5.39 | 2.7E-22 |
| *chil-23* | WBGene00011167 | protein_coding_gene | 2.4 | 5.39 | 1.8E-07 |
| *F13A7.11* | WBGene00008726 | protein_coding_gene | 2.4 | 5.38 | 2.7E-07 |
| *C43D7.8* | WBGene00008070 | protein_coding_gene | 2.4 | 5.38 | 4.3E-04 |
| *K08F9.1* | WBGene00010684 | protein_coding_gene | 2.4 | 5.37 | 7.5E-05 |
| *zag-1* | WBGene00006970 | protein_coding_gene | 2.4 | 5.36 | 2.3E-13 |
| *K01A6.7* | WBGene00045096 | protein_coding_gene | 2.4 | 5.35 | 8.7E-03 |

| | | | | | |
|---|---|---|---|---|---|
| *acn-1* | WBGene00000039 | protein_coding_gene | 2.4 | 5.33 | 1.4E-11 |
| *W10G11.4* | WBGene00021137 | protein_coding_gene | 2.4 | 5.32 | 1.5E-03 |
| *F13H6.5* | WBGene00017433 | protein_coding_gene | 2.4 | 5.30 | 5.4E-08 |
| *str-173* | WBGene00006217 | protein_coding_gene | 2.4 | 5.30 | 9.2E-03 |
| *T16G1.4* | WBGene00011798 | protein_coding_gene | 2.4 | 5.30 | 9.7E-09 |
| *tbb-4* | WBGene00006538 | protein_coding_gene | 2.4 | 5.29 | 1.0E-08 |
| *C32F10.4* | WBGene00016332 | protein_coding_gene | 2.4 | 5.28 | 1.4E-15 |
| *F14B4.1* | WBGene00008779 | protein_coding_gene | 2.4 | 5.26 | 3.5E-13 |
| *Y37F4.8* | WBGene00044289 | protein_coding_gene | 2.4 | 5.25 | 4.1E-05 |
| *E03A3.10* | WBGene00189944 | pseudogene | 2.4 | 5.24 | 5.4E-04 |
| *cki-1* | WBGene00000516 | protein_coding_gene | 2.4 | 5.24 | 6.4E-19 |
| *mlt-8* | WBGene00021095 | protein_coding_gene | 2.4 | 5.22 | 1.2E-12 |
| *K08C9.5* | WBGene00010652 | protein_coding_gene | 2.4 | 5.22 | 1.0E-02 |
| *F35D2.3* | WBGene00018041 | protein_coding_gene | 2.4 | 5.20 | 4.8E-14 |
| *F59E11.7* | WBGene00019113 | protein_coding_gene | 2.4 | 5.19 | 1.1E-06 |
| *asp-11* | WBGene00017677 | pseudogene | 2.4 | 5.18 | 1.8E-03 |
| *F23F12.8* | WBGene00017754 | protein_coding_gene | 2.4 | 5.16 | 1.7E-04 |
| *K01A2.4* | WBGene00019279 | protein_coding_gene | 2.4 | 5.16 | 1.9E-11 |
| *C01G6.9* | WBGene00007229 | protein_coding_gene | 2.4 | 5.15 | 4.6E-04 |
| *pros-1* | WBGene00000448 | protein_coding_gene | 2.4 | 5.13 | 1.9E-10 |
| *Y54G2A.36* | WBGene00021898 | protein_coding_gene | 2.4 | 5.12 | 2.0E-24 |
| *ZK512.1* | WBGene00013982 | protein_coding_gene | 2.4 | 5.12 | 3.9E-10 |
| *C13C12.2* | WBGene00007552 | protein_coding_gene | 2.4 | 5.12 | 3.2E-05 |
| *cnc-6* | WBGene00000560 | protein_coding_gene | 2.4 | 5.10 | 6.2E-03 |
| *clec-82* | WBGene00021873 | protein_coding_gene | 2.4 | 5.10 | 4.6E-03 |
| *C23G10.11* | WBGene00016017 | protein_coding_gene | 2.4 | 5.10 | 1.2E-03 |
| *W07G4.1* | WBGene00012335 | protein_coding_gene | 2.3 | 5.10 | 9.7E-03 |
| *Y70C5A.3* | WBGene00050904 | protein_coding_gene | 2.3 | 5.09 | 2.6E-08 |
| *C54F6.18* | WBGene00235093 | protein_coding_gene | 2.3 | 5.08 | 3.7E-03 |
| *F56H9.9* | WBGene00206460 | protein_coding_gene | 2.3 | 5.07 | 1.2E-02 |
| *ast-1* | WBGene00020368 | protein_coding_gene | 2.3 | 5.06 | 2.2E-06 |
| *syd-9* | WBGene00044068 | protein_coding_gene | 2.3 | 5.06 | 2.3E-17 |
| *pqn-16* | WBGene00004107 | protein_coding_gene | 2.3 | 5.06 | 1.2E-02 |
| *nstp-3* | WBGene00022577 | protein_coding_gene | 2.3 | 5.06 | 3.7E-03 |
| *C14A4.9* | WBGene00007560 | protein_coding_gene | 2.3 | 5.05 | 3.3E-04 |
| *F53B3.5* | WBGene00018743 | protein_coding_gene | 2.3 | 5.05 | 3.1E-17 |
| *hlh-14* | WBGene00001958 | protein_coding_gene | 2.3 | 5.02 | 1.1E-12 |
| *lin-32* | WBGene00003018 | protein_coding_gene | 2.3 | 5.02 | 6.4E-06 |
| *F31E3.12* | WBGene00219317 | protein_coding_gene | 2.3 | 5.02 | 3.6E-03 |
| *ZK1307.2* | WBGene00014245 | protein_coding_gene | 2.3 | 5.02 | 1.7E-04 |
| *K08D9.4* | WBGene00019525 | protein_coding_gene | 2.3 | 5.02 | 5.6E-06 |
| *F10E9.12* | WBGene00017361 | protein_coding_gene | 2.3 | 5.02 | 7.9E-18 |
| *C37A5.3* | WBGene00014698 | protein_coding_gene | 2.3 | 5.00 | 4.7E-03 |
| *fkb-7* | WBGene00001432 | protein_coding_gene | 2.3 | 4.99 | 3.8E-11 |
| *gcy-34* | WBGene00001554 | protein_coding_gene | 2.3 | 4.99 | 8.8E-03 |
| *Y44A6C.1* | WBGene00012850 | protein_coding_gene | 2.3 | 4.97 | 3.8E-24 |
| *K10D6.2* | WBGene00010742 | protein_coding_gene | 2.3 | 4.97 | 7.7E-11 |
| *Y53C12B.7* | WBGene00013146 | protein_coding_gene | 2.3 | 4.96 | 2.9E-09 |
| *hsp-12.1* | WBGene00011906 | protein_coding_gene | 2.3 | 4.96 | 8.3E-06 |
| *ZK813.6* | WBGene00044564 | protein_coding_gene | 2.3 | 4.96 | 1.0E-02 |
| *droe-4* | WBGene00010262 | protein_coding_gene | 2.3 | 4.95 | 1.2E-22 |
| *cutl-13* | WBGene00021025 | protein_coding_gene | 2.3 | 4.94 | 4.1E-03 |
| *tag-234* | WBGene00018874 | protein_coding_gene | 2.3 | 4.94 | 4.6E-19 |
| *F30H5.3* | WBGene00017937 | protein_coding_gene | 2.3 | 4.92 | 1.9E-07 |
| *F46F3.3* | WBGene00009791 | protein_coding_gene | 2.3 | 4.92 | 2.4E-03 |
| *C08F11.13* | WBGene00007460 | protein_coding_gene | 2.3 | 4.91 | 3.1E-20 |
| *C29F7.11* | WBGene00206531 | ncRNA_gene | 2.3 | 4.91 | 1.1E-02 |
| *F23H12.5* | WBGene00009094 | protein_coding_gene | 2.3 | 4.91 | 2.9E-05 |
| *unc-3* | WBGene00006743 | protein_coding_gene | 2.3 | 4.90 | 1.6E-05 |
| *nlp-28* | WBGene00003766 | protein_coding_gene | 2.3 | 4.89 | 3.0E-18 |
| *C08D8.1* | WBGene00015592 | protein_coding_gene | 2.3 | 4.89 | 3.8E-06 |
| *F46E10.2* | WBGene00018489 | protein_coding_gene | 2.3 | 4.87 | 2.7E-16 |
| *srz-56* | WBGene00016519 | protein_coding_gene | 2.3 | 4.86 | 8.0E-03 |
| *ddr-2* | WBGene00017381 | protein_coding_gene | 2.3 | 4.86 | 1.4E-19 |
| *F07H5.13* | WBGene00008563 | protein_coding_gene | 2.3 | 4.86 | 2.0E-18 |

| | | | | | |
|---|---|---|---|---|---|
| *T05C1.3* | WBGene00020250 | protein_coding_gene | 2.3 | 4.84 | 7.6E-03 |
| *F46G11.6* | WBGene00045269 | protein_coding_gene | 2.3 | 4.82 | 2.1E-03 |
| *C53A5.2* | WBGene00008263 | protein_coding_gene | 2.3 | 4.82 | 5.7E-04 |
| *skr-21* | WBGene00004827 | protein_coding_gene | 2.3 | 4.81 | 1.7E-22 |
| *ugt-43* | WBGene00008485 | protein_coding_gene | 2.3 | 4.81 | 9.6E-07 |
| *fbxa-105* | WBGene00015362 | protein_coding_gene | 2.3 | 4.81 | 1.5E-09 |
| *igcm-1* | WBGene00018215 | protein_coding_gene | 2.3 | 4.80 | 4.5E-14 |
| *cuti-1* | WBGene00022591 | protein_coding_gene | 2.3 | 4.80 | 1.1E-06 |
| *B0393.5* | WBGene00007170 | protein_coding_gene | 2.3 | 4.78 | 5.6E-08 |
| *K09C4.1* | WBGene00019547 | protein_coding_gene | 2.3 | 4.78 | 1.8E-07 |
| *nab-1* | WBGene00003516 | protein_coding_gene | 2.3 | 4.77 | 7.1E-12 |
| *R08C7.4* | WBGene00019948 | protein_coding_gene | 2.3 | 4.76 | 8.9E-03 |
| *F53A9.6* | WBGene00018729 | protein_coding_gene | 2.3 | 4.76 | 1.1E-23 |
| *F53F4.8* | WBGene00009991 | pseudogene | 2.2 | 4.76 | 1.6E-05 |
| *ZK971.1* | WBGene00014175 | protein_coding_gene | 2.2 | 4.75 | 1.5E-05 |
| *F48C1.11* | WBGene00195179 | protein_coding_gene | 2.2 | 4.75 | 1.3E-02 |
| *his-1* | WBGene00001875 | protein_coding_gene | 2.2 | 4.74 | 1.1E-02 |
| *Y43C5A.3* | WBGene00012783 | protein_coding_gene | 2.2 | 4.73 | 1.9E-30 |
| *dyf-7* | WBGene00001123 | protein_coding_gene | 2.2 | 4.73 | 2.2E-09 |
| *ZC443.4* | WBGene00013899 | protein_coding_gene | 2.2 | 4.71 | 2.1E-18 |
| *ttr-26* | WBGene00013079 | protein_coding_gene | 2.2 | 4.70 | 7.2E-10 |
| *oac-6* | WBGene00007833 | protein_coding_gene | 2.2 | 4.70 | 1.1E-04 |
| *Y87G2A.16* | WBGene00013602 | protein_coding_gene | 2.2 | 4.70 | 4.8E-05 |
| *ptr-9* | WBGene00004223 | protein_coding_gene | 2.2 | 4.70 | 1.6E-02 |
| *math-42* | WBGene00020532 | protein_coding_gene | 2.2 | 4.69 | 4.0E-03 |
| *F26F12.3* | WBGene00017834 | protein_coding_gene | 2.2 | 4.69 | 8.1E-27 |
| *C50B8.4* | WBGene00008226 | protein_coding_gene | 2.2 | 4.68 | 2.0E-06 |
| *cnc-7* | WBGene00010005 | protein_coding_gene | 2.2 | 4.68 | 5.4E-05 |
| *F53B7.7* | WBGene00009969 | protein_coding_gene | 2.2 | 4.68 | 3.3E-03 |
| *clec-66* | WBGene00009397 | protein_coding_gene | 2.2 | 4.66 | 8.0E-12 |
| *srbc-82* | WBGene00007832 | protein_coding_gene | 2.2 | 4.66 | 1.6E-02 |
| *nsy-4* | WBGene00021415 | protein_coding_gene | 2.2 | 4.66 | 9.2E-10 |
| *ncx-9* | WBGene00003574 | protein_coding_gene | 2.2 | 4.65 | 2.1E-03 |
| *grsp-3* | WBGene00016403 | protein_coding_gene | 2.2 | 4.64 | 6.6E-08 |
| *ikb-1* | WBGene00002069 | protein_coding_gene | 2.2 | 4.63 | 3.3E-18 |
| *F53H2.1* | WBGene00010004 | protein_coding_gene | 2.2 | 4.63 | 1.1E-02 |
| *C33A12.4* | WBGene00007883 | protein_coding_gene | 2.2 | 4.63 | 3.4E-09 |
| *mir-238* | WBGene00003331 | miRNA_gene | 2.2 | 4.62 | 7.9E-03 |
| *R160.10* | WBGene00219369 | protein_coding_gene | 2.2 | 4.62 | 1.4E-03 |
| *acd-1* | WBGene00016064 | protein_coding_gene | 2.2 | 4.62 | 3.1E-03 |
| *K01A2.10* | WBGene00019284 | protein_coding_gene | 2.2 | 4.61 | 1.4E-11 |
| *D1014.5* | WBGene00017018 | protein_coding_gene | 2.2 | 4.61 | 5.1E-04 |
| *C35C5.9* | WBGene00007958 | protein_coding_gene | 2.2 | 4.60 | 2.2E-17 |
| *F26F12.4* | WBGene00017835 | protein_coding_gene | 2.2 | 4.60 | 1.0E-03 |
| *nhr-99* | WBGene00003689 | protein_coding_gene | 2.2 | 4.59 | 2.7E-08 |
| *zig-4* | WBGene00006981 | protein_coding_gene | 2.2 | 4.58 | 1.2E-05 |
| *oac-51* | WBGene00020976 | protein_coding_gene | 2.2 | 4.56 | 4.1E-04 |
| *Y48A6B.7* | WBGene00012968 | protein_coding_gene | 2.2 | 4.55 | 2.5E-31 |
| *C54F6.6* | WBGene00016924 | protein_coding_gene | 2.2 | 4.55 | 9.9E-03 |
| *cdf-2* | WBGene00011821 | protein_coding_gene | 2.2 | 4.54 | 4.3E-15 |
| *Y38H8A.1* | WBGene00012635 | protein_coding_gene | 2.2 | 4.54 | 6.3E-03 |
| *B0462.5* | WBGene00194695 | protein_coding_gene | 2.2 | 4.54 | 3.7E-05 |
| *let-381* | WBGene00002601 | protein_coding_gene | 2.2 | 4.53 | 2.2E-04 |
| *cpr-4* | WBGene00000784 | protein_coding_gene | 2.2 | 4.53 | 1.3E-13 |
| *bath-25* | WBGene00020859 | protein_coding_gene | 2.2 | 4.53 | 1.5E-02 |
| *W03D2.9* | WBGene00020983 | protein_coding_gene | 2.2 | 4.53 | 9.5E-03 |
| *R10E8.6* | WBGene00011212 | protein_coding_gene | 2.2 | 4.53 | 5.2E-15 |
| *eva-1* | WBGene00009304 | protein_coding_gene | 2.2 | 4.53 | 9.5E-10 |
| *cbn-1* | WBGene00019285 | protein_coding_gene | 2.2 | 4.52 | 3.4E-04 |
| *ztf-30* | WBGene00015523 | protein_coding_gene | 2.2 | 4.51 | 8.0E-04 |
| *cnd-1* | WBGene00000561 | protein_coding_gene | 2.2 | 4.51 | 4.5E-07 |
| *glb-5* | WBGene00015964 | protein_coding_gene | 2.2 | 4.50 | 6.2E-03 |
| *W10G11.1* | WBGene00021134 | protein_coding_gene | 2.2 | 4.50 | 1.1E-02 |
| *cyp-14A3* | WBGene00010707 | protein_coding_gene | 2.2 | 4.50 | 9.0E-04 |
| *arl-6* | WBGene00000193 | protein_coding_gene | 2.2 | 4.49 | 8.6E-03 |

| | | | | | |
|---|---|---|---|---|---|
| *linc-103* | WBGene00219743 | lincRNA_gene | 2.2 | 4.49 | 2.3E-02 |
| *Y73B6BL.28* | WBGene00022249 | pseudogene | 2.2 | 4.49 | 2.0E-02 |
| *F38B6.3* | WBGene00018173 | protein_coding_gene | 2.2 | 4.49 | 2.0E-02 |
| *T09E11.11* | WBGene00011660 | pseudogene | 2.2 | 4.47 | 6.0E-05 |
| *F32H2.6* | WBGene00009343 | protein_coding_gene | 2.2 | 4.45 | 4.2E-04 |
| *dhs-16* | WBGene00000979 | protein_coding_gene | 2.2 | 4.44 | 4.9E-03 |
| *F54B11.10* | WBGene00010033 | protein_coding_gene | 2.1 | 4.43 | 8.6E-03 |
| *C44C1.6* | WBGene00016645 | protein_coding_gene | 2.1 | 4.43 | 5.0E-09 |
| *C39D10.5* | WBGene00016532 | protein_coding_gene | 2.1 | 4.42 | 1.3E-02 |
| *plx-2* | WBGene00004048 | protein_coding_gene | 2.1 | 4.41 | 8.4E-10 |
| *osm-8* | WBGene00003888 | protein_coding_gene | 2.1 | 4.40 | 3.7E-03 |
| *B0284.5* | WBGene00045188 | pseudogene | 2.1 | 4.39 | 2.4E-02 |
| *bigr-1* | WBGene00009512 | protein_coding_gene | 2.1 | 4.39 | 9.8E-03 |
| *fbxb-114* | WBGene00016254 | protein_coding_gene | 2.1 | 4.39 | 1.3E-02 |
| *T01B10.5* | WBGene00020137 | protein_coding_gene | 2.1 | 4.39 | 1.9E-04 |
| *ZK836.3* | WBGene00014099 | protein_coding_gene | 2.1 | 4.37 | 8.9E-06 |
| *spi-1* | WBGene00023416 | protein_coding_gene | 2.1 | 4.36 | 8.0E-04 |
| *F22F7.3* | WBGene00017721 | protein_coding_gene | 2.1 | 4.36 | 2.8E-03 |
| *clec-69* | WBGene00044642 | pseudogene | 2.1 | 4.36 | 1.2E-02 |
| *slc-17.9* | WBGene00011643 | protein_coding_gene | 2.1 | 4.35 | 6.4E-04 |
| *W02D7.8* | WBGene00020943 | protein_coding_gene | 2.1 | 4.35 | 2.7E-05 |
| *R160.4* | WBGene00020121 | protein_coding_gene | 2.1 | 4.34 | 1.6E-02 |
| *cut-6* | WBGene00000853 | protein_coding_gene | 2.1 | 4.34 | 2.5E-03 |
| *best-1* | WBGene00007203 | protein_coding_gene | 2.1 | 4.34 | 6.2E-05 |
| *F07A5.4* | WBGene00008543 | protein_coding_gene | 2.1 | 4.33 | 1.9E-03 |
| *Y34F4.3* | WBGene00021338 | protein_coding_gene | 2.1 | 4.31 | 8.0E-04 |
| *ZC84.1* | WBGene00013846 | protein_coding_gene | 2.1 | 4.31 | 4.9E-04 |
| *F21A3.11* | WBGene00219316 | protein_coding_gene | 2.1 | 4.31 | 2.0E-13 |
| *fipr-26* | WBGene00009963 | protein_coding_gene | 2.1 | 4.30 | 1.9E-02 |
| *lgc-27* | WBGene00022283 | protein_coding_gene | 2.1 | 4.30 | 9.0E-07 |
| *ham-2* | WBGene00001821 | protein_coding_gene | 2.1 | 4.30 | 5.1E-14 |
| *F36F2.2* | WBGene00009476 | protein_coding_gene | 2.1 | 4.29 | 8.4E-13 |
| *nhr-19* | WBGene00003618 | protein_coding_gene | 2.1 | 4.29 | 3.4E-15 |
| *F49E10.2* | WBGene00018640 | protein_coding_gene | 2.1 | 4.29 | 6.3E-04 |
| *F20C5.6* | WBGene00008971 | protein_coding_gene | 2.1 | 4.29 | 1.1E-10 |
| *H32K16.2* | WBGene00010422 | protein_coding_gene | 2.1 | 4.28 | 1.2E-05 |
| *ceh-48* | WBGene00015934 | protein_coding_gene | 2.1 | 4.27 | 3.4E-06 |
| *Y17D7B.2* | WBGene00012450 | protein_coding_gene | 2.1 | 4.27 | 2.2E-02 |
| *T10G3.1* | WBGene00011692 | protein_coding_gene | 2.1 | 4.27 | 1.1E-09 |
| *Y97E10AR.1* | WBGene00022396 | protein_coding_gene | 2.1 | 4.26 | 2.6E-10 |
| *mks-3* | WBGene00018042 | protein_coding_gene | 2.1 | 4.26 | 4.3E-03 |
| *Y55D5A.1* | WBGene00021915 | protein_coding_gene | 2.1 | 4.26 | 6.0E-07 |
| *C02F4.4* | WBGene00007264 | protein_coding_gene | 2.1 | 4.26 | 4.4E-03 |
| *F32D1.3* | WBGene00017983 | protein_coding_gene | 2.1 | 4.26 | 3.1E-09 |
| *rocf-1* | WBGene00011948 | protein_coding_gene | 2.1 | 4.25 | 9.3E-03 |
| *sups-1* | WBGene00017716 | protein_coding_gene | 2.1 | 4.25 | 5.9E-05 |
| *T28C12.15* | WBGene00271821 | protein_coding_gene | 2.1 | 4.24 | 3.0E-02 |
| *C54C8.12* | WBGene00044242 | protein_coding_gene | 2.1 | 4.23 | 2.7E-02 |
| *mks-2* | WBGene00194710 | protein_coding_gene | 2.1 | 4.23 | 2.5E-02 |
| *pag-3* | WBGene00003909 | protein_coding_gene | 2.1 | 4.23 | 2.0E-03 |
| *ceh-45* | WBGene00022837 | protein_coding_gene | 2.1 | 4.22 | 2.2E-03 |
| *C45B11.6* | WBGene00045379 | protein_coding_gene | 2.1 | 4.22 | 2.1E-04 |
| *clec-172* | WBGene00044719 | protein_coding_gene | 2.1 | 4.22 | 7.3E-04 |
| *K04H4.2* | WBGene00010573 | protein_coding_gene | 2.1 | 4.21 | 1.0E-06 |
| *Y65A5A.1* | WBGene00013418 | protein_coding_gene | 2.1 | 4.20 | 1.4E-13 |
| *Y57G7A.8* | WBGene00021971 | pseudogene | 2.1 | 4.20 | 6.5E-06 |
| *W05B2.2* | WBGene00012271 | protein_coding_gene | 2.1 | 4.20 | 8.4E-03 |
| *Y46E12BR.1* | WBGene00021598 | protein_coding_gene | 2.1 | 4.19 | 7.8E-03 |
| *D2024.10* | WBGene00044505 | pseudogene | 2.1 | 4.17 | 3.0E-02 |
| *F09F9.2* | WBGene00017307 | protein_coding_gene | 2.1 | 4.15 | 2.0E-03 |
| *C32D5.12* | WBGene00016319 | protein_coding_gene | 2.1 | 4.15 | 2.4E-06 |
| *C43H6.6* | WBGene00016619 | protein_coding_gene | 2.1 | 4.14 | 1.3E-02 |
| *E02C12.10* | WBGene00017095 | protein_coding_gene | 2.1 | 4.14 | 1.8E-02 |
| *R11F4.1* | WBGene00020007 | protein_coding_gene | 2.0 | 4.13 | 1.8E-13 |
| *dct-1* | WBGene00015776 | protein_coding_gene | 2.0 | 4.13 | 6.7E-24 |

| | | | | | |
|---|---|---|---|---|---|
| *C32B5.18* | WBGene00235351 | protein_coding_gene | 2.0 | 4.13 | 1.3E-02 |
| *F21C10.10* | WBGene00017659 | protein_coding_gene | 2.0 | 4.12 | 1.9E-03 |
| *F17C11.4* | WBGene00008915 | protein_coding_gene | 2.0 | 4.11 | 2.5E-16 |
| *EGAP1.1* | WBGene00017146 | protein_coding_gene | 2.0 | 4.11 | 1.3E-02 |
| *Y52B11A.12* | WBGene00077539 | protein_coding_gene | 2.0 | 4.11 | 5.5E-04 |
| *mam-5* | WBGene00011572 | protein_coding_gene | 2.0 | 4.10 | 1.4E-02 |
| *unc-86* | WBGene00006818 | protein_coding_gene | 2.0 | 4.10 | 2.2E-04 |
| *fbxa-175* | WBGene00045293 | pseudogene | 2.0 | 4.09 | 6.5E-06 |
| *ZC168.2* | WBGene00013856 | protein_coding_gene | 2.0 | 4.08 | 5.6E-03 |
| *K02E10.4* | WBGene00019318 | protein_coding_gene | 2.0 | 4.08 | 6.2E-06 |
| *T28F4.5* | WBGene00012140 | protein_coding_gene | 2.0 | 4.08 | 2.7E-20 |
| *gpa-9* | WBGene00001671 | protein_coding_gene | 2.0 | 4.08 | 5.5E-06 |
| *ceh-34* | WBGene00000455 | protein_coding_gene | 2.0 | 4.07 | 3.1E-07 |
| *M163.1* | WBGene00010935 | protein_coding_gene | 2.0 | 4.07 | 1.6E-42 |
| *ins-7* | WBGene00002090 | protein_coding_gene | 2.0 | 4.07 | 9.4E-06 |
| *aakg-4* | WBGene00020135 | protein_coding_gene | 2.0 | 4.07 | 2.3E-23 |
| *C27B7.9* | WBGene00007765 | protein_coding_gene | 2.0 | 4.06 | 1.1E-38 |
| *tbx-7* | WBGene00006544 | protein_coding_gene | 2.0 | 4.06 | 2.9E-03 |
| *R05G6.10* | WBGene00019902 | protein_coding_gene | 2.0 | 4.05 | 4.8E-22 |
| *alh-5* | WBGene00000111 | protein_coding_gene | 2.0 | 4.05 | 2.3E-19 |
| *M7.8* | WBGene00010884 | protein_coding_gene | 2.0 | 4.05 | 4.1E-10 |
| *C54F6.12* | WBGene00016929 | protein_coding_gene | 2.0 | 4.04 | 4.1E-03 |
| *F19G12.3* | WBGene00017611 | protein_coding_gene | 2.0 | 4.04 | 3.2E-02 |
| *ceh-30* | WBGene00000451 | protein_coding_gene | 2.0 | 4.04 | 1.1E-03 |
| *T16G1.7* | WBGene00011801 | protein_coding_gene | 2.0 | 4.03 | 5.8E-08 |
| *nlp-29* | WBGene00003767 | protein_coding_gene | 2.0 | 4.03 | 1.3E-09 |
| *fbxc-7* | WBGene00016302 | protein_coding_gene | 2.0 | 4.02 | 2.4E-02 |
| *H10D18.1* | WBGene00019177 | protein_coding_gene | 2.0 | 4.02 | 2.0E-02 |
| *C06G3.3* | WBGene00015549 | protein_coding_gene | 2.0 | 4.02 | 2.4E-03 |
| *C03A3.1* | WBGene00007266 | protein_coding_gene | 2.0 | 4.02 | 2.1E-05 |
| *F59F4.6* | WBGene00303371 | ncRNA_gene | 2.0 | 4.02 | 1.9E-02 |
| *arrd-4* | WBGene00014159 | protein_coding_gene | 2.0 | 4.02 | 9.9E-11 |
| *F26G1.15* | WBGene00270318 | protein_coding_gene | 2.0 | 4.02 | 3.7E-02 |
| *mec-3* | WBGene00003167 | protein_coding_gene | 2.0 | 4.01 | 2.0E-03 |
| *fax-1* | WBGene00001400 | protein_coding_gene | 2.0 | 4.01 | 1.1E-02 |
| *Y40H7A.7* | WBGene00012745 | pseudogene | 2.0 | 4.01 | 3.7E-03 |
| *F28G4.2* | WBGene00009227 | protein_coding_gene | 2.0 | 4.01 | 1.5E-02 |
| *F43C11.7* | WBGene00018384 | protein_coding_gene | 2.0 | 4.01 | 1.2E-13 |
| *his-64* | WBGene00001938 | protein_coding_gene | 2.0 | 4.00 | 6.1E-06 |
| *Y50E8A.5* | WBGene00013048 | protein_coding_gene | 2.0 | 4.00 | 7.8E-06 |
| *pef-1* | WBGene00003969 | protein_coding_gene | 2.0 | 4.00 | 4.7E-07 |
| *ceh-5* | WBGene00000430 | protein_coding_gene | 2.0 | 3.99 | 2.7E-04 |
| *C54F6.17* | WBGene00235092 | protein_coding_gene | 2.0 | 3.99 | 3.3E-02 |
| *C35A5.5* | WBGene00007949 | protein_coding_gene | 2.0 | 3.99 | 7.4E-03 |
| *F59B10.4* | WBGene00010320 | protein_coding_gene | 2.0 | 3.97 | 4.8E-08 |
| *osm-12* | WBGene00003892 | protein_coding_gene | 2.0 | 3.97 | 3.3E-03 |
| *D1086.2* | WBGene00008389 | protein_coding_gene | 2.0 | 3.97 | 3.5E-04 |
| *cyp-23A1* | WBGene00015135 | protein_coding_gene | 2.0 | 3.95 | 1.5E-02 |
| *Y5H2B.1* | WBGene00021164 | protein_coding_gene | 2.0 | 3.95 | 2.8E-02 |
| *C05D9.9* | WBGene00015476 | protein_coding_gene | 2.0 | 3.94 | 1.6E-13 |
| *C06A12.19* | WBGene00255702 | protein_coding_gene | 2.0 | 3.94 | 3.0E-02 |
| *rpr-1* | WBGene00045238 | ncRNA_gene | 2.0 | 3.93 | 1.6E-02 |
| *set-28* | WBGene00022224 | protein_coding_gene | 2.0 | 3.93 | 2.6E-02 |
| *F18C5.10* | WBGene00017563 | protein_coding_gene | 2.0 | 3.93 | 8.7E-40 |
| *rrn-1.2* | WBGene00004513 | rRNA_gene | 2.0 | 3.92 | 2.2E-02 |
| *nhr-150* | WBGene00007367 | protein_coding_gene | 2.0 | 3.92 | 8.9E-03 |
| *C36B1.13* | WBGene00044120 | protein_coding_gene | 2.0 | 3.91 | 2.1E-02 |
| *K10H10.13* | WBGene00194888 | protein_coding_gene | 2.0 | 3.91 | 3.9E-02 |
| *smf-3* | WBGene00004878 | protein_coding_gene | 2.0 | 3.90 | 9.7E-06 |
| *Y50E8A.1* | WBGene00013046 | protein_coding_gene | 2.0 | 3.90 | 1.1E-05 |
| *hst-3.2* | WBGene00044303 | protein_coding_gene | 2.0 | 3.89 | 2.0E-05 |
| *B0228.6* | WBGene00015063 | protein_coding_gene | 2.0 | 3.87 | 3.0E-08 |
| *fbxc-51* | WBGene00021576 | protein_coding_gene | 2.0 | 3.87 | 1.3E-22 |
| *nlp-66* | WBGene00011141 | protein_coding_gene | 2.0 | 3.87 | 5.5E-03 |
| *C07E3.3* | WBGene00007414 | protein_coding_gene | 1.9 | 3.86 | 3.8E-11 |

| | | | | | |
|---|---|---|---|---|---|
| *zip-7* | WBGene00013100 | protein_coding_gene | 1.9 | 3.86 | 1.0E-06 |
| *R102.1* | WBGene00011288 | protein_coding_gene | 1.9 | 3.86 | 3.2E-02 |
| *pqn-73* | WBGene00004155 | protein_coding_gene | 1.9 | 3.85 | 2.2E-02 |
| *F59B2.11* | WBGene00010313 | pseudogene | 1.9 | 3.85 | 5.9E-03 |
| *C17E4.11* | WBGene00044236 | protein_coding_gene | 1.9 | 3.85 | 4.2E-17 |
| *Y38H6C.16* | WBGene00012628 | protein_coding_gene | 1.9 | 3.84 | 1.2E-04 |
| *T24C2.5* | WBGene00011984 | protein_coding_gene | 1.9 | 3.84 | 1.1E-08 |
| *cutl-21* | WBGene00018861 | protein_coding_gene | 1.9 | 3.84 | 2.1E-04 |
| *F21D5.9* | WBGene00009014 | protein_coding_gene | 1.9 | 3.83 | 5.6E-03 |
| *F31D5.6* | WBGene00017949 | protein_coding_gene | 1.9 | 3.83 | 8.1E-04 |
| *egas-4* | WBGene00018906 | protein_coding_gene | 1.9 | 3.82 | 3.0E-03 |
| *rga-2* | WBGene00013135 | protein_coding_gene | 1.9 | 3.82 | 3.5E-07 |
| *C39B10.7* | WBGene00164973 | protein_coding_gene | 1.9 | 3.82 | 3.7E-03 |
| *nlp-2* | WBGene00003740 | protein_coding_gene | 1.9 | 3.81 | 2.6E-03 |
| *M01A8.1* | WBGene00010795 | protein_coding_gene | 1.9 | 3.81 | 4.1E-06 |
| *igdb-3* | WBGene00020691 | protein_coding_gene | 1.9 | 3.80 | 1.2E-03 |
| *chpf-2* | WBGene00019108 | protein_coding_gene | 1.9 | 3.80 | 1.3E-07 |
| *C10A4.3* | WBGene00015664 | protein_coding_gene | 1.9 | 3.80 | 6.5E-03 |
| *pgp-11* | WBGene00004005 | protein_coding_gene | 1.9 | 3.79 | 1.8E-10 |
| *Y45F10B.59* | WBGene00194917 | protein_coding_gene | 1.9 | 3.79 | 3.9E-02 |
| *aagr-1* | WBGene00017071 | protein_coding_gene | 1.9 | 3.78 | 8.8E-10 |
| *nspe-2* | WBGene00012604 | protein_coding_gene | 1.9 | 3.78 | 2.3E-03 |
| *C36A4.11* | WBGene00044167 | protein_coding_gene | 1.9 | 3.78 | 2.5E-03 |
| *linc-37* | WBGene00219709 | lincRNA_gene | 1.9 | 3.77 | 3.0E-02 |
| *K11H12.11* | WBGene00044723 | protein_coding_gene | 1.9 | 3.77 | 2.4E-02 |
| *K01F9.2* | WBGene00010475 | protein_coding_gene | 1.9 | 3.77 | 1.1E-06 |
| *klp-11* | WBGene00002222 | protein_coding_gene | 1.9 | 3.76 | 5.0E-07 |
| *C18H9.5* | WBGene00016003 | protein_coding_gene | 1.9 | 3.76 | 1.8E-07 |
| *K01D12.1* | WBGene00010461 | protein_coding_gene | 1.9 | 3.75 | 2.0E-02 |
| *cyp-13B1* | WBGene00008519 | protein_coding_gene | 1.9 | 3.75 | 3.8E-06 |
| *nhr-165* | WBGene00008158 | protein_coding_gene | 1.9 | 3.75 | 4.1E-02 |
| *dlg-1* | WBGene00001006 | protein_coding_gene | 1.9 | 3.75 | 1.0E-24 |
| *F15H9.8* | WBGene00194652 | pseudogene | 1.9 | 3.75 | 5.7E-03 |
| *clec-218* | WBGene00020938 | protein_coding_gene | 1.9 | 3.75 | 1.3E-03 |
| *C17E7.13* | WBGene00015906 | protein_coding_gene | 1.9 | 3.75 | 1.4E-02 |
| *W08A12.2* | WBGene00021079 | protein_coding_gene | 1.9 | 3.75 | 8.9E-08 |
| *W03B1.3* | WBGene00020973 | protein_coding_gene | 1.9 | 3.74 | 3.4E-05 |
| *fbxc-41* | WBGene00016304 | pseudogene | 1.9 | 3.74 | 2.9E-02 |
| *srz-99* | WBGene00012028 | protein_coding_gene | 1.9 | 3.74 | 4.0E-02 |
| *clec-40* | WBGene00011853 | protein_coding_gene | 1.9 | 3.74 | 6.3E-03 |
| *tts-2* | WBGene00006651 | ncRNA_gene | 1.9 | 3.72 | 1.3E-05 |
| *F49D11.6* | WBGene00018630 | protein_coding_gene | 1.9 | 3.72 | 1.6E-02 |
| *C35E7.12* | WBGene00271779 | protein_coding_gene | 1.9 | 3.72 | 4.1E-02 |
| *Y39G10AR.11* | WBGene00021469 | protein_coding_gene | 1.9 | 3.71 | 4.2E-37 |
| *Y69H2.10* | WBGene00013485 | protein_coding_gene | 1.9 | 3.71 | 7.4E-03 |
| *Y82E9BR.17* | WBGene00022349 | protein_coding_gene | 1.9 | 3.71 | 4.6E-25 |
| *egl-43* | WBGene00001207 | protein_coding_gene | 1.9 | 3.70 | 2.1E-07 |
| *C47E12.13* | WBGene00008156 | protein_coding_gene | 1.9 | 3.70 | 4.9E-02 |
| *F36H1.11* | WBGene00009496 | protein_coding_gene | 1.9 | 3.70 | 2.6E-02 |
| *cutl-8* | WBGene00013960 | protein_coding_gene | 1.9 | 3.69 | 2.4E-02 |
| *trx-3* | WBGene00019717 | protein_coding_gene | 1.9 | 3.69 | 1.3E-06 |
| *dex-1* | WBGene00017028 | protein_coding_gene | 1.9 | 3.68 | 2.3E-12 |
| *pho-6* | WBGene00018696 | protein_coding_gene | 1.9 | 3.68 | 2.6E-05 |
| *linc-43* | WBGene00219579 | lincRNA_gene | 1.9 | 3.68 | 3.0E-02 |
| *F42G4.6* | WBGene00009643 | protein_coding_gene | 1.9 | 3.68 | 9.7E-04 |
| *rom-2* | WBGene00004401 | protein_coding_gene | 1.9 | 3.67 | 1.1E-03 |
| *comt-4* | WBGene00021491 | protein_coding_gene | 1.9 | 3.67 | 4.2E-02 |
| *cest-1.2* | WBGene00011364 | protein_coding_gene | 1.9 | 3.67 | 1.3E-13 |
| *F58E6.11* | WBGene00010258 | protein_coding_gene | 1.9 | 3.67 | 7.6E-03 |
| *F22B7.1* | WBGene00017692 | protein_coding_gene | 1.9 | 3.67 | 5.5E-06 |
| *ins-23* | WBGene00002106 | protein_coding_gene | 1.9 | 3.67 | 7.0E-03 |
| *T21E8.6* | WBGene00164971 | protein_coding_gene | 1.9 | 3.67 | 3.9E-02 |
| *F21H7.12* | WBGene00009033 | protein_coding_gene | 1.9 | 3.67 | 2.0E-02 |
| *C26E1.2* | WBGene00007748 | protein_coding_gene | 1.9 | 3.67 | 2.0E-14 |
| *W08G11.6* | WBGene00023429 | snoRNA_gene | 1.9 | 3.66 | 1.3E-02 |

| | | | | | |
|---|---|---|---|---|---|
| *ceh-31* | WBGene00000452 | protein_coding_gene | 1.9 | 3.66 | 6.4E-03 |
| *F16G10.9* | WBGene00017524 | protein_coding_gene | 1.9 | 3.66 | 4.7E-02 |
| *ceh-2* | WBGene00000429 | protein_coding_gene | 1.9 | 3.66 | 1.8E-02 |
| *F53F10.1* | WBGene00018763 | protein_coding_gene | 1.9 | 3.66 | 8.6E-08 |
| *chil-18* | WBGene00011161 | protein_coding_gene | 1.9 | 3.66 | 2.9E-02 |
| *osr-1* | WBGene00016329 | protein_coding_gene | 1.9 | 3.66 | 9.3E-03 |
| *C37A5.6* | WBGene00014700 | protein_coding_gene | 1.9 | 3.66 | 4.0E-02 |
| *tth-1* | WBGene00006649 | protein_coding_gene | 1.9 | 3.66 | 1.1E-18 |
| *pha-2* | WBGene00004011 | protein_coding_gene | 1.9 | 3.66 | 7.0E-03 |
| *grdn-1* | WBGene00013082 | protein_coding_gene | 1.9 | 3.65 | 1.5E-18 |
| *clec-3* | WBGene00016577 | protein_coding_gene | 1.9 | 3.65 | 4.7E-04 |
| *D2030.2* | WBGene00008412 | protein_coding_gene | 1.9 | 3.65 | 5.2E-45 |
| *F23A7.1* | WBGene00009067 | protein_coding_gene | 1.9 | 3.65 | 3.2E-03 |
| *cdh-5* | WBGene00000397 | protein_coding_gene | 1.9 | 3.64 | 9.5E-05 |
| *slc-28.1* | WBGene00017867 | protein_coding_gene | 1.9 | 3.64 | 1.6E-05 |
| *clec-33* | WBGene00009859 | protein_coding_gene | 1.9 | 3.64 | 2.2E-02 |
| *ceh-27* | WBGene00000449 | protein_coding_gene | 1.9 | 3.63 | 2.2E-09 |
| *hlh-32* | WBGene00013665 | protein_coding_gene | 1.9 | 3.63 | 4.6E-02 |
| *Y15E3A.5* | WBGene00012444 | protein_coding_gene | 1.9 | 3.62 | 3.7E-12 |
| *F46F11.12* | WBGene00045082 | snoRNA_gene | 1.9 | 3.62 | 3.7E-02 |
| *magu-3* | WBGene00016841 | protein_coding_gene | 1.9 | 3.61 | 3.1E-04 |
| *fbxb-88* | WBGene00016359 | protein_coding_gene | 1.9 | 3.61 | 2.6E-08 |
| *maph-9* | WBGene00016397 | protein_coding_gene | 1.8 | 3.60 | 9.7E-03 |
| *mfb-1* | WBGene00008439 | protein_coding_gene | 1.8 | 3.60 | 4.7E-13 |
| *F22B7.9* | WBGene00017698 | protein_coding_gene | 1.8 | 3.60 | 1.2E-11 |
| *C07A4.3* | WBGene00007398 | protein_coding_gene | 1.8 | 3.59 | 4.5E-05 |
| *cyp-13A5* | WBGene00011672 | protein_coding_gene | 1.8 | 3.59 | 7.5E-11 |
| *W05H9.1* | WBGene00021048 | protein_coding_gene | 1.8 | 3.58 | 1.1E-45 |
| *ZK822.1* | WBGene00014089 | protein_coding_gene | 1.8 | 3.58 | 8.1E-05 |
| *lhfp-4* | WBGene00009155 | protein_coding_gene | 1.8 | 3.58 | 1.3E-02 |
| *spp-8* | WBGene00004993 | protein_coding_gene | 1.8 | 3.58 | 3.3E-12 |
| *marc-4* | WBGene00016903 | protein_coding_gene | 1.8 | 3.56 | 4.0E-04 |
| *B0478.3* | WBGene00015202 | protein_coding_gene | 1.8 | 3.56 | 1.1E-02 |
| *ccep-290* | WBGene00021643 | protein_coding_gene | 1.8 | 3.56 | 1.8E-04 |
| *F48E3.6* | WBGene00018606 | protein_coding_gene | 1.8 | 3.55 | 5.5E-12 |
| *F35G2.5* | WBGene00009438 | protein_coding_gene | 1.8 | 3.54 | 4.0E-02 |
| *C29F7.2* | WBGene00007811 | protein_coding_gene | 1.8 | 3.54 | 7.1E-25 |
| *fbxb-78* | WBGene00017116 | protein_coding_gene | 1.8 | 3.53 | 5.2E-04 |
| *F13G3.3* | WBGene00008763 | protein_coding_gene | 1.8 | 3.53 | 7.6E-03 |
| *ttr-17* | WBGene00012383 | protein_coding_gene | 1.8 | 3.53 | 1.6E-22 |
| *F21A9.2* | WBGene00017651 | protein_coding_gene | 1.8 | 3.53 | 5.0E-06 |
| *W03F8.6* | WBGene00020995 | protein_coding_gene | 1.8 | 3.53 | 2.1E-06 |
| *clc-2* | WBGene00000523 | protein_coding_gene | 1.8 | 3.52 | 1.2E-05 |
| *oac-31* | WBGene00018707 | protein_coding_gene | 1.8 | 3.52 | 4.8E-03 |
| *F35E12.9* | WBGene00009433 | protein_coding_gene | 1.8 | 3.52 | 1.2E-06 |
| *C53A3.1* | WBGene00016891 | protein_coding_gene | 1.8 | 3.52 | 4.2E-02 |
| *fbxa-127* | WBGene00012027 | protein_coding_gene | 1.8 | 3.51 | 3.0E-02 |
| *bbs-1* | WBGene00000241 | protein_coding_gene | 1.8 | 3.50 | 2.2E-02 |
| *srh-149* | WBGene00005365 | protein_coding_gene | 1.8 | 3.49 | 3.8E-02 |
| *C38C3.4* | WBGene00016513 | protein_coding_gene | 1.8 | 3.49 | 1.4E-11 |
| *W09G10.9* | WBGene00271830 | protein_coding_gene | 1.8 | 3.48 | 3.0E-02 |
| *mrpr-1* | WBGene00045095 | ncRNA_gene | 1.8 | 3.48 | 1.8E-02 |
| *C24A8.5* | WBGene00016039 | protein_coding_gene | 1.8 | 3.48 | 2.6E-02 |
| *Y95B8A.6* | WBGene00022386 | protein_coding_gene | 1.8 | 3.47 | 9.5E-12 |
| *M60.4* | WBGene00019780 | protein_coding_gene | 1.8 | 3.47 | 7.5E-20 |
| *T19C4.5* | WBGene00011839 | protein_coding_gene | 1.8 | 3.47 | 1.3E-07 |
| *ugt-4* | WBGene00013905 | protein_coding_gene | 1.8 | 3.47 | 7.8E-07 |
| *T14E8.4* | WBGene00043981 | protein_coding_gene | 1.8 | 3.46 | 6.7E-03 |
| *ceh-43* | WBGene00000463 | protein_coding_gene | 1.8 | 3.46 | 1.2E-18 |
| *F13C5.1* | WBGene00017422 | protein_coding_gene | 1.8 | 3.46 | 6.0E-09 |
| *Y45F10D.6* | WBGene00012886 | protein_coding_gene | 1.8 | 3.45 | 6.9E-05 |
| *clec-54* | WBGene00008596 | protein_coding_gene | 1.8 | 3.45 | 1.6E-16 |
| *F26A10.2* | WBGene00017814 | protein_coding_gene | 1.8 | 3.45 | 3.9E-07 |
| *F13A7.12* | WBGene00008727 | protein_coding_gene | 1.8 | 3.45 | 3.5E-02 |
| *F42C5.3* | WBGene00018345 | protein_coding_gene | 1.8 | 3.44 | 1.2E-02 |

| | | | | | |
|---|---|---|---|---|---|
| *glt-5* | WBGene00001623 | protein_coding_gene | 1.8 | 3.44 | 3.8E-08 |
| *R04B3.1* | WBGene00019866 | protein_coding_gene | 1.8 | 3.44 | 4.2E-02 |
| *C05A9.2* | WBGene00007317 | protein_coding_gene | 1.8 | 3.44 | 2.7E-10 |
| *F41C3.1* | WBGene00018267 | protein_coding_gene | 1.8 | 3.43 | 2.1E-02 |
| *Y87G2A.25* | WBGene00219288 | protein_coding_gene | 1.8 | 3.43 | 1.5E-03 |
| *F48E3.8* | WBGene00018607 | protein_coding_gene | 1.8 | 3.43 | 3.8E-07 |
| *twk-7* | WBGene00006662 | protein_coding_gene | 1.8 | 3.43 | 7.7E-09 |
| *ptr-15* | WBGene00004229 | protein_coding_gene | 1.8 | 3.43 | 1.3E-03 |
| *R11D1.12* | WBGene00045336 | protein_coding_gene | 1.8 | 3.43 | 2.5E-02 |
| *W07B8.4* | WBGene00021072 | protein_coding_gene | 1.8 | 3.42 | 1.5E-03 |
| *Y17D7C.4* | WBGene00050903 | pseudogene | 1.8 | 3.41 | 1.9E-02 |
| *C50F4.9* | WBGene00008234 | protein_coding_gene | 1.8 | 3.41 | 2.5E-04 |
| *Y102A11A.5* | WBGene00022415 | protein_coding_gene | 1.8 | 3.41 | 2.7E-03 |
| *C24H12.1* | WBGene00016071 | protein_coding_gene | 1.8 | 3.41 | 6.7E-06 |
| *F20D6.8* | WBGene00017637 | protein_coding_gene | 1.8 | 3.40 | 2.6E-03 |
| *Y51H4A.1* | WBGene00013098 | protein_coding_gene | 1.8 | 3.40 | 1.1E-02 |
| *nhr-42* | WBGene00003632 | protein_coding_gene | 1.8 | 3.40 | 1.0E-11 |
| *C04C11.25* | WBGene00284855 | protein_coding_gene | 1.8 | 3.39 | 7.3E-04 |
| *ham-1* | WBGene00001820 | protein_coding_gene | 1.8 | 3.39 | 2.4E-16 |
| *cutl-11* | WBGene00011443 | protein_coding_gene | 1.8 | 3.39 | 8.6E-03 |
| *unc-129* | WBGene00006852 | protein_coding_gene | 1.8 | 3.38 | 4.6E-08 |
| *T10H9.8* | WBGene00044779 | protein_coding_gene | 1.8 | 3.38 | 2.5E-09 |
| *F53C3.6* | WBGene00018750 | protein_coding_gene | 1.8 | 3.38 | 2.6E-02 |
| *ttr-50* | WBGene00010234 | protein_coding_gene | 1.8 | 3.37 | 2.3E-40 |
| *Y43F8C.3* | WBGene00012825 | protein_coding_gene | 1.8 | 3.37 | 2.5E-16 |
| *lam-3* | WBGene00002248 | protein_coding_gene | 1.8 | 3.37 | 3.1E-16 |
| *gad-3* | WBGene00015057 | protein_coding_gene | 1.8 | 3.37 | 5.7E-22 |
| *mnp-1* | WBGene00007139 | protein_coding_gene | 1.7 | 3.36 | 8.2E-25 |
| *clec-44* | WBGene00019928 | protein_coding_gene | 1.7 | 3.36 | 2.6E-02 |
| *Y71H2AM.15* | WBGene00022180 | protein_coding_gene | 1.7 | 3.36 | 1.4E-04 |
| *T04F8.9* | WBGene00011447 | protein_coding_gene | 1.7 | 3.35 | 5.0E-03 |
| *ZK593.2* | WBGene00014002 | protein_coding_gene | 1.7 | 3.35 | 2.4E-05 |
| *Y71F9AL.25* | WBGene00304792 | protein_coding_gene | 1.7 | 3.34 | 3.6E-04 |
| *F13E9.12* | WBGene00008761 | protein_coding_gene | 1.7 | 3.34 | 3.4E-02 |
| *F33H2.8* | WBGene00009371 | protein_coding_gene | 1.7 | 3.34 | 3.5E-03 |
| *K10G6.9* | WBGene00206374 | protein_coding_gene | 1.7 | 3.33 | 1.3E-02 |
| *bus-18* | WBGene00044631 | protein_coding_gene | 1.7 | 3.32 | 1.1E-02 |
| *C12D5.3* | WBGene00015706 | protein_coding_gene | 1.7 | 3.32 | 1.5E-02 |
| *T25B9.3* | WBGene00012009 | protein_coding_gene | 1.7 | 3.32 | 1.2E-02 |
| *C25F6.7* | WBGene00016106 | protein_coding_gene | 1.7 | 3.32 | 2.6E-04 |
| *jbts-14* | WBGene00018727 | protein_coding_gene | 1.7 | 3.32 | 2.2E-02 |
| *clec-107* | WBGene00012502 | protein_coding_gene | 1.7 | 3.32 | 4.5E-02 |
| *cpr-1* | WBGene00000781 | protein_coding_gene | 1.7 | 3.31 | 8.4E-27 |
| *Y73E7A.8* | WBGene00022274 | protein_coding_gene | 1.7 | 3.31 | 4.3E-03 |
| *C07A9.5* | WBGene00007401 | protein_coding_gene | 1.7 | 3.31 | 4.7E-02 |
| *B0284.3* | WBGene00007133 | protein_coding_gene | 1.7 | 3.30 | 8.6E-05 |
| *fbxa-69* | WBGene00010018 | protein_coding_gene | 1.7 | 3.30 | 2.9E-12 |
| *nlp-48* | WBGene00008002 | protein_coding_gene | 1.7 | 3.30 | 1.4E-02 |
| *Y43F8B.2* | WBGene00012813 | protein_coding_gene | 1.7 | 3.30 | 9.8E-27 |
| *pitr-3* | WBGene00015055 | protein_coding_gene | 1.7 | 3.30 | 5.2E-06 |
| *fkh-2* | WBGene00001434 | protein_coding_gene | 1.7 | 3.30 | 1.8E-05 |
| *fbxa-165* | WBGene00015600 | protein_coding_gene | 1.7 | 3.30 | 4.8E-02 |
| *C40H1.9* | WBGene00008044 | protein_coding_gene | 1.7 | 3.30 | 1.4E-02 |
| *Y75B8A.28* | WBGene00013559 | protein_coding_gene | 1.7 | 3.29 | 1.8E-05 |
| *W01C9.1* | WBGene00012175 | protein_coding_gene | 1.7 | 3.29 | 2.3E-02 |
| *cutl-29* | WBGene00017421 | protein_coding_gene | 1.7 | 3.29 | 1.8E-02 |
| *C47D12.4* | WBGene00008138 | protein_coding_gene | 1.7 | 3.29 | 1.0E-05 |
| *F13E9.13* | WBGene00044797 | protein_coding_gene | 1.7 | 3.29 | 5.2E-06 |
| *clec-196* | WBGene00009156 | protein_coding_gene | 1.7 | 3.29 | 2.1E-15 |
| *F18C5.5* | WBGene00017560 | protein_coding_gene | 1.7 | 3.28 | 1.0E-02 |
| *kal-1* | WBGene00002181 | protein_coding_gene | 1.7 | 3.28 | 1.0E-10 |
| *W01B11.6* | WBGene00020917 | protein_coding_gene | 1.7 | 3.28 | 1.3E-07 |
| *Y62F5A.10* | WBGene00013388 | protein_coding_gene | 1.7 | 3.28 | 1.6E-04 |
| *osm-6* | WBGene00003886 | protein_coding_gene | 1.7 | 3.28 | 1.1E-02 |
| *fbxc-53* | WBGene00016711 | protein_coding_gene | 1.7 | 3.28 | 3.7E-02 |

| | | | | | |
|---|---|---|---|---|---|
| *F57C12.6* | WBGene00194687 | protein_coding_gene | 1.7 | 3.28 | 1.9E-03 |
| *F52D1.2* | WBGene00018683 | protein_coding_gene | 1.7 | 3.28 | 7.1E-04 |
| *C26B9.6* | WBGene00016135 | protein_coding_gene | 1.7 | 3.27 | 5.6E-06 |
| *B0391.14* | WBGene00206462 | protein_coding_gene | 1.7 | 3.27 | 3.1E-02 |
| *T08A9.13* | WBGene00044442 | protein_coding_gene | 1.7 | 3.27 | 3.0E-02 |
| *D1054.18* | WBGene00077783 | protein_coding_gene | 1.7 | 3.26 | 3.8E-02 |
| *Y53H1B.4* | WBGene00013182 | protein_coding_gene | 1.7 | 3.26 | 3.6E-02 |
| *F07H5.4* | WBGene00014308 | pseudogene | 1.7 | 3.25 | 6.6E-21 |
| *tab-1* | WBGene00006380 | protein_coding_gene | 1.7 | 3.25 | 1.5E-02 |
| *ceh-32* | WBGene00000453 | protein_coding_gene | 1.7 | 3.25 | 3.2E-16 |
| *sptf-1* | WBGene00009743 | protein_coding_gene | 1.7 | 3.25 | 8.9E-08 |
| *Y51H7C.12* | WBGene00021790 | protein_coding_gene | 1.7 | 3.25 | 2.6E-02 |
| *C26B9.3* | WBGene00016133 | protein_coding_gene | 1.7 | 3.25 | 2.4E-04 |
| *C11G10.1* | WBGene00007525 | protein_coding_gene | 1.7 | 3.25 | 9.3E-05 |
| *fkh-7* | WBGene00001439 | protein_coding_gene | 1.7 | 3.24 | 1.2E-20 |
| *Y71G12B.31* | WBGene00044348 | protein_coding_gene | 1.7 | 3.24 | 3.6E-03 |
| *C13C4.6* | WBGene00007550 | protein_coding_gene | 1.7 | 3.23 | 3.0E-05 |
| *F17C8.3* | WBGene00008910 | protein_coding_gene | 1.7 | 3.23 | 2.2E-05 |
| *Y53F4B.20* | WBGene00013166 | protein_coding_gene | 1.7 | 3.23 | 1.9E-02 |
| *unk-1* | WBGene00016407 | protein_coding_gene | 1.7 | 3.22 | 3.5E-21 |
| *T18D3.7* | WBGene00011824 | protein_coding_gene | 1.7 | 3.22 | 6.0E-11 |
| *F13H8.1* | WBGene00017434 | protein_coding_gene | 1.7 | 3.21 | 1.1E-04 |
| *srpr-1.2* | WBGene00045168 | ncRNA_gene | 1.7 | 3.21 | 2.7E-02 |
| *F54B8.4* | WBGene00010019 | protein_coding_gene | 1.7 | 3.21 | 7.5E-06 |
| *Y77E11A.14* | WBGene00022314 | protein_coding_gene | 1.7 | 3.20 | 4.4E-03 |
| *F33D4.6* | WBGene00017998 | protein_coding_gene | 1.7 | 3.20 | 3.9E-04 |
| *slc-36.3* | WBGene00012629 | protein_coding_gene | 1.7 | 3.20 | 6.4E-04 |
| *frpr-11* | WBGene00019444 | protein_coding_gene | 1.7 | 3.19 | 1.1E-06 |
| *F08G2.5* | WBGene00008577 | protein_coding_gene | 1.7 | 3.19 | 8.4E-03 |
| *ZK930.5* | WBGene00014155 | protein_coding_gene | 1.7 | 3.19 | 1.0E-02 |
| *Y71F9AM.9* | WBGene00271781 | lincRNA_gene | 1.7 | 3.19 | 3.1E-02 |
| *tpra-1* | WBGene00008366 | protein_coding_gene | 1.7 | 3.19 | 2.5E-14 |
| *C53C11.5* | WBGene00044770 | protein_coding_gene | 1.7 | 3.19 | 1.4E-08 |
| *kpc-1* | WBGene00002232 | protein_coding_gene | 1.7 | 3.18 | 9.7E-30 |
| *fbxa-219* | WBGene00044403 | protein_coding_gene | 1.7 | 3.18 | 2.5E-05 |
| *asp-14* | WBGene00019619 | protein_coding_gene | 1.7 | 3.18 | 7.4E-14 |
| *F44F1.6* | WBGene00009698 | protein_coding_gene | 1.7 | 3.18 | 3.0E-10 |
| *F41E6.5* | WBGene00018286 | protein_coding_gene | 1.7 | 3.18 | 4.8E-09 |
| *nhr-18* | WBGene00003617 | protein_coding_gene | 1.7 | 3.17 | 8.1E-09 |
| *F45D3.3* | WBGene00009723 | protein_coding_gene | 1.7 | 3.17 | 1.1E-08 |
| *fut-2* | WBGene00001506 | protein_coding_gene | 1.7 | 3.17 | 1.3E-02 |
| *nphp-4* | WBGene00011261 | protein_coding_gene | 1.7 | 3.15 | 2.6E-03 |
| *clec-55* | WBGene00008597 | protein_coding_gene | 1.7 | 3.15 | 9.6E-04 |
| *lipl-6* | WBGene00021963 | protein_coding_gene | 1.7 | 3.15 | 6.5E-03 |
| *pha-4* | WBGene00004013 | protein_coding_gene | 1.7 | 3.15 | 1.6E-14 |
| *qua-1* | WBGene00004264 | protein_coding_gene | 1.7 | 3.15 | 2.3E-04 |
| *T09B4.5* | WBGene00020379 | protein_coding_gene | 1.7 | 3.14 | 4.5E-35 |
| *C31H2.3* | WBGene00016293 | protein_coding_gene | 1.7 | 3.14 | 3.9E-02 |
| *ins-4* | WBGene00002087 | protein_coding_gene | 1.7 | 3.14 | 1.3E-03 |
| *jmjd-3.2* | WBGene00009089 | protein_coding_gene | 1.7 | 3.14 | 3.5E-08 |
| *F46B6.9* | WBGene00009774 | protein_coding_gene | 1.6 | 3.13 | 7.6E-03 |
| *Y105E8A.34* | WBGene00077689 | pseudogene | 1.6 | 3.12 | 4.5E-02 |
| *Y54G2A.52* | WBGene00045397 | protein_coding_gene | 1.6 | 3.12 | 8.2E-10 |
| *C06B3.7* | WBGene00007366 | protein_coding_gene | 1.6 | 3.12 | 6.4E-03 |
| *daf-10* | WBGene00000906 | protein_coding_gene | 1.6 | 3.11 | 5.0E-03 |
| *fkh-8* | WBGene00001440 | protein_coding_gene | 1.6 | 3.11 | 1.9E-03 |
| *nep-14* | WBGene00018225 | pseudogene | 1.6 | 3.11 | 6.8E-06 |
| *ZK669.3* | WBGene00014053 | protein_coding_gene | 1.6 | 3.11 | 1.8E-19 |
| *K07H8.11* | WBGene00044510 | protein_coding_gene | 1.6 | 3.10 | 1.6E-02 |
| *efl-3* | WBGene00009899 | protein_coding_gene | 1.6 | 3.10 | 1.1E-18 |
| *egl-15* | WBGene00001184 | protein_coding_gene | 1.6 | 3.10 | 1.8E-14 |
| *lec-2* | WBGene00002265 | protein_coding_gene | 1.6 | 3.10 | 3.0E-08 |
| *unc-39* | WBGene00006775 | protein_coding_gene | 1.6 | 3.10 | 2.0E-03 |
| *F20D6.10* | WBGene00017639 | protein_coding_gene | 1.6 | 3.09 | 3.2E-04 |
| *C50D2.1* | WBGene00016805 | protein_coding_gene | 1.6 | 3.09 | 1.9E-02 |

| | | | | | |
|---|---|---|---|---|---|
| *T28B8.1* | WBGene00012113 | protein_coding_gene | 1.6 | 3.09 | 1.5E-22 |
| *dao-6* | WBGene00000932 | protein_coding_gene | 1.6 | 3.08 | 4.0E-02 |
| *C17C3.3* | WBGene00015889 | protein_coding_gene | 1.6 | 3.08 | 3.5E-02 |
| *D2005.6* | WBGene00008401 | protein_coding_gene | 1.6 | 3.08 | 1.5E-04 |
| *C08E8.4* | WBGene00007440 | protein_coding_gene | 1.6 | 3.08 | 7.8E-03 |
| *Y6D1A.2* | WBGene00012396 | protein_coding_gene | 1.6 | 3.07 | 1.5E-02 |
| *F38B6.6* | WBGene00018175 | protein_coding_gene | 1.6 | 3.07 | 4.8E-04 |
| *C13C4.7* | WBGene00007551 | protein_coding_gene | 1.6 | 3.07 | 5.9E-04 |
| *C05E11.3* | WBGene00015494 | protein_coding_gene | 1.6 | 3.07 | 1.4E-06 |
| *agr-1* | WBGene00018304 | protein_coding_gene | 1.6 | 3.07 | 7.1E-06 |
| *T16A1.10* | WBGene00269431 | protein_coding_gene | 1.6 | 3.07 | 2.9E-02 |
| *F52E1.5* | WBGene00018694 | protein_coding_gene | 1.6 | 3.06 | 3.2E-05 |
| *C16H3.3* | WBGene00015868 | protein_coding_gene | 1.6 | 3.06 | 4.3E-19 |
| *T01B6.1* | WBGene00020134 | protein_coding_gene | 1.6 | 3.06 | 1.3E-03 |
| *flp-32* | WBGene00010982 | protein_coding_gene | 1.6 | 3.06 | 2.8E-05 |
| *dyf-5* | WBGene00001121 | protein_coding_gene | 1.6 | 3.05 | 1.6E-02 |
| *Y116A8C.40* | WBGene00013812 | protein_coding_gene | 1.6 | 3.05 | 4.5E-02 |
| *grh-1* | WBGene00001707 | protein_coding_gene | 1.6 | 3.05 | 1.3E-05 |
| *C33D9.5* | WBGene00007899 | pseudogene | 1.6 | 3.05 | 9.1E-06 |
| *K09F5.6* | WBGene00044698 | protein_coding_gene | 1.6 | 3.05 | 1.3E-07 |
| *nhr-178* | WBGene00017510 | protein_coding_gene | 1.6 | 3.05 | 3.5E-02 |
| *nhr-91* | WBGene00003681 | protein_coding_gene | 1.6 | 3.05 | 2.8E-06 |
| *hmit-1.3* | WBGene00010811 | protein_coding_gene | 1.6 | 3.05 | 1.7E-13 |
| *C28G1.6* | WBGene00016191 | protein_coding_gene | 1.6 | 3.04 | 2.3E-05 |
| *C55A1.6* | WBGene00008323 | protein_coding_gene | 1.6 | 3.04 | 4.1E-02 |
| *fbxc-24* | WBGene00019927 | protein_coding_gene | 1.6 | 3.03 | 1.3E-03 |
| *eya-1* | WBGene00001377 | protein_coding_gene | 1.6 | 3.03 | 3.5E-11 |
| *M01H9.10* | WBGene00271788 | protein_coding_gene | 1.6 | 3.03 | 1.4E-02 |
| *ZK6.8* | WBGene00022643 | protein_coding_gene | 1.6 | 3.02 | 6.6E-04 |
| *cbp-2* | WBGene00010545 | protein_coding_gene | 1.6 | 3.01 | 3.5E-09 |
| *skr-19* | WBGene00004825 | protein_coding_gene | 1.6 | 3.01 | 1.8E-11 |
| *eyg-1* | WBGene00013147 | protein_coding_gene | 1.6 | 3.01 | 2.7E-02 |
| *F01D5.6* | WBGene00008496 | protein_coding_gene | 1.6 | 3.01 | 1.1E-02 |
| *fbxb-66* | WBGene00044066 | protein_coding_gene | 1.6 | 3.01 | 2.4E-06 |
| *npr-20* | WBGene00011578 | protein_coding_gene | 1.6 | 3.01 | 5.1E-05 |
| *mls-2* | WBGene00003377 | protein_coding_gene | 1.6 | 3.01 | 2.1E-03 |
| *cdh-9* | WBGene00000401 | protein_coding_gene | 1.6 | 3.01 | 4.2E-03 |
| *K08B5.2* | WBGene00019519 | protein_coding_gene | 1.6 | 3.00 | 1.0E-05 |
| *sym-2* | WBGene00006367 | protein_coding_gene | 1.6 | 3.00 | 2.1E-07 |
| *fbxb-60* | WBGene00010104 | protein_coding_gene | 1.6 | 3.00 | 2.9E-02 |
| *pat-9* | WBGene00003933 | protein_coding_gene | 1.6 | 2.99 | 1.1E-09 |
| *hhat-2* | WBGene00013312 | protein_coding_gene | 1.6 | 2.99 | 1.9E-03 |
| *fbxb-69* | WBGene00008613 | protein_coding_gene | 1.6 | 2.99 | 2.4E-02 |
| *C31H5.4* | WBGene00007855 | protein_coding_gene | 1.6 | 2.99 | 3.1E-17 |
| *cyp-13A3* | WBGene00011675 | protein_coding_gene | 1.6 | 2.98 | 7.3E-03 |
| *F53A2.1* | WBGene00009949 | protein_coding_gene | 1.6 | 2.98 | 4.3E-03 |
| *math-45* | WBGene00021772 | protein_coding_gene | 1.6 | 2.98 | 1.4E-03 |
| *F25B3.5* | WBGene00009102 | protein_coding_gene | 1.6 | 2.98 | 7.6E-04 |
| *lron-1* | WBGene00008093 | protein_coding_gene | 1.6 | 2.98 | 3.5E-02 |
| *Y41G9A.10* | WBGene00044901 | protein_coding_gene | 1.6 | 2.98 | 1.3E-04 |
| *ttr-37* | WBGene00017667 | protein_coding_gene | 1.6 | 2.97 | 7.3E-11 |
| *hlh-6* | WBGene00001952 | protein_coding_gene | 1.6 | 2.97 | 2.7E-02 |
| *unc-75* | WBGene00006807 | protein_coding_gene | 1.6 | 2.97 | 5.6E-03 |
| *cex-1* | WBGene00023407 | protein_coding_gene | 1.6 | 2.97 | 8.3E-11 |
| *Y75B8A.6* | WBGene00013543 | protein_coding_gene | 1.6 | 2.96 | 2.9E-15 |
| *mlt-9* | WBGene00008605 | protein_coding_gene | 1.6 | 2.96 | 2.0E-03 |
| *F02A9.1* | WBGene00008513 | protein_coding_gene | 1.6 | 2.96 | 1.3E-02 |
| *F40F9.10* | WBGene00009586 | protein_coding_gene | 1.6 | 2.96 | 2.1E-02 |
| *Y42A5A.3* | WBGene00012778 | protein_coding_gene | 1.6 | 2.96 | 1.7E-02 |
| *C11H1.7* | WBGene00007531 | protein_coding_gene | 1.6 | 2.96 | 3.7E-03 |
| *Y54G2A.11* | WBGene00021876 | protein_coding_gene | 1.6 | 2.95 | 9.0E-10 |
| *C06E7.4* | WBGene00015541 | protein_coding_gene | 1.6 | 2.95 | 3.5E-08 |
| *F56C4.4* | WBGene00044977 | protein_coding_gene | 1.6 | 2.95 | 4.9E-03 |
| *egl-38* | WBGene00001204 | protein_coding_gene | 1.6 | 2.94 | 1.2E-02 |
| *C46C11.4* | WBGene00044567 | protein_coding_gene | 1.6 | 2.94 | 8.2E-03 |

| | | | | | |
|---|---|---|---|---|---|
| *T19E7.6* | WBGene00044500 | protein_coding_gene | 1.6 | 2.94 | 1.3E-06 |
| *his-24* | WBGene00001898 | protein_coding_gene | 1.6 | 2.94 | 1.2E-39 |
| *igcm-3* | WBGene00020160 | protein_coding_gene | 1.6 | 2.94 | 7.4E-32 |
| *F52C9.5* | WBGene00018675 | protein_coding_gene | 1.6 | 2.93 | 5.0E-02 |
| *C42D4.1* | WBGene00016594 | protein_coding_gene | 1.6 | 2.93 | 2.0E-13 |
| *bkip-1* | WBGene00014569 | protein_coding_gene | 1.5 | 2.93 | 2.4E-02 |
| *che-2* | WBGene00000484 | protein_coding_gene | 1.5 | 2.92 | 4.0E-02 |
| *dyf-11* | WBGene00001127 | protein_coding_gene | 1.5 | 2.92 | 1.1E-02 |
| *sox-2* | WBGene00004949 | protein_coding_gene | 1.5 | 2.92 | 3.7E-12 |
| *col-76* | WBGene00000652 | protein_coding_gene | 1.5 | 2.91 | 1.5E-06 |
| *Y64G10A.7* | WBGene00013416 | protein_coding_gene | 1.5 | 2.91 | 2.3E-05 |
| *ugt-5* | WBGene00013906 | protein_coding_gene | 1.5 | 2.90 | 2.1E-09 |
| *K12B6.9* | WBGene00023432 | protein_coding_gene | 1.5 | 2.90 | 2.7E-06 |
| *T22F3.8* | WBGene00020698 | protein_coding_gene | 1.5 | 2.90 | 4.8E-02 |
| *gcy-22* | WBGene00001547 | protein_coding_gene | 1.5 | 2.90 | 2.1E-02 |
| *col-99* | WBGene00000674 | protein_coding_gene | 1.5 | 2.90 | 1.2E-06 |
| *moe-3* | WBGene00003388 | protein_coding_gene | 1.5 | 2.90 | 2.0E-08 |
| *ncam-1* | WBGene00017184 | protein_coding_gene | 1.5 | 2.90 | 9.3E-12 |
| *F08H9.3* | WBGene00008591 | protein_coding_gene | 1.5 | 2.89 | 1.8E-02 |
| *T16G1.6* | WBGene00011800 | protein_coding_gene | 1.5 | 2.89 | 2.9E-07 |
| *W04C9.8* | WBGene00235387 | ncRNA_gene | 1.5 | 2.89 | 2.1E-02 |
| *Y11D7A.19* | WBGene00077756 | protein_coding_gene | 1.5 | 2.89 | 4.3E-02 |
| *pdf-2* | WBGene00018590 | protein_coding_gene | 1.5 | 2.89 | 6.3E-07 |
| *C07B5.4* | WBGene00007409 | protein_coding_gene | 1.5 | 2.89 | 6.6E-05 |
| *EEED8.15* | WBGene00017143 | protein_coding_gene | 1.5 | 2.89 | 8.7E-15 |
| *C40H1.2* | WBGene00008038 | protein_coding_gene | 1.5 | 2.89 | 1.1E-04 |
| *F13D11.3* | WBGene00017428 | protein_coding_gene | 1.5 | 2.89 | 3.1E-07 |
| *C49H3.12* | WBGene00016795 | protein_coding_gene | 1.5 | 2.89 | 1.4E-02 |
| *cpn-2* | WBGene00000778 | protein_coding_gene | 1.5 | 2.89 | 7.3E-05 |
| *nhr-25* | WBGene00003623 | protein_coding_gene | 1.5 | 2.88 | 1.2E-10 |
| *gstk-2* | WBGene00017054 | protein_coding_gene | 1.5 | 2.88 | 4.4E-04 |
| *C35D10.3* | WBGene00016441 | protein_coding_gene | 1.5 | 2.88 | 3.9E-02 |
| *cest-17* | WBGene00019477 | protein_coding_gene | 1.5 | 2.88 | 2.0E-08 |
| *F59B2.9* | WBGene00010311 | protein_coding_gene | 1.5 | 2.87 | 1.0E-02 |
| *F46H6.5* | WBGene00023460 | protein_coding_gene | 1.5 | 2.87 | 3.5E-02 |
| *F08A7.1* | WBGene00045013 | protein_coding_gene | 1.5 | 2.87 | 1.3E-05 |
| *sto-1* | WBGene00006063 | protein_coding_gene | 1.5 | 2.87 | 1.4E-15 |
| *M03A1.8* | WBGene00077490 | protein_coding_gene | 1.5 | 2.86 | 1.0E-02 |
| *che-11* | WBGene00000490 | protein_coding_gene | 1.5 | 2.86 | 1.1E-02 |
| *ins-37* | WBGene00002120 | protein_coding_gene | 1.5 | 2.86 | 2.3E-02 |
| *D1086.5* | WBGene00008392 | protein_coding_gene | 1.5 | 2.86 | 1.3E-07 |
| *nhr-112* | WBGene00003702 | protein_coding_gene | 1.5 | 2.86 | 9.6E-04 |
| *nlp-30* | WBGene00003768 | protein_coding_gene | 1.5 | 2.86 | 6.8E-04 |
| *C32H11.4* | WBGene00007867 | protein_coding_gene | 1.5 | 2.86 | 4.0E-04 |
| *zip-8* | WBGene00017755 | protein_coding_gene | 1.5 | 2.86 | 2.7E-14 |
| *unc-130* | WBGene00006853 | protein_coding_gene | 1.5 | 2.85 | 1.7E-06 |
| *C08B6.2* | WBGene00007423 | protein_coding_gene | 1.5 | 2.85 | 4.0E-02 |
| *M02D8.6* | WBGene00019732 | protein_coding_gene | 1.5 | 2.85 | 7.8E-07 |
| *F42C5.4* | WBGene00018346 | protein_coding_gene | 1.5 | 2.85 | 5.9E-08 |
| *C16B8.4* | WBGene00015824 | protein_coding_gene | 1.5 | 2.85 | 2.9E-06 |
| *linc-46* | WBGene00219748 | lincRNA_gene | 1.5 | 2.85 | 1.2E-02 |
| *vab-3* | WBGene00006870 | protein_coding_gene | 1.5 | 2.84 | 3.2E-07 |
| *fbxb-18* | WBGene00019039 | protein_coding_gene | 1.5 | 2.84 | 7.5E-03 |
| *swip-10* | WBGene00018738 | protein_coding_gene | 1.5 | 2.84 | 4.2E-03 |
| *F31F7.1* | WBGene00017964 | protein_coding_gene | 1.5 | 2.84 | 8.9E-38 |
| *C35E7.4* | WBGene00016456 | protein_coding_gene | 1.5 | 2.83 | 6.8E-04 |
| *ZK757.2* | WBGene00014074 | protein_coding_gene | 1.5 | 2.83 | 1.5E-02 |
| *icl-1* | WBGene00001564 | protein_coding_gene | 1.5 | 2.82 | 2.1E-03 |
| *unc-120* | WBGene00006844 | protein_coding_gene | 1.5 | 2.82 | 2.0E-07 |
| *K12G11.6* | WBGene00010792 | protein_coding_gene | 1.5 | 2.82 | 4.5E-02 |
| *phlp-2* | WBGene00009647 | protein_coding_gene | 1.5 | 2.82 | 1.4E-02 |
| *gem-4* | WBGene00001577 | protein_coding_gene | 1.5 | 2.82 | 1.6E-11 |
| *pxd-1* | WBGene00007982 | protein_coding_gene | 1.5 | 2.82 | 2.2E-18 |
| *ceh-22* | WBGene00000445 | protein_coding_gene | 1.5 | 2.82 | 8.4E-03 |
| *Y67D8C.9* | WBGene00022073 | protein_coding_gene | 1.5 | 2.82 | 1.1E-08 |

| | | | | | |
|---|---|---|---|---|---|
| *hil-3* | WBGene00001854 | protein_coding_gene | 1.5 | 2.81 | 8.5E-27 |
| *igcm-4* | WBGene00022418 | protein_coding_gene | 1.5 | 2.81 | 2.2E-12 |
| *F45C12.17* | WBGene00269388 | protein_coding_gene | 1.5 | 2.81 | 1.9E-04 |
| *fbxb-19* | WBGene00019040 | protein_coding_gene | 1.5 | 2.81 | 1.0E-02 |
| *K10D3.6* | WBGene00010740 | protein_coding_gene | 1.5 | 2.81 | 1.5E-06 |
| *M01H9.4* | WBGene00019720 | protein_coding_gene | 1.5 | 2.81 | 6.5E-08 |
| *K04A8.20* | WBGene00235332 | protein_coding_gene | 1.5 | 2.81 | 1.0E-02 |
| *pqn-42* | WBGene00004129 | protein_coding_gene | 1.5 | 2.81 | 2.4E-03 |
| *cyp-42A1* | WBGene00013585 | protein_coding_gene | 1.5 | 2.80 | 7.9E-08 |
| *ngn-1* | WBGene00003595 | protein_coding_gene | 1.5 | 2.80 | 1.5E-03 |
| *fbxb-77* | WBGene00021204 | protein_coding_gene | 1.5 | 2.80 | 4.5E-02 |
| *W10G11.2* | WBGene00021135 | protein_coding_gene | 1.5 | 2.80 | 5.8E-03 |
| *F09F3.8* | WBGene00014732 | pseudogene | 1.5 | 2.79 | 3.7E-02 |
| *osm-5* | WBGene00003885 | protein_coding_gene | 1.5 | 2.79 | 1.9E-02 |
| *F46C8.3* | WBGene00018485 | protein_coding_gene | 1.5 | 2.79 | 2.3E-06 |
| *D1053.3* | WBGene00008368 | protein_coding_gene | 1.5 | 2.79 | 7.7E-04 |
| *T01D3.6* | WBGene00011330 | protein_coding_gene | 1.5 | 2.79 | 1.1E-09 |
| *ref-2* | WBGene00004335 | protein_coding_gene | 1.5 | 2.78 | 4.2E-04 |
| *F33E2.5* | WBGene00009361 | protein_coding_gene | 1.5 | 2.78 | 3.3E-13 |
| *plpr-1* | WBGene00011524 | protein_coding_gene | 1.5 | 2.78 | 4.9E-03 |
| *dep-1* | WBGene00009717 | protein_coding_gene | 1.5 | 2.78 | 1.9E-08 |
| *clec-266* | WBGene00016088 | protein_coding_gene | 1.5 | 2.78 | 6.0E-15 |
| *hmg-11* | WBGene00001976 | protein_coding_gene | 1.5 | 2.78 | 5.0E-12 |
| *C33D9.6* | WBGene00007900 | protein_coding_gene | 1.5 | 2.78 | 4.8E-06 |
| *dlc-5* | WBGene00043067 | protein_coding_gene | 1.5 | 2.78 | 1.4E-02 |
| *F57C12.2* | WBGene00019014 | protein_coding_gene | 1.5 | 2.78 | 4.2E-04 |
| *lgg-1* | WBGene00002980 | protein_coding_gene | 1.5 | 2.78 | 3.5E-19 |
| *cyp-34A2* | WBGene00011699 | protein_coding_gene | 1.5 | 2.77 | 1.8E-03 |
| *ptr-19* | WBGene00004233 | protein_coding_gene | 1.5 | 2.77 | 2.7E-02 |
| *F20C5.5* | WBGene00008970 | protein_coding_gene | 1.5 | 2.77 | 3.3E-03 |
| *K12C11.6* | WBGene00045052 | protein_coding_gene | 1.5 | 2.76 | 6.1E-06 |
| *W03G11.4* | WBGene00012226 | protein_coding_gene | 1.5 | 2.76 | 8.9E-09 |
| *ace-3* | WBGene00000037 | protein_coding_gene | 1.5 | 2.76 | 1.7E-03 |
| *Y54G2A.28* | WBGene00021892 | protein_coding_gene | 1.5 | 2.76 | 3.6E-02 |
| *W02B8.1* | WBGene00012197 | protein_coding_gene | 1.5 | 2.76 | 1.4E-05 |
| *T16G1.2* | WBGene00011796 | protein_coding_gene | 1.5 | 2.76 | 2.0E-02 |
| *bath-47* | WBGene00006464 | protein_coding_gene | 1.5 | 2.75 | 9.6E-03 |
| *nhr-67* | WBGene00003657 | protein_coding_gene | 1.5 | 2.75 | 5.0E-03 |
| *mks-6* | WBGene00010642 | protein_coding_gene | 1.5 | 2.75 | 2.9E-02 |
| *lips-10* | WBGene00008803 | protein_coding_gene | 1.5 | 2.75 | 3.0E-05 |
| *W05G11.8* | WBGene00271636 | ncRNA_gene | 1.5 | 2.75 | 3.9E-02 |
| *ugt-33* | WBGene00007946 | protein_coding_gene | 1.5 | 2.75 | 2.2E-03 |
| *W05H9.3* | WBGene00021050 | ncRNA_gene | 1.5 | 2.75 | 2.9E-12 |
| *K02D7.1* | WBGene00019298 | protein_coding_gene | 1.5 | 2.74 | 1.3E-10 |
| *ltd-1* | WBGene00003089 | protein_coding_gene | 1.5 | 2.74 | 3.4E-05 |
| *F45E10.2* | WBGene00009729 | protein_coding_gene | 1.5 | 2.74 | 8.7E-06 |
| *srpr-2.1* | WBGene00195016 | ncRNA_gene | 1.5 | 2.74 | 3.5E-02 |
| *sulp-8* | WBGene00013963 | protein_coding_gene | 1.5 | 2.74 | 1.1E-05 |
| *ZK1128.7* | WBGene00014233 | protein_coding_gene | 1.5 | 2.74 | 4.9E-12 |
| *Y58A7A.3* | WBGene00021977 | protein_coding_gene | 1.5 | 2.74 | 2.1E-05 |
| *daao-1* | WBGene00022076 | protein_coding_gene | 1.5 | 2.74 | 4.6E-03 |
| *linc-91* | WBGene00219746 | lincRNA_gene | 1.5 | 2.73 | 5.0E-02 |
| *F38E9.1* | WBGene00018184 | protein_coding_gene | 1.5 | 2.73 | 2.7E-13 |
| *Y51H4A.8* | WBGene00013104 | protein_coding_gene | 1.5 | 2.73 | 7.8E-06 |
| *fbxb-100* | WBGene00013403 | protein_coding_gene | 1.4 | 2.73 | 7.1E-03 |
| *C36E8.4* | WBGene00007983 | protein_coding_gene | 1.4 | 2.73 | 1.9E-05 |
| *Y38F1A.8* | WBGene00012610 | protein_coding_gene | 1.4 | 2.72 | 1.2E-03 |
| *cht-1* | WBGene00000503 | protein_coding_gene | 1.4 | 2.72 | 1.6E-17 |
| *F58H1.5* | WBGene00010288 | protein_coding_gene | 1.4 | 2.72 | 1.0E-07 |
| *dhs-15* | WBGene00000978 | protein_coding_gene | 1.4 | 2.72 | 1.5E-02 |
| *T21D12.14* | WBGene00045177 | protein_coding_gene | 1.4 | 2.71 | 4.9E-02 |
| *nhr-32* | WBGene00003626 | protein_coding_gene | 1.4 | 2.71 | 8.2E-12 |
| *F21D12.3* | WBGene00017663 | protein_coding_gene | 1.4 | 2.71 | 2.3E-08 |
| *capa-1* | WBGene00021268 | protein_coding_gene | 1.4 | 2.70 | 2.9E-04 |
| *Y27F2A.8* | WBGene00021299 | protein_coding_gene | 1.4 | 2.70 | 3.1E-03 |

| | | | | | |
|---|---|---|---|---|---|
| *linc-6* | WBGene00219700 | lincRNA_gene | 1.4 | 2.70 | 5.4E-04 |
| *T04C12.8* | WBGene00011430 | protein_coding_gene | 1.4 | 2.70 | 2.4E-03 |
| *T24B8.3* | WBGene00011977 | protein_coding_gene | 1.4 | 2.70 | 3.3E-24 |
| *fbxl-1* | WBGene00015350 | protein_coding_gene | 1.4 | 2.70 | 1.1E-16 |
| *dhs-31* | WBGene00011424 | protein_coding_gene | 1.4 | 2.70 | 6.5E-06 |
| *F53F1.6* | WBGene00009984 | protein_coding_gene | 1.4 | 2.70 | 9.8E-04 |
| *vab-7* | WBGene00006873 | protein_coding_gene | 1.4 | 2.69 | 7.7E-03 |
| *Y22D7AL.15* | WBGene00021253 | protein_coding_gene | 1.4 | 2.69 | 1.1E-07 |
| *sta-2* | WBGene00010251 | protein_coding_gene | 1.4 | 2.69 | 6.9E-08 |
| *C54D10.13* | WBGene00045400 | protein_coding_gene | 1.4 | 2.69 | 4.8E-03 |
| *ttx-1* | WBGene00006652 | protein_coding_gene | 1.4 | 2.69 | 1.4E-03 |
| *hsp-12.6* | WBGene00002013 | protein_coding_gene | 1.4 | 2.68 | 6.1E-03 |
| *crb-1* | WBGene00000792 | protein_coding_gene | 1.4 | 2.68 | 2.5E-08 |
| *faah-2* | WBGene00015048 | protein_coding_gene | 1.4 | 2.68 | 7.1E-05 |
| *aff-1* | WBGene00016625 | protein_coding_gene | 1.4 | 2.68 | 1.6E-02 |
| *ugt-14* | WBGene00019233 | protein_coding_gene | 1.4 | 2.68 | 4.3E-03 |
| *osg-1* | WBGene00019832 | protein_coding_gene | 1.4 | 2.68 | 2.3E-04 |
| *cbp-3* | WBGene00009595 | protein_coding_gene | 1.4 | 2.67 | 1.1E-04 |
| *poml-4* | WBGene00044697 | protein_coding_gene | 1.4 | 2.67 | 4.0E-02 |
| *cpg-7* | WBGene00010723 | protein_coding_gene | 1.4 | 2.67 | 1.5E-03 |
| *C30G4.4* | WBGene00016269 | protein_coding_gene | 1.4 | 2.67 | 2.4E-08 |
| *Y57G11B.2* | WBGene00013295 | protein_coding_gene | 1.4 | 2.67 | 6.0E-05 |
| *ttr-21* | WBGene00008628 | protein_coding_gene | 1.4 | 2.67 | 7.7E-08 |
| *ins-34* | WBGene00002117 | protein_coding_gene | 1.4 | 2.67 | 1.1E-04 |
| *xtr-1* | WBGene00006965 | protein_coding_gene | 1.4 | 2.67 | 4.1E-02 |
| *F43C9.1* | WBGene00018375 | protein_coding_gene | 1.4 | 2.67 | 4.4E-03 |
| *F21D9.4* | WBGene00009018 | protein_coding_gene | 1.4 | 2.67 | 2.9E-02 |
| *pax-3* | WBGene00003939 | protein_coding_gene | 1.4 | 2.67 | 1.2E-03 |
| *F13E9.15* | WBGene00045208 | protein_coding_gene | 1.4 | 2.67 | 9.4E-07 |
| *T16H12.9* | WBGene00011817 | protein_coding_gene | 1.4 | 2.67 | 5.5E-03 |
| *T23F11.6* | WBGene00011957 | protein_coding_gene | 1.4 | 2.66 | 1.6E-02 |
| *dpy-1* | WBGene00001063 | protein_coding_gene | 1.4 | 2.66 | 1.6E-02 |
| *K03H1.5* | WBGene00010540 | protein_coding_gene | 1.4 | 2.66 | 9.2E-10 |
| *psa-3* | WBGene00009560 | protein_coding_gene | 1.4 | 2.66 | 2.2E-03 |
| *nhr-21* | WBGene00003620 | protein_coding_gene | 1.4 | 2.66 | 8.8E-08 |
| *lgc-22* | WBGene00017483 | protein_coding_gene | 1.4 | 2.65 | 6.2E-14 |
| *C47E12.9* | WBGene00008152 | protein_coding_gene | 1.4 | 2.65 | 1.3E-02 |
| *T01D1.8* | WBGene00044641 | protein_coding_gene | 1.4 | 2.65 | 3.6E-02 |
| *M162.5* | WBGene00010931 | protein_coding_gene | 1.4 | 2.64 | 4.2E-04 |
| *F47B8.2* | WBGene00009803 | protein_coding_gene | 1.4 | 2.64 | 3.8E-06 |
| *F32D1.11* | WBGene00044530 | protein_coding_gene | 1.4 | 2.64 | 1.1E-02 |
| *W08F4.13* | WBGene00044733 | protein_coding_gene | 1.4 | 2.63 | 2.2E-02 |
| *aqp-3* | WBGene00000171 | protein_coding_gene | 1.4 | 2.63 | 4.0E-02 |
| *cex-2* | WBGene00023408 | protein_coding_gene | 1.4 | 2.63 | 1.3E-07 |
| *dhrs-4* | WBGene00010063 | protein_coding_gene | 1.4 | 2.63 | 8.2E-14 |
| *T05B11.7* | WBGene00044342 | protein_coding_gene | 1.4 | 2.62 | 2.9E-02 |
| *ncr-2* | WBGene00003562 | protein_coding_gene | 1.4 | 2.62 | 7.5E-04 |
| *Y57A10A.26* | WBGene00013266 | protein_coding_gene | 1.4 | 2.62 | 2.6E-12 |
| *K08E7.6* | WBGene00010674 | protein_coding_gene | 1.4 | 2.62 | 4.5E-02 |
| *C05G5.5* | WBGene00007351 | protein_coding_gene | 1.4 | 2.62 | 2.4E-02 |
| *Y92H12BR.4* | WBGene00022370 | pseudogene | 1.4 | 2.61 | 4.0E-03 |
| *msp-42* | WBGene00003437 | pseudogene | 1.4 | 2.61 | 4.0E-02 |
| *madd-2* | WBGene00016539 | protein_coding_gene | 1.4 | 2.61 | 5.5E-10 |
| *F18A11.2* | WBGene00008929 | protein_coding_gene | 1.4 | 2.61 | 3.3E-02 |
| *C18H9.6* | WBGene00016004 | protein_coding_gene | 1.4 | 2.61 | 1.3E-05 |
| *dhs-2* | WBGene00000966 | protein_coding_gene | 1.4 | 2.61 | 4.1E-07 |
| *memo-1* | WBGene00016500 | protein_coding_gene | 1.4 | 2.60 | 1.7E-14 |
| *tos-1* | WBGene00019466 | protein_coding_gene | 1.4 | 2.60 | 2.0E-46 |
| *tag-243* | WBGene00044078 | protein_coding_gene | 1.4 | 2.60 | 3.5E-04 |
| *best-3* | WBGene00015286 | protein_coding_gene | 1.4 | 2.59 | 2.0E-05 |
| *T14B1.1* | WBGene00011763 | protein_coding_gene | 1.4 | 2.59 | 6.5E-05 |
| *F44A6.5* | WBGene00009677 | protein_coding_gene | 1.4 | 2.59 | 3.6E-07 |
| *Y50D4A.10* | WBGene00235315 | protein_coding_gene | 1.4 | 2.59 | 6.6E-03 |
| *F56H11.2* | WBGene00010175 | protein_coding_gene | 1.4 | 2.59 | 5.7E-04 |
| *K10D6.4* | WBGene00010744 | protein_coding_gene | 1.4 | 2.59 | 2.0E-09 |

| | | | | | |
|---|---|---|---|---|---|
| *F47B8.3* | WBGene00009804 | protein_coding_gene | 1.4 | 2.58 | 1.1E-05 |
| *C08G5.7* | WBGene00044418 | protein_coding_gene | 1.4 | 2.58 | 3.0E-03 |
| *vap-2* | WBGene00011462 | protein_coding_gene | 1.4 | 2.58 | 7.8E-04 |
| *C24H12.11* | WBGene00016080 | pseudogene | 1.4 | 2.58 | 5.0E-03 |
| *cav-1* | WBGene00000301 | protein_coding_gene | 1.4 | 2.57 | 1.6E-20 |
| *sma-10* | WBGene00020649 | protein_coding_gene | 1.4 | 2.57 | 4.6E-14 |
| *nhr-117* | WBGene00003707 | protein_coding_gene | 1.4 | 2.57 | 2.2E-04 |
| *Y55F3BR.11* | WBGene00044757 | protein_coding_gene | 1.4 | 2.56 | 4.3E-03 |
| *ugt-15* | WBGene00008097 | protein_coding_gene | 1.4 | 2.56 | 3.1E-02 |
| *mod-5* | WBGene00003387 | protein_coding_gene | 1.4 | 2.56 | 2.0E-03 |
| *tag-89* | WBGene00006459 | protein_coding_gene | 1.4 | 2.56 | 7.4E-03 |
| *Y52B11A.4* | WBGene00013124 | protein_coding_gene | 1.4 | 2.56 | 3.3E-03 |
| *cfi-1* | WBGene00000476 | protein_coding_gene | 1.3 | 2.55 | 1.0E-03 |
| *Y71F9AL.7* | WBGene00022112 | protein_coding_gene | 1.3 | 2.54 | 7.8E-04 |
| *ceh-10* | WBGene00000435 | protein_coding_gene | 1.3 | 2.54 | 3.3E-02 |
| *cwn-1* | WBGene00000857 | protein_coding_gene | 1.3 | 2.54 | 5.1E-07 |
| *ncx-3* | WBGene00003568 | protein_coding_gene | 1.3 | 2.54 | 1.6E-06 |
| *ZK384.7* | WBGene00269383 | ncRNA_gene | 1.3 | 2.54 | 1.4E-03 |
| *M60.7* | WBGene00019782 | protein_coding_gene | 1.3 | 2.54 | 5.8E-03 |
| *insc-1* | WBGene00018392 | protein_coding_gene | 1.3 | 2.54 | 1.6E-05 |
| *F41G4.8* | WBGene00018311 | protein_coding_gene | 1.3 | 2.54 | 6.4E-04 |
| *F59F5.3* | WBGene00010342 | protein_coding_gene | 1.3 | 2.54 | 8.5E-03 |
| *nlp-71* | WBGene00022254 | protein_coding_gene | 1.3 | 2.54 | 9.9E-04 |
| *Y17G9B.8* | WBGene00021205 | protein_coding_gene | 1.3 | 2.54 | 2.1E-12 |
| *slt-1* | WBGene00004854 | protein_coding_gene | 1.3 | 2.53 | 4.6E-04 |
| *F54D1.6* | WBGene00010047 | protein_coding_gene | 1.3 | 2.53 | 1.1E-16 |
| *Y73F4A.3* | WBGene00013516 | protein_coding_gene | 1.3 | 2.53 | 9.0E-04 |
| *mltn-12* | WBGene00008277 | protein_coding_gene | 1.3 | 2.53 | 4.7E-02 |
| *nmad-1* | WBGene00017304 | protein_coding_gene | 1.3 | 2.53 | 2.0E-06 |
| *cest-27* | WBGene00018958 | protein_coding_gene | 1.3 | 2.53 | 2.3E-07 |
| *spp-1* | WBGene00004986 | protein_coding_gene | 1.3 | 2.53 | 2.2E-02 |
| *inx-2* | WBGene00002124 | protein_coding_gene | 1.3 | 2.52 | 3.1E-13 |
| *ceh-20* | WBGene00000443 | protein_coding_gene | 1.3 | 2.52 | 2.7E-12 |
| *Y65B4BL.7* | WBGene00022039 | protein_coding_gene | 1.3 | 2.52 | 1.4E-02 |
| *C03A7.13* | WBGene00015371 | protein_coding_gene | 1.3 | 2.52 | 2.2E-03 |
| *T13C2.3* | WBGene00020479 | protein_coding_gene | 1.3 | 2.52 | 2.6E-03 |
| *R148.5* | WBGene00020104 | protein_coding_gene | 1.3 | 2.52 | 7.3E-11 |
| *ser-4* | WBGene00004779 | protein_coding_gene | 1.3 | 2.51 | 2.6E-03 |
| *F10G8.8* | WBGene00008666 | protein_coding_gene | 1.3 | 2.51 | 1.6E-07 |
| *unc-5* | WBGene00006745 | protein_coding_gene | 1.3 | 2.51 | 2.1E-10 |
| *K04F10.2* | WBGene00019395 | protein_coding_gene | 1.3 | 2.51 | 3.8E-02 |
| *C29H12.6* | WBGene00016237 | protein_coding_gene | 1.3 | 2.51 | 5.2E-04 |
| *F36H5.13* | WBGene00206519 | protein_coding_gene | 1.3 | 2.51 | 1.2E-02 |
| *mdl-1* | WBGene00003163 | protein_coding_gene | 1.3 | 2.51 | 1.8E-05 |
| *lad-2* | WBGene00002243 | protein_coding_gene | 1.3 | 2.51 | 4.5E-03 |
| *C52E2.4* | WBGene00016883 | protein_coding_gene | 1.3 | 2.51 | 1.7E-02 |
| *flp-8* | WBGene00001451 | protein_coding_gene | 1.3 | 2.51 | 3.0E-07 |
| *cpna-5* | WBGene00007044 | protein_coding_gene | 1.3 | 2.50 | 4.5E-08 |
| *C18A11.2* | WBGene00015948 | protein_coding_gene | 1.3 | 2.50 | 4.5E-02 |
| *F43G6.8* | WBGene00009660 | protein_coding_gene | 1.3 | 2.50 | 3.0E-12 |
| *acs-17* | WBGene00016716 | protein_coding_gene | 1.3 | 2.50 | 1.4E-09 |
| *mig-13* | WBGene00003245 | protein_coding_gene | 1.3 | 2.50 | 2.5E-06 |
| *T25D10.4* | WBGene00020801 | protein_coding_gene | 1.3 | 2.50 | 4.0E-07 |
| *C44H9.5* | WBGene00008100 | protein_coding_gene | 1.3 | 2.50 | 8.5E-09 |
| *fkb-4* | WBGene00001429 | protein_coding_gene | 1.3 | 2.50 | 2.2E-02 |
| *Y37D8A.16* | WBGene00012554 | protein_coding_gene | 1.3 | 2.50 | 5.5E-06 |
| *nhr-206* | WBGene00011097 | protein_coding_gene | 1.3 | 2.50 | 9.5E-03 |
| *F32B4.5* | WBGene00009315 | protein_coding_gene | 1.3 | 2.50 | 2.2E-02 |
| *T26H2.10* | WBGene00044976 | pseudogene | 1.3 | 2.49 | 1.5E-02 |
| *C54E10.6* | WBGene00008310 | protein_coding_gene | 1.3 | 2.49 | 1.7E-03 |
| *T04A6.1* | WBGene00020200 | protein_coding_gene | 1.3 | 2.49 | 8.3E-09 |
| *fat-3* | WBGene00001395 | protein_coding_gene | 1.3 | 2.48 | 1.1E-02 |
| *C06G3.5* | WBGene00015551 | protein_coding_gene | 1.3 | 2.48 | 1.2E-07 |
| *slc-28.2* | WBGene00017868 | protein_coding_gene | 1.3 | 2.48 | 1.8E-02 |
| *tsp-6* | WBGene00006632 | protein_coding_gene | 1.3 | 2.48 | 2.8E-02 |

| | | | | | |
|---|---|---|---|---|---|
| *Y75B7AL.2* | WBGene00022285 | protein_coding_gene | 1.3 | 2.48 | 2.1E-02 |
| *T08B6.9* | WBGene00020356 | protein_coding_gene | 1.3 | 2.47 | 3.2E-02 |
| *BE0003N10.3* | WBGene00015268 | protein_coding_gene | 1.3 | 2.47 | 2.9E-04 |
| *ZC455.1* | WBGene00013902 | protein_coding_gene | 1.3 | 2.46 | 2.5E-05 |
| *C34B4.3* | WBGene00007908 | protein_coding_gene | 1.3 | 2.46 | 3.0E-02 |
| *C24H12.7* | WBGene00016076 | pseudogene | 1.3 | 2.46 | 2.7E-02 |
| *cest-24* | WBGene00020891 | protein_coding_gene | 1.3 | 2.46 | 3.5E-04 |
| *epg-2* | WBGene00021468 | protein_coding_gene | 1.3 | 2.46 | 2.9E-12 |
| *Y71F9AL.6* | WBGene00022111 | protein_coding_gene | 1.3 | 2.46 | 1.2E-02 |
| *F12A10.1* | WBGene00017390 | protein_coding_gene | 1.3 | 2.45 | 2.6E-05 |
| *F48E3.9* | WBGene00018608 | protein_coding_gene | 1.3 | 2.45 | 3.4E-02 |
| *M01G12.14* | WBGene00010825 | protein_coding_gene | 1.3 | 2.45 | 3.7E-02 |
| *glb-15* | WBGene00009387 | protein_coding_gene | 1.3 | 2.45 | 3.6E-02 |
| *fbxc-18* | WBGene00016307 | protein_coding_gene | 1.3 | 2.44 | 2.7E-04 |
| *vab-2* | WBGene00006869 | protein_coding_gene | 1.3 | 2.44 | 6.9E-04 |
| *C06G3.6* | WBGene00015552 | protein_coding_gene | 1.3 | 2.44 | 1.8E-07 |
| *vab-15* | WBGene00006881 | protein_coding_gene | 1.3 | 2.44 | 5.3E-03 |
| *ZC487.1* | WBGene00022623 | protein_coding_gene | 1.3 | 2.44 | 2.9E-02 |
| *F39E9.28* | WBGene00271716 | pseudogene | 1.3 | 2.44 | 8.5E-04 |
| *ttr-44* | WBGene00008341 | protein_coding_gene | 1.3 | 2.44 | 6.0E-04 |
| *fbxb-107* | WBGene00017912 | protein_coding_gene | 1.3 | 2.44 | 2.3E-02 |
| *T28H10.1* | WBGene00012142 | protein_coding_gene | 1.3 | 2.44 | 6.2E-05 |
| *C35A5.6* | WBGene00007950 | protein_coding_gene | 1.3 | 2.44 | 7.7E-06 |
| *zmp-2* | WBGene00019212 | protein_coding_gene | 1.3 | 2.43 | 3.1E-02 |
| *flp-21* | WBGene00001464 | protein_coding_gene | 1.3 | 2.43 | 9.9E-04 |
| *MTCE.7* | WBGene00014454 | rRNA_gene | 1.3 | 2.43 | 3.2E-04 |
| *Y47G6A.5* | WBGene00021635 | protein_coding_gene | 1.3 | 2.43 | 2.7E-09 |
| *flp-12* | WBGene00001455 | protein_coding_gene | 1.3 | 2.43 | 7.7E-06 |
| *F19C7.2* | WBGene00017592 | protein_coding_gene | 1.3 | 2.43 | 1.1E-03 |
| *cyn-6* | WBGene00000882 | protein_coding_gene | 1.3 | 2.43 | 7.5E-10 |
| *T19A5.3* | WBGene00020554 | protein_coding_gene | 1.3 | 2.43 | 1.7E-02 |
| *Y51A2D.13* | WBGene00013080 | protein_coding_gene | 1.3 | 2.43 | 4.6E-14 |
| *ZK1290.13* | WBGene00022893 | protein_coding_gene | 1.3 | 2.42 | 2.4E-05 |
| *lact-5* | WBGene00016355 | protein_coding_gene | 1.3 | 2.42 | 3.0E-06 |
| *ttm-2* | WBGene00017840 | protein_coding_gene | 1.3 | 2.42 | 4.7E-04 |
| *atz-1* | WBGene00009212 | protein_coding_gene | 1.3 | 2.42 | 4.6E-11 |
| *F36A4.2* | WBGene00018081 | protein_coding_gene | 1.3 | 2.42 | 2.0E-02 |
| *sqst-1* | WBGene00011737 | protein_coding_gene | 1.3 | 2.42 | 1.3E-07 |
| *W02D7.4* | WBGene00020940 | protein_coding_gene | 1.3 | 2.42 | 2.6E-07 |
| *R13A5.15* | WBGene00194726 | protein_coding_gene | 1.3 | 2.42 | 2.7E-02 |
| *R11G1.2* | WBGene00020011 | protein_coding_gene | 1.3 | 2.42 | 2.4E-02 |
| *C18A11.1* | WBGene00015947 | protein_coding_gene | 1.3 | 2.41 | 2.1E-04 |
| *nspc-16* | WBGene00008357 | protein_coding_gene | 1.3 | 2.41 | 3.3E-02 |
| *syg-2* | WBGene00007750 | protein_coding_gene | 1.3 | 2.41 | 4.4E-05 |
| *tsp-16* | WBGene00006642 | protein_coding_gene | 1.3 | 2.41 | 9.4E-03 |
| *Y75B12B.3* | WBGene00013568 | protein_coding_gene | 1.3 | 2.41 | 2.5E-02 |
| *mab-7* | WBGene00003104 | protein_coding_gene | 1.3 | 2.41 | 3.7E-02 |
| *let-522* | WBGene00010870 | protein_coding_gene | 1.3 | 2.41 | 1.1E-06 |
| *clec-86* | WBGene00016912 | protein_coding_gene | 1.3 | 2.41 | 8.2E-05 |
| *kel-1* | WBGene00002184 | protein_coding_gene | 1.3 | 2.41 | 7.3E-03 |
| *C07D10.5* | WBGene00015569 | protein_coding_gene | 1.3 | 2.40 | 3.6E-11 |
| *hlh-1* | WBGene00001948 | protein_coding_gene | 1.3 | 2.40 | 2.9E-04 |
| *sulp-3* | WBGene00018283 | protein_coding_gene | 1.3 | 2.39 | 3.0E-02 |
| *R03E9.2* | WBGene00019843 | protein_coding_gene | 1.3 | 2.39 | 3.4E-04 |
| *pho-4* | WBGene00004023 | protein_coding_gene | 1.3 | 2.39 | 2.9E-06 |
| *nhr-43* | WBGene00003633 | protein_coding_gene | 1.3 | 2.39 | 8.0E-04 |
| *let-653* | WBGene00002827 | protein_coding_gene | 1.3 | 2.39 | 3.2E-06 |
| *F59F5.7* | WBGene00010345 | protein_coding_gene | 1.3 | 2.39 | 2.4E-02 |
| *Y77E11A.23* | WBGene00235267 | protein_coding_gene | 1.3 | 2.39 | 4.0E-03 |
| *tyr-1* | WBGene00015332 | protein_coding_gene | 1.3 | 2.38 | 4.4E-05 |
| *F26G1.2* | WBGene00017838 | protein_coding_gene | 1.3 | 2.38 | 3.5E-04 |
| *lect-2* | WBGene00019407 | protein_coding_gene | 1.2 | 2.38 | 3.2E-07 |
| *ztf-11* | WBGene00009939 | protein_coding_gene | 1.2 | 2.37 | 1.2E-10 |
| *fbxb-80* | WBGene00017117 | pseudogene | 1.2 | 2.37 | 3.9E-02 |
| *T05E11.8* | WBGene00011484 | protein_coding_gene | 1.2 | 2.37 | 3.3E-03 |

| | | | | | |
|---|---|---|---|---|---|
| *srm-1* | WBGene00005638 | protein_coding_gene | 1.2 | 2.37 | 4.9E-02 |
| *Y62F5A.9* | WBGene00013387 | protein_coding_gene | 1.2 | 2.37 | 4.7E-02 |
| *madf-10* | WBGene00013717 | protein_coding_gene | 1.2 | 2.37 | 7.1E-03 |
| *K12C11.7* | WBGene00045053 | protein_coding_gene | 1.2 | 2.37 | 9.5E-04 |
| *C35E7.5* | WBGene00016457 | protein_coding_gene | 1.2 | 2.37 | 2.0E-08 |
| *zig-8* | WBGene00006985 | protein_coding_gene | 1.2 | 2.37 | 4.9E-02 |
| *B0507.1* | WBGene00015218 | protein_coding_gene | 1.2 | 2.36 | 3.1E-03 |
| *F20C5.11* | WBGene00303015 | protein_coding_gene | 1.2 | 2.36 | 2.9E-03 |
| *Y70G10A.3* | WBGene00013499 | protein_coding_gene | 1.2 | 2.36 | 1.1E-05 |
| *Y73B6A.1* | WBGene00022228 | protein_coding_gene | 1.2 | 2.36 | 3.0E-02 |
| *K02E10.1* | WBGene00019317 | protein_coding_gene | 1.2 | 2.36 | 1.2E-05 |
| *C34F11.8* | WBGene00016417 | protein_coding_gene | 1.2 | 2.36 | 2.8E-05 |
| *ugt-8* | WBGene00019234 | protein_coding_gene | 1.2 | 2.36 | 1.6E-09 |
| *F35B3.7* | WBGene00018033 | protein_coding_gene | 1.2 | 2.36 | 4.8E-03 |
| *pqn-13* | WBGene00004104 | protein_coding_gene | 1.2 | 2.35 | 1.9E-02 |
| *daam-1* | WBGene00018976 | protein_coding_gene | 1.2 | 2.35 | 5.6E-03 |
| *Y38C1AA.6* | WBGene00021397 | protein_coding_gene | 1.2 | 2.35 | 2.1E-02 |
| *nlr-1* | WBGene00003772 | protein_coding_gene | 1.2 | 2.35 | 7.6E-05 |
| *C05B5.5* | WBGene00007322 | protein_coding_gene | 1.2 | 2.34 | 2.6E-02 |
| *fozi-1* | WBGene00010453 | protein_coding_gene | 1.2 | 2.34 | 2.9E-04 |
| *bus-17* | WBGene00044630 | protein_coding_gene | 1.2 | 2.34 | 4.2E-02 |
| *sup-26* | WBGene00006331 | protein_coding_gene | 1.2 | 2.34 | 6.9E-22 |
| *R11G1.7* | WBGene00044725 | protein_coding_gene | 1.2 | 2.34 | 2.5E-03 |
| *T09A12.1* | WBGene00020372 | protein_coding_gene | 1.2 | 2.34 | 2.9E-02 |
| *epg-9* | WBGene00022078 | protein_coding_gene | 1.2 | 2.34 | 1.1E-08 |
| *cwn-2* | WBGene00000858 | protein_coding_gene | 1.2 | 2.33 | 8.0E-05 |
| *C01A2.1* | WBGene00007213 | protein_coding_gene | 1.2 | 2.33 | 4.1E-02 |
| *C06B8.2* | WBGene00007368 | protein_coding_gene | 1.2 | 2.33 | 2.1E-04 |
| *Y42G9A.1* | WBGene00021531 | protein_coding_gene | 1.2 | 2.33 | 7.0E-04 |
| *fbxb-116* | WBGene00012066 | protein_coding_gene | 1.2 | 2.33 | 1.9E-02 |
| *cal-2* | WBGene00000286 | protein_coding_gene | 1.2 | 2.32 | 3.5E-04 |
| *Y50D4A.5* | WBGene00021739 | protein_coding_gene | 1.2 | 2.32 | 1.1E-04 |
| *T27A10.6* | WBGene00020846 | protein_coding_gene | 1.2 | 2.32 | 3.0E-04 |
| *lgg-2* | WBGene00002981 | protein_coding_gene | 1.2 | 2.32 | 1.7E-15 |
| *nhr-55* | WBGene00003645 | protein_coding_gene | 1.2 | 2.32 | 2.9E-05 |
| *sfrp-1* | WBGene00022242 | protein_coding_gene | 1.2 | 2.32 | 3.5E-03 |
| *ptr-3* | WBGene00004218 | protein_coding_gene | 1.2 | 2.32 | 2.4E-02 |
| *Y17G9B.2* | WBGene00021199 | protein_coding_gene | 1.2 | 2.32 | 1.4E-02 |
| *Y44E3A.1* | WBGene00021547 | protein_coding_gene | 1.2 | 2.32 | 3.4E-03 |
| *grl-7* | WBGene00001716 | protein_coding_gene | 1.2 | 2.31 | 3.6E-02 |
| *clec-180* | WBGene00017991 | protein_coding_gene | 1.2 | 2.31 | 4.7E-02 |
| *cest-16* | WBGene00020330 | protein_coding_gene | 1.2 | 2.31 | 2.9E-02 |
| *ptp-4* | WBGene00010558 | protein_coding_gene | 1.2 | 2.31 | 4.9E-03 |
| *trh-1* | WBGene00007827 | protein_coding_gene | 1.2 | 2.31 | 2.6E-02 |
| *ZK550.2* | WBGene00013996 | protein_coding_gene | 1.2 | 2.31 | 1.6E-03 |
| *nhr-110* | WBGene00003700 | protein_coding_gene | 1.2 | 2.30 | 2.0E-06 |
| *strm-1* | WBGene00019198 | protein_coding_gene | 1.2 | 2.30 | 1.7E-02 |
| *C05C12.4* | WBGene00007336 | protein_coding_gene | 1.2 | 2.30 | 3.8E-12 |
| *T04F8.7* | WBGene00011445 | protein_coding_gene | 1.2 | 2.30 | 1.4E-05 |
| *C34D1.4* | WBGene00007931 | protein_coding_gene | 1.2 | 2.30 | 1.3E-03 |
| *fbxb-93* | WBGene00013408 | protein_coding_gene | 1.2 | 2.30 | 1.7E-02 |
| *F13C5.2* | WBGene00017423 | protein_coding_gene | 1.2 | 2.29 | 5.0E-15 |
| *C13E3.1* | WBGene00015740 | protein_coding_gene | 1.2 | 2.29 | 5.3E-03 |
| *T19H5.4* | WBGene00011849 | protein_coding_gene | 1.2 | 2.29 | 1.7E-13 |
| *tbx-8* | WBGene00006545 | protein_coding_gene | 1.2 | 2.29 | 1.8E-02 |
| *Y41D4A.3* | WBGene00021505 | protein_coding_gene | 1.2 | 2.29 | 2.2E-03 |
| *C49F5.7* | WBGene00008211 | protein_coding_gene | 1.2 | 2.29 | 6.0E-07 |
| *ZK863.8* | WBGene00014125 | protein_coding_gene | 1.2 | 2.29 | 4.8E-05 |
| *syg-1* | WBGene00006365 | protein_coding_gene | 1.2 | 2.29 | 4.6E-04 |
| *R07G3.8* | WBGene00019944 | protein_coding_gene | 1.2 | 2.29 | 9.1E-05 |
| *T28A11.22* | WBGene00044526 | protein_coding_gene | 1.2 | 2.29 | 1.3E-02 |
| *F08C6.5* | WBGene00017243 | protein_coding_gene | 1.2 | 2.28 | 3.7E-02 |
| *nhr-152* | WBGene00015705 | protein_coding_gene | 1.2 | 2.28 | 3.5E-03 |
| *inx-6* | WBGene00002128 | protein_coding_gene | 1.2 | 2.28 | 5.9E-03 |
| *mig-18* | WBGene00003249 | protein_coding_gene | 1.2 | 2.27 | 3.2E-05 |

| | | | | | |
|---|---|---|---|---|---|
| *ptp-1* | WBGene00004213 | protein_coding_gene | 1.2 | 2.27 | 2.0E-05 |
| *F58H1.7* | WBGene00010290 | protein_coding_gene | 1.2 | 2.27 | 9.6E-04 |
| *Y13C8A.4* | WBGene00219303 | pseudogene | 1.2 | 2.27 | 7.7E-03 |
| *ins-18* | WBGene00002101 | protein_coding_gene | 1.2 | 2.27 | 5.0E-02 |
| *E02H4.7* | WBGene00077757 | protein_coding_gene | 1.2 | 2.27 | 1.0E-07 |
| *tag-275* | WBGene00016423 | protein_coding_gene | 1.2 | 2.26 | 2.4E-03 |
| *fbxa-211* | WBGene00012566 | protein_coding_gene | 1.2 | 2.26 | 2.1E-07 |
| *sdz-14* | WBGene00008782 | protein_coding_gene | 1.2 | 2.26 | 4.7E-04 |
| *F58A3.3* | WBGene00010223 | protein_coding_gene | 1.2 | 2.26 | 1.6E-02 |
| *fbxc-36* | WBGene00019918 | protein_coding_gene | 1.2 | 2.26 | 6.1E-07 |
| *elt-6* | WBGene00001253 | protein_coding_gene | 1.2 | 2.26 | 5.9E-03 |
| *R10E9.3* | WBGene00011215 | protein_coding_gene | 1.2 | 2.26 | 4.5E-03 |
| *vab-23* | WBGene00014153 | protein_coding_gene | 1.2 | 2.25 | 3.7E-03 |
| *ist-1* | WBGene00002163 | protein_coding_gene | 1.2 | 2.25 | 1.6E-05 |
| *DH11.2* | WBGene00008436 | protein_coding_gene | 1.2 | 2.25 | 7.3E-11 |
| *ceh-6* | WBGene00000431 | protein_coding_gene | 1.2 | 2.24 | 3.3E-02 |
| *C27B7.7* | WBGene00007764 | protein_coding_gene | 1.2 | 2.24 | 3.8E-06 |
| *cdh-8* | WBGene00017576 | protein_coding_gene | 1.2 | 2.24 | 4.6E-03 |
| *K11D12.12* | WBGene00019651 | protein_coding_gene | 1.2 | 2.24 | 7.4E-07 |
| *ZK154.1* | WBGene00022665 | protein_coding_gene | 1.2 | 2.24 | 4.8E-02 |
| *tag-273* | WBGene00013289 | protein_coding_gene | 1.2 | 2.24 | 2.9E-04 |
| *Y43B11AR.3* | WBGene00021544 | protein_coding_gene | 1.2 | 2.24 | 1.8E-05 |
| *clec-264* | WBGene00009291 | protein_coding_gene | 1.2 | 2.23 | 2.0E-03 |
| *spp-22* | WBGene00044283 | protein_coding_gene | 1.2 | 2.23 | 4.3E-04 |
| *nspc-9* | WBGene00009634 | protein_coding_gene | 1.2 | 2.23 | 2.2E-07 |
| *tmed-13* | WBGene00022255 | protein_coding_gene | 1.2 | 2.23 | 3.7E-02 |
| *R04B5.5* | WBGene00011003 | protein_coding_gene | 1.2 | 2.23 | 3.2E-06 |
| *ceh-37* | WBGene00000458 | protein_coding_gene | 1.2 | 2.23 | 4.7E-06 |
| *acs-3* | WBGene00020343 | protein_coding_gene | 1.2 | 2.23 | 6.4E-06 |
| *pudl-1* | WBGene00044489 | protein_coding_gene | 1.2 | 2.23 | 4.1E-04 |
| *C18D4.6* | WBGene00007675 | protein_coding_gene | 1.2 | 2.23 | 1.5E-07 |
| *jmjd-3.1* | WBGene00017571 | protein_coding_gene | 1.2 | 2.23 | 9.7E-07 |
| *T01E8.1* | WBGene00011331 | protein_coding_gene | 1.2 | 2.22 | 3.0E-05 |
| *mec-6* | WBGene00003170 | protein_coding_gene | 1.2 | 2.22 | 4.6E-02 |
| *nhr-94* | WBGene00003684 | protein_coding_gene | 1.2 | 2.22 | 7.5E-03 |
| *efn-4* | WBGene00001165 | protein_coding_gene | 1.2 | 2.22 | 1.6E-08 |
| *Y49A10A.2* | WBGene00044949 | protein_coding_gene | 1.1 | 2.22 | 6.0E-07 |
| *flp-34* | WBGene00019975 | protein_coding_gene | 1.1 | 2.22 | 2.8E-03 |
| *F56C9.8* | WBGene00018952 | protein_coding_gene | 1.1 | 2.22 | 1.3E-05 |
| *F19C7.3* | WBGene00017593 | protein_coding_gene | 1.1 | 2.21 | 3.0E-02 |
| *F58B4.6* | WBGene00010239 | protein_coding_gene | 1.1 | 2.21 | 2.3E-02 |
| *C10C5.7* | WBGene00007511 | protein_coding_gene | 1.1 | 2.21 | 4.8E-02 |
| *max-1* | WBGene00003143 | protein_coding_gene | 1.1 | 2.21 | 6.5E-04 |
| *rhgf-2* | WBGene00006476 | protein_coding_gene | 1.1 | 2.21 | 2.5E-04 |
| *nhr-35* | WBGene00003628 | protein_coding_gene | 1.1 | 2.21 | 6.3E-07 |
| *cdh-12* | WBGene00022103 | protein_coding_gene | 1.1 | 2.21 | 2.7E-03 |
| *F57A8.1* | WBGene00010177 | protein_coding_gene | 1.1 | 2.20 | 9.6E-07 |
| *lron-15* | WBGene00016974 | protein_coding_gene | 1.1 | 2.20 | 1.1E-02 |
| *T03F1.11* | WBGene00020192 | protein_coding_gene | 1.1 | 2.20 | 1.3E-04 |
| *F13B12.2* | WBGene00008730 | protein_coding_gene | 1.1 | 2.20 | 2.2E-03 |
| *clec-31* | WBGene00009854 | protein_coding_gene | 1.1 | 2.20 | 9.1E-03 |
| *ZK250.13* | WBGene00235352 | protein_coding_gene | 1.1 | 2.19 | 1.4E-02 |
| *lev-8* | WBGene00002975 | protein_coding_gene | 1.1 | 2.19 | 4.6E-02 |
| *F54C9.3* | WBGene00010041 | protein_coding_gene | 1.1 | 2.19 | 1.2E-12 |
| *C35E7.6* | WBGene00016458 | protein_coding_gene | 1.1 | 2.19 | 2.2E-11 |
| *linc-15* | WBGene00219571 | lincRNA_gene | 1.1 | 2.18 | 1.0E-03 |
| *C44E12.1* | WBGene00016657 | protein_coding_gene | 1.1 | 2.18 | 9.8E-06 |
| *magu-2* | WBGene00006467 | protein_coding_gene | 1.1 | 2.18 | 6.6E-06 |
| *C33G3.4* | WBGene00007904 | protein_coding_gene | 1.1 | 2.18 | 3.1E-11 |
| *che-12* | WBGene00000491 | protein_coding_gene | 1.1 | 2.18 | 3.5E-02 |
| *chil-22* | WBGene00011166 | protein_coding_gene | 1.1 | 2.18 | 1.9E-03 |
| *ptr-8* | WBGene00004222 | protein_coding_gene | 1.1 | 2.18 | 3.4E-03 |
| *nep-21* | WBGene00011794 | protein_coding_gene | 1.1 | 2.18 | 8.7E-04 |
| *T05E11.2* | WBGene00011479 | protein_coding_gene | 1.1 | 2.17 | 3.4E-02 |
| *R01H10.4* | WBGene00010972 | protein_coding_gene | 1.1 | 2.17 | 5.3E-05 |

| | | | | | |
|---|---|---|---|---|---|
| *jmjd-3.3* | WBGene00007813 | protein_coding_gene | 1.1 | 2.17 | 2.9E-02 |
| *C06B8.7* | WBGene00007372 | protein_coding_gene | 1.1 | 2.17 | 6.5E-12 |
| *D2063.1* | WBGene00017060 | protein_coding_gene | 1.1 | 2.17 | 2.2E-03 |
| *nspc-18* | WBGene00008358 | protein_coding_gene | 1.1 | 2.17 | 2.8E-03 |
| *ccch-2* | WBGene00009537 | protein_coding_gene | 1.1 | 2.17 | 3.4E-02 |
| *dcar-1* | WBGene00007395 | protein_coding_gene | 1.1 | 2.16 | 3.3E-08 |
| *glb-11* | WBGene00007981 | protein_coding_gene | 1.1 | 2.16 | 1.4E-02 |
| *unc-30* | WBGene00006766 | protein_coding_gene | 1.1 | 2.16 | 3.0E-02 |
| *Y71A12B.12* | WBGene00013509 | protein_coding_gene | 1.1 | 2.16 | 1.9E-03 |
| *clc-5* | WBGene00000526 | protein_coding_gene | 1.1 | 2.15 | 1.6E-02 |
| *Y73F8A.5* | WBGene00013518 | protein_coding_gene | 1.1 | 2.15 | 2.4E-02 |
| *exc-5* | WBGene00001366 | protein_coding_gene | 1.1 | 2.15 | 1.5E-08 |
| *siah-1* | WBGene00021369 | protein_coding_gene | 1.1 | 2.15 | 1.1E-10 |
| *W04B5.2* | WBGene00021019 | protein_coding_gene | 1.1 | 2.15 | 6.9E-03 |
| *Y45G5AM.5* | WBGene00021557 | protein_coding_gene | 1.1 | 2.15 | 2.4E-02 |
| *nhr-103* | WBGene00003693 | protein_coding_gene | 1.1 | 2.15 | 3.5E-03 |
| *F49C12.7* | WBGene00009877 | protein_coding_gene | 1.1 | 2.14 | 1.4E-03 |
| *R11.4* | WBGene00011229 | protein_coding_gene | 1.1 | 2.14 | 6.1E-03 |
| *flp-27* | WBGene00016114 | protein_coding_gene | 1.1 | 2.14 | 7.0E-03 |
| *cyp-25A3* | WBGene00014697 | protein_coding_gene | 1.1 | 2.14 | 5.0E-04 |
| *nhr-120* | WBGene00003710 | protein_coding_gene | 1.1 | 2.14 | 1.3E-03 |
| *dhp-2* | WBGene00000964 | protein_coding_gene | 1.1 | 2.14 | 4.2E-04 |
| *R10E8.8* | WBGene00011213 | protein_coding_gene | 1.1 | 2.14 | 1.1E-03 |
| *F08B4.3* | WBGene00017235 | pseudogene | 1.1 | 2.13 | 9.2E-06 |
| *EGAP9.3* | WBGene00017151 | protein_coding_gene | 1.1 | 2.13 | 3.0E-02 |
| *C06E1.1* | WBGene00015518 | protein_coding_gene | 1.1 | 2.13 | 3.6E-06 |
| *nlp-42* | WBGene00013588 | protein_coding_gene | 1.1 | 2.13 | 6.2E-03 |
| *alr-1* | WBGene00044330 | protein_coding_gene | 1.1 | 2.13 | 9.6E-04 |
| *H04M03.12* | WBGene00019156 | protein_coding_gene | 1.1 | 2.13 | 1.9E-02 |
| *atg-2* | WBGene00019748 | protein_coding_gene | 1.1 | 2.13 | 3.7E-10 |
| *cutl-20* | WBGene00018787 | protein_coding_gene | 1.1 | 2.13 | 2.0E-02 |
| *hlh-3* | WBGene00001950 | protein_coding_gene | 1.1 | 2.13 | 6.2E-03 |
| *M03F4.6* | WBGene00019759 | protein_coding_gene | 1.1 | 2.12 | 2.1E-02 |
| *fipr-1* | WBGene00009096 | protein_coding_gene | 1.1 | 2.12 | 6.7E-05 |
| *fbxa-33* | WBGene00013841 | pseudogene | 1.1 | 2.12 | 4.2E-02 |
| *nfm-1* | WBGene00003593 | protein_coding_gene | 1.1 | 2.12 | 5.6E-07 |
| *ZK550.6* | WBGene00014000 | protein_coding_gene | 1.1 | 2.12 | 5.8E-10 |
| *nhr-88* | WBGene00003678 | protein_coding_gene | 1.1 | 2.12 | 2.4E-04 |
| *nhr-136* | WBGene00003726 | protein_coding_gene | 1.1 | 2.12 | 1.2E-03 |
| *elt-7* | WBGene00015981 | protein_coding_gene | 1.1 | 2.12 | 4.8E-05 |
| *R02D5.1* | WBGene00010976 | protein_coding_gene | 1.1 | 2.12 | 2.4E-04 |
| *lido-15* | WBGene00018804 | protein_coding_gene | 1.1 | 2.12 | 1.0E-02 |
| *ilys-4* | WBGene00016958 | protein_coding_gene | 1.1 | 2.11 | 2.7E-03 |
| *F22E5.20* | WBGene00017713 | protein_coding_gene | 1.1 | 2.11 | 4.2E-02 |
| *R07D5.2* | WBGene00011101 | protein_coding_gene | 1.1 | 2.11 | 4.3E-02 |
| *aat-1* | WBGene00000002 | protein_coding_gene | 1.1 | 2.11 | 1.1E-03 |
| *pqm-1* | WBGene00004096 | protein_coding_gene | 1.1 | 2.11 | 9.1E-07 |
| *T05E11.9* | WBGene00014505 | ncRNA_gene | 1.1 | 2.11 | 3.7E-07 |
| *shn-1* | WBGene00006444 | protein_coding_gene | 1.1 | 2.10 | 2.7E-05 |
| *irx-1* | WBGene00007984 | protein_coding_gene | 1.1 | 2.10 | 1.3E-05 |
| *H29C22.1* | WBGene00045261 | protein_coding_gene | 1.1 | 2.10 | 1.7E-06 |
| *ZK1010.8* | WBGene00014181 | protein_coding_gene | 1.1 | 2.10 | 3.2E-02 |
| *sdz-5* | WBGene00016578 | protein_coding_gene | 1.1 | 2.10 | 2.8E-02 |
| *Y69A2AL.2* | WBGene00044468 | protein_coding_gene | 1.1 | 2.10 | 4.5E-03 |
| *C28G1.10* | WBGene00219376 | protein_coding_gene | 1.1 | 2.10 | 3.8E-03 |
| *myo-3* | WBGene00003515 | protein_coding_gene | 1.1 | 2.10 | 1.4E-08 |
| *lin-12* | WBGene00003001 | protein_coding_gene | 1.1 | 2.09 | 1.9E-06 |
| *hot-6* | WBGene00001991 | protein_coding_gene | 1.1 | 2.09 | 4.9E-03 |
| *F59E11.5* | WBGene00019111 | protein_coding_gene | 1.1 | 2.09 | 5.8E-05 |
| *C32E12.4* | WBGene00016330 | protein_coding_gene | 1.1 | 2.09 | 6.1E-08 |
| *T23B3.2* | WBGene00020710 | protein_coding_gene | 1.1 | 2.09 | 1.4E-06 |
| *tag-278* | WBGene00015356 | protein_coding_gene | 1.1 | 2.09 | 3.9E-04 |
| *F42G2.2* | WBGene00018351 | protein_coding_gene | 1.1 | 2.09 | 1.2E-05 |
| *C33D9.13* | WBGene00194651 | protein_coding_gene | 1.1 | 2.08 | 7.1E-04 |
| *crtc-1* | WBGene00021237 | protein_coding_gene | 1.1 | 2.08 | 2.5E-06 |

| | | | | | |
|---|---|---|---|---|---|
| *M02H5.8* | WBGene00019744 | protein_coding_gene | 1.1 | 2.08 | 2.3E-02 |
| *akt-2* | WBGene00000103 | protein_coding_gene | 1.1 | 2.08 | 4.1E-06 |
| *tcl-2* | WBGene00006561 | protein_coding_gene | 1.1 | 2.07 | 4.5E-03 |
| *eri-6* | WBGene00016561 | protein_coding_gene | 1.1 | 2.07 | 3.5E-06 |
| *fbxa-212* | WBGene00012568 | pseudogene | 1.1 | 2.07 | 4.8E-04 |
| *nac-2* | WBGene00003518 | protein_coding_gene | 1.0 | 2.07 | 1.1E-03 |
| *lin-17* | WBGene00003006 | protein_coding_gene | 1.0 | 2.07 | 4.6E-04 |
| *nex-2* | WBGene00003589 | protein_coding_gene | 1.0 | 2.07 | 1.4E-10 |
| *C40H1.7* | WBGene00008042 | protein_coding_gene | 1.0 | 2.07 | 6.1E-05 |
| *twk-31* | WBGene00006683 | protein_coding_gene | 1.0 | 2.07 | 2.8E-03 |
| *endu-2* | WBGene00019779 | protein_coding_gene | 1.0 | 2.06 | 1.5E-05 |
| *unc-6* | WBGene00006746 | protein_coding_gene | 1.0 | 2.06 | 1.9E-04 |
| *Y34D9A.8* | WBGene00021333 | protein_coding_gene | 1.0 | 2.06 | 6.3E-03 |
| *MTCE.33* | WBGene00014472 | rRNA_gene | 1.0 | 2.06 | 9.4E-04 |
| *cdh-3* | WBGene00000395 | protein_coding_gene | 1.0 | 2.06 | 9.3E-05 |
| *T22C8.6* | WBGene00011927 | protein_coding_gene | 1.0 | 2.06 | 2.7E-02 |
| *F09C3.2* | WBGene00008610 | protein_coding_gene | 1.0 | 2.05 | 1.9E-03 |
| *C34C12.7* | WBGene00007926 | protein_coding_gene | 1.0 | 2.05 | 4.7E-03 |
| *Y54G2A.32* | WBGene00021894 | protein_coding_gene | 1.0 | 2.05 | 2.7E-03 |
| *nhr-44* | WBGene00003634 | protein_coding_gene | 1.0 | 2.05 | 2.6E-02 |
| *R10D12.1* | WBGene00011185 | protein_coding_gene | 1.0 | 2.05 | 4.5E-02 |
| *ugt-51* | WBGene00015369 | protein_coding_gene | 1.0 | 2.05 | 8.7E-03 |
| *Y73B6BL.44* | WBGene00044350 | protein_coding_gene | 1.0 | 2.05 | 2.3E-03 |
| *dhs-27* | WBGene00000990 | protein_coding_gene | 1.0 | 2.05 | 1.4E-04 |
| *R10H1.1* | WBGene00019998 | protein_coding_gene | 1.0 | 2.05 | 6.1E-04 |
| *oig-3* | WBGene00003861 | protein_coding_gene | 1.0 | 2.05 | 1.9E-04 |
| *mtl-2* | WBGene00003474 | protein_coding_gene | 1.0 | 2.05 | 3.7E-05 |
| *D2045.8* | WBGene00008425 | protein_coding_gene | 1.0 | 2.05 | 7.0E-04 |
| *F14F9.5* | WBGene00017468 | protein_coding_gene | 1.0 | 2.05 | 1.1E-02 |
| *T21B4.21* | WBGene00220076 | ncRNA_gene | 1.0 | 2.05 | 8.8E-03 |
| *valv-1* | WBGene00015216 | protein_coding_gene | 1.0 | 2.04 | 1.3E-02 |
| *C13C4.4* | WBGene00007548 | protein_coding_gene | 1.0 | 2.04 | 7.6E-04 |
| *Y51B9A.9* | WBGene00013091 | protein_coding_gene | 1.0 | 2.04 | 5.1E-03 |
| *K02A6.3* | WBGene00019294 | protein_coding_gene | 1.0 | 2.04 | 1.6E-04 |
| *exc-9* | WBGene00017644 | protein_coding_gene | 1.0 | 2.04 | 7.1E-04 |
| *F52B10.3* | WBGene00044304 | protein_coding_gene | 1.0 | 2.04 | 5.0E-02 |
| *C33H5.13* | WBGene00016379 | protein_coding_gene | 1.0 | 2.04 | 3.0E-04 |
| *Y32F6B.1* | WBGene00012531 | protein_coding_gene | 1.0 | 2.04 | 2.0E-07 |
| *srj-14* | WBGene00005602 | protein_coding_gene | 1.0 | 2.04 | 8.1E-03 |
| *mig-10* | WBGene00003243 | protein_coding_gene | 1.0 | 2.04 | 1.2E-07 |
| *ZK1053.4* | WBGene00014199 | protein_coding_gene | 1.0 | 2.04 | 1.8E-09 |
| *clec-151* | WBGene00008659 | protein_coding_gene | 1.0 | 2.04 | 4.6E-02 |
| *Y105E8B.9* | WBGene00013693 | protein_coding_gene | 1.0 | 2.03 | 1.4E-03 |
| *ins-17* | WBGene00002100 | protein_coding_gene | 1.0 | 2.03 | 2.3E-03 |
| *sulp-2* | WBGene00017464 | protein_coding_gene | 1.0 | 2.03 | 1.2E-02 |
| *Y57A10A.14* | WBGene00013257 | protein_coding_gene | 1.0 | 2.03 | 6.7E-03 |
| *hnd-1* | WBGene00001981 | protein_coding_gene | 1.0 | 2.03 | 3.8E-03 |
| *nspd-6* | WBGene00043147 | protein_coding_gene | 1.0 | 2.03 | 2.6E-02 |
| *nhr-119* | WBGene00003709 | protein_coding_gene | 1.0 | 2.03 | 3.7E-05 |
| *C41G7.8* | WBGene00077525 | protein_coding_gene | 1.0 | 2.03 | 1.7E-02 |
| *F56H9.2* | WBGene00010174 | protein_coding_gene | 1.0 | 2.03 | 3.5E-04 |
| *sul-2* | WBGene00006309 | protein_coding_gene | 1.0 | 2.02 | 1.6E-02 |
| *Y2H9A.4* | WBGene00012381 | protein_coding_gene | 1.0 | 2.02 | 1.3E-02 |
| *ric-3* | WBGene00004363 | protein_coding_gene | 1.0 | 2.02 | 1.1E-02 |
| *nhr-107* | WBGene00003697 | protein_coding_gene | 1.0 | 2.02 | 2.9E-03 |
| *F46G10.2* | WBGene00009797 | protein_coding_gene | 1.0 | 2.02 | 3.7E-05 |
| *nob-1* | WBGene00003779 | protein_coding_gene | 1.0 | 2.01 | 2.0E-02 |
| *C30G7.4* | WBGene00044178 | protein_coding_gene | 1.0 | 2.01 | 3.4E-02 |
| *zig-7* | WBGene00006984 | protein_coding_gene | 1.0 | 2.01 | 2.9E-07 |
| *R02D5.3* | WBGene00010977 | protein_coding_gene | 1.0 | 2.01 | 4.5E-03 |
| *C34E11.4* | WBGene00007937 | protein_coding_gene | 1.0 | 2.01 | 4.9E-02 |
| *mab-21* | WBGene00003112 | protein_coding_gene | 1.0 | 2.01 | 8.8E-03 |
| *cpz-2* | WBGene00000789 | protein_coding_gene | 1.0 | 2.01 | 2.0E-03 |
| *srap-1* | WBGene00011522 | protein_coding_gene | 1.0 | 2.01 | 2.3E-06 |
| *asah-2* | WBGene00009192 | protein_coding_gene | 1.0 | 2.01 | 6.3E-04 |

| | | | | | |
|---|---|---|---|---|---|
| *C35D10.8* | WBGene00016445 | protein_coding_gene | 1.0 | 2.01 | 4.8E-03 |
| *F53F8.4* | WBGene00010001 | protein_coding_gene | 1.0 | 2.01 | 4.6E-03 |
| *glo-3* | WBGene00010341 | protein_coding_gene | 1.0 | 2.00 | 1.7E-02 |
| *T19D7.6* | WBGene00020574 | protein_coding_gene | 1.0 | 2.00 | 2.8E-02 |
| *T05H4.15* | WBGene00020276 | protein_coding_gene | 1.0 | 2.00 | 3.5E-02 |
| *skr-20* | WBGene00004826 | protein_coding_gene | 1.0 | 2.00 | 1.9E-06 |
| *tsp-20* | WBGene00006646 | protein_coding_gene | 1.0 | 2.00 | 6.6E-03 |
| *lgc-32* | WBGene00020569 | protein_coding_gene | -1.0 | 0.50 | 4.6E-03 |
| *trap-3* | WBGene00021420 | protein_coding_gene | -1.0 | 0.50 | 1.6E-10 |
| *ifd-1* | WBGene00002057 | protein_coding_gene | -1.0 | 0.50 | 4.3E-03 |
| *M01G5.1* | WBGene00019715 | protein_coding_gene | -1.0 | 0.50 | 3.3E-14 |
| *cdr-7* | WBGene00010472 | protein_coding_gene | -1.0 | 0.50 | 2.0E-03 |
| *Y102A11A.3* | WBGene00022413 | protein_coding_gene | -1.0 | 0.50 | 1.7E-04 |
| *pid-5* | WBGene00021555 | protein_coding_gene | -1.0 | 0.50 | 8.2E-12 |
| *let-805* | WBGene00002915 | protein_coding_gene | -1.0 | 0.50 | 4.3E-06 |
| *col-150* | WBGene00000723 | protein_coding_gene | -1.0 | 0.50 | 5.8E-03 |
| *cyp-33E2* | WBGene00018334 | protein_coding_gene | -1.0 | 0.50 | 4.9E-03 |
| *cpn-4* | WBGene00000780 | protein_coding_gene | -1.0 | 0.50 | 3.7E-06 |
| *F54D5.12* | WBGene00010055 | protein_coding_gene | -1.0 | 0.50 | 1.3E-04 |
| *Y75B8A.24* | WBGene00013557 | protein_coding_gene | -1.0 | 0.50 | 2.5E-10 |
| *set-9* | WBGene00017482 | protein_coding_gene | -1.0 | 0.50 | 2.2E-13 |
| *tkt-1* | WBGene00008506 | protein_coding_gene | -1.0 | 0.49 | 6.4E-14 |
| *lsy-27* | WBGene00009834 | protein_coding_gene | -1.0 | 0.49 | 2.0E-04 |
| *rhy-1* | WBGene00012324 | protein_coding_gene | -1.0 | 0.49 | 6.6E-05 |
| *Y119D3B.21* | WBGene00022497 | protein_coding_gene | -1.0 | 0.49 | 6.3E-05 |
| *F53C11.1* | WBGene00009971 | protein_coding_gene | -1.0 | 0.49 | 3.6E-03 |
| *elo-1* | WBGene00001239 | protein_coding_gene | -1.0 | 0.49 | 6.2E-11 |
| *F55H12.4* | WBGene00010135 | protein_coding_gene | -1.0 | 0.49 | 1.3E-04 |
| *F35H10.10* | WBGene00018073 | protein_coding_gene | -1.0 | 0.49 | 3.6E-05 |
| *pfas-1* | WBGene00008654 | protein_coding_gene | -1.0 | 0.49 | 5.4E-06 |
| *mua-3* | WBGene00003482 | protein_coding_gene | -1.0 | 0.49 | 1.1E-05 |
| *memi-2* | WBGene00013380 | protein_coding_gene | -1.0 | 0.49 | 1.1E-05 |
| *C05D2.8* | WBGene00015469 | protein_coding_gene | -1.0 | 0.49 | 9.0E-06 |
| *C17C3.1* | WBGene00015887 | protein_coding_gene | -1.0 | 0.49 | 7.6E-04 |
| *vha-8* | WBGene00006917 | protein_coding_gene | -1.0 | 0.49 | 2.4E-11 |
| *C17H12.8* | WBGene00015933 | protein_coding_gene | -1.0 | 0.49 | 1.8E-02 |
| *tfg-1* | WBGene00006565 | protein_coding_gene | -1.0 | 0.49 | 1.4E-09 |
| *vha-6* | WBGene00006915 | protein_coding_gene | -1.0 | 0.49 | 1.2E-05 |
| *bckd-1B* | WBGene00006518 | protein_coding_gene | -1.0 | 0.49 | 9.3E-13 |
| *vha-19* | WBGene00021952 | protein_coding_gene | -1.0 | 0.49 | 5.2E-11 |
| *Y102E9.5* | WBGene00044452 | protein_coding_gene | -1.0 | 0.49 | 2.0E-02 |
| *T13C5.6* | WBGene00020486 | protein_coding_gene | -1.0 | 0.49 | 1.4E-04 |
| *ZK909.6* | WBGene00014150 | protein_coding_gene | -1.0 | 0.49 | 3.4E-04 |
| *papl-1* | WBGene00008936 | protein_coding_gene | -1.0 | 0.48 | 7.3E-07 |
| *glt-1* | WBGene00001620 | protein_coding_gene | -1.0 | 0.48 | 1.8E-10 |
| *T16H12.2* | WBGene00011812 | protein_coding_gene | -1.0 | 0.48 | 3.1E-02 |
| *oatr-1* | WBGene00015814 | protein_coding_gene | -1.0 | 0.48 | 2.3E-15 |
| *R06C1.4* | WBGene00011059 | protein_coding_gene | -1.0 | 0.48 | 4.8E-06 |
| *C11E4.7* | WBGene00007521 | protein_coding_gene | -1.0 | 0.48 | 1.1E-03 |
| *F54F7.2* | WBGene00010064 | protein_coding_gene | -1.1 | 0.48 | 4.1E-04 |
| *B0511.11* | WBGene00015236 | protein_coding_gene | -1.1 | 0.48 | 8.0E-03 |
| *C06A1.4* | WBGene00014667 | pseudogene | -1.1 | 0.48 | 4.7E-07 |
| *C52G5.2* | WBGene00008260 | protein_coding_gene | -1.1 | 0.48 | 2.1E-03 |
| *ccm-2* | WBGene00019462 | protein_coding_gene | -1.1 | 0.48 | 2.2E-02 |
| *F14H12.3* | WBGene00017471 | protein_coding_gene | -1.1 | 0.48 | 1.3E-02 |
| *F56F11.2* | WBGene00018989 | protein_coding_gene | -1.1 | 0.48 | 2.5E-02 |
| *ctsa-1.2* | WBGene00008741 | protein_coding_gene | -1.1 | 0.48 | 6.3E-05 |
| *C14B1.7* | WBGene00007578 | pseudogene | -1.1 | 0.48 | 4.4E-11 |
| *C14C11.2* | WBGene00015766 | protein_coding_gene | -1.1 | 0.48 | 1.7E-07 |
| *cal-5* | WBGene00006861 | protein_coding_gene | -1.1 | 0.48 | 3.5E-05 |
| *egg-5* | WBGene00020035 | protein_coding_gene | -1.1 | 0.48 | 1.5E-08 |
| *C14A11.6* | WBGene00015751 | protein_coding_gene | -1.1 | 0.48 | 4.2E-02 |
| *F17C8.9* | WBGene00185014 | protein_coding_gene | -1.1 | 0.48 | 2.4E-04 |
| *F47F2.1* | WBGene00018569 | protein_coding_gene | -1.1 | 0.48 | 7.3E-04 |
| *C10G8.8* | WBGene00015684 | protein_coding_gene | -1.1 | 0.48 | 1.4E-11 |

| | | | | | |
|---|---|---|---|---|---|
| *nas-20* | WBGene00003539 | protein_coding_gene | -1.1 | 0.48 | 4.7E-03 |
| *C55A6.12* | WBGene00044019 | protein_coding_gene | -1.1 | 0.48 | 4.2E-06 |
| *B0303.11* | WBGene00015131 | protein_coding_gene | -1.1 | 0.48 | 7.0E-10 |
| *del-6* | WBGene00011891 | protein_coding_gene | -1.1 | 0.48 | 2.4E-07 |
| *odc-1* | WBGene00003844 | protein_coding_gene | -1.1 | 0.47 | 2.4E-09 |
| *ltah-1.2* | WBGene00022610 | protein_coding_gene | -1.1 | 0.47 | 1.8E-11 |
| *T04F3.1* | WBGene00011436 | protein_coding_gene | -1.1 | 0.47 | 1.0E-06 |
| *C01G10.9* | WBGene00007236 | protein_coding_gene | -1.1 | 0.47 | 4.6E-05 |
| *Y53C10A.5* | WBGene00013136 | protein_coding_gene | -1.1 | 0.47 | 3.9E-06 |
| *H10E21.1* | WBGene00019182 | protein_coding_gene | -1.1 | 0.47 | 3.6E-02 |
| *ZK652.8* | WBGene00022786 | protein_coding_gene | -1.1 | 0.47 | 4.4E-03 |
| *nceh-1* | WBGene00012810 | protein_coding_gene | -1.1 | 0.47 | 3.6E-02 |
| *vrp-1* | WBGene00021869 | protein_coding_gene | -1.1 | 0.47 | 9.2E-18 |
| *C48B4.7* | WBGene00008170 | protein_coding_gene | -1.1 | 0.47 | 8.3E-06 |
| *pmp-4* | WBGene00004061 | protein_coding_gene | -1.1 | 0.47 | 6.6E-06 |
| *srd-14* | WBGene00005092 | pseudogene | -1.1 | 0.47 | 1.6E-02 |
| *ZK105.1* | WBGene00022653 | protein_coding_gene | -1.1 | 0.47 | 6.5E-06 |
| *cest-9.1* | WBGene00007692 | protein_coding_gene | -1.1 | 0.47 | 2.1E-04 |
| *C26B9.5* | WBGene00016134 | protein_coding_gene | -1.1 | 0.47 | 8.2E-09 |
| *nduo-6* | WBGene00010957 | protein_coding_gene | -1.1 | 0.47 | 1.5E-03 |
| *Y111B2A.1* | WBGene00013727 | protein_coding_gene | -1.1 | 0.47 | 5.4E-04 |
| *ldh-1* | WBGene00002262 | protein_coding_gene | -1.1 | 0.47 | 9.4E-11 |
| *pgph-3* | WBGene00016892 | protein_coding_gene | -1.1 | 0.47 | 9.9E-03 |
| *K08C7.1* | WBGene00010646 | protein_coding_gene | -1.1 | 0.47 | 6.1E-05 |
| *cyp-25A2* | WBGene00007964 | protein_coding_gene | -1.1 | 0.46 | 1.1E-05 |
| *twk-17* | WBGene00006671 | protein_coding_gene | -1.1 | 0.46 | 2.4E-02 |
| *trpp-12* | WBGene00016614 | protein_coding_gene | -1.1 | 0.46 | 2.1E-06 |
| *slc-17.3* | WBGene00015333 | protein_coding_gene | -1.1 | 0.46 | 4.8E-03 |
| *coq-1* | WBGene00000761 | protein_coding_gene | -1.1 | 0.46 | 4.3E-11 |
| *B0222.5* | WBGene00015056 | protein_coding_gene | -1.1 | 0.46 | 8.2E-08 |
| *mct-4* | WBGene00003986 | protein_coding_gene | -1.1 | 0.46 | 7.9E-08 |
| *acs-5* | WBGene00013575 | protein_coding_gene | -1.1 | 0.46 | 1.2E-10 |
| *F35F10.1* | WBGene00018052 | protein_coding_gene | -1.1 | 0.46 | 3.0E-02 |
| *ptr-16* | WBGene00004230 | protein_coding_gene | -1.1 | 0.46 | 2.2E-02 |
| *T01C8.2* | WBGene00020143 | protein_coding_gene | -1.1 | 0.46 | 4.4E-07 |
| *col-106* | WBGene00000680 | protein_coding_gene | -1.1 | 0.46 | 1.9E-11 |
| *W04B5.3* | WBGene00021020 | protein_coding_gene | -1.1 | 0.46 | 5.4E-04 |
| *mpc-1* | WBGene00011119 | protein_coding_gene | -1.1 | 0.46 | 3.3E-14 |
| *fah-1* | WBGene00019620 | protein_coding_gene | -1.1 | 0.46 | 7.7E-13 |
| *cest-13* | WBGene00010116 | protein_coding_gene | -1.1 | 0.46 | 9.8E-04 |
| *C31H2.4* | WBGene00016294 | protein_coding_gene | -1.1 | 0.46 | 3.4E-04 |
| *fbxc-44* | WBGene00015819 | protein_coding_gene | -1.1 | 0.46 | 4.9E-06 |
| *F31D4.8* | WBGene00009293 | protein_coding_gene | -1.1 | 0.46 | 2.6E-03 |
| *ugt-26* | WBGene00015695 | protein_coding_gene | -1.1 | 0.46 | 7.5E-04 |
| *C16E9.1* | WBGene00015865 | protein_coding_gene | -1.1 | 0.46 | 3.2E-02 |
| *Y43F11A.6* | WBGene00014576 | pseudogene | -1.1 | 0.46 | 2.7E-06 |
| *Y46H3C.7* | WBGene00021607 | protein_coding_gene | -1.1 | 0.46 | 1.3E-02 |
| *tatn-1* | WBGene00009628 | protein_coding_gene | -1.1 | 0.46 | 1.0E-05 |
| *F13H8.8* | WBGene00017441 | protein_coding_gene | -1.1 | 0.46 | 1.1E-02 |
| *Y54E10A.17* | WBGene00021835 | protein_coding_gene | -1.1 | 0.45 | 7.7E-05 |
| *pat-12* | WBGene00003936 | protein_coding_gene | -1.1 | 0.45 | 2.9E-08 |
| *tald-1* | WBGene00021286 | protein_coding_gene | -1.1 | 0.45 | 4.0E-13 |
| *K10C2.12* | WBGene00206386 | protein_coding_gene | -1.1 | 0.45 | 5.0E-02 |
| *spds-1* | WBGene00012909 | protein_coding_gene | -1.1 | 0.45 | 3.6E-07 |
| *C23H3.2* | WBGene00016018 | protein_coding_gene | -1.1 | 0.45 | 2.6E-08 |
| *mpc-2* | WBGene00018765 | protein_coding_gene | -1.1 | 0.45 | 1.1E-08 |
| *R102.4* | WBGene00011291 | protein_coding_gene | -1.1 | 0.45 | 1.8E-11 |
| *him-4* | WBGene00001863 | protein_coding_gene | -1.1 | 0.45 | 7.4E-06 |
| *H43E16.1* | WBGene00019273 | protein_coding_gene | -1.1 | 0.45 | 3.2E-04 |
| *lbp-6* | WBGene00002258 | protein_coding_gene | -1.1 | 0.45 | 1.2E-12 |
| *Y14H12B.2* | WBGene00021192 | protein_coding_gene | -1.1 | 0.45 | 9.0E-10 |
| *F21H7.5* | WBGene00009031 | protein_coding_gene | -1.1 | 0.45 | 4.0E-02 |
| *gly-8* | WBGene00001633 | protein_coding_gene | -1.1 | 0.45 | 7.0E-08 |
| *DC2.5* | WBGene00017082 | protein_coding_gene | -1.2 | 0.45 | 2.1E-04 |
| *dhs-14* | WBGene00000977 | protein_coding_gene | -1.2 | 0.45 | 4.3E-09 |

| | | | | | |
|---|---|---|---|---|---|
| *Y46H3C.5* | WBGene00021605 | protein_coding_gene | -1.2 | 0.45 | 3.5E-03 |
| *C26C6.9* | WBGene00007743 | protein_coding_gene | -1.2 | 0.45 | 6.9E-07 |
| *R17.3* | WBGene00011269 | protein_coding_gene | -1.2 | 0.45 | 3.6E-03 |
| *clec-41* | WBGene00007153 | protein_coding_gene | -1.2 | 0.45 | 4.0E-07 |
| *C06A1.12* | WBGene00219803 | ncRNA_gene | -1.2 | 0.45 | 1.1E-03 |
| *T11F9.12* | WBGene00011713 | protein_coding_gene | -1.2 | 0.45 | 4.5E-08 |
| *scav-5* | WBGene00011076 | protein_coding_gene | -1.2 | 0.45 | 5.2E-03 |
| *H03A11.2* | WBGene00010357 | protein_coding_gene | -1.2 | 0.45 | 5.2E-05 |
| *acdh-9* | WBGene00017874 | protein_coding_gene | -1.2 | 0.44 | 1.2E-06 |
| *gale-1* | WBGene00008132 | protein_coding_gene | -1.2 | 0.44 | 5.0E-18 |
| *Y62H9A.4* | WBGene00013392 | protein_coding_gene | -1.2 | 0.44 | 2.6E-07 |
| *sec-61.A* | WBGene00013311 | protein_coding_gene | -1.2 | 0.44 | 3.1E-20 |
| *trap-1* | WBGene00022122 | protein_coding_gene | -1.2 | 0.44 | 1.7E-15 |
| *T22F7.4* | WBGene00020703 | protein_coding_gene | -1.2 | 0.44 | 3.3E-02 |
| *cysl-3* | WBGene00019962 | protein_coding_gene | -1.2 | 0.44 | 2.3E-04 |
| *vha-13* | WBGene00013025 | protein_coding_gene | -1.2 | 0.44 | 6.3E-17 |
| *pqn-72* | WBGene00004154 | protein_coding_gene | -1.2 | 0.44 | 2.0E-02 |
| *lec-8* | WBGene00002271 | protein_coding_gene | -1.2 | 0.44 | 8.8E-05 |
| *gon-1* | WBGene00001650 | protein_coding_gene | -1.2 | 0.44 | 7.9E-12 |
| *cytb-5.1* | WBGene00007848 | protein_coding_gene | -1.2 | 0.44 | 1.9E-09 |
| *vha-5* | WBGene00006914 | protein_coding_gene | -1.2 | 0.44 | 1.2E-11 |
| *drd-5* | WBGene00018885 | protein_coding_gene | -1.2 | 0.44 | 7.2E-04 |
| *ZK1193.2* | WBGene00022858 | protein_coding_gene | -1.2 | 0.43 | 4.8E-05 |
| *F19C7.1* | WBGene00017591 | protein_coding_gene | -1.2 | 0.43 | 4.6E-04 |
| *C48D1.5* | WBGene00008178 | protein_coding_gene | -1.2 | 0.43 | 8.0E-03 |
| *mct-3* | WBGene00010834 | protein_coding_gene | -1.2 | 0.43 | 1.4E-09 |
| *Y26E6A.3* | WBGene00012514 | protein_coding_gene | -1.2 | 0.43 | 2.2E-05 |
| *K03B8.8* | WBGene00010525 | protein_coding_gene | -1.2 | 0.43 | 4.3E-03 |
| *C52B11.5* | WBGene00016874 | protein_coding_gene | -1.2 | 0.43 | 1.7E-07 |
| *idh-1* | WBGene00010317 | protein_coding_gene | -1.2 | 0.43 | 3.5E-21 |
| *cnnm-3* | WBGene00016343 | protein_coding_gene | -1.2 | 0.43 | 8.6E-07 |
| *F20G2.2* | WBGene00008986 | protein_coding_gene | -1.2 | 0.43 | 3.5E-10 |
| *R09H10.5* | WBGene00011182 | protein_coding_gene | -1.2 | 0.43 | 9.0E-05 |
| *eri-7* | WBGene00016566 | protein_coding_gene | -1.2 | 0.43 | 1.1E-10 |
| *C05C10.3* | WBGene00007330 | protein_coding_gene | -1.2 | 0.43 | 1.1E-10 |
| *gly-13* | WBGene00001638 | protein_coding_gene | -1.2 | 0.42 | 5.8E-12 |
| *C18B2.3* | WBGene00015954 | protein_coding_gene | -1.2 | 0.42 | 1.6E-03 |
| *pcyt-1* | WBGene00017241 | protein_coding_gene | -1.2 | 0.42 | 6.0E-15 |
| *F36G9.3* | WBGene00009485 | protein_coding_gene | -1.2 | 0.42 | 9.1E-03 |
| *fbp-1* | WBGene00001404 | protein_coding_gene | -1.2 | 0.42 | 2.0E-21 |
| *gba-4* | WBGene00021160 | protein_coding_gene | -1.2 | 0.42 | 5.1E-04 |
| *cyp-36A1* | WBGene00007913 | protein_coding_gene | -1.2 | 0.42 | 4.1E-05 |
| *suro-1* | WBGene00011235 | protein_coding_gene | -1.2 | 0.42 | 1.7E-02 |
| *toh-1* | WBGene00006591 | protein_coding_gene | -1.2 | 0.42 | 7.6E-05 |
| *add-1* | WBGene00000072 | protein_coding_gene | -1.2 | 0.42 | 1.3E-06 |
| *pxn-1* | WBGene00004256 | protein_coding_gene | -1.2 | 0.42 | 7.3E-08 |
| *Y73B3A.18* | WBGene00022219 | pseudogene | -1.2 | 0.42 | 3.1E-03 |
| *ugt-47* | WBGene00011006 | protein_coding_gene | -1.2 | 0.42 | 8.2E-14 |
| *adt-3* | WBGene00020567 | protein_coding_gene | -1.3 | 0.42 | 3.1E-03 |
| *asna-2* | WBGene00018577 | pseudogene | -1.3 | 0.42 | 4.1E-03 |
| *F56F10.1* | WBGene00018984 | protein_coding_gene | -1.3 | 0.42 | 1.3E-04 |
| *col-96* | WBGene00000671 | protein_coding_gene | -1.3 | 0.42 | 1.2E-03 |
| *nkb-3* | WBGene00010117 | protein_coding_gene | -1.3 | 0.42 | 1.9E-11 |
| *kat-1* | WBGene00002183 | protein_coding_gene | -1.3 | 0.42 | 5.9E-08 |
| *ech-6* | WBGene00001155 | protein_coding_gene | -1.3 | 0.42 | 1.2E-05 |
| *let-767* | WBGene00002891 | protein_coding_gene | -1.3 | 0.42 | 3.1E-15 |
| *pmt-2* | WBGene00018811 | protein_coding_gene | -1.3 | 0.42 | 3.6E-09 |
| *zmp-3* | WBGene00016283 | protein_coding_gene | -1.3 | 0.42 | 3.0E-05 |
| *C14E2.12* | WBGene00219816 | snoRNA_gene | -1.3 | 0.42 | 2.0E-02 |
| *D1054.8* | WBGene00008375 | protein_coding_gene | -1.3 | 0.41 | 8.8E-11 |
| *T01C8.3* | WBGene00020144 | protein_coding_gene | -1.3 | 0.41 | 3.3E-02 |
| *B0403.3* | WBGene00015167 | protein_coding_gene | -1.3 | 0.41 | 2.2E-03 |
| *F28H7.8* | WBGene00009241 | protein_coding_gene | -1.3 | 0.41 | 8.9E-04 |
| *cld-9* | WBGene00009432 | protein_coding_gene | -1.3 | 0.41 | 8.0E-03 |
| *R04A9.9* | WBGene00195183 | protein_coding_gene | -1.3 | 0.41 | 1.2E-04 |

| | | | | | |
|---|---|---|---|---|---|
| *sth-1* | WBGene00006060 | protein_coding_gene | -1.3 | 0.41 | 5.5E-03 |
| *ser-5* | WBGene00008890 | protein_coding_gene | -1.3 | 0.41 | 2.8E-02 |
| *dpf-6* | WBGene00001059 | protein_coding_gene | -1.3 | 0.41 | 6.4E-03 |
| *C06E2.1* | WBGene00015527 | protein_coding_gene | -1.3 | 0.41 | 2.0E-02 |
| *T08B1.1* | WBGene00020340 | protein_coding_gene | -1.3 | 0.41 | 1.1E-03 |
| *T08D2.7* | WBGene00011612 | protein_coding_gene | -1.3 | 0.41 | 4.6E-02 |
| *C44B7.7* | WBGene00016629 | protein_coding_gene | -1.3 | 0.41 | 8.2E-05 |
| *cth-2* | WBGene00022856 | protein_coding_gene | -1.3 | 0.41 | 7.5E-26 |
| *hprt-1* | WBGene00013690 | protein_coding_gene | -1.3 | 0.41 | 9.0E-20 |
| *his-38* | WBGene00001912 | protein_coding_gene | -1.3 | 0.41 | 4.0E-02 |
| *mans-2* | WBGene00008258 | protein_coding_gene | -1.3 | 0.41 | 4.8E-14 |
| *C18E9.5* | WBGene00007685 | protein_coding_gene | -1.3 | 0.41 | 4.1E-10 |
| *bcmo-2* | WBGene00018755 | protein_coding_gene | -1.3 | 0.41 | 6.9E-05 |
| *pmp-1* | WBGene00004058 | protein_coding_gene | -1.3 | 0.41 | 4.1E-11 |
| *afmd-1* | WBGene00017051 | protein_coding_gene | -1.3 | 0.41 | 1.7E-03 |
| *D2096.6* | WBGene00017073 | protein_coding_gene | -1.3 | 0.41 | 3.5E-02 |
| *ctb-1* | WBGene00000829 | protein_coding_gene | -1.3 | 0.40 | 8.6E-12 |
| *ckc-1* | WBGene00000515 | protein_coding_gene | -1.3 | 0.40 | 6.5E-06 |
| *gst-36* | WBGene00001784 | protein_coding_gene | -1.3 | 0.40 | 4.5E-07 |
| *klu-2* | WBGene00022592 | protein_coding_gene | -1.3 | 0.40 | 5.4E-05 |
| *bas-1* | WBGene00000239 | protein_coding_gene | -1.3 | 0.40 | 1.6E-07 |
| *C18A11.3* | WBGene00015949 | protein_coding_gene | -1.3 | 0.40 | 1.2E-02 |
| *Y37H2A.14* | WBGene00045416 | protein_coding_gene | -1.3 | 0.40 | 4.3E-02 |
| *grsp-1* | WBGene00012840 | protein_coding_gene | -1.3 | 0.40 | 5.4E-04 |
| *ZC449.7* | WBGene00045365 | protein_coding_gene | -1.3 | 0.40 | 1.5E-02 |
| *K08D8.3* | WBGene00010657 | protein_coding_gene | -1.3 | 0.40 | 8.4E-06 |
| *R08F11.4* | WBGene00019968 | protein_coding_gene | -1.3 | 0.40 | 4.3E-02 |
| *col-113* | WBGene00000687 | protein_coding_gene | -1.3 | 0.40 | 1.6E-02 |
| *C01B10.3* | WBGene00015278 | protein_coding_gene | -1.3 | 0.40 | 4.2E-08 |
| *ttll-15* | WBGene00010630 | protein_coding_gene | -1.3 | 0.40 | 3.4E-04 |
| *ZK512.7* | WBGene00013986 | protein_coding_gene | -1.3 | 0.40 | 2.0E-02 |
| *slc-17.5* | WBGene00017681 | protein_coding_gene | -1.3 | 0.40 | 1.7E-05 |
| *ptr-24* | WBGene00004238 | protein_coding_gene | -1.3 | 0.40 | 1.3E-12 |
| *nspc-1* | WBGene00023477 | protein_coding_gene | -1.3 | 0.40 | 1.9E-02 |
| *nas-7* | WBGene00003526 | protein_coding_gene | -1.3 | 0.40 | 2.4E-03 |
| *ahcy-1* | WBGene00019322 | protein_coding_gene | -1.3 | 0.40 | 3.0E-17 |
| *aass-1* | WBGene00019819 | protein_coding_gene | -1.3 | 0.40 | 7.5E-12 |
| *ule-2* | WBGene00045038 | protein_coding_gene | -1.3 | 0.40 | 1.0E-13 |
| *qdpr-1* | WBGene00011398 | protein_coding_gene | -1.3 | 0.40 | 7.3E-21 |
| *daf-28* | WBGene00000920 | protein_coding_gene | -1.3 | 0.40 | 7.0E-03 |
| *tmem-135* | WBGene00019345 | protein_coding_gene | -1.3 | 0.39 | 3.7E-07 |
| *col-62* | WBGene00000638 | protein_coding_gene | -1.3 | 0.39 | 1.5E-02 |
| *Y49E10.29* | WBGene00045239 | protein_coding_gene | -1.3 | 0.39 | 1.8E-04 |
| *F23D12.11* | WBGene00194674 | protein_coding_gene | -1.3 | 0.39 | 5.6E-05 |
| *cest-2.2* | WBGene00013874 | protein_coding_gene | -1.3 | 0.39 | 2.8E-10 |
| *sptl-1* | WBGene00016020 | protein_coding_gene | -1.3 | 0.39 | 8.2E-17 |
| *gpx-5* | WBGene00007516 | protein_coding_gene | -1.4 | 0.39 | 2.0E-10 |
| *linc-7* | WBGene00219609 | lincRNA_gene | -1.4 | 0.39 | 2.5E-05 |
| *pgph-2* | WBGene00018424 | protein_coding_gene | -1.4 | 0.39 | 2.3E-06 |
| *F41D9.2* | WBGene00018282 | protein_coding_gene | -1.4 | 0.39 | 5.2E-04 |
| *T06A4.3* | WBGene00020283 | protein_coding_gene | -1.4 | 0.39 | 2.1E-02 |
| *F22F4.5* | WBGene00017718 | protein_coding_gene | -1.4 | 0.39 | 4.1E-03 |
| *dhc-4* | WBGene00012272 | protein_coding_gene | -1.4 | 0.39 | 5.2E-03 |
| *PDB1.1* | WBGene00019803 | protein_coding_gene | -1.4 | 0.39 | 4.9E-09 |
| *unc-52* | WBGene00006787 | protein_coding_gene | -1.4 | 0.39 | 4.0E-09 |
| *lgc-4* | WBGene00017580 | protein_coding_gene | -1.4 | 0.39 | 1.2E-02 |
| *twk-26* | WBGene00006679 | protein_coding_gene | -1.4 | 0.39 | 1.1E-03 |
| *F56D6.16* | WBGene00219304 | pseudogene | -1.4 | 0.39 | 2.8E-02 |
| *pqn-22* | WBGene00004112 | protein_coding_gene | -1.4 | 0.39 | 2.6E-07 |
| *gpd-2* | WBGene00001684 | protein_coding_gene | -1.4 | 0.38 | 3.2E-12 |
| *Y14H12A.1* | WBGene00021190 | protein_coding_gene | -1.4 | 0.38 | 2.1E-05 |
| *F26C11.1* | WBGene00009146 | protein_coding_gene | -1.4 | 0.38 | 1.3E-02 |
| *col-48* | WBGene00000625 | protein_coding_gene | -1.4 | 0.38 | 2.8E-04 |
| *nhr-31* | WBGene00003625 | protein_coding_gene | -1.4 | 0.38 | 4.0E-08 |
| *C50D2.6* | WBGene00016809 | protein_coding_gene | -1.4 | 0.38 | 7.8E-05 |

| | | | | | |
|---|---|---|---|---|---|
| *pmp-5* | WBGene00004062 | protein_coding_gene | -1.4 | 0.38 | 3.7E-09 |
| *tbc-19* | WBGene00007849 | protein_coding_gene | -1.4 | 0.38 | 9.2E-08 |
| *C17F4.7* | WBGene00015913 | protein_coding_gene | -1.4 | 0.38 | 6.0E-07 |
| *pqn-36* | WBGene00004123 | protein_coding_gene | -1.4 | 0.38 | 3.6E-03 |
| *C06H5.6* | WBGene00007394 | protein_coding_gene | -1.4 | 0.38 | 2.5E-10 |
| *aqp-4* | WBGene00000172 | protein_coding_gene | -1.4 | 0.38 | 1.3E-07 |
| *F42A10.7* | WBGene00018343 | protein_coding_gene | -1.4 | 0.38 | 8.3E-05 |
| *chk-2* | WBGene00000499 | protein_coding_gene | -1.4 | 0.38 | 1.5E-07 |
| *C32E8.9* | WBGene00016325 | protein_coding_gene | -1.4 | 0.38 | 2.5E-09 |
| *R06B10.1* | WBGene00019912 | pseudogene | -1.4 | 0.38 | 2.8E-02 |
| *sec-61.B* | WBGene00021427 | protein_coding_gene | -1.4 | 0.38 | 5.9E-21 |
| *math-18* | WBGene00016544 | protein_coding_gene | -1.4 | 0.38 | 6.2E-03 |
| *Y39F10A.1* | WBGene00021452 | protein_coding_gene | -1.4 | 0.38 | 4.7E-02 |
| *cest-10* | WBGene00016595 | protein_coding_gene | -1.4 | 0.38 | 1.9E-08 |
| *W04E12.7* | WBGene00012252 | protein_coding_gene | -1.4 | 0.38 | 1.2E-03 |
| *C09B8.4* | WBGene00015623 | protein_coding_gene | -1.4 | 0.38 | 7.5E-05 |
| *cpt-5* | WBGene00008629 | protein_coding_gene | -1.4 | 0.38 | 1.7E-06 |
| *aqp-7* | WBGene00000175 | protein_coding_gene | -1.4 | 0.38 | 2.6E-16 |
| *C14F11.4* | WBGene00015780 | protein_coding_gene | -1.4 | 0.38 | 8.5E-08 |
| *F43C1.5* | WBGene00009649 | protein_coding_gene | -1.4 | 0.38 | 1.7E-02 |
| *C06E8.5* | WBGene00015544 | protein_coding_gene | -1.4 | 0.38 | 2.7E-02 |
| *Y73E7A.3* | WBGene00022270 | protein_coding_gene | -1.4 | 0.37 | 3.5E-05 |
| *T03D3.5* | WBGene00020183 | protein_coding_gene | -1.4 | 0.37 | 1.2E-10 |
| *dod-19* | WBGene00022644 | protein_coding_gene | -1.4 | 0.37 | 2.0E-02 |
| *clec-205* | WBGene00021602 | protein_coding_gene | -1.4 | 0.37 | 8.3E-03 |
| *cah-5* | WBGene00000283 | protein_coding_gene | -1.4 | 0.37 | 2.5E-08 |
| *flr-4* | WBGene00001468 | protein_coding_gene | -1.4 | 0.37 | 5.2E-03 |
| *plep-1* | WBGene00021797 | protein_coding_gene | -1.4 | 0.37 | 1.7E-02 |
| *col-101* | WBGene00000675 | protein_coding_gene | -1.4 | 0.37 | 1.2E-02 |
| *art-1* | WBGene00000198 | protein_coding_gene | -1.4 | 0.37 | 5.3E-17 |
| *hmit-1.2* | WBGene00013074 | protein_coding_gene | -1.4 | 0.37 | 1.6E-04 |
| *pes-9* | WBGene00003982 | protein_coding_gene | -1.4 | 0.37 | 2.2E-26 |
| *K10C2.7* | WBGene00019622 | protein_coding_gene | -1.4 | 0.37 | 4.5E-03 |
| *ptps-1* | WBGene00015010 | protein_coding_gene | -1.4 | 0.37 | 3.4E-07 |
| *wrt-4* | WBGene00006950 | protein_coding_gene | -1.4 | 0.37 | 1.2E-02 |
| *arrd-10* | WBGene00010268 | protein_coding_gene | -1.4 | 0.37 | 1.2E-03 |
| *F08F3.4* | WBGene00017262 | protein_coding_gene | -1.4 | 0.37 | 1.6E-18 |
| *nlp-26* | WBGene00003764 | protein_coding_gene | -1.4 | 0.37 | 1.2E-04 |
| *F23F12.3* | WBGene00017751 | protein_coding_gene | -1.4 | 0.37 | 5.6E-03 |
| *col-80* | WBGene00000656 | protein_coding_gene | -1.4 | 0.37 | 4.2E-06 |
| *C15H9.9* | WBGene00015803 | protein_coding_gene | -1.4 | 0.37 | 7.1E-22 |
| *acl-1* | WBGene00010339 | protein_coding_gene | -1.4 | 0.37 | 5.5E-21 |
| *bca-1* | WBGene00000245 | protein_coding_gene | -1.4 | 0.37 | 5.8E-08 |
| *gst-24* | WBGene00001772 | protein_coding_gene | -1.4 | 0.37 | 2.8E-03 |
| *F22E5.1* | WBGene00017703 | protein_coding_gene | -1.4 | 0.37 | 1.1E-02 |
| *F52H2.6* | WBGene00018718 | protein_coding_gene | -1.4 | 0.37 | 1.4E-17 |
| *ZK218.1* | WBGene00013935 | protein_coding_gene | -1.5 | 0.37 | 1.6E-03 |
| *R08E5.3* | WBGene00019963 | protein_coding_gene | -1.5 | 0.37 | 1.3E-07 |
| *vha-12* | WBGene00006921 | protein_coding_gene | -1.5 | 0.36 | 9.9E-23 |
| *R13A5.10* | WBGene00020052 | protein_coding_gene | -1.5 | 0.36 | 9.1E-04 |
| *W06A7.4* | WBGene00012293 | protein_coding_gene | -1.5 | 0.36 | 4.3E-05 |
| *piit-1* | WBGene00009384 | protein_coding_gene | -1.5 | 0.36 | 6.1E-04 |
| *C52D10.3* | WBGene00016877 | protein_coding_gene | -1.5 | 0.36 | 3.2E-05 |
| *mtp-18* | WBGene00077690 | protein_coding_gene | -1.5 | 0.36 | 6.0E-06 |
| *cnp-2* | WBGene00012901 | protein_coding_gene | -1.5 | 0.36 | 9.0E-04 |
| *ZK185.3* | WBGene00022683 | protein_coding_gene | -1.5 | 0.36 | 9.5E-07 |
| *F37B12.1* | WBGene00009504 | protein_coding_gene | -1.5 | 0.36 | 2.3E-03 |
| *col-166* | WBGene00000739 | protein_coding_gene | -1.5 | 0.36 | 5.3E-05 |
| *clec-187* | WBGene00014137 | protein_coding_gene | -1.5 | 0.36 | 6.3E-09 |
| *folt-1* | WBGene00007388 | protein_coding_gene | -1.5 | 0.36 | 1.6E-07 |
| *F42G10.1* | WBGene00009645 | protein_coding_gene | -1.5 | 0.36 | 5.5E-05 |
| *cyp-33C8* | WBGene00019967 | protein_coding_gene | -1.5 | 0.36 | 4.6E-03 |
| *clec-57* | WBGene00008594 | protein_coding_gene | -1.5 | 0.36 | 1.2E-04 |
| *T25B9.9* | WBGene00012015 | protein_coding_gene | -1.5 | 0.36 | 3.1E-20 |
| *alh-9* | WBGene00000115 | protein_coding_gene | -1.5 | 0.36 | 5.6E-21 |

| | | | | | |
|---|---|---|---|---|---|
| *T25C12.3* | WBGene00012018 | protein_coding_gene | -1.5 | 0.35 | 8.0E-03 |
| *B0454.5* | WBGene00015196 | protein_coding_gene | -1.5 | 0.35 | 1.6E-05 |
| *C12D12.1* | WBGene00015713 | protein_coding_gene | -1.5 | 0.35 | 3.7E-06 |
| *ZC302.3* | WBGene00013863 | protein_coding_gene | -1.5 | 0.35 | 2.8E-02 |
| *ugt-48* | WBGene00015965 | protein_coding_gene | -1.5 | 0.35 | 4.8E-07 |
| *gldc-1* | WBGene00020022 | protein_coding_gene | -1.5 | 0.35 | 2.2E-25 |
| *Y43F8C.13* | WBGene00012834 | protein_coding_gene | -1.5 | 0.35 | 4.0E-17 |
| *slc-17.4* | WBGene00012130 | protein_coding_gene | -1.5 | 0.35 | 8.0E-09 |
| *dct-16* | WBGene00012615 | protein_coding_gene | -1.5 | 0.35 | 2.5E-03 |
| *oat-1* | WBGene00003837 | protein_coding_gene | -1.5 | 0.35 | 4.1E-06 |
| *dsc-4* | WBGene00001099 | protein_coding_gene | -1.5 | 0.35 | 6.2E-17 |
| *sprr-2* | WBGene00009629 | protein_coding_gene | -1.5 | 0.35 | 3.0E-03 |
| *acox-1.3* | WBGene00008566 | protein_coding_gene | -1.5 | 0.35 | 6.0E-04 |
| *pks-1* | WBGene00016558 | protein_coding_gene | -1.5 | 0.35 | 3.2E-07 |
| *Y65B4A.9* | WBGene00022032 | protein_coding_gene | -1.5 | 0.35 | 2.0E-02 |
| *dhs-3* | WBGene00000967 | protein_coding_gene | -1.5 | 0.35 | 1.2E-19 |
| *fut-4* | WBGene00001507 | protein_coding_gene | -1.5 | 0.35 | 2.1E-02 |
| *Y67H2A.9* | WBGene00014941 | pseudogene | -1.5 | 0.35 | 1.7E-13 |
| *ugt-60* | WBGene00007402 | protein_coding_gene | -1.5 | 0.35 | 4.6E-05 |
| *C39D10.8* | WBGene00016535 | protein_coding_gene | -1.5 | 0.35 | 1.8E-07 |
| *acr-16* | WBGene00000055 | protein_coding_gene | -1.5 | 0.34 | 7.1E-04 |
| *clec-56* | WBGene00008595 | protein_coding_gene | -1.5 | 0.34 | 1.5E-13 |
| *F20D6.5* | WBGene00017635 | protein_coding_gene | -1.5 | 0.34 | 2.6E-03 |
| *lgc-54* | WBGene00020528 | protein_coding_gene | -1.6 | 0.34 | 1.8E-04 |
| *mec-19* | WBGene00008216 | protein_coding_gene | -1.6 | 0.34 | 1.6E-02 |
| *cln-3.1* | WBGene00000539 | protein_coding_gene | -1.6 | 0.34 | 8.3E-04 |
| *K08H10.6* | WBGene00010698 | protein_coding_gene | -1.6 | 0.34 | 3.8E-02 |
| *poml-2* | WBGene00017089 | protein_coding_gene | -1.6 | 0.34 | 2.6E-05 |
| *wrt-8* | WBGene00006954 | protein_coding_gene | -1.6 | 0.34 | 4.2E-02 |
| *hacd-1* | WBGene00019978 | protein_coding_gene | -1.6 | 0.34 | 3.5E-07 |
| *col-110* | WBGene00000684 | protein_coding_gene | -1.6 | 0.34 | 2.0E-02 |
| *C01H6.4* | WBGene00007254 | protein_coding_gene | -1.6 | 0.34 | 2.3E-05 |
| *C45E5.1* | WBGene00016664 | protein_coding_gene | -1.6 | 0.34 | 8.0E-09 |
| *K11H12.3* | WBGene00019659 | protein_coding_gene | -1.6 | 0.33 | 4.7E-02 |
| *K01G12.3* | WBGene00010483 | protein_coding_gene | -1.6 | 0.33 | 7.3E-04 |
| *cyp-13A12* | WBGene00008810 | protein_coding_gene | -1.6 | 0.33 | 3.0E-02 |
| *F29G6.1* | WBGene00009257 | protein_coding_gene | -1.6 | 0.33 | 3.0E-02 |
| *ttr-18* | WBGene00007142 | protein_coding_gene | -1.6 | 0.33 | 9.0E-12 |
| *T24E12.11* | WBGene00020778 | protein_coding_gene | -1.6 | 0.33 | 3.4E-04 |
| *ctsa-4.1* | WBGene00019605 | protein_coding_gene | -1.6 | 0.33 | 4.3E-08 |
| *Y22D7AL.11* | WBGene00021249 | protein_coding_gene | -1.6 | 0.33 | 5.7E-04 |
| *F13H8.3* | WBGene00017436 | protein_coding_gene | -1.6 | 0.33 | 1.1E-13 |
| *twk-22* | WBGene00006675 | protein_coding_gene | -1.6 | 0.33 | 2.4E-06 |
| *nmat-1* | WBGene00012295 | protein_coding_gene | -1.6 | 0.33 | 1.1E-04 |
| *ttr-14* | WBGene00011460 | protein_coding_gene | -1.6 | 0.33 | 9.9E-06 |
| *zip-3* | WBGene00012330 | protein_coding_gene | -1.6 | 0.33 | 1.5E-13 |
| *lgc-45* | WBGene00021146 | protein_coding_gene | -1.6 | 0.33 | 5.2E-11 |
| *efn-3* | WBGene00001164 | protein_coding_gene | -1.6 | 0.33 | 4.3E-04 |
| *ctsa-2* | WBGene00019617 | protein_coding_gene | -1.6 | 0.33 | 7.5E-03 |
| *F33E2.4* | WBGene00009360 | protein_coding_gene | -1.6 | 0.33 | 1.0E-02 |
| *nlp-77* | WBGene00015514 | protein_coding_gene | -1.6 | 0.33 | 4.6E-27 |
| *fmo-3* | WBGene00001478 | protein_coding_gene | -1.6 | 0.33 | 7.6E-06 |
| *Y66C5A.1* | WBGene00044156 | protein_coding_gene | -1.6 | 0.33 | 6.2E-03 |
| *pgp-1* | WBGene00003995 | protein_coding_gene | -1.6 | 0.32 | 1.7E-02 |
| *H23N18.5* | WBGene00019236 | protein_coding_gene | -1.6 | 0.32 | 3.9E-03 |
| *sptl-3* | WBGene00011932 | protein_coding_gene | -1.6 | 0.32 | 9.5E-29 |
| *T25E4.1* | WBGene00020803 | protein_coding_gene | -1.6 | 0.32 | 1.7E-02 |
| *F26G1.5* | WBGene00017841 | protein_coding_gene | -1.6 | 0.32 | 1.6E-04 |
| *let-383* | WBGene00044898 | protein_coding_gene | -1.6 | 0.32 | 8.8E-05 |
| *col-155* | WBGene00000728 | protein_coding_gene | -1.6 | 0.32 | 1.1E-05 |
| *F31A3.3* | WBGene00017938 | protein_coding_gene | -1.6 | 0.32 | 6.7E-03 |
| *ZK742.3* | WBGene00022813 | protein_coding_gene | -1.6 | 0.32 | 1.2E-05 |
| *ZC53.1* | WBGene00022508 | protein_coding_gene | -1.6 | 0.32 | 3.9E-06 |
| *F10A3.4* | WBGene00008634 | protein_coding_gene | -1.7 | 0.32 | 1.2E-03 |
| *F10D11.6* | WBGene00008652 | protein_coding_gene | -1.7 | 0.32 | 8.3E-05 |

| | | | | | |
|---|---|---|---|---|---|
| *hpd-1* | WBGene00001993 | protein_coding_gene | -1.7 | 0.32 | 1.1E-12 |
| *ZK228.3* | WBGene00013946 | protein_coding_gene | -1.7 | 0.32 | 1.4E-09 |
| *ctsa-3.1* | WBGene00017969 | protein_coding_gene | -1.7 | 0.32 | 1.9E-02 |
| *C26B2.8* | WBGene00016130 | protein_coding_gene | -1.7 | 0.32 | 2.2E-02 |
| *T25B9.1* | WBGene00012007 | protein_coding_gene | -1.7 | 0.32 | 1.1E-08 |
| *col-39* | WBGene00000616 | protein_coding_gene | -1.7 | 0.31 | 1.3E-02 |
| *lbp-8* | WBGene00002260 | protein_coding_gene | -1.7 | 0.31 | 4.9E-02 |
| *sdz-12* | WBGene00017406 | protein_coding_gene | -1.7 | 0.31 | 4.1E-02 |
| *B0244.7* | WBGene00015082 | protein_coding_gene | -1.7 | 0.31 | 4.5E-02 |
| *vha-11* | WBGene00006920 | protein_coding_gene | -1.7 | 0.31 | 5.3E-21 |
| *comt-3* | WBGene00021487 | protein_coding_gene | -1.7 | 0.31 | 3.8E-06 |
| *Y55F3BR.10* | WBGene00044742 | protein_coding_gene | -1.7 | 0.31 | 3.8E-04 |
| *T10B5.10* | WBGene00020396 | protein_coding_gene | -1.7 | 0.31 | 2.1E-02 |
| *nlp-24* | WBGene00003762 | protein_coding_gene | -1.7 | 0.31 | 1.7E-02 |
| *Y60A3A.11* | WBGene00014933 | pseudogene | -1.7 | 0.31 | 3.9E-02 |
| *ets-5* | WBGene00016600 | protein_coding_gene | -1.7 | 0.31 | 4.6E-02 |
| *Y39B6A.27* | WBGene00012686 | protein_coding_gene | -1.7 | 0.31 | 8.7E-05 |
| *nhr-93* | WBGene00003683 | protein_coding_gene | -1.7 | 0.31 | 3.5E-02 |
| *Y102A11A.7* | WBGene00022417 | protein_coding_gene | -1.7 | 0.31 | 4.4E-02 |
| *F07G6.10* | WBGene00045271 | protein_coding_gene | -1.7 | 0.31 | 3.4E-03 |
| *T01D1.3* | WBGene00020148 | protein_coding_gene | -1.7 | 0.31 | 3.5E-08 |
| *elo-9* | WBGene00001247 | protein_coding_gene | -1.7 | 0.31 | 3.1E-09 |
| *srb-14* | WBGene00016338 | pseudogene | -1.7 | 0.30 | 4.4E-02 |
| *cima-1* | WBGene00018473 | protein_coding_gene | -1.7 | 0.30 | 2.3E-06 |
| *lgc-40* | WBGene00020767 | protein_coding_gene | -1.7 | 0.30 | 3.2E-03 |
| *Y59E9AL.4* | WBGene00021994 | protein_coding_gene | -1.7 | 0.30 | 4.7E-06 |
| *T20D4.19* | WBGene00020624 | protein_coding_gene | -1.7 | 0.30 | 4.6E-02 |
| *col-8* | WBGene00000597 | protein_coding_gene | -1.7 | 0.30 | 1.3E-10 |
| *VB0395L.1* | WBGene00044150 | protein_coding_gene | -1.7 | 0.30 | 3.4E-04 |
| *col-178* | WBGene00000751 | protein_coding_gene | -1.7 | 0.30 | 4.1E-37 |
| *skpo-3* | WBGene00017968 | protein_coding_gene | -1.7 | 0.30 | 1.5E-09 |
| *C27A2.5* | WBGene00016153 | protein_coding_gene | -1.7 | 0.30 | 4.7E-02 |
| *mup-4* | WBGene00003497 | protein_coding_gene | -1.7 | 0.30 | 1.1E-20 |
| *haao-1* | WBGene00010595 | protein_coding_gene | -1.7 | 0.30 | 8.9E-20 |
| *amt-1* | WBGene00000133 | protein_coding_gene | -1.7 | 0.30 | 1.1E-03 |
| *dao-4* | WBGene00000930 | protein_coding_gene | -1.7 | 0.30 | 4.2E-02 |
| *grd-5* | WBGene00001694 | protein_coding_gene | -1.7 | 0.30 | 2.3E-06 |
| *ugt-46* | WBGene00015141 | protein_coding_gene | -1.7 | 0.30 | 3.4E-13 |
| *EGAP4.1* | WBGene00017149 | protein_coding_gene | -1.7 | 0.30 | 3.2E-02 |
| *nas-38* | WBGene00003554 | protein_coding_gene | -1.8 | 0.30 | 1.5E-02 |
| *E04D5.5* | WBGene00008484 | protein_coding_gene | -1.8 | 0.30 | 1.1E-04 |
| *str-148* | WBGene00006195 | protein_coding_gene | -1.8 | 0.30 | 4.4E-02 |
| *nduo-3* | WBGene00010966 | protein_coding_gene | -1.8 | 0.30 | 6.5E-04 |
| *ugt-21* | WBGene00007885 | protein_coding_gene | -1.8 | 0.30 | 3.8E-05 |
| *pho-13* | WBGene00008560 | protein_coding_gene | -1.8 | 0.29 | 2.0E-02 |
| *nep-10* | WBGene00017555 | protein_coding_gene | -1.8 | 0.29 | 2.5E-02 |
| *fbxb-21* | WBGene00022558 | protein_coding_gene | -1.8 | 0.29 | 4.1E-02 |
| *ttr-31* | WBGene00010225 | protein_coding_gene | -1.8 | 0.29 | 8.0E-08 |
| *linc-130* | WBGene00219651 | lincRNA_gene | -1.8 | 0.29 | 4.1E-02 |
| *nas-36* | WBGene00003552 | protein_coding_gene | -1.8 | 0.29 | 5.8E-03 |
| *nas-1* | WBGene00003520 | protein_coding_gene | -1.8 | 0.29 | 4.3E-02 |
| *C04G6.2* | WBGene00015453 | protein_coding_gene | -1.8 | 0.29 | 3.5E-02 |
| *K06B9.4* | WBGene00019441 | pseudogene | -1.8 | 0.29 | 3.7E-08 |
| *col-81* | WBGene00000657 | protein_coding_gene | -1.8 | 0.29 | 1.1E-02 |
| *cbl-1* | WBGene00007533 | protein_coding_gene | -1.8 | 0.29 | 9.1E-13 |
| *clec-204* | WBGene00021603 | protein_coding_gene | -1.8 | 0.29 | 4.6E-04 |
| *col-58* | WBGene00000634 | protein_coding_gene | -1.8 | 0.29 | 4.8E-03 |
| *F58B4.5* | WBGene00010238 | protein_coding_gene | -1.8 | 0.29 | 2.2E-10 |
| *col-169* | WBGene00000742 | protein_coding_gene | -1.8 | 0.29 | 1.4E-02 |
| *C46F2.1* | WBGene00016714 | protein_coding_gene | -1.8 | 0.29 | 1.2E-07 |
| *pqn-2* | WBGene00004098 | protein_coding_gene | -1.8 | 0.29 | 4.9E-02 |
| *oac-37* | WBGene00010391 | protein_coding_gene | -1.8 | 0.29 | 3.9E-02 |
| *twk-10* | WBGene00006665 | protein_coding_gene | -1.8 | 0.29 | 1.1E-02 |
| *F28H7.3* | WBGene00009237 | protein_coding_gene | -1.8 | 0.29 | 4.0E-07 |
| *Y57G11C.23* | WBGene00013318 | protein_coding_gene | -1.8 | 0.29 | 1.9E-02 |

| | | | | | |
|---|---|---|---|---|---|
| *C55A6.4* | WBGene00008333 | protein_coding_gene | -1.8 | 0.29 | 8.4E-10 |
| *cpr-5* | WBGene00000785 | protein_coding_gene | -1.8 | 0.28 | 1.4E-07 |
| *K10C2.8* | WBGene00045265 | protein_coding_gene | -1.8 | 0.28 | 7.6E-10 |
| *pcp-3* | WBGene00003958 | protein_coding_gene | -1.8 | 0.28 | 3.3E-10 |
| *wago-2* | WBGene00018862 | pseudogene | -1.8 | 0.28 | 1.3E-18 |
| *pept-1* | WBGene00003877 | protein_coding_gene | -1.8 | 0.28 | 7.5E-15 |
| *T02G6.11* | WBGene00086562 | protein_coding_gene | -1.8 | 0.28 | 3.5E-02 |
| *str-227* | WBGene00006258 | protein_coding_gene | -1.8 | 0.28 | 3.5E-02 |
| *F11E6.3* | WBGene00008707 | protein_coding_gene | -1.8 | 0.28 | 6.2E-11 |
| *ZK662.2* | WBGene00014039 | protein_coding_gene | -1.8 | 0.28 | 3.8E-03 |
| *C18A11.4* | WBGene00015950 | protein_coding_gene | -1.8 | 0.28 | 1.2E-02 |
| *mth-1* | WBGene00017948 | protein_coding_gene | -1.8 | 0.28 | 5.6E-07 |
| *vit-4* | WBGene00006928 | protein_coding_gene | -1.8 | 0.28 | 2.9E-02 |
| *C15F1.1* | WBGene00015794 | protein_coding_gene | -1.8 | 0.28 | 1.7E-10 |
| *F57F4.4* | WBGene00019017 | protein_coding_gene | -1.8 | 0.28 | 1.5E-02 |
| *ZK228.4* | WBGene00013947 | protein_coding_gene | -1.8 | 0.28 | 1.5E-13 |
| *F22F1.2* | WBGene00017714 | protein_coding_gene | -1.8 | 0.28 | 3.7E-02 |
| *mpst-4* | WBGene00017387 | protein_coding_gene | -1.8 | 0.28 | 3.9E-03 |
| *sams-1* | WBGene00008205 | protein_coding_gene | -1.8 | 0.28 | 2.4E-45 |
| *cut-4* | WBGene00008482 | protein_coding_gene | -1.8 | 0.28 | 1.0E-03 |
| *R12E2.6* | WBGene00020032 | protein_coding_gene | -1.9 | 0.28 | 3.3E-02 |
| *ldp-1* | WBGene00017719 | protein_coding_gene | -1.9 | 0.28 | 3.5E-39 |
| *C05C9.3* | WBGene00007327 | protein_coding_gene | -1.9 | 0.28 | 9.0E-07 |
| *lrg-1* | WBGene00003067 | pseudogene | -1.9 | 0.28 | 9.4E-12 |
| *Y48B6A.10* | WBGene00012981 | protein_coding_gene | -1.9 | 0.28 | 5.1E-03 |
| *pqn-11* | WBGene00004103 | pseudogene | -1.9 | 0.28 | 3.2E-02 |
| *C41H7.1* | WBGene00016571 | protein_coding_gene | -1.9 | 0.28 | 1.6E-03 |
| *ZC449.8* | WBGene00195189 | protein_coding_gene | -1.9 | 0.28 | 7.0E-06 |
| *C17H12.6* | WBGene00015932 | protein_coding_gene | -1.9 | 0.28 | 4.4E-02 |
| *doxa-1* | WBGene00015519 | protein_coding_gene | -1.9 | 0.27 | 4.8E-06 |
| *pqn-54* | WBGene00004139 | protein_coding_gene | -1.9 | 0.27 | 1.1E-02 |
| *C50A2.3* | WBGene00016797 | protein_coding_gene | -1.9 | 0.27 | 3.9E-09 |
| *C53B7.6* | WBGene00016895 | protein_coding_gene | -1.9 | 0.27 | 2.8E-02 |
| *pgp-9* | WBGene00004003 | protein_coding_gene | -1.9 | 0.27 | 5.2E-03 |
| *F56D5.3* | WBGene00010148 | protein_coding_gene | -1.9 | 0.27 | 1.0E-06 |
| *F35B3.4* | WBGene00018031 | protein_coding_gene | -1.9 | 0.27 | 7.0E-03 |
| *clec-206* | WBGene00019089 | protein_coding_gene | -1.9 | 0.27 | 3.2E-02 |
| *F35E12.2* | WBGene00009426 | protein_coding_gene | -1.9 | 0.27 | 3.8E-02 |
| *F44E5.16* | WBGene00219949 | ncRNA_gene | -1.9 | 0.27 | 2.9E-02 |
| *ZC434.9* | WBGene00013895 | protein_coding_gene | -1.9 | 0.27 | 5.4E-09 |
| *abu-7* | WBGene00000030 | protein_coding_gene | -1.9 | 0.27 | 3.7E-02 |
| *Y47G6A.33* | WBGene00045399 | protein_coding_gene | -1.9 | 0.27 | 8.2E-14 |
| *ncx-7* | WBGene00003572 | protein_coding_gene | -1.9 | 0.27 | 2.9E-19 |
| *C02E7.7* | WBGene00015340 | protein_coding_gene | -1.9 | 0.27 | 4.7E-04 |
| *Y32B12C.3* | WBGene00012526 | pseudogene | -1.9 | 0.27 | 2.6E-02 |
| *nas-30* | WBGene00003548 | protein_coding_gene | -1.9 | 0.27 | 1.1E-02 |
| *ZK6.11* | WBGene00022645 | protein_coding_gene | -1.9 | 0.27 | 6.5E-03 |
| *grd-10* | WBGene00001699 | protein_coding_gene | -1.9 | 0.27 | 1.2E-03 |
| *clec-118* | WBGene00021032 | protein_coding_gene | -1.9 | 0.27 | 2.7E-02 |
| *dct-18* | WBGene00010266 | protein_coding_gene | -1.9 | 0.27 | 1.0E-28 |
| *C45B11.2* | WBGene00008105 | protein_coding_gene | -1.9 | 0.27 | 2.7E-02 |
| *col-143* | WBGene00000716 | protein_coding_gene | -1.9 | 0.27 | 8.3E-11 |
| *C35A5.3* | WBGene00007947 | protein_coding_gene | -1.9 | 0.27 | 6.1E-03 |
| *R03H10.12* | WBGene00303363 | pseudogene | -1.9 | 0.27 | 2.4E-02 |
| *T25G12.6* | WBGene00020813 | protein_coding_gene | -1.9 | 0.26 | 7.6E-08 |
| *gsto-1* | WBGene00016204 | protein_coding_gene | -1.9 | 0.26 | 2.5E-07 |
| *F52C6.14* | WBGene00018671 | protein_coding_gene | -1.9 | 0.26 | 2.4E-02 |
| *pgp-8* | WBGene00004002 | protein_coding_gene | -1.9 | 0.26 | 1.8E-03 |
| *C49F5.8* | WBGene00008212 | protein_coding_gene | -1.9 | 0.26 | 4.2E-02 |
| *Y57G11C.42* | WBGene00013331 | protein_coding_gene | -2.0 | 0.26 | 9.4E-04 |
| *T10E9.3* | WBGene00020413 | protein_coding_gene | -2.0 | 0.26 | 3.3E-04 |
| *F11F1.1* | WBGene00008714 | protein_coding_gene | -2.0 | 0.26 | 2.1E-09 |
| *asns-1* | WBGene00003816 | protein_coding_gene | -2.0 | 0.26 | 5.9E-04 |
| *H06A10.1* | WBGene00010368 | protein_coding_gene | -2.0 | 0.26 | 3.5E-06 |
| *T23F6.1* | WBGene00011949 | protein_coding_gene | -2.0 | 0.26 | 1.1E-02 |

| | | | | | |
|---|---|---|---|---|---|
| *ncx-8* | WBGene00003573 | protein_coding_gene | -2.0 | 0.26 | 2.3E-04 |
| *act-5* | WBGene00000067 | protein_coding_gene | -2.0 | 0.25 | 1.4E-03 |
| *W02H3.1* | WBGene00012217 | protein_coding_gene | -2.0 | 0.25 | 2.6E-05 |
| *C49A9.4* | WBGene00016758 | protein_coding_gene | -2.0 | 0.25 | 1.2E-07 |
| *T07A5.1* | WBGene00011554 | protein_coding_gene | -2.0 | 0.25 | 5.0E-04 |
| *T06E4.8* | WBGene00011535 | protein_coding_gene | -2.0 | 0.25 | 2.2E-02 |
| *K05F6.12* | WBGene00045486 | protein_coding_gene | -2.0 | 0.25 | 3.1E-02 |
| *Y43C5A.7* | WBGene00044922 | protein_coding_gene | -2.0 | 0.25 | 1.1E-05 |
| *C16C4.17* | WBGene00044359 | protein_coding_gene | -2.0 | 0.25 | 2.1E-02 |
| *F58G6.9* | WBGene00044107 | protein_coding_gene | -2.0 | 0.25 | 9.1E-03 |
| *F41E6.11* | WBGene00018292 | protein_coding_gene | -2.0 | 0.25 | 1.1E-02 |
| *cpg-9* | WBGene00022072 | protein_coding_gene | -2.0 | 0.25 | 1.6E-07 |
| *comt-5* | WBGene00021492 | protein_coding_gene | -2.0 | 0.25 | 2.3E-03 |
| *abu-6* | WBGene00000029 | protein_coding_gene | -2.0 | 0.25 | 2.5E-02 |
| *C39B5.5* | WBGene00016523 | protein_coding_gene | -2.0 | 0.25 | 9.4E-05 |
| *lips-14* | WBGene00019208 | pseudogene | -2.0 | 0.25 | 1.7E-02 |
| *col-160* | WBGene00000733 | protein_coding_gene | -2.0 | 0.25 | 3.7E-16 |
| *fbxa-72* | WBGene00017962 | protein_coding_gene | -2.0 | 0.25 | 1.9E-07 |
| *Y47H9C.14* | WBGene00012956 | pseudogene | -2.0 | 0.25 | 2.0E-02 |
| *Y105C5B.15* | WBGene00013654 | protein_coding_gene | -2.0 | 0.24 | 8.2E-04 |
| *T21C9.9* | WBGene00011895 | protein_coding_gene | -2.0 | 0.24 | 1.9E-03 |
| *W10G11.3* | WBGene00021136 | protein_coding_gene | -2.0 | 0.24 | 5.0E-05 |
| *C04C3.14* | WBGene00219410 | protein_coding_gene | -2.0 | 0.24 | 2.4E-02 |
| *T15B7.14* | WBGene00020526 | protein_coding_gene | -2.1 | 0.24 | 4.1E-04 |
| *adt-2* | WBGene00000083 | protein_coding_gene | -2.1 | 0.24 | 2.7E-21 |
| *lact-1* | WBGene00018523 | protein_coding_gene | -2.1 | 0.24 | 3.0E-08 |
| *Y73B6BL.31* | WBGene00022252 | protein_coding_gene | -2.1 | 0.24 | 4.0E-18 |
| *arf-1.1* | WBGene00000190 | protein_coding_gene | -2.1 | 0.24 | 7.0E-06 |
| *F01D5.2* | WBGene00008493 | protein_coding_gene | -2.1 | 0.24 | 3.2E-02 |
| *Y55F3AM.14* | WBGene00021931 | protein_coding_gene | -2.1 | 0.24 | 1.2E-02 |
| *cyp-14A1* | WBGene00010705 | protein_coding_gene | -2.1 | 0.24 | 2.9E-06 |
| *cept-1* | WBGene00009057 | protein_coding_gene | -2.1 | 0.24 | 7.1E-45 |
| *tba-4* | WBGene00006530 | protein_coding_gene | -2.1 | 0.24 | 3.9E-19 |
| *C45B2.2* | WBGene00016659 | protein_coding_gene | -2.1 | 0.24 | 1.4E-03 |
| *col-42* | WBGene00013489 | protein_coding_gene | -2.1 | 0.24 | 9.4E-21 |
| *nspb-12* | WBGene00017305 | protein_coding_gene | -2.1 | 0.24 | 1.8E-02 |
| *grl-21* | WBGene00001730 | protein_coding_gene | -2.1 | 0.23 | 2.3E-03 |
| *T06E4.14* | WBGene00077691 | protein_coding_gene | -2.1 | 0.23 | 5.2E-03 |
| *nas-27* | WBGene00003545 | protein_coding_gene | -2.1 | 0.23 | 1.1E-02 |
| *rol-8* | WBGene00004398 | protein_coding_gene | -2.1 | 0.23 | 1.7E-04 |
| *twk-42* | WBGene00006692 | protein_coding_gene | -2.1 | 0.23 | 8.6E-08 |
| *col-43* | WBGene00000620 | protein_coding_gene | -2.1 | 0.23 | 4.3E-03 |
| *mxl-3* | WBGene00003511 | protein_coding_gene | -2.1 | 0.23 | 3.7E-21 |
| *col-79* | WBGene00000655 | protein_coding_gene | -2.1 | 0.23 | 1.1E-02 |
| *col-90* | WBGene00000665 | protein_coding_gene | -2.1 | 0.23 | 6.1E-04 |
| *vit-1* | WBGene00006925 | protein_coding_gene | -2.1 | 0.23 | 9.6E-03 |
| *Y47D3B.6* | WBGene00012942 | protein_coding_gene | -2.1 | 0.23 | 7.4E-04 |
| *clec-1* | WBGene00017772 | protein_coding_gene | -2.1 | 0.23 | 4.4E-16 |
| *C30G12.10* | WBGene00284859 | ncRNA_gene | -2.1 | 0.23 | 1.4E-02 |
| *C33D3.3* | WBGene00007894 | protein_coding_gene | -2.1 | 0.23 | 1.6E-03 |
| *col-104* | WBGene00000678 | protein_coding_gene | -2.1 | 0.23 | 8.1E-08 |
| *Y51A2D.18* | WBGene00013083 | protein_coding_gene | -2.1 | 0.23 | 1.0E-18 |
| *daf-36* | WBGene00007536 | protein_coding_gene | -2.1 | 0.23 | 7.3E-19 |
| *linc-21* | WBGene00219688 | lincRNA_gene | -2.1 | 0.23 | 1.5E-02 |
| *pcp-2* | WBGene00003957 | protein_coding_gene | -2.1 | 0.23 | 2.0E-03 |
| *tyr-2* | WBGene00010661 | protein_coding_gene | -2.2 | 0.22 | 1.3E-09 |
| *pes-8* | WBGene00003981 | protein_coding_gene | -2.2 | 0.22 | 3.5E-06 |
| *F21A3.3* | WBGene00008993 | protein_coding_gene | -2.2 | 0.22 | 5.1E-08 |
| *nspc-4* | WBGene00044339 | protein_coding_gene | -2.2 | 0.22 | 6.7E-17 |
| *aat-7* | WBGene00000008 | protein_coding_gene | -2.2 | 0.22 | 1.0E-03 |
| *C30H6.5* | WBGene00007823 | protein_coding_gene | -2.2 | 0.22 | 6.2E-03 |
| *col-107* | WBGene00000681 | protein_coding_gene | -2.2 | 0.22 | 6.2E-07 |
| *cya-2* | WBGene00000864 | protein_coding_gene | -2.2 | 0.22 | 1.7E-02 |
| *scl-24* | WBGene00008575 | protein_coding_gene | -2.2 | 0.22 | 1.6E-02 |
| *sqt-2* | WBGene00005017 | protein_coding_gene | -2.2 | 0.22 | 1.1E-05 |

| | | | | | |
|---|---|---|---|---|---|
| *sdz-16* | WBGene00009297 | protein_coding_gene | -2.2 | 0.22 | 5.8E-05 |
| *oac-32* | WBGene00018708 | protein_coding_gene | -2.2 | 0.22 | 1.6E-02 |
| *pqn-74* | WBGene00004156 | protein_coding_gene | -2.2 | 0.22 | 5.8E-06 |
| *cpr-8* | WBGene00021070 | protein_coding_gene | -2.2 | 0.22 | 2.9E-07 |
| *ugt-65* | WBGene00008583 | protein_coding_gene | -2.2 | 0.21 | 6.8E-07 |
| *kynu-1* | WBGene00015802 | protein_coding_gene | -2.2 | 0.21 | 2.8E-41 |
| *F01D5.3* | WBGene00008494 | protein_coding_gene | -2.2 | 0.21 | 2.1E-03 |
| *Y16B4A.2* | WBGene00012445 | protein_coding_gene | -2.2 | 0.21 | 1.0E-19 |
| *cdr-2* | WBGene00008296 | protein_coding_gene | -2.2 | 0.21 | 1.6E-06 |
| *abu-15* | WBGene00004099 | protein_coding_gene | -2.2 | 0.21 | 5.1E-03 |
| *fbxa-15* | WBGene00022327 | protein_coding_gene | -2.2 | 0.21 | 1.9E-02 |
| *atic-1* | WBGene00016957 | protein_coding_gene | -2.2 | 0.21 | 4.3E-22 |
| *gfi-1* | WBGene00001581 | protein_coding_gene | -2.2 | 0.21 | 1.7E-03 |
| *T17H7.1* | WBGene00020550 | protein_coding_gene | -2.3 | 0.21 | 7.0E-05 |
| *F45B8.5* | WBGene00194794 | protein_coding_gene | -2.3 | 0.21 | 1.6E-02 |
| *acdh-2* | WBGene00015894 | protein_coding_gene | -2.3 | 0.21 | 1.8E-10 |
| *rft-1* | WBGene00044637 | protein_coding_gene | -2.3 | 0.21 | 1.8E-10 |
| *nep-22* | WBGene00020788 | protein_coding_gene | -2.3 | 0.21 | 4.2E-34 |
| *F58G6.7* | WBGene00010277 | protein_coding_gene | -2.3 | 0.21 | 6.5E-08 |
| *sqt-1* | WBGene00005016 | protein_coding_gene | -2.3 | 0.21 | 1.4E-04 |
| *F31D5.1* | WBGene00017944 | protein_coding_gene | -2.3 | 0.21 | 1.9E-03 |
| *pho-7* | WBGene00011879 | protein_coding_gene | -2.3 | 0.21 | 6.8E-09 |
| *ppat-1* | WBGene00011407 | protein_coding_gene | -2.3 | 0.21 | 9.7E-23 |
| *B0391.10* | WBGene00007165 | pseudogene | -2.3 | 0.21 | 1.9E-17 |
| *ugt-62* | WBGene00010904 | protein_coding_gene | -2.3 | 0.20 | 1.2E-04 |
| *Y58A7A.1* | WBGene00021975 | protein_coding_gene | -2.3 | 0.20 | 9.7E-19 |
| *Y41C4A.32* | WBGene00270321 | protein_coding_gene | -2.3 | 0.20 | 1.1E-03 |
| *vit-3* | WBGene00006927 | protein_coding_gene | -2.3 | 0.20 | 4.5E-03 |
| *hrg-7* | WBGene00007605 | protein_coding_gene | -2.3 | 0.20 | 2.7E-04 |
| *C05D9.12* | WBGene00219800 | ncRNA_gene | -2.3 | 0.20 | 1.4E-04 |
| *ugt-22* | WBGene00007455 | protein_coding_gene | -2.3 | 0.20 | 1.4E-05 |
| *pqn-37* | WBGene00004124 | protein_coding_gene | -2.3 | 0.20 | 1.3E-07 |
| *nspb-10* | WBGene00007246 | protein_coding_gene | -2.3 | 0.20 | 1.4E-02 |
| *F07C6.6* | WBGene00045067 | protein_coding_gene | -2.3 | 0.20 | 1.2E-02 |
| *abu-8* | WBGene00000031 | protein_coding_gene | -2.3 | 0.20 | 2.0E-04 |
| *oac-52* | WBGene00020977 | protein_coding_gene | -2.3 | 0.20 | 5.2E-03 |
| *hsp-17* | WBGene00002021 | protein_coding_gene | -2.3 | 0.20 | 1.9E-17 |
| *smd-1* | WBGene00004875 | protein_coding_gene | -2.3 | 0.20 | 2.0E-21 |
| *F54D8.10* | WBGene00284847 | protein_coding_gene | -2.3 | 0.20 | 1.4E-05 |
| *mec-5* | WBGene00003169 | protein_coding_gene | -2.3 | 0.20 | 9.3E-21 |
| *col-73* | WBGene00000649 | protein_coding_gene | -2.4 | 0.20 | 8.0E-04 |
| *ins-30* | WBGene00002113 | protein_coding_gene | -2.4 | 0.20 | 3.2E-03 |
| *col-176* | WBGene00000749 | protein_coding_gene | -2.4 | 0.20 | 3.3E-03 |
| *col-109* | WBGene00000683 | protein_coding_gene | -2.4 | 0.20 | 1.3E-08 |
| *rol-6* | WBGene00004397 | protein_coding_gene | -2.4 | 0.20 | 2.4E-05 |
| *paic-1* | WBGene00015116 | protein_coding_gene | -2.4 | 0.19 | 9.1E-32 |
| *K02F2.11* | WBGene00220000 | ncRNA_gene | -2.4 | 0.19 | 6.0E-03 |
| *Y46G5A.29* | WBGene00012917 | protein_coding_gene | -2.4 | 0.19 | 7.1E-04 |
| *Y95B8A.2* | WBGene00022383 | protein_coding_gene | -2.4 | 0.19 | 2.2E-05 |
| *ugt-30* | WBGene00011340 | protein_coding_gene | -2.4 | 0.19 | 7.0E-08 |
| *str-146* | WBGene00006193 | protein_coding_gene | -2.4 | 0.19 | 2.7E-03 |
| *nhr-114* | WBGene00003704 | protein_coding_gene | -2.4 | 0.19 | 3.0E-30 |
| *C56A3.10* | WBGene00077568 | pseudogene | -2.4 | 0.19 | 5.2E-03 |
| *abu-10* | WBGene00000033 | protein_coding_gene | -2.4 | 0.19 | 1.3E-03 |
| *hil-6* | WBGene00001857 | protein_coding_gene | -2.4 | 0.19 | 5.1E-03 |
| *col-147* | WBGene00000720 | protein_coding_gene | -2.4 | 0.19 | 1.5E-03 |
| *H25K10.1* | WBGene00010411 | protein_coding_gene | -2.4 | 0.19 | 3.2E-04 |
| *F35F10.5* | WBGene00018055 | protein_coding_gene | -2.4 | 0.19 | 4.8E-03 |
| *B0334.13* | WBGene00007149 | protein_coding_gene | -2.4 | 0.19 | 2.8E-03 |
| *grl-4* | WBGene00001713 | protein_coding_gene | -2.4 | 0.19 | 6.6E-18 |
| *ugt-19* | WBGene00019515 | protein_coding_gene | -2.4 | 0.18 | 3.6E-04 |
| *col-173* | WBGene00000746 | protein_coding_gene | -2.4 | 0.18 | 8.8E-04 |
| *AC3.5* | WBGene00007071 | protein_coding_gene | -2.4 | 0.18 | 3.3E-27 |
| *scav-4* | WBGene00008692 | protein_coding_gene | -2.4 | 0.18 | 2.8E-10 |
| *col-17* | WBGene00000606 | protein_coding_gene | -2.4 | 0.18 | 1.6E-05 |

| | | | | | |
|---|---|---|---|---|---|
| *lon-1* | WBGene00003055 | protein_coding_gene | -2.4 | 0.18 | 6.5E-44 |
| *col-103* | WBGene00000677 | protein_coding_gene | -2.4 | 0.18 | 4.6E-49 |
| *ZK6.6* | WBGene00022641 | protein_coding_gene | -2.5 | 0.18 | 3.2E-05 |
| *ZK402.3* | WBGene00022730 | protein_coding_gene | -2.5 | 0.18 | 4.2E-03 |
| *F57B1.9* | WBGene00045488 | protein_coding_gene | -2.5 | 0.18 | 7.5E-07 |
| *col-156* | WBGene00000729 | protein_coding_gene | -2.5 | 0.18 | 5.3E-05 |
| *F09B12.3* | WBGene00008607 | protein_coding_gene | -2.5 | 0.18 | 5.6E-30 |
| *C42D4.3* | WBGene00016596 | protein_coding_gene | -2.5 | 0.18 | 2.0E-05 |
| *col-137* | WBGene00000710 | protein_coding_gene | -2.5 | 0.18 | 8.2E-04 |
| *K10H10.4* | WBGene00010760 | protein_coding_gene | -2.5 | 0.18 | 1.6E-03 |
| *gst-10* | WBGene00001758 | protein_coding_gene | -2.5 | 0.18 | 9.8E-18 |
| *R07E3.4* | WBGene00011105 | protein_coding_gene | -2.5 | 0.17 | 1.2E-15 |
| *C35A11.4* | WBGene00016431 | protein_coding_gene | -2.5 | 0.17 | 1.5E-19 |
| *cest-7* | WBGene00007693 | protein_coding_gene | -2.5 | 0.17 | 1.5E-03 |
| *Y47D7A.13* | WBGene00021625 | protein_coding_gene | -2.5 | 0.17 | 1.4E-04 |
| *T06E4.9* | WBGene00011536 | protein_coding_gene | -2.5 | 0.17 | 3.4E-03 |
| *Y82E9BL.18* | WBGene00044402 | protein_coding_gene | -2.5 | 0.17 | 3.0E-03 |
| *col-60* | WBGene00000636 | protein_coding_gene | -2.5 | 0.17 | 4.8E-07 |
| *T16G12.1* | WBGene00011803 | protein_coding_gene | -2.6 | 0.17 | 8.8E-21 |
| *T06E4.12* | WBGene00044011 | protein_coding_gene | -2.6 | 0.17 | 2.3E-03 |
| *Y47H9C.9* | WBGene00012952 | protein_coding_gene | -2.6 | 0.17 | 2.8E-03 |
| *E01G4.6* | WBGene00008448 | protein_coding_gene | -2.6 | 0.17 | 5.9E-08 |
| *cyp-35A4* | WBGene00016786 | protein_coding_gene | -2.6 | 0.17 | 1.6E-03 |
| *twk-16* | WBGene00006670 | protein_coding_gene | -2.6 | 0.17 | 2.8E-08 |
| *col-125* | WBGene00000699 | protein_coding_gene | -2.6 | 0.17 | 1.4E-04 |
| *ZK84.1* | WBGene00022649 | protein_coding_gene | -2.6 | 0.17 | 4.1E-09 |
| *pqn-63* | WBGene00004146 | protein_coding_gene | -2.6 | 0.17 | 3.3E-03 |
| *nspc-5* | WBGene00044340 | protein_coding_gene | -2.6 | 0.16 | 5.1E-03 |
| *col-14* | WBGene00000603 | protein_coding_gene | -2.6 | 0.16 | 1.6E-10 |
| *R12E2.7* | WBGene00020033 | protein_coding_gene | -2.6 | 0.16 | 9.8E-07 |
| *F19G12.t1* | WBGene00023271 | pseudogene | -2.6 | 0.16 | 2.2E-03 |
| *ZC239.15* | WBGene00022573 | protein_coding_gene | -2.6 | 0.16 | 3.8E-04 |
| *gst-26* | WBGene00001774 | protein_coding_gene | -2.6 | 0.16 | 4.1E-38 |
| *tag-10* | WBGene00006404 | protein_coding_gene | -2.6 | 0.16 | 2.2E-25 |
| *clec-245* | WBGene00007851 | protein_coding_gene | -2.6 | 0.16 | 1.2E-03 |
| *col-158* | WBGene00000731 | protein_coding_gene | -2.7 | 0.16 | 1.6E-05 |
| *gst-4* | WBGene00001752 | protein_coding_gene | -2.7 | 0.16 | 2.8E-21 |
| *col-71* | WBGene00000647 | protein_coding_gene | -2.7 | 0.15 | 9.9E-10 |
| *Y47G6A.19* | WBGene00021645 | protein_coding_gene | -2.7 | 0.15 | 4.9E-18 |
| *mpst-7* | WBGene00011307 | protein_coding_gene | -2.7 | 0.15 | 3.8E-09 |
| *col-159* | WBGene00000732 | protein_coding_gene | -2.7 | 0.15 | 1.2E-12 |
| *T26E3.11* | WBGene00044905 | pseudogene | -2.7 | 0.15 | 1.5E-03 |
| *F38B6.4* | WBGene00018174 | protein_coding_gene | -2.7 | 0.15 | 3.2E-25 |
| *Y50D4B.6* | WBGene00021745 | protein_coding_gene | -2.7 | 0.15 | 2.3E-05 |
| *gpdh-1* | WBGene00009824 | protein_coding_gene | -2.7 | 0.15 | 1.1E-07 |
| *M03B6.1* | WBGene00010833 | protein_coding_gene | -2.7 | 0.15 | 6.5E-11 |
| *col-98* | WBGene00000673 | protein_coding_gene | -2.7 | 0.15 | 3.1E-51 |
| *acp-6* | WBGene00022245 | protein_coding_gene | -2.8 | 0.15 | 3.9E-42 |
| *acs-1* | WBGene00018488 | protein_coding_gene | -2.8 | 0.15 | 1.7E-92 |
| *F22H10.6* | WBGene00017730 | protein_coding_gene | -2.8 | 0.14 | 5.8E-11 |
| *col-120* | WBGene00000694 | protein_coding_gene | -2.8 | 0.14 | 2.8E-08 |
| *kin-15* | WBGene00002199 | protein_coding_gene | -2.8 | 0.14 | 2.0E-15 |
| *cest-28* | WBGene00019654 | protein_coding_gene | -2.8 | 0.14 | 1.4E-05 |
| *C05E7.2* | WBGene00007344 | protein_coding_gene | -2.8 | 0.14 | 1.2E-06 |
| *col-133* | WBGene00000707 | protein_coding_gene | -2.8 | 0.14 | 5.1E-05 |
| *ZC250.4* | WBGene00045305 | protein_coding_gene | -2.8 | 0.14 | 5.9E-25 |
| *grd-13* | WBGene00001702 | protein_coding_gene | -2.8 | 0.14 | 8.3E-07 |
| *C05C8.8* | WBGene00015465 | protein_coding_gene | -2.8 | 0.14 | 6.9E-05 |
| *mua-6* | WBGene00003485 | protein_coding_gene | -2.8 | 0.14 | 1.5E-31 |
| *col-161* | WBGene00000734 | protein_coding_gene | -2.8 | 0.14 | 1.1E-04 |
| *nduo-2* | WBGene00010961 | protein_coding_gene | -2.9 | 0.14 | 1.9E-07 |
| *col-63* | WBGene00000639 | protein_coding_gene | -2.9 | 0.14 | 1.9E-11 |
| *spp-23* | WBGene00021335 | protein_coding_gene | -2.9 | 0.14 | 3.0E-08 |
| *col-130* | WBGene00000704 | protein_coding_gene | -2.9 | 0.14 | 1.1E-08 |
| *col-91* | WBGene00000666 | protein_coding_gene | -2.9 | 0.13 | 2.1E-07 |

| | | | | | |
|---|---|---|---|---|---|
| *abu-11* | WBGene00000034 | protein_coding_gene | -2.9 | 0.13 | 4.5E-05 |
| *Y51H7C.1* | WBGene00021779 | protein_coding_gene | -2.9 | 0.13 | 7.0E-10 |
| *zipt-2.3* | WBGene00013207 | protein_coding_gene | -2.9 | 0.13 | 1.4E-06 |
| *col-77* | WBGene00000653 | protein_coding_gene | -2.9 | 0.13 | 2.0E-13 |
| *T28D6.3* | WBGene00012123 | protein_coding_gene | -2.9 | 0.13 | 4.0E-27 |
| *Y50D4B.4* | WBGene00021743 | protein_coding_gene | -2.9 | 0.13 | 1.2E-32 |
| *col-146* | WBGene00000719 | protein_coding_gene | -2.9 | 0.13 | 1.0E-07 |
| *bli-6* | WBGene00000256 | protein_coding_gene | -2.9 | 0.13 | 6.0E-12 |
| *drd-1* | WBGene00009902 | protein_coding_gene | -3.0 | 0.13 | 3.6E-58 |
| *aat-4* | WBGene00000005 | protein_coding_gene | -3.0 | 0.13 | 7.8E-19 |
| *dhs-25* | WBGene00000988 | protein_coding_gene | -3.0 | 0.13 | 8.3E-08 |
| *dpy-13* | WBGene00001074 | protein_coding_gene | -3.0 | 0.12 | 7.0E-12 |
| *T20D4.3* | WBGene00020609 | protein_coding_gene | -3.1 | 0.12 | 3.4E-04 |
| *hsd-2* | WBGene00022498 | protein_coding_gene | -3.1 | 0.12 | 8.8E-04 |
| *elo-6* | WBGene00001244 | protein_coding_gene | -3.1 | 0.12 | 1.1E-28 |
| *F18E3.13* | WBGene00045276 | protein_coding_gene | -3.1 | 0.12 | 9.4E-11 |
| *pho-1* | WBGene00004020 | protein_coding_gene | -3.1 | 0.12 | 1.6E-07 |
| *sptl-2* | WBGene00018398 | protein_coding_gene | -3.1 | 0.12 | 2.1E-43 |
| *elo-2* | WBGene00001240 | protein_coding_gene | -3.1 | 0.11 | 7.0E-79 |
| *T10G3.3* | WBGene00011694 | protein_coding_gene | -3.1 | 0.11 | 1.9E-30 |
| *F16H6.4* | WBGene00008894 | protein_coding_gene | -3.2 | 0.11 | 2.3E-04 |
| *F40H3.2* | WBGene00018250 | protein_coding_gene | -3.2 | 0.11 | 8.5E-16 |
| *grd-6* | WBGene00001695 | protein_coding_gene | -3.2 | 0.11 | 2.5E-07 |
| *F53B3.6* | WBGene00018744 | protein_coding_gene | -3.2 | 0.11 | 8.3E-11 |
| *col-97* | WBGene00000672 | protein_coding_gene | -3.2 | 0.11 | 1.3E-15 |
| *ech-7* | WBGene00001156 | protein_coding_gene | -3.2 | 0.11 | 6.9E-33 |
| *grd-14* | WBGene00001703 | protein_coding_gene | -3.3 | 0.10 | 2.6E-13 |
| *dhs-26* | WBGene00000989 | protein_coding_gene | -3.3 | 0.10 | 1.8E-10 |
| *elo-5* | WBGene00001243 | protein_coding_gene | -3.3 | 0.10 | 1.9E-108 |
| *col-38* | WBGene00000615 | protein_coding_gene | -3.3 | 0.10 | 8.0E-25 |
| *col-162* | WBGene00000735 | protein_coding_gene | -3.3 | 0.10 | 7.2E-06 |
| *acox-1.4* | WBGene00008567 | protein_coding_gene | -3.3 | 0.10 | 2.3E-28 |
| *F53F1.4* | WBGene00009982 | protein_coding_gene | -3.3 | 0.10 | 9.3E-24 |
| *col-175* | WBGene00000748 | protein_coding_gene | -3.3 | 0.10 | 4.5E-15 |
| *dpy-4* | WBGene00001066 | protein_coding_gene | -3.3 | 0.10 | 5.5E-14 |
| *D1086.3* | WBGene00008390 | protein_coding_gene | -3.4 | 0.10 | 7.1E-15 |
| *C04E12.5* | WBGene00015435 | protein_coding_gene | -3.4 | 0.09 | 7.1E-05 |
| *ZK180.6* | WBGene00022680 | protein_coding_gene | -3.4 | 0.09 | 1.6E-12 |
| *C04E12.10* | WBGene00015439 | pseudogene | -3.4 | 0.09 | 6.1E-05 |
| *Y47G6A.15* | WBGene00021641 | protein_coding_gene | -3.4 | 0.09 | 1.3E-14 |
| *dpy-6* | WBGene00001068 | protein_coding_gene | -3.5 | 0.09 | 1.8E-09 |
| *col-138* | WBGene00000711 | protein_coding_gene | -3.5 | 0.09 | 2.5E-15 |
| *col-157* | WBGene00000730 | protein_coding_gene | -3.5 | 0.09 | 1.6E-07 |
| *col-88* | WBGene00000663 | protein_coding_gene | -3.5 | 0.09 | 4.2E-13 |
| *col-49* | WBGene00000626 | protein_coding_gene | -3.5 | 0.09 | 3.1E-11 |
| *F41E6.12* | WBGene00018293 | protein_coding_gene | -3.5 | 0.09 | 1.6E-23 |
| *F46F2.5* | WBGene00009789 | protein_coding_gene | -3.5 | 0.09 | 2.2E-12 |
| *dpy-5* | WBGene00001067 | protein_coding_gene | -3.6 | 0.08 | 6.9E-17 |
| *Y51H7C.13* | WBGene00021791 | protein_coding_gene | -3.6 | 0.08 | 7.5E-13 |
| *nhr-74* | WBGene00003664 | protein_coding_gene | -3.6 | 0.08 | 7.4E-05 |
| *sur-5* | WBGene00006351 | protein_coding_gene | -3.6 | 0.08 | 6.7E-203 |
| *F58G6.3* | WBGene00010274 | protein_coding_gene | -3.7 | 0.08 | 7.0E-26 |
| *col-180* | WBGene00000753 | protein_coding_gene | -3.7 | 0.08 | 2.7E-11 |
| *C07G1.7* | WBGene00015573 | protein_coding_gene | -3.7 | 0.08 | 1.1E-06 |
| *hrg-4* | WBGene00009493 | protein_coding_gene | -3.8 | 0.07 | 2.3E-10 |
| *K02E11.10* | WBGene00044109 | protein_coding_gene | -3.8 | 0.07 | 6.1E-08 |
| *Y48G8AL.12* | WBGene00021690 | protein_coding_gene | -3.8 | 0.07 | 2.9E-07 |
| *col-167* | WBGene00000740 | protein_coding_gene | -3.8 | 0.07 | 2.4E-11 |
| *R03G8.4* | WBGene00010999 | pseudogene | -3.8 | 0.07 | 3.0E-05 |
| *col-149* | WBGene00000722 | protein_coding_gene | -3.9 | 0.07 | 4.8E-13 |
| *gcsh-1* | WBGene00008354 | protein_coding_gene | -3.9 | 0.07 | 7.2E-37 |
| *F18E3.11* | WBGene00017567 | protein_coding_gene | -3.9 | 0.07 | 1.5E-20 |
| *rol-1* | WBGene00004394 | protein_coding_gene | -3.9 | 0.07 | 9.9E-17 |
| *K09C4.5* | WBGene00019550 | protein_coding_gene | -4.0 | 0.06 | 3.5E-32 |
| *col-145* | WBGene00000718 | protein_coding_gene | -4.0 | 0.06 | 1.6E-09 |

| | | | | | |
|---|---|---|---|---|---|
| *Y47D7A.15* | WBGene00021627 | protein_coding_gene | -4.0 | 0.06 | 1.6E-10 |
| *F42A8.1* | WBGene00009626 | protein_coding_gene | -4.0 | 0.06 | 1.9E-43 |
| *grd-4* | WBGene00001693 | protein_coding_gene | -4.1 | 0.06 | 4.5E-06 |
| *nas-3* | WBGene00003522 | protein_coding_gene | -4.3 | 0.05 | 3.2E-16 |
| *abhd-3.2* | WBGene00016644 | protein_coding_gene | -4.3 | 0.05 | 1.9E-73 |
| *R07E5.4* | WBGene00011112 | protein_coding_gene | -4.5 | 0.04 | 1.4E-27 |
| *F18E3.12* | WBGene00045275 | protein_coding_gene | -4.5 | 0.04 | 6.8E-19 |
| *R193.2* | WBGene00020128 | protein_coding_gene | -4.6 | 0.04 | 1.4E-44 |
| *F46F2.3* | WBGene00009787 | protein_coding_gene | -4.8 | 0.04 | 1.6E-17 |

Genes expressed in the same direction in CB4856 (vs. N2) fed *E. coli* OP50 or *S. epidermidis*

| | | | *E. coli* OP50 | | | *S. epidermidis* | | |
|---|---|---|---|---|---|---|---|---|
| Gene Name | Wormbase Gene ID | Type | $log_2$FC | FC | Adj P-value | $log_2$FC | FC | Adj P-value |
| *nspa-1* | WBGene00010377 | protein_coding_gene | 8.1 | 271.084 | 8.9E-20 | 6.4 | 85.47 | 5.9E-18 |
| *clec-77* | WBGene00044343 | pseudogene | 7.8 | 219.595 | 2.3E-25 | 5.5 | 43.91 | 5.5E-136 |
| *cyp-37B1* | WBGene00009226 | protein_coding_gene | 7.7 | 206.507 | 4.8E-60 | 2.6 | 5.86 | 3.6E-14 |
| *pud-2.2* | WBGene00017500 | protein_coding_gene | 7.3 | 153.739 | 1.0E-16 | 1.3 | 2.45 | 1.4E-09 |
| *K02F6.4* | WBGene00019338 | protein_coding_gene | 7.2 | 151.450 | 1.6E-16 | 6.0 | 64.68 | 7.4E-28 |
| *clec-110* | WBGene00008909 | protein_coding_gene | 7.1 | 141.223 | 3.8E-16 | 6.5 | 89.28 | 5.6E-37 |
| *clec-208* | WBGene00018097 | protein_coding_gene | 7.1 | 140.507 | 3.7E-36 | 5.5 | 44.46 | 1.6E-124 |
| *fipr-18* | WBGene00044450 | protein_coding_gene | 6.9 | 117.843 | 2.3E-20 | 5.1 | 35.15 | 5.4E-79 |
| *C39B5.10* | WBGene00016528 | protein_coding_gene | 6.8 | 107.731 | 2.2E-34 | 4.3 | 19.22 | 7.1E-10 |
| *T23G4.5* | WBGene00011961 | protein_coding_gene | 6.6 | 96.062 | 4.0E-14 | 5.5 | 45.69 | 2.5E-69 |
| *clec-124* | WBGene00015909 | protein_coding_gene | 6.4 | 85.625 | 2.2E-13 | 6.2 | 73.80 | 2.2E-29 |
| *M7.10* | WBGene00010886 | protein_coding_gene | 6.4 | 83.926 | 2.1E-13 | 4.0 | 16.42 | 6.1E-41 |
| *fipr-17* | WBGene00012965 | protein_coding_gene | 6.3 | 80.244 | 1.8E-31 | 4.8 | 27.59 | 2.5E-78 |
| *T10D4.1* | WBGene00020406 | protein_coding_gene | 6.3 | 77.877 | 2.0E-11 | 4.4 | 21.61 | 1.6E-12 |
| *ZK1290.1* | WBGene00022886 | protein_coding_gene | 6.2 | 74.969 | 1.2E-12 | 4.8 | 27.53 | 6.5E-70 |
| *lipl-2* | WBGene00009773 | protein_coding_gene | 6.2 | 72.425 | 3.9E-45 | 1.0 | 2.03 | 1.5E-02 |
| *clec-158* | WBGene00020067 | protein_coding_gene | 6.2 | 71.598 | 4.3E-16 | 5.0 | 31.13 | 1.5E-70 |
| *M176.10* | WBGene00010946 | protein_coding_gene | 6.2 | 71.438 | 2.0E-16 | 4.5 | 22.37 | 1.4E-75 |
| *Y110A2AL.7* | WBGene00022444 | protein_coding_gene | 6.1 | 70.435 | 2.2E-16 | 4.3 | 19.12 | 1.5E-61 |
| *fipr-20* | WBGene00044449 | protein_coding_gene | 6.1 | 70.085 | 1.2E-19 | 4.8 | 27.55 | 2.0E-79 |
| *Y43F8C.17* | WBGene00012838 | protein_coding_gene | 6.1 | 67.164 | 8.4E-11 | 5.8 | 55.60 | 5.2E-38 |
| *T10B10.9* | WBGene00011685 | protein_coding_gene | 6.0 | 66.234 | 9.5E-11 | 4.6 | 24.85 | 6.1E-37 |
| *F46B6.13* | WBGene00044192 | protein_coding_gene | 6.0 | 64.373 | 7.3E-12 | 4.1 | 17.69 | 6.9E-45 |
| *R03H10.6* | WBGene00019858 | protein_coding_gene | 6.0 | 63.165 | 2.7E-57 | 6.3 | 81.21 | 6.3E-160 |
| *scl-7* | WBGene00008029 | protein_coding_gene | 6.0 | 62.814 | 1.1E-11 | 4.9 | 30.18 | 2.2E-52 |
| *clec-141* | WBGene00017139 | protein_coding_gene | 5.9 | 61.679 | 1.0E-11 | 5.2 | 36.11 | 1.2E-45 |
| *hpo-2* | WBGene00012500 | protein_coding_gene | 5.9 | 60.038 | 7.3E-32 | 4.6 | 23.71 | 7.5E-81 |
| *Y25C1A.2* | WBGene00021289 | protein_coding_gene | 5.9 | 59.343 | 1.9E-11 | 4.7 | 26.55 | 9.9E-36 |
| *ZK1053.3* | WBGene00014198 | protein_coding_gene | 5.9 | 59.049 | 2.9E-46 | 4.7 | 26.78 | 2.7E-60 |
| *E02H4.4* | WBGene00008462 | protein_coding_gene | 5.8 | 55.944 | 1.1E-51 | 2.7 | 6.71 | 5.1E-109 |
| *clec-232* | WBGene00009487 | protein_coding_gene | 5.8 | 53.938 | 3.9E-14 | 4.4 | 21.80 | 5.4E-52 |
| *clec-161* | WBGene00010228 | protein_coding_gene | 5.7 | 53.636 | 5.1E-20 | 4.6 | 23.83 | 5.4E-64 |
| *clec-133* | WBGene00021142 | protein_coding_gene | 5.7 | 53.234 | 7.3E-10 | 4.5 | 22.14 | 3.4E-34 |
| *C49C3.11* | WBGene00008201 | protein_coding_gene | 5.7 | 52.668 | 2.4E-62 | 5.1 | 34.28 | 6.5E-151 |
| *B0250.3* | WBGene00007120 | pseudogene | 5.7 | 52.173 | 6.8E-104 | 5.8 | 55.68 | 9.7E-111 |
| *fipr-28* | WBGene00013120 | protein_coding_gene | 5.7 | 51.238 | 2.1E-22 | 4.6 | 24.97 | 1.2E-62 |
| *F46A8.5* | WBGene00009748 | protein_coding_gene | 5.7 | 50.381 | 1.2E-09 | 4.7 | 26.46 | 7.0E-39 |
| *clec-132* | WBGene00021143 | protein_coding_gene | 5.6 | 49.716 | 6.6E-11 | 5.3 | 40.17 | 5.7E-15 |
| *scl-9* | WBGene00009890 | protein_coding_gene | 5.6 | 47.603 | 1.7E-13 | 4.2 | 18.59 | 4.3E-39 |
| *clec-24* | WBGene00009856 | protein_coding_gene | 5.5 | 45.709 | 1.9E-18 | 2.3 | 4.90 | 1.6E-07 |
| *Y6G8.15* | WBGene00189937 | protein_coding_gene | 5.5 | 45.586 | 4.4E-10 | 5.6 | 47.60 | 9.3E-16 |
| *F49F1.15* | WBGene00235365 | protein_coding_gene | 5.5 | 44.924 | 3.2E-10 | 4.4 | 21.33 | 2.1E-44 |
| *C17E4.19* | WBGene00219310 | protein_coding_gene | 5.5 | 44.778 | 3.3E-32 | 3.6 | 12.21 | 4.1E-46 |
| *F07G6.8* | WBGene00017227 | protein_coding_gene | 5.4 | 42.889 | 5.3E-09 | 4.1 | 17.51 | 1.8E-21 |
| *pqn-84* | WBGene00004165 | protein_coding_gene | 5.4 | 42.343 | 2.7E-42 | 4.8 | 28.30 | 6.7E-131 |
| *Y59H11AM.1* | WBGene00022005 | protein_coding_gene | 5.4 | 42.329 | 1.6E-12 | 5.4 | 41.59 | 1.9E-13 |
| *R53.8* | WBGene00011277 | protein_coding_gene | 5.4 | 42.279 | 7.2E-20 | 4.6 | 24.46 | 1.7E-56 |
| *ZK1290.11* | WBGene00022892 | protein_coding_gene | 5.4 | 41.770 | 1.6E-19 | 4.4 | 20.89 | 1.4E-49 |
| *Y82E9BL.9* | WBGene00023395 | pseudogene | 5.4 | 41.268 | 1.0E-18 | 4.7 | 25.14 | 5.3E-25 |
| *tag-329* | WBGene00008231 | protein_coding_gene | 5.4 | 40.930 | 1.6E-37 | 4.5 | 21.87 | 2.1E-97 |
| *nspc-12* | WBGene00023493 | protein_coding_gene | 5.3 | 40.560 | 5.3E-14 | 4.5 | 22.97 | 4.5E-25 |
| *R09E10.8* | WBGene00011177 | protein_coding_gene | 5.3 | 40.200 | 3.2E-12 | 4.7 | 25.20 | 1.6E-34 |

| | | | | | | | | |
|---|---|---|---|---|---|---|---|---|
| *F43C11.2* | WBGene00018379 | protein_coding_gene | 5.3 | 39.272 | 1.4E-14 | 4.8 | 28.46 | 7.1E-56 |
| *F42A6.2* | WBGene00018324 | protein_coding_gene | 5.3 | 39.242 | 7.3E-10 | 4.0 | 16.55 | 4.3E-19 |
| *D1022.2* | WBGene00017021 | protein_coding_gene | 5.3 | 39.237 | 1.2E-08 | 4.3 | 19.97 | 1.1E-27 |
| *clec-138* | WBGene00018049 | protein_coding_gene | 5.3 | 38.891 | 4.1E-12 | 4.3 | 19.72 | 1.1E-44 |
| *F16G10.10* | WBGene00017525 | protein_coding_gene | 5.2 | 37.650 | 2.1E-09 | 4.7 | 25.54 | 4.7E-25 |
| *C45G9.10* | WBGene00016681 | protein_coding_gene | 5.2 | 37.248 | 3.3E-14 | 4.3 | 19.73 | 2.3E-46 |
| *F47C12.12* | WBGene00219377 | pseudogene | 5.2 | 37.076 | 3.1E-09 | 4.8 | 27.37 | 5.3E-16 |
| *F11A6.16* | WBGene00271780 | protein_coding_gene | 5.1 | 35.397 | 2.1E-09 | 4.1 | 17.30 | 1.2E-19 |
| *Y6G8.14* | WBGene00138715 | protein_coding_gene | 5.1 | 35.388 | 2.9E-08 | 4.2 | 17.99 | 3.8E-26 |
| *clec-125* | WBGene00021118 | protein_coding_gene | 5.1 | 35.338 | 6.1E-51 | 5.3 | 40.12 | 4.0E-117 |
| *T05A7.11* | WBGene00044383 | protein_coding_gene | 5.1 | 35.157 | 3.5E-11 | 1.2 | 2.36 | 2.4E-04 |
| *B0222.10* | WBGene00015058 | ncRNA_gene | 5.1 | 35.104 | 7.3E-28 | 4.9 | 29.03 | 5.3E-90 |
| *T26A8.3* | WBGene00020826 | protein_coding_gene | 5.1 | 34.850 | 2.2E-17 | 4.3 | 20.19 | 1.2E-49 |
| *plg-1* | WBGene00004041 | pseudogene | 5.1 | 34.769 | 3.9E-51 | 5.6 | 49.38 | 1.3E-108 |
| *F13E9.10* | WBGene00008759 | protein_coding_gene | 5.1 | 34.109 | 2.0E-13 | 4.8 | 28.57 | 2.8E-37 |
| *C28C12.3* | WBGene00016179 | protein_coding_gene | 5.1 | 34.032 | 1.2E-25 | 4.2 | 18.29 | 6.4E-73 |
| *nas-24* | WBGene00003543 | protein_coding_gene | 5.1 | 33.855 | 1.8E-26 | 4.4 | 21.11 | 5.3E-80 |
| *C35B1.3* | WBGene00016433 | protein_coding_gene | 5.1 | 33.628 | 3.5E-09 | 4.1 | 17.37 | 3.1E-18 |
| *B0513.7* | WBGene00007199 | pseudogene | 5.0 | 33.051 | 4.0E-09 | 2.9 | 7.52 | 2.1E-06 |
| *ins-20* | WBGene00002103 | protein_coding_gene | 5.0 | 32.770 | 6.1E-08 | 2.4 | 5.32 | 8.8E-08 |
| *C47A10.12* | WBGene00008129 | protein_coding_gene | 5.0 | 32.703 | 9.4E-11 | 5.1 | 33.62 | 2.0E-41 |
| *C16C8.7* | WBGene00015845 | protein_coding_gene | 5.0 | 32.556 | 5.2E-13 | 4.5 | 22.88 | 4.3E-34 |
| *T16G12.10* | WBGene00044222 | protein_coding_gene | 5.0 | 31.742 | 2.0E-30 | 4.7 | 25.85 | 1.6E-88 |
| *F22B3.5* | WBGene00009036 | protein_coding_gene | 5.0 | 31.698 | 6.1E-09 | 4.0 | 15.47 | 1.1E-09 |
| *tag-138* | WBGene00006484 | protein_coding_gene | 5.0 | 31.628 | 9.4E-11 | 1.6 | 3.05 | 1.6E-06 |
| *clec-130* | WBGene00021144 | protein_coding_gene | 5.0 | 31.389 | 6.8E-09 | 2.5 | 5.58 | 1.9E-09 |
| *F47C12.8* | WBGene00018551 | protein_coding_gene | 5.0 | 31.263 | 1.0E-10 | 5.2 | 35.53 | 1.5E-12 |
| *H25K10.4* | WBGene00010413 | protein_coding_gene | 4.9 | 30.684 | 2.0E-12 | 1.8 | 3.54 | 2.4E-05 |
| *clec-32* | WBGene00009860 | protein_coding_gene | 4.9 | 30.573 | 8.6E-09 | 1.5 | 2.78 | 3.2E-03 |
| *R05H10.7* | WBGene00011046 | protein_coding_gene | 4.9 | 30.025 | 4.3E-17 | 5.3 | 38.52 | 1.2E-68 |
| *F26C11.4* | WBGene00044159 | protein_coding_gene | 4.9 | 29.900 | 1.8E-10 | 4.6 | 24.28 | 8.2E-18 |
| *W02D7.12* | WBGene00044356 | protein_coding_gene | 4.9 | 29.092 | 1.5E-07 | 3.6 | 12.00 | 2.4E-19 |
| *try-8* | WBGene00017791 | protein_coding_gene | 4.9 | 29.007 | 2.5E-36 | 4.9 | 30.29 | 1.5E-155 |
| *Y40H7A.147* | WBGene00077755 | pseudogene | 4.8 | 28.812 | 1.5E-08 | 4.0 | 16.43 | 2.4E-09 |
| *F01D5.4* | WBGene00014730 | pseudogene | 4.8 | 28.475 | 2.0E-07 | 2.8 | 6.86 | 3.8E-07 |
| *Y110A2AL.6* | WBGene00022443 | protein_coding_gene | 4.8 | 28.010 | 2.0E-08 | 4.0 | 15.87 | 2.6E-21 |
| *F38E1.10* | WBGene00018182 | protein_coding_gene | 4.8 | 27.880 | 2.1E-08 | 3.8 | 13.49 | 1.2E-21 |
| *T14G12.11* | WBGene00219411 | protein_coding_gene | 4.8 | 27.640 | 4.7E-08 | 5.0 | 31.05 | 5.3E-12 |
| *C14F11.7* | WBGene00015782 | protein_coding_gene | 4.8 | 27.543 | 4.2E-22 | 4.8 | 27.30 | 1.3E-55 |
| *B0207.5* | WBGene00015028 | protein_coding_gene | 4.8 | 27.449 | 7.4E-32 | 4.9 | 29.44 | 3.0E-118 |
| *C09G12.5* | WBGene00015655 | protein_coding_gene | 4.8 | 27.190 | 7.1E-17 | 4.1 | 16.71 | 2.9E-55 |
| *fipr-16* | WBGene00015521 | protein_coding_gene | 4.7 | 26.796 | 7.7E-18 | 4.3 | 19.97 | 2.2E-34 |
| *clec-181* | WBGene00044511 | protein_coding_gene | 4.7 | 26.754 | 3.1E-08 | 3.9 | 15.15 | 7.4E-11 |
| *Y43F8C.15* | WBGene00012836 | protein_coding_gene | 4.7 | 26.675 | 3.2E-08 | 4.3 | 19.98 | 2.2E-12 |
| *F26H11.8* | WBGene00195149 | protein_coding_gene | 4.7 | 26.576 | 2.9E-11 | 5.0 | 32.27 | 3.1E-13 |
| *F02H6.6* | WBGene00008539 | protein_coding_gene | 4.7 | 26.017 | 4.0E-07 | 4.1 | 17.06 | 5.9E-21 |
| *F46A8.3* | WBGene00009746 | protein_coding_gene | 4.7 | 25.860 | 5.7E-11 | 4.1 | 16.72 | 2.5E-29 |
| *M151.10* | WBGene00303365 | protein_coding_gene | 4.7 | 25.680 | 5.0E-27 | 5.1 | 33.83 | 2.7E-71 |
| *K09C8.2* | WBGene00010715 | protein_coding_gene | 4.7 | 25.670 | 4.6E-07 | 3.9 | 14.61 | 3.0E-16 |
| *Y39A1A.27* | WBGene00194731 | protein_coding_gene | 4.7 | 25.454 | 4.8E-07 | 3.6 | 12.42 | 2.5E-18 |
| *K03B8.11* | WBGene00010526 | protein_coding_gene | 4.7 | 25.194 | 5.3E-07 | 4.0 | 15.64 | 7.9E-14 |
| *clec-157* | WBGene00017810 | protein_coding_gene | 4.6 | 24.948 | 6.2E-30 | 4.9 | 30.72 | 5.5E-82 |
| *E02H9.1* | WBGene00017099 | protein_coding_gene | 4.6 | 24.853 | 2.2E-12 | 3.7 | 12.60 | 2.9E-23 |
| *clec-92* | WBGene00013125 | protein_coding_gene | 4.6 | 24.592 | 1.6E-14 | 4.2 | 18.74 | 1.2E-52 |
| *W02B12.16* | WBGene00195148 | protein_coding_gene | 4.6 | 24.496 | 3.8E-09 | 3.6 | 11.89 | 8.3E-19 |
| *srh-254* | WBGene00005460 | pseudogene | 4.6 | 23.879 | 8.1E-07 | 5.0 | 32.87 | 2.4E-13 |
| *clec-263* | WBGene00009171 | protein_coding_gene | 4.6 | 23.745 | 8.5E-07 | 3.1 | 8.75 | 1.8E-19 |
| *F49H6.5* | WBGene00009908 | protein_coding_gene | 4.6 | 23.736 | 1.4E-15 | 1.9 | 3.80 | 8.9E-11 |
| *linc-1* | WBGene00194659 | lincRNA_gene | 4.5 | 23.277 | 8.0E-25 | 4.0 | 15.47 | 8.5E-45 |
| *T12A2.5* | WBGene00020439 | protein_coding_gene | 4.5 | 23.272 | 1.5E-13 | 4.7 | 26.30 | 5.8E-55 |
| *scl-8* | WBGene00008030 | protein_coding_gene | 4.5 | 23.004 | 2.4E-10 | 2.8 | 6.96 | 2.8E-21 |
| *ptr-22* | WBGene00004236 | protein_coding_gene | 4.5 | 22.995 | 1.3E-44 | 1.8 | 3.56 | 1.5E-12 |
| *C16C8.19* | WBGene00015856 | protein_coding_gene | 4.5 | 22.862 | 1.1E-12 | 4.2 | 17.91 | 5.9E-33 |
| *F17E9.3* | WBGene00017540 | protein_coding_gene | 4.5 | 22.826 | 6.9E-09 | 3.6 | 12.40 | 2.6E-21 |
| *clec-219* | WBGene00020945 | protein_coding_gene | 4.5 | 22.305 | 3.7E-10 | 5.7 | 50.62 | 7.2E-185 |

| | | | | | | | | |
|---|---|---|---|---|---|---|---|---|
| *fbxa-58* | WBGene00016225 | pseudogene | 4.5 | 22.251 | 1.6E-08 | 5.0 | 32.96 | 3.4E-12 |
| *F28B1.9* | WBGene00045381 | protein_coding_gene | 4.5 | 22.211 | 1.3E-06 | 3.7 | 13.13 | 8.7E-14 |
| *nspf-3* | WBGene00044366 | protein_coding_gene | 4.5 | 22.196 | 1.2E-11 | 3.2 | 9.14 | 8.7E-22 |
| *F46A8.8* | WBGene00009751 | protein_coding_gene | 4.4 | 21.564 | 6.1E-07 | 3.5 | 11.41 | 5.0E-17 |
| *Y51A2D.1* | WBGene00013072 | protein_coding_gene | 4.4 | 21.509 | 2.3E-07 | 4.1 | 17.59 | 7.4E-15 |
| *Y49F6B.6* | WBGene00021718 | protein_coding_gene | 4.4 | 21.347 | 8.4E-10 | 4.1 | 17.29 | 6.6E-24 |
| *Y43F8B.19* | WBGene00077688 | protein_coding_gene | 4.4 | 20.939 | 2.9E-07 | 4.2 | 17.81 | 1.3E-09 |
| *C06E2.9* | WBGene00044579 | protein_coding_gene | 4.4 | 20.926 | 6.2E-07 | 3.2 | 9.00 | 3.1E-13 |
| *ZK1225.10* | WBGene00044714 | pseudogene | 4.4 | 20.864 | 3.0E-07 | 4.0 | 16.42 | 3.1E-11 |
| *Y18D10A.2* | WBGene00012475 | protein_coding_gene | 4.4 | 20.779 | 9.4E-10 | 3.9 | 14.75 | 3.6E-30 |
| *F49F1.5* | WBGene00018645 | protein_coding_gene | 4.4 | 20.765 | 2.1E-21 | 2.5 | 5.51 | 2.9E-28 |
| *F25E5.3* | WBGene00017784 | protein_coding_gene | 4.3 | 20.371 | 2.9E-24 | 4.8 | 27.35 | 8.5E-71 |
| *clec-126* | WBGene00021117 | protein_coding_gene | 4.3 | 20.127 | 7.6E-22 | 3.7 | 13.22 | 7.2E-41 |
| *hpo-7* | WBGene00016548 | protein_coding_gene | 4.3 | 20.062 | 4.2E-07 | 3.1 | 8.47 | 2.3E-08 |
| *cwp-1* | WBGene00000859 | protein_coding_gene | 4.3 | 19.967 | 2.8E-06 | 3.8 | 13.87 | 3.8E-20 |
| *F30A10.11* | WBGene00009268 | protein_coding_gene | 4.3 | 19.870 | 7.2E-20 | 2.3 | 5.01 | 1.9E-08 |
| *clec-105* | WBGene00012488 | protein_coding_gene | 4.3 | 19.844 | 4.7E-07 | 4.4 | 20.78 | 3.4E-10 |
| *clec-135* | WBGene00015821 | protein_coding_gene | 4.3 | 19.381 | 4.9E-12 | 4.3 | 20.26 | 3.0E-41 |
| *R13.5* | WBGene00014825 | pseudogene | 4.3 | 19.337 | 5.8E-07 | 3.8 | 14.18 | 1.1E-08 |
| *clec-261* | WBGene00013834 | protein_coding_gene | 4.3 | 19.313 | 4.6E-08 | 3.8 | 13.82 | 5.3E-18 |
| *dsl-6* | WBGene00001108 | protein_coding_gene | 4.3 | 19.195 | 3.5E-09 | 1.5 | 2.76 | 1.3E-09 |
| *T22C1.12* | WBGene00011921 | protein_coding_gene | 4.2 | 18.854 | 1.9E-06 | 4.1 | 17.03 | 1.8E-09 |
| *F26D2.16* | WBGene00009154 | protein_coding_gene | 4.2 | 18.835 | 1.6E-09 | 5.4 | 43.22 | 8.8E-158 |
| *T12A7.10* | WBGene00194933 | protein_coding_gene | 4.2 | 18.470 | 5.0E-06 | 3.5 | 11.45 | 2.4E-08 |
| *C41D11.10* | WBGene00195180 | protein_coding_gene | 4.2 | 18.465 | 5.0E-06 | 3.3 | 9.62 | 4.9E-11 |
| *msp-79* | WBGene00003466 | protein_coding_gene | 4.2 | 18.312 | 1.4E-07 | 3.6 | 11.81 | 5.2E-15 |
| *F13E9.19* | WBGene00194922 | protein_coding_gene | 4.2 | 17.909 | 2.8E-06 | 4.1 | 17.44 | 4.4E-10 |
| *nspa-4* | WBGene00044272 | protein_coding_gene | 4.1 | 17.738 | 6.8E-10 | 3.5 | 10.94 | 3.8E-17 |
| *clec-106* | WBGene00012482 | protein_coding_gene | 4.1 | 17.721 | 1.1E-10 | 4.6 | 24.21 | 1.9E-24 |
| *B0513.90* | WBGene00235301 | protein_coding_gene | 4.1 | 17.356 | 7.8E-06 | 3.3 | 9.65 | 1.9E-16 |
| *F13E9.9* | WBGene00008758 | protein_coding_gene | 4.1 | 17.195 | 1.6E-06 | 2.5 | 5.76 | 2.9E-08 |
| *K08C9.6* | WBGene00010653 | protein_coding_gene | 4.1 | 17.035 | 1.7E-06 | 3.6 | 12.11 | 2.2E-08 |
| *T23E7.6* | WBGene00044772 | protein_coding_gene | 4.1 | 16.801 | 7.4E-20 | 1.2 | 2.23 | 9.6E-03 |
| *fbxa-161* | WBGene00015596 | pseudogene | 4.0 | 16.560 | 1.3E-05 | 1.8 | 3.45 | 1.8E-04 |
| *fbxc-20* | WBGene00018352 | protein_coding_gene | 4.0 | 16.520 | 3.4E-14 | 1.6 | 2.94 | 2.4E-08 |
| *K01A6.4* | WBGene00010446 | protein_coding_gene | 4.0 | 16.480 | 2.2E-06 | 1.4 | 2.64 | 5.3E-03 |
| *F56D6.11* | WBGene00044473 | protein_coding_gene | 4.0 | 16.443 | 1.3E-05 | 3.9 | 14.55 | 9.4E-11 |
| *F59A6.11* | WBGene00044664 | protein_coding_gene | 4.0 | 16.434 | 2.3E-06 | 3.6 | 12.17 | 4.1E-13 |
| *srp-5* | WBGene00005646 | pseudogene | 4.0 | 16.285 | 6.4E-06 | 3.7 | 12.56 | 5.0E-10 |
| *ZC328.5* | WBGene00022594 | protein_coding_gene | 4.0 | 16.165 | 8.2E-16 | 4.1 | 17.25 | 2.4E-39 |
| *Y43F8C.24* | WBGene00045345 | pseudogene | 4.0 | 16.142 | 2.6E-06 | 3.5 | 11.13 | 1.4E-09 |
| *F40G9.7* | WBGene00018243 | protein_coding_gene | 4.0 | 16.020 | 1.4E-05 | 3.7 | 12.67 | 1.2E-14 |
| *K12H6.5* | WBGene00019685 | protein_coding_gene | 4.0 | 15.883 | 1.5E-05 | 3.5 | 11.20 | 6.8E-21 |
| *C17E4.12* | WBGene00044769 | pseudogene | 4.0 | 15.700 | 5.6E-09 | 3.1 | 8.55 | 6.0E-07 |
| *fbxa-163* | WBGene00015598 | protein_coding_gene | 4.0 | 15.636 | 1.6E-10 | 2.2 | 4.49 | 2.9E-06 |
| *nspc-17* | WBGene00008359 | protein_coding_gene | 4.0 | 15.470 | 3.3E-50 | 3.0 | 7.92 | 9.3E-74 |
| *E02H9.9* | WBGene00017106 | protein_coding_gene | 3.9 | 15.378 | 1.5E-18 | 3.9 | 14.81 | 1.3E-40 |
| *K04H8.3* | WBGene00010578 | protein_coding_gene | 3.9 | 15.327 | 1.3E-11 | 4.2 | 18.83 | 1.7E-26 |
| *Y71A12B.19* | WBGene00077443 | pseudogene | 3.9 | 15.269 | 5.8E-22 | 1.3 | 2.47 | 4.1E-04 |
| *Y17G9A.4* | WBGene00021197 | protein_coding_gene | 3.9 | 15.114 | 1.9E-24 | 1.2 | 2.29 | 6.7E-05 |
| *F47C12.6* | WBGene00018549 | protein_coding_gene | 3.9 | 14.951 | 2.2E-05 | 3.1 | 8.43 | 2.5E-08 |
| *flp-23* | WBGene00017693 | protein_coding_gene | 3.9 | 14.863 | 5.2E-06 | 3.3 | 9.59 | 7.2E-12 |
| *T05E12.3* | WBGene00011486 | protein_coding_gene | 3.9 | 14.821 | 3.6E-75 | 1.5 | 2.85 | 1.9E-30 |
| *msp-71* | WBGene00003460 | pseudogene | 3.9 | 14.643 | 7.1E-27 | 4.4 | 20.75 | 5.1E-64 |
| *clec-108* | WBGene00012504 | protein_coding_gene | 3.9 | 14.523 | 1.9E-05 | 3.5 | 11.55 | 2.2E-15 |
| *Y20C6A.4* | WBGene00077495 | protein_coding_gene | 3.9 | 14.502 | 3.4E-08 | 2.1 | 4.25 | 8.6E-05 |
| *plp-2* | WBGene00008976 | protein_coding_gene | 3.9 | 14.464 | 1.8E-20 | 3.6 | 11.99 | 1.5E-26 |
| *M01G12.10* | WBGene00010823 | protein_coding_gene | 3.9 | 14.433 | 6.5E-06 | 3.5 | 11.31 | 1.4E-09 |
| *F56D2.8* | WBGene00018968 | protein_coding_gene | 3.8 | 14.384 | 2.0E-07 | 3.7 | 12.81 | 5.9E-17 |
| *nspd-8* | WBGene00021280 | protein_coding_gene | 3.8 | 14.369 | 6.8E-06 | 3.3 | 10.04 | 4.4E-09 |
| *F58F9.8* | WBGene00044306 | protein_coding_gene | 3.8 | 14.096 | 3.2E-07 | 4.0 | 15.79 | 5.6E-20 |
| *F28C6.10* | WBGene00009209 | protein_coding_gene | 3.8 | 14.085 | 1.4E-22 | 3.7 | 12.72 | 2.9E-55 |
| *3R5.2* | WBGene00007066 | pseudogene | 3.8 | 13.968 | 8.4E-06 | 2.2 | 4.46 | 8.2E-06 |
| *K12H6.9* | WBGene00019689 | protein_coding_gene | 3.8 | 13.786 | 4.0E-07 | 4.5 | 22.53 | 1.1E-87 |
| *C16C8.9* | WBGene00015847 | protein_coding_gene | 3.8 | 13.720 | 5.3E-07 | 4.8 | 27.29 | 2.3E-97 |

| | | | | | | | | |
|---|---|---|---|---|---|---|---|---|
| *clec-137* | WBGene00018048 | protein_coding_gene | 3.7 | 13.367 | 5.3E-07 | 5.2 | 35.66 | 1.1E-113 |
| *Y49F6B.13* | WBGene00044377 | protein_coding_gene | 3.7 | 13.362 | 3.1E-05 | 3.2 | 9.08 | 5.2E-07 |
| *F56C3.8* | WBGene00018945 | protein_coding_gene | 3.7 | 13.346 | 3.8E-13 | 2.9 | 7.46 | 1.3E-19 |
| *Y46D2A.3* | WBGene00021591 | protein_coding_gene | 3.7 | 13.336 | 3.2E-05 | 3.9 | 15.23 | 1.0E-11 |
| *F35C5.3* | WBGene00009391 | protein_coding_gene | 3.7 | 13.321 | 3.1E-05 | 3.3 | 10.12 | 8.3E-14 |
| *C35E7.7* | WBGene00016459 | protein_coding_gene | 3.7 | 13.040 | 5.0E-10 | 3.4 | 10.64 | 4.7E-22 |
| *F59A1.6* | WBGene00010292 | protein_coding_gene | 3.7 | 12.959 | 1.5E-05 | 2.8 | 6.79 | 3.9E-07 |
| *C49G7.13* | WBGene00269425 | protein_coding_gene | 3.7 | 12.900 | 1.7E-06 | 3.4 | 10.57 | 2.6E-09 |
| *ZC443.2* | WBGene00013897 | protein_coding_gene | 3.7 | 12.766 | 7.8E-06 | 1.1 | 2.10 | 1.9E-02 |
| *clec-221* | WBGene00008916 | protein_coding_gene | 3.7 | 12.764 | 2.1E-07 | 4.2 | 18.76 | 8.4E-127 |
| *F58A4.1* | WBGene00010226 | protein_coding_gene | 3.7 | 12.705 | 5.5E-16 | 4.0 | 16.32 | 1.1E-59 |
| *D2096.5* | WBGene00017072 | protein_coding_gene | 3.7 | 12.694 | 9.5E-11 | 3.7 | 13.38 | 1.6E-33 |
| *M142.3* | WBGene00014809 | pseudogene | 3.6 | 12.508 | 8.8E-10 | 3.4 | 10.30 | 6.0E-23 |
| *C36C5.14* | WBGene00016483 | protein_coding_gene | 3.6 | 12.367 | 1.6E-06 | 3.4 | 10.21 | 3.7E-09 |
| *W07G1.7* | WBGene00012334 | protein_coding_gene | 3.6 | 12.340 | 2.2E-05 | 2.3 | 4.76 | 3.2E-06 |
| *scl-26* | WBGene00016821 | protein_coding_gene | 3.6 | 12.118 | 6.4E-05 | 4.0 | 15.67 | 2.7E-09 |
| *B0462.1* | WBGene00007181 | pseudogene | 3.6 | 11.855 | 4.5E-06 | 2.2 | 4.62 | 4.1E-08 |
| *C01G5.7* | WBGene00015309 | protein_coding_gene | 3.6 | 11.842 | 1.2E-07 | 2.0 | 3.98 | 1.9E-07 |
| *arrd-11* | WBGene00011052 | protein_coding_gene | 3.6 | 11.839 | 1.6E-07 | 2.7 | 6.71 | 1.9E-10 |
| *fip-3* | WBGene00015520 | protein_coding_gene | 3.6 | 11.776 | 8.0E-05 | 2.5 | 5.80 | 1.2E-07 |
| *F25E5.7* | WBGene00017788 | protein_coding_gene | 3.6 | 11.738 | 1.3E-06 | 3.9 | 14.64 | 1.6E-82 |
| *F45D11.1* | WBGene00018448 | pseudogene | 3.6 | 11.731 | 1.6E-07 | 1.7 | 3.20 | 1.5E-07 |
| *Y37E11AL.4* | WBGene00021360 | pseudogene | 3.5 | 11.708 | 1.1E-13 | 4.5 | 22.68 | 5.6E-32 |
| *Y6E2A.4* | WBGene00012398 | protein_coding_gene | 3.5 | 11.684 | 2.0E-13 | 1.1 | 2.12 | 2.2E-03 |
| *F16G10.15* | WBGene00017529 | pseudogene | 3.5 | 11.550 | 5.6E-15 | 1.3 | 2.43 | 2.7E-04 |
| *clc-4* | WBGene00000525 | protein_coding_gene | 3.5 | 11.545 | 1.4E-10 | 3.5 | 11.45 | 9.8E-22 |
| *T02H6.9* | WBGene00020179 | protein_coding_gene | 3.5 | 11.516 | 5.6E-10 | 3.5 | 11.66 | 1.1E-23 |
| *fbxa-100* | WBGene00008826 | protein_coding_gene | 3.5 | 11.449 | 3.9E-05 | 1.8 | 3.38 | 1.3E-03 |
| *glf-1* | WBGene00019154 | protein_coding_gene | 3.5 | 11.429 | 2.1E-12 | 1.3 | 2.47 | 2.5E-08 |
| *T19H5.6* | WBGene00044807 | protein_coding_gene | 3.5 | 11.144 | 2.1E-10 | 1.0 | 2.04 | 2.5E-03 |
| *Y73F8A.10* | WBGene00013519 | protein_coding_gene | 3.5 | 11.128 | 1.6E-04 | 3.2 | 8.92 | 1.5E-09 |
| *Y102A5C.4* | WBGene00013613 | protein_coding_gene | 3.4 | 10.868 | 8.6E-10 | 4.0 | 16.50 | 1.3E-20 |
| *F41C3.2* | WBGene00018268 | protein_coding_gene | 3.4 | 10.828 | 2.4E-16 | 1.0 | 2.04 | 1.4E-03 |
| *Y75B12A.1* | WBGene00014949 | pseudogene | 3.4 | 10.749 | 1.8E-04 | 2.2 | 4.75 | 4.8E-06 |
| *Y64G10A.2* | WBGene00013412 | protein_coding_gene | 3.4 | 10.719 | 6.3E-05 | 3.2 | 9.00 | 3.8E-07 |
| *clec-136* | WBGene00018047 | protein_coding_gene | 3.4 | 10.672 | 1.4E-09 | 2.7 | 6.49 | 1.6E-18 |
| *W09G12.6* | WBGene00021120 | protein_coding_gene | 3.4 | 10.595 | 6.9E-05 | 1.6 | 3.09 | 7.2E-04 |
| *F46A8.4* | WBGene00009747 | protein_coding_gene | 3.4 | 10.473 | 2.2E-04 | 2.8 | 7.00 | 1.4E-07 |
| *K03B8.14* | WBGene00194783 | protein_coding_gene | 3.4 | 10.417 | 3.8E-09 | 4.0 | 16.32 | 3.3E-14 |
| *ZC178.2* | WBGene00022536 | protein_coding_gene | 3.4 | 10.320 | 8.3E-05 | 3.1 | 8.51 | 5.3E-07 |
| *Y116A8C.44* | WBGene00013815 | protein_coding_gene | 3.4 | 10.275 | 2.8E-07 | 3.7 | 12.81 | 1.1E-16 |
| *fbxa-30* | WBGene00013837 | protein_coding_gene | 3.4 | 10.214 | 1.0E-08 | 1.5 | 2.84 | 1.1E-04 |
| *abf-2* | WBGene00000013 | protein_coding_gene | 3.3 | 10.137 | 1.0E-08 | 1.1 | 2.12 | 3.1E-03 |
| *Y71A12B.24* | WBGene00195228 | protein_coding_gene | 3.3 | 10.131 | 9.3E-05 | 3.0 | 8.21 | 2.7E-08 |
| *Y57E12AL.4* | WBGene00021958 | protein_coding_gene | 3.3 | 10.122 | 9.2E-05 | 2.3 | 5.08 | 1.1E-06 |
| *F17E9.19* | WBGene00249825 | pseudogene | 3.3 | 9.954 | 3.5E-04 | 3.4 | 10.62 | 2.6E-09 |
| *Y82E9BR.8* | WBGene00022341 | protein_coding_gene | 3.3 | 9.940 | 1.1E-04 | 3.3 | 10.01 | 3.7E-09 |
| *pkd-2* | WBGene00004035 | protein_coding_gene | 3.3 | 9.934 | 2.2E-04 | 3.7 | 13.11 | 2.7E-16 |
| *F43G6.17* | WBGene00219378 | protein_coding_gene | 3.3 | 9.930 | 2.7E-04 | 2.6 | 6.02 | 1.3E-05 |
| *oac-4* | WBGene00015873 | protein_coding_gene | 3.3 | 9.917 | 2.2E-05 | 3.0 | 8.19 | 1.2E-06 |
| *fbxa-136* | WBGene00007672 | protein_coding_gene | 3.3 | 9.571 | 3.8E-04 | 2.6 | 5.87 | 3.7E-11 |
| *F21D9.5* | WBGene00009019 | pseudogene | 3.2 | 9.463 | 7.0E-16 | 1.7 | 3.33 | 5.0E-07 |
| *F22E5.7* | WBGene00017706 | protein_coding_gene | 3.2 | 9.442 | 4.5E-04 | 2.5 | 5.78 | 1.0E-06 |
| *R05G6.3* | WBGene00043466 | protein_coding_gene | 3.2 | 9.293 | 1.7E-04 | 2.5 | 5.67 | 2.3E-05 |
| *nspf-1* | WBGene00044364 | protein_coding_gene | 3.2 | 9.240 | 1.8E-04 | 2.0 | 4.06 | 4.8E-05 |
| *W04E12.9* | WBGene00012254 | protein_coding_gene | 3.2 | 9.236 | 3.6E-04 | 1.3 | 2.39 | 6.5E-03 |
| *Y74C10AR.2* | WBGene00022282 | protein_coding_gene | 3.2 | 9.222 | 1.2E-05 | 5.1 | 34.43 | 1.0E-118 |
| *F11C7.6* | WBGene00017378 | protein_coding_gene | 3.2 | 9.210 | 5.0E-06 | 1.8 | 3.56 | 4.3E-16 |
| *Y49F6B.20* | WBGene00303369 | protein_coding_gene | 3.2 | 9.203 | 1.4E-05 | 3.5 | 11.55 | 2.5E-20 |
| *try-5* | WBGene00006623 | protein_coding_gene | 3.2 | 9.108 | 8.4E-09 | 2.9 | 7.21 | 2.8E-17 |
| *F13B6.1* | WBGene00017416 | protein_coding_gene | 3.2 | 9.065 | 1.2E-08 | 3.0 | 8.21 | 3.0E-31 |
| *fbxa-103* | WBGene00008612 | protein_coding_gene | 3.2 | 9.015 | 3.4E-14 | 2.0 | 4.01 | 1.2E-09 |
| *Y37F4.3* | WBGene00021384 | protein_coding_gene | 3.2 | 8.953 | 2.1E-04 | 2.9 | 7.47 | 2.0E-06 |
| *Y18H1A.8* | WBGene00021212 | protein_coding_gene | 3.2 | 8.911 | 9.2E-08 | 3.7 | 13.41 | 1.4E-17 |
| *C17E4.20* | WBGene00219313 | protein_coding_gene | 3.1 | 8.835 | 1.8E-114 | 2.3 | 4.83 | 2.6E-34 |

| | | | | | | | | |
|---|---|---|---|---|---|---|---|---|
| *ZK218.8* | WBGene00013942 | pseudogene | 3.1 | 8.772 | 2.5E-04 | 2.2 | 4.58 | 6.5E-06 |
| *T02E9.6* | WBGene00044203 | protein_coding_gene | 3.1 | 8.767 | 2.5E-04 | 2.8 | 6.74 | 3.1E-08 |
| *C25A6.1* | WBGene00016081 | protein_coding_gene | 3.1 | 8.692 | 1.1E-07 | 1.4 | 2.67 | 4.5E-05 |
| *Y43F8C.16* | WBGene00012837 | protein_coding_gene | 3.1 | 8.677 | 8.0E-04 | 3.1 | 8.68 | 6.7E-11 |
| *C36C5.15* | WBGene00016484 | protein_coding_gene | 3.1 | 8.668 | 7.7E-04 | 2.0 | 3.95 | 2.5E-05 |
| *nep-25* | WBGene00022874 | protein_coding_gene | 3.1 | 8.649 | 1.1E-11 | 4.1 | 16.61 | 2.1E-26 |
| *K12H6.4* | WBGene00019684 | protein_coding_gene | 3.1 | 8.645 | 3.1E-05 | 4.4 | 21.32 | 1.9E-85 |
| *fipr-19* | WBGene00044451 | protein_coding_gene | 3.1 | 8.434 | 9.3E-04 | 1.9 | 3.85 | 2.3E-05 |
| *clec-99* | WBGene00013933 | protein_coding_gene | 3.1 | 8.315 | 7.6E-05 | 5.0 | 31.48 | 7.6E-104 |
| *T11F9.22* | WBGene00194784 | protein_coding_gene | 3.1 | 8.302 | 9.9E-04 | 1.9 | 3.79 | 1.8E-05 |
| *ZC204.6* | WBGene00022556 | protein_coding_gene | 3.1 | 8.291 | 3.6E-04 | 2.3 | 4.78 | 1.0E-04 |
| *ttr-29* | WBGene00011286 | protein_coding_gene | 3.0 | 8.202 | 3.8E-10 | 1.0 | 2.05 | 1.7E-03 |
| *skr-11* | WBGene00004817 | pseudogene | 3.0 | 8.196 | 1.1E-36 | 3.2 | 9.10 | 4.1E-63 |
| *clec-162* | WBGene00007399 | protein_coding_gene | 3.0 | 8.172 | 3.6E-04 | 2.6 | 5.87 | 7.2E-10 |
| *Y116F11A.3* | WBGene00013817 | protein_coding_gene | 3.0 | 8.011 | 4.6E-04 | 2.9 | 7.58 | 1.7E-06 |
| *C08E3.14* | WBGene00044415 | protein_coding_gene | 3.0 | 8.010 | 4.6E-04 | 2.1 | 4.27 | 2.4E-04 |
| *C16C8.8* | WBGene00015846 | protein_coding_gene | 3.0 | 7.956 | 3.6E-04 | 3.1 | 8.51 | 3.1E-08 |
| *T16H12.1* | WBGene00011811 | protein_coding_gene | 3.0 | 7.949 | 4.8E-57 | 2.8 | 6.76 | 7.2E-46 |
| *2RSSE.3* | WBGene00044951 | pseudogene | 3.0 | 7.932 | 4.8E-04 | 1.4 | 2.56 | 1.1E-03 |
| *Y105C5B.7* | WBGene00013648 | pseudogene | 3.0 | 7.884 | 4.7E-04 | 2.2 | 4.45 | 2.0E-08 |
| *Y59E9AR.1* | WBGene00021997 | protein_coding_gene | 3.0 | 7.839 | 5.0E-04 | 2.1 | 4.41 | 4.0E-06 |
| *Y39A3A.4* | WBGene00021435 | protein_coding_gene | 3.0 | 7.836 | 3.9E-07 | 1.9 | 3.81 | 1.1E-07 |
| *dct-10* | WBGene00012618 | pseudogene | 3.0 | 7.818 | 2.7E-04 | 1.7 | 3.36 | 8.4E-04 |
| *F17E9.18* | WBGene00235282 | protein_coding_gene | 3.0 | 7.806 | 5.2E-04 | 2.2 | 4.53 | 1.4E-04 |
| *Y66H1A.8* | WBGene00235263 | protein_coding_gene | 3.0 | 7.785 | 1.5E-05 | 1.8 | 3.51 | 2.8E-04 |
| *W06G6.11* | WBGene00012313 | protein_coding_gene | 3.0 | 7.737 | 5.4E-04 | 1.7 | 3.22 | 1.4E-04 |
| *nspf-2* | WBGene00044365 | protein_coding_gene | 2.9 | 7.716 | 1.1E-03 | 2.3 | 5.05 | 1.2E-06 |
| *nlp-45* | WBGene00020136 | protein_coding_gene | 2.9 | 7.706 | 7.0E-06 | 2.2 | 4.53 | 7.7E-11 |
| *W01G7.2* | WBGene00014851 | pseudogene | 2.9 | 7.703 | 5.9E-04 | 2.9 | 7.51 | 6.1E-08 |
| *Y17G7B.23* | WBGene00044030 | protein_coding_gene | 2.9 | 7.689 | 2.8E-04 | 2.8 | 7.10 | 3.7E-10 |
| *K07A1.6* | WBGene00010613 | protein_coding_gene | 2.9 | 7.631 | 1.5E-82 | 1.8 | 3.49 | 5.3E-54 |
| *try-9* | WBGene00023425 | protein_coding_gene | 2.9 | 7.631 | 4.4E-08 | 1.6 | 3.03 | 4.7E-09 |
| *F46F5.1* | WBGene00018493 | protein_coding_gene | 2.9 | 7.594 | 4.3E-05 | 2.0 | 4.05 | 4.8E-08 |
| *clec-207* | WBGene00018096 | protein_coding_gene | 2.9 | 7.582 | 7.0E-05 | 4.8 | 27.57 | 2.5E-101 |
| *F26C11.3* | WBGene00009147 | protein_coding_gene | 2.9 | 7.569 | 6.3E-04 | 2.4 | 5.40 | 1.7E-07 |
| *K12H6.8* | WBGene00019688 | protein_coding_gene | 2.9 | 7.567 | 5.5E-05 | 3.0 | 7.87 | 1.5E-17 |
| *F32B5.3* | WBGene00017977 | protein_coding_gene | 2.9 | 7.536 | 7.1E-05 | 2.8 | 6.99 | 3.8E-09 |
| *clec-217* | WBGene00017824 | protein_coding_gene | 2.9 | 7.399 | 2.6E-04 | 4.6 | 24.98 | 3.7E-51 |
| *F49F1.2* | WBGene00235363 | protein_coding_gene | 2.9 | 7.397 | 6.7E-04 | 2.5 | 5.64 | 8.4E-07 |
| *M01D1.11* | WBGene00220026 | snoRNA_gene | 2.9 | 7.363 | 2.0E-03 | 1.1 | 2.11 | 3.3E-02 |
| *srab-25* | WBGene00022807 | protein_coding_gene | 2.9 | 7.340 | 8.0E-04 | 1.5 | 2.86 | 4.3E-03 |
| *C41G7.15* | WBGene00219284 | protein_coding_gene | 2.9 | 7.329 | 8.0E-04 | 2.4 | 5.30 | 4.5E-05 |
| *str-163* | WBGene00006208 | protein_coding_gene | 2.9 | 7.262 | 1.4E-04 | 1.1 | 2.09 | 3.2E-02 |
| *Y51A2D.8* | WBGene00013076 | protein_coding_gene | 2.9 | 7.236 | 2.6E-04 | 4.8 | 27.08 | 4.3E-156 |
| *F55B11.5* | WBGene00010087 | protein_coding_gene | 2.9 | 7.223 | 5.6E-07 | 1.0 | 2.01 | 9.9E-03 |
| *F42G2.7* | WBGene00044362 | protein_coding_gene | 2.8 | 7.170 | 4.4E-04 | 2.6 | 6.17 | 3.7E-08 |
| *F21D9.11* | WBGene00219452 | protein_coding_gene | 2.8 | 7.157 | 3.9E-09 | 2.0 | 4.06 | 1.3E-07 |
| *F25E5.4* | WBGene00017785 | protein_coding_gene | 2.8 | 7.123 | 2.2E-31 | 2.4 | 5.26 | 2.2E-35 |
| *fip-7* | WBGene00013105 | protein_coding_gene | 2.8 | 6.997 | 2.2E-03 | 2.6 | 5.95 | 9.0E-06 |
| *C04F12.6* | WBGene00007300 | pseudogene | 2.8 | 6.883 | 2.6E-06 | 3.5 | 11.04 | 6.2E-19 |
| *clec-96* | WBGene00013930 | protein_coding_gene | 2.8 | 6.862 | 3.1E-04 | 5.0 | 32.27 | 8.2E-38 |
| *T02D1.7* | WBGene00011373 | protein_coding_gene | 2.8 | 6.800 | 1.2E-03 | 2.9 | 7.22 | 4.3E-08 |
| *F25D7.5* | WBGene00009114 | protein_coding_gene | 2.8 | 6.792 | 2.9E-03 | 2.4 | 5.15 | 1.7E-08 |
| *M01G12.7* | WBGene00010821 | protein_coding_gene | 2.8 | 6.768 | 1.2E-03 | 2.4 | 5.44 | 1.4E-06 |
| *abf-1* | WBGene00000012 | protein_coding_gene | 2.8 | 6.743 | 5.5E-07 | 3.0 | 8.08 | 3.4E-18 |
| *C16C8.10* | WBGene00015848 | protein_coding_gene | 2.8 | 6.738 | 2.6E-03 | 1.7 | 3.27 | 6.8E-04 |
| *irg-6* | WBGene00007864 | protein_coding_gene | 2.8 | 6.730 | 2.7E-04 | 2.4 | 5.15 | 9.0E-07 |
| *clec-260* | WBGene00013357 | protein_coding_gene | 2.7 | 6.715 | 5.6E-06 | 2.8 | 6.88 | 2.3E-14 |
| *F28A10.5* | WBGene00017873 | protein_coding_gene | 2.7 | 6.647 | 5.8E-05 | 4.1 | 16.81 | 2.1E-11 |
| *B0310.3* | WBGene00015139 | protein_coding_gene | 2.7 | 6.595 | 7.0E-09 | 1.2 | 2.31 | 7.2E-04 |
| *C17E4.1* | WBGene00007641 | protein_coding_gene | 2.7 | 6.542 | 3.6E-10 | 2.2 | 4.74 | 5.7E-20 |
| *clpr-1* | WBGene00012233 | protein_coding_gene | 2.7 | 6.525 | 3.9E-03 | 2.5 | 5.79 | 1.8E-05 |
| *fbxa-217* | WBGene00012954 | pseudogene | 2.7 | 6.511 | 1.2E-31 | 2.1 | 4.26 | 2.4E-26 |
| *clec-42* | WBGene00008891 | protein_coding_gene | 2.7 | 6.486 | 2.3E-03 | 1.6 | 3.02 | 4.8E-05 |
| *F16H6.9* | WBGene00008899 | protein_coding_gene | 2.7 | 6.444 | 1.6E-06 | 3.3 | 9.85 | 7.5E-23 |

| | | | | | | | | |
|---|---|---|---|---|---|---|---|---|
| *cwp-2* | WBGene00000860 | protein_coding_gene | 2.7 | 6.441 | 1.7E-03 | 2.6 | 5.94 | 2.3E-06 |
| *R09D1.9* | WBGene00011165 | pseudogene | 2.7 | 6.428 | 8.5E-12 | 2.5 | 5.80 | 1.4E-13 |
| *K10C2.14* | WBGene00206388 | protein_coding_gene | 2.7 | 6.401 | 1.0E-12 | 2.5 | 5.72 | 4.8E-26 |
| *clec-129* | WBGene00021145 | protein_coding_gene | 2.7 | 6.392 | 4.0E-03 | 1.6 | 2.97 | 1.3E-03 |
| *Y41C4A.21* | WBGene00194733 | protein_coding_gene | 2.7 | 6.367 | 1.3E-03 | 2.2 | 4.48 | 2.1E-05 |
| *clec-100* | WBGene00013179 | protein_coding_gene | 2.7 | 6.289 | 2.1E-03 | 4.6 | 23.95 | 5.3E-13 |
| *dmsr-12* | WBGene00019262 | protein_coding_gene | 2.6 | 6.218 | 1.6E-06 | 1.3 | 2.40 | 4.3E-05 |
| *C35B1.8* | WBGene00016437 | protein_coding_gene | 2.6 | 6.149 | 1.5E-05 | 2.6 | 6.26 | 5.9E-16 |
| *F10D2.8* | WBGene00017334 | protein_coding_gene | 2.6 | 6.149 | 3.8E-06 | 2.7 | 6.62 | 2.2E-27 |
| *Y57G11C.1132* | WBGene00194666 | antisense_lncRNA_gene | 2.6 | 6.122 | 2.2E-03 | 2.4 | 5.40 | 1.5E-06 |
| *C10G8.2* | WBGene00015681 | protein_coding_gene | 2.6 | 6.101 | 2.4E-03 | 1.9 | 3.77 | 3.8E-04 |
| *C31B8.12* | WBGene00016285 | protein_coding_gene | 2.6 | 6.086 | 6.1E-04 | 3.4 | 10.51 | 1.1E-16 |
| *C40A11.3* | WBGene00016546 | protein_coding_gene | 2.6 | 6.062 | 2.2E-08 | 1.2 | 2.36 | 6.2E-05 |
| *mboa-1* | WBGene00007174 | protein_coding_gene | 2.6 | 6.060 | 1.7E-06 | 1.0 | 2.06 | 5.4E-03 |
| *Y82E9BR.1* | WBGene00022334 | protein_coding_gene | 2.6 | 6.049 | 3.5E-31 | 1.8 | 3.54 | 2.0E-22 |
| *C16C8.17* | WBGene00015854 | protein_coding_gene | 2.6 | 6.047 | 2.6E-03 | 2.4 | 5.28 | 7.1E-07 |
| *ttr-22* | WBGene00011450 | protein_coding_gene | 2.6 | 6.046 | 4.3E-03 | 1.4 | 2.73 | 4.6E-03 |
| *F13E9.4* | WBGene00008753 | protein_coding_gene | 2.6 | 6.039 | 2.5E-03 | 2.5 | 5.59 | 2.6E-05 |
| *C17B7.2* | WBGene00015876 | pseudogene | 2.6 | 6.017 | 5.3E-03 | 1.0 | 2.00 | 4.1E-02 |
| *Y54G2A.24* | WBGene00021889 | pseudogene | 2.6 | 6.014 | 7.3E-09 | 3.1 | 8.46 | 4.4E-20 |
| *K11D12.6* | WBGene00019646 | protein_coding_gene | 2.6 | 5.972 | 6.8E-04 | 4.8 | 28.18 | 1.4E-85 |
| *ppt-1* | WBGene00004092 | protein_coding_gene | 2.6 | 5.918 | 7.2E-27 | 2.0 | 3.93 | 6.1E-23 |
| *scl-18* | WBGene00011724 | protein_coding_gene | 2.6 | 5.910 | 2.7E-03 | 2.1 | 4.27 | 2.5E-04 |
| *F59A1.16* | WBGene00010301 | protein_coding_gene | 2.6 | 5.866 | 5.1E-03 | 2.6 | 5.93 | 1.3E-05 |
| *C06E1.11* | WBGene00015526 | protein_coding_gene | 2.6 | 5.863 | 1.1E-13 | 1.9 | 3.70 | 6.2E-11 |
| *math-47* | WBGene00022688 | pseudogene | 2.5 | 5.840 | 1.2E-03 | 2.7 | 6.62 | 1.7E-09 |
| *lov-1* | WBGene00003058 | protein_coding_gene | 2.5 | 5.836 | 5.3E-03 | 2.9 | 7.55 | 1.4E-07 |
| *fbxc-49* | WBGene00020534 | protein_coding_gene | 2.5 | 5.820 | 2.6E-04 | 2.2 | 4.69 | 2.1E-09 |
| *R09A8.9* | WBGene00219278 | protein_coding_gene | 2.5 | 5.818 | 2.3E-06 | 2.0 | 4.03 | 1.2E-08 |
| *linc-140* | WBGene00219596 | lincRNA_gene | 2.5 | 5.815 | 6.1E-03 | 1.7 | 3.21 | 3.4E-04 |
| *Y47D7A.12* | WBGene00021624 | protein_coding_gene | 2.5 | 5.743 | 3.7E-03 | 1.9 | 3.67 | 9.3E-07 |
| *R05H10.1* | WBGene00011042 | protein_coding_gene | 2.5 | 5.733 | 1.9E-13 | 1.7 | 3.24 | 7.8E-11 |
| *F25B3.2* | WBGene00009099 | protein_coding_gene | 2.5 | 5.728 | 8.1E-04 | 2.0 | 3.92 | 5.4E-05 |
| *tyr-5* | WBGene00021553 | protein_coding_gene | 2.5 | 5.726 | 1.9E-10 | 3.3 | 9.99 | 1.4E-39 |
| *C04F12.12* | WBGene00077685 | protein_coding_gene | 2.5 | 5.691 | 6.5E-11 | 2.7 | 6.37 | 5.0E-21 |
| *ZK783.3* | WBGene00022818 | pseudogene | 2.5 | 5.683 | 2.0E-03 | 2.0 | 3.97 | 2.8E-06 |
| *wrt-1* | WBGene00006947 | protein_coding_gene | 2.5 | 5.631 | 1.7E-05 | 1.1 | 2.13 | 9.7E-07 |
| *R10D12.18* | WBGene00194678 | ncRNA_gene | 2.5 | 5.610 | 2.0E-04 | 2.6 | 6.11 | 2.6E-11 |
| *W03D8.10* | WBGene00020991 | pseudogene | 2.5 | 5.604 | 2.2E-08 | 3.1 | 8.37 | 1.5E-27 |
| *bath-10* | WBGene00021726 | protein_coding_gene | 2.5 | 5.593 | 1.1E-19 | 1.3 | 2.53 | 7.2E-07 |
| *Y47D7A.11* | WBGene00021623 | protein_coding_gene | 2.5 | 5.586 | 7.8E-03 | 2.1 | 4.26 | 1.0E-05 |
| *fbxa-113* | WBGene00013757 | protein_coding_gene | 2.5 | 5.569 | 1.1E-05 | 2.0 | 3.92 | 1.1E-07 |
| *Y6G8.16* | WBGene00194801 | protein_coding_gene | 2.5 | 5.536 | 3.0E-03 | 2.0 | 3.90 | 1.9E-07 |
| *F54D12.9* | WBGene00018822 | protein_coding_gene | 2.5 | 5.496 | 4.9E-04 | 5.1 | 33.24 | 7.9E-76 |
| *C44B12.9* | WBGene00044462 | protein_coding_gene | 2.4 | 5.463 | 7.8E-07 | 3.6 | 11.83 | 8.1E-20 |
| *fbxa-172* | WBGene00008014 | protein_coding_gene | 2.4 | 5.448 | 1.3E-04 | 1.5 | 2.91 | 1.1E-07 |
| *C05B5.11* | WBGene00045242 | protein_coding_gene | 2.4 | 5.433 | 1.7E-03 | 4.6 | 24.62 | 2.5E-69 |
| *C06A12.8* | WBGene00007359 | protein_coding_gene | 2.4 | 5.393 | 7.8E-03 | 2.1 | 4.41 | 1.2E-04 |
| *M7.9* | WBGene00010885 | protein_coding_gene | 2.4 | 5.372 | 1.5E-03 | 5.2 | 36.79 | 9.9E-134 |
| *F46B3.15* | WBGene00009766 | protein_coding_gene | 2.4 | 5.354 | 4.8E-03 | 2.9 | 7.30 | 2.4E-08 |
| *F58F9.9* | WBGene00044307 | protein_coding_gene | 2.4 | 5.321 | 5.0E-03 | 3.5 | 11.18 | 1.4E-09 |
| *Y71F9B.9* | WBGene00022130 | protein_coding_gene | 2.4 | 5.280 | 7.6E-12 | 1.1 | 2.09 | 2.3E-06 |
| *clec-242* | WBGene00013471 | protein_coding_gene | 2.4 | 5.249 | 1.0E-02 | 2.3 | 4.89 | 3.7E-06 |
| *nas-17* | WBGene00003536 | protein_coding_gene | 2.4 | 5.225 | 2.7E-03 | 5.2 | 36.67 | 1.7E-33 |
| *C33G3.5* | WBGene00007905 | protein_coding_gene | 2.4 | 5.219 | 9.9E-04 | 5.2 | 35.98 | 2.0E-63 |
| *C43D7.11* | WBGene00077664 | pseudogene | 2.4 | 5.217 | 1.0E-05 | 1.6 | 3.13 | 1.9E-04 |
| *T26F2.2* | WBGene00012058 | pseudogene | 2.4 | 5.197 | 5.6E-13 | 2.4 | 5.44 | 3.9E-27 |
| *sknr-1* | WBGene00020961 | protein_coding_gene | 2.4 | 5.161 | 1.1E-02 | 1.8 | 3.45 | 5.0E-04 |
| *F46B3.14* | WBGene00009765 | protein_coding_gene | 2.4 | 5.142 | 6.0E-03 | 2.8 | 7.07 | 8.6E-09 |
| *trpl-5* | WBGene00004149 | protein_coding_gene | 2.4 | 5.109 | 7.6E-04 | 3.0 | 7.73 | 1.1E-07 |
| *C44B12.3* | WBGene00016637 | protein_coding_gene | 2.3 | 5.054 | 5.9E-06 | 2.6 | 6.17 | 1.0E-14 |
| *F57A8.7* | WBGene00010182 | protein_coding_gene | 2.3 | 5.005 | 9.5E-03 | 1.0 | 2.06 | 2.1E-02 |
| *T22B7.3* | WBGene00020672 | protein_coding_gene | 2.3 | 4.984 | 4.8E-06 | 1.2 | 2.28 | 3.1E-03 |
| *K10G4.5* | WBGene00010754 | protein_coding_gene | 2.3 | 4.984 | 1.5E-02 | 1.2 | 2.32 | 1.3E-02 |
| *F18E9.7* | WBGene00017573 | protein_coding_gene | 2.3 | 4.979 | 1.7E-03 | 5.1 | 33.84 | 3.5E-92 |

| | | | | | | | | |
|---|---|---|---|---|---|---|---|---|
| *ssp-19* | WBGene00006047 | protein_coding_gene | 2.3 | 4.975 | 2.2E-06 | 3.4 | 10.50 | 1.7E-42 |
| *scl-5* | WBGene00008027 | protein_coding_gene | 2.3 | 4.954 | 5.6E-09 | 1.2 | 2.34 | 2.2E-12 |
| *F30A10.2* | WBGene00009261 | protein_coding_gene | 2.3 | 4.940 | 3.0E-04 | 1.1 | 2.07 | 2.2E-03 |
| *Y6E2A.1* | WBGene00012397 | protein_coding_gene | 2.3 | 4.934 | 7.0E-03 | 1.0 | 2.04 | 7.2E-03 |
| *C35D6.5* | WBGene00007962 | pseudogene | 2.3 | 4.926 | 3.0E-03 | 1.1 | 2.15 | 1.1E-03 |
| *Y46D2A.1* | WBGene00021589 | protein_coding_gene | 2.3 | 4.921 | 8.2E-04 | 2.0 | 4.08 | 4.9E-07 |
| *nhr-246* | WBGene00014192 | protein_coding_gene | 2.3 | 4.876 | 2.4E-03 | 1.2 | 2.30 | 1.7E-02 |
| *C50F2.5* | WBGene00016839 | protein_coding_gene | 2.3 | 4.861 | 3.5E-05 | 1.5 | 2.92 | 3.1E-06 |
| *Y82E9BL.19* | WBGene00194735 | protein_coding_gene | 2.3 | 4.834 | 1.1E-02 | 1.3 | 2.54 | 1.0E-02 |
| *W09C3.1* | WBGene00021109 | protein_coding_gene | 2.3 | 4.820 | 1.1E-02 | 1.4 | 2.64 | 4.1E-03 |
| *ZK666.14* | WBGene00194952 | protein_coding_gene | 2.3 | 4.777 | 1.6E-02 | 2.2 | 4.63 | 4.8E-06 |
| *clec-95* | WBGene00013927 | protein_coding_gene | 2.3 | 4.762 | 3.1E-04 | 5.2 | 37.63 | 1.3E-114 |
| *ins-31* | WBGene00002114 | protein_coding_gene | 2.2 | 4.701 | 7.9E-04 | 5.1 | 34.29 | 2.8E-194 |
| *Y39B6A.21* | WBGene00012680 | pseudogene | 2.2 | 4.689 | 2.1E-03 | 3.4 | 10.85 | 3.9E-82 |
| *W02A2.4* | WBGene00012189 | pseudogene | 2.2 | 4.670 | 4.8E-04 | 2.2 | 4.44 | 1.5E-07 |
| *R11E3.19* | WBGene00235374 | protein_coding_gene | 2.2 | 4.659 | 8.4E-04 | 1.6 | 3.04 | 3.0E-04 |
| *F20B6.1* | WBGene00017627 | pseudogene | 2.2 | 4.642 | 1.0E-02 | 1.2 | 2.37 | 7.4E-03 |
| *B0348.2* | WBGene00015152 | protein_coding_gene | 2.2 | 4.641 | 6.2E-03 | 2.0 | 4.07 | 6.3E-07 |
| *gasr-8* | WBGene00008144 | protein_coding_gene | 2.2 | 4.636 | 4.8E-03 | 1.3 | 2.47 | 4.8E-03 |
| *pals-7* | WBGene00007661 | protein_coding_gene | 2.2 | 4.613 | 1.0E-02 | 1.1 | 2.11 | 3.3E-02 |
| *F59A7.2* | WBGene00019090 | protein_coding_gene | 2.2 | 4.575 | 2.0E-07 | 1.1 | 2.19 | 4.0E-03 |
| *W05F2.8* | WBGene00195045 | protein_coding_gene | 2.2 | 4.536 | 7.5E-03 | 1.7 | 3.19 | 7.8E-04 |
| *clec-170* | WBGene00009515 | protein_coding_gene | 2.2 | 4.522 | 9.0E-12 | 3.2 | 9.16 | 1.8E-69 |
| *K07A1.3* | WBGene00010610 | protein_coding_gene | 2.2 | 4.509 | 4.5E-16 | 1.4 | 2.63 | 2.0E-14 |
| *fbxa-176* | WBGene00008017 | protein_coding_gene | 2.2 | 4.480 | 1.1E-03 | 1.5 | 2.76 | 4.7E-05 |
| *C17H12.37* | WBGene00236786 | protein_coding_gene | 2.1 | 4.391 | 2.4E-02 | 2.8 | 6.76 | 4.6E-06 |
| *clec-98* | WBGene00013932 | protein_coding_gene | 2.1 | 4.386 | 1.7E-02 | 2.4 | 5.31 | 1.5E-06 |
| *fbxa-87* | WBGene00044660 | pseudogene | 2.1 | 4.377 | 5.9E-07 | 1.2 | 2.26 | 8.4E-05 |
| *W03G1.5* | WBGene00021011 | protein_coding_gene | 2.1 | 4.369 | 6.3E-04 | 1.4 | 2.61 | 1.8E-05 |
| *F10E7.3* | WBGene00017345 | pseudogene | 2.1 | 4.367 | 2.4E-02 | 1.7 | 3.30 | 9.4E-04 |
| *eol-1* | WBGene00045401 | protein_coding_gene | 2.1 | 4.362 | 2.0E-03 | 1.7 | 3.16 | 3.3E-07 |
| *R155.4* | WBGene00020118 | protein_coding_gene | 2.1 | 4.347 | 9.8E-03 | 2.0 | 4.05 | 8.0E-07 |
| *clec-28* | WBGene00009857 | protein_coding_gene | 2.1 | 4.344 | 2.5E-03 | 1.5 | 2.83 | 2.6E-04 |
| *T10D4.7* | WBGene00020409 | protein_coding_gene | 2.1 | 4.342 | 2.0E-02 | 1.4 | 2.59 | 6.0E-03 |
| *C17E4.14* | WBGene00195884 | ncRNA_gene | 2.1 | 4.336 | 2.4E-02 | 1.2 | 2.31 | 1.0E-02 |
| *F58A6.9* | WBGene00019026 | protein_coding_gene | 2.1 | 4.330 | 7.1E-08 | 2.1 | 4.28 | 4.4E-09 |
| *fbxa-160* | WBGene00015604 | protein_coding_gene | 2.1 | 4.323 | 1.4E-02 | 1.3 | 2.42 | 1.8E-03 |
| *W08G11.1* | WBGene00012346 | protein_coding_gene | 2.1 | 4.319 | 7.0E-03 | 1.1 | 2.08 | 1.3E-02 |
| *pph-2* | WBGene00004084 | protein_coding_gene | 2.1 | 4.318 | 1.4E-03 | 1.8 | 3.52 | 8.5E-08 |
| *F57G9.6* | WBGene00044041 | pseudogene | 2.1 | 4.291 | 1.7E-05 | 1.6 | 3.11 | 2.2E-07 |
| *F53C3.8* | WBGene00018752 | protein_coding_gene | 2.1 | 4.291 | 1.9E-03 | 1.3 | 2.45 | 1.7E-03 |
| *F22B7.3* | WBGene00017694 | protein_coding_gene | 2.1 | 4.278 | 1.5E-02 | 1.1 | 2.12 | 5.1E-03 |
| *F59A6.3* | WBGene00019085 | protein_coding_gene | 2.1 | 4.278 | 1.5E-02 | 2.8 | 6.89 | 4.8E-07 |
| *F44E5.3* | WBGene00009690 | protein_coding_gene | 2.1 | 4.265 | 4.4E-04 | 1.5 | 2.80 | 3.4E-04 |
| *T04B2.1* | WBGene00011421 | pseudogene | 2.1 | 4.241 | 2.3E-02 | 1.7 | 3.27 | 1.2E-03 |
| *Y47D7A.7* | WBGene00021620 | protein_coding_gene | 2.1 | 4.234 | 6.3E-03 | 2.2 | 4.57 | 6.3E-09 |
| *C46A5.4* | WBGene00016700 | protein_coding_gene | 2.1 | 4.209 | 3.5E-03 | 1.1 | 2.20 | 2.2E-04 |
| *F57A10.2* | WBGene00010184 | protein_coding_gene | 2.1 | 4.186 | 3.1E-03 | 2.2 | 4.55 | 1.1E-06 |
| *T05H10.3* | WBGene00011508 | protein_coding_gene | 2.1 | 4.162 | 1.4E-02 | 1.3 | 2.40 | 2.5E-10 |
| *C49C8.6* | WBGene00016770 | protein_coding_gene | 2.1 | 4.143 | 1.7E-02 | 1.8 | 3.54 | 1.2E-04 |
| *clec-183* | WBGene00011855 | protein_coding_gene | 2.0 | 4.140 | 2.2E-02 | 1.6 | 3.14 | 1.9E-03 |
| *clec-197* | WBGene00008202 | protein_coding_gene | 2.0 | 4.138 | 3.9E-03 | 4.8 | 28.56 | 6.3E-169 |
| *cal-6* | WBGene00022214 | pseudogene | 2.0 | 4.134 | 2.9E-02 | 2.2 | 4.50 | 5.0E-05 |
| *math-2* | WBGene00015606 | protein_coding_gene | 2.0 | 4.122 | 2.3E-02 | 3.0 | 7.92 | 1.3E-06 |
| *B0228.8* | WBGene00015065 | protein_coding_gene | 2.0 | 4.122 | 2.9E-02 | 2.1 | 4.17 | 2.9E-04 |
| *F20A1.8* | WBGene00017619 | protein_coding_gene | 2.0 | 4.118 | 2.8E-02 | 1.9 | 3.70 | 3.7E-04 |
| *F21D9.1* | WBGene00009015 | protein_coding_gene | 2.0 | 4.078 | 9.9E-04 | 1.3 | 2.41 | 6.4E-03 |
| *lips-16* | WBGene00012588 | pseudogene | 2.0 | 4.060 | 1.2E-10 | 1.6 | 3.02 | 5.7E-12 |
| *K11H12.5* | WBGene00019661 | protein_coding_gene | 2.0 | 4.024 | 3.2E-06 | 1.5 | 2.86 | 5.7E-09 |
| *mec-8* | WBGene00003172 | protein_coding_gene | 2.0 | 4.002 | 3.9E-41 | 1.2 | 2.36 | 6.9E-16 |
| *C31B8.9* | WBGene00016284 | protein_coding_gene | 2.0 | 4.002 | 3.2E-02 | 1.7 | 3.29 | 1.8E-03 |
| *F28B1.2* | WBGene00009196 | protein_coding_gene | 2.0 | 3.999 | 1.8E-02 | 1.7 | 3.36 | 1.7E-04 |
| *Y62H9A.16* | WBGene00304791 | protein_coding_gene | 2.0 | 3.999 | 4.3E-03 | 1.3 | 2.44 | 3.6E-03 |
| *F59B2.12* | WBGene00010314 | protein_coding_gene | 2.0 | 3.992 | 2.7E-03 | 5.1 | 33.41 | 0.0E+00 |
| *F26F2.11* | WBGene00195158 | protein_coding_gene | 2.0 | 3.986 | 3.4E-02 | 1.3 | 2.46 | 7.8E-03 |

| | | | | | | | | |
|---|---|---|---|---|---|---|---|---|
| *F28B1.3* | WBGene00009197 | pseudogene | 2.0 | 3.980 | 2.1E-02 | 1.6 | 2.95 | 2.4E-03 |
| *F49C5.10* | WBGene00044162 | protein_coding_gene | 2.0 | 3.978 | 2.0E-02 | 1.5 | 2.78 | 3.3E-03 |
| *Y57E12B.10* | WBGene00202501 | protein_coding_gene | 2.0 | 3.962 | 2.6E-02 | 1.2 | 2.26 | 2.0E-02 |
| *R13A1.1* | WBGene00020041 | protein_coding_gene | 2.0 | 3.948 | 2.2E-02 | 1.7 | 3.18 | 1.0E-03 |
| *Y20C6A.1* | WBGene00012491 | protein_coding_gene | 2.0 | 3.948 | 2.4E-11 | 1.0 | 2.03 | 9.2E-08 |
| *Y39B6A.31* | WBGene00012690 | pseudogene | 2.0 | 3.942 | 4.3E-04 | 3.4 | 10.44 | 3.1E-30 |
| *ZK402.5* | WBGene00022731 | protein_coding_gene | 2.0 | 3.924 | 2.7E-02 | 3.2 | 9.30 | 2.8E-07 |
| *C27A2.8* | WBGene00044387 | protein_coding_gene | 2.0 | 3.906 | 3.1E-02 | 1.2 | 2.25 | 1.3E-02 |
| *K08C9.10* | WBGene00086564 | protein_coding_gene | 2.0 | 3.891 | 3.6E-02 | 2.3 | 4.83 | 9.3E-05 |
| *B0334.10* | WBGene00007148 | protein_coding_gene | 1.9 | 3.863 | 2.9E-02 | 2.3 | 5.02 | 1.5E-05 |
| *M195.4* | WBGene00010949 | protein_coding_gene | 1.9 | 3.842 | 2.5E-02 | 2.2 | 4.73 | 5.3E-05 |
| *R11G10.4* | WBGene00050968 | protein_coding_gene | 1.9 | 3.832 | 1.5E-02 | 2.3 | 5.04 | 2.6E-06 |
| *tig-3* | WBGene00021594 | protein_coding_gene | 1.9 | 3.824 | 1.3E-03 | 1.4 | 2.58 | 3.1E-03 |
| *K08E5.1* | WBGene00014801 | protein_coding_gene | 1.9 | 3.809 | 1.0E-07 | 2.1 | 4.19 | 8.9E-13 |
| *oac-43* | WBGene00011653 | protein_coding_gene | 1.9 | 3.752 | 4.2E-02 | 1.2 | 2.22 | 1.4E-02 |
| *B0238.13* | WBGene00015077 | pseudogene | 1.9 | 3.746 | 4.1E-02 | 1.0 | 2.00 | 4.6E-02 |
| *T12A2.6* | WBGene00020440 | protein_coding_gene | 1.9 | 3.739 | 2.8E-02 | 1.4 | 2.57 | 9.1E-03 |
| *T22G5.1* | WBGene00011929 | protein_coding_gene | 1.9 | 3.723 | 6.2E-03 | 4.9 | 28.99 | 3.2E-57 |
| *C08G5.1* | WBGene00015614 | pseudogene | 1.9 | 3.714 | 7.2E-03 | 3.3 | 9.96 | 2.2E-10 |
| *Y47H9C.12* | WBGene00012955 | protein_coding_gene | 1.9 | 3.694 | 2.8E-02 | 1.4 | 2.73 | 4.3E-03 |
| *T27A1.2* | WBGene00020834 | protein_coding_gene | 1.9 | 3.670 | 1.3E-07 | 1.3 | 2.48 | 1.4E-09 |
| *Y71G12B.5* | WBGene00022145 | protein_coding_gene | 1.9 | 3.665 | 8.0E-03 | 2.5 | 5.62 | 7.8E-11 |
| *R10D12.10* | WBGene00011191 | protein_coding_gene | 1.9 | 3.645 | 1.9E-05 | 2.0 | 3.90 | 1.0E-14 |
| *F20A1.6* | WBGene00017618 | protein_coding_gene | 1.9 | 3.633 | 3.1E-09 | 1.5 | 2.87 | 2.8E-15 |
| *T21E12.5* | WBGene00020656 | protein_coding_gene | 1.9 | 3.611 | 1.5E-02 | 1.4 | 2.68 | 1.6E-03 |
| *dod-23* | WBGene00009898 | protein_coding_gene | 1.9 | 3.606 | 2.1E-13 | 1.2 | 2.22 | 5.0E-30 |
| *fbxa-182* | WBGene00018353 | protein_coding_gene | 1.8 | 3.603 | 2.9E-07 | 2.4 | 5.24 | 1.7E-33 |
| *Y75B12B.13* | WBGene00045402 | protein_coding_gene | 1.8 | 3.603 | 3.3E-02 | 1.2 | 2.34 | 9.2E-03 |
| *F46F5.4* | WBGene00018495 | pseudogene | 1.8 | 3.598 | 7.6E-03 | 2.6 | 6.09 | 2.3E-08 |
| *F19H6.5* | WBGene00008960 | protein_coding_gene | 1.8 | 3.597 | 3.2E-02 | 2.4 | 5.41 | 3.2E-09 |
| *Y52D5A.2* | WBGene00021793 | protein_coding_gene | 1.8 | 3.586 | 1.1E-02 | 1.2 | 2.36 | 3.8E-03 |
| *B0024.4* | WBGene00007097 | protein_coding_gene | 1.8 | 3.581 | 1.2E-04 | 1.2 | 2.30 | 1.2E-02 |
| *ugt-66* | WBGene00016013 | protein_coding_gene | 1.8 | 3.576 | 1.6E-06 | 2.1 | 4.43 | 5.4E-15 |
| *xtr-2* | WBGene00006966 | protein_coding_gene | 1.8 | 3.566 | 3.0E-02 | 1.3 | 2.52 | 8.5E-03 |
| *K07A1.15* | WBGene00010619 | protein_coding_gene | 1.8 | 3.563 | 1.8E-09 | 1.2 | 2.32 | 4.0E-07 |
| *msrp-1* | WBGene00017978 | protein_coding_gene | 1.8 | 3.557 | 4.8E-06 | 1.9 | 3.72 | 3.9E-07 |
| *Y53F4B.8* | WBGene00013155 | pseudogene | 1.8 | 3.556 | 1.5E-03 | 1.1 | 2.19 | 7.3E-03 |
| *folr-1* | WBGene00007648 | protein_coding_gene | 1.8 | 3.555 | 4.8E-02 | 1.7 | 3.29 | 7.0E-04 |
| *K07A1.13* | WBGene00010617 | protein_coding_gene | 1.8 | 3.528 | 4.7E-26 | 1.5 | 2.92 | 9.1E-32 |
| *clec-74* | WBGene00021585 | protein_coding_gene | 1.8 | 3.522 | 3.3E-03 | 1.0 | 2.06 | 4.2E-02 |
| *fbxa-168* | WBGene00007838 | pseudogene | 1.8 | 3.519 | 1.5E-07 | 2.0 | 3.97 | 4.3E-12 |
| *nep-24* | WBGene00013786 | protein_coding_gene | 1.8 | 3.518 | 7.6E-03 | 1.8 | 3.45 | 2.3E-06 |
| *rrf-2* | WBGene00004509 | protein_coding_gene | 1.8 | 3.475 | 1.3E-17 | 2.0 | 4.07 | 4.0E-41 |
| *C08B6.4* | WBGene00007425 | protein_coding_gene | 1.8 | 3.472 | 2.4E-02 | 1.5 | 2.89 | 1.2E-09 |
| *Y6E2A.5* | WBGene00012399 | protein_coding_gene | 1.8 | 3.451 | 1.2E-03 | 1.1 | 2.11 | 6.7E-03 |
| *Y64G10A.10* | WBGene00045144 | protein_coding_gene | 1.8 | 3.444 | 5.0E-02 | 2.3 | 5.06 | 2.6E-06 |
| *math-37* | WBGene00020080 | protein_coding_gene | 1.8 | 3.440 | 9.1E-04 | 1.0 | 2.06 | 1.3E-04 |
| *F10A3.1* | WBGene00008631 | protein_coding_gene | 1.8 | 3.428 | 3.4E-02 | 1.0 | 2.04 | 7.2E-03 |
| *C02B10.6* | WBGene00015331 | protein_coding_gene | 1.8 | 3.416 | 4.1E-02 | 1.6 | 2.98 | 9.3E-04 |
| *R05G9R.1* | WBGene00019906 | protein_coding_gene | 1.8 | 3.407 | 4.6E-03 | 1.0 | 2.05 | 5.0E-04 |
| *C36F7.5* | WBGene00007986 | protein_coding_gene | 1.8 | 3.399 | 1.3E-04 | 1.0 | 2.06 | 4.1E-04 |
| *bus-8* | WBGene00044623 | protein_coding_gene | 1.8 | 3.395 | 2.2E-03 | 1.0 | 2.03 | 6.8E-05 |
| *Y37H2A.7* | WBGene00012567 | protein_coding_gene | 1.8 | 3.373 | 2.1E-07 | 1.9 | 3.62 | 1.7E-10 |
| *C38C3.8* | WBGene00016516 | pseudogene | 1.7 | 3.361 | 6.3E-03 | 1.4 | 2.59 | 6.3E-04 |
| *C48B4.12* | WBGene00008175 | protein_coding_gene | 1.7 | 3.348 | 5.6E-03 | 4.6 | 25.07 | 4.0E-93 |
| *Y13C8A.1* | WBGene00021188 | protein_coding_gene | 1.7 | 3.345 | 2.1E-04 | 1.6 | 2.94 | 5.4E-06 |
| *AC3.9* | WBGene00044022 | pseudogene | 1.7 | 3.315 | 4.3E-02 | 1.0 | 2.03 | 4.7E-02 |
| *Y43C5B.2* | WBGene00012785 | protein_coding_gene | 1.7 | 3.294 | 5.1E-03 | 1.9 | 3.82 | 2.0E-09 |
| *trf-1* | WBGene00006612 | protein_coding_gene | 1.7 | 3.290 | 4.7E-02 | 1.8 | 3.38 | 1.1E-04 |
| *C14E2.4* | WBGene00015773 | protein_coding_gene | 1.7 | 3.285 | 4.9E-02 | 1.8 | 3.56 | 9.3E-04 |
| *her-1* | WBGene00001842 | protein_coding_gene | 1.7 | 3.280 | 2.1E-04 | 1.7 | 3.18 | 1.0E-05 |
| *Y1A5A.1* | WBGene00012379 | protein_coding_gene | 1.7 | 3.265 | 1.3E-03 | 2.1 | 4.35 | 3.7E-12 |
| *ZK546.7* | WBGene00022763 | pseudogene | 1.7 | 3.263 | 9.3E-03 | 1.9 | 3.75 | 1.1E-06 |
| *F13H6.4* | WBGene00017432 | pseudogene | 1.7 | 3.225 | 2.1E-03 | 1.3 | 2.42 | 6.6E-05 |
| *K06A4.8* | WBGene00044015 | pseudogene | 1.7 | 3.191 | 3.8E-03 | 1.8 | 3.52 | 2.3E-09 |

| | | | | | | | | |
|---|---|---|---|---|---|---|---|---|
| *F16B3.2* | WBGene00017502 | protein_coding_gene | 1.7 | 3.185 | 3.6E-02 | 1.1 | 2.18 | 1.0E-02 |
| *T06G6.11* | WBGene00011550 | pseudogene | 1.6 | 3.110 | 3.0E-02 | 1.4 | 2.56 | 6.9E-03 |
| *W08F4.7* | WBGene00021096 | protein_coding_gene | 1.6 | 3.106 | 1.3E-03 | 2.5 | 5.77 | 1.3E-10 |
| *B0393.4* | WBGene00007169 | protein_coding_gene | 1.6 | 3.068 | 4.1E-04 | 2.4 | 5.14 | 1.4E-18 |
| *T24C4.4* | WBGene00020760 | protein_coding_gene | 1.6 | 3.052 | 5.8E-03 | 1.4 | 2.60 | 1.3E-04 |
| *cri-1* | WBGene00010614 | protein_coding_gene | 1.6 | 3.037 | 6.3E-03 | 1.6 | 3.04 | 1.6E-06 |
| *math-48* | WBGene00022689 | protein_coding_gene | 1.6 | 3.035 | 2.2E-02 | 2.0 | 3.96 | 7.9E-07 |
| *ZK896.4* | WBGene00014135 | protein_coding_gene | 1.6 | 3.031 | 3.2E-03 | 1.4 | 2.65 | 1.8E-04 |
| *nspc-10* | WBGene00009640 | protein_coding_gene | 1.6 | 3.021 | 1.1E-11 | 1.1 | 2.15 | 1.2E-12 |
| *Y40A1A.6* | WBGene00304815 | ncRNA_gene | 1.6 | 2.982 | 4.7E-15 | 1.4 | 2.71 | 4.1E-14 |
| *exc-13* | WBGene00015393 | protein_coding_gene | 1.6 | 2.962 | 3.6E-03 | 1.2 | 2.25 | 1.3E-04 |
| *skr-12* | WBGene00004818 | protein_coding_gene | 1.6 | 2.958 | 1.3E-08 | 1.5 | 2.90 | 2.5E-11 |
| *C17E4.10* | WBGene00007647 | protein_coding_gene | 1.6 | 2.940 | 1.8E-21 | 1.4 | 2.64 | 4.0E-50 |
| *F59A1.13* | WBGene00010299 | protein_coding_gene | 1.6 | 2.929 | 5.1E-03 | 1.7 | 3.30 | 8.6E-08 |
| *ugt-9* | WBGene00020587 | protein_coding_gene | 1.5 | 2.919 | 7.7E-09 | 1.4 | 2.68 | 6.3E-05 |
| *clec-36* | WBGene00044332 | protein_coding_gene | 1.5 | 2.913 | 1.2E-03 | 1.4 | 2.60 | 3.0E-06 |
| *fbxb-57* | WBGene00010101 | protein_coding_gene | 1.5 | 2.902 | 2.6E-03 | 1.2 | 2.23 | 8.5E-03 |
| *msp-55* | WBGene00003448 | protein_coding_gene | 1.5 | 2.896 | 2.2E-02 | 1.4 | 2.67 | 2.0E-04 |
| *C17E4.2* | WBGene00007642 | protein_coding_gene | 1.5 | 2.894 | 5.7E-10 | 1.5 | 2.81 | 3.1E-12 |
| *acc-3* | WBGene00018880 | protein_coding_gene | 1.5 | 2.890 | 3.4E-02 | 1.5 | 2.74 | 1.8E-04 |
| *msp-57* | WBGene00003450 | protein_coding_gene | 1.5 | 2.889 | 3.6E-04 | 1.3 | 2.54 | 1.5E-03 |
| *fbxc-48* | WBGene00015615 | protein_coding_gene | 1.5 | 2.883 | 2.9E-02 | 1.8 | 3.48 | 1.5E-05 |
| *B0391.8* | WBGene00014643 | pseudogene | 1.5 | 2.874 | 1.2E-06 | 1.2 | 2.28 | 2.9E-06 |
| *Y47G6A.26* | WBGene00021650 | protein_coding_gene | 1.5 | 2.862 | 1.3E-03 | 1.8 | 3.56 | 4.5E-10 |
| *lys-6* | WBGene00003095 | protein_coding_gene | 1.5 | 2.857 | 8.4E-04 | 1.2 | 2.35 | 8.9E-06 |
| *F43E2.6* | WBGene00018394 | protein_coding_gene | 1.5 | 2.845 | 1.9E-03 | 1.3 | 2.47 | 6.7E-05 |
| *nspd-9* | WBGene00017808 | protein_coding_gene | 1.5 | 2.840 | 2.0E-03 | 1.5 | 2.88 | 5.8E-11 |
| *C09B9.4* | WBGene00015629 | protein_coding_gene | 1.5 | 2.826 | 1.7E-03 | 2.1 | 4.17 | 2.7E-16 |
| *F17C8.7* | WBGene00008912 | protein_coding_gene | 1.5 | 2.813 | 8.8E-04 | 1.8 | 3.53 | 4.8E-10 |
| *Y71G12B.35* | WBGene00185098 | protein_coding_gene | 1.5 | 2.804 | 1.8E-04 | 1.9 | 3.75 | 5.8E-14 |
| *pals-33* | WBGene00021081 | protein_coding_gene | 1.5 | 2.797 | 4.5E-02 | 1.2 | 2.24 | 2.2E-04 |
| *C36C5.12* | WBGene00016481 | protein_coding_gene | 1.5 | 2.787 | 3.7E-03 | 2.5 | 5.47 | 1.0E-13 |
| *nep-19* | WBGene00020292 | protein_coding_gene | 1.5 | 2.774 | 4.0E-02 | 1.8 | 3.55 | 7.3E-06 |
| *F55A12.6* | WBGene00018865 | protein_coding_gene | 1.5 | 2.772 | 2.0E-02 | 1.0 | 2.03 | 9.0E-03 |
| *H37A05.2* | WBGene00010426 | protein_coding_gene | 1.5 | 2.772 | 3.5E-05 | 1.2 | 2.29 | 1.2E-04 |
| *C38D9.8* | WBGene00008016 | protein_coding_gene | 1.5 | 2.761 | 2.4E-03 | 1.6 | 2.95 | 3.7E-05 |
| *F11C7.2* | WBGene00017376 | protein_coding_gene | 1.5 | 2.760 | 1.7E-04 | 1.0 | 2.02 | 3.7E-07 |
| *T10E9.6* | WBGene00020416 | protein_coding_gene | 1.5 | 2.737 | 1.9E-02 | 1.2 | 2.25 | 3.2E-03 |
| *F35E2.10* | WBGene00044270 | pseudogene | 1.5 | 2.734 | 3.6E-02 | 1.5 | 2.87 | 1.2E-04 |
| *T02B11.4* | WBGene00020156 | protein_coding_gene | 1.4 | 2.732 | 3.1E-03 | 1.2 | 2.28 | 8.7E-05 |
| *Y116A8A.10* | WBGene00014975 | pseudogene | 1.4 | 2.726 | 5.4E-03 | 1.7 | 3.23 | 1.7E-05 |
| *ZK896.5* | WBGene00014136 | protein_coding_gene | 1.4 | 2.718 | 1.7E-02 | 1.9 | 3.83 | 1.6E-12 |
| *sst-20* | WBGene00006058 | protein_coding_gene | 1.4 | 2.700 | 9.0E-03 | 1.4 | 2.56 | 5.9E-06 |
| *nhr-54* | WBGene00003644 | protein_coding_gene | 1.4 | 2.690 | 6.3E-03 | 1.4 | 2.67 | 6.4E-06 |
| *K08C7.4* | WBGene00010647 | protein_coding_gene | 1.4 | 2.673 | 2.6E-02 | 1.6 | 2.94 | 3.0E-05 |
| *Y55F3AM.21* | WBGene00219399 | protein_coding_gene | 1.4 | 2.670 | 3.3E-03 | 2.2 | 4.54 | 5.1E-10 |
| *F26D10.13* | WBGene00044244 | protein_coding_gene | 1.4 | 2.641 | 2.7E-02 | 5.7 | 51.99 | 7.0E-90 |
| *F02H6.4* | WBGene00008537 | pseudogene | 1.4 | 2.638 | 9.6E-12 | 2.3 | 5.04 | 3.9E-50 |
| *C35A11.2* | WBGene00016429 | protein_coding_gene | 1.4 | 2.604 | 4.1E-02 | 1.0 | 2.04 | 1.9E-02 |
| *Y39F10A.3* | WBGene00021454 | protein_coding_gene | 1.4 | 2.573 | 4.4E-02 | 1.7 | 3.17 | 3.4E-05 |
| *Y50D4C.2* | WBGene00021747 | pseudogene | 1.3 | 2.547 | 1.0E-08 | 1.5 | 2.75 | 3.7E-11 |
| *F52F12.9* | WBGene00009942 | protein_coding_gene | 1.3 | 2.546 | 2.3E-02 | 1.8 | 3.49 | 6.1E-07 |
| *ZK809.9* | WBGene00194934 | protein_coding_gene | 1.3 | 2.545 | 1.6E-03 | 1.4 | 2.57 | 1.0E-04 |
| *F54H5.5* | WBGene00018841 | protein_coding_gene | 1.3 | 2.513 | 2.7E-02 | 1.5 | 2.84 | 1.6E-07 |
| *C31H1.1* | WBGene00016286 | protein_coding_gene | 1.3 | 2.500 | 1.1E-02 | 1.7 | 3.27 | 5.1E-09 |
| *F55C5.2* | WBGene00010092 | pseudogene | 1.3 | 2.494 | 1.3E-03 | 1.7 | 3.14 | 2.6E-08 |
| *R05G6.1* | WBGene00019896 | protein_coding_gene | 1.3 | 2.488 | 3.6E-02 | 1.9 | 3.82 | 4.4E-07 |
| *clec-79* | WBGene00018548 | protein_coding_gene | 1.3 | 2.482 | 7.8E-03 | 1.6 | 3.08 | 3.9E-13 |
| *B0507.6* | WBGene00015223 | protein_coding_gene | 1.3 | 2.456 | 1.8E-02 | 1.3 | 2.40 | 6.4E-05 |
| *gst-33* | WBGene00001781 | protein_coding_gene | 1.3 | 2.455 | 2.5E-02 | 1.0 | 2.03 | 2.5E-03 |
| *R08A2.1* | WBGene00011132 | protein_coding_gene | 1.3 | 2.446 | 2.4E-03 | 1.1 | 2.16 | 2.1E-05 |
| *msp-3* | WBGene00003424 | protein_coding_gene | 1.3 | 2.413 | 1.2E-03 | 1.8 | 3.37 | 3.6E-13 |
| *C08F11.12* | WBGene00007459 | protein_coding_gene | 1.3 | 2.407 | 2.8E-04 | 1.2 | 2.23 | 1.3E-05 |
| *vap-1* | WBGene00006886 | protein_coding_gene | 1.3 | 2.399 | 6.2E-07 | 1.0 | 2.04 | 2.3E-11 |
| *aqp-9* | WBGene00000177 | protein_coding_gene | 1.3 | 2.379 | 2.3E-07 | 1.6 | 2.96 | 3.2E-21 |

| | | | | | | | | |
|---|---|---|---|---|---|---|---|---|
| *Y53F4B.11* | WBGene00013158 | protein_coding_gene | 1.2 | 2.368 | 1.2E-02 | 2.1 | 4.30 | 1.8E-13 |
| *T23F11.2* | WBGene00011954 | protein_coding_gene | 1.2 | 2.368 | 6.6E-03 | 1.3 | 2.49 | 3.8E-03 |
| *Y41C4A.8* | WBGene00012755 | protein_coding_gene | 1.2 | 2.354 | 5.1E-05 | 1.6 | 3.08 | 1.3E-12 |
| *dhhc-13* | WBGene00019257 | protein_coding_gene | 1.2 | 2.337 | 3.6E-02 | 1.4 | 2.57 | 1.4E-04 |
| *ssp-16* | WBGene00006044 | protein_coding_gene | 1.2 | 2.324 | 2.1E-02 | 1.1 | 2.14 | 1.6E-02 |
| *Y50E8A.12* | WBGene00013055 | protein_coding_gene | 1.2 | 2.315 | 2.3E-02 | 1.6 | 2.93 | 1.5E-06 |
| *C47A4.1* | WBGene00008122 | protein_coding_gene | 1.2 | 2.303 | 4.2E-02 | 1.2 | 2.35 | 4.3E-03 |
| *F37A8.1* | WBGene00009501 | protein_coding_gene | 1.2 | 2.299 | 2.9E-02 | 1.6 | 2.96 | 1.7E-07 |
| *F55B12.11* | WBGene00077570 | pseudogene | 1.2 | 2.299 | 4.0E-02 | 1.4 | 2.64 | 1.3E-03 |
| *M7.7* | WBGene00010883 | protein_coding_gene | 1.2 | 2.298 | 4.8E-02 | 1.3 | 2.38 | 1.2E-05 |
| *zig-6* | WBGene00006983 | protein_coding_gene | 1.2 | 2.289 | 3.6E-03 | 1.3 | 2.47 | 3.5E-06 |
| *BE10.1* | WBGene00007209 | protein_coding_gene | 1.2 | 2.281 | 1.9E-02 | 1.0 | 2.01 | 3.2E-03 |
| *C50C3.2* | WBGene00016800 | protein_coding_gene | 1.2 | 2.278 | 2.2E-02 | 1.7 | 3.30 | 1.4E-13 |
| *C18E9.8* | WBGene00007688 | protein_coding_gene | 1.2 | 2.275 | 2.2E-02 | 1.7 | 3.15 | 2.0E-09 |
| *Y41C4A.18* | WBGene00012761 | protein_coding_gene | 1.2 | 2.253 | 3.4E-02 | 1.1 | 2.08 | 3.0E-03 |
| *C06A8.6* | WBGene00015516 | protein_coding_gene | 1.2 | 2.244 | 3.4E-02 | 1.5 | 2.75 | 9.1E-09 |
| *gad-2* | WBGene00015666 | protein_coding_gene | 1.2 | 2.236 | 4.8E-02 | 1.5 | 2.83 | 1.9E-05 |
| *nspc-3* | WBGene00044338 | protein_coding_gene | 1.2 | 2.232 | 4.4E-02 | 1.2 | 2.30 | 1.5E-03 |
| *F55G11.8* | WBGene00010128 | protein_coding_gene | 1.1 | 2.214 | 3.2E-02 | 1.2 | 2.23 | 1.7E-03 |
| *F12E12.6* | WBGene00017407 | protein_coding_gene | 1.1 | 2.211 | 1.9E-02 | 1.1 | 2.16 | 1.0E-02 |
| *ZK418.2* | WBGene00022732 | protein_coding_gene | 1.1 | 2.204 | 3.3E-02 | 2.3 | 4.86 | 7.6E-27 |
| *lgc-47* | WBGene00009799 | protein_coding_gene | 1.1 | 2.204 | 3.0E-02 | 1.1 | 2.17 | 2.1E-05 |
| *Y57A10B.6* | WBGene00013278 | protein_coding_gene | 1.1 | 2.201 | 8.1E-04 | 1.3 | 2.42 | 2.0E-05 |
| *srp-3* | WBGene00005644 | protein_coding_gene | 1.1 | 2.196 | 2.2E-02 | 1.6 | 3.04 | 2.9E-08 |
| *ant-1.4* | WBGene00020140 | protein_coding_gene | 1.1 | 2.193 | 1.7E-02 | 1.6 | 3.09 | 1.0E-04 |
| *msp-65* | WBGene00003458 | protein_coding_gene | 1.1 | 2.185 | 4.0E-02 | 1.3 | 2.50 | 7.5E-05 |
| *T28C6.5* | WBGene00012120 | protein_coding_gene | 1.1 | 2.163 | 2.5E-02 | 1.4 | 2.58 | 3.4E-05 |
| *C34D4.10* | WBGene00016402 | protein_coding_gene | 1.1 | 2.154 | 7.5E-08 | 1.5 | 2.78 | 6.6E-21 |
| *fbxb-17* | WBGene00019037 | protein_coding_gene | 1.1 | 2.118 | 1.6E-02 | 1.2 | 2.31 | 1.9E-04 |
| *ZK84.2* | WBGene00022650 | protein_coding_gene | 1.1 | 2.106 | 1.2E-02 | 1.1 | 2.15 | 1.4E-04 |
| *sdz-30* | WBGene00011432 | protein_coding_gene | 1.0 | 2.061 | 7.3E-04 | 1.1 | 2.18 | 5.6E-05 |
| *ate-1* | WBGene00010615 | protein_coding_gene | 1.0 | 2.057 | 5.9E-14 | 1.0 | 2.02 | 1.2E-25 |
| *C15H7.3* | WBGene00007610 | protein_coding_gene | 1.0 | 2.040 | 5.0E-02 | 1.2 | 2.26 | 1.7E-05 |
| *T05G5.4* | WBGene00011499 | protein_coding_gene | 1.0 | 2.019 | 2.9E-02 | 1.1 | 2.15 | 3.3E-03 |
| *F42G4.2* | WBGene00009641 | protein_coding_gene | 1.0 | 2.017 | 4.2E-02 | 1.2 | 2.30 | 3.5E-05 |
| *clec-190* | WBGene00010953 | protein_coding_gene | 1.0 | 2.014 | 2.9E-02 | 3.6 | 11.87 | 1.1E-24 |
| *C06A5.12* | WBGene00044414 | protein_coding_gene | -1.0 | 0.493 | 3.1E-02 | -1.3 | 0.40 | 2.5E-04 |
| *acs-21* | WBGene00016849 | protein_coding_gene | -1.0 | 0.485 | 1.3E-02 | -1.1 | 0.48 | 1.0E-04 |
| *M02B7.2* | WBGene00019724 | protein_coding_gene | -1.0 | 0.483 | 5.6E-06 | -1.2 | 0.44 | 5.3E-08 |
| *tbc-10* | WBGene00019915 | protein_coding_gene | -1.1 | 0.471 | 3.9E-04 | -1.3 | 0.40 | 7.1E-11 |
| *nhr-132* | WBGene00003722 | protein_coding_gene | -1.1 | 0.461 | 9.7E-03 | -1.2 | 0.45 | 6.1E-05 |
| *F58D5.2* | WBGene00010242 | protein_coding_gene | -1.1 | 0.459 | 2.9E-02 | -1.7 | 0.30 | 2.1E-07 |
| *nhr-80* | WBGene00003670 | protein_coding_gene | -1.2 | 0.440 | 5.7E-05 | -1.2 | 0.44 | 2.8E-13 |
| *R06B10.2* | WBGene00019913 | protein_coding_gene | -1.2 | 0.438 | 3.6E-02 | -1.6 | 0.34 | 3.6E-04 |
| *sea-1* | WBGene00004750 | protein_coding_gene | -1.2 | 0.437 | 9.4E-05 | -1.5 | 0.35 | 8.4E-18 |
| *clec-52* | WBGene00015052 | protein_coding_gene | -1.2 | 0.436 | 1.2E-03 | -1.7 | 0.31 | 6.8E-23 |
| *F39B2.5* | WBGene00009556 | protein_coding_gene | -1.2 | 0.433 | 8.7E-08 | -1.5 | 0.35 | 3.6E-13 |
| *nduo-1* | WBGene00010959 | protein_coding_gene | -1.2 | 0.431 | 5.4E-08 | -1.4 | 0.38 | 6.1E-19 |
| *irg-5* | WBGene00009429 | protein_coding_gene | -1.2 | 0.429 | 2.1E-02 | -1.7 | 0.31 | 8.2E-05 |
| *lsl-1* | WBGene00009937 | protein_coding_gene | -1.2 | 0.426 | 3.6E-18 | -1.1 | 0.46 | 1.5E-16 |
| *fbxb-84* | WBGene00020544 | protein_coding_gene | -1.2 | 0.423 | 1.1E-02 | -1.0 | 0.49 | 1.8E-02 |
| *D2096.11* | WBGene00017078 | protein_coding_gene | -1.3 | 0.418 | 3.2E-02 | -1.4 | 0.38 | 2.5E-04 |
| *D1054.1* | WBGene00008370 | protein_coding_gene | -1.3 | 0.416 | 5.6E-08 | -1.2 | 0.43 | 7.0E-10 |
| *asp-12* | WBGene00017678 | protein_coding_gene | -1.3 | 0.415 | 3.3E-03 | -1.0 | 0.49 | 1.9E-03 |
| *sdz-28* | WBGene00020073 | protein_coding_gene | -1.3 | 0.405 | 8.7E-04 | -1.3 | 0.40 | 1.1E-05 |
| *Y49F6C.8* | WBGene00021730 | protein_coding_gene | -1.3 | 0.400 | 3.7E-06 | -1.5 | 0.36 | 7.3E-10 |
| *F25D1.5* | WBGene00009110 | protein_coding_gene | -1.3 | 0.399 | 1.5E-04 | -1.3 | 0.41 | 7.3E-08 |
| *F42A9.6* | WBGene00018335 | protein_coding_gene | -1.3 | 0.398 | 9.2E-11 | -1.3 | 0.41 | 2.1E-19 |
| *K02F6.3* | WBGene00019337 | protein_coding_gene | -1.4 | 0.387 | 4.5E-03 | -1.1 | 0.48 | 7.8E-05 |
| *Y82E9BR.22* | WBGene00022354 | protein_coding_gene | -1.4 | 0.386 | 2.6E-07 | -2.2 | 0.21 | 1.3E-32 |
| *Y82E9BR.14* | WBGene00022347 | protein_coding_gene | -1.4 | 0.384 | 2.4E-19 | -1.2 | 0.42 | 1.5E-19 |
| *Y116A8C.33* | WBGene00013809 | protein_coding_gene | -1.4 | 0.383 | 1.4E-02 | -1.4 | 0.38 | 2.6E-06 |
| *fbxa-14* | WBGene00022326 | protein_coding_gene | -1.4 | 0.381 | 8.5E-03 | -1.7 | 0.31 | 1.1E-05 |
| *ZC239.16* | WBGene00022574 | protein_coding_gene | -1.4 | 0.381 | 4.2E-02 | -2.2 | 0.21 | 2.6E-10 |
| *fbxb-108* | WBGene00017249 | protein_coding_gene | -1.4 | 0.378 | 3.3E-02 | -1.1 | 0.48 | 3.7E-02 |

| | | | | | | | | |
|---|---|---|---|---|---|---|---|---|
| *F31F4.1* | WBGene00017957 | protein_coding_gene | -1.4 | 0.373 | 1.9E-05 | -1.3 | 0.41 | 2.5E-08 |
| *Y32B12C.1* | WBGene00012525 | protein_coding_gene | -1.4 | 0.372 | 4.1E-02 | -2.3 | 0.20 | 7.8E-10 |
| *his-60* | WBGene00001934 | protein_coding_gene | -1.4 | 0.370 | 6.8E-03 | -2.0 | 0.24 | 3.1E-08 |
| *C08F1.6* | WBGene00015610 | protein_coding_gene | -1.4 | 0.367 | 7.9E-04 | -2.2 | 0.22 | 7.1E-12 |
| *fbxb-35* | WBGene00017208 | protein_coding_gene | -1.5 | 0.366 | 3.2E-02 | -1.7 | 0.31 | 1.7E-04 |
| *Y57G11C.51* | WBGene00044258 | protein_coding_gene | -1.5 | 0.366 | 7.1E-08 | -1.0 | 0.49 | 8.4E-06 |
| *T03E6.8* | WBGene00011397 | protein_coding_gene | -1.5 | 0.365 | 1.7E-03 | -3.1 | 0.12 | 1.7E-29 |
| *gpx-1* | WBGene00009165 | protein_coding_gene | -1.5 | 0.360 | 9.5E-11 | -1.5 | 0.35 | 7.7E-16 |
| *C04G2.14* | WBGene00255426 | ncRNA_gene | -1.5 | 0.360 | 9.1E-03 | -1.1 | 0.45 | 1.4E-03 |
| *pals-18* | WBGene00022353 | protein_coding_gene | -1.5 | 0.358 | 1.5E-02 | -2.4 | 0.18 | 1.4E-14 |
| *str-176* | WBGene00006220 | protein_coding_gene | -1.5 | 0.355 | 2.4E-10 | -1.2 | 0.44 | 4.4E-13 |
| *ncx-6* | WBGene00003571 | protein_coding_gene | -1.5 | 0.352 | 1.8E-04 | -1.0 | 0.49 | 7.5E-06 |
| *pzf-1* | WBGene00011505 | protein_coding_gene | -1.5 | 0.351 | 5.3E-08 | -1.4 | 0.37 | 3.0E-08 |
| *K12H6.7* | WBGene00019687 | protein_coding_gene | -1.5 | 0.349 | 1.3E-02 | -1.2 | 0.45 | 9.5E-03 |
| *bath-4* | WBGene00018665 | protein_coding_gene | -1.5 | 0.346 | 1.6E-05 | -1.4 | 0.39 | 1.6E-05 |
| *Y110A2AL.9* | WBGene00022445 | protein_coding_gene | -1.5 | 0.344 | 1.4E-02 | -2.5 | 0.18 | 2.5E-16 |
| *twk-34* | WBGene00006686 | protein_coding_gene | -1.6 | 0.340 | 5.7E-09 | -1.1 | 0.46 | 4.1E-08 |
| *F40D4.13* | WBGene00009572 | protein_coding_gene | -1.6 | 0.335 | 2.3E-02 | -2.3 | 0.20 | 5.0E-24 |
| *Y54G11A.19* | WBGene00235256 | protein_coding_gene | -1.6 | 0.327 | 4.8E-04 | -1.1 | 0.47 | 6.9E-03 |
| *col-135* | WBGene00000708 | protein_coding_gene | -1.6 | 0.326 | 5.0E-22 | -1.7 | 0.31 | 6.5E-26 |
| *pals-17* | WBGene00022346 | protein_coding_gene | -1.6 | 0.322 | 1.3E-04 | -1.0 | 0.49 | 9.9E-05 |
| *fip-5* | WBGene00009620 | protein_coding_gene | -1.6 | 0.319 | 7.7E-07 | -1.0 | 0.49 | 1.9E-03 |
| *Y24F12A.4* | WBGene00012499 | pseudogene | -1.6 | 0.319 | 7.6E-03 | -1.2 | 0.45 | 1.3E-02 |
| *clp-3* | WBGene00000544 | protein_coding_gene | -1.7 | 0.317 | 5.1E-03 | -1.4 | 0.39 | 3.8E-05 |
| *nhr-65* | WBGene00003655 | protein_coding_gene | -1.7 | 0.316 | 1.1E-05 | -1.0 | 0.49 | 1.1E-03 |
| *linc-24* | WBGene00219771 | lincRNA_gene | -1.7 | 0.314 | 4.5E-02 | -1.7 | 0.31 | 1.7E-04 |
| *F20D1.22* | WBGene00304908 | protein_coding_gene | -1.7 | 0.308 | 1.9E-03 | -1.4 | 0.39 | 4.7E-04 |
| *Y34F4.6* | WBGene00189956 | protein_coding_gene | -1.7 | 0.307 | 1.6E-02 | -1.9 | 0.26 | 1.2E-06 |
| *ZK1240.9* | WBGene00022873 | protein_coding_gene | -1.7 | 0.306 | 1.6E-04 | -2.0 | 0.25 | 5.7E-12 |
| *fbxa-138* | WBGene00022345 | protein_coding_gene | -1.7 | 0.305 | 4.7E-02 | -2.6 | 0.16 | 4.9E-07 |
| *mir-8199* | WBGene00016553 | miRNA_gene | -1.7 | 0.305 | 4.9E-02 | -1.7 | 0.32 | 1.8E-04 |
| *fbxb-83* | WBGene00020543 | protein_coding_gene | -1.7 | 0.305 | 3.4E-03 | -1.1 | 0.45 | 9.4E-03 |
| *skr-5* | WBGene00004811 | protein_coding_gene | -1.7 | 0.304 | 4.5E-02 | -2.0 | 0.25 | 1.6E-05 |
| *fbxb-90* | WBGene00044694 | protein_coding_gene | -1.7 | 0.302 | 1.7E-02 | -1.5 | 0.35 | 2.4E-03 |
| *drd-10* | WBGene00016361 | protein_coding_gene | -1.7 | 0.301 | 1.0E-04 | -1.8 | 0.28 | 2.6E-09 |
| *fat-7* | WBGene00001399 | protein_coding_gene | -1.7 | 0.300 | 2.1E-03 | -1.3 | 0.42 | 2.5E-15 |
| *K06G5.1* | WBGene00010605 | protein_coding_gene | -1.7 | 0.299 | 4.4E-14 | -1.8 | 0.29 | 6.3E-93 |
| *Y48D7A.1* | WBGene00021656 | protein_coding_gene | -1.8 | 0.295 | 4.1E-02 | -1.1 | 0.48 | 6.5E-03 |
| *Y17D7C.3* | WBGene00045366 | pseudogene | -1.8 | 0.294 | 1.3E-06 | -2.2 | 0.22 | 2.8E-19 |
| *Y43F8B.23* | WBGene00194815 | protein_coding_gene | -1.8 | 0.294 | 4.3E-02 | -2.4 | 0.19 | 3.4E-05 |
| *Y38H6C.23* | WBGene00012634 | protein_coding_gene | -1.8 | 0.294 | 1.0E-05 | -3.3 | 0.10 | 6.5E-29 |
| *T04A8.18* | WBGene00119203 | protein_coding_gene | -1.8 | 0.294 | 1.5E-02 | -1.2 | 0.43 | 1.5E-03 |
| *F26H11.4* | WBGene00009182 | protein_coding_gene | -1.8 | 0.293 | 3.2E-06 | -1.4 | 0.37 | 8.1E-07 |
| *rgs-11* | WBGene00004354 | protein_coding_gene | -1.8 | 0.292 | 3.2E-07 | -1.9 | 0.26 | 3.7E-16 |
| *H16D19.5* | WBGene00195253 | pseudogene | -1.8 | 0.291 | 1.8E-02 | -1.6 | 0.33 | 6.3E-04 |
| *Y35H6.3* | WBGene00021342 | pseudogene | -1.8 | 0.290 | 9.0E-07 | -1.5 | 0.35 | 8.7E-06 |
| *F49F1.12* | WBGene00018652 | protein_coding_gene | -1.8 | 0.290 | 2.4E-02 | -1.3 | 0.40 | 1.1E-03 |
| *F01D5.1* | WBGene00008492 | protein_coding_gene | -1.8 | 0.288 | 1.9E-04 | -1.4 | 0.37 | 8.3E-04 |
| *C50E3.11* | WBGene00016822 | protein_coding_gene | -1.8 | 0.286 | 3.1E-13 | -2.0 | 0.25 | 2.1E-25 |
| *ugt-31* | WBGene00021464 | protein_coding_gene | -1.8 | 0.285 | 1.4E-03 | -1.2 | 0.43 | 3.2E-04 |
| *math-10* | WBGene00015839 | protein_coding_gene | -1.8 | 0.283 | 3.2E-02 | -3.5 | 0.09 | 1.3E-09 |
| *clec-80* | WBGene00043062 | protein_coding_gene | -1.8 | 0.281 | 1.5E-06 | -1.2 | 0.44 | 5.1E-23 |
| *C01C10.2* | WBGene00015294 | protein_coding_gene | -1.9 | 0.274 | 1.9E-08 | -1.7 | 0.31 | 6.3E-11 |
| *ZK218.5* | WBGene00013939 | protein_coding_gene | -1.9 | 0.274 | 3.5E-02 | -1.8 | 0.28 | 1.8E-04 |
| *K02E7.10* | WBGene00019314 | protein_coding_gene | -1.9 | 0.274 | 3.0E-02 | -2.9 | 0.13 | 1.6E-06 |
| *sdz-29* | WBGene00020081 | pseudogene | -1.9 | 0.273 | 8.1E-05 | -1.7 | 0.30 | 1.2E-04 |
| *T02G5.4* | WBGene00020164 | pseudogene | -1.9 | 0.272 | 1.3E-11 | -1.9 | 0.28 | 1.3E-25 |
| *fbxa-199* | WBGene00005931 | protein_coding_gene | -1.9 | 0.271 | 4.0E-02 | -1.1 | 0.46 | 4.1E-03 |
| *rmd-6* | WBGene00020070 | protein_coding_gene | -1.9 | 0.271 | 1.7E-02 | -1.8 | 0.29 | 1.0E-04 |
| *F35F10.6* | WBGene00018056 | protein_coding_gene | -1.9 | 0.271 | 8.0E-03 | -1.2 | 0.44 | 9.6E-03 |
| *drh-2* | WBGene00001091 | pseudogene | -1.9 | 0.268 | 3.3E-27 | -1.5 | 0.36 | 4.8E-05 |
| *K10G4.10* | WBGene00044202 | protein_coding_gene | -1.9 | 0.267 | 2.9E-02 | -1.0 | 0.50 | 8.8E-03 |
| *ppw-1* | WBGene00004093 | protein_coding_gene | -1.9 | 0.267 | 1.7E-18 | -1.5 | 0.35 | 5.5E-26 |
| *F10C1.8* | WBGene00017327 | protein_coding_gene | -1.9 | 0.266 | 5.2E-03 | -1.1 | 0.47 | 1.8E-03 |
| *ptp-5.3* | WBGene00021207 | protein_coding_gene | -1.9 | 0.266 | 3.7E-03 | -2.0 | 0.25 | 4.2E-09 |

| | | | | | | | | |
|---|---|---|---|---|---|---|---|---|
| *Y37H2A.12* | WBGene00044758 | protein_coding_gene | -1.9 | 0.266 | 1.8E-10 | -2.4 | 0.19 | 3.1E-15 |
| *osm-3* | WBGene00003884 | protein_coding_gene | -1.9 | 0.266 | 8.9E-05 | -1.3 | 0.39 | 5.6E-05 |
| *fbxa-55* | WBGene00020453 | protein_coding_gene | -1.9 | 0.264 | 7.3E-06 | -1.9 | 0.27 | 6.9E-11 |
| *R11A8.1* | WBGene00011236 | protein_coding_gene | -1.9 | 0.263 | 2.3E-20 | -1.0 | 0.49 | 5.4E-13 |
| *R12C12.7* | WBGene00020027 | protein_coding_gene | -1.9 | 0.262 | 7.0E-25 | -1.7 | 0.31 | 2.4E-45 |
| *egas-3* | WBGene00013480 | protein_coding_gene | -1.9 | 0.261 | 3.6E-02 | -1.6 | 0.33 | 1.6E-03 |
| *K06B4.15* | WBGene00077450 | protein_coding_gene | -1.9 | 0.260 | 2.8E-02 | -2.0 | 0.24 | 7.7E-05 |
| *nhr-226* | WBGene00020850 | protein_coding_gene | -1.9 | 0.259 | 4.6E-03 | -1.8 | 0.29 | 1.5E-06 |
| *fbxa-59* | WBGene00020455 | protein_coding_gene | -2.0 | 0.254 | 3.9E-06 | -2.0 | 0.24 | 2.9E-22 |
| *Y37H2B.1* | WBGene00012572 | pseudogene | -2.0 | 0.254 | 2.3E-02 | -1.8 | 0.29 | 7.8E-04 |
| *fbxc-43* | WBGene00045297 | protein_coding_gene | -2.0 | 0.249 | 5.9E-05 | -1.7 | 0.30 | 3.7E-07 |
| *ugt-29* | WBGene00021709 | protein_coding_gene | -2.0 | 0.249 | 2.6E-19 | -2.6 | 0.17 | 1.1E-22 |
| *F10G2.4* | WBGene00017365 | protein_coding_gene | -2.0 | 0.248 | 3.2E-03 | -1.4 | 0.38 | 4.1E-03 |
| *sid-1* | WBGene00004795 | protein_coding_gene | -2.0 | 0.248 | 9.9E-18 | -1.3 | 0.41 | 1.5E-23 |
| *nep-8* | WBGene00017553 | protein_coding_gene | -2.0 | 0.247 | 1.7E-03 | -1.4 | 0.38 | 5.2E-05 |
| *gst-41* | WBGene00001789 | protein_coding_gene | -2.0 | 0.247 | 4.2E-04 | -1.4 | 0.37 | 1.3E-08 |
| *fbxb-81* | WBGene00020541 | protein_coding_gene | -2.0 | 0.247 | 9.5E-03 | -1.3 | 0.41 | 9.0E-03 |
| *timm-17B.2* | WBGene00017069 | protein_coding_gene | -2.0 | 0.247 | 5.2E-15 | -1.4 | 0.38 | 1.5E-09 |
| *fbxa-3* | WBGene00020363 | protein_coding_gene | -2.0 | 0.247 | 3.5E-02 | -2.3 | 0.20 | 8.9E-05 |
| *fbxb-36* | WBGene00017211 | protein_coding_gene | -2.0 | 0.244 | 1.2E-02 | -1.7 | 0.30 | 3.1E-04 |
| *K08D12.6* | WBGene00019540 | protein_coding_gene | -2.0 | 0.242 | 1.4E-04 | -1.4 | 0.38 | 6.9E-20 |
| *clec-233* | WBGene00009030 | protein_coding_gene | -2.1 | 0.241 | 1.6E-02 | -2.0 | 0.25 | 2.2E-05 |
| *C28D4.5* | WBGene00007793 | protein_coding_gene | -2.1 | 0.241 | 5.2E-03 | -2.0 | 0.26 | 5.6E-10 |
| *K02E7.6* | WBGene00019311 | pseudogene | -2.1 | 0.240 | 2.4E-05 | -2.2 | 0.22 | 5.7E-13 |
| *F54A3.2* | WBGene00018781 | protein_coding_gene | -2.1 | 0.239 | 4.9E-24 | -1.7 | 0.31 | 1.6E-17 |
| *nhr-183* | WBGene00018266 | protein_coding_gene | -2.1 | 0.238 | 6.3E-04 | -2.3 | 0.20 | 1.5E-08 |
| *fbxa-210* | WBGene00012565 | protein_coding_gene | -2.1 | 0.238 | 2.0E-34 | -2.2 | 0.21 | 1.8E-46 |
| *C16C4.18* | WBGene00194998 | protein_coding_gene | -2.1 | 0.237 | 1.5E-04 | -2.3 | 0.20 | 6.9E-09 |
| *fbxa-51* | WBGene00020448 | protein_coding_gene | -2.1 | 0.235 | 2.2E-06 | -1.3 | 0.42 | 8.5E-05 |
| *nhr-263* | WBGene00017367 | protein_coding_gene | -2.1 | 0.233 | 2.7E-02 | -1.5 | 0.36 | 2.6E-03 |
| *R03H10.7* | WBGene00019859 | protein_coding_gene | -2.1 | 0.233 | 1.5E-02 | -2.0 | 0.26 | 3.1E-05 |
| *F59H6.3* | WBGene00019134 | pseudogene | -2.1 | 0.231 | 1.4E-02 | -1.1 | 0.47 | 4.6E-03 |
| *trp-2* | WBGene00006615 | protein_coding_gene | -2.1 | 0.230 | 1.9E-05 | -1.9 | 0.26 | 9.1E-10 |
| *clec-47* | WBGene00011668 | protein_coding_gene | -2.1 | 0.229 | 2.0E-20 | -2.2 | 0.21 | 3.6E-08 |
| *ZK637.6* | WBGene00014987 | pseudogene | -2.1 | 0.228 | 2.3E-20 | -1.3 | 0.39 | 3.2E-11 |
| *T05F1.5* | WBGene00011491 | protein_coding_gene | -2.1 | 0.227 | 4.1E-03 | -1.6 | 0.33 | 1.0E-06 |
| *fbxa-95* | WBGene00009222 | protein_coding_gene | -2.2 | 0.225 | 7.2E-09 | -2.9 | 0.14 | 1.2E-17 |
| *M162.12* | WBGene00050946 | pseudogene | -2.2 | 0.225 | 1.9E-02 | -1.1 | 0.46 | 4.7E-03 |
| *ctl-1* | WBGene00000830 | protein_coding_gene | -2.2 | 0.223 | 1.9E-02 | -2.4 | 0.19 | 2.5E-06 |
| *H04D03.3* | WBGene00010364 | protein_coding_gene | -2.2 | 0.222 | 1.9E-03 | -1.3 | 0.41 | 3.0E-03 |
| *CE7X_3.1* | WBGene00008349 | pseudogene | -2.2 | 0.220 | 3.8E-04 | -1.9 | 0.27 | 1.6E-04 |
| *irg-4* | WBGene00008584 | protein_coding_gene | -2.2 | 0.218 | 1.6E-08 | -2.1 | 0.24 | 2.3E-08 |
| *rgs-9* | WBGene00004352 | protein_coding_gene | -2.2 | 0.217 | 1.7E-15 | -2.2 | 0.22 | 4.2E-20 |
| *enu-3.5* | WBGene00012552 | protein_coding_gene | -2.2 | 0.216 | 2.7E-05 | -1.4 | 0.38 | 6.7E-04 |
| *fbxa-187* | WBGene00009838 | pseudogene | -2.2 | 0.215 | 2.0E-02 | -2.1 | 0.23 | 1.3E-04 |
| *F31E9.6* | WBGene00009299 | protein_coding_gene | -2.2 | 0.214 | 4.8E-06 | -1.7 | 0.30 | 2.7E-06 |
| *W04E12.5* | WBGene00012250 | protein_coding_gene | -2.2 | 0.213 | 1.4E-02 | -1.6 | 0.34 | 2.6E-03 |
| *fkb-8* | WBGene00001433 | protein_coding_gene | -2.2 | 0.212 | 7.3E-06 | -2.2 | 0.21 | 7.6E-12 |
| *C29F9.2* | WBGene00016217 | protein_coding_gene | -2.2 | 0.211 | 3.9E-03 | -1.4 | 0.37 | 2.1E-03 |
| *Y17D7B.7* | WBGene00014562 | pseudogene | -2.2 | 0.211 | 6.3E-05 | -2.0 | 0.26 | 2.0E-12 |
| *T08E11.8* | WBGene00020364 | protein_coding_gene | -2.2 | 0.211 | 7.3E-03 | -1.0 | 0.50 | 7.8E-03 |
| *C13G5.2* | WBGene00015747 | protein_coding_gene | -2.3 | 0.207 | 7.5E-40 | -2.2 | 0.21 | 3.2E-49 |
| *sqst-5* | WBGene00269434 | protein_coding_gene | -2.3 | 0.206 | 1.4E-03 | -1.8 | 0.29 | 1.4E-06 |
| *F11A5.18* | WBGene00045511 | protein_coding_gene | -2.3 | 0.202 | 6.5E-03 | -1.4 | 0.39 | 1.1E-03 |
| *nhr-227* | WBGene00020851 | protein_coding_gene | -2.3 | 0.201 | 7.4E-03 | -2.1 | 0.24 | 1.6E-05 |
| *ZC266.1* | WBGene00022584 | protein_coding_gene | -2.3 | 0.200 | 7.6E-10 | -1.5 | 0.36 | 6.8E-06 |
| *clec-73* | WBGene00021579 | protein_coding_gene | -2.3 | 0.198 | 2.8E-05 | -2.4 | 0.18 | 2.8E-20 |
| *K08H2.3* | WBGene00010690 | protein_coding_gene | -2.3 | 0.197 | 1.6E-13 | -1.1 | 0.45 | 7.0E-05 |
| *Y38H8A.8* | WBGene00045062 | pseudogene | -2.3 | 0.196 | 1.2E-02 | -2.6 | 0.17 | 1.4E-05 |
| *F08D12.2* | WBGene00017246 | protein_coding_gene | -2.4 | 0.195 | 2.8E-12 | -2.7 | 0.15 | 2.6E-31 |
| *C30G12.2* | WBGene00016274 | protein_coding_gene | -2.4 | 0.195 | 1.6E-04 | -1.3 | 0.39 | 6.2E-11 |
| *fbxa-64* | WBGene00020883 | protein_coding_gene | -2.4 | 0.194 | 1.6E-10 | -2.2 | 0.23 | 2.4E-14 |
| *E01G4.7* | WBGene00045410 | pseudogene | -2.4 | 0.192 | 3.6E-12 | -4.2 | 0.05 | 2.7E-50 |
| *K09F6.5* | WBGene00019588 | protein_coding_gene | -2.4 | 0.191 | 1.2E-02 | -2.1 | 0.24 | 1.8E-04 |
| *peel-1* | WBGene00077563 | protein_coding_gene | -2.4 | 0.191 | 4.9E-03 | -1.5 | 0.36 | 3.8E-04 |

| | | | | | | | | |
|---|---|---|---|---|---|---|---|---|
| *fbxa-192* | WBGene00010212 | protein_coding_gene | -2.4 | 0.191 | 2.3E-23 | -2.2 | 0.21 | 2.0E-23 |
| *bath-13* | WBGene00018223 | protein_coding_gene | -2.4 | 0.190 | 5.6E-04 | -1.9 | 0.26 | 1.3E-07 |
| *C23H5.8* | WBGene00016027 | protein_coding_gene | -2.4 | 0.190 | 8.8E-06 | -1.5 | 0.34 | 8.4E-24 |
| *H11L12.1* | WBGene00019189 | protein_coding_gene | -2.4 | 0.190 | 1.4E-05 | -1.7 | 0.30 | 1.9E-07 |
| *R10E8.3* | WBGene00011209 | protein_coding_gene | -2.4 | 0.189 | 4.0E-03 | -2.2 | 0.23 | 1.1E-04 |
| *C25F9.6* | WBGene00007726 | protein_coding_gene | -2.4 | 0.189 | 5.1E-03 | -1.2 | 0.44 | 1.2E-02 |
| *C46G7.5* | WBGene00044502 | protein_coding_gene | -2.4 | 0.186 | 4.2E-05 | -2.3 | 0.20 | 5.4E-11 |
| *fbxc-50* | WBGene00015912 | protein_coding_gene | -2.4 | 0.185 | 4.2E-27 | -2.3 | 0.21 | 3.0E-42 |
| *his-48* | WBGene00001922 | protein_coding_gene | -2.4 | 0.184 | 7.4E-03 | -1.0 | 0.49 | 7.5E-03 |
| *bath-21* | WBGene00019138 | protein_coding_gene | -2.5 | 0.183 | 8.5E-06 | -1.7 | 0.32 | 3.3E-05 |
| *ZC239.13* | WBGene00022571 | protein_coding_gene | -2.5 | 0.183 | 3.2E-03 | -1.3 | 0.41 | 1.7E-03 |
| *clec-147* | WBGene00007313 | protein_coding_gene | -2.5 | 0.181 | 1.4E-07 | -3.2 | 0.11 | 5.4E-37 |
| *fbxa-54* | WBGene00020449 | protein_coding_gene | -2.5 | 0.180 | 1.6E-06 | -2.0 | 0.26 | 5.7E-12 |
| *Y47G6A.14* | WBGene00021640 | protein_coding_gene | -2.5 | 0.180 | 1.9E-27 | -3.2 | 0.11 | 2.1E-45 |
| *math-49* | WBGene00022690 | protein_coding_gene | -2.5 | 0.179 | 8.3E-03 | -1.8 | 0.28 | 7.8E-04 |
| *F55A3.7* | WBGene00018852 | pseudogene | -2.5 | 0.178 | 5.0E-07 | -1.3 | 0.41 | 3.3E-06 |
| *F23D12.3* | WBGene00009087 | protein_coding_gene | -2.5 | 0.178 | 2.2E-04 | -1.9 | 0.26 | 2.8E-08 |
| *T28B8.4* | WBGene00012115 | protein_coding_gene | -2.5 | 0.177 | 1.6E-04 | -1.1 | 0.48 | 3.7E-03 |
| *F52C6.13* | WBGene00018670 | protein_coding_gene | -2.5 | 0.177 | 3.5E-03 | -1.3 | 0.41 | 6.4E-03 |
| *cdr-4* | WBGene00010470 | protein_coding_gene | -2.5 | 0.176 | 2.0E-05 | -1.1 | 0.47 | 1.9E-12 |
| *linc-72* | WBGene00219590 | lincRNA_gene | -2.5 | 0.173 | 3.1E-03 | -1.1 | 0.48 | 3.2E-02 |
| *F52C6.2* | WBGene00018659 | protein_coding_gene | -2.5 | 0.171 | 6.1E-30 | -1.6 | 0.32 | 6.1E-14 |
| *srw-85* | WBGene00005832 | protein_coding_gene | -2.6 | 0.171 | 3.0E-03 | -2.4 | 0.19 | 4.5E-05 |
| *C32H11.7* | WBGene00007870 | pseudogene | -2.6 | 0.169 | 2.7E-03 | -1.8 | 0.29 | 1.0E-04 |
| *fbxb-53* | WBGene00018104 | protein_coding_gene | -2.6 | 0.168 | 9.1E-04 | -1.7 | 0.30 | 9.8E-04 |
| *fbxb-12* | WBGene00018109 | pseudogene | -2.6 | 0.168 | 2.7E-03 | -1.5 | 0.36 | 5.3E-04 |
| *K05F6.15* | WBGene00269419 | pseudogene | -2.6 | 0.166 | 3.1E-04 | -2.3 | 0.21 | 2.3E-05 |
| *bath-20* | WBGene00015012 | protein_coding_gene | -2.6 | 0.164 | 2.3E-03 | -2.0 | 0.25 | 2.1E-05 |
| *B0281.5* | WBGene00015112 | protein_coding_gene | -2.6 | 0.164 | 1.5E-18 | -2.6 | 0.17 | 7.3E-17 |
| *nhr-220* | WBGene00020591 | pseudogene | -2.6 | 0.164 | 1.4E-03 | -1.9 | 0.26 | 2.5E-04 |
| *fbxb-51* | WBGene00019414 | protein_coding_gene | -2.6 | 0.162 | 2.2E-03 | -1.7 | 0.30 | 1.2E-04 |
| *Y23H5A.8* | WBGene00023172 | protein_coding_gene | -2.6 | 0.161 | 1.0E-03 | -2.1 | 0.24 | 2.0E-04 |
| *Y53C10A.1* | WBGene00014915 | pseudogene | -2.7 | 0.158 | 1.1E-23 | -1.2 | 0.45 | 9.8E-07 |
| *F59E12.15* | WBGene00235269 | protein_coding_gene | -2.7 | 0.158 | 1.9E-03 | -1.5 | 0.35 | 4.5E-04 |
| *R03D7.2* | WBGene00010989 | protein_coding_gene | -2.7 | 0.157 | 9.6E-22 | -1.6 | 0.33 | 1.5E-07 |
| *Y40C5A.4* | WBGene00021497 | protein_coding_gene | -2.7 | 0.156 | 7.3E-11 | -1.9 | 0.26 | 3.3E-17 |
| *nspc-19* | WBGene00008360 | protein_coding_gene | -2.7 | 0.154 | 1.3E-04 | -3.1 | 0.11 | 3.6E-21 |
| *Y69A2AR.12* | WBGene00022083 | protein_coding_gene | -2.7 | 0.154 | 3.8E-03 | -1.7 | 0.30 | 5.7E-04 |
| *F21D9.2* | WBGene00009016 | protein_coding_gene | -2.7 | 0.154 | 2.1E-04 | -2.0 | 0.26 | 4.8E-07 |
| *K09F6.13* | WBGene00235164 | protein_coding_gene | -2.7 | 0.153 | 1.6E-03 | -3.1 | 0.12 | 1.8E-08 |
| *npl-4.1* | WBGene00019120 | protein_coding_gene | -2.7 | 0.151 | 1.1E-22 | -2.5 | 0.17 | 2.3E-25 |
| *scrm-3* | WBGene00015437 | protein_coding_gene | -2.7 | 0.151 | 1.2E-03 | -1.9 | 0.26 | 3.0E-05 |
| *acox-1.2* | WBGene00008565 | protein_coding_gene | -2.7 | 0.150 | 5.1E-16 | -1.2 | 0.45 | 4.8E-05 |
| *F36H5.17* | WBGene00270293 | protein_coding_gene | -2.7 | 0.150 | 1.4E-03 | -1.5 | 0.35 | 4.5E-04 |
| *bath-30* | WBGene00017460 | protein_coding_gene | -2.7 | 0.150 | 5.7E-37 | -2.7 | 0.15 | 4.5E-49 |
| *F11A5.15* | WBGene00008680 | protein_coding_gene | -2.8 | 0.149 | 1.3E-03 | -2.2 | 0.21 | 6.5E-06 |
| *F12E12.11* | WBGene00044811 | protein_coding_gene | -2.8 | 0.149 | 2.6E-09 | -2.0 | 0.25 | 3.8E-09 |
| *Y102A5C.35* | WBGene00044220 | pseudogene | -2.8 | 0.148 | 1.3E-03 | -1.5 | 0.36 | 3.8E-03 |
| *Y38H6C.19* | WBGene00012631 | protein_coding_gene | -2.8 | 0.146 | 6.7E-06 | -3.2 | 0.11 | 7.2E-35 |
| *sdz-18* | WBGene00018442 | protein_coding_gene | -2.8 | 0.145 | 1.1E-03 | -1.6 | 0.32 | 2.5E-04 |
| *bath-6* | WBGene00018666 | pseudogene | -2.8 | 0.143 | 1.5E-06 | -1.5 | 0.36 | 1.2E-03 |
| *fbxb-43* | WBGene00021774 | protein_coding_gene | -2.8 | 0.142 | 4.9E-04 | -1.8 | 0.28 | 5.7E-04 |
| *arrd-20* | WBGene00015440 | pseudogene | -2.8 | 0.142 | 1.0E-03 | -2.2 | 0.22 | 1.3E-04 |
| *C25F9.9* | WBGene00007728 | protein_coding_gene | -2.8 | 0.141 | 9.8E-04 | -3.3 | 0.10 | 9.5E-09 |
| *F16H6.7* | WBGene00008897 | protein_coding_gene | -2.8 | 0.141 | 9.1E-04 | -1.7 | 0.31 | 1.3E-04 |
| *CE7X_3.2* | WBGene00008350 | pseudogene | -2.8 | 0.140 | 2.3E-03 | -2.0 | 0.26 | 2.1E-05 |
| *ZC239.20* | WBGene00044371 | protein_coding_gene | -2.8 | 0.140 | 1.1E-04 | -2.1 | 0.23 | 9.2E-06 |
| *F13B6.3* | WBGene00017418 | protein_coding_gene | -2.9 | 0.139 | 8.8E-12 | -2.4 | 0.19 | 4.4E-26 |
| *ugt-28* | WBGene00015693 | protein_coding_gene | -2.9 | 0.138 | 3.0E-21 | -2.0 | 0.25 | 3.3E-13 |
| *C17B7.5* | WBGene00015879 | protein_coding_gene | -2.9 | 0.138 | 8.2E-04 | -2.2 | 0.22 | 9.9E-06 |
| *D1022.17* | WBGene00271809 | ncRNA_gene | -2.9 | 0.136 | 2.7E-05 | -1.9 | 0.26 | 1.4E-04 |
| *scrm-4* | WBGene00008681 | protein_coding_gene | -2.9 | 0.136 | 2.2E-22 | -2.0 | 0.24 | 2.7E-26 |
| *folt-2* | WBGene00018138 | protein_coding_gene | -2.9 | 0.135 | 2.3E-04 | -1.9 | 0.27 | 3.1E-35 |
| *fbxa-57* | WBGene00016224 | pseudogene | -2.9 | 0.135 | 7.7E-04 | -2.4 | 0.19 | 5.2E-05 |
| *C40A11.4* | WBGene00016547 | protein_coding_gene | -2.9 | 0.135 | 1.7E-03 | -2.3 | 0.20 | 1.4E-05 |

| | | | | | | | | |
|---|---|---|---|---|---|---|---|---|
| *fbxb-52* | WBGene00019417 | protein_coding_gene | -2.9 | 0.135 | 7.3E-04 | -1.4 | 0.39 | 1.1E-03 |
| *F57E7.1* | WBGene00010202 | protein_coding_gene | -2.9 | 0.134 | 7.1E-04 | -3.0 | 0.13 | 6.5E-08 |
| *fbxc-19* | WBGene00020361 | protein_coding_gene | -2.9 | 0.133 | 6.5E-04 | -2.1 | 0.23 | 1.7E-04 |
| *fbxb-13* | WBGene00015607 | protein_coding_gene | -2.9 | 0.133 | 2.9E-04 | -2.0 | 0.25 | 1.8E-04 |
| *Y43F8B.14* | WBGene00012824 | protein_coding_gene | -2.9 | 0.131 | 6.2E-04 | -1.4 | 0.38 | 8.8E-04 |
| *fbxb-44* | WBGene00019416 | protein_coding_gene | -2.9 | 0.130 | 5.9E-04 | -2.0 | 0.26 | 3.3E-05 |
| *clec-119* | WBGene00018575 | protein_coding_gene | -2.9 | 0.130 | 9.4E-06 | -2.5 | 0.18 | 1.0E-09 |
| *Y45F10C.6* | WBGene00044256 | protein_coding_gene | -3.0 | 0.129 | 5.4E-04 | -1.8 | 0.30 | 8.6E-05 |
| *col-141* | WBGene00000714 | protein_coding_gene | -3.0 | 0.128 | 3.3E-05 | -1.7 | 0.32 | 1.2E-04 |
| *F40G9.6* | WBGene00018242 | protein_coding_gene | -3.0 | 0.126 | 4.7E-04 | -2.2 | 0.22 | 1.1E-04 |
| *fbxb-97* | WBGene00016885 | protein_coding_gene | -3.0 | 0.125 | 4.5E-04 | -1.4 | 0.39 | 1.1E-03 |
| *F21D9.6* | WBGene00009020 | pseudogene | -3.0 | 0.125 | 4.6E-04 | -2.1 | 0.23 | 2.5E-04 |
| *R10E8.1* | WBGene00011207 | protein_coding_gene | -3.0 | 0.124 | 2.7E-14 | -3.2 | 0.11 | 1.5E-27 |
| *C50E3.13* | WBGene00016824 | protein_coding_gene | -3.0 | 0.123 | 8.6E-30 | -3.0 | 0.13 | 4.7E-39 |
| *msp-53* | WBGene00003446 | protein_coding_gene | -3.0 | 0.123 | 2.2E-04 | -2.6 | 0.16 | 8.0E-07 |
| *ZK1240.11* | WBGene00206413 | protein_coding_gene | -3.0 | 0.123 | 4.2E-04 | -1.7 | 0.30 | 1.1E-04 |
| *M01G12.9* | WBGene00010822 | protein_coding_gene | -3.0 | 0.122 | 4.0E-09 | -2.9 | 0.13 | 1.3E-18 |
| *M04C3.5* | WBGene00206418 | protein_coding_gene | -3.0 | 0.122 | 3.9E-04 | -4.0 | 0.06 | 3.9E-11 |
| *C40A11.8* | WBGene00016551 | protein_coding_gene | -3.0 | 0.121 | 9.3E-09 | -1.7 | 0.30 | 2.5E-07 |
| *Y39G10AR.16* | WBGene00021472 | protein_coding_gene | -3.1 | 0.120 | 1.6E-05 | -3.1 | 0.11 | 5.0E-20 |
| *K08D10.9* | WBGene00019532 | pseudogene | -3.1 | 0.118 | 3.0E-04 | -2.1 | 0.23 | 1.2E-05 |
| *linc-11* | WBGene00219758 | lincRNA_gene | -3.1 | 0.115 | 7.6E-05 | -3.4 | 0.09 | 1.5E-07 |
| *C33C12.7* | WBGene00016339 | protein_coding_gene | -3.1 | 0.113 | 2.3E-04 | -2.4 | 0.19 | 2.1E-05 |
| *ZC239.22* | WBGene00194717 | protein_coding_gene | -3.1 | 0.113 | 6.9E-04 | -3.5 | 0.09 | 9.7E-10 |
| *clp-6* | WBGene00000546 | protein_coding_gene | -3.1 | 0.113 | 2.4E-06 | -2.3 | 0.20 | 1.7E-14 |
| *col-12* | WBGene00000601 | protein_coding_gene | -3.2 | 0.113 | 1.5E-06 | -1.6 | 0.32 | 6.2E-16 |
| *fbxa-156* | WBGene00007392 | protein_coding_gene | -3.2 | 0.112 | 5.9E-05 | -2.9 | 0.14 | 8.2E-10 |
| *T20D4.11* | WBGene00020617 | protein_coding_gene | -3.2 | 0.111 | 6.5E-04 | -2.1 | 0.24 | 2.5E-04 |
| *fbxb-98* | WBGene00015830 | protein_coding_gene | -3.2 | 0.110 | 1.9E-04 | -1.6 | 0.34 | 3.2E-04 |
| *T24E12.5* | WBGene00020774 | protein_coding_gene | -3.2 | 0.110 | 1.7E-04 | -4.3 | 0.05 | 4.1E-10 |
| *F47H4.2* | WBGene00009835 | protein_coding_gene | -3.2 | 0.109 | 5.6E-04 | -4.9 | 0.03 | 2.6E-14 |
| *C50E3.12* | WBGene00016823 | protein_coding_gene | -3.2 | 0.108 | 2.1E-19 | -1.8 | 0.30 | 1.2E-08 |
| *F09C6.12* | WBGene00044182 | protein_coding_gene | -3.2 | 0.108 | 5.1E-04 | -3.8 | 0.07 | 1.9E-08 |
| *Y113G7A.13* | WBGene00013750 | protein_coding_gene | -3.2 | 0.108 | 1.7E-04 | -2.1 | 0.23 | 1.7E-04 |
| *ZK1025.3* | WBGene00014183 | protein_coding_gene | -3.2 | 0.108 | 1.5E-04 | -1.9 | 0.28 | 5.1E-04 |
| *C52E2.5* | WBGene00016884 | protein_coding_gene | -3.2 | 0.108 | 4.7E-04 | -3.2 | 0.11 | 5.4E-07 |
| *nspa-5* | WBGene00013887 | protein_coding_gene | -3.2 | 0.106 | 1.1E-04 | -3.7 | 0.08 | 2.3E-08 |
| *gst-38* | WBGene00001786 | protein_coding_gene | -3.2 | 0.106 | 1.5E-04 | -1.4 | 0.38 | 4.6E-03 |
| *W02A11.6* | WBGene00014852 | pseudogene | -3.2 | 0.106 | 1.5E-04 | -1.4 | 0.37 | 7.0E-04 |
| *B0281.4* | WBGene00015111 | protein_coding_gene | -3.2 | 0.106 | 1.4E-04 | -1.5 | 0.36 | 4.9E-04 |
| *duxl-1* | WBGene00022554 | protein_coding_gene | -3.3 | 0.105 | 2.6E-26 | -3.8 | 0.07 | 4.8E-48 |
| *W05G11.2* | WBGene00021040 | protein_coding_gene | -3.3 | 0.104 | 9.0E-05 | -2.6 | 0.17 | 1.5E-05 |
| *Y46B2A.4* | WBGene00284850 | protein_coding_gene | -3.3 | 0.104 | 1.1E-04 | -2.1 | 0.23 | 1.9E-04 |
| *C08F11.6* | WBGene00014672 | pseudogene | -3.3 | 0.103 | 2.7E-06 | -2.8 | 0.14 | 1.4E-09 |
| *kin-21* | WBGene00002204 | protein_coding_gene | -3.3 | 0.103 | 1.2E-04 | -2.6 | 0.16 | 9.2E-06 |
| *F59H6.15* | WBGene00269394 | protein_coding_gene | -3.3 | 0.102 | 1.1E-04 | -2.6 | 0.17 | 1.3E-05 |
| *Y47H10A.6* | WBGene00077537 | pseudogene | -3.3 | 0.100 | 9.9E-05 | -1.8 | 0.29 | 7.5E-05 |
| *fbxb-50* | WBGene00019413 | protein_coding_gene | -3.3 | 0.100 | 9.9E-05 | -1.9 | 0.28 | 4.3E-05 |
| *C40A11.6* | WBGene00016549 | protein_coding_gene | -3.3 | 0.099 | 9.0E-05 | -2.7 | 0.15 | 2.3E-07 |
| *Y19D10B.6* | WBGene00021235 | protein_coding_gene | -3.3 | 0.099 | 3.9E-06 | -2.3 | 0.21 | 1.1E-06 |
| *Y49E10.37* | WBGene00304219 | ncRNA_gene | -3.3 | 0.098 | 9.7E-16 | -2.1 | 0.23 | 2.9E-11 |
| *T02G6.10* | WBGene00077772 | pseudogene | -3.3 | 0.098 | 1.1E-05 | -2.5 | 0.17 | 1.8E-05 |
| *F16H6.10* | WBGene00008900 | protein_coding_gene | -3.4 | 0.097 | 9.5E-15 | -3.5 | 0.09 | 3.0E-22 |
| *Y47G7B.2* | WBGene00021654 | protein_coding_gene | -3.4 | 0.096 | 4.4E-05 | -1.2 | 0.44 | 2.0E-02 |
| *arrd-22* | WBGene00020612 | protein_coding_gene | -3.4 | 0.095 | 1.2E-05 | -2.0 | 0.25 | 1.3E-06 |
| *fbxb-42* | WBGene00021773 | protein_coding_gene | -3.4 | 0.095 | 2.2E-04 | -2.3 | 0.20 | 2.7E-06 |
| *fbxb-15* | WBGene00022557 | protein_coding_gene | -3.4 | 0.095 | 5.3E-06 | -2.6 | 0.17 | 1.8E-05 |
| *npr-30* | WBGene00019183 | protein_coding_gene | -3.4 | 0.095 | 2.5E-04 | -2.0 | 0.25 | 1.7E-06 |
| *sdz-11* | WBGene00017408 | protein_coding_gene | -3.4 | 0.094 | 6.4E-05 | -1.6 | 0.33 | 2.4E-04 |
| *btb-17* | WBGene00018200 | protein_coding_gene | -3.4 | 0.094 | 2.4E-06 | -4.4 | 0.05 | 1.3E-16 |
| *W07A12.8* | WBGene00012325 | protein_coding_gene | -3.4 | 0.093 | 1.4E-06 | -1.3 | 0.40 | 1.5E-03 |
| *stdh-2* | WBGene00008678 | protein_coding_gene | -3.4 | 0.093 | 5.7E-05 | -2.4 | 0.19 | 2.4E-06 |
| *Y60A3A.25* | WBGene00044211 | protein_coding_gene | -3.4 | 0.092 | 3.2E-05 | -2.0 | 0.24 | 3.7E-07 |
| *Y43F8B.24* | WBGene00195176 | protein_coding_gene | -3.5 | 0.091 | 4.9E-05 | -3.2 | 0.11 | 3.8E-07 |
| *phat-3* | WBGene00016782 | protein_coding_gene | -3.5 | 0.090 | 6.6E-17 | -4.0 | 0.06 | 2.6E-60 |

| | | | | | | | | |
|---|---|---|---|---|---|---|---|---|
| *C17F4.8* | WBGene00015914 | protein_coding_gene | -3.5 | 0.089 | 4.6E-05 | -2.2 | 0.21 | 5.9E-06 |
| *F26A3.5* | WBGene00009143 | protein_coding_gene | -3.5 | 0.089 | 1.4E-06 | -3.4 | 0.09 | 8.9E-16 |
| *F14D2.5* | WBGene00017453 | protein_coding_gene | -3.5 | 0.089 | 4.6E-05 | -1.3 | 0.40 | 1.4E-03 |
| *F36H5.14* | WBGene00219271 | protein_coding_gene | -3.5 | 0.089 | 4.4E-05 | -3.1 | 0.12 | 2.1E-08 |
| *sdz-24* | WBGene00019495 | protein_coding_gene | -3.5 | 0.089 | 2.2E-05 | -5.7 | 0.02 | 2.2E-101 |
| *F19B2.6* | WBGene00008945 | protein_coding_gene | -3.5 | 0.089 | 8.2E-07 | -2.1 | 0.23 | 1.8E-08 |
| *fbxb-96* | WBGene00016886 | protein_coding_gene | -3.5 | 0.088 | 3.9E-05 | -2.2 | 0.22 | 7.5E-06 |
| *bath-3* | WBGene00019140 | protein_coding_gene | -3.5 | 0.086 | 2.8E-13 | -2.5 | 0.18 | 6.8E-10 |
| *asp-17* | WBGene00012683 | protein_coding_gene | -3.6 | 0.085 | 4.0E-06 | -3.0 | 0.13 | 9.0E-17 |
| *F39F10.3* | WBGene00018203 | protein_coding_gene | -3.6 | 0.085 | 7.7E-10 | -2.9 | 0.13 | 5.0E-11 |
| *fbxa-188* | WBGene00009839 | protein_coding_gene | -3.6 | 0.085 | 3.0E-05 | -3.2 | 0.11 | 1.4E-08 |
| *Y73B6BL.14* | WBGene00022237 | protein_coding_gene | -3.6 | 0.083 | 1.1E-04 | -1.2 | 0.44 | 3.4E-03 |
| *ZC204.12* | WBGene00022562 | protein_coding_gene | -3.6 | 0.082 | 1.7E-07 | -5.6 | 0.02 | 3.1E-42 |
| *ZK1240.5* | WBGene00022870 | protein_coding_gene | -3.6 | 0.081 | 2.2E-05 | -2.8 | 0.14 | 1.6E-07 |
| *scrm-8* | WBGene00019530 | protein_coding_gene | -3.6 | 0.081 | 1.9E-05 | -2.6 | 0.16 | 9.1E-06 |
| *T20D4.5* | WBGene00020611 | protein_coding_gene | -3.6 | 0.080 | 2.0E-05 | -2.3 | 0.20 | 3.1E-06 |
| *C04G6.6* | WBGene00015456 | protein_coding_gene | -3.7 | 0.079 | 1.7E-18 | -2.1 | 0.23 | 4.8E-15 |
| *F54E2.1* | WBGene00018823 | protein_coding_gene | -3.7 | 0.076 | 5.1E-08 | -3.5 | 0.09 | 1.2E-26 |
| *ZC239.21* | WBGene00044372 | protein_coding_gene | -3.7 | 0.076 | 1.3E-05 | -1.6 | 0.33 | 2.3E-04 |
| *F14D2.15* | WBGene00023427 | protein_coding_gene | -3.7 | 0.075 | 1.0E-20 | -3.6 | 0.08 | 9.7E-24 |
| *msp-64* | WBGene00003457 | protein_coding_gene | -3.7 | 0.075 | 7.6E-07 | -3.8 | 0.07 | 1.3E-34 |
| *bath-27* | WBGene00017450 | protein_coding_gene | -3.8 | 0.074 | 4.6E-16 | -3.5 | 0.09 | 3.9E-17 |
| *btb-3* | WBGene00015013 | pseudogene | -3.8 | 0.073 | 1.0E-05 | -2.3 | 0.21 | 8.7E-05 |
| *F19B10.2* | WBGene00017584 | protein_coding_gene | -3.8 | 0.072 | 4.5E-22 | -3.7 | 0.08 | 2.4E-30 |
| *F59H6.2* | WBGene00019133 | pseudogene | -3.8 | 0.072 | 8.7E-06 | -1.6 | 0.34 | 3.0E-04 |
| *math-20* | WBGene00016555 | protein_coding_gene | -3.8 | 0.072 | 2.3E-10 | -3.5 | 0.09 | 7.2E-24 |
| *fbxa-117* | WBGene00014907 | pseudogene | -3.8 | 0.072 | 8.3E-06 | -2.2 | 0.22 | 6.8E-06 |
| *scrm-5* | WBGene00019531 | protein_coding_gene | -3.8 | 0.071 | 6.4E-06 | -3.5 | 0.09 | 1.4E-09 |
| *T19C9.8* | WBGene00011844 | protein_coding_gene | -3.8 | 0.071 | 6.3E-06 | -4.4 | 0.05 | 2.5E-10 |
| *fbxa-10* | WBGene00020450 | protein_coding_gene | -3.8 | 0.070 | 7.1E-06 | -3.2 | 0.11 | 4.7E-07 |
| *fbxb-49* | WBGene00019412 | protein_coding_gene | -3.8 | 0.070 | 6.5E-06 | -1.6 | 0.34 | 2.9E-03 |
| *nhr-115* | WBGene00003705 | protein_coding_gene | -3.8 | 0.070 | 4.4E-10 | -2.0 | 0.25 | 5.4E-07 |
| *F52C6.4* | WBGene00018661 | protein_coding_gene | -3.9 | 0.068 | 2.7E-05 | -2.6 | 0.17 | 2.5E-06 |
| *F10E7.1* | WBGene00017343 | protein_coding_gene | -3.9 | 0.068 | 5.6E-06 | -2.2 | 0.22 | 6.8E-06 |
| *K09H9.7* | WBGene00019598 | protein_coding_gene | -3.9 | 0.068 | 6.2E-28 | -2.9 | 0.13 | 3.8E-27 |
| *fat-5* | WBGene00001397 | protein_coding_gene | -3.9 | 0.067 | 3.0E-36 | -1.4 | 0.38 | 2.6E-23 |
| *T12B5.14* | WBGene00050913 | protein_coding_gene | -3.9 | 0.066 | 4.4E-06 | -3.0 | 0.12 | 3.0E-08 |
| *T26E3.8* | WBGene00012041 | protein_coding_gene | -3.9 | 0.066 | 4.4E-06 | -2.2 | 0.21 | 8.7E-05 |
| *nspd-7* | WBGene00012781 | protein_coding_gene | -4.0 | 0.063 | 8.5E-08 | -3.8 | 0.07 | 1.7E-40 |
| *fbxa-11* | WBGene00020451 | protein_coding_gene | -4.0 | 0.063 | 2.9E-06 | -3.0 | 0.13 | 5.5E-08 |
| *math-40* | WBGene00020359 | protein_coding_gene | -4.0 | 0.062 | 4.0E-08 | -3.4 | 0.10 | 7.3E-13 |
| *F14D2.19* | WBGene00206522 | protein_coding_gene | -4.0 | 0.060 | 2.1E-06 | -3.9 | 0.07 | 9.7E-11 |
| *fbxa-6* | WBGene00008352 | protein_coding_gene | -4.0 | 0.060 | 2.1E-06 | -3.6 | 0.08 | 2.3E-08 |
| *Y57G11C.40* | WBGene00013329 | protein_coding_gene | -4.1 | 0.060 | 2.1E-06 | -2.3 | 0.21 | 4.3E-06 |
| *Y9D1A.1* | WBGene00021186 | protein_coding_gene | -4.1 | 0.060 | 7.8E-16 | -2.9 | 0.14 | 3.7E-15 |
| *pud-3* | WBGene00017501 | protein_coding_gene | -4.1 | 0.059 | 1.2E-06 | -4.4 | 0.05 | 1.6E-11 |
| *K05F6.4* | WBGene00019415 | protein_coding_gene | -4.1 | 0.059 | 5.8E-09 | -2.8 | 0.15 | 3.9E-08 |
| *F42G2.5* | WBGene00018354 | protein_coding_gene | -4.1 | 0.058 | 1.6E-06 | -4.4 | 0.05 | 3.8E-11 |
| *C02G6.1* | WBGene00015359 | pseudogene | -4.1 | 0.057 | 2.1E-07 | -1.0 | 0.49 | 5.3E-03 |
| *F31E9.11* | WBGene00195233 | ncRNA_gene | -4.1 | 0.057 | 7.1E-06 | -3.6 | 0.09 | 1.1E-09 |
| *math-24* | WBGene00016719 | protein_coding_gene | -4.2 | 0.056 | 1.9E-09 | -4.2 | 0.05 | 6.7E-22 |
| *F26D2.10* | WBGene00009149 | protein_coding_gene | -4.2 | 0.056 | 9.9E-07 | -2.9 | 0.14 | 1.1E-09 |
| *fbxb-46* | WBGene00019420 | protein_coding_gene | -4.2 | 0.056 | 1.1E-06 | -1.9 | 0.26 | 3.2E-05 |
| *ugt-25* | WBGene00015692 | protein_coding_gene | -4.2 | 0.055 | 2.0E-25 | -4.0 | 0.06 | 6.5E-37 |
| *clec-72* | WBGene00021583 | protein_coding_gene | -4.2 | 0.055 | 9.8E-07 | -3.1 | 0.12 | 2.6E-08 |
| *fbxb-20* | WBGene00022559 | protein_coding_gene | -4.2 | 0.054 | 8.6E-07 | -2.0 | 0.26 | 1.9E-04 |
| *fbxa-38* | WBGene00016522 | protein_coding_gene | -4.2 | 0.054 | 8.5E-07 | -3.5 | 0.09 | 1.6E-09 |
| *D1065.6* | WBGene00268212 | ncRNA_gene | -4.2 | 0.053 | 6.9E-08 | -2.8 | 0.14 | 5.0E-06 |
| *fbxb-95* | WBGene00016880 | protein_coding_gene | -4.3 | 0.053 | 6.3E-07 | -2.5 | 0.18 | 1.4E-06 |
| *F52C6.12* | WBGene00018669 | protein_coding_gene | -4.3 | 0.052 | 3.8E-06 | -3.4 | 0.09 | 3.3E-09 |
| *Y75B8A.32* | WBGene00013562 | protein_coding_gene | -4.3 | 0.051 | 5.0E-07 | -3.7 | 0.08 | 4.1E-10 |
| *F19B10.1* | WBGene00017583 | protein_coding_gene | -4.3 | 0.051 | 3.6E-14 | -3.5 | 0.09 | 5.8E-19 |
| *fbxa-60* | WBGene00020456 | protein_coding_gene | -4.3 | 0.050 | 1.4E-28 | -3.4 | 0.09 | 2.0E-65 |
| *fbxb-54* | WBGene00019418 | protein_coding_gene | -4.4 | 0.048 | 2.9E-07 | -2.8 | 0.14 | 1.3E-07 |
| *btb-14* | WBGene00013275 | protein_coding_gene | -4.4 | 0.048 | 6.0E-25 | -3.1 | 0.12 | 2.1E-29 |

| | | | | | | | | |
|---|---|---|---|---|---|---|---|---|
| *F52C6.3* | WBGene00018660 | protein_coding_gene | -4.4 | 0.047 | 2.6E-07 | -3.2 | 0.11 | 3.4E-07 |
| *grsp-4* | WBGene00017201 | protein_coding_gene | -4.4 | 0.047 | 3.4E-47 | -2.2 | 0.22 | 1.3E-20 |
| *M04C3.2* | WBGene00010851 | protein_coding_gene | -4.4 | 0.047 | 2.4E-07 | -4.4 | 0.05 | 5.9E-11 |
| *ctsa-4.2* | WBGene00021503 | protein_coding_gene | -4.4 | 0.047 | 7.5E-44 | -3.2 | 0.11 | 1.8E-84 |
| *K08D10.14* | WBGene00219216 | protein_coding_gene | -4.5 | 0.045 | 1.6E-07 | -4.0 | 0.06 | 1.4E-09 |
| *Y73B6BL.288* | WBGene00219218 | protein_coding_gene | -4.5 | 0.045 | 3.1E-15 | -2.2 | 0.21 | 7.6E-06 |
| *zeel-1* | WBGene00021463 | protein_coding_gene | -4.5 | 0.045 | 1.5E-07 | -3.4 | 0.10 | 1.3E-09 |
| *C04E12.2* | WBGene00015433 | protein_coding_gene | -4.5 | 0.044 | 1.1E-06 | -4.0 | 0.06 | 6.5E-11 |
| *fbxa-180* | WBGene00009295 | pseudogene | -4.5 | 0.043 | 9.0E-07 | -3.9 | 0.07 | 2.0E-09 |
| *R07E5.15* | WBGene00011120 | protein_coding_gene | -4.6 | 0.042 | 3.5E-07 | -3.5 | 0.09 | 1.9E-14 |
| *F49F1.7* | WBGene00018647 | protein_coding_gene | -4.6 | 0.041 | 7.1E-07 | -1.1 | 0.47 | 2.0E-02 |
| *math-39* | WBGene00020358 | protein_coding_gene | -4.6 | 0.040 | 5.0E-07 | -3.5 | 0.09 | 6.8E-15 |
| *F14D2.14* | WBGene00017462 | protein_coding_gene | -4.7 | 0.039 | 4.3E-08 | -3.1 | 0.12 | 2.5E-08 |
| *bath-11* | WBGene00018664 | pseudogene | -4.7 | 0.039 | 4.0E-08 | -2.6 | 0.16 | 1.3E-07 |
| *Y54G2A.45* | WBGene00044488 | protein_coding_gene | -4.7 | 0.038 | 3.9E-27 | -2.5 | 0.17 | 1.3E-30 |
| *math-41* | WBGene00020360 | protein_coding_gene | -4.7 | 0.038 | 2.4E-11 | -3.3 | 0.10 | 1.6E-12 |
| *T22B7.7* | WBGene00020674 | protein_coding_gene | -4.7 | 0.038 | 1.9E-30 | -2.2 | 0.22 | 7.2E-20 |
| *T28A11.2* | WBGene00020869 | protein_coding_gene | -4.7 | 0.038 | 8.6E-15 | -2.9 | 0.13 | 5.0E-11 |
| *F14D2.11* | WBGene00017459 | protein_coding_gene | -4.7 | 0.037 | 3.0E-08 | -3.3 | 0.10 | 1.6E-07 |
| *fbxb-91* | WBGene00044692 | protein_coding_gene | -4.8 | 0.037 | 2.5E-08 | -2.8 | 0.14 | 2.0E-06 |
| *Y82E9BR.5* | WBGene00022338 | protein_coding_gene | -4.8 | 0.037 | 2.4E-08 | -3.8 | 0.07 | 1.5E-08 |
| *Y46G5A.7* | WBGene00012899 | protein_coding_gene | -4.8 | 0.035 | 1.7E-08 | -3.9 | 0.07 | 6.0E-09 |
| *nspc-14* | WBGene00009638 | protein_coding_gene | -4.8 | 0.035 | 6.1E-31 | -5.2 | 0.03 | 1.0E-105 |
| *fbxa-216* | WBGene00012953 | protein_coding_gene | -4.9 | 0.034 | 4.1E-10 | -3.3 | 0.10 | 6.0E-09 |
| *T16G12.3* | WBGene00011804 | protein_coding_gene | -4.9 | 0.034 | 1.4E-40 | -3.4 | 0.10 | 1.4E-38 |
| *Y7A9C.1* | WBGene00012418 | pseudogene | -4.9 | 0.033 | 9.2E-09 | -4.7 | 0.04 | 5.3E-12 |
| *C25F9.10* | WBGene00044291 | protein_coding_gene | -4.9 | 0.033 | 9.1E-09 | -4.0 | 0.06 | 7.3E-11 |
| *fbxa-65* | WBGene00015885 | protein_coding_gene | -5.0 | 0.032 | 1.3E-12 | -3.6 | 0.08 | 2.2E-16 |
| *C25F9.14* | WBGene00045461 | protein_coding_gene | -5.0 | 0.032 | 6.5E-09 | -4.0 | 0.06 | 4.4E-11 |
| *M162.9* | WBGene00014811 | pseudogene | -5.0 | 0.031 | 5.2E-09 | -3.2 | 0.11 | 8.0E-09 |
| *btb-10* | WBGene00019313 | protein_coding_gene | -5.0 | 0.030 | 4.2E-09 | -3.5 | 0.09 | 6.9E-08 |
| *C08F11.7* | WBGene00007454 | protein_coding_gene | -5.0 | 0.030 | 3.6E-17 | -3.5 | 0.09 | 4.9E-19 |
| *R10E8.7* | WBGene00014821 | pseudogene | -5.1 | 0.030 | 6.6E-11 | -4.0 | 0.06 | 2.7E-12 |
| *F14D2.8* | WBGene00017456 | protein_coding_gene | -5.1 | 0.030 | 3.6E-09 | -3.2 | 0.11 | 6.6E-07 |
| *K09F6.15* | WBGene00271744 | protein_coding_gene | -5.1 | 0.030 | 3.6E-09 | -4.1 | 0.06 | 3.0E-10 |
| *bath-7* | WBGene00018667 | protein_coding_gene | -5.1 | 0.030 | 7.4E-09 | -3.2 | 0.11 | 5.6E-11 |
| *bath-22* | WBGene00018658 | pseudogene | -5.1 | 0.029 | 2.9E-09 | -2.8 | 0.14 | 3.4E-06 |
| *bath-14* | WBGene00015016 | pseudogene | -5.1 | 0.029 | 2.7E-09 | -2.4 | 0.19 | 2.6E-05 |
| *Y46G5A.8* | WBGene00012900 | protein_coding_gene | -5.2 | 0.028 | 1.7E-09 | -2.8 | 0.15 | 3.2E-06 |
| *bath-24* | WBGene00015014 | protein_coding_gene | -5.2 | 0.028 | 1.7E-09 | -3.7 | 0.07 | 5.8E-09 |
| *K09F6.3* | WBGene00019586 | protein_coding_gene | -5.2 | 0.028 | 9.9E-09 | -4.3 | 0.05 | 2.0E-27 |
| *fbxa-189* | WBGene00009840 | protein_coding_gene | -5.2 | 0.027 | 1.6E-09 | -4.2 | 0.05 | 9.5E-12 |
| *ZC239.6* | WBGene00022569 | protein_coding_gene | -5.2 | 0.027 | 1.4E-09 | -3.2 | 0.11 | 7.9E-09 |
| *R08E5.1* | WBGene00019961 | protein_coding_gene | -5.2 | 0.027 | 1.3E-09 | -3.4 | 0.09 | 6.3E-08 |
| *K09F6.7* | WBGene00019590 | protein_coding_gene | -5.3 | 0.026 | 9.0E-10 | -4.3 | 0.05 | 4.7E-14 |
| *K09F6.6* | WBGene00019589 | protein_coding_gene | -5.3 | 0.026 | 7.1E-10 | -4.3 | 0.05 | 4.0E-14 |
| *C25F9.4* | WBGene00007724 | protein_coding_gene | -5.3 | 0.025 | 6.4E-10 | -4.8 | 0.03 | 3.7E-16 |
| *math-26* | WBGene00018100 | protein_coding_gene | -5.3 | 0.025 | 1.2E-21 | -4.3 | 0.05 | 7.4E-42 |
| *K05C4.9* | WBGene00010585 | protein_coding_gene | -5.3 | 0.025 | 5.5E-10 | -4.0 | 0.06 | 3.4E-09 |
| *K09F6.4* | WBGene00019587 | protein_coding_gene | -5.3 | 0.025 | 8.3E-09 | -3.6 | 0.08 | 2.7E-10 |
| *C38D9.13* | WBGene00235090 | ncRNA_gene | -5.4 | 0.024 | 2.1E-12 | -3.2 | 0.11 | 5.2E-20 |
| *skr-4* | WBGene00004810 | protein_coding_gene | -5.4 | 0.024 | 1.2E-23 | -4.9 | 0.03 | 8.5E-43 |
| *Y46D2A.2* | WBGene00021590 | protein_coding_gene | -5.4 | 0.024 | 3.8E-10 | -1.7 | 0.30 | 1.2E-04 |
| *K09F6.10* | WBGene00019593 | protein_coding_gene | -5.4 | 0.023 | 2.9E-10 | -4.8 | 0.04 | 1.3E-11 |
| *msp-19* | WBGene00003426 | protein_coding_gene | -5.4 | 0.023 | 7.5E-10 | -4.6 | 0.04 | 1.3E-20 |
| *fbxa-207* | WBGene00013612 | pseudogene | -5.5 | 0.022 | 1.8E-10 | -3.5 | 0.09 | 5.7E-10 |
| *math-14* | WBGene00015828 | protein_coding_gene | -5.5 | 0.022 | 1.5E-10 | -5.7 | 0.02 | 2.3E-25 |
| *nspa-8* | WBGene00012294 | protein_coding_gene | -5.5 | 0.022 | 1.7E-10 | -4.4 | 0.05 | 2.4E-10 |
| *math-27* | WBGene00018101 | pseudogene | -5.5 | 0.022 | 1.7E-18 | -3.8 | 0.07 | 2.8E-20 |
| *math-3* | WBGene00015608 | protein_coding_gene | -5.5 | 0.022 | 5.3E-26 | -4.3 | 0.05 | 6.4E-28 |
| *F59E12.6* | WBGene00019122 | protein_coding_gene | -5.6 | 0.021 | 9.5E-11 | -3.6 | 0.08 | 7.5E-10 |
| *K09F6.9* | WBGene00019592 | protein_coding_gene | -5.6 | 0.021 | 2.0E-09 | -3.4 | 0.10 | 1.7E-15 |
| *gst-22* | WBGene00001770 | protein_coding_gene | -5.6 | 0.021 | 8.2E-11 | -4.2 | 0.05 | 1.3E-11 |
| *E01G4.5* | WBGene00008447 | protein_coding_gene | -5.6 | 0.021 | 1.8E-10 | -4.5 | 0.04 | 2.4E-17 |
| *fbxa-129* | WBGene00010294 | pseudogene | -5.6 | 0.020 | 7.0E-11 | -3.5 | 0.09 | 1.8E-09 |

| | | | | | | | | |
|---|---|---|---|---|---|---|---|---|
| *C29G2.2* | WBGene00016230 | pseudogene | -5.6 | 0.020 | 6.8E-11 | -3.6 | 0.08 | 3.8E-08 |
| *fbxa-67* | WBGene00020457 | protein_coding_gene | -5.7 | 0.020 | 5.2E-11 | -4.6 | 0.04 | 7.9E-11 |
| *T07D3.4* | WBGene00020307 | protein_coding_gene | -5.7 | 0.019 | 9.5E-10 | -4.3 | 0.05 | 1.1E-10 |
| *Y102A5C.36* | WBGene00044213 | protein_coding_gene | -5.8 | 0.018 | 2.4E-11 | -4.4 | 0.05 | 2.3E-12 |
| *pals-23* | WBGene00016218 | protein_coding_gene | -5.9 | 0.017 | 1.1E-11 | -3.5 | 0.09 | 7.8E-16 |
| *Y48E1B.8* | WBGene00013007 | protein_coding_gene | -5.9 | 0.017 | 5.7E-31 | -1.7 | 0.31 | 4.3E-04 |
| *bath-1* | WBGene00019139 | protein_coding_gene | -5.9 | 0.017 | 8.5E-12 | -4.5 | 0.04 | 1.5E-10 |
| *ZK1240.2* | WBGene00022867 | protein_coding_gene | -5.9 | 0.016 | 7.0E-12 | -4.8 | 0.04 | 2.1E-12 |
| *bath-19* | WBGene00019132 | protein_coding_gene | -5.9 | 0.016 | 6.7E-12 | -4.0 | 0.06 | 5.6E-11 |
| *ZC204.14* | WBGene00022564 | protein_coding_gene | -5.9 | 0.016 | 6.5E-12 | -4.3 | 0.05 | 6.8E-10 |
| *C25F9.5* | WBGene00007725 | protein_coding_gene | -6.0 | 0.016 | 4.6E-12 | -5.1 | 0.03 | 1.1E-14 |
| *msp-33* | WBGene00003431 | protein_coding_gene | -6.1 | 0.015 | 7.8E-17 | -1.6 | 0.33 | 9.2E-07 |
| *btb-4* | WBGene00019142 | protein_coding_gene | -6.1 | 0.015 | 2.2E-12 | -4.4 | 0.05 | 4.5E-14 |
| *B0281.3* | WBGene00015110 | protein_coding_gene | -6.1 | 0.015 | 1.7E-12 | -5.2 | 0.03 | 6.1E-14 |
| *pals-24* | WBGene00016219 | protein_coding_gene | -6.2 | 0.014 | 1.3E-12 | -5.3 | 0.02 | 1.1E-22 |
| *Y113G7B.12* | WBGene00013762 | protein_coding_gene | -6.2 | 0.014 | 1.1E-12 | -4.7 | 0.04 | 3.6E-11 |
| *fbxa-193* | WBGene00010295 | pseudogene | -6.2 | 0.014 | 8.5E-13 | -3.8 | 0.07 | 5.0E-14 |
| *F10E7.2* | WBGene00017344 | protein_coding_gene | -6.2 | 0.014 | 8.5E-13 | -4.9 | 0.03 | 1.8E-25 |
| *Y38E10A.14* | WBGene00012592 | protein_coding_gene | -6.2 | 0.013 | 1.2E-12 | -4.9 | 0.03 | 1.4E-22 |
| *R07C12.2* | WBGene00019932 | pseudogene | -6.3 | 0.013 | 4.1E-13 | -4.9 | 0.03 | 2.0E-22 |
| *C04E12.4* | WBGene00015434 | protein_coding_gene | -6.3 | 0.013 | 3.3E-13 | -5.6 | 0.02 | 2.4E-35 |
| *Y113G7B.14* | WBGene00013763 | pseudogene | -6.4 | 0.012 | 2.6E-13 | -4.9 | 0.03 | 7.3E-12 |
| *bath-2* | WBGene00018668 | protein_coding_gene | -6.4 | 0.011 | 9.3E-18 | -4.3 | 0.05 | 7.8E-33 |
| *Y102A5C.2* | WBGene00013611 | protein_coding_gene | -6.5 | 0.011 | 7.1E-14 | -4.4 | 0.05 | 1.3E-16 |
| *math-22* | WBGene00016717 | protein_coding_gene | -6.6 | 0.011 | 4.2E-14 | -4.0 | 0.06 | 3.5E-20 |
| *C38D9.2* | WBGene00008010 | protein_coding_gene | -6.6 | 0.010 | 1.9E-182 | -4.6 | 0.04 | 2.1E-193 |
| *bath-28* | WBGene00017461 | protein_coding_gene | -6.9 | 0.009 | 4.1E-15 | -5.0 | 0.03 | 2.9E-28 |
| *math-4* | WBGene00015609 | protein_coding_gene | -6.9 | 0.008 | 2.0E-15 | -5.7 | 0.02 | 1.5E-36 |
| *gst-27* | WBGene00001775 | protein_coding_gene | -7.1 | 0.007 | 2.8E-45 | -4.7 | 0.04 | 1.7E-112 |
| *ZK1240.3* | WBGene00022868 | protein_coding_gene | -7.1 | 0.007 | 4.7E-16 | -5.5 | 0.02 | 3.1E-31 |
| *F54D10.7* | WBGene00018808 | protein_coding_gene | -7.1 | 0.007 | 3.7E-16 | -5.1 | 0.03 | 2.3E-12 |
| *clec-10* | WBGene00015403 | protein_coding_gene | -7.1 | 0.007 | 3.6E-16 | -5.1 | 0.03 | 1.9E-12 |
| *F59A7.8* | WBGene00019095 | pseudogene | -7.2 | 0.007 | 2.1E-16 | -5.5 | 0.02 | 9.7E-20 |
| *Y102A5C.5* | WBGene00014954 | pseudogene | -7.2 | 0.007 | 1.9E-16 | -5.7 | 0.02 | 5.0E-16 |
| *F55B11.6* | WBGene00044249 | protein_coding_gene | -7.3 | 0.006 | 7.1E-17 | -5.9 | 0.02 | 1.0E-39 |
| *bath-5* | WBGene00019141 | protein_coding_gene | -7.4 | 0.006 | 4.6E-17 | -5.3 | 0.03 | 7.0E-26 |
| *T12B5.15* | WBGene00050914 | protein_coding_gene | -7.5 | 0.006 | 2.0E-17 | -5.7 | 0.02 | 3.5E-39 |
| *F19B10.10* | WBGene00017589 | protein_coding_gene | -7.5 | 0.005 | 1.6E-15 | -4.9 | 0.03 | 1.3E-48 |
| *fbxa-206* | WBGene00013610 | protein_coding_gene | -7.6 | 0.005 | 5.8E-18 | -4.9 | 0.03 | 1.0E-11 |
| *C27C7.1* | WBGene00007766 | protein_coding_gene | -7.7 | 0.005 | 1.3E-18 | -6.0 | 0.02 | 8.3E-60 |
| *M04C3.1* | WBGene00010850 | protein_coding_gene | -8.0 | 0.004 | 2.4E-19 | -6.4 | 0.01 | 1.9E-55 |
| *F19B2.5* | WBGene00008944 | protein_coding_gene | -8.1 | 0.004 | 5.6E-20 | -5.6 | 0.02 | 2.6E-24 |
| *sri-40* | WBGene00005552 | protein_coding_gene | -8.2 | 0.003 | 3.9E-18 | -5.3 | 0.02 | 2.9E-29 |
| *Y102A5C.6* | WBGene00014955 | pseudogene | -8.3 | 0.003 | 6.9E-21 | -6.4 | 0.01 | 9.4E-55 |
| *Y94H6A.10* | WBGene00022382 | protein_coding_gene | -8.6 | 0.002 | 3.6E-22 | -5.4 | 0.02 | 3.6E-23 |

DEGs unique to *S. epidermidis* condition

| Gene Name | Wormbase Gene ID | Type | $log_2FC$ | FC | Adj P-value |
|---|---|---|---|---|---|
| *F25C8.1* | WBGene00009104 | pseudogene | 3.3 | 9.96 | 3.9E-13 |
| *gmd-2* | WBGene00010166 | protein_coding_gene | 3.0 | 8.02 | 3.7E-16 |
| *col-182* | WBGene00000755 | pseudogene | 2.9 | 7.71 | 2.9E-23 |
| *C50E3.9* | WBGene00016820 | protein_coding_gene | 2.7 | 6.71 | 5.8E-06 |
| *F56H6.7* | WBGene00010168 | protein_coding_gene | 2.7 | 6.52 | 3.8E-10 |
| *F29A7.3* | WBGene00017913 | protein_coding_gene | 2.6 | 6.03 | 3.9E-07 |
| *clec-169* | WBGene00009526 | protein_coding_gene | 2.6 | 5.96 | 1.6E-23 |
| *mab-3* | WBGene00003100 | protein_coding_gene | 2.5 | 5.72 | 1.3E-18 |
| *F19H6.6* | WBGene00008961 | protein_coding_gene | 2.4 | 5.27 | 1.3E-07 |
| *far-3* | WBGene00001387 | protein_coding_gene | 2.4 | 5.22 | 1.4E-24 |
| *oac-35* | WBGene00010171 | protein_coding_gene | 2.4 | 5.10 | 5.7E-05 |
| *bgnt-1.6* | WBGene00010167 | protein_coding_gene | 2.3 | 4.99 | 6.5E-06 |
| *F45E1.4* | WBGene00018465 | protein_coding_gene | 2.3 | 4.91 | 1.5E-24 |
| *Y39E4A.1* | WBGene00012711 | protein_coding_gene | 2.2 | 4.59 | 1.3E-07 |
| *H10D18.6* | WBGene00019181 | pseudogene | 2.2 | 4.52 | 3.3E-07 |
| *cyp-35A5* | WBGene00019473 | protein_coding_gene | 2.1 | 4.42 | 2.1E-20 |
| *C47F8.6* | WBGene00008162 | protein_coding_gene | 2.1 | 4.41 | 1.2E-04 |

| | | | | | |
|---|---|---|---|---|---|
| *F35F10.13* | WBGene00018062 | protein_coding_gene | 2.1 | 4.26 | 2.4E-04 |
| *F40F8.12* | WBGene00044917 | protein_coding_gene | 2.1 | 4.25 | 5.0E-05 |
| *ZK849.1* | WBGene00014100 | protein_coding_gene | 2.1 | 4.15 | 3.3E-11 |
| *clec-93* | WBGene00013929 | protein_coding_gene | 2.1 | 4.14 | 9.9E-06 |
| *Y43F8C.23* | WBGene00045344 | protein_coding_gene | 2.0 | 4.07 | 1.9E-05 |
| *F44G3.10* | WBGene00009710 | protein_coding_gene | 2.0 | 4.04 | 2.1E-06 |
| *F27E5.3* | WBGene00009193 | protein_coding_gene | 2.0 | 4.01 | 3.0E-10 |
| *Y73C8C.10* | WBGene00022267 | protein_coding_gene | 2.0 | 4.00 | 6.2E-07 |
| *C10F3.7* | WBGene00015679 | protein_coding_gene | 2.0 | 3.95 | 5.0E-09 |
| *C09B9.85* | WBGene00220250 | protein_coding_gene | 2.0 | 3.95 | 1.0E-05 |
| *C05B5.1* | WBGene00007319 | protein_coding_gene | 2.0 | 3.93 | 2.7E-04 |
| *ant-1.3* | WBGene00010485 | protein_coding_gene | 1.9 | 3.82 | 5.9E-18 |
| *F17E9.15* | WBGene00194833 | protein_coding_gene | 1.9 | 3.79 | 4.8E-04 |
| *C34D4.2* | WBGene00016398 | protein_coding_gene | 1.9 | 3.70 | 3.2E-11 |
| *oac-23* | WBGene00018143 | protein_coding_gene | 1.9 | 3.70 | 3.4E-07 |
| *C07G3.10* | WBGene00015578 | protein_coding_gene | 1.9 | 3.68 | 1.1E-04 |
| *F56H6.9* | WBGene00010170 | protein_coding_gene | 1.9 | 3.68 | 2.3E-04 |
| *F46F5.11* | WBGene00018502 | protein_coding_gene | 1.9 | 3.67 | 6.5E-08 |
| *T10B11.4* | WBGene00020400 | protein_coding_gene | 1.9 | 3.66 | 9.4E-05 |
| *gmeb-3* | WBGene00008092 | protein_coding_gene | 1.9 | 3.65 | 3.8E-07 |
| *F32B5.1* | WBGene00017975 | protein_coding_gene | 1.9 | 3.65 | 1.5E-11 |
| *T07H8.11* | WBGene00304201 | protein_coding_gene | 1.9 | 3.63 | 2.2E-19 |
| *K09H11.4* | WBGene00019601 | protein_coding_gene | 1.9 | 3.63 | 4.7E-05 |
| *C17G10.10* | WBGene00195182 | protein_coding_gene | 1.9 | 3.63 | 4.4E-05 |
| *F28A10.1* | WBGene00017869 | protein_coding_gene | 1.9 | 3.61 | 2.4E-08 |
| *clec-210* | WBGene00022261 | protein_coding_gene | 1.8 | 3.57 | 2.4E-04 |
| *ZK1290.7* | WBGene00022889 | protein_coding_gene | 1.8 | 3.56 | 1.9E-04 |
| *C01G10.17* | WBGene00044658 | protein_coding_gene | 1.8 | 3.55 | 8.0E-05 |
| *F46F5.14* | WBGene00018505 | protein_coding_gene | 1.8 | 3.54 | 1.7E-06 |
| *math-23* | WBGene00016718 | protein_coding_gene | 1.8 | 3.53 | 2.5E-04 |
| *ugt-32* | WBGene00018543 | protein_coding_gene | 1.8 | 3.49 | 1.5E-06 |
| *Y50D4B.7* | WBGene00021746 | pseudogene | 1.8 | 3.39 | 3.1E-04 |
| *fbxa-73* | WBGene00020640 | protein_coding_gene | 1.8 | 3.36 | 6.7E-05 |
| *fer-1* | WBGene00001414 | protein_coding_gene | 1.7 | 3.36 | 3.8E-13 |
| *K07E8.1* | WBGene00019494 | pseudogene | 1.7 | 3.35 | 1.5E-03 |
| *F33E2.6* | WBGene00009362 | protein_coding_gene | 1.7 | 3.34 | 4.5E-04 |
| *F56D5.2* | WBGene00010147 | protein_coding_gene | 1.7 | 3.33 | 1.9E-05 |
| *ZC513.3* | WBGene00022628 | protein_coding_gene | 1.7 | 3.32 | 8.0E-07 |
| *pcp-4* | WBGene00003959 | protein_coding_gene | 1.7 | 3.30 | 6.8E-15 |
| *Y62H9A.8* | WBGene00013396 | protein_coding_gene | 1.7 | 3.27 | 1.2E-04 |
| *F07D3.3* | WBGene00008556 | protein_coding_gene | 1.7 | 3.23 | 8.1E-05 |
| *H20J04.1* | WBGene00019216 | protein_coding_gene | 1.7 | 3.22 | 3.0E-09 |
| *nkb-1* | WBGene00007646 | protein_coding_gene | 1.7 | 3.22 | 1.6E-79 |
| *F55G11.1* | WBGene00010122 | protein_coding_gene | 1.7 | 3.19 | 7.1E-05 |
| *C29F9.5* | WBGene00016220 | protein_coding_gene | 1.7 | 3.17 | 1.2E-03 |
| *clec-165* | WBGene00009523 | protein_coding_gene | 1.7 | 3.15 | 1.8E-08 |
| *F55C9.12* | WBGene00044196 | pseudogene | 1.7 | 3.14 | 1.0E-03 |
| *Y69E1A.8* | WBGene00013478 | protein_coding_gene | 1.6 | 3.13 | 3.0E-08 |
| *rpl-42* | WBGene00004455 | protein_coding_gene | 1.6 | 3.12 | 7.9E-06 |
| *C08F11.14* | WBGene00007461 | pseudogene | 1.6 | 3.09 | 5.1E-06 |
| *F58A6.5* | WBGene00019024 | protein_coding_gene | 1.6 | 3.09 | 1.5E-09 |
| *Y69A2AR.19* | WBGene00022090 | protein_coding_gene | 1.6 | 3.09 | 5.4E-19 |
| *lips-6* | WBGene00014009 | protein_coding_gene | 1.6 | 3.09 | 3.8E-09 |
| *W05B2.3* | WBGene00014854 | pseudogene | 1.6 | 3.08 | 7.4E-04 |
| *R12E2.14* | WBGene00020039 | protein_coding_gene | 1.6 | 3.08 | 1.2E-08 |
| *msp-50* | WBGene00003443 | protein_coding_gene | 1.6 | 3.06 | 6.6E-17 |
| *ZC412.5* | WBGene00013886 | protein_coding_gene | 1.6 | 3.05 | 5.7E-07 |
| *C44B9.2* | WBGene00008079 | protein_coding_gene | 1.6 | 3.05 | 1.5E-10 |
| *cyc-2.2* | WBGene00013854 | protein_coding_gene | 1.6 | 3.05 | 1.2E-12 |
| *Y67D8C.7* | WBGene00022071 | protein_coding_gene | 1.6 | 3.05 | 1.4E-03 |
| *Y82E9BL.12* | WBGene00022328 | protein_coding_gene | 1.6 | 3.05 | 1.6E-03 |
| *Y7A5A.9* | WBGene00012415 | protein_coding_gene | 1.6 | 3.03 | 2.0E-03 |
| *wago-10* | WBGene00020707 | protein_coding_gene | 1.6 | 3.03 | 4.8E-12 |
| *haf-8* | WBGene00001818 | protein_coding_gene | 1.6 | 3.00 | 2.7E-04 |
| *F18F11.4* | WBGene00017577 | protein_coding_gene | 1.6 | 2.99 | 6.5E-04 |

| | | | | | |
|---|---|---|---|---|---|
| *cest-35.2* | WBGene00016863 | protein_coding_gene | 1.6 | 2.99 | 9.5E-08 |
| *gln-4* | WBGene00001605 | pseudogene | 1.6 | 2.98 | 4.7E-05 |
| *clec-195* | WBGene00013798 | protein_coding_gene | 1.6 | 2.98 | 3.0E-04 |
| *F49C5.12* | WBGene00184956 | protein_coding_gene | 1.6 | 2.96 | 9.4E-04 |
| *nhr-219* | WBGene00020555 | protein_coding_gene | 1.6 | 2.96 | 2.4E-03 |
| *C02G6.2* | WBGene00015360 | protein_coding_gene | 1.6 | 2.95 | 2.4E-04 |
| *T09E11.12* | WBGene00045103 | protein_coding_gene | 1.6 | 2.95 | 3.1E-04 |
| *fut-5* | WBGene00043986 | protein_coding_gene | 1.6 | 2.94 | 8.5E-04 |
| *W02B3.7* | WBGene00020927 | protein_coding_gene | 1.6 | 2.94 | 2.6E-03 |
| *cut-3* | WBGene00009041 | protein_coding_gene | 1.6 | 2.94 | 5.1E-13 |
| *nep-7* | WBGene00017552 | protein_coding_gene | 1.6 | 2.94 | 1.8E-03 |
| *R155.5* | WBGene00189953 | protein_coding_gene | 1.6 | 2.93 | 2.1E-03 |
| *W03F8.2* | WBGene00020992 | pseudogene | 1.5 | 2.93 | 1.9E-05 |
| *pax-2* | WBGene00003938 | protein_coding_gene | 1.5 | 2.93 | 1.2E-03 |
| *oac-45* | WBGene00011656 | protein_coding_gene | 1.5 | 2.92 | 8.2E-04 |
| *Y39E4B.11* | WBGene00012720 | protein_coding_gene | 1.5 | 2.90 | 5.4E-06 |
| *C53B4.2* | WBGene00008272 | protein_coding_gene | 1.5 | 2.89 | 6.1E-05 |
| *M05B5.3* | WBGene00010871 | protein_coding_gene | 1.5 | 2.89 | 4.9E-08 |
| *K07E12.2* | WBGene00019501 | protein_coding_gene | 1.5 | 2.88 | 4.9E-07 |
| *Y53F4B.16* | WBGene00013162 | protein_coding_gene | 1.5 | 2.88 | 3.4E-04 |
| *F55F10.3* | WBGene00050916 | protein_coding_gene | 1.5 | 2.88 | 1.1E-04 |
| *C16C10.9* | WBGene00007629 | protein_coding_gene | 1.5 | 2.87 | 2.0E-03 |
| *T06D8.10* | WBGene00011530 | protein_coding_gene | 1.5 | 2.87 | 1.2E-07 |
| *F22B3.10* | WBGene00045122 | protein_coding_gene | 1.5 | 2.86 | 4.1E-04 |
| *C06A6.7* | WBGene00015511 | protein_coding_gene | 1.5 | 2.85 | 7.1E-07 |
| *pals-6* | WBGene00007660 | protein_coding_gene | 1.5 | 2.85 | 2.3E-05 |
| *T25D1.2* | WBGene00020794 | protein_coding_gene | 1.5 | 2.84 | 1.9E-03 |
| *C47F8.5* | WBGene00008161 | protein_coding_gene | 1.5 | 2.84 | 3.1E-03 |
| *sre-6* | WBGene00009242 | protein_coding_gene | 1.5 | 2.84 | 5.5E-04 |
| *wht-5* | WBGene00008950 | protein_coding_gene | 1.5 | 2.83 | 2.2E-08 |
| *decr-1.3* | WBGene00011467 | protein_coding_gene | 1.5 | 2.83 | 5.3E-08 |
| *D1081.5* | WBGene00008383 | protein_coding_gene | 1.5 | 2.83 | 4.9E-07 |
| *F58D5.6* | WBGene00010245 | protein_coding_gene | 1.5 | 2.83 | 1.9E-04 |
| *T09D3.8* | WBGene00044538 | protein_coding_gene | 1.5 | 2.82 | 3.9E-03 |
| *msp-51* | WBGene00003444 | protein_coding_gene | 1.5 | 2.82 | 8.4E-11 |
| *K07A1.4* | WBGene00010611 | protein_coding_gene | 1.5 | 2.81 | 7.4E-10 |
| *F02H6.7* | WBGene00008540 | protein_coding_gene | 1.5 | 2.81 | 4.5E-03 |
| *col-74* | WBGene00000650 | protein_coding_gene | 1.5 | 2.81 | 3.6E-10 |
| *R102.3* | WBGene00011290 | protein_coding_gene | 1.5 | 2.81 | 1.7E-04 |
| *ZK1251.3* | WBGene00014241 | protein_coding_gene | 1.5 | 2.81 | 2.3E-04 |
| *C35E7.10* | WBGene00016462 | protein_coding_gene | 1.5 | 2.79 | 1.5E-11 |
| *C09H10.9* | WBGene00007503 | protein_coding_gene | 1.5 | 2.78 | 3.0E-06 |
| *K07F5.6* | WBGene00010634 | protein_coding_gene | 1.5 | 2.77 | 7.5E-07 |
| *Y18D10A.26* | WBGene00023459 | protein_coding_gene | 1.5 | 2.77 | 5.3E-04 |
| *T25B9.6* | WBGene00012012 | protein_coding_gene | 1.5 | 2.75 | 1.7E-09 |
| *Y57A10A.1* | WBGene00013246 | protein_coding_gene | 1.5 | 2.75 | 3.3E-22 |
| *msp-45* | WBGene00003438 | protein_coding_gene | 1.5 | 2.74 | 2.2E-12 |
| *inpp-1* | WBGene00012016 | protein_coding_gene | 1.5 | 2.74 | 5.6E-11 |
| *F08H9.4* | WBGene00008592 | protein_coding_gene | 1.4 | 2.73 | 3.0E-03 |
| *acdh-8* | WBGene00019406 | protein_coding_gene | 1.4 | 2.73 | 1.4E-11 |
| *T07G12.3* | WBGene00011594 | protein_coding_gene | 1.4 | 2.72 | 4.7E-05 |
| *C09D4.3* | WBGene00015634 | protein_coding_gene | 1.4 | 2.71 | 9.5E-09 |
| *his-70* | WBGene00001944 | protein_coding_gene | 1.4 | 2.71 | 8.0E-06 |
| *F40H6.5* | WBGene00018255 | protein_coding_gene | 1.4 | 2.70 | 4.3E-03 |
| *T11F8.4* | WBGene00020435 | protein_coding_gene | 1.4 | 2.70 | 1.6E-07 |
| *C25H3.16* | WBGene00044392 | protein_coding_gene | 1.4 | 2.69 | 2.1E-03 |
| *dhhc-12* | WBGene00010323 | protein_coding_gene | 1.4 | 2.69 | 2.0E-06 |
| *ZK484.7* | WBGene00022753 | protein_coding_gene | 1.4 | 2.69 | 1.8E-06 |
| *oac-44* | WBGene00011654 | protein_coding_gene | 1.4 | 2.69 | 3.2E-03 |
| *F44F1.1* | WBGene00009693 | pseudogene | 1.4 | 2.69 | 2.5E-03 |
| *col-165* | WBGene00000738 | protein_coding_gene | 1.4 | 2.68 | 9.7E-12 |
| *F21F3.2* | WBGene00017672 | protein_coding_gene | 1.4 | 2.68 | 1.7E-05 |
| *F36H12.9* | WBGene00018123 | protein_coding_gene | 1.4 | 2.67 | 3.6E-03 |
| *clec-67* | WBGene00018971 | protein_coding_gene | 1.4 | 2.66 | 3.4E-09 |
| *F09E10.1* | WBGene00017294 | protein_coding_gene | 1.4 | 2.66 | 4.7E-13 |

| | | | | | |
|---|---|---|---|---|---|
| *Y38F1A.7* | WBGene00012609 | protein_coding_gene | 1.4 | 2.66 | 1.1E-03 |
| *Y47D7A.2* | WBGene00021616 | protein_coding_gene | 1.4 | 2.66 | 4.1E-03 |
| *C35A11.3* | WBGene00016430 | protein_coding_gene | 1.4 | 2.66 | 4.1E-03 |
| *trf-2* | WBGene00022454 | protein_coding_gene | 1.4 | 2.66 | 1.2E-03 |
| *F01G10.5* | WBGene00008508 | protein_coding_gene | 1.4 | 2.65 | 2.2E-09 |
| *spe-47* | WBGene00021271 | protein_coding_gene | 1.4 | 2.65 | 2.3E-05 |
| *F36H1.3* | WBGene00009492 | protein_coding_gene | 1.4 | 2.64 | 1.0E-06 |
| *str-7* | WBGene00006075 | protein_coding_gene | 1.4 | 2.64 | 5.1E-03 |
| *kin-24* | WBGene00002206 | protein_coding_gene | 1.4 | 2.64 | 5.6E-06 |
| *Y2H9A.6* | WBGene00086568 | protein_coding_gene | 1.4 | 2.64 | 1.9E-05 |
| *F49E11.7* | WBGene00009893 | protein_coding_gene | 1.4 | 2.63 | 2.7E-05 |
| *Y48B6A.5* | WBGene00012979 | protein_coding_gene | 1.4 | 2.63 | 2.0E-07 |
| *F54C8.1* | WBGene00010035 | protein_coding_gene | 1.4 | 2.63 | 6.1E-08 |
| *acdh-5* | WBGene00016491 | protein_coding_gene | 1.4 | 2.63 | 5.5E-08 |
| *T27F6.1* | WBGene00012102 | protein_coding_gene | 1.4 | 2.63 | 2.8E-04 |
| *T19H12.15* | WBGene00271827 | protein_coding_gene | 1.4 | 2.62 | 4.7E-18 |
| *clec-134* | WBGene00021141 | protein_coding_gene | 1.4 | 2.62 | 8.0E-03 |
| *spe-42* | WBGene00007117 | protein_coding_gene | 1.4 | 2.62 | 2.0E-04 |
| *T15D6.10* | WBGene00011784 | protein_coding_gene | 1.4 | 2.61 | 5.6E-03 |
| *F36H12.14* | WBGene00018127 | pseudogene | 1.4 | 2.61 | 4.6E-05 |
| *tsp-19* | WBGene00006645 | protein_coding_gene | 1.4 | 2.61 | 1.8E-04 |
| *T04A6.3* | WBGene00020202 | protein_coding_gene | 1.4 | 2.60 | 1.6E-03 |
| *T25B9.5* | WBGene00012011 | protein_coding_gene | 1.4 | 2.60 | 7.1E-05 |
| *C55C3.7* | WBGene00043065 | protein_coding_gene | 1.4 | 2.60 | 4.4E-03 |
| *F21A3.4* | WBGene00008994 | protein_coding_gene | 1.4 | 2.60 | 3.4E-04 |
| *K08D8.5* | WBGene00010659 | protein_coding_gene | 1.4 | 2.60 | 7.7E-07 |
| *M05D6.1* | WBGene00010874 | protein_coding_gene | 1.4 | 2.59 | 2.7E-03 |
| *Y47D7A.18* | WBGene00235165 | protein_coding_gene | 1.4 | 2.59 | 9.8E-04 |
| *K07A1.10* | WBGene00010616 | protein_coding_gene | 1.4 | 2.59 | 5.3E-36 |
| *F38A1.9* | WBGene00009522 | protein_coding_gene | 1.4 | 2.59 | 1.2E-06 |
| *C16C8.18* | WBGene00015855 | protein_coding_gene | 1.4 | 2.58 | 5.7E-07 |
| *nhr-81* | WBGene00003671 | protein_coding_gene | 1.4 | 2.58 | 4.6E-03 |
| *Y66A7A.4* | WBGene00013424 | protein_coding_gene | 1.4 | 2.58 | 2.4E-04 |
| *ZK1010.6* | WBGene00014180 | pseudogene | 1.4 | 2.58 | 9.4E-03 |
| *F15D4.5* | WBGene00008862 | protein_coding_gene | 1.4 | 2.58 | 3.0E-05 |
| *F23B12.1* | WBGene00009079 | protein_coding_gene | 1.4 | 2.57 | 2.2E-05 |
| *F10C1.3* | WBGene00017325 | protein_coding_gene | 1.4 | 2.57 | 2.9E-06 |
| *B0207.7* | WBGene00015030 | protein_coding_gene | 1.4 | 2.57 | 3.8E-05 |
| *C47F8.3* | WBGene00008159 | protein_coding_gene | 1.4 | 2.57 | 8.9E-03 |
| *C39H7.1* | WBGene00016541 | protein_coding_gene | 1.4 | 2.56 | 2.1E-11 |
| *ZK892.4* | WBGene00014128 | protein_coding_gene | 1.4 | 2.56 | 3.3E-03 |
| *W03A5.1* | WBGene00020966 | protein_coding_gene | 1.4 | 2.56 | 4.0E-06 |
| *T20B6.2* | WBGene00020598 | protein_coding_gene | 1.4 | 2.56 | 2.4E-03 |
| *C27D6.15* | WBGene00303362 | protein_coding_gene | 1.4 | 2.56 | 9.7E-04 |
| *Y39B6A.25* | WBGene00012684 | protein_coding_gene | 1.4 | 2.56 | 1.1E-04 |
| *mboa-4* | WBGene00007446 | protein_coding_gene | 1.4 | 2.56 | 7.5E-07 |
| *ugt-63* | WBGene00015449 | protein_coding_gene | 1.4 | 2.56 | 3.3E-14 |
| *T05H4.7* | WBGene00020270 | protein_coding_gene | 1.3 | 2.55 | 1.4E-03 |
| *tag-293* | WBGene00015398 | protein_coding_gene | 1.3 | 2.55 | 8.9E-03 |
| *ZK354.2* | WBGene00022705 | protein_coding_gene | 1.3 | 2.54 | 6.5E-11 |
| *F36A2.14* | WBGene00023490 | protein_coding_gene | 1.3 | 2.54 | 6.4E-09 |
| *spch-2* | WBGene00015689 | protein_coding_gene | 1.3 | 2.54 | 6.5E-05 |
| *Y50D4B.9* | WBGene00268189 | pseudogene | 1.3 | 2.54 | 6.7E-03 |
| *Y38A10A.2* | WBGene00021389 | protein_coding_gene | 1.3 | 2.54 | 3.3E-06 |
| *F16C3.4* | WBGene00044243 | protein_coding_gene | 1.3 | 2.54 | 3.7E-04 |
| *irld-14* | WBGene00011322 | protein_coding_gene | 1.3 | 2.53 | 5.2E-05 |
| *Y53F4B.27* | WBGene00013173 | protein_coding_gene | 1.3 | 2.53 | 4.5E-05 |
| *LLC1.121* | WBGene00235121 | pseudogene | 1.3 | 2.53 | 7.4E-03 |
| *oac-48* | WBGene00011960 | protein_coding_gene | 1.3 | 2.53 | 2.2E-03 |
| *4R79.2* | WBGene00007067 | protein_coding_gene | 1.3 | 2.52 | 1.4E-04 |
| *F15H9.7* | WBGene00077535 | pseudogene | 1.3 | 2.51 | 3.6E-03 |
| *T27A8.7* | WBGene00206527 | protein_coding_gene | 1.3 | 2.51 | 9.4E-03 |
| *F16G10.2* | WBGene00017517 | protein_coding_gene | 1.3 | 2.51 | 1.3E-03 |
| *C36H8.1* | WBGene00007987 | protein_coding_gene | 1.3 | 2.50 | 6.0E-08 |
| *M199.9* | WBGene00044998 | protein_coding_gene | 1.3 | 2.50 | 1.1E-03 |

| | | | | | |
|---|---|---|---|---|---|
| *C49A1.5* | WBGene00008187 | protein_coding_gene | 1.3 | 2.50 | 8.6E-03 |
| *C17H12.3* | WBGene00015929 | protein_coding_gene | 1.3 | 2.50 | 1.2E-07 |
| *spe-45* | WBGene00009217 | protein_coding_gene | 1.3 | 2.50 | 7.0E-04 |
| *msrp-6* | WBGene00017058 | protein_coding_gene | 1.3 | 2.50 | 1.4E-06 |
| *snb-5* | WBGene00004899 | protein_coding_gene | 1.3 | 2.50 | 6.9E-03 |
| *F54H12.8* | WBGene00045247 | protein_coding_gene | 1.3 | 2.49 | 1.0E-04 |
| *C54E10.10* | WBGene00206523 | protein_coding_gene | 1.3 | 2.49 | 1.3E-03 |
| *irld-6* | WBGene00009239 | protein_coding_gene | 1.3 | 2.49 | 6.0E-05 |
| *ugt-16* | WBGene00013901 | protein_coding_gene | 1.3 | 2.49 | 6.4E-11 |
| *K03H1.12* | WBGene00010547 | protein_coding_gene | 1.3 | 2.49 | 2.3E-05 |
| *spp-17* | WBGene00005002 | protein_coding_gene | 1.3 | 2.49 | 6.1E-04 |
| *R02F11.10* | WBGene00202497 | protein_coding_gene | 1.3 | 2.49 | 1.4E-03 |
| *nspd-5* | WBGene00017386 | protein_coding_gene | 1.3 | 2.49 | 7.2E-04 |
| *cyp-25A6* | WBGene00019438 | protein_coding_gene | 1.3 | 2.48 | 8.4E-03 |
| *oac-36* | WBGene00010172 | protein_coding_gene | 1.3 | 2.46 | 8.8E-03 |
| *C16D9.5* | WBGene00015860 | protein_coding_gene | 1.3 | 2.46 | 7.3E-04 |
| *T15D6.8* | WBGene00011782 | protein_coding_gene | 1.3 | 2.46 | 8.2E-03 |
| *F32H2.7* | WBGene00009344 | protein_coding_gene | 1.3 | 2.46 | 3.3E-12 |
| *uggt-2* | WBGene00009178 | protein_coding_gene | 1.3 | 2.46 | 2.8E-06 |
| *F46A9.2* | WBGene00009755 | protein_coding_gene | 1.3 | 2.46 | 1.7E-06 |
| *K08E7.4* | WBGene00010672 | protein_coding_gene | 1.3 | 2.46 | 7.5E-03 |
| *nlp-16* | WBGene00003754 | protein_coding_gene | 1.3 | 2.45 | 4.2E-23 |
| *oac-34* | WBGene00010157 | protein_coding_gene | 1.3 | 2.45 | 8.0E-03 |
| *F37A4.5* | WBGene00018135 | protein_coding_gene | 1.3 | 2.45 | 2.0E-07 |
| *C14A4.8* | WBGene00007559 | protein_coding_gene | 1.3 | 2.44 | 1.3E-03 |
| *Y40B10A.4* | WBGene00021489 | pseudogene | 1.3 | 2.44 | 4.4E-03 |
| *T08G3.13* | WBGene00011621 | protein_coding_gene | 1.3 | 2.44 | 1.2E-02 |
| *C48D1.9* | WBGene00194643 | protein_coding_gene | 1.3 | 2.44 | 4.7E-03 |
| *gst-39* | WBGene00001787 | protein_coding_gene | 1.3 | 2.44 | 1.3E-08 |
| *F39B3.7* | WBGene00304237 | protein_coding_gene | 1.3 | 2.44 | 9.7E-03 |
| *rsef-1* | WBGene00016344 | protein_coding_gene | 1.3 | 2.43 | 2.6E-05 |
| *C33F10.8* | WBGene00016356 | protein_coding_gene | 1.3 | 2.43 | 5.2E-06 |
| *scrm-6* | WBGene00009753 | protein_coding_gene | 1.3 | 2.43 | 4.7E-05 |
| *M28.9* | WBGene00010900 | protein_coding_gene | 1.3 | 2.43 | 1.6E-09 |
| *elo-7* | WBGene00001245 | protein_coding_gene | 1.3 | 2.43 | 3.8E-04 |
| *nas-19* | WBGene00003538 | protein_coding_gene | 1.3 | 2.42 | 9.3E-03 |
| *C33F10.12* | WBGene00016358 | protein_coding_gene | 1.3 | 2.42 | 6.9E-08 |
| *C08F11.1* | WBGene00007450 | protein_coding_gene | 1.3 | 2.42 | 5.9E-04 |
| *C31B8.7* | WBGene00016282 | protein_coding_gene | 1.3 | 2.41 | 5.8E-06 |
| *Y47D3B.12* | WBGene00185055 | protein_coding_gene | 1.3 | 2.41 | 3.2E-04 |
| *F13D12.9* | WBGene00008743 | protein_coding_gene | 1.3 | 2.41 | 2.0E-07 |
| *qui-1* | WBGene00004265 | protein_coding_gene | 1.3 | 2.41 | 1.0E-08 |
| *C29E4.14* | WBGene00044634 | protein_coding_gene | 1.3 | 2.41 | 1.2E-03 |
| *clec-164* | WBGene00021949 | protein_coding_gene | 1.3 | 2.41 | 1.4E-02 |
| *F36D3.8* | WBGene00009473 | protein_coding_gene | 1.3 | 2.40 | 1.9E-03 |
| *F58E6.5* | WBGene00010254 | protein_coding_gene | 1.3 | 2.40 | 5.1E-09 |
| *clec-193* | WBGene00013774 | protein_coding_gene | 1.3 | 2.40 | 1.2E-02 |
| *E03H4.11* | WBGene00008478 | protein_coding_gene | 1.3 | 2.40 | 7.4E-03 |
| *AC7.3* | WBGene00014997 | protein_coding_gene | 1.3 | 2.39 | 1.8E-03 |
| *arrd-8* | WBGene00012467 | protein_coding_gene | 1.3 | 2.39 | 6.3E-03 |
| *C01G5.3* | WBGene00015305 | protein_coding_gene | 1.3 | 2.39 | 6.3E-04 |
| *W06H8.4* | WBGene00021067 | pseudogene | 1.3 | 2.39 | 1.0E-02 |
| *Y26G10.3* | WBGene00050973 | pseudogene | 1.3 | 2.39 | 8.1E-03 |
| *R02D1.2* | WBGene00235310 | protein_coding_gene | 1.3 | 2.39 | 1.7E-03 |
| *ipla-5* | WBGene00019229 | protein_coding_gene | 1.3 | 2.39 | 2.3E-03 |
| *K07A3.3* | WBGene00019461 | protein_coding_gene | 1.3 | 2.38 | 1.4E-07 |
| *F10G7.12* | WBGene00044744 | protein_coding_gene | 1.3 | 2.38 | 1.9E-03 |
| *lips-9* | WBGene00010267 | protein_coding_gene | 1.3 | 2.38 | 1.7E-03 |
| *nhr-169* | WBGene00008289 | protein_coding_gene | 1.3 | 2.38 | 8.6E-03 |
| *csp-1* | WBGene00000819 | protein_coding_gene | 1.2 | 2.37 | 2.4E-03 |
| *F57G8.7* | WBGene00010216 | protein_coding_gene | 1.2 | 2.37 | 1.5E-02 |
| *C35A5.4* | WBGene00007948 | protein_coding_gene | 1.2 | 2.36 | 1.2E-04 |
| *ptp-5.1* | WBGene00016053 | protein_coding_gene | 1.2 | 2.36 | 1.1E-06 |
| *msp-31* | WBGene00003429 | protein_coding_gene | 1.2 | 2.36 | 6.2E-11 |
| *ZK616.61* | WBGene00194816 | protein_coding_gene | 1.2 | 2.36 | 1.8E-06 |

| | | | | | |
|---|---|---|---|---|---|
| *spch-1* | WBGene00007307 | protein_coding_gene | 1.2 | 2.36 | 2.2E-03 |
| *C47E8.1* | WBGene00008141 | protein_coding_gene | 1.2 | 2.36 | 1.1E-03 |
| *irld-7* | WBGene00018168 | protein_coding_gene | 1.2 | 2.36 | 3.6E-03 |
| *C06A1.3* | WBGene00007354 | protein_coding_gene | 1.2 | 2.36 | 3.9E-07 |
| *Y38H8A.4* | WBGene00012638 | protein_coding_gene | 1.2 | 2.36 | 3.6E-06 |
| *F35E2.5* | WBGene00009412 | protein_coding_gene | 1.2 | 2.36 | 3.7E-04 |
| *ZK1248.5* | WBGene00022876 | protein_coding_gene | 1.2 | 2.36 | 5.7E-05 |
| *C54E4.5* | WBGene00016920 | protein_coding_gene | 1.2 | 2.35 | 1.2E-13 |
| *T10H4.13* | WBGene00045339 | protein_coding_gene | 1.2 | 2.35 | 6.1E-03 |
| *best-6* | WBGene00015628 | protein_coding_gene | 1.2 | 2.35 | 1.2E-02 |
| *best-10* | WBGene00008185 | protein_coding_gene | 1.2 | 2.35 | 3.7E-03 |
| *T05F1.8* | WBGene00011493 | protein_coding_gene | 1.2 | 2.34 | 8.0E-07 |
| *C09B9.7* | WBGene00015630 | protein_coding_gene | 1.2 | 2.34 | 6.7E-04 |
| *fbxb-64* | WBGene00008071 | protein_coding_gene | 1.2 | 2.34 | 1.3E-02 |
| *F58G1.3* | WBGene00010265 | protein_coding_gene | 1.2 | 2.34 | 8.5E-06 |
| *bgnt-1.8* | WBGene00008290 | protein_coding_gene | 1.2 | 2.34 | 1.5E-02 |
| *ssp-10* | WBGene00006039 | protein_coding_gene | 1.2 | 2.33 | 1.1E-03 |
| *ent-3* | WBGene00010510 | protein_coding_gene | 1.2 | 2.33 | 2.0E-06 |
| *paf-1* | WBGene00003906 | protein_coding_gene | 1.2 | 2.33 | 1.2E-03 |
| *T14A8.2* | WBGene00020497 | protein_coding_gene | 1.2 | 2.33 | 3.5E-06 |
| *W03D8.1* | WBGene00020984 | protein_coding_gene | 1.2 | 2.33 | 5.0E-03 |
| *ssq-2* | WBGene00006051 | protein_coding_gene | 1.2 | 2.32 | 1.5E-17 |
| *C30B5.6* | WBGene00016247 | protein_coding_gene | 1.2 | 2.32 | 1.2E-03 |
| *Y50E8A.6* | WBGene00013049 | protein_coding_gene | 1.2 | 2.32 | 1.1E-02 |
| *msp-81* | WBGene00003467 | protein_coding_gene | 1.2 | 2.32 | 4.2E-09 |
| *F44G4.5* | WBGene00009714 | protein_coding_gene | 1.2 | 2.32 | 5.1E-04 |
| *T27A3.5* | WBGene00020841 | protein_coding_gene | 1.2 | 2.32 | 3.3E-06 |
| *F56F3.4* | WBGene00010155 | protein_coding_gene | 1.2 | 2.31 | 2.0E-08 |
| *M163.22* | WBGene00304195 | protein_coding_gene | 1.2 | 2.31 | 2.4E-03 |
| *ZK616.1* | WBGene00022771 | protein_coding_gene | 1.2 | 2.31 | 3.9E-03 |
| *Y49G5A.1* | WBGene00021731 | protein_coding_gene | 1.2 | 2.30 | 1.5E-02 |
| *F35C5.4* | WBGene00009392 | protein_coding_gene | 1.2 | 2.30 | 2.3E-03 |
| *cdc-25.4* | WBGene00000389 | protein_coding_gene | 1.2 | 2.30 | 1.3E-03 |
| *Y32G9A.3* | WBGene00021302 | protein_coding_gene | 1.2 | 2.30 | 1.5E-02 |
| *nas-22* | WBGene00003541 | protein_coding_gene | 1.2 | 2.30 | 4.3E-03 |
| *wht-4* | WBGene00017179 | protein_coding_gene | 1.2 | 2.30 | 4.0E-04 |
| *K06A1.2* | WBGene00019425 | protein_coding_gene | 1.2 | 2.29 | 2.0E-03 |
| *try-7* | WBGene00006625 | protein_coding_gene | 1.2 | 2.29 | 4.2E-05 |
| *str-11* | WBGene00006079 | pseudogene | 1.2 | 2.29 | 1.8E-02 |
| *Y67A6A.1* | WBGene00013449 | protein_coding_gene | 1.2 | 2.29 | 5.9E-05 |
| *gly-1* | WBGene00001626 | protein_coding_gene | 1.2 | 2.29 | 1.8E-02 |
| *F58D5.7* | WBGene00010246 | protein_coding_gene | 1.2 | 2.29 | 5.4E-07 |
| *B0244.9* | WBGene00015084 | protein_coding_gene | 1.2 | 2.29 | 3.1E-04 |
| *F36G3.3* | WBGene00009484 | protein_coding_gene | 1.2 | 2.29 | 1.6E-02 |
| *grl-15* | WBGene00001724 | protein_coding_gene | 1.2 | 2.29 | 1.6E-04 |
| *F38C2.4* | WBGene00009536 | protein_coding_gene | 1.2 | 2.28 | 1.5E-02 |
| *K08F4.12* | WBGene00010680 | protein_coding_gene | 1.2 | 2.28 | 1.4E-02 |
| *F36D3.5* | WBGene00009471 | protein_coding_gene | 1.2 | 2.28 | 8.5E-05 |
| *F13H8.6* | WBGene00017439 | protein_coding_gene | 1.2 | 2.28 | 1.3E-02 |
| *dhhc-5* | WBGene00020066 | protein_coding_gene | 1.2 | 2.28 | 9.4E-03 |
| *R13H9.5* | WBGene00020071 | protein_coding_gene | 1.2 | 2.28 | 1.2E-03 |
| *C29F5.3* | WBGene00016212 | protein_coding_gene | 1.2 | 2.27 | 6.6E-04 |
| *C36C9.4* | WBGene00016487 | protein_coding_gene | 1.2 | 2.27 | 1.7E-02 |
| *C32B5.13* | WBGene00016306 | protein_coding_gene | 1.2 | 2.27 | 1.9E-02 |
| *msp-142* | WBGene00003469 | protein_coding_gene | 1.2 | 2.27 | 2.5E-08 |
| *ZC262.4* | WBGene00022581 | protein_coding_gene | 1.2 | 2.27 | 1.8E-02 |
| *T21G5.1* | WBGene00020659 | protein_coding_gene | 1.2 | 2.27 | 1.5E-05 |
| *Y45F10B.3* | WBGene00012871 | protein_coding_gene | 1.2 | 2.27 | 9.2E-04 |
| *Y39B6A.9* | WBGene00012671 | protein_coding_gene | 1.2 | 2.27 | 8.3E-03 |
| *ZC581.2* | WBGene00022632 | protein_coding_gene | 1.2 | 2.26 | 1.3E-04 |
| *Y116F11B.11* | WBGene00013827 | pseudogene | 1.2 | 2.26 | 1.1E-02 |
| *Y37H2A.13* | WBGene00045417 | protein_coding_gene | 1.2 | 2.26 | 4.4E-04 |
| *glct-4* | WBGene00008160 | protein_coding_gene | 1.2 | 2.26 | 1.8E-02 |
| *C47E12.12* | WBGene00008155 | protein_coding_gene | 1.2 | 2.26 | 5.3E-03 |
| *msp-40* | WBGene00003435 | protein_coding_gene | 1.2 | 2.26 | 5.1E-03 |

| | | | | | |
|---|---|---|---|---|---|
| *Y23H5B.12* | WBGene00194701 | protein_coding_gene | 1.2 | 2.26 | 1.8E-02 |
| *F56F4.4* | WBGene00018981 | protein_coding_gene | 1.2 | 2.26 | 1.4E-02 |
| *F28E10.4* | WBGene00017902 | protein_coding_gene | 1.2 | 2.26 | 1.7E-03 |
| *C34C6.3* | WBGene00007916 | protein_coding_gene | 1.2 | 2.26 | 1.3E-06 |
| *F35C11.2* | WBGene00009401 | protein_coding_gene | 1.2 | 2.26 | 4.2E-03 |
| *B0496.1* | WBGene00015210 | protein_coding_gene | 1.2 | 2.26 | 4.4E-03 |
| *bgnt-1.4* | WBGene00015982 | protein_coding_gene | 1.2 | 2.25 | 1.3E-02 |
| *gst-40* | WBGene00001788 | protein_coding_gene | 1.2 | 2.25 | 1.1E-02 |
| *Y45F10B.15* | WBGene00012876 | pseudogene | 1.2 | 2.25 | 2.0E-02 |
| *F10G2.11* | WBGene00271802 | pseudogene | 1.2 | 2.25 | 2.0E-02 |
| *smz-2* | WBGene00020661 | protein_coding_gene | 1.2 | 2.25 | 1.5E-07 |
| *wht-8* | WBGene00012925 | protein_coding_gene | 1.2 | 2.25 | 5.0E-06 |
| *D1081.12* | WBGene00086554 | protein_coding_gene | 1.2 | 2.25 | 1.4E-05 |
| *C06C6.6* | WBGene00007380 | pseudogene | 1.2 | 2.25 | 9.6E-03 |
| *F44B9.10* | WBGene00018411 | protein_coding_gene | 1.2 | 2.25 | 1.2E-02 |
| *C07B5.3* | WBGene00007408 | protein_coding_gene | 1.2 | 2.24 | 6.6E-03 |
| *C15B12.8* | WBGene00015788 | protein_coding_gene | 1.2 | 2.24 | 3.4E-05 |
| *F38H4.6* | WBGene00009550 | protein_coding_gene | 1.2 | 2.24 | 1.9E-03 |
| *K08E7.5* | WBGene00010673 | protein_coding_gene | 1.2 | 2.24 | 2.8E-03 |
| *F41G3.5* | WBGene00018301 | protein_coding_gene | 1.2 | 2.24 | 7.2E-05 |
| *C28H8.7* | WBGene00016198 | protein_coding_gene | 1.2 | 2.24 | 2.2E-02 |
| *frk-1* | WBGene00001487 | protein_coding_gene | 1.2 | 2.24 | 1.9E-06 |
| *F59C6.3* | WBGene00010324 | protein_coding_gene | 1.2 | 2.24 | 7.5E-04 |
| *F36G9.17* | WBGene00077605 | pseudogene | 1.2 | 2.23 | 1.2E-02 |
| *F33D11.1* | WBGene00018000 | protein_coding_gene | 1.2 | 2.23 | 2.1E-02 |
| *T23G11.10* | WBGene00011973 | protein_coding_gene | 1.2 | 2.23 | 1.7E-02 |
| *tag-191* | WBGene00007049 | protein_coding_gene | 1.2 | 2.23 | 5.3E-04 |
| *F46F5.16* | WBGene00018507 | protein_coding_gene | 1.2 | 2.23 | 2.1E-02 |
| *T02E1.7* | WBGene00011379 | protein_coding_gene | 1.2 | 2.23 | 5.9E-05 |
| *F53B2.5* | WBGene00009956 | protein_coding_gene | 1.2 | 2.22 | 8.0E-06 |
| *gsp-3* | WBGene00021113 | protein_coding_gene | 1.2 | 2.22 | 1.1E-07 |
| *C03C10.2* | WBGene00007269 | protein_coding_gene | 1.2 | 2.22 | 1.2E-03 |
| *Y105C5B.11* | WBGene00013651 | protein_coding_gene | 1.2 | 2.22 | 6.8E-04 |
| *F17C11.11* | WBGene00008922 | protein_coding_gene | 1.2 | 2.22 | 2.8E-15 |
| *F22G12.8* | WBGene00044712 | pseudogene | 1.2 | 2.22 | 7.4E-06 |
| *F32A11.3* | WBGene00009308 | protein_coding_gene | 1.2 | 2.22 | 3.8E-10 |
| *C46H11.6* | WBGene00016730 | protein_coding_gene | 1.1 | 2.22 | 2.0E-03 |
| *F18G5.5* | WBGene00017581 | pseudogene | 1.1 | 2.21 | 2.4E-02 |
| *F32B4.2* | WBGene00009313 | protein_coding_gene | 1.1 | 2.21 | 2.7E-06 |
| *M7.12* | WBGene00010887 | protein_coding_gene | 1.1 | 2.21 | 4.1E-07 |
| *W03D8.3* | WBGene00020986 | protein_coding_gene | 1.1 | 2.21 | 8.3E-03 |
| *R08C7.5* | WBGene00019949 | protein_coding_gene | 1.1 | 2.21 | 9.5E-05 |
| *R03D7.5* | WBGene00010991 | protein_coding_gene | 1.1 | 2.21 | 1.5E-03 |
| *F28A10.4* | WBGene00017872 | protein_coding_gene | 1.1 | 2.21 | 1.6E-02 |
| *ZK930.7* | WBGene00014157 | protein_coding_gene | 1.1 | 2.21 | 1.9E-03 |
| *ptp-5.2* | WBGene00013304 | protein_coding_gene | 1.1 | 2.20 | 4.0E-05 |
| *F09C12.8* | WBGene00017279 | protein_coding_gene | 1.1 | 2.20 | 3.8E-06 |
| *R05C11.2* | WBGene00019874 | protein_coding_gene | 1.1 | 2.20 | 2.4E-02 |
| *F19B6.3* | WBGene00008949 | protein_coding_gene | 1.1 | 2.20 | 1.8E-02 |
| *grd-16* | WBGene00001705 | protein_coding_gene | 1.1 | 2.20 | 1.3E-02 |
| *F31E9.8* | WBGene00045408 | pseudogene | 1.1 | 2.20 | 3.7E-03 |
| *F26F12.2* | WBGene00017833 | protein_coding_gene | 1.1 | 2.20 | 2.1E-02 |
| *F25B3.4* | WBGene00009101 | protein_coding_gene | 1.1 | 2.20 | 1.6E-03 |
| *F44D12.8* | WBGene00009685 | protein_coding_gene | 1.1 | 2.20 | 2.6E-04 |
| *T09F5.12* | WBGene00014836 | lincRNA_gene | 1.1 | 2.20 | 1.1E-05 |
| *msrp-4* | WBGene00018500 | protein_coding_gene | 1.1 | 2.20 | 1.3E-02 |
| *comp-1* | WBGene00018158 | protein_coding_gene | 1.1 | 2.19 | 3.4E-05 |
| *K07A1.20* | WBGene00219285 | protein_coding_gene | 1.1 | 2.19 | 2.4E-02 |
| *R01H2.4* | WBGene00019812 | protein_coding_gene | 1.1 | 2.19 | 2.9E-03 |
| *clec-156* | WBGene00017809 | protein_coding_gene | 1.1 | 2.18 | 3.8E-03 |
| *C06A5.2* | WBGene00015500 | protein_coding_gene | 1.1 | 2.18 | 9.6E-04 |
| *Y62H9A.13* | WBGene00013401 | protein_coding_gene | 1.1 | 2.18 | 2.5E-02 |
| *clec-68* | WBGene00018970 | protein_coding_gene | 1.1 | 2.18 | 2.5E-02 |
| *ugt-41* | WBGene00017336 | protein_coding_gene | 1.1 | 2.18 | 1.3E-07 |
| *cut-2* | WBGene00009983 | protein_coding_gene | 1.1 | 2.18 | 4.4E-07 |

| | | | | | |
|---|---|---|---|---|---|
| *Y57G7A.6* | WBGene00021969 | protein_coding_gene | 1.1 | 2.17 | 4.8E-06 |
| *tbx-34* | WBGene00006553 | protein_coding_gene | 1.1 | 2.17 | 2.4E-04 |
| *alg-4* | WBGene00006449 | protein_coding_gene | 1.1 | 2.17 | 1.9E-03 |
| *C25F9.8* | WBGene00007727 | protein_coding_gene | 1.1 | 2.17 | 4.3E-03 |
| *ssp-9* | WBGene00006038 | protein_coding_gene | 1.1 | 2.17 | 3.5E-03 |
| *C10C6.3* | WBGene00007512 | protein_coding_gene | 1.1 | 2.17 | 3.3E-03 |
| *F37A8.2* | WBGene00009502 | protein_coding_gene | 1.1 | 2.17 | 1.2E-03 |
| *ZC477.2* | WBGene00022617 | protein_coding_gene | 1.1 | 2.17 | 2.3E-03 |
| *C27D6.11* | WBGene00044388 | protein_coding_gene | 1.1 | 2.17 | 3.9E-05 |
| *K10D2.8* | WBGene00189952 | protein_coding_gene | 1.1 | 2.17 | 6.5E-04 |
| *Y7A5A.11* | WBGene00044155 | pseudogene | 1.1 | 2.17 | 3.7E-03 |
| *H32C10.1* | WBGene00019255 | protein_coding_gene | 1.1 | 2.16 | 7.9E-06 |
| *Y57G11A.2* | WBGene00013290 | protein_coding_gene | 1.1 | 2.16 | 9.0E-04 |
| *cwp-3* | WBGene00000861 | protein_coding_gene | 1.1 | 2.16 | 1.5E-02 |
| *F42G8.8* | WBGene00018359 | protein_coding_gene | 1.1 | 2.16 | 4.5E-06 |
| *ttbk-4* | WBGene00012169 | protein_coding_gene | 1.1 | 2.16 | 2.5E-06 |
| *ZK1248.20* | WBGene00044393 | protein_coding_gene | 1.1 | 2.16 | 6.5E-03 |
| *C27D8.1* | WBGene00007777 | protein_coding_gene | 1.1 | 2.16 | 2.9E-07 |
| *Y37D8A.8* | WBGene00012549 | protein_coding_gene | 1.1 | 2.16 | 5.3E-04 |
| *C17C3.9* | WBGene00015892 | protein_coding_gene | 1.1 | 2.16 | 2.3E-02 |
| *B0280.2* | WBGene00015100 | protein_coding_gene | 1.1 | 2.16 | 2.9E-03 |
| *ZC581.10* | WBGene00195178 | protein_coding_gene | 1.1 | 2.16 | 1.5E-03 |
| *T16G12.7* | WBGene00011808 | protein_coding_gene | 1.1 | 2.15 | 6.9E-04 |
| *glct-3* | WBGene00011781 | protein_coding_gene | 1.1 | 2.15 | 2.7E-02 |
| *C47E12.11* | WBGene00008154 | protein_coding_gene | 1.1 | 2.15 | 1.4E-04 |
| *msp-152* | WBGene00003470 | protein_coding_gene | 1.1 | 2.15 | 5.6E-03 |
| *skpo-2* | WBGene00015841 | protein_coding_gene | 1.1 | 2.15 | 4.1E-04 |
| *gsp-4* | WBGene00020187 | protein_coding_gene | 1.1 | 2.15 | 6.1E-09 |
| *ZK1128.3* | WBGene00014229 | protein_coding_gene | 1.1 | 2.15 | 1.9E-04 |
| *ssp-11* | WBGene00006040 | protein_coding_gene | 1.1 | 2.15 | 8.2E-03 |
| *C30A5.10* | WBGene00016242 | protein_coding_gene | 1.1 | 2.15 | 2.0E-02 |
| *clec-123* | WBGene00021288 | protein_coding_gene | 1.1 | 2.15 | 2.5E-06 |
| *argk-1* | WBGene00009706 | protein_coding_gene | 1.1 | 2.15 | 1.9E-02 |
| *K08F8.5* | WBGene00010682 | protein_coding_gene | 1.1 | 2.15 | 4.5E-04 |
| *C14A6.10* | WBGene00045348 | pseudogene | 1.1 | 2.14 | 2.5E-02 |
| *msp-78* | WBGene00003465 | protein_coding_gene | 1.1 | 2.14 | 3.7E-05 |
| *Y71G12B.33* | WBGene00044358 | protein_coding_gene | 1.1 | 2.14 | 3.1E-02 |
| *C18F10.2* | WBGene00015977 | protein_coding_gene | 1.1 | 2.14 | 4.1E-04 |
| *F46F5.7* | WBGene00018498 | protein_coding_gene | 1.1 | 2.14 | 5.5E-03 |
| *F38E1.3* | WBGene00018178 | protein_coding_gene | 1.1 | 2.14 | 1.1E-03 |
| *srr-3* | WBGene00005654 | protein_coding_gene | 1.1 | 2.14 | 2.8E-02 |
| *C17H12.5* | WBGene00015931 | protein_coding_gene | 1.1 | 2.14 | 7.5E-06 |
| *ZK105.14* | WBGene00271707 | protein_coding_gene | 1.1 | 2.13 | 4.8E-03 |
| *F13A7.1* | WBGene00008724 | protein_coding_gene | 1.1 | 2.13 | 5.7E-06 |
| *W05B10.3* | WBGene00012278 | protein_coding_gene | 1.1 | 2.13 | 2.9E-02 |
| *C15H7.2* | WBGene00014678 | pseudogene | 1.1 | 2.13 | 2.7E-02 |
| *alg-3* | WBGene00011910 | protein_coding_gene | 1.1 | 2.13 | 2.1E-03 |
| *msd-4* | WBGene00016447 | protein_coding_gene | 1.1 | 2.13 | 8.8E-04 |
| *C53D6.5* | WBGene00008282 | pseudogene | 1.1 | 2.13 | 3.2E-02 |
| *F27C8.5* | WBGene00009185 | protein_coding_gene | 1.1 | 2.13 | 1.4E-04 |
| *dmsr-3* | WBGene00012993 | protein_coding_gene | 1.1 | 2.13 | 9.2E-03 |
| *cnnm-5* | WBGene00011260 | protein_coding_gene | 1.1 | 2.13 | 5.5E-03 |
| *cwp-5* | WBGene00009844 | protein_coding_gene | 1.1 | 2.13 | 3.4E-03 |
| *Y65B4A.7* | WBGene00022030 | protein_coding_gene | 1.1 | 2.13 | 1.3E-02 |
| *C55B7.10* | WBGene00016946 | protein_coding_gene | 1.1 | 2.13 | 2.0E-03 |
| *F57F4.1* | WBGene00019015 | protein_coding_gene | 1.1 | 2.13 | 6.0E-06 |
| *ttbk-3* | WBGene00009324 | protein_coding_gene | 1.1 | 2.12 | 2.0E-04 |
| *B0218.5* | WBGene00015049 | protein_coding_gene | 1.1 | 2.12 | 8.1E-04 |
| *C09H5.7* | WBGene00015661 | protein_coding_gene | 1.1 | 2.12 | 4.9E-03 |
| *Y57G11B.3* | WBGene00013296 | protein_coding_gene | 1.1 | 2.12 | 5.0E-03 |
| *srb-3* | WBGene00005068 | protein_coding_gene | 1.1 | 2.12 | 2.9E-02 |
| *K11C4.1* | WBGene00019642 | protein_coding_gene | 1.1 | 2.12 | 1.9E-04 |
| *clec-236* | WBGene00013495 | pseudogene | 1.1 | 2.12 | 1.3E-02 |
| *Y23H5B.2* | WBGene00021273 | pseudogene | 1.1 | 2.12 | 2.0E-02 |
| *spe-11* | WBGene00004965 | protein_coding_gene | 1.1 | 2.12 | 1.2E-06 |

| | | | | | |
|---|---|---|---|---|---|
| *R07H5.9* | WBGene00011129 | protein_coding_gene | 1.1 | 2.12 | 3.5E-03 |
| *K05F1.1* | WBGene00019405 | protein_coding_gene | 1.1 | 2.12 | 1.3E-02 |
| *nhr-175* | WBGene00008630 | protein_coding_gene | 1.1 | 2.12 | 3.3E-02 |
| *pck-3* | WBGene00019151 | protein_coding_gene | 1.1 | 2.11 | 1.4E-05 |
| *cpb-2* | WBGene00000771 | protein_coding_gene | 1.1 | 2.11 | 5.5E-05 |
| *F58H1.6* | WBGene00010289 | protein_coding_gene | 1.1 | 2.11 | 8.4E-06 |
| *F07F6.1* | WBGene00017215 | protein_coding_gene | 1.1 | 2.11 | 1.0E-03 |
| *msp-59* | WBGene00003452 | protein_coding_gene | 1.1 | 2.11 | 1.3E-06 |
| *C55C3.4* | WBGene00016954 | protein_coding_gene | 1.1 | 2.11 | 2.6E-04 |
| *linc-23* | WBGene00219692 | lincRNA_gene | 1.1 | 2.11 | 1.8E-02 |
| *ZC581.7* | WBGene00022634 | protein_coding_gene | 1.1 | 2.11 | 1.6E-03 |
| *C38C10.3* | WBGene00008001 | protein_coding_gene | 1.1 | 2.11 | 9.1E-04 |
| *swt-5* | WBGene00010594 | protein_coding_gene | 1.1 | 2.11 | 2.6E-02 |
| *nkb-2* | WBGene00008074 | protein_coding_gene | 1.1 | 2.11 | 1.2E-05 |
| *msp-76* | WBGene00003463 | protein_coding_gene | 1.1 | 2.11 | 1.5E-07 |
| *W07A12.4* | WBGene00012322 | protein_coding_gene | 1.1 | 2.11 | 7.7E-04 |
| *hdl-1* | WBGene00001839 | protein_coding_gene | 1.1 | 2.10 | 9.6E-05 |
| *C15A11.2* | WBGene00007597 | protein_coding_gene | 1.1 | 2.10 | 1.2E-03 |
| *acs-6* | WBGene00022849 | protein_coding_gene | 1.1 | 2.10 | 2.1E-05 |
| *rab-3* | WBGene00004267 | protein_coding_gene | 1.1 | 2.10 | 3.7E-13 |
| *C17G10.3* | WBGene00015917 | protein_coding_gene | 1.1 | 2.10 | 1.1E-03 |
| *snf-2* | WBGene00004901 | protein_coding_gene | 1.1 | 2.10 | 1.4E-04 |
| *Y53F4B.24* | WBGene00013170 | protein_coding_gene | 1.1 | 2.10 | 6.7E-03 |
| *F23C8.8* | WBGene00017737 | protein_coding_gene | 1.1 | 2.10 | 2.8E-03 |
| *F23F12.12* | WBGene00017756 | protein_coding_gene | 1.1 | 2.10 | 2.5E-02 |
| *T22B3.3* | WBGene00011911 | protein_coding_gene | 1.1 | 2.10 | 1.9E-07 |
| *R03D7.8* | WBGene00010992 | protein_coding_gene | 1.1 | 2.10 | 1.4E-03 |
| *F31E8.5* | WBGene00017955 | protein_coding_gene | 1.1 | 2.10 | 2.9E-05 |
| *Y7A5A.21* | WBGene00219352 | pseudogene | 1.1 | 2.10 | 3.6E-02 |
| *clp-10* | WBGene00021041 | protein_coding_gene | 1.1 | 2.10 | 3.3E-02 |
| *fbxb-68* | WBGene00008611 | protein_coding_gene | 1.1 | 2.10 | 1.9E-02 |
| *F21C10.9* | WBGene00017658 | protein_coding_gene | 1.1 | 2.09 | 9.5E-12 |
| *phf-33* | WBGene00010708 | protein_coding_gene | 1.1 | 2.09 | 2.9E-02 |
| *K04F1.8* | WBGene00019387 | protein_coding_gene | 1.1 | 2.09 | 5.8E-03 |
| *C45G9.4* | WBGene00016675 | protein_coding_gene | 1.1 | 2.09 | 2.1E-03 |
| *F13E9.5* | WBGene00008754 | protein_coding_gene | 1.1 | 2.09 | 2.8E-04 |
| *C32D5.4* | WBGene00016312 | protein_coding_gene | 1.1 | 2.09 | 1.2E-03 |
| *mam-3* | WBGene00010714 | protein_coding_gene | 1.1 | 2.09 | 6.9E-03 |
| *oac-9* | WBGene00016753 | protein_coding_gene | 1.1 | 2.09 | 2.3E-05 |
| *K03H1.9* | WBGene00010544 | protein_coding_gene | 1.1 | 2.09 | 6.9E-03 |
| *C18E3.1* | WBGene00015970 | protein_coding_gene | 1.1 | 2.09 | 1.9E-03 |
| *Y38E10A.3* | WBGene00012581 | protein_coding_gene | 1.1 | 2.08 | 8.0E-03 |
| *ttr-56* | WBGene00050896 | protein_coding_gene | 1.1 | 2.08 | 2.0E-02 |
| *msrp-5* | WBGene00017059 | protein_coding_gene | 1.1 | 2.08 | 1.0E-04 |
| *acs-18* | WBGene00011173 | protein_coding_gene | 1.1 | 2.08 | 4.1E-06 |
| *Y4C6A.1* | WBGene00021151 | pseudogene | 1.1 | 2.08 | 1.4E-02 |
| *T04F3.3* | WBGene00011438 | protein_coding_gene | 1.1 | 2.08 | 4.0E-04 |
| *kin-5* | WBGene00002193 | protein_coding_gene | 1.1 | 2.08 | 1.8E-04 |
| *spe-49* | WBGene00019667 | protein_coding_gene | 1.1 | 2.08 | 5.4E-03 |
| *kvs-5* | WBGene00021948 | protein_coding_gene | 1.1 | 2.08 | 1.7E-06 |
| *C49C8.1* | WBGene00016765 | protein_coding_gene | 1.1 | 2.08 | 4.2E-03 |
| *R04B5.11* | WBGene00011008 | protein_coding_gene | 1.1 | 2.08 | 2.8E-02 |
| *twk-6* | WBGene00006661 | protein_coding_gene | 1.1 | 2.07 | 2.0E-02 |
| *F45C12.9* | WBGene00018440 | protein_coding_gene | 1.1 | 2.07 | 2.1E-02 |
| *M163.8* | WBGene00010940 | protein_coding_gene | 1.1 | 2.07 | 1.8E-06 |
| *T05A8.7* | WBGene00020233 | protein_coding_gene | 1.1 | 2.07 | 3.8E-02 |
| *klp-17* | WBGene00002227 | protein_coding_gene | 1.1 | 2.07 | 2.4E-03 |
| *best-11* | WBGene00008186 | protein_coding_gene | 1.1 | 2.07 | 1.4E-02 |
| *sfxn-1.2* | WBGene00009513 | protein_coding_gene | 1.1 | 2.07 | 3.9E-03 |
| *Y106G6A.4* | WBGene00013696 | protein_coding_gene | 1.1 | 2.07 | 2.3E-06 |
| *F41H10.5* | WBGene00018318 | protein_coding_gene | 1.0 | 2.07 | 3.3E-03 |
| *Y59E9AL.3* | WBGene00021993 | protein_coding_gene | 1.0 | 2.07 | 7.3E-04 |
| *F42C5.5* | WBGene00018347 | protein_coding_gene | 1.0 | 2.07 | 3.3E-04 |
| *Y40H4A.2* | WBGene00012741 | protein_coding_gene | 1.0 | 2.07 | 1.4E-03 |
| *W06D12.6* | WBGene00012305 | protein_coding_gene | 1.0 | 2.07 | 6.5E-03 |

| | | | | | |
|---|---|---|---|---|---|
| *Y73F8A.20* | WBGene00013526 | protein_coding_gene | 1.0 | 2.07 | 4.6E-04 |
| *ttr-9* | WBGene00007888 | protein_coding_gene | 1.0 | 2.07 | 1.3E-02 |
| *ugt-42* | WBGene00017959 | protein_coding_gene | 1.0 | 2.06 | 1.8E-02 |
| *K09E4.1* | WBGene00010719 | protein_coding_gene | 1.0 | 2.06 | 5.4E-04 |
| *Y24D9B.1* | WBGene00021287 | protein_coding_gene | 1.0 | 2.06 | 2.8E-03 |
| *F33C8.4* | WBGene00009358 | protein_coding_gene | 1.0 | 2.06 | 2.3E-08 |
| *C26C6.6* | WBGene00007740 | protein_coding_gene | 1.0 | 2.06 | 1.7E-02 |
| *T09B4.3* | WBGene00020377 | protein_coding_gene | 1.0 | 2.06 | 3.5E-03 |
| *wht-6* | WBGene00007060 | protein_coding_gene | 1.0 | 2.06 | 4.7E-05 |
| *F58D5.8* | WBGene00010247 | protein_coding_gene | 1.0 | 2.06 | 1.3E-02 |
| *clec-97* | WBGene00013931 | protein_coding_gene | 1.0 | 2.06 | 7.6E-07 |
| *C28A5.6* | WBGene00007791 | protein_coding_gene | 1.0 | 2.06 | 3.4E-08 |
| *C24H12.2* | WBGene00016072 | protein_coding_gene | 1.0 | 2.06 | 3.8E-02 |
| *rnh-1.1* | WBGene00022888 | protein_coding_gene | 1.0 | 2.06 | 3.2E-04 |
| *ZK970.8* | WBGene00014174 | protein_coding_gene | 1.0 | 2.06 | 1.4E-03 |
| *Y37E11AL.2* | WBGene00021358 | protein_coding_gene | 1.0 | 2.06 | 3.9E-05 |
| *F59A1.2* | WBGene00014779 | pseudogene | 1.0 | 2.06 | 6.1E-03 |
| *spe-10* | WBGene00004964 | protein_coding_gene | 1.0 | 2.06 | 2.1E-03 |
| *M05B5.1* | WBGene00010869 | protein_coding_gene | 1.0 | 2.05 | 3.1E-04 |
| *spe-9* | WBGene00004963 | protein_coding_gene | 1.0 | 2.05 | 6.5E-03 |
| *Y80D3A.8* | WBGene00013586 | protein_coding_gene | 1.0 | 2.05 | 1.6E-02 |
| *ZK616.65* | WBGene00235383 | protein_coding_gene | 1.0 | 2.05 | 3.6E-03 |
| *ssp-33* | WBGene00011134 | protein_coding_gene | 1.0 | 2.05 | 4.2E-03 |
| *Y73F8A.15* | WBGene00013524 | protein_coding_gene | 1.0 | 2.05 | 9.0E-03 |
| *nhr-198* | WBGene00010603 | protein_coding_gene | 1.0 | 2.05 | 2.2E-02 |
| *fbxa-42* | WBGene00044676 | protein_coding_gene | 1.0 | 2.05 | 1.0E-02 |
| *C05C12.1* | WBGene00007335 | protein_coding_gene | 1.0 | 2.05 | 3.7E-04 |
| *fbxa-88* | WBGene00008632 | protein_coding_gene | 1.0 | 2.05 | 3.5E-02 |
| *H05L14.1* | WBGene00010366 | protein_coding_gene | 1.0 | 2.05 | 3.1E-07 |
| *Y71G12B.3* | WBGene00022143 | protein_coding_gene | 1.0 | 2.05 | 1.3E-03 |
| *C28D4.8* | WBGene00007795 | protein_coding_gene | 1.0 | 2.05 | 1.2E-02 |
| *msp-38* | WBGene00003434 | protein_coding_gene | 1.0 | 2.04 | 1.8E-04 |
| *C32E12.1* | WBGene00016327 | protein_coding_gene | 1.0 | 2.04 | 1.1E-05 |
| *tsp-18* | WBGene00006644 | protein_coding_gene | 1.0 | 2.04 | 2.0E-02 |
| *Y53C12A.8* | WBGene00044158 | protein_coding_gene | 1.0 | 2.04 | 2.1E-02 |
| *ZK795.2* | WBGene00014082 | protein_coding_gene | 1.0 | 2.04 | 1.9E-05 |
| *C18A3.7* | WBGene00015944 | protein_coding_gene | 1.0 | 2.04 | 2.4E-03 |
| *fbxa-166* | WBGene00015601 | protein_coding_gene | 1.0 | 2.04 | 3.9E-02 |
| *F40H6.1* | WBGene00018253 | protein_coding_gene | 1.0 | 2.04 | 5.0E-05 |
| *nhr-251* | WBGene00022756 | protein_coding_gene | 1.0 | 2.04 | 2.2E-02 |
| *oac-56* | WBGene00013450 | protein_coding_gene | 1.0 | 2.04 | 4.2E-02 |
| *R02E4.3* | WBGene00044724 | protein_coding_gene | 1.0 | 2.04 | 1.0E-02 |
| *kin-26* | WBGene00002208 | protein_coding_gene | 1.0 | 2.03 | 1.4E-03 |
| *Y53F4B.17* | WBGene00013163 | protein_coding_gene | 1.0 | 2.03 | 6.7E-03 |
| *F57A8.6* | WBGene00010181 | protein_coding_gene | 1.0 | 2.03 | 3.7E-03 |
| *T15H9.4* | WBGene00011789 | protein_coding_gene | 1.0 | 2.03 | 2.4E-02 |
| *F59A6.4* | WBGene00019086 | protein_coding_gene | 1.0 | 2.03 | 8.7E-05 |
| *Y102A5C.38* | WBGene00050883 | pseudogene | 1.0 | 2.03 | 7.6E-03 |
| *T16A1.2* | WBGene00020533 | protein_coding_gene | 1.0 | 2.03 | 1.5E-02 |
| *T16A9.5* | WBGene00011795 | protein_coding_gene | 1.0 | 2.03 | 1.6E-05 |
| *ZK945.6* | WBGene00014168 | protein_coding_gene | 1.0 | 2.03 | 1.6E-03 |
| *gst-32* | WBGene00001780 | protein_coding_gene | 1.0 | 2.03 | 4.6E-02 |
| *clec-109* | WBGene00008907 | protein_coding_gene | 1.0 | 2.03 | 4.5E-02 |
| *nep-23* | WBGene00013785 | protein_coding_gene | 1.0 | 2.03 | 2.2E-04 |
| *Y41E3.461* | WBGene00219444 | ncRNA_gene | 1.0 | 2.03 | 4.6E-02 |
| *cbs-2* | WBGene00018783 | protein_coding_gene | 1.0 | 2.02 | 2.0E-03 |
| *M04G7.2* | WBGene00019773 | protein_coding_gene | 1.0 | 2.02 | 2.3E-02 |
| *snf-5* | WBGene00004904 | protein_coding_gene | 1.0 | 2.02 | 6.3E-03 |
| *Y119C1B.1* | WBGene00022467 | protein_coding_gene | 1.0 | 2.02 | 3.0E-03 |
| *Y39A1A.2* | WBGene00012642 | protein_coding_gene | 1.0 | 2.02 | 8.2E-03 |
| *glna-1* | WBGene00007480 | protein_coding_gene | 1.0 | 2.02 | 4.9E-04 |
| *C09D4.9* | WBGene00206357 | protein_coding_gene | 1.0 | 2.02 | 1.6E-02 |
| *Y7A5A.20* | WBGene00219351 | protein_coding_gene | 1.0 | 2.02 | 4.1E-02 |
| *Y67D8B.5* | WBGene00022064 | protein_coding_gene | 1.0 | 2.02 | 4.5E-02 |
| *snf-4* | WBGene00004903 | pseudogene | 1.0 | 2.01 | 1.5E-02 |

| | | | | | |
|---|---|---|---|---|---|
| *clec-6* | WBGene00016451 | protein_coding_gene | 1.0 | 2.01 | 4.4E-02 |
| *T03F6.6* | WBGene00011401 | protein_coding_gene | 1.0 | 2.01 | 4.1E-05 |
| *R13H4.8* | WBGene00011266 | protein_coding_gene | 1.0 | 2.01 | 4.7E-02 |
| *oac-20* | WBGene00009488 | protein_coding_gene | 1.0 | 2.01 | 3.1E-02 |
| *tor-2* | WBGene00006598 | protein_coding_gene | 1.0 | 2.01 | 4.5E-02 |
| *F13A2.2* | WBGene00017411 | protein_coding_gene | 1.0 | 2.01 | 4.5E-02 |
| *oig-2* | WBGene00003860 | protein_coding_gene | 1.0 | 2.01 | 5.5E-05 |
| *Y71G12B.18* | WBGene00022156 | protein_coding_gene | 1.0 | 2.01 | 4.3E-02 |
| *F13H8.5* | WBGene00017438 | protein_coding_gene | 1.0 | 2.01 | 4.3E-04 |
| *T16G1.13* | WBGene00194787 | protein_coding_gene | 1.0 | 2.01 | 1.3E-02 |
| *Y54G2A.20* | WBGene00021885 | pseudogene | 1.0 | 2.01 | 4.4E-02 |
| *nhr-113* | WBGene00003703 | protein_coding_gene | 1.0 | 2.01 | 4.4E-02 |
| *F35E2.9* | WBGene00009416 | protein_coding_gene | 1.0 | 2.01 | 5.7E-05 |
| *abu-12* | WBGene00004127 | protein_coding_gene | 1.0 | 2.01 | 1.5E-03 |
| *Y116A8C.24* | WBGene00013801 | protein_coding_gene | 1.0 | 2.00 | 2.6E-02 |
| *T28B8.6* | WBGene00044122 | protein_coding_gene | 1.0 | 2.00 | 2.3E-03 |
| *F36A2.11* | WBGene00009458 | protein_coding_gene | 1.0 | 2.00 | 4.1E-03 |
| *F58E6.13* | WBGene00077697 | protein_coding_gene | 1.0 | 2.00 | 4.8E-04 |
| *fbxa-4* | WBGene00044672 | protein_coding_gene | -1.0 | 0.49 | 2.7E-03 |
| *skr-13* | WBGene00004819 | protein_coding_gene | -1.0 | 0.49 | 5.8E-04 |
| *Y49F6A.5* | WBGene00021713 | protein_coding_gene | -1.0 | 0.49 | 4.8E-02 |
| *acs-2* | WBGene00009221 | protein_coding_gene | -1.0 | 0.49 | 3.1E-03 |
| *Y69A2AR.8* | WBGene00022079 | pseudogene | -1.0 | 0.49 | 1.7E-02 |
| *fbxa-93* | WBGene00012368 | pseudogene | -1.0 | 0.49 | 6.0E-04 |
| *trm-2A* | WBGene00019241 | protein_coding_gene | -1.0 | 0.49 | 1.8E-12 |
| *C33C12.11* | WBGene00044360 | protein_coding_gene | -1.0 | 0.49 | 2.3E-02 |
| *fbxa-185* | WBGene00009836 | protein_coding_gene | -1.0 | 0.49 | 7.7E-03 |
| *irld-27* | WBGene00077751 | protein_coding_gene | -1.0 | 0.49 | 7.6E-03 |
| *T08E11.1* | WBGene00020357 | protein_coding_gene | -1.0 | 0.48 | 3.7E-02 |
| *csn-4* | WBGene00000816 | protein_coding_gene | -1.0 | 0.48 | 2.9E-16 |
| *F41B5.1* | WBGene00023284 | protein_coding_gene | -1.1 | 0.48 | 2.4E-04 |
| *Y37A1B.5* | WBGene00012538 | protein_coding_gene | -1.1 | 0.48 | 5.6E-18 |
| *hil-2* | WBGene00001853 | protein_coding_gene | -1.1 | 0.48 | 2.3E-09 |
| *fbxa-29* | WBGene00013836 | protein_coding_gene | -1.1 | 0.48 | 1.9E-03 |
| *F57F4.2* | WBGene00019016 | protein_coding_gene | -1.1 | 0.48 | 3.4E-02 |
| *Y73B3A.1* | WBGene00022203 | protein_coding_gene | -1.1 | 0.48 | 5.7E-11 |
| *eri-5* | WBGene00021419 | protein_coding_gene | -1.1 | 0.48 | 5.0E-10 |
| *T09F5.10* | WBGene00011669 | protein_coding_gene | -1.1 | 0.48 | 2.0E-02 |
| *fbxa-108* | WBGene00010293 | protein_coding_gene | -1.1 | 0.48 | 2.7E-08 |
| *nhr-123* | WBGene00003713 | protein_coding_gene | -1.1 | 0.47 | 2.8E-02 |
| *cest-1.1* | WBGene00011362 | protein_coding_gene | -1.1 | 0.47 | 3.4E-10 |
| *ctc-1* | WBGene00010964 | protein_coding_gene | -1.1 | 0.47 | 9.4E-21 |
| *D1044.7* | WBGene00017032 | protein_coding_gene | -1.1 | 0.47 | 9.9E-03 |
| *clec-225* | WBGene00007152 | protein_coding_gene | -1.1 | 0.47 | 2.2E-02 |
| *Y18D10A.22* | WBGene00014864 | protein_coding_gene | -1.1 | 0.47 | 1.8E-03 |
| *Y105C5B.25* | WBGene00013662 | protein_coding_gene | -1.1 | 0.47 | 3.1E-02 |
| *oac-54* | WBGene00012323 | protein_coding_gene | -1.1 | 0.47 | 2.0E-02 |
| *C29F3.7* | WBGene00007807 | protein_coding_gene | -1.1 | 0.47 | 5.7E-11 |
| *B0250.12* | WBGene00050927 | pseudogene | -1.1 | 0.47 | 3.5E-03 |
| *pals-22* | WBGene00016216 | protein_coding_gene | -1.1 | 0.46 | 2.5E-03 |
| *zipt-1* | WBGene00009286 | protein_coding_gene | -1.1 | 0.46 | 1.5E-34 |
| *nape-1* | WBGene00021371 | protein_coding_gene | -1.1 | 0.46 | 7.9E-07 |
| *K03D3.2* | WBGene00010528 | protein_coding_gene | -1.1 | 0.46 | 4.1E-03 |
| *fbxc-40* | WBGene00016295 | protein_coding_gene | -1.1 | 0.46 | 1.4E-03 |
| *Y69A2AR.47* | WBGene00235294 | protein_coding_gene | -1.1 | 0.46 | 2.3E-02 |
| *bath-35* | WBGene00012196 | protein_coding_gene | -1.1 | 0.46 | 4.7E-03 |
| *Y56A3A.19* | WBGene00013237 | protein_coding_gene | -1.1 | 0.45 | 1.6E-25 |
| *fbxc-39* | WBGene00018456 | protein_coding_gene | -1.2 | 0.45 | 1.2E-02 |
| *oac-18* | WBGene00009230 | pseudogene | -1.2 | 0.45 | 1.5E-02 |
| *C49C8.2* | WBGene00016766 | protein_coding_gene | -1.2 | 0.45 | 6.1E-03 |
| *F58E1.13* | WBGene00019044 | protein_coding_gene | -1.2 | 0.45 | 3.8E-03 |
| *ver-1* | WBGene00006894 | protein_coding_gene | -1.2 | 0.45 | 2.0E-02 |
| *cdr-1* | WBGene00000412 | protein_coding_gene | -1.2 | 0.45 | 1.4E-02 |
| *T20D4.13* | WBGene00020619 | pseudogene | -1.2 | 0.44 | 2.0E-02 |
| *fbxb-45* | WBGene00019705 | protein_coding_gene | -1.2 | 0.44 | 5.9E-03 |

| | | | | | |
|---|---|---|---|---|---|
| *btb-2* | WBGene00020802 | protein_coding_gene | -1.2 | 0.44 | 4.1E-03 |
| *C33G8.2* | WBGene00016360 | protein_coding_gene | -1.2 | 0.44 | 6.4E-15 |
| *W04B5.1* | WBGene00021018 | pseudogene | -1.2 | 0.44 | 1.6E-07 |
| *Y47H9C.1* | WBGene00012947 | protein_coding_gene | -1.2 | 0.44 | 4.6E-03 |
| *F47G3.4* | WBGene00194641 | protein_coding_gene | -1.2 | 0.44 | 4.2E-06 |
| *T07D3.6* | WBGene00020309 | pseudogene | -1.2 | 0.44 | 1.7E-02 |
| *his-72* | WBGene00001946 | protein_coding_gene | -1.2 | 0.44 | 1.8E-28 |
| *T07D3.9* | WBGene00020311 | protein_coding_gene | -1.2 | 0.44 | 1.4E-11 |
| *Y57G11C.1143* | WBGene00219342 | pseudogene | -1.2 | 0.44 | 3.0E-03 |
| *T24A6.1* | WBGene00020744 | protein_coding_gene | -1.2 | 0.43 | 9.4E-03 |
| *F21H7.2* | WBGene00009028 | protein_coding_gene | -1.2 | 0.43 | 5.7E-06 |
| *nhr-72* | WBGene00003662 | protein_coding_gene | -1.2 | 0.43 | 1.8E-02 |
| *ttr-12* | WBGene00009759 | protein_coding_gene | -1.2 | 0.43 | 2.7E-03 |
| *Y43F8B.13* | WBGene00012823 | protein_coding_gene | -1.2 | 0.43 | 2.4E-03 |
| *Y43F8B.15* | WBGene00045415 | protein_coding_gene | -1.2 | 0.43 | 2.6E-03 |
| *ubc-19* | WBGene00006714 | protein_coding_gene | -1.2 | 0.43 | 1.6E-04 |
| *M116.2* | WBGene00019789 | pseudogene | -1.2 | 0.43 | 1.3E-10 |
| *C28C12.4* | WBGene00016180 | protein_coding_gene | -1.2 | 0.43 | 3.0E-03 |
| *mxl-2* | WBGene00003510 | protein_coding_gene | -1.2 | 0.43 | 7.7E-15 |
| *C02B8.6* | WBGene00015324 | protein_coding_gene | -1.2 | 0.43 | 1.8E-10 |
| *fbxa-107* | WBGene00012564 | protein_coding_gene | -1.2 | 0.43 | 4.4E-11 |
| *H16D19.6* | WBGene00195254 | pseudogene | -1.2 | 0.42 | 1.5E-02 |
| *W06D12.8* | WBGene00198082 | ncRNA_gene | -1.3 | 0.42 | 1.2E-02 |
| *Y6G8.9* | WBGene00050974 | pseudogene | -1.3 | 0.42 | 1.2E-02 |
| *T02G5.11* | WBGene00020167 | pseudogene | -1.3 | 0.42 | 6.2E-04 |
| *F47B10.3* | WBGene00009814 | protein_coding_gene | -1.3 | 0.42 | 8.2E-03 |
| *ZC239.5* | WBGene00022568 | protein_coding_gene | -1.3 | 0.41 | 1.2E-02 |
| *F09C6.3* | WBGene00008617 | protein_coding_gene | -1.3 | 0.41 | 1.3E-02 |
| *Y116F11B.7* | WBGene00013823 | protein_coding_gene | -1.3 | 0.41 | 1.1E-02 |
| *nhr-148* | WBGene00015396 | protein_coding_gene | -1.3 | 0.41 | 1.1E-02 |
| *C31C9.7* | WBGene00007841 | protein_coding_gene | -1.3 | 0.41 | 1.2E-05 |
| *Y111B2A.21* | WBGene00013741 | protein_coding_gene | -1.3 | 0.40 | 5.3E-04 |
| *F01D5.5* | WBGene00008495 | protein_coding_gene | -1.3 | 0.40 | 3.8E-05 |
| *F58D5.3* | WBGene00010243 | protein_coding_gene | -1.3 | 0.39 | 1.1E-03 |
| *M01H9.2* | WBGene00019718 | protein_coding_gene | -1.3 | 0.39 | 2.0E-03 |
| *Y113G7A.14* | WBGene00013751 | protein_coding_gene | -1.4 | 0.39 | 6.1E-03 |
| *Y51A2B.5* | WBGene00013069 | protein_coding_gene | -1.4 | 0.39 | 2.4E-03 |
| *fbxc-34* | WBGene00019921 | protein_coding_gene | -1.4 | 0.39 | 4.6E-03 |
| *C14A6.16* | WBGene00206536 | pseudogene | -1.4 | 0.38 | 3.8E-03 |
| *ZK250.2* | WBGene00022685 | protein_coding_gene | -1.4 | 0.38 | 9.5E-04 |
| *fbxb-10* | WBGene00020362 | protein_coding_gene | -1.4 | 0.38 | 2.1E-03 |
| *F11A5.17* | WBGene00045510 | protein_coding_gene | -1.4 | 0.38 | 7.9E-05 |
| *Y80D4G.1* | WBGene00206459 | pseudogene | -1.4 | 0.38 | 1.5E-11 |
| *Y51A2B.6* | WBGene00013070 | protein_coding_gene | -1.4 | 0.38 | 2.5E-03 |
| *ZC239.17* | WBGene00022575 | protein_coding_gene | -1.4 | 0.38 | 8.6E-04 |
| *T15B7.10* | WBGene00020522 | pseudogene | -1.4 | 0.37 | 1.2E-05 |
| *C25F9.11* | WBGene00045411 | protein_coding_gene | -1.4 | 0.37 | 5.5E-04 |
| *linc-16* | WBGene00219728 | lincRNA_gene | -1.4 | 0.37 | 5.4E-04 |
| *C25F9.15* | WBGene00050967 | protein_coding_gene | -1.4 | 0.37 | 6.3E-04 |
| *dhs-23* | WBGene00000986 | protein_coding_gene | -1.4 | 0.37 | 8.5E-10 |
| *C40A11.2* | WBGene00016545 | protein_coding_gene | -1.5 | 0.36 | 2.1E-03 |
| *Y110A2AL.3* | WBGene00022440 | protein_coding_gene | -1.5 | 0.36 | 2.0E-05 |
| *R07C12.1* | WBGene00019931 | protein_coding_gene | -1.5 | 0.36 | 2.9E-04 |
| *K10G4.3* | WBGene00010752 | protein_coding_gene | -1.5 | 0.36 | 2.4E-03 |
| *fbxa-26* | WBGene00022324 | protein_coding_gene | -1.5 | 0.36 | 1.0E-06 |
| *M02E1.3* | WBGene00023458 | protein_coding_gene | -1.5 | 0.36 | 3.3E-03 |
| *T03E6.9* | WBGene00045356 | protein_coding_gene | -1.5 | 0.36 | 5.3E-03 |
| *Y57G11C.52* | WBGene00044741 | protein_coding_gene | -1.5 | 0.36 | 3.2E-03 |
| *clec-2* | WBGene00015198 | protein_coding_gene | -1.5 | 0.36 | 1.2E-03 |
| *F55C12.19* | WBGene00271817 | protein_coding_gene | -1.5 | 0.35 | 1.4E-03 |
| *ZK686.1* | WBGene00022791 | protein_coding_gene | -1.5 | 0.35 | 3.2E-29 |
| *F49F1.18* | WBGene00235368 | protein_coding_gene | -1.5 | 0.35 | 3.9E-04 |
| *F59A7.7* | WBGene00019094 | protein_coding_gene | -1.5 | 0.35 | 3.5E-03 |
| *Y73B6A.6* | WBGene00044503 | pseudogene | -1.5 | 0.34 | 2.4E-03 |
| *F09C6.14* | WBGene00044278 | protein_coding_gene | -1.5 | 0.34 | 2.3E-03 |

| | | | | | |
|---|---|---|---|---|---|
| *cpr-2* | WBGene00000782 | protein_coding_gene | -1.5 | 0.34 | 1.2E-03 |
| *C17B7.4* | WBGene00015878 | protein_coding_gene | -1.5 | 0.34 | 3.6E-04 |
| *C14C6.2* | WBGene00015756 | protein_coding_gene | -1.5 | 0.34 | 1.8E-32 |
| *F31E9.3* | WBGene00009296 | protein_coding_gene | -1.6 | 0.34 | 5.8E-04 |
| *srh-70* | WBGene00005291 | protein_coding_gene | -1.6 | 0.34 | 3.2E-03 |
| *fbxa-114* | WBGene00013758 | protein_coding_gene | -1.6 | 0.34 | 7.2E-16 |
| *F07E5.5* | WBGene00017210 | protein_coding_gene | -1.6 | 0.34 | 1.1E-25 |
| *ZK1240.1* | WBGene00022866 | protein_coding_gene | -1.6 | 0.33 | 1.0E-03 |
| *fbxa-85* | WBGene00044654 | protein_coding_gene | -1.6 | 0.33 | 1.9E-03 |
| *F26F2.1* | WBGene00009166 | protein_coding_gene | -1.6 | 0.33 | 2.8E-04 |
| *arrd-21* | WBGene00015441 | pseudogene | -1.6 | 0.33 | 8.2E-04 |
| *Y71H2AM.14* | WBGene00022179 | protein_coding_gene | -1.6 | 0.32 | 7.9E-05 |
| *pals-37* | WBGene00138721 | protein_coding_gene | -1.6 | 0.32 | 5.0E-04 |
| *F53C3.3* | WBGene00018747 | protein_coding_gene | -1.6 | 0.32 | 1.2E-03 |
| *F15B9.6* | WBGene00008850 | protein_coding_gene | -1.6 | 0.32 | 1.5E-05 |
| *Y47H10A.3* | WBGene00012959 | protein_coding_gene | -1.6 | 0.32 | 4.9E-04 |
| *C08F1.11* | WBGene00044417 | protein_coding_gene | -1.6 | 0.32 | 9.3E-05 |
| *F41D3.9* | WBGene00009614 | protein_coding_gene | -1.7 | 0.32 | 2.0E-04 |
| *nhr-58* | WBGene00003648 | protein_coding_gene | -1.7 | 0.32 | 3.0E-06 |
| *F09C6.10* | WBGene00008620 | protein_coding_gene | -1.7 | 0.32 | 1.6E-04 |
| *Y32B12C.5* | WBGene00194769 | pseudogene | -1.7 | 0.31 | 5.1E-11 |
| *Y48G9A.7* | WBGene00021701 | protein_coding_gene | -1.7 | 0.31 | 1.1E-03 |
| *F07C4.6* | WBGene00017200 | protein_coding_gene | -1.7 | 0.31 | 4.1E-06 |
| *nhr-225* | WBGene00020849 | protein_coding_gene | -1.7 | 0.30 | 9.9E-04 |
| *Y46C8AM.1* | WBGene00271795 | protein_coding_gene | -1.8 | 0.29 | 2.1E-07 |
| *H16D19.4* | WBGene00010402 | pseudogene | -1.8 | 0.29 | 1.3E-03 |
| *Y6G8.2* | WBGene00012404 | protein_coding_gene | -1.8 | 0.29 | 2.7E-15 |
| *dach-1* | WBGene00020386 | protein_coding_gene | -1.8 | 0.29 | 8.8E-13 |
| *clec-37* | WBGene00013609 | protein_coding_gene | -1.8 | 0.28 | 1.2E-03 |
| *Y82E9BL.3* | WBGene00022320 | protein_coding_gene | -1.8 | 0.28 | 2.8E-04 |
| *pals-19* | WBGene00044761 | protein_coding_gene | -1.9 | 0.28 | 4.7E-08 |
| *cyp-35B3* | WBGene00019470 | protein_coding_gene | -1.9 | 0.27 | 5.6E-04 |
| *F57E7.2* | WBGene00010203 | protein_coding_gene | -1.9 | 0.27 | 5.0E-05 |
| *nspe-4* | WBGene00012589 | protein_coding_gene | -1.9 | 0.27 | 6.9E-07 |
| *gcy-15* | WBGene00001541 | protein_coding_gene | -1.9 | 0.26 | 1.3E-06 |
| *K03D3.5* | WBGene00010531 | protein_coding_gene | -2.0 | 0.26 | 3.4E-05 |
| *C08F1.10* | WBGene00015613 | protein_coding_gene | -2.0 | 0.25 | 1.2E-13 |
| *K02E7.5* | WBGene00019310 | pseudogene | -2.0 | 0.25 | 4.9E-05 |
| *C14C6.8* | WBGene00015762 | protein_coding_gene | -2.0 | 0.24 | 1.3E-04 |
| *C08F11.3* | WBGene00007452 | protein_coding_gene | -2.1 | 0.23 | 6.2E-11 |
| *K05F6.10* | WBGene00019421 | protein_coding_gene | -2.2 | 0.22 | 5.8E-11 |
| *nhr-116* | WBGene00003706 | protein_coding_gene | -2.2 | 0.22 | 8.3E-09 |
| *C52E12.11* | WBGene00304189 | pseudogene | -2.3 | 0.20 | 2.0E-08 |
| *F53C3.4* | WBGene00018748 | protein_coding_gene | -2.4 | 0.19 | 7.4E-07 |
| *F09C6.13* | WBGene00044183 | protein_coding_gene | -2.4 | 0.19 | 5.6E-11 |
| *F55G11.4* | WBGene00010124 | protein_coding_gene | -2.4 | 0.19 | 3.7E-13 |
| *F09C6.11* | WBGene00044181 | protein_coding_gene | -2.5 | 0.18 | 1.4E-06 |
| *F57E7.4* | WBGene00077616 | protein_coding_gene | -2.5 | 0.18 | 9.9E-07 |
| *Y43F8B.9* | WBGene00012819 | protein_coding_gene | -2.7 | 0.16 | 3.3E-17 |
| *ilys-2* | WBGene00016669 | protein_coding_gene | -2.7 | 0.16 | 9.7E-12 |
| *F14D7.10* | WBGene00008797 | protein_coding_gene | -2.8 | 0.15 | 9.8E-08 |
| *pals-39* | WBGene00008301 | protein_coding_gene | -2.8 | 0.14 | 4.2E-09 |
| *C14C6.5* | WBGene00015759 | protein_coding_gene | -2.9 | 0.13 | 3.0E-16 |
| *Y68A4A.13* | WBGene00044212 | protein_coding_gene | -3.1 | 0.12 | 1.7E-13 |
| *btb-16* | WBGene00018195 | protein_coding_gene | -3.4 | 0.10 | 1.5E-59 |
| *clec-76* | WBGene00021587 | protein_coding_gene | -5.4 | 0.02 | 4.0E-24 |

Genes expressed in opposing directions in CB4856 (vs. N2) fed *E. coli* OP50 or *S. epidermidis*

| | | | *E. coli* OP50 | | | *S. epidermidis* | | |
|---|---|---|---|---|---|---|---|---|
| Gene Name | Wormbase Gene ID | Type | $log_2FC$ | FC | Adj P-value | $log_2FC$ | FC | Adj P-value |
| *C50F7.5* | WBGene00016845 | protein_coding_gene | 7.3 | 156.3 | 2.2E-70 | -1.1 | 0.5 | 6.5E-07 |
| *cest-29* | WBGene00017205 | pseudogene | 5.8 | 55.1 | 5.6E-10 | -1.6 | 0.3 | 2.6E-10 |
| *math-38* | WBGene00020326 | protein_coding_gene | 3.6 | 12.1 | 7.3E-26 | -1.2 | 0.4 | 7.0E-13 |
| *oac-14* | WBGene00008602 | protein_coding_gene | 3.5 | 11.5 | 2.1E-18 | -1.2 | 0.4 | 1.5E-04 |
| *thn-1* | WBGene00009213 | protein_coding_gene | 3.1 | 8.5 | 3.5E-05 | -3.1 | 0.1 | 2.3E-65 |

| | | | | | | | | |
|---|---|---|---|---|---|---|---|---|
| *clec-75* | WBGene00021586 | protein_coding_gene | 2.4 | 5.3 | 1.0E-03 | -1.1 | 0.5 | 1.1E-02 |
| *thn-2* | WBGene00009215 | protein_coding_gene | 1.8 | 3.6 | 4.5E-11 | -1.1 | 0.5 | 3.9E-17 |
| *glc-1* | WBGene00001591 | protein_coding_gene | 1.8 | 3.4 | 8.2E-16 | -1.0 | 0.5 | 3.1E-12 |
| *srh-2* | WBGene00005228 | protein_coding_gene | 1.7 | 3.3 | 1.8E-02 | -1.3 | 0.4 | 4.0E-04 |
| *Y8A9A.2* | WBGene00021171 | protein_coding_gene | 1.6 | 3.1 | 9.3E-08 | -1.1 | 0.5 | 6.1E-04 |
| *clec-7* | WBGene00017364 | protein_coding_gene | 1.6 | 3.1 | 3.0E-10 | -1.5 | 0.4 | 3.7E-25 |
| *lys-10* | WBGene00003099 | protein_coding_gene | 1.6 | 3.0 | 2.1E-02 | -1.6 | 0.3 | 7.2E-06 |
| *nlp-31* | WBGene00003769 | protein_coding_gene | 1.6 | 3.0 | 2.1E-07 | -1.0 | 0.5 | 6.8E-10 |
| *Y110A2AL.4* | WBGene00022441 | protein_coding_gene | 1.4 | 2.6 | 4.2E-02 | -1.4 | 0.4 | 2.2E-09 |
| *W04A8.4* | WBGene00012239 | protein_coding_gene | 1.3 | 2.5 | 1.9E-05 | -1.3 | 0.4 | 1.1E-13 |
| *col-65* | WBGene00000641 | protein_coding_gene | -1.1 | 0.5 | 4.4E-02 | 1.4 | 2.7 | 2.7E-16 |
| *phat-4* | WBGene00020237 | protein_coding_gene | -1.4 | 0.4 | 2.2E-02 | 1.0 | 2.1 | 7.6E-06 |
| *H10E21.4* | WBGene00019184 | protein_coding_gene | -1.4 | 0.4 | 1.5E-02 | 1.1 | 2.1 | 7.5E-04 |
| *ZK180.5* | WBGene00022679 | protein_coding_gene | -1.5 | 0.4 | 3.0E-03 | 1.1 | 2.1 | 3.8E-10 |
| *col-144* | WBGene00000717 | protein_coding_gene | -1.5 | 0.4 | 4.0E-02 | 1.1 | 2.1 | 1.6E-02 |
| *K01D12.9* | WBGene00010468 | protein_coding_gene | -1.5 | 0.4 | 1.6E-02 | 1.2 | 2.4 | 2.4E-04 |
| *R12E2.15* | WBGene00020040 | protein_coding_gene | -1.5 | 0.3 | 5.1E-03 | 1.1 | 2.1 | 2.2E-05 |
| *col-154* | WBGene00000727 | protein_coding_gene | -1.6 | 0.3 | 9.5E-03 | 1.9 | 3.8 | 3.3E-07 |
| *col-115* | WBGene00000689 | protein_coding_gene | -1.7 | 0.3 | 2.3E-02 | 1.2 | 2.3 | 3.3E-03 |
| *col-170* | WBGene00000743 | protein_coding_gene | -1.8 | 0.3 | 2.0E-02 | 1.3 | 2.4 | 5.7E-03 |
| *grl-16* | WBGene00001725 | protein_coding_gene | -2.0 | 0.3 | 3.5E-03 | 1.2 | 2.3 | 1.1E-10 |
| *C05D12.3* | WBGene00014666 | protein_coding_gene | -2.0 | 0.2 | 1.1E-23 | 1.3 | 2.5 | 2.4E-21 |
| *pgp-12* | WBGene00004006 | protein_coding_gene | -2.1 | 0.2 | 5.9E-07 | 1.1 | 2.2 | 4.4E-03 |
| *ram-2* | WBGene00004300 | protein_coding_gene | -2.2 | 0.2 | 6.0E-08 | 1.3 | 2.4 | 1.9E-15 |
| *B0205.13* | WBGene00044644 | protein_coding_gene | -2.4 | 0.2 | 1.3E-03 | 1.1 | 2.2 | 1.7E-02 |
| *col-168* | WBGene00000741 | protein_coding_gene | -2.4 | 0.2 | 2.6E-04 | 1.1 | 2.1 | 1.5E-02 |
| *scl-6* | WBGene00008028 | protein_coding_gene | -3.2 | 0.1 | 1.8E-06 | 1.3 | 2.5 | 2.5E-08 |

**Supplemental Table 15**

**GO terms enriched in genesets from Table 14 comparing N2, CB4856 after infection by *S. epidermidis* for genes that were initially asymmetrically expressed on *E. coli***

**Supplemental Table 15 - Gene Ontology Terms Differentiating *E. coli* and *S. epidermidis* Conditions**

DEGs in both genotypes, but regulated in opposing directions

| Category | Number of DEGs | Genes in Geneset | Downregulated Genes in Geneset | Upregulated Genes in Geneset | p-value | FDR-adjusted p-value | Differentially Expressed Genes within a Given Geneset |
|---|---|---|---|---|---|---|---|
| Extracellular material: collagen | 7 | 185 | 7 (OP50); 0 (*S. epidermidis* ) | 0 (OP50); 7 (*S. epidermidis* ) | 3.5E-09 | 4.5E-08 | WBGene00000641,WBGene00000689,WBGene00000717,WBGene00000727,WBGene00000741,WBGene00000743,WBGene00004300 |
| Stress response: pathogen | 4 | 177 | 0 (OP50); 4 (*S. epidermidis* ) | 4 (OP50); 0 (*S. epidermidis* ) | 5.2E-05 | 6.8E-04 | WBGene00003769,WBGene00009213,WBGene00009215,WBGene00020326 |

DEGs identified in *S. epidermidis* condition only

| Category | Number of DEGs | Genes in Geneset | Downregulated Genes in Geneset | Upregulated Genes in Geneset | p-value | FDR-adjusted p-value | Differentially Expressed Genes within a Given Geneset |
|---|---|---|---|---|---|---|---|
| Unknown | 364 | 8160 | 79 | 285 | 4.1E-18 | 3.7E-16 | WBGene00000861,WBGene00003860,WBGene00004127,WBGene00005002,WBGene00006038,WBGene00006039,WBGene00006040,WBGene00006051,WBGene00007319,WBGene00007408,WBGene00007450,WBGene00007452,WBGene00007503,WBGene00007512,WBGene00007559,WBGene00007597,WBGene00007629,WBGene00007660,WBGene00007727,WBGene00007795,WBGene00007841,WBGene00007888,WBGene00007987,WBGene00008079,WBGene00008141,WBGene00008154,WBGene00008155,WBGene00008187,WBGene00008272,WBGene00008301,WBGene00008383,WBGene00008508,WBGene00008540,WBGene00008617,WBGene00008620,WBGene00008724,WBGene00008743,WBGene00008754,WBGene00008797,WBGene00008850,WBGene00008862,WBGene00008949,WBGene00008961,WBGene00008994,WBGene00009028,WBGene00009166,WBGene00009185,WBGene00009193,WBGene00009217,WBGene00009239,WBGene00009308,WBGene00009344,WBGene00009358,WBGene00009362,WBGene00009392,WBGene00009412,WBGene00009416,WBGene00009458,WBGene00009471,WBGene00009473,WBGene00009484,WBGene00009502,WBGene00009522,WBGene00009550,WBGene00009614,WBGene00009685,WBGene00009714,WBGene00009755,WBGene00009759,WBGene00009814,WBGene00009956,WBGene00010122,WBGene00010147,WBGene00010168,WBGene00010170,WBGene00010181,WBGene00010203,WBGene00010216,WBGene00010243,WBGene00010245,WBGene00010246,WBGene00010247,WBGene00010254,WBGene00010289,WBGene00010324,WBGene00010528,WBGene00010531,WBGene00010544,WBGene00010547,WBGene00010611,WBGene00010616,WBGene00010672,WBGene00010673,WBGene00010680,WBGene00010682,WBGene00010708,WBGene00010714,WBGene00010752,WBGene00010871,WBGene00010887,WBGene00010900,WBGene00010940,WBGene00011008,WBGene00011129,WBGene00011134,WBGene00011266,WBGene00011290,WBGene00011322,WBGene00011362,WBGene00018440,WBGene00018498,WBGene00018500,WBGene00018502,WBGene00018505,WBGene00018507,WBGene00018747,WBGene00018748,WBGene00018981,WBGene00019016,WBGene00019024,WBGene00019044,WBGene00019216,WBGene00019255,WBGene00019387,WBGene00019405,WBGene00019421,WBGene00019425,WBGene00019501,WBGene00019601,WBGene00019667,WBGene00019718,WBGene00019812,WBGene00019874,WBGene00019931,WBGene00019949,WBGene00020039,WBGene00020202,WBGene00020233,WBGene00020377,WBGene00020400,WBGene00020533,WBGene00020598,WBGene00020744,WBGene00020794,WBGene00020802,WBGene00020927,WBGene00020984,WBGene00020986,WBGene00021271,WBGene00021302,WBGene00021358,WBGene00021389,WBGene00021616,WBGene00011379,WBGene00011401,WBGene00011530,WBGene00011594,WBGene00011669,WBGene00011782,<br>WBGene00011784,WBGene00011789,WBGene00011795,WBGene00011911,WBGene00011973,WBGene00012012,WBGene00012196,WBGene00012278,WBGene00012305,WBGene00012322,WBGene00012404,WBGene00012415,WBGene00012538,WBGene00012549,WBGene00012581,WBGene00012589,WBGene00012609,WBGene00012642,WBGene00012671,WBGene00012711,WBGene00012720,WBGene00012819,WBGene00012823,WBGene00012871,WBGene00012947,WBGene00012959,WBGene00012979,WBGene00013069,WBGene00013070,WBGene00013162,WBGene00013163,WBGene00013170,WBGene00013173,WBGene00013246,WBGene00013296,WBGene00013396,WBGene00013401,WBGene00013424,WBGene00013449,WBGene00013478,WBGene00013524,WBGene00013526,WBGene00013651,WBGene00013662,WBGene00013696,WBGene00013741,WBGene00013751,WBGene00013823,WBGene00013886,WBGene00014082,WBGene00014100,WBGene00014157,WBGene00014168,WBGene00014229,WBGene00014836,WBGene00014997,WBGene00015084,WBGene00015210,WBGene00015305,WBGene00015500,WBGene00015511,WBGene00015613,WBGene00015630,WBGene00015679,WBGene00015756,WBGene00015762,WBGene00015788,WBGene00015841,WBGene00015860,WBGene00015878,WBGene00015892,WBGene00015917,WBGene00015944,WBGene00015970,WBGene00015977,WBGene00016072,WBGene00016180,WBGene00016198,WBGene00016216,WBGene00016247,WBGene00016312,WBGene00016430,WBGene00016447,WBGene00016487,WBGene00016675,WBGene00016718,WBGene00016730,WBGene00016766,WBGene00016820,WBGene00016863,WBGene00017032,WBGene00017058,WBGene00017059,WBGene00017200,WBGene00017215,WBGene00017294,WBGene00017325,WBGene00017386,WBGene00017411,WBGene00017438,WBGene00017439,WBGene00017517,WBGene00017577,WBGene00017581,WBGene00017658,WBGene00017756,WBGene00017833,WBGene00017872,WBGene00017902,WBGene00017913,WBGene00017955,WBGene00018062,WBGene00018158,WBGene00018168,WBGene00018195,WBGene00018253,WBGene00018318,WBGene00021701,WBGene00021713,WBGene00021969,WBGene00021993,WBGene00022030,WBGene00022064,WBGene00022071,WBGene00022143,WBGene00022156,WBGene00022267,WBGene00022320,WBGene00022328,WBGene00022440,WBGene00022467,WBGene00022581,WBGene00022617,WBGene00022628,WBGene00022685,WBGene00022791,WBGene00022876,WBGene00022889,WBGene00023458,WBGene00023459,WBGene00023490,WBGene00043065,WBGene00044122,WBGene00044158,WBGene00044181,WBGene00044183,WBGene00044212,WBGene00044243,WBGene00044278,WBGene00044358,WBGene00044360,WBGene00044392,WBGene00044393,WBGene00044417,WBGene00044538<br>WBGene00044658,WBGene00044724,WBGene00044741,WBGene00044744,WBGene00044761,WBGene00044917,WBGene00044998,WBGene00045122,WBGene00045247,WBGene00045344,WBGene00045356,WBGene00045411,WBGene00045415,WBGene00045417,WBGene00045510,WBGene00050896,WBGene00050916,WBGene00050967,WBGene00077616,WBGene00077697,WBGene00077751,WBGene00086554,WBGene00086568,WBGene00184956,WBGene00185055,WBGene00189952,WBGene00189953,WBGene00194641,WBGene00194643,WBGene00194701,WBGene00194787,WBGene00194816,WBGene00194833,WBGene00195178,WBGene00195182,WBGene00202497,WBGene00206357,WBGene00206523,WBGene00206527,WBGene00219285,WBGene00219342,WBGene00219351,WBGene00219444,WBGene00235165,WBGene00235294,WBGene00235310,WBGene00271707,WBGene00271795,WBGene00271817,WBGene00271827 |
| Signaling: phosphatase | 33 | 197 | 0 | 33 | 8.2E-16 | 7.3E-14 | WBGene00000389,WBGene00007354,WBGene00009079,WBGene00009101,WBGene00009401,WBGene00009492,WBGene00009893,WBGene00010265,WBGene00010634,WBGene00010869,WBGene00010992,WBGene00011438,WBGene00011808,WBGene00012741,WBGene00013304,WBGene00013586,WBGene00015661,WBGene00015929,WBGene00015931,WBGene00016053,WBGene00016242,WBGene00016356,WBGene00016398,WBGene00017279,WBGene00018347,WBGene00018359,WBGene00019773,WBGene00020187,WBGene00020841,WBGene00021113,WBGene00022090,WBGene00022753,WBGene00235383 |
| Cytoskeleton: microtubule | 24 | 128 | 0 | 24 | 7.0E-13 | 6.3E-11 | WBGene00007049,WBGene00007269,WBGene00007335,WBGene00007777,WBGene00009324,WBGene00010366,WBGene00010874,WBGene00012169,WBGene00015030,WBGene00015049,WBGene00015634,WBGene00015855,WBGene00016541,WBGene00016765,WBGene00016946,WBGene00018123,WBGene00018178,WBGene00018301,WBGene00019086,WBGene00019642,WBGene00020071,WBGene00020435,WBGene00022632,WBGene00022705 |
| Major sperm protein | 12 | 31 | 0 | 12 | 3.7E-10 | 3.3E-08 | WBGene00003429,WBGene00003434,WBGene00003435,WBGene00003438,WBGene00003443,WBGene00003444,WBGene00003452,WBGene00003463,WBGene00003465,WBGene00003467,WBGene00003469,WBGene00003470 |
| Signaling: Tyrosine kinase | 12 | 75 | 1 | 11 | 1.6E-06 | 1.4E-04 | WBGene00001487,WBGene00002193,WBGene00002206,WBGene00002208,WBGene00006894,WBGene00012011,WBGene00013801,WBGene00016462,WBGene00016954,WBGene00020659,WBGene00022454,WBGene00022634 |
| Protein modification: carbohydrate | 15 | 127 | 1 | 14 | 2.6E-06 | 2.4E-04 | WBGene00001626,WBGene00008159,WBGene00008160,WBGene00008161,WBGene00008162,WBGene00008290,WBGene00008478,WBGene00009178,WBGene00010167,WBGene00011781,WBGene00012102,WBGene00015982,WBGene00022179,WBGene00043986,WBGene00045103 |
| Stress response: C-type lectin | 20 | 257 | 4 | 16 | 2.3E-05 | 2.1E-03 | WBGene00007152,WBGene00008907,WBGene00009523,WBGene00009526,WBGene00009844,WBGene00013609,WBGene00013774,WBGene00013798,WBGene00013929,WBGene00013931,WBGene00015198,WBGene00016451,WBGene00017809,WBGene00018970,WBGene00018971,WBGene00021141,WBGene00021288,WBGene00021587,WBGene00021949,WBGene00022261 |
